# Single-cell transcriptomics, scRNA-Seq and C1 CAGE discovered distinct phases of pluripotency during naïve-to-primed conversion in mice

**DOI:** 10.1101/2020.09.25.313239

**Authors:** Michael Böttcher, Yuhki Tada, Jonathan Moody, Masayo Kondo, Hiroki Ura, Imad Abugessaisa, Takeya Kasukawa, Chung-Chau Hon, Koji Nagao, Piero Carninci, Kuniya Abe

**Affiliations:** RIKEN Center for Integrative Medical Sciences (IMS), 1-7-22 Suehiro-cho, Tsurumi-ku, Yokohama, Kanagawa, 230-0045 Japan; RIKEN BioResource Research Center, 3-1-1 Koyadai, Tsukuba, Ibaraki, 305-0074 Japan; Department of Biological Sciences, Graduate School of Science, Osaka University, 1-1 Machikaneyama-cho, Toyonaka, Osaka 560-0043 Japan; Animal Developmental Genetics, Graduate School of Life and Environmental Sciences, University of Tsukuba, 1-1-1, Tennodai, Tsukuba, Ibaraki, 305-8577 Japan

**Keywords:** Single cell transcriptomics, Pluripotent stem cells, Cell differentiation, Naïve pluripotency, Primed pluripotency, Developmental transition, enhancer RNA, non-coding RNA, X chromosome inactivation

## Abstract

**Background:** Two types of mammalian pluripotent stem cells (PSC), i.e. naïve and primed possess distinct cellular characteristics. It is largely unknown how these differences are generated during naïve-to-primed transition process. We have established a robust *in vitro* transition system using a Wnt inhibitor for the first time and analyzed dynamic changes in cellular status via single-cell RNA-sequencing and C1 CAGE analyses.

**Results:** Analysis of known marker genes suggested that the cell transition process progresses as expected. However, cluster analyses revealed a sudden increase in expression profile diversities three and four days after induction of the transition. These expression diversities can be reconciled by the presence of two subpopulations with distinct transcription profiles emerging at these time points. One of the subpopulations appears transiently, and surprisingly these cells showed a global downregulation of gene expression. Moreover, initiation of random X chromosome inactivation (XCI) coincides with the appearance of these transient cells. The other subpopulation can be maintained as a stem cell line and possesses expression profiles more similar to those of primed epiblast stem cells (EpiSC) than embryonic stem cells (ESC). However, there are important differences in gene expression related to epithelial-mesenchymal transition (EMT), suggesting that this subpopulation may represent a novel pluripotent state that has an intermediate cellular phenotype between ESC and EpiSC.

**Conclusions:** These findings should contribute to our understanding of the establishment and maintenance of distinct differentiation statuses of mammalian PSCs and provide new insights into the pluripotency spectrum in general.

## Introduction

Pluripotency of cells becomes restricted during development. Cells are undergoing differentiation and acquire distinct functions required for each cell type and cell lineage. In mammals, there exists cell lineage maintaining pluripotency in the early stage of development, and cultured stem cell lines which can be propagated indefinitely *in vitro* while retaining pluripotency have been derived from these pluripotent cells. Currently, at least two types of PSCs are known in mammals, i.e. naïve and primed. Mouse ESCs correspond to naïve PSCs, while mouse EpiSCs, human ESCs and human induced pluripotent stem cells (iPSCs) are classified as primed PSCs. The mouse ESCs are derived from preimplantation blastocysts, while EpiSCs are derivative of epiblast cells of mouse postimplantation embryos. Naïve and primed PSCs, both have capacities to differentiate into multiple cell types from the three germ layers, although they are different in various aspects. For example, there are differences between mouse ESCs and EpiSCs in their epigenetic status, e.g. DNA methylation [1], enhancer usage [2, 3], expression of naïve pluripotent markers [4], cell adhesion properties [5], nuclear architecture/replication timing [6], and metabolism [7]. Furthermore, in female cells X chromosome inactivation (XCI) takes place in EpiSCs, whereas mESCs show no XCI [8]. These differences were revealed by comparisons between mouse ESCs and EpiSCs, but it is still largely unknown how these differences are generated during the transition process from naïve to primed status or how cells exit from the naïve state to gain primed pluripotency. On the other hand, it has been suggested that mammalian PSCs may have greater diversities than previously thought [9; 10]. For example, it was reported that EpiSC-like cells may be present in the mES cell population or vice versa [11, 12]. Recently, “formative state”, a hypothetical state representing the intermediate state between naïve and primed states has been proposed [13, 14]. However, such an intermediate state between naïve and primed has previously not been clearly defined. This is probably due to the lack of an experimental model system that recapitulates the naïve-to-primed transition reproducibly *in vitro*. Mouse ESCs can be converted to primed PSCs by changing the culture medium, but massive cell death occurs, which hampers a precise analysis of the transition process [15, 16]. Epiblast-like cells (EpiLC) possess cellular characteristics similar to the primed EpiSCs, but these cells appear only transiently after induction from mESCs and cannot be maintained as a stem cell line [17]. We recently reported a robust method to efficiently establish EpiSC cell lines by using an Wnt inhibitor [18]. Using a modified culture condition with the Wnt inhibitor we succeeded to establish an *in vitro* system, in which we could efficiently and reproducibly convert ESC to primed PSC-like cells for the first time. The primed PSC-like cells generated in this way show cellular morphologies highly similar to those of the existing EpiSC lines and can be maintained *in vitro* for at least 20 passages (this work) without losing the primed PSC characteristics. As a preliminary experiment, we have converted mES cells carrying a fluorescence reporter specific to the naïve state and found that the transition process proceeds asynchronously, and that cells with distinct cellular states were intermingled within a colony. Therefore, we applied two methods of single-cell RNA sequencing; as the Fluidigm single-cell RNA-Seq (scRNA-Seq) [19] and single-cell C1 Cap Analysis of Gene Expression (C1 CAGE) [20] to elucidate dynamic changes in cellular status during the naïve-to-primed transition process at single-cell resolution for the first time. CAGE detects 5’-end of coding mRNA as well as non-coding RNA including enhancer or antisense RNAs [21]. Thus, this technique may provide insights into the enhancer/promoter interplay or non-coding RNA functions, which drives hierarchical regulations of gene expression during development.

Single-cell transcriptome data revealed distinct cell clusters in addition to the clusters mainly composed of ESCs or EpiSCs. The temporal order of emergence of these intermediary clusters was estimated by pseudotime analysis. Surprisingly, thousands of genes are globally downregulated in one of the intermediary clusters. Moreover, initiation of XCI coincides with the appearance of this cell cluster. The other subpopulation represents self-renewing stem cells exhibiting distinct expression profiles from the EpiSC cells, suggesting that this subpopulation may represent novel stem cells that have an intermediate cellular phenotype between mESC and EpiSC.

These findings should contribute to our understanding of the establishment and maintenance of distinct differentiation statuses of mammalian PSCs and provide new insights into the pluripotency spectrum in general.

## Materials and Methods

### Cell line

ESCs used in this study were established from female F1 inter-subspecific hybrid embryos (MB3), a cross between C57BL/6J (B6) and MSM/Ms (MSM) (RIKEN RBC No. RBRC00209). MSM is an inbred mouse strain derived from the Japanese wild mouse Mus musculus molossinus. We also used female EpiSCs, 129Ba2, a 129xB6N F1 hybrid line [18]. In addition, we sampled the primed PSC-like cells at Day 22 (P10) and a clonal cell line isolated from the primed PSC-like cells sampled at passage 20 (Clone 1E). All animal experiments were approved by the Institution Animal Experiment Committee of RIKEN Tsukuba Institute.

### ES cell culture

Mouse ESCs were cultured in ES medium composed of Glasgow-Minimal Essential Medium (GMEM) (Sigma-Aldrich) supplemented with 14% knockout serum replacement (KSR) (Life Technologies), 1% ES culture grade fetal calf serum (FCS) (Life Technologies), 1x non-essential amino acid (NEAA) (Life Technologies), 1000 units/mL LIF, 100 µM 2-mercaptoethanol and penicillin/streptomycin. Mouse ESCs were maintained on mitomycin C (Sigma-Aldrich) treated mouse embryonic fibroblast (MEF) feeder cells [22].

### Naïve-to-primed conversion

Mouse ESCs were seeded onto MEF feeders at a density of 1-3 x 10^5^ cells per 3 cm dish and cultured in the ES medium over night at 37°C. For conversion of ES cells to EpiSC-like cells, ES cell medium was replaced with EpiSC medium (DMEM/F12 plus glutamax (Gibco), 1xNEAA (Life Technologies), 15% KSR (Life Technologies), 5 ng/mL of basic FGF (Reprocell), 10 ng/mL of Activin A (Wako) and 2 µM IWP-2 (Stemgent) and the cells were incubated at 37°C overnight. The day of the medium change was set as Day 0. On the next day (Day 1), cells were passaged using CTKCa dissociation buffer (phosphate buffered saline containing 0.25% trypsin (BD Diagnostic Systems), 1 mg/ml of collagenase (Life Technologies), 20% KSR (Life Technologies), 1 mM CaCl_2_) essentially as described by Sugimoto et al. [18]. The medium was changed every day and cells were passaged every other day. For harvesting primed PSC-like cells, cells were dissociated by 0.25% Trypsin, 1 mM EDTA and the single cell suspension was used for single-cell capture or plate purification was done to remove feeder cells before harvesting.

### Single-cell capture, RT and cDNA synthesis

For each sample 3,000 cells were loaded in a C1 single-cell Auto Prep array (Fluidigm, 100-5760) for mRNA-sequencing (10–17 μm). We processed samples of all time points following the Fluidigm manufacturer’s instructions and recommended reagents (PN 100-7168 l1) as well as the C1 CAGE protocol (https://www.fluidigm.com/c1openapp/scripthub/script/2015-07/c1-cage-1436761405138-3) [23]. After priming the C1 array and loading of the cell mix we added a Calcein AM/ Ethidium homodimer-1 staining mix (LIVE/DEAD kit, Life Technologies). Both protocols follow the manufacturer guide to perform the cell mix loading, staining, loading of reagent mixes for lysis, reverse transcription, PCR amplification and cDNA harvest. We used External RNA Controls Consortium (ERCC) spike Mix 1 (Thermo Fisher, 4456740) [24] instead of ArrayControl RNA spikes.

### Single-cell capture imaging

Imaging of the cell capture chambers was done in brightfield, green filter and red filter mode. Due to the different sample acquisition time points for both Fluidigm scRNA-Seq protocol and C1 CAGE two different imaging systems have been used. The first device was Cellomics ArrayScan VTI High Content Analysis Reader (Thermo Scientific) and it was applied as described elsewhere [25]. The main difference between the Cellomics platform and the follow up IN Cell Analyzer 6000 system (GE Healthcare) is the eased use in automated C1 array scans and the capability of the IN Cell Analyzer to take z-stacked images, which show a vertical cross section of the capture chamber. All images from the two platforms are available from SCPortalen at (http://single-cell.clst.riken.jp/riken_data/mES2EpiSC_summary_view.php) [26]

### Library preparation and sequencing

The optimal concentration range for harvested single-cell cDNA is between 0.1 to 0.3 ng/µL. In case of the Fluidigm scRNA-Seq protocol 2 µL of each cell have been diluted in appropriate amounts of harvest dilution buffer based on prior picogreen (Thermo Fisher, P11496) cDNA concentration measurements for each cDNA cell sample. The workflow for the library preparation equally follows the Fluidigm manufacturer instructions and used reagents from Illumina (FC-131-1096, FC-131-1002). In brief, after cDNA sample dilution comes the tagmentation reaction, followed by an enzyme deactivation step and finally an indexing PCR for multiplexing samples. Fluidigm scRNA-Seq utilizes the Nextera XT index primer kit with 96 indices, whereas C1 CAGE uses a custom primer set [20](Invitrogen) instead of the kit’s S index primer set. All samples are pooled after the index PCR and the pooled mix is purified using Agencourt AMPure XP magnetic beads as described in the Fluidigm manual. Prior to sequencing on Illumina HiSeq2500 we quantified all libraries (KAPA Library Quantification kit, KK4835) and adjusted the library concentration for loading on the flow cell to 9 pM. Library quality has been checked with Agilent High Sensitivity DNA kit (5067-4626) prior to loading on the flow cell. Fluidigm scRNA-Seq protocol samples were sequenced in high-output mode, paired end, 100 bases and C1 CAGE in high output mode, paired end, 50 bases.

### Fluidigm scRNA-Seq data processing

All FASTQ files from Fluidigm scRNA-Seq runs where mapped using STAR v2.4.1d [27] against the GRCm38p4 reference genome and Gencode M8 as annotation reference. The mapping output was used for upload to ZENBU. We used Tagdust v2.13 [28] to remove library primer and adapter sequence artifacts, rRNA sequences, Spike sequences, and other non-desirable sequences before RNA-seq quantification. Estimates of RNA expression were generated with Kallisto v0.44.0 [29, 30] using Gencode M8 transcript IDs as reference. We combined the resulting single-cell expression matrices into two comprehensive matrices with single cells in columns and rows with gene level expression values as estimated counts and TPM values respectively.

### C1 CAGE sequence data processing

Two different C1 CAGE data processing workflows have been applied. For the first, C1 CAGE FASTQ files have been processed using the Moirai software platform [31] (https://github.com/Population-Transcriptomics/C1%20CAGE-preview/blob/master/OP-WORKFLOW-CAGEscan-short-reads-v2.0.ipynb). The Moirai pipeline creates BED12 files for all C1 CAGE samples, which are used to make a CAGEexp object with the CAGEr R Bioconductor package [32] (https://rdrr.io/bioc/CAGEr/). We made a custom BED file for annotating expressed TSS in order to make a C1 CAGE gene expression matrix. The annotation BED file from refTSS [33] combines annotations from DRA000914 [34], the FANTOM5 mouse promotor and enhancer atlas (https://fantom.gsc.riken.jp/data/) and the Eukaryotic Promotor Database EPDnew mouse promotors (https://epd.vital-it.ch/EPDnew_database.php), as well as Gencode M8. The gene expression matrix was generated with the CAGEr function CTSStoGenes. The resulting expression matrix was used to perform DEG analysis and k-means clustering analog to how it was done on Fluidigm scRNA-Seq data. This was done for direct comparison of Fluidigm scRNA-Seq and C1 CAGE data (Figure 1D, S2A, 4A)

**Figure 1:**
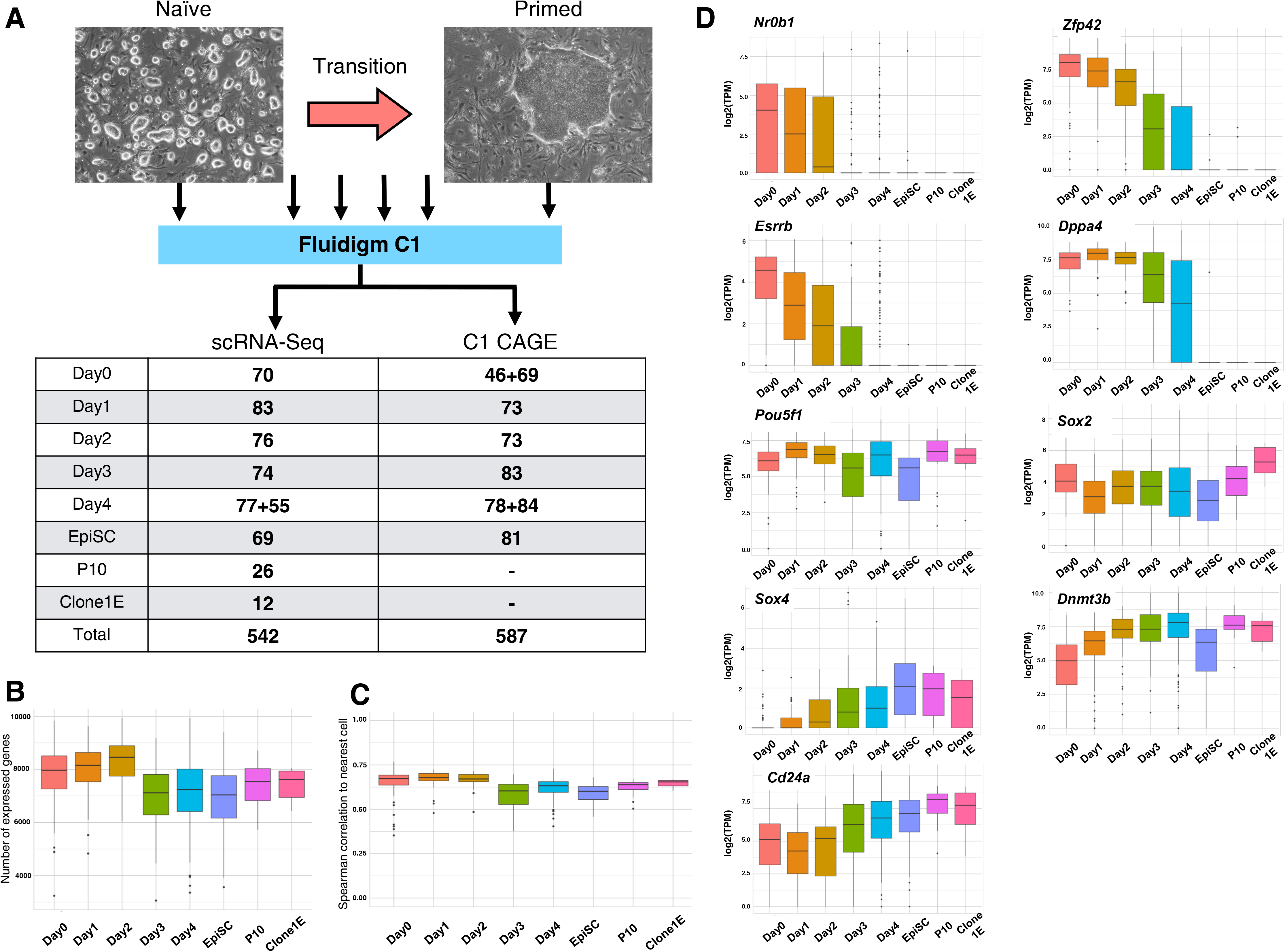
Single-cell transcriptome profiling of a time course of mouse embryonic stem cells undergoing naïve to primed transition. A) Outline of the experimental setup showing the number of cells passing initial quality filtering for each time point for both Fluidigm scRNA-Seq and C1 CAGE data. B) Distribution of the number of expressed genes per time point of the scRNA-Seq data. Only genes expressed in more than 10 cells with a TPM > 1 are considered. C) Quality assessment via neighboring cell similarities. D) Expression profiles of selected pluripotency related marker genes. Box plots represent medians (center lines) with lower and upper quartiles. Whiskers represent 1.5x the interquartile range. Outliers are represented as dots.

### Expression data analysis

All expression data analysis was done on the respective gene expression matrices for Fluidigm scRNA-Seq and C1 CAGE after removing cells that fail quality controls and have been tagged for removal in the affiliated experimental metadata tables. Quality was assessed from various sources such as capture images, cDNA concentration or sequencing reads. Based on t-Distributed Stochastic Neighbor Embedding (t-SNE) k-means clusters we performed differential gene expression analysis between all clusters using the SCDE v2.10.1 R package [35]. Pseudotime analysis was done with TSCAN v1.20.0 [36] using the set of differentially expressed genes between the Day 0 cells and the EpiSC cells and the differentially expressed genes between t-SNE k-means cluster 1 and 5 in case of pseudotime sorting of C1 CAGE samples. Hierarchical clustering heatmaps have been created with the pheatmap v1.0.12 R package [37]. Gene ontology analysis was done with the Enrichr web tool [38, 39]. Cell cycle assignment was done using a set of orthologous mouse genes based on the set from Whitfield et al. [40] with the phase scoring method described in [41]. All sample BAM files of the STAR alignment output and C1 CAGE BED12 files have been uploaded to the ZENBU browser for expression visualization and data exploration [42] (Figure S1J).

### Promotor/ enhancer analysis

A promotor/ enhancer expression matrix was constructed intersecting read 5’ ends with FANTOM5 promotor/enhancer annotation using a second C1 CAGE data processing workflow (https://fantom.gsc.riken.jp/data/). The data were processed using Seurat [43] v3.1.1, excluding features detected in fewer than 3 cells and cells tagged for removal in metadata, and normalized with Seurat NormalizeData (normalization.method = “LogNormalize”, scale.factor = 10000). Differential expression testing was performed with Seurat FindAllMarkers (min.pct = 0.05, logfc.threshold = 0.25, using a Wilcoxon Rank Sum test). Pseudotime analysis was performed with Slingshot v1.4.0, tradeSeq v1.1.03 and clusterExperiment v2.6.1: PCA1-30 of the top 10000 promotors/enhancers were clustered using Seurat FindClusters (algorithm = 4 (Leiden), resolution = 0.7). Pseudotime curve was generated with Slingshot getLineages using the previous PCA embeddings specifying the start and end cluster. NB-GAM model fit with Slingshot fitGAM (nknots=7) to the top 20% of features by variance across cells (4334 promotors and 341 enhancers). Consensus clustering of the expression patterns was performed with tradeSeq clusterExpressionPatterns (minSizes = 50) and merged with mergeClusters(mergeMethod=”adjP”,DEMethod=”limma”,cutoff=0.95) from into 5 enhancer/promotor clusters.

### RNA-FISH and immunostaining

RNA-FISH analysis of *Xist* RNA using strand-specific DNA probe and immunofluorescence analysis of H3K27me3 histone modifications were performed as described in Shiura and Abe [44].

### Allelic expression preprocessing

The single nucleotide polymorphisms (SNPs) data for MSM/Ms was downloaded from NIG Mouse Genome Database (MSMv4HQ, http://molossinus.lab.nig.ac.jp/msmdb/index.jsp). We used X chromosome SNPs of the coding region and filtered out multi allelic SNPs. The information about indels was also filtered out. The SNPs lifted over from the mm10 genome to the mm9 genome with CrossMap-0.2.6 [45]. MSM/Ms mouse genome was reconstructed from mm9 using the SNPs with bigBedToBed (http://hgdownload.soe.ucsc.edu/admin/exe/macOSX.x86_64/) and SeqKit v0.7.0 [46].

### Allelic expression analysis

For allelic expression analysis, we aligned all reads to both B6 mouse genome (mm9) and MSM/Ms mouse genome independently using STAR-2.5.3a. We sorted and merged reads from both B6 and MSM using SAMtools version 1.5 [47]. Variant calling was performed using the Genome Analysis Toolkit (GATK) version 3.7-0-gcfedb67 [48]. Variant annotation was performed using SnpEff [49] /SnpSift [50] 4.3r (build 2017-09-06 16:41). To identify high-confidence SNPs, we considered only heterozygous bases present in dbSNP (build 128) and MSMv4HQ reference database. SNPs detected from B6 and MSM genome were collected.

The samtools mpileup command (pileup2base_no_strand.pl, https://github.com/riverlee/pileup2base) was used to count the reads at each SNPs genomic position from the merged reads from both B6 and MSM.

We classified the reads with SNPs as biallelic, B6 monoallelic or MSM monoallelic. Allelic expression was measured as the total number of reads mapped on the B6 genome divided by the total number of reads for each SNP: Allelic-percentage = (B6 reads/(B6 + MSM) reads) * 100 [%].

biallelic: allelic-percentage ≧10 or ≦90 [%]

B6 monoallelic: allelic-percentage > 90 [%]

MSM monoallelic: allelic-percentage <10 [%]

Not detected: The reads were less than 10

We used two criteria to define the XCI state of each cell: one is biallelic expression ratio and the other is B6 and MSM monoallelic expression ratio. In clone 1E cells, which are supposed to complete XCI, the biallelic expression ratio of each cell was found to be 11% or less. Therefore, cells with a biallelic ratio of 11% or less are defined as ‘XCI’, while the rest of the cells are defined as ‘XC_Active’. We also used the MSM and B6 monoallelic expression ratio for defining XCI state. The clone 1E cells, in which B6 chromosome X is inactivated, showed MSM monoallelic expression ratio of ≧72%.

Thus, we defined the cells with B6 or MSM monoallelic expression ratio of more than 72% as cells undergone rXCI. When both criteria were fulfilled, a cell was defined as either ‘XCI’ or ‘XCI_active’. If the two criteria are not fulfilled, a cell was classified as ‘XCI_Intermediate’. Cells with less than 50 variants were labeled as ‘No_definition’.

## Results

### Transition from naïve to primed pluripotency

Naïve state to primed state transition was initiated by replacing ES cell culture medium with EpiSC medium containing an Wnt inhibitor, IWP-2, and the day of the medium change was set as Day 0. Cells at Day 0 showed typical morphologies of naïve ESCs, i.e. round and dome-shaped compact colonies (Figure S1A). These dome-shaped colonies were observed until Day 2 (Figure S1B, C) but larger and flatter colonies appeared from Day 3 on (Figure S1D, E). Morphologies of these flat colonies are similar to those of EpiSCs directly derived from post-implantation embryos (Figure S1F), indicating that primed PSC-like cells appear to form after Day 3. These primed PSC-like cells can be propagated stably for more than 12 passages (∼22 days after the initiation of transition). From the primed PSC-like cells, clonal cell lines can be obtained. Those clones were also morphologically stable even after 20 passages. Addition of IWP-2 to the medium is highly effective for transition to primed type stem cells. Cells cultured in the medium containing IWP-2 were converted efficiently to the primed type cells, whereas high mortality was observed in cell culture without the Wnt inhibitor (Figure S1G, H). In this study, we used a female ES cell line derived from intersubspecific hybrid embryos, which can be used for XCI analysis. Taking advantage of numerous SNPs existing between the two subspecies, it is possible to perform allele-specific gene expression analysis. We also used a female EpiSC line as a reference primed PSCs [18]. In addition, we sampled the primed PSC-like cells at Day 22 (P10) and a clonal cell line isolated from the primed PSC-like cells (Clone 1E), which underwent >20 passages.

### Single cell transcriptome analyses of the transition process using scRNA-Seq and C1 CAGE

We used scRNA-Seq on a time-course of pluripotent mESCs triggered to undergo the transition from a naïve to primed pluripotent state. In total we obtained 579 single cell transcriptome profiles via the Fluidigm scRNA-Seq protocol and 587 cells via C1 CAGE (Figure 1A). These cells passed stringent quality screenings before applying computational analysis and represent sampling time points from a transition stage between these two pluripotent states. They have been deeply sequenced with average 3.1 million sequencing reads per cell for scRNA-Seq and 1 million reads for C1 CAGE respectively.

We observed a reduction of the median number of expressed genes within each group of time points after Day 2 from more than 8500 expressed genes to less than 8000 genes (Figure 1B). Furthermore, the variability of expressed genes in individual cells was larger in cells from the Day 3, Day 4 and EpiSC group compared to earlier time points. Plotting the Spearman correlation of nearest cells [51] also shows a more variable distribution for the same groups (Figure 1C), thus indicating a global change in cellular expression profiles during the transition process from naïve to primed stem cells.

We also checked known marker genes of the naïve state (shown here *Esrrb, Nr0b1*, *Dppa4*, *Zfp42*), pluripotency markers (*Pou5f1*, *Sox2*) and primed state markers (*Sox4*, *Cd24a*, *Dnmt3b*) and could validate our data by matching the expression of these known markers with our time point samples (Figure 1D).

We performed differential gene expression analysis between the cells from the Day 0 mES group and the EpiSC group. This resulted in 950 significantly differentially expressed (DE) genes (p adjust < 0.01) between these groups (File S2) which allowed us to visualize our data via hierarchical cluster analysis (Figure 2A). Many genes appear to be specifically downregulated in the cluster 3 group (Figure 2A, Figure S7G, Figure S14A and S14B, File S3). Principal component analysis (PCA) demonstrates that PC1 and PC2 separate the cells depending on their developmental progression from naïve to primed (Figure 2B). The Day 0 to Day 2 samples form a dense cluster of cells, whereas after Day 2 cells start to show larger expression heterogeneity and thus distribute more widespread in the PCA plot. This observation is consistent with the wider distribution seen in Figure 1B and C. EpiSC cells are clustered together on the opposite side of the naïve cells, i.e. Day 0 (Figure 2B), and the Day 3 and Day 4 samples are mapped in between Day 0 and EpiSC, indicating that these cells are in transition states. Next, we used t-SNE based on the same set of differentially expressed genes and applied a k-means clustering with 5 clusters to organize our cells into comparable groups (Figure 2C and 2D, Figure S4A). These cluster results were obtained after removing a group of 37 cells that formed a distinct sixth cluster via t-SNE (Figure S2A). These cells were found to be contaminating feeder cells due to their expression of Y chromosome genes and the expression of the fibroblast marker *Vimentin* as well as their lack of *Pou5f1* expression (Figure S2B-F).

**Figure 2:**
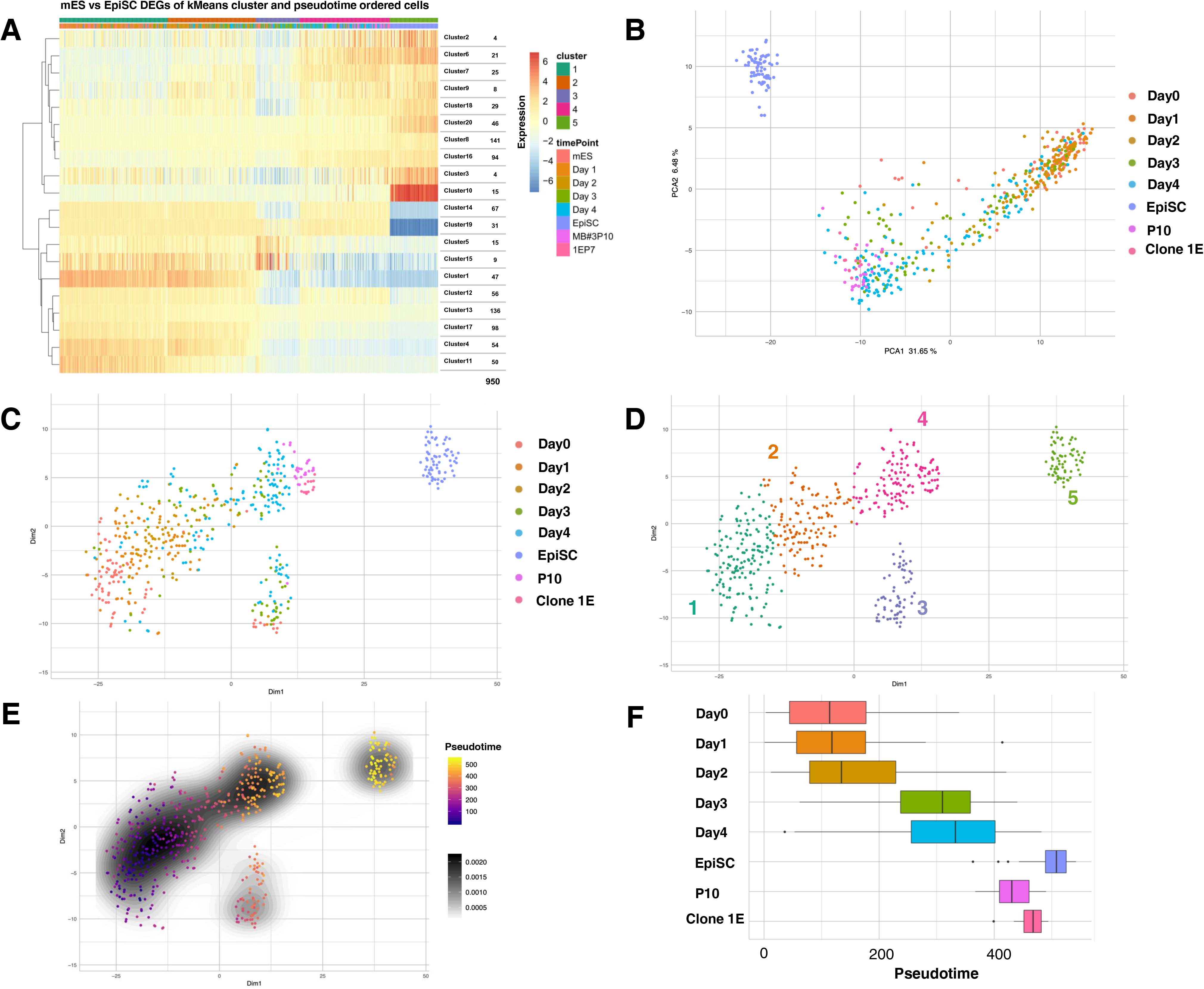
Clustering and pseudotime sorting of scRNA-Seq data based on 950 DE genes (p-adjusted < 0.01) between the mES and EpiSC time point samples. A) Heatmap with cells sorted by t-SNE k-means cluster groups and pseudotime. Twenty k-means gene clusters formed via hierarchical clustering. Expression scale log_2_(TPM+1) - rowMeans(log_2_(TPM+1)). B) PCA and C) t-SNE plot of all cells. D) Five k-means cluster groups based on t-SNE data. E) Color coded pseudotime of all cells within the t-SNE visualization. F) Pseudotime ordered cells grouped by sampling time points and sample origin. Box plots represent medians (center lines) with lower and upper quartiles. Whiskers represent 1.5x the interquartile range. Outliers are represented as dots.

In order to rule out confounding effects contributed due to the cell cycle phase of cells we performed a cell cycle phase assignment based on the expression of known phase marker genes [43; 52]. The cell cycle distribution among the cells (Figure S3A and S3B) indicates that cell cycle did not contribute to the results obtained through pseudotime analysis.

We also used pseudotime analysis to determine the temporal order of cell samples from transitioning time points and overlaid the t-SNE plot with the pseudotime order of cells (Figure 2E, Figure S4B). This pseudotime sorting enabled us to determine the developmental trajectory of samples within the five k-means cluster groups. The pseudotime order reflects the actual time points of cell sampling and serves as a validation of temporal developmental order purely based on cellular gene expression profiles (Figure 2F).

Following the trajectory indicated by the pseudotime sorting, the developmental order of the clusters is 1, 2, 3, 4 and 5. Cluster 1 is mainly composed of Day 0 and Day 1 cells, representing mostly naïve pluripotent cells. The Day 2 cells are contained in both cluster 1 and cluster 2, indicating that the Day 2 cells are heterogenous and a fraction of the cells start transitioning their pluripotency state. Part of cluster 2 is composed of Day 3 and Day 4 cells. All the cells belonging to cluster 5 correspond to EpiSCs. Surprisingly, we found two intermediary clusters (3 and 4) between the naïve and the primed state. Cluster 3 contains mainly Day 3 and 4 cells, while cluster 4 includes Day 3 and 4 as well as the primed PSC-like cells which have gone through 10∼20 more passages compared to Day 3 and 4 cells, i.e. P10 and Clone 1E (Figure 2C and 2D). It should be noted that morphologies of P10 and Clone 1E cells are highly similar to those of EpiSCs, but the cluster 4 clearly demonstrates distinct expression profiles from those of cluster 5 according to the t-SNE results.

### Characterization of t-SNE clusters based on single cell gene expression profiles

After grouping cells into five clusters, we performed differential gene expression analysis between the clusters (File S2). As shown in Figure 3A, there is a large increase in the number of significant DE genes between cluster 2 and 3, as well as 3 and 4, suggesting that cluster 3 exhibits distinct expression profiles compared to other clusters. Expression of each DE gene can be visualized at single-cell resolution by overlaying single-cell expression levels onto the t-SNE map (Figure 3B, Figure S5). By manually examining such visualizations for 1044 selected DE genes, we identified genes specific to each cluster, as well as genes enriched in multiple clusters, or absent from all but one cluster. Based on these DE genes expression patterns, we can outline characteristics of each cluster.

**Figure 3:**
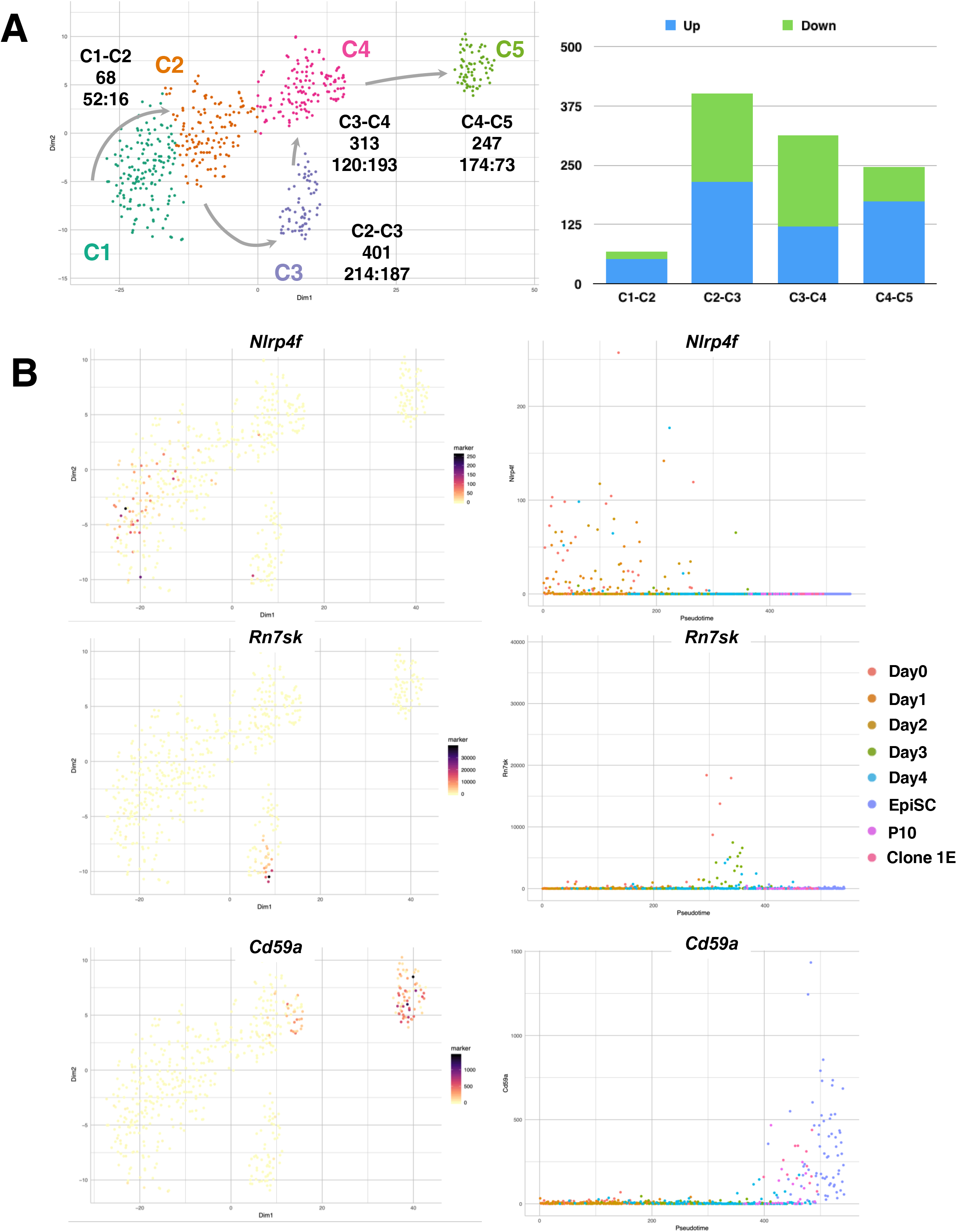
Differential gene expression between t-SNE k-means clusters for marker gene identification. A) Number of up and downregulated DE genes (p-adjusted < 0.01) between clusters. B) Selected cluster specific genes for the naïve (*Nlrp4f*), transition phase (*Rn7sk*) and primed state (*Cd59a*) shown as overlay of the t-SNE plot and the expression plotted against the pseudotime scale.

Cluster 1 is enriched with naïve pluripotency genes such as *Esrrb* or *Zfp42*. Expression of these genes is also detected in cluster 2, thus they are not very specific to cluster 1. There are some genes highly enriched in cluster 1, e.g. *Nlrp4f* and *Arl14epl*, whose expressions are detected predominantly in oocytes and preimplantation embryos [53].

Most of the DE genes in cluster 2 are expressed in other clusters as well. Many naïve pluripotency genes are heterogeneously expressed in this cluster and are downregulated as cell differentiation progresses. There are some genes, e.g. *Tmem59l* or *Car4*, whose expression is initiated in cluster 2 on and continued to be expressed until later stages, indicating naïve to primed conversion already commenced from this cluster. There are only a few genes exhibiting cluster 2-specific expression, e.g. *Wnt8a*.

The intermediary cluster 3 is characterized by specific downregulation of thousands of genes; approximately one third of the transcriptome shows downregulation in this cluster (Figure 2A, Figure S14A and S14B, File S3). Therefore, there are many examples for genes specifically downregulated in cluster 3 such as *Tmem263*, *Trp53* or *Ccnb2* (Figure S5, File S4). On the other hand, there is also a group of genes exhibiting specific upregulation only in this cluster, e.g. *H1fx*, *Itga7*, *Ccdc36* and *Rpph1*. Along this line, it is interesting to find cluster 3-specific expression of *Rn7sk*, which is a small nuclear RNA known to act as a transcriptional regulator in embryonic stem cells by decreasing the rate of RNA PolII elongation and inhibiting the CDK9/Cyclin T complex [54, 55]. This observation can be an indicator that gene regulatory networks are re-configured in this transient state in order to prepare cells for later lineage commitment. Besides these genes unique to cluster 3, the cells in cluster 3 show residual expression of naïve pluripotency genes and initial expression of primed marker genes same as cluster 2 cells.

In cluster 4 known primed marker genes are expressed, while naïve pluripotency gene expression has been almost diminished, suggesting their primed identity. In fact, known primed marker genes like *Fgf5* or *Pou3f1* are positives for cluster 4 as well as cluster 5, which is solely composed of EpiSCs. However, there are several genes expressed in clusters 2, 3 and 4, but greatly reduced in cluster 5. In particular, the cell adhesion molecule E-cadherin (*Cdh1*) is known to be expressed in naïve type ESCs, but not in primed EpiSCs [56]. *Cdh1* is clearly expressed in cluster 4, while downregulated in cluster 5. Other genes like *Cyp24a1* or *Krt18* demonstrate cluster 4 specific expression as well, suggesting that cluster 4 cells have distinct expression profiles compared to those of cluster 5.

Cluster 5 is composed of only EpiSCs, therefore express primed PSC markers, many of which are shared by cluster 4 cells. However, there are genes whose expressions are specific to cluster 5, but not to cluster 4 cells. For example, expression of *Cdh2* which encodes N-cadherin or *Vim* which encodes vimentin are detected only in cluster 5. *Cdh2* and *Vim* are known to be involved in EMT, and the results suggest that cluster 5 cells have completed EMT, whereas cluster 4 cells have not. This is significant, because EMT is one of the hallmarks of naïve-to-primed transition [57]. In other words, this finding indicates that cluster 4 cells have not completed EMT, representing a novel, intermediate pluripotency state between naïve and primed pluripotency. In addition, we manually identified 54 cluster 5-specific genes (File S4); one of which is *Cd59a* representing a novel, highly specific EpiSC marker (Figure 3B, Figure S5).

Based on the significant DE genes we performed gene set enrichment analysis with the web-based Enrichr tool [39, 40]. We identified DE genes enriched in KEGG pathways (Figure S6). In the differences between cluster 1 and 2 we find genes linked to pluripotency maintenance, whereas cluster 3 vs 4 show many DE genes belonging to metabolic pathways and in the cluster 4 vs 5 differences we can see striking changes in genes linked to cell adhesion molecules, which suggests that cell surface properties of the cluster 4 and 5 are different.

### C1 CAGE revealed dynamic changes in promoter/enhancer activities during the transition process

Like the procedure used to cluster the Fluidigm scRNA-Seq derived data, we generated a t-SNE plot for the C1 CAGE data using 635 genes differentially expressed between Day 0 mES and EpiSC samples. Strikingly, we can independently validate our cluster results from the Fluidigm scRNA-Seq protocol with the C1 CAGE data. There are also two naïve k-means clusters (1 and 2), two transition stage clusters (cluster 3 and 4), as well as an EpiSC specific cluster (5) and a small cluster comprising of feeder or differentiated cells (6) (Figure 4A, S7G). Unlike the Fluidigm scRNA-Seq protocol C1 CAGE allows the detection of both non-poly adenylated transcripts and poly(A)+ RNA. Cluster 7 in the heat map consists of 48 histone gene transcripts, most of which show upregulated expression in the k-means cluster 4 and 5 (Figure 4B, File S3). Such a histone cluster upregulation is not detected by the Fluidigm scRNA-Seq, as they are mostly non-poly adenylated [58]. Due to the different priming strategies and thus in RNA capture between these protocols, there is a larger variability with regards to which expressed genes have been detected. Nevertheless, we could observe that marker genes are expressed appropriately in the clusters (Figure S7A-F). According to the results, the k-means clusters 1, 2, 3, 4, 5 generated from the C1 CAGE data correspond to the clusters 1, 2, 3, 4, 5 of the scRNA-Seq analysis, respectively (Fig. 2D, Fig. 4A).

**Figure 4:**
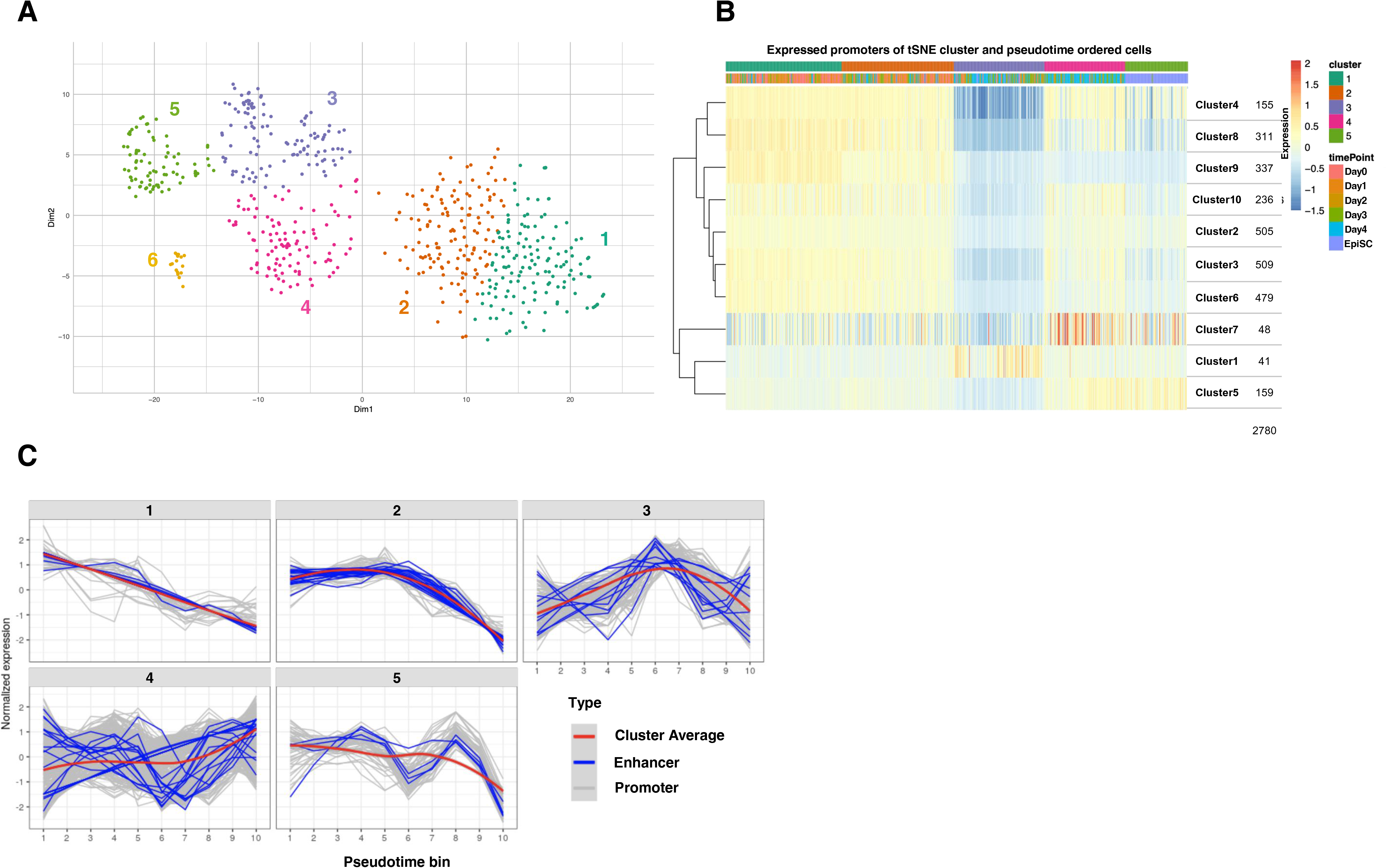
Clustering of the C1 CAGE data. A) t-SNE based on 635 DE genes (p-adjusted < 0.01) between the mES and EpiSC time point samples. B) Changes in promotor/enhancer expression detected by C1 CAGE during the time course. Heatmap with cells sorted by the t-SNE k-means cluster groups and Slingshot pseudotime. 10 k-means gene clusters formed via hierarchical clustering. C) Five expression modules of promotors and enhancers from C1 CAGE data. Cells pooled into 10 bins along a pseudotime axis generated with Slingshot. Promotors and enhancers are clustered with tradeSeq/clusterExperiment.

NASTs are a class of short, low abundance non-coding RNA expressed specifically in naïve ESCs [34]. We found that a number of NAST genes are expressed during the naïve-to-primed transition process, and some of them appear to be naïve state-specific downregulated upon entering the primed state, e.g. heatmap row cluster 1 (Figure S7G). We can also observe a decrease in expression of many NASTs during the naïve to primed transition phase cluster 3 (Figure S7G). It is not well studied to what extent annotated NASTs may be part of annotated genes from other annotation sources and thus to what degree individual NASTs are genuinely unique genes.

With the C1 CAGE data we could demonstrate dynamic changes in single-cell promotor and enhancer usage during the naïve-to-primed conversion process (Figure 4B, Figure S8A-B). There are enhancer RNAs (eRNAs) that show specificity for the naïve state, the transition period, and the primed state (Figure S8C).

In accordance with the findings from the scRNA-Seq data, C1 CAGE data also shows that TSS level (Transcription Start Site) expression is reduced in cluster 3 (Figure 4B). Figure S9A shows the promotors exhibiting great reduction of expression only in the cluster 3 (Figure S9A), while nine promotors show specific upregulation in the cluster 3 (Figure S9B). We also identified 10 non-coding eRNAs downregulated in k-means cluster 3 and two eRNAs that are upregulated (Figure S9C), suggesting that the enhancer activities are also altered in the cluster 3 cells. Figure S10 shows the top nine differentially expressed promotors and enhancers for each C1 CAGE k-means cluster group.

We calculated pseudotimes for all cells based on TSS expression, using Slingshot [58] for C1 CAGE pseudotime analysis. We divided the Slingshot pseudotime scale into 10 bins. By comparing Slingshot pseudotime bins with k-means clusters and sampling time points (Figure S11A-C), we could show that the scRNA-Seq k-means cluster 3 corresponds to the Slingshot pseudotime bin 6 or [19.9-23.8](Figure S11D). Here we identified 5 modules of promotors and enhancers with similar expression patterns depicted along the Slingshot pseudotime bin (Figure 4C, Figure S12). Modules 1 is active in the naïve state and the activities is decreased progressively as differentiation proceeds. Module 2 is constant until the bin 6 and declines thereafter. Other modules 3, 4 and 5 also show changes in their activities at around the bin 6, suggesting it corresponds to the transition point. These expression pattern modules of promotors and enhancers might correspond to gene regulatory networks interactions involved in establishment and maintenance of pluripotency states.

### X chromosome inactivation initiated at Day 3 as revealed by RNA-FISH and scRNA-Seq

As described before, the period between Day 2 and Day 3 corresponds to the transition point, where cells exit from a naïve state to a more differentiated state. To support this notion, we analyzed XCI status of cells, since XCI is one of the most reliable indicators of cell differentiation [59, 60] Random X chromosome inactivation (rXCI) is a phenomenon in which one of the two X chromosomes is randomly inactivated in a female mammalian cell during development [61]. It results in chromosome-wide silencing of either the maternal or paternal X chromosome. Once established, the XCI pattern of individual cells will be clonally inherited to the daughter cells. The large non-coding RNA *Xist* is known to be involved in the initiation of XCI, leading to silencing of most X-linked genes except for escapees, genes known to be exempted from XCI. XCI is thought to occur as the cells exit from the naïve state, though precise timing of the XCI initiation has not been determined [44]. Since we obtained global expression profiles of single cells transitioning from naïve to primed, we reasoned that we could delineate progression of the XCI process, taking advantage of our *in vitro* transition system.

First, we conducted RNA-FISH analysis of *Xist* RNA expression (Figure 5A). The result indicates that *Xist* RNA clouds can increasingly be observed within nucleus of each cell from Day 3. Analysis of H3K27me3 deposits, another landmark of inactive X, also showed the same trend (Figure 5B). Next, we calculated and compared X chromosome/autosome (X/A) expression ratios in each single cell (Figure 5C). The ratio is close to 2 at Day 0, Day 1 and Day 2, whereas it decreased to about 1 after Day 3. This indicates that total expression levels of the X-linked genes are reduced to about half at Day 3 compared to Day 0, Day 1 and Day 2. These results suggest that XCI initiates between Day 2 and Day 3.

**Figure 5:**
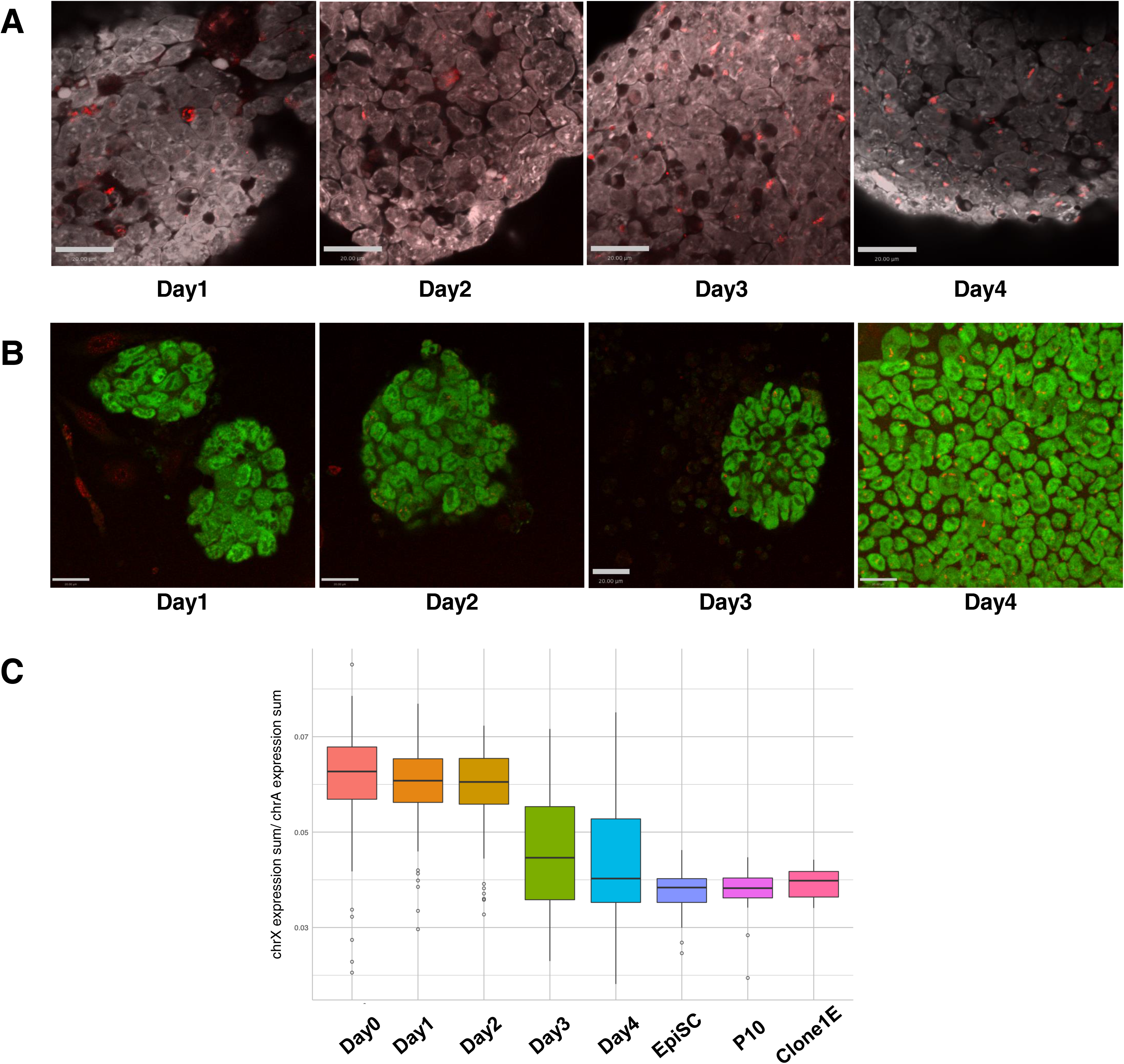
RNA-FISH, immunostaining and dosage analysis of the X-linked genes suggest that XCI initiates between Day 2 and Day 3 in our cell conversion system. A) RNA-FISH of *Xist* RNA. Red signals were found only in intercellular space in Day 1, indicating these were artifacts. Day 2 cells were mostly negative for the signal. In Day 3, *Xist*-positive cells appeared and increased in Day 4. B) Immunostaining for H3K27me3 (red) and OCT4 (green)). Day 1 and Day 2 cells were negative for the staining. Approximately 40% of nuclei in the Day 3 colony were positive for the H3K27me3 signal, while majority of the nuclei were positive in the Day 4 colony. C) Differences in ratios of X-chromosome expression levels to autosomal expression levels, from mESCs to EpiSCs. Box plots represent medians (center lines) with lower and upper quartiles. Whiskers represent 1.5x the interquartile range. Outliers are represented as dots.

### Allele-specific expression analysis of X-linked genes during the transition process

To analyze allele specific gene expression, we developed a rXCI pipeline based on the detected variants. We detected 1570 SNPs in the transcripts and focused on the 137 informative SNPs with reads > 10 expressed in at least 50% of the cells. As shown in Figure 6A, we colored allelic expression status for each gene; blue for maternal (MSM strain) allele, red for paternal (B6 strain) allele, green for biallelic expression and gray for not detected. We observed a trend that biallelic expression of each X-linked gene continues until Day 2, while mono-allelic expression of X-linked genes appears to increase from Day 3 onwards. At Day 4, more than half of the cells underwent rXCI. These findings demonstrate that rXCI begins at Day 3, thus supporting the RNA-FISH results. At Day 3 and Day 4, there are cells still showing biallelic expression (green), but P10 cells which have undergone 12 passages show much less biallelic expression, suggesting that rXCI may be completed in these cells. Analysis of the clone 1E sample indicates that all the single cells derived from the same clone show the same allelic expression pattern of the X-linked genes as expected (Figure 6A).

**Figure 6:**
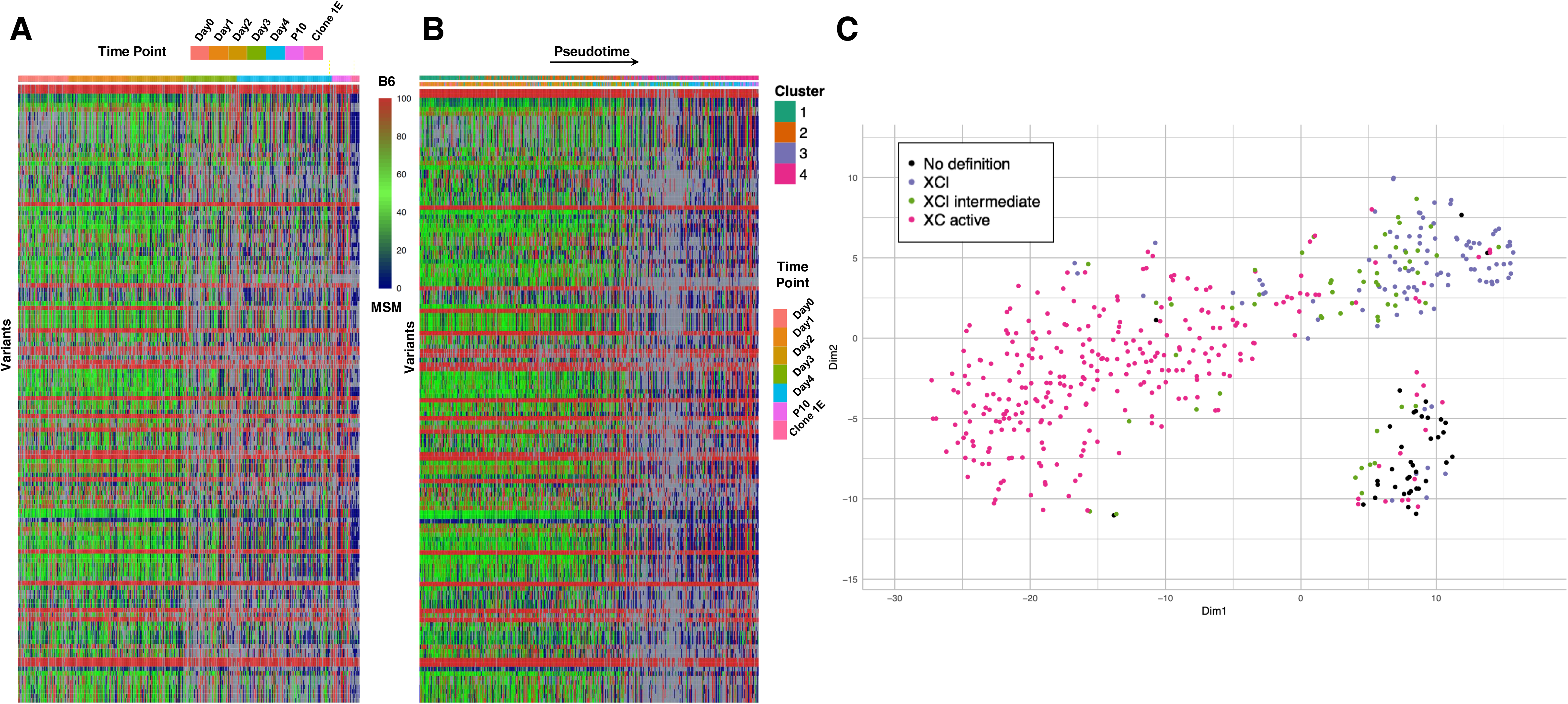
Allele specific expression analysis at the single-cell level revealed heterogeneity of XCI status among cells. A) Heatmap representing allele-specific expression from mESCs to ESC-derived primed PSC-like cells of X-linked genes. Red: specifically expressed from B6 allele (allelic percentage > 90%); Green: biallelically expressed (allelic percentage <= 90%, >= 10%); Blue: specifically expressed from MSM allele (allelic percentage < 10%). Gray colors were shown for data not available (less than 10 reads). SNPs are ordered based on genomic position. N = 137 informative SNPs. B) Pseudotime-ordered heatmap representing allele-specific expression which indicates the onset of rXCI. C) XCI status plotted onto the t-SNE clustering reveals coordinated XCI during stem cell conversion process.

### Identification of known and novel escape genes

As just described, XCI is completed in P10 and clone 1E cells. However, several cells showing biallelic expression were detected in these cells and we noticed that most of the genes are known escape genes. Variants showing biallelic expression in at least two cells from P10 and 1E clone were identified as escape genes, and among them we found known escapees such as *Ddx3x*, *Eif2s3x*, *Kdm5c*, *Kdm6a* (Figure S13A). These results confirmed that our computational pipeline is appropriate for the analysis of XCI status. We also identified some genes (*Slc7a3*, *Hnrnpa1* or *Cetn2*) as potential novel escapee candidates. Furthermore, regardless of the cell type or the differentiation stage, genes expressed specifically from the B6 or MSM allele in almost all the single cells were detected. These are also considered to be escape genes, but their expressions are biased strongly to one of the two alleles. To validate our findings, we performed Sanger sequencing of several candidate genes and confirmed that *Cetn2*, *Slc7a3* and *Hnrnpa1* are novel escape genes whose expression is biased to one of the two alleles.

### rXCI analysis and pseudotime estimation suggests that rXCI initiation coincides with global downregulation of gene expression

Based on the bioinformatics analysis of the scRNA-Seq data, each cell was ordered along the pseudotime axis to identify the starting time of rXCI on the pseudotime axis (Figure 6B). Surprisingly, we observed a transient downregulation of many X-linked genes at a specific period during the transition. Such transient downregulations do not seem to be X-linked gene specific. A heatmap visualization of 21,777 autosomal genes shows that many of the genes are downregulated during this period (Figure S14A), while 965 X-linked genes show similar results (Figure S14B).

During the downregulation period, it is not possible to assess XCI status. However, cells undergone XCI begin to emerge after this downregulation period, implying that cells might have to go through the downregulation period to attain XCI.

To visualize the XCI state of each single cell in a different way, we first categorized the cells into four groups based on X chromosome states; i.e. XCI, XCI_Intermediates, XC_Active, No_definition, according to the definition criteria described in the method section. The assigned XCI status of each cell was overlaid onto the t-SNE map (Figure 6C). Almost all the cluster 1 and 2 cells are XC active. On the other hand, all four categories of cells, especially considerable number of XCI cells, were identified in the cluster 3. It is interesting to find that *Xist* RNA expression is upregulated in some of the cluster 3 cells, whereas *Tsix*, antisense partner of *Xist* with repressive function on *Xist* expression, is being downregulated in the same cluster (Fig. S13B). There are more cells undergone XCI in the cluster 4 than in the cluster 3, while number of XCI_Intermediate cells is similar to that of the XCI cells in the cluster 4. In the cluster 4, cells corresponded to P10 or clone 1E (Figure 2C) represent mainly XCI cells, indicating that XCI is completed at later stages of the development. All the above results indicate that cells in the cluster 3 just exited from the naïve state begin to undergo XCI accompanied by a transient downregulation of gene expression not just limited to the X-chromosome, and that XCI process is more advanced in the cluster 4 and almost completed in P10 and clone 1E cells.

## Discussion

In this study, transcription dynamics of the naïve-to-primed transition process have been explored for the first time by using two different single-cell transcriptomics techniques, i.e. scRNA-Seq and C1 CAGE. The data obtained could thus generate a comprehensive catalog of genes exhibiting characteristic changes during the transition. Differential gene expression analysis identified known and novel marker genes that should be extremely useful for functional characterization of this developmental transition process. Interestingly, cluster analyses revealed intermediary subpopulations of cells in addition to the naïve and the primed PSCs. The presence of such subpopulations cannot be discovered by bulk expression analysis, emphasizing the merits of the single-cell technologies. Here we used female ESCs from intersubspecific hybrid embryos. Taking advantage of existing SNPs between the two subspecies of mice [62], we could perform allele-specific expression analysis at the single-cell level and adopted this technique for the analysis of the random X chromosome inactivation phenomenon.

### Discovery of transient global downregulation of gene expression in the transition stage

One of the most intriguing findings of this study is that approximately one third of the transcriptome (∼6000 genes) is downregulated transiently and specifically in cells classified as the cluster 3 (Figures 2A, 4B, 6B). Both autosomal and X-linked genes showed this transient gene repression. The cluster 3 cells exhibited expression profiles highly divergent from those in cells of other identified clusters. This is probably due to the global gene repression occurring in those cells. Heterogeneities and high variation in expression profiles among cluster 3 cells may also be explained by different degrees of gene repression at the time of sample collection. Such a subpopulation of cells, i.e. cluster 3, was detected reproducibly in three different batches (two Day4 samples for scRNA-Seq and one C1 CAGE Day4) of samples by using two different single-cell technologies. Although the cluster 3 cells exhibited very distinct expression profiles, pseudotime analysis estimated the cluster 3 emerged just after cells exited from the naïve state. In fact, the cluster 3 cells express some of the naïve genes as well as early markers for the primed state, suggesting that the cluster 3 cells position at an intermediate step between naïve and primed. Cell cycle assignment analysis indicated that the cluster 3 cells do not correspond to any specific cell cycle phase. There are many genes specifically downregulated in cluster 3, whereas those genes are highly expressed in other clusters. On the other hand, there is a set of genes exhibiting transient upregulation only in the cluster 3, which may provide clues to the global gene repression phenomenon in this cluster. One interesting example is *Rn7sk*, which encodes a small non-coding RNA involved in transcription repression. *Rn7sk* is an RNA component of a small nuclear ribonucleoprotein complex (snRNP) and known to inhibits the cyclin dependent kinase activity of the positive transcription elongation factor P-TEFb [54], acting as a gene-specific transcription repressor in ESCs [55] Therefore, it is possible that *Rn7sk* may contribute to the global gene repression occurring in the cluster 3. Experimental tests of this hypothesis are currently underway.

### The cluster 4 represents the third pluripotent stem cells with intermediate characteristics between naïve and primed

The second unexpected finding in this study is the discovery of the cluster 4 (Figure 2D). Cells in this cluster show morphologies similar to the primed PSCs and express a number of the primed state marker genes. However, bioinformatical analysis classified these cells to the cluster distinct from the EpiSC cluster, i.e. cluster 5, and the pathway analysis suggested that genes involved in cell adhesion are expressed differentially between the cluster 4 and 5. We noticed that the cluster 4 cells express *Cdh1* (E-cadherin) but do not express *Cdh2* (N-cadherin)(Figure S5). It is known that naïve PSCs undergo epithelial-mesenchymal transition (EMT) process, in which *Cdh1* expression of the naïve PSCs is replaced with *Cdh2* expression that is specific to the primed PSCs [64]. The absence of *Cdh2* expression in the cluster 4 cells suggests that the EMT may not be complete in these cells. Absence of *vimentin* expression in the cluster 4 supports this notion (Figure S2, S5). Since the completion of EMT is one of the criteria defining the EpiSCs, the cluster 4 cells stay at the stage prior to the EpiSC state and self-renew this cellular state. In other words, the cluster 4 cells may represent novel pluripotent stem cells in mice besides ESCs and EpiSCs, exhibiting an intermediate state between ESCs and EpiSCs. A third pluripotency state called “formative” has previously been proposed [13]. The formative state is thought to be an intermediate state between naïve and primed, although the formative PSCs have not been established in mice. Whereas EpiLC [17] is suggested to be in the formative state, it is a transient cell type and not self-renewing stem cell unlike our cluster 4 cells. Our preliminary analysis suggests that EpiLC is more to the naïve state compared to the cluster 4 cells. Although stem cells with intermediary pluripotency states had been reported [65, 66], relationships of these cells with the formative state remain elusive.

Recently, it was reported that human naïve PSCs can acquire novel pluripotency comparable to the formative state, if the naïve cells are cultured in medium containing Wnt signaling inhibitor [14]. It is thus possible that our cluster 4 cells represent a mouse counter part of their formative state cells. Formative state PSCs or PSCs cultured in the presence of Wnt inhibitor seem to have greater capacities for multi-lineage differentiation compared to the existing naïve or primed PSCs [18, 67, 68] and therefore those new versions of PSCs have a potential to replace the naïve or primed PSCs in stem cell sciences. However, research on those novel PSCs is still in its infancy and further studies must be conducted to elucidate its full potential. Comparison of the putative formative-like PSCs between human and mice should contribute to the understanding of this novel pluripotent state, and the cluster 4 cells of this study provide a good reference for these comparisons.

### Initiation of XCI coincides with emergence of the cluster 3

In our *in vitro* experimental system, random XCI happens between the time points Day 2 and Day 3. This was confirmed by RNA-FISH, immunostaining and allele-specific gene expression analysis at single cell resolution (Figure 5A-B). Allele-specific expression analysis enabled to classify each single cell arbitrarily into three categories, i.e. biallelic, intermediate and inactivated. Detailed analysis of these three categories of cells should yield important information about initiation and progression of this epigenetic reprogramming event. Moreover, the analysis could detect known and novel escaped genes as well as monoallelic expressed genes showing genetic-origin-dependency. Combined, random XCI appears to be initiated in cells of the cluster 3 and more advanced in the cluster 4 cells. As described above, gene repression takes place in the cluster 3. Currently, we do not know whether this is just a coincidence or indicative of mechanistic relationships between the two phenomena. Perturbation experiments for either one of the phenomena could help to infer whether these two are interdependent or not. There are precedents of the global gene repression: XCI in mammalian female embryos, meiotic chromosome inactivation during male spermatogenesis or global epigenomic changes in primordial germ cells [69, 70, 71, 72]. Failures in these global repression phenomena lead to various abnormalities such as embryonic lethality and infertility, clearly indicating the biological importance of the global repression. Common feature of these phenomena is that they occur when cells undergo major epigenetic reprogramming events. Therefore, the cluster 3 cells should be analyzed with regards to epigenetic changes. In any case, our experimental system should provide unprecedented opportunity for the studies of global gene repression and epigenetic reprogramming.

### C1 CAGE: a single cell transcriptome profiling beyond scRNA-Seq

In this study, we tried to use two different single cell expression profiling techniques and compared the results. Basically, the results from the two methods are highly consistent. In addition, as C1 CAGE can detect non-polyadenylated RNA, we were able to observe expression dynamics of eRNAs, histone mRNAs and NASTs during the transition process for the first time. Interestingly, some NASTs seem to show specificity only to the naïve pluripotency states. Perturbation experiments on the specific NASTs might help to shed light on the regulatory role of this class of non-coding RNA in naïve states. It is known that usage of enhancers changes during the naïve-primed transition [2, 3]. For example, it is well known that *Pou5f1* gene has both distal and proximal enhancers, of which proximal enhancer drives the primed state-specific expression [73]. In this particular case eRNA expression was not observed in our analysis. This may be due to either very low level or no expression of eRNAs in this locus, because even a bulk analysis using hundreds of cells conducted at the same time as the C1 CAGE analysis could not detect CAGE counts in this region. Thus, identification of enhancer should not rely on single parameter/technique alone. Nevertheless, the present C1 CAGE analysis could detect novel RNA expression at a number of enhancer regions annotated by FANTOM5 atlas, and some of which show specificities to either naïve or primed state, confirming the previous notion [2, 3]. Therefore, we consider the C1 CAGE data of this study a valuable resource for further studies on the regulatory roles of diverse classes of expressed non-coding RNAs including eRNAs in the early mammalian developmental process.

## Abbreviations

PSC: pluripotent stem cell
XCI: X chromosome inactivation
EpiSC: epiblast stem cell
eRNA: enhancer RNA
ESC: embryonic stem cell
EMT: epithelial-mesenchymal transition
iPSC: induced pluripotent stem cell
EpiLC: epiblast-like cell
scRNA-Seq: single-cell RNA-Seq
C1 CAGE: single-cell Cap Analysis of Gene Expression
GMEM: Glasgow-Minimal Essential Medium
KSR: knockout serum replacement
FCS: fetal calf serum
NEAA: non-essential amino acid
MEF: mouse embryonic fibroblast
ERCC: External RNA Controls Consortium
t-SNE: t-Distributed Stochastic Neighbor Embedding
SNP: single nucleotide polymorphism
GATK: Genome Analysis Toolkit
NAST: non-annotated stem cell transcript
rXCI: random X chromosome inactivation
snRNP: small nuclear ribonucleoprotein

## Availability of data and materials

All raw FASTQ sequencing files can be downloaded from DDBJ with the accession numbers DRA010828 and DRA010829. All C1 capture array images as well as additional files affiliated with the samples are available on SCPortalen [26] (http://single-cell.clst.riken.jp/riken_data/mES2EpiSC_summary_view.php). ZENBU exploratory tracks can be found here after sign in:

Fluidigm scRNA-Seq: https://fantom.gsc.riken.jp/zenbu/gLyphs/#config=1qUudPWiDNTgcknv0TJkp;loc=mm10::chr12:86353254..86594021+

C1 CAGE: https://fantom.gsc.riken.jp/zenbu/gLyphs/#config=bYYvK4ICElFj8aWmkAJ7z;loc=mm10::chr8:106586626..106686971+

## Authors’ contributions

KA and PC conceived the project. MK and HU maintained cell cultures. MB performed all single-cell experiments. MB and IA managed the data. MB, JM and YT did bioinformatics analysis, and IA, TK, CCH and KN helped with some parts. PC and KA supervised the project. MB, YT, JM and KA wrote the manuscript. All authors read and approved the final manuscript.

## Acknowledgements

We thank Dr. M. Furuno for coordinating the project. We also thank Super Computer Facilities of National Institute of Genetics, Mishima, Japan, as computations were partially performed on the NIG Supercomputer. This work was supported by a grant from the RIKEN Single Cell Project grant and in part by grants to KA from the Ministry of Education, Culture, Sports and Technology of Japan. The project was also supported in part by the RIKEN institutional budget to the RIKEN Center for Integrative Medical Sciences and for the former Center for Life Science Technologies. MB was supported by RIKEN as an International Program Associate.

## Additional files

Figure S1: Microscopic images of the cell culture at each time point. A) Day 0, B) Day 1, C) Day 2, D) Day 3, E) Day 4, F) EpiSC derived from embryos. Morphologies of cells transitioned with (G) and without IWP-2 (H). Photos were taken at Day 4. I) Cellular morphologies of clone 1E cells. J) Screen capture of Zenbu browser expression histograms of *Pou5f1* locus.

Figure S2: A) Initial t-SNE clustering of scRNA-Seq data based on 916 DE genes (p-adjusted < 0.01) between the mES and EpiSC time point samples. B - F) Expression of selected genes plotted onto the t-SNE clustering. B) and C) are Y-linked genes. A cluster of cells marked by dotted circle likely corresponds to contaminated feeder cells.

Figure S3: Cell cycle analysis of Fluidigm scRNA-Seq data. Cell cycle scoring based on 176 phase marker genes [40]. A) Each cell’s estimated cycle phase plotted onto the t-SNE clustering. B) Pie charts showing cell cycle distribution per t-SNE k-means cluster.

Figure S4: Alternative PCA visualizations. A) t-SNE k-means cluster groups overlaid onto PCA plot. B) Color coded pseudotime of all cells within the PCA plot.

Figure S5: Expression of selected DE genes between all t-SNE k-means clusters plotted onto the t-SNE clustering. Shown are genes that are either specific to a k-means cluster or absent from a cluster. A) and B) enriched in cluster 1, i.e. naïve-specific. C) specific to cluster 2. D) an example of gene upregulated from cluster 2 on except for cluster 3. E) and F) examples of genes expressed in all the clusters except for cluster 3. G) and H) examples of genes enriched in cluster 3 but not in other clusters. I) and J) genes known for their specificity to primed PSCs. K), L), M) and N) genes related to EMT. O) and P) examples of genes with specificity to cluster 4.

Figure S6: Enrichr gene set enrichment analysis based on DE genes from t-SNE k-means cluster comparisons. A) KEGG Pathways enriched in DE genes between cluster 1 and cluster 2. B) Pathways enriched in DE genes between cluster 2 and cluster 3. C) Pathways enriched in DE genes between cluster 3 and cluster 4. D) Pathways enriched in DE genes between cluster 4 and cluster 5.

Figure S7: Clustering and expression visualization of C1 CAGE data. A - F) Expression of selected genes between k-means clusters 1-5 plotted onto the t-SNE clustering. G) Heatmap of DE NASTs between the mES and EpiSC time point samples. Cells sorted by t-SNE k-means cluster groups and pseudotime. Twenty k-means NAST clusters formed via hierarchical clustering. Expression scale log_2_(count+1) - rowMeans(log_2_(count+1)).

Figure S8: Promotors and enhancers differentially expressed at sample time points A) Dotplot of gene promotors with significantly upregulated (Wilcoxon Rank Sum test, Bonferroni adjusted p < 0.05) expression in one time point. B) Dotplot of enhancer loci with significantly upregulated (Wilcoxon Rank Sum test, Bonferroni adjusted p < 0.05) expression in one time point. C) Expression of selected enhancers from B) left: smoothed expression along the pseudotime, right: percentage of cells where the enhancer was detected in each time point.

Figure S9: Promotors and enhancers differentially expressed in C1 CAGE k-means cluster 3. A) Dotplot of the top 12 differentially expressed promotors during the time course, all are downregulated in k-means cluster 3. B) Dotplot of significantly upregulated gene promotors in k-means cluster 3. (Wilcoxon Rank Sum test, Bonferroni adjusted p < 0.05). C) Dotplot of all differentially expressed enhancers when comparing k-means clusters (Wilcoxon Rank Sum test, Bonferroni adjusted p < 0.05).

Figure S10: Promotors and enhancers differentially expressed during the Slingshot pseudotime. A - E) The top 9 differentially expressed promotors or enhancers from each k-means cluster group plotted across the Slingshot pseudotime.

Figure S11: Relationship between time point, C1 CAGE k-means clusters, and Slingshot pseudotime. A) Barplot where cells from each k-means cluster appear on the Slingshot pseudotime. B) Barplot where cells from each time point appear on the Slingshot pseudotime. C) Barplot where cells from each pseudotime bin appear on the Slingshot pseudotime. D) Number of cells from each k-means cluster appearing in each pseudotime bin.

Figure S12: Enhancers differentially expressed during the Slingshot pseudotime. All differentially expressed enhancers from each expression module of Fig. 4C plotted across the Slingshot pseudotime.

Figure S13: Single-cell allelic expression analysis detected escape genes. A) In this bar graph: black shows known escape genes, red shows novel biased escape genes and false-positive results are shown in green. Each line indicates the position on the X chromosome. B) Expression of *Tsix* and *Xist* plotted onto the t-SNE clustering. The cluster 3 cells are marked by the dotted circle.

Figure S14: Global downregulation of genes in Fluidigm scRNA-Seq t-SNE k-means cluster 3. Heatmaps with A) autosomal genes and B) X linked genes. Cells sorted by t-SNE k-means cluster groups and pseudotime. Twenty k-means gene clusters formed via hierarchical clustering. Expression scale log_2_(TPM+1) - rowMeans(log_2_(TPM+1)).

Table S1: This 2-column table contains the cell_id and the cell cycle phase assigned to each cell_id.

Table S2: Gene information parsed from the M8 Gencode GTF reference file. This table was used to filter genes by chromosomes.

File S1: This zip file contains all scRNA-Seq and C1 CAGE metadata files, expression tables and tables containing t-SNE dimensions and k-means clusters that have been used to create figures. The metadata file discard column can be used to remove all cells that fail quality criteria. These cells are tagged as TRUE. All analysis was done on the subset that is tagged as discard FALSE.

File S2: Zip file containing all tables for differential gene expression results.

File S3: Zip file containing heatmap related tables for Figure 2A, 4B, S7G and S14A-B. These tables list all genes for each of the heatmap k-means clusters.

File S4: All t-SNE visualizations overlaid with expression of selected genes. Examples are shown in Figure S5.

File S5: Tables providing variant position, allelic expression status and other information related to Figure 6A-C.

File S6: Various source code files.

