## Supplementary Figures for "Single-cell transcriptomics, scRNA-Seq and C1 CAGE discovered distinct phases of pluripotency during naïve-to-primed conversion in mice"

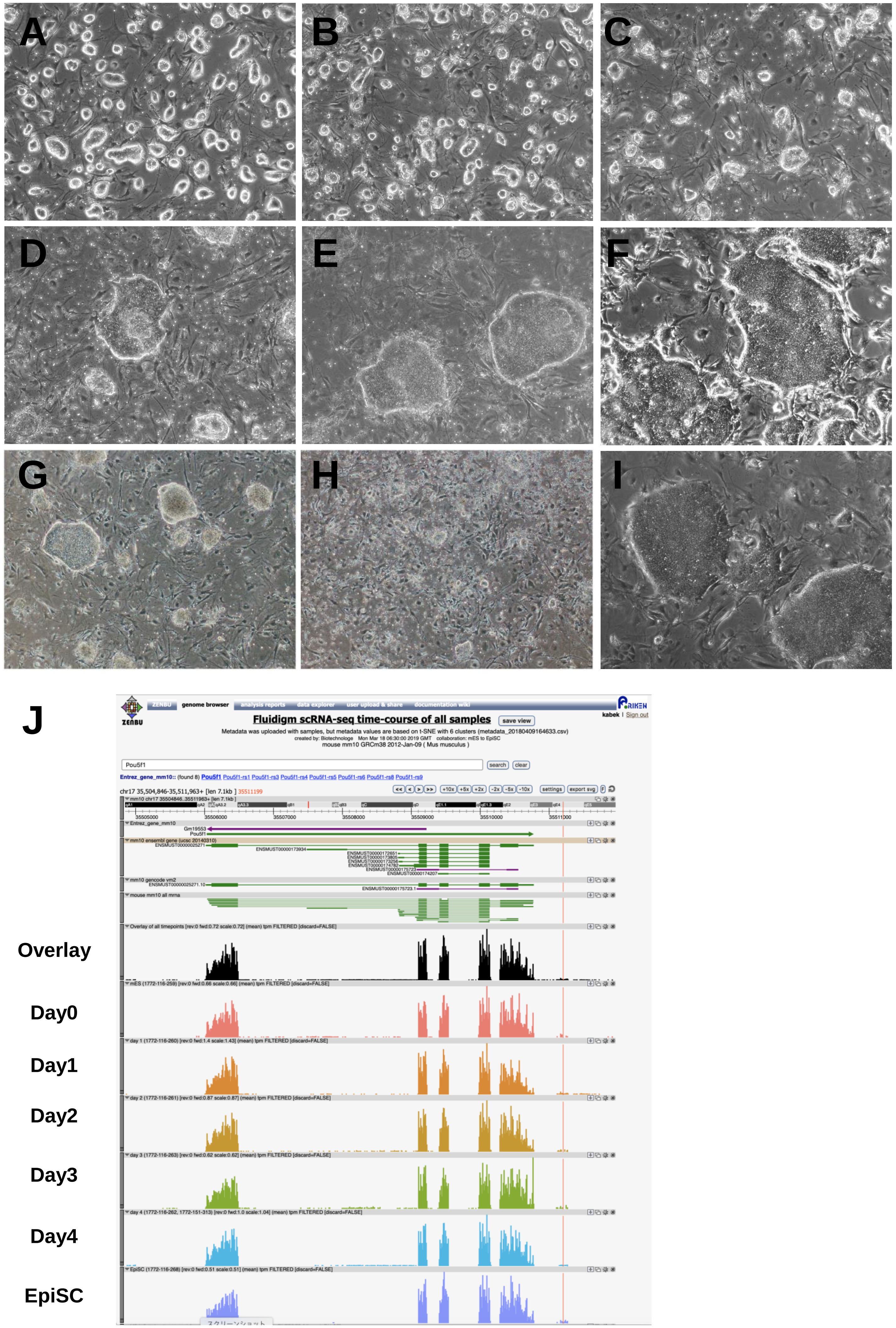
A
B
C
D
E
F
G
H
I
J
Fluidigm scRNA-seq time-course of all samples
Overlay
Day0
Day1
Day2
Day3
Day4
EpiSC

Fig. S1

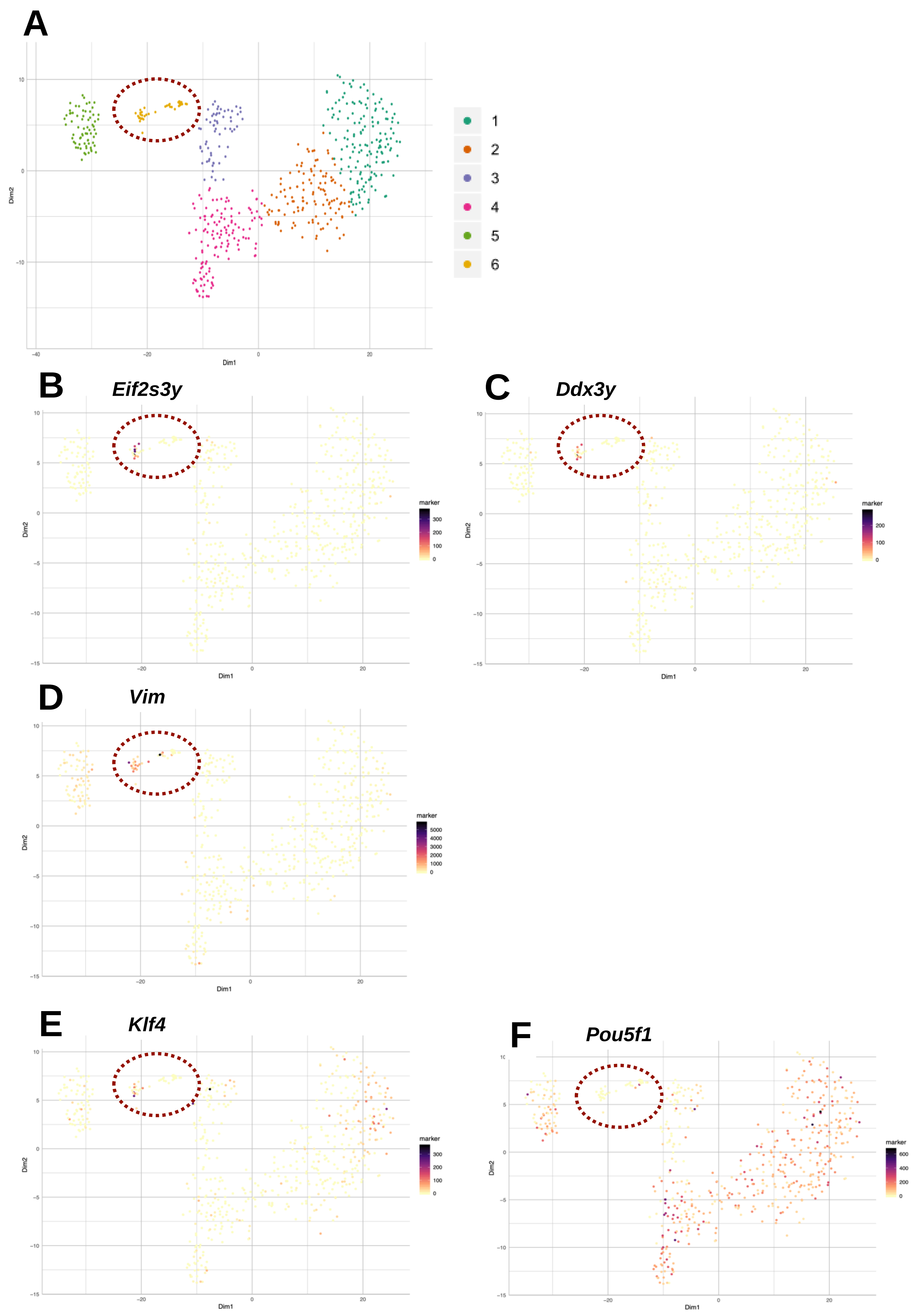
A
Dim1
Dim2
1
2
3
4
5
6
B
Eif2s3y
C
Ddx3y
D
Vim
E
Klf4
F
Pou5f1
marker

Fig. S2

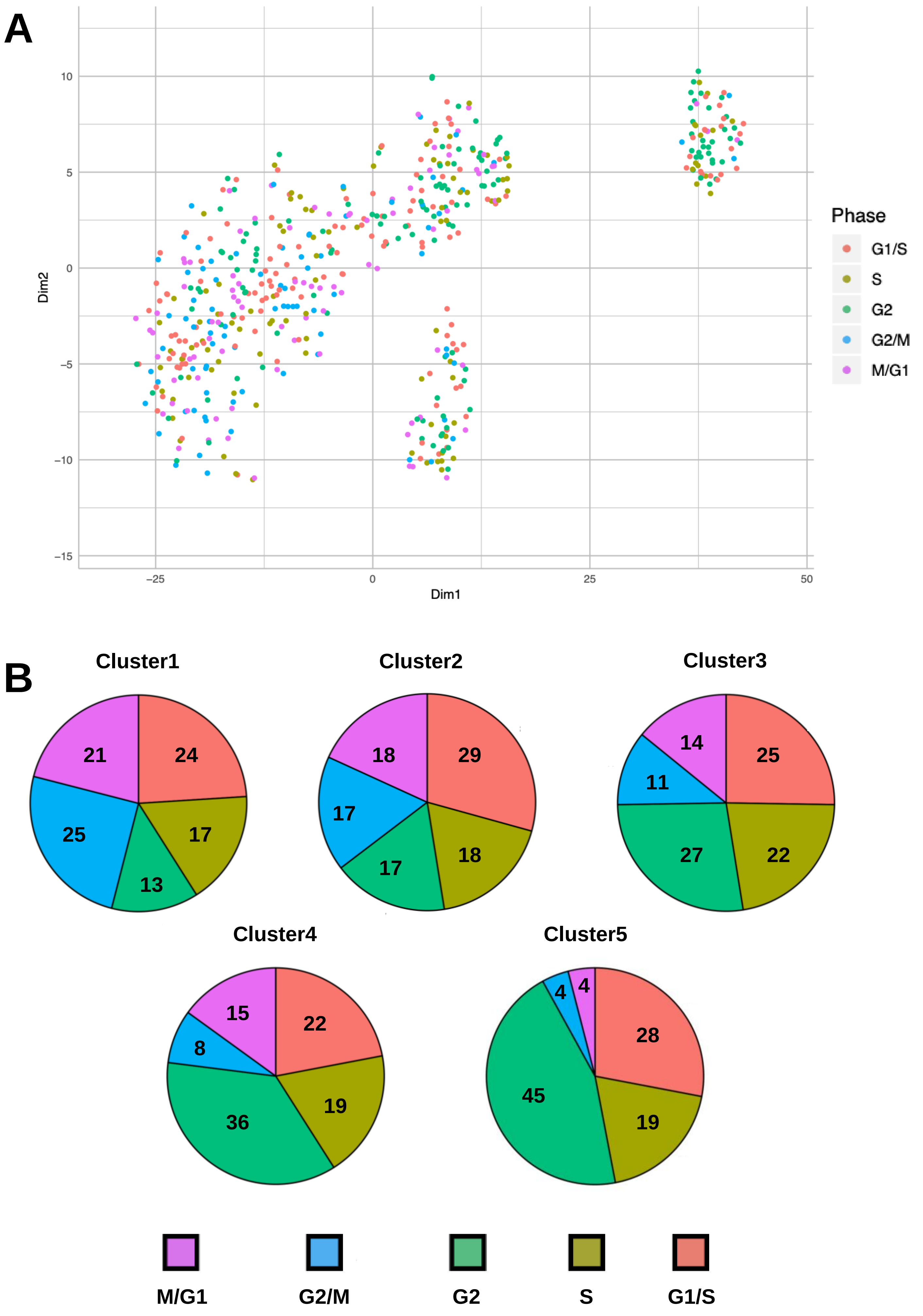
A
10
5
0
−5
−10
−15
Dim2
−25
0
25
50
Dim1
Phase
G1/S
S
G2
G2/M
M/G1
B
Cluster1
21
24
25
17
13
Cluster2
18
29
17
18
17
Cluster3
14
25
11
27
22
Cluster4
15
22
8
19
36
Cluster5
4
4
28
45
19
M/G1
G2/M
G2
S
G1/S

Fig. S3

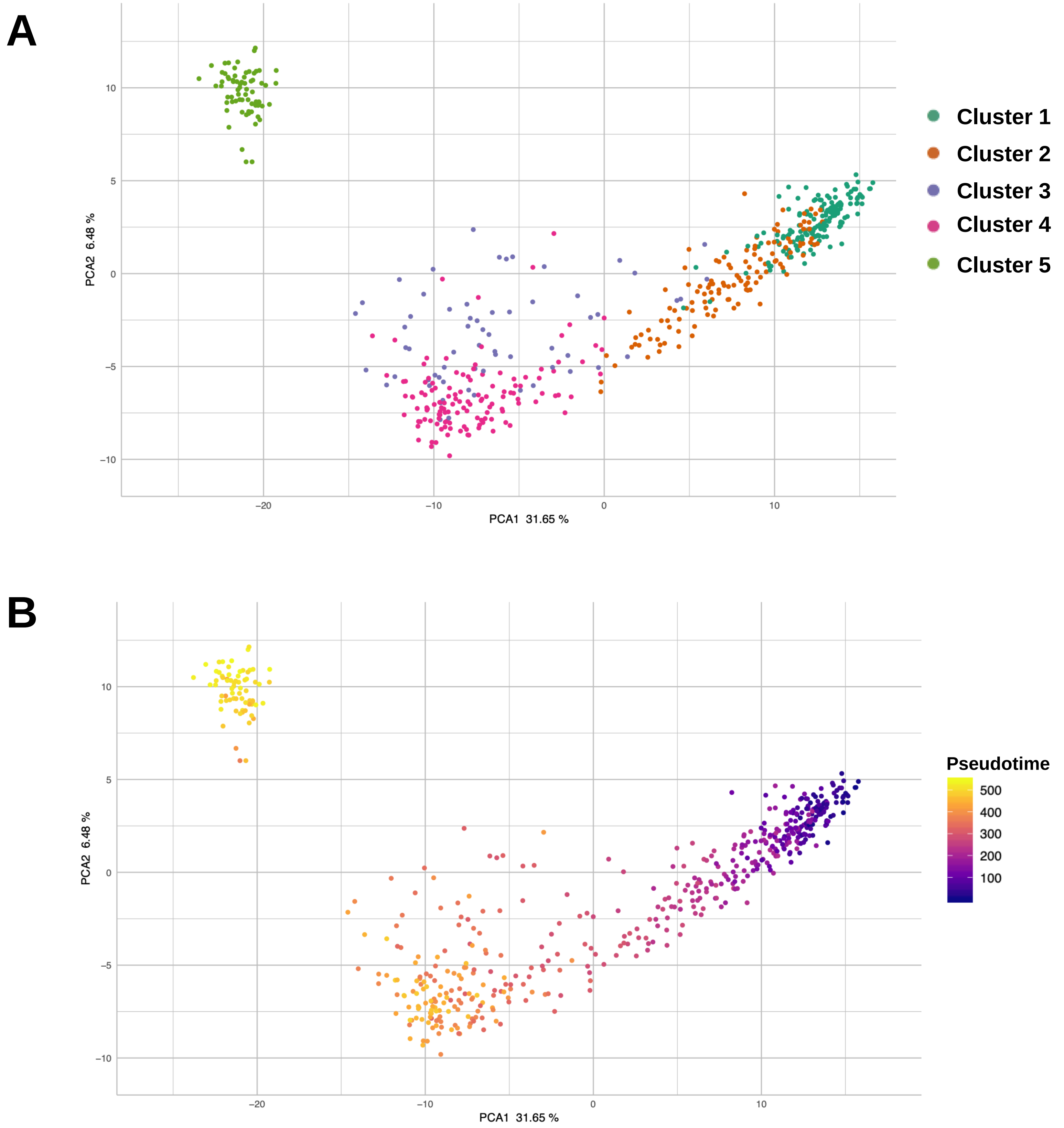
A
Cluster 1
Cluster 2
Cluster 3
Cluster 4
Cluster 5
PCA2 6.48 %
PCA1 31.65 %
10
5
0
−5
−10
−20
−10
0
10
B
Pseudotime
500
400
300
200
100
PCA2 6.48 %
PCA1 31.65 %
10
5
0
−5
−10
−20
−10
0
10

Fig. S4

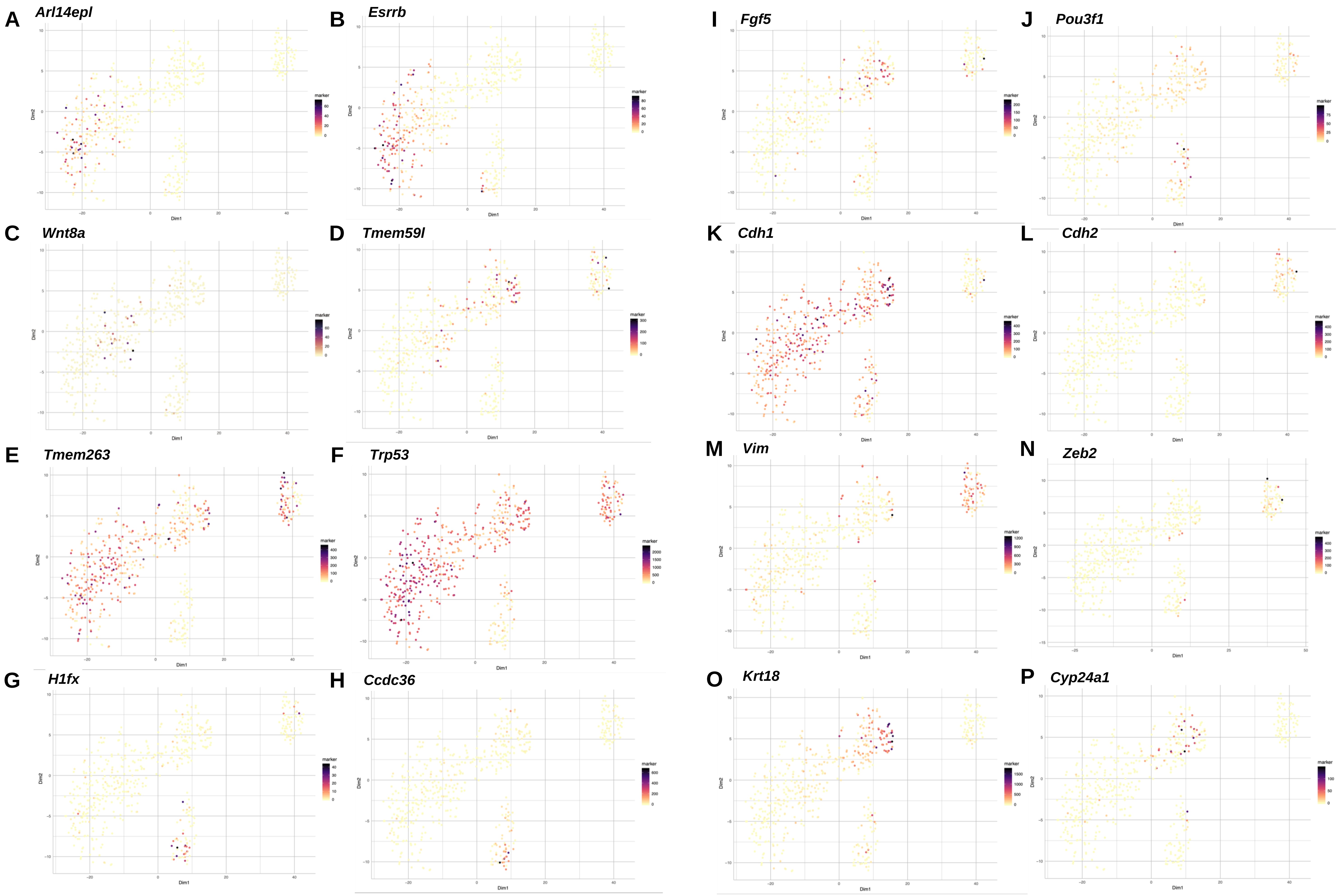
A Arl14epl
B Esrrb
C Wnt8a
D Tmem59l
E Tmem263
F Trp53
G H1fx
H Ccdc36
I Fgf5
J Pou3f1
K Cdh1
L Cdh2
M Vim
N Zeb2
O Krt18
P Cyp24a1

Fig. S5

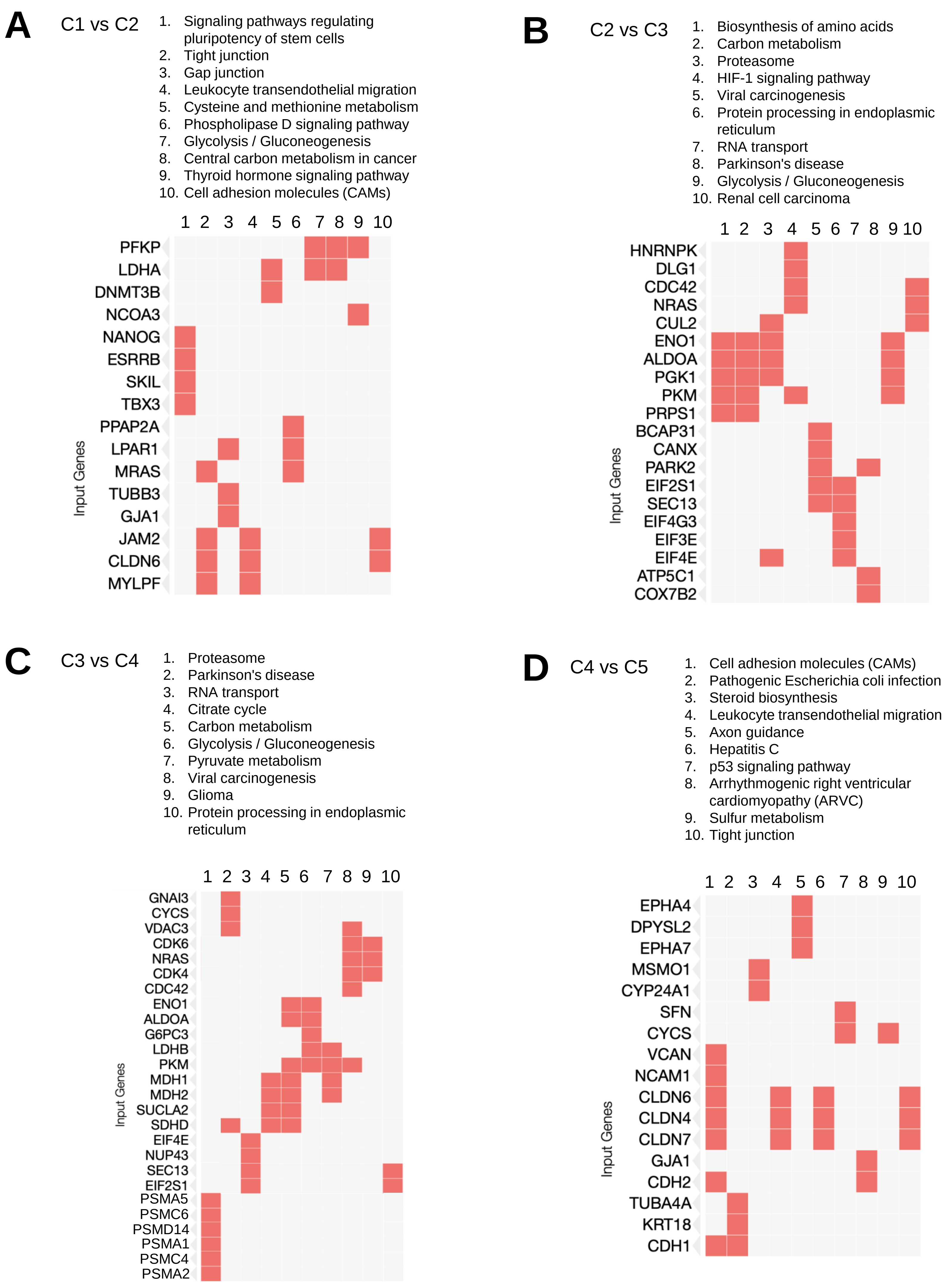
A
C1 vs C2
1. Signaling pathways regulating pluripotency of stem cells
2. Tight junction
3. Gap junction
4. Leukocyte transendothelial migration
5. Cysteine and methionine metabolism
6. Phospholipase D signaling pathway
7. Glycolysis / Gluconeogenesis
8. Central carbon metabolism in cancer
9. Thyroid hormone signaling pathway
10. Cell adhesion molecules (CAMs)
1 2 3 4 5 6 7 8 9 10
PFKP
LDHA
DNMT3B
NCOA3
NANOG
ESRRB
SKIL
TBX3
PPAP2A
LPAR1
MRAS
TUBB3
GJA1
JAM2
CLDN6
MYLPF
Input Genes
B
C2 vs C3
1. Biosynthesis of amino acids
2. Carbon metabolism
3. Proteasome
4. HIF-1 signaling pathway
5. Viral carcinogenesis
6. Protein processing in endoplasmic reticulum
7. RNA transport
8. Parkinson's disease
9. Glycolysis / Gluconeogenesis
10. Renal cell carcinoma
1 2 3 4 5 6 7 8 9 10
HNRNPK
DLG1
CDC42
NRAS
CUL2
ENO1
ALDOA
PGK1
PKM
PRPS1
BCAP31
CANX
PARK2
EIF2S1
SEC13
EIF4G3
EIF3E
EIF4E
ATP5C1
COX7B2
Input Genes
C
C3 vs C4
1. Proteasome
2. Parkinson's disease
3. RNA transport
4. Citrate cycle
5. Carbon metabolism
6. Glycolysis / Gluconeogenesis
7. Pyruvate metabolism
8. Viral carcinogenesis
9. Glioma
10. Protein processing in endoplasmic reticulum
1 2 3 4 5 6 7 8 9 10
GNAI3
CYCS
VDAC3
CDK6
NRAS
CDK4
CDC42
ENO1
ALDOA
G6PC3
LDHB
PKM
MDH1
MDH2
SUCLA2
SDHD
EIF4E
NUP43
SEC13
EIF2S1
PSMA5
PSMC6
PSMD14
PSMA1
PSMC4
PSMA2
Input Genes
D
C4 vs C5
1. Cell adhesion molecules (CAMs)
2. Pathogenic Escherichia coli infection
3. Steroid biosynthesis
4. Leukocyte transendothelial migration
5. Axon guidance
6. Hepatitis C
7. p53 signaling pathway
8. Arrhythmogenic right ventricular cardiomyopathy (ARVC)
9. Sulfur metabolism
10. Tight junction
1 2 3 4 5 6 7 8 9 10
EPHA4
DPYSL2
EPHA7
MSMO1
CYP24A1
SFN
CYCS
VCAN
NCAM1
CLDN6
CLDN4
CLDN7
GJA1
CDH2
TUBA4A
KRT18
CDH1
Input Genes

Fig. S6

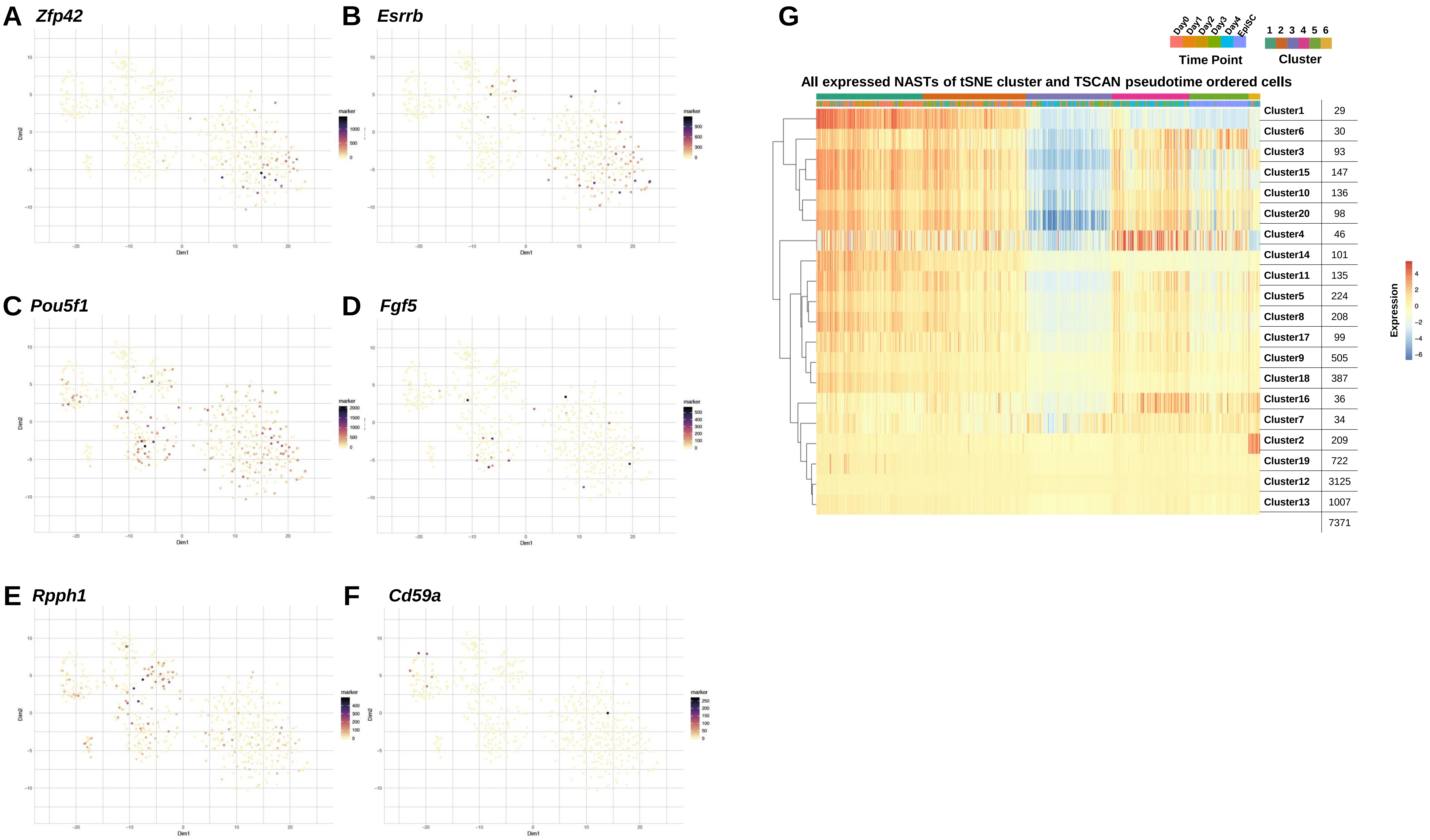

A Zfp42
B Esrrb
C Pou5f1
D Fgf5
E Rpph1
F Cd59a
Dim1
Dim2
marker
G
Day0
Day1
Day2
Day3
Day4
EpiSC
Time Point
1 2 3 4 5 6
Cluster
All expressed NASTs of tSNE cluster and TSCAN pseudotime ordered cells
Cluster1 29
Cluster6 30
Cluster3 93
Cluster15 147
Cluster10 136
Cluster20 98
Cluster4 46
Cluster14 101
Cluster11 135
Cluster5 224
Cluster8 208
Cluster17 99
Cluster9 505
Cluster18 387
Cluster16 36
Cluster7 34
Cluster2 209
Cluster19 722
Cluster12 3125
Cluster13 1007
7371
Expression


Fig. S7

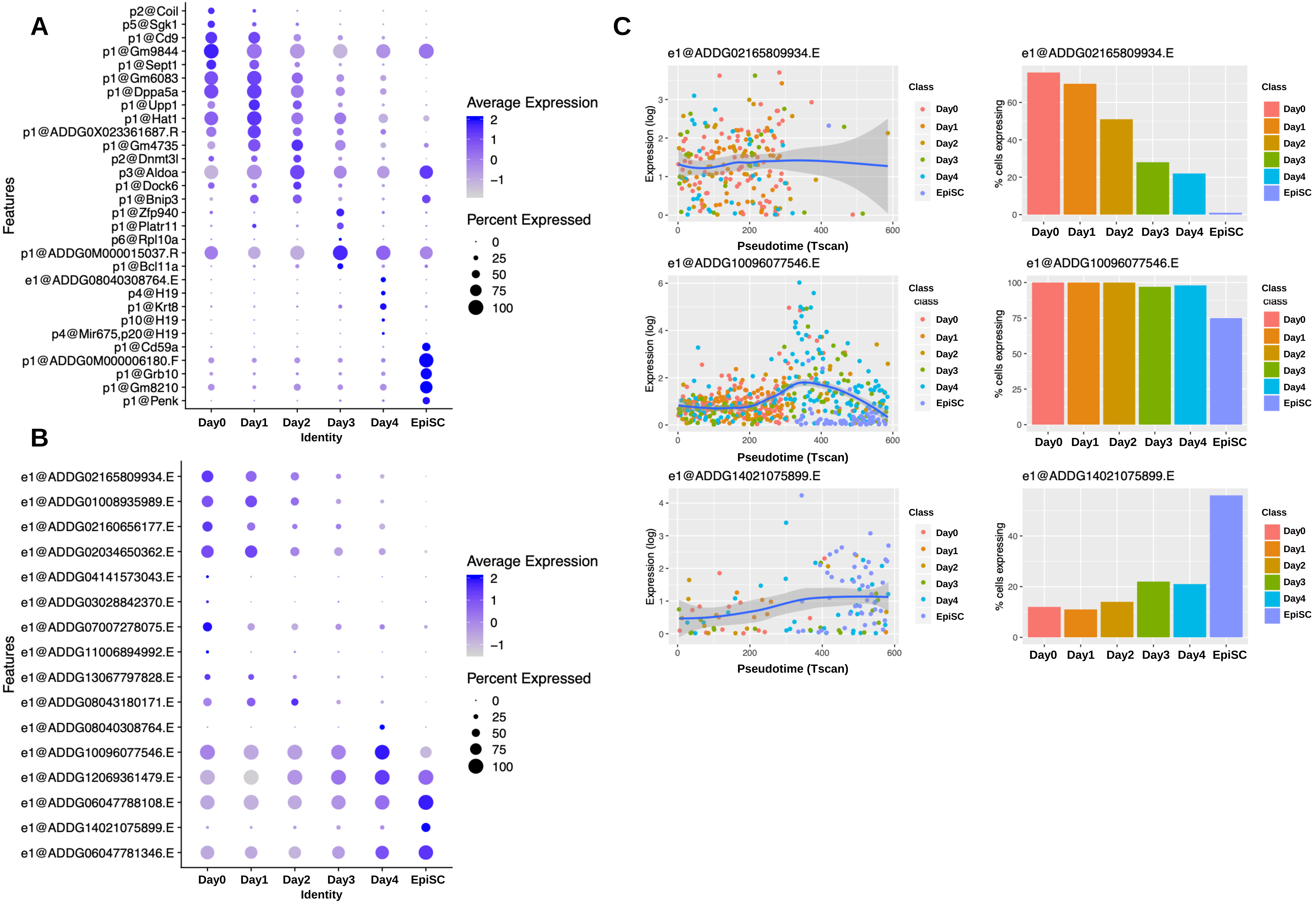
A
p2@Coil
p5@Sgk1
p1@Cd9
p1@Gm9844
p1@Sept1
p1@Gm6083
p1@Dppa5a
p1@Upp1
p1@Hat1
p1@ADDG0X023361687.R
p1@Gm4735
p2@Dnmt3l
p3@Aldoa
p1@Dock6
p1@Bnip3
p1@Zfp940
p1@Platr11
p6@Rpl10a
p1@ADDG0M000015037.R
p1@Bcl11a
e1@ADDG08040308764.E
p4@H19
p1@Krt8
p10@H19
p4@Mir675,p20@H19
p1@Cd59a
p1@ADDG0M000006180.F
p1@Grb10
p1@Gm8210
p1@Penk
Features
Day0
Day1
Day2
Day3
Day4
EpiSC
Identity
Average Expression
2
1
0
−1
Percent Expressed
0
25
50
75
100
B
e1@ADDG02165809934.E
e1@ADDG01008935989.E
e1@ADDG02160656177.E
e1@ADDG02034650362.E
e1@ADDG04141573043.E
e1@ADDG03028842370.E
e1@ADDG07007278075.E
e1@ADDG11006894992.E
e1@ADDG13067797828.E
e1@ADDG08043180171.E
e1@ADDG08040308764.E
e1@ADDG10096077546.E
e1@ADDG12069361479.E
e1@ADDG06047788108.E
e1@ADDG14021075899.E
e1@ADDG06047781346.E
Features
Day0
Day1
Day2
Day3
Day4
EpiSC
Identity
Average Expression
2
1
0
−1
Percent Expressed
0
25
50
75
100
C
e1@ADDG02165809934.E
Expression (log)
Pseudotime (Tscan)
Class
Day0
Day1
Day2
Day3
Day4
EpiSC
% cells expressing
e1@ADDG10096077546.E
e1@ADDG14021075899.E

Fig. S8

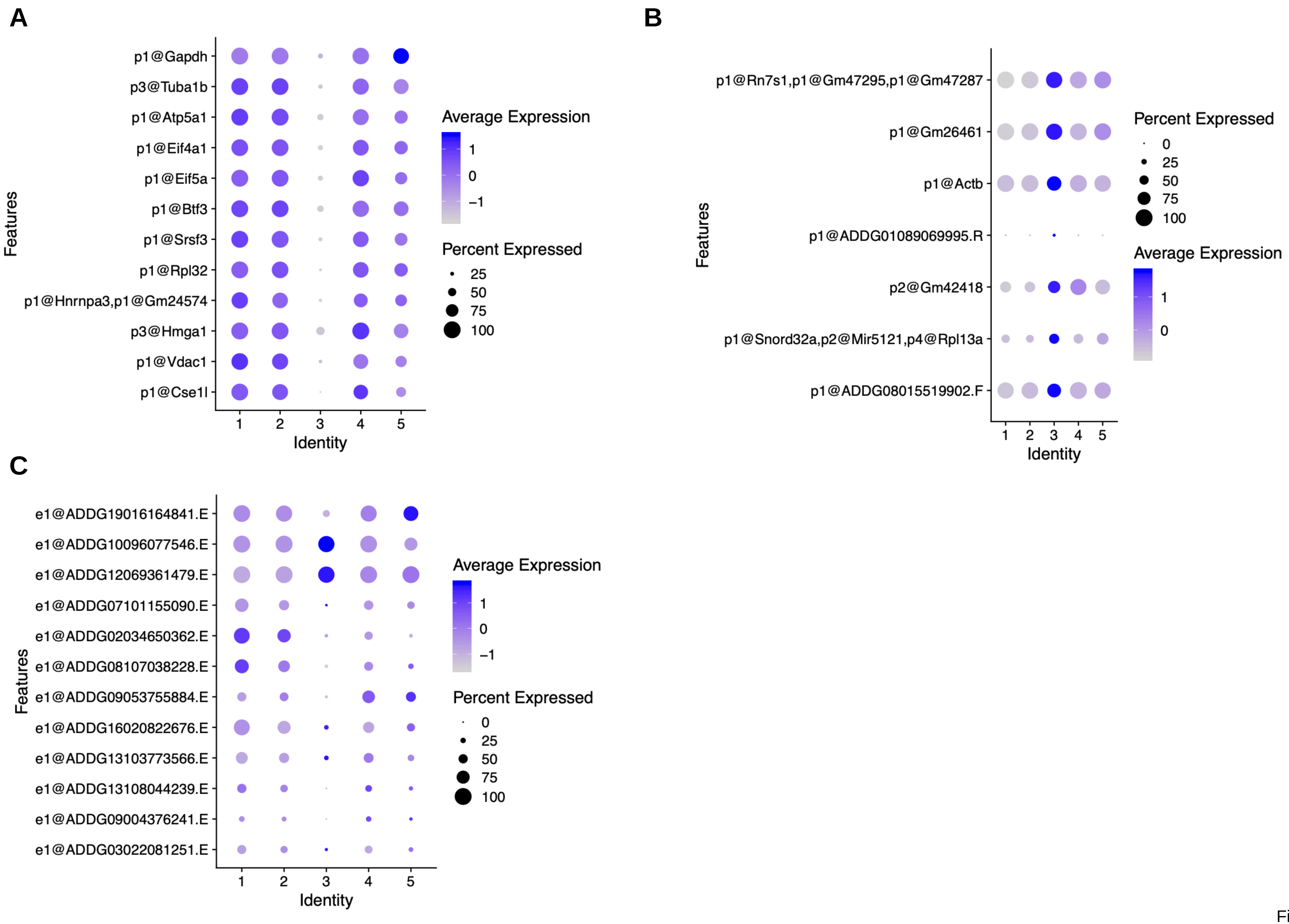

A
p1@Gapdh
p3@Tuba1b
p1@Atp5a1
p1@Eif4a1
p1@Eif5a
p1@Btf3
p1@Srsf3
p1@Rpl32
p1@Hnrnpa3,p1@Gm24574
p3@Hmga1
p1@Vdac1
p1@Cse1l
Features
1
2
3
4
5
Identity
Average Expression
1
0
−1
Percent Expressed
25
50
75
100
B
p1@Rn7s1,p1@Gm47295,p1@Gm47287
p1@Gm26461
p1@Actb
p1@ADDG01089069995.R
p2@Gm42418
p1@Snord32a,p2@Mir5121,p4@Rpl13a
p1@ADDG08015519902.F
Features
1
2
3
4
5
Identity
Percent Expressed
0
25
50
75
100
Average Expression
1
0
C
e1@ADDG19016164841.E
e1@ADDG10096077546.E
e1@ADDG12069361479.E
e1@ADDG07101155090.E
e1@ADDG02034650362.E
e1@ADDG08107038228.E
e1@ADDG09053755884.E
e1@ADDG16020822676.E
e1@ADDG13103773566.E
e1@ADDG13108044239.E
e1@ADDG09004376241.E
e1@ADDG03022081251.E
Features
1
2
3
4
5
Identity
Average Expression
1
0
−1
Percent Expressed
0
25
50
75
100


Fig. S9

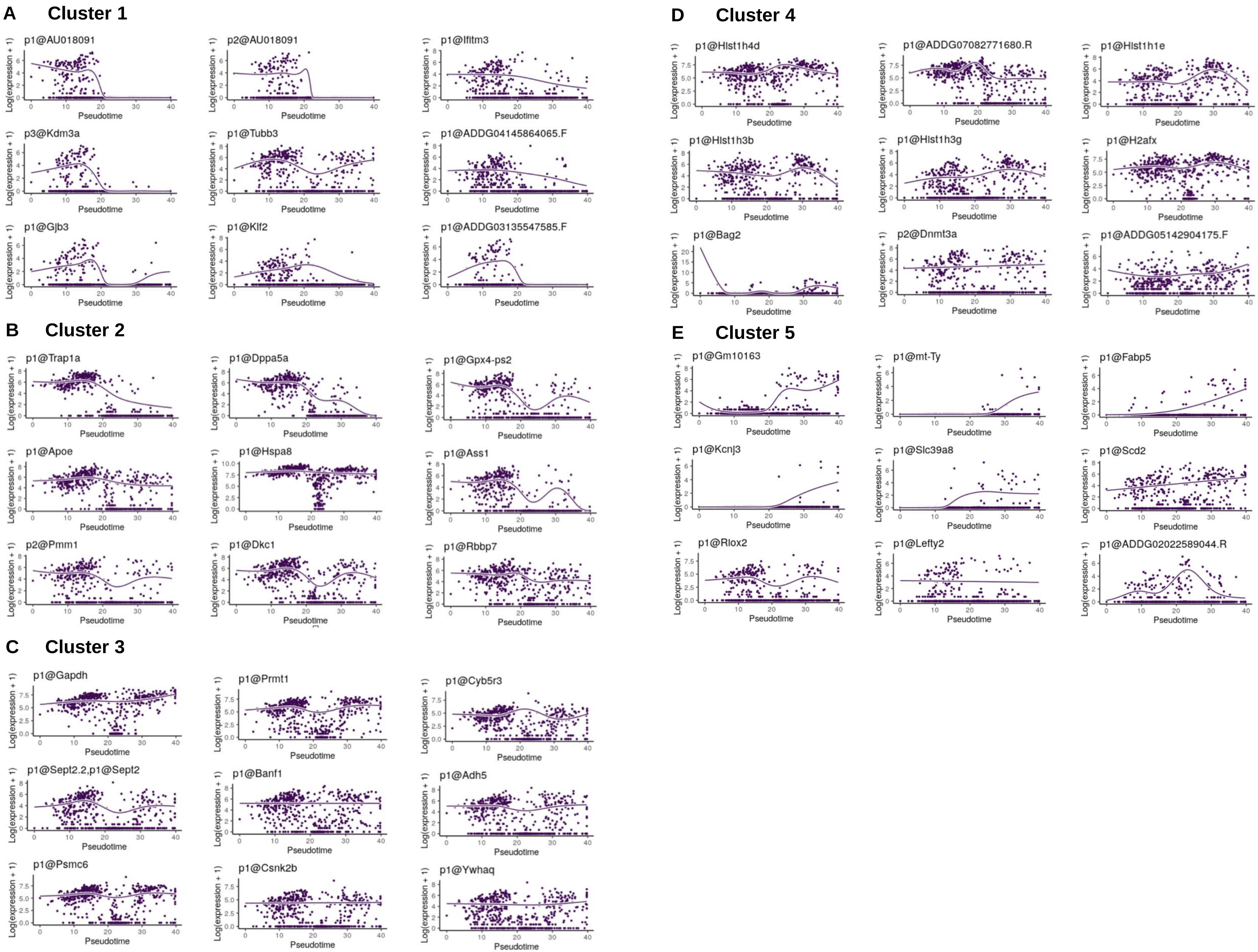
A Cluster 1
p1@AU018091
p2@AU018091
p1@Ifitm3
p3@Kdm3a
p1@Tubb3
p1@ADDG04145864065.F
p1@Gjb3
p1@Klf2
p1@ADDG03135547585.F
B Cluster 2
p1@Trap1a
p1@Dppa5a
p1@Gpx4-ps2
p1@Apoe
p1@Hspa8
p1@Ass1
p2@Pmm1
p1@Dkc1
p1@Rbbp7
C Cluster 3
p1@Gapdh
p1@Prmt1
p1@Cyb5r3
p1@Sept2.2,p1@Sept2
p1@Banf1
p1@Adh5
p1@Psmc6
p1@Csnk2b
p1@Ywhaq
D Cluster 4
p1@Hist1h4d
p1@ADDG07082771680.R
p1@Hist1h1e
p1@Hist1h3b
p1@Hist1h3g
p1@H2afx
p1@Bag2
p2@Dnmt3a
p1@ADDG05142904175.F
E Cluster 5
p1@Gm10163
p1@mt-Ty
p1@Fabp5
p1@Kcnj3
p1@Slc39a8
p1@Scd2
p1@Rlox2
p1@Lefty2
p1@ADDG02022589044.R
Log(expression + 1)
Pseudotime

Fig. S10

A

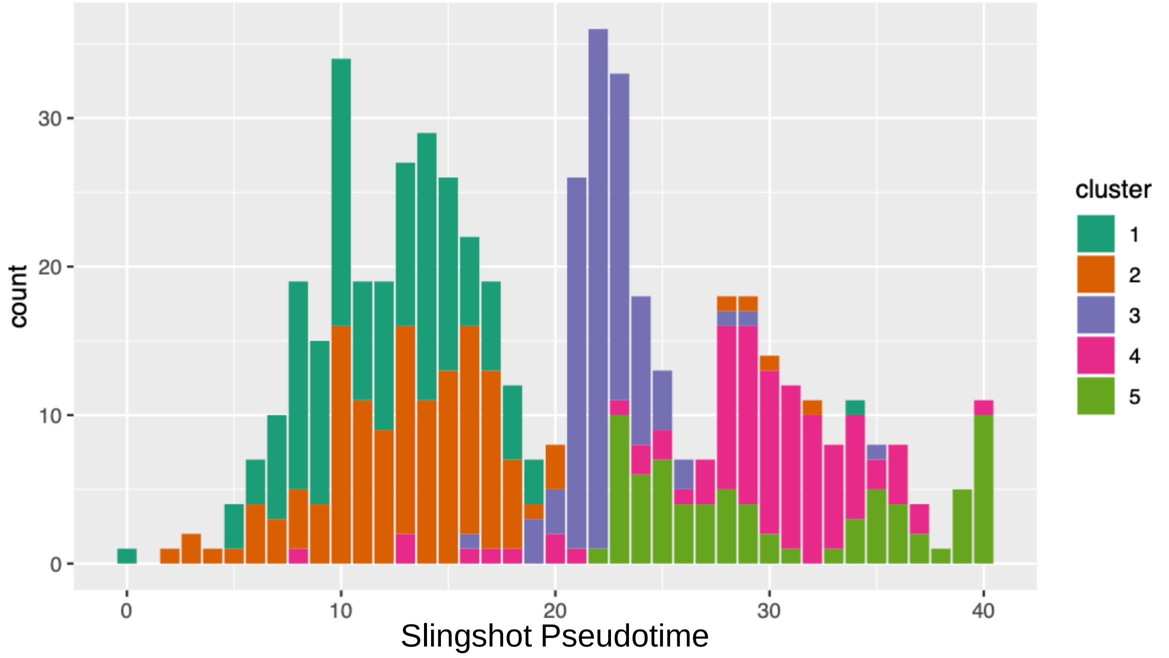


B

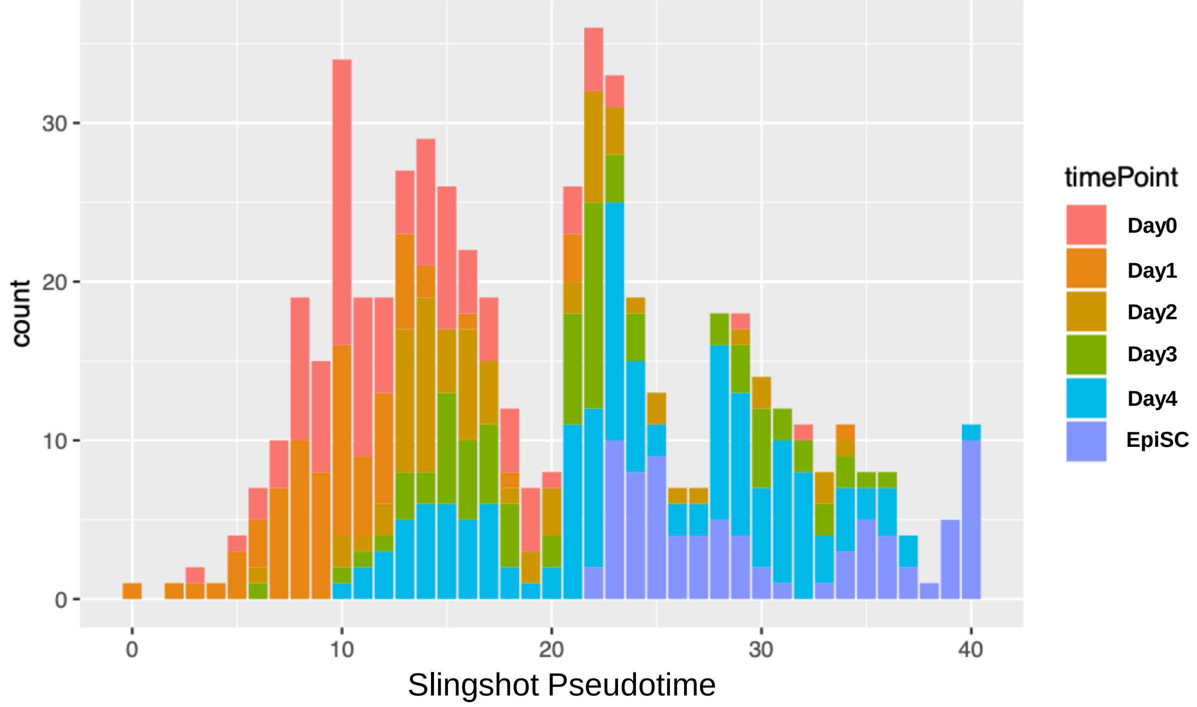


C

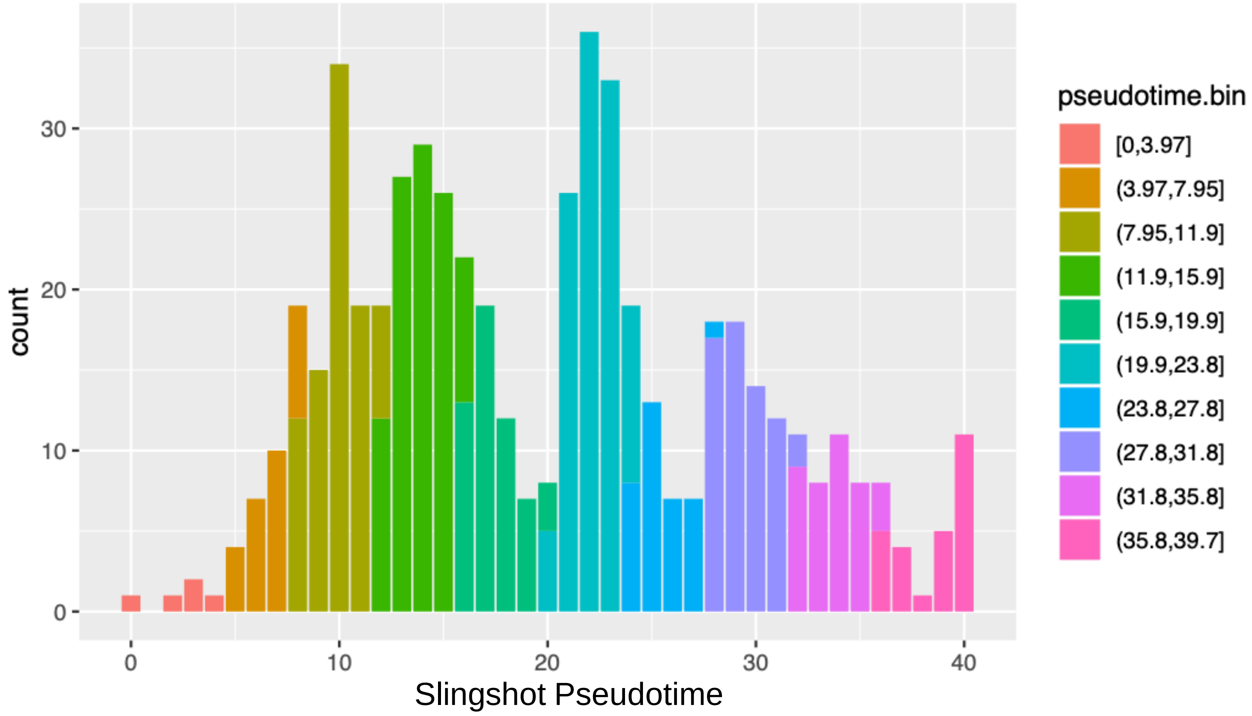


D

**Cluster**

| bin [Slingshot Pseudotime] | 1 | 2 | 3 | 4 | 5 |
|---|---|---|---|---|---|
| 1 [0,3.97] | 1 | 4 | 0 | 0 | 0 |
| 2 [3.97,7.95] | 18 | 10 | 0 | 0 | 0 |
| 3 [7.95,11.9] | 51 | 35 | 0 | 1 | 0 |
| 4 [11.9,15.9] | 51 | 50 | 0 | 2 | 0 |
| 5 [15.9,19.9] | 16 | 29 | 5 | 4 | 0 |
| 6 [19.9,23.8] | 0 | 2 | 91 | 3 | 14 |
| 7 [23.8,27.8] | 0 | 0 | 9 | 9 | 18 |
| 8 [27.8,31.8] | 0 | 4 | 2 | 45 | 12 |
| 9 [31.8,35.8] | 1 | 0 | 1 | 27 | 10 |
| 10 [35.8,39.7] | 0 | 0 | 0 | 5 | 21 |
| Total cell number | 138 | 134 | 108 | 96 | 75 |

Fig. S11

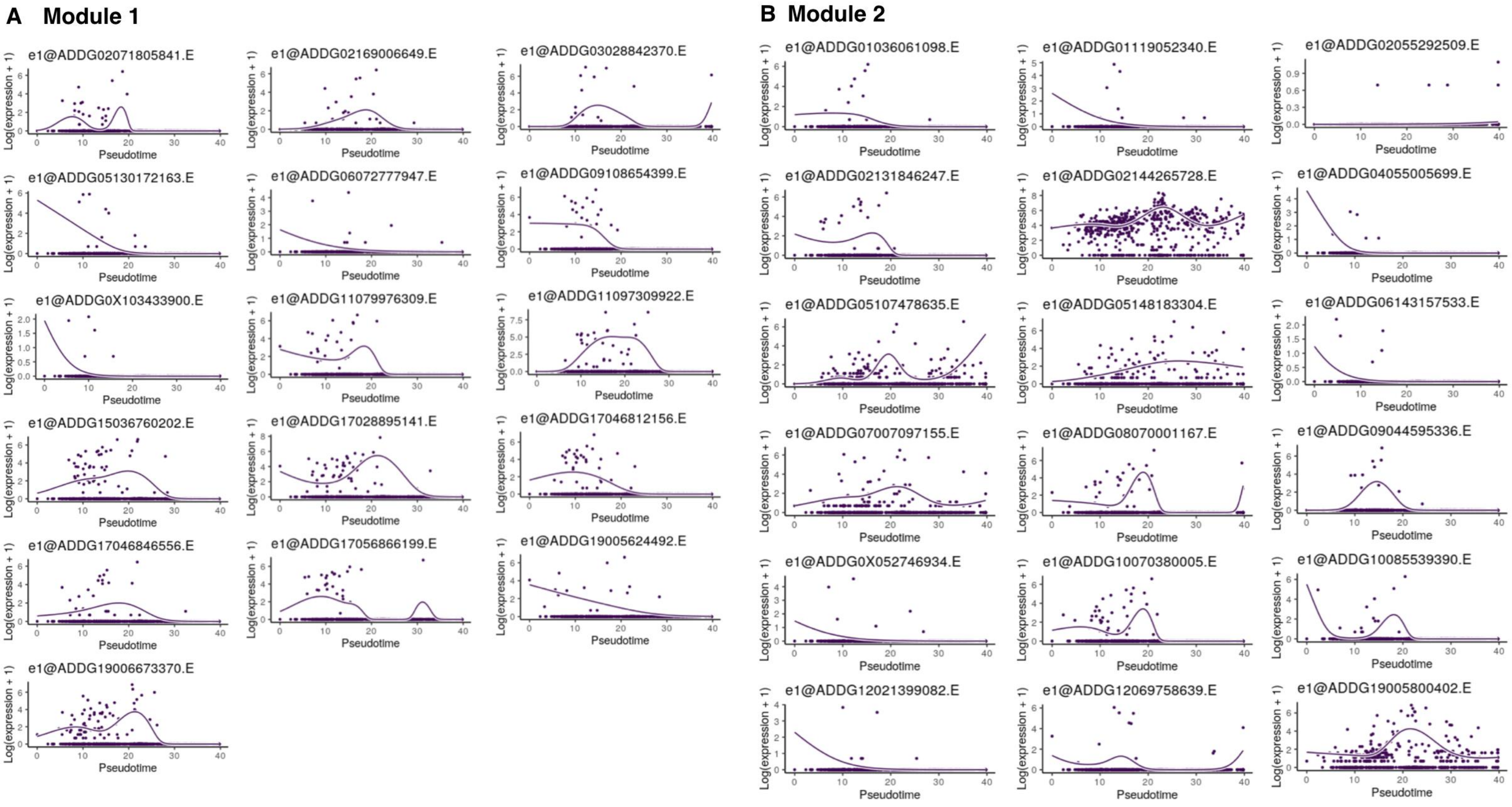
A Module 1
e1@ADDG02071805841.E
e1@ADDG02169006649.E
e1@ADDG03028842370.E
e1@ADDG05130172163.E
e1@ADDG06072777947.E
e1@ADDG09108654399.E
e1@ADDG0X103433900.E
e1@ADDG11079976309.E
e1@ADDG11097309922.E
e1@ADDG15036760202.E
e1@ADDG17028895141.E
e1@ADDG17046812156.E
e1@ADDG17046846556.E
e1@ADDG17056866199.E
e1@ADDG19005624492.E
e1@ADDG19006673370.E
B Module 2
e1@ADDG01036061098.E
e1@ADDG01119052340.E
e1@ADDG02055292509.E
e1@ADDG02131846247.E
e1@ADDG02144265728.E
e1@ADDG04055005699.E
e1@ADDG05107478635.E
e1@ADDG05148183304.E
e1@ADDG06143157533.E
e1@ADDG07007097155.E
e1@ADDG08070001167.E
e1@ADDG09044595336.E
e1@ADDG0X052746934.E
e1@ADDG10070380005.E
e1@ADDG10085539390.E
e1@ADDG12021399082.E
e1@ADDG12069758639.E
e1@ADDG19005800402.E
Log(expression + 1)
Pseudotime

Fig. S12

**C Module 3**

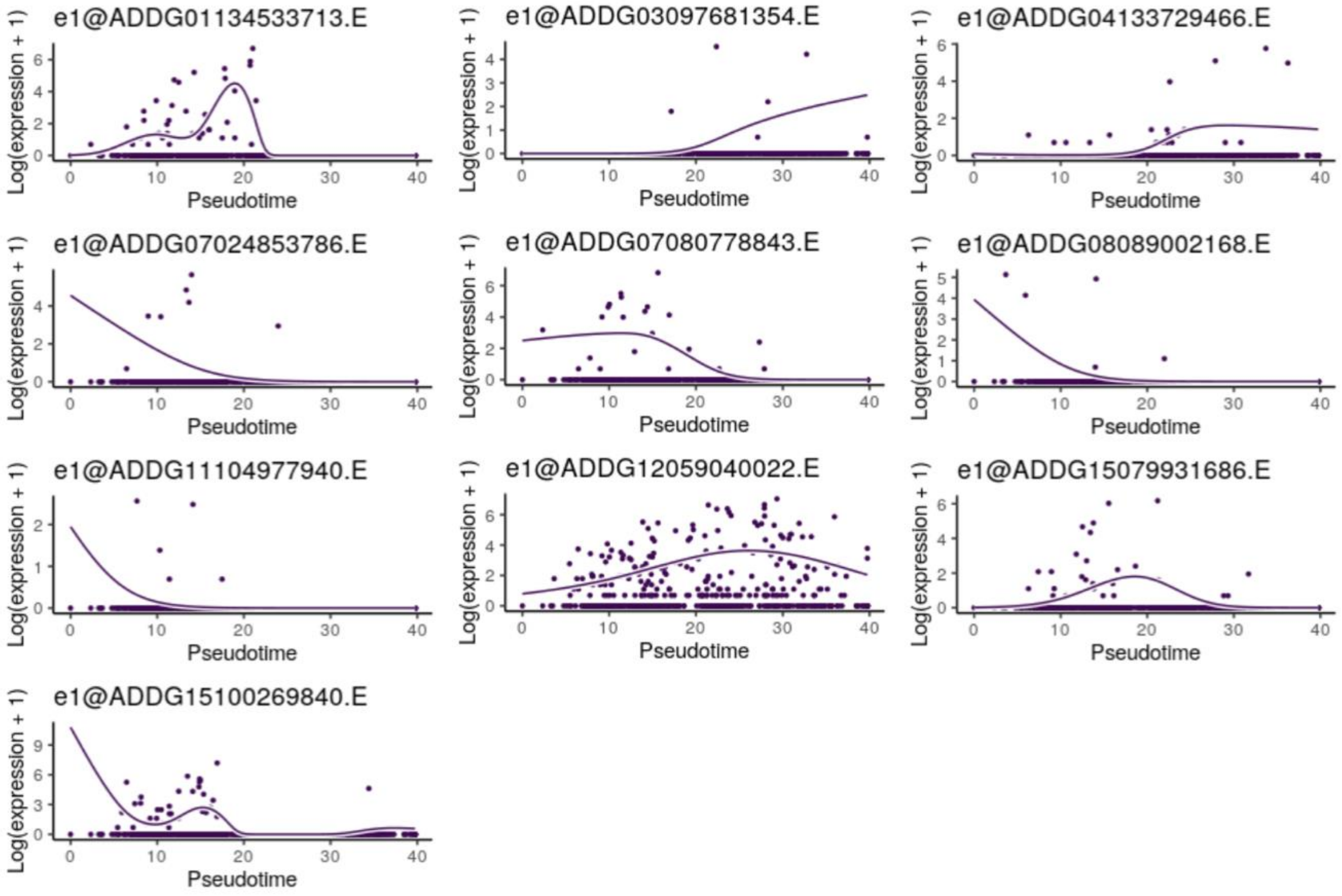


**D Module 4**

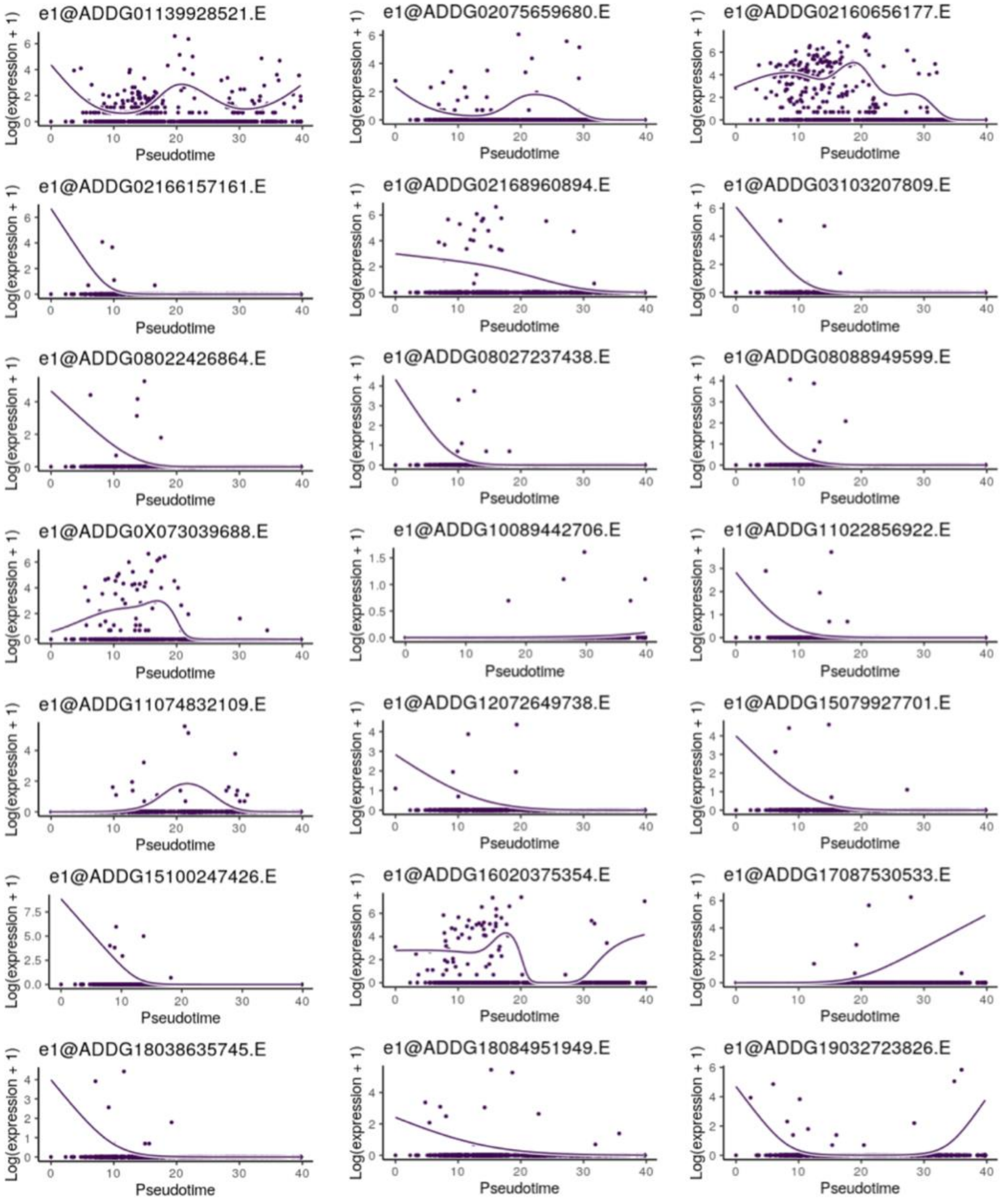


**E Module 5**

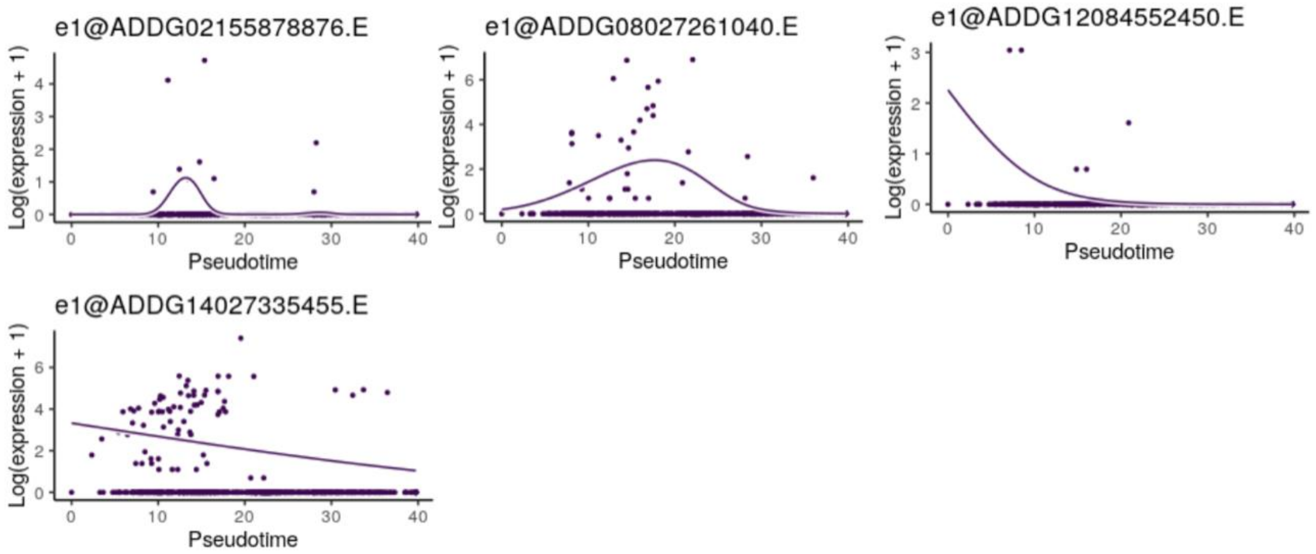


Fig. S12

**A**

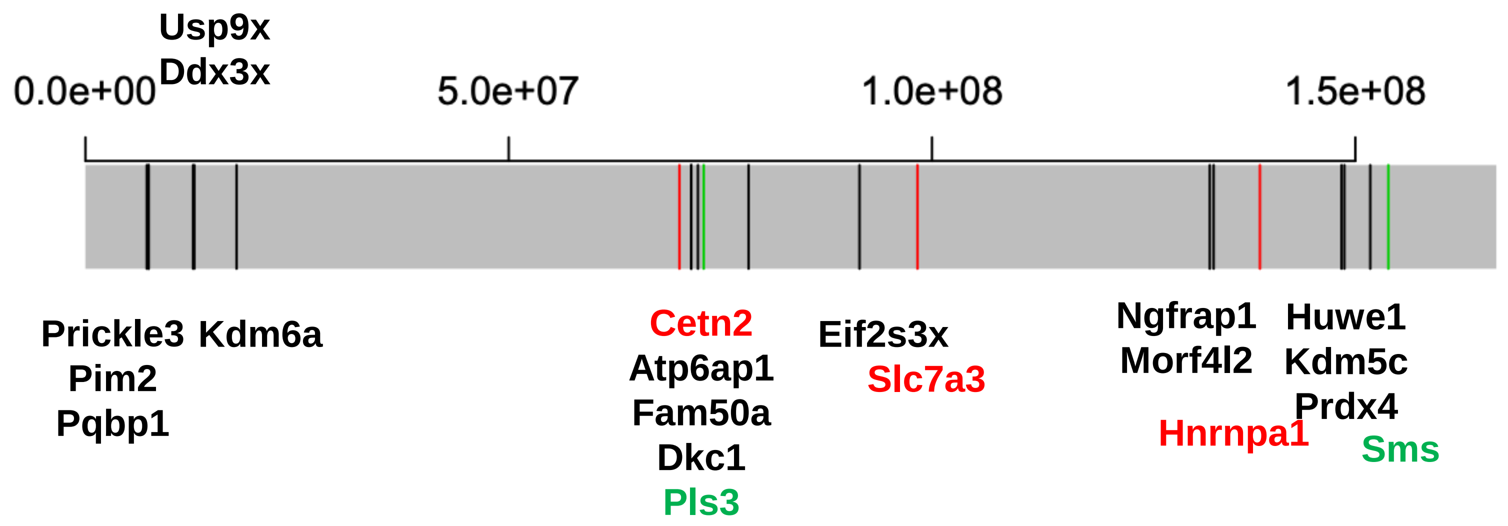

Usp9x
Ddx3x
0.0e+00
5.0e+07
1.0e+08
1.5e+08
Prickle3
Kdm6a
Pim2
Pqbp1
Cetn2
Atp6ap1
Fam50a
Dkc1
Pls3
Eif2s3x
Slc7a3
Ngfrap1
Morf4l2
Hnrnpa1
Huwe1
Kdm5c
Prdx4
Sms


**B**

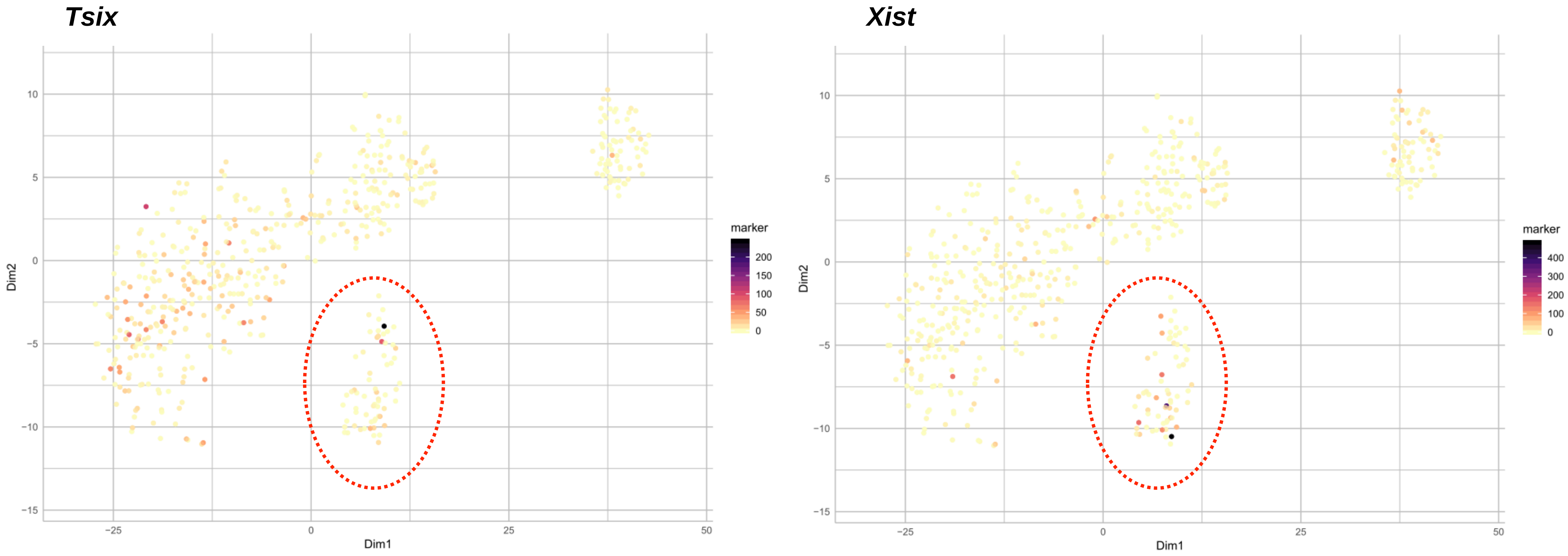

Tsix
Xist
Dim1
Dim2
marker


Fig. S13

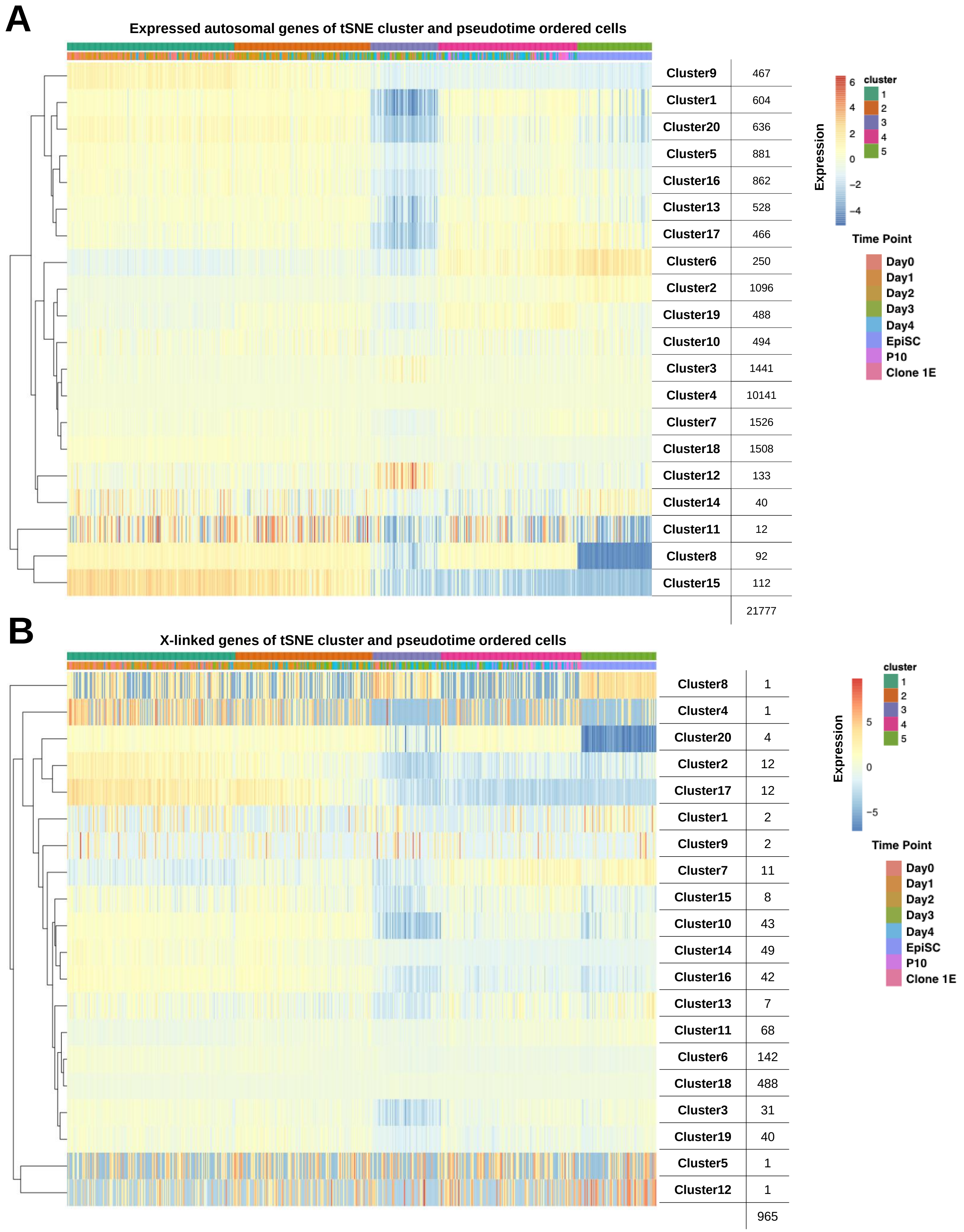

A
Expressed autosomal genes of tSNE cluster and pseudotime ordered cells
Cluster9 467
Cluster1 604
Cluster20 636
Cluster5 881
Cluster16 862
Cluster13 528
Cluster17 466
Cluster6 250
Cluster2 1096
Cluster19 488
Cluster10 494
Cluster3 1441
Cluster4 10141
Cluster7 1526
Cluster18 1508
Cluster12 133
Cluster14 40
Cluster11 12
Cluster8 92
Cluster15 112
21777
Expression
6
4
2
0
−2
−4
cluster
1
2
3
4
5
Time Point
Day0
Day1
Day2
Day3
Day4
EpiSC
P10
Clone 1E
B
X-linked genes of tSNE cluster and pseudotime ordered cells
Cluster8 1
Cluster4 1
Cluster20 4
Cluster2 12
Cluster17 12
Cluster1 2
Cluster9 2
Cluster7 11
Cluster15 8
Cluster10 43
Cluster14 49
Cluster16 42
Cluster13 7
Cluster11 68
Cluster6 142
Cluster18 488
Cluster3 31
Cluster19 40
Cluster5 1
Cluster12 1
965
Expression
5
0
−5
cluster
1
2
3
4
5
Time Point
Day0
Day1
Day2
Day3
Day4
EpiSC
P10
Clone 1E


Fig. S14
