## Supplementary material for "Single-cell transcriptomics, scRNA-Seq and C1 CAGE discovered distinct phases of pluripotency during naïve-to-primed conversion in mice": tSNE_heatscaleCustomGenesC1CAGE_20190317042758.pdf

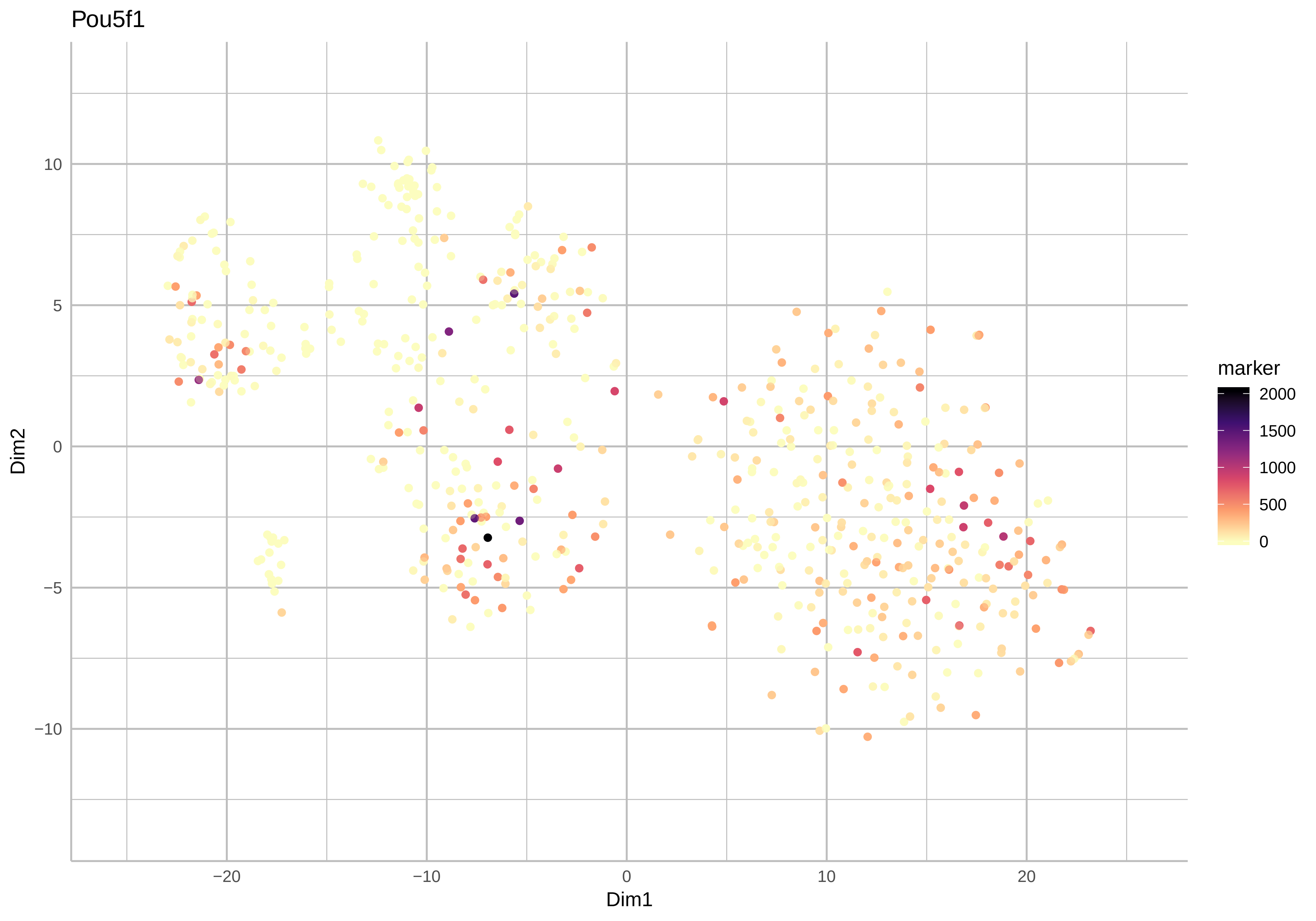

Pou5f1
Dim2
Dim1
10
5
0
−5
−10
−20
−10
0
10
20
marker
2000
1500
1000
500
0

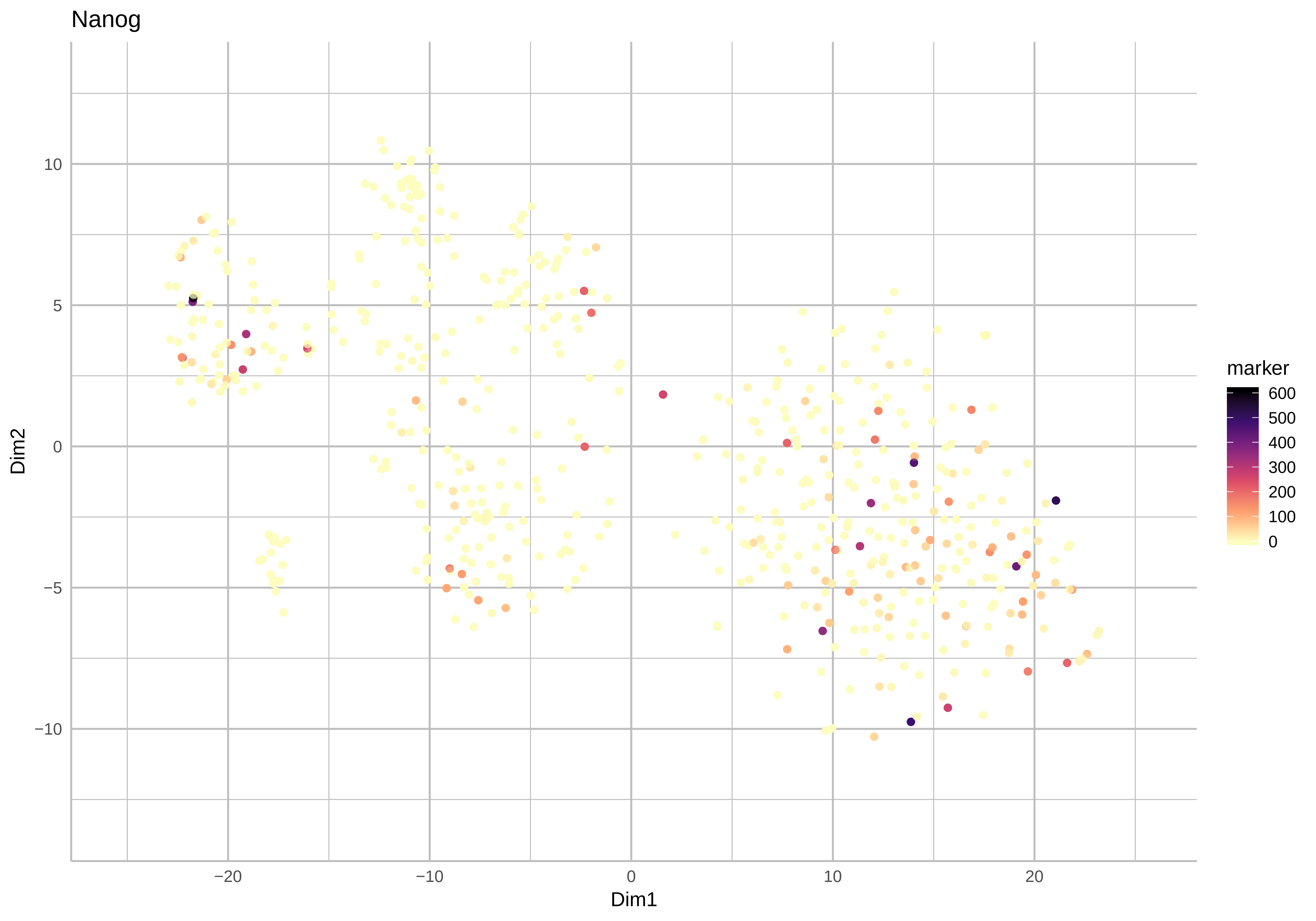

Nanog
Dim2
Dim1
10
5
0
−5
−10
−20
−10
0
10
20
marker
600
500
400
300
200
100
0

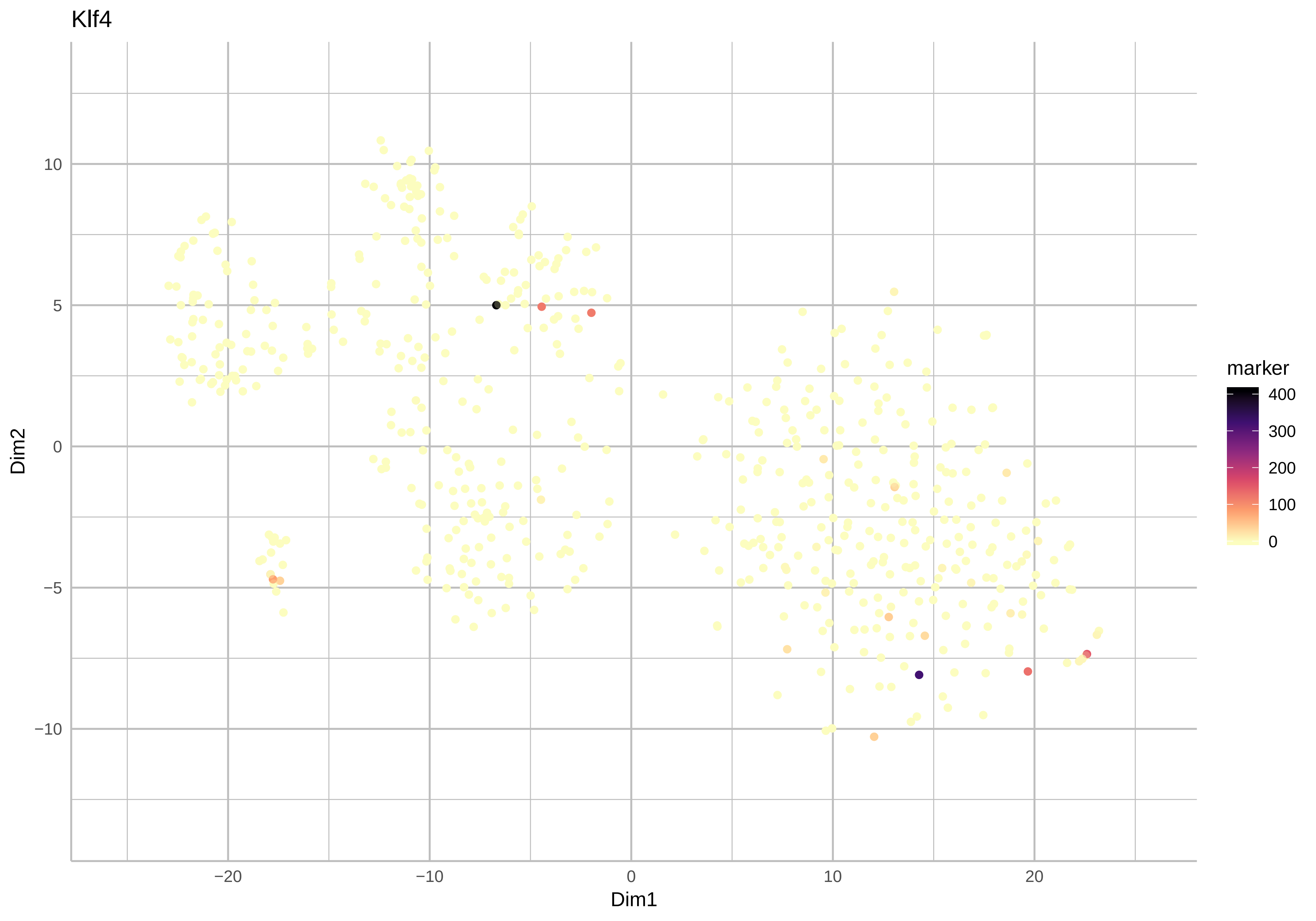
Klf4
Dim2
Dim1
marker
400
300
200
100
0
10
5
0
−5
−10
−20
−10
0
10
20

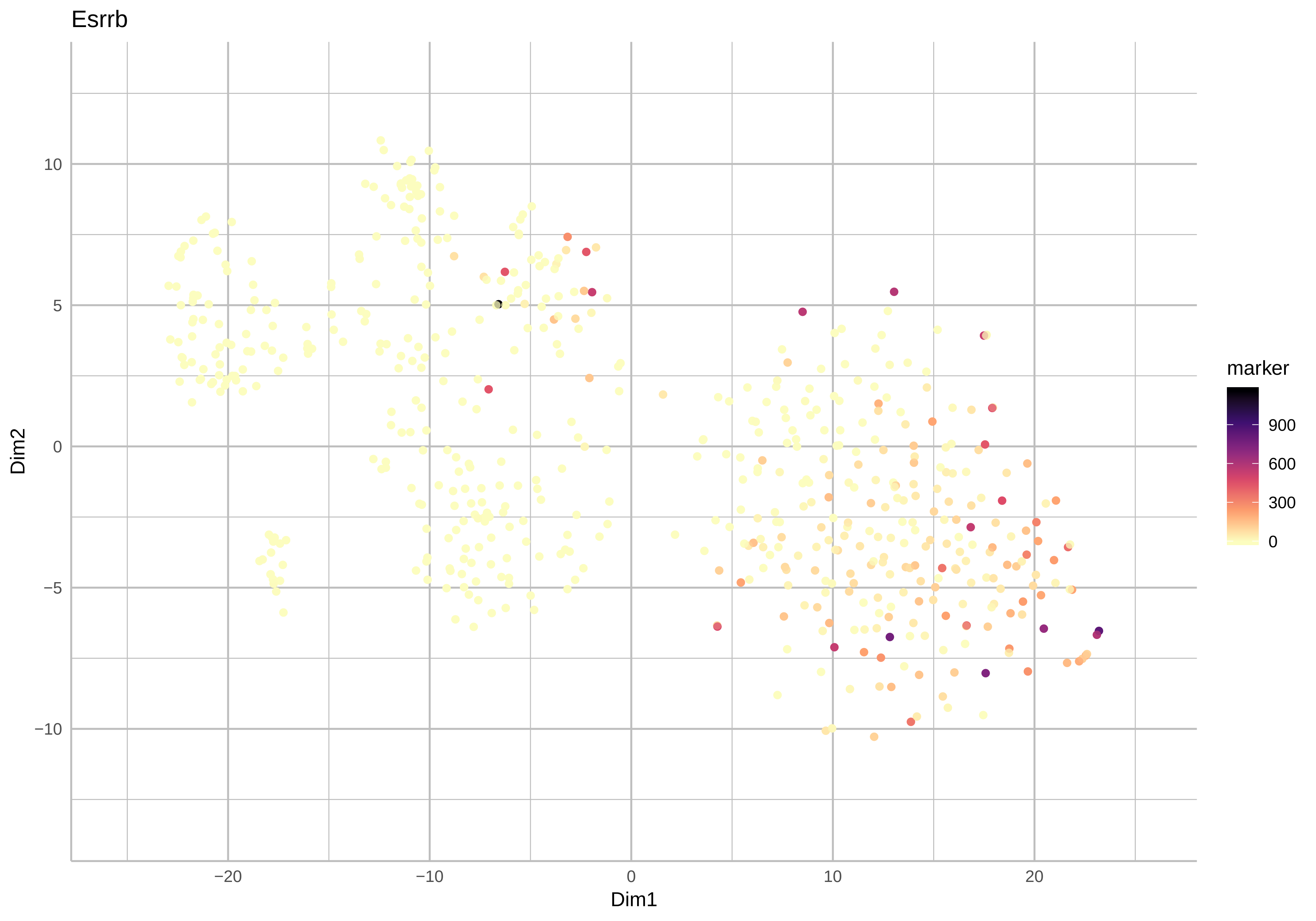
Esrrb
Dim1
Dim2
marker
900
600
300
0
−20
−10
0
10
20
10
5
0
−5
−10

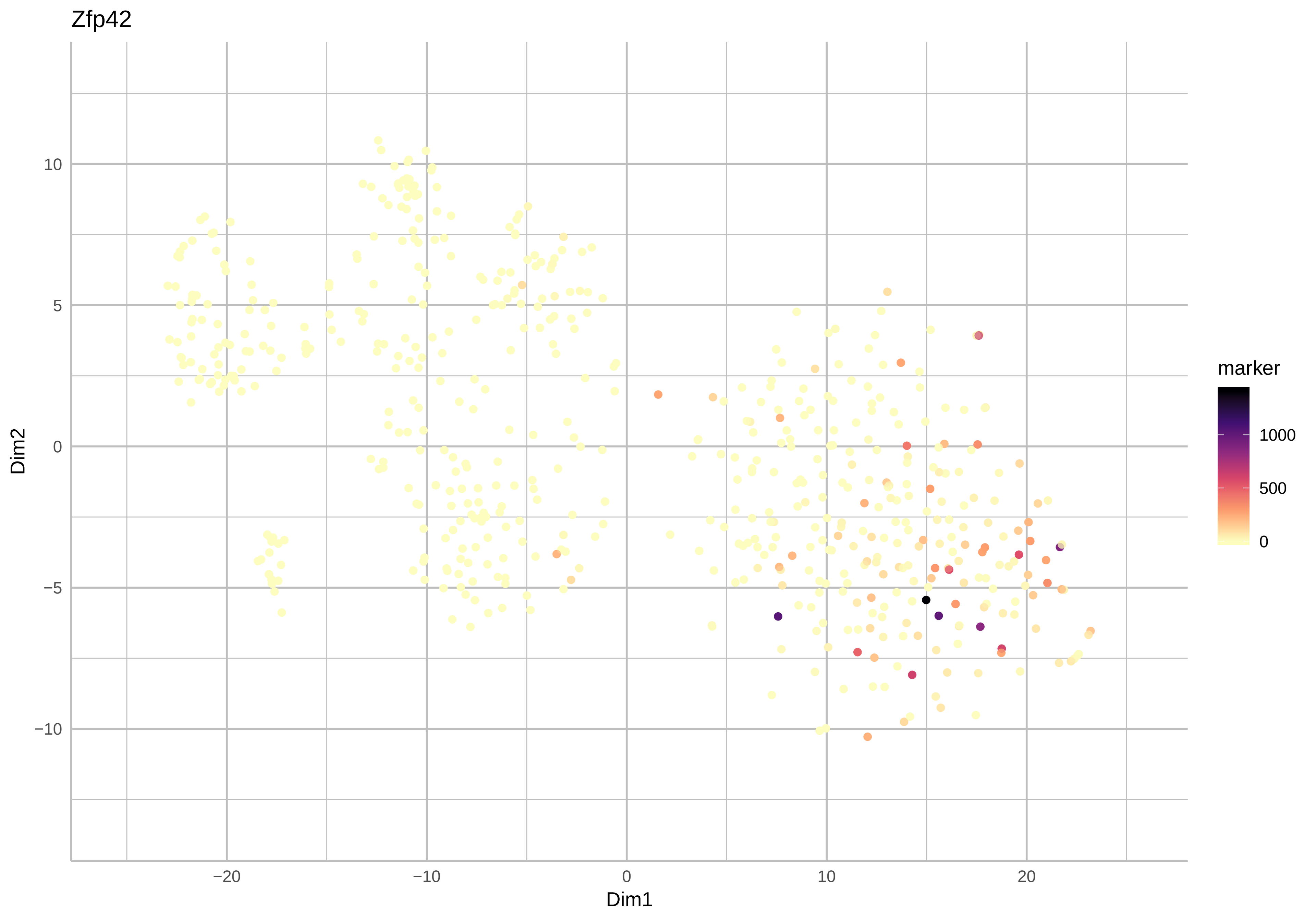

Zfp42
Dim2
Dim1
10
5
0
−5
−10
−20
−10
0
10
20
marker
1000
500
0

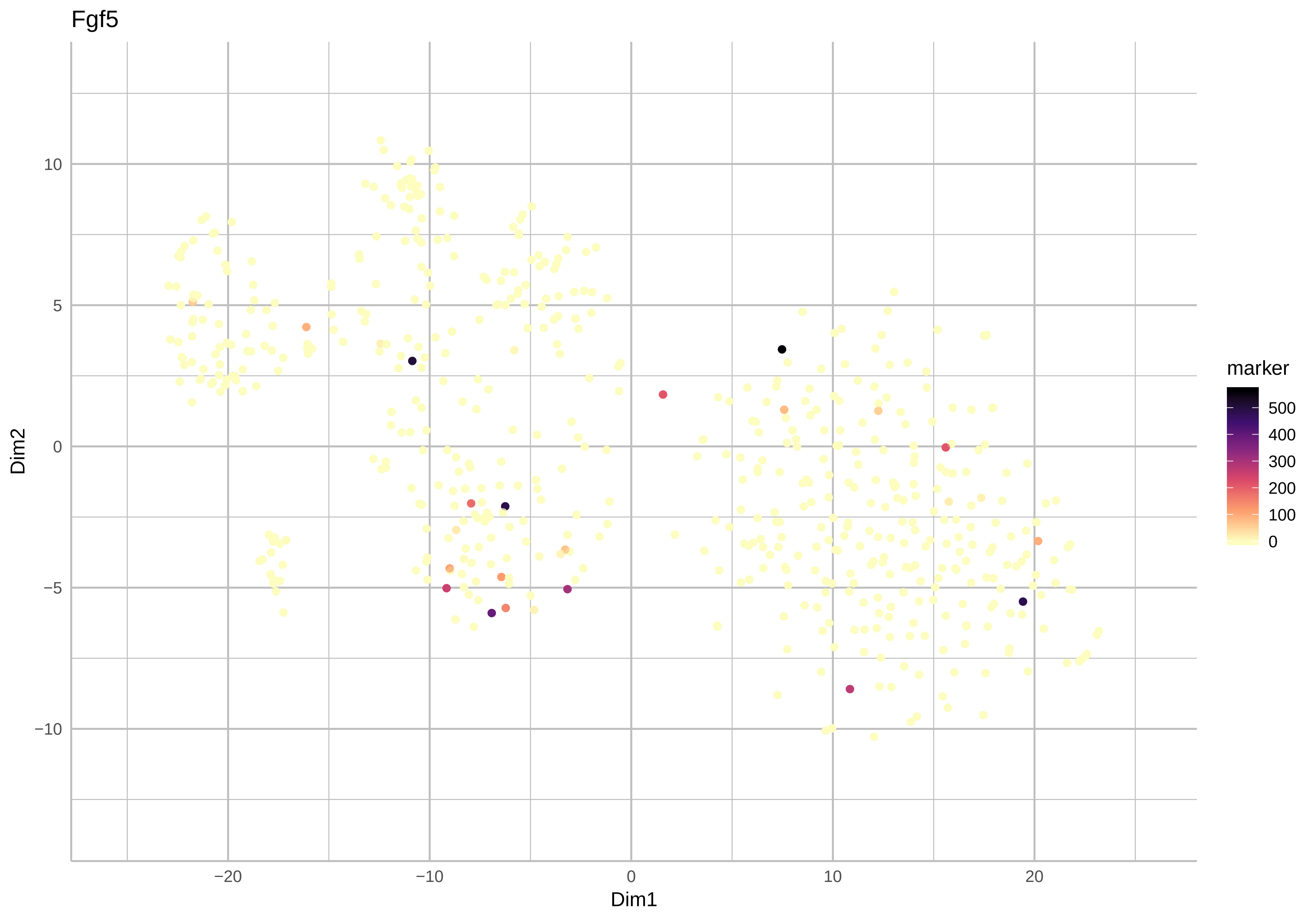

Fgf5
Dim2
Dim1
10
5
0
−5
−10
−20
−10
0
10
20
marker
500
400
300
200
100
0

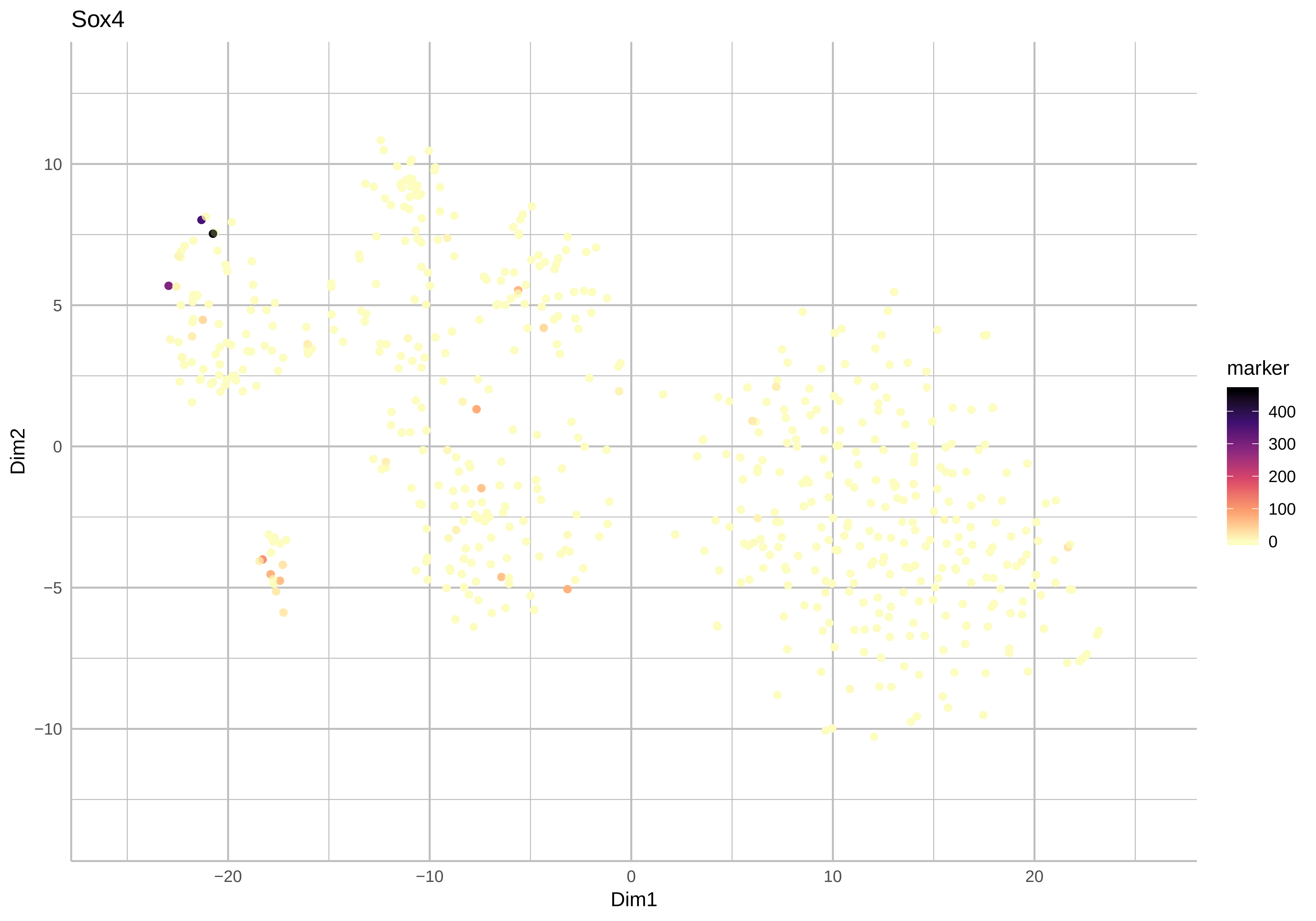
Sox4
Dim2
Dim1
marker
400
300
200
100
0
10
5
0
−5
−10
−20
−10
0
10
20

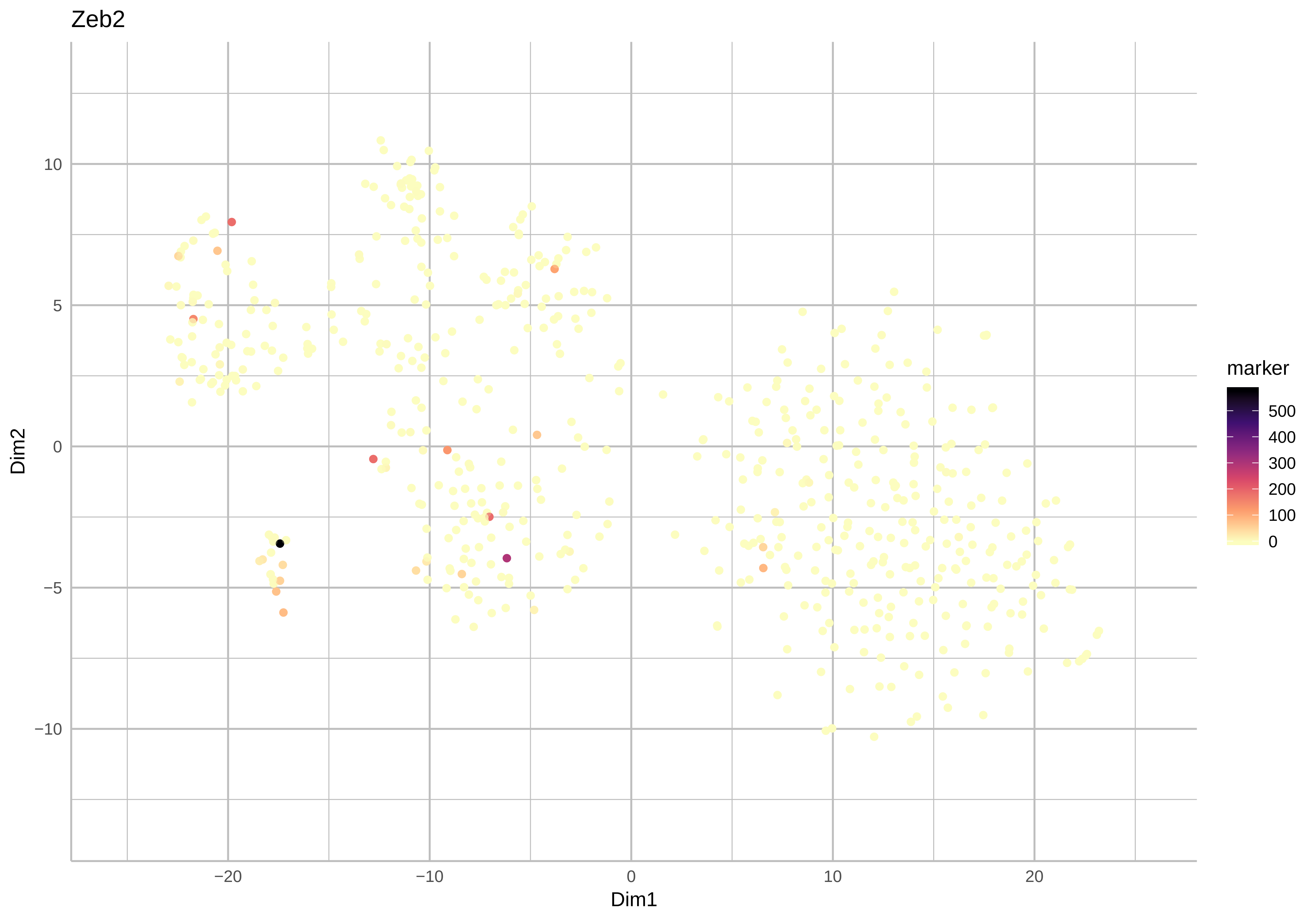
Zeb2
Dim1
Dim2
−20
−10
0
10
20
−10
−5
0
5
10
marker
500
400
300
200
100
0

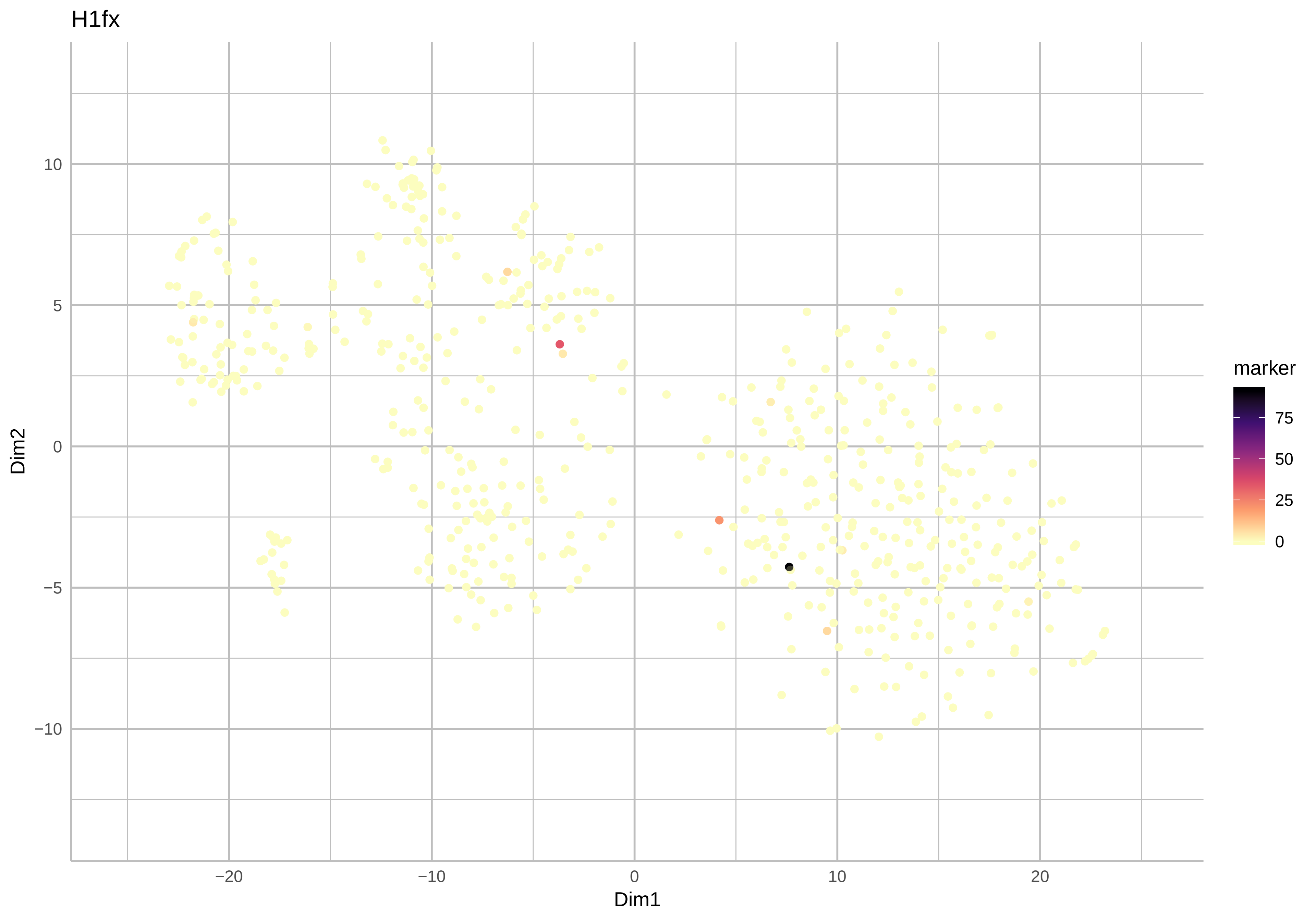
H1fx
Dim2
Dim1
marker
75
50
25
0
10
5
0
−5
−10
−20
−10
0
10
20

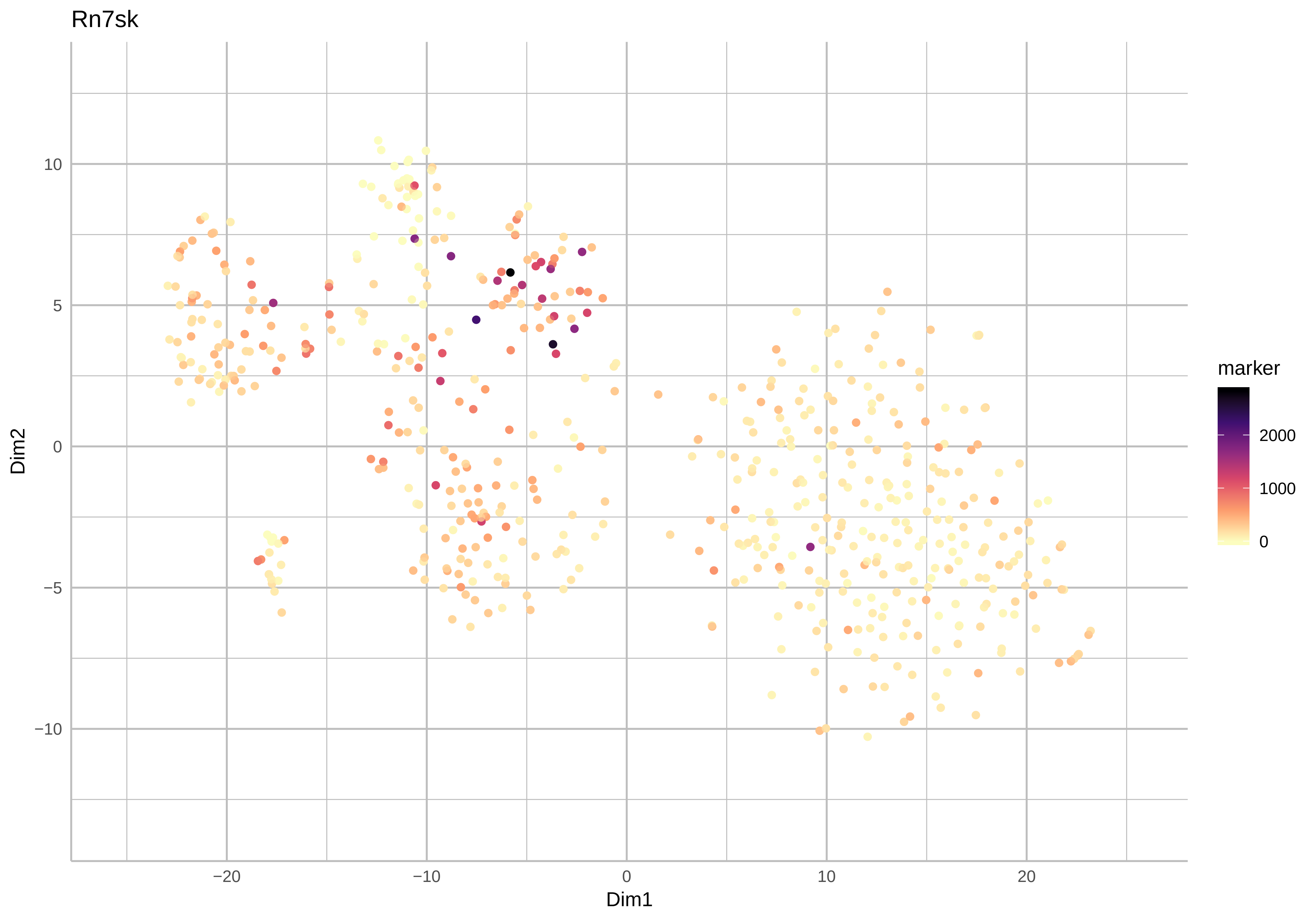

Rn7sk
Dim2
Dim1
10
5
0
−5
−10
−20
−10
0
10
20
marker
2000
1000
0

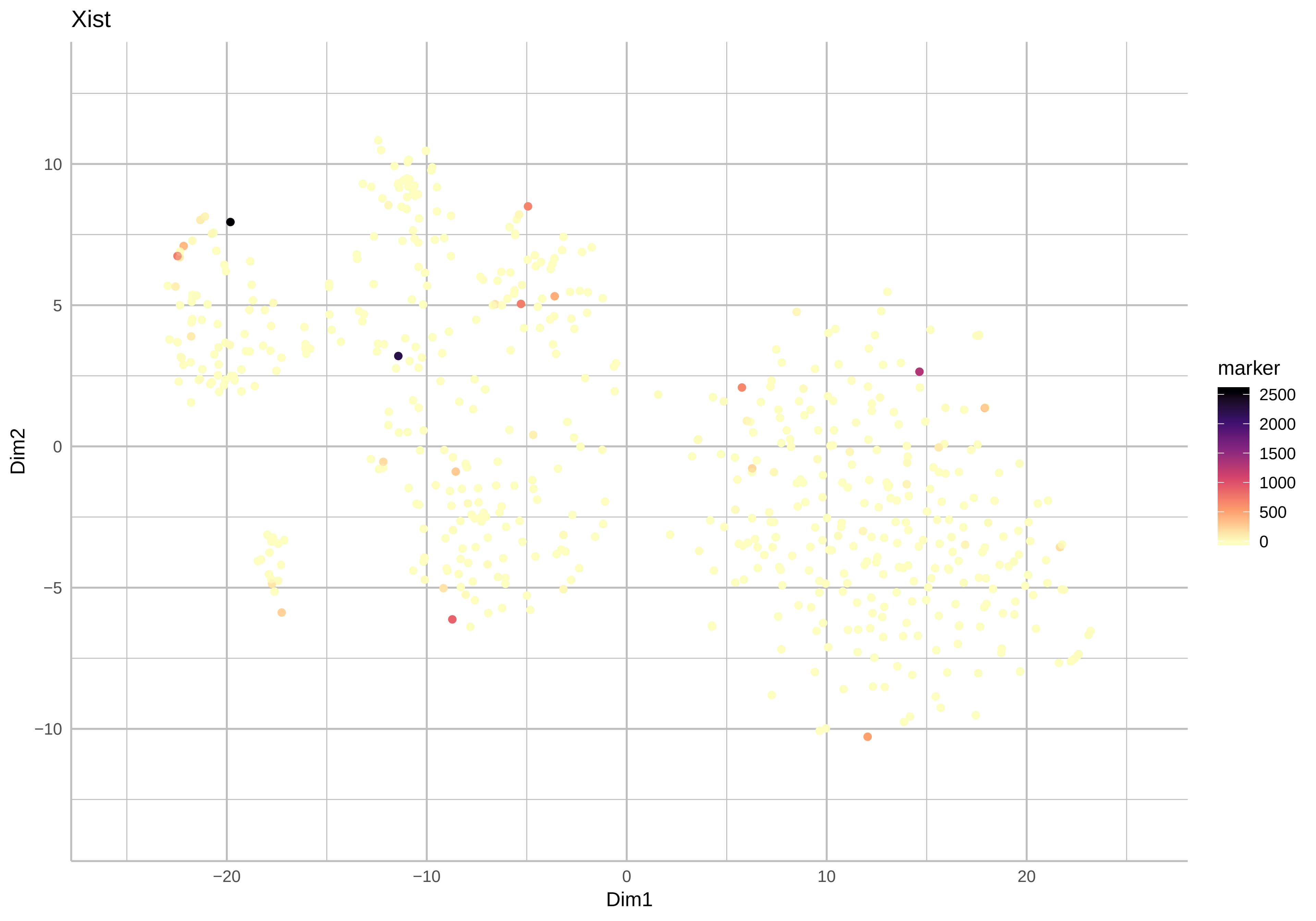
Xist
Dim2
Dim1
marker
2500
2000
1500
1000
500
0
10
5
0
−5
−10
−20
−10
0
10
20

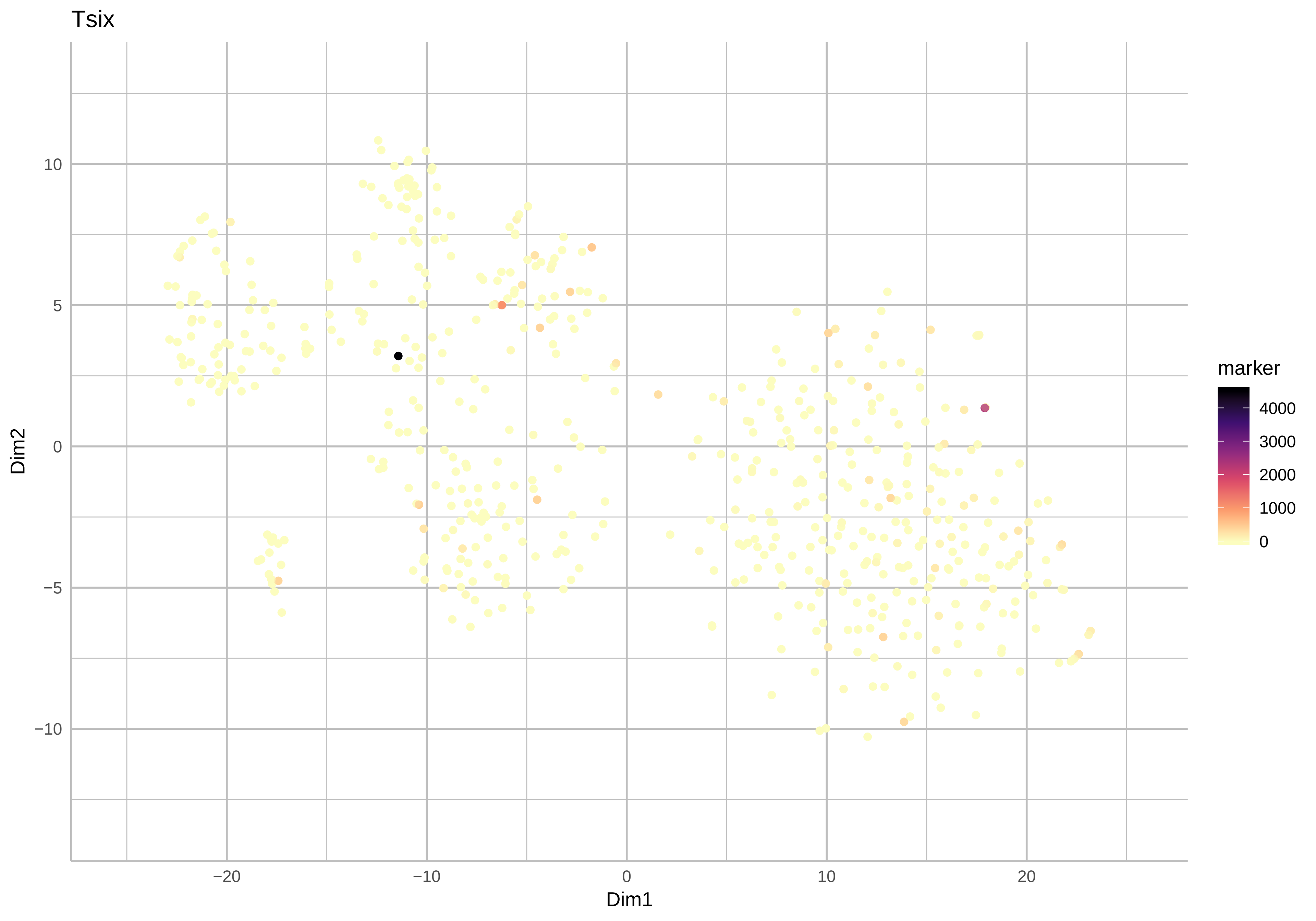

Tsix
Dim2
Dim1
marker
4000
3000
2000
1000
0
10
5
0
−5
−10
−20
−10
0
10
20

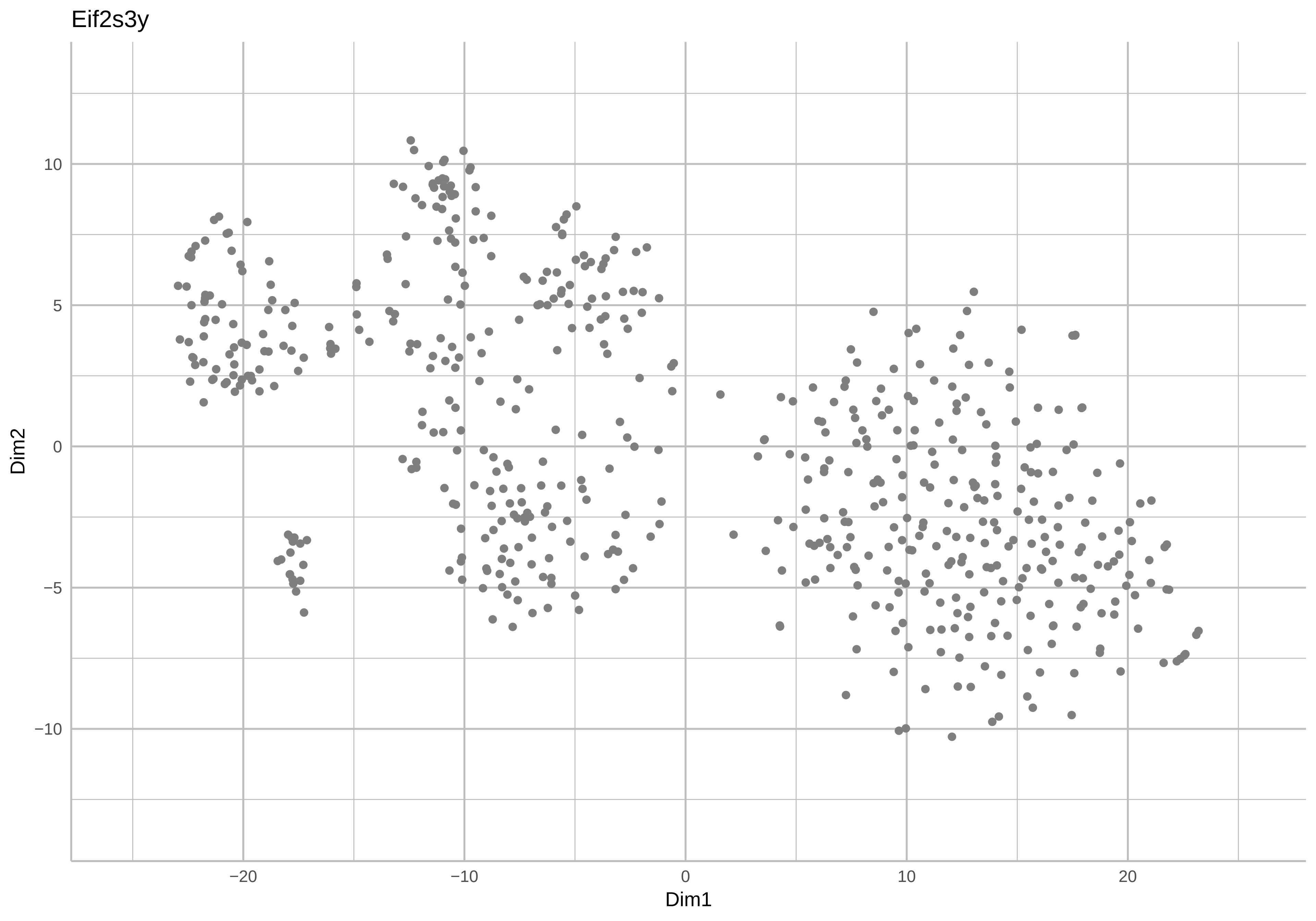
Eif2s3y
Dim2
10
5
0
−5
−10
−20
−10
0
10
20
Dim1

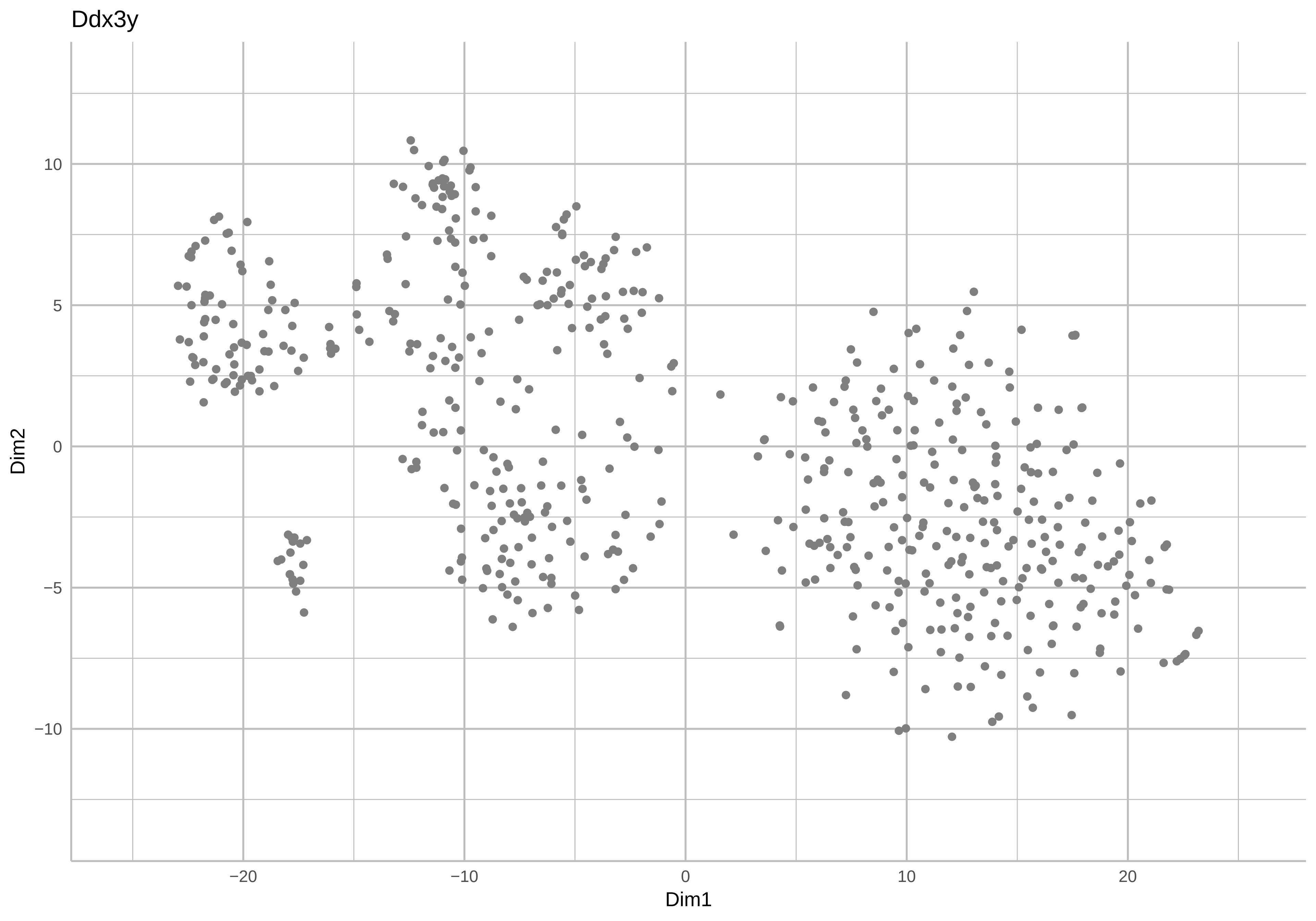
Ddx3y
10
5
0
−5
−10
Dim2
−20
−10
0
10
20
Dim1

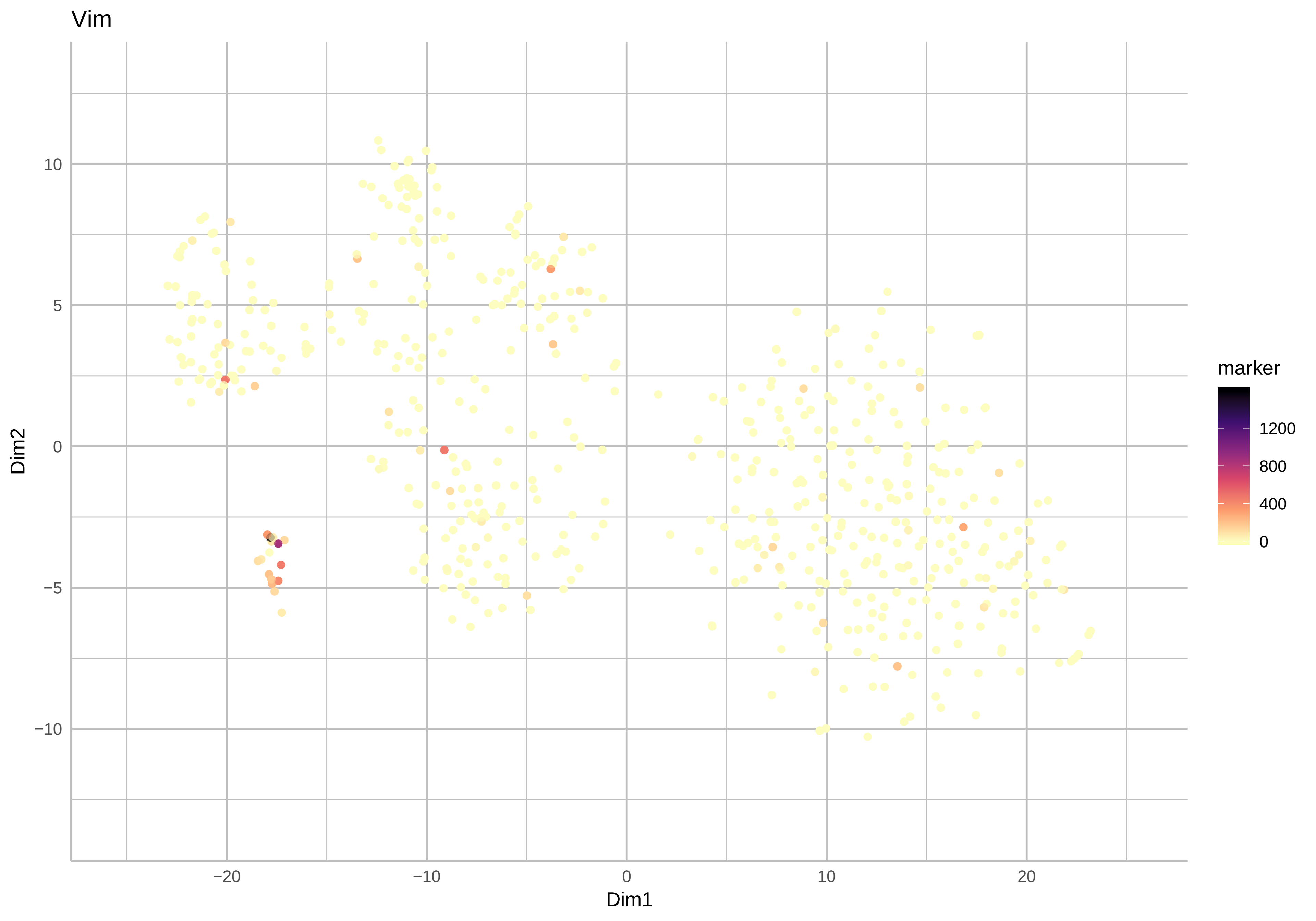
Vim
Dim2
Dim1
10
5
0
−5
−10
−20
−10
0
10
20
marker
1200
800
400
0

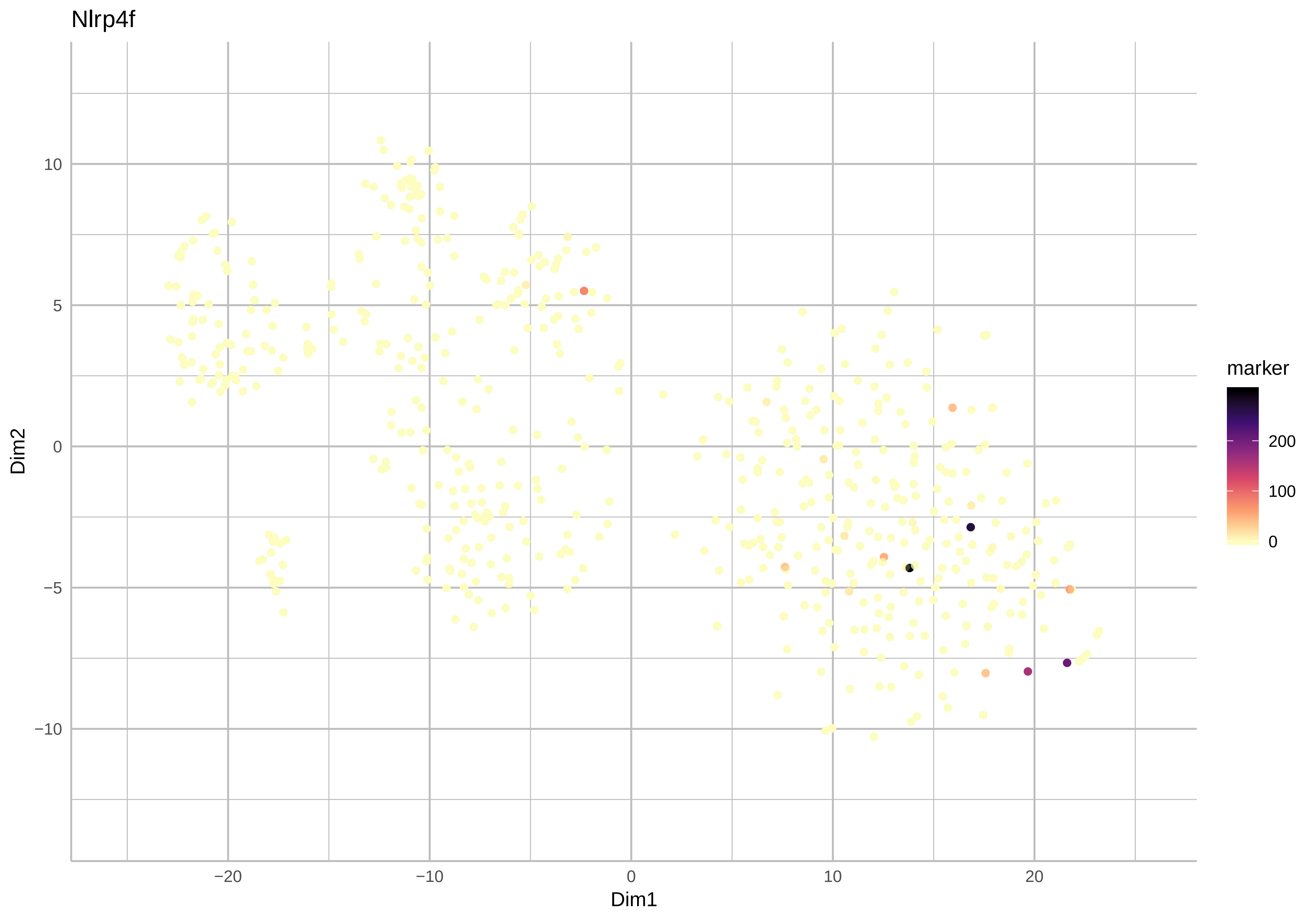

Nlrp4f
Dim2
Dim1
10
5
0
−5
−10
−20
−10
0
10
20
marker
200
100
0

Cd59a

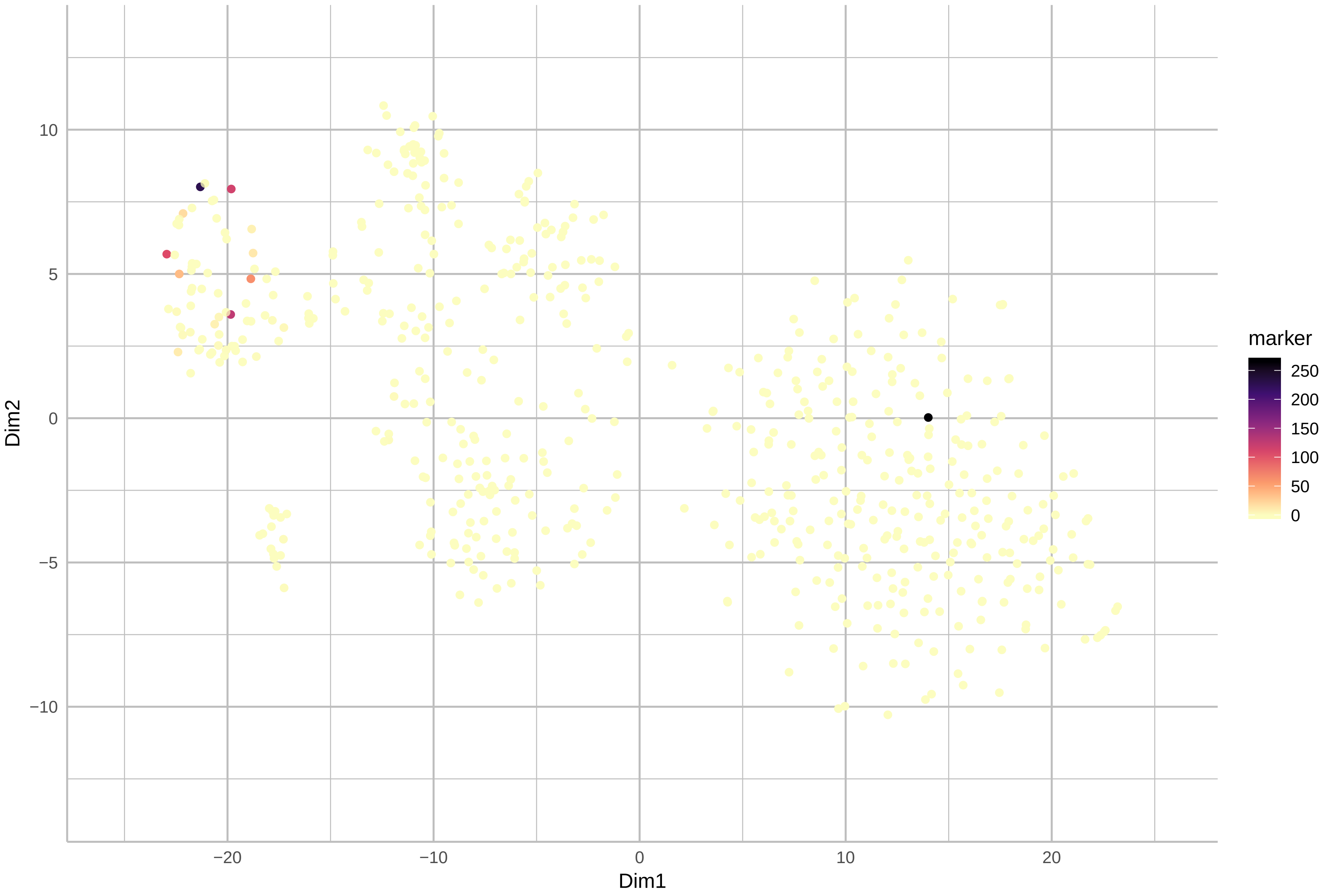

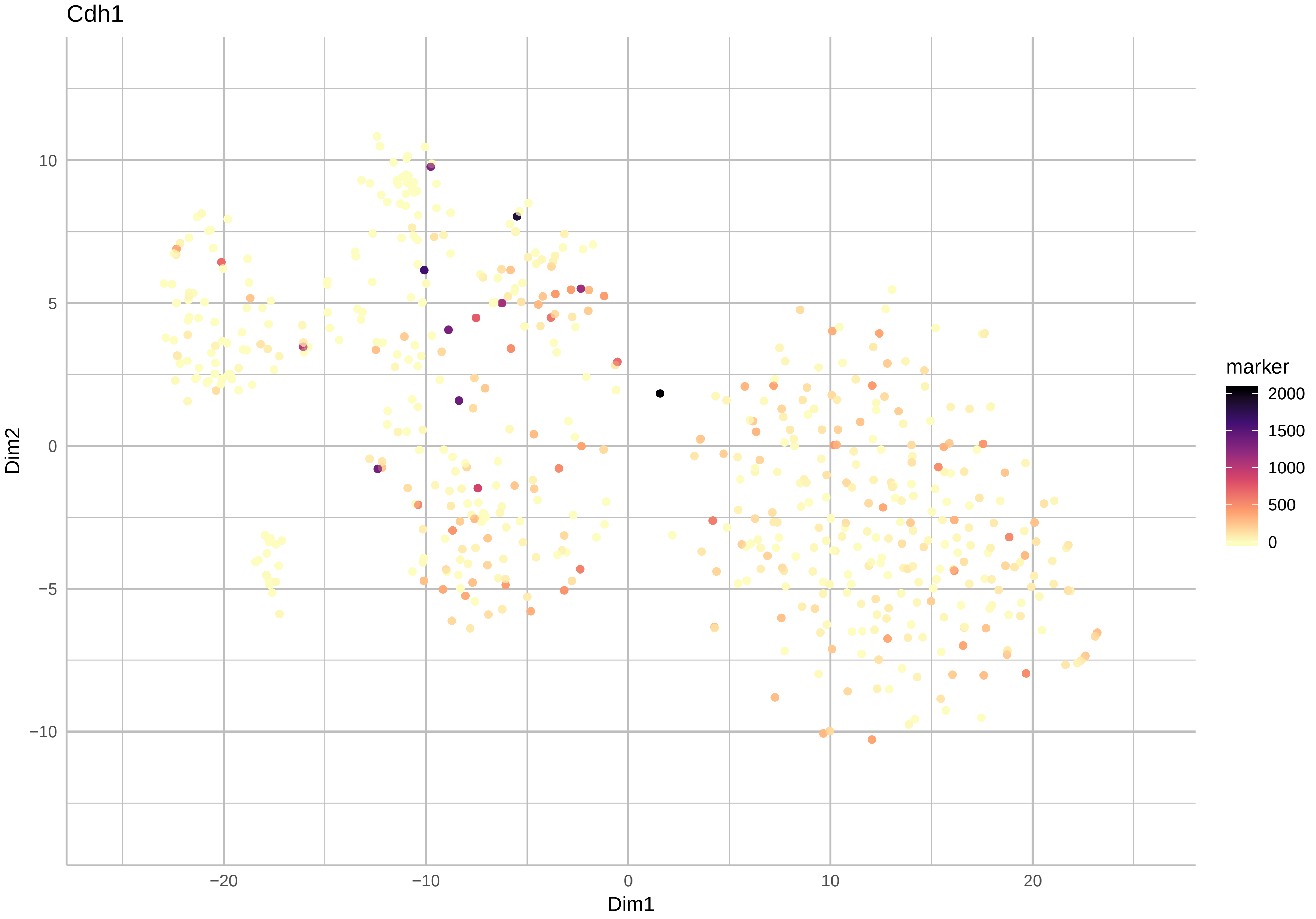

Cdh1
10
5
0
−5
−10
−20
−10
0
10
20
Dim1
Dim2
marker
2000
1500
1000
500
0

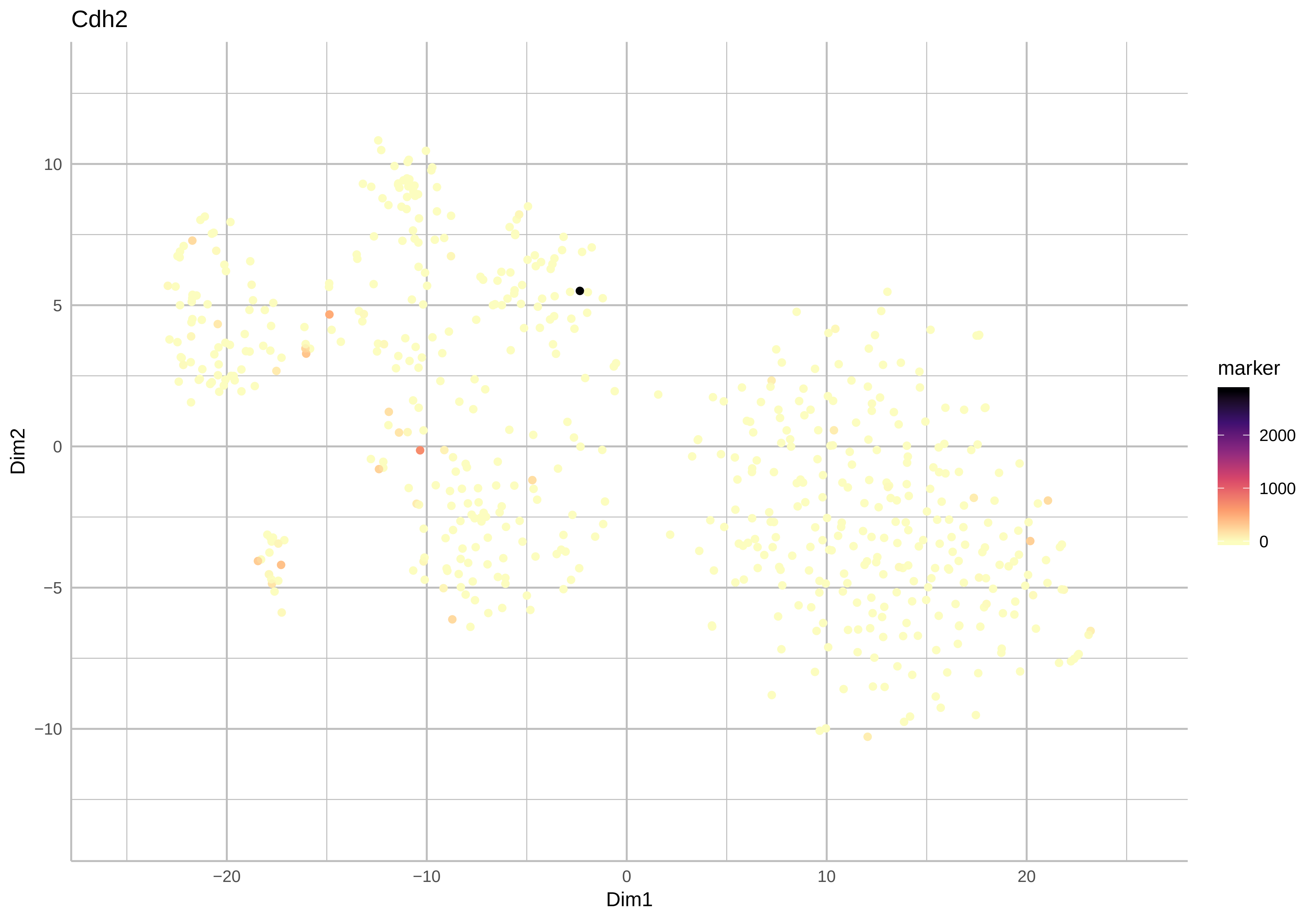

Cdh2
Dim2
Dim1
10
5
0
−5
−10
−20
−10
0
10
20
marker
2000
1000
0

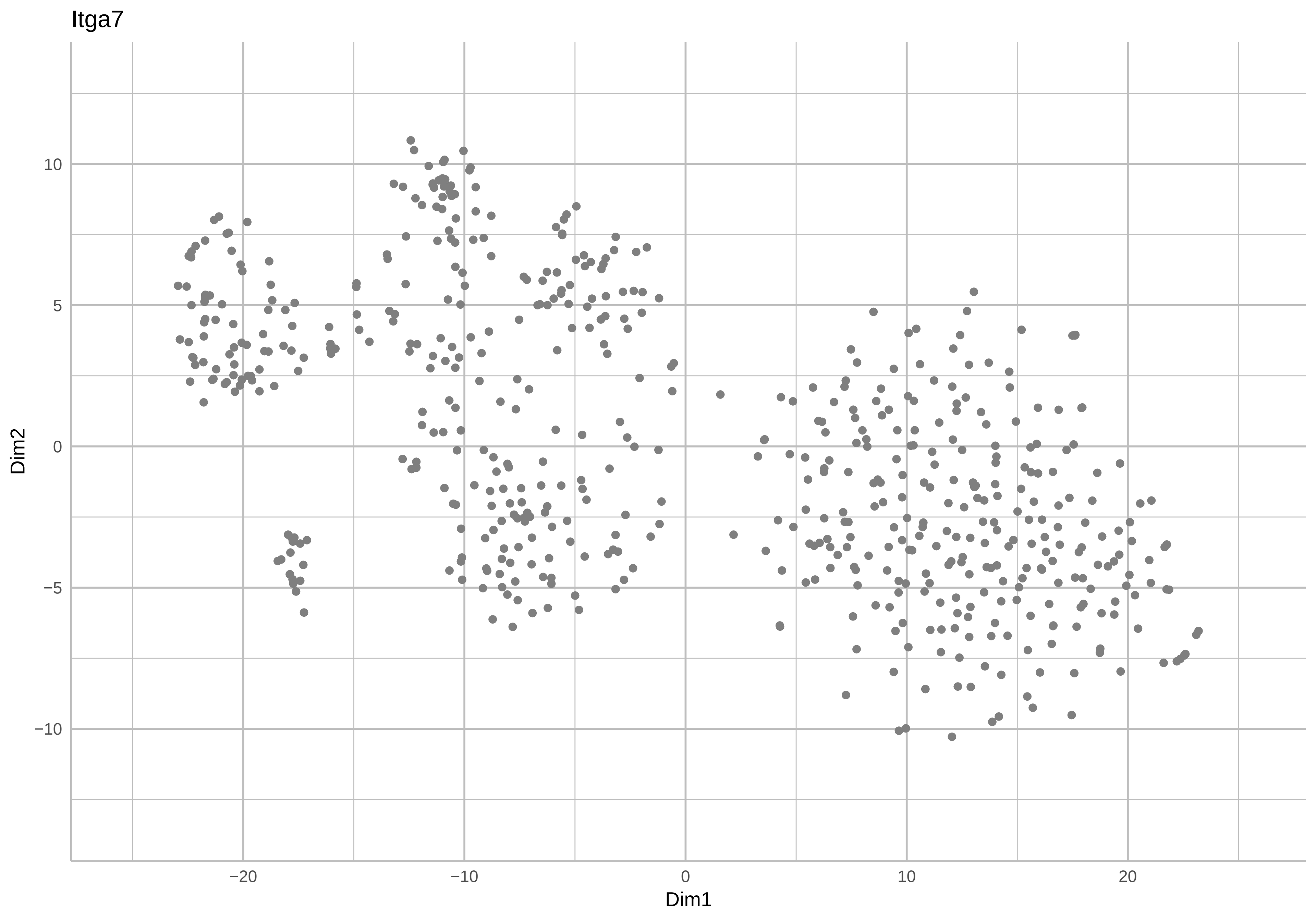
Itga7
Dim2
Dim1
10
5
0
−5
−10
−20
−10
0
10
20

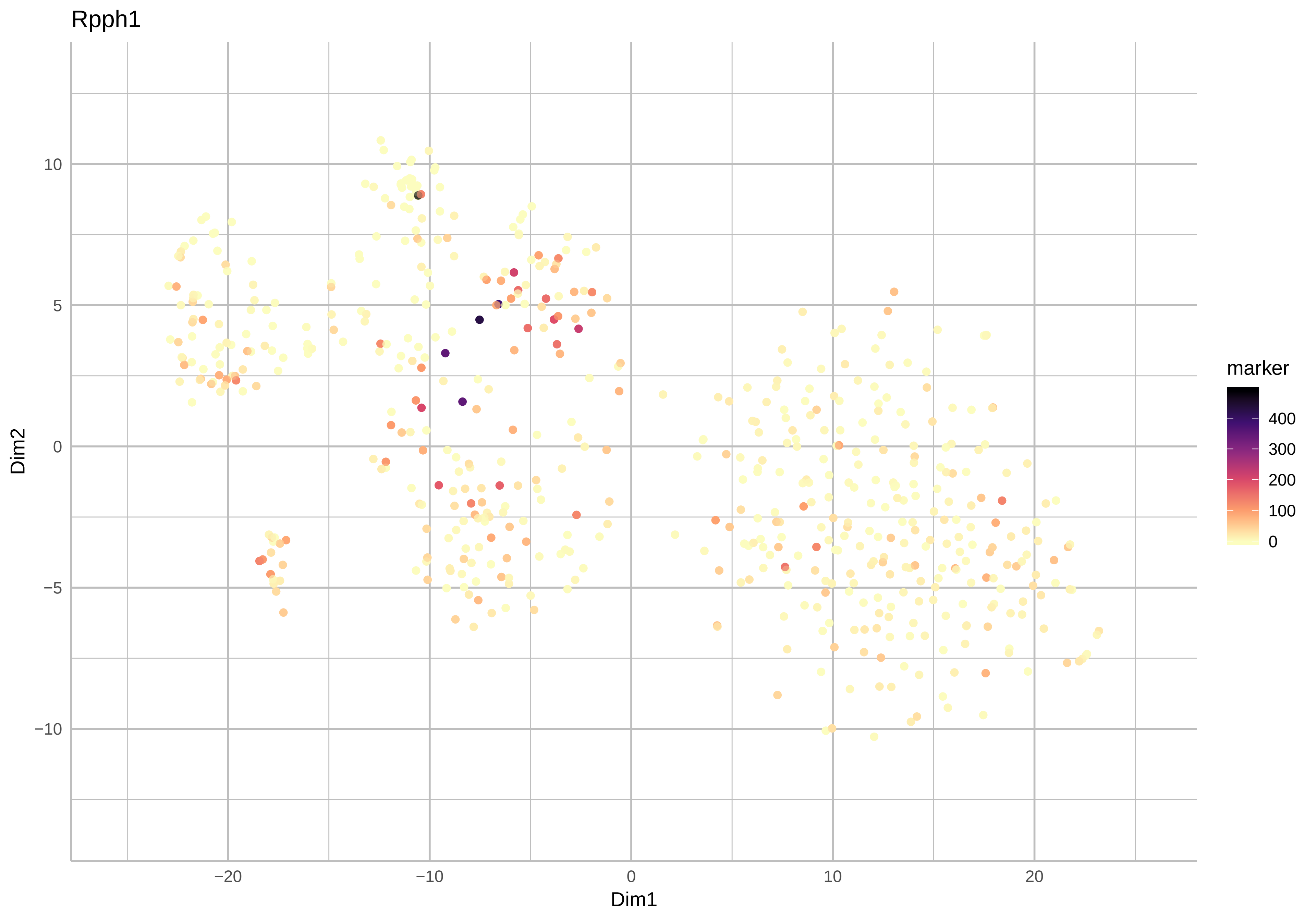
Rpph1
Dim2
Dim1
marker
400
300
200
100
0
10
5
0
−5
−10
−20
−10
0
10
20

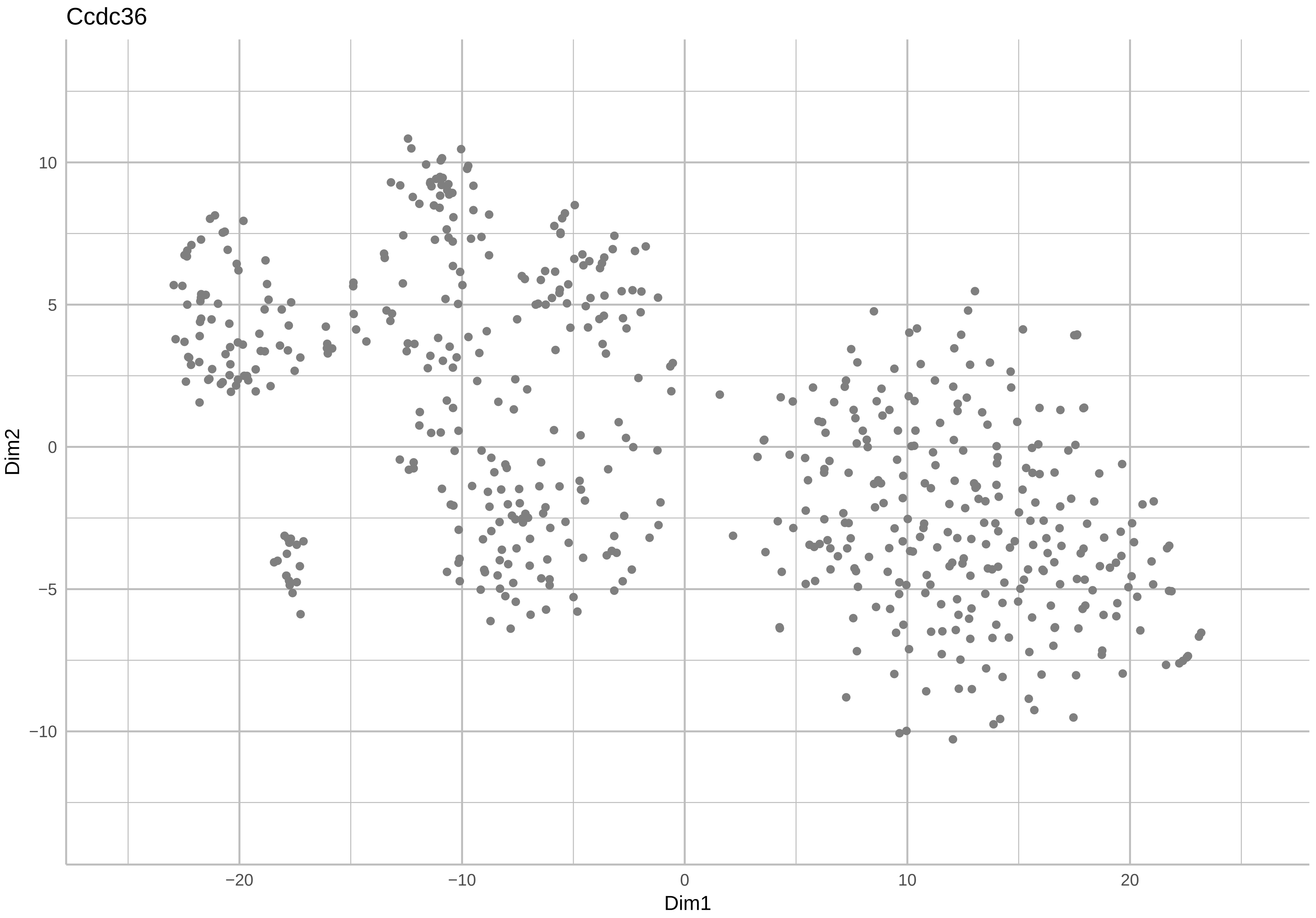
Ccdc36
Dim2
Dim1
10
5
0
−5
−10
−20
−10
0
10
20

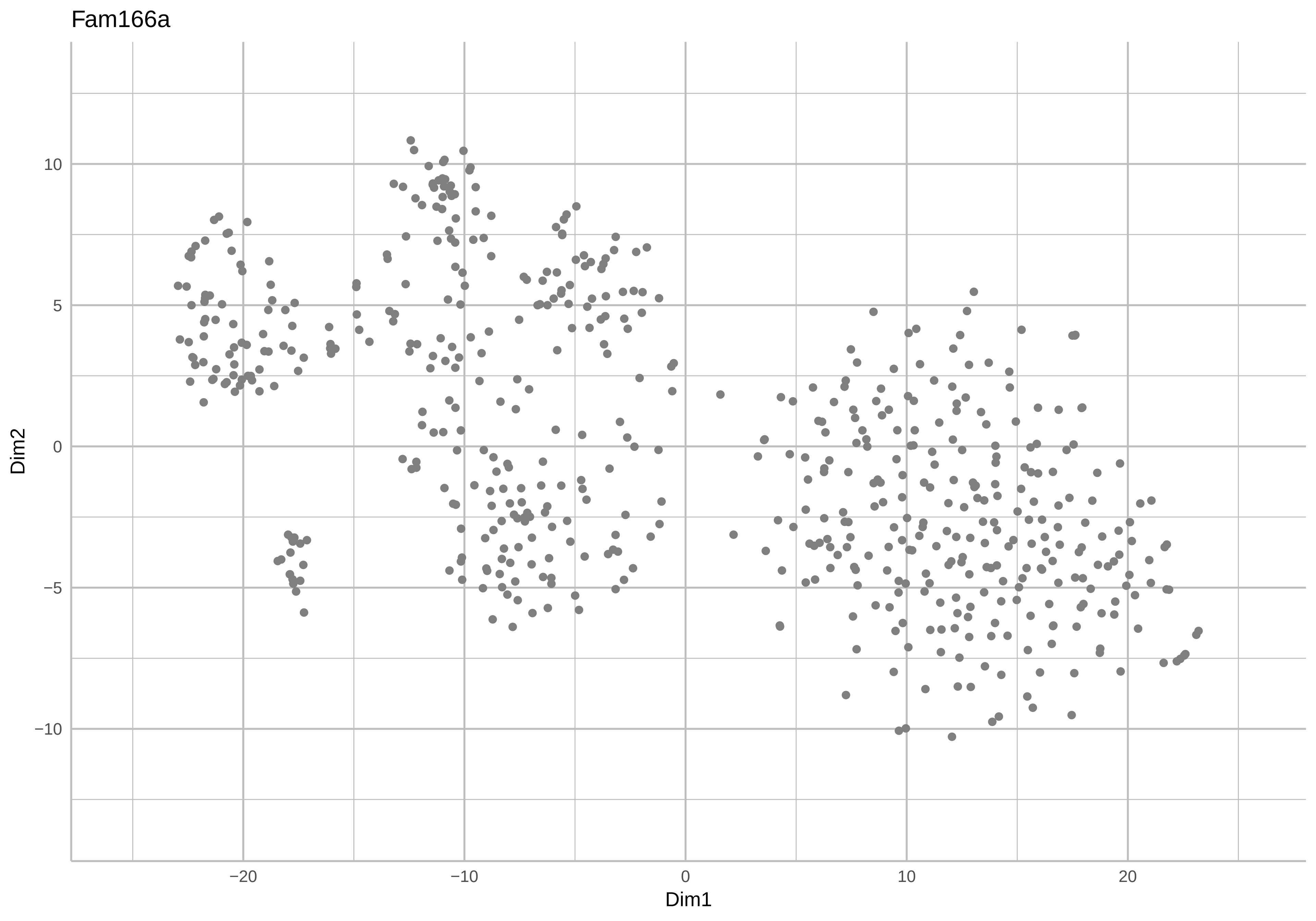
Fam166a
Dim2
10
5
0
−5
−10
−20
−10
0
10
20
Dim1

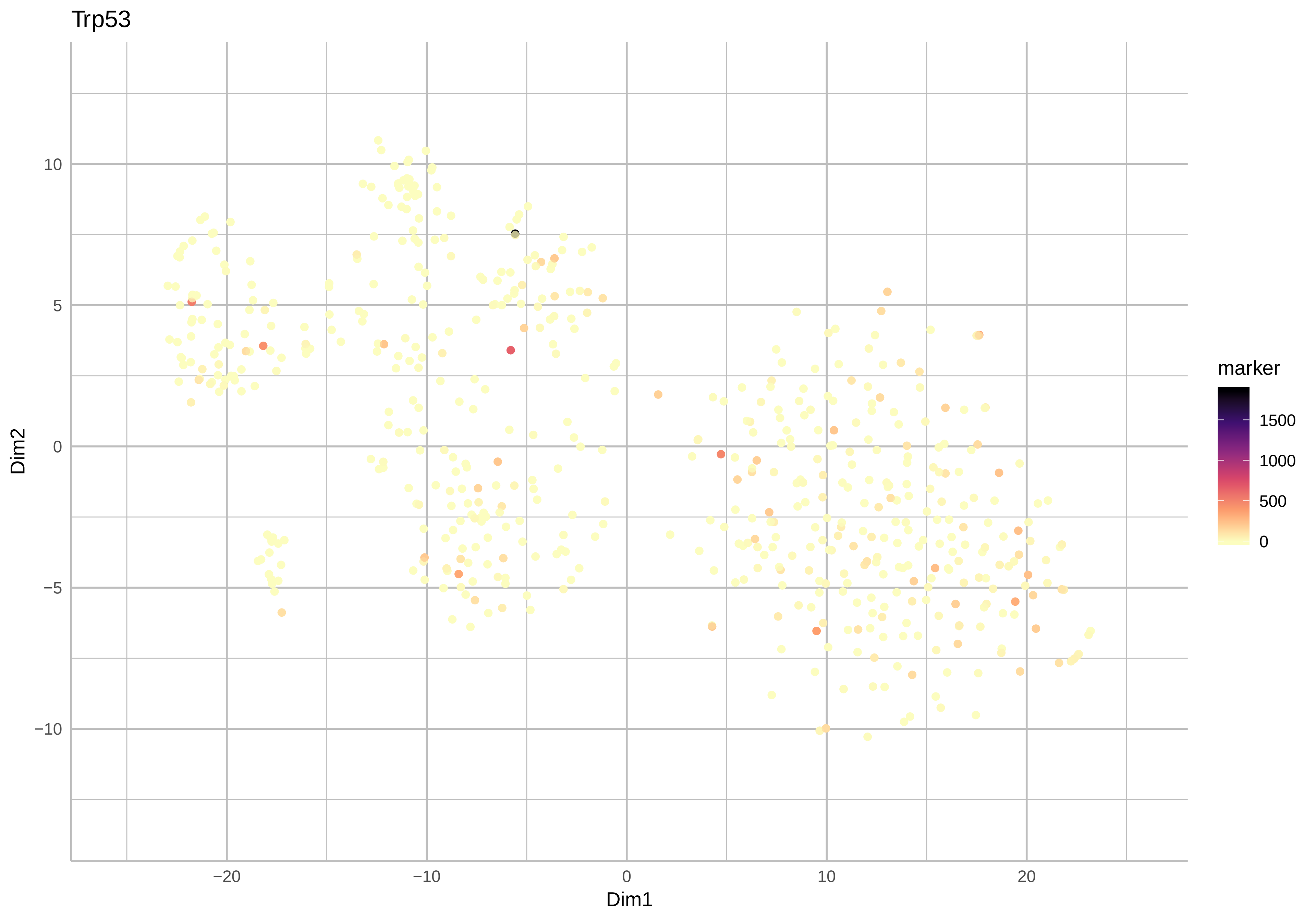
Trp53
10
5
0
−5
−10
−20
−10
0
10
20
Dim1
Dim2
marker
1500
1000
500
0

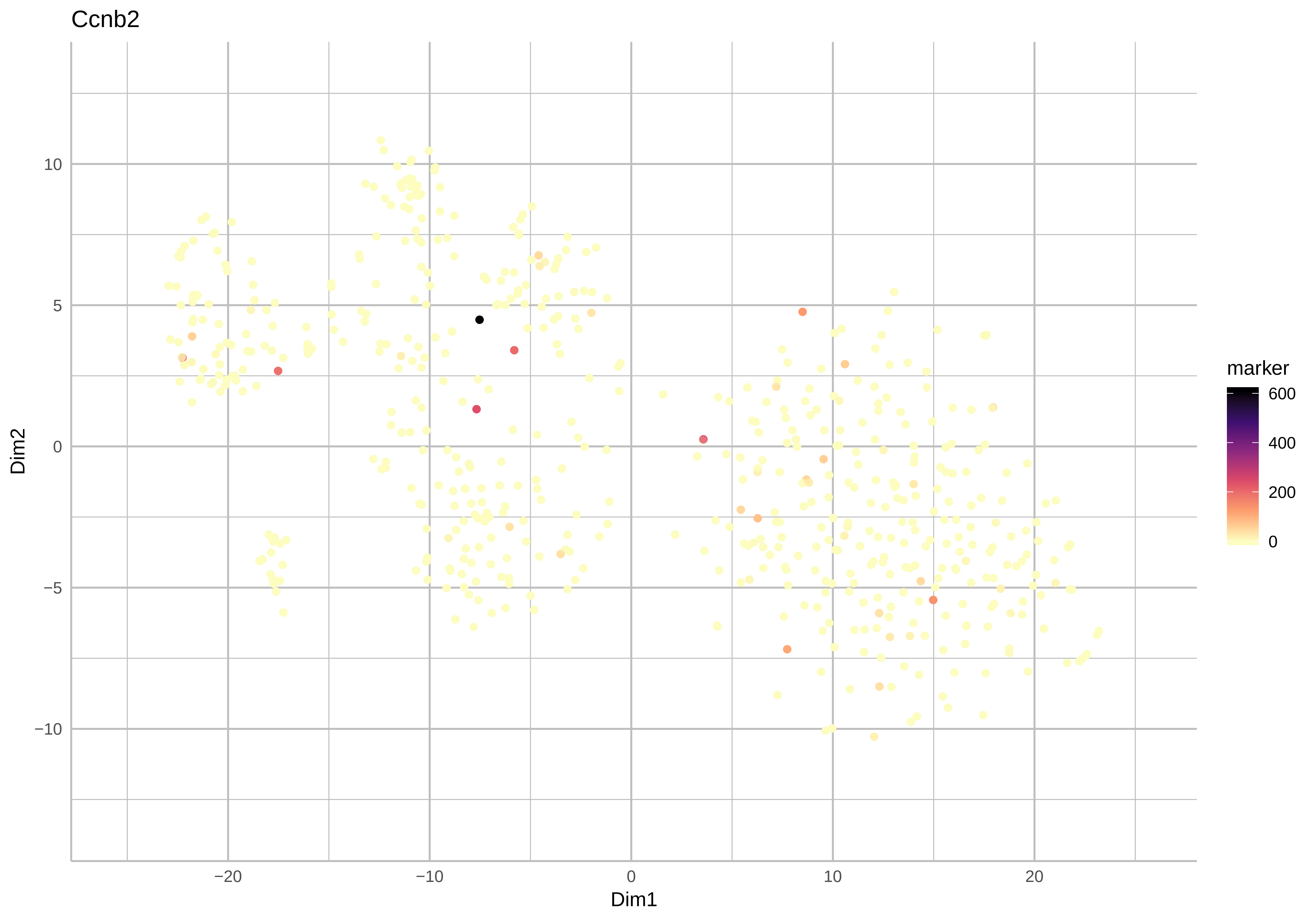
Ccnb2
Dim2
10
5
0
−5
−10
−20
−10
0
10
20
Dim1
marker
600
400
200
0

Tmem263

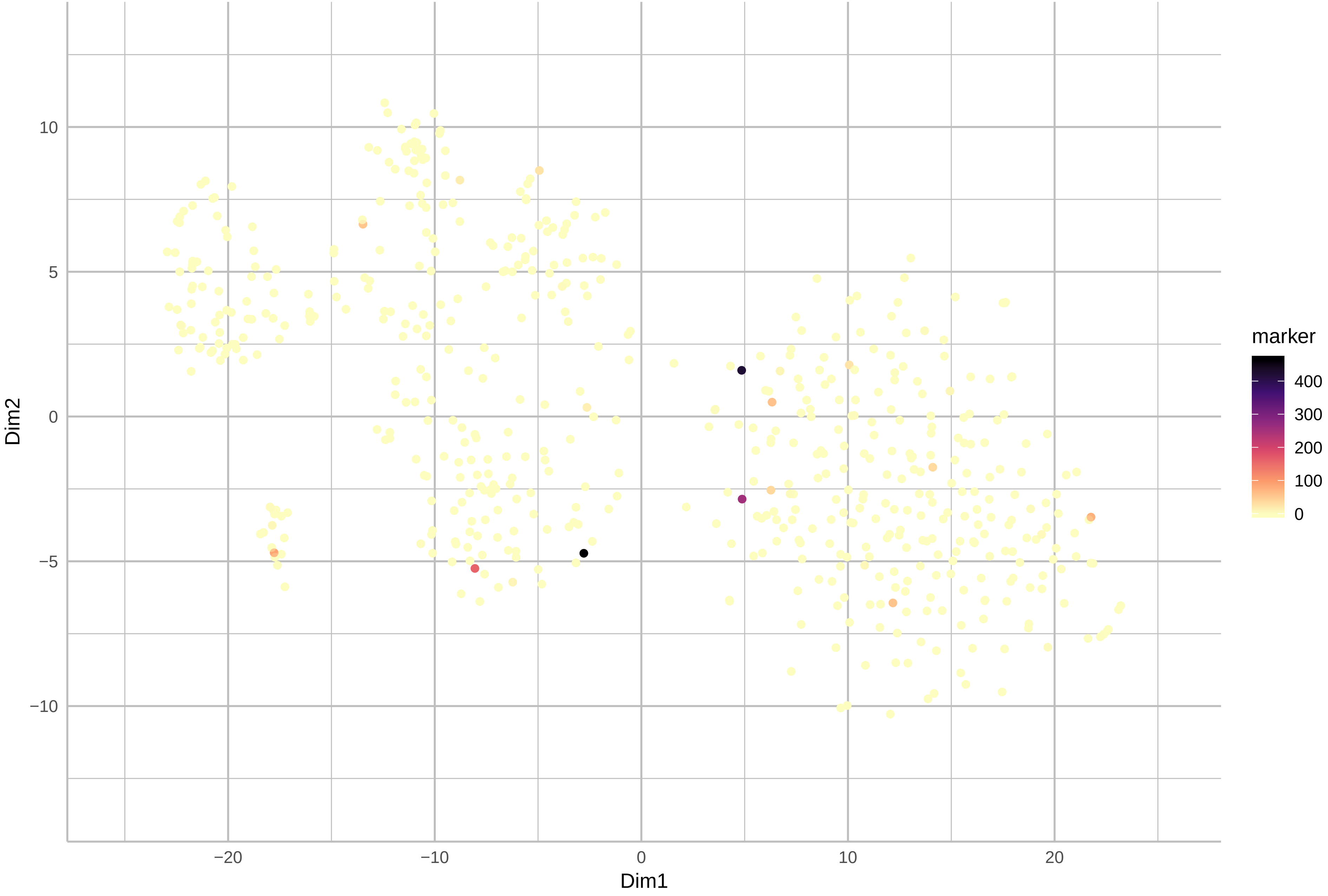
