## Supplementary material for "Single-cell transcriptomics, scRNA-Seq and C1 CAGE discovered distinct phases of pluripotency during naïve-to-primed conversion in mice": tSNEheatscaleCustomGenesKallisto_20190317200828.pdf

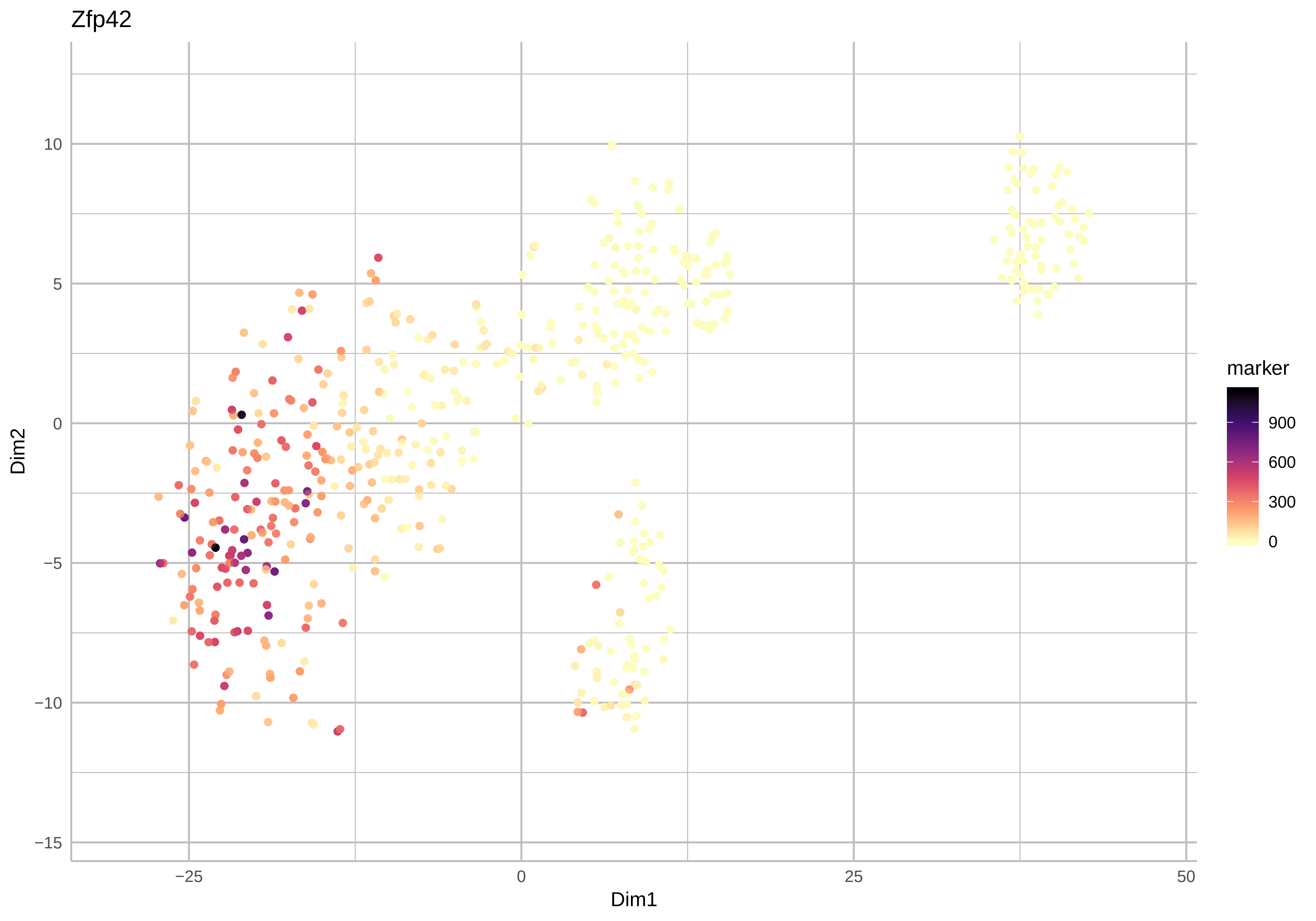

Zfp42
Dim2
Dim1
10
5
0
−5
−10
−15
−25
0
25
50
marker
900
600
300
0

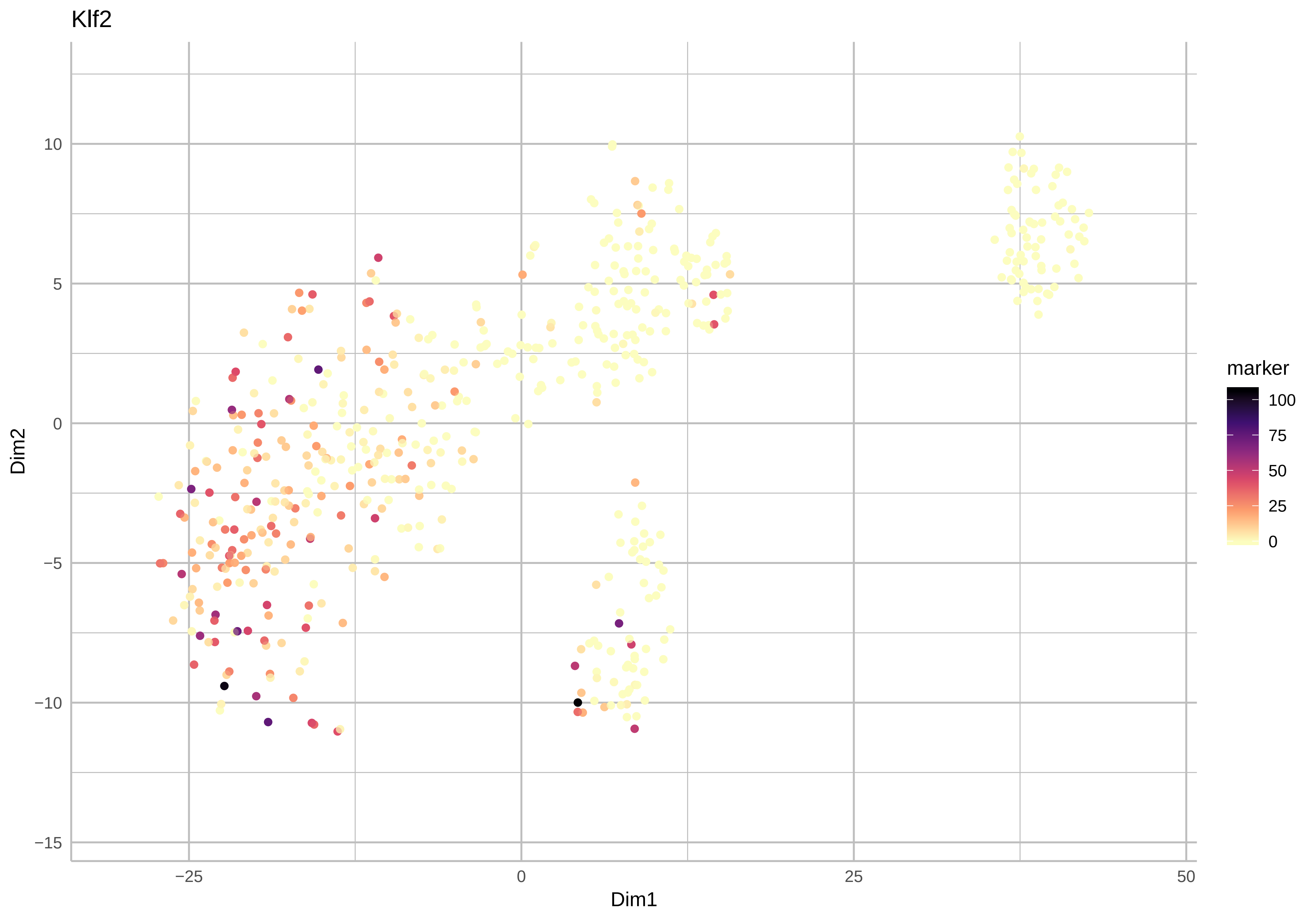

Klf2
Dim2
Dim1
10
5
0
−5
−10
−15
−25
0
25
50
marker
100
75
50
25
0

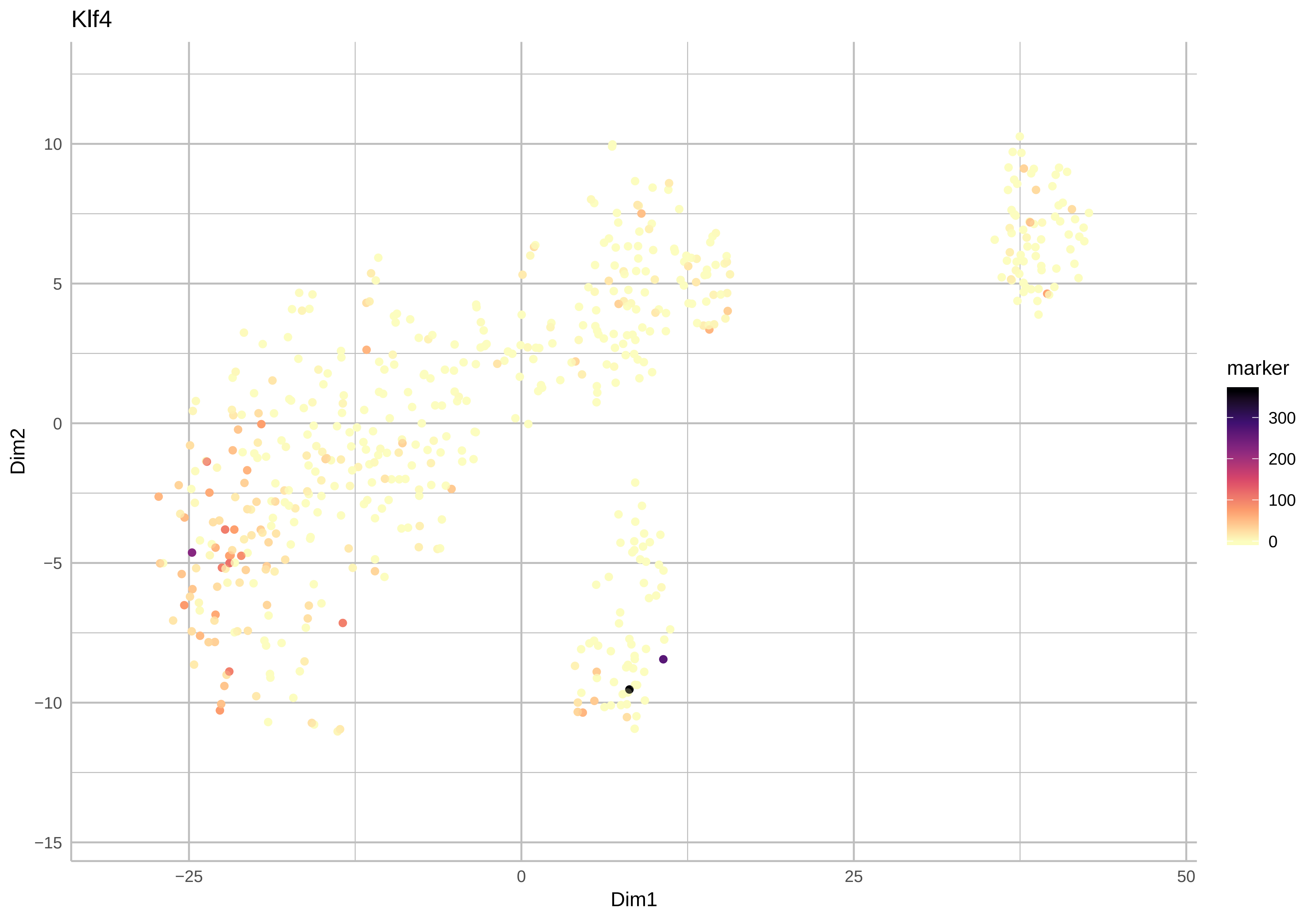
Klf4
Dim2
Dim1
10
5
0
−5
−10
−15
−25
0
25
50
marker
300
200
100
0

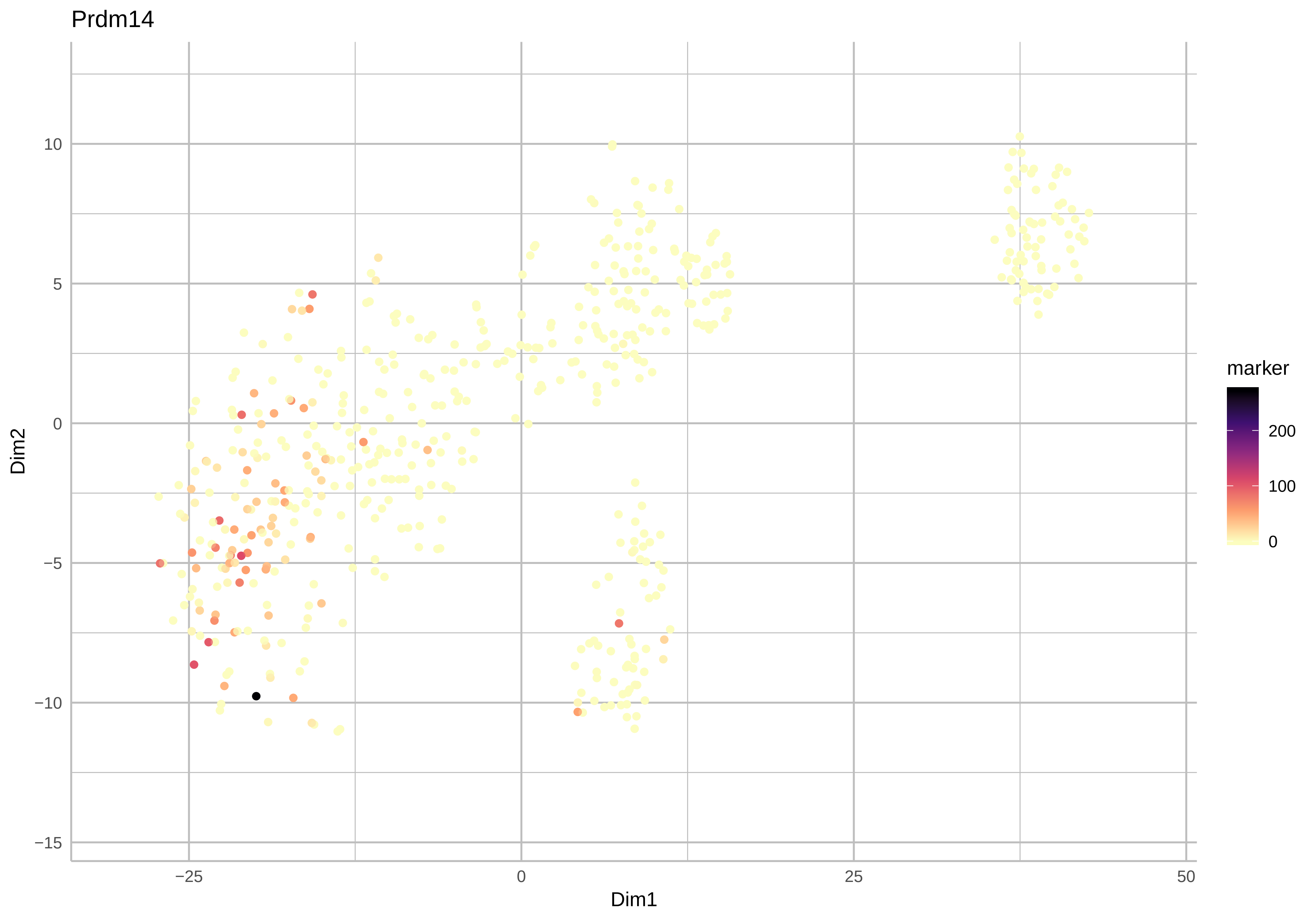
Prdm14
marker
200
100
0
Dim2
10
5
0
−5
−10
−15
Dim1
−25
0
25
50

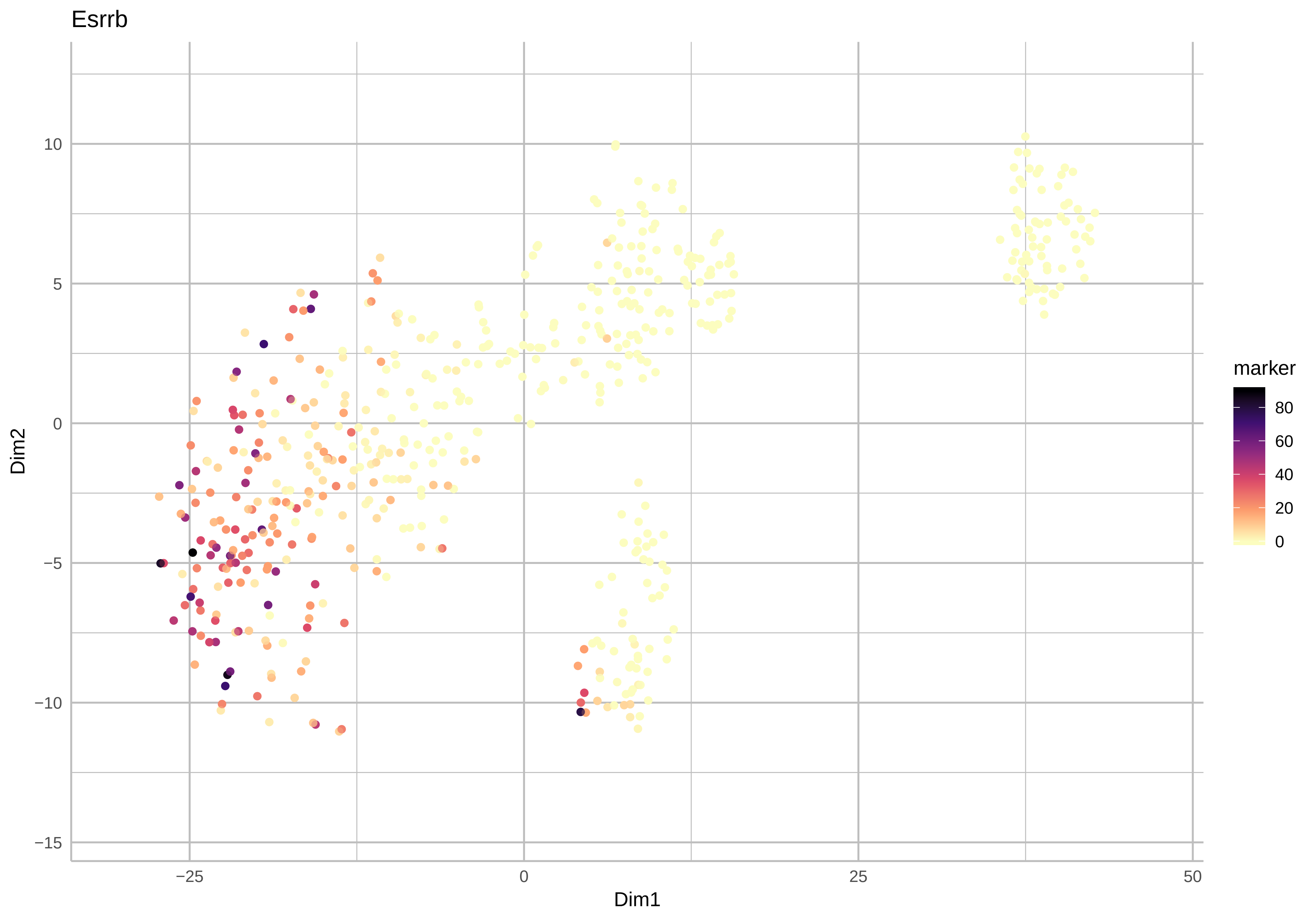
Esrrb
Dim2
Dim1
marker
80
60
40
20
0
10
5
0
−5
−10
−15
−25
0
25
50

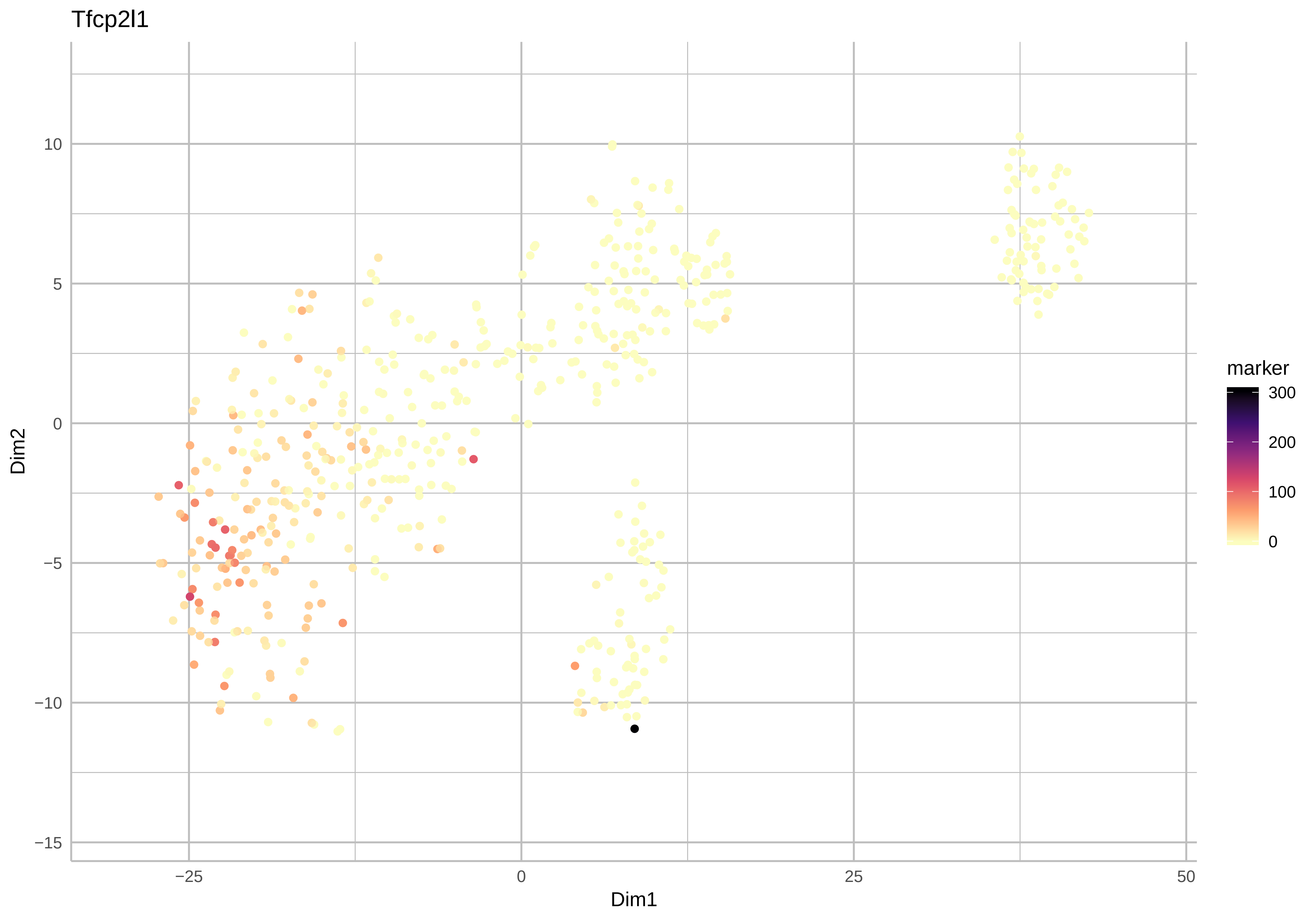
Tfcp2l1
Dim2
Dim1
10
5
0
−5
−10
−15
−25
0
25
50
marker
300
200
100
0

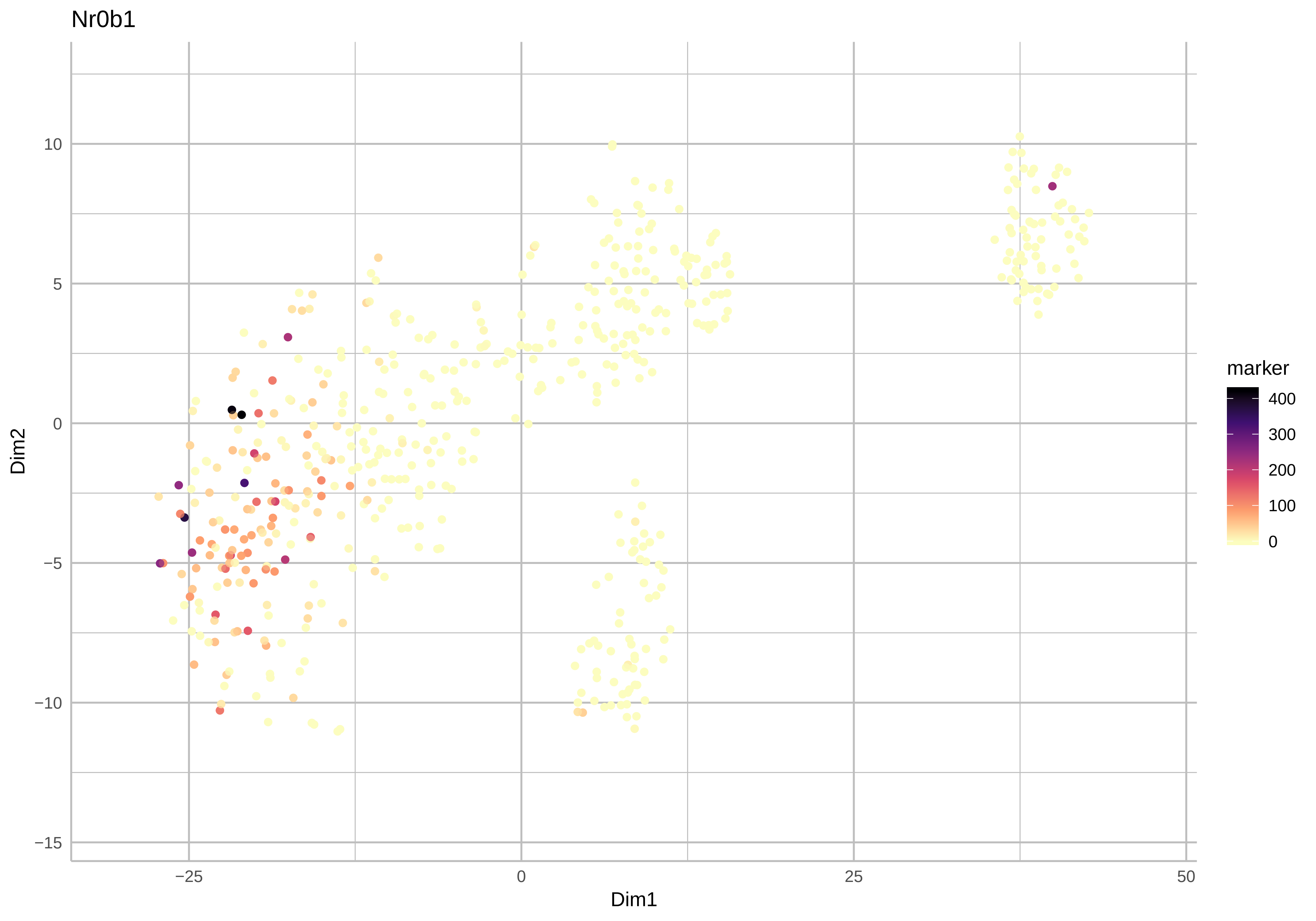
Nr0b1
Dim2
Dim1
10
5
0
−5
−10
−15
−25
0
25
50
marker
400
300
200
100
0

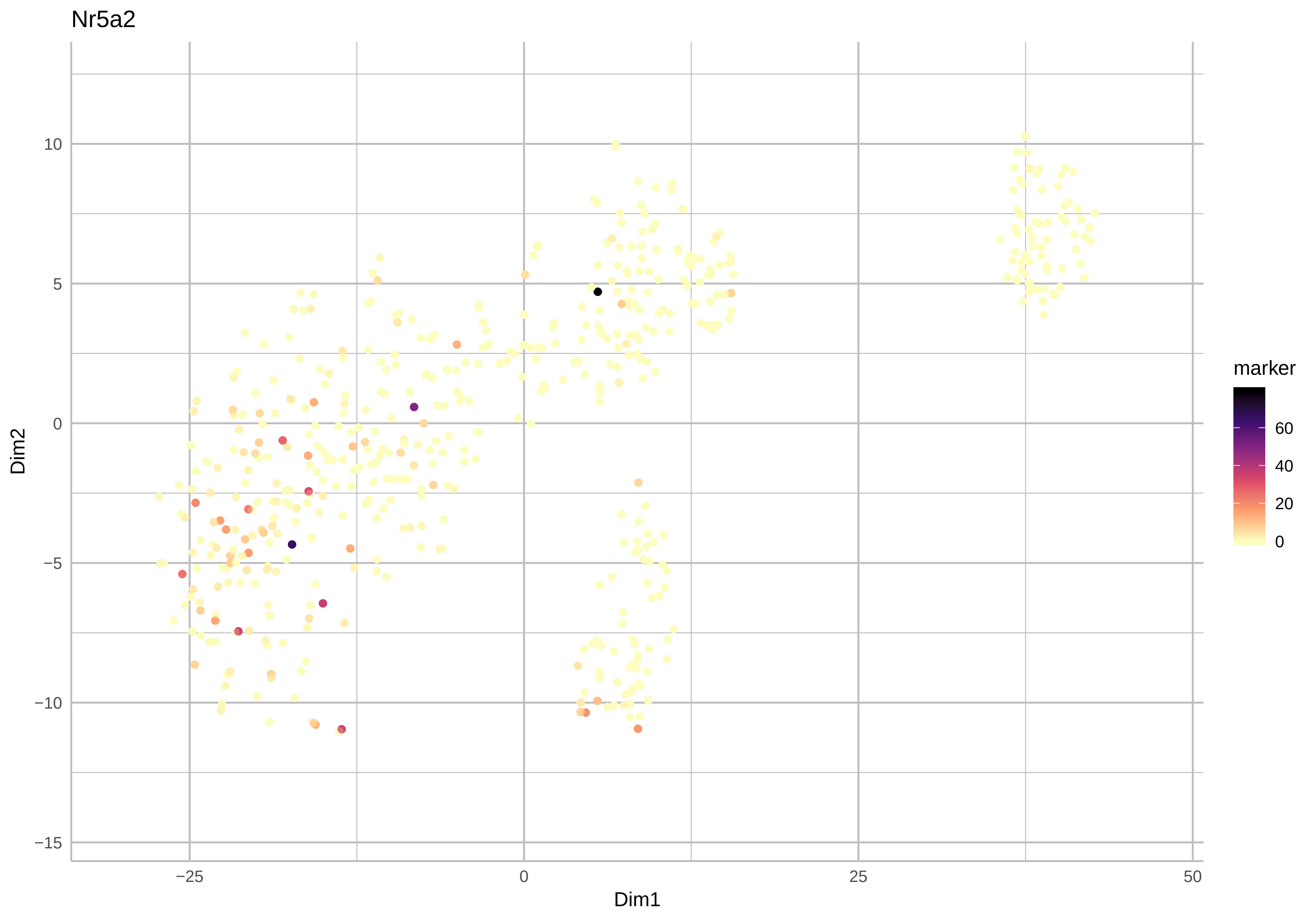
Nr5a2
Dim2
10
5
0
−5
−10
−15
−25
0
25
50
Dim1
marker
60
40
20
0

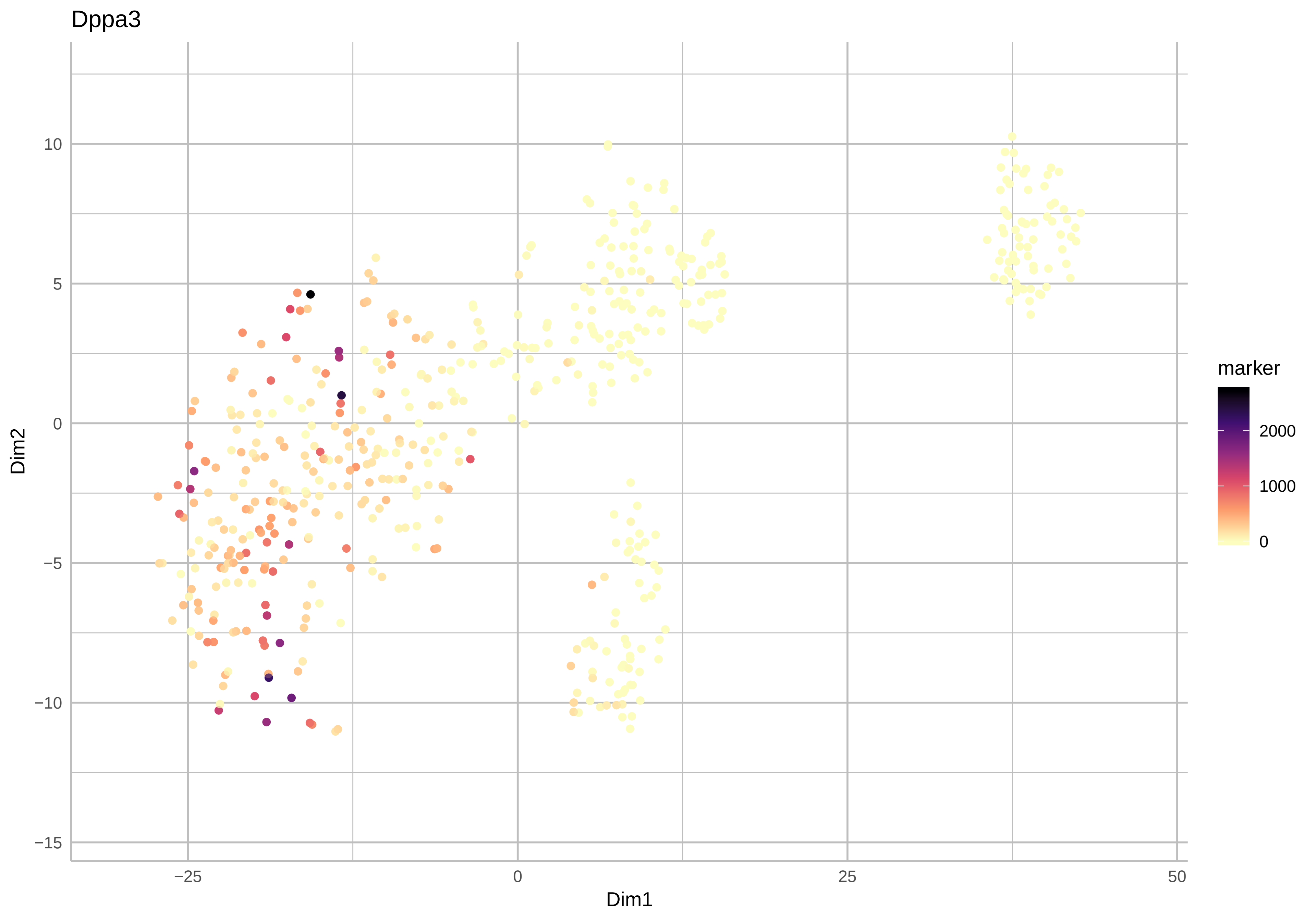
Dppa3
Dim2
Dim1
10
5
0
−5
−10
−15
−25
0
25
50
marker
2000
1000
0

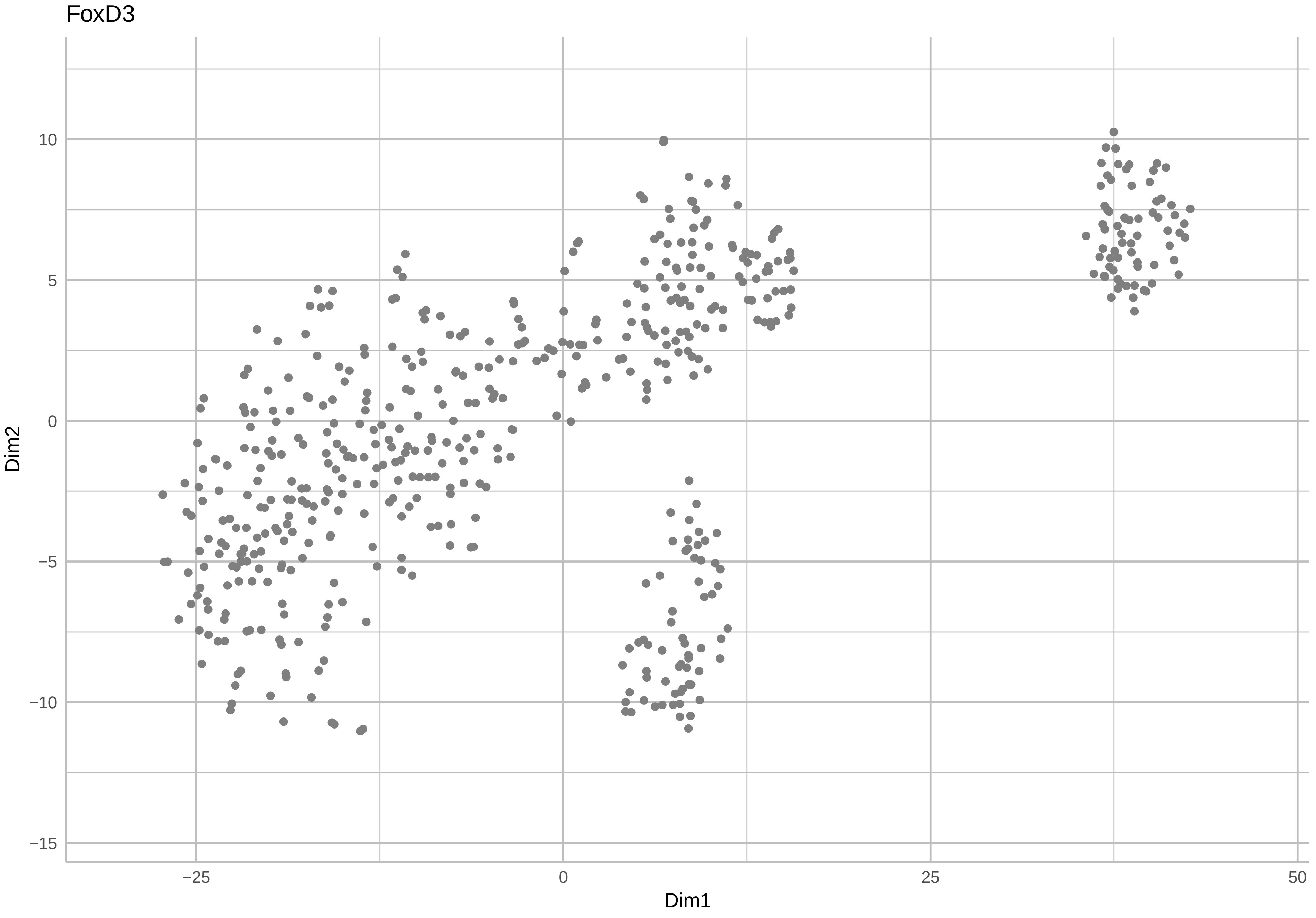
FoxD3
Dim2
Dim1
10
5
0
−5
−10
−15
−25
0
25
50

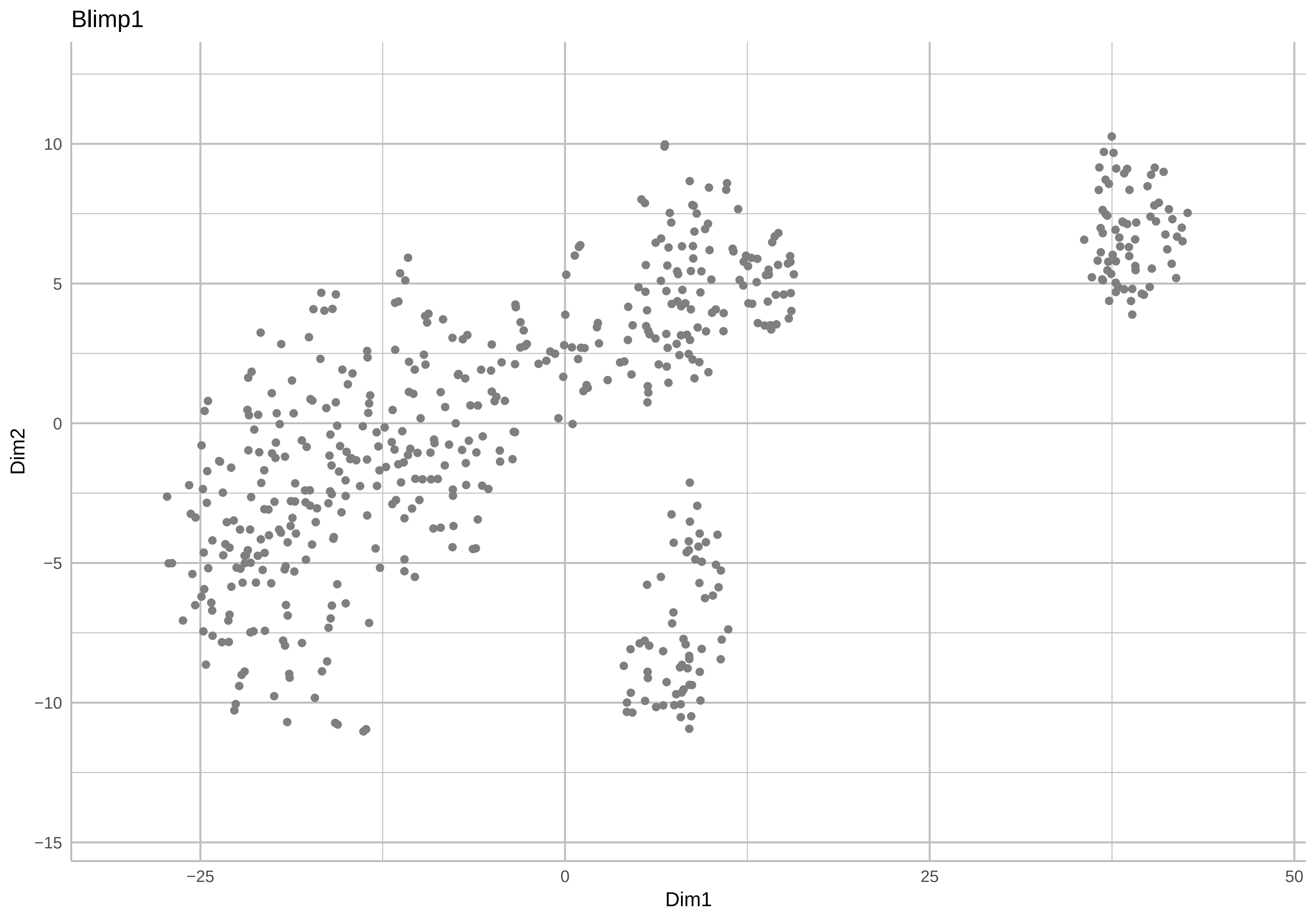
Blimp1
Dim2
Dim1
10
5
0
−5
−10
−15
−25
0
25
50

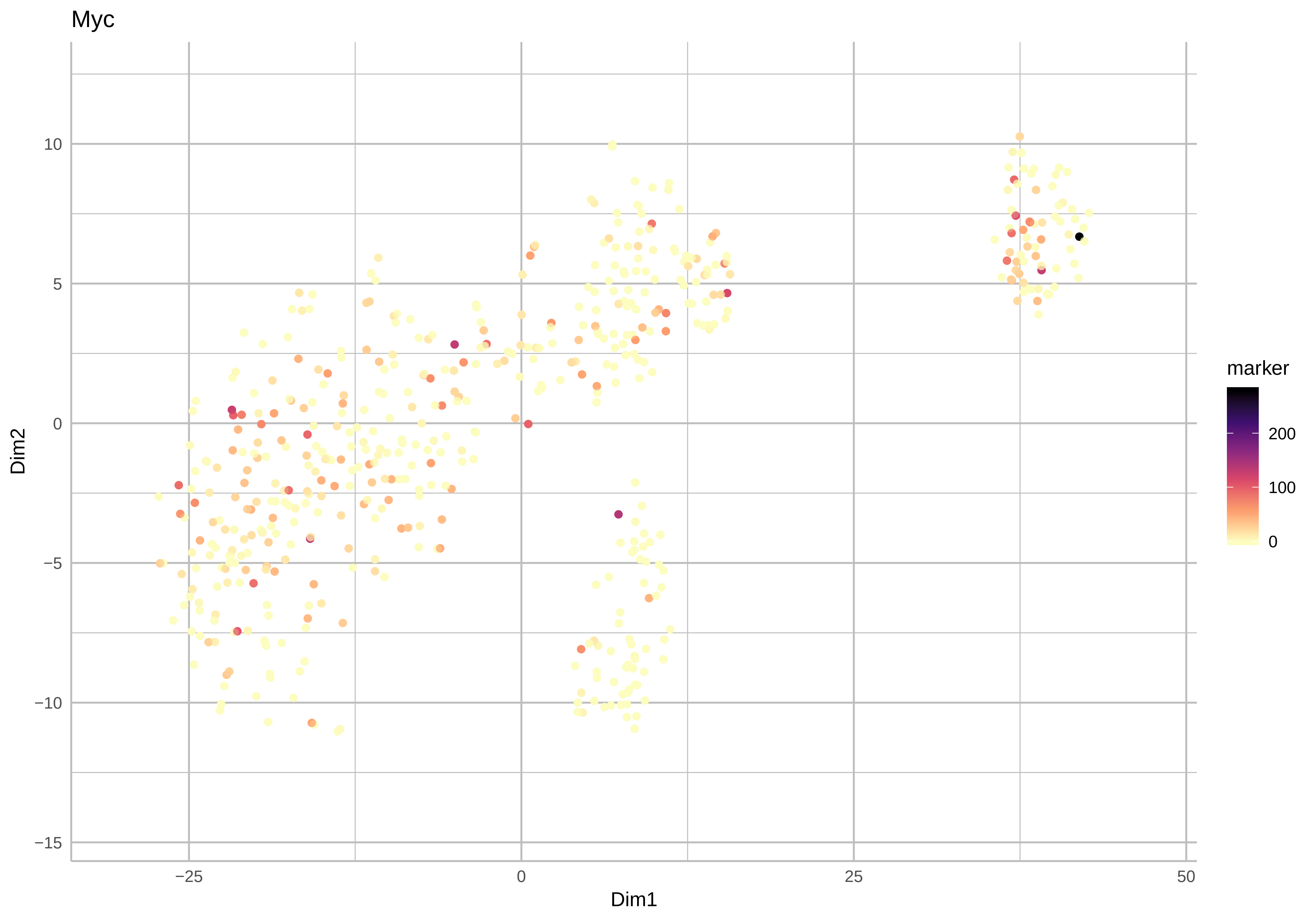

Myc
Dim2
Dim1
marker
200
100
0
10
5
0
−5
−10
−15
−25
0
25
50

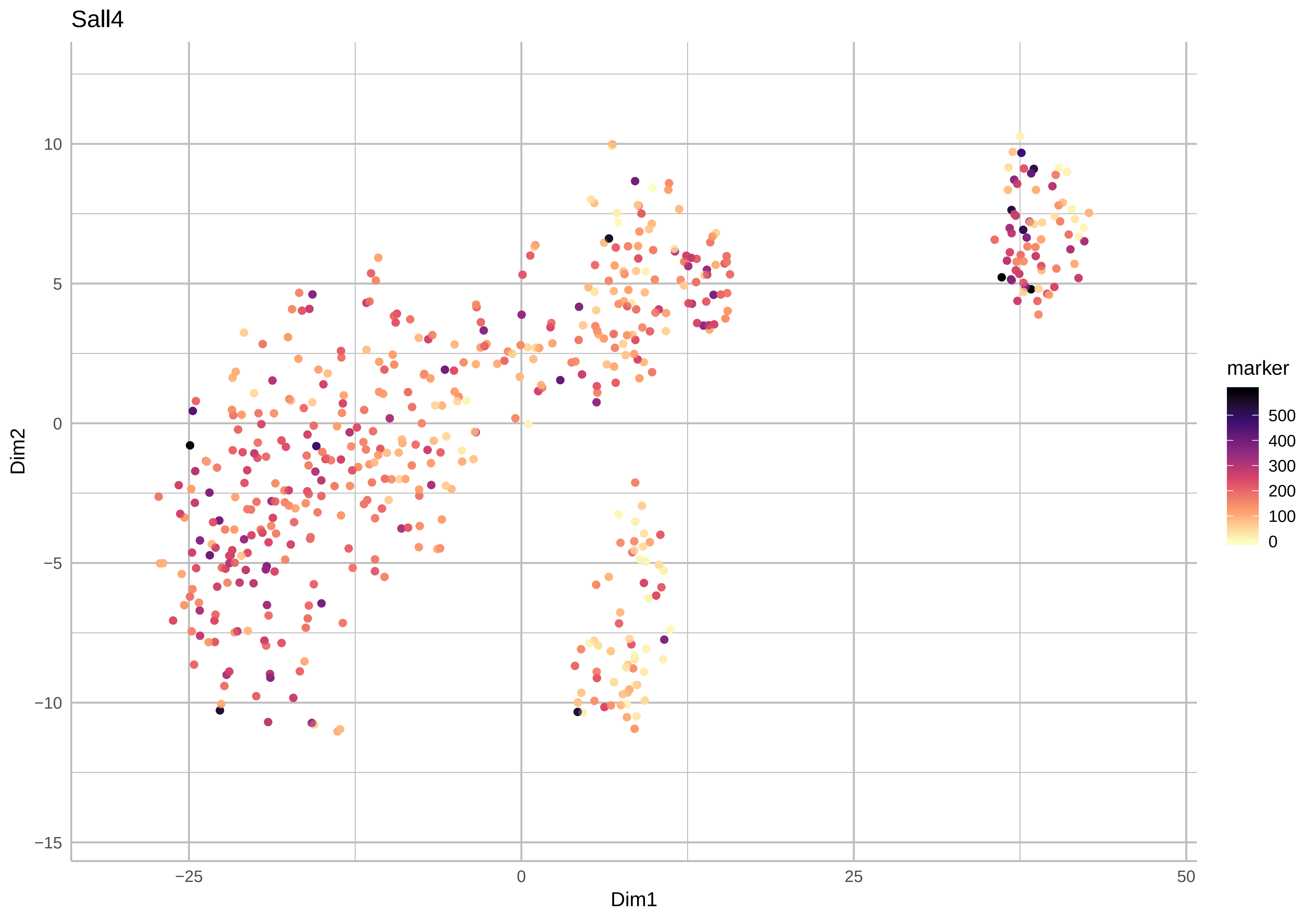

Sall4
10
5
0
−5
−10
−15
Dim2
−25
0
25
50
Dim1
marker
500
400
300
200
100
0

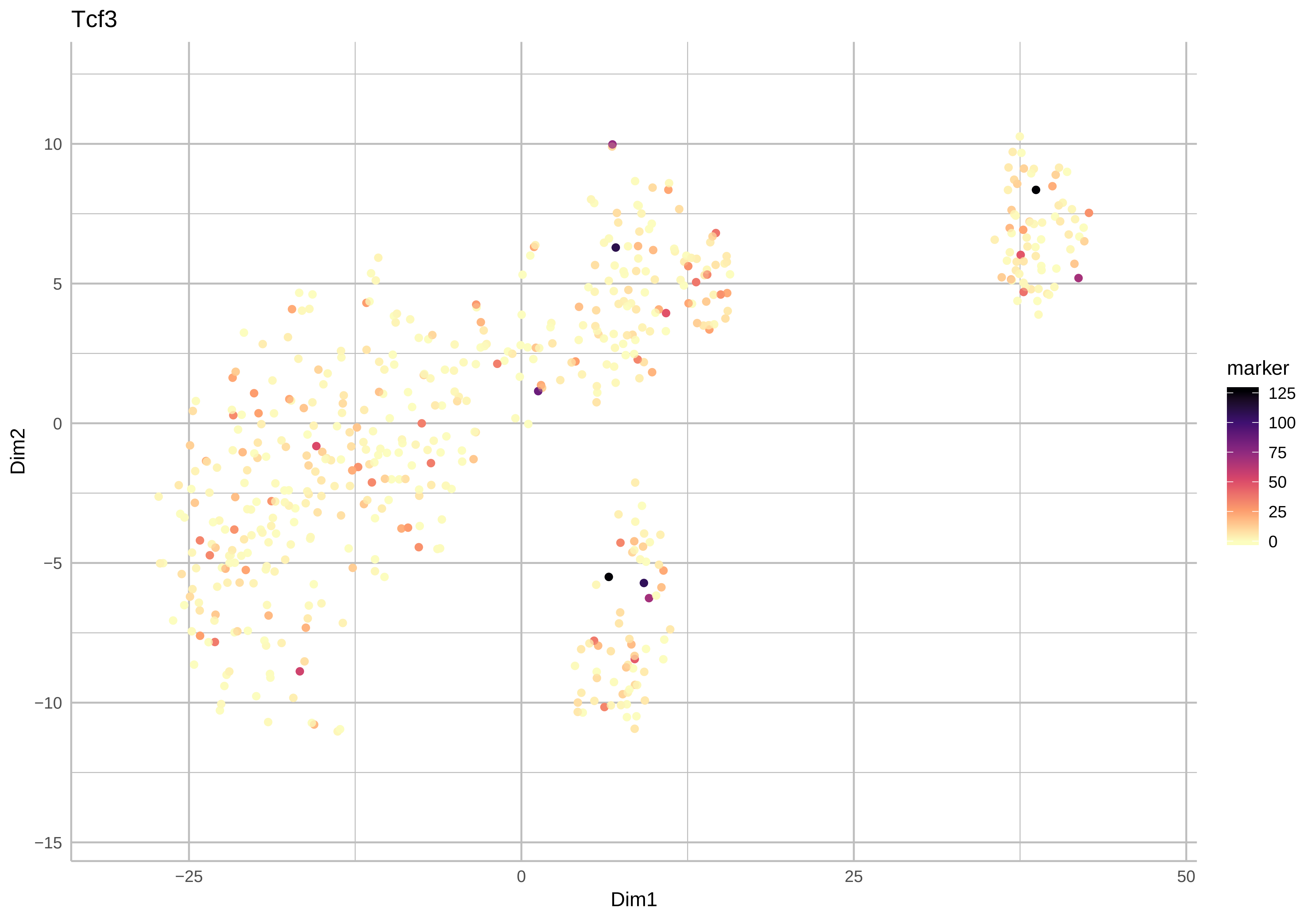
Tcf3
Dim2
Dim1
10
5
0
−5
−10
−15
−25
0
25
50
marker
125
100
75
50
25
0

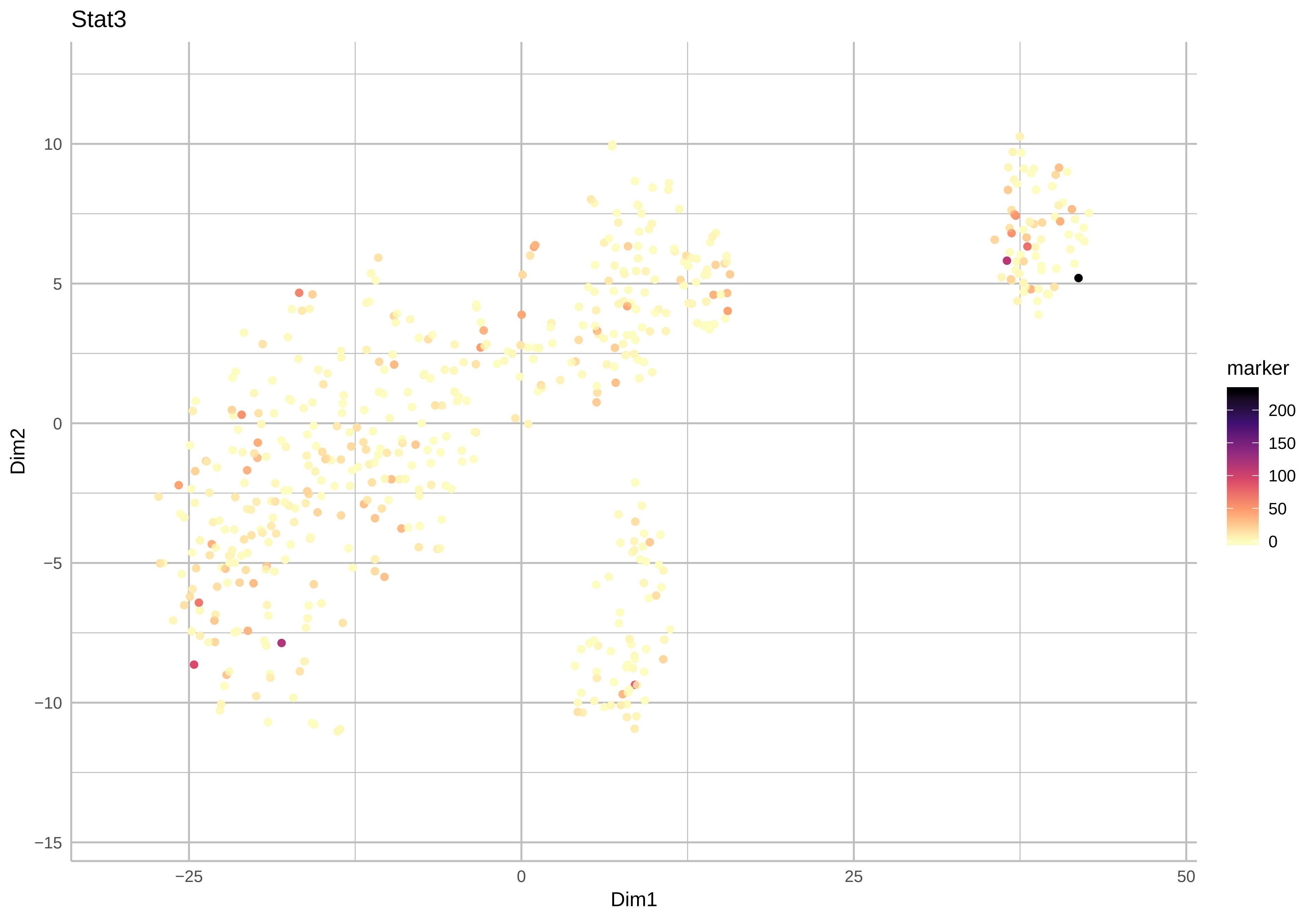

Stat3
10
5
0
−5
−10
−15
Dim2
−25
0
25
50
Dim1
marker
200
150
100
50
0

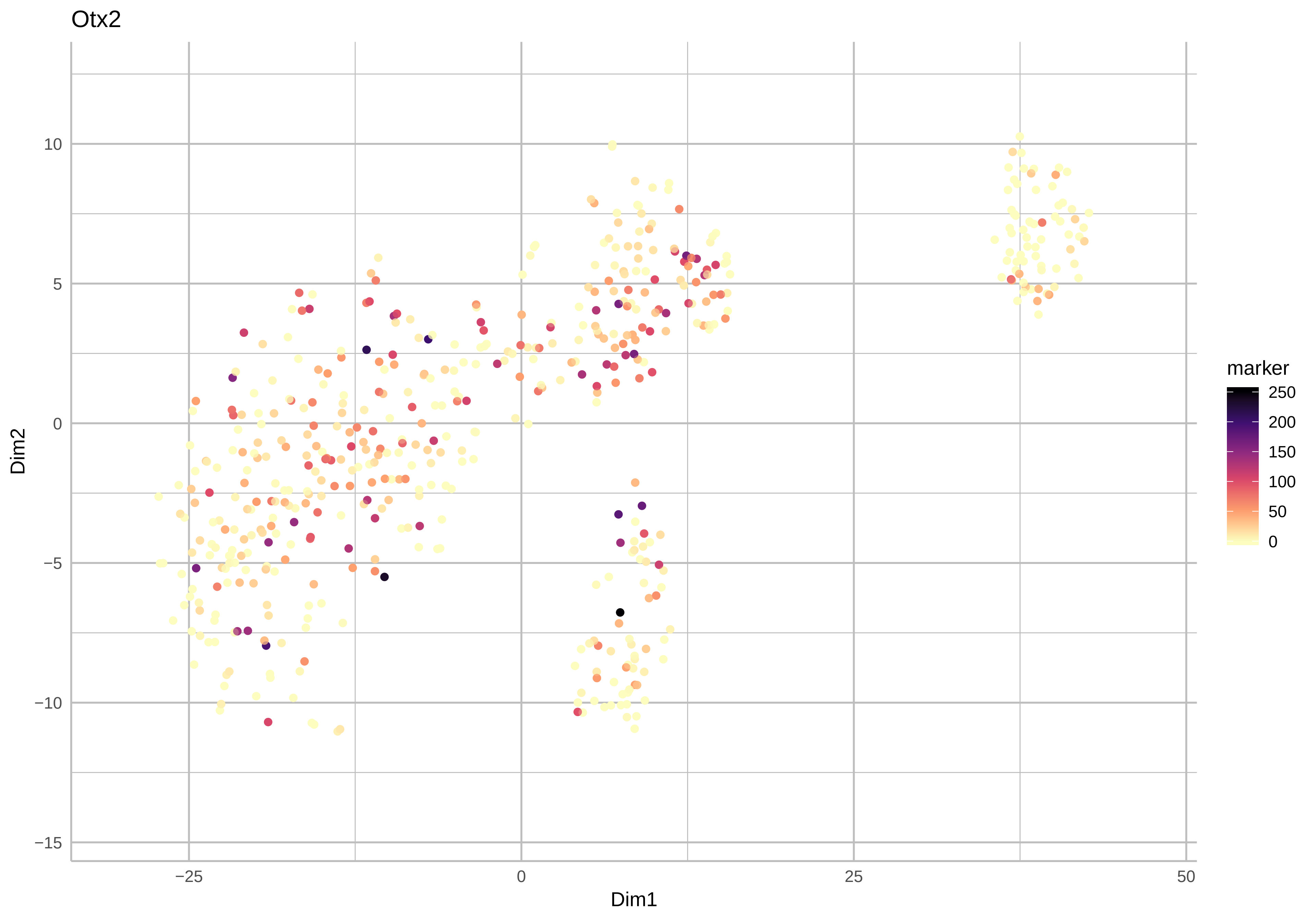

Otx2
Dim2
Dim1
10
5
0
−5
−10
−15
−25
0
25
50
marker
250
200
150
100
50
0

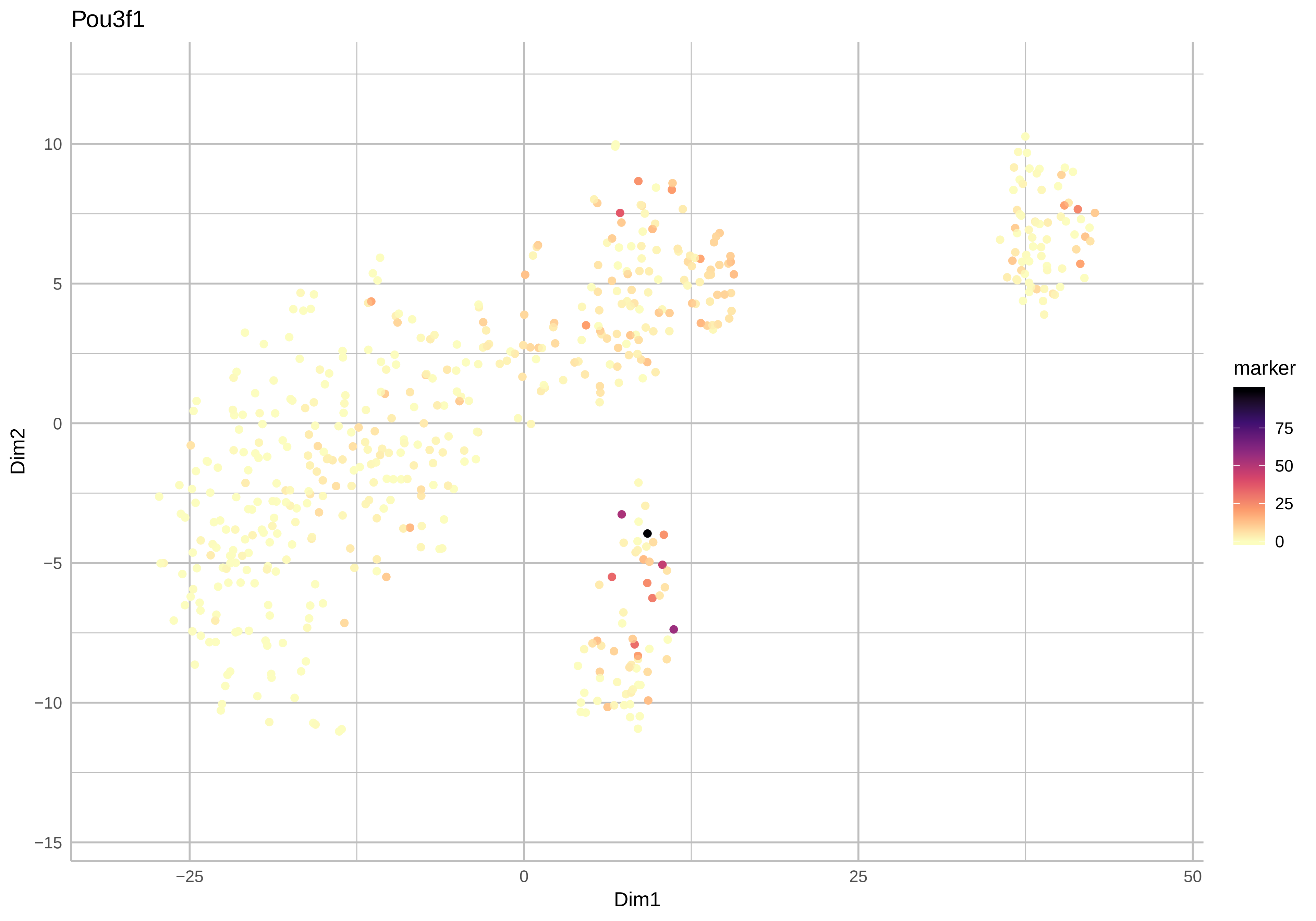

Pou3f1
Dim2
Dim1
marker
10
5
0
−5
−10
−15
−25
0
25
50
75
50
25
0

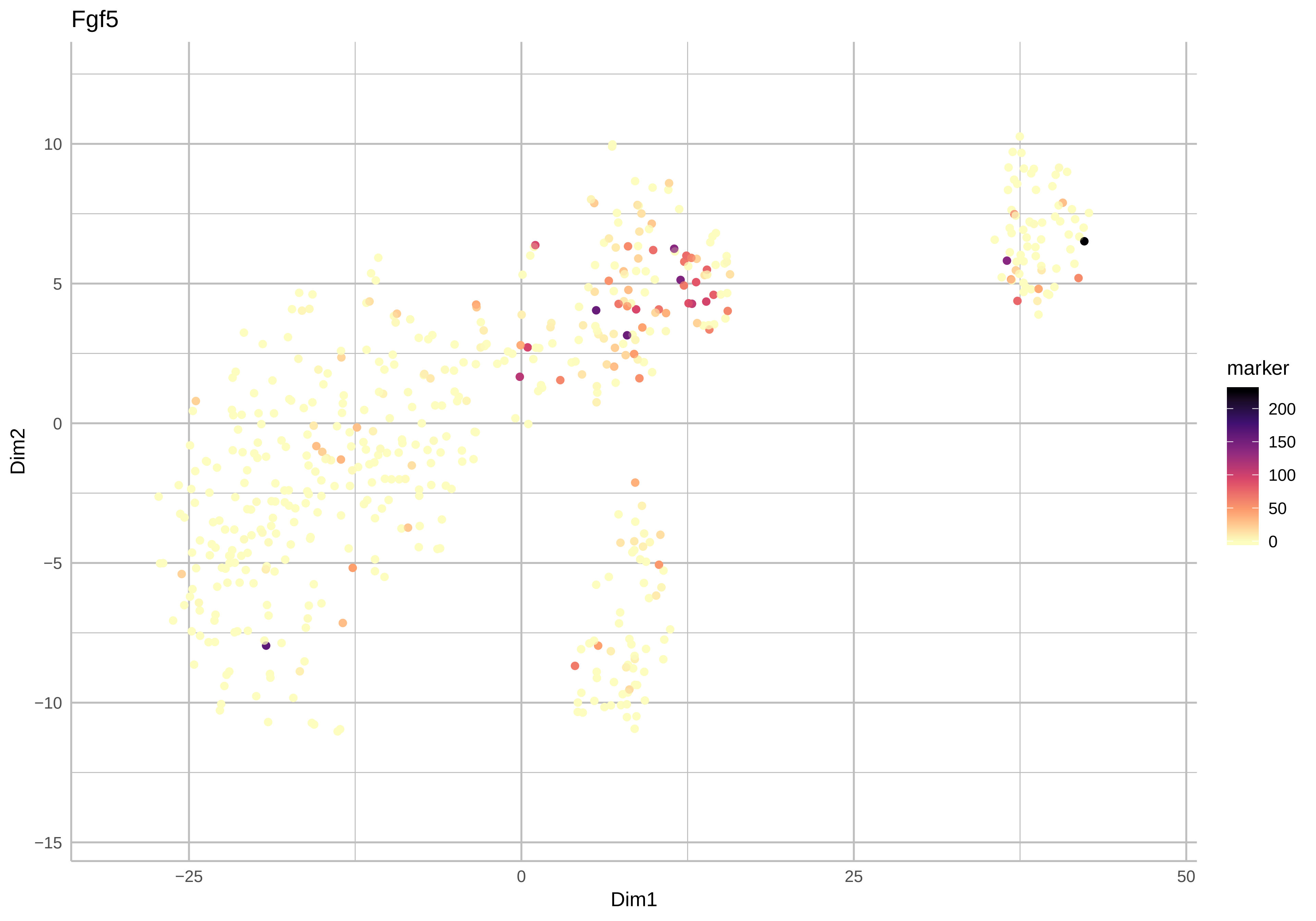
Fgf5
Dim2
Dim1
marker
10
5
0
−5
−10
−15
−25
0
25
50
200
150
100
50
0

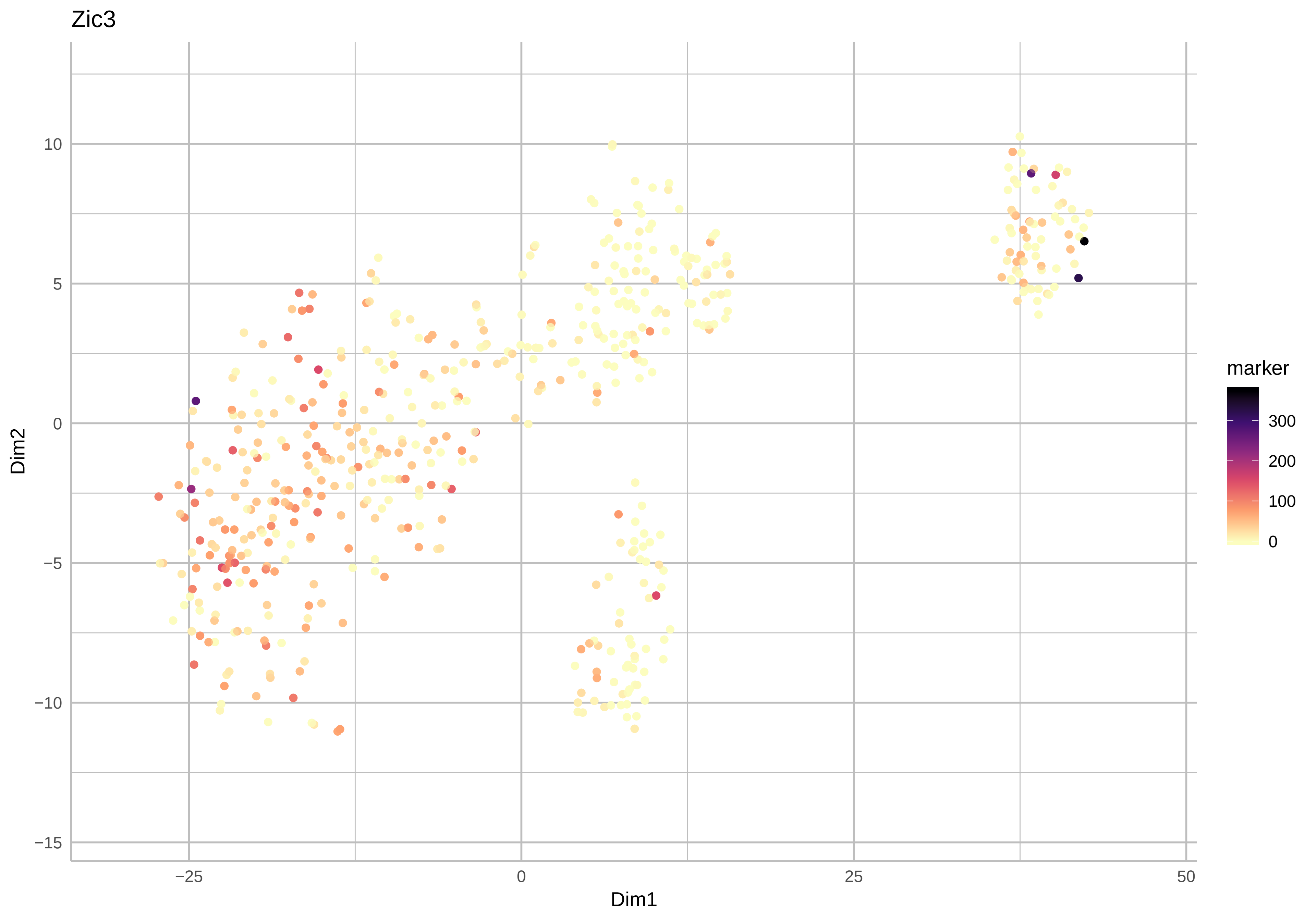
Zic3
Dim2
Dim1
10
5
0
−5
−10
−15
−25
0
25
50
marker
300
200
100
0

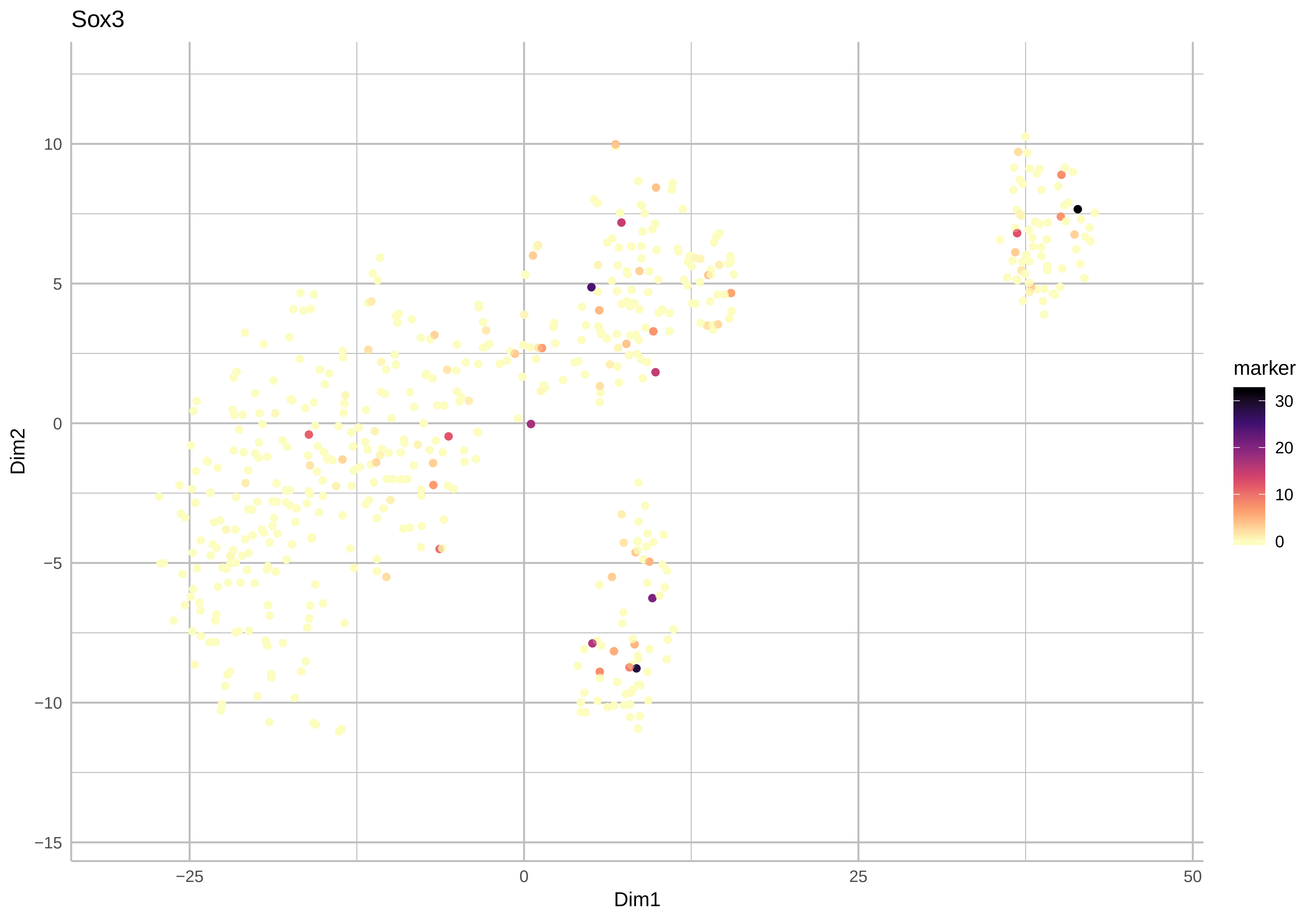
Sox3
Dim2
Dim1
10
5
0
−5
−10
−15
−25
0
25
50
marker
30
20
10
0

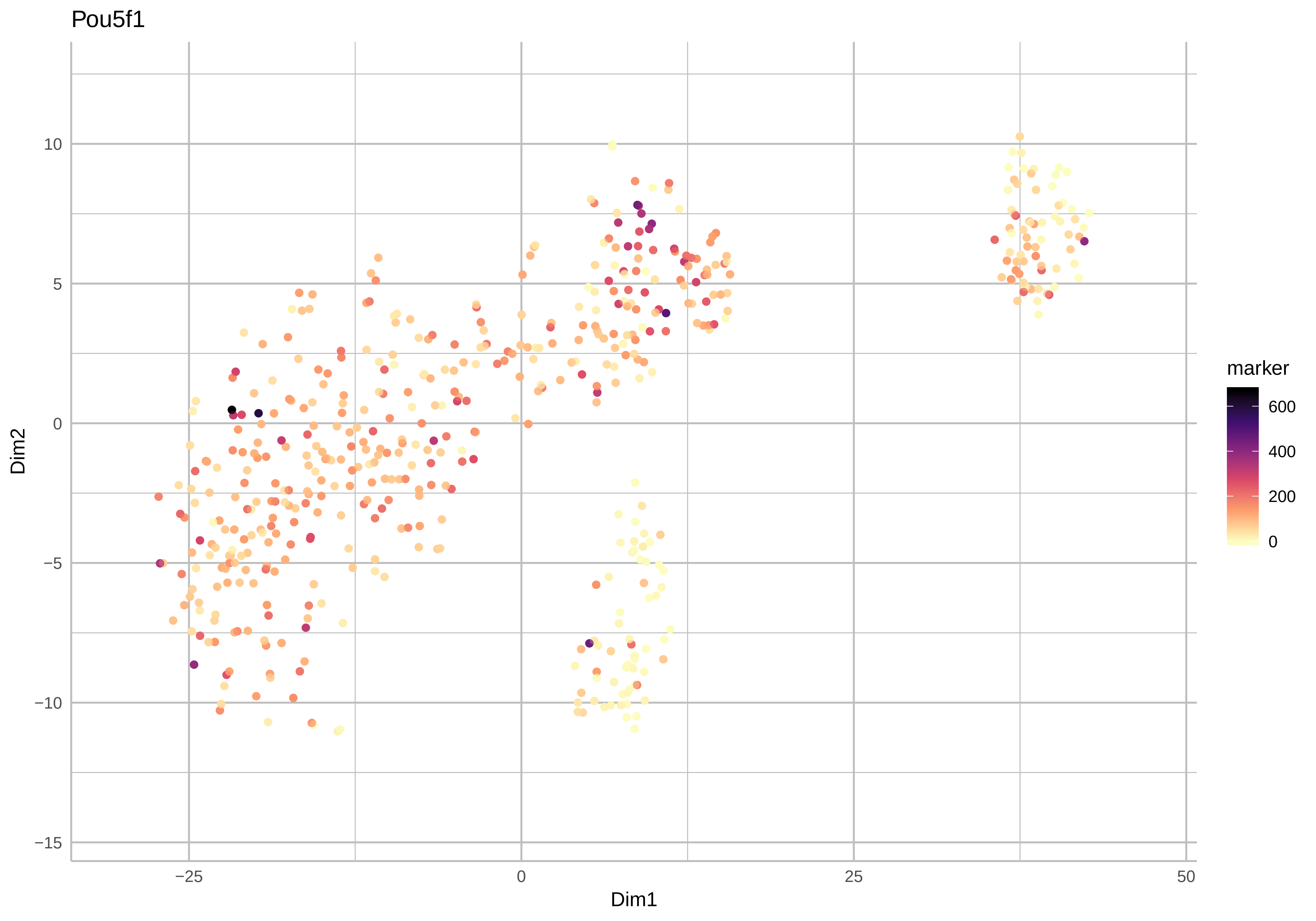

Pou5f1
Dim2
Dim1
10
5
0
−5
−10
−15
−25
0
25
50
marker
600
400
200
0

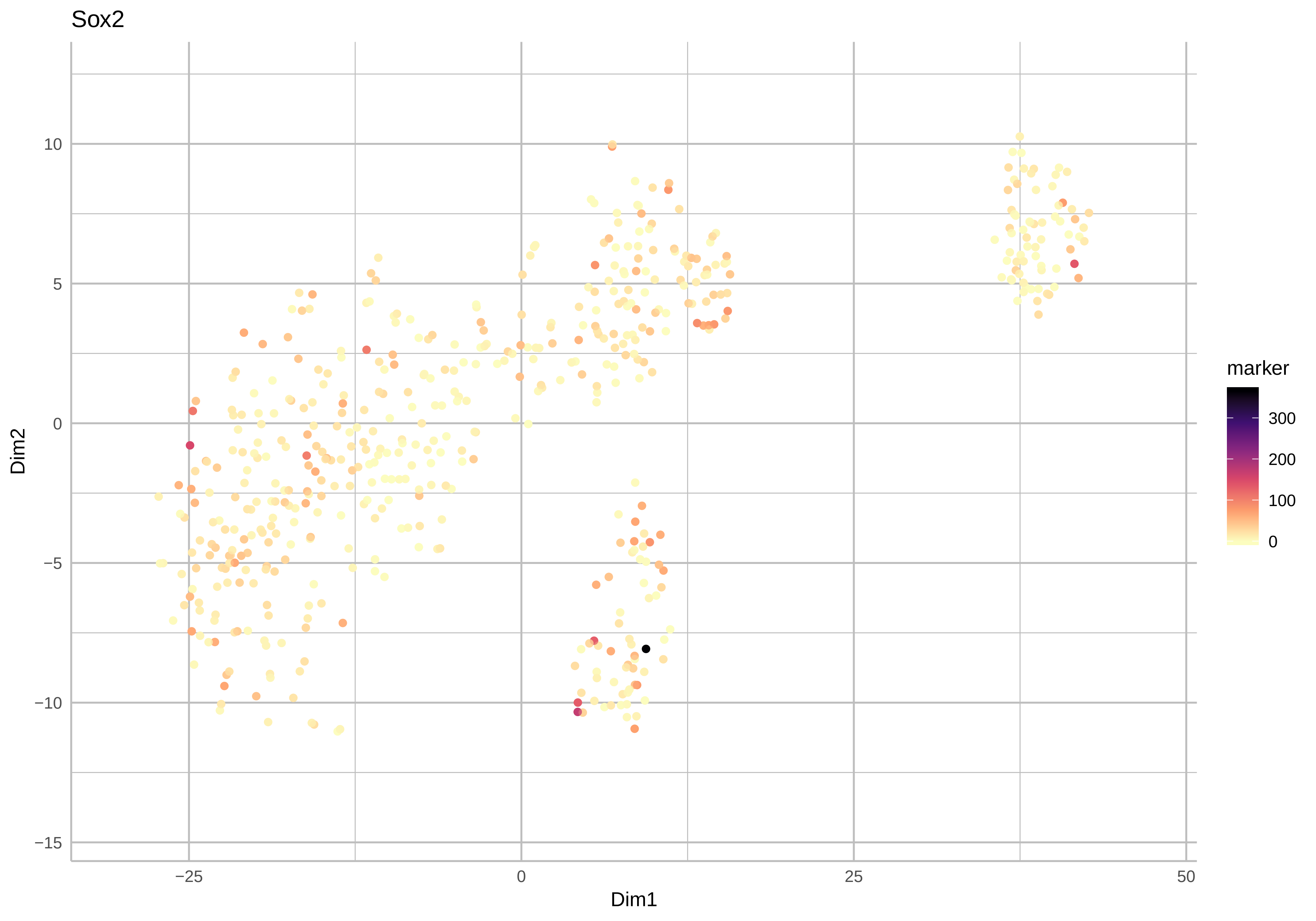

Sox2
Dim2
Dim1
10
5
0
−5
−10
−15
−25
0
25
50
marker
300
200
100
0

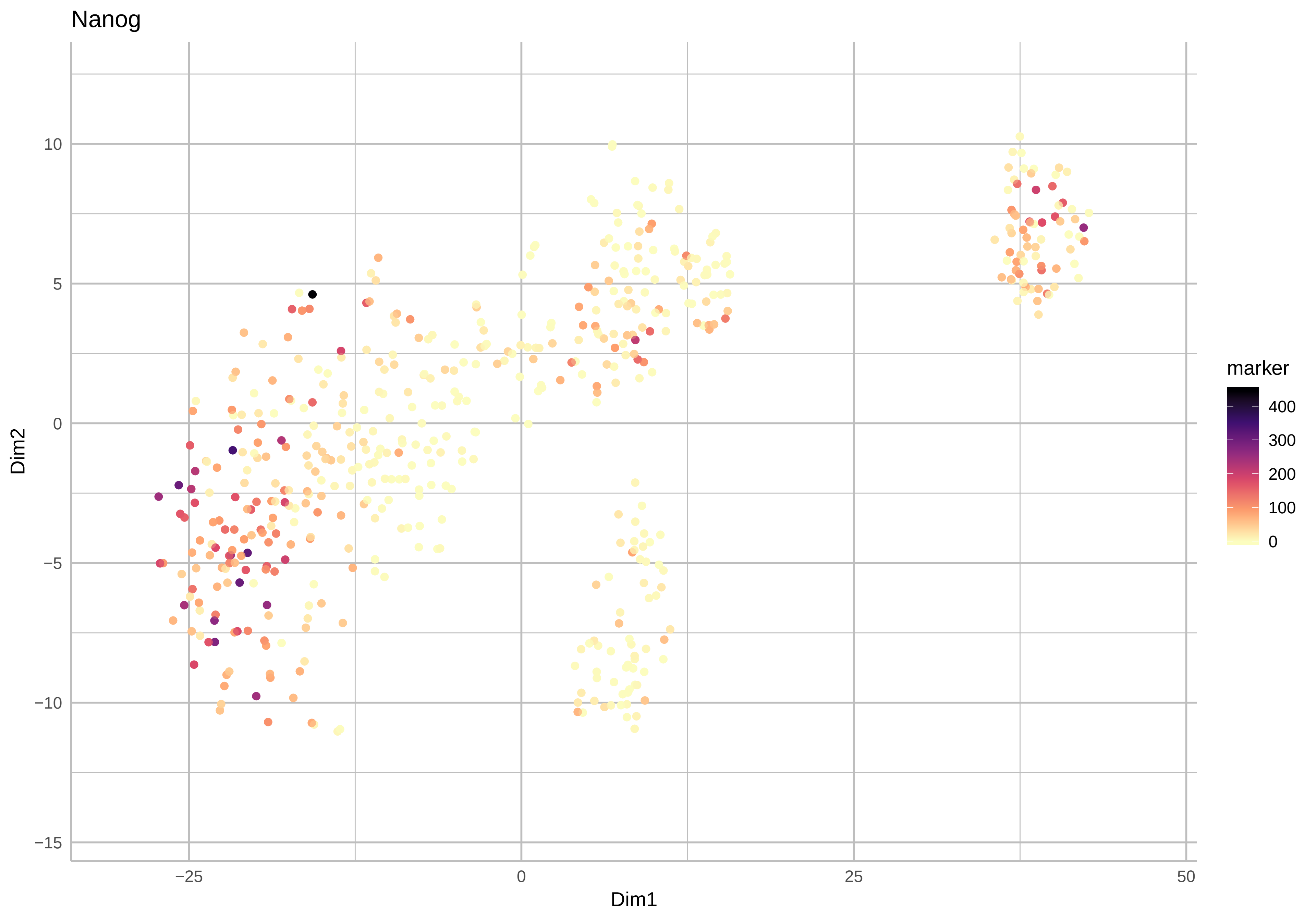

Nanog
Dim2
Dim1
10
5
0
−5
−10
−15
−25
0
25
50
marker
400
300
200
100
0

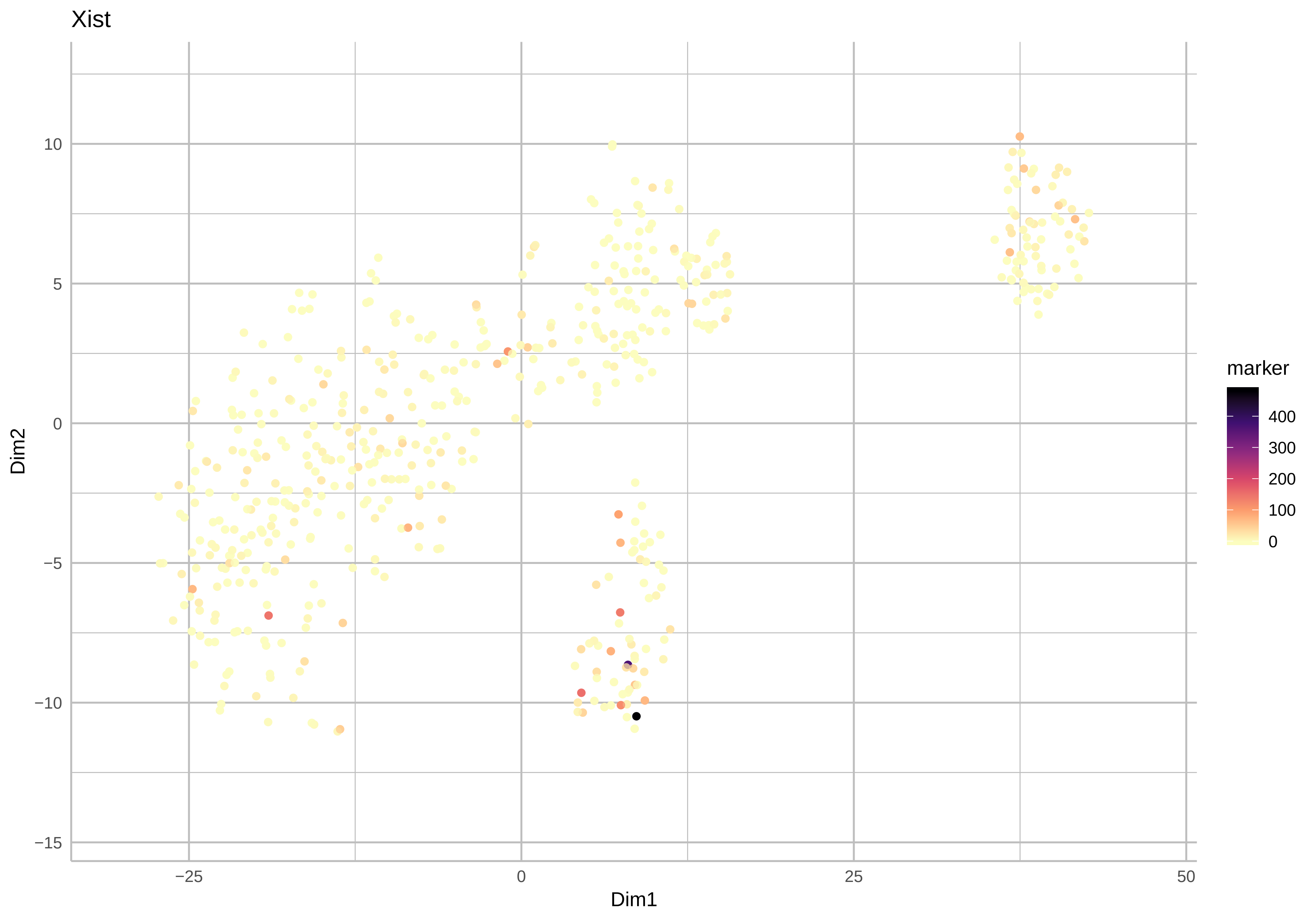

Xist
Dim2
Dim1
10
5
0
−5
−10
−15
−25
0
25
50
marker
400
300
200
100
0

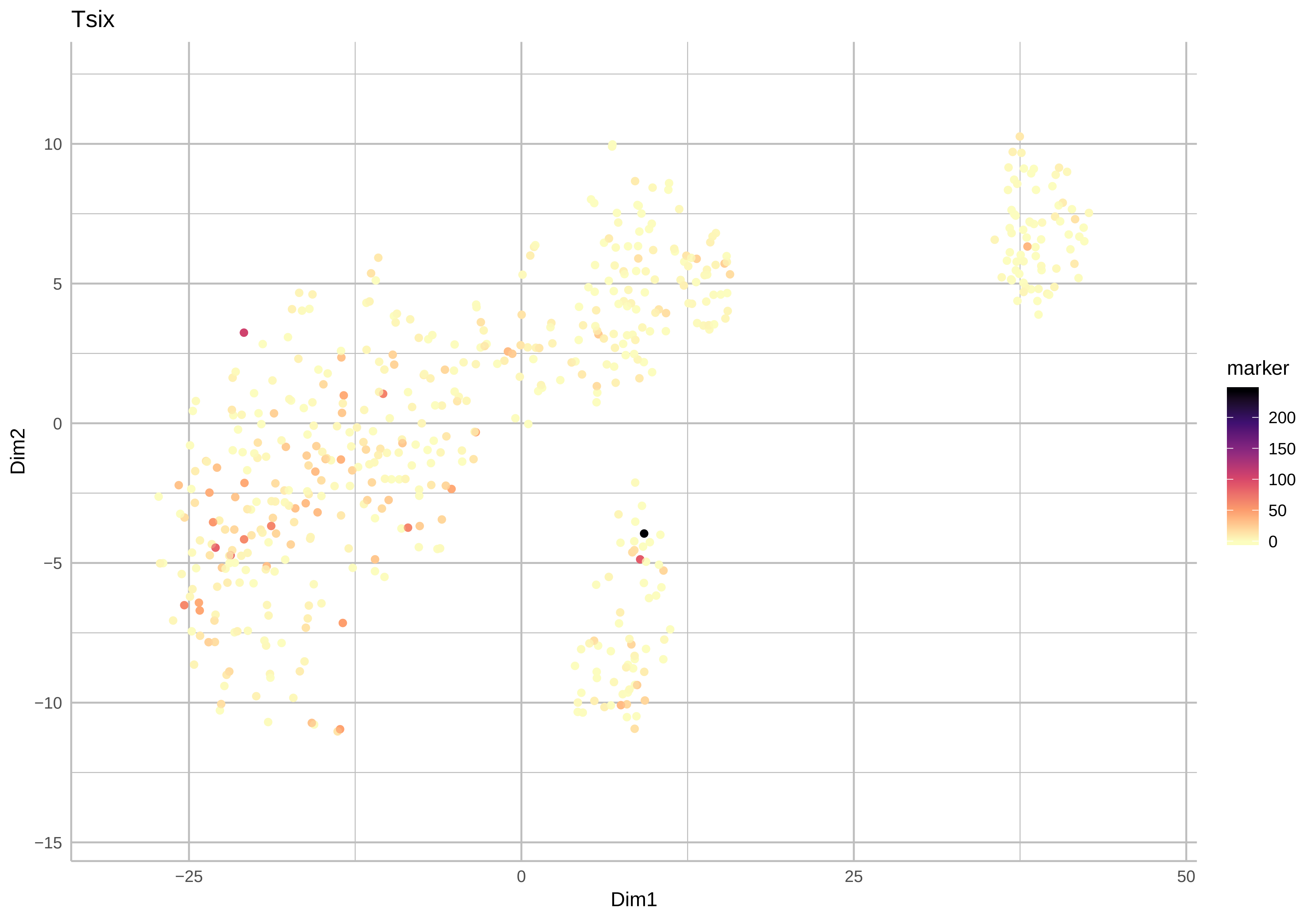
Tsix
Dim2
Dim1
10
5
0
−5
−10
−15
−25
0
25
50
marker
200
150
100
50
0

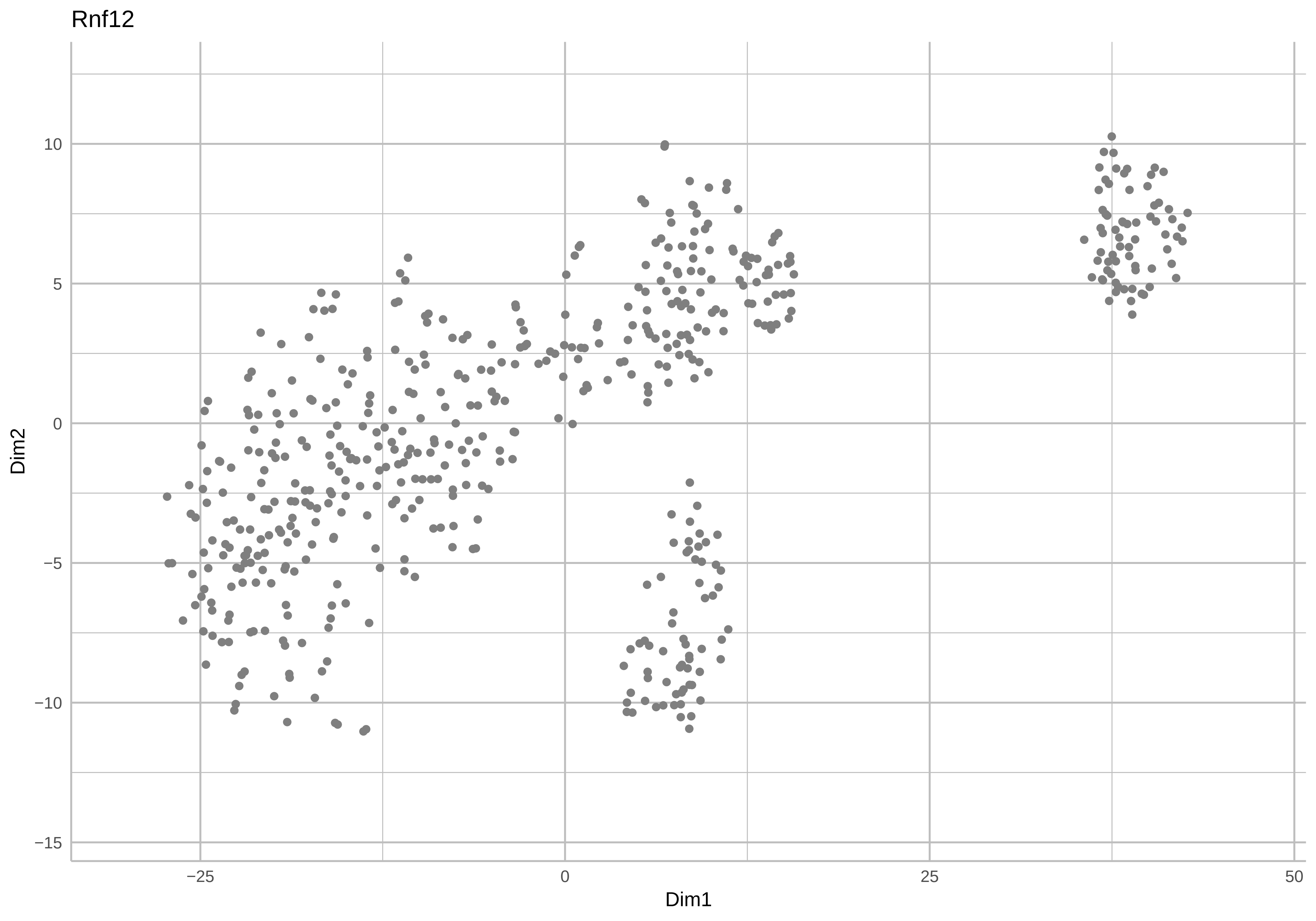
Rnf12
Dim2
Dim1
10
5
0
−5
−10
−15
−25
0
25
50

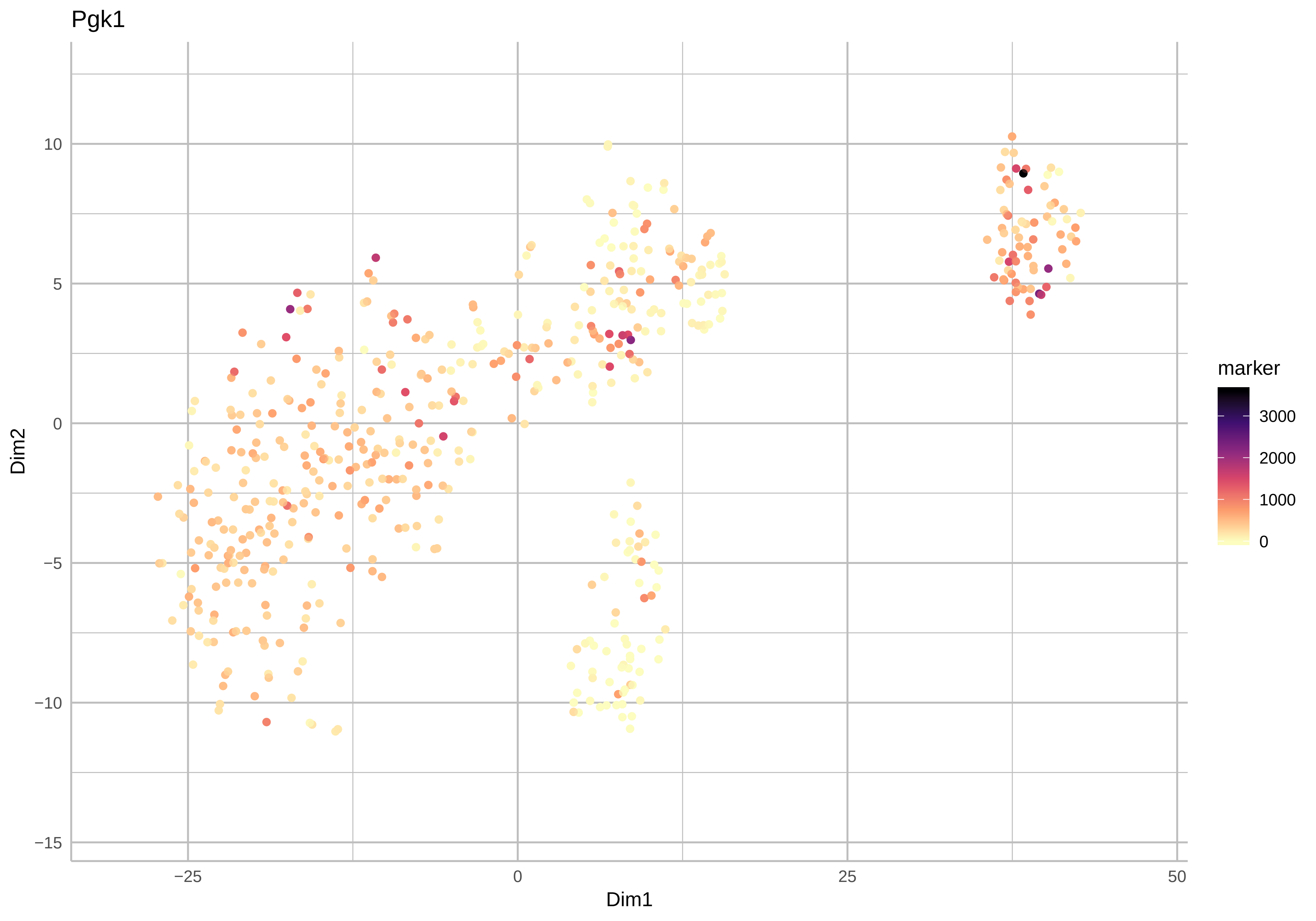

Pgk1
Dim2
Dim1
10
5
0
−5
−10
−15
−25
0
25
50
marker
3000
2000
1000
0

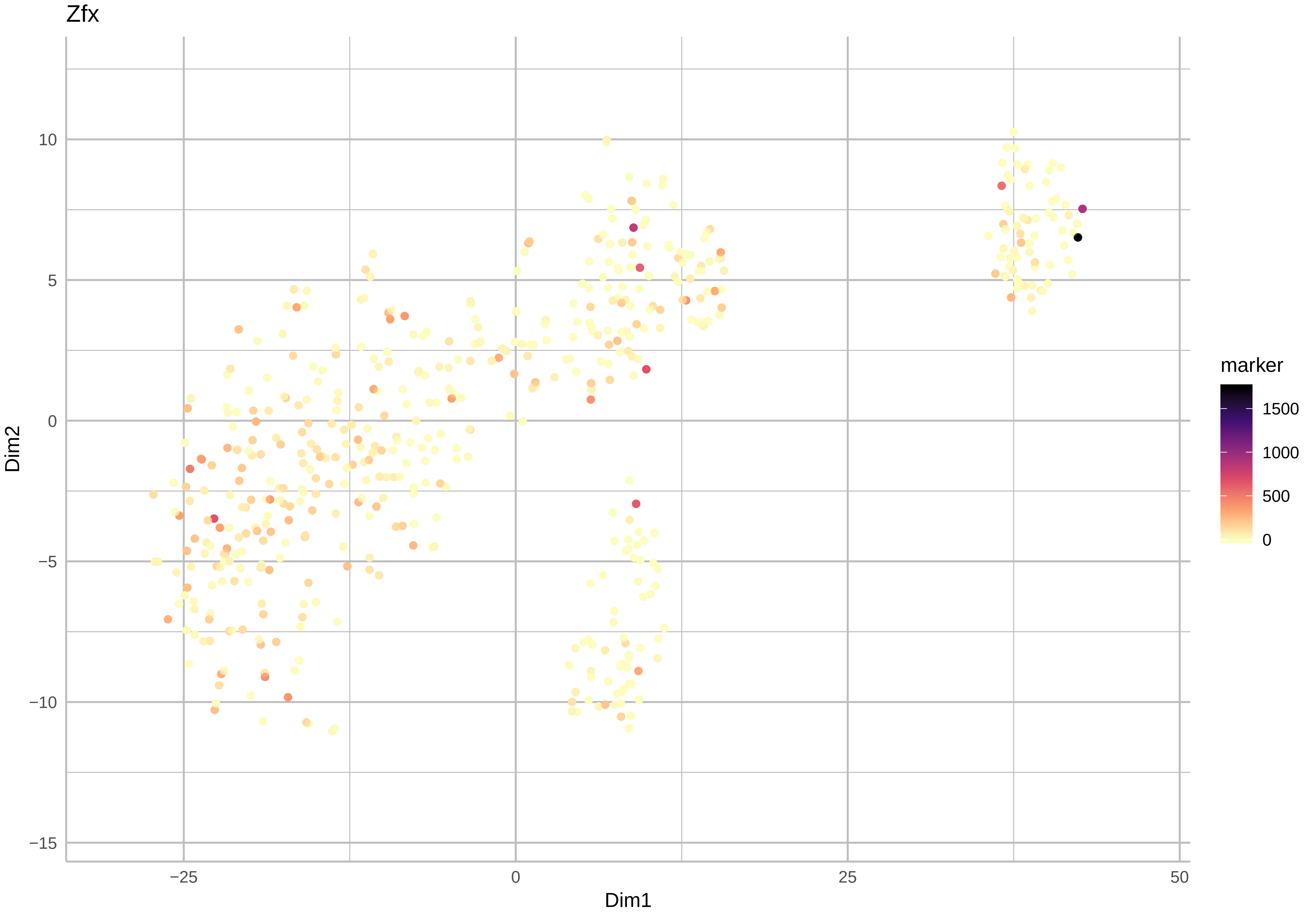

Zfx
10
5
0
−5
−10
−15
−25
0
25
50
Dim1
Dim2
marker
1500
1000
500
0

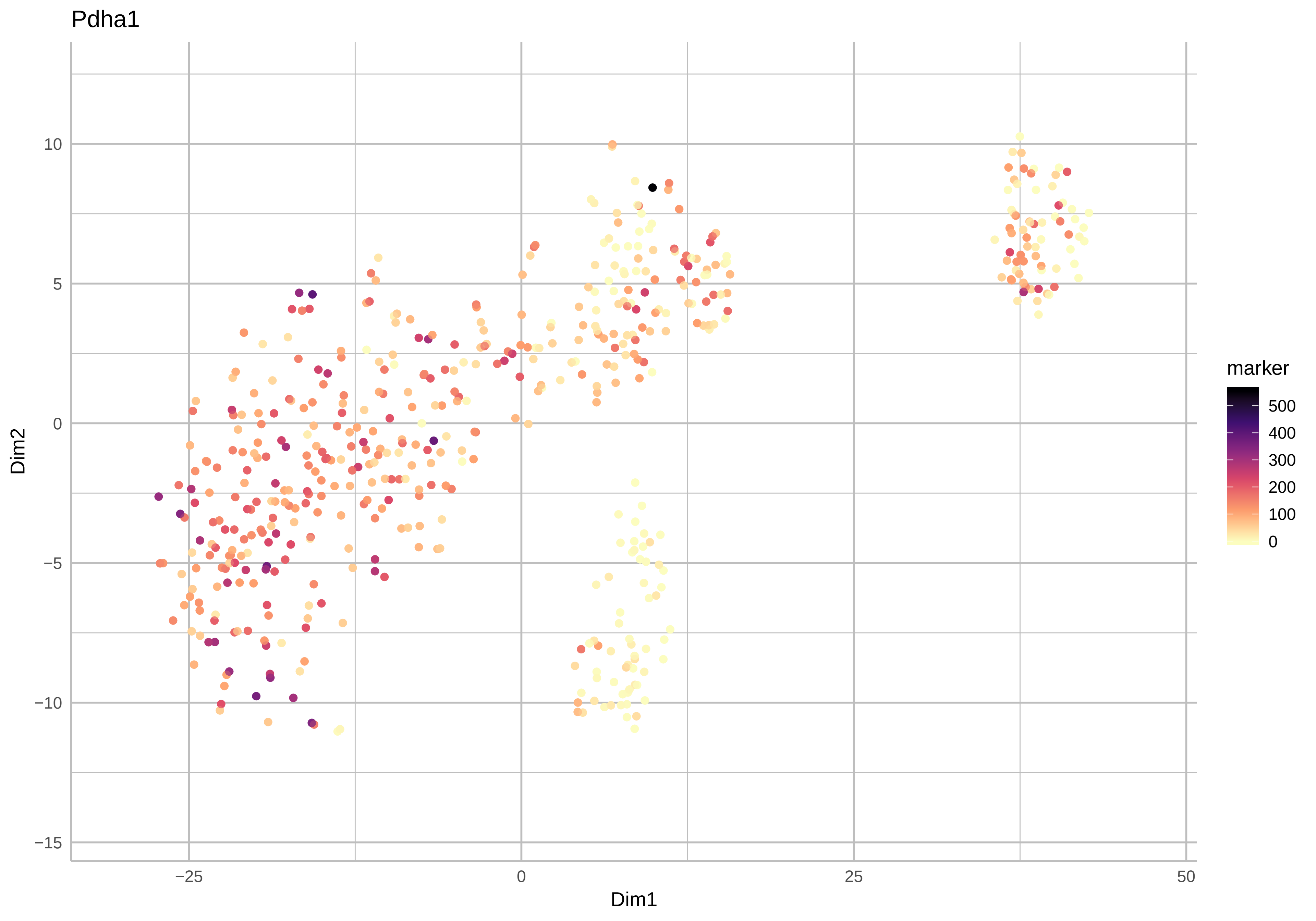

Pdha1
Dim2
Dim1
10
5
0
−5
−10
−15
−25
0
25
50
marker
500
400
300
200
100
0

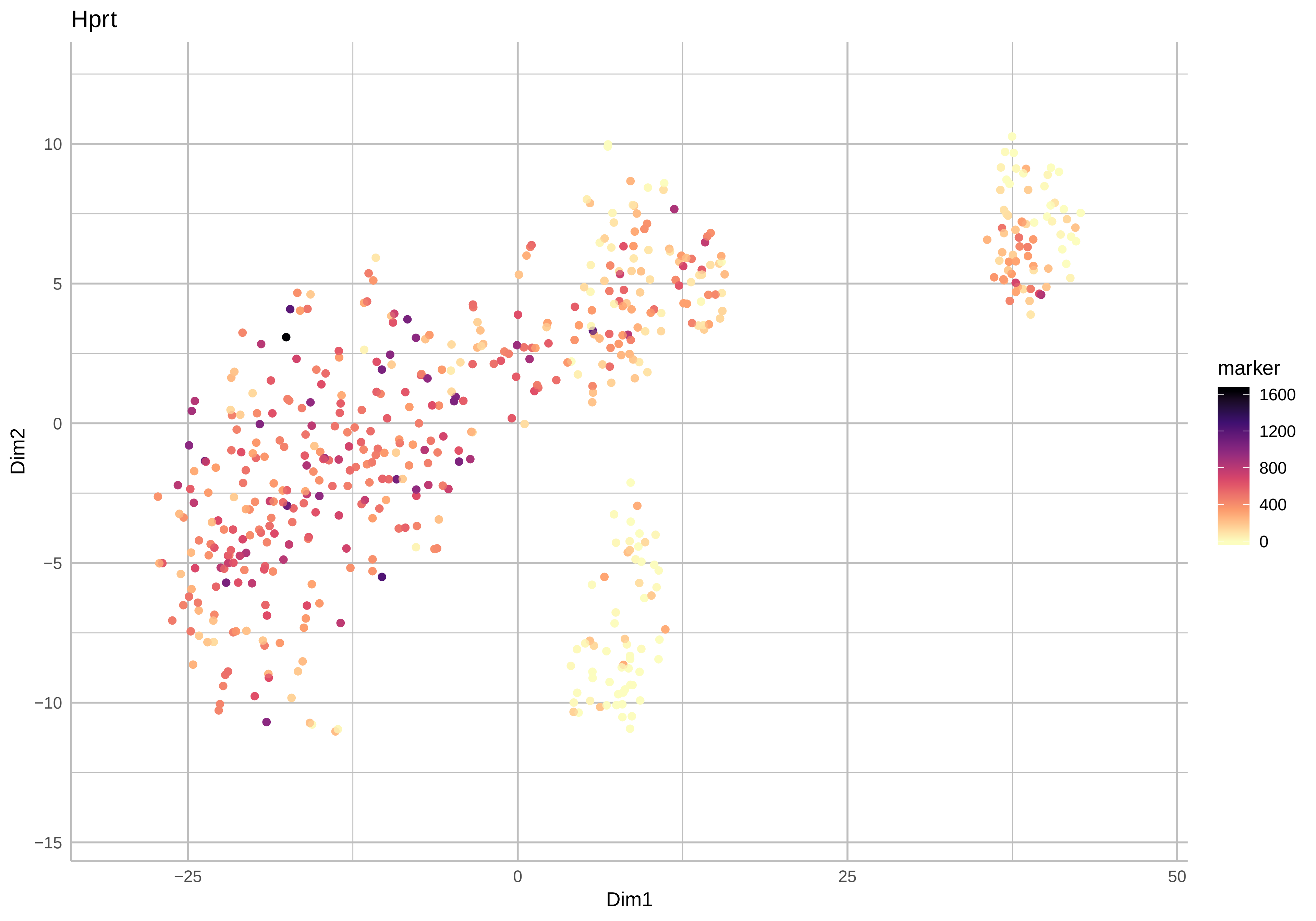
Hprt
Dim2
Dim1
marker
1600
1200
800
400
0
10
5
0
−5
−10
−15
−25
0
25
50

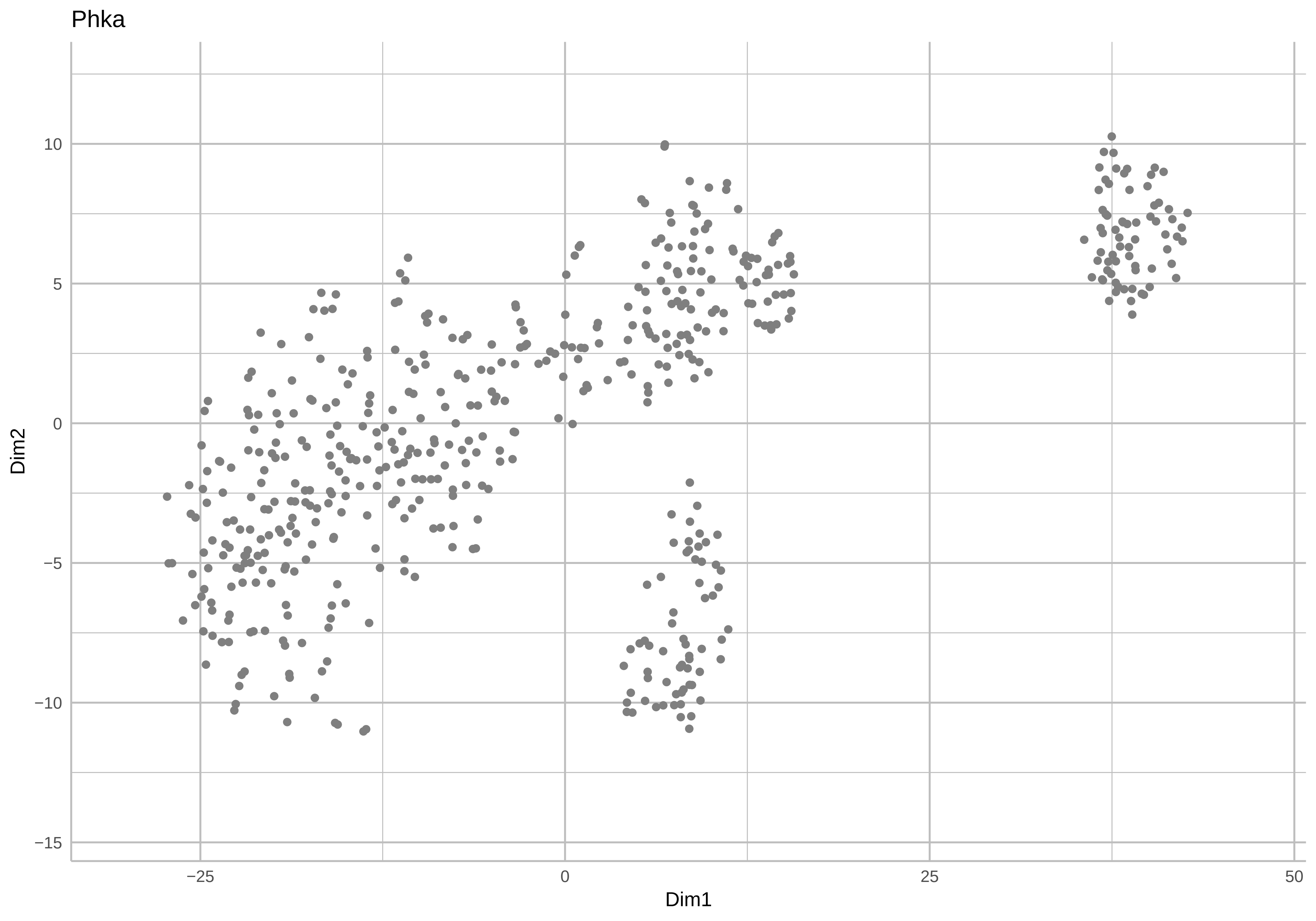

Phka
Dim2
10
5
0
−5
−10
−15
−25
0
25
50
Dim1

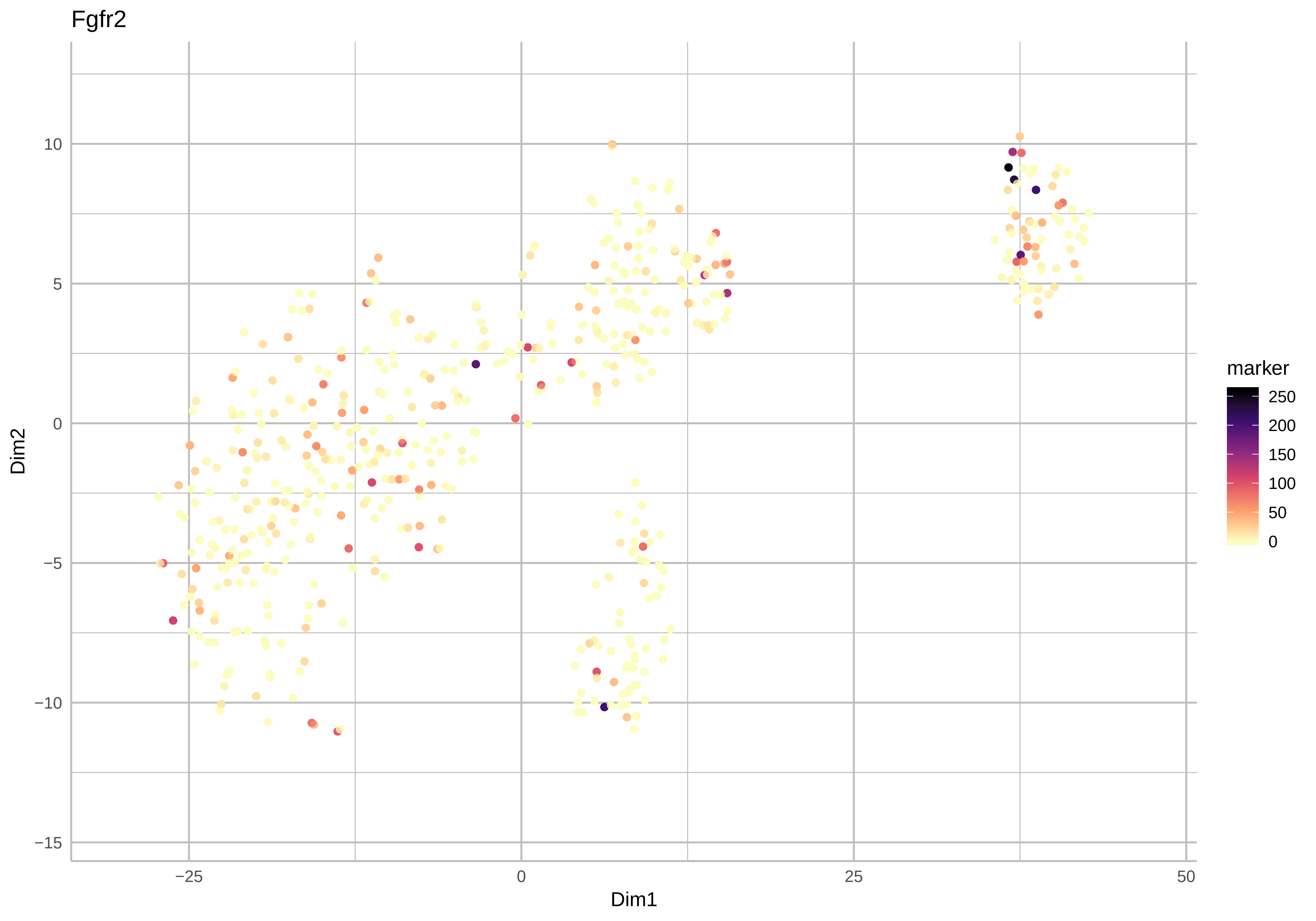
Fgfr2
Dim2
Dim1
10
5
0
−5
−10
−15
−25
0
25
50
marker
250
200
150
100
50
0

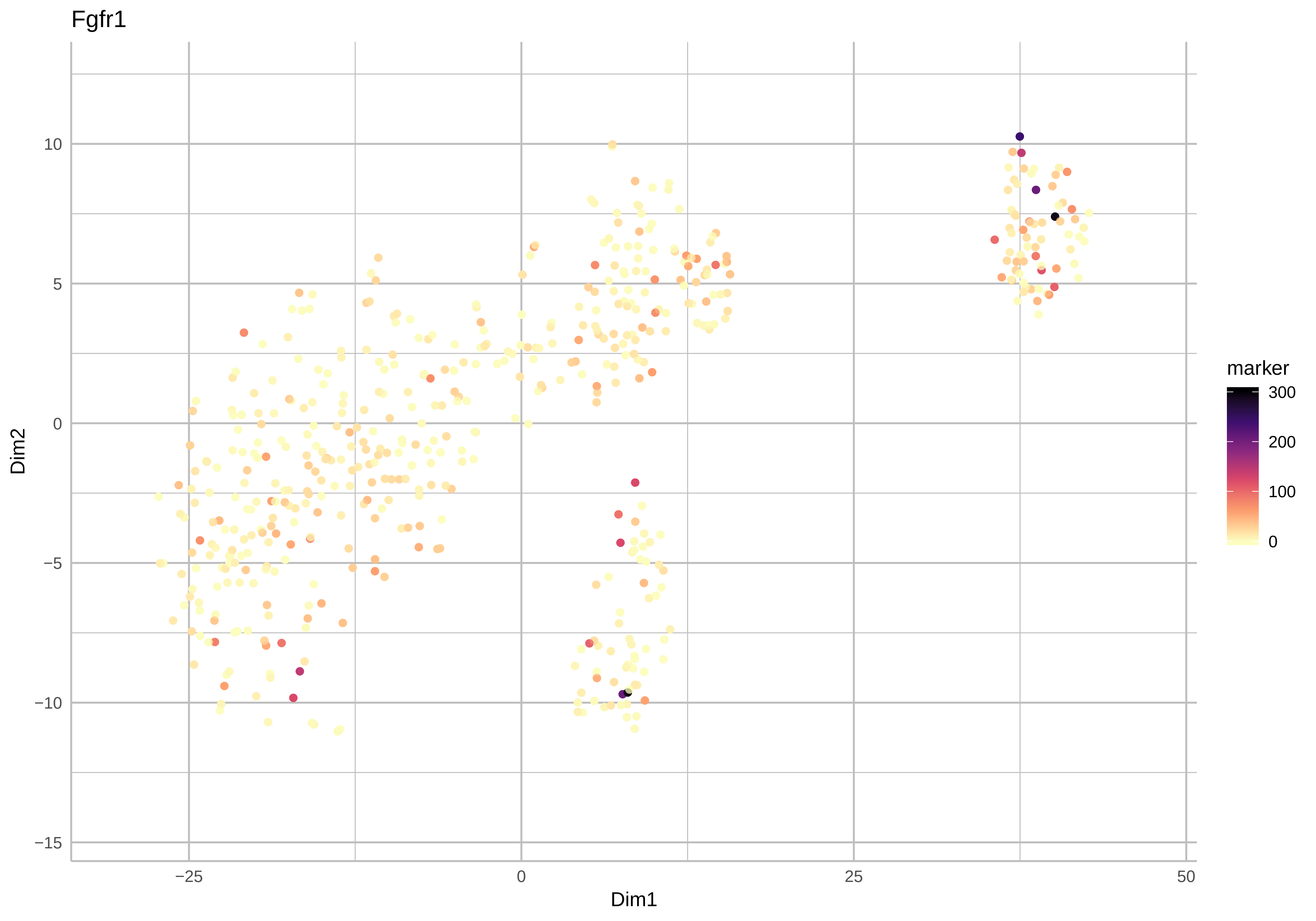
Fgfr1
Dim2
Dim1
10
5
0
−5
−10
−15
−25
0
25
50
marker
300
200
100
0

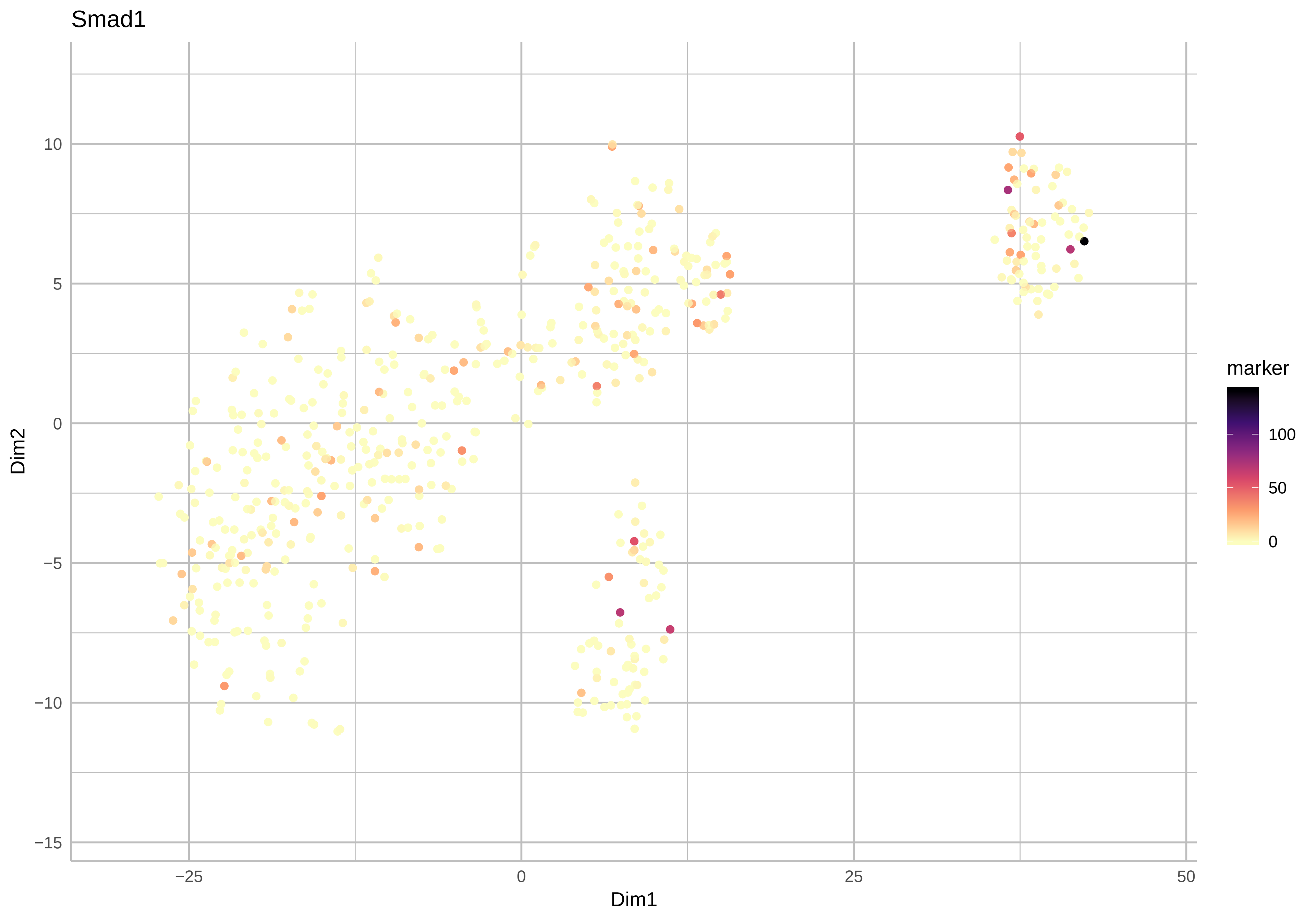

Smad1
Dim2
Dim1
marker
100
50
0
10
5
0
−5
−10
−15
−25
0
25
50

Smad5

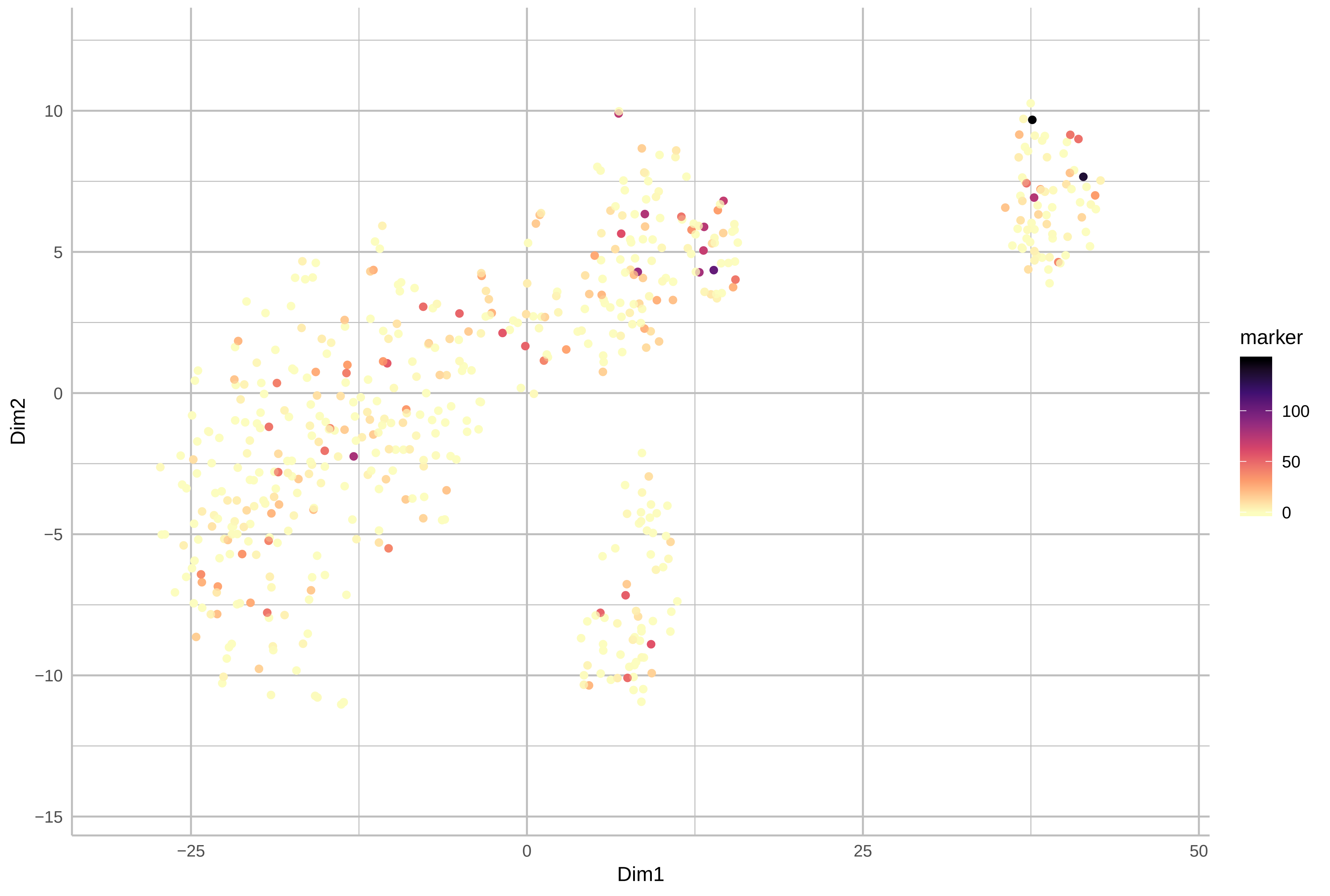

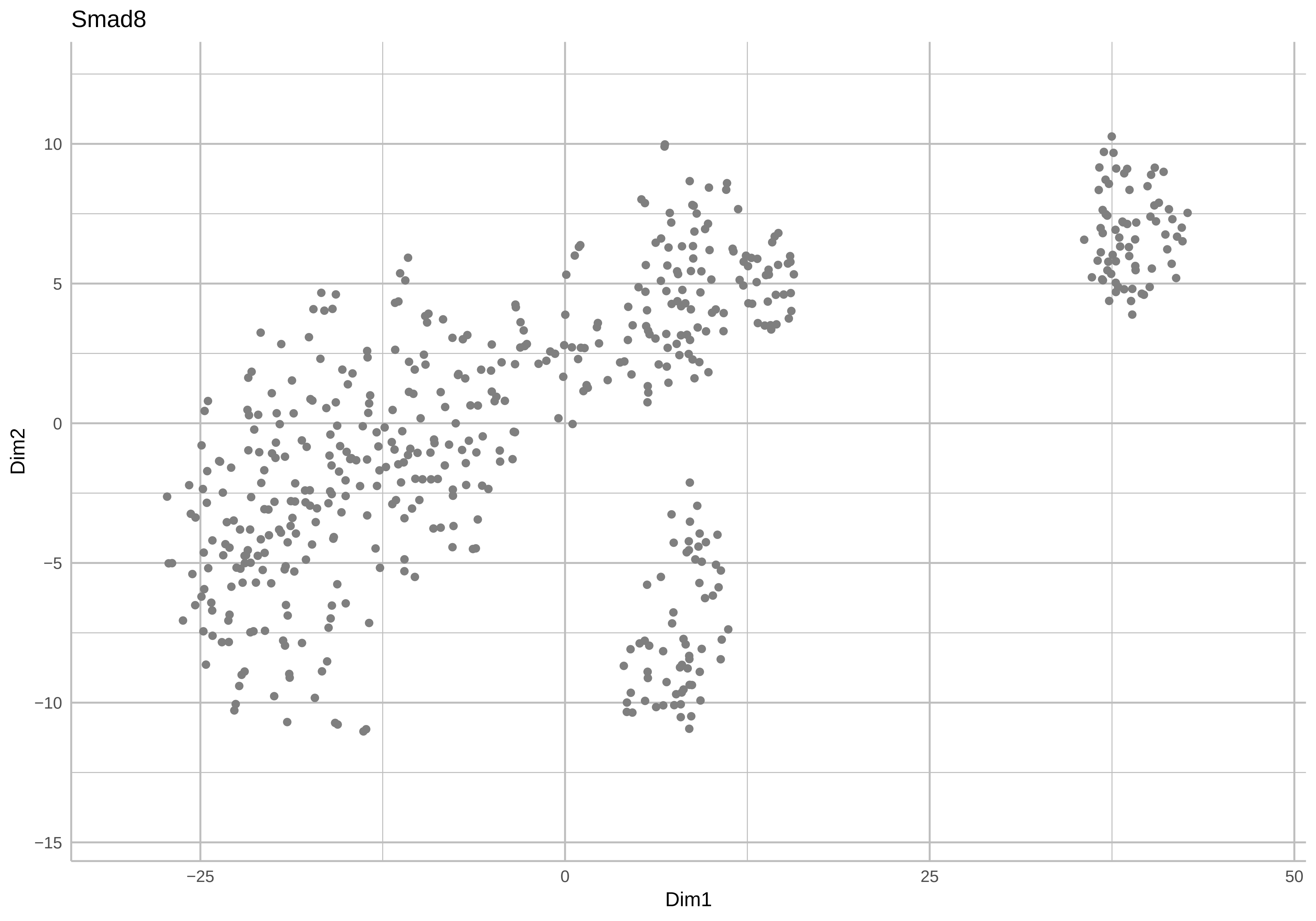
Smad8
Dim2
Dim1
10
5
0
−5
−10
−15
−25
0
25
50

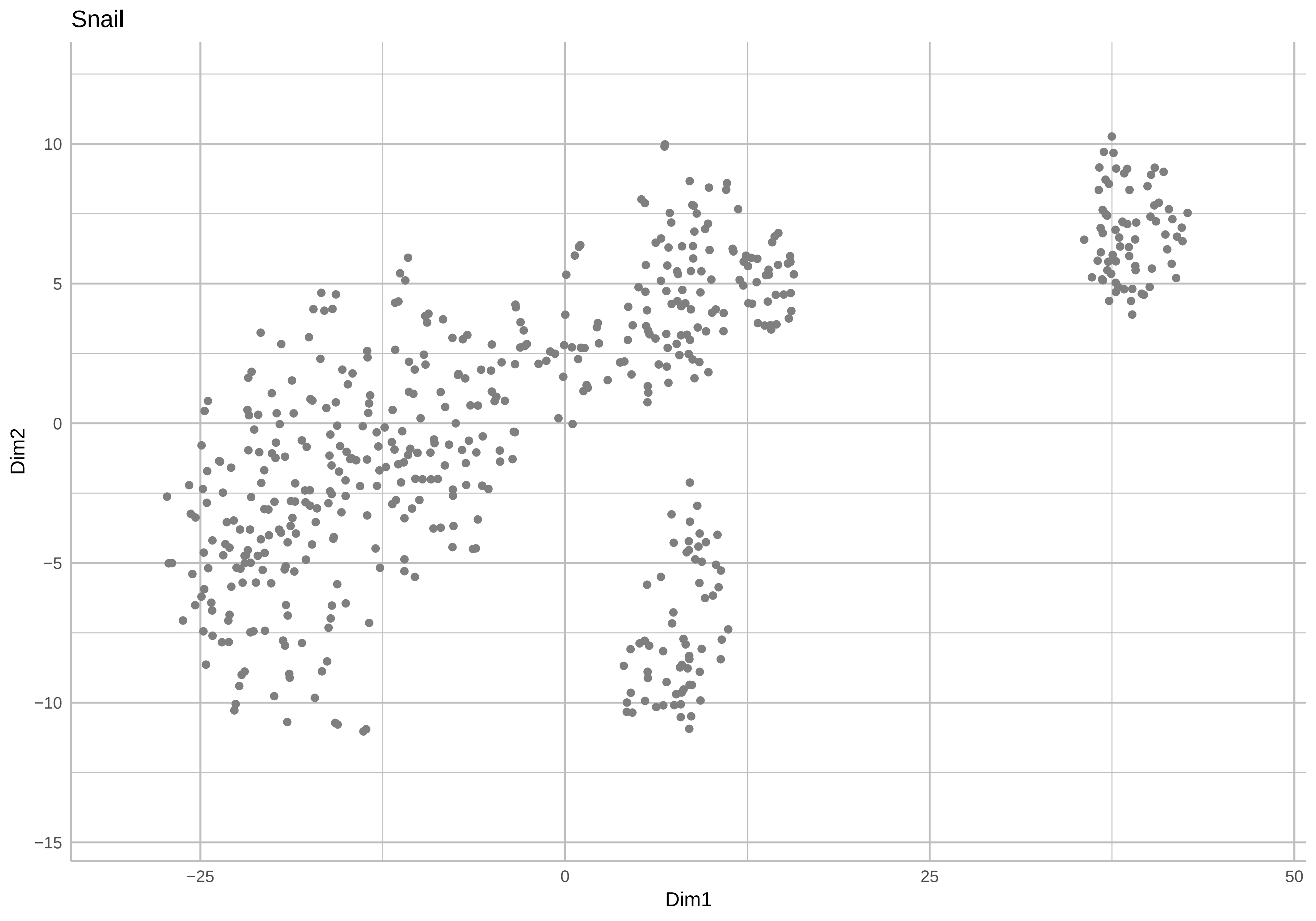
Snail
Dim2
10
5
0
−5
−10
−15
−25
0
25
50
Dim1

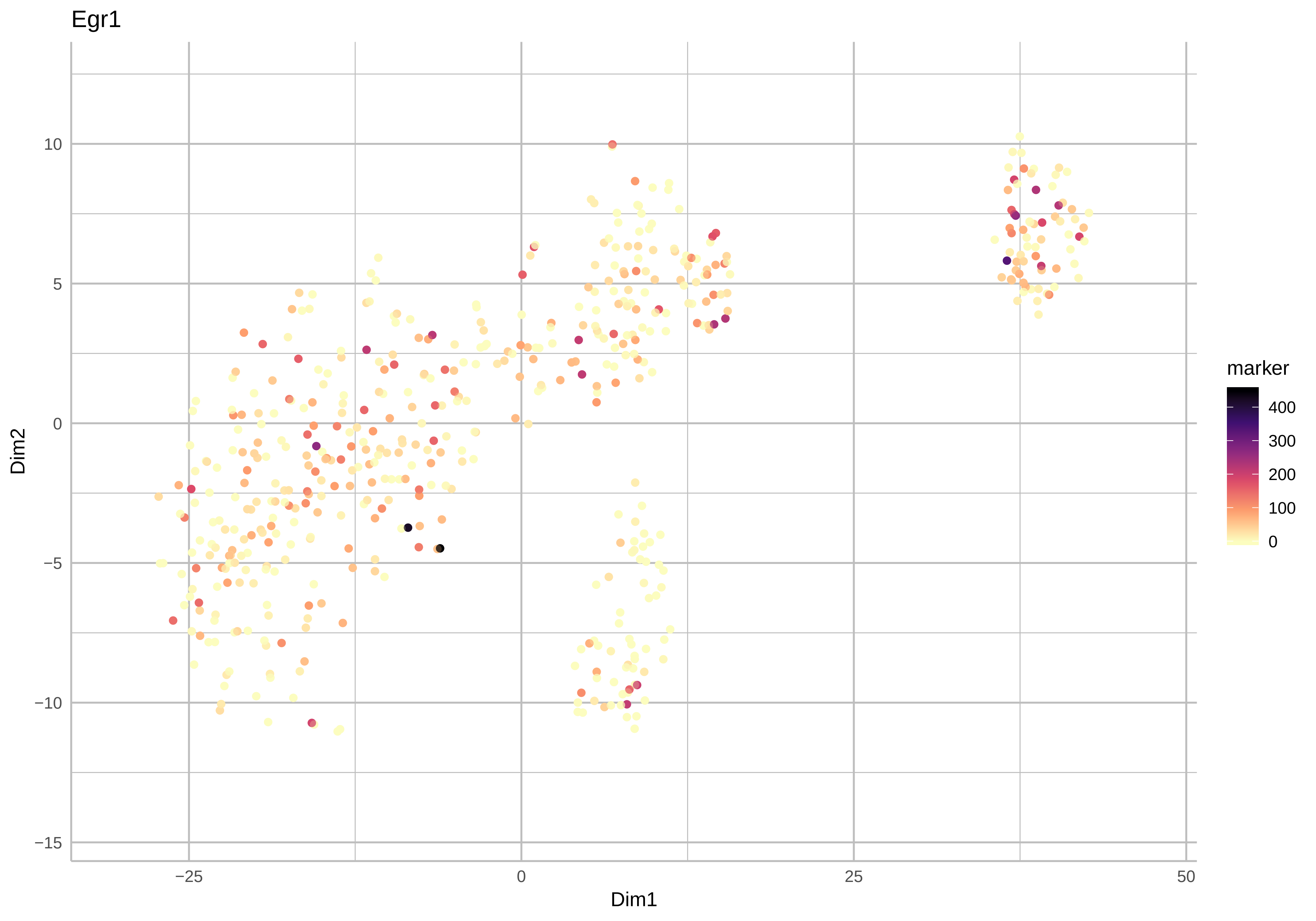

Egr1
Dim2
Dim1
10
5
0
−5
−10
−15
−25
0
25
50
marker
400
300
200
100
0

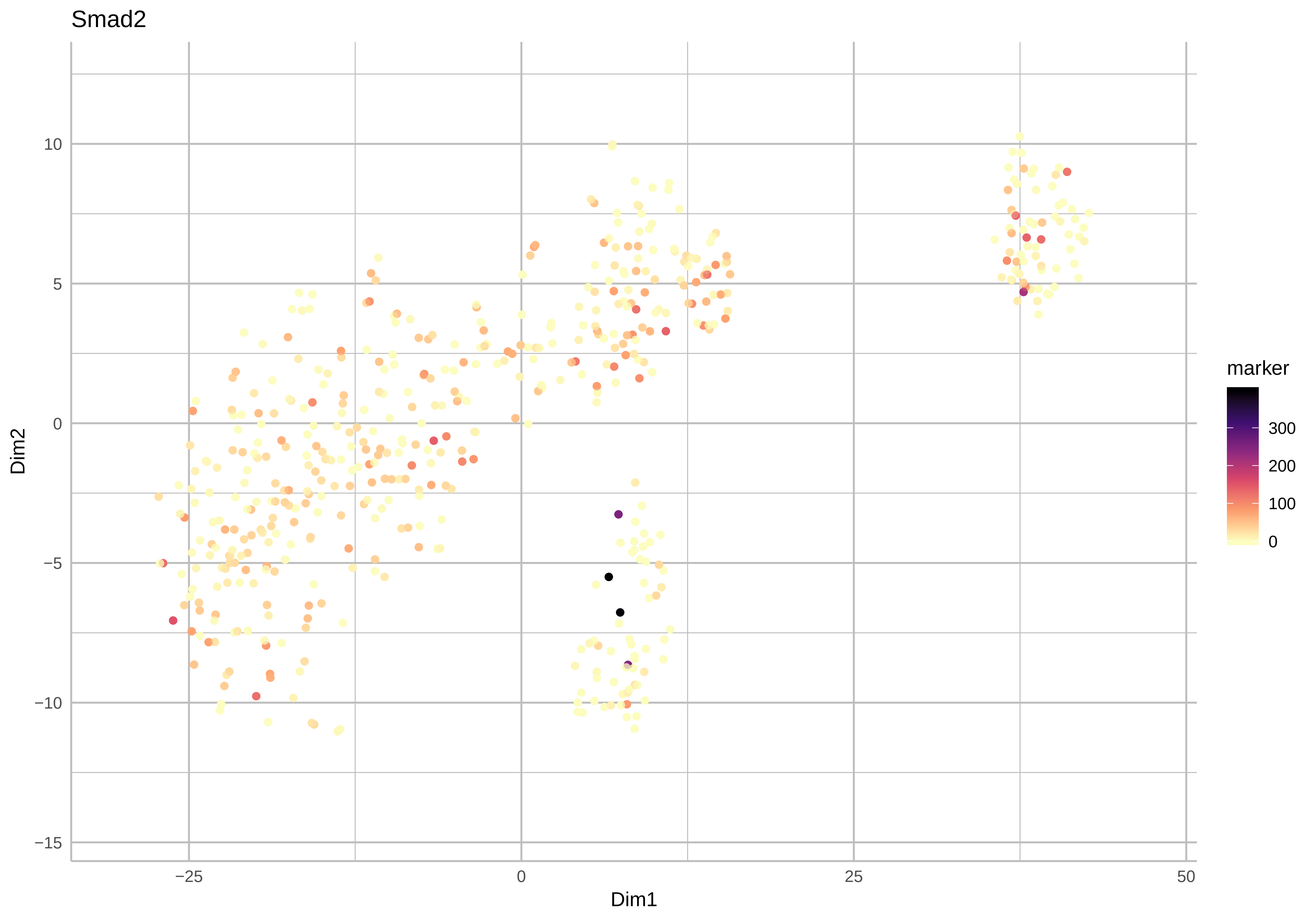

Smad2
10
5
0
−5
−10
−15
−25
0
25
50
Dim1
Dim2
marker
300
200
100
0

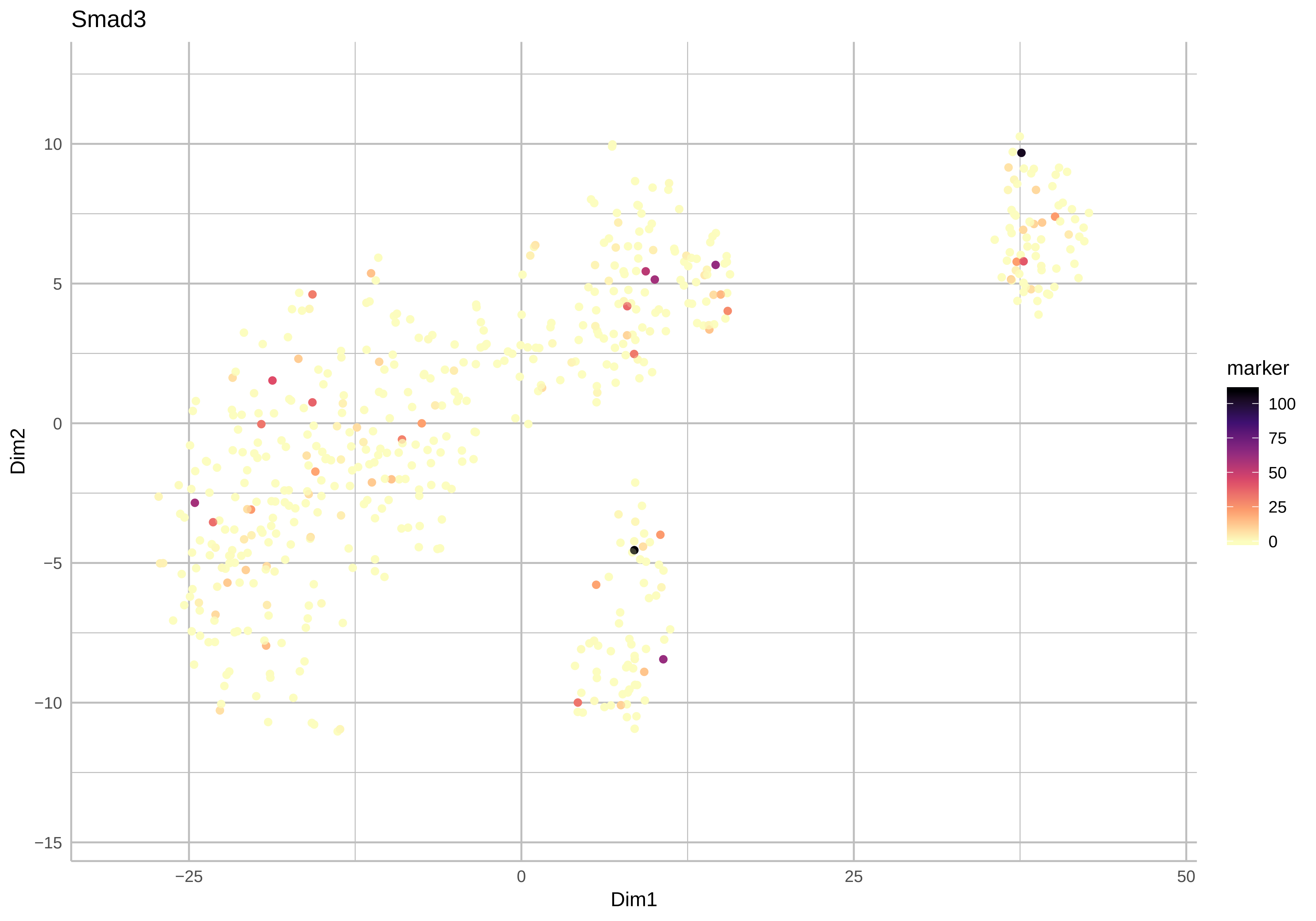
Smad3
Dim1
Dim2
marker
100
75
50
25
0
−25
0
25
50
10
5
0
−5
−10
−15

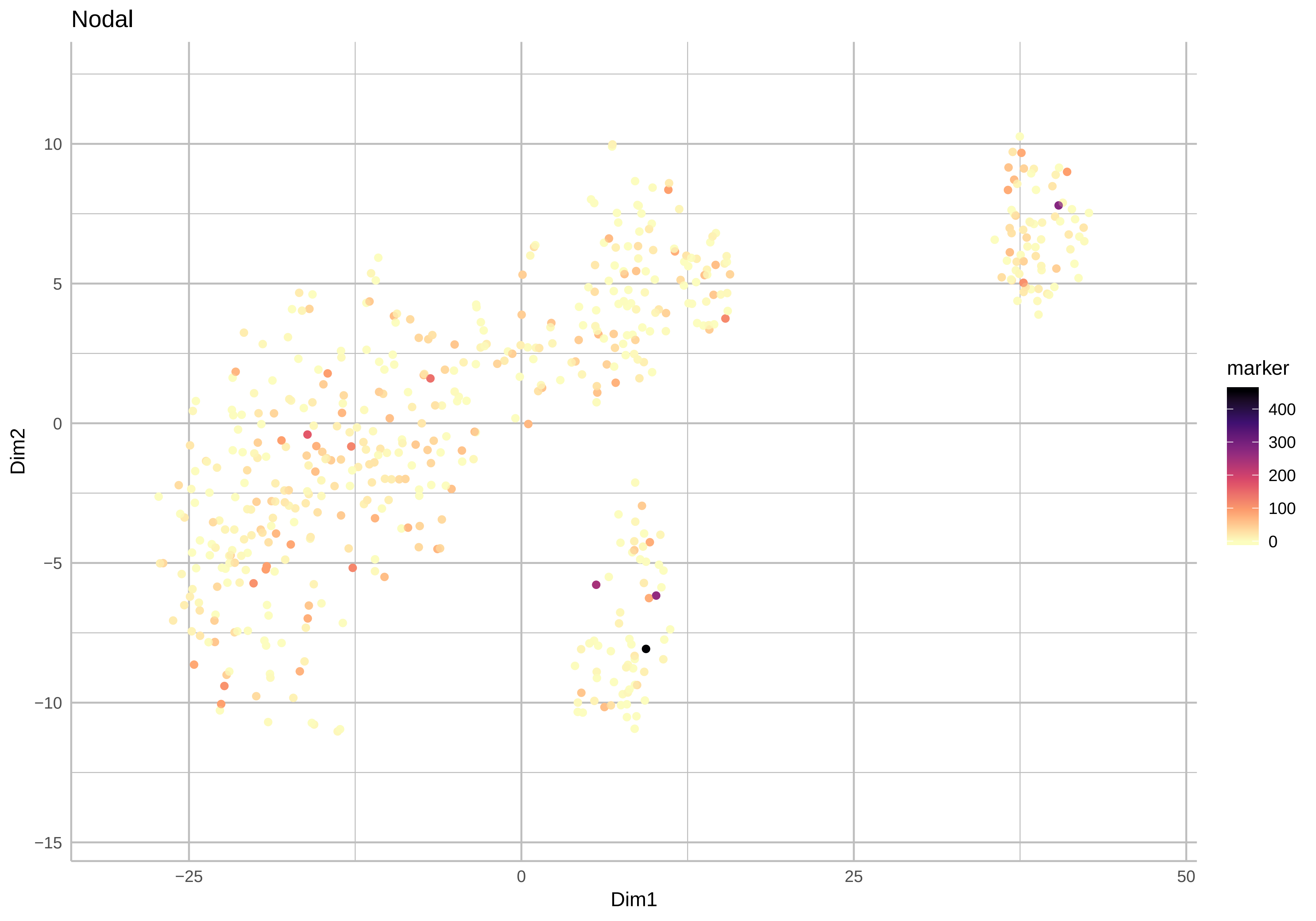

Nodal
Dim2
Dim1
10
5
0
−5
−10
−15
−25
0
25
50
marker
400
300
200
100
0

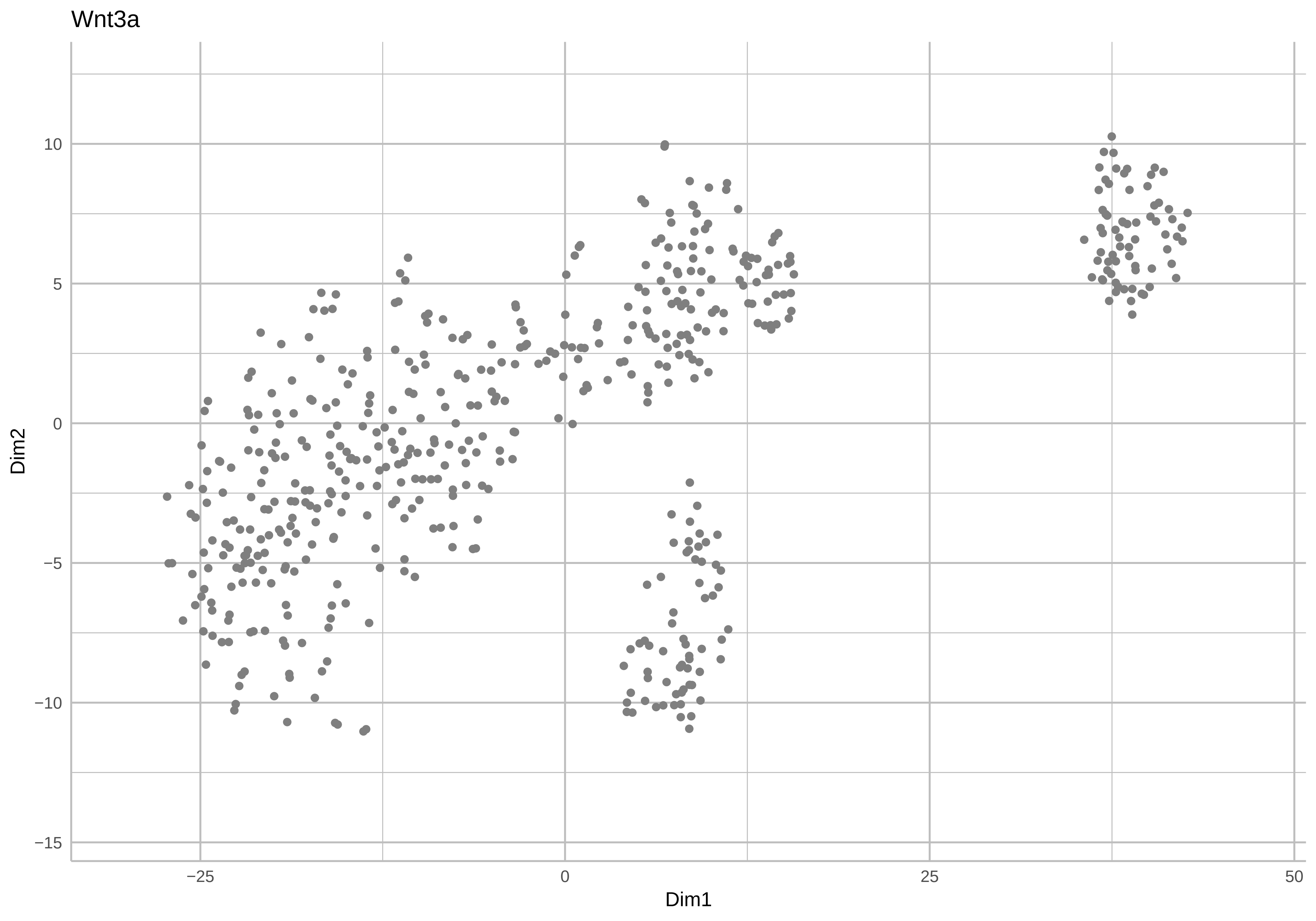
Wnt3a
Dim2
10
5
0
−5
−10
−15
−25
0
25
50
Dim1

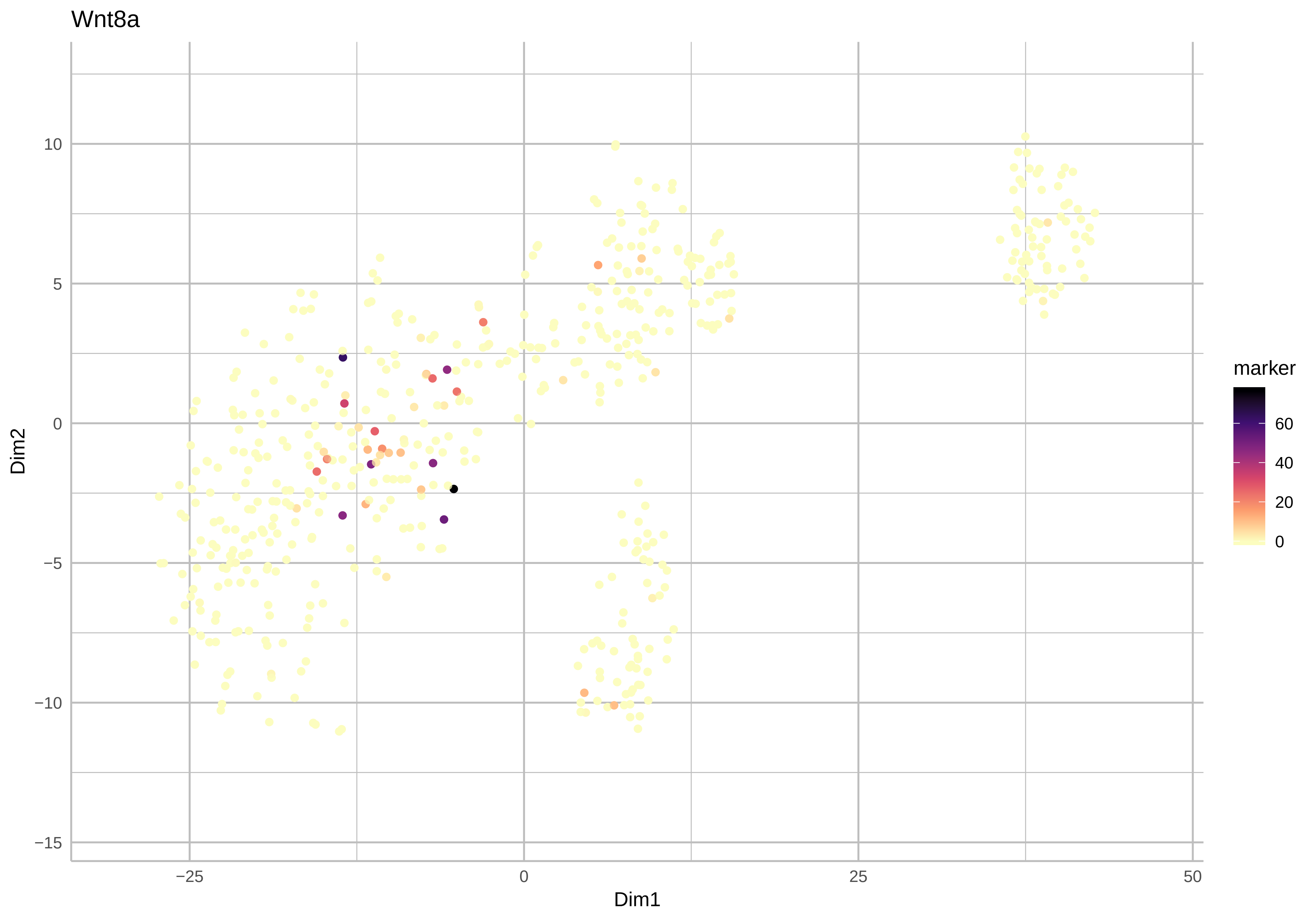
Wnt8a
Dim2
10
5
0
−5
−10
−15
Dim1
−25
0
25
50
marker
60
40
20
0

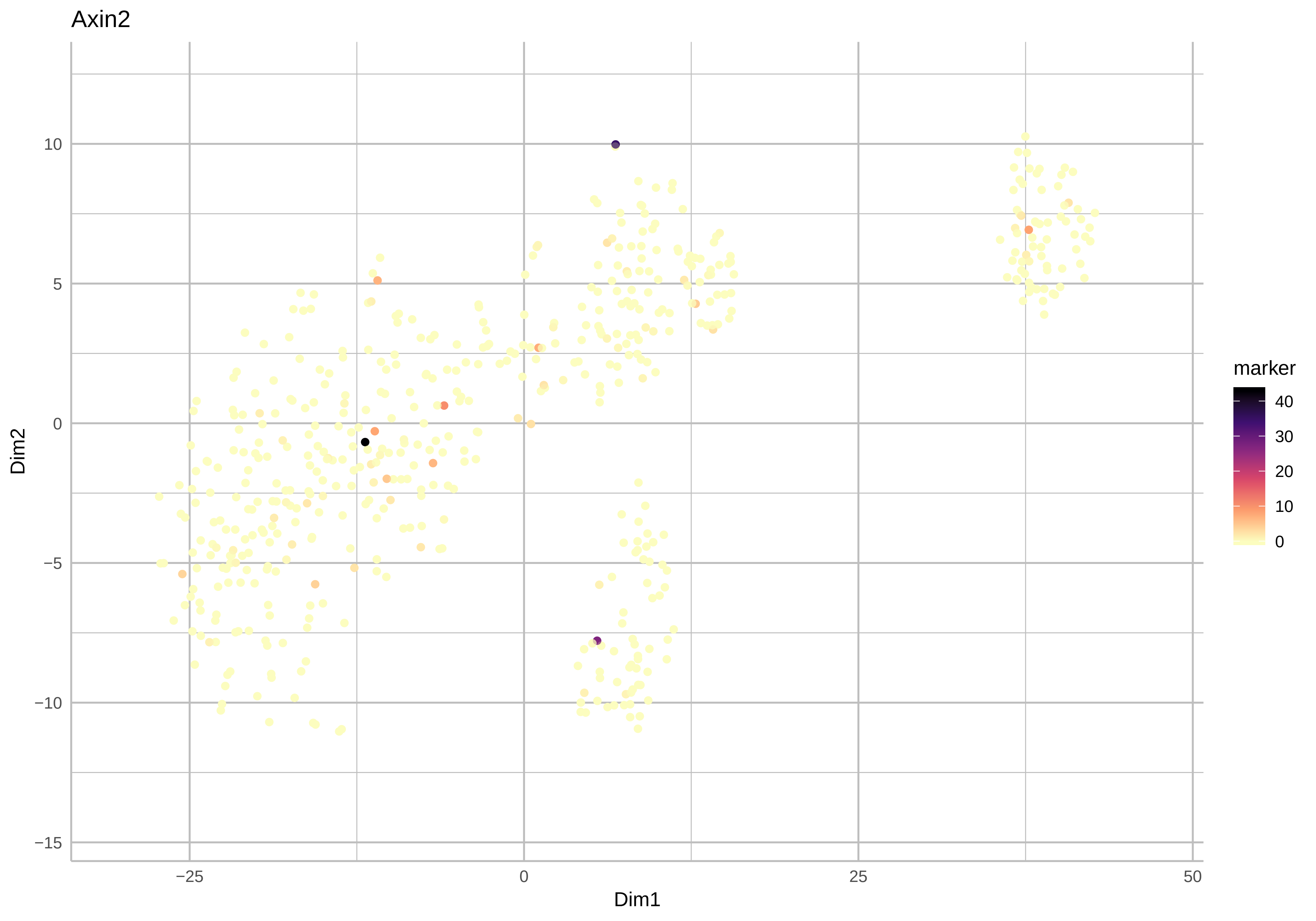
Axin2
Dim2
Dim1
10
5
0
−5
−10
−15
−25
0
25
50
marker
40
30
20
10
0

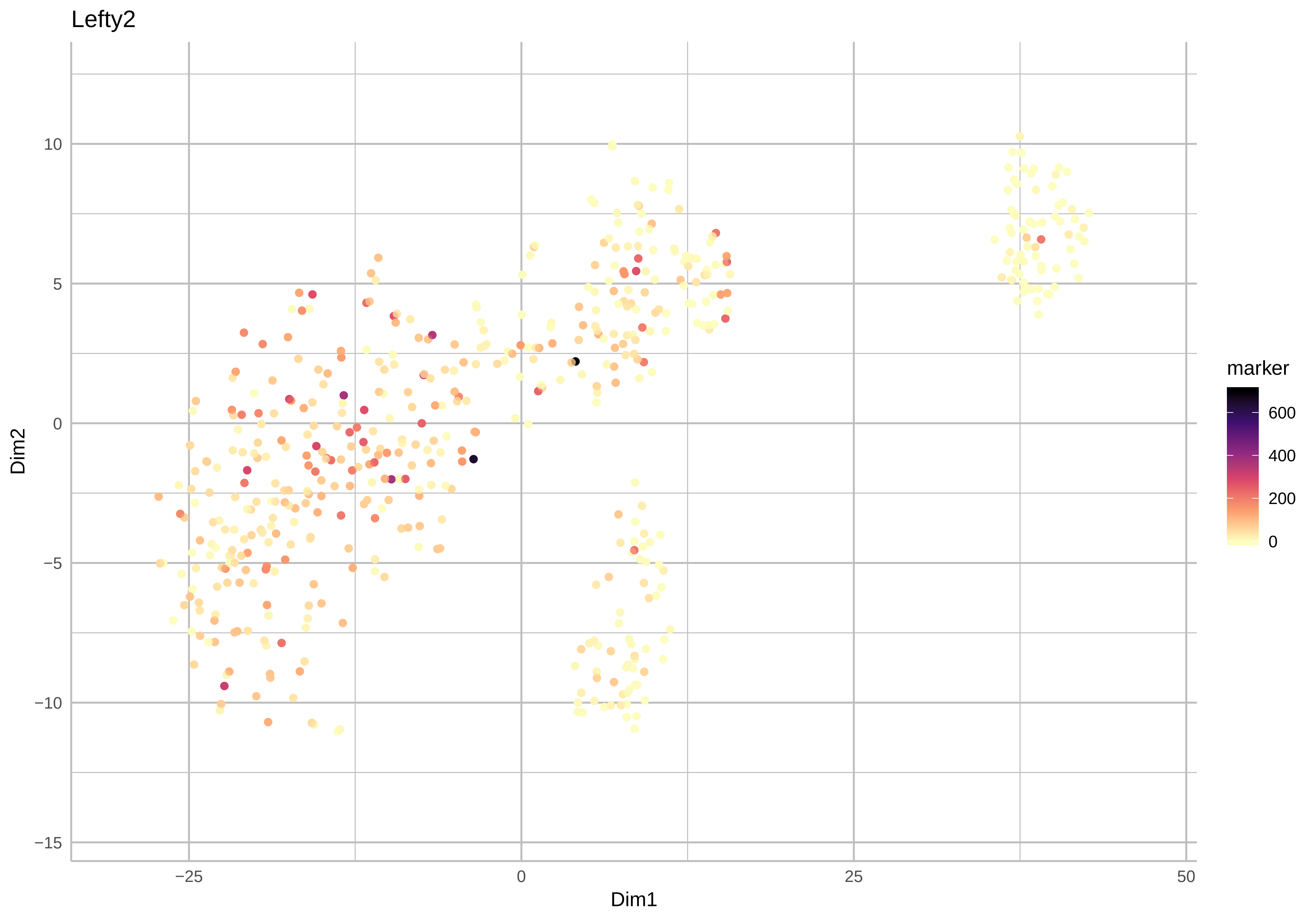

Lefty2
10
5
0
−5
−10
−15
Dim2
−25
0
25
50
Dim1
marker
600
400
200
0

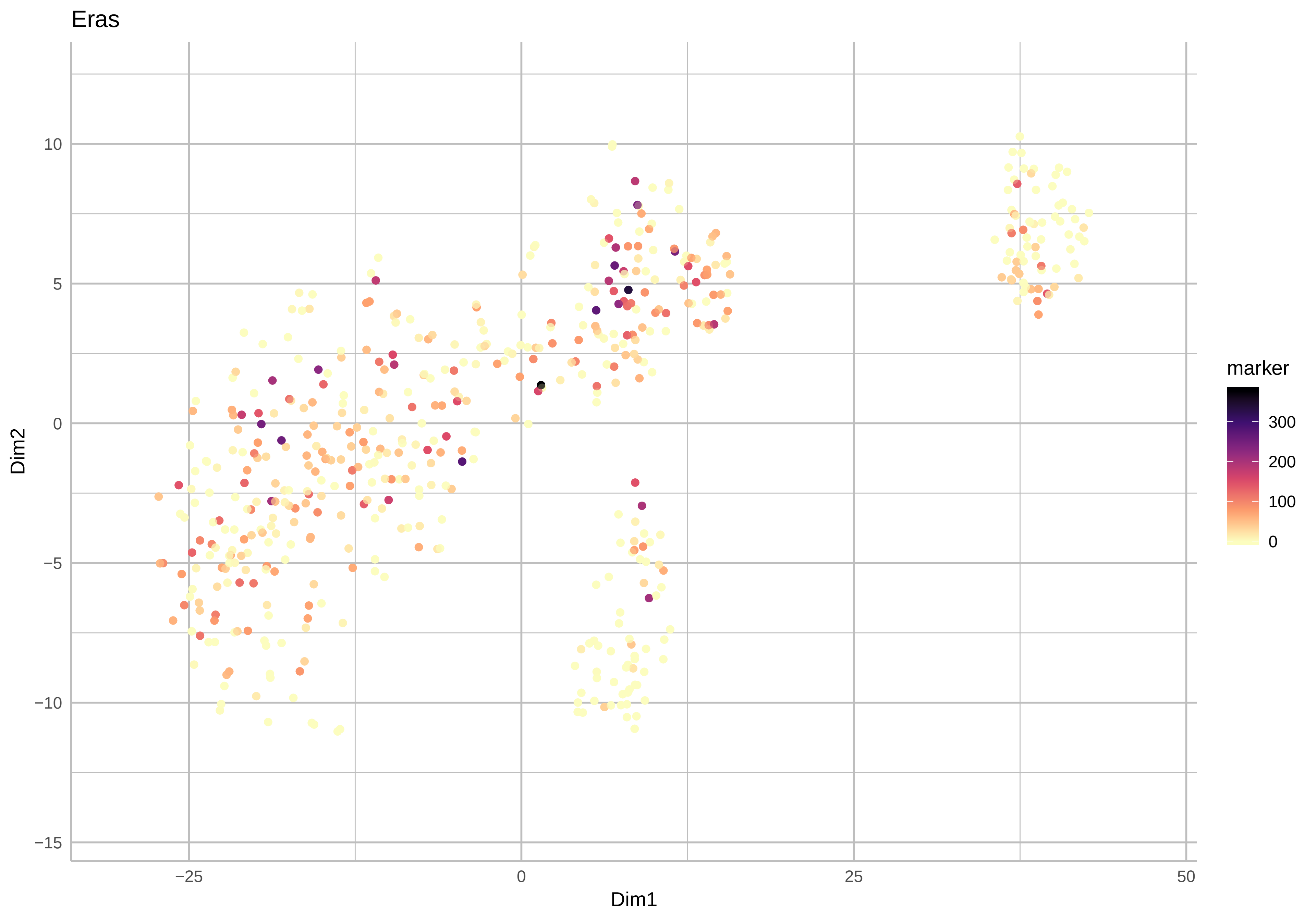

Eras
Dim2
Dim1
10
5
0
−5
−10
−15
−25
0
25
50
marker
300
200
100
0

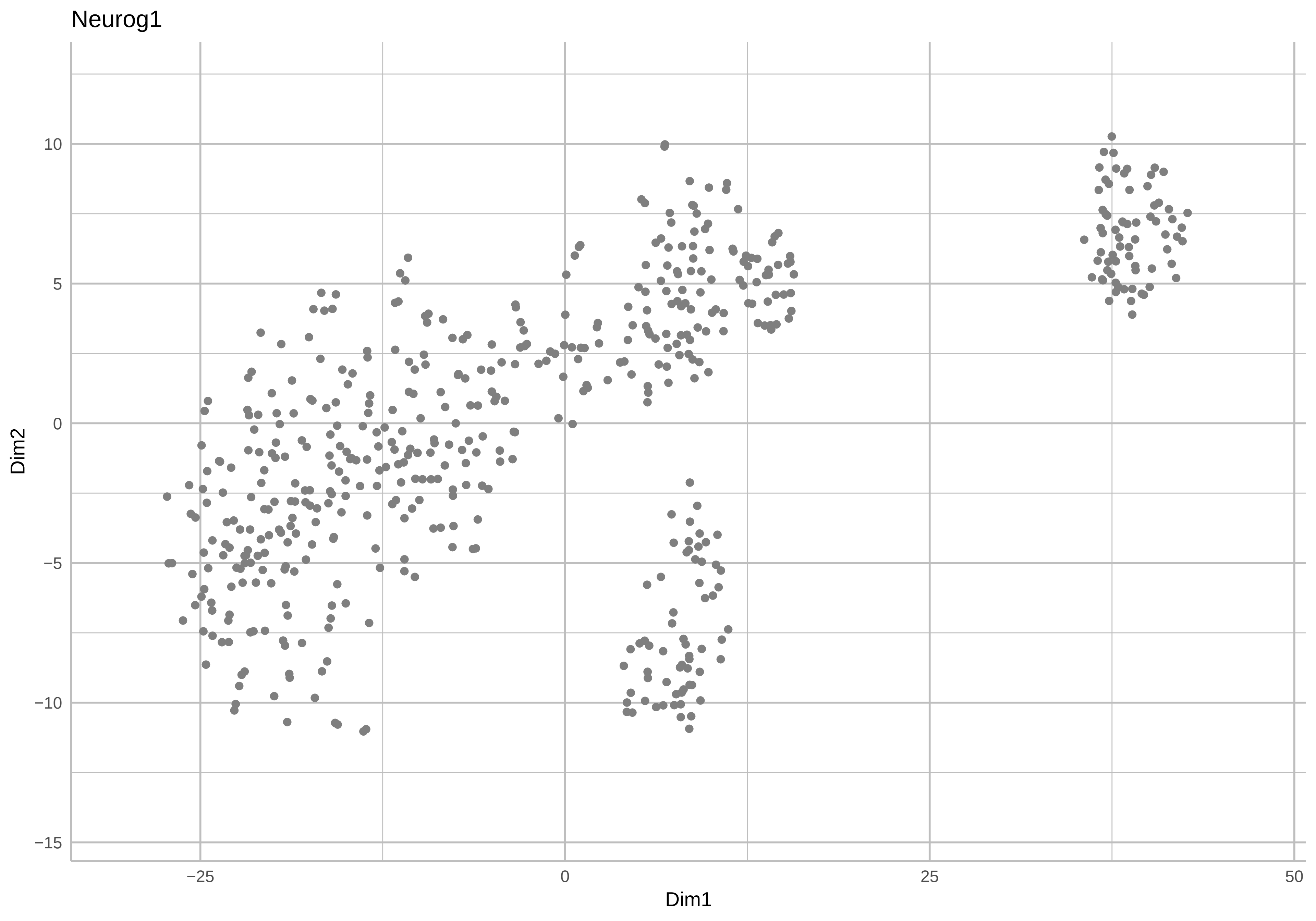
Neurog1
10
5
0
−5
−10
−15
Dim2
−25
0
25
50
Dim1

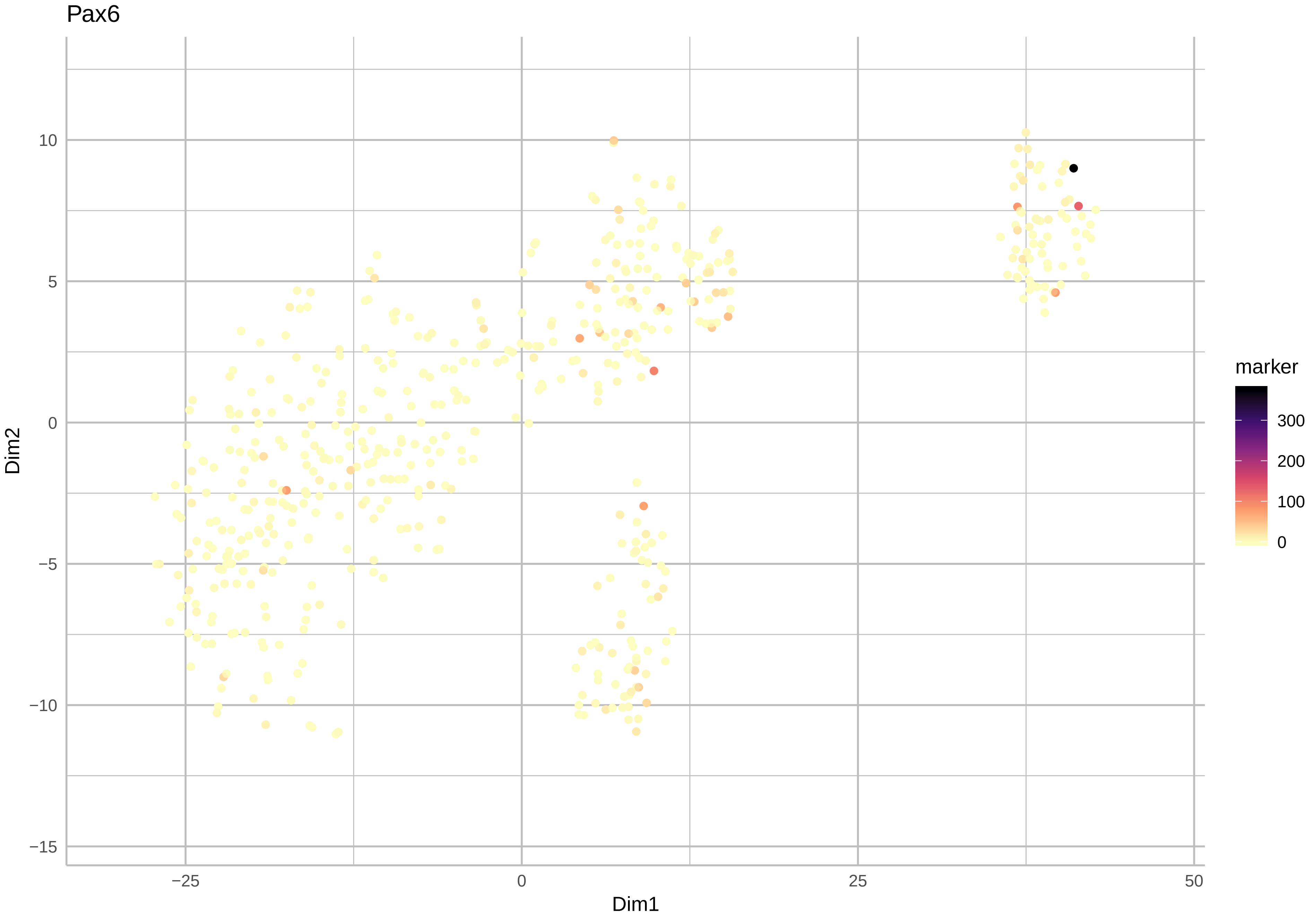

Pax6
Dim2
Dim1
10
5
0
−5
−10
−15
−25
0
25
50
marker
300
200
100
0

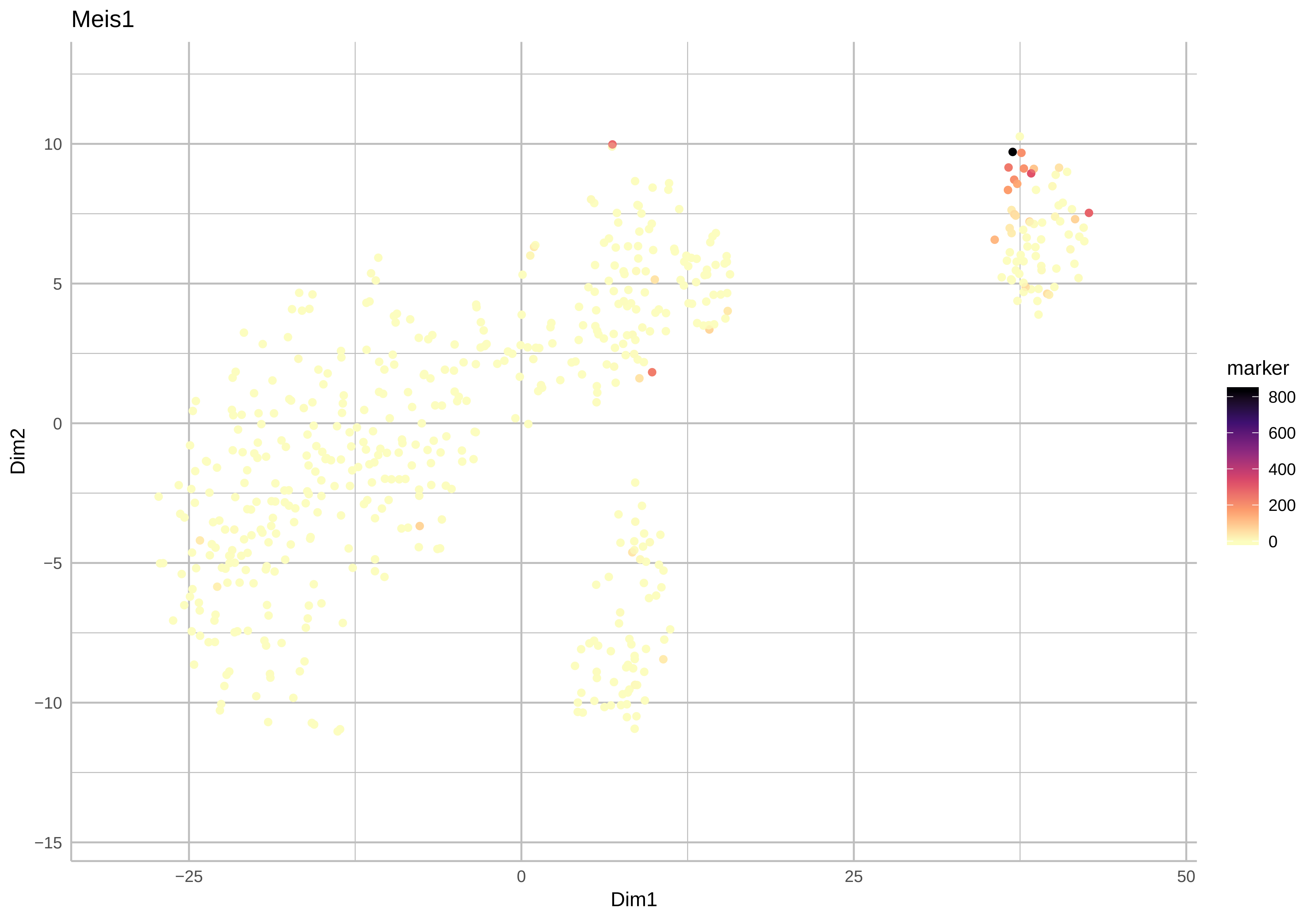
Meis1
Dim2
Dim1
10
5
0
−5
−10
−15
−25
0
25
50
marker
800
600
400
200
0

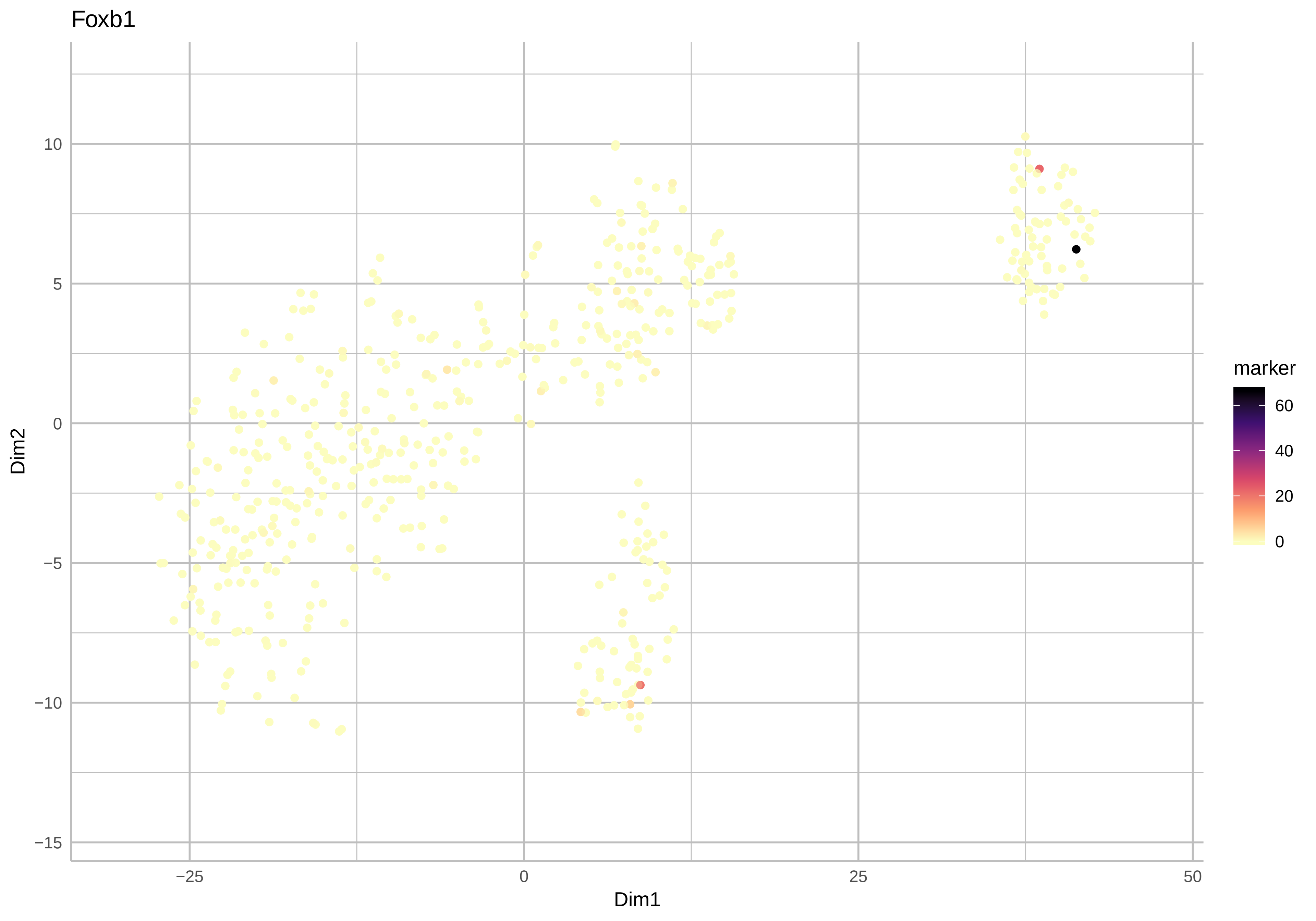

Foxb1
Dim2
Dim1
10
5
0
−5
−10
−15
−25
0
25
50
marker
60
40
20
0

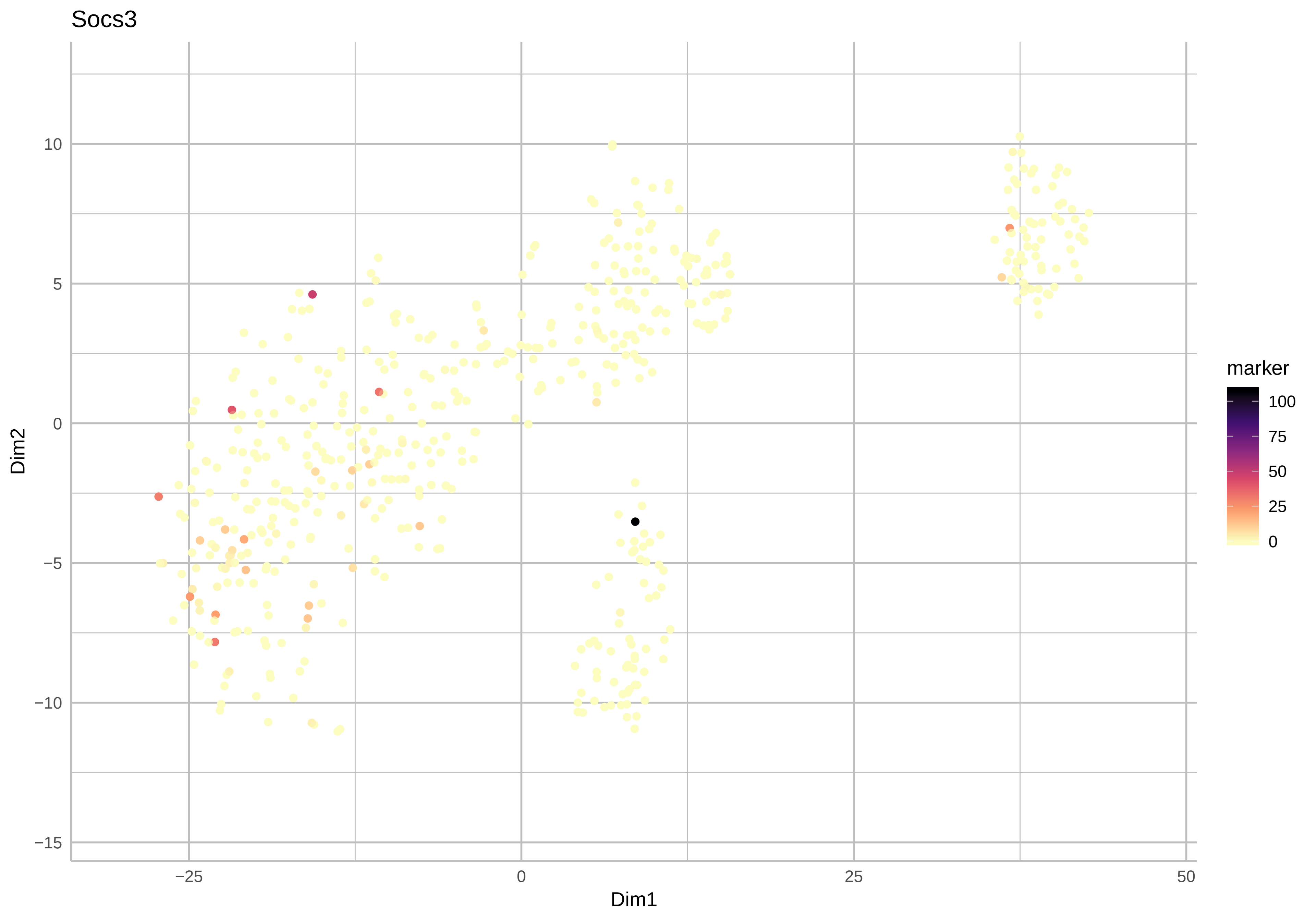
Socs3
Dim2
Dim1
10
5
0
−5
−10
−15
−25
0
25
50
marker
100
75
50
25
0

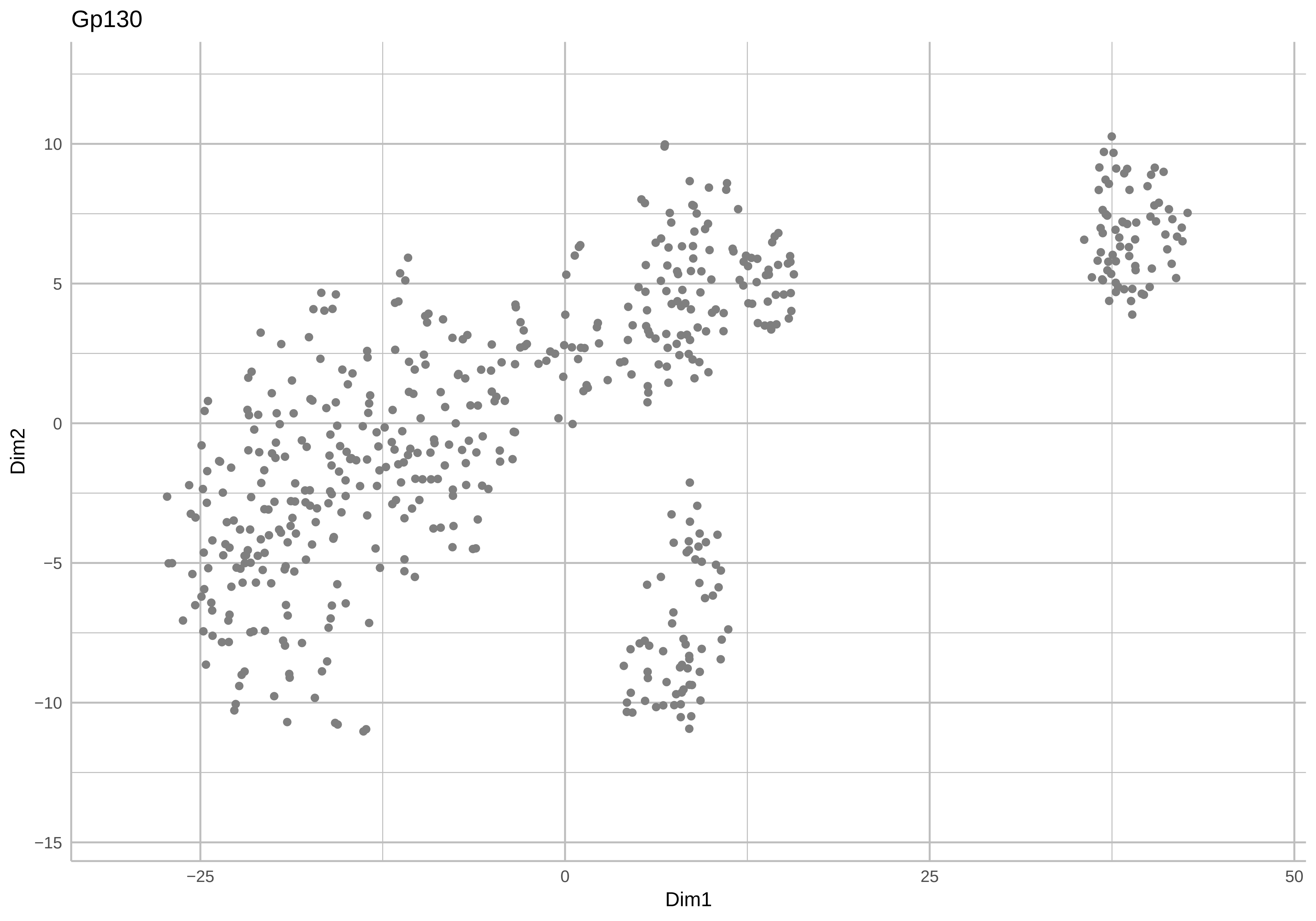

Gp130
Dim2
Dim1
10
5
0
−5
−10
−15
−25
0
25
50

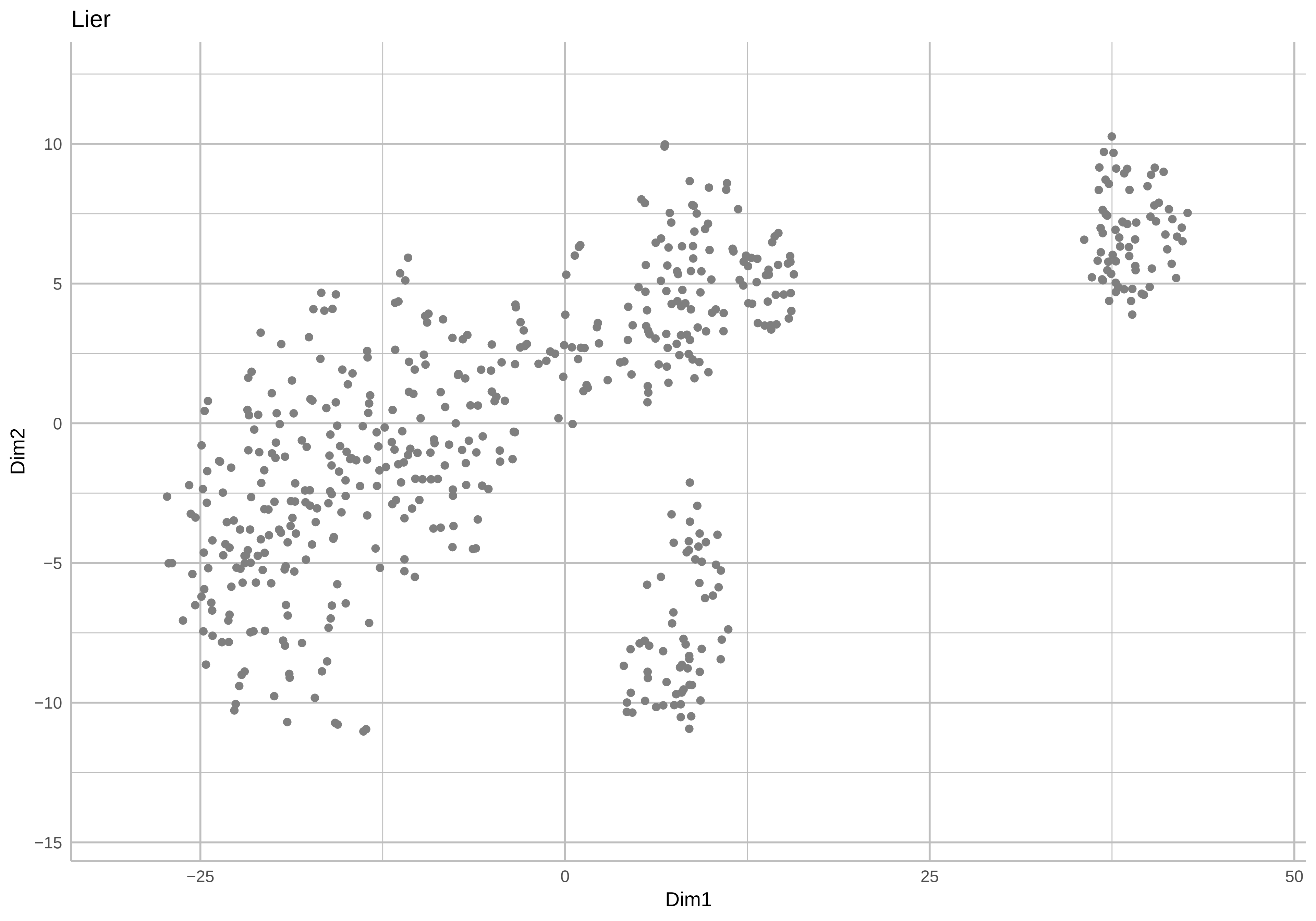

Lier
Dim2
Dim1
10
5
0
−5
−10
−15
−25
0
25
50

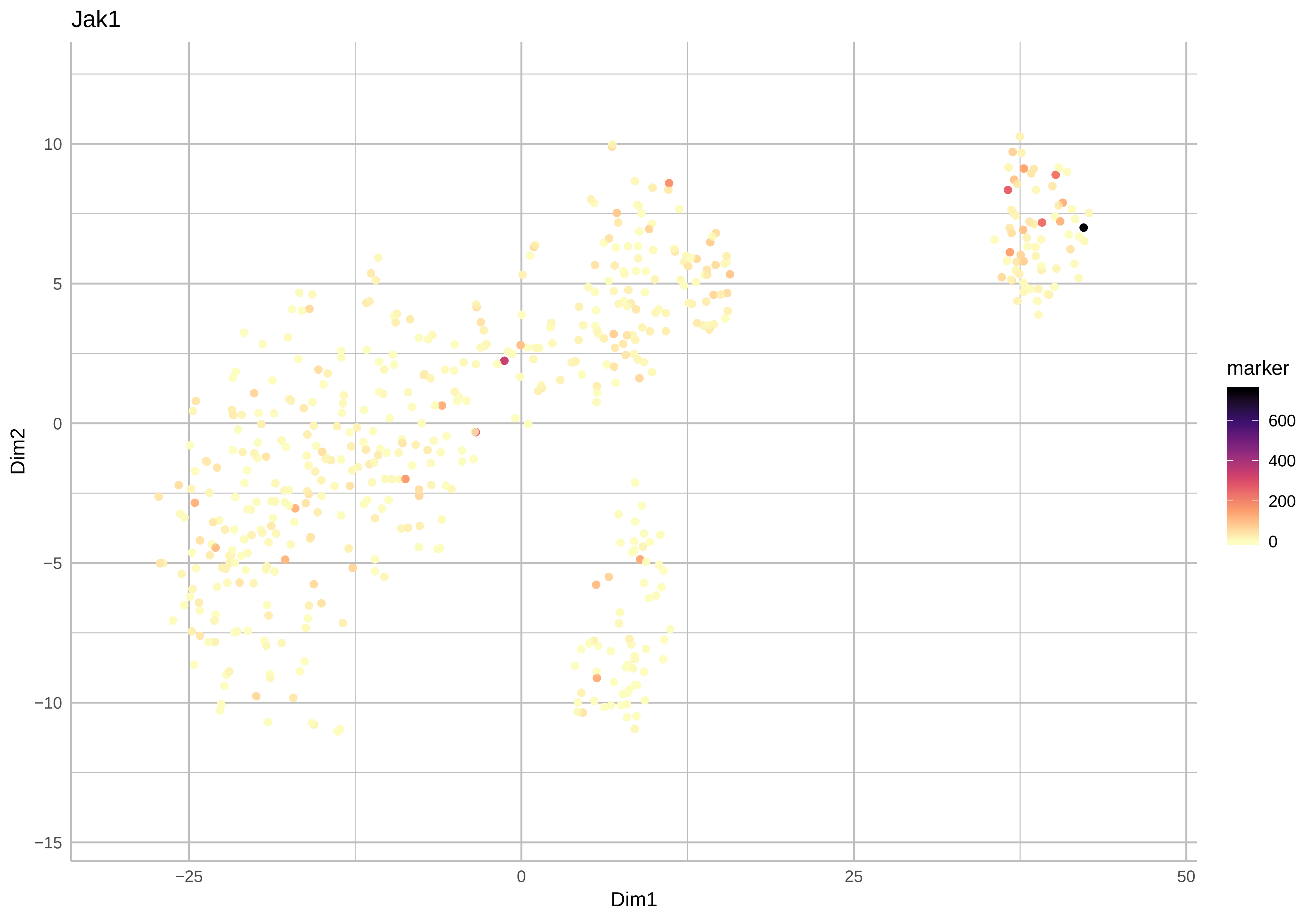
Jak1
Dim2
Dim1
10
5
0
−5
−10
−15
−25
0
25
50
marker
600
400
200
0

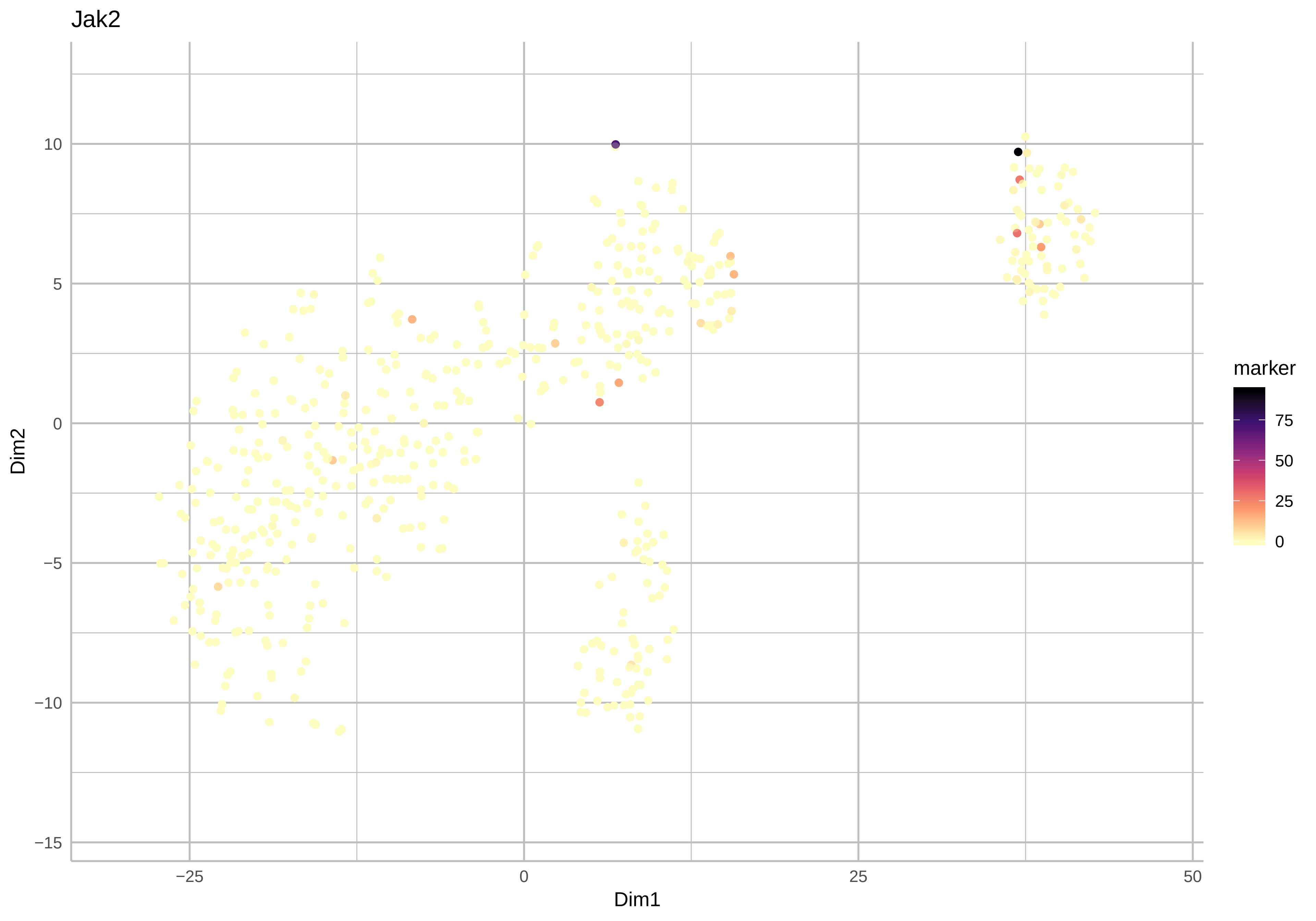

Jak2
Dim2
Dim1
10
5
0
−5
−10
−15
−25
0
25
50
marker
75
50
25
0

Tyk2

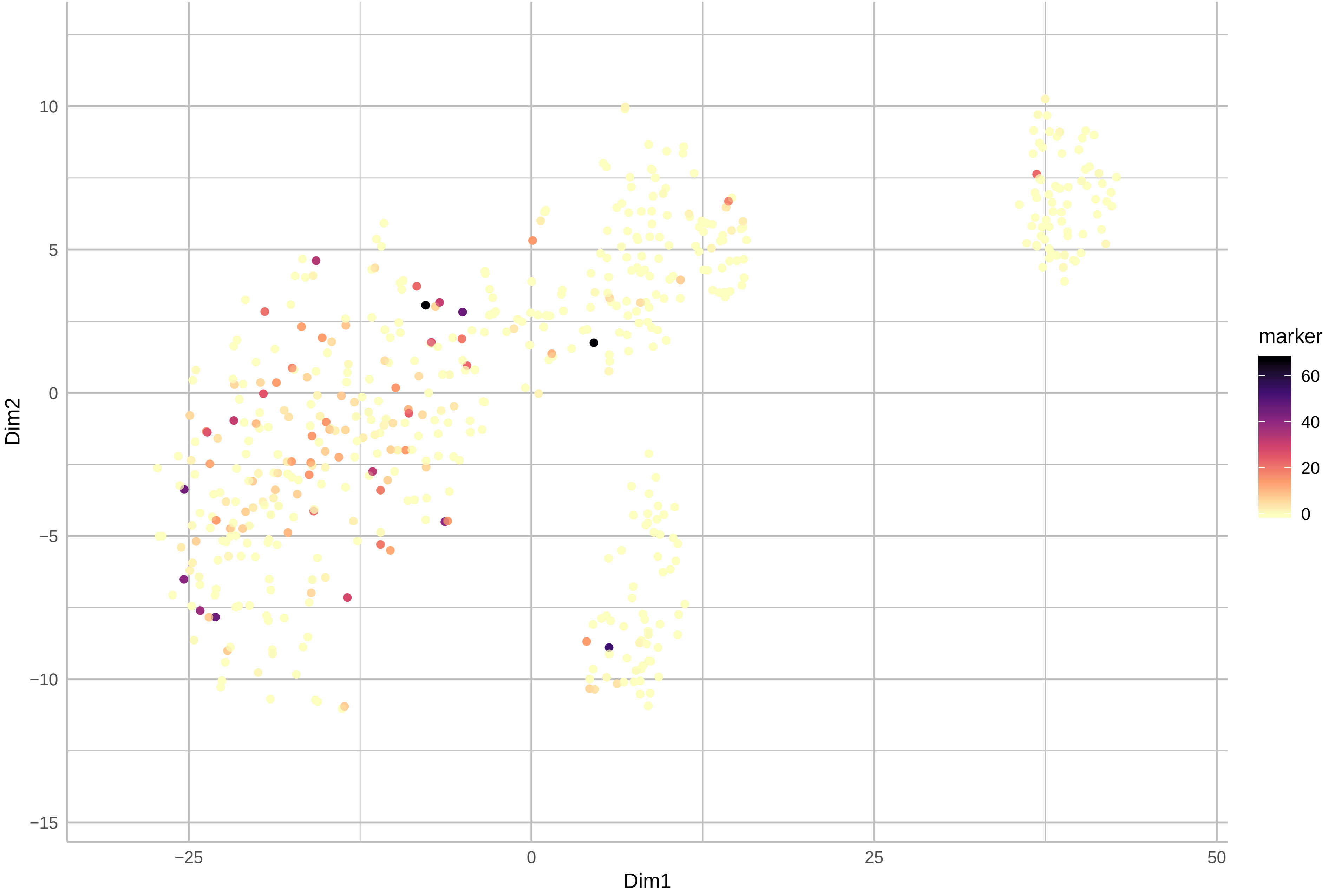

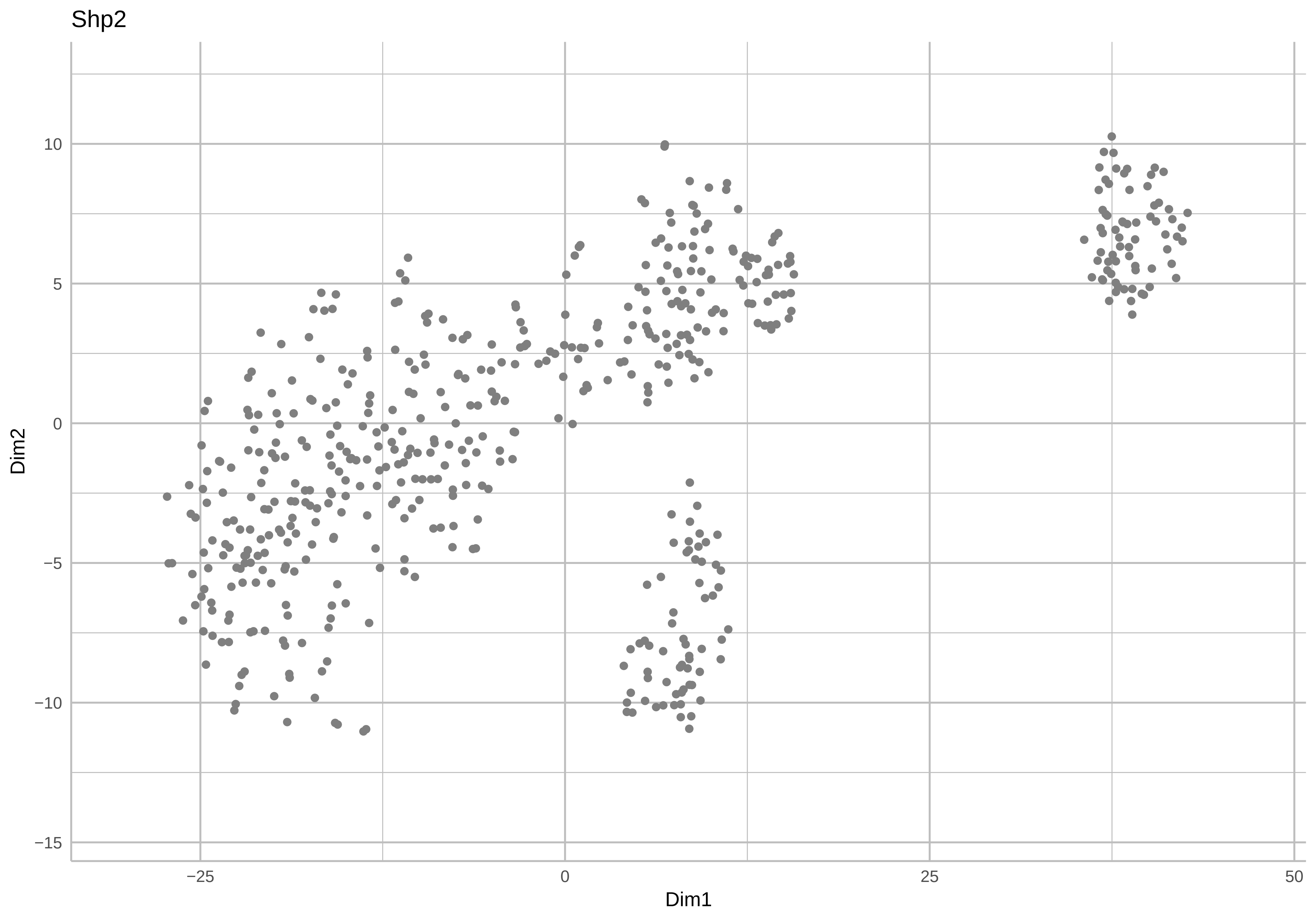

Shp2
Dim2
Dim1
10
5
0
−5
−10
−15
−25
0
25
50

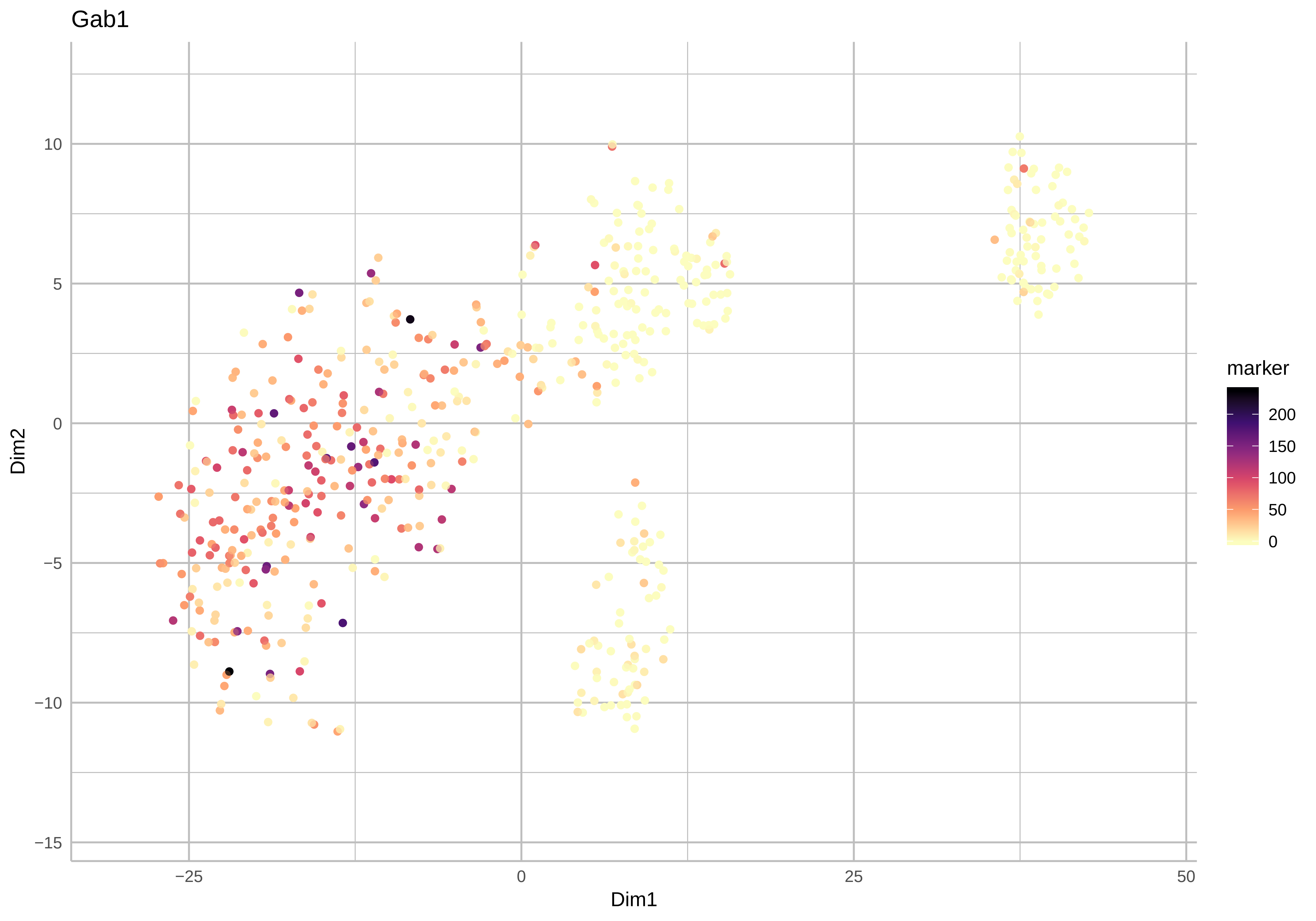

Gab1
10
5
0
−5
−10
−15
Dim2
−25
0
25
50
Dim1
marker
200
150
100
50
0

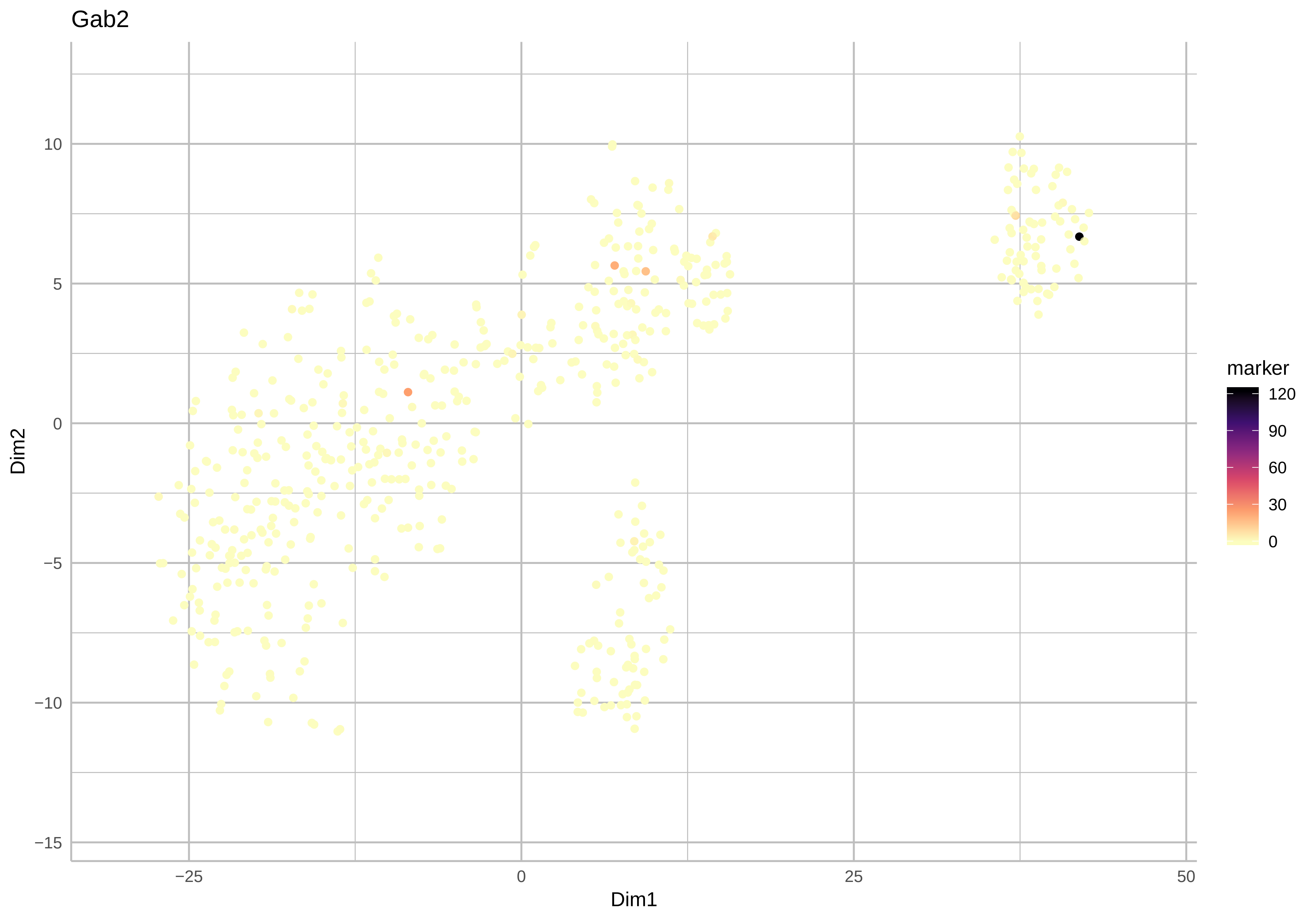

Gab2
Dim2
Dim1
marker
120
90
60
30
0
10
5
0
−5
−10
−15
−25
0
25
50

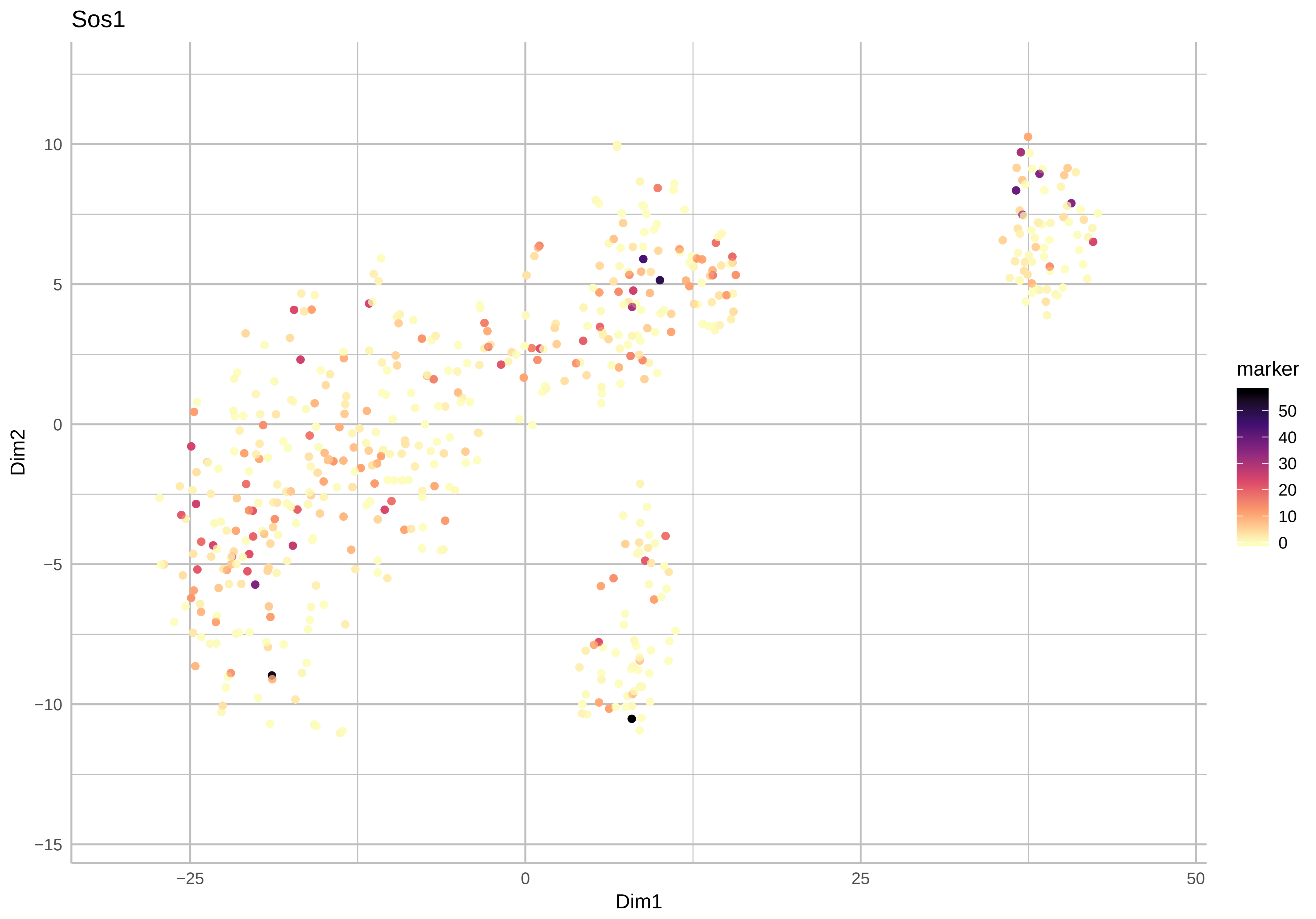

Sos1
Dim2
Dim1
10
5
0
−5
−10
−15
−25
0
25
50
marker
50
40
30
20
10
0

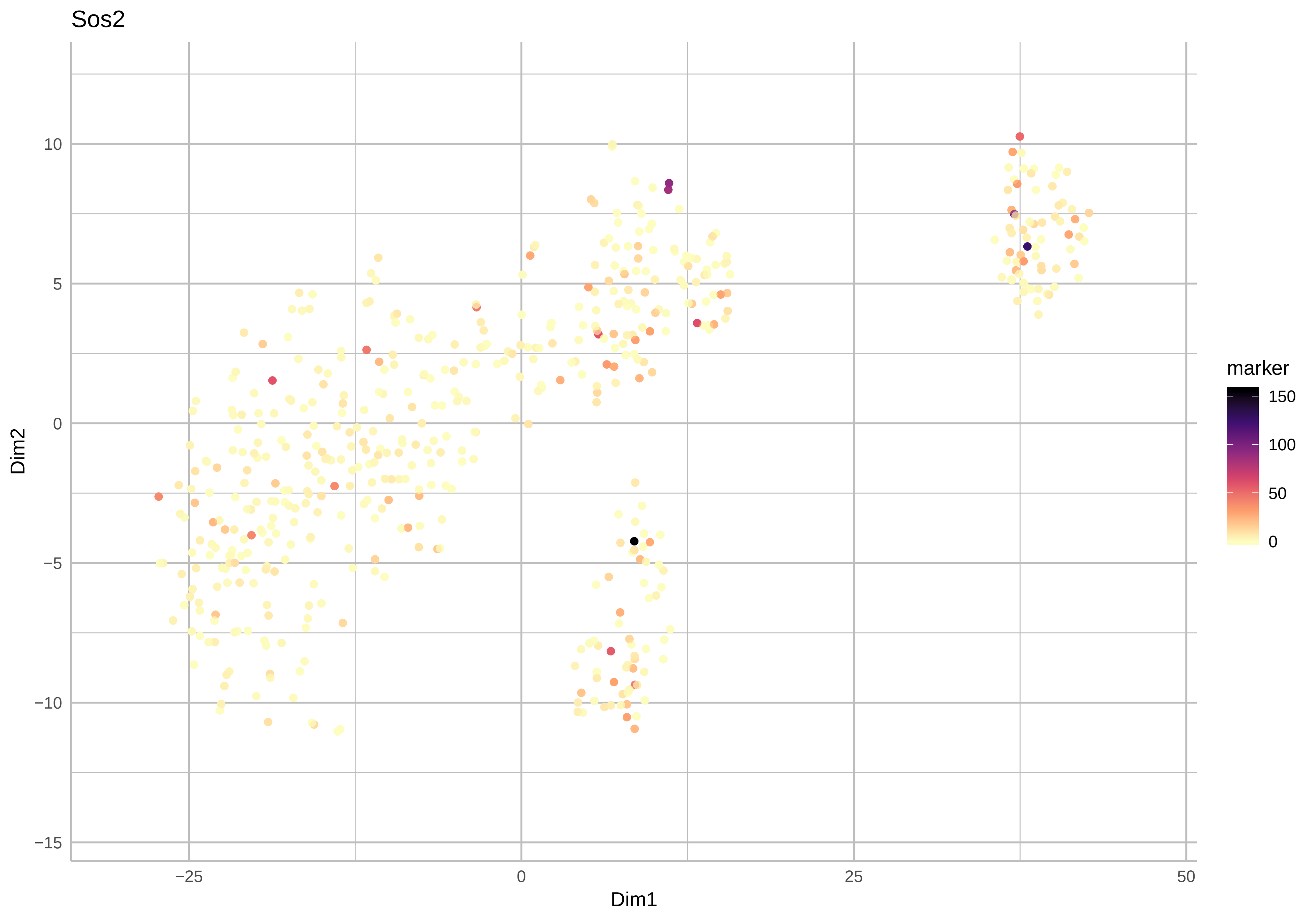
Sos2
Dim2
Dim1
10
5
0
−5
−10
−15
−25
0
25
50
marker
150
100
50
0

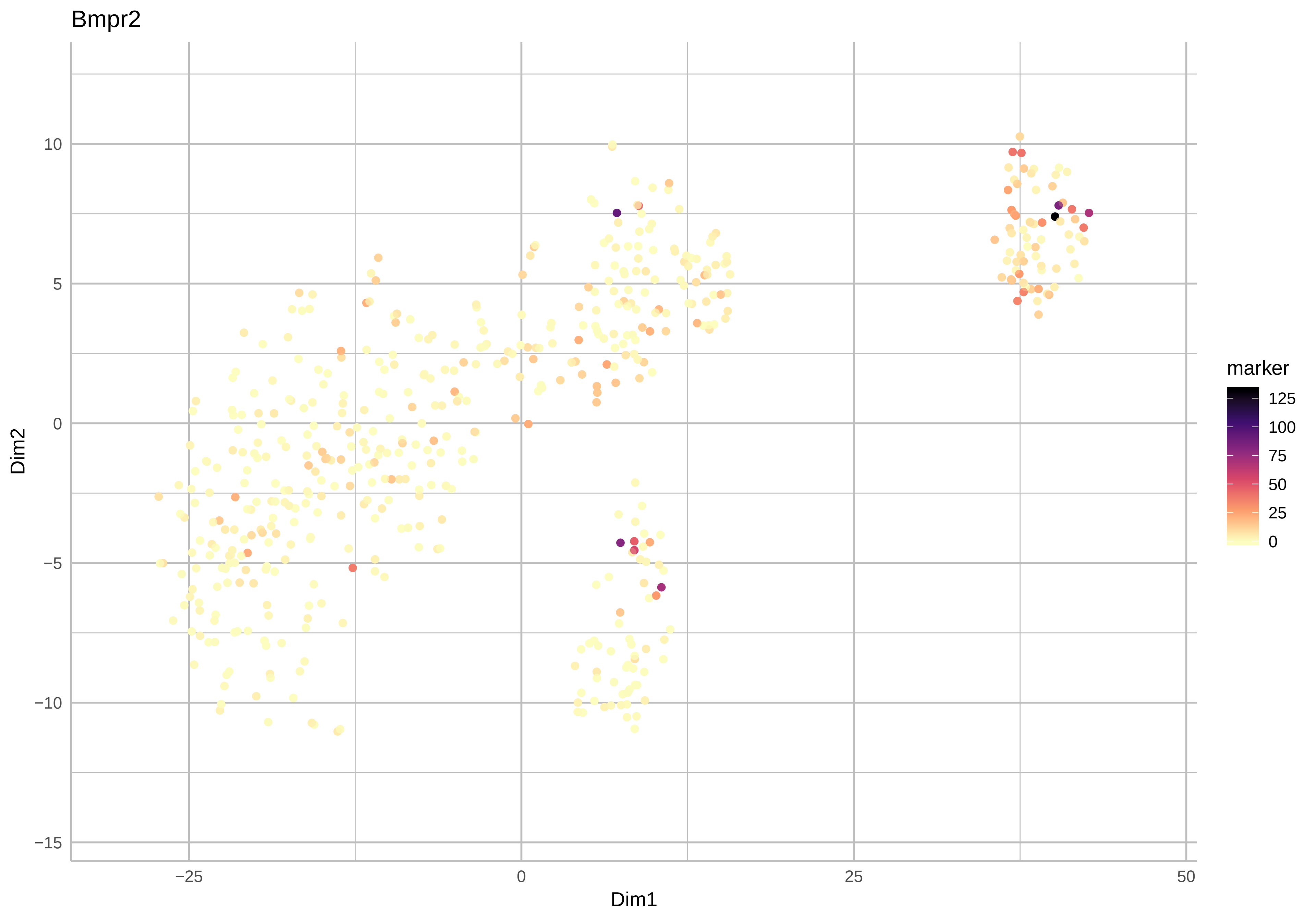
Bmpr2
10
5
0
−5
−10
−15
−25
0
25
50
Dim1
Dim2
marker
125
100
75
50
25
0

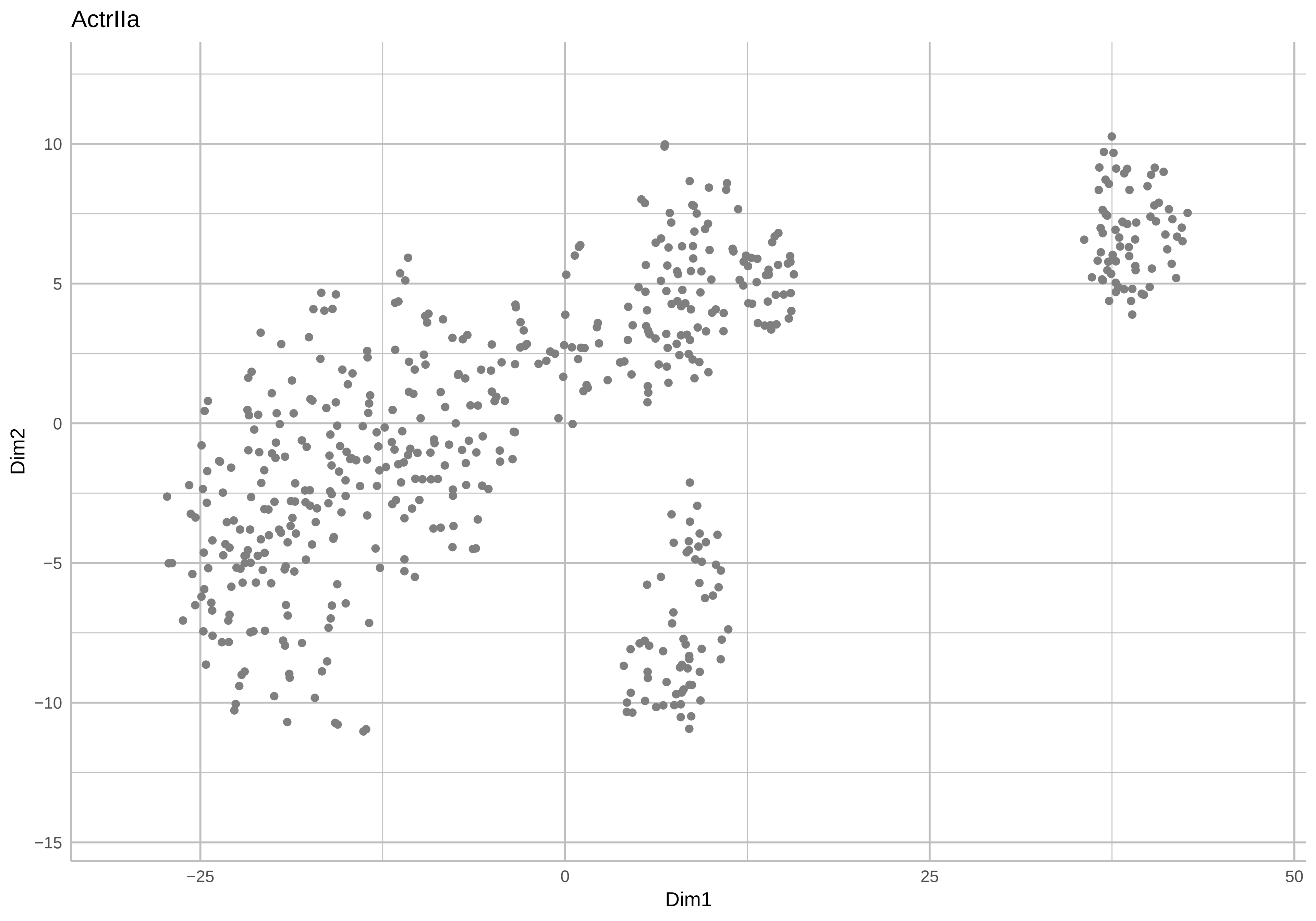
ActrIIa
Dim2
Dim1
10
5
0
−5
−10
−15
−25
0
25
50

ActrIIb

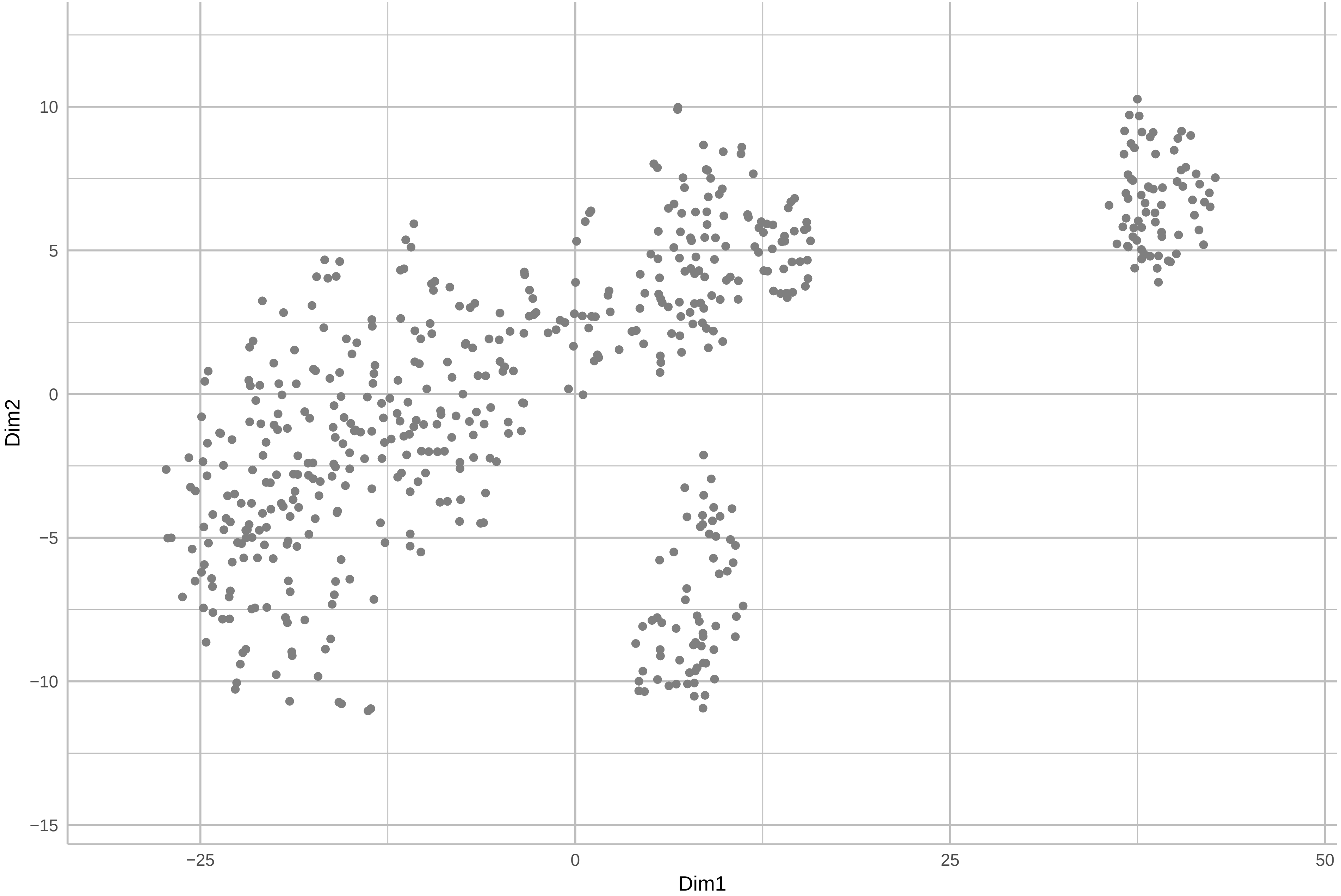

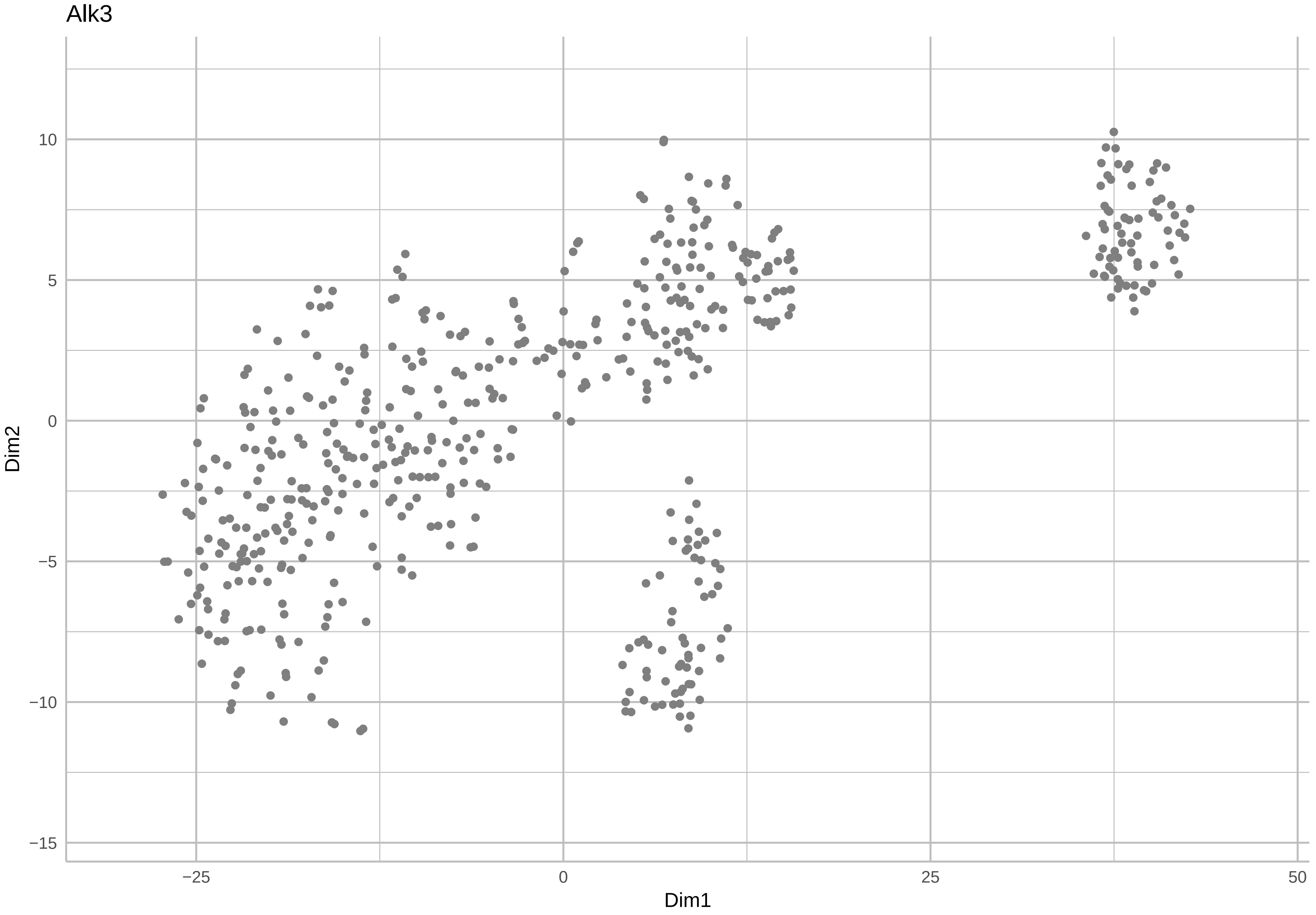

Alk3
10
5
0
−5
−10
−15
Dim2
−25
0
25
50
Dim1

Alk6

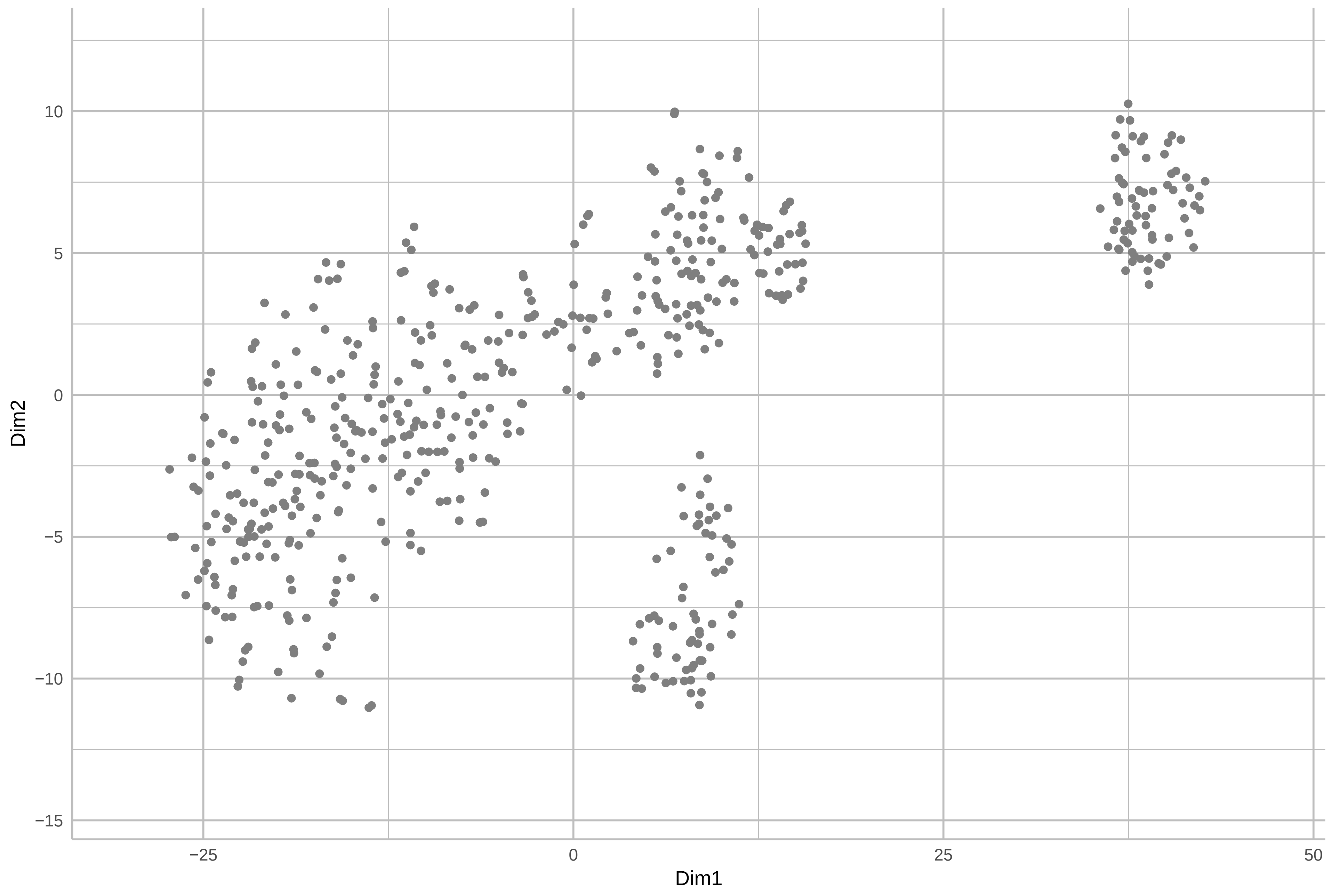

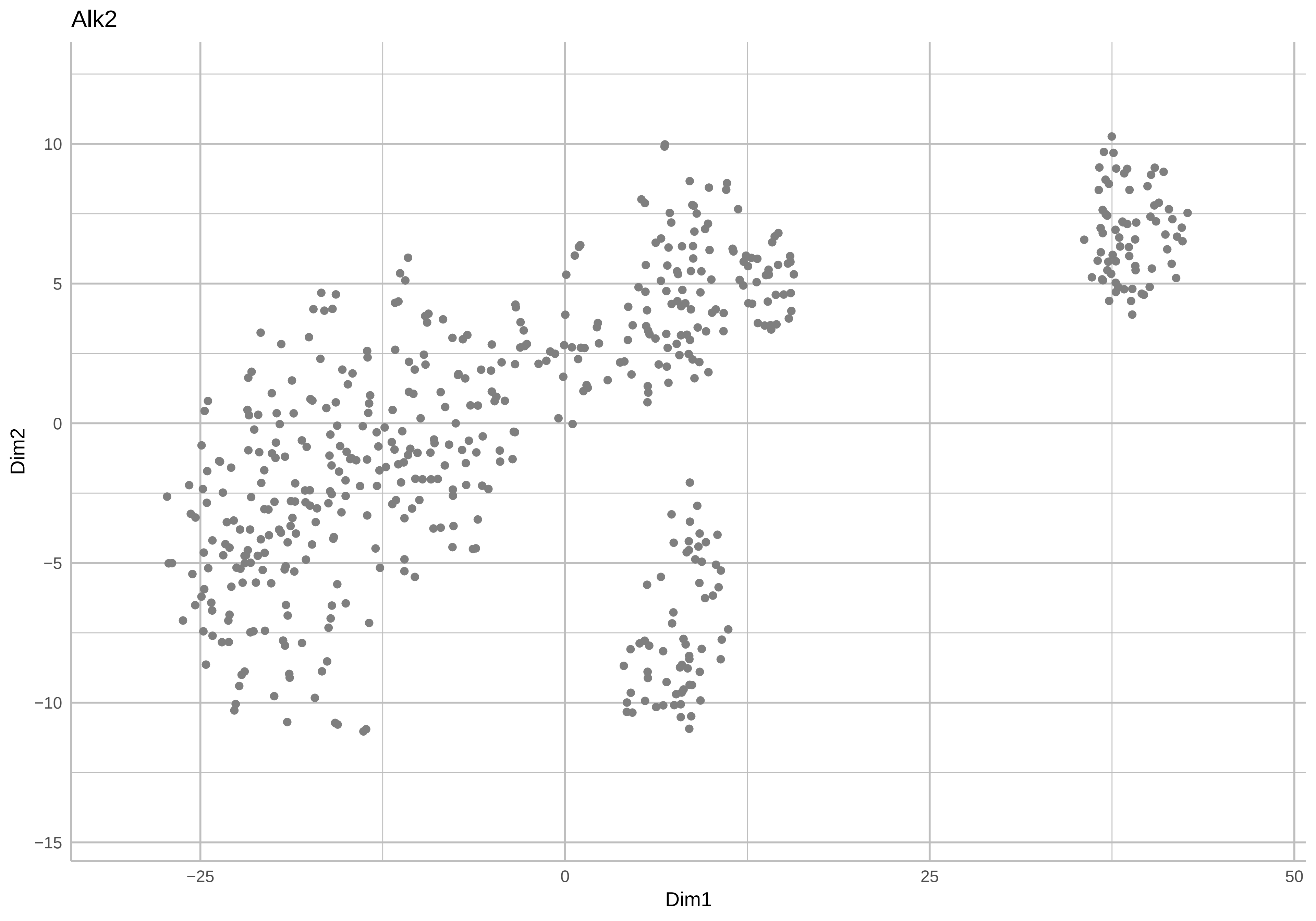
Alk2
Dim2
Dim1
10
5
0
−5
−10
−15
−25
0
25
50

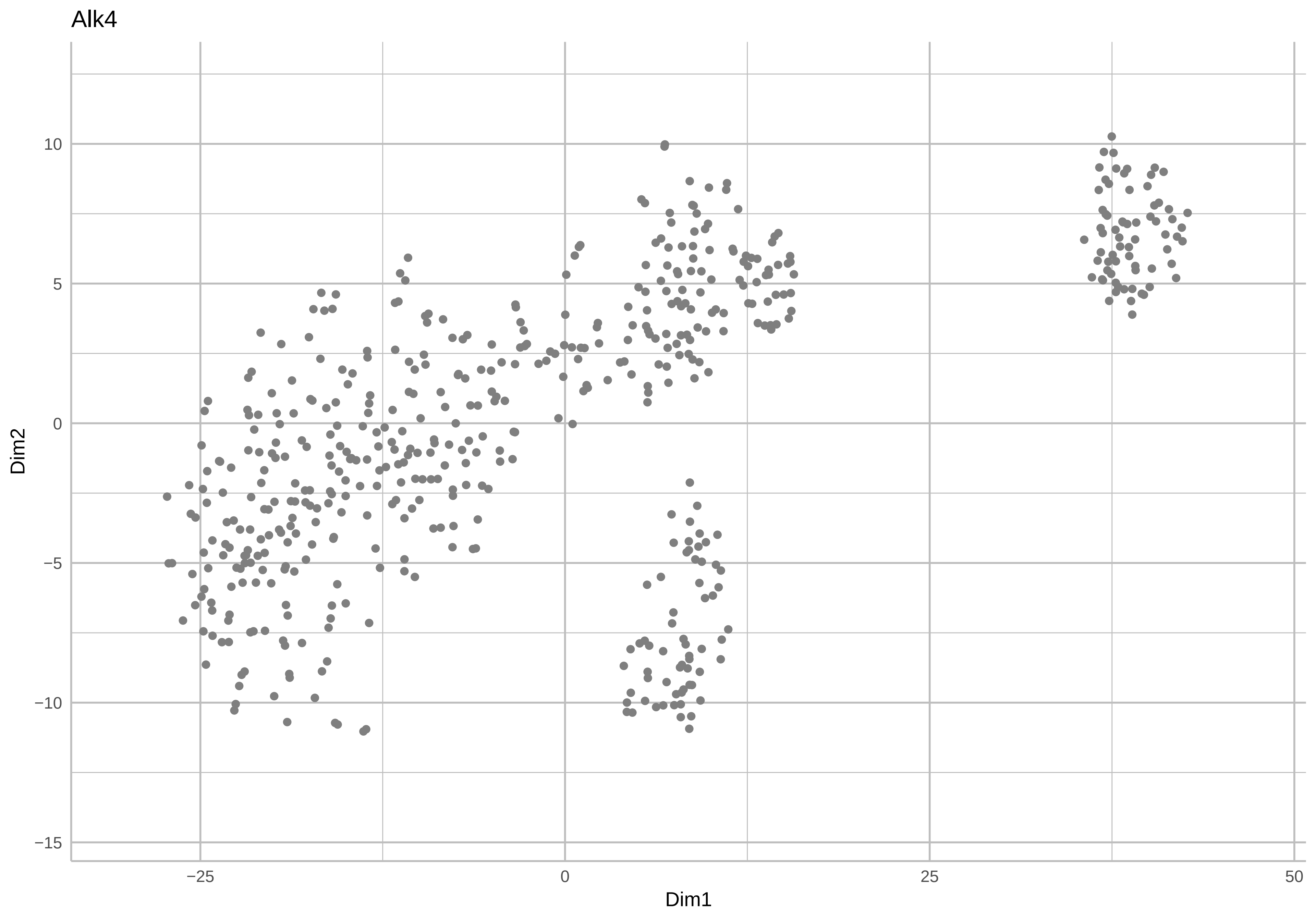
Alk4
Dim2
Dim1
10
5
0
−5
−10
−15
−25
0
25
50

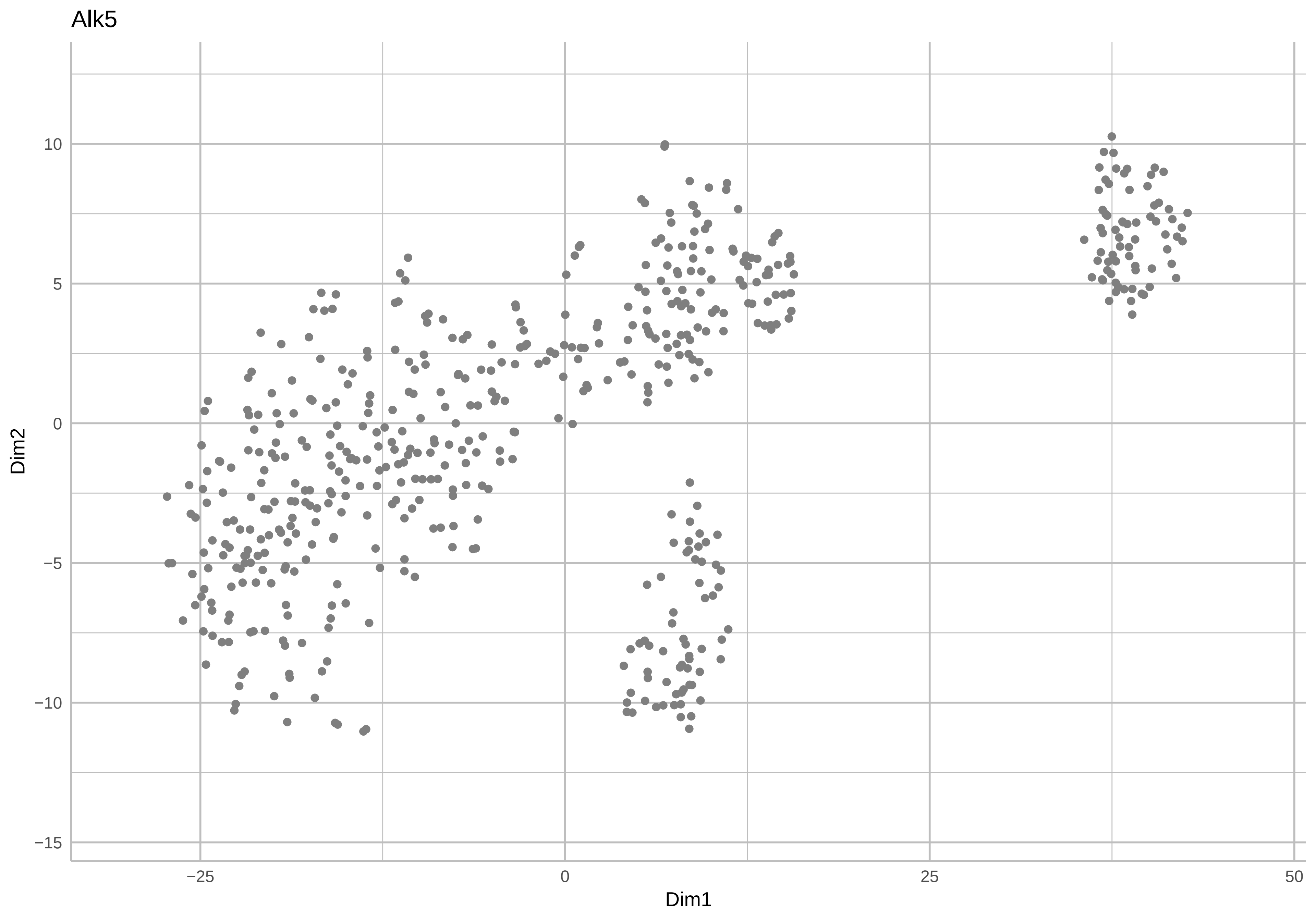
Alk5
Dim2
10
5
0
−5
−10
−15
−25
0
25
50
Dim1

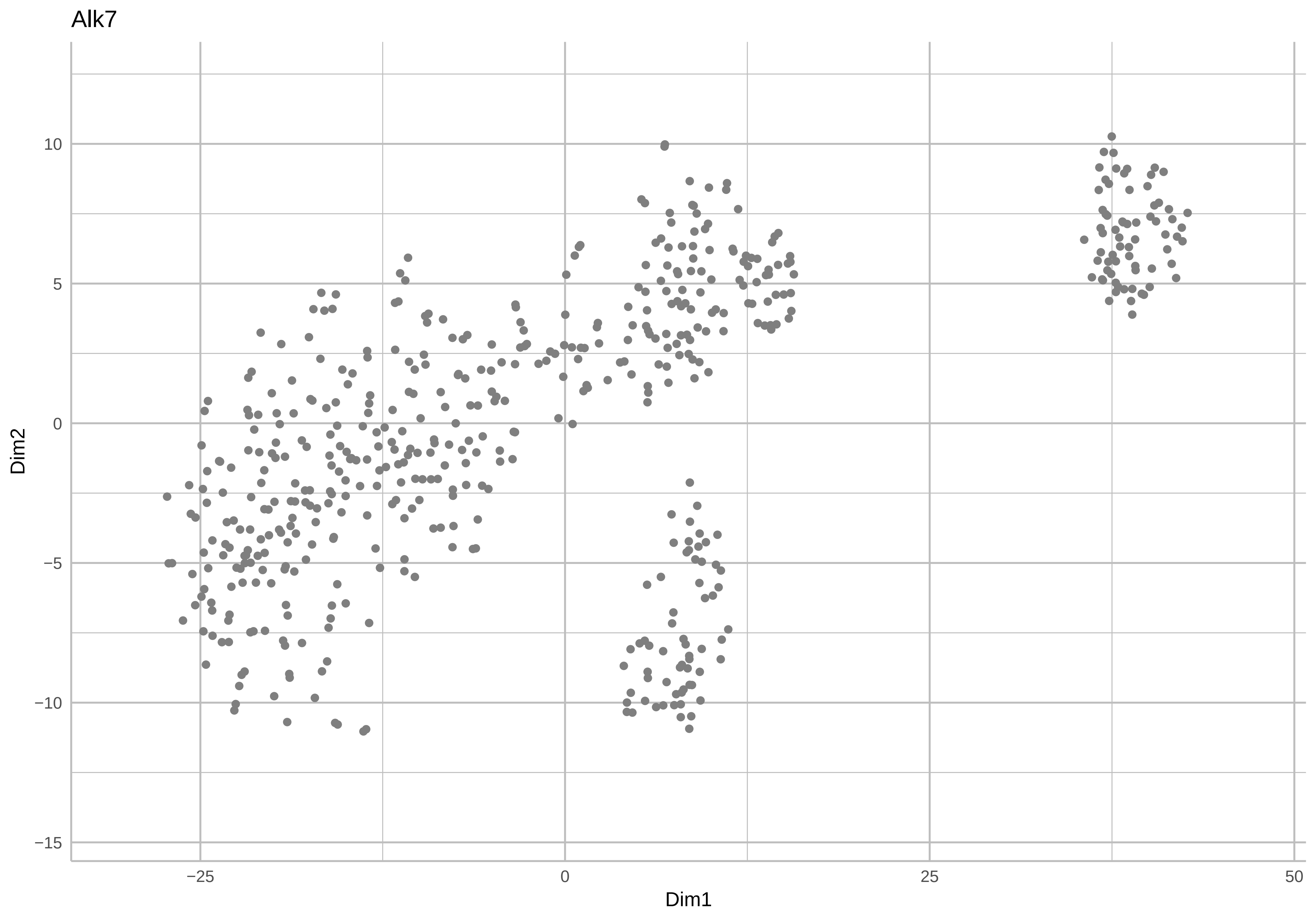
Alk7
Dim2
Dim1
10
5
0
−5
−10
−15
−25
0
25
50

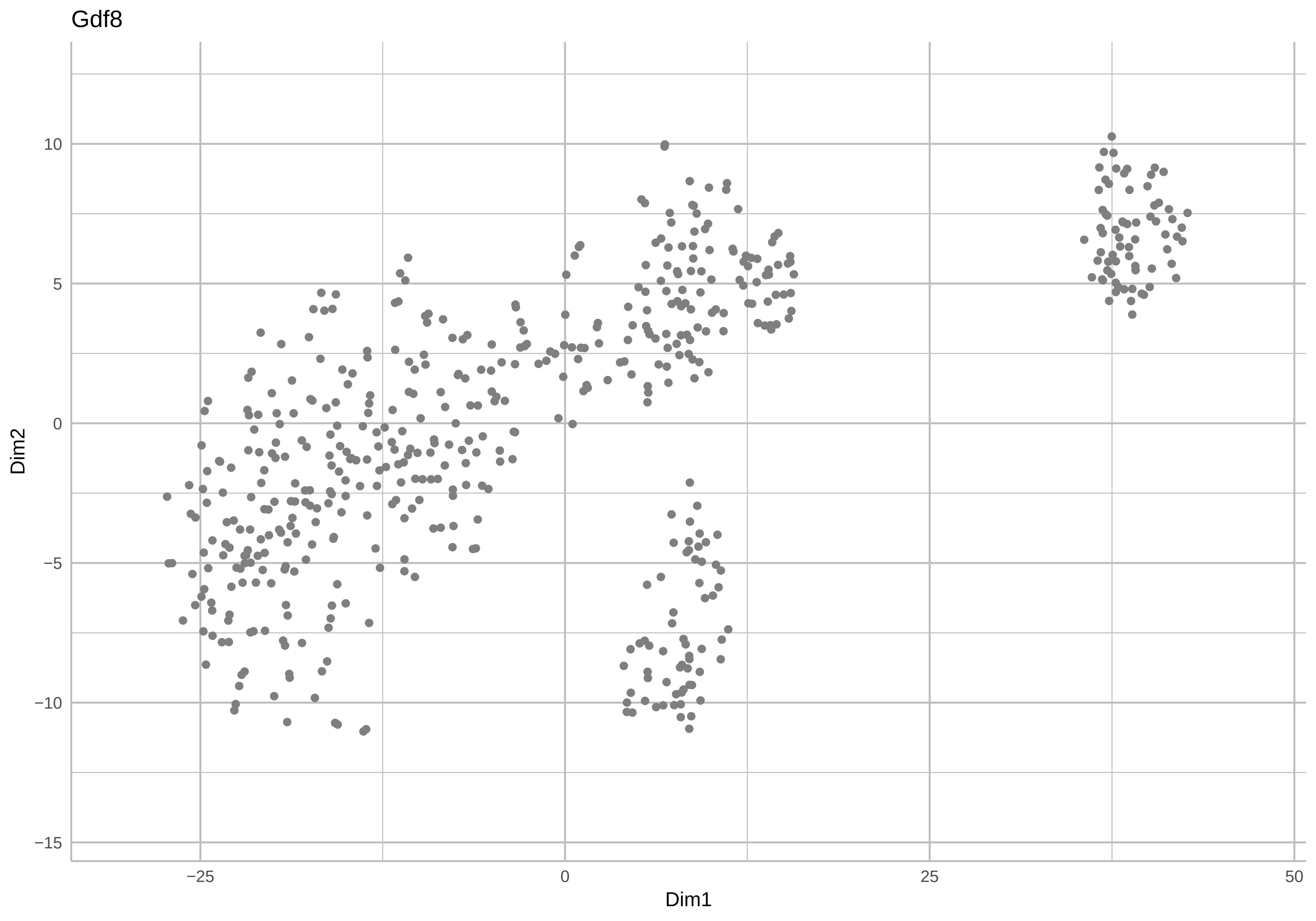
Gdf8
Dim2
10
5
0
−5
−10
−15
−25
0
25
50
Dim1

Gata3

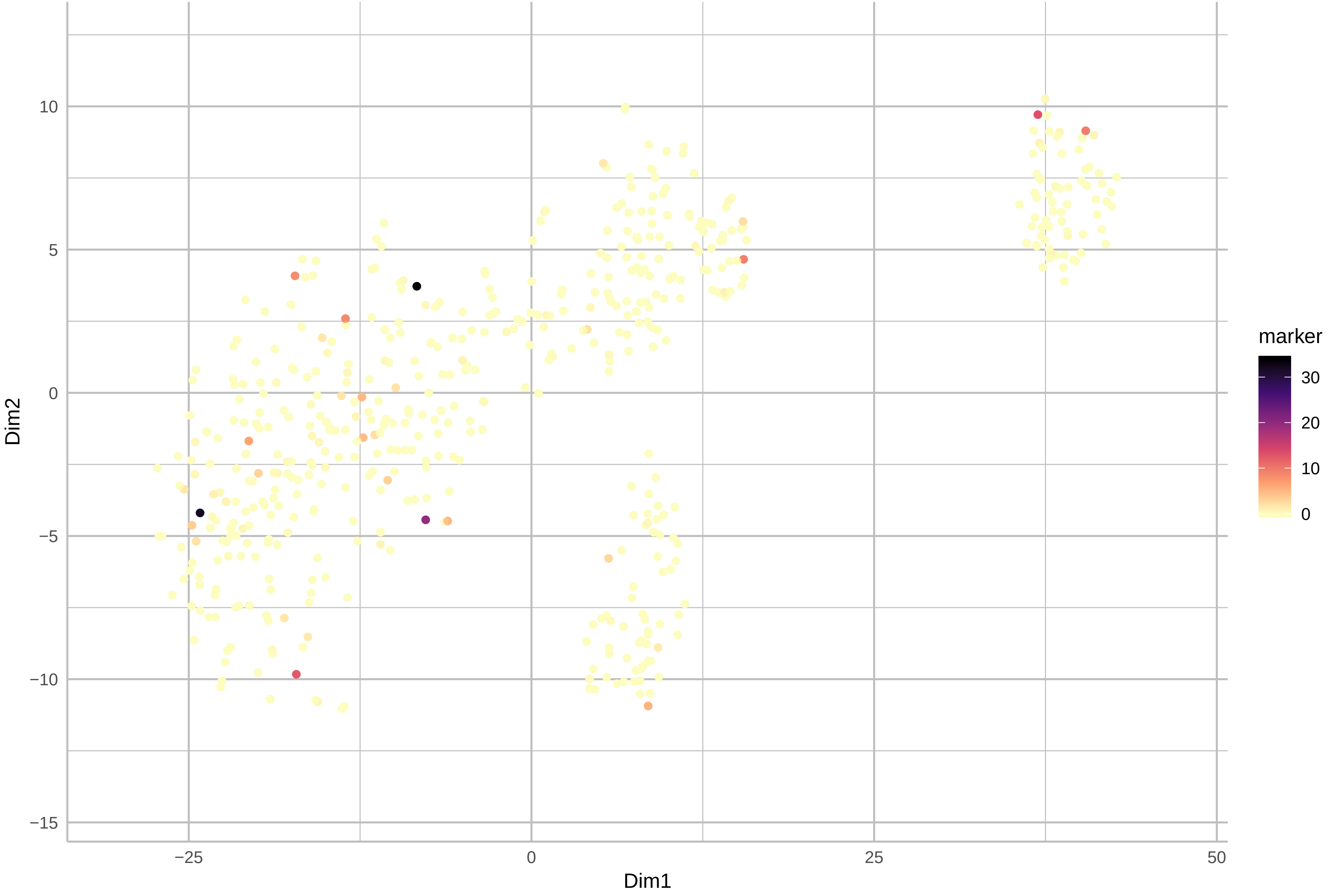

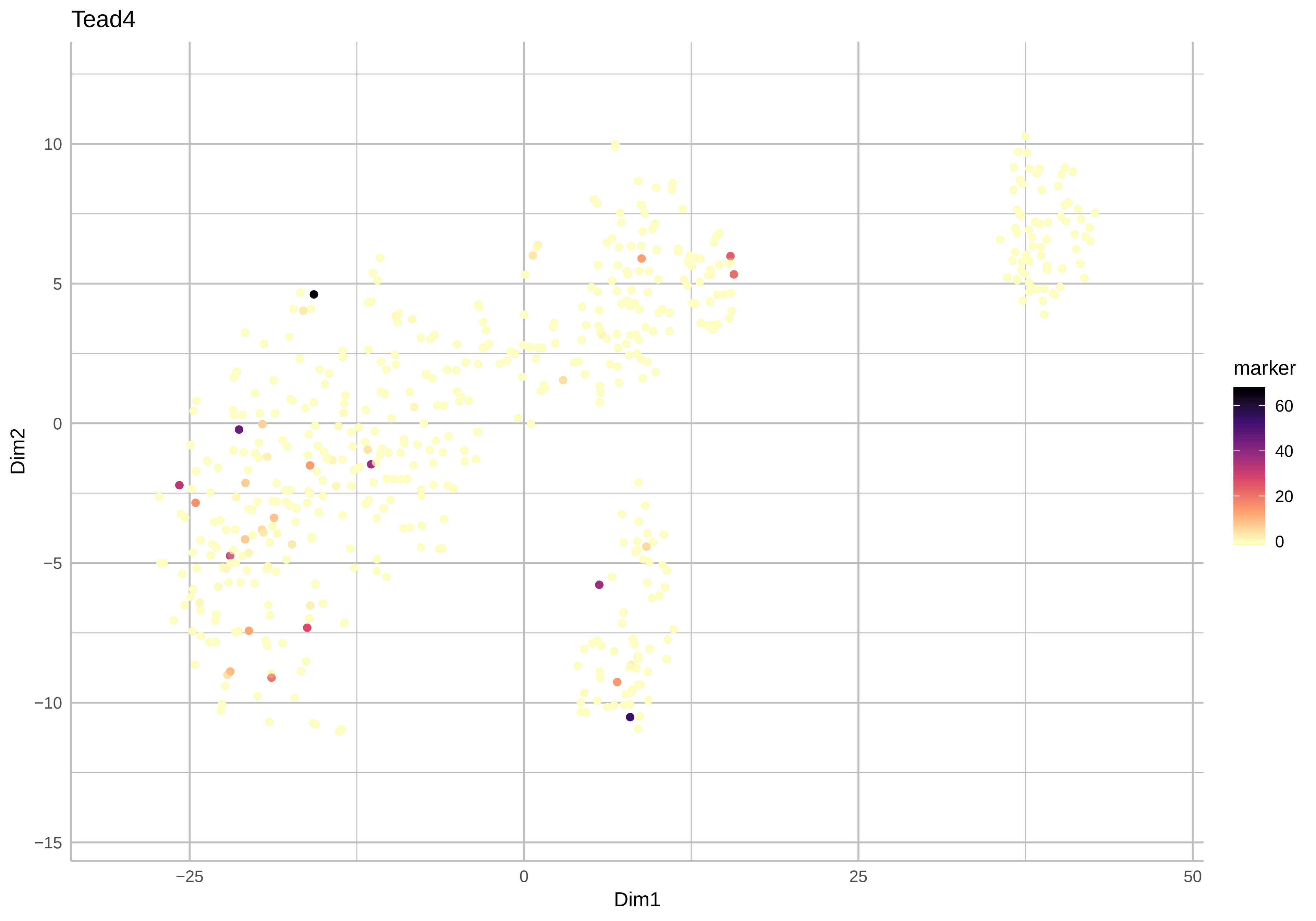
Tead4
Dim1
Dim2
marker
60
40
20
0
−25
0
25
50
10
5
0
−5
−10
−15

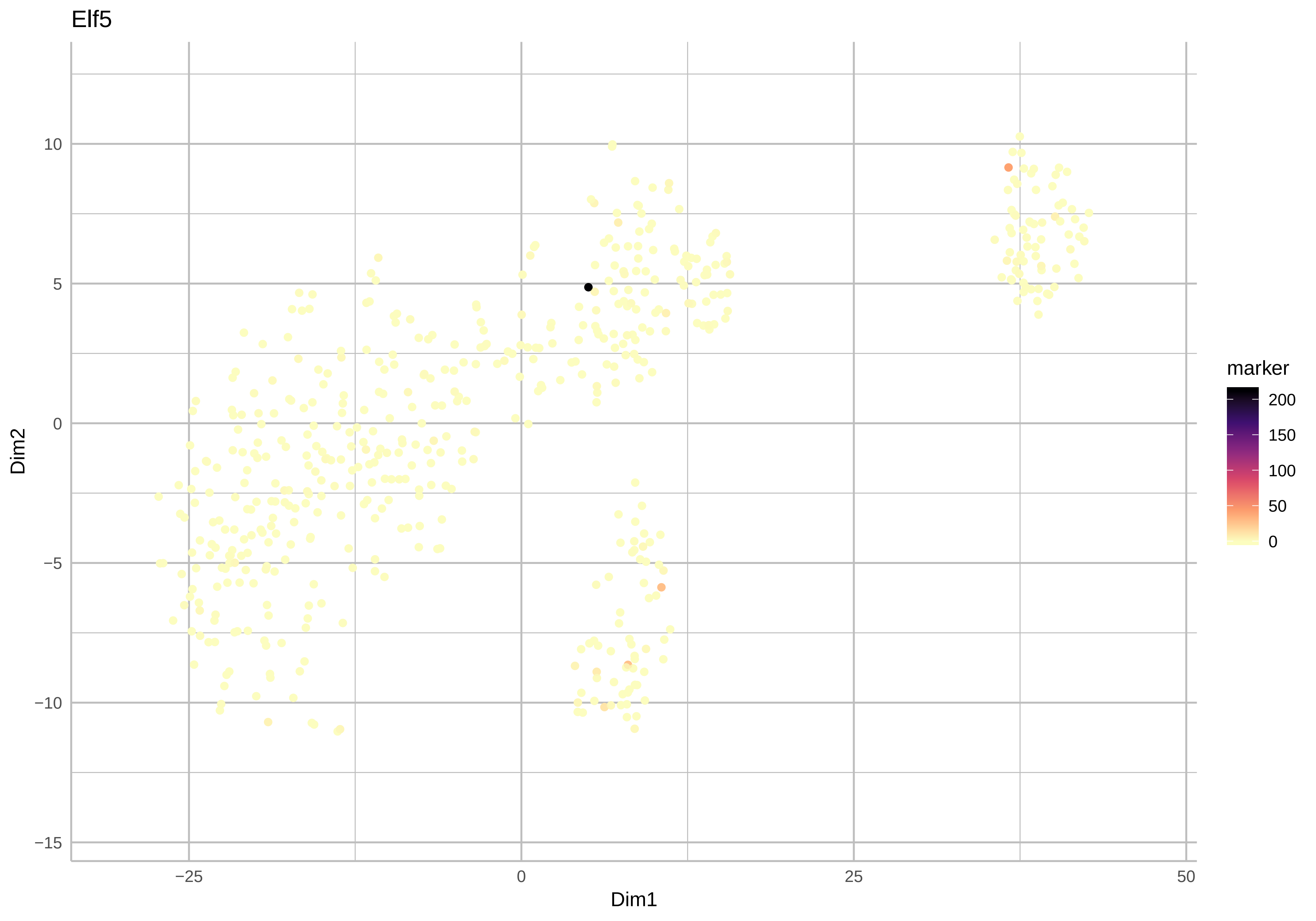

Elf5
Dim2
Dim1
marker
200
150
100
50
0
10
5
0
−5
−10
−15
−25
0
25
50

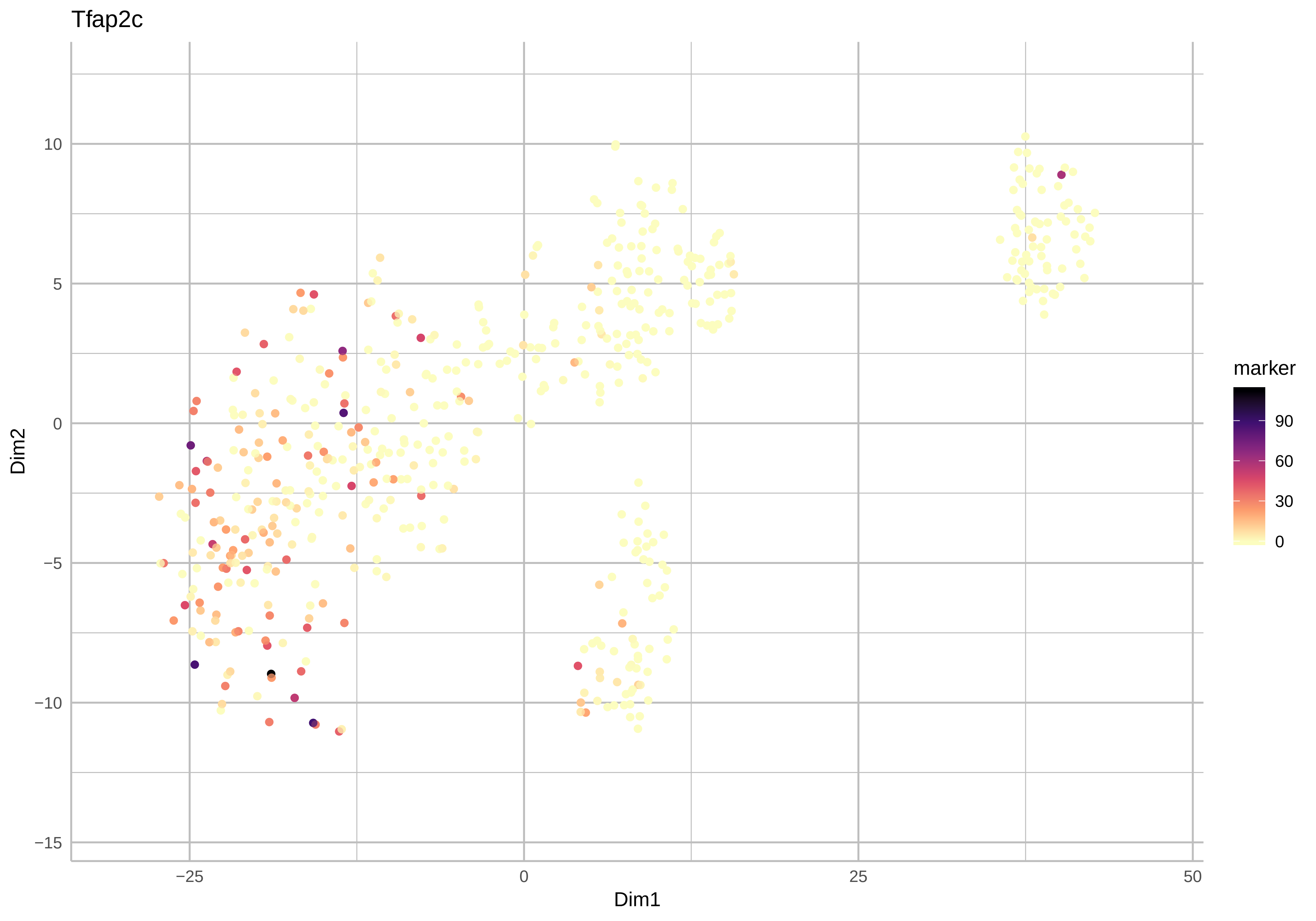
Tfap2c
Dim2
10
5
0
−5
−10
−15
−25
0
25
50
Dim1
marker
90
60
30
0

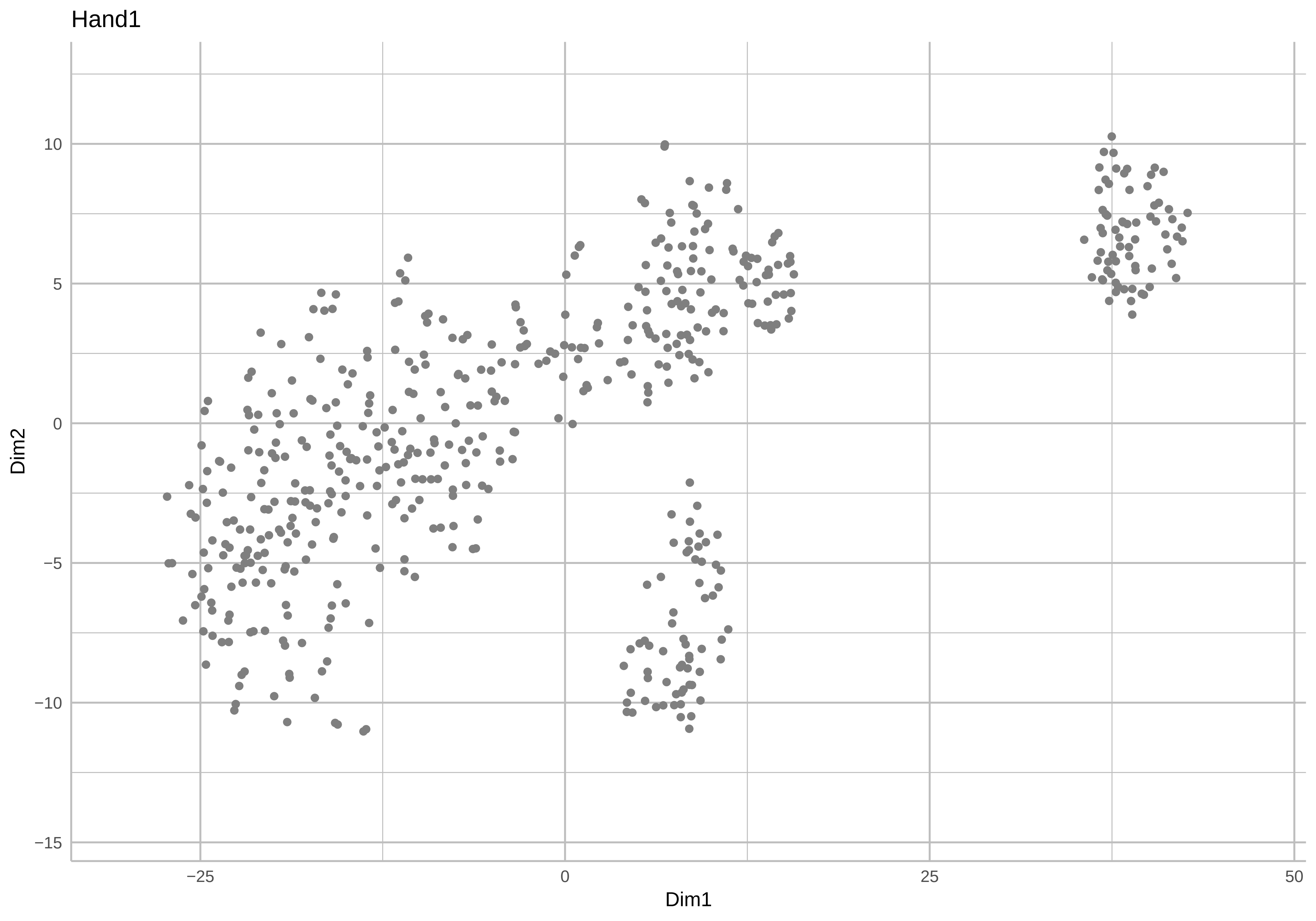
Hand1
Dim2
10
5
0
−5
−10
−15
−25
0
25
50
Dim1

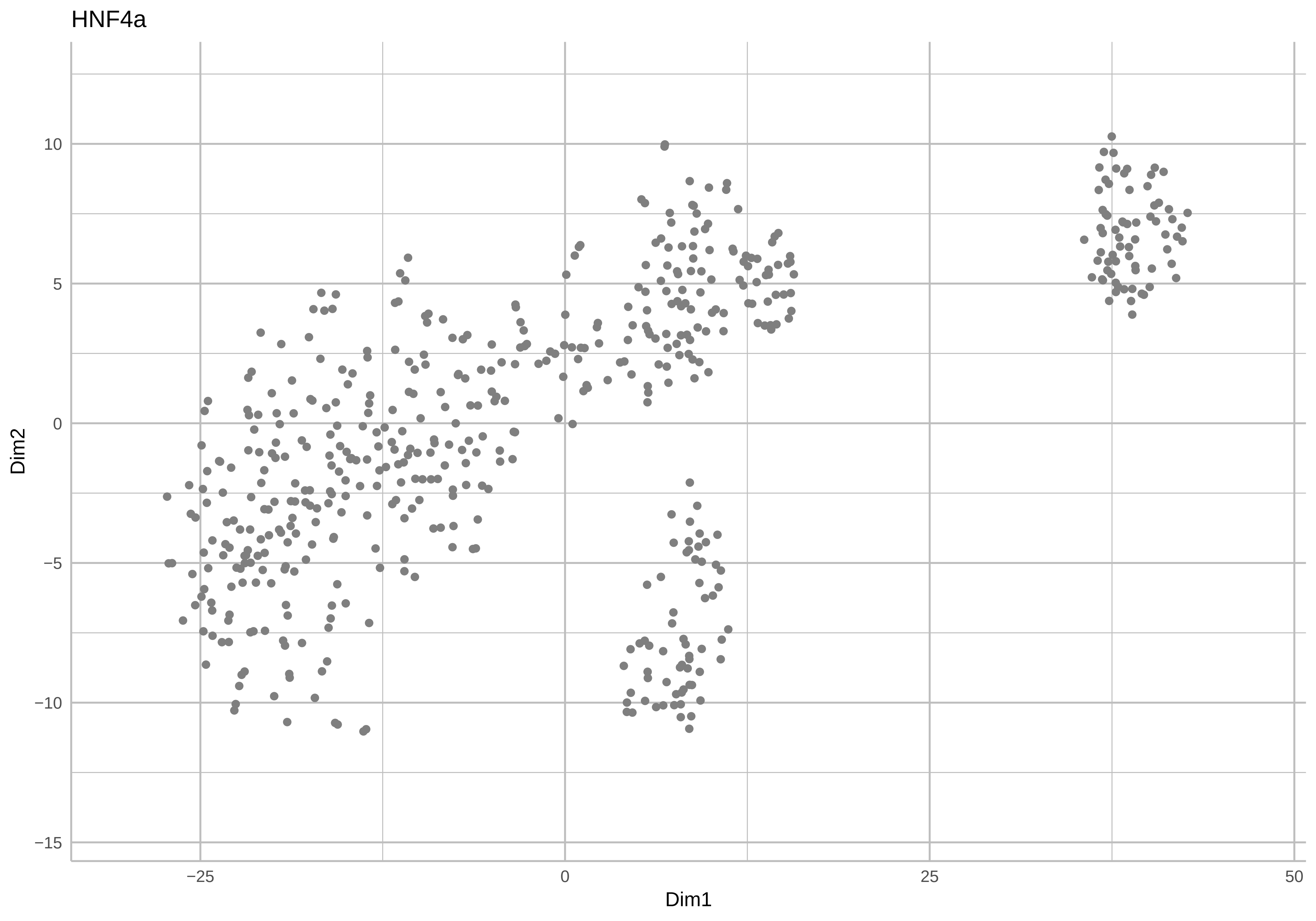
HNF4a
Dim2
Dim1
10
5
0
−5
−10
−15
−25
0
25
50

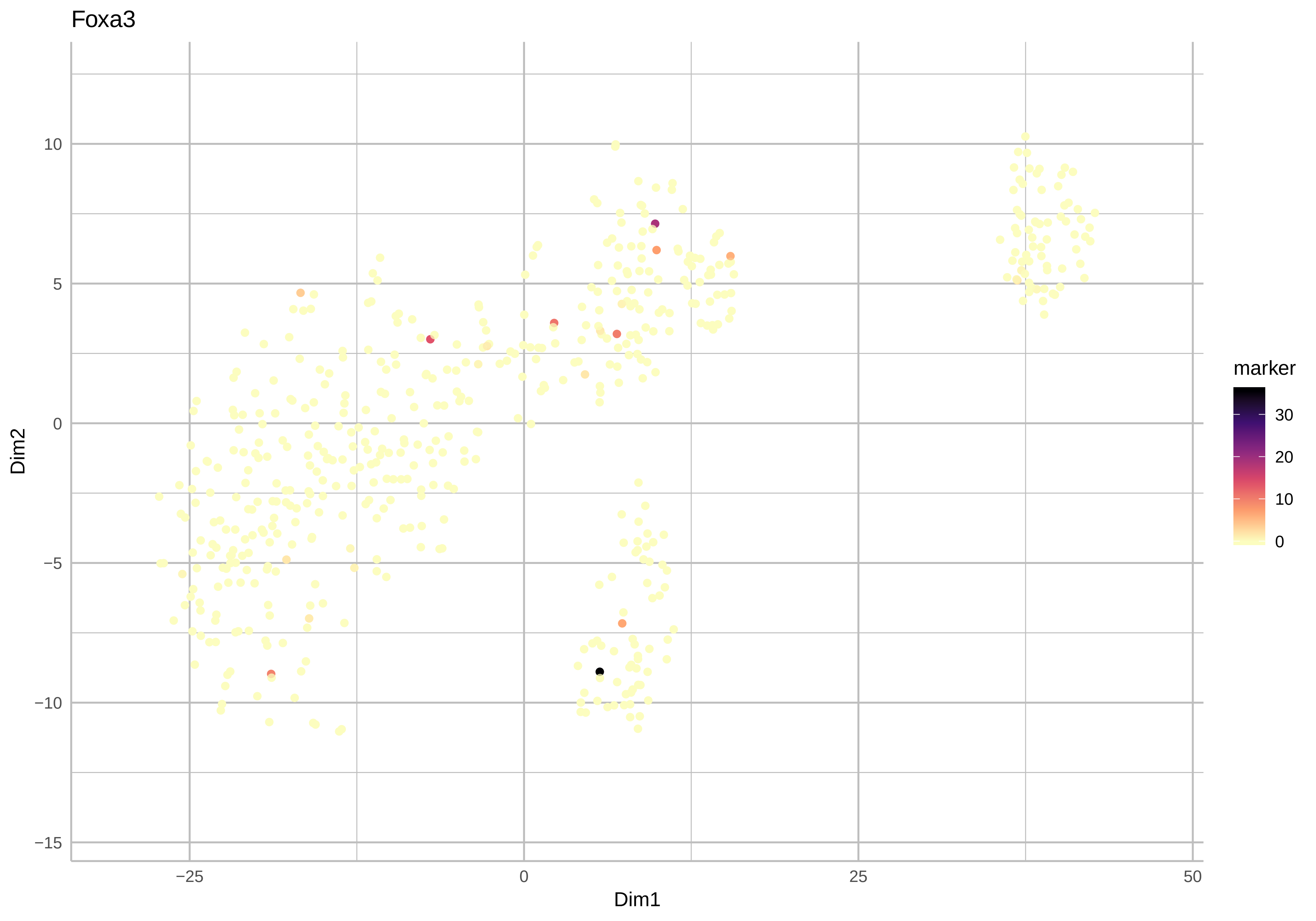

Foxa3
10
5
0
−5
−10
−15
Dim2
−25
0
25
50
Dim1
marker
30
20
10
0

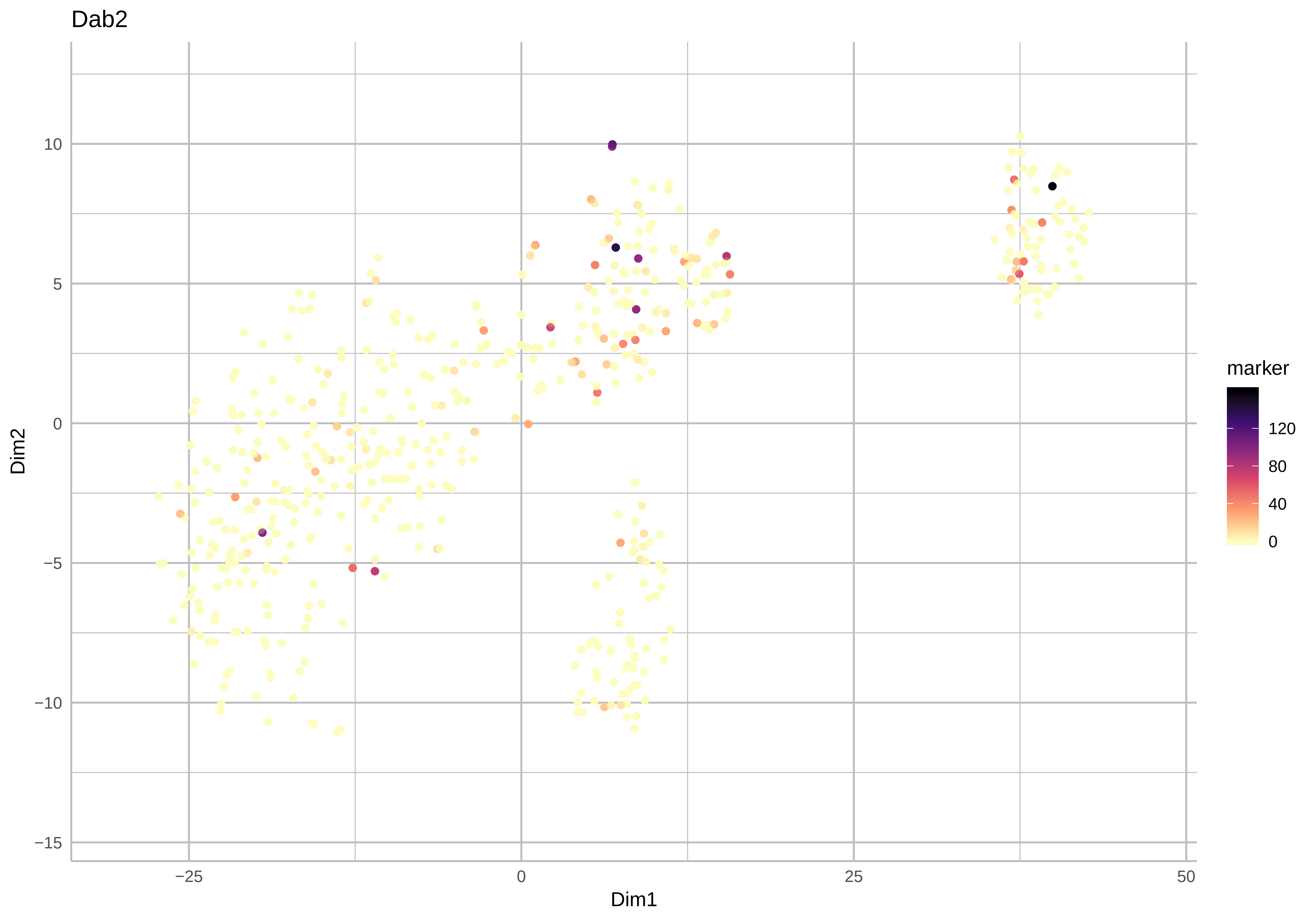
Dab2
10
5
0
−5
−10
−15
Dim2
−25
0
25
50
Dim1
marker
120
80
40
0

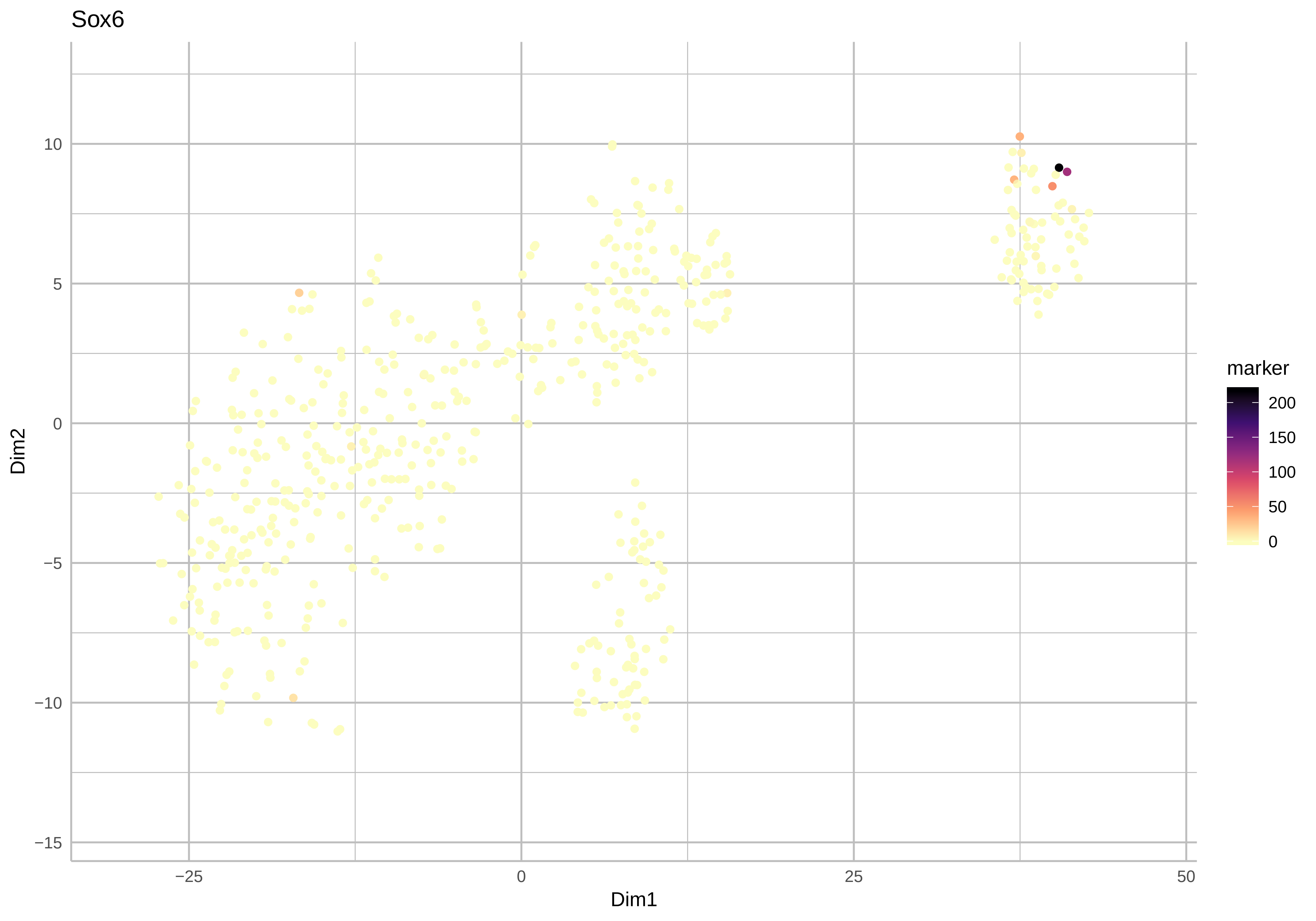
Sox6
Dim2
Dim1
10
5
0
−5
−10
−15
−25
0
25
50
marker
200
150
100
50
0

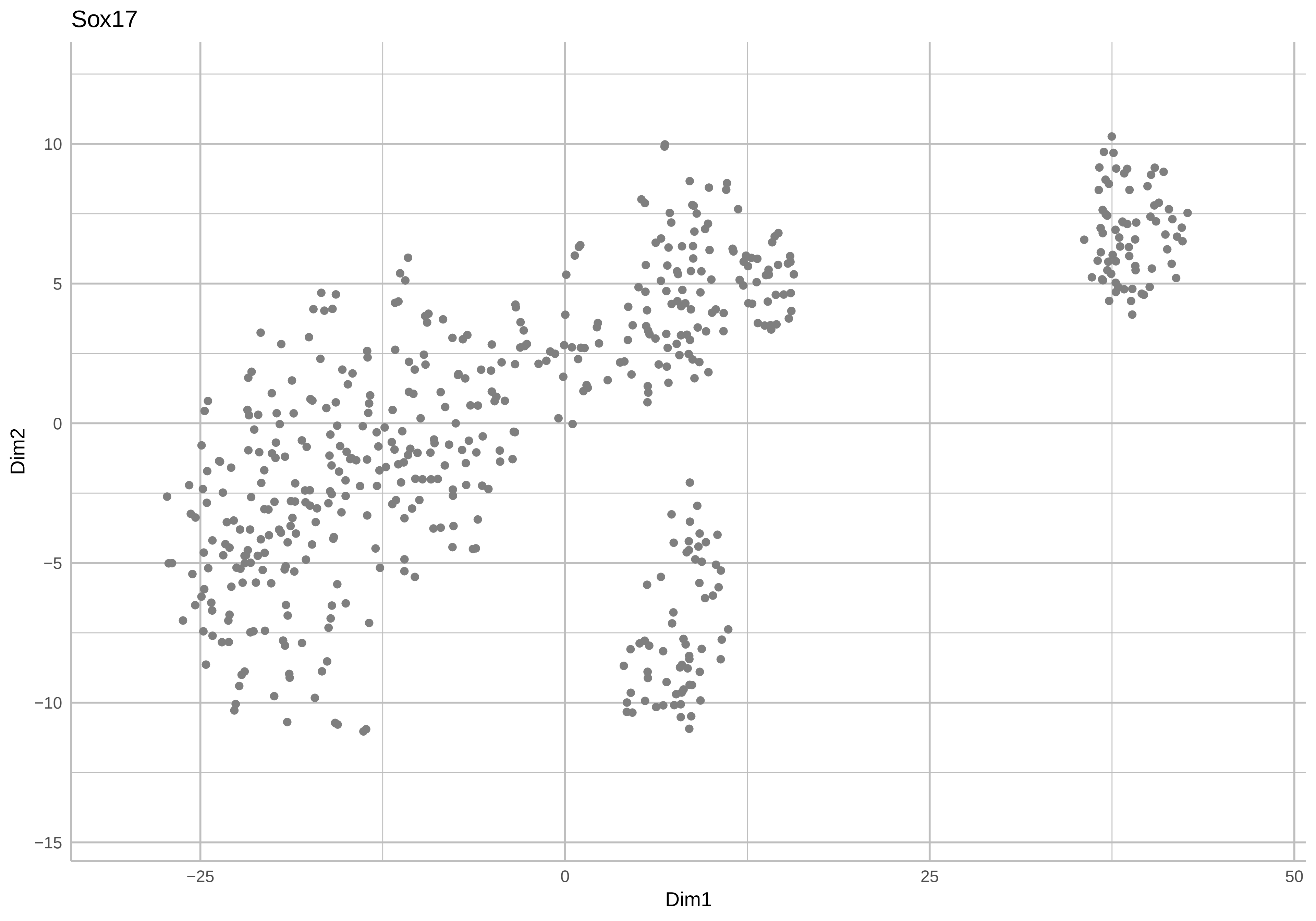
Sox17
Dim2
Dim1
10
5
0
−5
−10
−15
−25
0
25
50

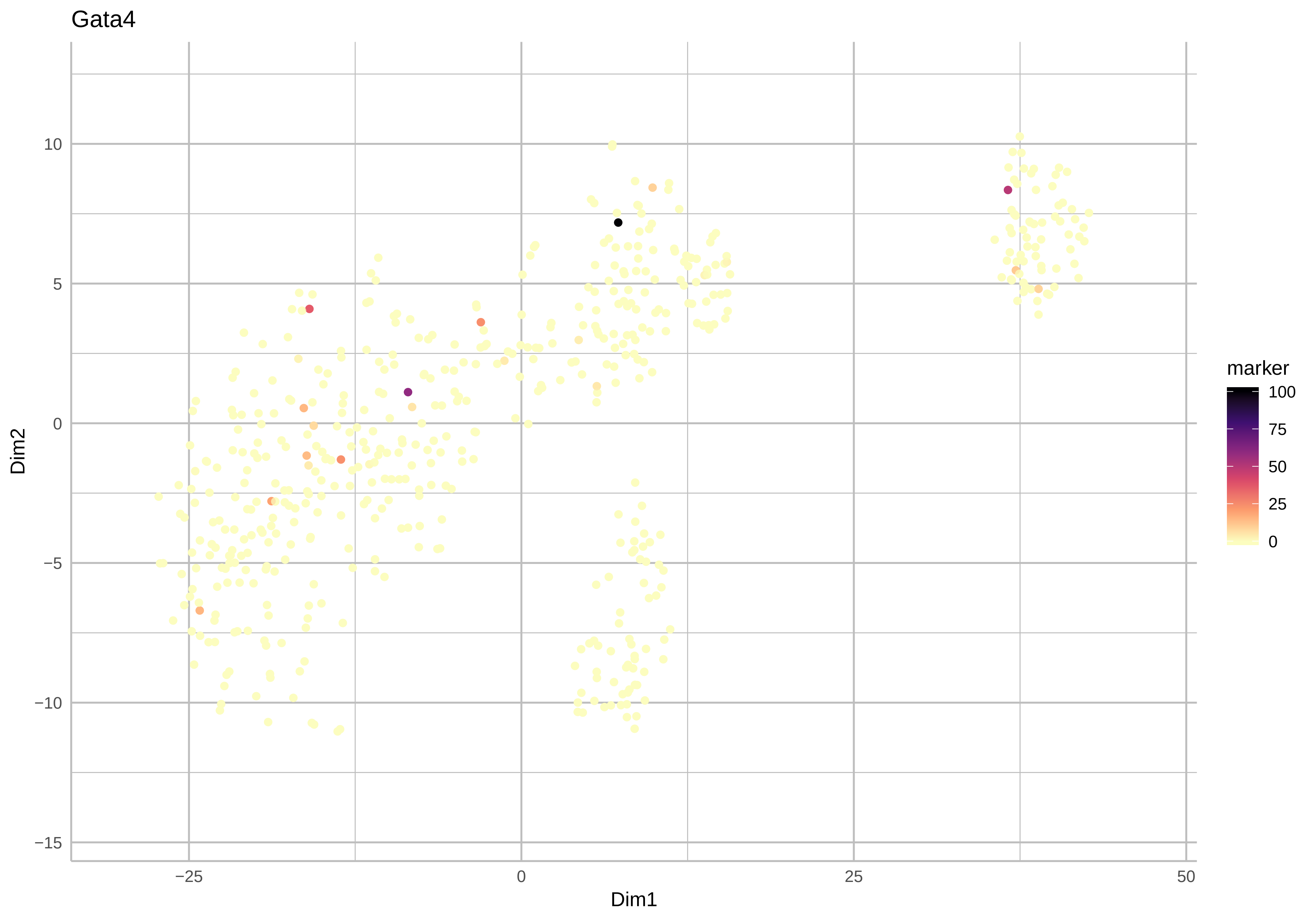
Gata4
Dim2
10
5
0
−5
−10
−15
−25
0
25
50
Dim1
marker
100
75
50
25
0

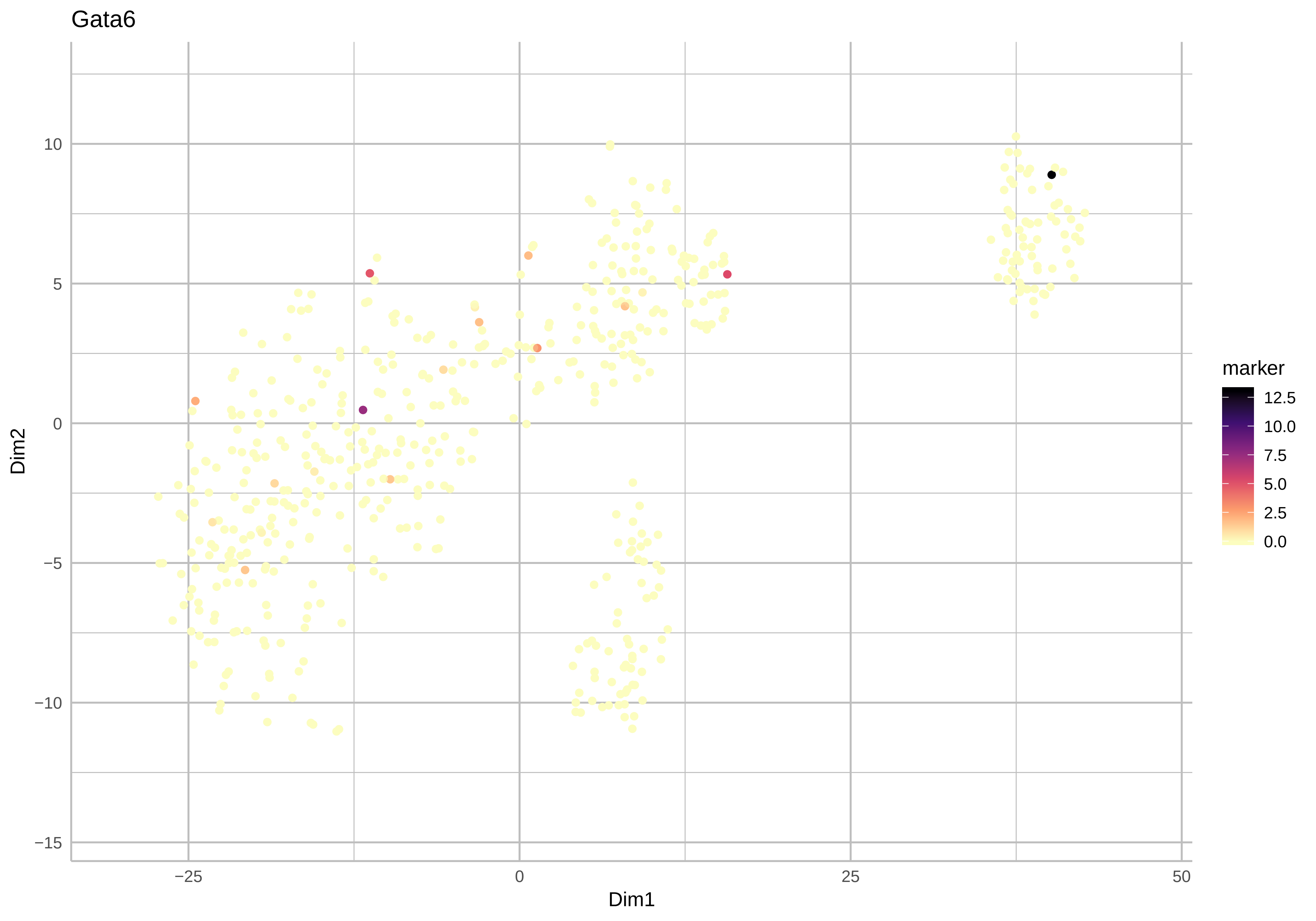

Gata6
Dim2
Dim1
10
5
0
−5
−10
−15
−25
0
25
50
marker
12.5
10.0
7.5
5.0
2.5
0.0

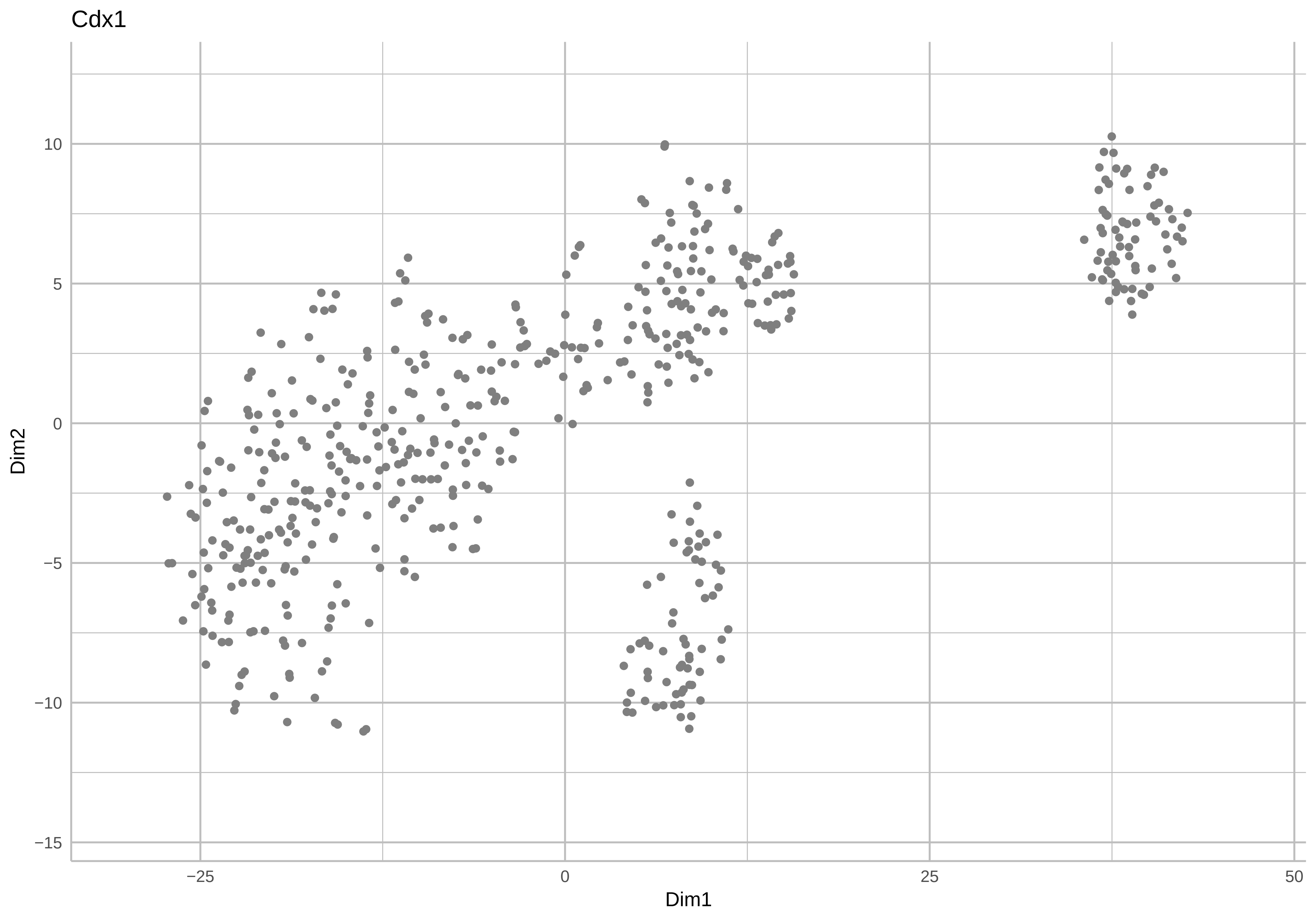
Cdx1
Dim2
Dim1
10
5
0
−5
−10
−15
−25
0
25
50

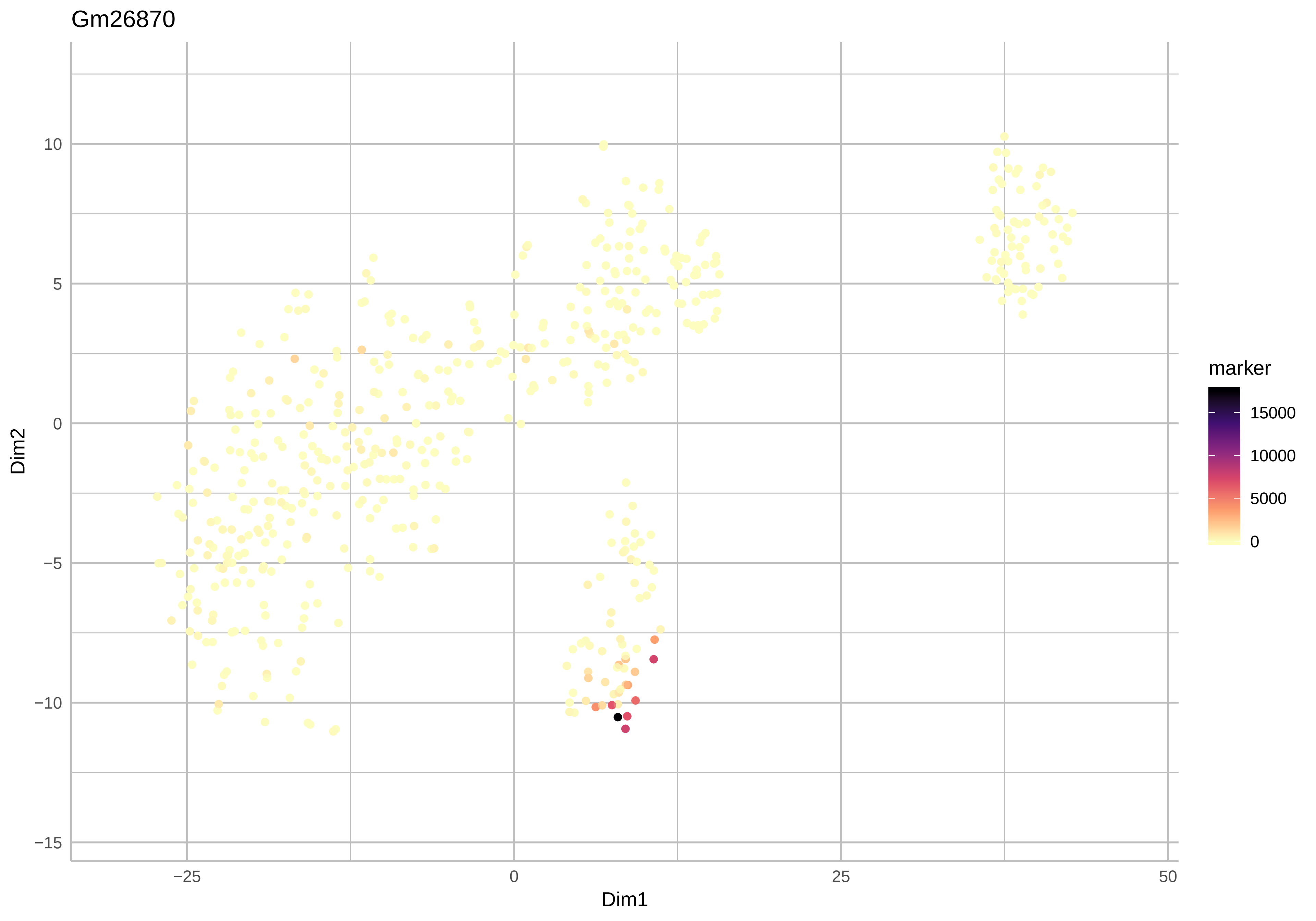
Gm26870
Dim2
Dim1
marker
15000
10000
5000
0
10
5
0
−5
−10
−15
−25
0
25
50

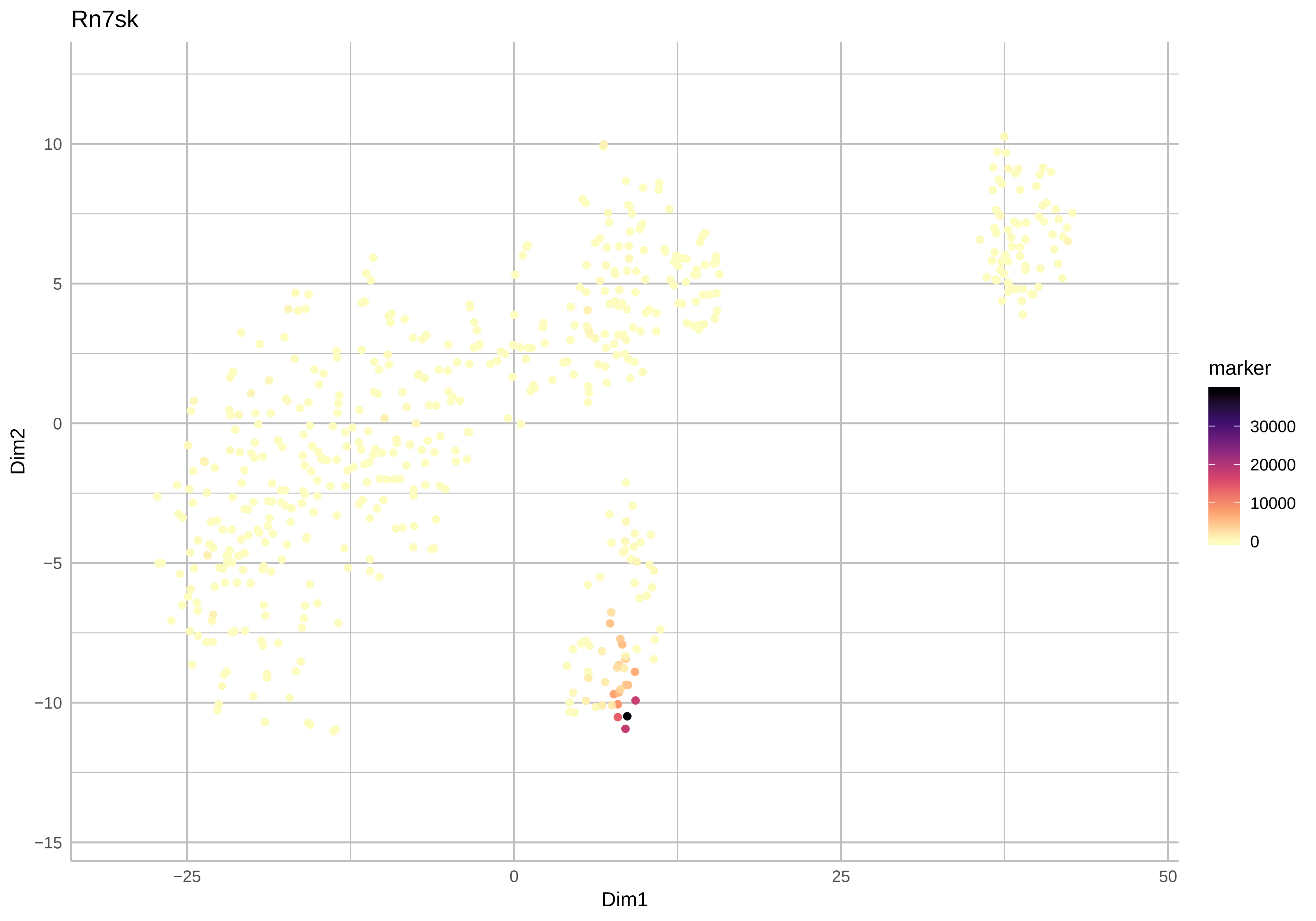
Rn7sk
Dim2
Dim1
marker
30000
20000
10000
0
10
5
0
−5
−10
−15
−25
0
25
50

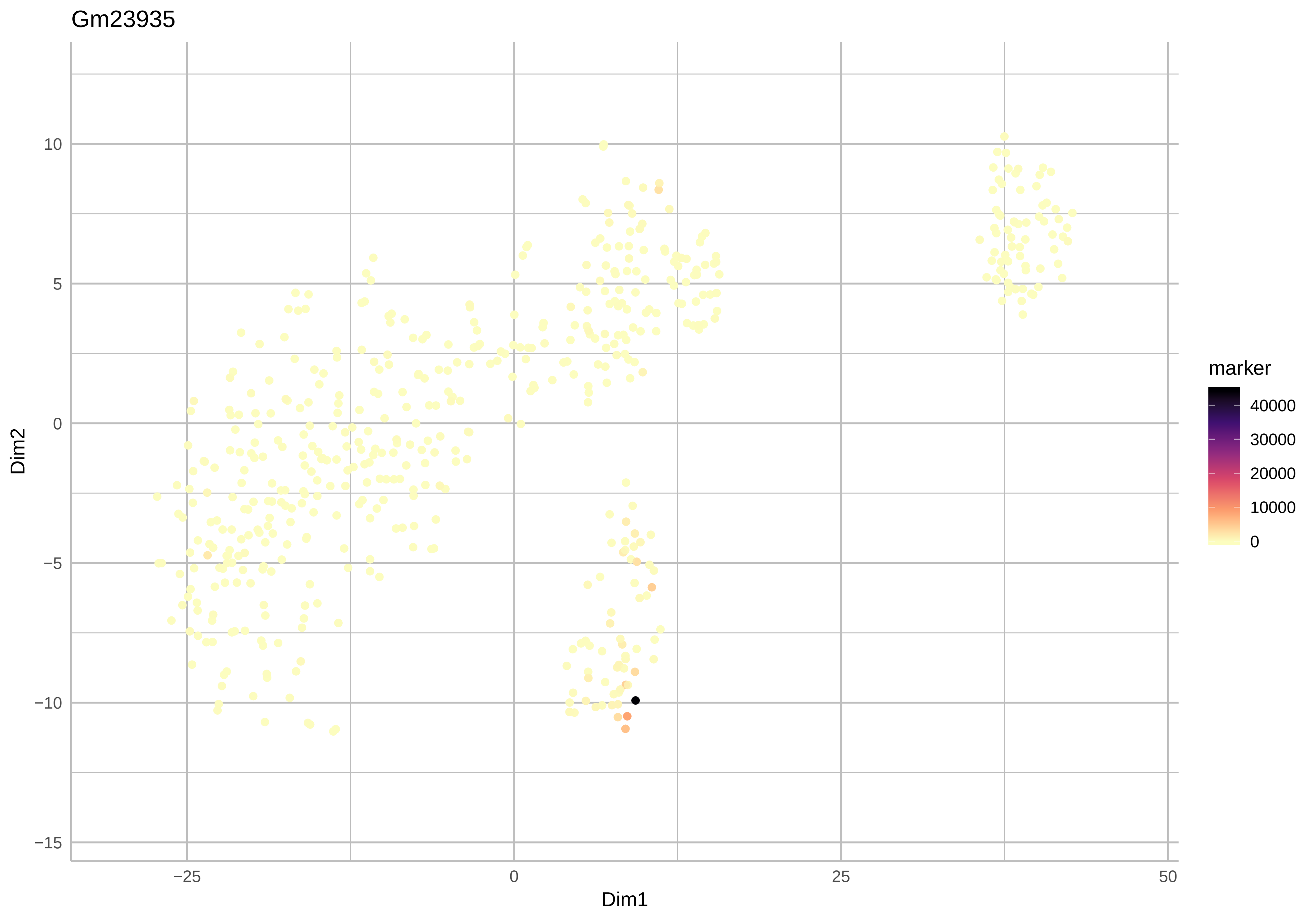

Gm23935
Dim2
Dim1
10
5
0
−5
−10
−15
−25
0
25
50
marker
40000
30000
20000
10000
0

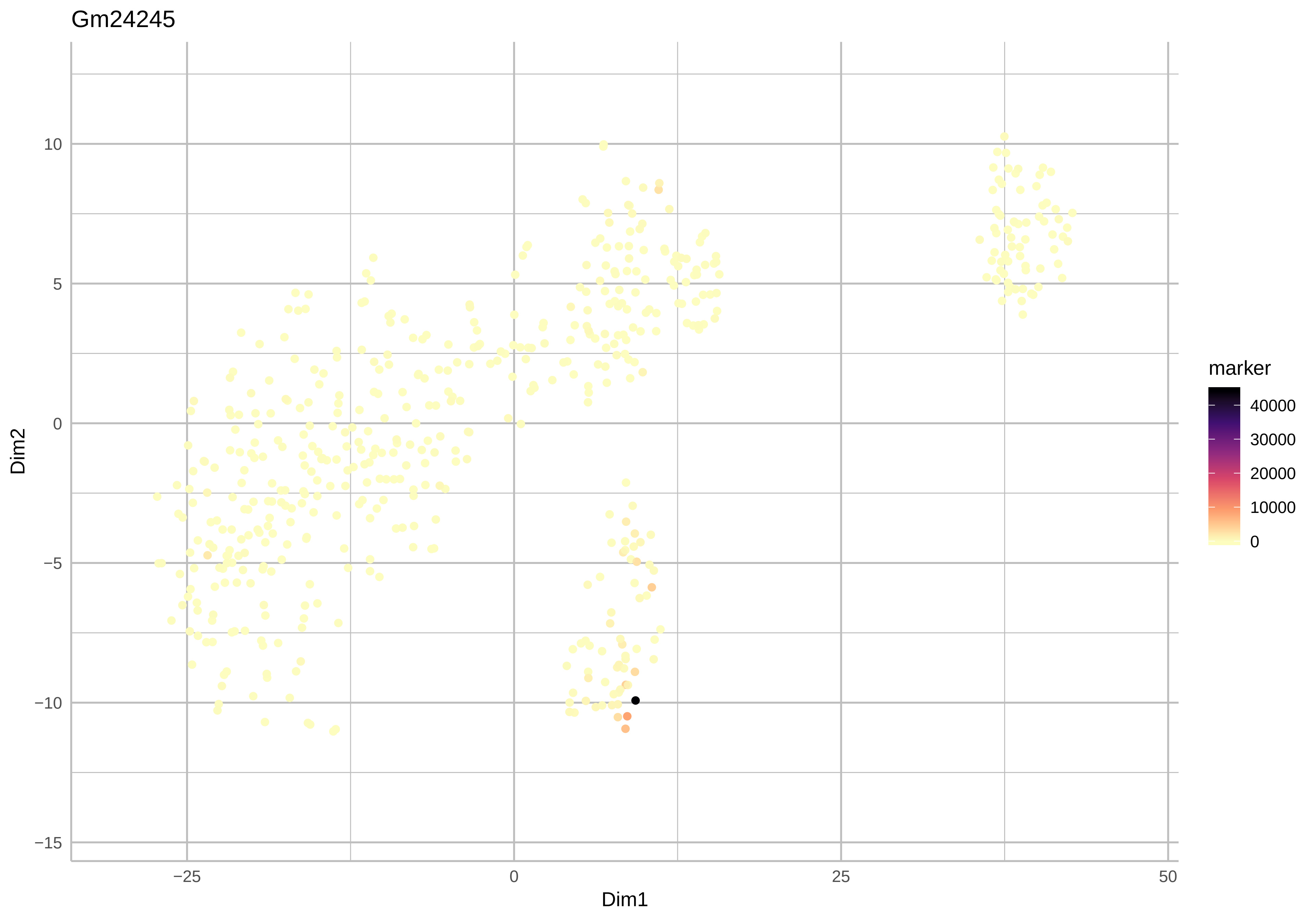
Gm24245
marker
40000
30000
20000
10000
0
10
5
0
−5
−10
−15
Dim2
−25
0
25
50
Dim1

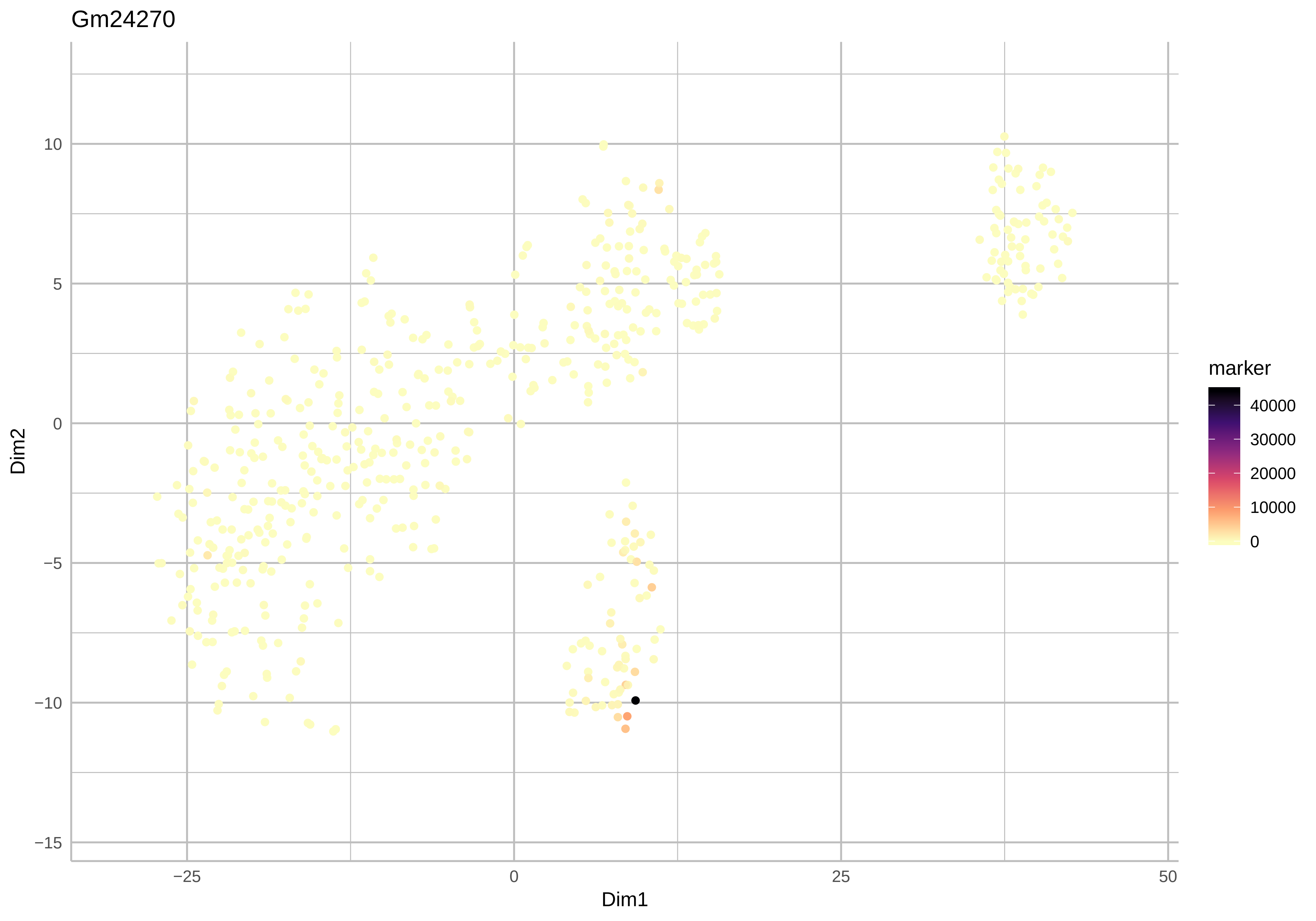
Gm24270
Dim2
Dim1
10
5
0
−5
−10
−15
−25
0
25
50
marker
40000
30000
20000
10000
0

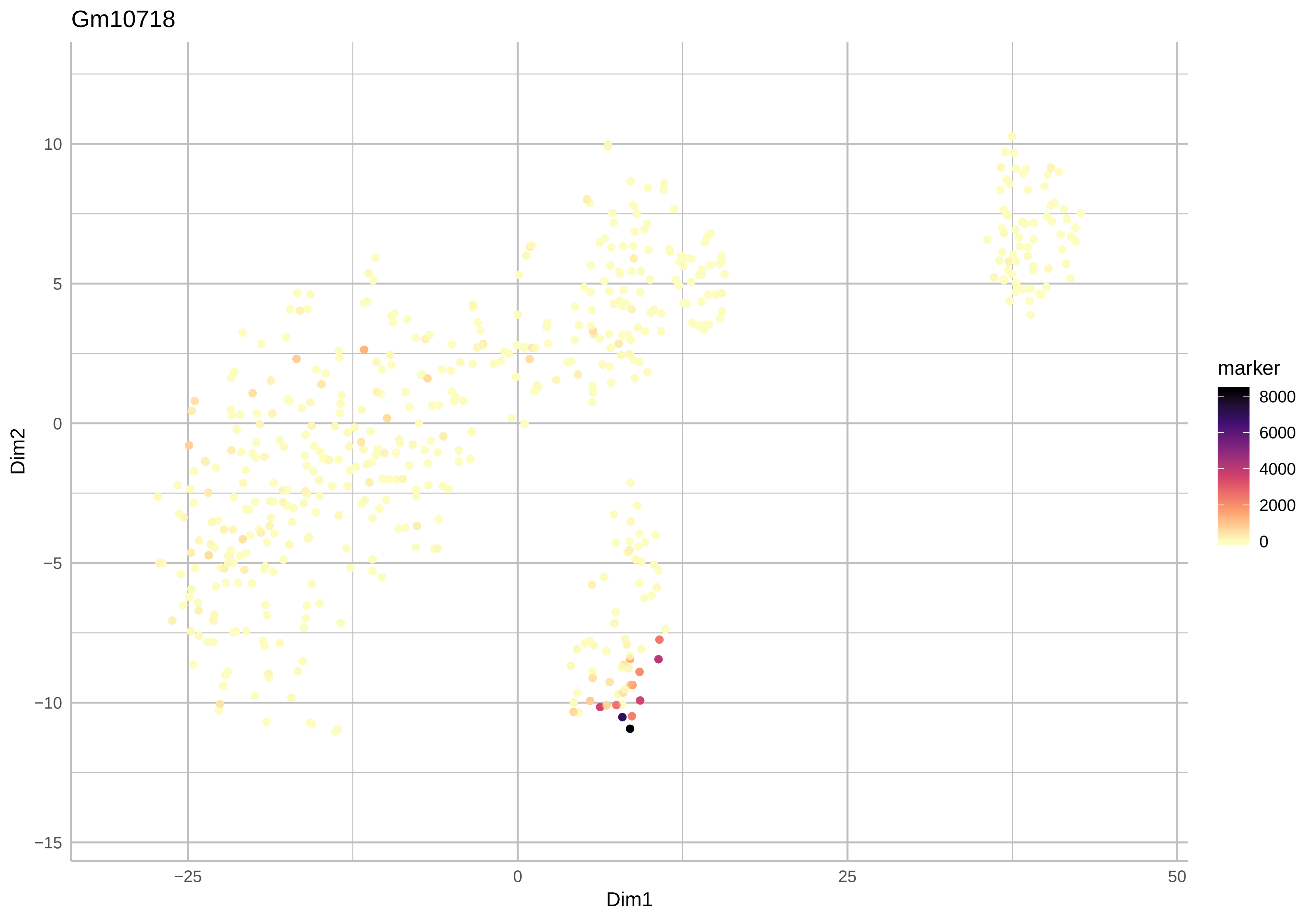
Gm10718
Dim2
10
5
0
−5
−10
−15
−25
0
25
50
Dim1
marker
8000
6000
4000
2000
0

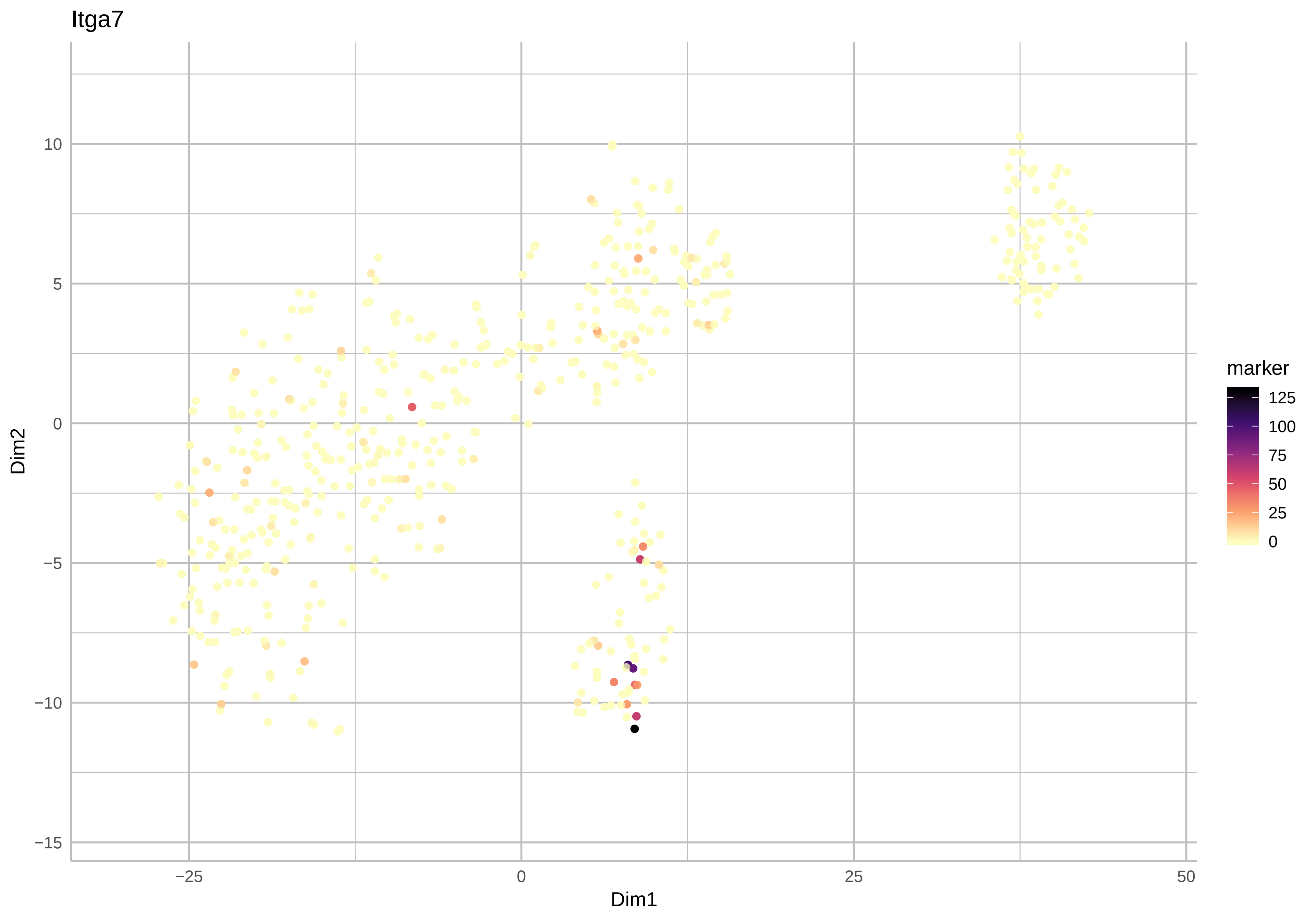

Itga7
10
5
0
−5
−10
−15
Dim2
−25
0
25
50
Dim1
marker
125
100
75
50
25
0

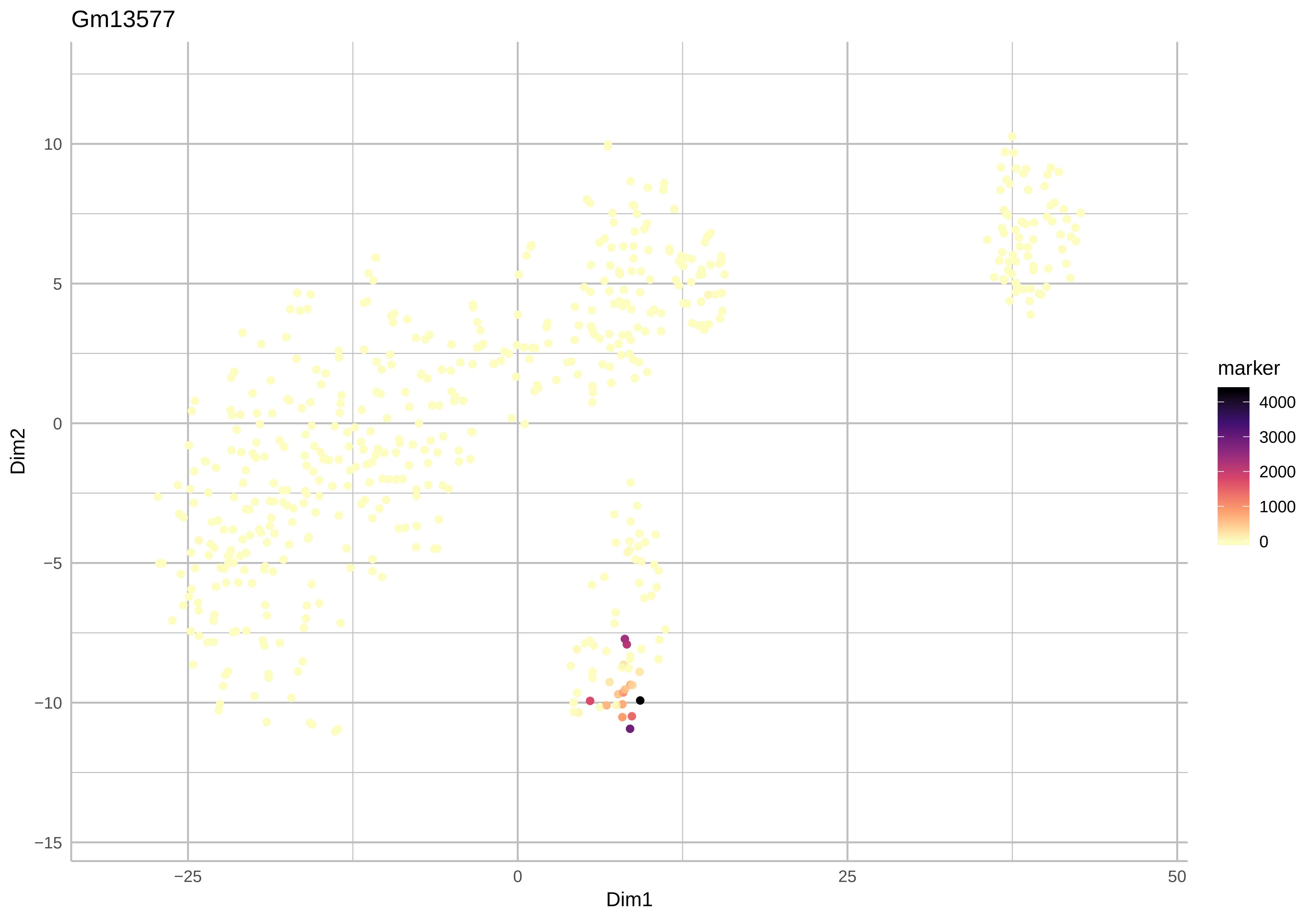

Gm13577
Dim2
Dim1
10
5
0
−5
−10
−15
−25
0
25
50
marker
4000
3000
2000
1000
0

Rpph1

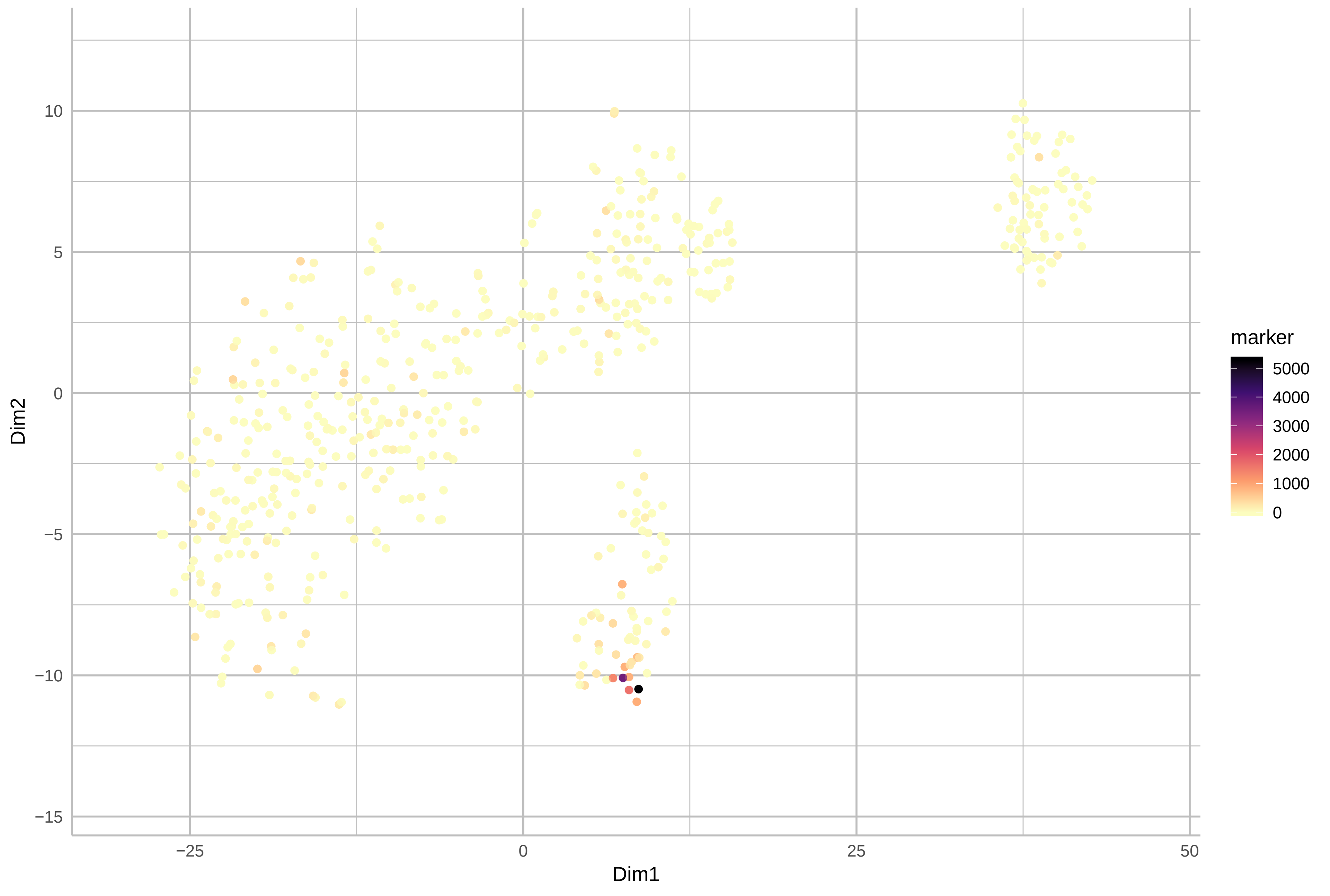

10
5
0
−5
−10
−15
Dim2
−25
0
25
50
Dim1
marker
5000
4000
3000
2000
1000
0

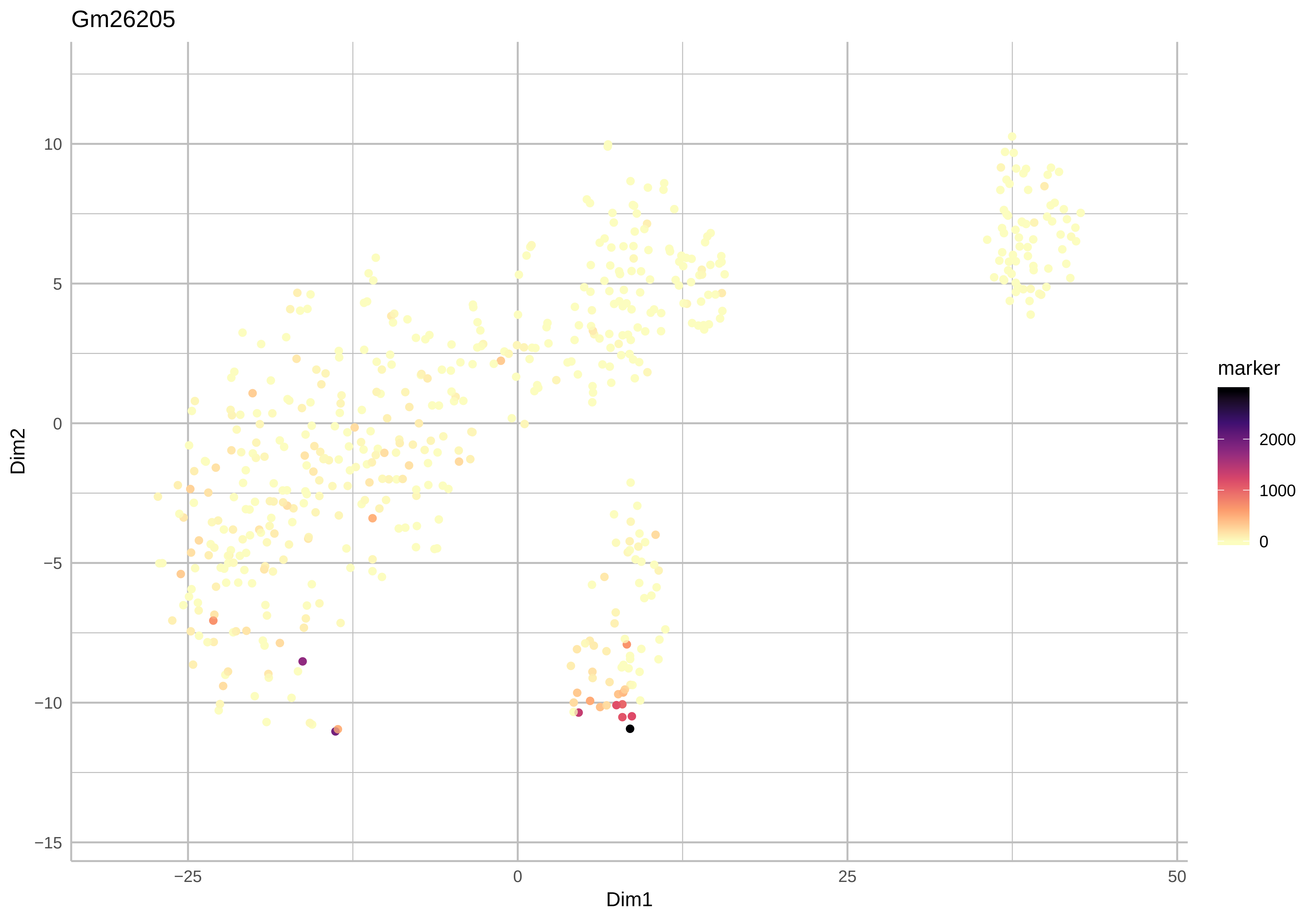
Gm26205
Dim2
Dim1
marker
2000
1000
0
10
5
0
−5
−10
−15
−25
0
25
50

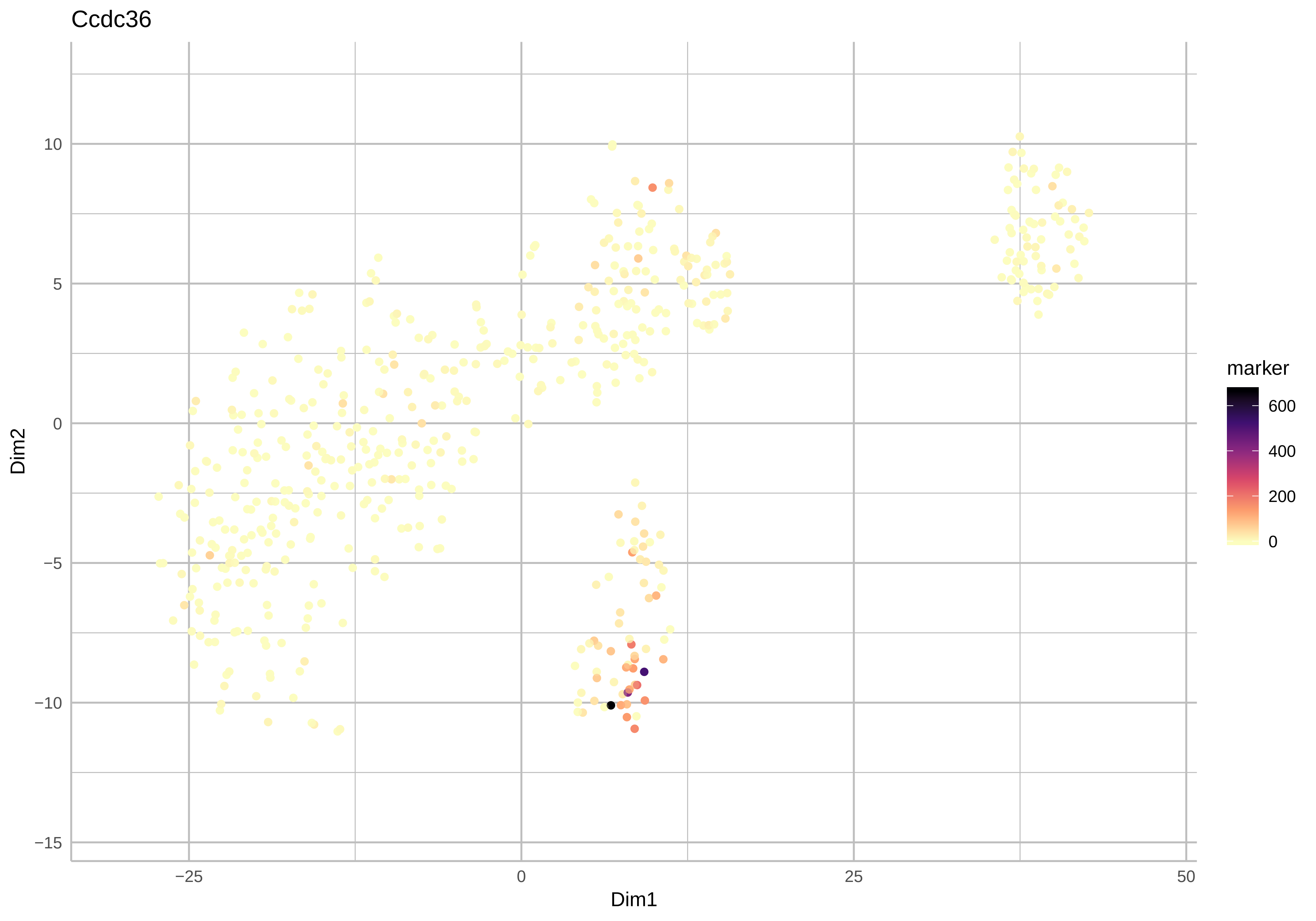
Ccdc36
Dim1
Dim2
−25
0
25
50
10
5
0
−5
−10
−15
marker
600
400
200
0

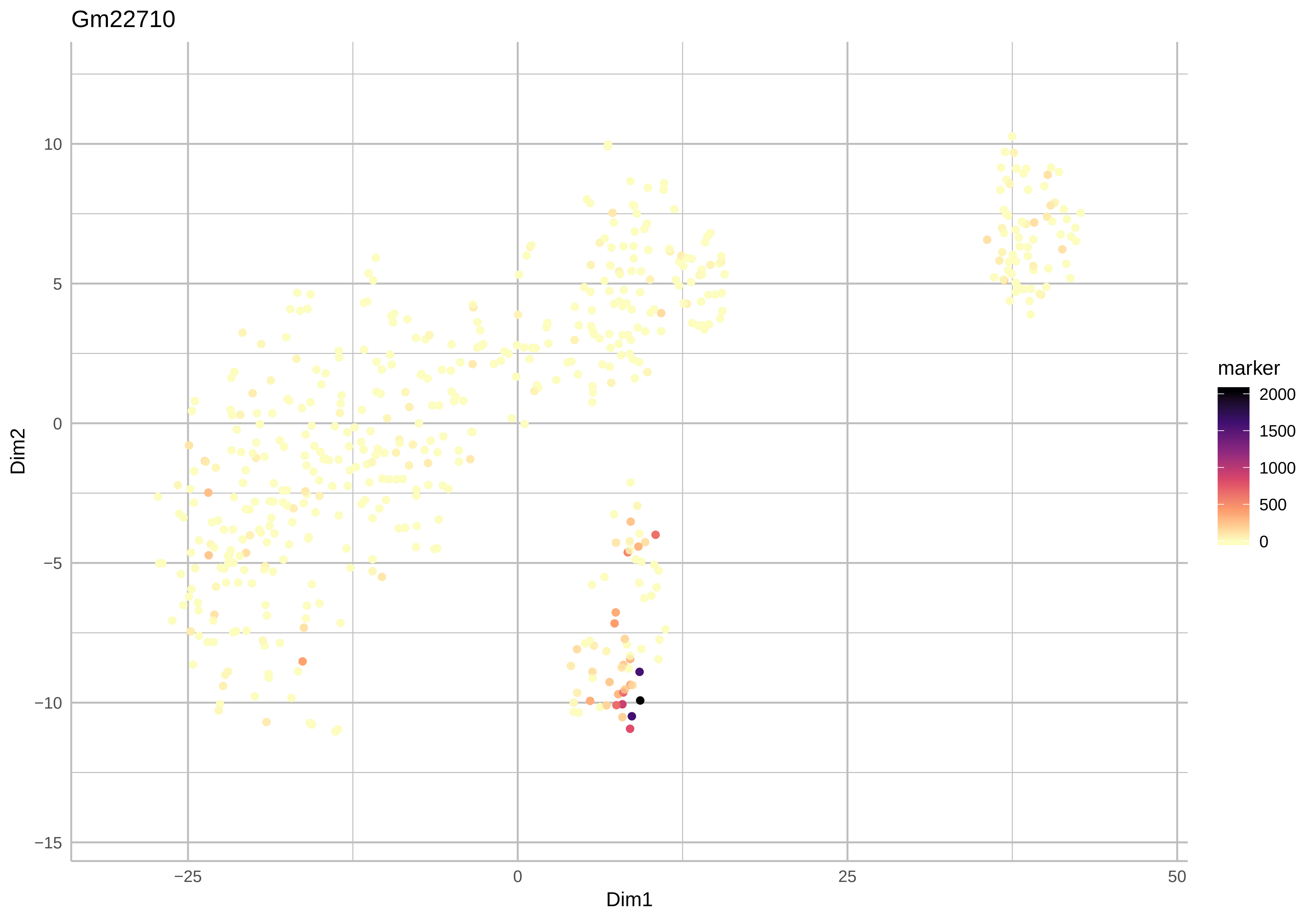
Gm22710
Dim2
Dim1
10
5
0
−5
−10
−15
−25
0
25
50
marker
2000
1500
1000
500
0

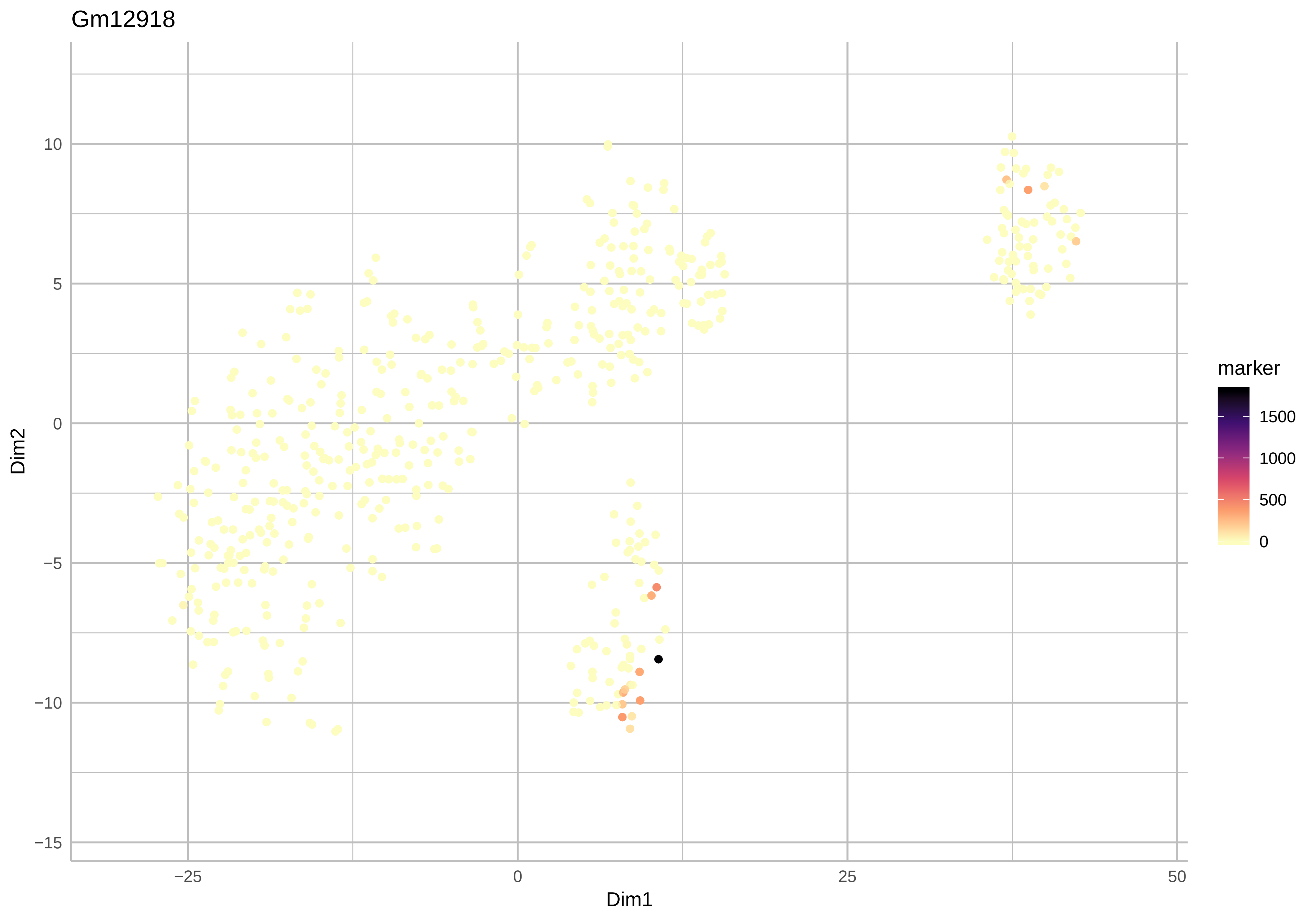
Gm12918
10
5
0
−5
−10
−15
Dim2
−25
0
25
50
Dim1
marker
1500
1000
500
0

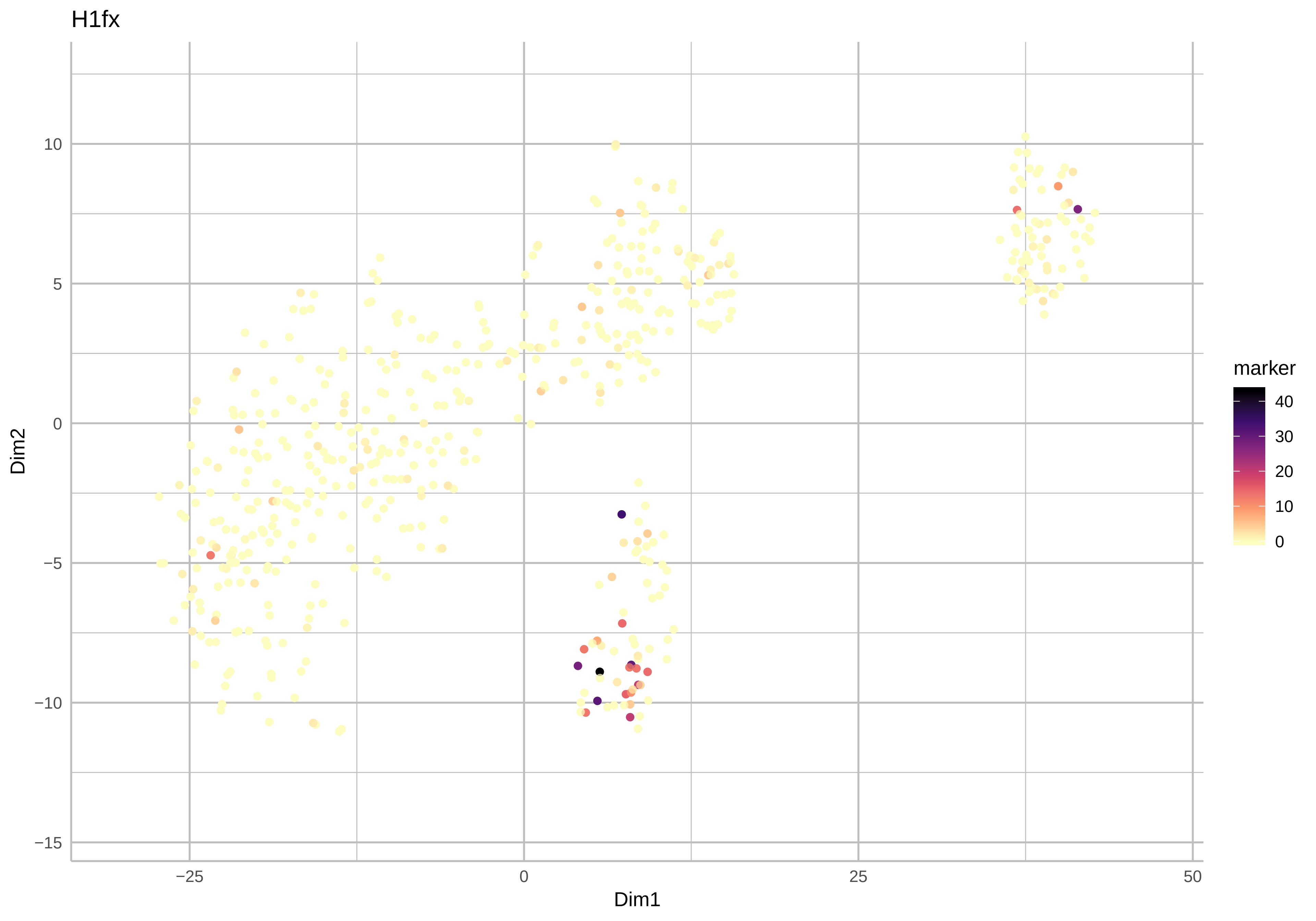

H1fx
Dim2
Dim1
marker
40
30
20
10
0
10
5
0
−5
−10
−15
−25
0
25
50

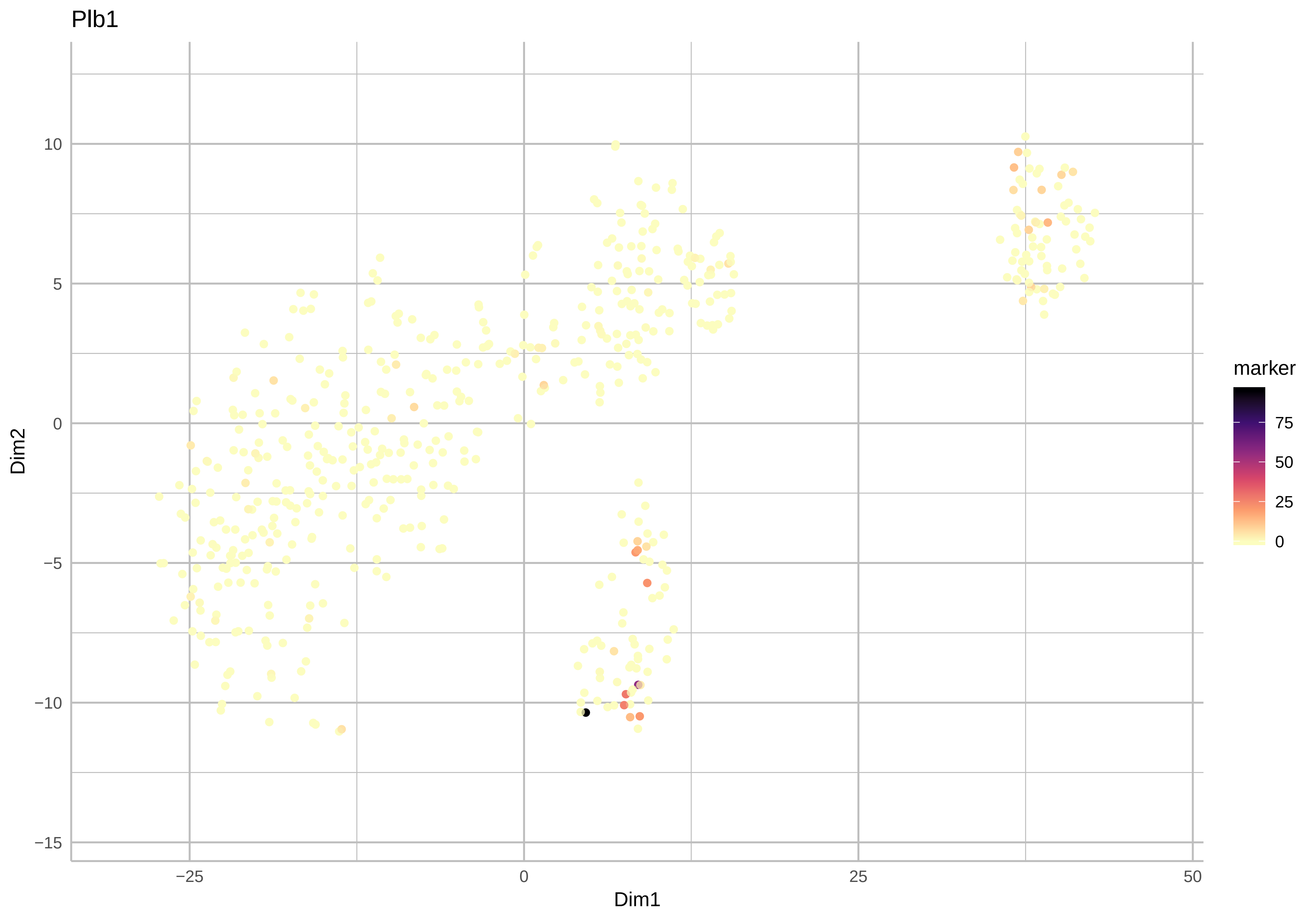
Plb1
Dim2
Dim1
10
5
0
−5
−10
−15
−25
0
25
50
marker
75
50
25
0

Gm21847

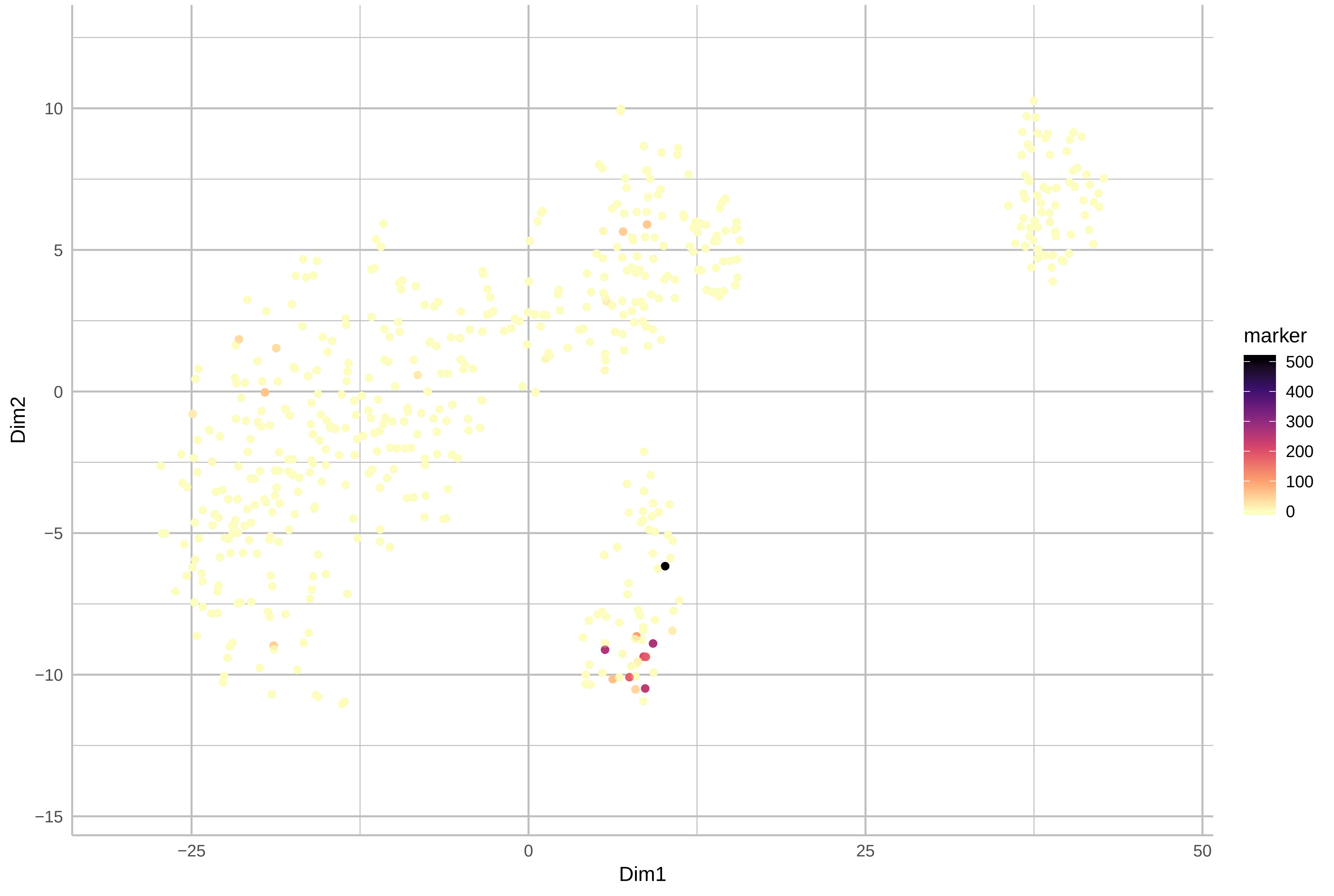

Rnu1b6

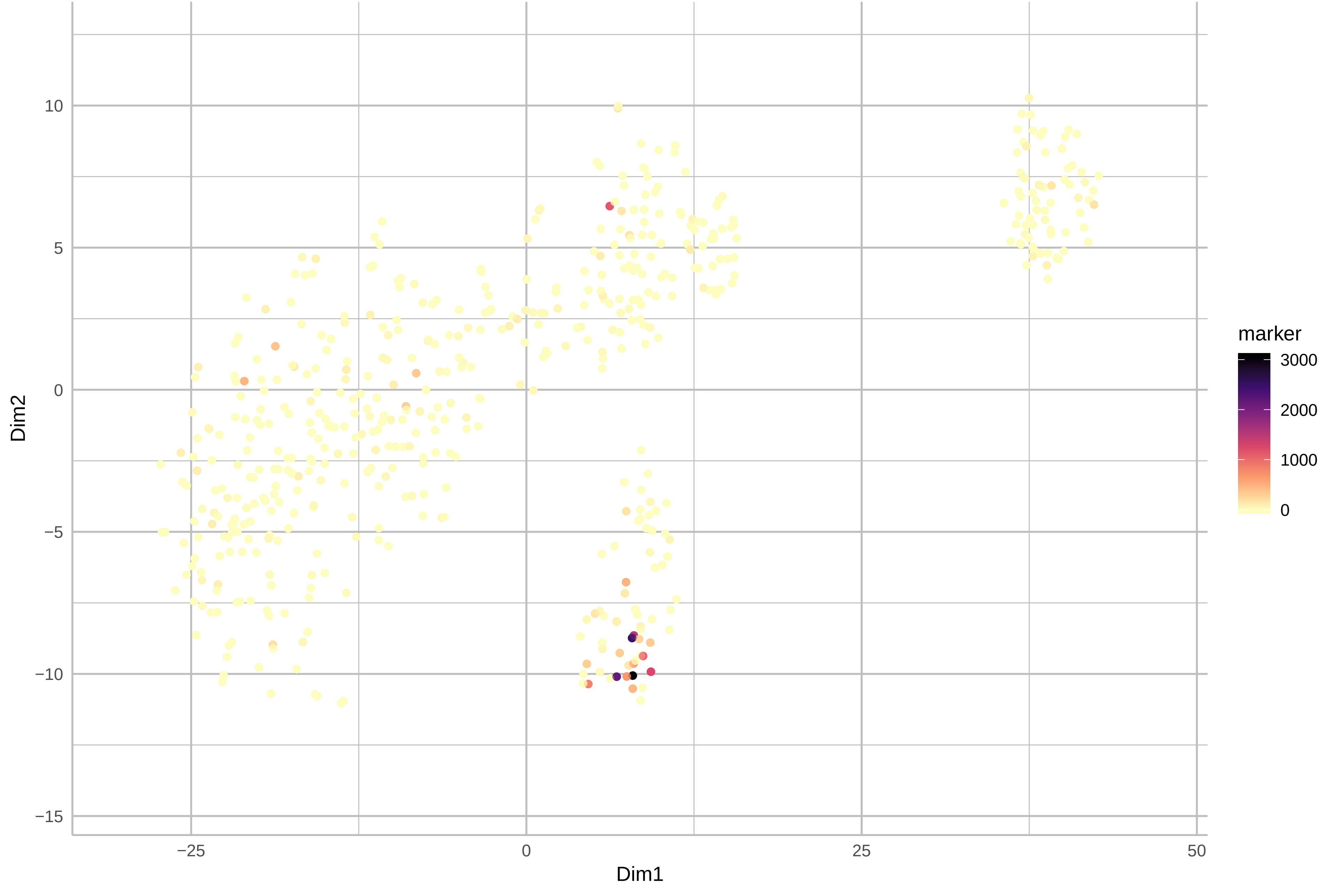

Gm10801

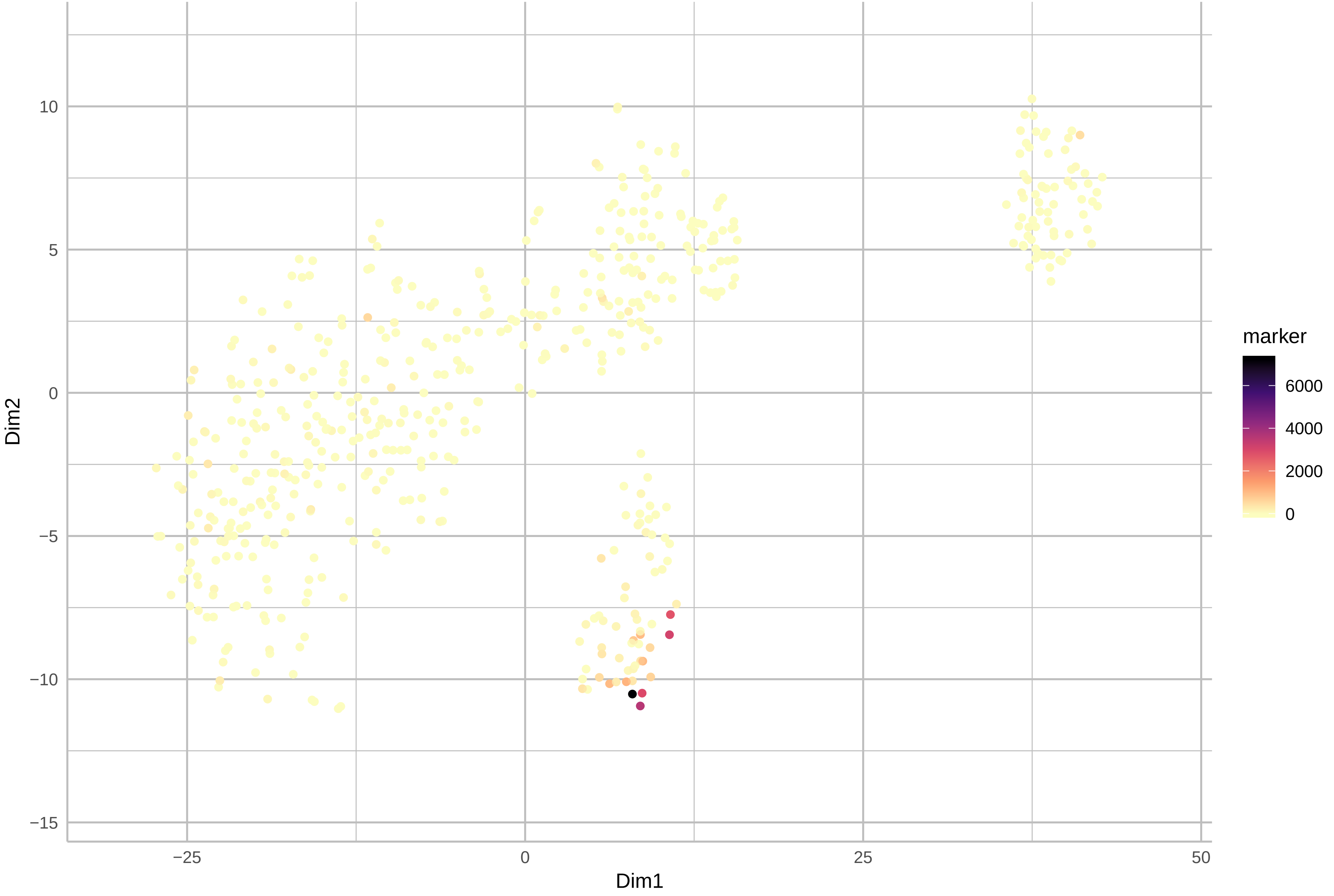

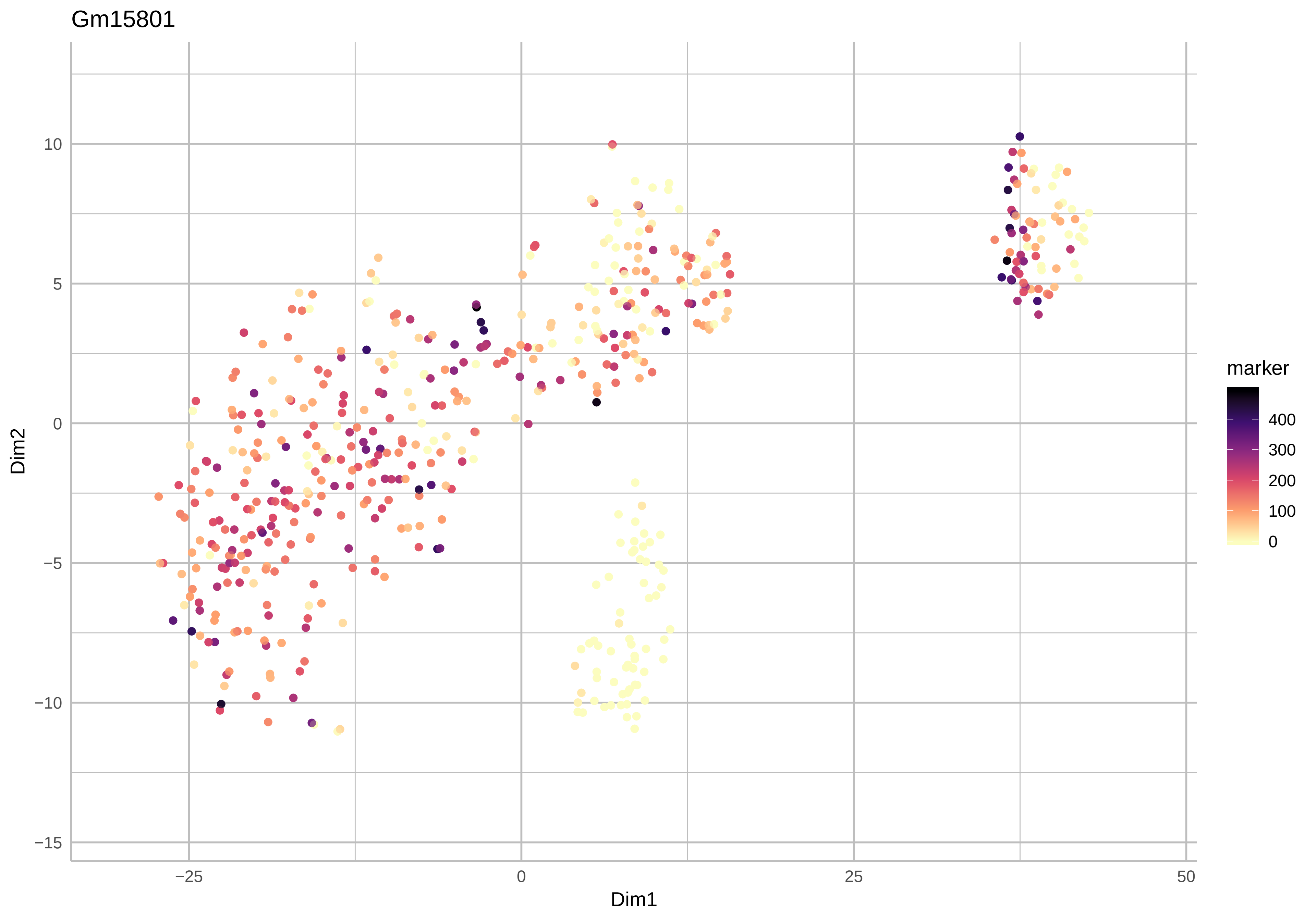
Gm15801
Dim2
Dim1
marker
400
300
200
100
0
10
5
0
−5
−10
−15
−25
0
25
50

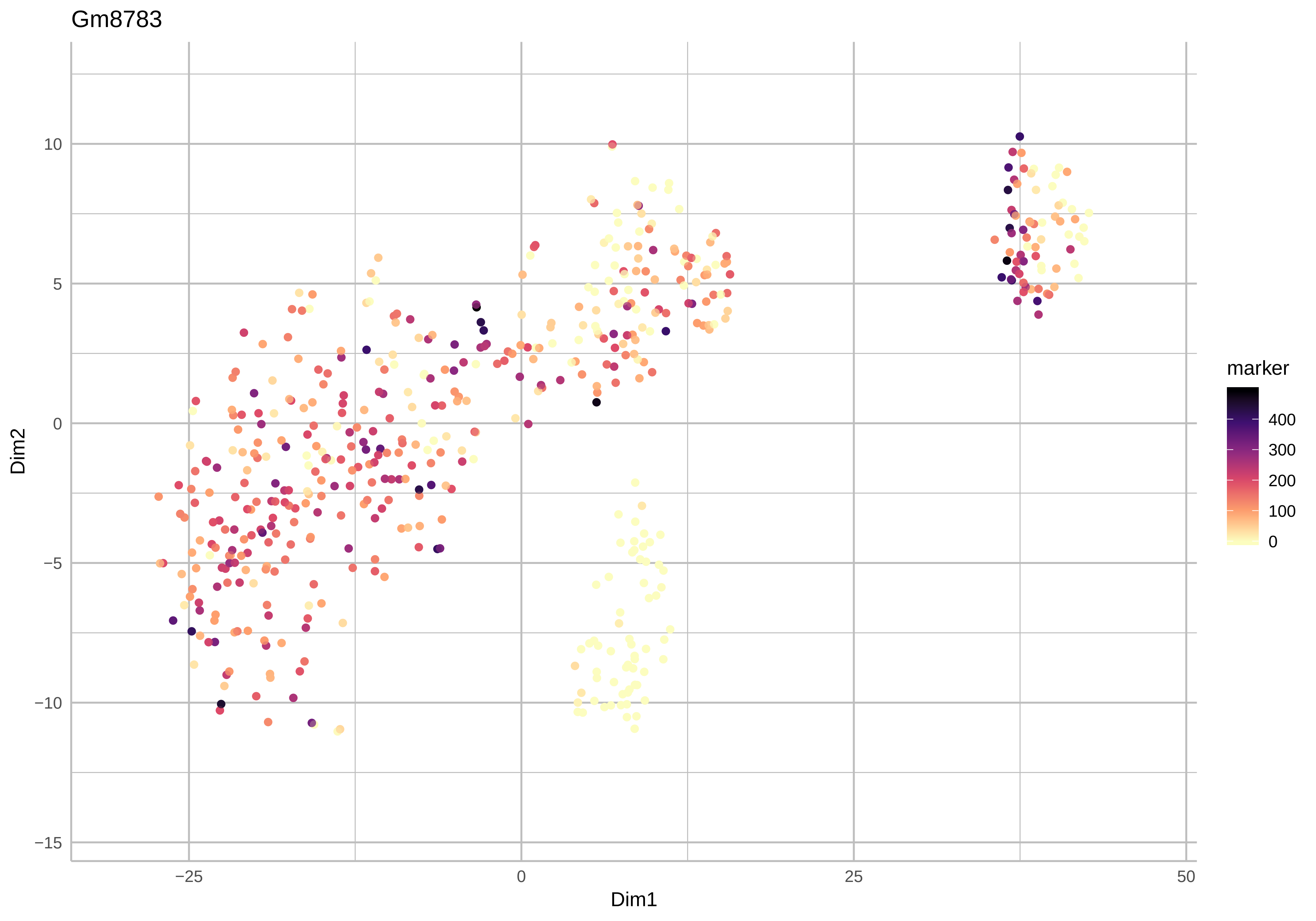
Gm8783
Dim2
Dim1
10
5
0
−5
−10
−15
−25
0
25
50
marker
400
300
200
100
0

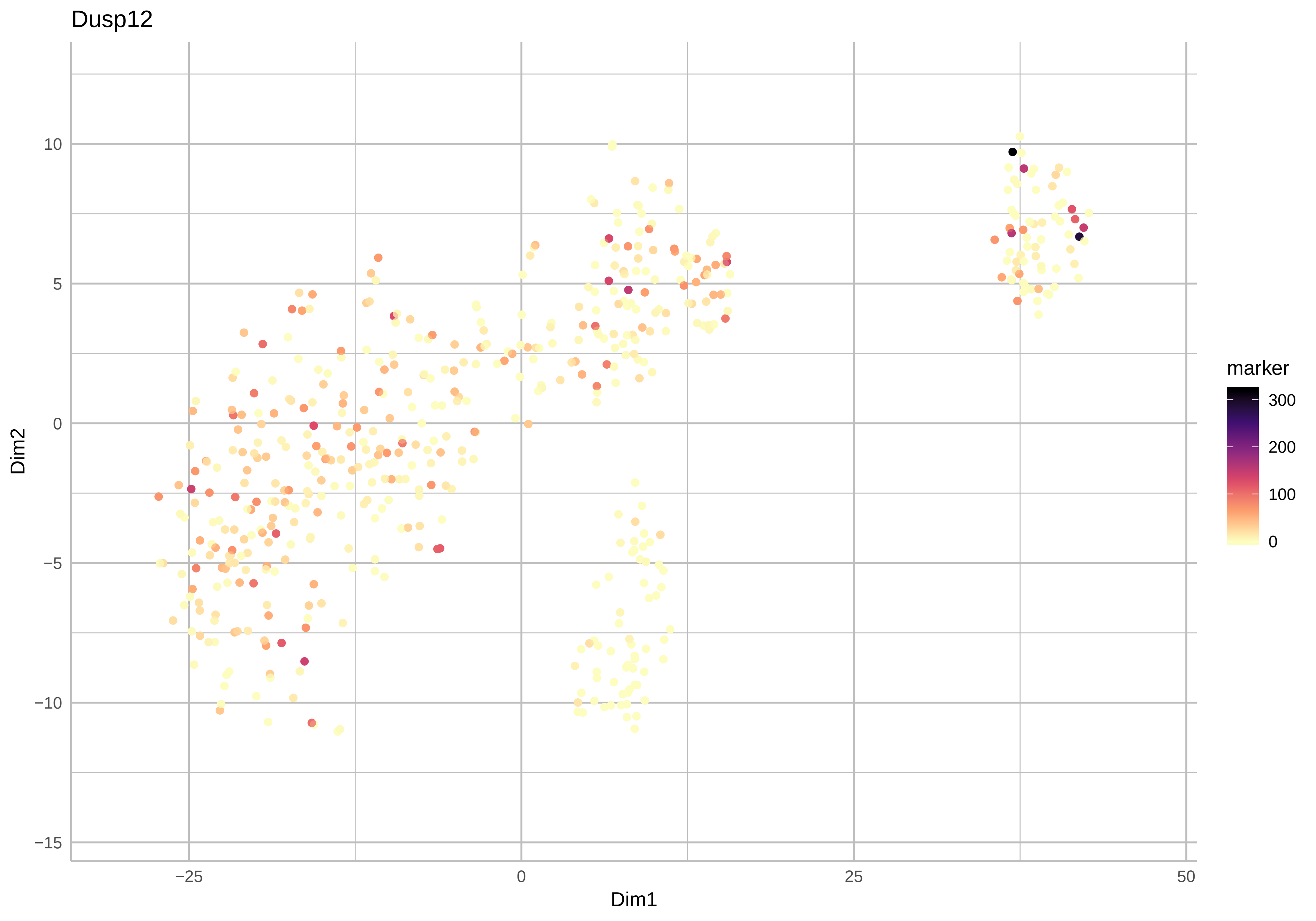

Dusp12
Dim2
Dim1
10
5
0
−5
−10
−15
−25
0
25
50
marker
300
200
100
0

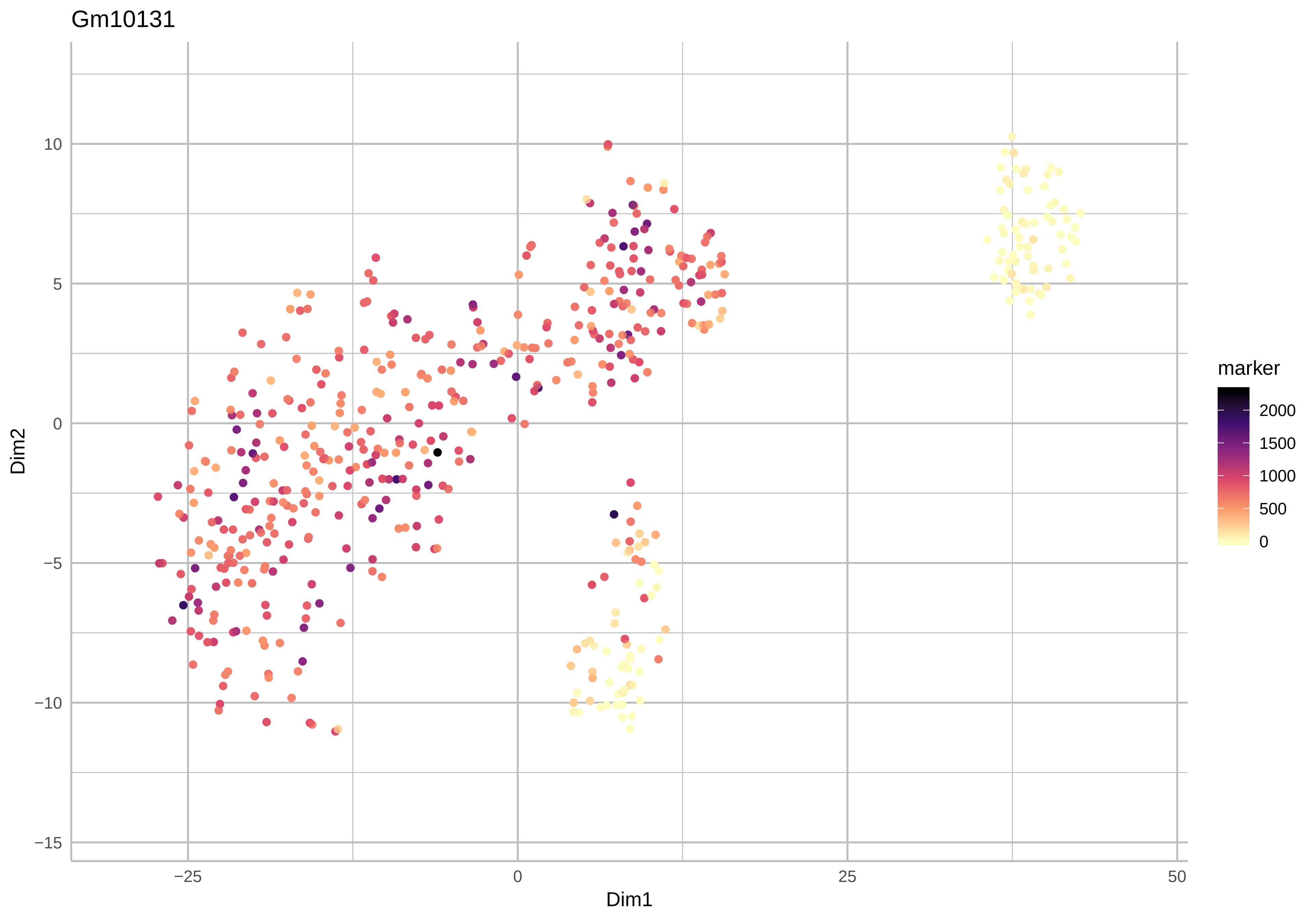
Gm10131
Dim2
Dim1
marker
2000
1500
1000
500
0
10
5
0
−5
−10
−15
−25
0
25
50

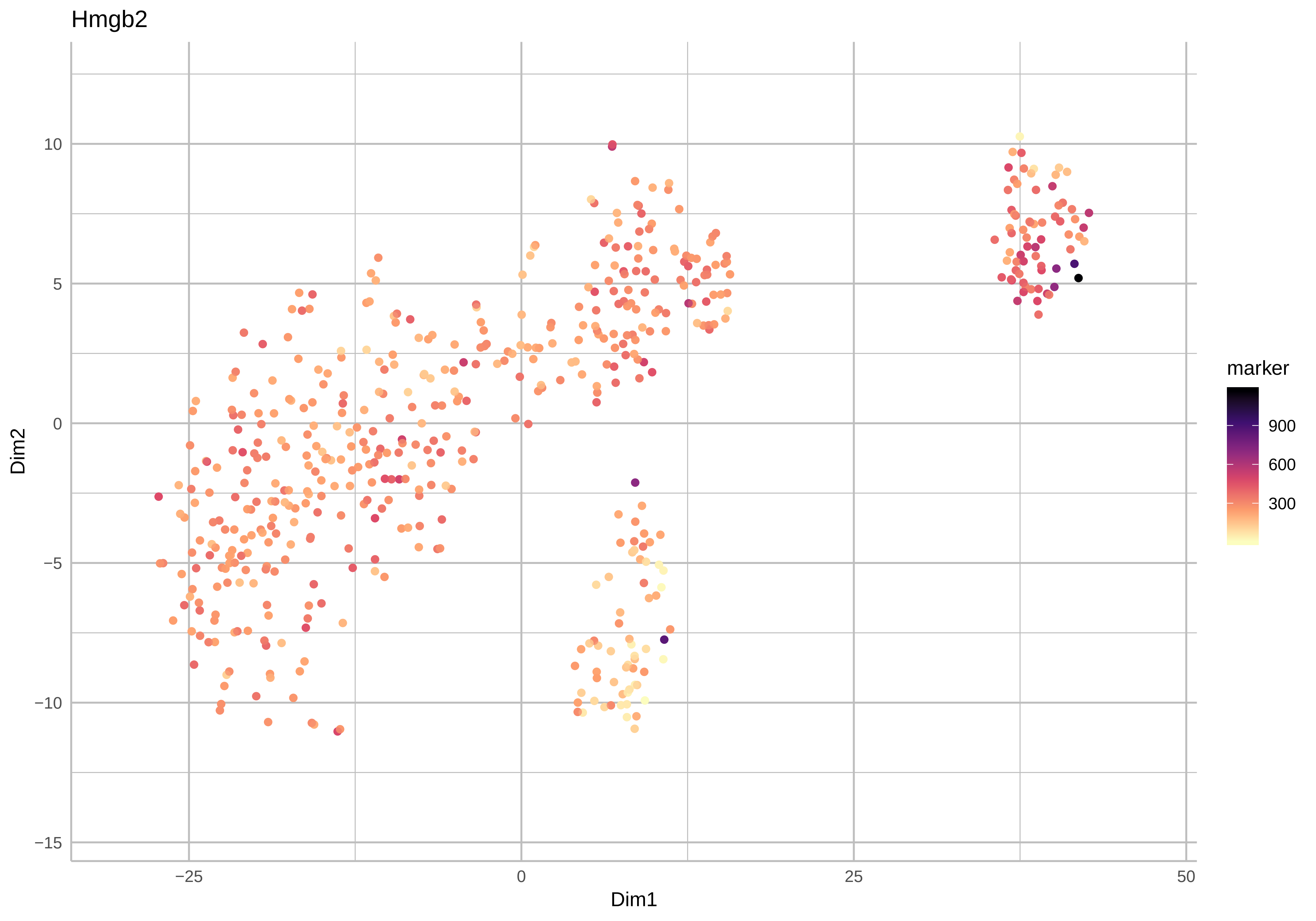
Hmgb2
Dim2
Dim1
10
5
0
−5
−10
−15
−25
0
25
50
marker
900
600
300

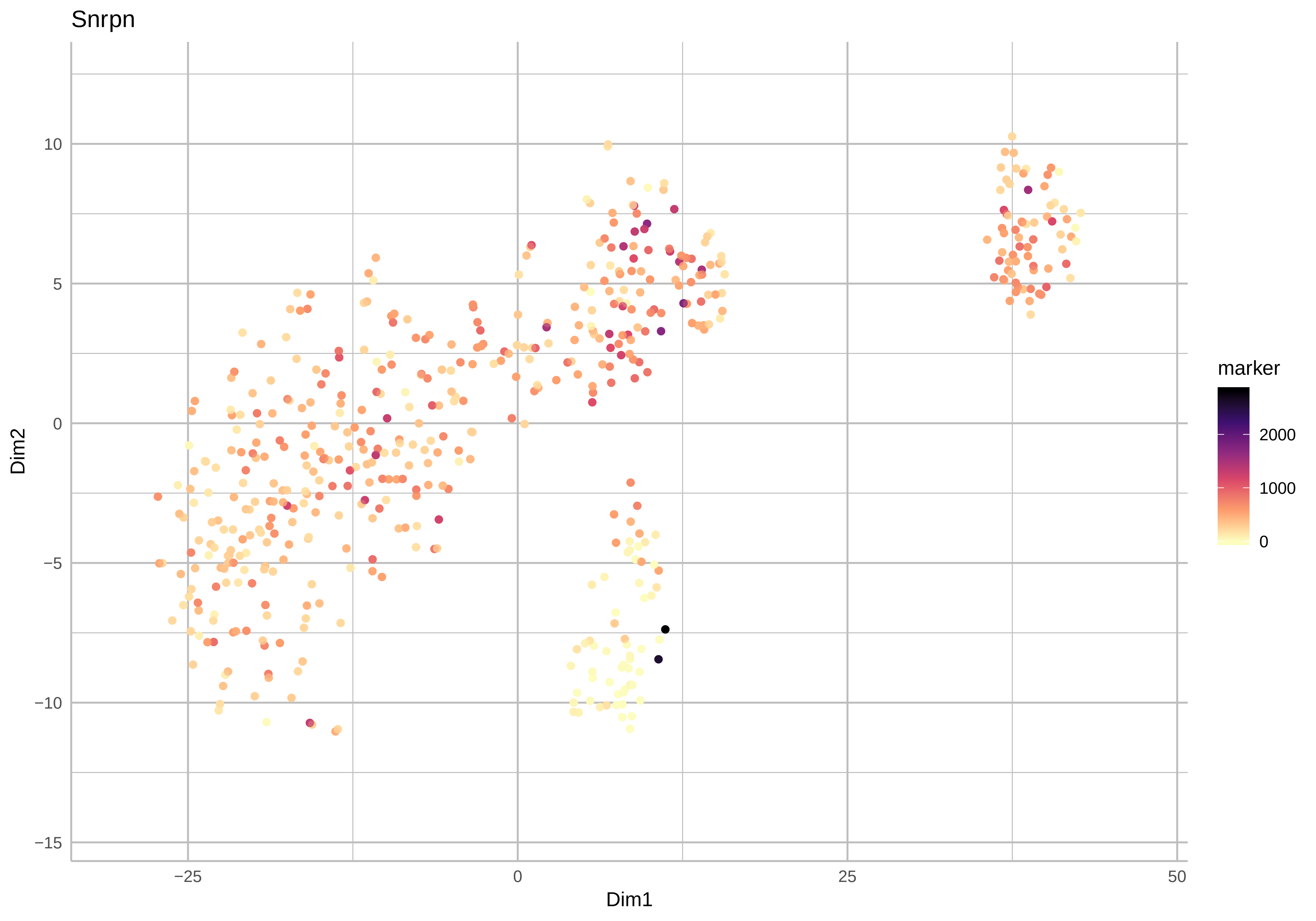
Snrpn
10
5
0
−5
−10
−15
Dim2
−25
0
25
50
Dim1
marker
2000
1000
0

Uhrf1

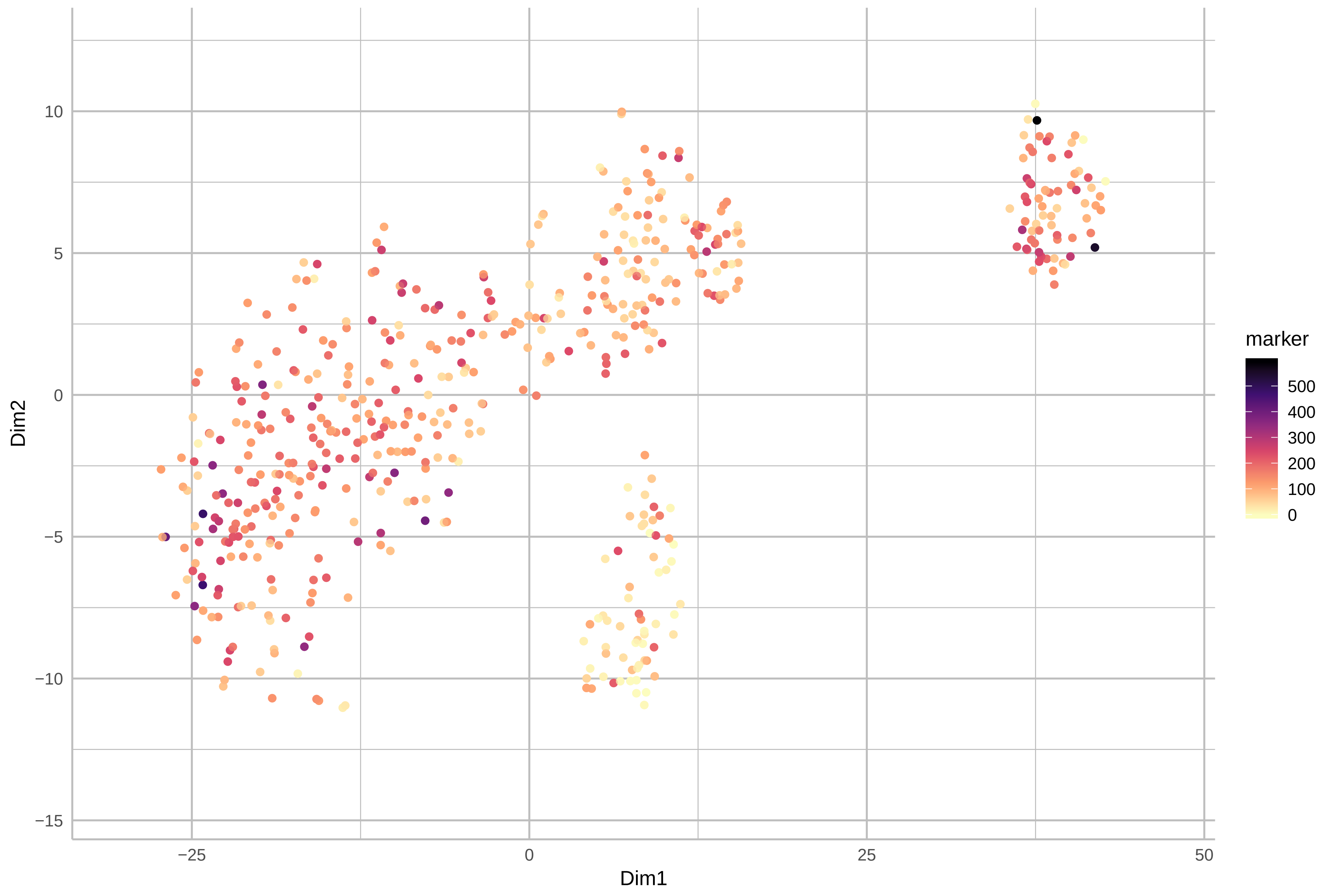

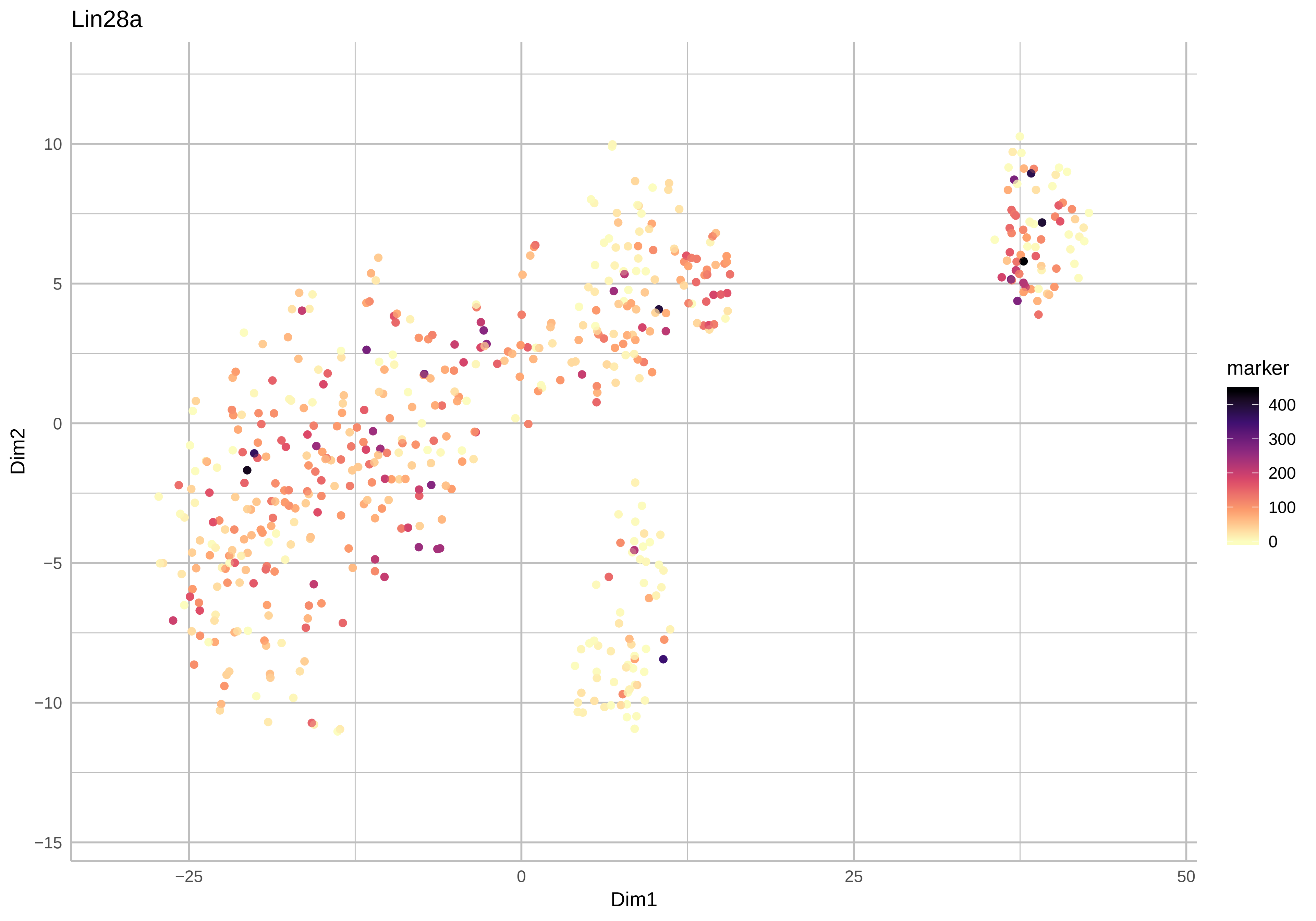

Lin28a
10
5
0
−5
−10
−15
Dim2
−25
0
25
50
Dim1
marker
400
300
200
100
0

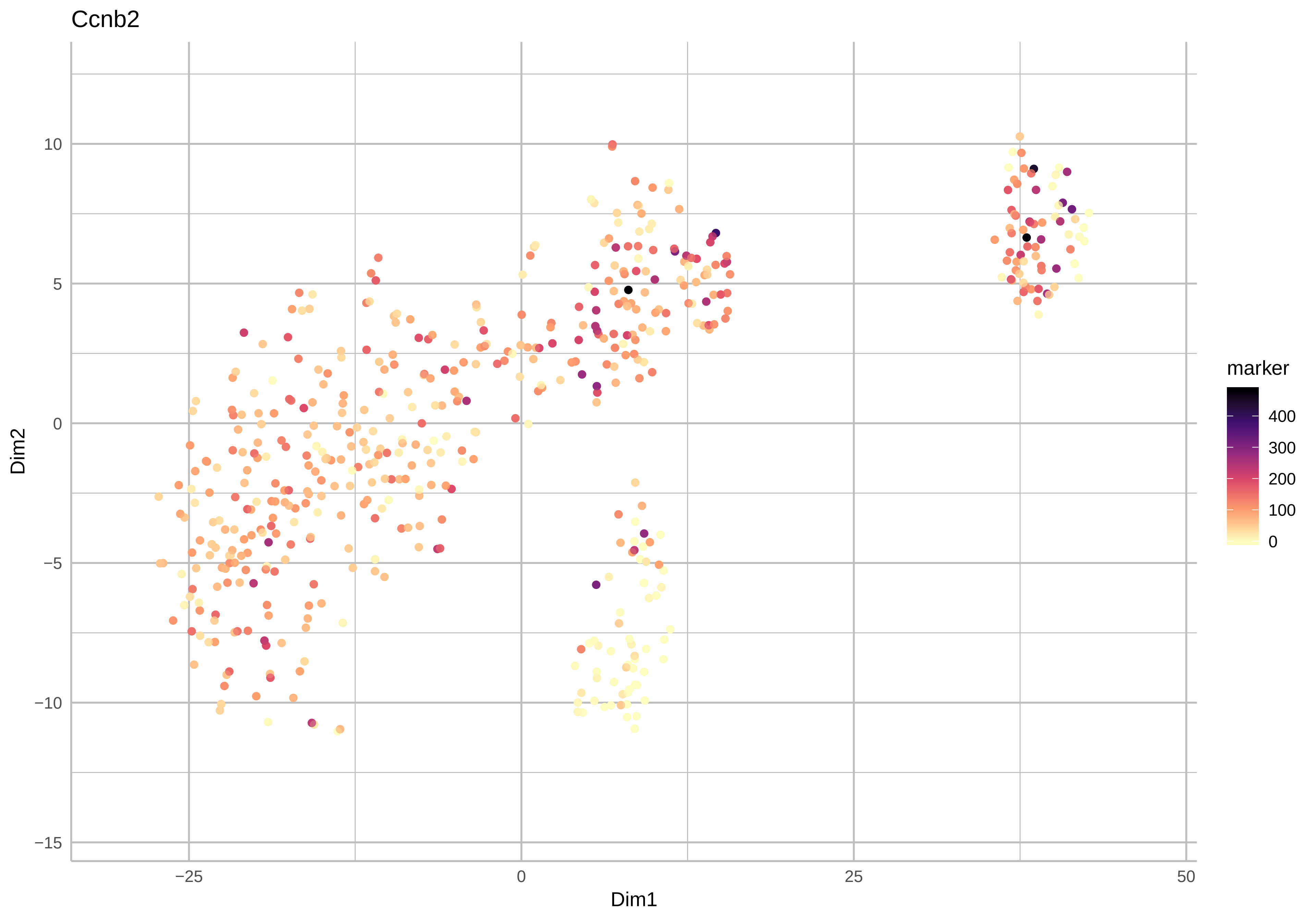

Ccnb2
Dim2
Dim1
marker
400
300
200
100
0
10
5
0
−5
−10
−15
−25
0
25
50

Cdca3

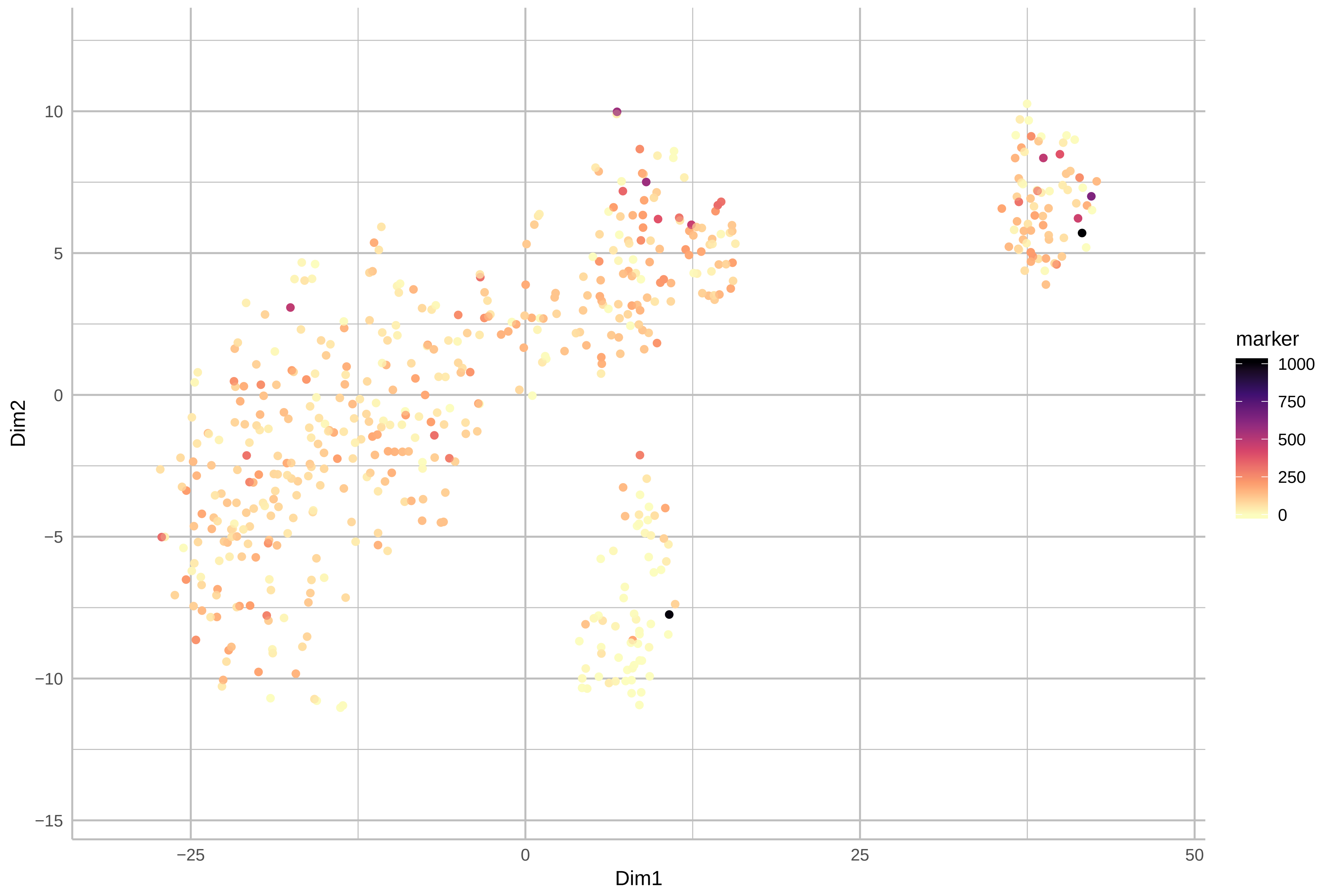

Mki67

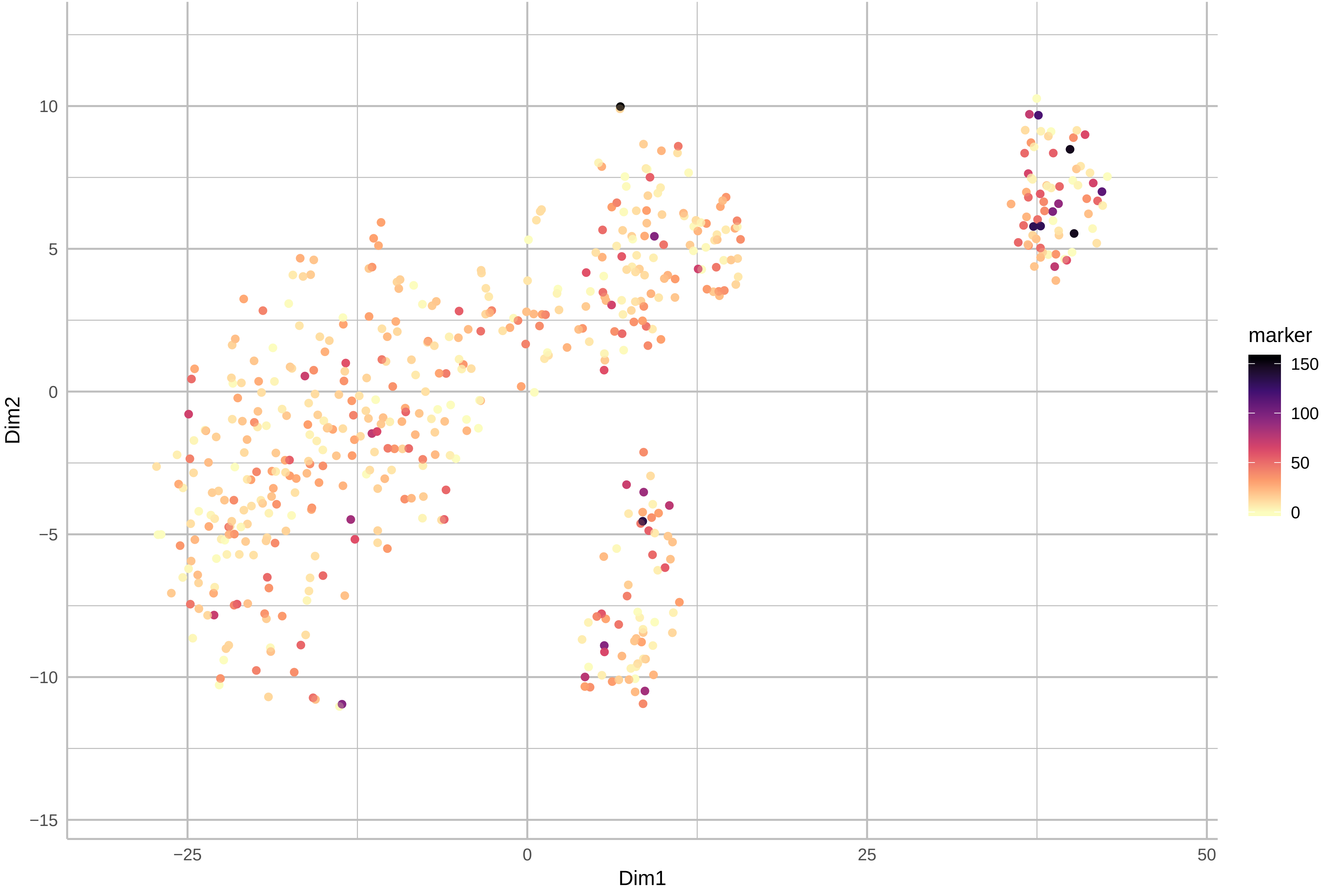

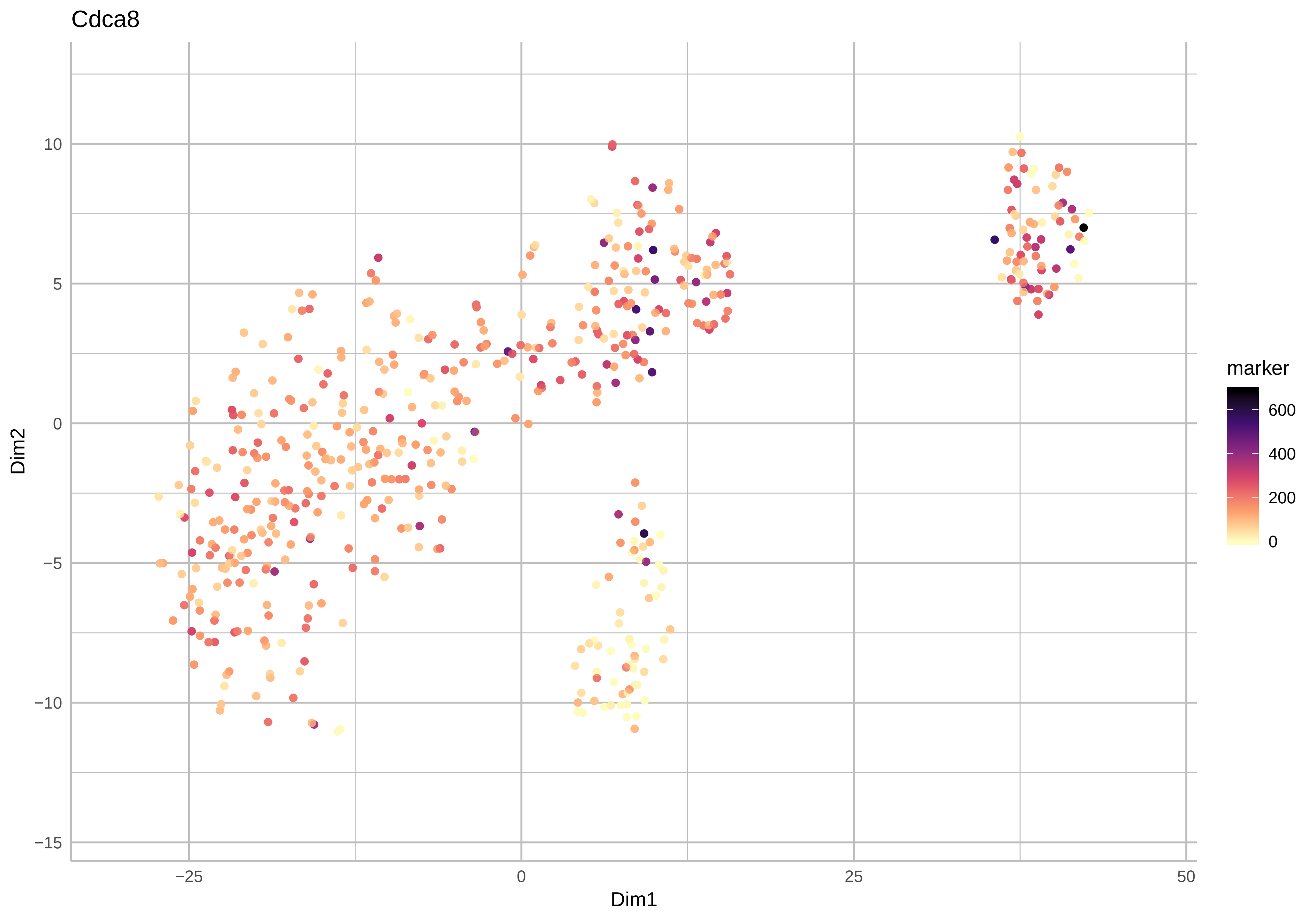

Cdca8
Dim2
Dim1
10
5
0
−5
−10
−15
−25
0
25
50
marker
600
400
200
0

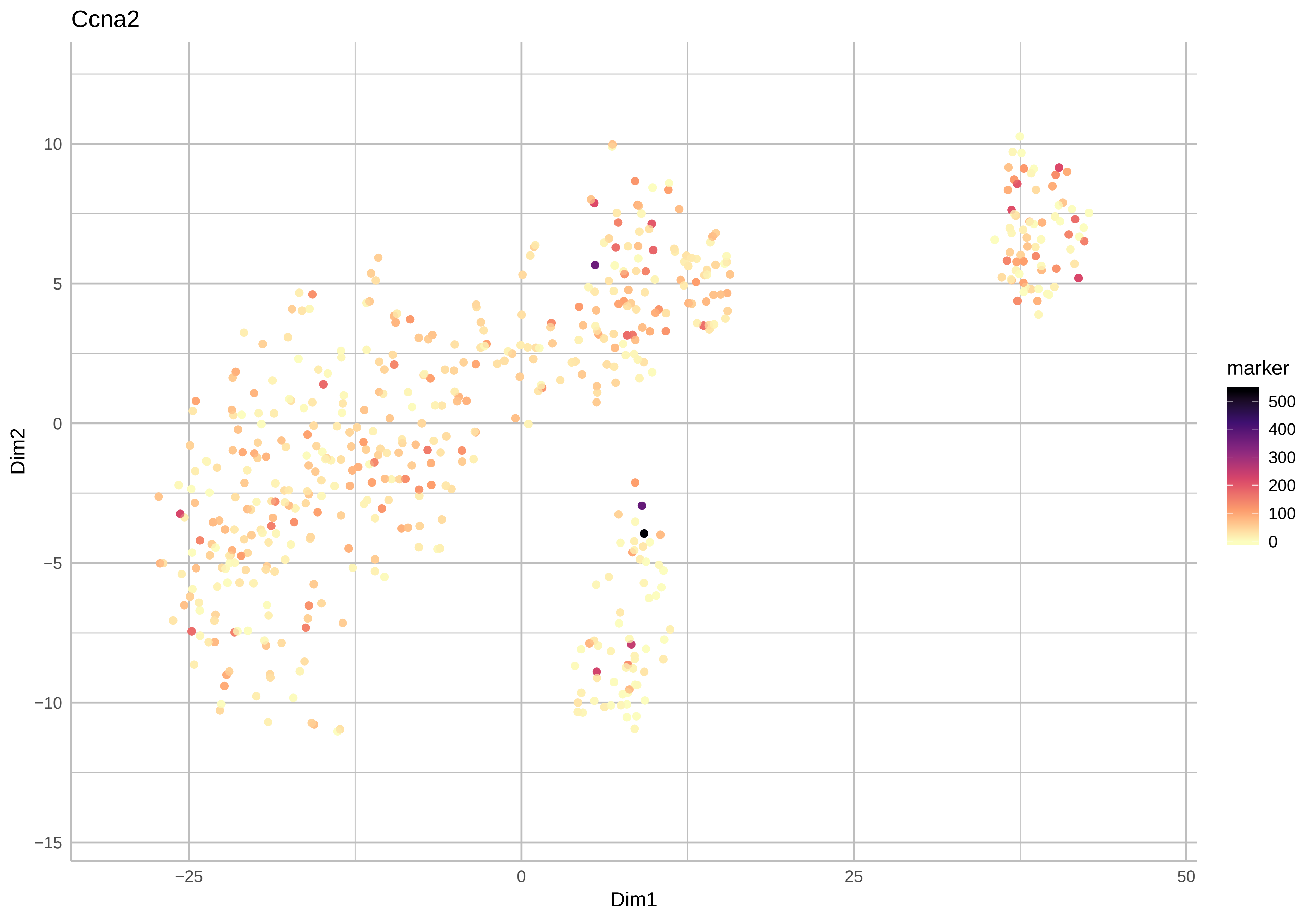
Ccna2
Dim2
Dim1
10
5
0
−5
−10
−15
−25
0
25
50
marker
500
400
300
200
100
0

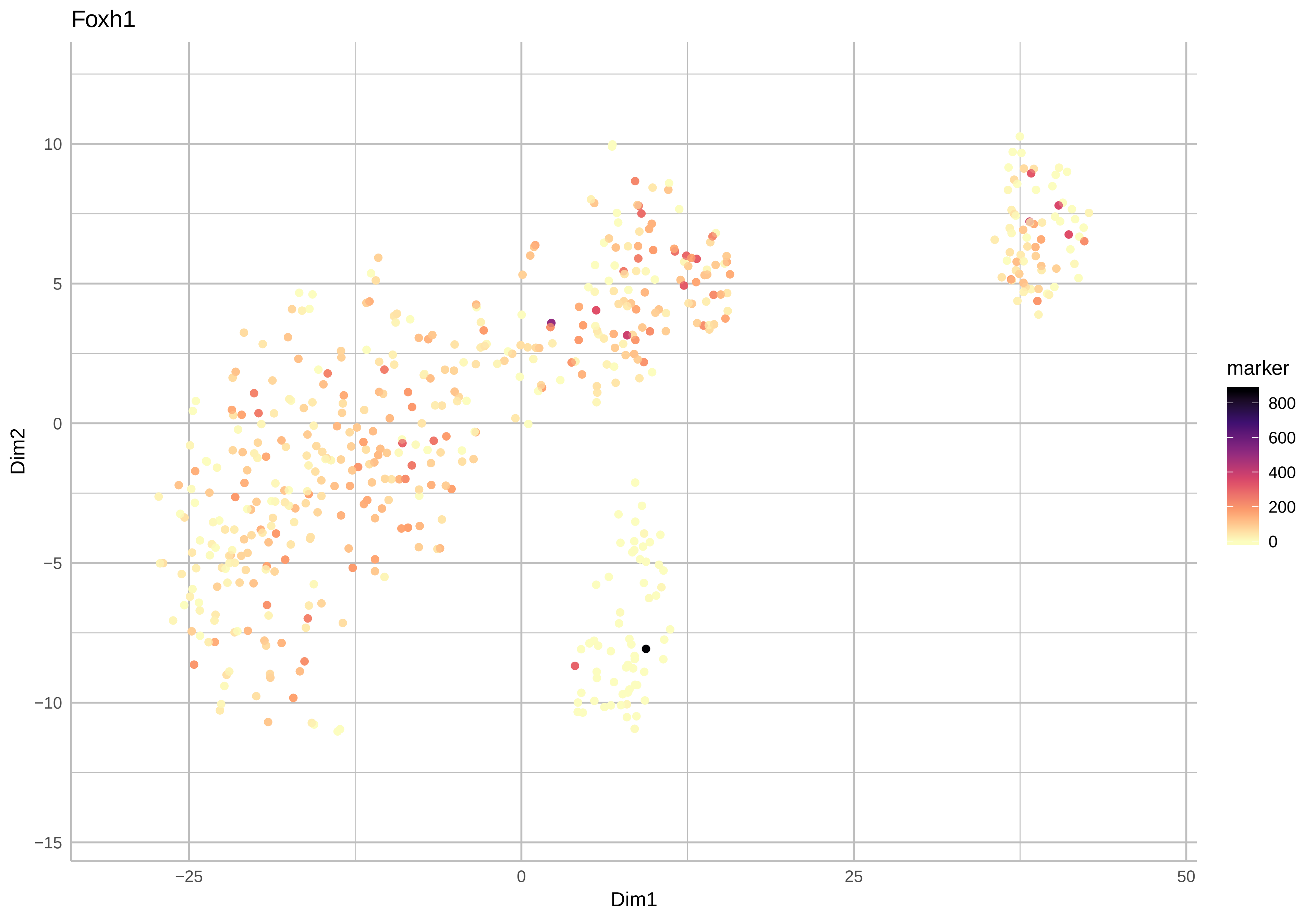

Foxh1
Dim2
Dim1
10
5
0
−5
−10
−15
−25
0
25
50
marker
800
600
400
200
0

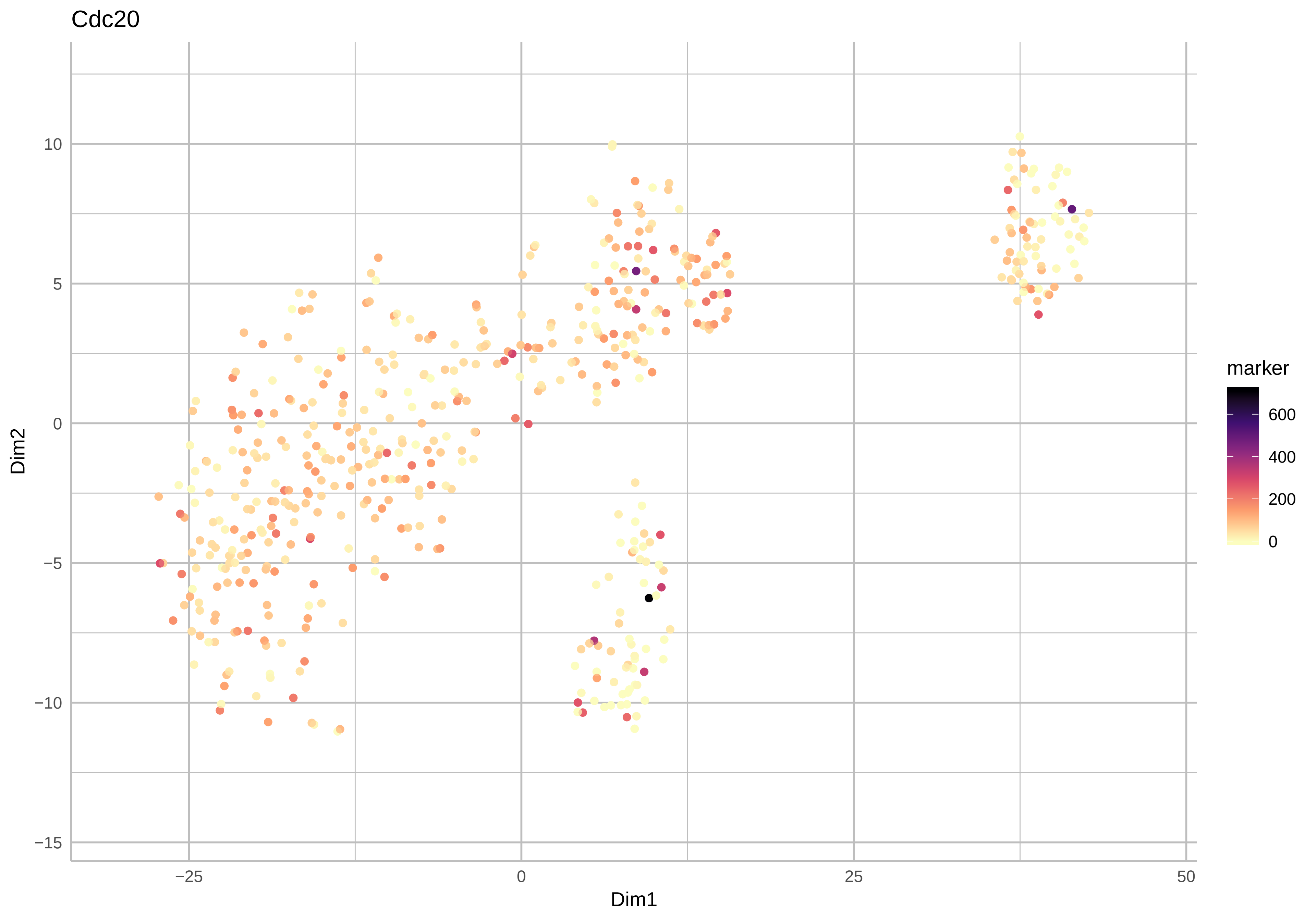
Cdc20
Dim2
Dim1
10
5
0
−5
−10
−15
−25
0
25
50
marker
600
400
200
0

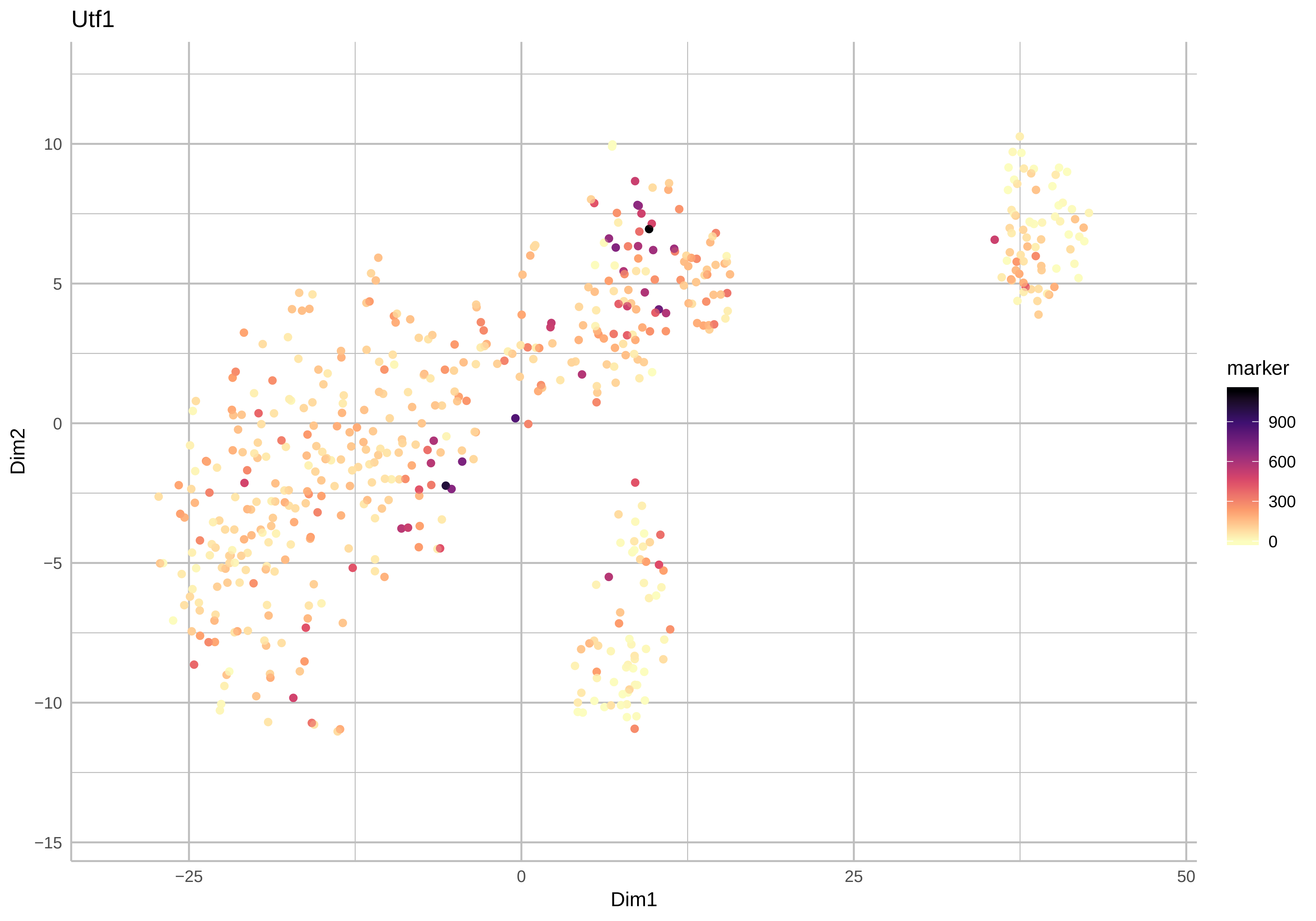

Utf1
10
5
0
−5
−10
−15
Dim2
−25
0
25
50
Dim1
marker
900
600
300
0

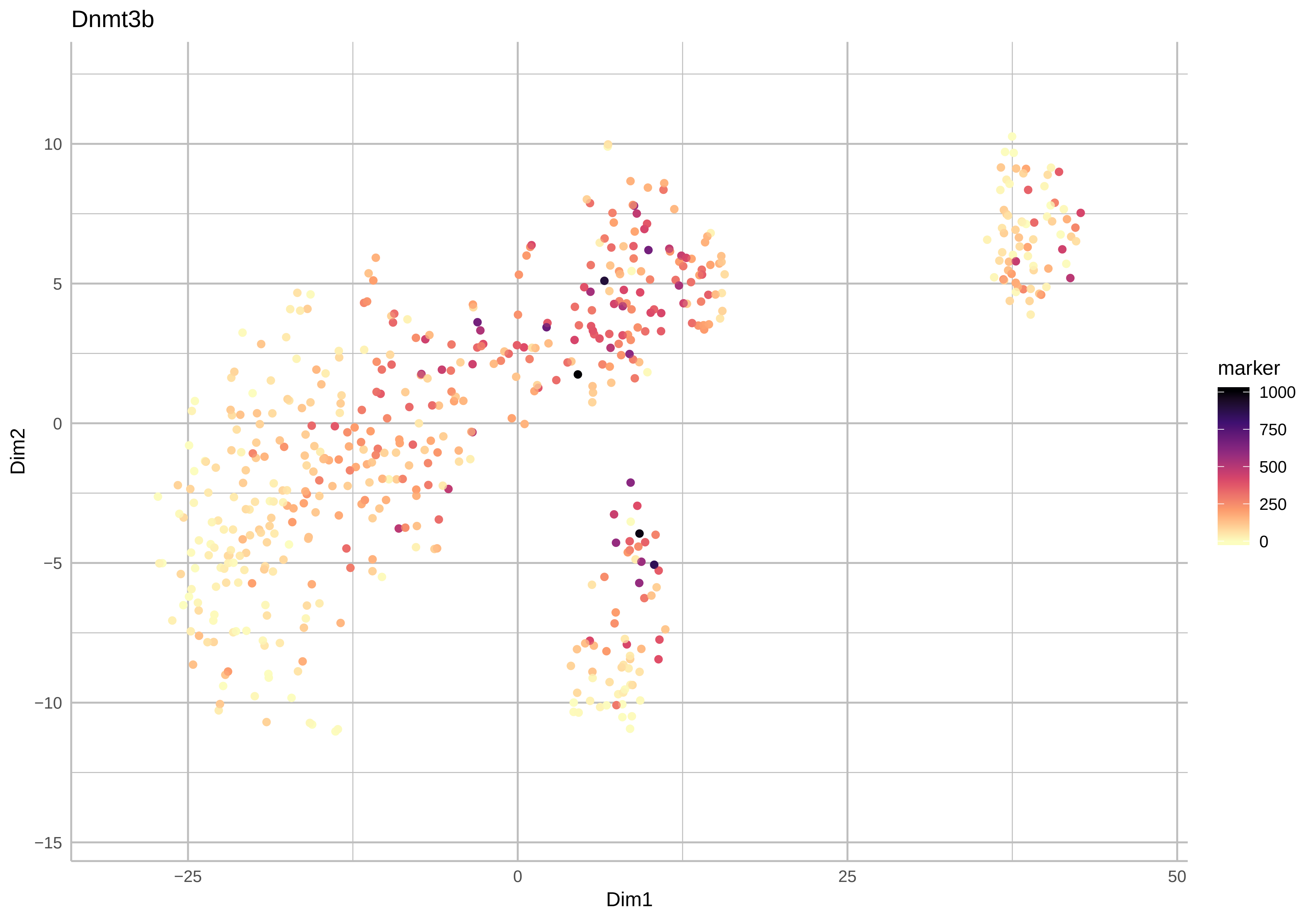

Dnmt3b
Dim2
Dim1
10
5
0
−5
−10
−15
−25
0
25
50
marker
1000
750
500
250
0

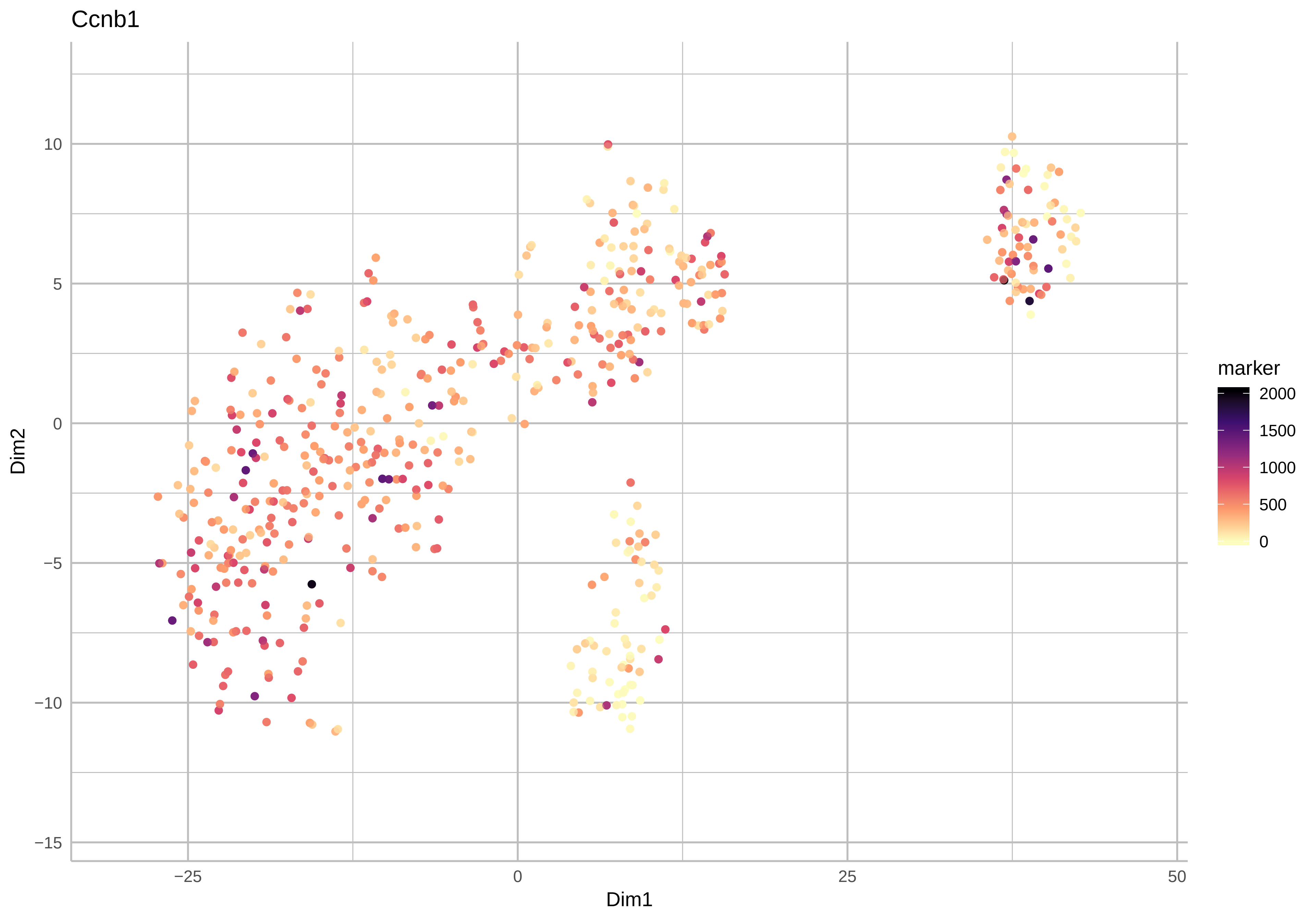

Ccnb1
Dim2
Dim1
10
5
0
−5
−10
−15
−25
0
25
50
marker
2000
1500
1000
500
0

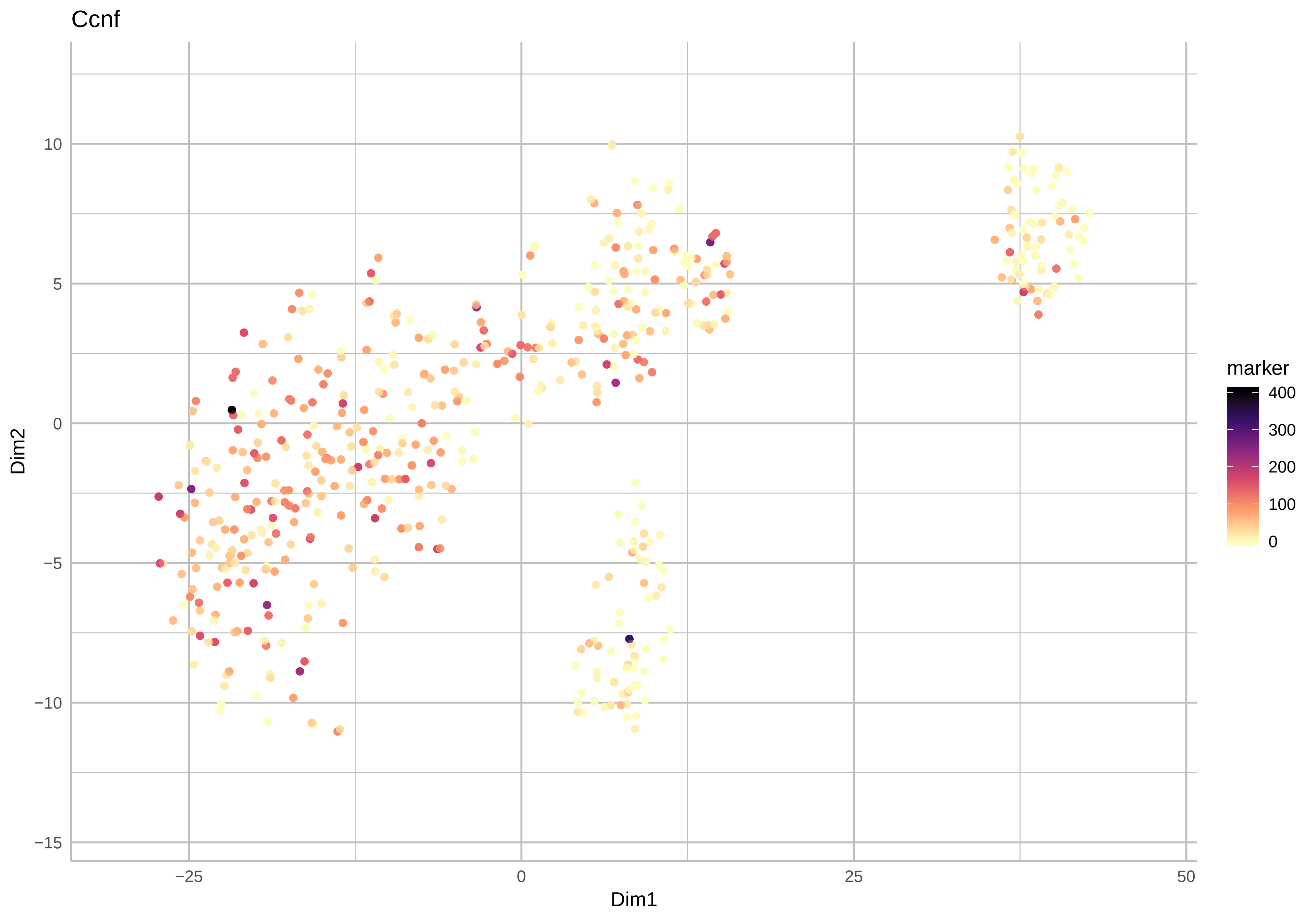

Ccnf
Dim2
Dim1
10
5
0
−5
−10
−15
−25
0
25
50
marker
400
300
200
100
0

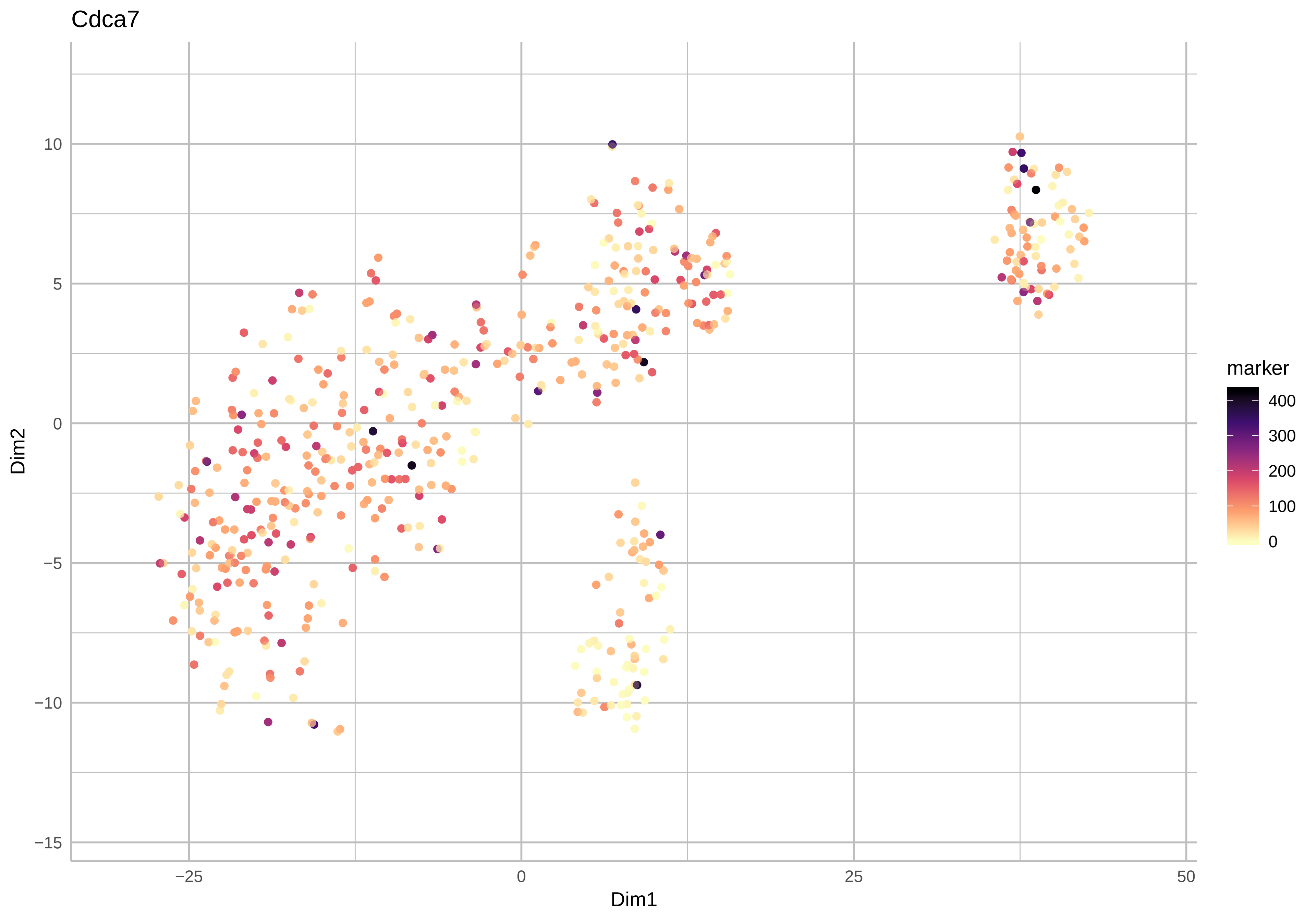

Cdca7
Dim2
Dim1
10
5
0
−5
−10
−15
−25
0
25
50
marker
400
300
200
100
0

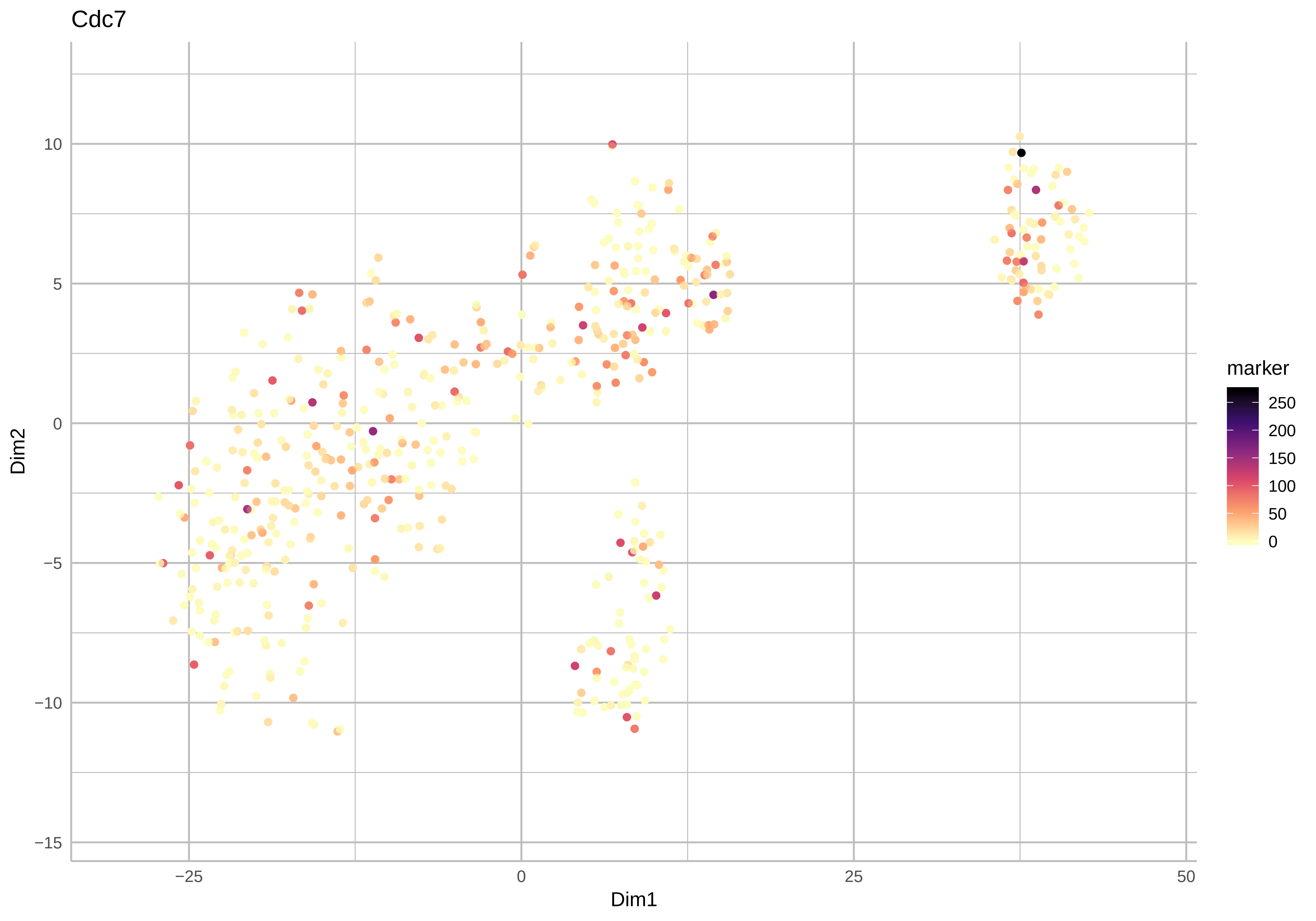
Cdc7
Dim2
Dim1
marker
250
200
150
100
50
0
10
5
0
−5
−10
−15
−25
0
25
50

Larp7

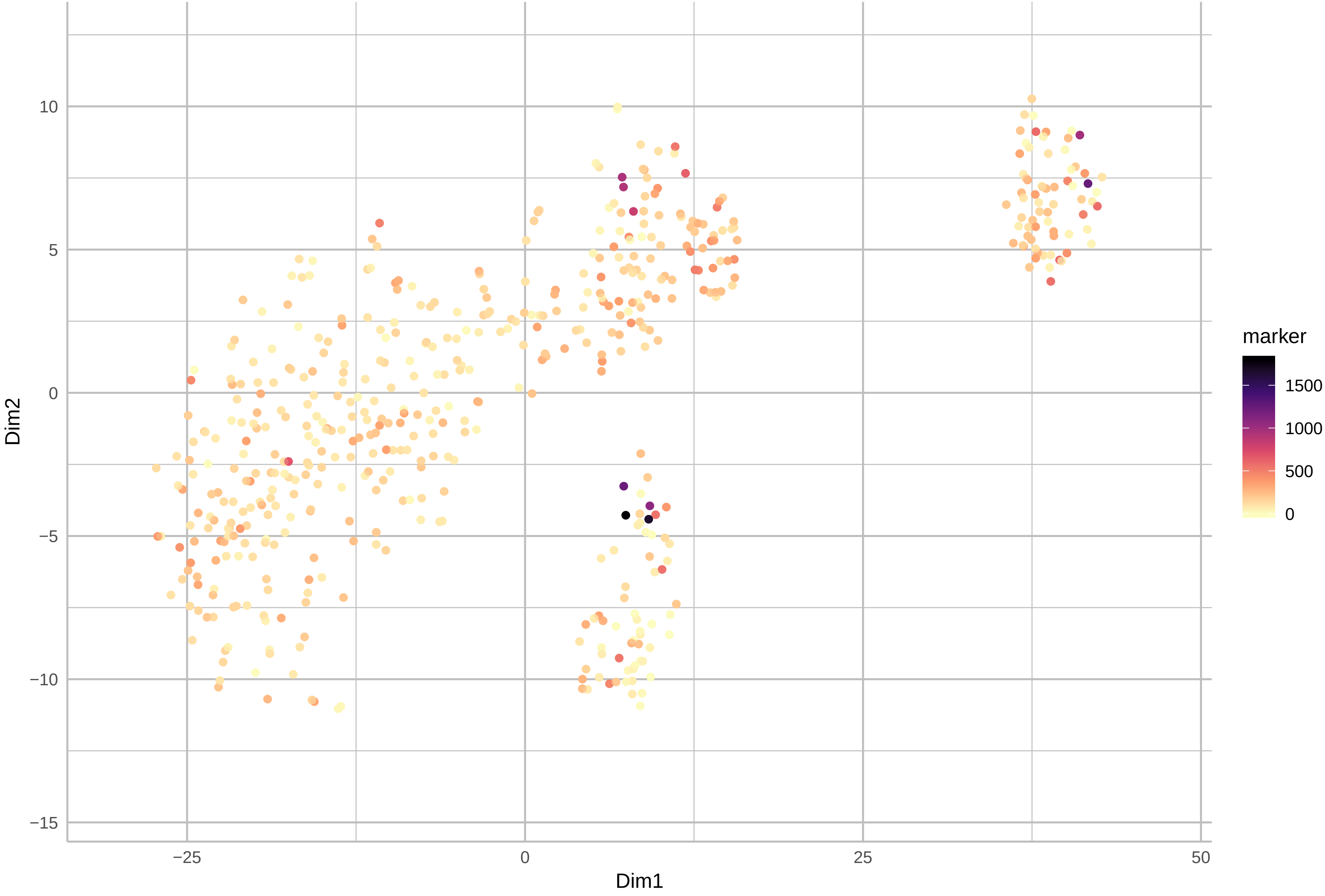

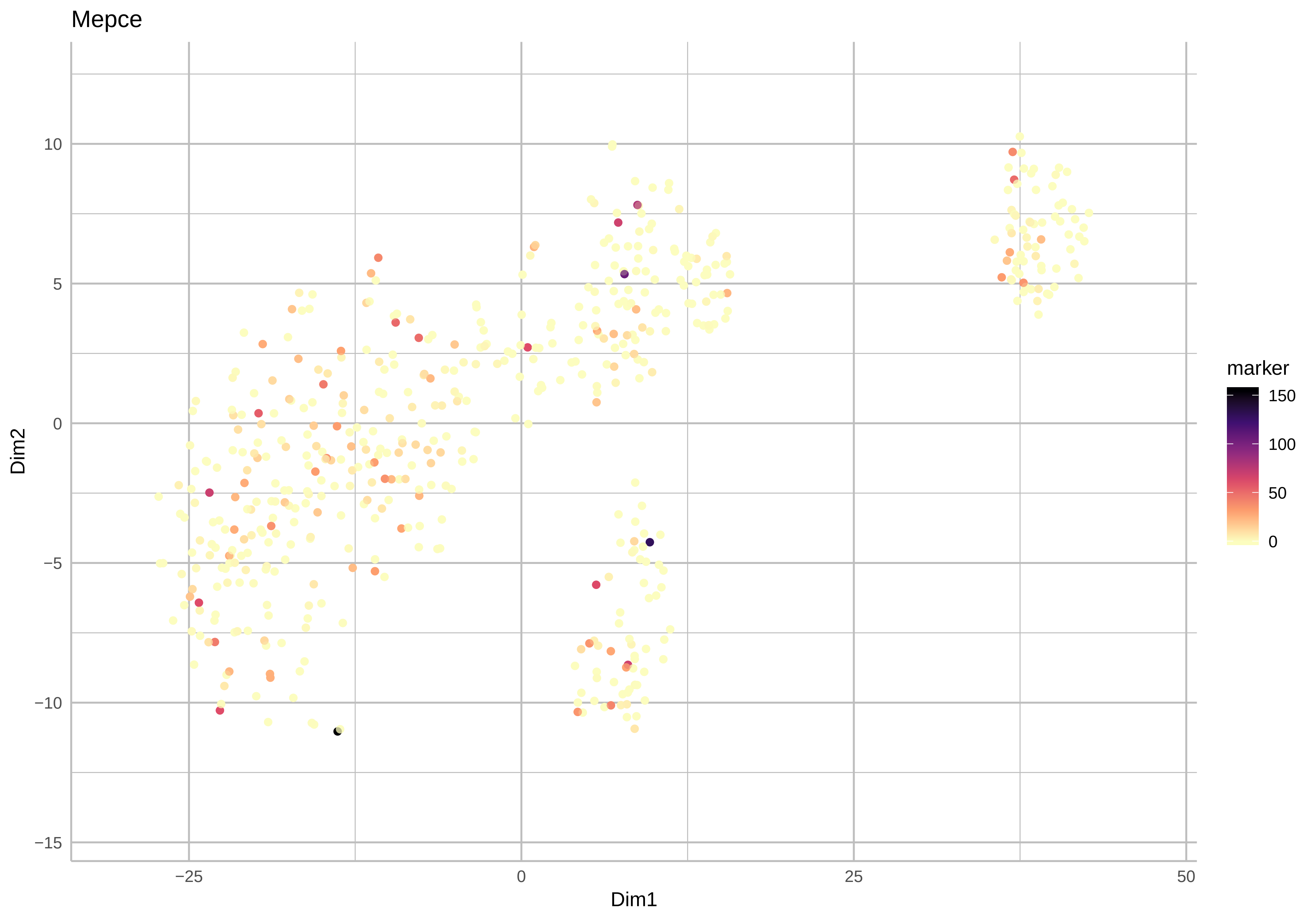

Mepce
Dim1
Dim2
marker
150
100
50
0
−25
0
25
50
10
5
0
−5
−10
−15

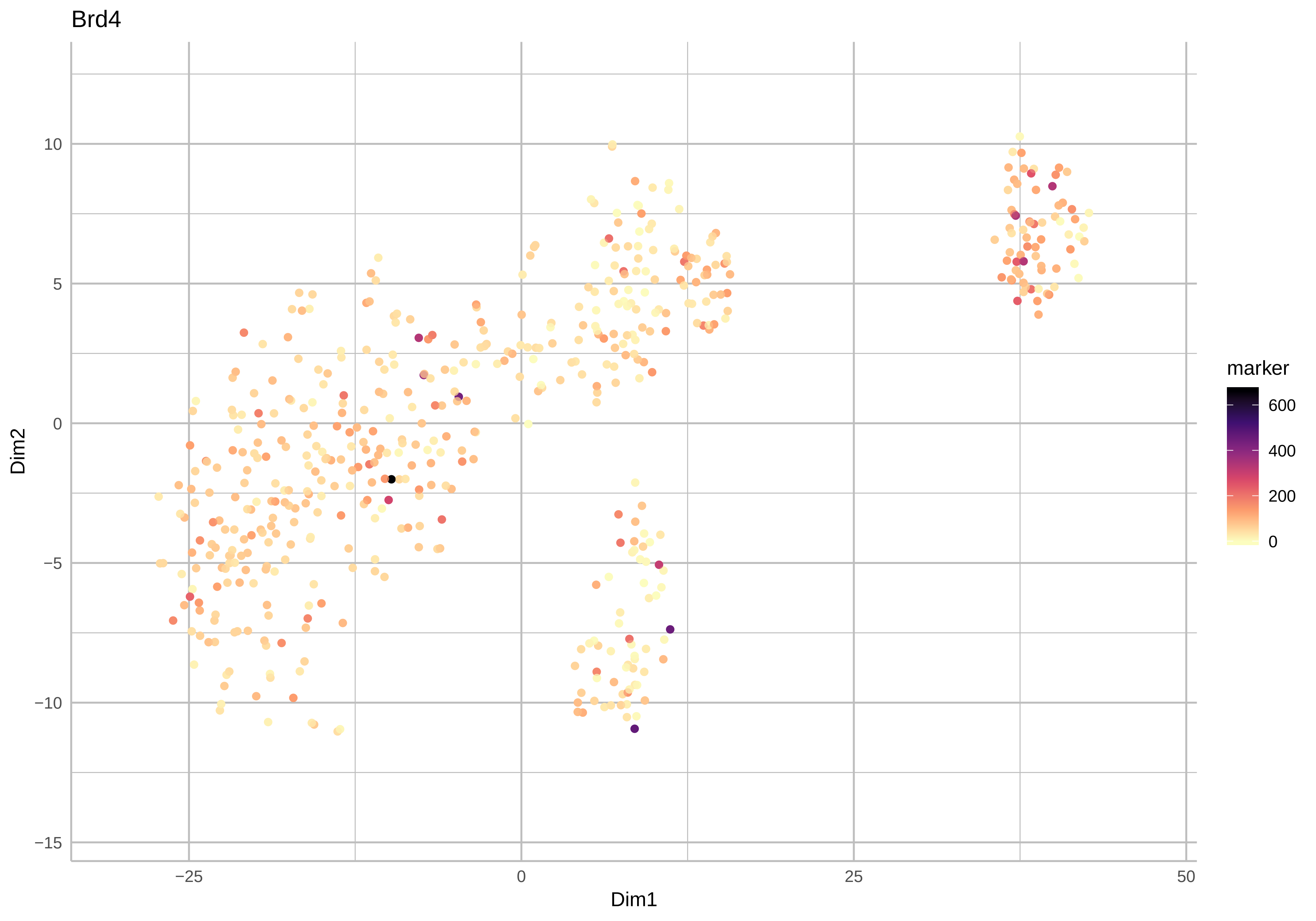
Brd4
Dim1
Dim2
marker
600
400
200
0
−25
0
25
50
10
5
0
−5
−10
−15

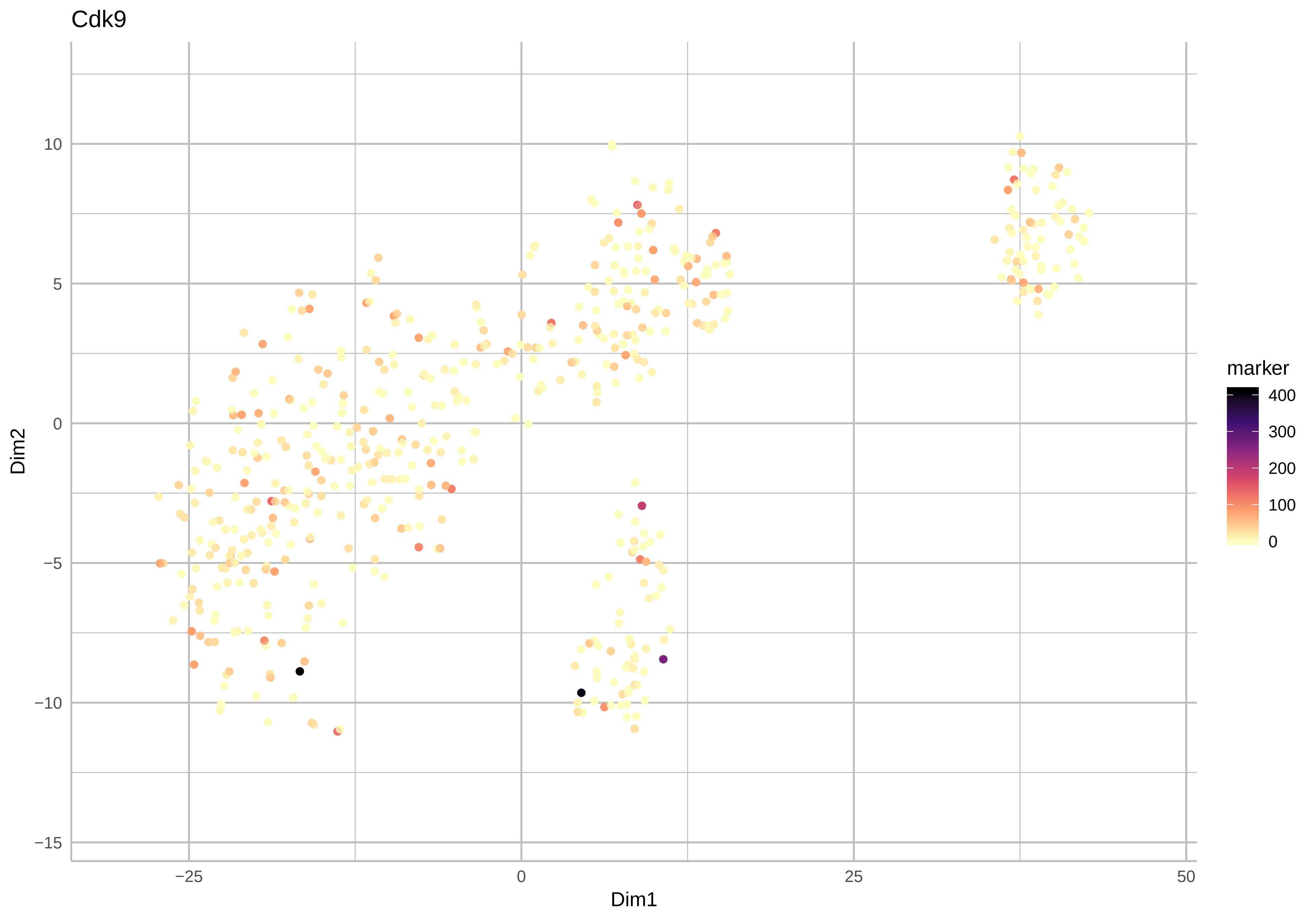
Cdk9
Dim2
Dim1
10
5
0
−5
−10
−15
−25
0
25
50
marker
400
300
200
100
0

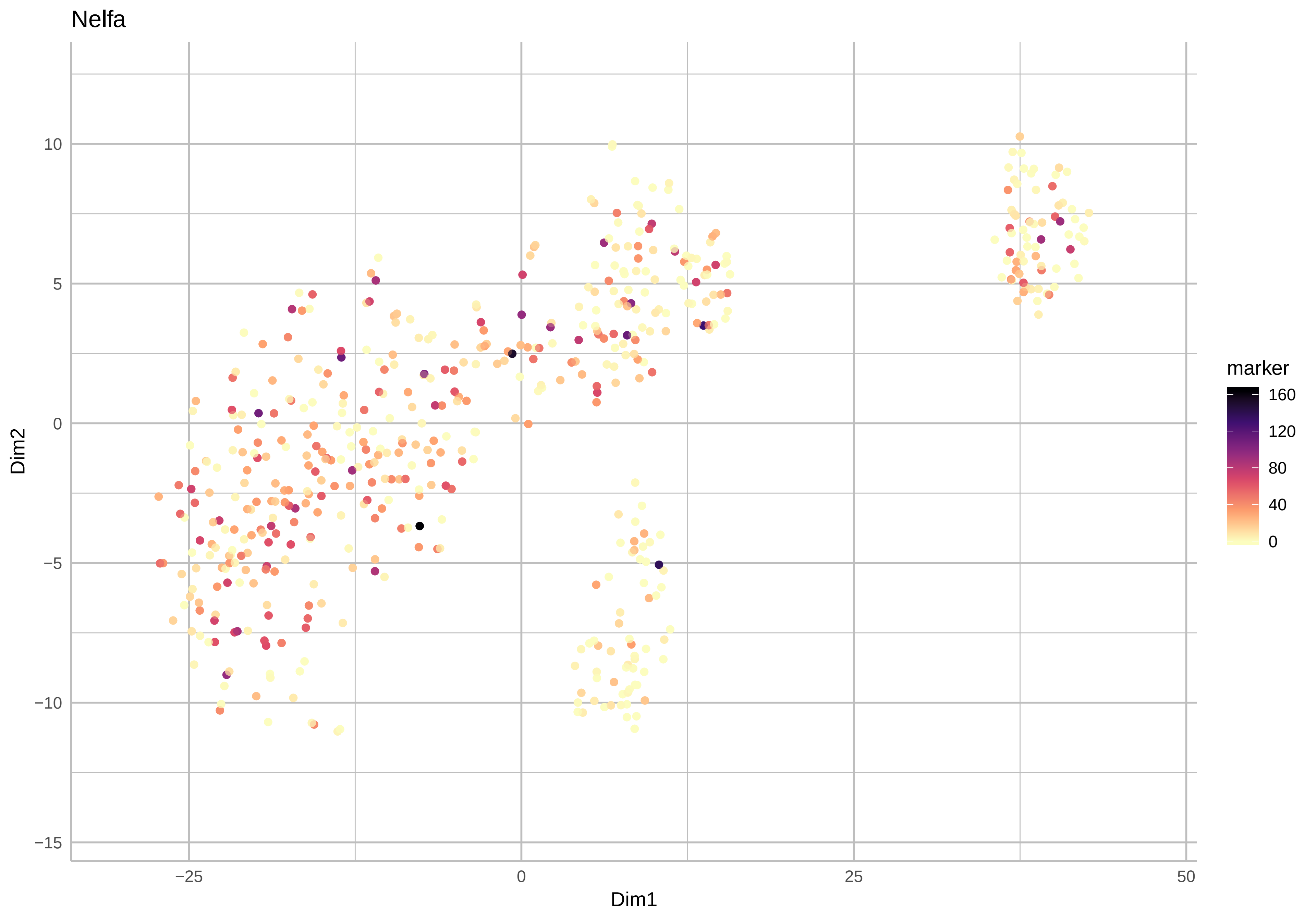

Nelfa
Dim2
Dim1
marker
160
120
80
40
0
10
5
0
−5
−10
−15
−25
0
25
50

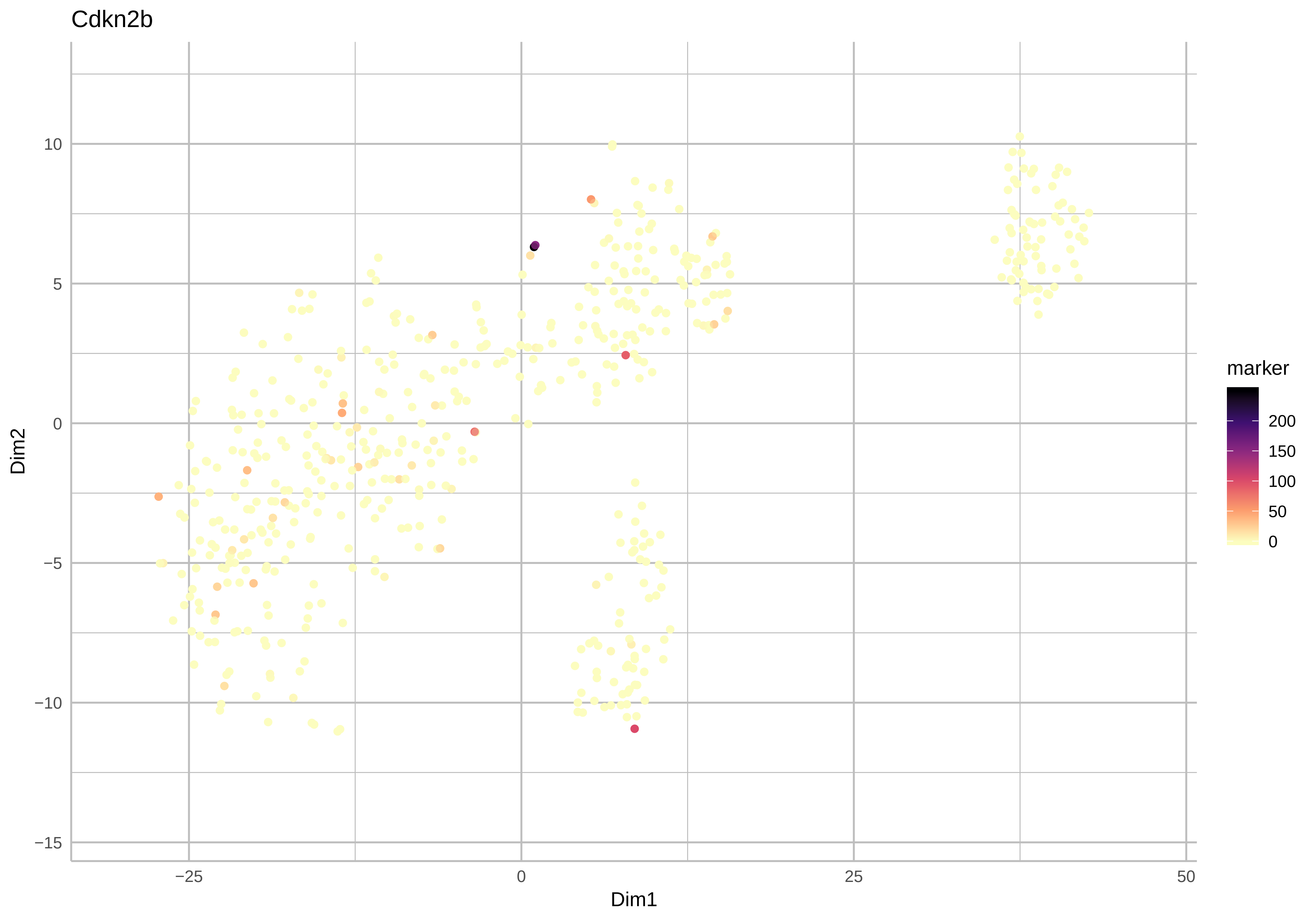
Cdkn2b
Dim1
Dim2
−25
0
25
50
10
5
0
−5
−10
−15
marker
200
150
100
50
0

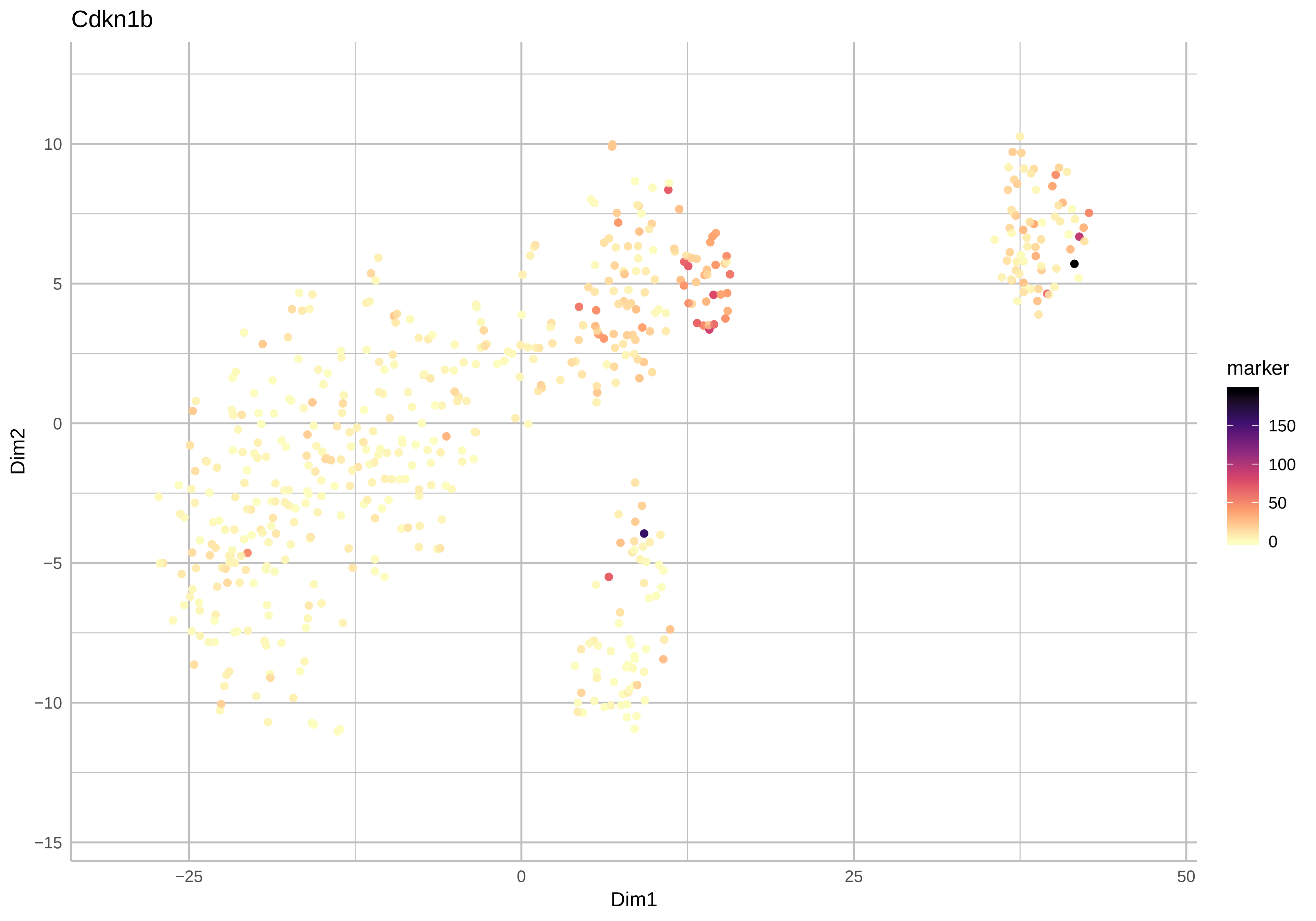
Cdkn1b
Dim2
Dim1
10
5
0
−5
−10
−15
−25
0
25
50
marker
150
100
50
0

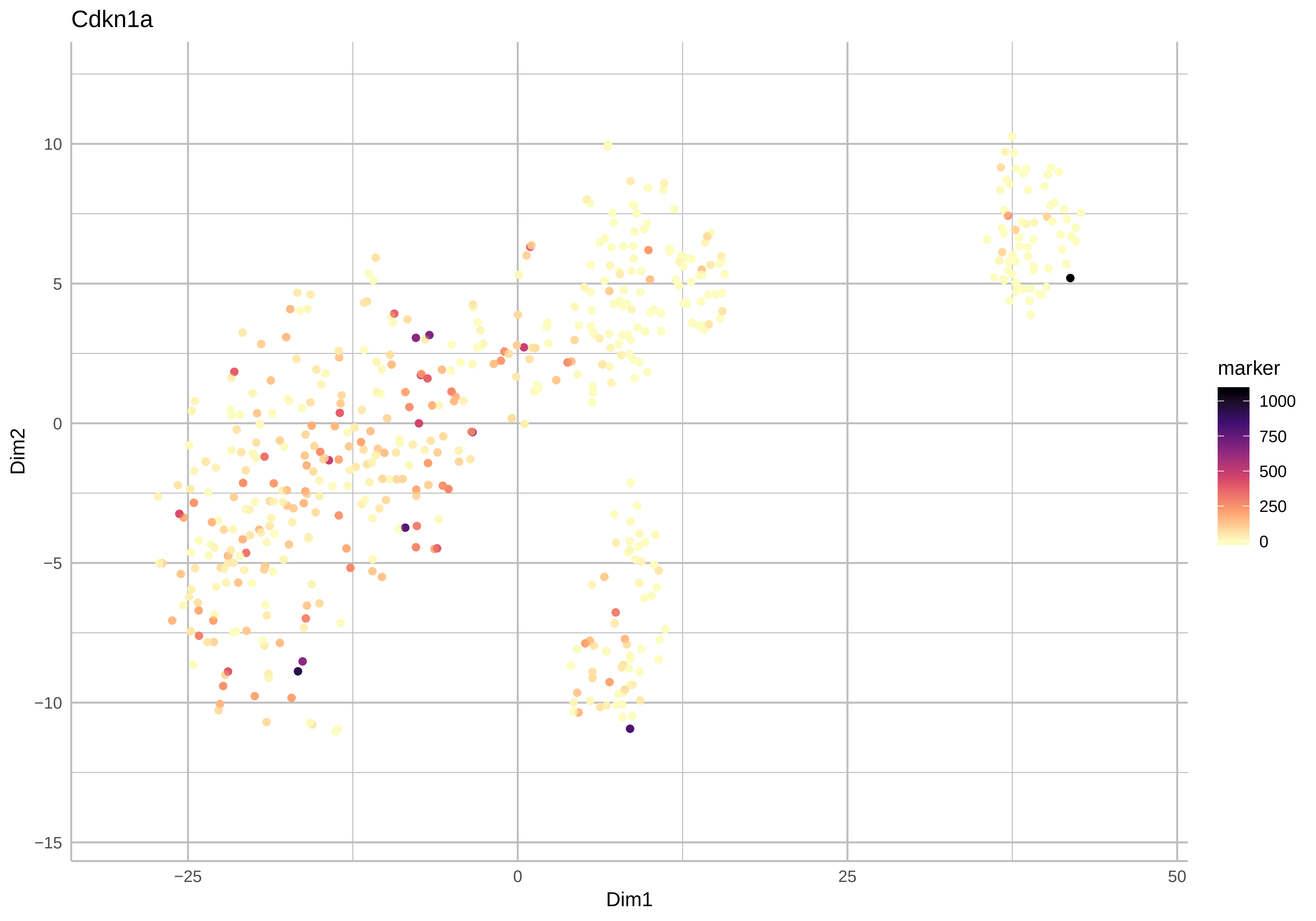
Cdkn1a
Dim2
Dim1
10
5
0
−5
−10
−15
−25
0
25
50
marker
1000
750
500
250
0

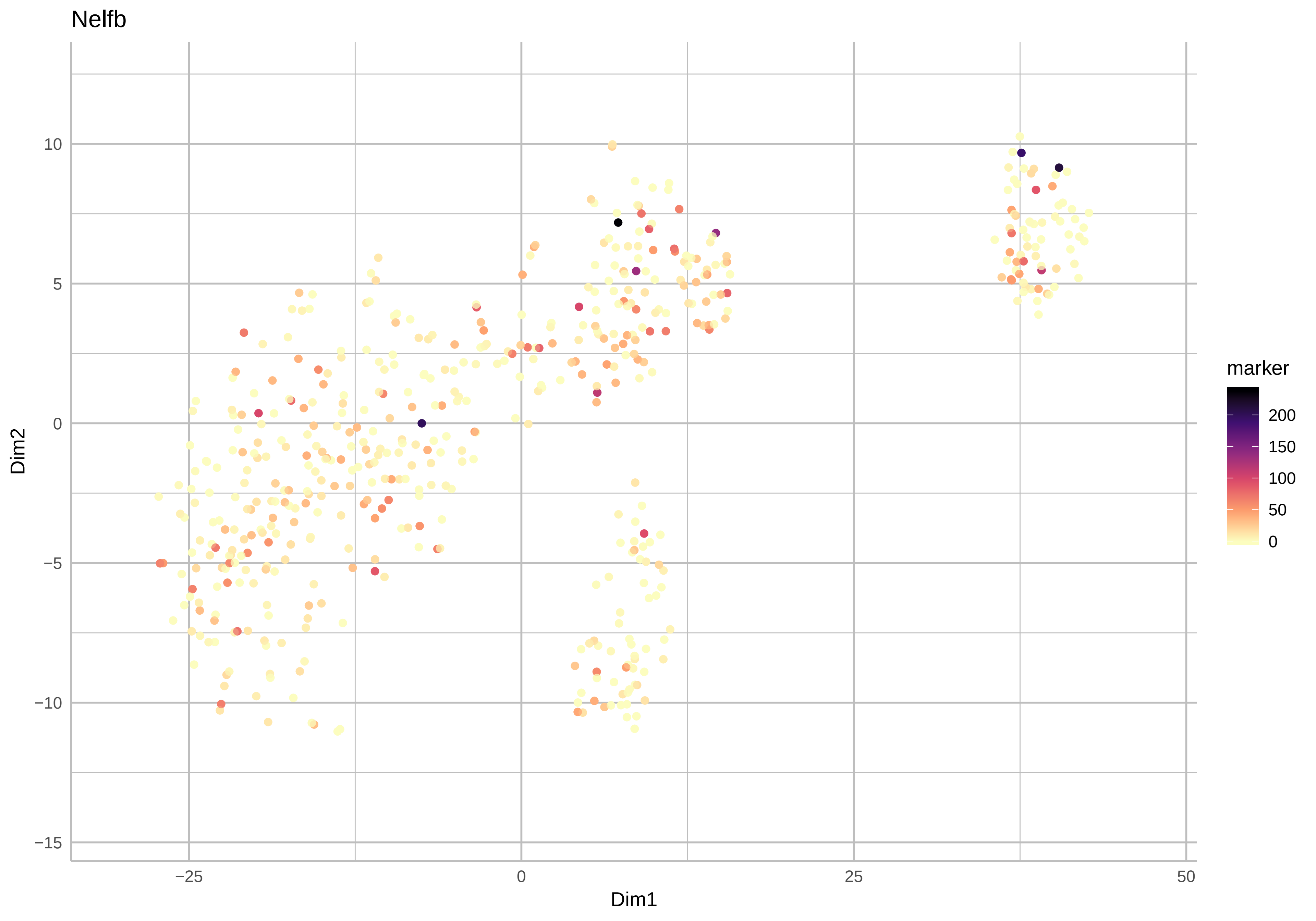
Nelfb
10
5
0
−5
−10
−15
Dim2
−25
0
25
50
Dim1
marker
200
150
100
50
0

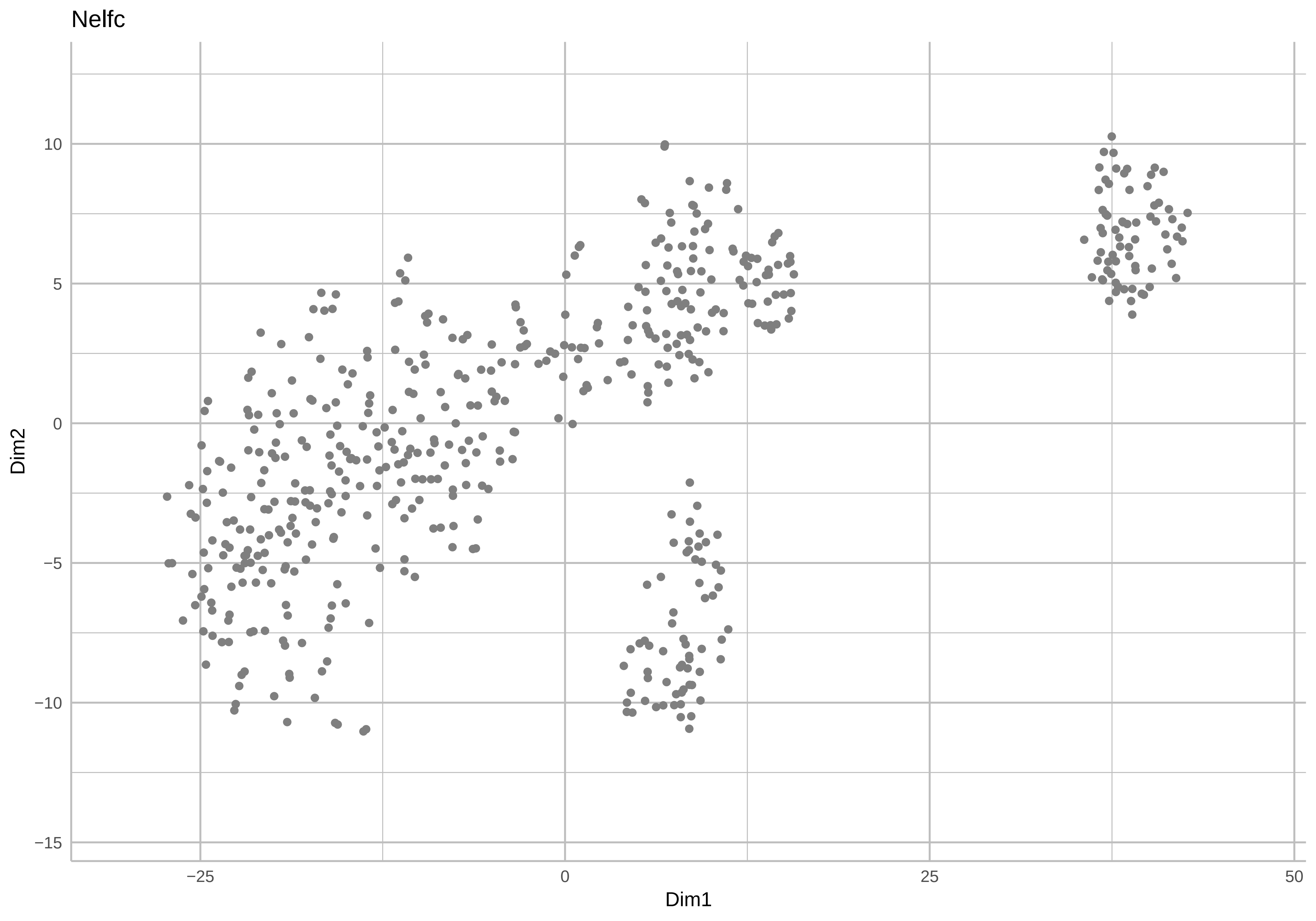

Nelfc
Dim2
Dim1
10
5
0
−5
−10
−15
−25
0
25
50

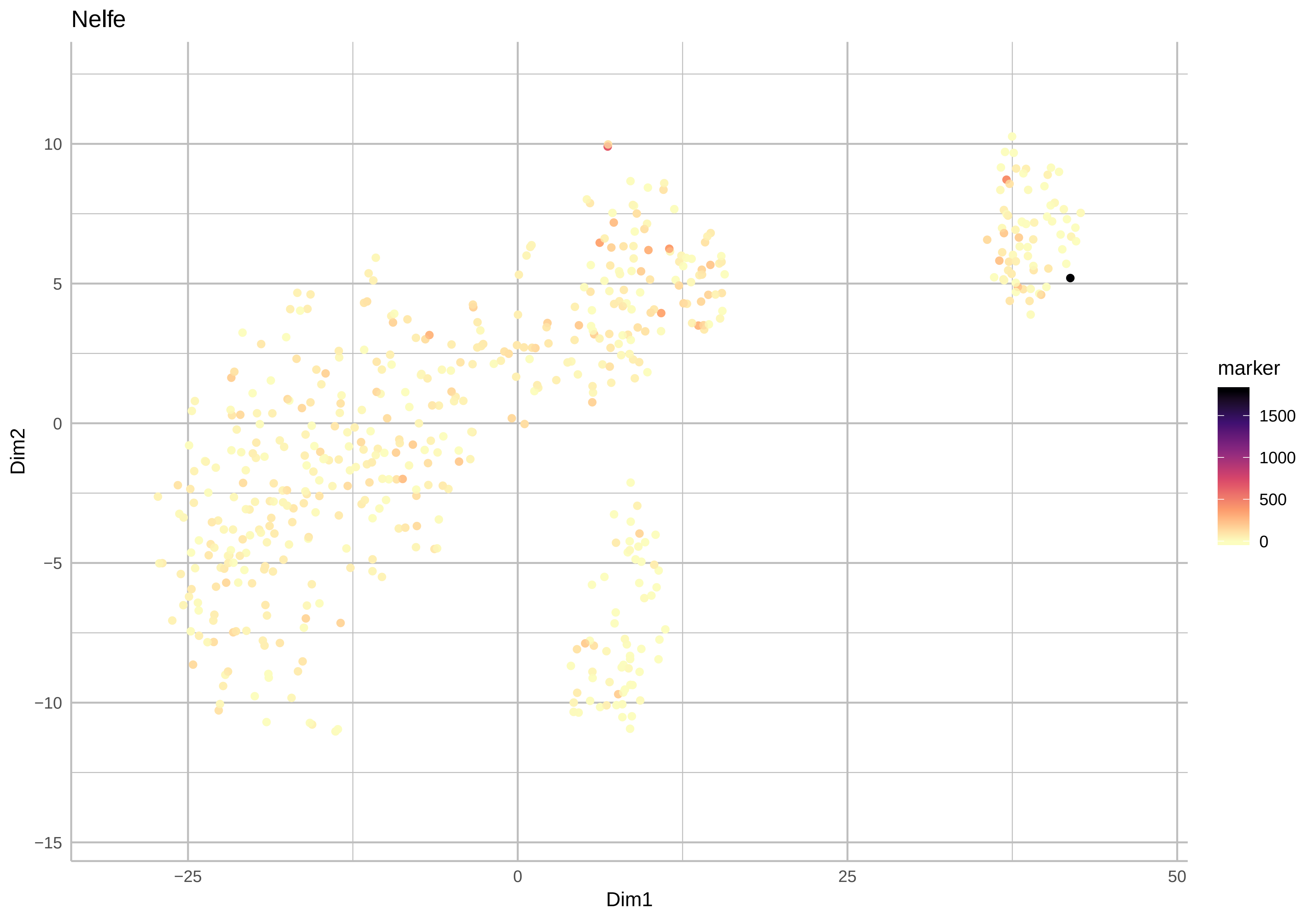

Nelfe
Dim2
Dim1
10
5
0
−5
−10
−15
−25
0
25
50
marker
1500
1000
500
0

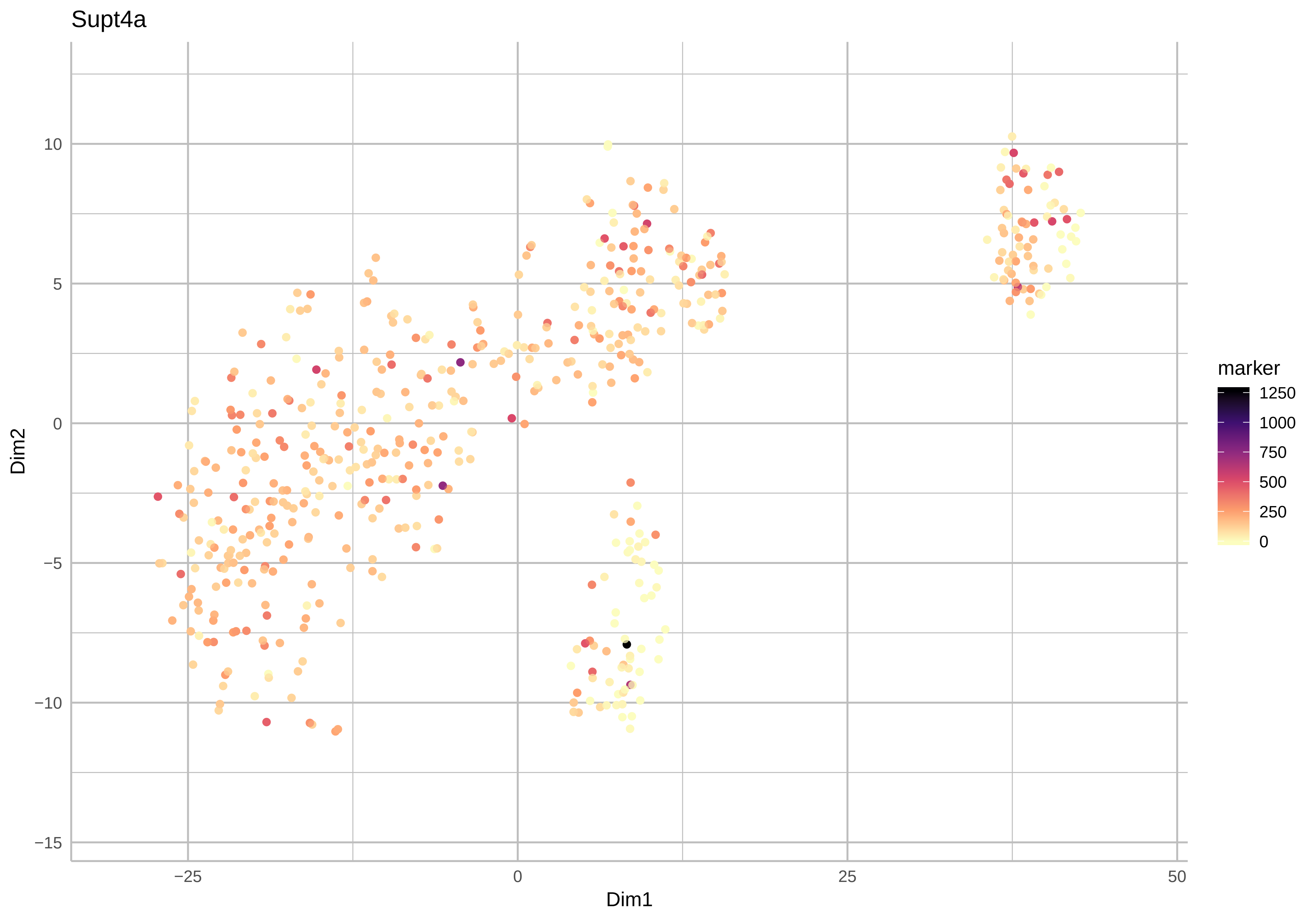

Supt4a
Dim2
Dim1
10
5
0
−5
−10
−15
−25
0
25
50
marker
1250
1000
750
500
250
0

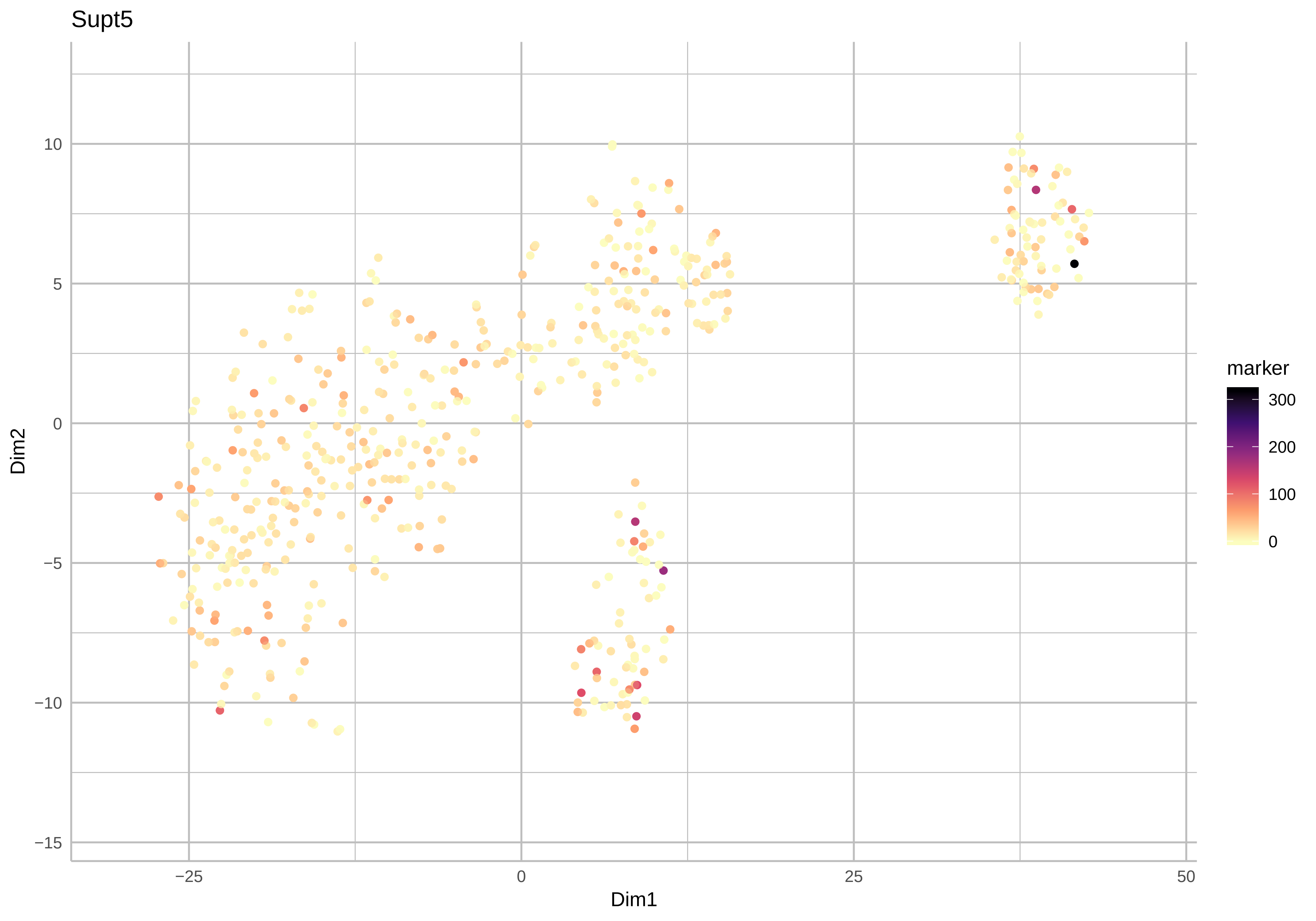

Supt5
Dim2
Dim1
10
5
0
−5
−10
−15
−25
0
25
50
marker
300
200
100
0

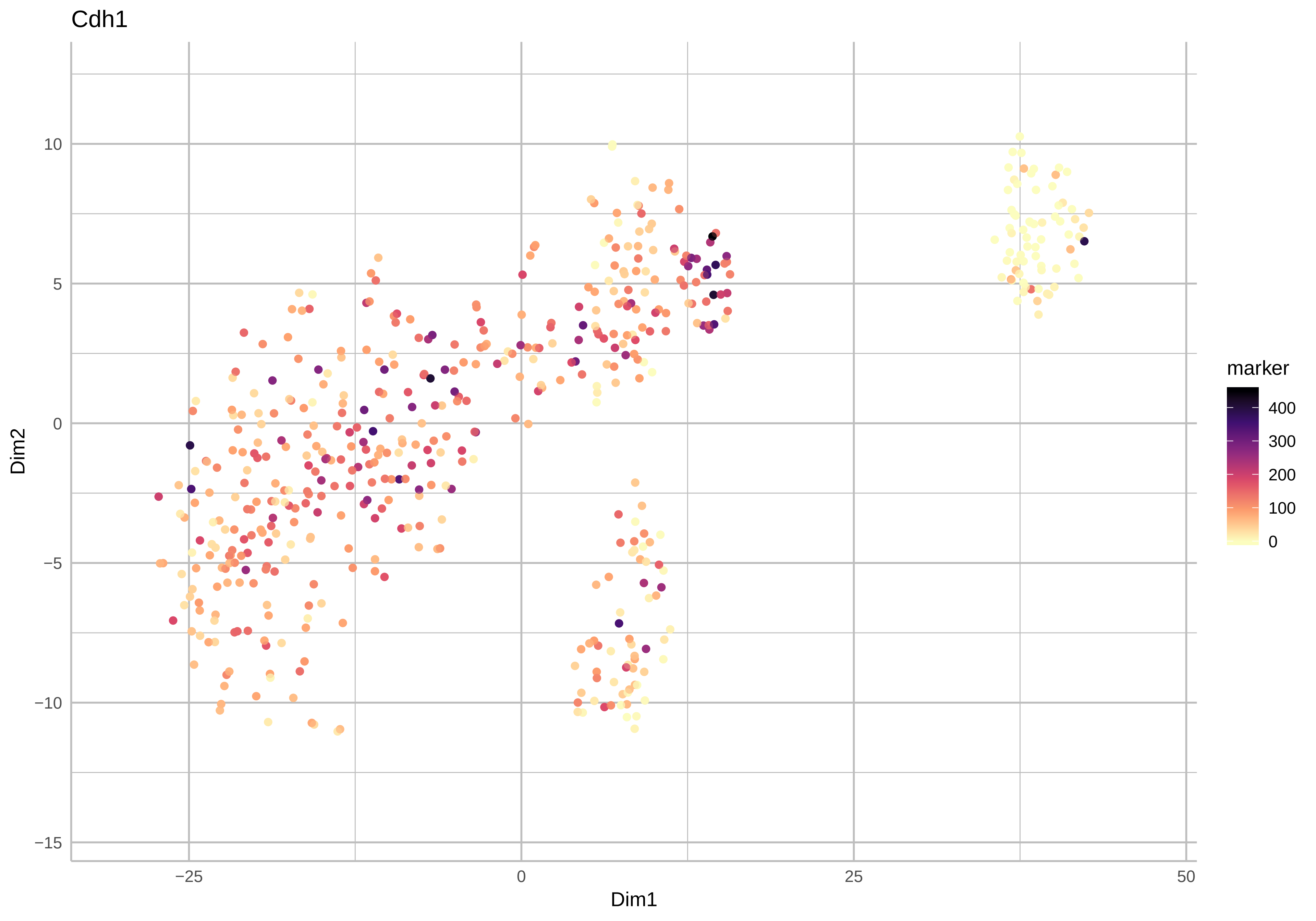
Cdh1
Dim2
Dim1
marker
400
300
200
100
0
10
5
0
−5
−10
−15
−25
0
25
50

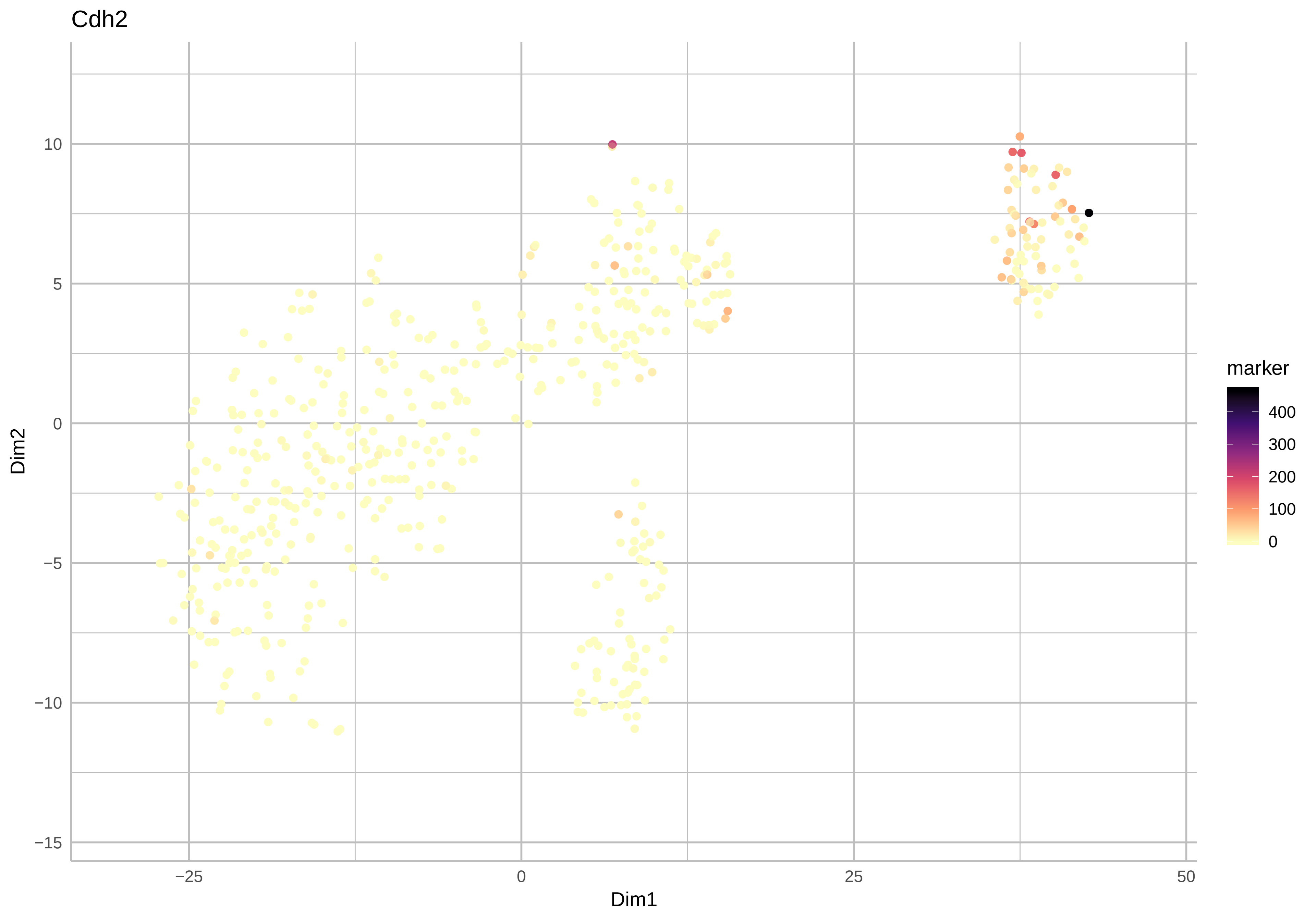
Cdh2
Dim2
Dim1
−25
0
25
50
10
5
0
−5
−10
−15
marker
400
300
200
100
0
