## Supplementary material for "Single-cell transcriptomics, scRNA-Seq and C1 CAGE discovered distinct phases of pluripotency during naïve-to-primed conversion in mice": tSNEheatscaleMarkersKallisto_20190317195223.pdf

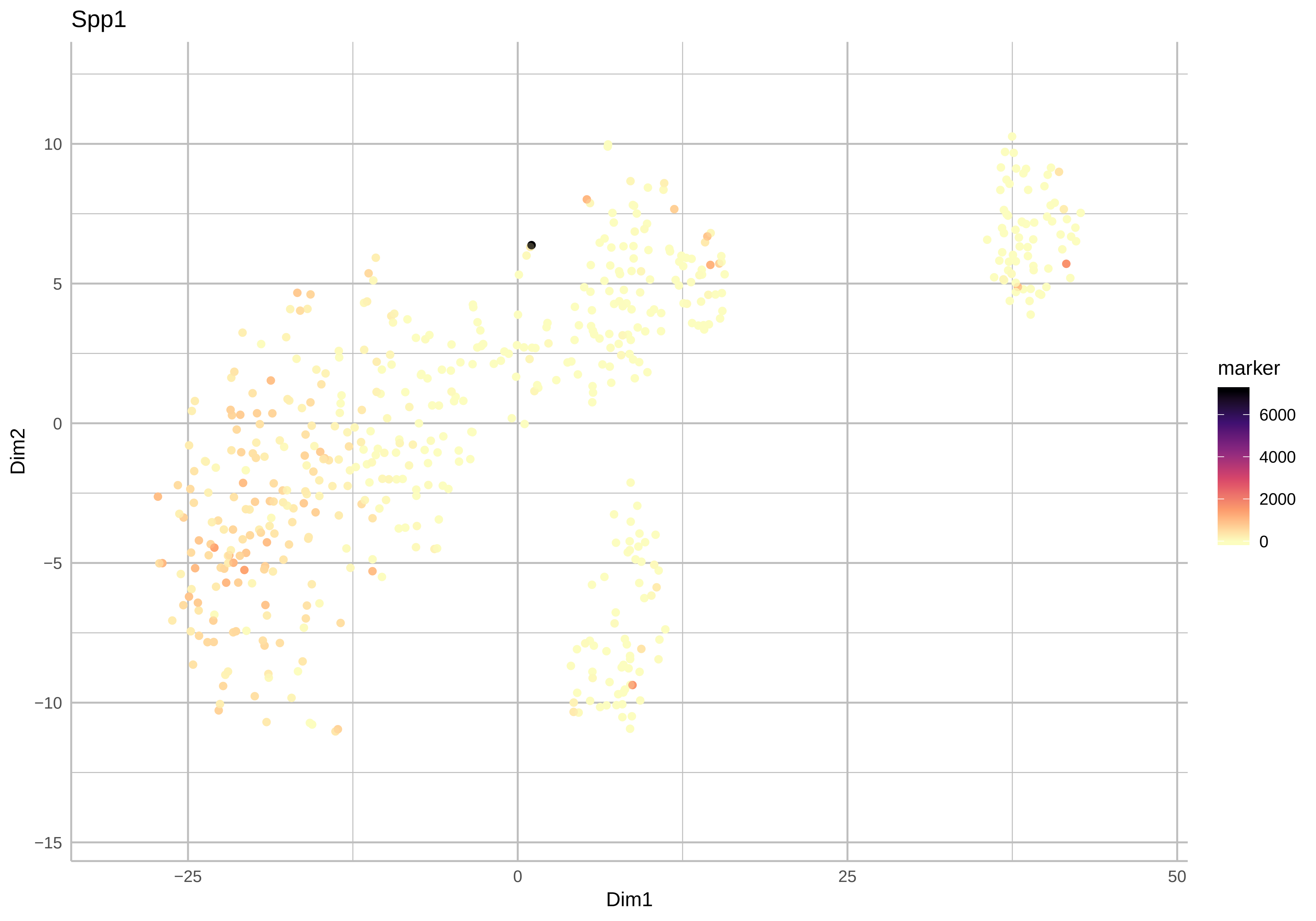
Spp1
10
5
0
−5
−10
−15
−25
0
25
50
Dim1
Dim2
marker
6000
4000
2000
0

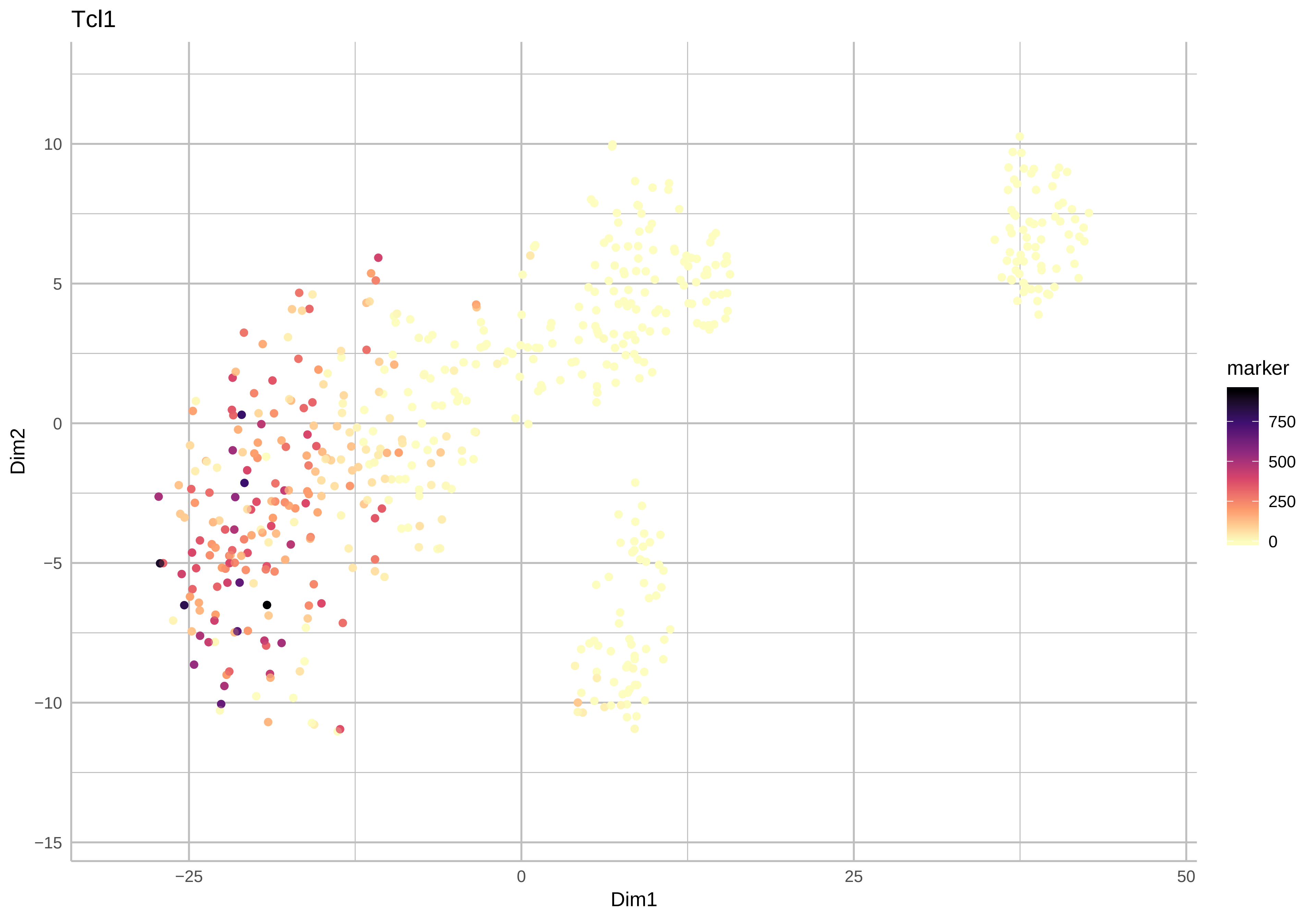

Tcl1
10
5
0
−5
−10
−15
−25
0
25
50
Dim1
Dim2
marker
750
500
250
0

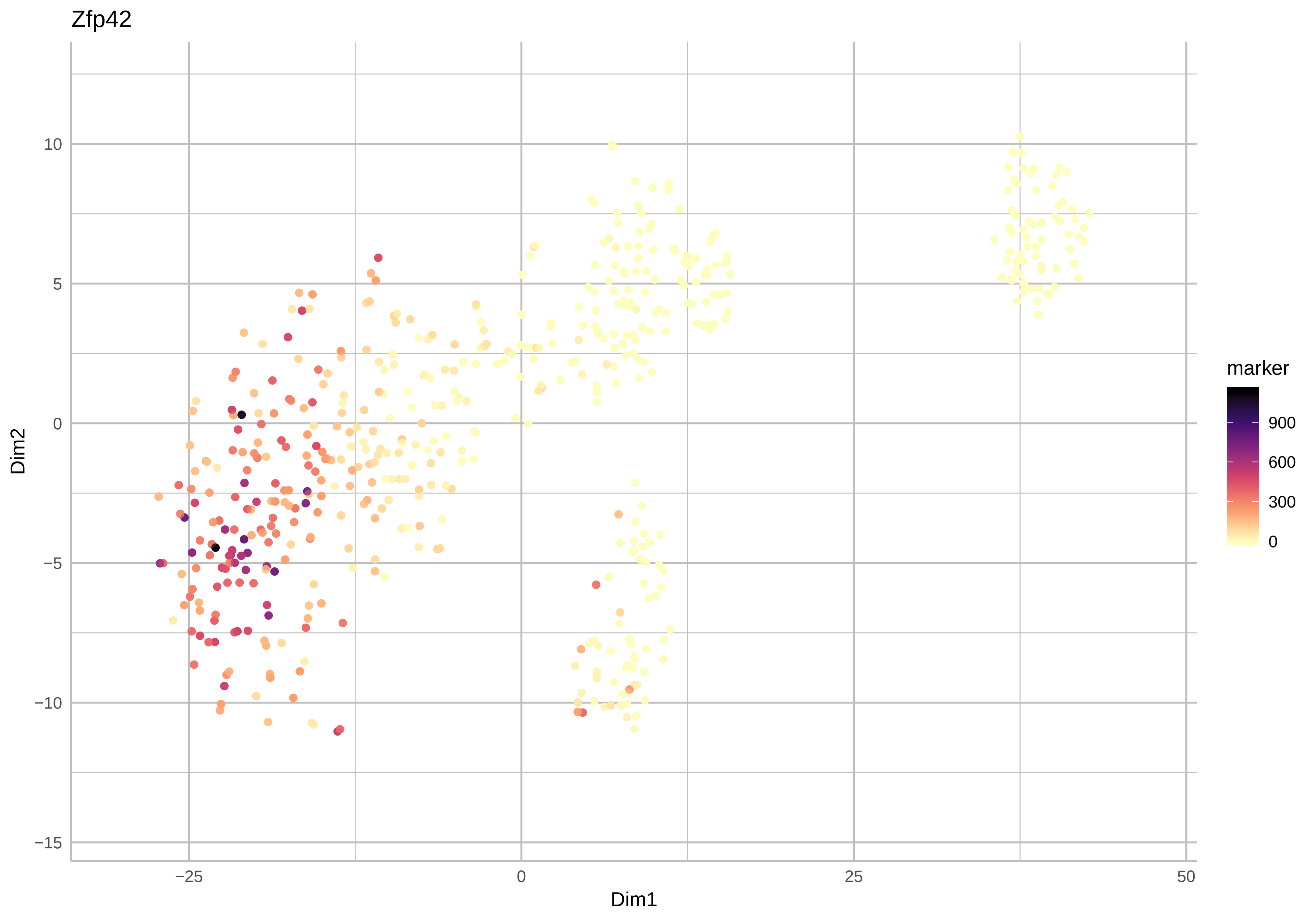
Zfp42
Dim2
Dim1
marker
900
600
300
0
10
5
0
−5
−10
−15
−25
0
25
50

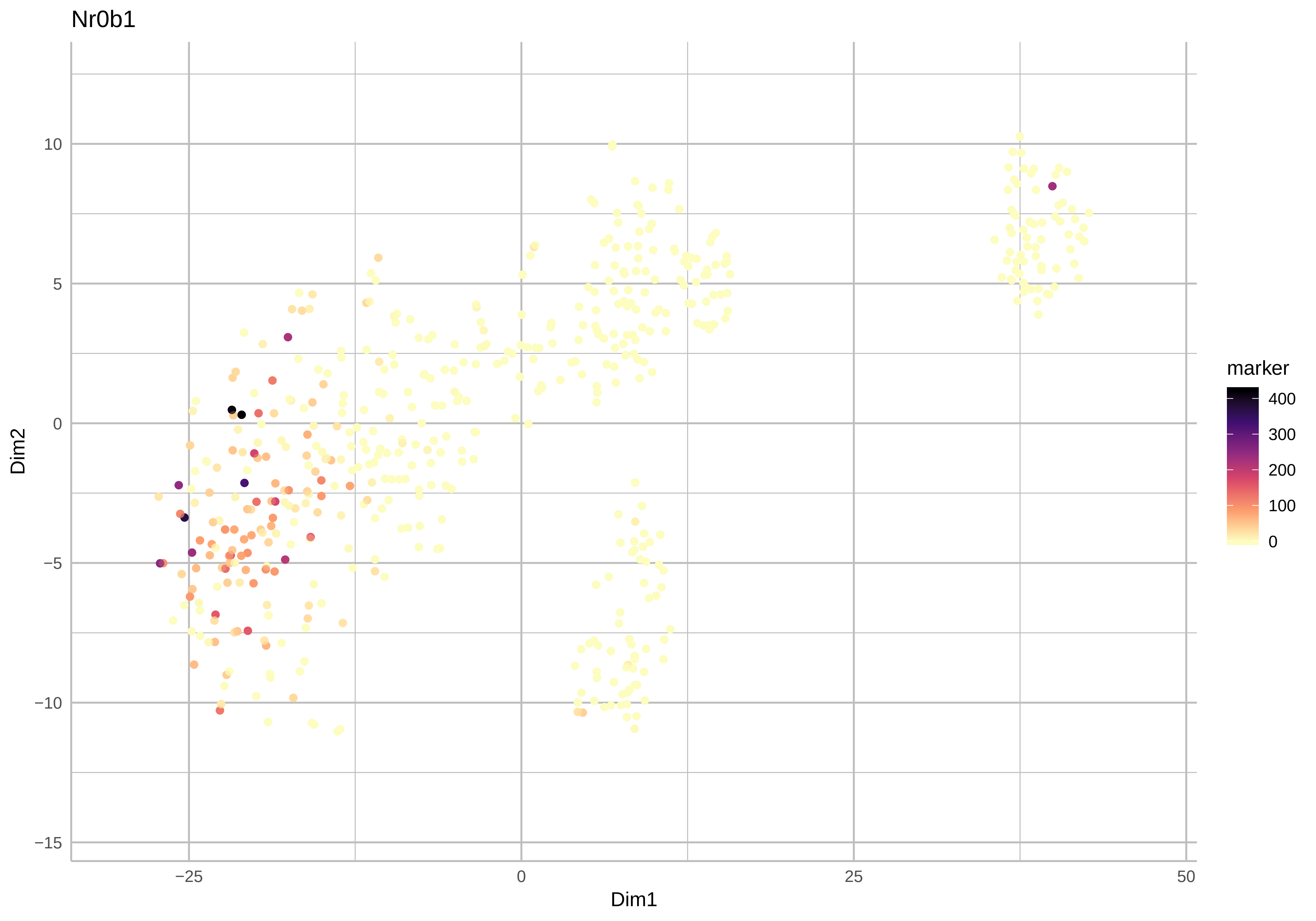

Nr0b1
10
5
0
−5
−10
−15
Dim2
−25
0
25
50
Dim1
marker
400
300
200
100
0

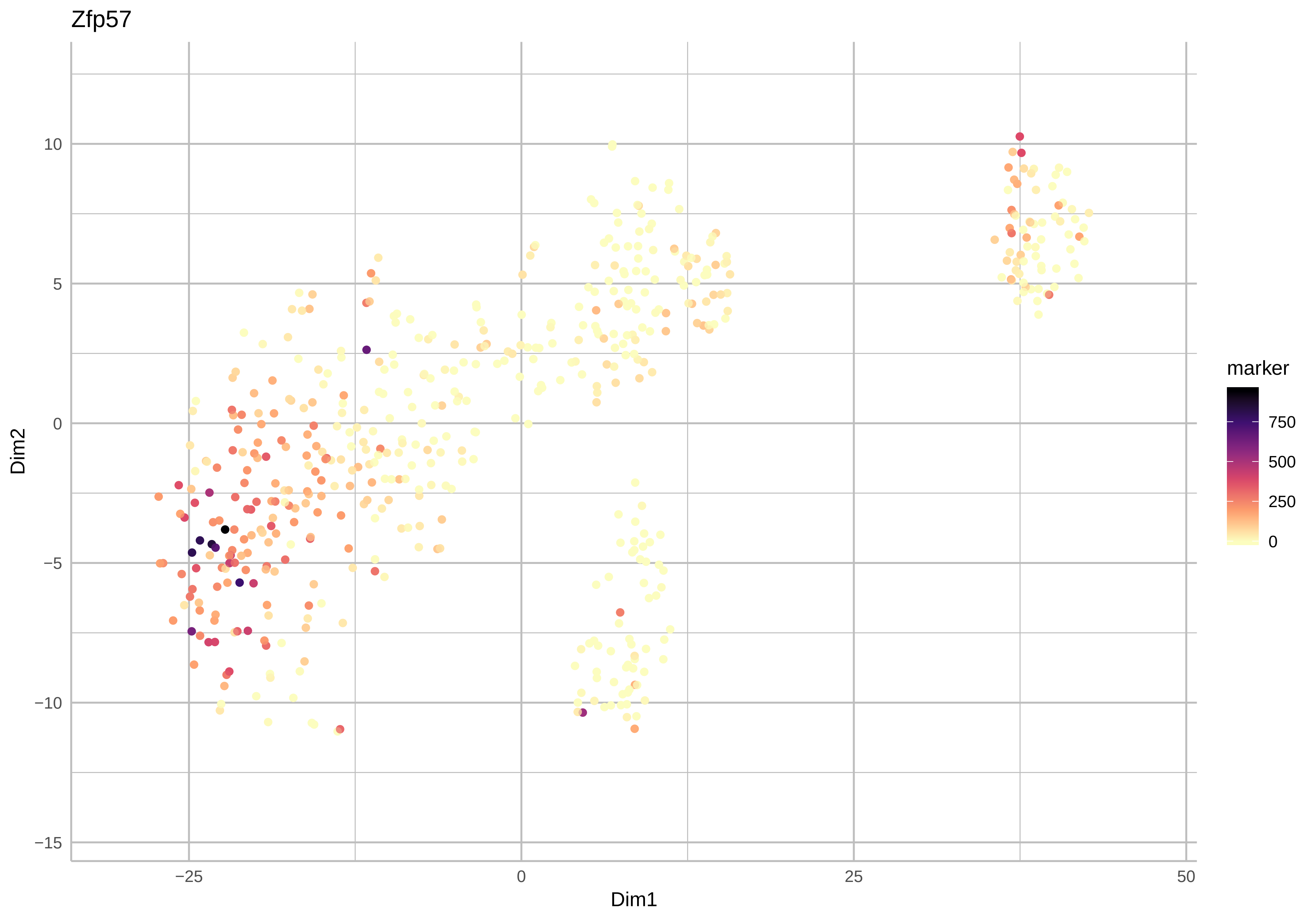

Zfp57
Dim2
Dim1
marker
750
500
250
0
10
5
0
−5
−10
−15
−25
0
25
50

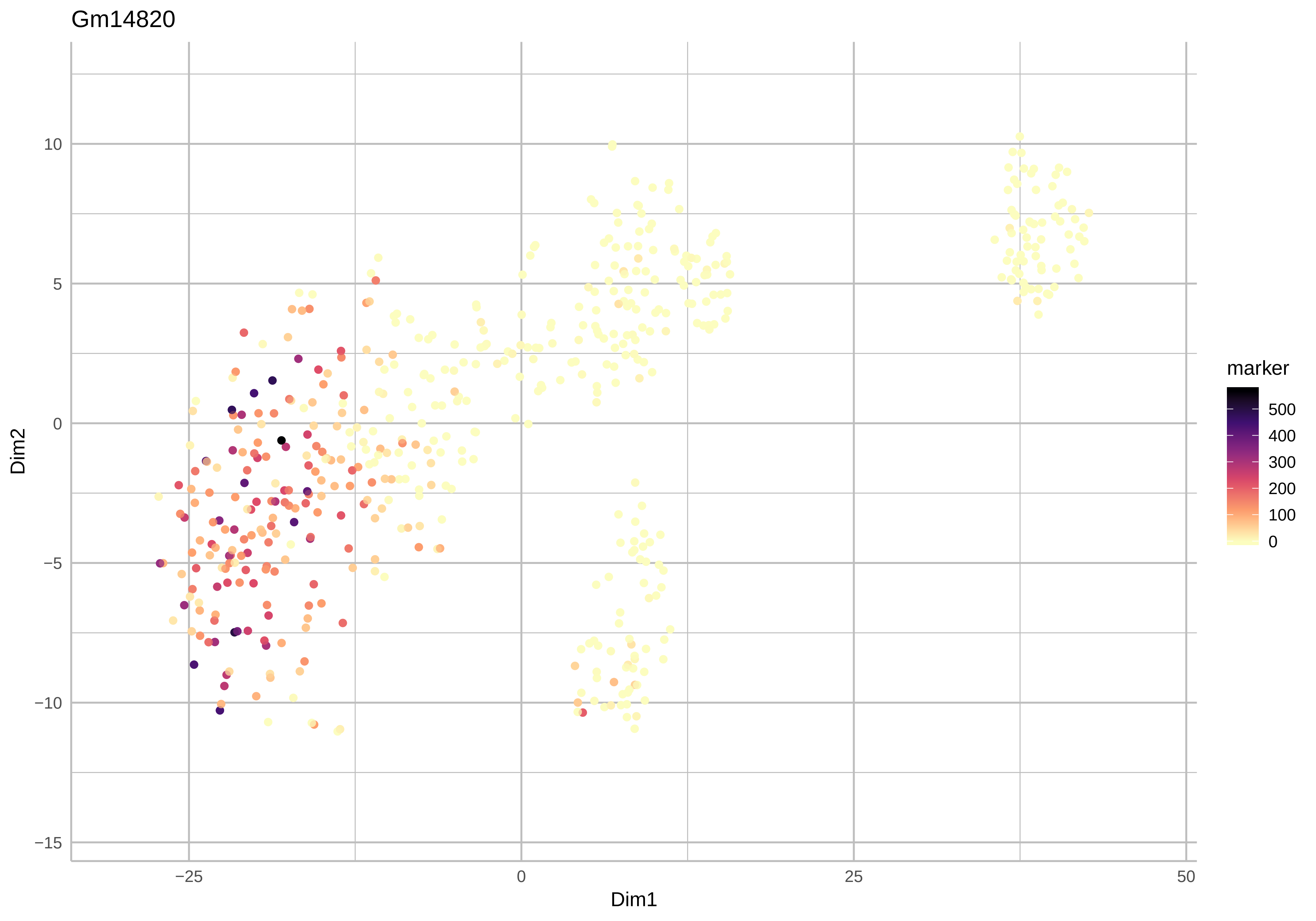
Gm14820
Dim2
Dim1
10
5
0
−5
−10
−15
−25
0
25
50
marker
500
400
300
200
100
0

Mras

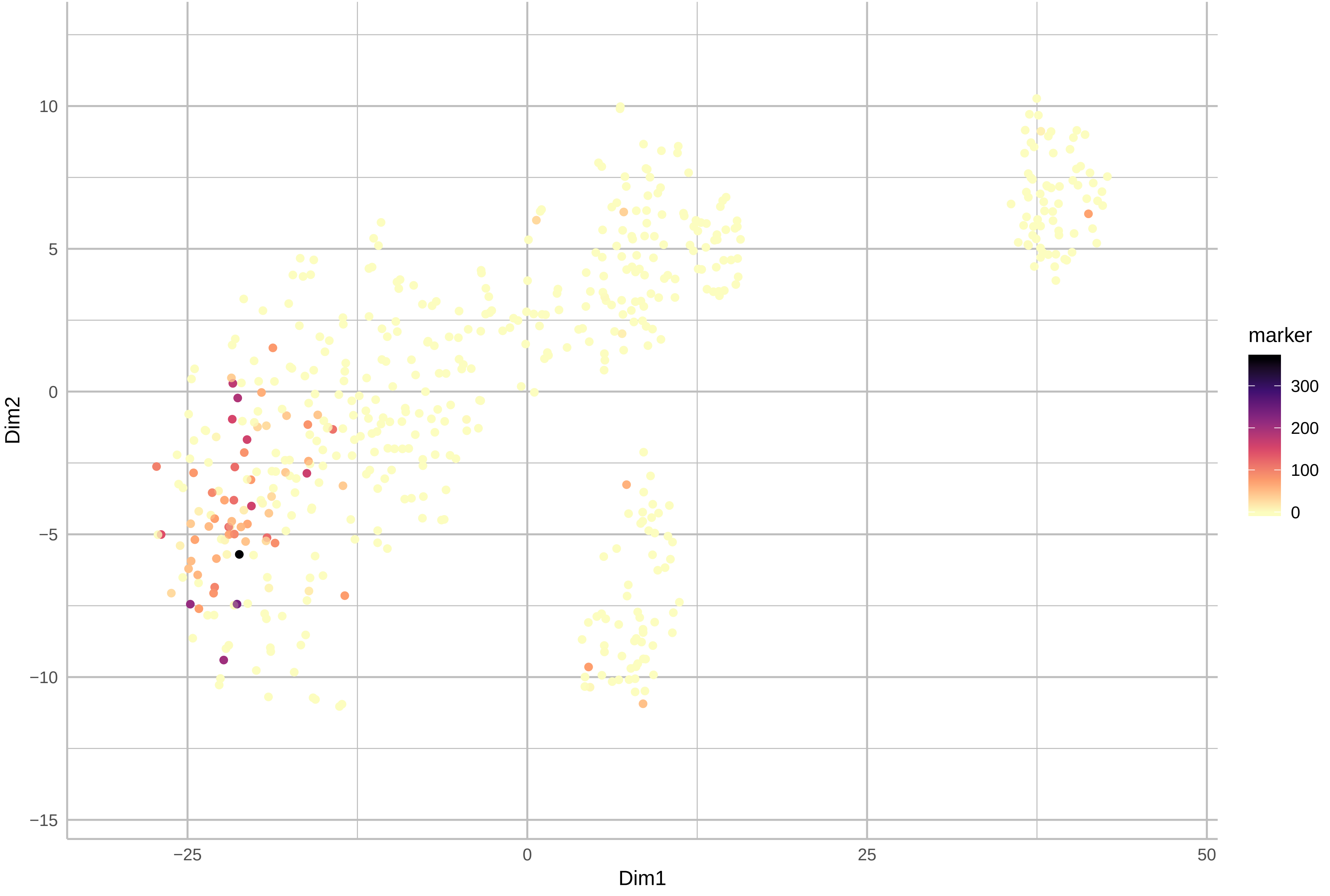

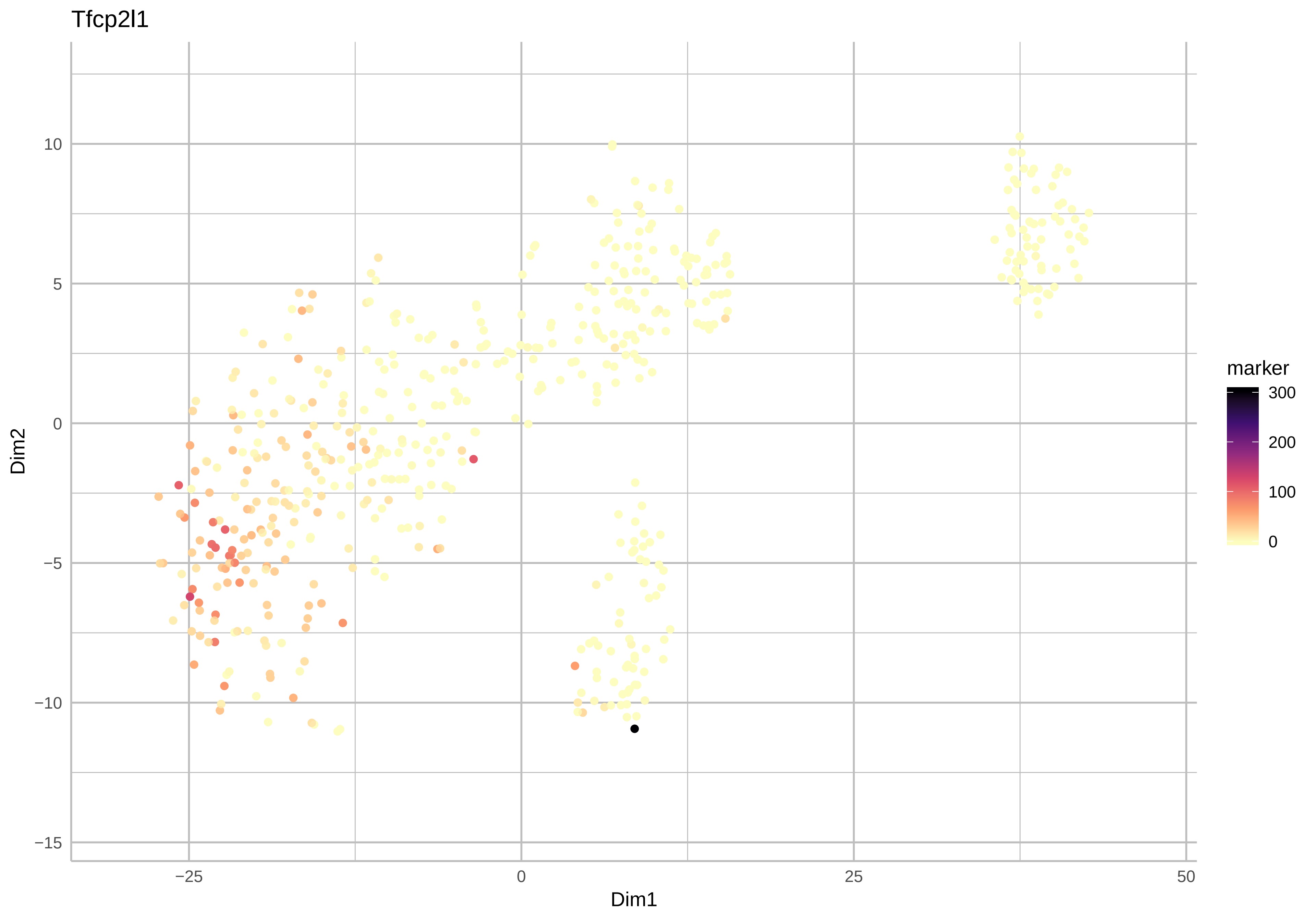
Tfcp2l1
Dim2
Dim1
10
5
0
−5
−10
−15
−25
0
25
50
marker
300
200
100
0

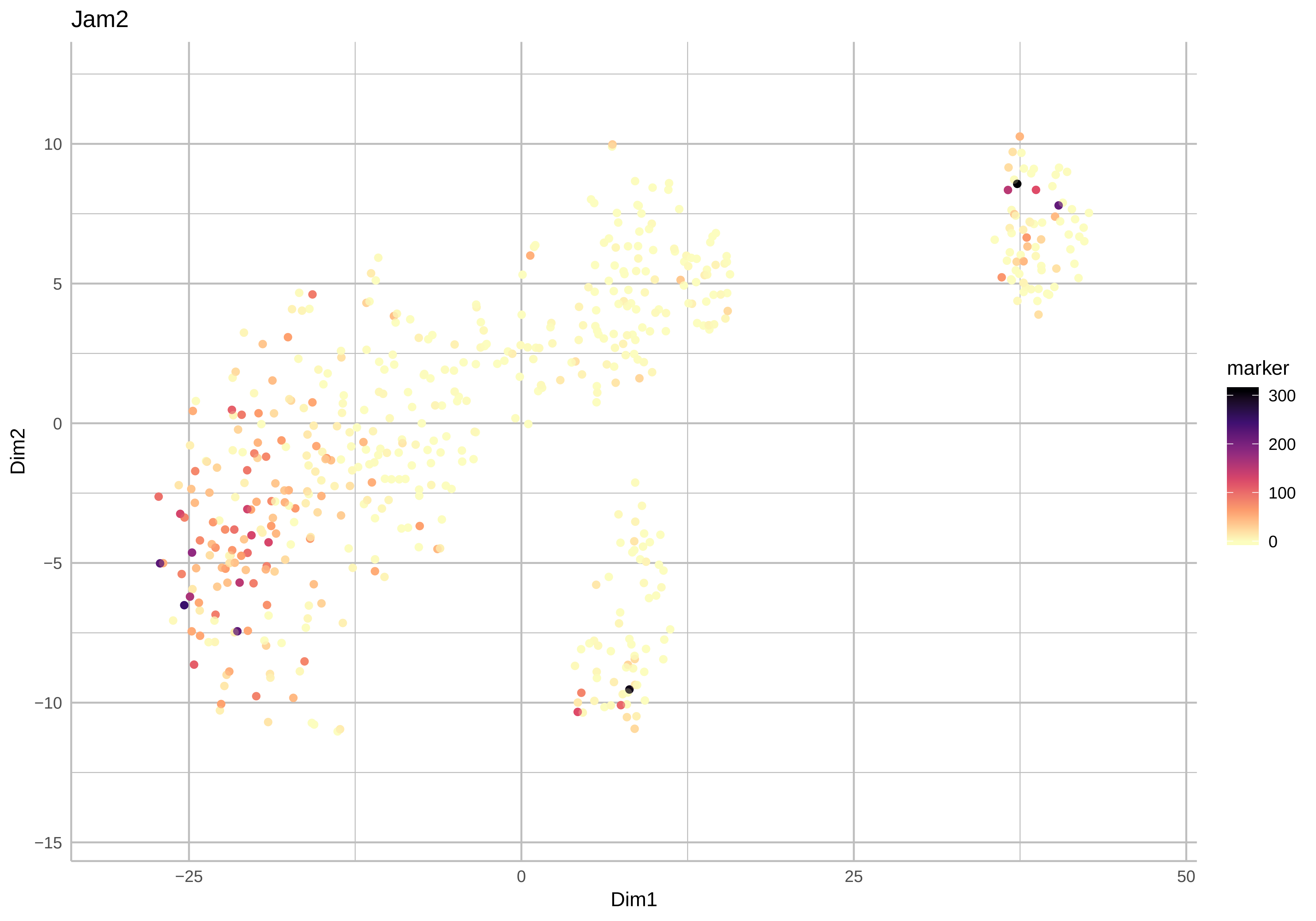

Jam2
Dim2
Dim1
10
5
0
−5
−10
−15
−25
0
25
50
marker
300
200
100
0

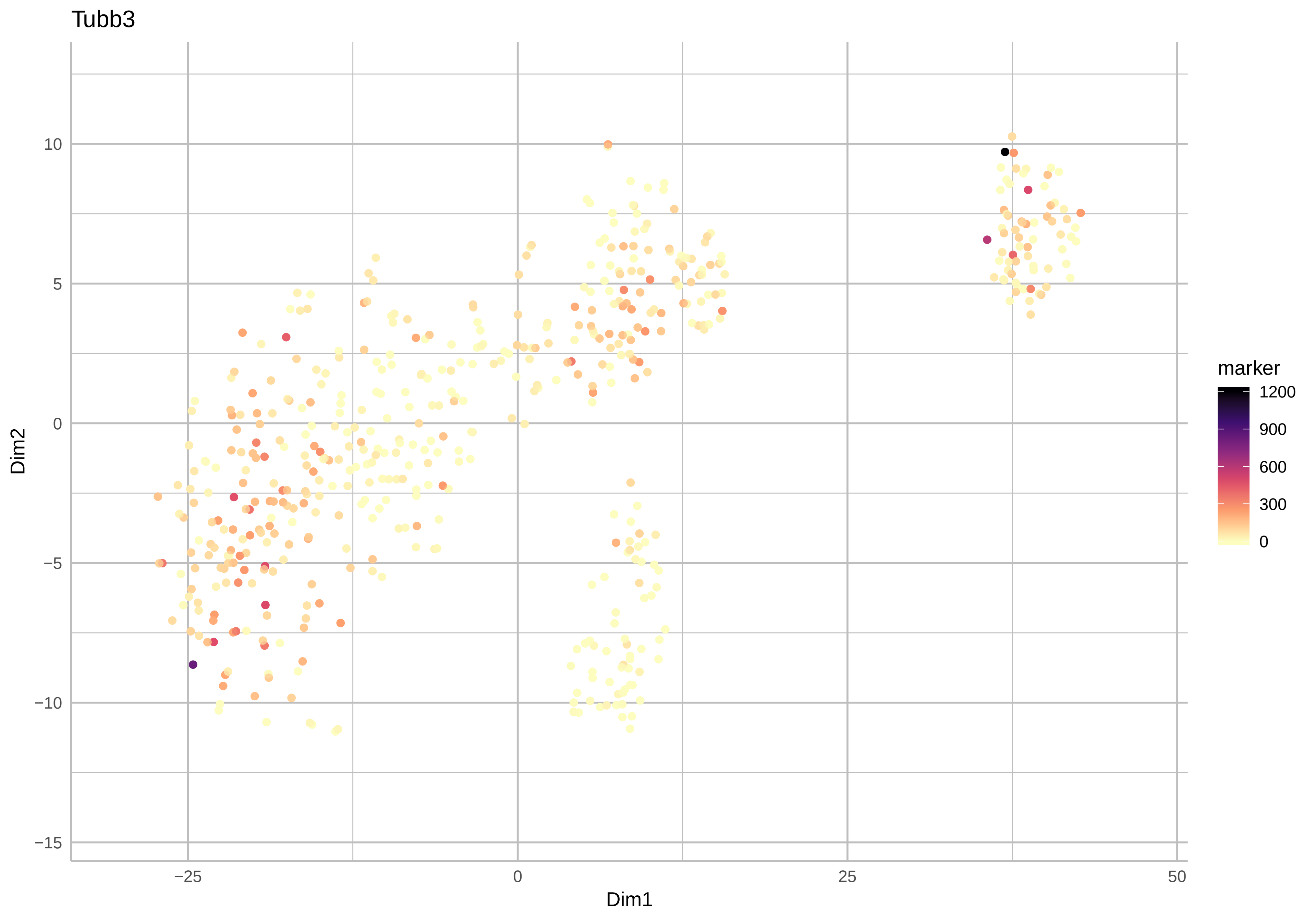
Tubb3
Dim2
Dim1
10
5
0
−5
−10
−15
−25
0
25
50
marker
1200
900
600
300
0

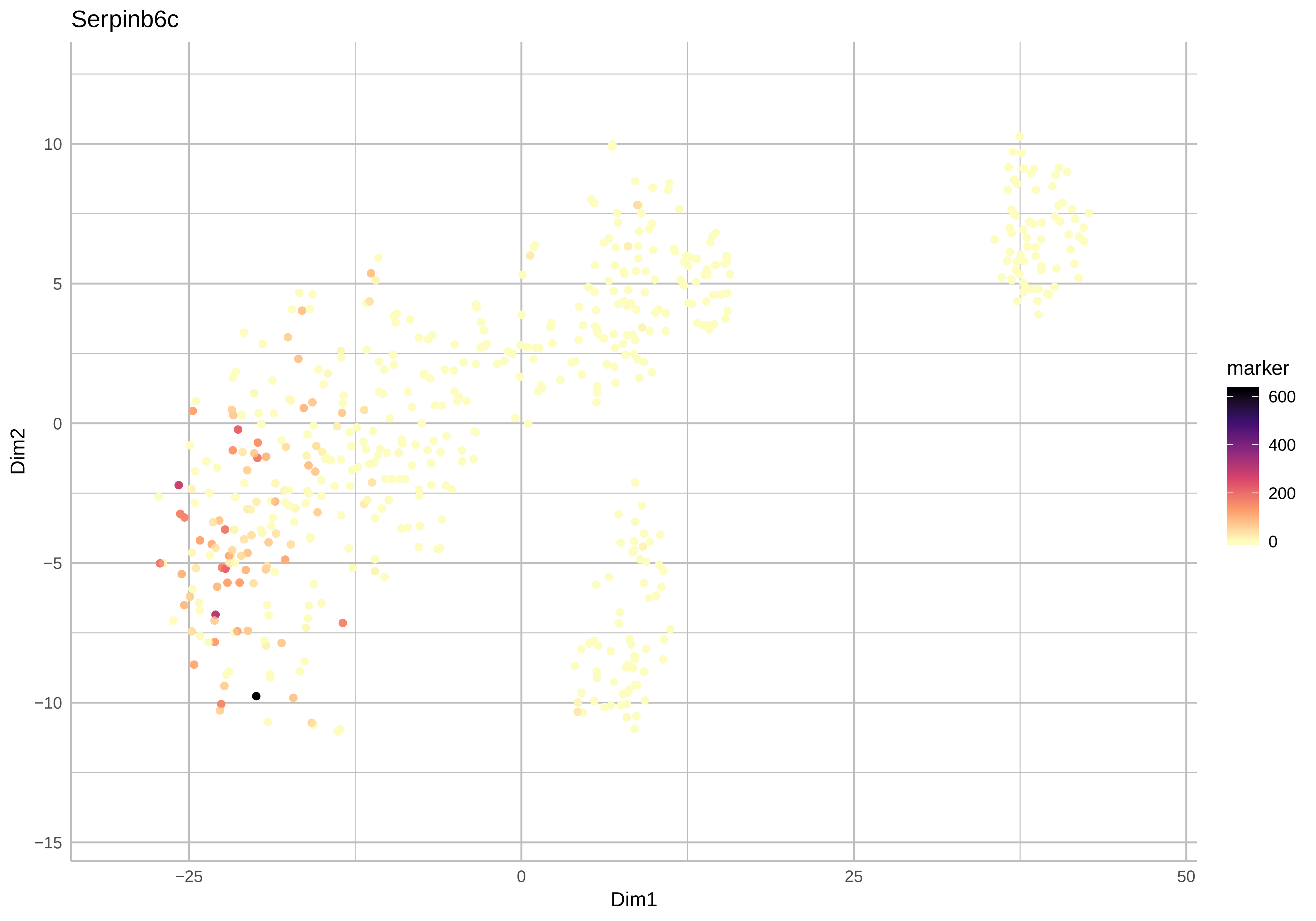
Serpinb6c
Dim2
Dim1
10
5
0
−5
−10
−15
−25
0
25
50
marker
600
400
200
0

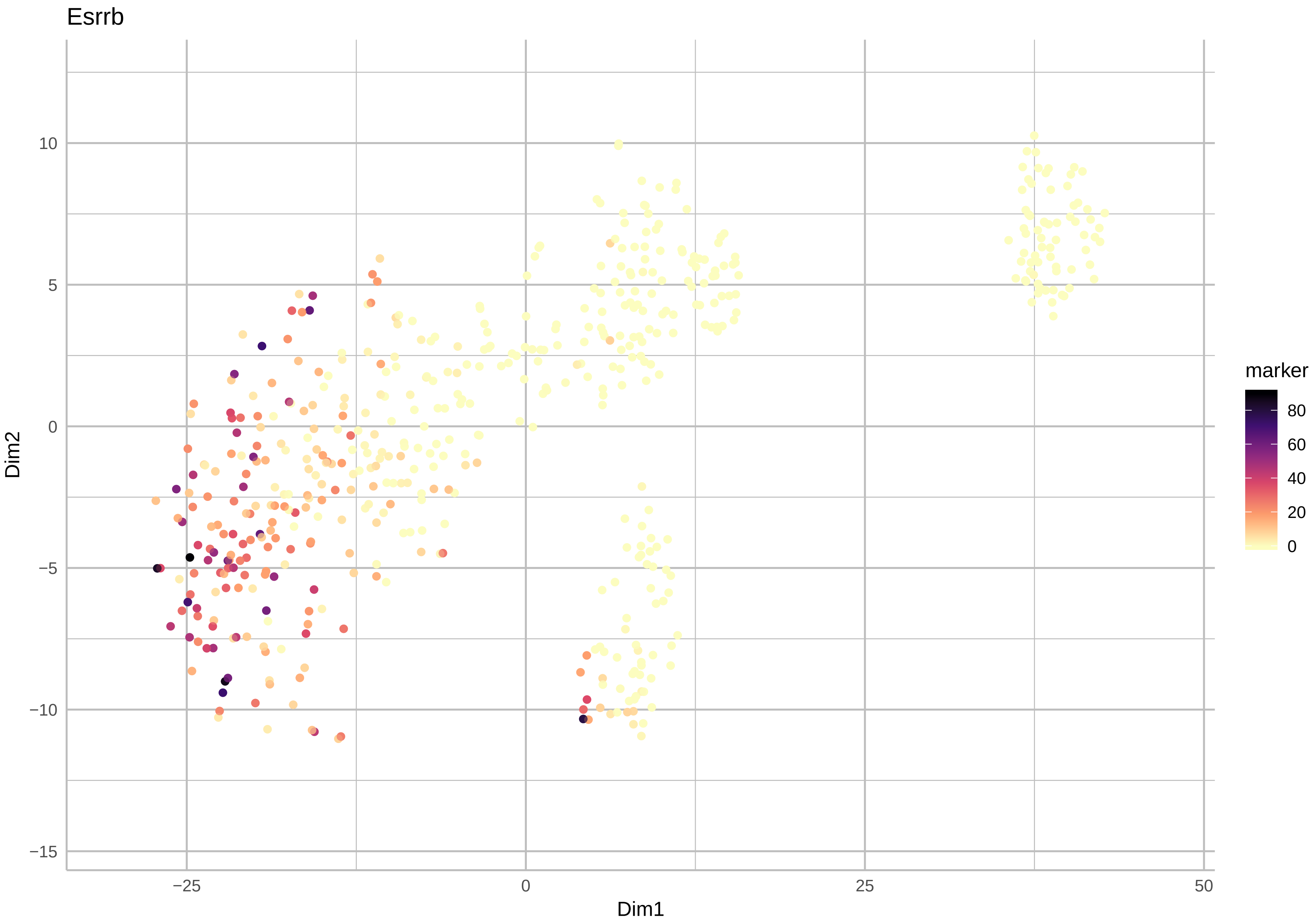

Esrrb
Dim2
Dim1
marker
80
60
40
20
0
10
5
0
−5
−10
−15
−25
0
25
50

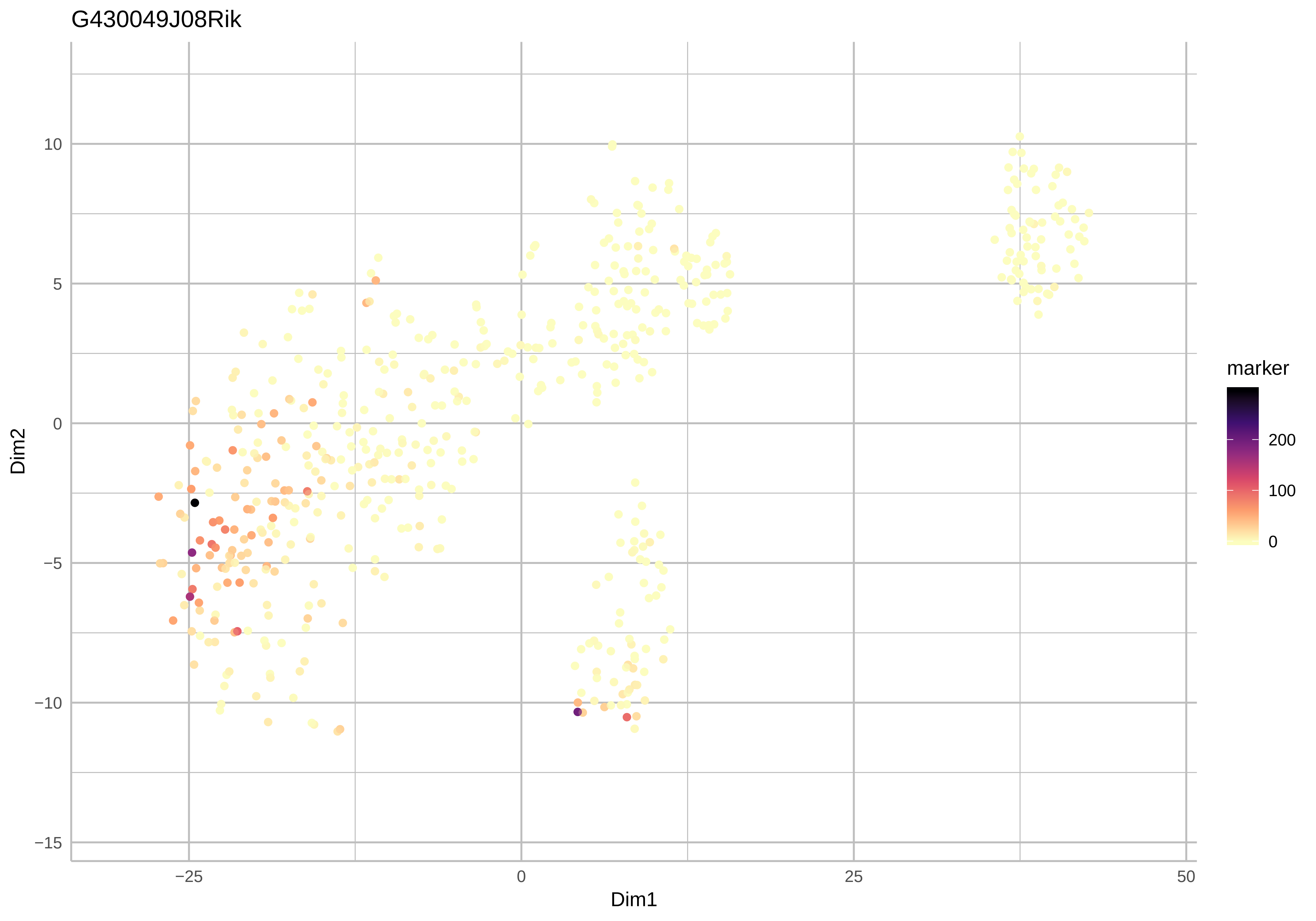

G430049J08Rik
Dim2
Dim1
10
5
0
−5
−10
−15
−25
0
25
50
marker
200
100
0

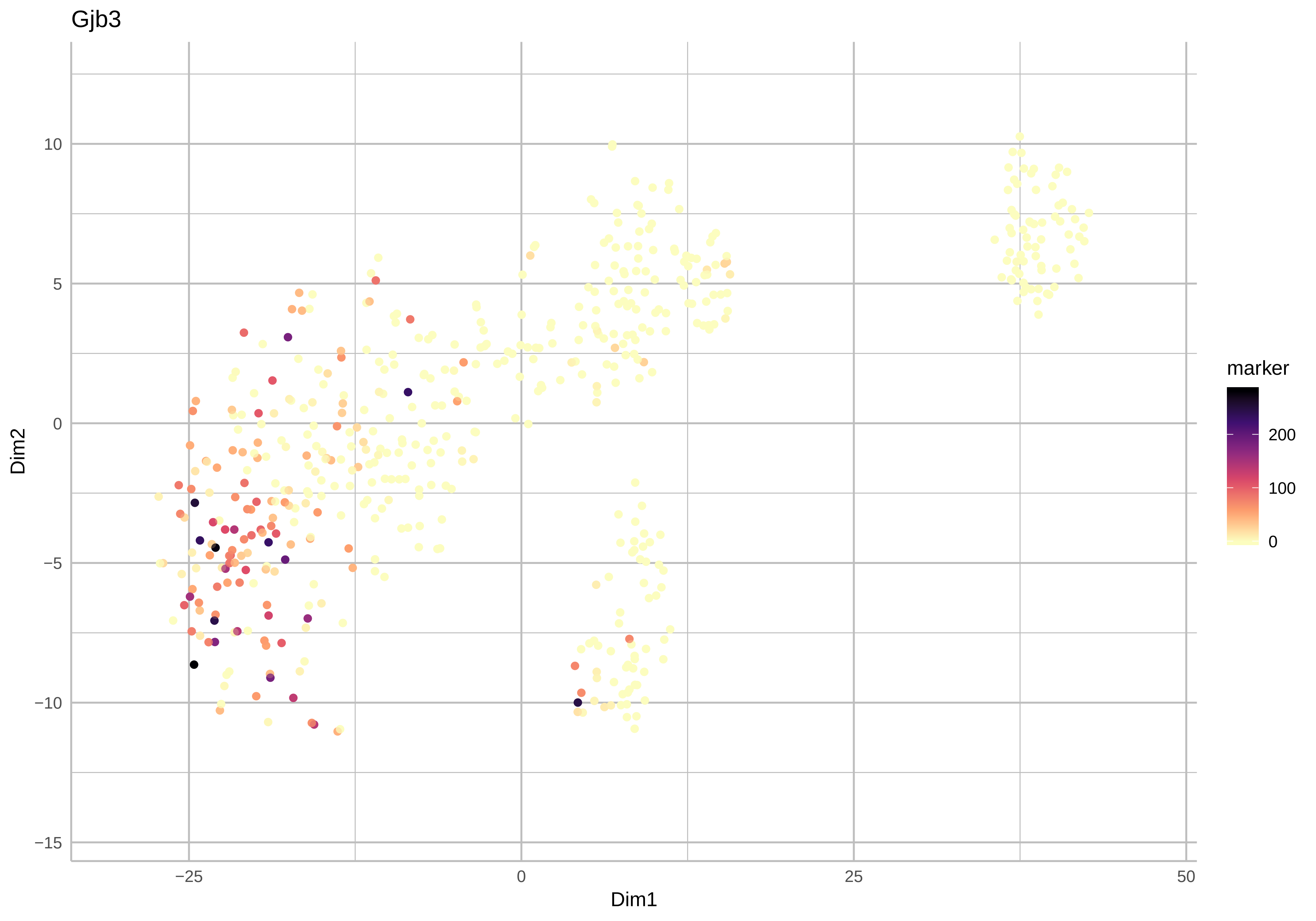
Gjb3
Dim2
Dim1
10
5
0
−5
−10
−15
−25
0
25
50
marker
200
100
0

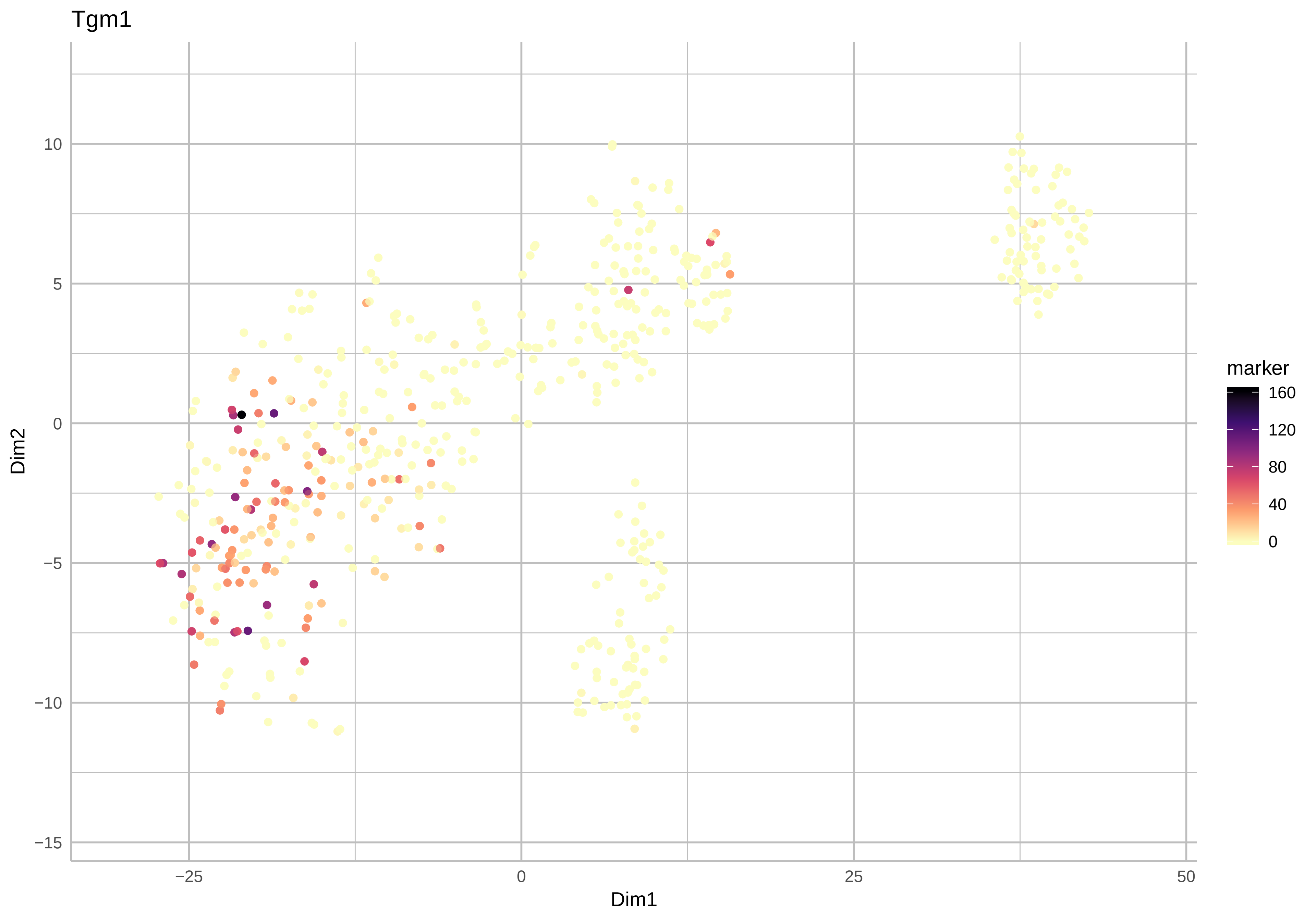

Tgm1
10
5
0
−5
−10
−15
−25
0
25
50
Dim1
Dim2
marker
160
120
80
40
0

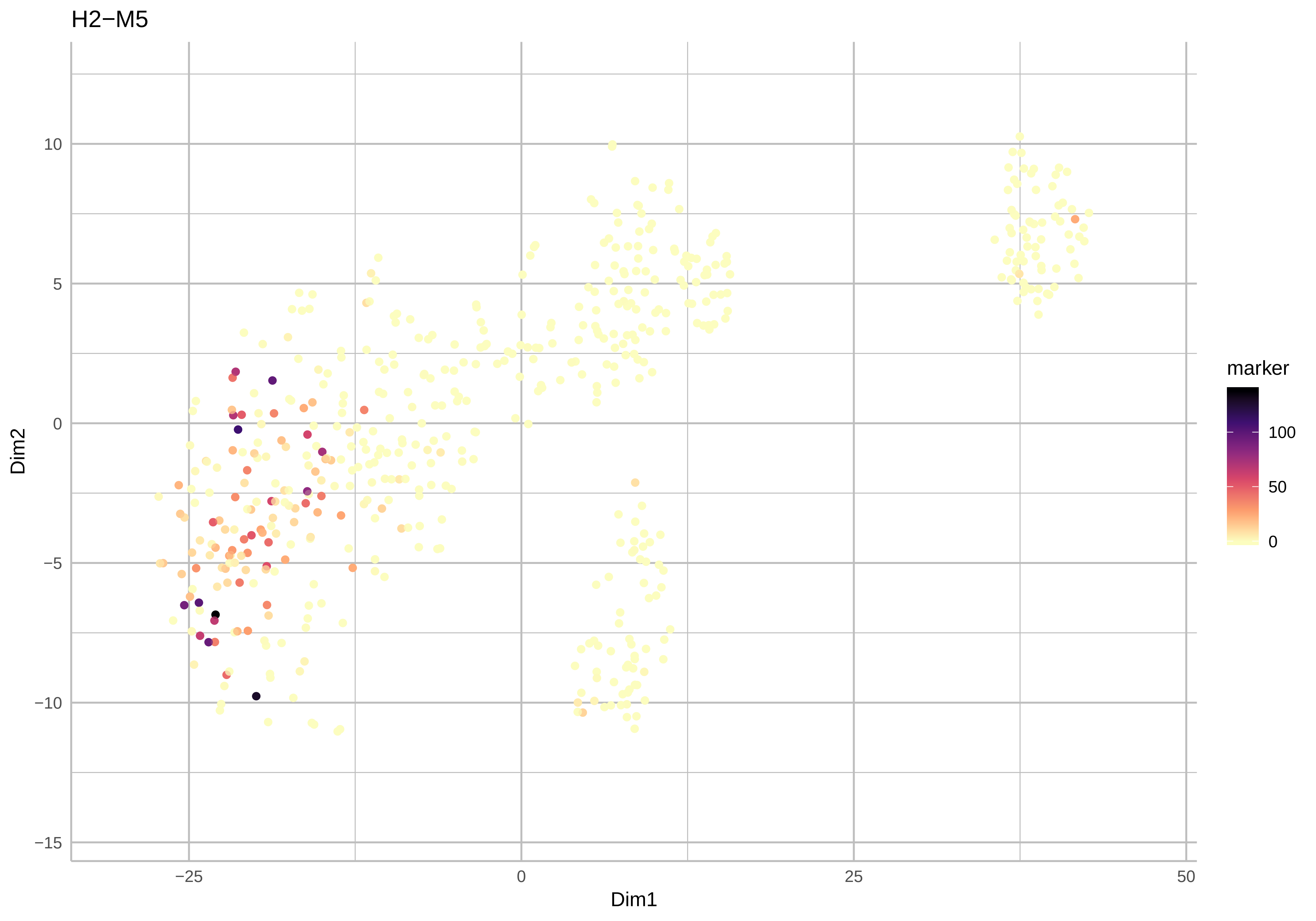

H2−M5
10
5
0
−5
−10
−15
−25
0
25
50
Dim1
Dim2
marker
100
50
0

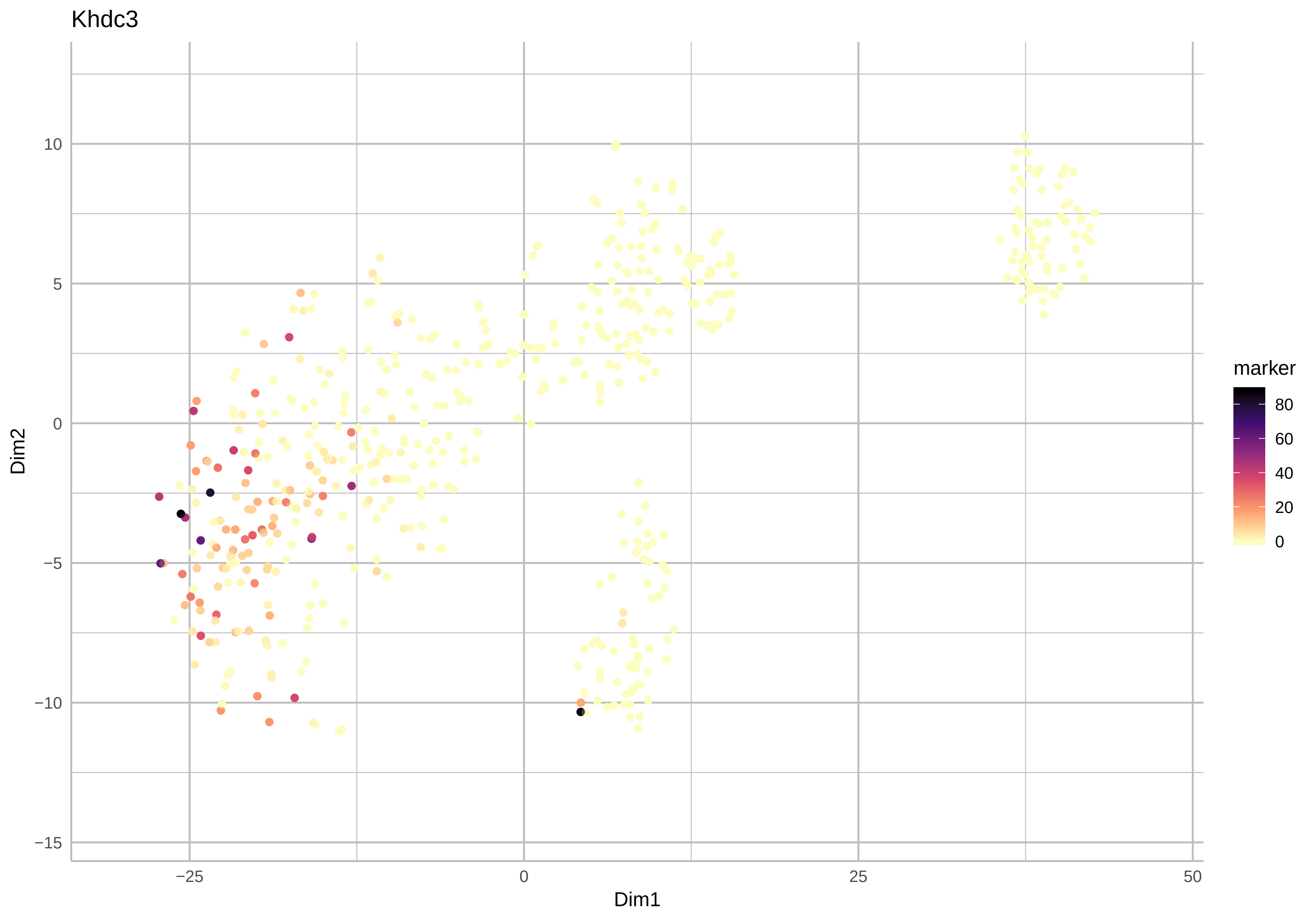

Khdc3
Dim2
Dim1
10
5
0
−5
−10
−15
−25
0
25
50
marker
80
60
40
20
0

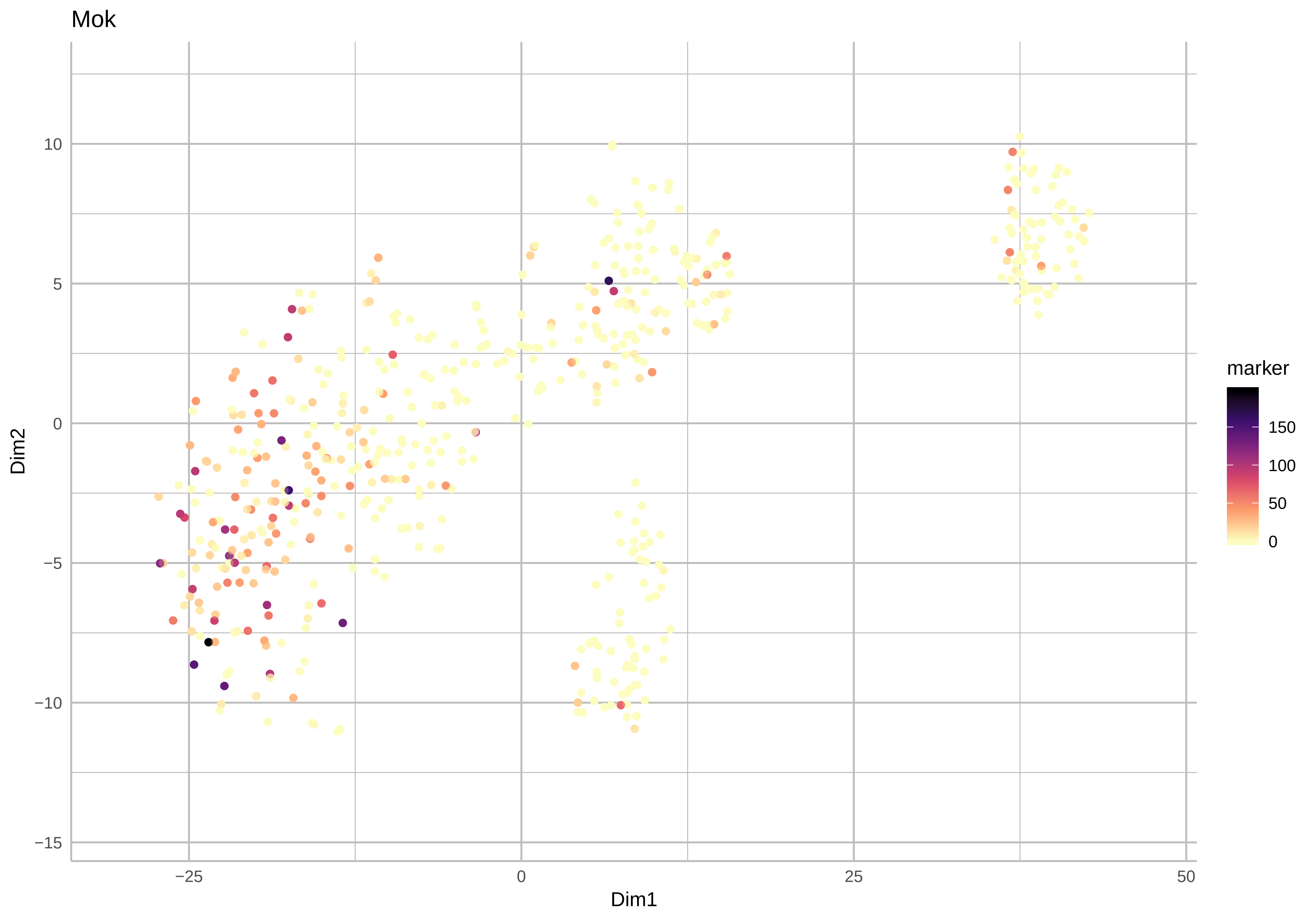

Mok
Dim2
Dim1
10
5
0
−5
−10
−15
−25
0
25
50
marker
150
100
50
0

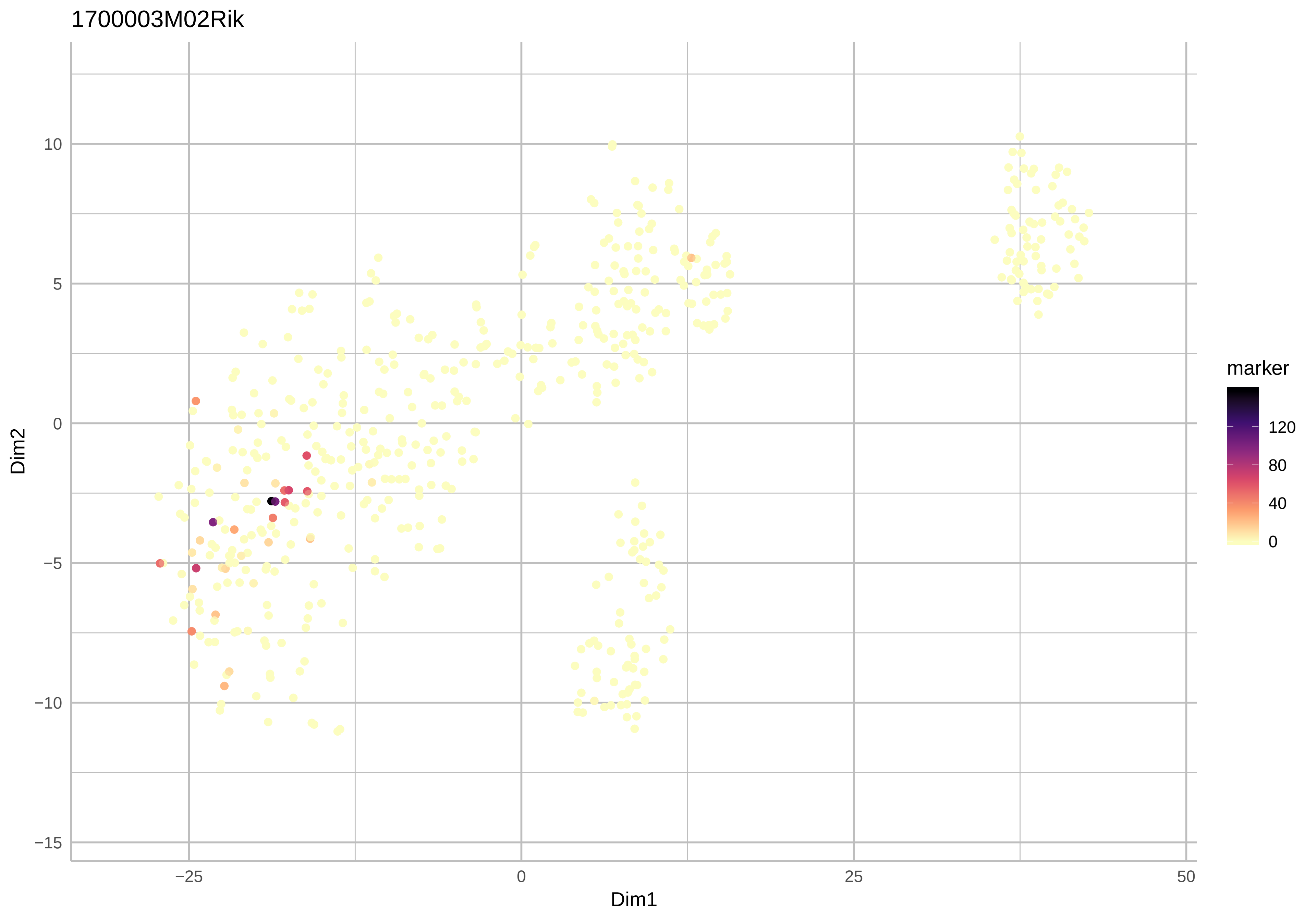

1700003M02Rik
Dim2
Dim1
10
5
0
−5
−10
−15
−25
0
25
50
marker
120
80
40
0

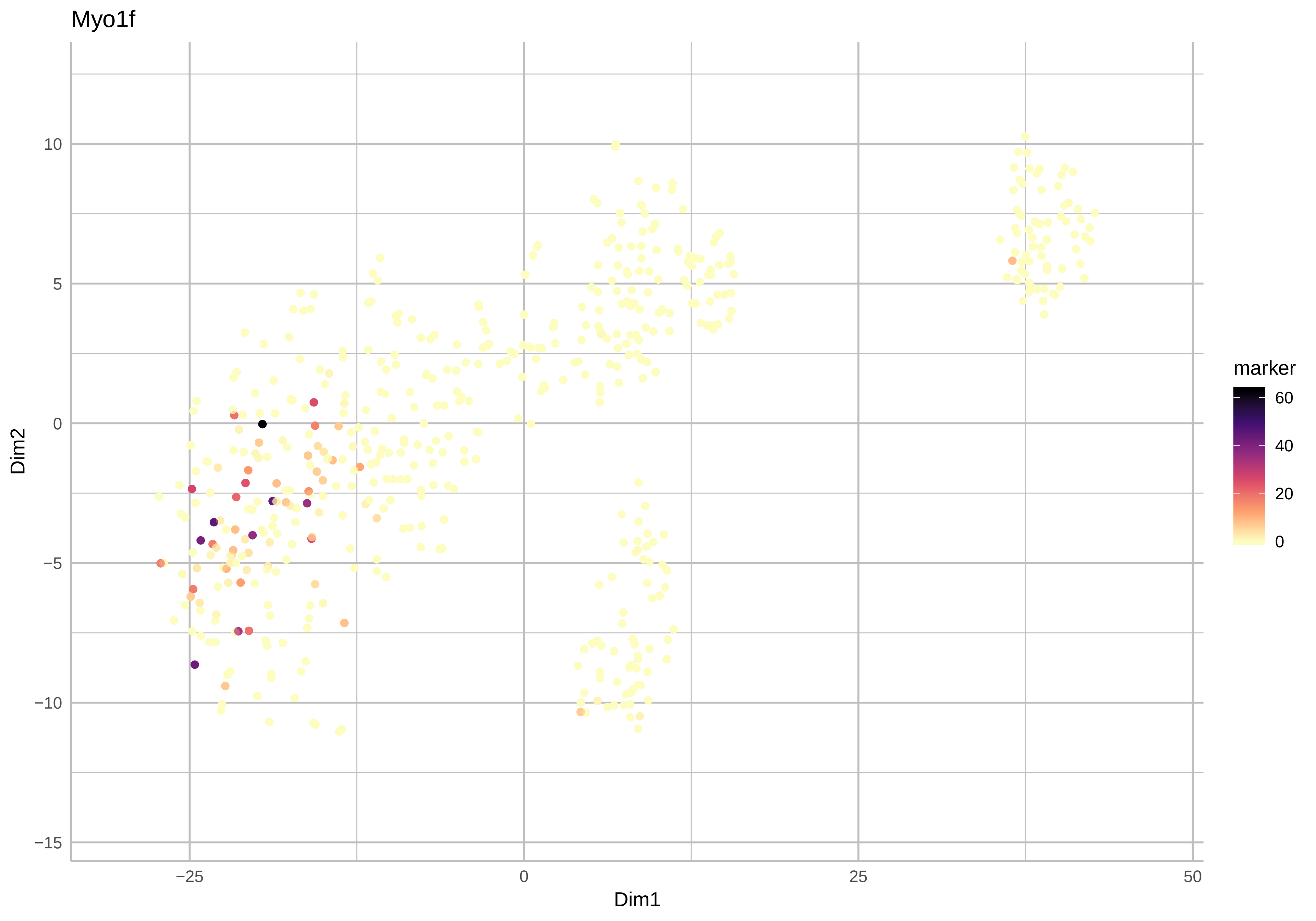
Myo1f
Dim2
Dim1
10
5
0
−5
−10
−15
−25
0
25
50
marker
60
40
20
0

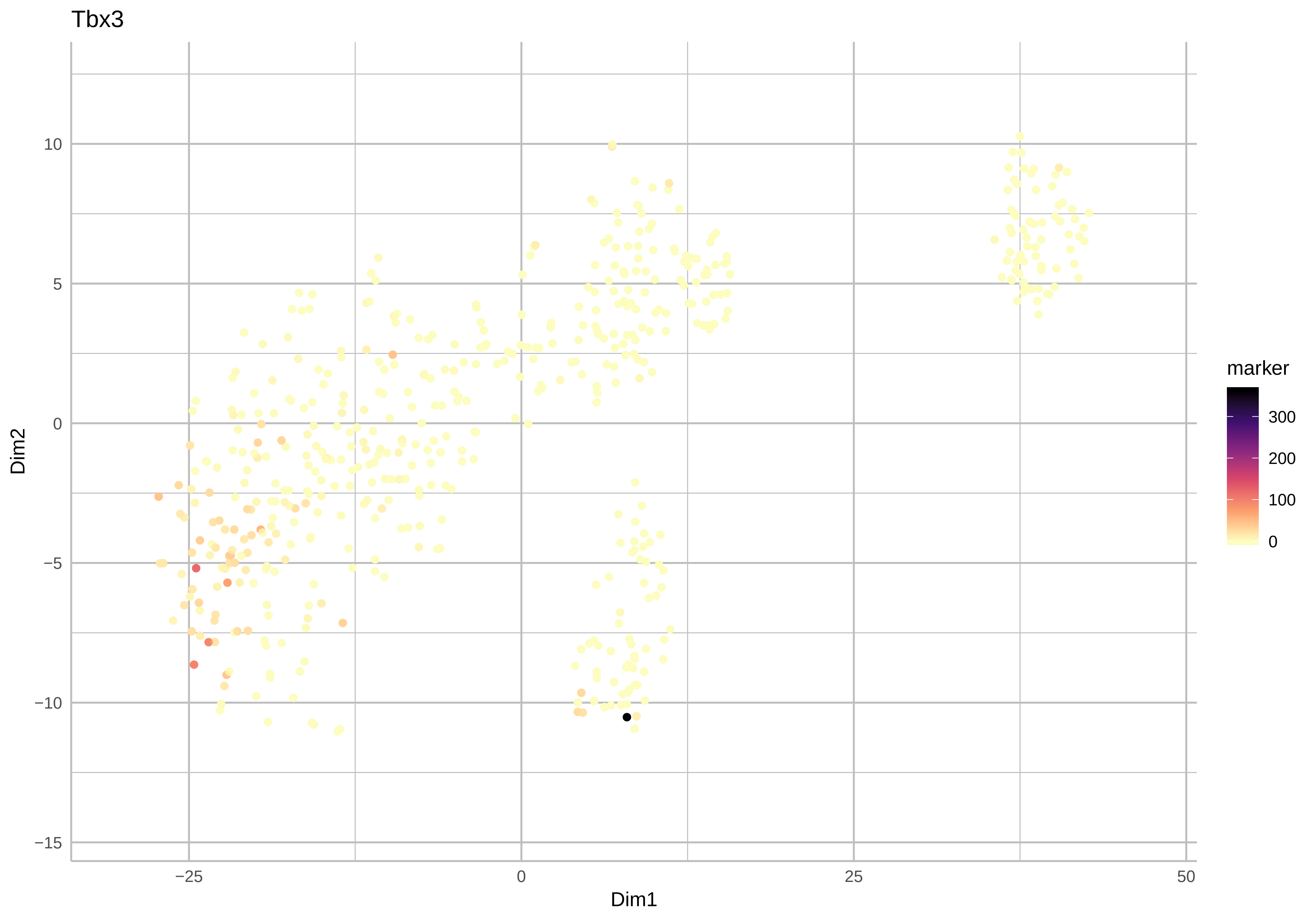
Tbx3
Dim2
Dim1
10
5
0
−5
−10
−15
−25
0
25
50
marker
300
200
100
0

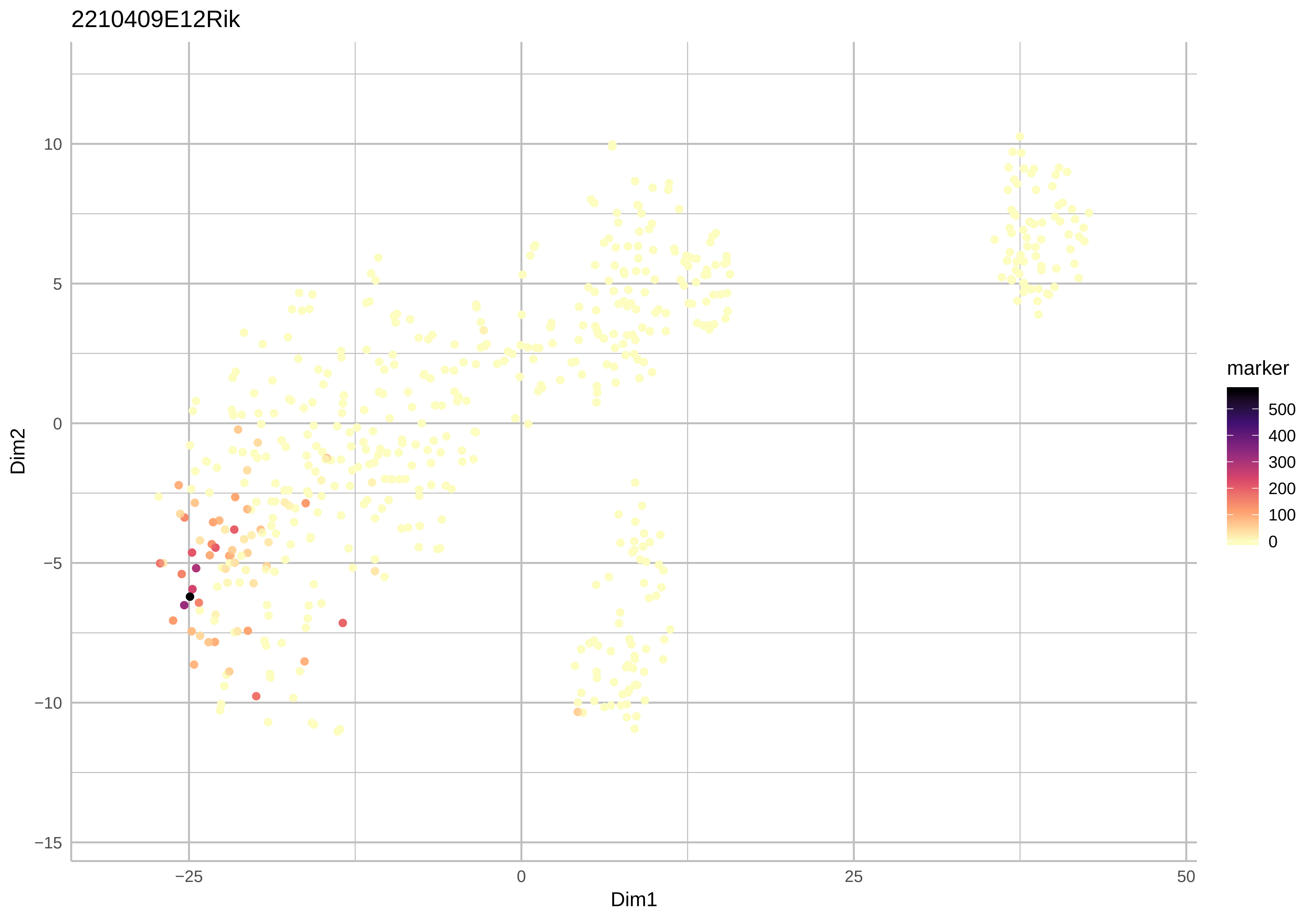
2210409E12Rik
Dim2
10
5
0
−5
−10
−15
−25
0
25
50
Dim1
marker
500
400
300
200
100
0

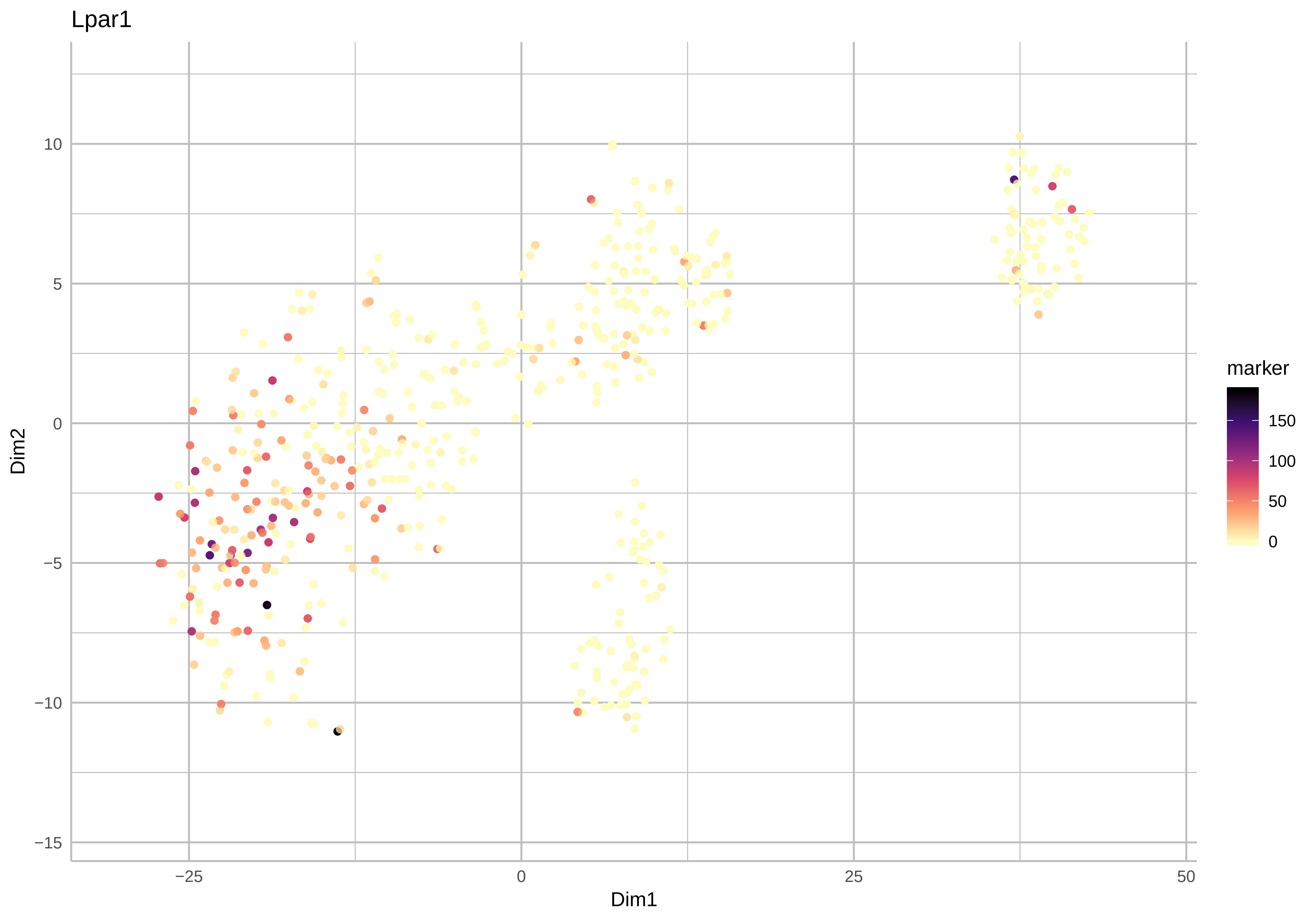

Lpar1
10
5
0
−5
−10
−15
Dim2
−25
0
25
50
Dim1
marker
150
100
50
0

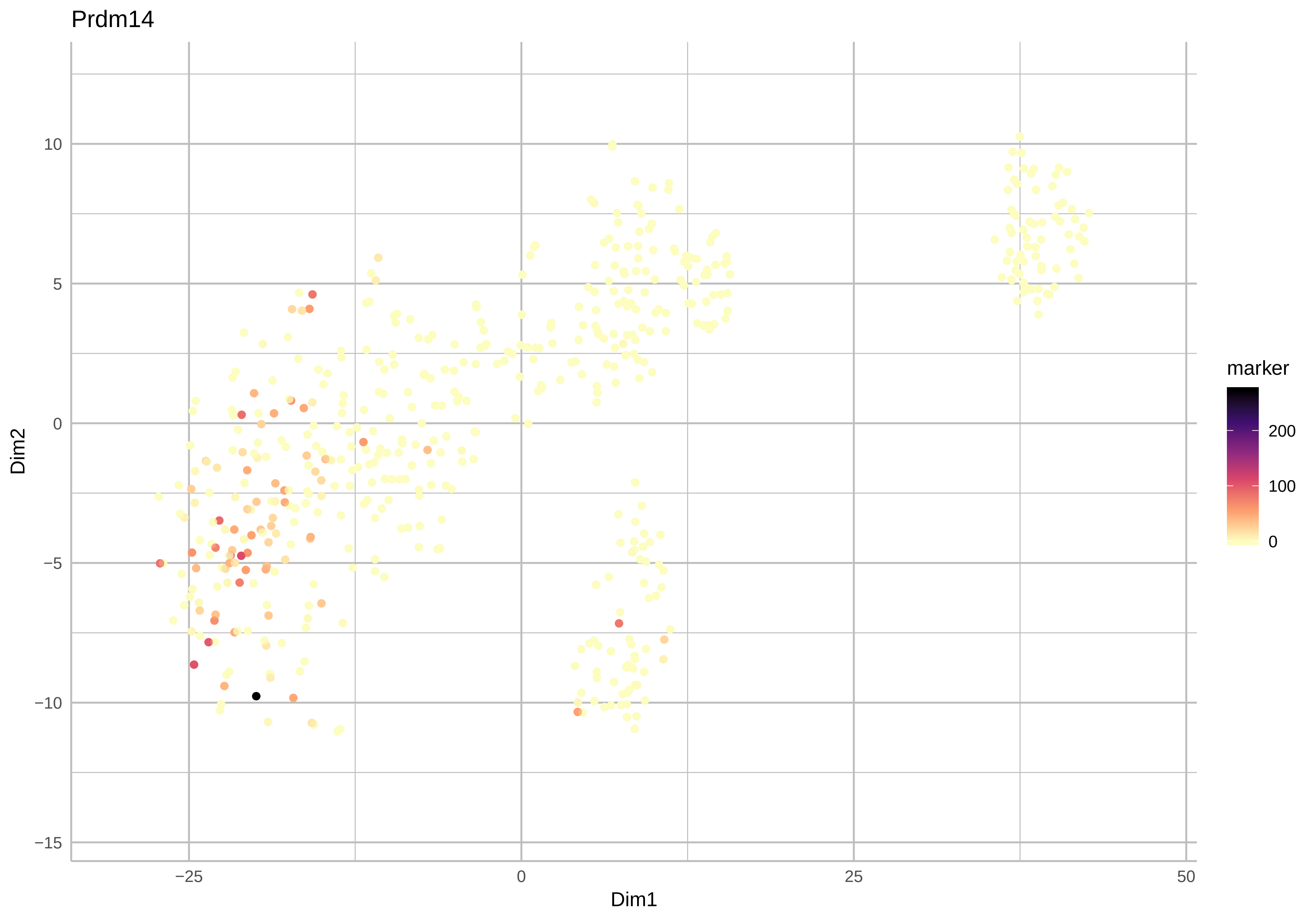
Prdm14
Dim2
Dim1
marker
200
100
0
10
5
0
−5
−10
−15
−25
0
25
50

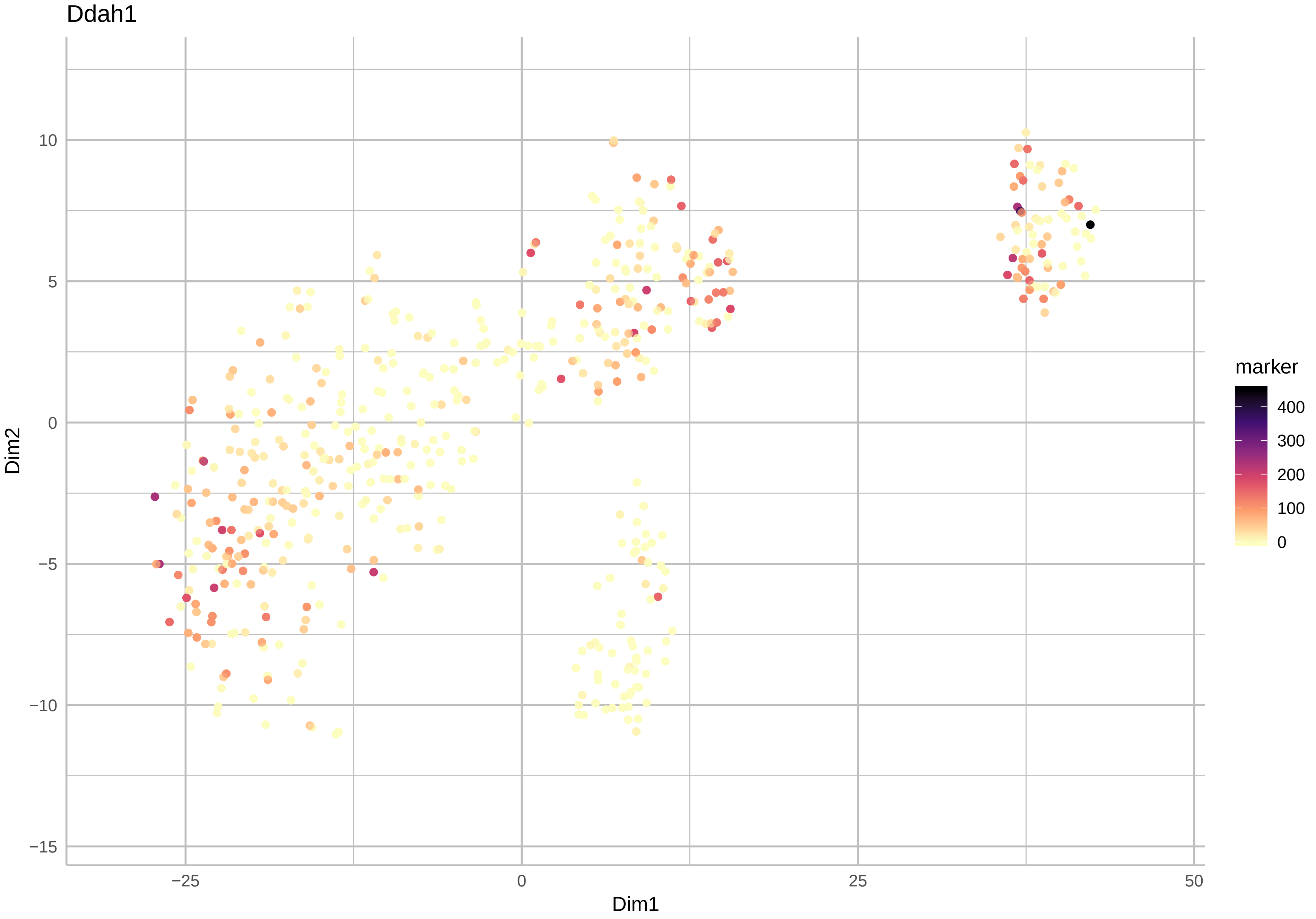
Ddah1
Dim2
Dim1
10
5
0
−5
−10
−15
−25
0
25
50
marker
400
300
200
100
0

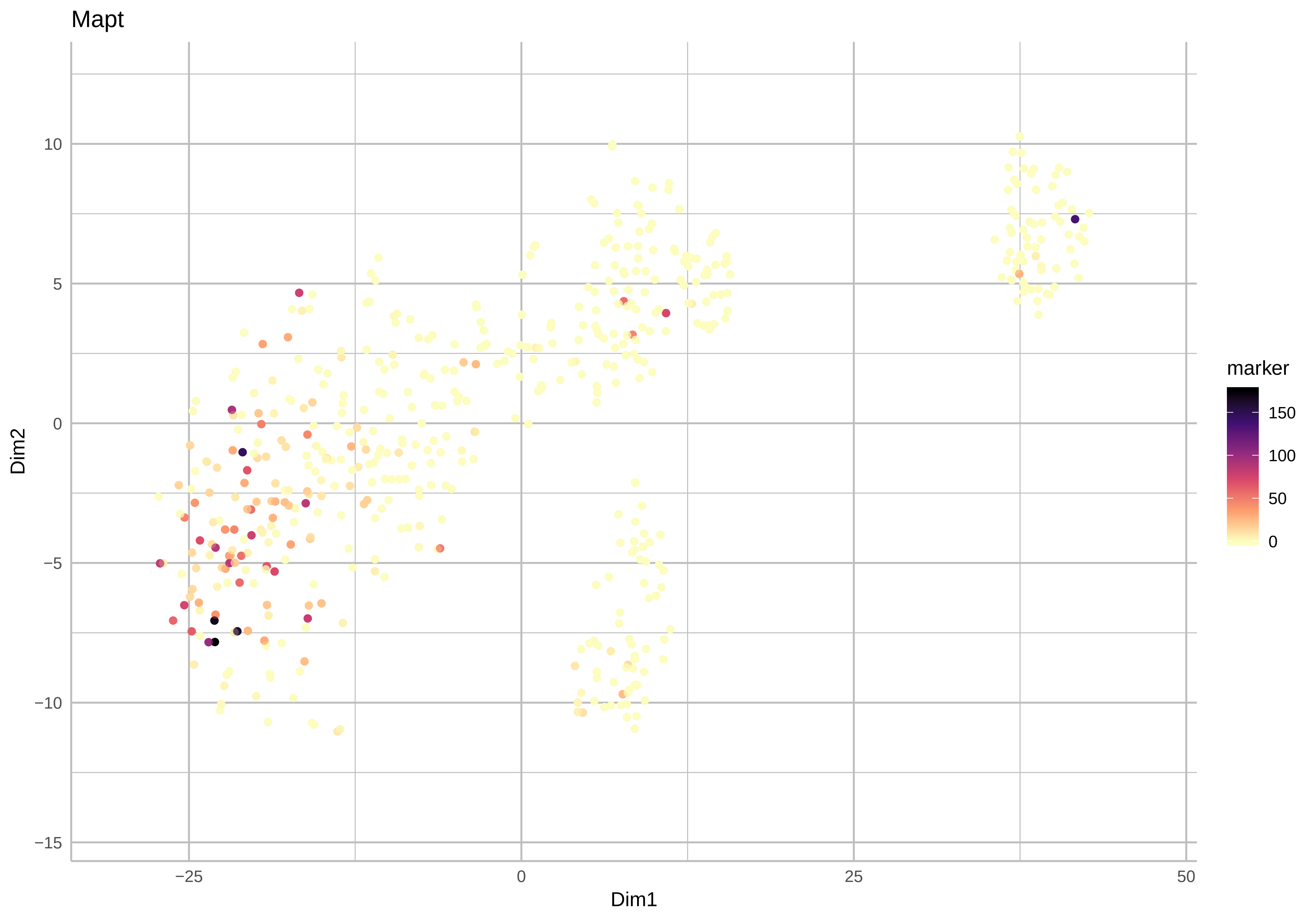

Mapt
10
5
0
−5
−10
−15
Dim2
−25
0
25
50
Dim1
marker
150
100
50
0

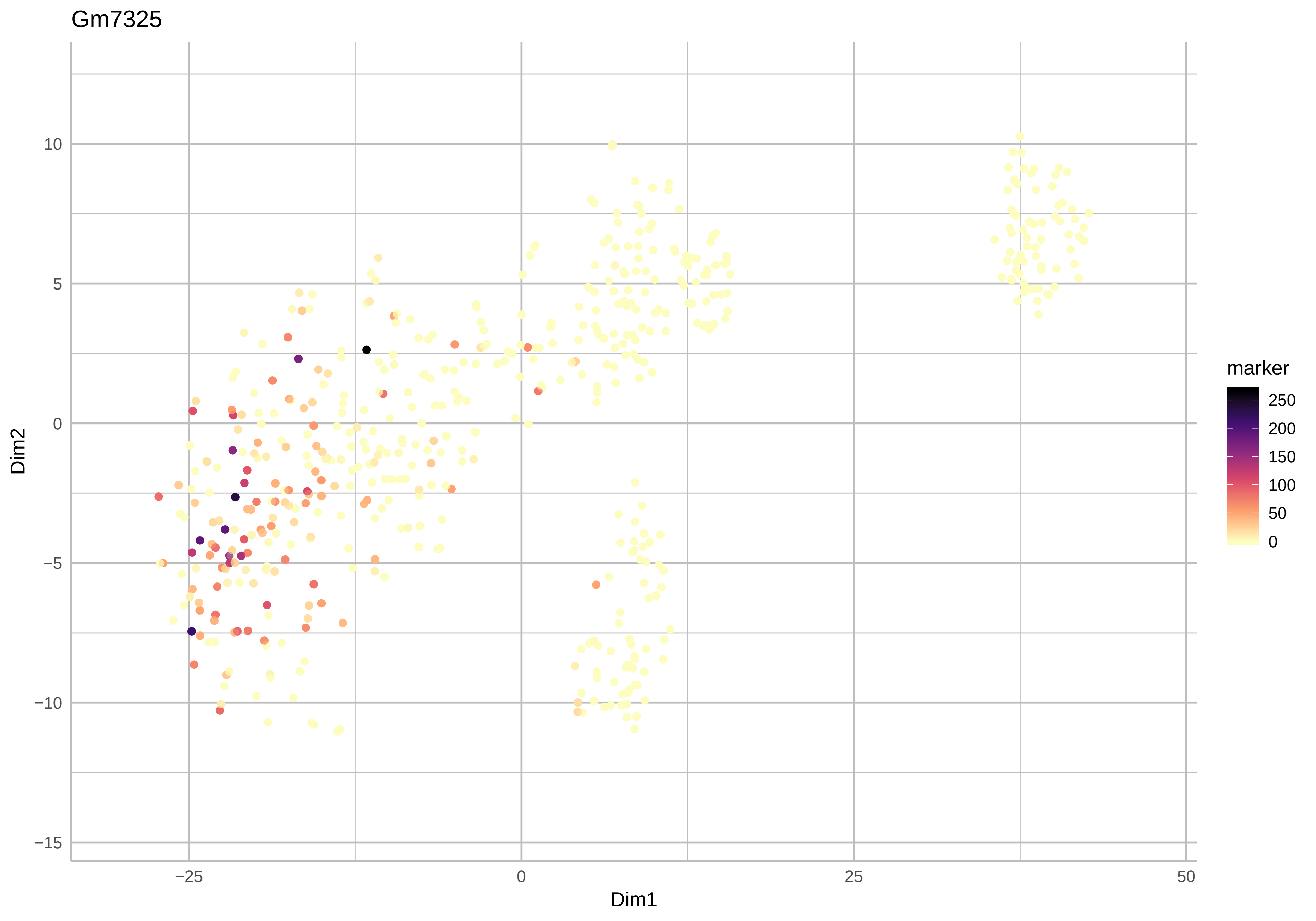
Gm7325
Dim2
Dim1
10
5
0
−5
−10
−15
−25
0
25
50
marker
250
200
150
100
50
0

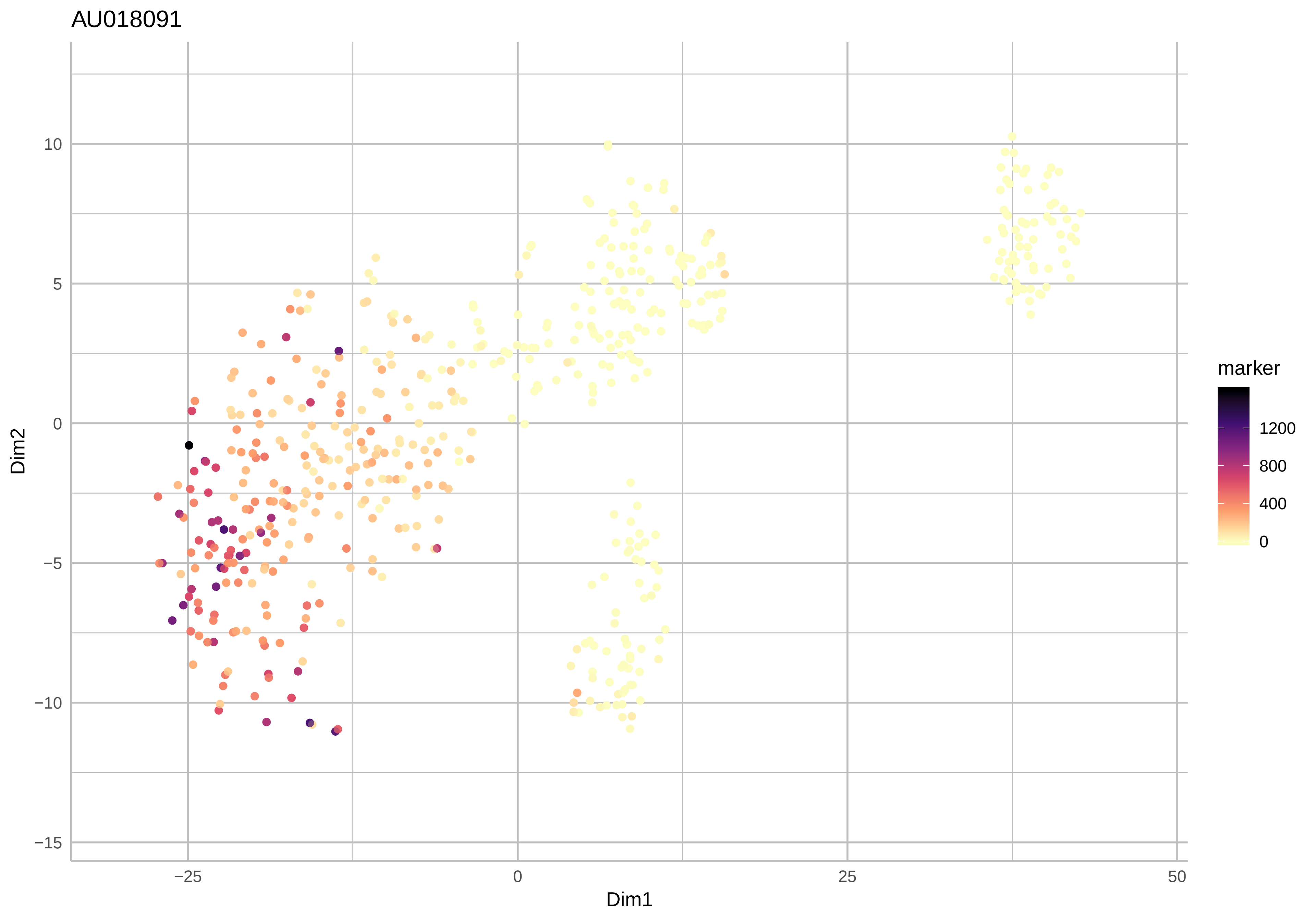
AU018091
Dim2
Dim1
10
5
0
−5
−10
−15
−25
0
25
50
marker
1200
800
400
0

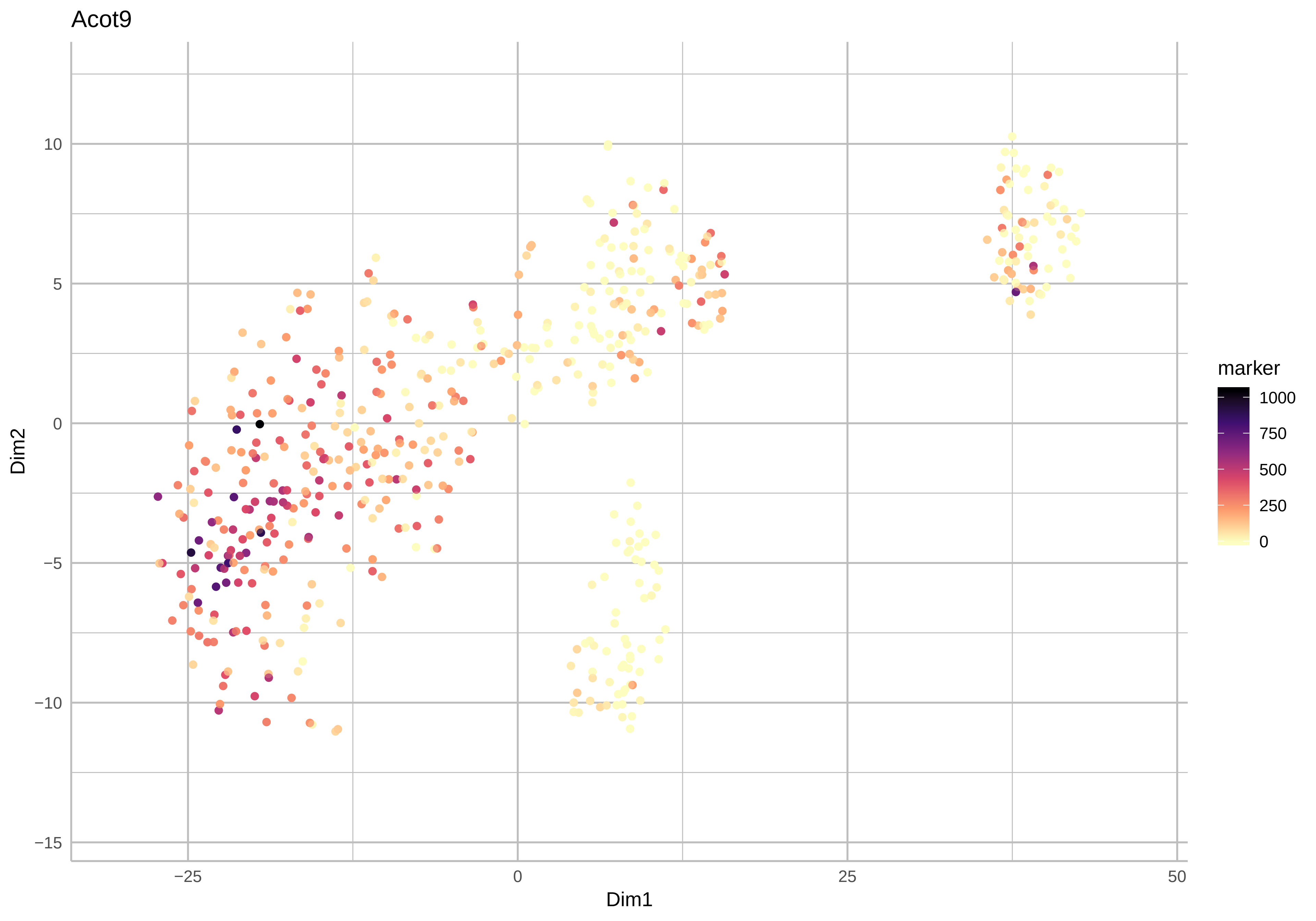
Acot9
10
5
0
−5
−10
−15
Dim2
−25
0
25
50
Dim1
marker
1000
750
500
250
0

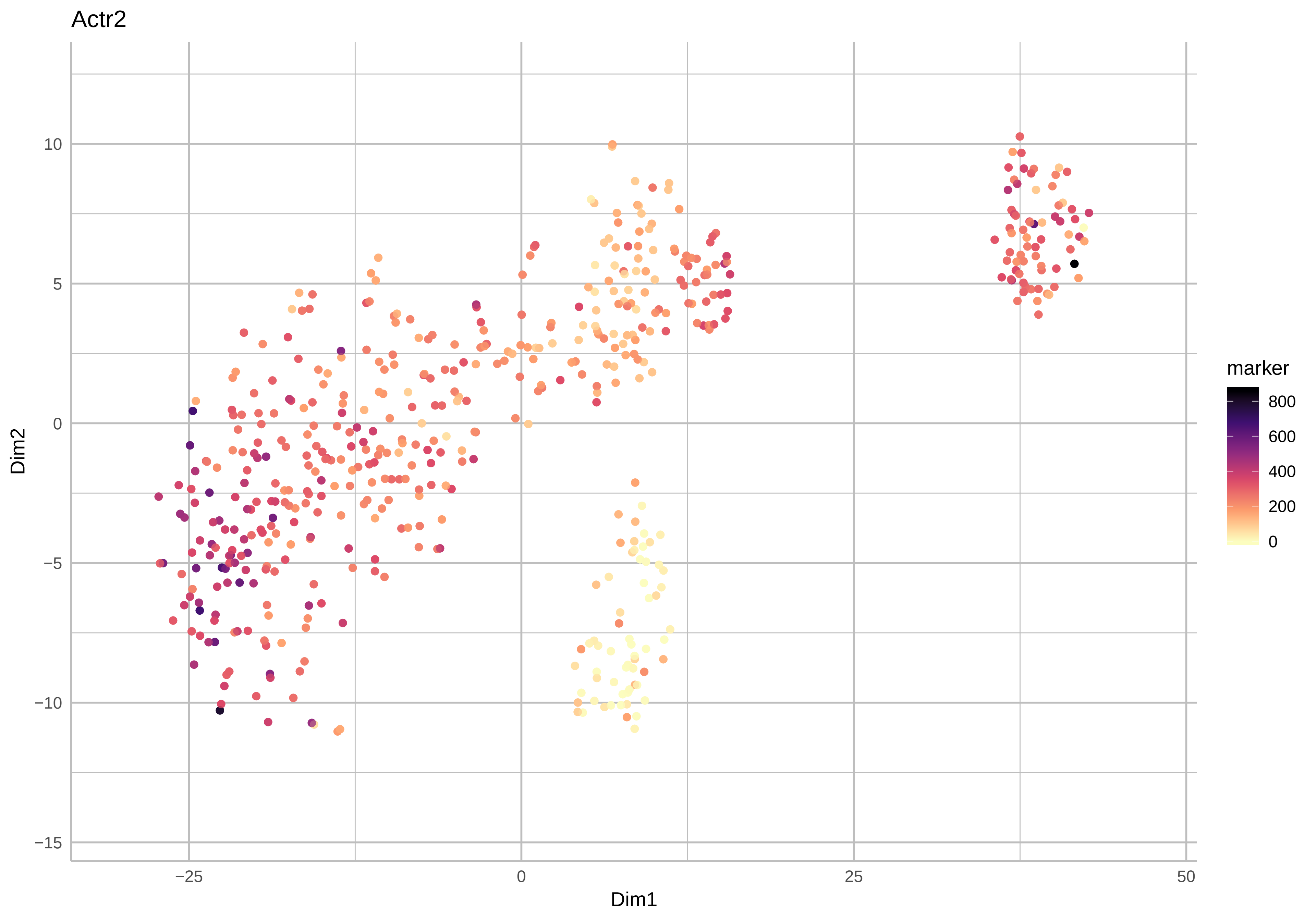

Actr2
Dim2
Dim1
10
5
0
−5
−10
−15
−25
0
25
50
marker
800
600
400
200
0

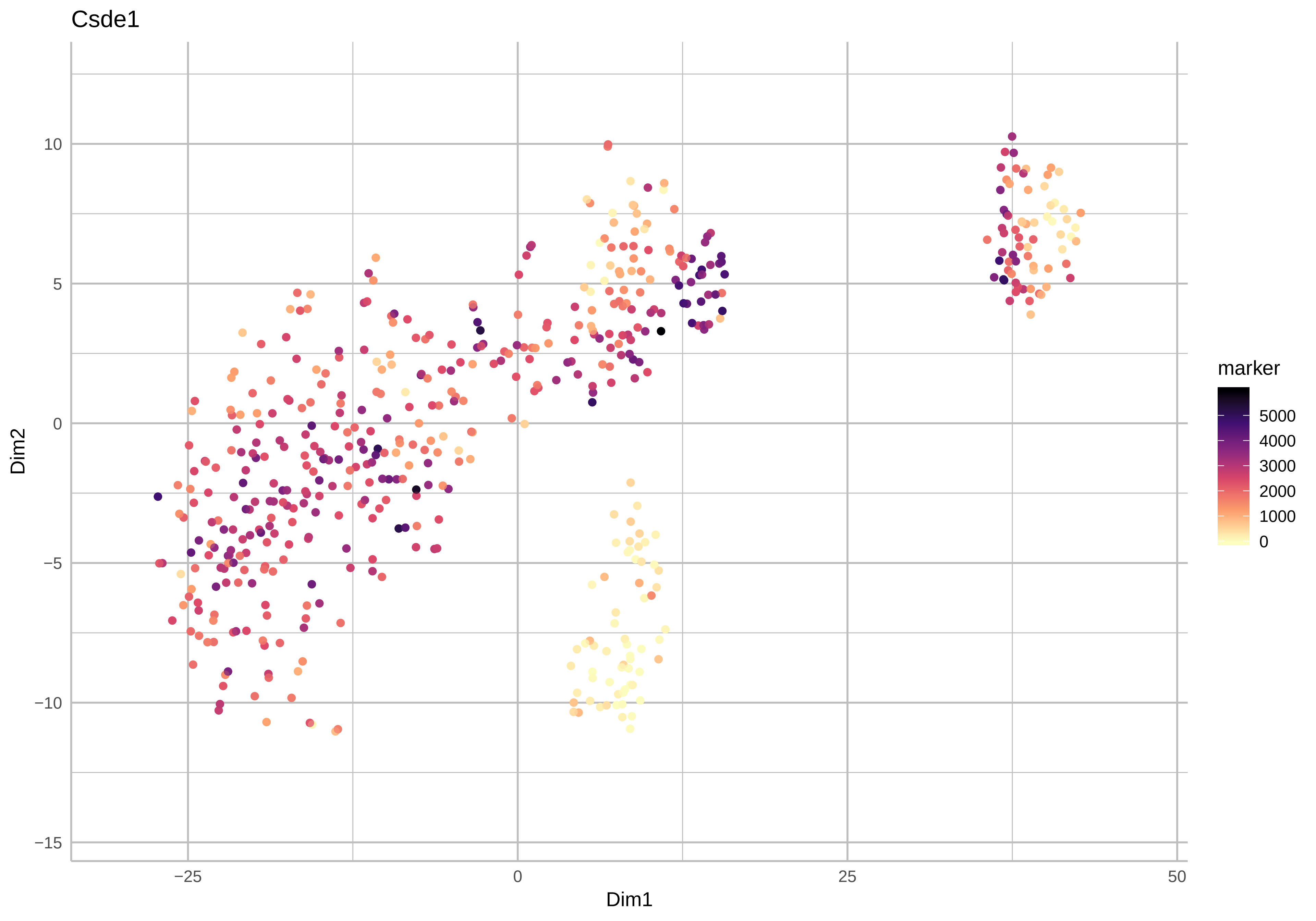

Csde1
Dim2
Dim1
10
5
0
−5
−10
−15
−25
0
25
50
marker
5000
4000
3000
2000
1000
0

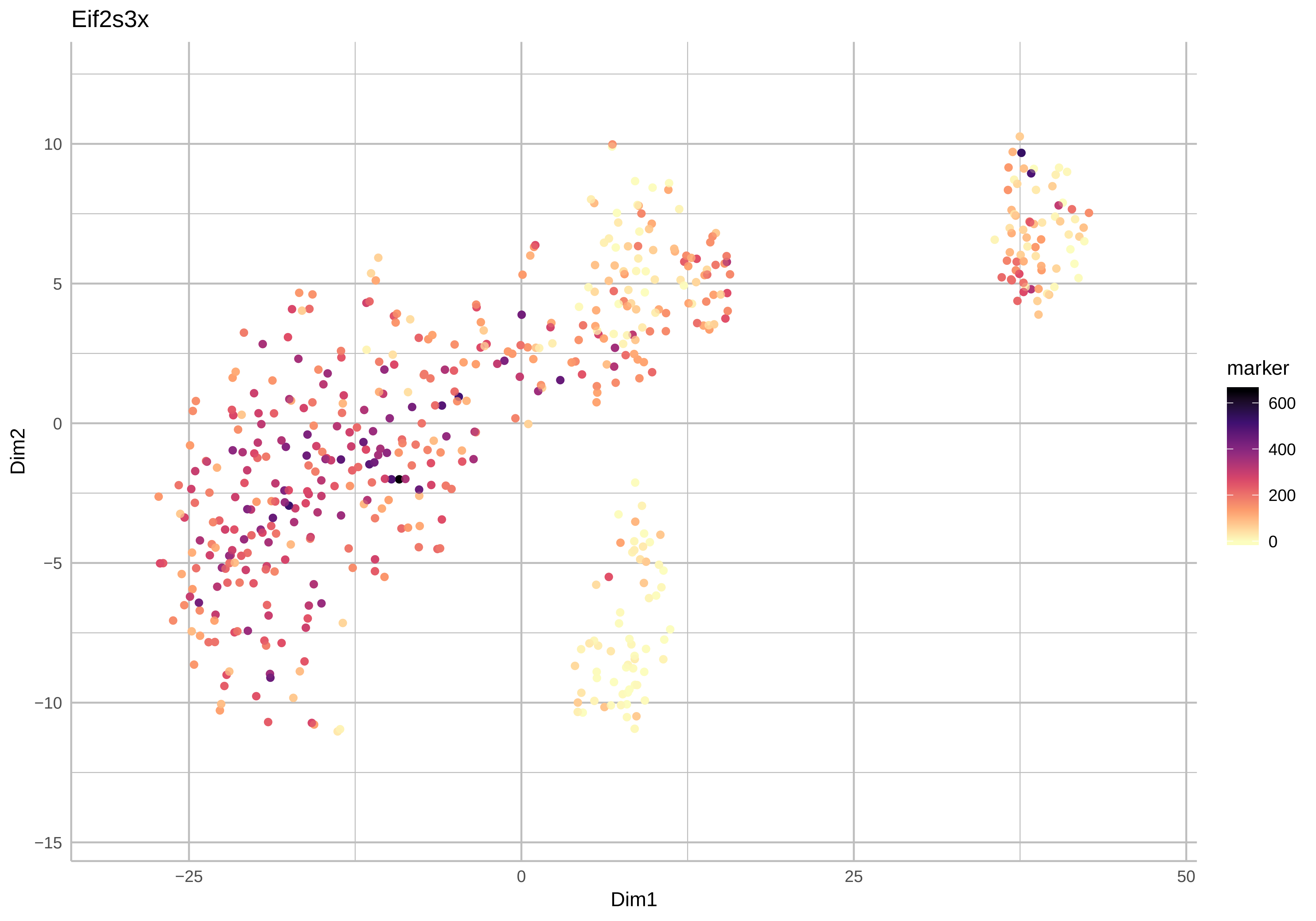
Eif2s3x
10
5
0
−5
−10
−15
−25
0
25
50
Dim1
Dim2
marker
600
400
200
0

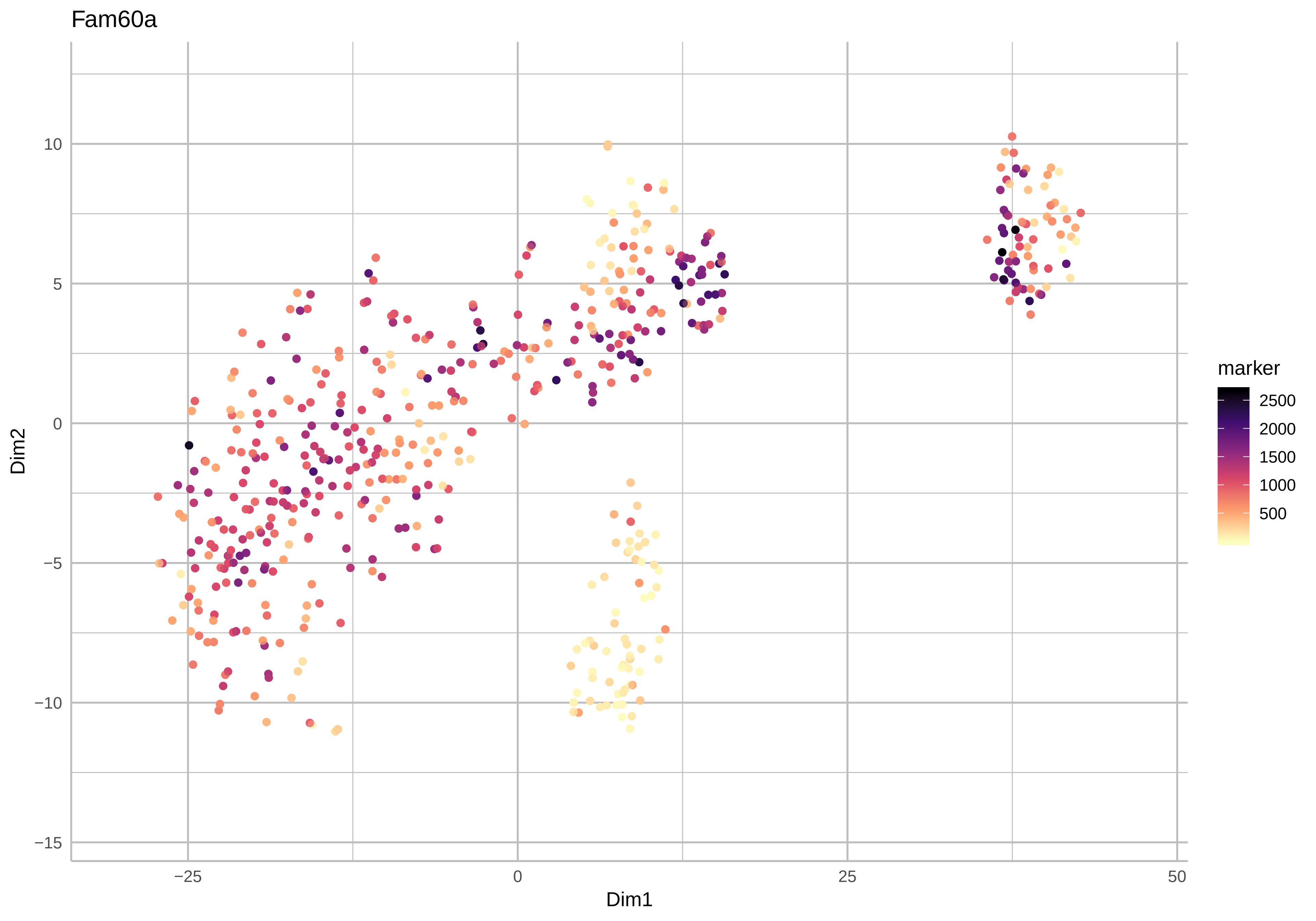

Fam60a
Dim2
Dim1
10
5
0
−5
−10
−15
−25
0
25
50
marker
2500
2000
1500
1000
500

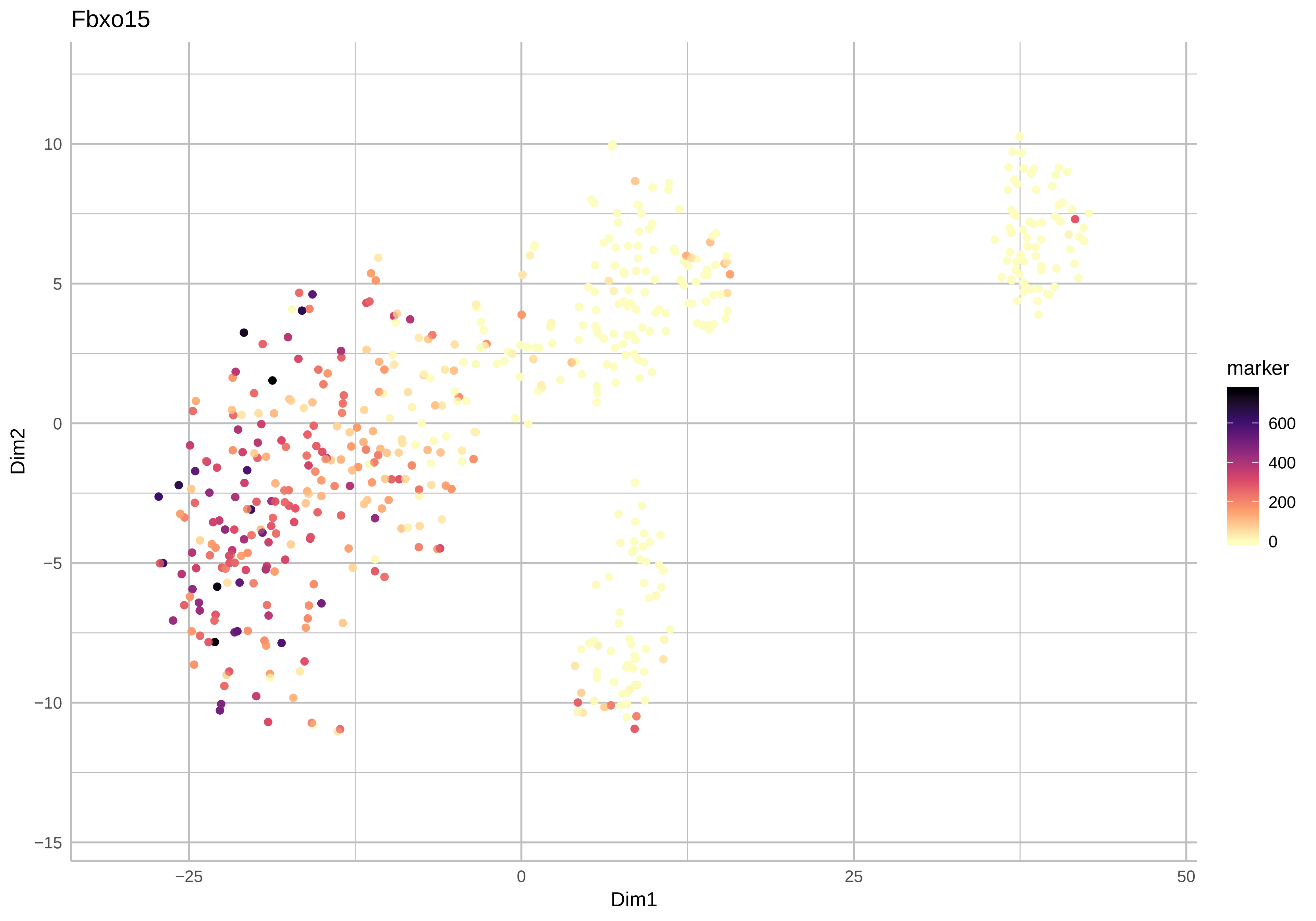
Fbxo15
10
5
0
−5
−10
−15
−25
0
25
50
Dim1
Dim2
marker
600
400
200
0

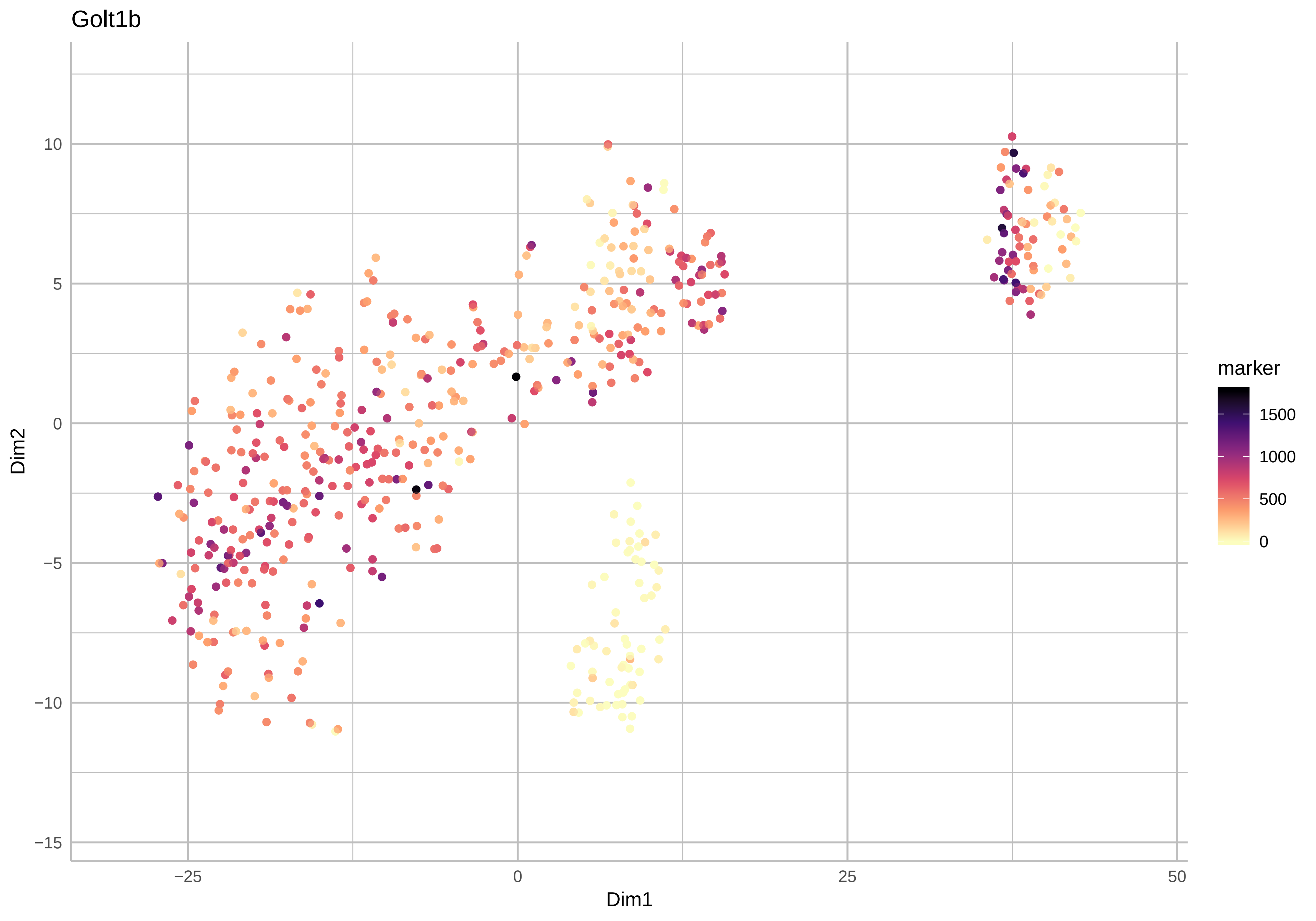
Golt1b
Dim2
Dim1
10
5
0
−5
−10
−15
−25
0
25
50
marker
1500
1000
500
0

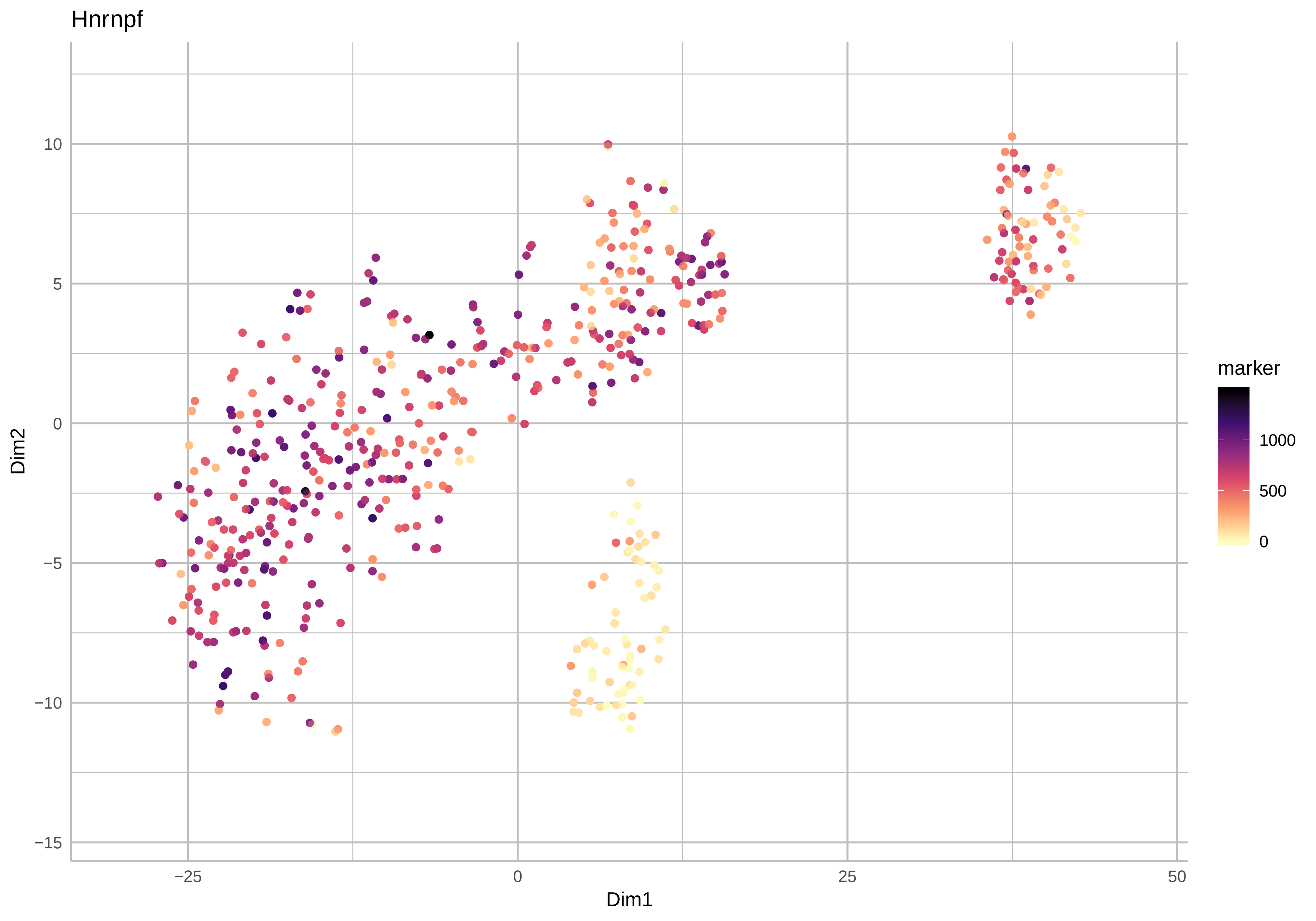

Hnrnpf
10
5
0
−5
−10
−15
Dim2
−25
0
25
50
Dim1
marker
1000
500
0

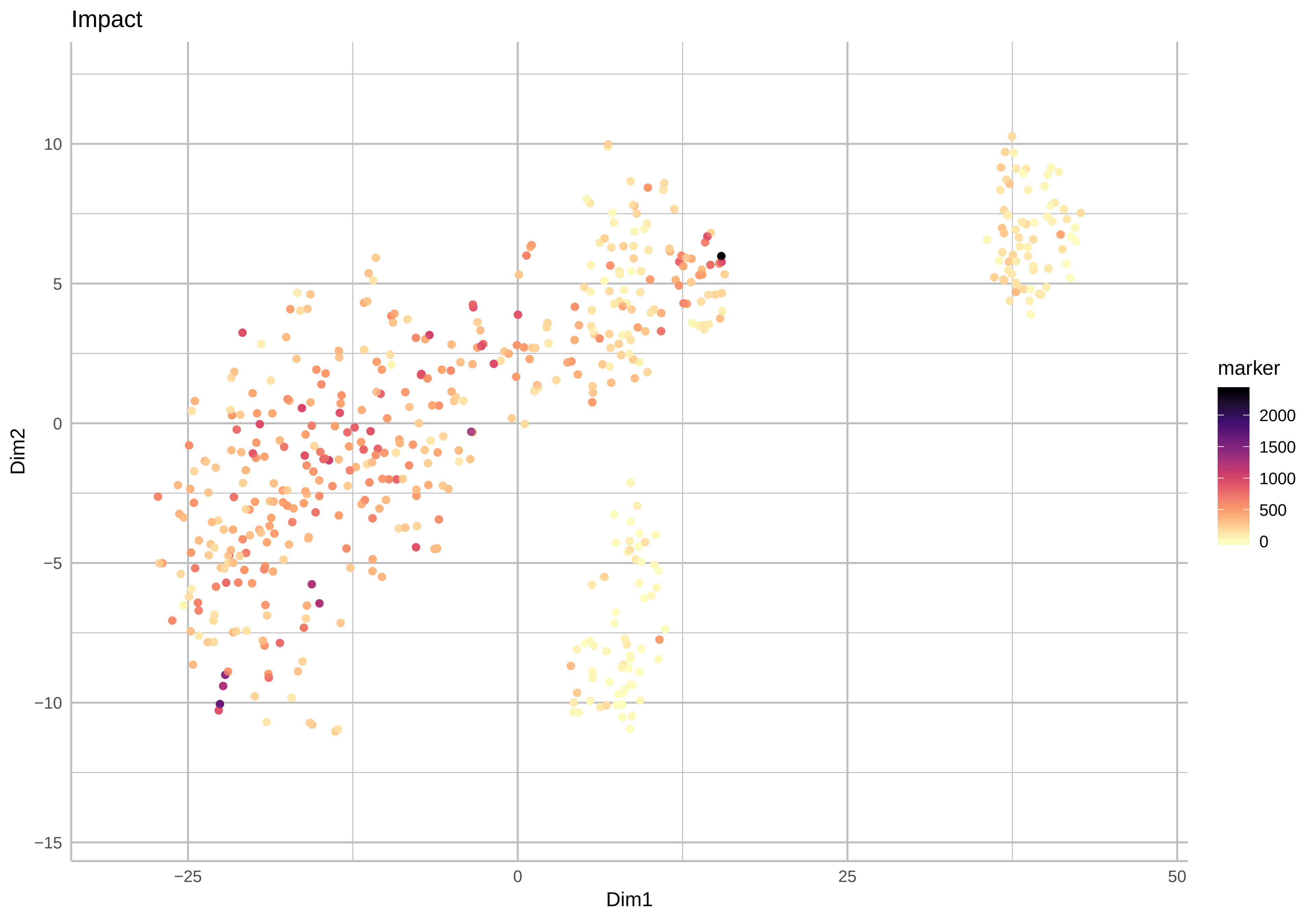
Impact
Dim2
Dim1
10
5
0
−5
−10
−15
−25
0
25
50
marker
2000
1500
1000
500
0

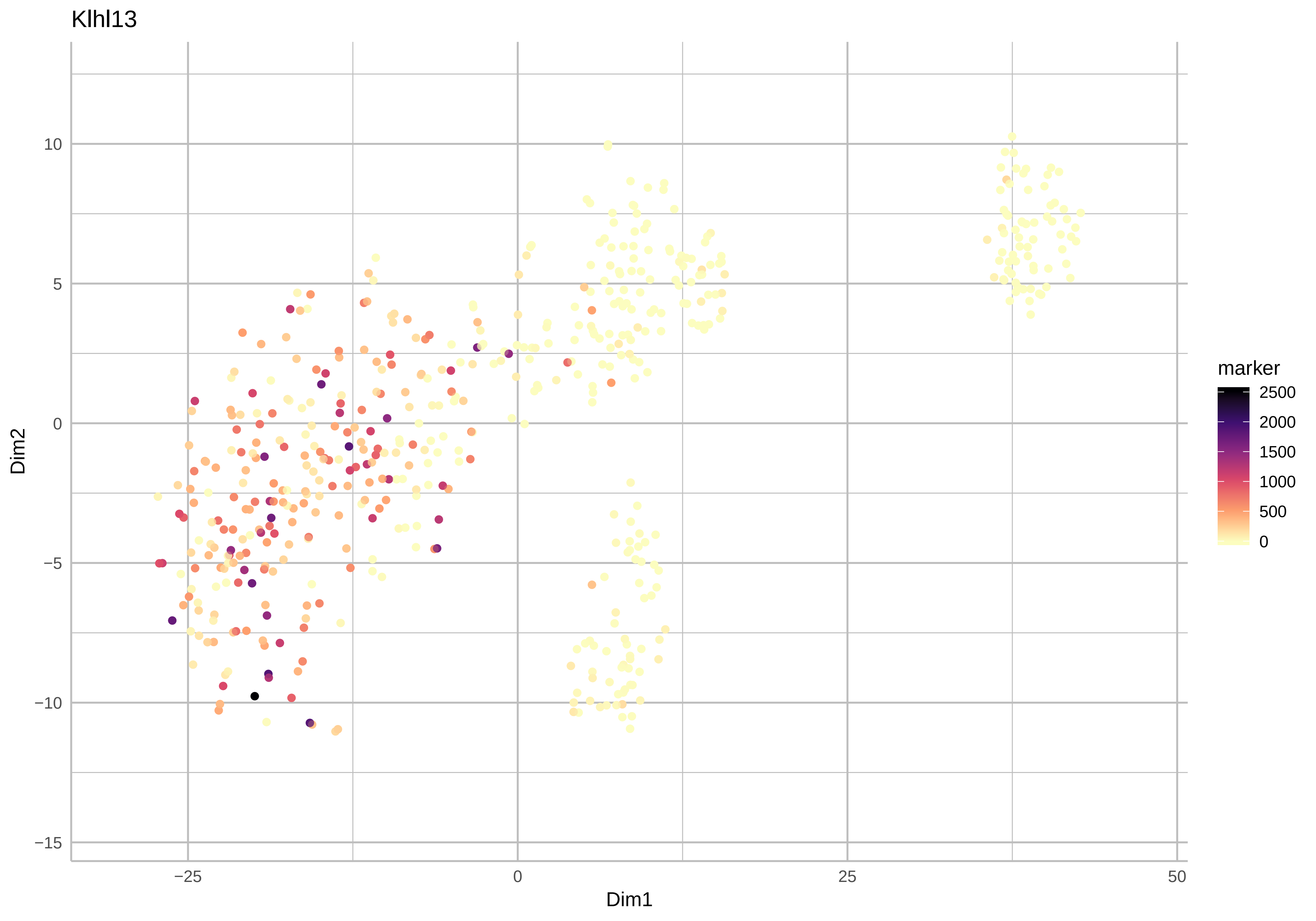

Klhl13
marker
2500
2000
1500
1000
500
0
Dim2
10
5
0
−5
−10
−15
Dim1
−25
0
25
50

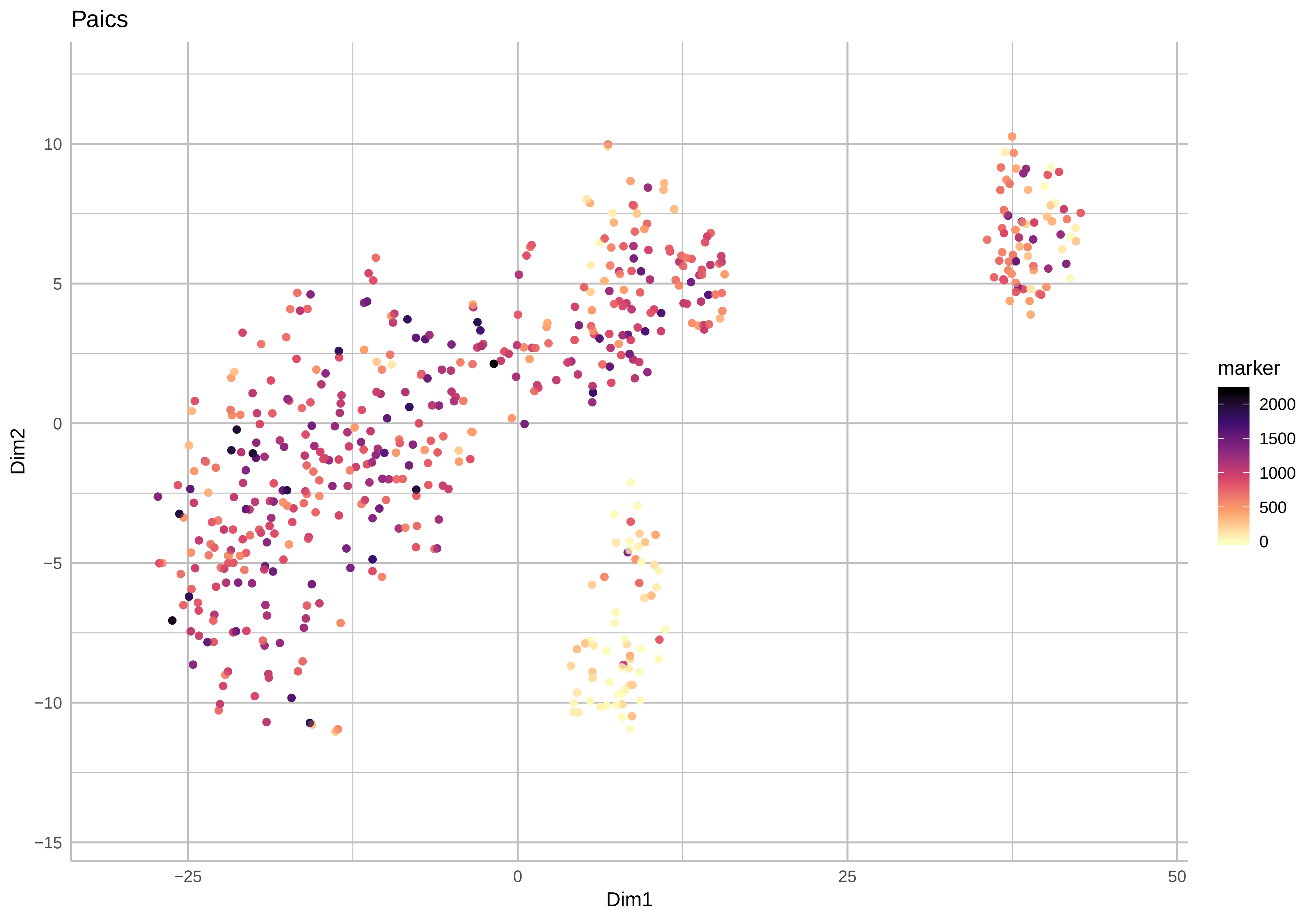
Paics
Dim2
10
5
0
−5
−10
−15
−25
0
25
50
Dim1
marker
2000
1500
1000
500
0

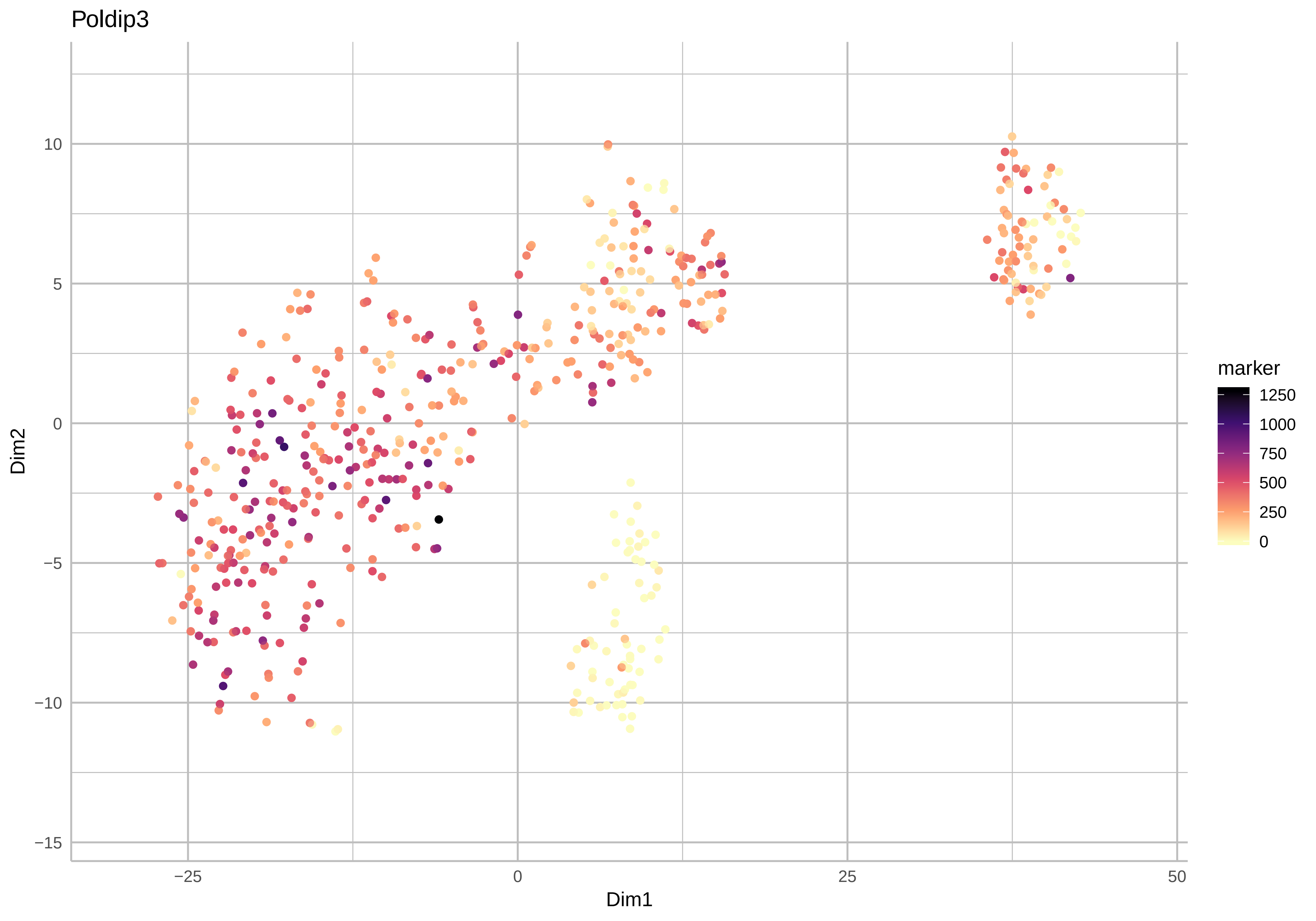
Poldip3
Dim2
Dim1
10
5
0
−5
−10
−15
−25
0
25
50
marker
1250
1000
750
500
250
0

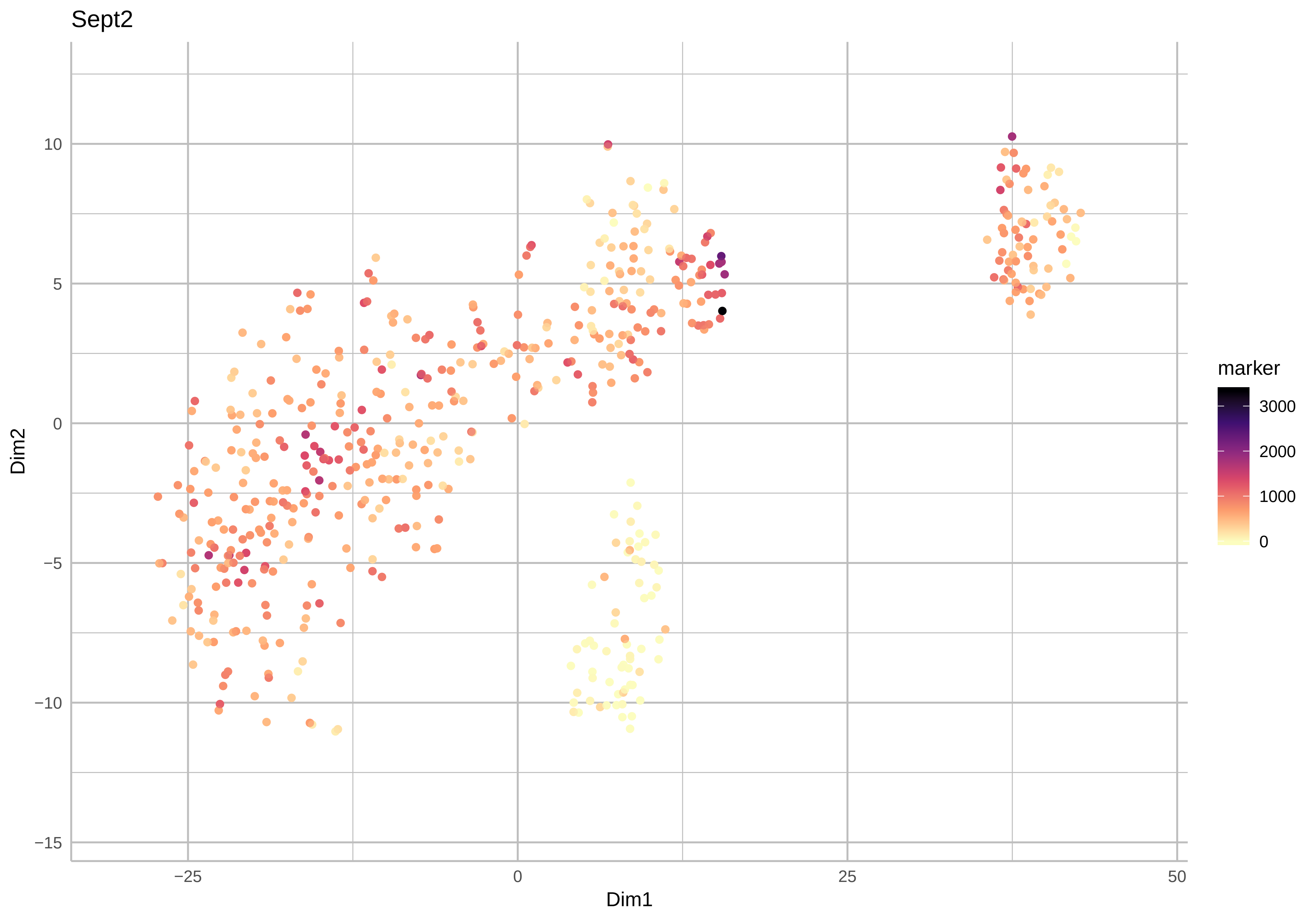
Sept2
Dim2
Dim1
marker
3000
2000
1000
0
10
5
0
−5
−10
−15
−25
0
25
50

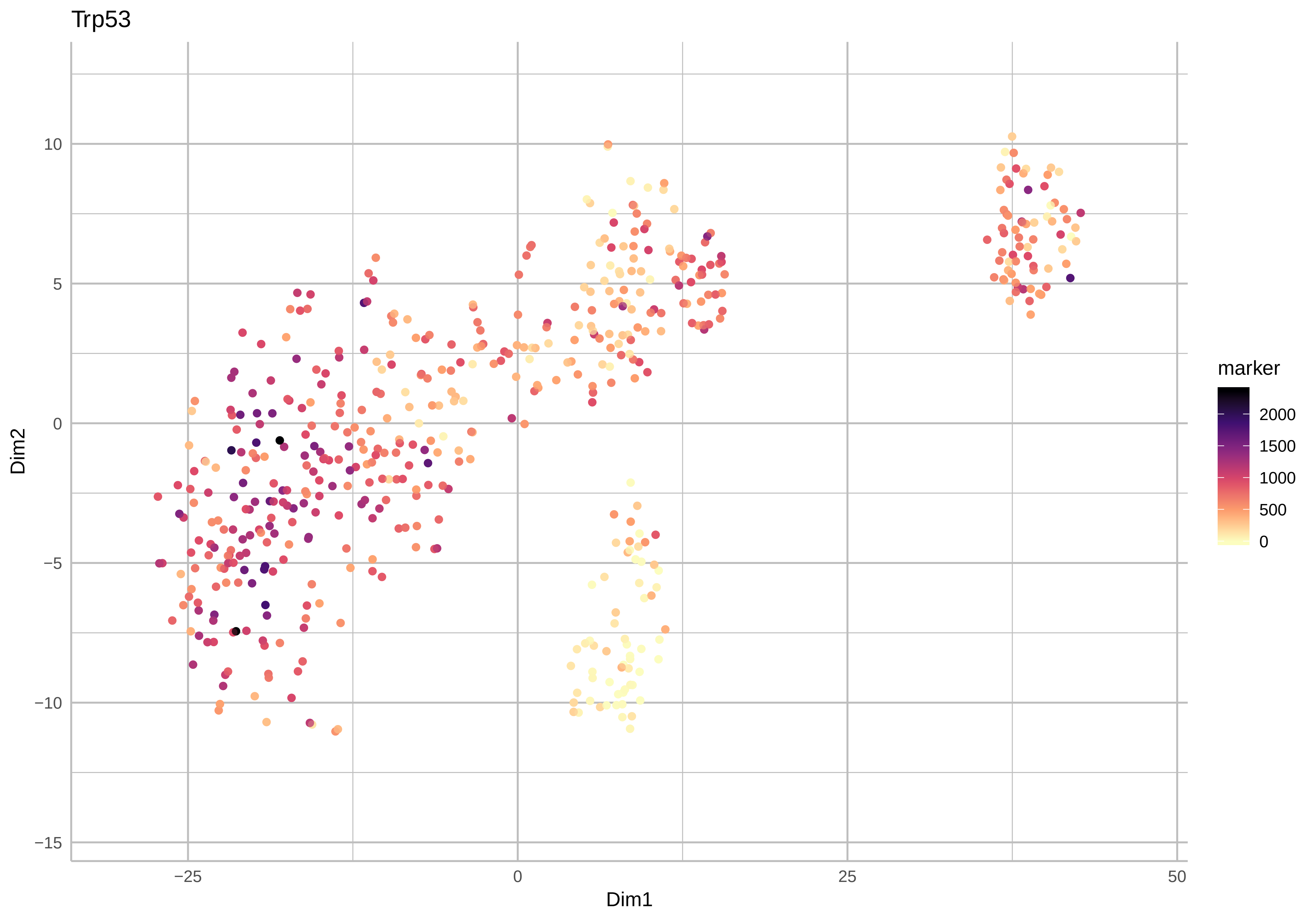
Trp53
Dim2
Dim1
10
5
0
−5
−10
−15
−25
0
25
50
marker
2000
1500
1000
500
0

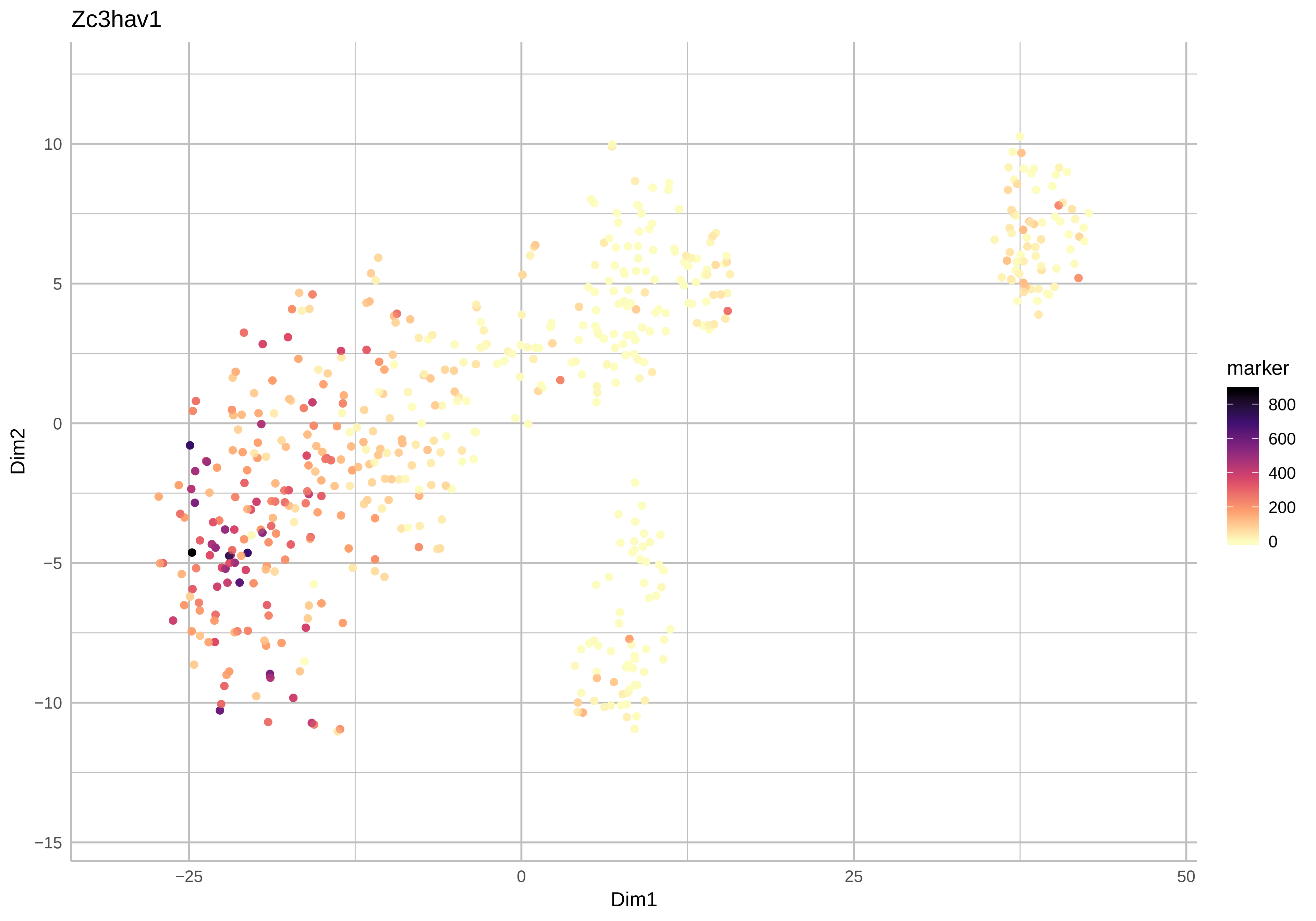

Zc3hav1
10
5
0
−5
−10
−15
Dim2
−25
0
25
50
Dim1
marker
800
600
400
200
0

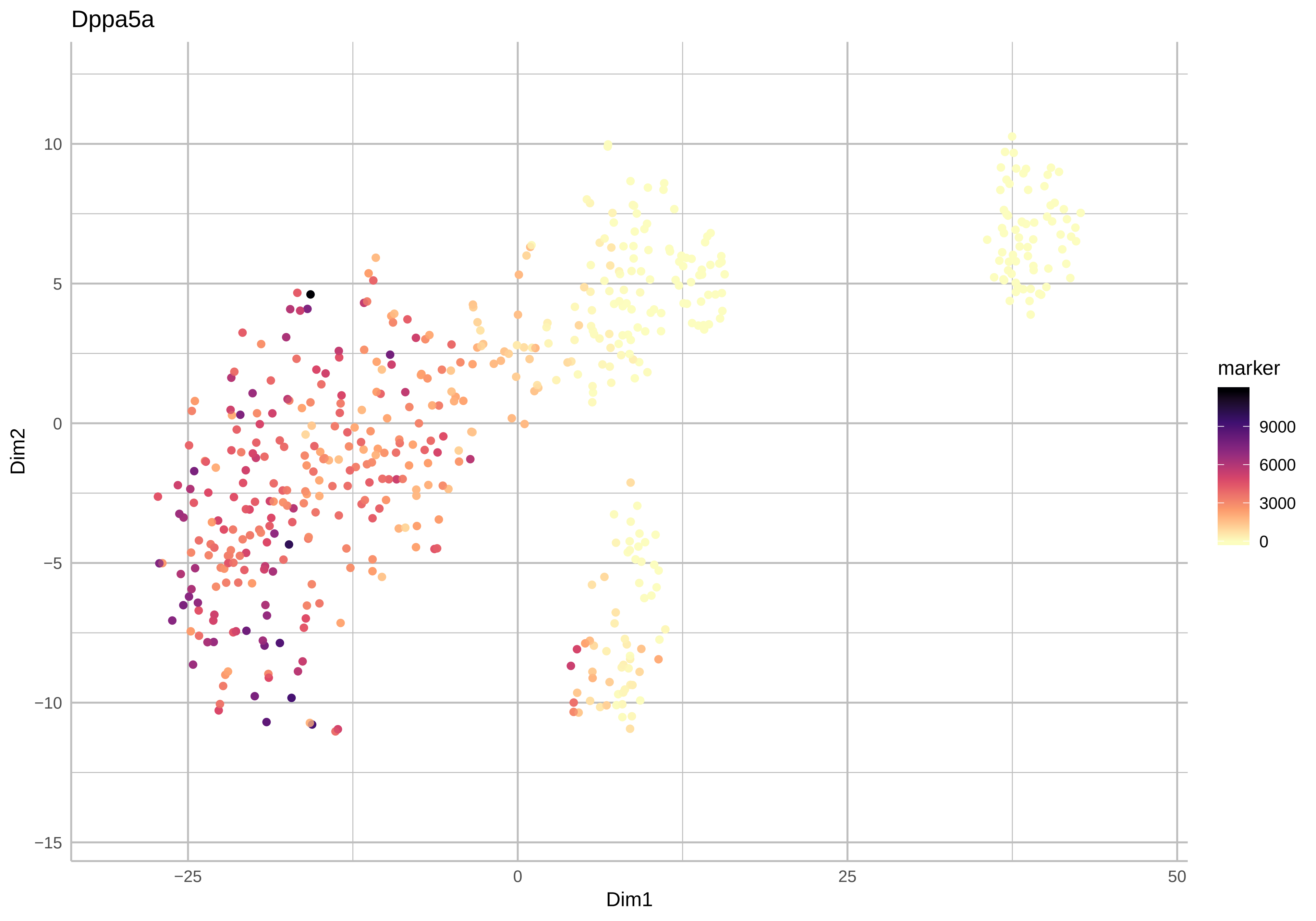

Dppa5a
Dim2
Dim1
10
5
0
−5
−10
−15
−25
0
25
50
marker
9000
6000
3000
0

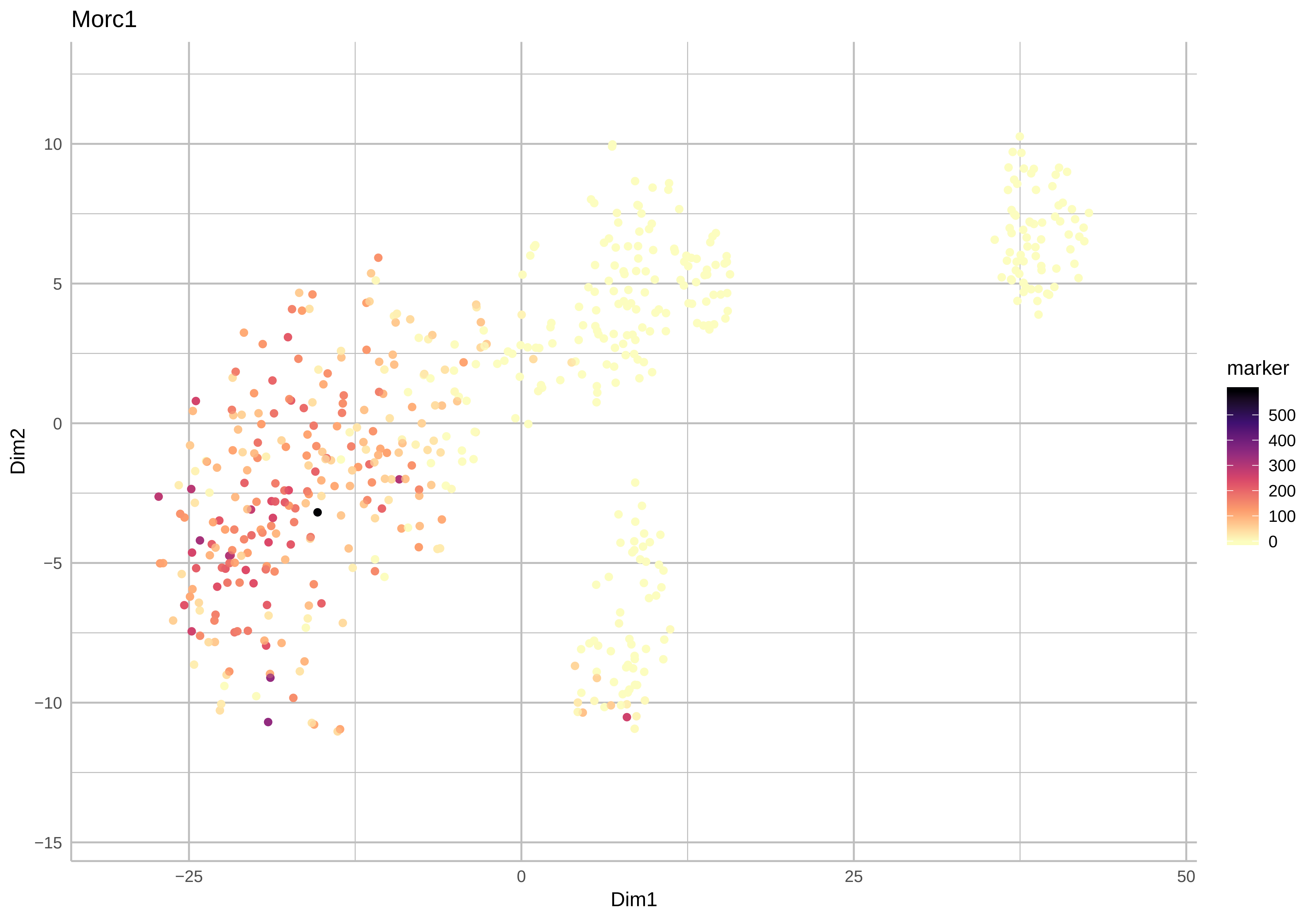

Morc1
Dim2
Dim1
10
5
0
−5
−10
−15
−25
0
25
50
marker
500
400
300
200
100
0

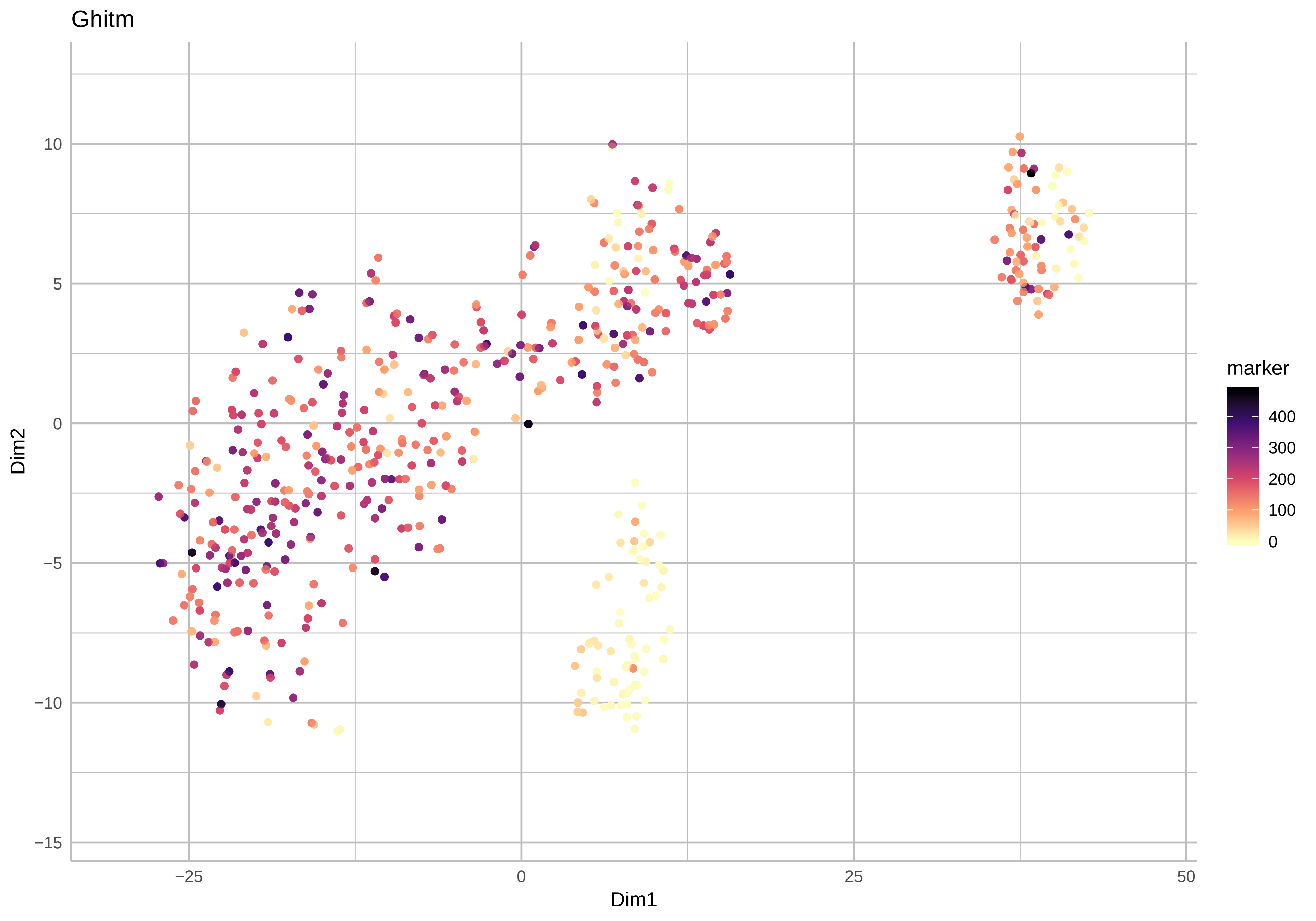

Ghitm
10
5
0
−5
−10
−15
−25
0
25
50
Dim1
Dim2
marker
400
300
200
100
0

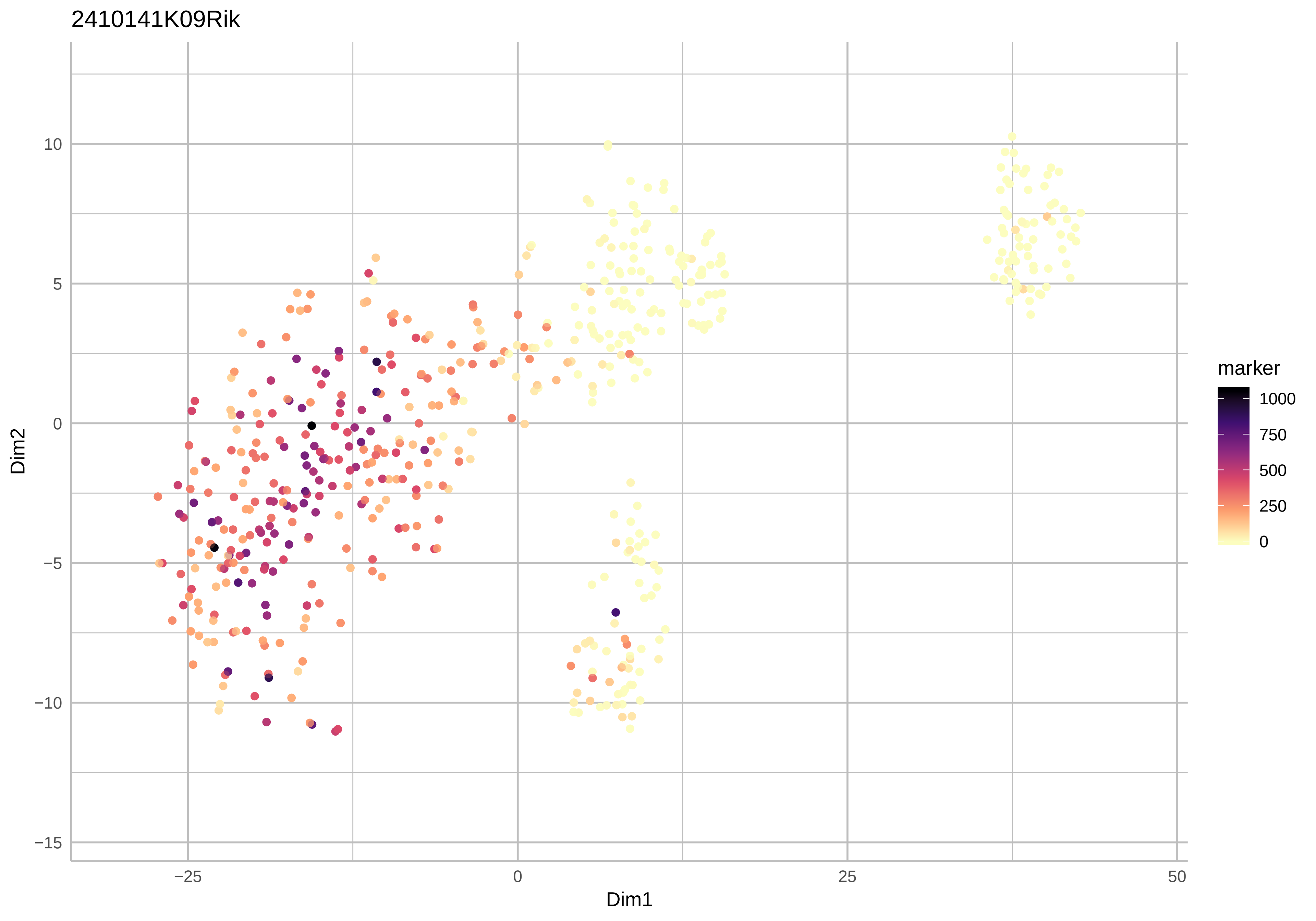

2410141K09Rik
Dim2
Dim1
10
5
0
−5
−10
−15
−25
0
25
50
marker
1000
750
500
250
0

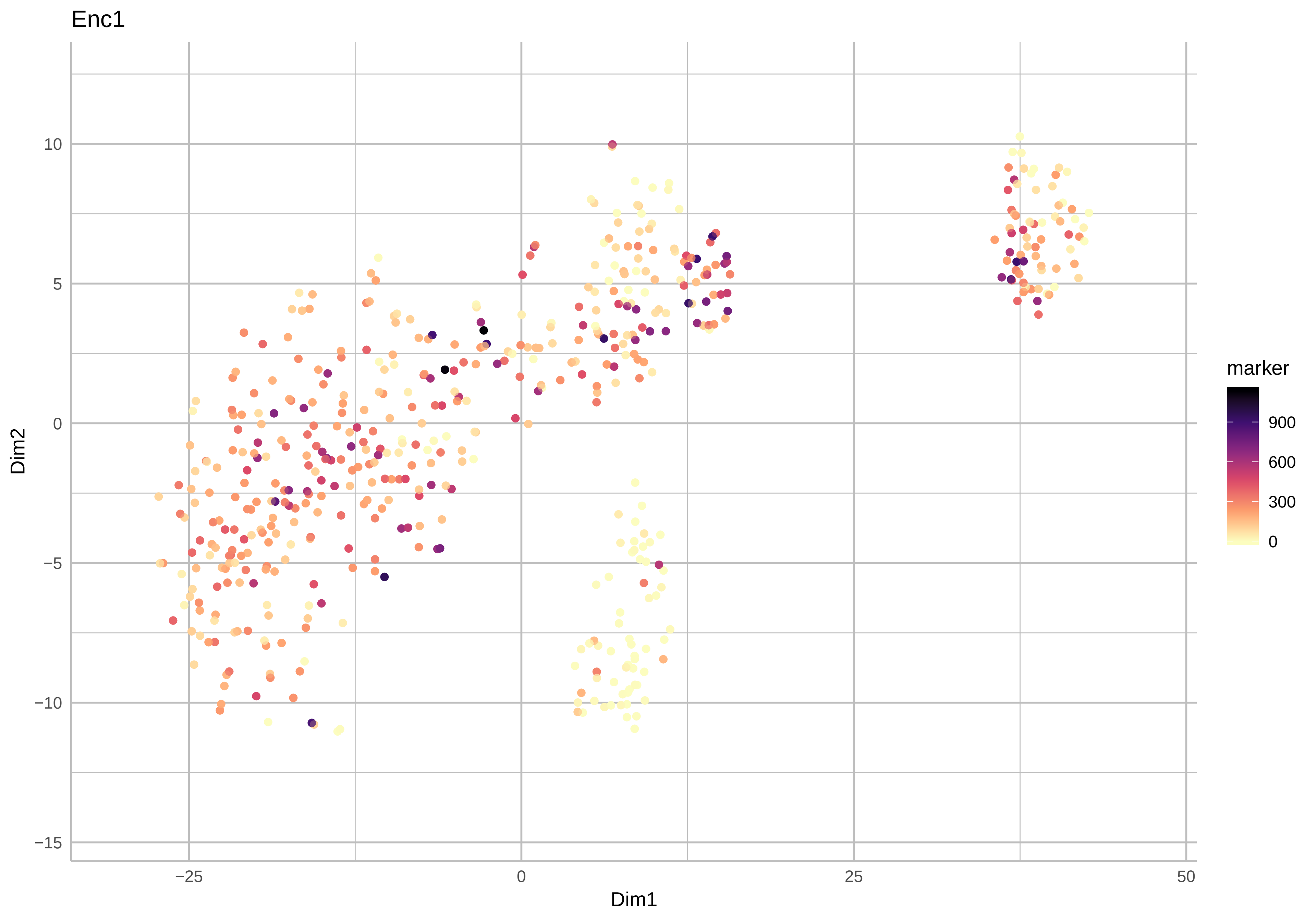

Enc1
Dim2
Dim1
10
5
0
−5
−10
−15
−25
0
25
50
marker
900
600
300
0

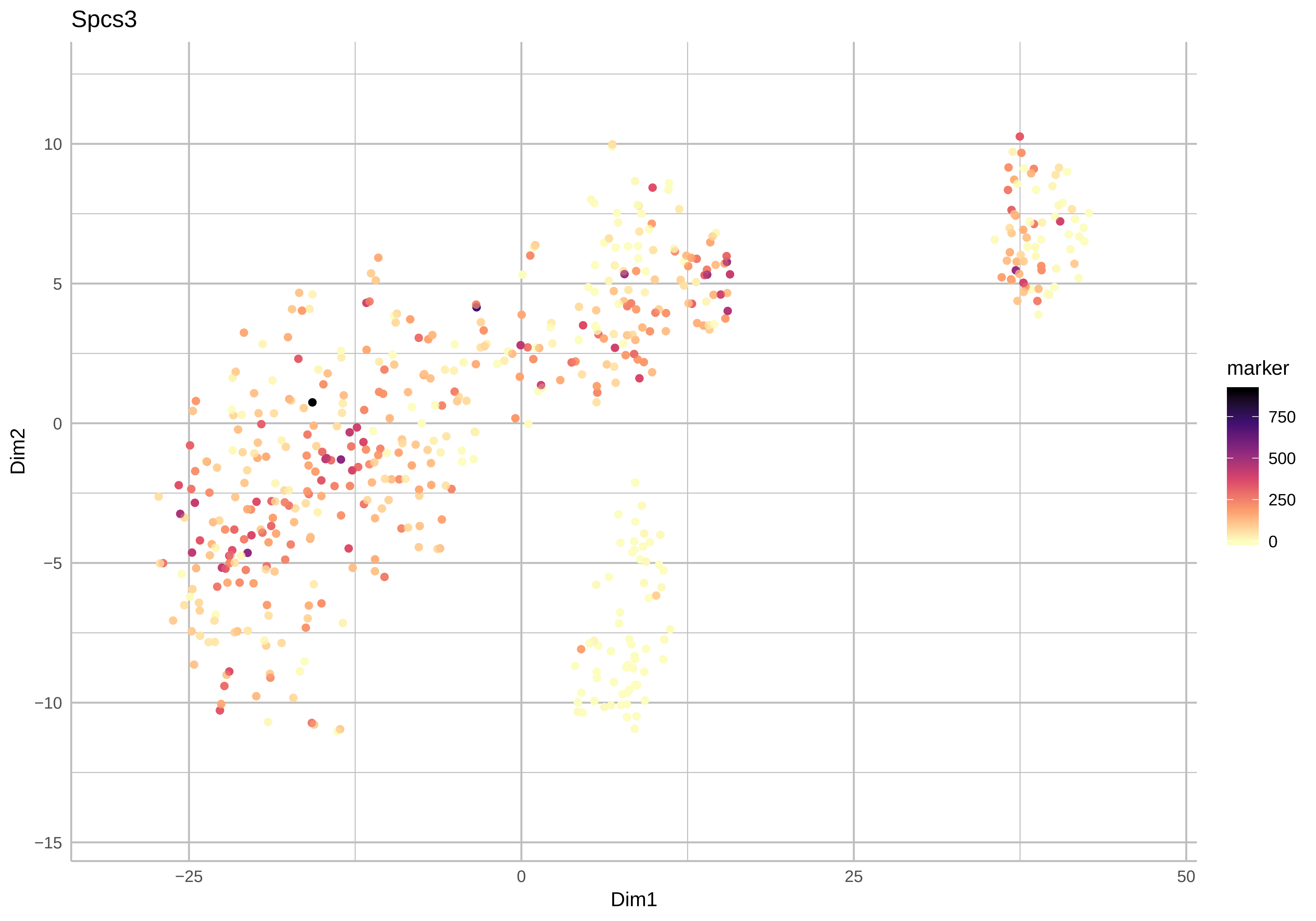
Spcs3
10
5
0
−5
−10
−15
−25
0
25
50
Dim1
Dim2
marker
750
500
250
0

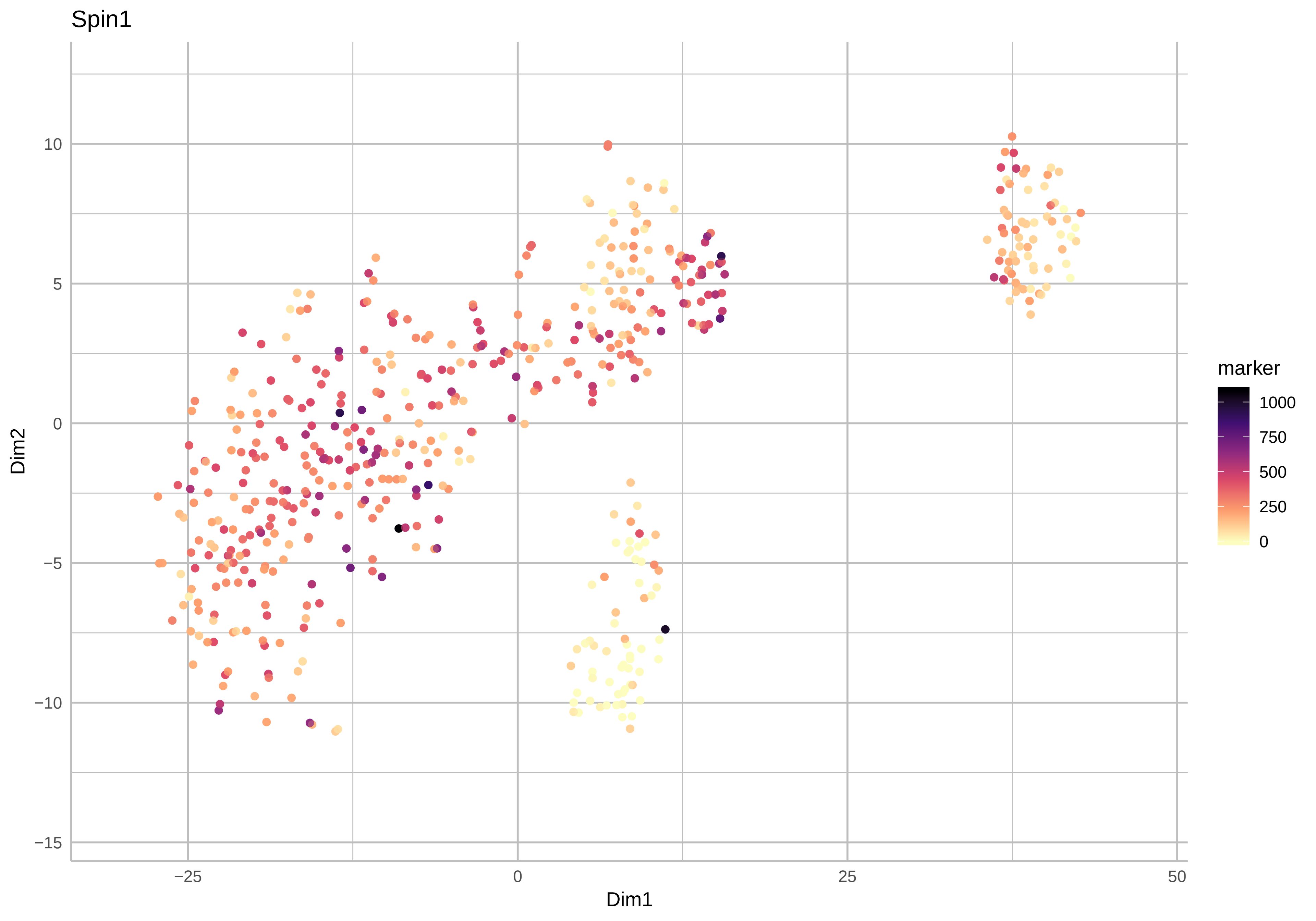

Spin1
Dim2
Dim1
10
5
0
−5
−10
−15
−25
0
25
50
marker
1000
750
500
250
0

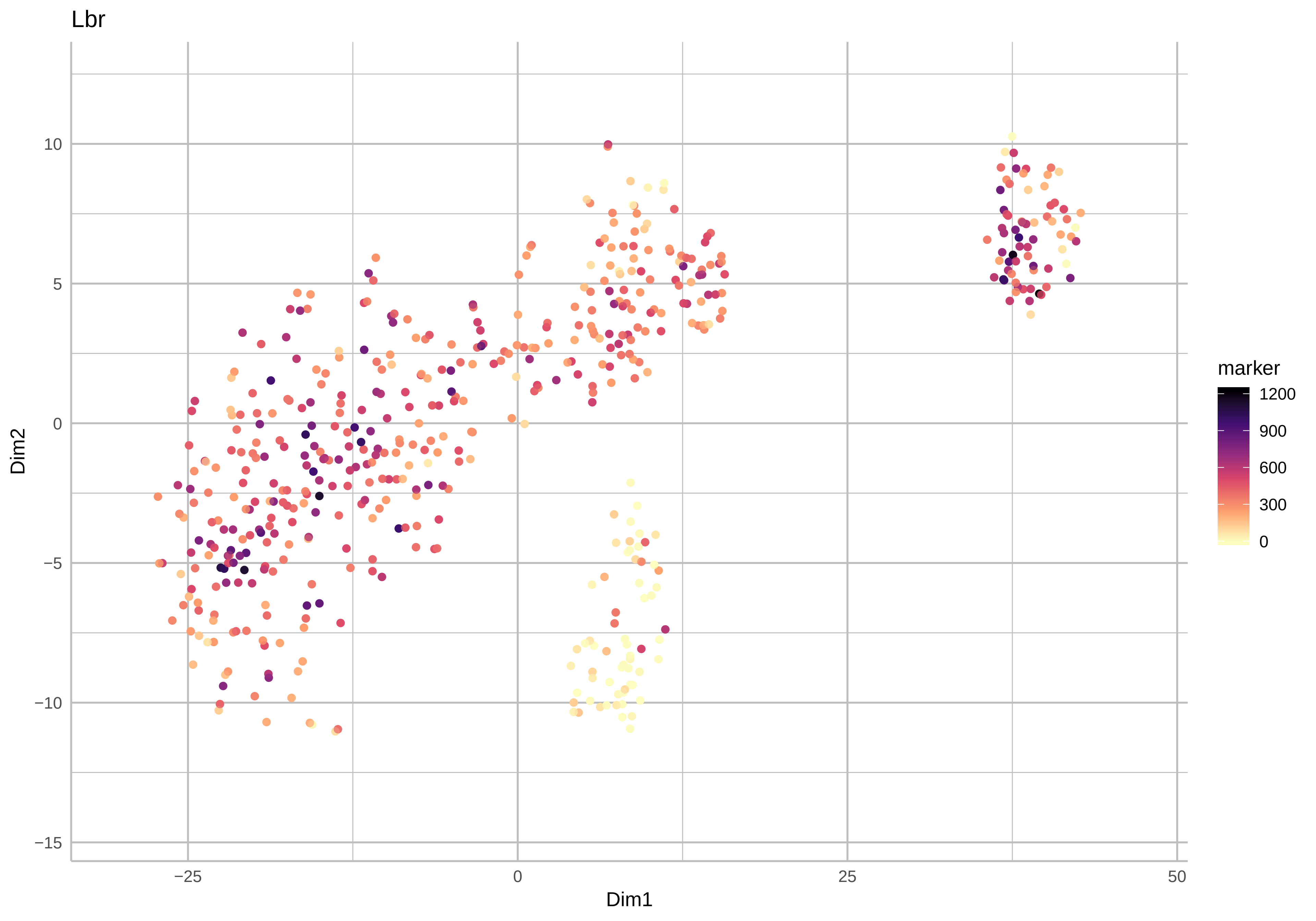
Lbr
Dim2
Dim1
10
5
0
−5
−10
−15
−25
0
25
50
marker
1200
900
600
300
0

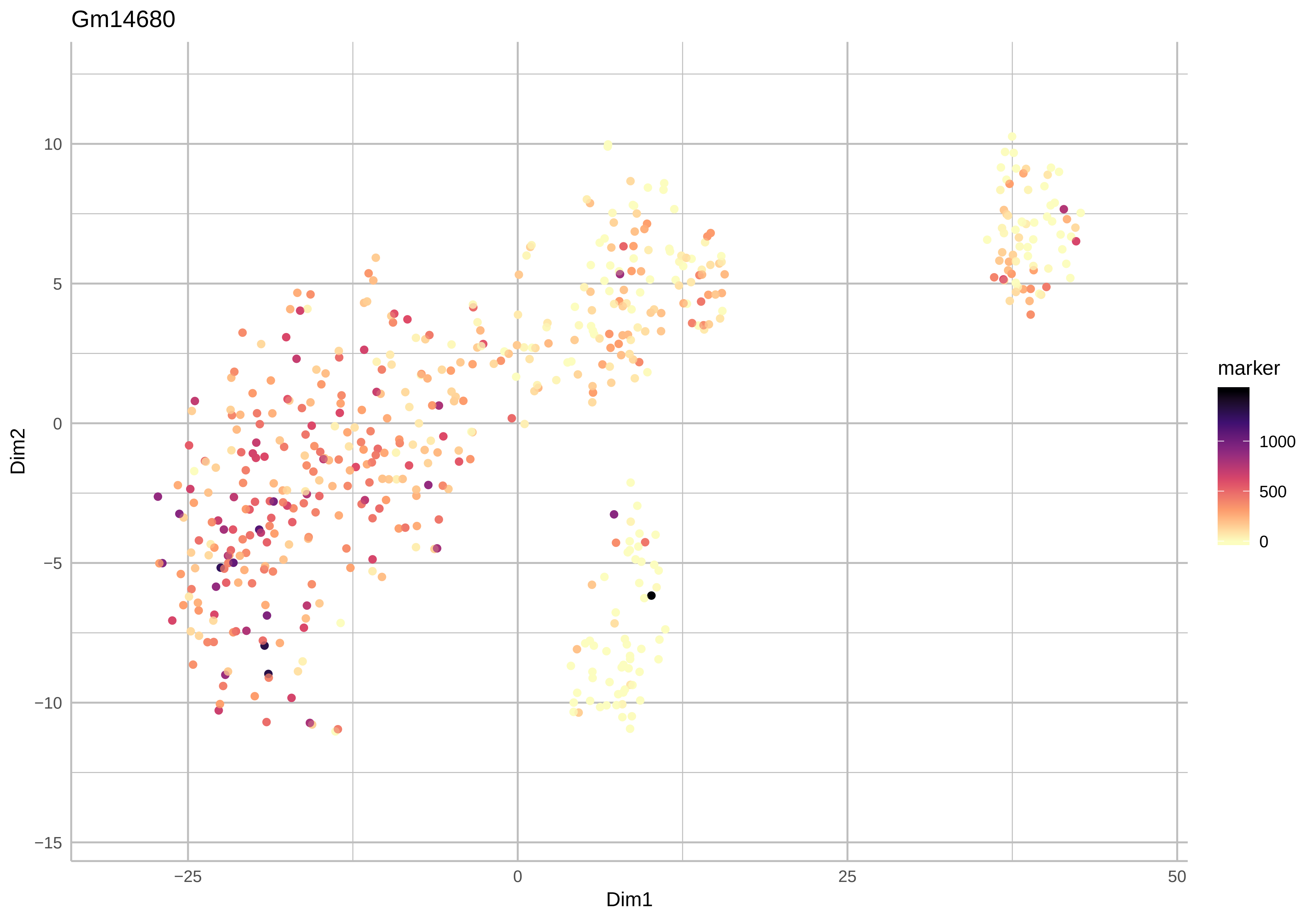
Gm14680
10
5
0
−5
−10
−15
Dim2
−25
0
25
50
Dim1
marker
1000
500
0

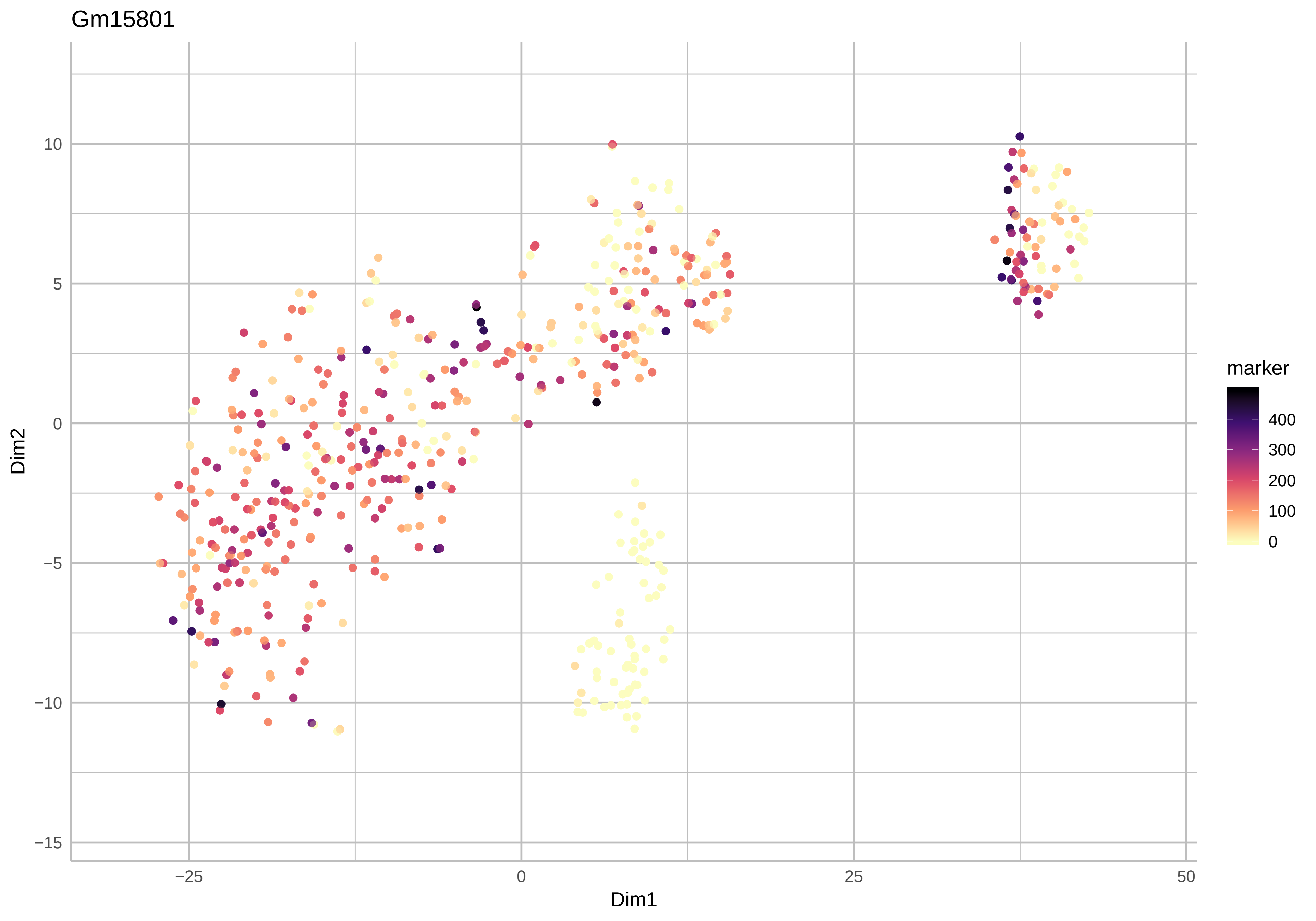
Gm15801
Dim2
Dim1
marker
400
300
200
100
0
10
5
0
−5
−10
−15
−25
0
25
50

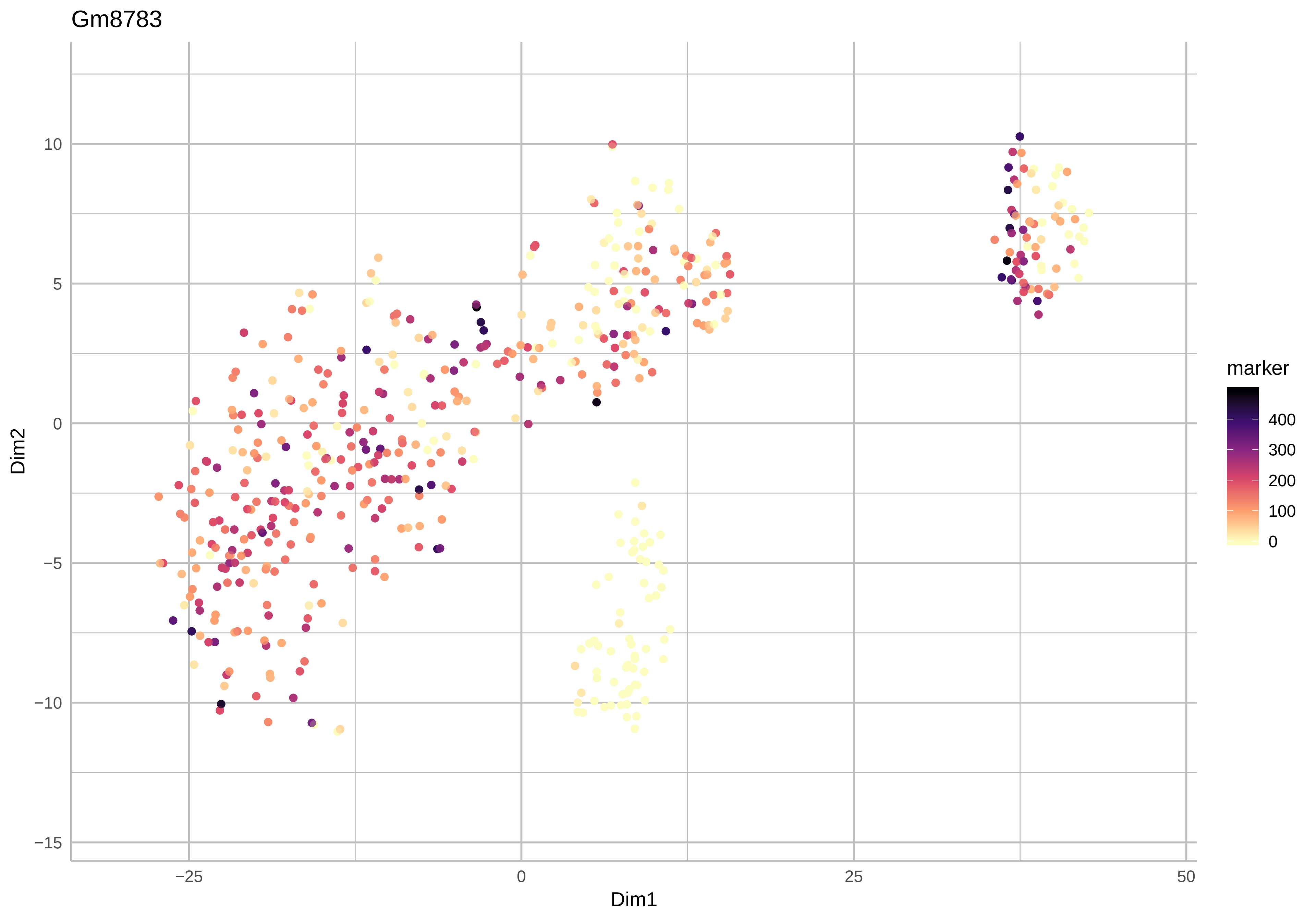

Gm8783
10
5
0
−5
−10
−15
Dim2
−25
0
25
50
Dim1
marker
400
300
200
100
0

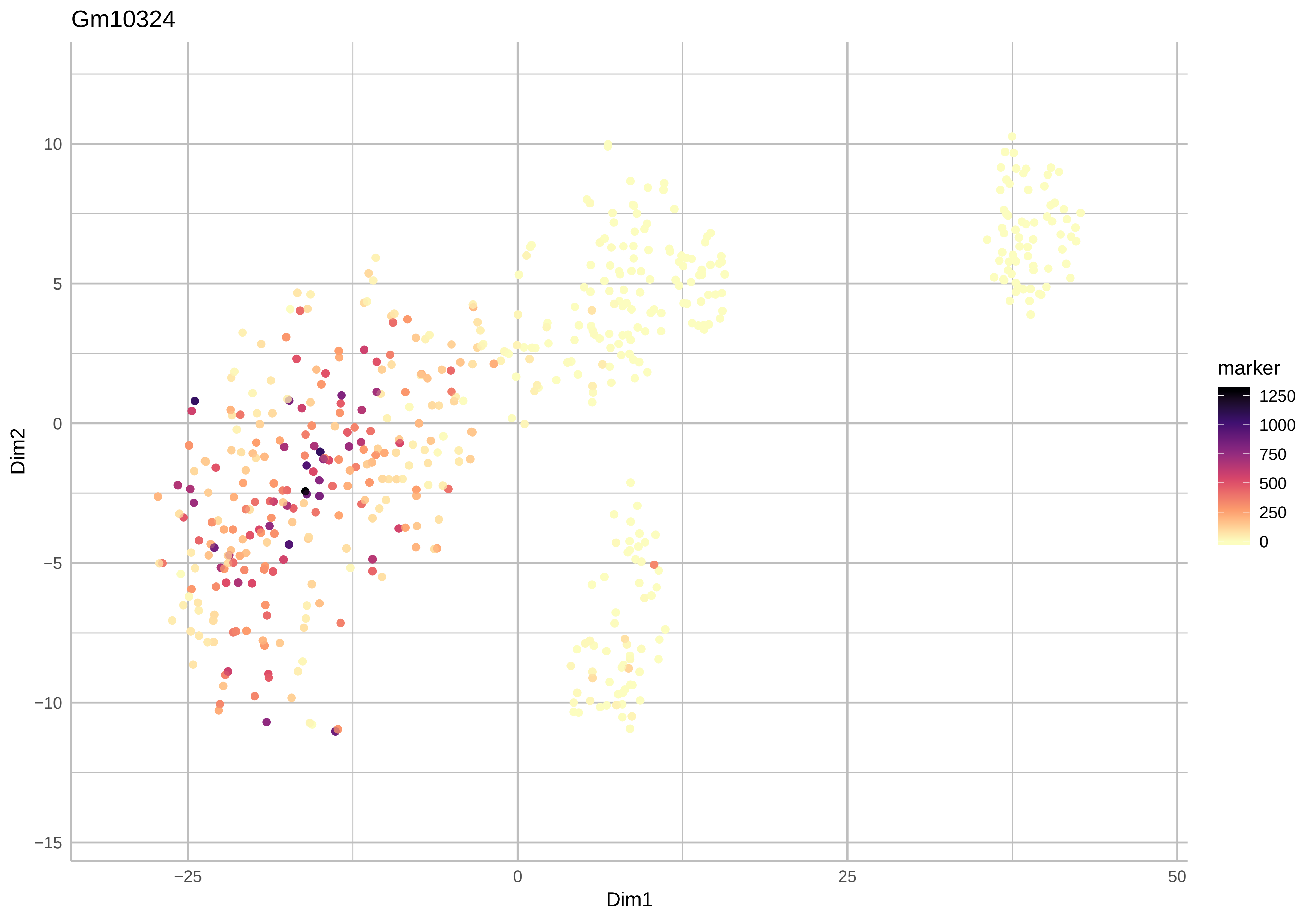

Gm10324
10
5
0
−5
−10
−15
Dim2
−25
0
25
50
Dim1
marker
1250
1000
750
500
250
0

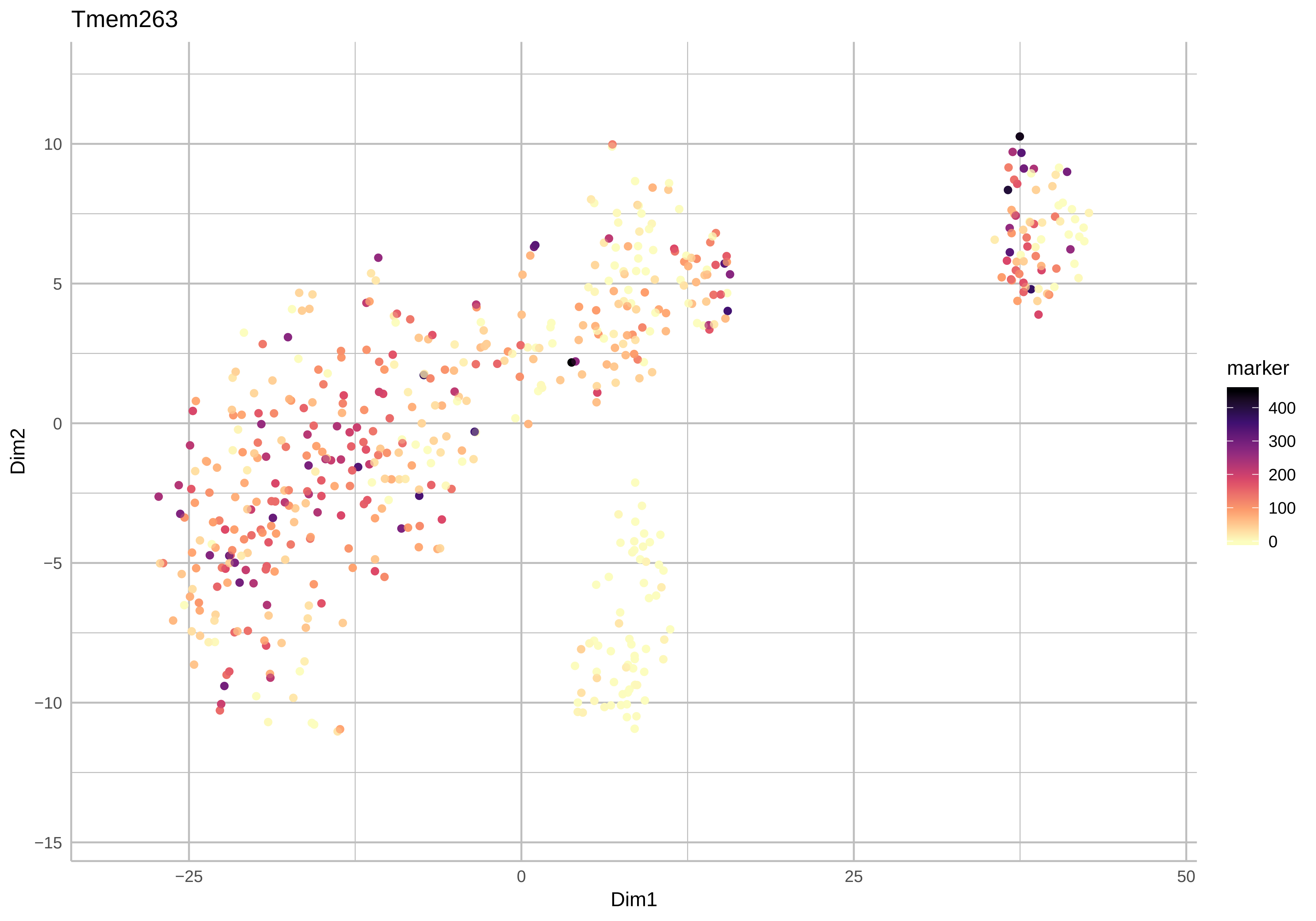
Tmem263
Dim1
Dim2
−25
0
25
50
10
5
0
−5
−10
−15
marker
400
300
200
100
0

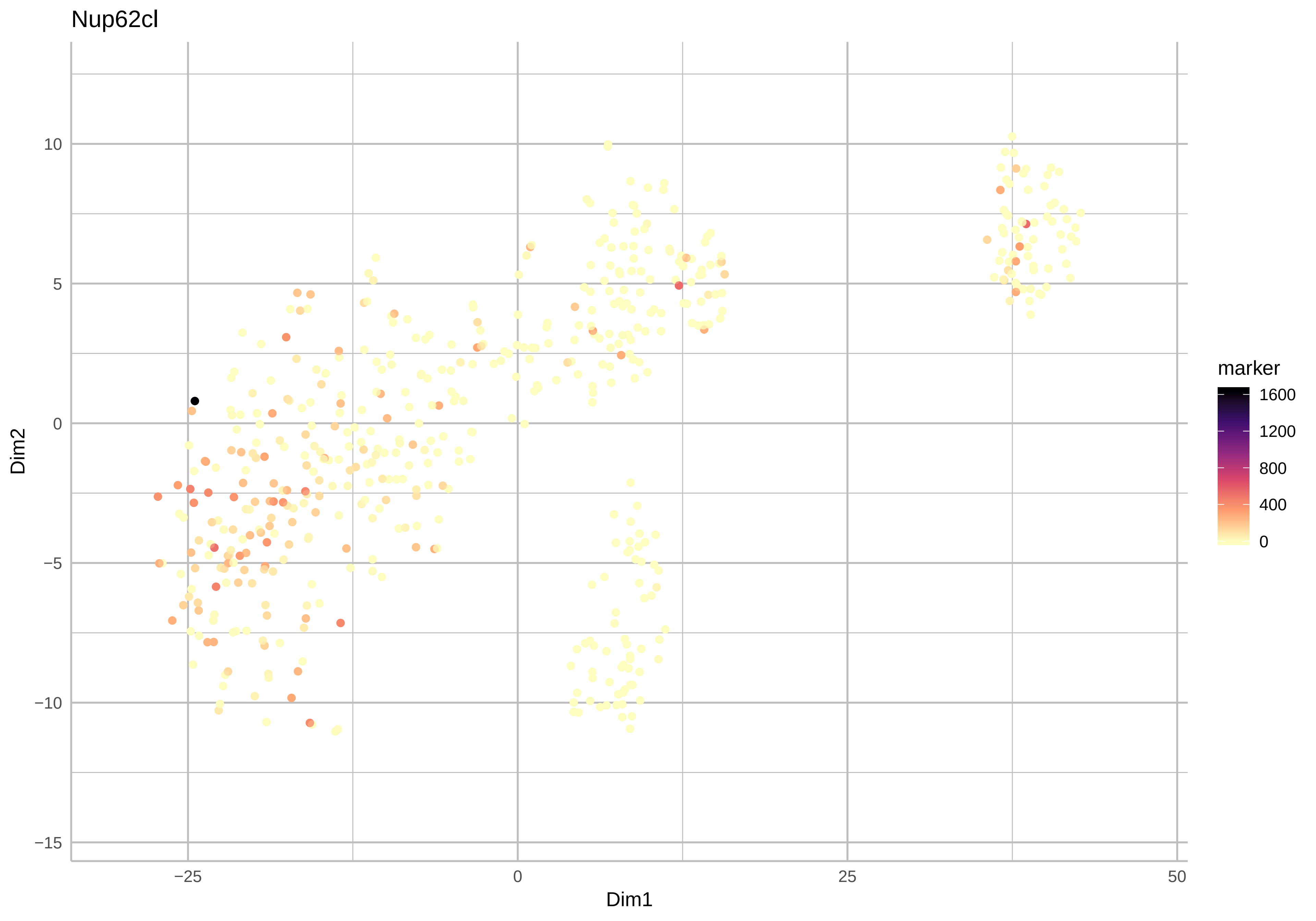
Nup62cl
Dim2
Dim1
10
5
0
−5
−10
−15
−25
0
25
50
marker
1600
1200
800
400
0

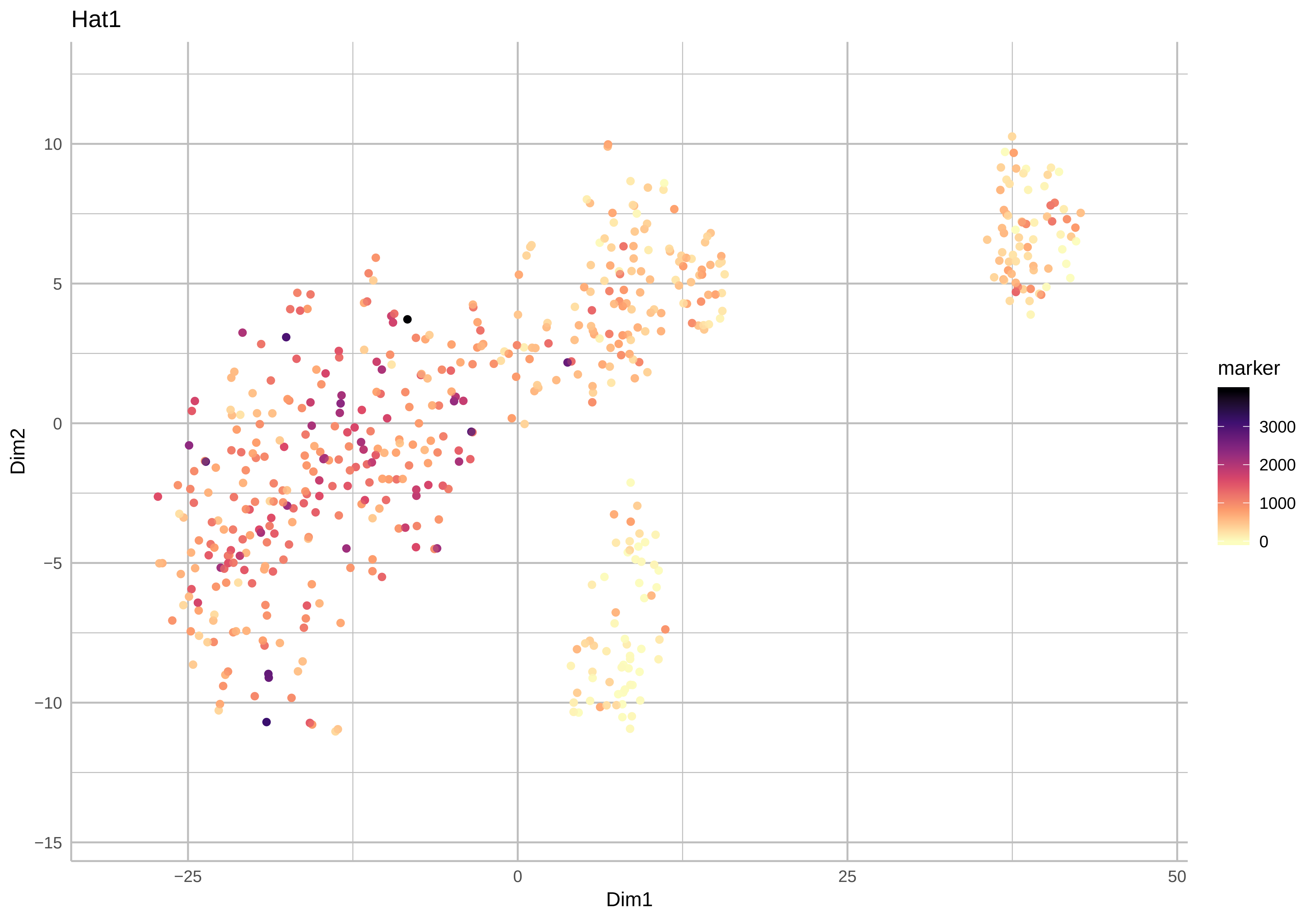
Hat1
Dim2
Dim1
marker
3000
2000
1000
0
10
5
0
−5
−10
−15
−25
0
25
50

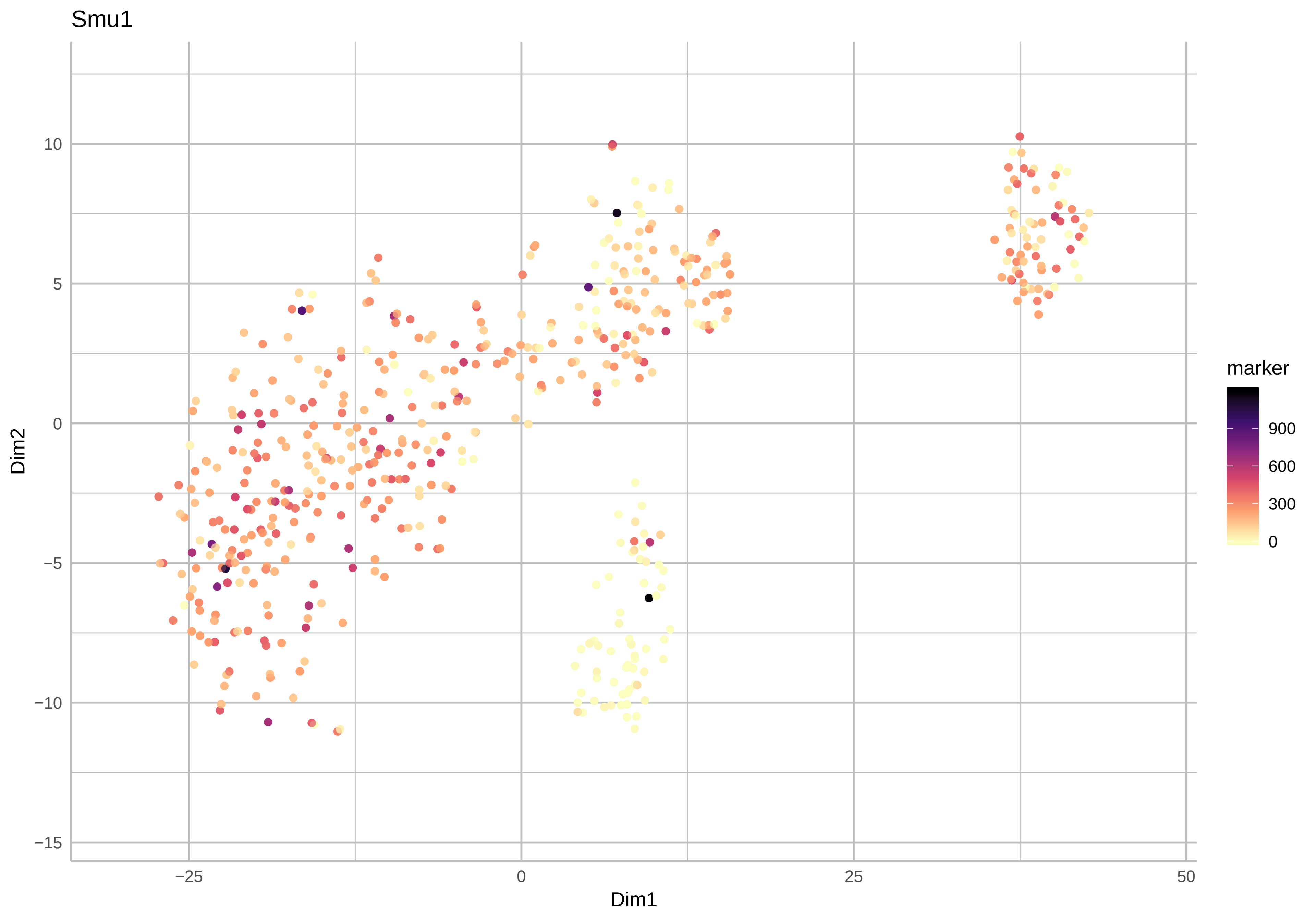
Smu1
Dim2
Dim1
10
5
0
−5
−10
−15
−25
0
25
50
marker
900
600
300
0

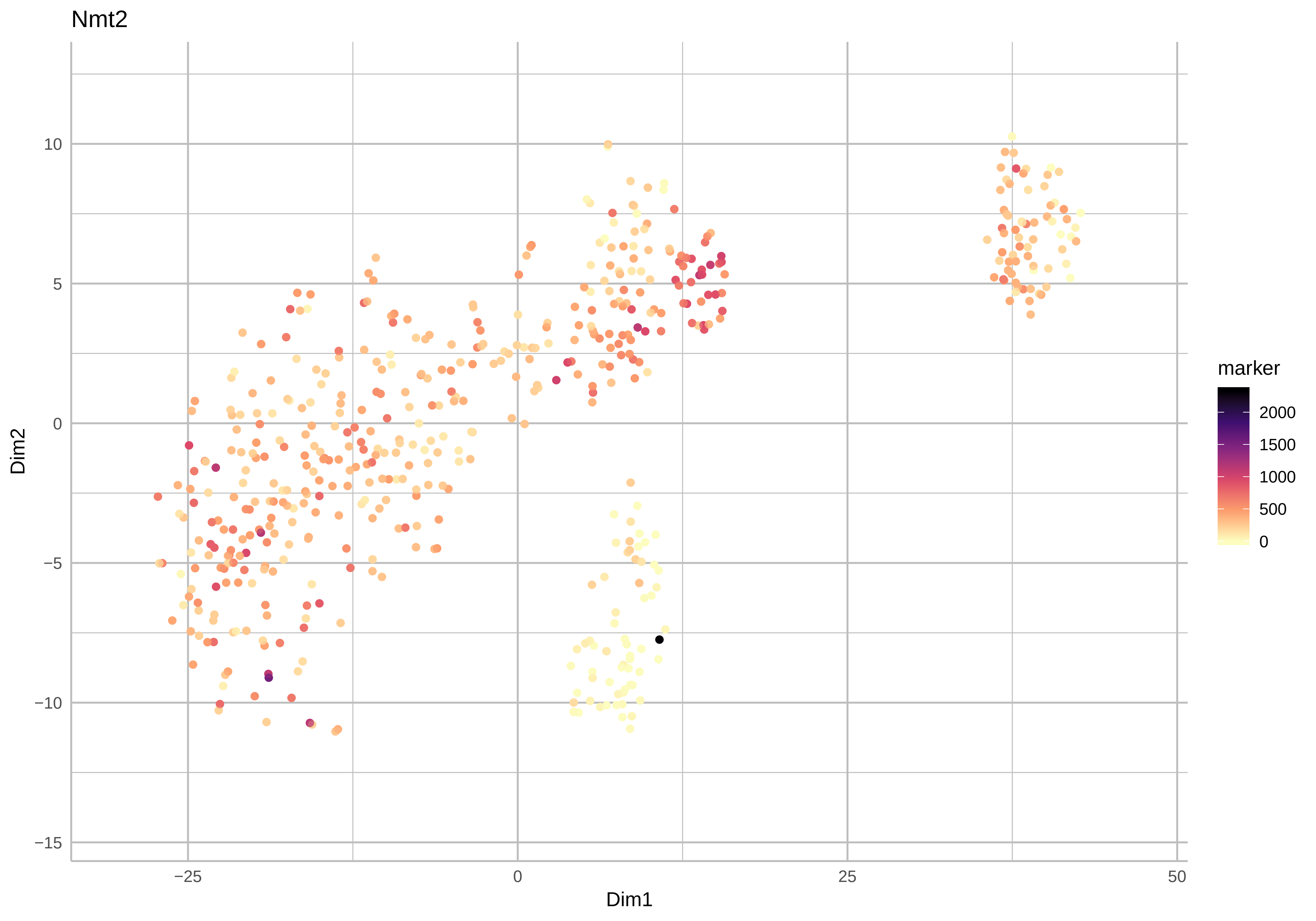
Nmt2
10
5
0
−5
−10
−15
Dim2
−25
0
25
50
Dim1
marker
2000
1500
1000
500
0

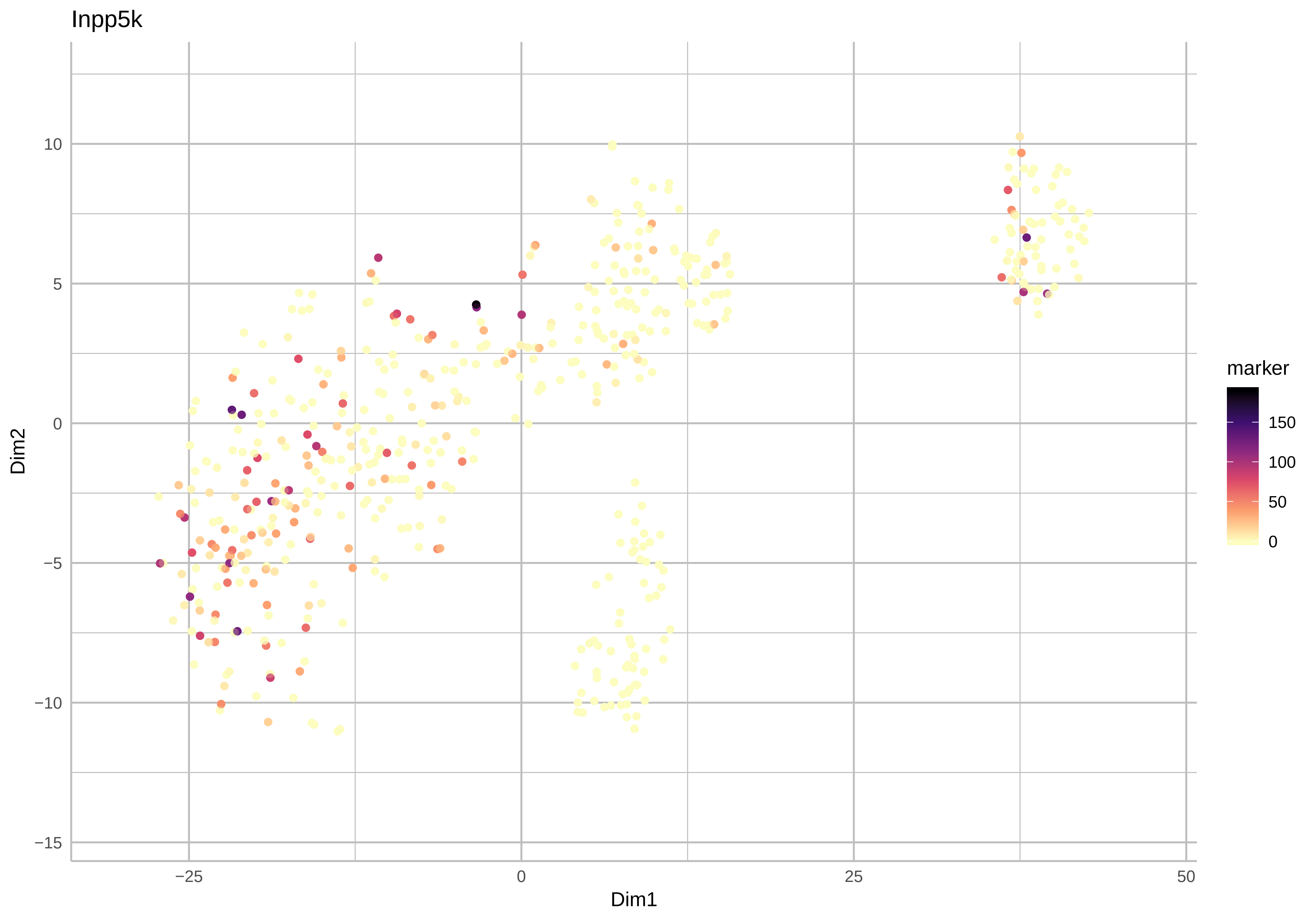

Inpp5k
Dim2
10
5
0
−5
−10
−15
−25
0
25
50
Dim1
marker
150
100
50
0

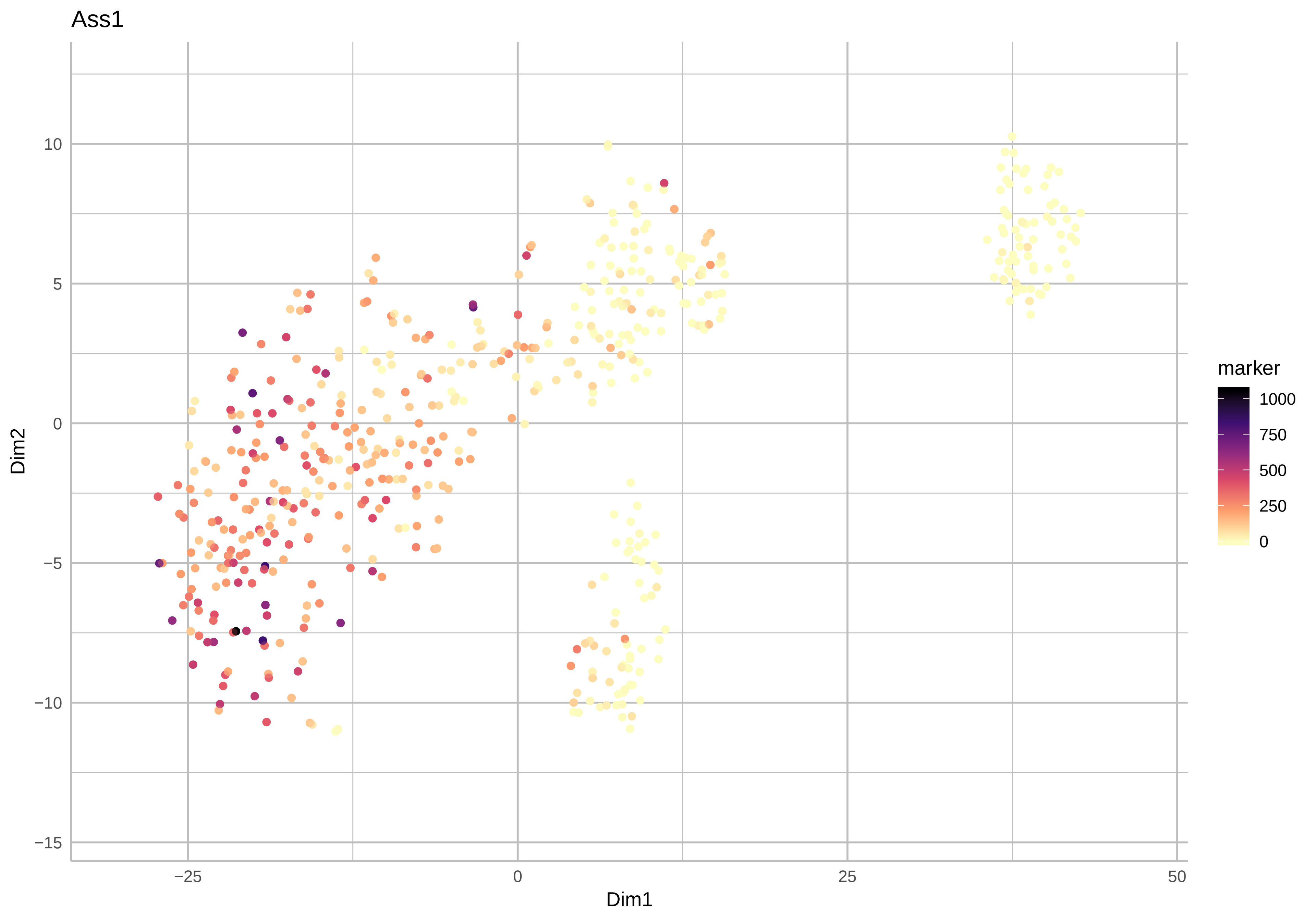

Ass1
Dim2
Dim1
10
5
0
−5
−10
−15
−25
0
25
50
marker
1000
750
500
250
0

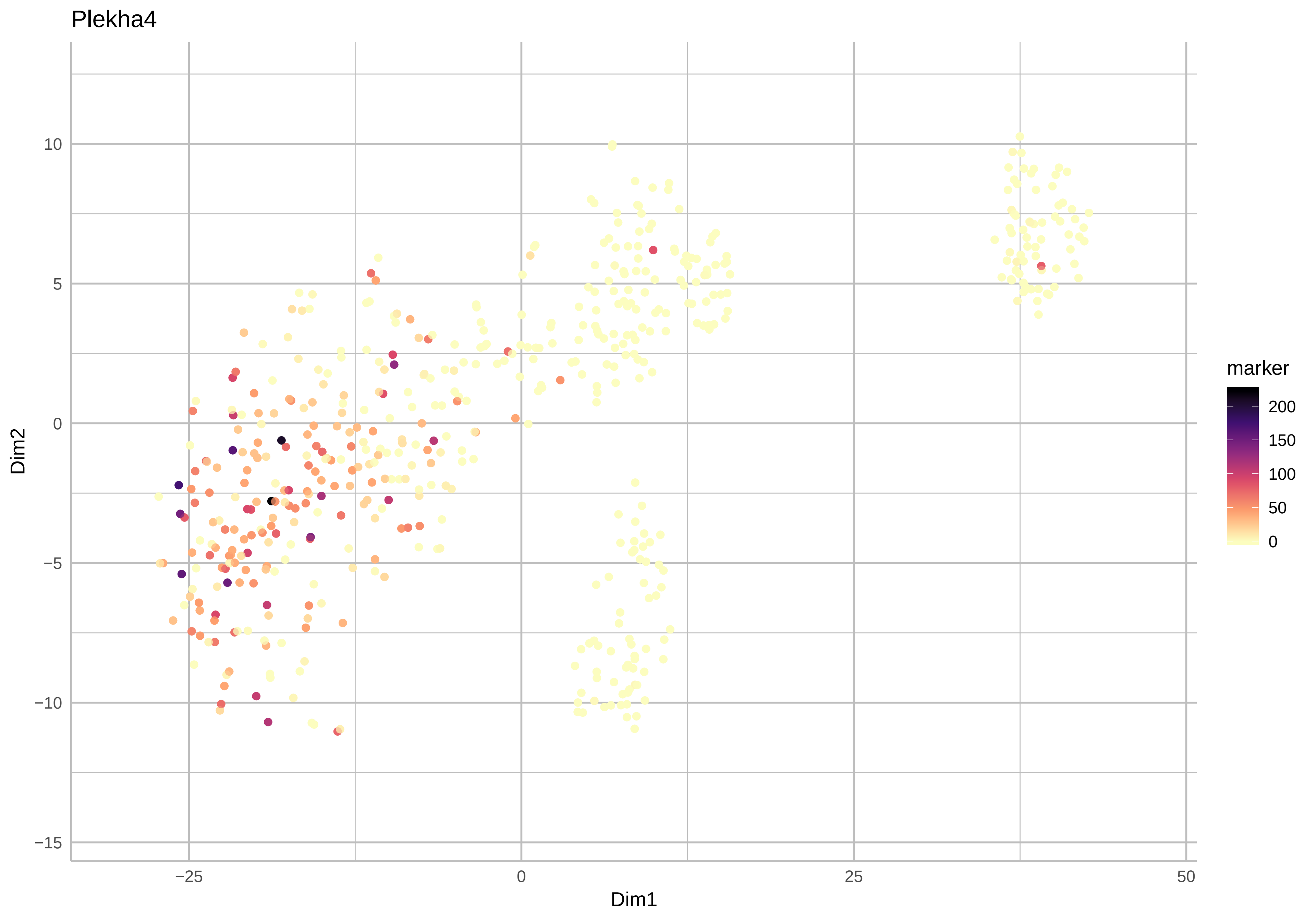

Plekha4
Dim2
Dim1
10
5
0
−5
−10
−15
−25
0
25
50
marker
200
150
100
50
0

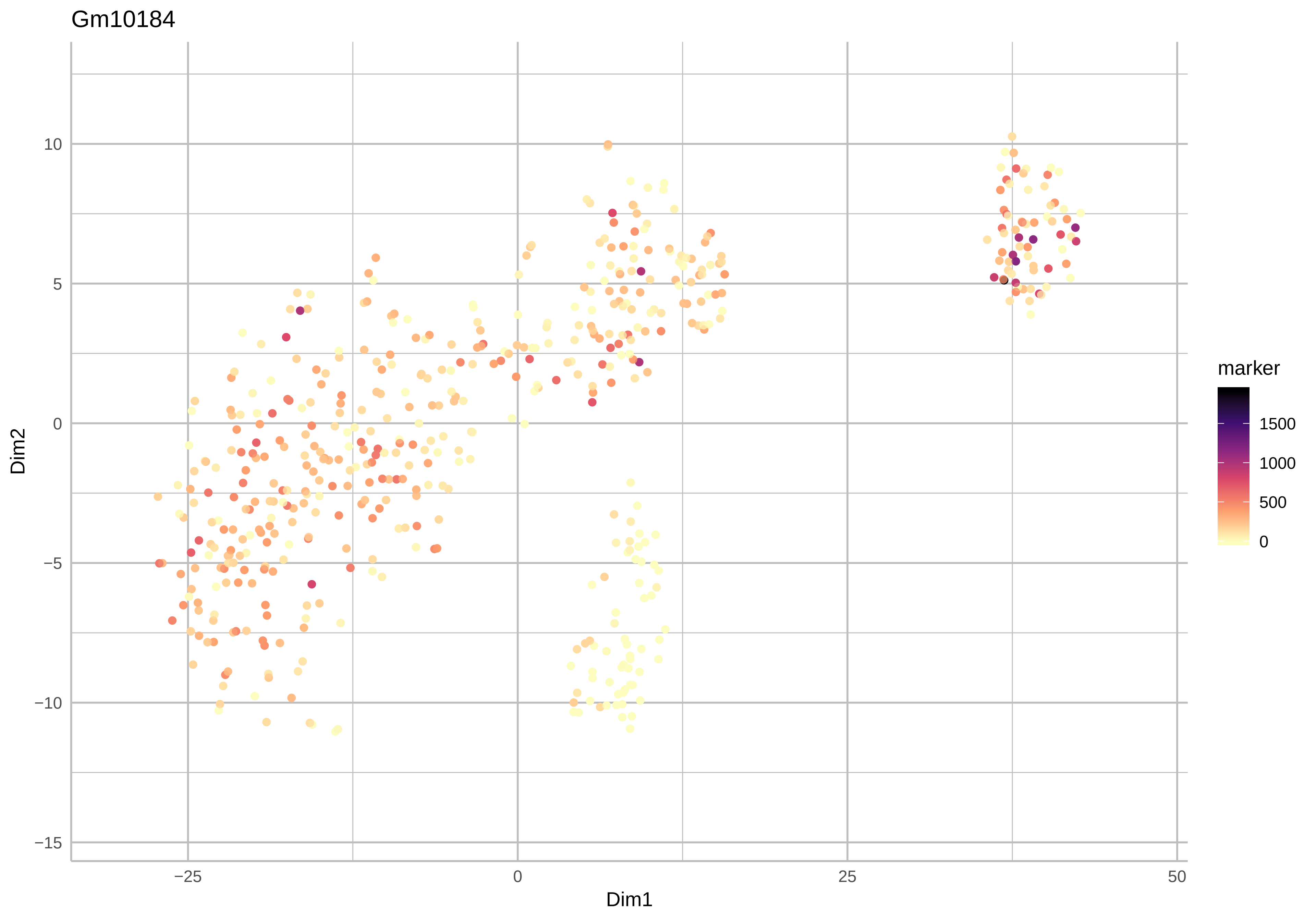
Gm10184
Dim2
Dim1
10
5
0
−5
−10
−15
−25
0
25
50
marker
1500
1000
500
0

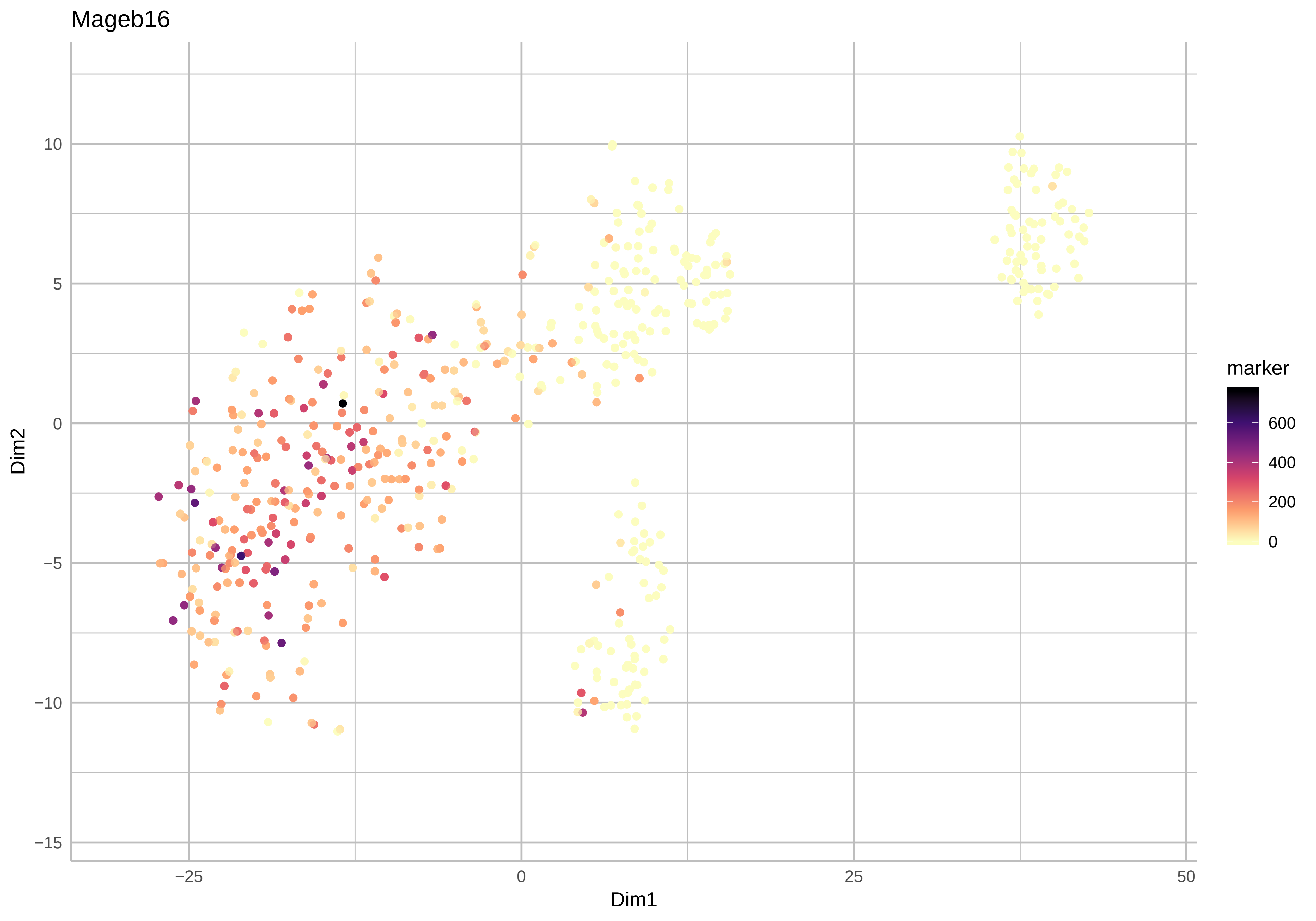
Mageb16
marker
600
400
200
0
Dim2
10
5
0
−5
−10
−15
−25
0
25
50
Dim1

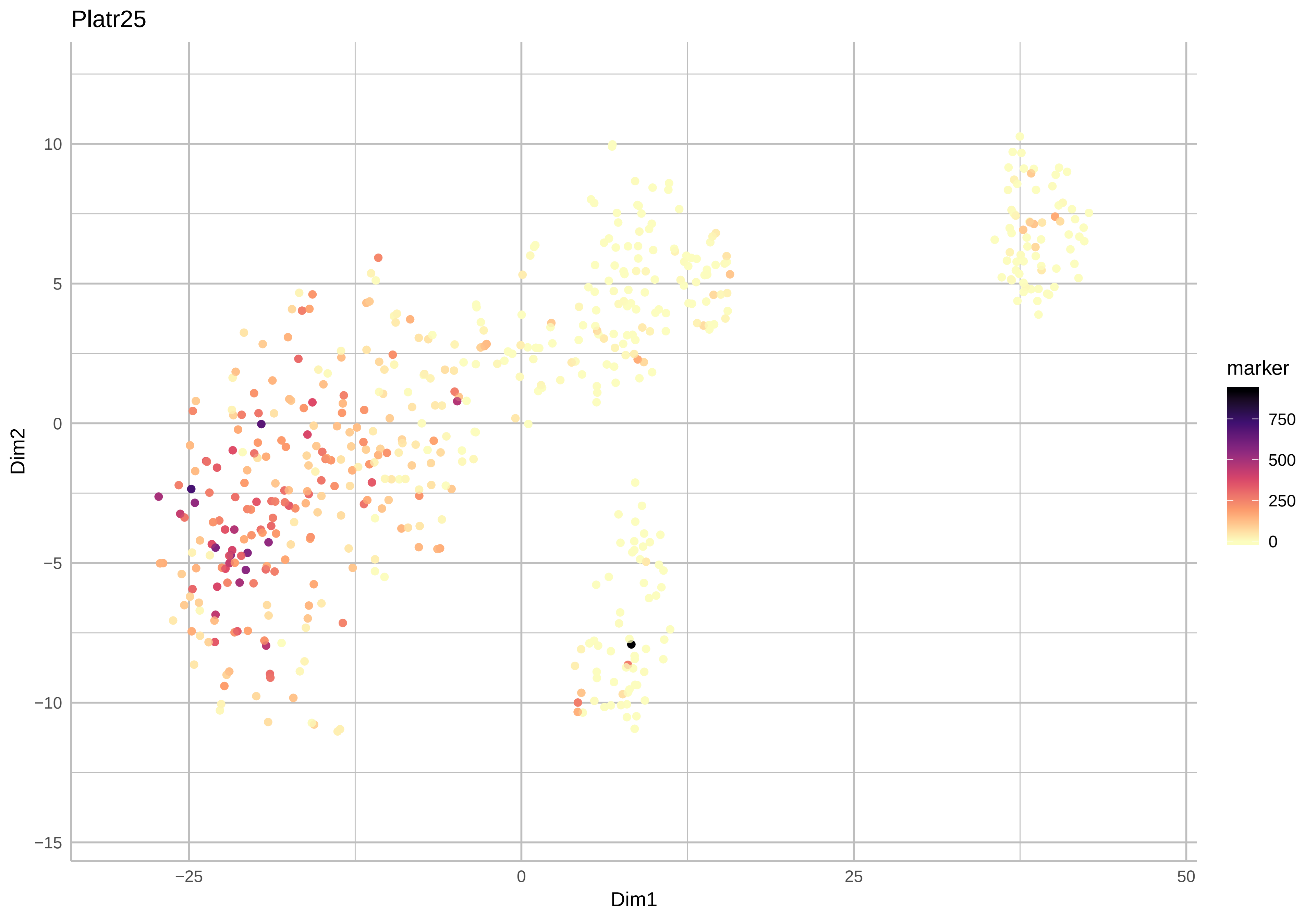
Platr25
Dim2
10
5
0
−5
−10
−15
Dim1
−25
0
25
50
marker
750
500
250
0

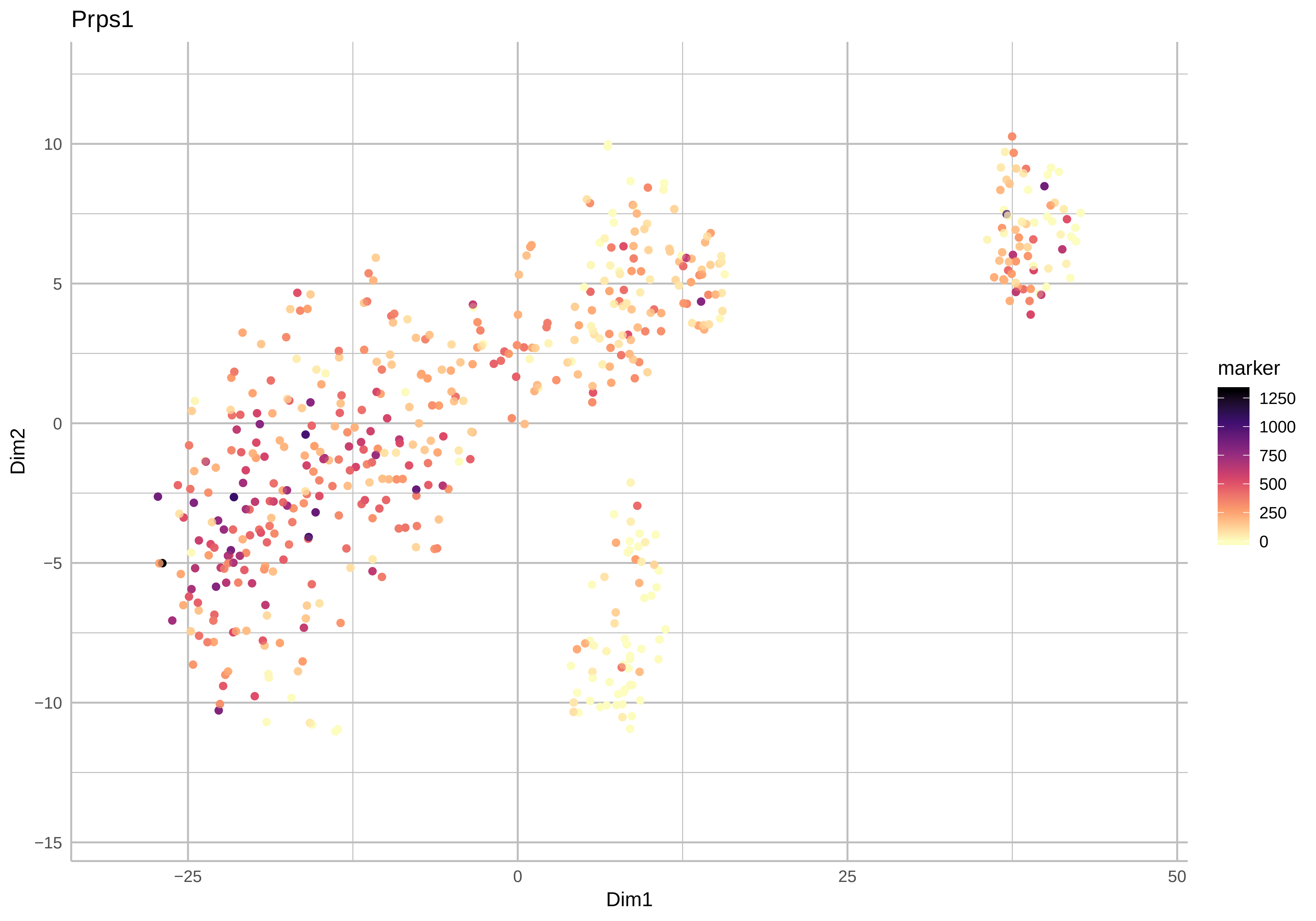

Prps1
Dim2
Dim1
10
5
0
−5
−10
−15
−25
0
25
50
marker
1250
1000
750
500
250
0

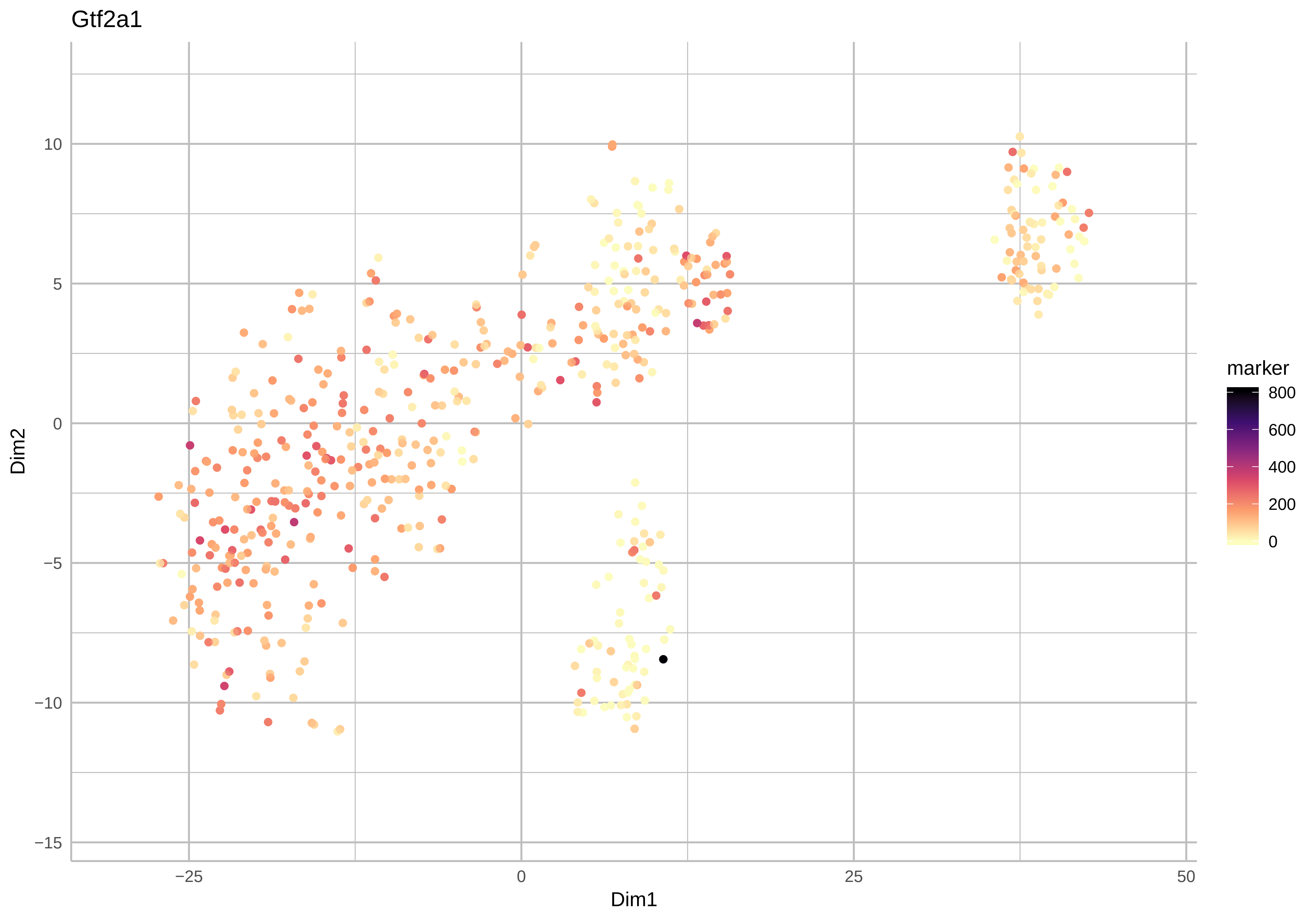
Gtf2a1
Dim2
Dim1
marker
800
600
400
200
0
10
5
0
−5
−10
−15
−25
0
25
50

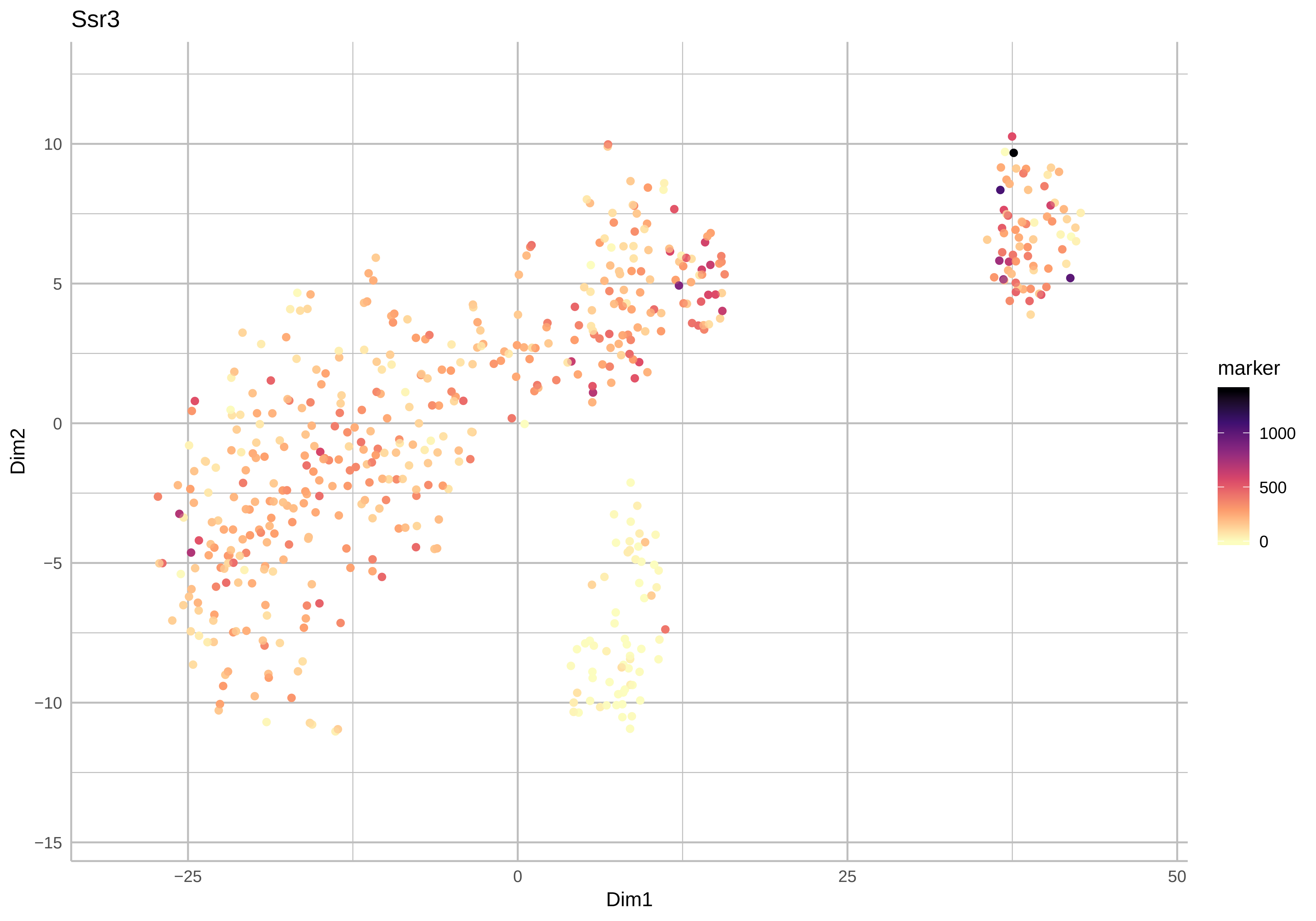
Ssr3
Dim2
Dim1
marker
1000
500
0
10
5
0
−5
−10
−15
−25
0
25
50

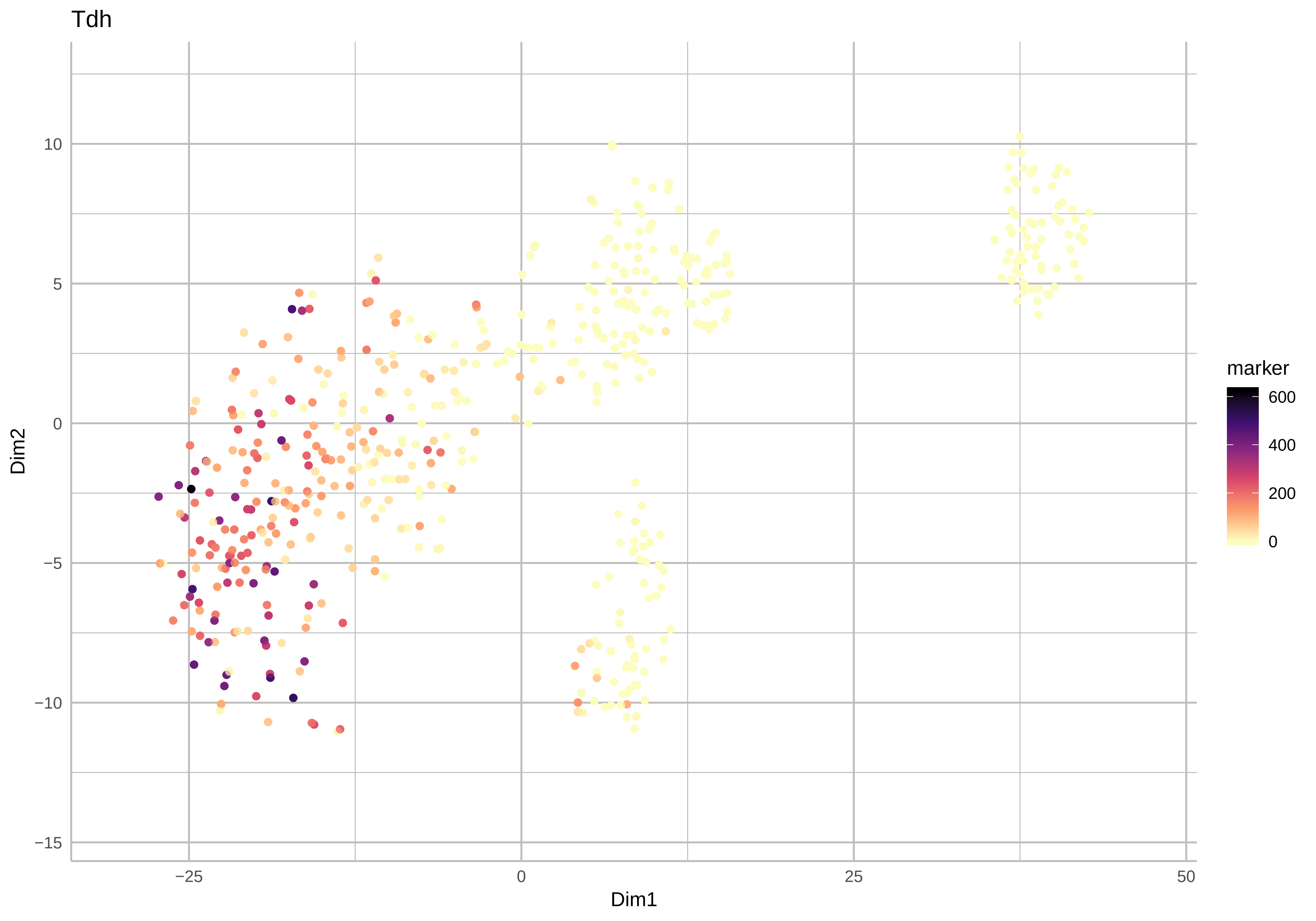

Tdh
10
5
0
−5
−10
−15
Dim2
−25
0
25
50
Dim1
marker
600
400
200
0

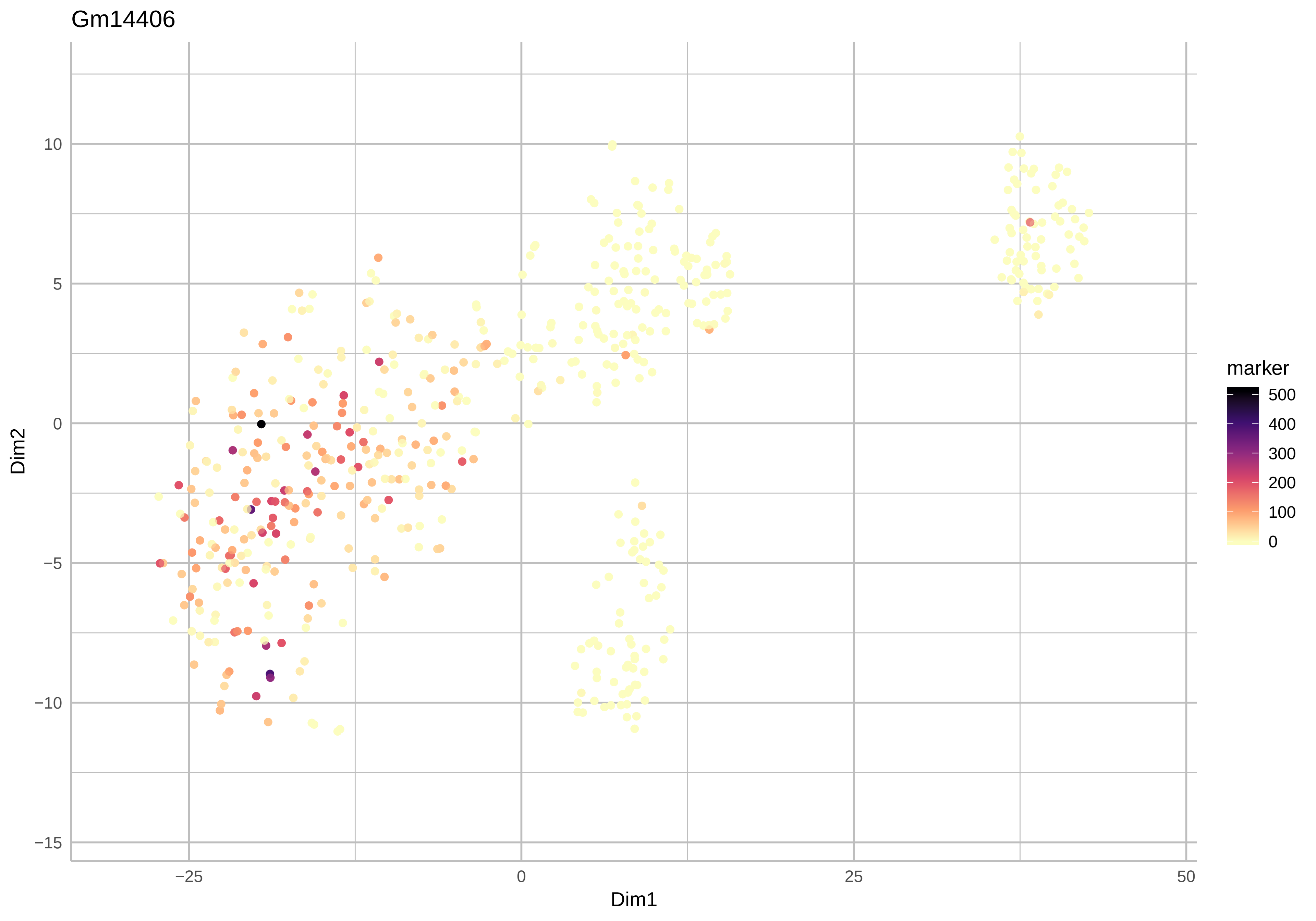
Gm14406
Dim2
Dim1
marker
500
400
300
200
100
0
10
5
0
−5
−10
−15
−25
0
25
50

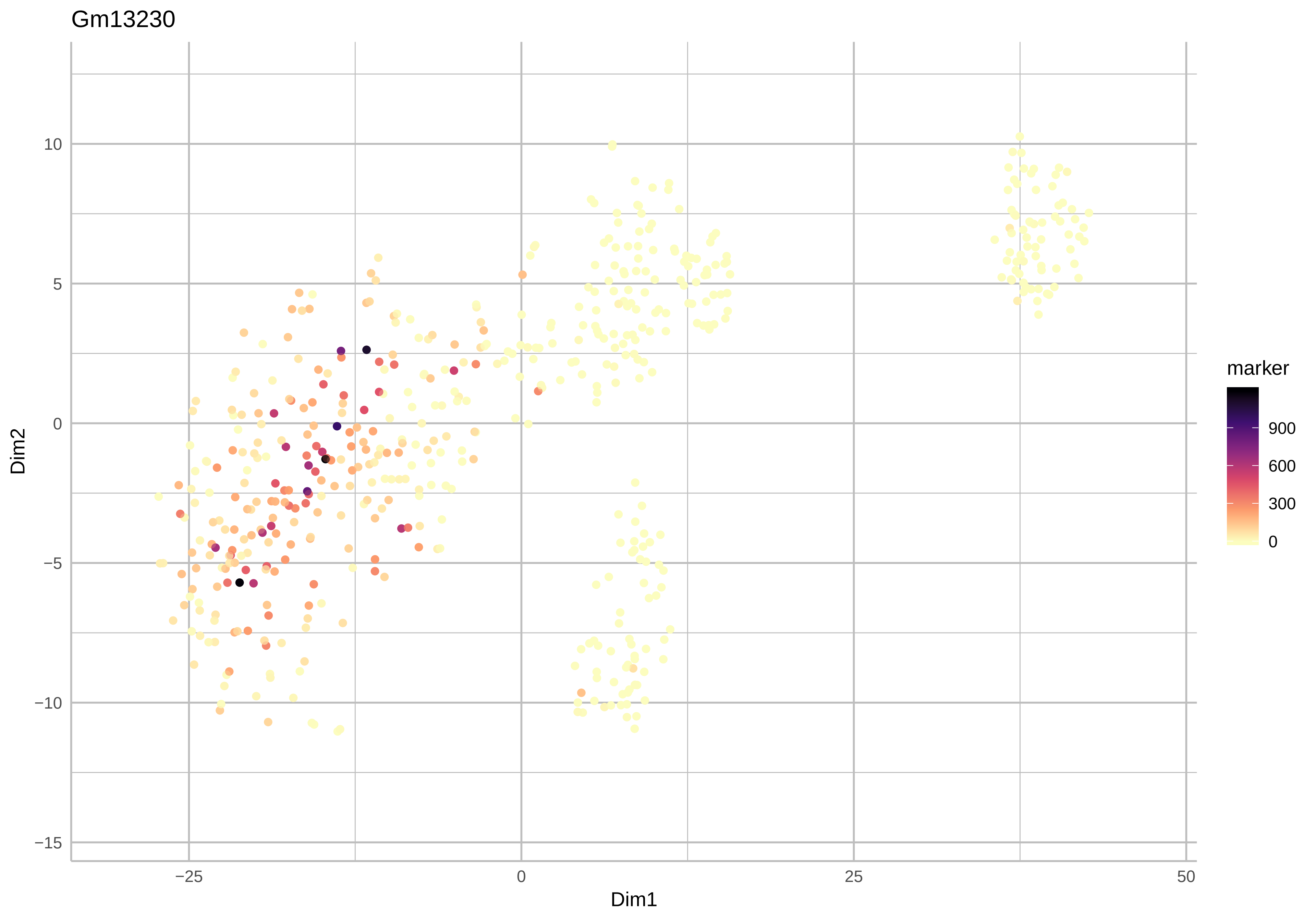
Gm13230
Dim2
Dim1
10
5
0
−5
−10
−15
−25
0
25
50
marker
900
600
300
0

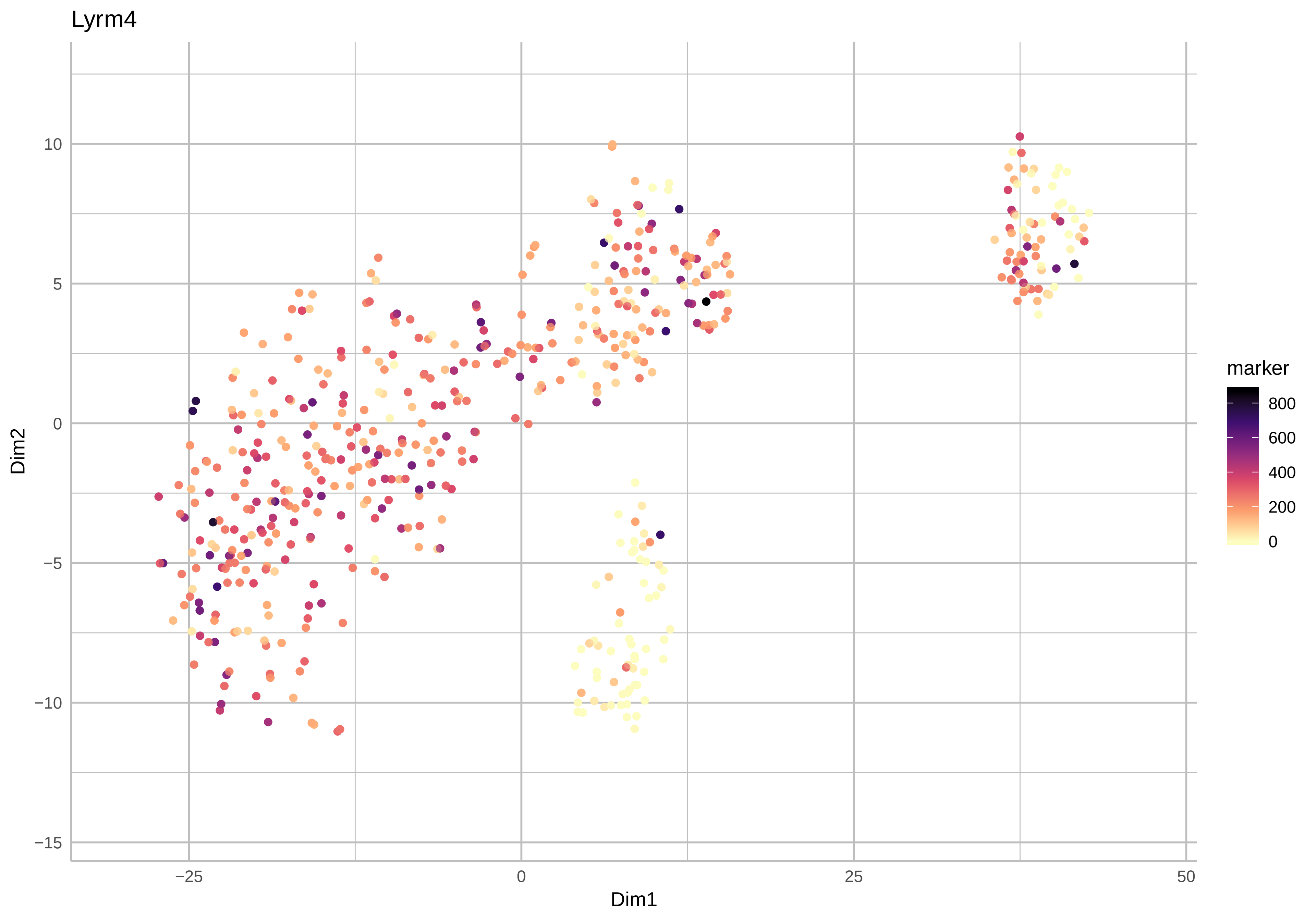

Lyrm4
10
5
0
−5
−10
−15
Dim2
−25
0
25
50
Dim1
marker
800
600
400
200
0

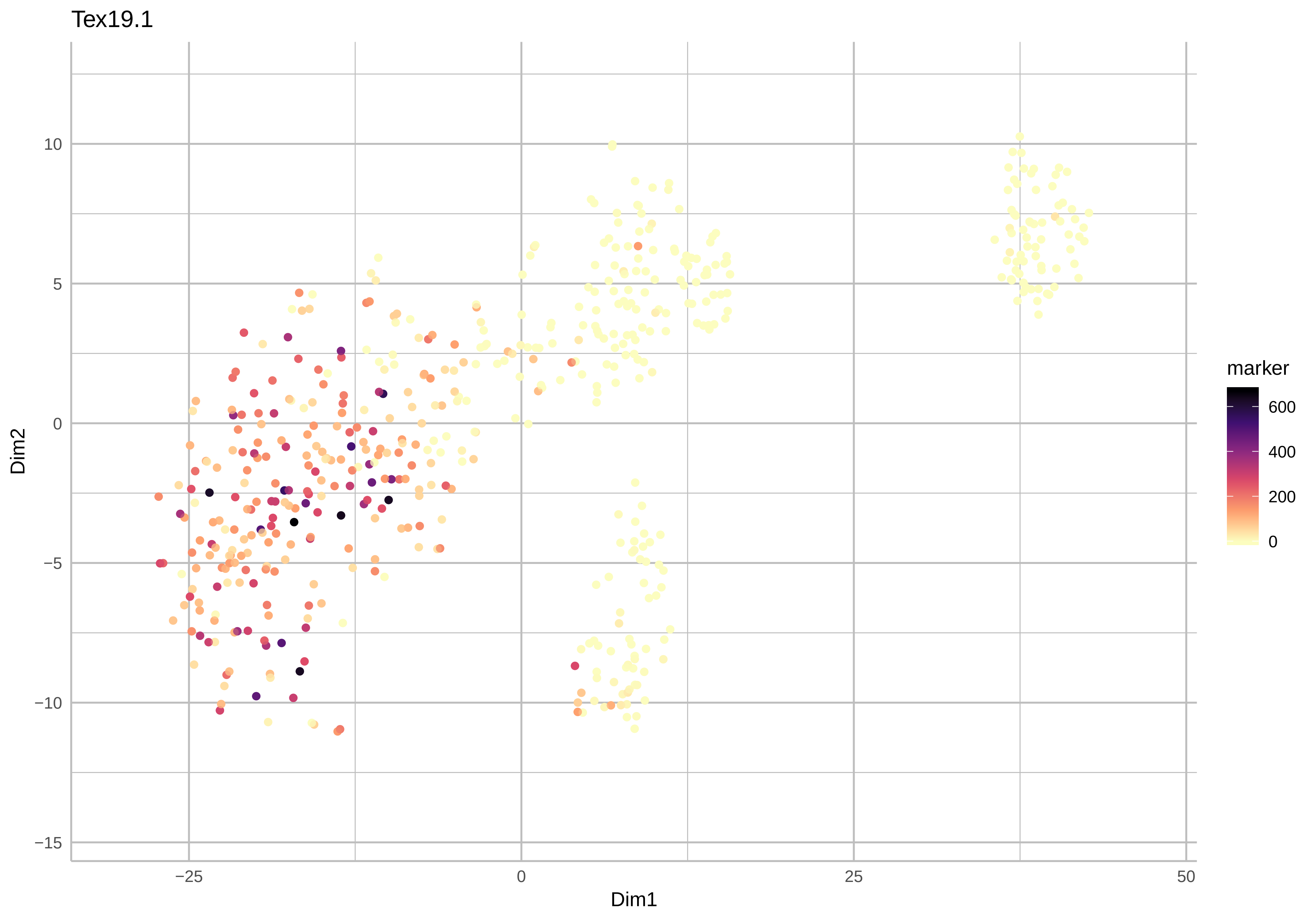

Tex19.1
Dim2
Dim1
10
5
0
−5
−10
−15
−25
0
25
50
marker
600
400
200
0

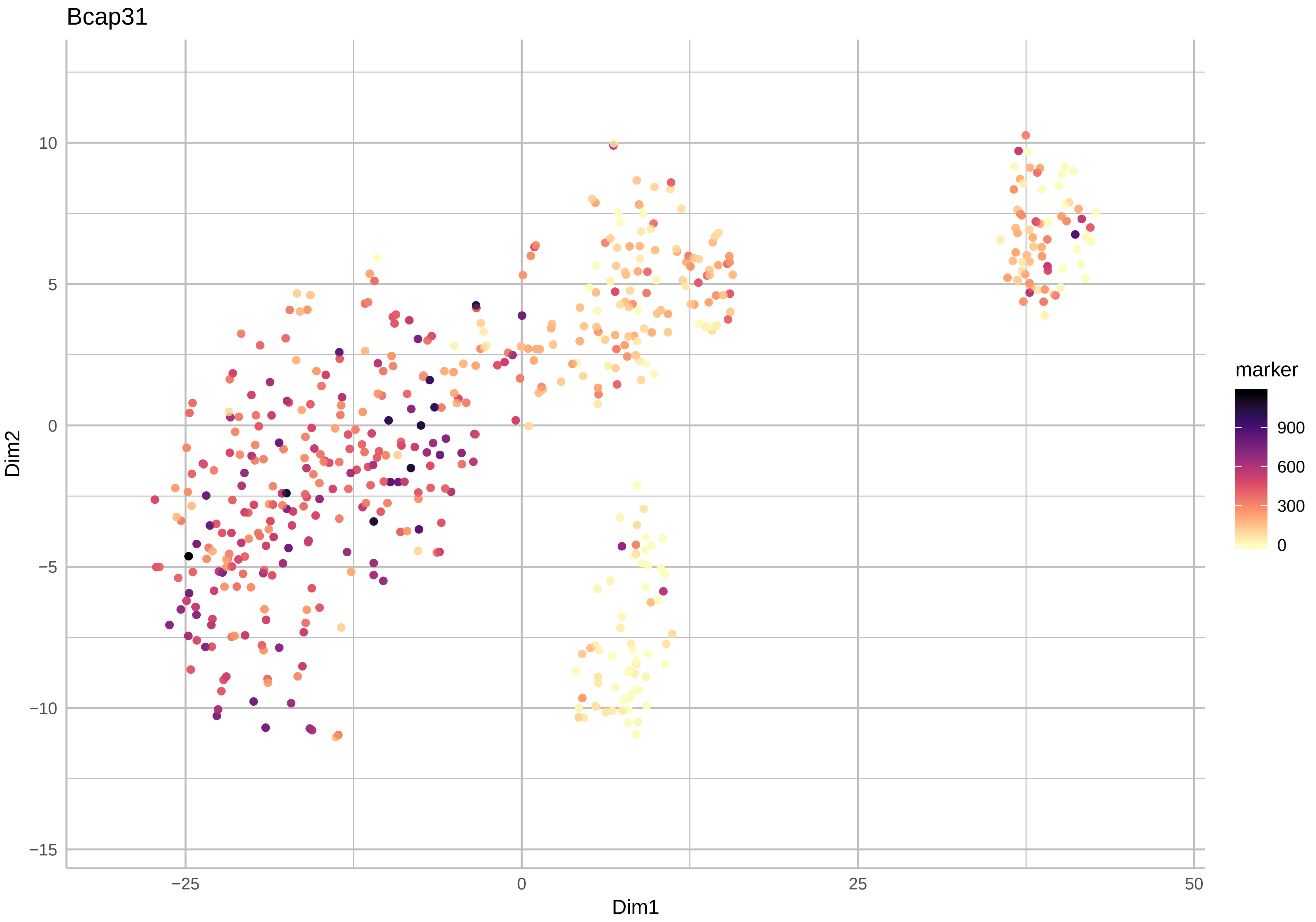

Bcap31
Dim2
Dim1
marker
900
600
300
0
10
5
0
−5
−10
−15
−25
0
25
50

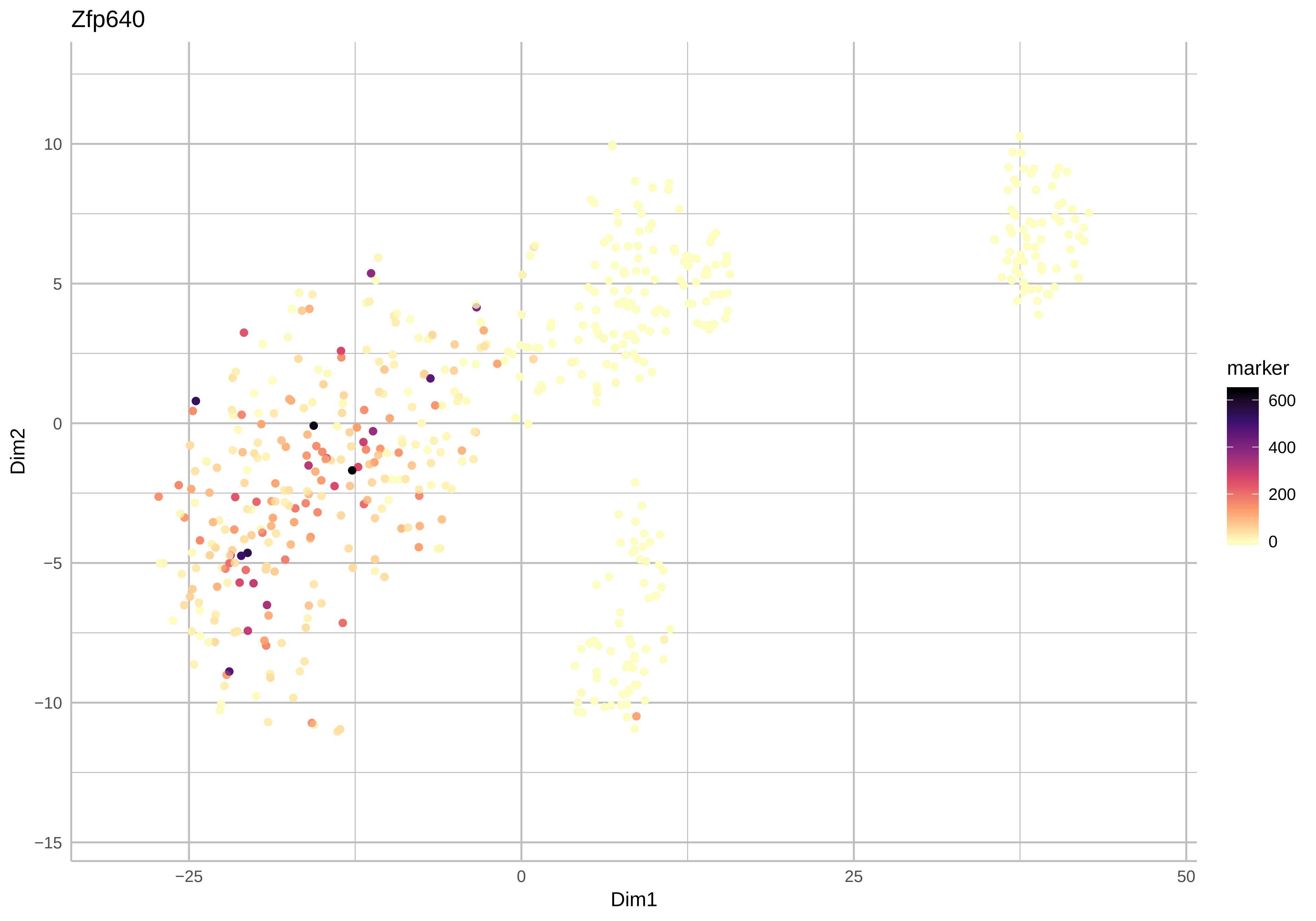

Zfp640
Dim2
Dim1
marker
600
400
200
0
10
5
0
−5
−10
−15
−25
0
25
50

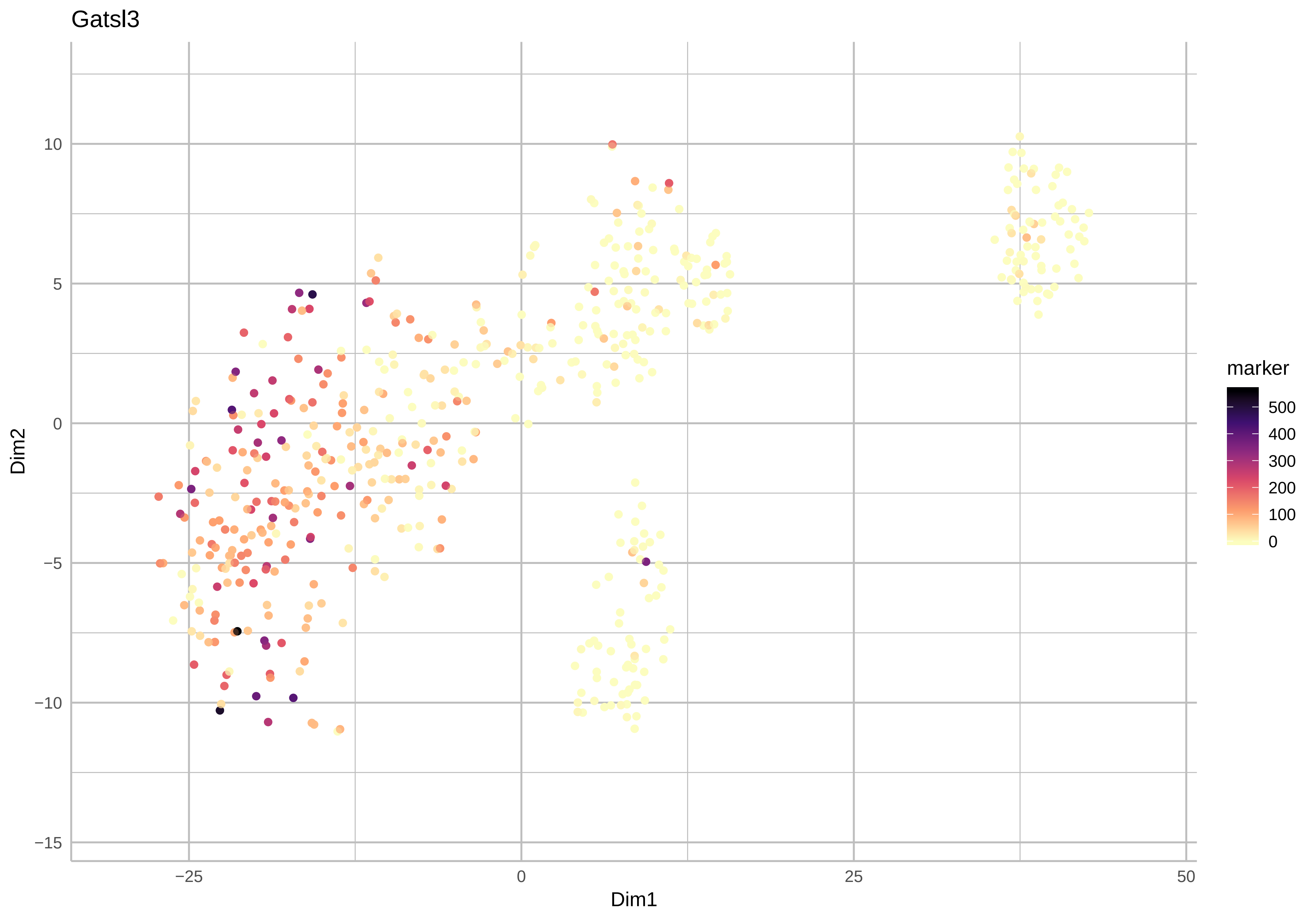

Gatsl3
10
5
0
−5
−10
−15
Dim2
−25
0
25
50
Dim1
marker
500
400
300
200
100
0

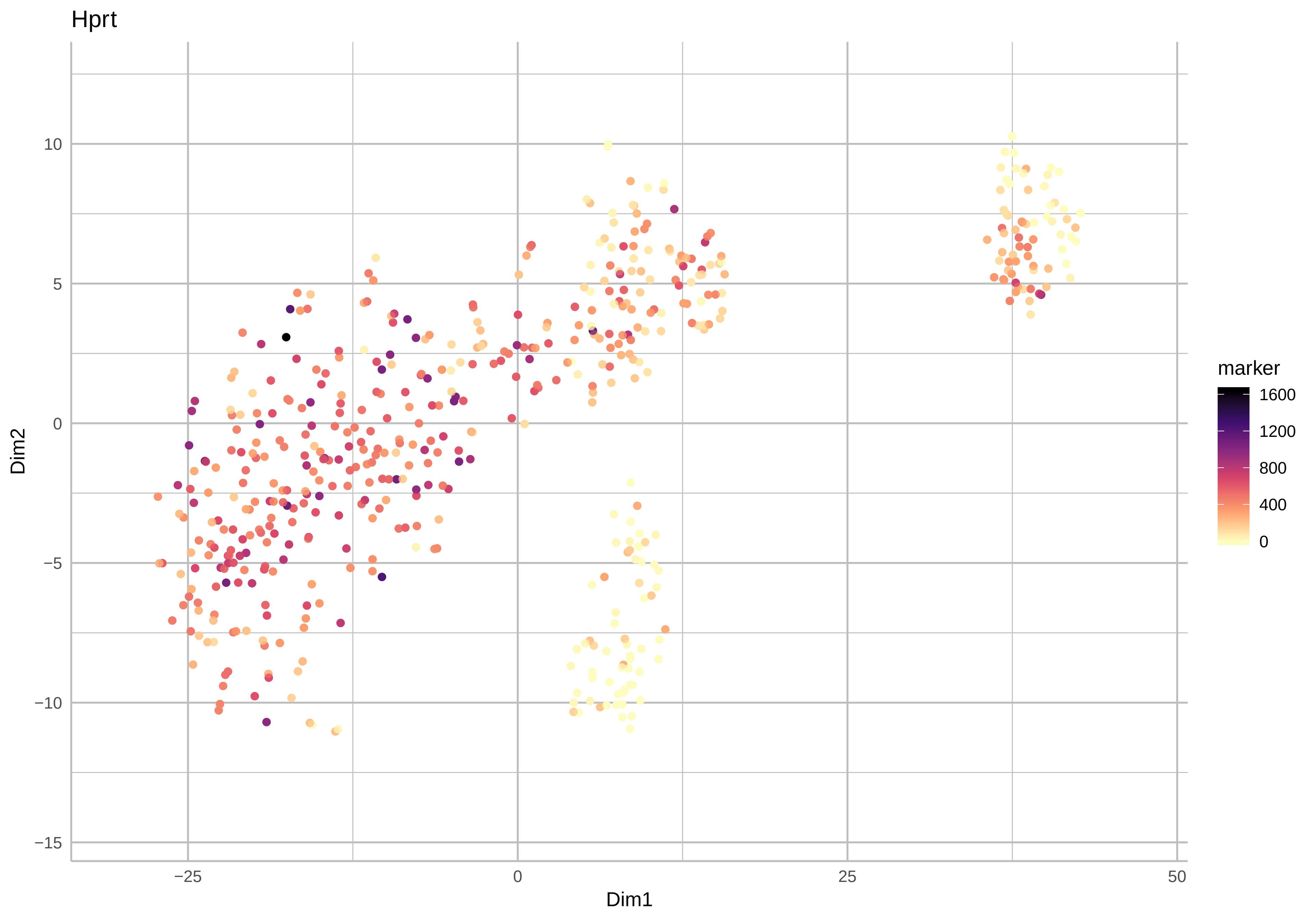

Hprt
Dim2
Dim1
10
5
0
−5
−10
−15
−25
0
25
50
marker
1600
1200
800
400
0

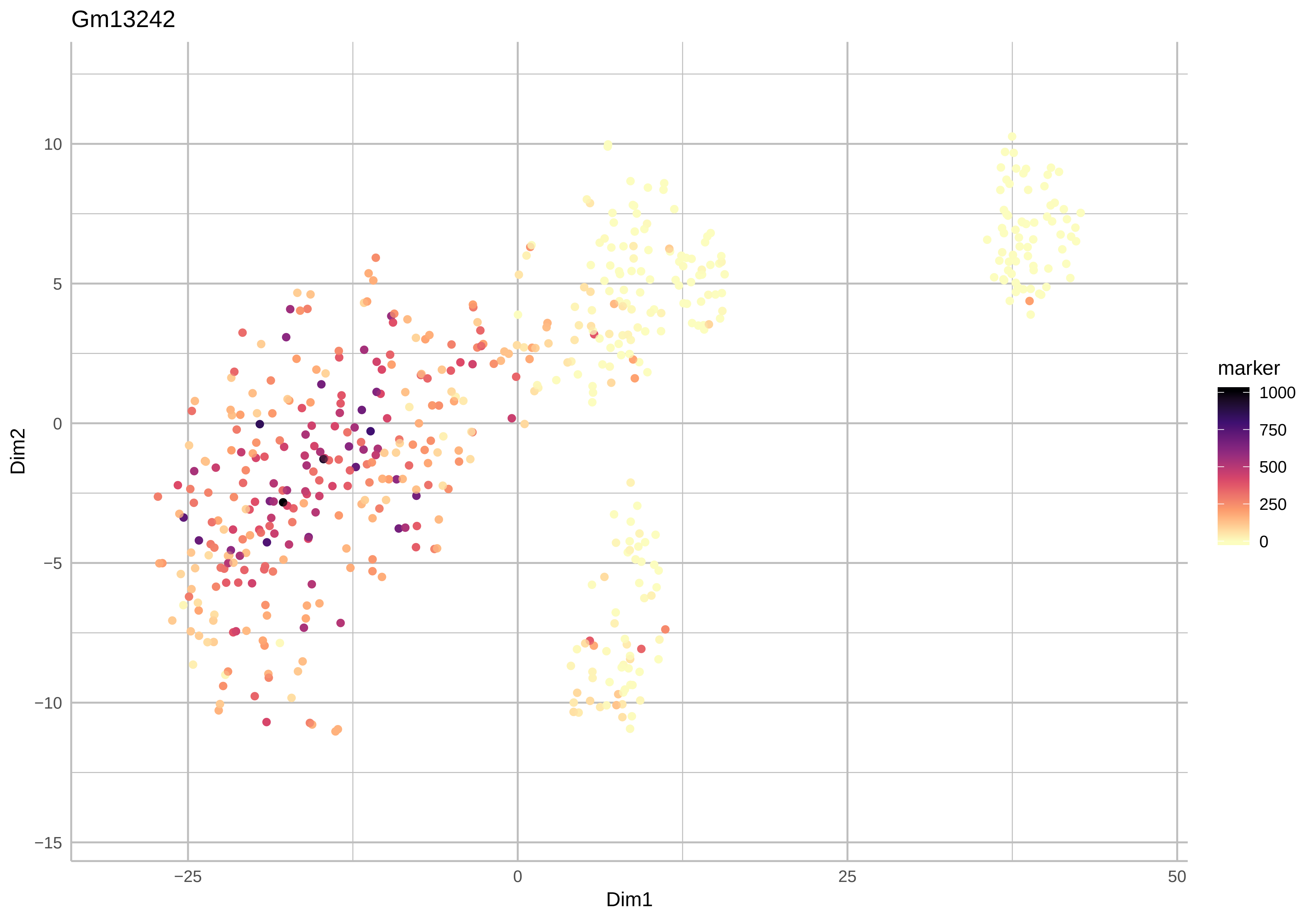
Gm13242
Dim2
10
5
0
−5
−10
−15
Dim1
−25
0
25
50
marker
1000
750
500
250
0

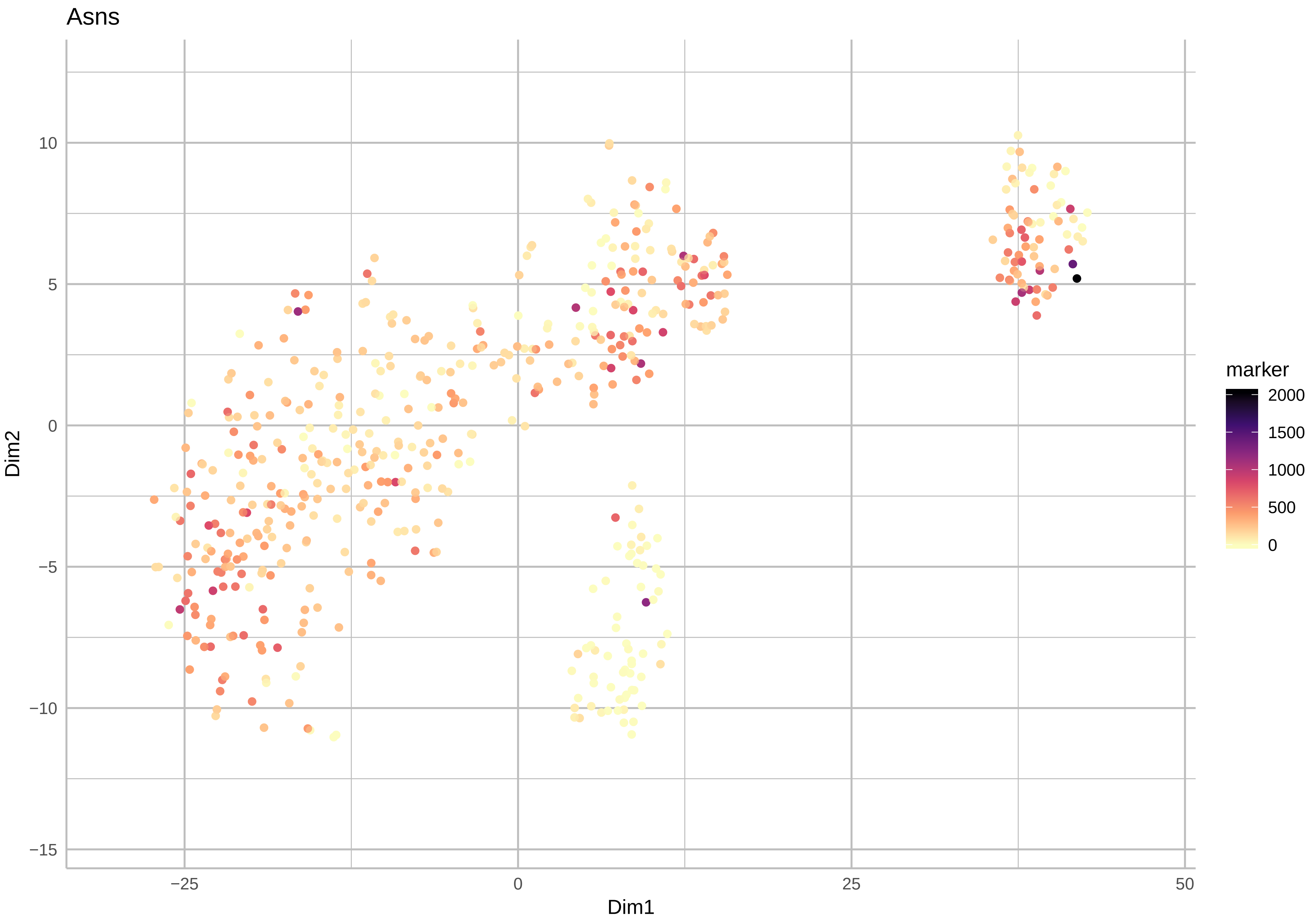

Asns
10
5
0
−5
−10
−15
Dim2
−25
0
25
50
Dim1
marker
2000
1500
1000
500
0

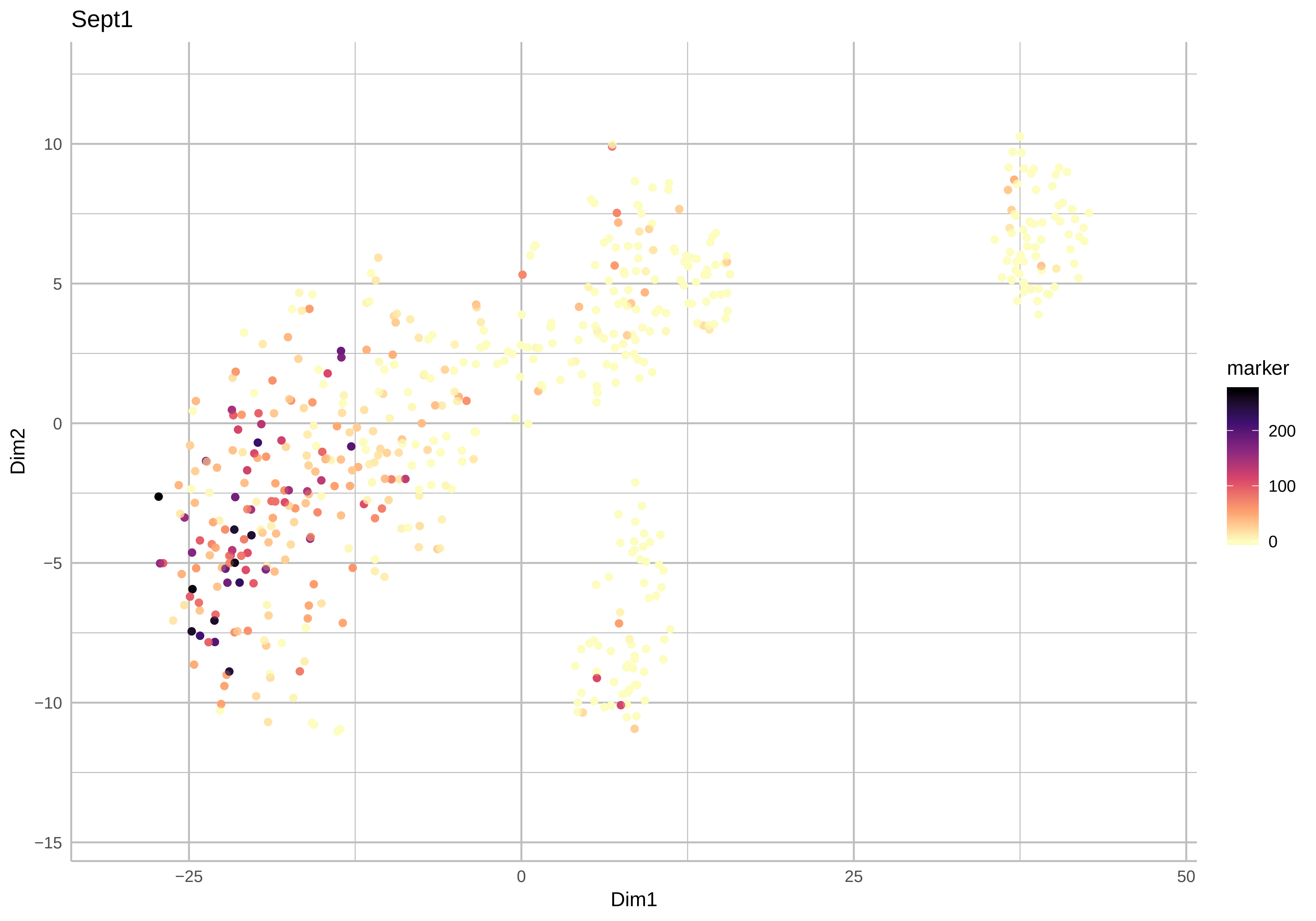

Sept1
10
5
0
−5
−10
−15
Dim2
−25
0
25
50
Dim1
marker
200
100
0

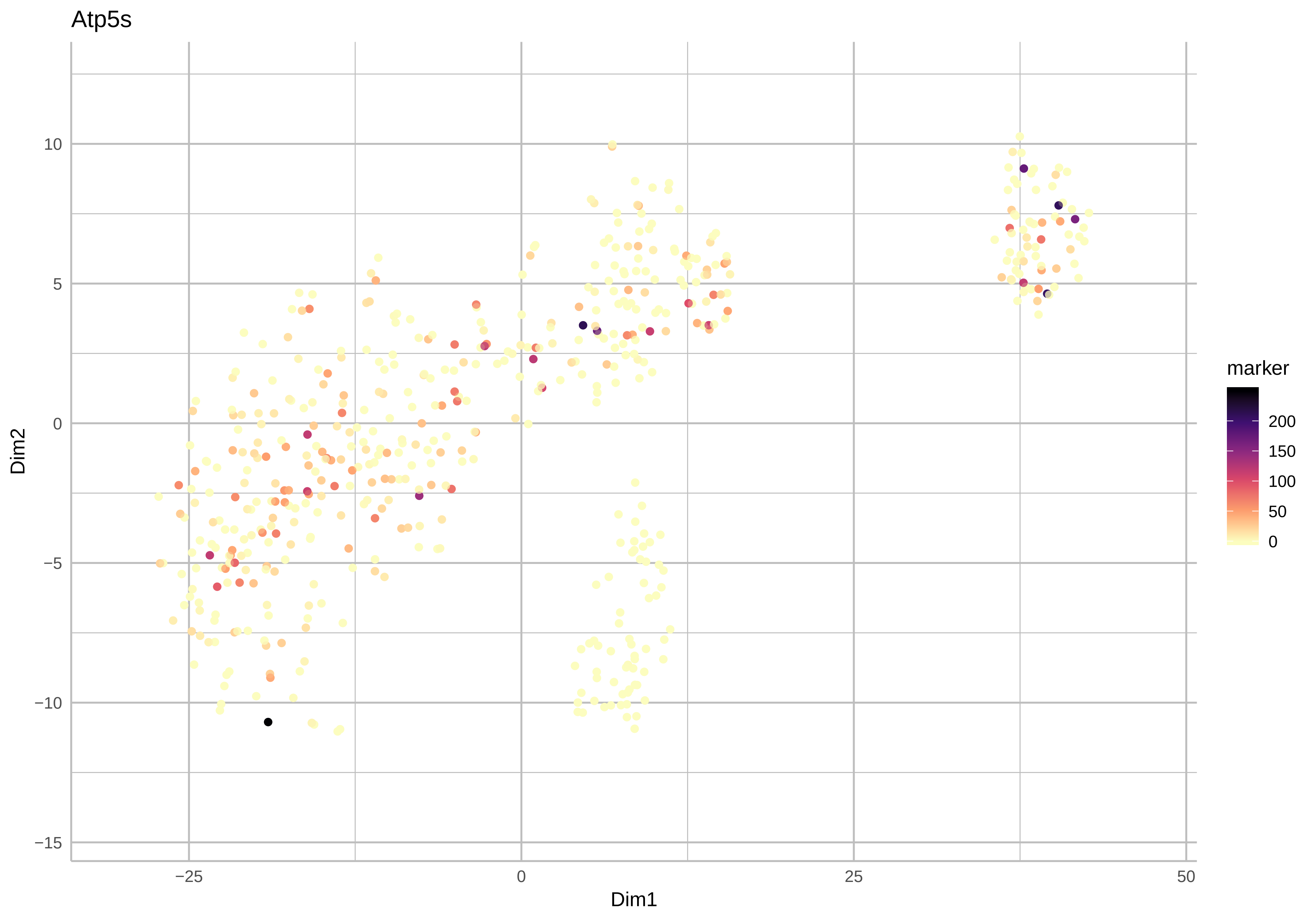

Atp5s
Dim2
Dim1
10
5
0
−5
−10
−15
−25
0
25
50
marker
200
150
100
50
0

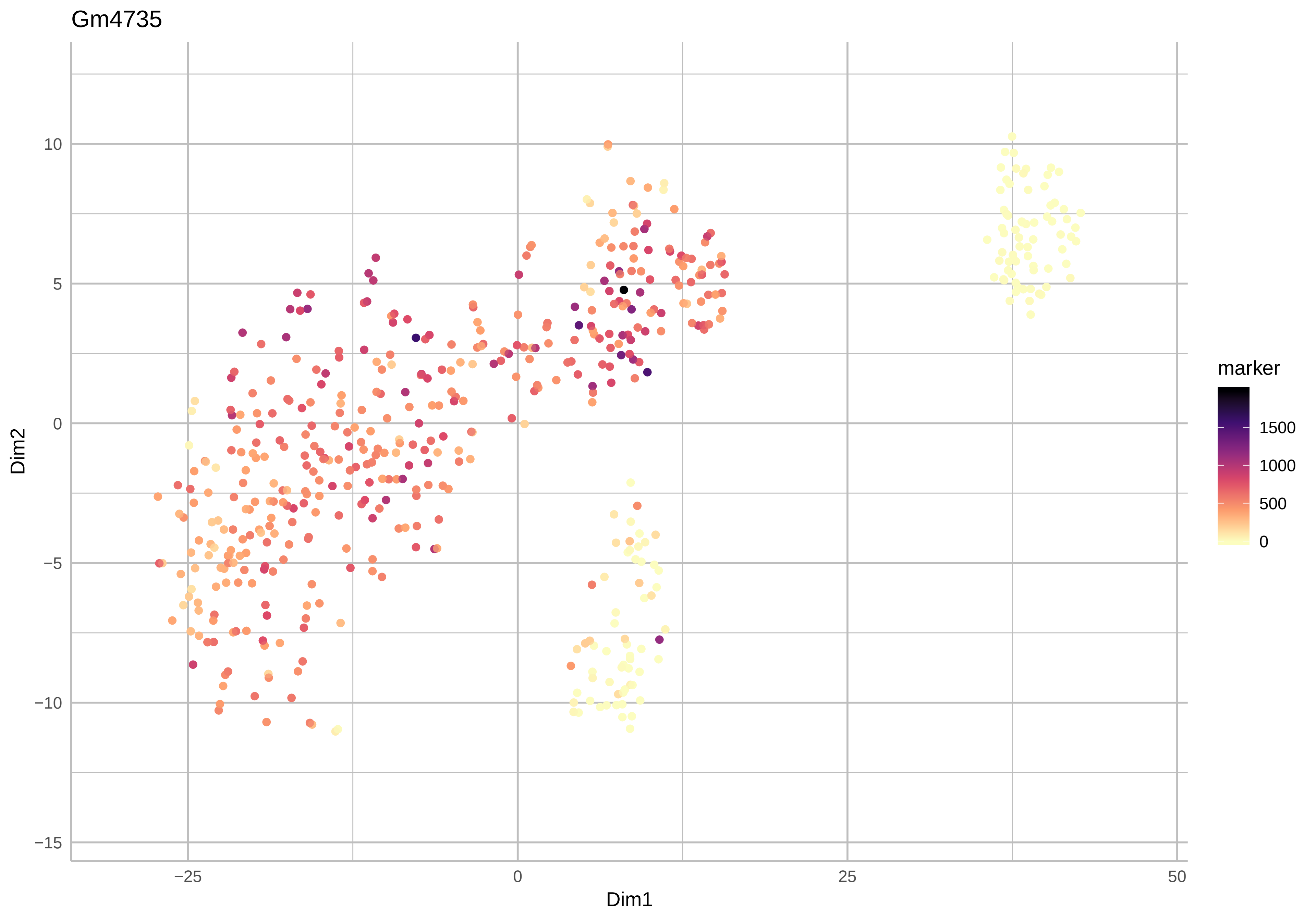
Gm4735
Dim2
Dim1
10
5
0
−5
−10
−15
−25
0
25
50
marker
1500
1000
500
0

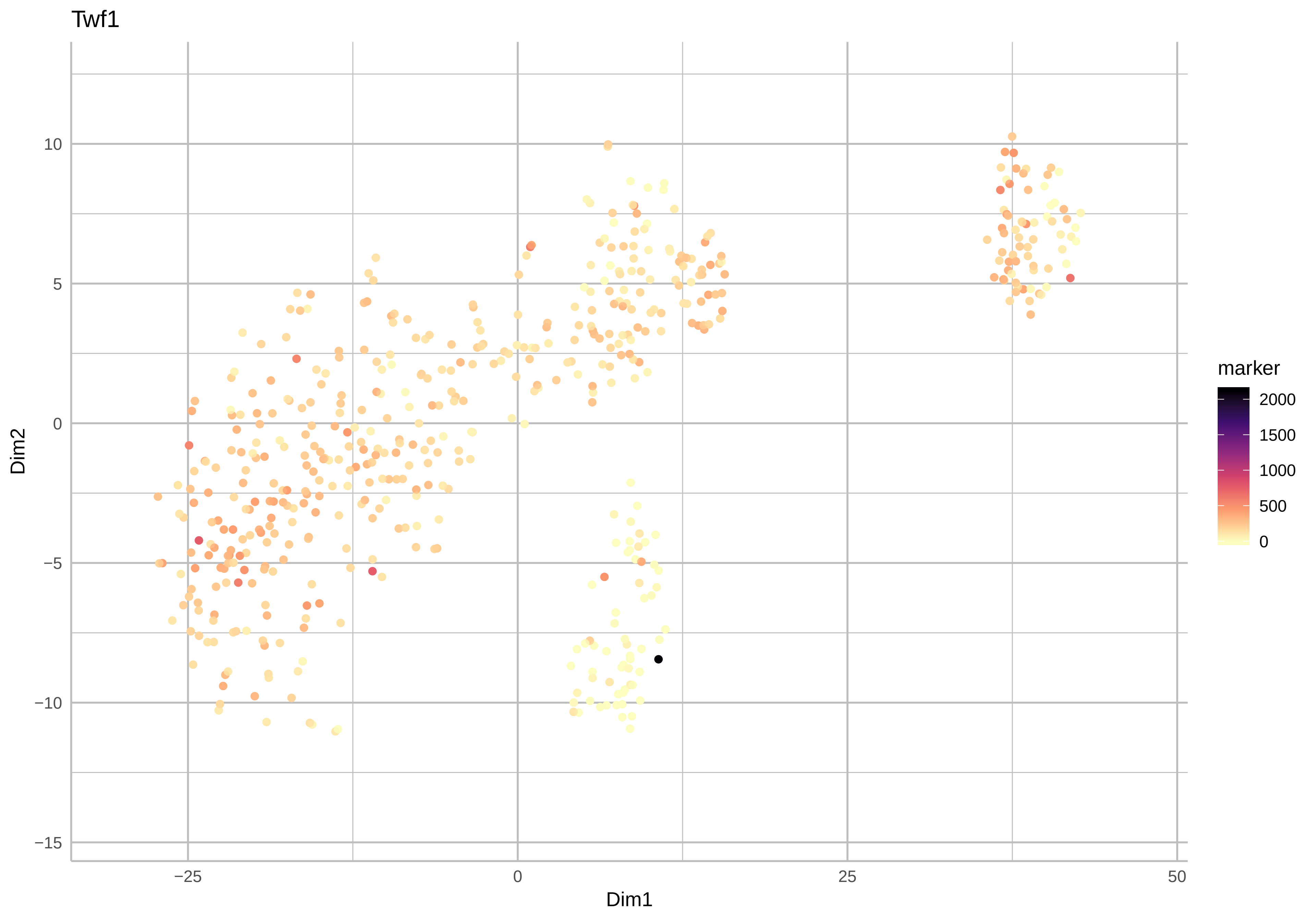

Twf1
Dim2
Dim1
10
5
0
−5
−10
−15
−25
0
25
50
marker
2000
1500
1000
500
0

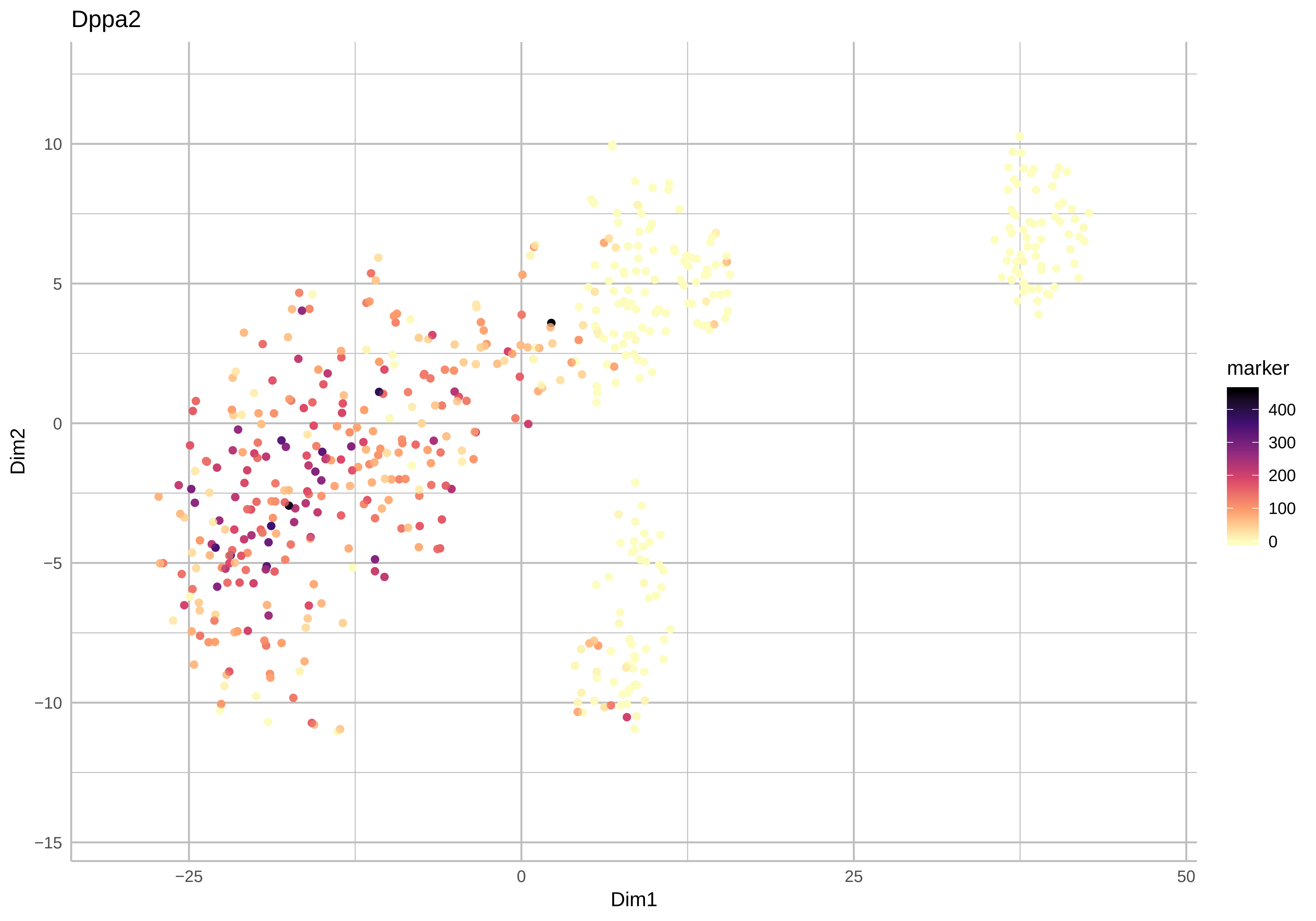

Dppa2
10
5
0
−5
−10
−15
Dim2
−25
0
25
50
Dim1
marker
400
300
200
100
0

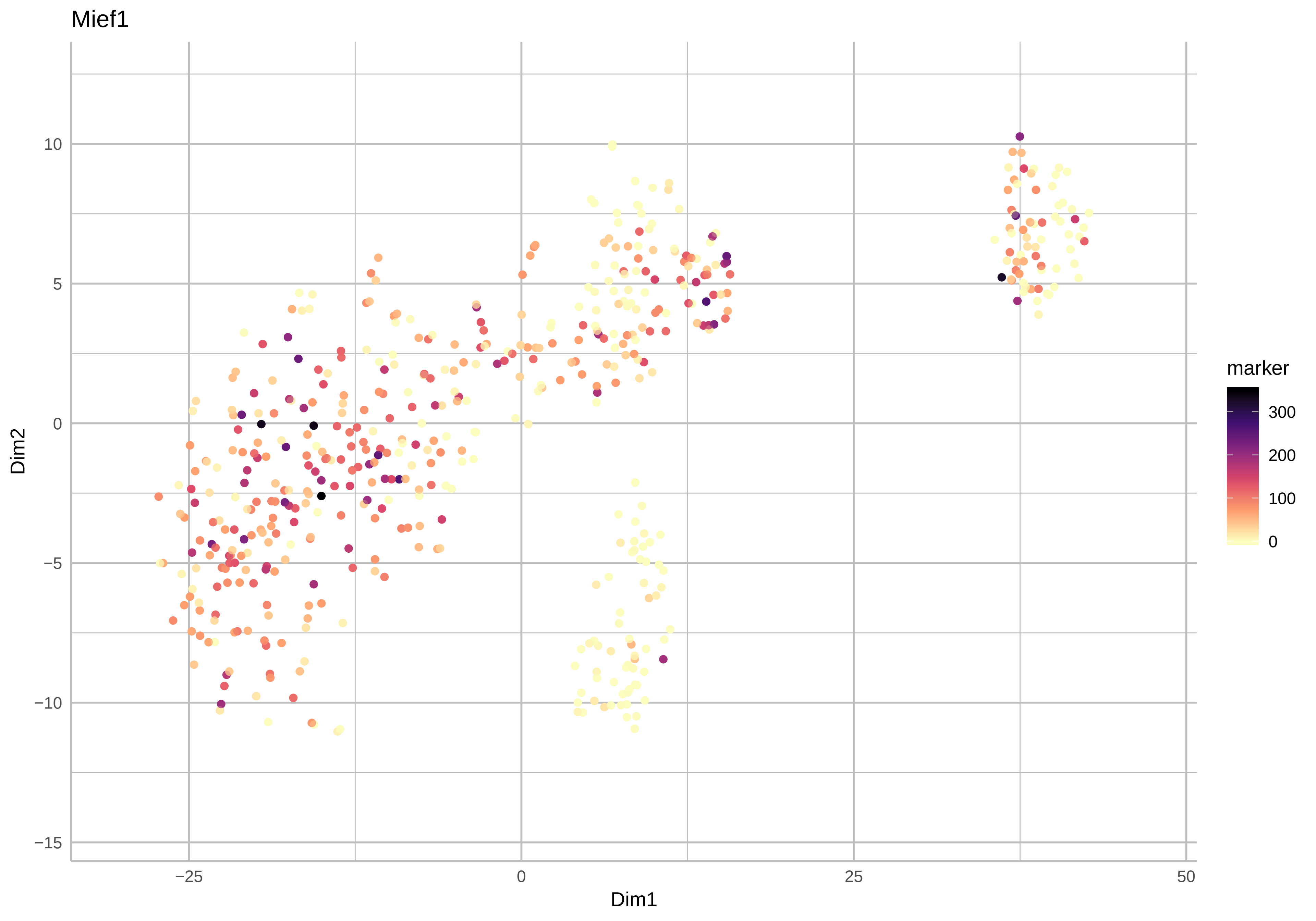
Mief1
Dim2
Dim1
10
5
0
−5
−10
−15
−25
0
25
50
marker
300
200
100
0

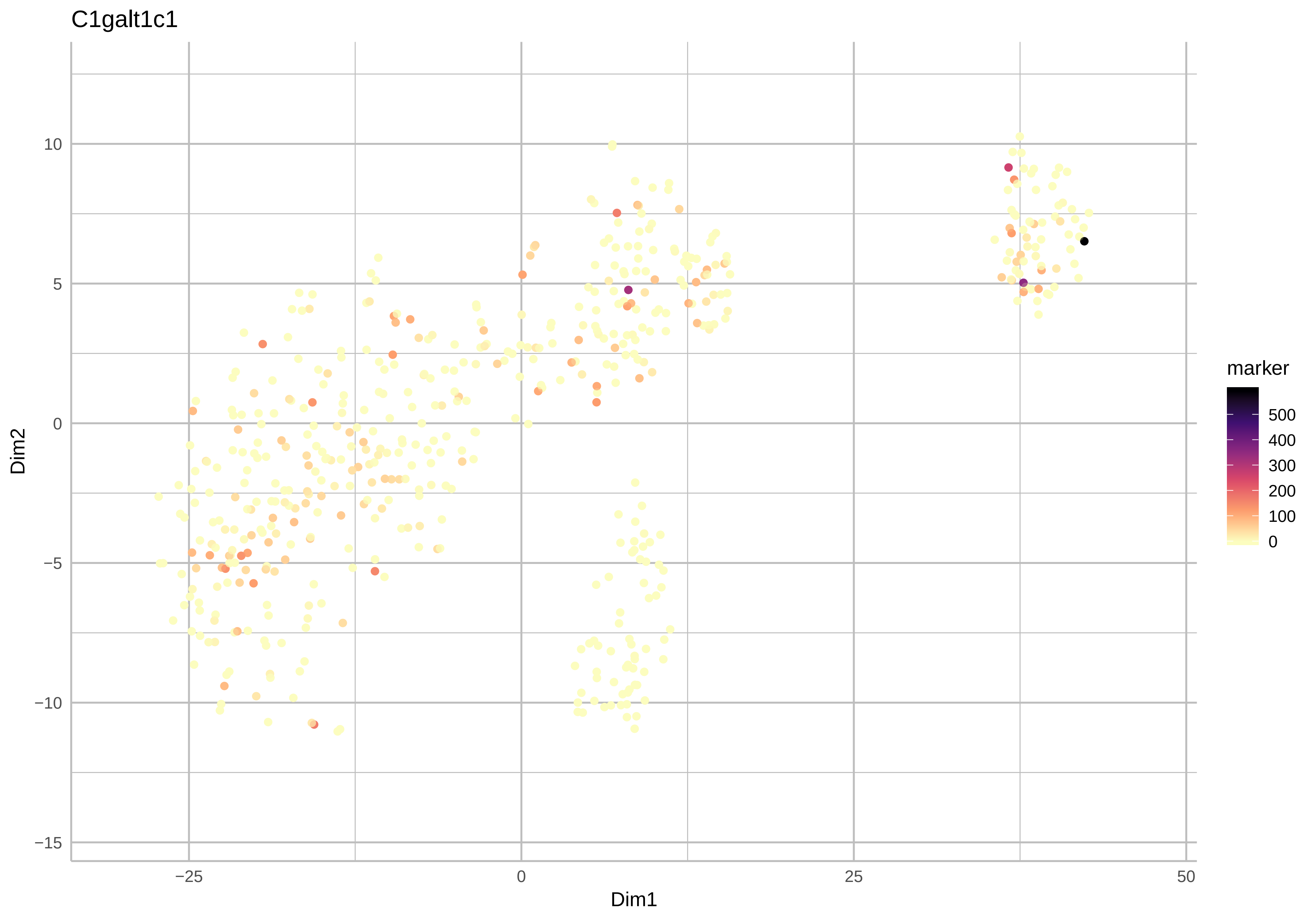
C1galt1c1
Dim2
10
5
0
−5
−10
−15
−25
0
25
50
Dim1
marker
500
400
300
200
100
0

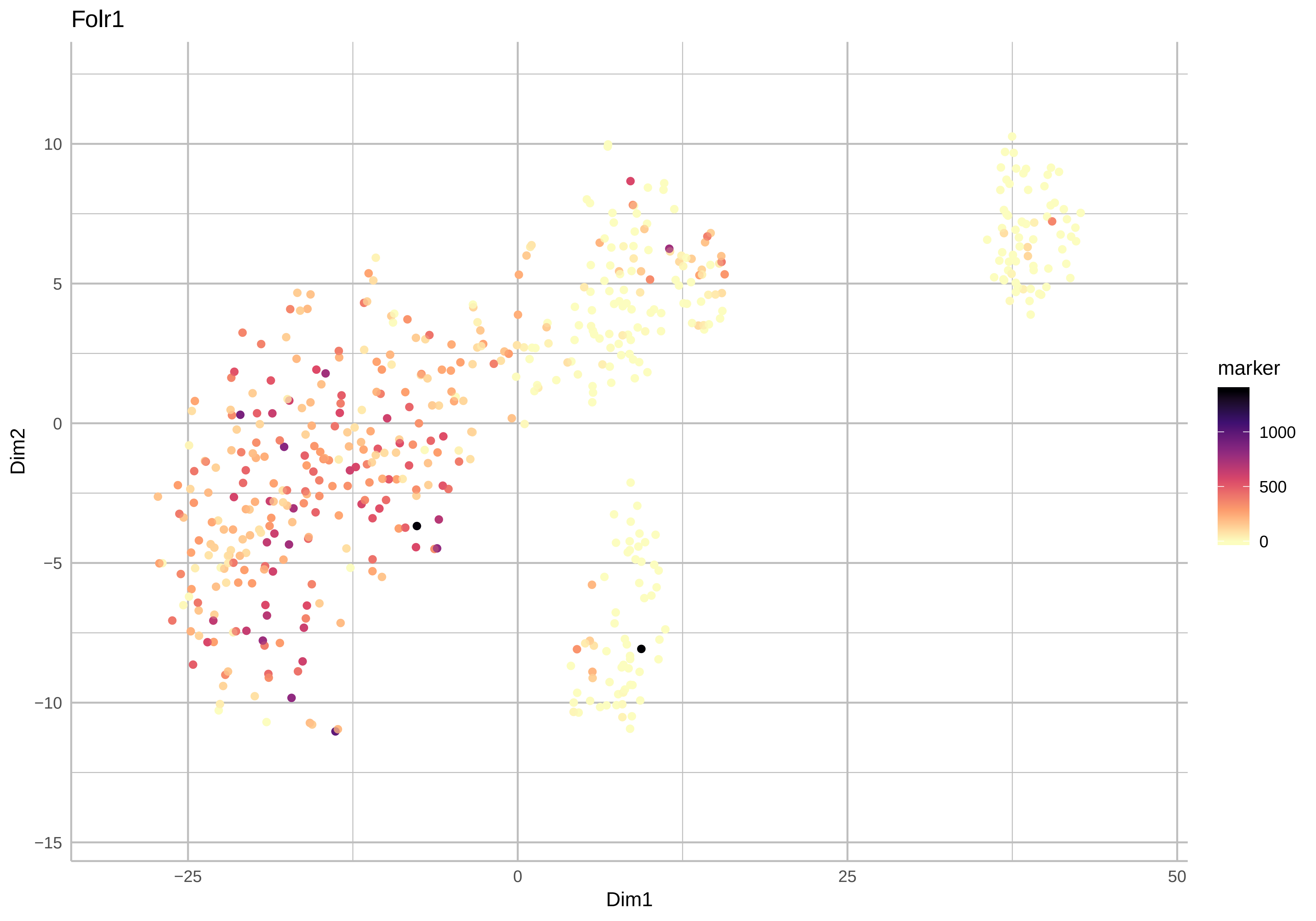
Folr1
Dim2
Dim1
marker
1000
500
0
10
5
0
−5
−10
−15
−25
0
25
50

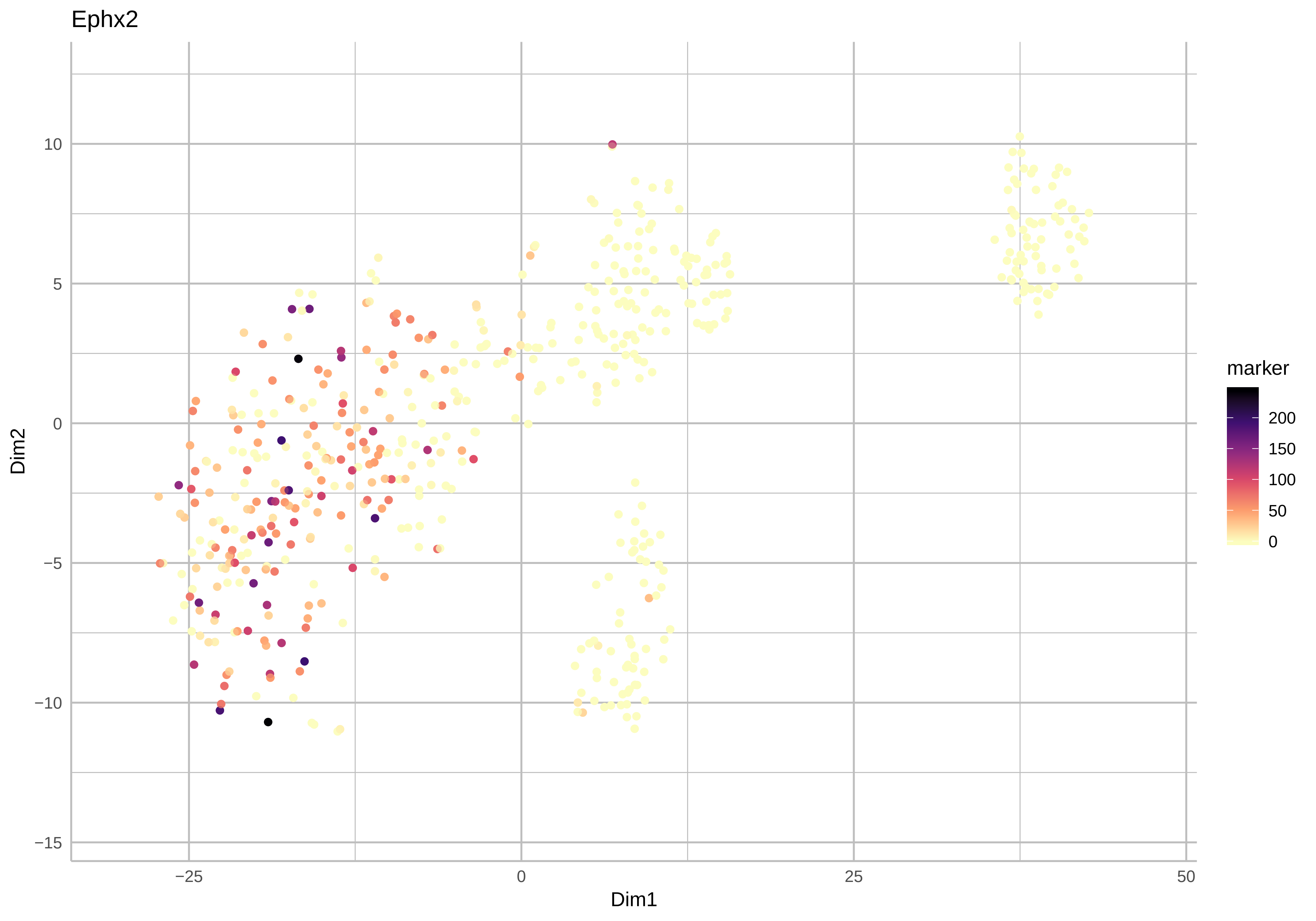

Ephx2
10
5
0
−5
−10
−15
Dim2
−25
0
25
50
Dim1
marker
200
150
100
50
0

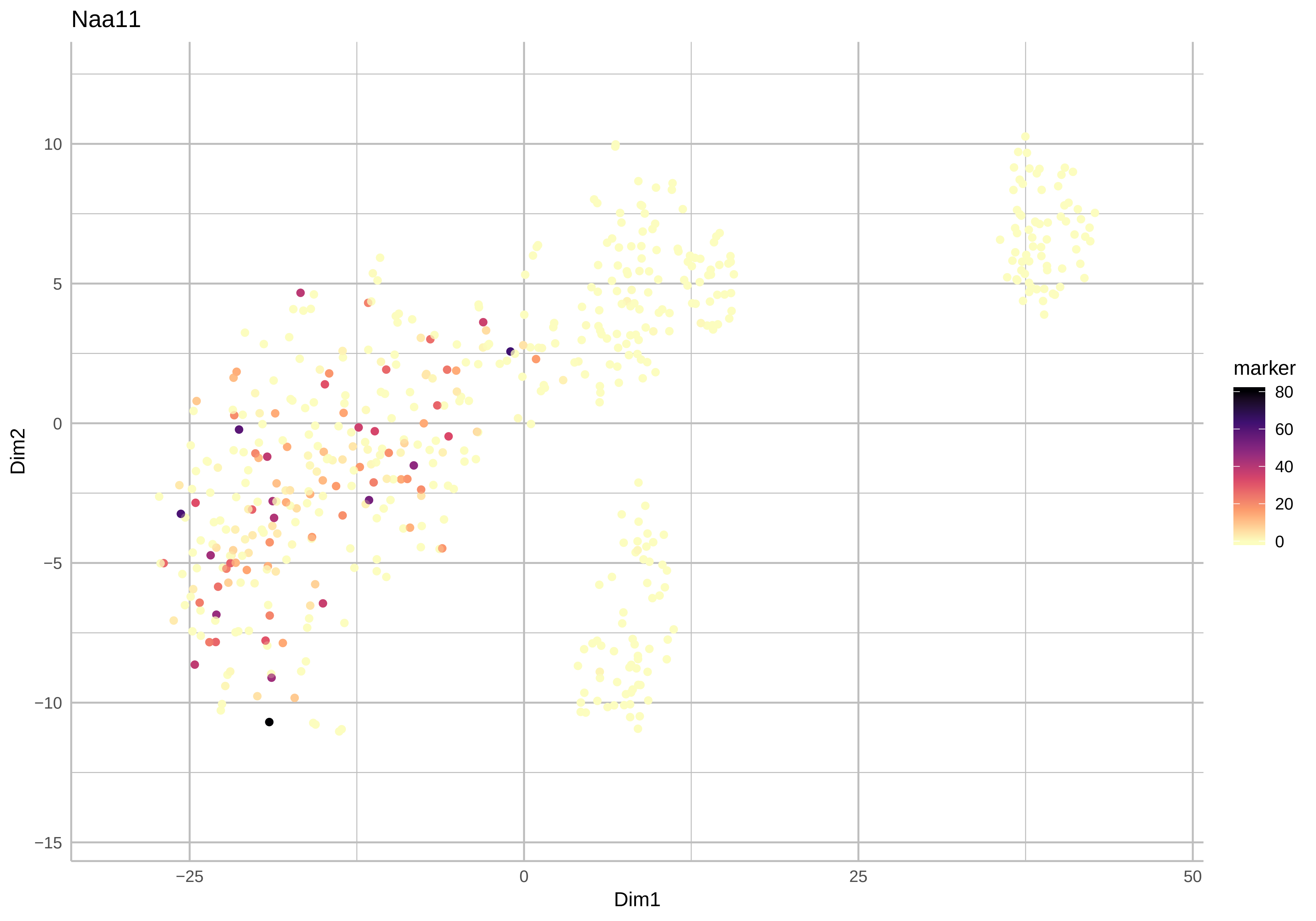

Naa11
10
5
0
−5
−10
−15
Dim2
−25
0
25
50
Dim1
marker
80
60
40
20
0

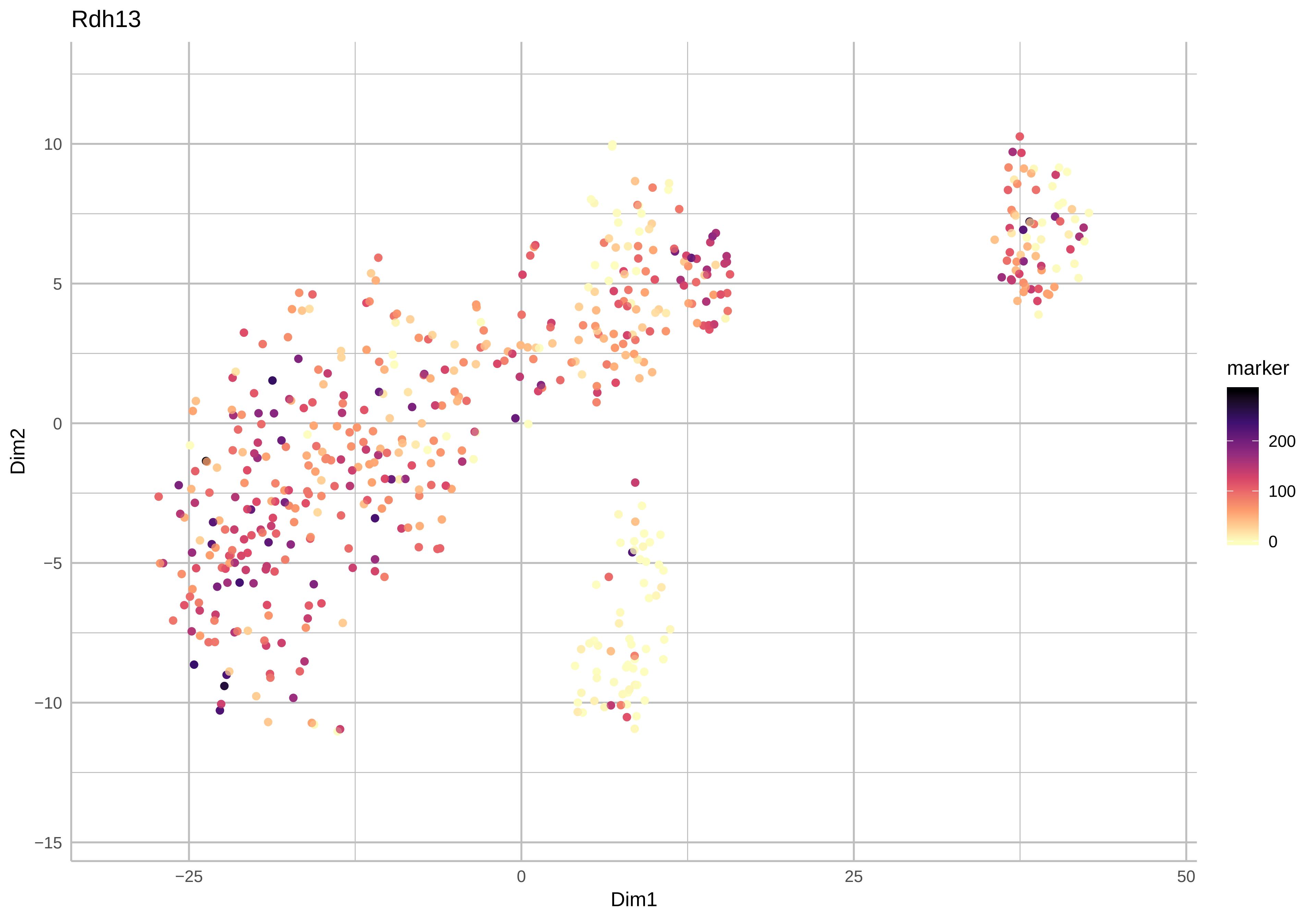

Rdh13
10
5
0
−5
−10
−15
Dim2
−25
0
25
50
Dim1
marker
200
100
0

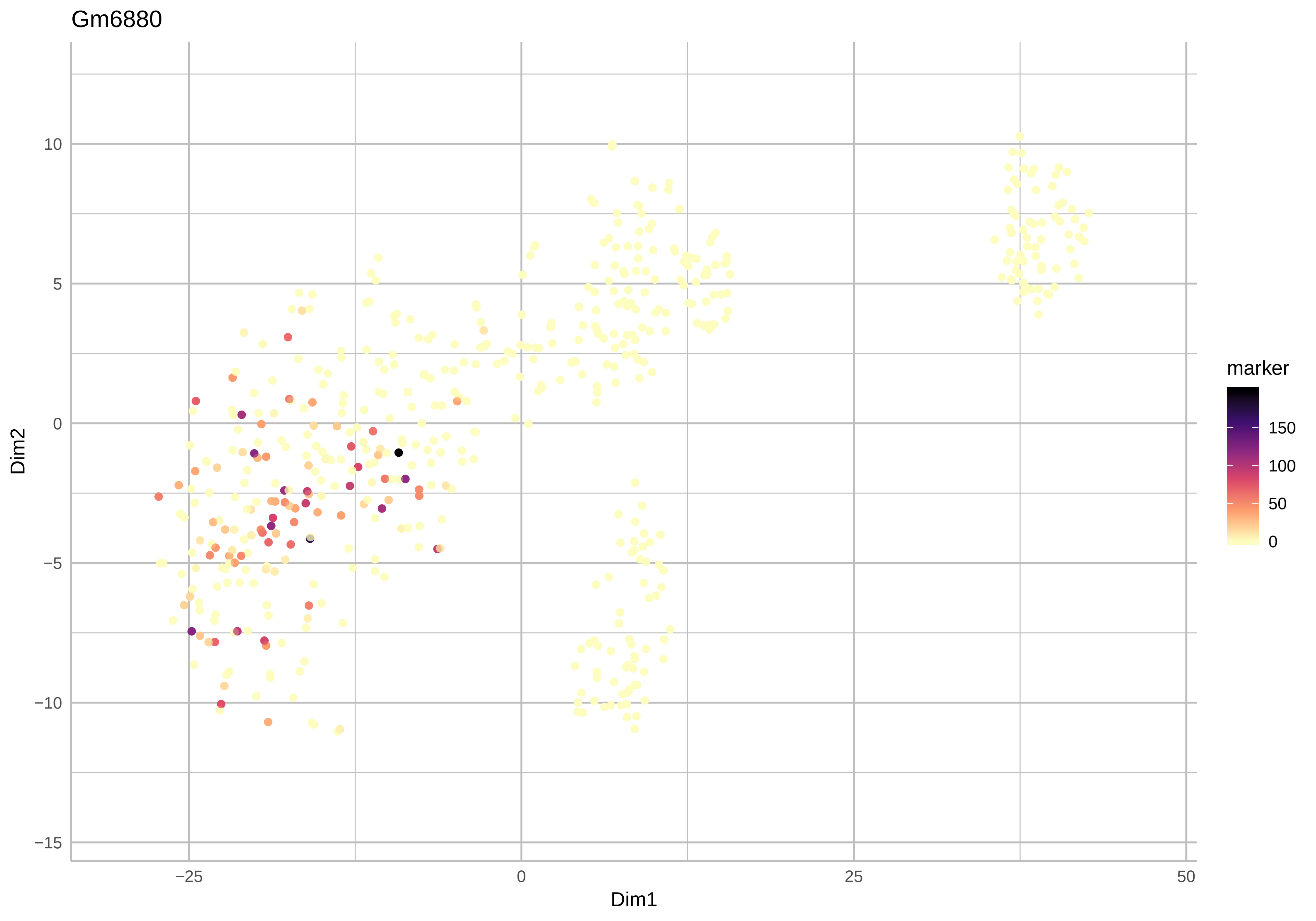
Gm6880
Dim2
Dim1
10
5
0
−5
−10
−15
−25
0
25
50
marker
150
100
50
0

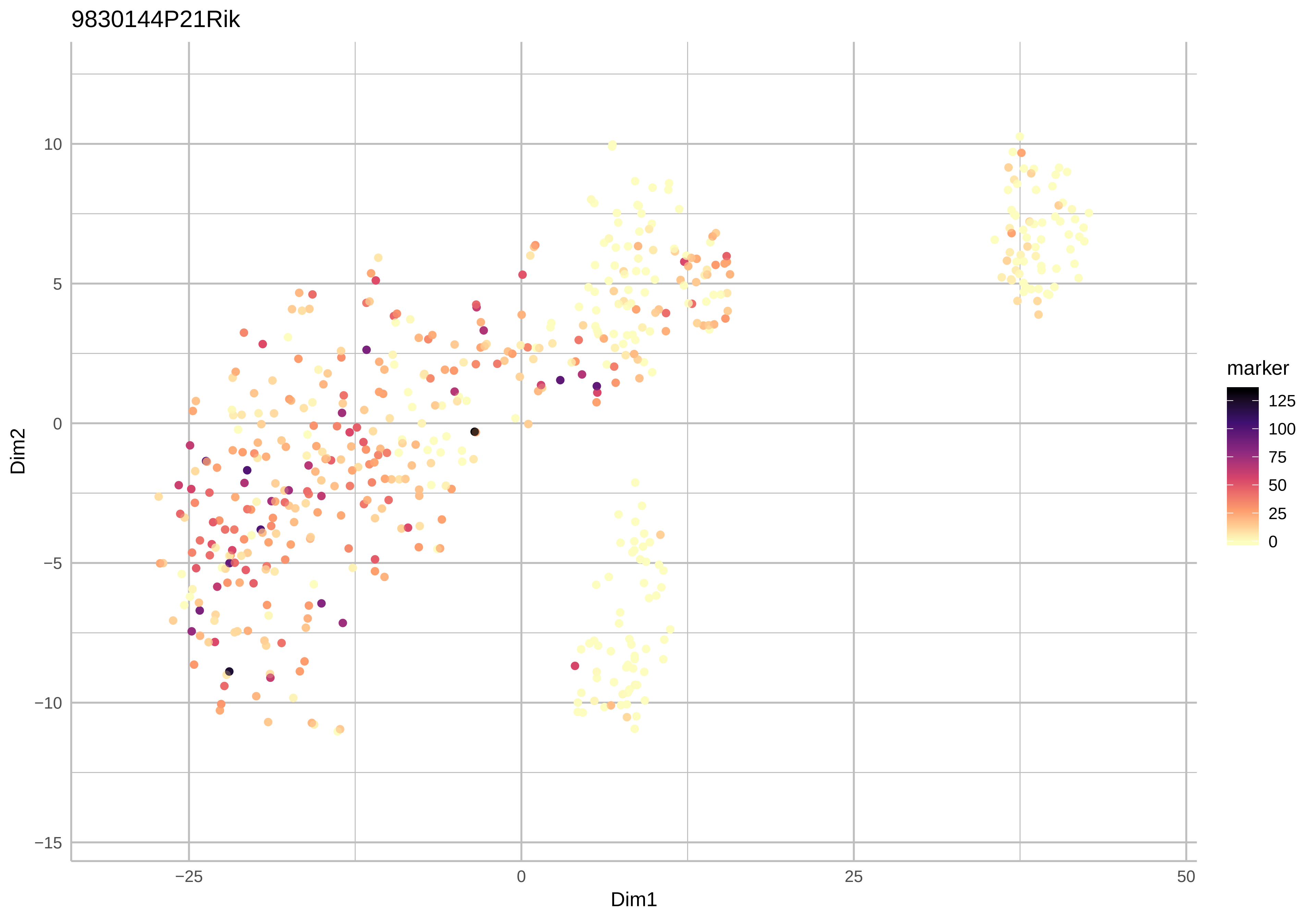

9830144P21Rik
Dim2
Dim1
10
5
0
−5
−10
−15
−25
0
25
50
marker
125
100
75
50
25
0

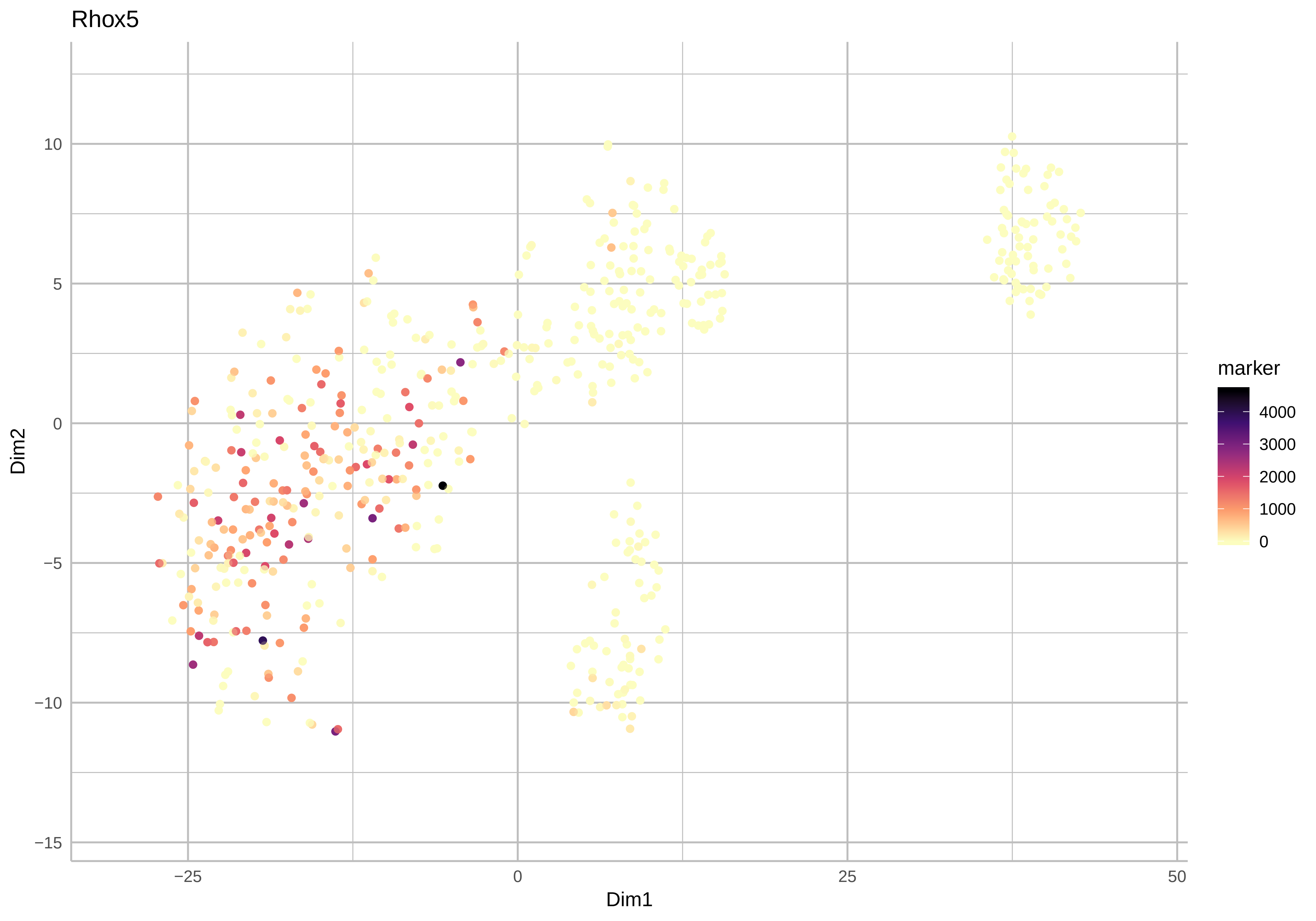

Rhox5
Dim2
Dim1
10
5
0
−5
−10
−15
−25
0
25
50
marker
4000
3000
2000
1000
0

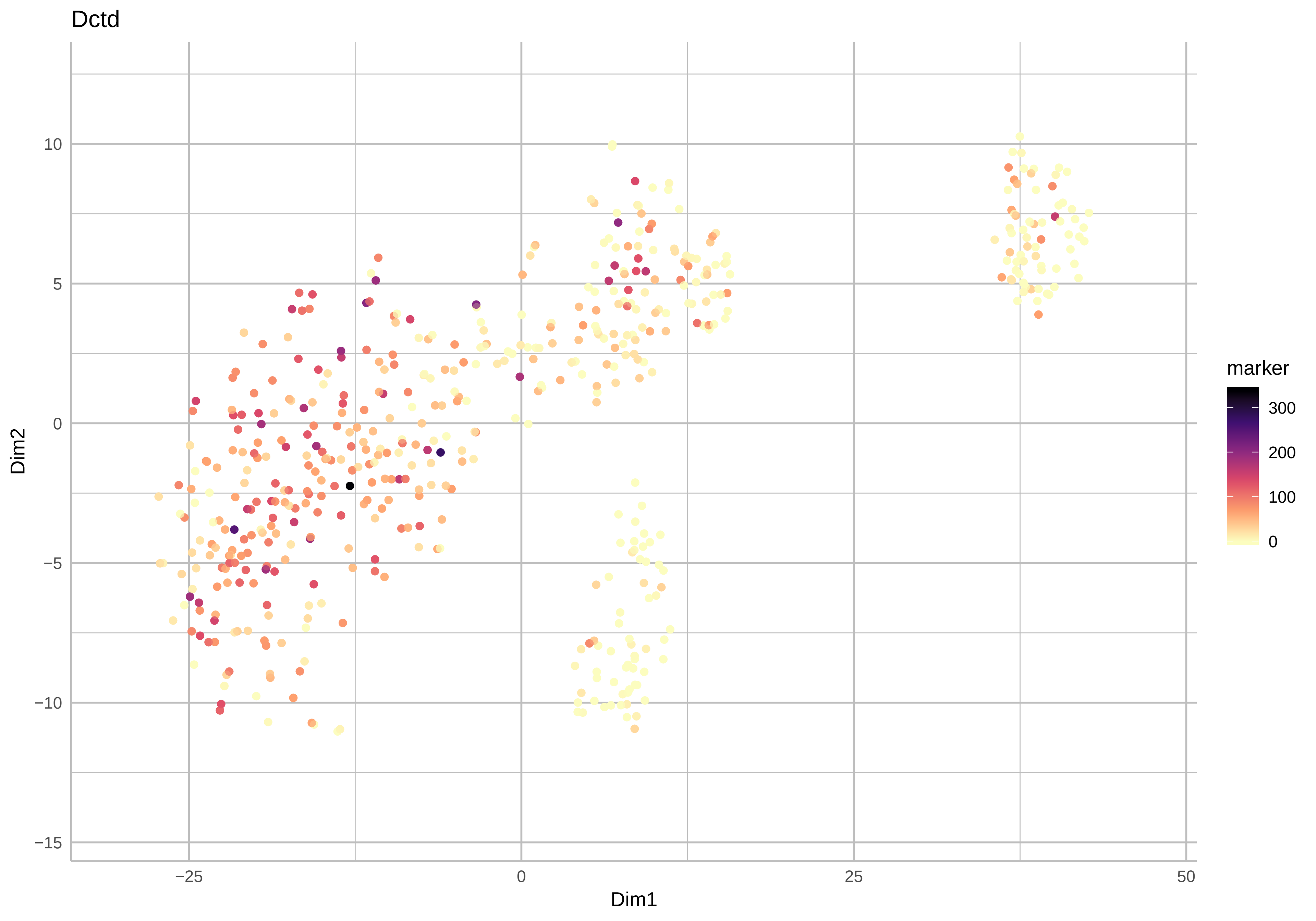
Dctd
Dim2
Dim1
marker
300
200
100
0
10
5
0
−5
−10
−15
−25
0
25
50

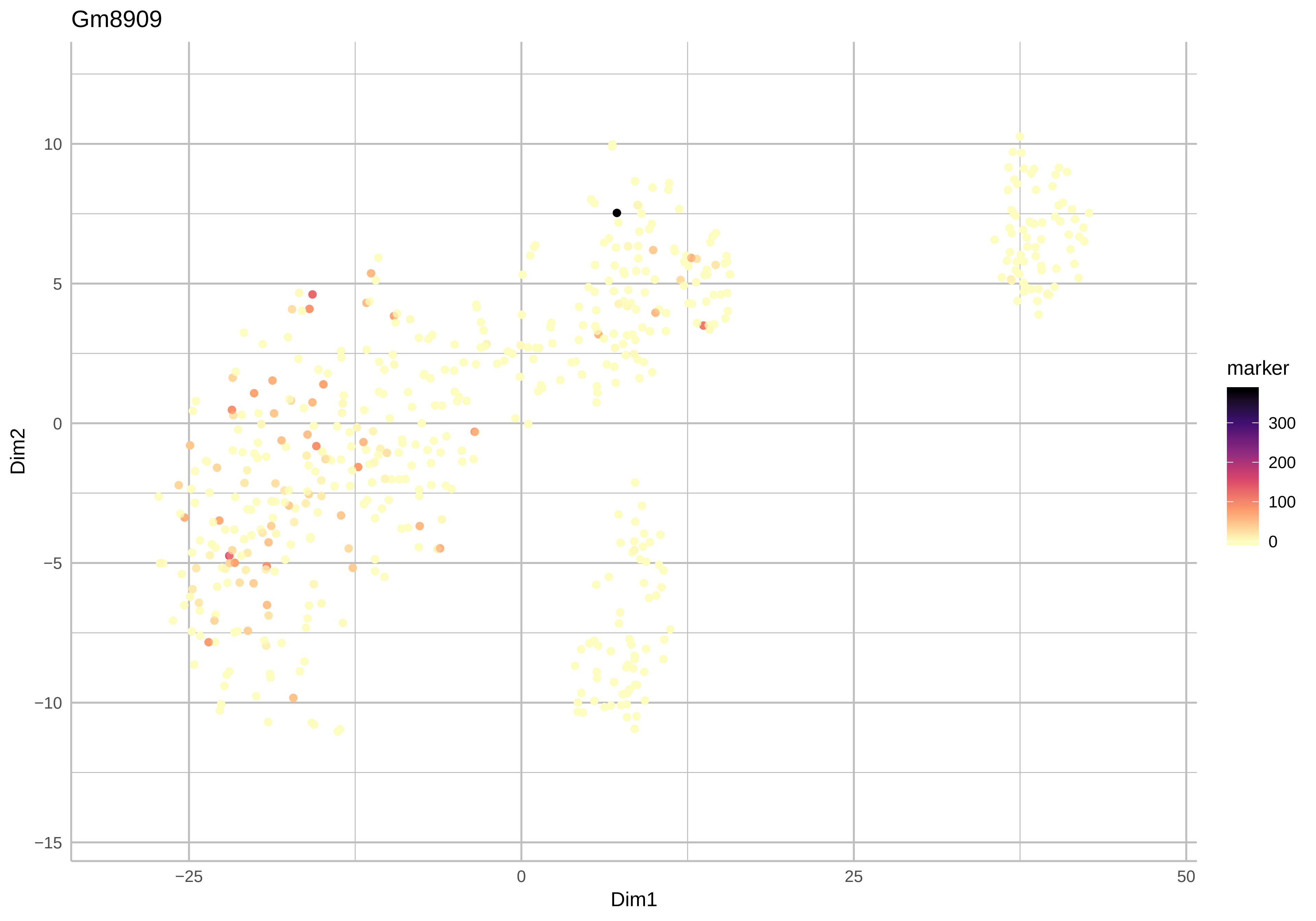
Gm8909
Dim1
Dim2
−25
0
25
50
10
5
0
−5
−10
−15
marker
300
200
100
0

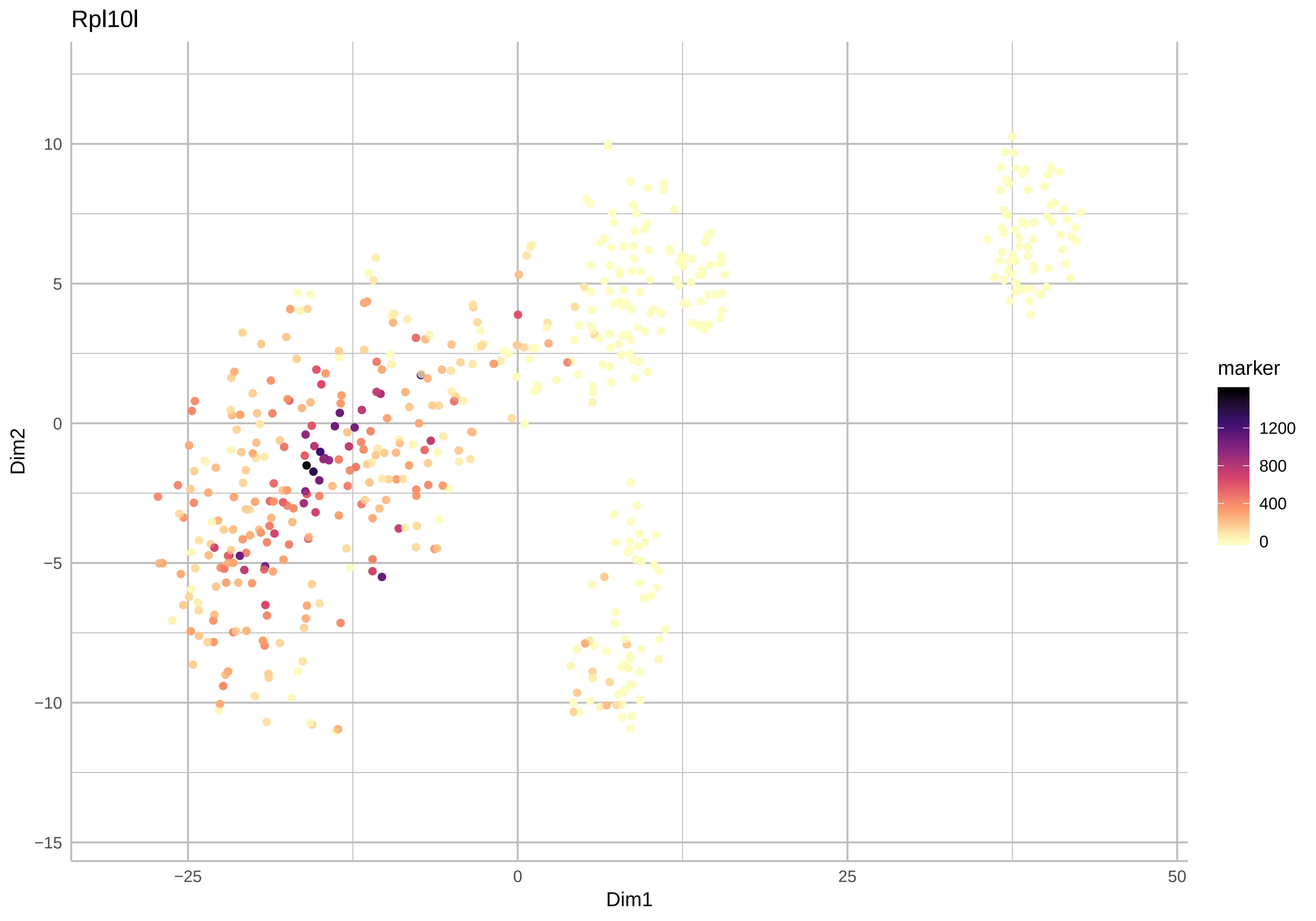
Rpl10l
Dim2
Dim1
marker
1200
800
400
0
10
5
0
−5
−10
−15
−25
0
25
50

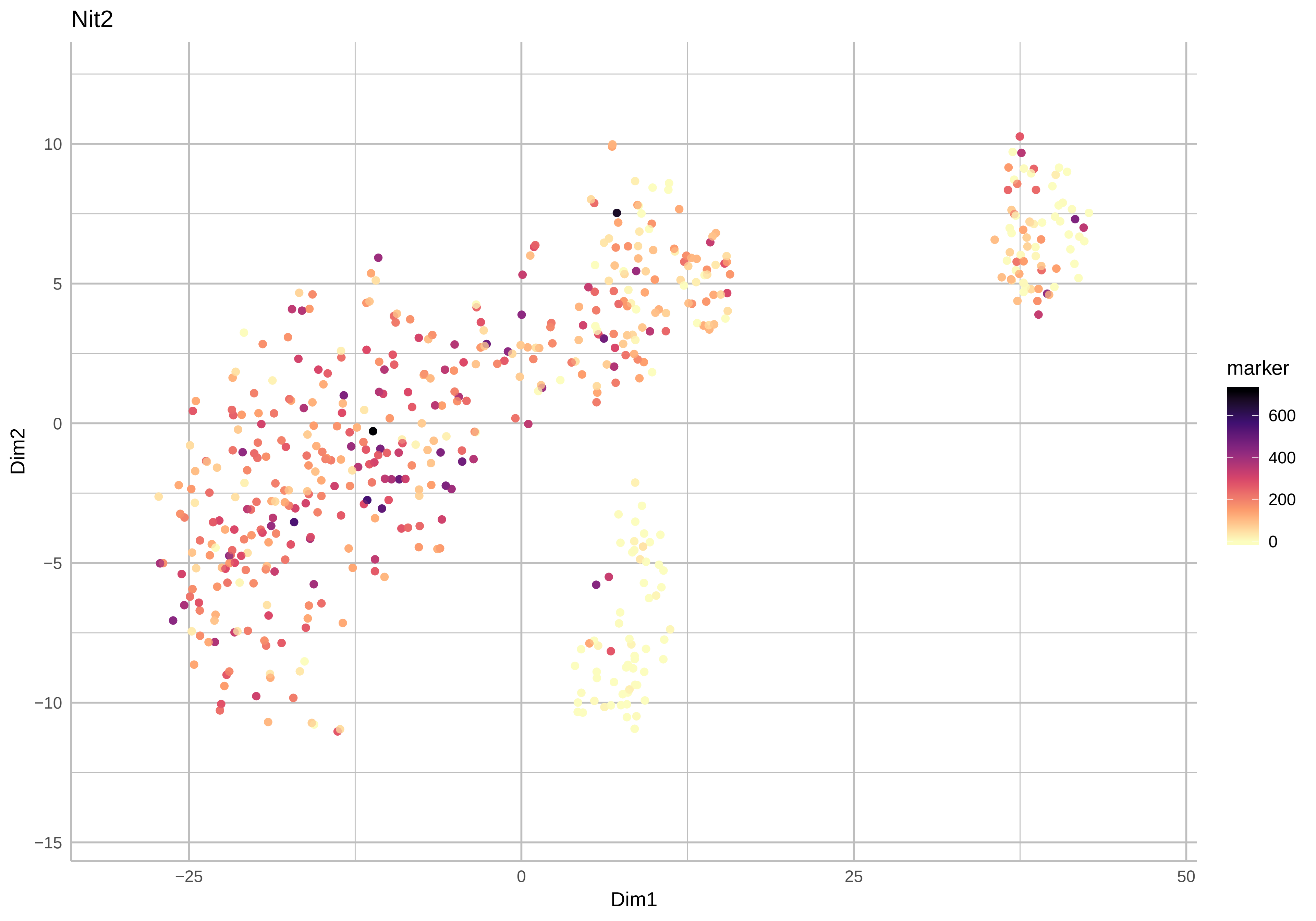
Nit2
Dim2
Dim1
10
5
0
−5
−10
−15
−25
0
25
50
marker
600
400
200
0

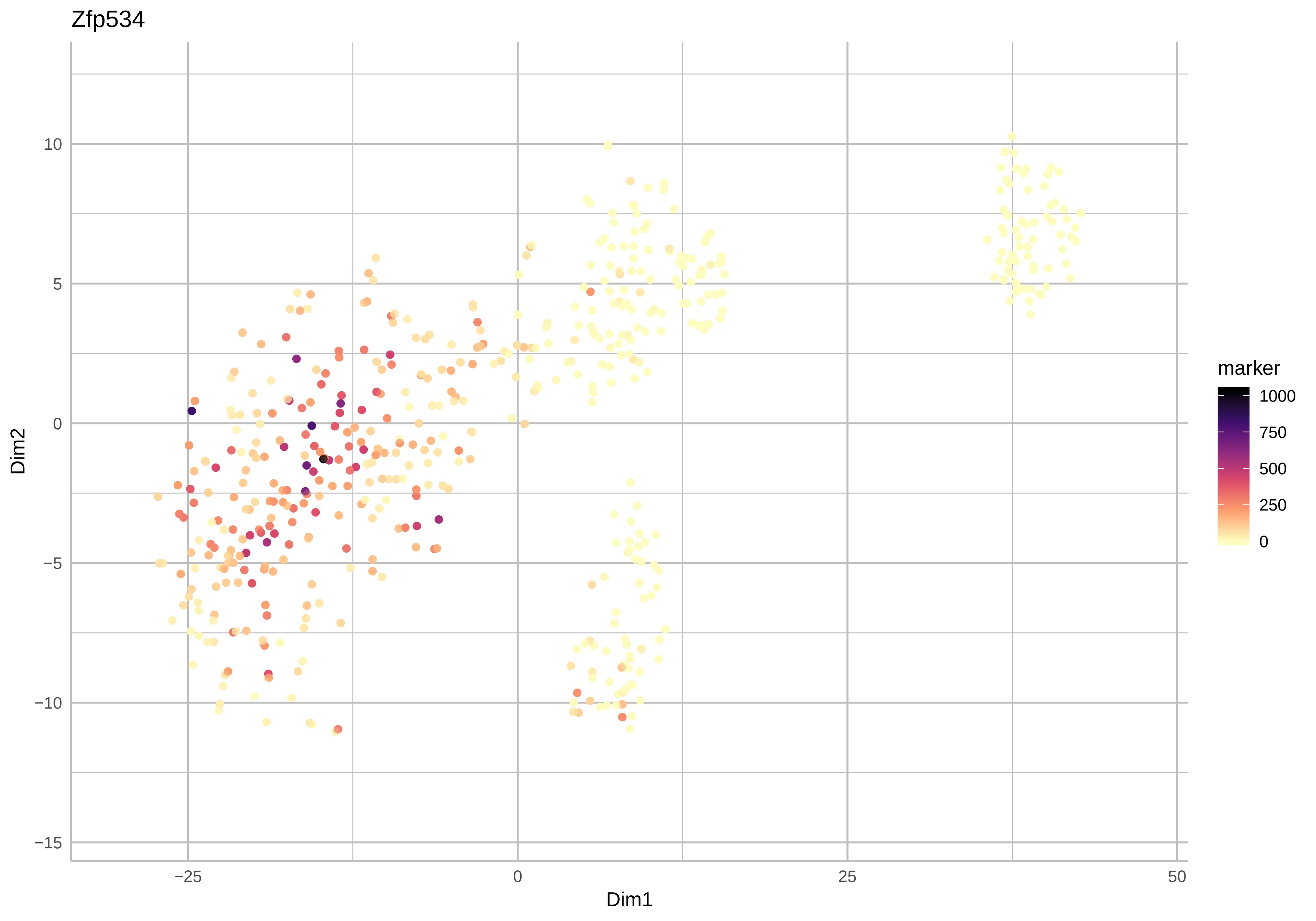

Zfp534
10
5
0
−5
−10
−15
−25
0
25
50
Dim1
Dim2
marker
1000
750
500
250
0

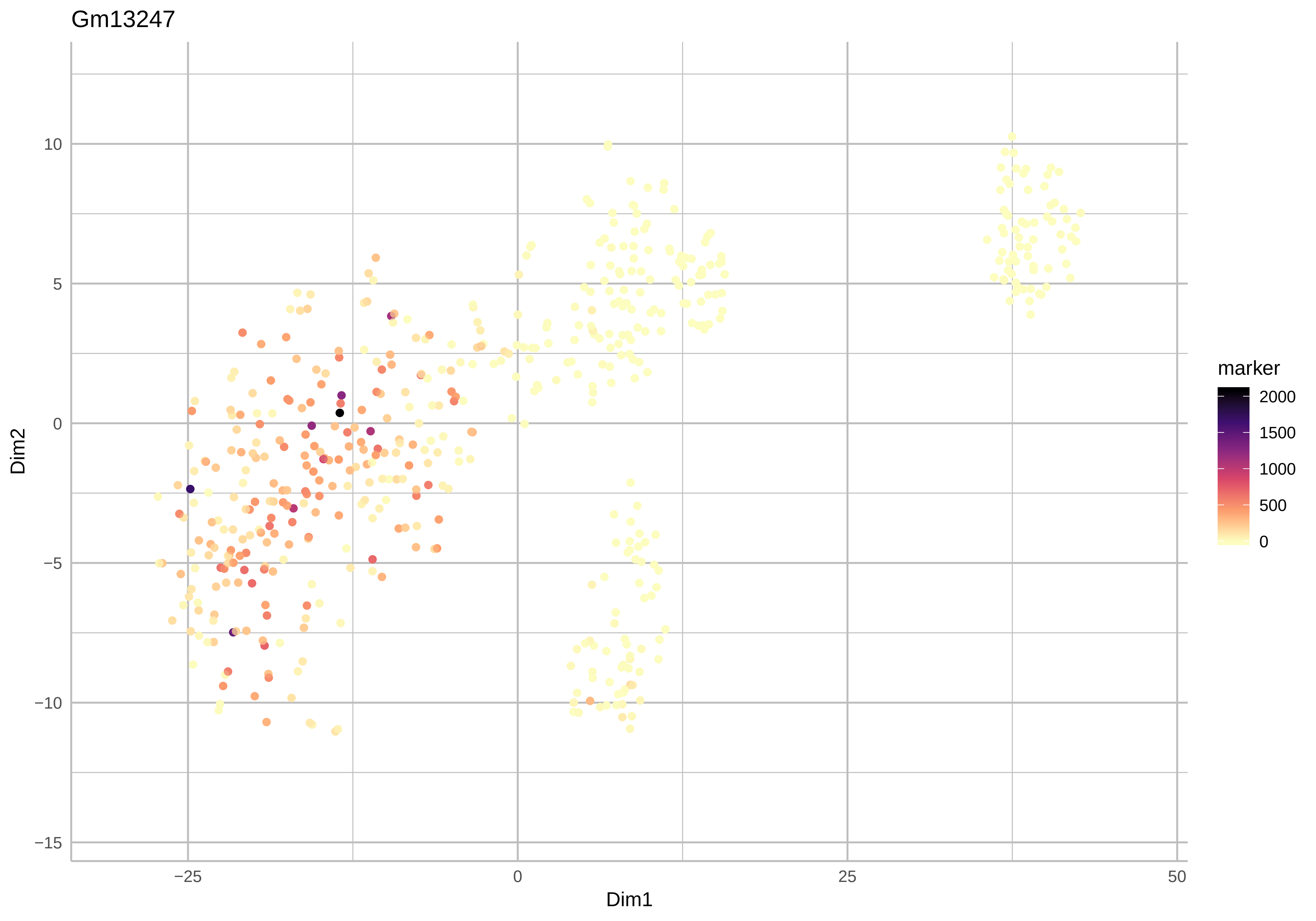

Gm13247
10
5
0
−5
−10
−15
Dim2
−25
0
25
50
Dim1
marker
2000
1500
1000
500
0

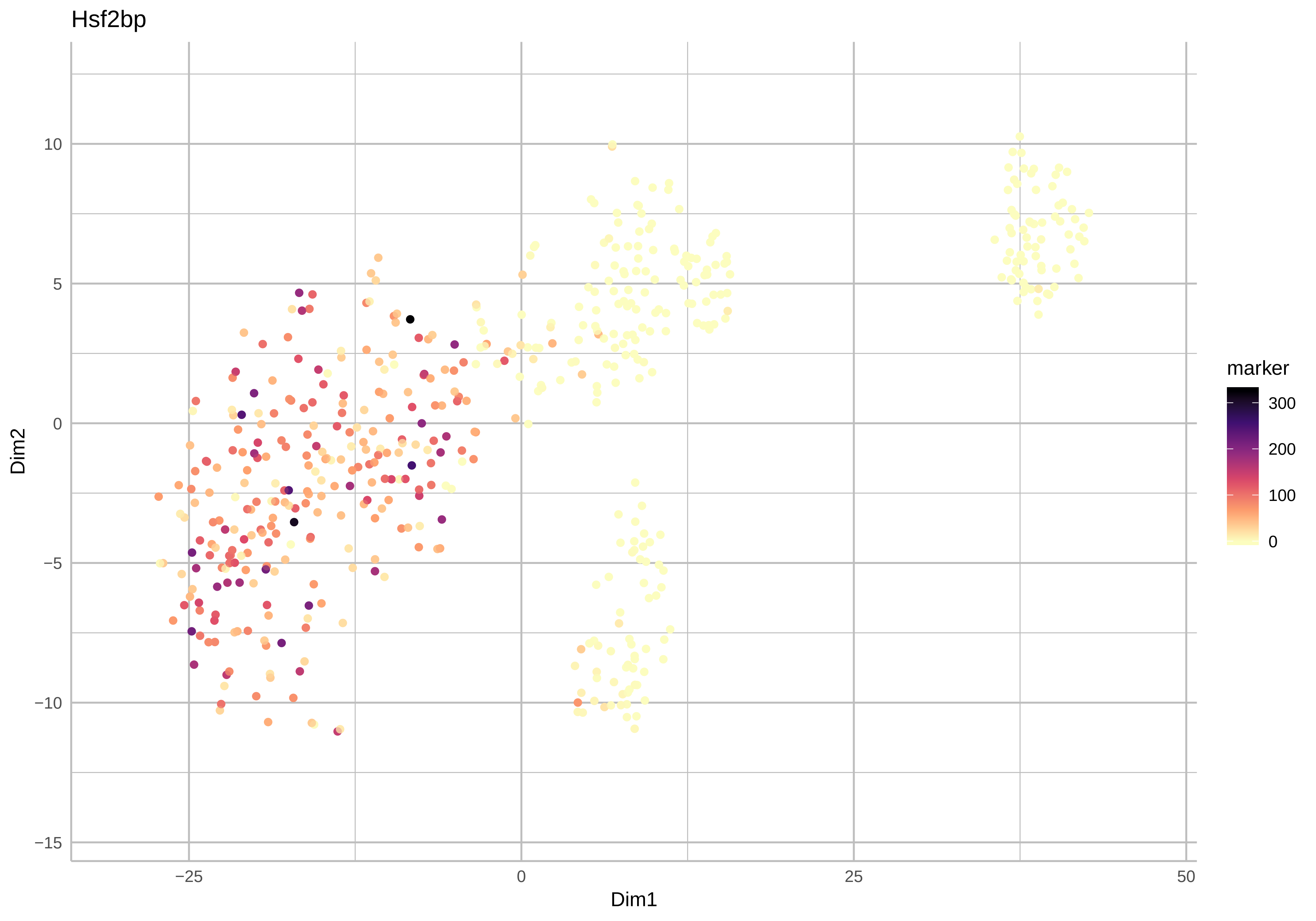

Hsf2bp
Dim2
Dim1
10
5
0
−5
−10
−15
−25
0
25
50
marker
300
200
100
0

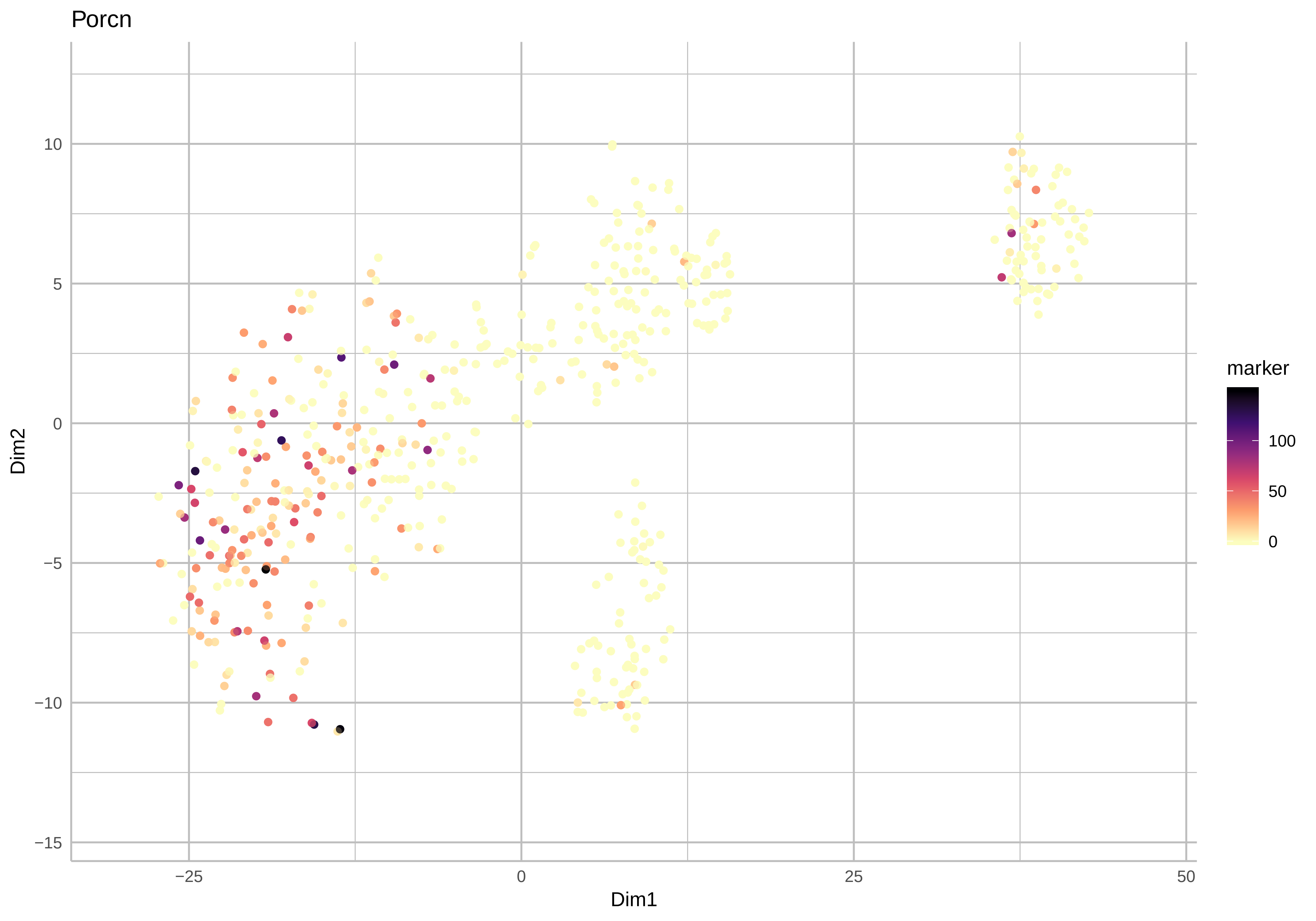

Porcn
Dim2
Dim1
10
5
0
−5
−10
−15
−25
0
25
50
marker
100
50
0

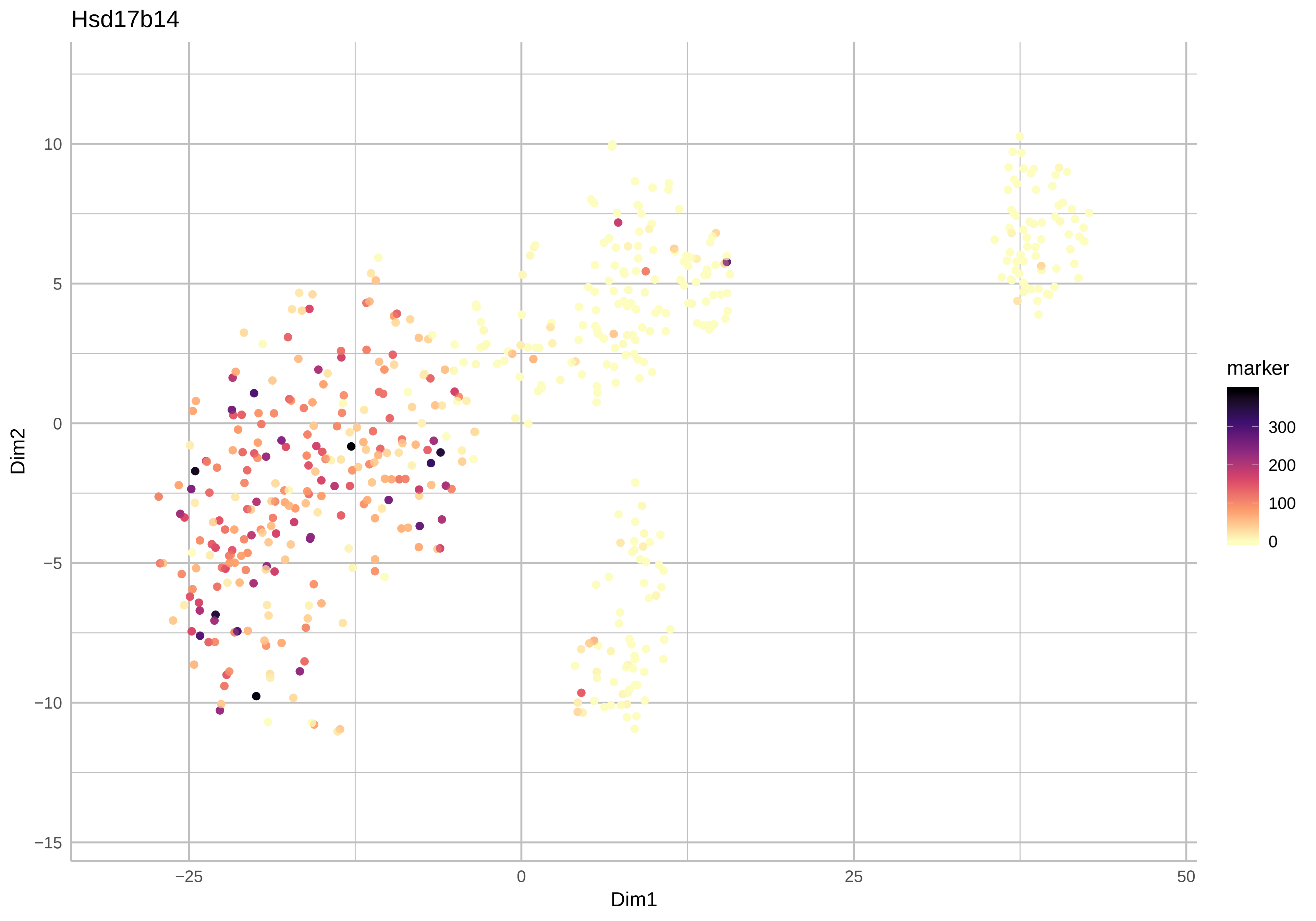
Hsd17b14
10
5
0
−5
−10
−15
−25
0
25
50
Dim1
Dim2
marker
300
200
100
0

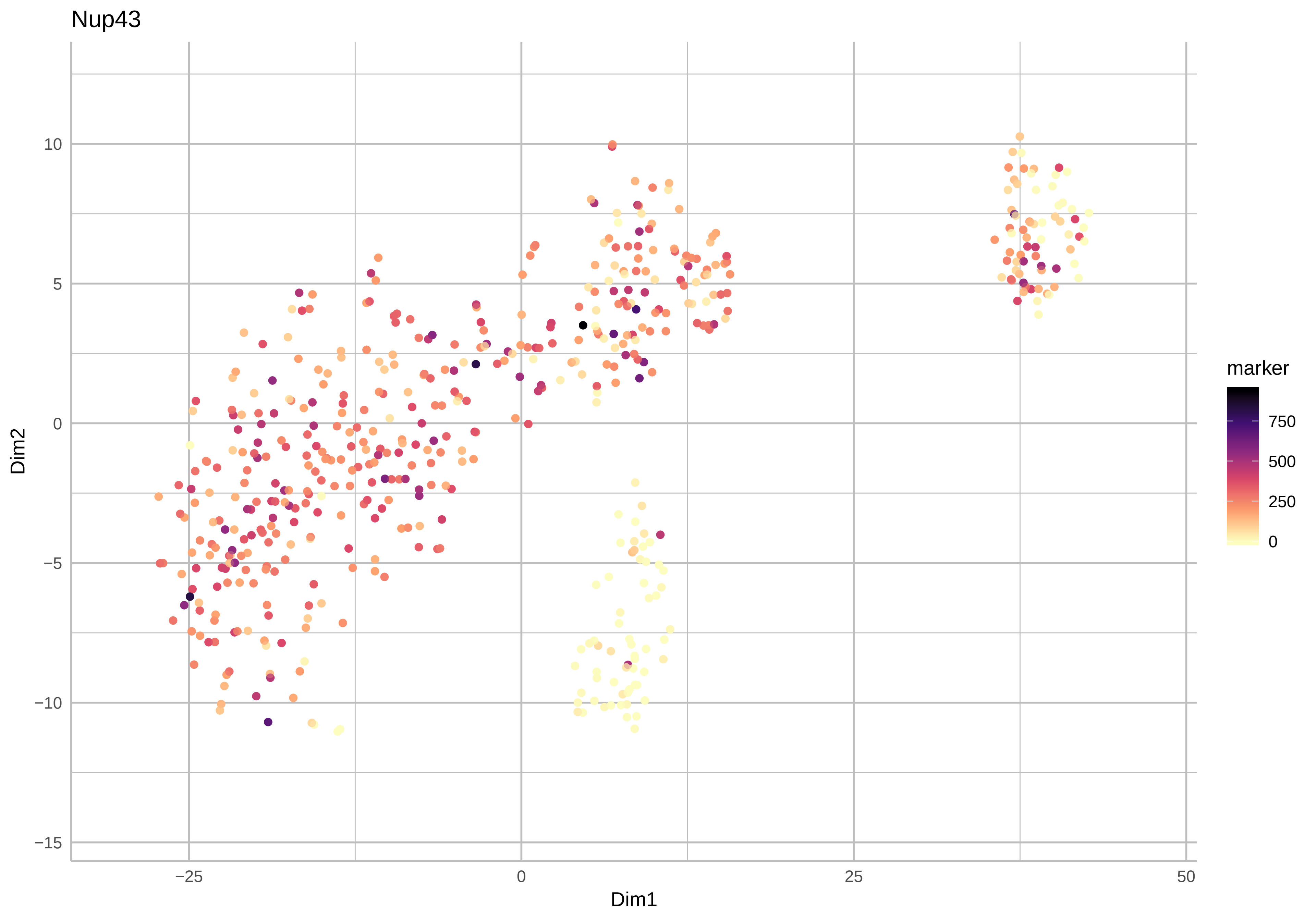
Nup43
Dim2
Dim1
10
5
0
−5
−10
−15
−25
0
25
50
marker
750
500
250
0

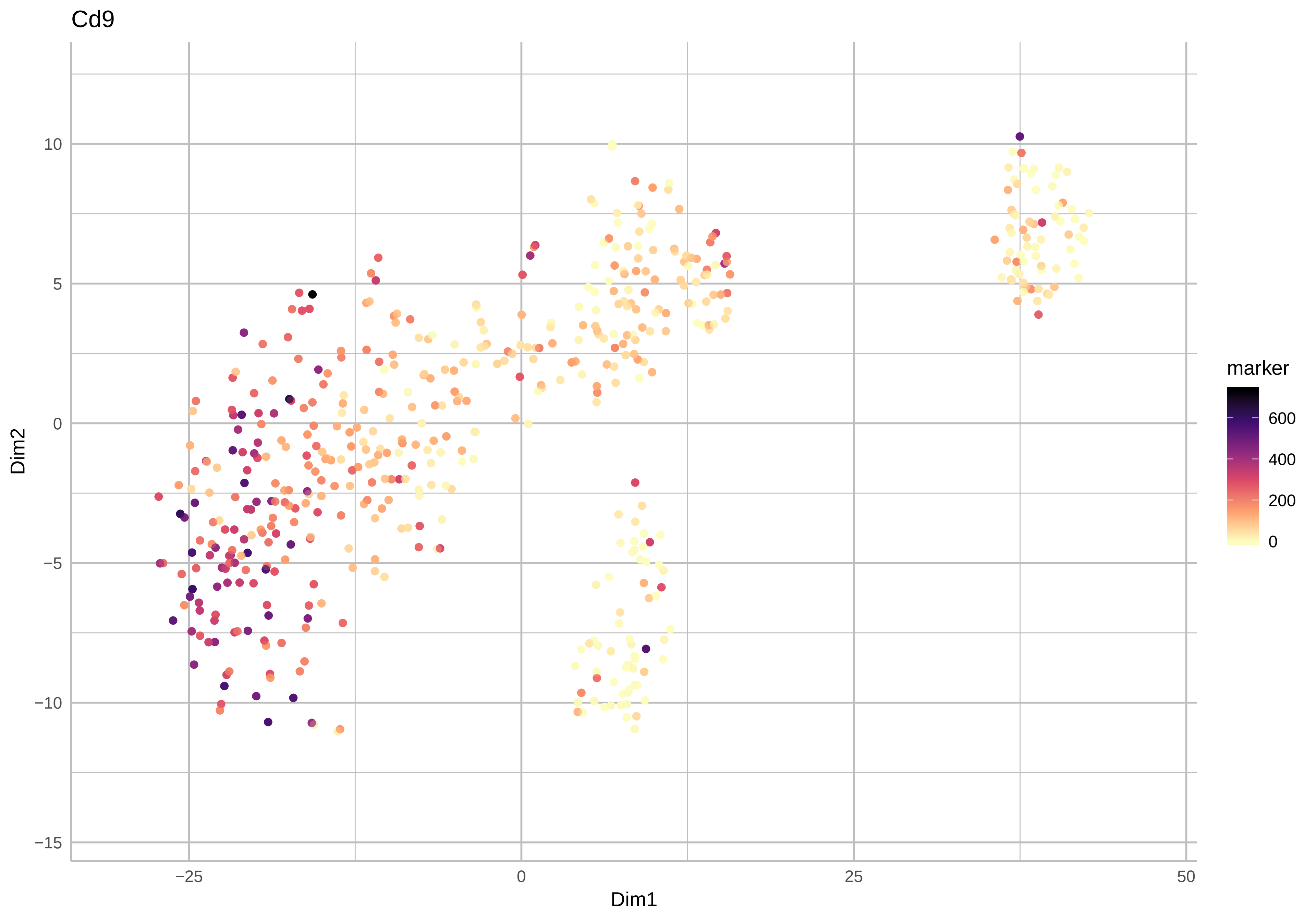
Cd9
Dim2
Dim1
10
5
0
−5
−10
−15
−25
0
25
50
marker
600
400
200
0

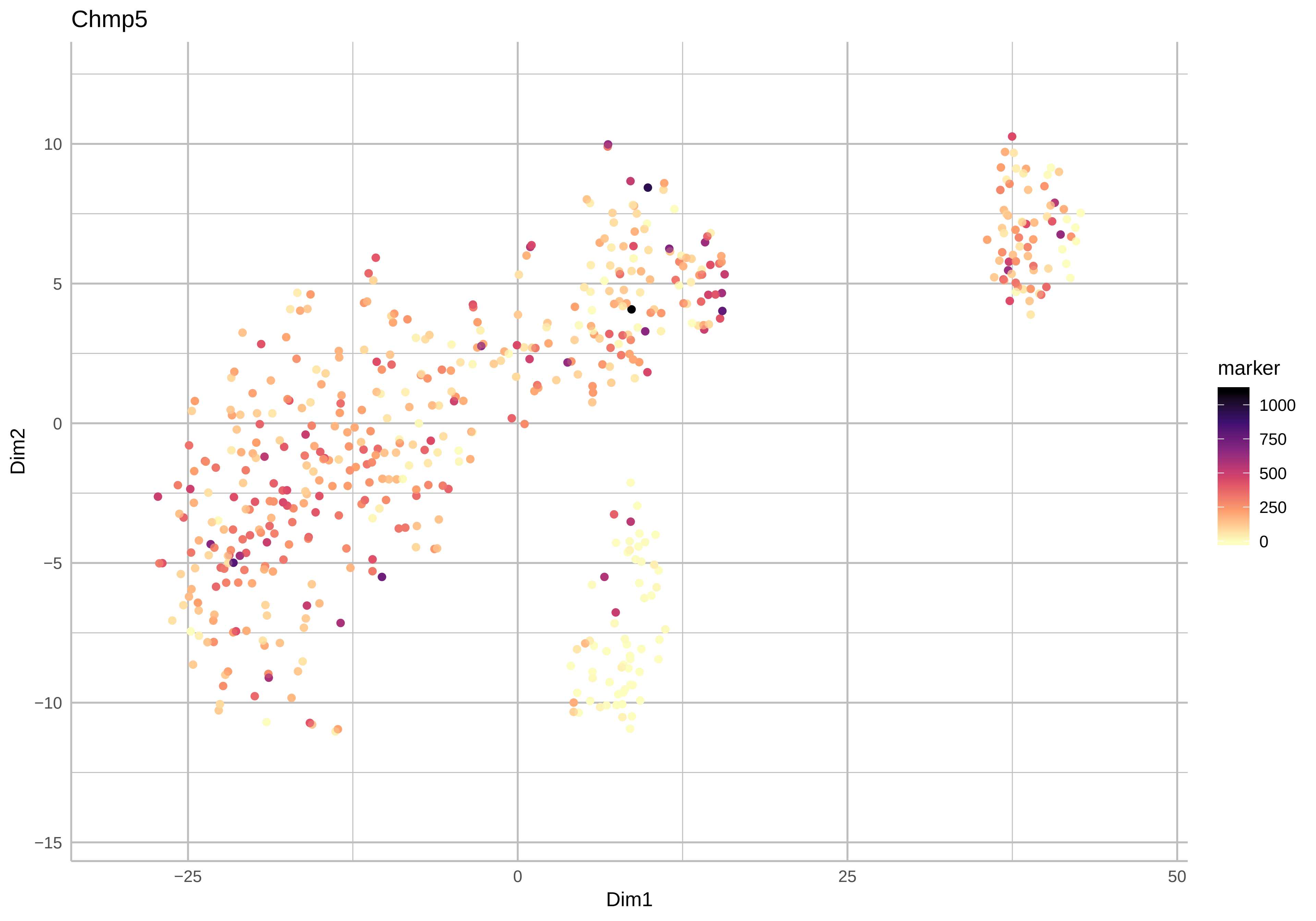
Chmp5
Dim2
Dim1
10
5
0
−5
−10
−15
−25
0
25
50
marker
1000
750
500
250
0

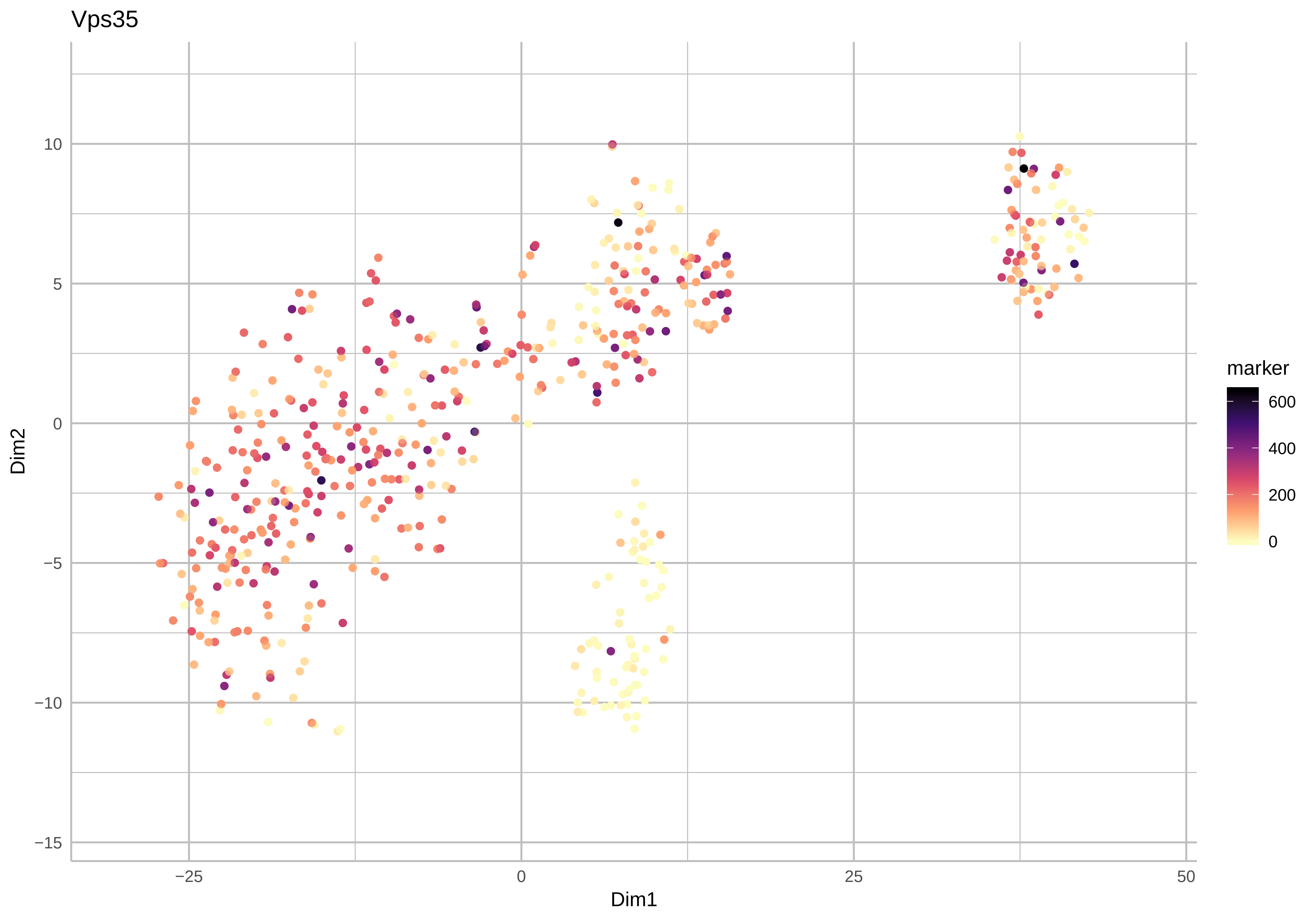

Vps35
Dim2
Dim1
10
5
0
−5
−10
−15
−25
0
25
50
marker
600
400
200
0

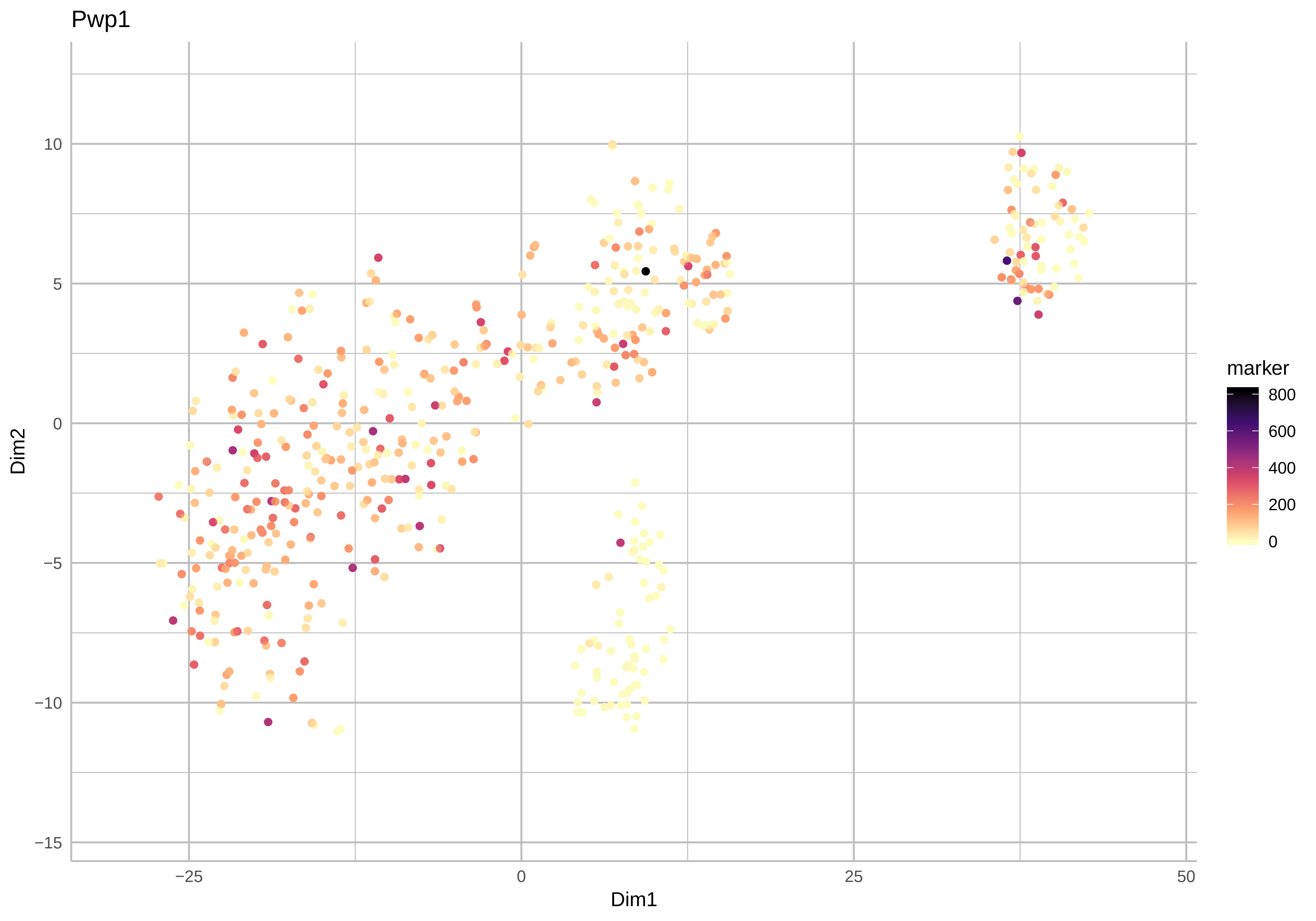

Pwp1
Dim2
Dim1
10
5
0
−5
−10
−15
−25
0
25
50
marker
800
600
400
200
0

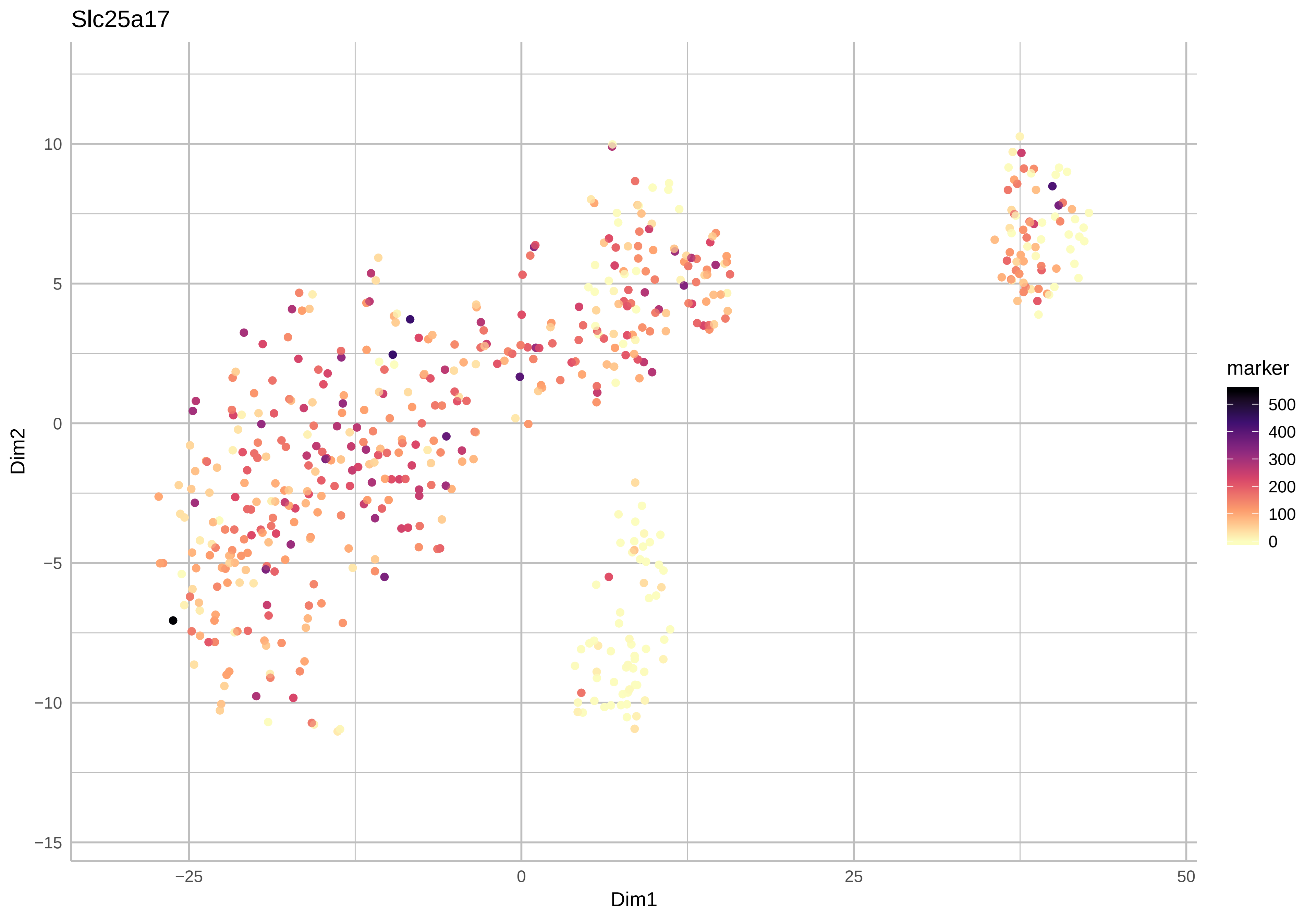

Slc25a17
10
5
0
−5
−10
−15
Dim2
−25
0
25
50
Dim1
marker
500
400
300
200
100
0

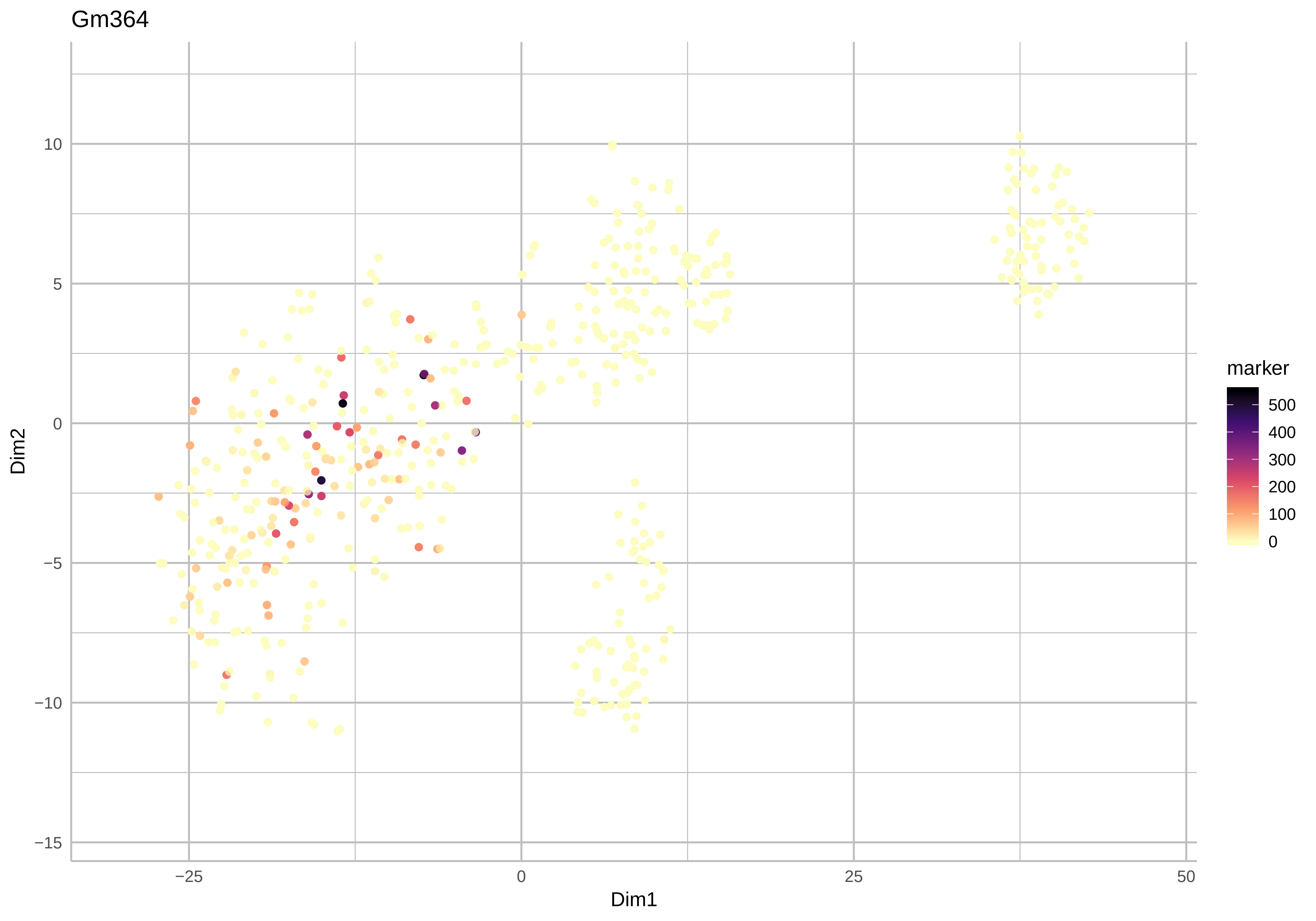
Gm364
Dim2
Dim1
10
5
0
−5
−10
−15
−25
0
25
50
marker
500
400
300
200
100
0

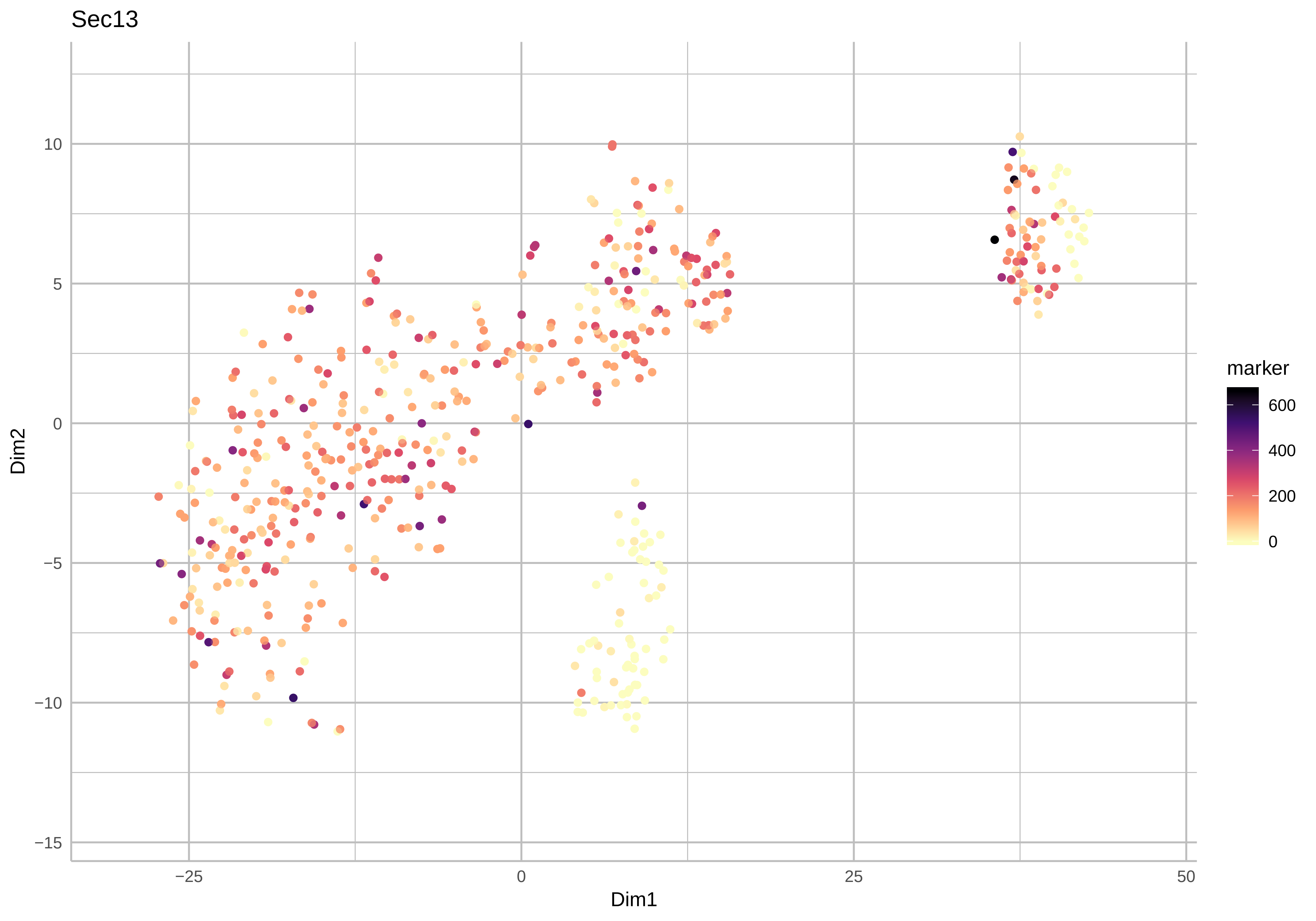
Sec13
Dim2
Dim1
10
5
0
−5
−10
−15
−25
0
25
50
marker
600
400
200
0

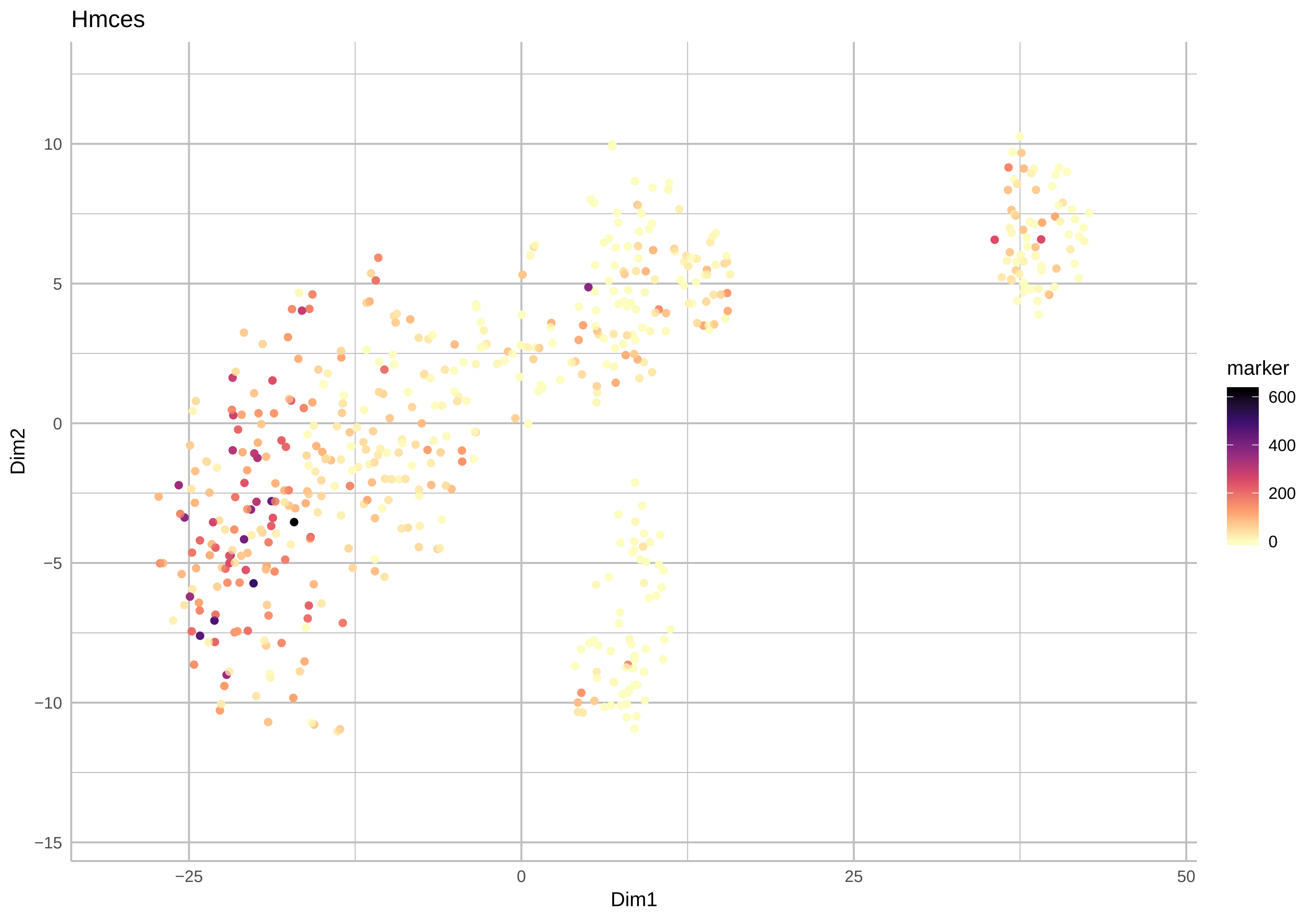
Hmces
Dim2
Dim1
marker
600
400
200
0
10
5
0
−5
−10
−15
−25
0
25
50

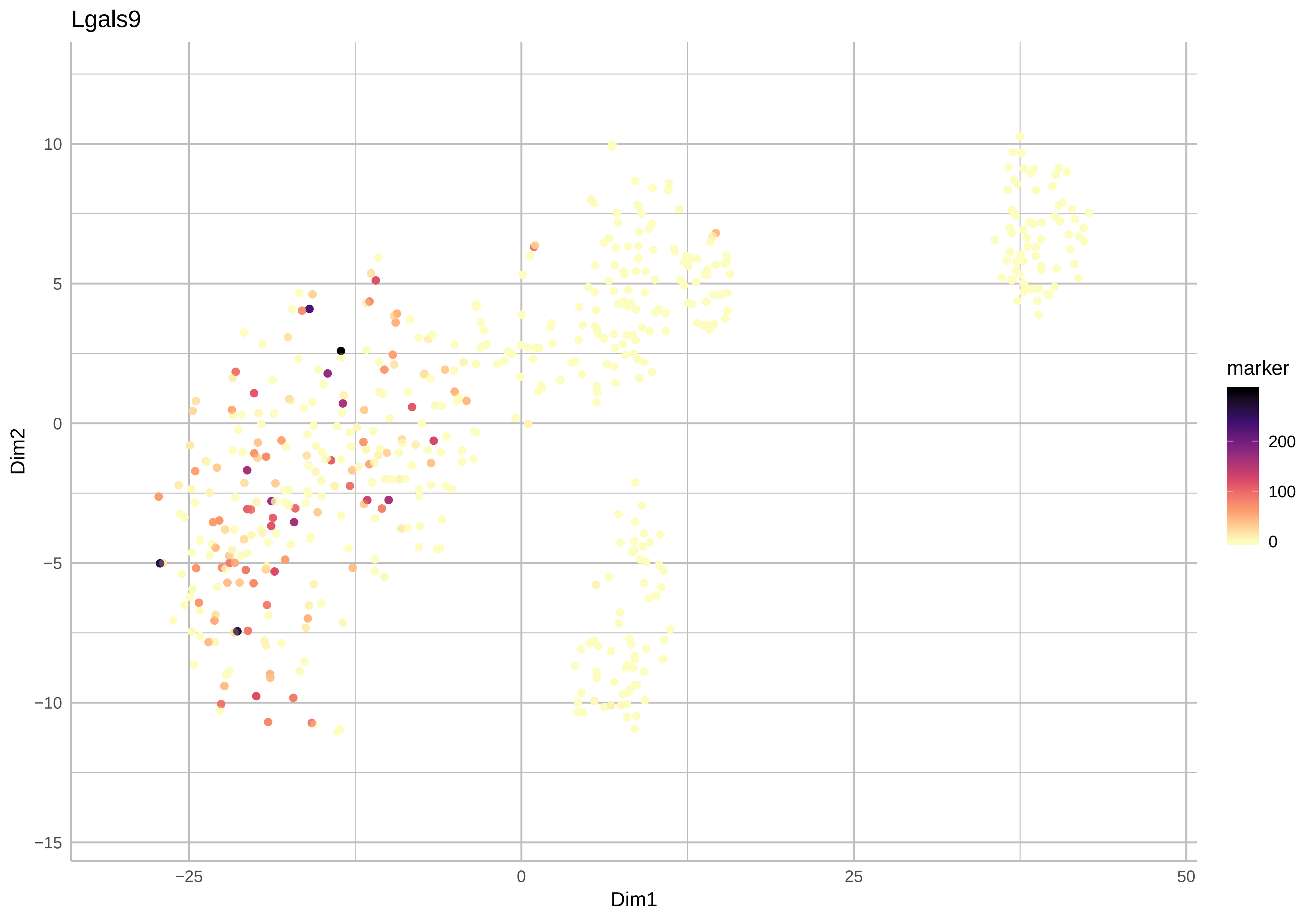

Lgals9
10
5
0
−5
−10
−15
Dim2
−25
0
25
50
Dim1
marker
200
100
0

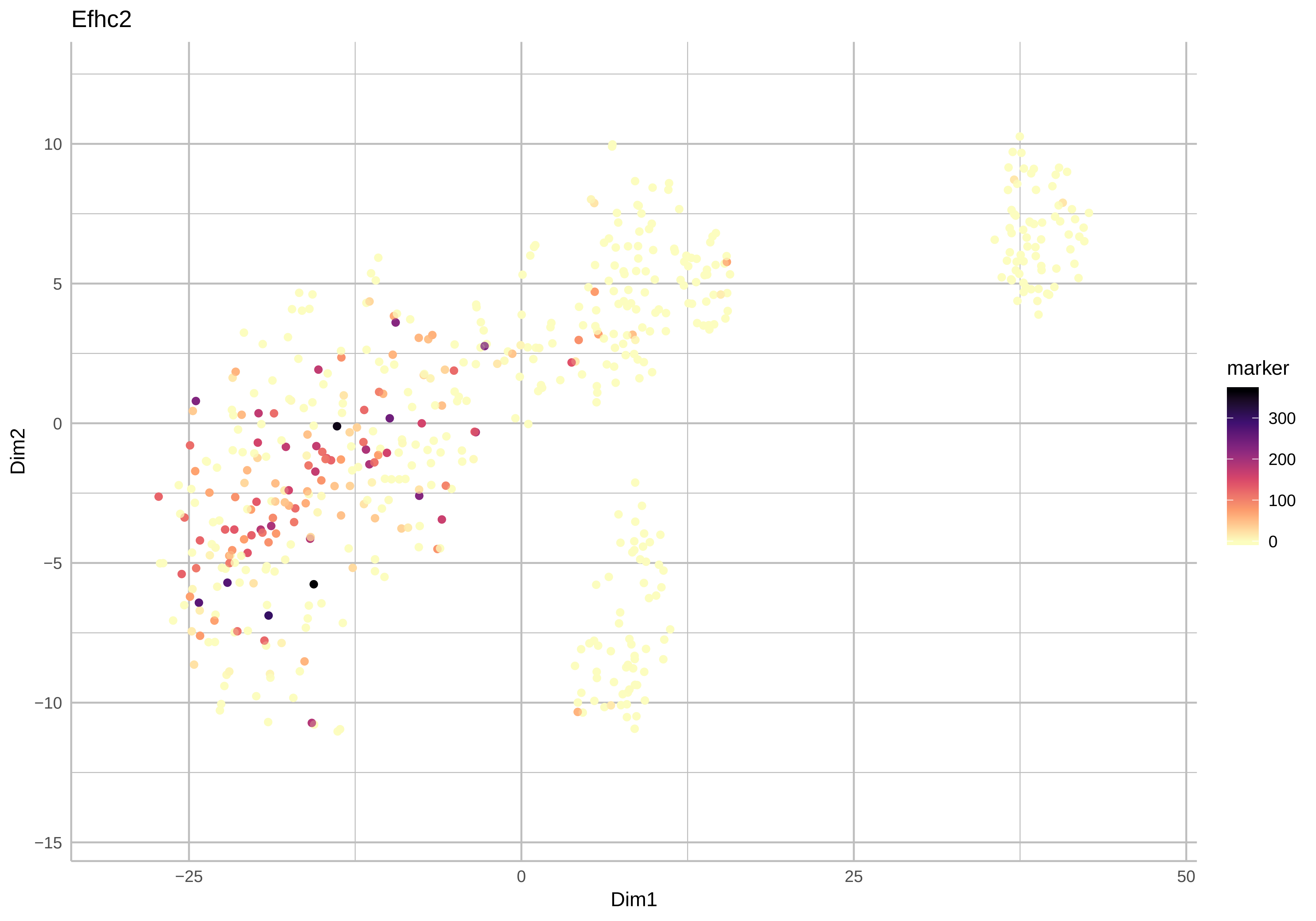

Efhc2
Dim2
Dim1
marker
300
200
100
0
10
5
0
−5
−10
−15
−25
0
25
50

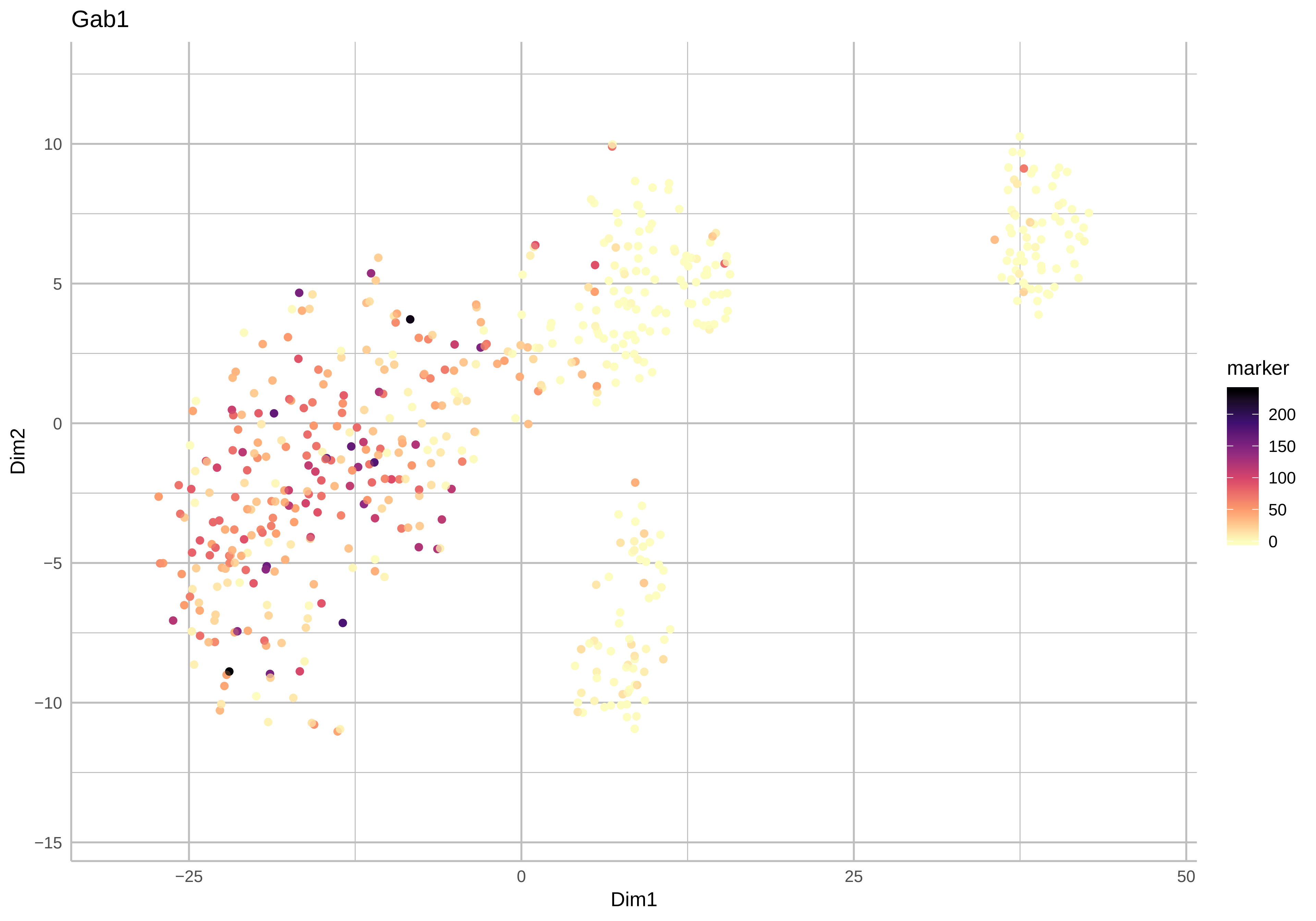
Gab1
Dim2
Dim1
10
5
0
−5
−10
−15
−25
0
25
50
marker
200
150
100
50
0

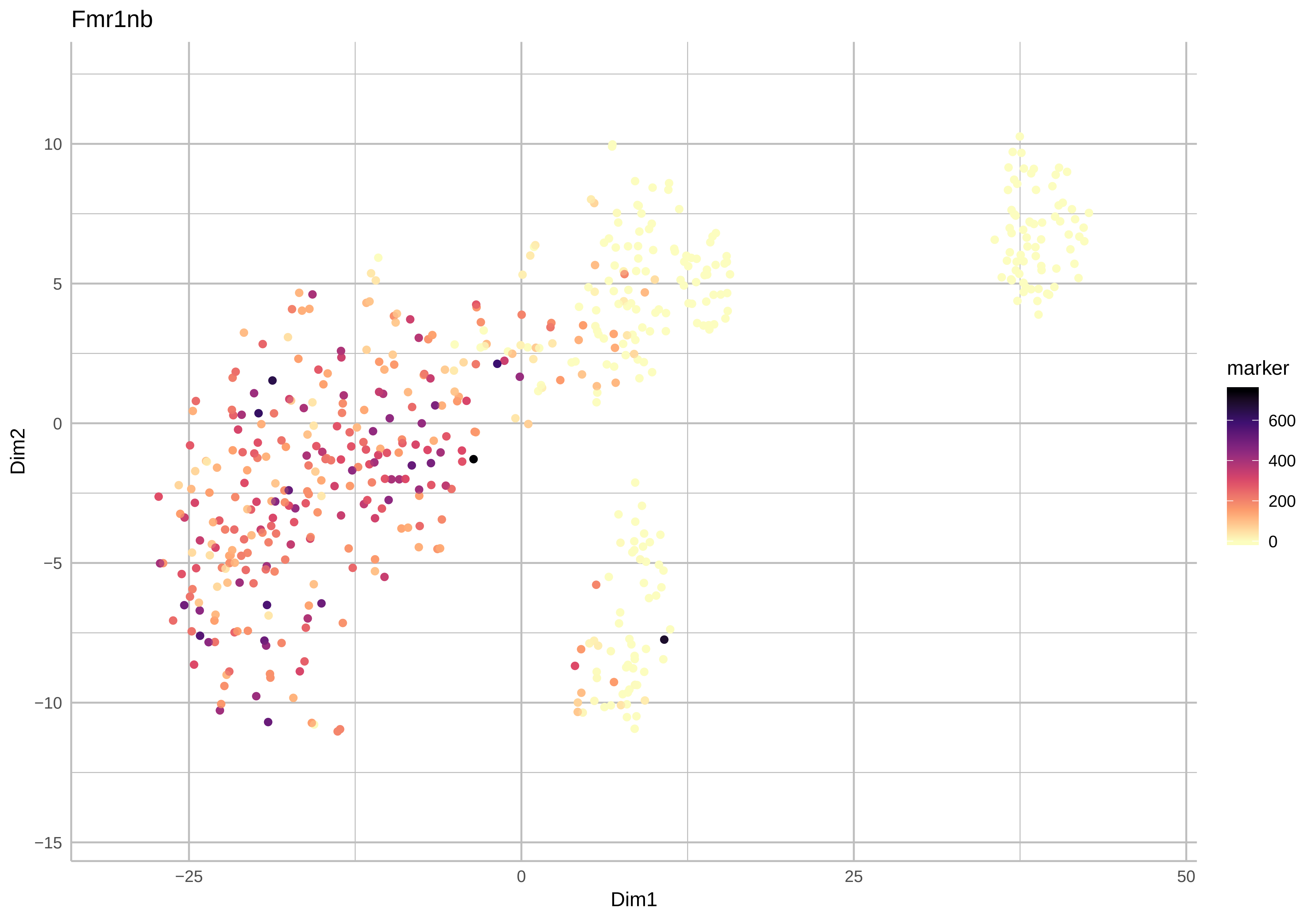

Fmr1nb
10
5
0
−5
−10
−15
Dim2
−25
0
25
50
Dim1
marker
600
400
200
0

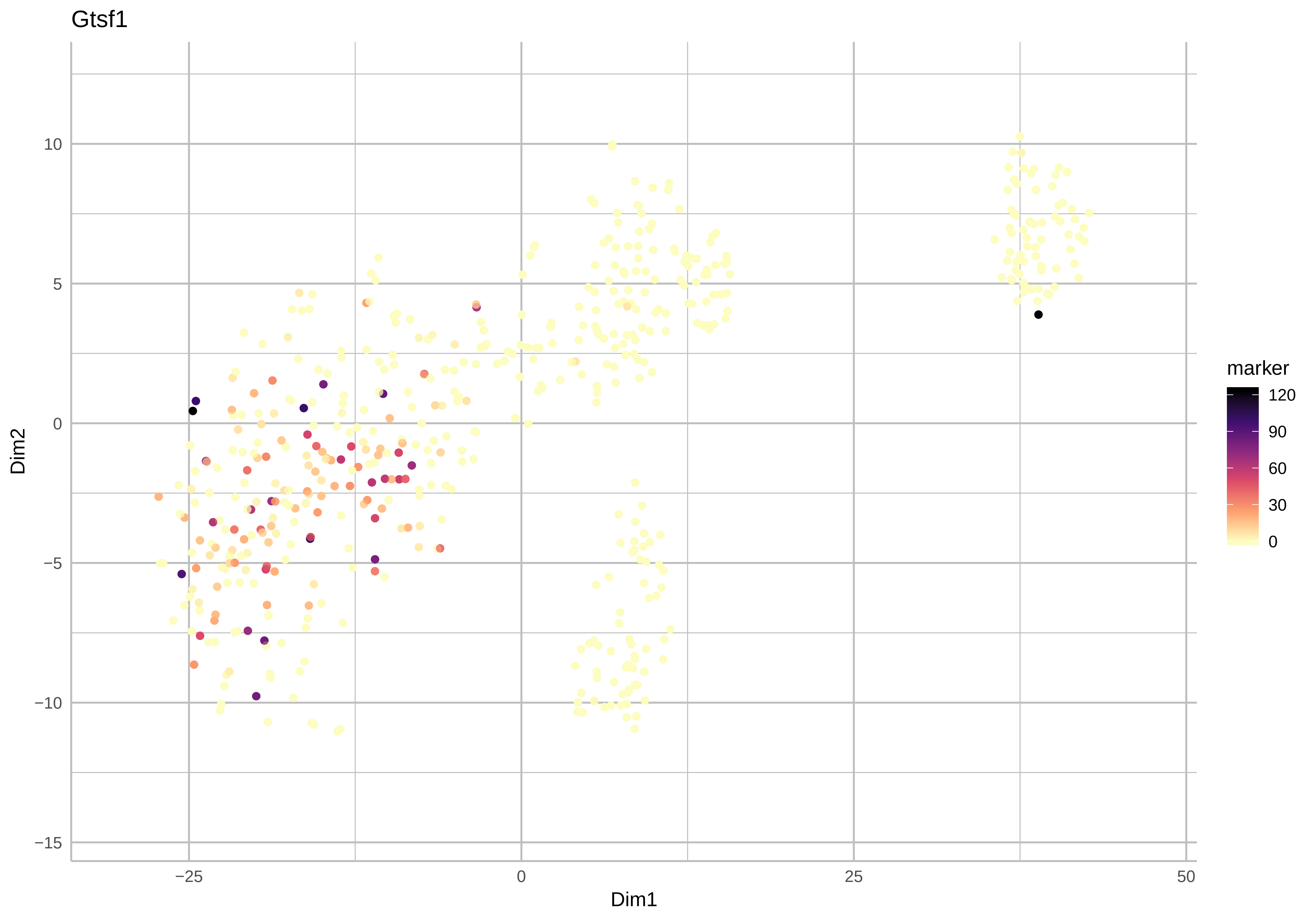

Gtsf1
Dim2
Dim1
marker
120
90
60
30
0
10
5
0
−5
−10
−15
−25
0
25
50

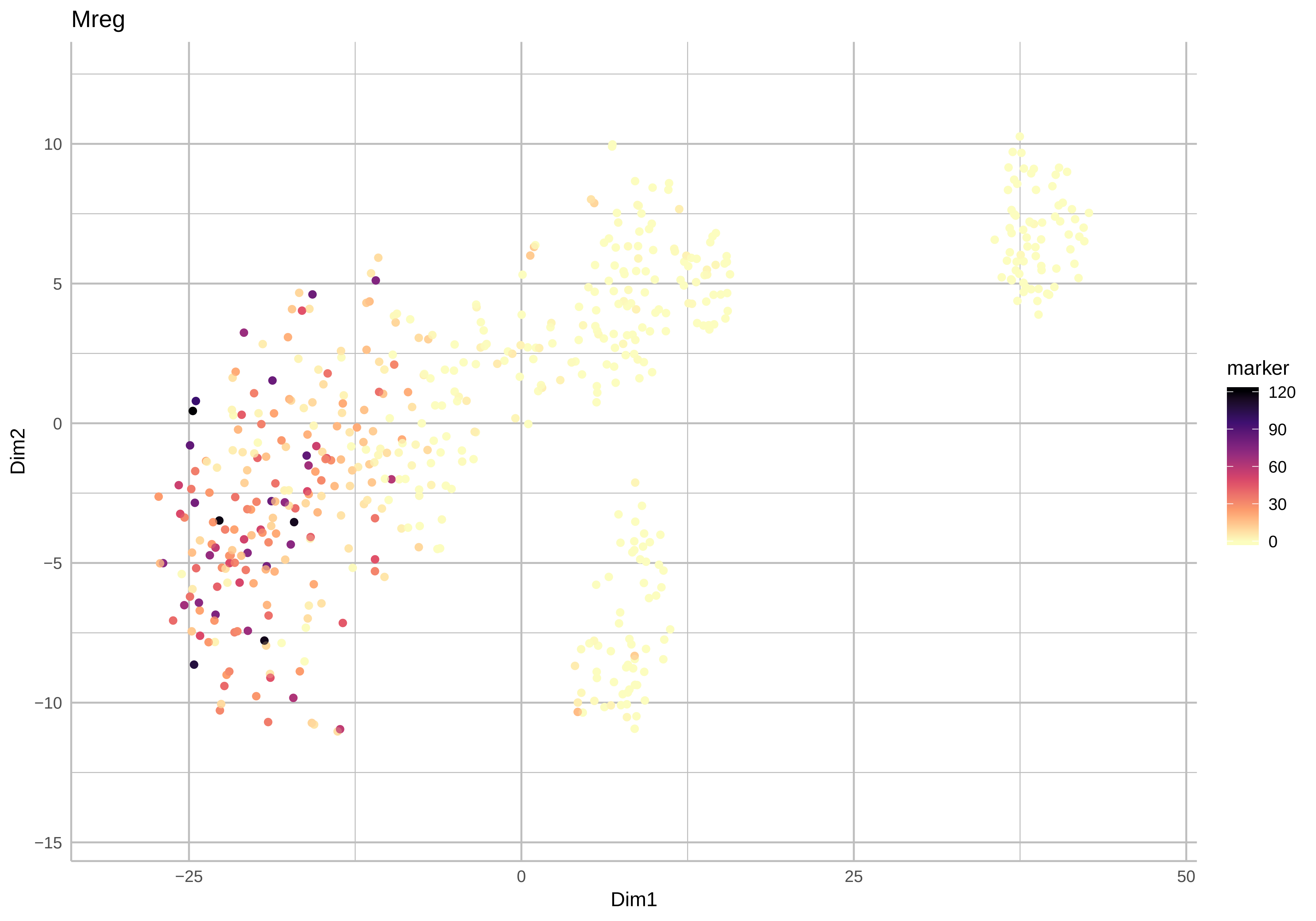
Mreg
Dim2
Dim1
10
5
0
−5
−10
−15
−25
0
25
50
marker
120
90
60
30
0

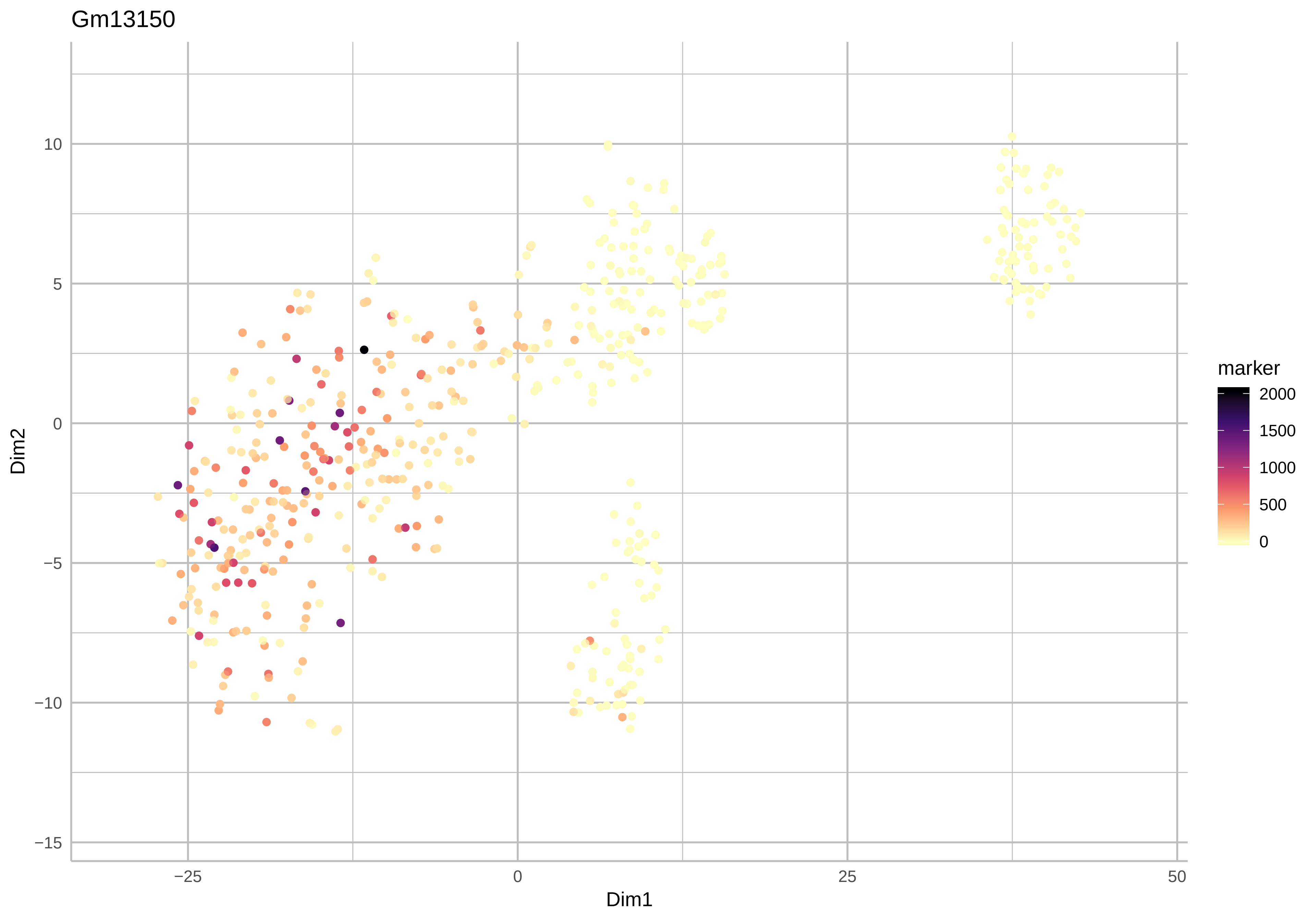

Gm13150
Dim2
10
5
0
−5
−10
−15
Dim1
−25
0
25
50
marker
2000
1500
1000
500
0

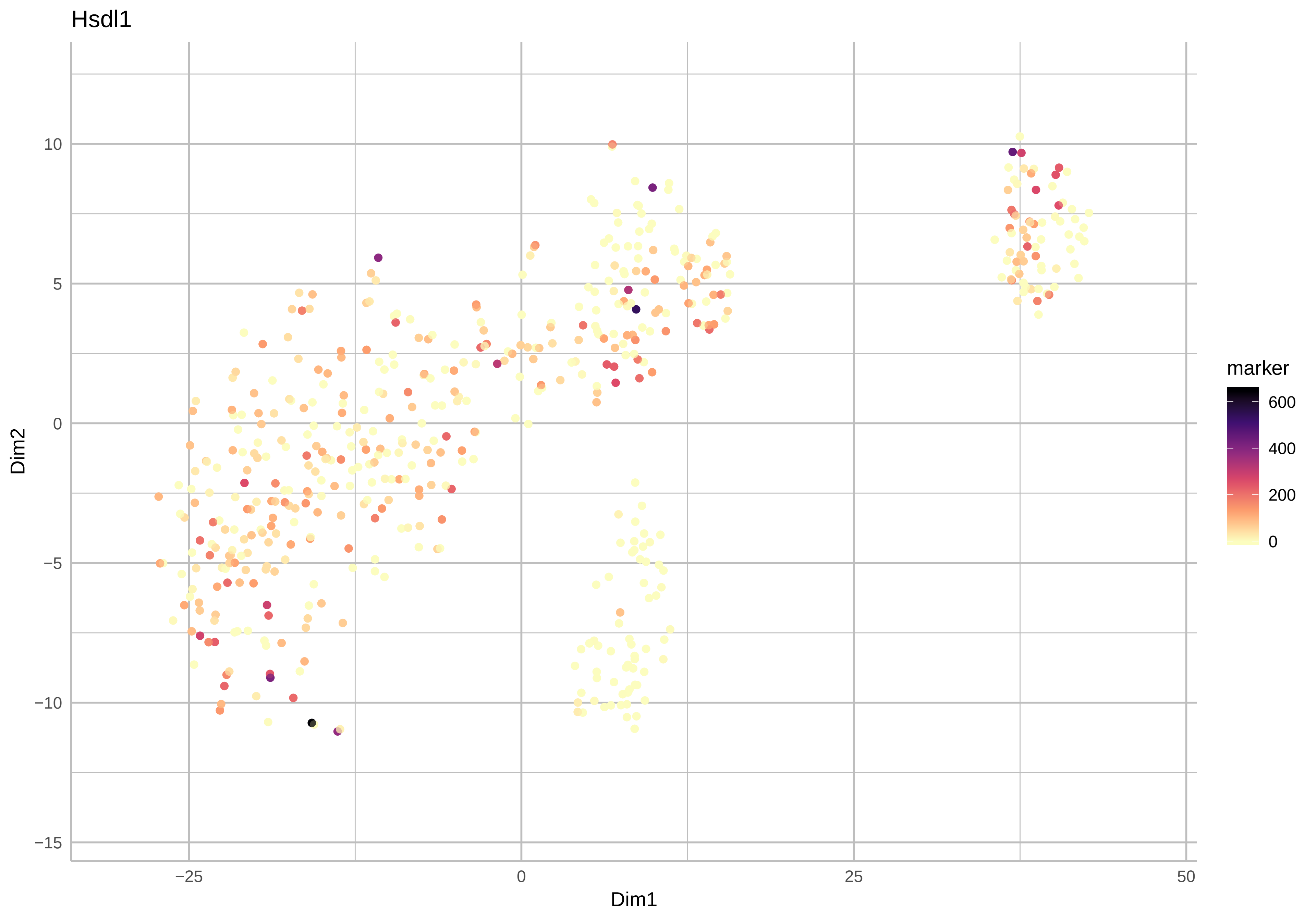
Hsdl1
Dim2
Dim1
10
5
0
−5
−10
−15
−25
0
25
50
marker
600
400
200
0

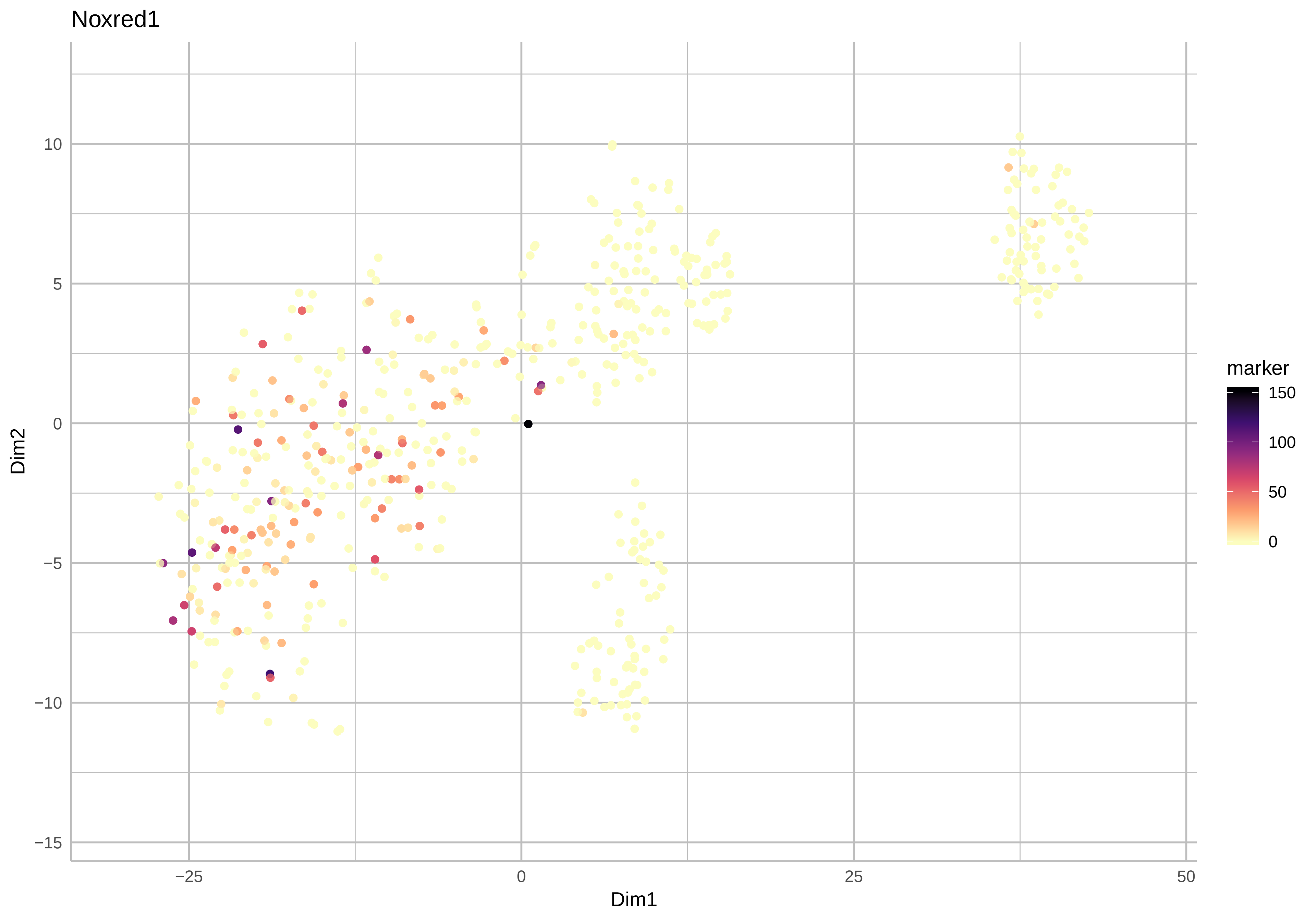
Noxred1
Dim2
Dim1
10
5
0
−5
−10
−15
−25
0
25
50
marker
150
100
50
0

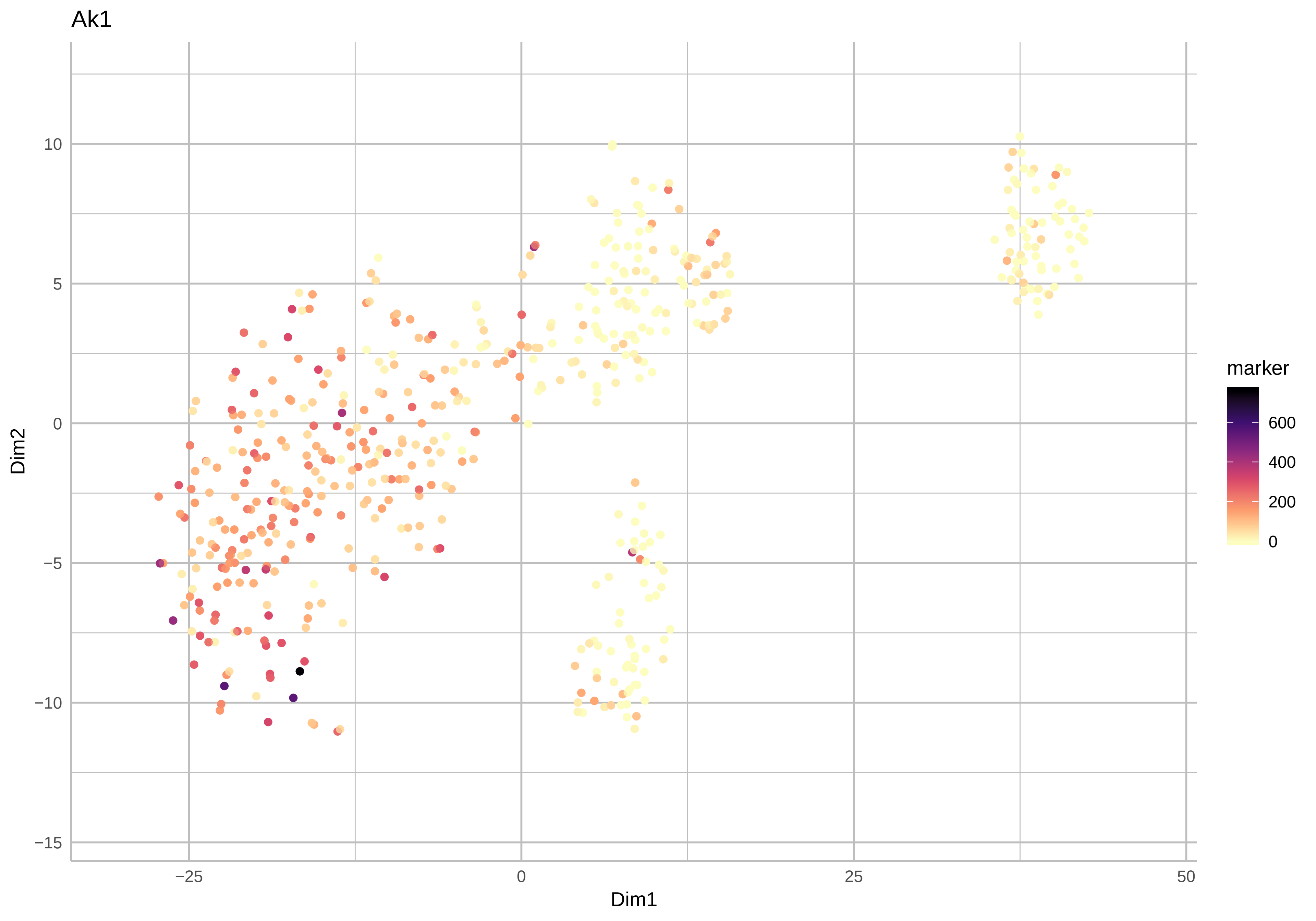

Ak1
10
5
0
−5
−10
−15
Dim2
−25
0
25
50
Dim1
marker
600
400
200
0

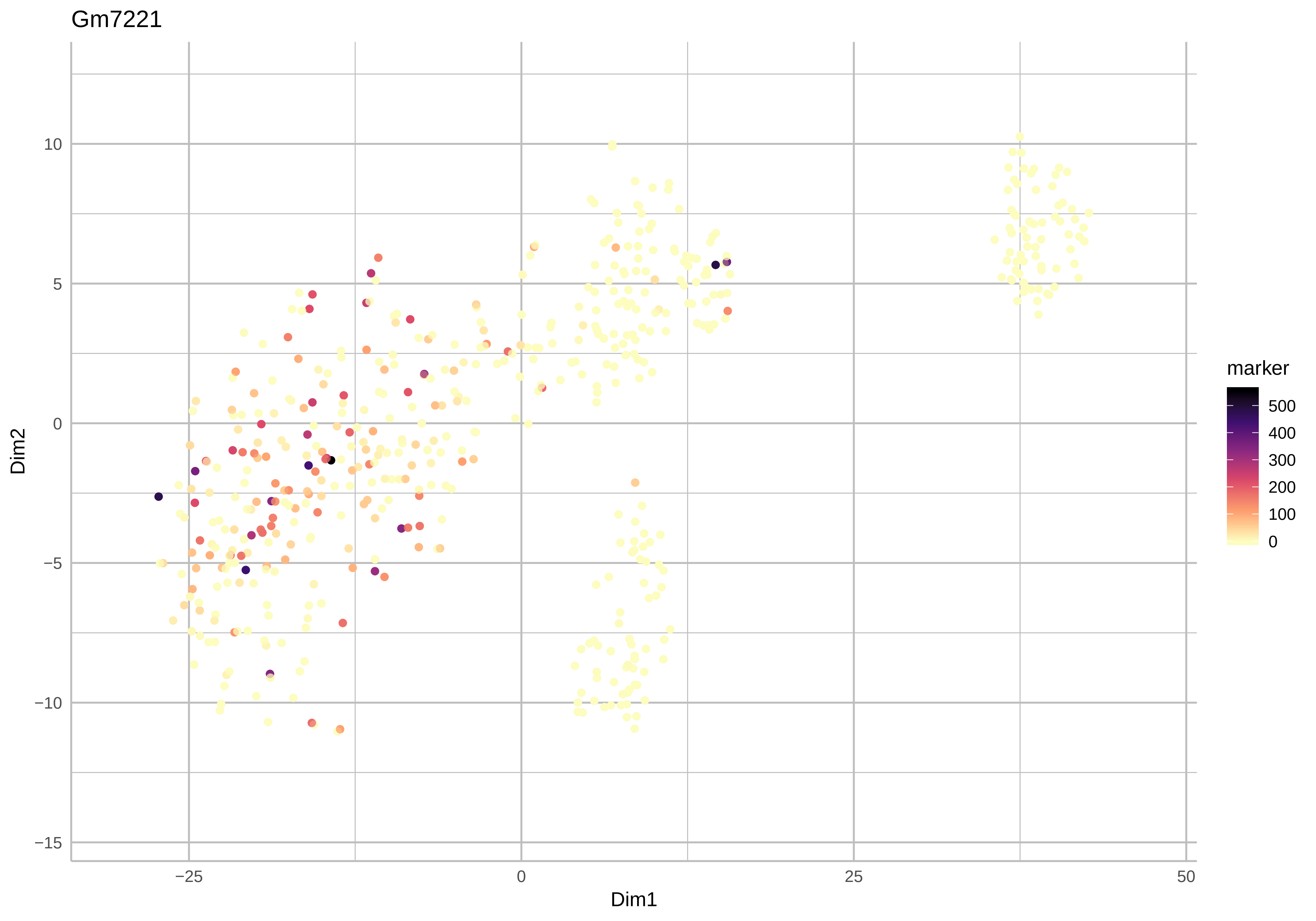
Gm7221
Dim2
Dim1
10
5
0
−5
−10
−15
−25
0
25
50
marker
500
400
300
200
100
0

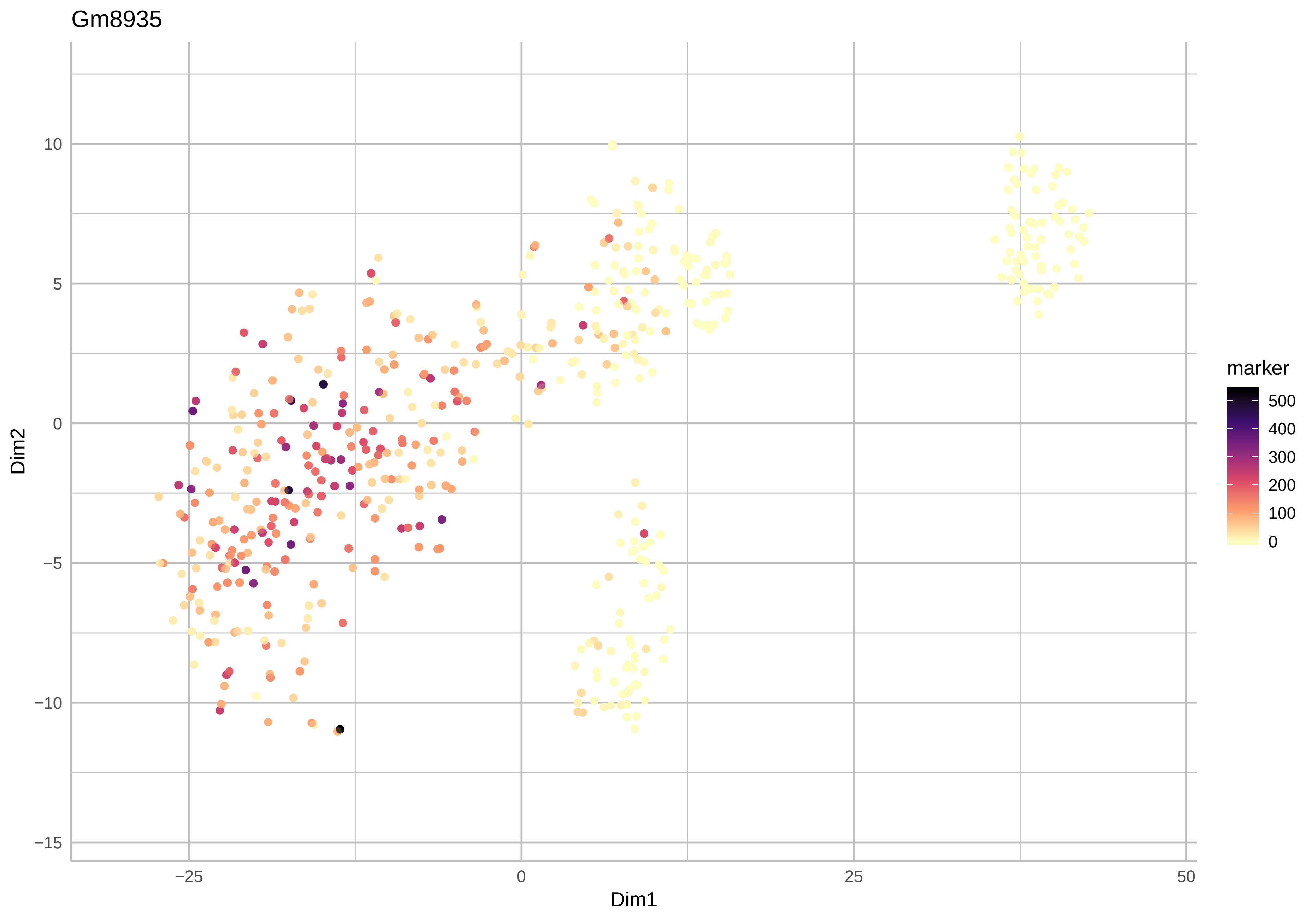

Gm8935
10
5
0
−5
−10
−15
−25
0
25
50
Dim1
Dim2
marker
500
400
300
200
100
0

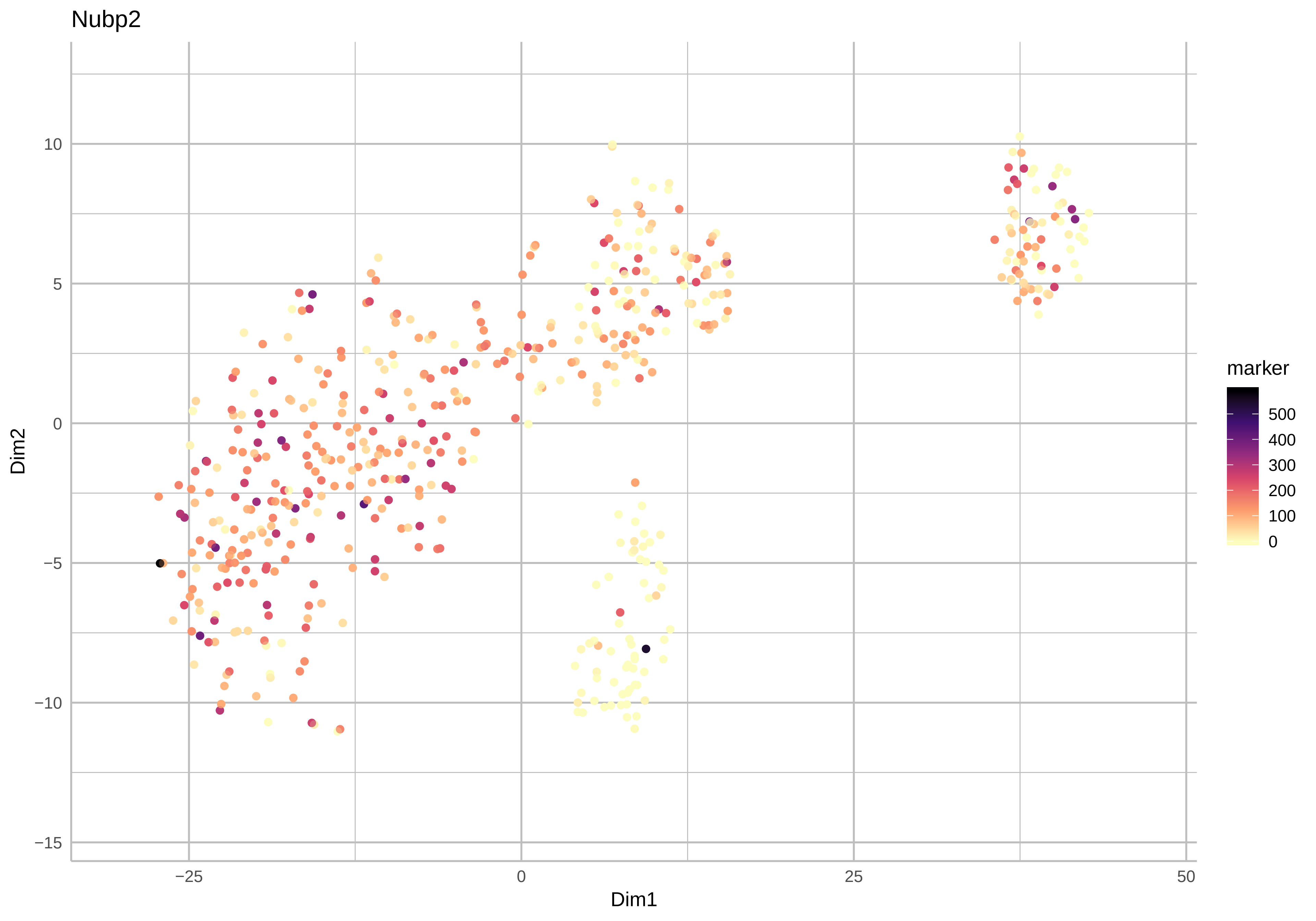

Nubp2
Dim2
Dim1
10
5
0
−5
−10
−15
−25
0
25
50
marker
500
400
300
200
100
0

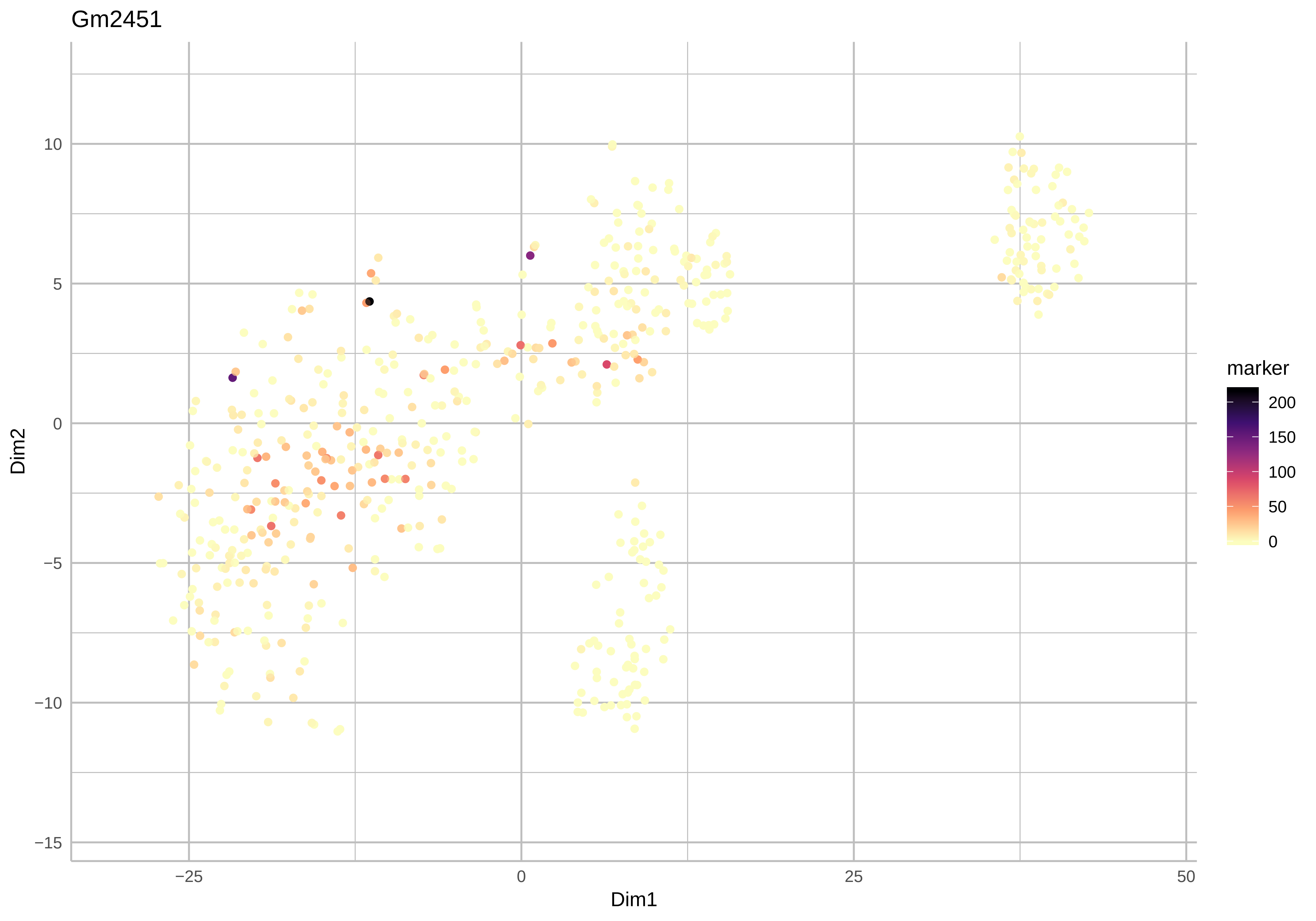

Gm2451
10
5
0
−5
−10
−15
Dim2
−25
0
25
50
Dim1
marker
200
150
100
50
0

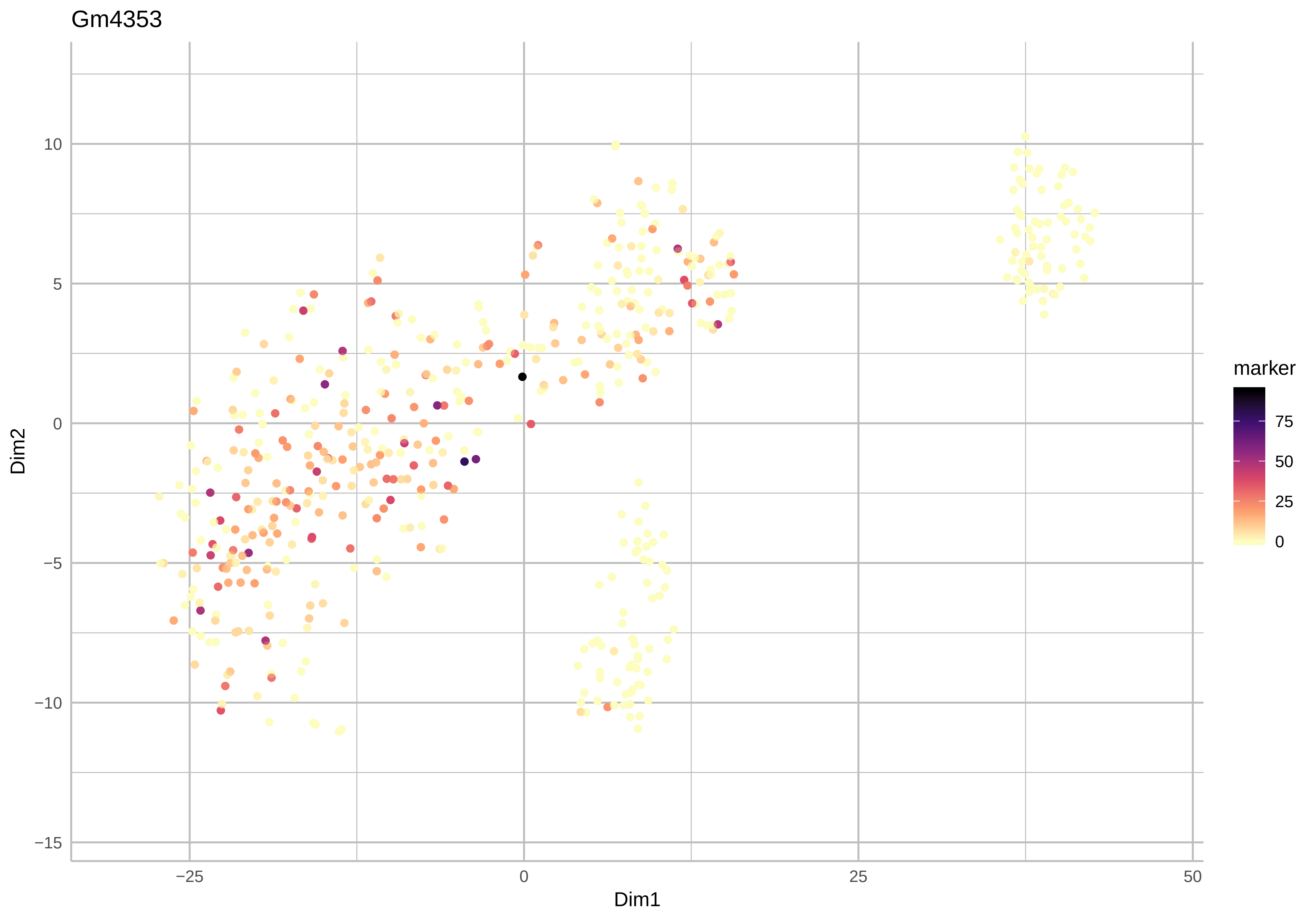
Gm4353
Dim2
Dim1
10
5
0
−5
−10
−15
−25
0
25
50
marker
75
50
25
0

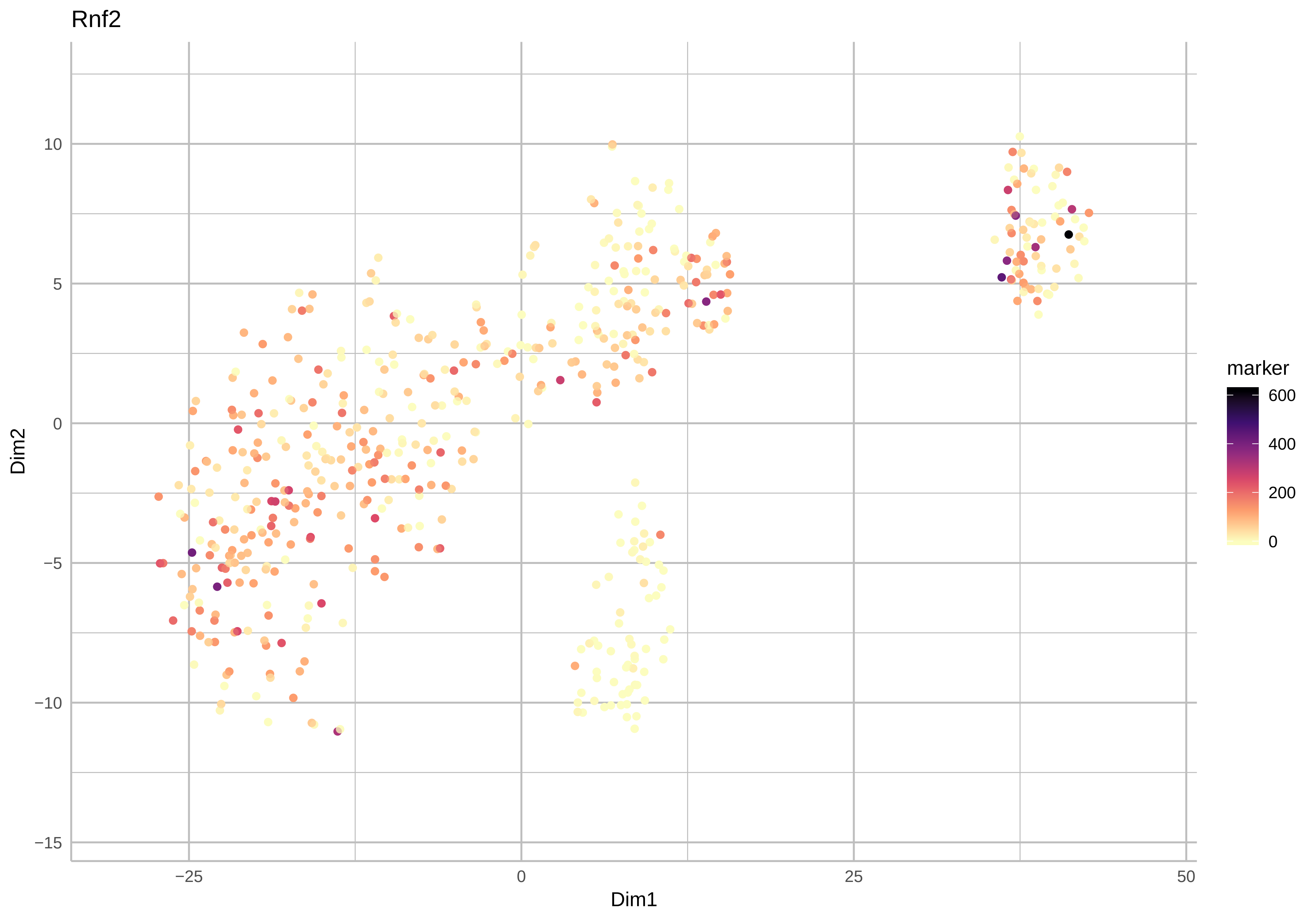

Rnf2
Dim2
Dim1
10
5
0
−5
−10
−15
−25
0
25
50
marker
600
400
200
0

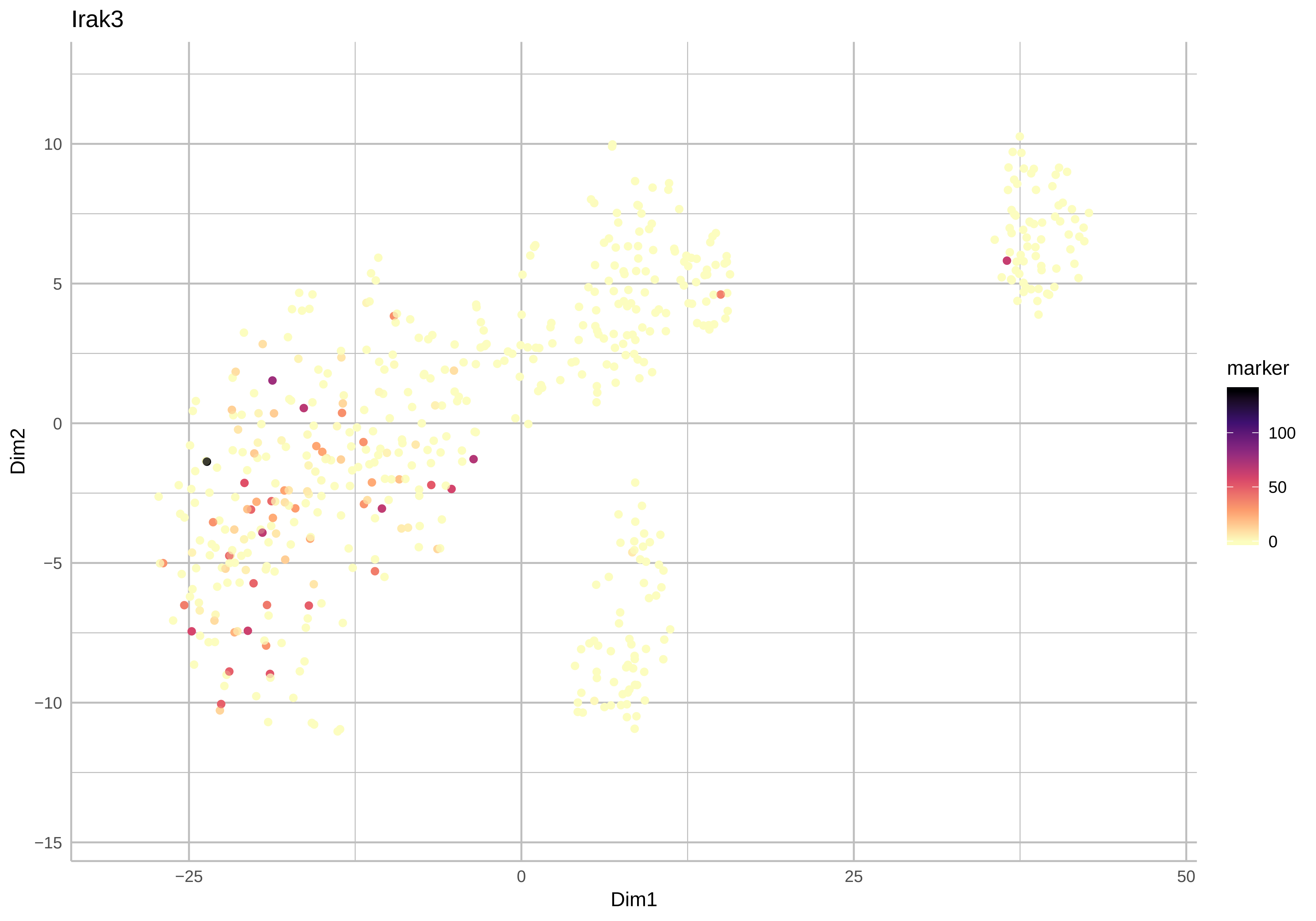
Irak3
Dim2
Dim1
10
5
0
−5
−10
−15
−25
0
25
50
marker
100
50
0

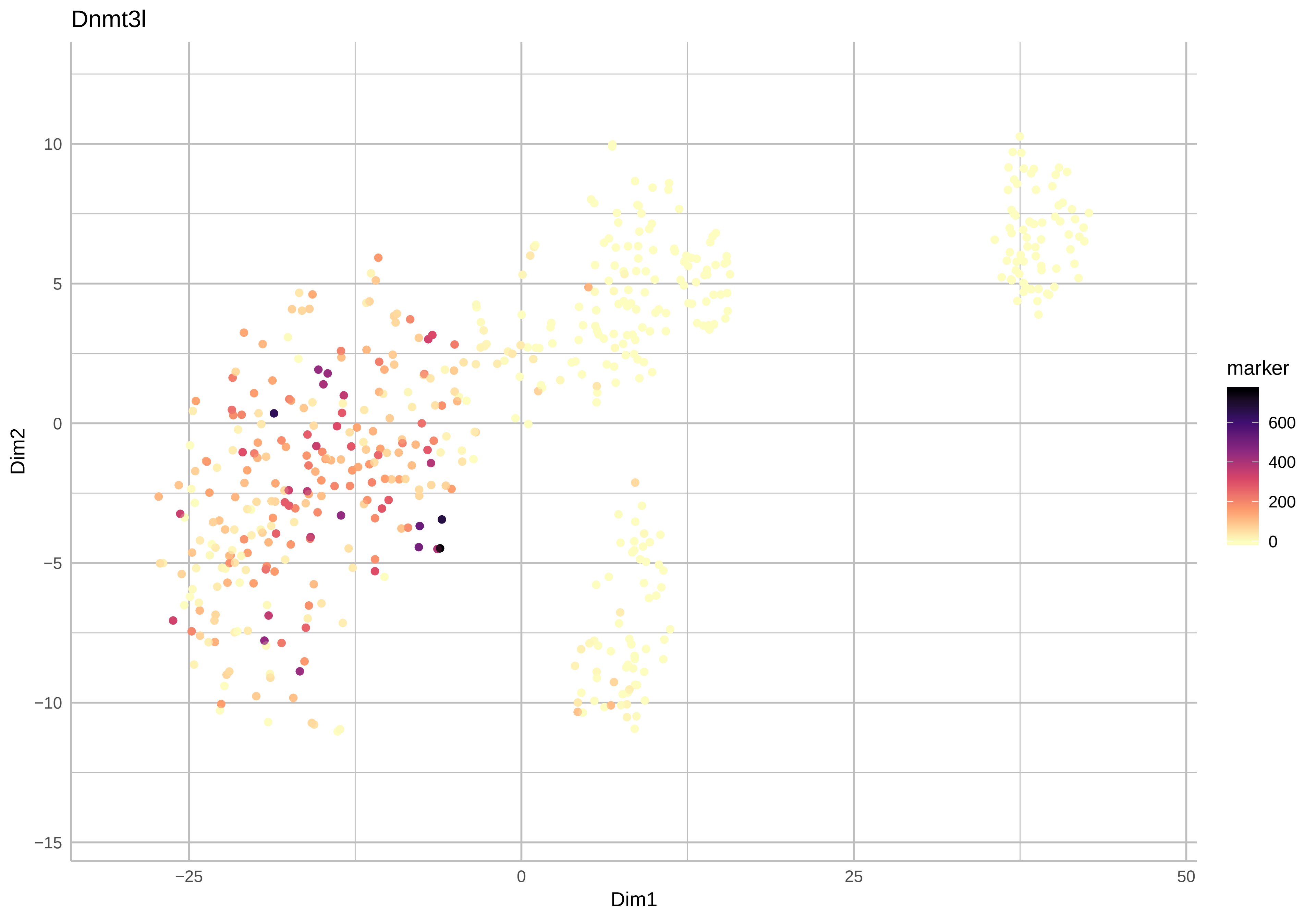

Dnmt3l
Dim2
Dim1
10
5
0
−5
−10
−15
−25
0
25
50
marker
600
400
200
0

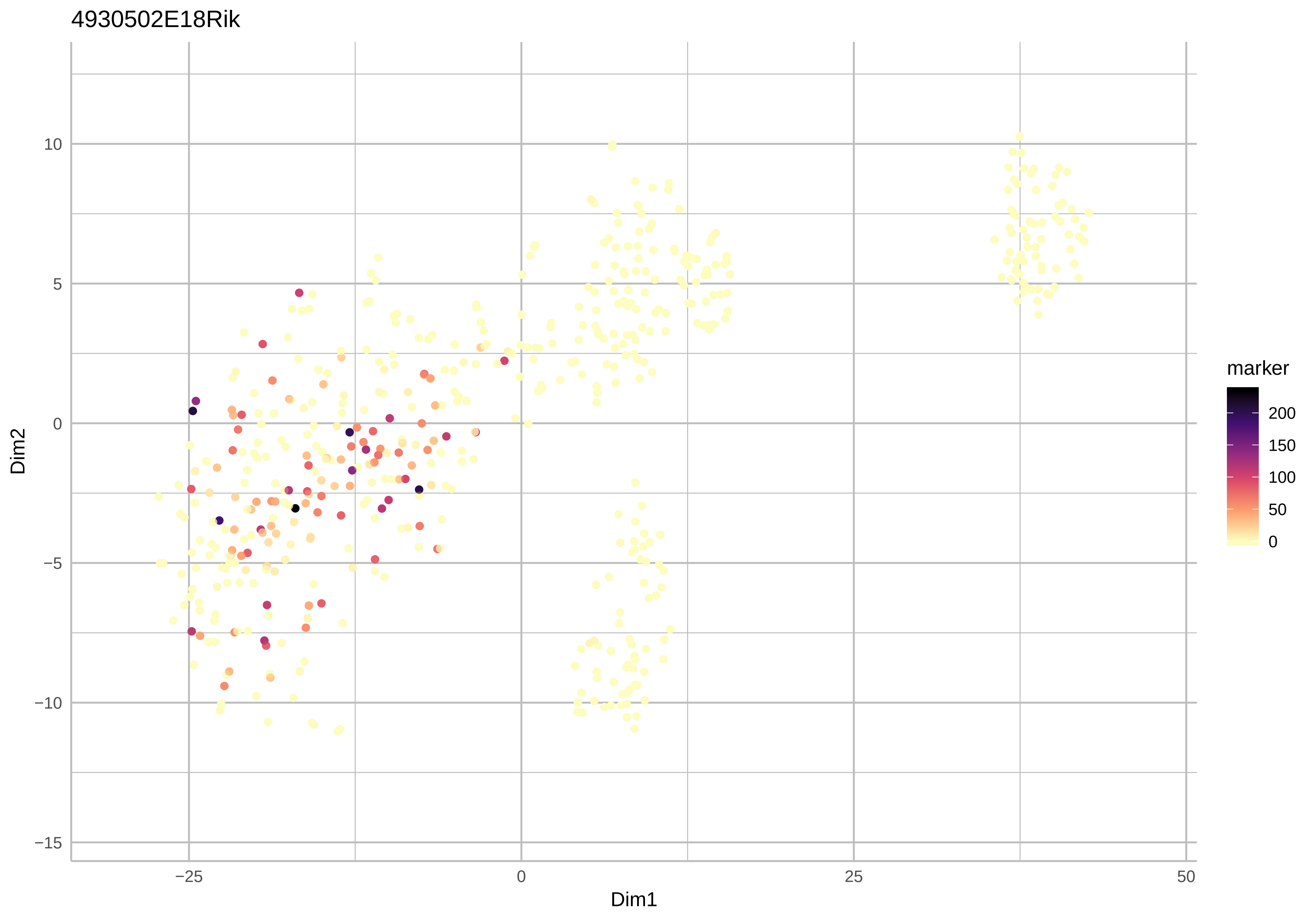

4930502E18Rik
Dim2
Dim1
marker
200
150
100
50
0
10
5
0
−5
−10
−15
−25
0
25
50

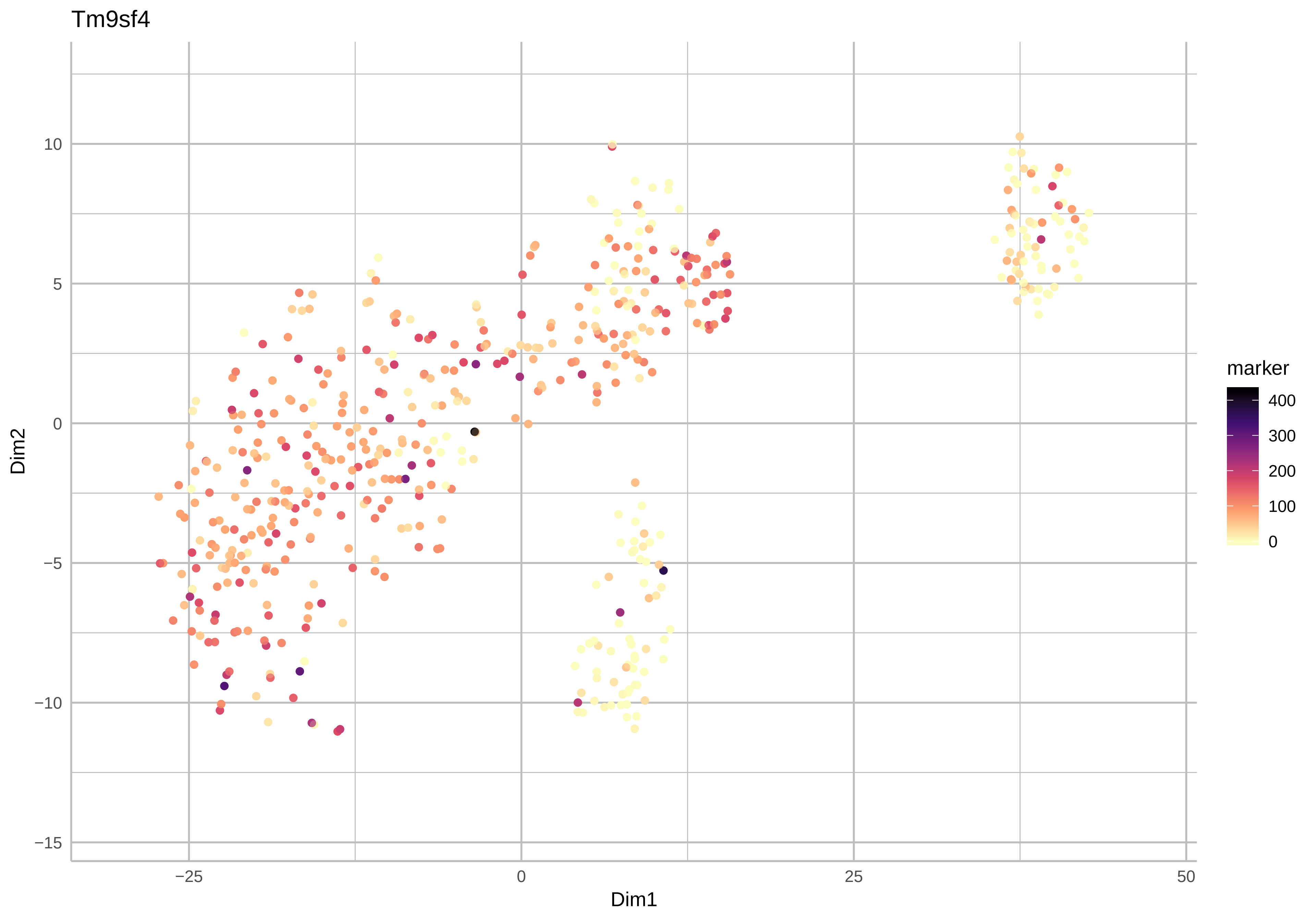
Tm9sf4
Dim2
Dim1
10
5
0
−5
−10
−15
−25
0
25
50
marker
400
300
200
100
0

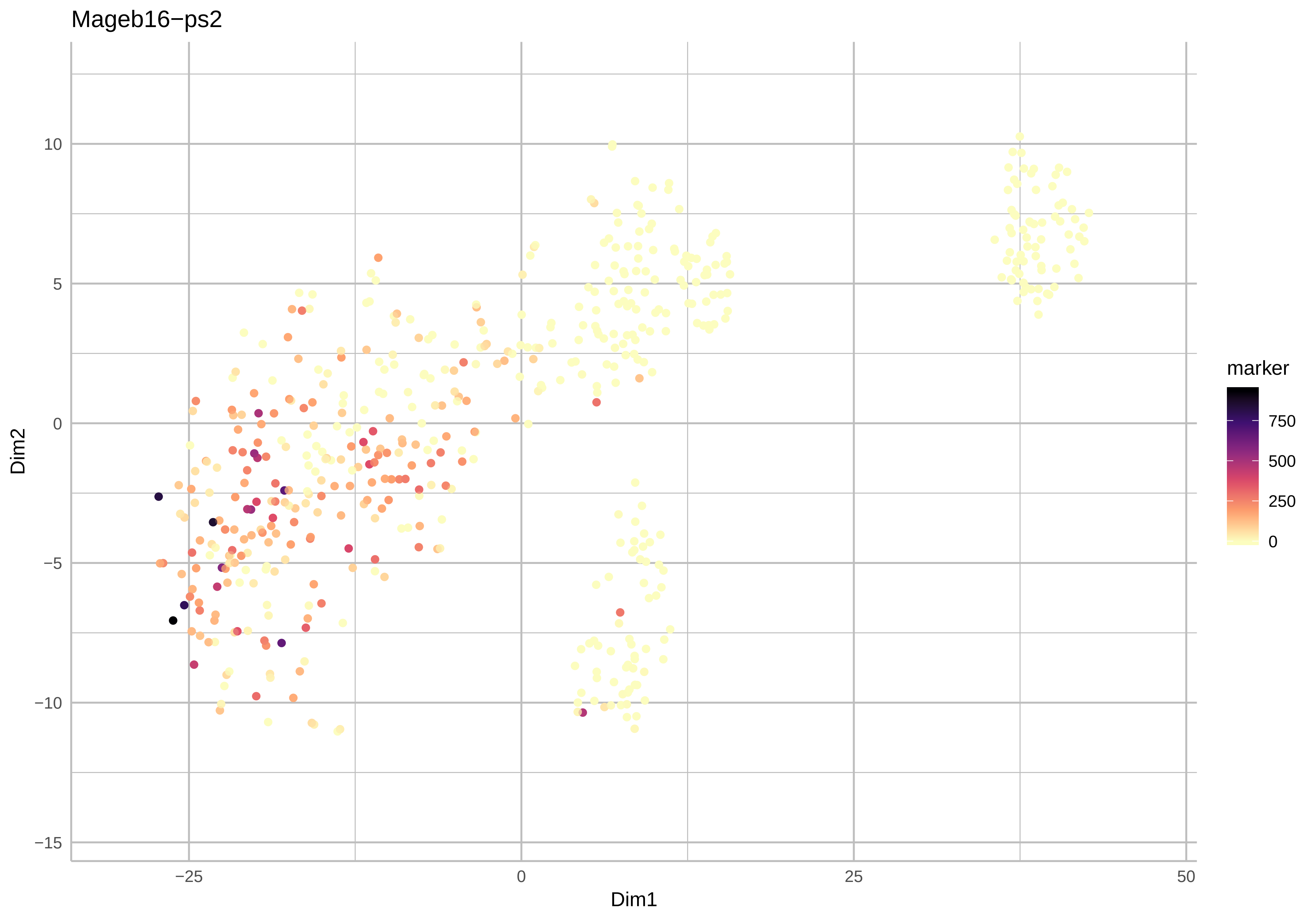
Mageb16−ps2
Dim2
Dim1
10
5
0
−5
−10
−15
−25
0
25
50
marker
750
500
250
0

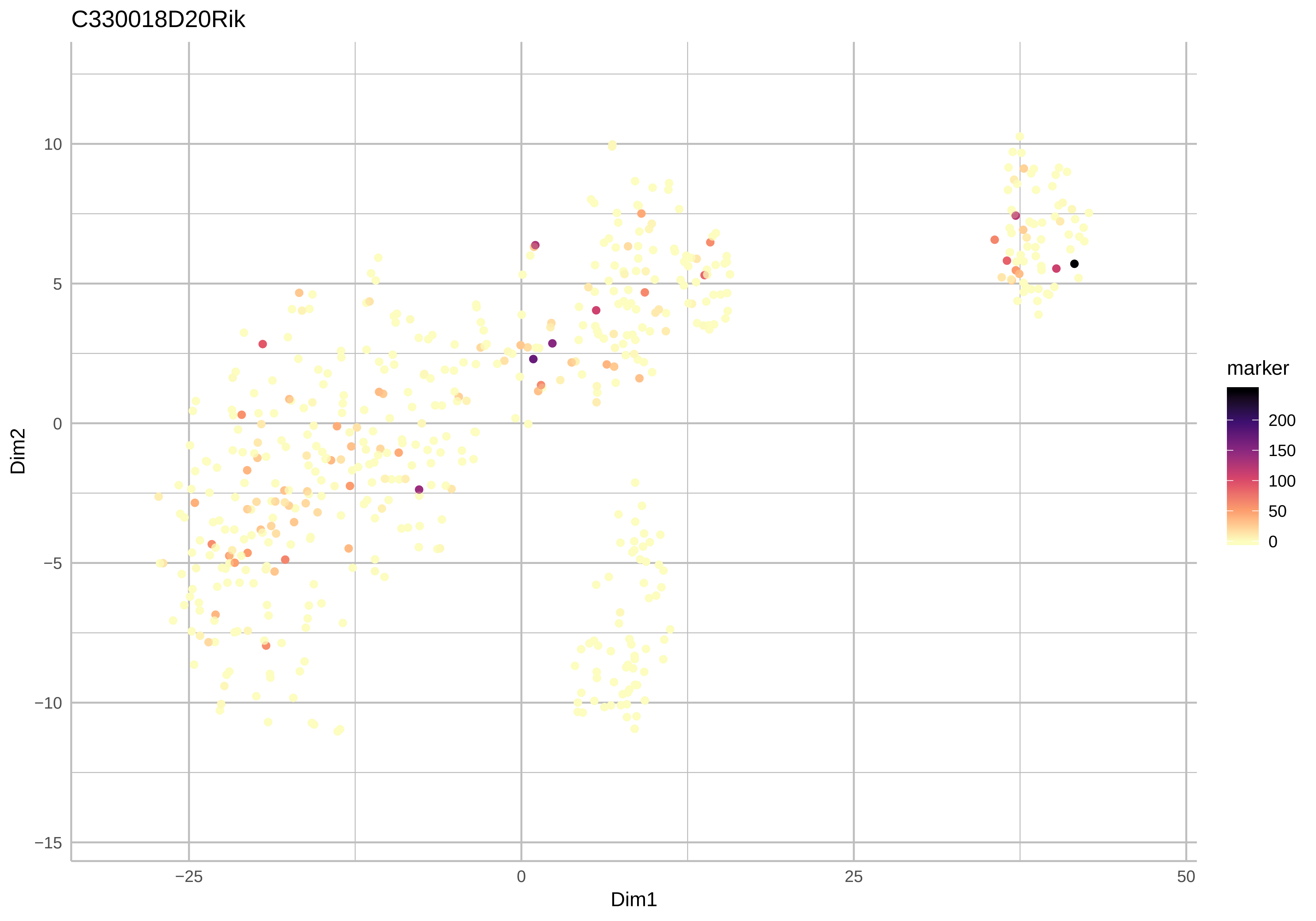

C330018D20Rik
10
5
0
−5
−10
−15
Dim2
−25
0
25
50
Dim1
marker
200
150
100
50
0

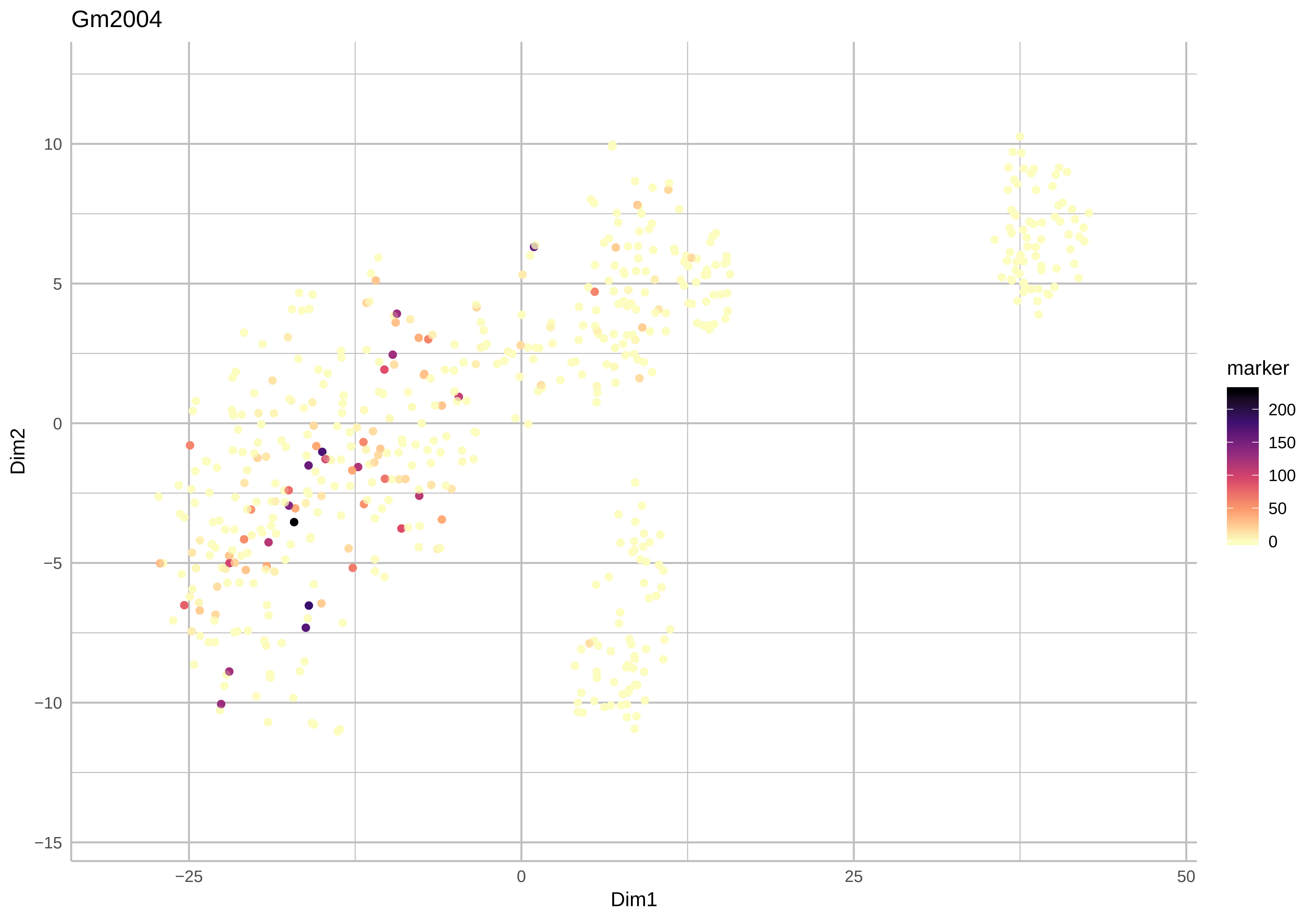
Gm2004
10
5
0
−5
−10
−15
Dim2
−25
0
25
50
Dim1
marker
200
150
100
50
0

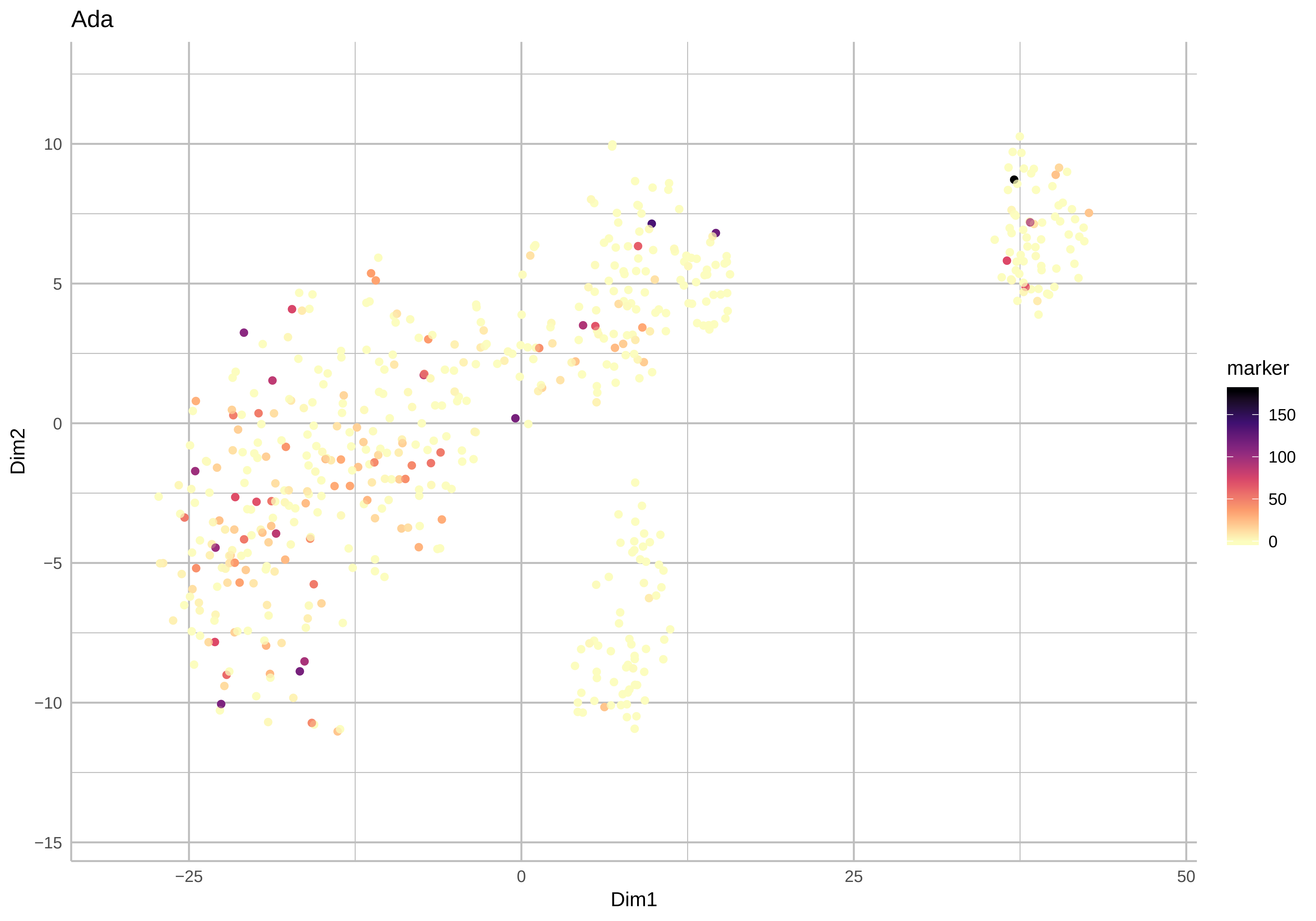

Ada
10
5
0
−5
−10
−15
Dim2
−25
0
25
50
Dim1
marker
150
100
50
0

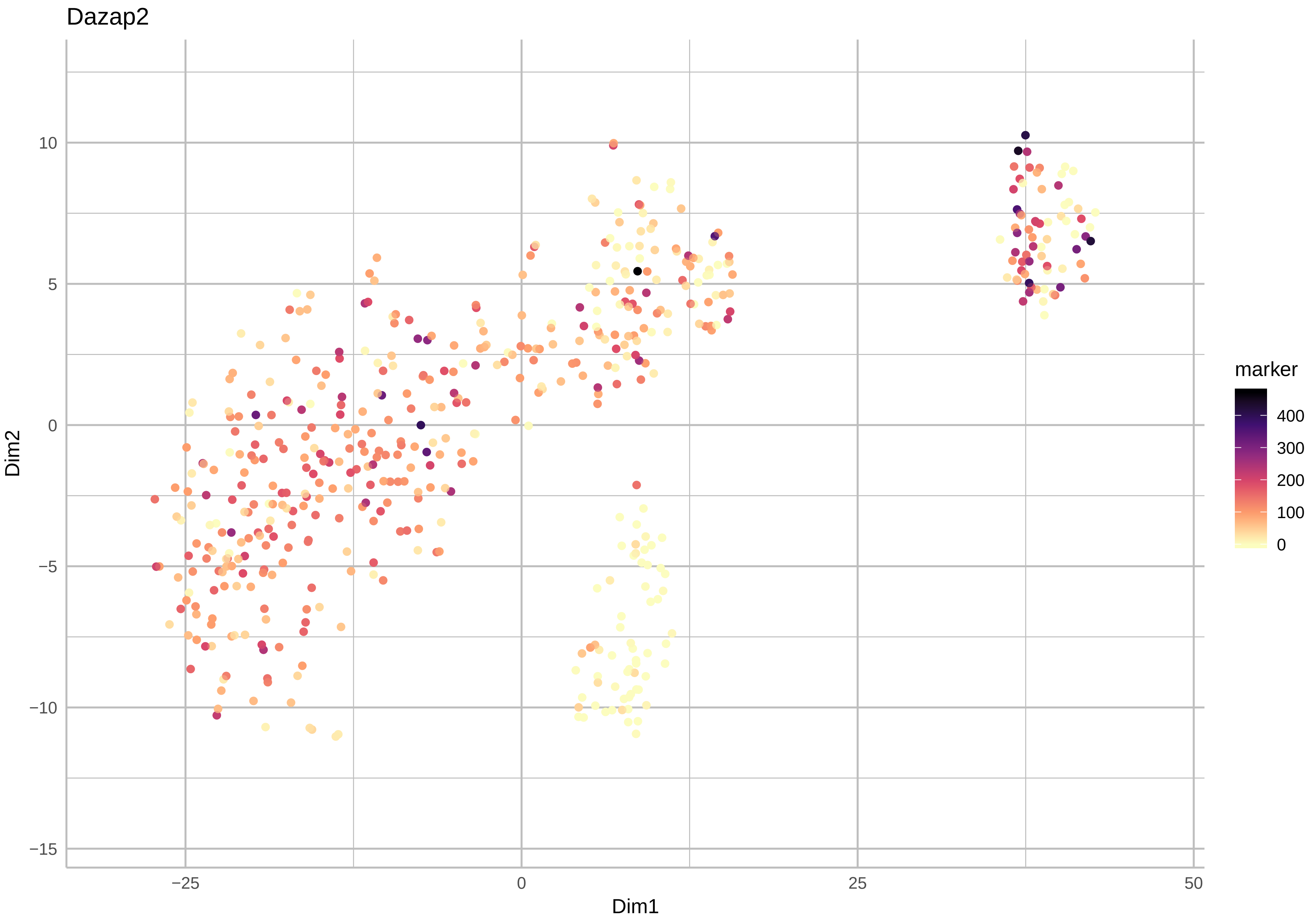
Dazap2
Dim2
Dim1
10
5
0
−5
−10
−15
−25
0
25
50
marker
400
300
200
100
0

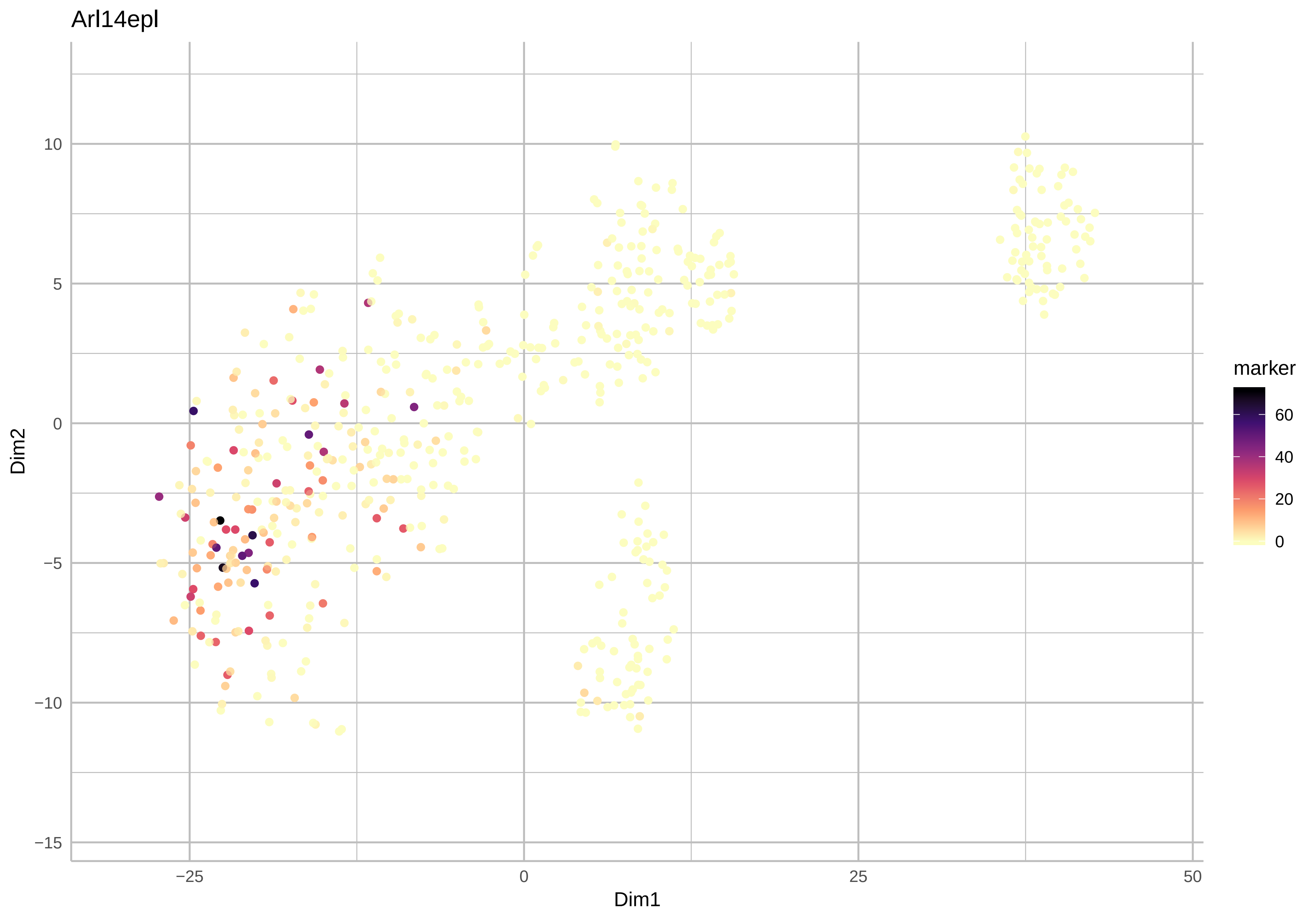
Arl14epl
Dim2
Dim1
10
5
0
−5
−10
−15
−25
0
25
50
marker
60
40
20
0

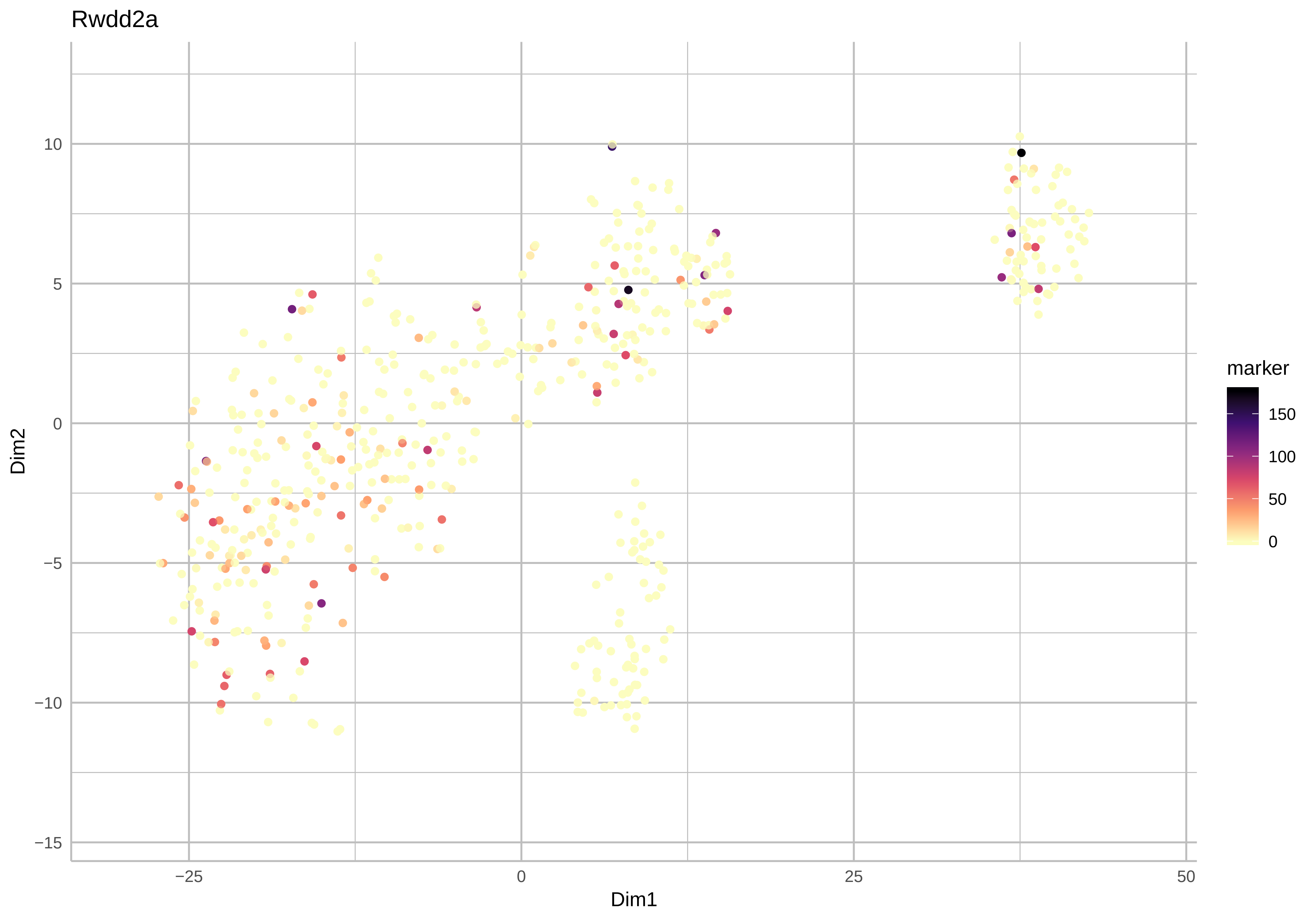

Rwdd2a
Dim2
Dim1
10
5
0
−5
−10
−15
−25
0
25
50
marker
150
100
50
0

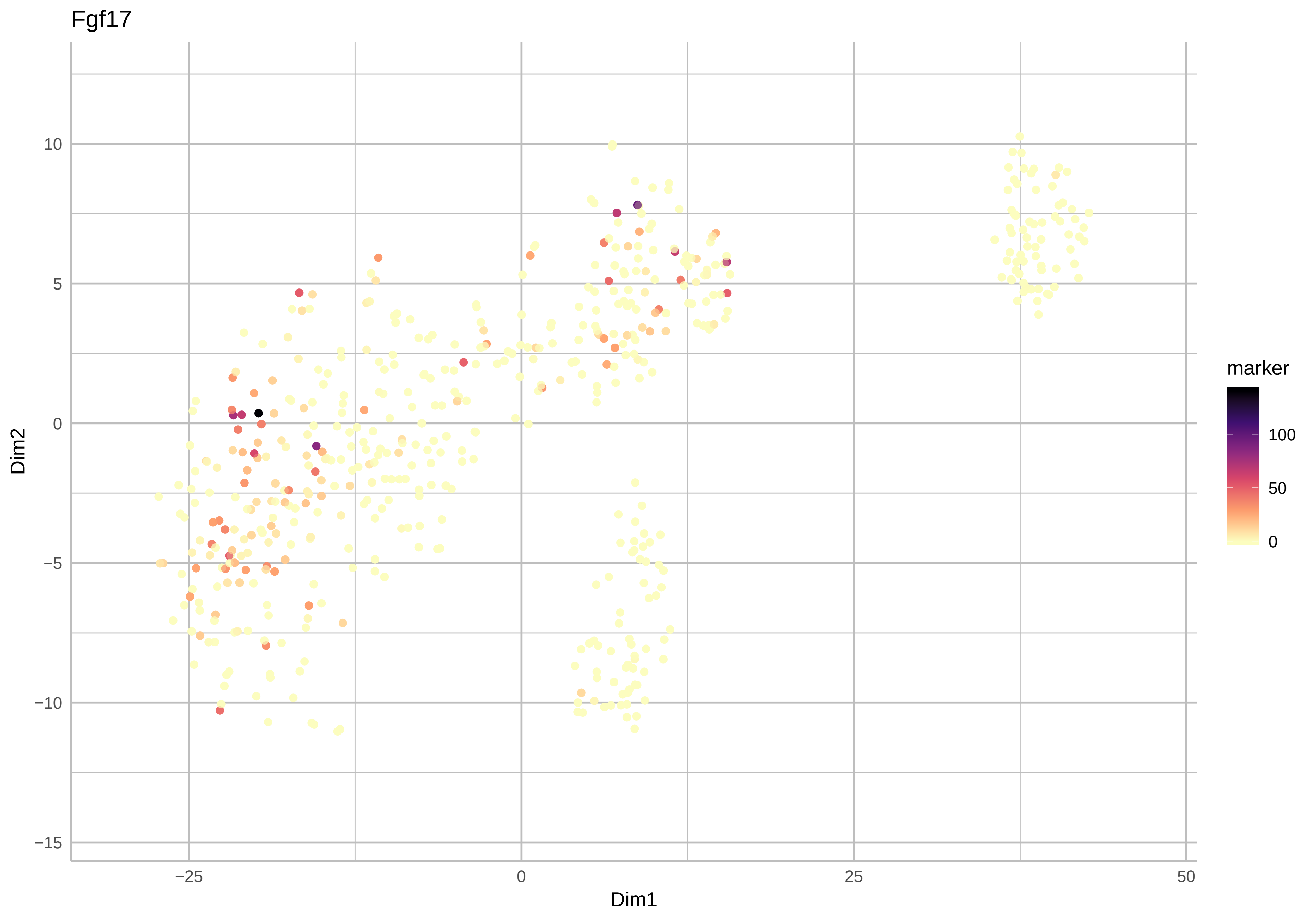

Fgf17
Dim2
Dim1
10
5
0
−5
−10
−15
−25
0
25
50
marker
100
50
0

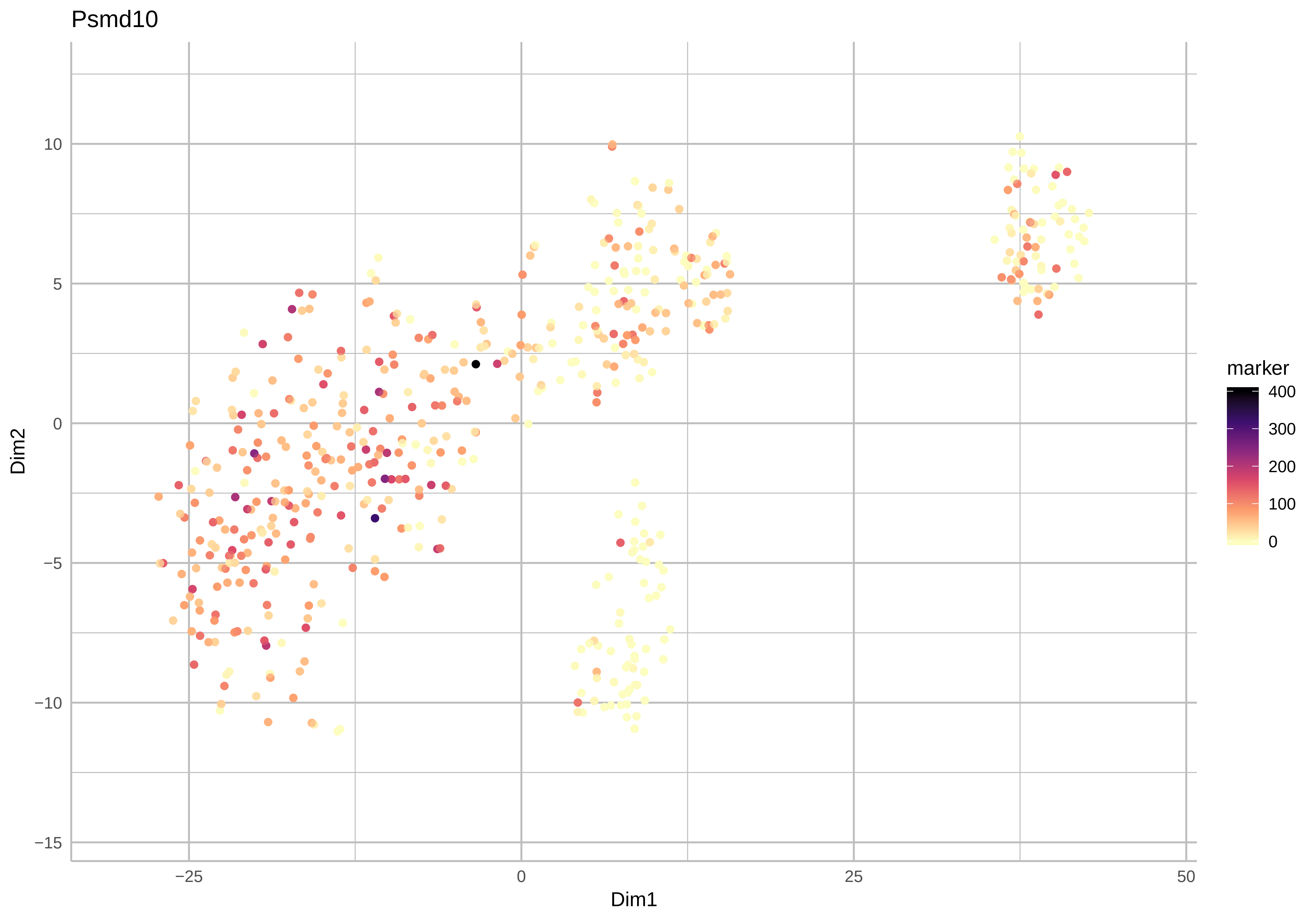
Psmd10
Dim2
Dim1
10
5
0
−5
−10
−15
−25
0
25
50
marker
400
300
200
100
0

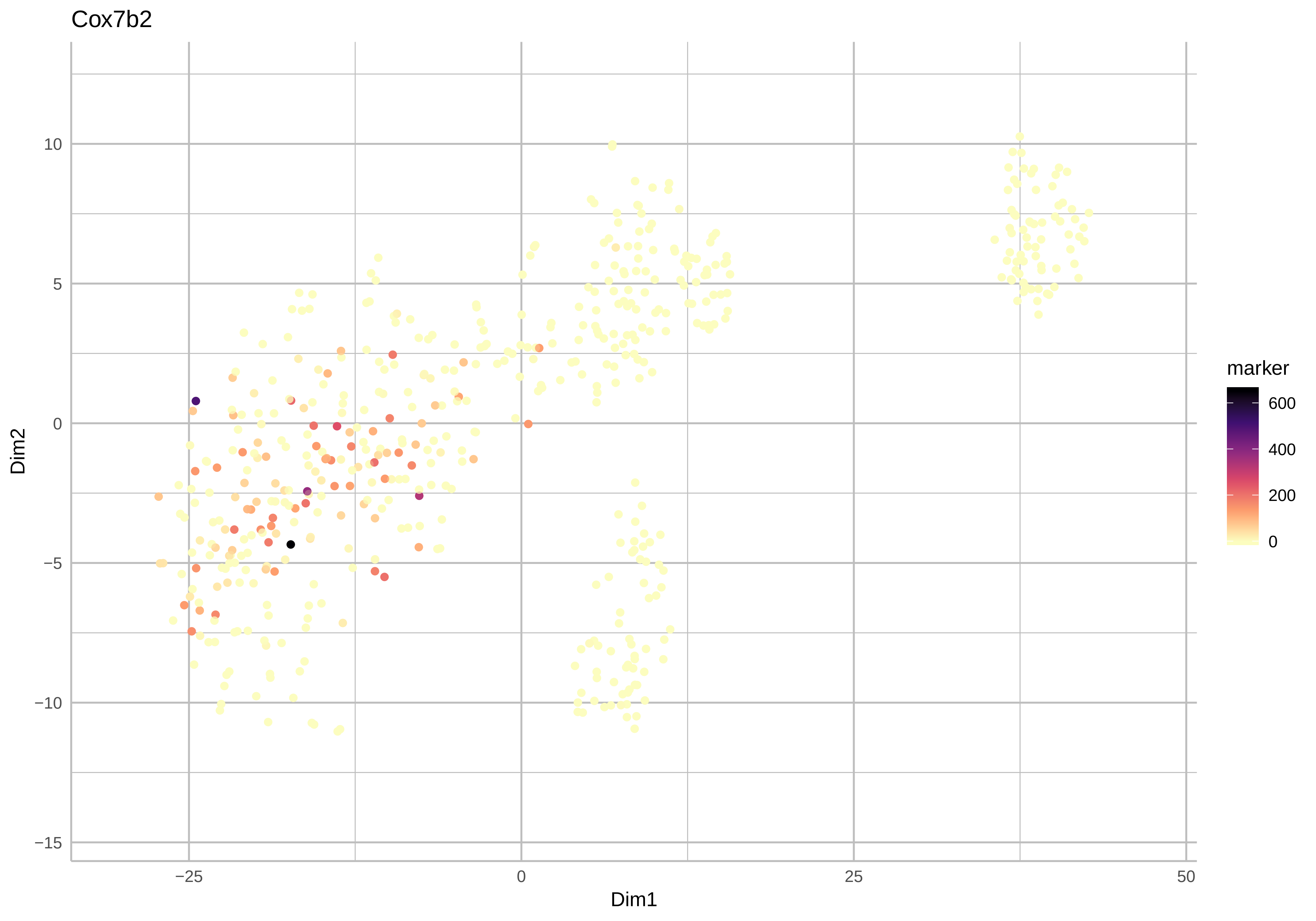

Cox7b2
Dim2
Dim1
10
5
0
−5
−10
−15
−25
0
25
50
marker
600
400
200
0

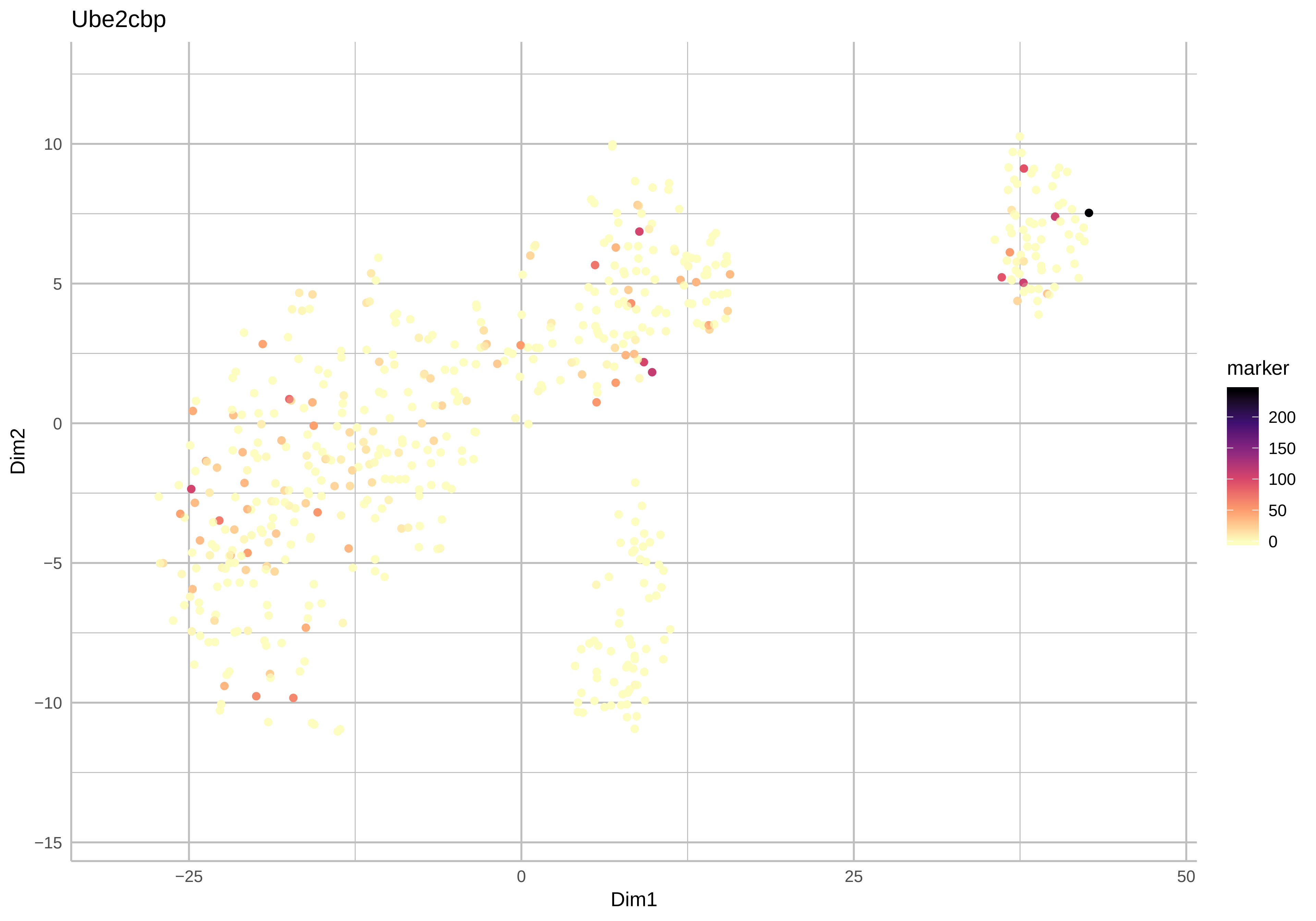

Ube2cbp
10
5
0
−5
−10
−15
Dim2
−25
0
25
50
Dim1
marker
200
150
100
50
0

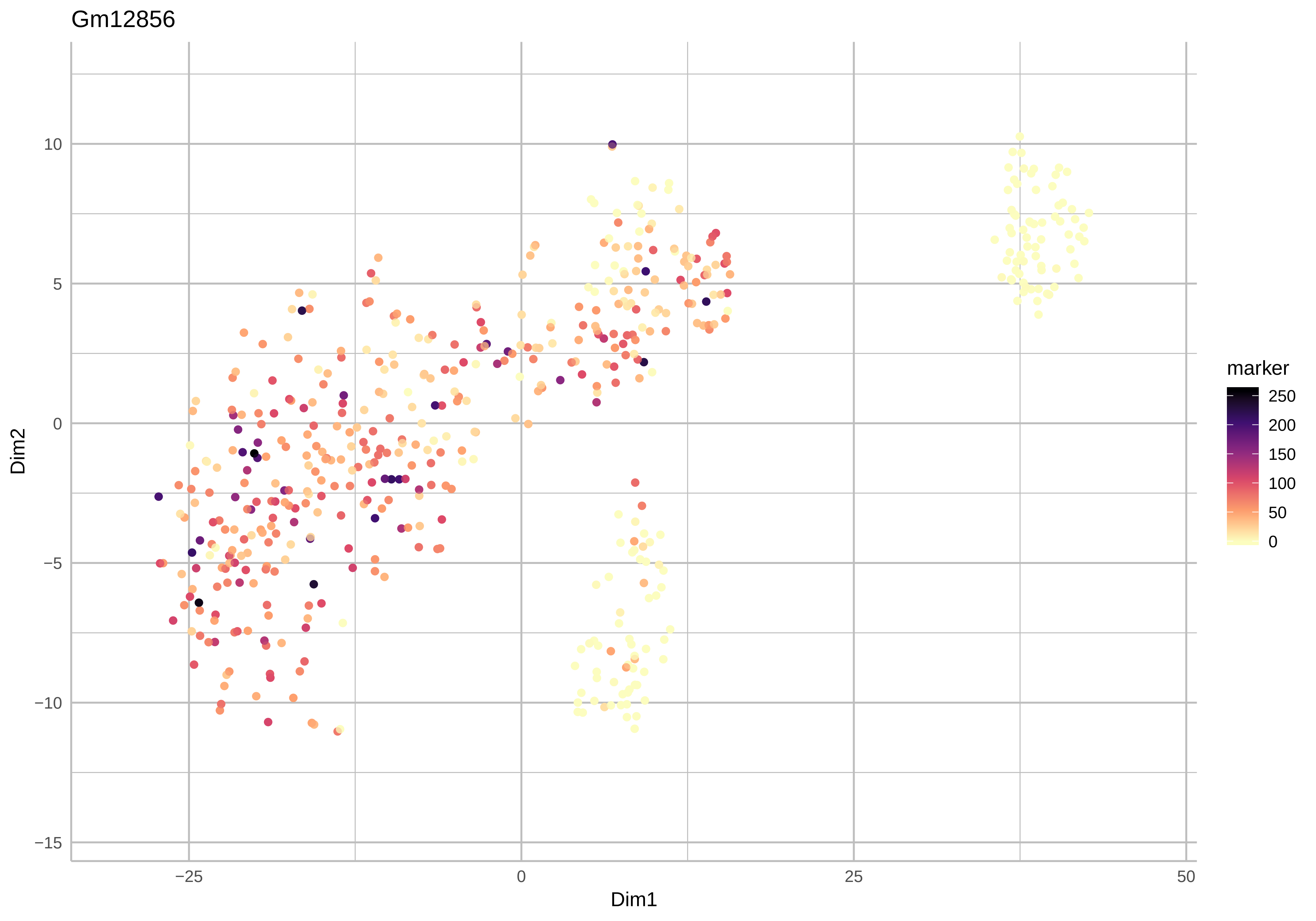
Gm12856
Dim2
10
5
0
−5
−10
−15
−25
0
25
50
Dim1
marker
250
200
150
100
50
0

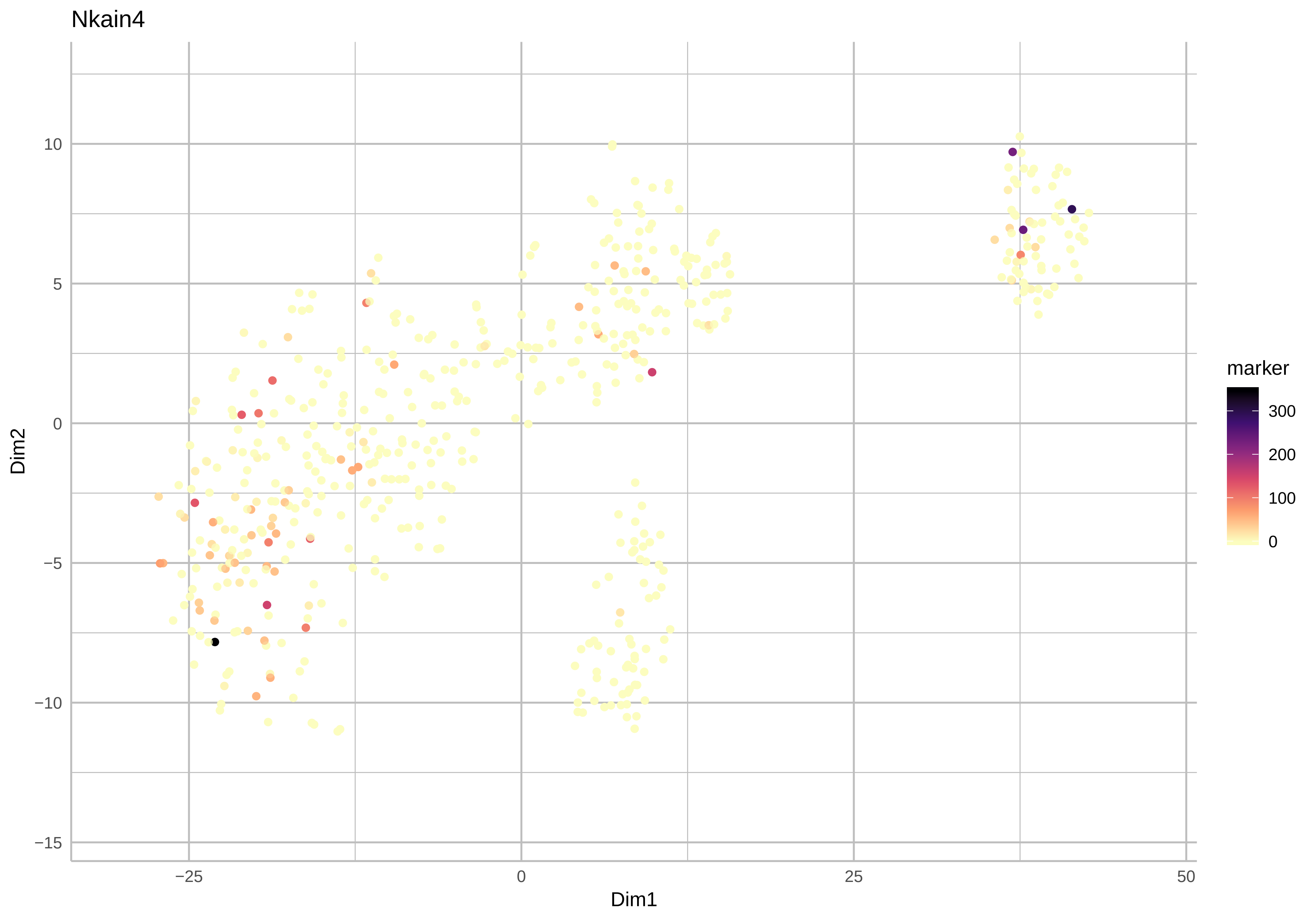

Nkain4
10
5
0
−5
−10
−15
Dim2
−25
0
25
50
Dim1
marker
300
200
100
0

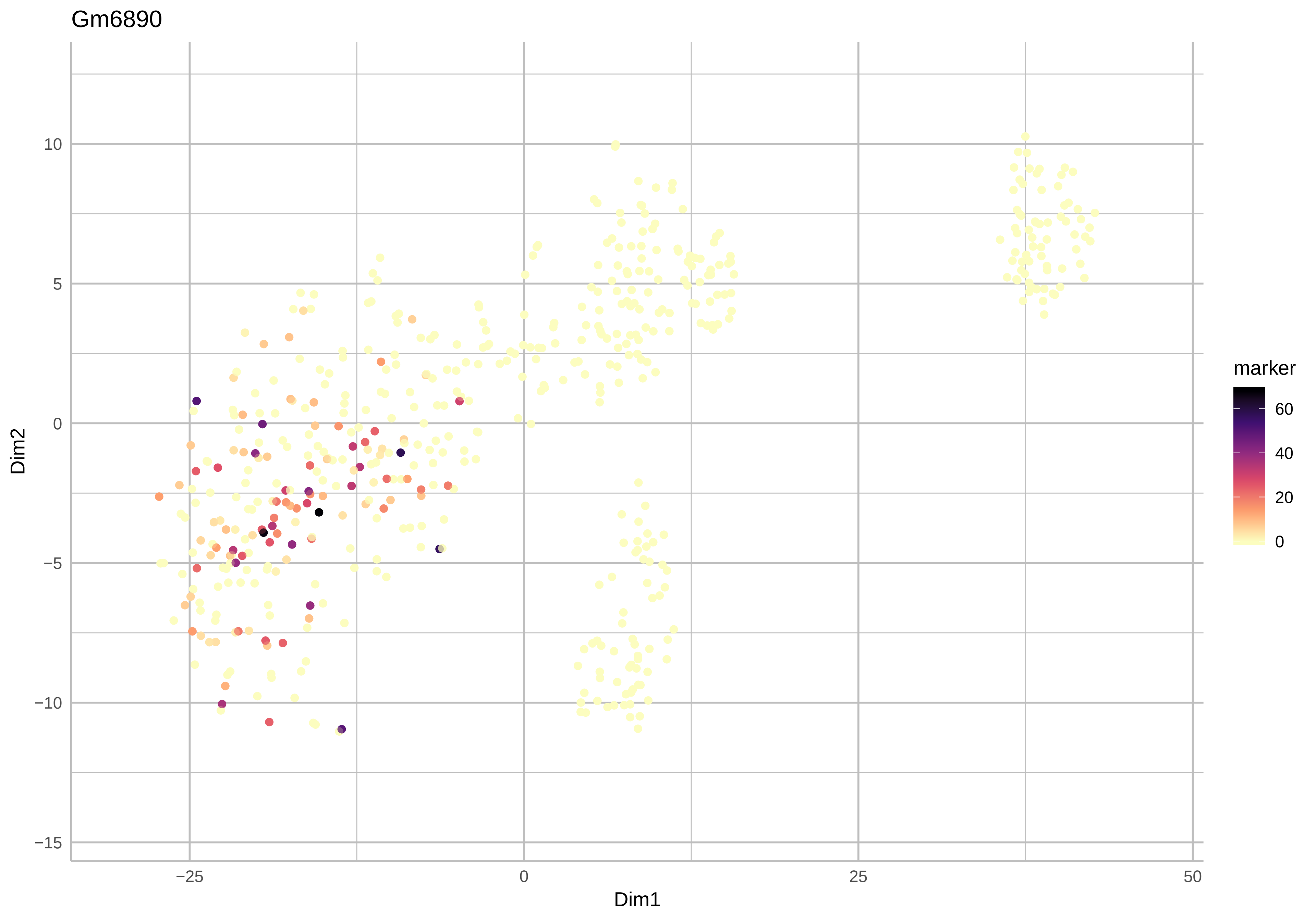

Gm6890
Dim2
Dim1
marker
10
5
0
−5
−10
−15
−25
0
25
50
60
40
20
0

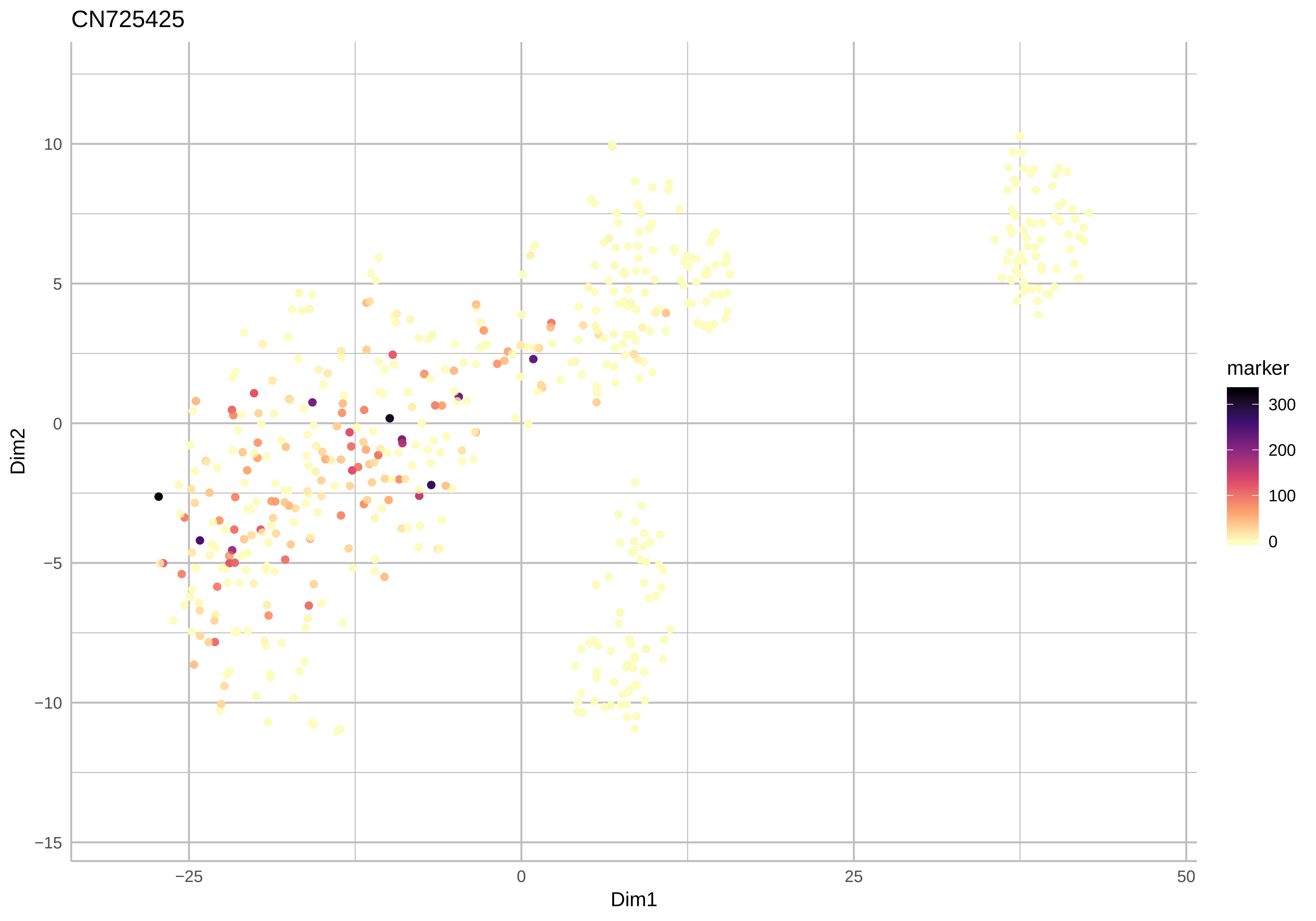

CN725425
Dim2
Dim1
10
5
0
−5
−10
−15
−25
0
25
50
marker
300
200
100
0

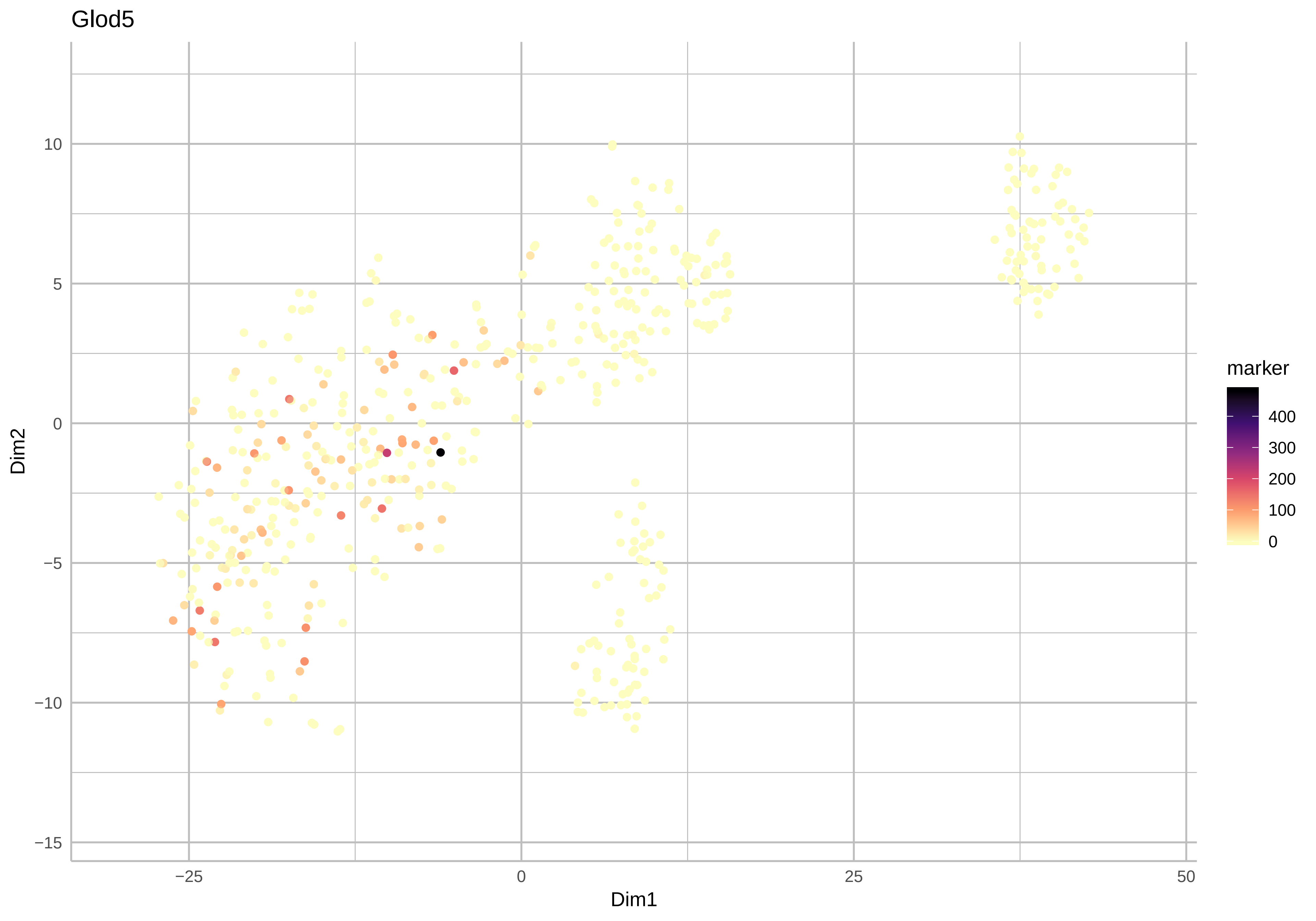
Glod5
Dim2
Dim1
10
5
0
−5
−10
−15
−25
0
25
50
marker
400
300
200
100
0

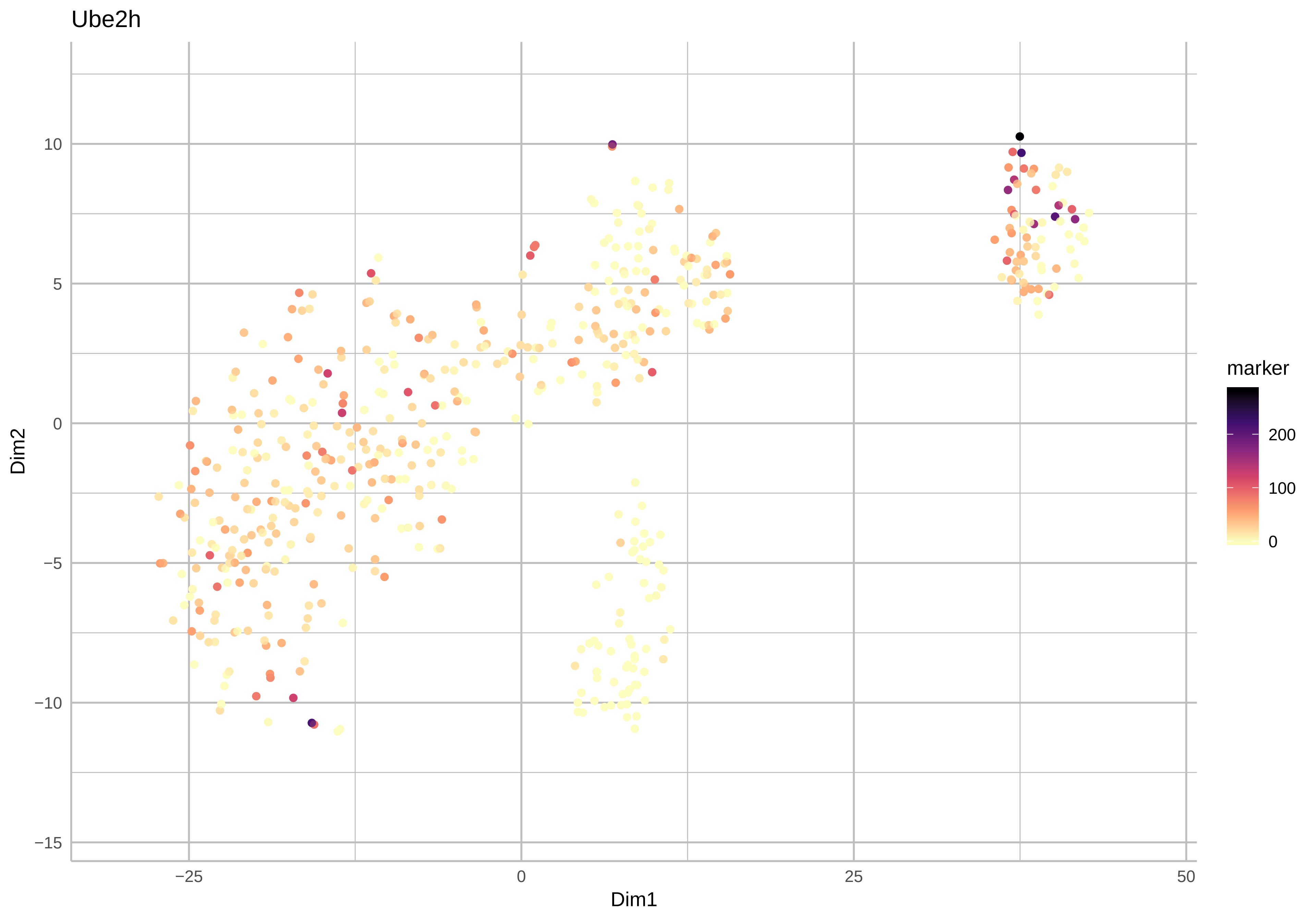
Ube2h
10
5
0
−5
−10
−15
Dim2
−25
0
25
50
Dim1
marker
200
100
0

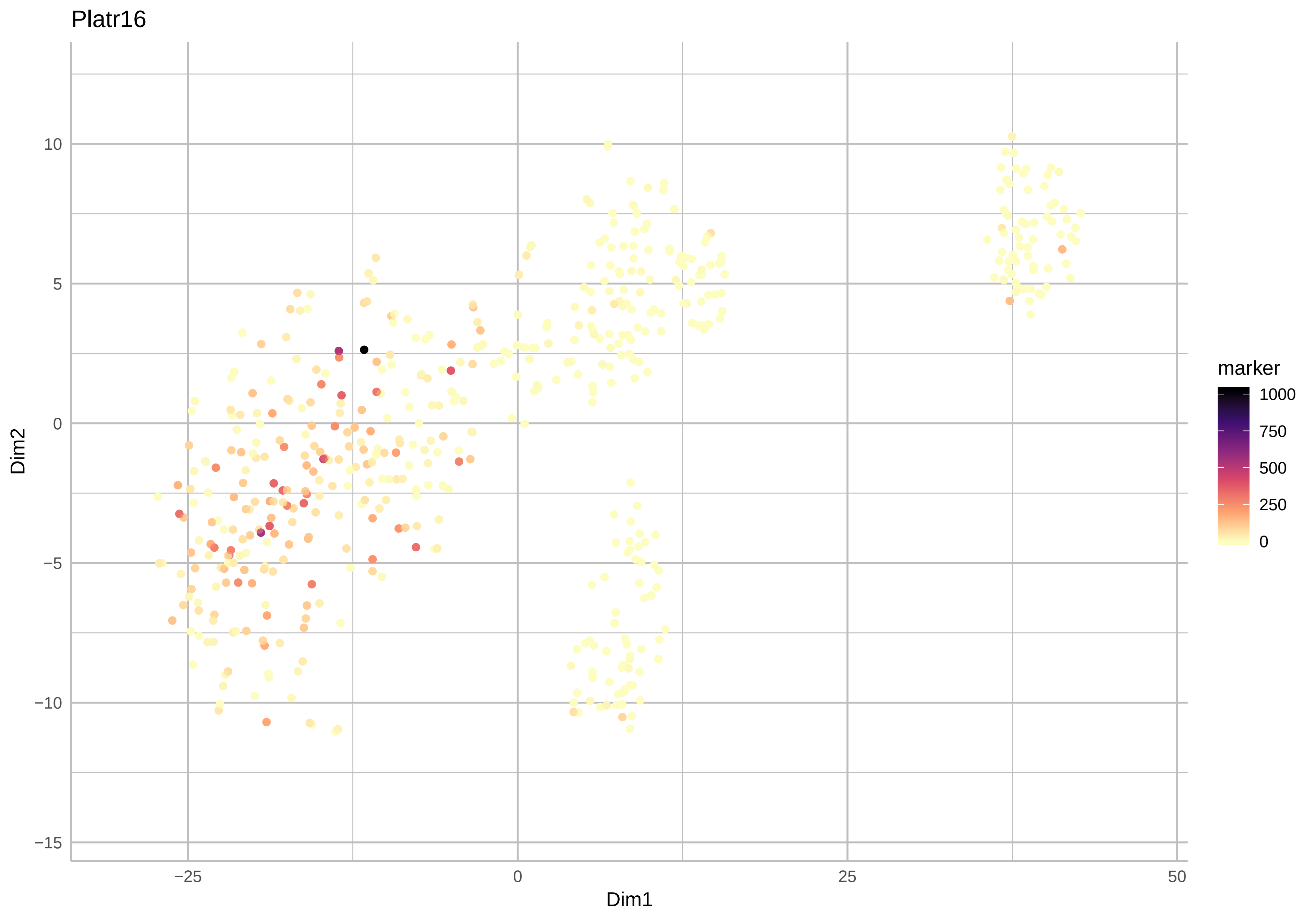
Platr16
10
5
0
−5
−10
−15
Dim2
−25
0
25
50
Dim1
marker
1000
750
500
250
0

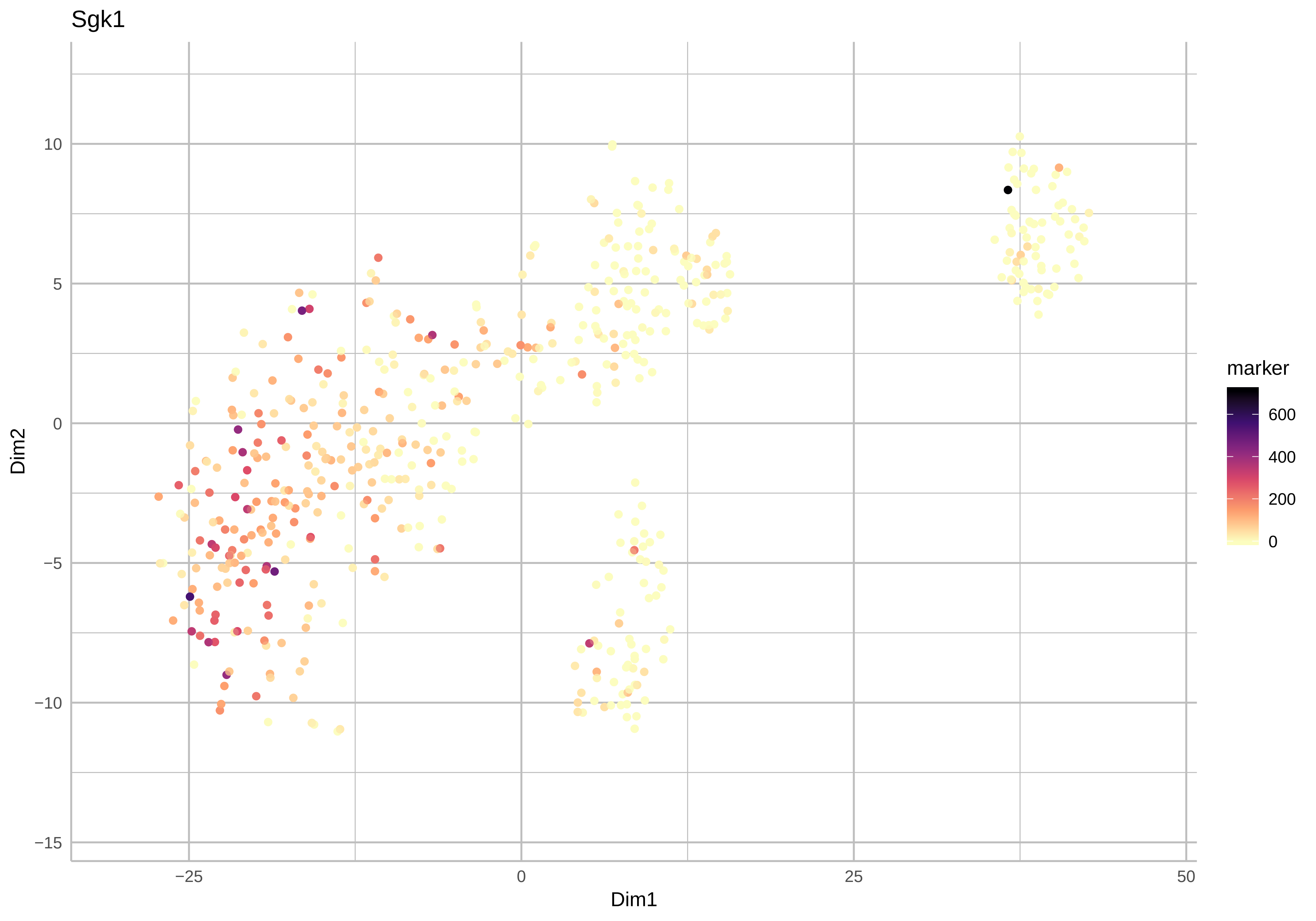

Sgk1
Dim2
Dim1
10
5
0
−5
−10
−15
−25
0
25
50
marker
600
400
200
0

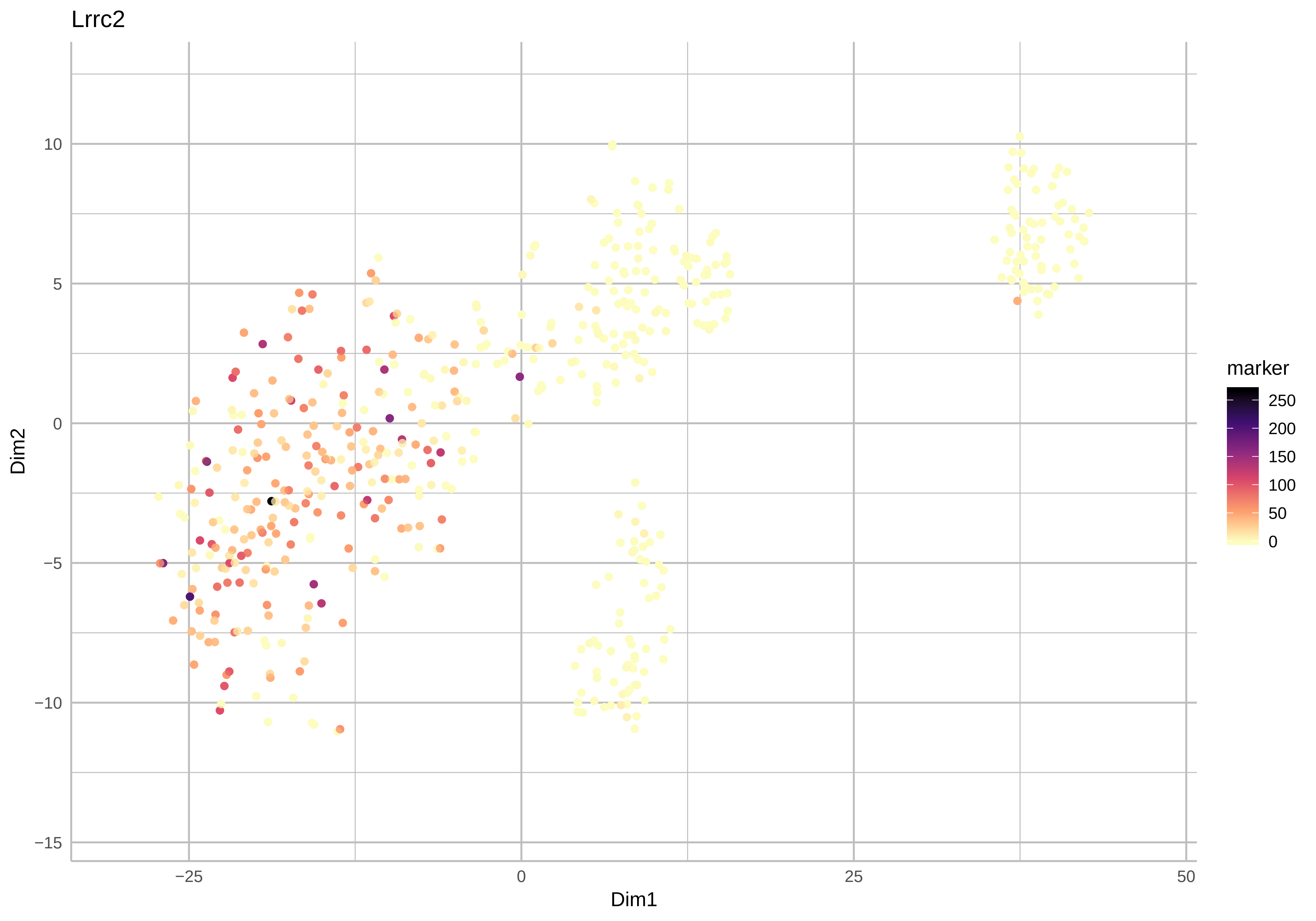

Lrrc2
Dim2
Dim1
marker
250
200
150
100
50
0
10
5
0
−5
−10
−15
−25
0
25
50

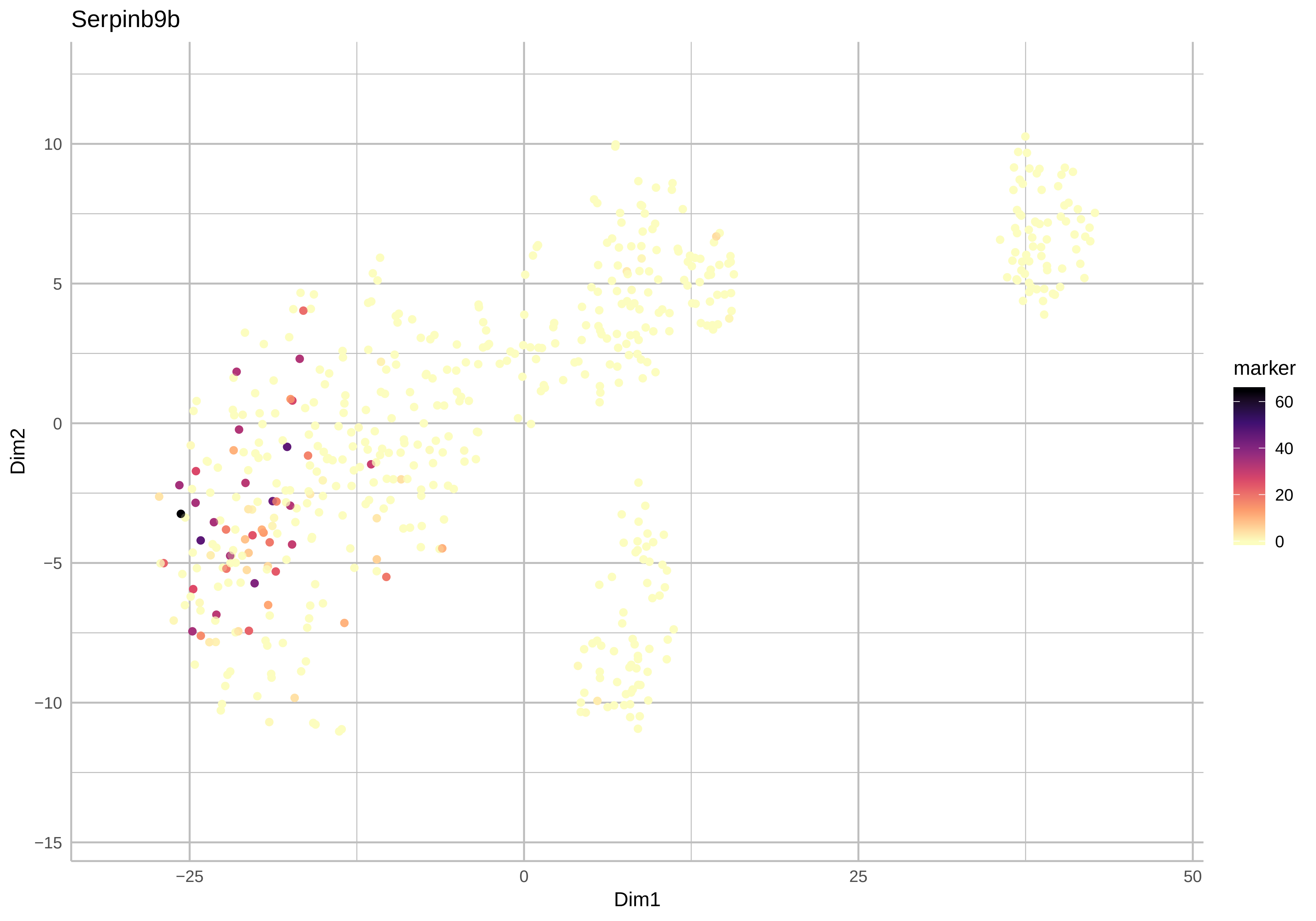

Serpinb9b
Dim2
Dim1
10
5
0
−5
−10
−15
−25
0
25
50
marker
60
40
20
0

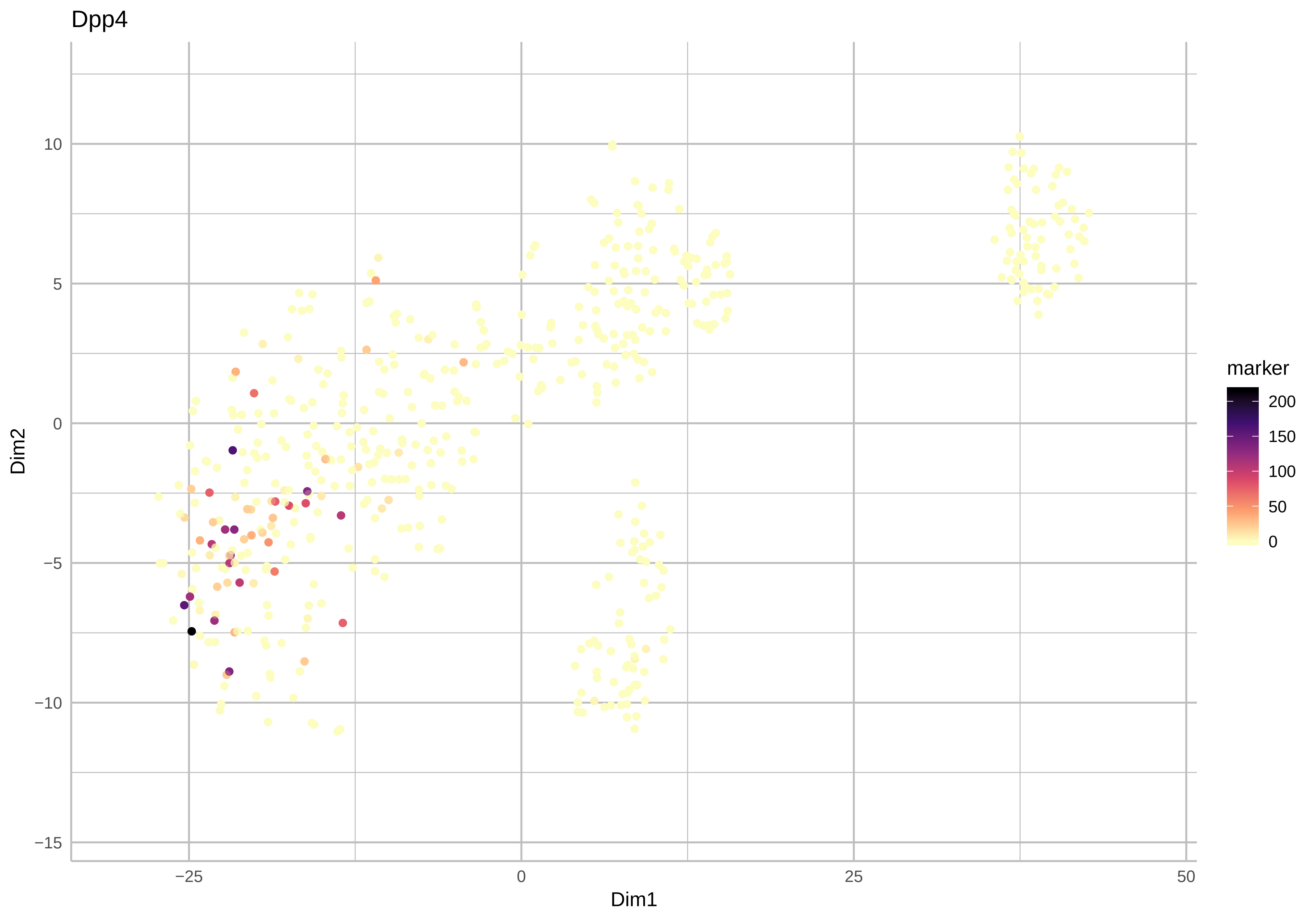
Dpp4
10
5
0
−5
−10
−15
Dim2
−25
0
25
50
Dim1
marker
200
150
100
50
0

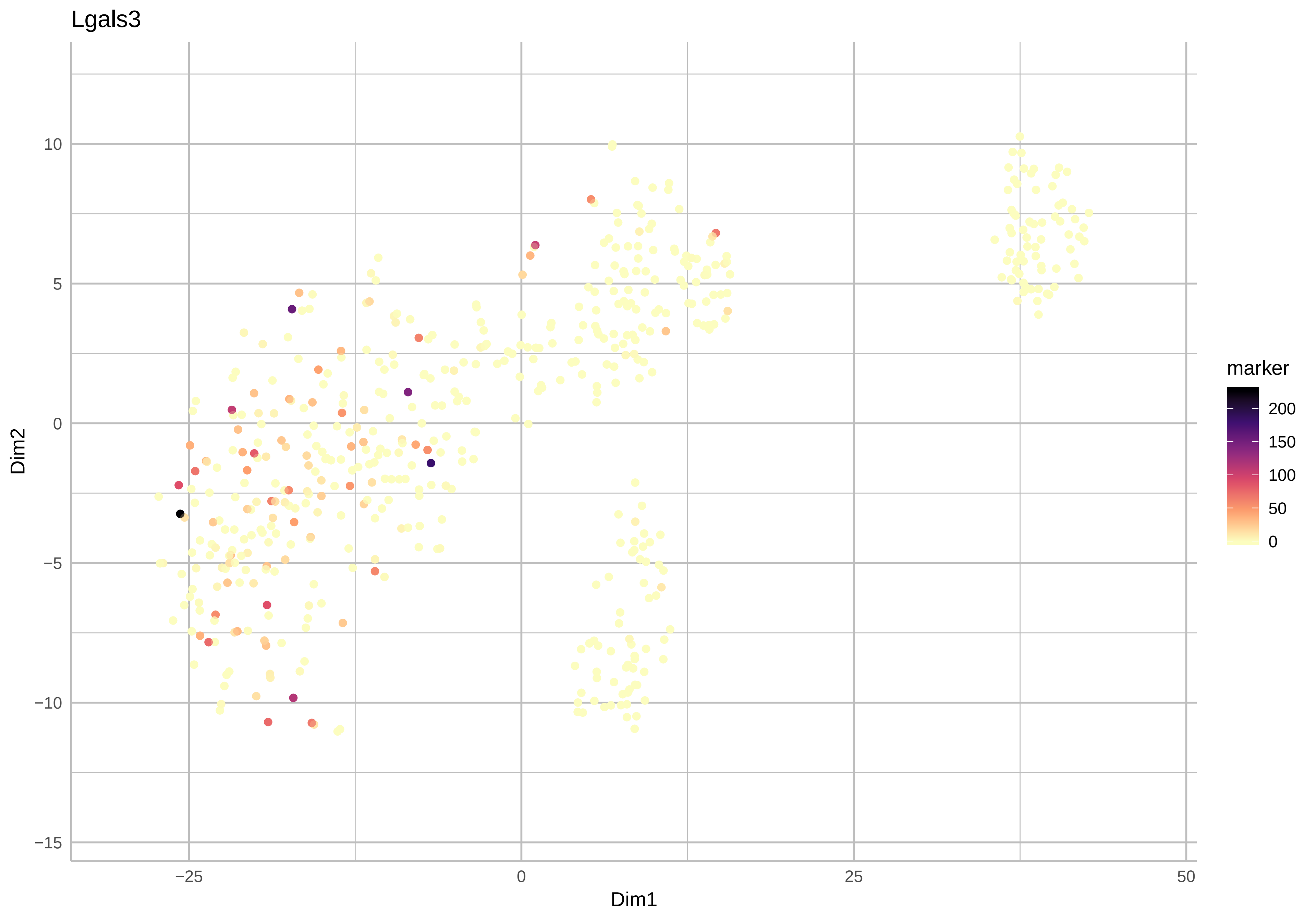
Lgals3
Dim2
Dim1
10
5
0
−5
−10
−15
−25
0
25
50
marker
200
150
100
50
0

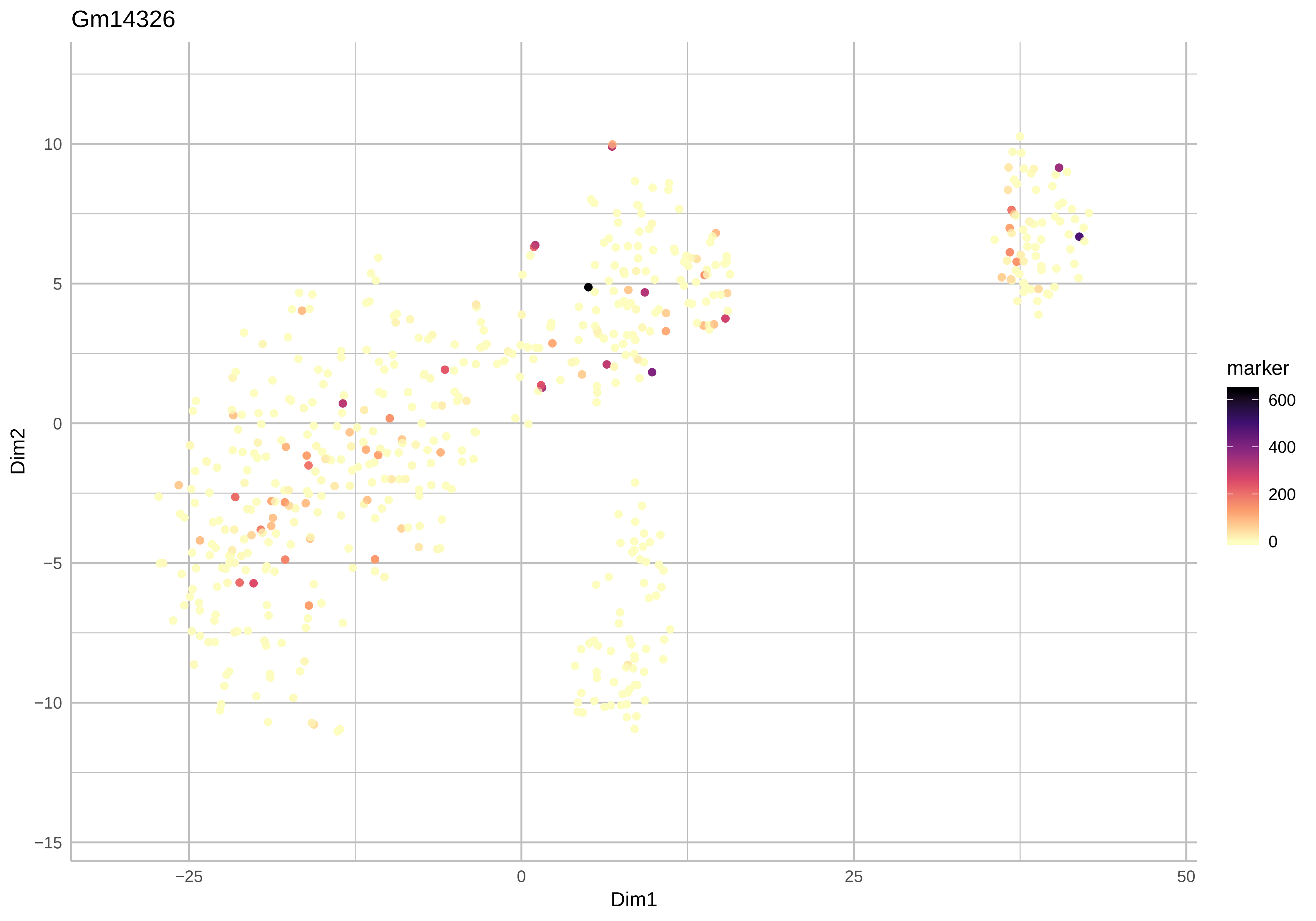
Gm14326
Dim2
10
5
0
−5
−10
−15
Dim1
−25
0
25
50
marker
600
400
200
0

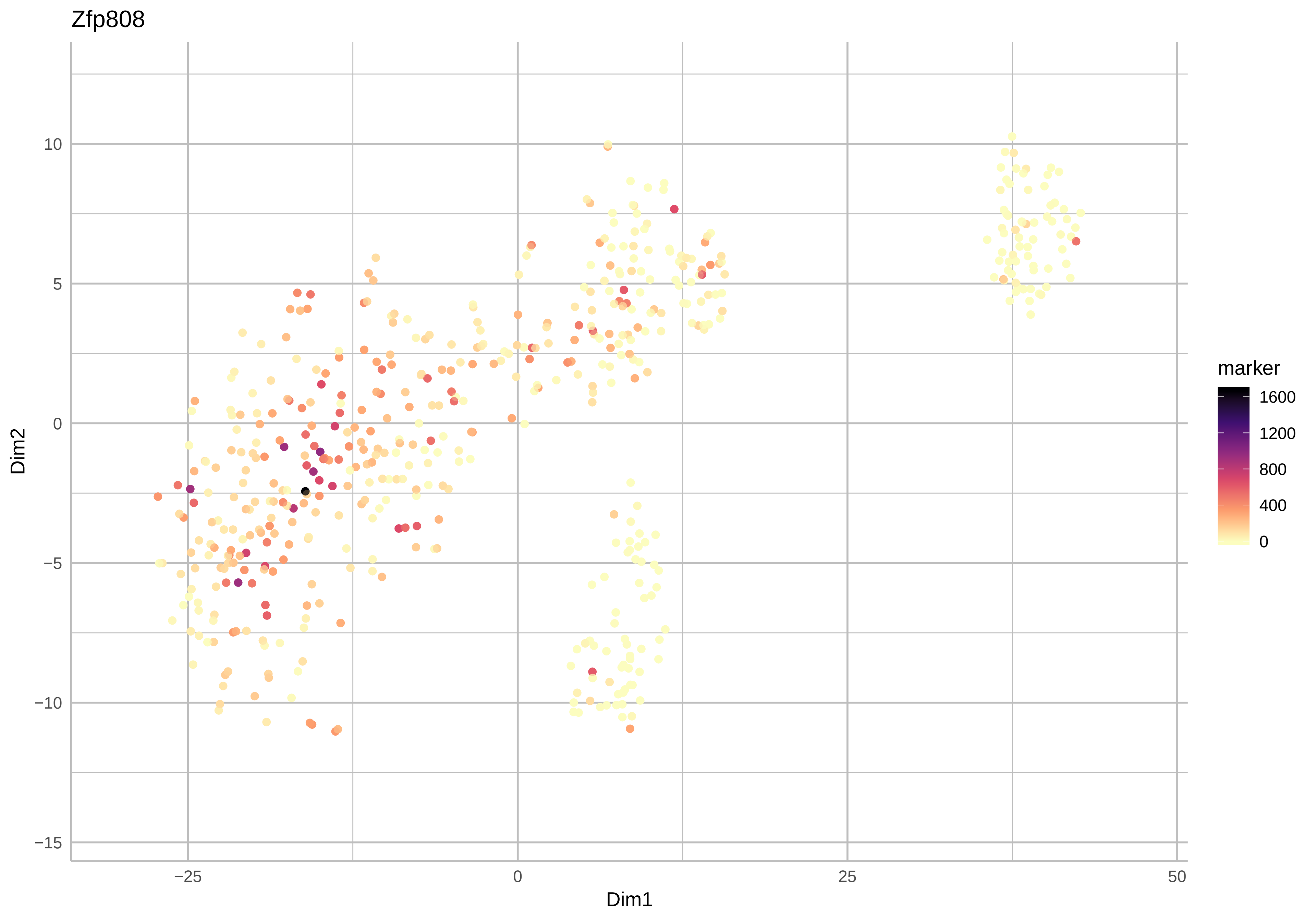

Zfp808
Dim2
Dim1
10
5
0
−5
−10
−15
−25
0
25
50
marker
1600
1200
800
400
0

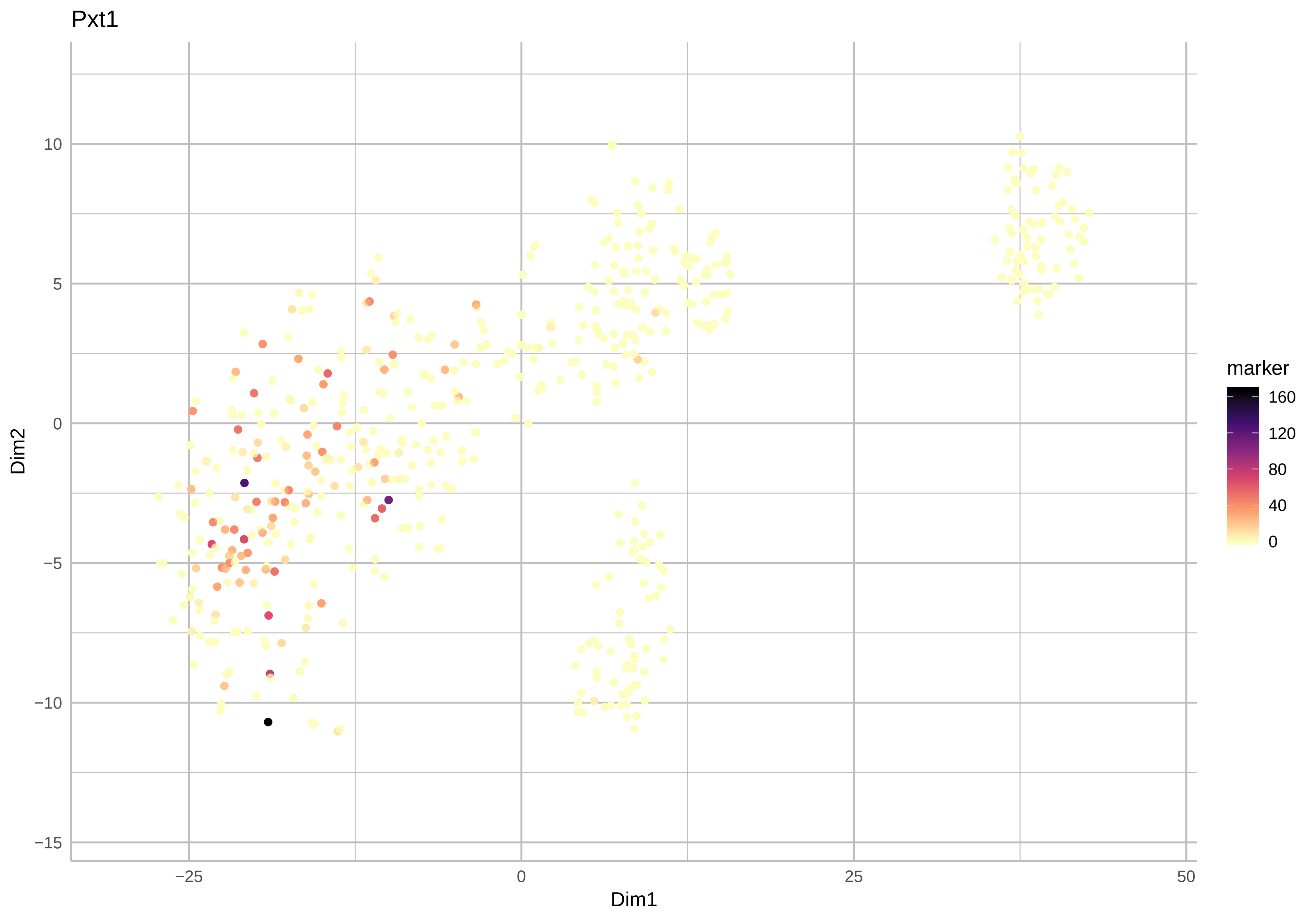
Pxt1
Dim2
Dim1
10
5
0
−5
−10
−15
−25
0
25
50
marker
160
120
80
40
0

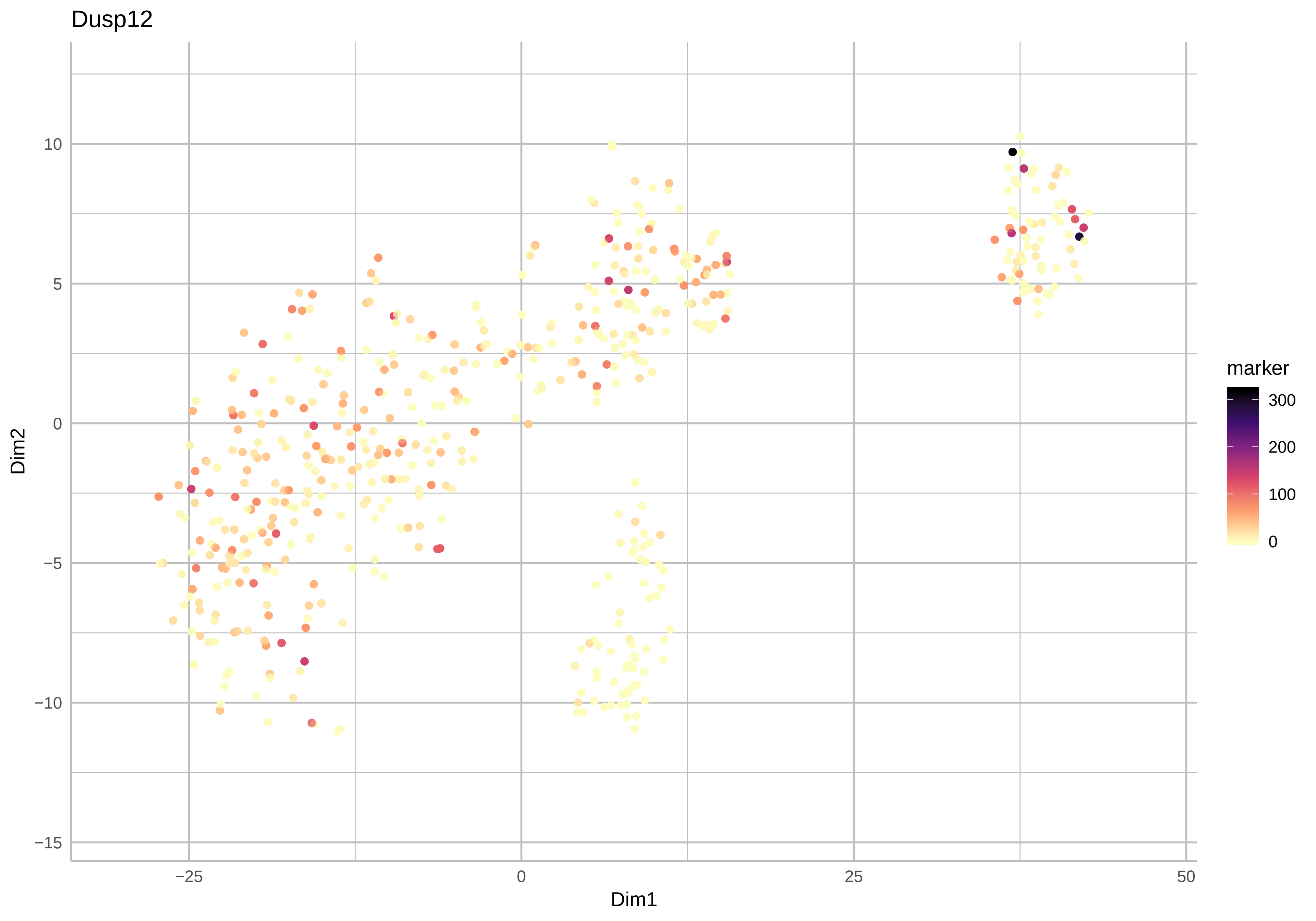

Dusp12
10
5
0
−5
−10
−15
Dim2
−25
0
25
50
Dim1
marker
300
200
100
0

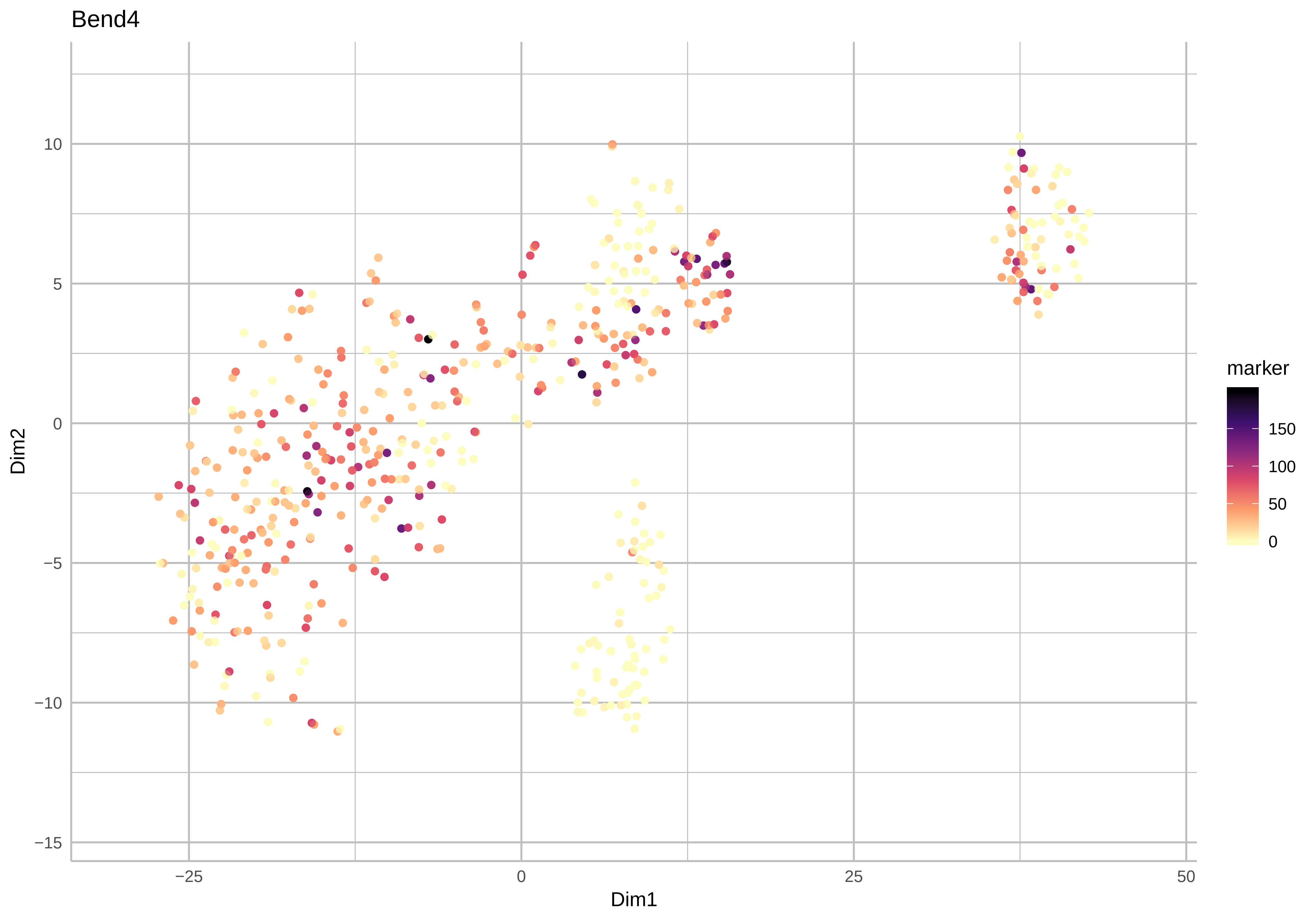
Bend4
Dim2
Dim1
10
5
0
−5
−10
−15
−25
0
25
50
marker
150
100
50
0

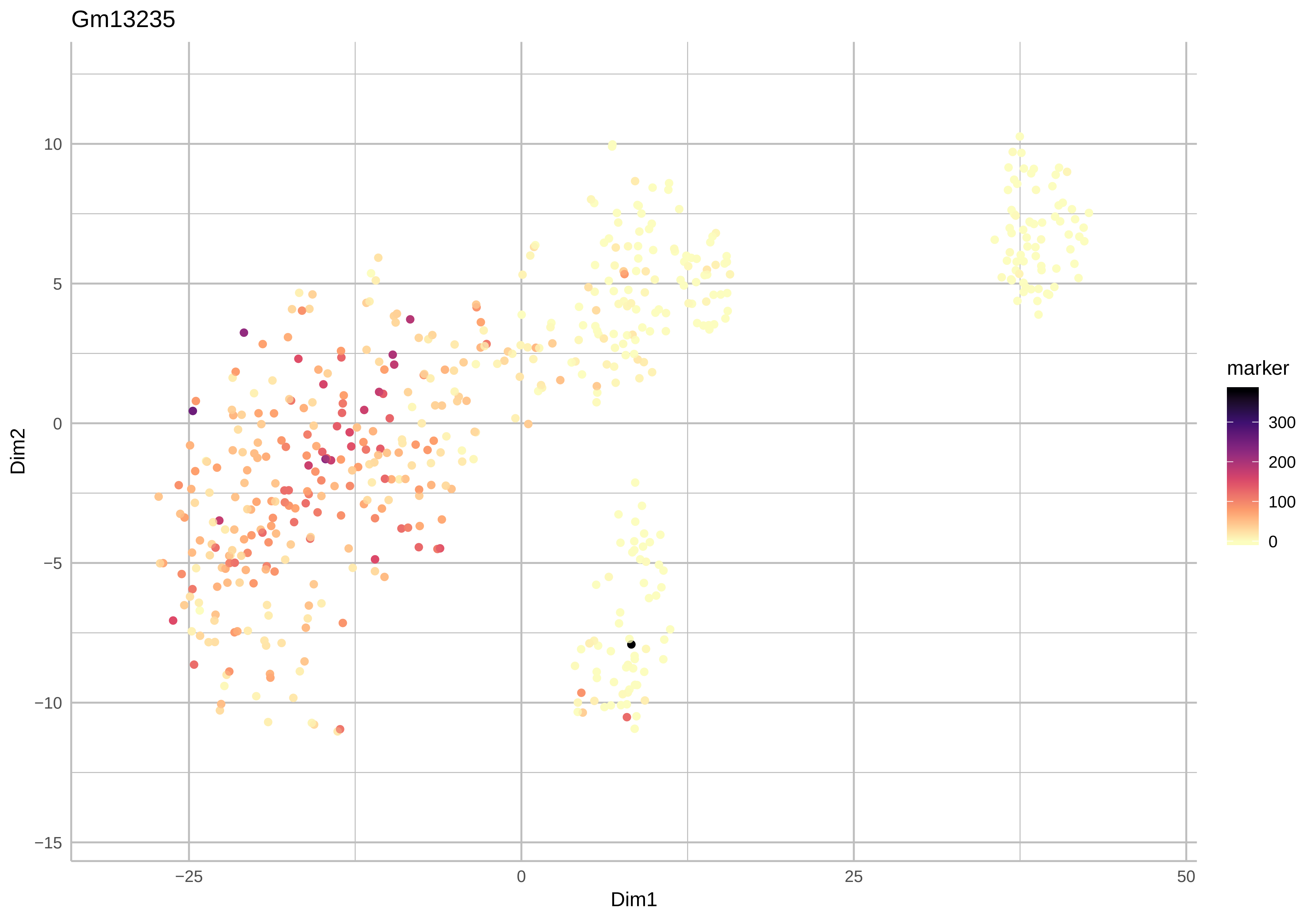

Gm13235
10
5
0
−5
−10
−15
Dim2
−25
0
25
50
Dim1
marker
300
200
100
0

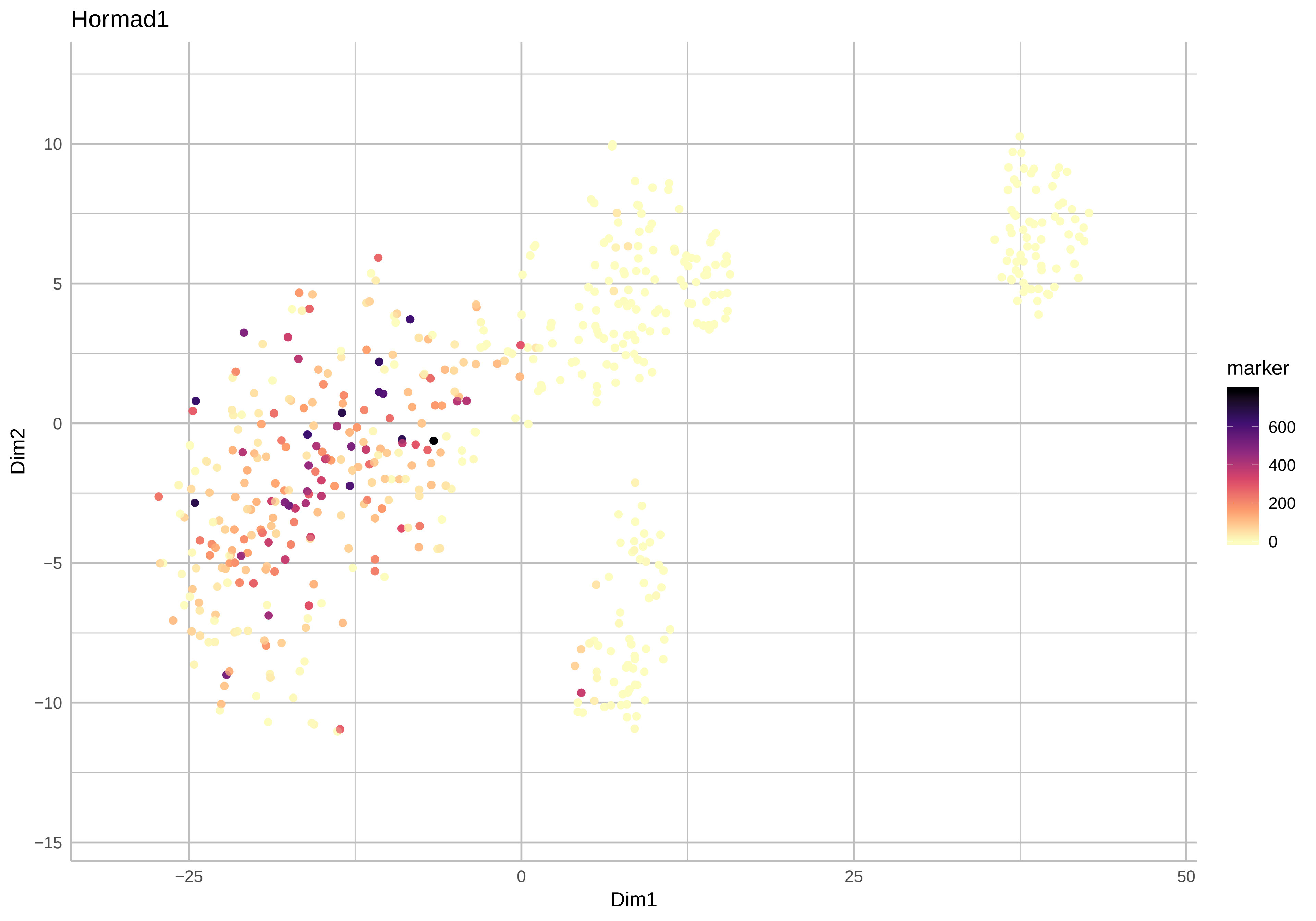
Hormad1
Dim2
Dim1
10
5
0
−5
−10
−15
−25
0
25
50
marker
600
400
200
0

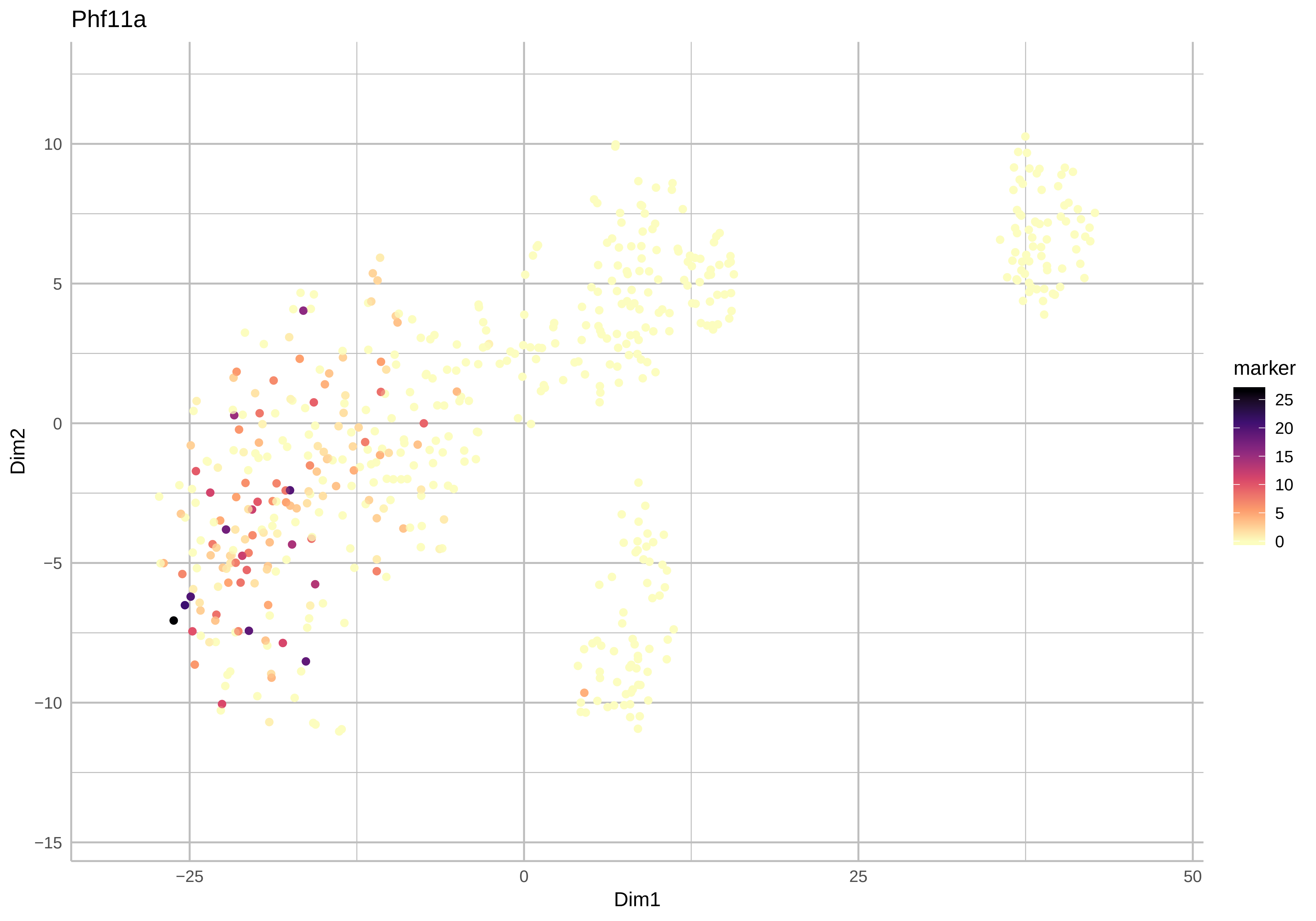
Phf11a
Dim2
Dim1
marker
25
20
15
10
5
0
10
5
0
−5
−10
−15
−25
0
25
50

Sucla2

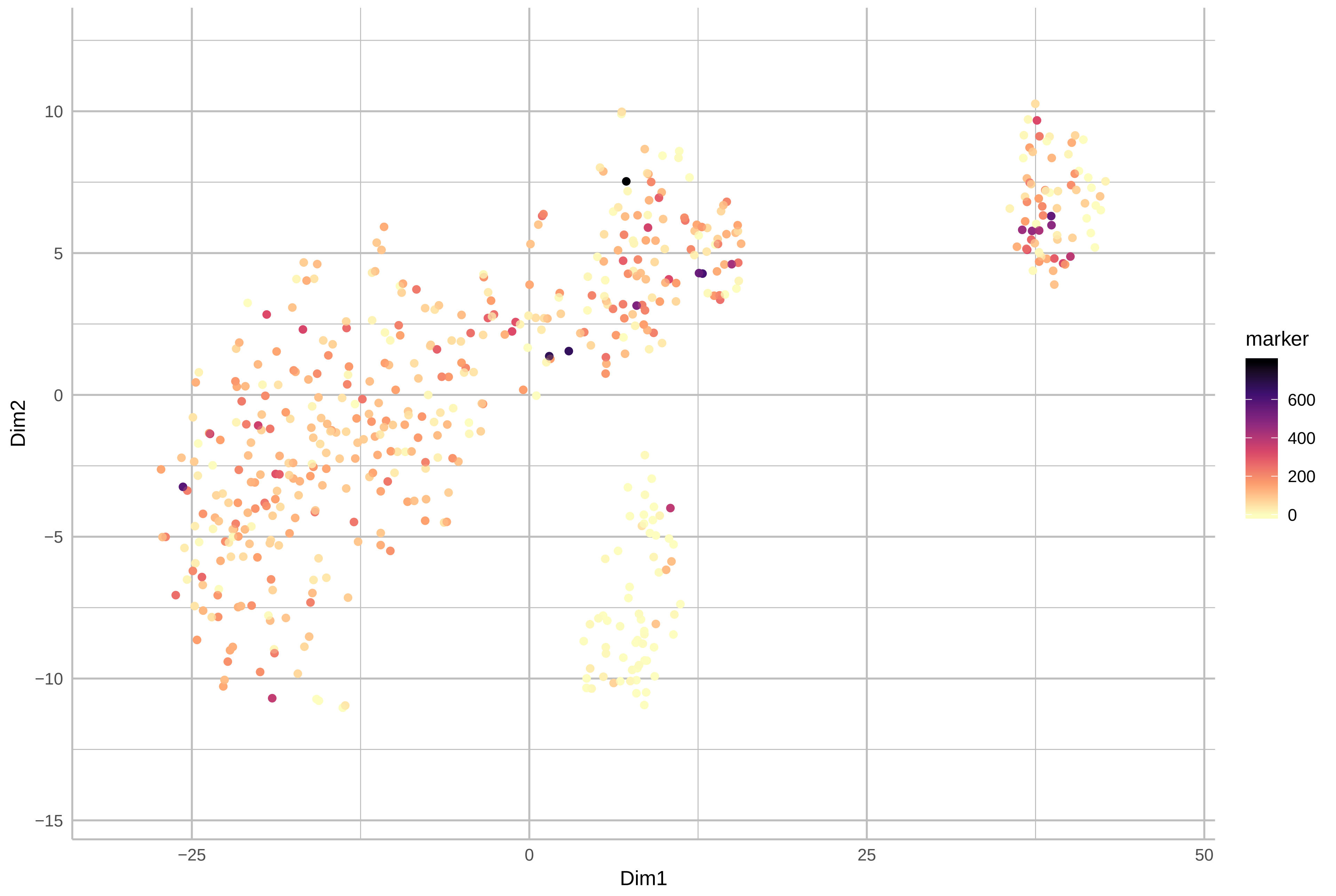

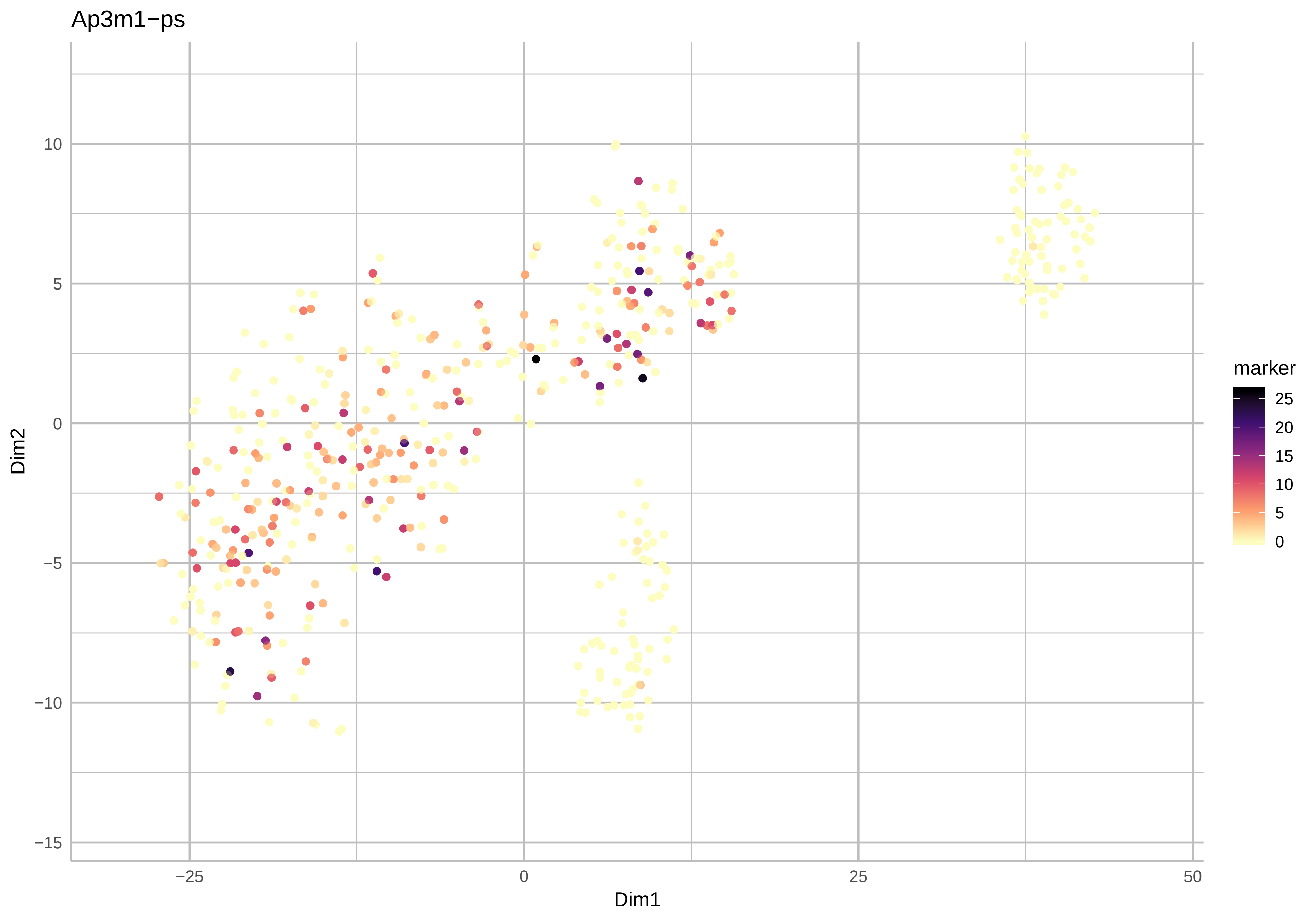
Ap3m1−ps
Dim2
Dim1
10
5
0
−5
−10
−15
−25
0
25
50
marker
25
20
15
10
5
0

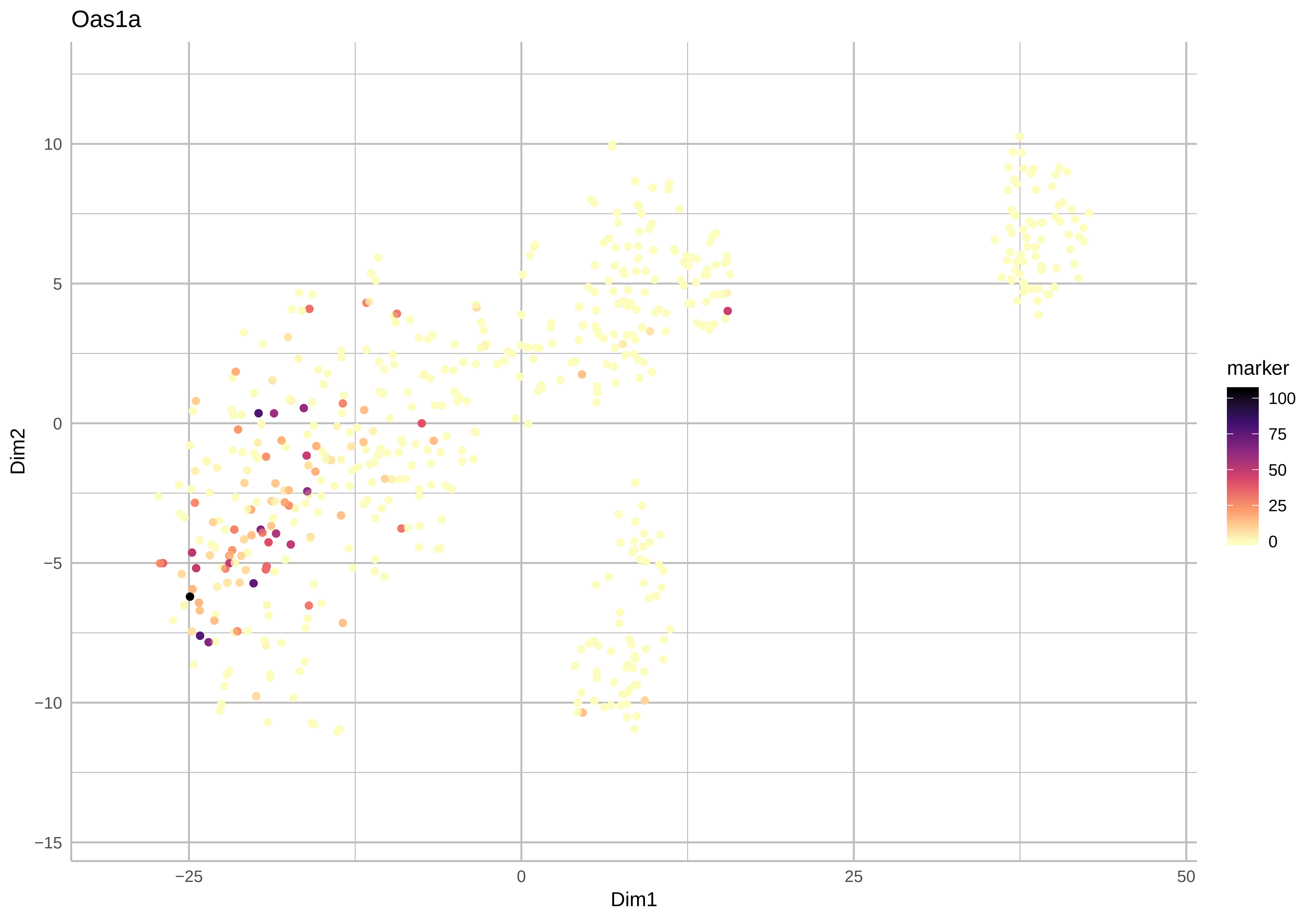

Oas1a
Dim1
Dim2
−25
0
25
50
10
5
0
−5
−10
−15
marker
100
75
50
25
0

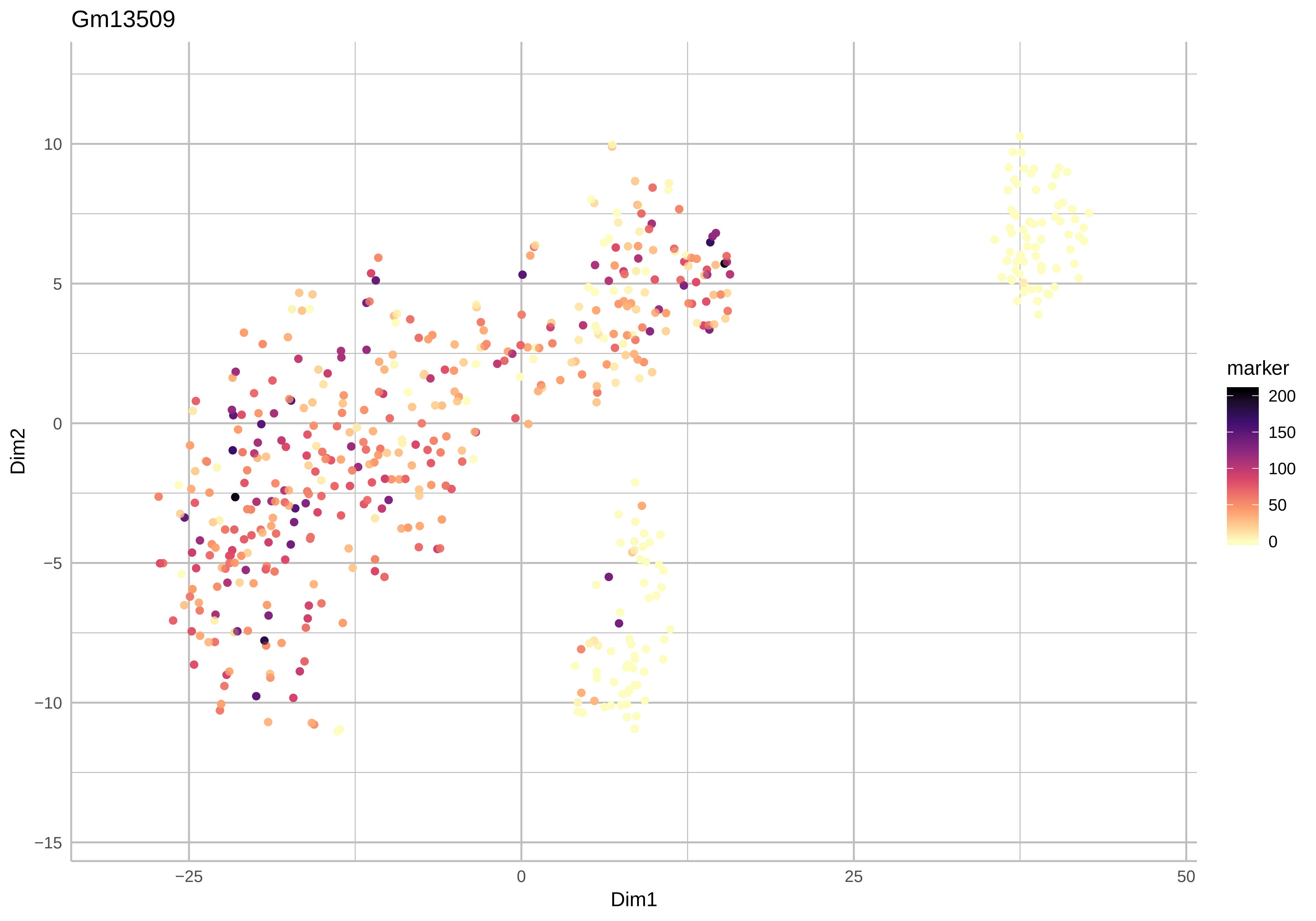
Gm13509
Dim2
Dim1
10
5
0
−5
−10
−15
−25
0
25
50
marker
200
150
100
50
0

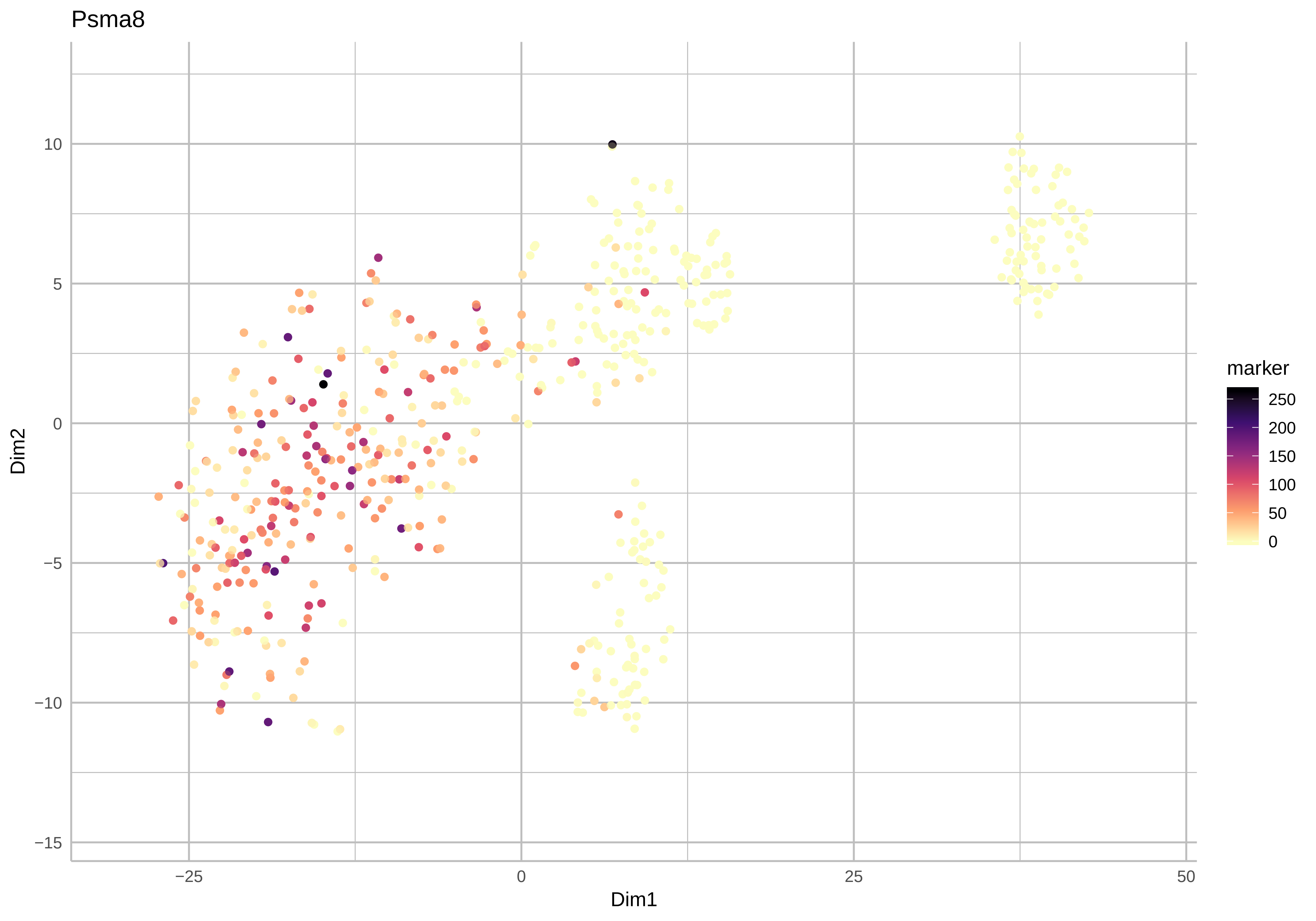

Psma8
Dim2
Dim1
10
5
0
−5
−10
−15
−25
0
25
50
marker
250
200
150
100
50
0

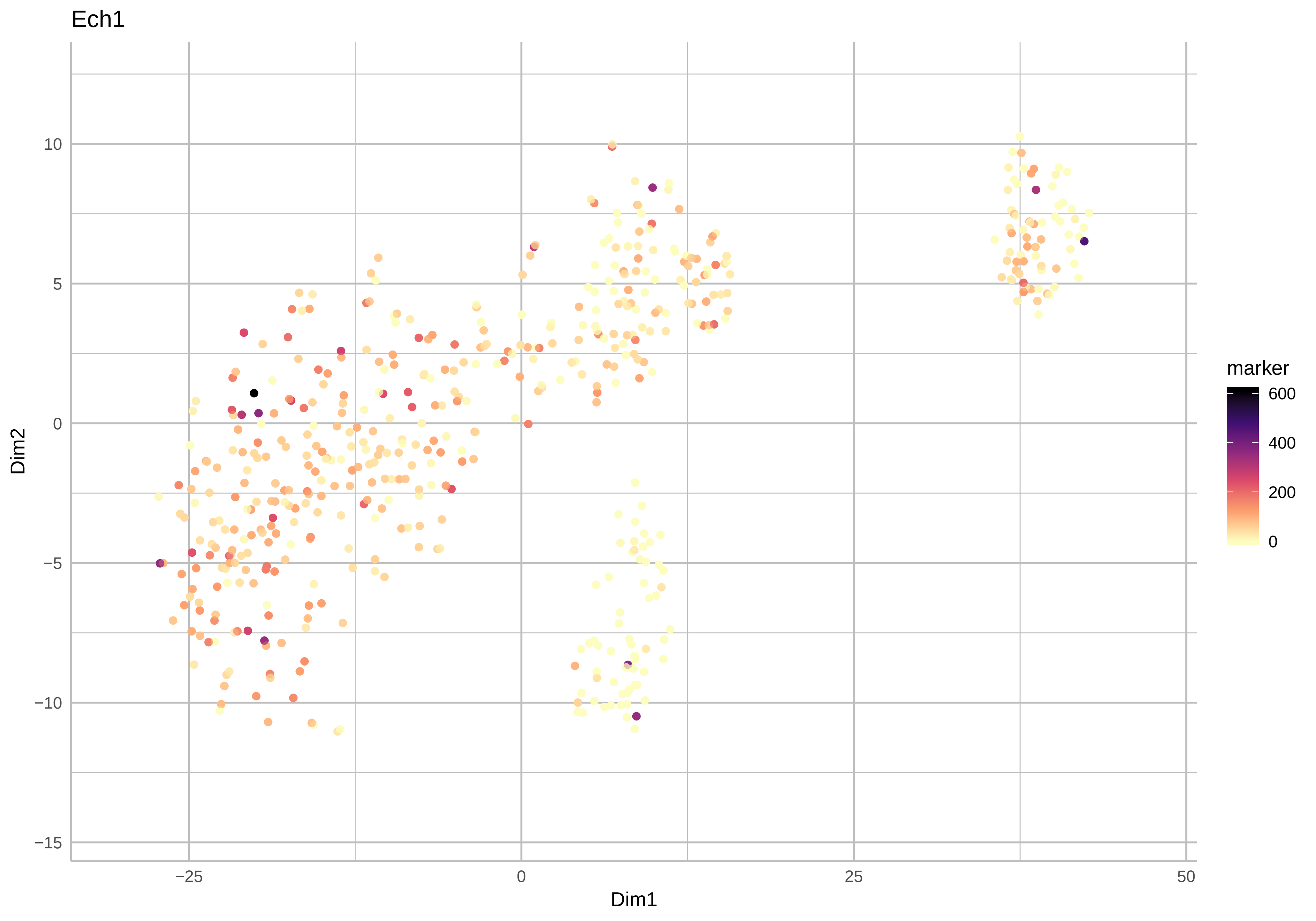

Ech1
10
5
0
−5
−10
−15
Dim2
−25
0
25
50
Dim1
marker
600
400
200
0

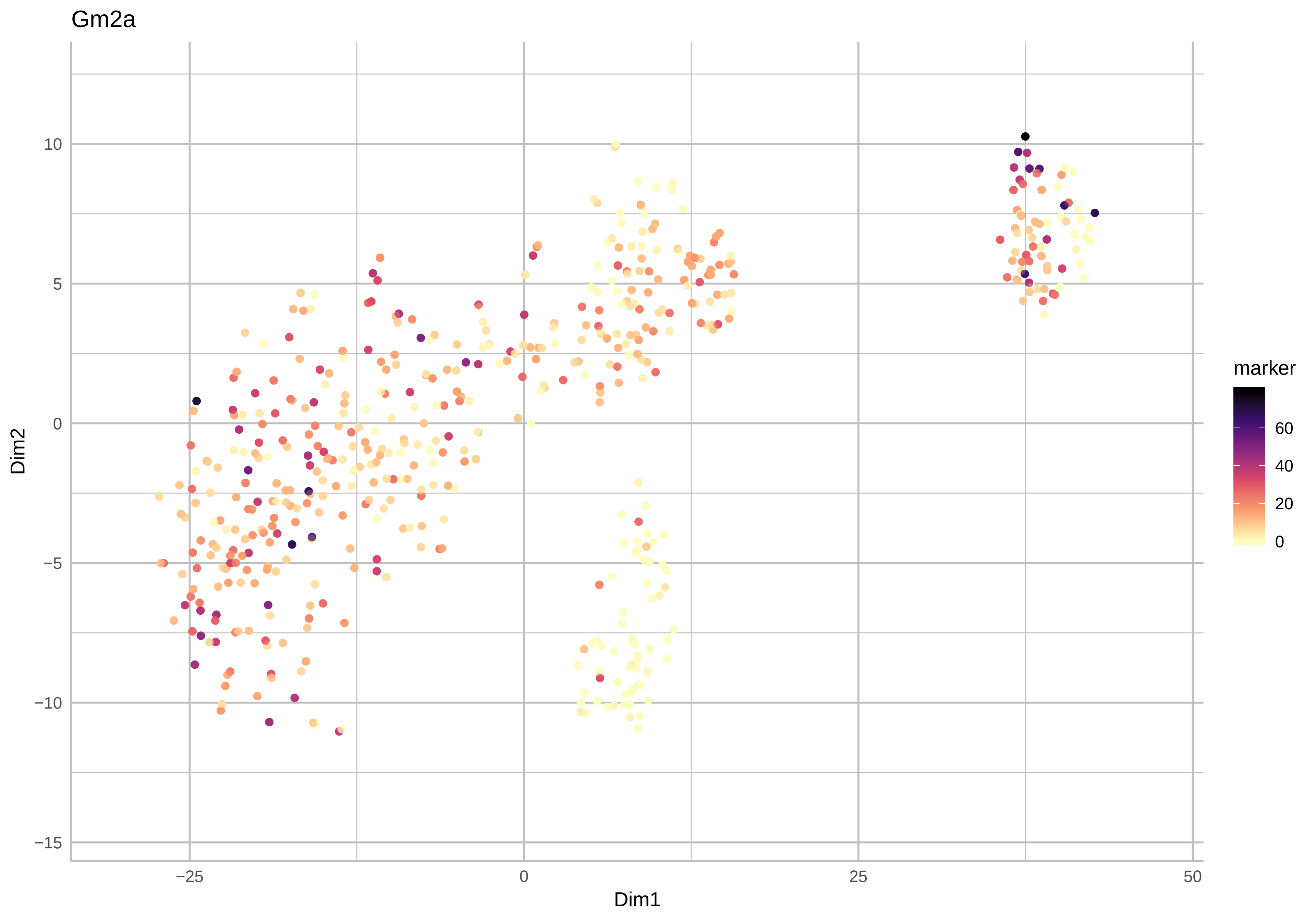

Gm2a
Dim2
Dim1
marker
60
40
20
0
10
5
0
−5
−10
−15
−25
0
25
50

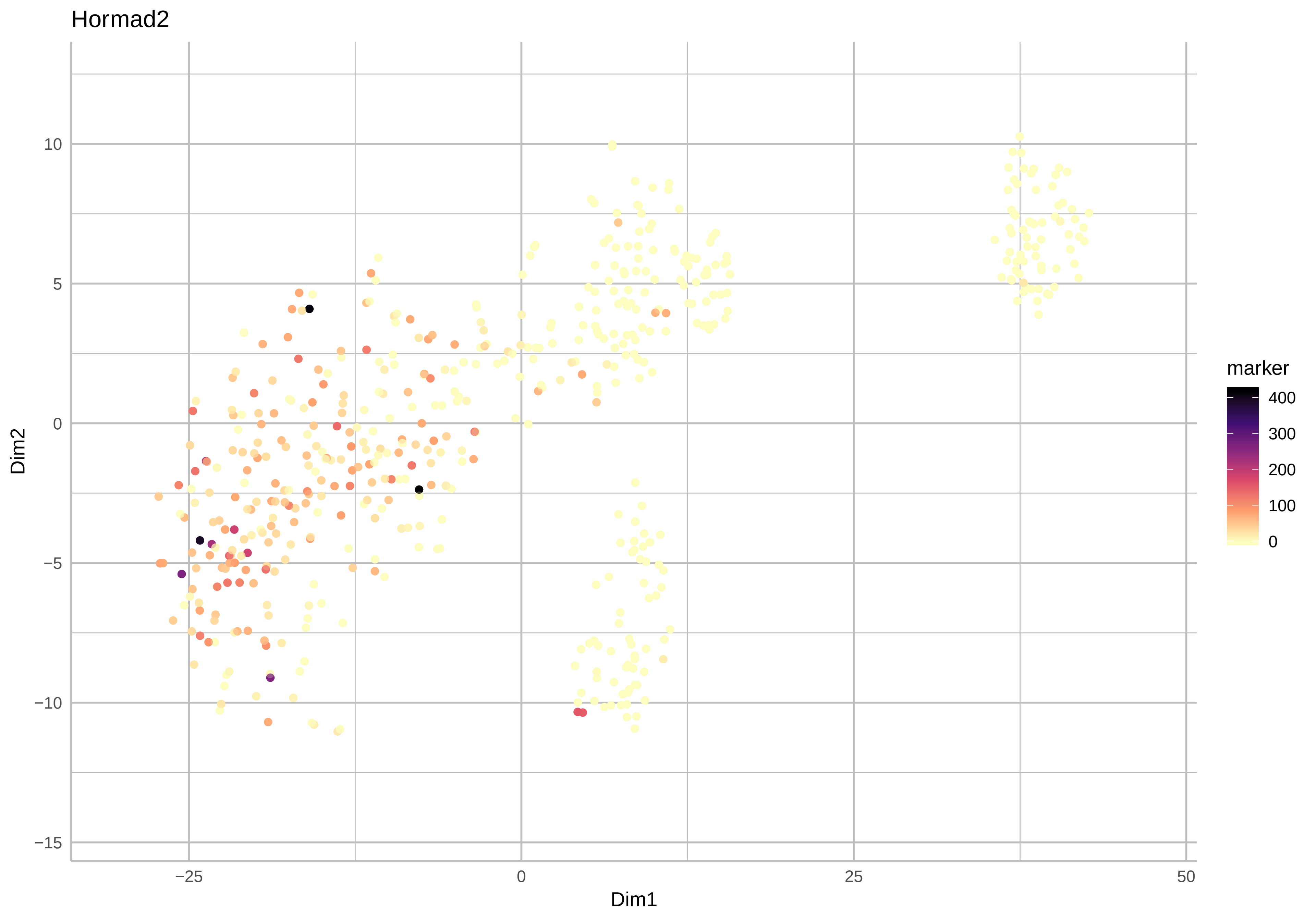

Hormad2
Dim2
Dim1
10
5
0
−5
−10
−15
−25
0
25
50
marker
400
300
200
100
0

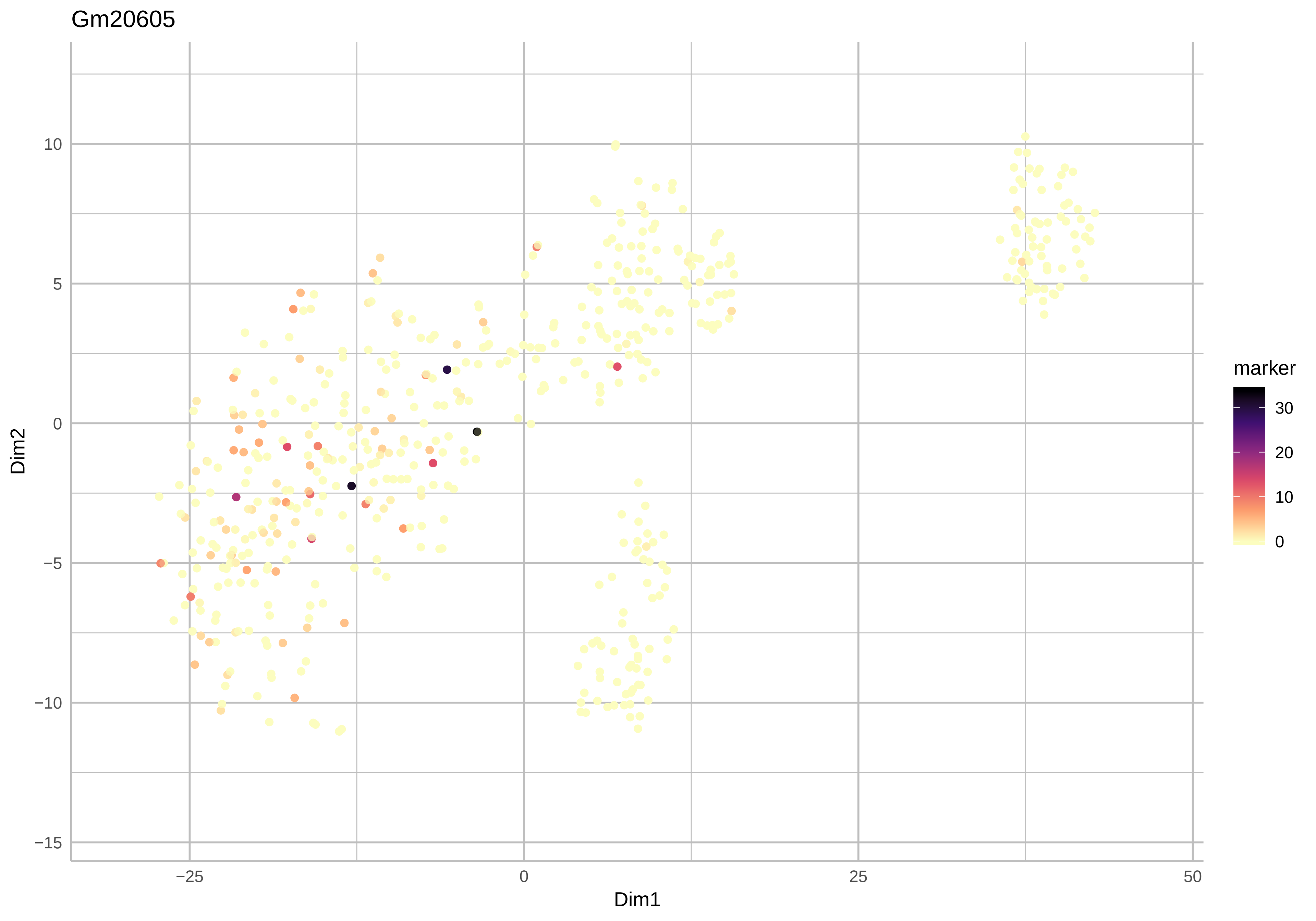

Gm20605
Dim2
Dim1
10
5
0
−5
−10
−15
−25
0
25
50
marker
30
20
10
0

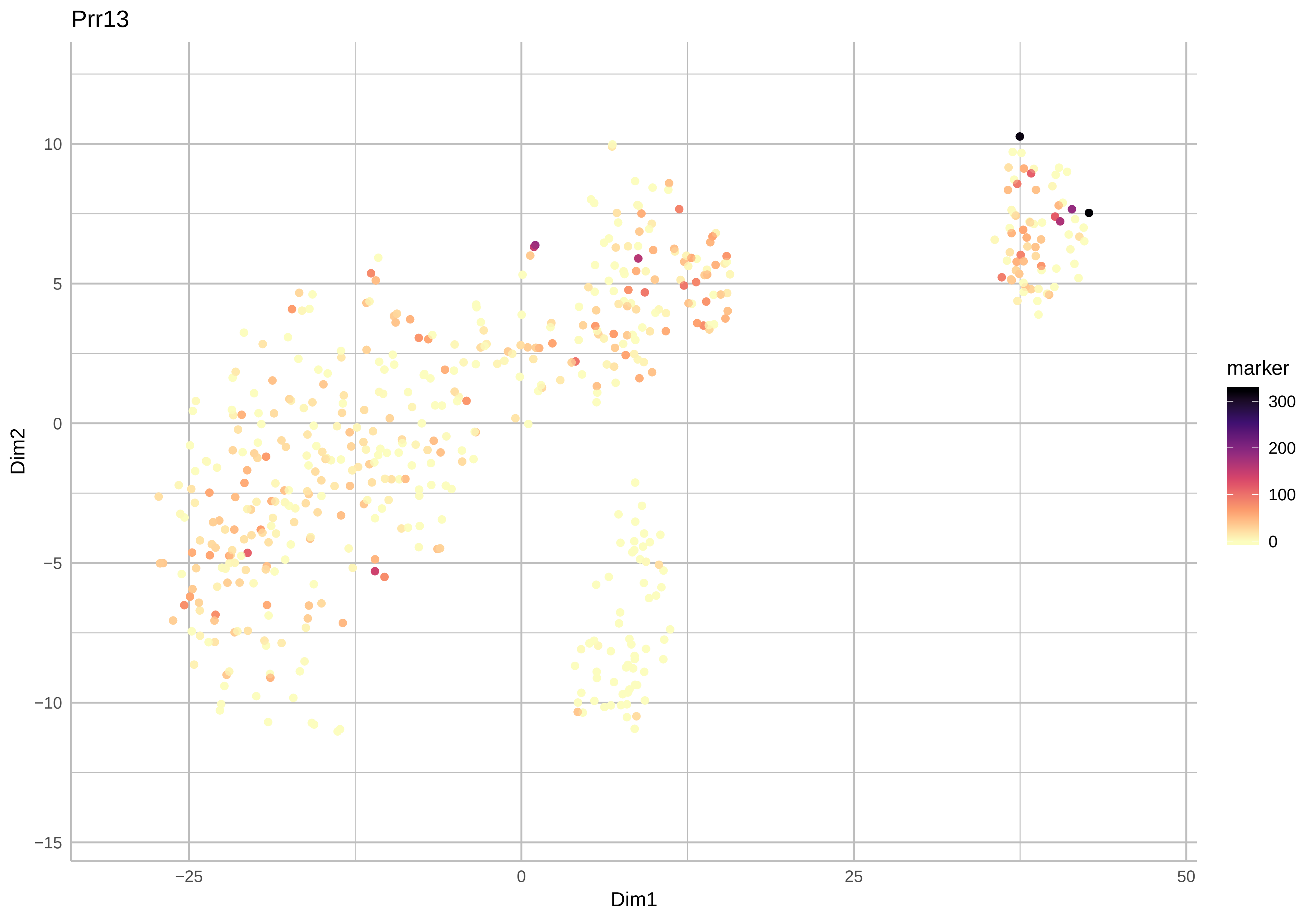

Prr13
Dim2
Dim1
10
5
0
−5
−10
−15
−25
0
25
50
marker
300
200
100
0

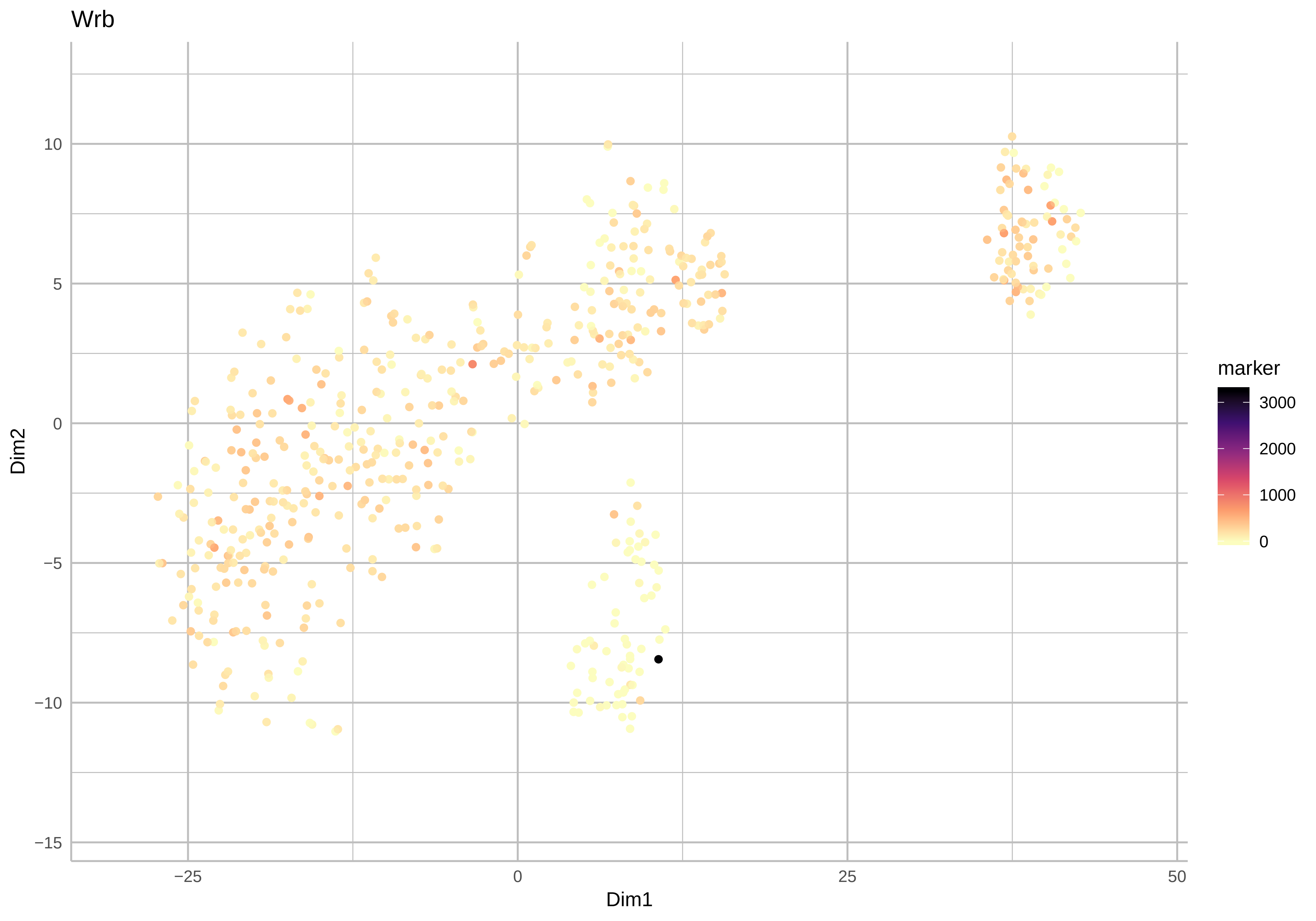
Wrb
10
5
0
−5
−10
−15
Dim2
−25
0
25
50
Dim1
marker
3000
2000
1000
0

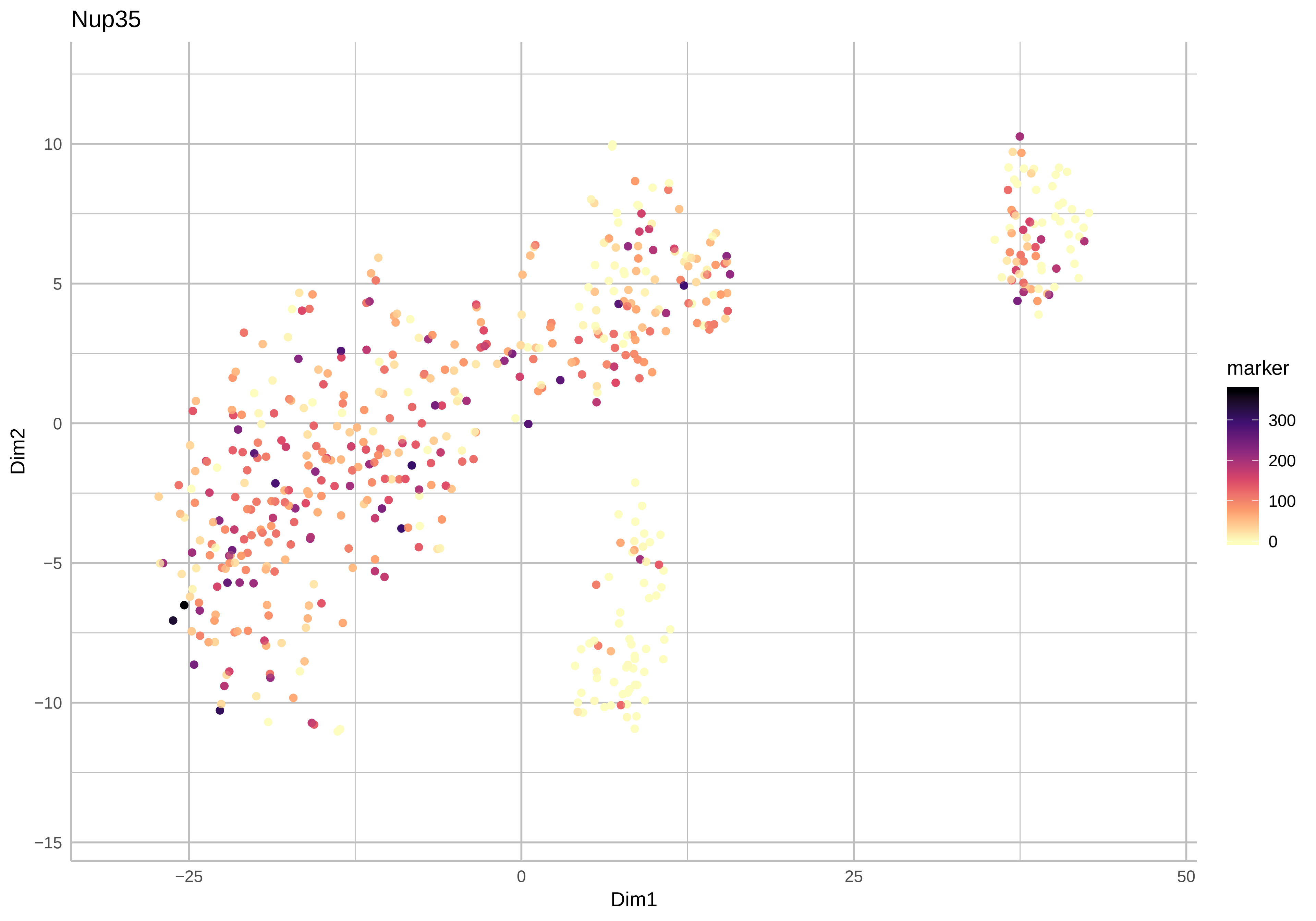

Nup35
Dim2
Dim1
10
5
0
−5
−10
−15
−25
0
25
50
marker
300
200
100
0

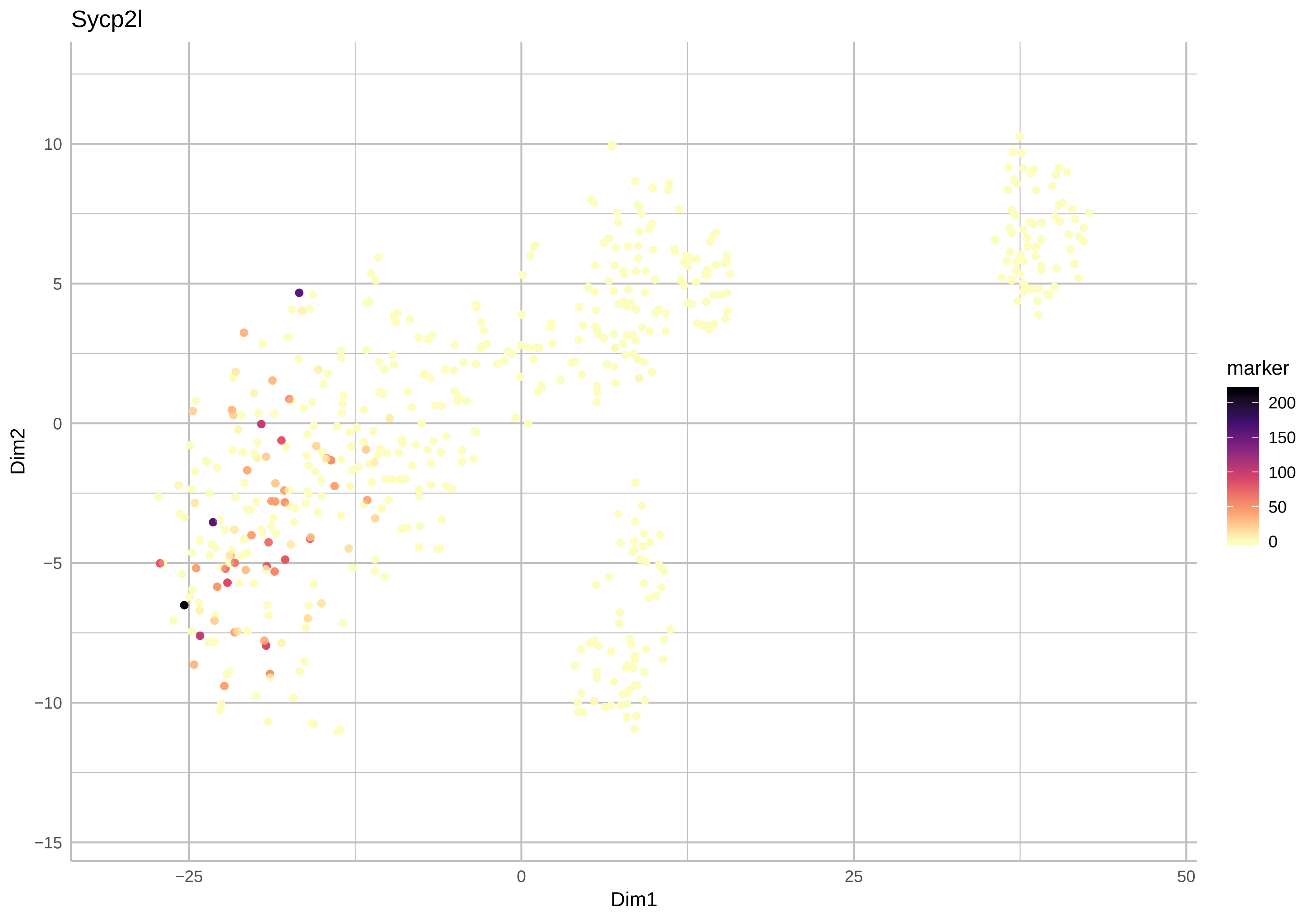

Sycp2l
10
5
0
−5
−10
−15
−25
0
25
50
Dim1
Dim2
marker
200
150
100
50
0

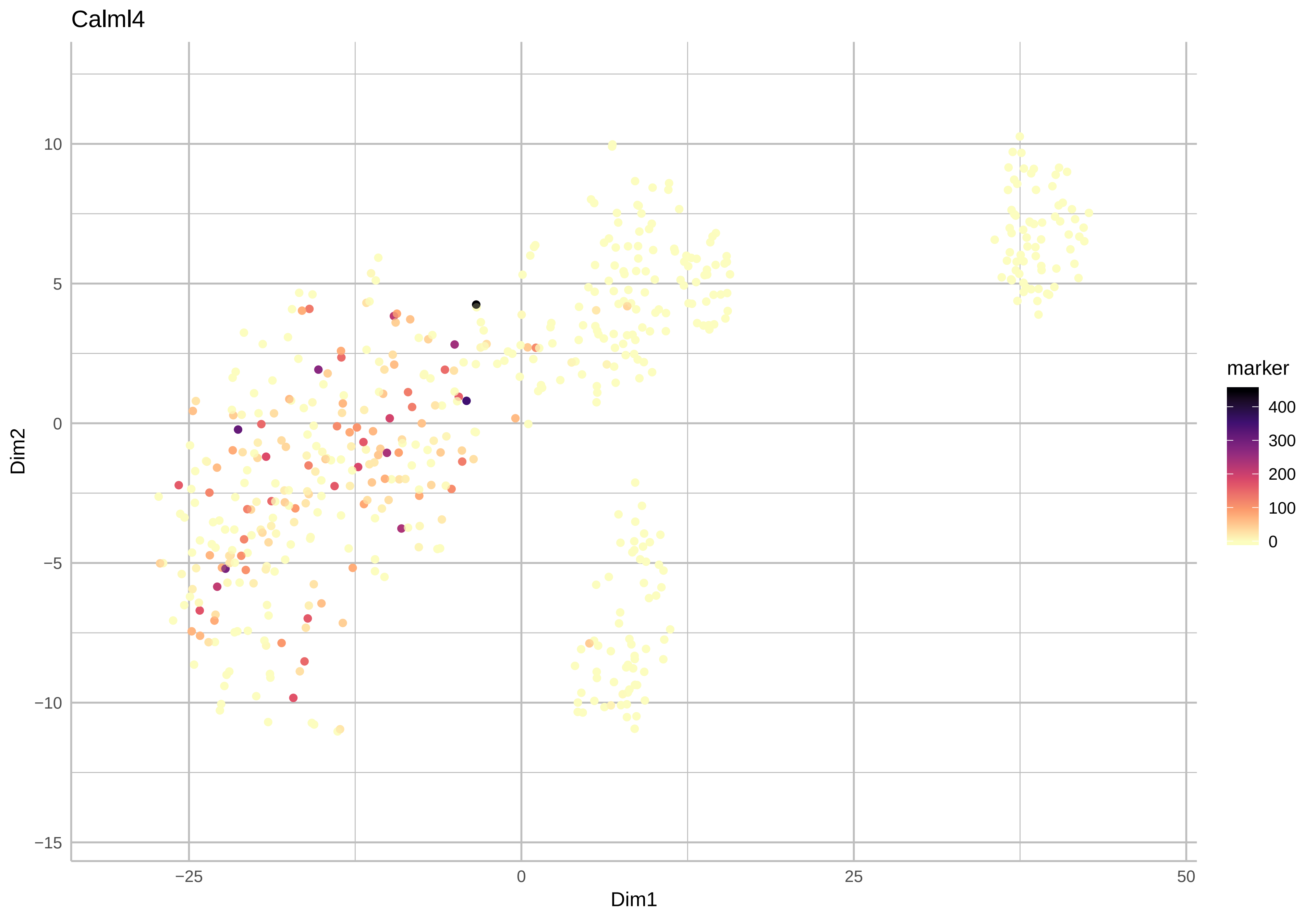

Calml4
10
5
0
−5
−10
−15
Dim2
−25
0
25
50
Dim1
marker
400
300
200
100
0

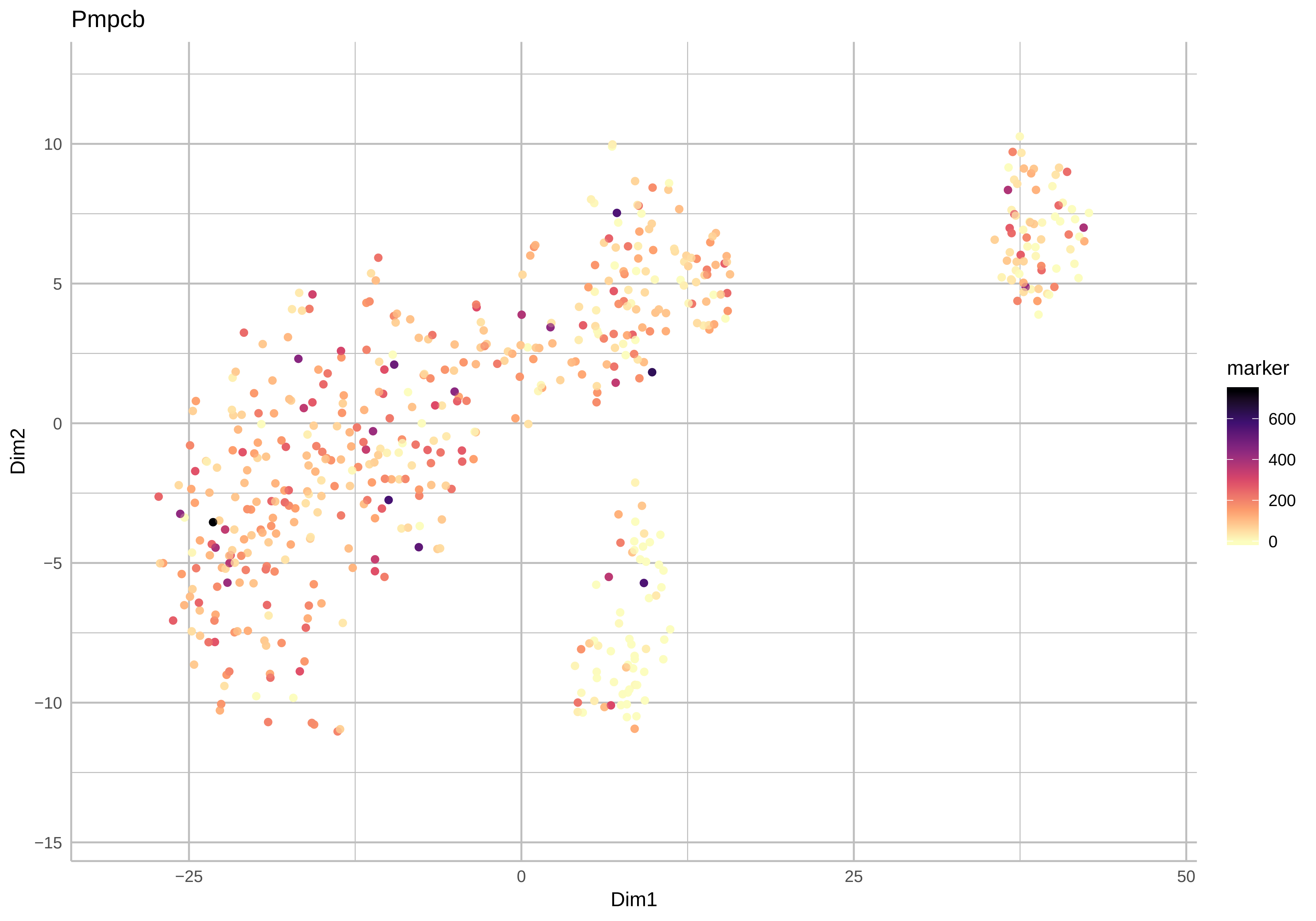
Pmpcb
Dim2
Dim1
10
5
0
−5
−10
−15
−25
0
25
50
marker
600
400
200
0

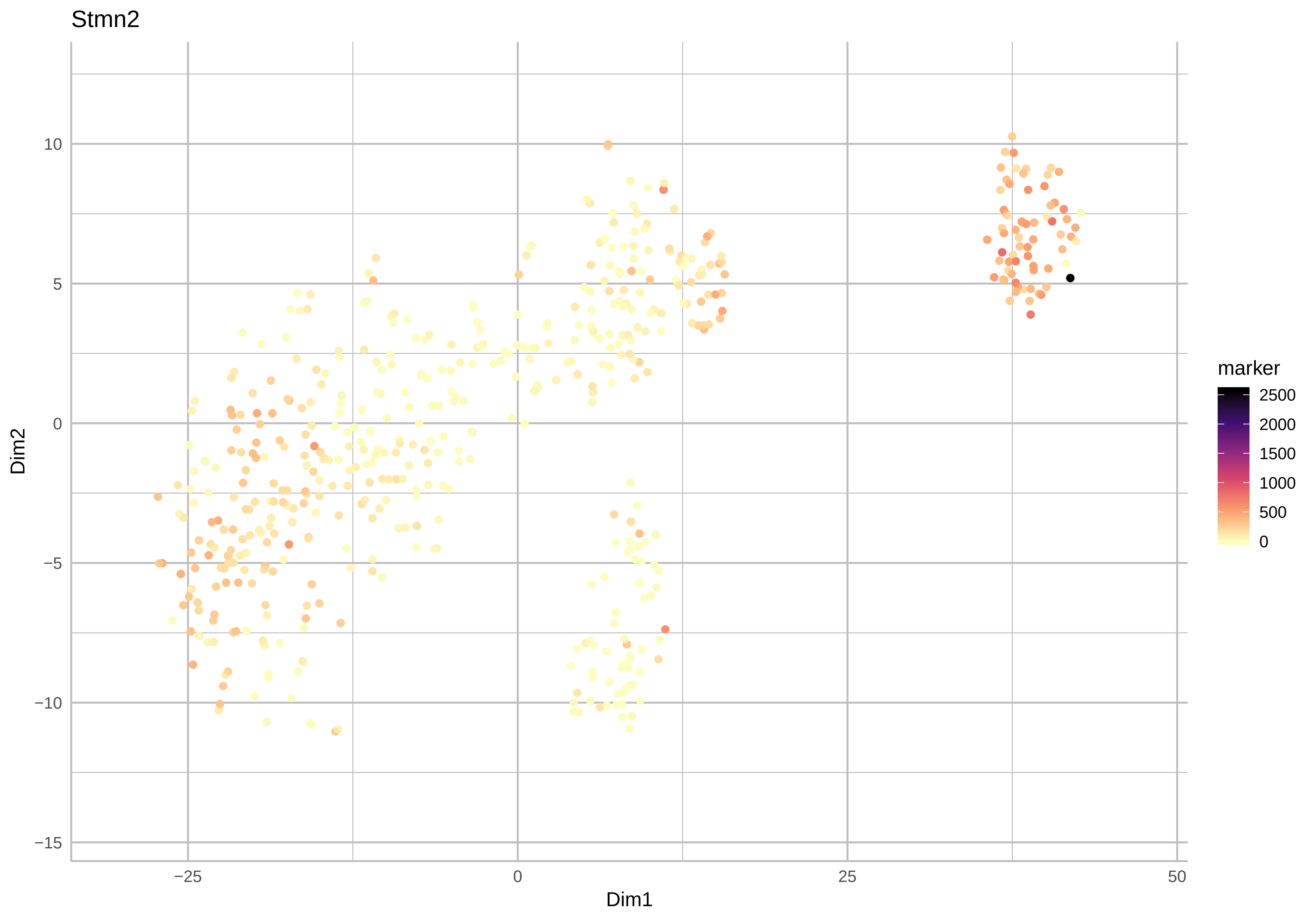
Stmn2
Dim2
Dim1
10
5
0
−5
−10
−15
−25
0
25
50
marker
2500
2000
1500
1000
500
0

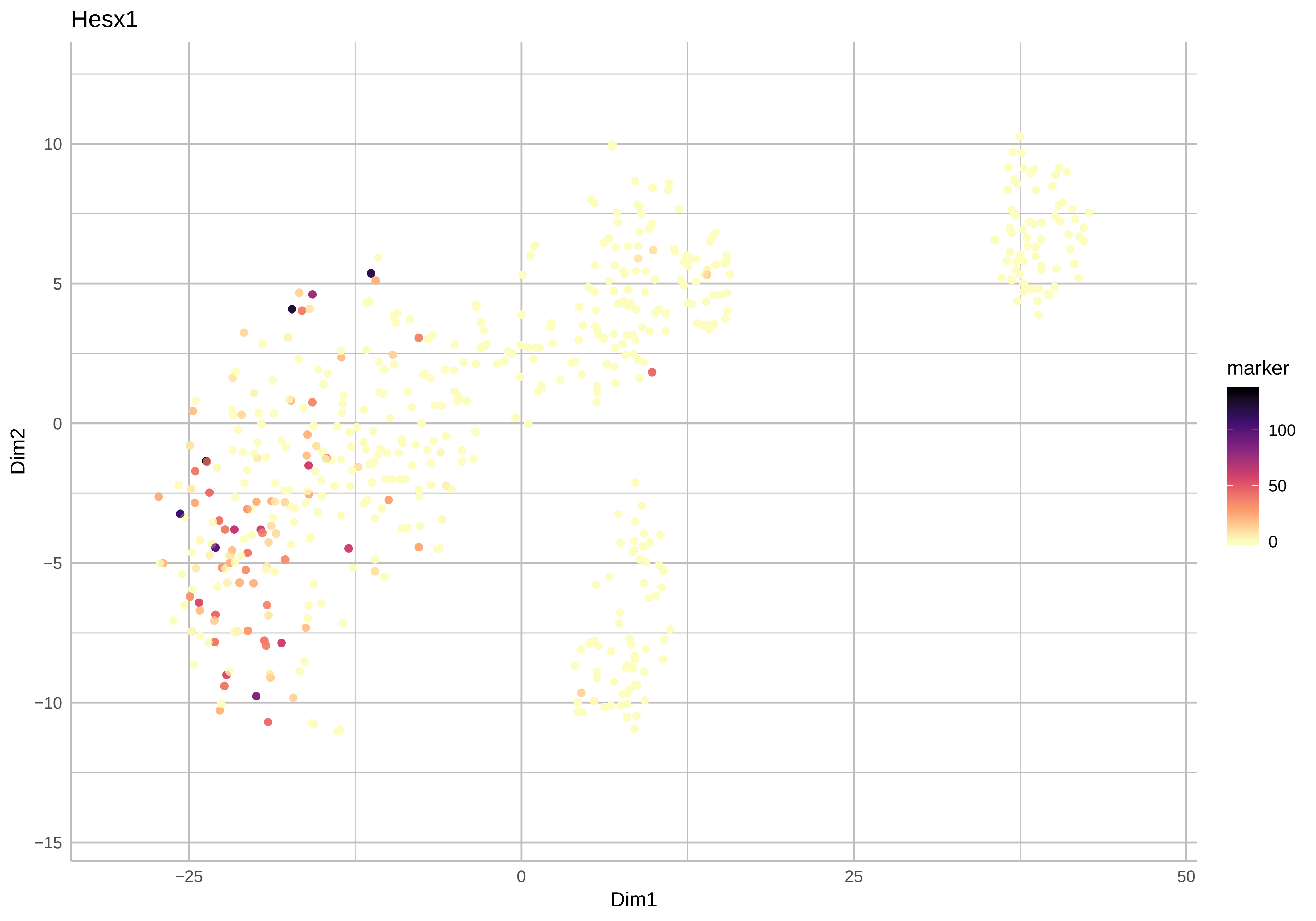
Hesx1
Dim2
Dim1
10
5
0
−5
−10
−15
−25
0
25
50
marker
100
50
0

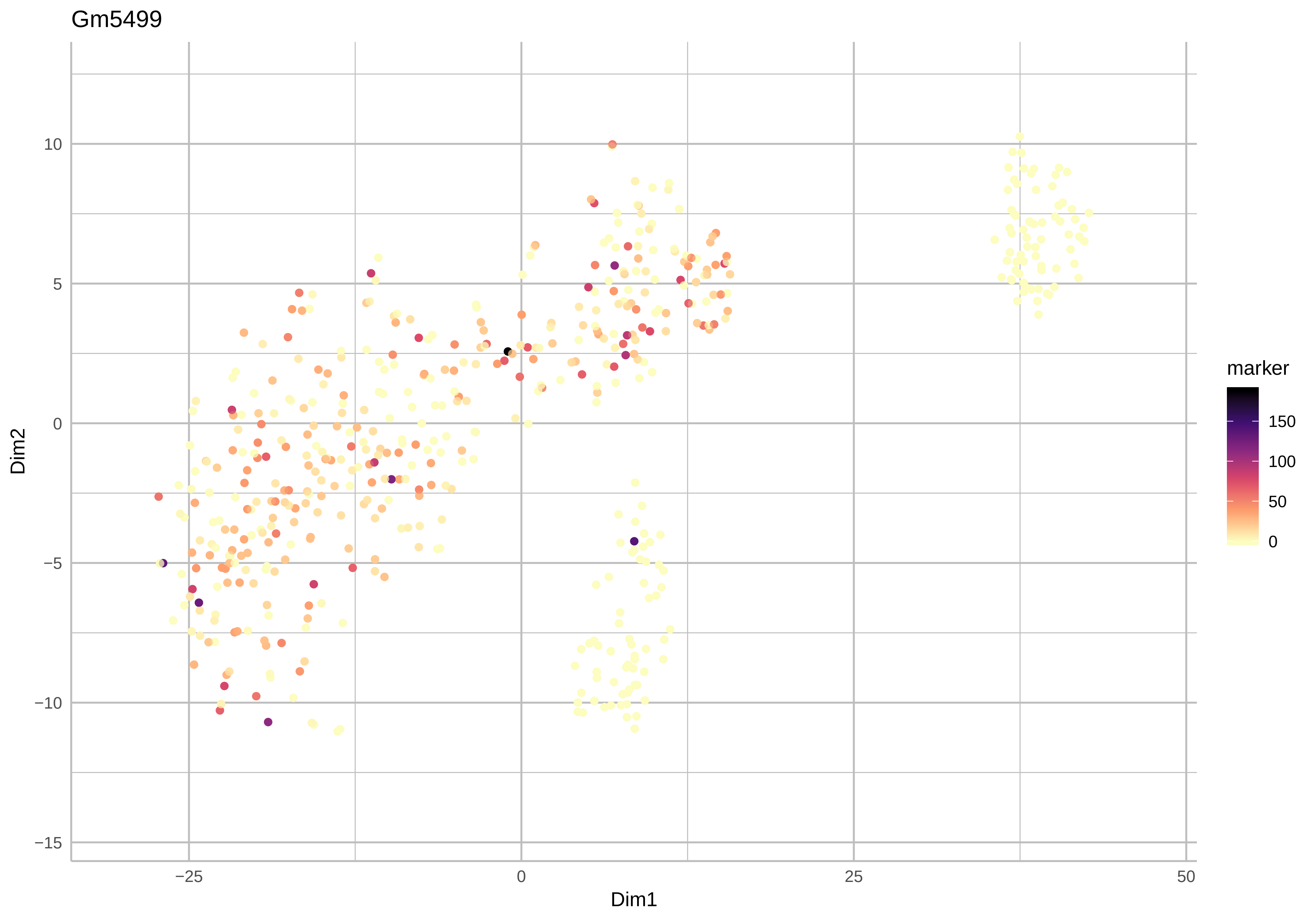

Gm5499
marker
150
100
50
0
Dim2
Dim1
10
5
0
−5
−10
−15
−25
0
25
50

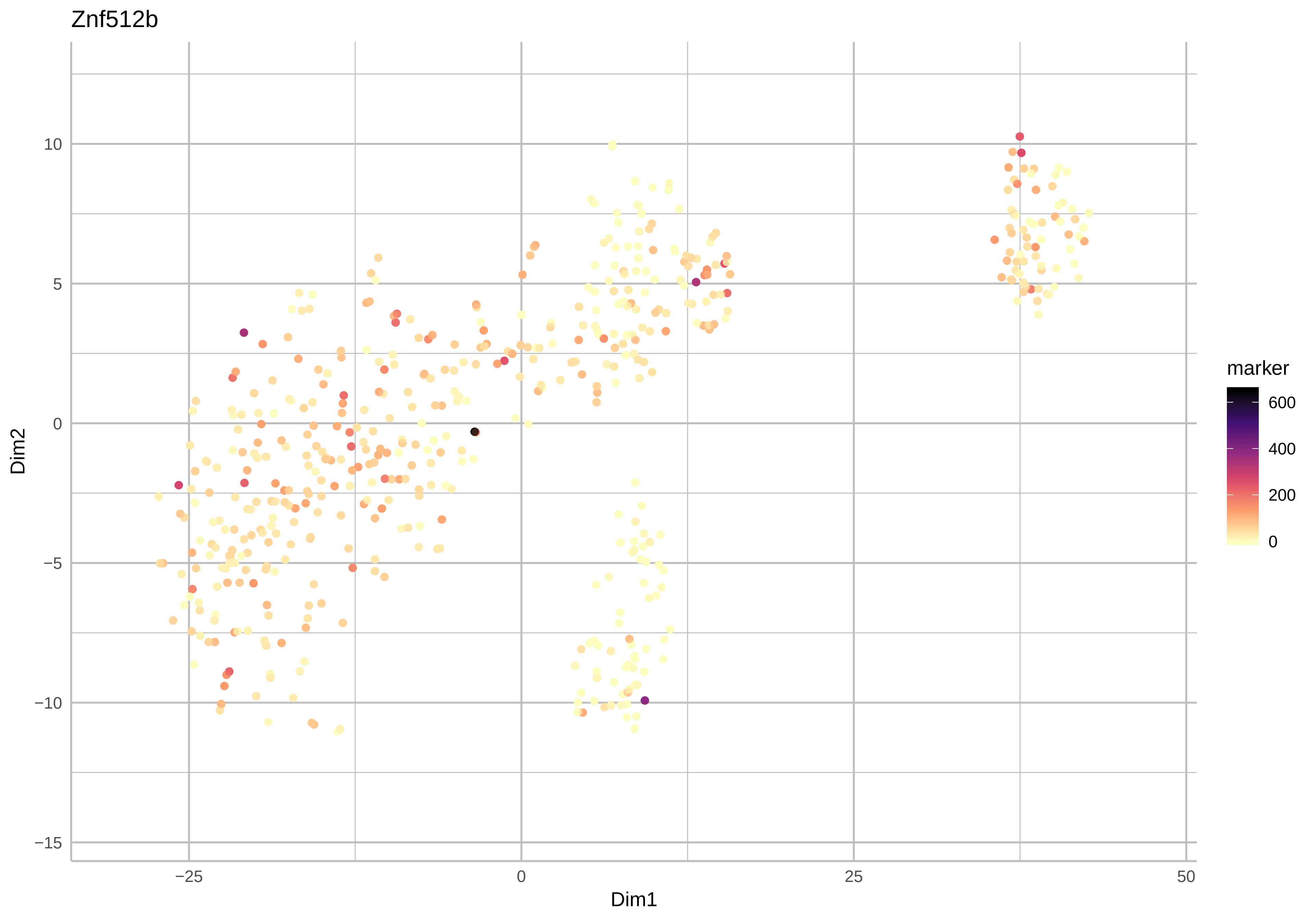

Znf512b
10
5
0
−5
−10
−15
Dim2
−25
0
25
50
Dim1
marker
600
400
200
0

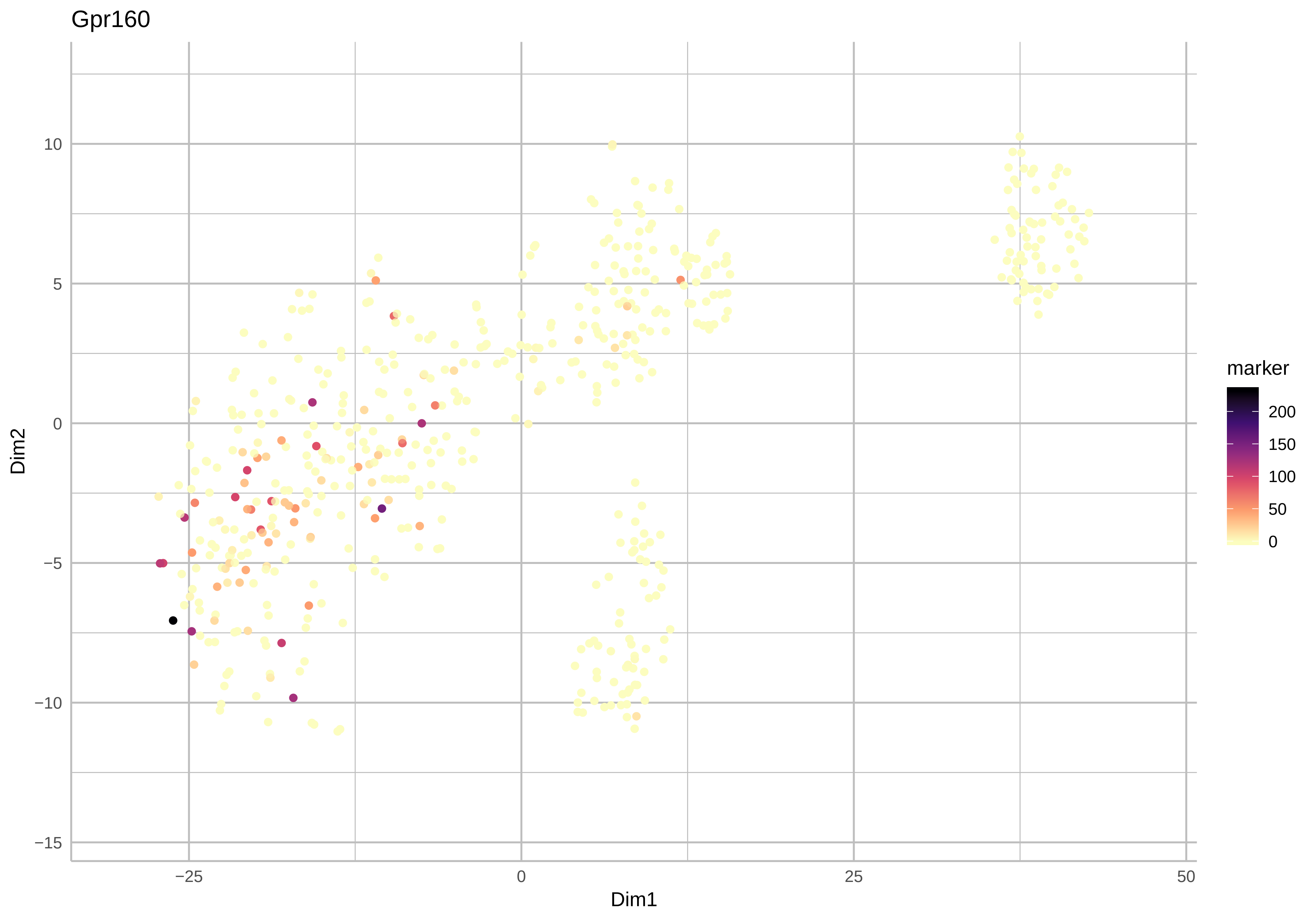
Gpr160
Dim2
Dim1
10
5
0
−5
−10
−15
−25
0
25
50
marker
200
150
100
50
0

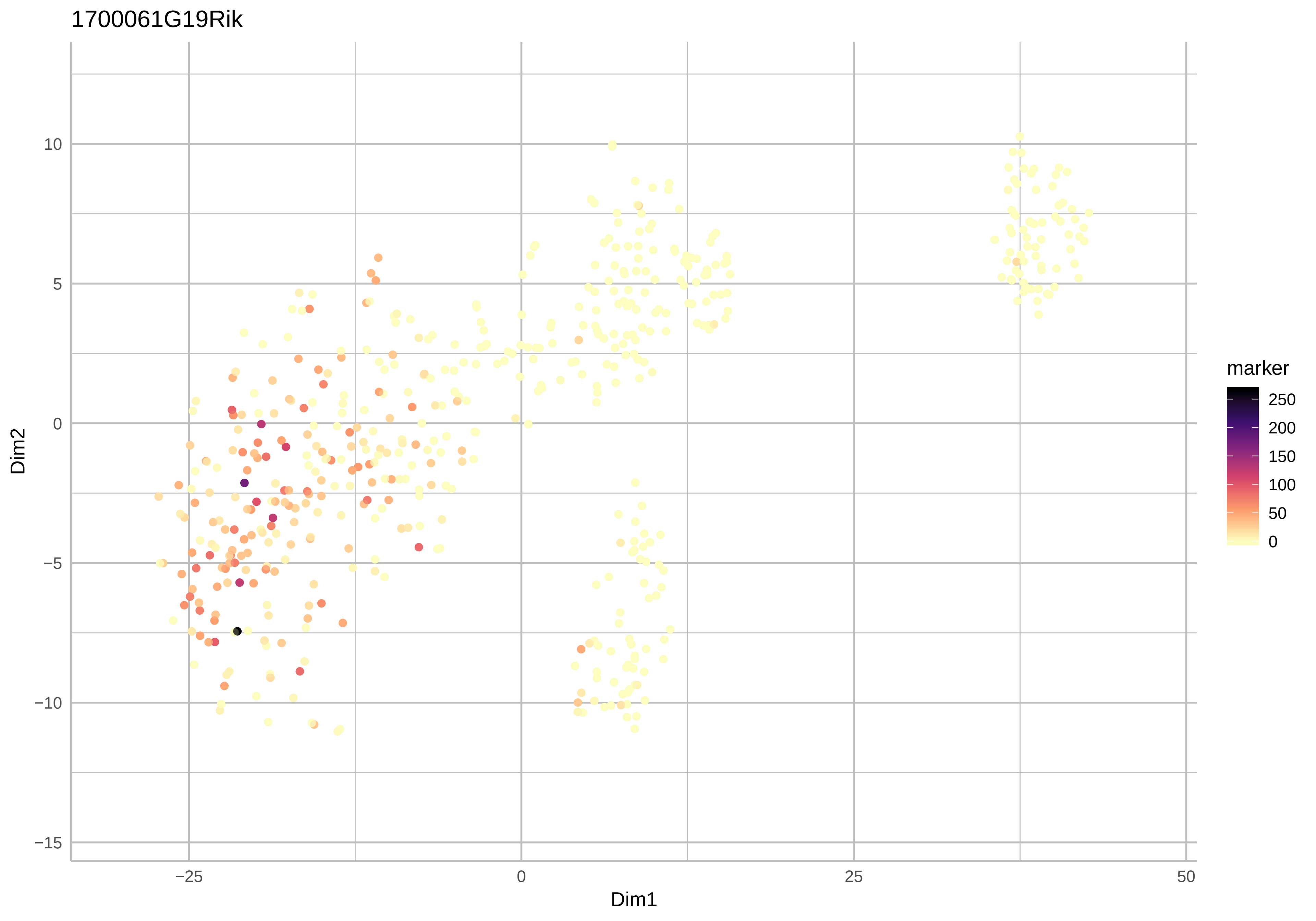
1700061G19Rik
Dim2
10
5
0
−5
−10
−15
−25
0
25
50
Dim1
marker
250
200
150
100
50
0

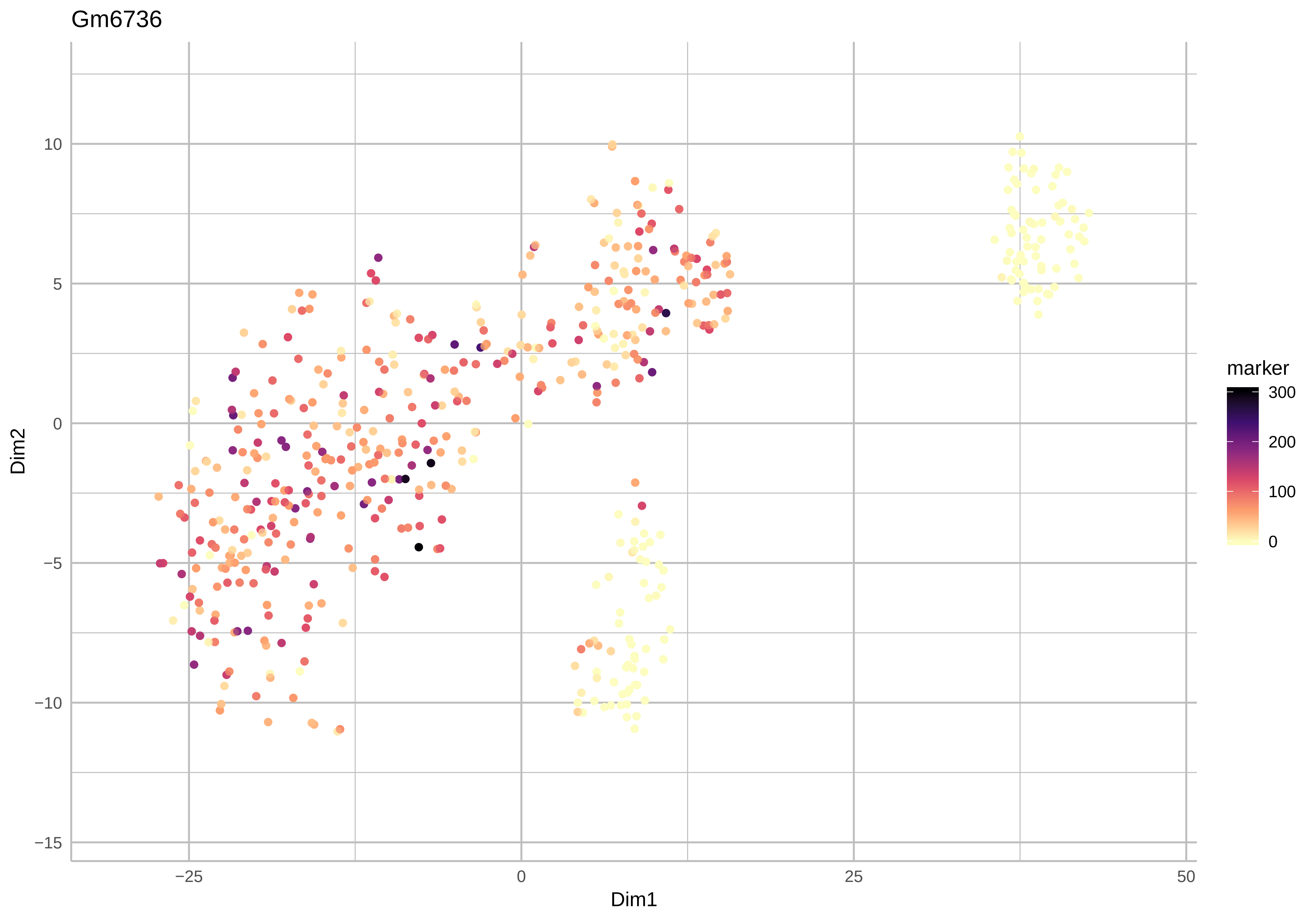

Gm6736
Dim2
Dim1
10
5
0
−5
−10
−15
−25
0
25
50
marker
300
200
100
0

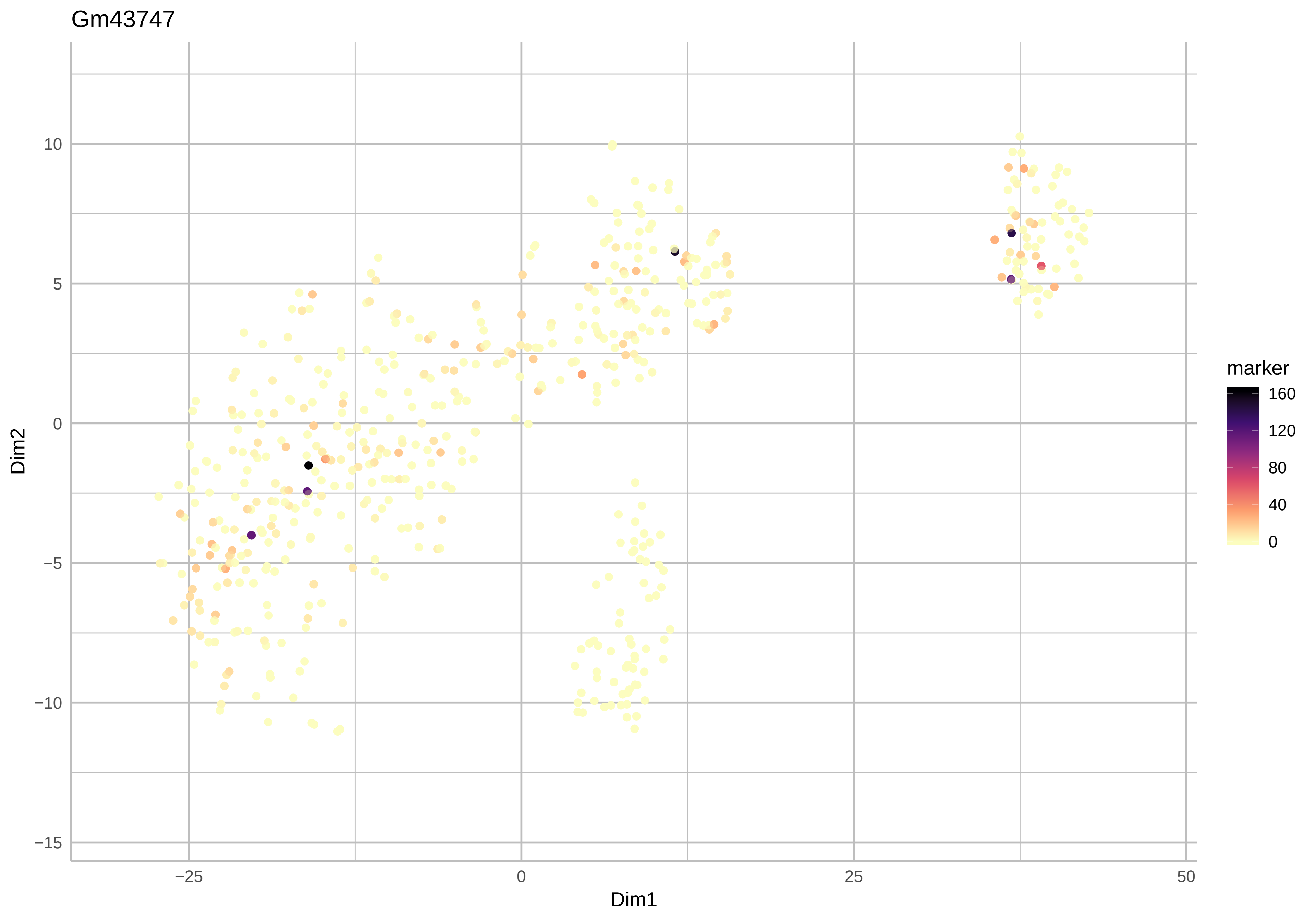

Gm43747
10
5
0
−5
−10
−15
Dim2
−25
0
25
50
Dim1
marker
160
120
80
40
0

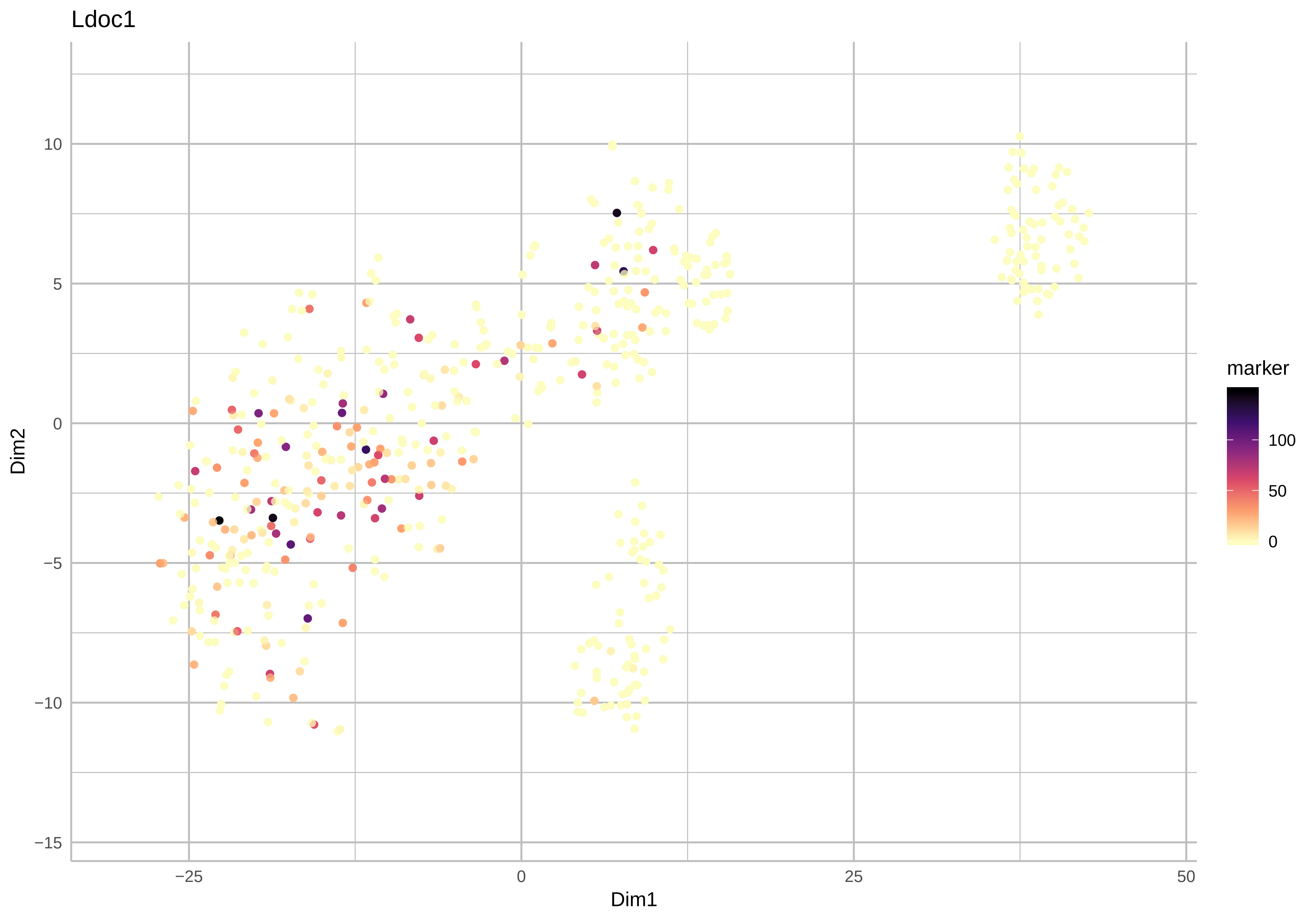

Ldoc1
Dim2
Dim1
10
5
0
−5
−10
−15
−25
0
25
50
marker
100
50
0

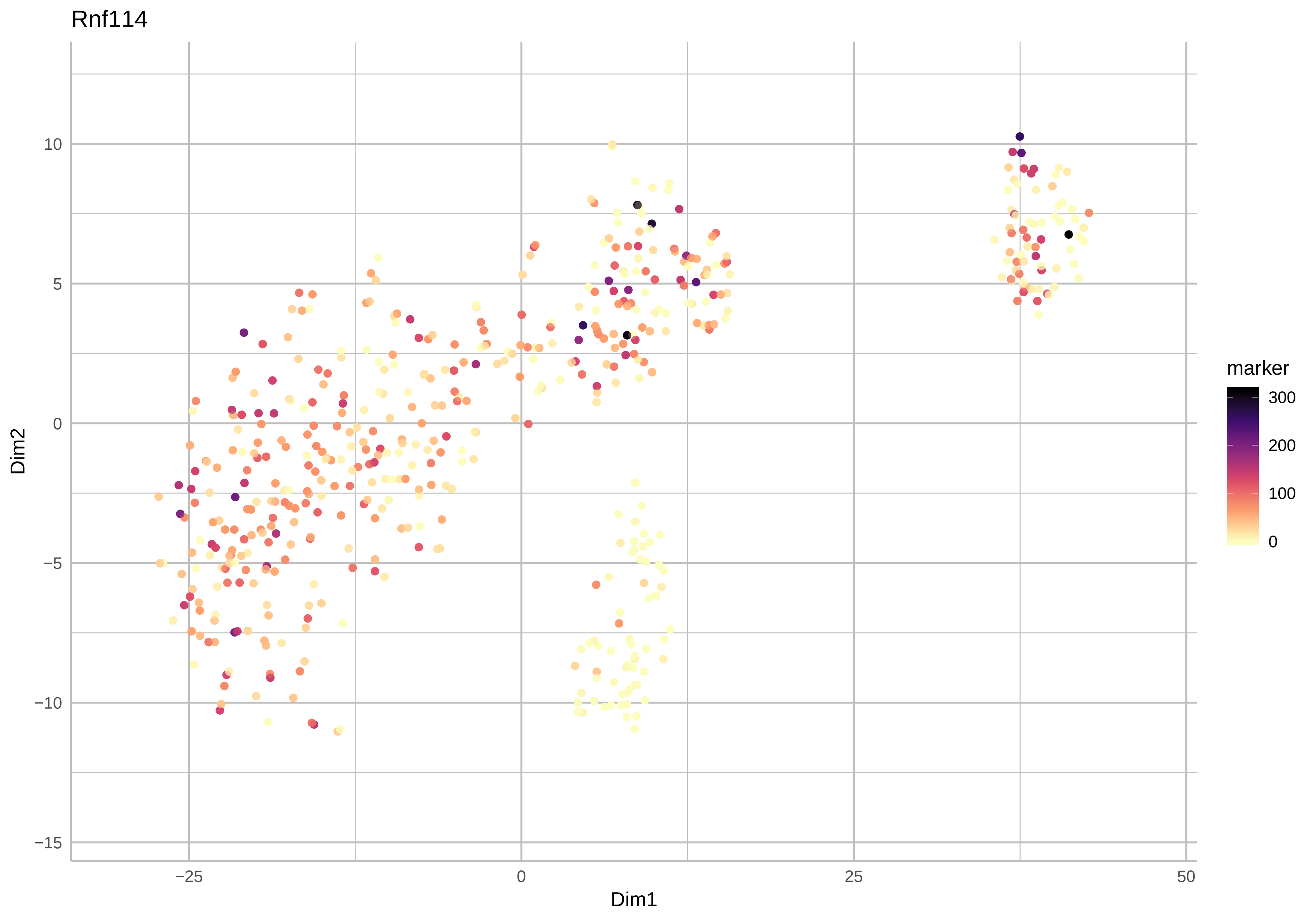

Rnf114
marker
300
200
100
0
10
5
0
−5
−10
−15
Dim2
−25
0
25
50
Dim1

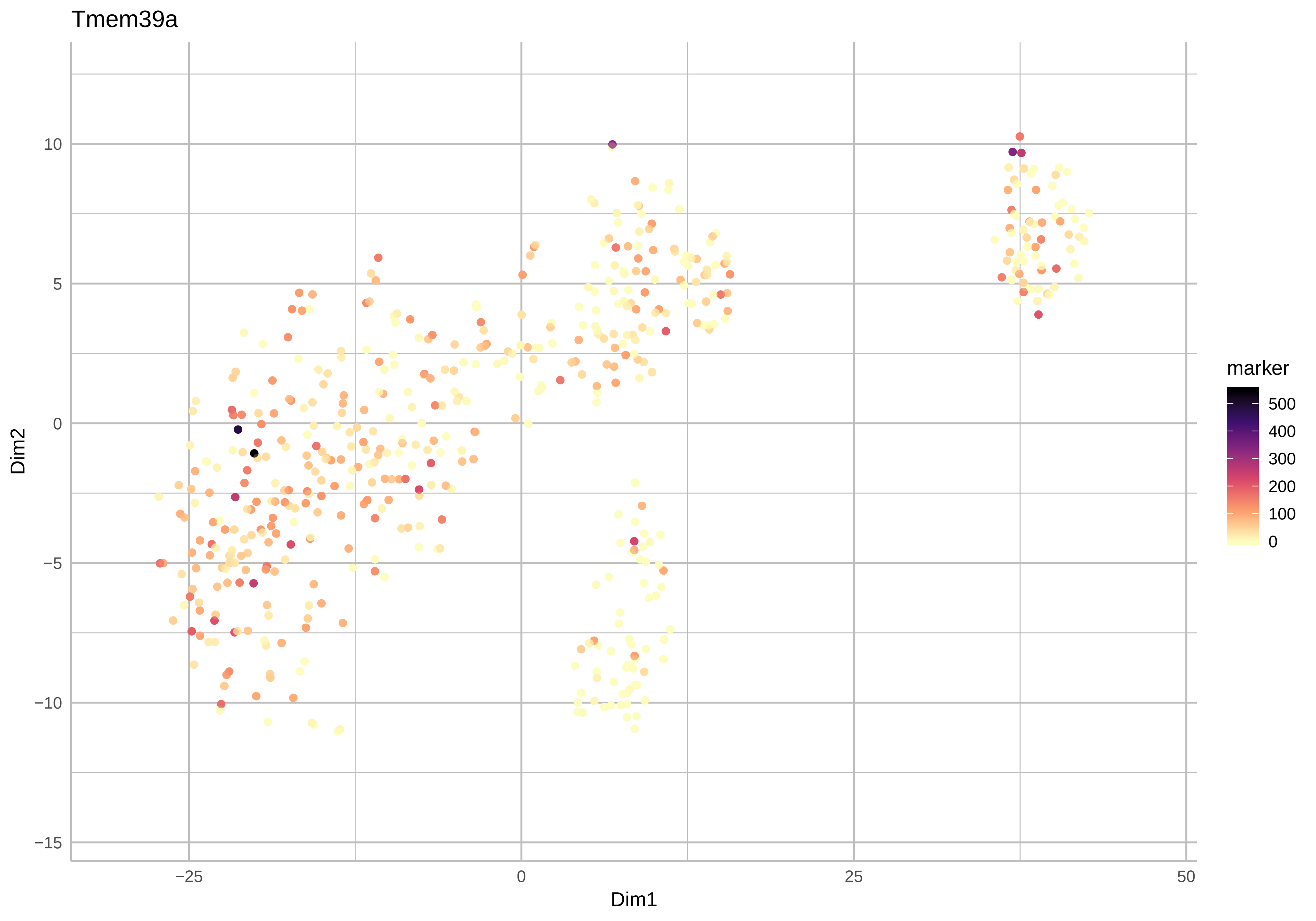

Tmem39a
10
5
0
−5
−10
−15
−25
0
25
50
Dim1
Dim2
marker
500
400
300
200
100
0

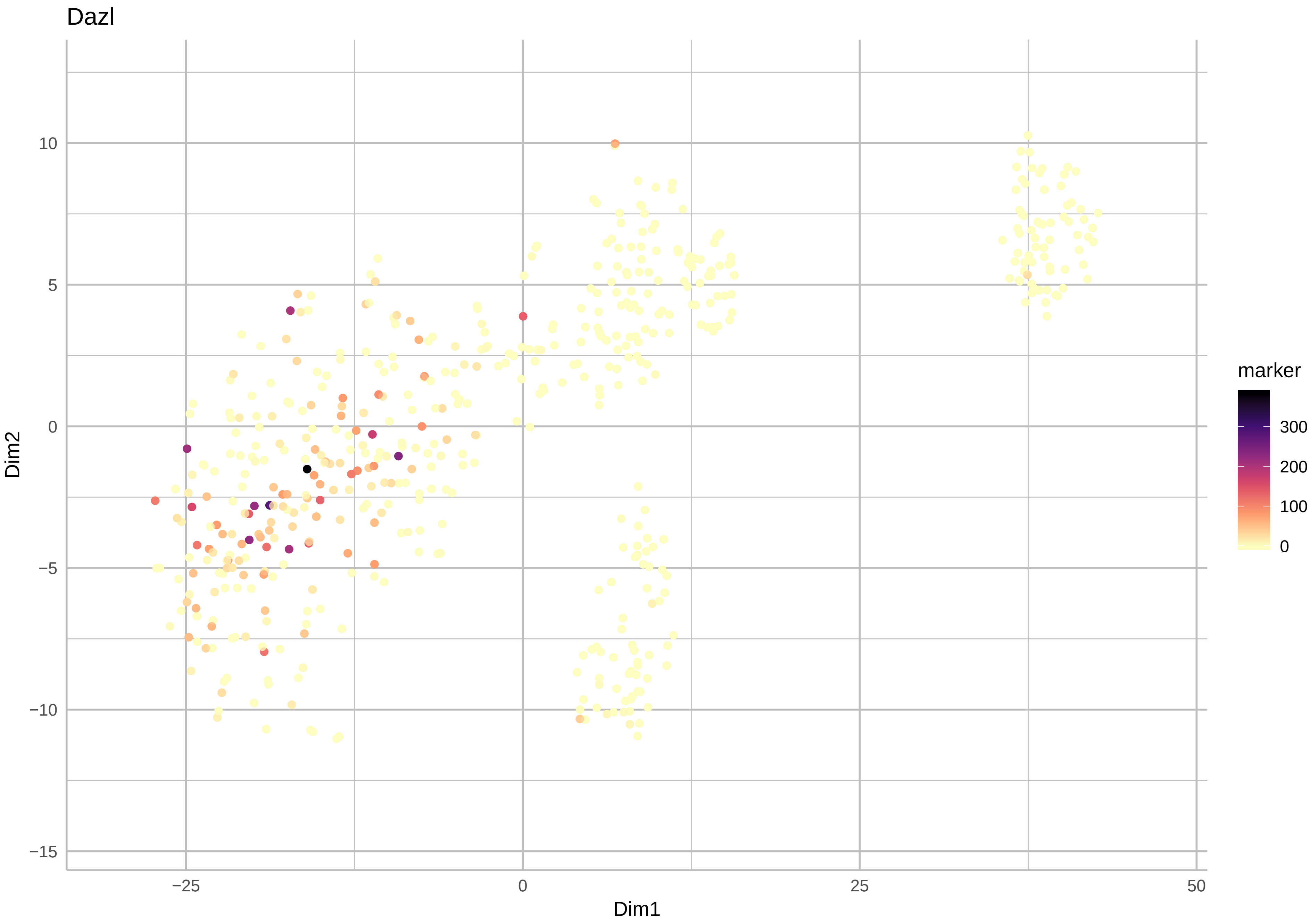

Dazl
Dim2
Dim1
marker
300
200
100
0
10
5
0
−5
−10
−15
−25
0
25
50

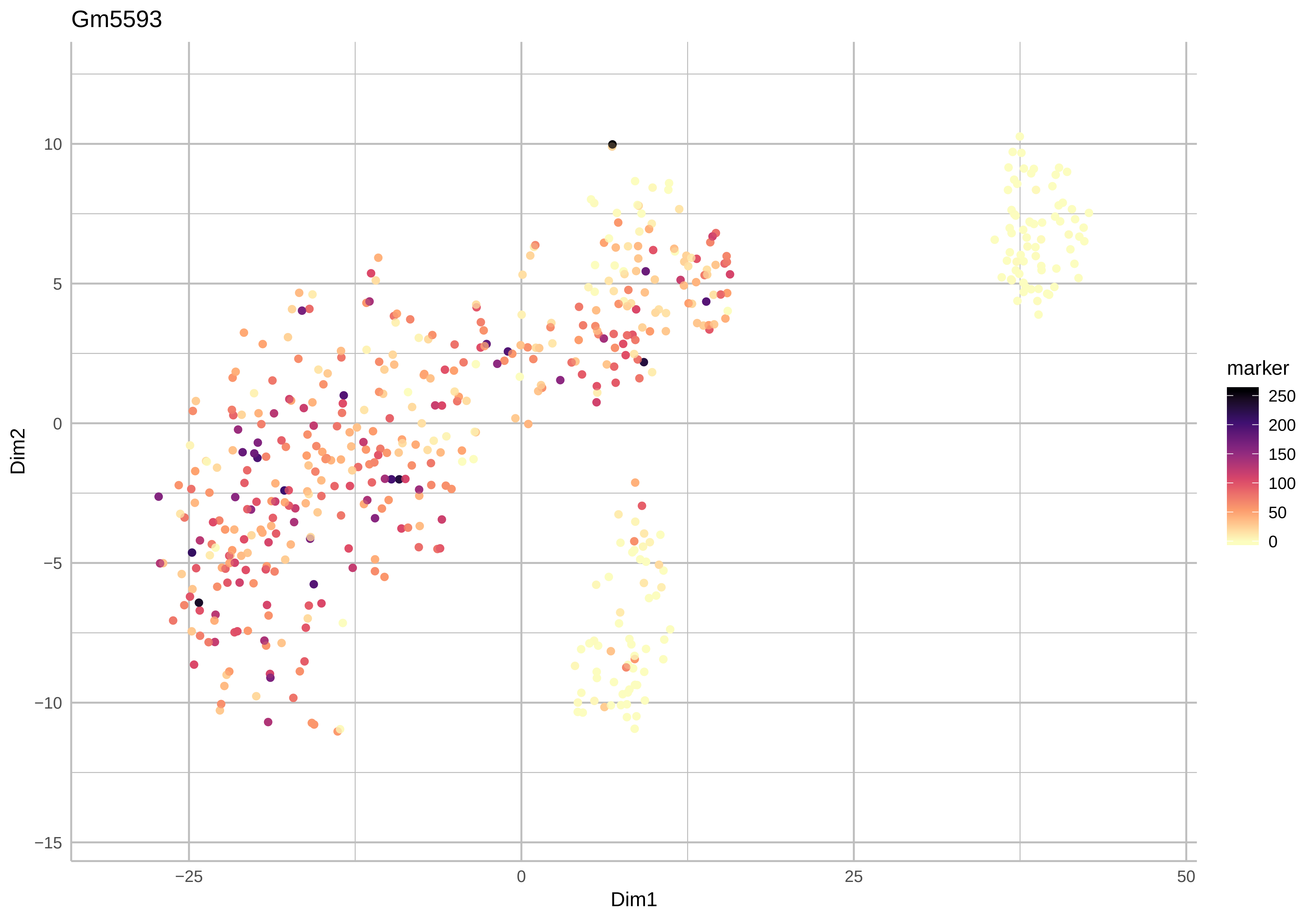

Gm5593
10
5
0
−5
−10
−15
−25
0
25
50
Dim1
Dim2
marker
250
200
150
100
50
0

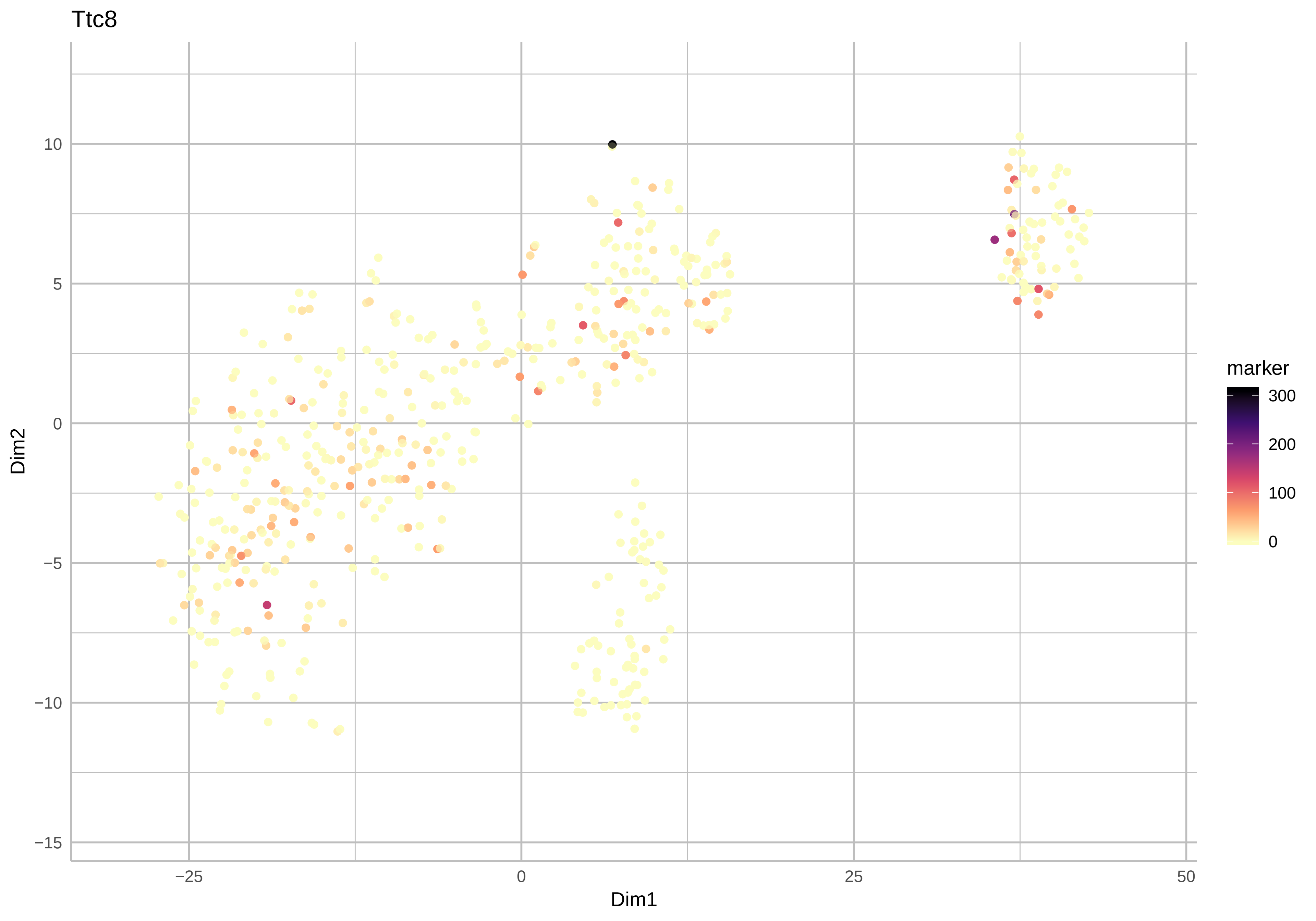

Ttc8
Dim2
Dim1
10
5
0
−5
−10
−15
−25
0
25
50
marker
300
200
100
0

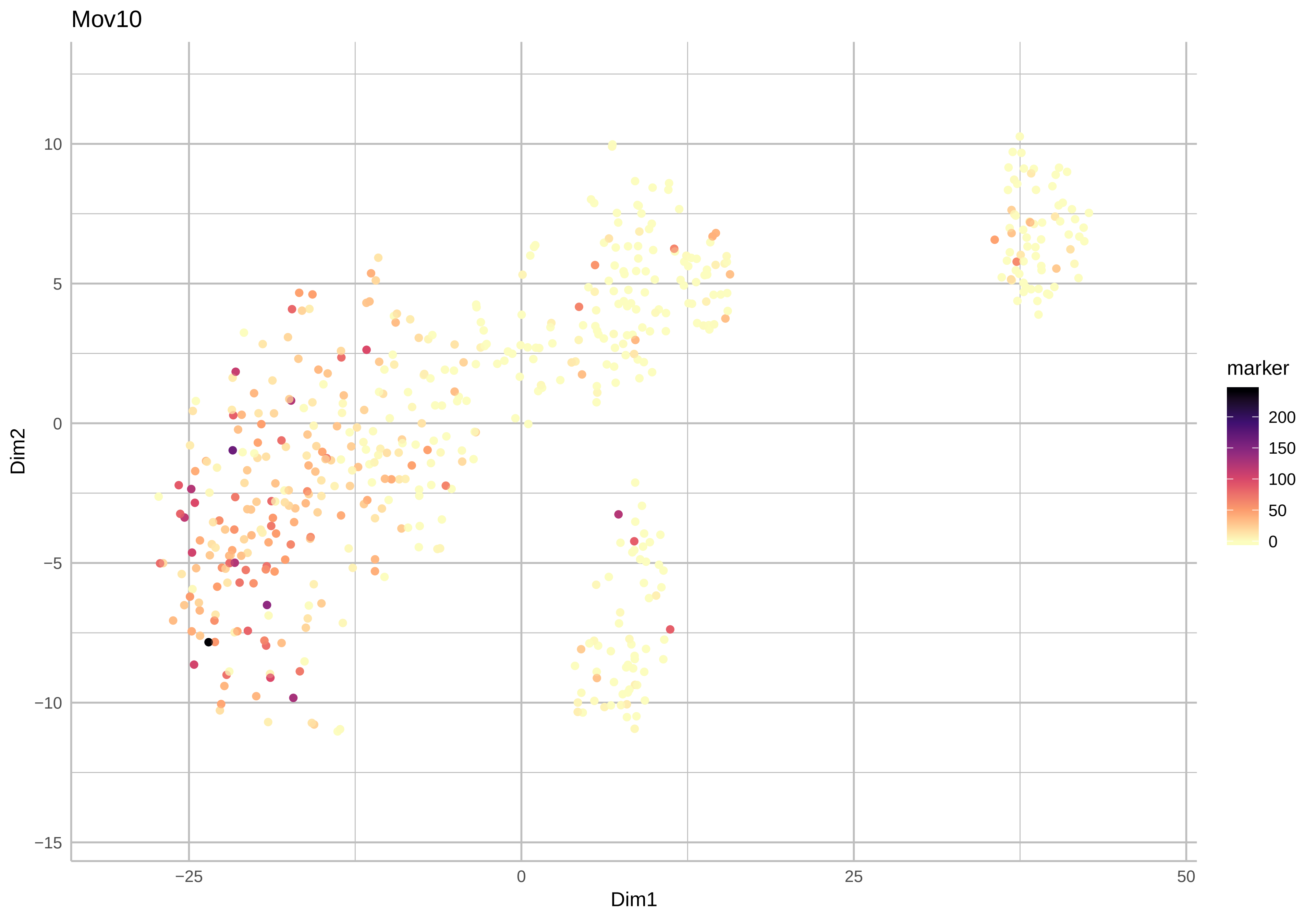
Mov10
Dim2
Dim1
10
5
0
−5
−10
−15
−25
0
25
50
marker
200
150
100
50
0

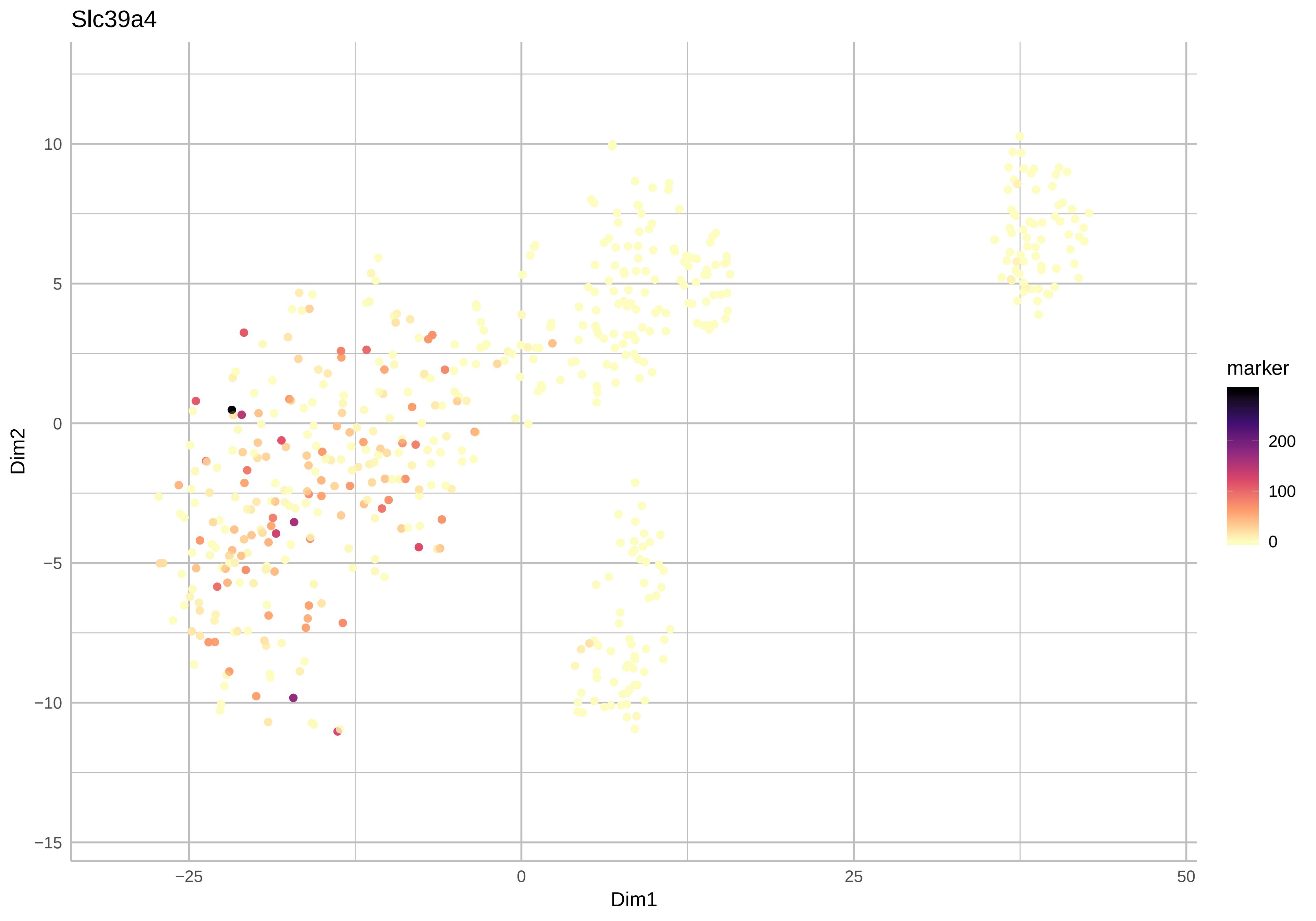
Slc39a4
Dim2
Dim1
marker
200
100
0
10
5
0
−5
−10
−15
−25
0
25
50

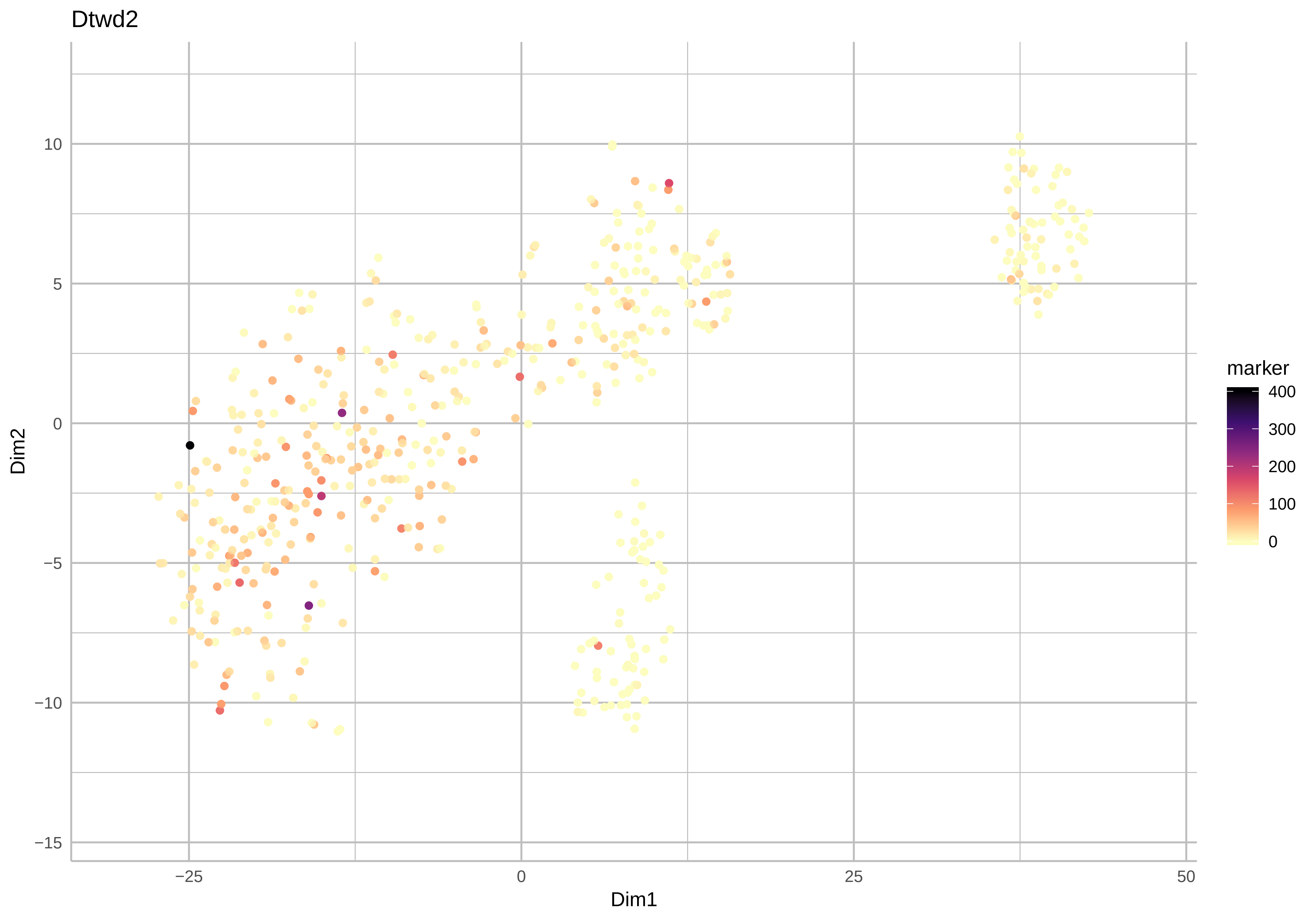
Dtwd2
Dim2
Dim1
10
5
0
−5
−10
−15
−25
0
25
50
marker
400
300
200
100
0

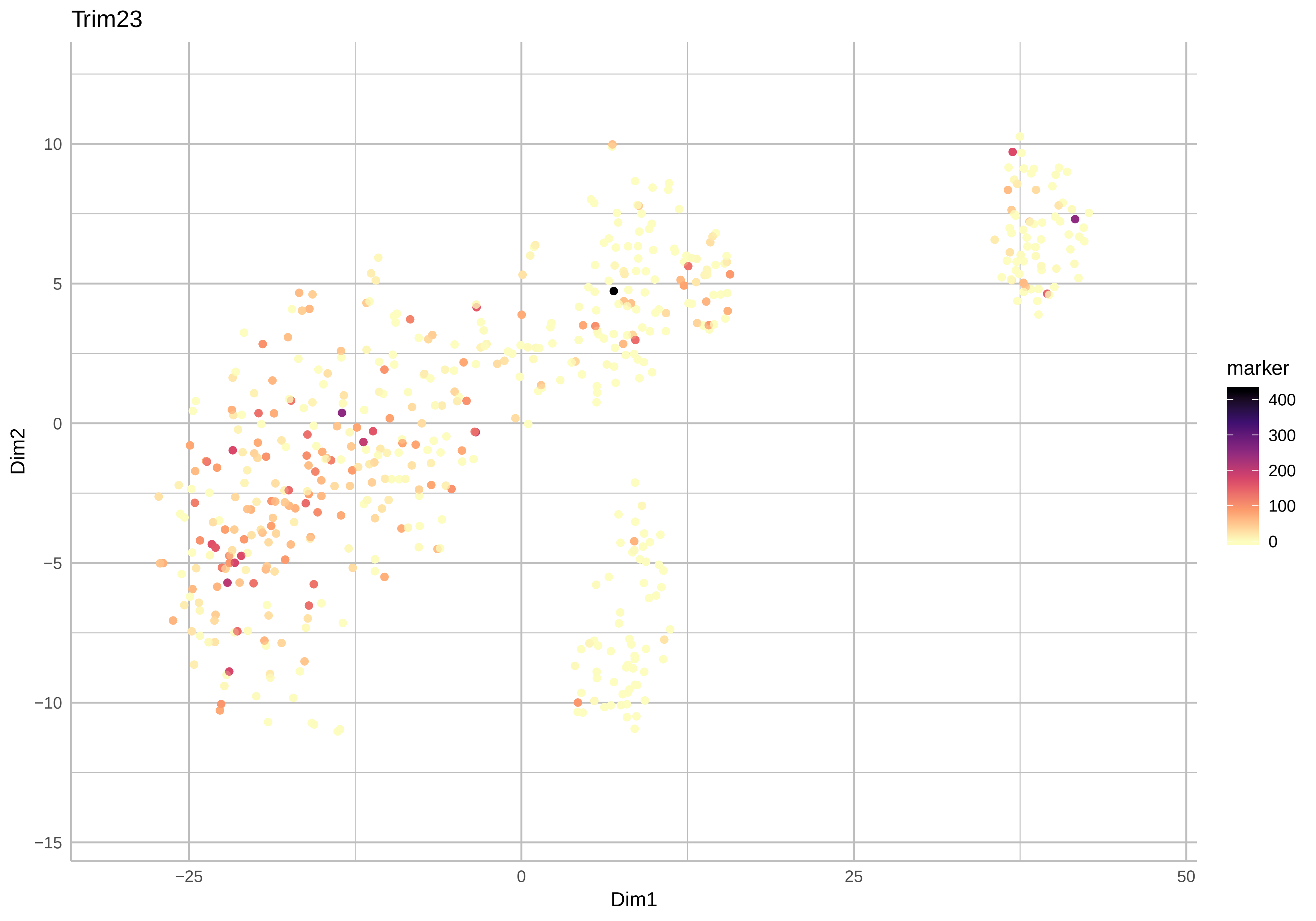

Trim23
Dim2
10
5
0
−5
−10
−15
−25
0
25
50
Dim1
marker
400
300
200
100
0

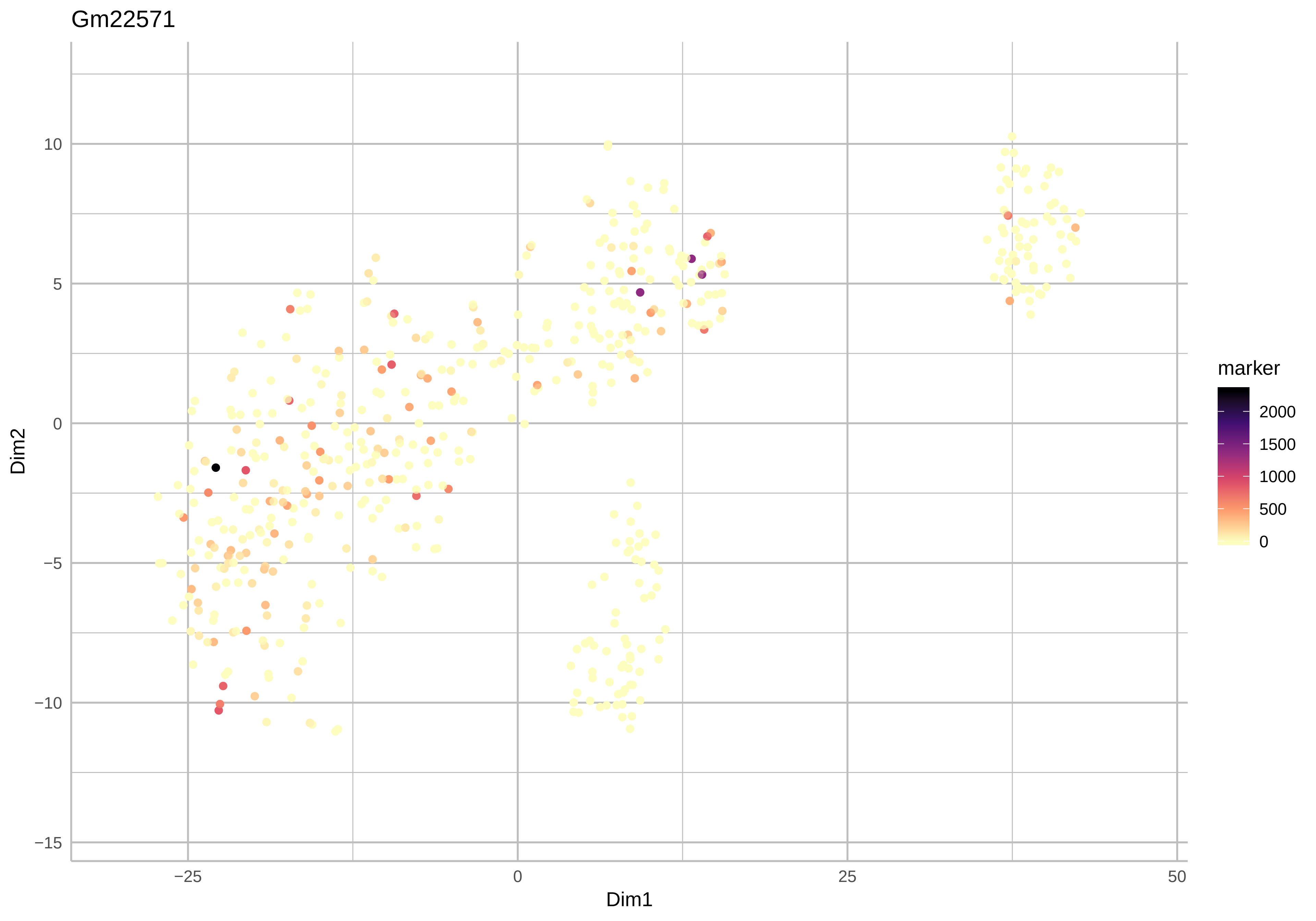
Gm22571
Dim2
Dim1
10
5
0
−5
−10
−15
−25
0
25
50
marker
2000
1500
1000
500
0

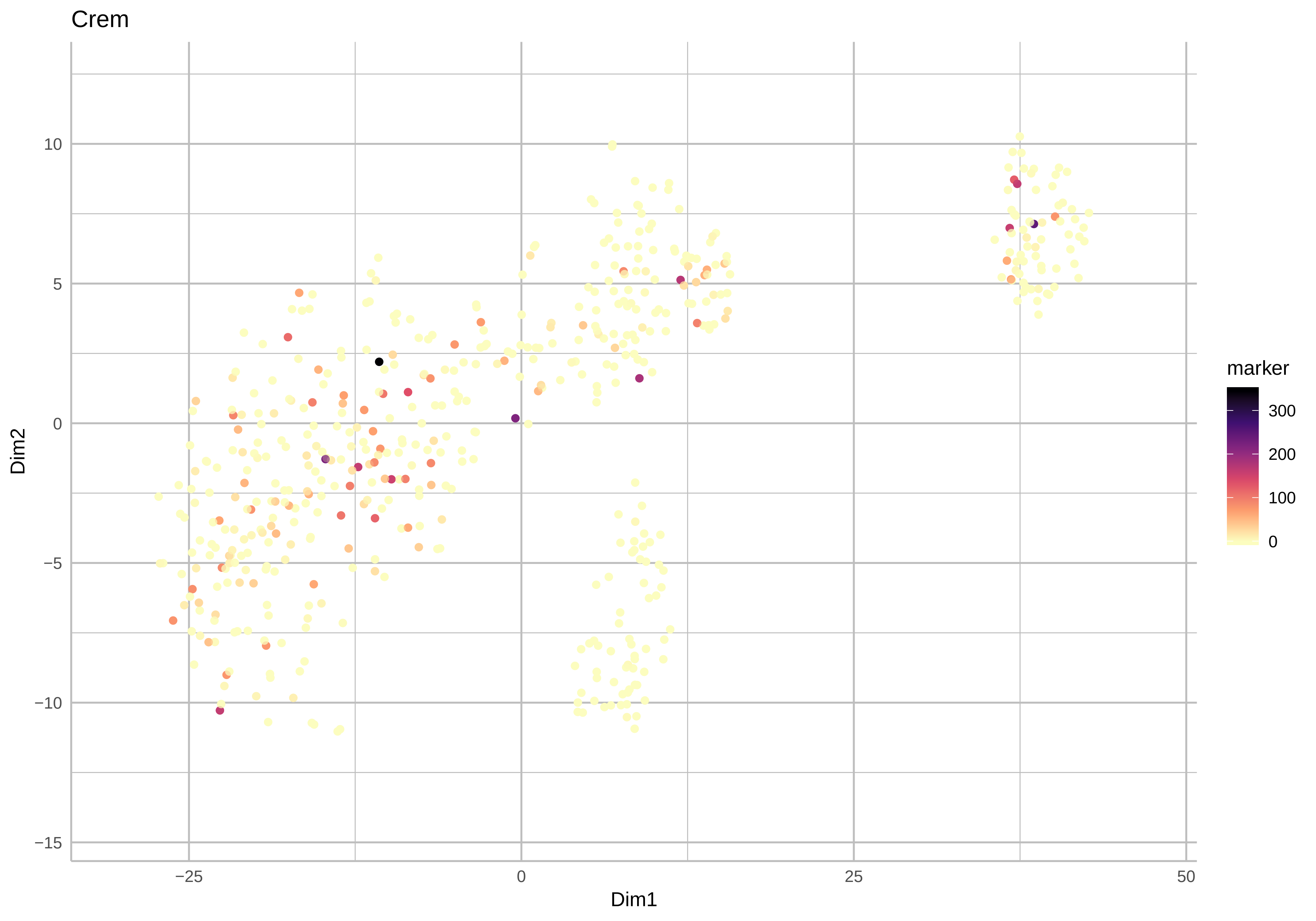
Crem
Dim2
Dim1
10
5
0
−5
−10
−15
−25
0
25
50
marker
300
200
100
0

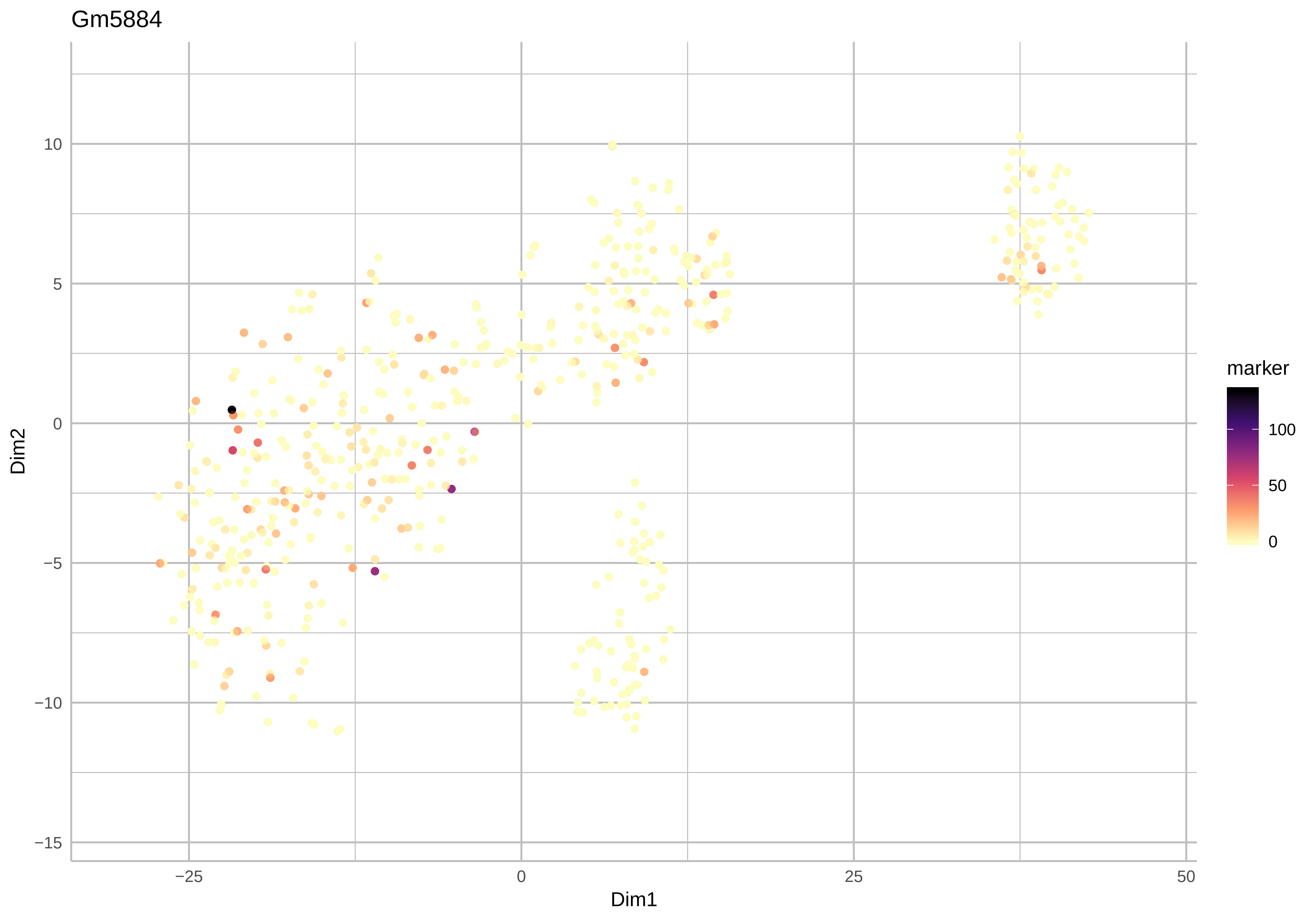

Gm5884
Dim2
Dim1
10
5
0
−5
−10
−15
−25
0
25
50
marker
100
50
0

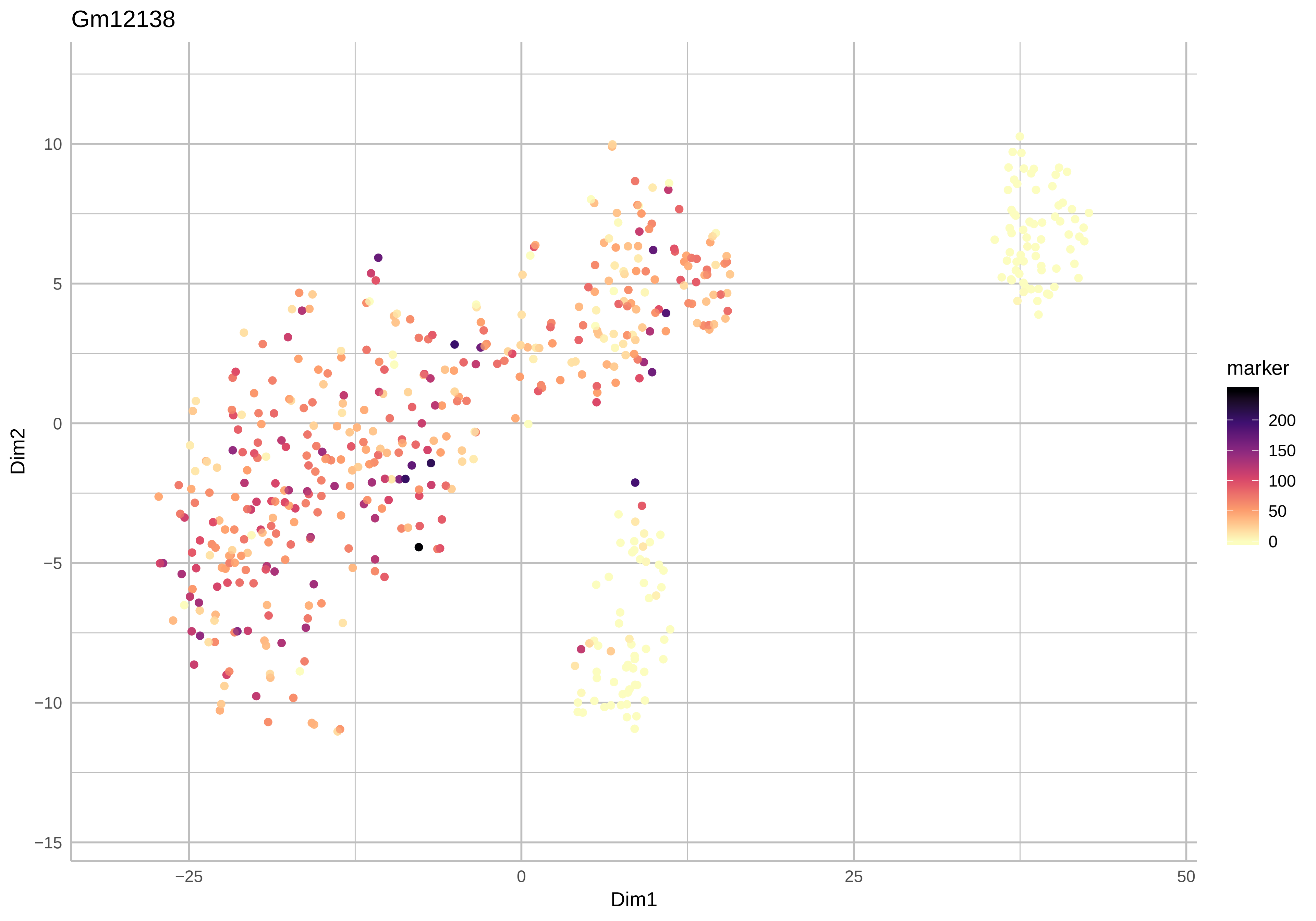

Gm12138
Dim2
Dim1
10
5
0
−5
−10
−15
−25
0
25
50
marker
200
150
100
50
0

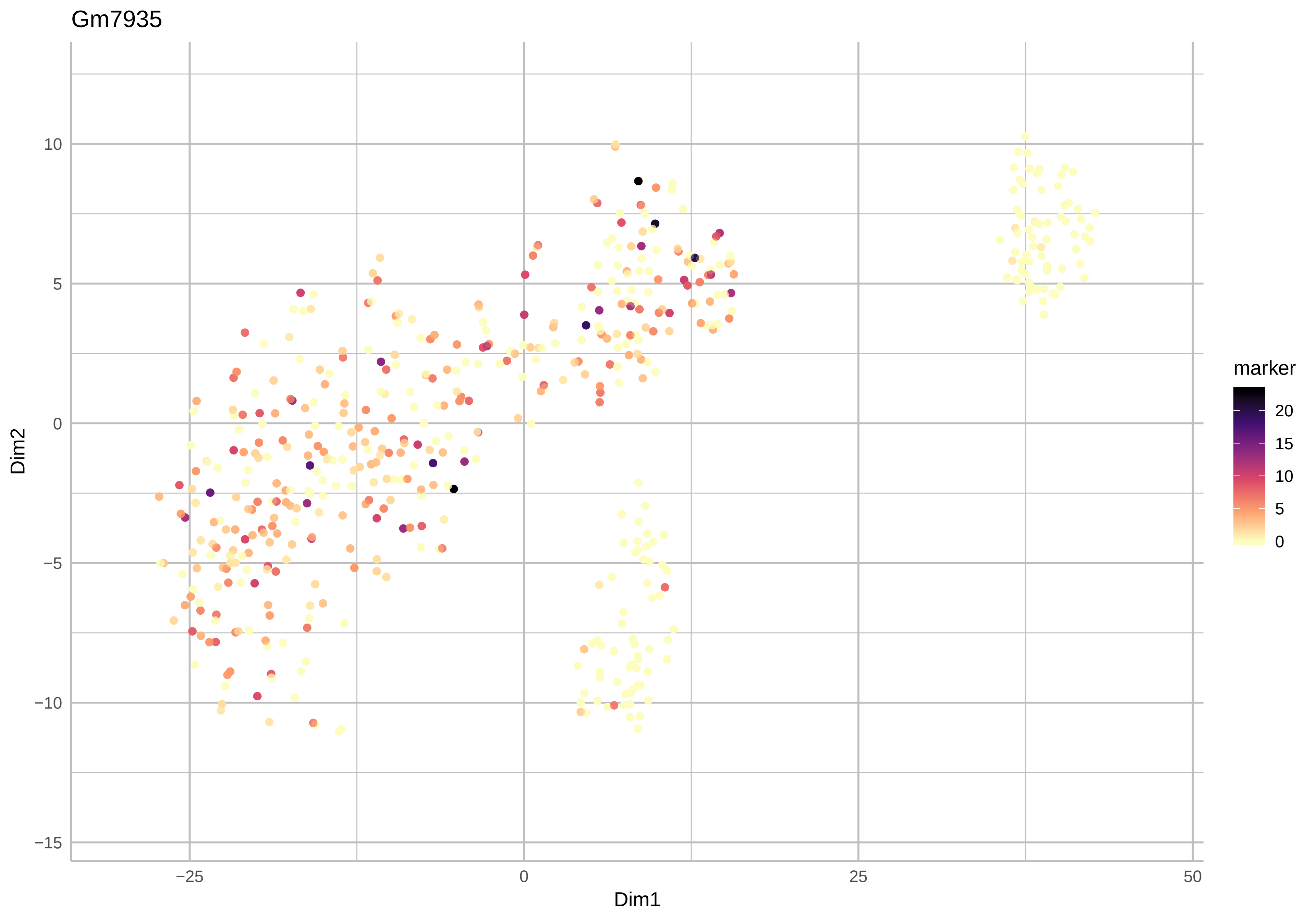
Gm7935
10
5
0
−5
−10
−15
Dim2
−25
0
25
50
Dim1
marker
20
15
10
5
0

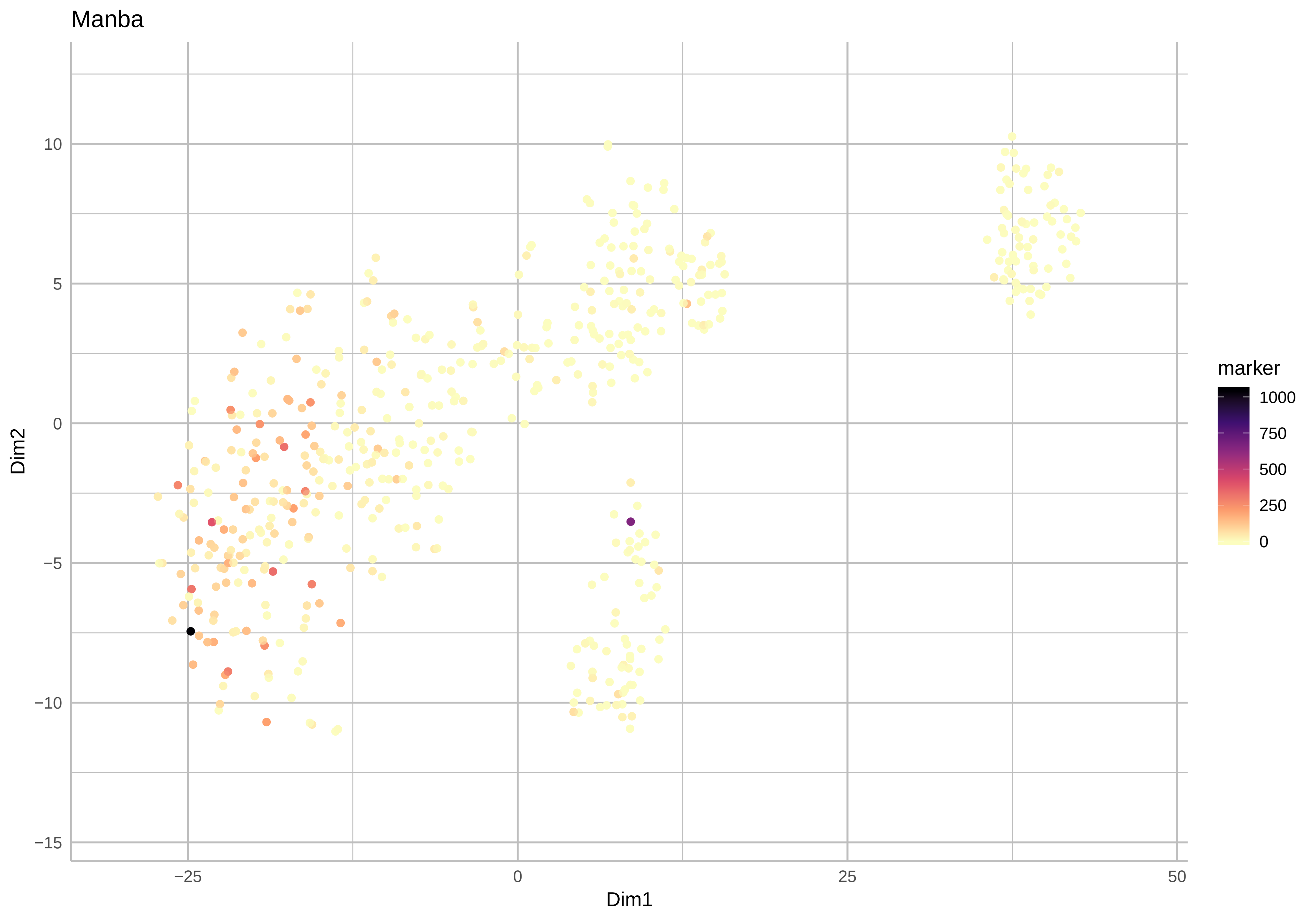
Manba
Dim2
Dim1
10
5
0
−5
−10
−15
−25
0
25
50
marker
1000
750
500
250
0

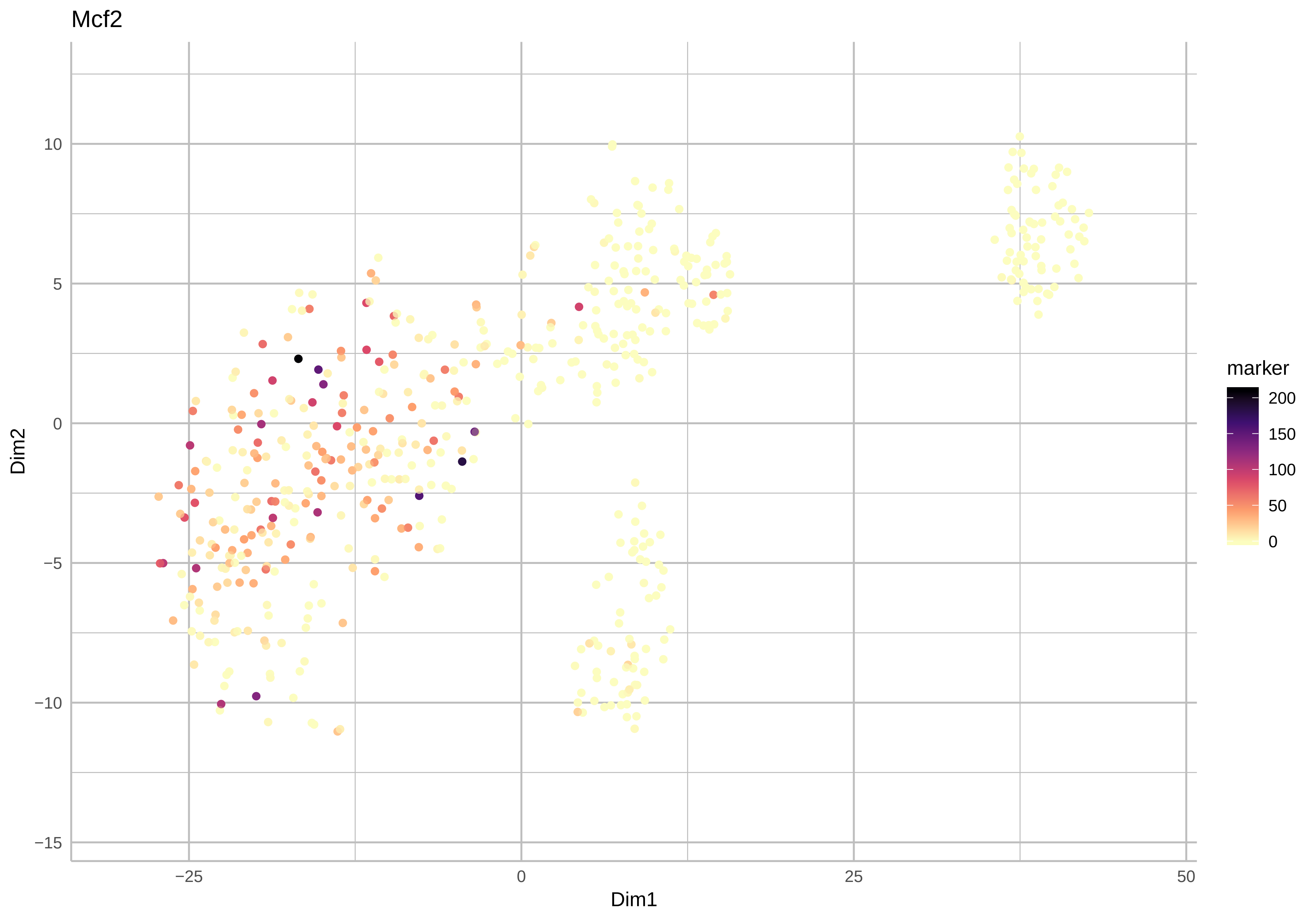

Mcf2
Dim2
Dim1
10
5
0
−5
−10
−15
−25
0
25
50
marker
200
150
100
50
0

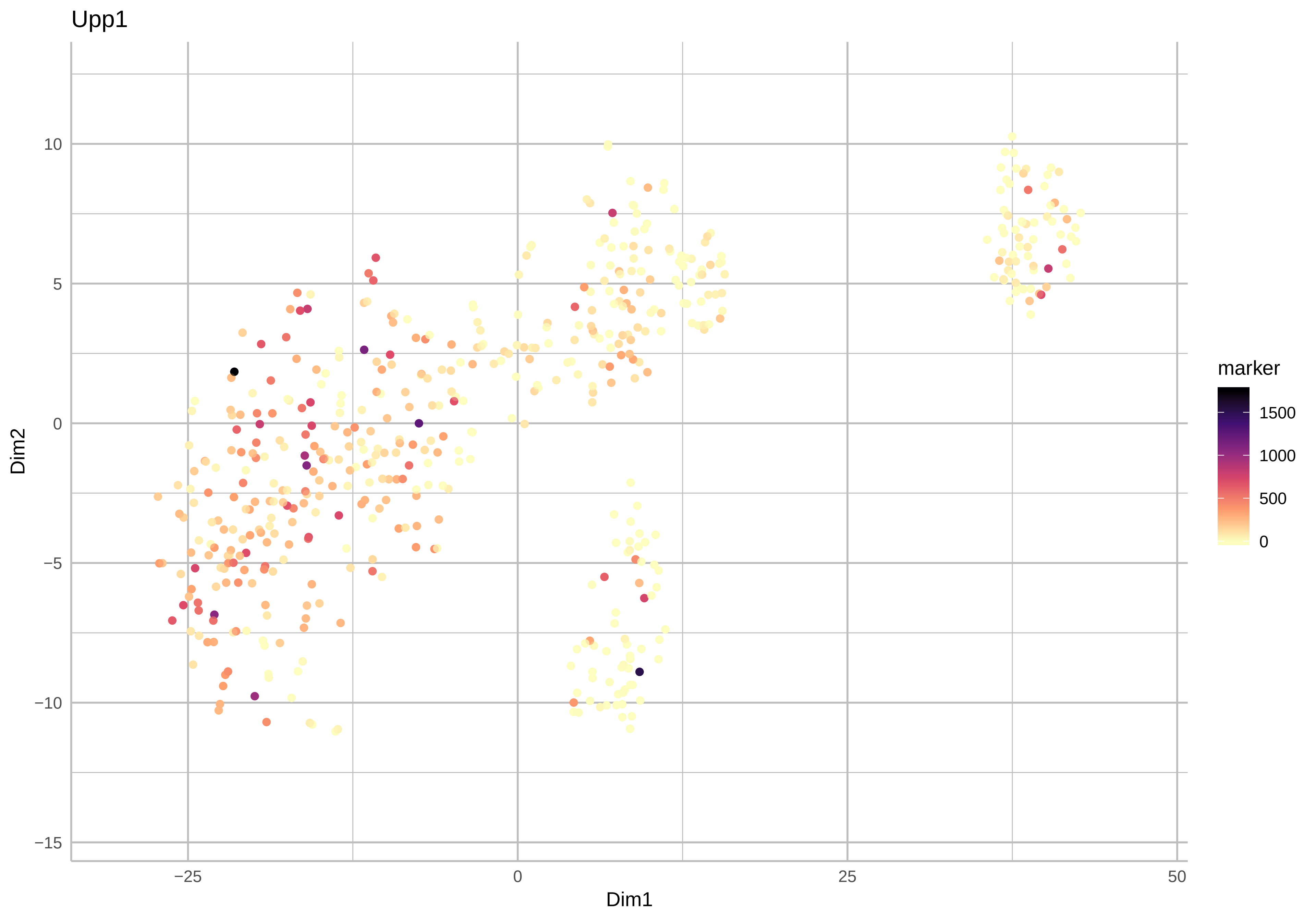
Upp1
Dim2
Dim1
10
5
0
−5
−10
−15
−25
0
25
50
marker
1500
1000
500
0

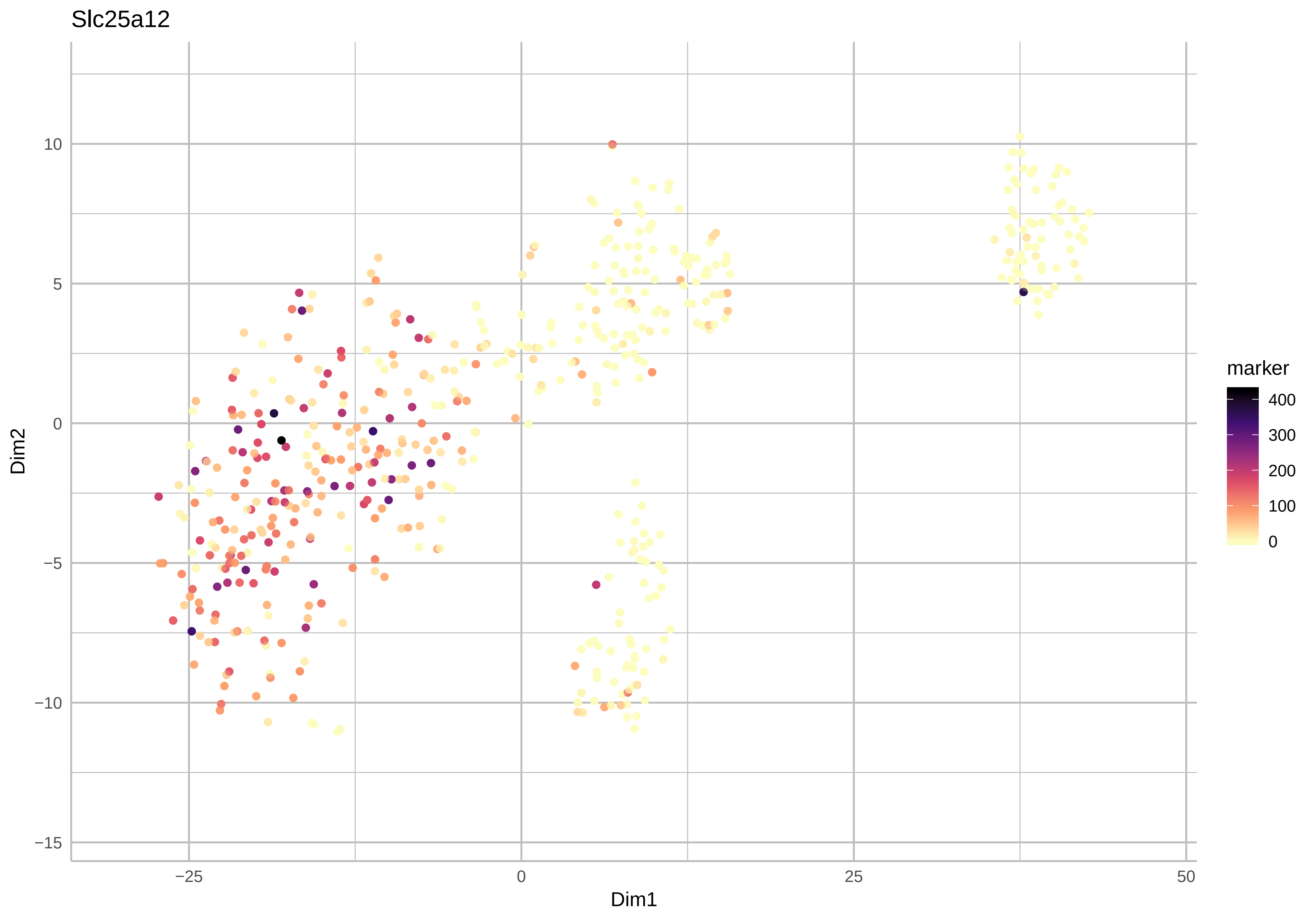

Slc25a12
Dim2
Dim1
10
5
0
−5
−10
−15
−25
0
25
50
marker
400
300
200
100
0

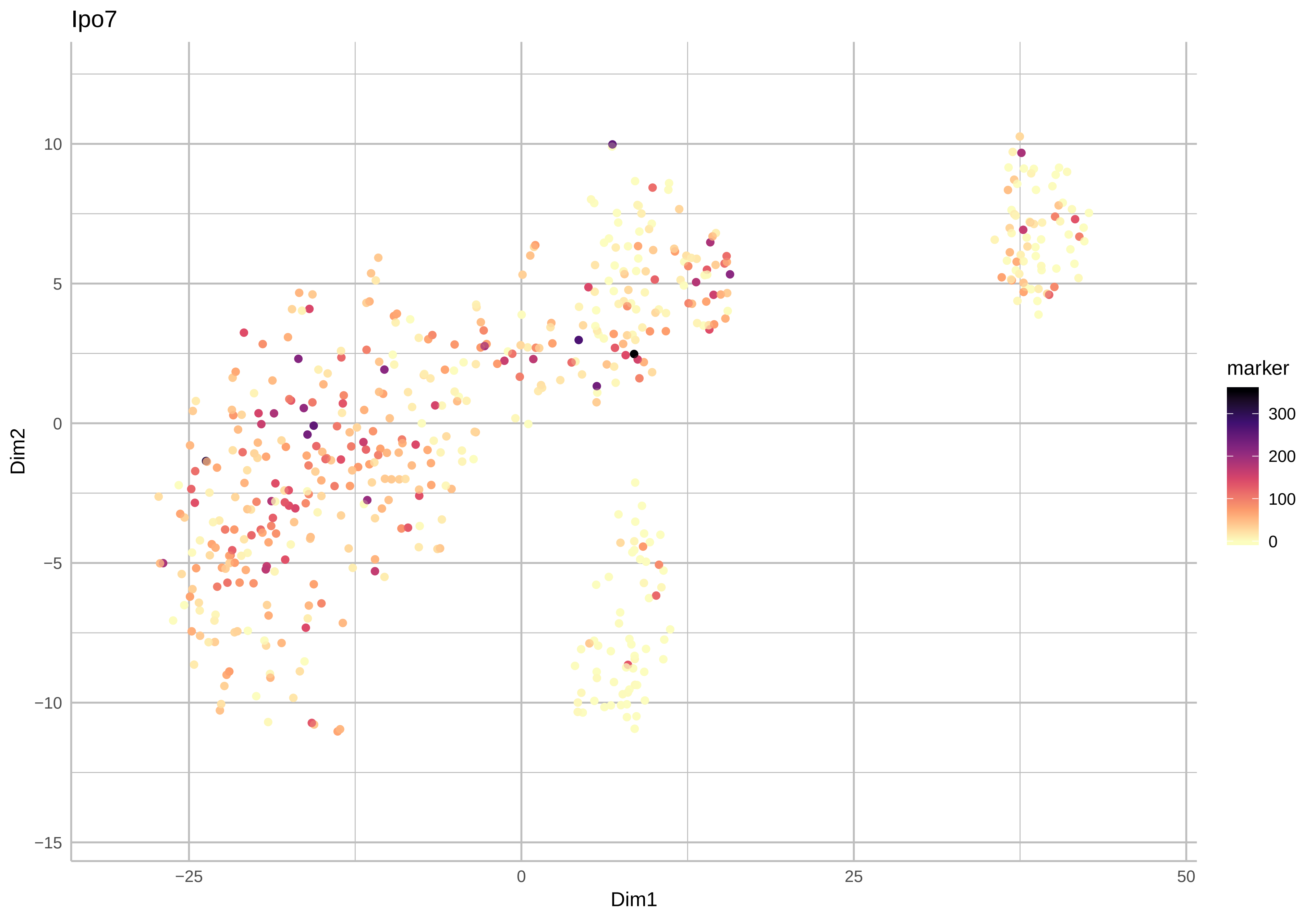

Ipo7
10
5
0
−5
−10
−15
−25
0
25
50
Dim1
Dim2
marker
300
200
100
0

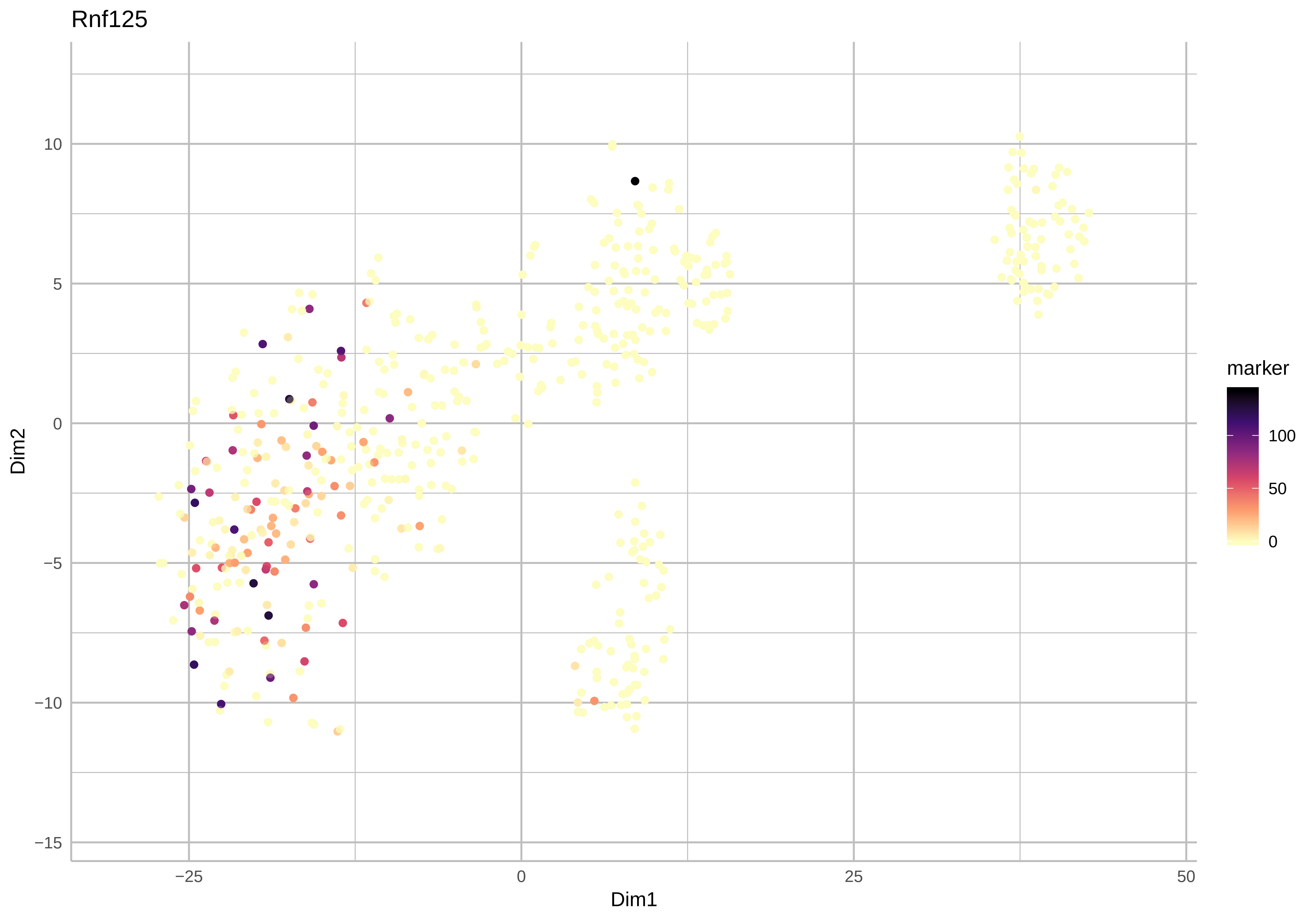
Rnf125
Dim2
Dim1
10
5
0
−5
−10
−15
−25
0
25
50
marker
100
50
0

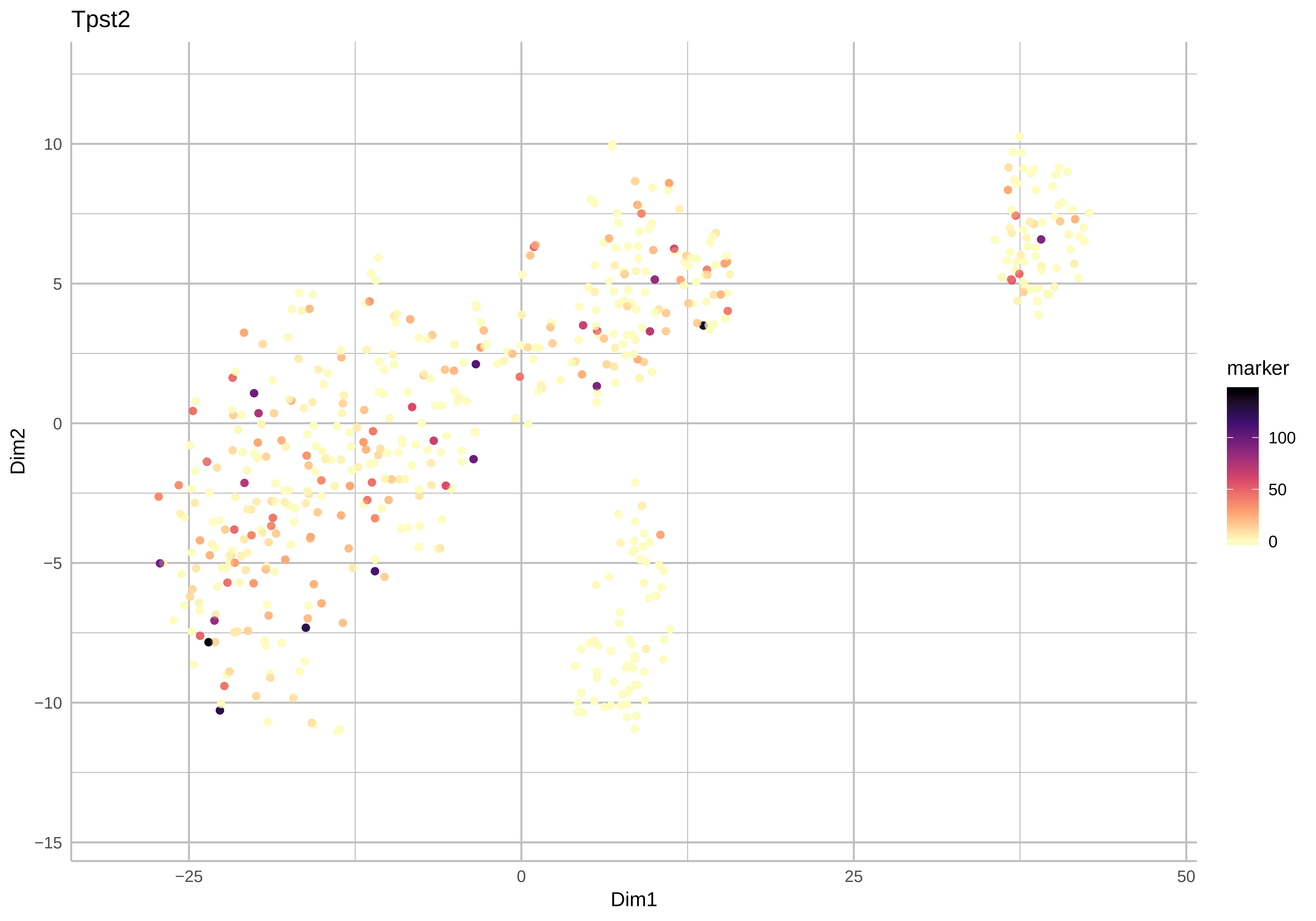

Tpst2
Dim2
Dim1
10
5
0
−5
−10
−15
−25
0
25
50
marker
100
50
0

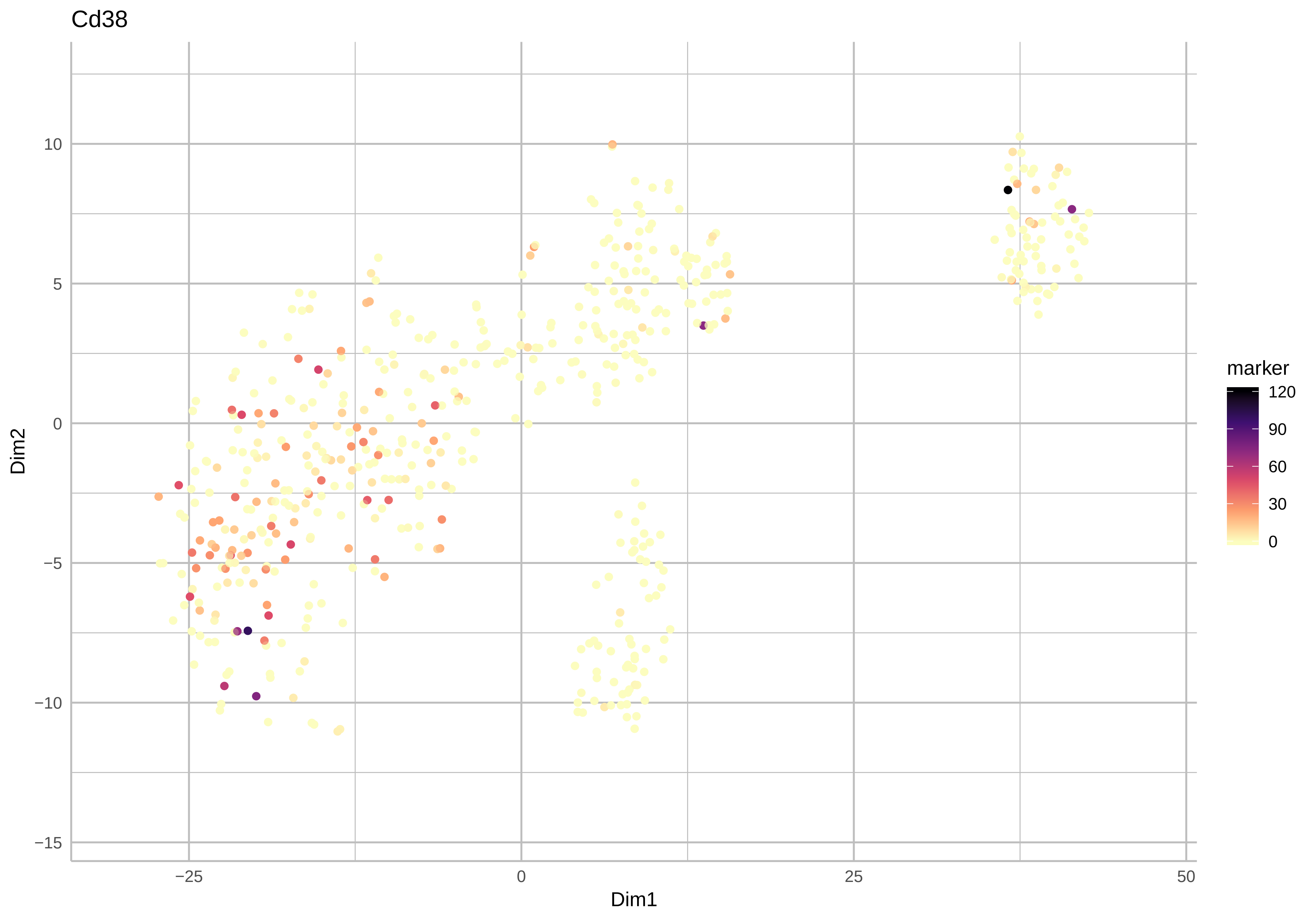
Cd38
Dim2
Dim1
10
5
0
−5
−10
−15
−25
0
25
50
marker
120
90
60
30
0

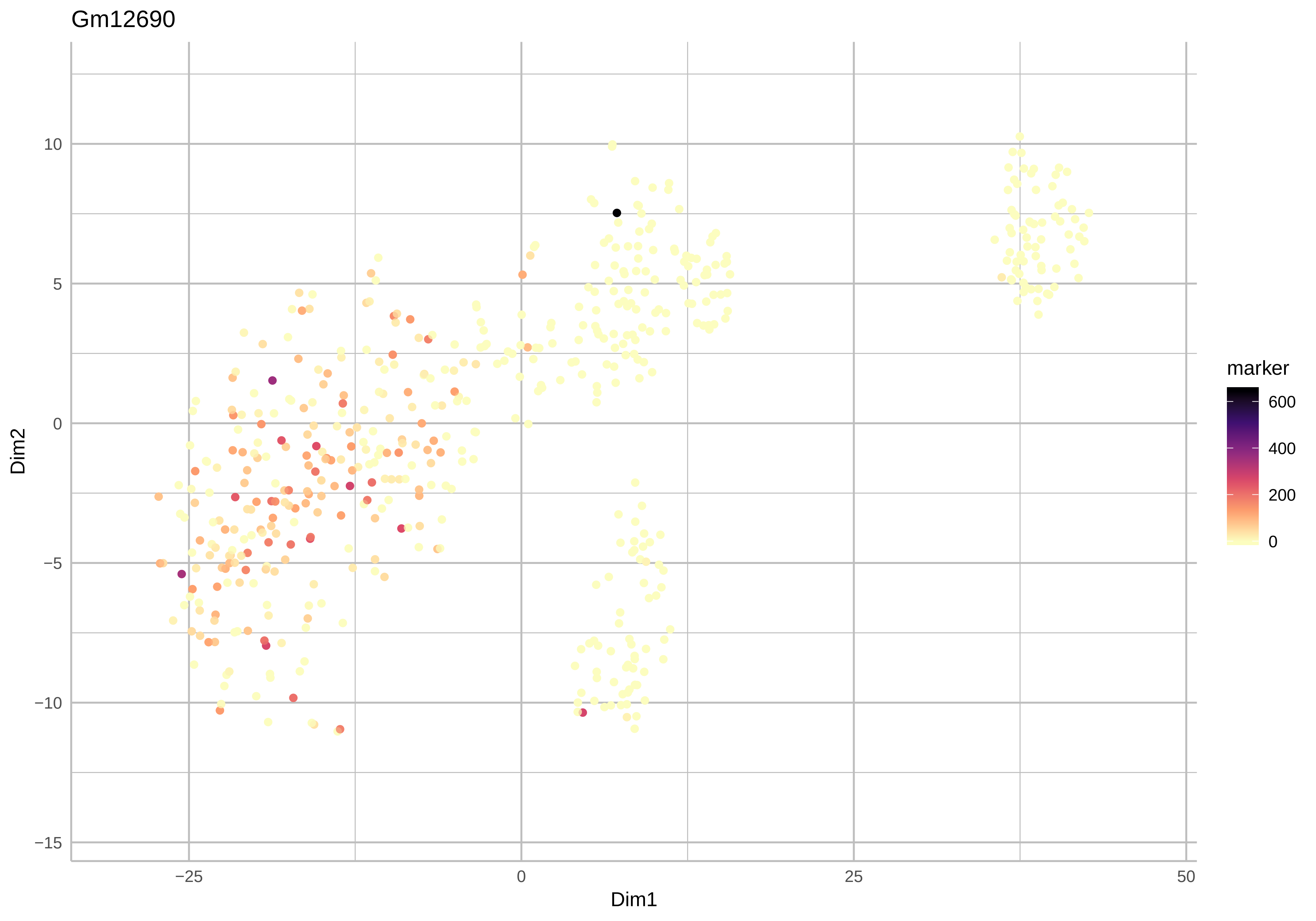
Gm12690
Dim2
Dim1
10
5
0
−5
−10
−15
−25
0
25
50
marker
600
400
200
0

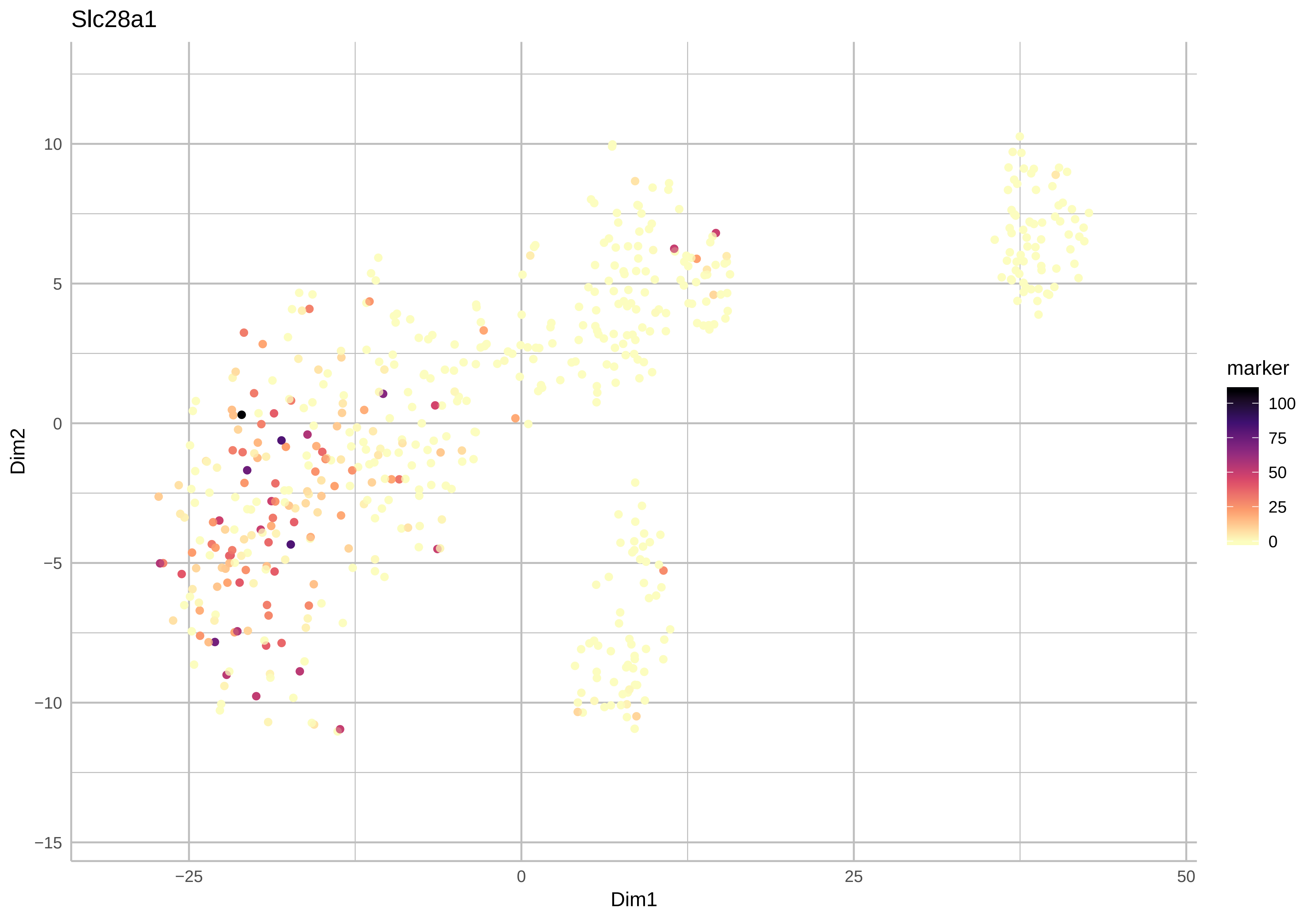

Slc28a1
Dim2
Dim1
10
5
0
−5
−10
−15
−25
0
25
50
marker
100
75
50
25
0

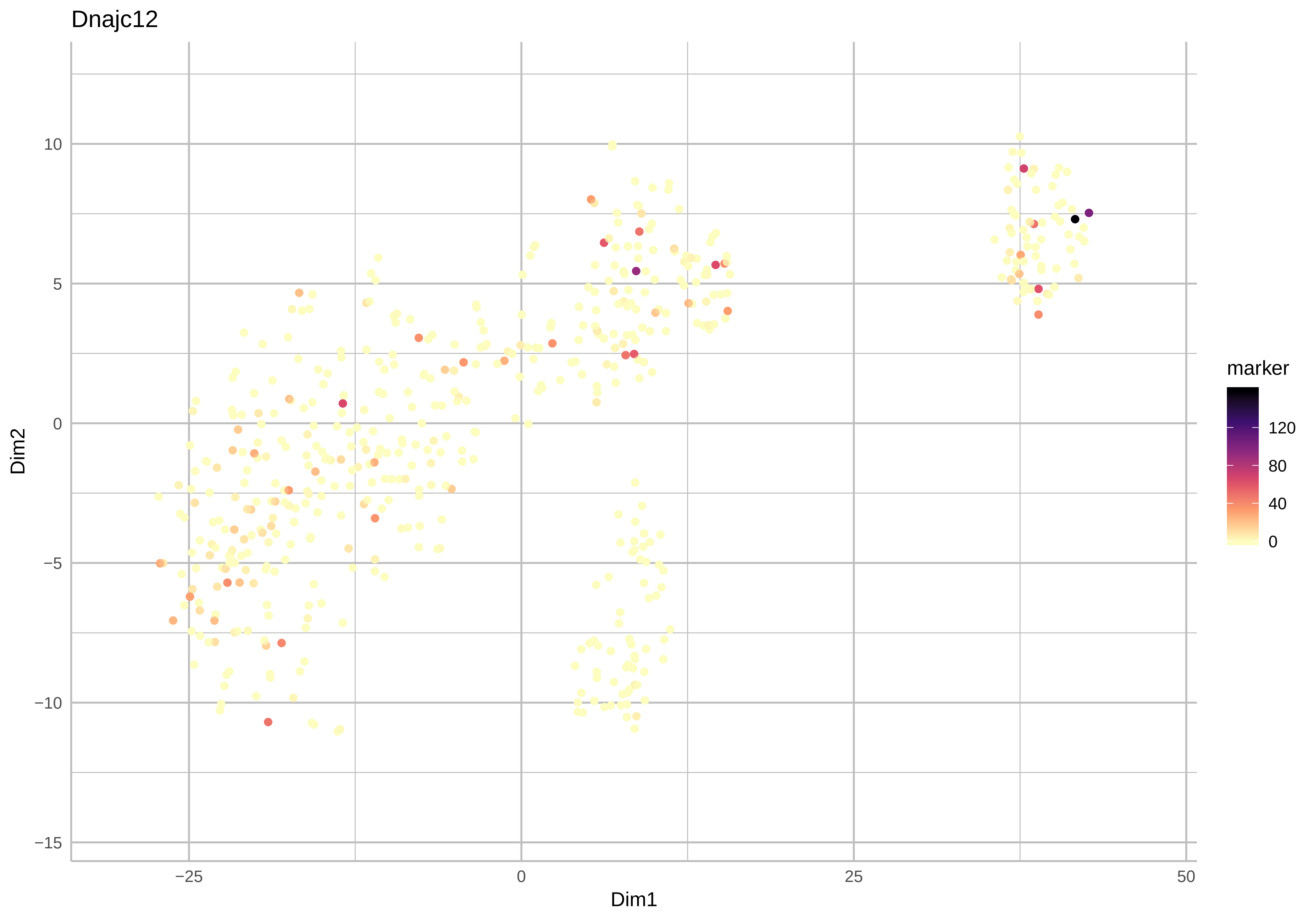

Dnajc12
Dim2
Dim1
10
5
0
−5
−10
−15
−25
0
25
50
marker
120
80
40
0

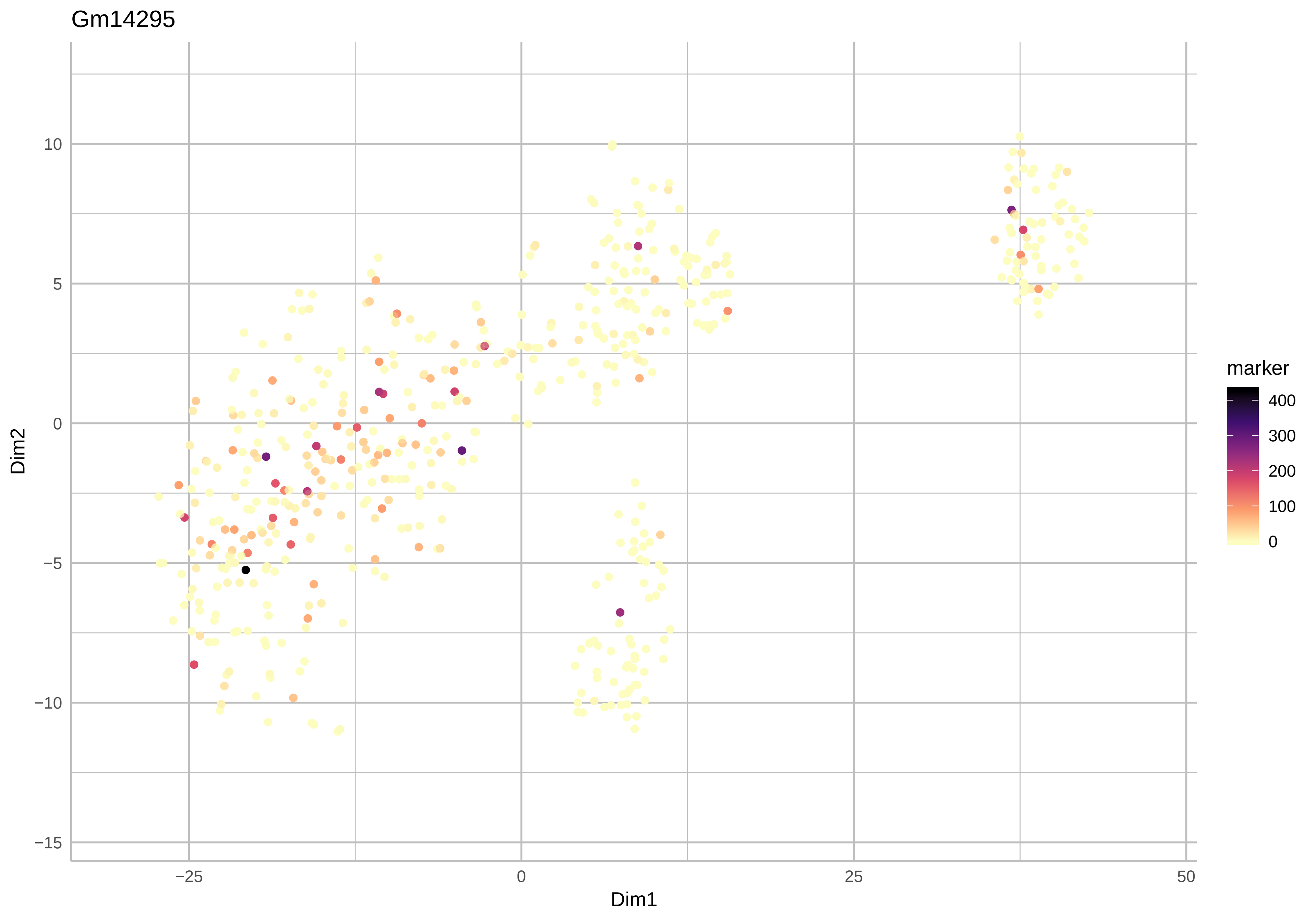
Gm14295
Dim2
Dim1
10
5
0
−5
−10
−15
−25
0
25
50
marker
400
300
200
100
0

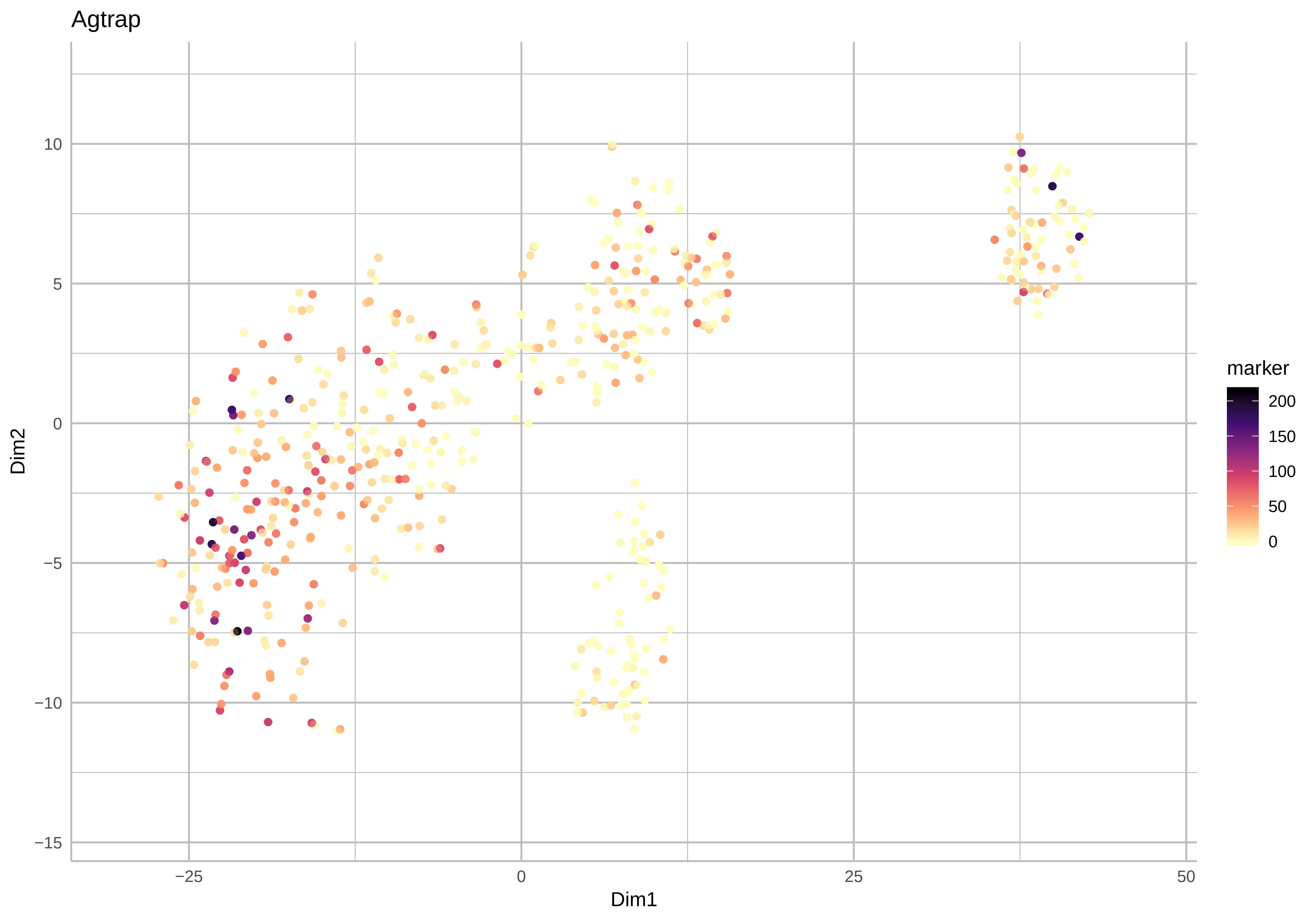
Agtrap
Dim2
Dim1
marker
200
150
100
50
0
10
5
0
−5
−10
−15
−25
0
25
50

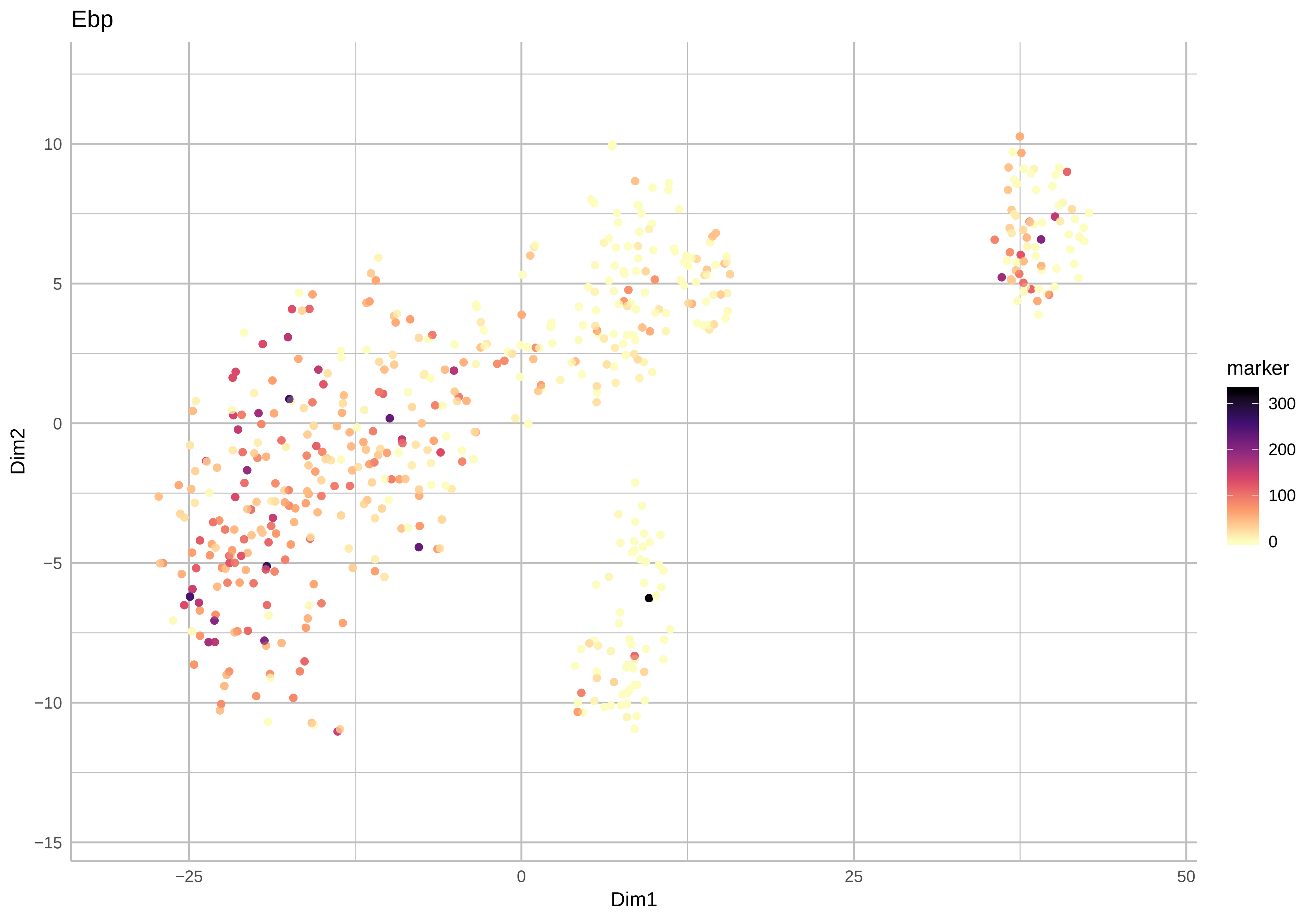
Ebp
10
5
0
−5
−10
−15
Dim2
−25
0
25
50
Dim1
marker
300
200
100
0

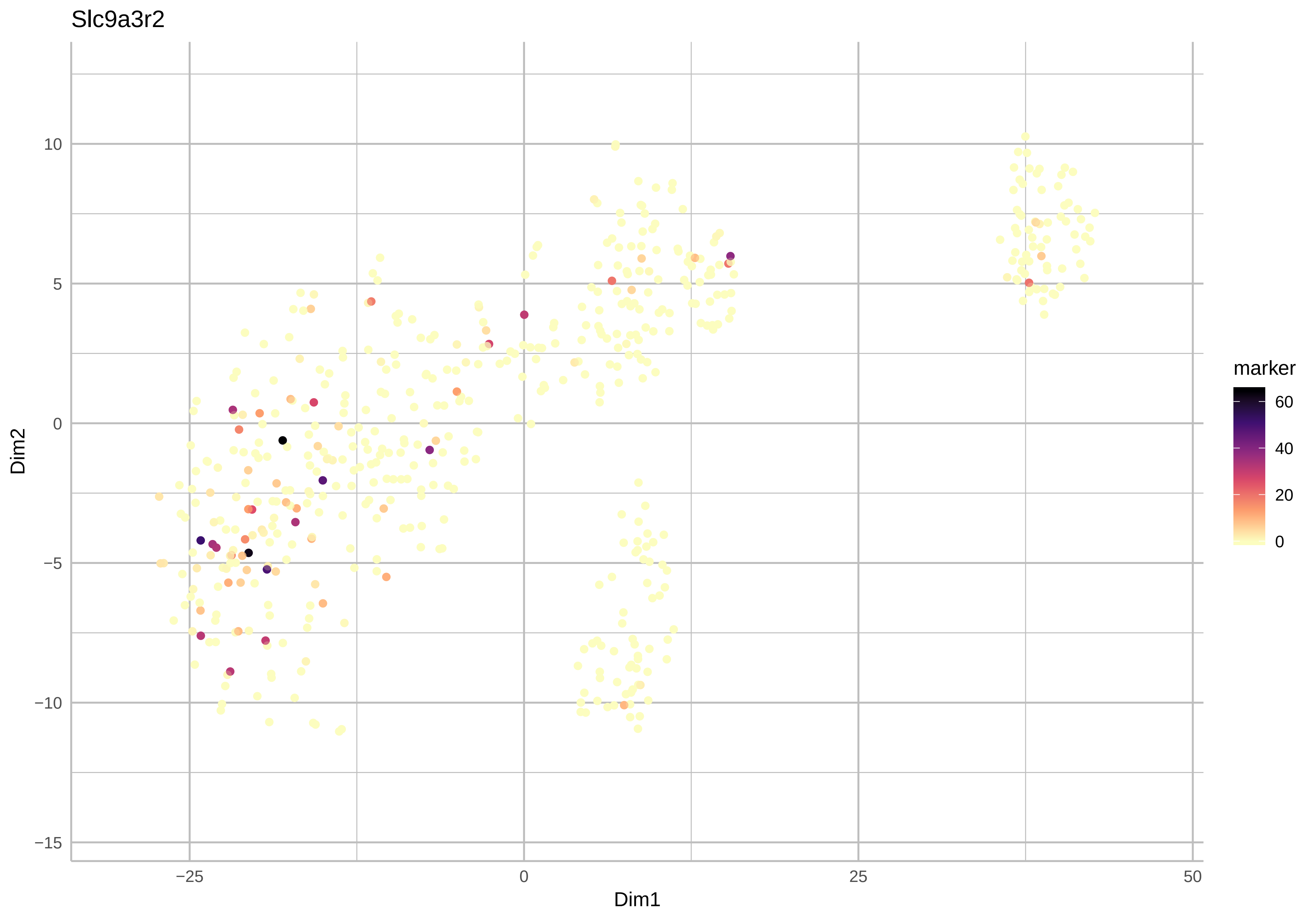
Slc9a3r2
Dim1
Dim2
−25
0
25
50
10
5
0
−5
−10
−15
marker
60
40
20
0

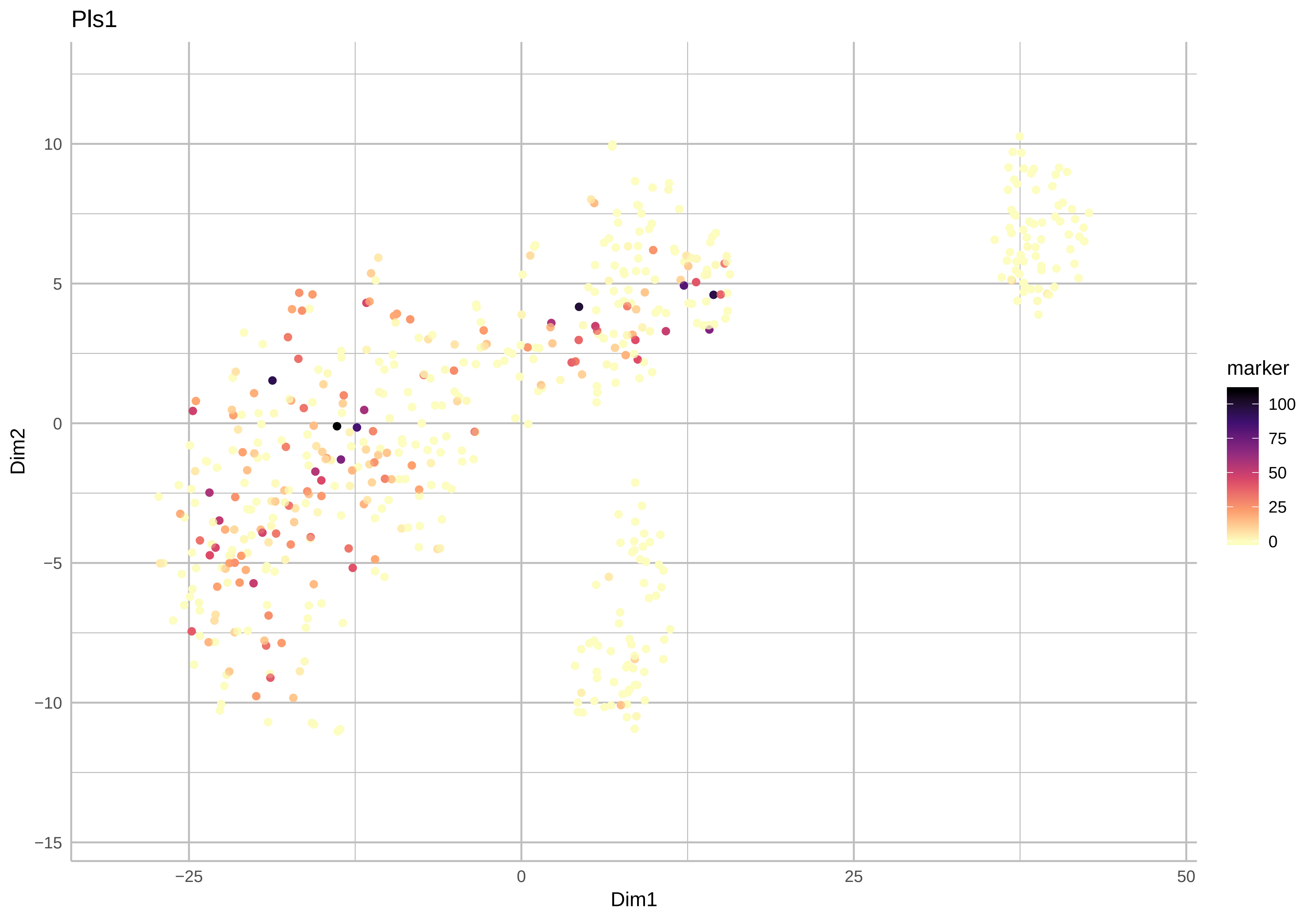

Pls1
10
5
0
−5
−10
−15
Dim2
−25
0
25
50
Dim1
marker
100
75
50
25
0

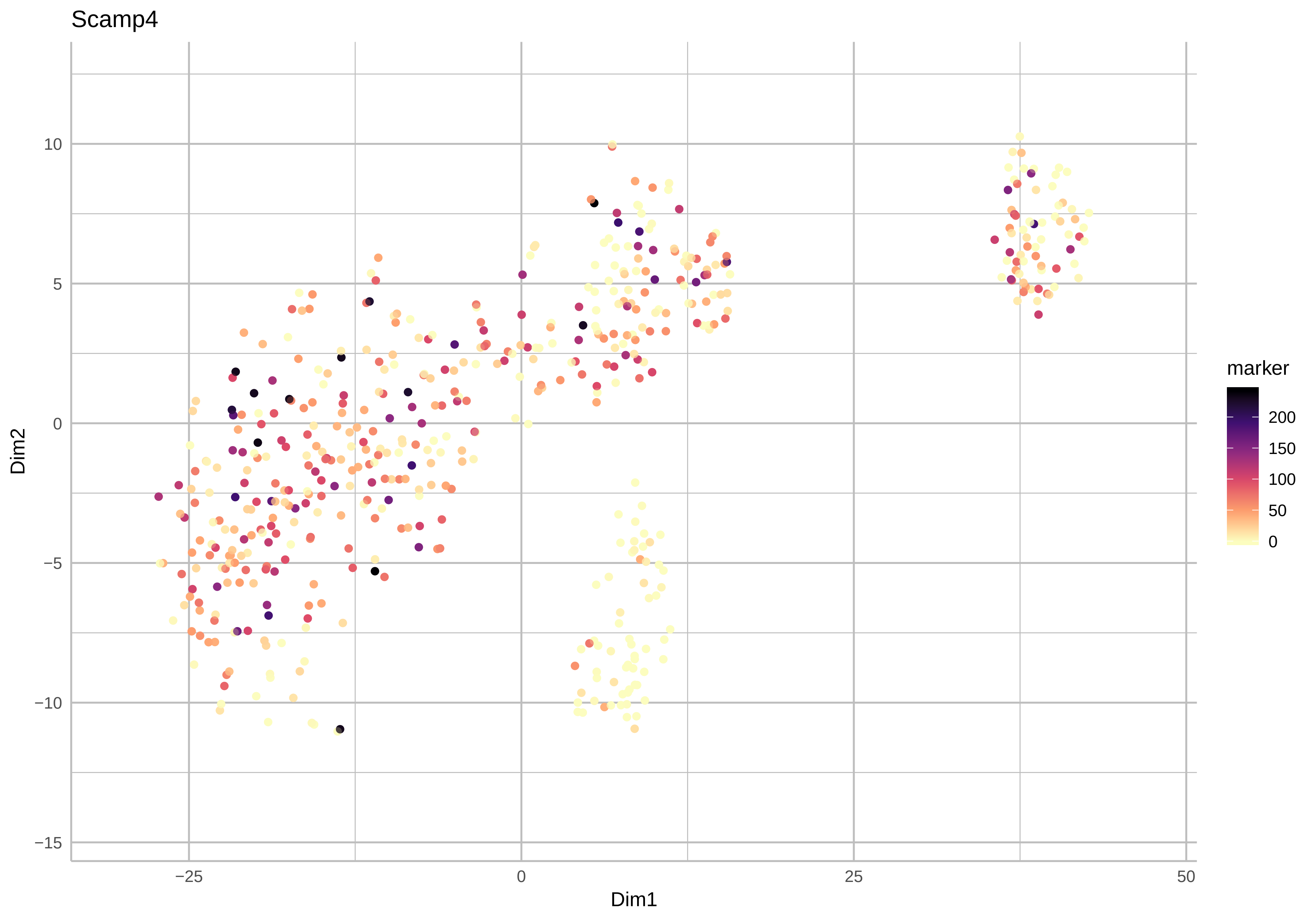
Scamp4
Dim2
Dim1
10
5
0
−5
−10
−15
−25
0
25
50
marker
200
150
100
50
0

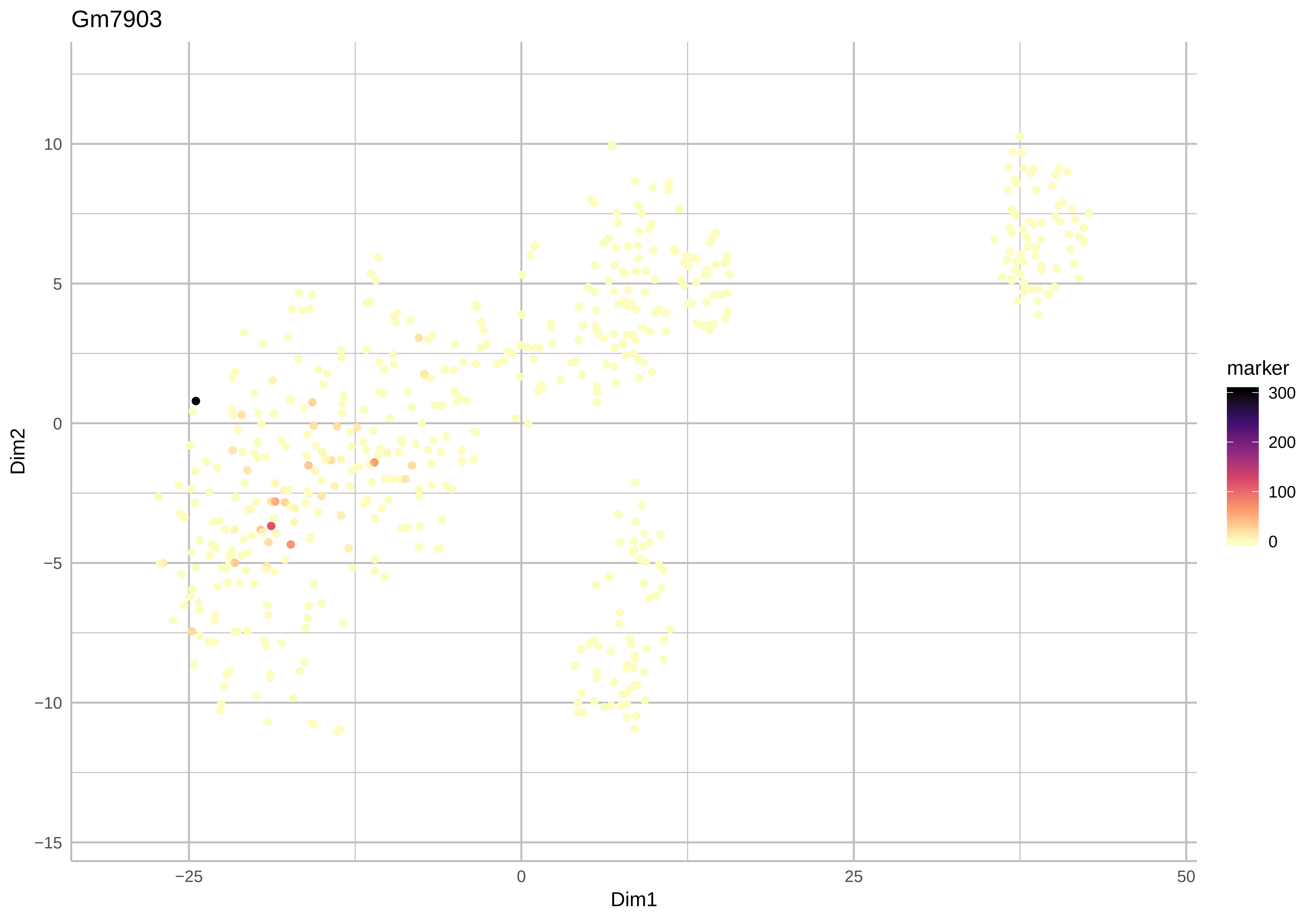
Gm7903
10
5
0
−5
−10
−15
Dim2
−25
0
25
50
Dim1
marker
300
200
100
0

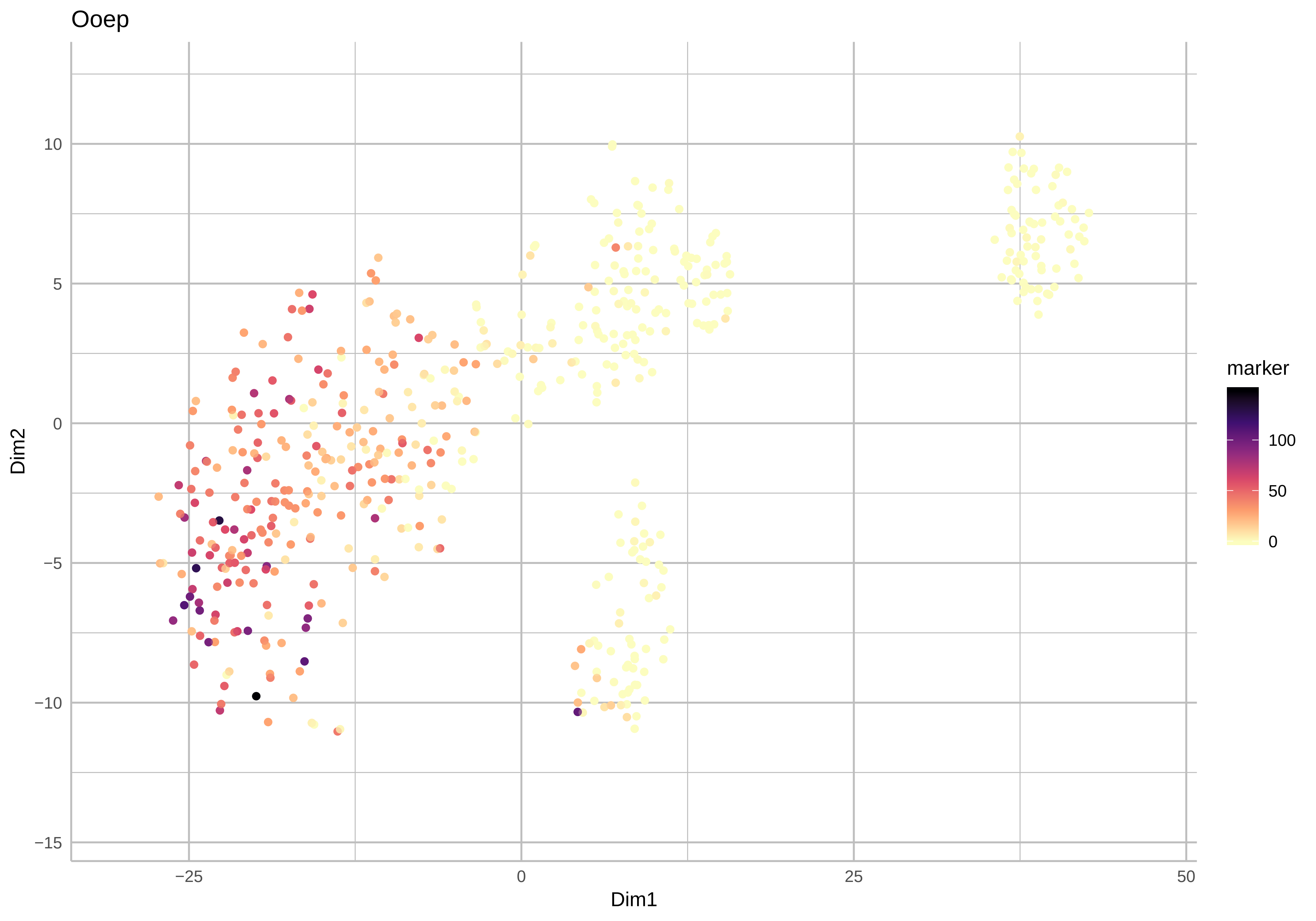
Ooep
Dim2
Dim1
10
5
0
−5
−10
−15
−25
0
25
50
marker
100
50
0

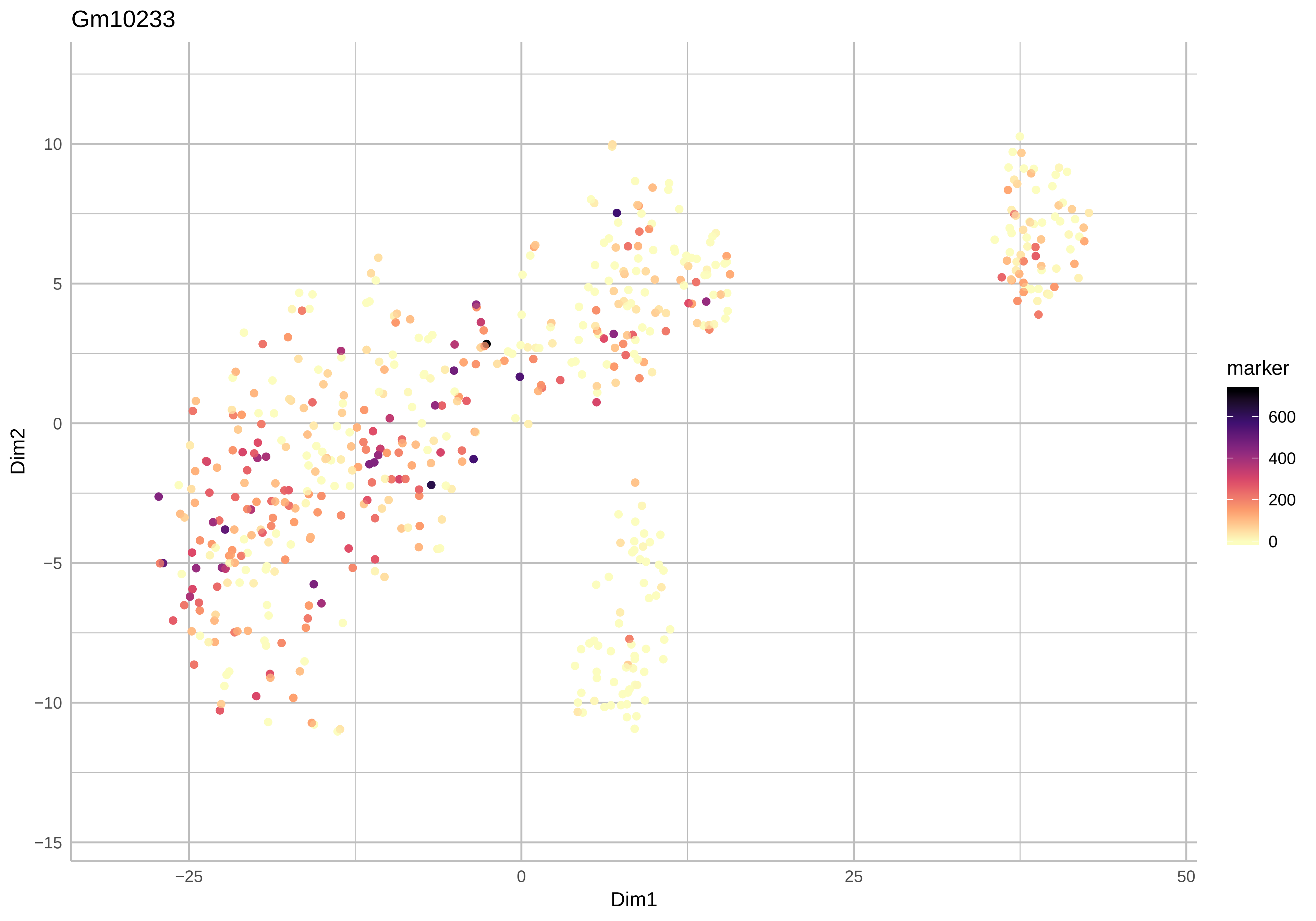

Gm10233
10
5
0
−5
−10
−15
Dim2
−25
0
25
50
Dim1
marker
600
400
200
0

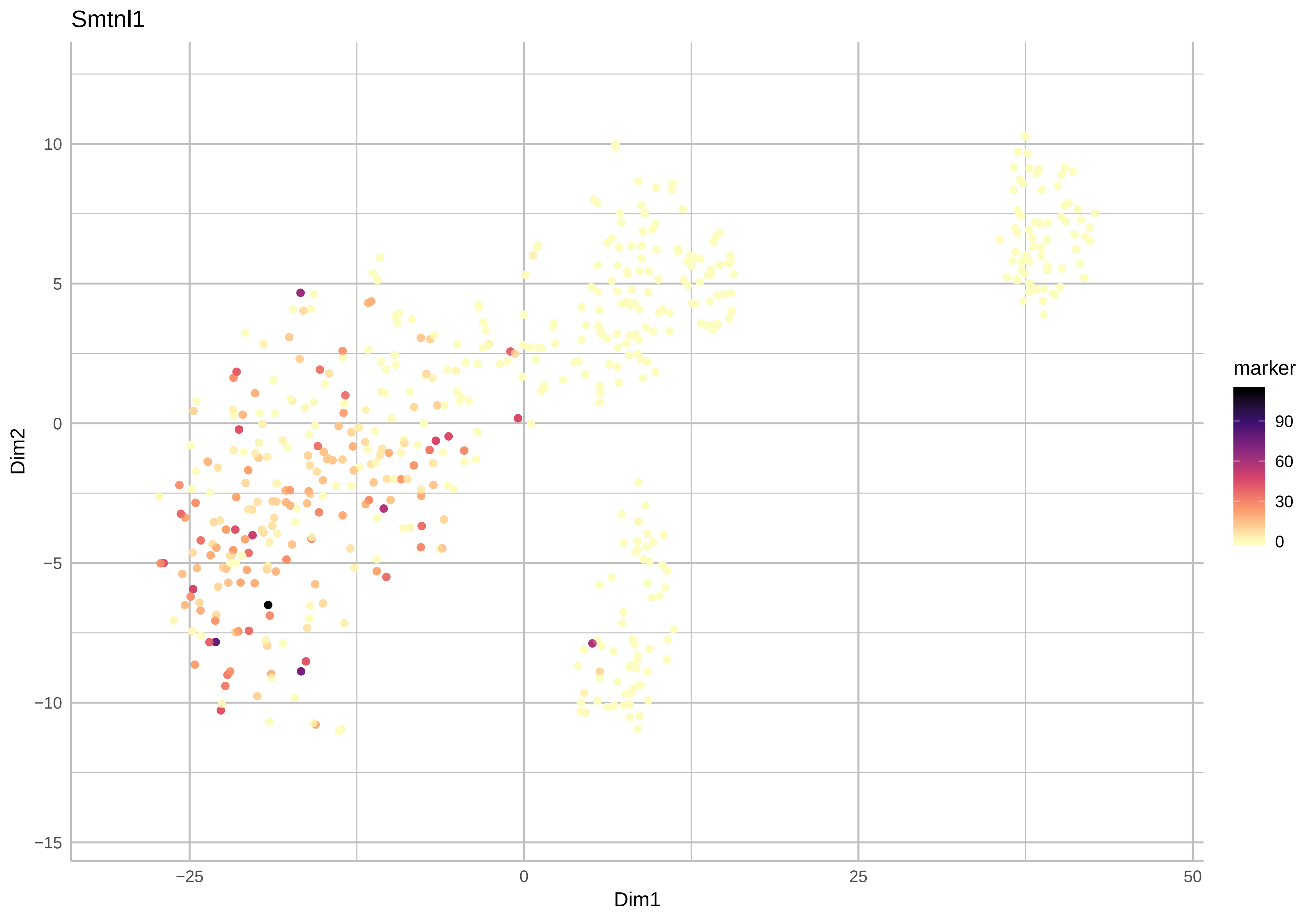

Smtnl1
10
5
0
−5
−10
−15
−25
0
25
50
Dim1
Dim2
marker
90
60
30
0

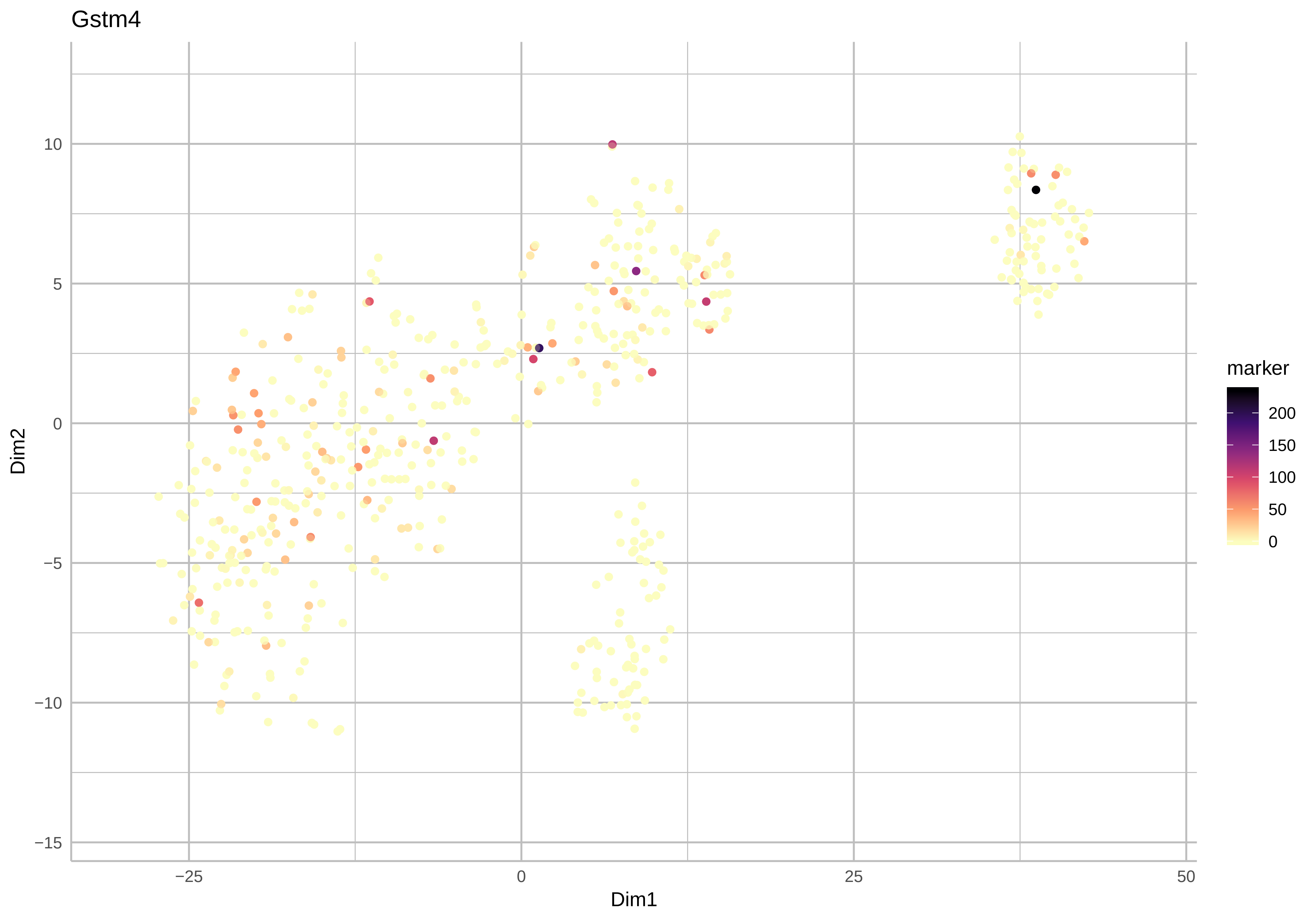

Gstm4
Dim2
Dim1
10
5
0
−5
−10
−15
−25
0
25
50
marker
200
150
100
50
0

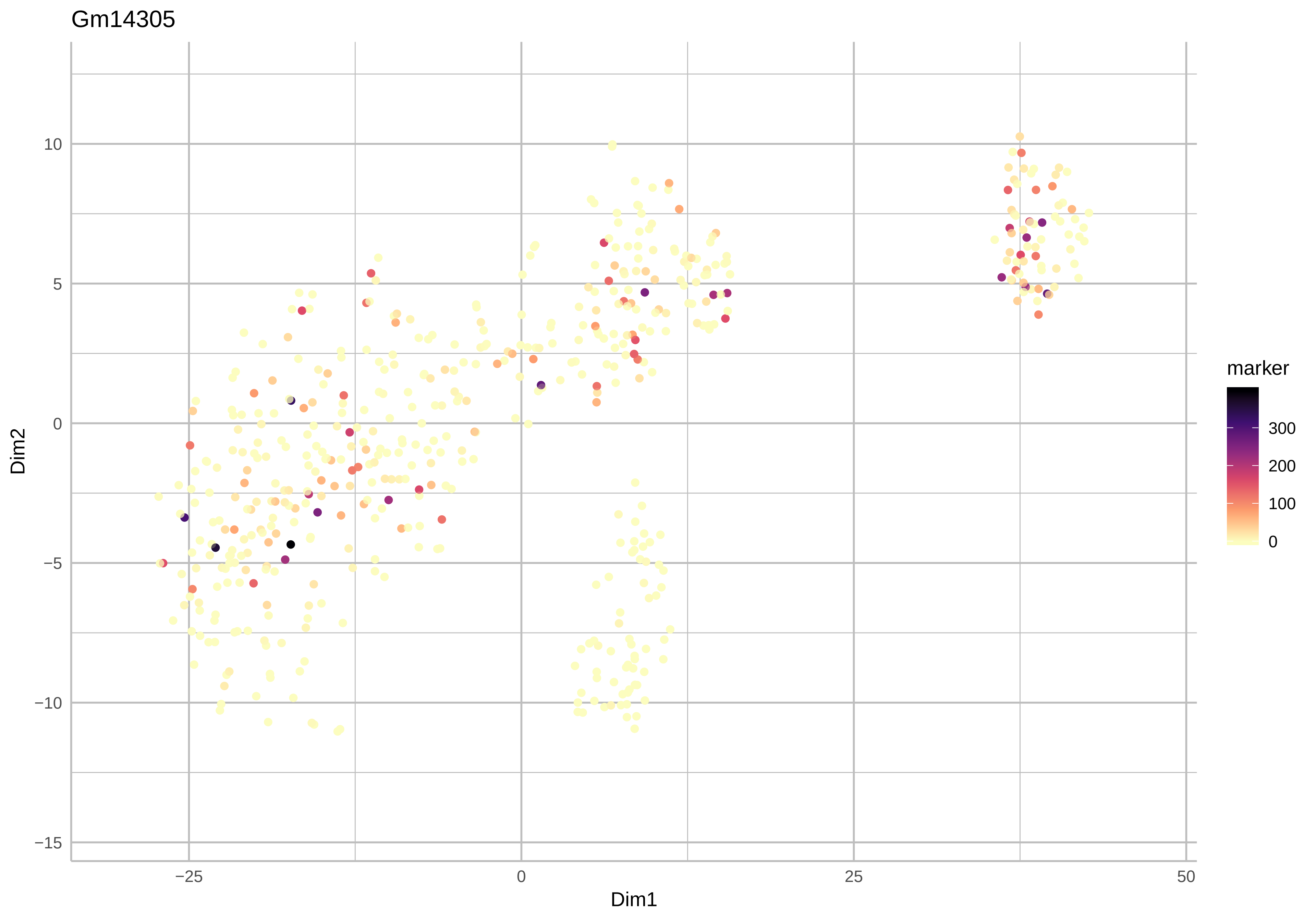

Gm14305
Dim2
Dim1
10
5
0
−5
−10
−15
−25
0
25
50
marker
300
200
100
0

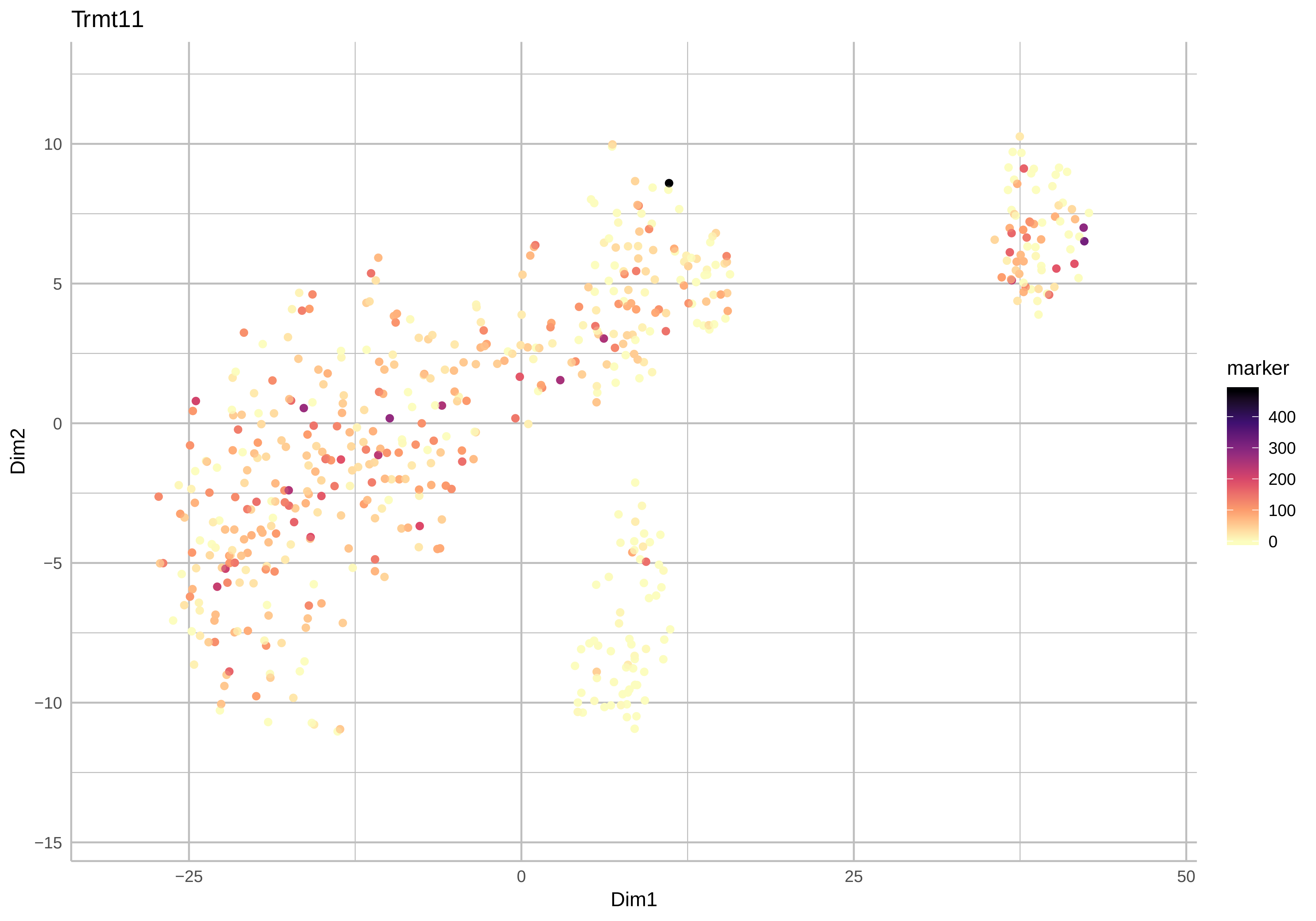
Trmt11
Dim2
Dim1
marker
400
300
200
100
0
10
5
0
−5
−10
−15
−25
0
25
50

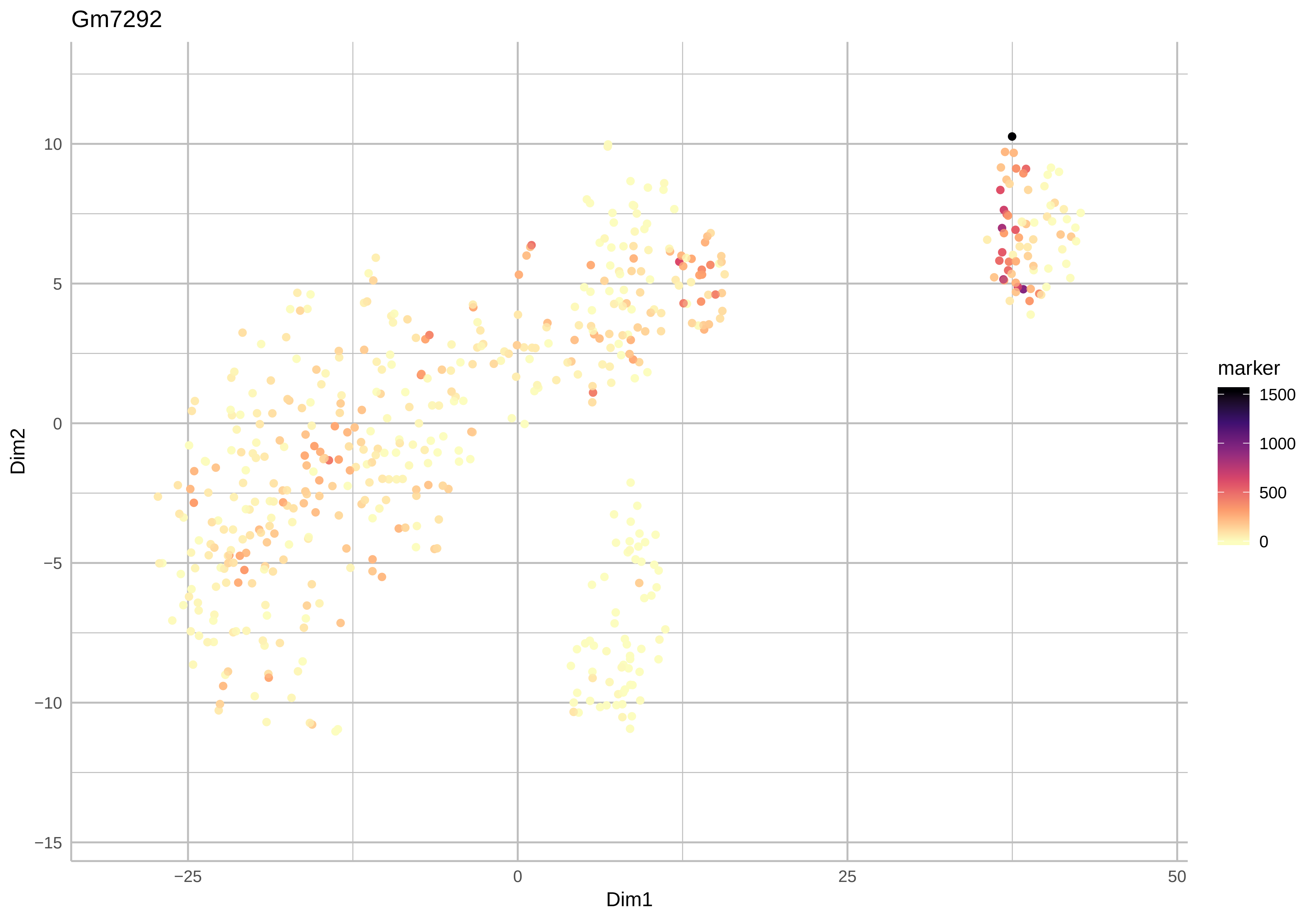

Gm7292
10
5
0
−5
−10
−15
Dim2
−25
0
25
50
Dim1
marker
1500
1000
500
0

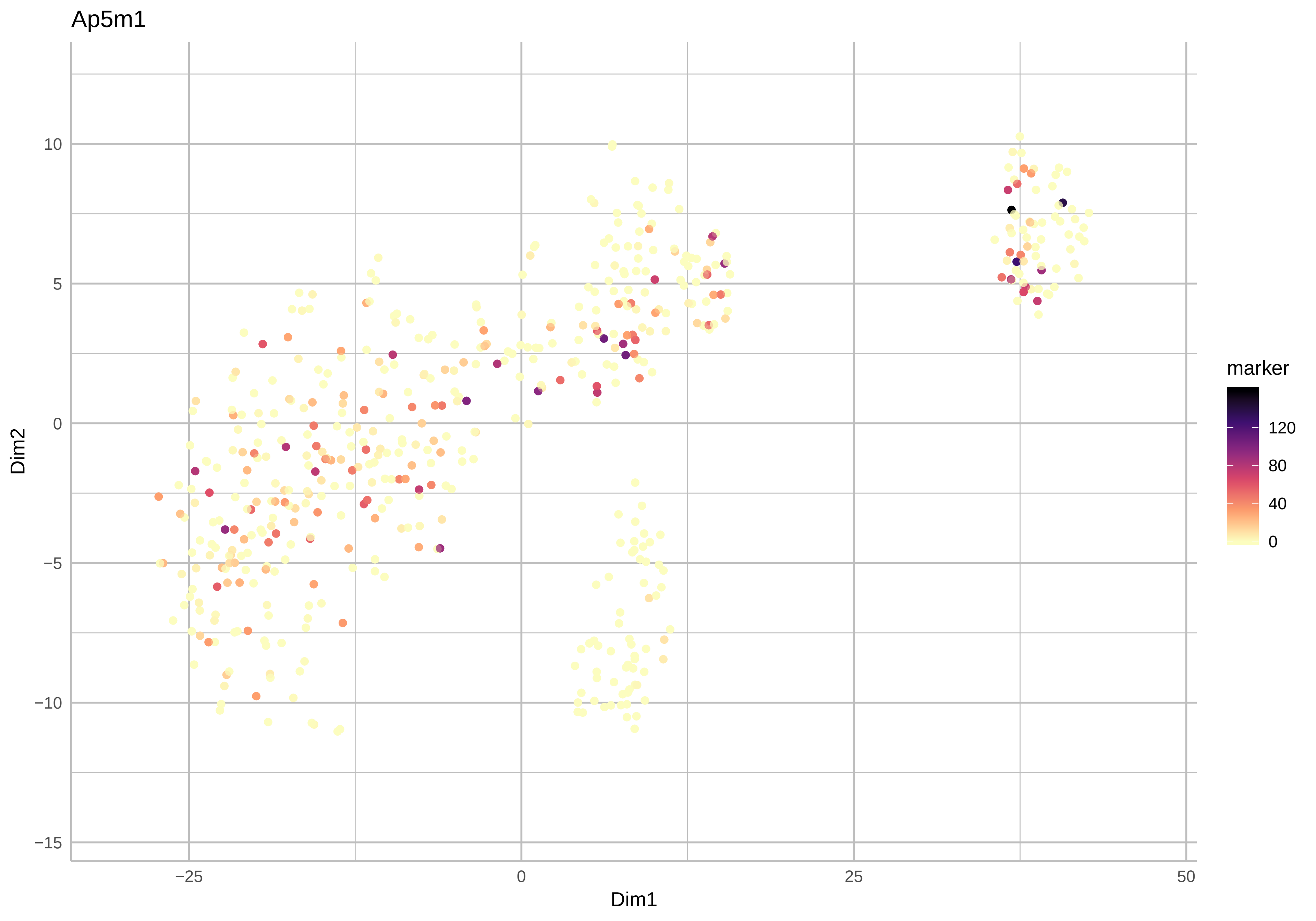

Ap5m1
Dim2
Dim1
10
5
0
−5
−10
−15
−25
0
25
50
marker
120
80
40
0

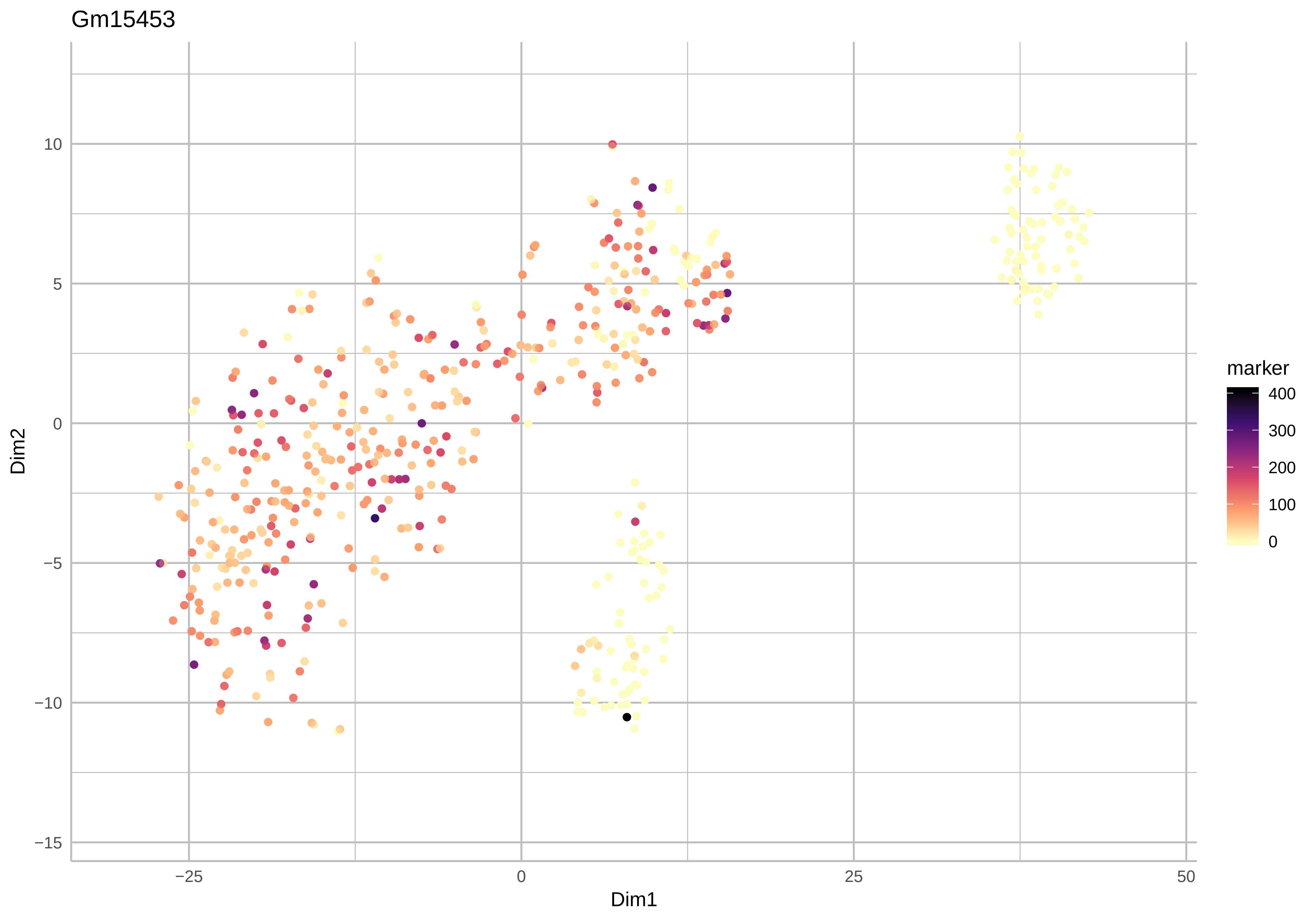

Gm15453
Dim2
Dim1
10
5
0
−5
−10
−15
−25
0
25
50
marker
400
300
200
100
0

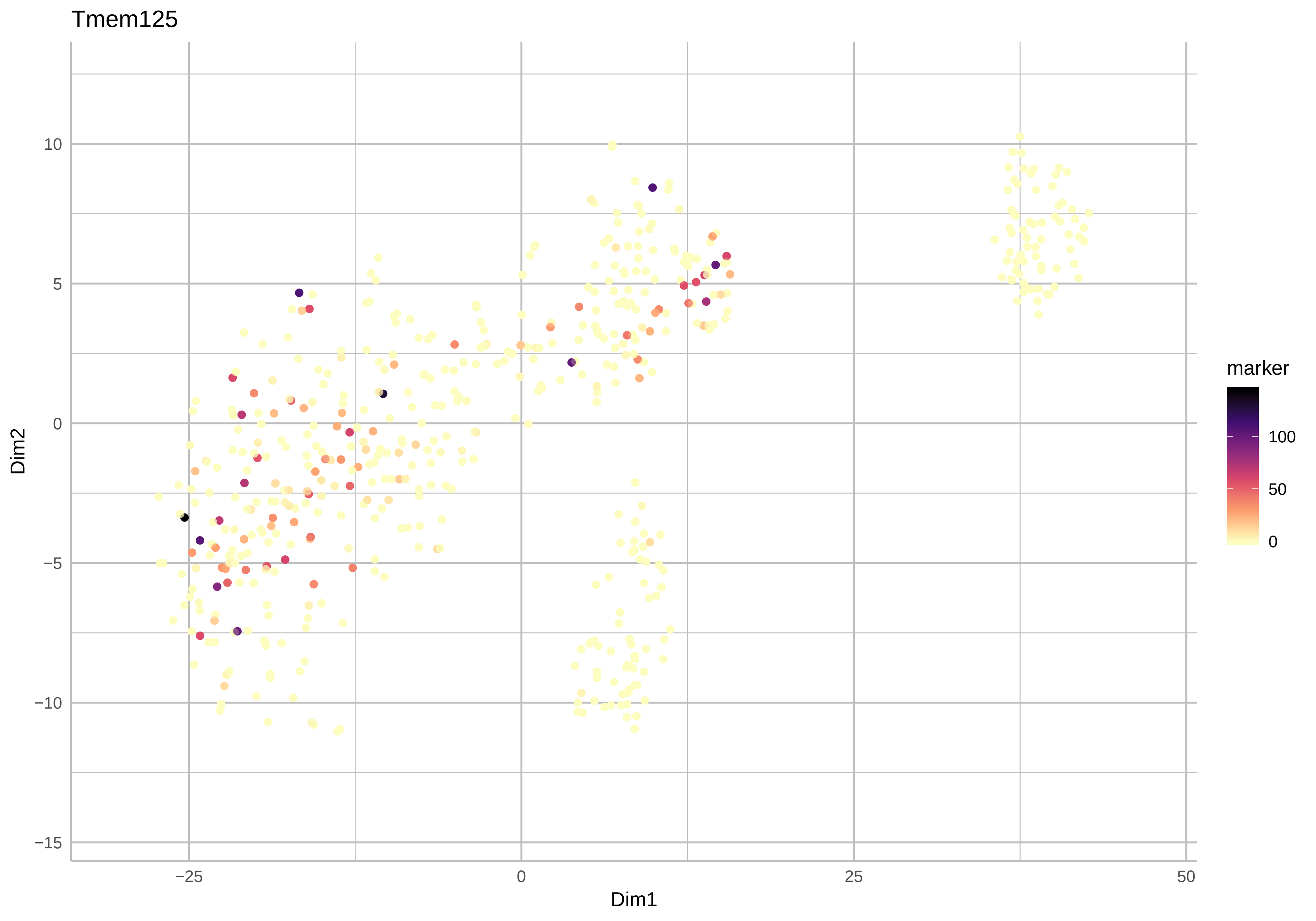
Tmem125
Dim2
Dim1
10
5
0
−5
−10
−15
−25
0
25
50
marker
100
50
0

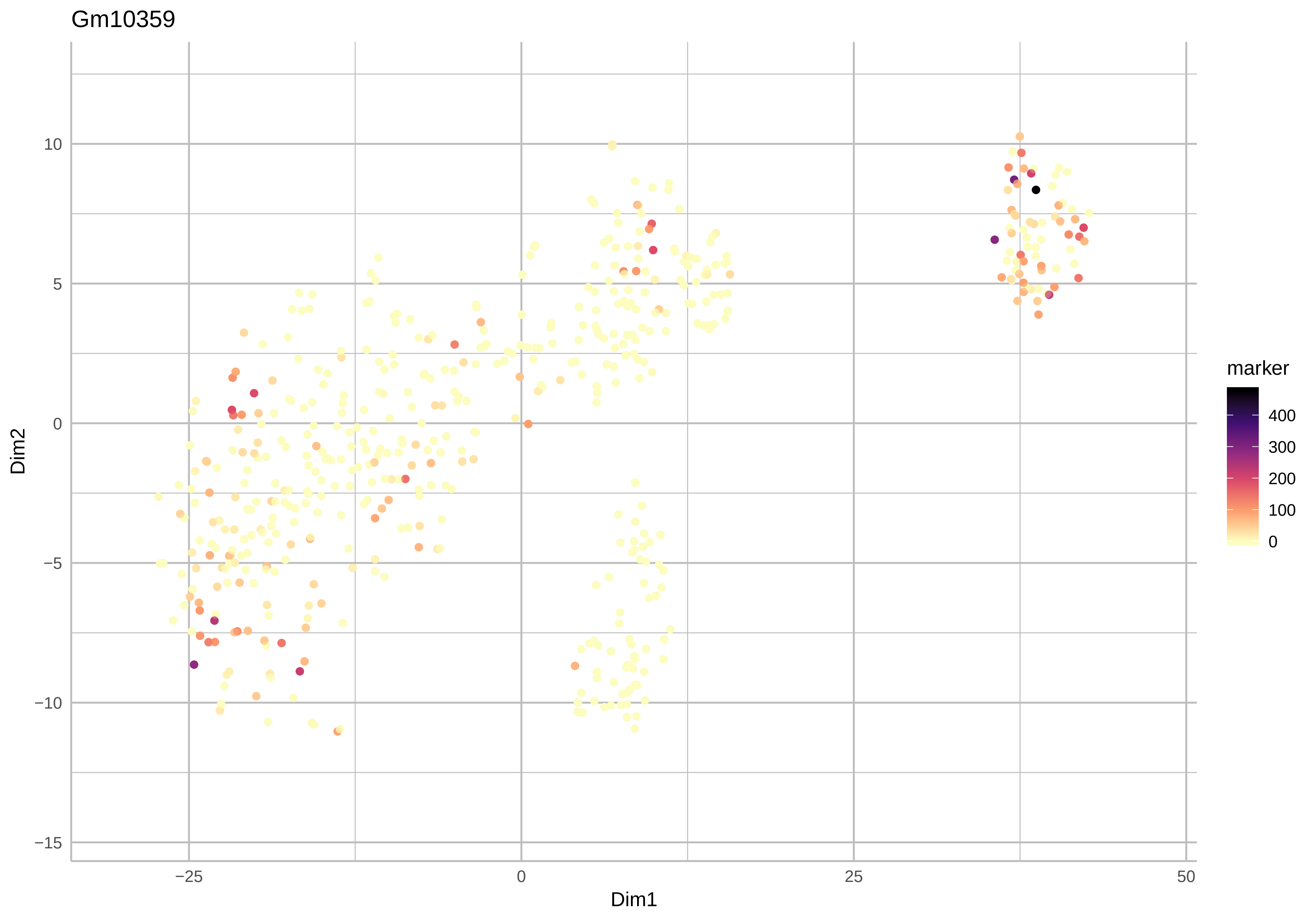

Gm10359
Dim2
Dim1
10
5
0
−5
−10
−15
−25
0
25
50
marker
400
300
200
100
0

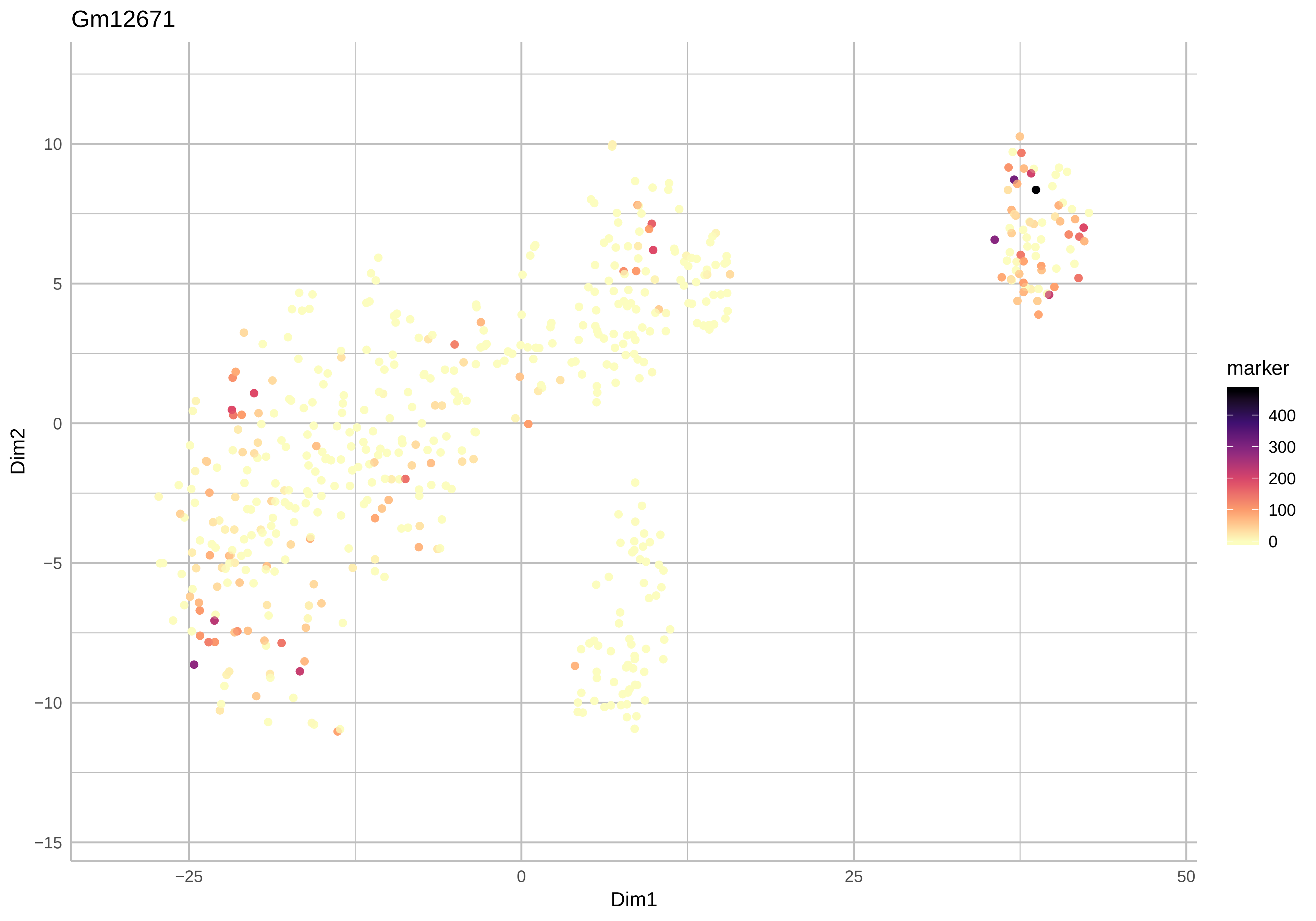

Gm12671
Dim2
Dim1
10
5
0
−5
−10
−15
−25
0
25
50
marker
400
300
200
100
0

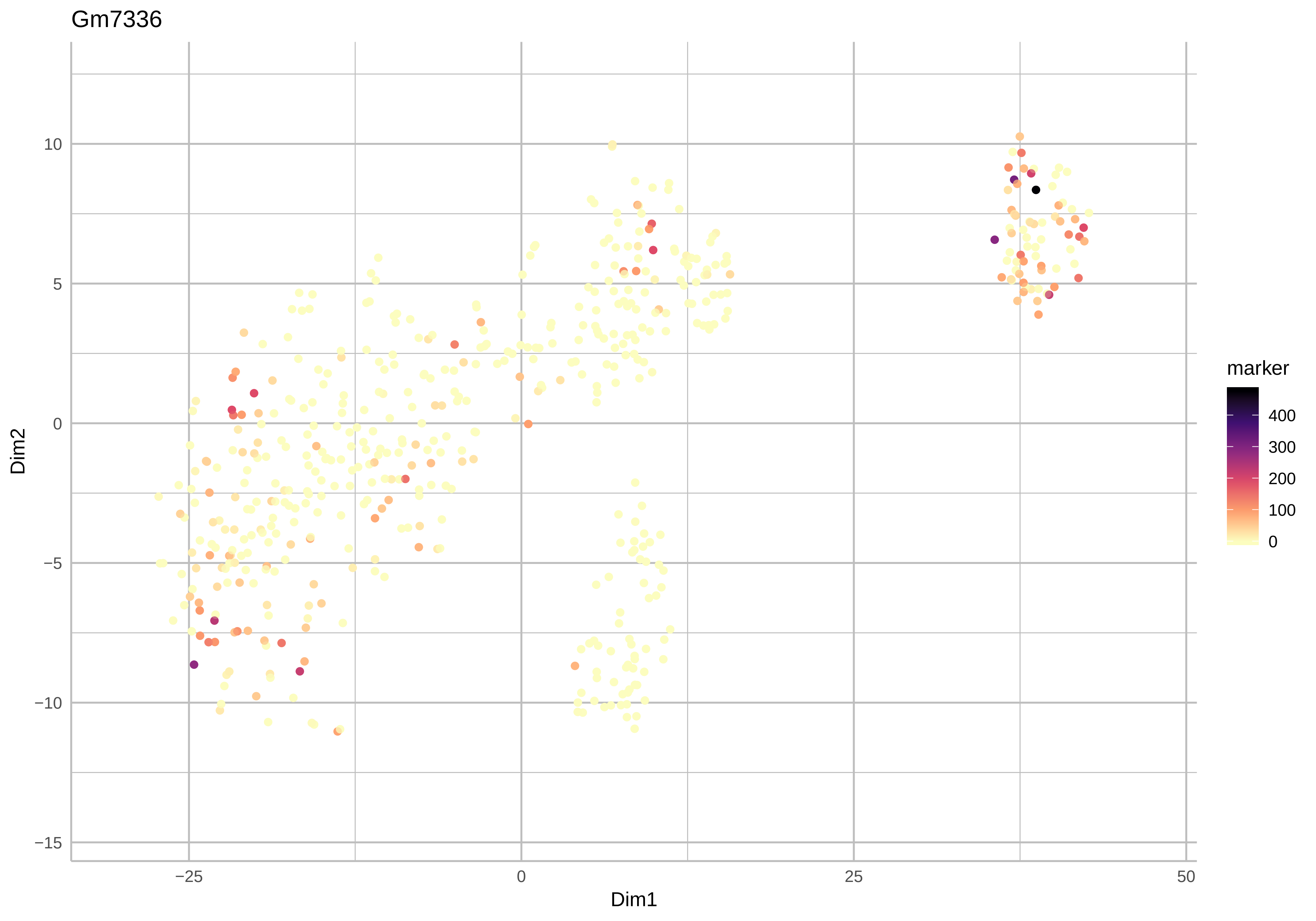

Gm7336
Dim2
Dim1
marker
400
300
200
100
0
10
5
0
−5
−10
−15
−25
0
25
50

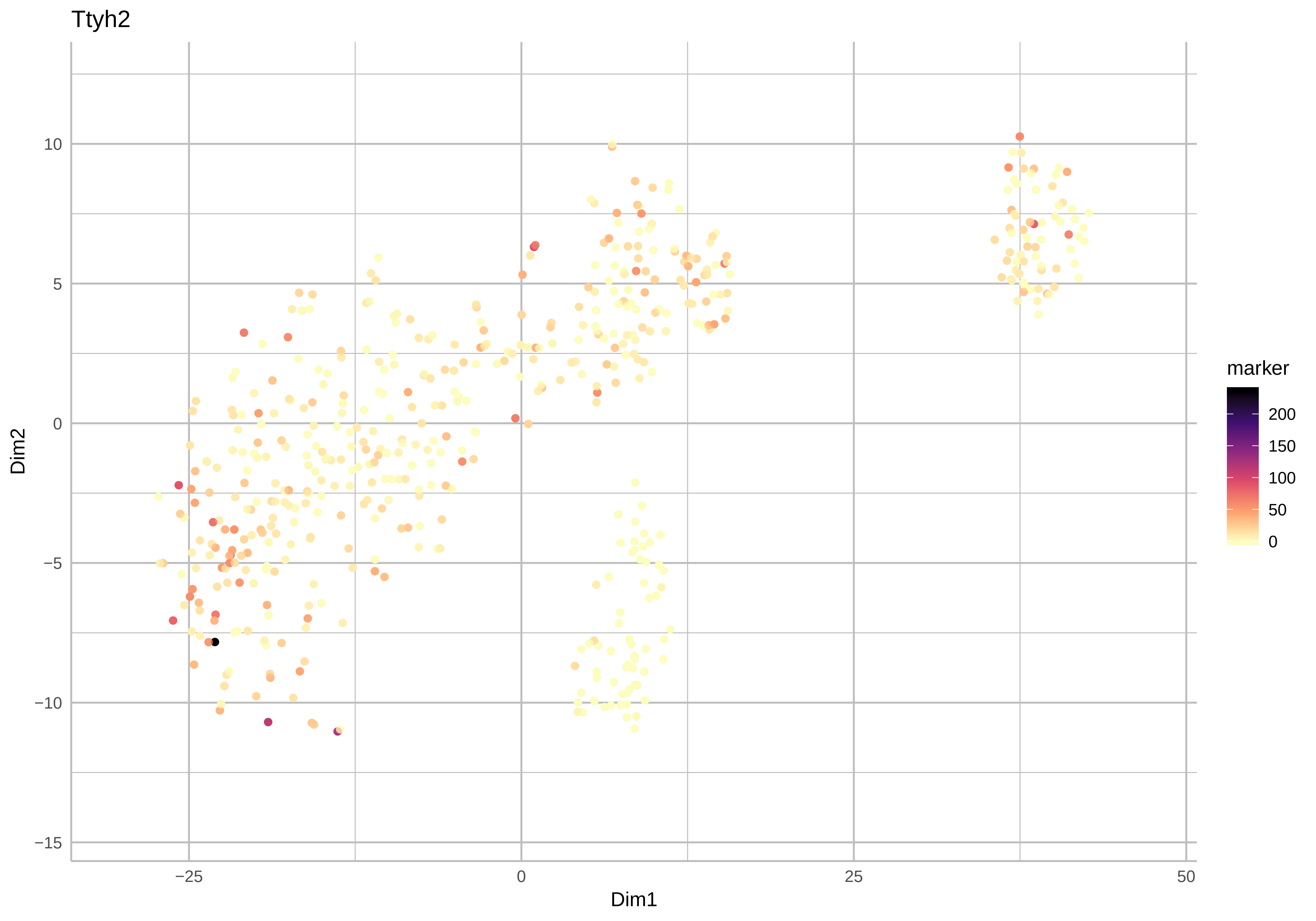

Ttyh2
Dim2
Dim1
marker
200
150
100
50
0
10
5
0
−5
−10
−15
−25
0
25
50

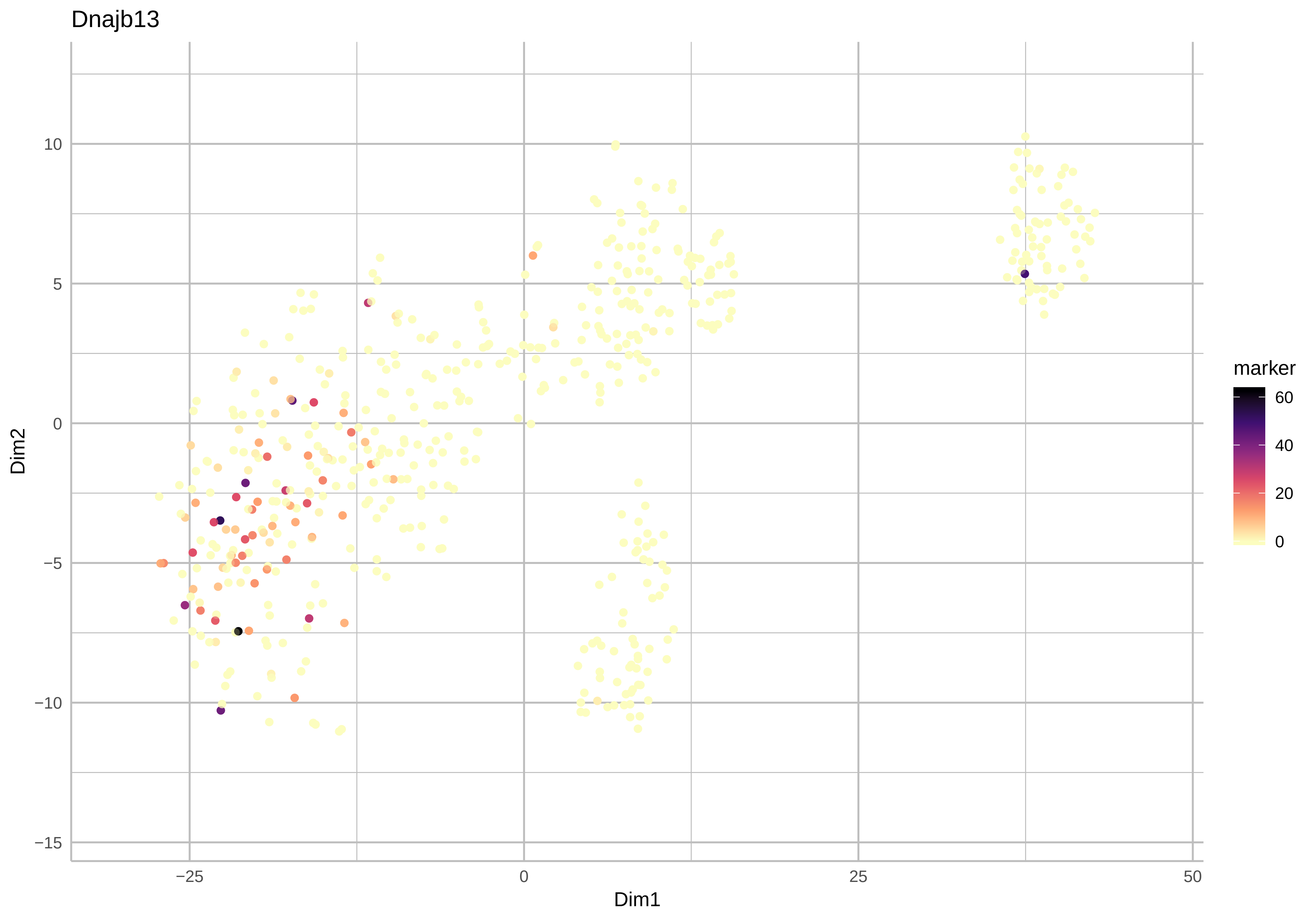

Dnajb13
Dim2
Dim1
10
5
0
−5
−10
−15
−25
0
25
50
marker
60
40
20
0

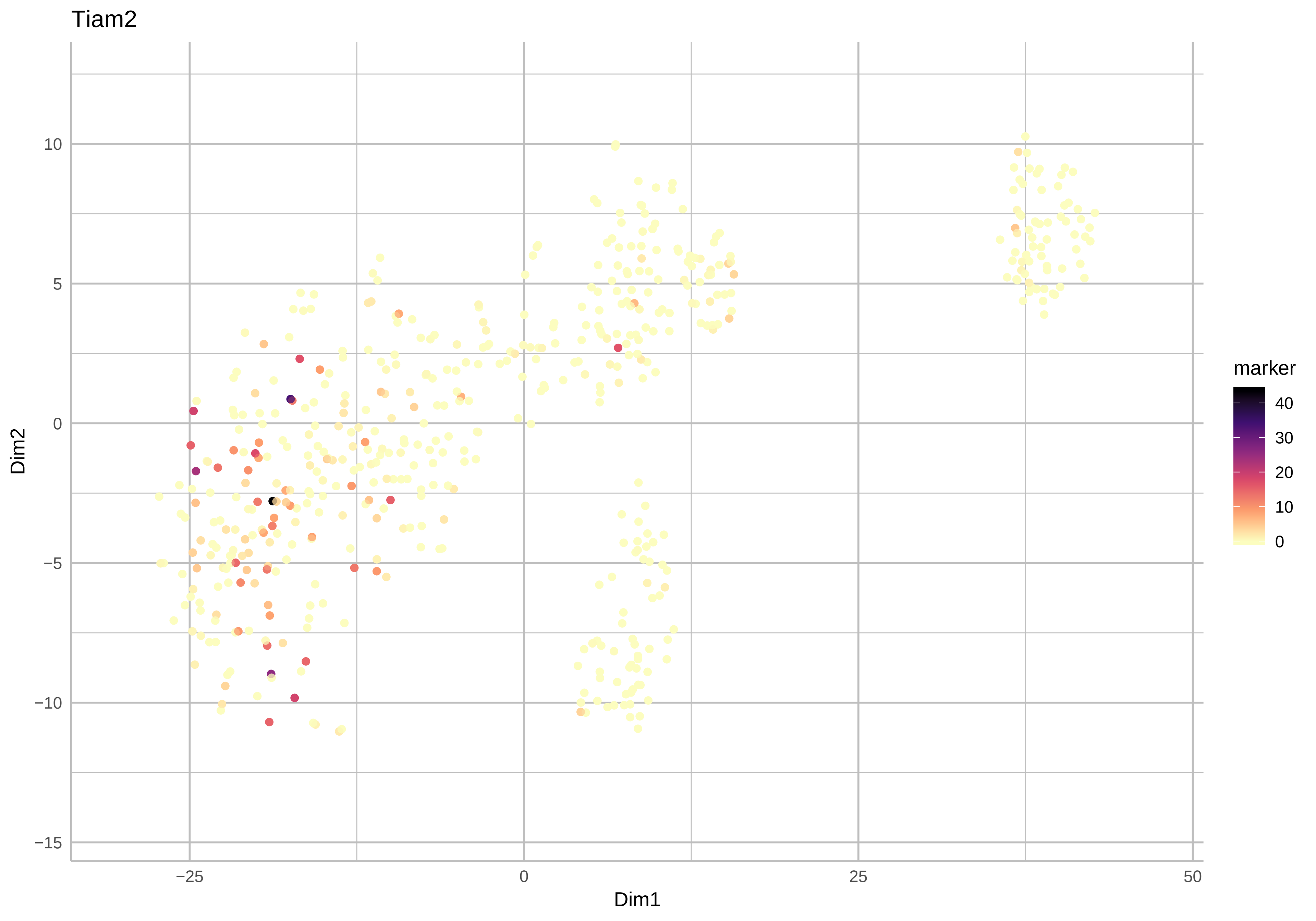

Tiam2
Dim2
Dim1
10
5
0
−5
−10
−15
−25
0
25
50
marker
40
30
20
10
0

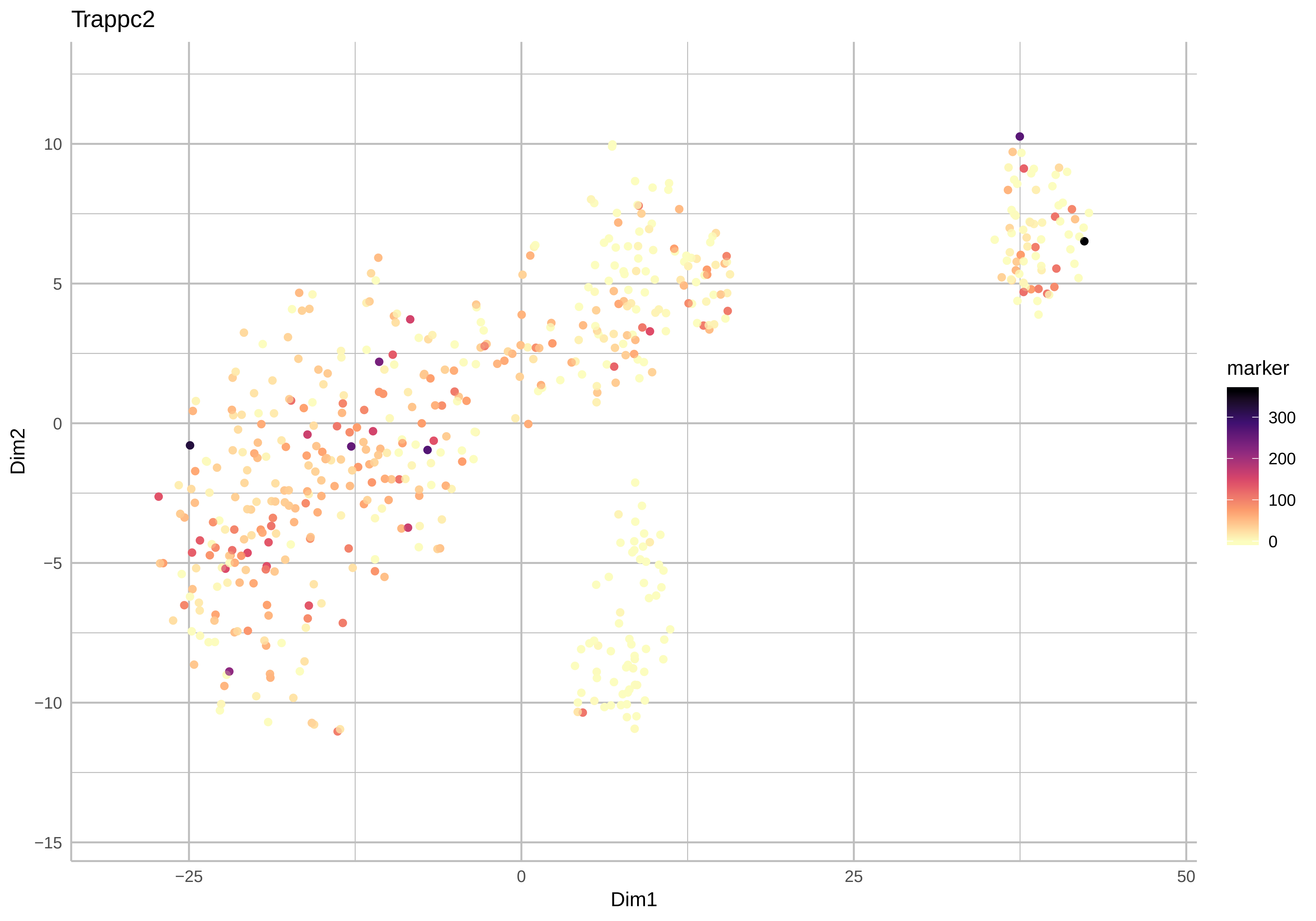

Trappc2
10
5
0
−5
−10
−15
Dim2
−25
0
25
50
Dim1
marker
300
200
100
0

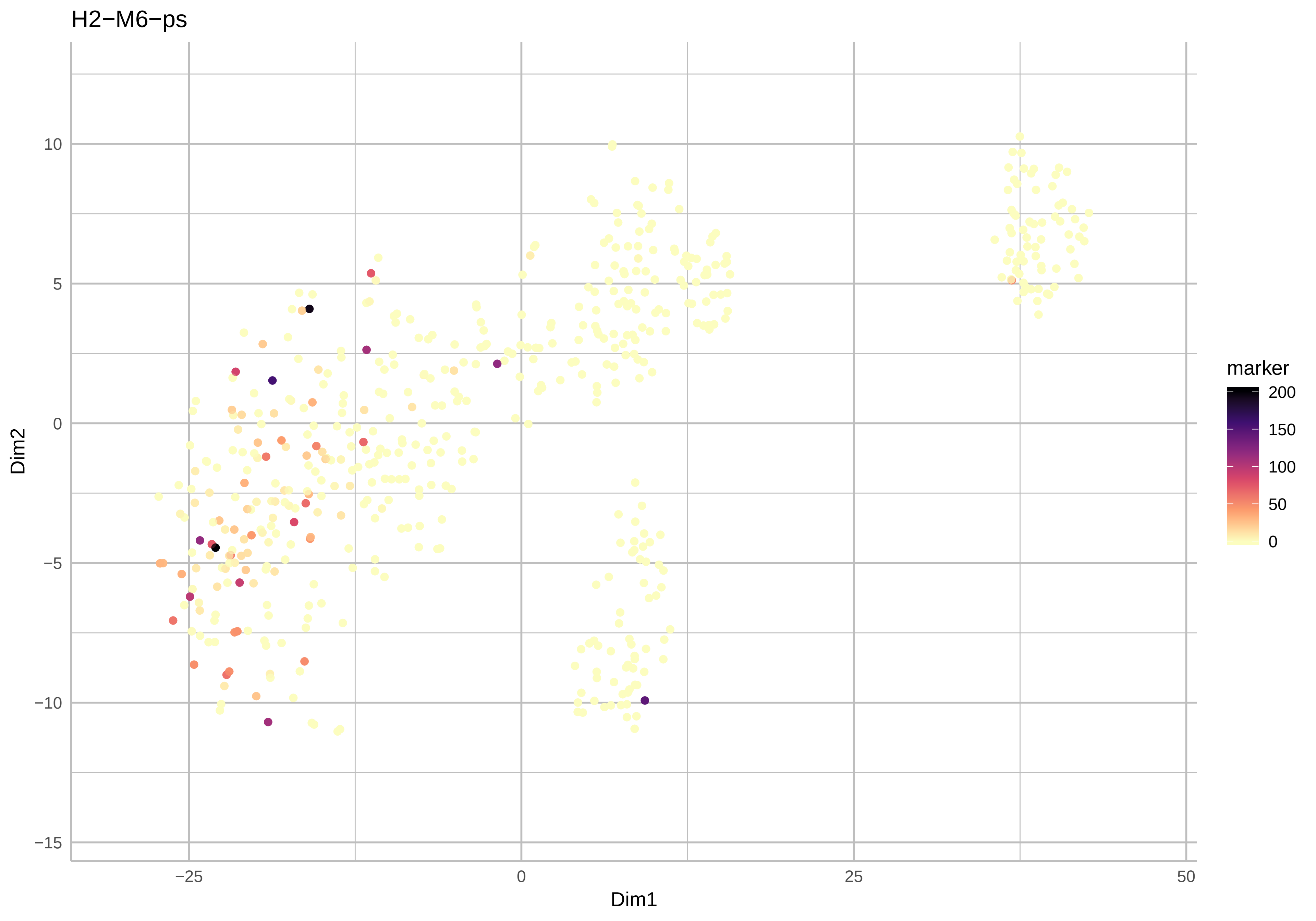
H2−M6−ps
Dim2
Dim1
10
5
0
−5
−10
−15
−25
0
25
50
marker
200
150
100
50
0

Gm7488

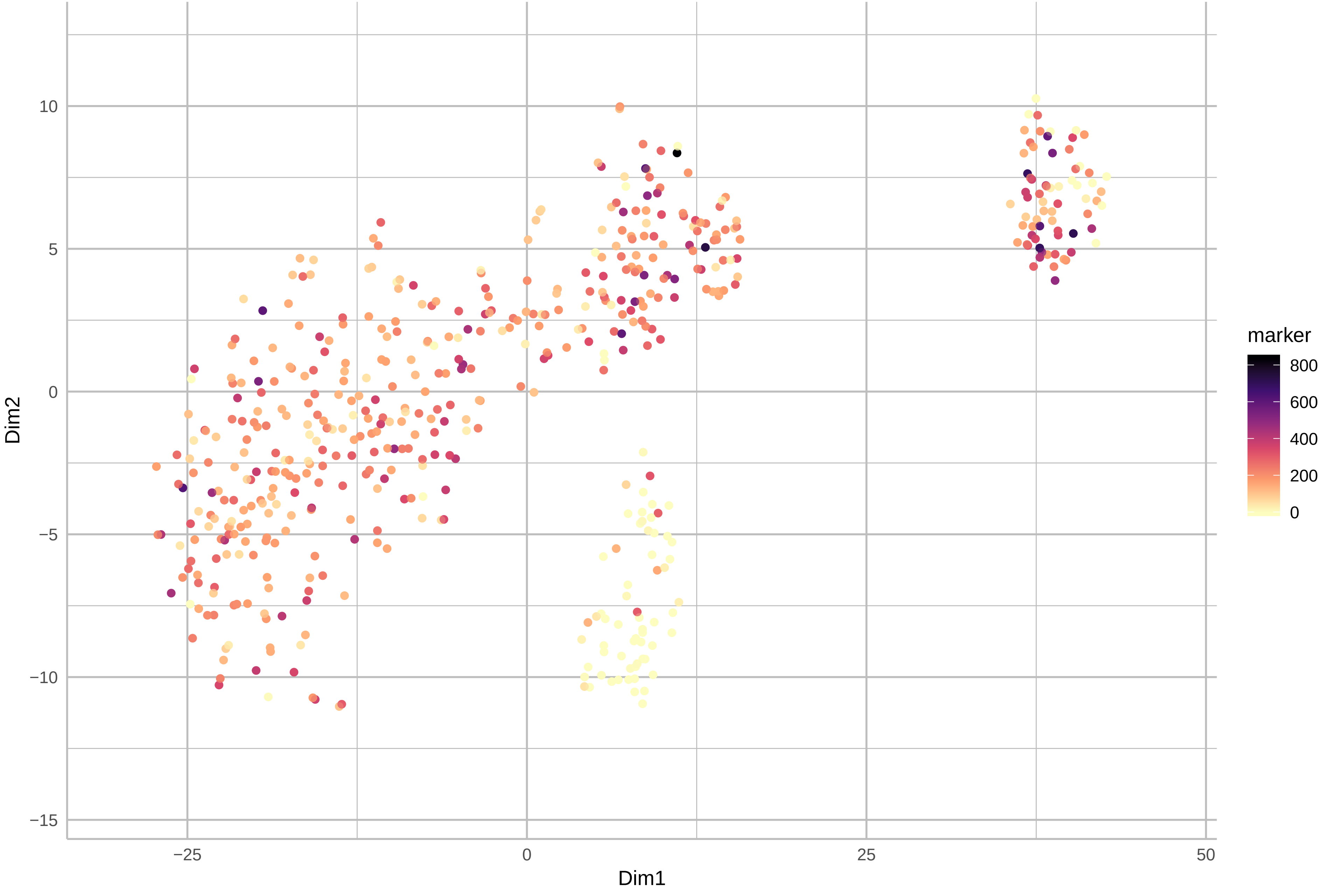

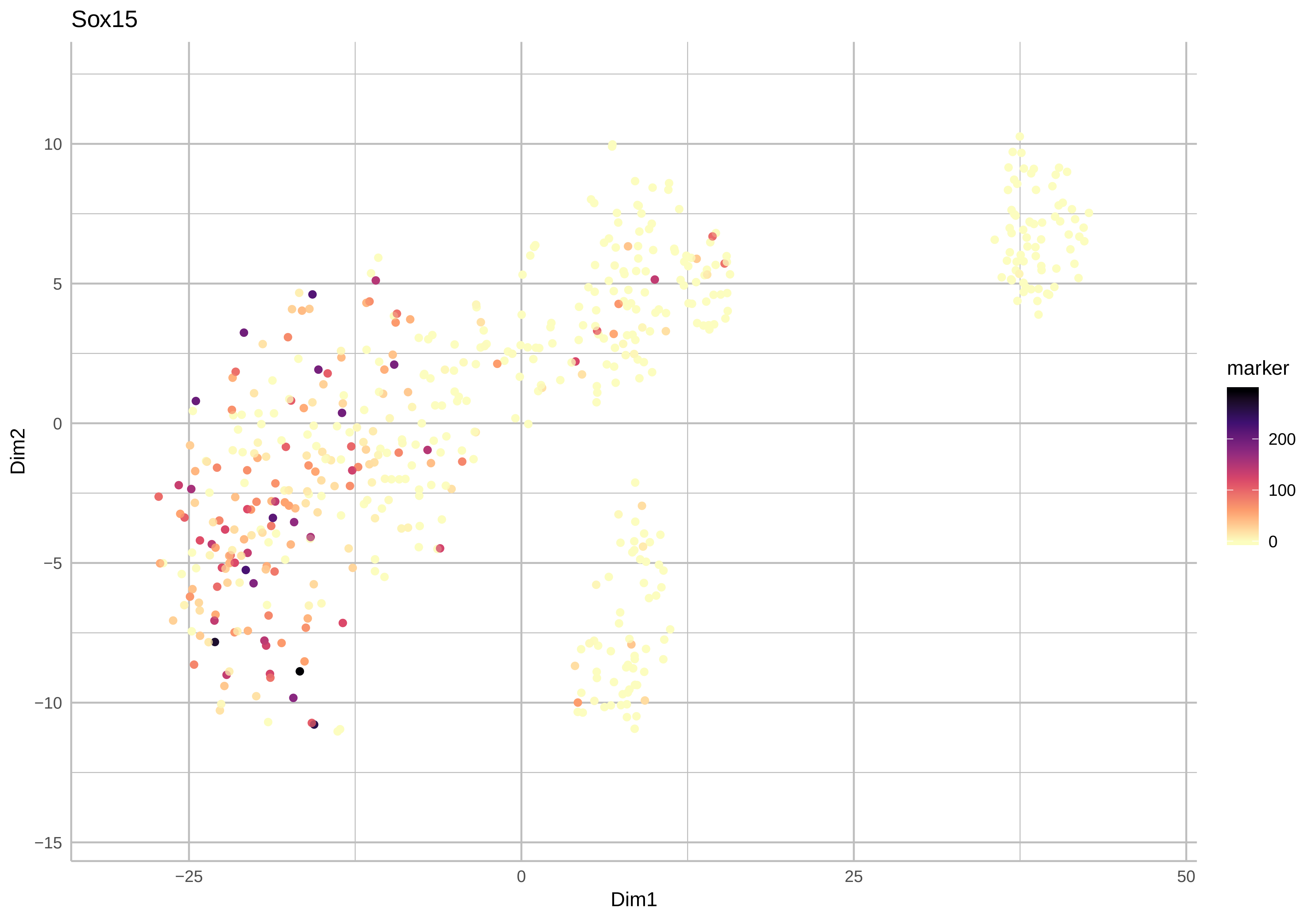

Sox15
Dim2
Dim1
10
5
0
−5
−10
−15
−25
0
25
50
marker
200
100
0

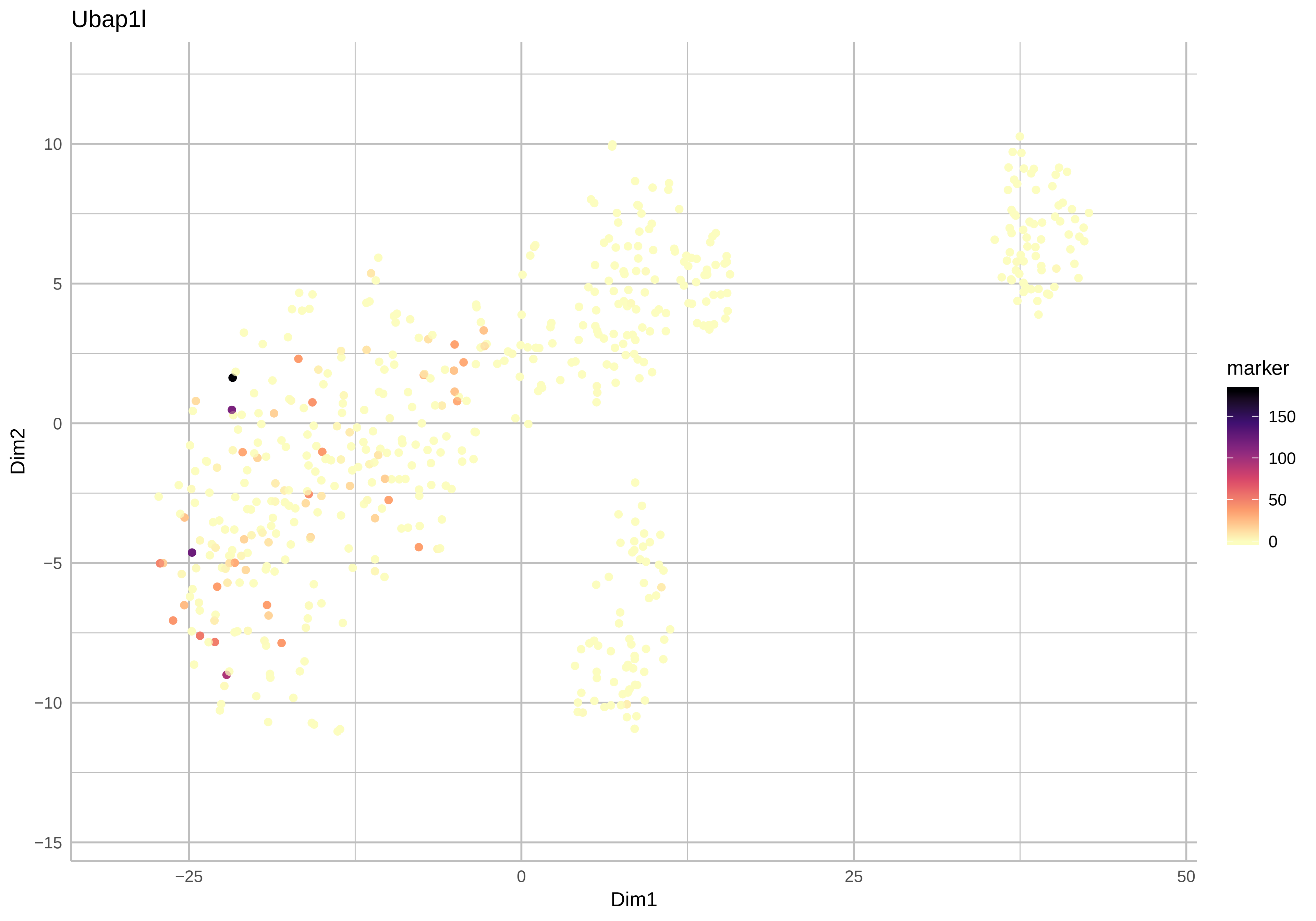

Ubap1l
Dim2
Dim1
10
5
0
−5
−10
−15
−25
0
25
50
marker
150
100
50
0

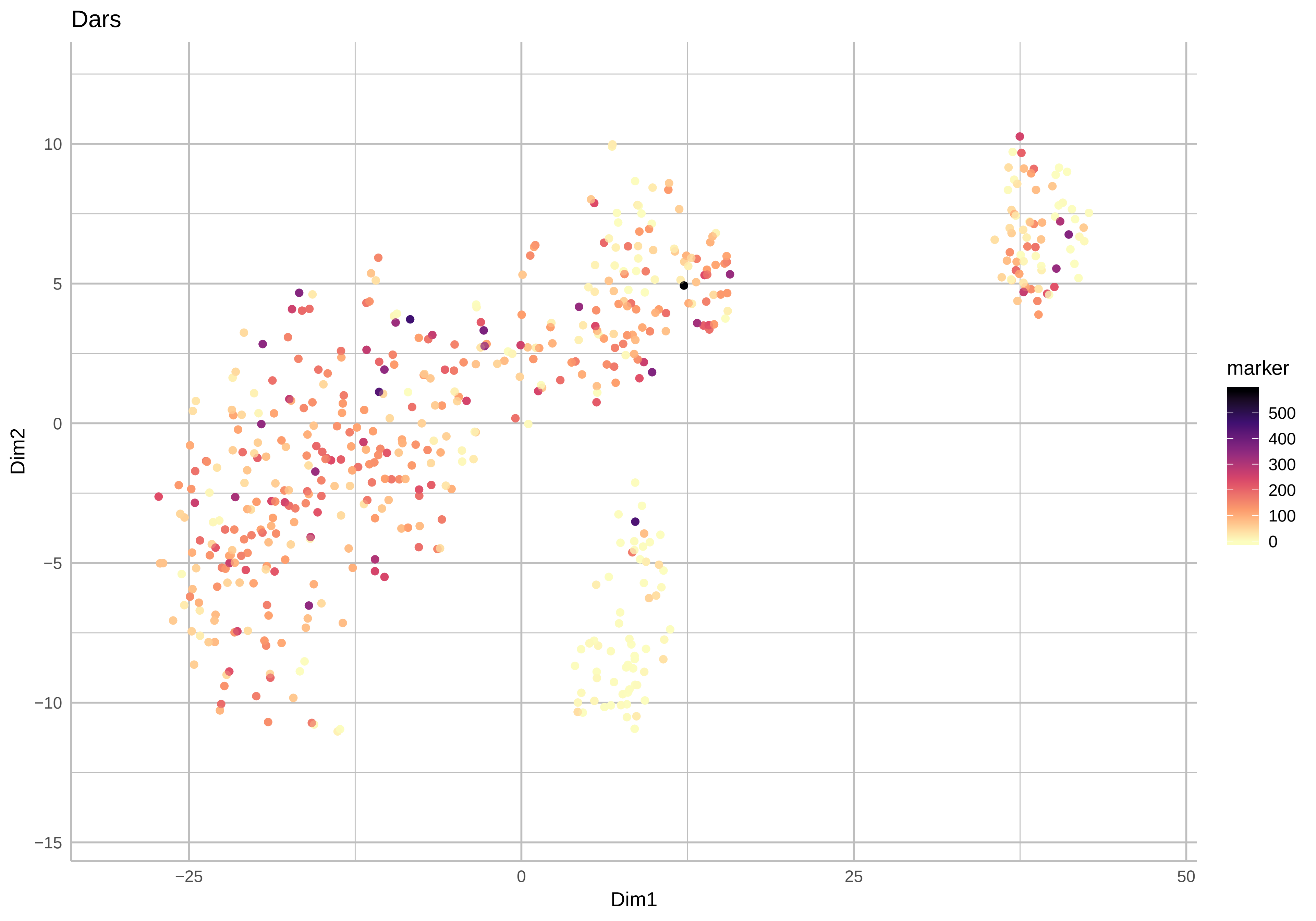

Dars
10
5
0
−5
−10
−15
Dim2
−25
0
25
50
Dim1
marker
500
400
300
200
100
0

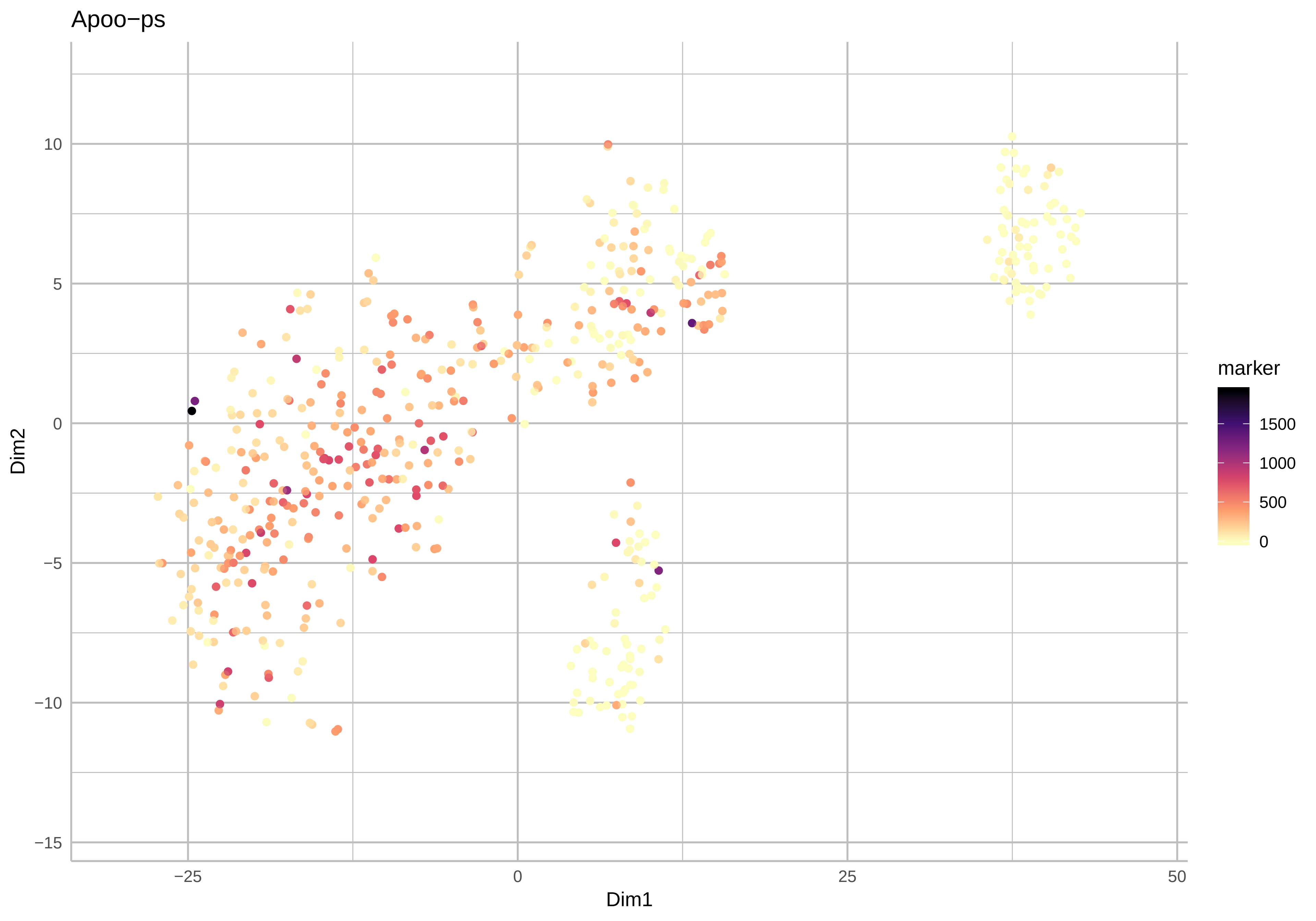

Apoo−ps
Dim2
Dim1
10
5
0
−5
−10
−15
−25
0
25
50
marker
1500
1000
500
0

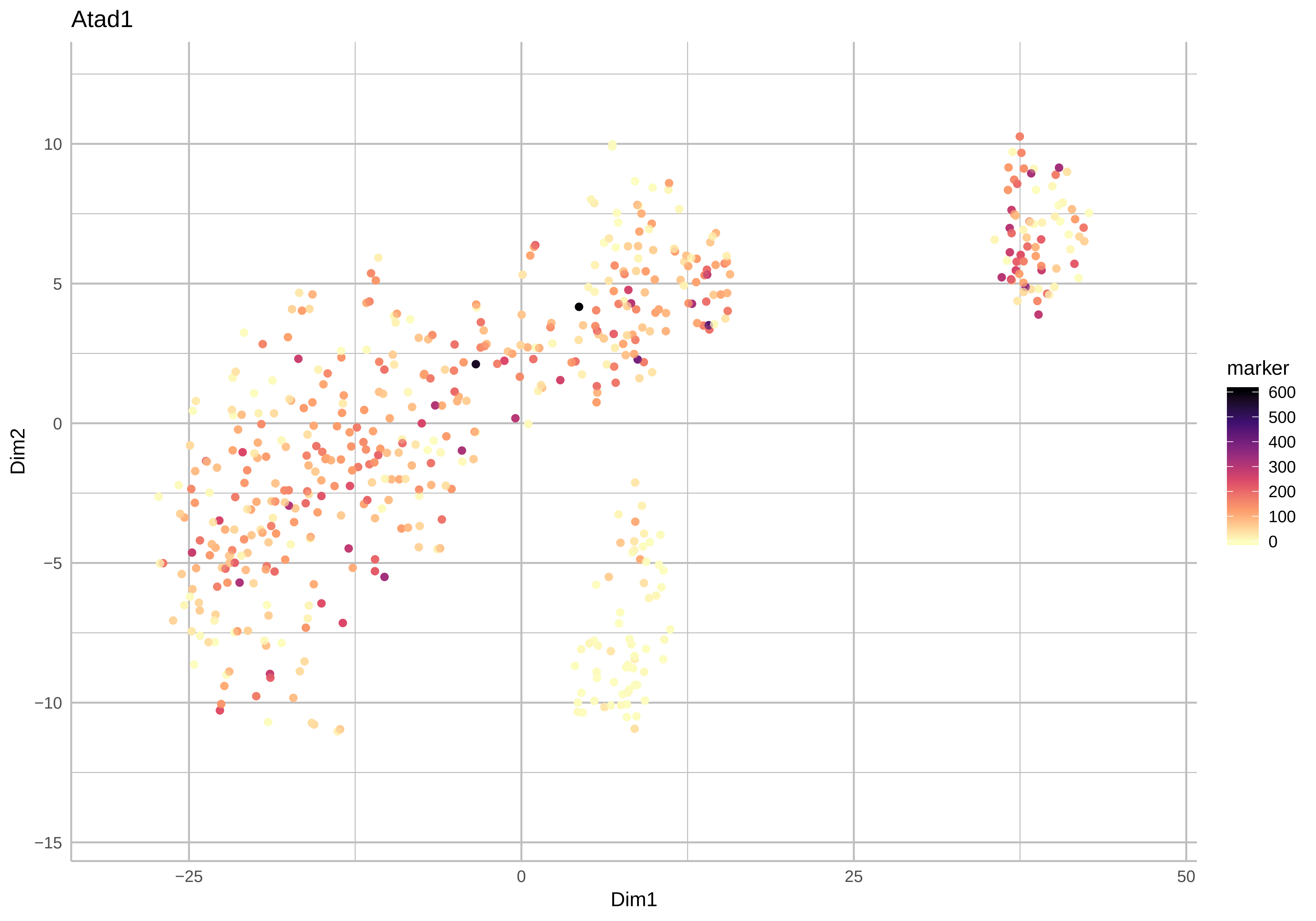

Atad1
10
5
0
−5
−10
−15
Dim2
−25
0
25
50
Dim1
marker
600
500
400
300
200
100
0

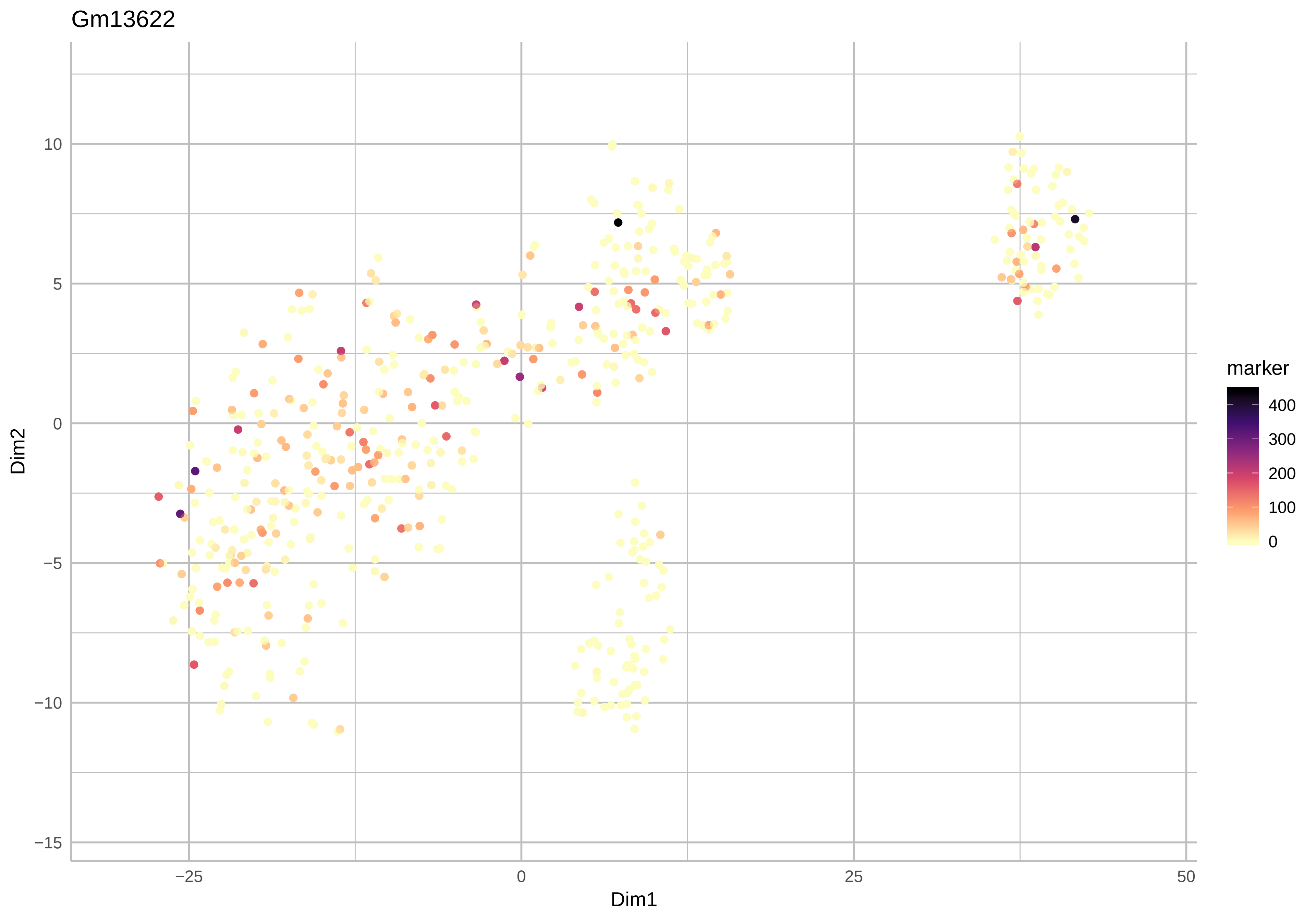

Gm13622
Dim2
Dim1
10
5
0
−5
−10
−15
−25
0
25
50
marker
400
300
200
100
0

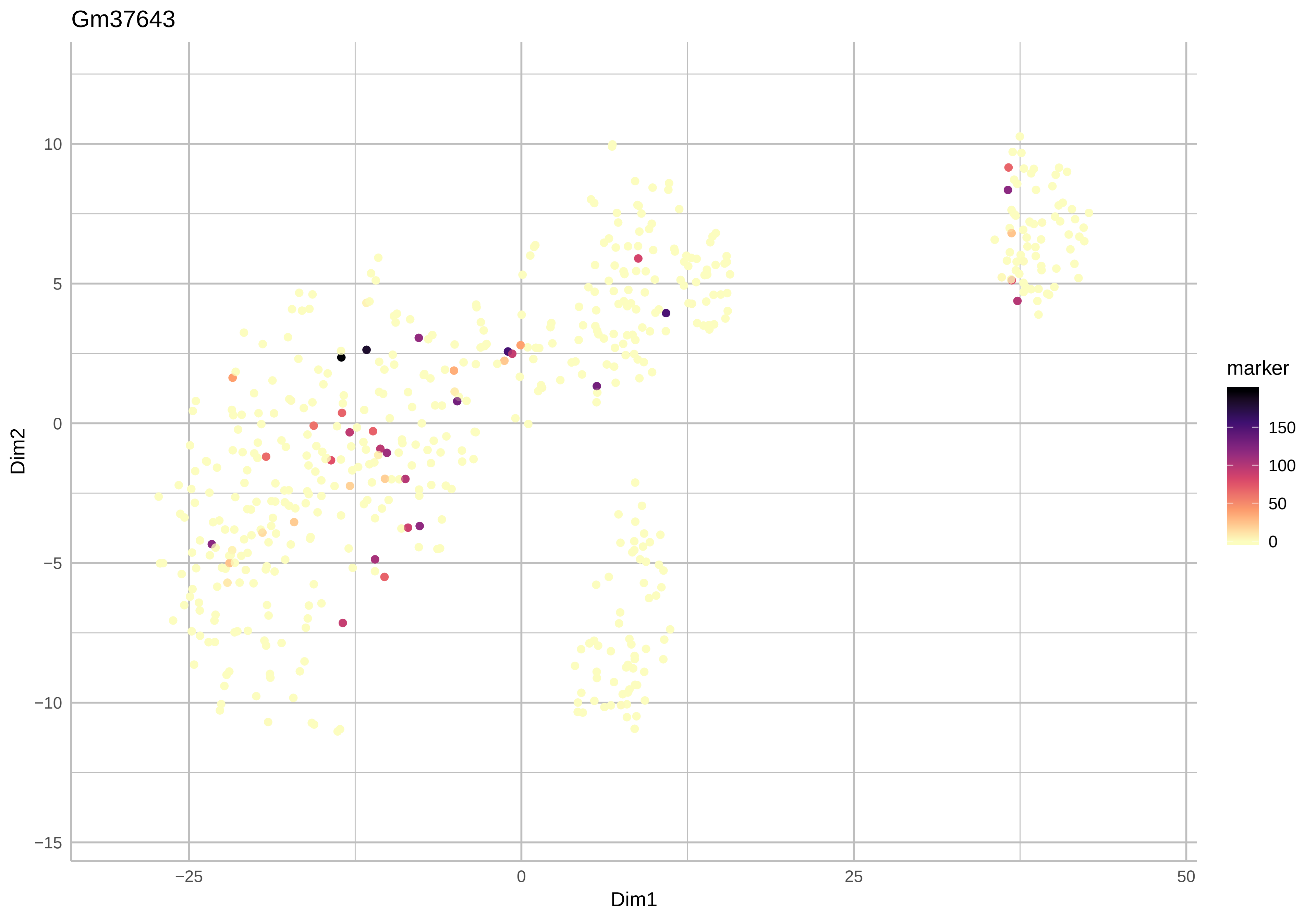

Gm37643
Dim2
Dim1
marker
150
100
50
0
10
5
0
−5
−10
−15
−25
0
25
50

Gm13051

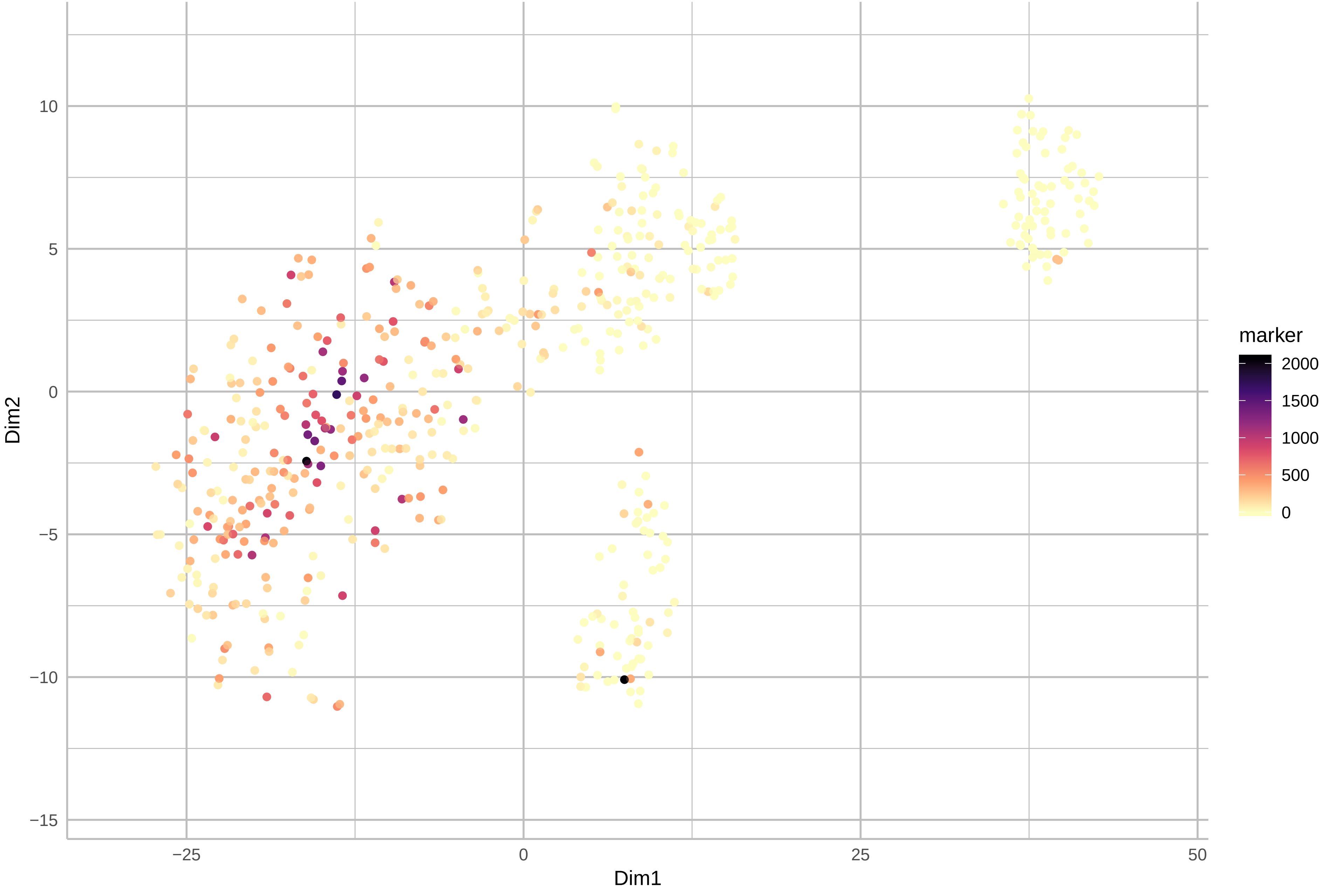

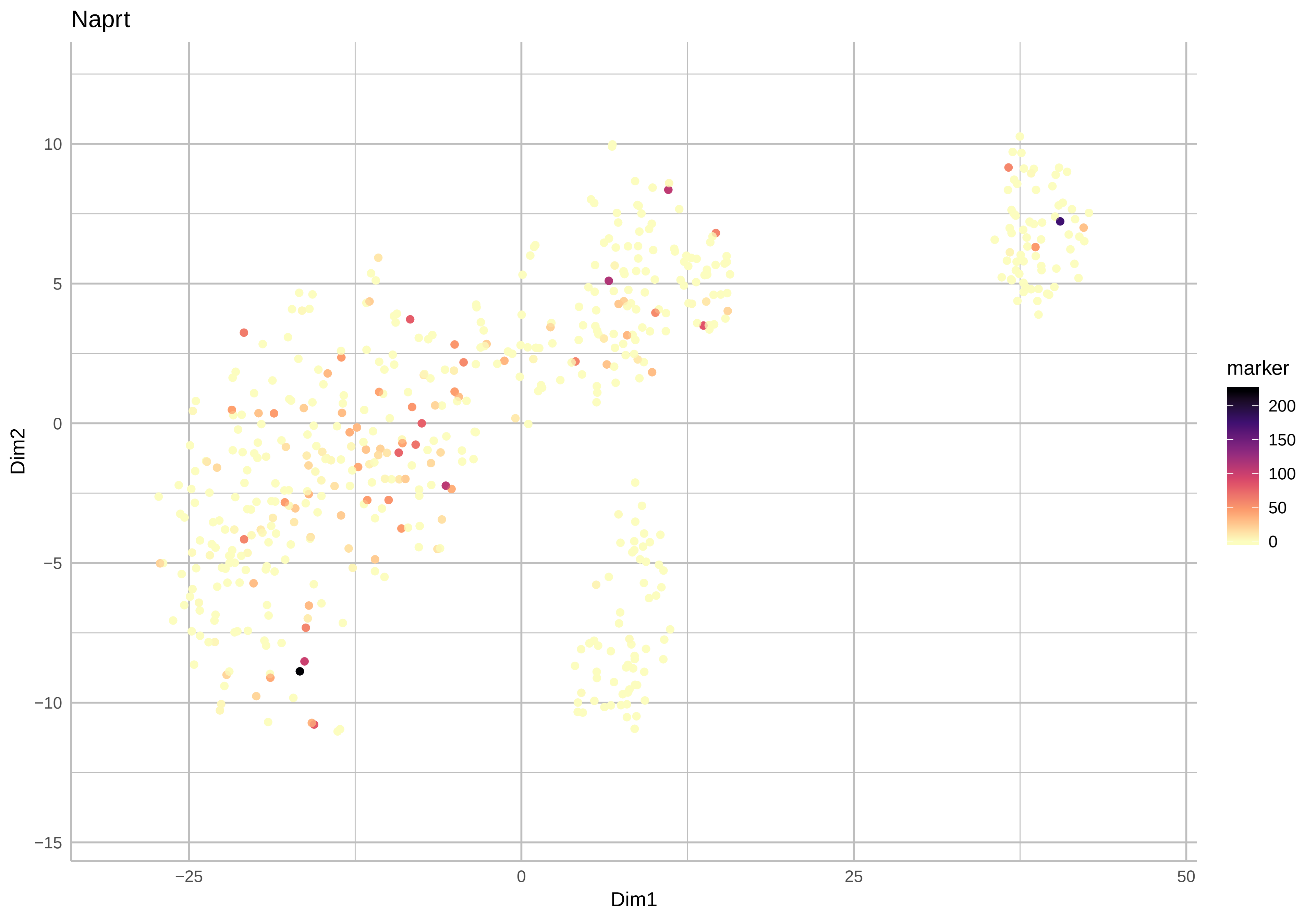
Naprt
Dim2
Dim1
10
5
0
−5
−10
−15
−25
0
25
50
marker
200
150
100
50
0

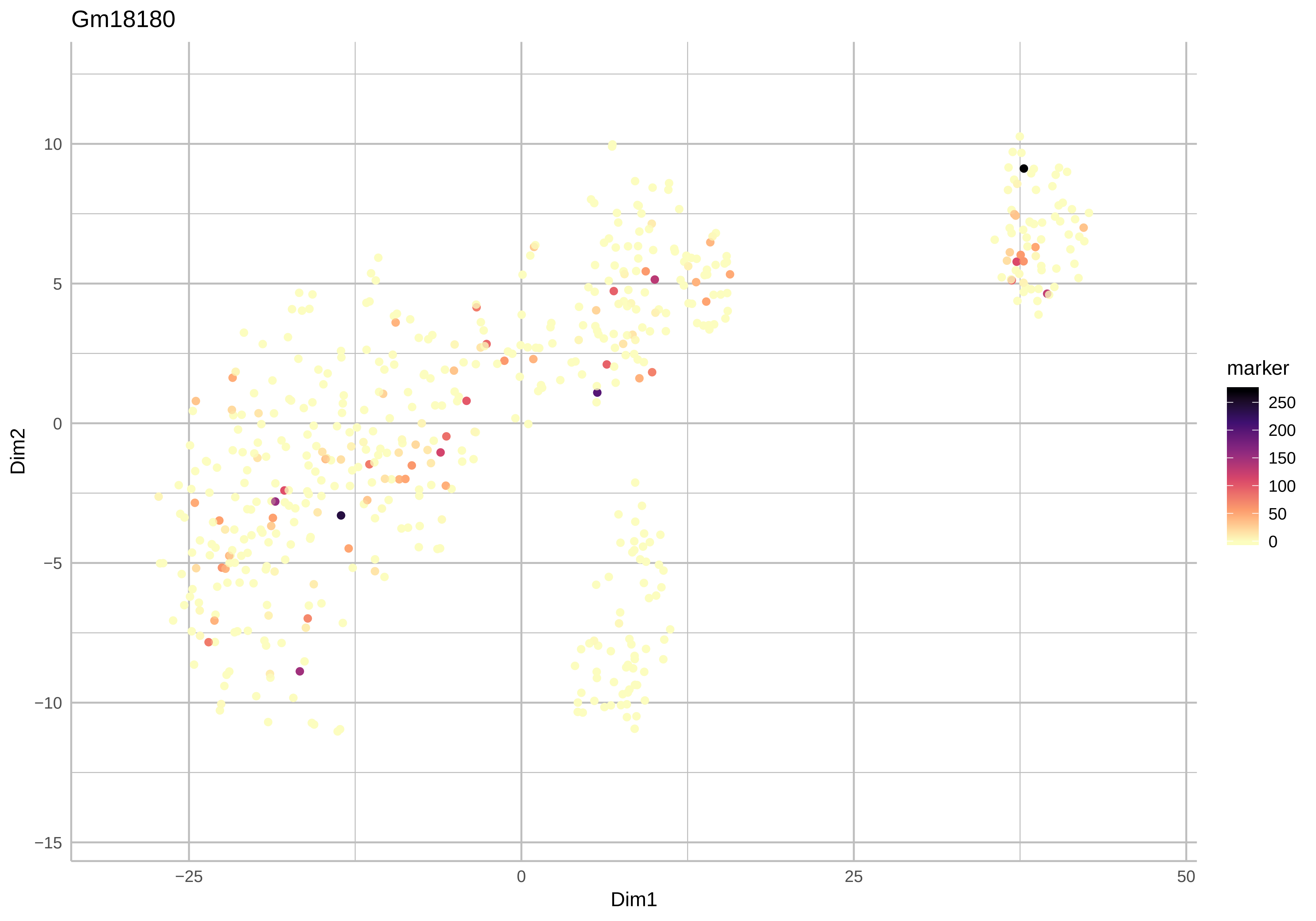

Gm18180
Dim2
Dim1
10
5
0
−5
−10
−15
−25
0
25
50
marker
250
200
150
100
50
0

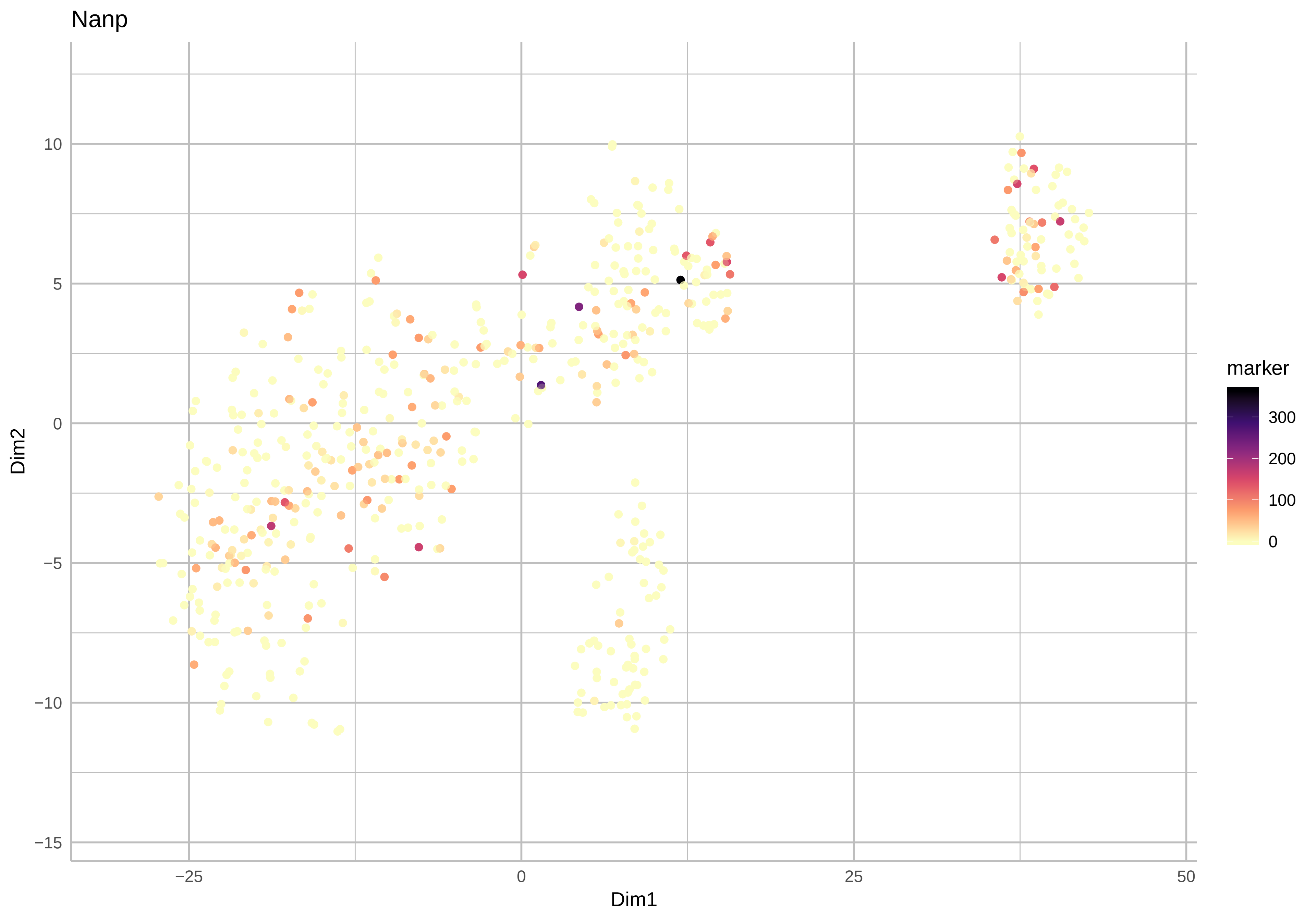

Nanp
Dim2
Dim1
10
5
0
−5
−10
−15
−25
0
25
50
marker
300
200
100
0

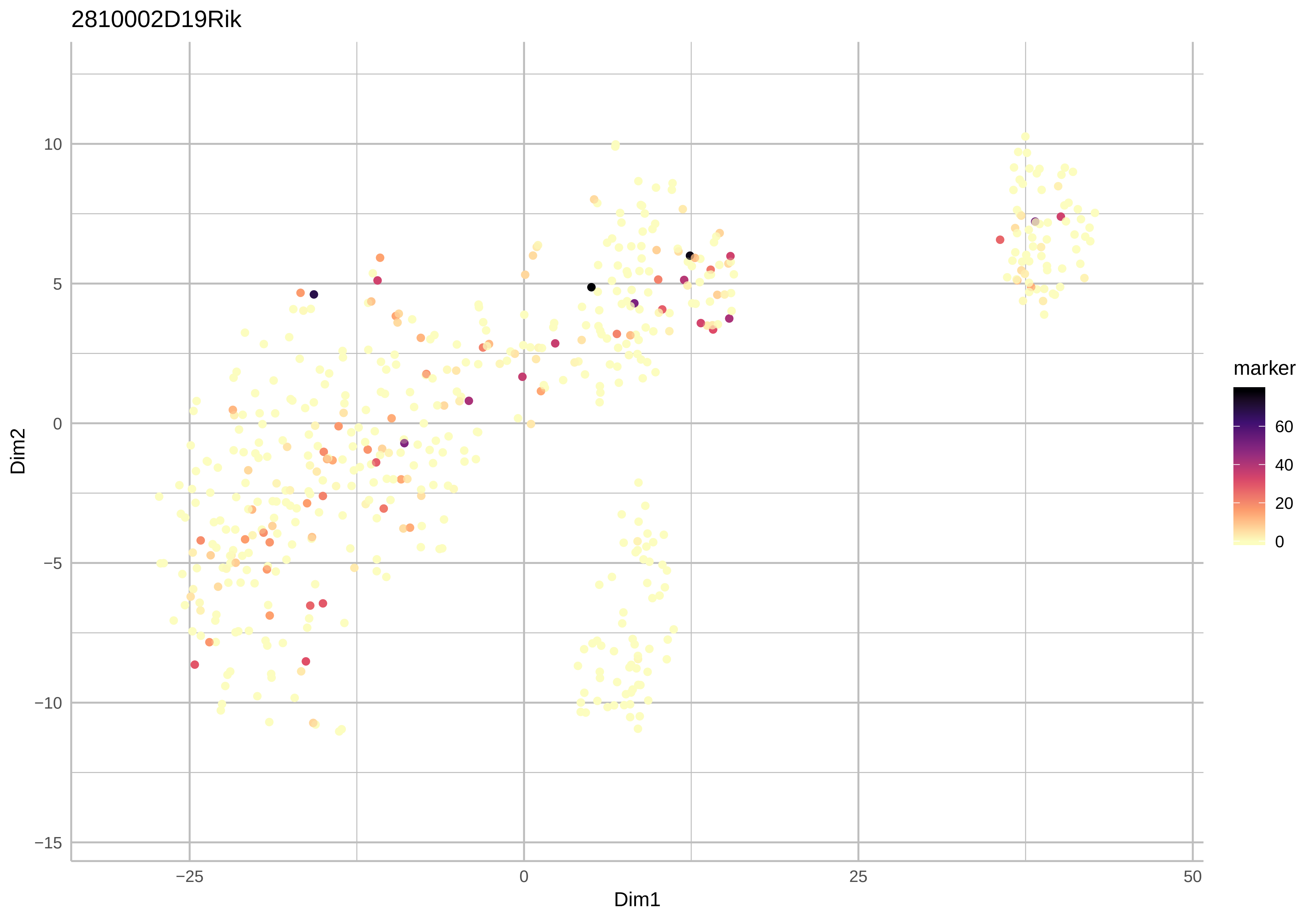
2810002D19Rik
Dim2
Dim1
marker
60
40
20
0
10
5
0
−5
−10
−15
−25
0
25
50

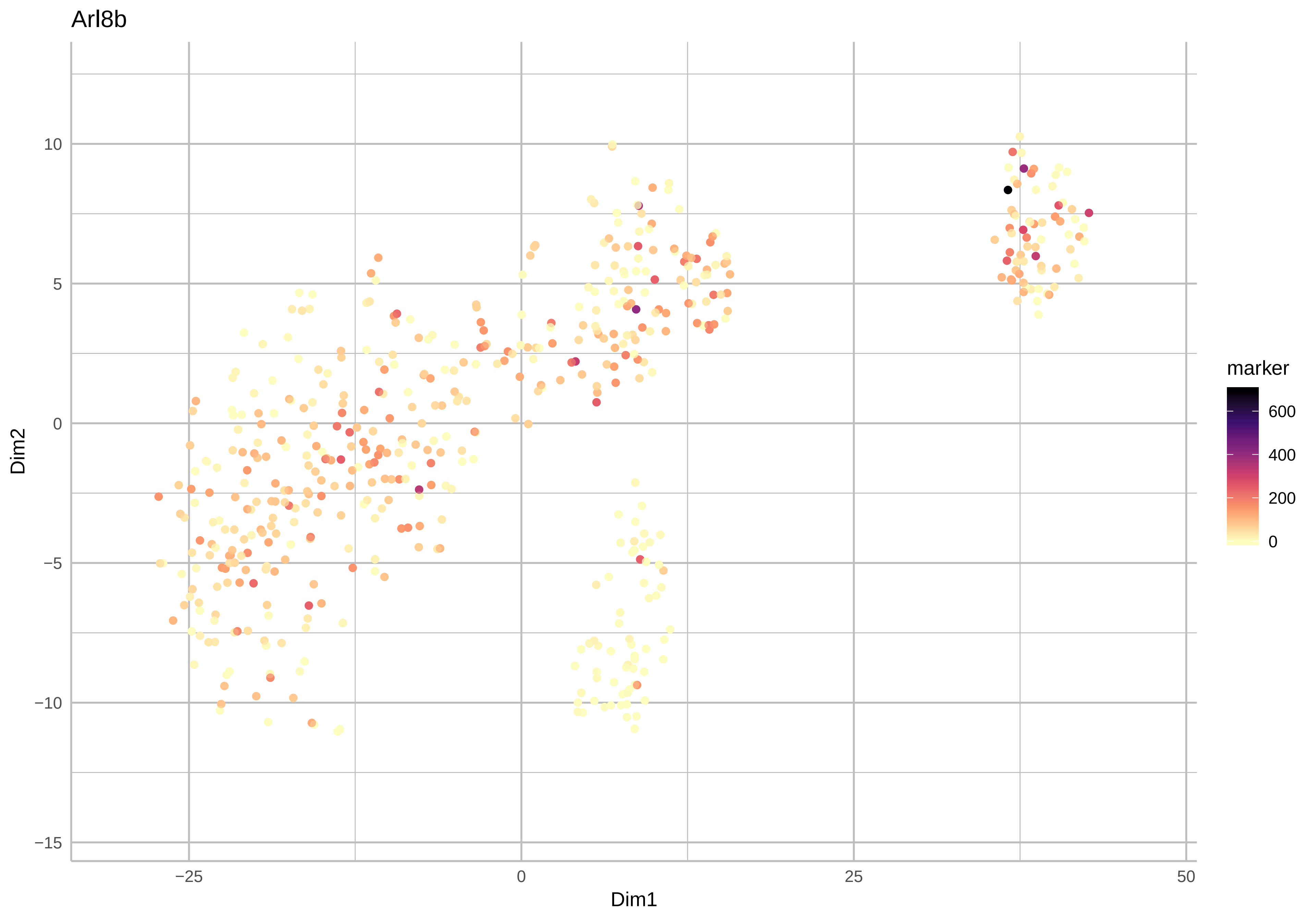
Arl8b
10
5
0
−5
−10
−15
−25
0
25
50
Dim1
Dim2
marker
600
400
200
0

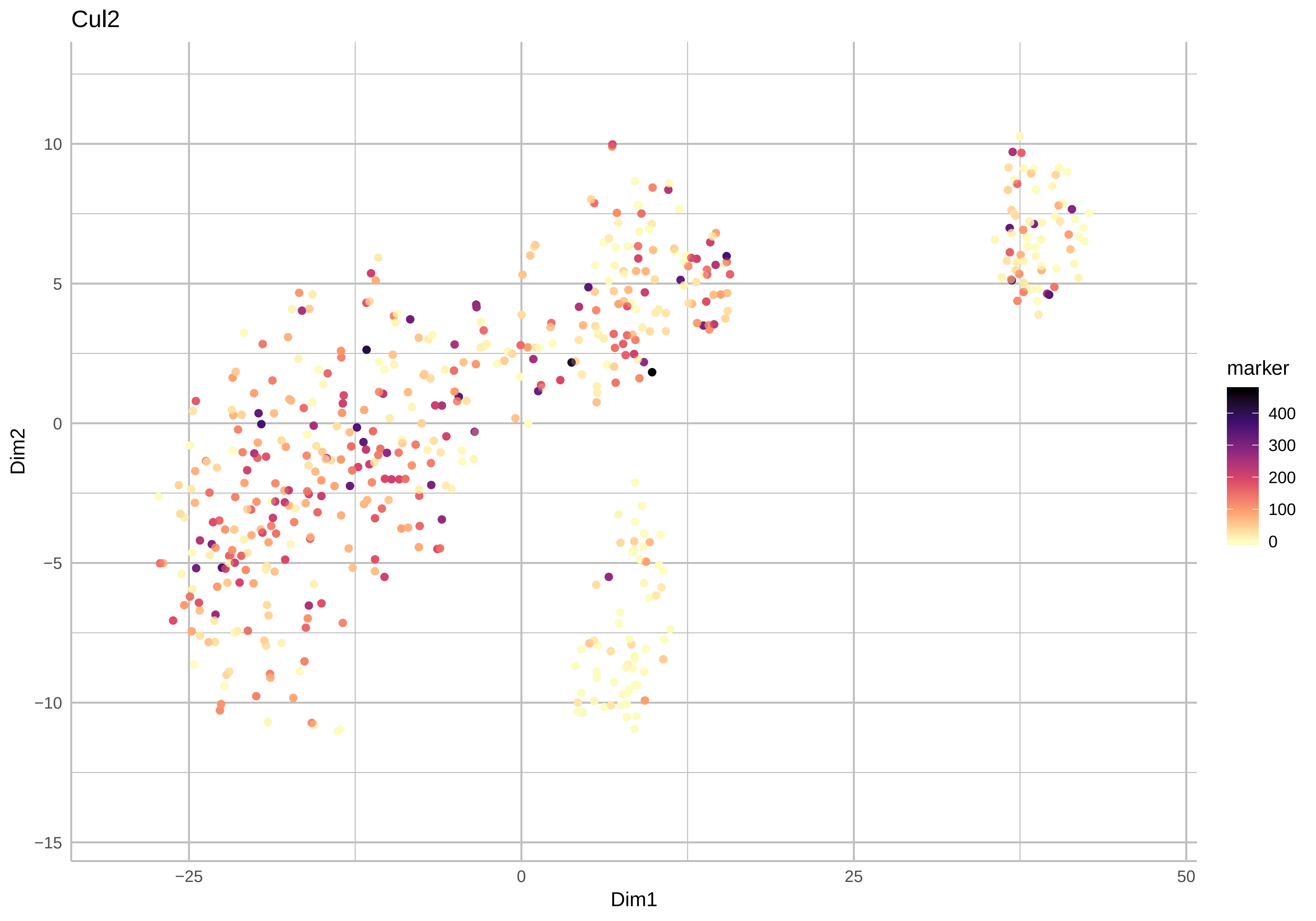
Cul2
Dim2
Dim1
10
5
0
−5
−10
−15
−25
0
25
50
marker
400
300
200
100
0

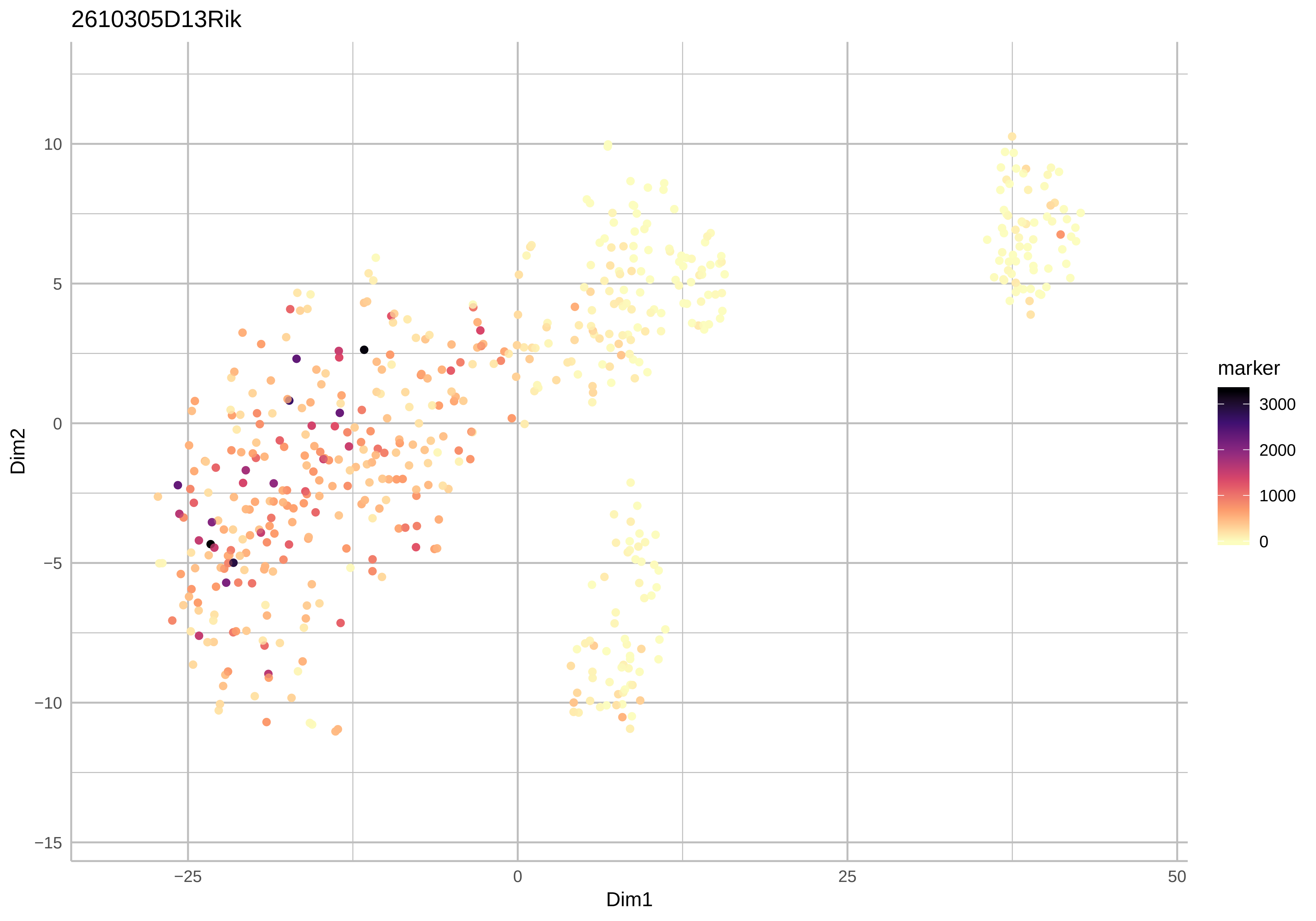
2610305D13Rik
Dim2
Dim1
10
5
0
−5
−10
−15
−25
0
25
50
marker
3000
2000
1000
0

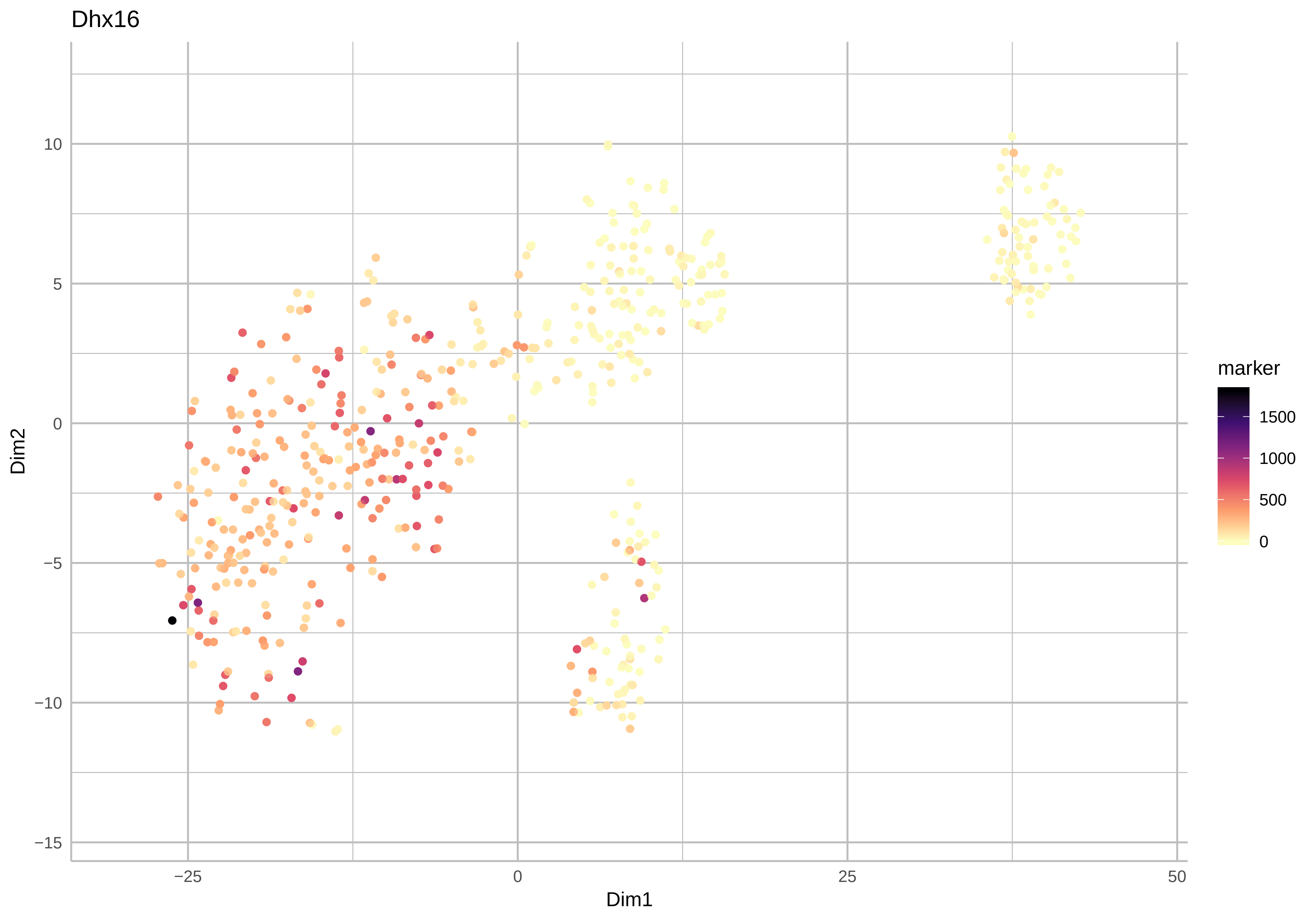
Dhx16
Dim2
Dim1
10
5
0
−5
−10
−15
−25
0
25
50
marker
1500
1000
500
0

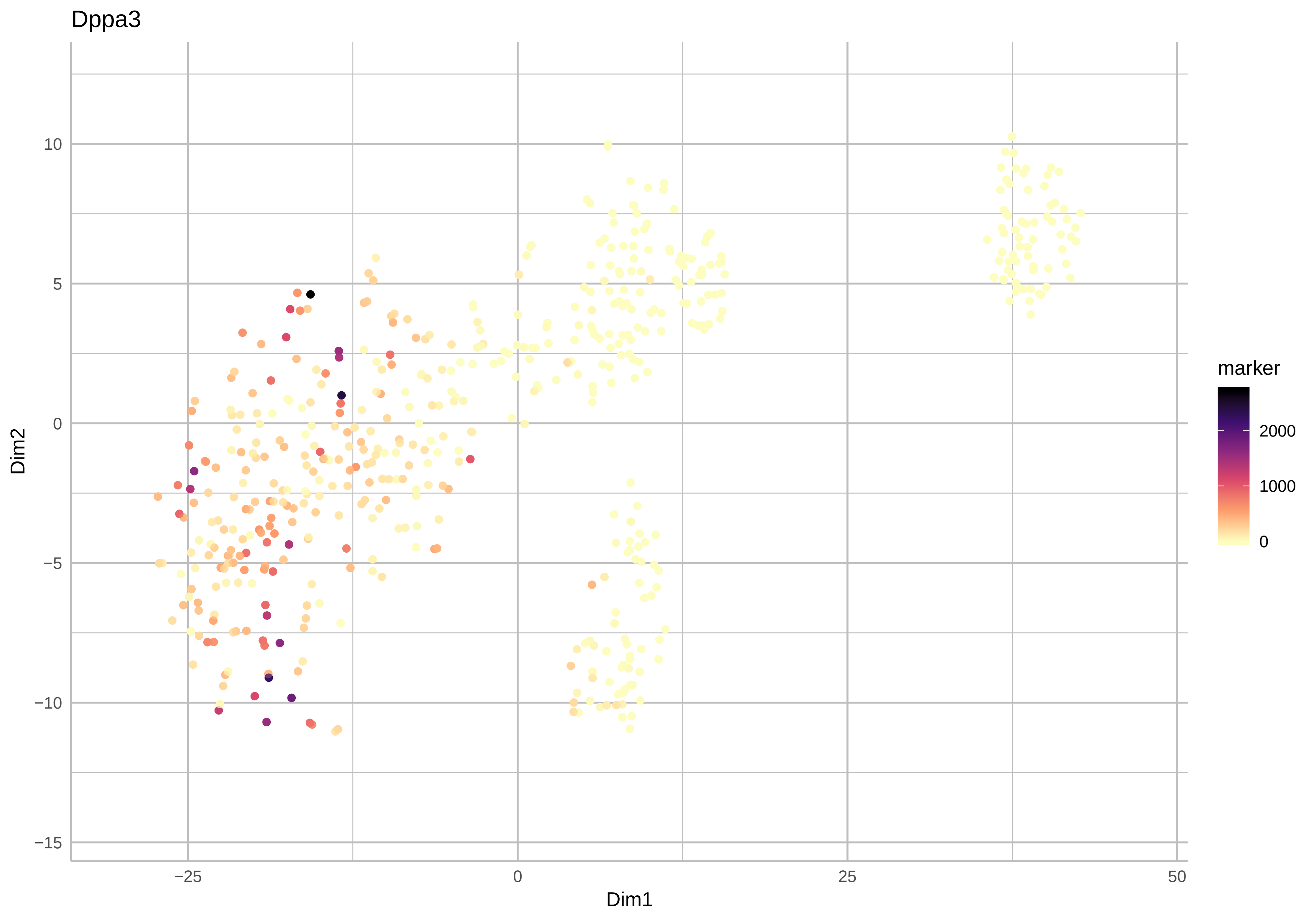
Dppa3
Dim2
Dim1
marker
2000
1000
0
10
5
0
−5
−10
−15
−25
0
25
50

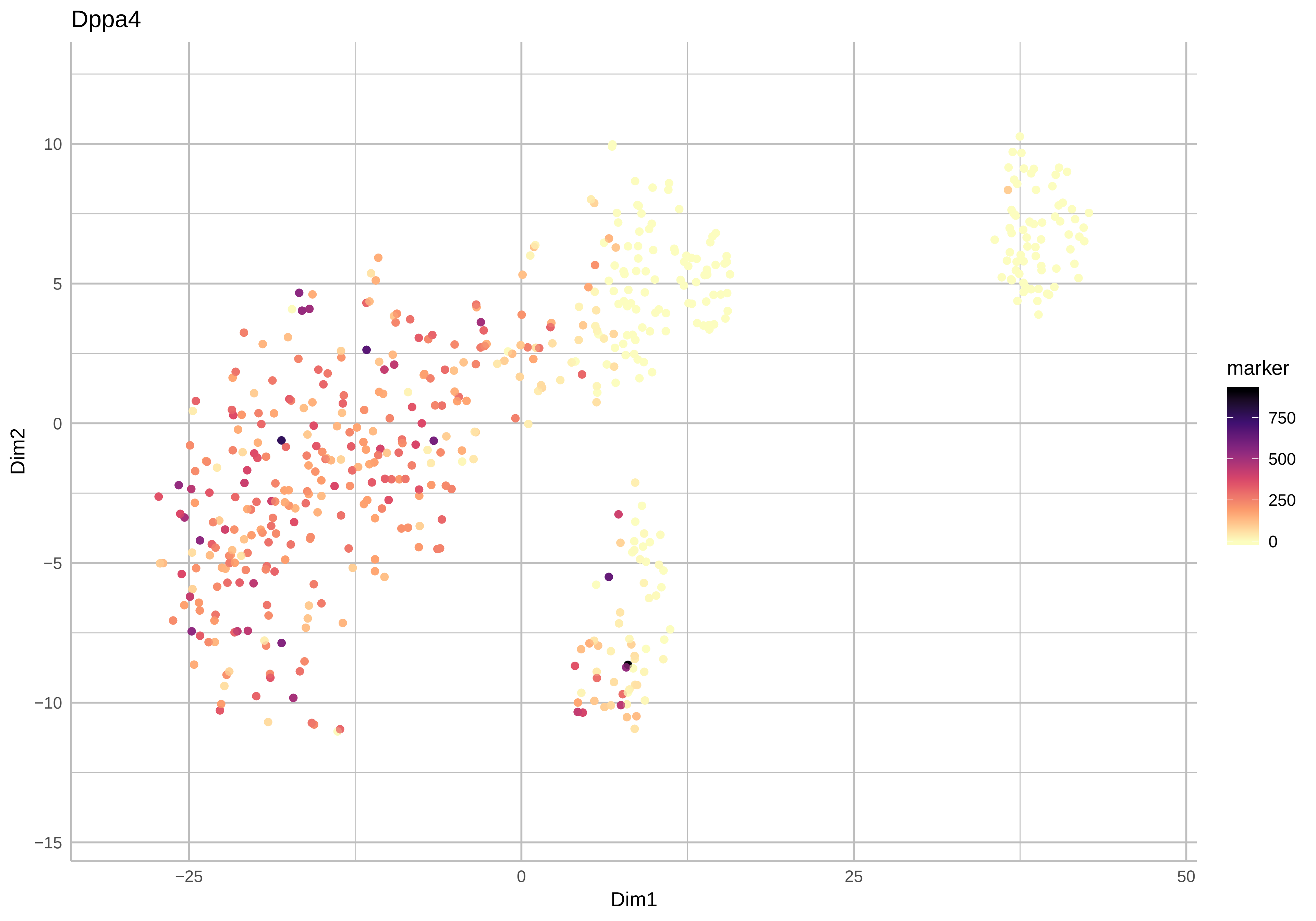

Dppa4
10
5
0
−5
−10
−15
Dim2
−25
0
25
50
Dim1
marker
750
500
250
0

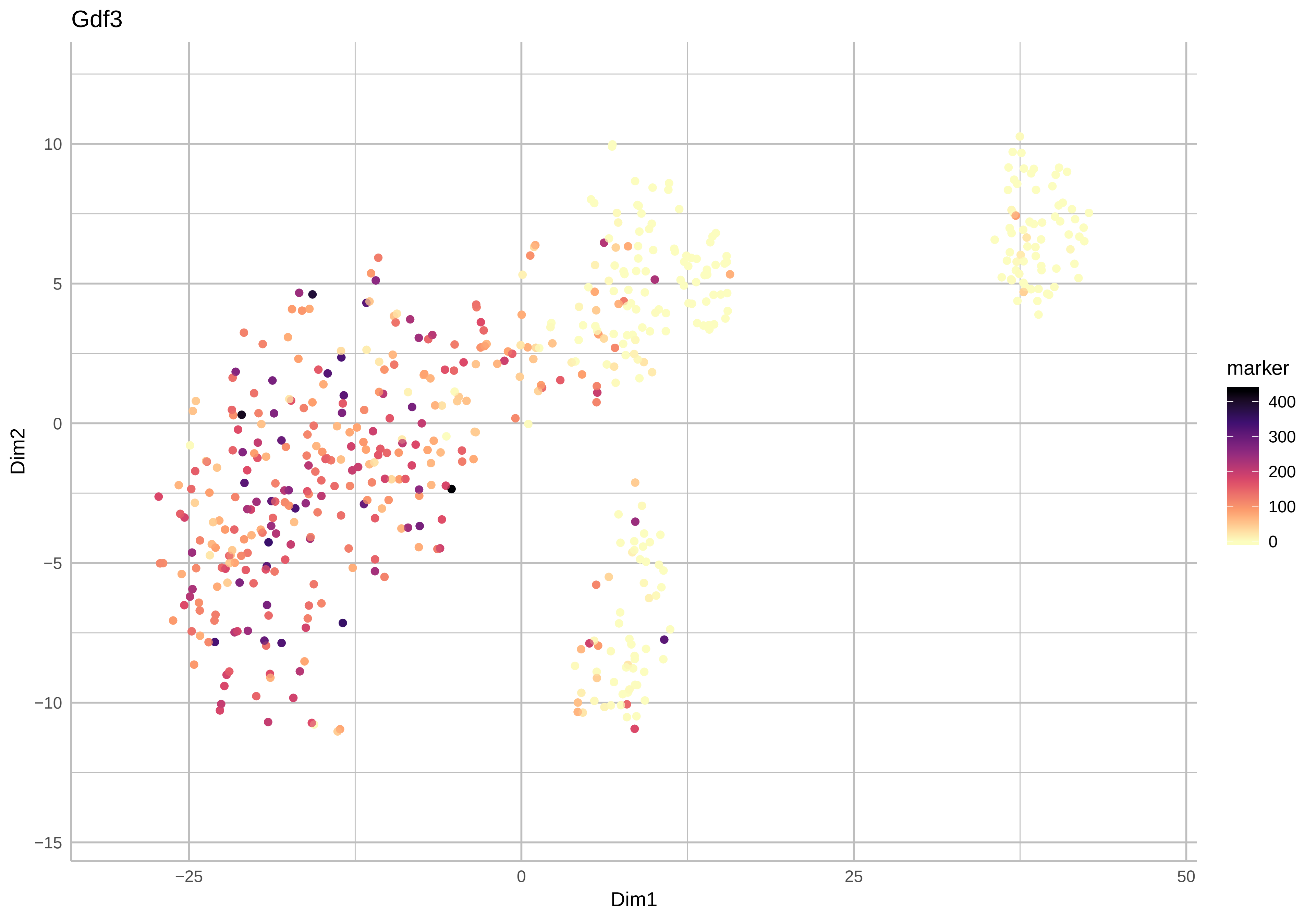

Gdf3
Dim2
Dim1
10
5
0
−5
−10
−15
−25
0
25
50
marker
400
300
200
100
0

Gm13145

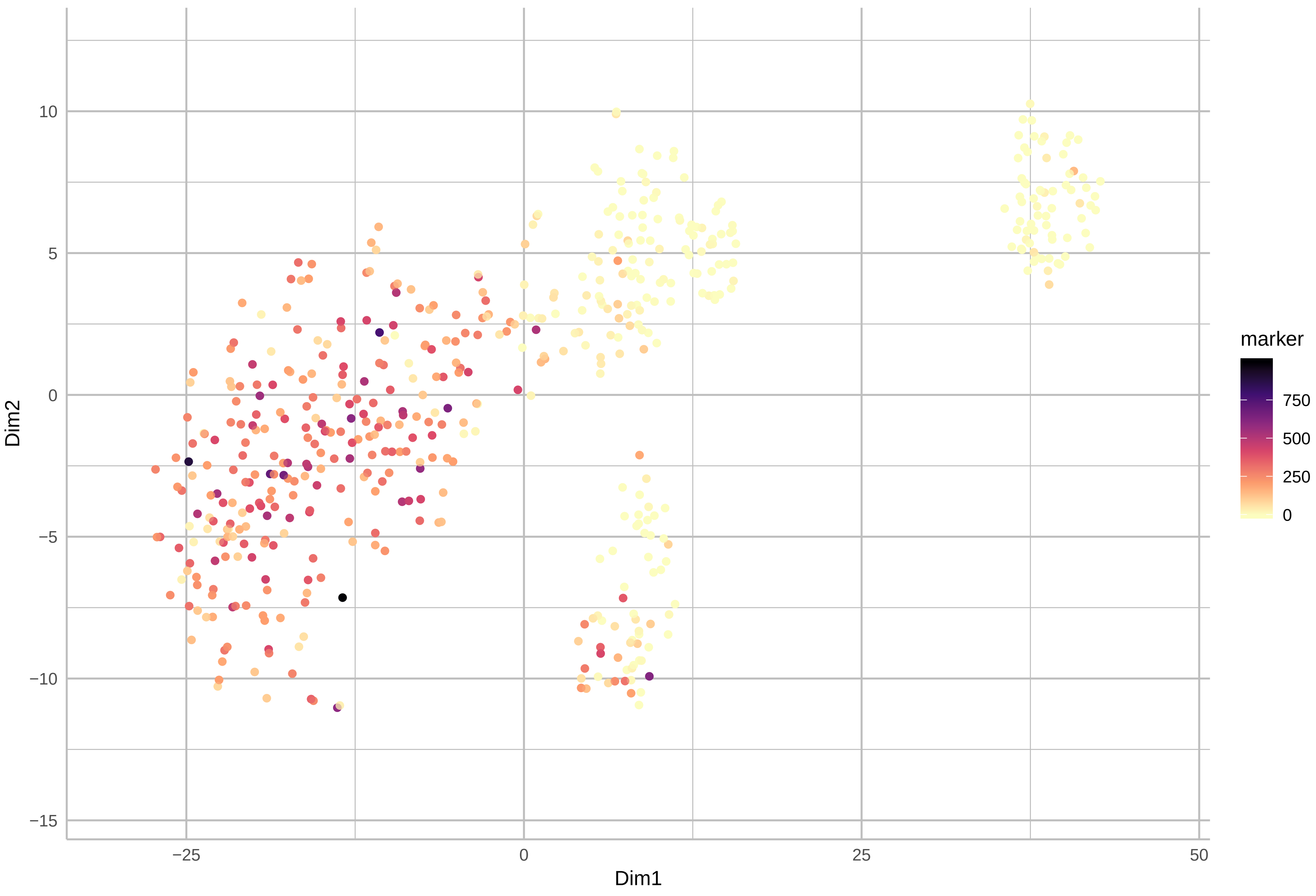

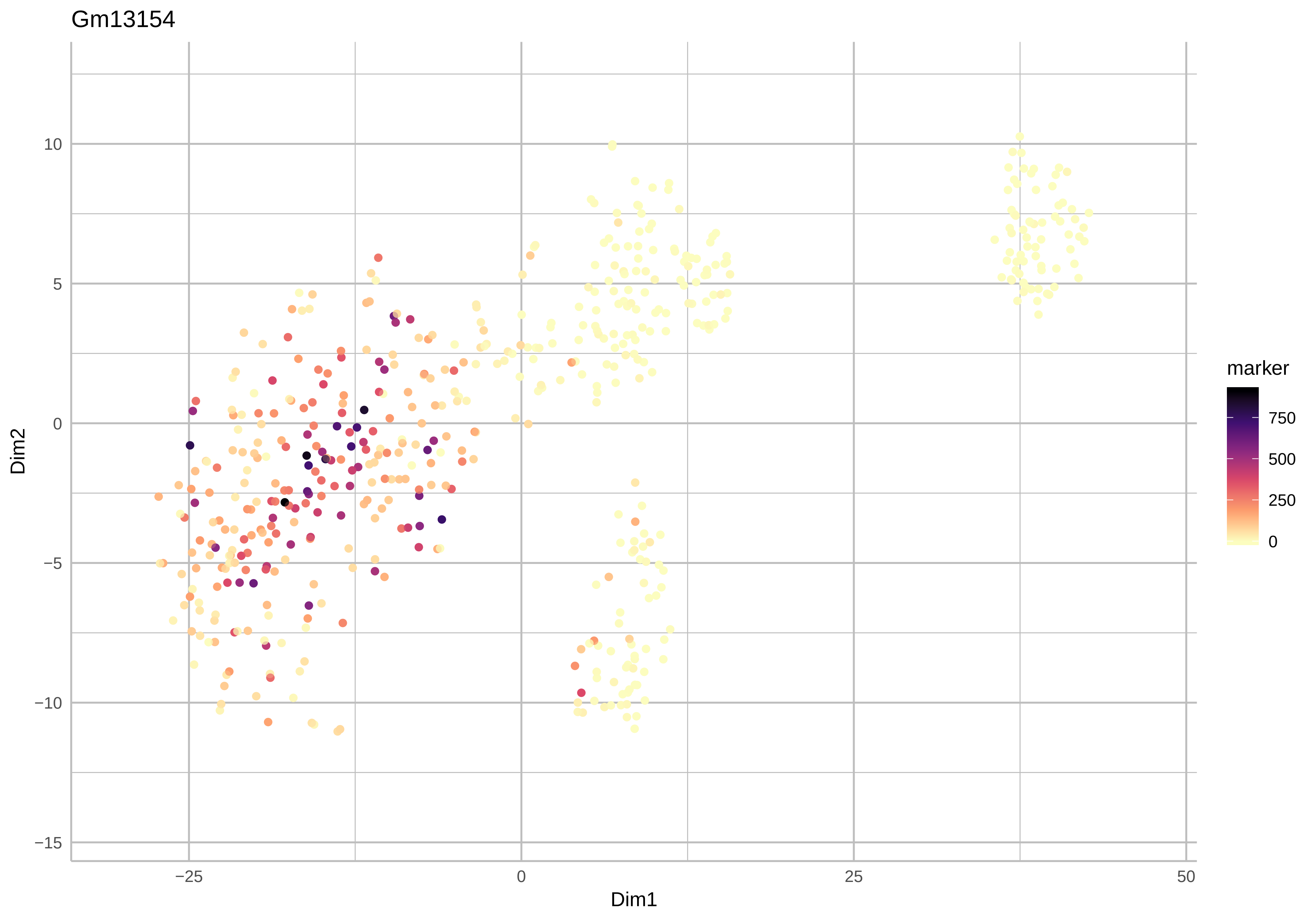

Gm13154
Dim2
Dim1
10
5
0
−5
−10
−15
−25
0
25
50
marker
750
500
250
0

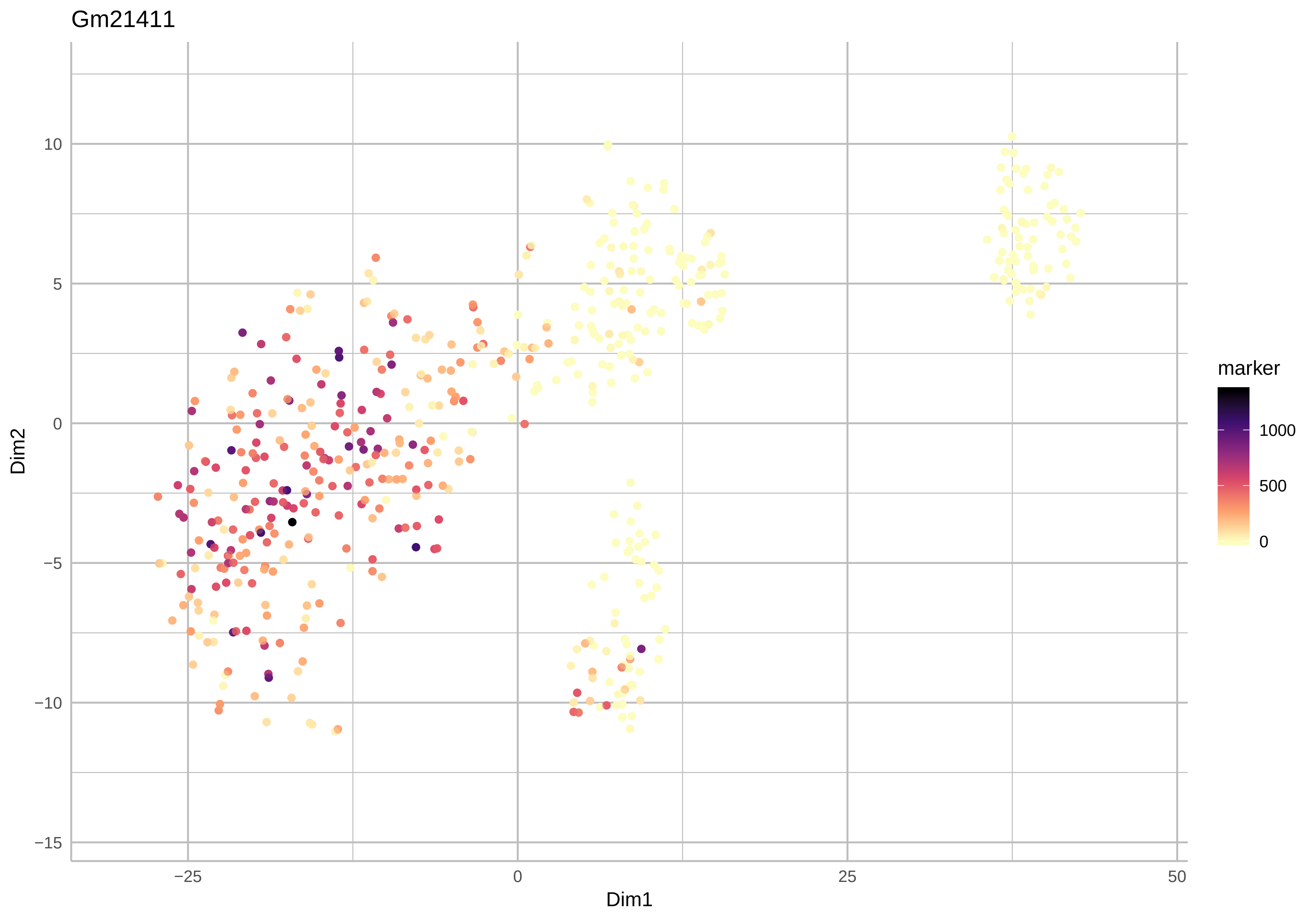

Gm21411
Dim2
Dim1
marker
1000
500
0
10
5
0
−5
−10
−15
−25
0
25
50

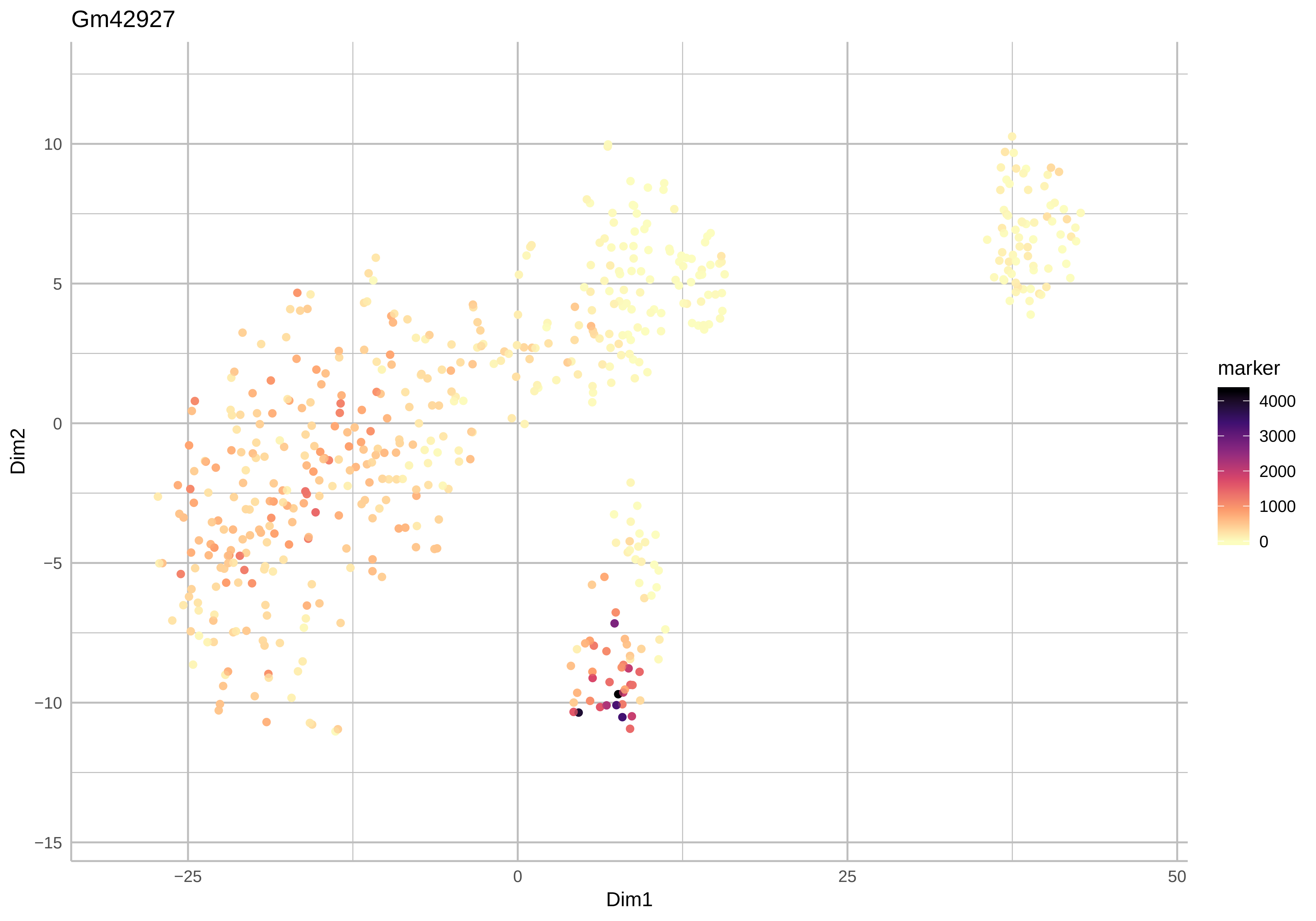

Gm42927
10
5
0
−5
−10
−15
Dim2
−25
0
25
50
Dim1
marker
4000
3000
2000
1000
0

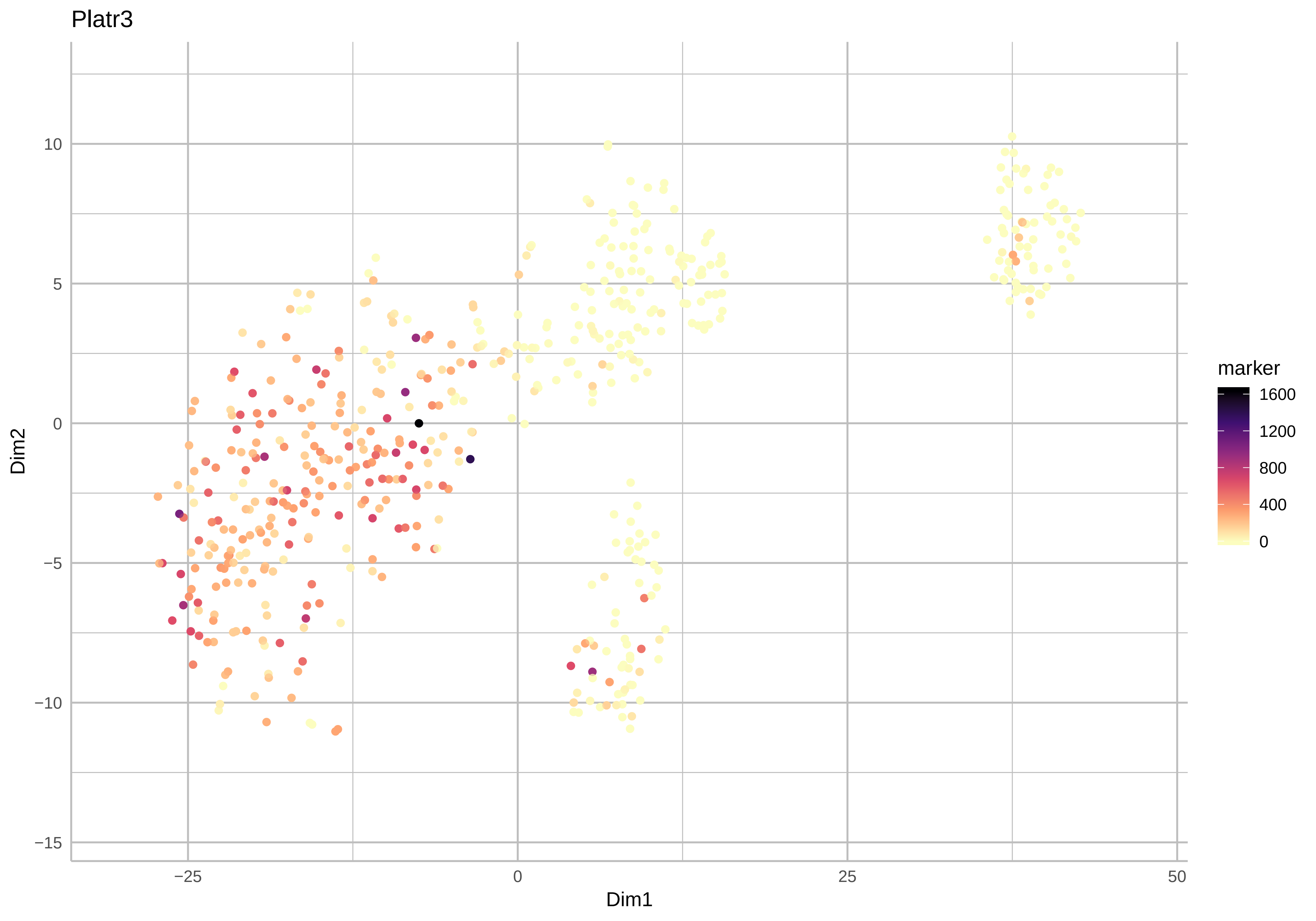

Platr3
Dim2
Dim1
marker
1600
1200
800
400
0
10
5
0
−5
−10
−15
−25
0
25
50

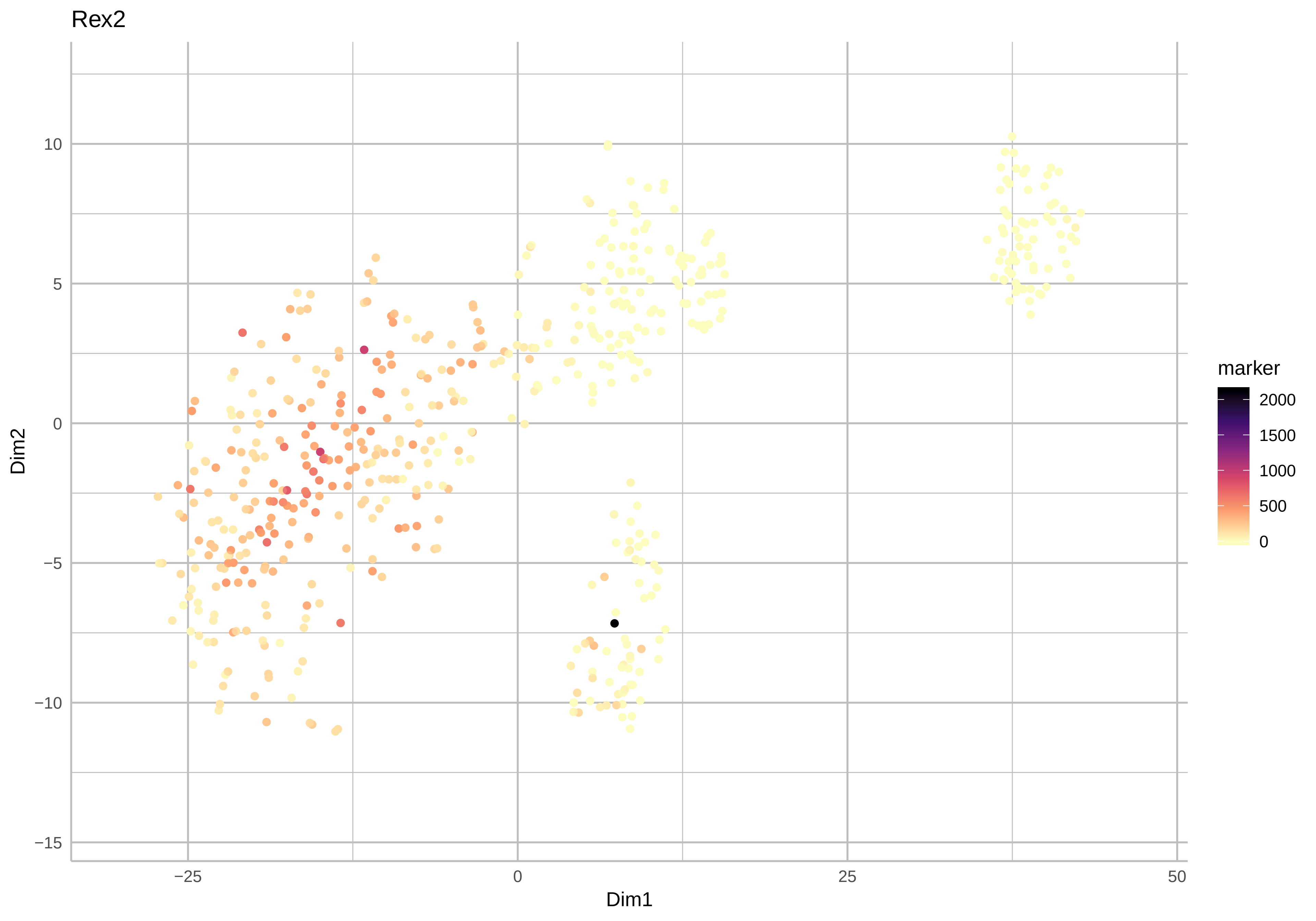
Rex2
Dim2
Dim1
10
5
0
−5
−10
−15
−25
0
25
50
marker
2000
1500
1000
500
0

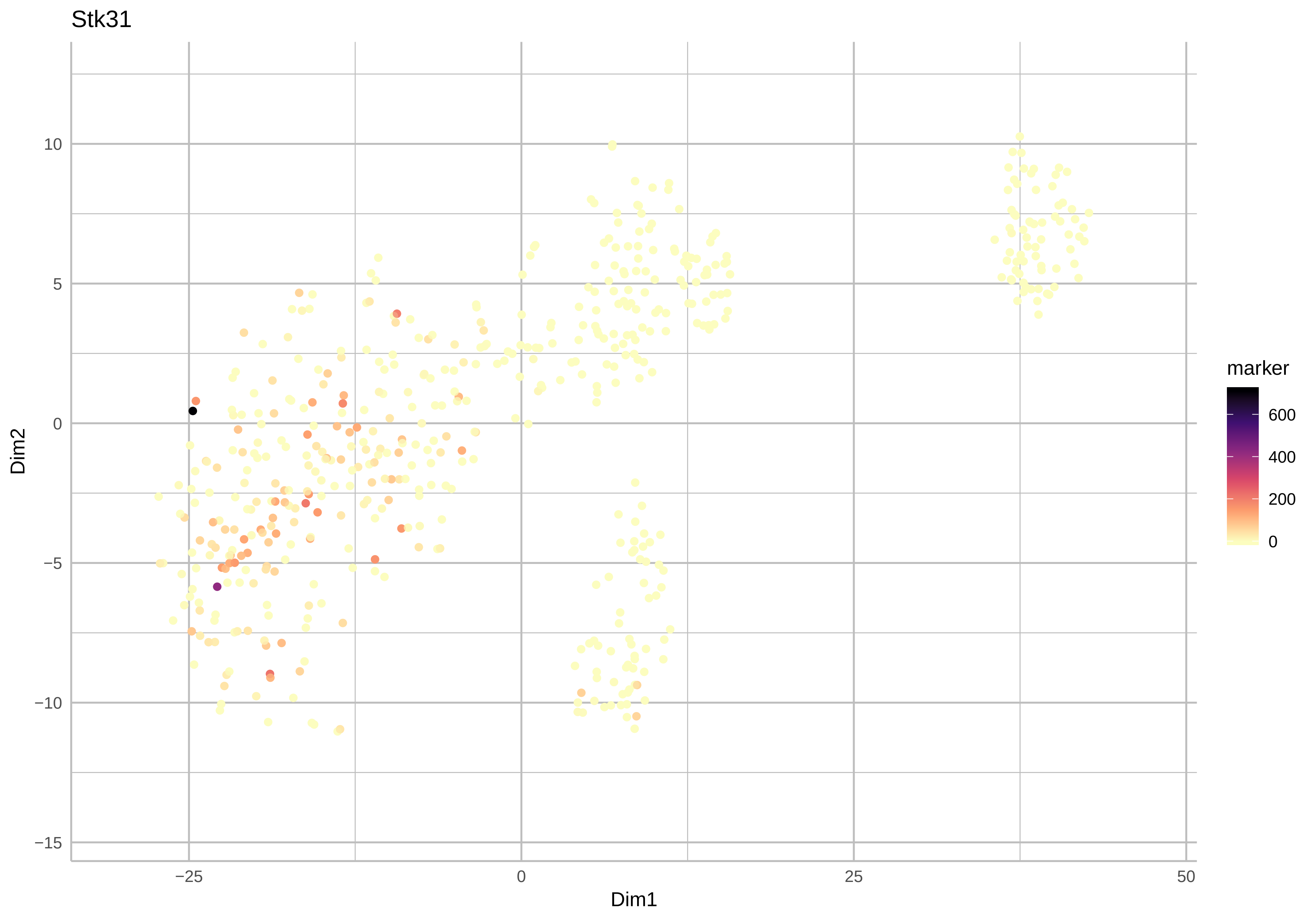
Stk31
Dim2
Dim1
10
5
0
−5
−10
−15
−25
0
25
50
marker
600
400
200
0

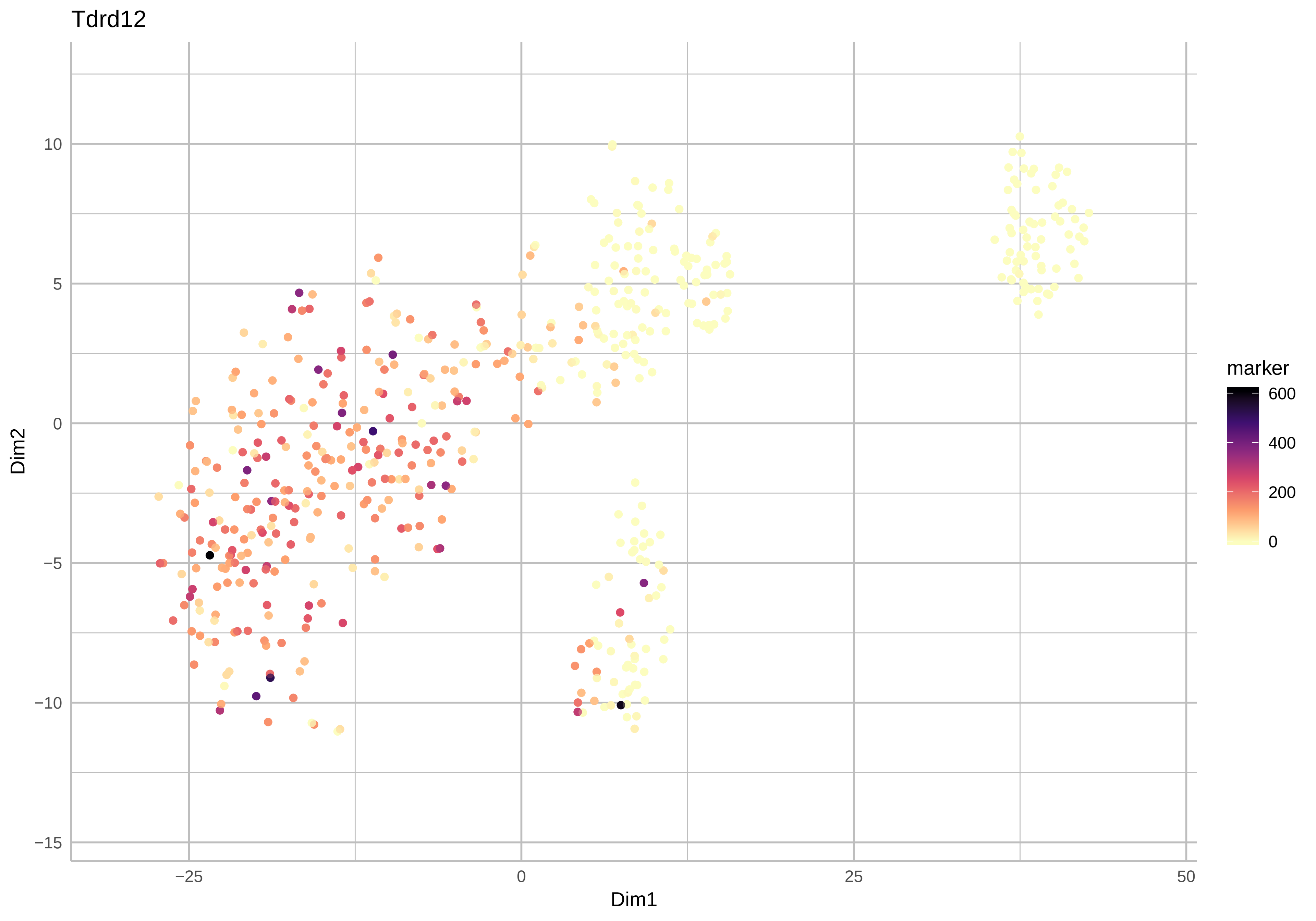

Tdrd12
Dim2
Dim1
marker
600
400
200
0
10
5
0
−5
−10
−15
−25
0
25
50

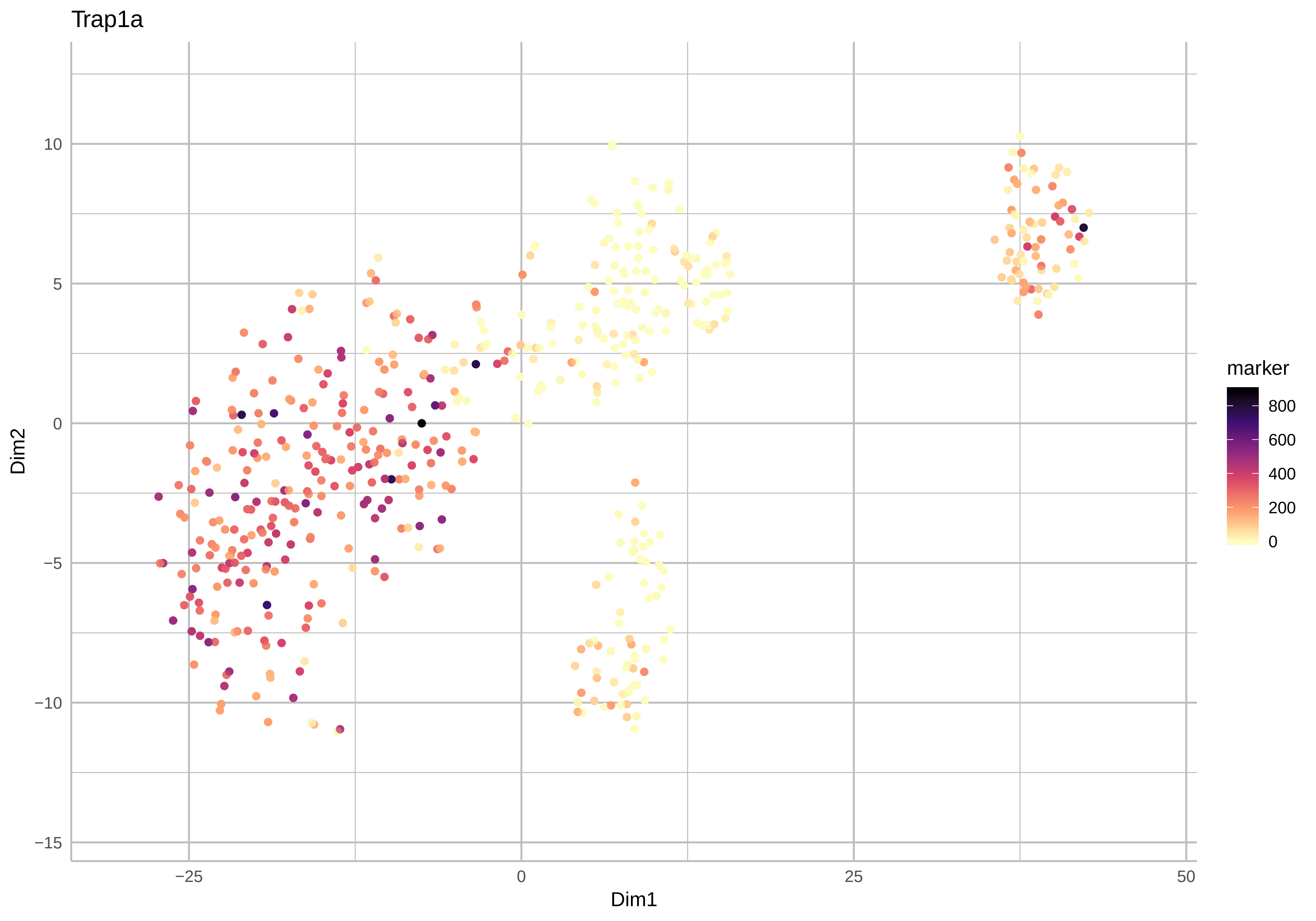

Trap1a
10
5
0
−5
−10
−15
Dim2
−25
0
25
50
Dim1
marker
800
600
400
200
0

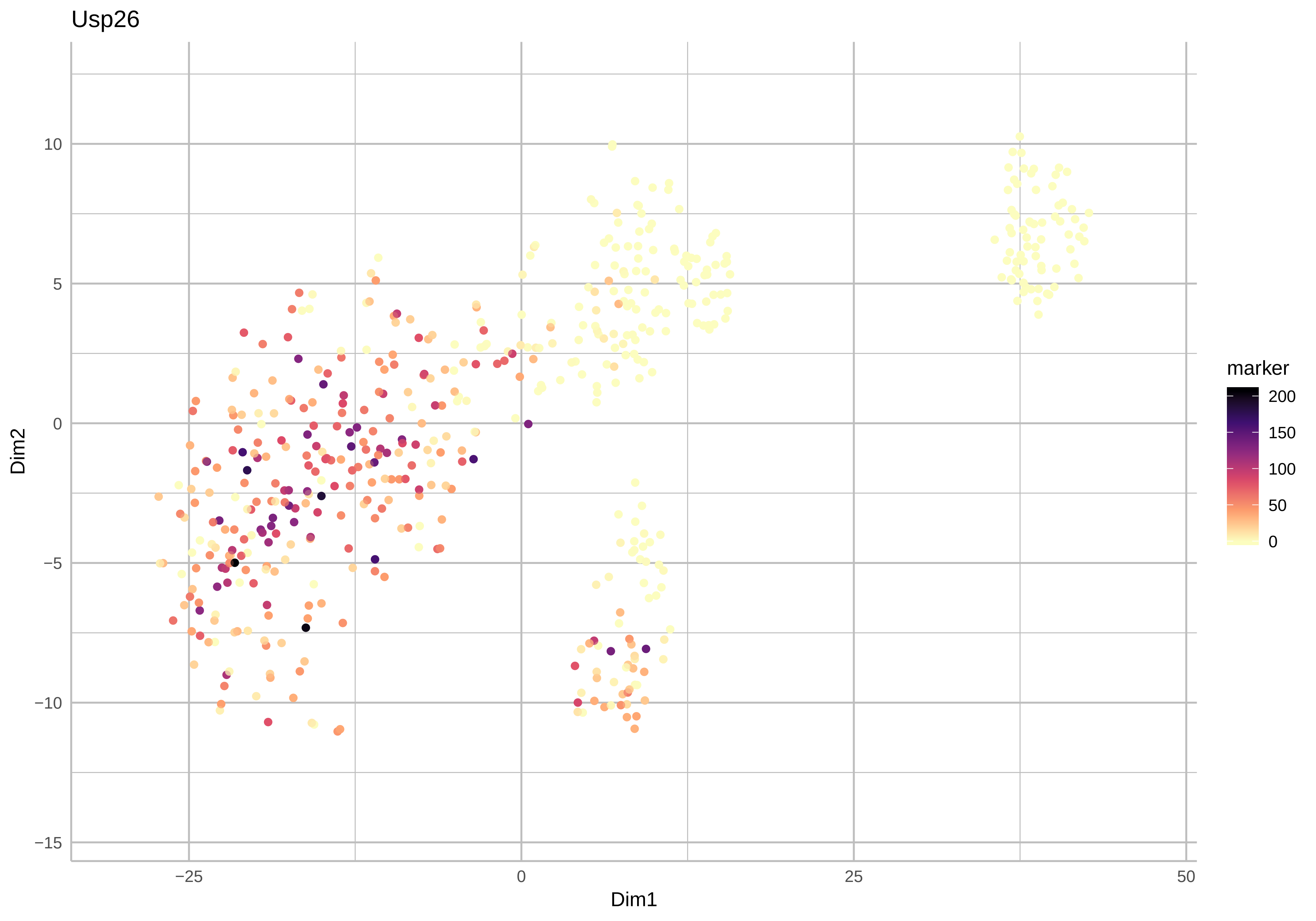

Usp26
Dim2
Dim1
10
5
0
−5
−10
−15
−25
0
25
50
marker
200
150
100
50
0

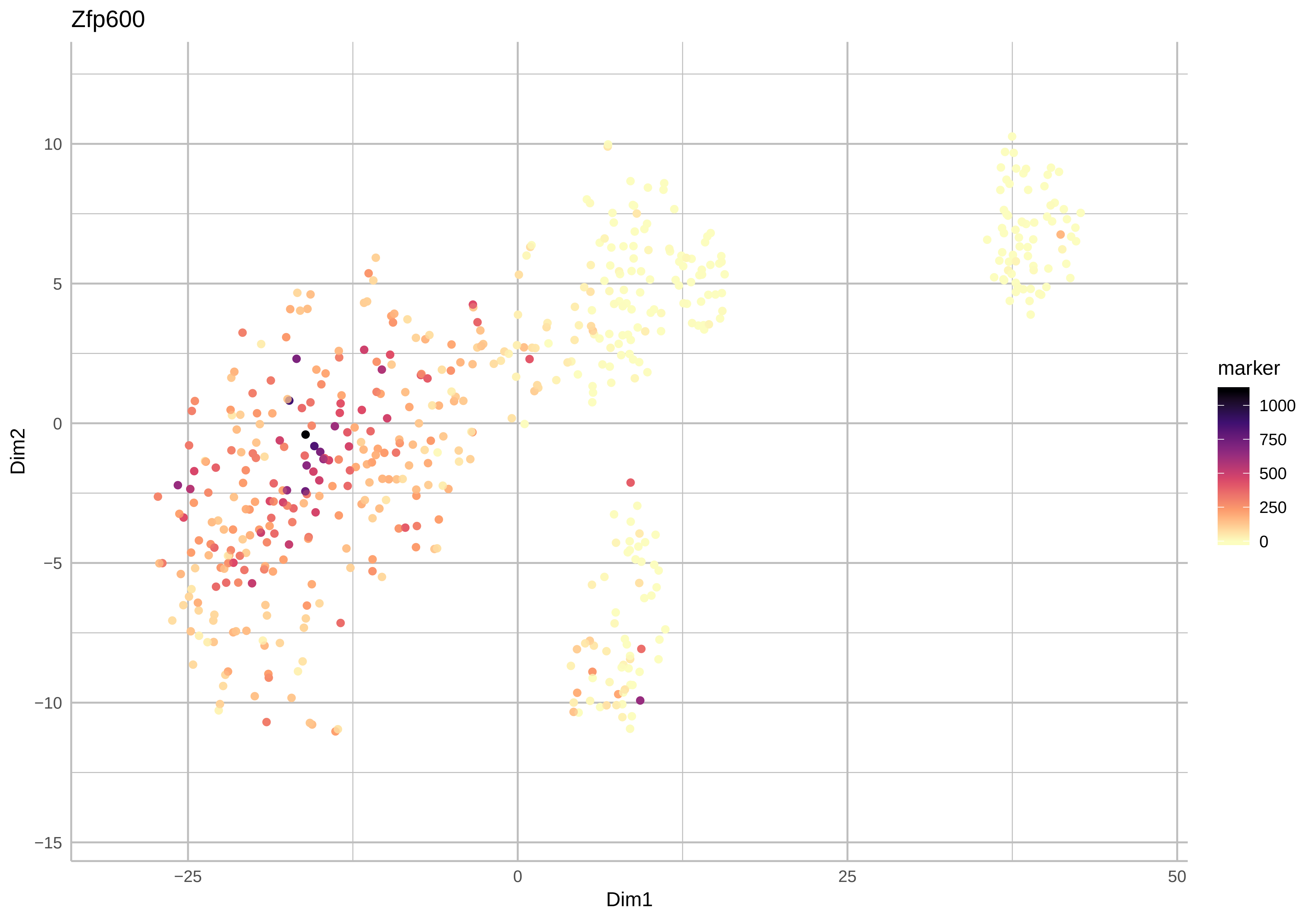

Zfp600
10
5
0
−5
−10
−15
Dim2
−25
0
25
50
Dim1
marker
1000
750
500
250
0

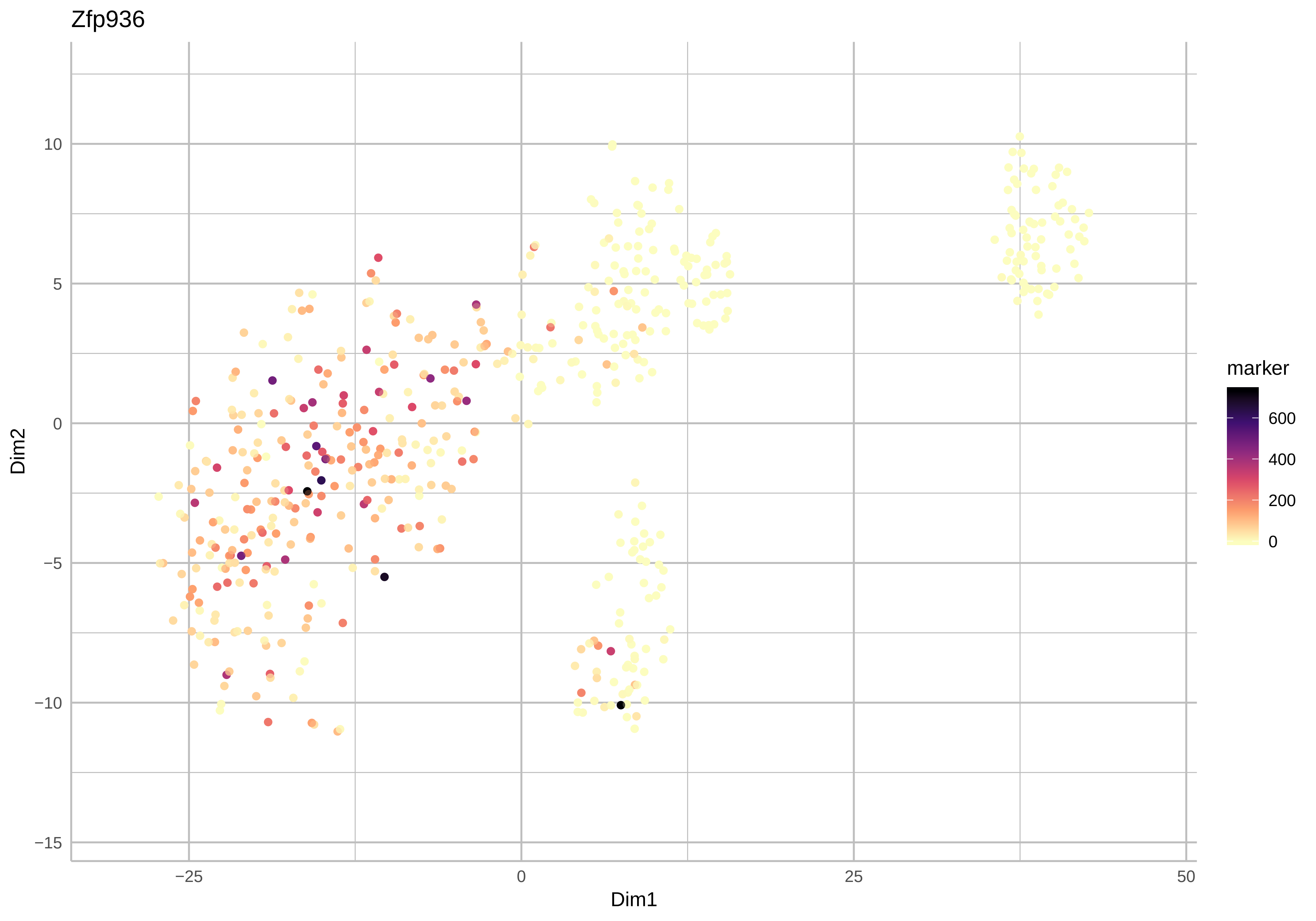

Zfp936
Dim2
10
5
0
−5
−10
−15
Dim1
−25
0
25
50
marker
600
400
200
0

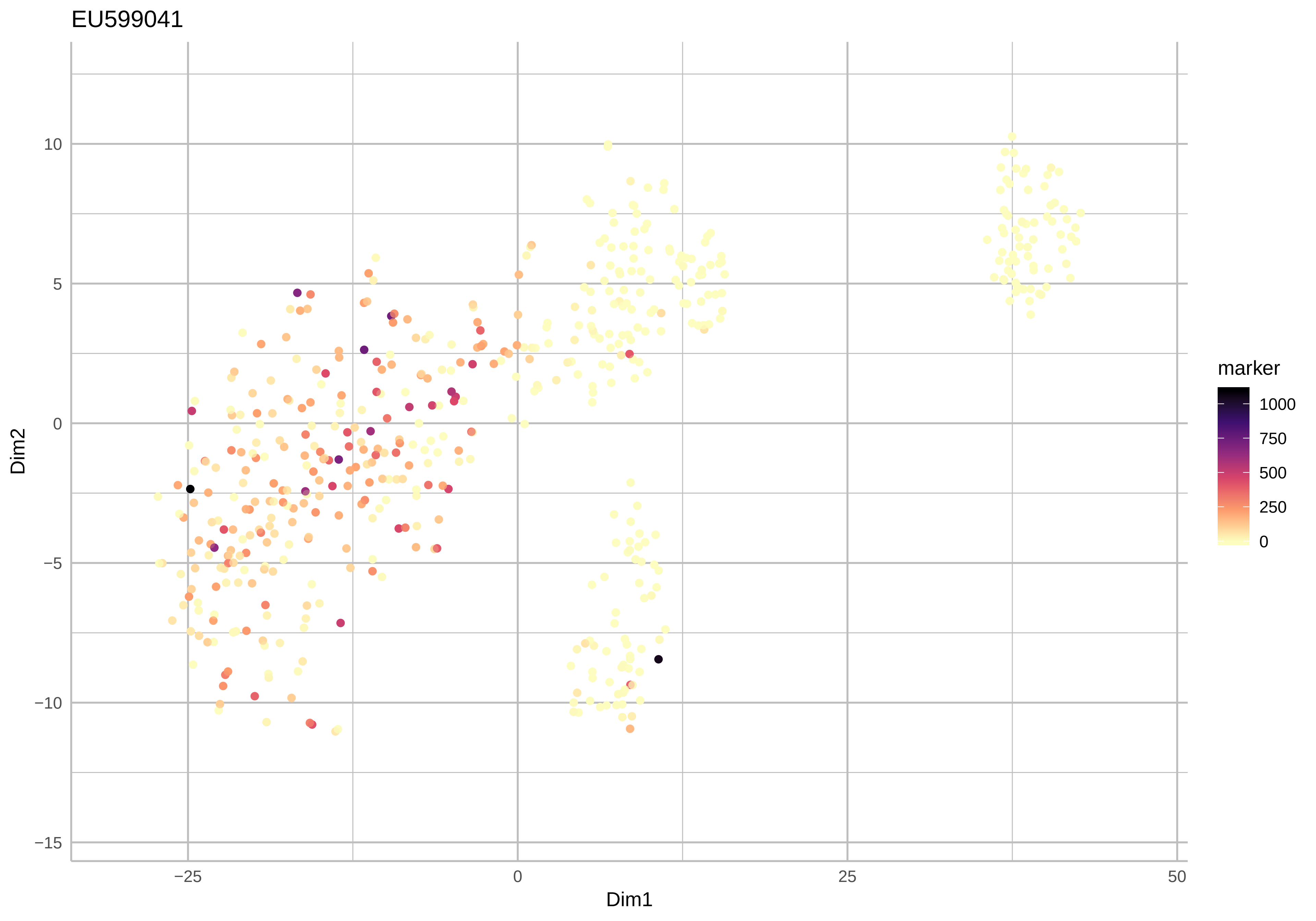
EU599041
Dim2
Dim1
10
5
0
−5
−10
−15
−25
0
25
50
marker
1000
750
500
250
0

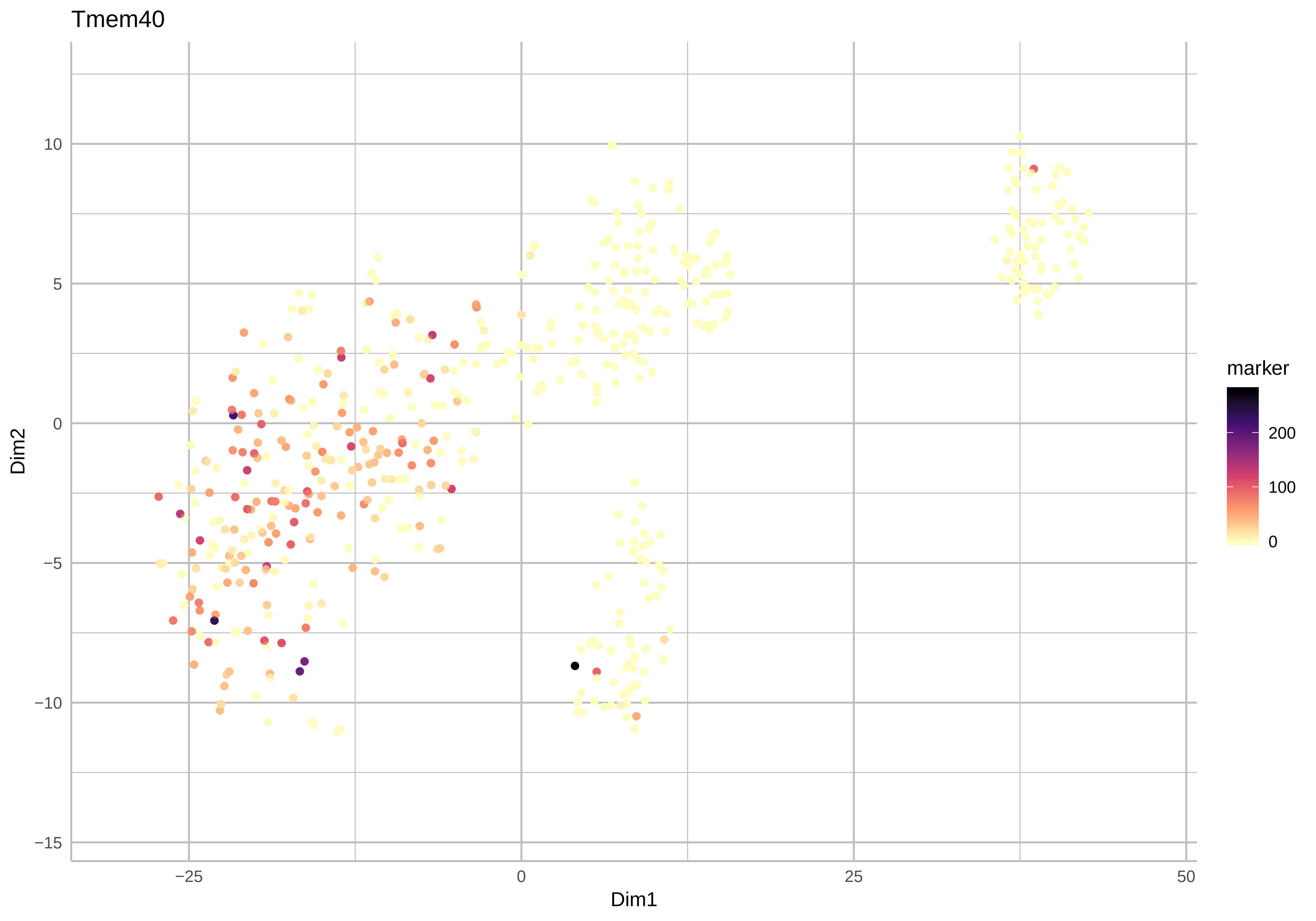

Tmem40
Dim2
10
5
0
−5
−10
−15
Dim1
−25
0
25
50
marker
200
100
0

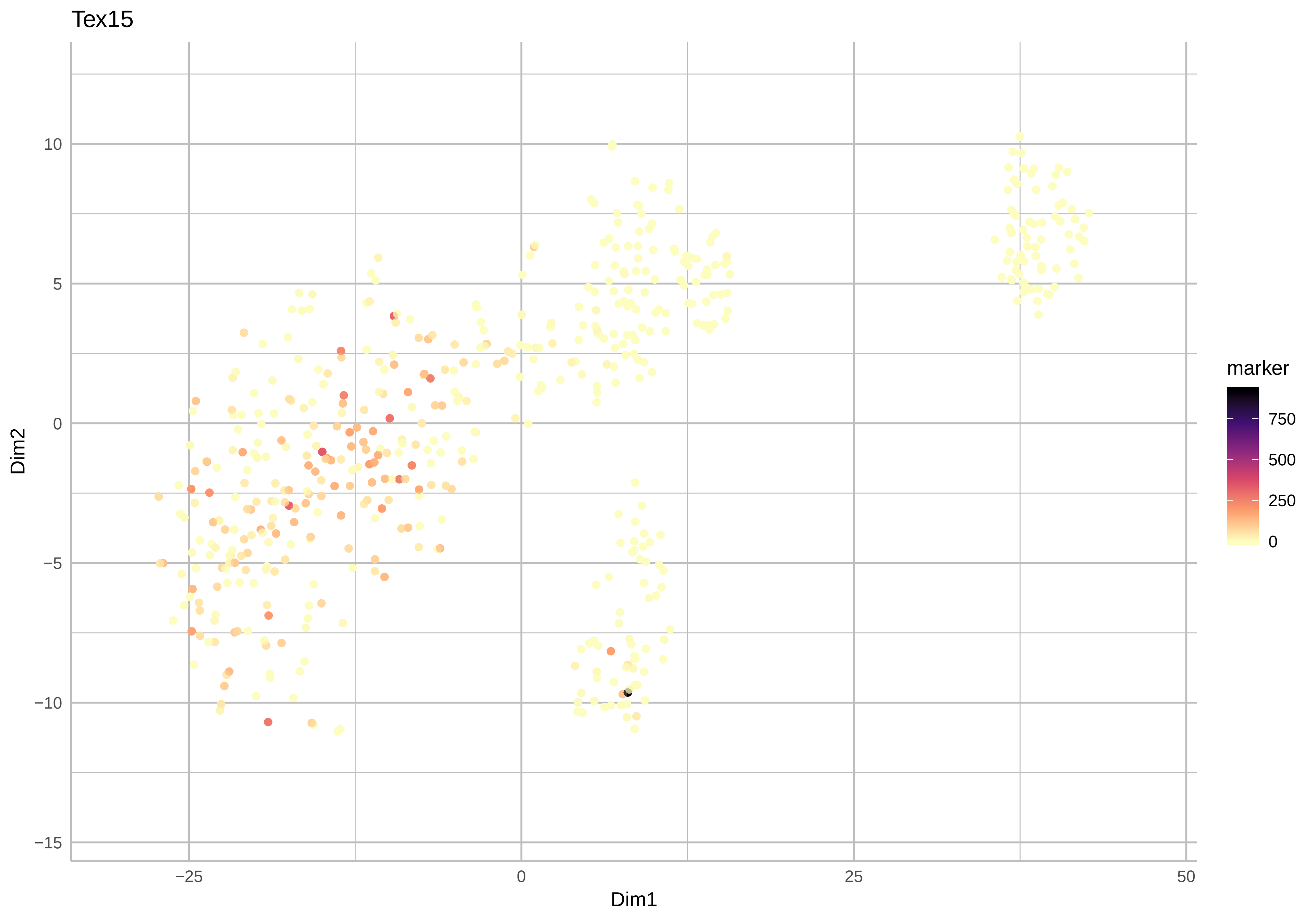

Tex15
Dim2
Dim1
10
5
0
−5
−10
−15
−25
0
25
50
marker
750
500
250
0

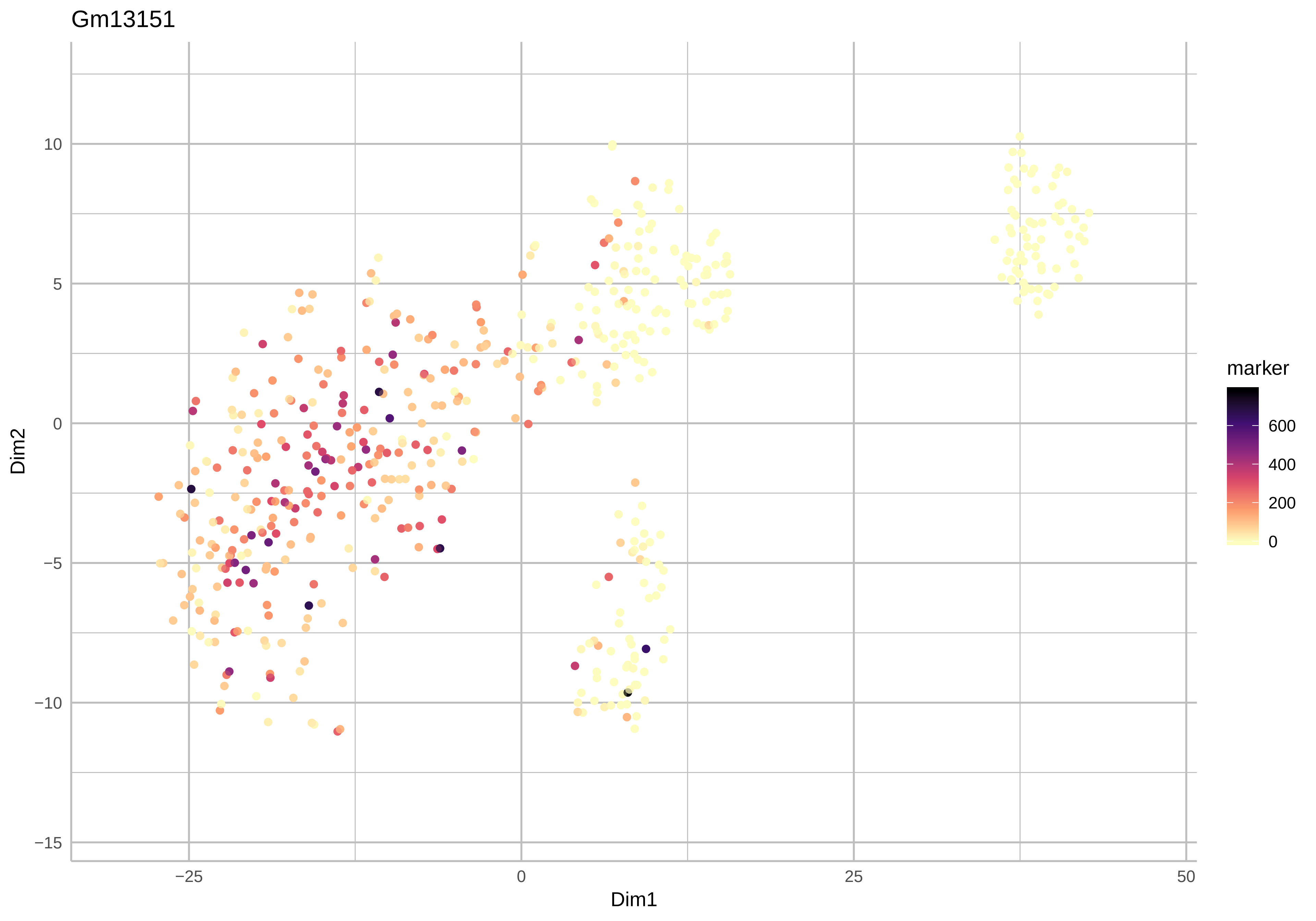
Gm13151
Dim2
Dim1
10
5
0
−5
−10
−15
−25
0
25
50
marker
600
400
200
0

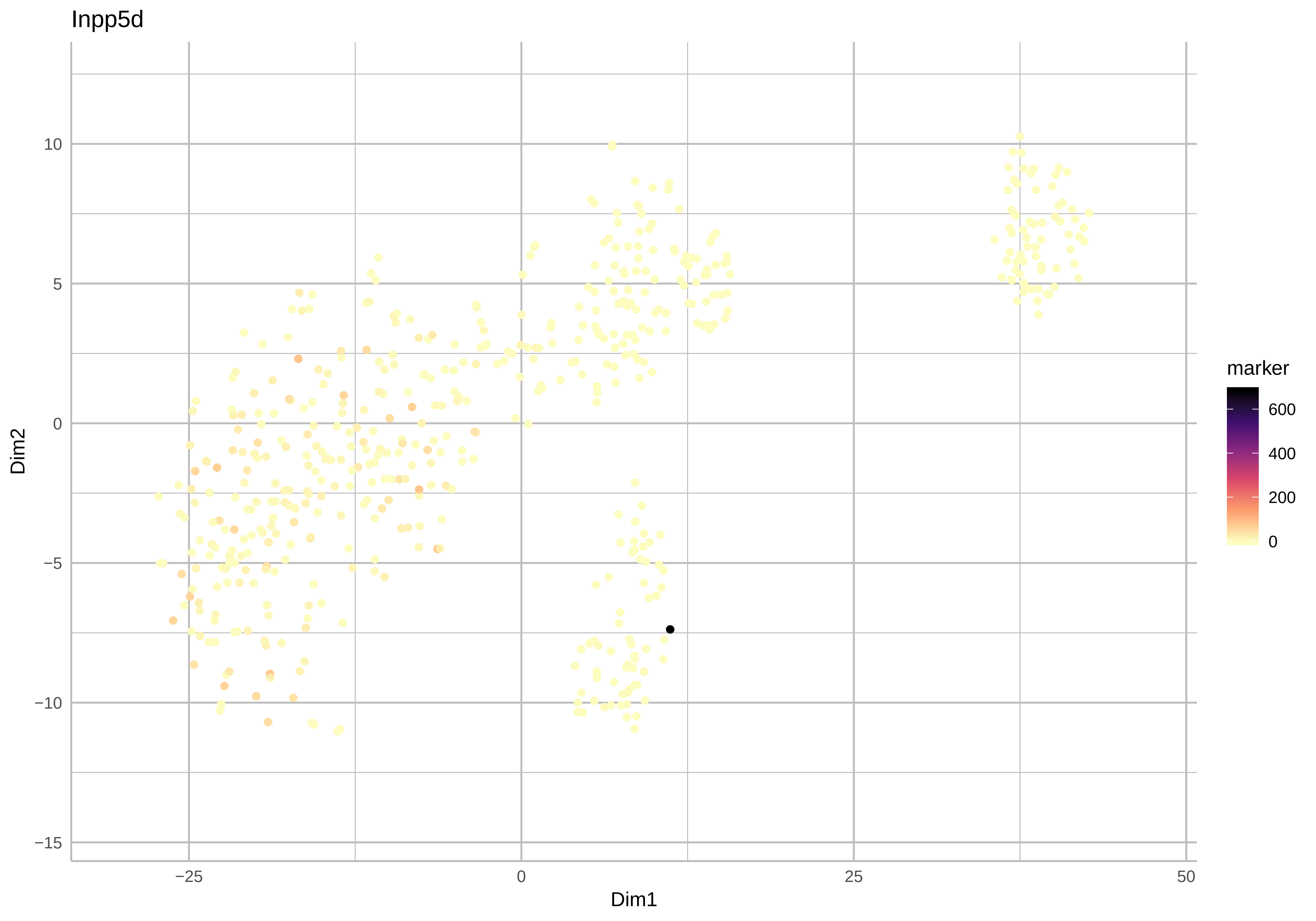
Inpp5d
Dim2
Dim1
10
5
0
−5
−10
−15
−25
0
25
50
marker
600
400
200
0

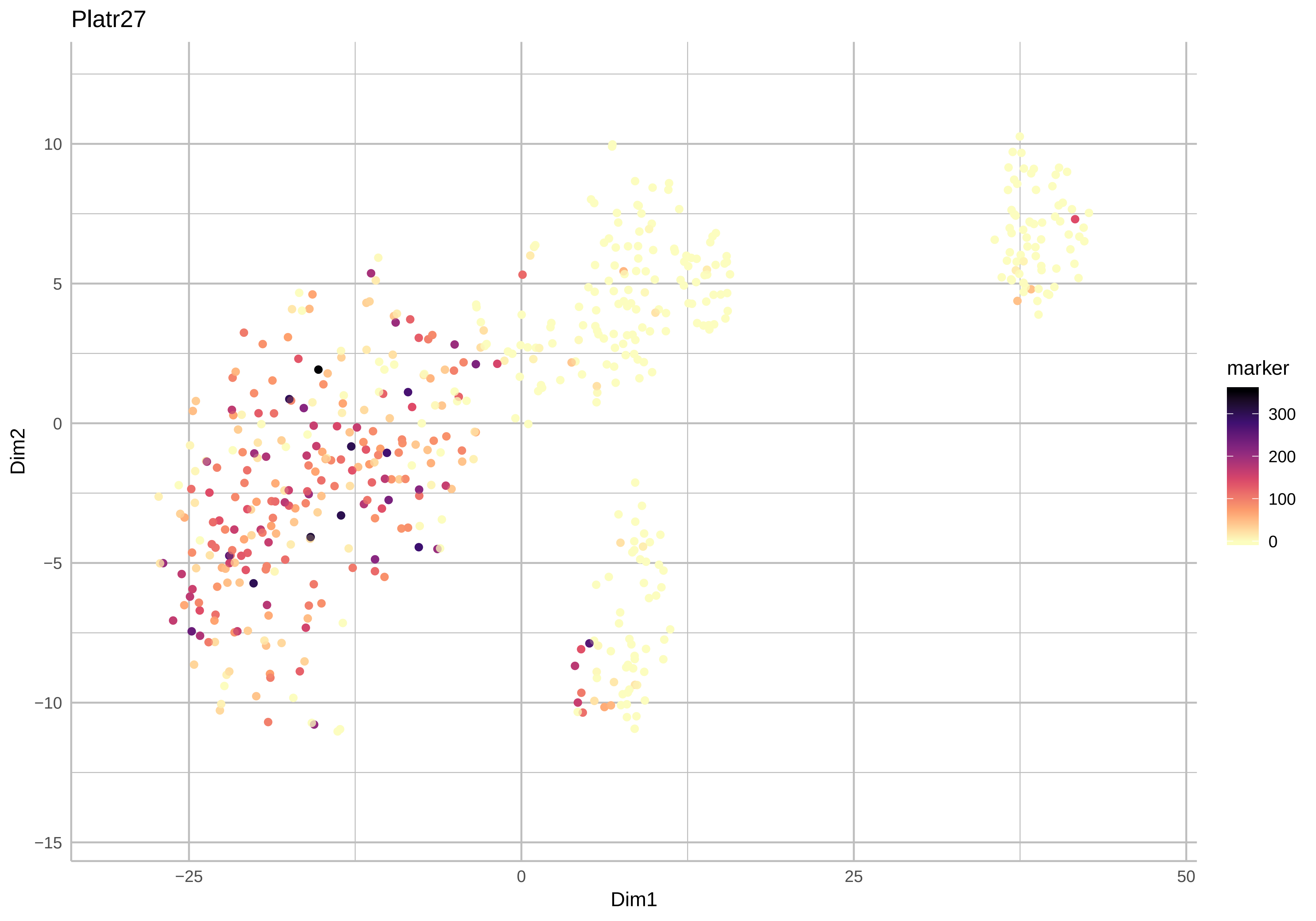

Platr27
10
5
0
−5
−10
−15
Dim2
−25
0
25
50
Dim1
marker
300
200
100
0

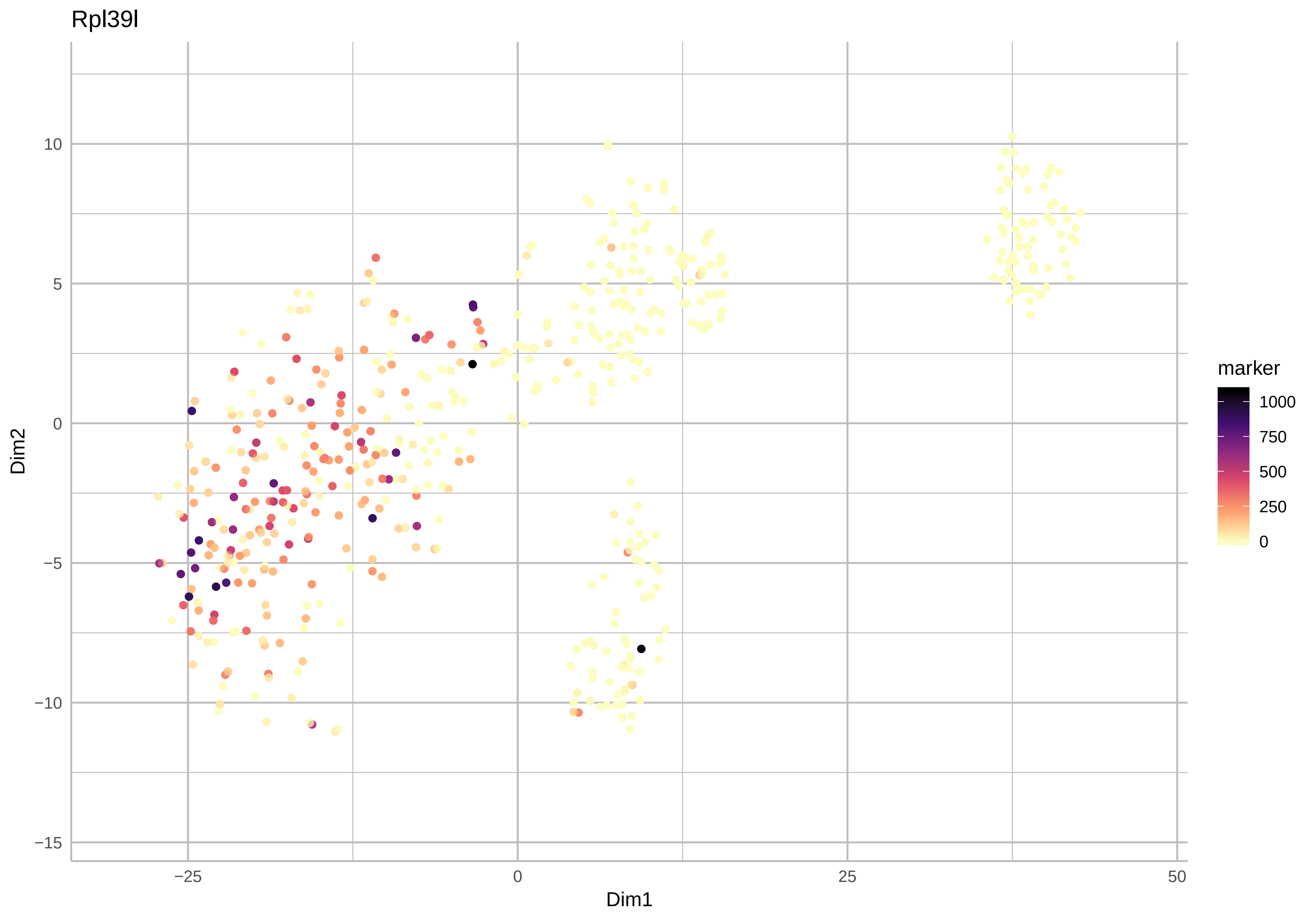

Rpl39l
10
5
0
−5
−10
−15
−25
0
25
50
Dim1
Dim2
marker
1000
750
500
250
0

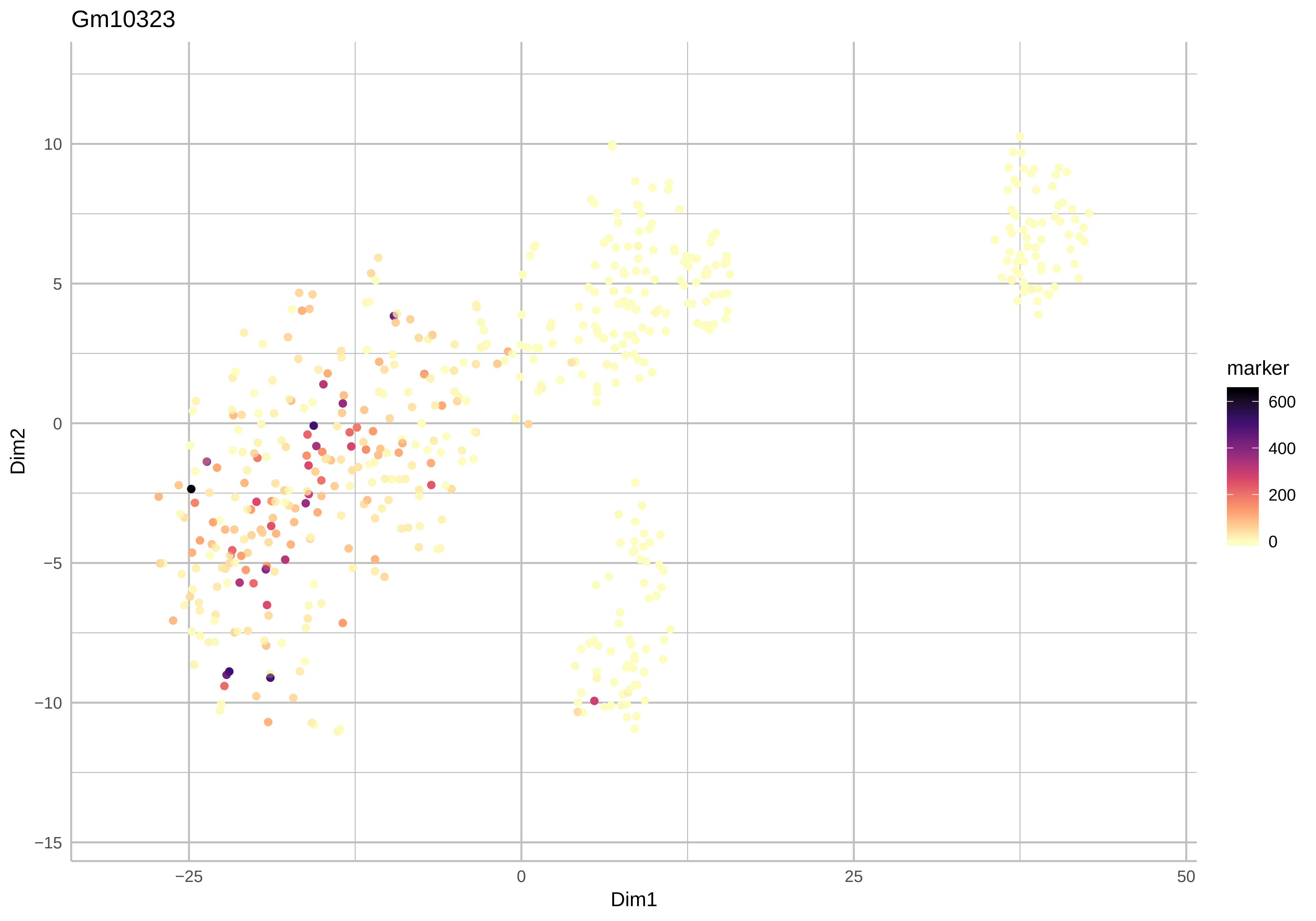

Gm10323
10
5
0
−5
−10
−15
Dim2
−25
0
25
50
Dim1
marker
600
400
200
0

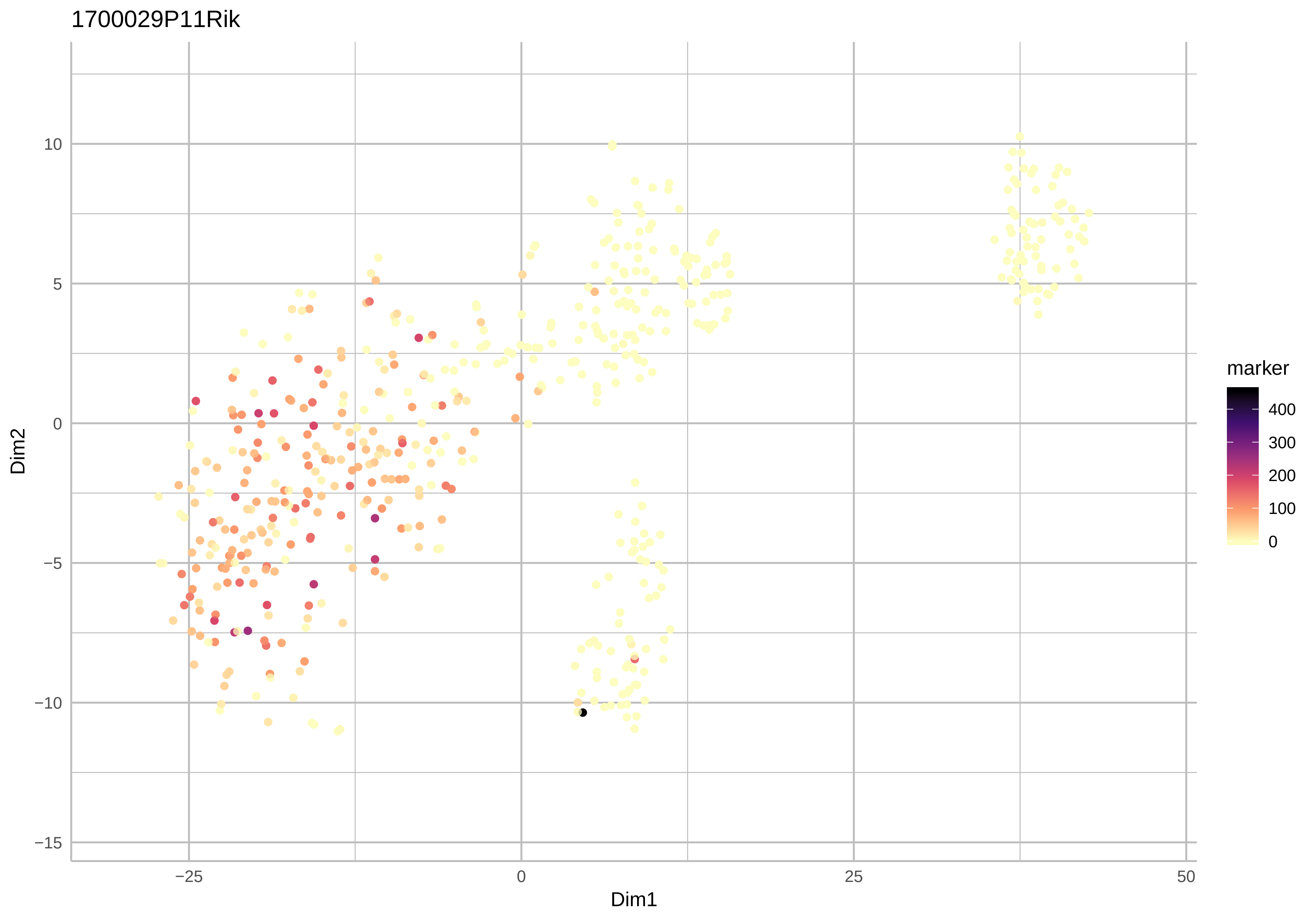
1700029P11Rik
marker
400
300
200
100
0
Dim2
10
5
0
−5
−10
−15
Dim1
−25
0
25
50

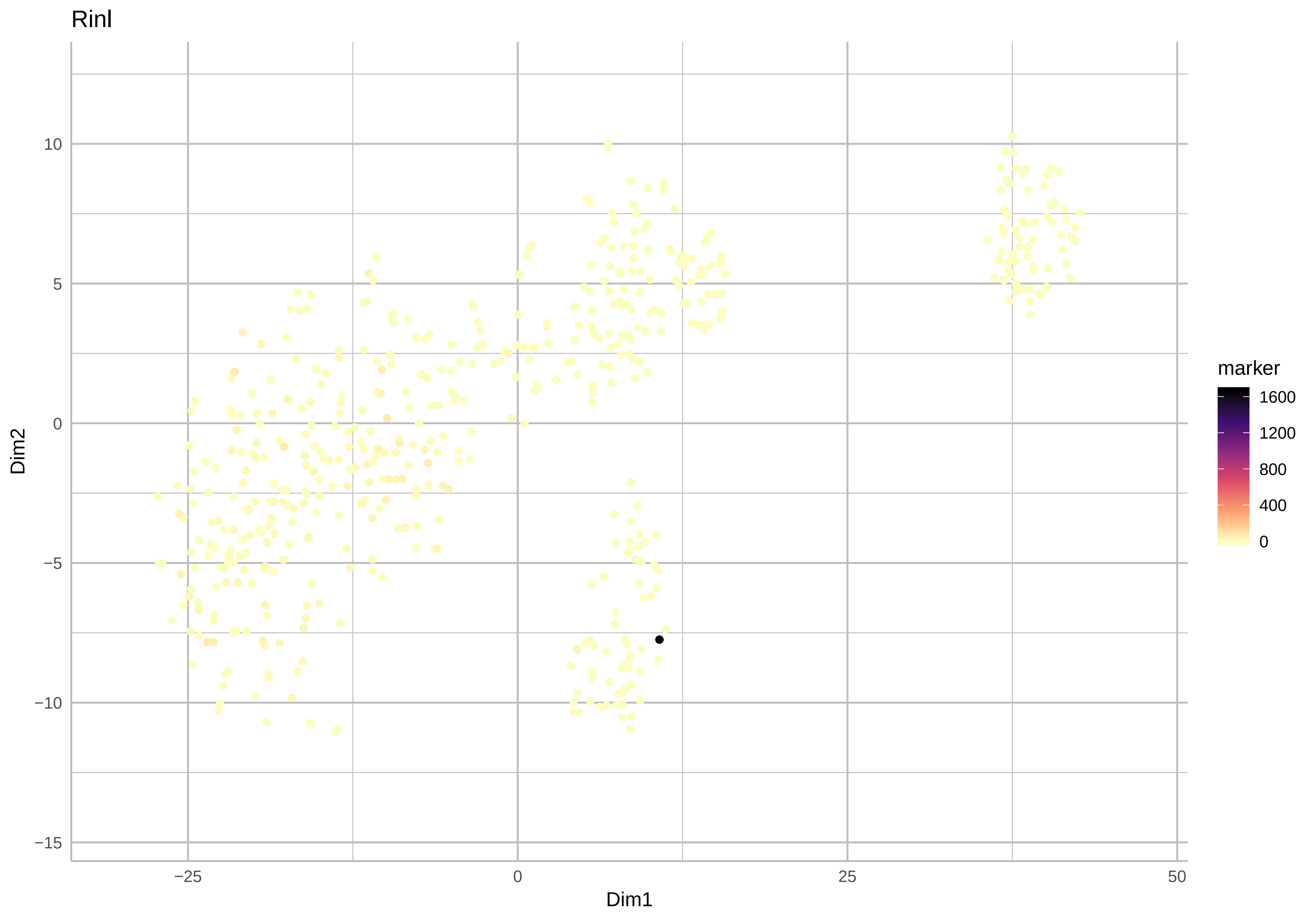

Rinl
Dim2
Dim1
10
5
0
−5
−10
−15
−25
0
25
50
marker
1600
1200
800
400
0

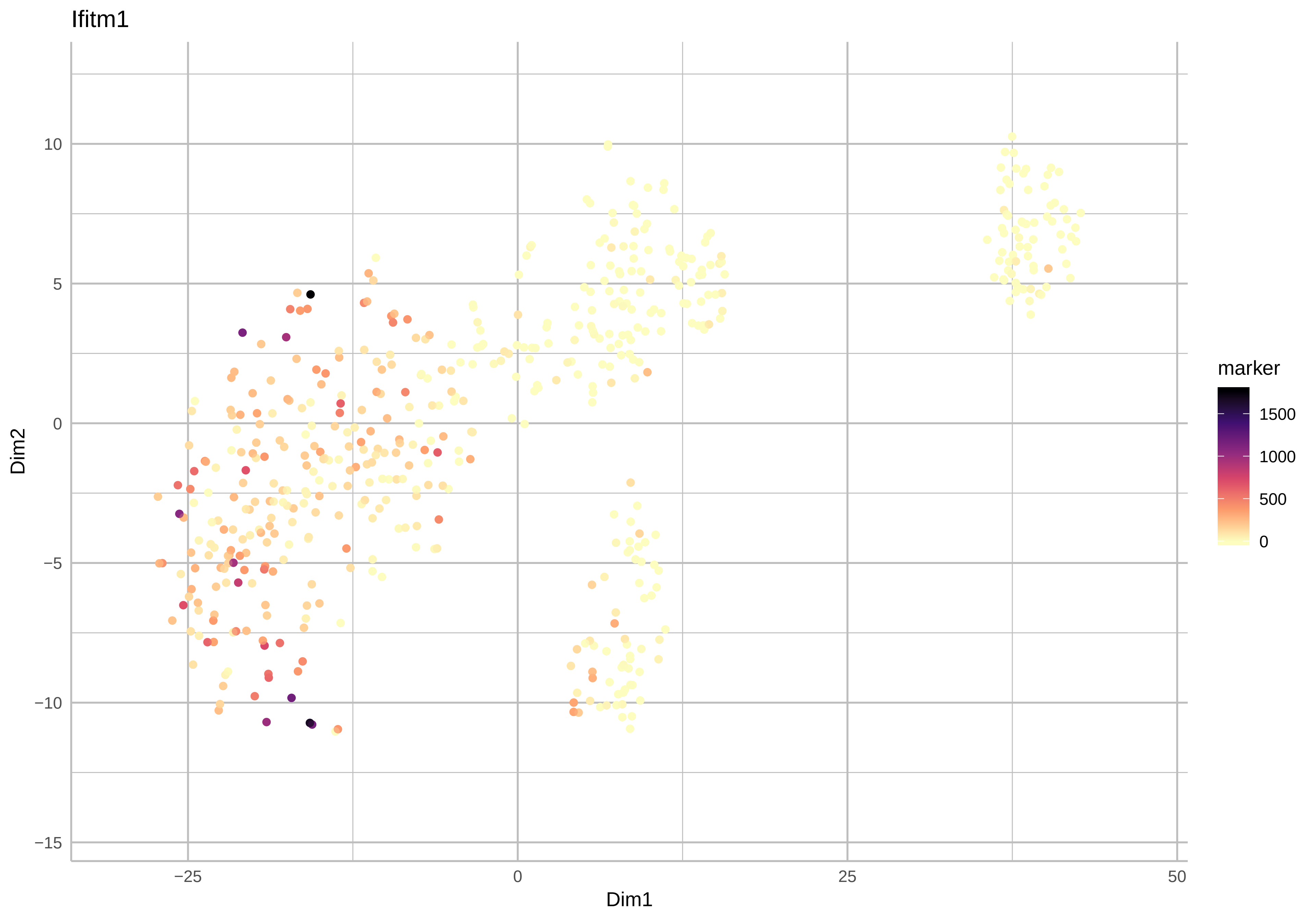
Ifitm1
Dim2
Dim1
10
5
0
−5
−10
−15
−25
0
25
50
marker
1500
1000
500
0

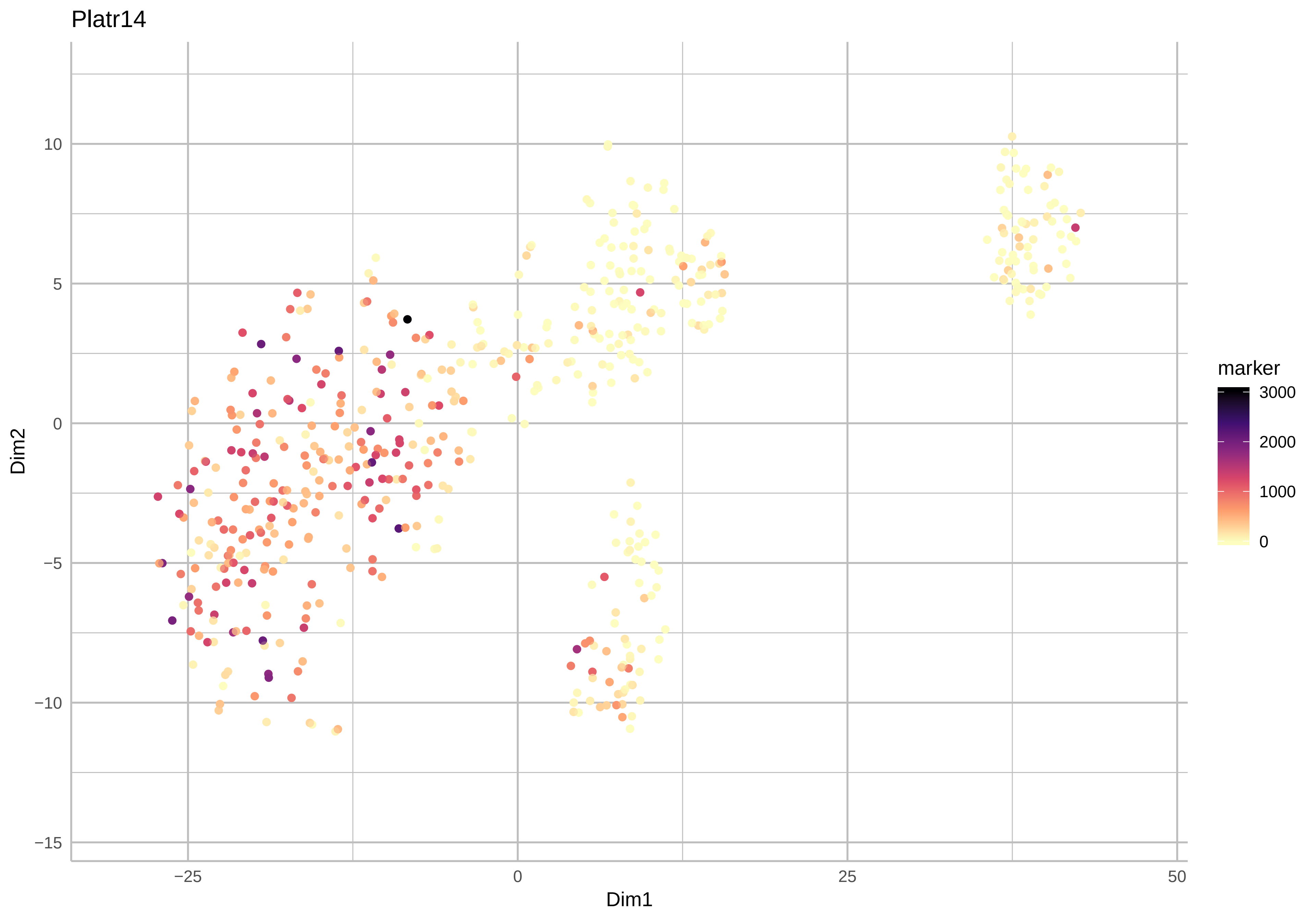

Platr14
Dim2
Dim1
10
5
0
−5
−10
−15
−25
0
25
50
marker
3000
2000
1000
0

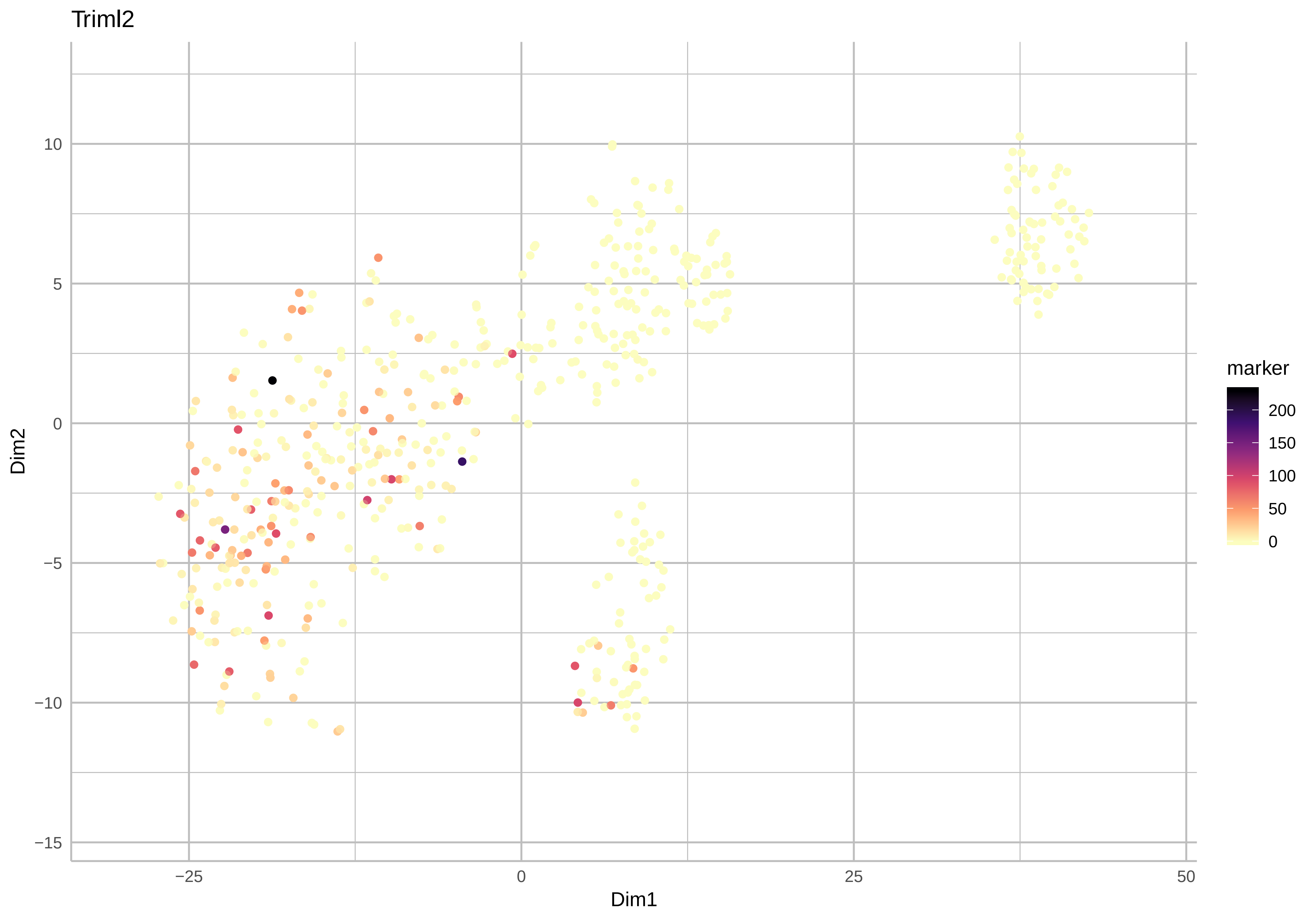
Triml2
Dim2
Dim1
marker
200
150
100
50
0
10
5
0
−5
−10
−15
−25
0
25
50

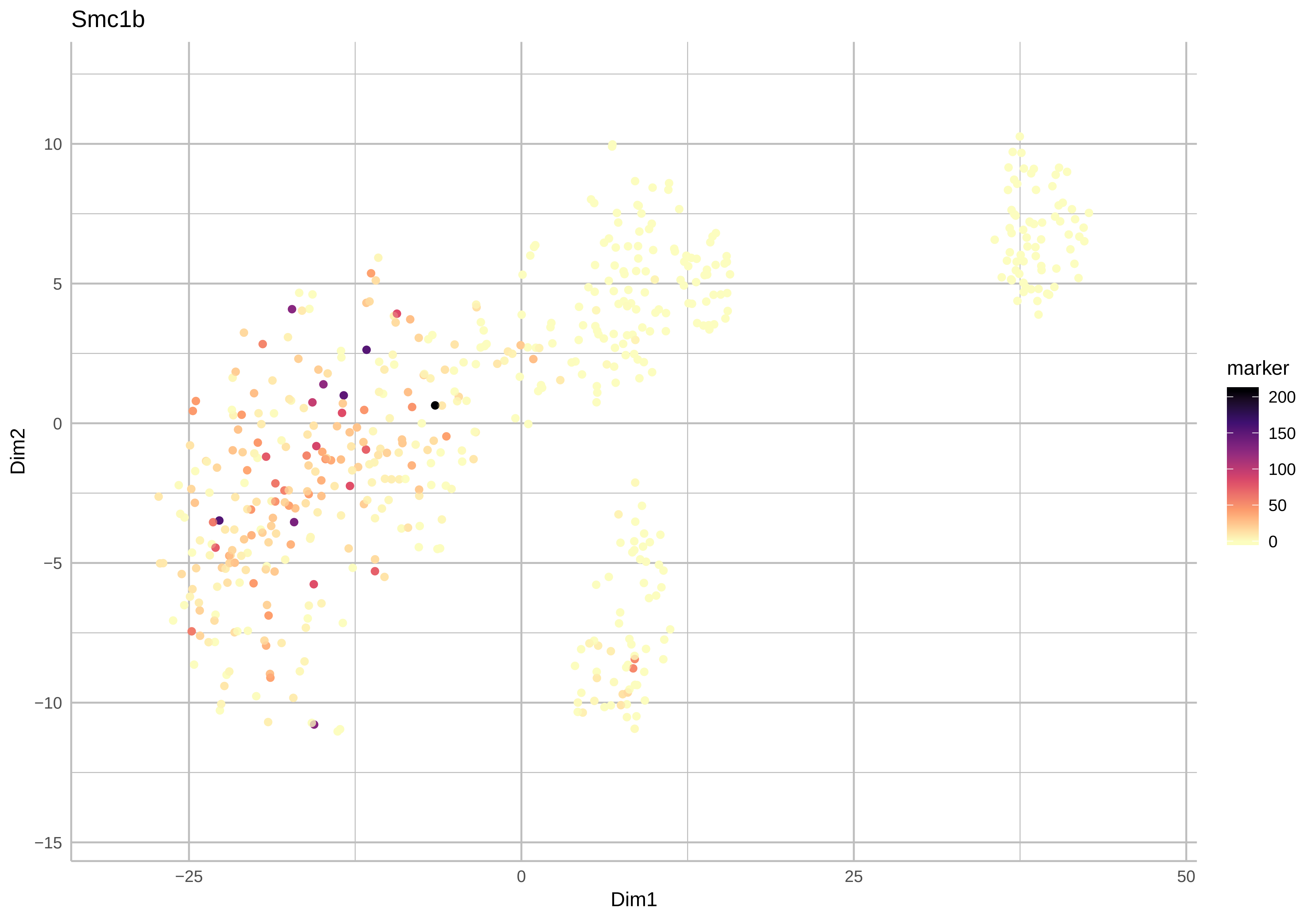
Smc1b
Dim2
Dim1
10
5
0
−5
−10
−15
−25
0
25
50
marker
200
150
100
50
0

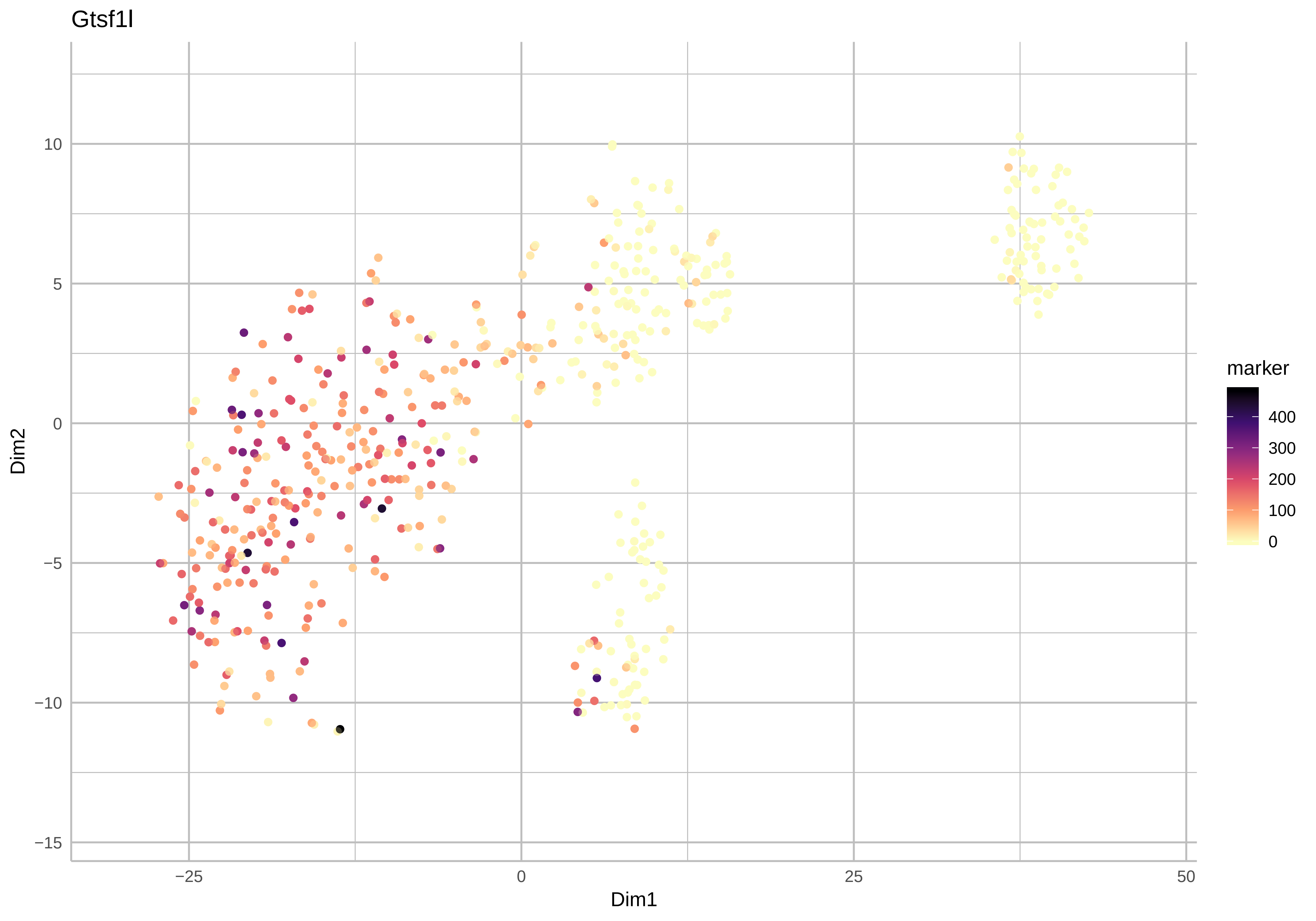

Gtsf1l
Dim2
Dim1
10
5
0
−5
−10
−15
−25
0
25
50
marker
400
300
200
100
0

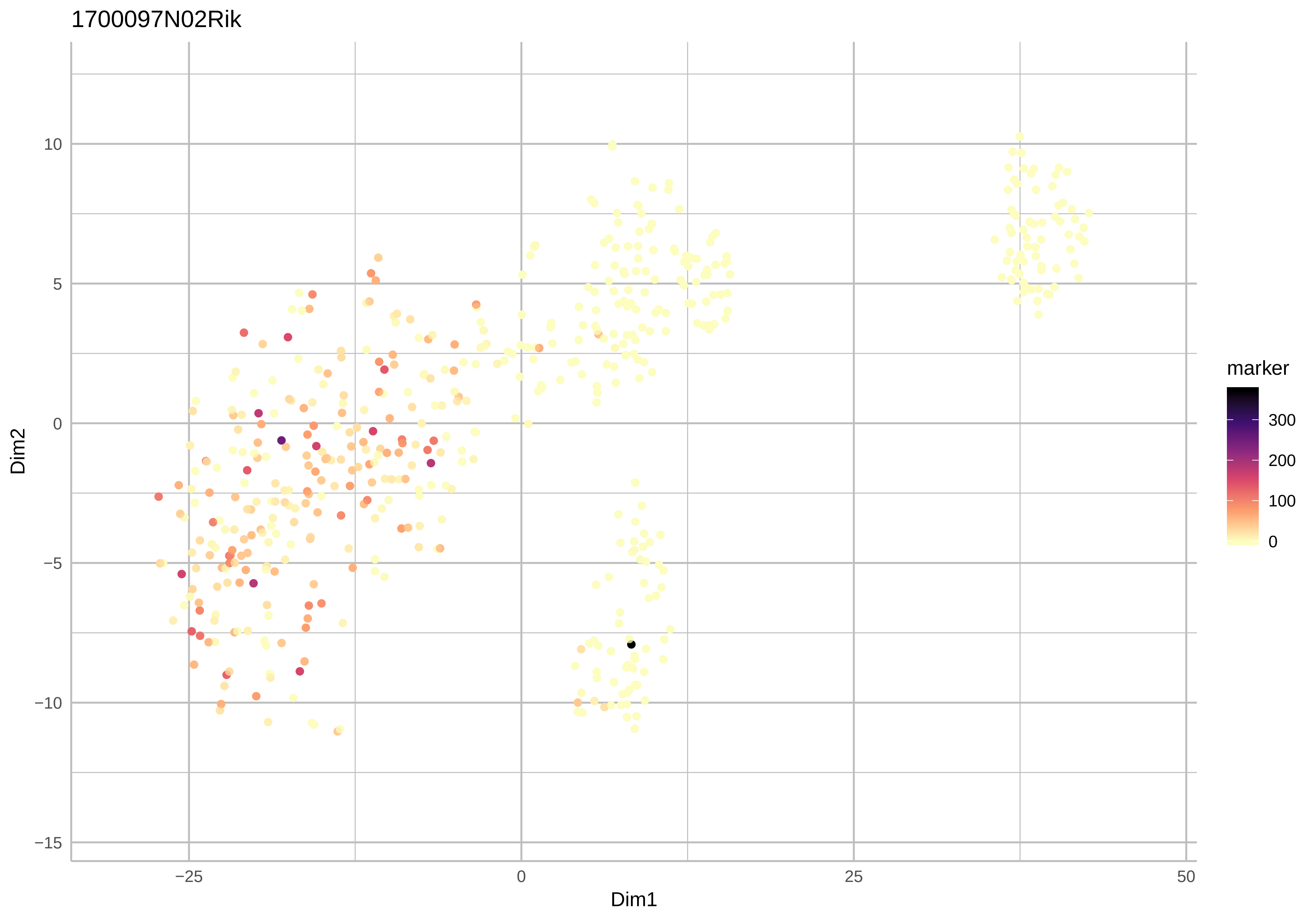

1700097N02Rik
Dim2
Dim1
10
5
0
−5
−10
−15
−25
0
25
50
marker
300
200
100
0

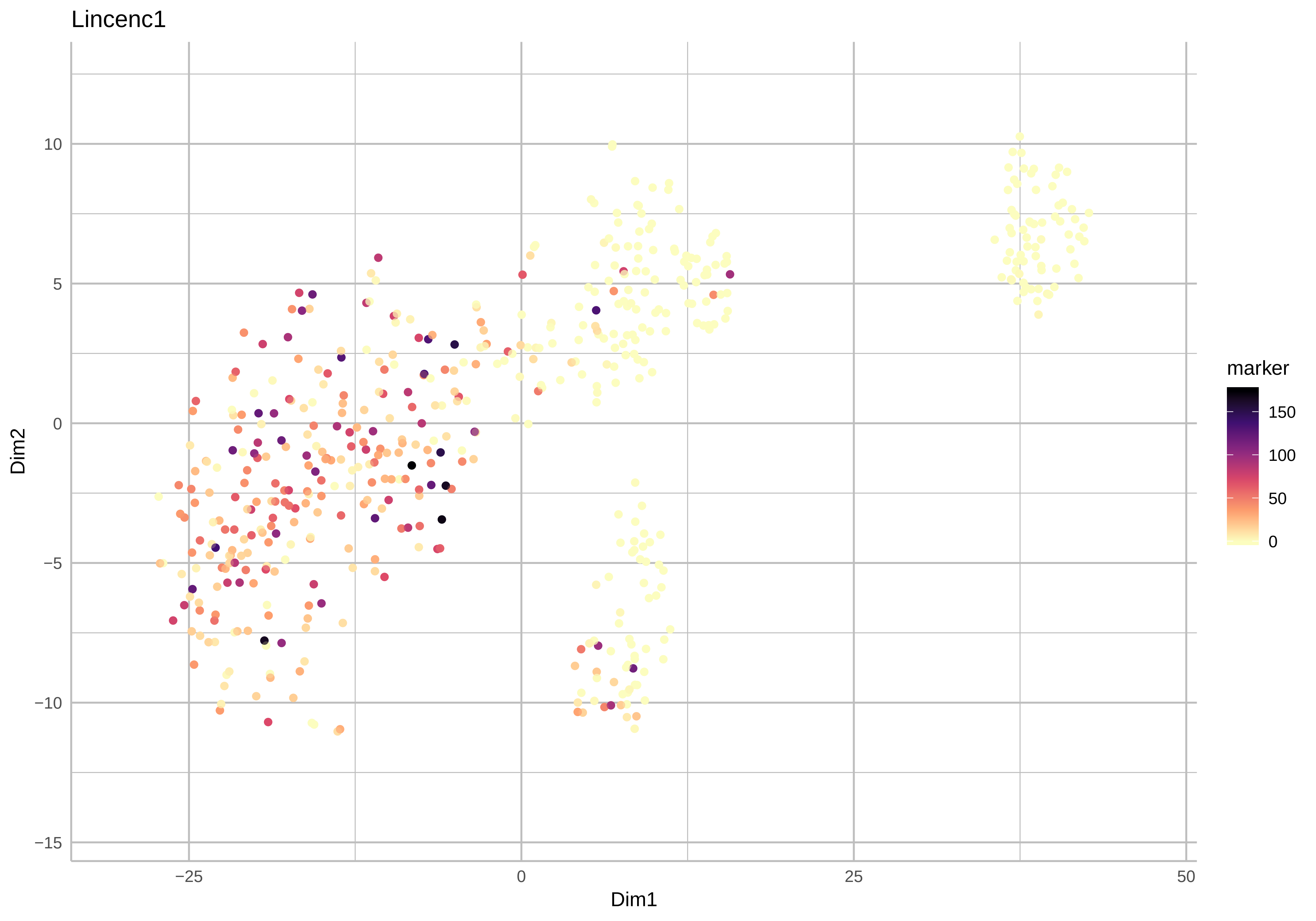
Lincenc1
Dim2
Dim1
10
5
0
−5
−10
−15
−25
0
25
50
marker
150
100
50
0

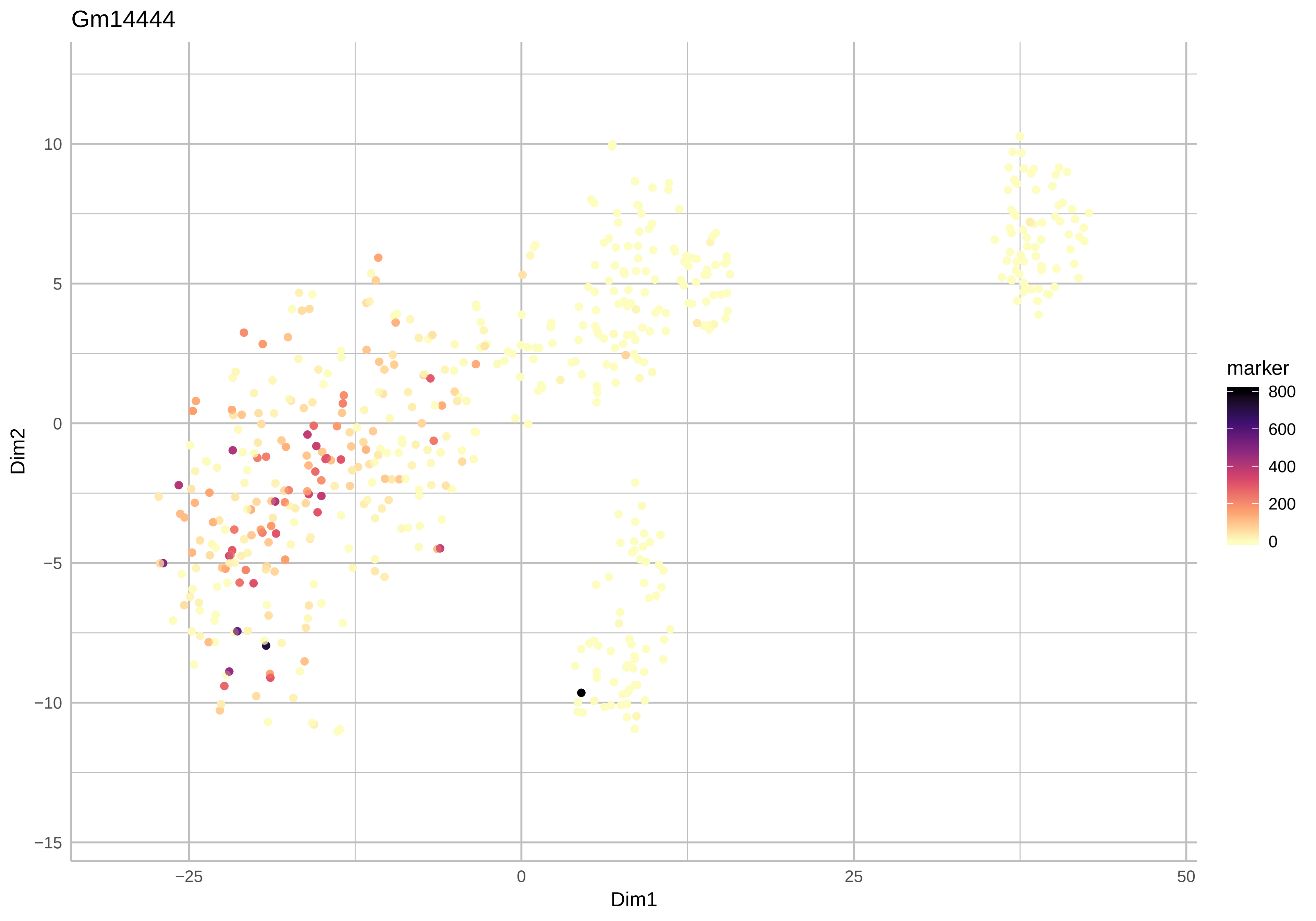
Gm14444
Dim2
Dim1
10
5
0
−5
−10
−15
−25
0
25
50
marker
800
600
400
200
0

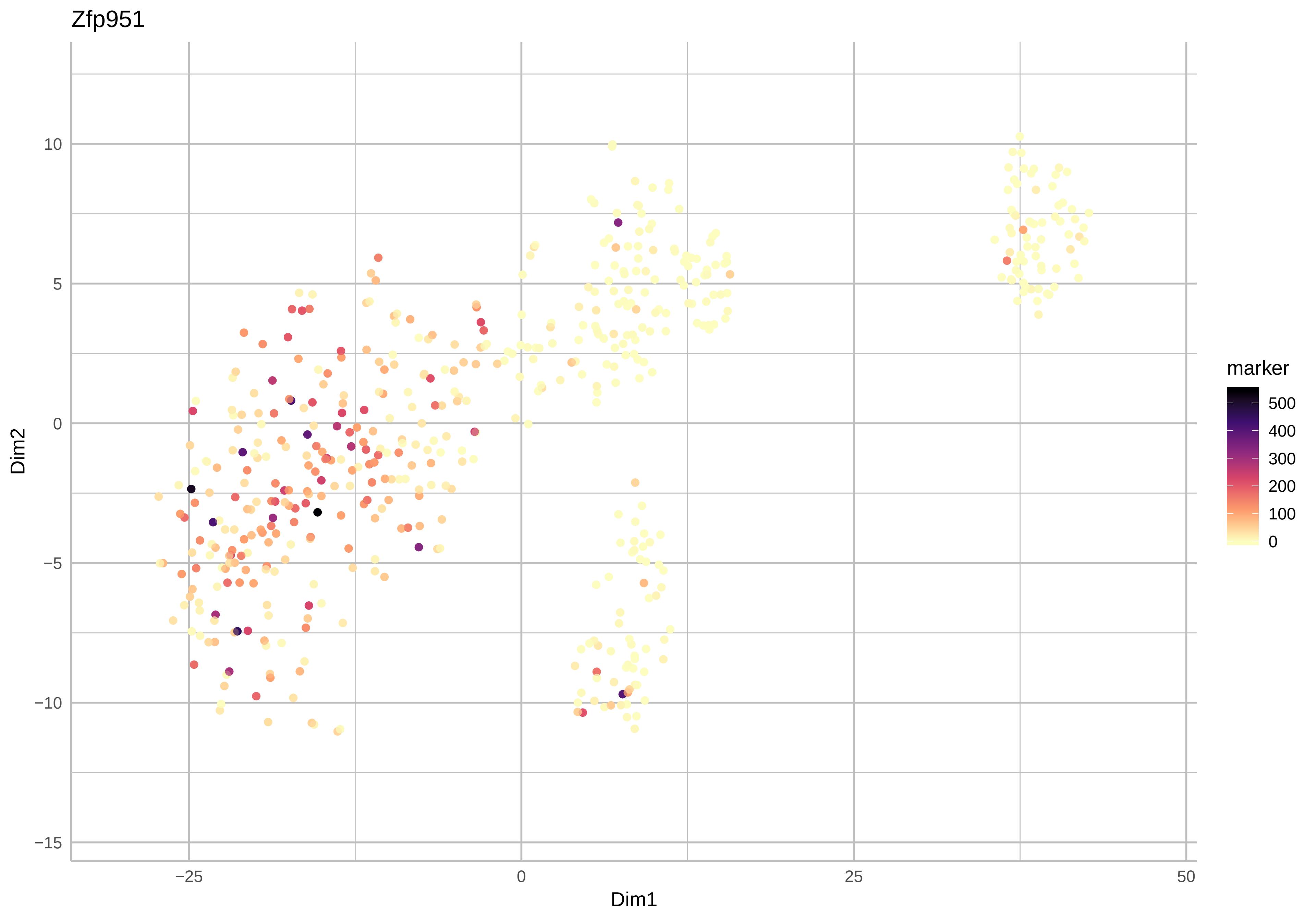
Zfp951
Dim2
Dim1
10
5
0
−5
−10
−15
−25
0
25
50
marker
500
400
300
200
100
0

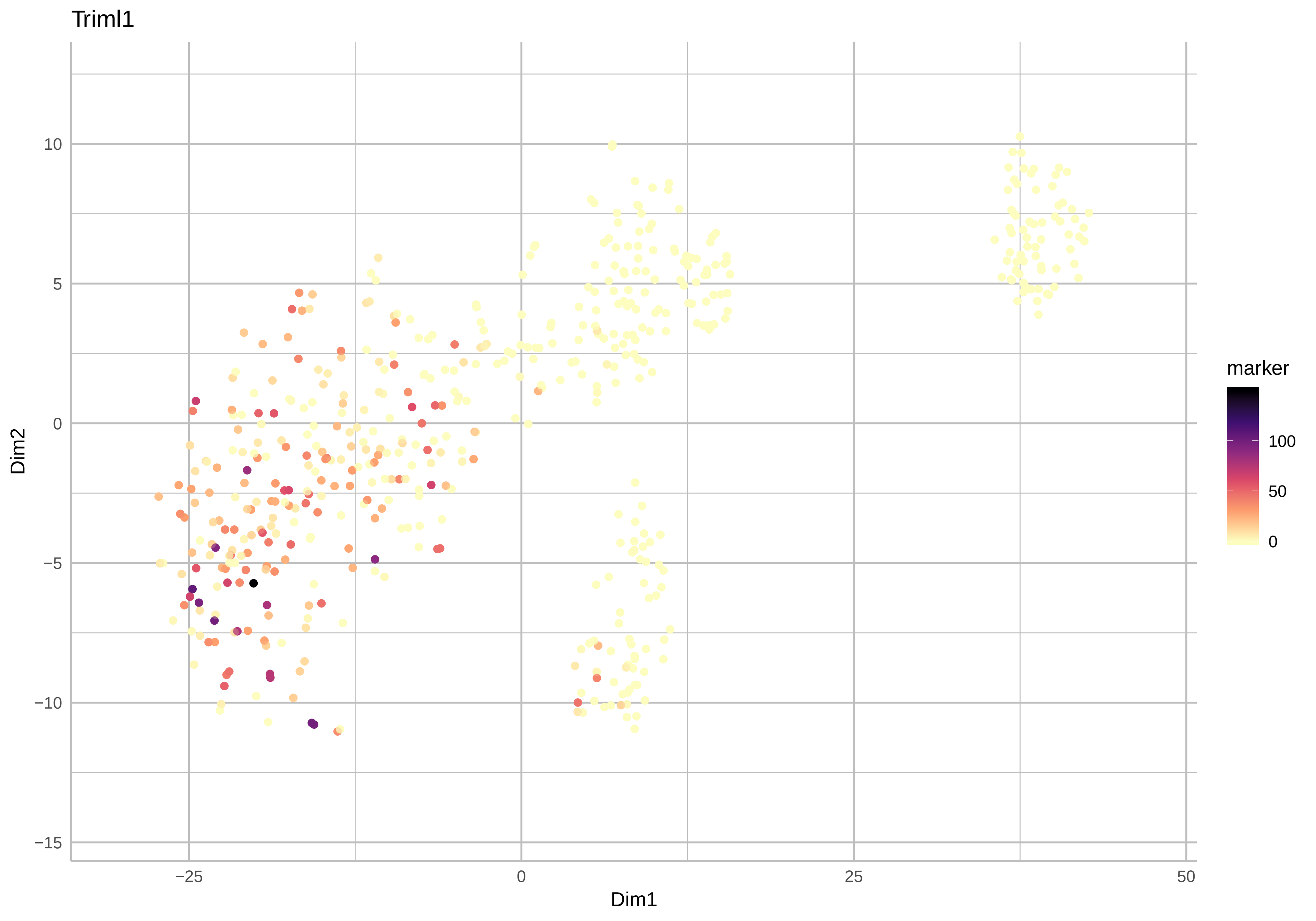
Triml1
Dim2
Dim1
10
5
0
−5
−10
−15
−25
0
25
50
marker
100
50
0

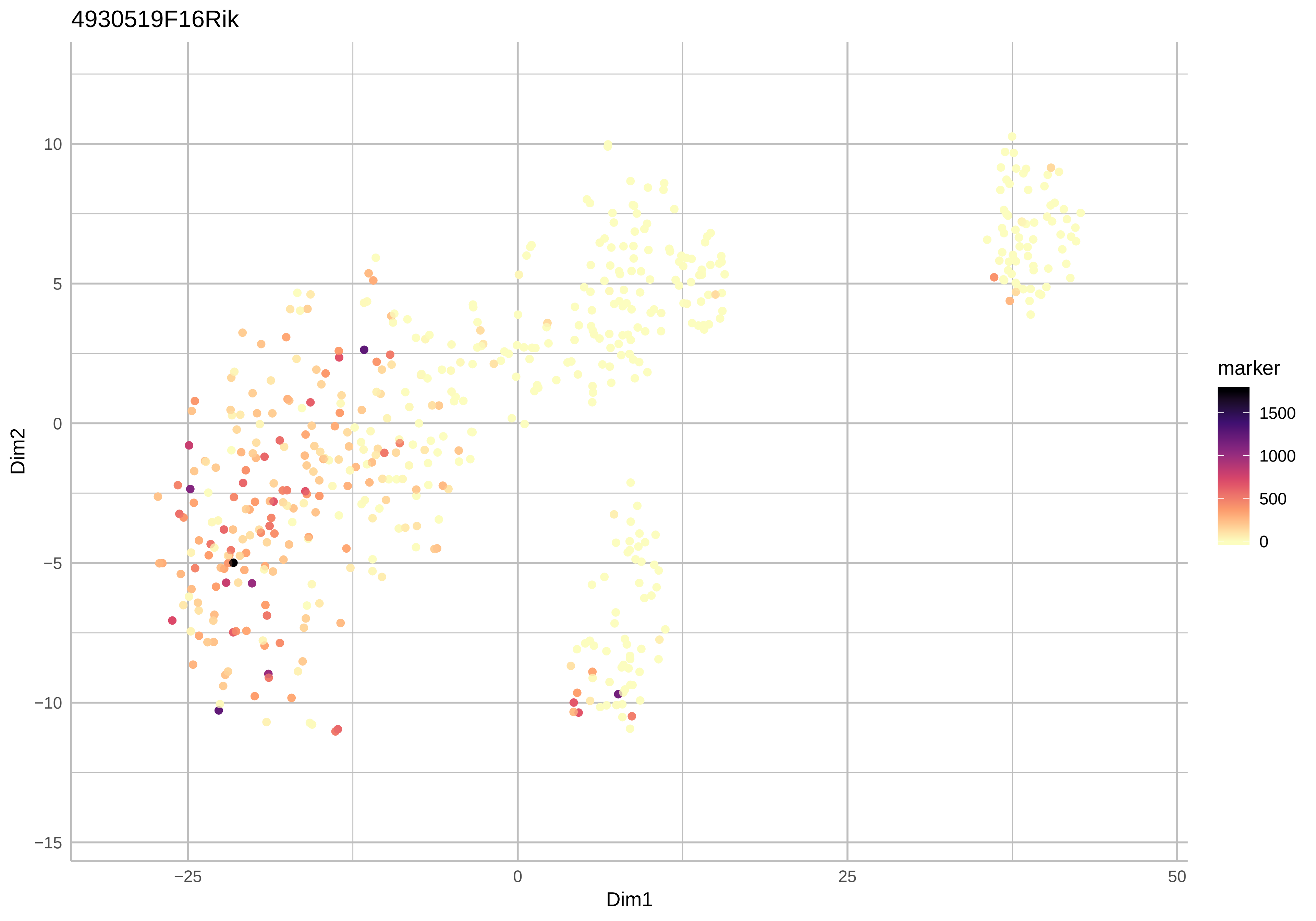
4930519F16Rik
Dim2
Dim1
10
5
0
−5
−10
−15
−25
0
25
50
marker
1500
1000
500
0

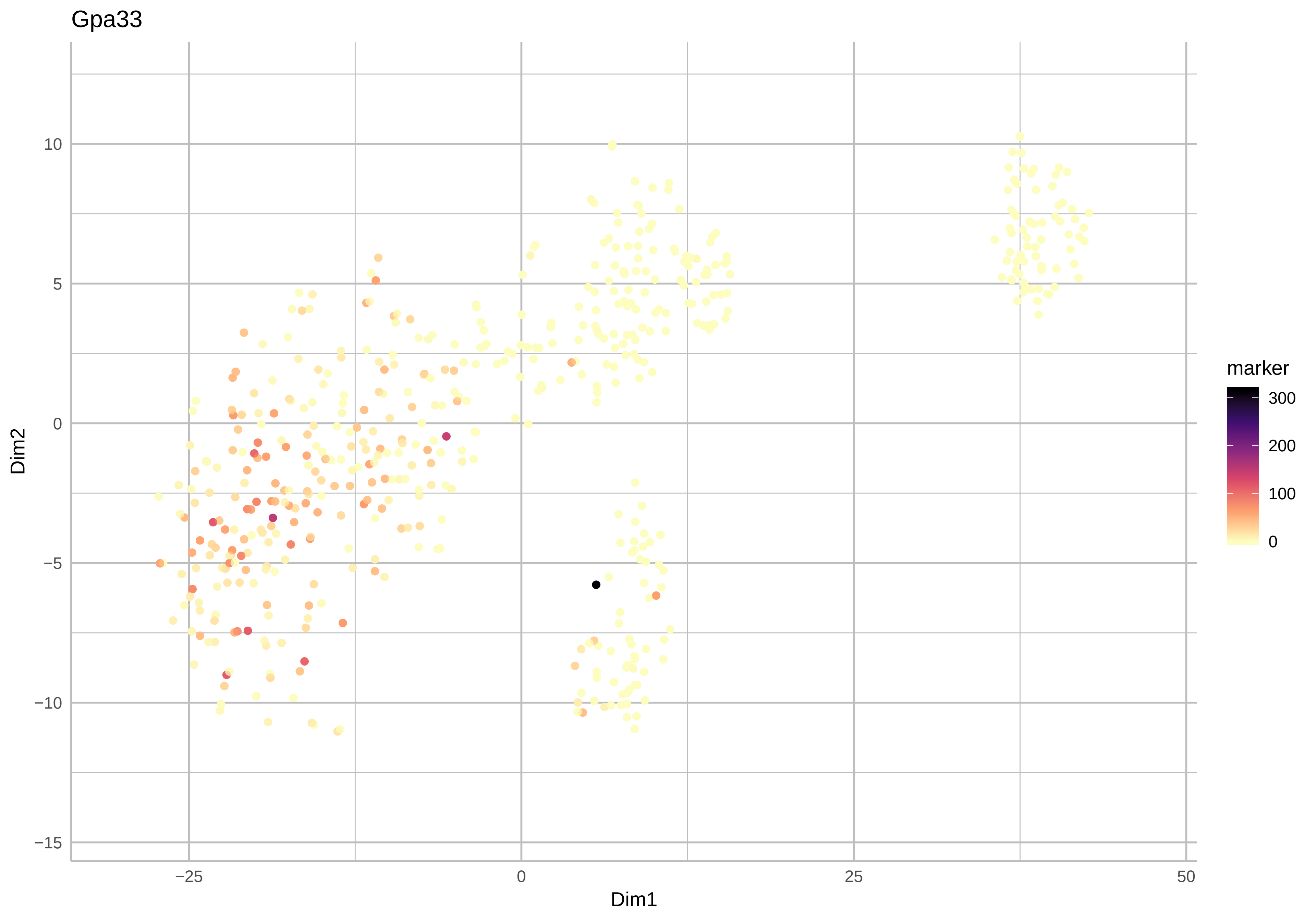

Gpa33
Dim2
Dim1
marker
300
200
100
0
10
5
0
−5
−10
−15
−25
0
25
50

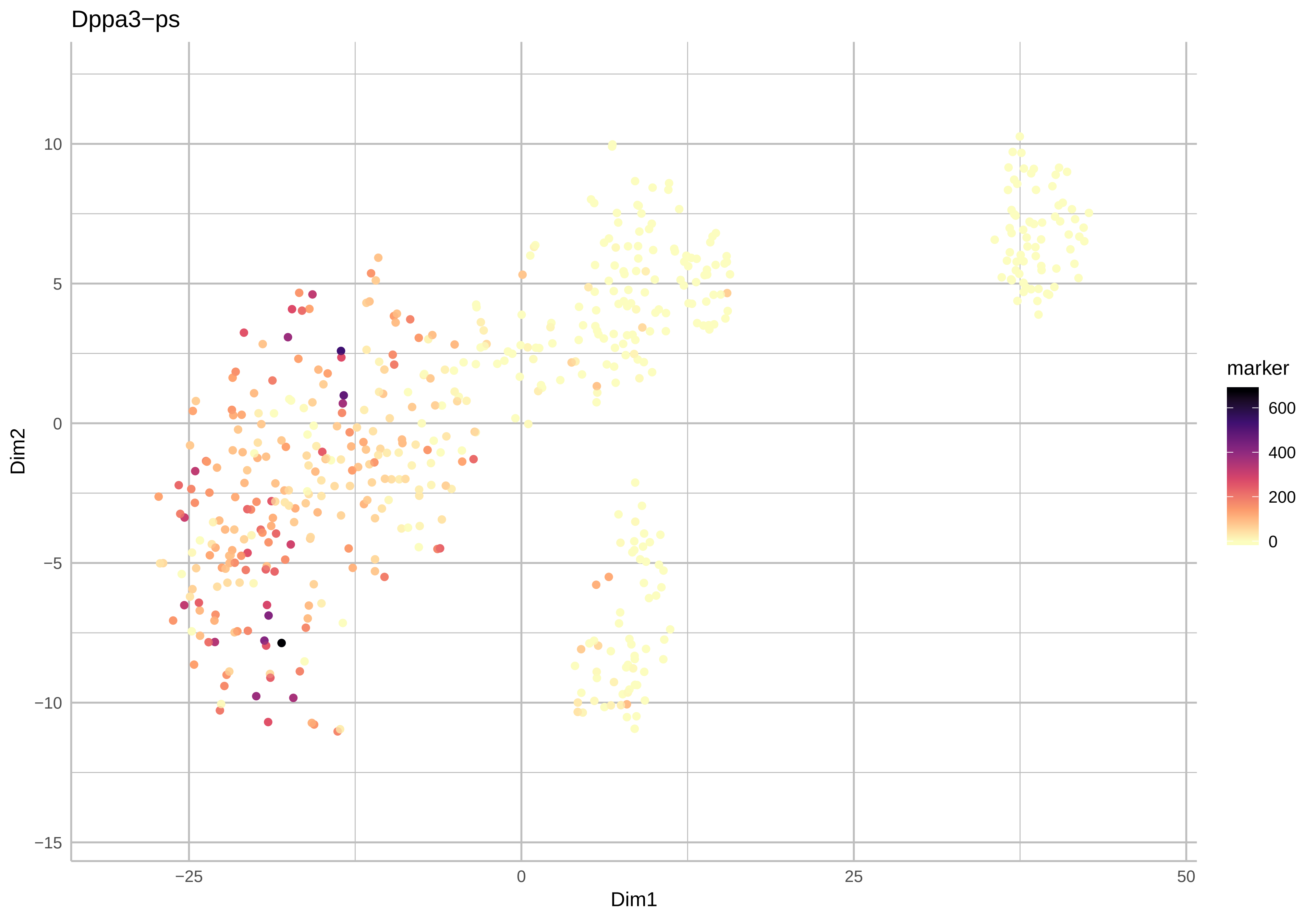

Dppa3−ps
Dim2
Dim1
10
5
0
−5
−10
−15
−25
0
25
50
marker
600
400
200
0

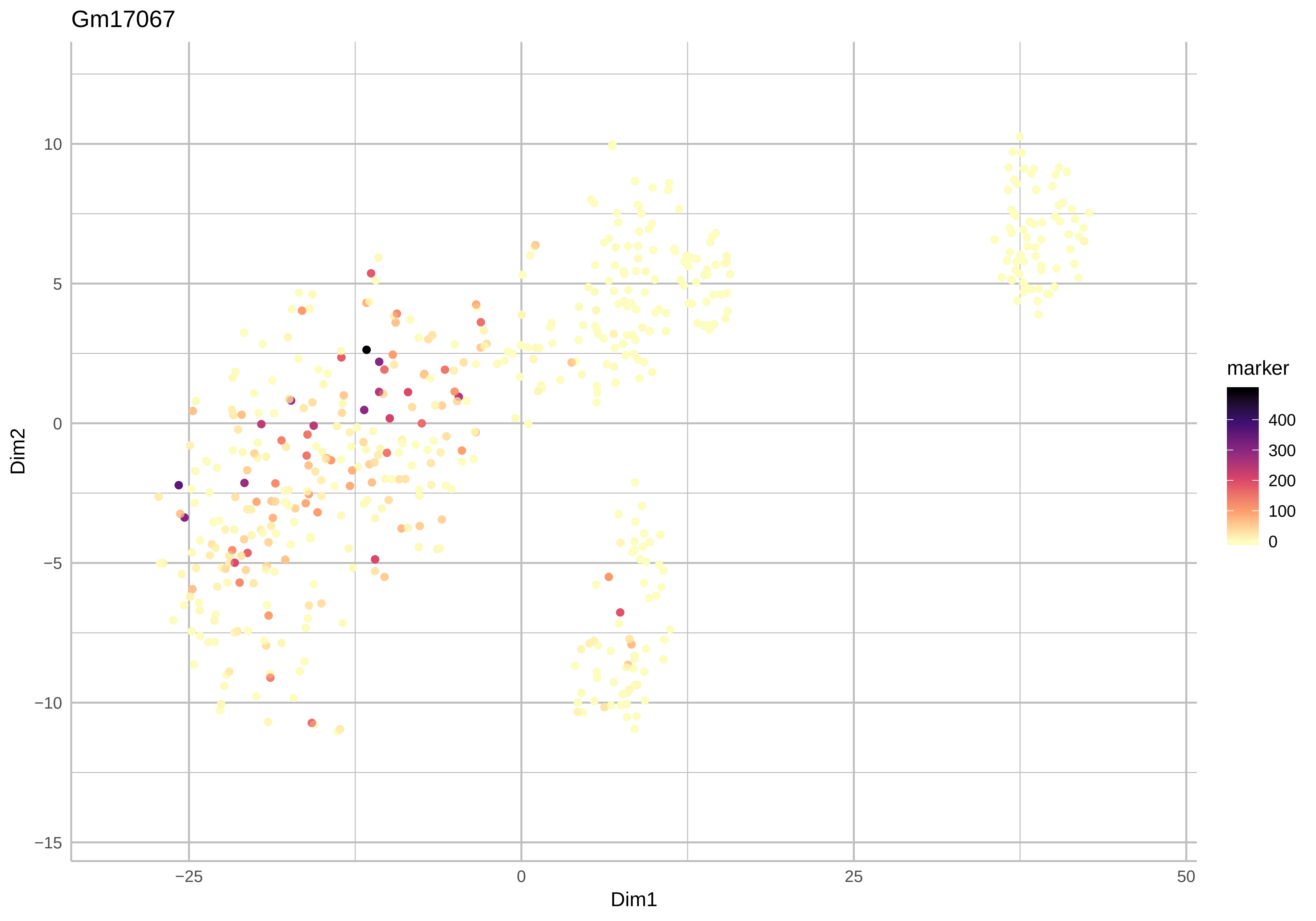

Gm17067
Dim2
Dim1
10
5
0
−5
−10
−15
−25
0
25
50
marker
400
300
200
100
0

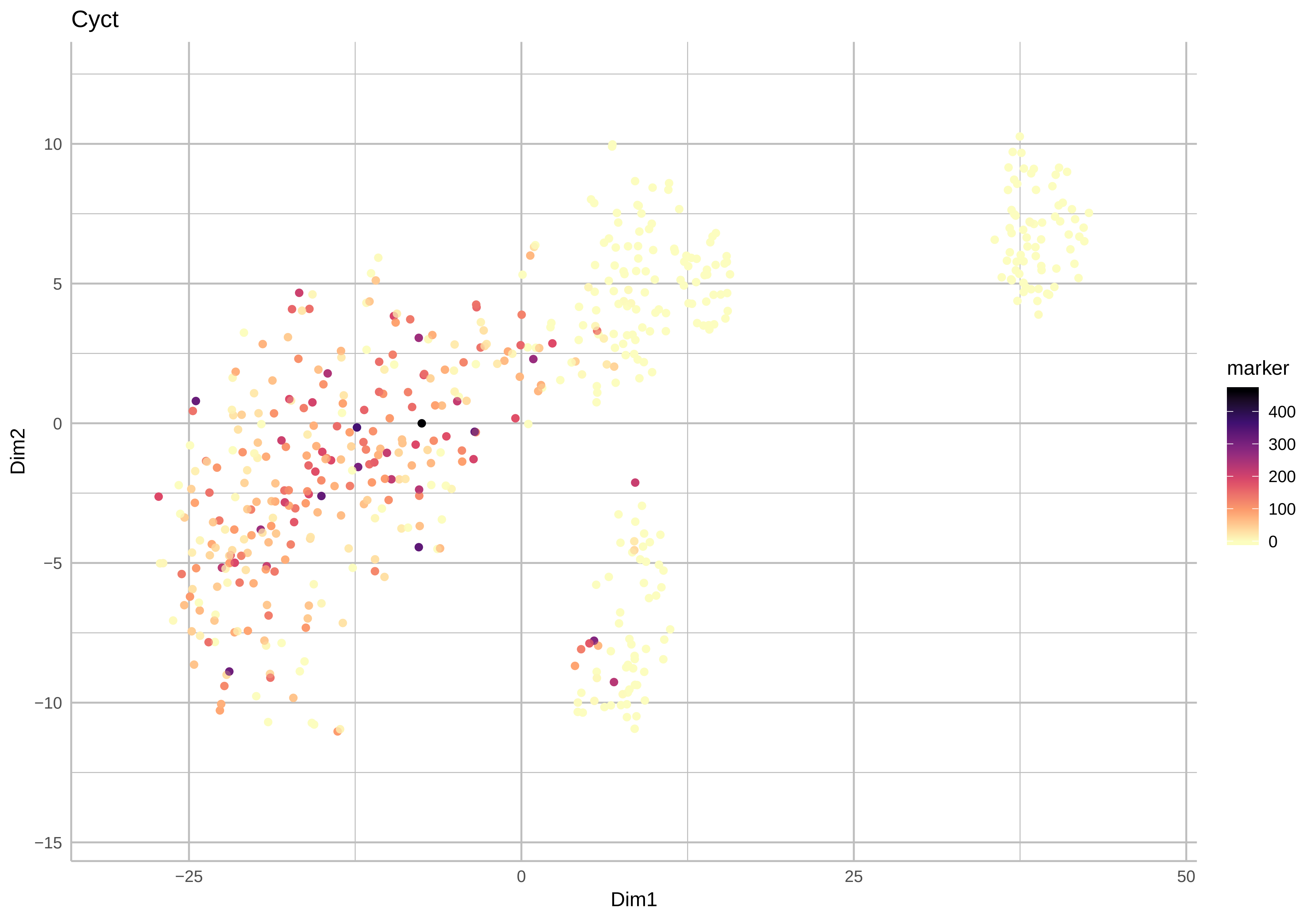
Cyct
Dim2
Dim1
10
5
0
−5
−10
−15
−25
0
25
50
marker
400
300
200
100
0

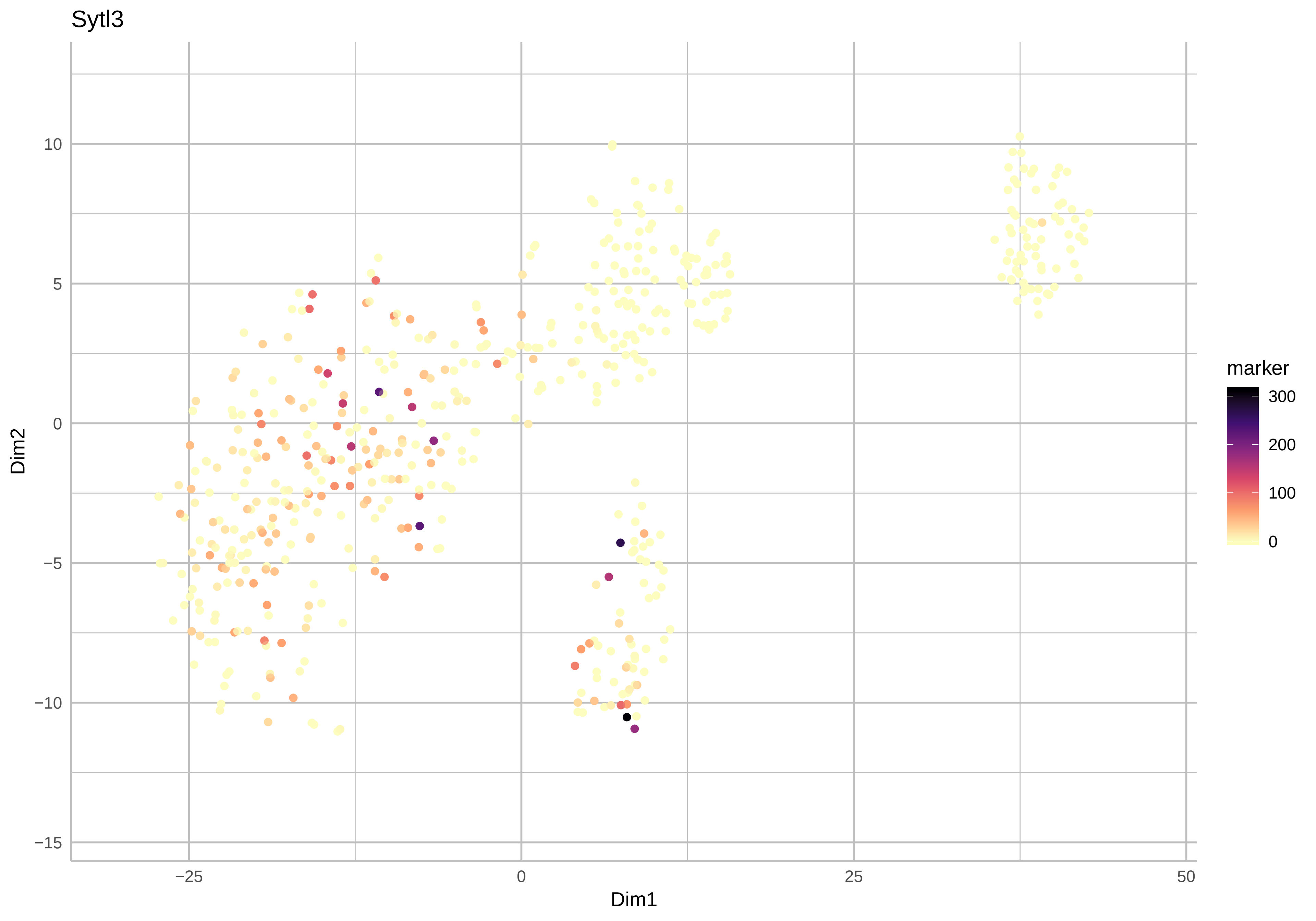

Sytl3
Dim2
Dim1
marker
300
200
100
0
10
5
0
−5
−10
−15
−25
0
25
50

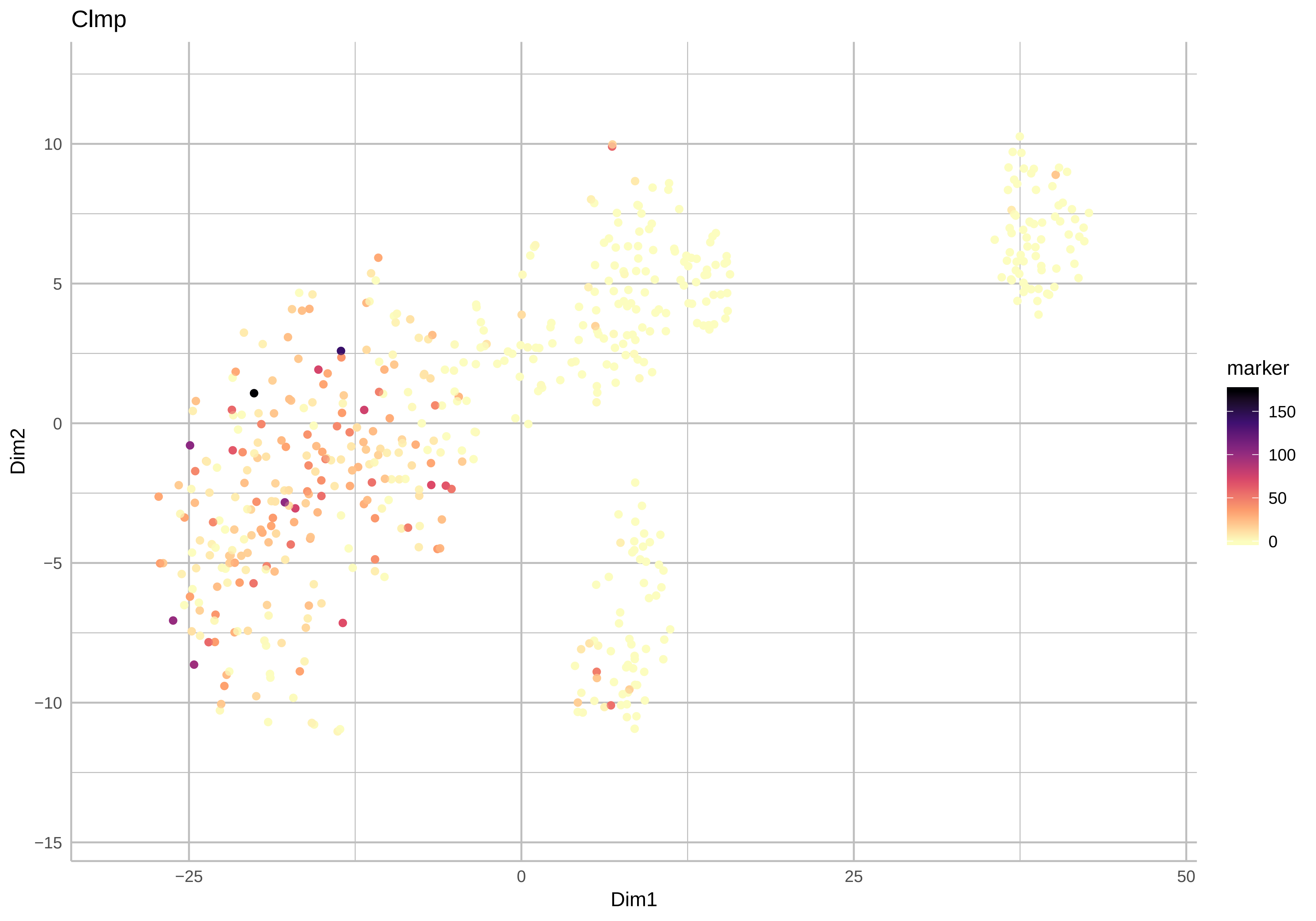

Clmp
10
5
0
−5
−10
−15
−25
0
25
50
Dim1
Dim2
marker
150
100
50
0

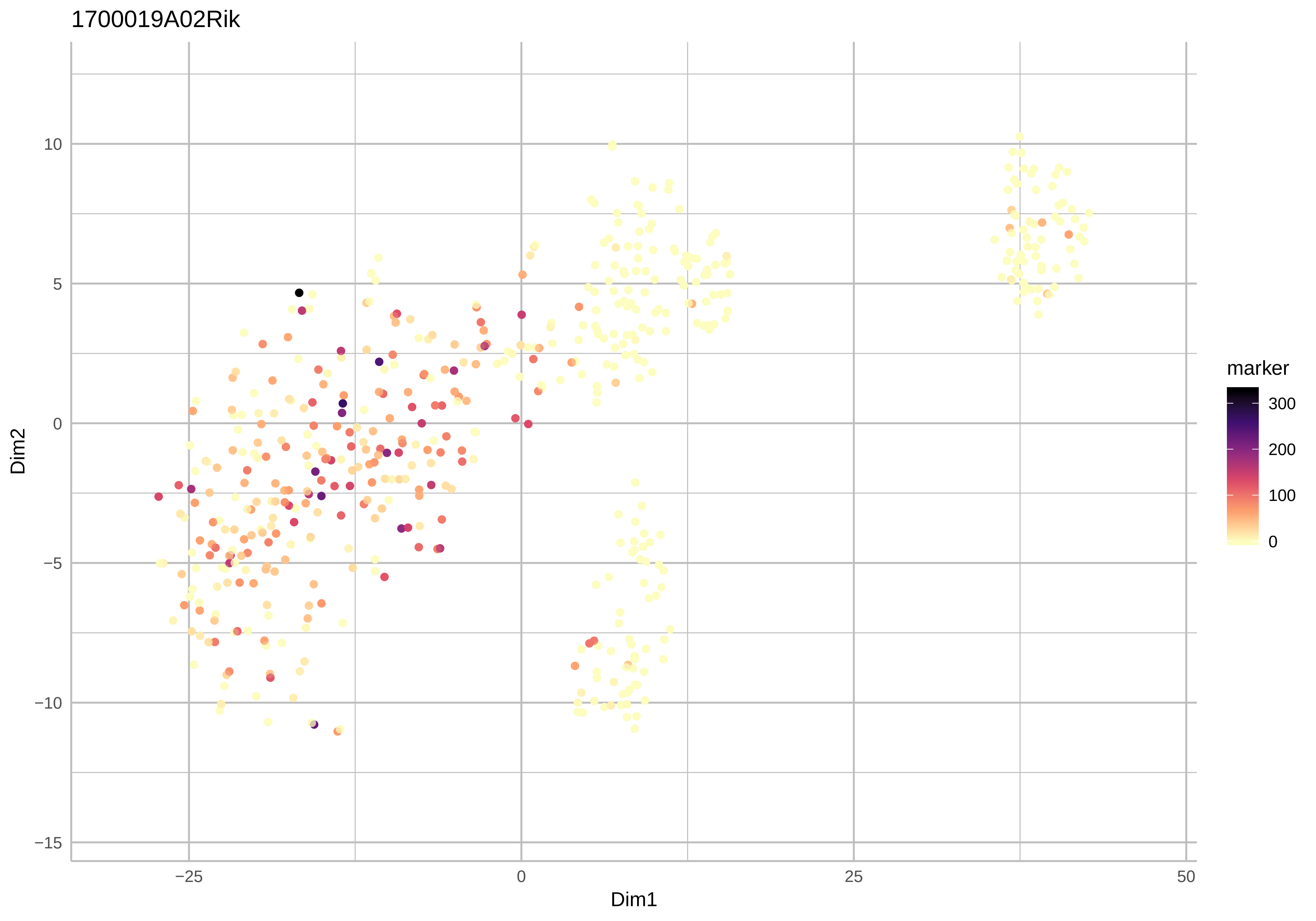

1700019A02Rik
Dim2
Dim1
10
5
0
−5
−10
−15
−25
0
25
50
marker
300
200
100
0

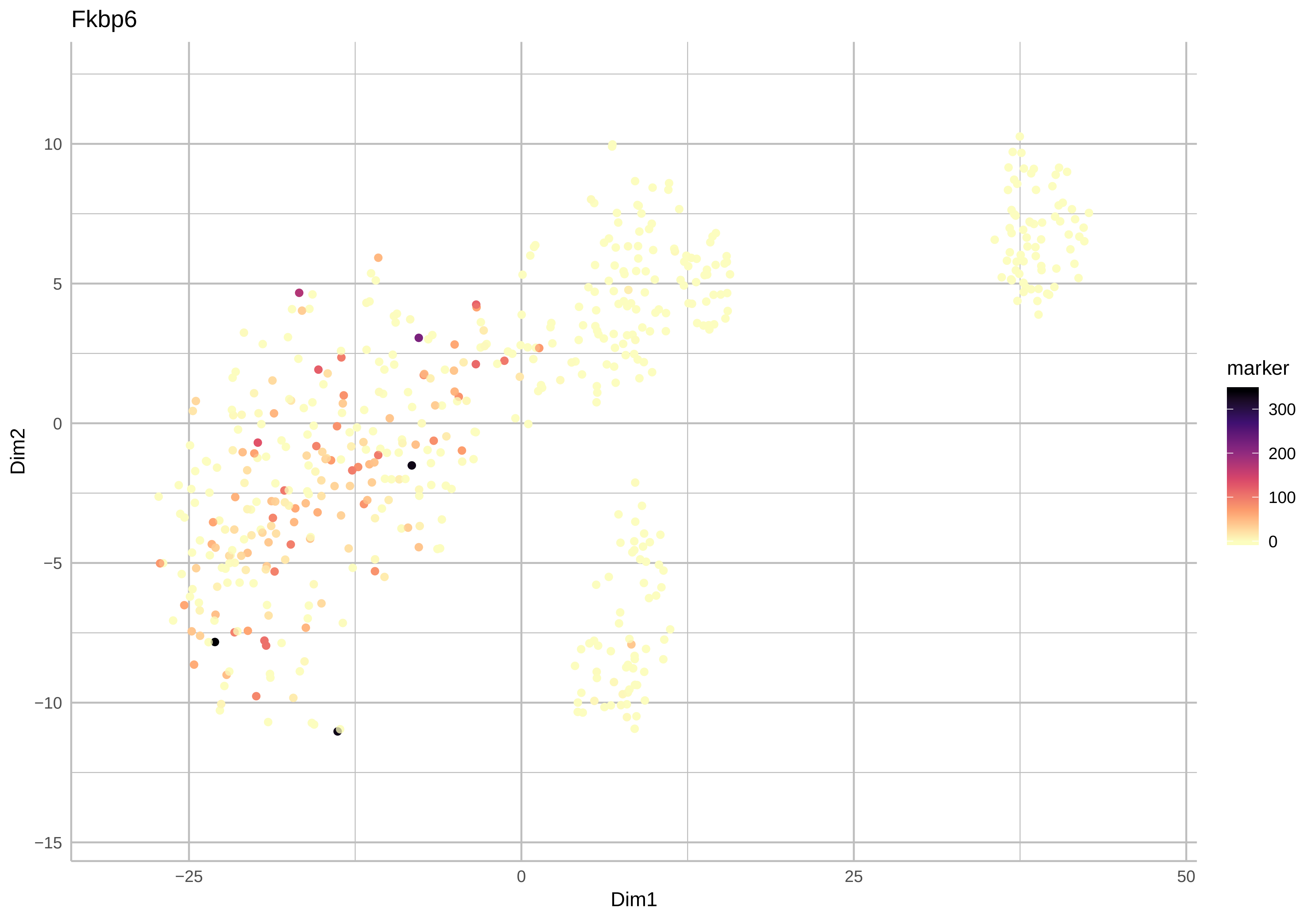

Fkbp6
Dim2
Dim1
10
5
0
−5
−10
−15
−25
0
25
50
marker
300
200
100
0

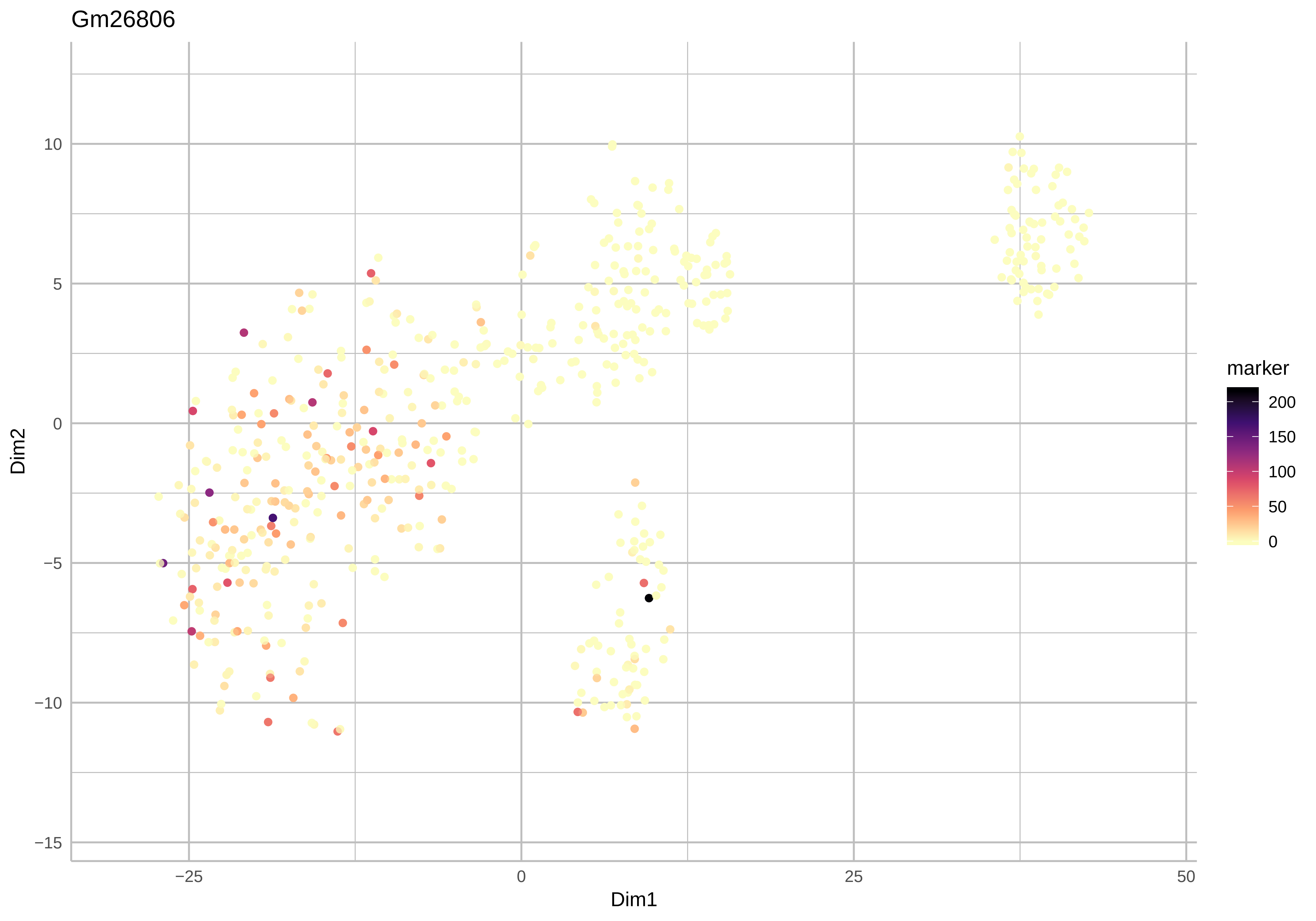
Gm26806
Dim2
Dim1
marker
200
150
100
50
0
10
5
0
−5
−10
−15
−25
0
25
50

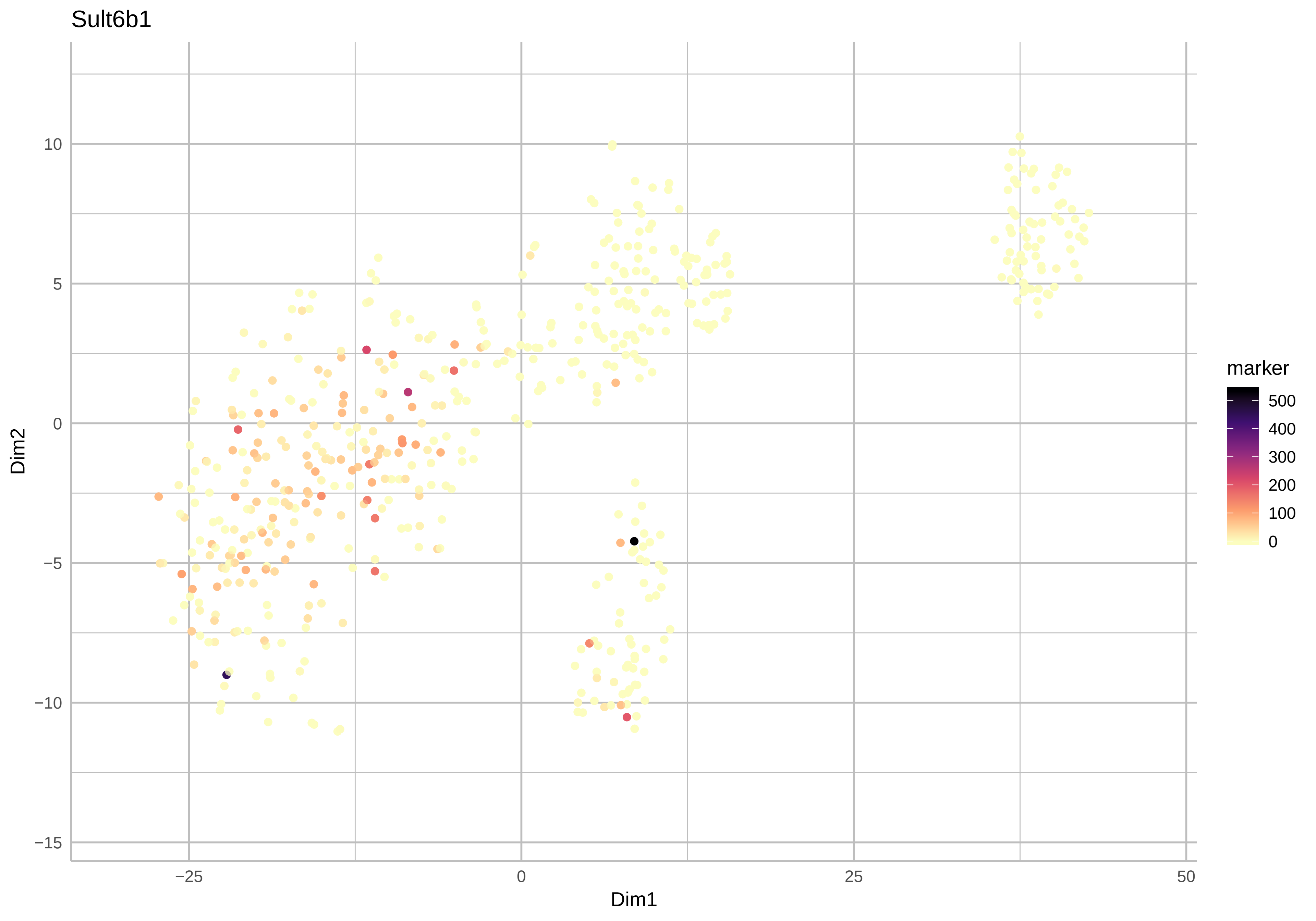

Sult6b1
Dim2
Dim1
10
5
0
−5
−10
−15
−25
0
25
50
marker
500
400
300
200
100
0

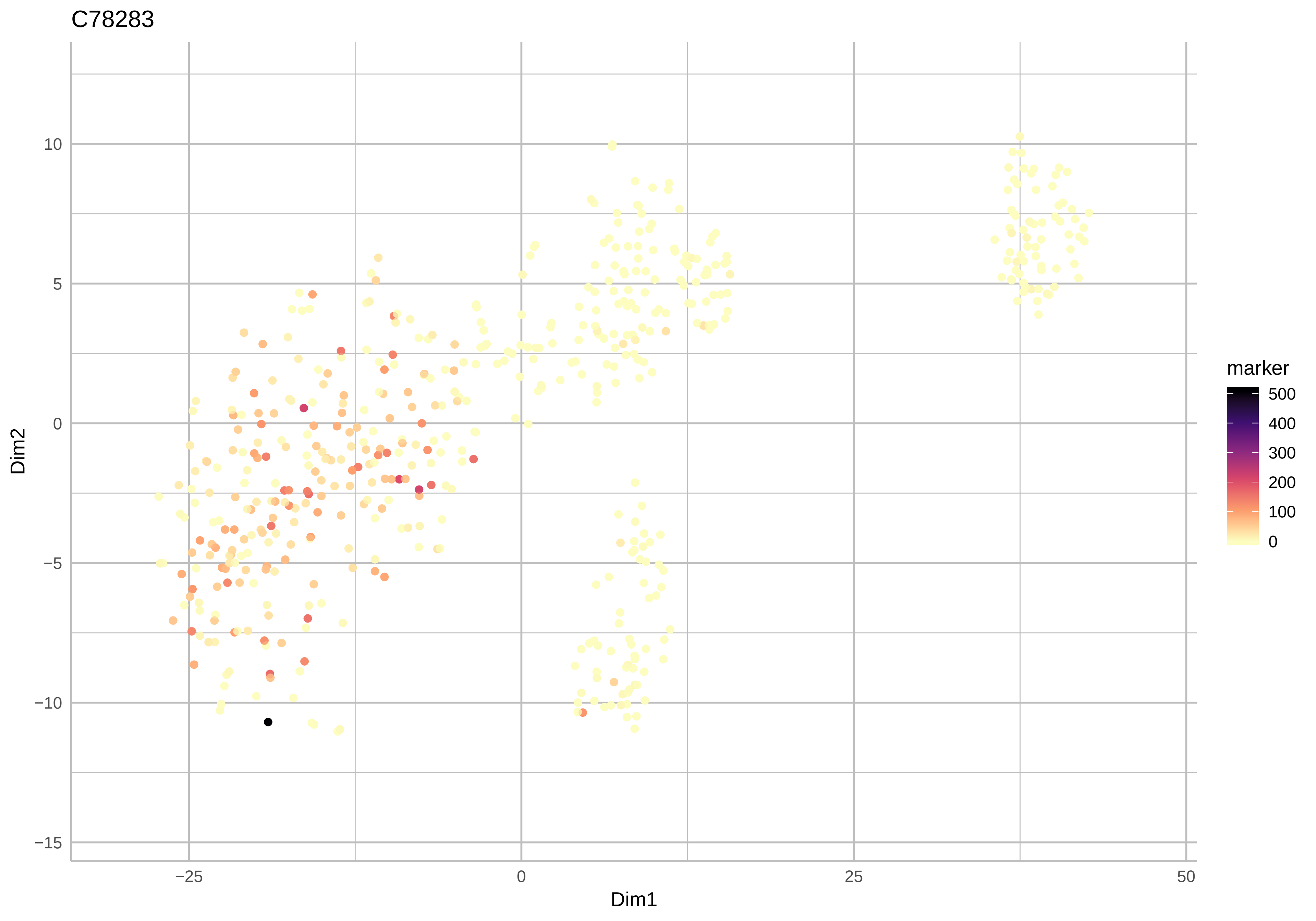
C78283
Dim2
Dim1
10
5
0
−5
−10
−15
−25
0
25
50
marker
500
400
300
200
100
0

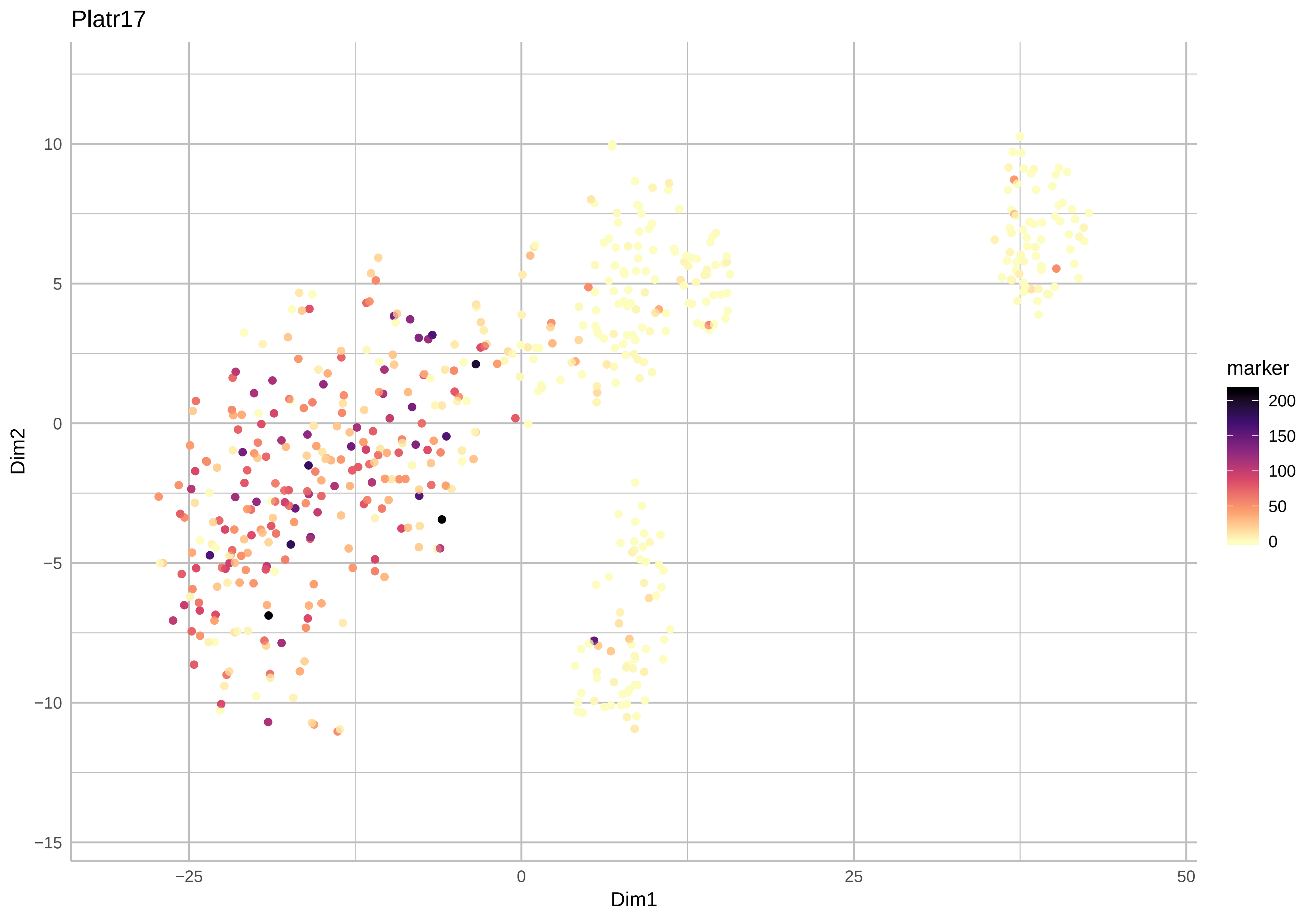
Platr17
Dim2
Dim1
10
5
0
−5
−10
−15
−25
0
25
50
marker
200
150
100
50
0

Platr21

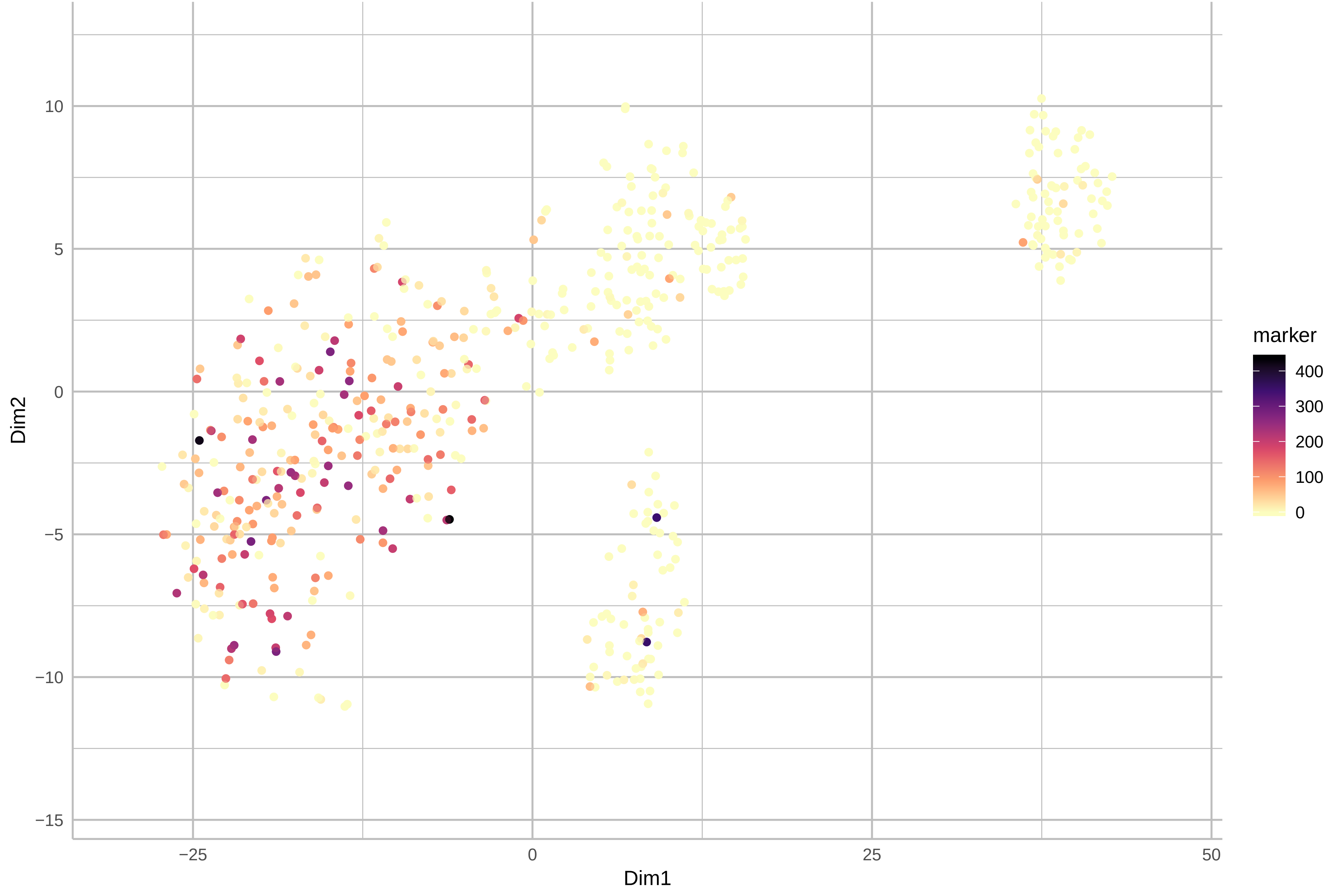

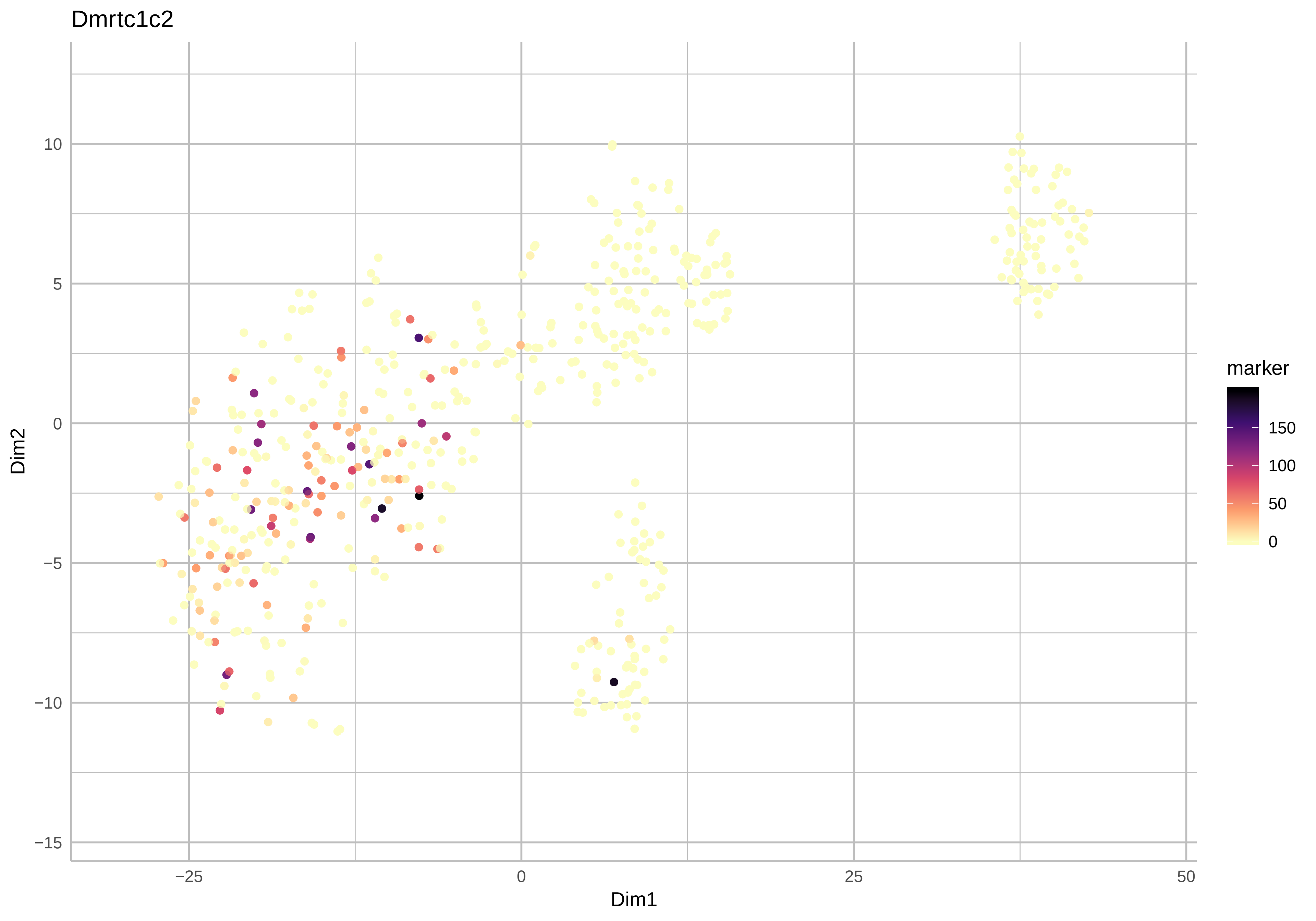
Dmrtc1c2
Dim2
Dim1
marker
150
100
50
0
10
5
0
−5
−10
−15
−25
0
25
50

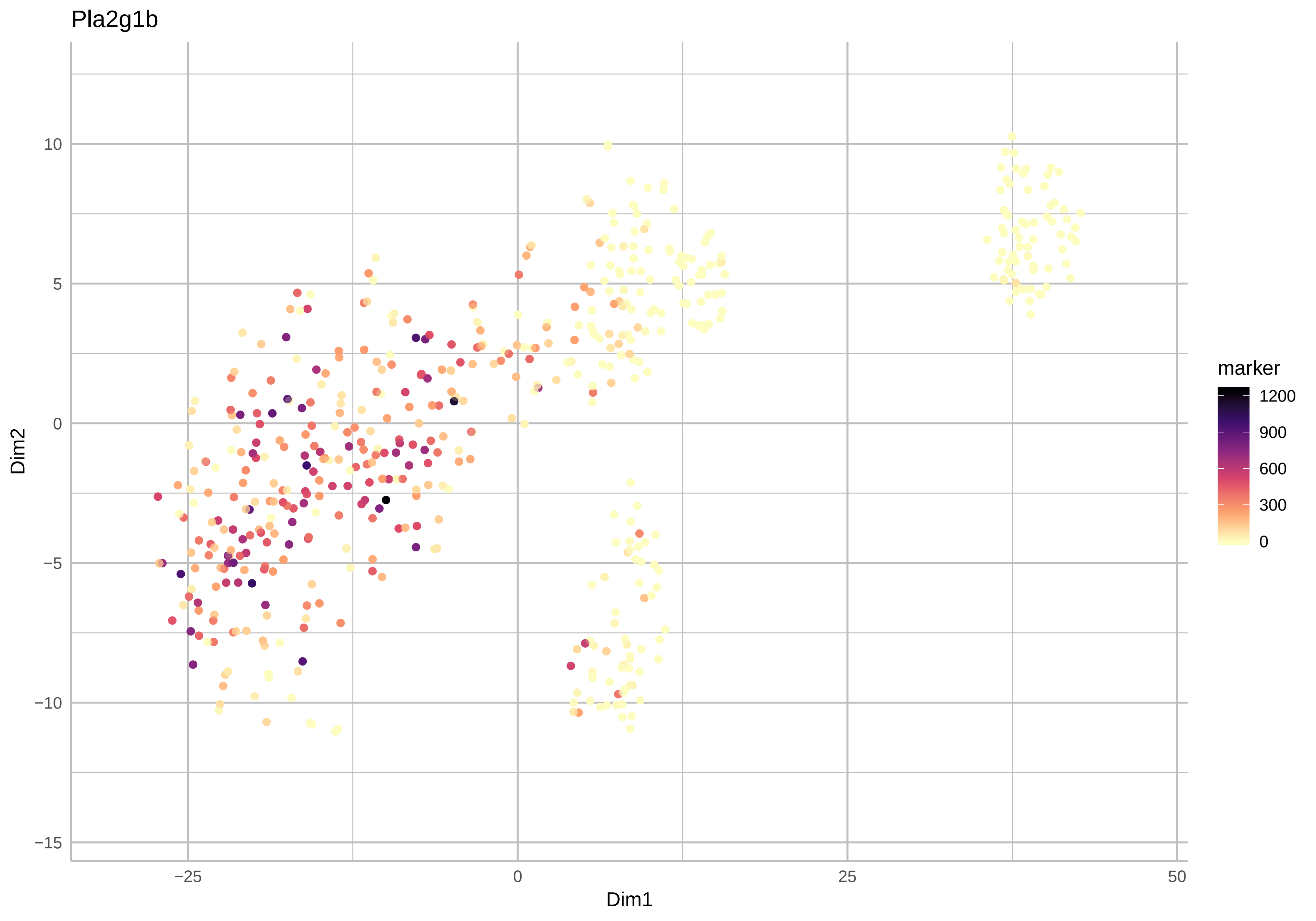
Pla2g1b
Dim2
Dim1
10
5
0
−5
−10
−15
−25
0
25
50
marker
1200
900
600
300
0

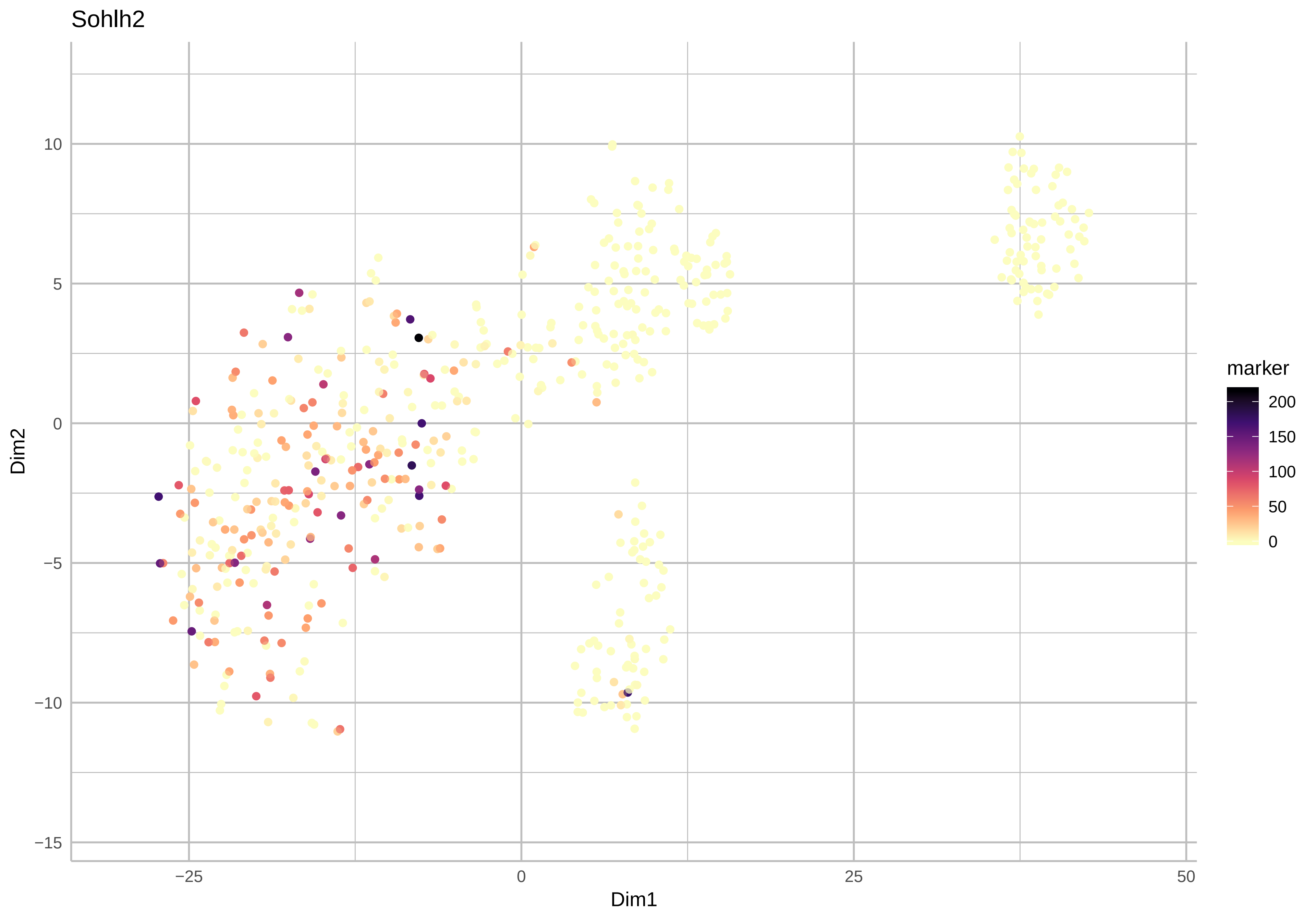

Sohlh2
Dim2
Dim1
marker
200
150
100
50
0
10
5
0
−5
−10
−15
−25
0
25
50

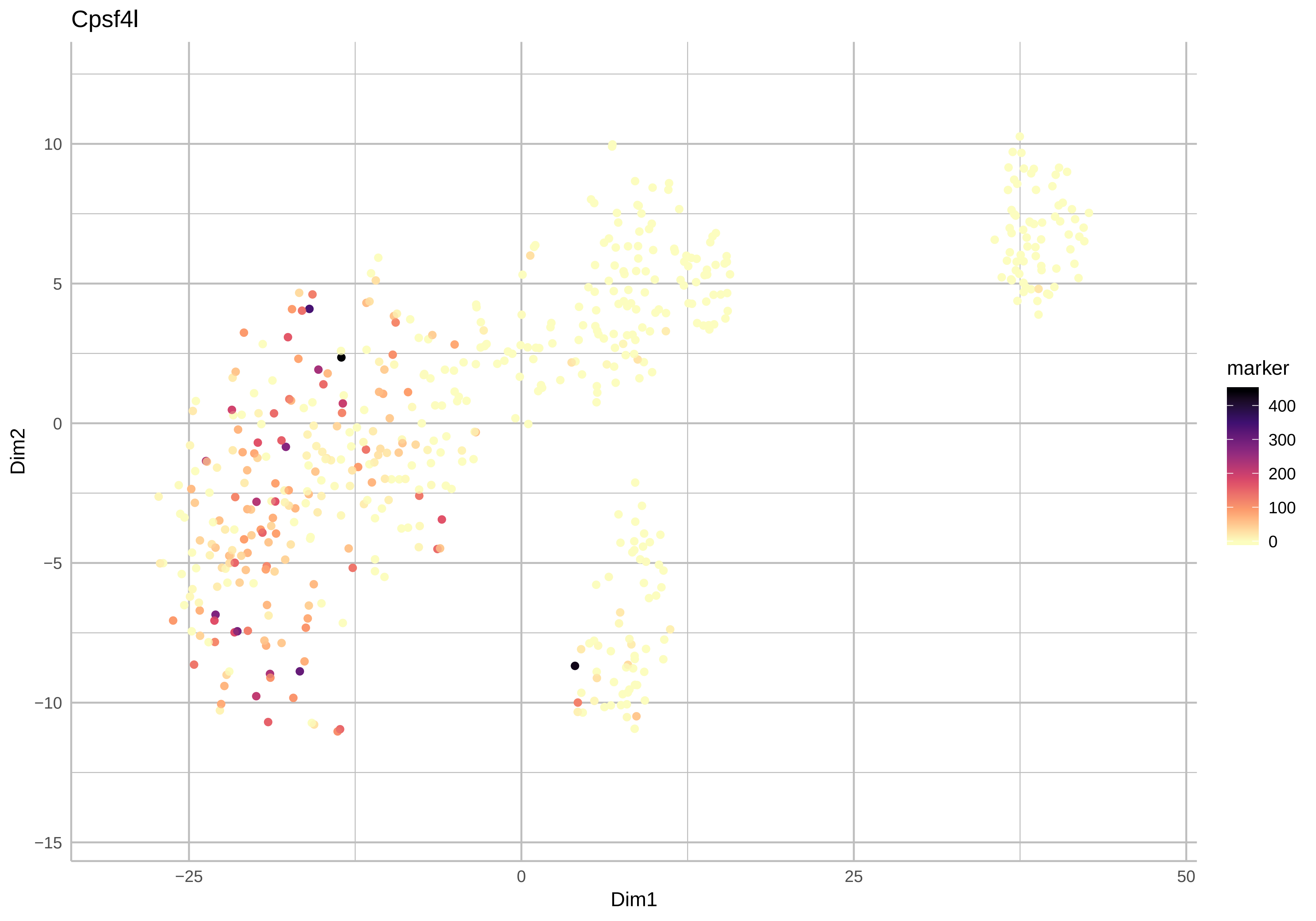
Cpsf4l
Dim2
Dim1
marker
400
300
200
100
0
10
5
0
−5
−10
−15
−25
0
25
50

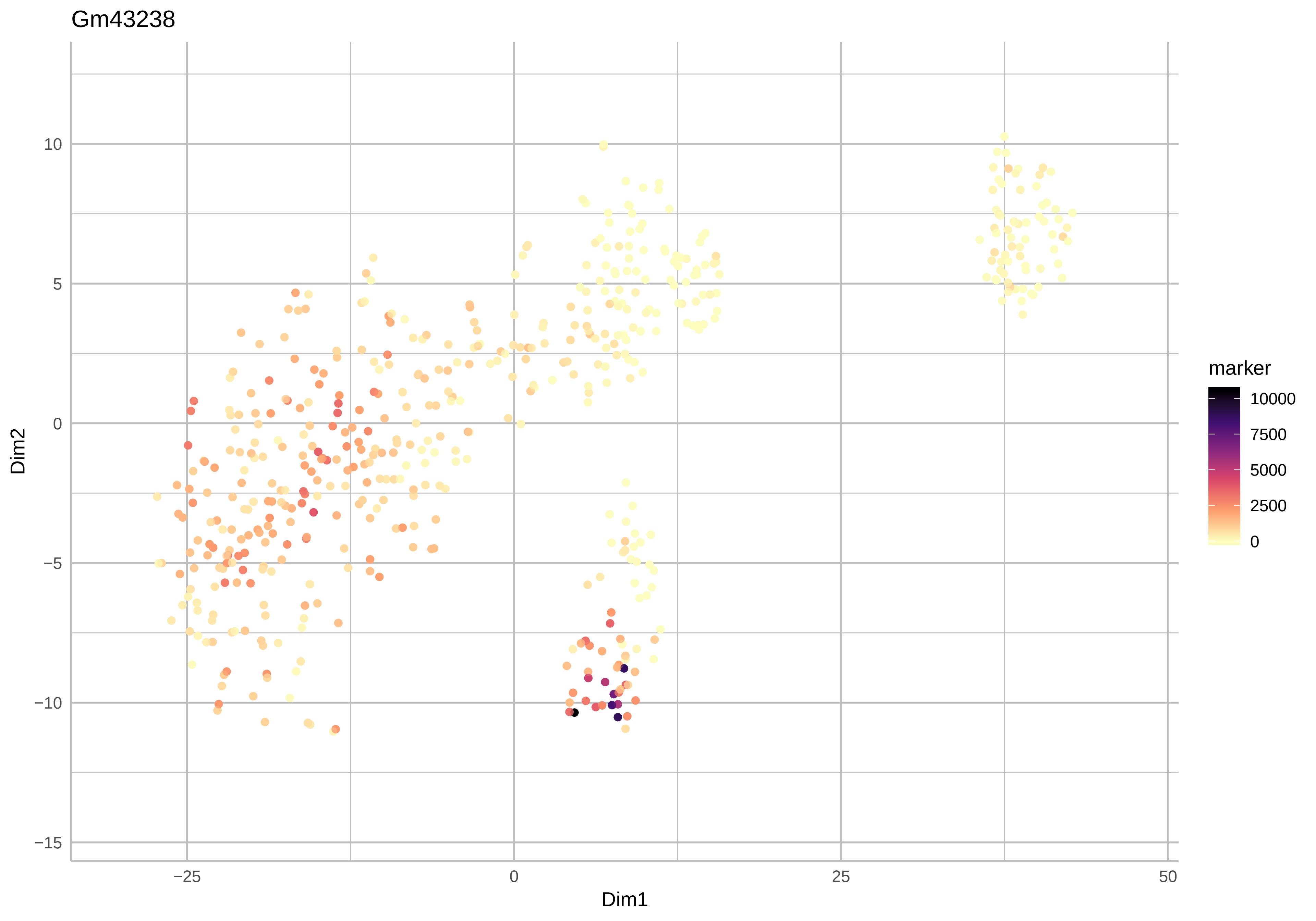

Gm43238
10
5
0
−5
−10
−15
Dim2
−25
0
25
50
Dim1
marker
10000
7500
5000
2500
0

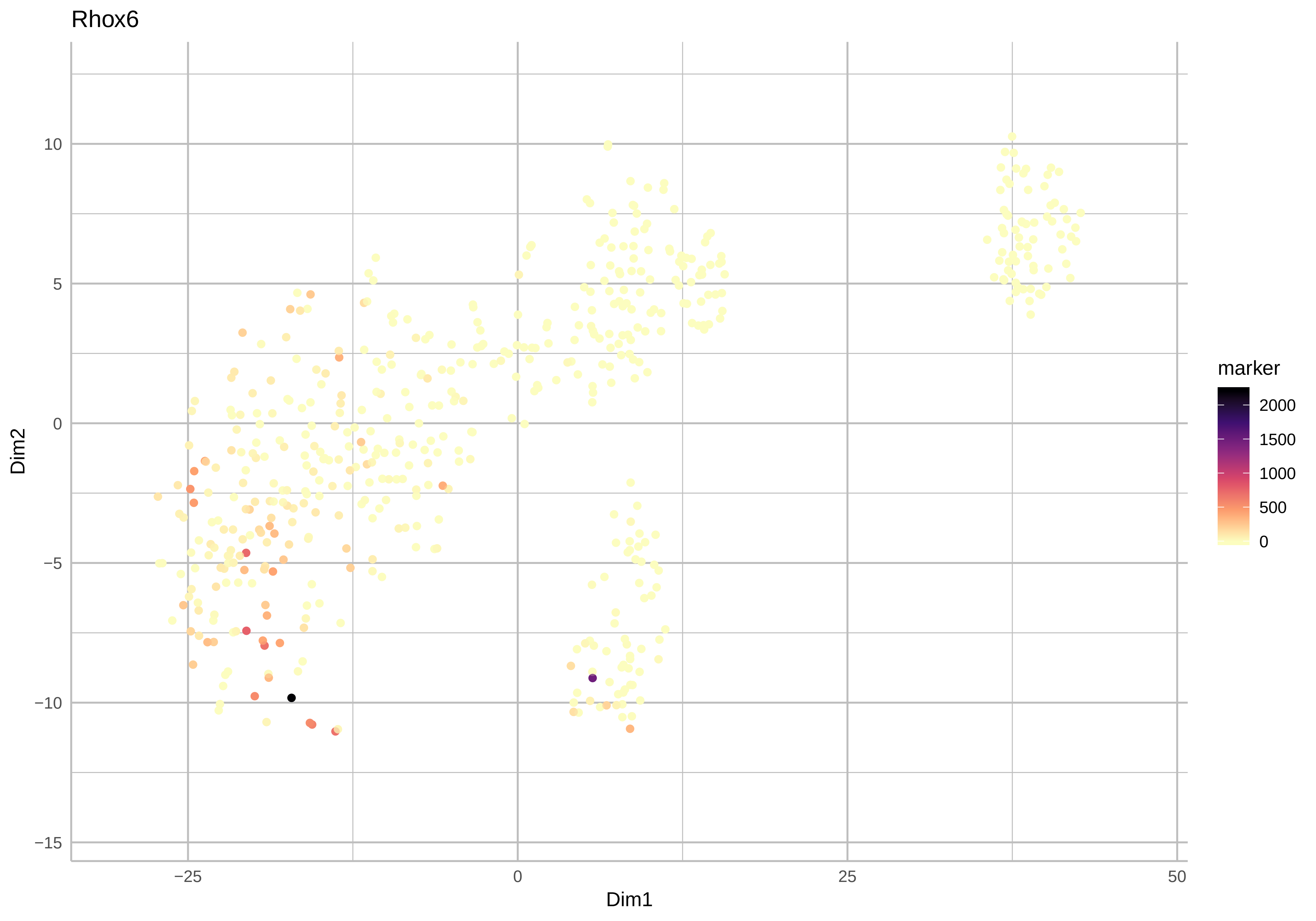

Rhox6
Dim2
Dim1
marker
2000
1500
1000
500
0
10
5
0
−5
−10
−15
−25
0
25
50

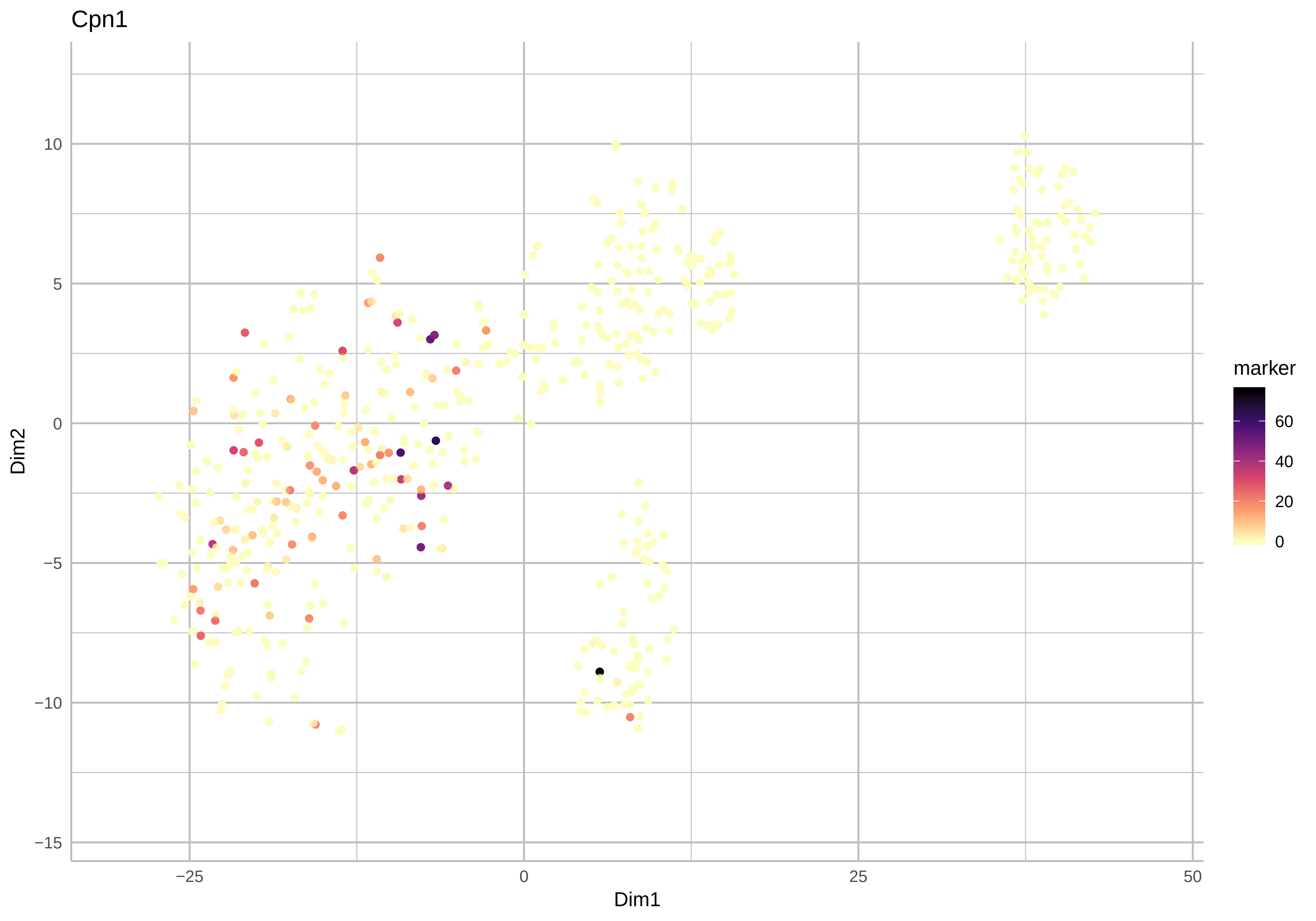

Cpn1
Dim2
Dim1
marker
60
40
20
0
10
5
0
−5
−10
−15
−25
0
25
50

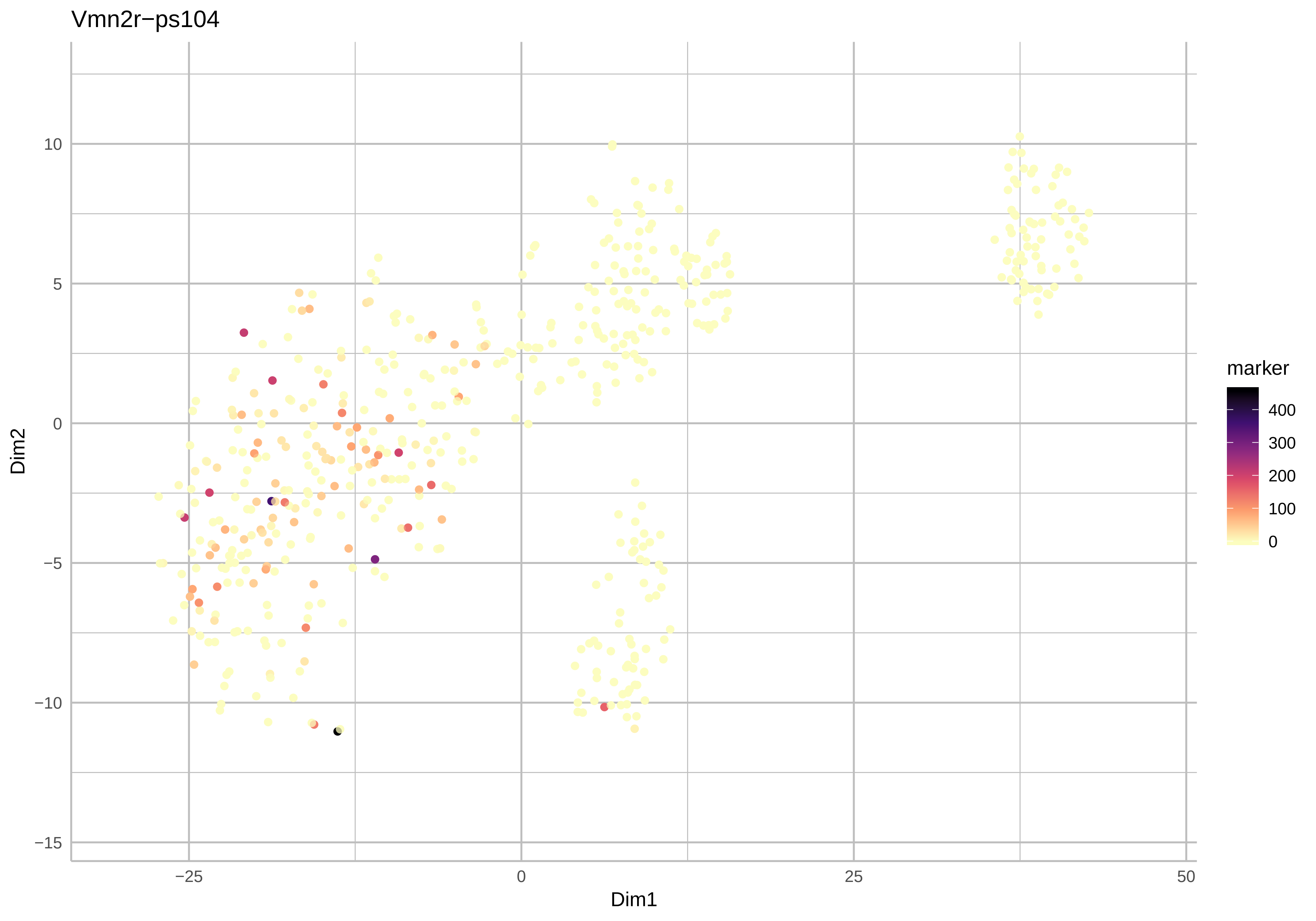

Vmn2r−ps104
Dim2
Dim1
marker
400
300
200
100
0
10
5
0
−5
−10
−15
−25
0
25
50

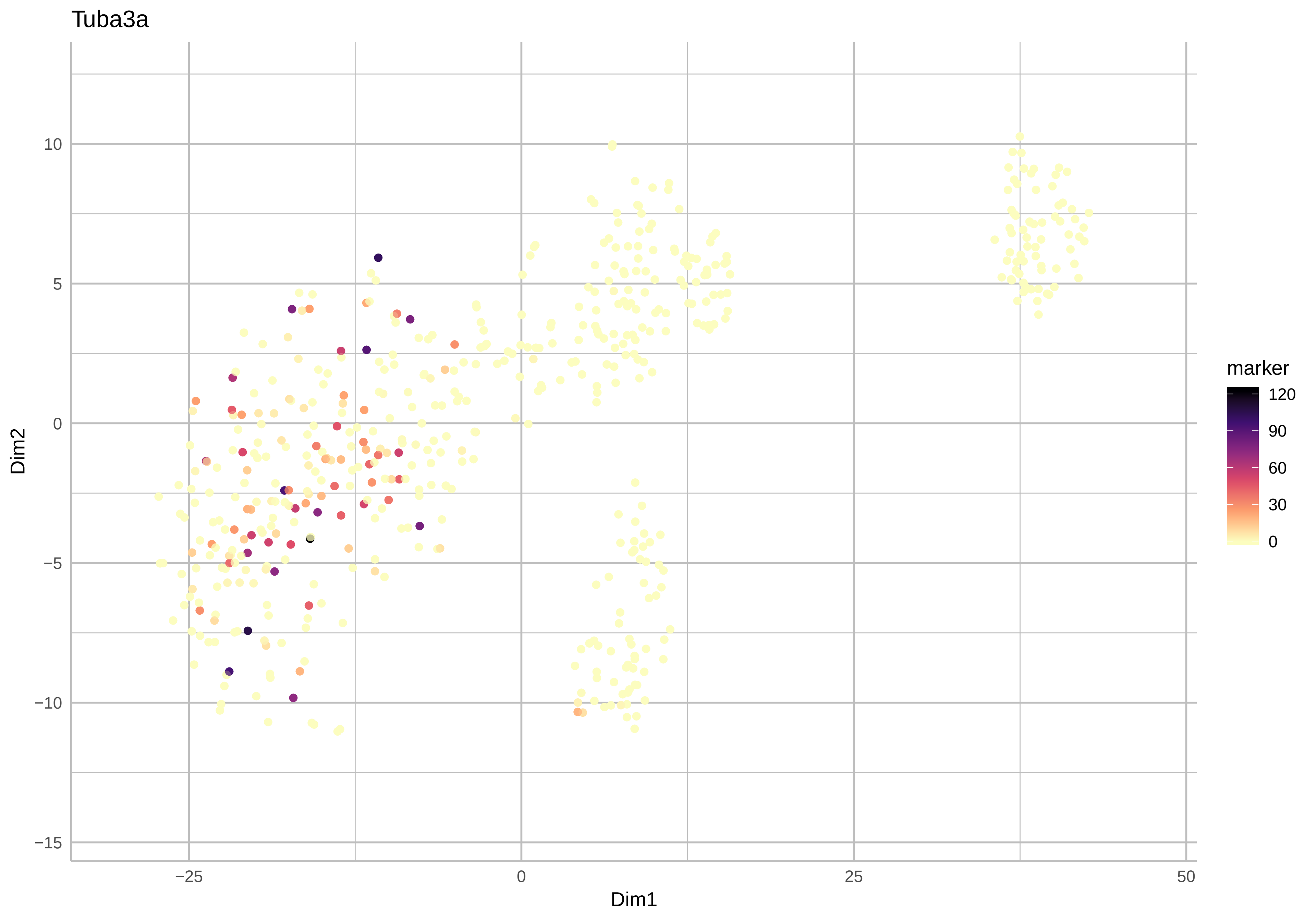

Tuba3a
10
5
0
−5
−10
−15
Dim2
−25
0
25
50
Dim1
marker
120
90
60
30
0

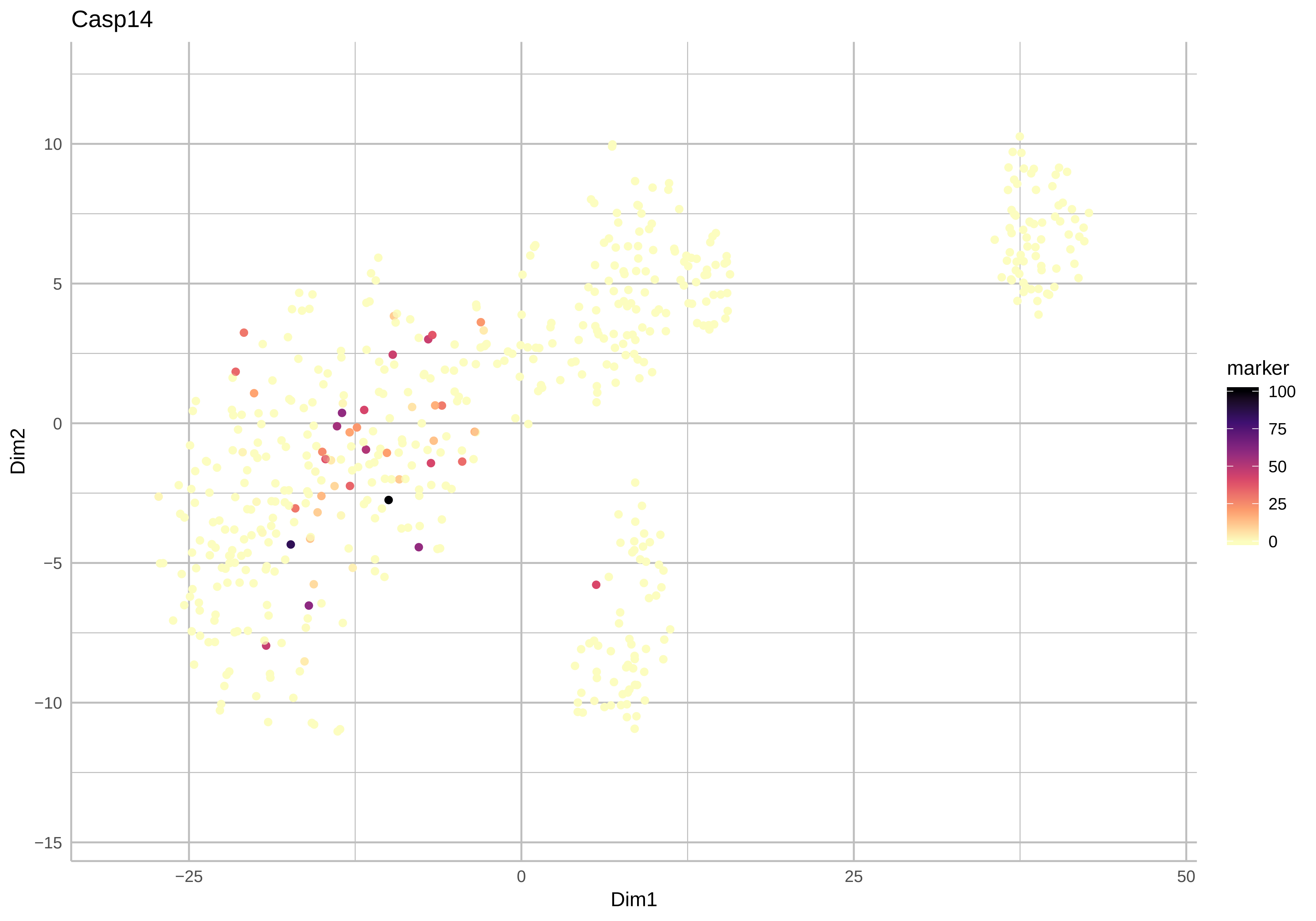

Casp14
Dim2
Dim1
10
5
0
−5
−10
−15
−25
0
25
50
marker
100
75
50
25
0

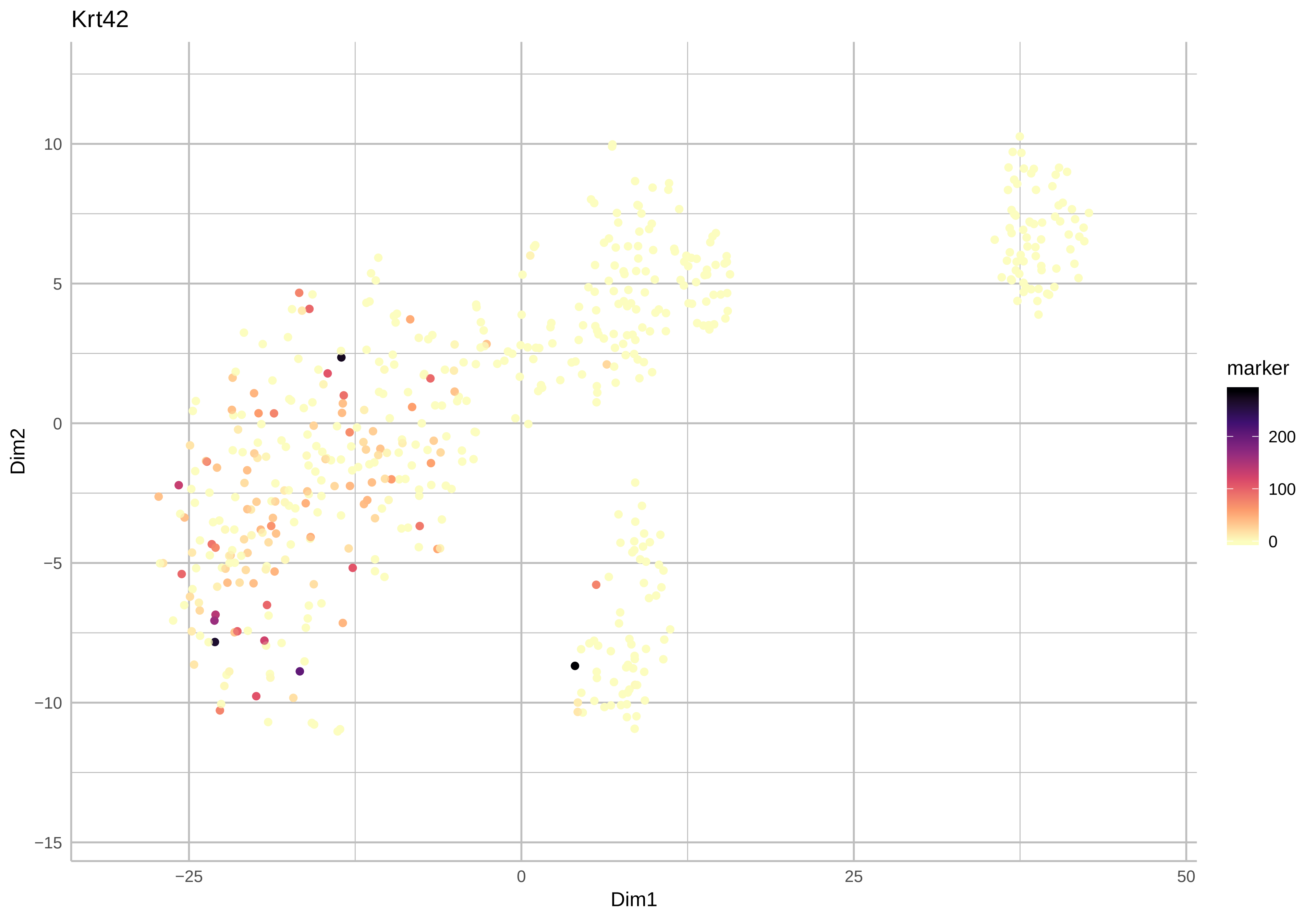

Krt42
10
5
0
−5
−10
−15
Dim2
−25
0
25
50
Dim1
marker
200
100
0

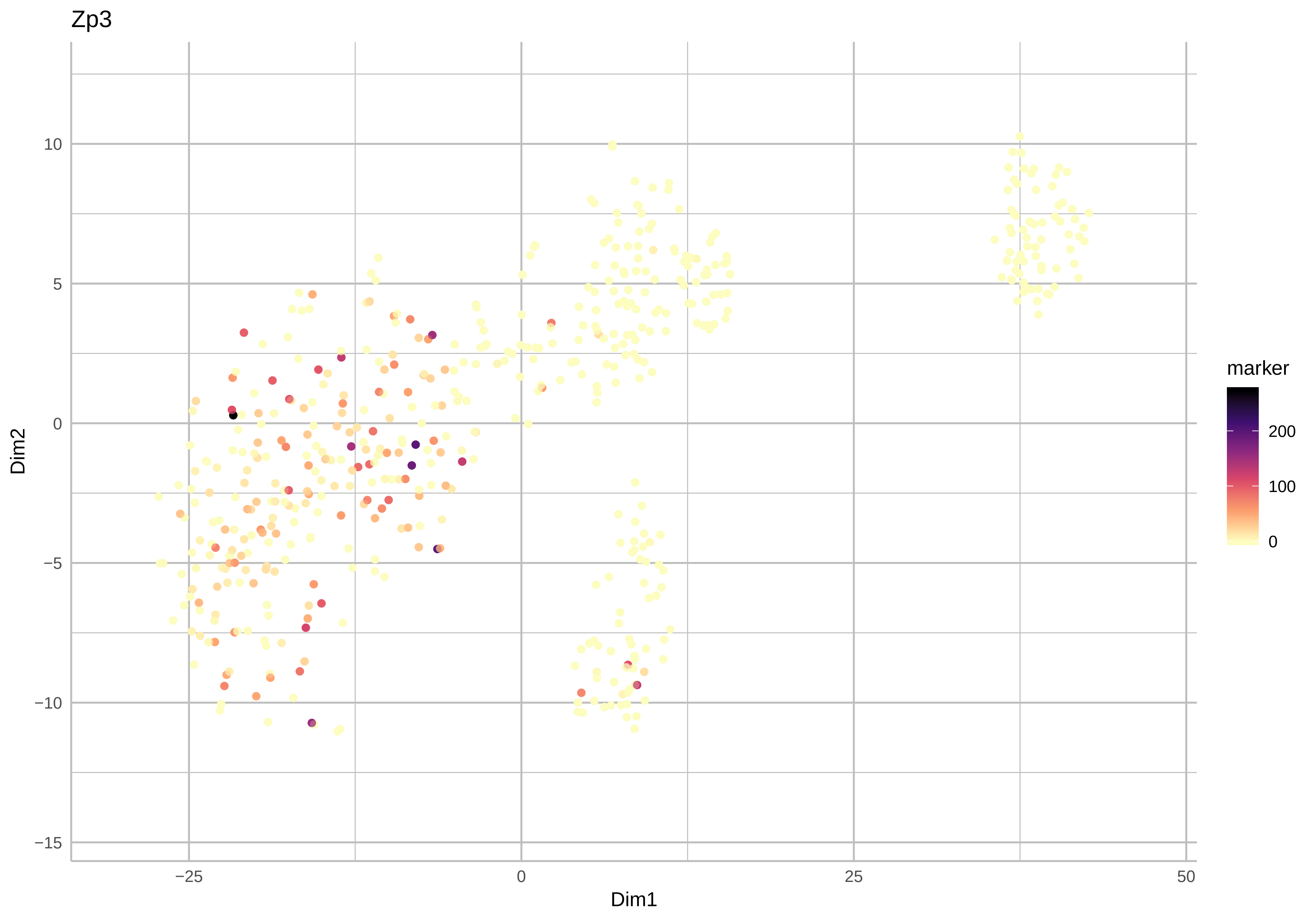
Zp3
Dim1
Dim2
marker
200
100
0
10
5
0
−5
−10
−15
−25
0
25
50

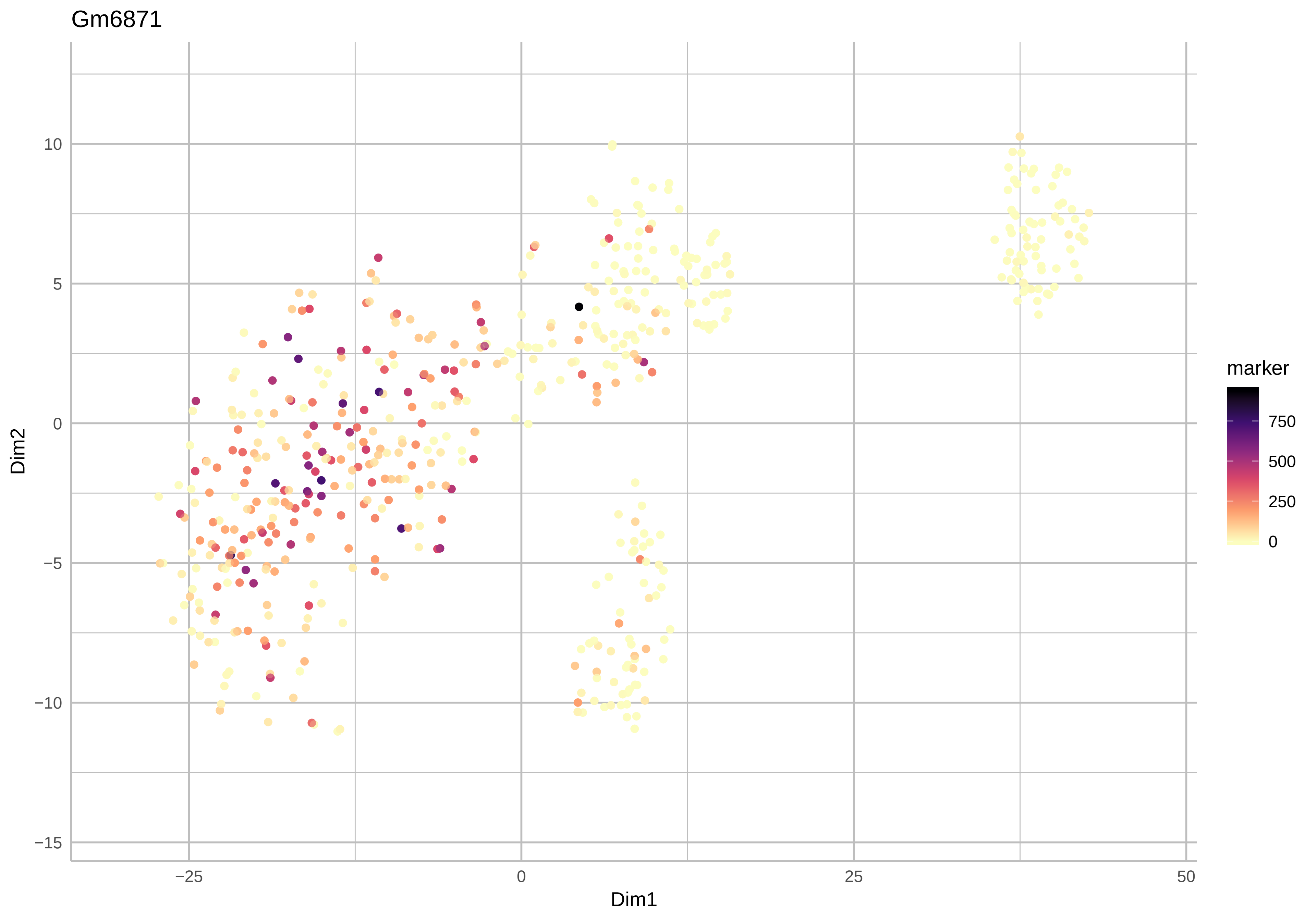

Gm6871
Dim2
Dim1
marker
750
500
250
0
10
5
0
−5
−10
−15
−25
0
25
50

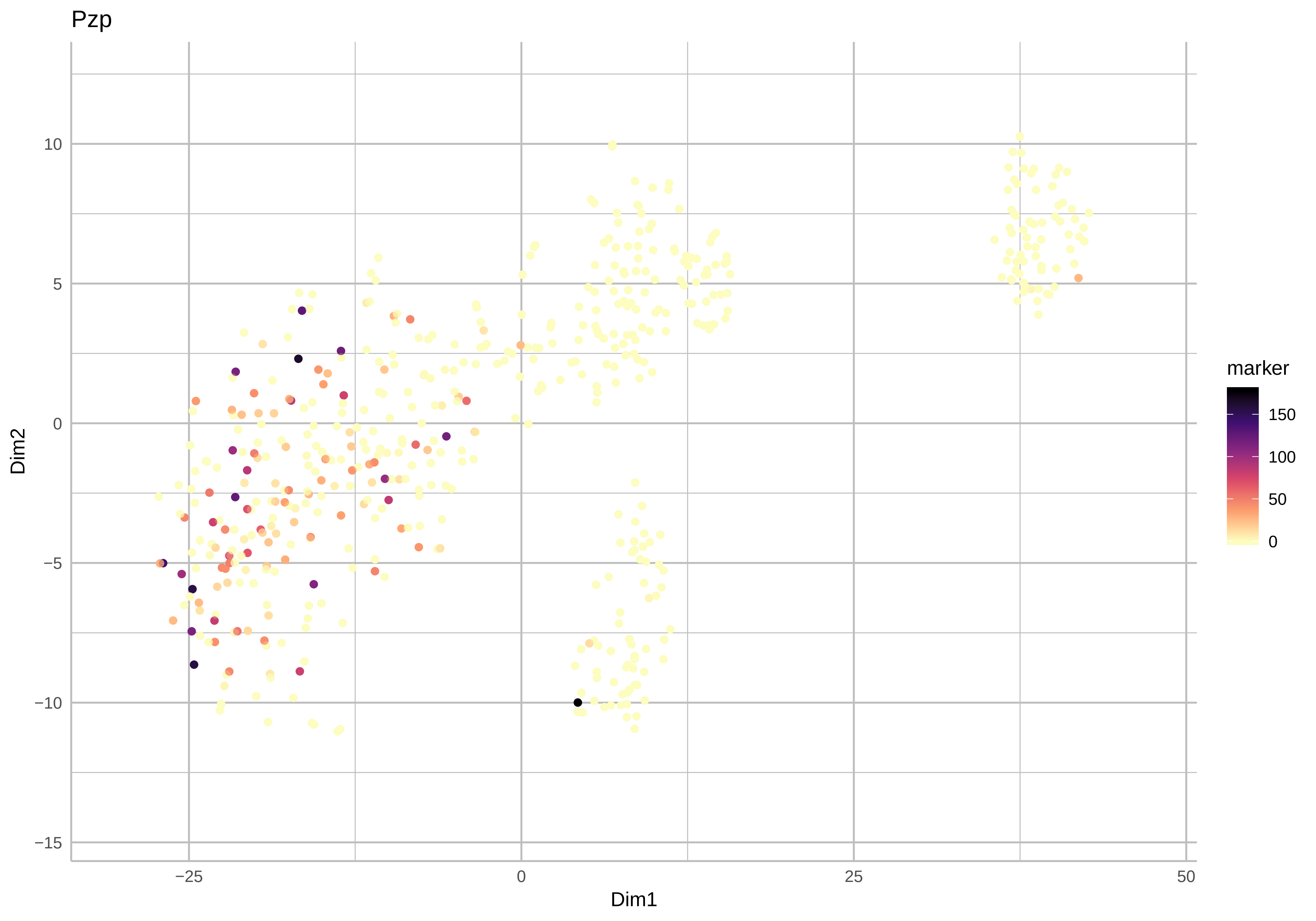

Pzp
Dim2
Dim1
10
5
0
−5
−10
−15
−25
0
25
50
marker
150
100
50
0

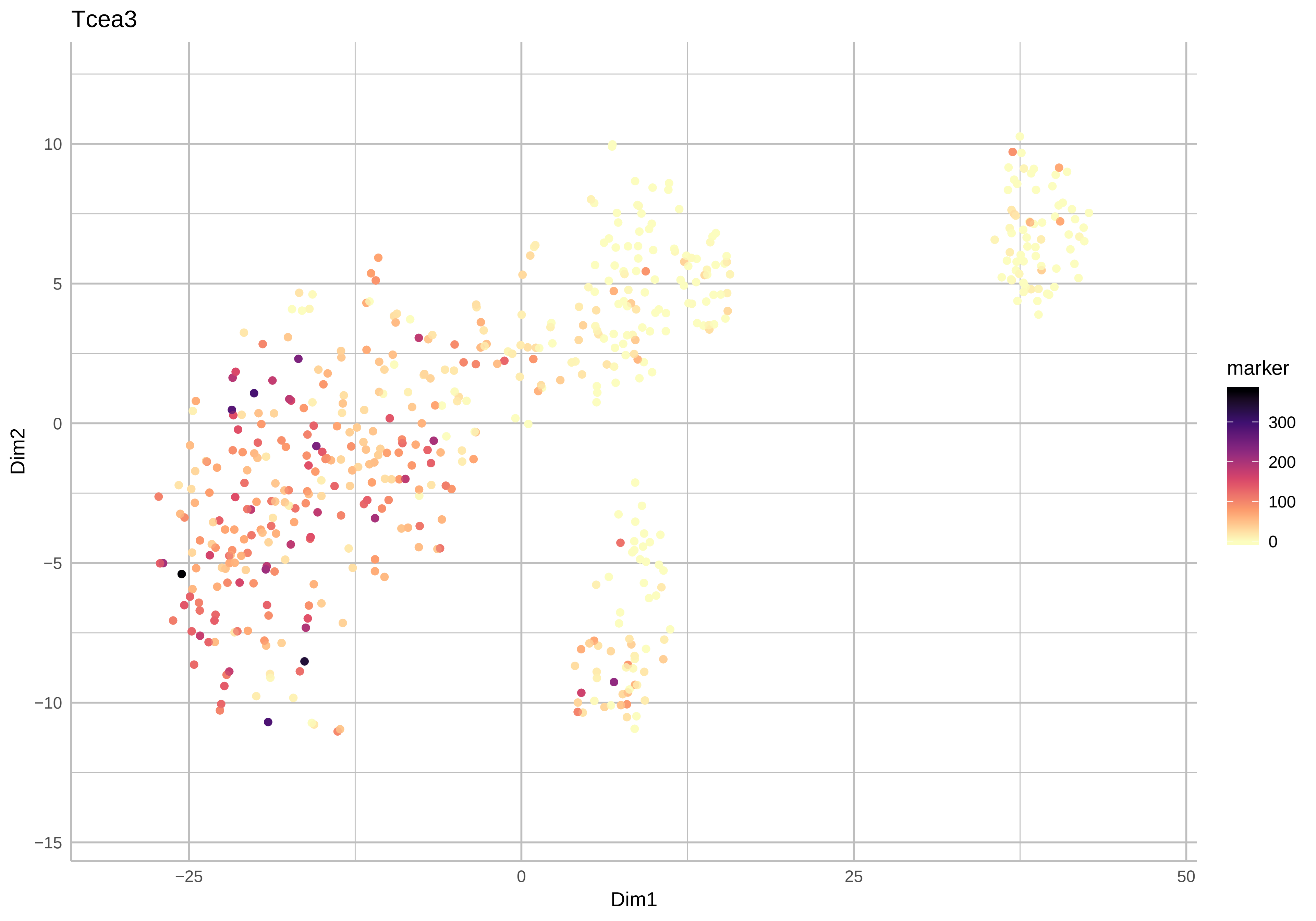
Tcea3
Dim1
Dim2
−25
0
25
50
−15
−10
−5
0
5
10
marker
300
200
100
0

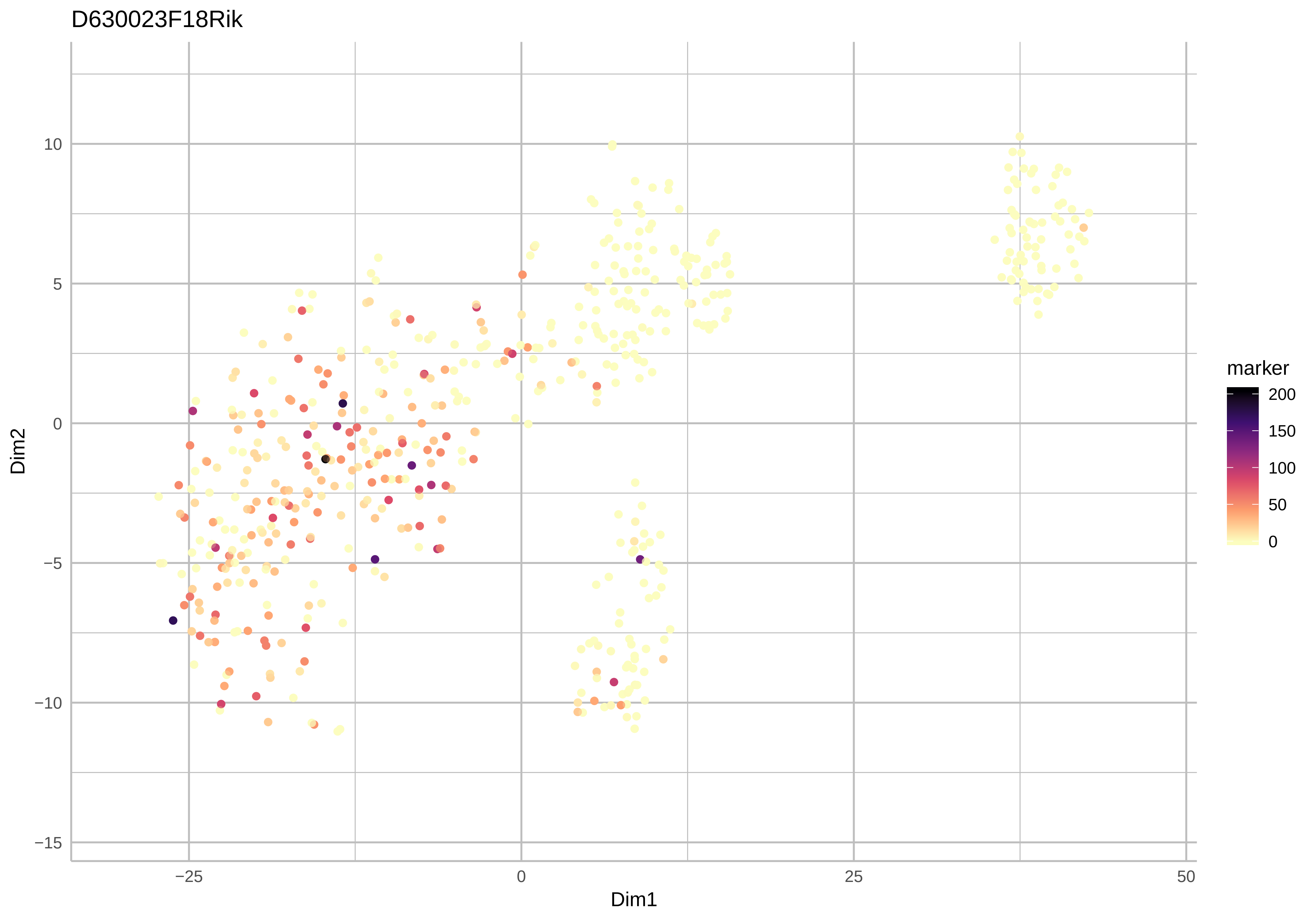
D630023F18Rik
Dim1
Dim2
marker
200
150
100
50
0
−25
0
25
50
10
5
0
−5
−10
−15

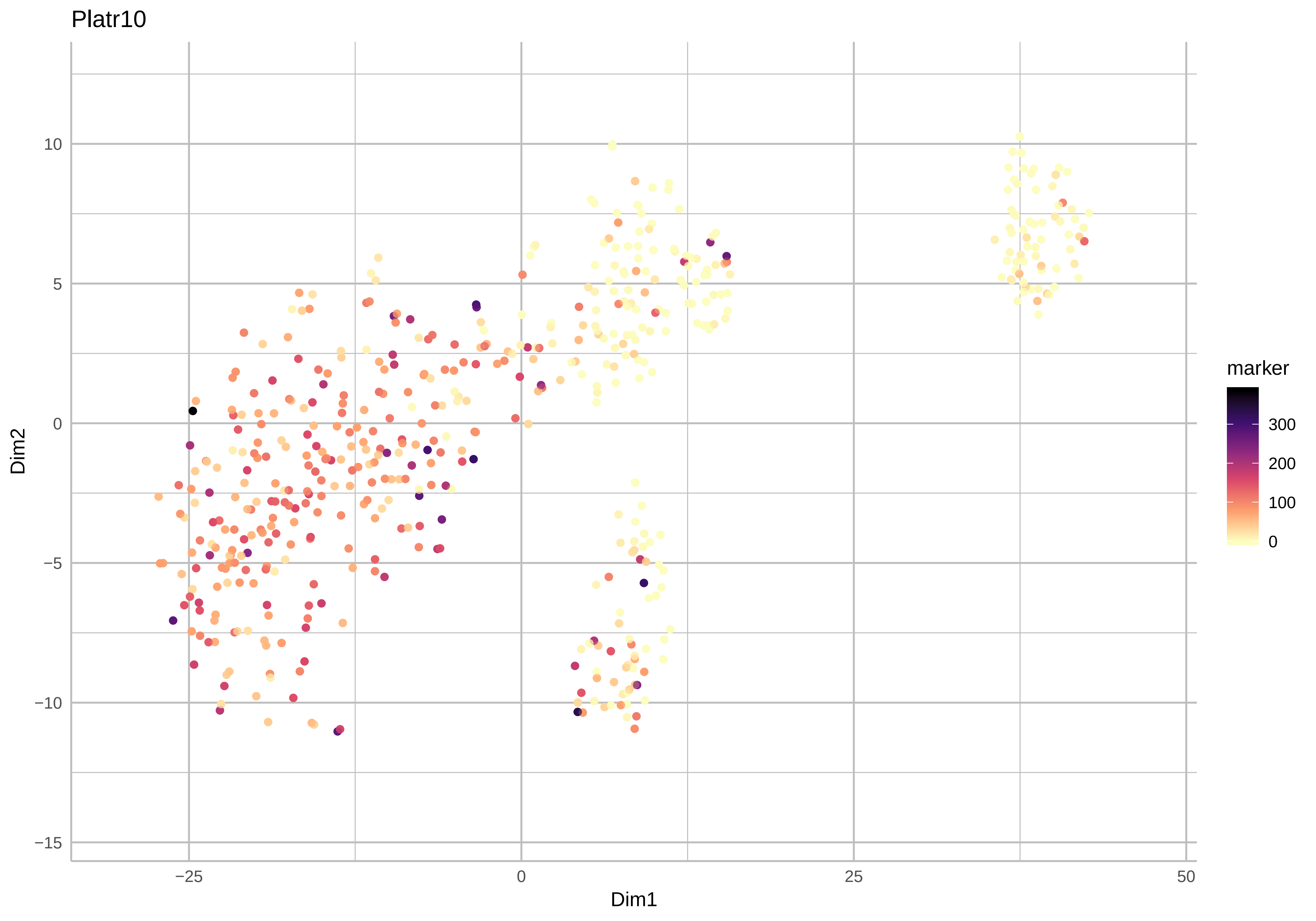
Platr10
Dim2
Dim1
10
5
0
−5
−10
−15
−25
0
25
50
marker
300
200
100
0

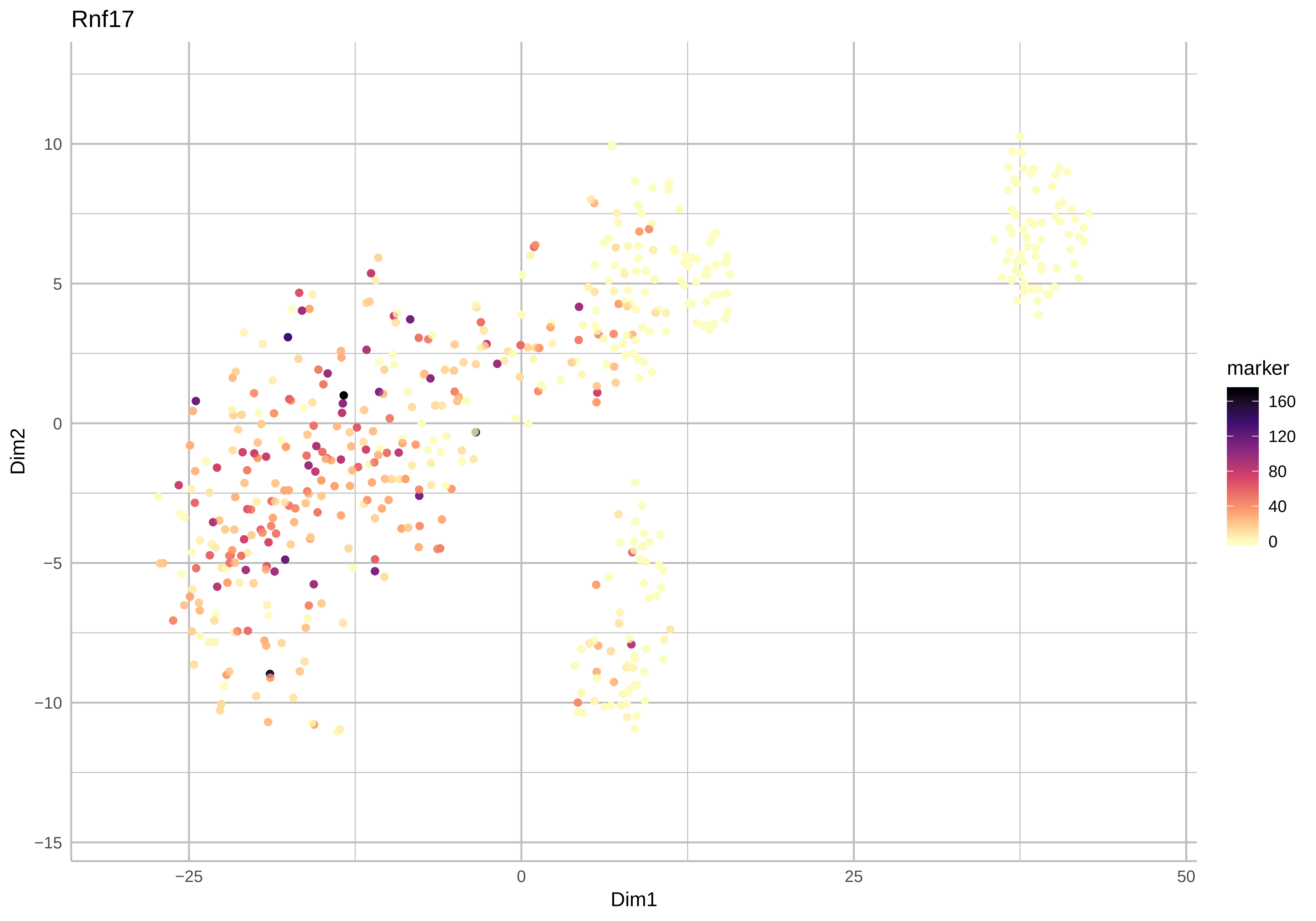

Rnf17
Dim2
Dim1
10
5
0
−5
−10
−15
−25
0
25
50
marker
160
120
80
40
0

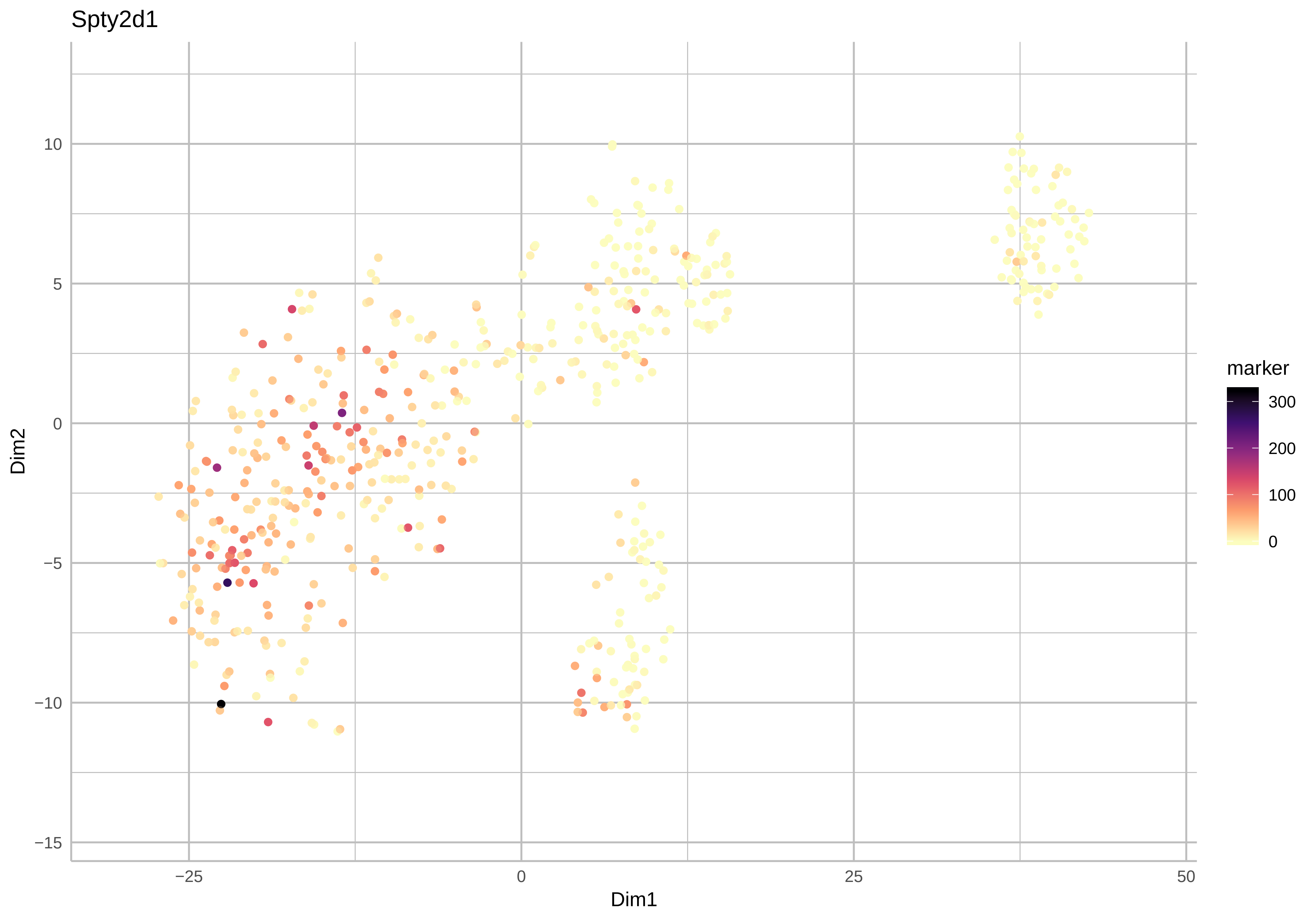
Spty2d1
Dim2
Dim1
10
5
0
−5
−10
−15
−25
0
25
50
marker
300
200
100
0

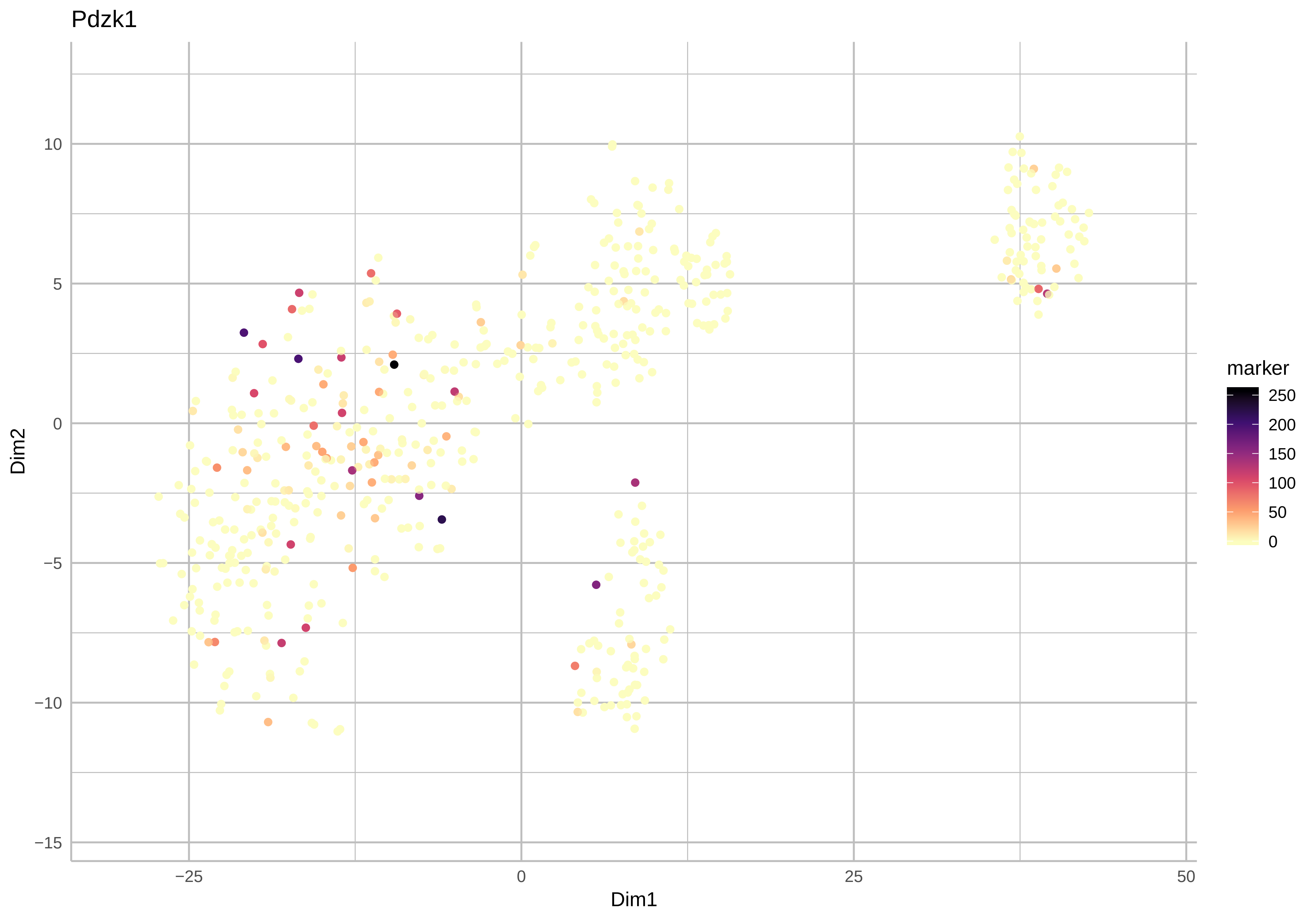
Pdzk1
Dim2
Dim1
10
5
0
−5
−10
−15
−25
0
25
50
marker
250
200
150
100
50
0

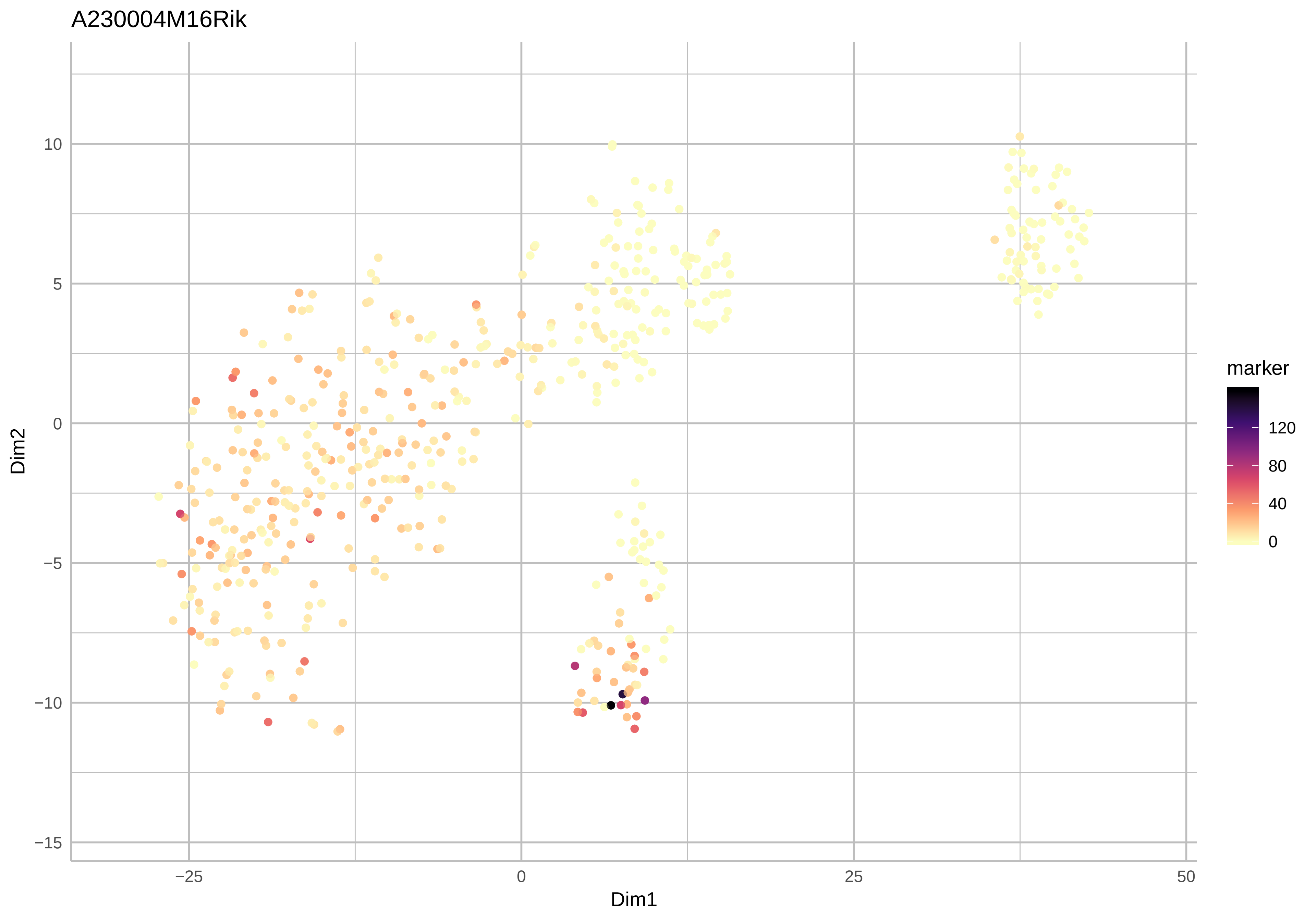
A230004M16Rik
Dim2
Dim1
10
5
0
−5
−10
−15
−25
0
25
50
marker
120
80
40
0

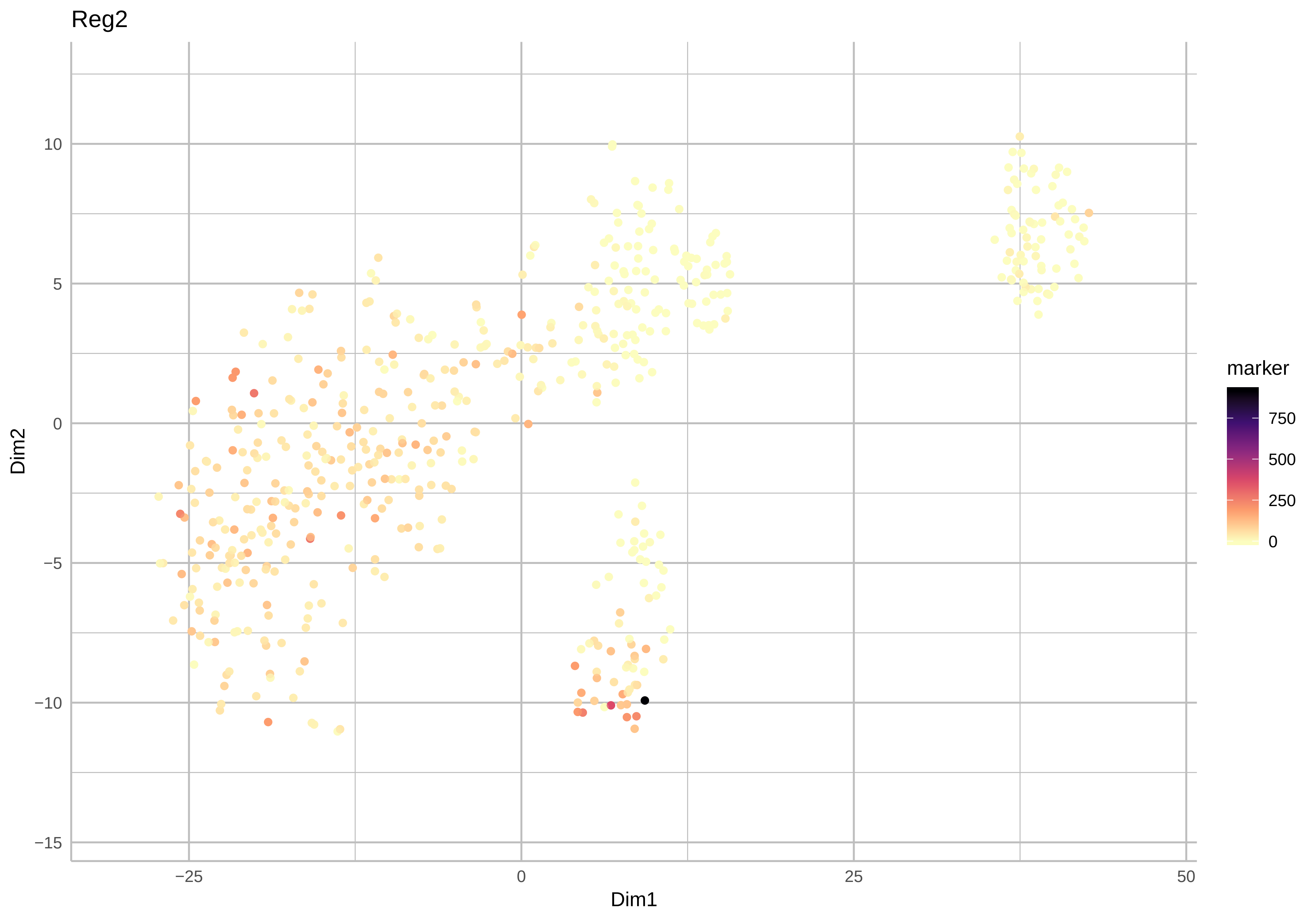
Reg2
Dim2
Dim1
10
5
0
−5
−10
−15
−25
0
25
50
marker
750
500
250
0

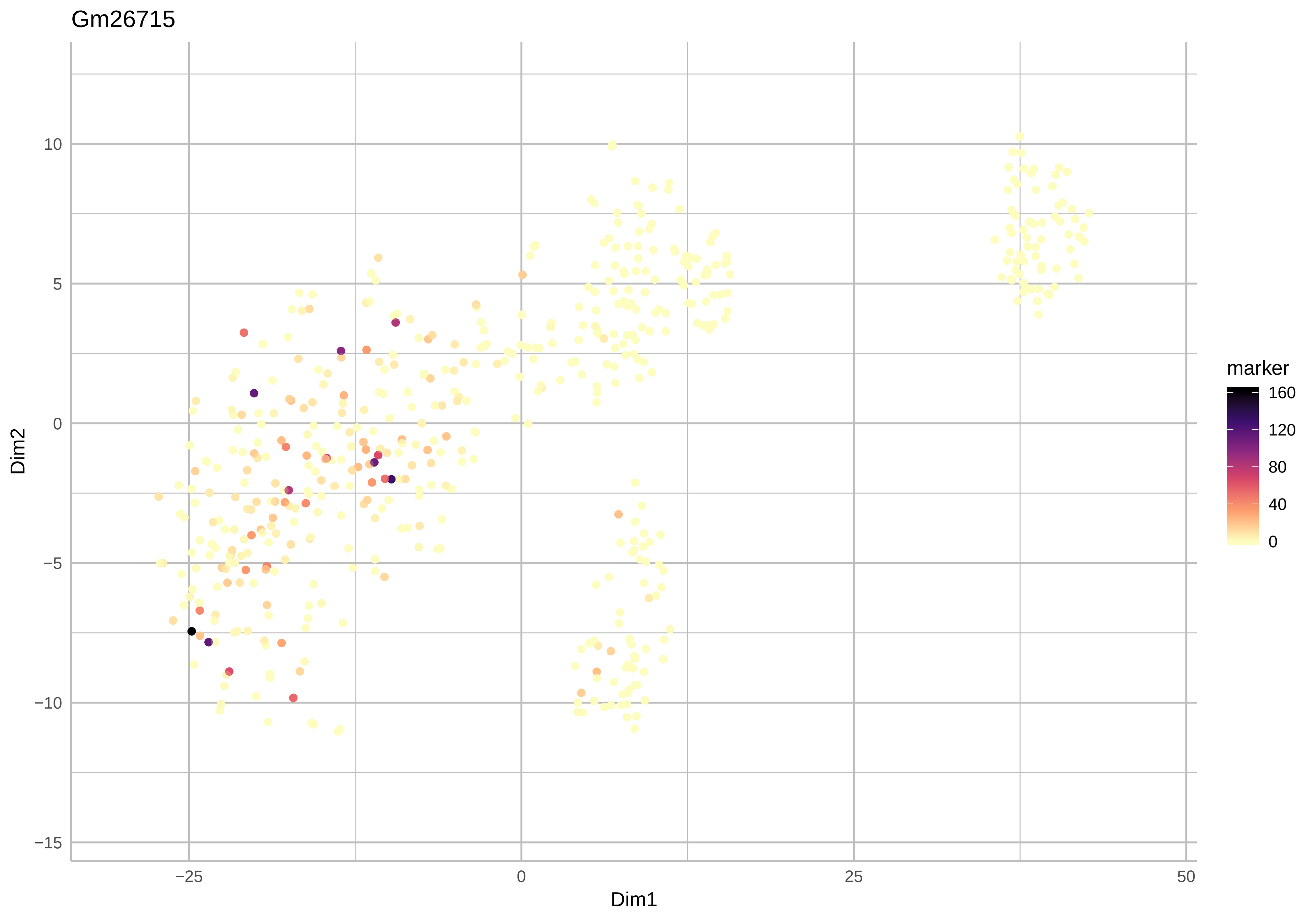

Gm26715
10
5
0
−5
−10
−15
−25
0
25
50
Dim1
Dim2
marker
160
120
80
40
0

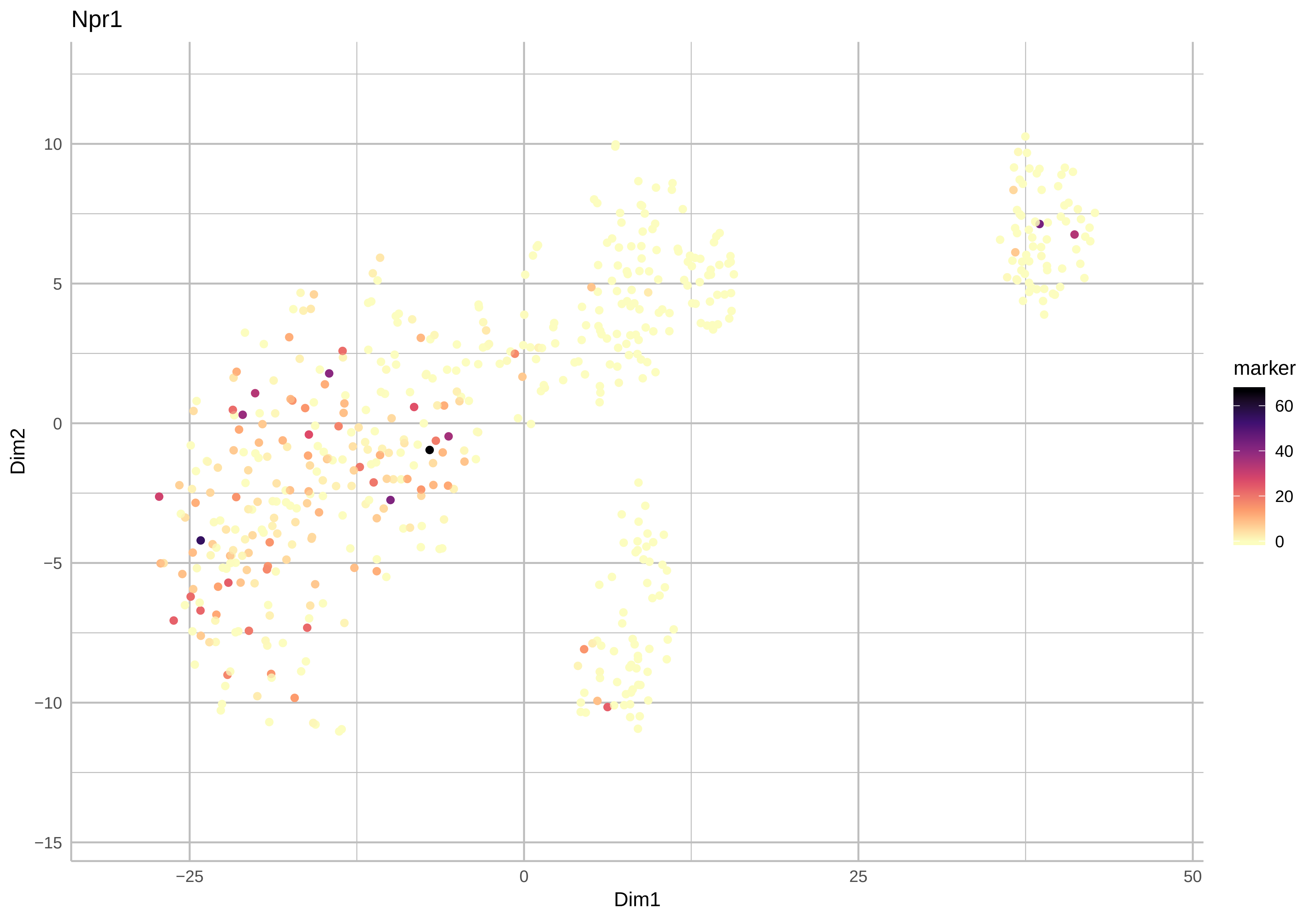

Npr1
Dim2
Dim1
10
5
0
−5
−10
−15
−25
0
25
50
marker
60
40
20
0

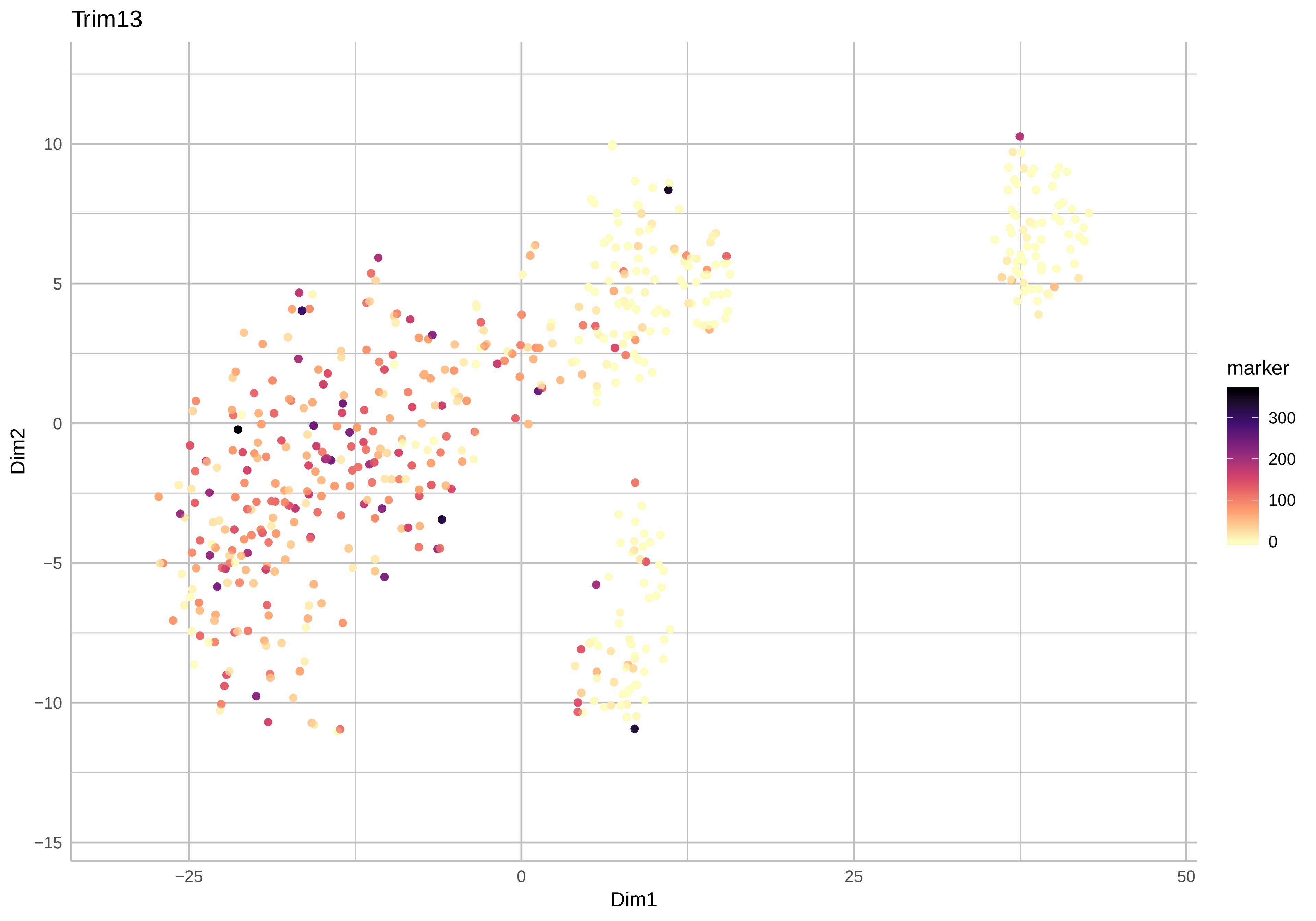

Trim13
Dim2
Dim1
10
5
0
−5
−10
−15
−25
0
25
50
marker
300
200
100
0

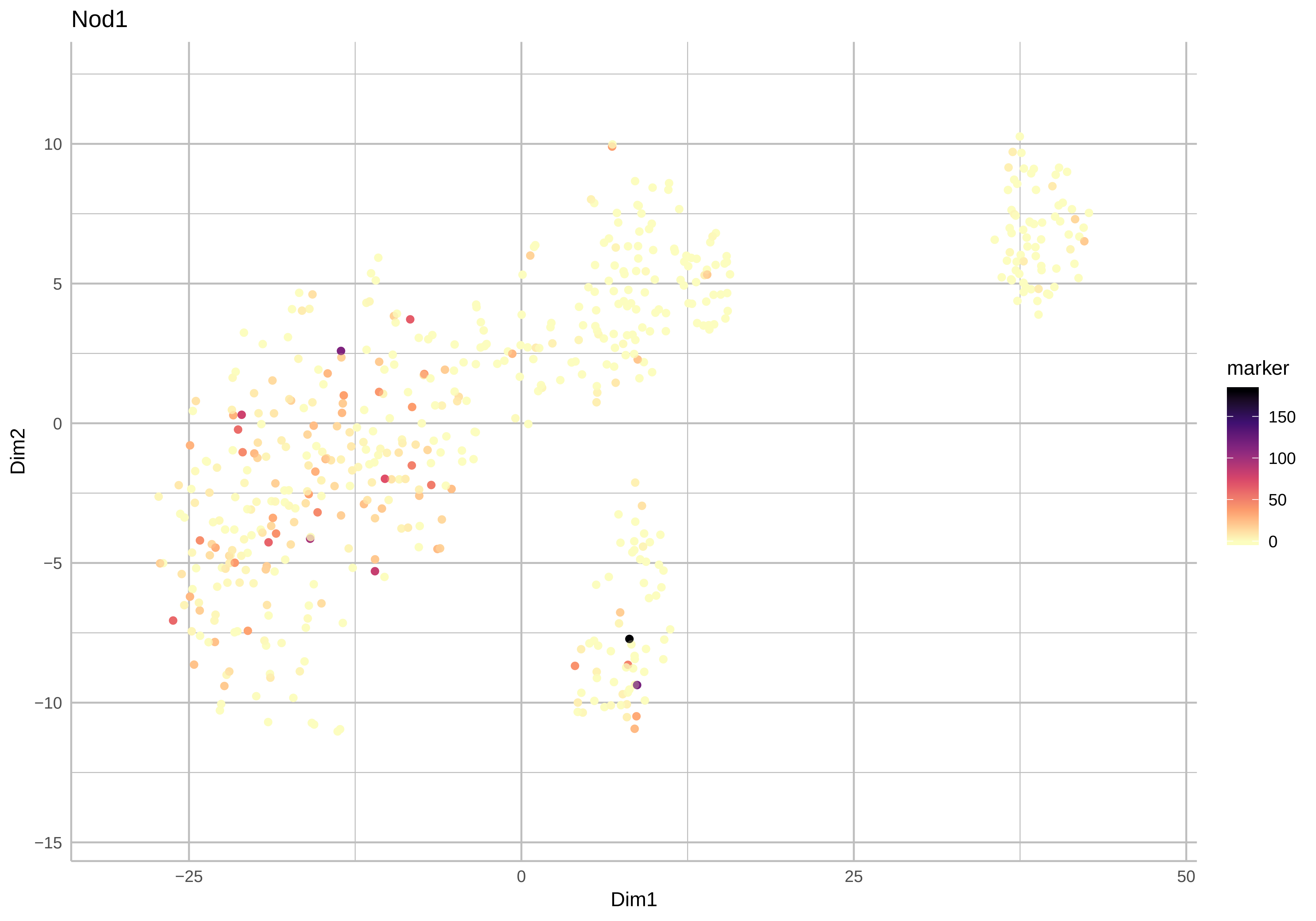

Nod1
Dim2
Dim1
10
5
0
−5
−10
−15
−25
0
25
50
marker
150
100
50
0

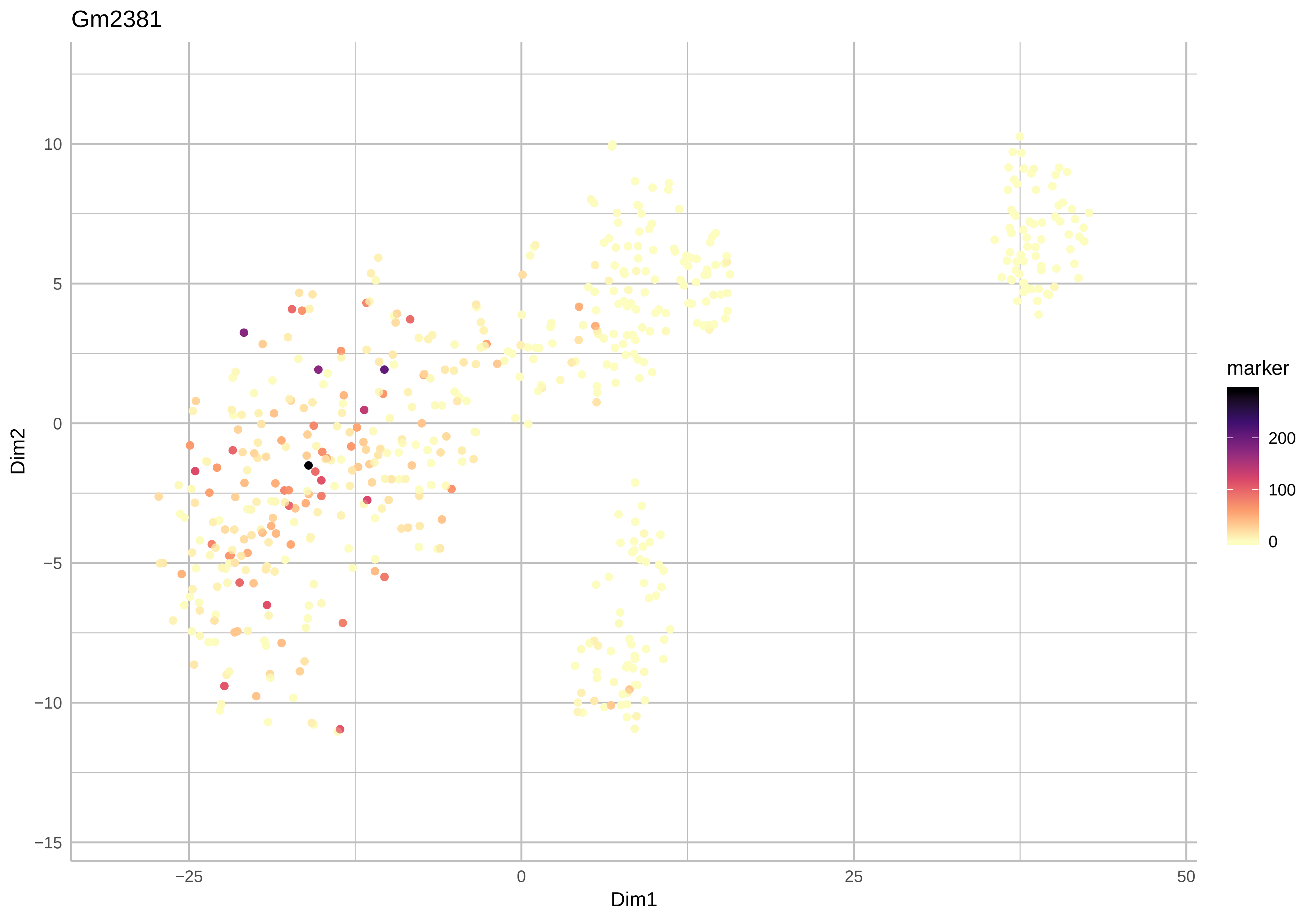

Gm2381
Dim2
Dim1
marker
200
100
0
10
5
0
−5
−10
−15
−25
0
25
50

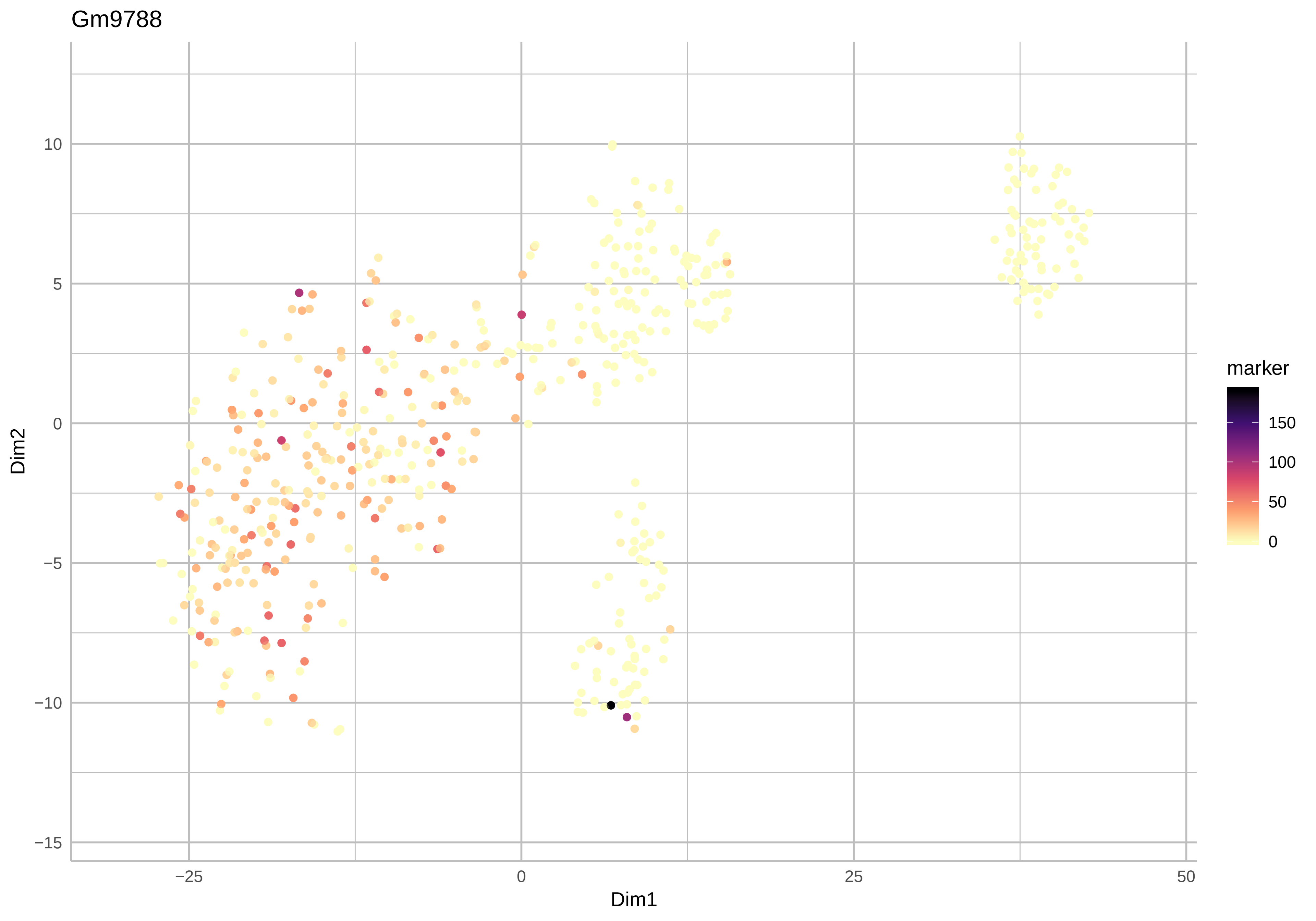
Gm9788
Dim2
Dim1
10
5
0
−5
−10
−15
−25
0
25
50
marker
150
100
50
0

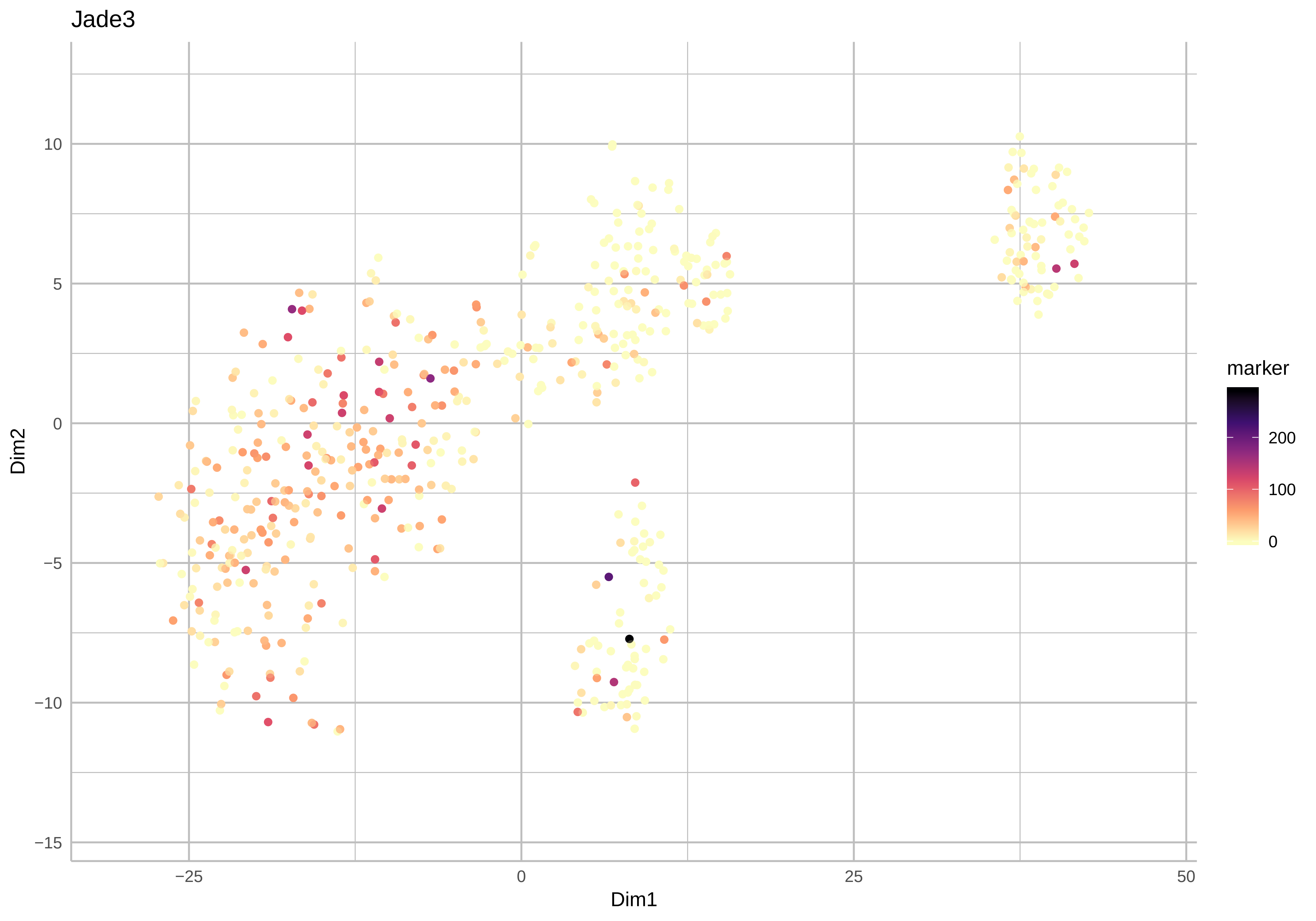

Jade3
10
5
0
−5
−10
−15
Dim2
−25
0
25
50
Dim1
marker
200
100
0

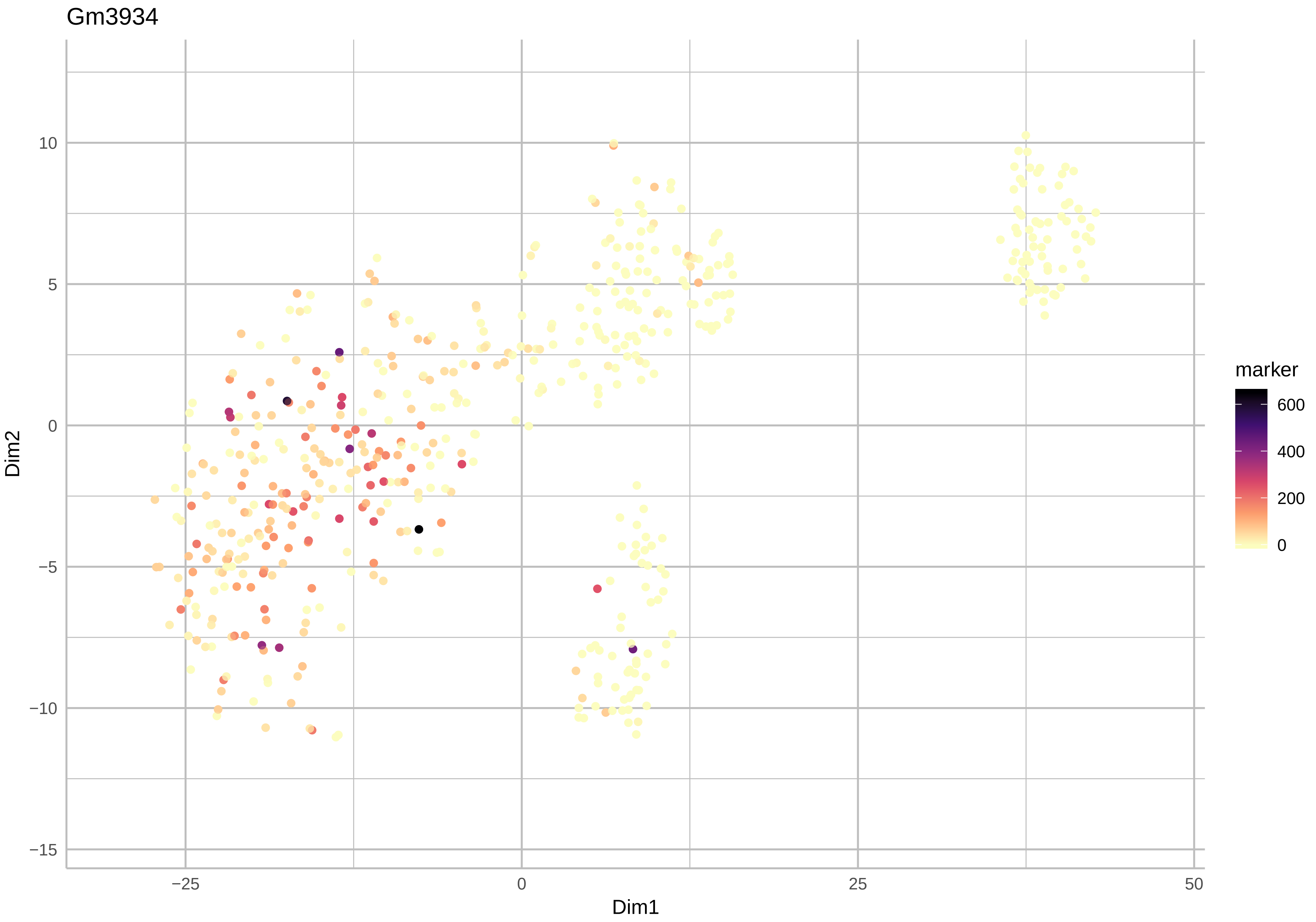

Gm3934
Dim2
Dim1
10
5
0
−5
−10
−15
−25
0
25
50
marker
600
400
200
0

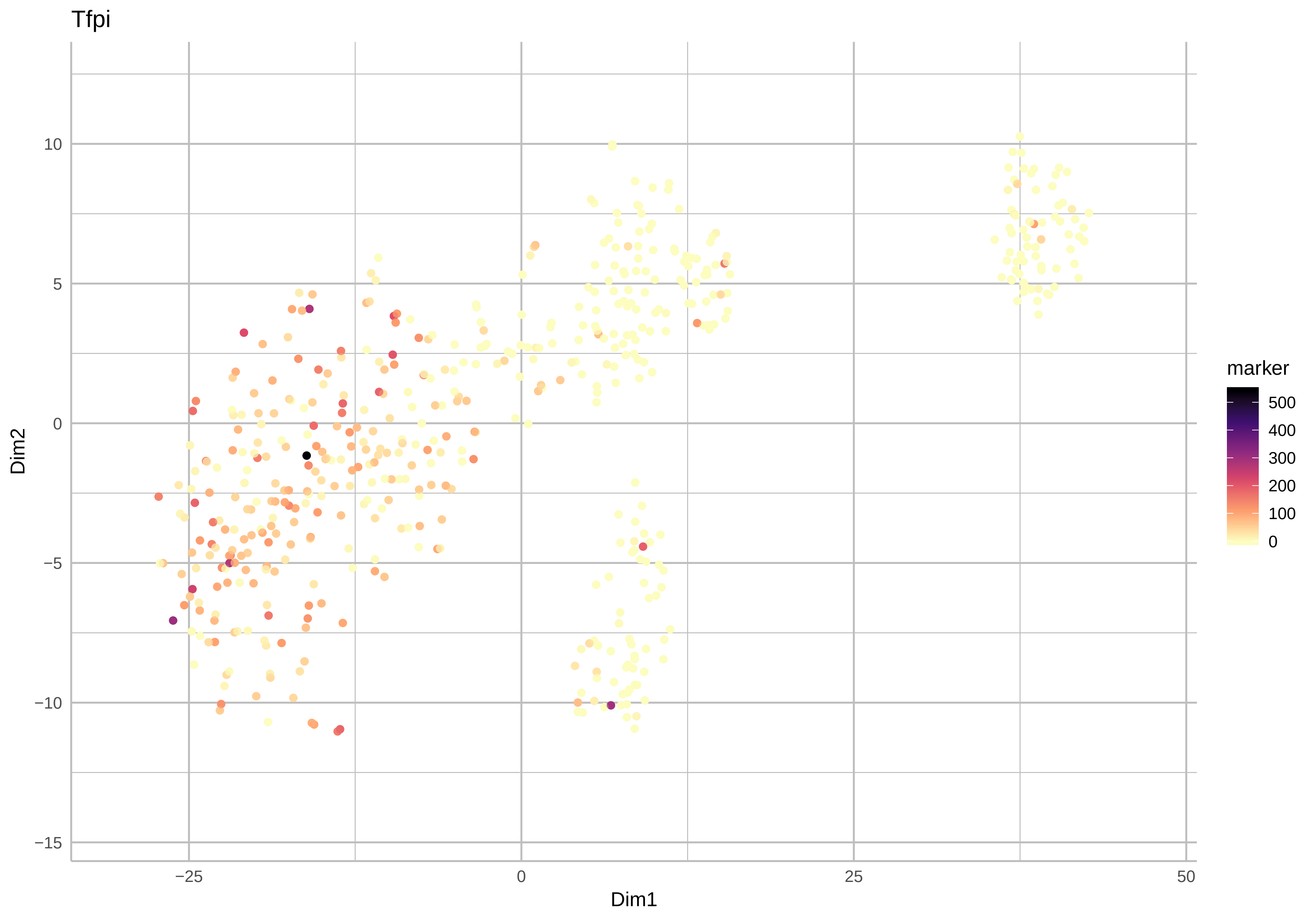
Tfpi
Dim2
Dim1
10
5
0
−5
−10
−15
−25
0
25
50
marker
500
400
300
200
100
0

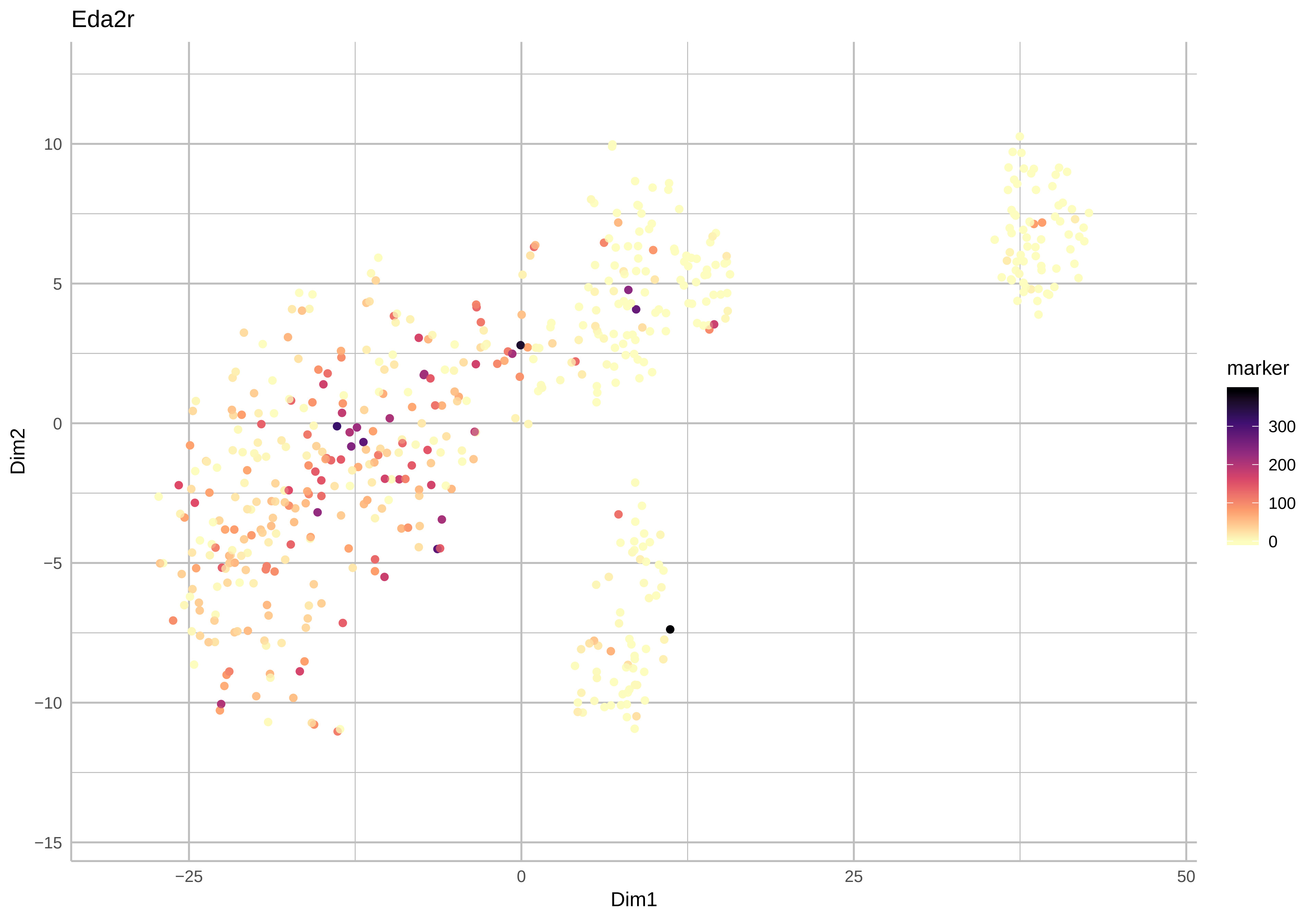
Eda2r
Dim2
Dim1
10
5
0
−5
−10
−15
−25
0
25
50
marker
300
200
100
0

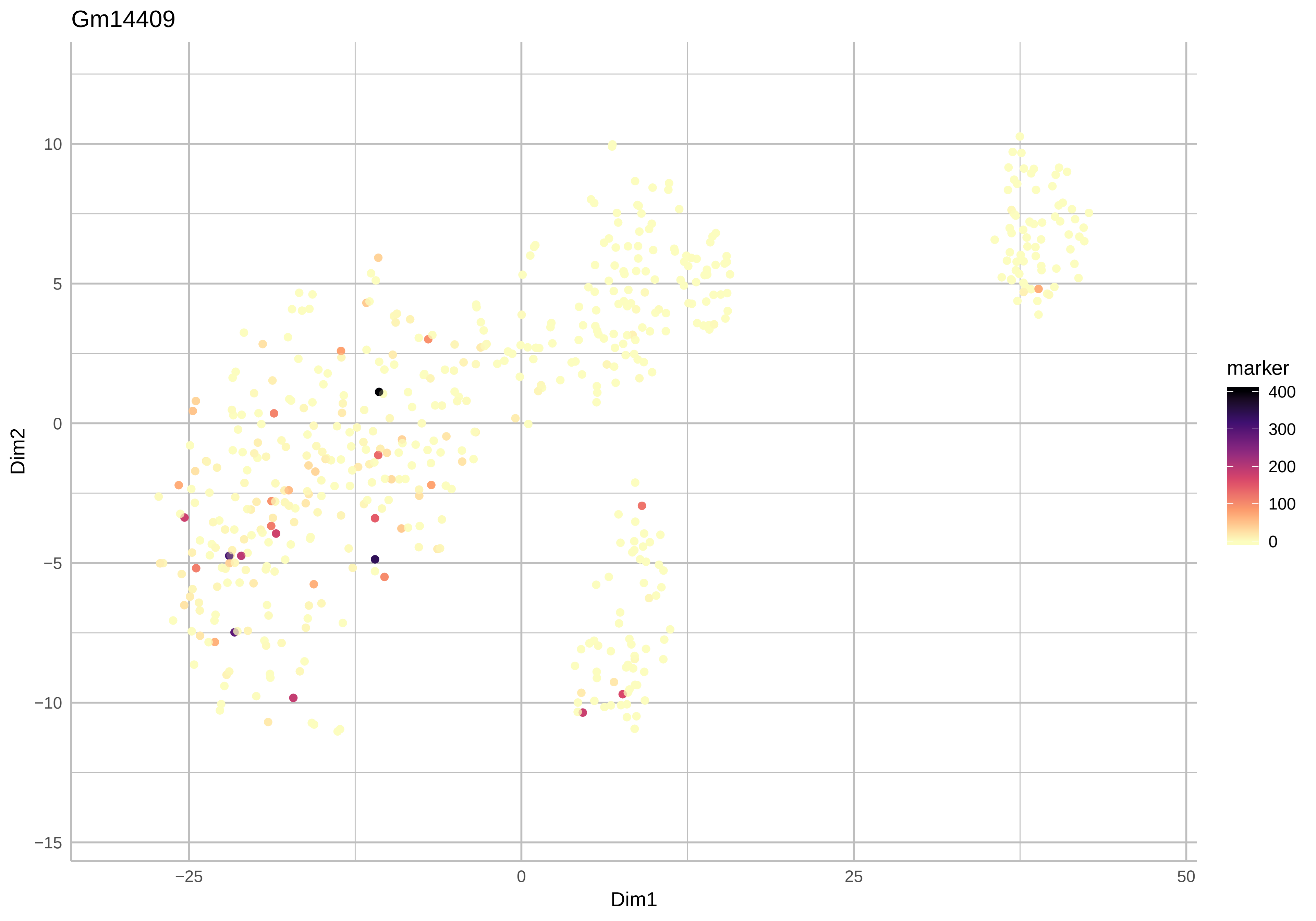
Gm14409
Dim2
10
5
0
−5
−10
−15
−25
0
25
50
Dim1
marker
400
300
200
100
0

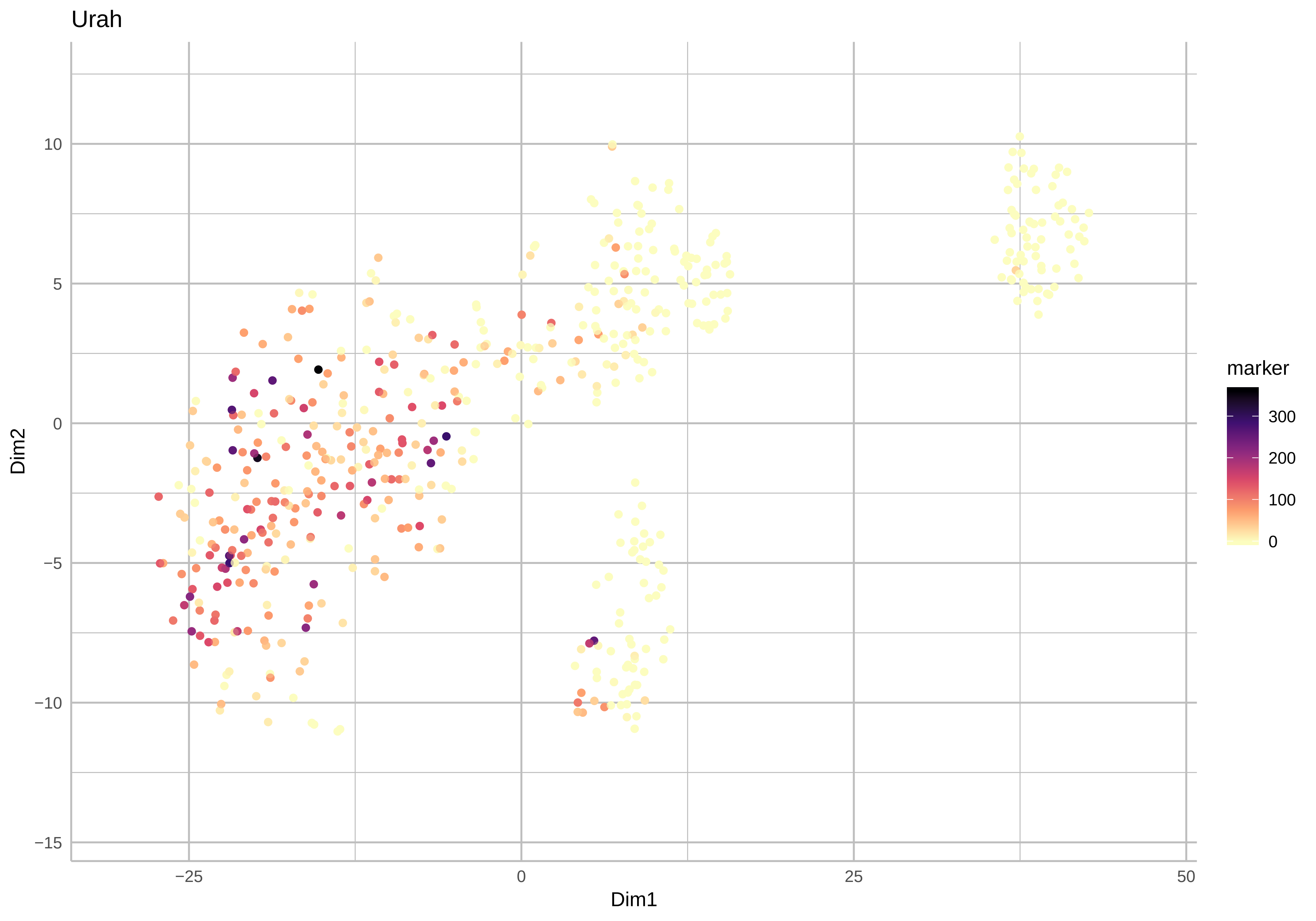
Urah
Dim2
Dim1
10
5
0
−5
−10
−15
−25
0
25
50
marker
300
200
100
0

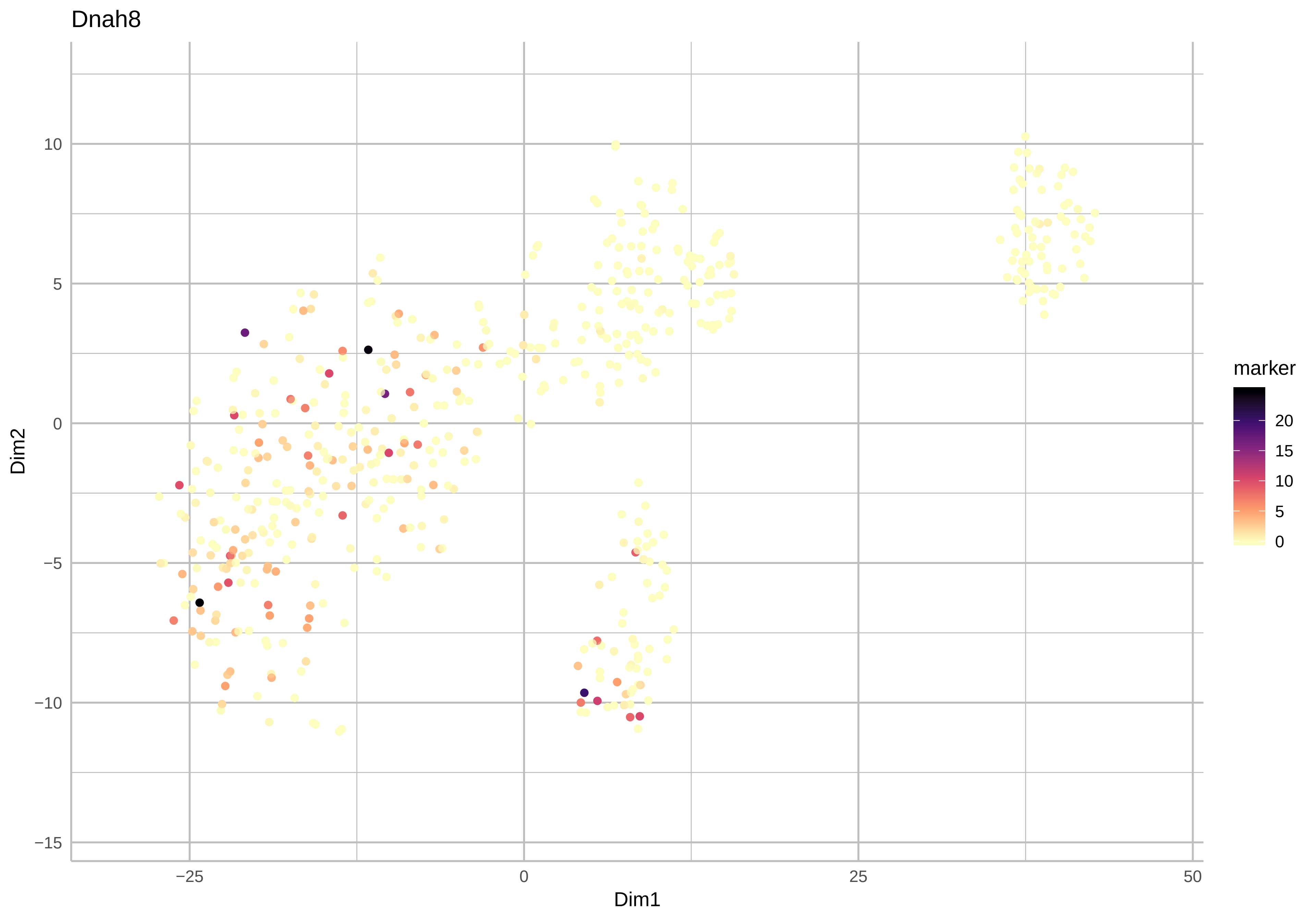
Dnah8
Dim2
Dim1
marker
10
5
0
−5
−10
−15
−25
0
25
50
20
15
10
5
0

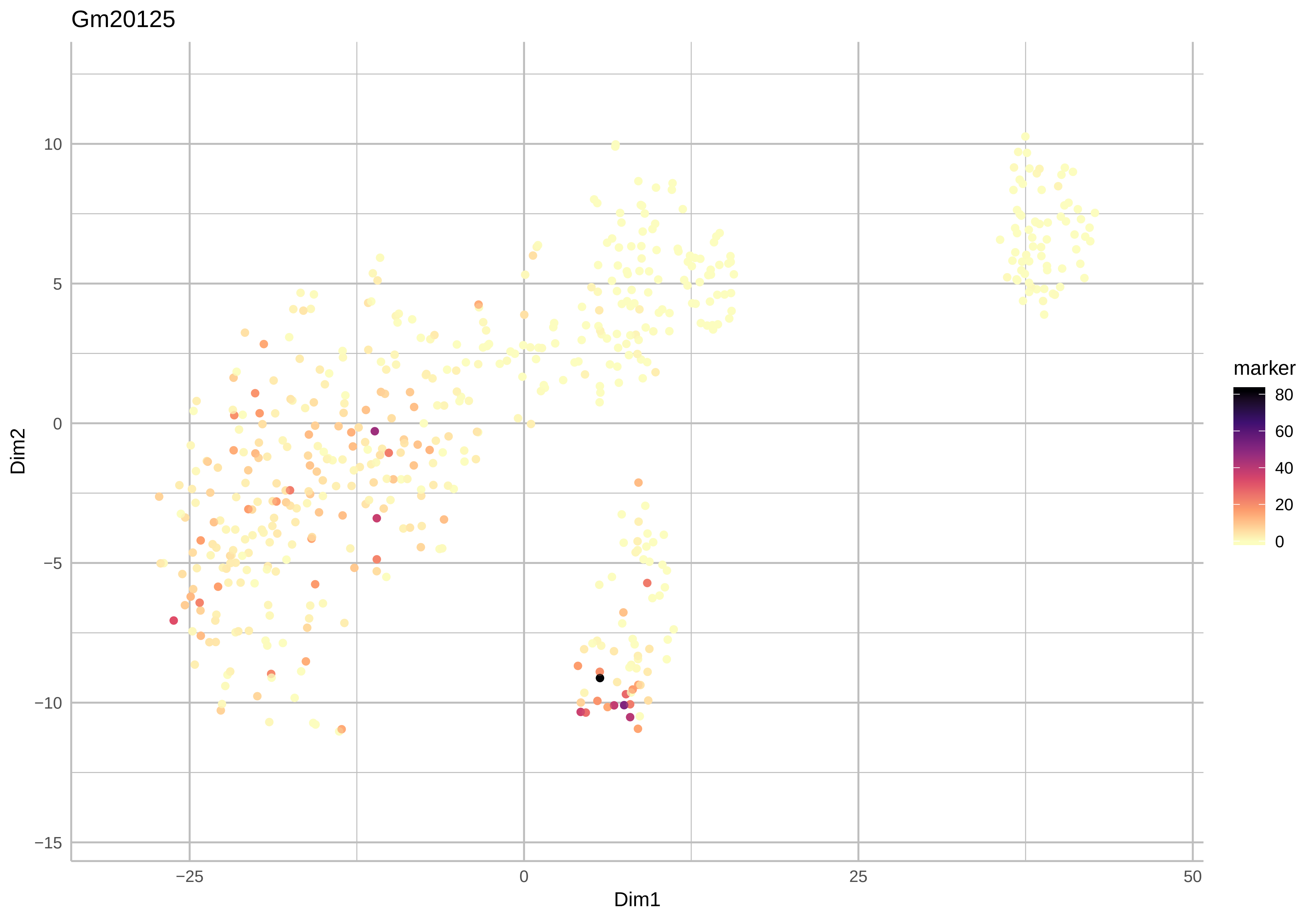

Gm20125
Dim2
Dim1
10
5
0
−5
−10
−15
−25
0
25
50
marker
80
60
40
20
0

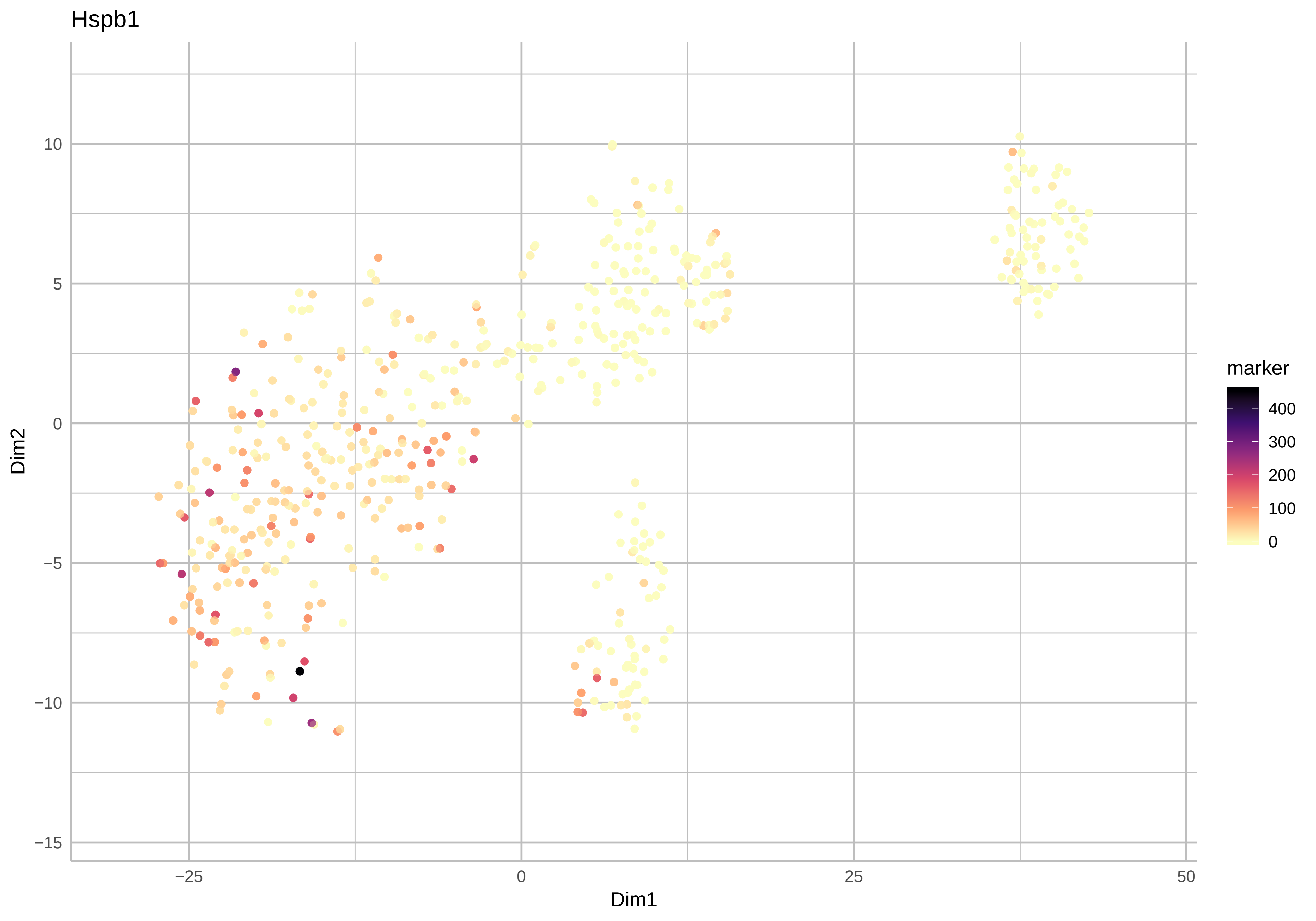
Hspb1
Dim2
Dim1
marker
400
300
200
100
0
10
5
0
−5
−10
−15
−25
0
25
50

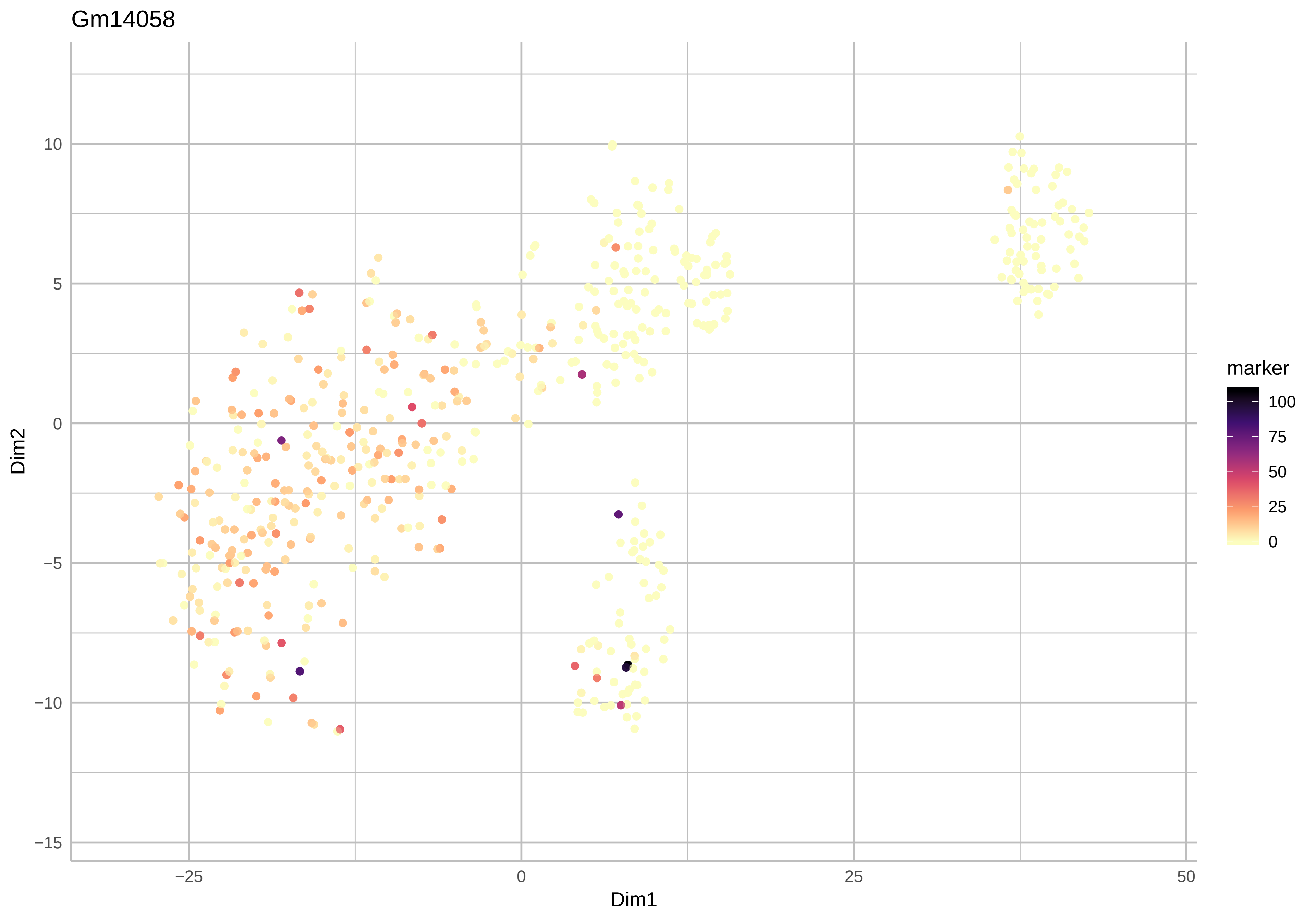

Gm14058
10
5
0
−5
−10
−15
−25
0
25
50
Dim1
Dim2
marker
100
75
50
25
0

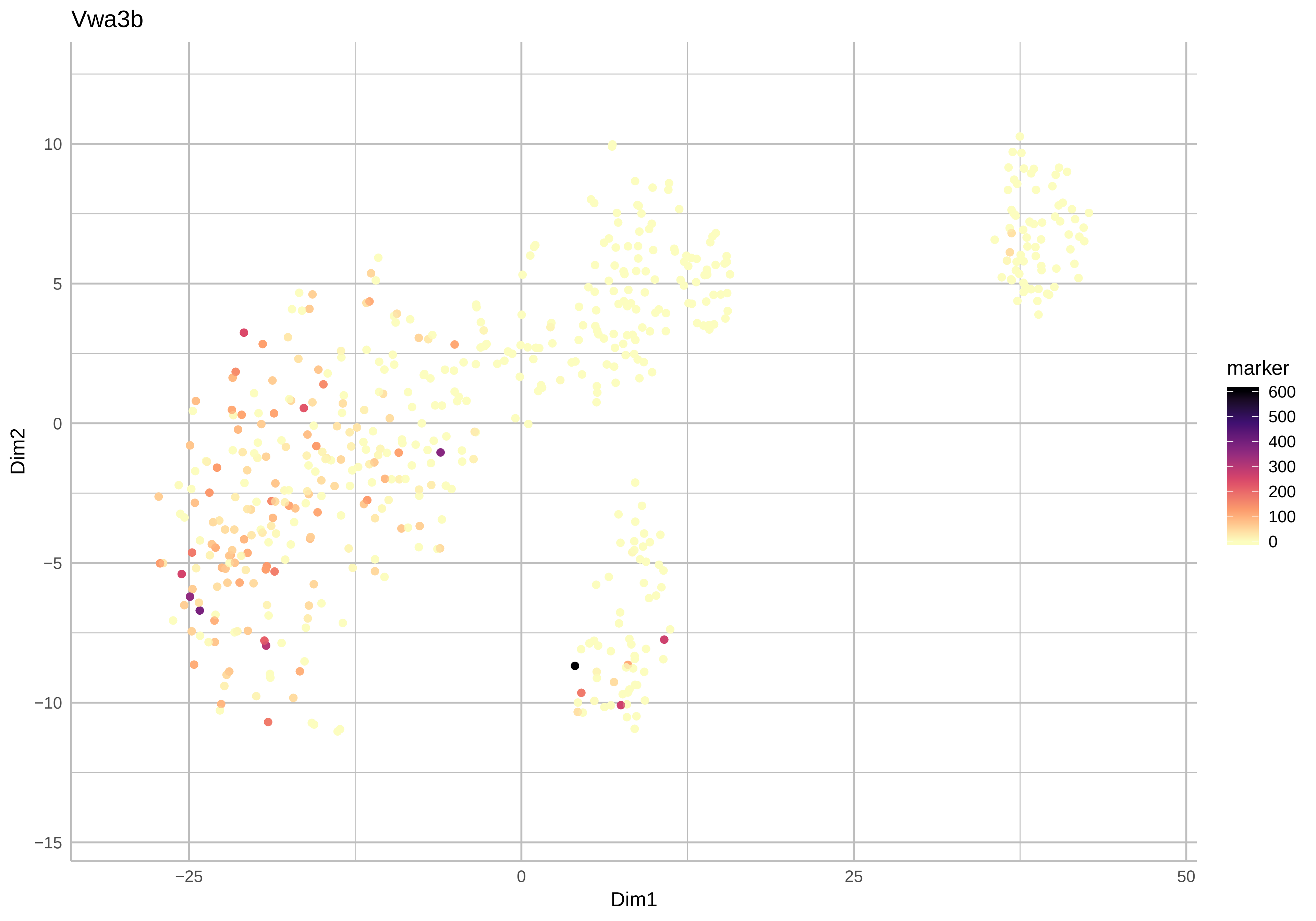

Vwa3b
Dim2
Dim1
10
5
0
−5
−10
−15
−25
0
25
50
marker
600
500
400
300
200
100
0

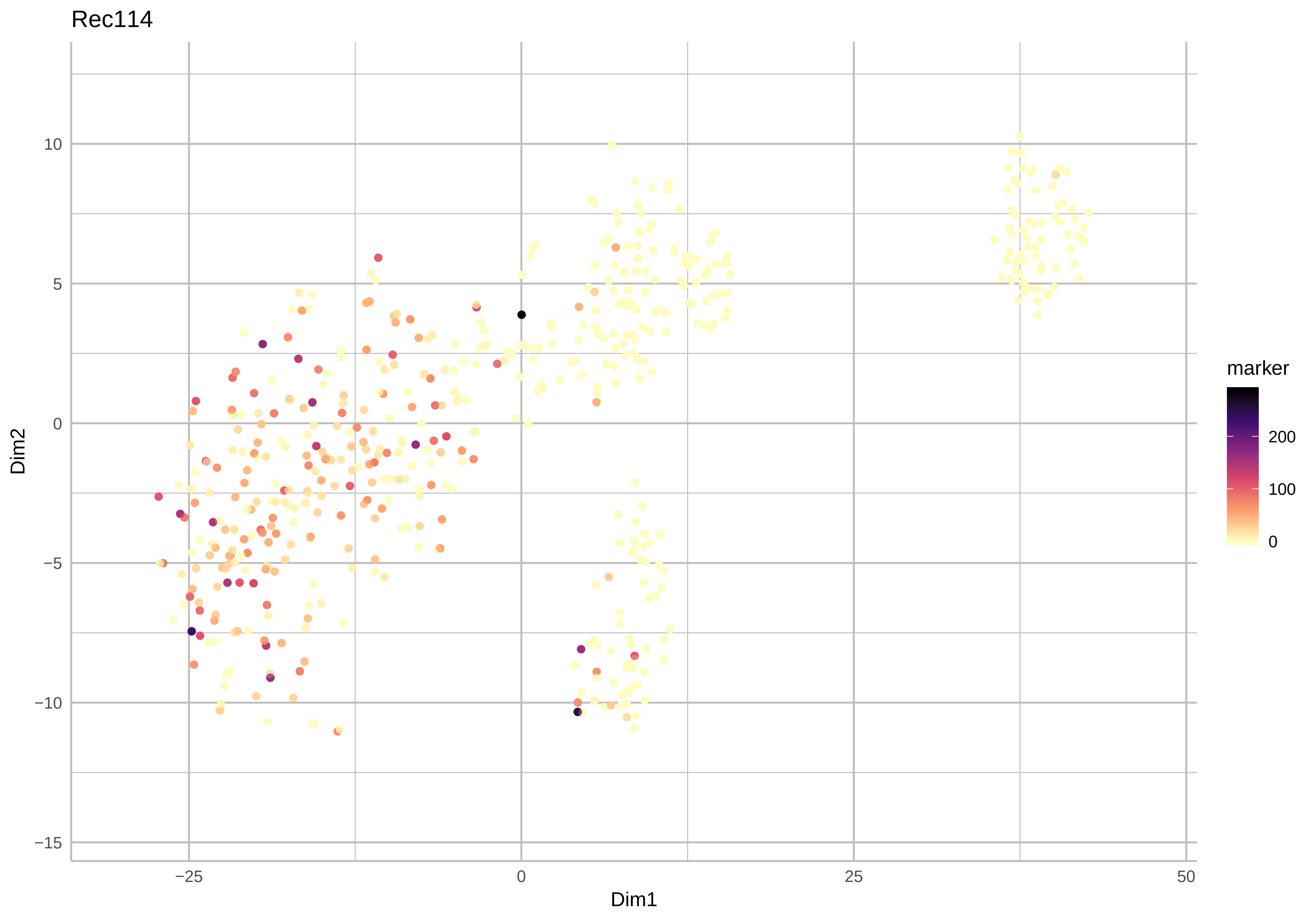
Rec114
10
5
0
−5
−10
−15
−25
0
25
50
Dim1
Dim2
marker
200
100
0

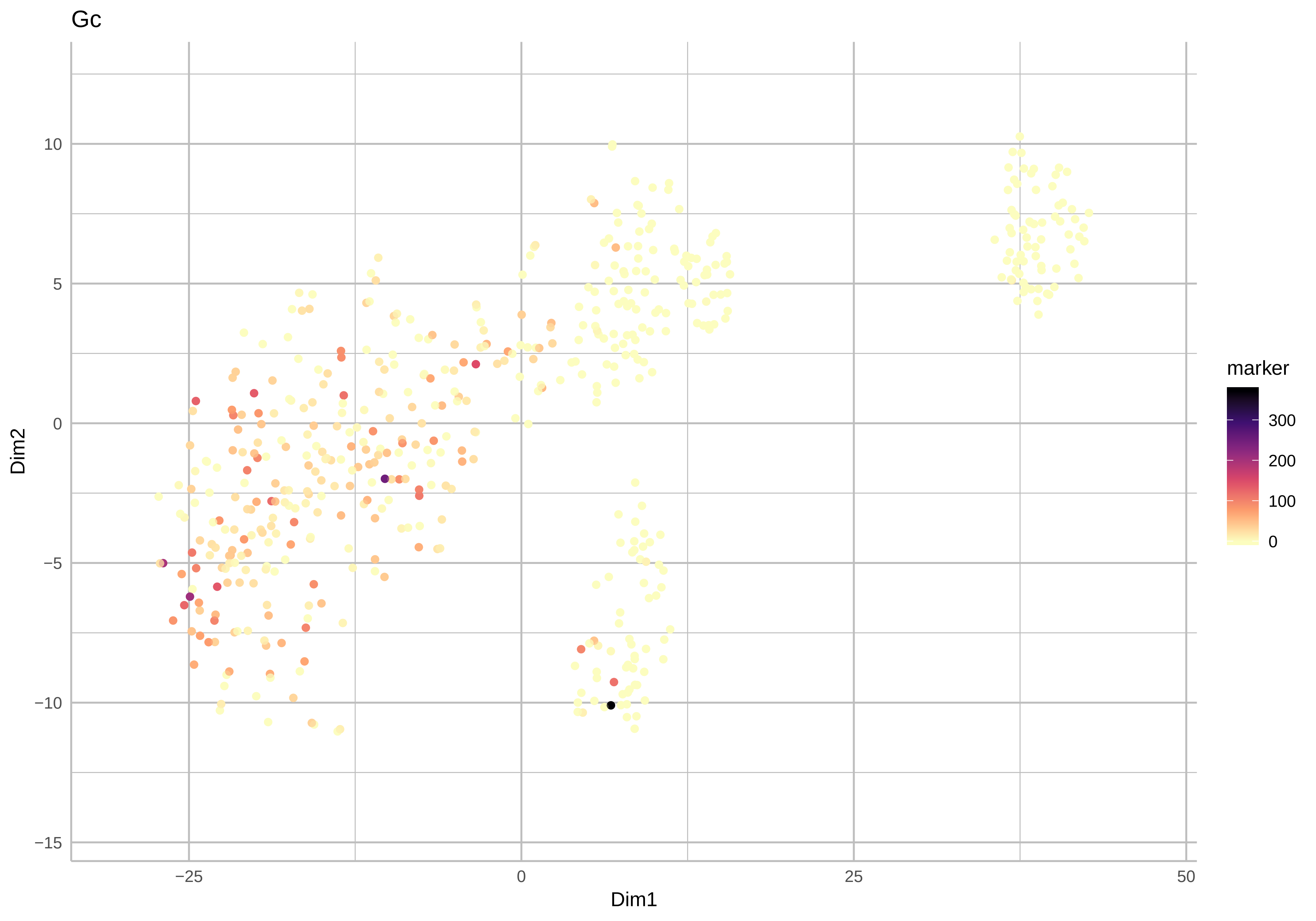
Gc
Dim2
10
5
0
−5
−10
−15
Dim1
−25
0
25
50
marker
300
200
100
0

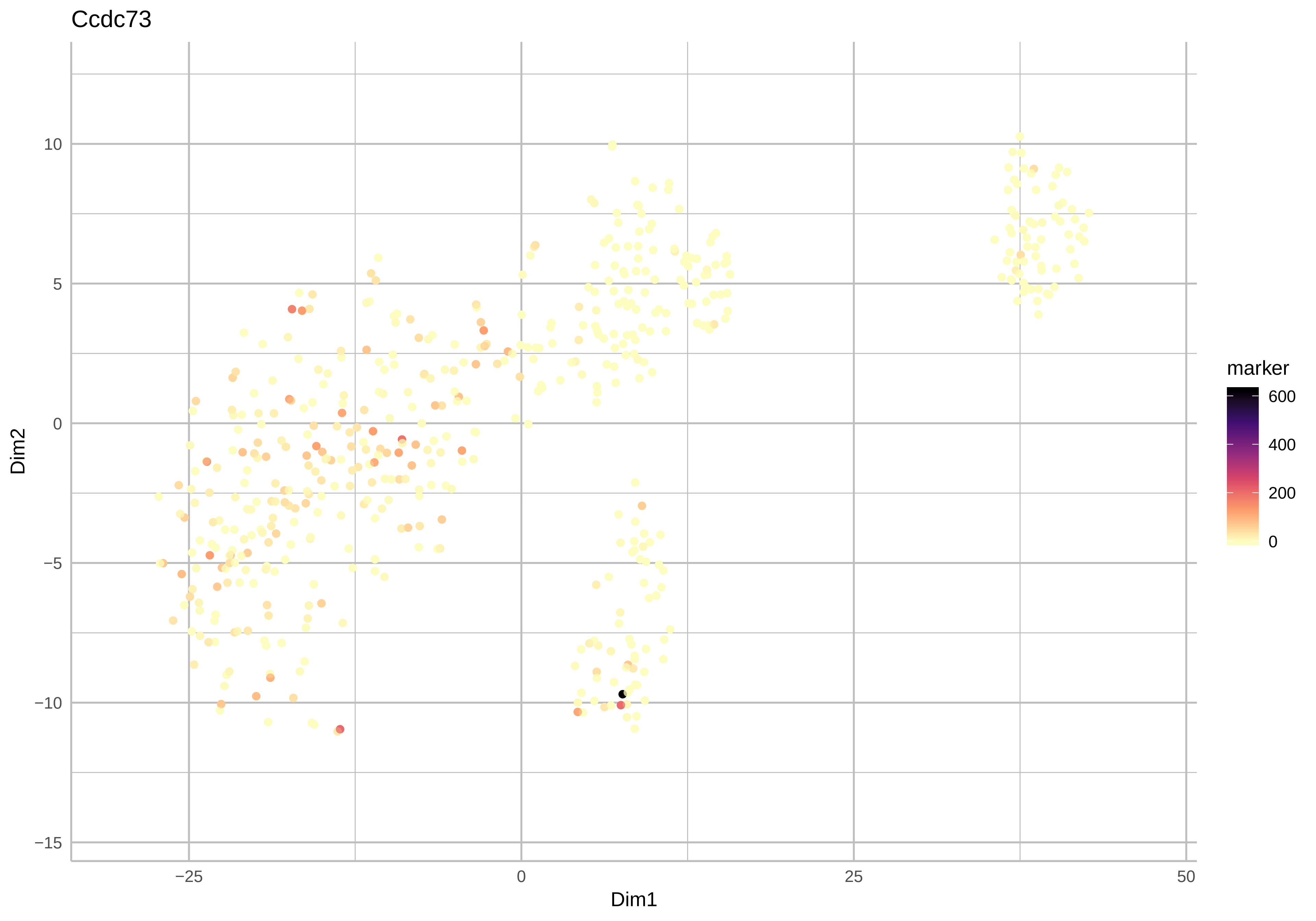

Ccdc73
Dim2
10
5
0
−5
−10
−15
−25
0
25
50
Dim1
marker
600
400
200
0

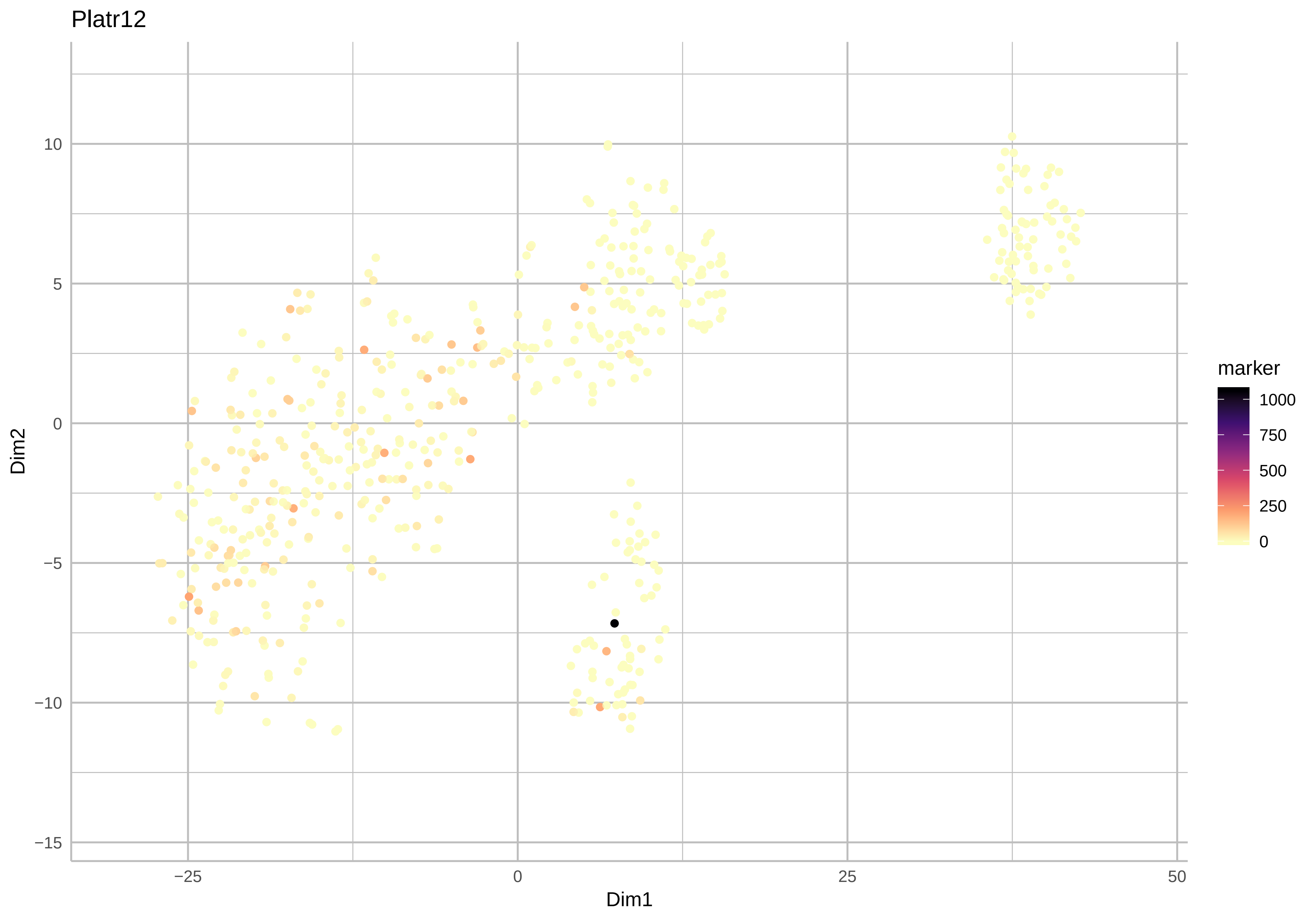

Platr12
Dim2
10
5
0
−5
−10
−15
−25
0
25
50
Dim1
marker
1000
750
500
250
0

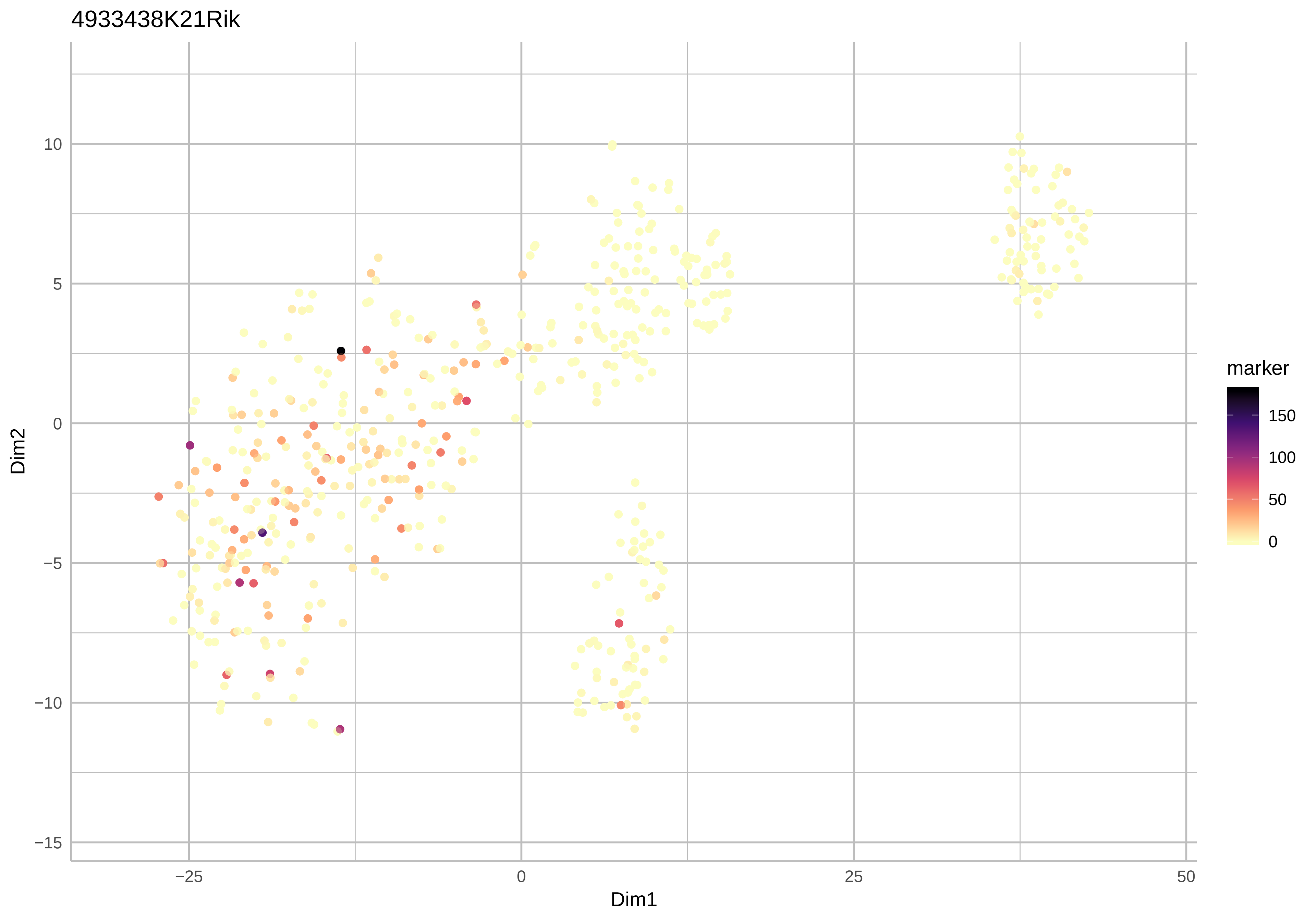
4933438K21Rik
Dim2
Dim1
marker
150
100
50
0
10
5
0
−5
−10
−15
−25
0
25
50

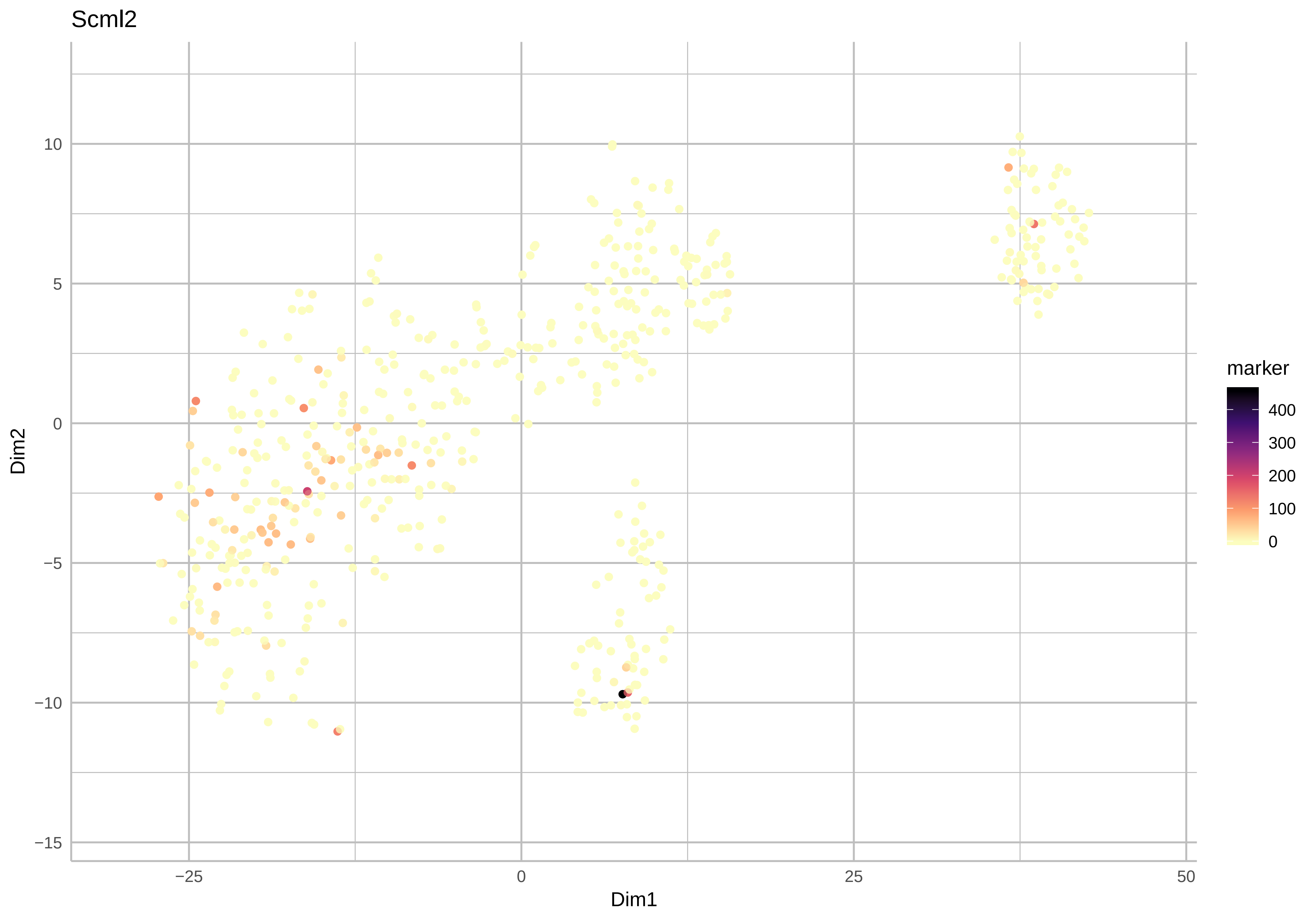

Scml2
Dim2
Dim1
marker
400
300
200
100
0
10
5
0
−5
−10
−15
−25
0
25
50

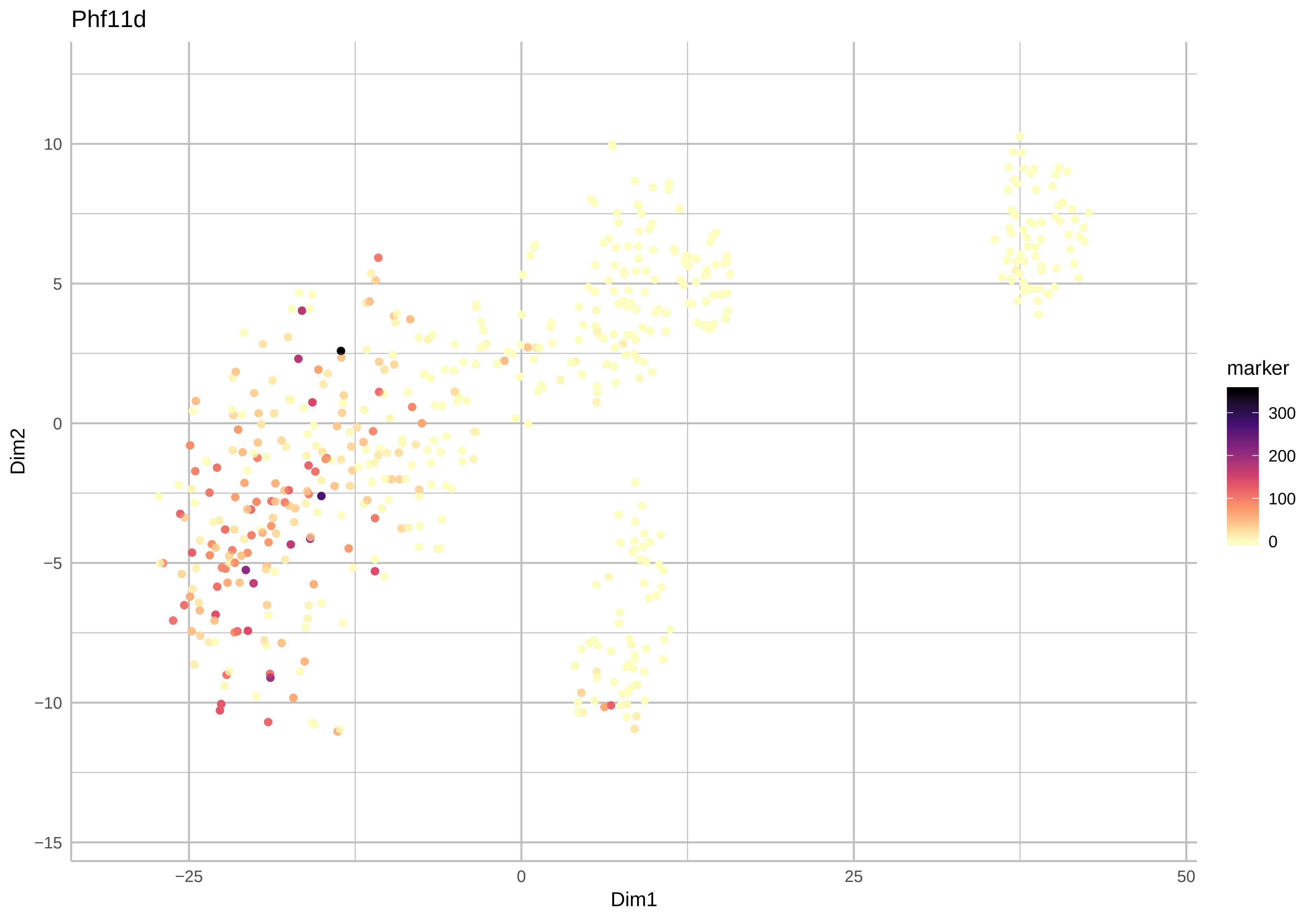
Phf11d
Dim2
Dim1
10
5
0
−5
−10
−15
−25
0
25
50
marker
300
200
100
0

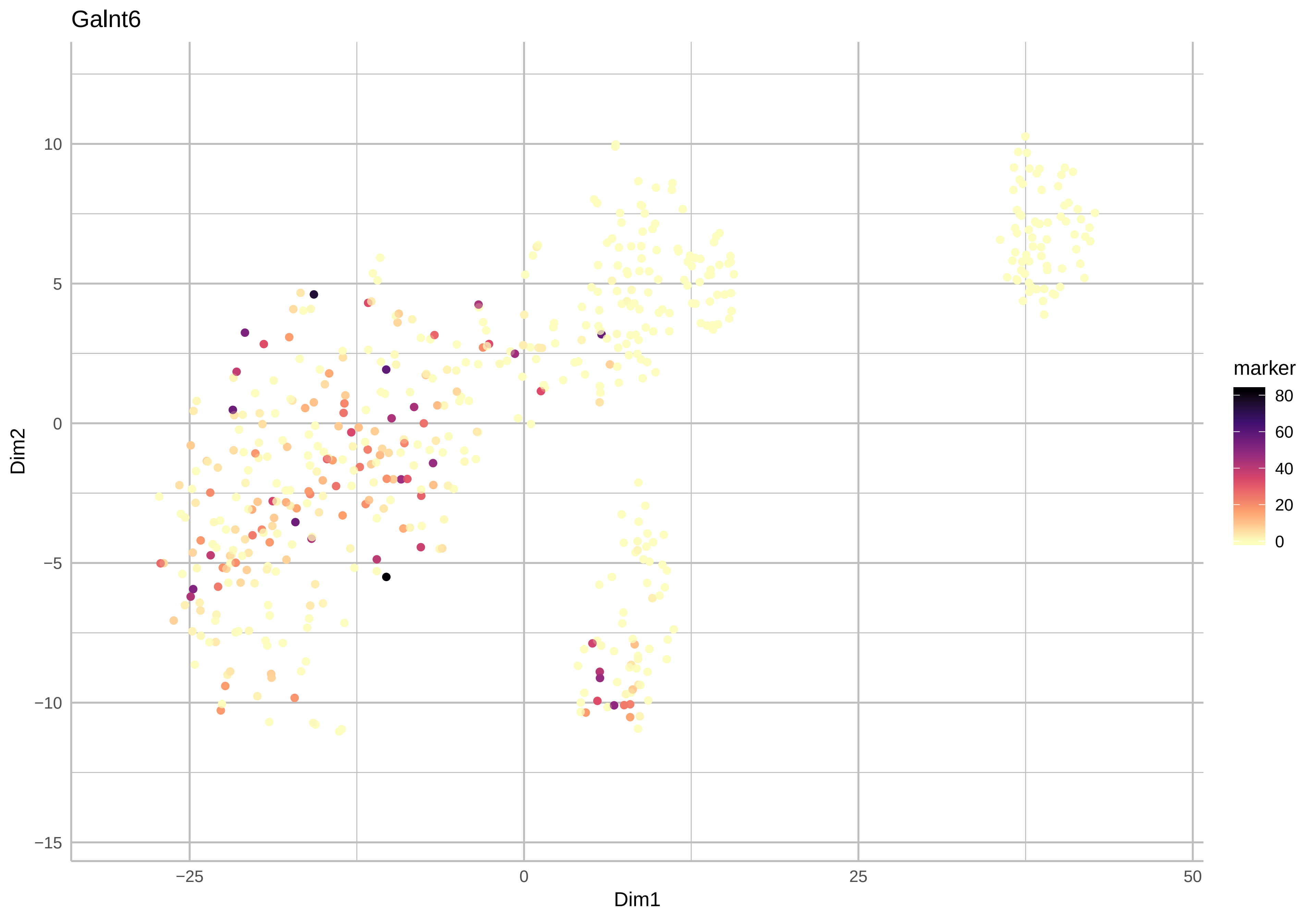
Galnt6
Dim2
Dim1
10
5
0
−5
−10
−15
−25
0
25
50
marker
80
60
40
20
0

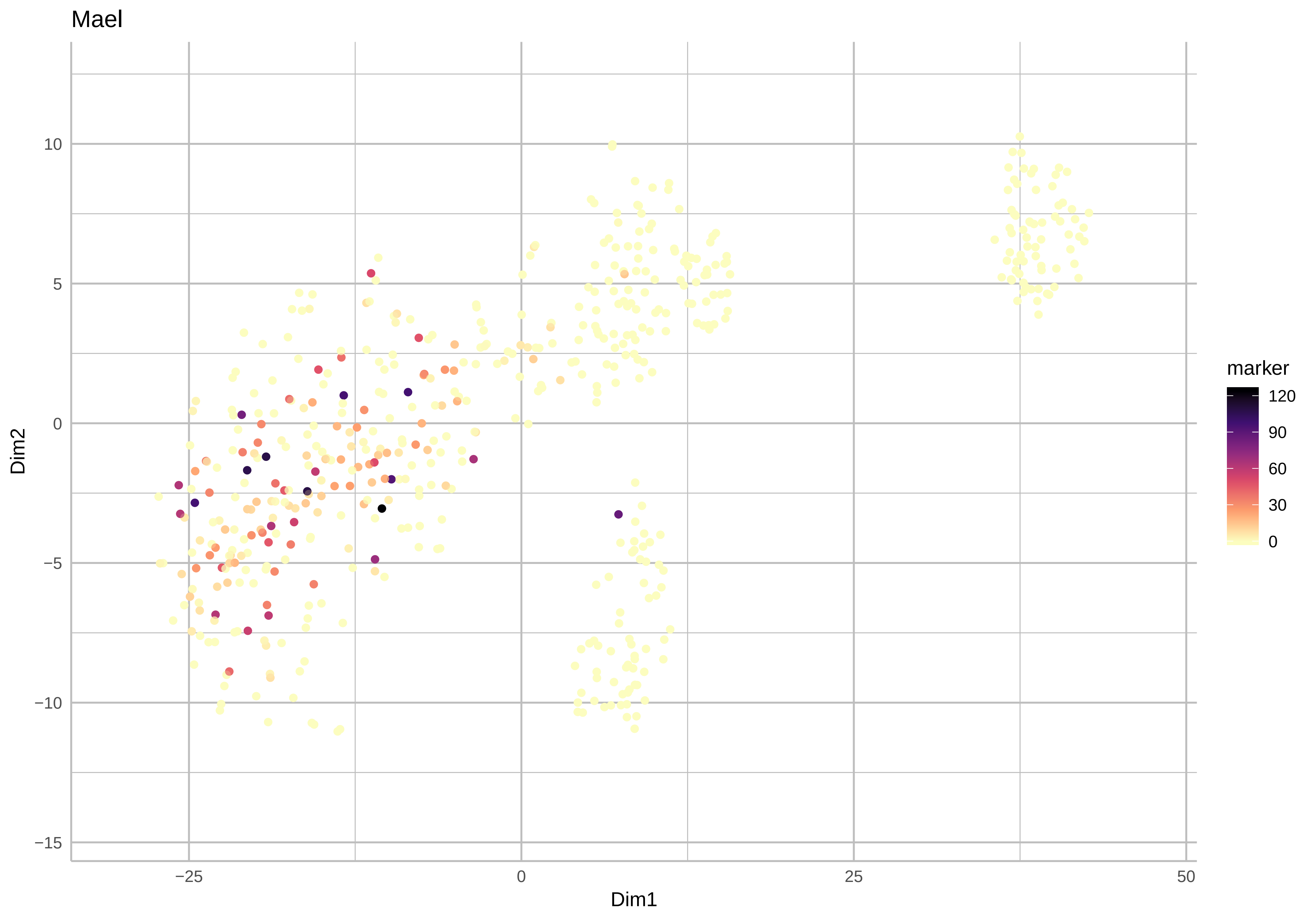
Mael
Dim2
Dim1
10
5
0
−5
−10
−15
−25
0
25
50
marker
120
90
60
30
0

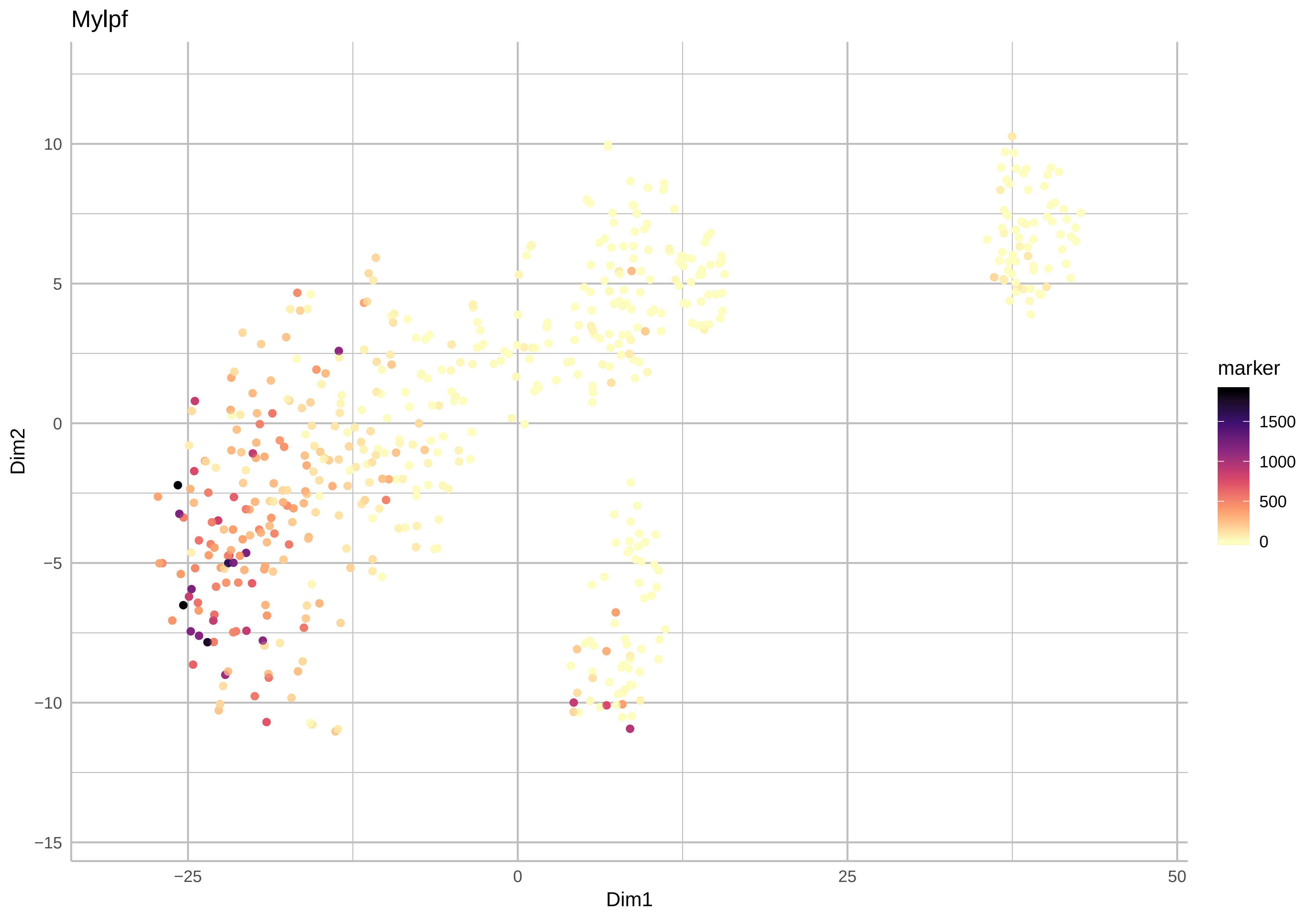

Mylpf
Dim2
Dim1
10
5
0
−5
−10
−15
−25
0
25
50
marker
1500
1000
500
0

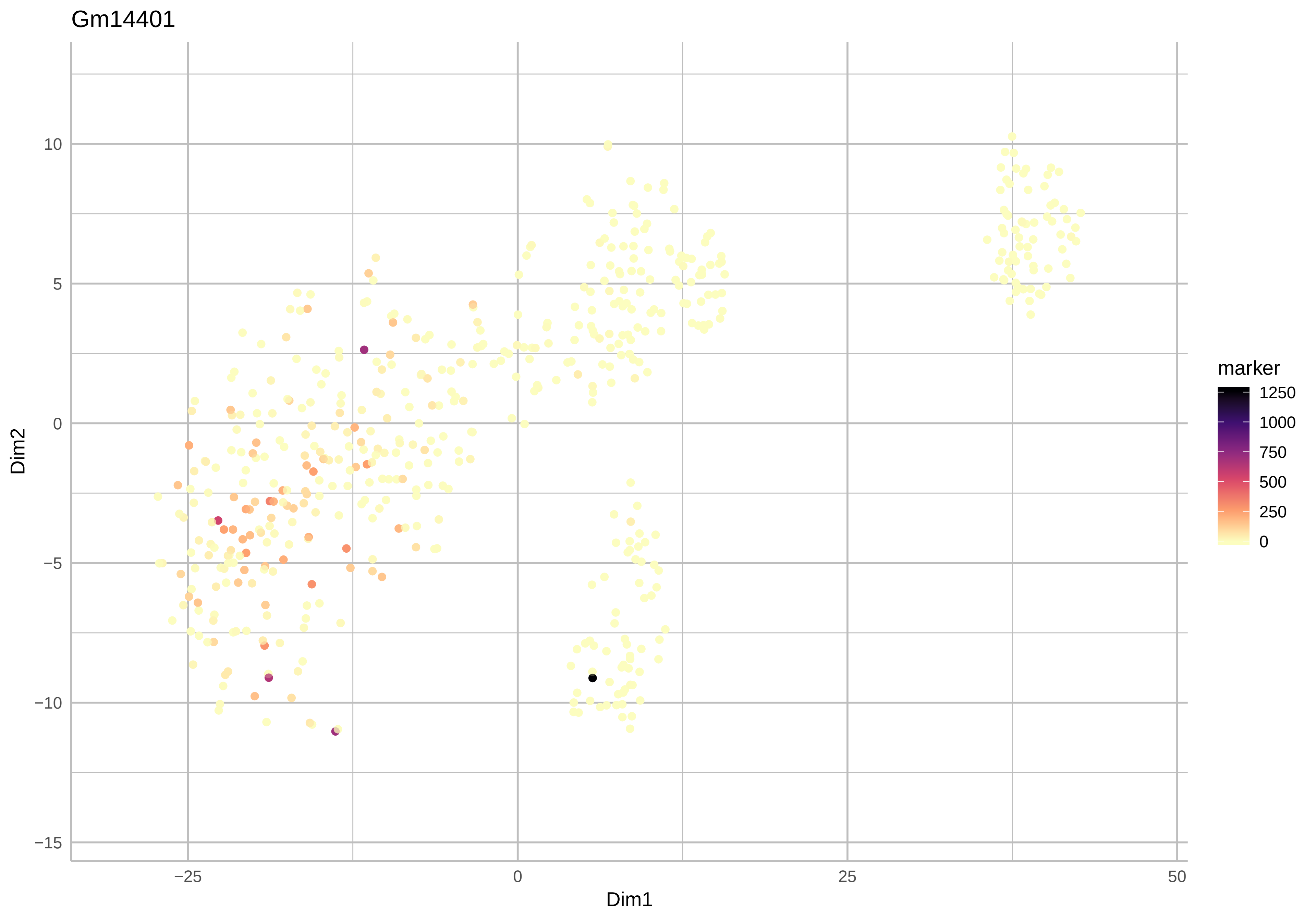

Gm14401
Dim2
10
5
0
−5
−10
−15
−25
0
25
50
Dim1
marker
1250
1000
750
500
250
0

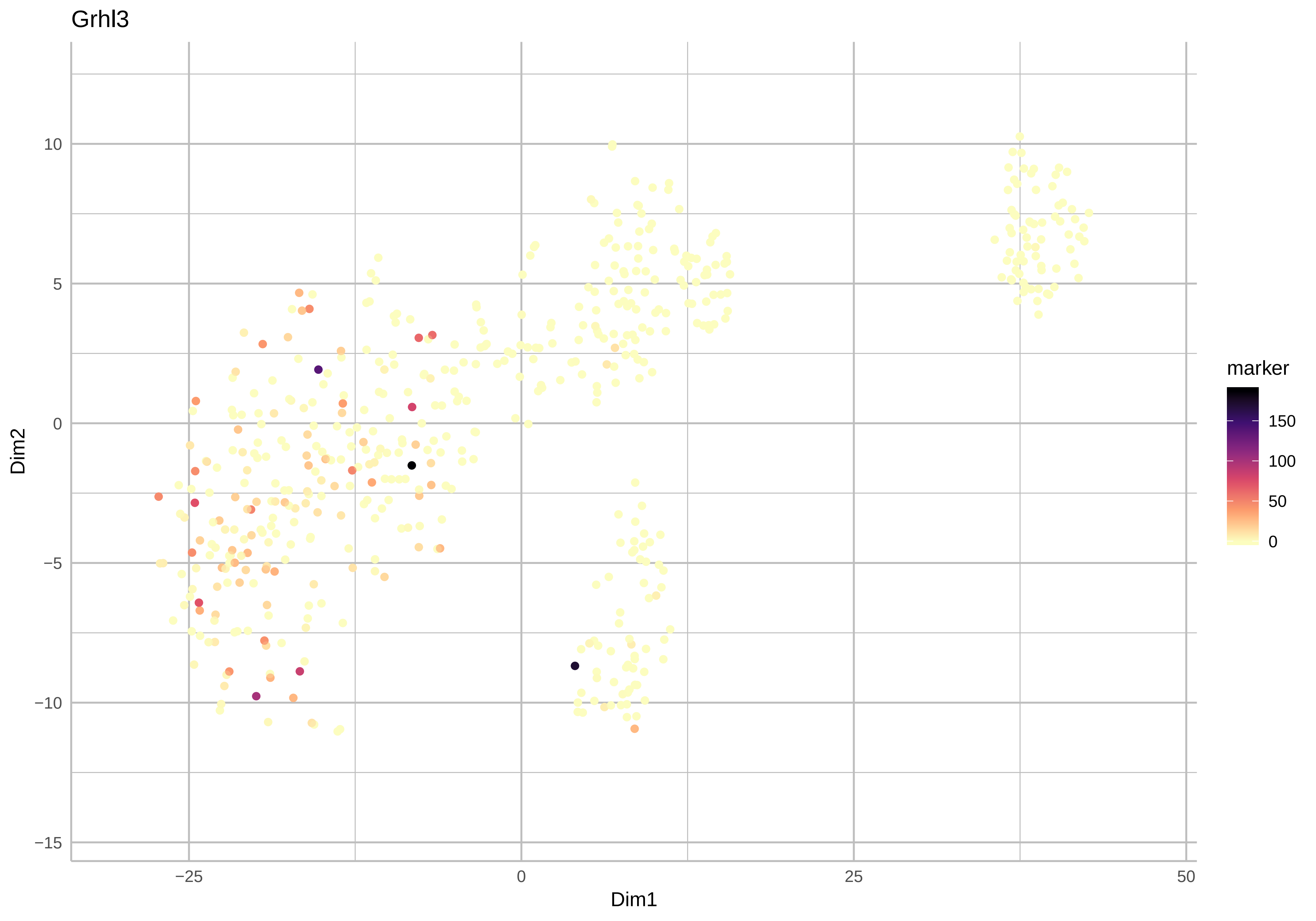
Grhl3
Dim2
Dim1
10
5
0
−5
−10
−15
−25
0
25
50
marker
150
100
50
0

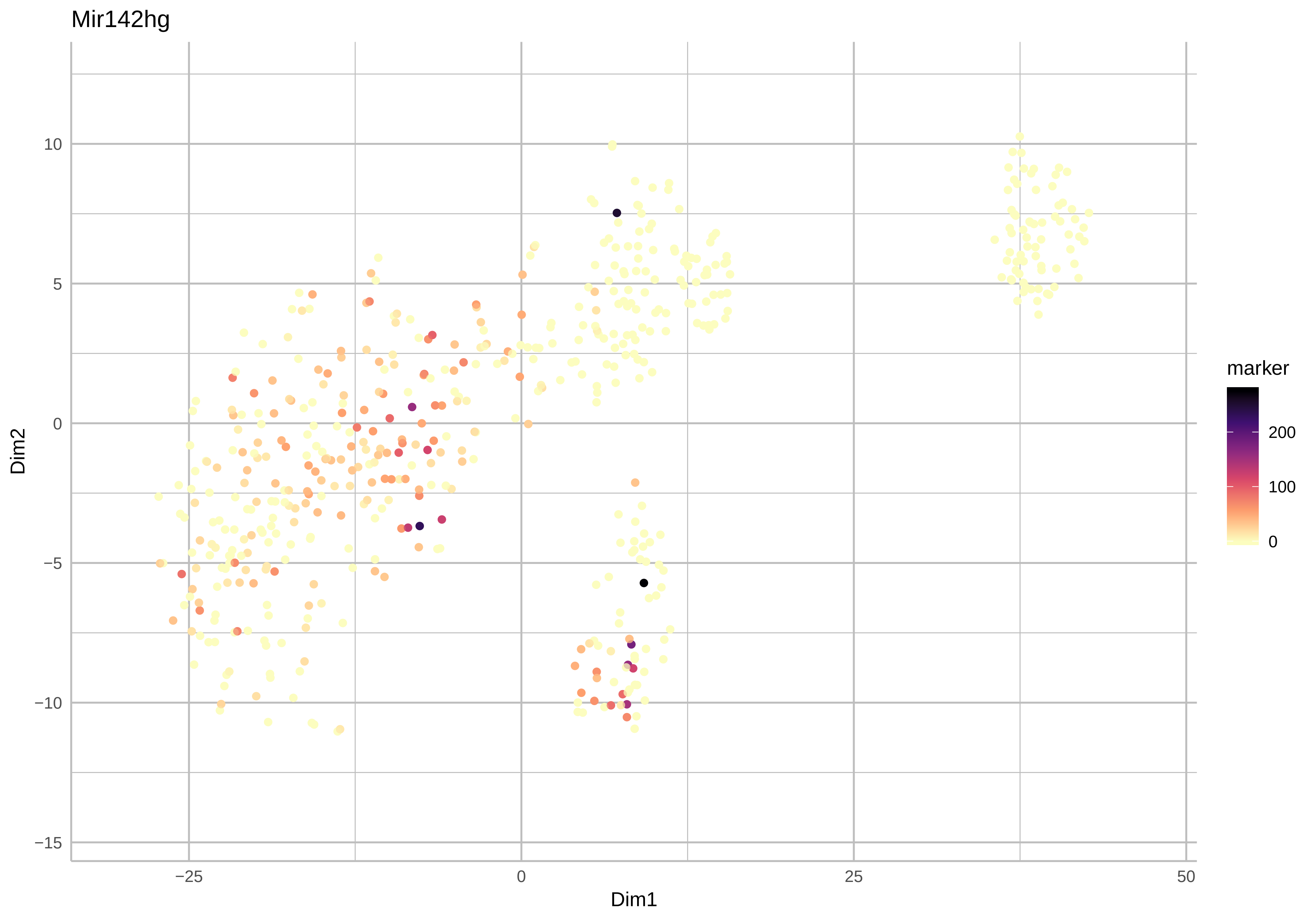

Mir142hg
Dim2
Dim1
10
5
0
−5
−10
−15
−25
0
25
50
marker
200
100
0

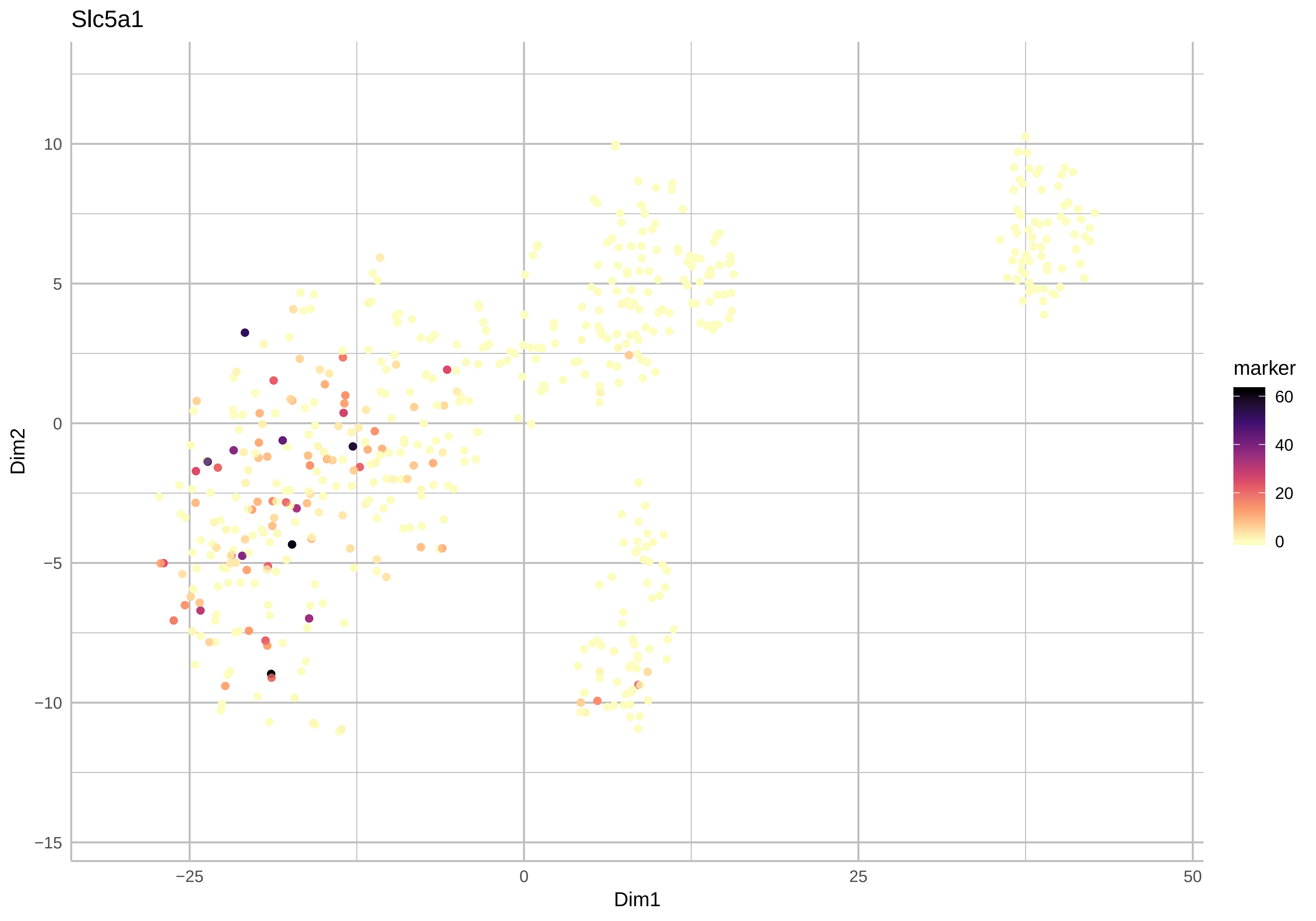
Slc5a1
Dim2
Dim1
marker
60
40
20
0
10
5
0
−5
−10
−15
−25
0
25
50

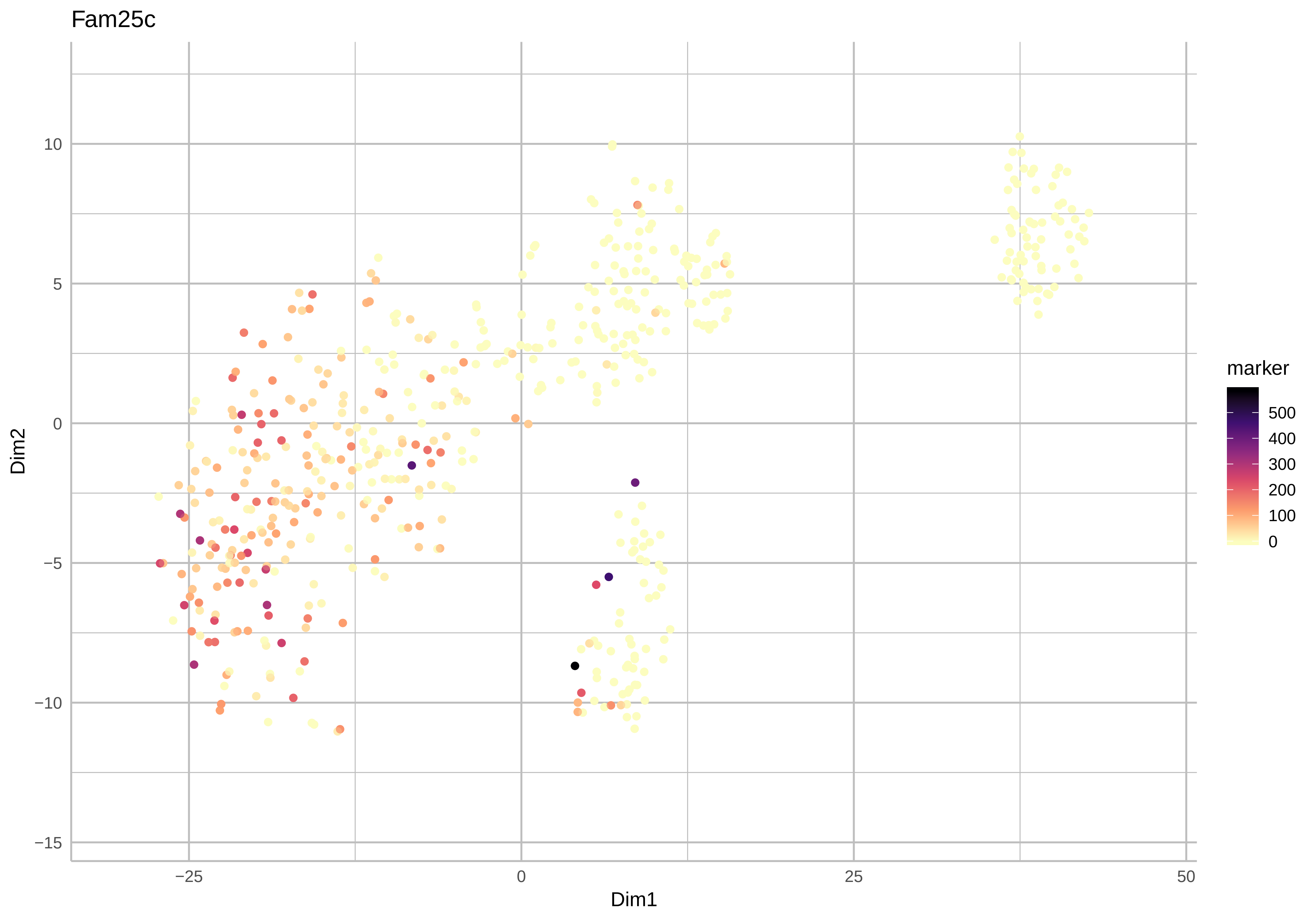

Fam25c
Dim1
Dim2
−25
0
25
50
10
5
0
−5
−10
−15
marker
500
400
300
200
100
0

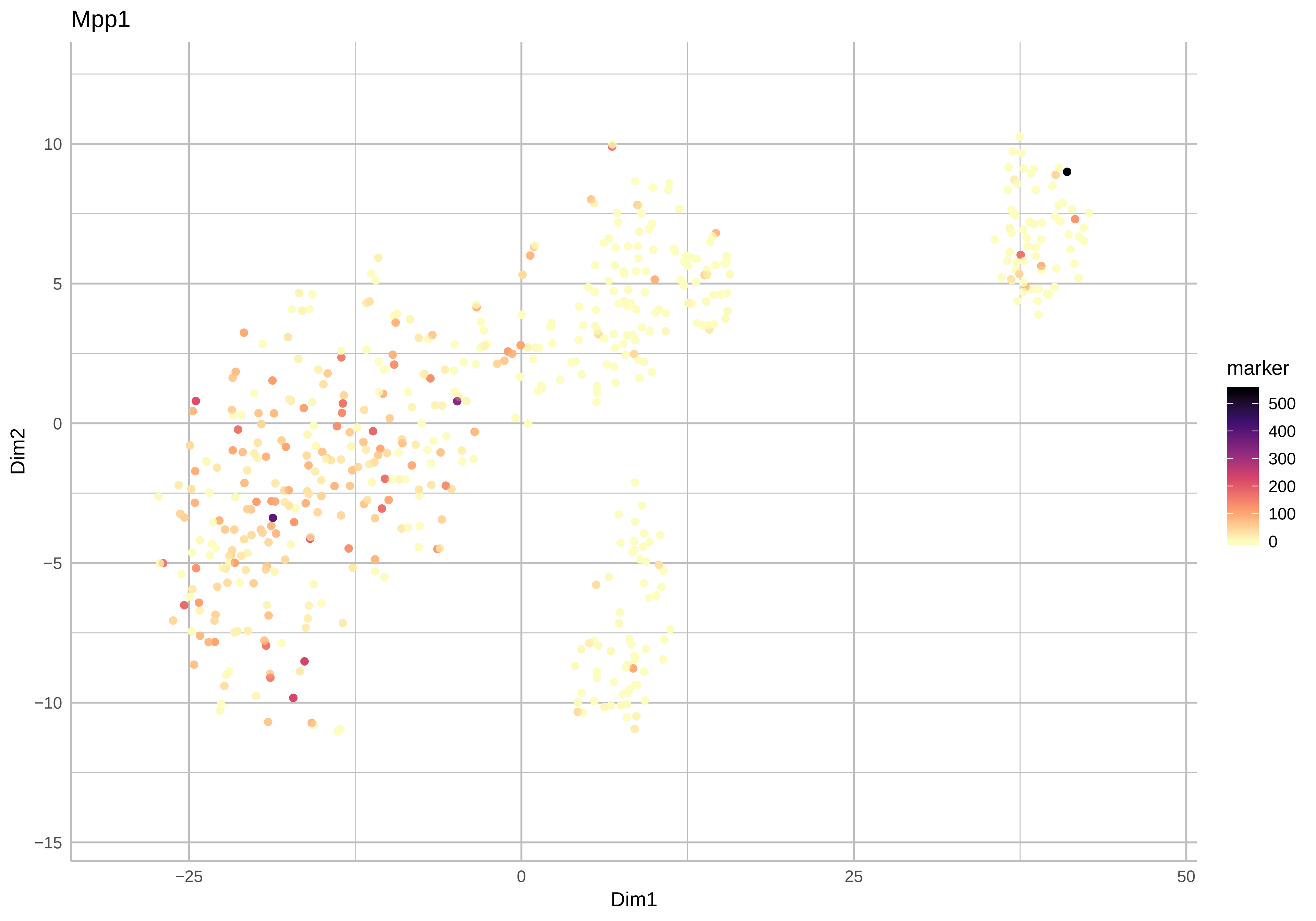

Mpp1
Dim2
Dim1
10
5
0
−5
−10
−15
−25
0
25
50
marker
500
400
300
200
100
0

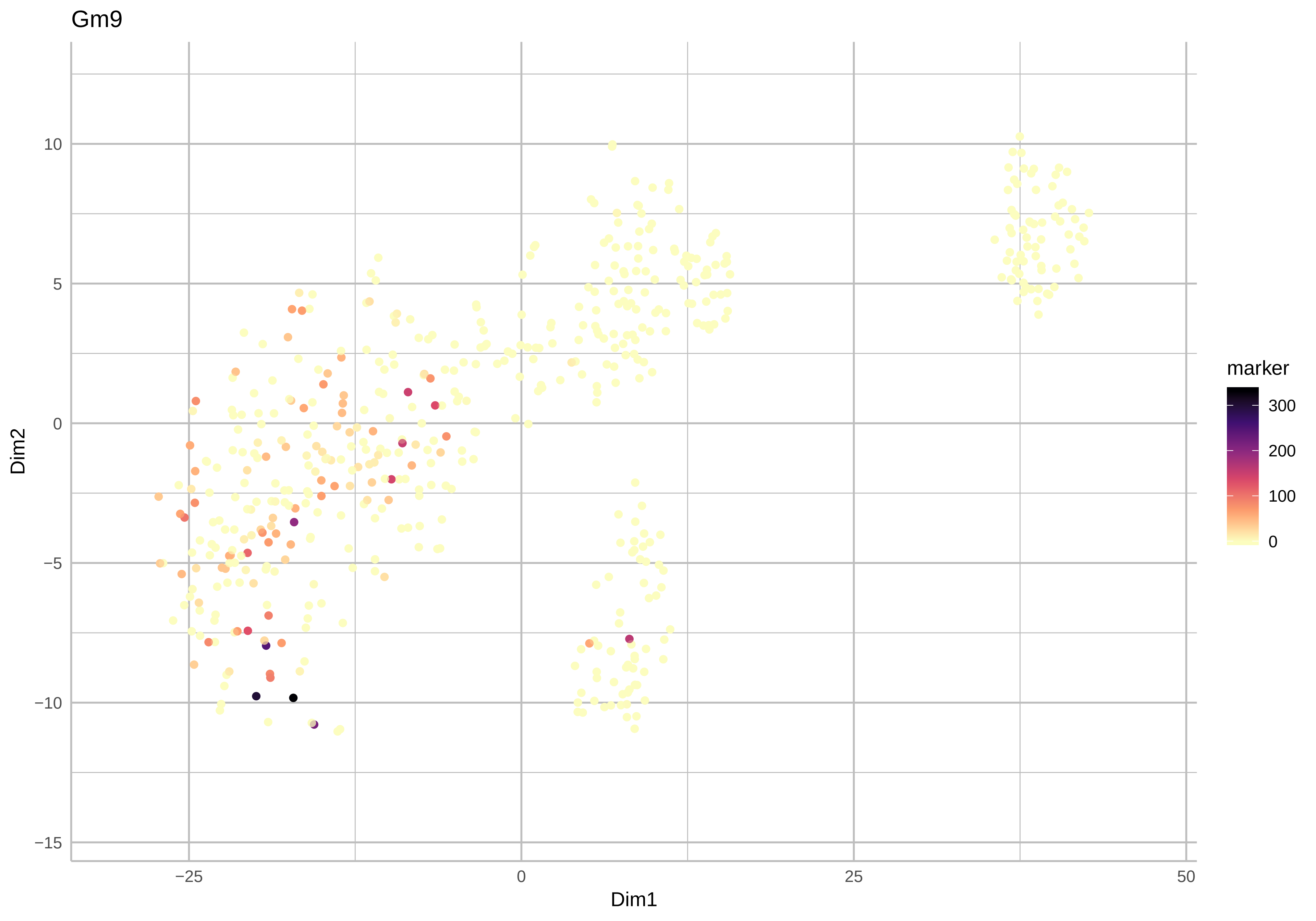
Gm9
10
5
0
−5
−10
−15
Dim2
−25
0
25
50
Dim1
marker
300
200
100
0

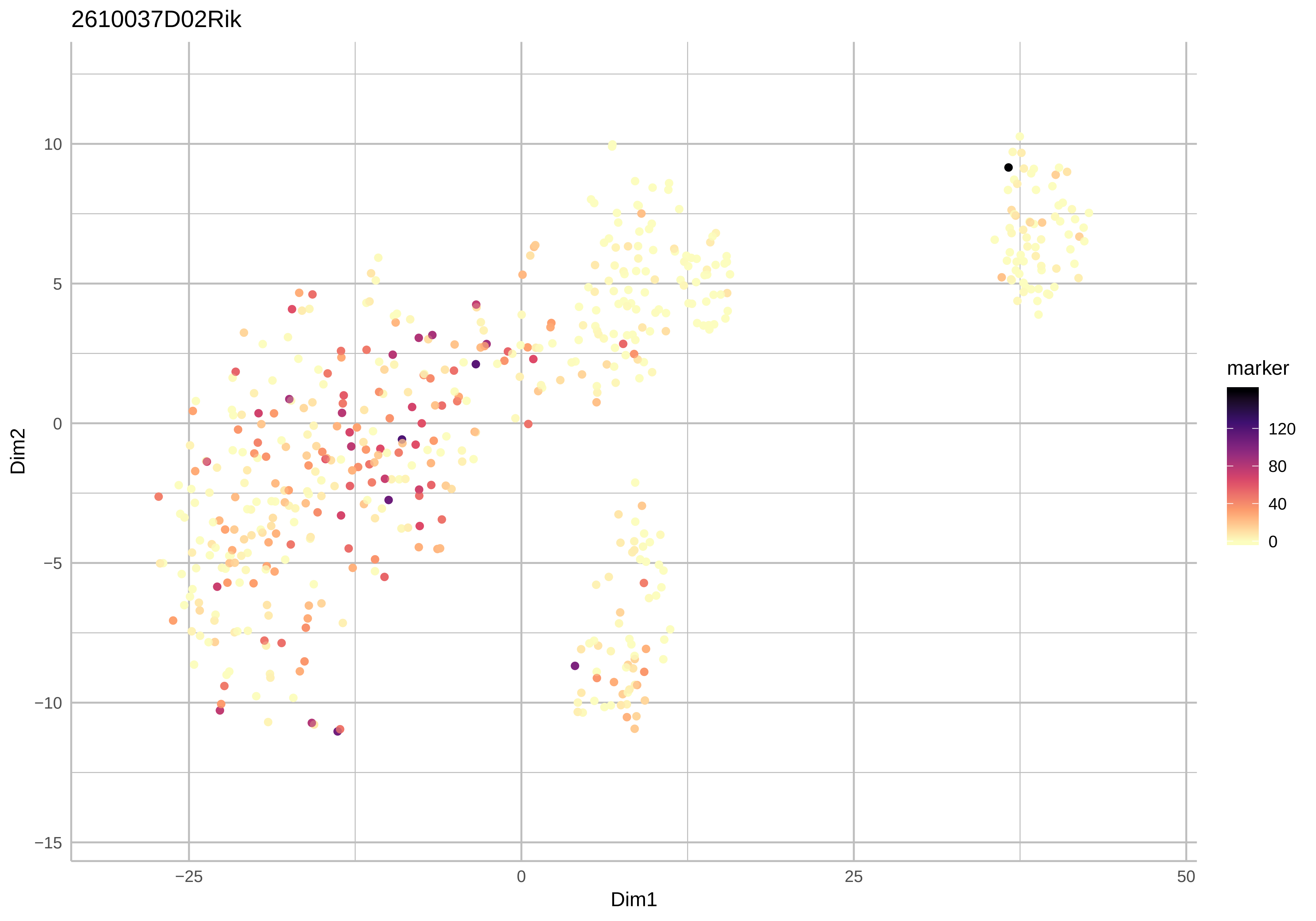
2610037D02Rik
Dim2
Dim1
10
5
0
−5
−10
−15
−25
0
25
50
marker
120
80
40
0

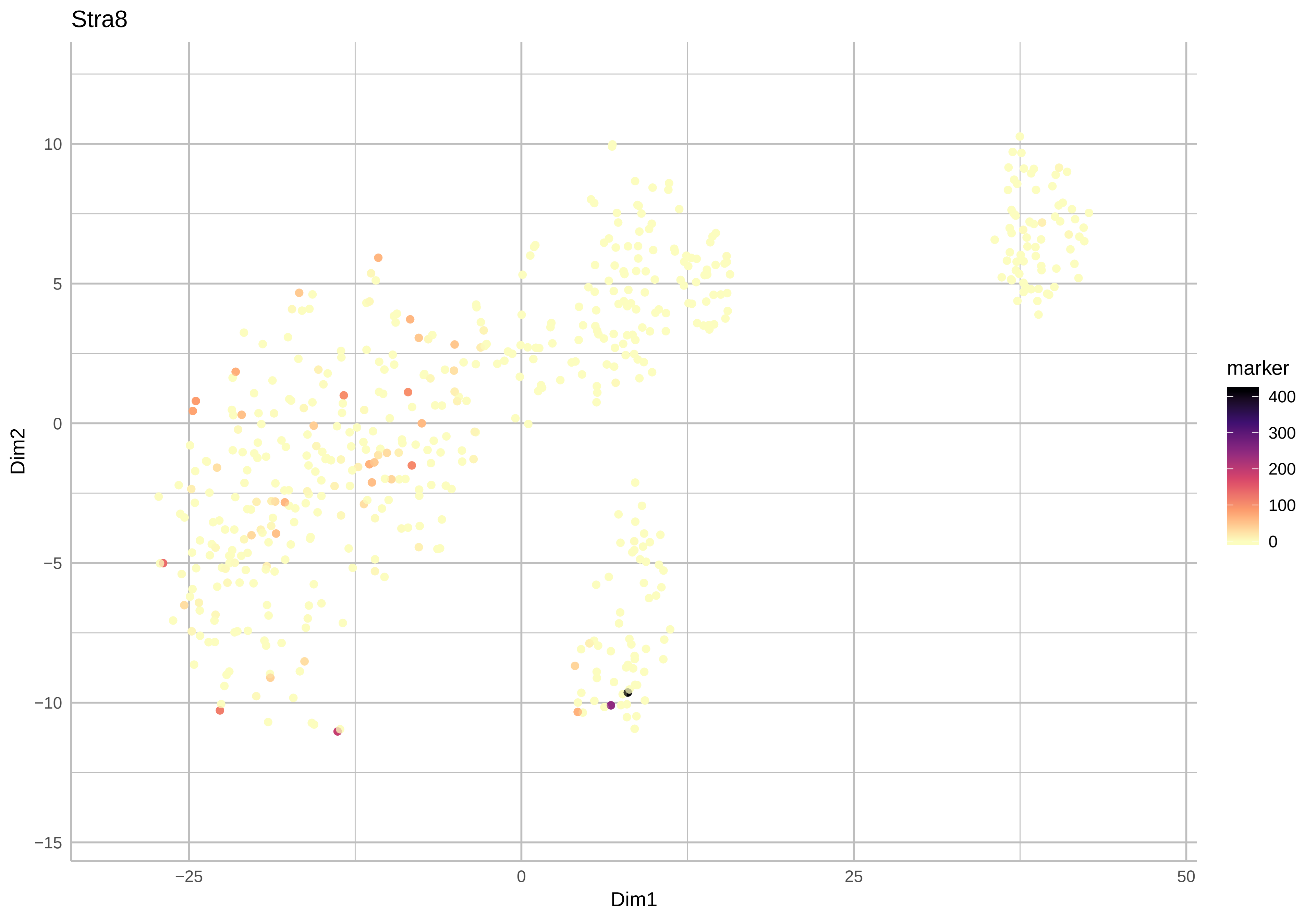

Stra8
Dim2
Dim1
10
5
0
−5
−10
−15
−25
0
25
50
marker
400
300
200
100
0

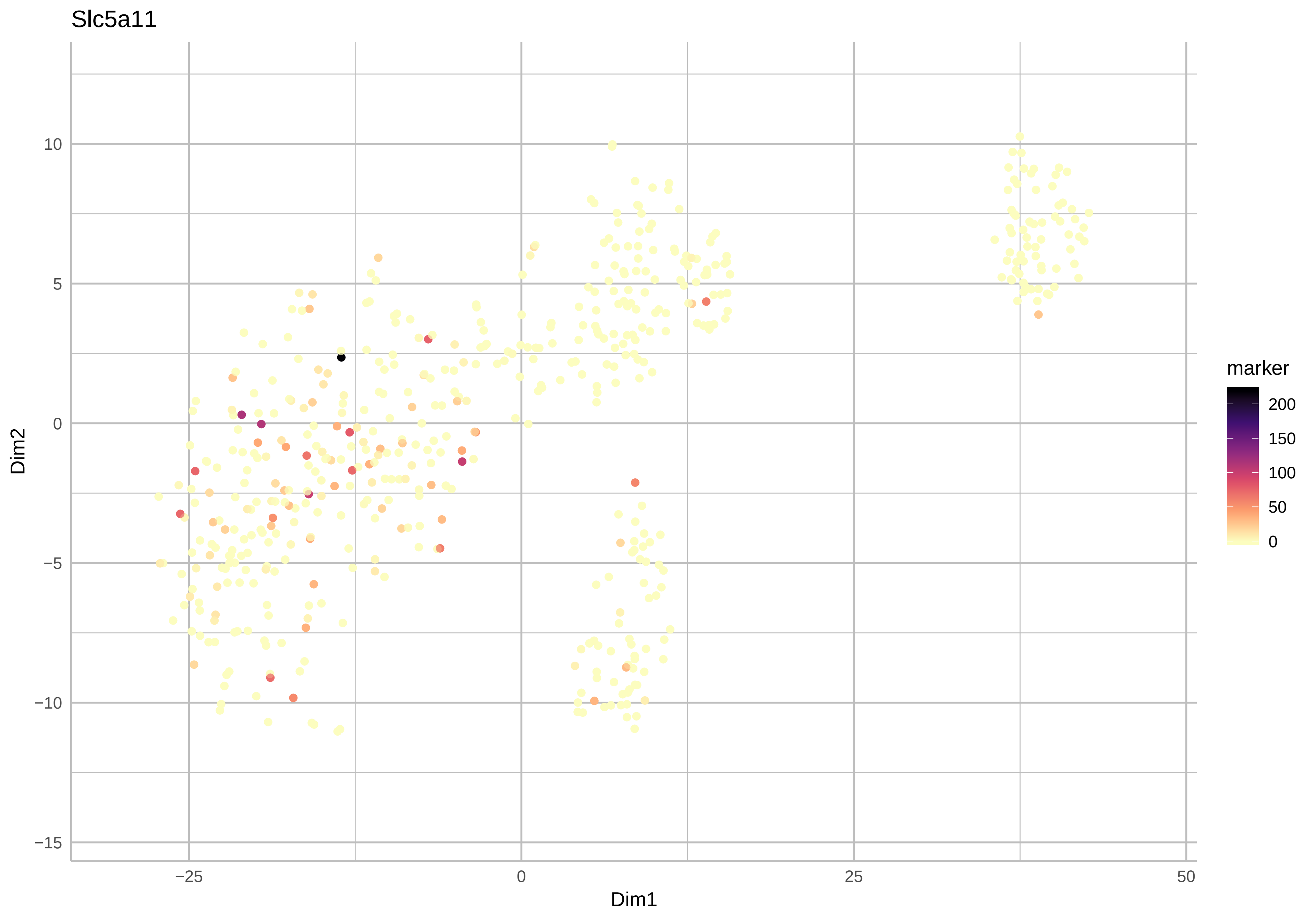
Slc5a11
Dim2
Dim1
10
5
0
−5
−10
−15
−25
0
25
50
marker
200
150
100
50
0

Fam178b

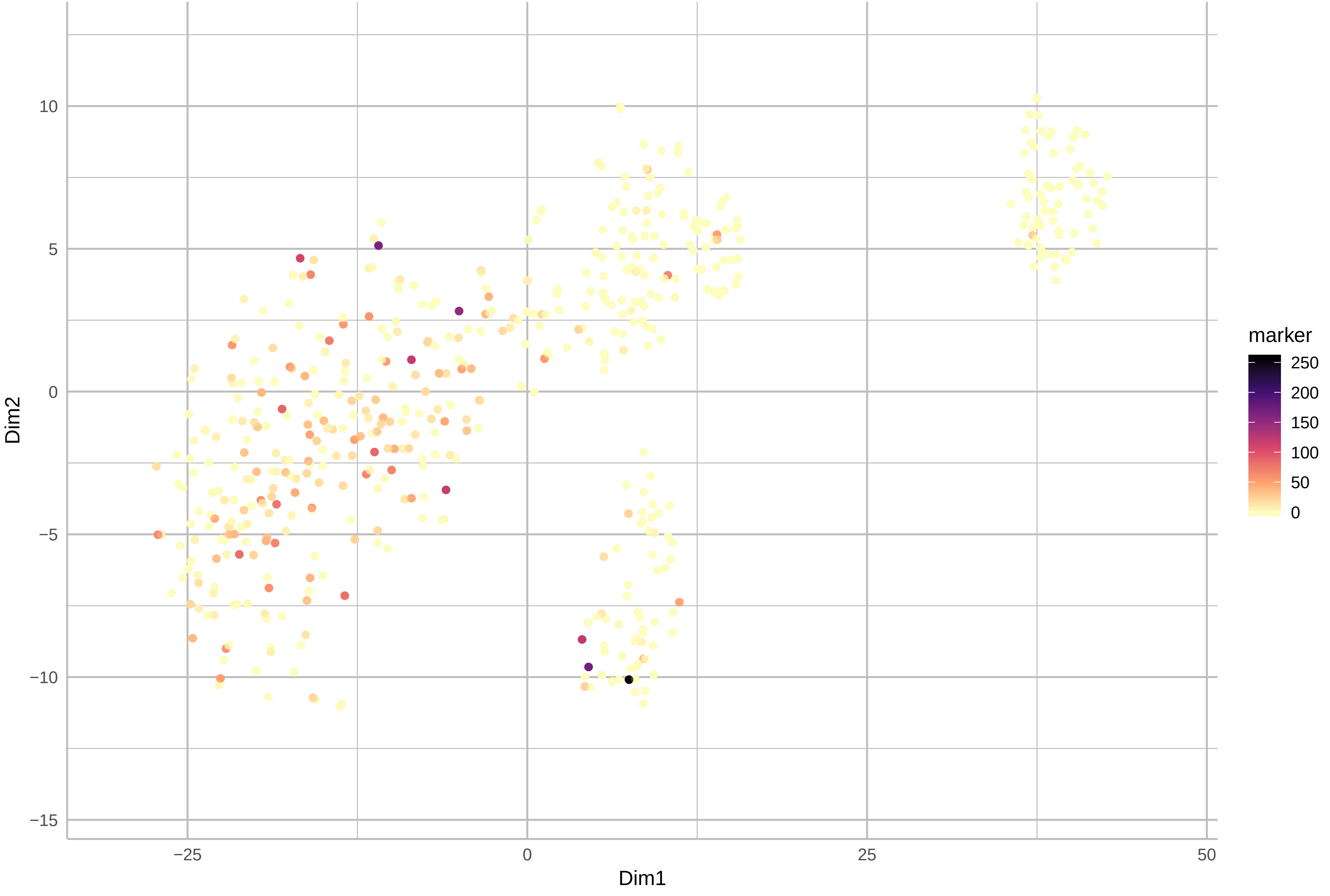

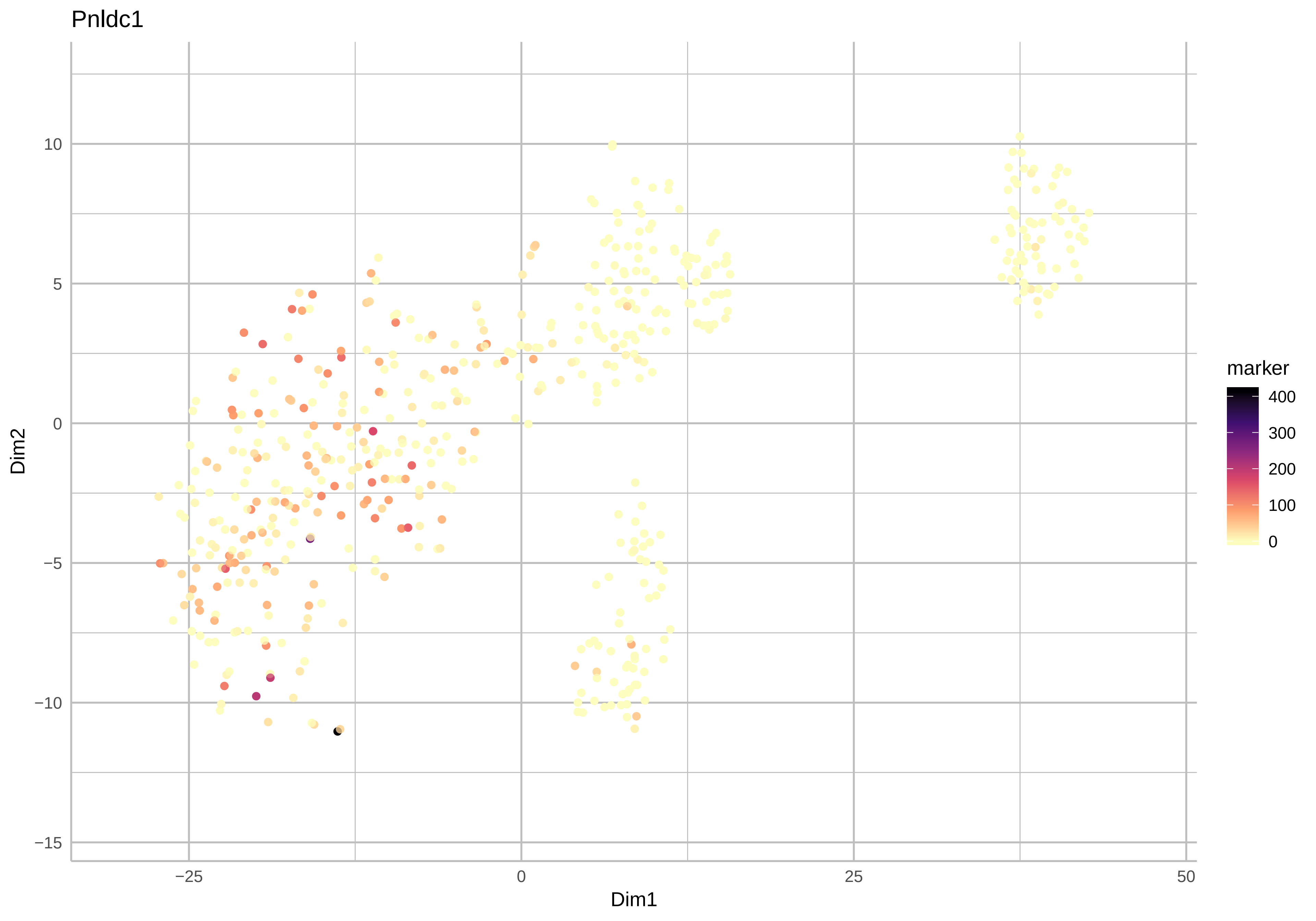

Pnldc1
Dim1
Dim2
−25
0
25
50
−15
−10
−5
0
5
10
marker
400
300
200
100
0

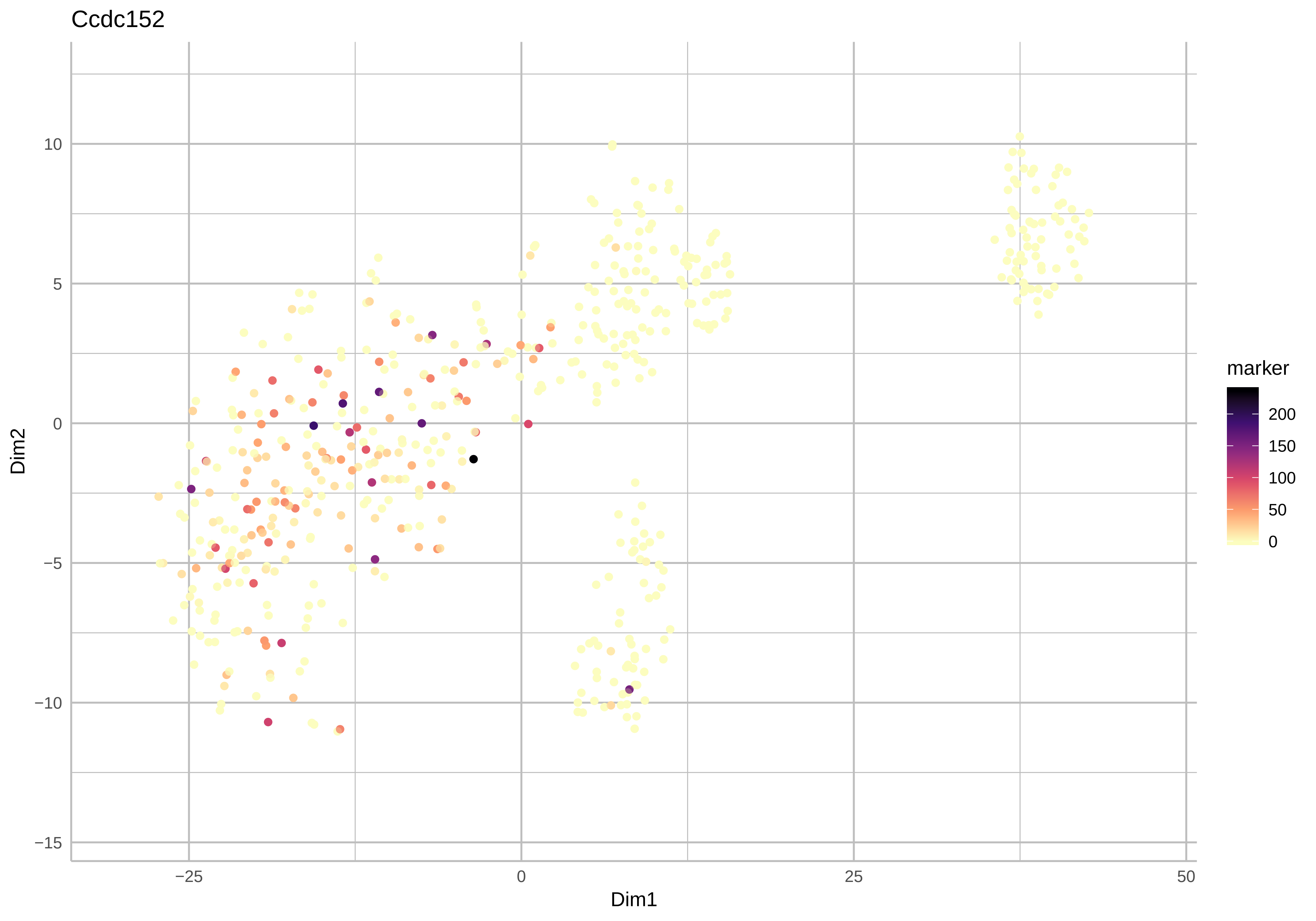
Ccdc152
Dim2
Dim1
10
5
0
−5
−10
−15
−25
0
25
50
marker
200
150
100
50
0

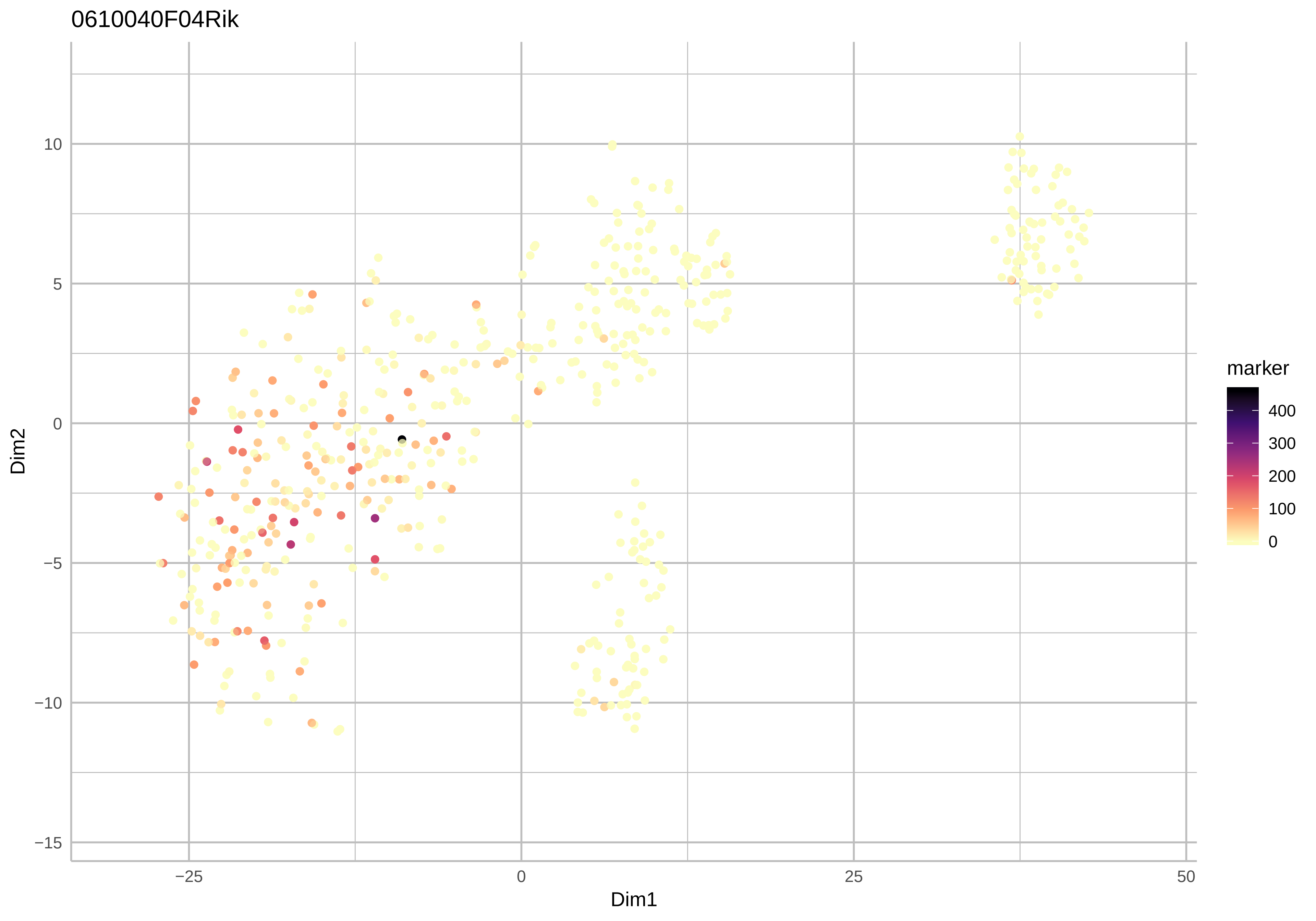
0610040F04Rik
Dim2
Dim1
10
5
0
−5
−10
−15
−25
0
25
50
marker
400
300
200
100
0

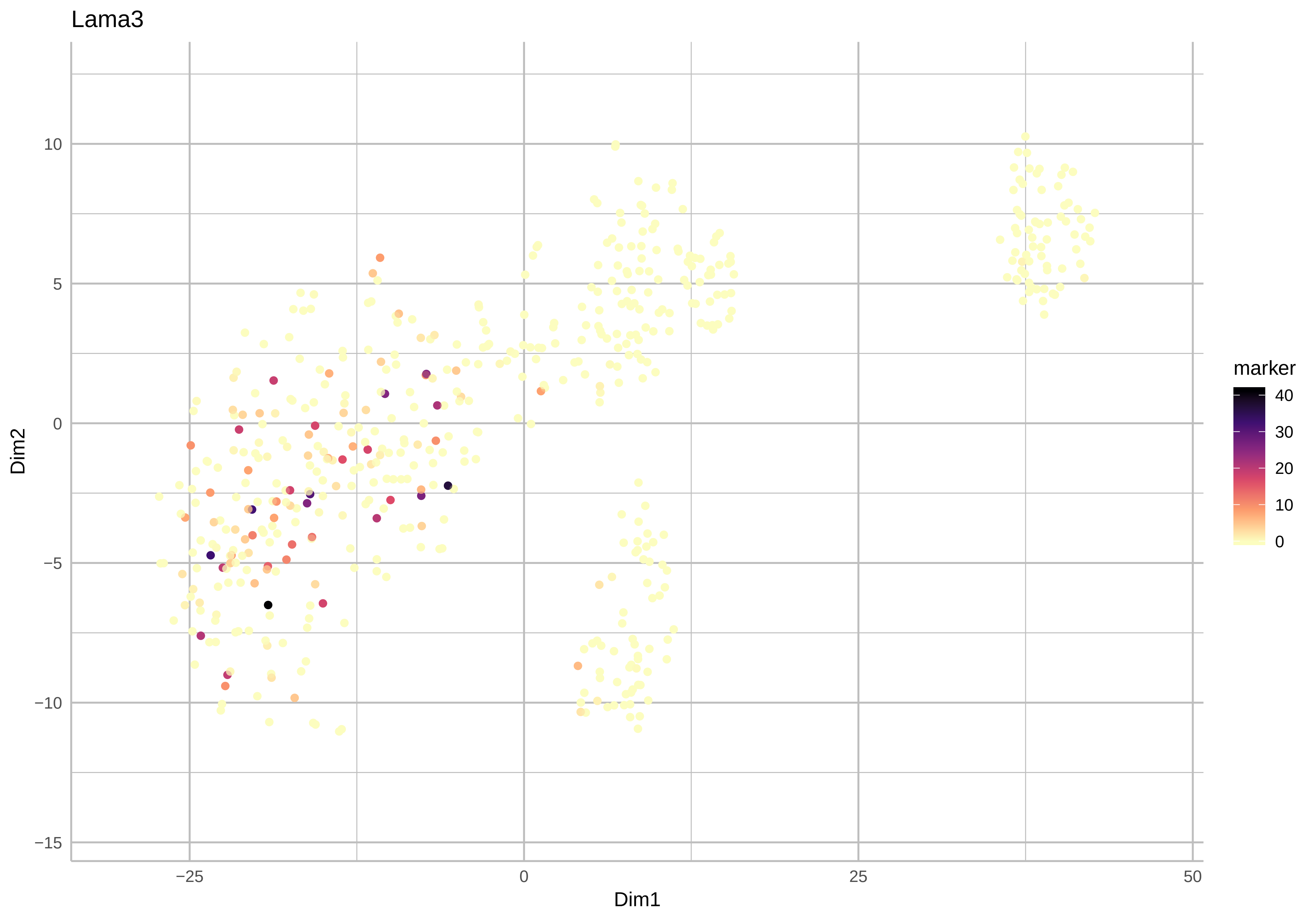
Lama3
10
5
0
−5
−10
−15
Dim2
−25
0
25
50
Dim1
marker
40
30
20
10
0

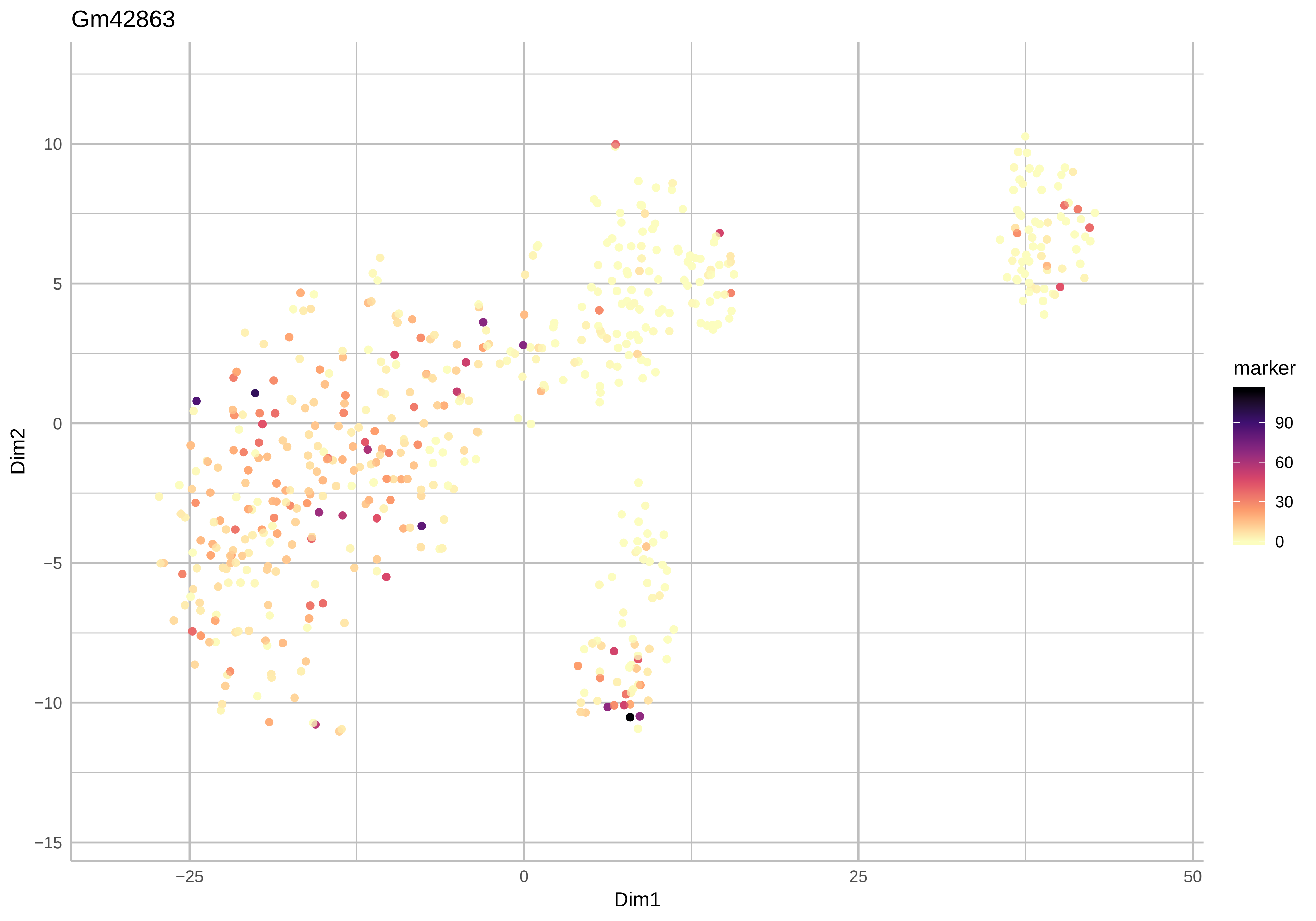
Gm42863
Dim2
Dim1
marker
90
60
30
0
10
5
0
−5
−10
−15
−25
0
25
50

Slc17a9

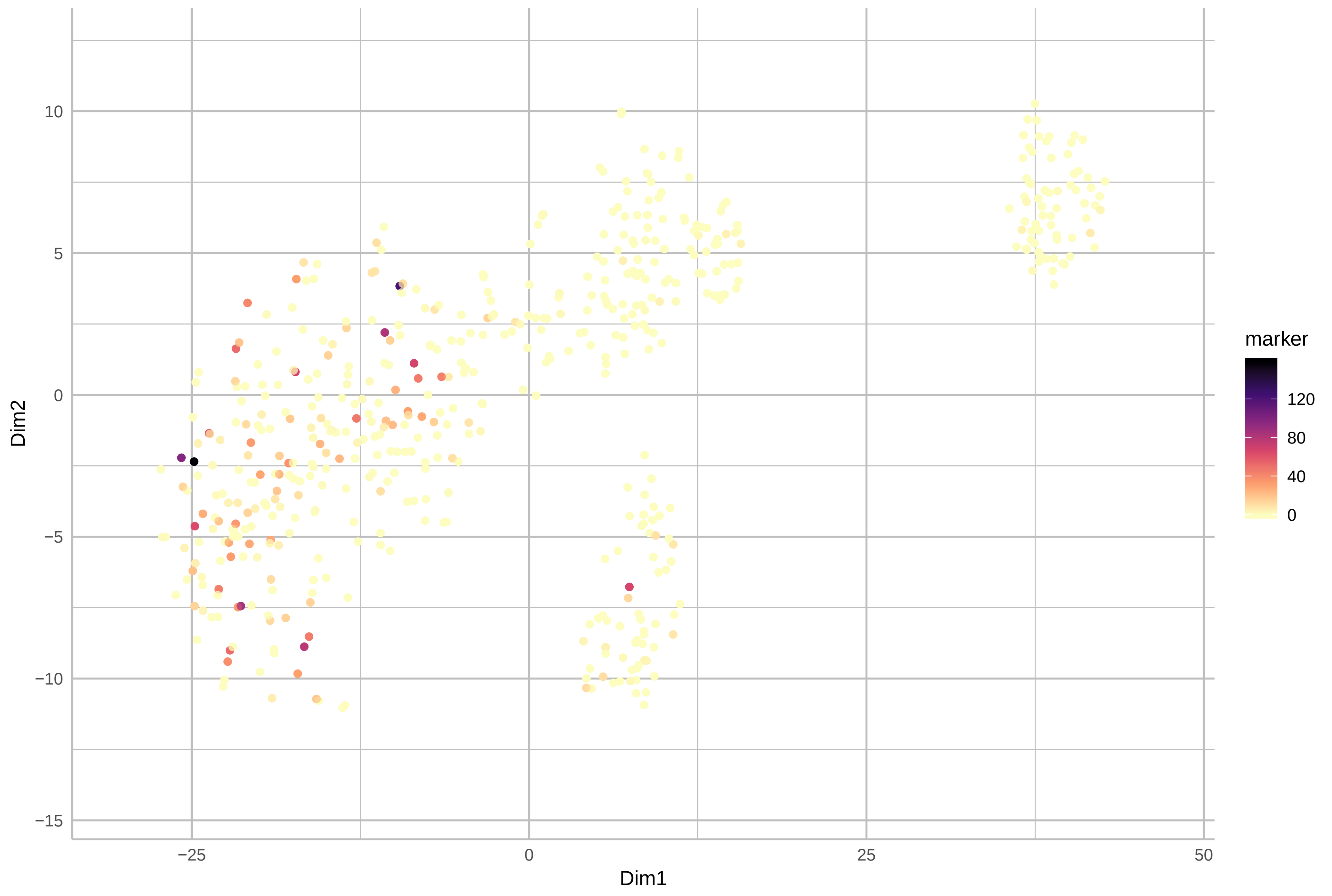

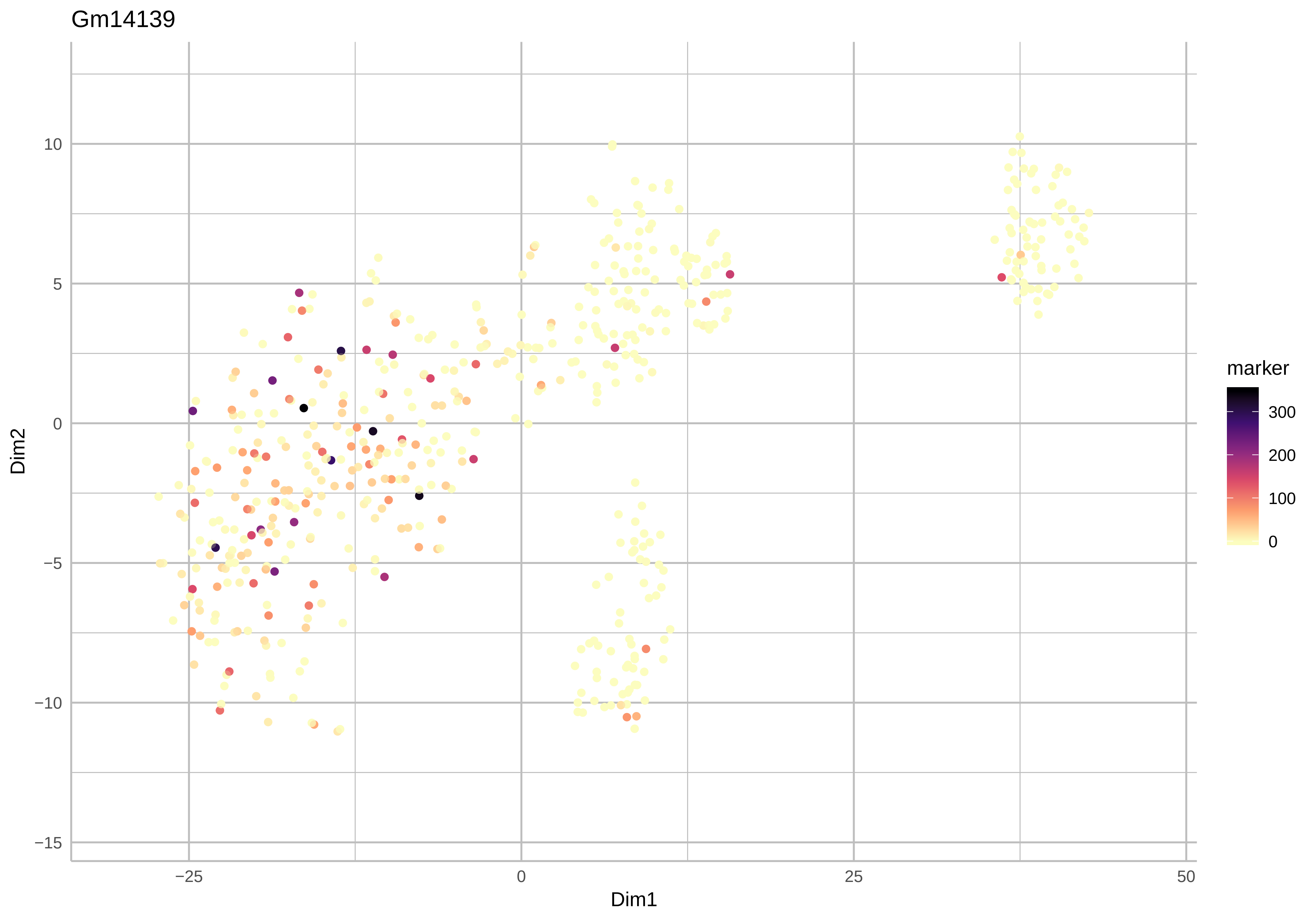
Gm14139
Dim2
Dim1
10
5
0
−5
−10
−15
−25
0
25
50
marker
300
200
100
0

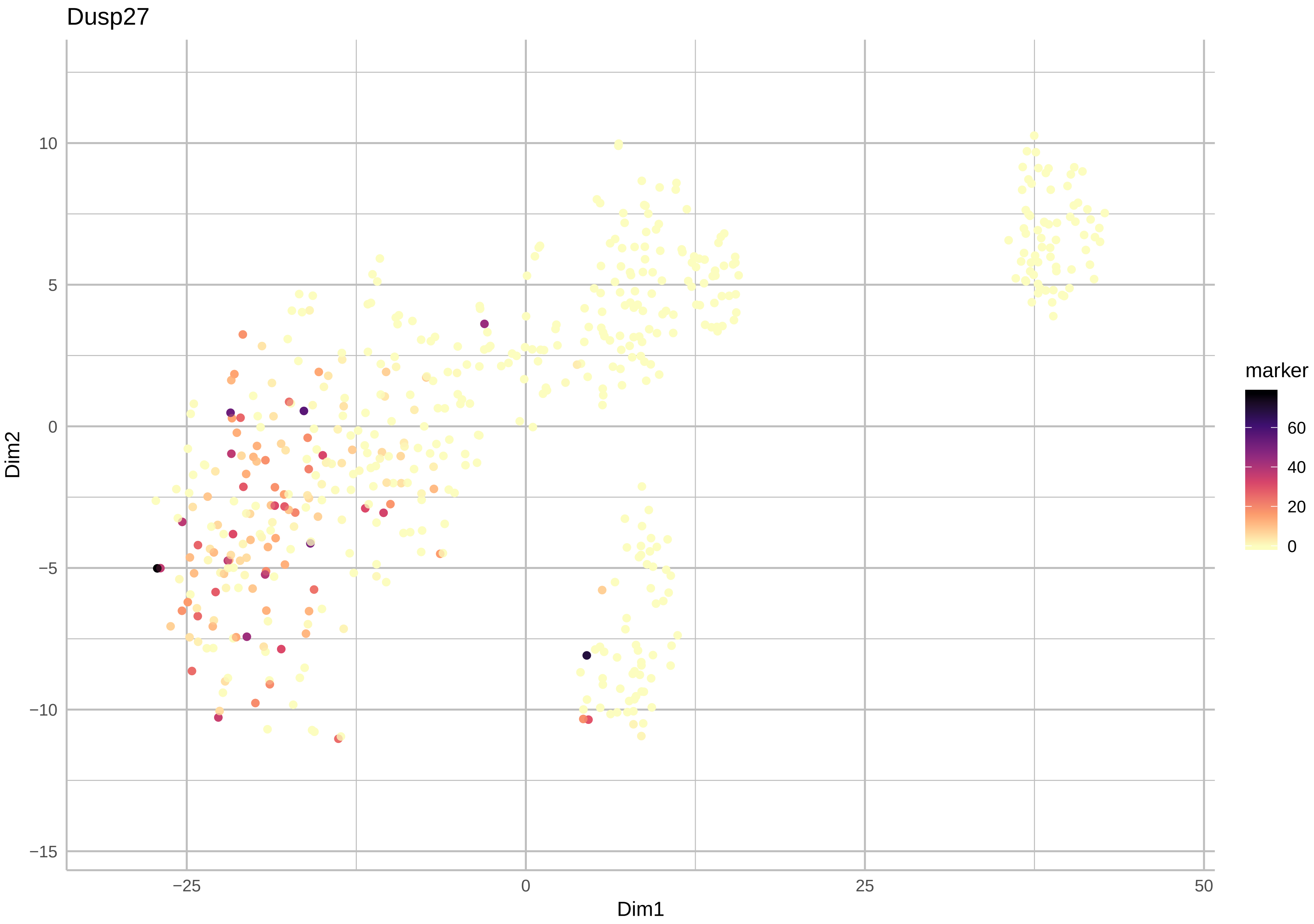

Dusp27
10
5
0
−5
−10
−15
−25
0
25
50
Dim1
Dim2
marker
60
40
20
0

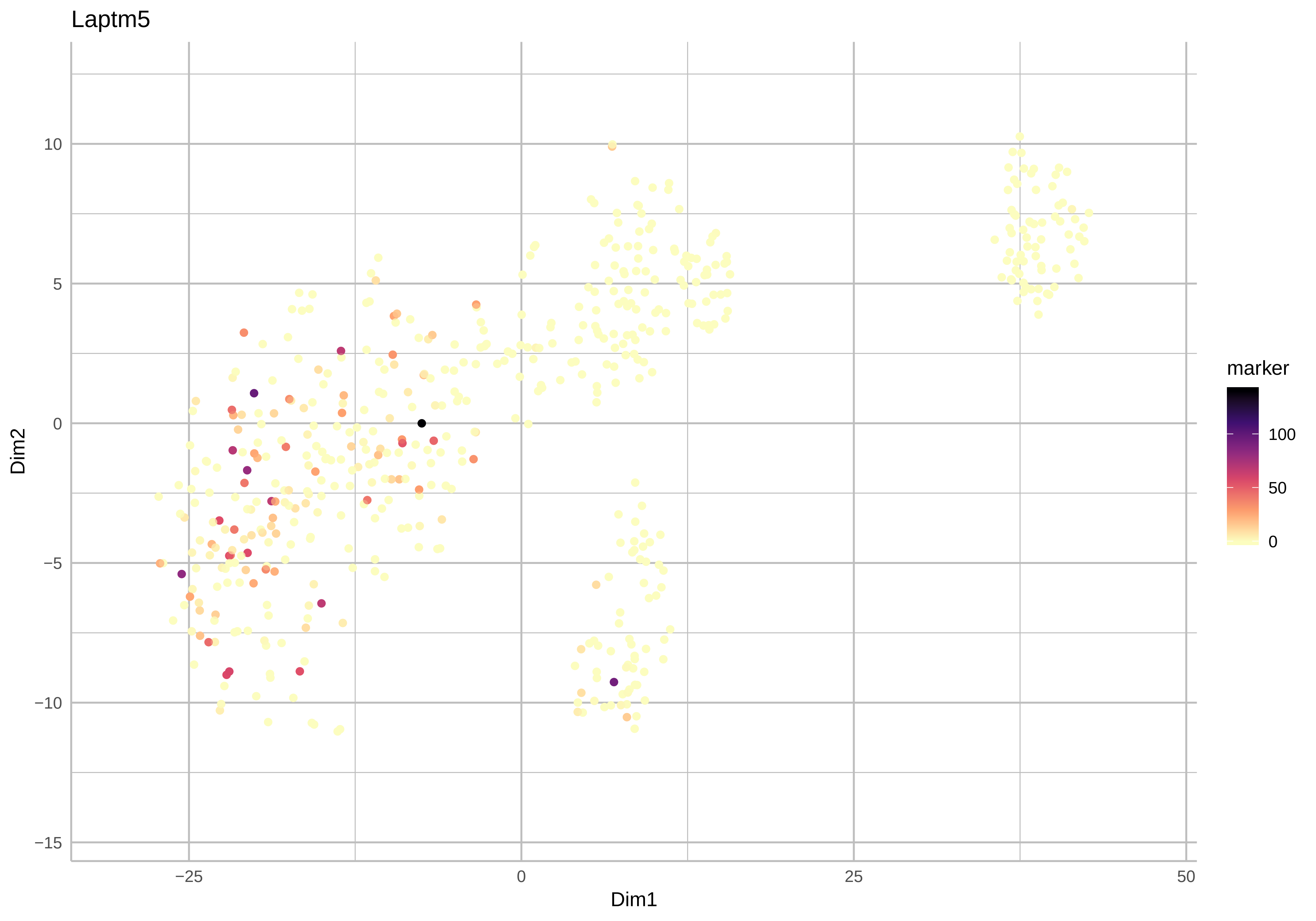

Laptm5
10
5
0
−5
−10
−15
−25
0
25
50
Dim1
Dim2
marker
100
50
0

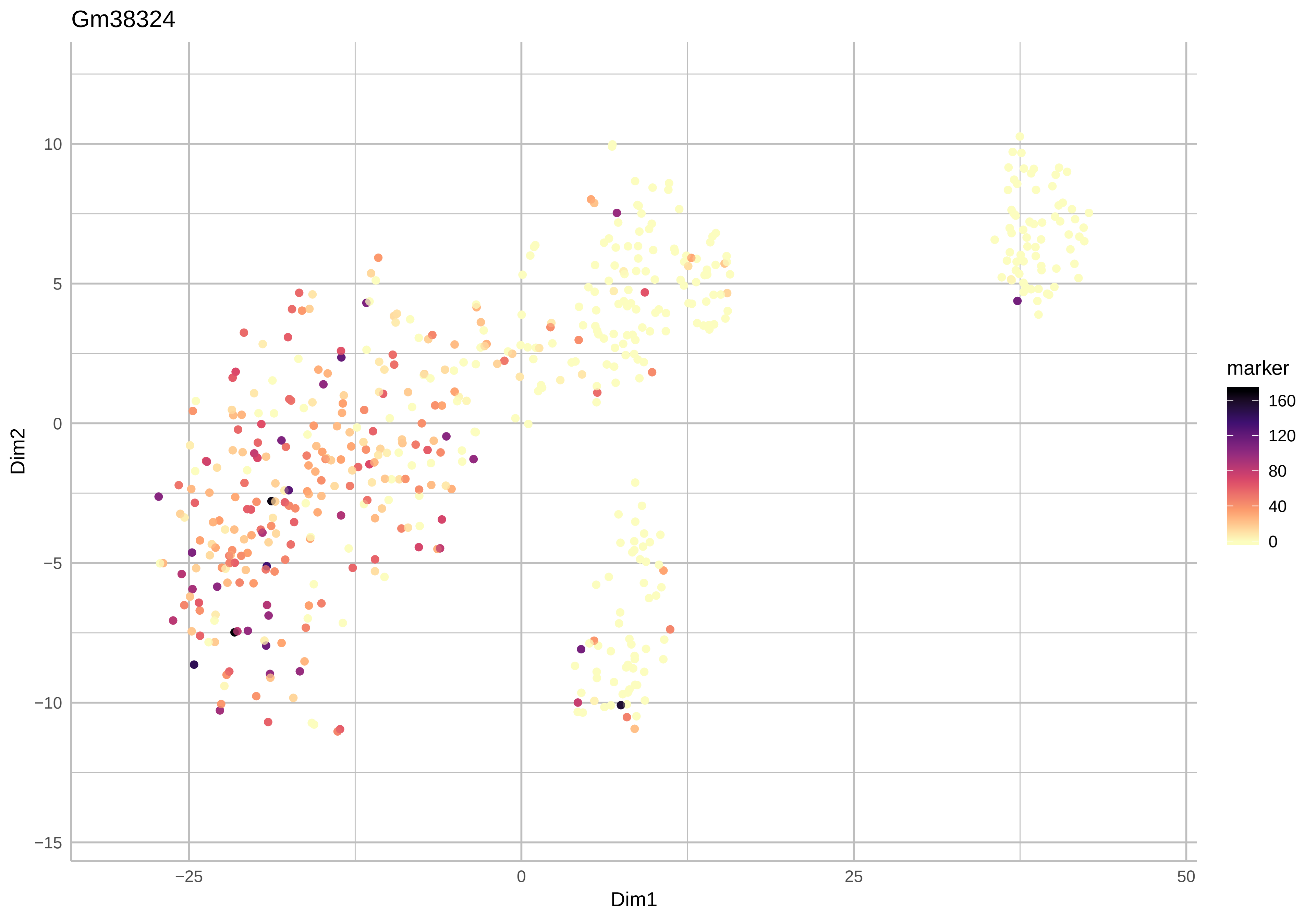
Gm38324
Dim2
Dim1
10
5
0
−5
−10
−15
−25
0
25
50
marker
160
120
80
40
0

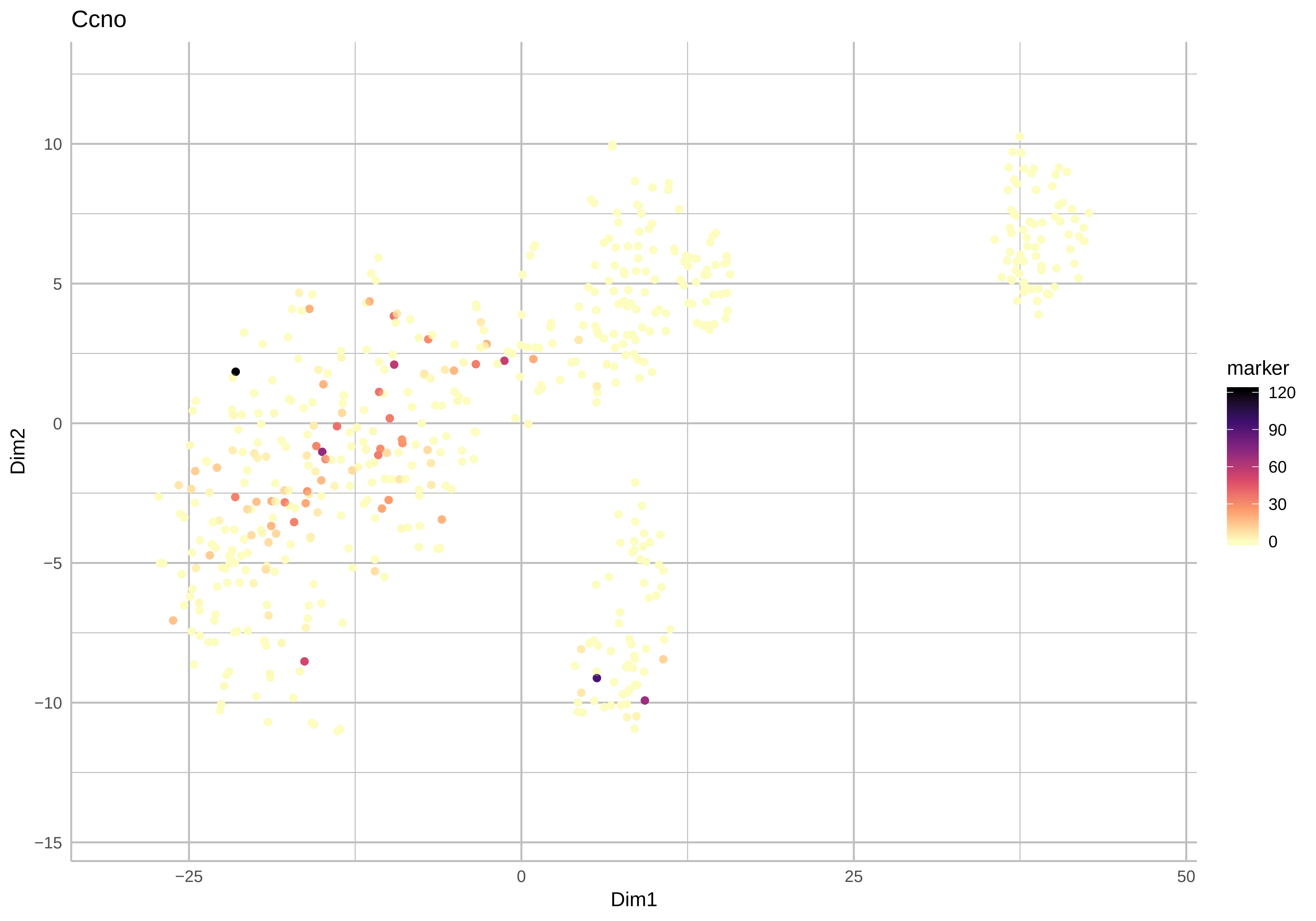
Ccno
Dim2
Dim1
marker
120
90
60
30
0
10
5
0
−5
−10
−15
−25
0
25
50

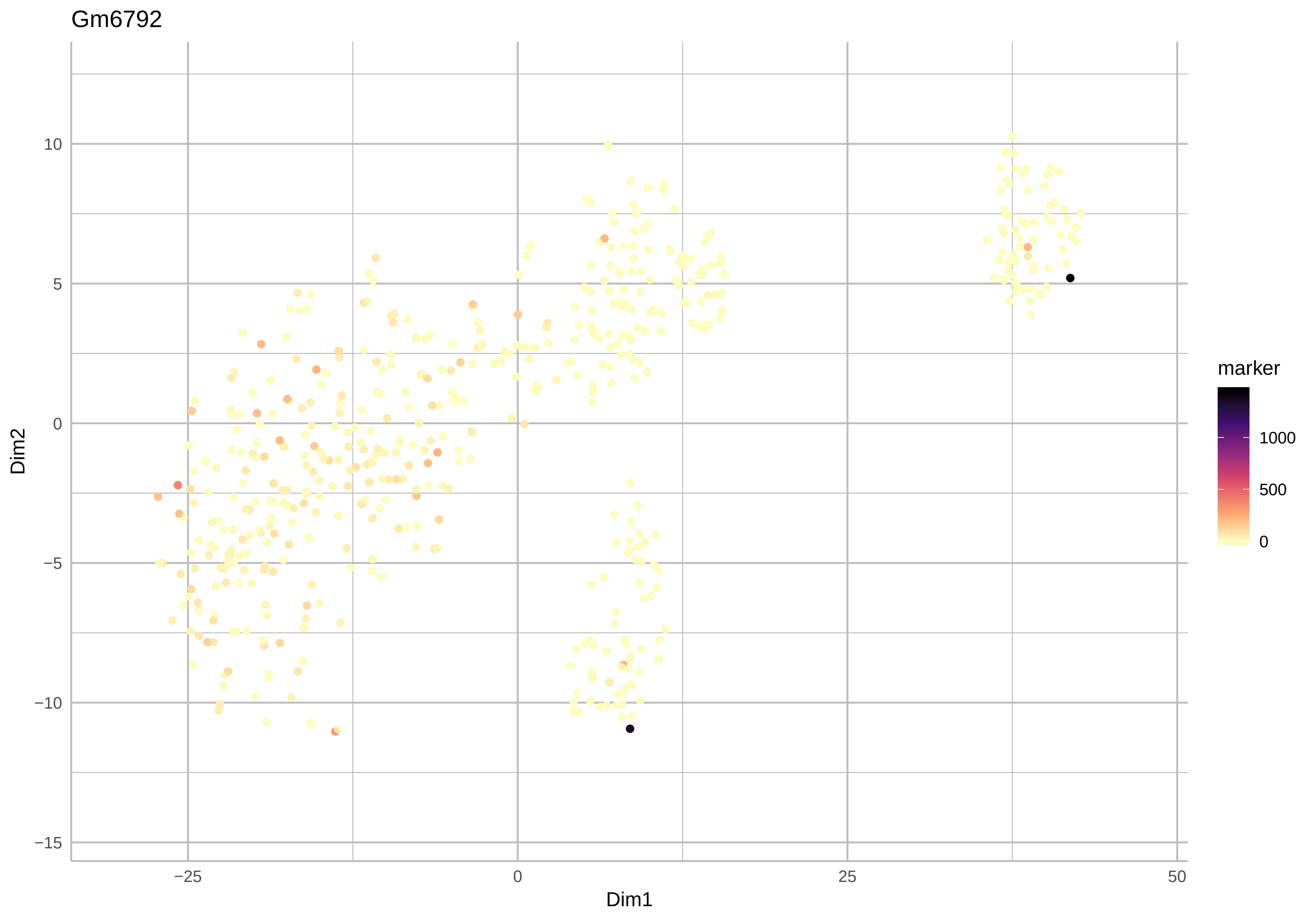

Gm6792
Dim2
Dim1
10
5
0
−5
−10
−15
−25
0
25
50
marker
1000
500
0

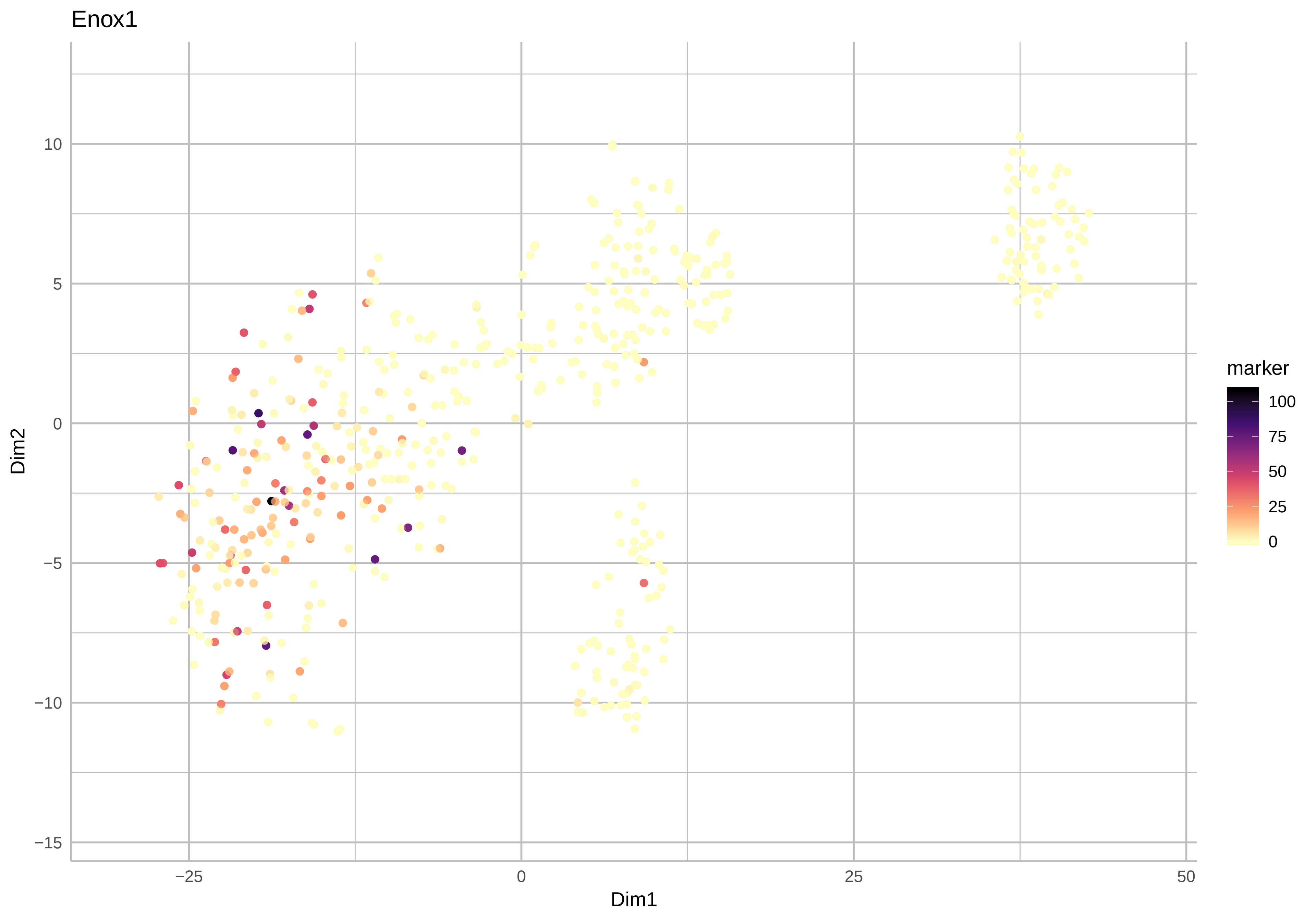
Enox1
Dim2
Dim1
10
5
0
−5
−10
−15
−25
0
25
50
marker
100
75
50
25
0

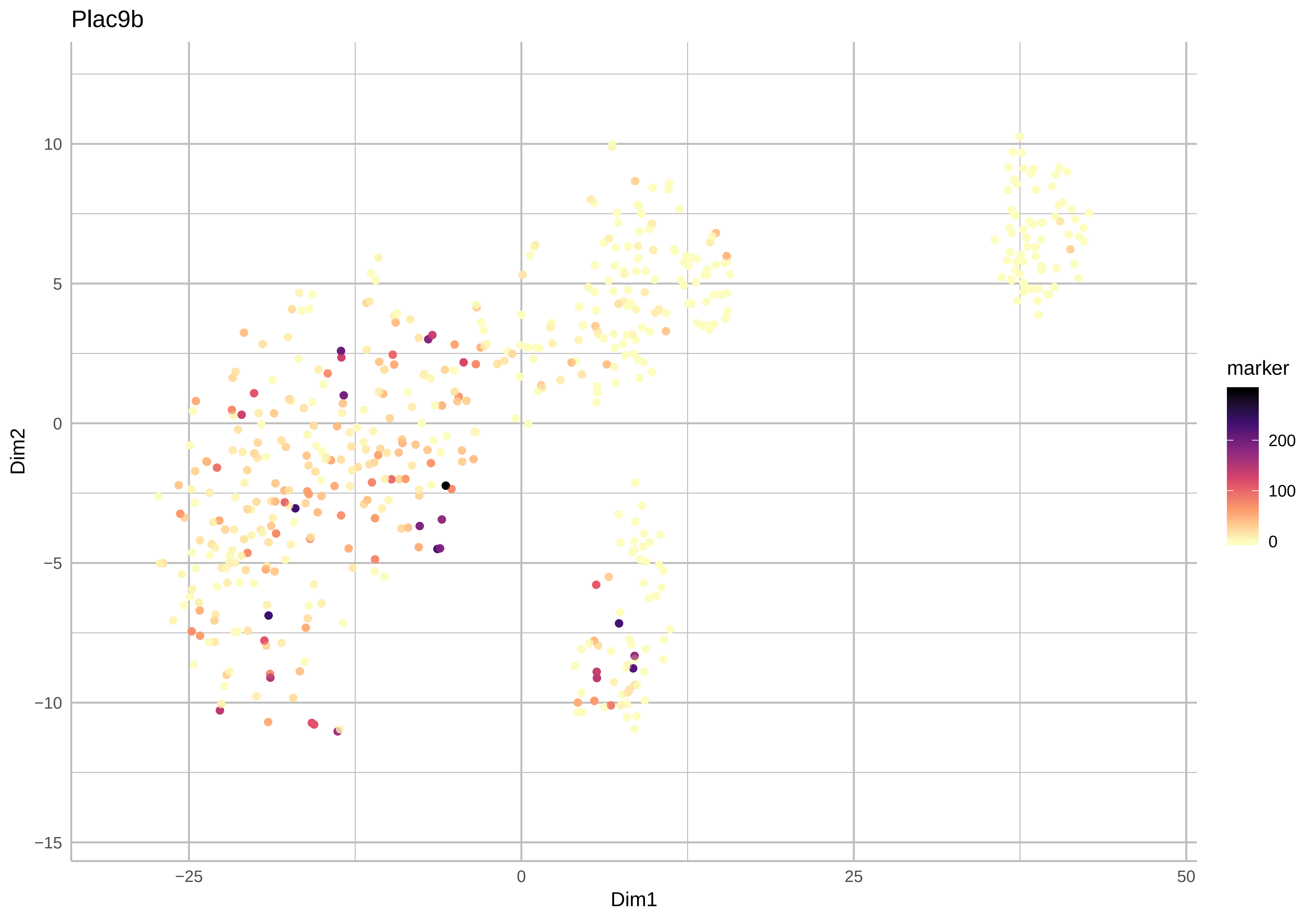
Plac9b
Dim2
Dim1
marker
200
100
0
10
5
0
−5
−10
−15
−25
0
25
50

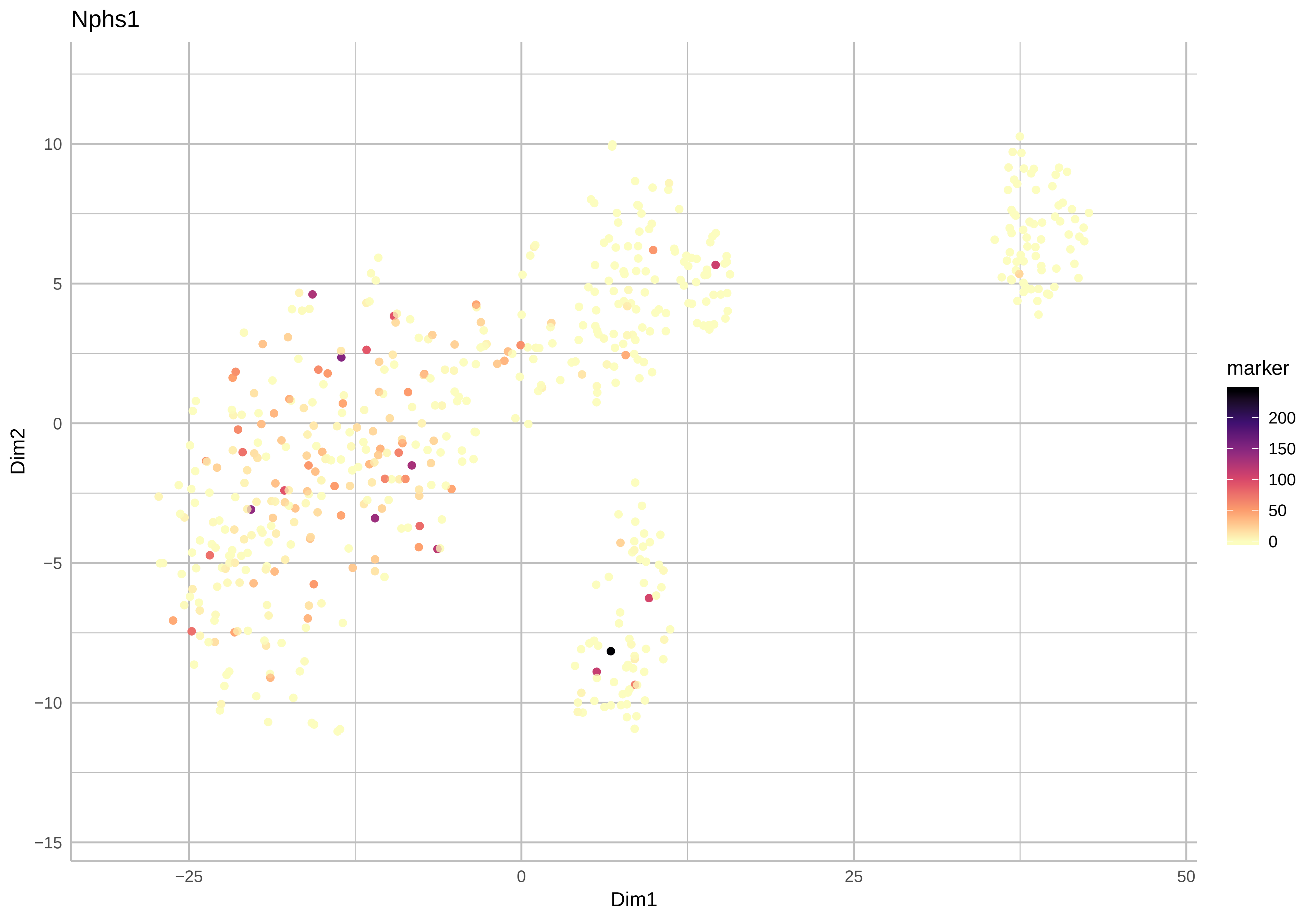

Nphs1
Dim2
Dim1
marker
200
150
100
50
0
10
5
0
−5
−10
−15
−25
0
25
50

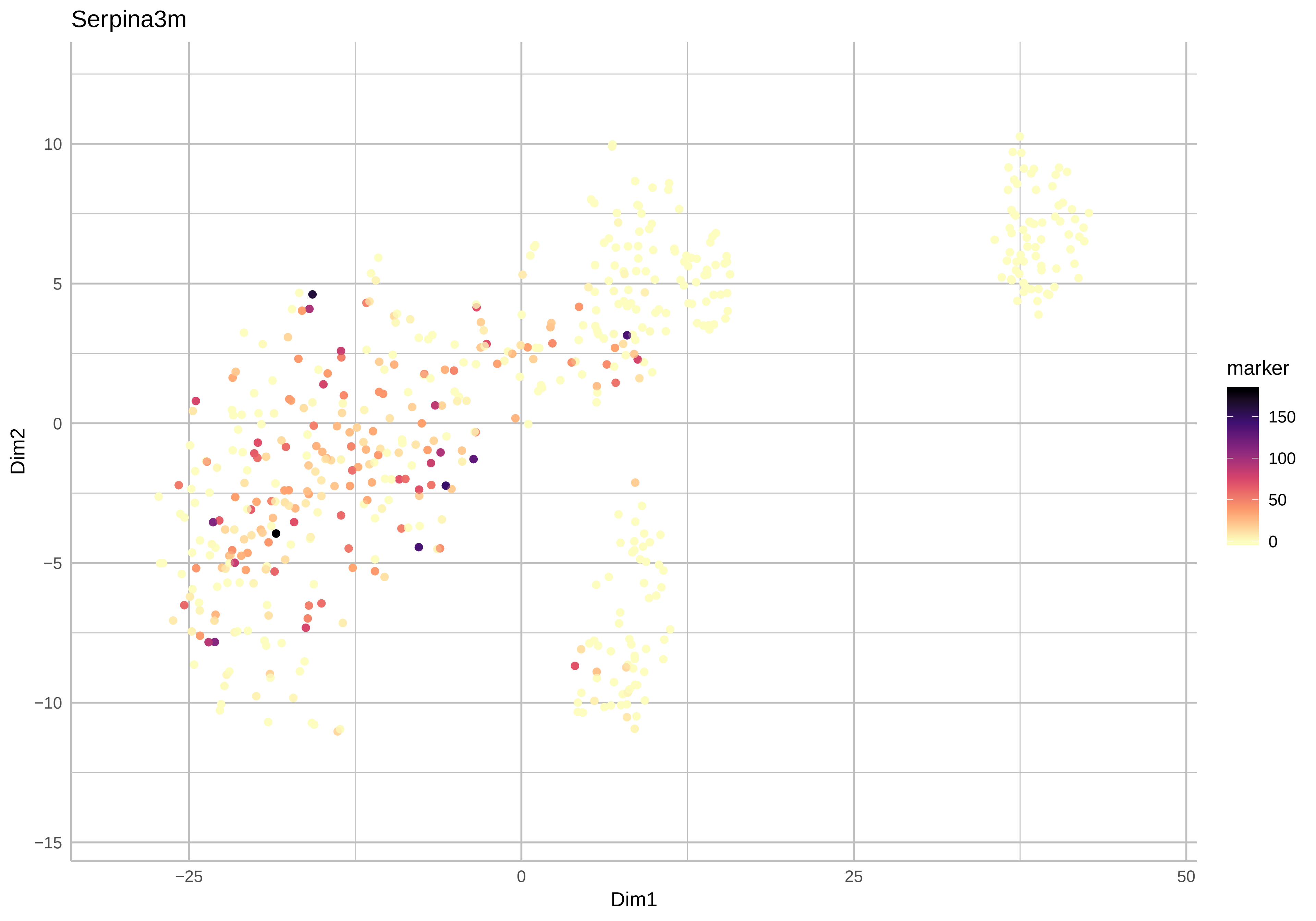
Serpina3m
Dim2
Dim1
marker
150
100
50
0
10
5
0
−5
−10
−15
−25
0
25
50

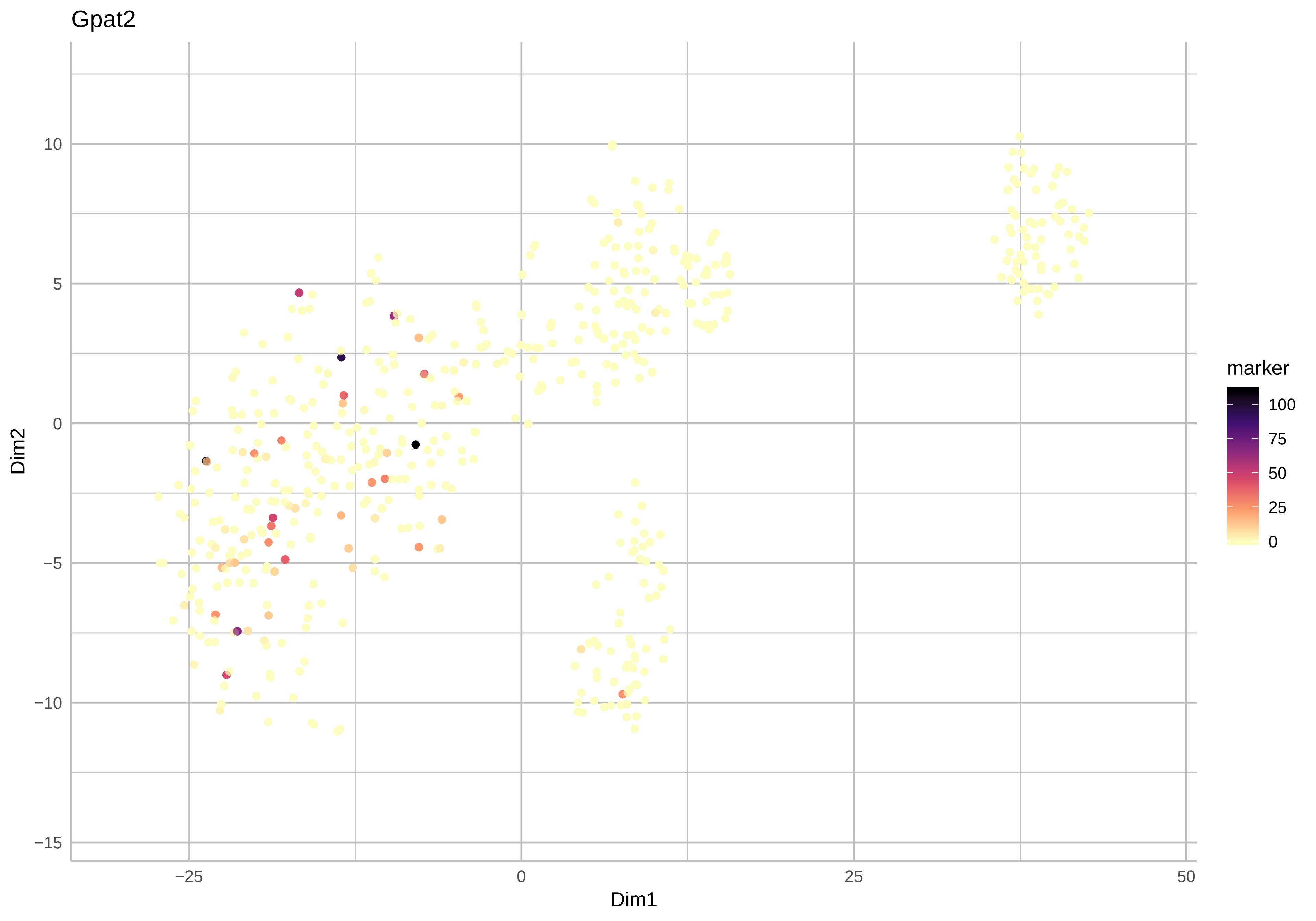
Gpat2
Dim2
Dim1
marker
100
75
50
25
0
10
5
0
−5
−10
−15
−25
0
25
50

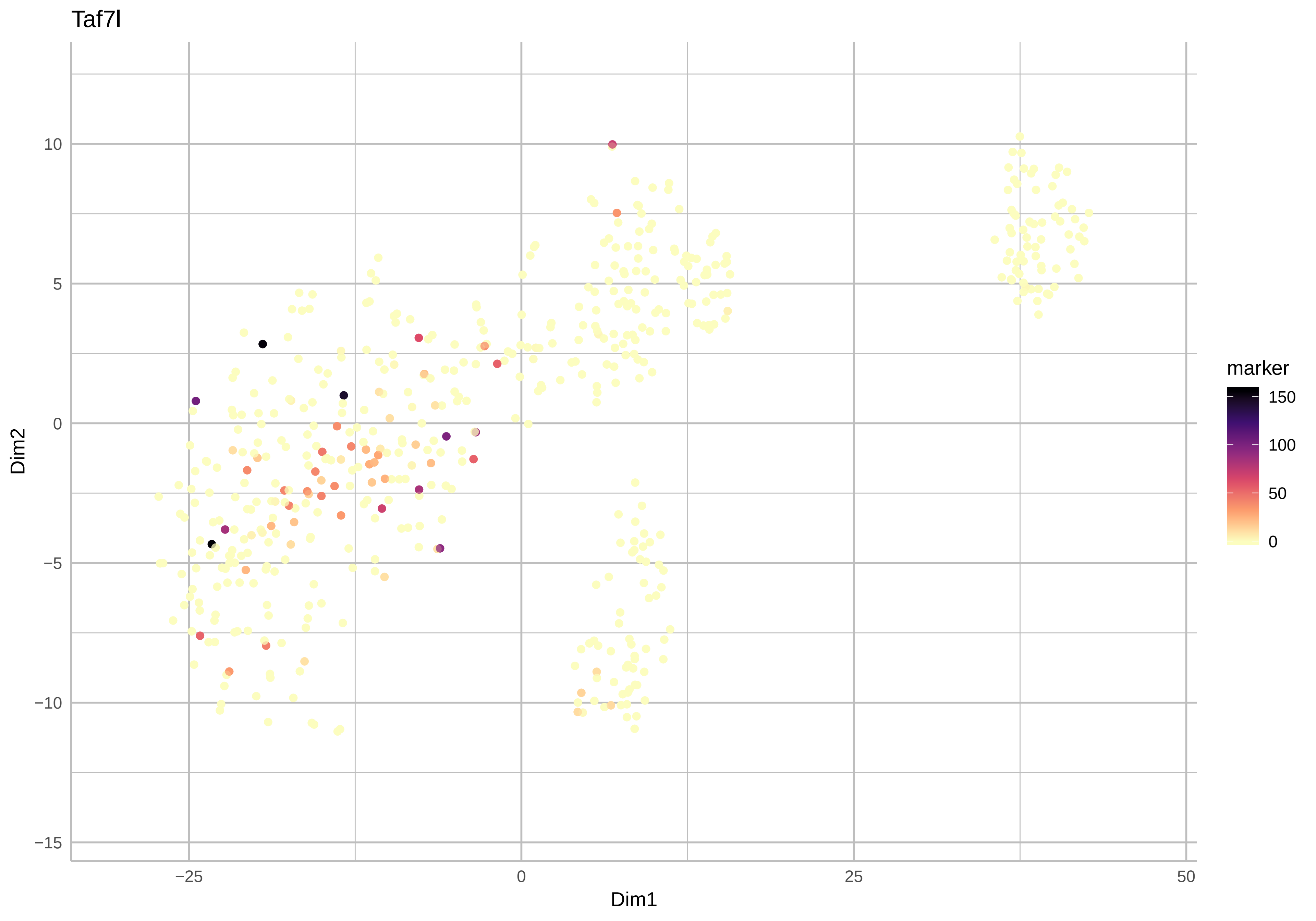

Taf7l
Dim2
Dim1
10
5
0
−5
−10
−15
−25
0
25
50
marker
150
100
50
0

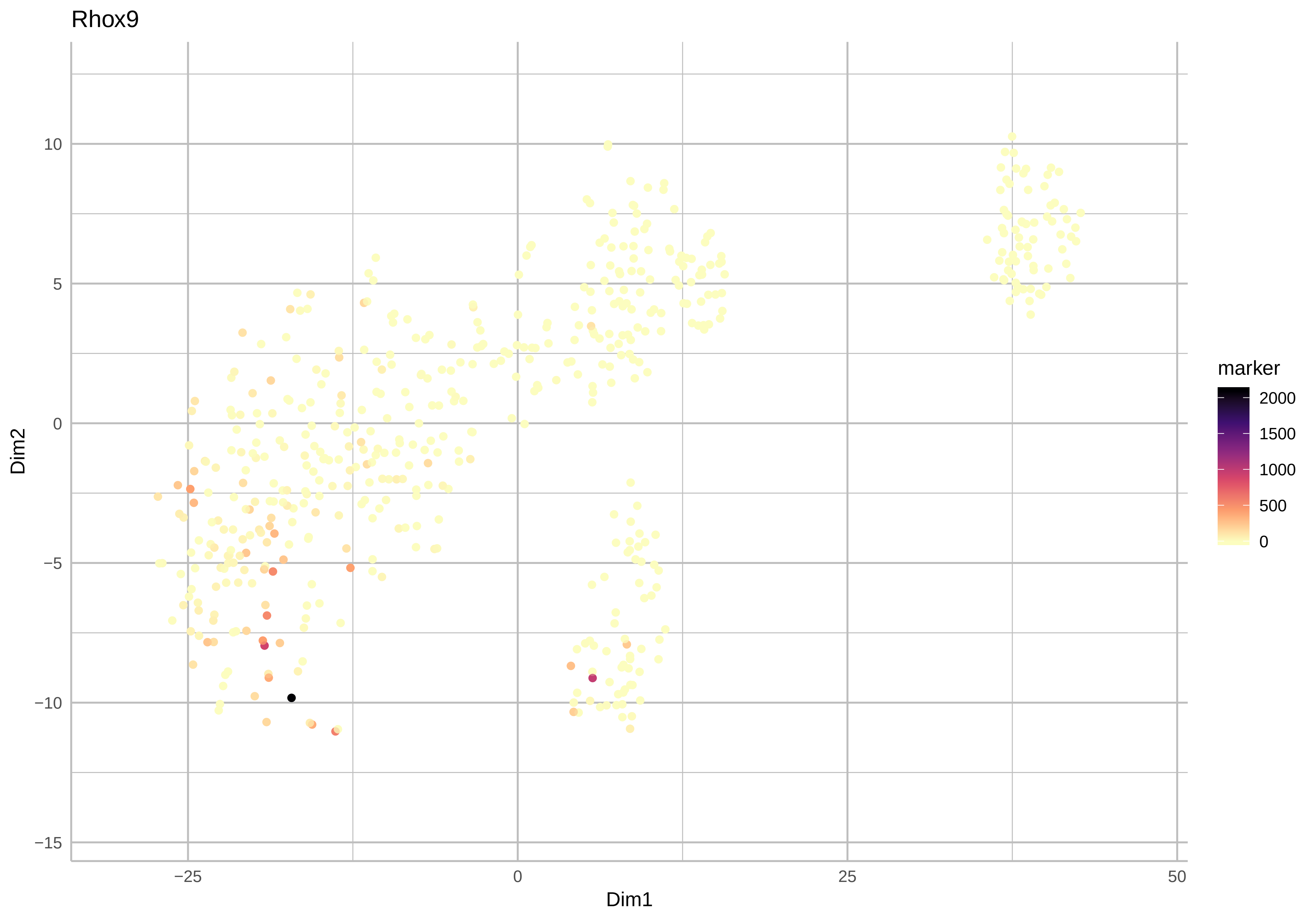
Rhox9
Dim2
10
5
0
−5
−10
−15
Dim1
−25
0
25
50
marker
2000
1500
1000
500
0

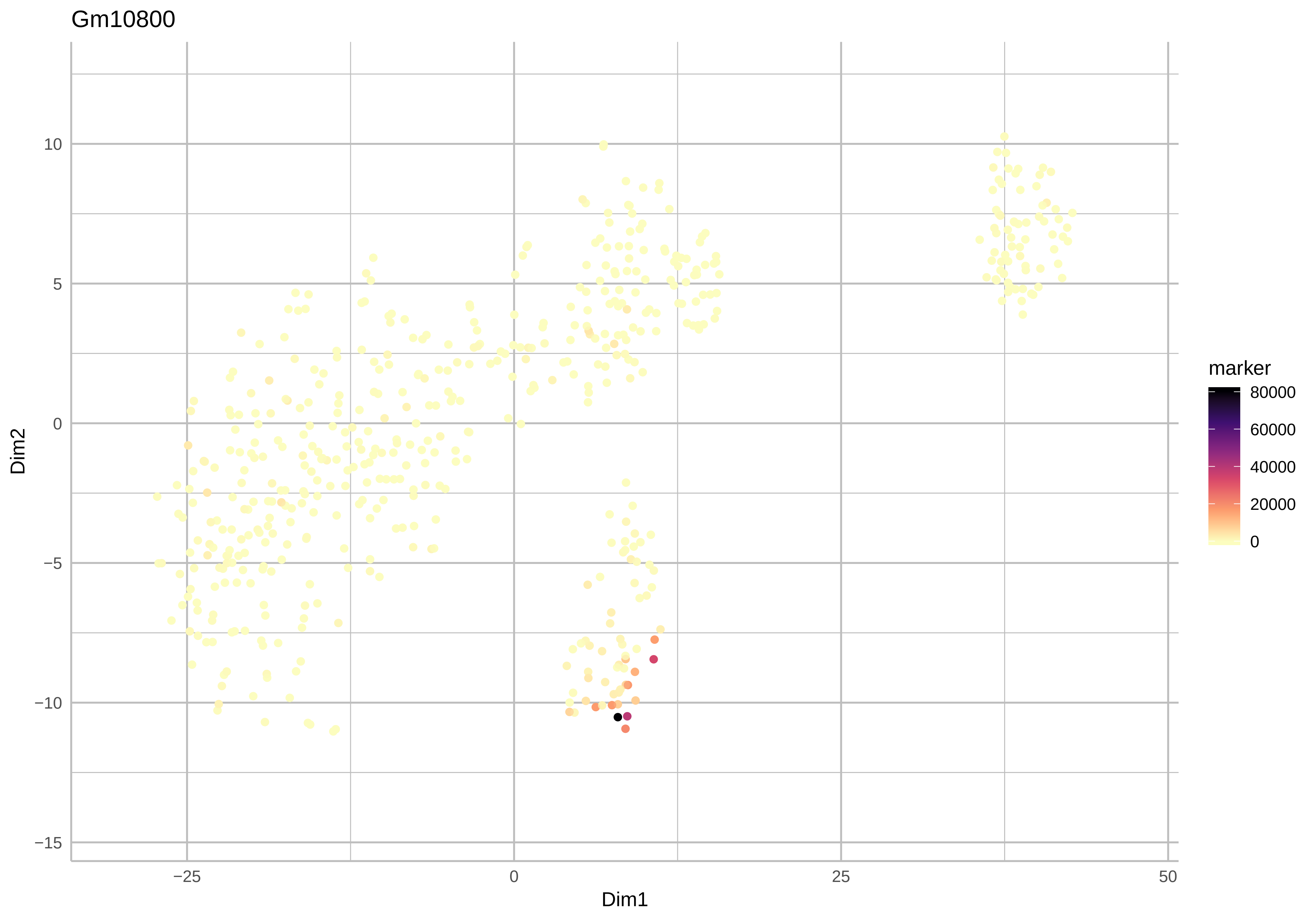
Gm10800
Dim2
Dim1
10
5
0
−5
−10
−15
−25
0
25
50
marker
80000
60000
40000
20000
0

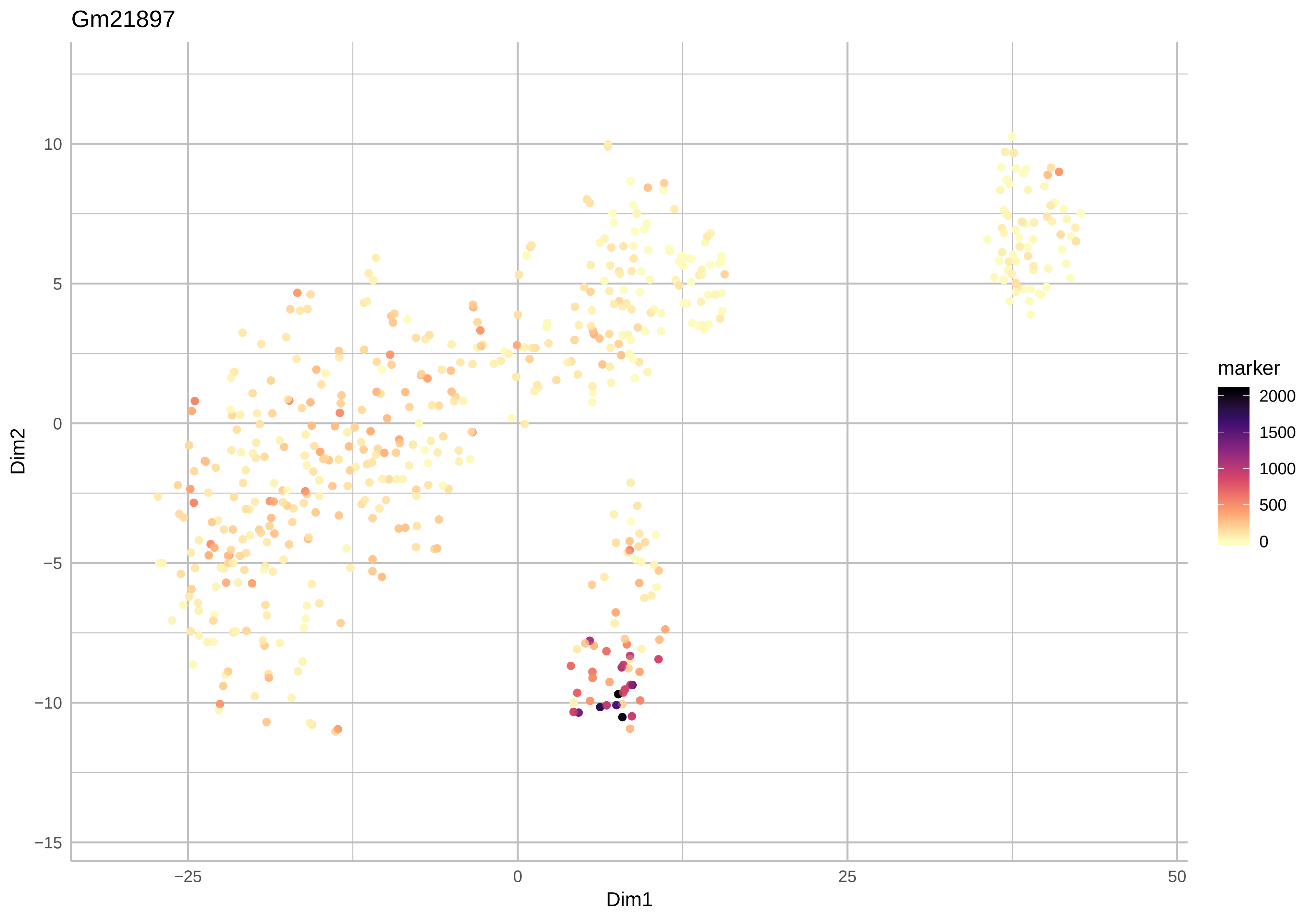
Gm21897
Dim2
Dim1
10
5
0
−5
−10
−15
−25
0
25
50
marker
2000
1500
1000
500
0

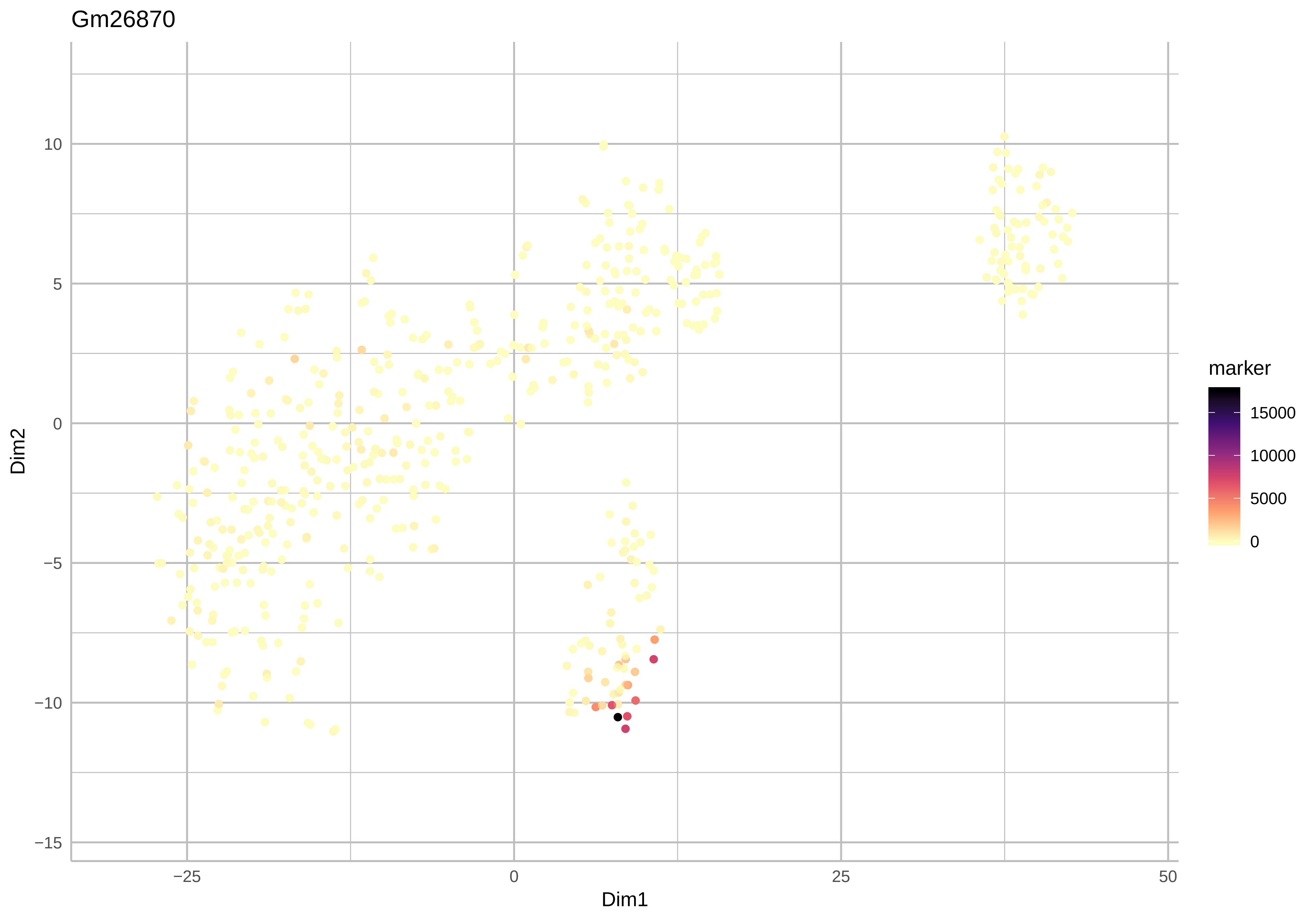

Gm26870
Dim2
Dim1
marker
15000
10000
5000
0
10
5
0
−5
−10
−15
−25
0
25
50

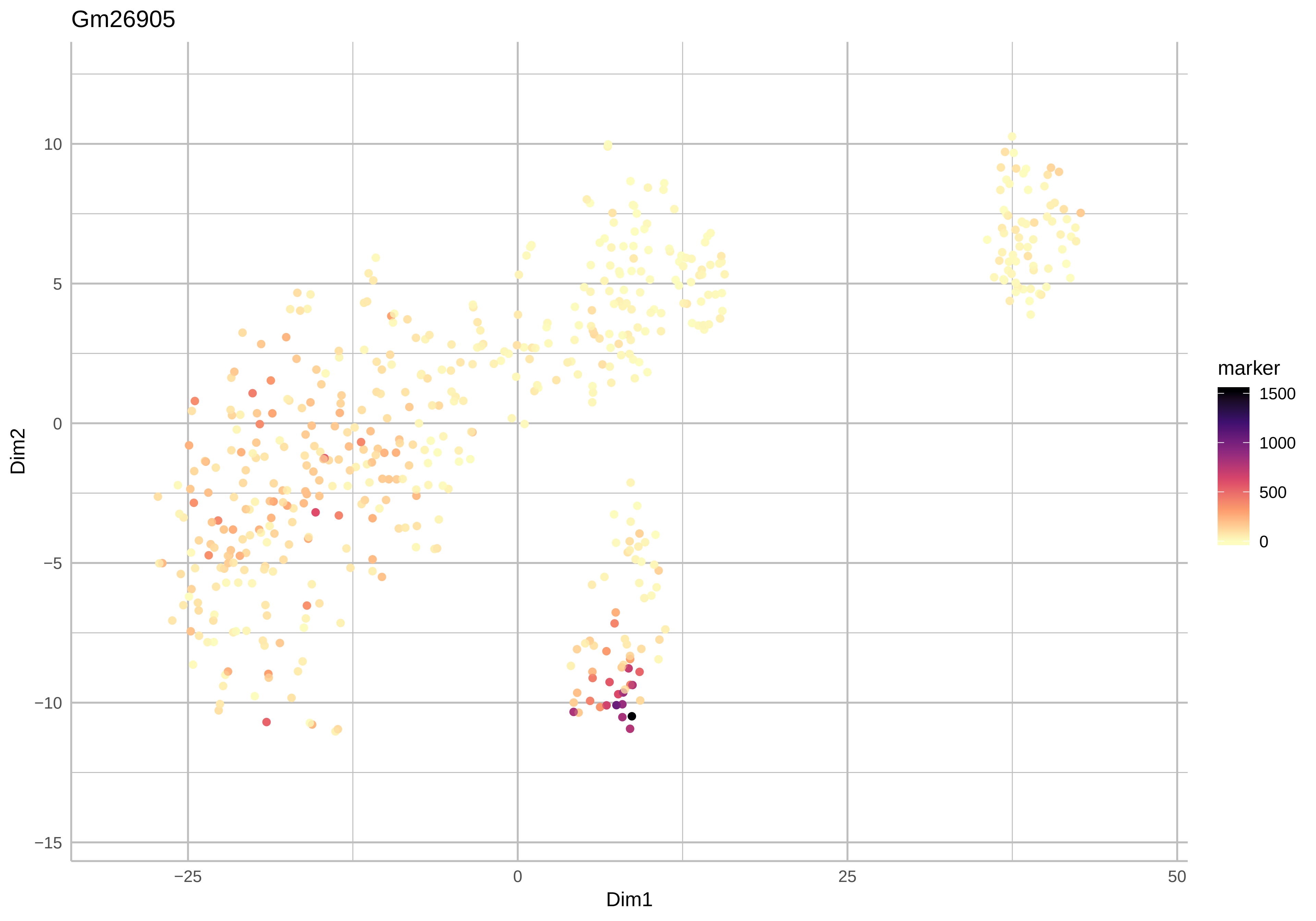
Gm26905
Dim2
Dim1
10
5
0
−5
−10
−15
−25
0
25
50
marker
1500
1000
500
0

Gm28268

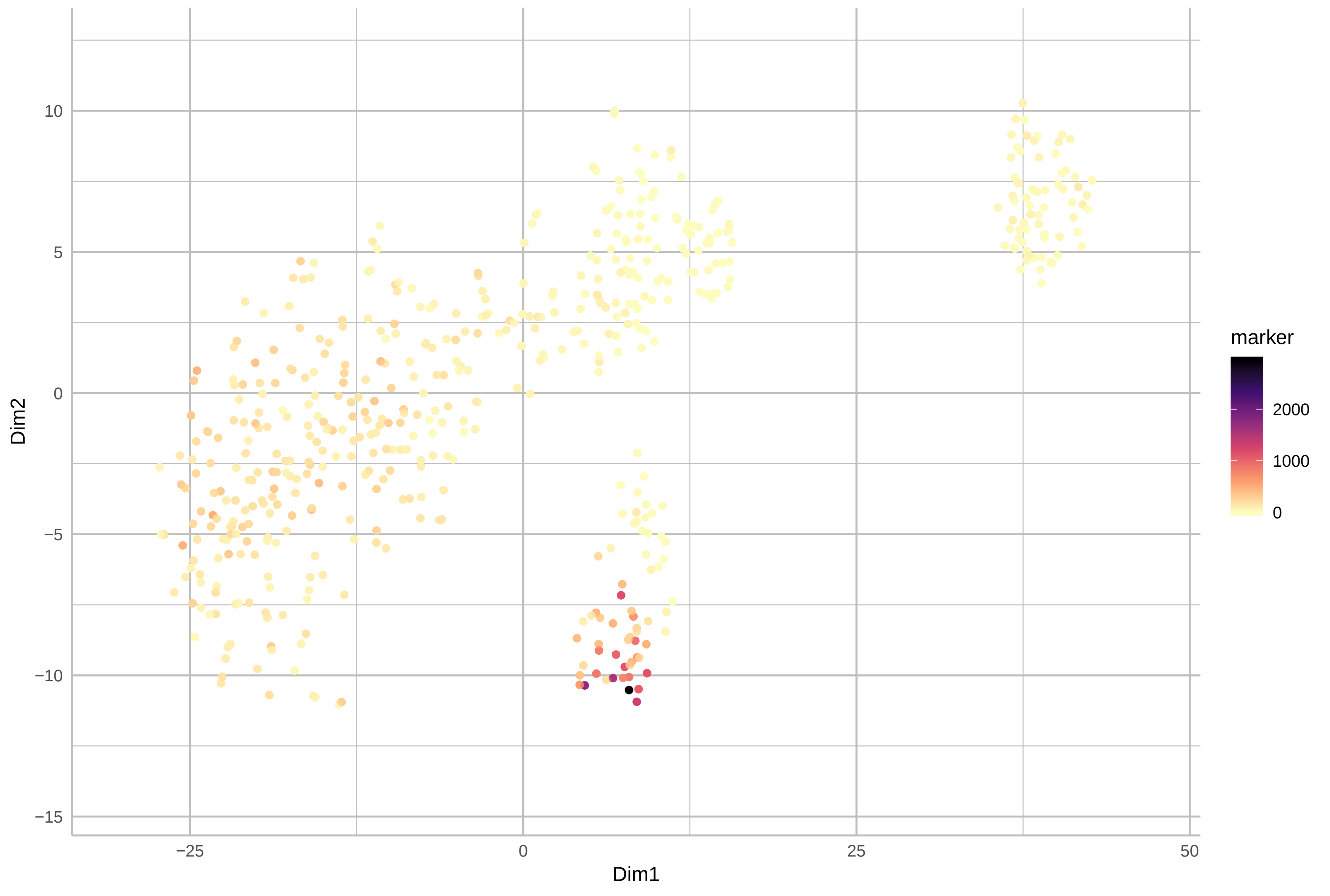

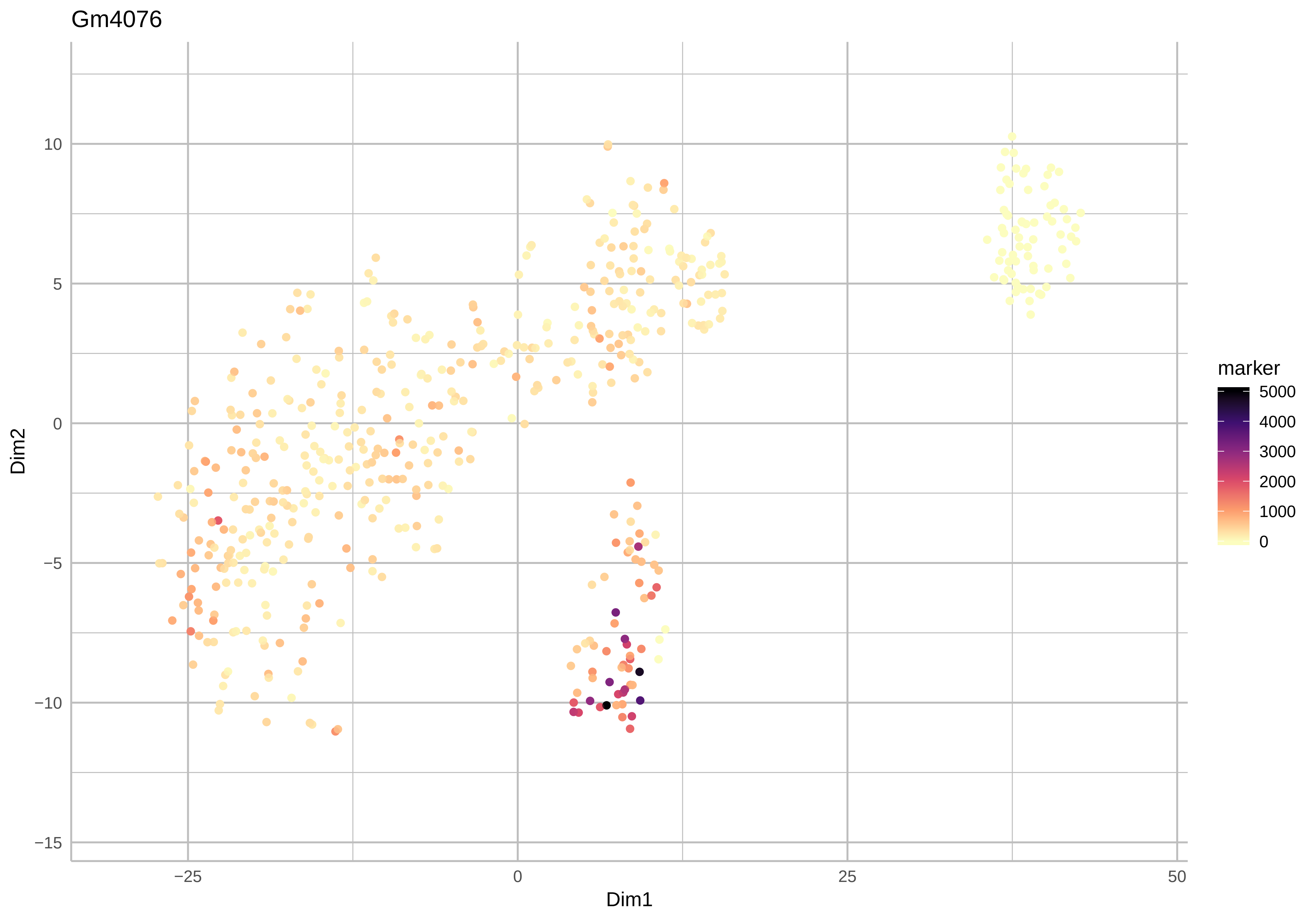
Gm4076
Dim2
10
5
0
−5
−10
−15
−25
0
25
50
Dim1
marker
5000
4000
3000
2000
1000
0

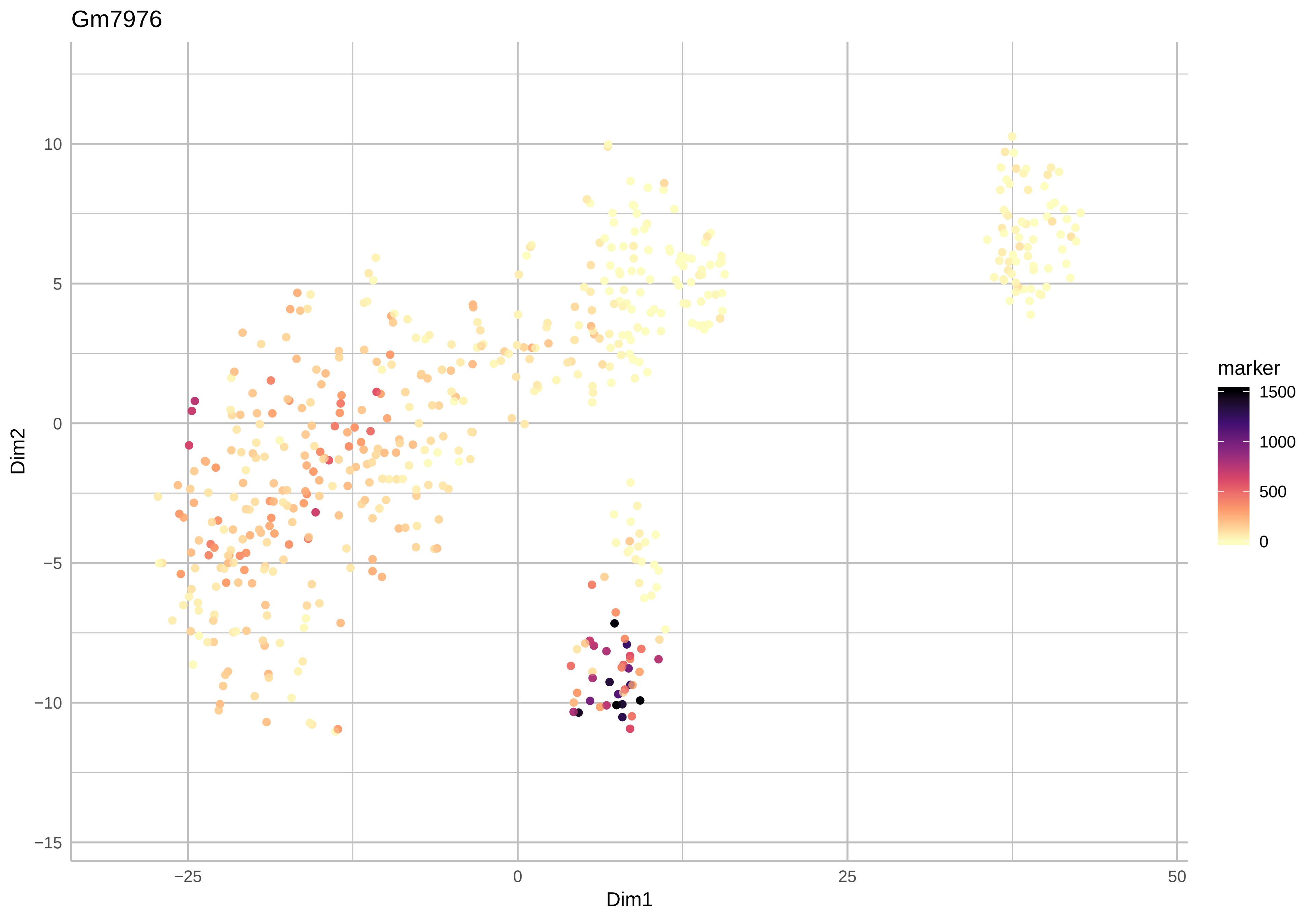
Gm7976
Dim2
Dim1
10
5
0
−5
−10
−15
−25
0
25
50
marker
1500
1000
500
0

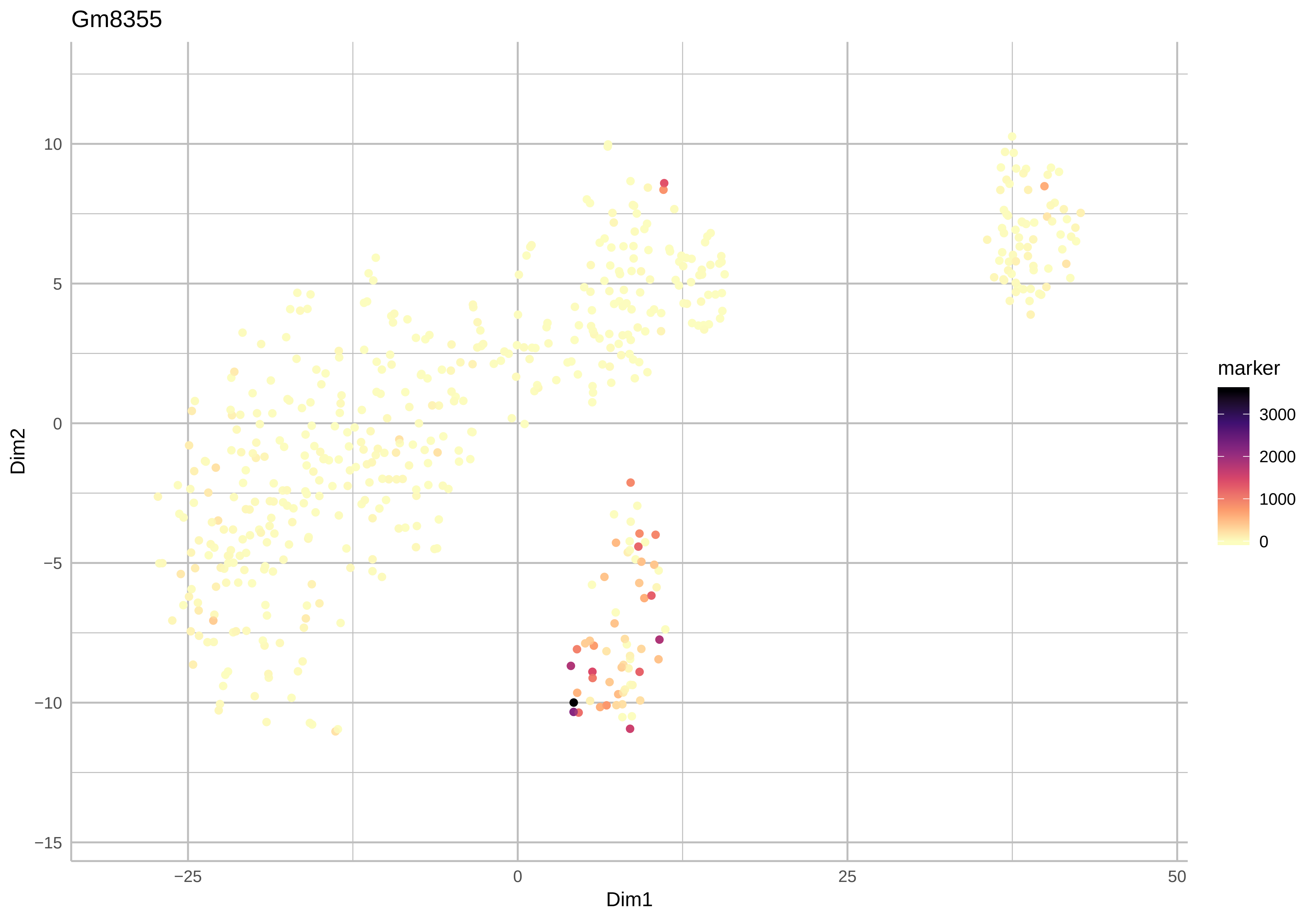

Gm8355
Dim2
Dim1
marker
3000
2000
1000
0
10
5
0
−5
−10
−15
−25
0
25
50

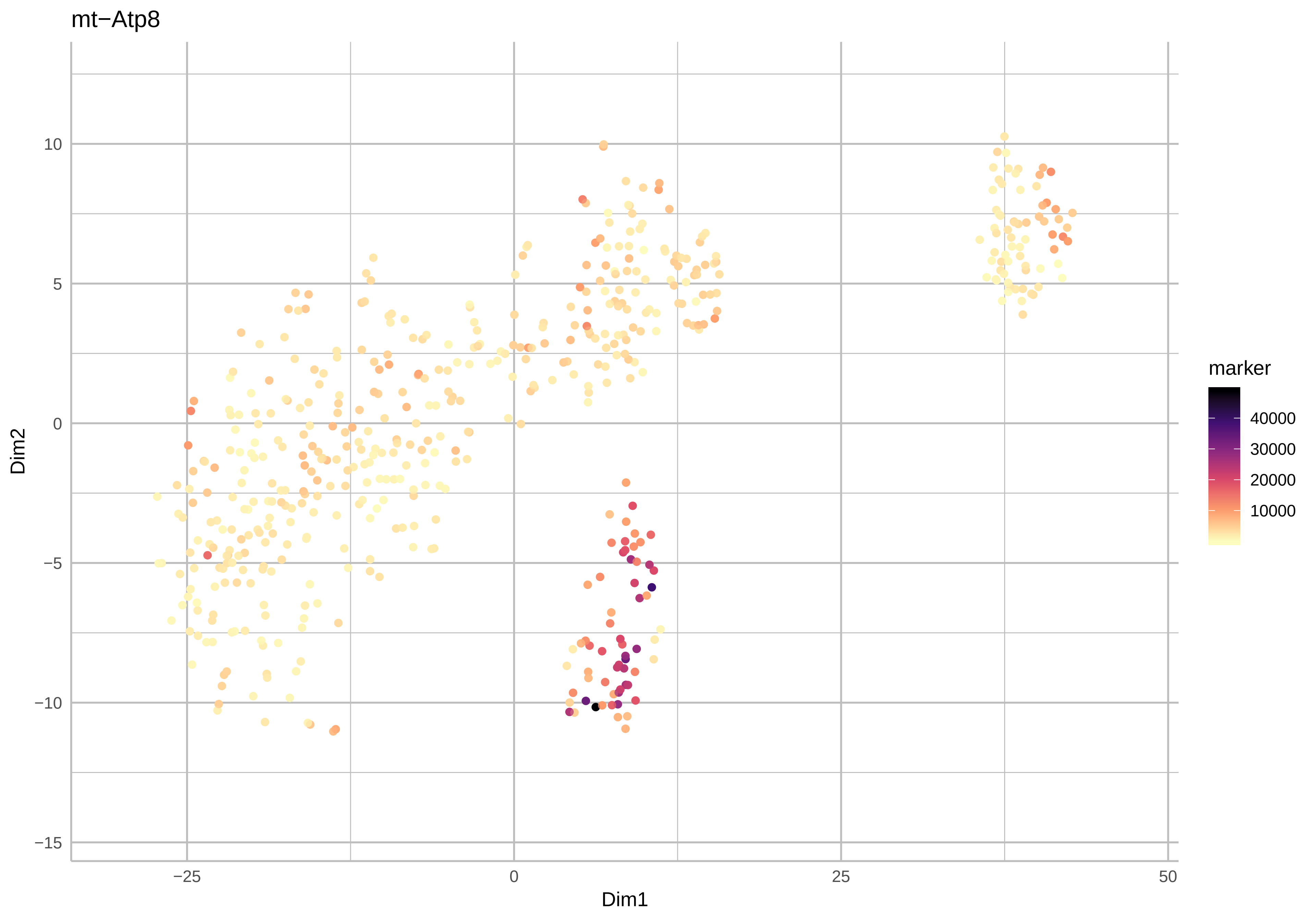

mt−Atp8
Dim2
Dim1
10
5
0
−5
−10
−15
−25
0
25
50
marker
40000
30000
20000
10000

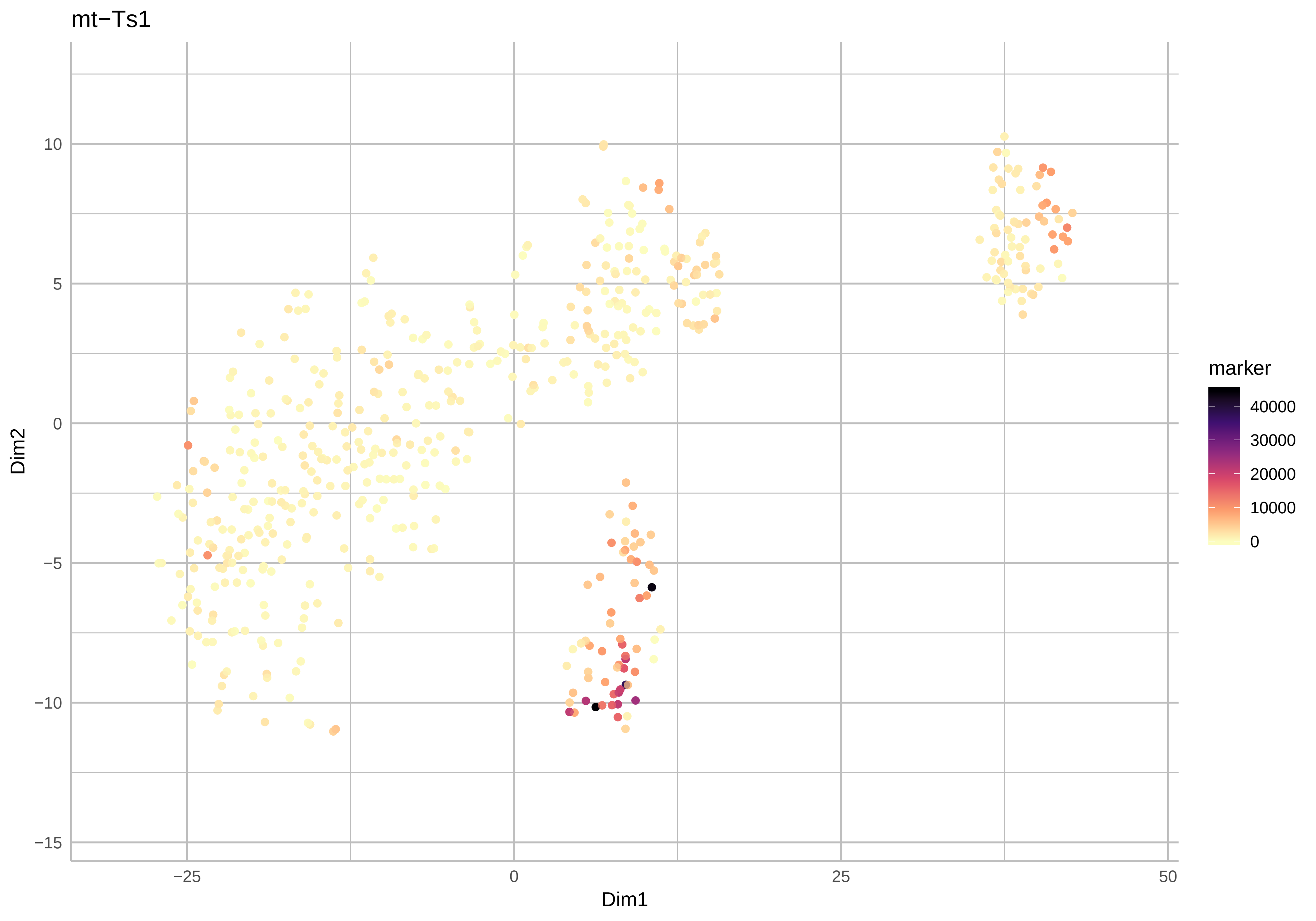

mt−Ts1
Dim2
Dim1
marker
40000
30000
20000
10000
0
10
5
0
−5
−10
−15
−25
0
25
50

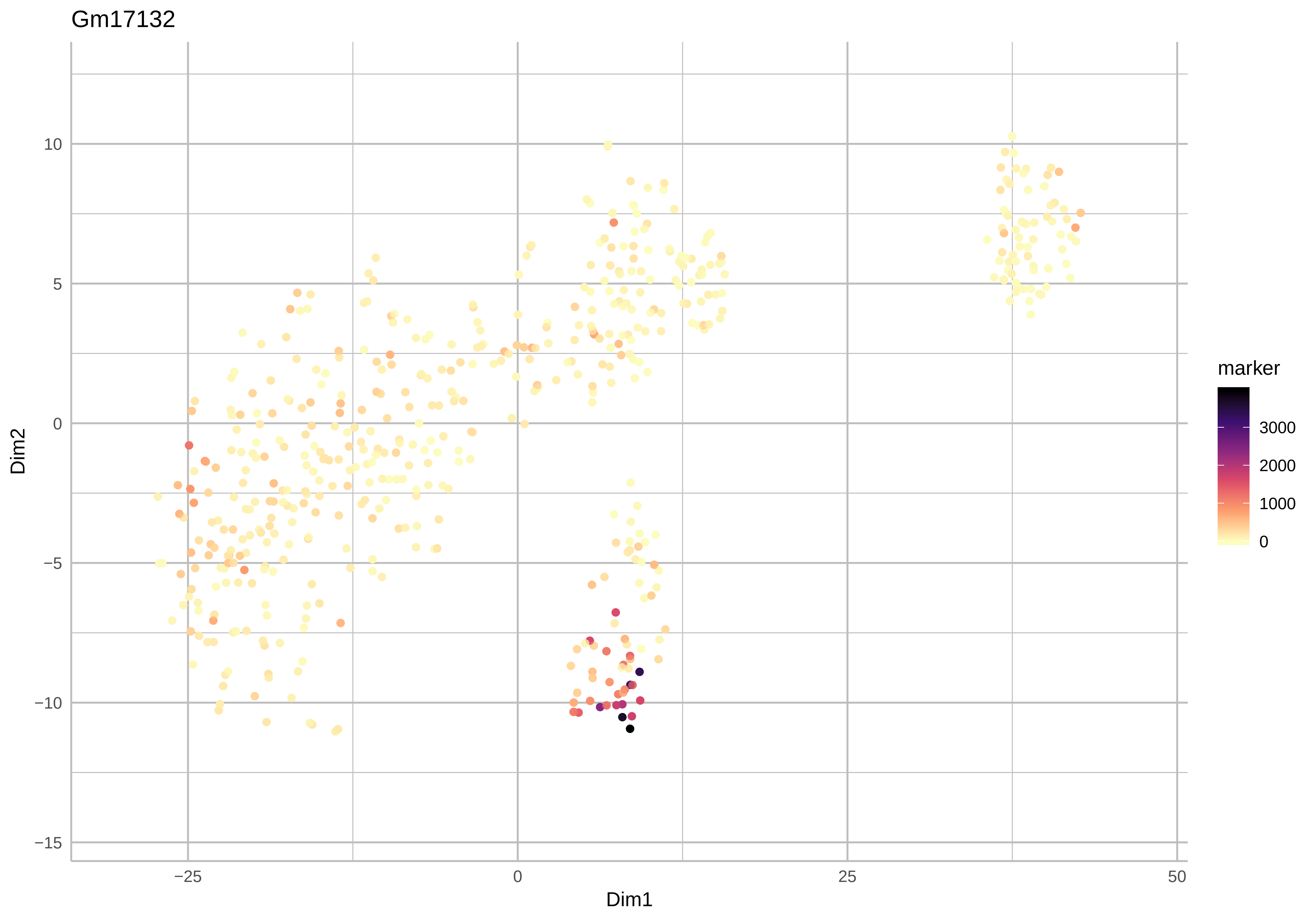
Gm17132
Dim2
Dim1
marker
3000
2000
1000
0
10
5
0
−5
−10
−15
−25
0
25
50

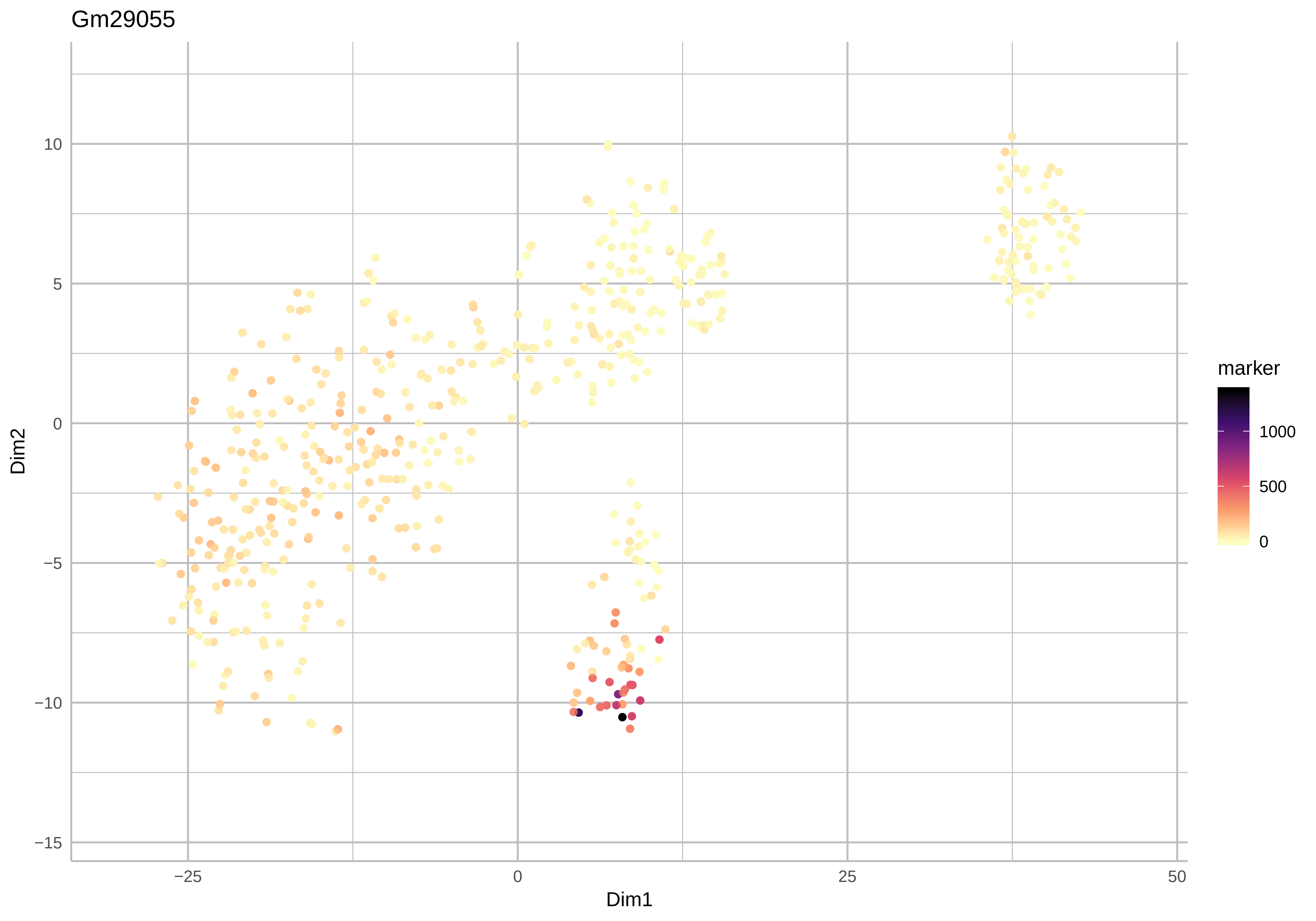

Gm29055
Dim2
10
5
0
−5
−10
−15
Dim1
−25
0
25
50
marker
1000
500
0

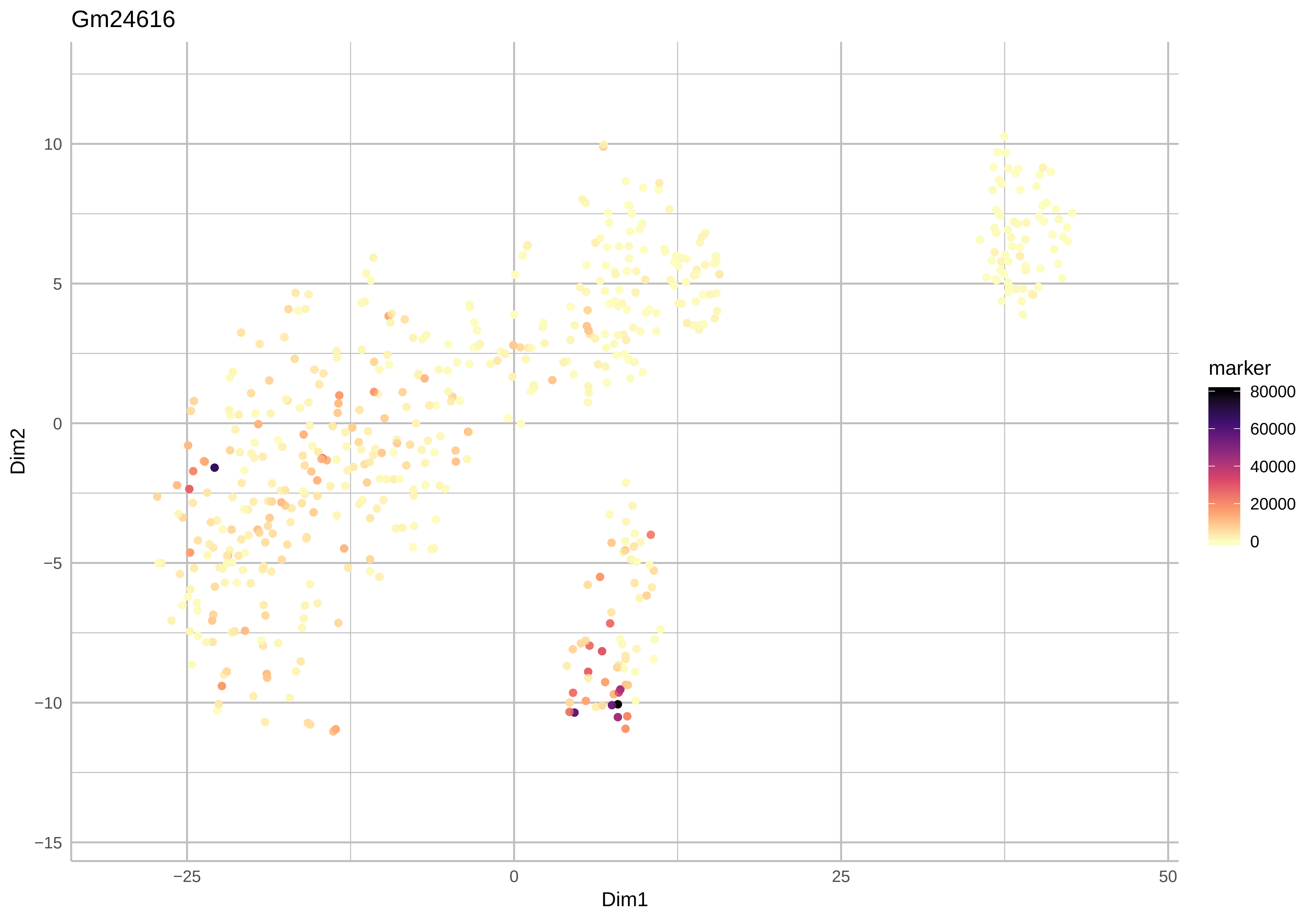

Gm24616
Dim2
Dim1
10
5
0
−5
−10
−15
−25
0
25
50
marker
80000
60000
40000
20000
0

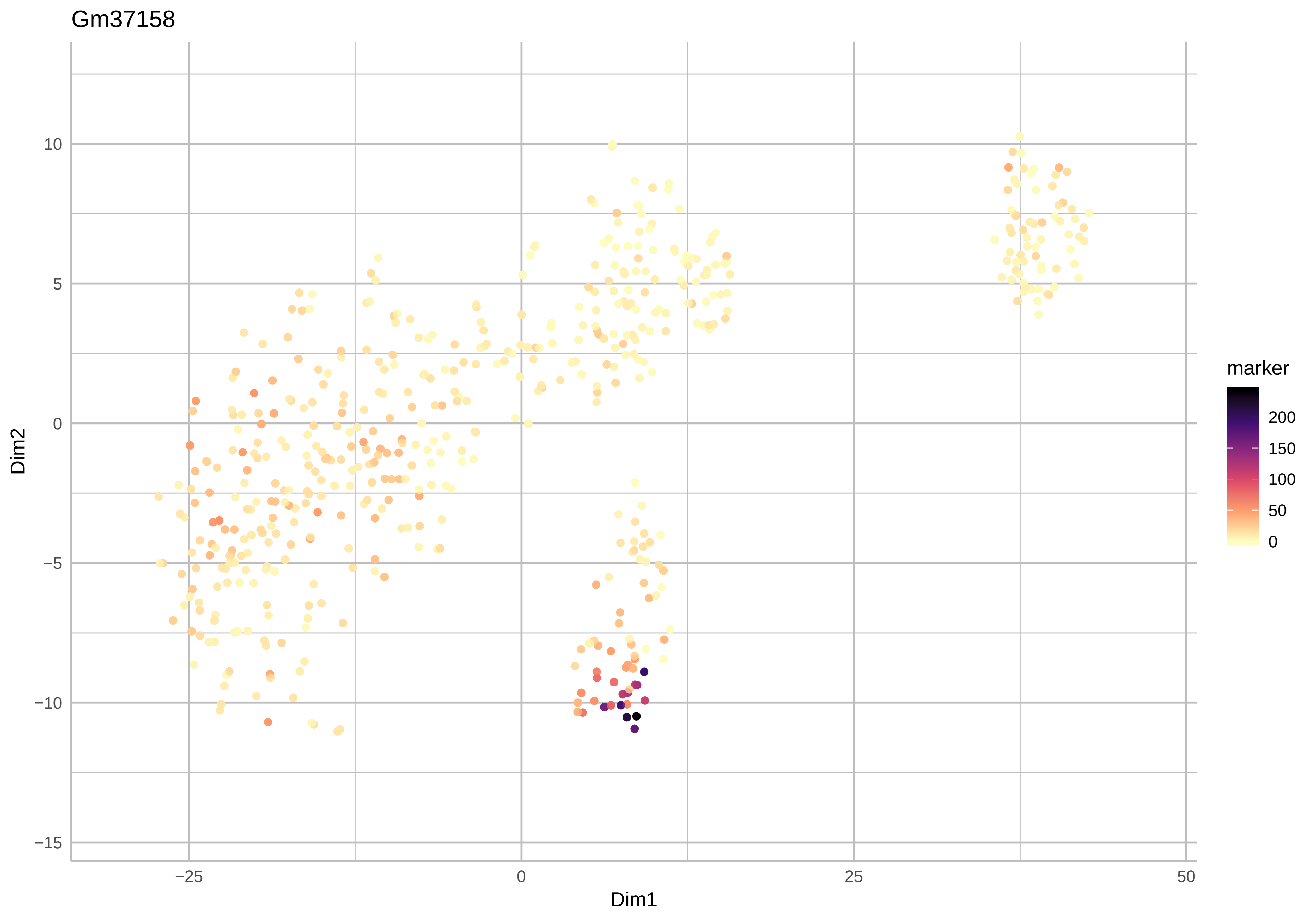
Gm37158
Dim2
Dim1
10
5
0
−5
−10
−15
−25
0
25
50
marker
200
150
100
50
0

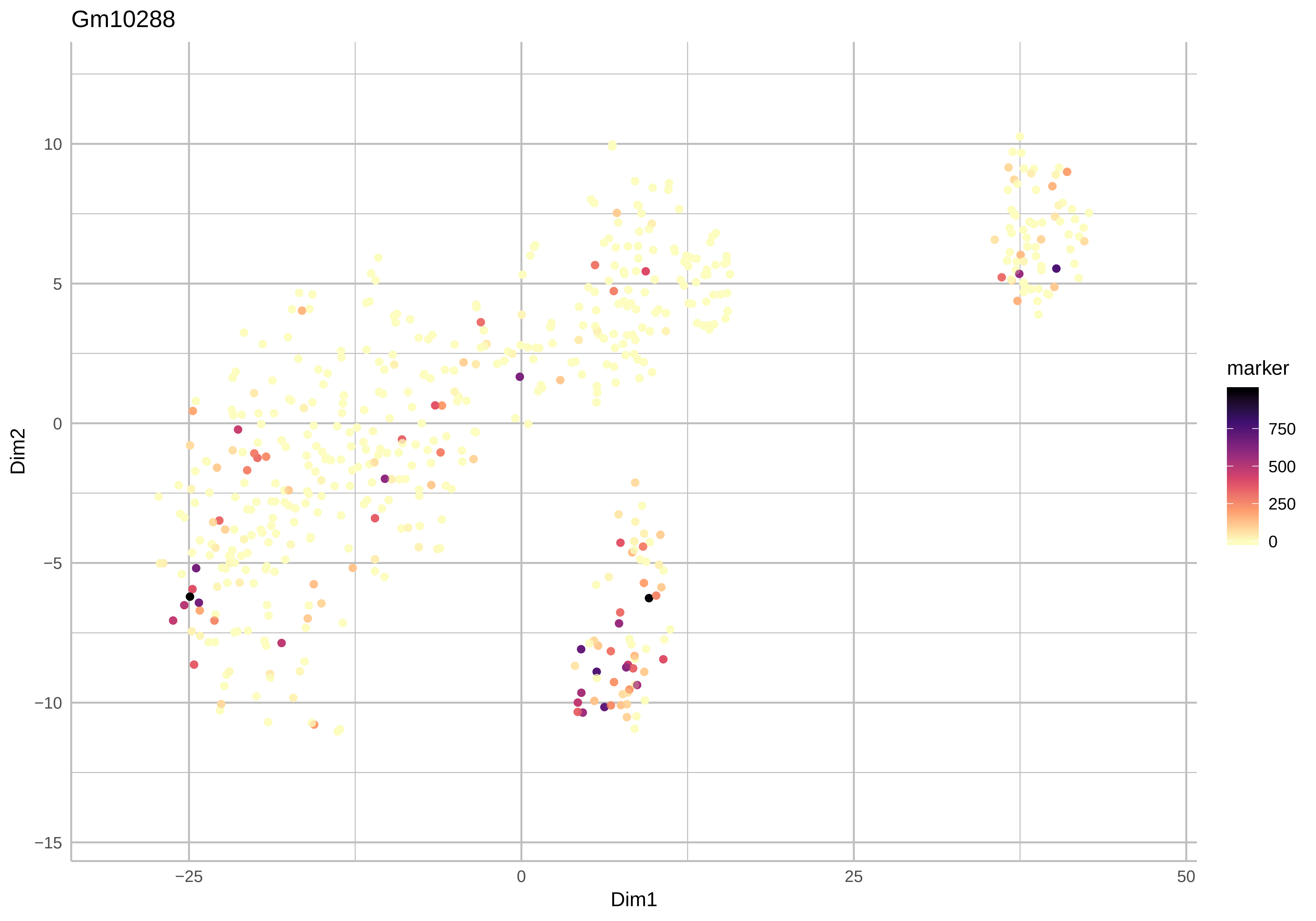

Gm10288
Dim2
Dim1
10
5
0
−5
−10
−15
−25
0
25
50
marker
750
500
250
0

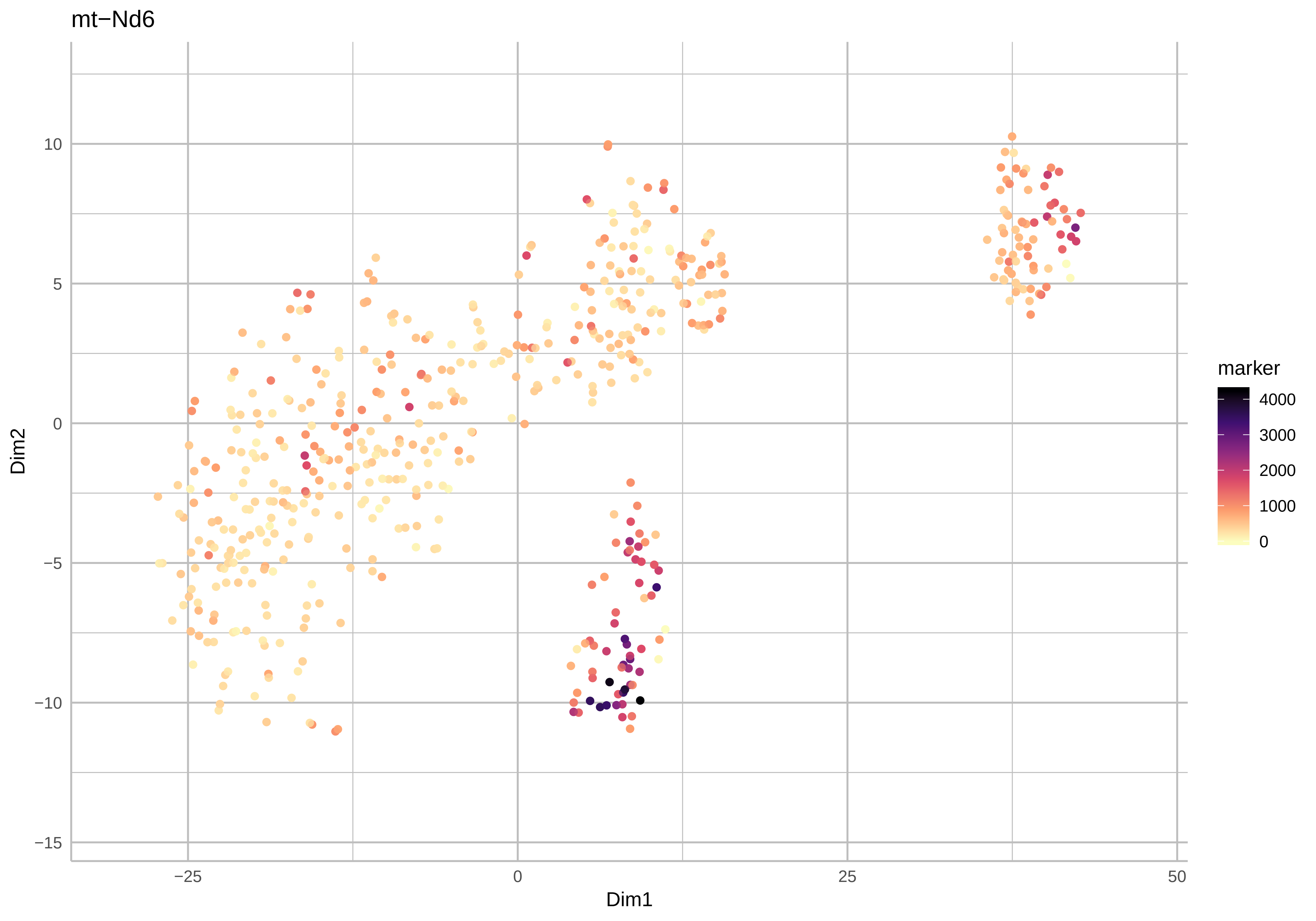

mt−Nd6
Dim2
Dim1
10
5
0
−5
−10
−15
−25
0
25
50
marker
4000
3000
2000
1000
0

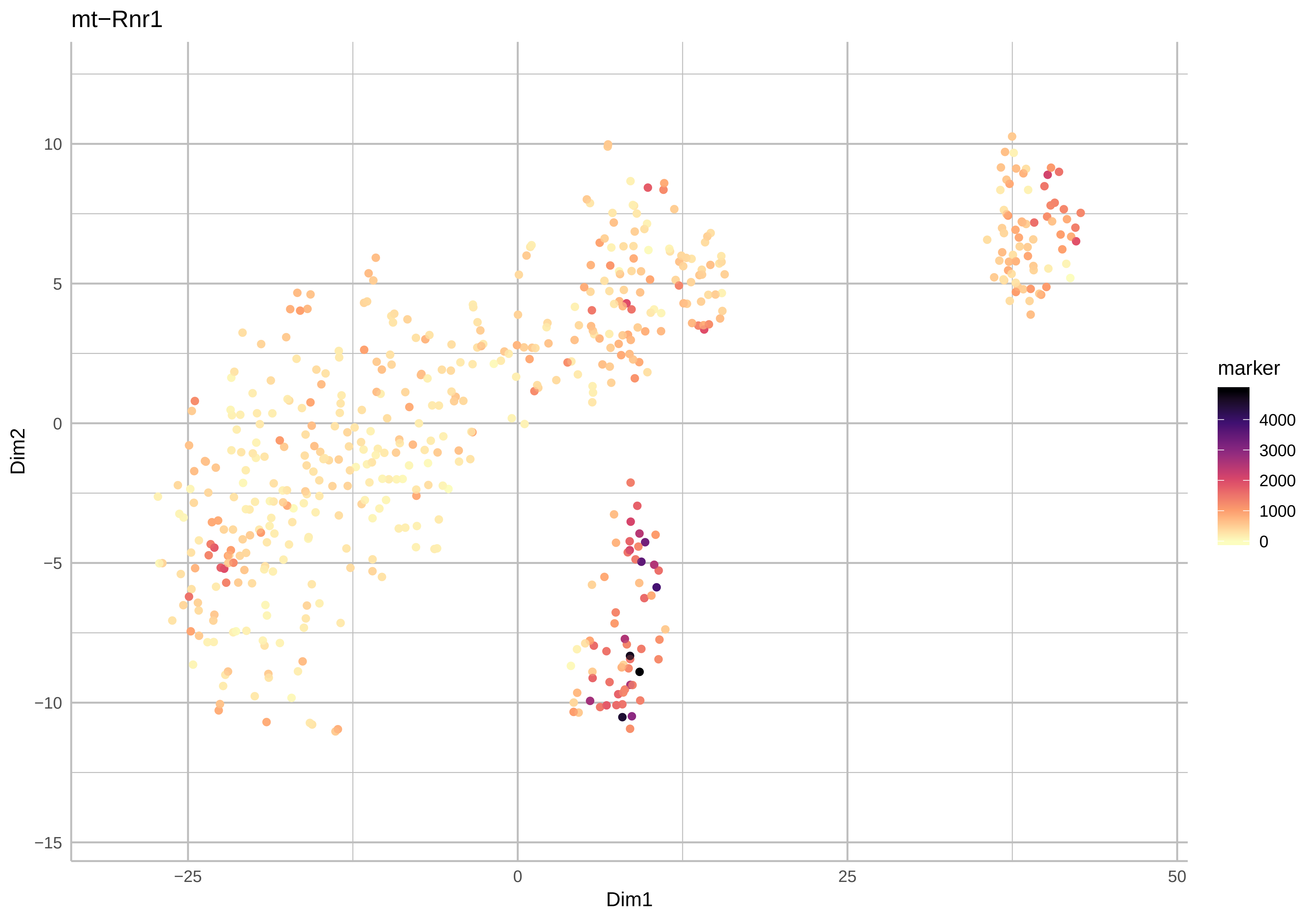
mt−Rnr1
10
5
0
−5
−10
−15
Dim2
−25
0
25
50
Dim1
marker
4000
3000
2000
1000
0

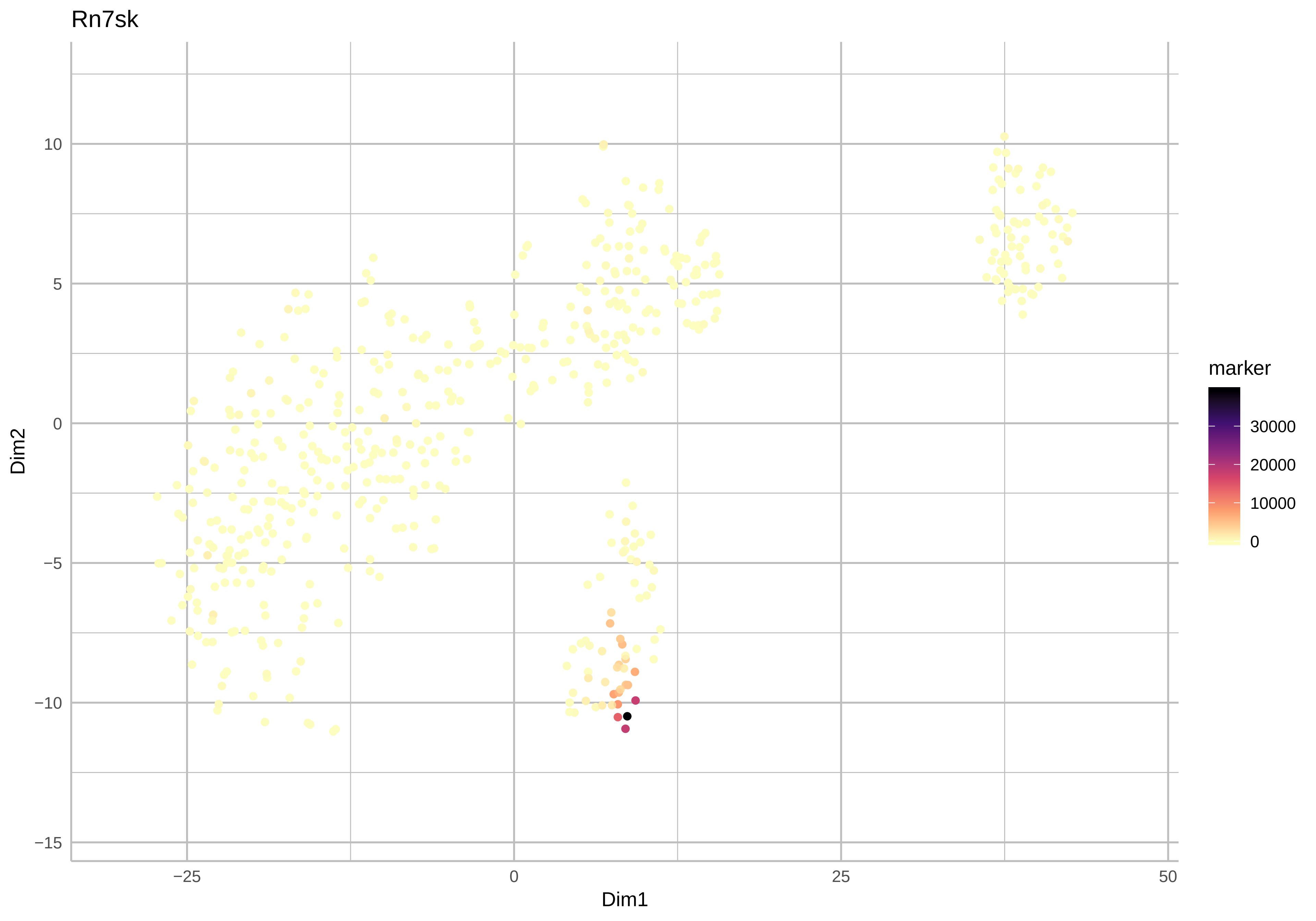
Rn7sk
Dim2
Dim1
10
5
0
−5
−10
−15
−25
0
25
50
marker
30000
20000
10000
0

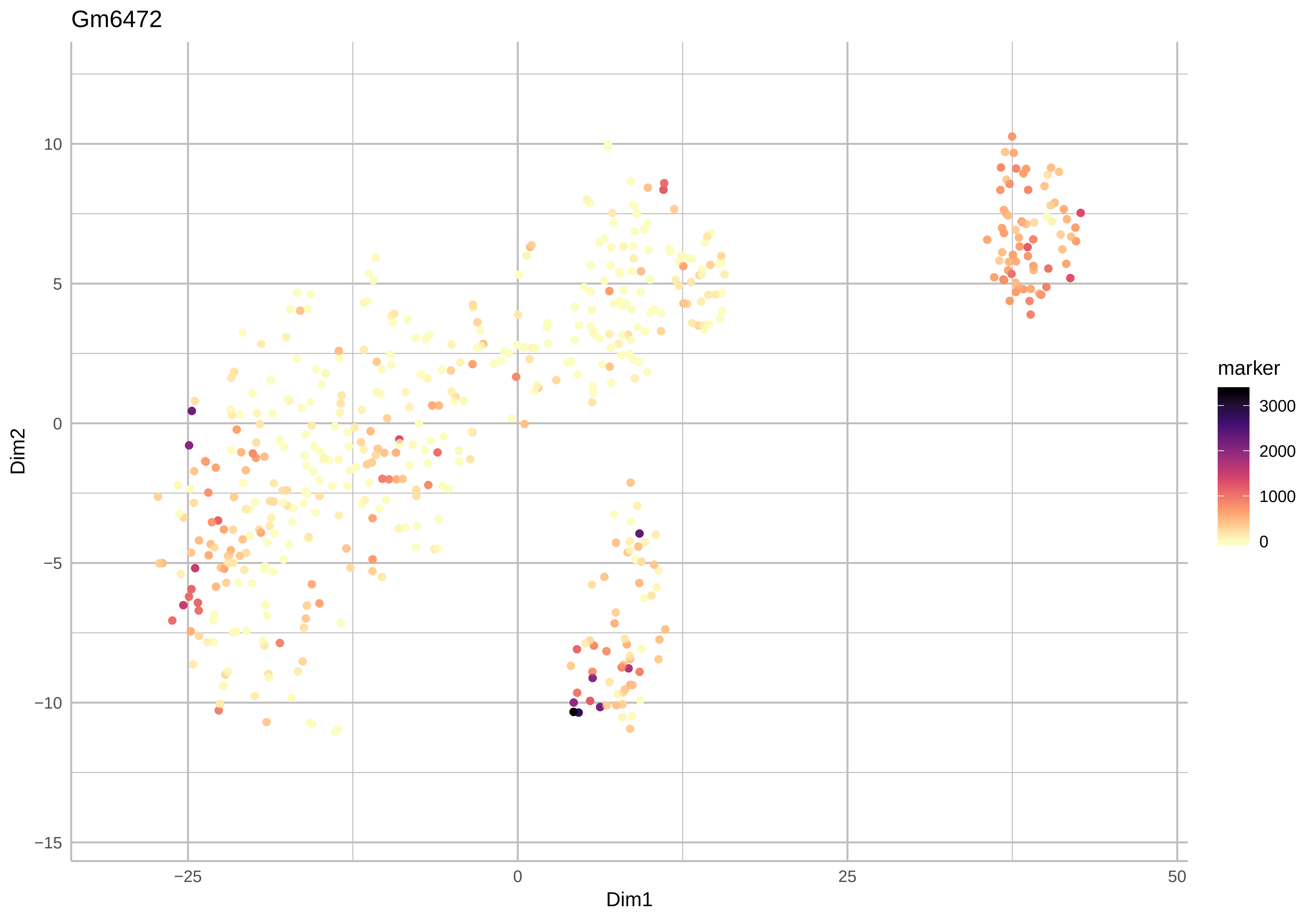

Gm6472
Dim2
Dim1
marker
3000
2000
1000
0
10
5
0
−5
−10
−15
−25
0
25
50

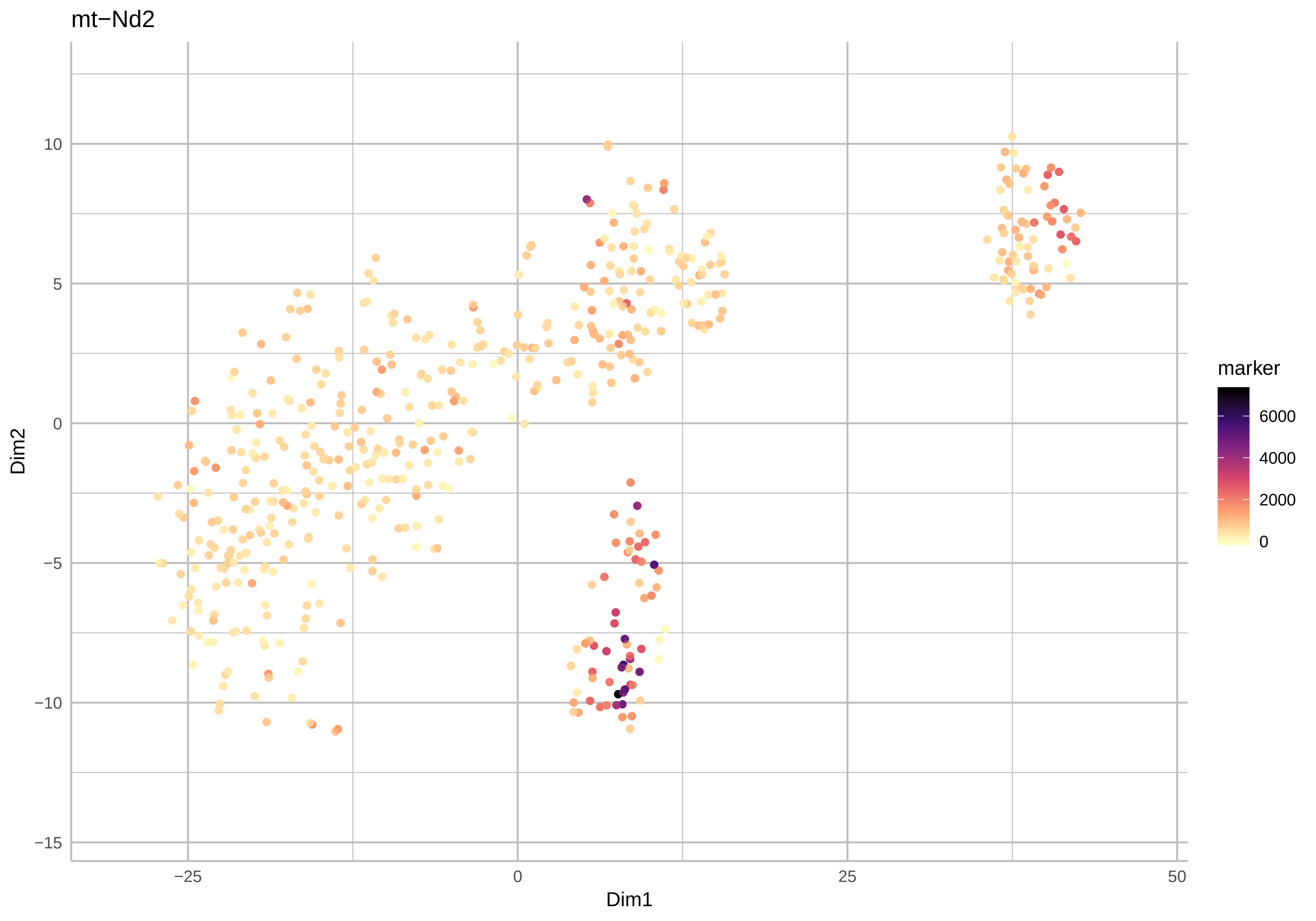
mt−Nd2
Dim2
Dim1
10
5
0
−5
−10
−15
−25
0
25
50
marker
6000
4000
2000
0

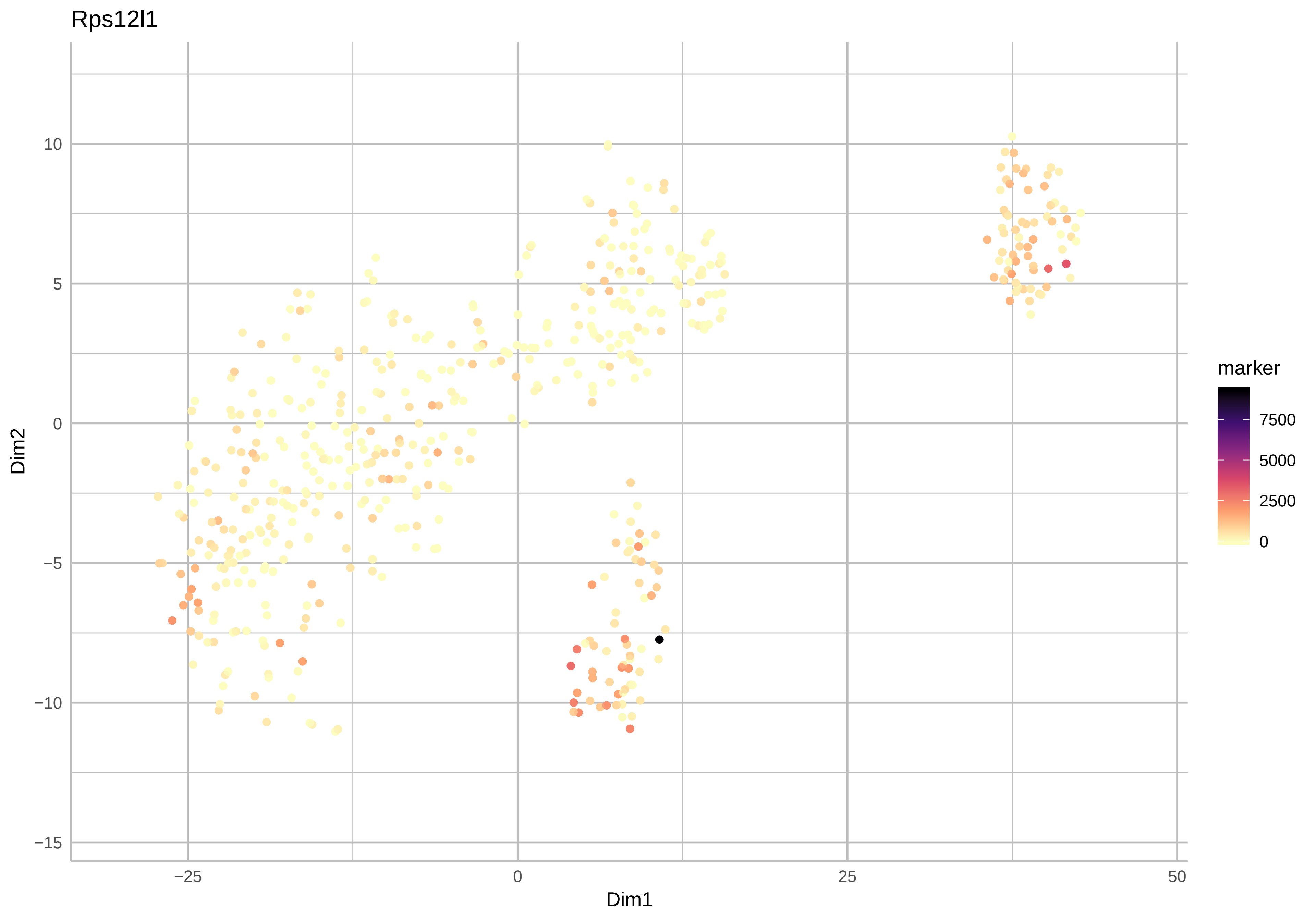
Rps12l1
Dim2
Dim1
10
5
0
−5
−10
−15
−25
0
25
50
marker
7500
5000
2500
0

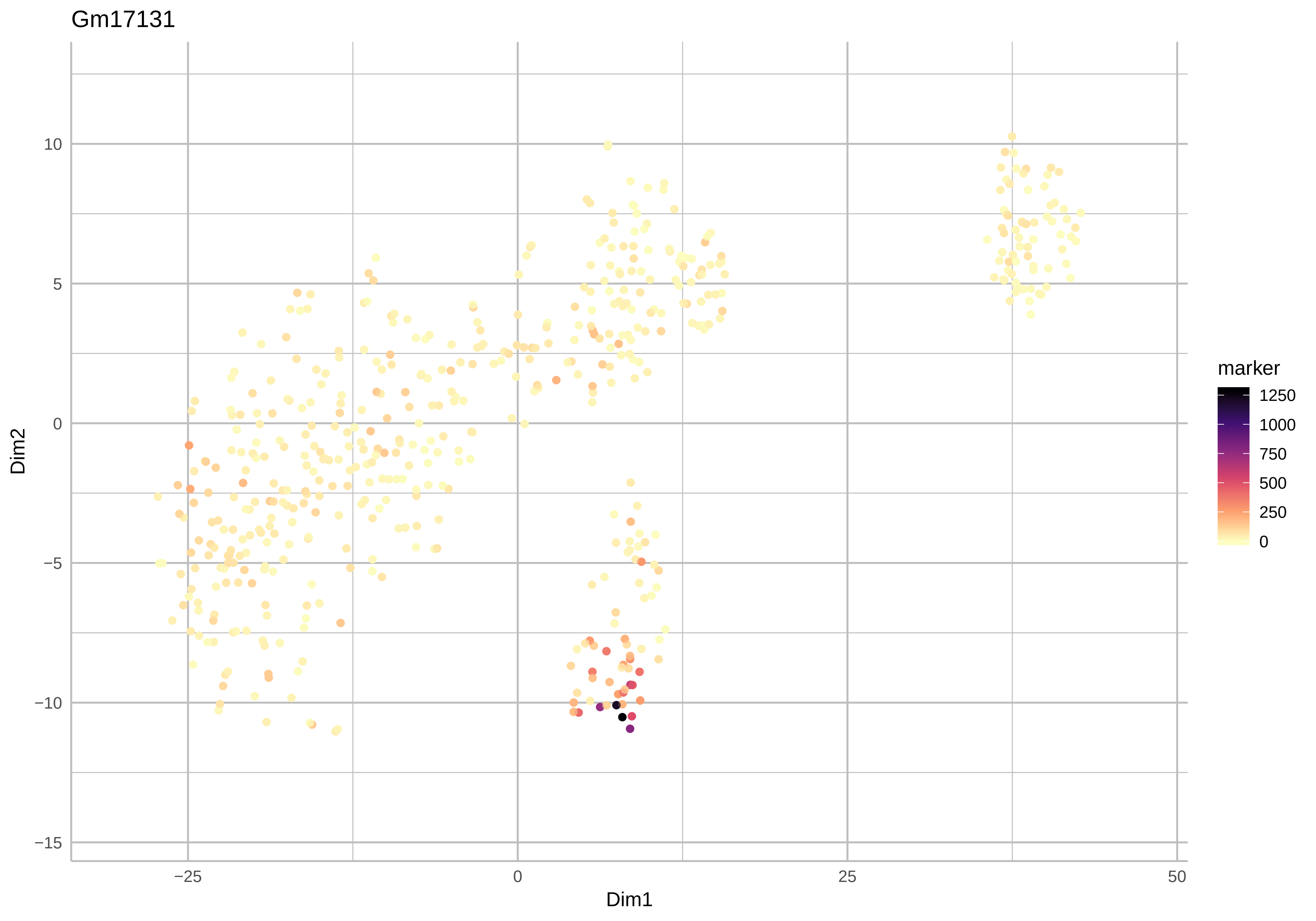

Gm17131
Dim2
Dim1
10
5
0
−5
−10
−15
−25
0
25
50
marker
1250
1000
750
500
250
0

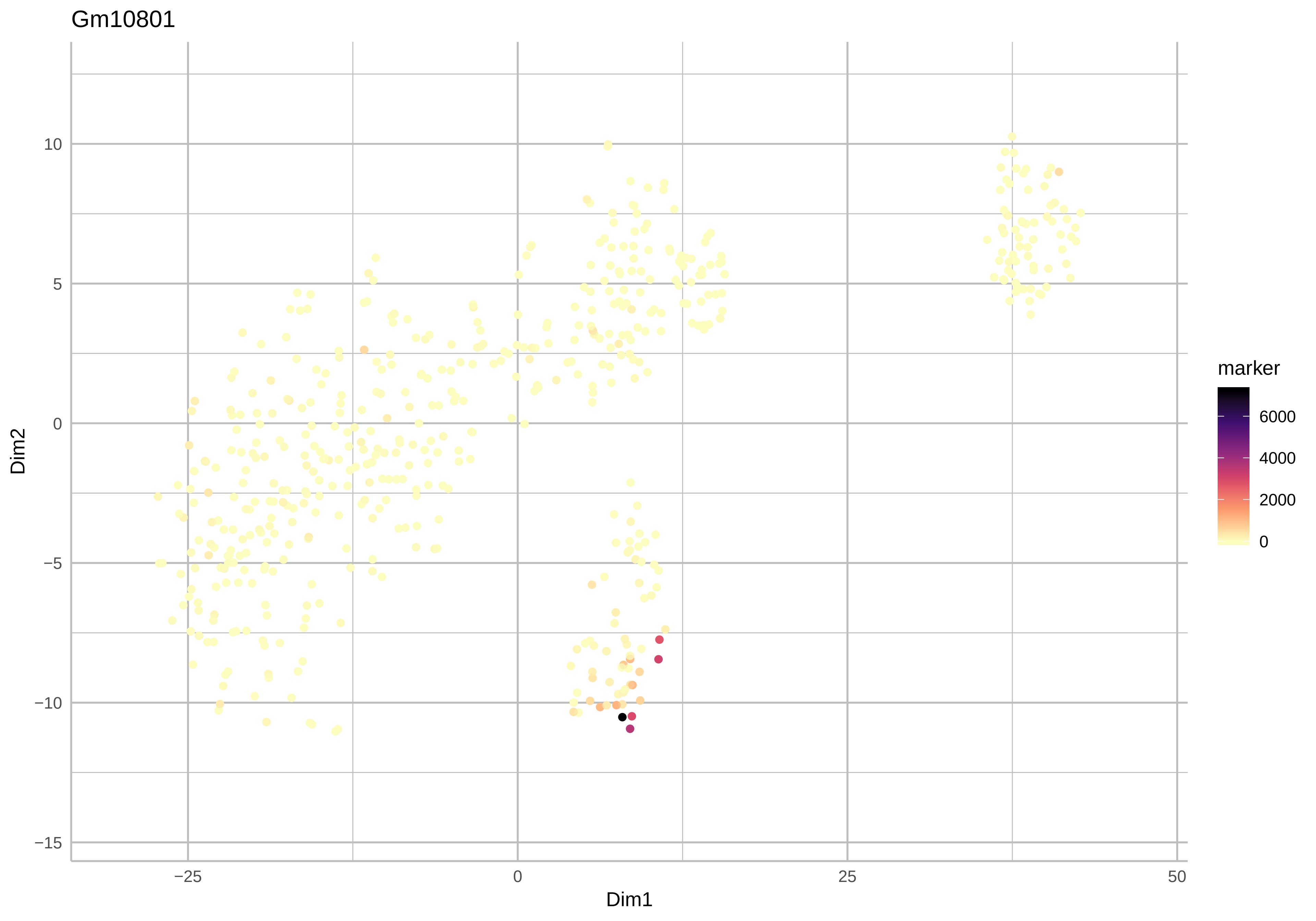

Gm10801
Dim2
Dim1
10
5
0
−5
−10
−15
−25
0
25
50
marker
6000
4000
2000
0

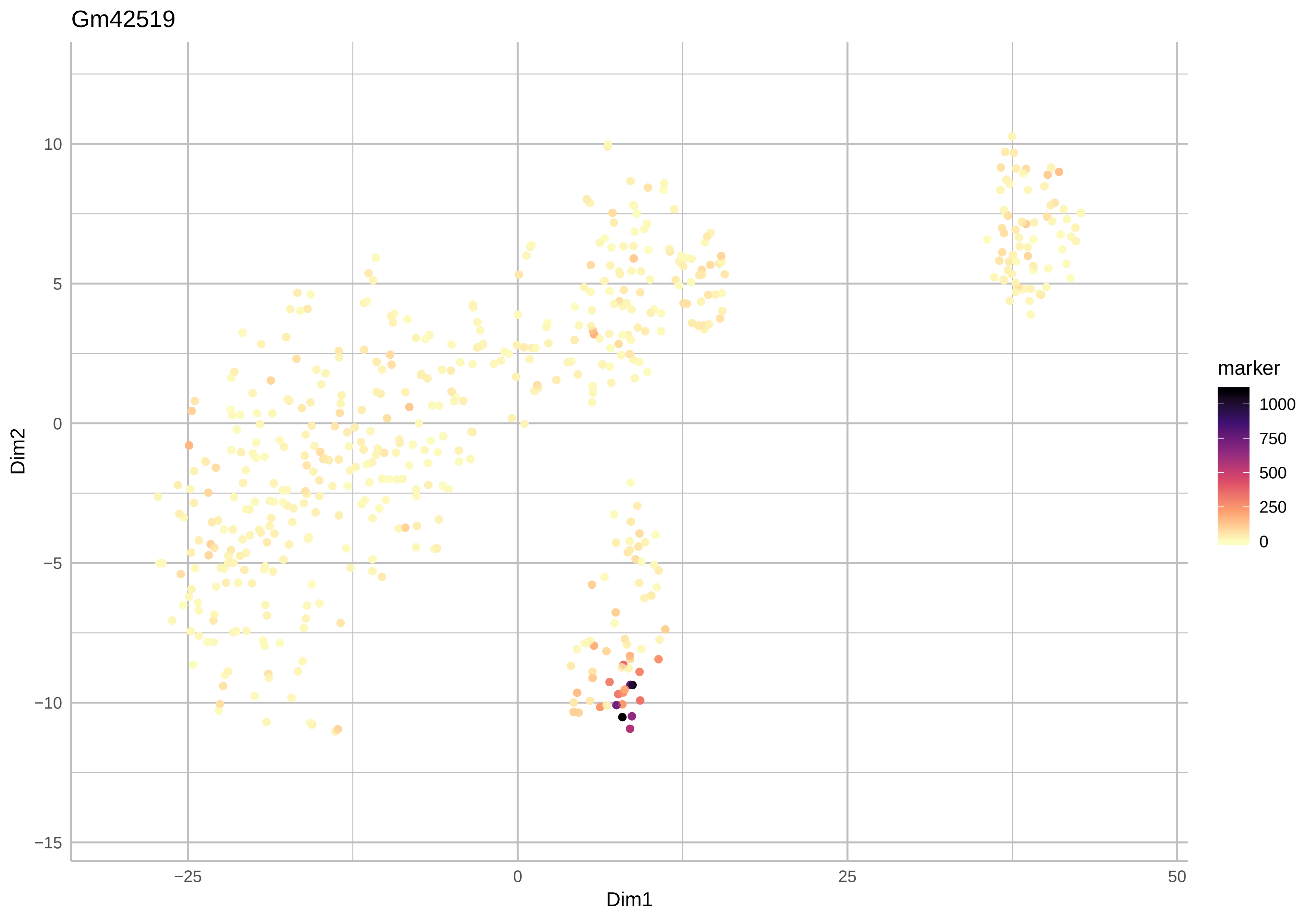

Gm42519
Dim2
Dim1
10
5
0
−5
−10
−15
−25
0
25
50
marker
1000
750
500
250
0

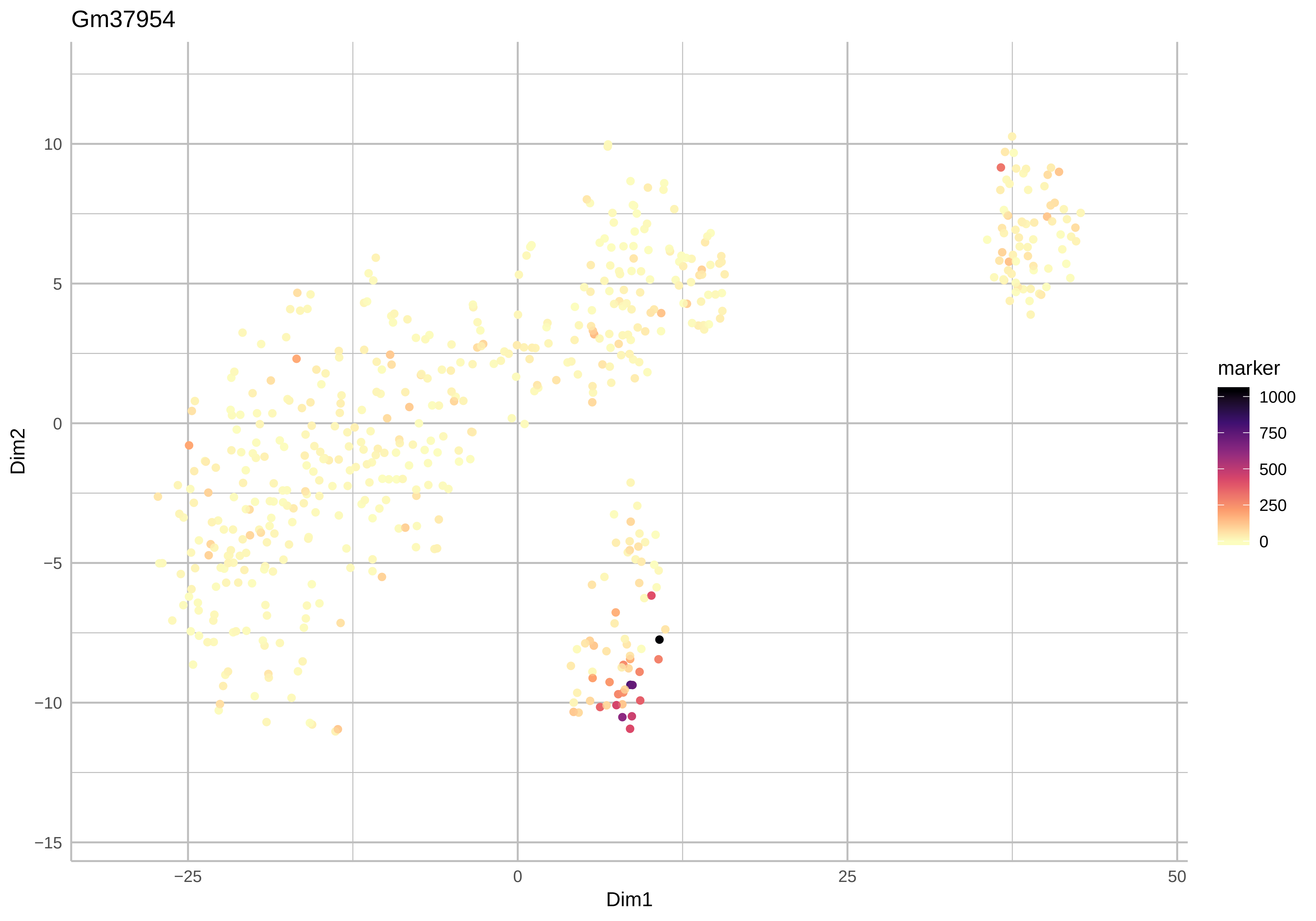

Gm37954
Dim2
Dim1
10
5
0
−5
−10
−15
−25
0
25
50
marker
1000
750
500
250
0

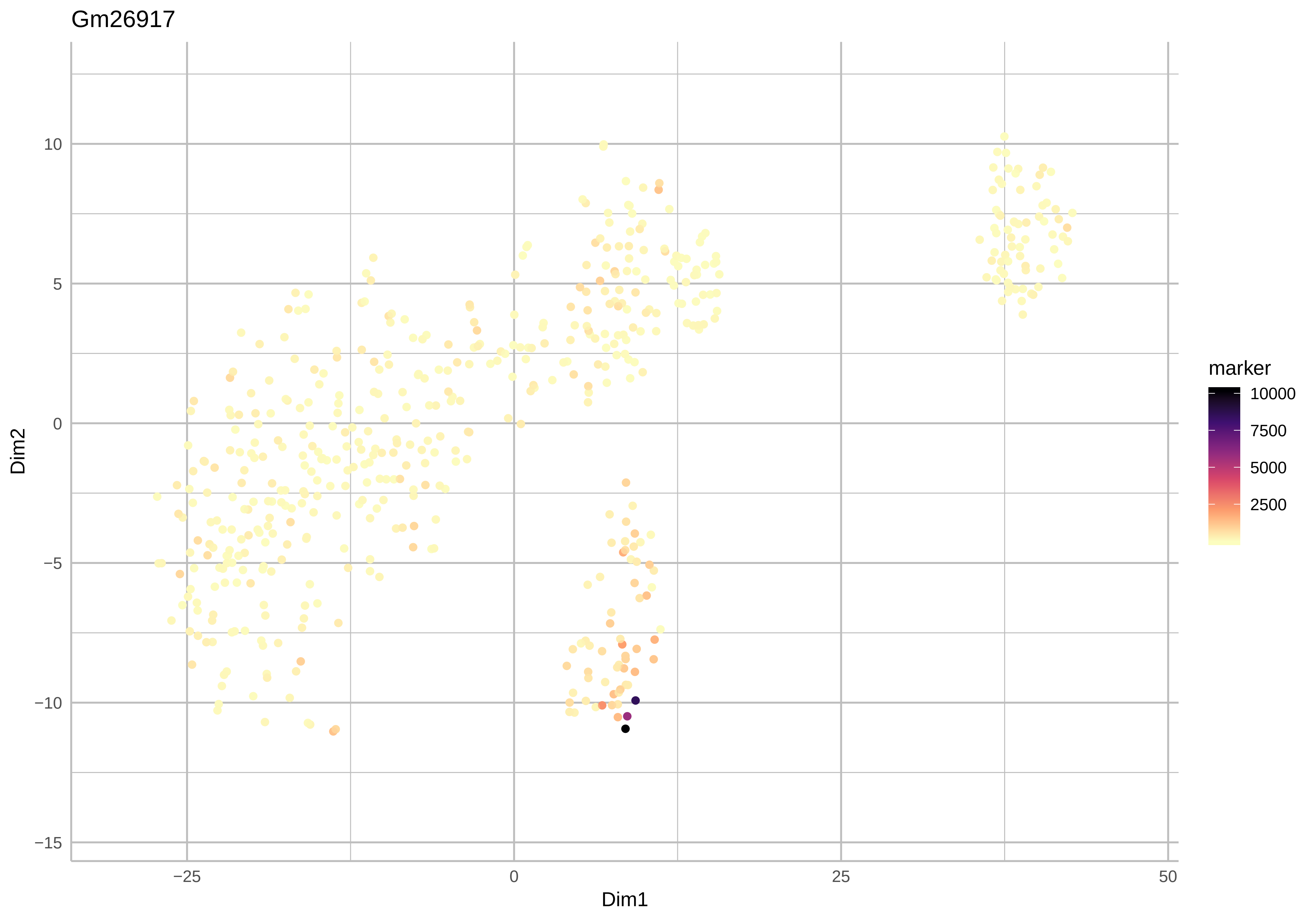

Gm26917
Dim2
Dim1
marker
10000
7500
5000
2500
10
5
0
−5
−10
−15
−25
0
25
50

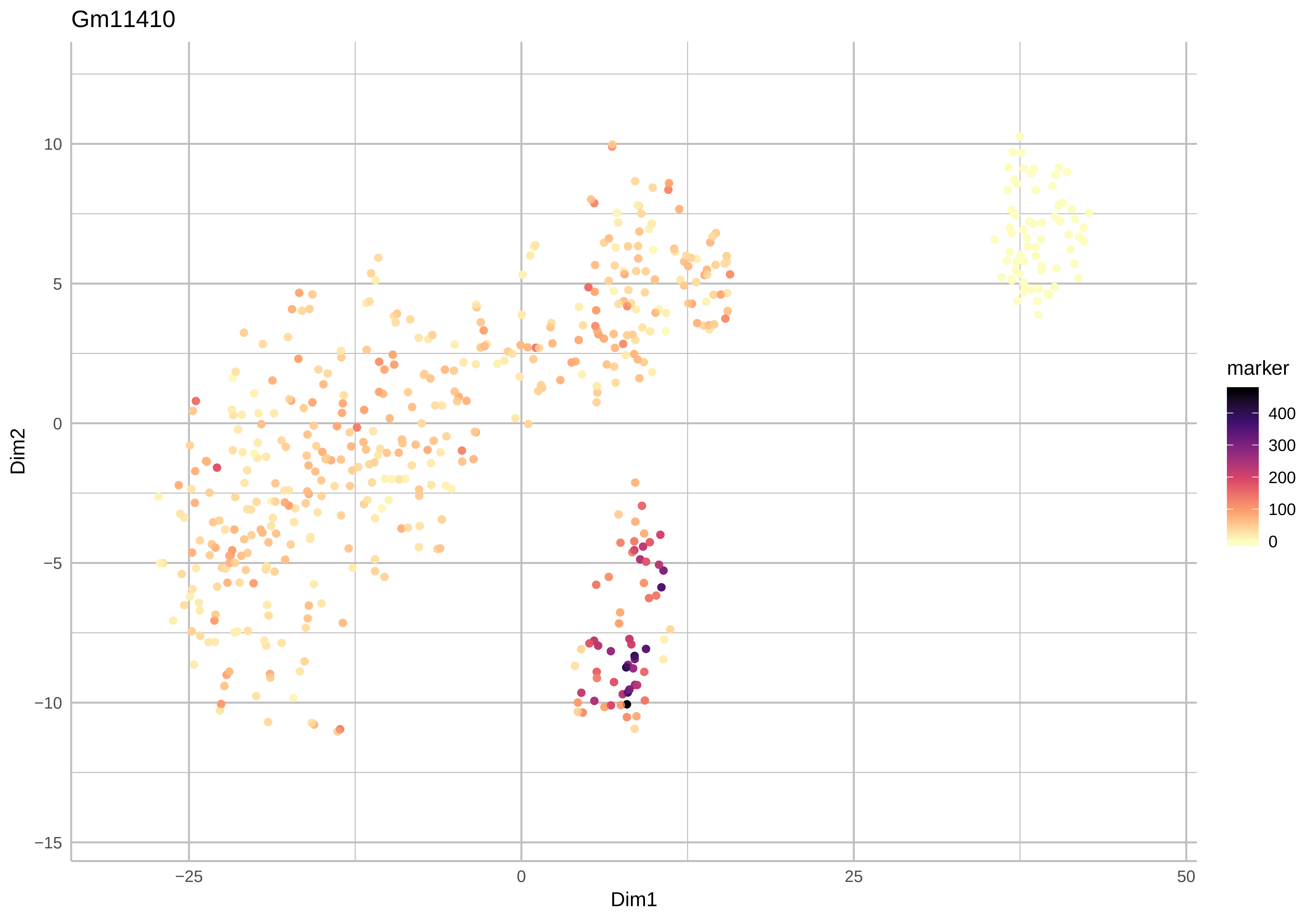
Gm11410
Dim2
Dim1
10
5
0
−5
−10
−15
−25
0
25
50
marker
400
300
200
100
0

Gm21962

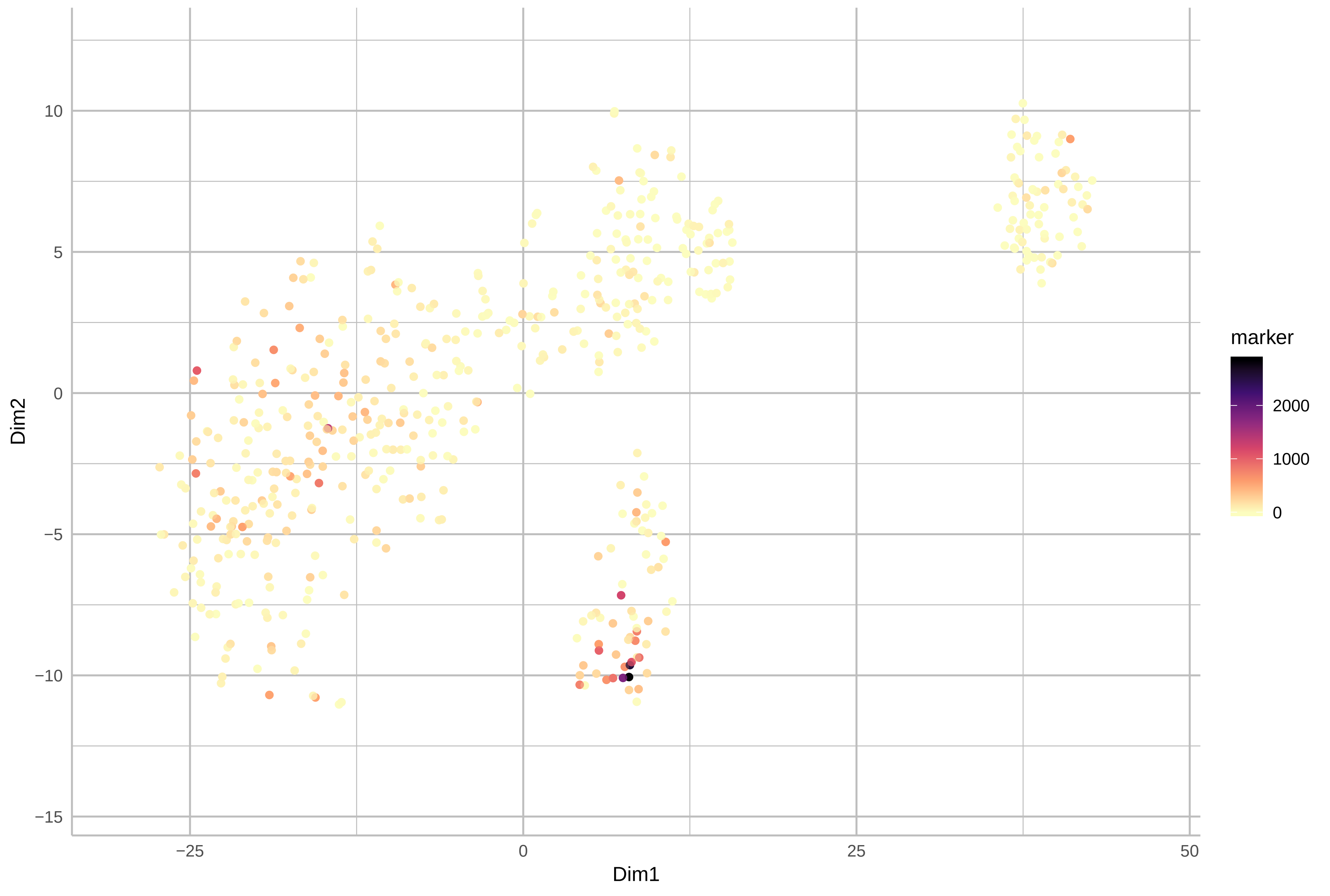

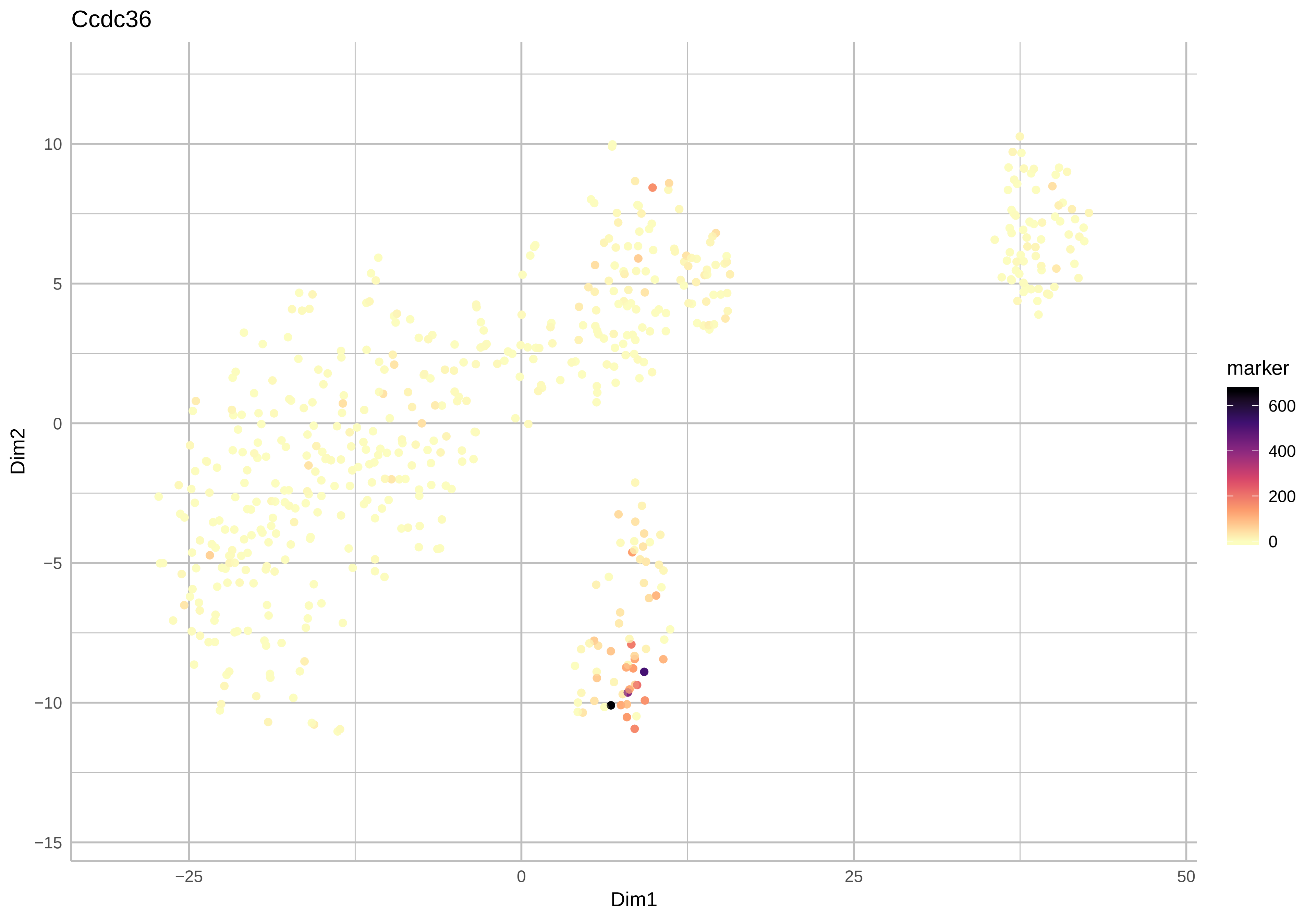
Ccdc36
Dim2
Dim1
10
5
0
−5
−10
−15
−25
0
25
50
marker
600
400
200
0

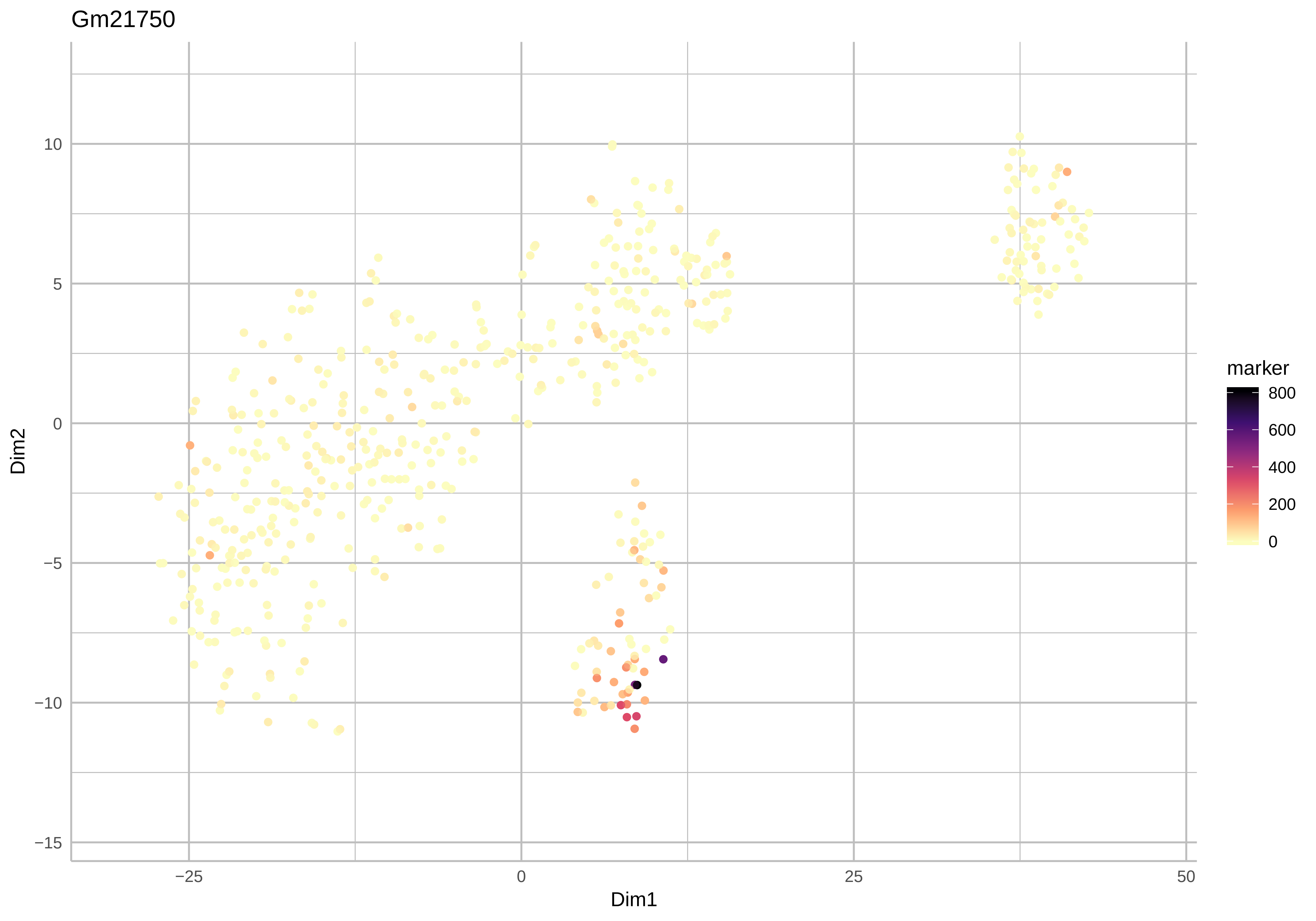

Gm21750
Dim2
Dim1
marker
800
600
400
200
0
10
5
0
−5
−10
−15
−25
0
25
50

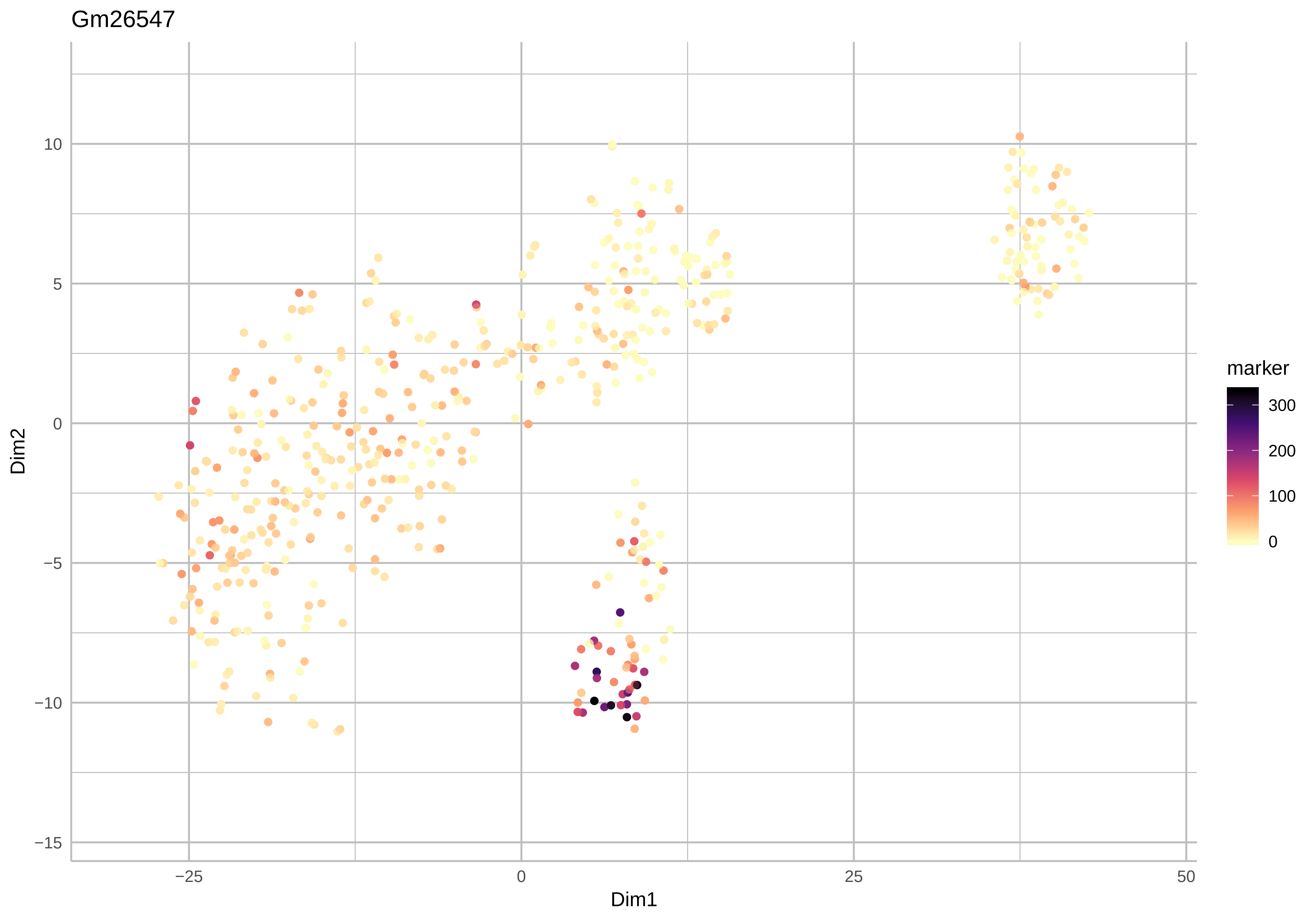

Gm26547
Dim2
Dim1
10
5
0
−5
−10
−15
−25
0
25
50
marker
300
200
100
0

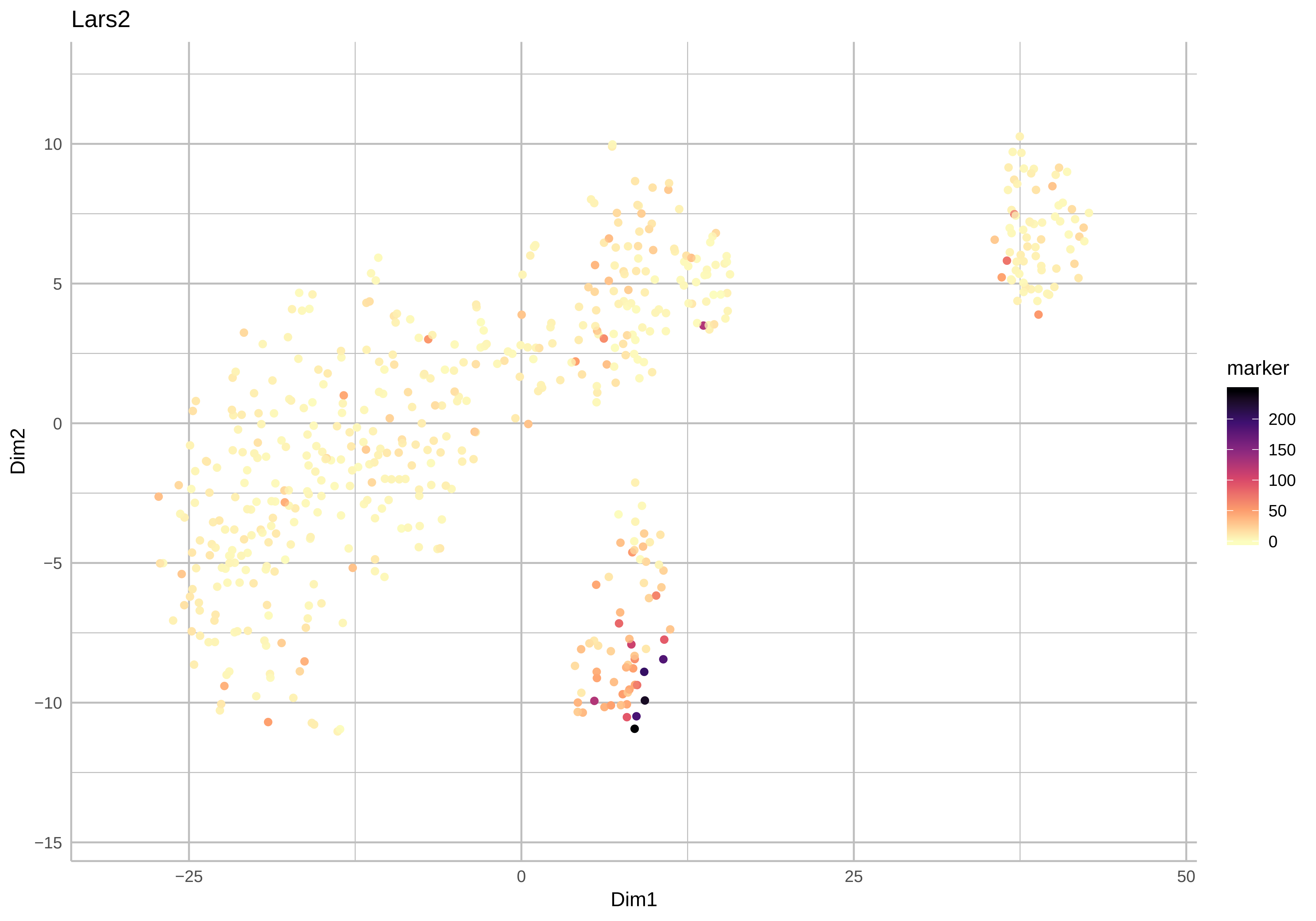

Lars2
Dim2
Dim1
10
5
0
−5
−10
−15
−25
0
25
50
marker
200
150
100
50
0

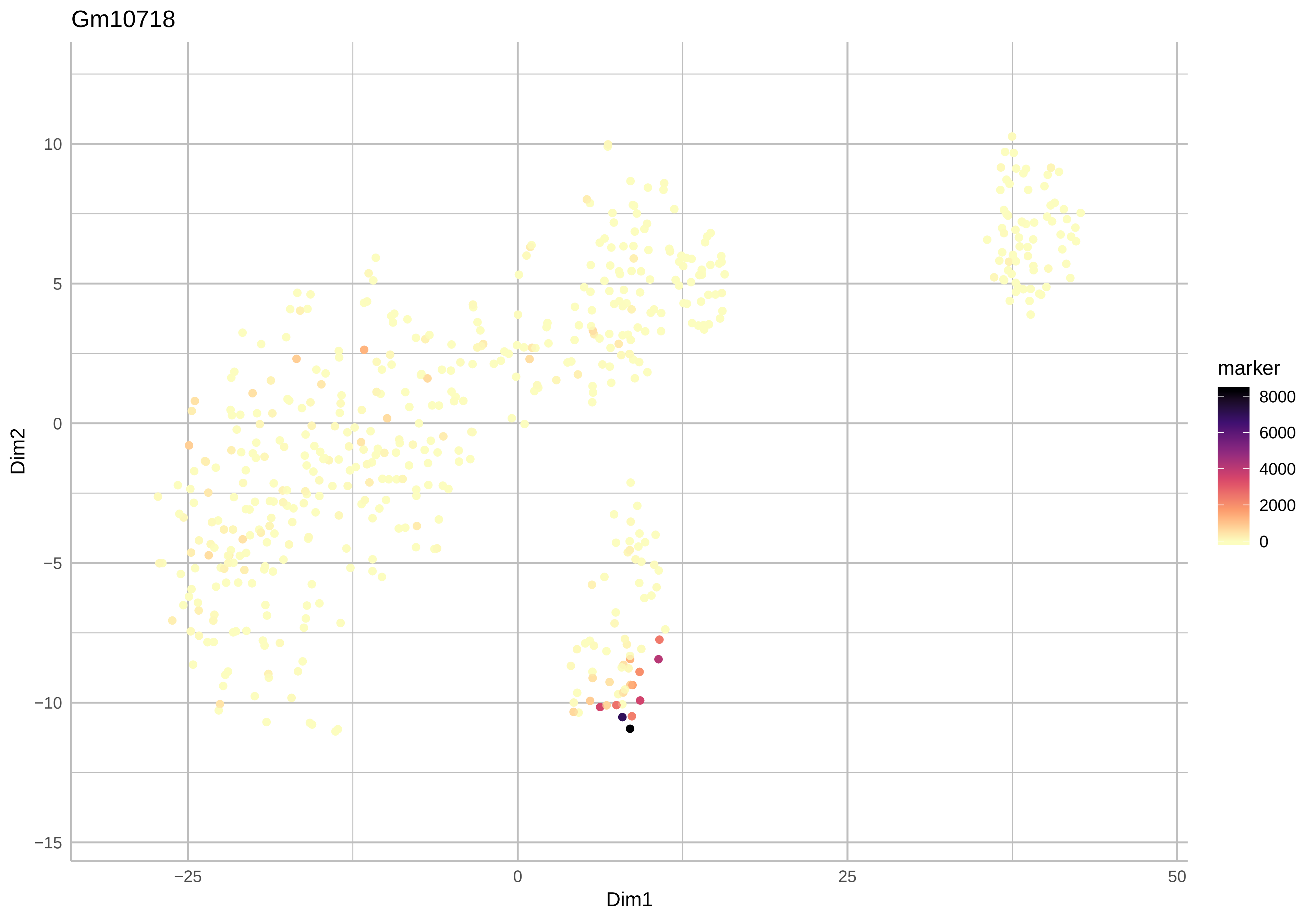
Gm10718
Dim2
10
5
0
−5
−10
−15
−25
0
25
50
Dim1
marker
8000
6000
4000
2000
0

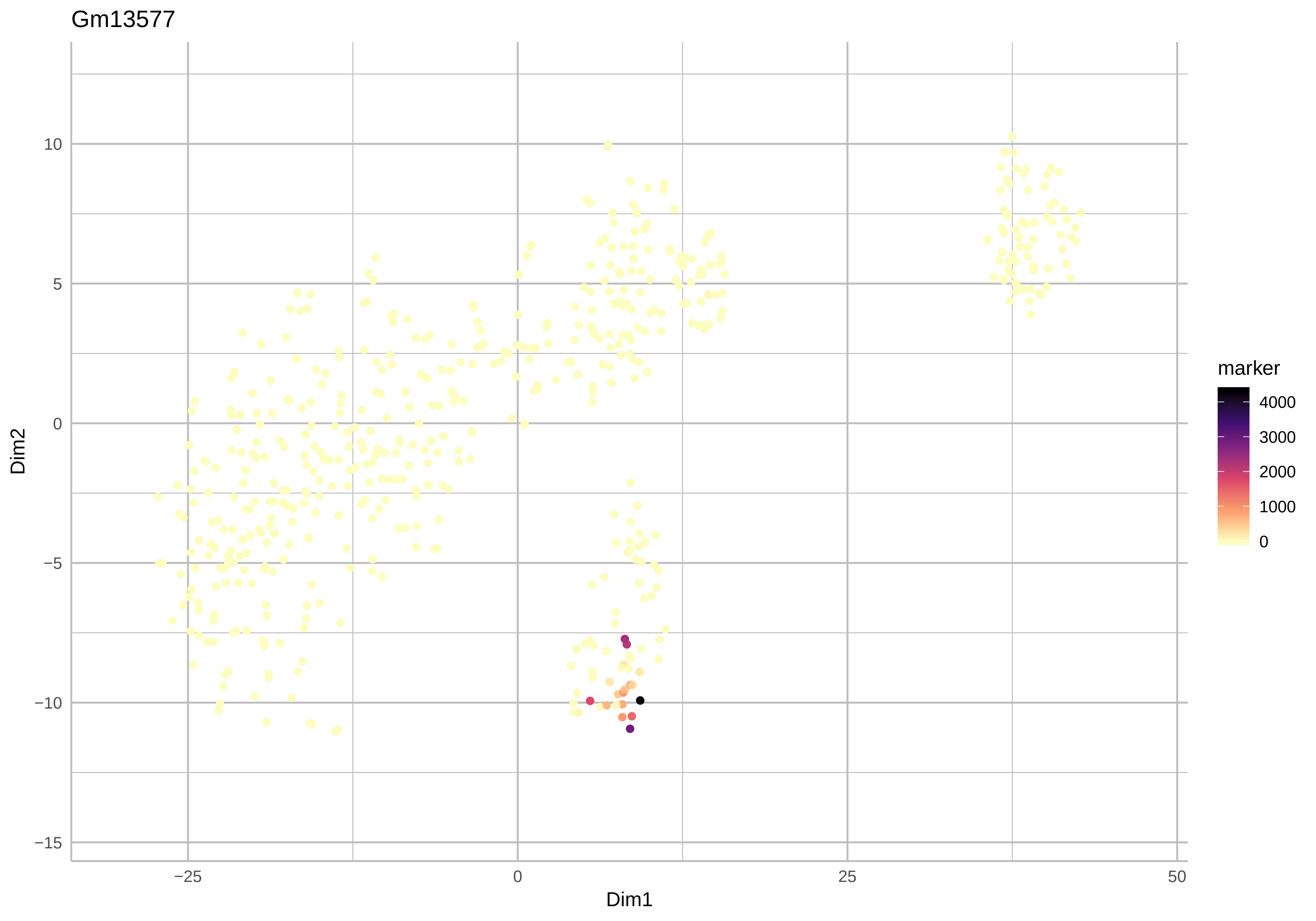

Gm13577
Dim2
Dim1
10
5
0
−5
−10
−15
−25
0
25
50
marker
4000
3000
2000
1000
0

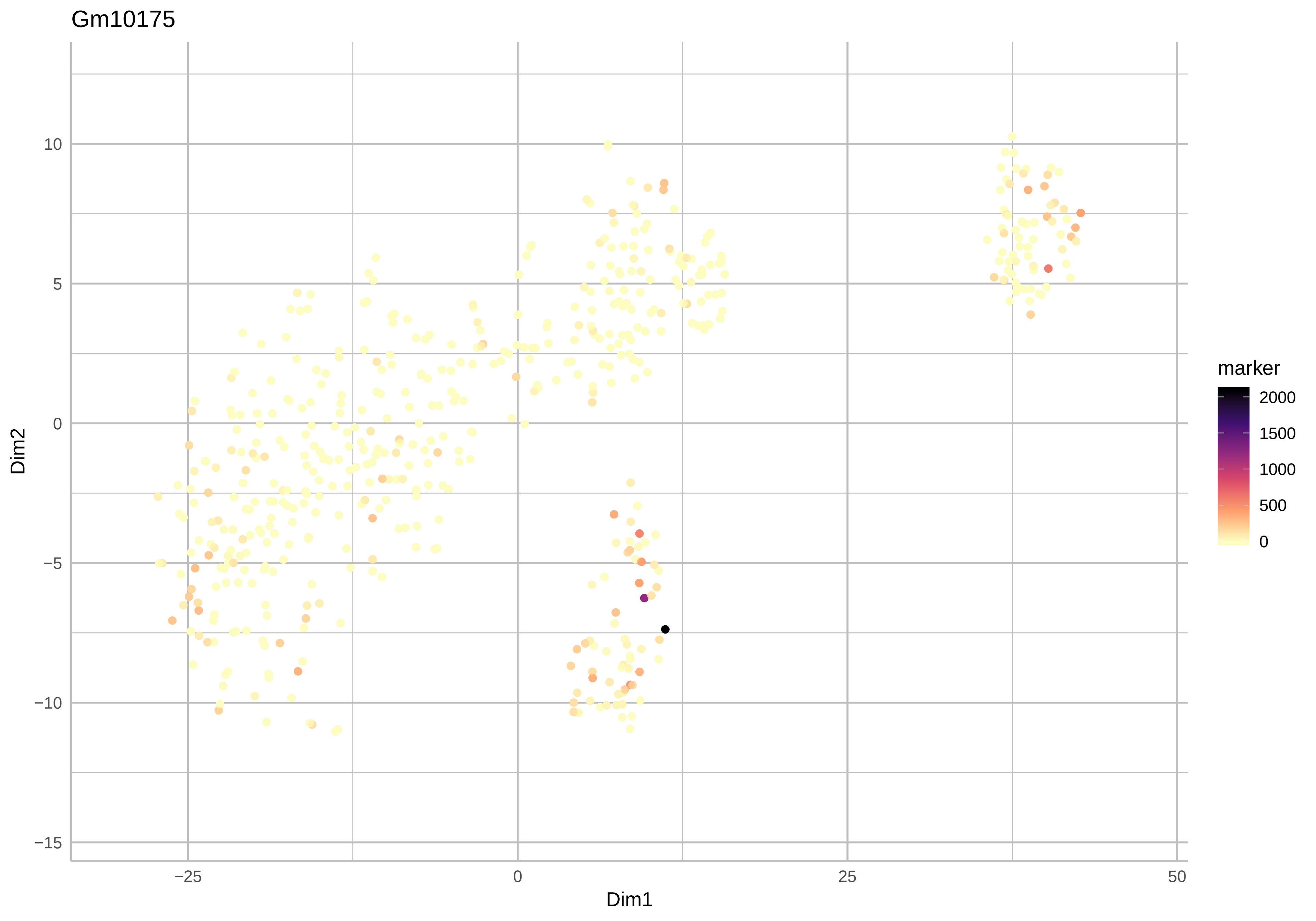
Gm10175
Dim2
Dim1
10
5
0
−5
−10
−15
−25
0
25
50
marker
2000
1500
1000
500
0

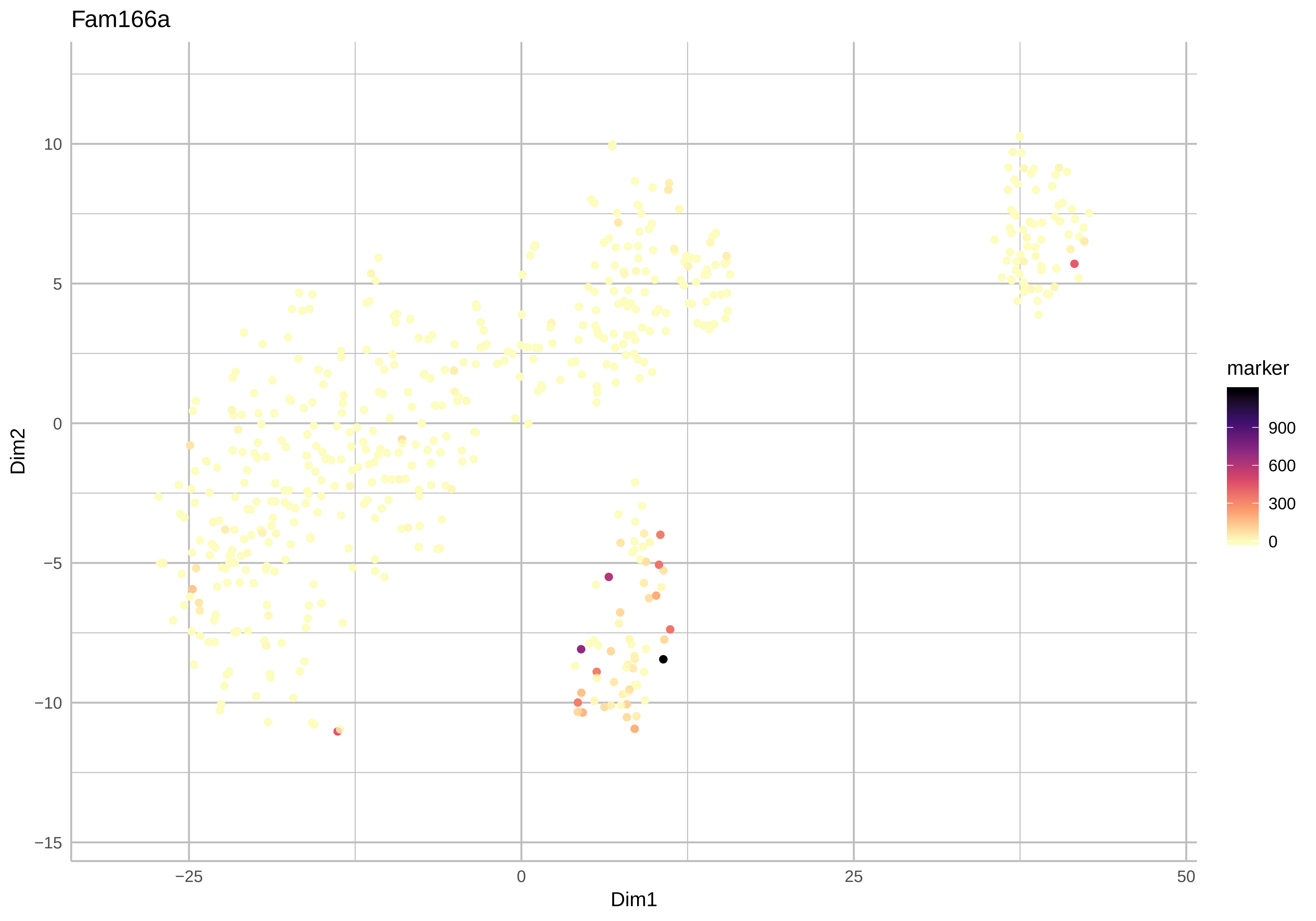

Fam166a
Dim2
Dim1
marker
900
600
300
0
10
5
0
−5
−10
−15
−25
0
25
50

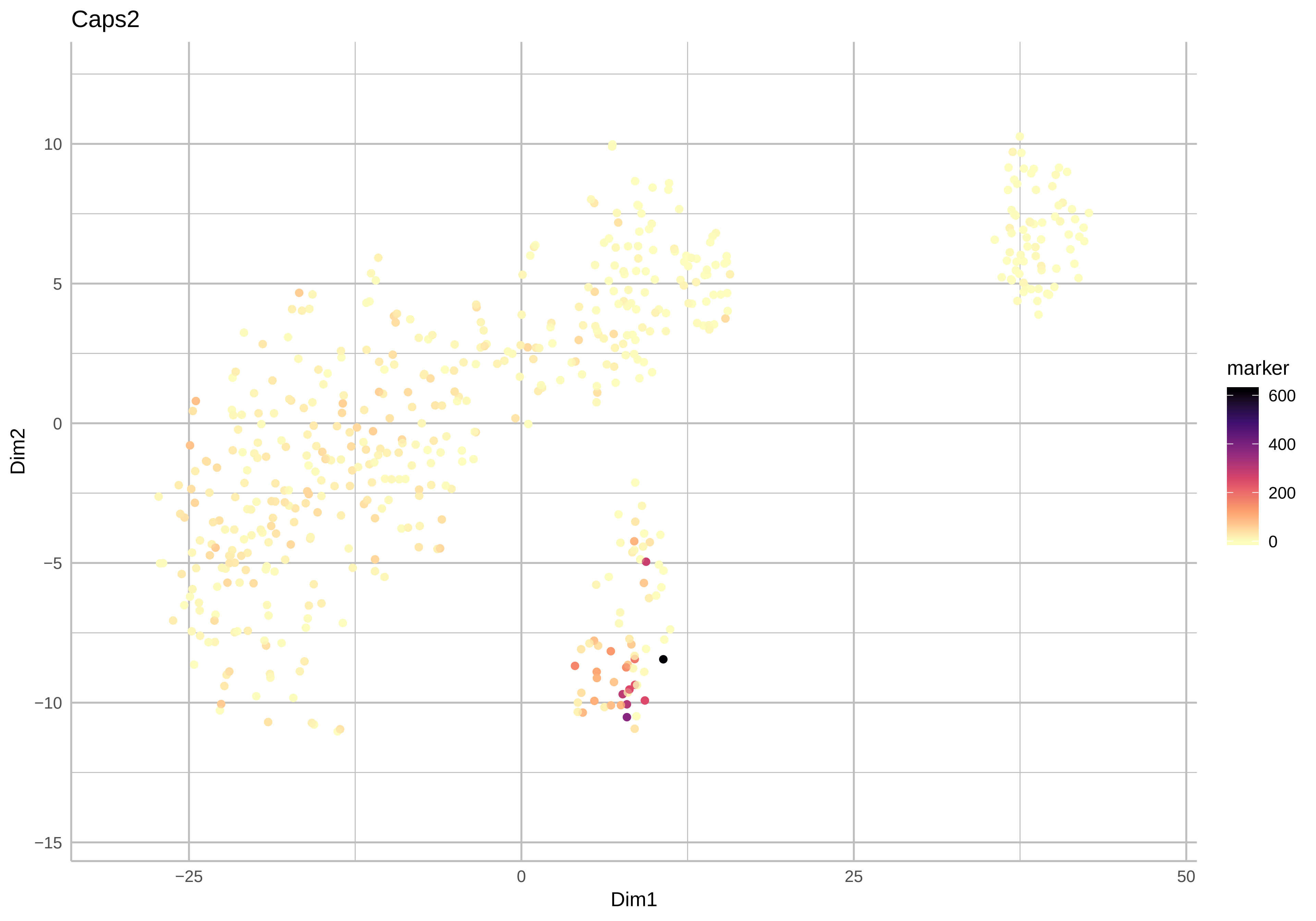
Caps2
Dim2
Dim1
marker
600
400
200
0
10
5
0
−5
−10
−15
−25
0
25
50

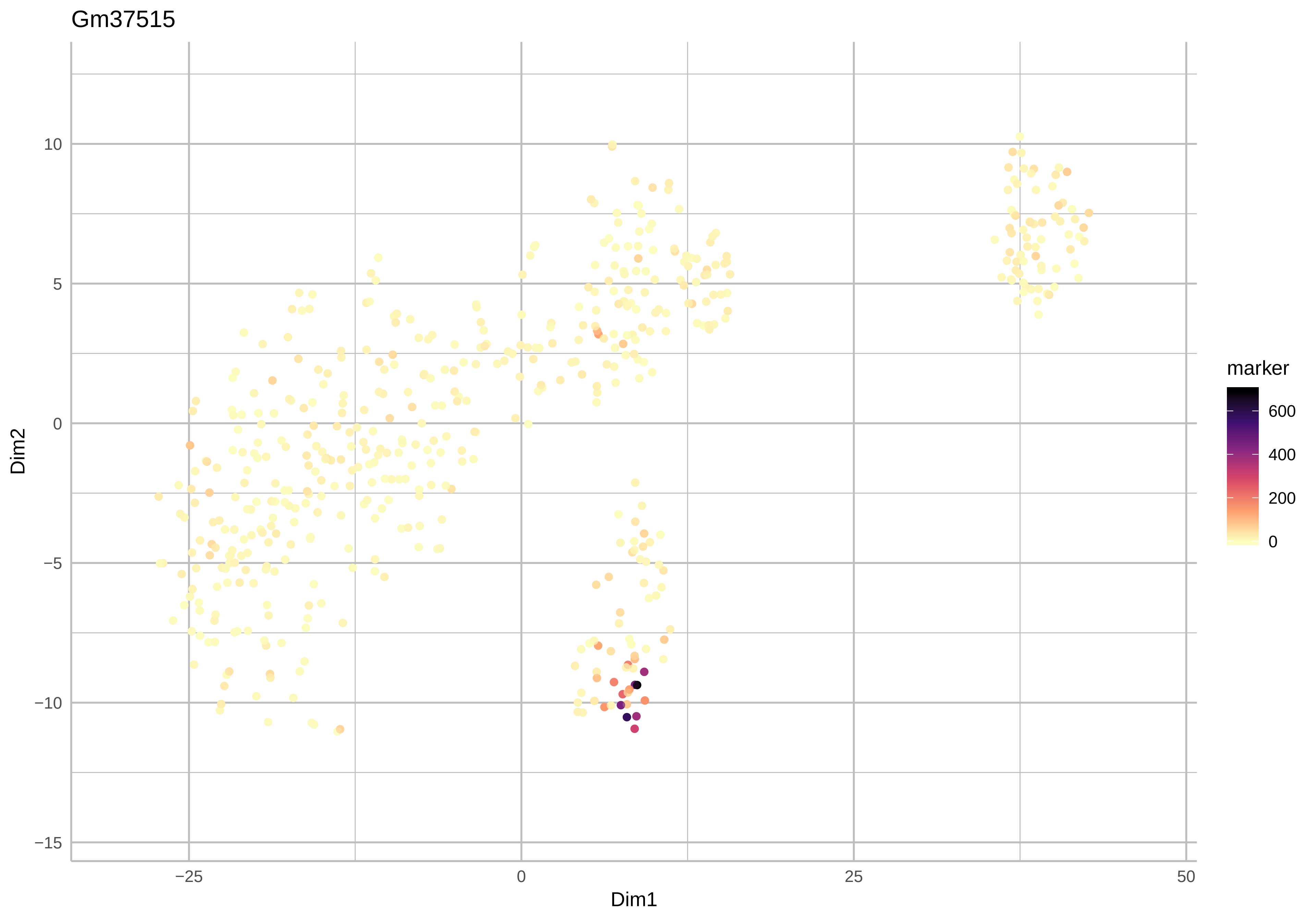

Gm37515
Dim2
Dim1
10
5
0
−5
−10
−15
−25
0
25
50
marker
600
400
200
0

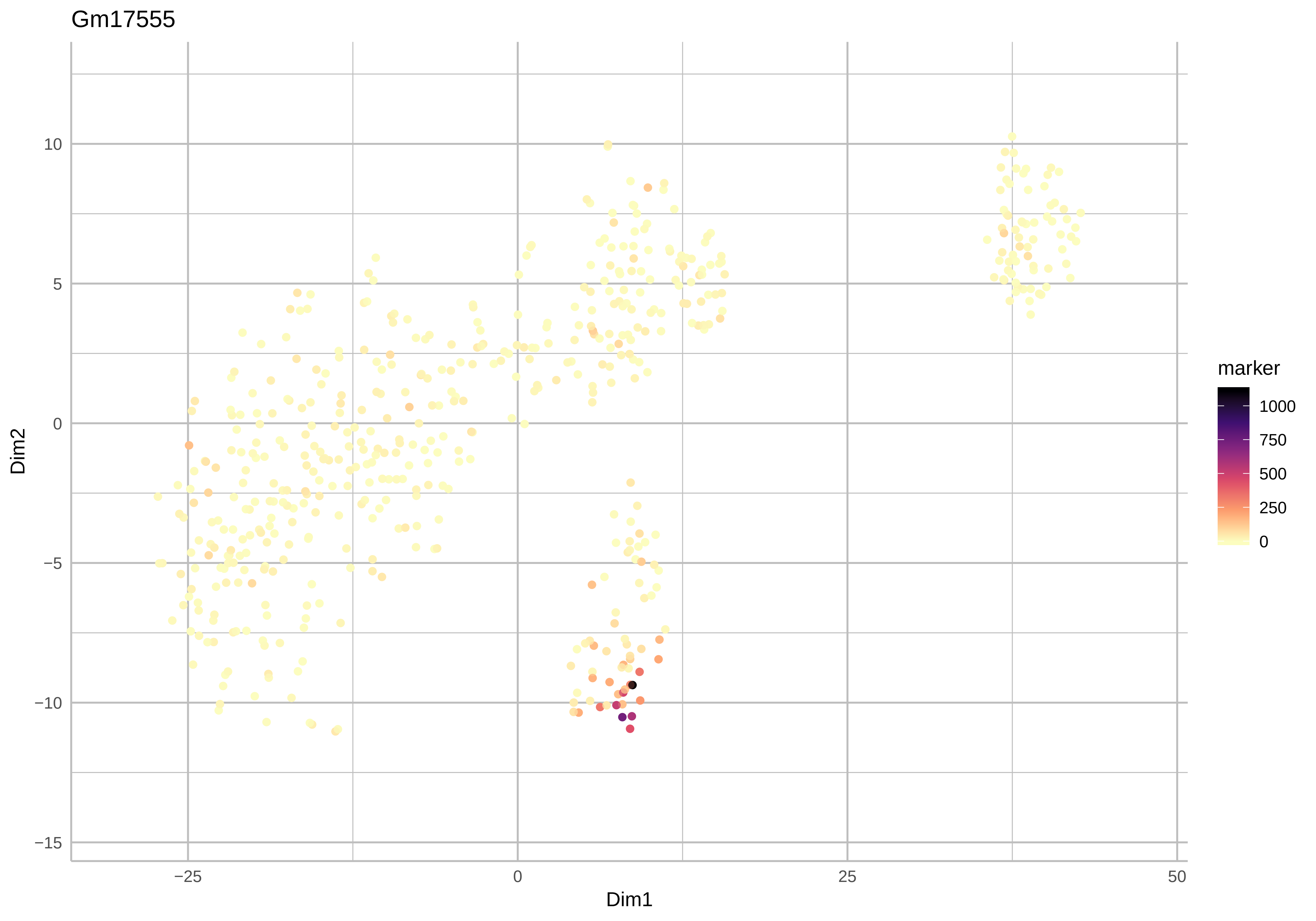
Gm17555
Dim2
Dim1
10
5
0
−5
−10
−15
−25
0
25
50
marker
1000
750
500
250
0

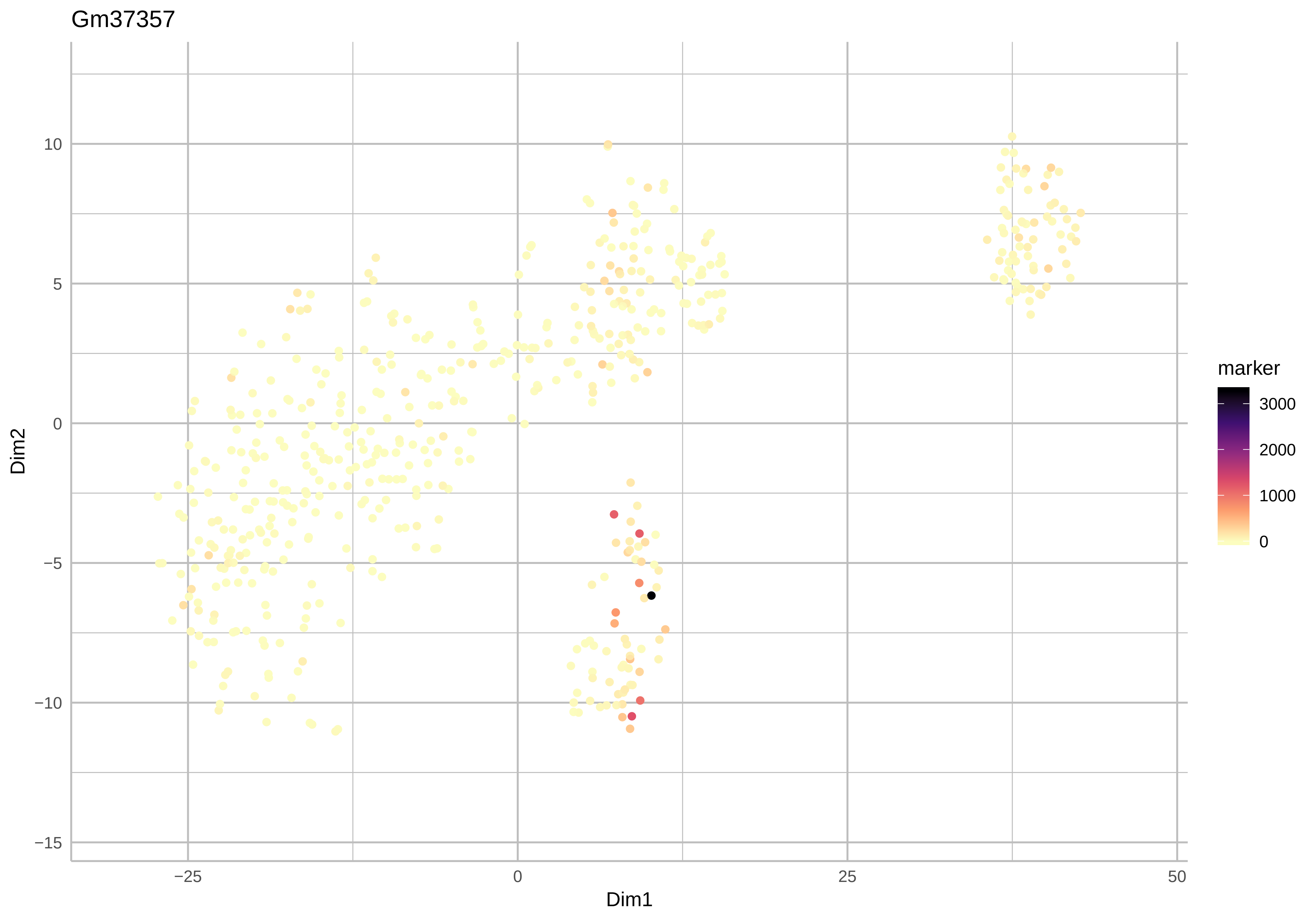

Gm37357
10
5
0
−5
−10
−15
Dim2
−25
0
25
50
Dim1
marker
3000
2000
1000
0

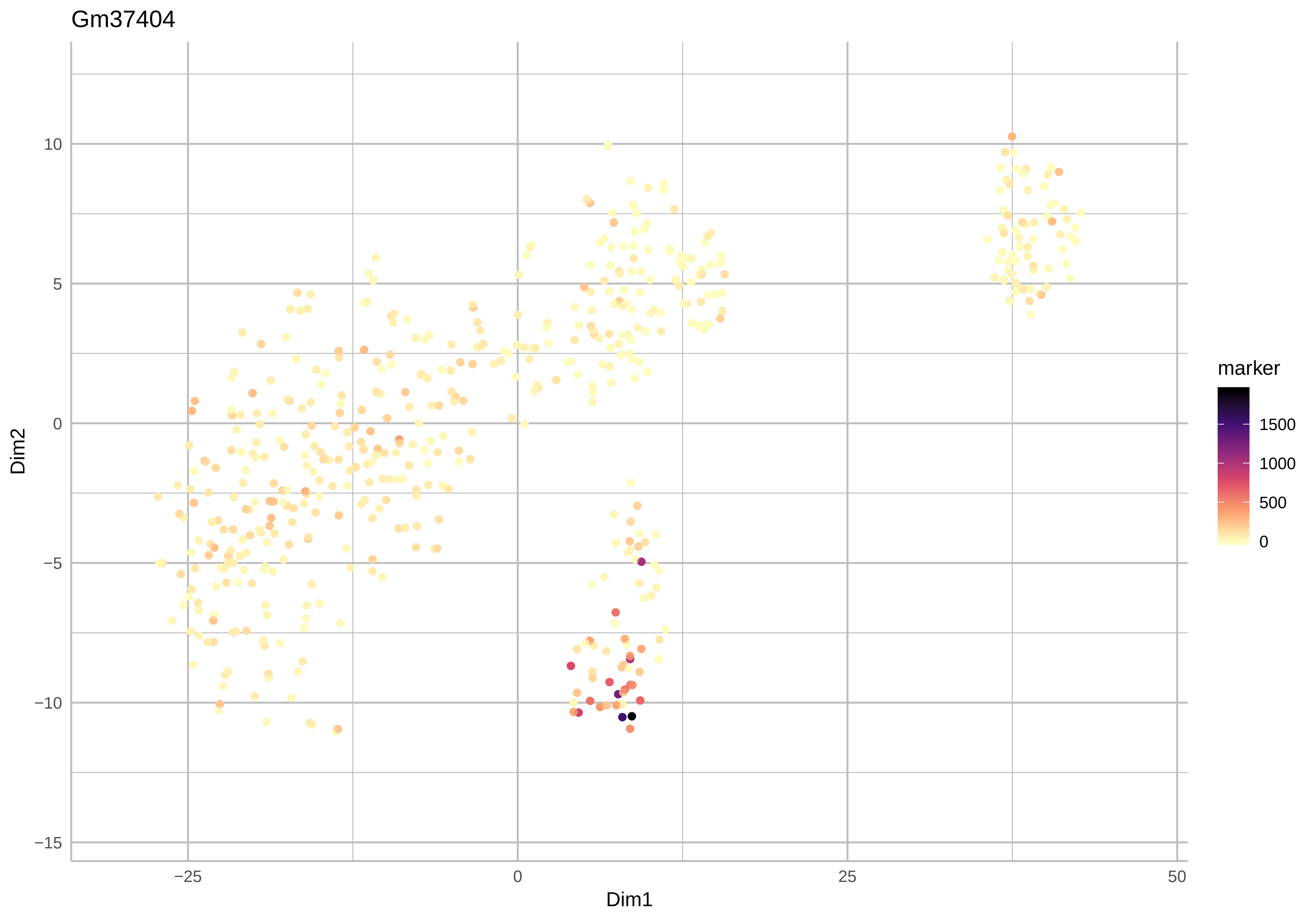

Gm37404
Dim2
Dim1
10
5
0
−5
−10
−15
−25
0
25
50
marker
1500
1000
500
0

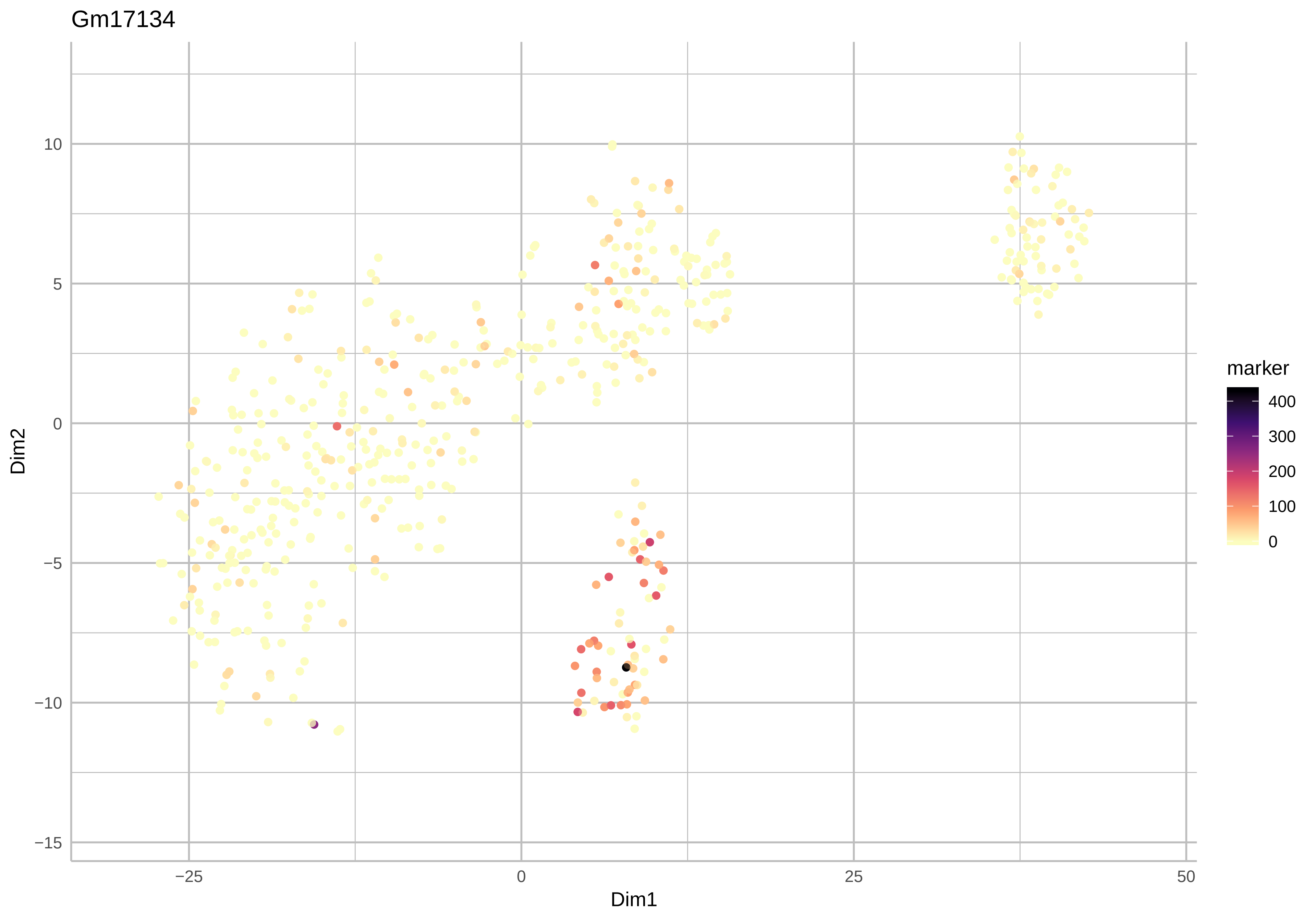
Gm17134
Dim2
Dim1
10
5
0
−5
−10
−15
−25
0
25
50
marker
400
300
200
100
0

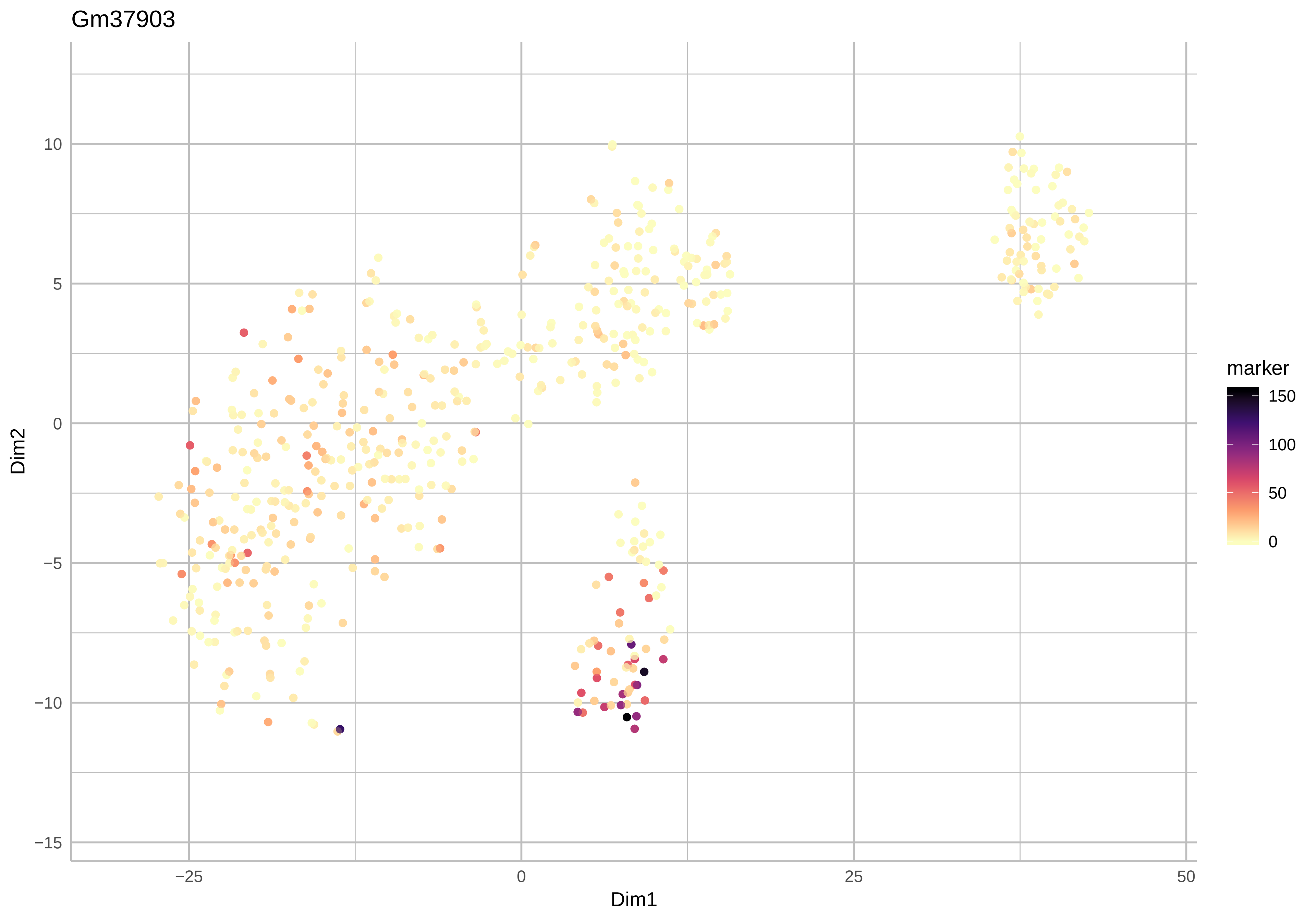

Gm37903
Dim2
Dim1
10
5
0
−5
−10
−15
−25
0
25
50
marker
150
100
50
0

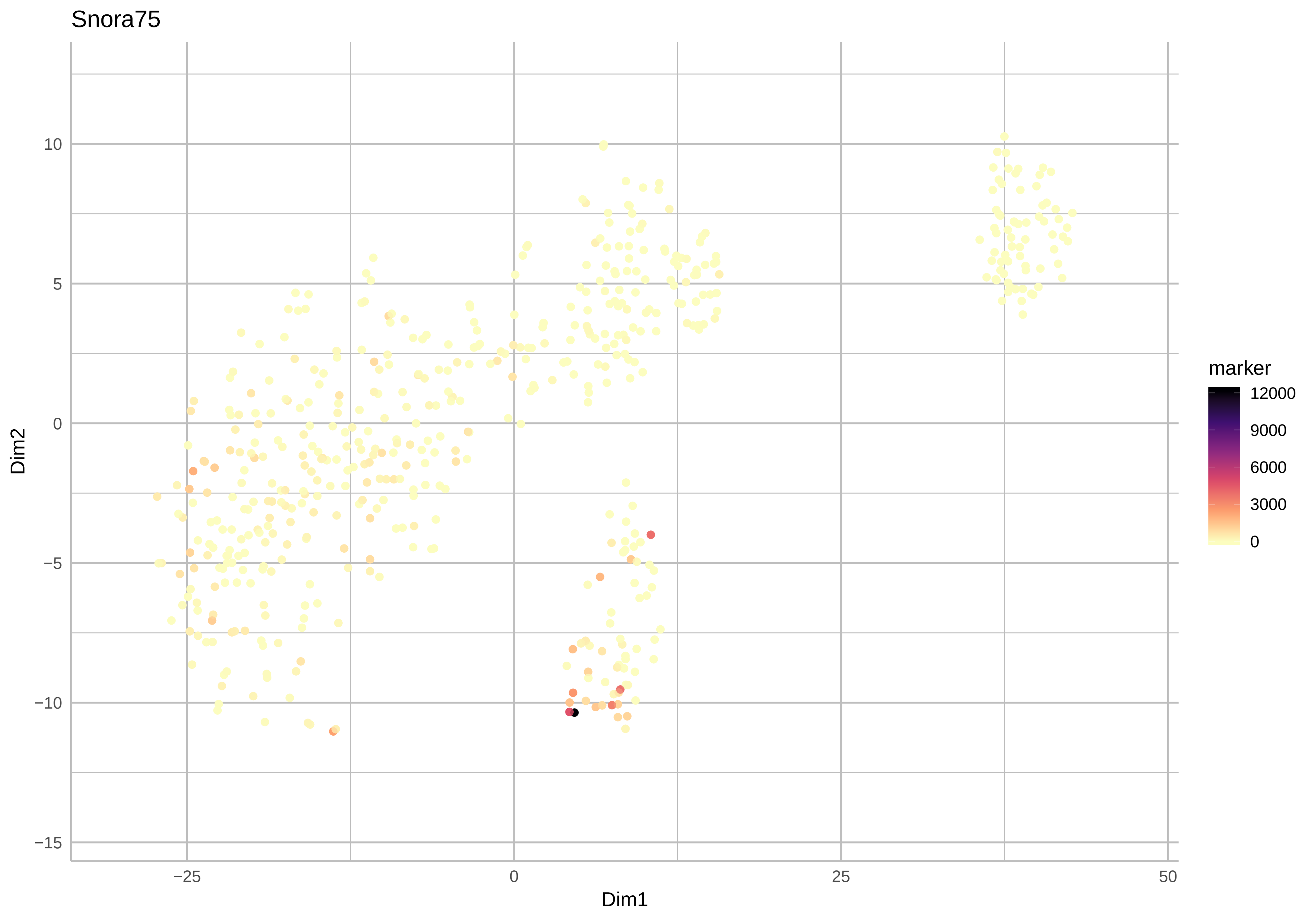
Snora75
Dim2
Dim1
marker
12000
9000
6000
3000
0
10
5
0
−5
−10
−15
−25
0
25
50

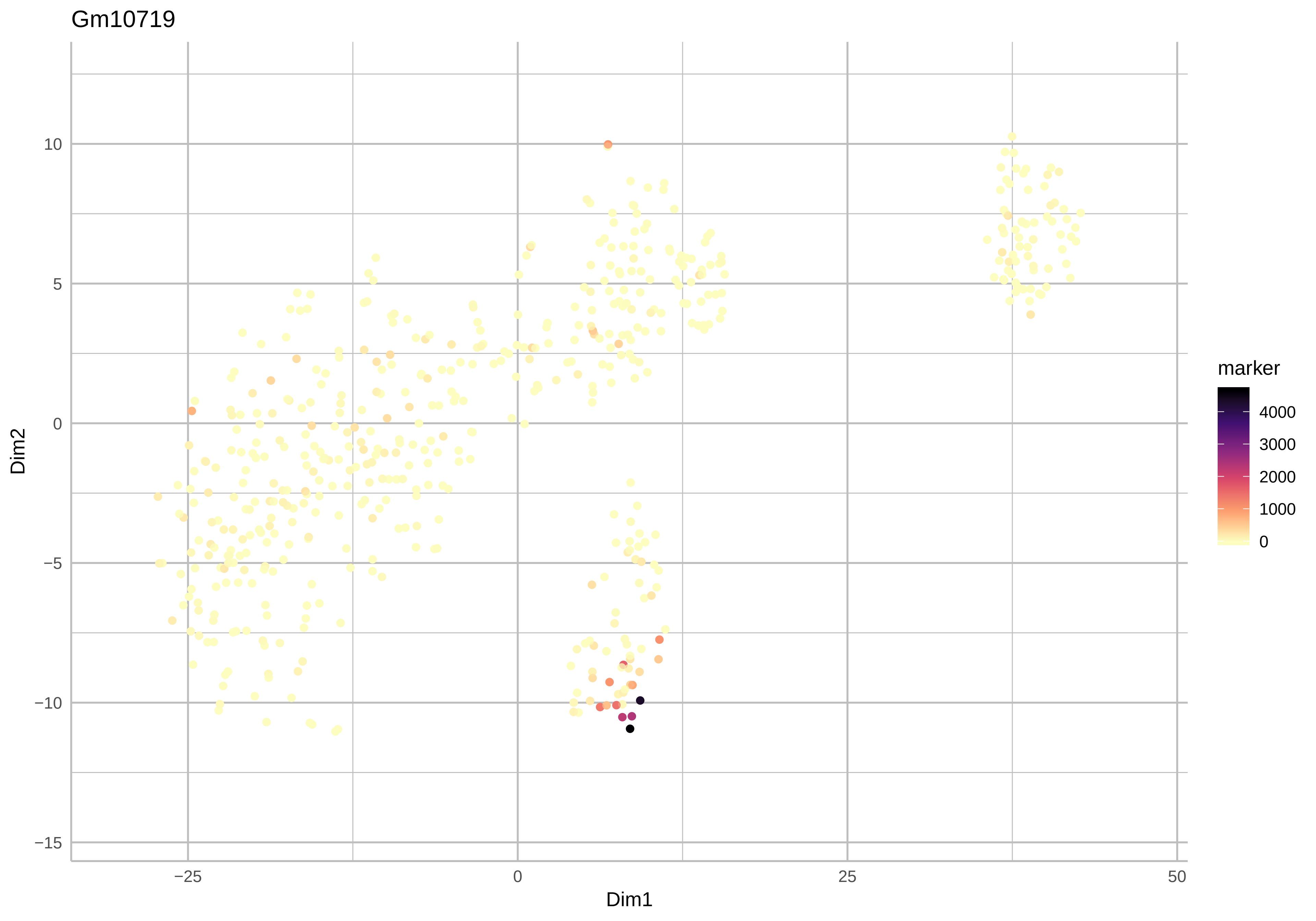

Gm10719
Dim2
10
5
0
−5
−10
−15
−25
0
25
50
Dim1
marker
4000
3000
2000
1000
0

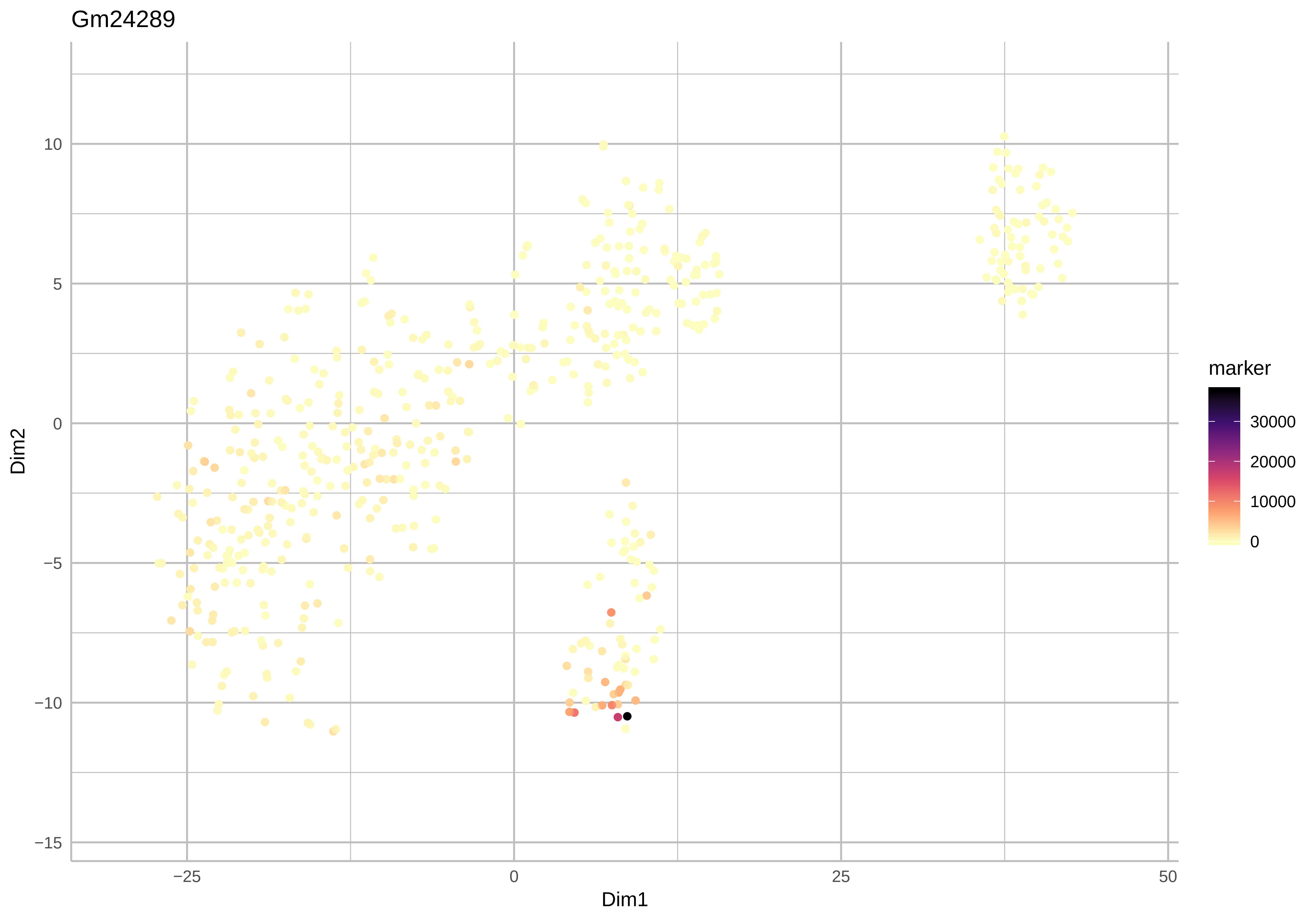
Gm24289
Dim2
Dim1
10
5
0
−5
−10
−15
−25
0
25
50
marker
30000
20000
10000
0

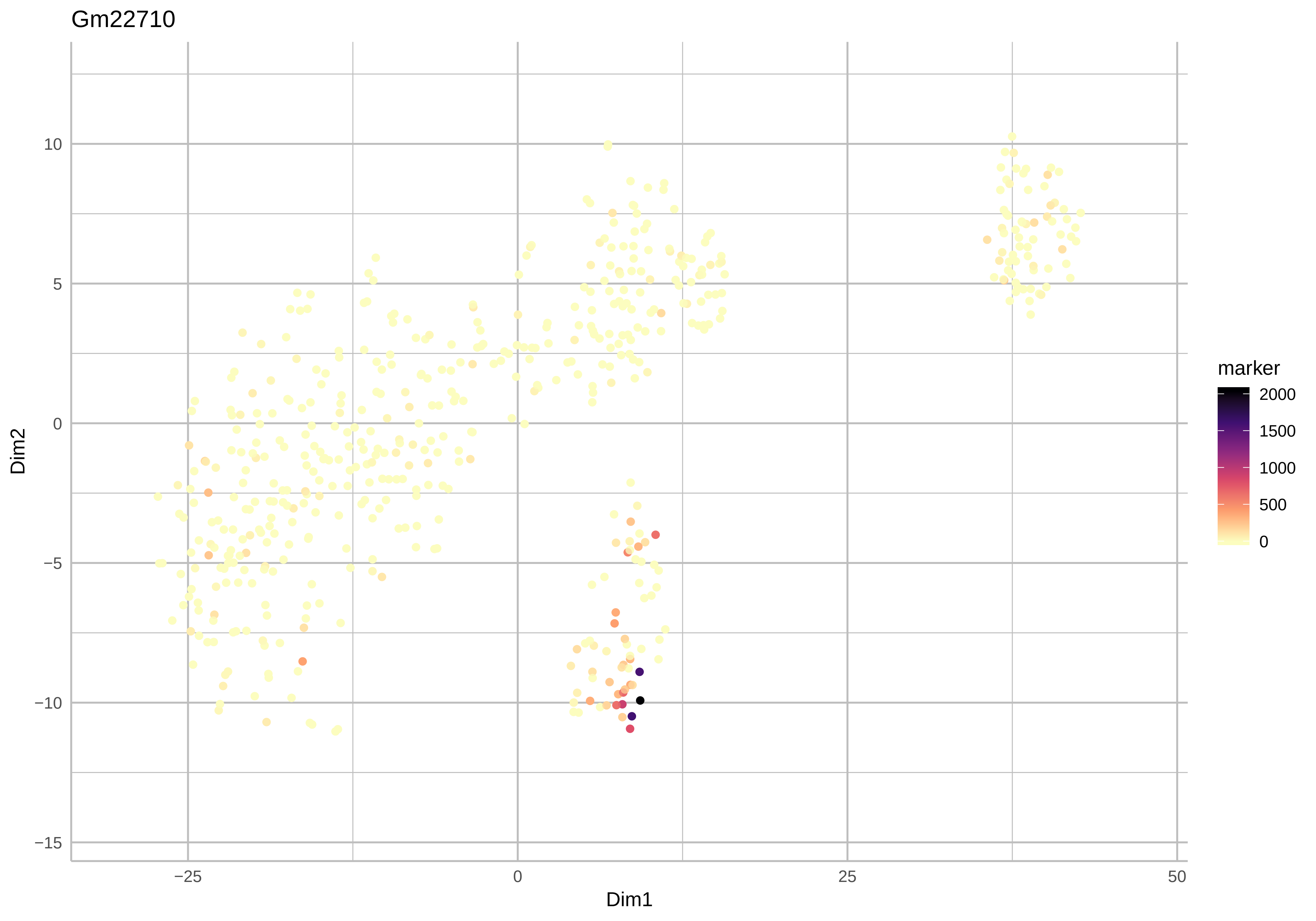

Gm22710
marker
2000
1500
1000
500
0
Dim2
Dim1
10
5
0
−5
−10
−15
−25
0
25
50

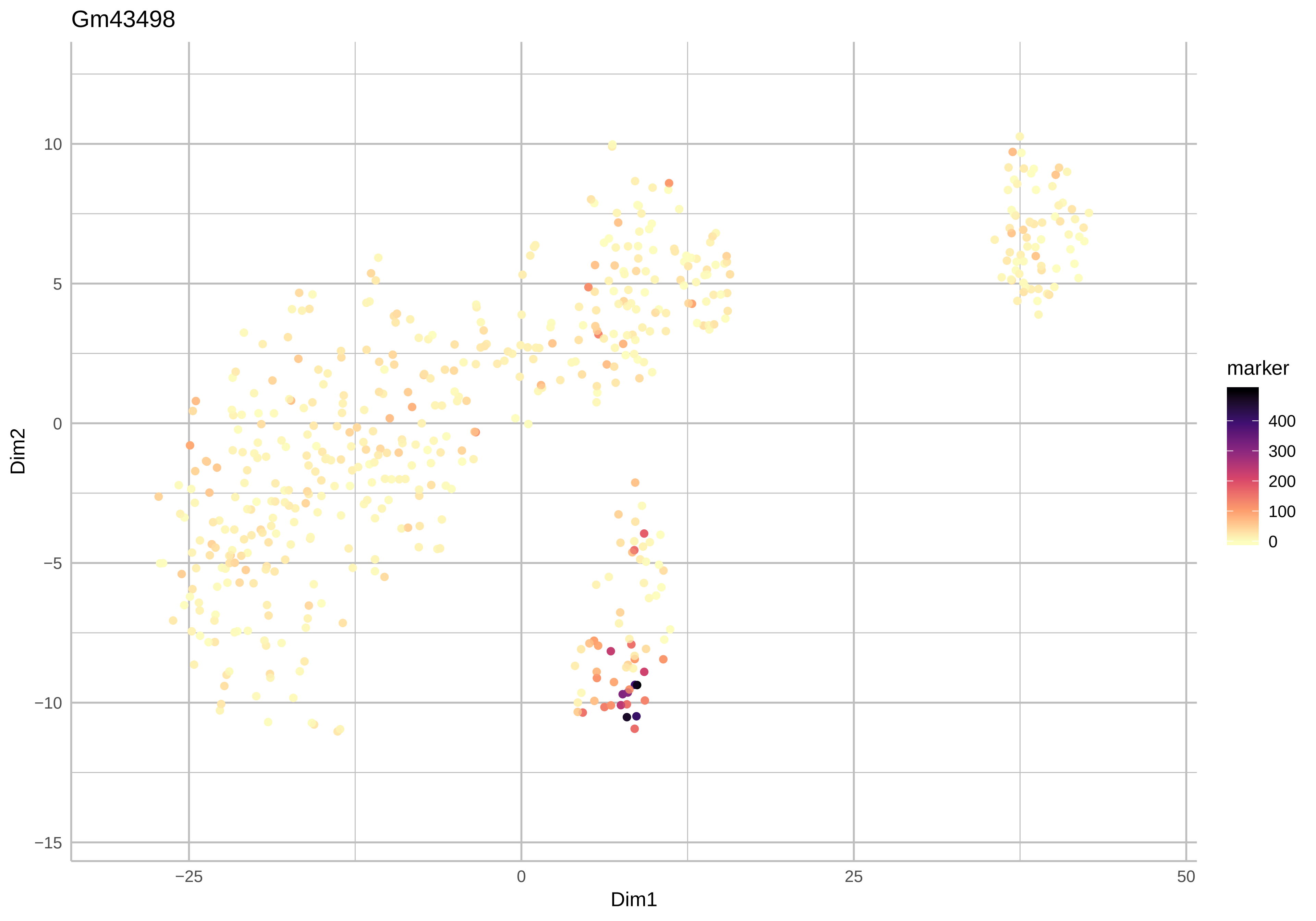

Gm43498
10
5
0
−5
−10
−15
Dim2
−25
0
25
50
Dim1
marker
400
300
200
100
0

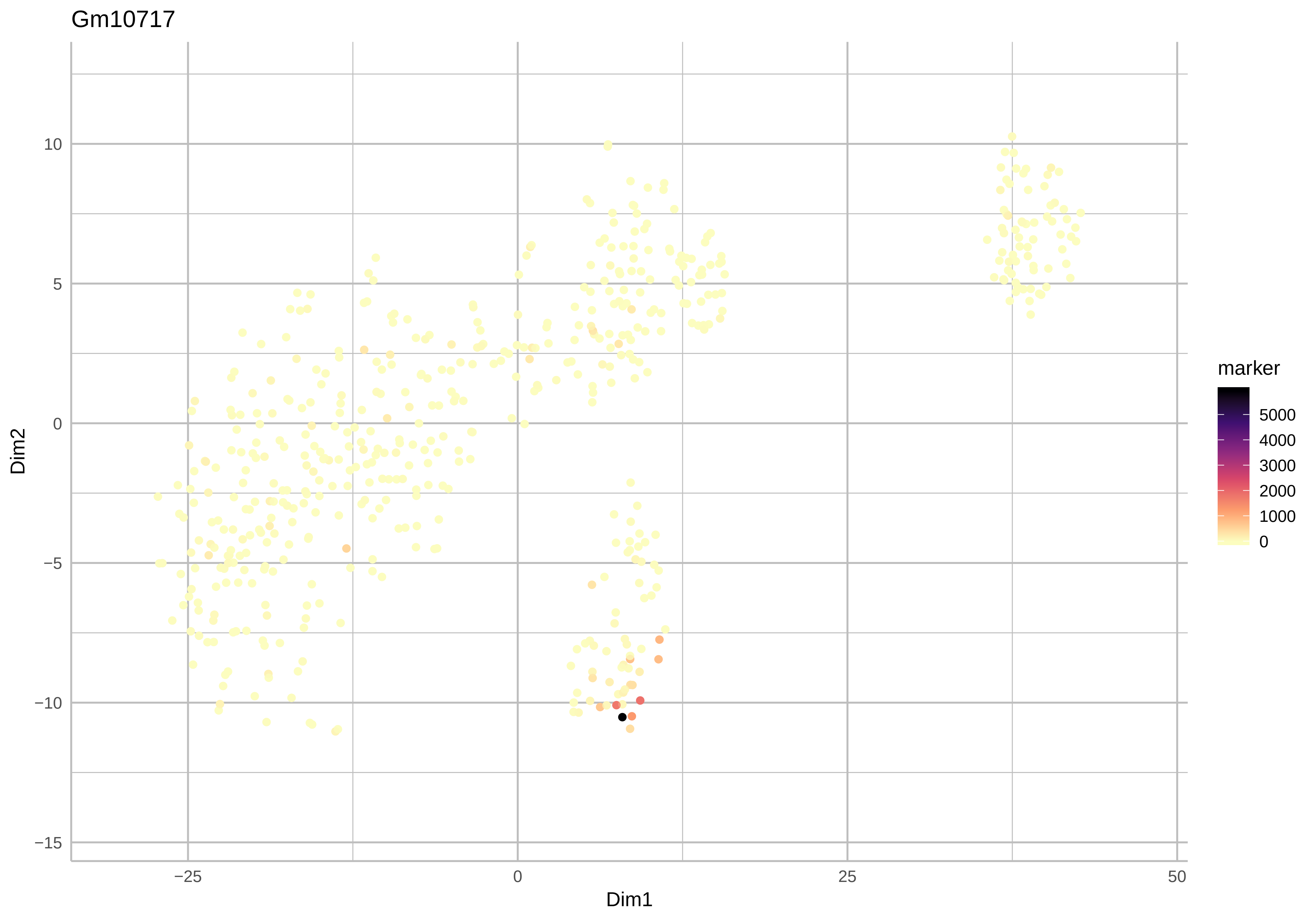

Gm10717
Dim2
Dim1
marker
5000
4000
3000
2000
1000
0
10
5
0
−5
−10
−15
−25
0
25
50

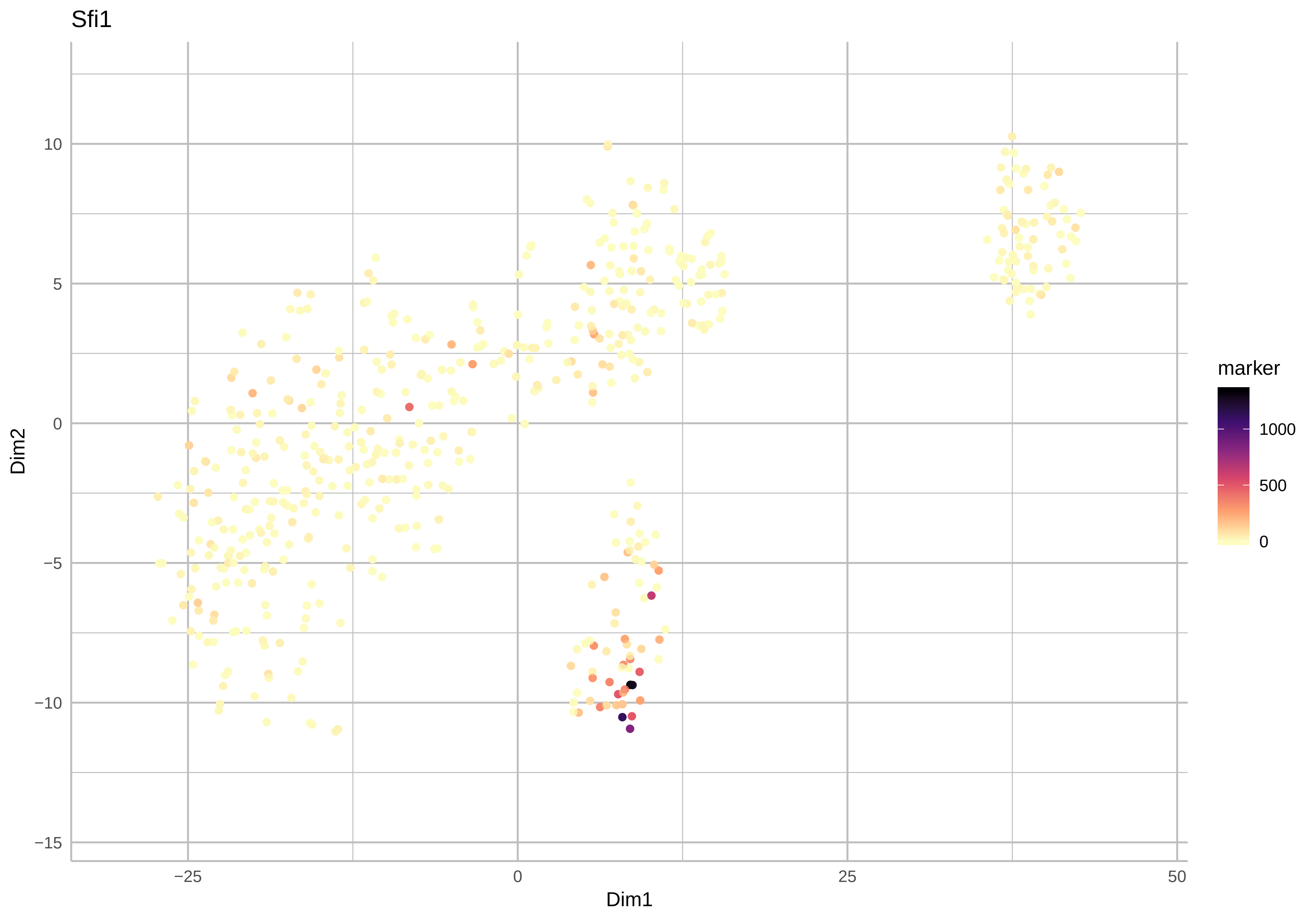
Sfi1
Dim2
Dim1
10
5
0
−5
−10
−15
−25
0
25
50
marker
1000
500
0

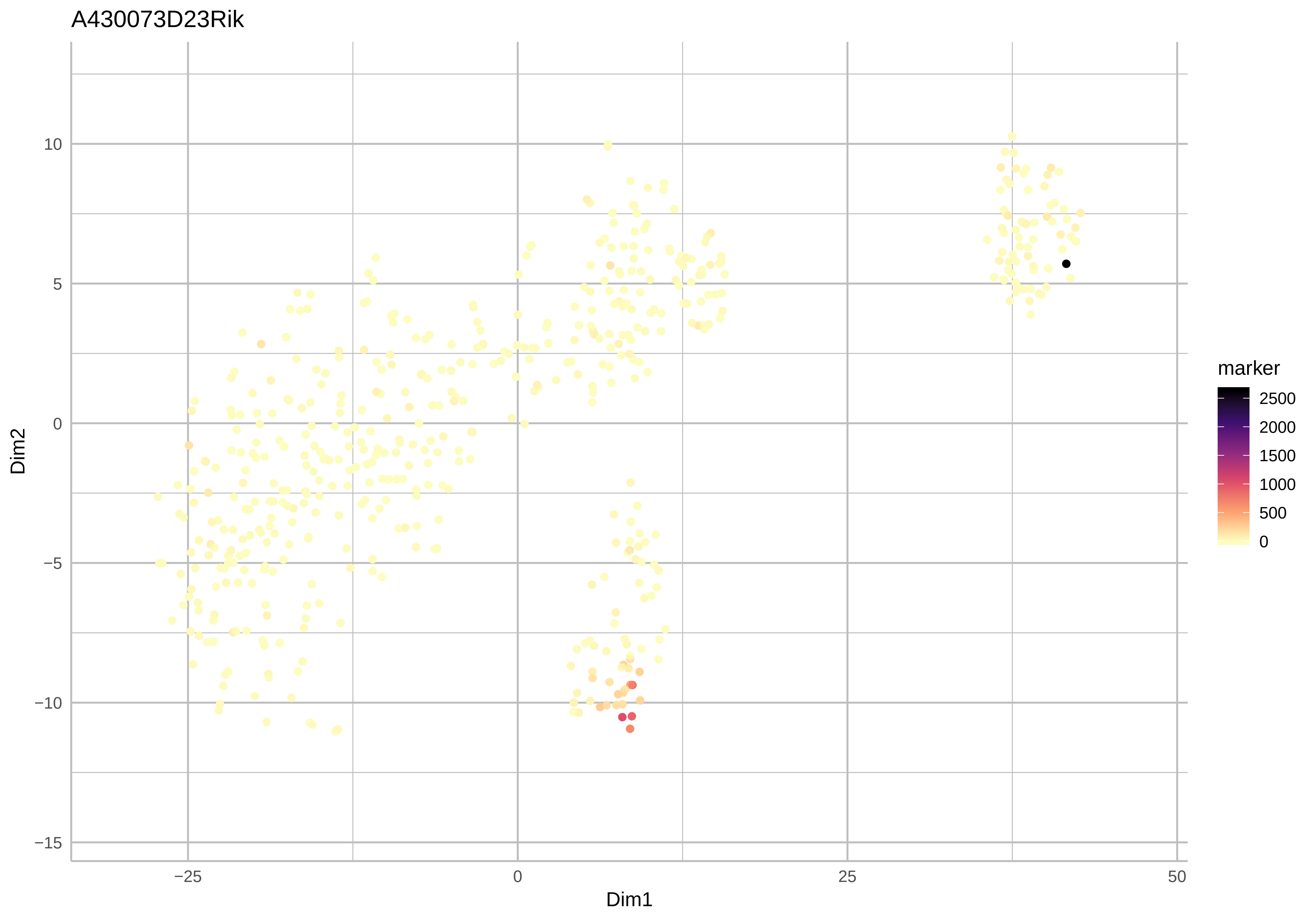
A430073D23Rik
Dim2
Dim1
marker
2500
2000
1500
1000
500
0
10
5
0
−5
−10
−15
−25
0
25
50

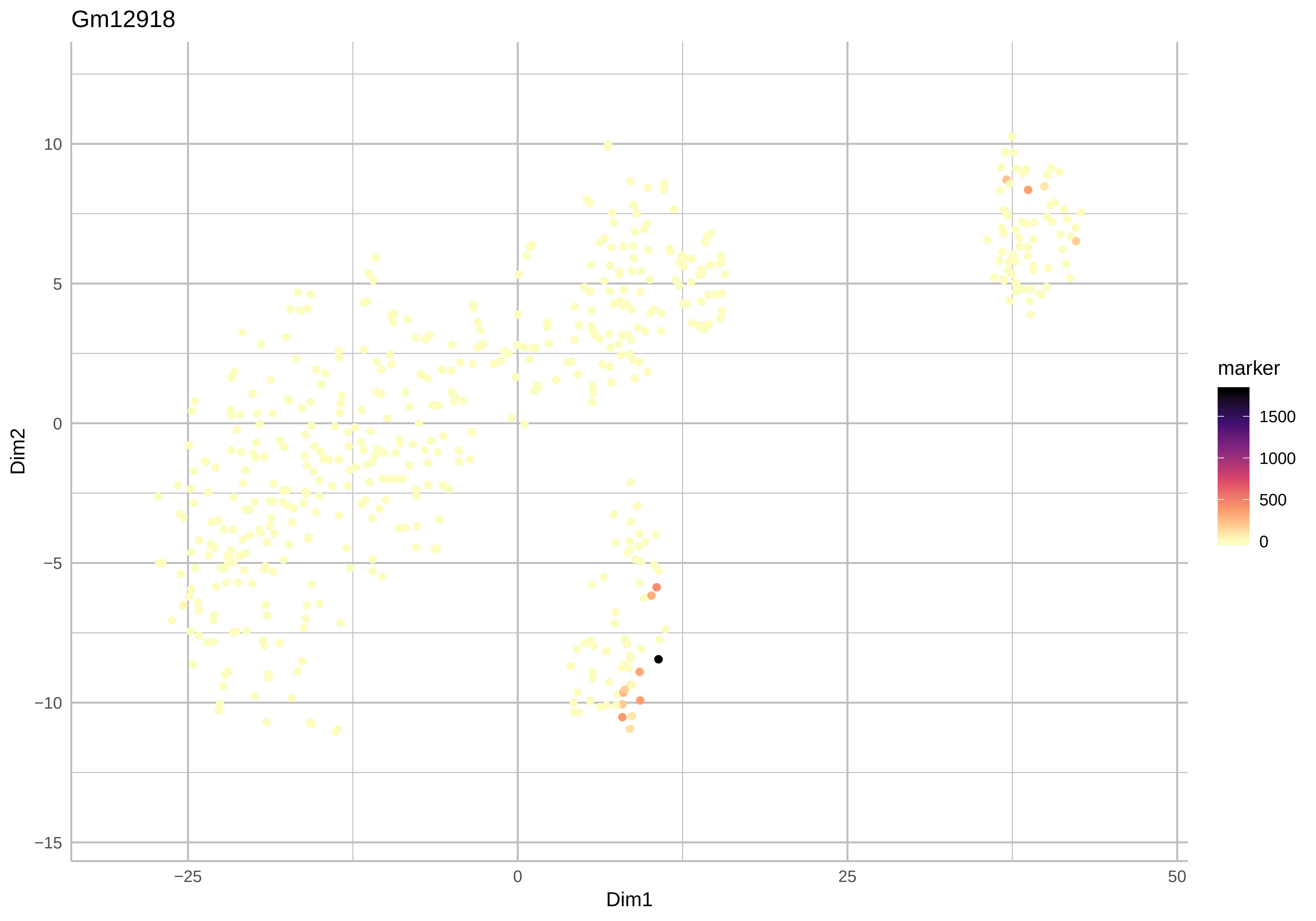

Gm12918
10
5
0
−5
−10
−15
Dim2
−25
0
25
50
Dim1
marker
1500
1000
500
0

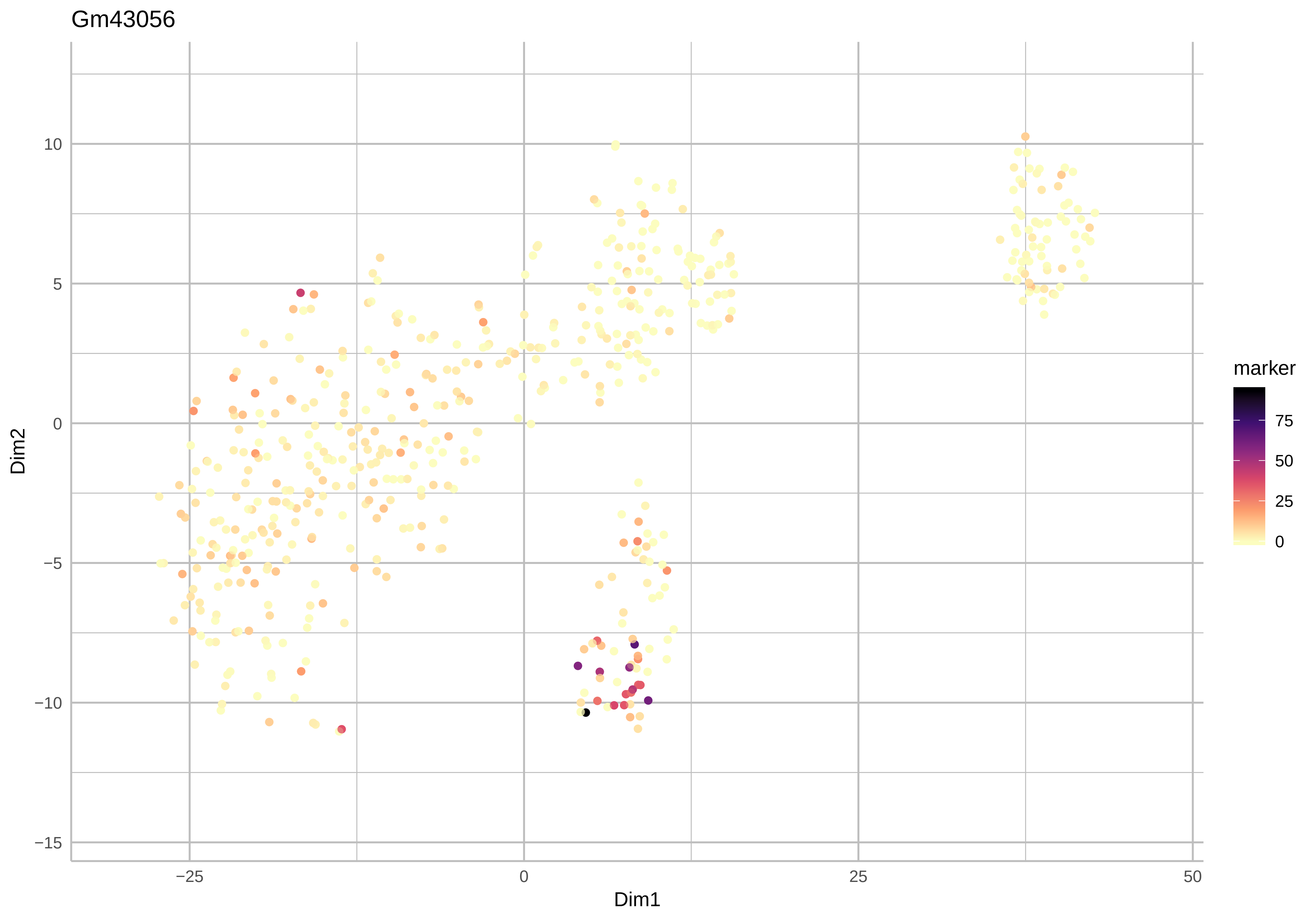
Gm43056
Dim2
Dim1
marker
75
50
25
0
10
5
0
−5
−10
−15
−25
0
25
50

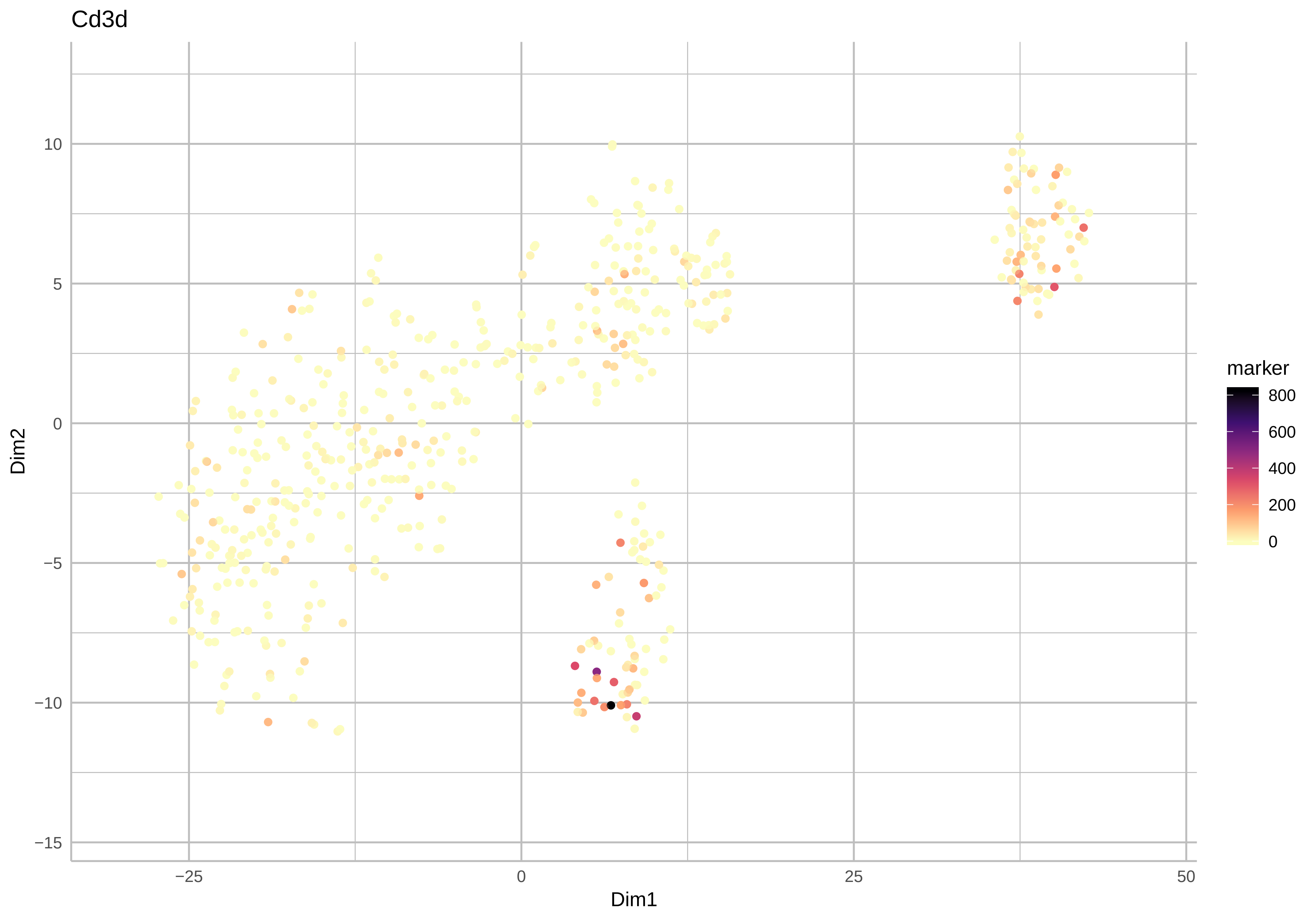
Cd3d
Dim2
Dim1
marker
800
600
400
200
0
10
5
0
−5
−10
−15
−25
0
25
50

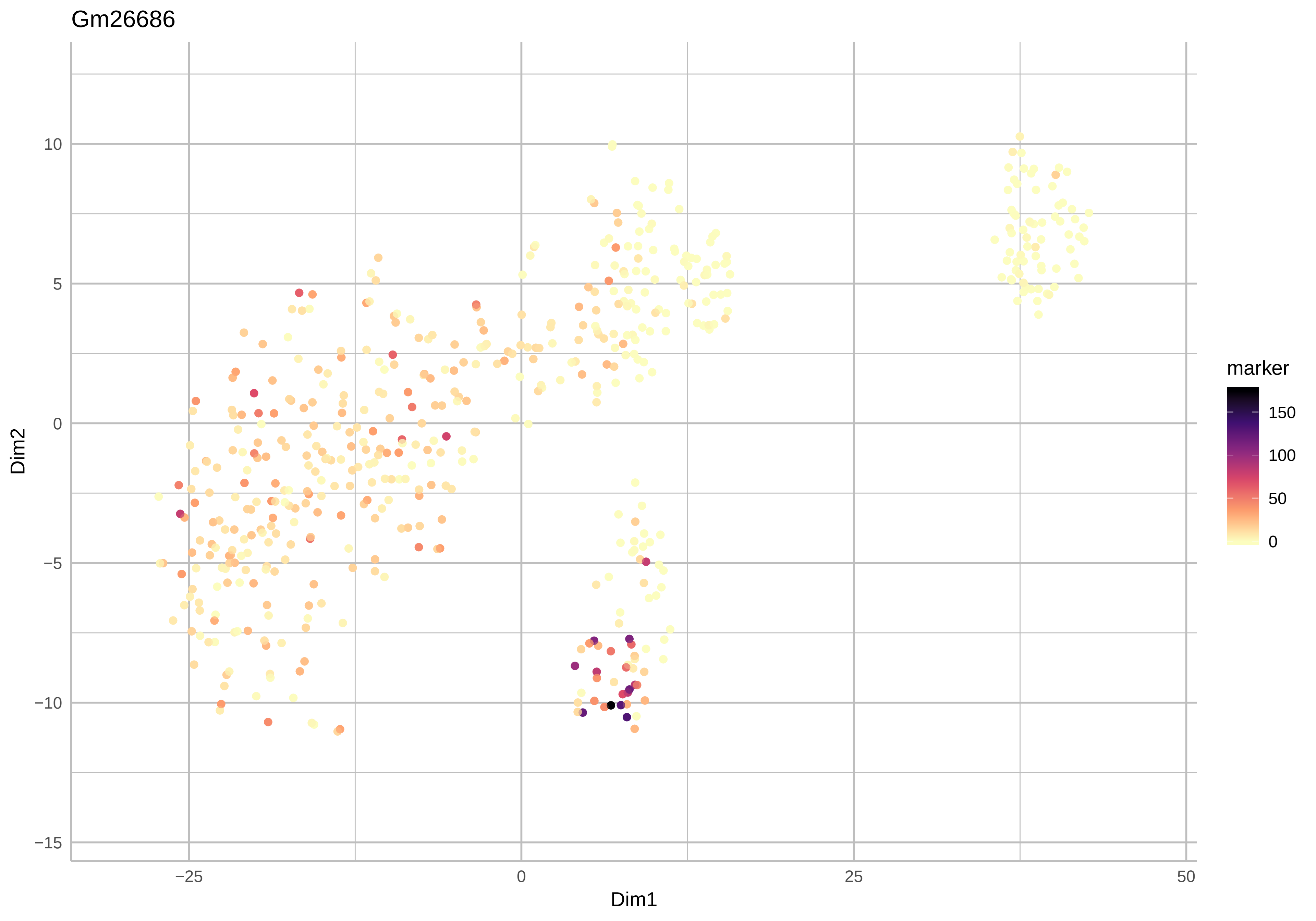

Gm26686
10
5
0
−5
−10
−15
Dim2
−25
0
25
50
Dim1
marker
150
100
50
0

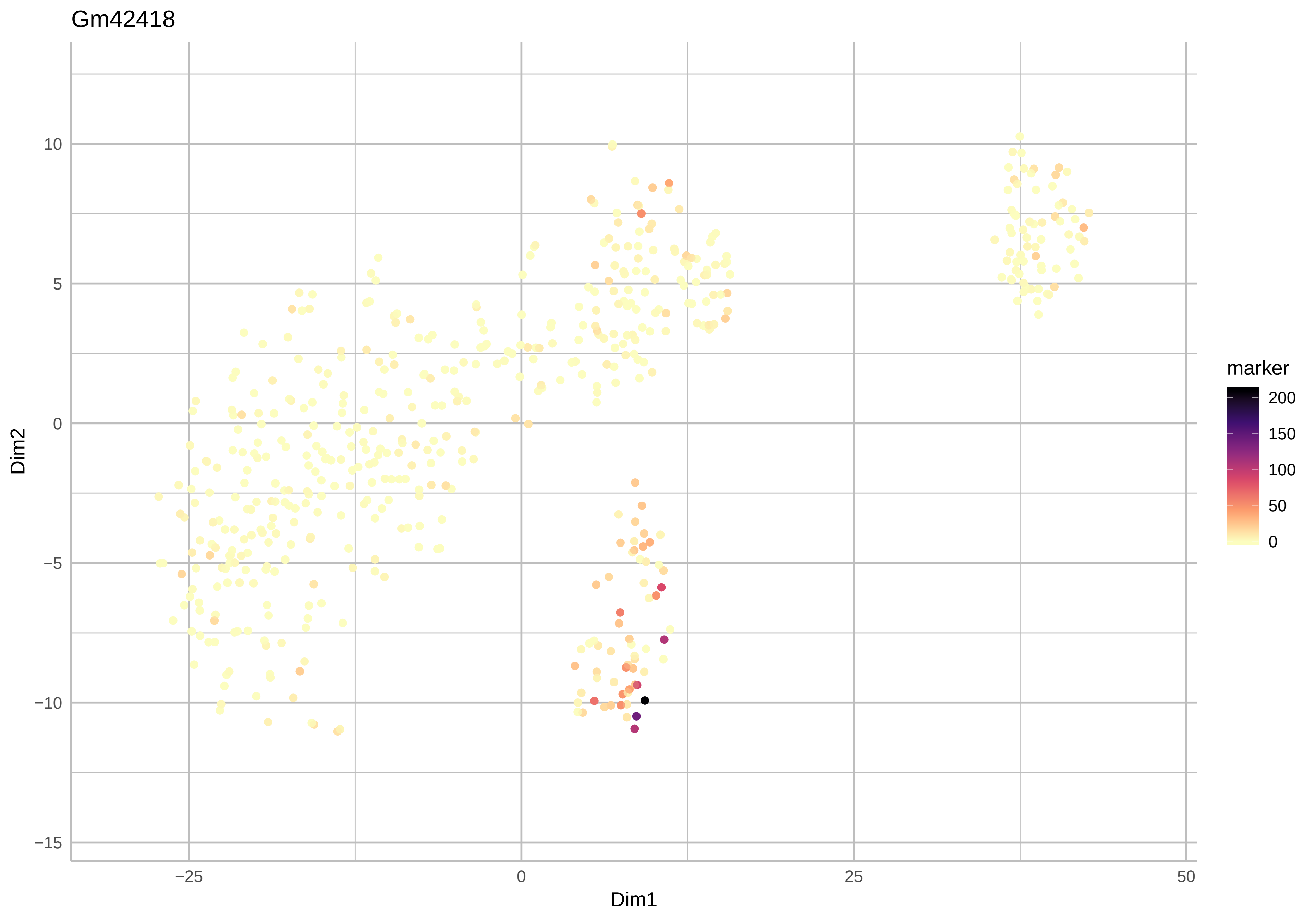
Gm42418
Dim2
Dim1
10
5
0
−5
−10
−15
−25
0
25
50
marker
200
150
100
50
0

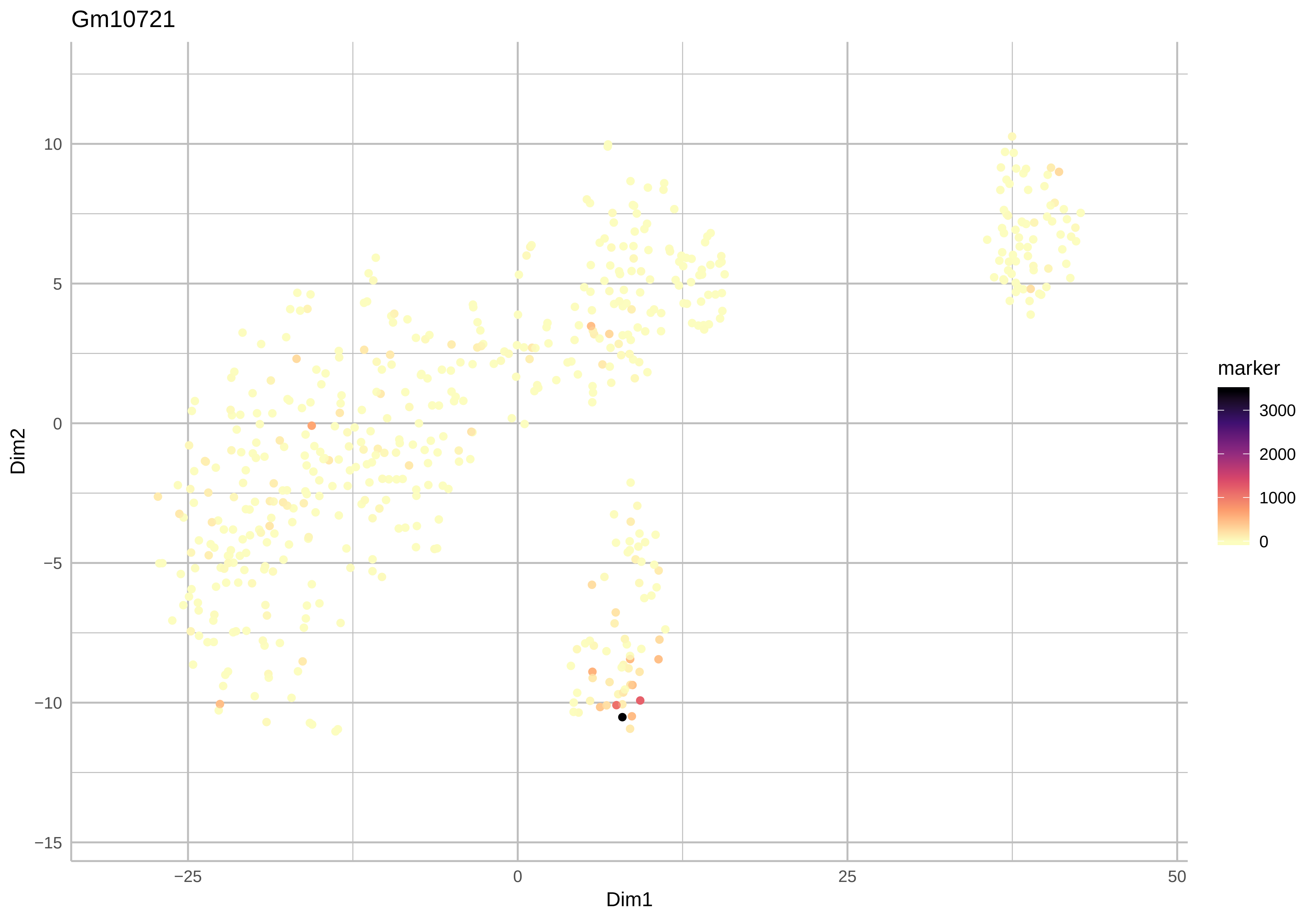

Gm10721
Dim2
Dim1
marker
3000
2000
1000
0
10
5
0
−5
−10
−15
−25
0
25
50

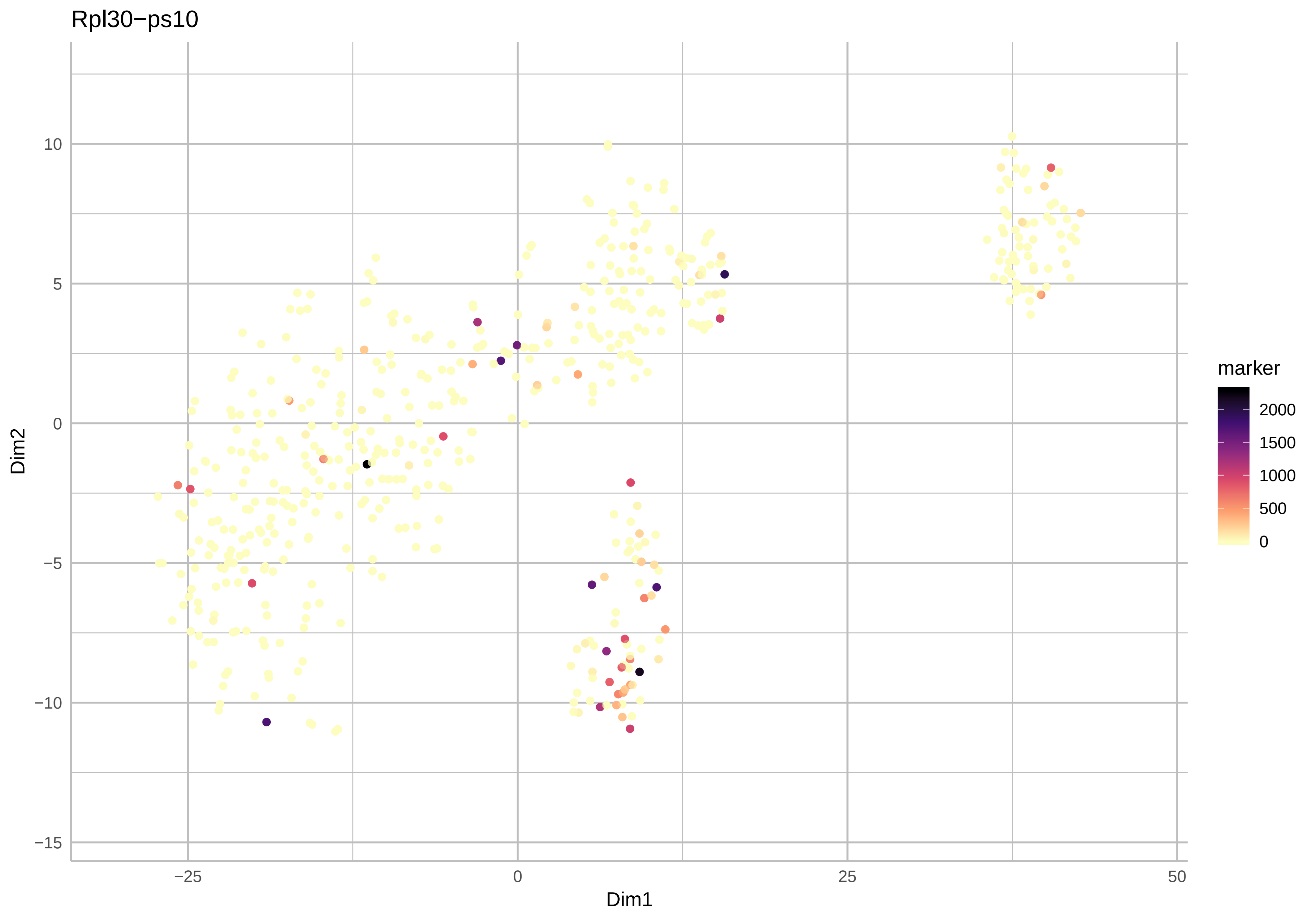
Rpl30−ps10
Dim2
Dim1
10
5
0
−5
−10
−15
−25
0
25
50
marker
2000
1500
1000
500
0

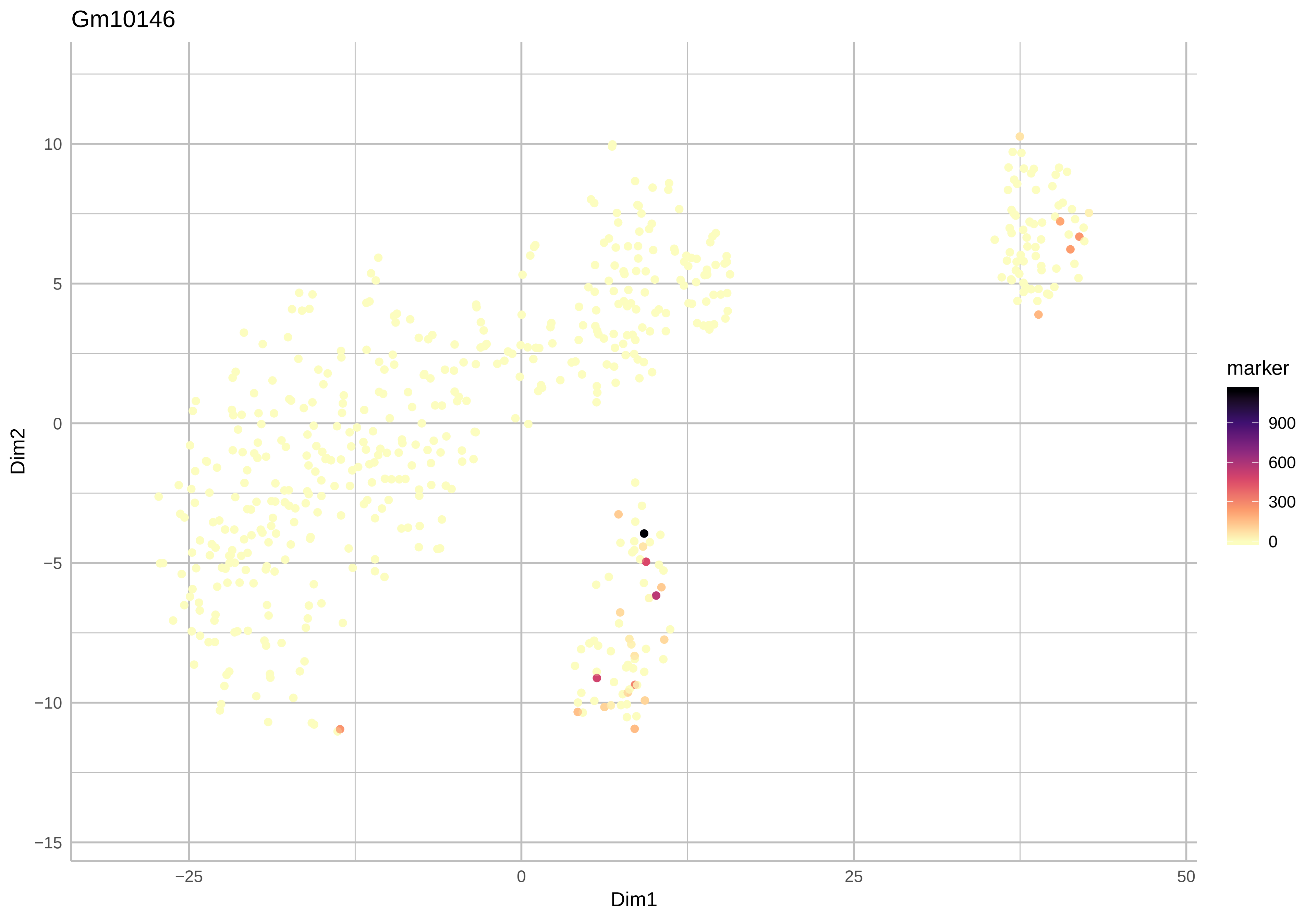
Gm10146
Dim2
Dim1
10
5
0
−5
−10
−15
−25
0
25
50
marker
900
600
300
0

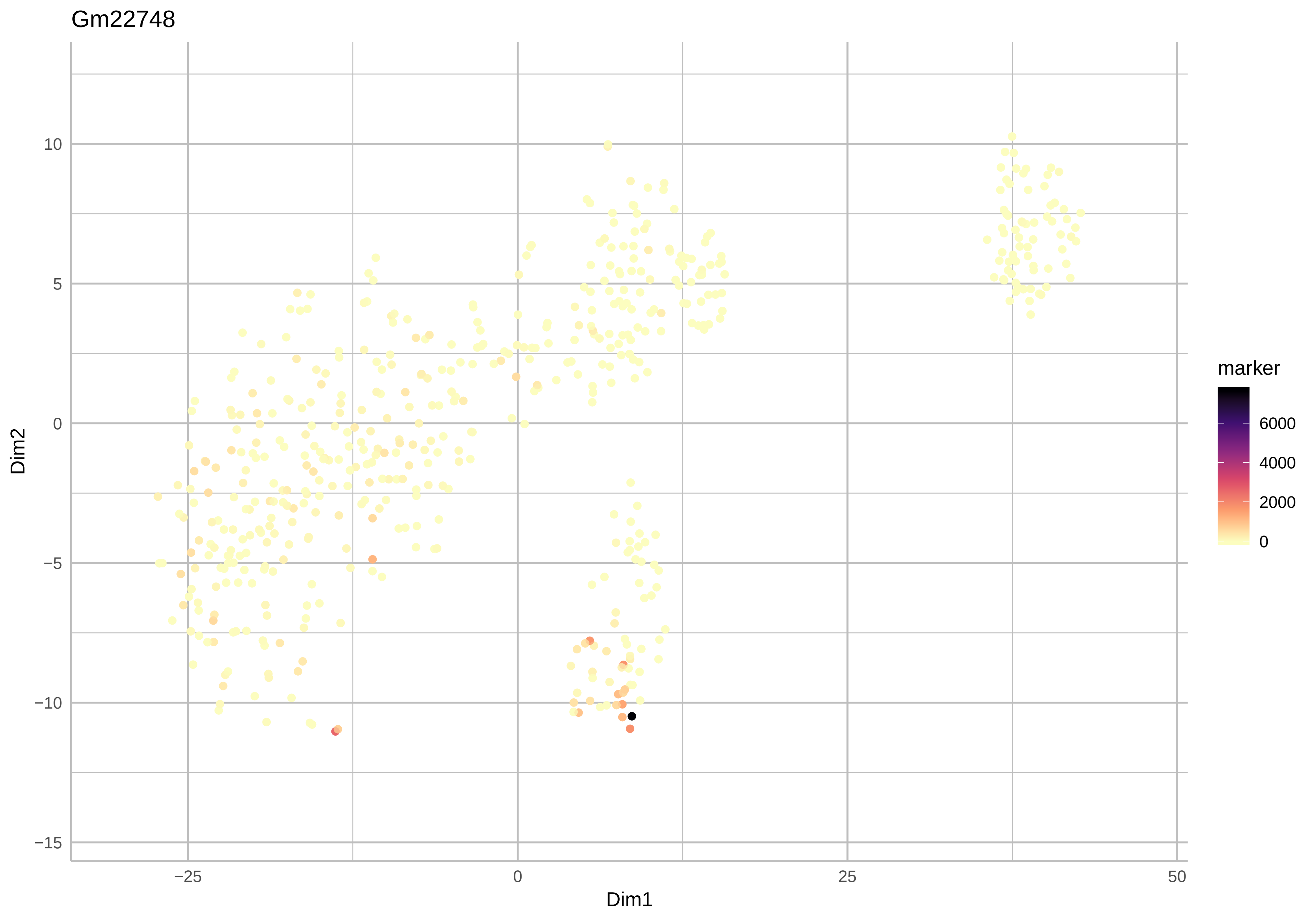

Gm22748
Dim2
Dim1
10
5
0
−5
−10
−15
−25
0
25
50
marker
6000
4000
2000
0

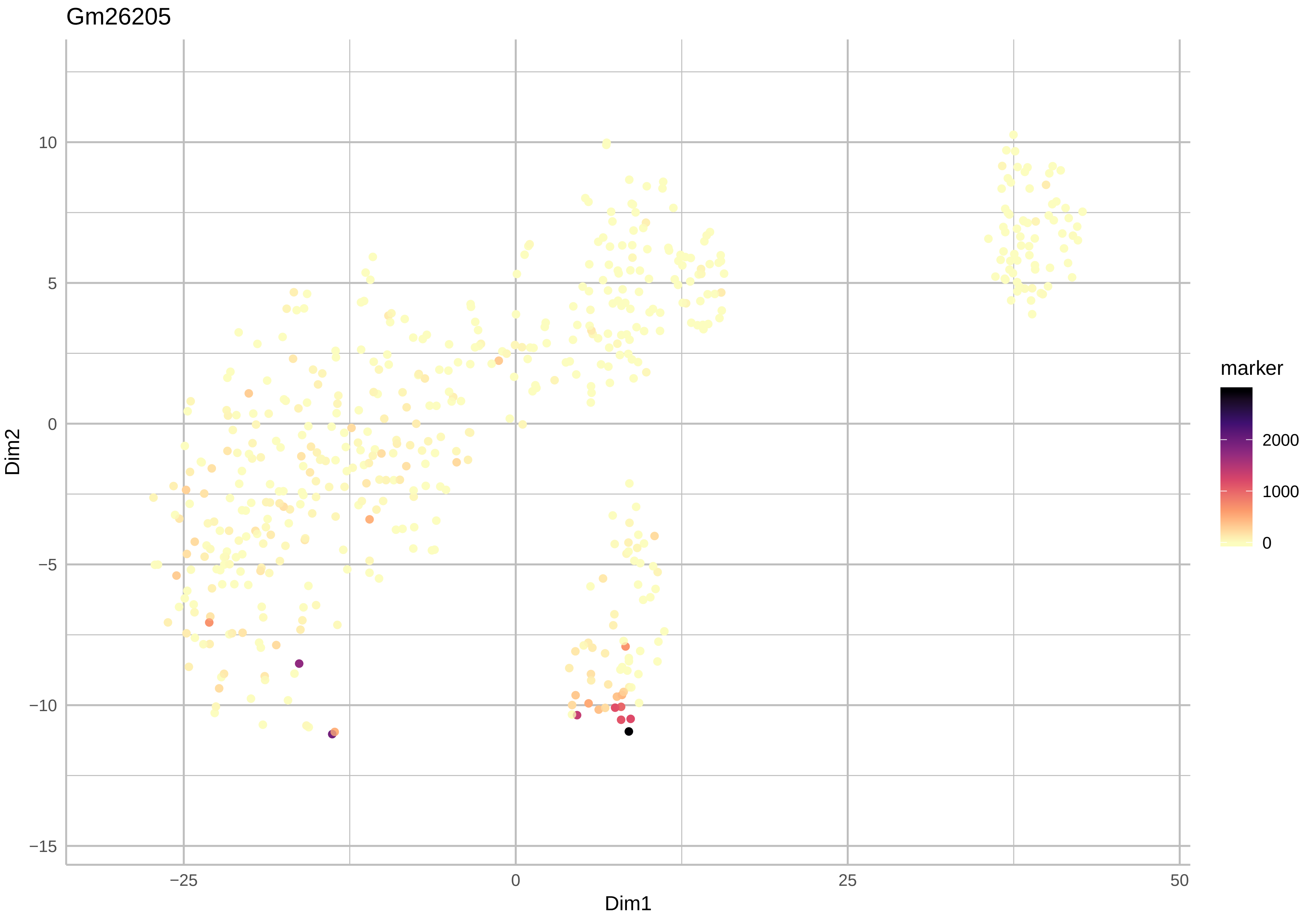

Gm26205
Dim2
Dim1
marker
2000
1000
0
10
5
0
−5
−10
−15
−25
0
25
50

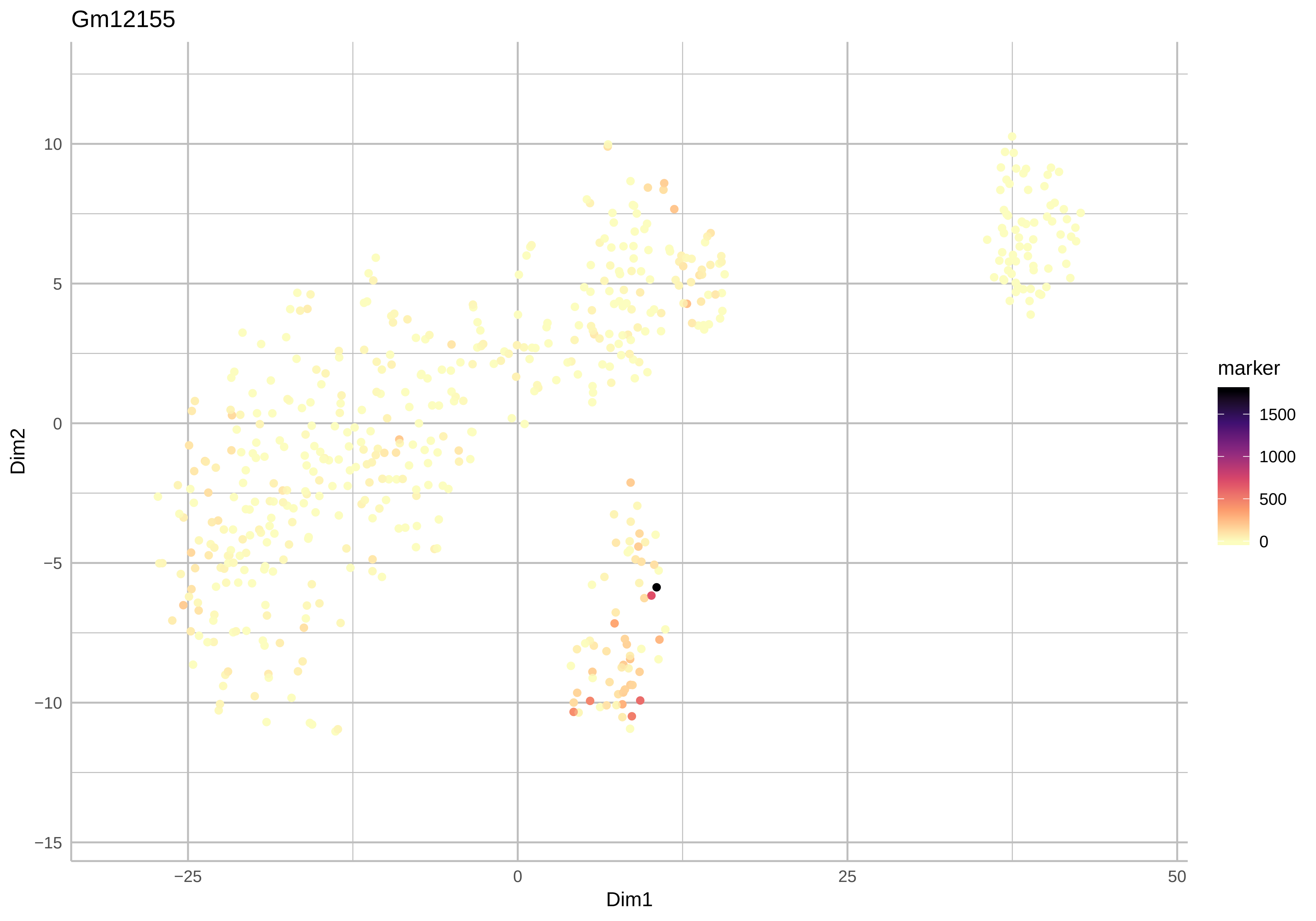
Gm12155
Dim2
Dim1
10
5
0
−5
−10
−15
−25
0
25
50
marker
1500
1000
500
0

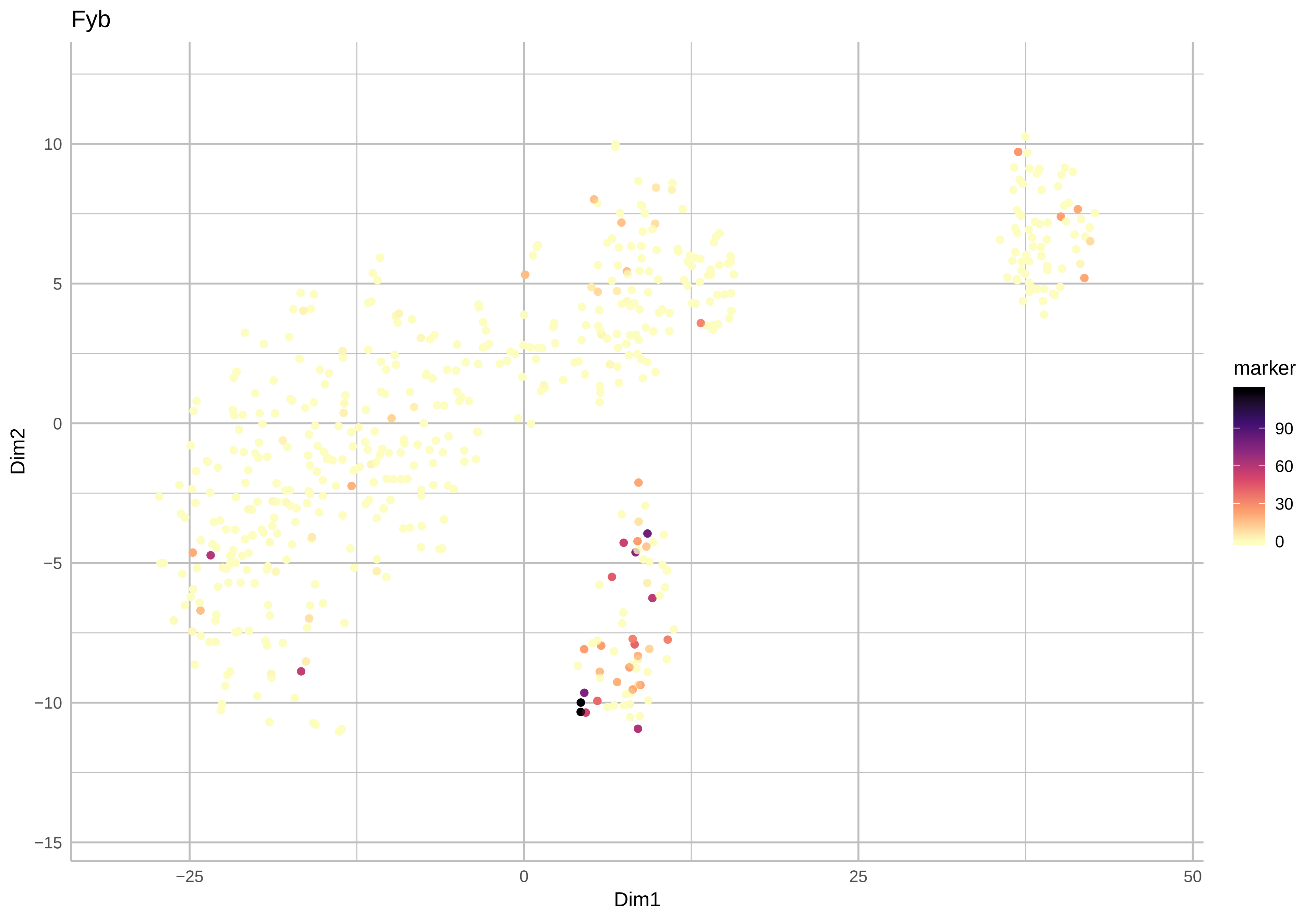

Fyb
Dim2
Dim1
10
5
0
−5
−10
−15
−25
0
25
50
marker
90
60
30
0

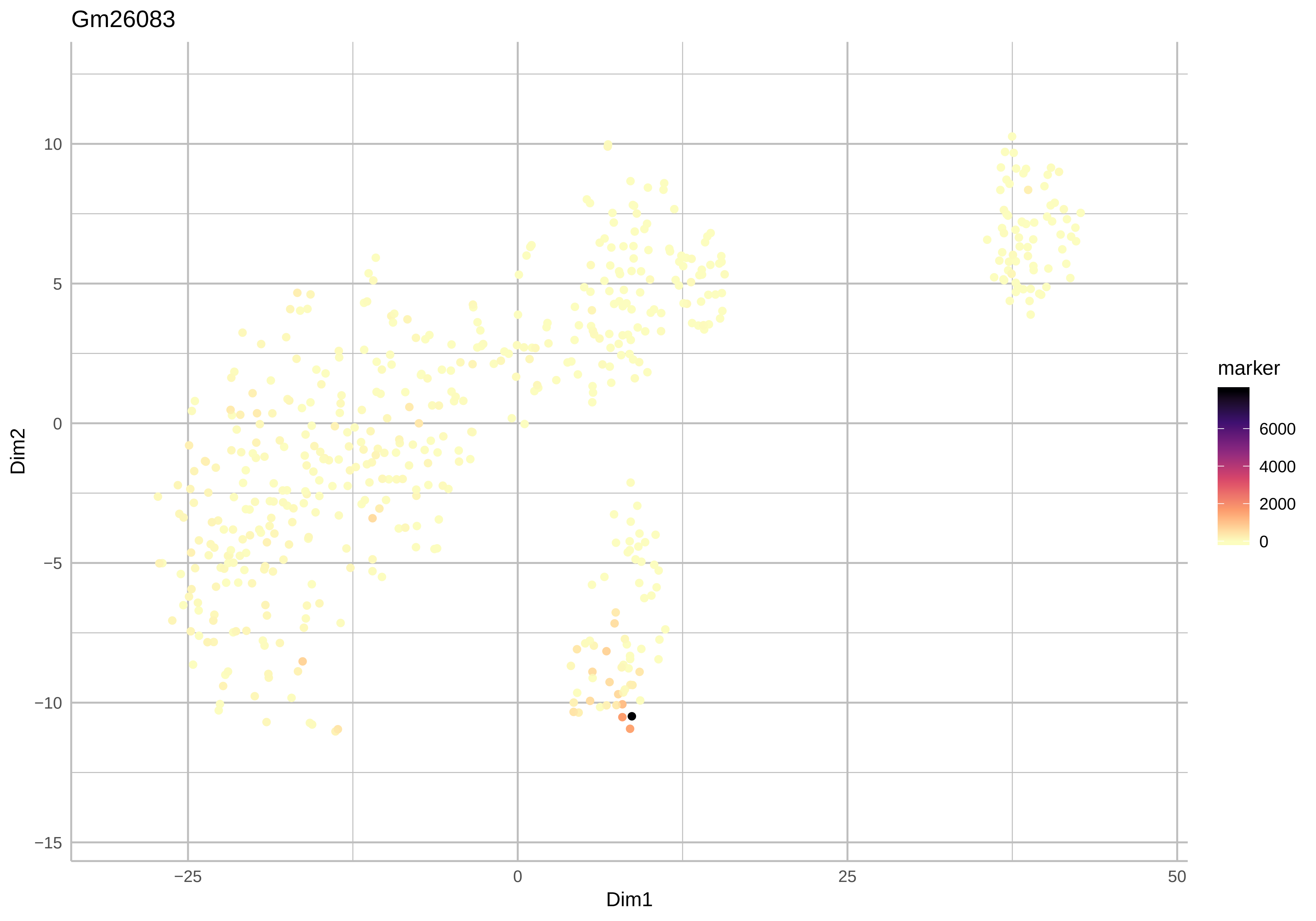
Gm26083
Dim2
Dim1
10
5
0
−5
−10
−15
−25
0
25
50
marker
6000
4000
2000
0

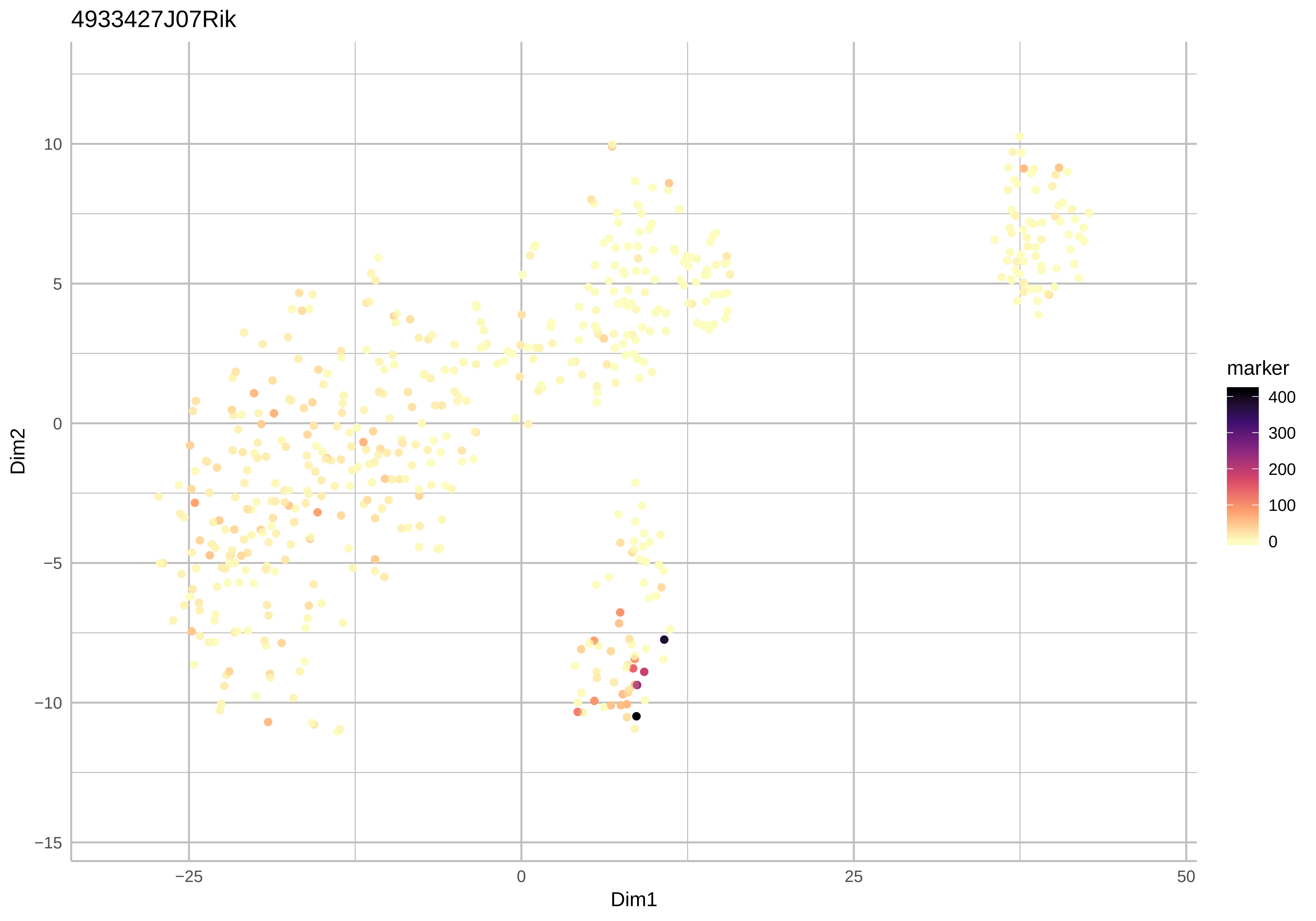
4933427J07Rik
Dim2
Dim1
10
5
0
−5
−10
−15
−25
0
25
50
marker
400
300
200
100
0

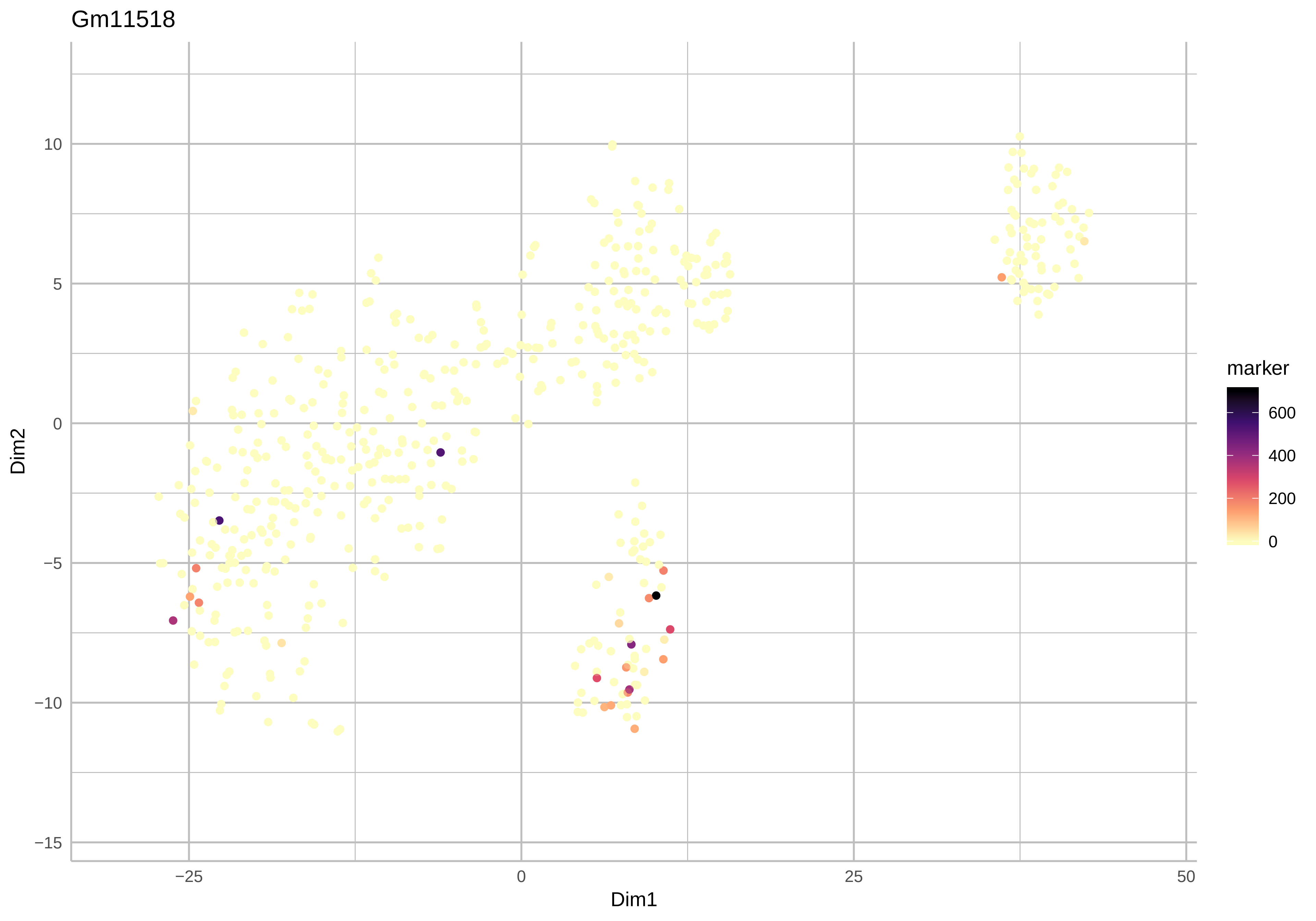

Gm11518
10
5
0
−5
−10
−15
Dim2
−25
0
25
50
Dim1
marker
600
400
200
0

Gm11400

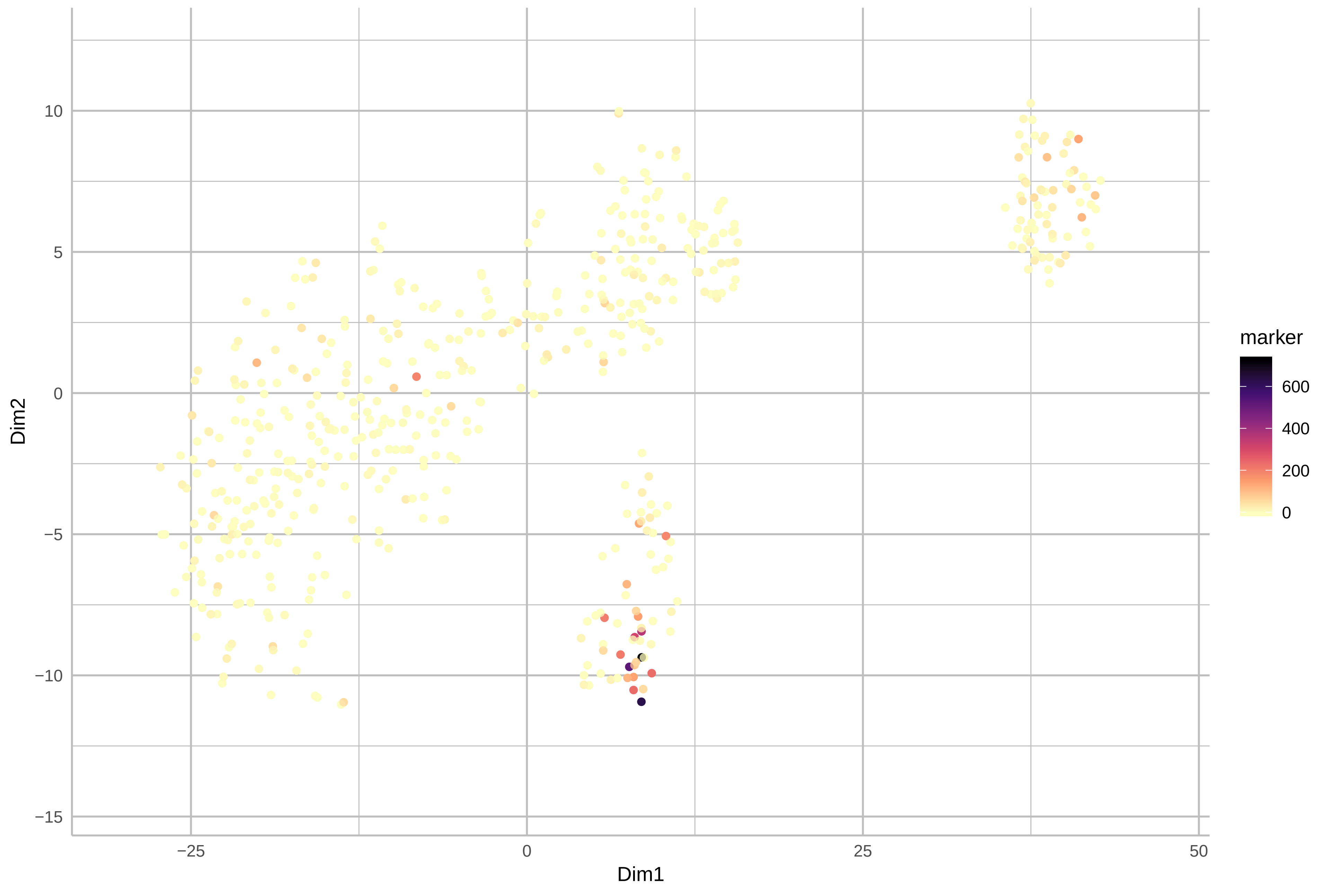

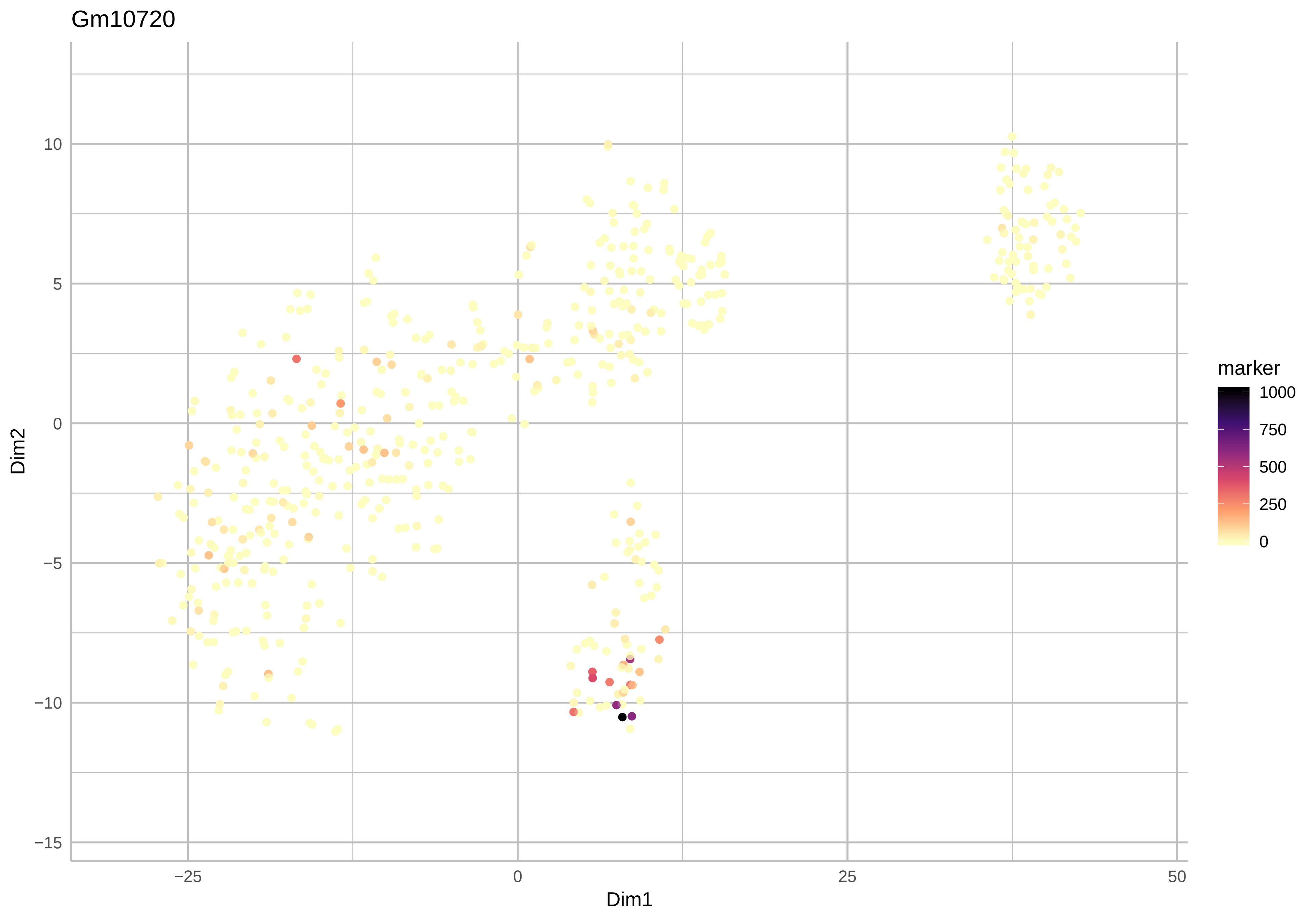
Gm10720
Dim2
Dim1
10
5
0
−5
−10
−15
−25
0
25
50
marker
1000
750
500
250
0

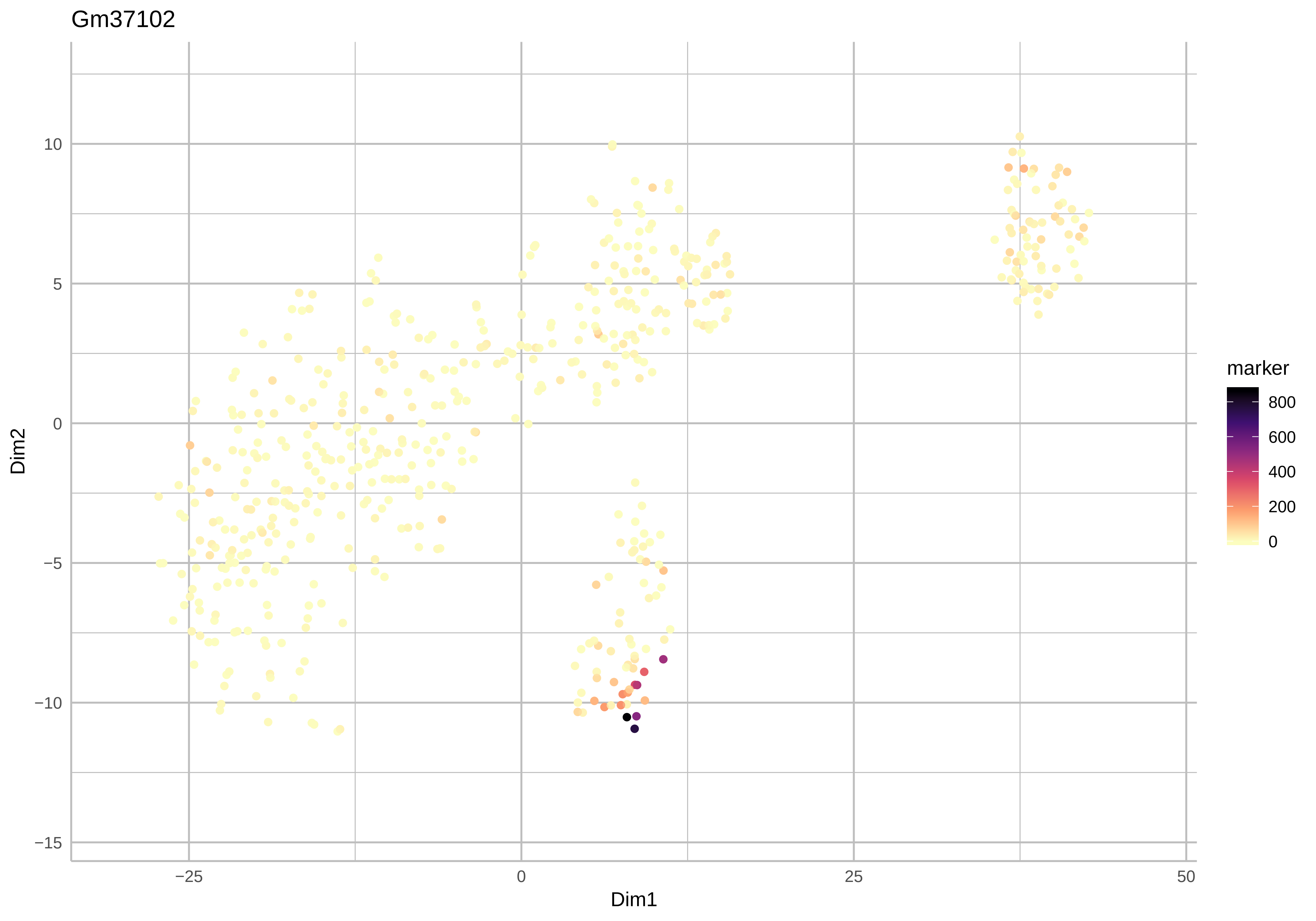

Gm37102
10
5
0
−5
−10
−15
−25
0
25
50
Dim1
Dim2
marker
800
600
400
200
0

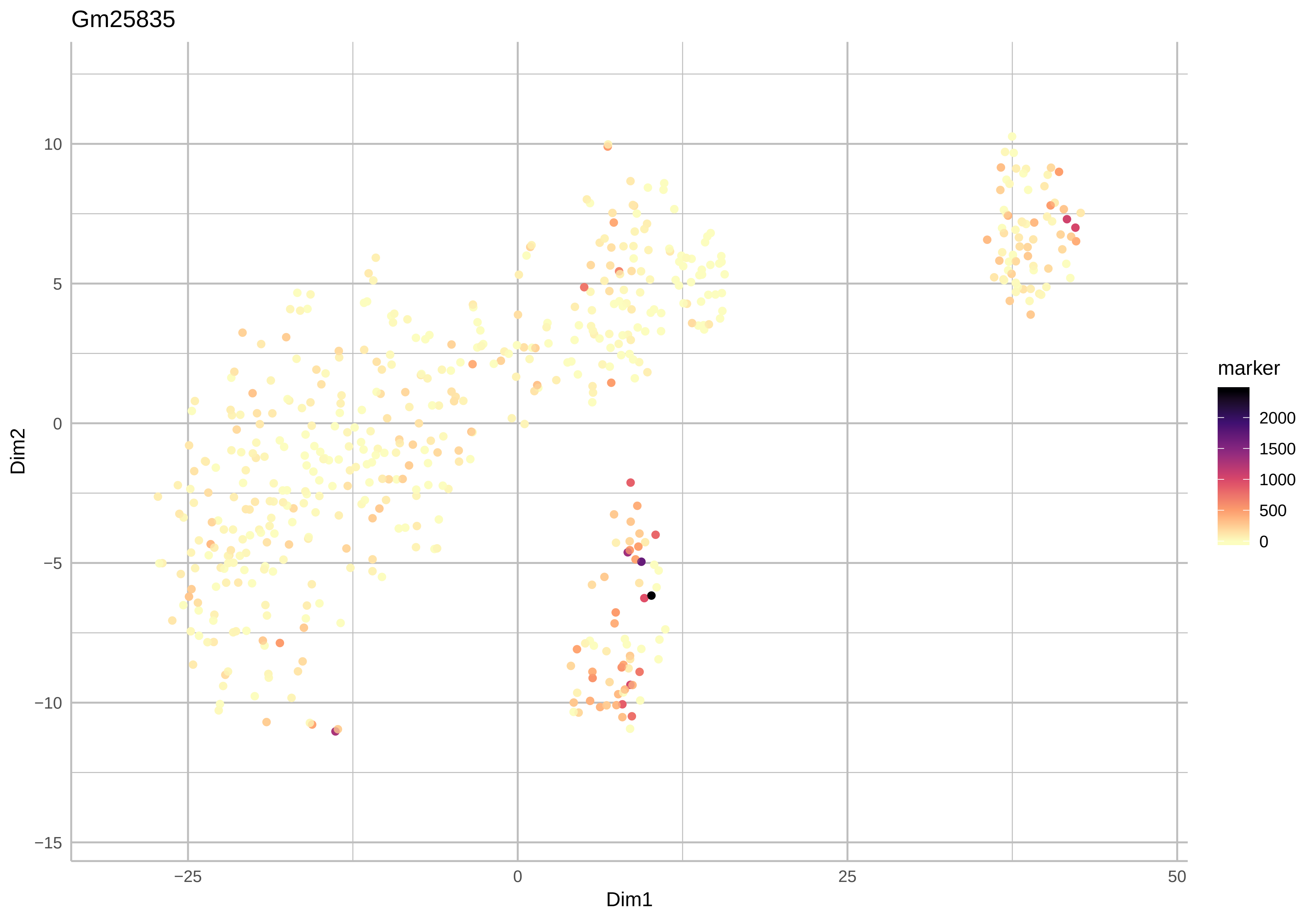

Gm25835
10
5
0
−5
−10
−15
Dim2
−25
0
25
50
Dim1
marker
2000
1500
1000
500
0

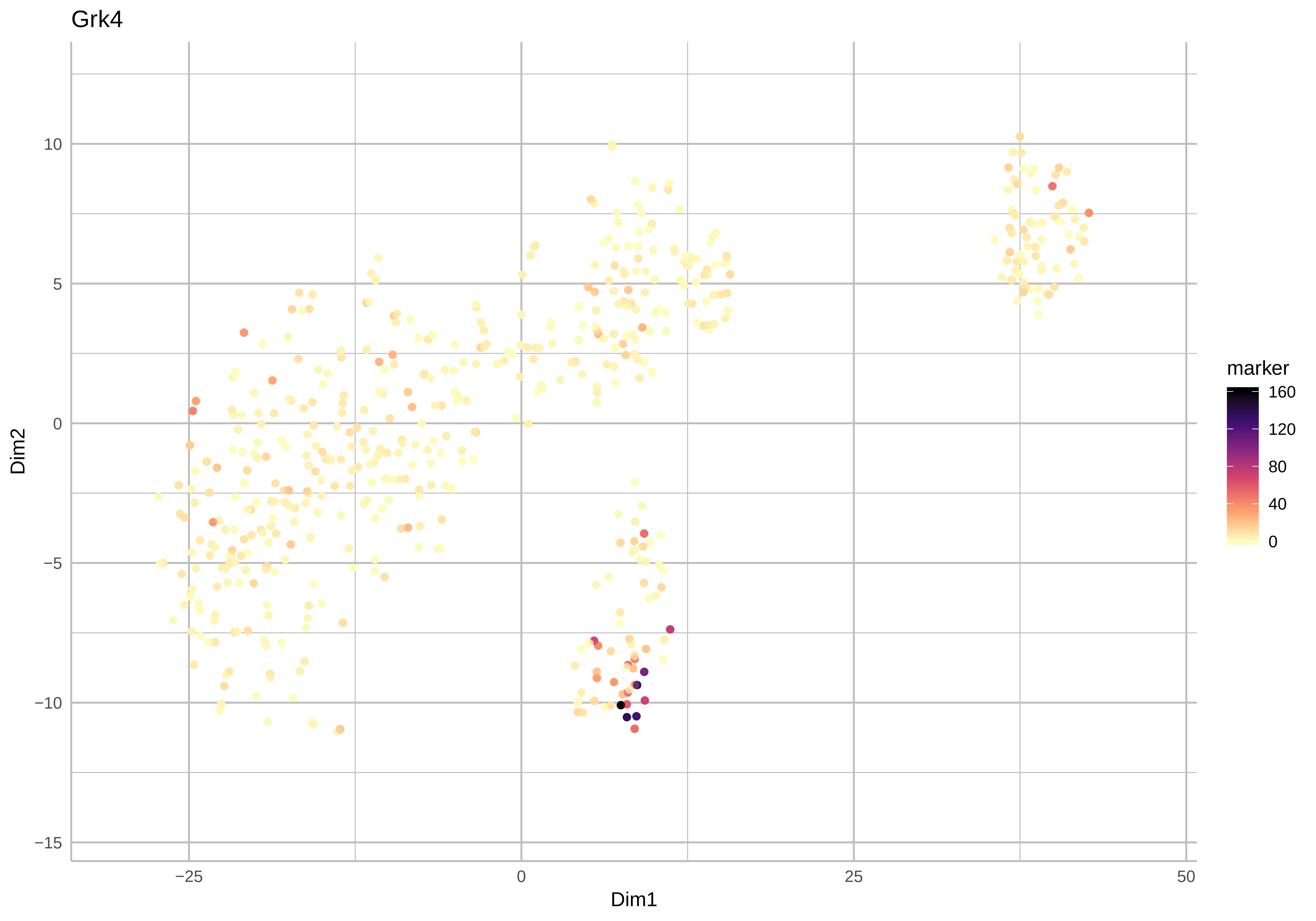

Grk4
Dim2
Dim1
marker
160
120
80
40
0
10
5
0
−5
−10
−15
−25
0
25
50

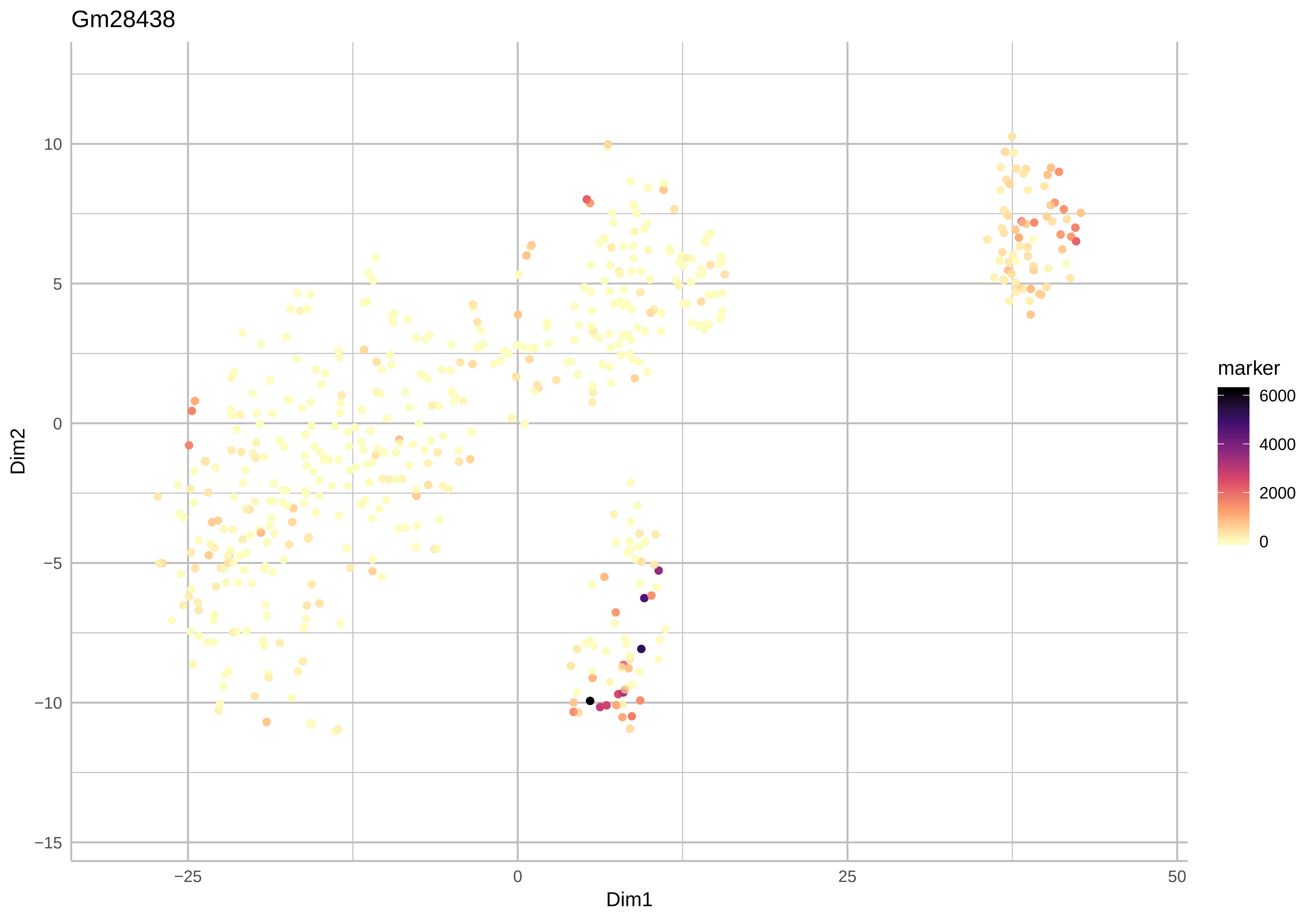

Gm28438
10
5
0
−5
−10
−15
Dim2
−25
0
25
50
Dim1
marker
6000
4000
2000
0

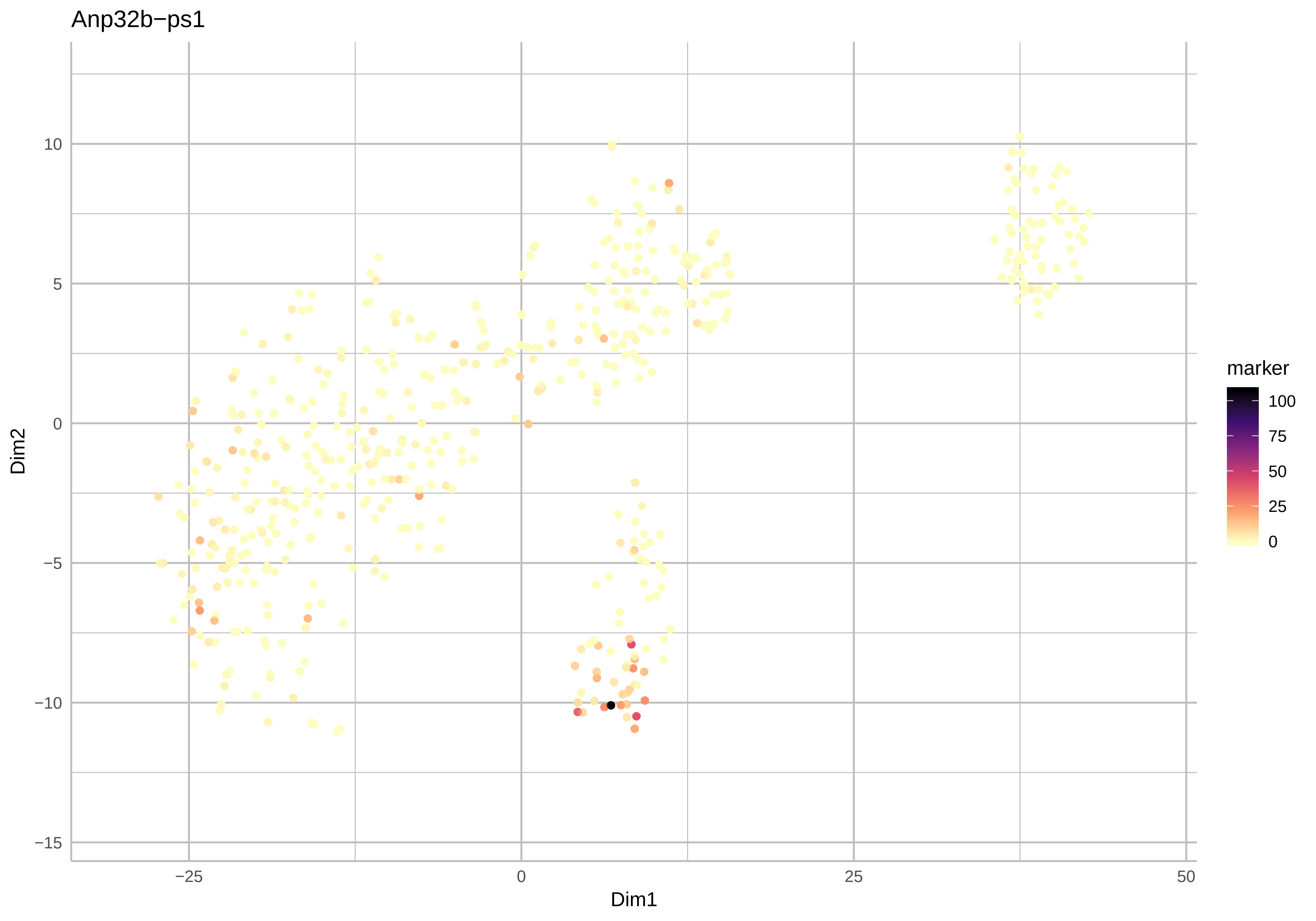
Anp32b−ps1
Dim2
Dim1
10
5
0
−5
−10
−15
−25
0
25
50
marker
100
75
50
25
0

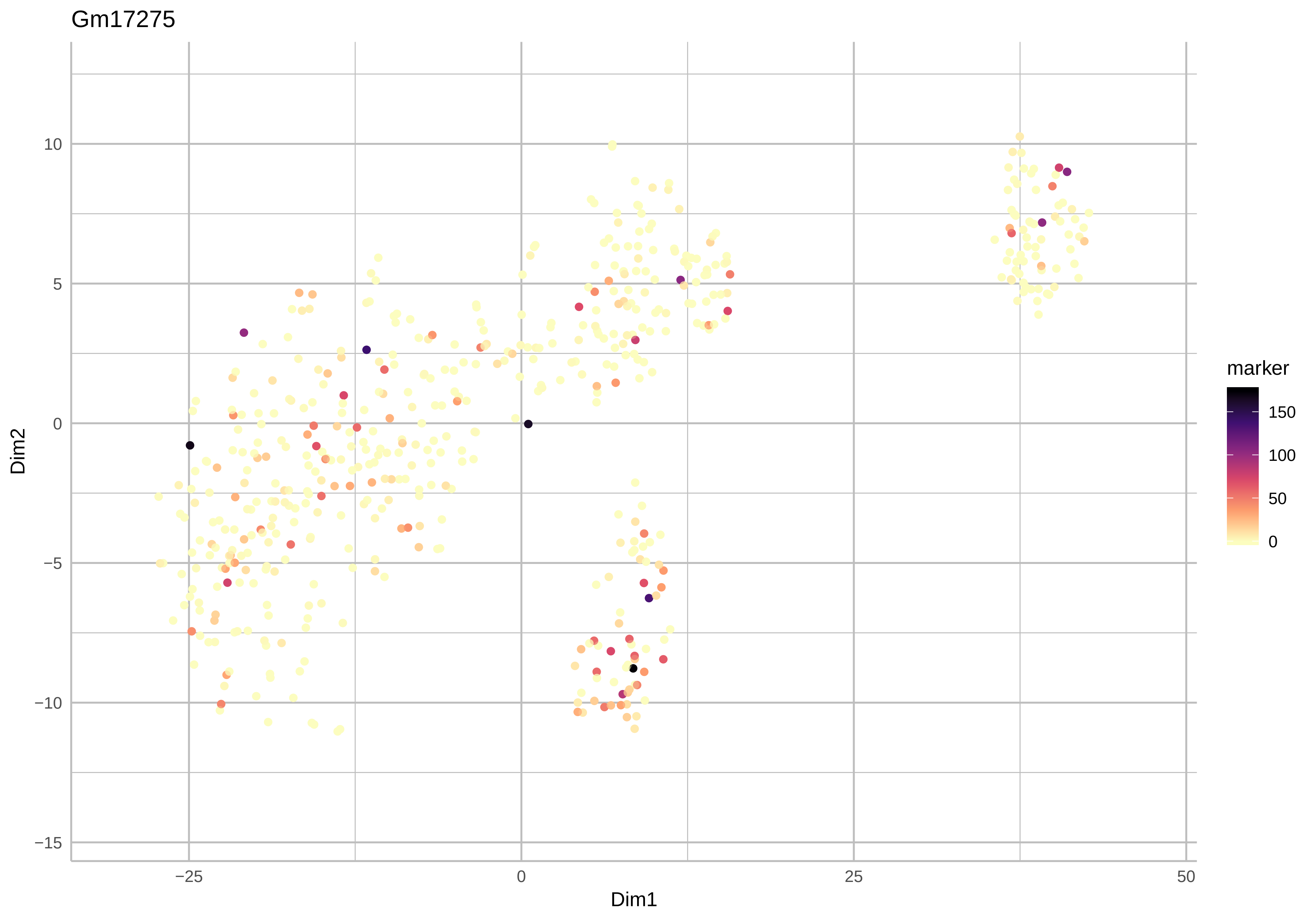
Gm17275
10
5
0
−5
−10
−15
−25
0
25
50
Dim1
Dim2
marker
150
100
50
0

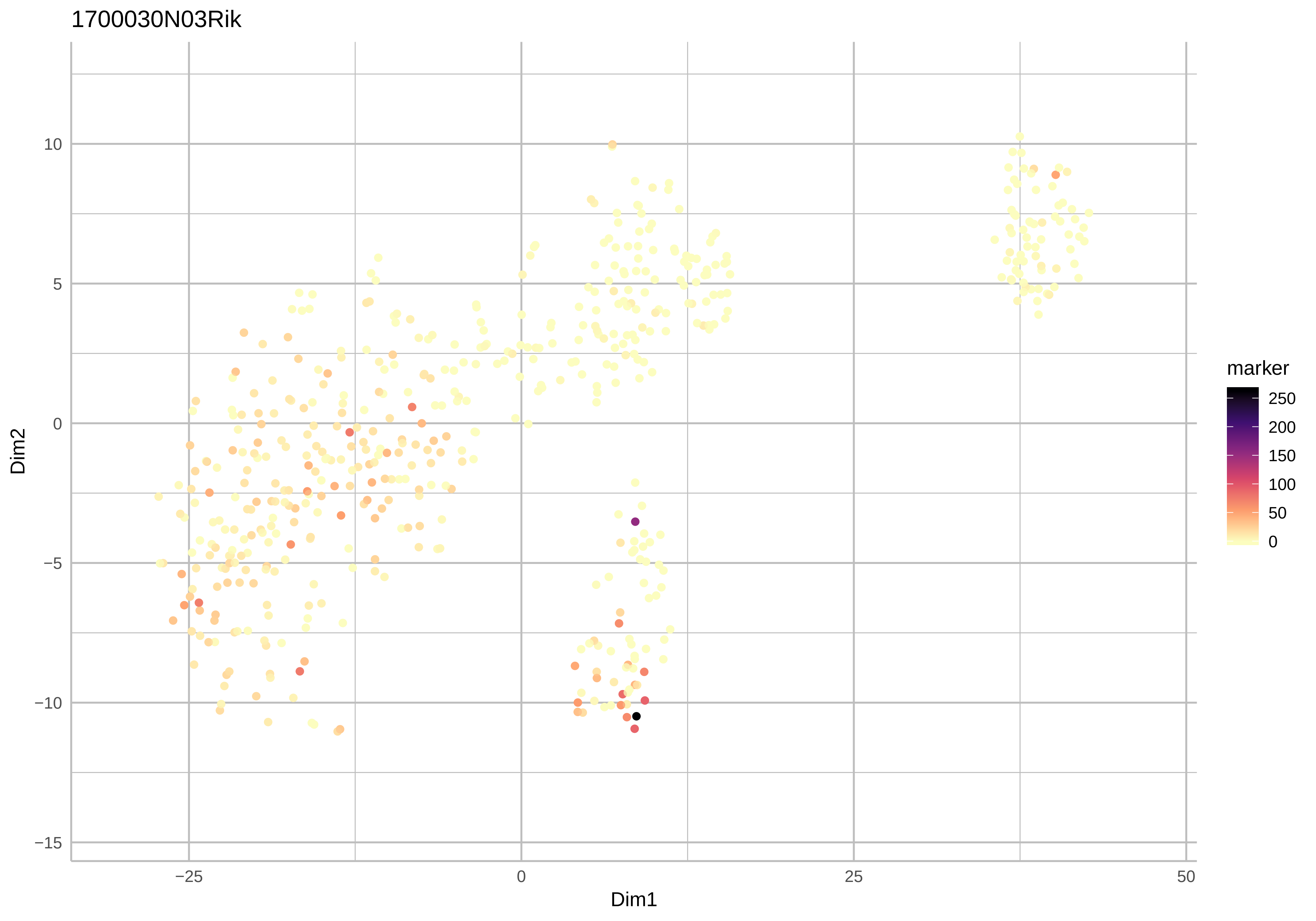

1700030N03Rik
Dim2
Dim1
10
5
0
−5
−10
−15
−25
0
25
50
marker
250
200
150
100
50
0

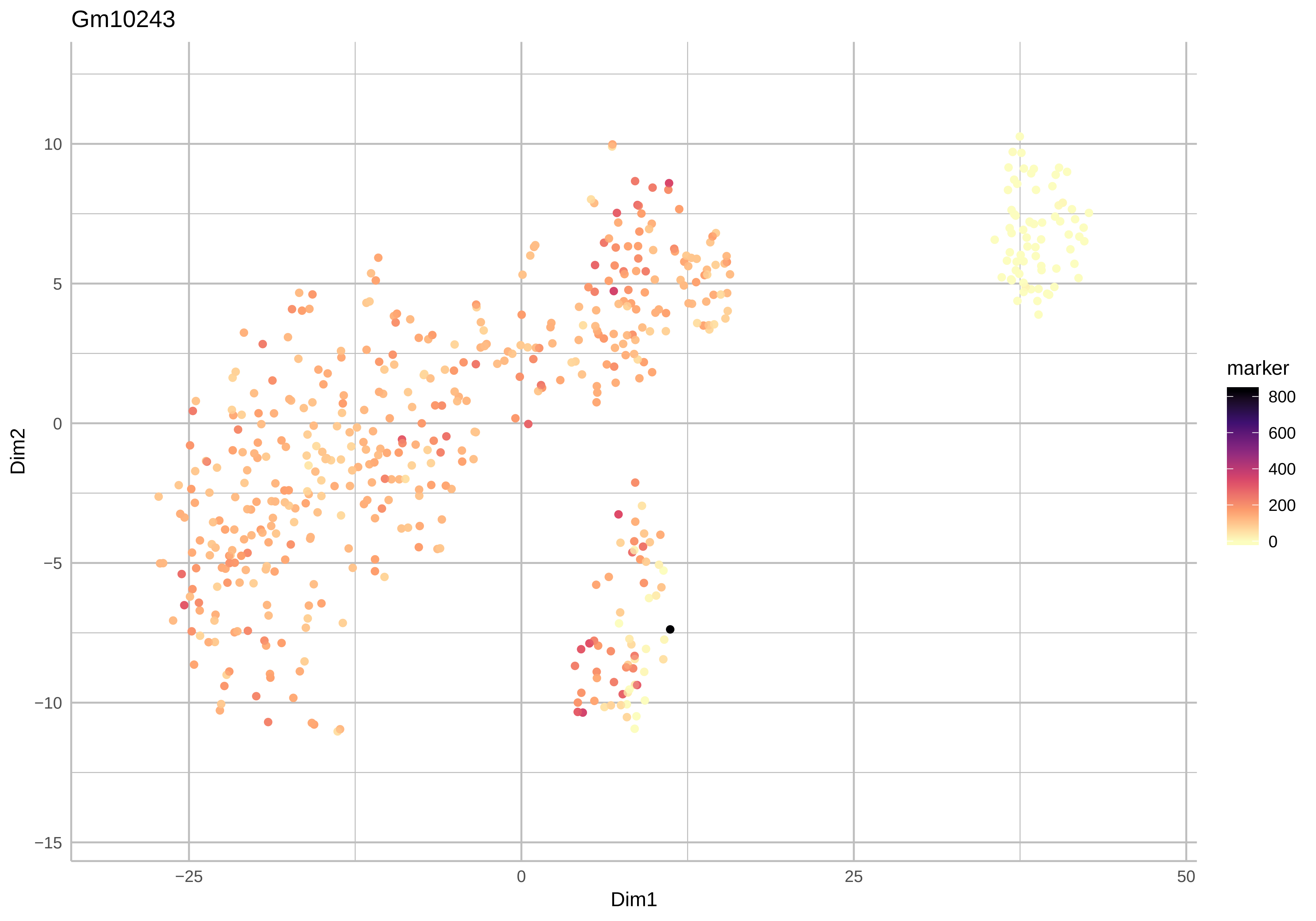
Gm10243
Dim2
Dim1
10
5
0
−5
−10
−15
−25
0
25
50
marker
800
600
400
200
0

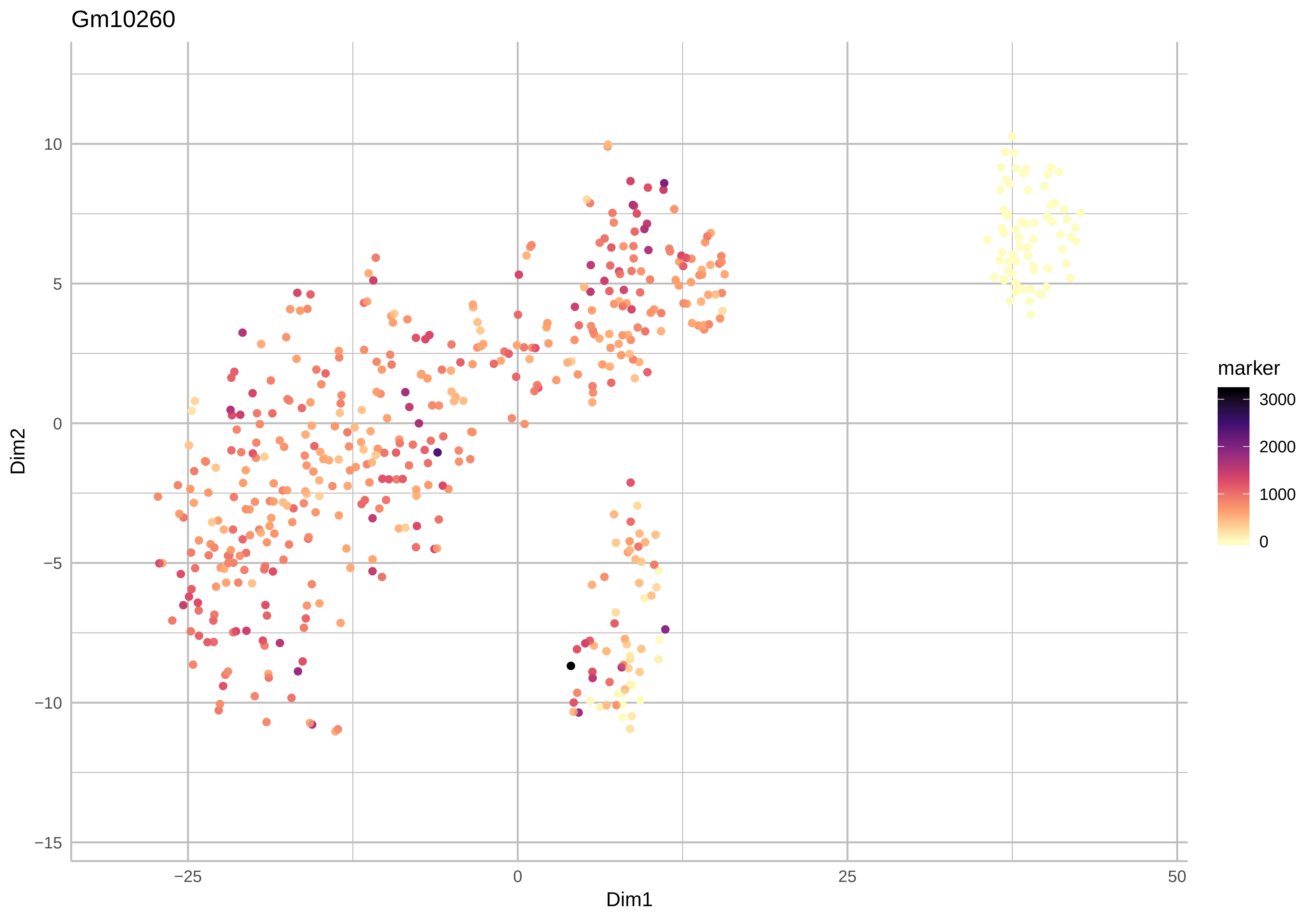
Gm10260
Dim2
Dim1
10
5
0
−5
−10
−15
−25
0
25
50
marker
3000
2000
1000
0

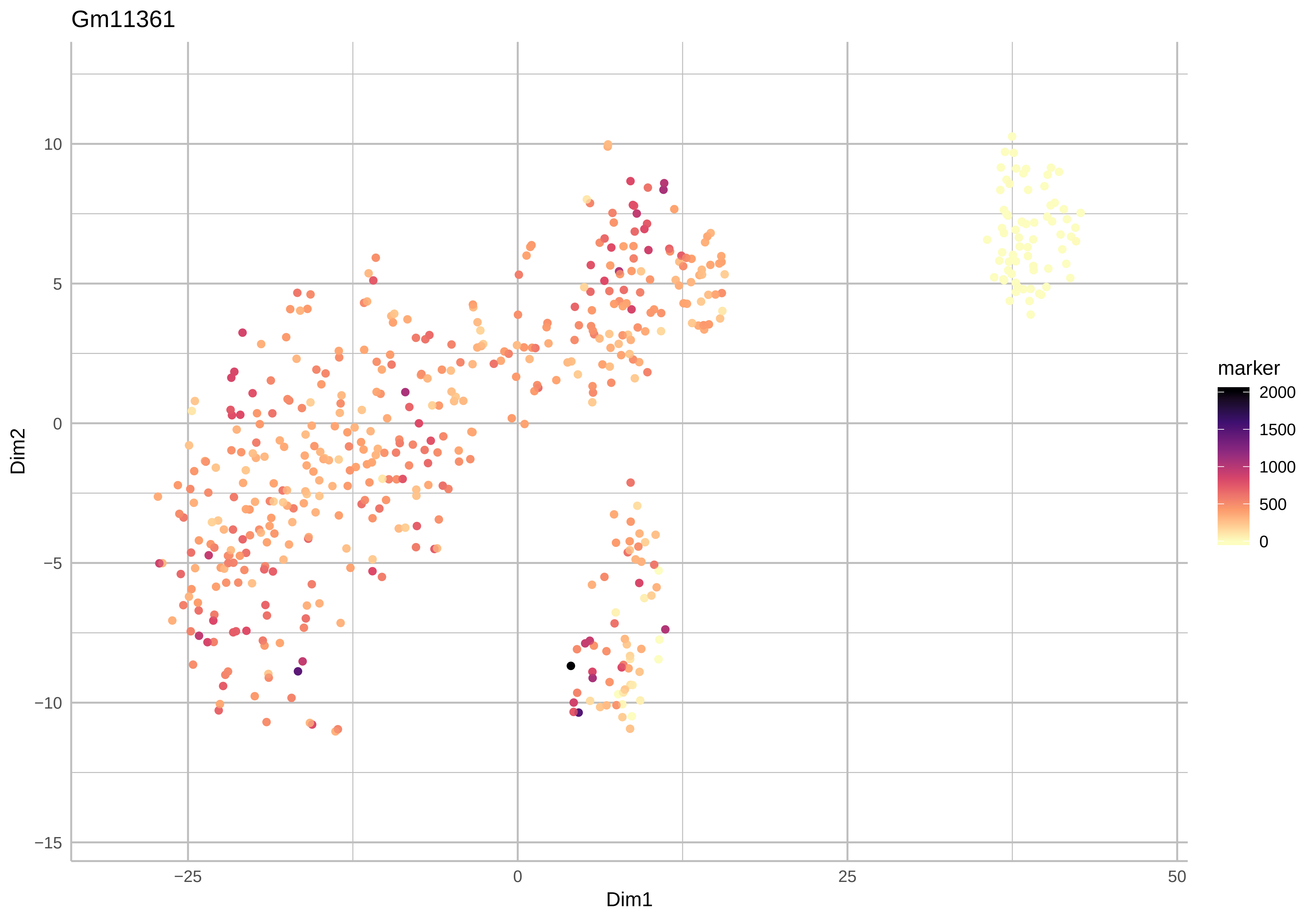
Gm11361
Dim2
Dim1
marker
2000
1500
1000
500
0
10
5
0
−5
−10
−15
−25
0
25
50

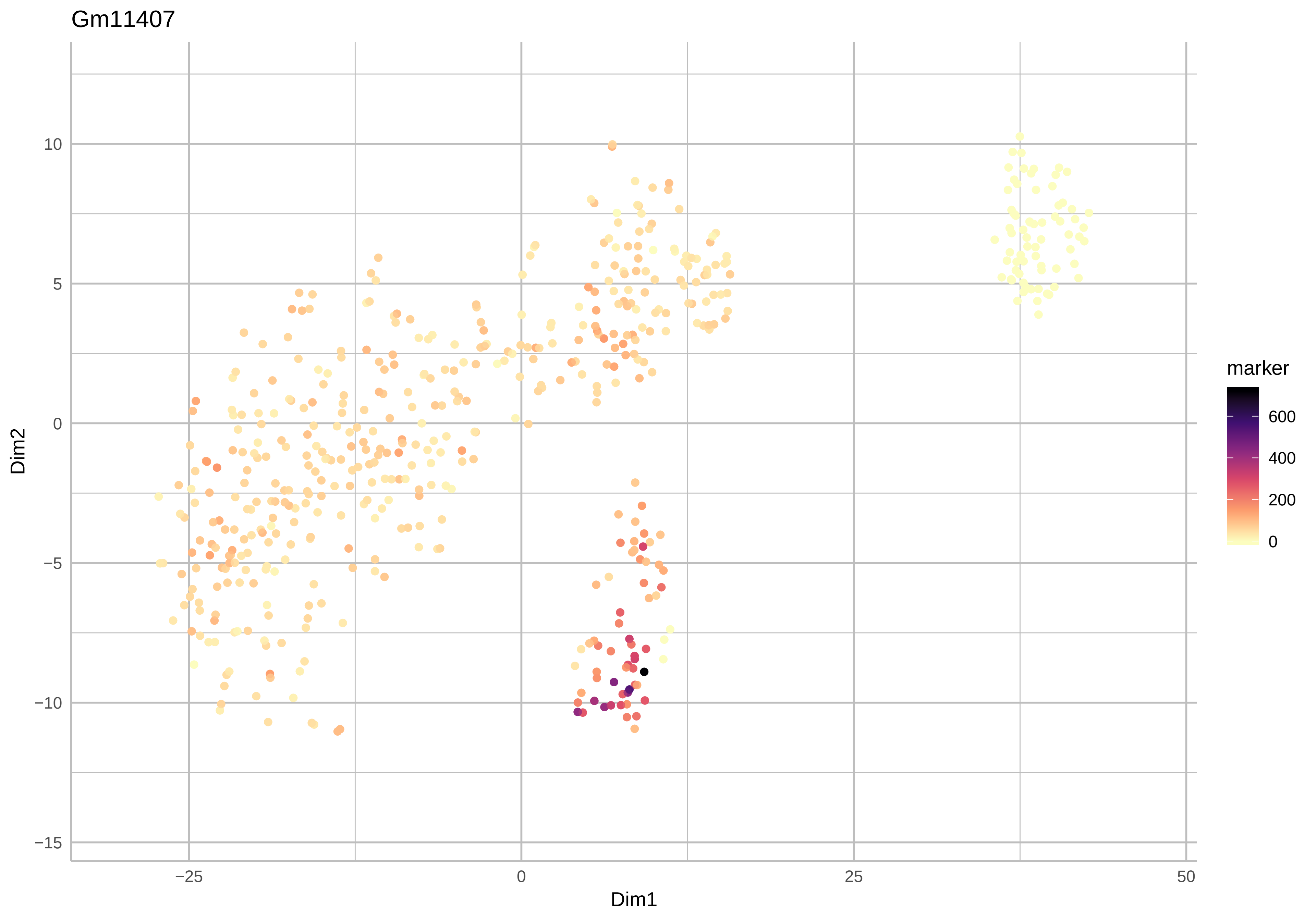

Gm11407
Dim2
Dim1
−25
0
25
50
10
5
0
−5
−10
−15
marker
600
400
200
0

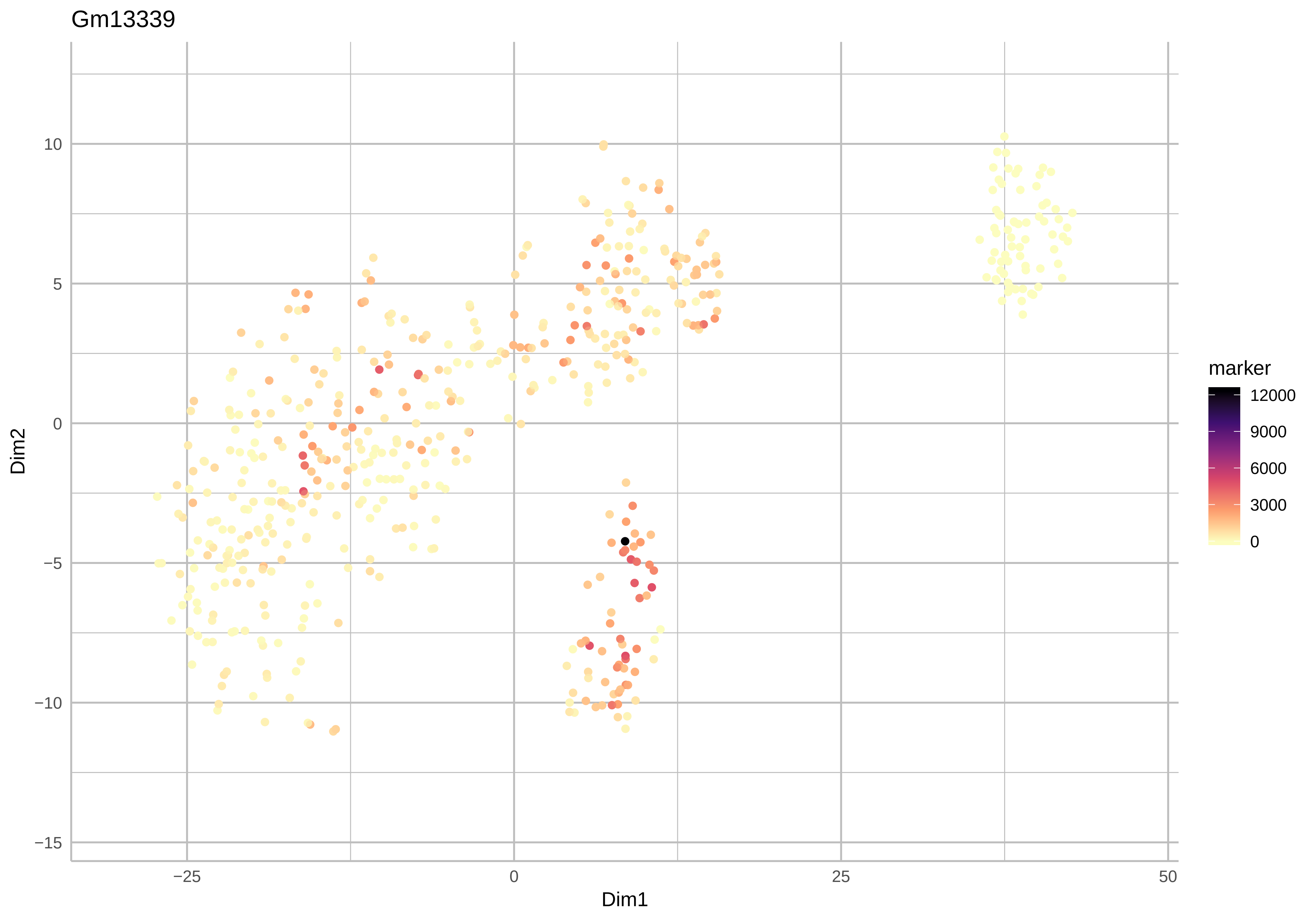
Gm13339
Dim2
Dim1
marker
12000
9000
6000
3000
0
10
5
0
−5
−10
−15
−25
0
25
50

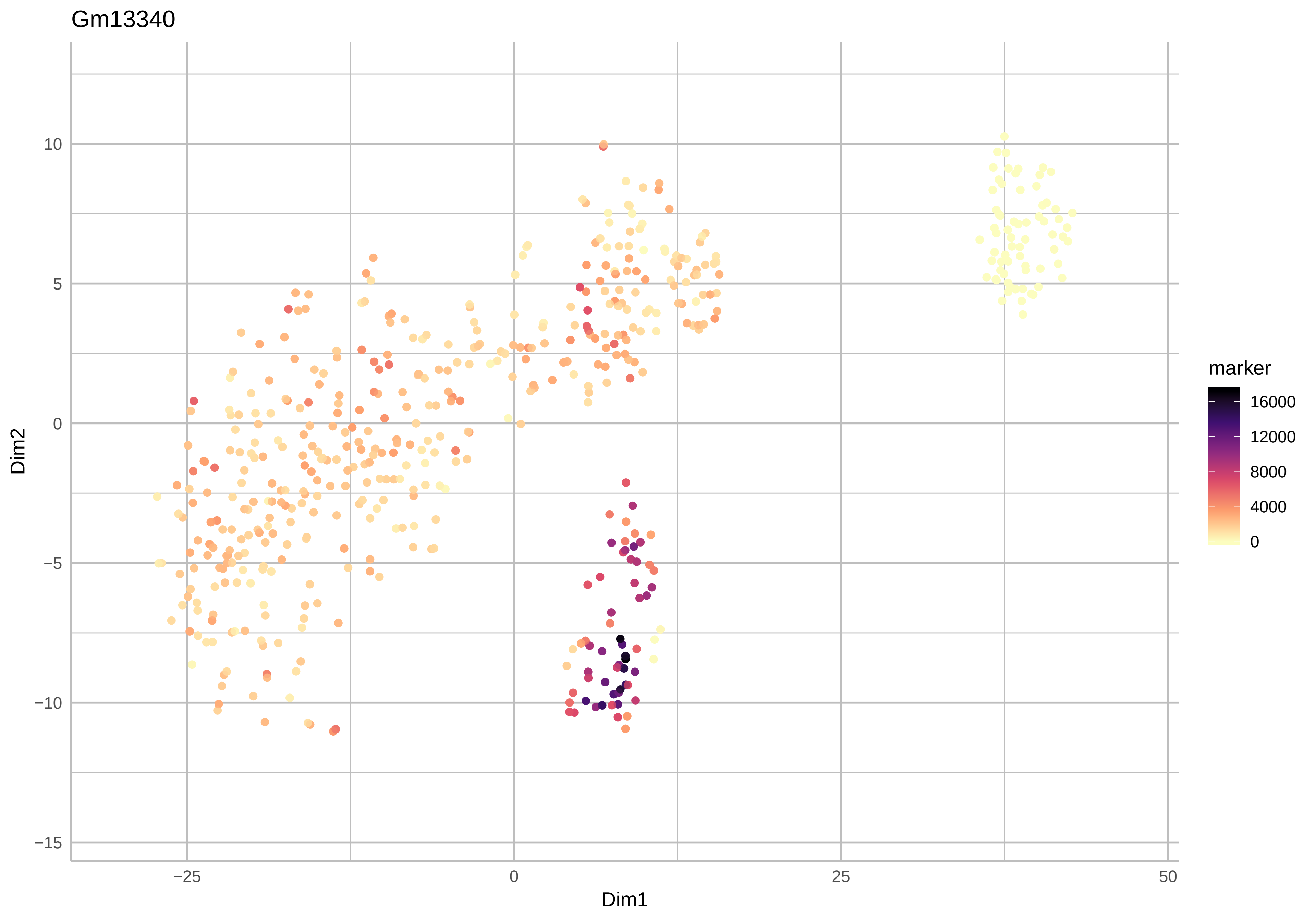

Gm13340
10
5
0
−5
−10
−15
Dim2
−25
0
25
50
Dim1
marker
16000
12000
8000
4000
0

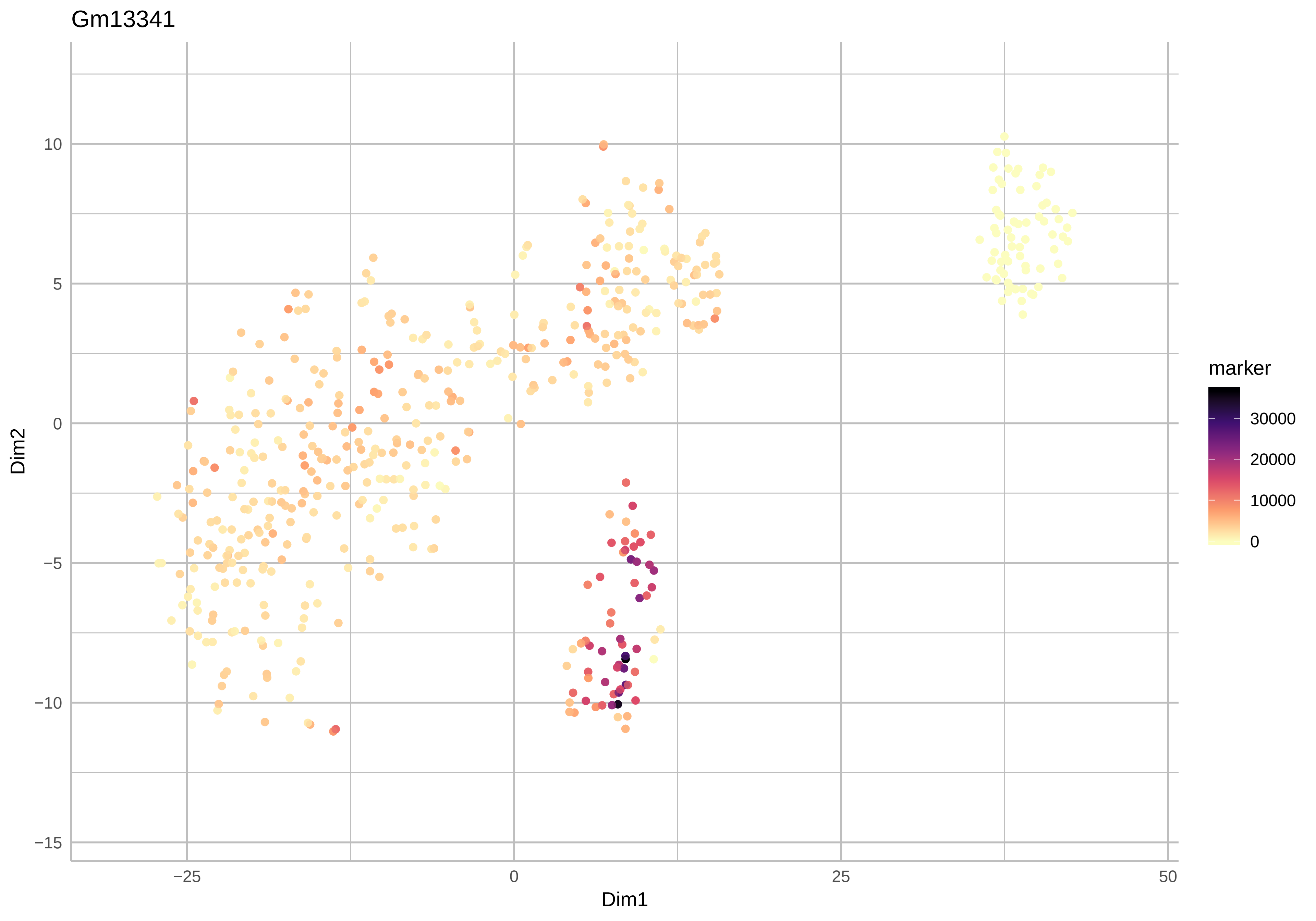

Gm13341
Dim2
10
5
0
−5
−10
−15
Dim1
−25
0
25
50
marker
30000
20000
10000
0

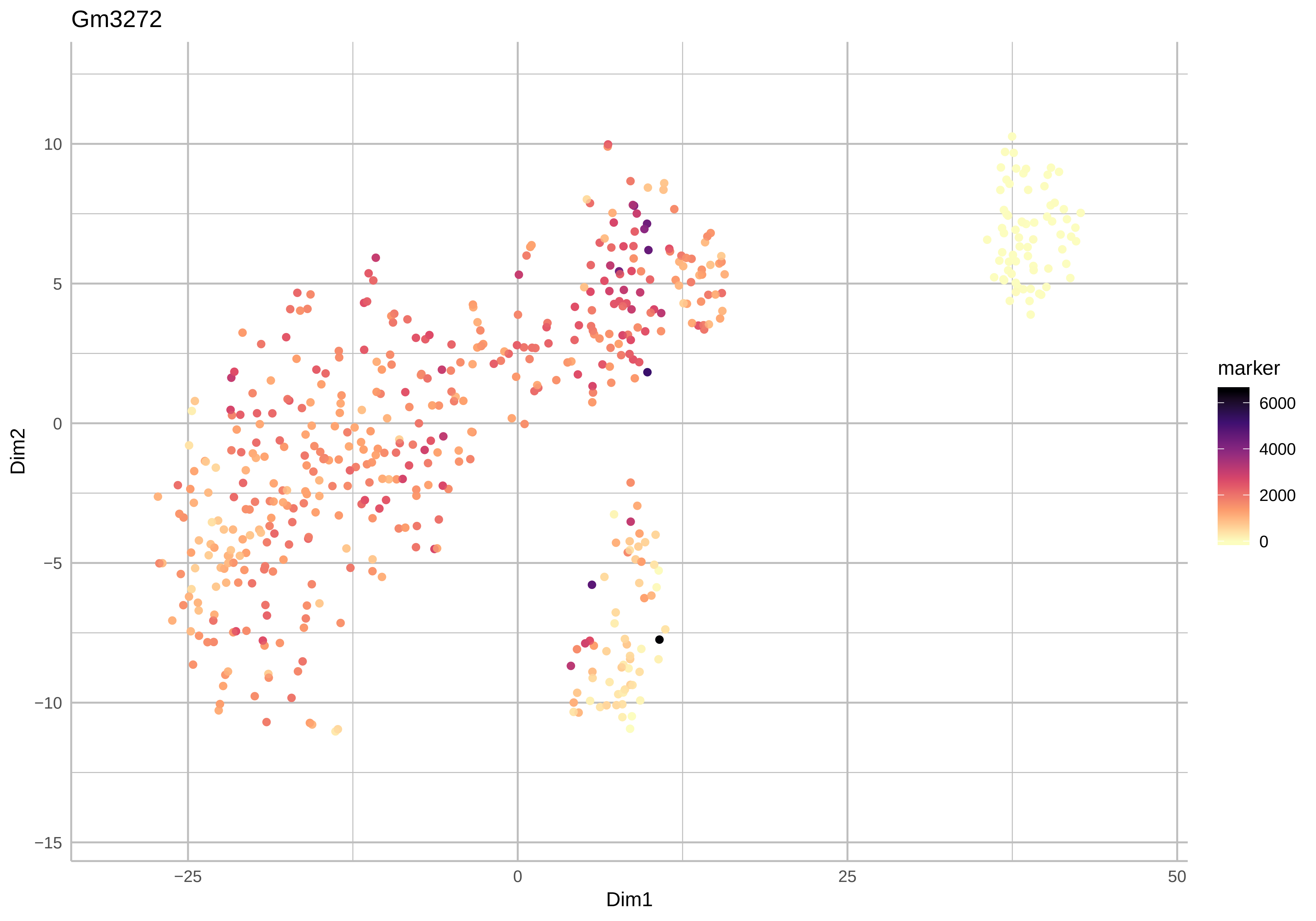

Gm3272
Dim2
Dim1
10
5
0
−5
−10
−15
−25
0
25
50
marker
6000
4000
2000
0

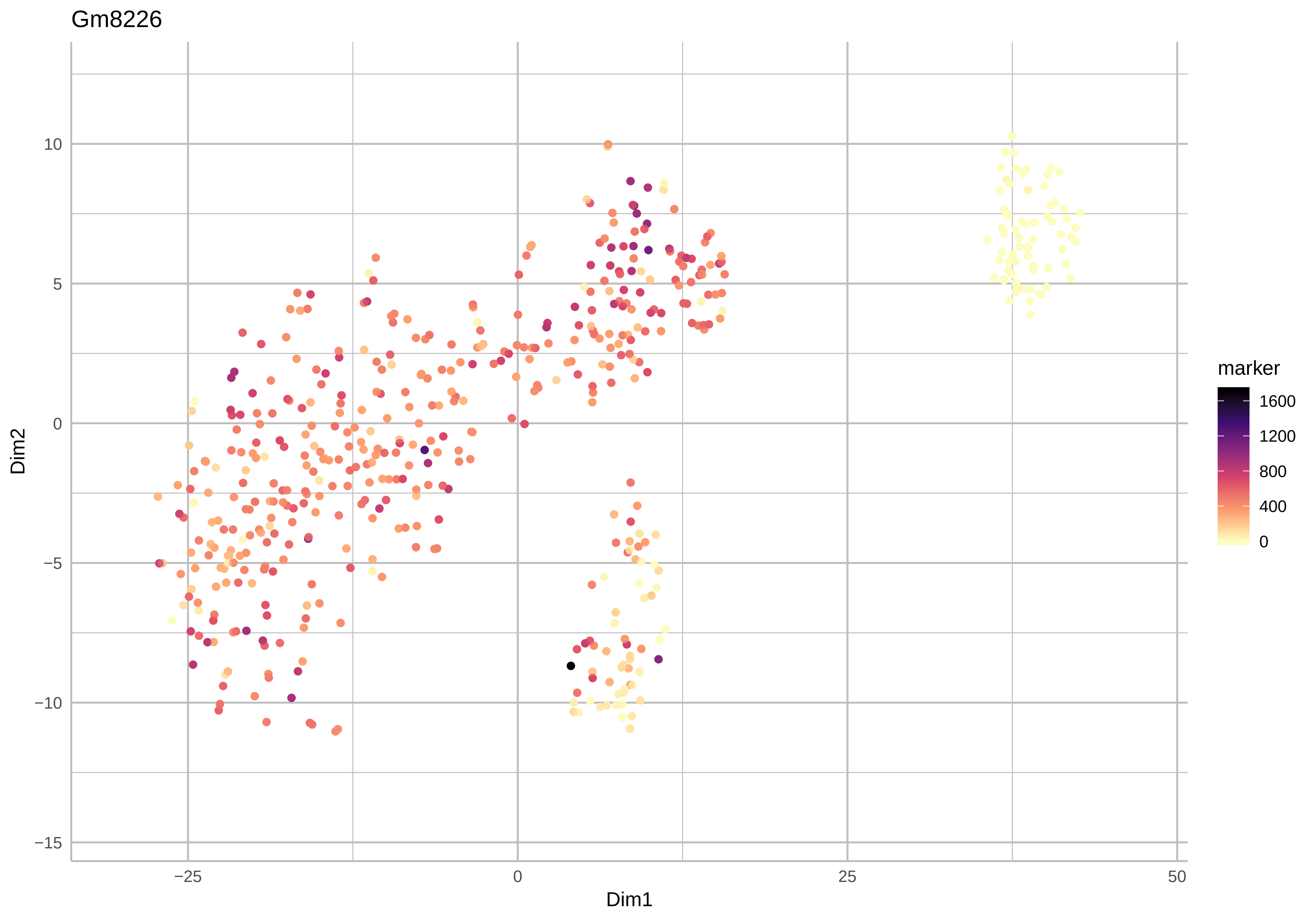
Gm8226
Dim2
Dim1
10
5
0
−5
−10
−15
−25
0
25
50
marker
1600
1200
800
400
0

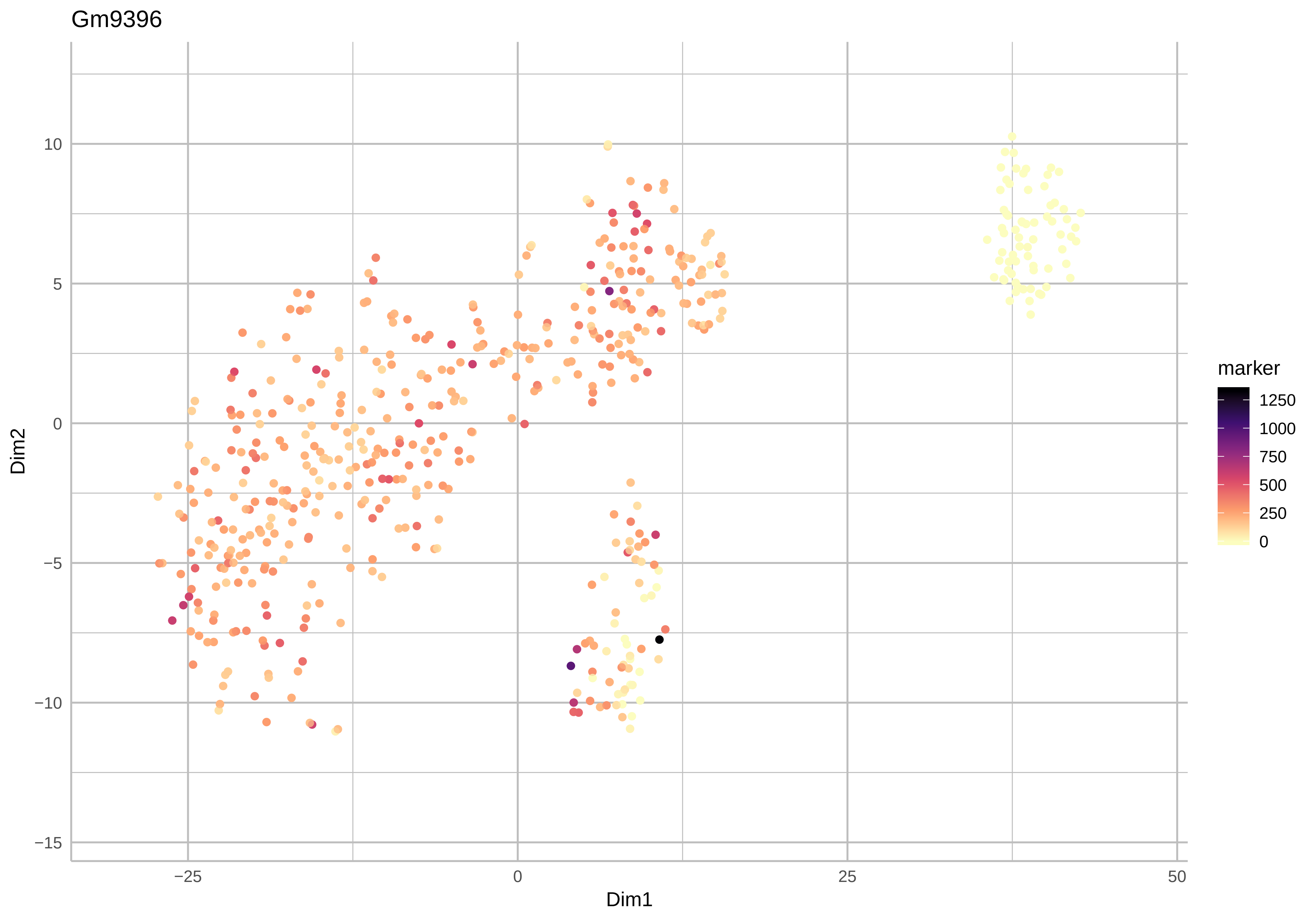
Gm9396
Dim2
Dim1
10
5
0
−5
−10
−15
−25
0
25
50
marker
1250
1000
750
500
250
0

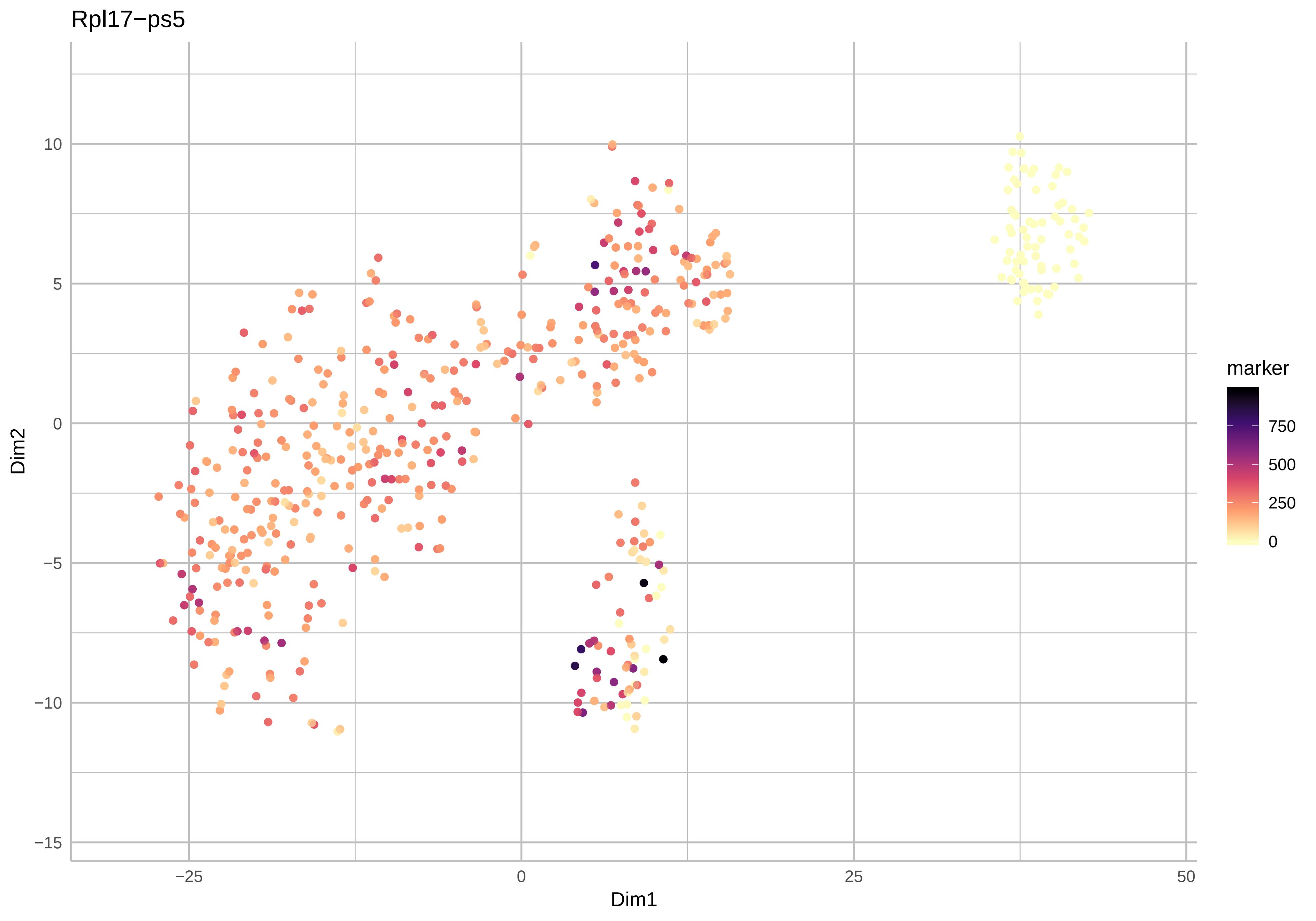

Rpl17−ps5
10
5
0
−5
−10
−15
Dim2
−25
0
25
50
Dim1
marker
750
500
250
0

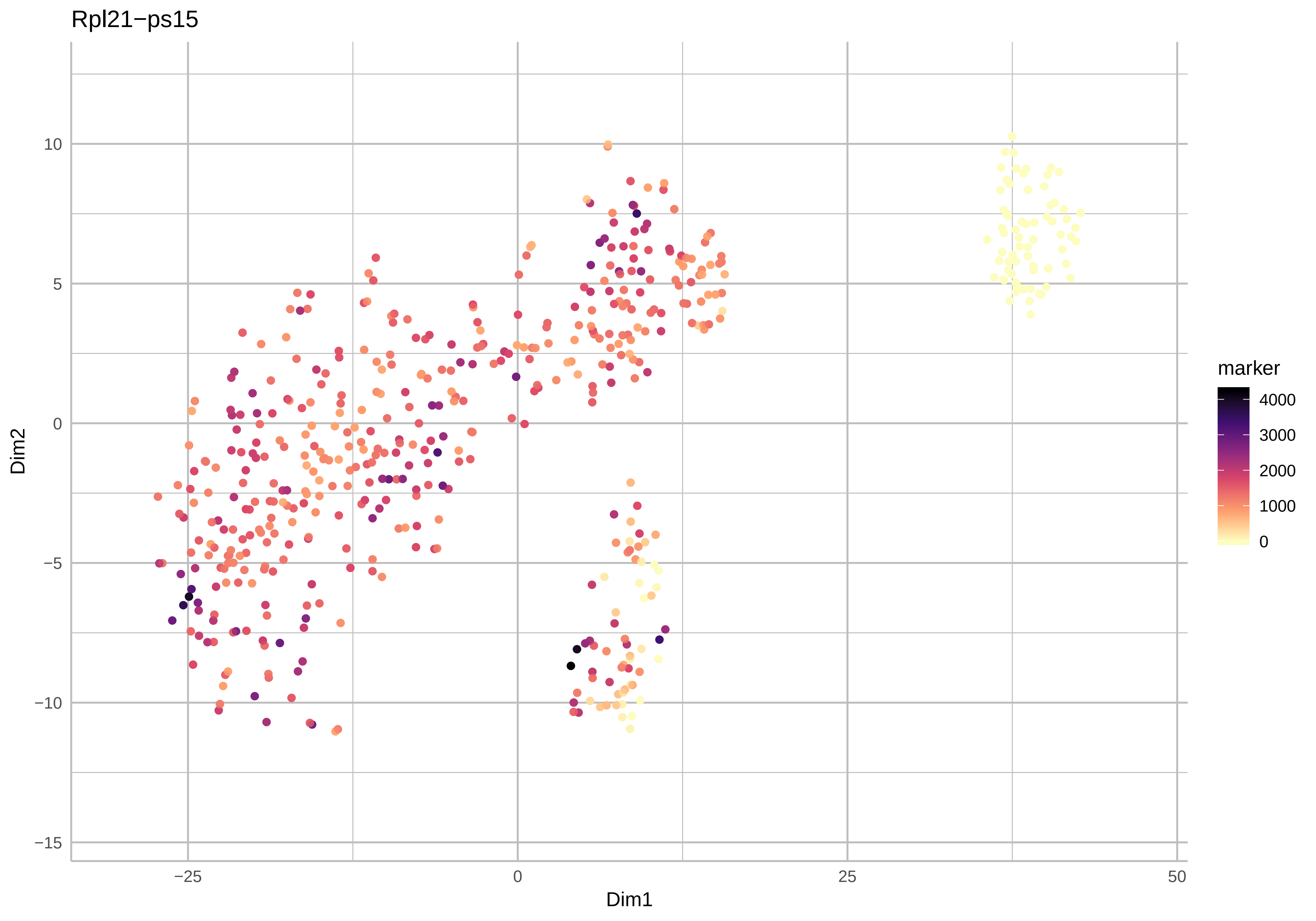

Rpl21−ps15
Dim2
Dim1
10
5
0
−5
−10
−15
−25
0
25
50
marker
4000
3000
2000
1000
0

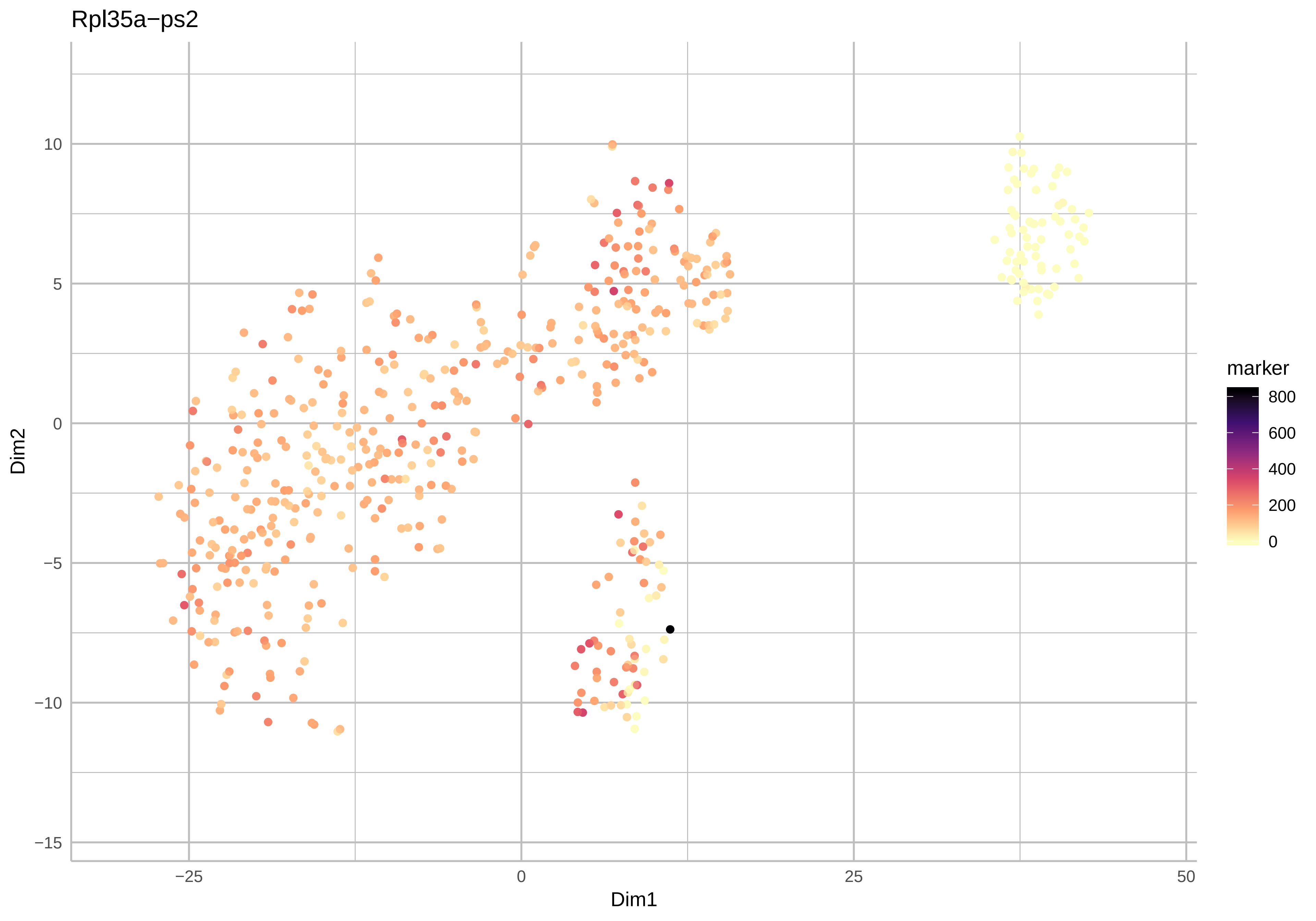

Rpl35a−ps2
10
5
0
−5
−10
−15
Dim2
−25
0
25
50
Dim1
marker
800
600
400
200
0

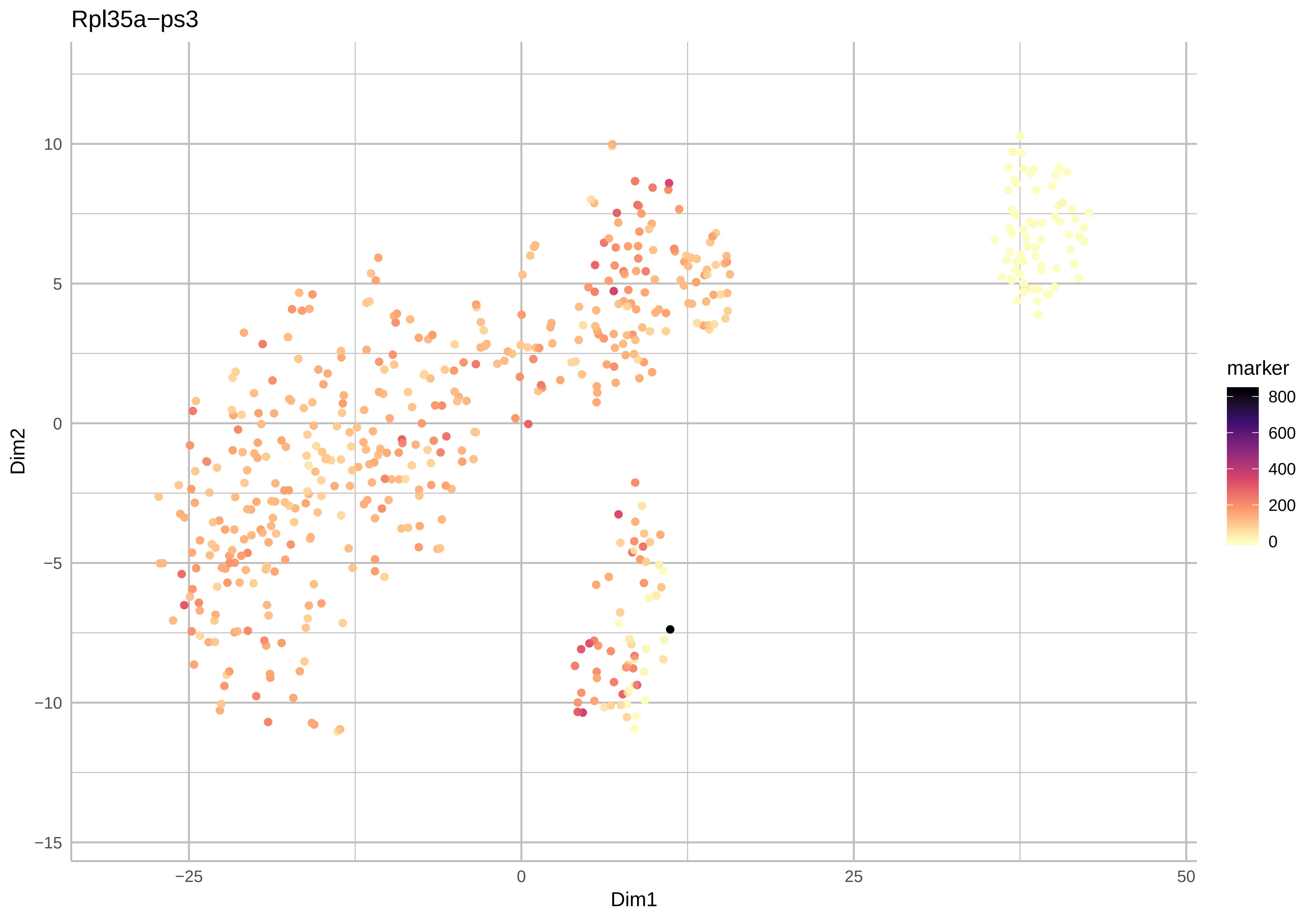
Rpl35a−ps3
Dim2
Dim1
10
5
0
−5
−10
−15
−25
0
25
50
marker
800
600
400
200
0

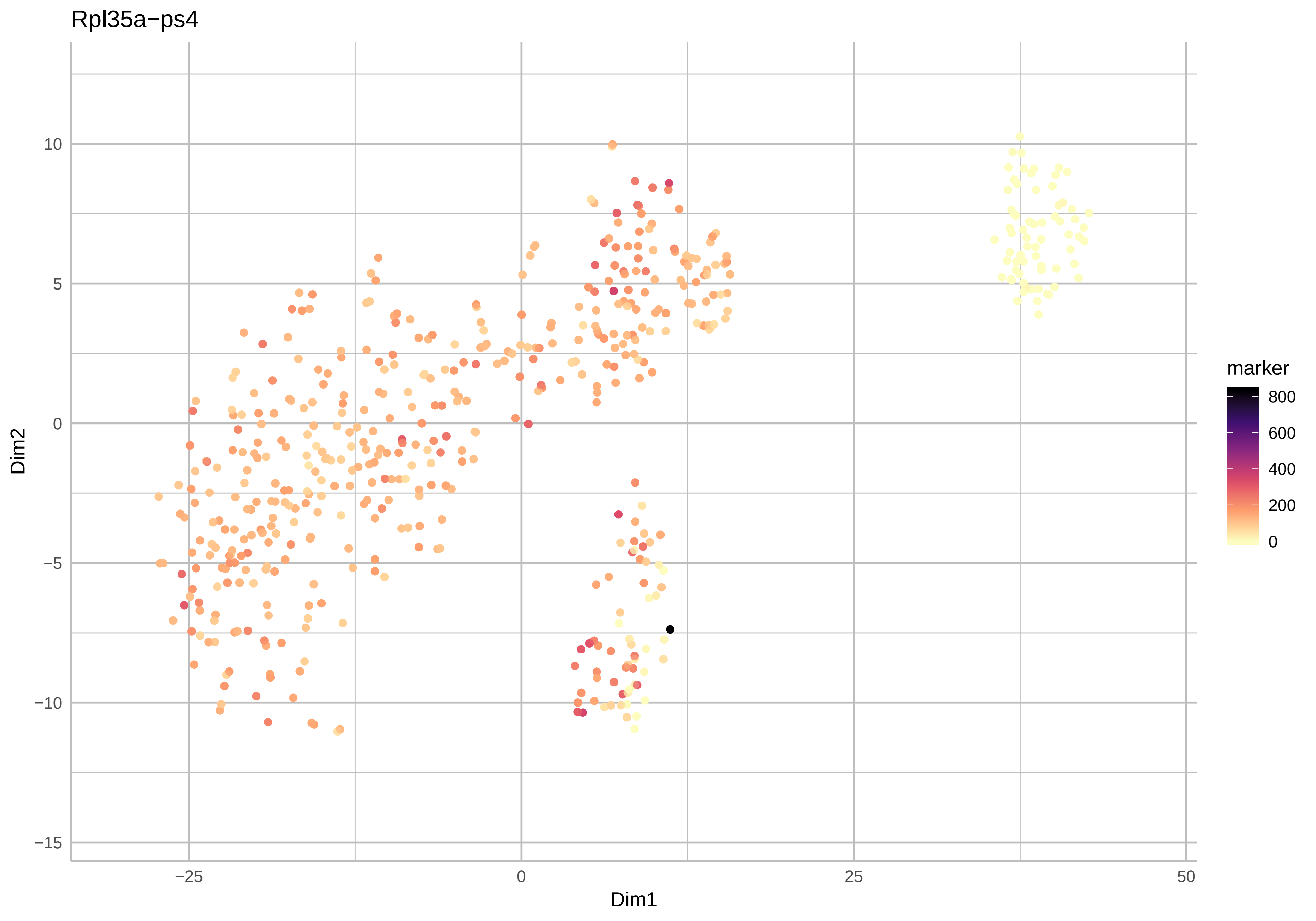

Rpl35a−ps4
Dim2
Dim1
10
5
0
−5
−10
−15
−25
0
25
50
marker
800
600
400
200
0

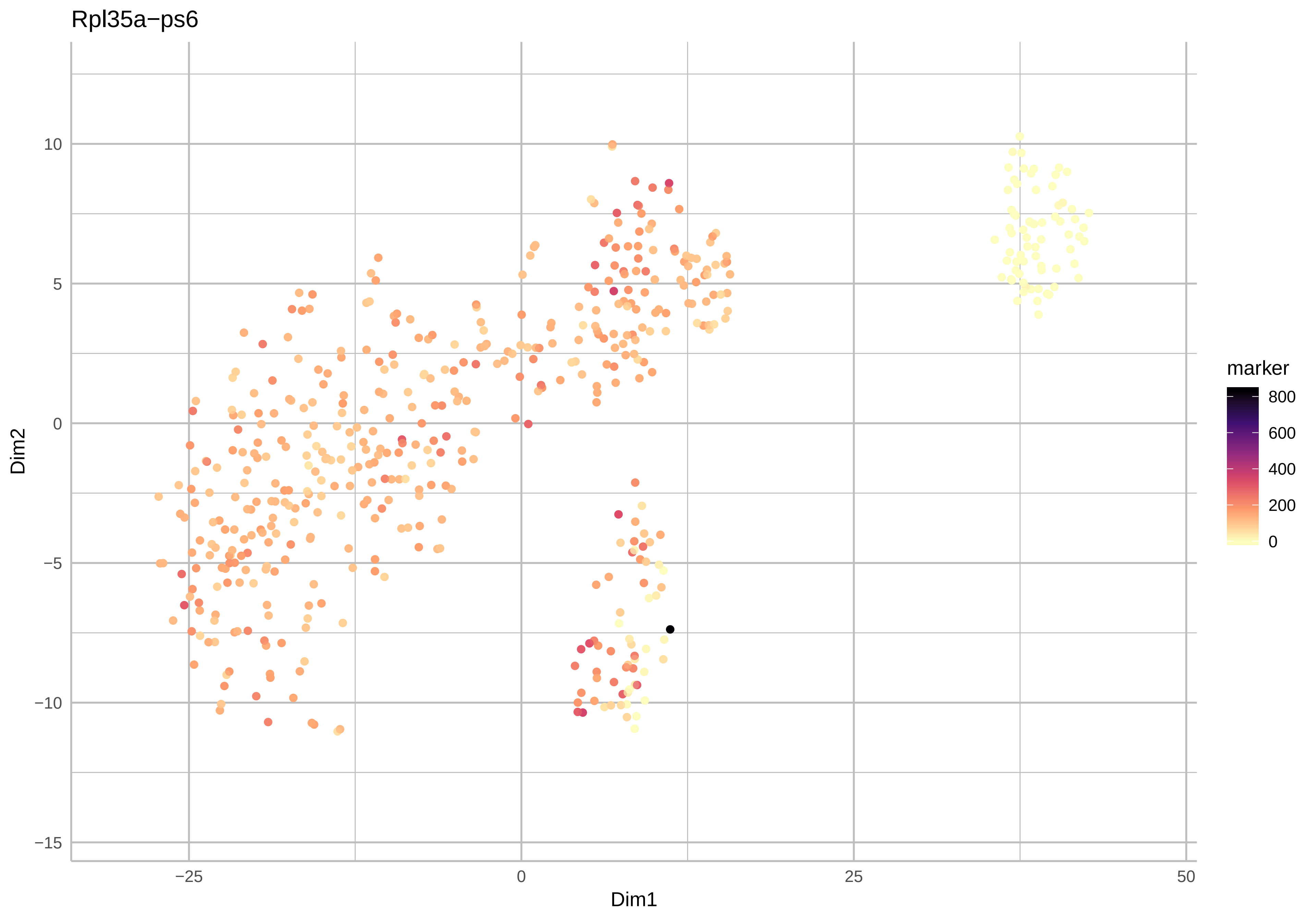
Rpl35a−ps6
Dim1
Dim2
marker
800
600
400
200
0
10
5
0
−5
−10
−15
−25
0
25
50

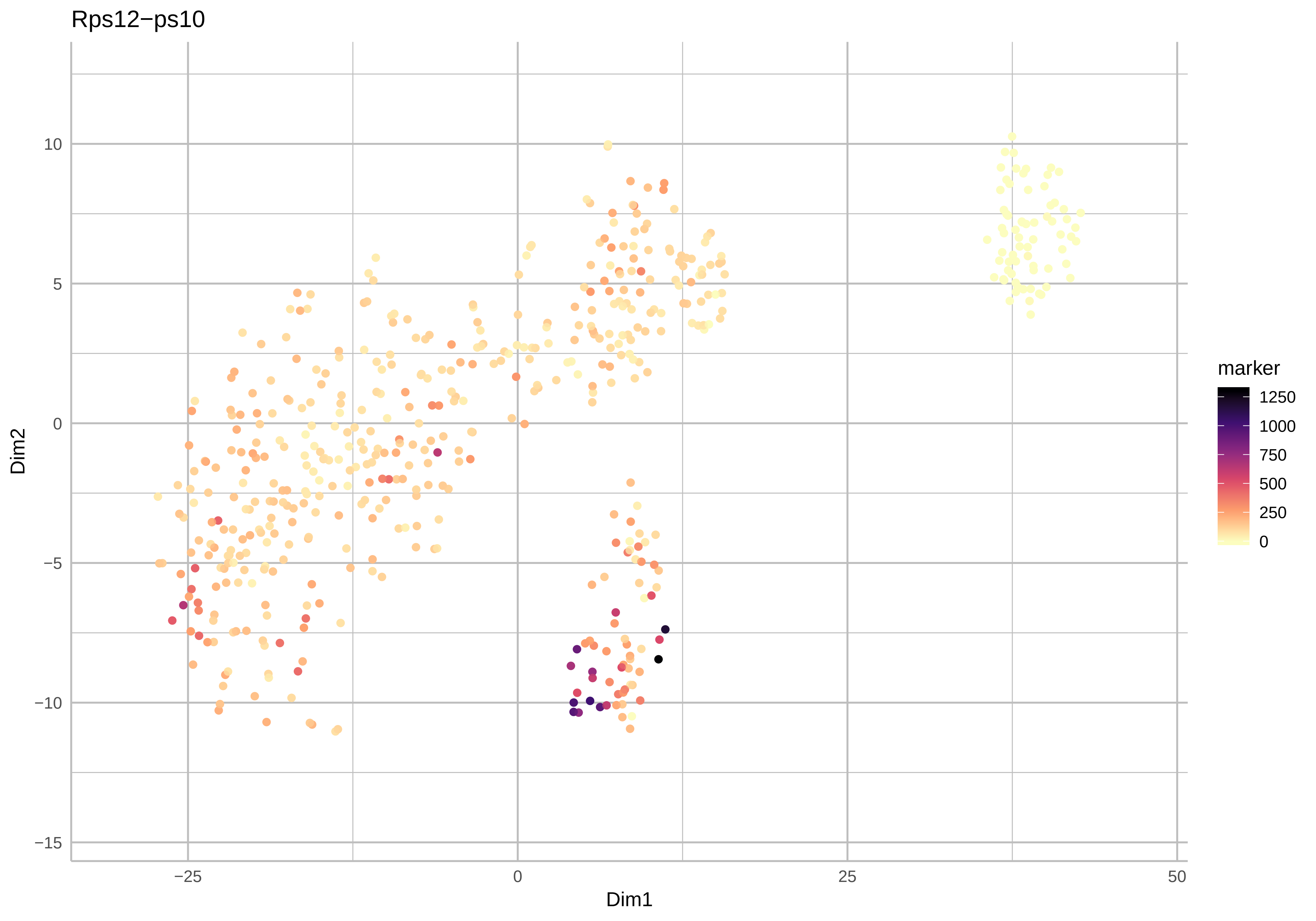

Rps12−ps10
Dim2
10
5
0
−5
−10
−15
Dim1
−25
0
25
50
marker
1250
1000
750
500
250
0

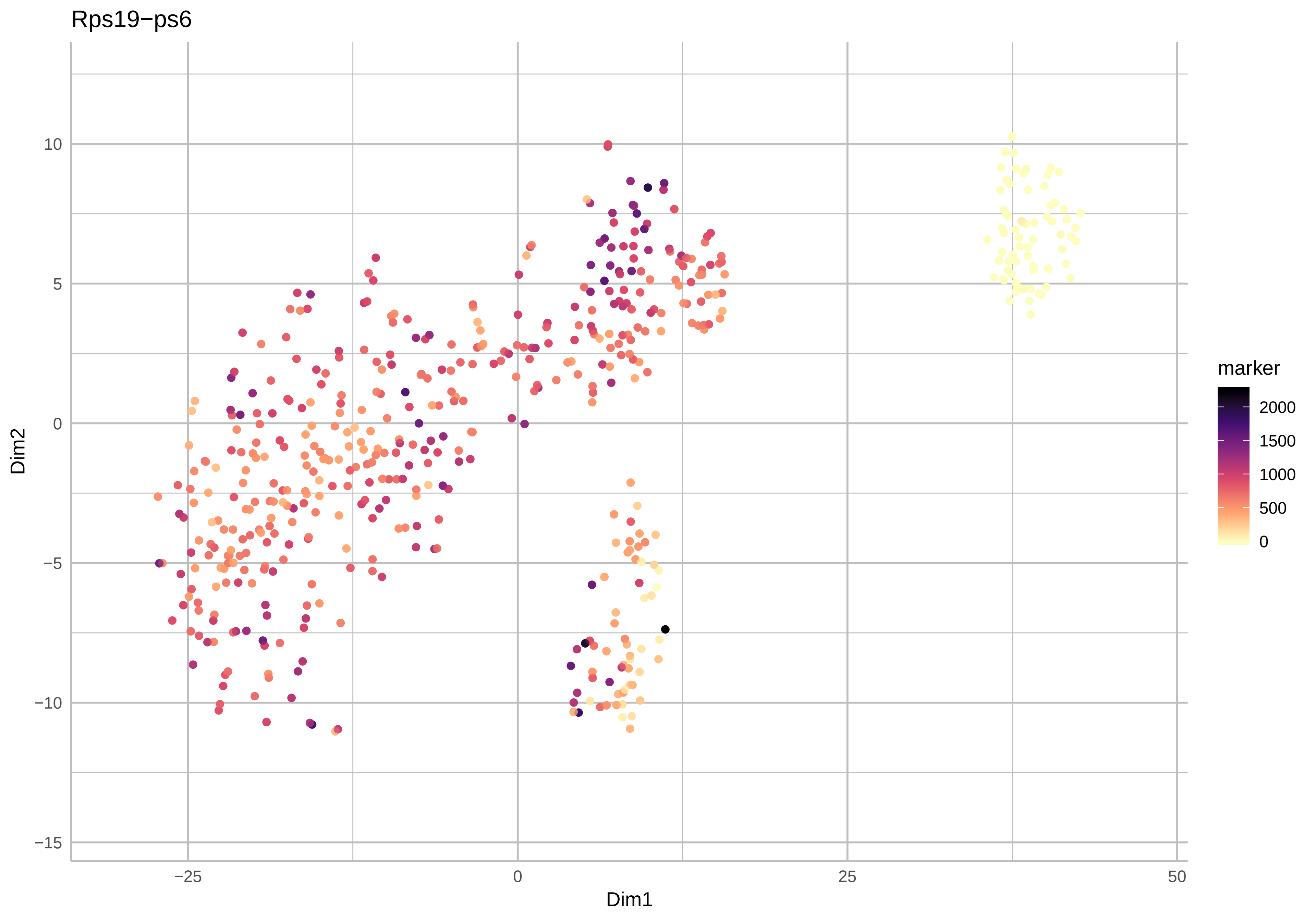
Rps19−ps6
Dim2
Dim1
10
5
0
−5
−10
−15
−25
0
25
50
marker
2000
1500
1000
500
0

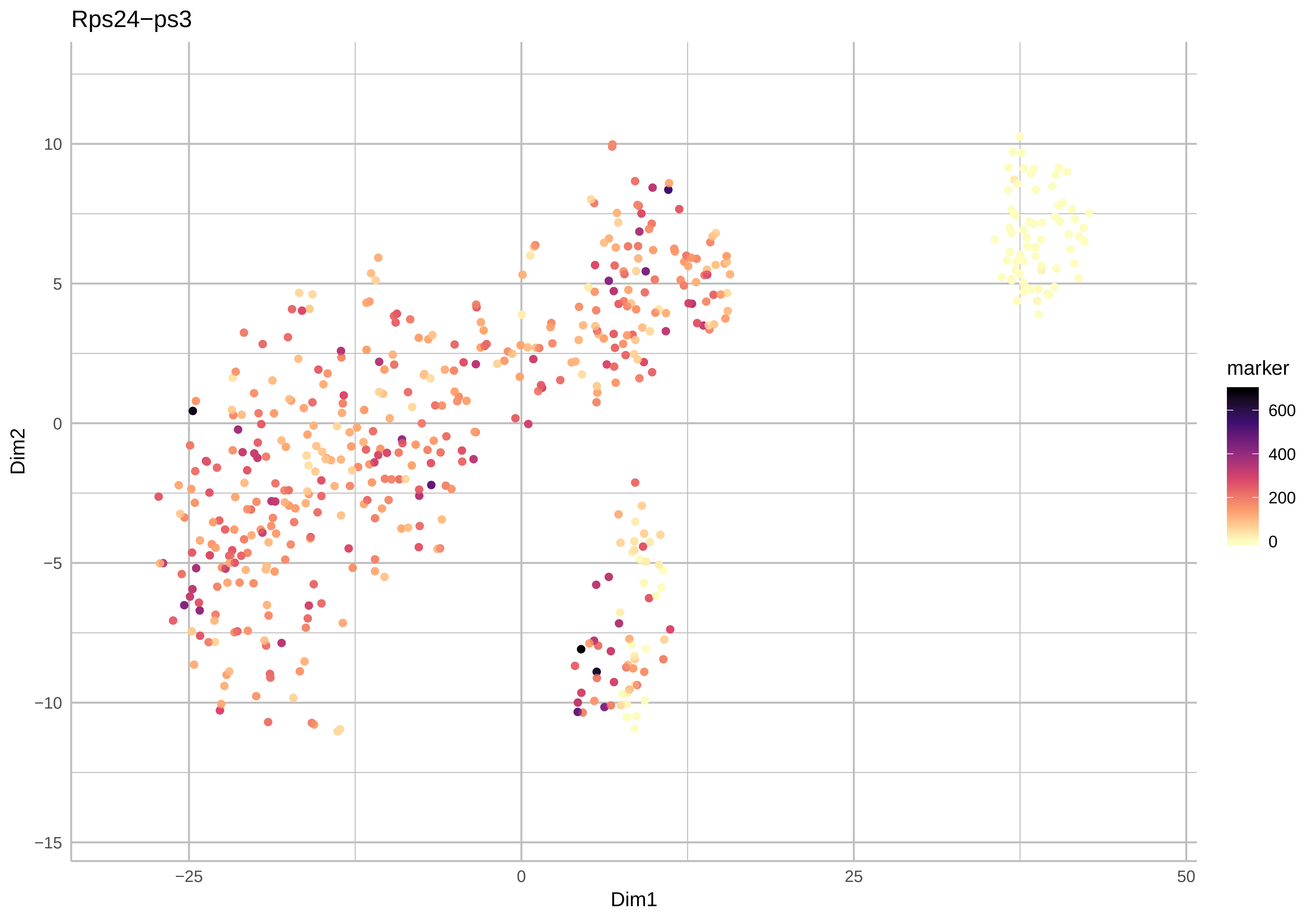

Rps24−ps3
Dim2
Dim1
10
5
0
−5
−10
−15
−25
0
25
50
marker
600
400
200
0

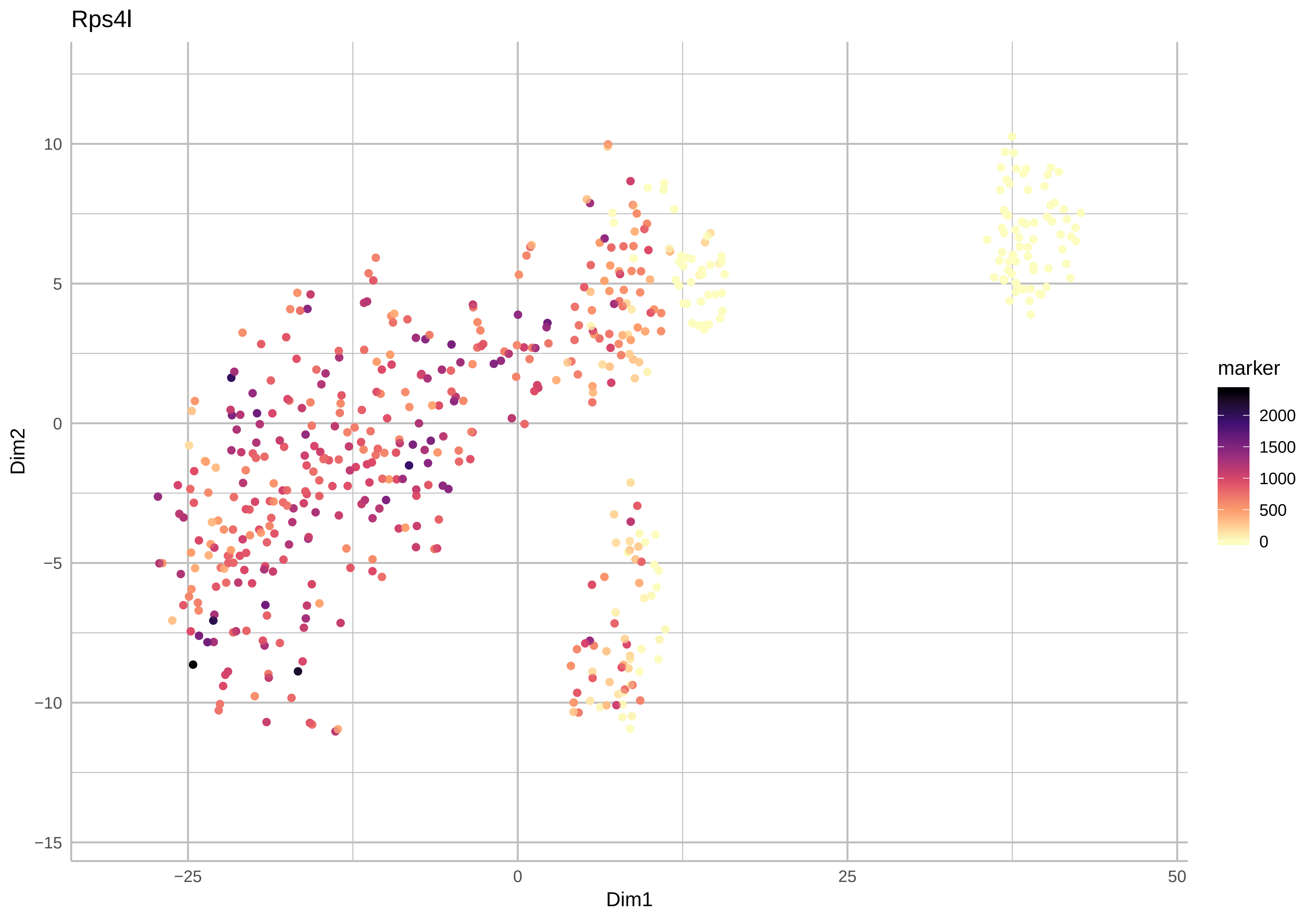
Rps4l
Dim2
Dim1
marker
2000
1500
1000
500
0
10
5
0
−5
−10
−15
−25
0
25
50

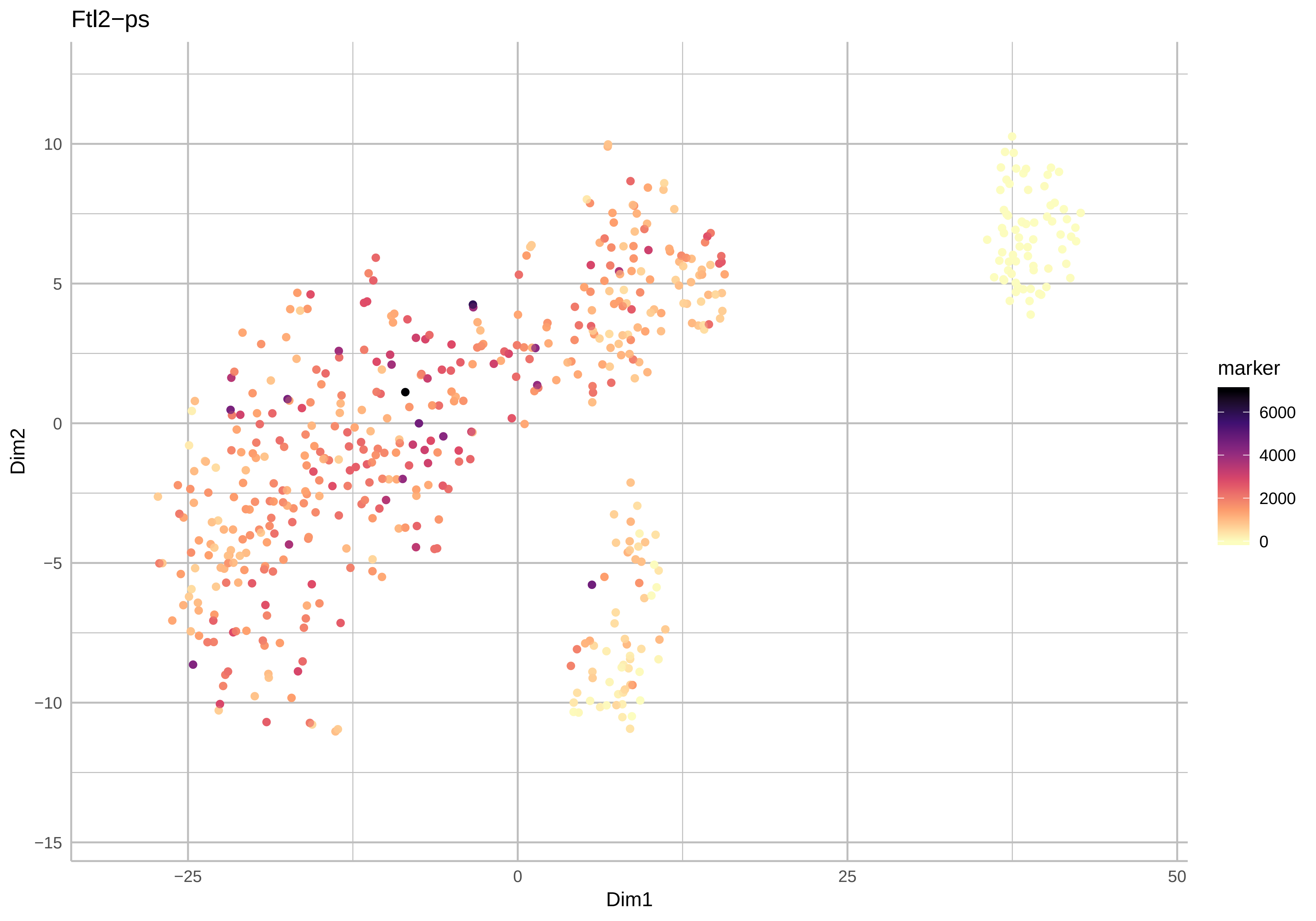
Ftl2−ps
Dim2
Dim1
10
5
0
−5
−10
−15
−25
0
25
50
marker
6000
4000
2000
0

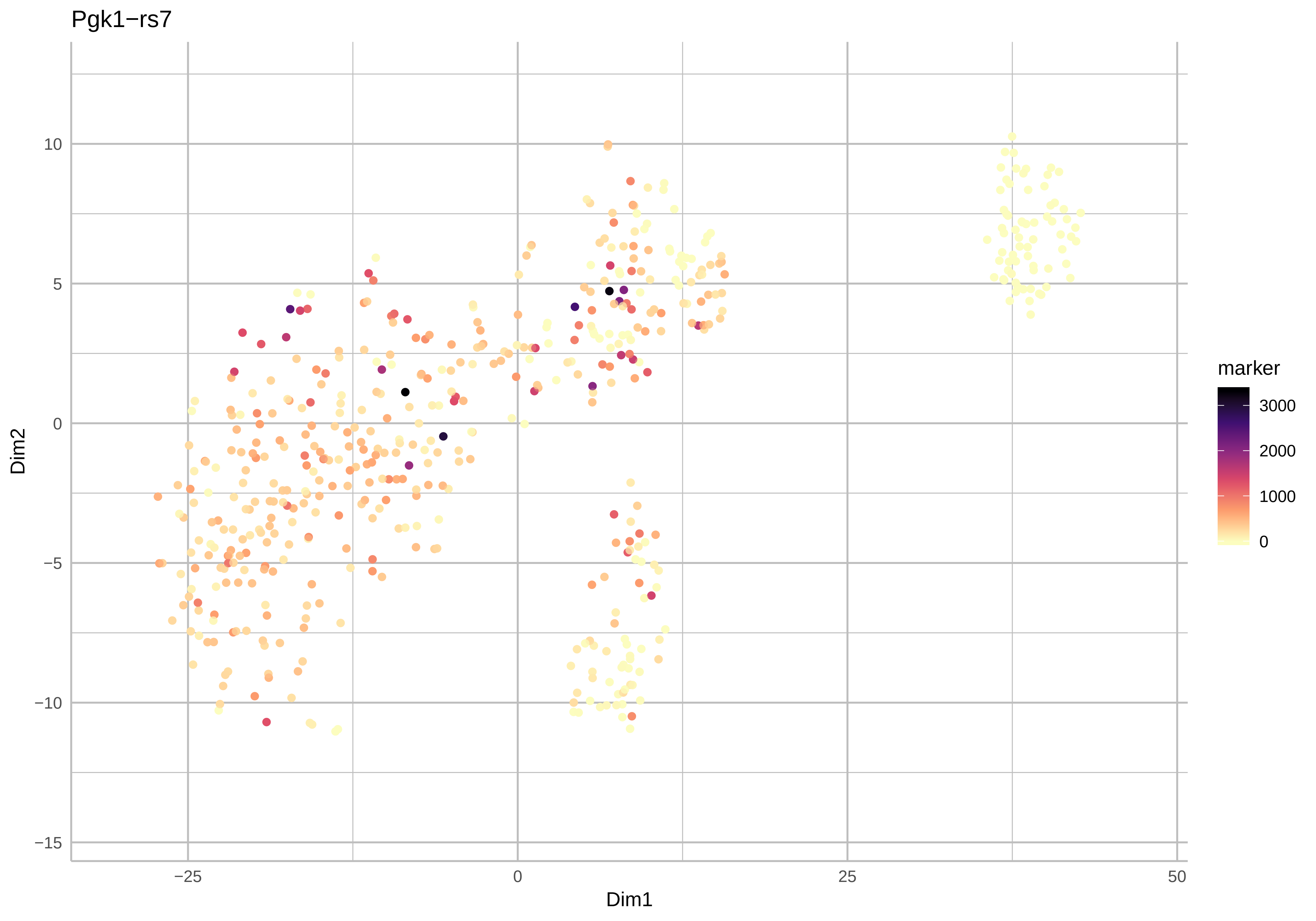

Pgk1−rs7
Dim2
Dim1
10
5
0
−5
−10
−15
−25
0
25
50
marker
3000
2000
1000
0

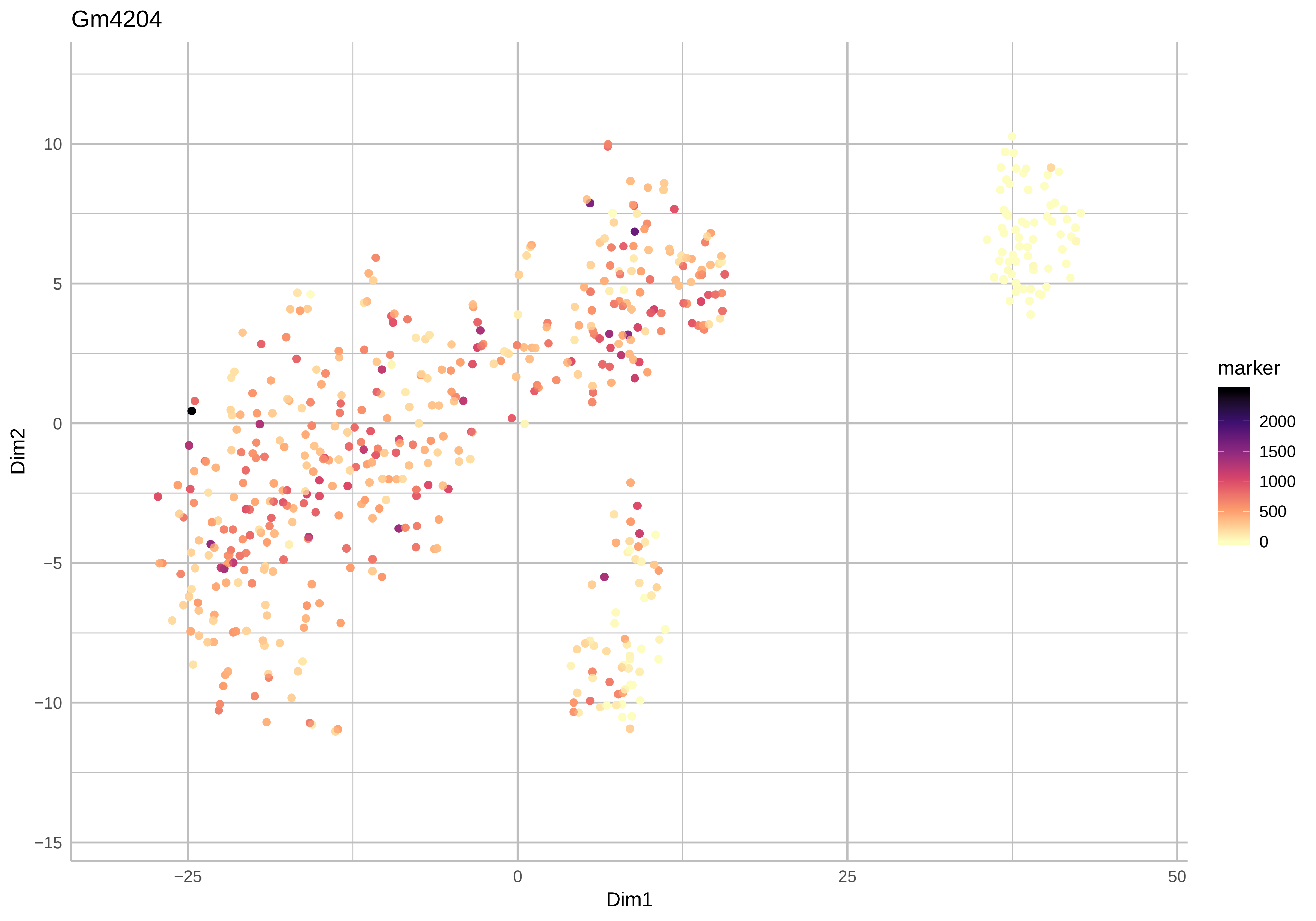

Gm4204
Dim2
Dim1
10
5
0
−5
−10
−15
−25
0
25
50
marker
2000
1500
1000
500
0

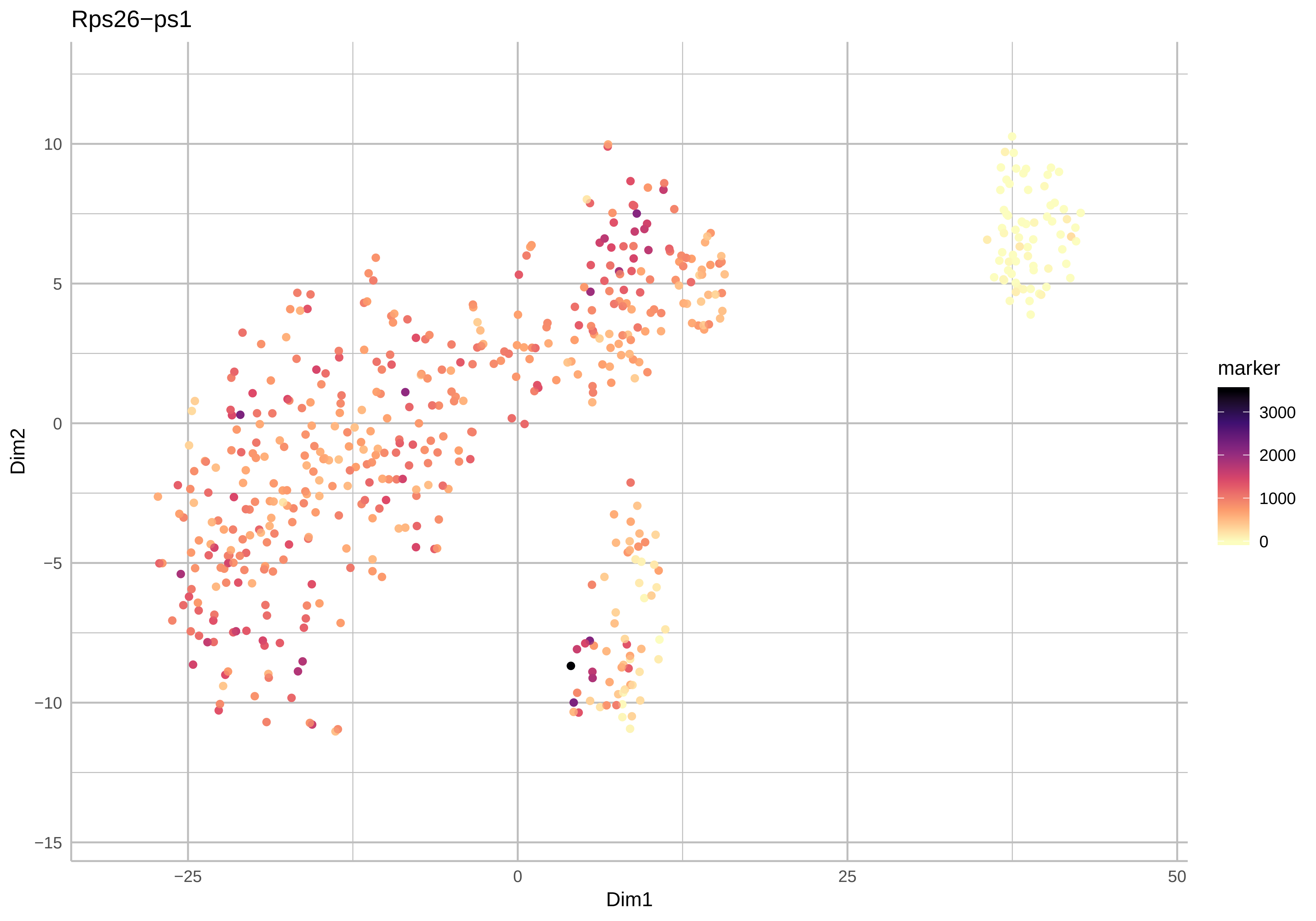
Rps26−ps1
Dim2
Dim1
10
5
0
−5
−10
−15
−25
0
25
50
marker
3000
2000
1000
0

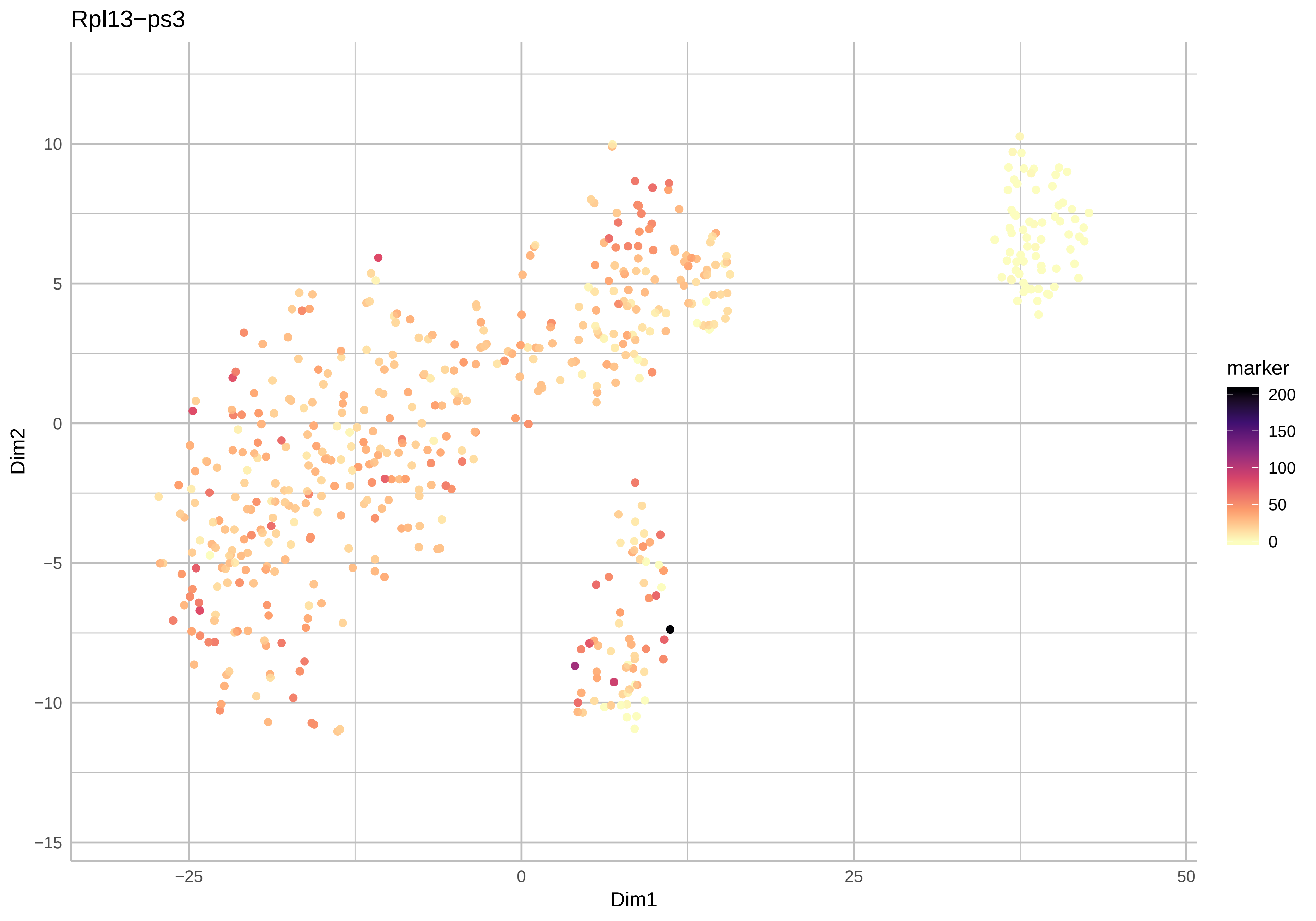
Rpl13−ps3
Dim2
Dim1
10
5
0
−5
−10
−15
−25
0
25
50
marker
200
150
100
50
0

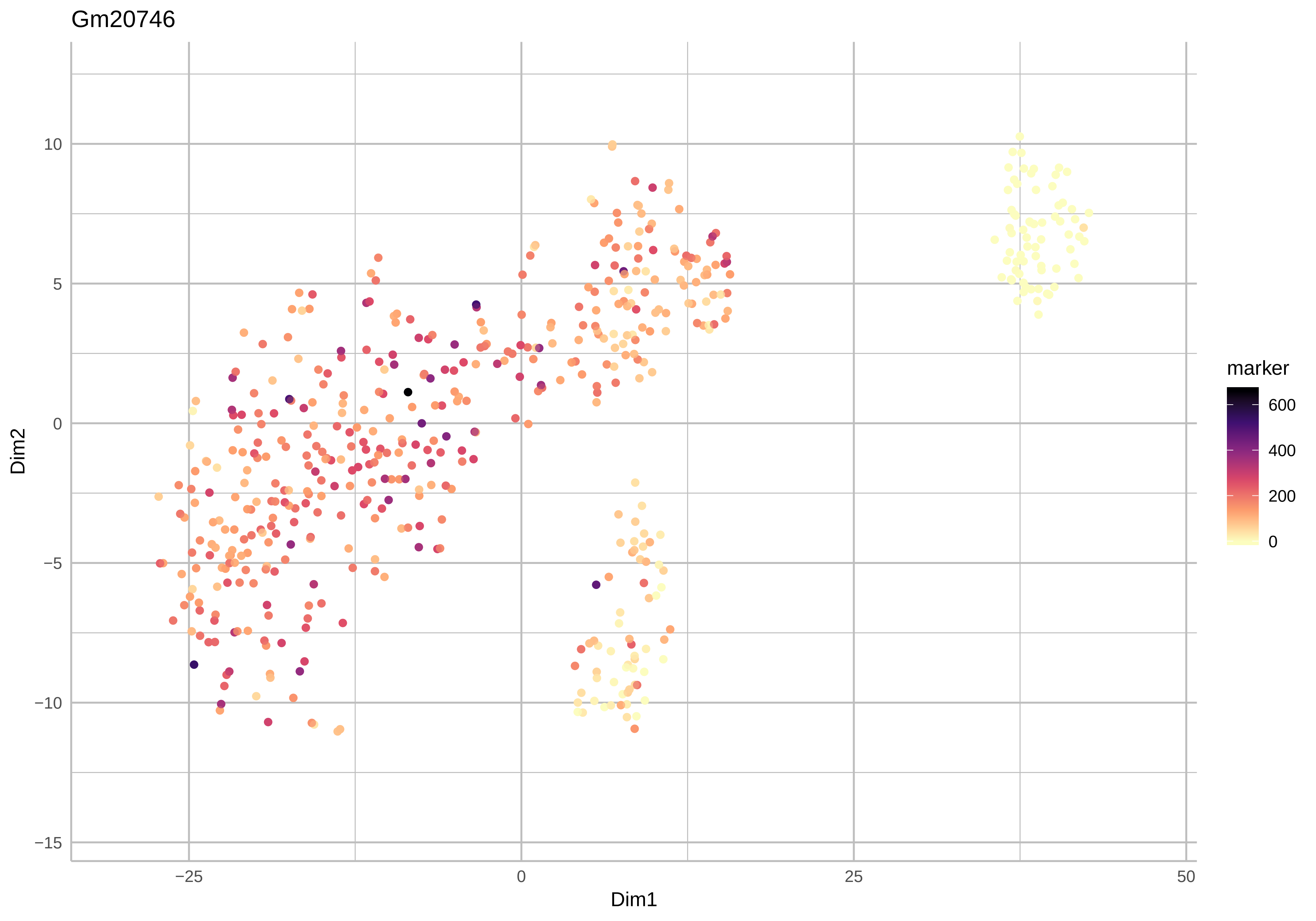

Gm20746
Dim2
Dim1
10
5
0
−5
−10
−15
−25
0
25
50
marker
600
400
200
0

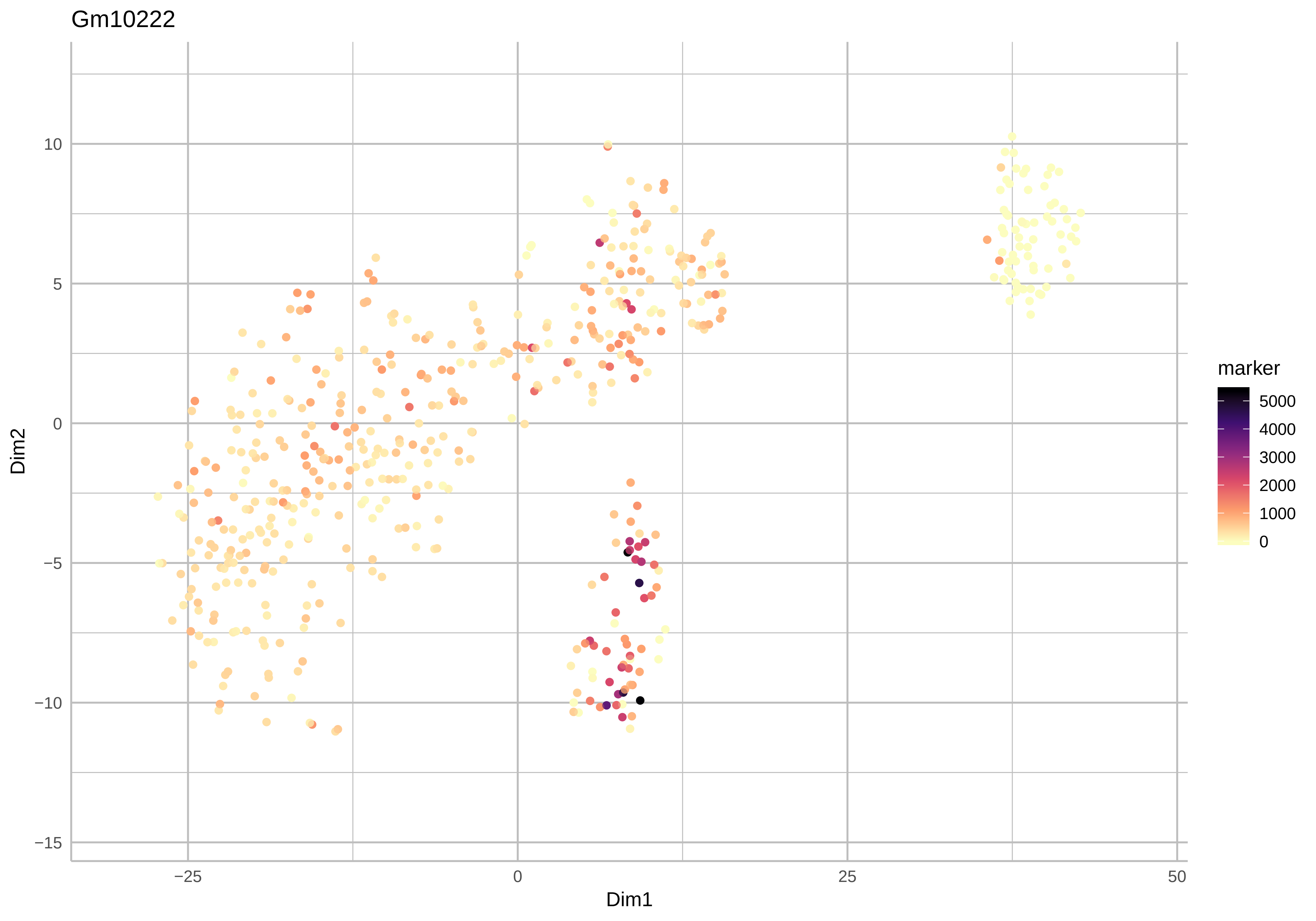

Gm10222
Dim2
Dim1
10
5
0
−5
−10
−15
−25
0
25
50
marker
5000
4000
3000
2000
1000
0

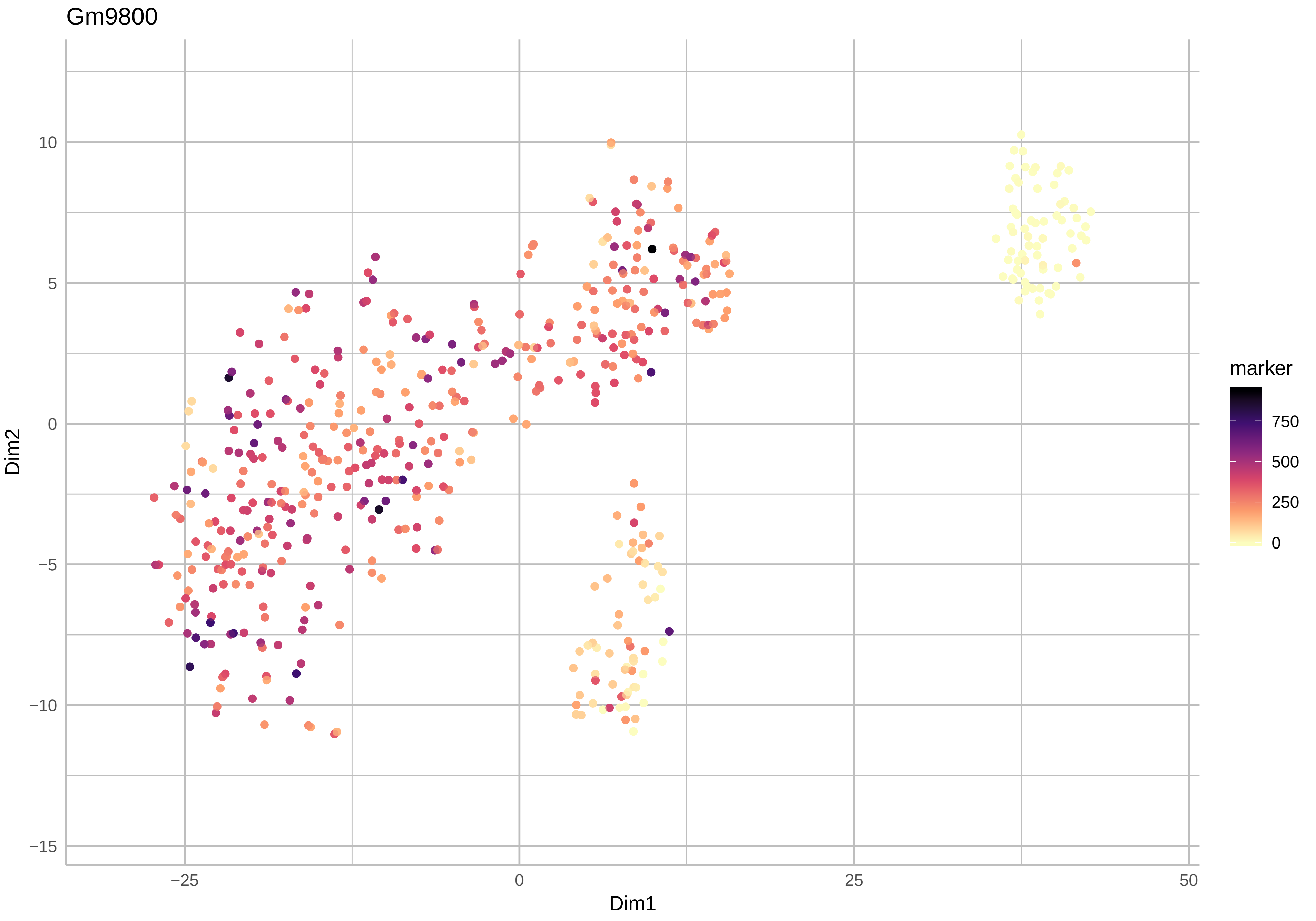

Gm9800
Dim2
Dim1
10
5
0
−5
−10
−15
−25
0
25
50
marker
750
500
250
0

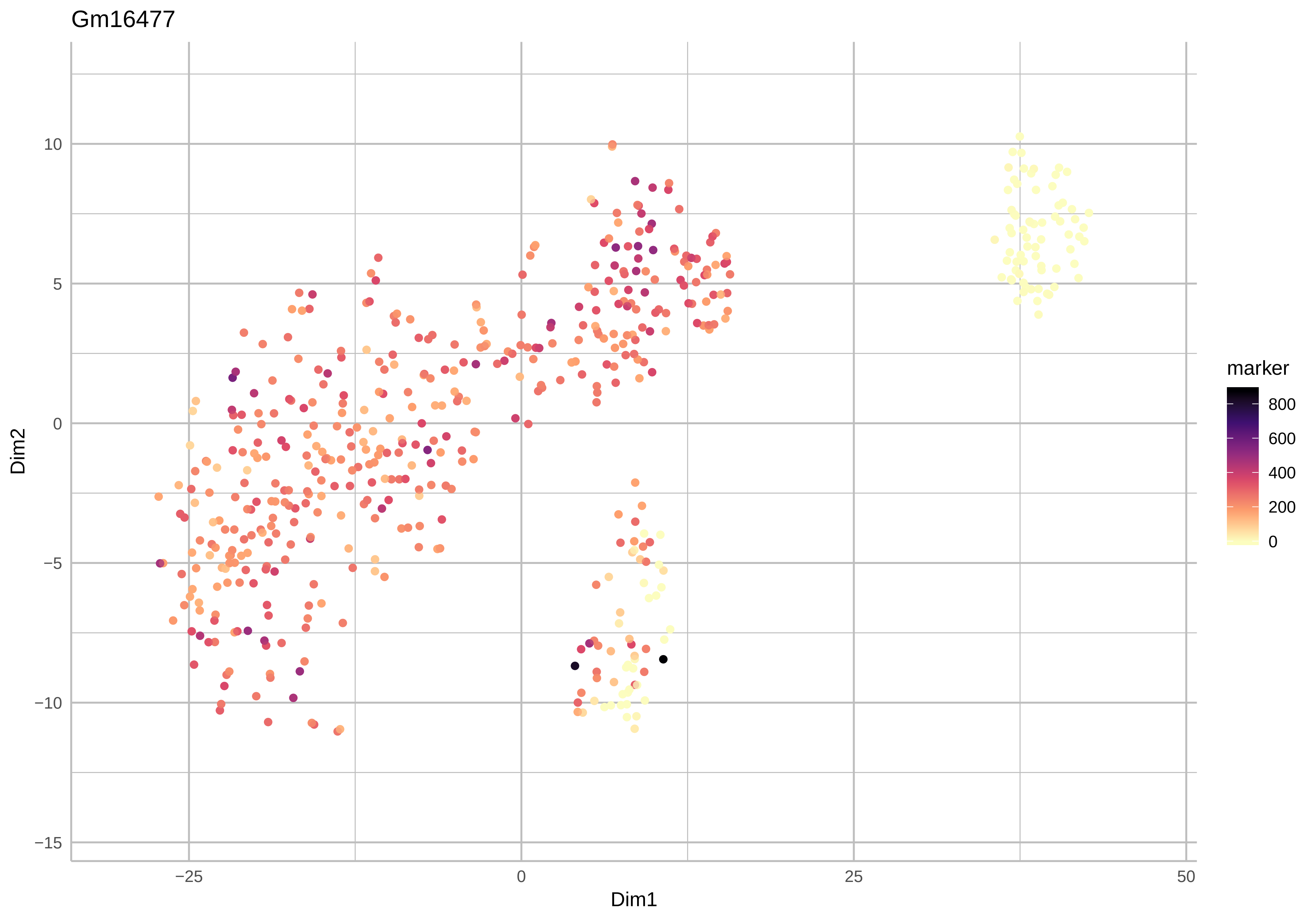
Gm16477
Dim2
Dim1
10
5
0
−5
−10
−15
−25
0
25
50
marker
800
600
400
200
0

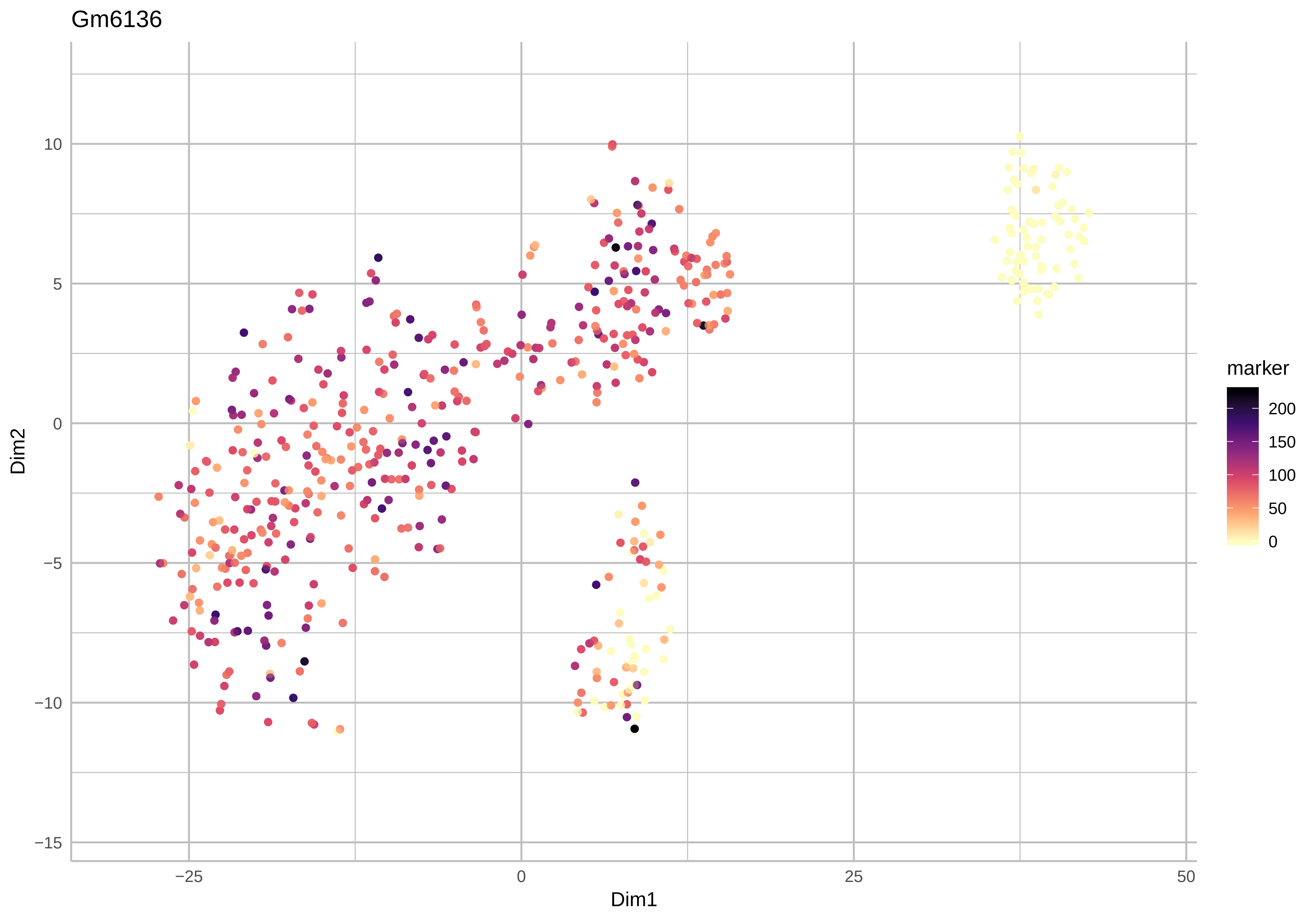

Gm6136
Dim2
Dim1
10
5
0
−5
−10
−15
−25
0
25
50
marker
200
150
100
50
0

Gm14681

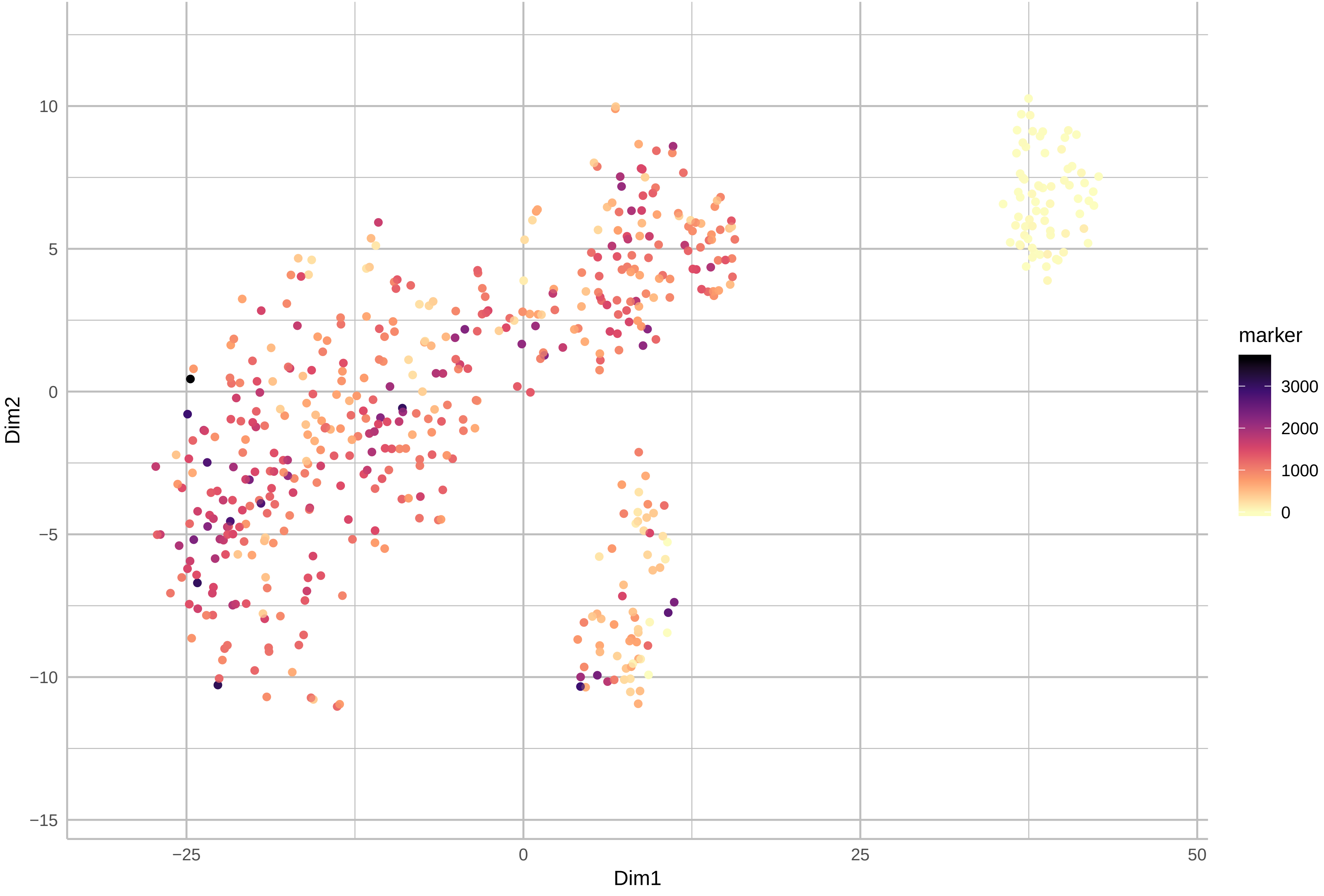

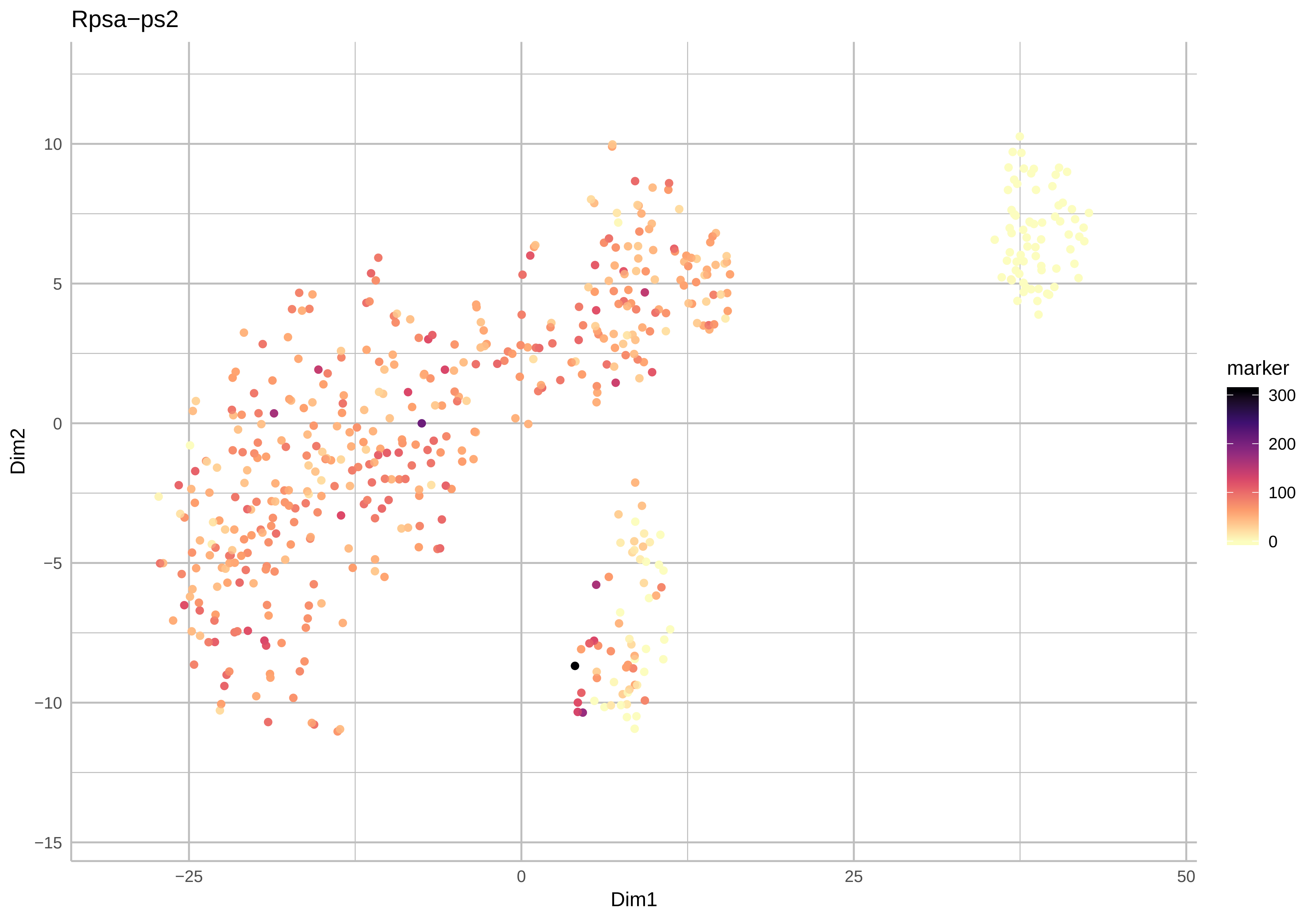

Rpsa−ps2
Dim2
Dim1
10
5
0
−5
−10
−15
−25
0
25
50
marker
300
200
100
0

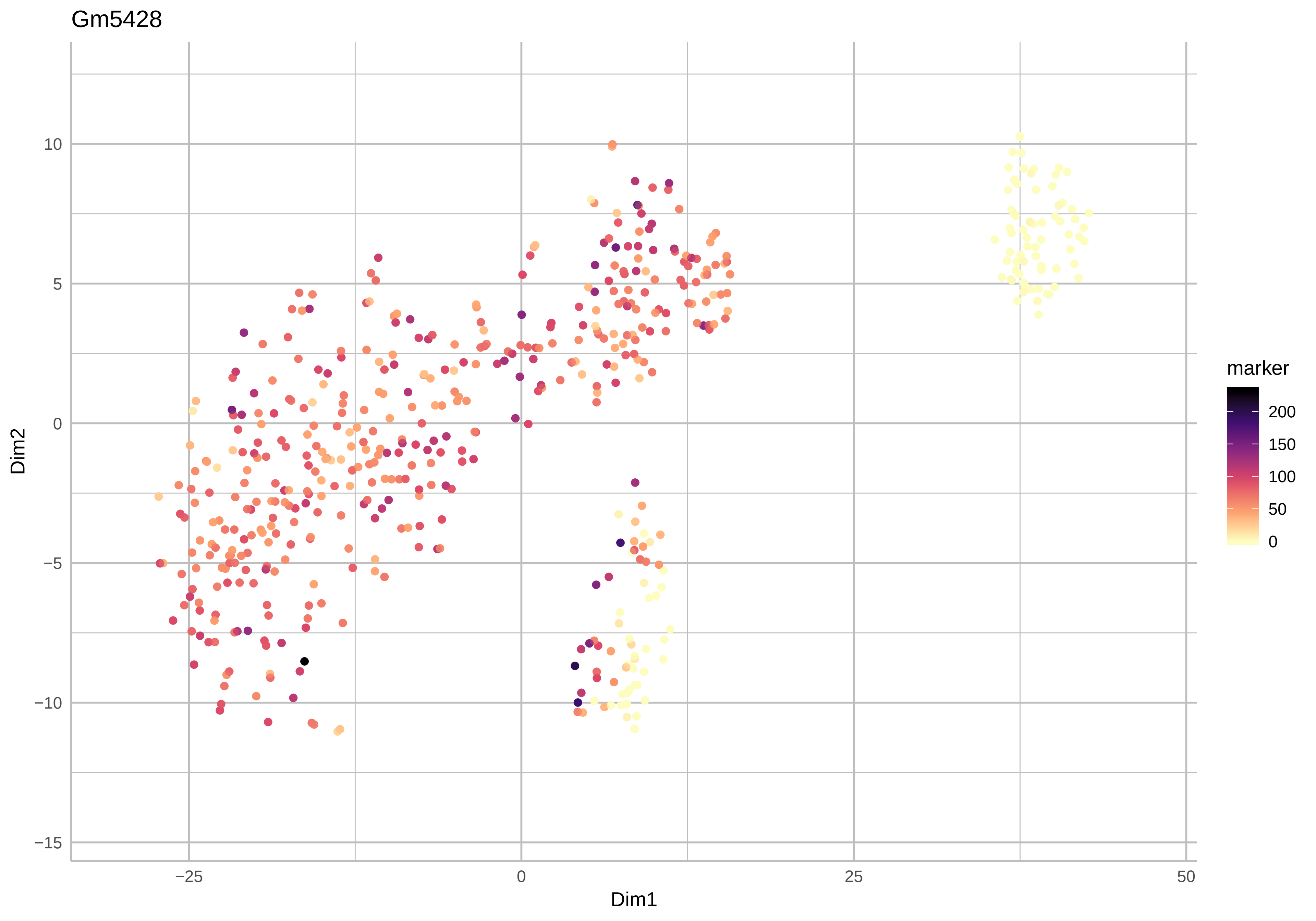
Gm5428
Dim2
Dim1
−25
0
25
50
10
5
0
−5
−10
−15
marker
200
150
100
50
0

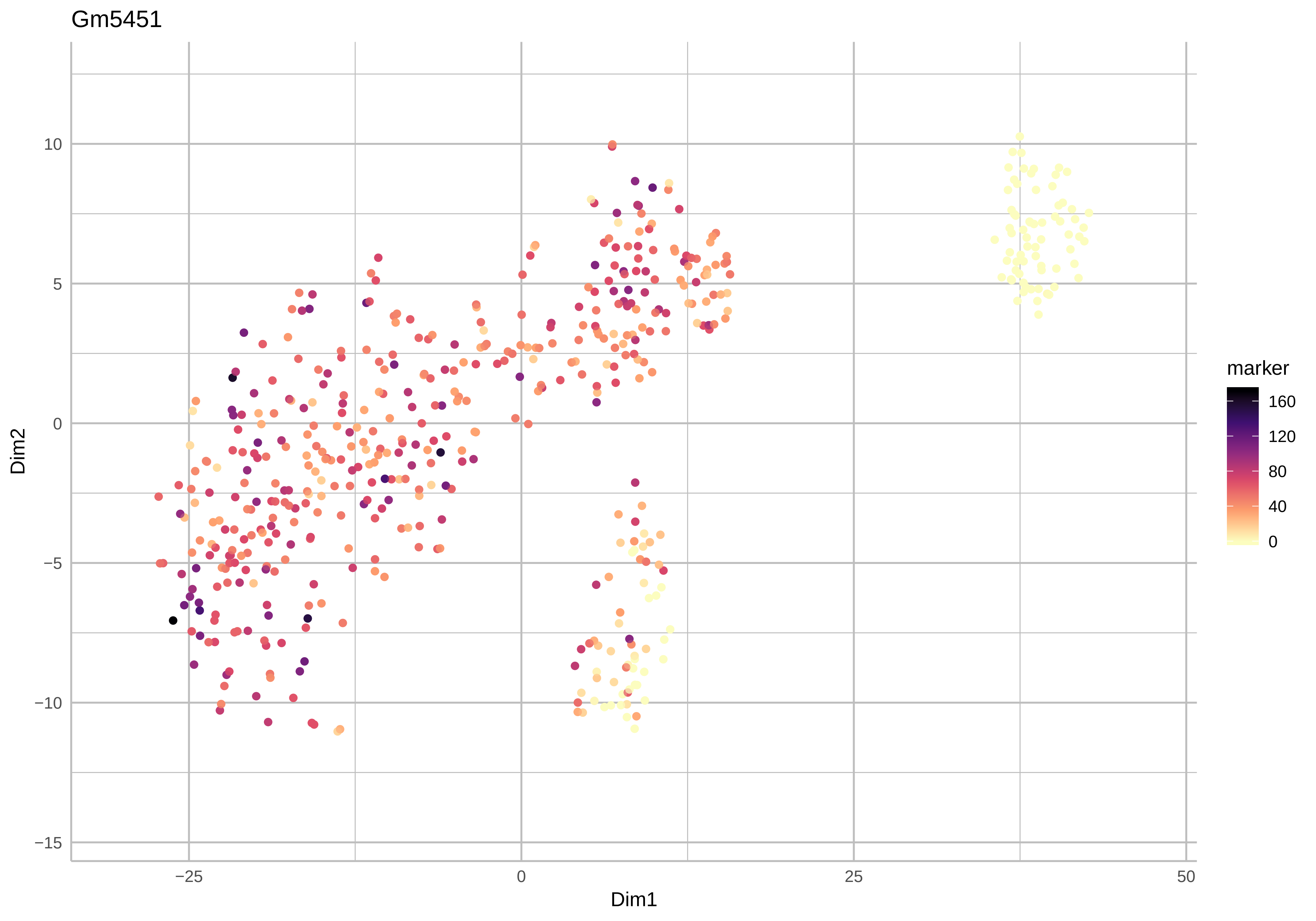

Gm5451
Dim2
Dim1
marker
160
120
80
40
0
10
5
0
−5
−10
−15
−25
0
25
50

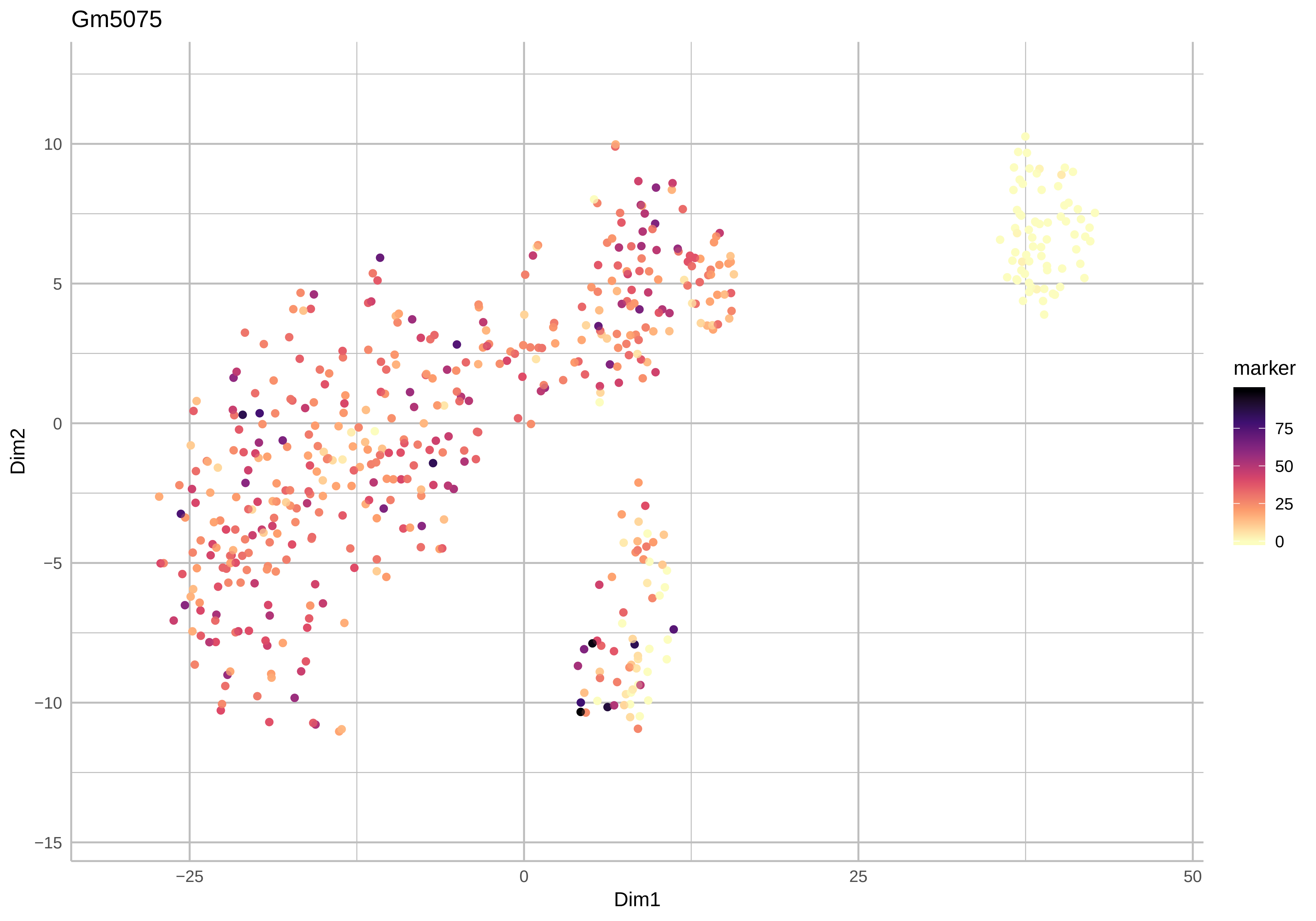
Gm5075
Dim2
Dim1
10
5
0
−5
−10
−15
−25
0
25
50
marker
75
50
25
0

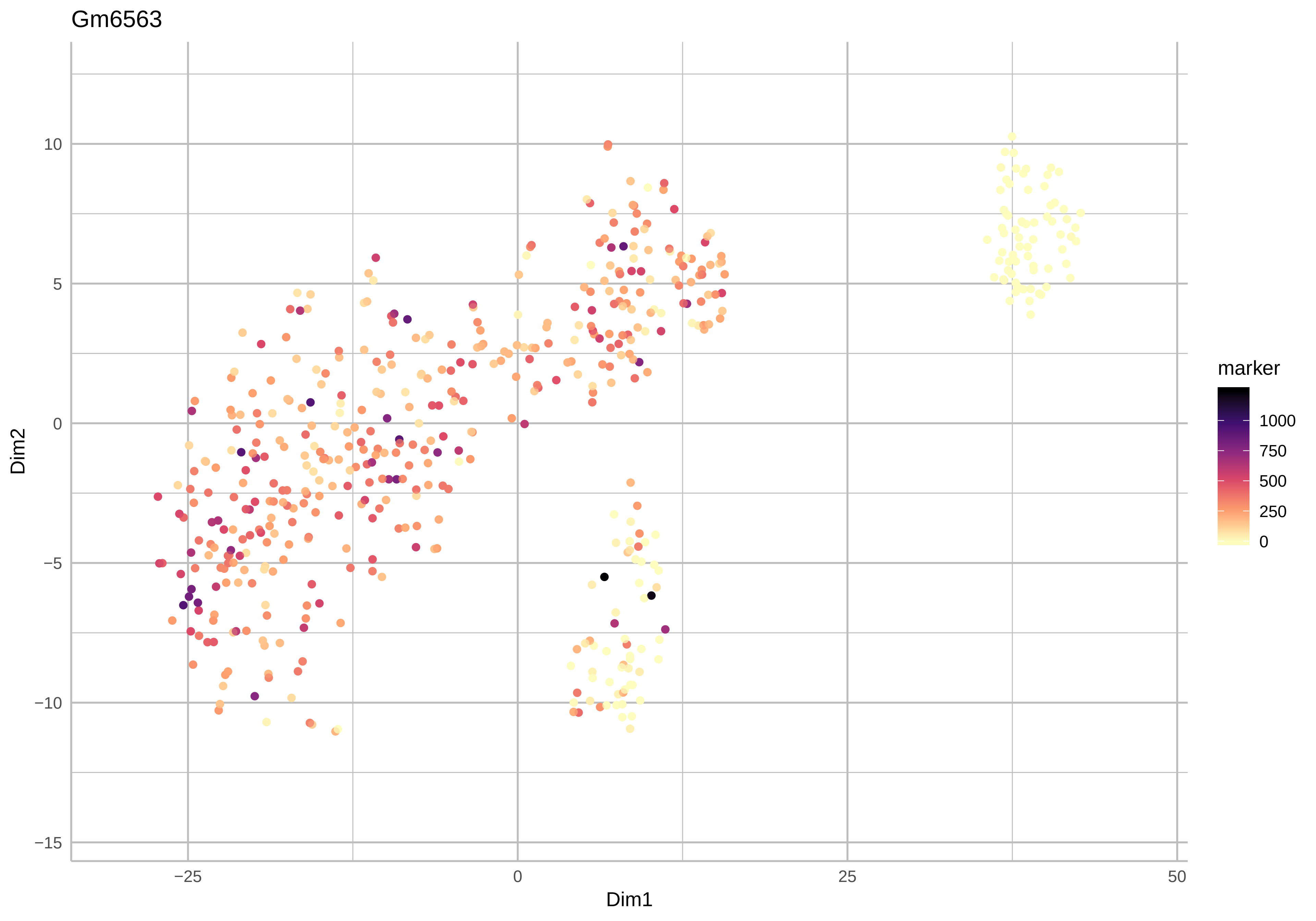
Gm6563
Dim2
Dim1
marker
1000
750
500
250
0
10
5
0
−5
−10
−15
−25
0
25
50

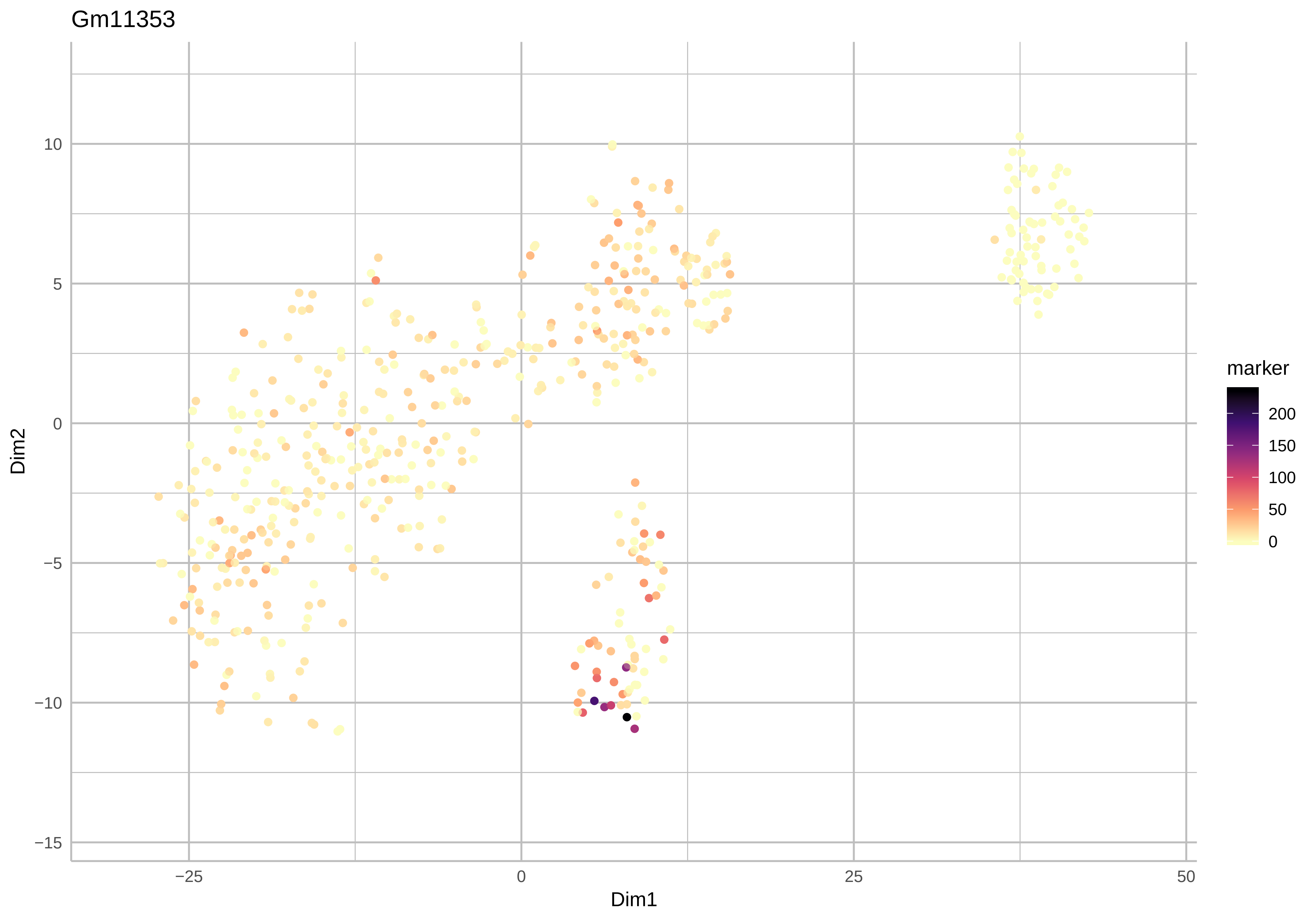

Gm11353
Dim2
Dim1
10
5
0
−5
−10
−15
−25
0
25
50
marker
200
150
100
50
0

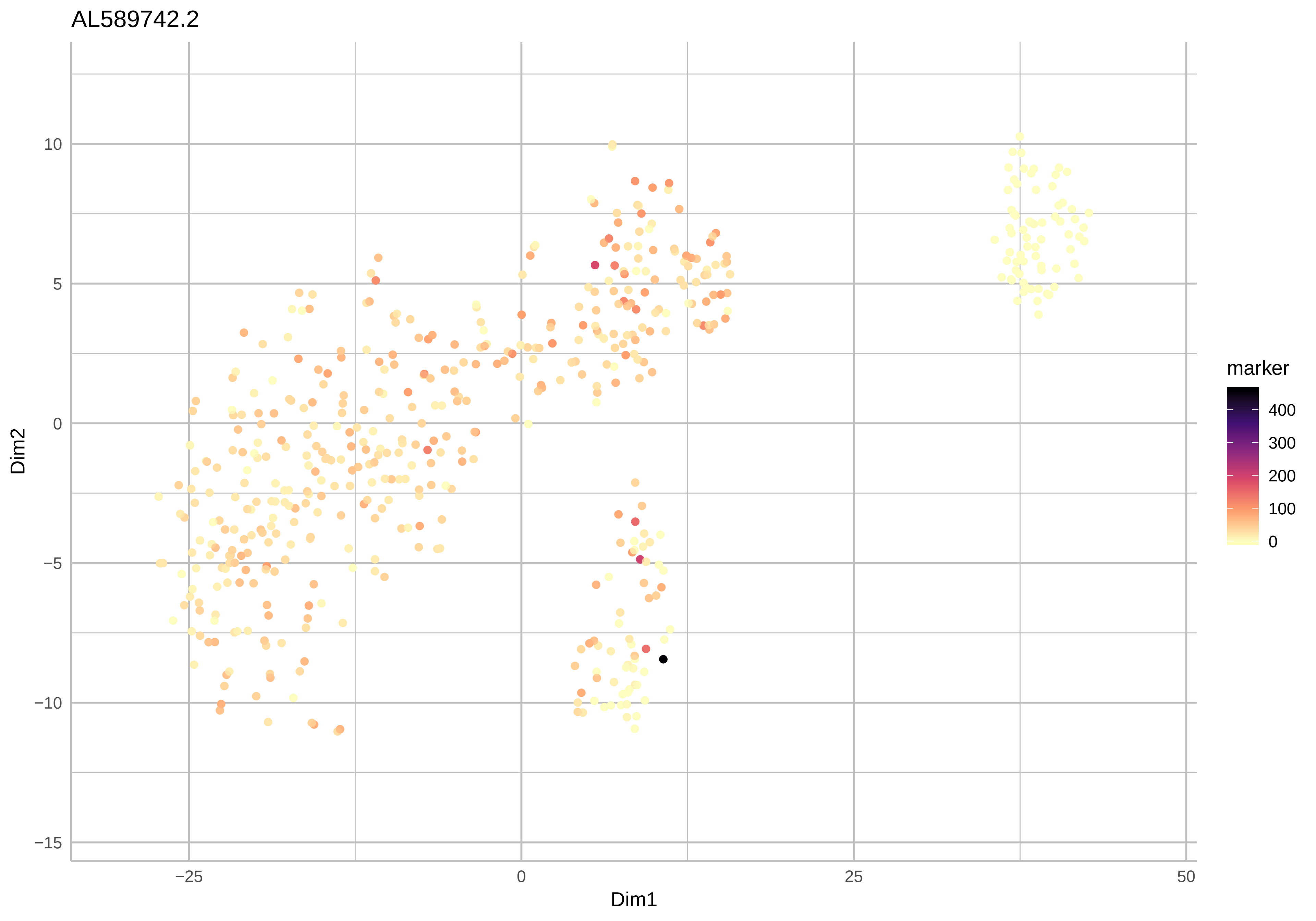
AL589742.2
Dim2
Dim1
10
5
0
−5
−10
−15
−25
0
25
50
marker
400
300
200
100
0

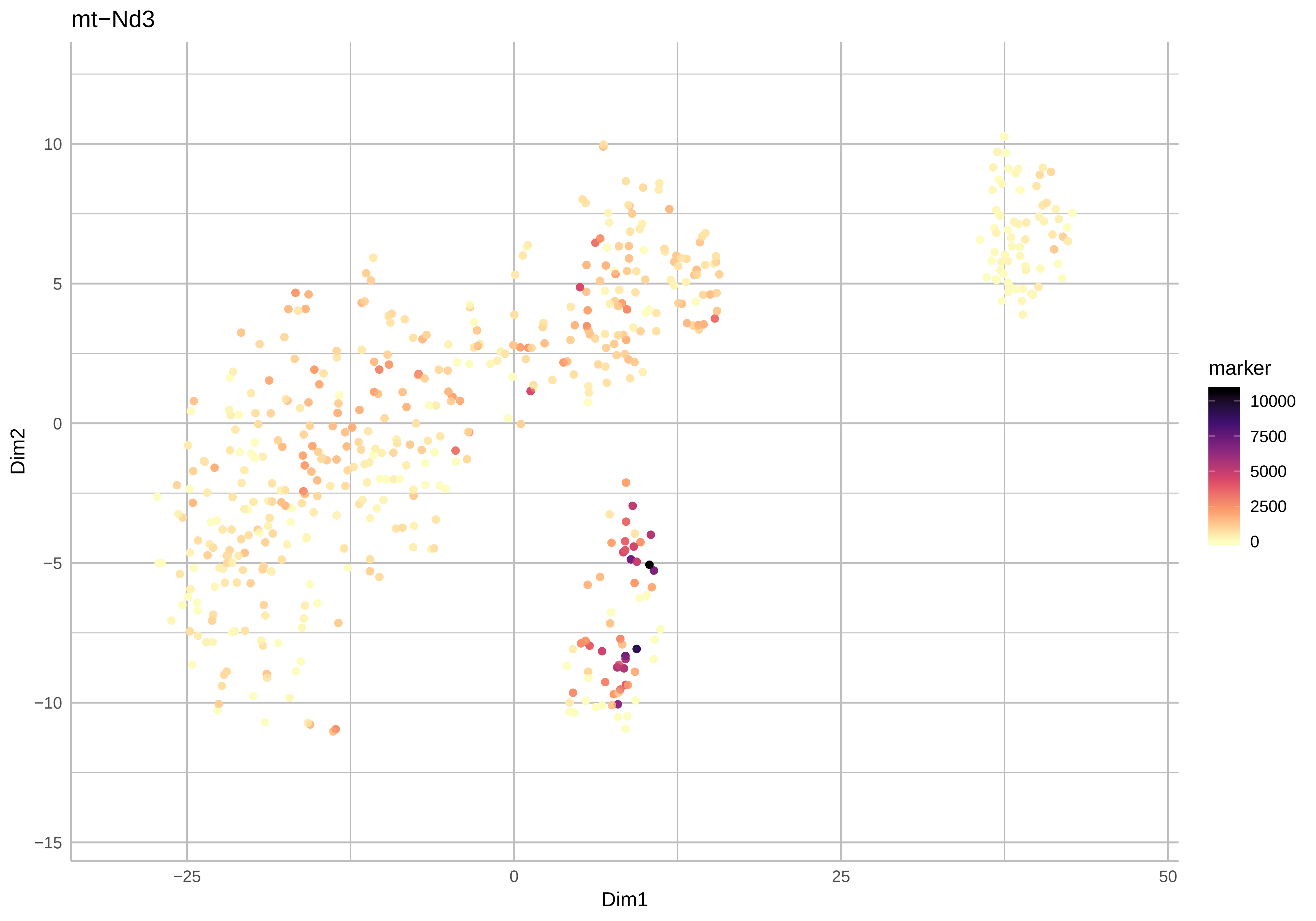
mt−Nd3
Dim2
Dim1
10
5
0
−5
−10
−15
−25
0
25
50
marker
10000
7500
5000
2500
0

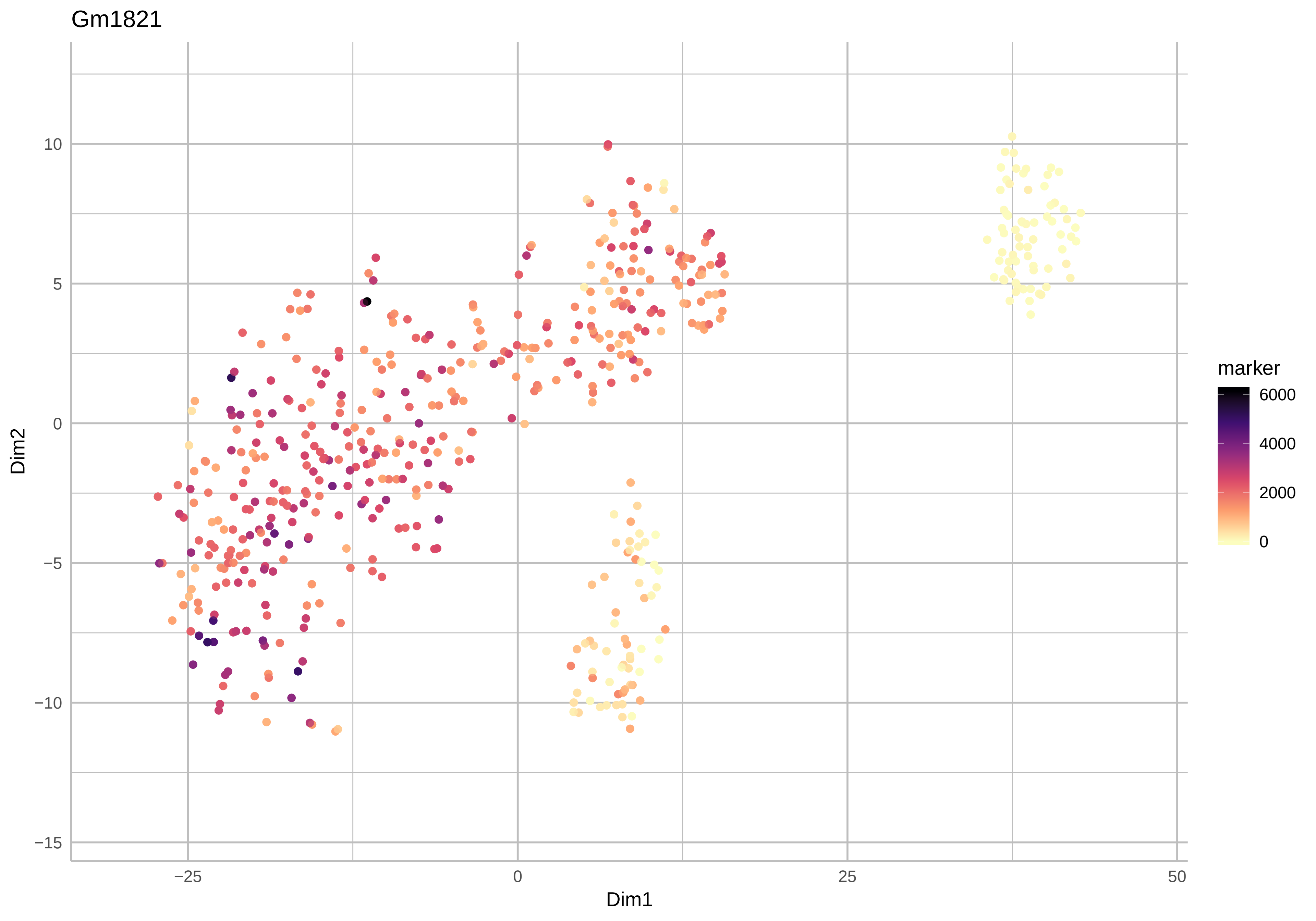

Gm1821
Dim2
Dim1
10
5
0
−5
−10
−15
−25
0
25
50
marker
6000
4000
2000
0

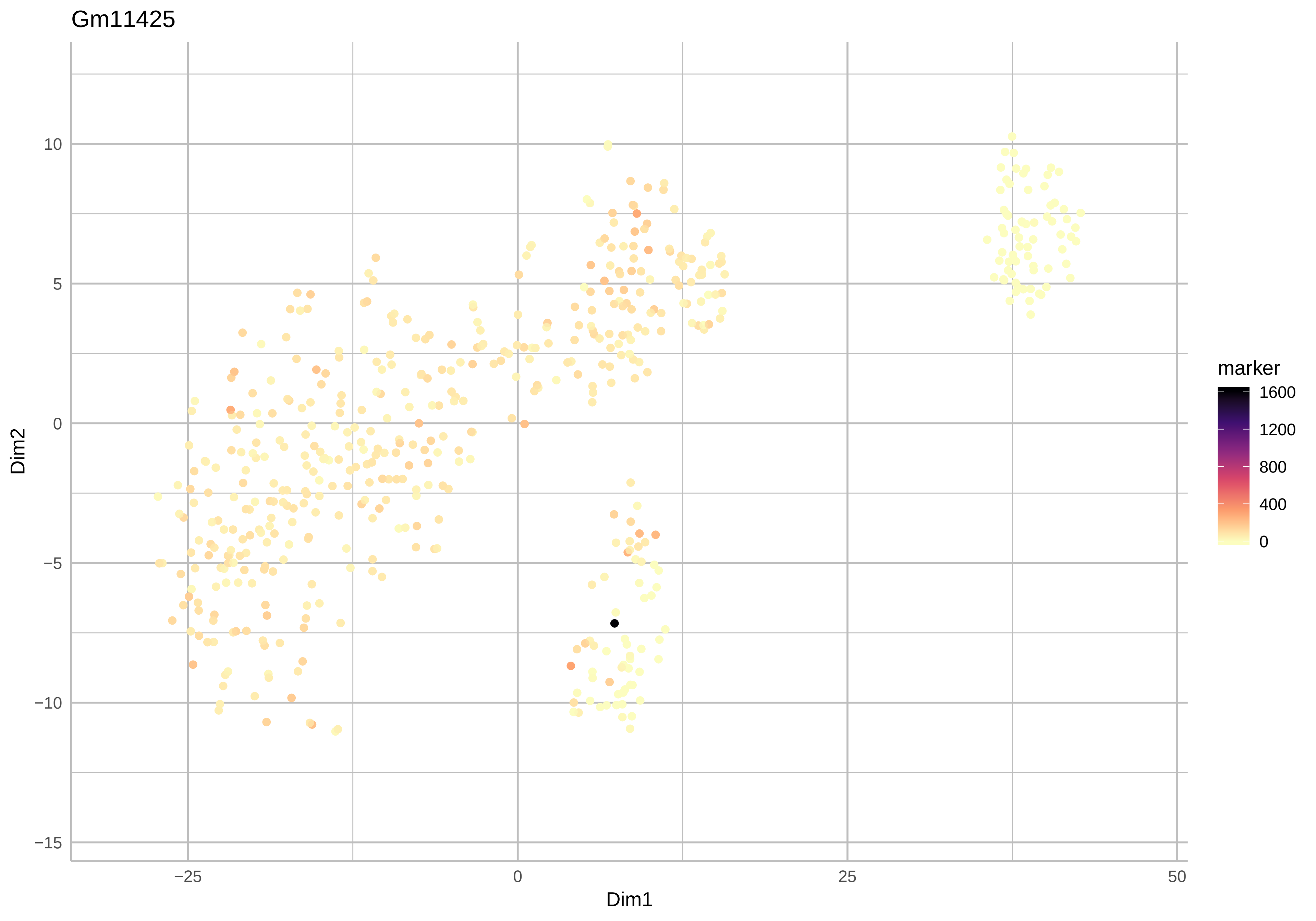
Gm11425
Dim2
Dim1
10
5
0
−5
−10
−15
−25
0
25
50
marker
1600
1200
800
400
0

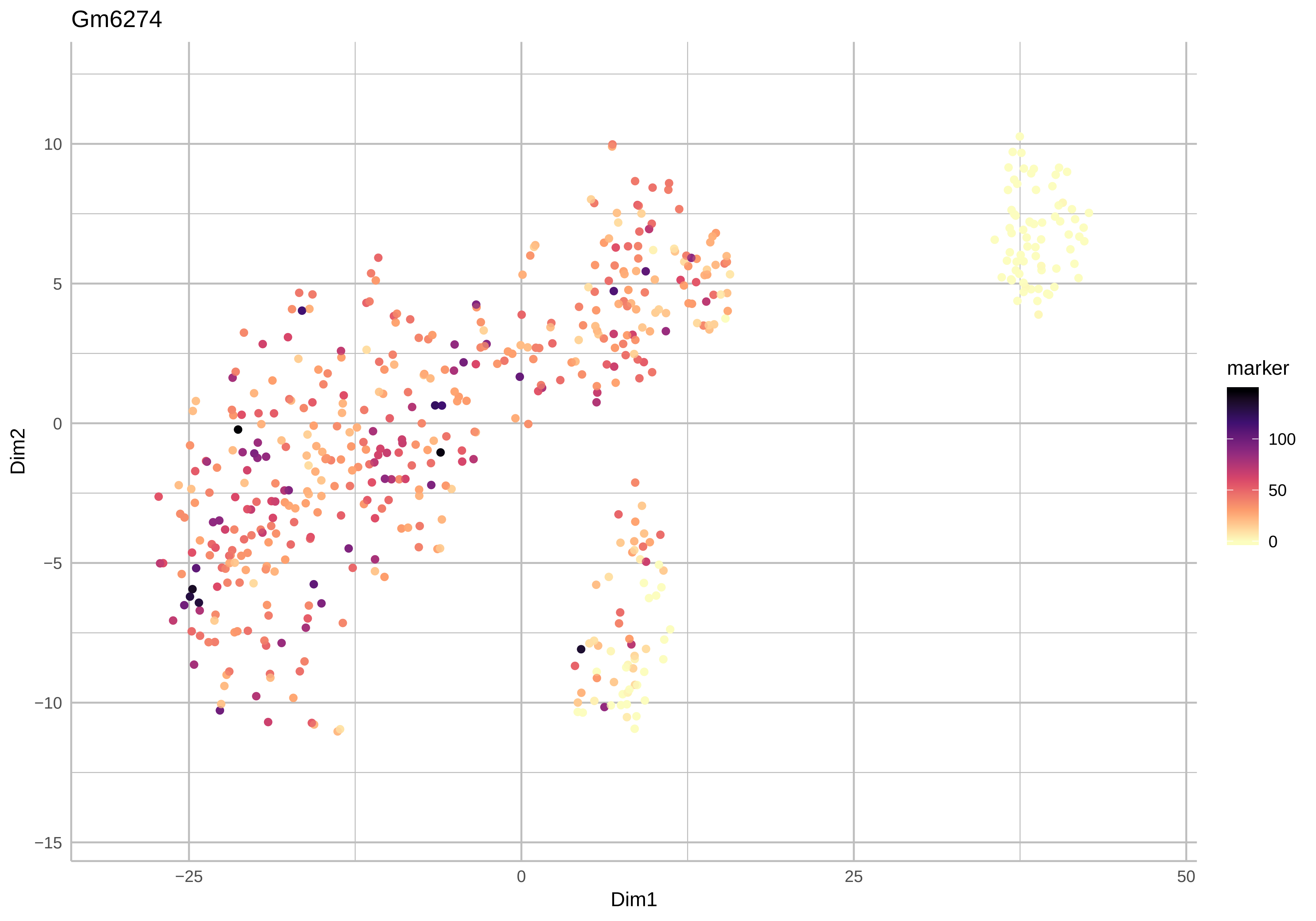

Gm6274
10
5
0
−5
−10
−15
−25
0
25
50
Dim1
Dim2
marker
100
50
0

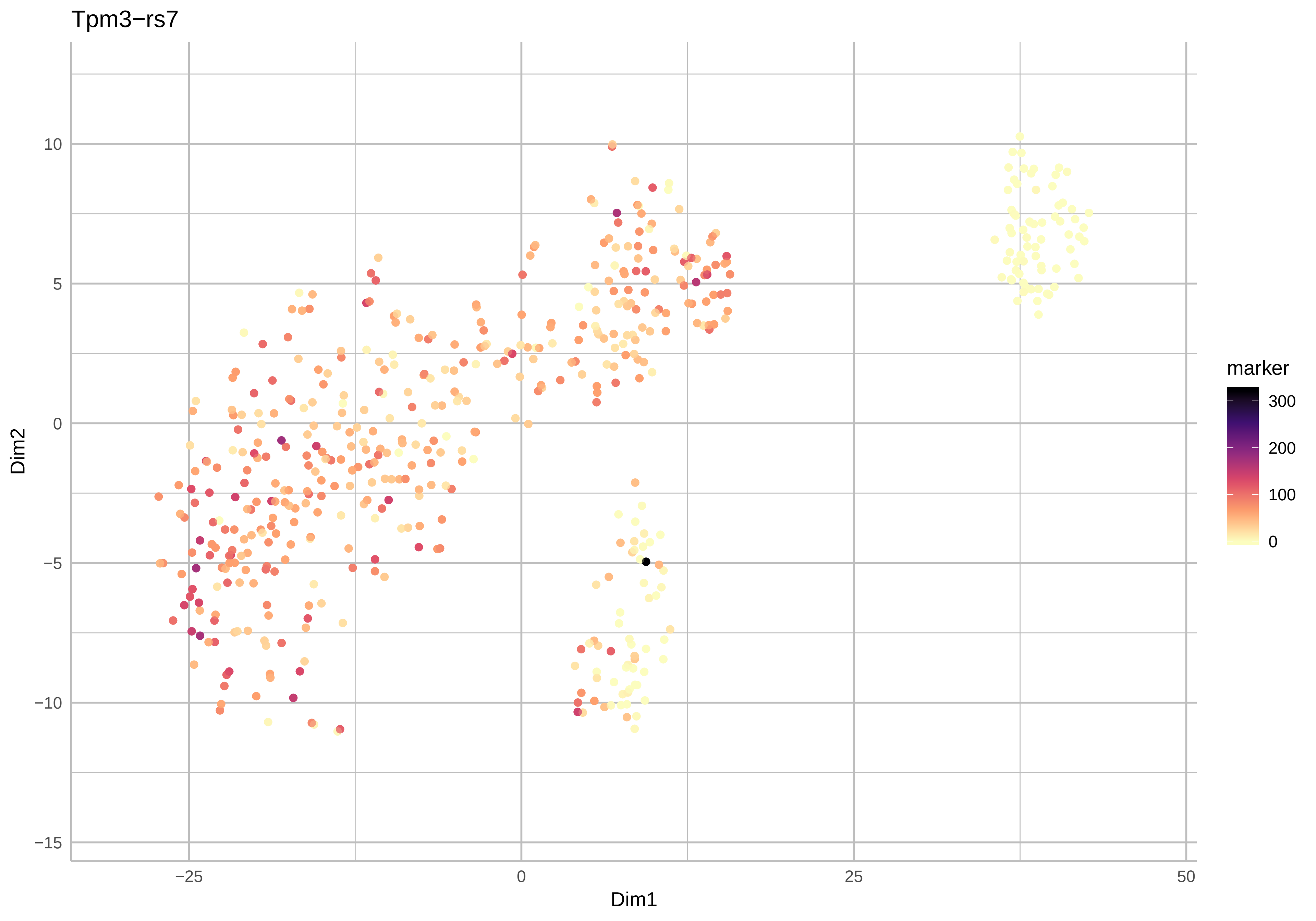
Tpm3−rs7
Dim2
Dim1
marker
300
200
100
0
10
5
0
−5
−10
−15
−25
0
25
50

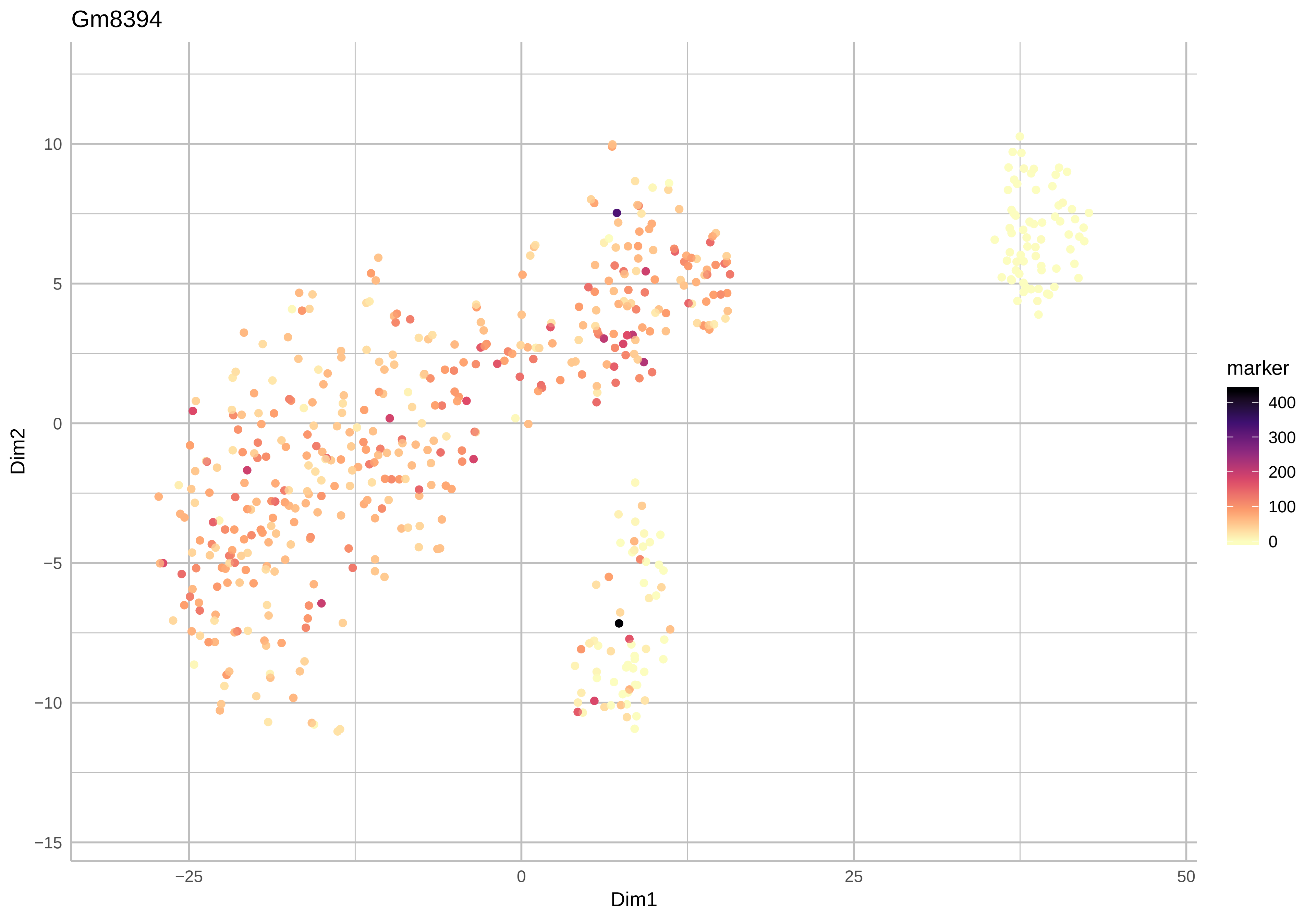

Gm8394
Dim2
Dim1
10
5
0
−5
−10
−15
−25
0
25
50
marker
400
300
200
100
0

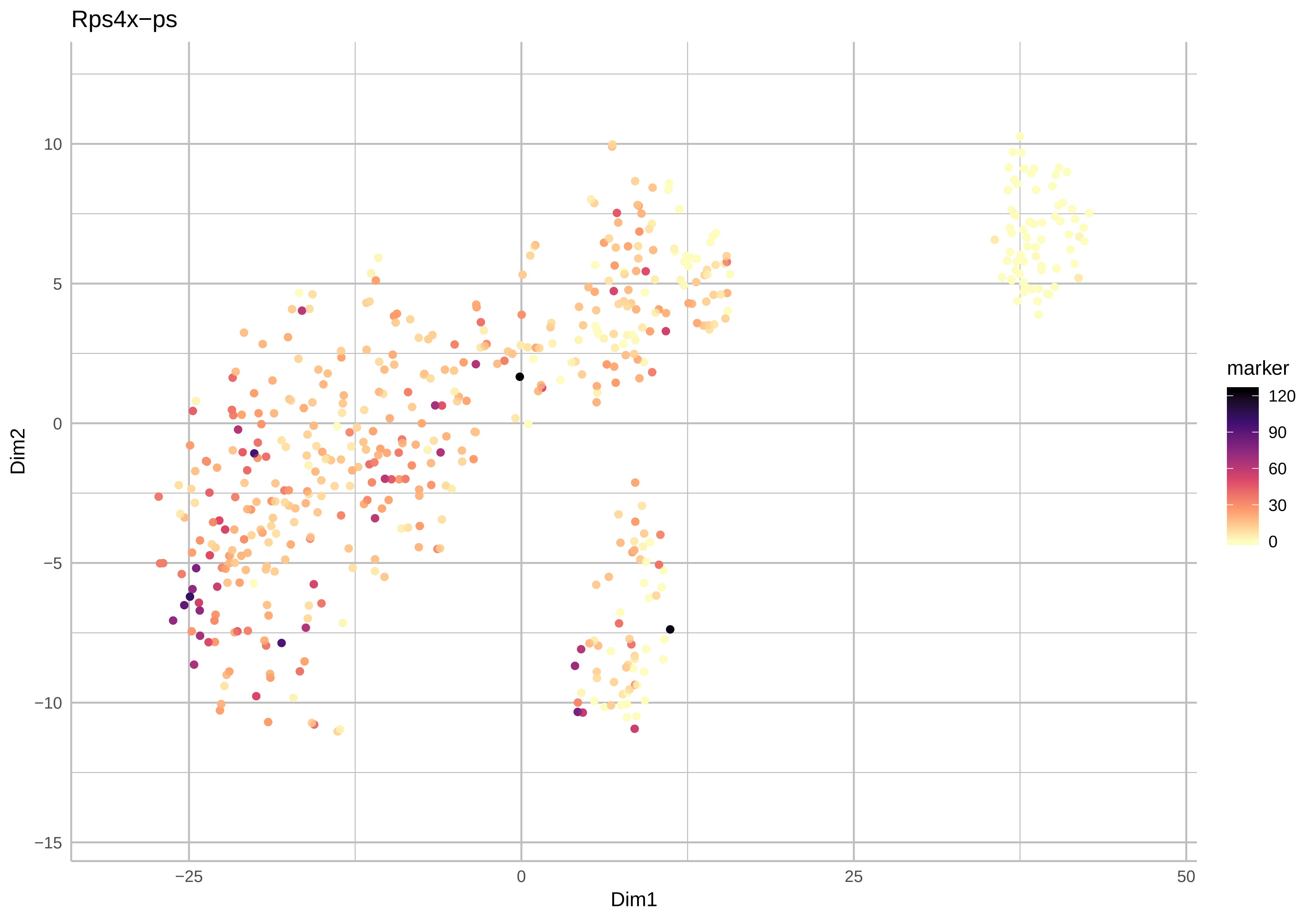
Rps4x−ps
Dim2
Dim1
10
5
0
−5
−10
−15
−25
0
25
50
marker
120
90
60
30
0

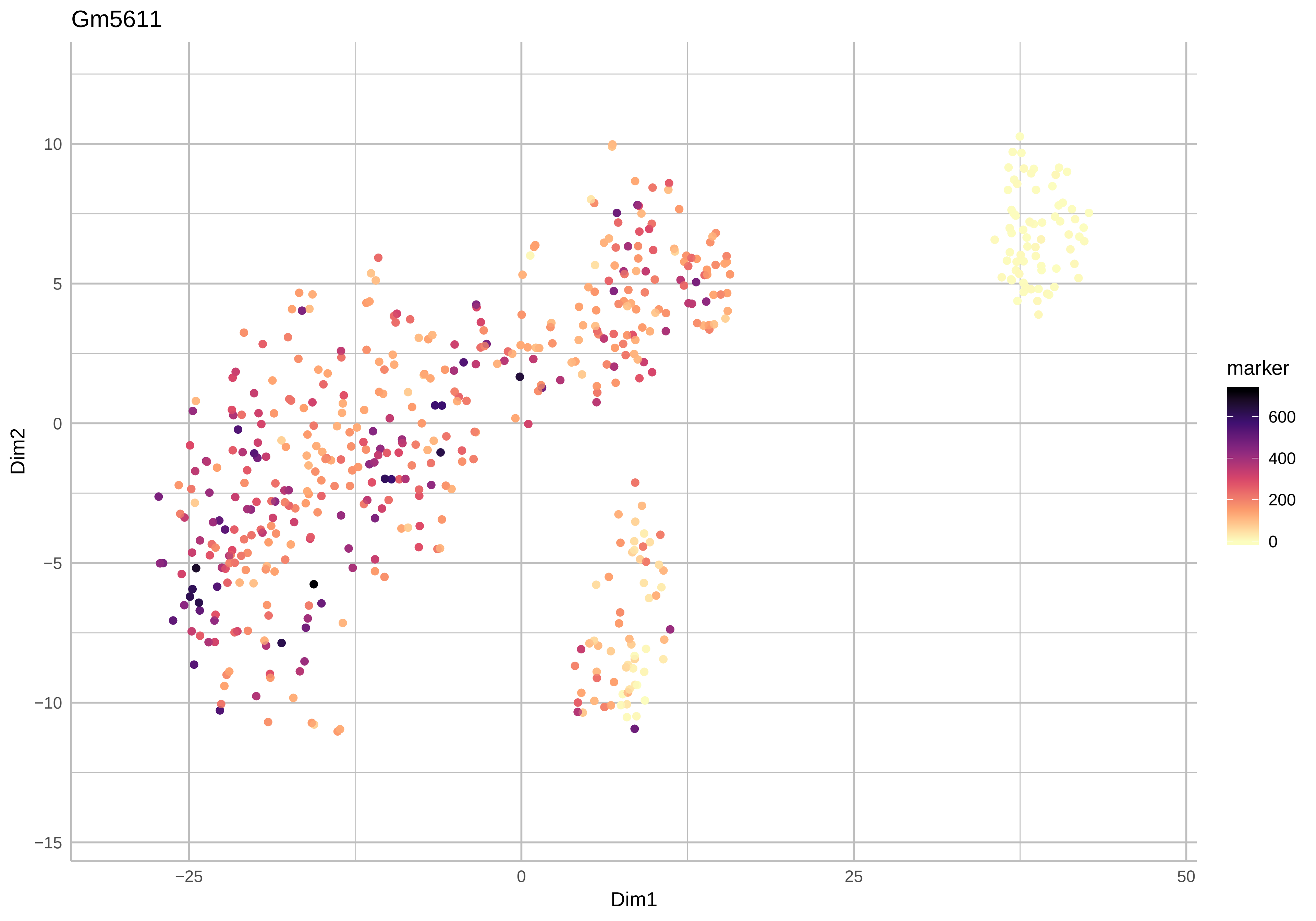

Gm5611
10
5
0
−5
−10
−15
−25
0
25
50
Dim1
Dim2
marker
600
400
200
0

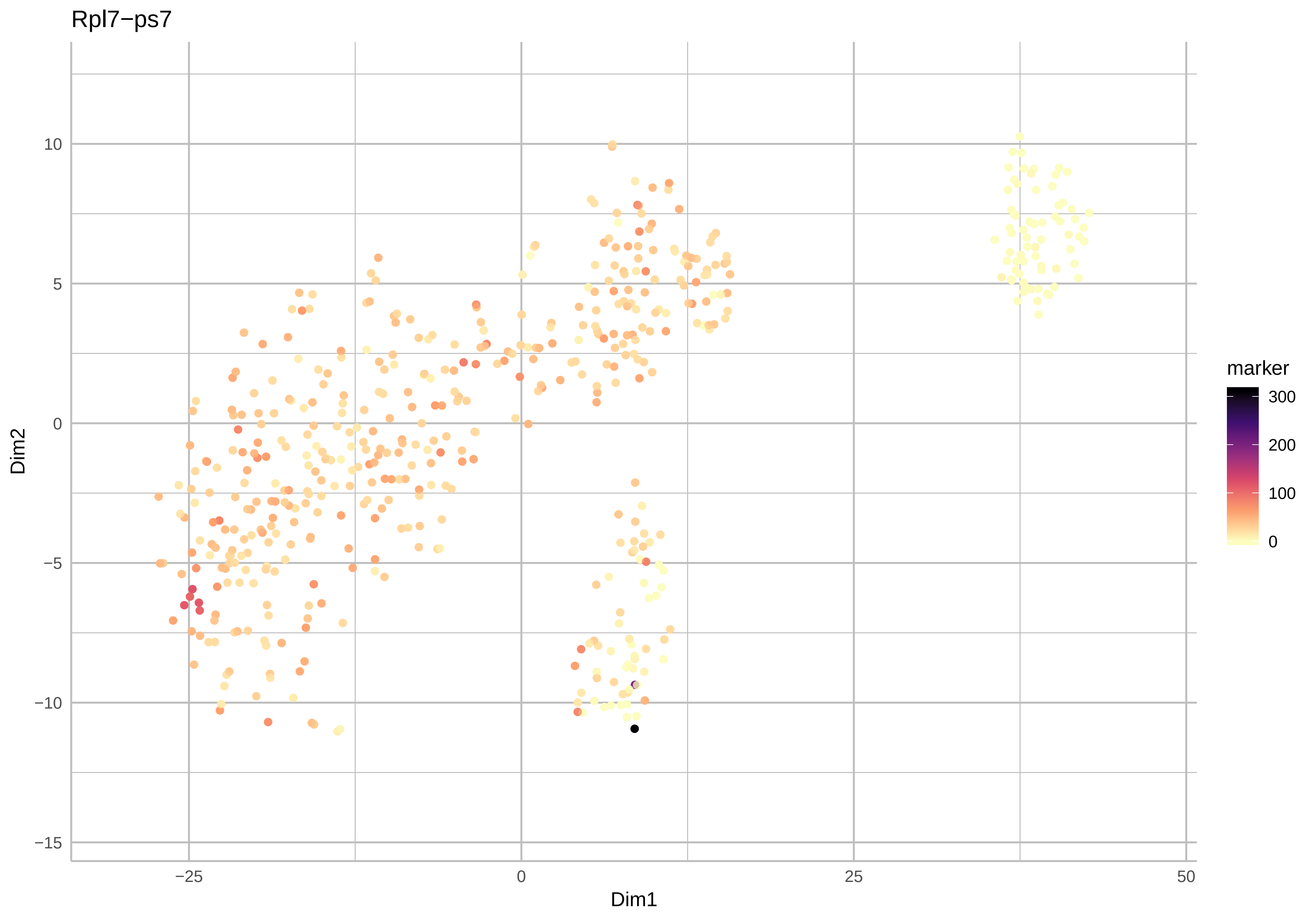

Rpl7−ps7
Dim2
Dim1
10
5
0
−5
−10
−15
−25
0
25
50
marker
300
200
100
0

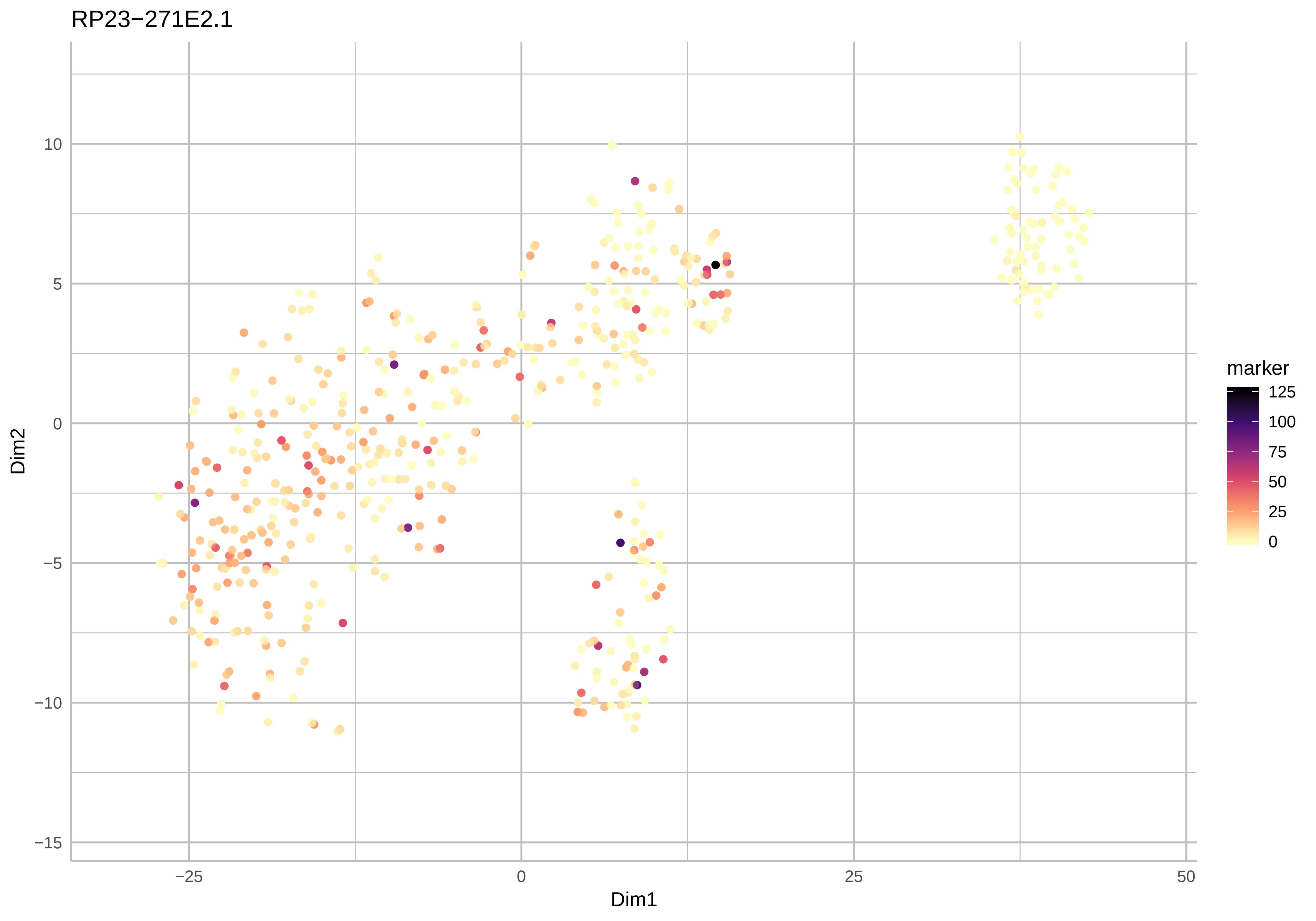

RP23−271E2.1
Dim2
Dim1
10
5
0
−5
−10
−15
−25
0
25
50
marker
125
100
75
50
25
0

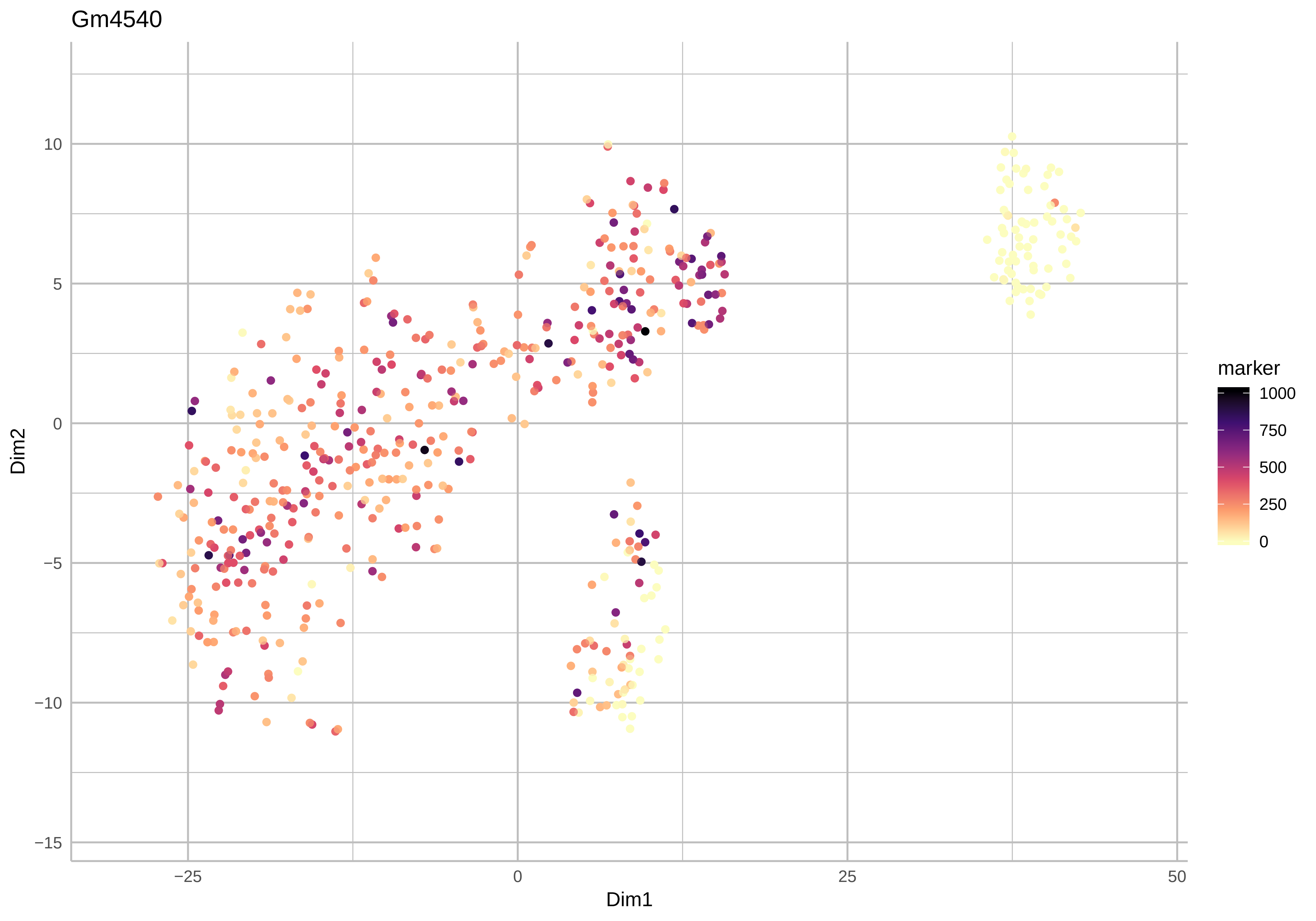
Gm4540
Dim2
Dim1
10
5
0
−5
−10
−15
−25
0
25
50
marker
1000
750
500
250
0

Gm21887

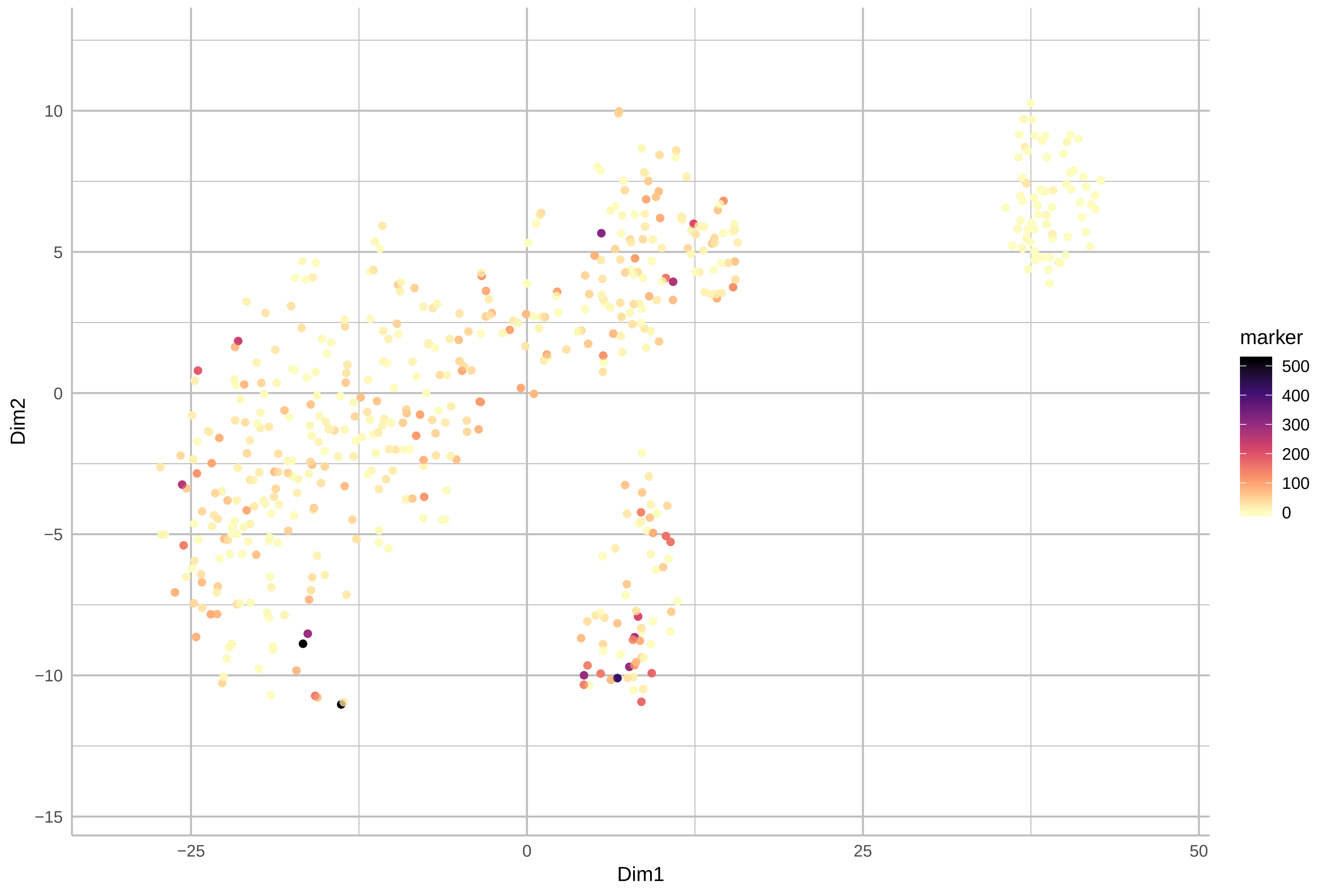

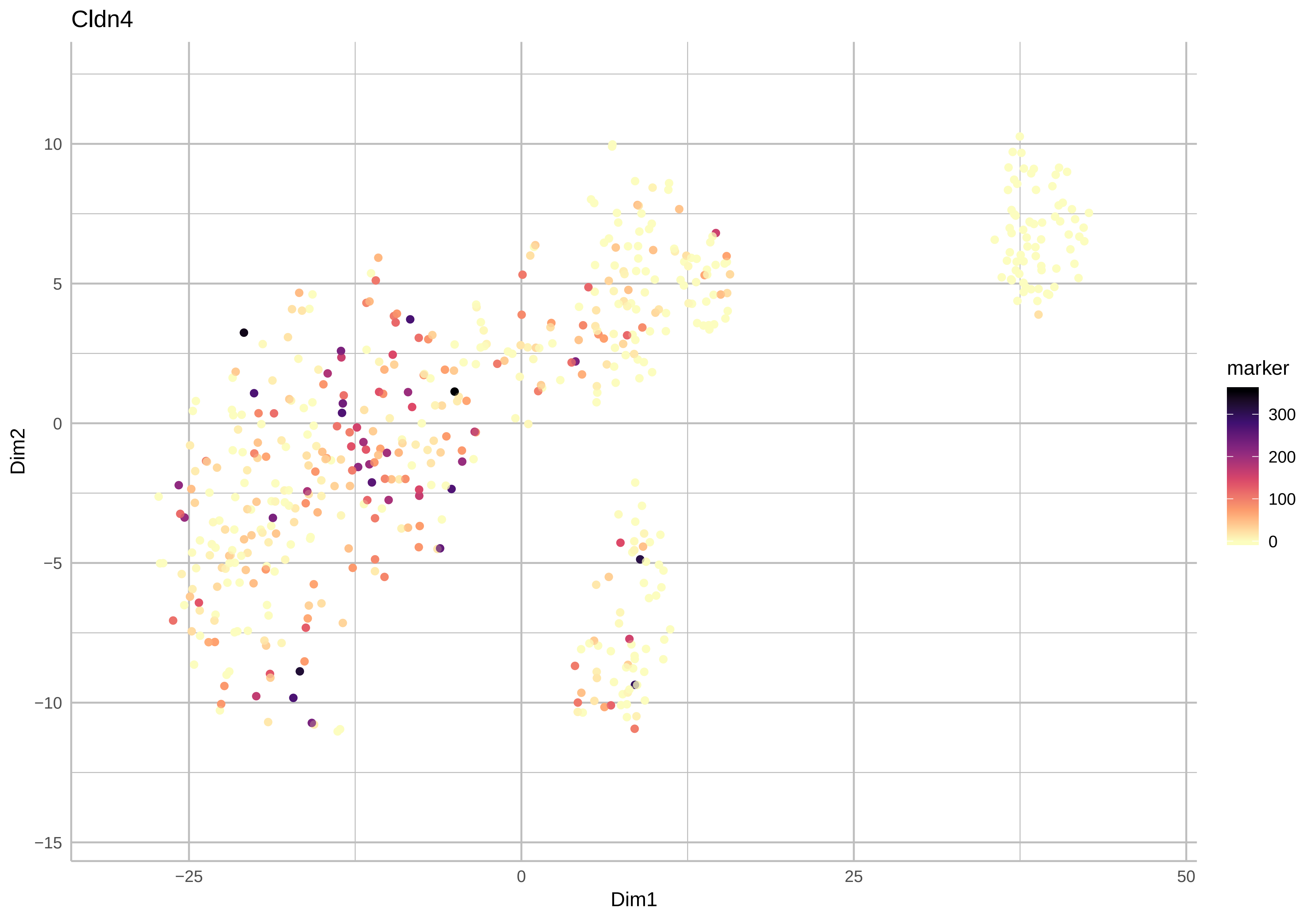

Cldn4
10
5
0
−5
−10
−15
−25
0
25
50
Dim1
Dim2
marker
300
200
100
0

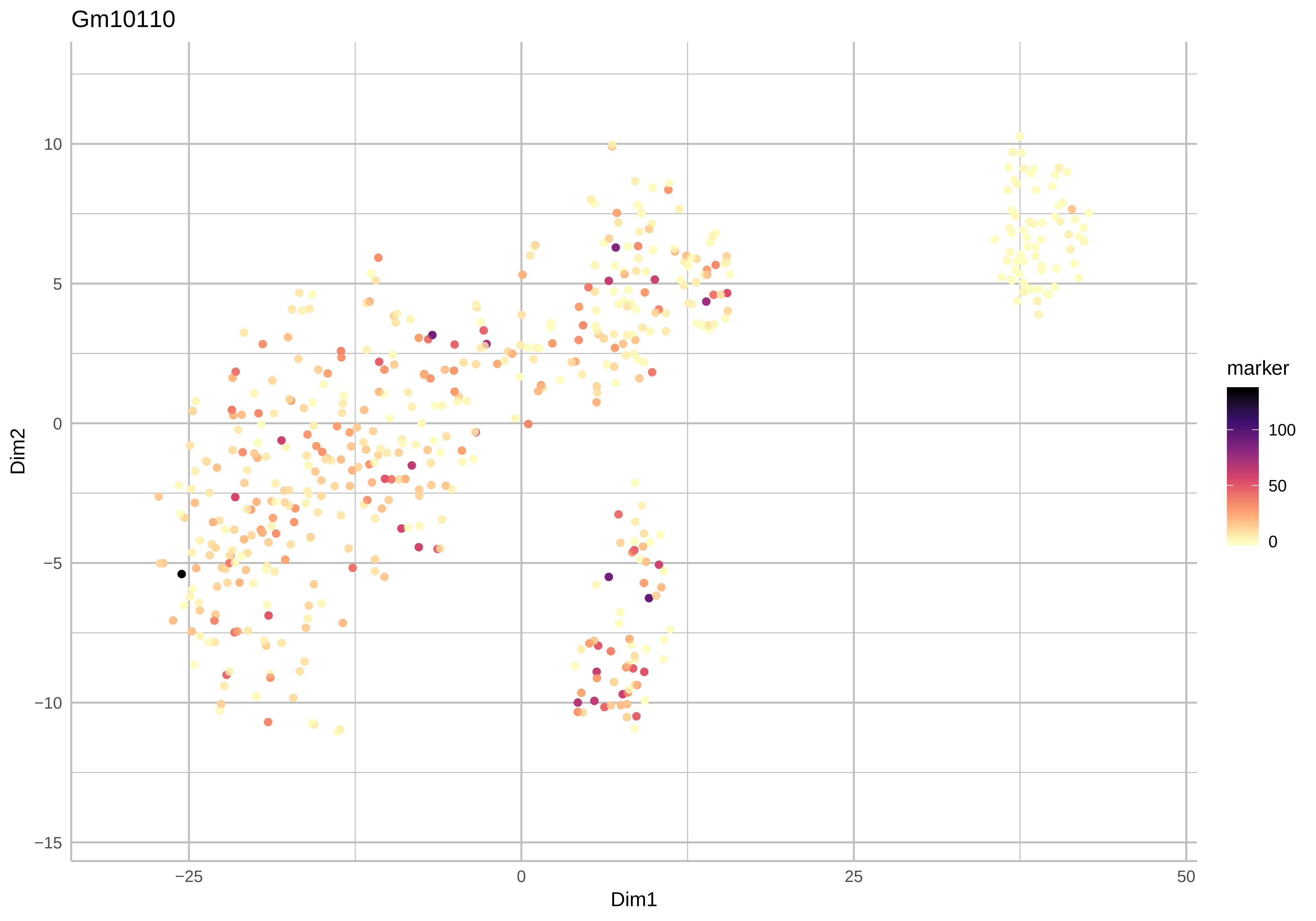
Gm10110
Dim2
Dim1
marker
100
50
0
10
5
0
−5
−10
−15
−25
0
25
50

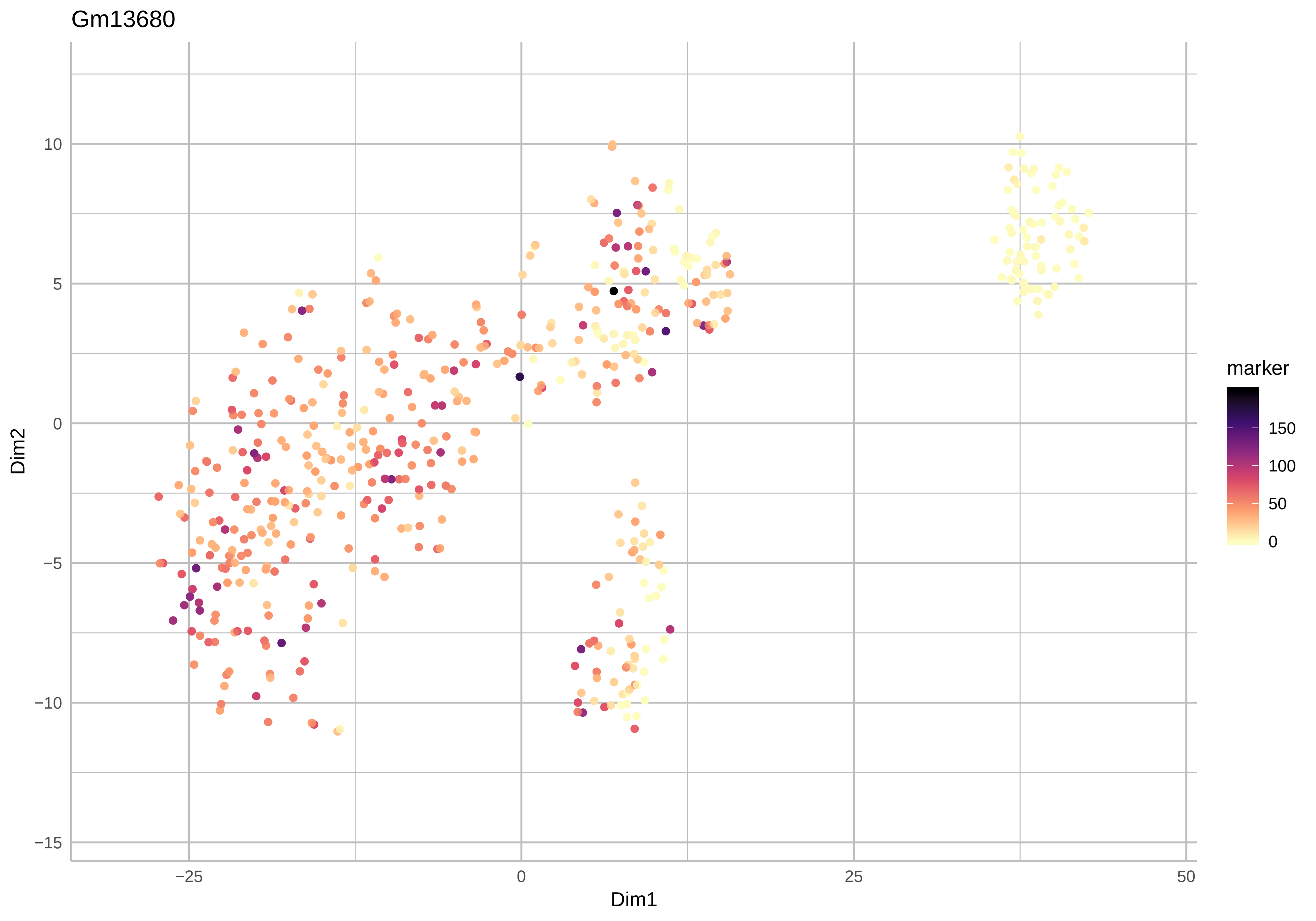

Gm13680
Dim2
Dim1
10
5
0
−5
−10
−15
−25
0
25
50
marker
150
100
50
0

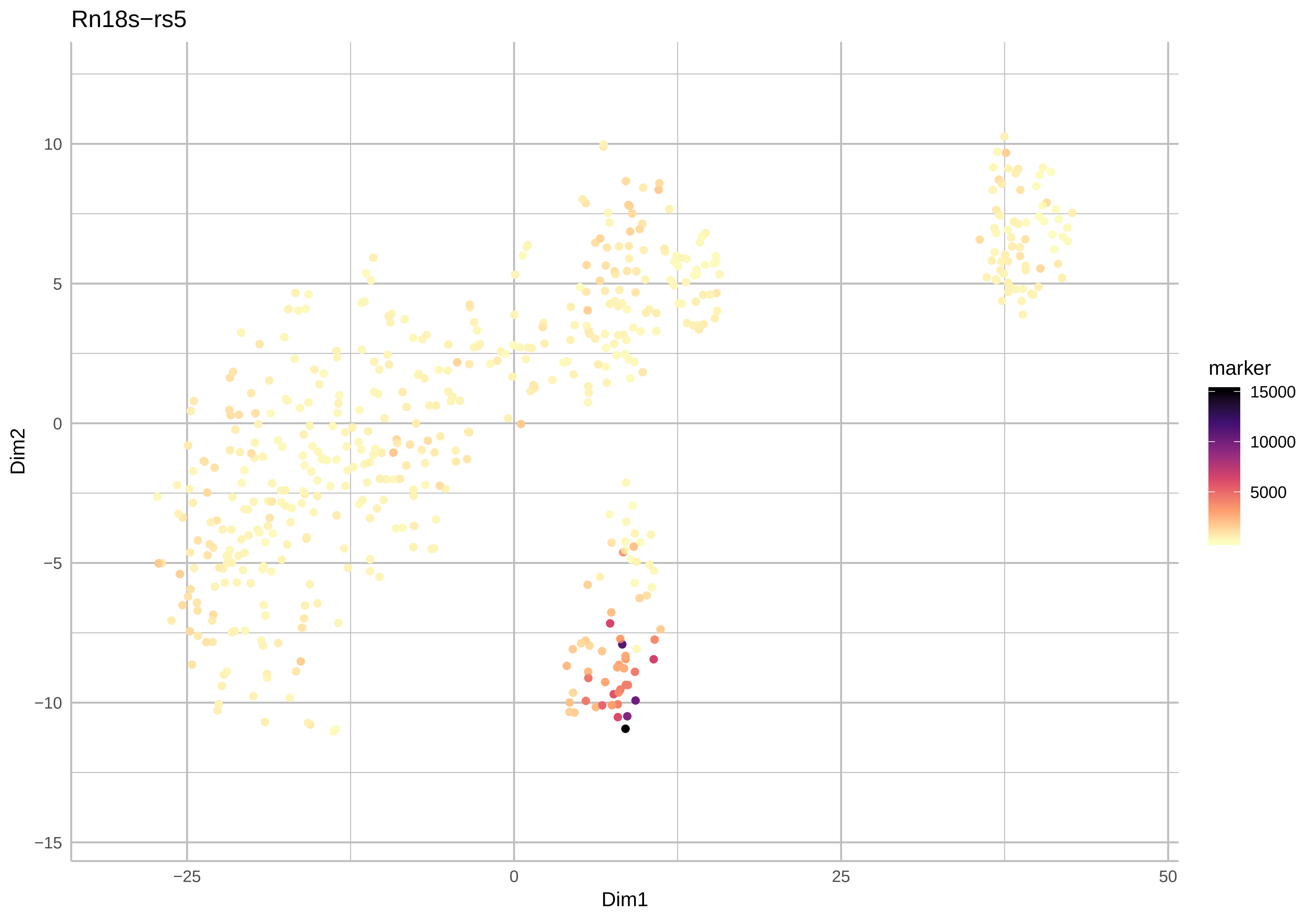
Rn18s−rs5
Dim2
Dim1
10
5
0
−5
−10
−15
−25
0
25
50
marker
15000
10000
5000

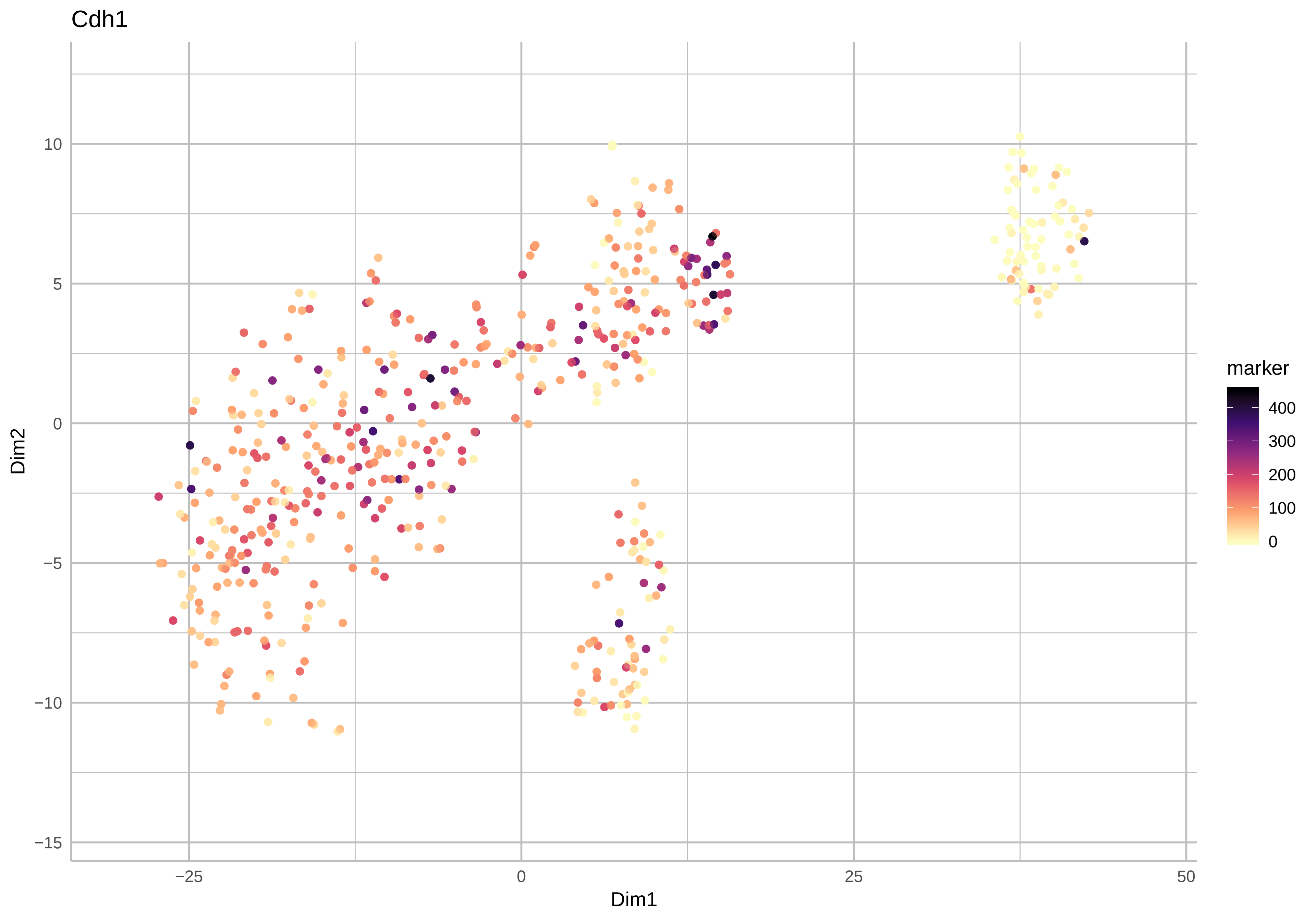
Cdh1
Dim2
Dim1
10
5
0
−5
−10
−15
−25
0
25
50
marker
400
300
200
100
0

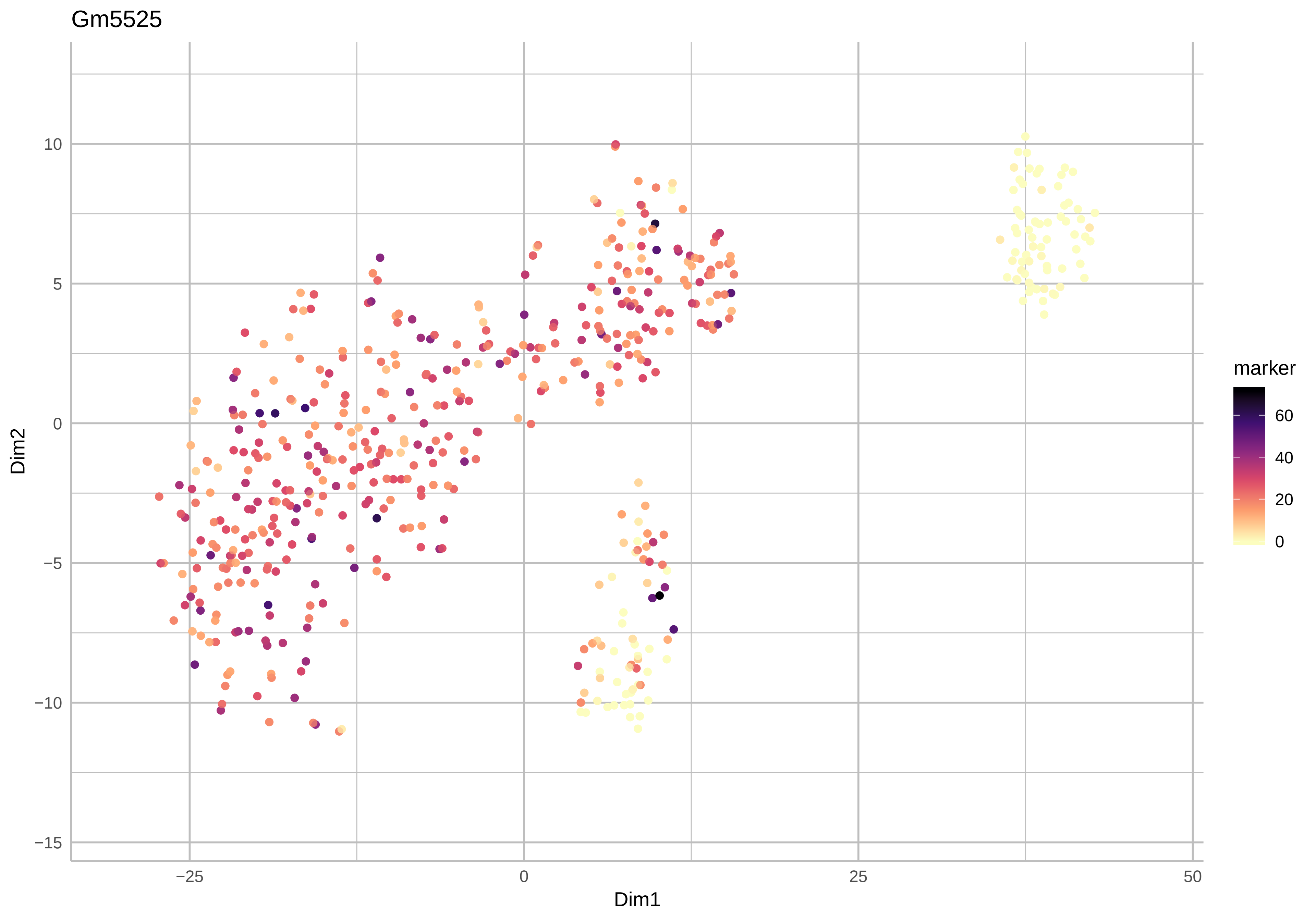
Gm5525
Dim2
Dim1
10
5
0
−5
−10
−15
−25
0
25
50
marker
60
40
20
0

Gm23935

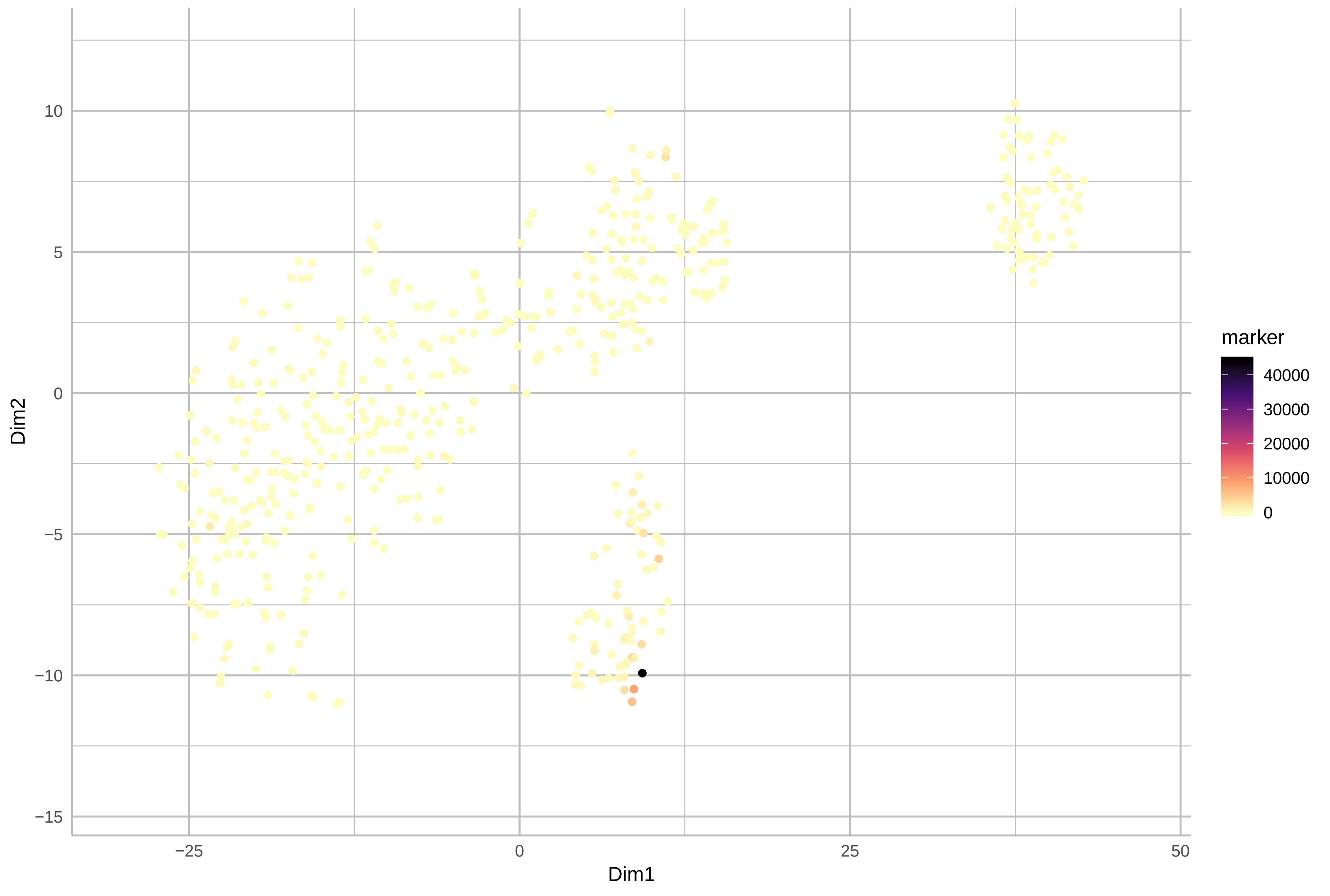

Dim2
10
5
0
−5
−10
−15
−25
0
25
50
Dim1
marker
40000
30000
20000
10000
0

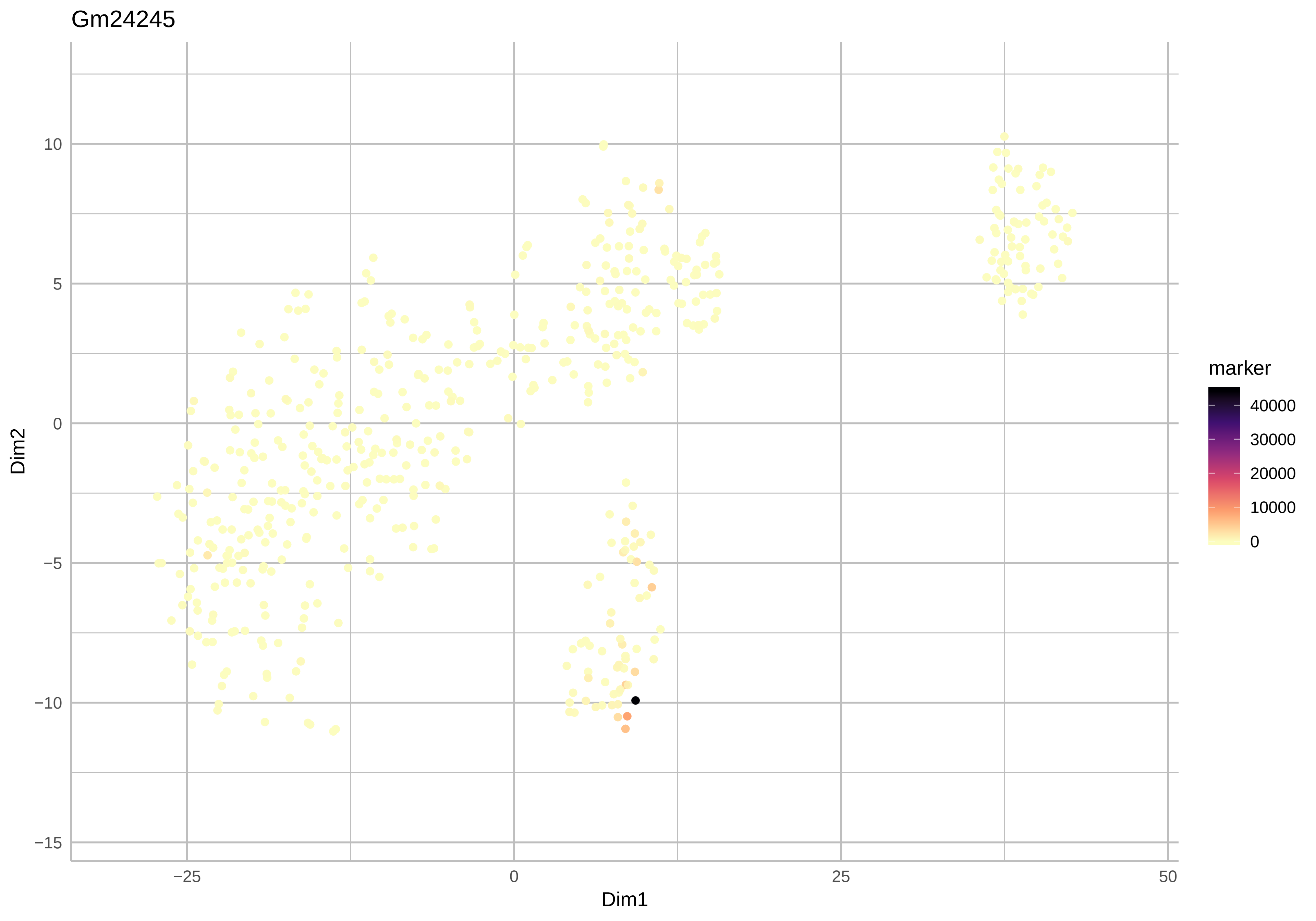
Gm24245
Dim2
Dim1
10
5
0
−5
−10
−15
−25
0
25
50
marker
40000
30000
20000
10000
0

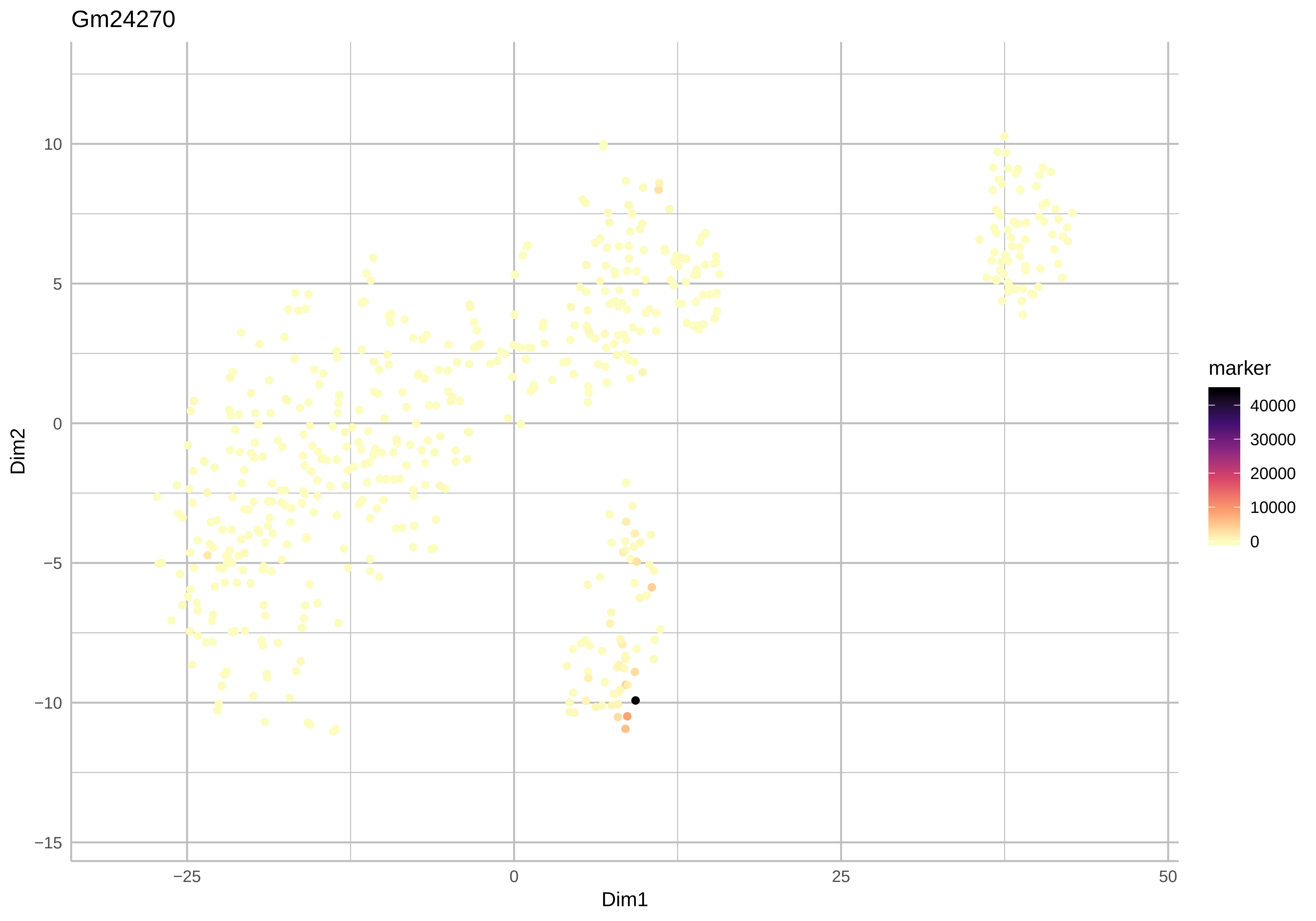

Gm24270
Dim2
Dim1
marker
40000
30000
20000
10000
0
10
5
0
−5
−10
−15
−25
0
25
50

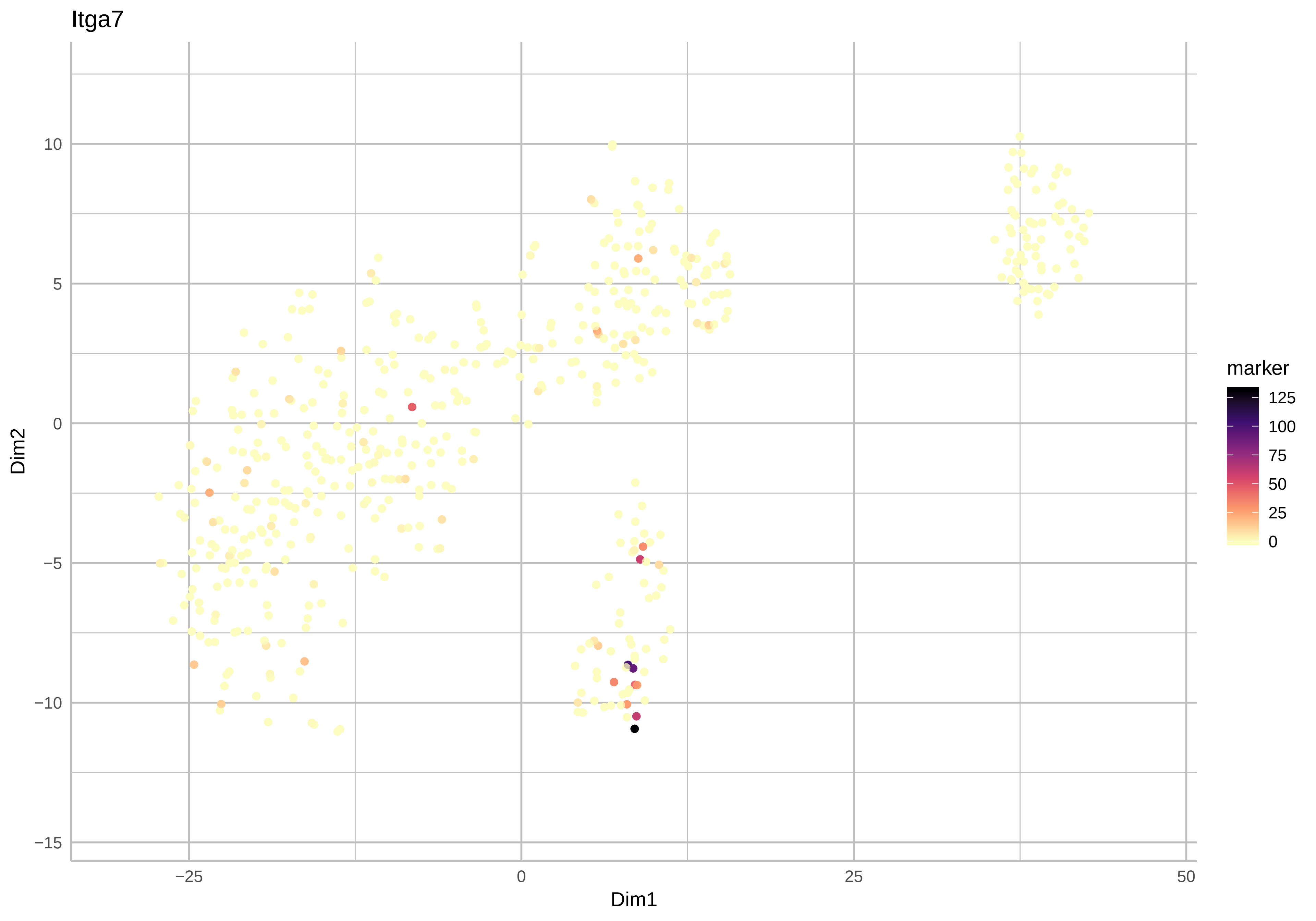
Itga7
Dim2
Dim1
10
5
0
−5
−10
−15
−25
0
25
50
marker
125
100
75
50
25
0

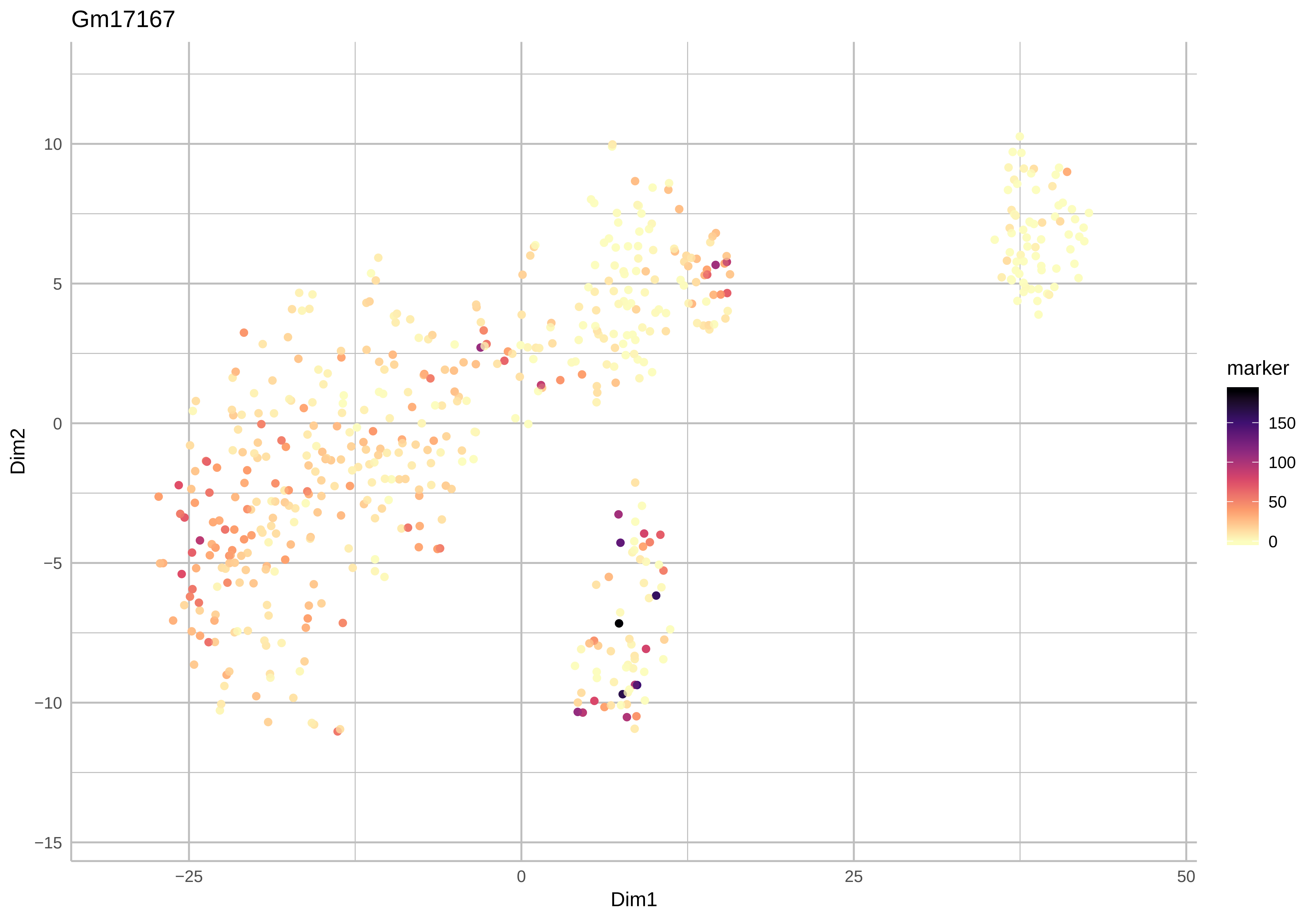

Gm17167
Dim2
Dim1
marker
150
100
50
0
10
5
0
−5
−10
−15
−25
0
25
50

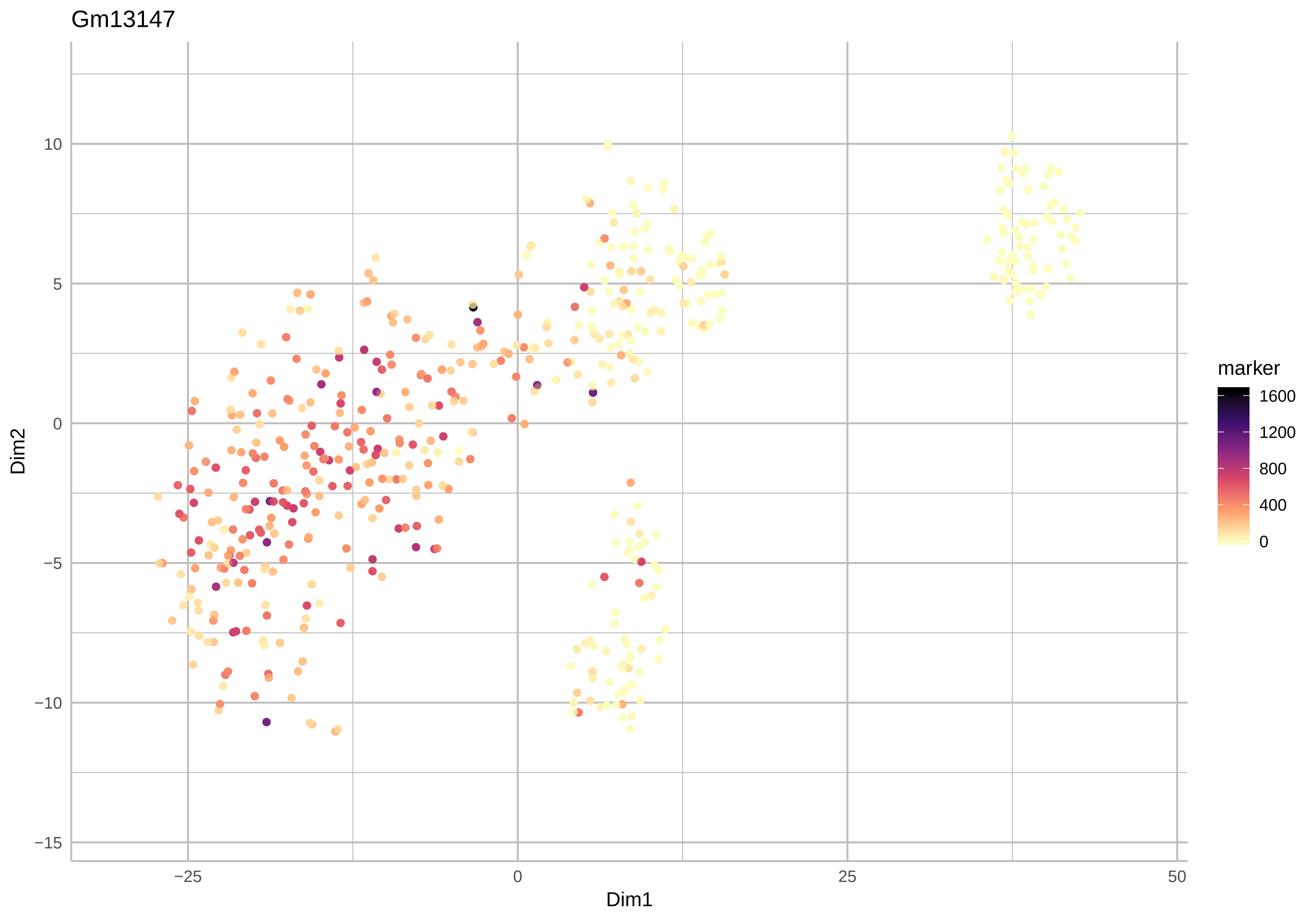
Gm13147
Dim2
10
5
0
−5
−10
−15
Dim1
−25
0
25
50
marker
1600
1200
800
400
0

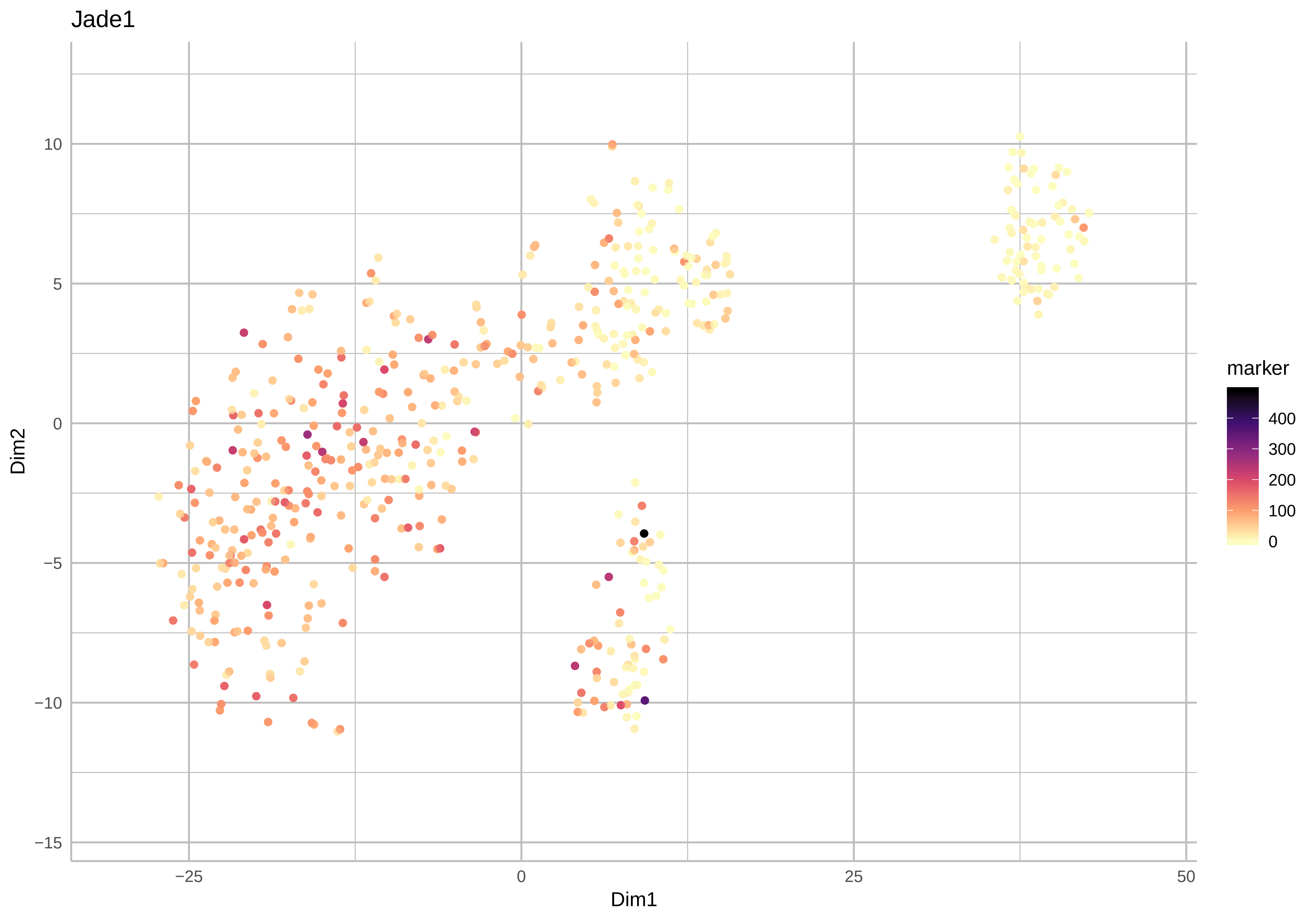

Jade1
Dim2
Dim1
10
5
0
−5
−10
−15
−25
0
25
50
marker
400
300
200
100
0

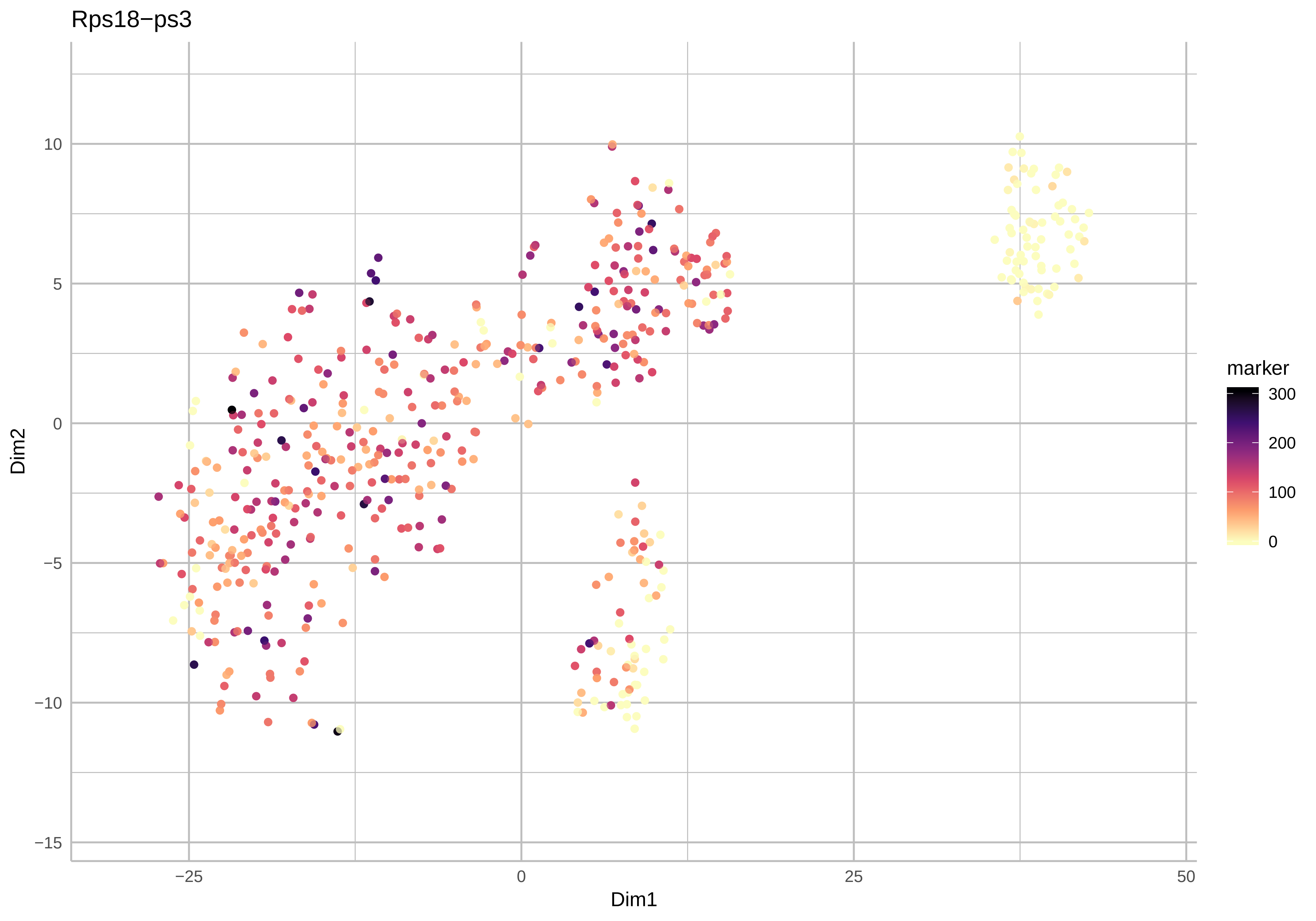
Rps18−ps3
10
5
0
−5
−10
−15
Dim2
−25
0
25
50
Dim1
marker
300
200
100
0

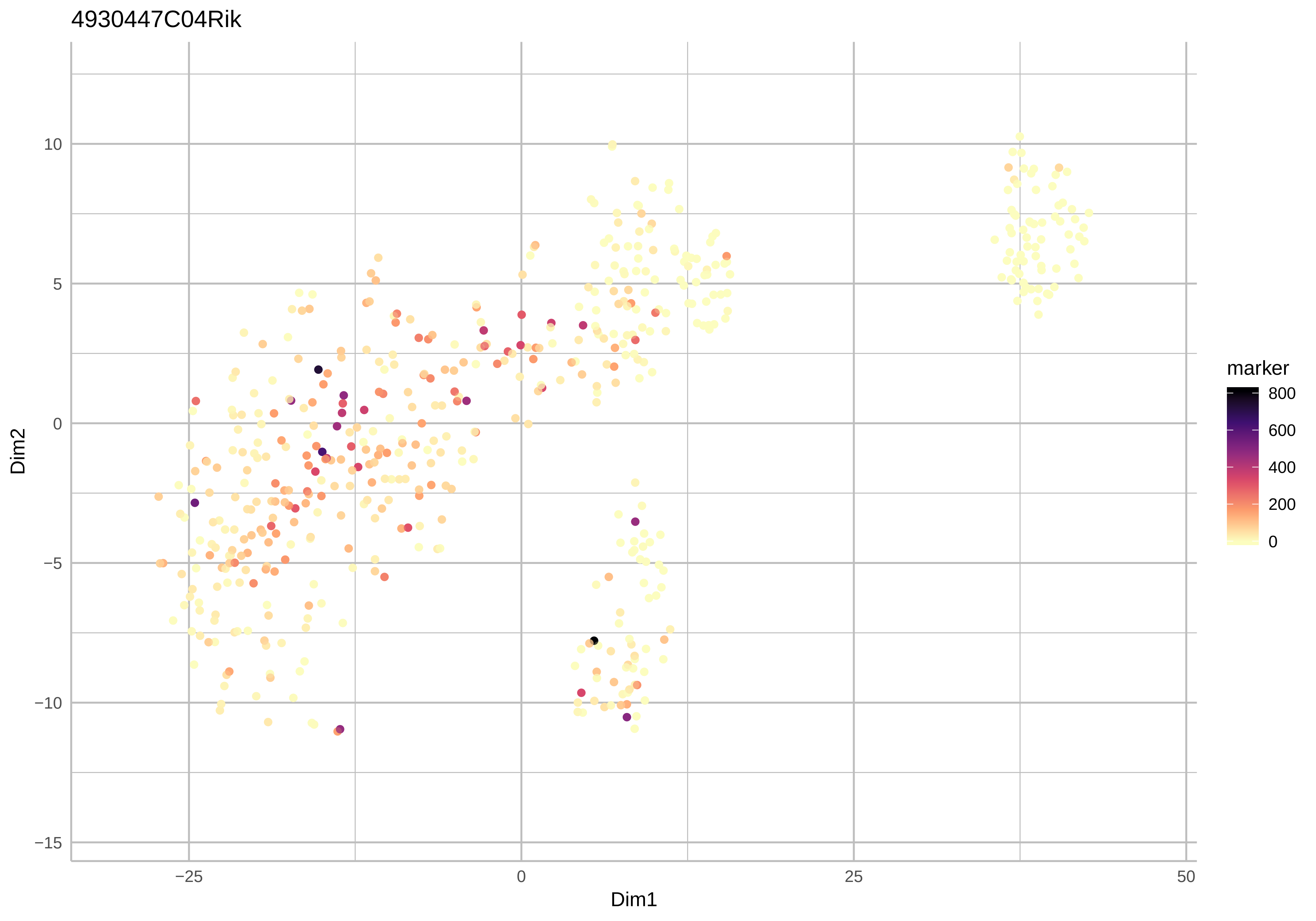
4930447C04Rik
Dim2
Dim1
10
5
0
−5
−10
−15
−25
0
25
50
marker
800
600
400
200
0

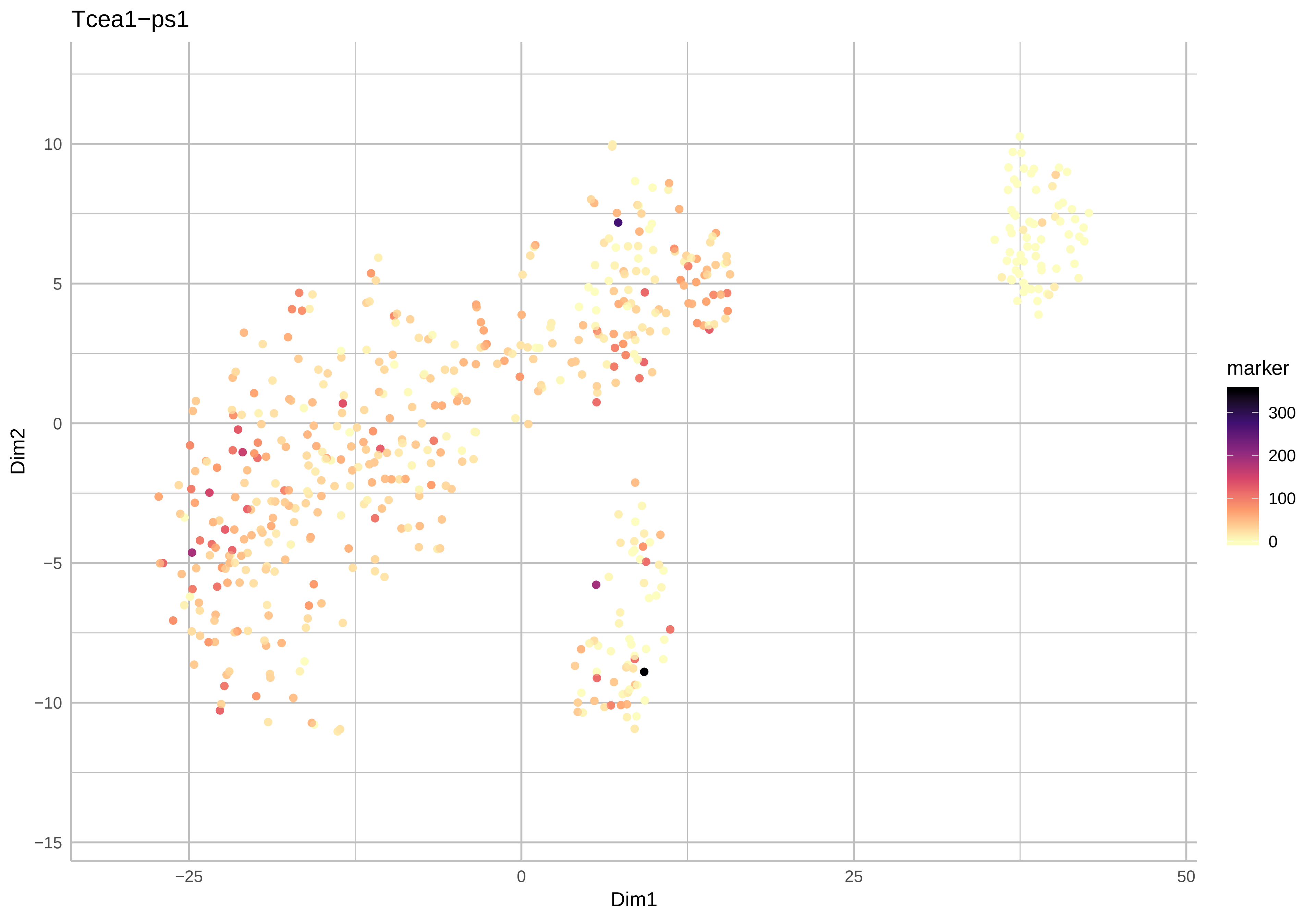

Tcea1−ps1
Dim2
Dim1
10
5
0
−5
−10
−15
−25
0
25
50
marker
300
200
100
0

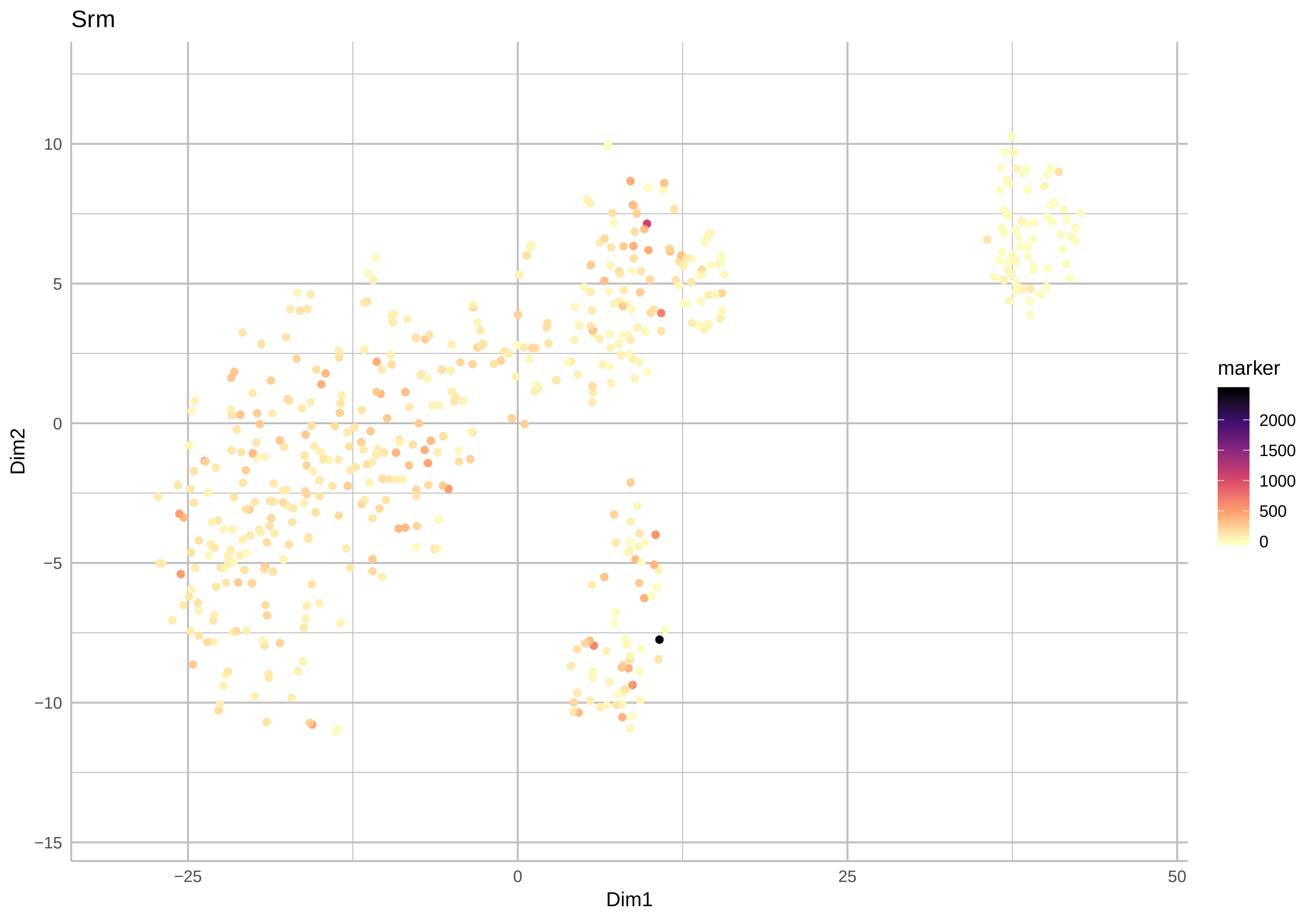

Srm
Dim2
Dim1
10
5
0
−5
−10
−15
−25
0
25
50
marker
2000
1500
1000
500
0

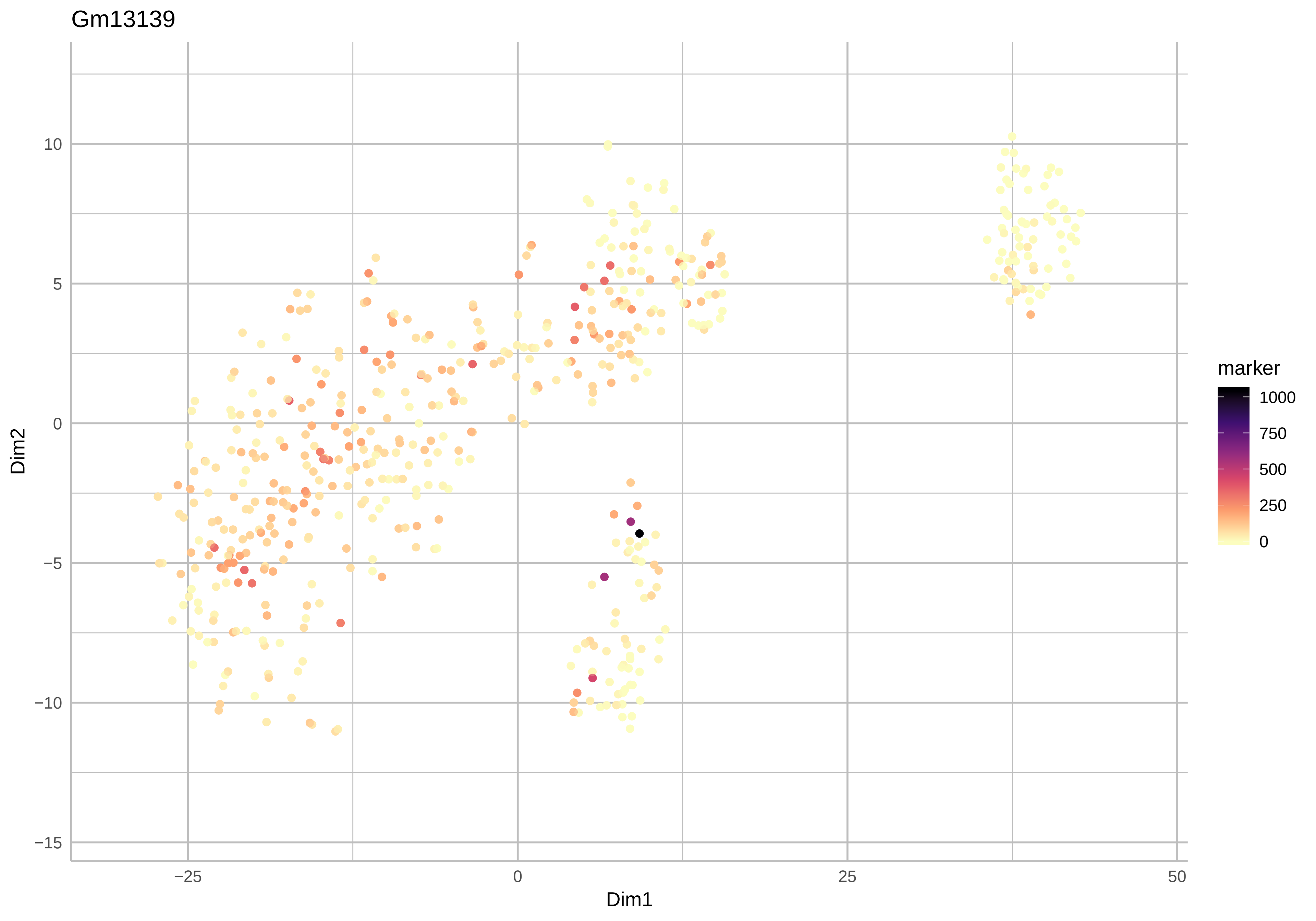
Gm13139
Dim2
10
5
0
−5
−10
−15
−25
0
25
50
Dim1
marker
1000
750
500
250
0

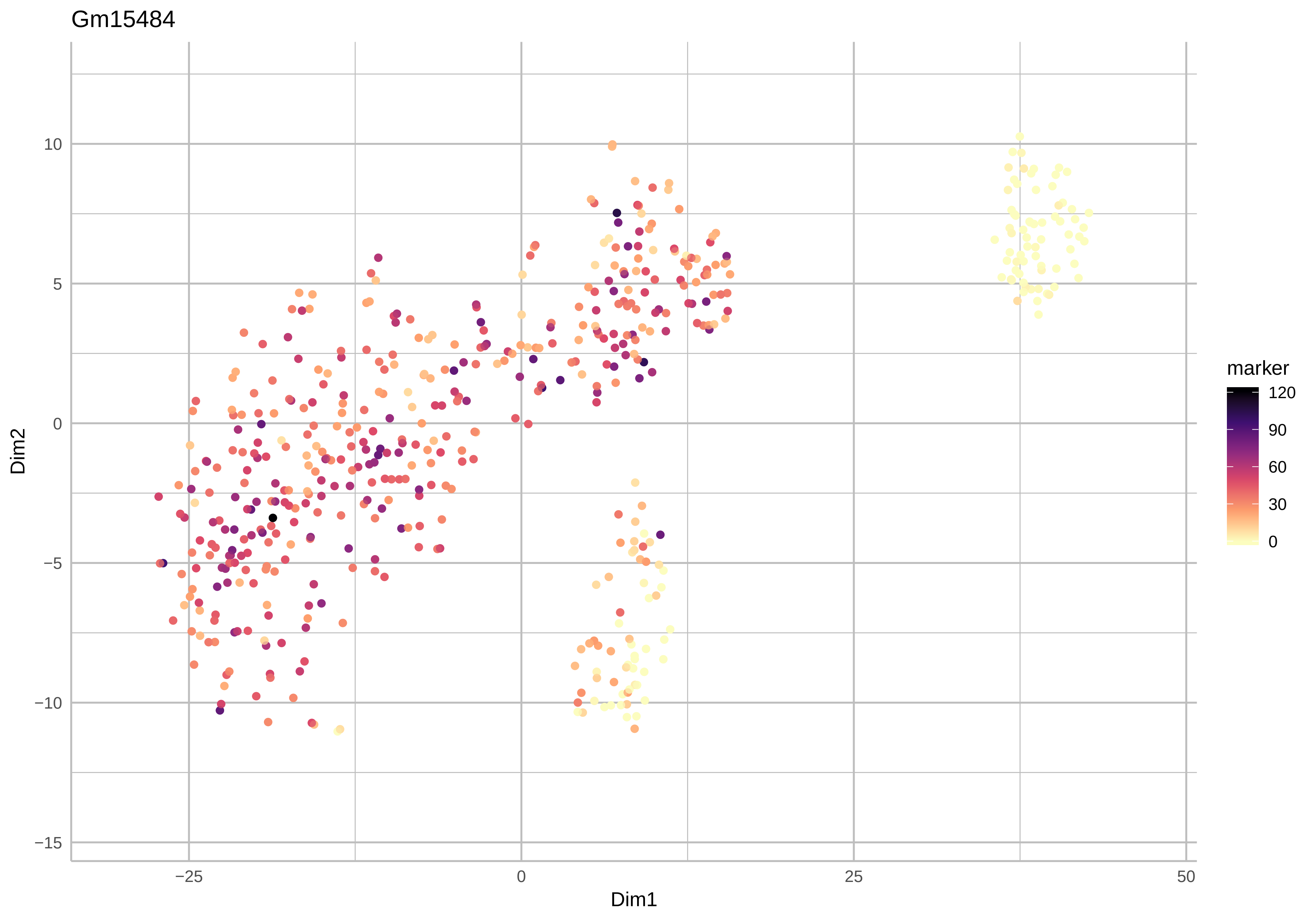
Gm15484
Dim2
Dim1
marker
120
90
60
30
0
10
5
0
−5
−10
−15
−25
0
25
50

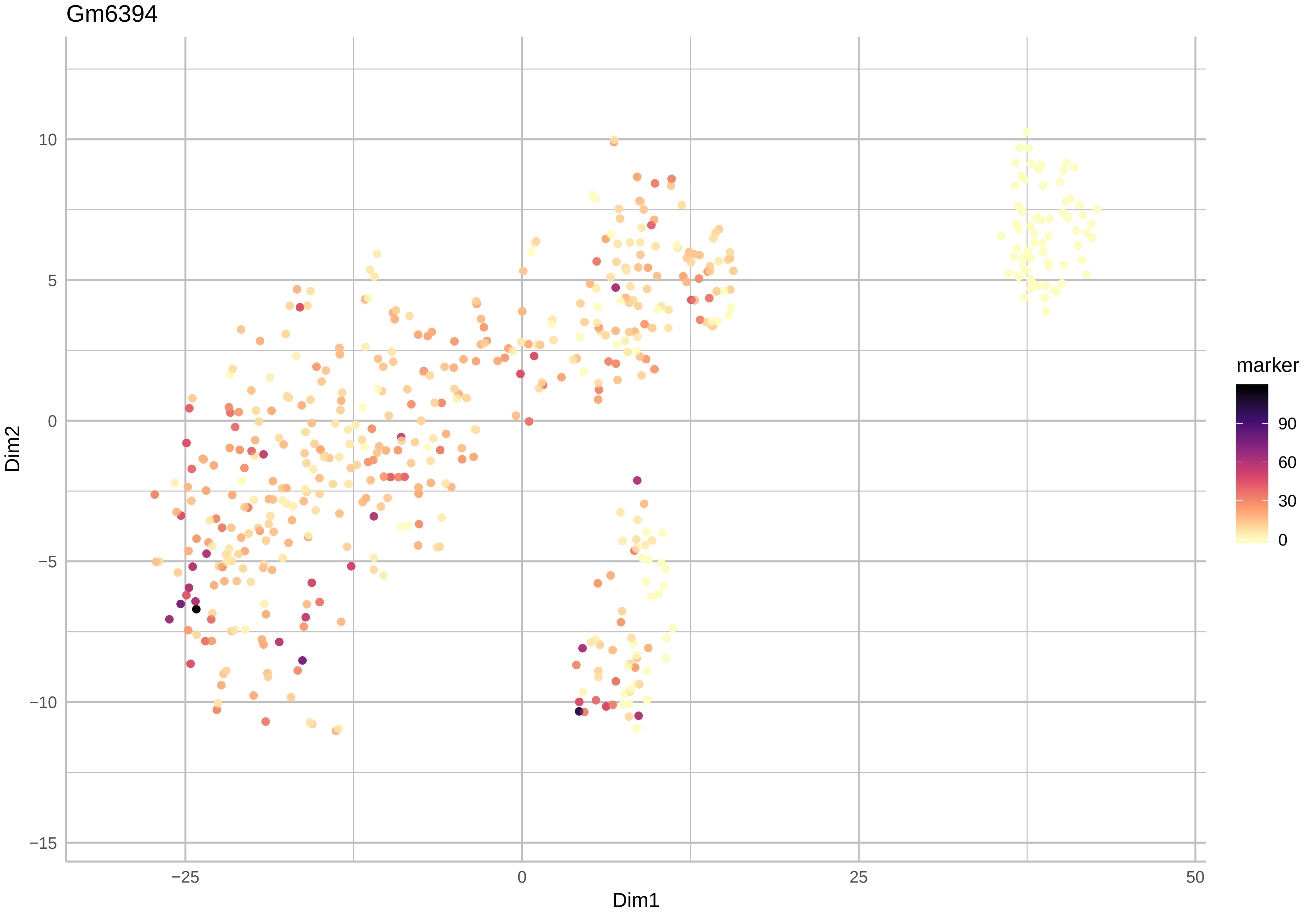

Gm6394
10
5
0
−5
−10
−15
Dim2
−25
0
25
50
Dim1
marker
90
60
30
0

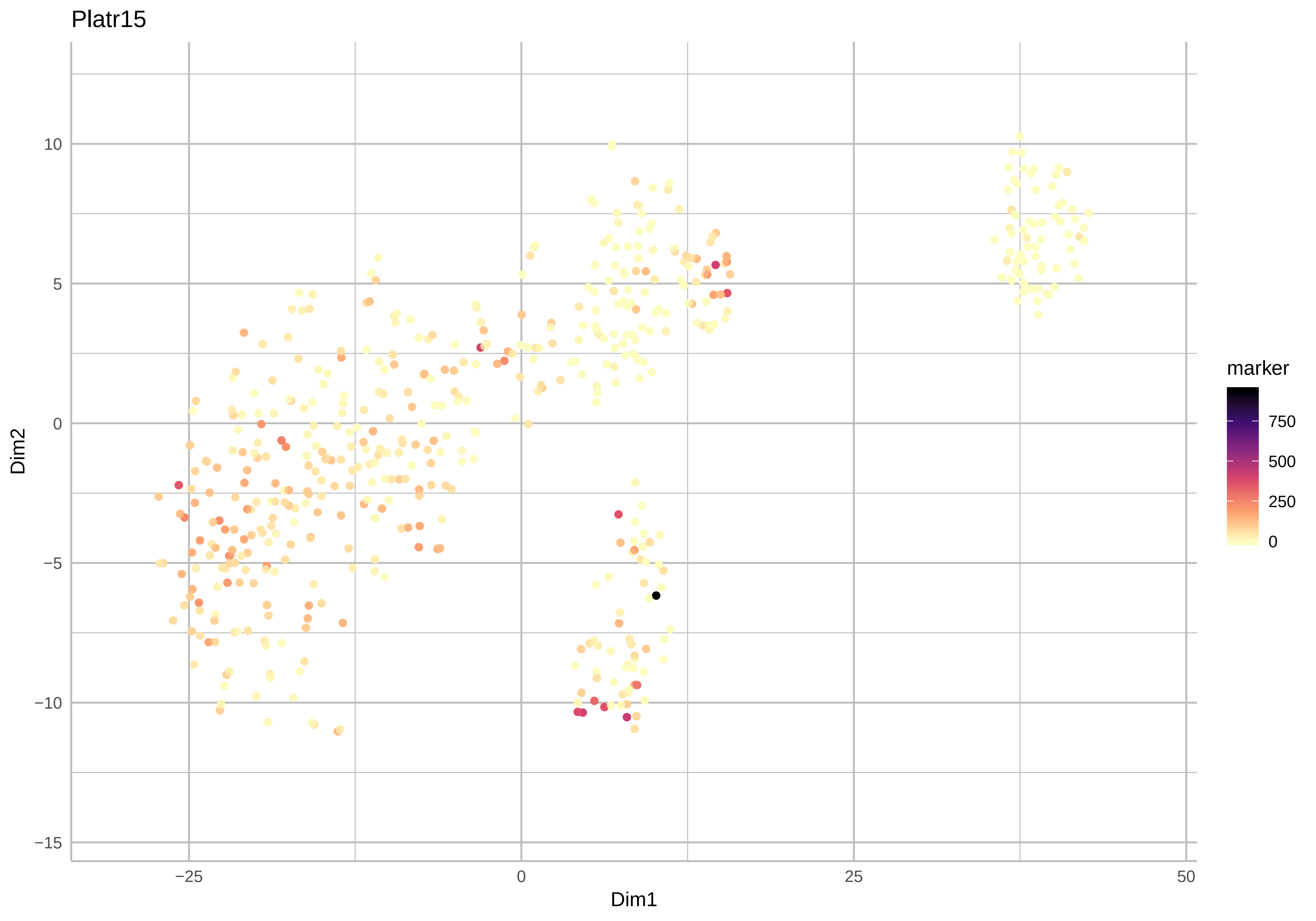
Platr15
Dim2
Dim1
10
5
0
−5
−10
−15
−25
0
25
50
marker
750
500
250
0

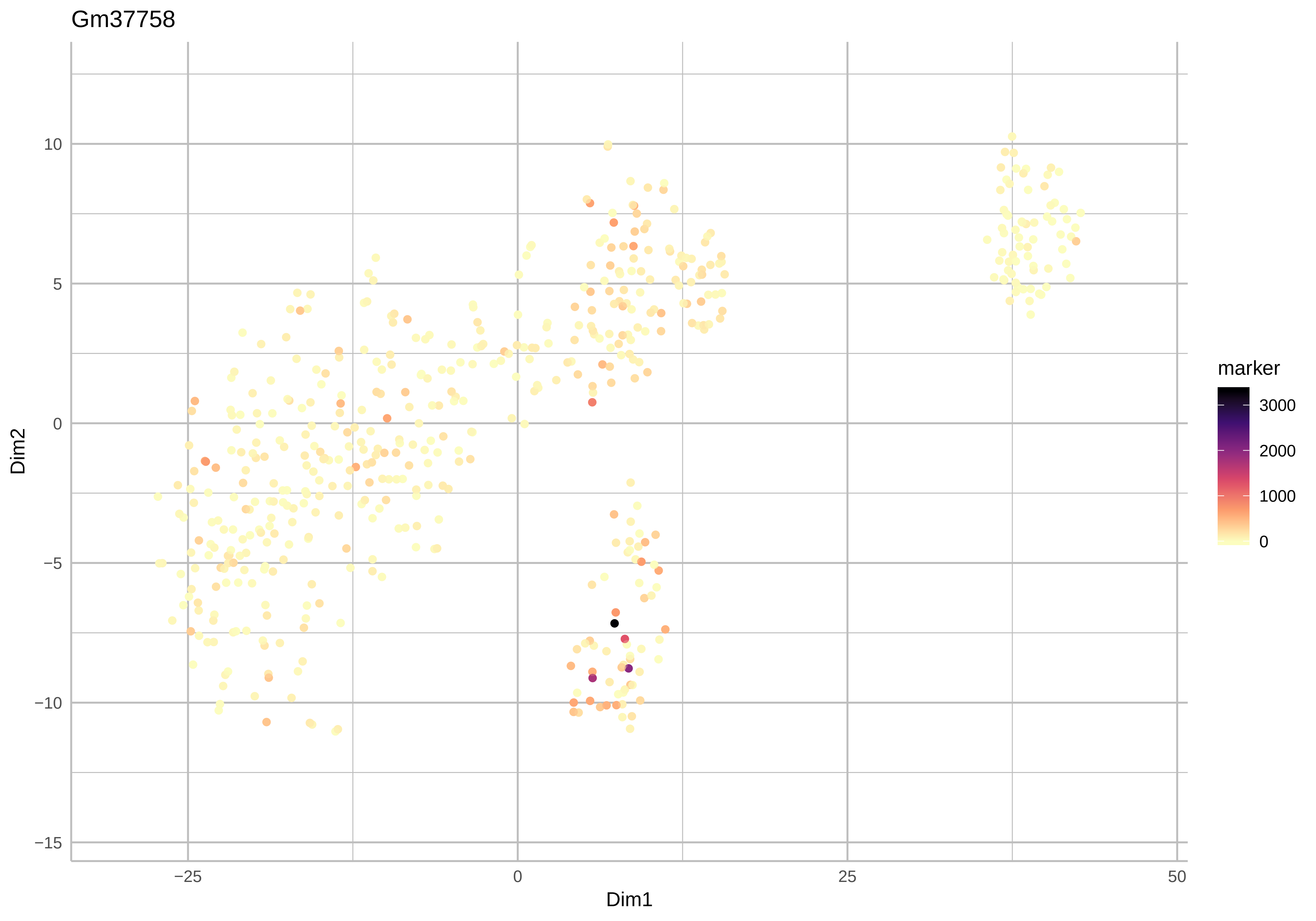

Gm37758
Dim2
Dim1
10
5
0
−5
−10
−15
−25
0
25
50
marker
3000
2000
1000
0

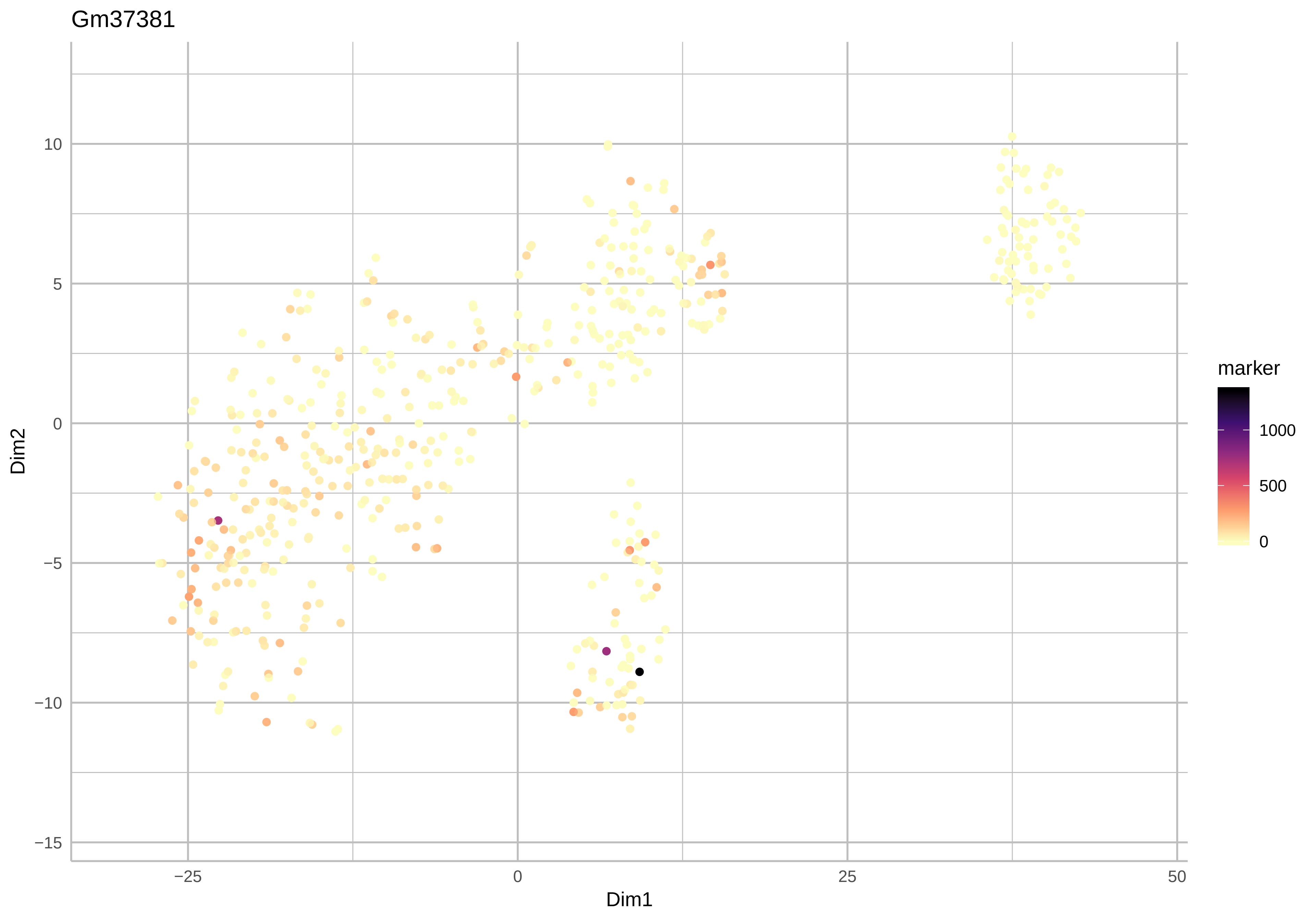
Gm37381
Dim2
Dim1
10
5
0
−5
−10
−15
−25
0
25
50
marker
1000
500
0

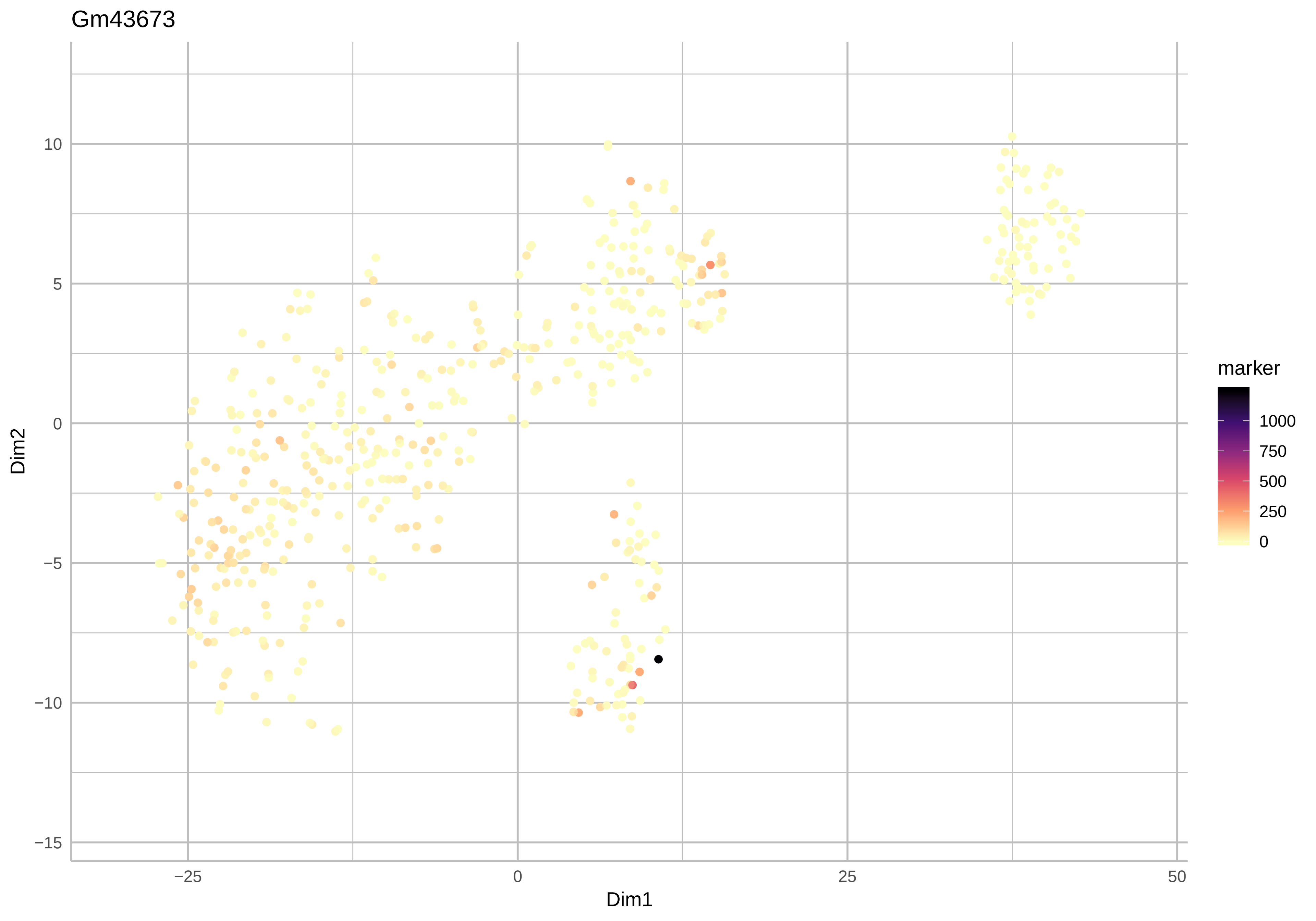

Gm43673
Dim2
Dim1
10
5
0
−5
−10
−15
−25
0
25
50
marker
1000
750
500
250
0

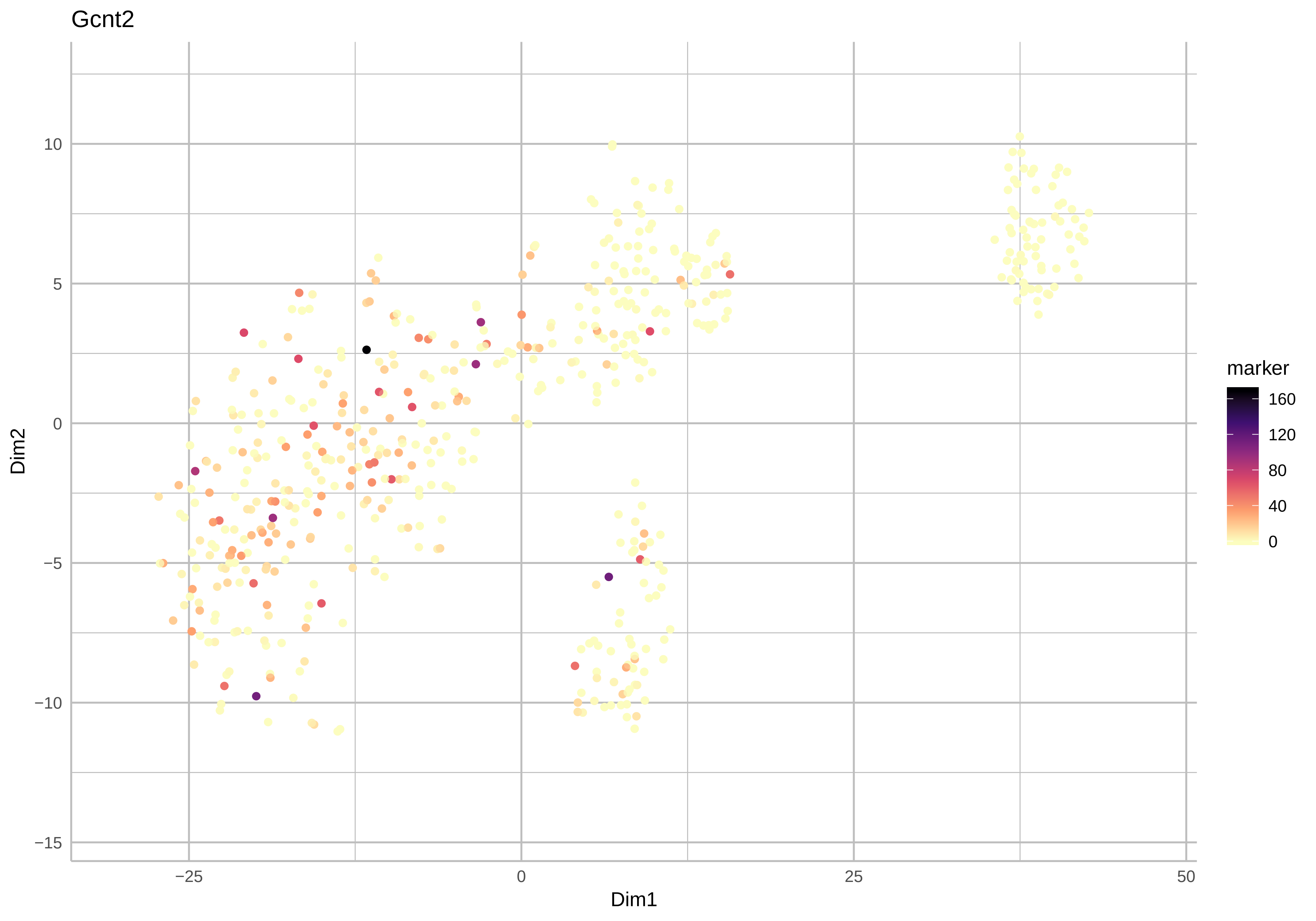
Gcnt2
Dim2
Dim1
10
5
0
−5
−10
−15
−25
0
25
50
marker
160
120
80
40
0

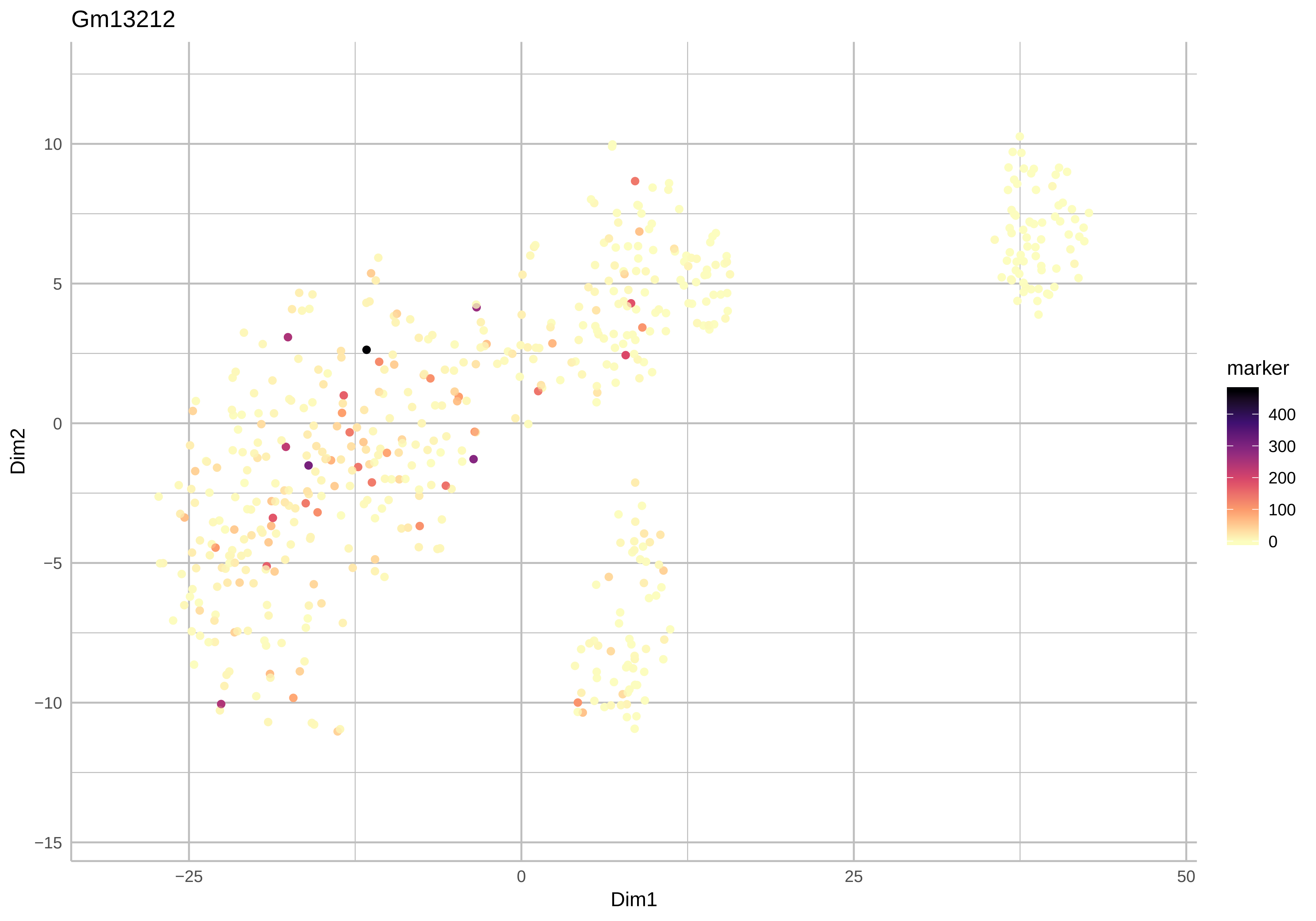

Gm13212
Dim2
Dim1
10
5
0
−5
−10
−15
−25
0
25
50
marker
400
300
200
100
0

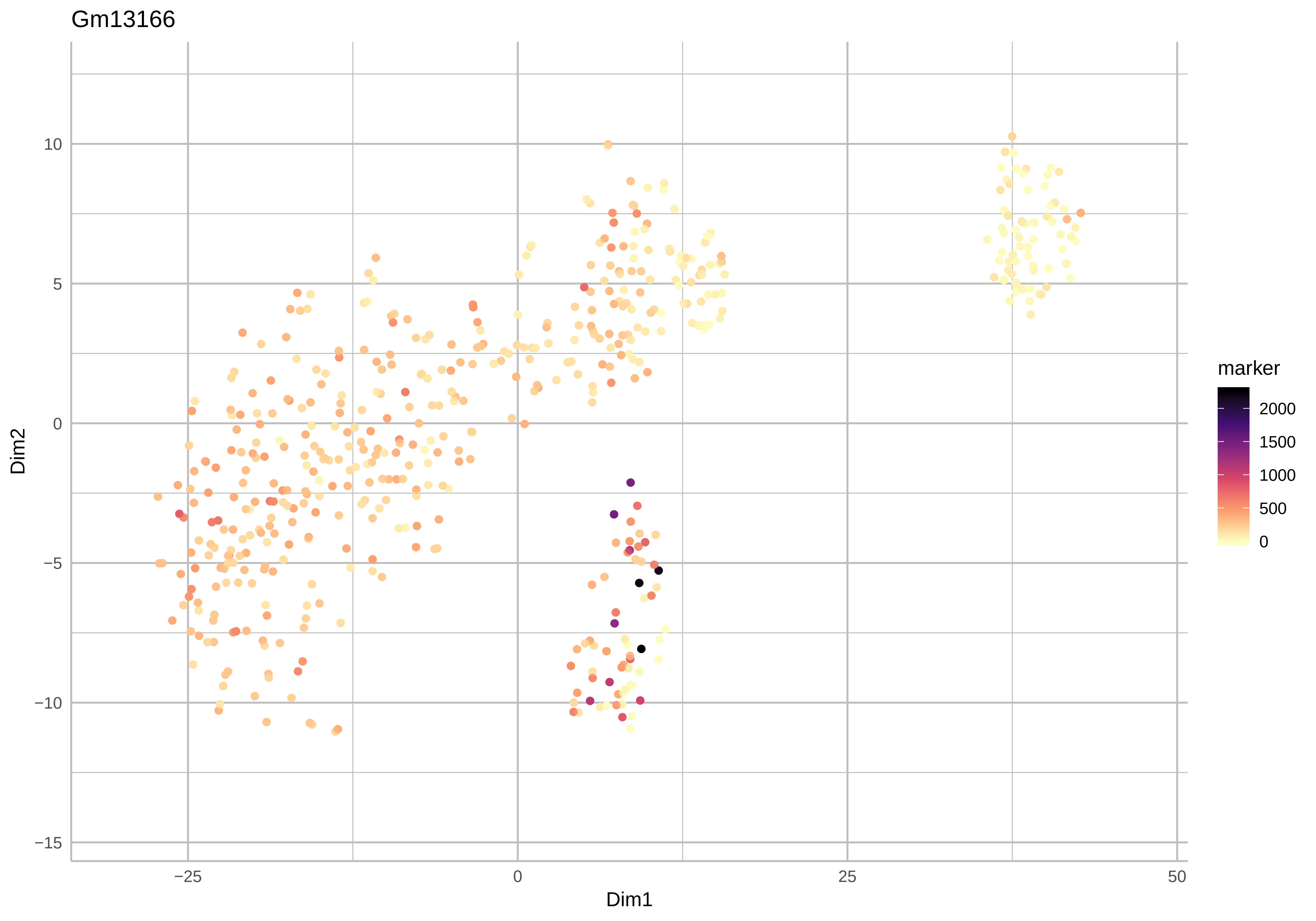

Gm13166
Dim2
Dim1
10
5
0
−5
−10
−15
−25
0
25
50
marker
2000
1500
1000
500
0

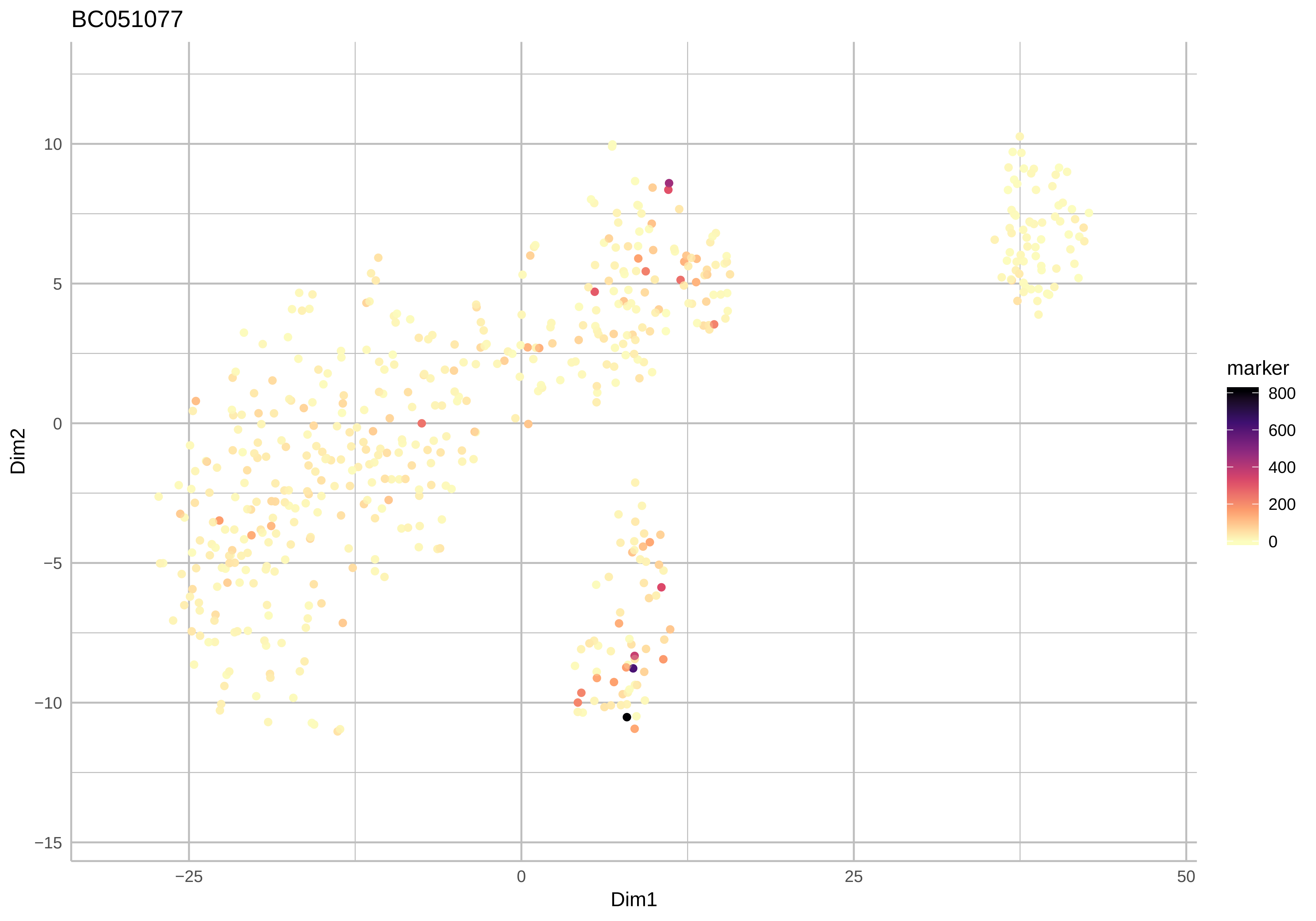

BC051077
Dim2
Dim1
10
5
0
−5
−10
−15
−25
0
25
50
marker
800
600
400
200
0

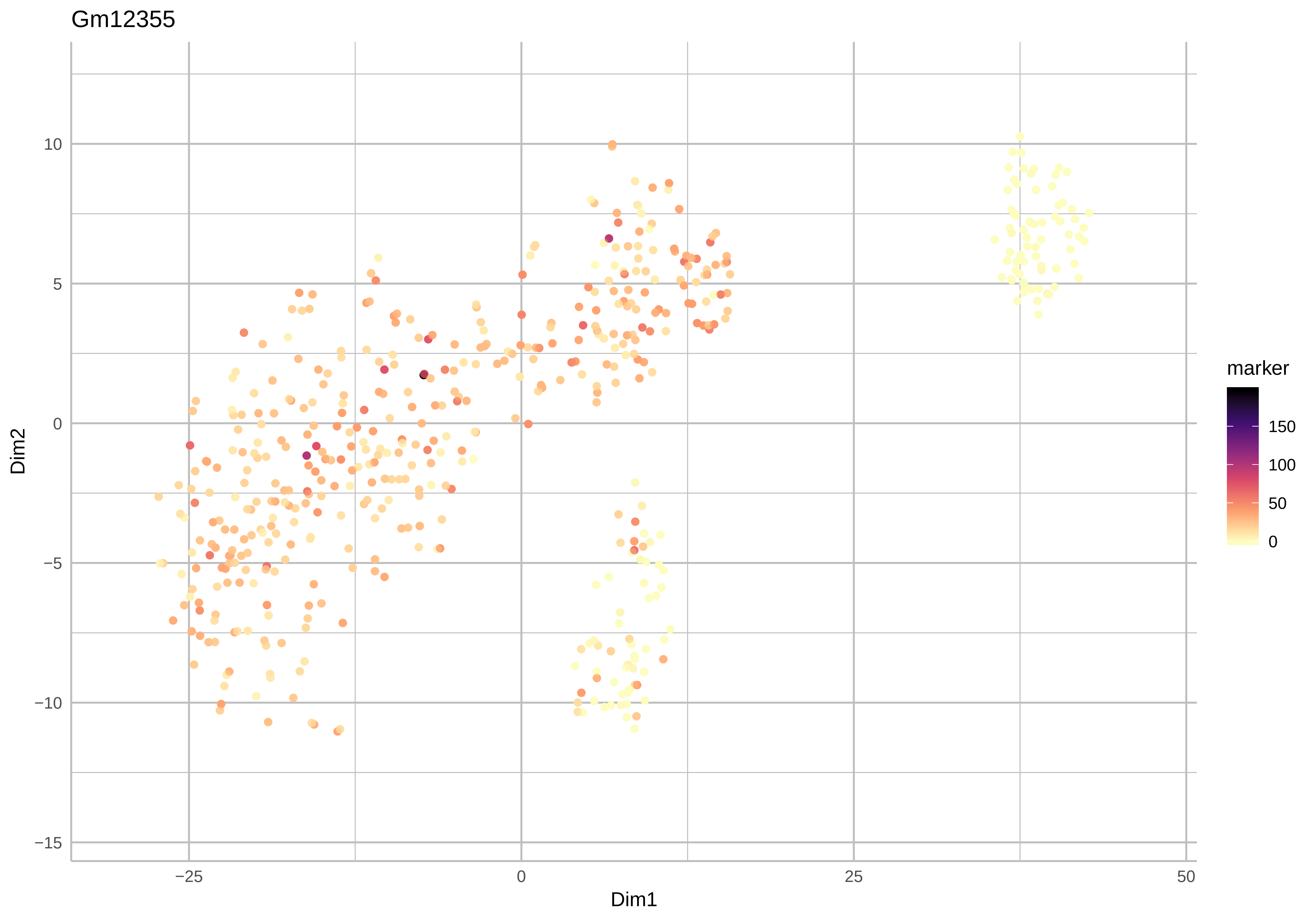

Gm12355
Dim2
Dim1
10
5
0
−5
−10
−15
−25
0
25
50
marker
150
100
50
0

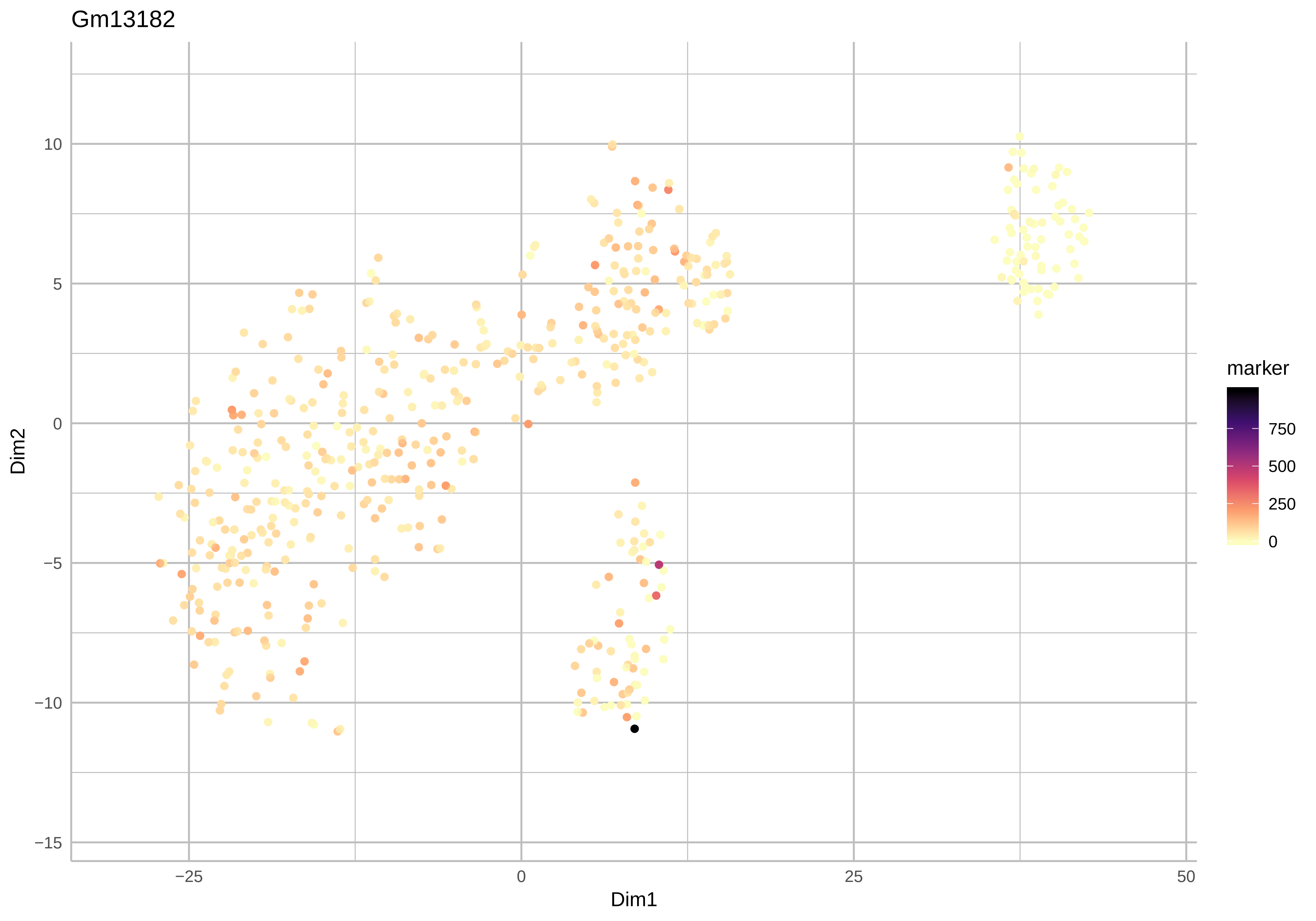

Gm13182
Dim2
Dim1
marker
750
500
250
0
10
5
0
−5
−10
−15
−25
0
25
50

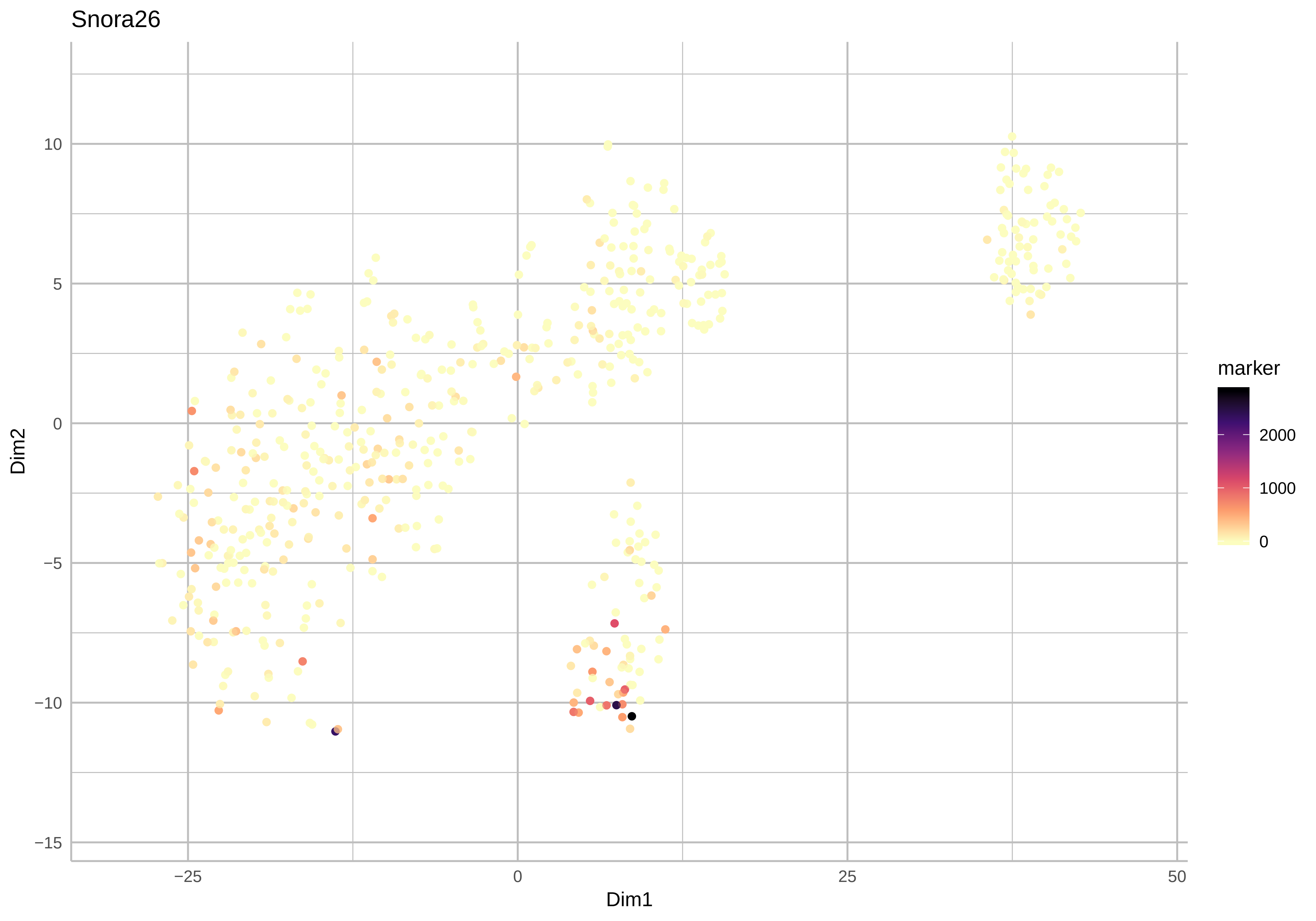
Snora26
Dim2
Dim1
10
5
0
−5
−10
−15
−25
0
25
50
marker
2000
1000
0

Gm13487

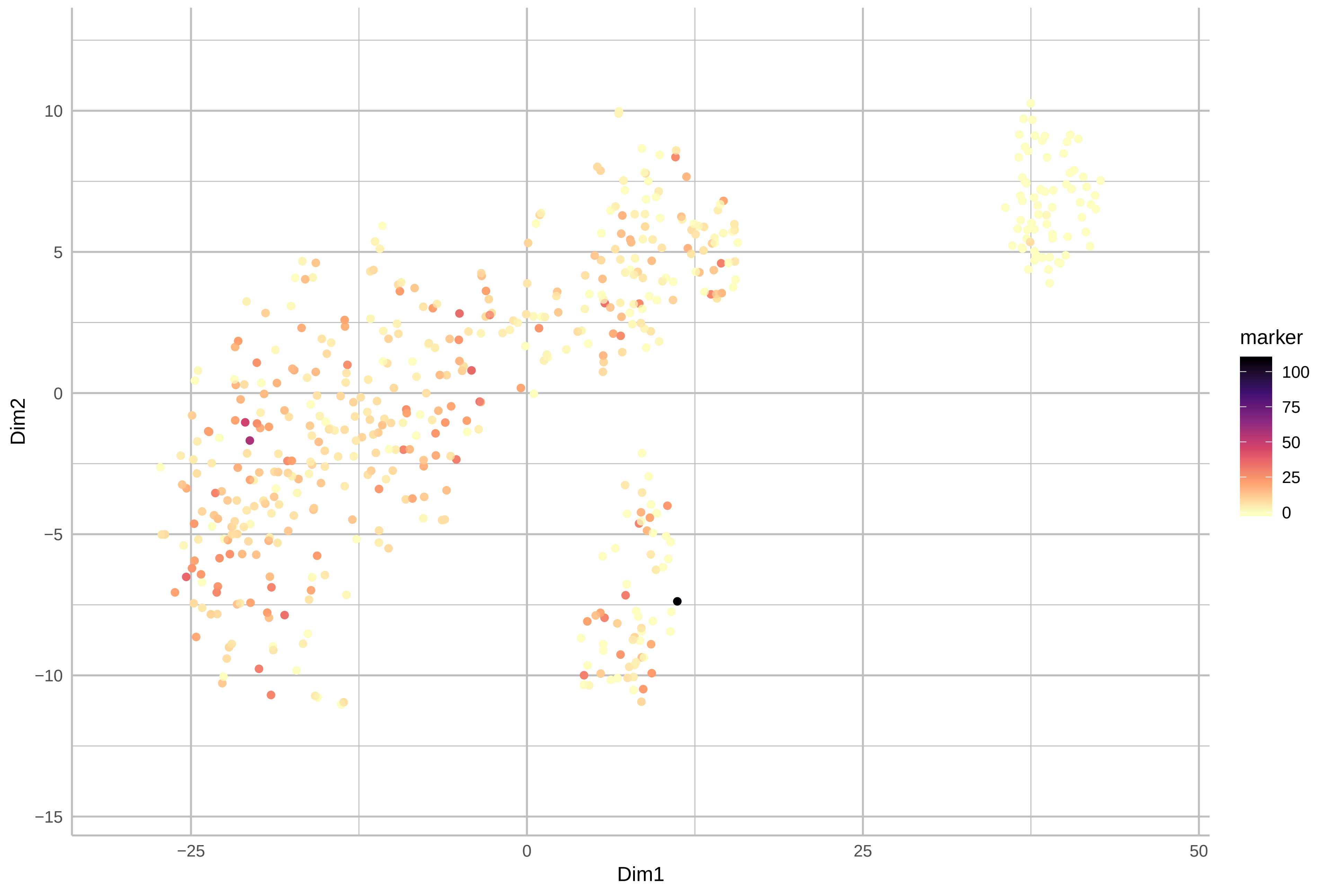

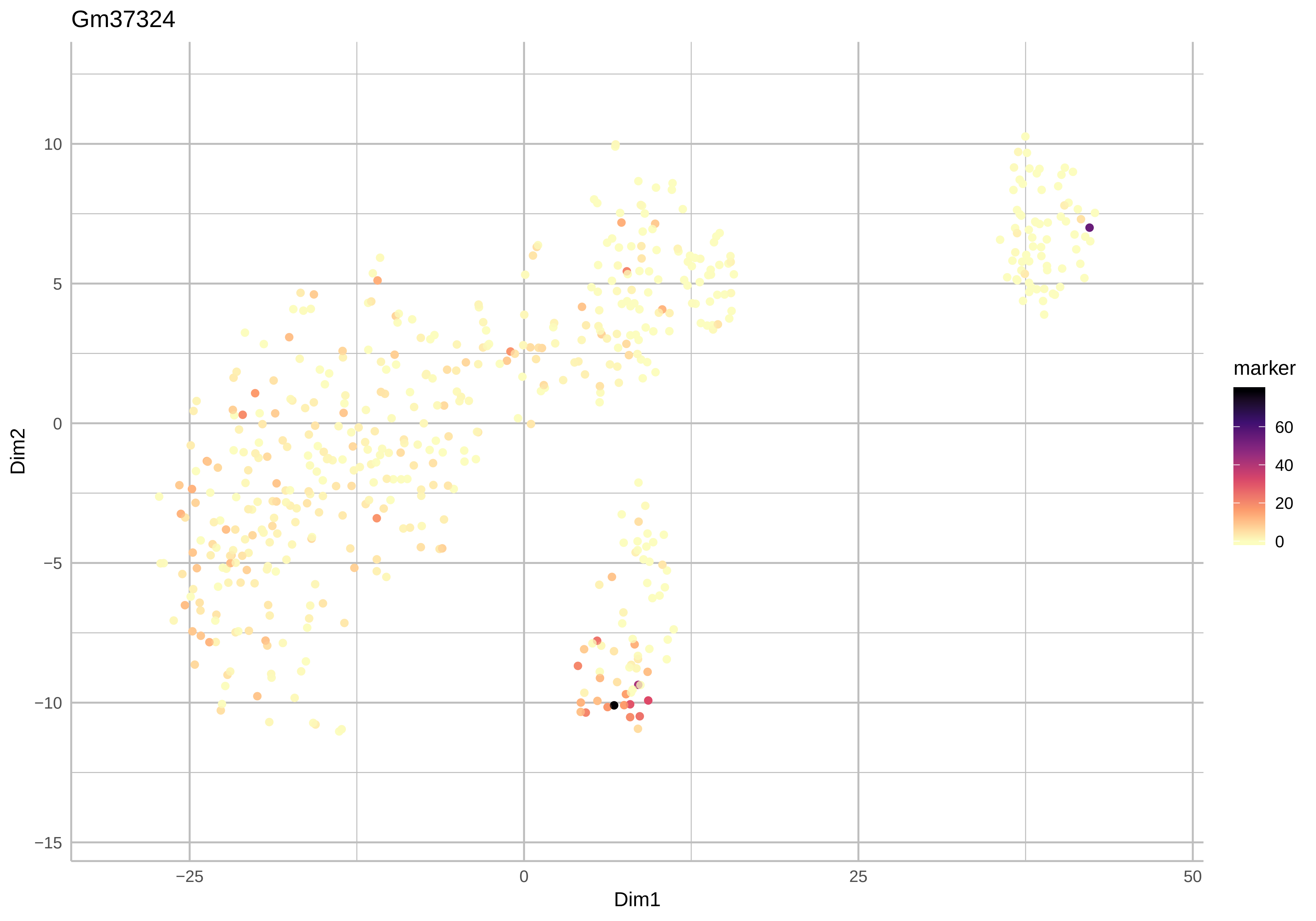

Gm37324
Dim2
Dim1
10
5
0
−5
−10
−15
−25
0
25
50
marker
60
40
20
0

Dancr

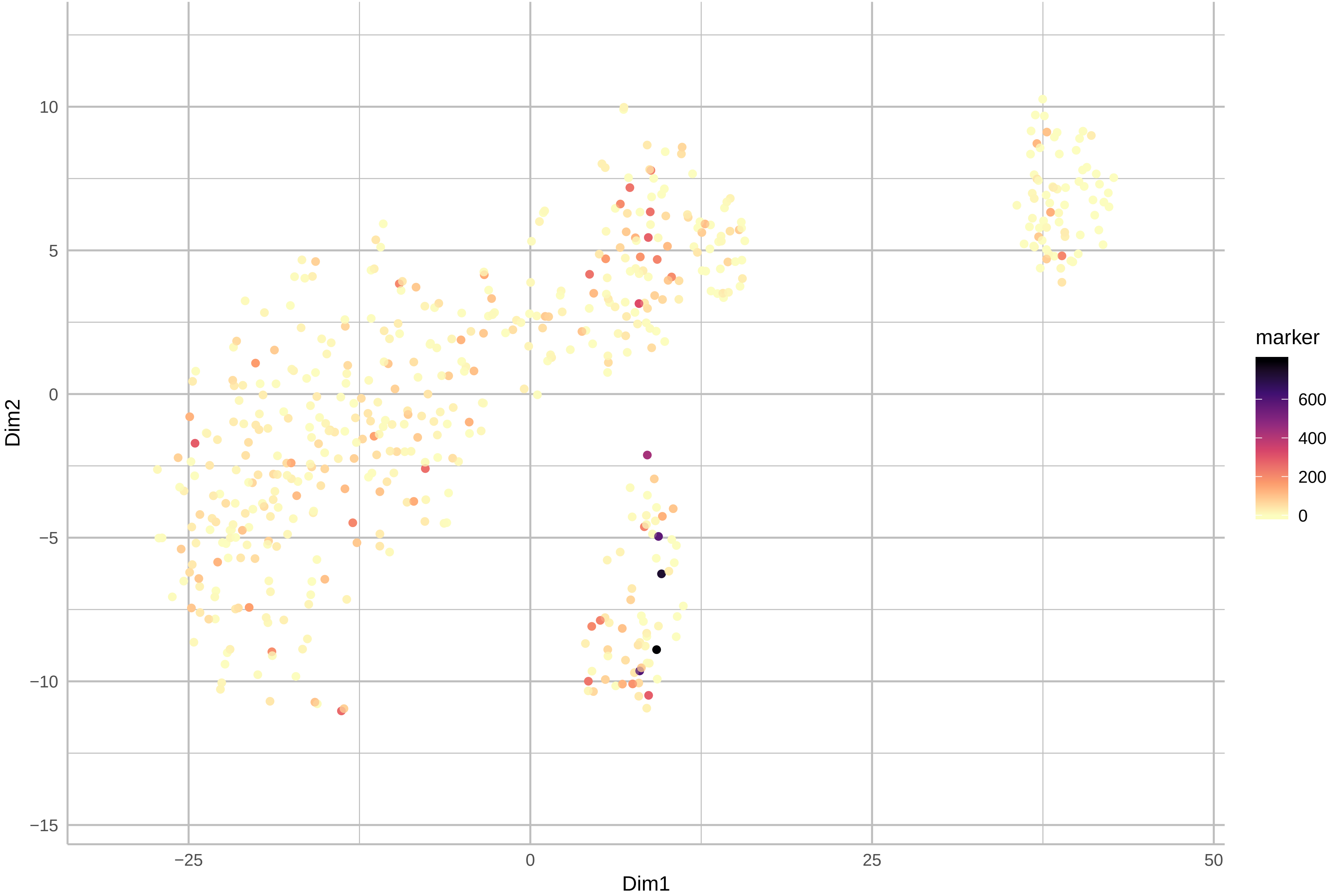

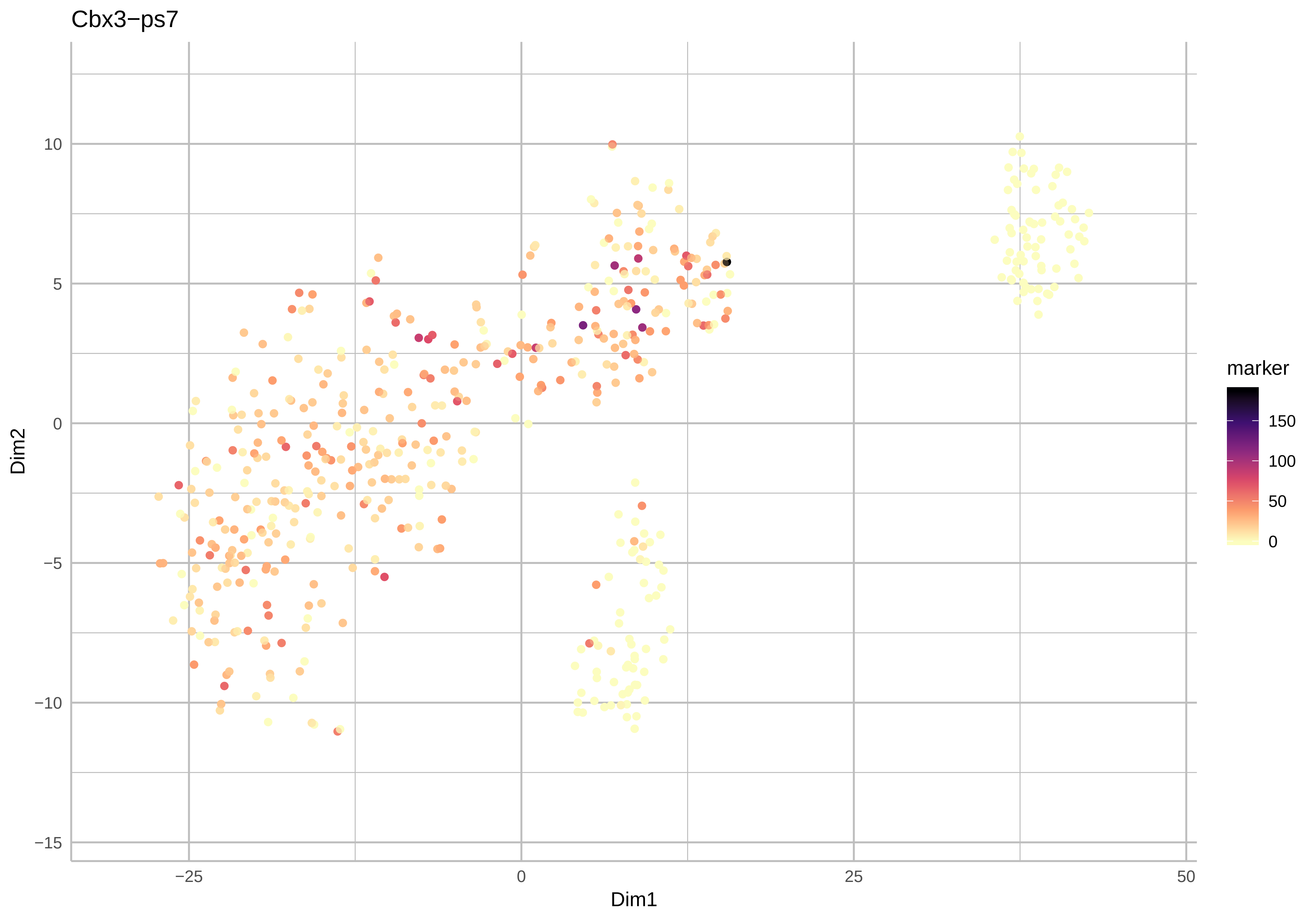

Cbx3−ps7
10
5
0
−5
−10
−15
−25
0
25
50
Dim1
Dim2
marker
150
100
50
0

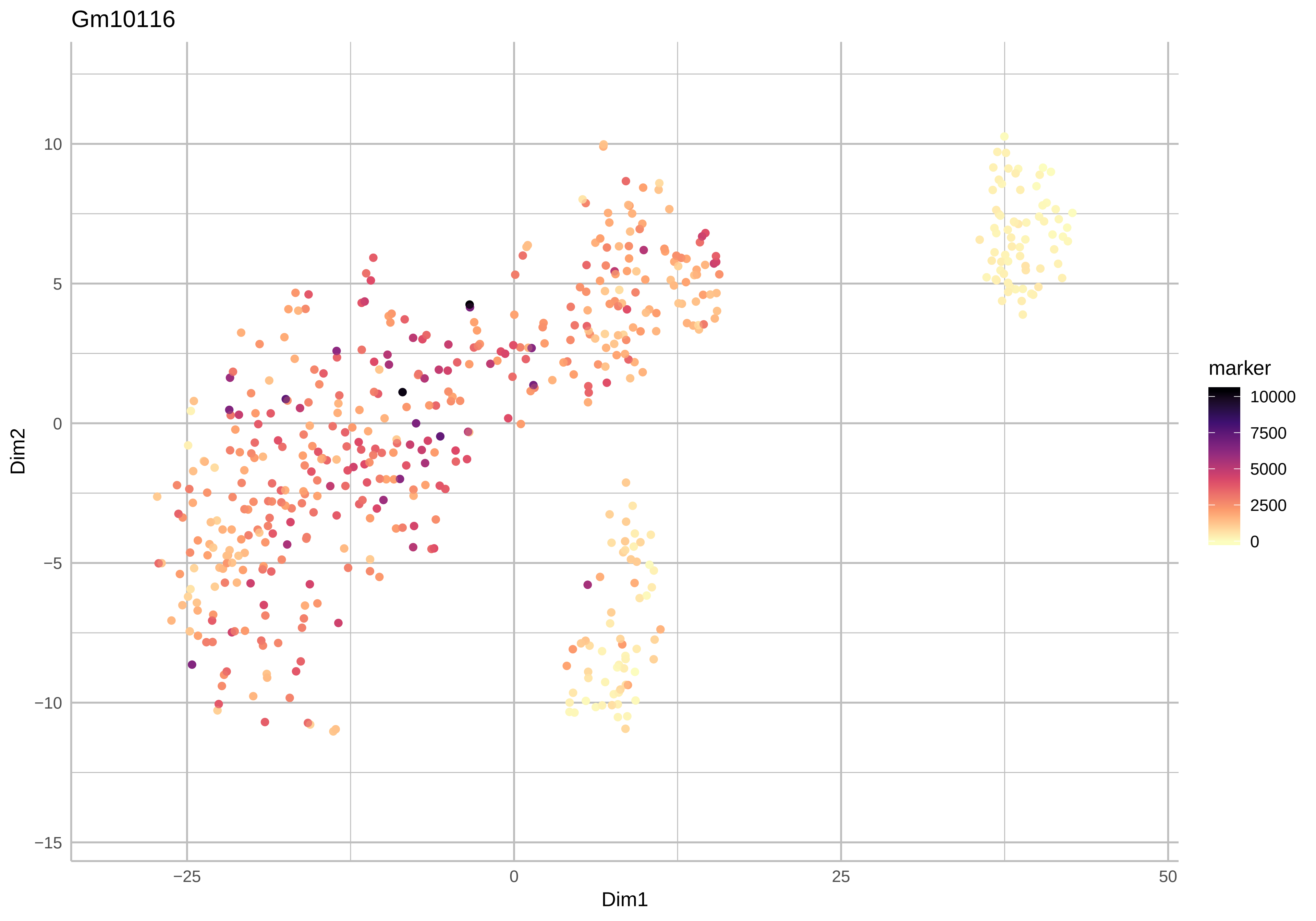

Gm10116
Dim2
Dim1
10
5
0
−5
−10
−15
−25
0
25
50
marker
10000
7500
5000
2500
0

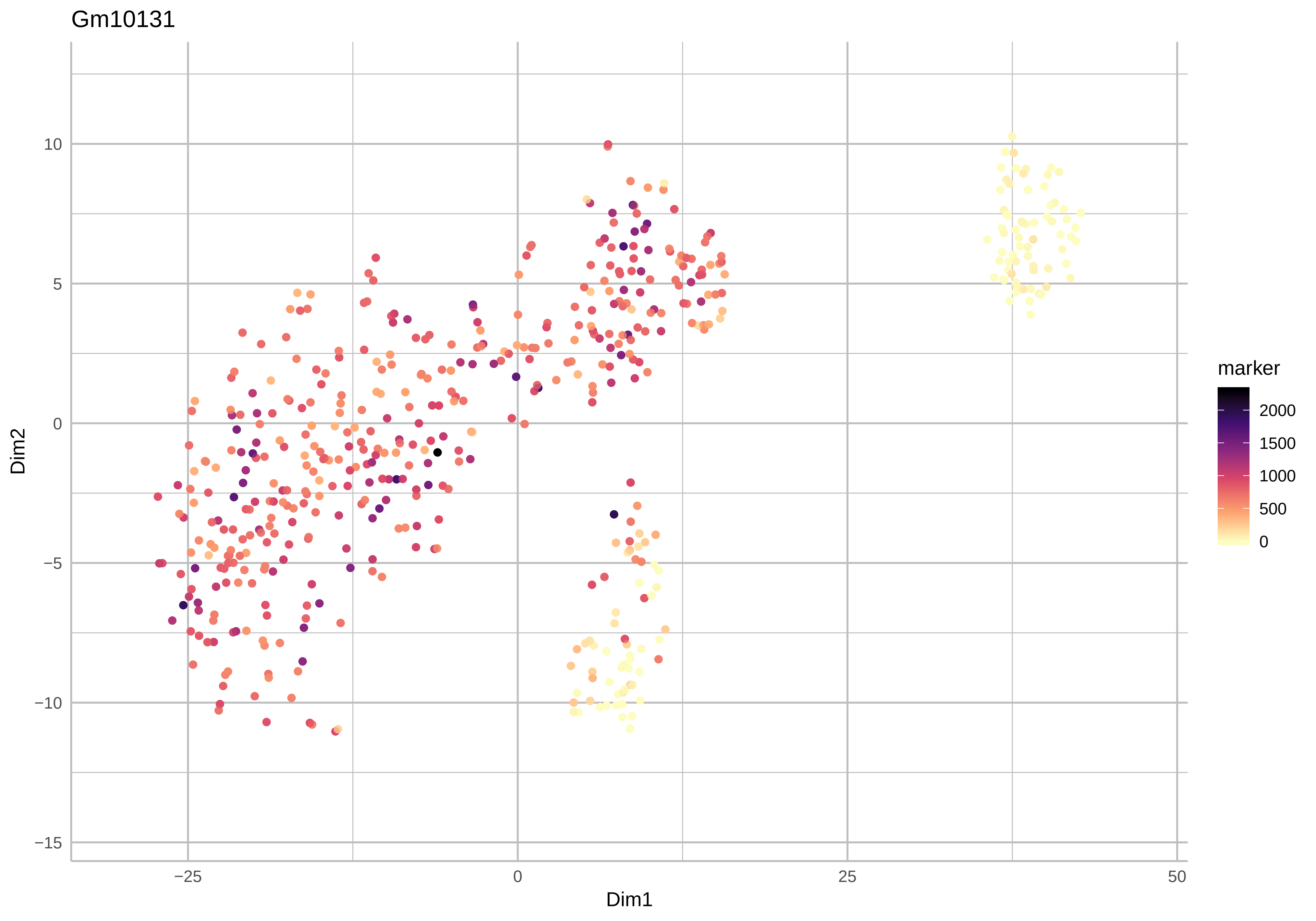
Gm10131
Dim2
Dim1
10
5
0
−5
−10
−15
−25
0
25
50
marker
2000
1500
1000
500
0

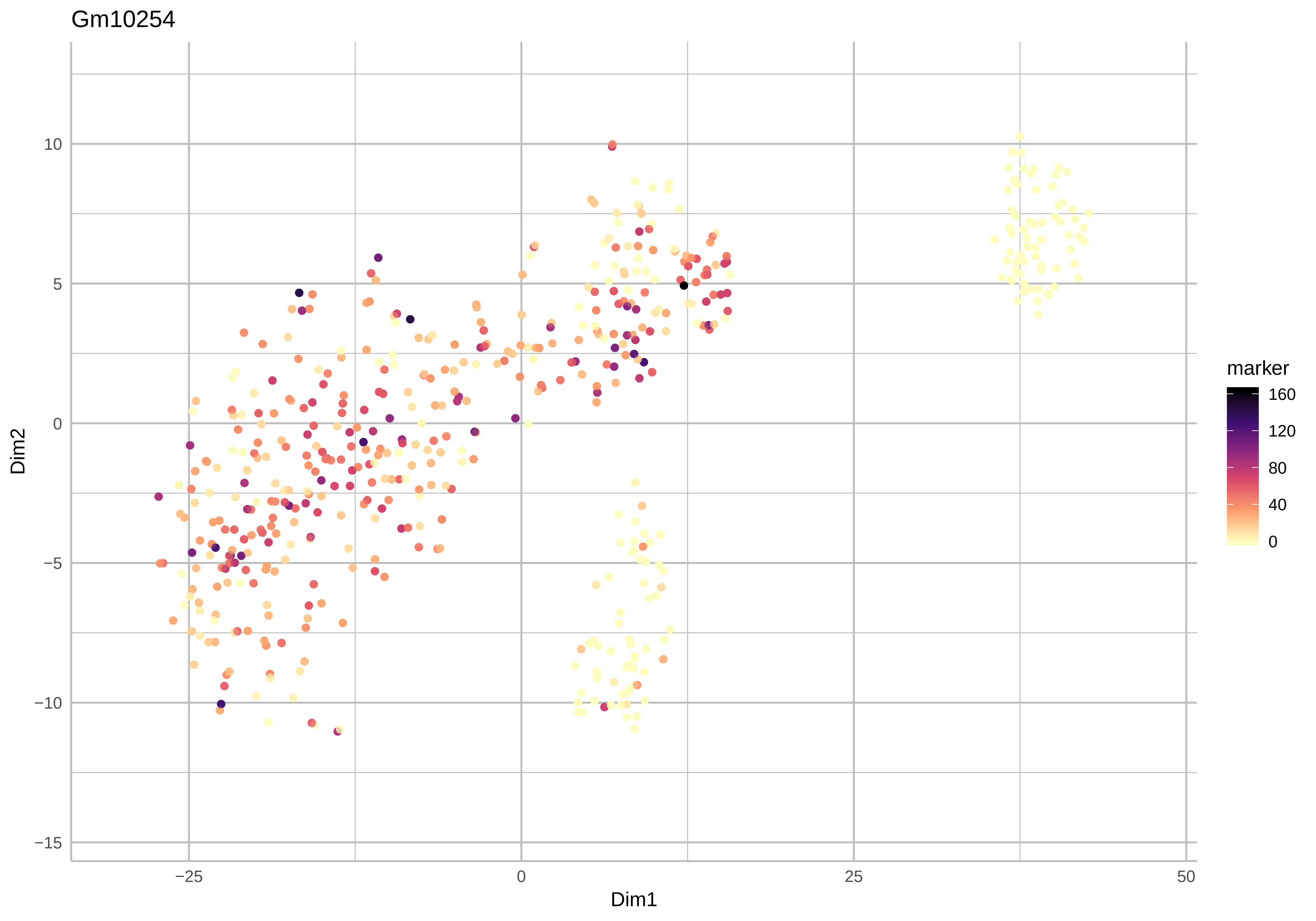

Gm10254
10
5
0
−5
−10
−15
Dim2
−25
0
25
50
Dim1
marker
160
120
80
40
0

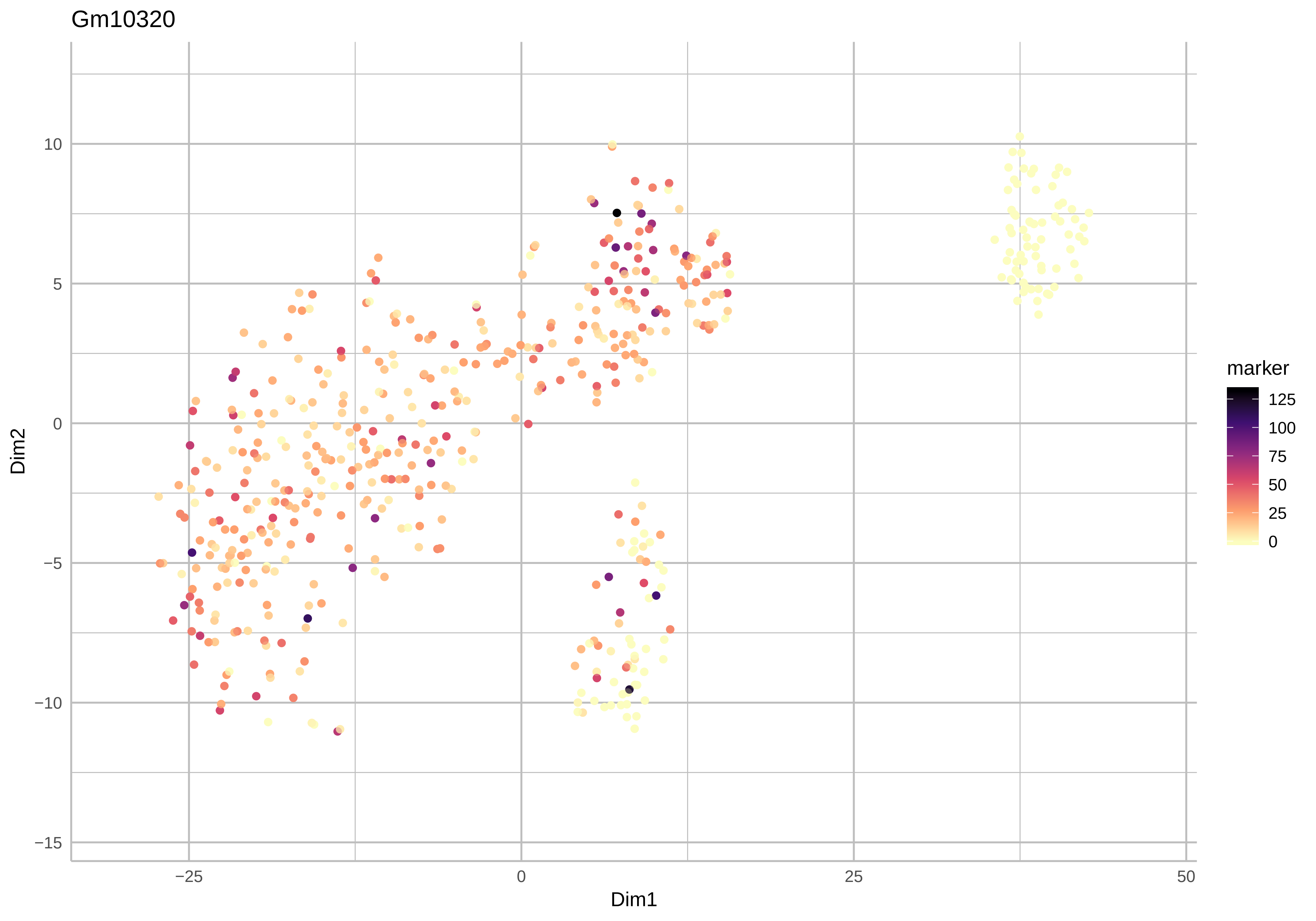
Gm10320
Dim2
Dim1
10
5
0
−5
−10
−15
−25
0
25
50
marker
125
100
75
50
25
0

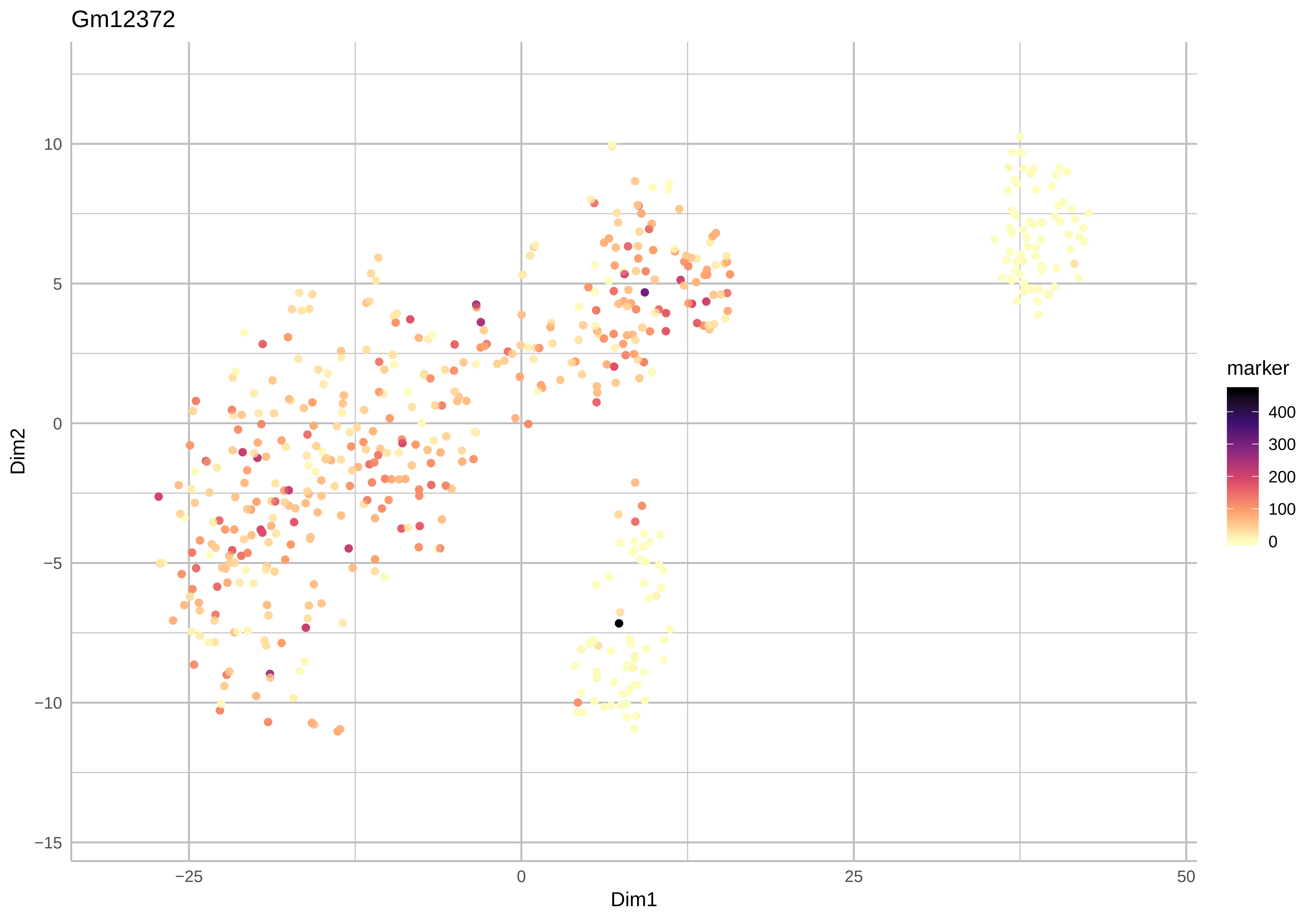
Gm12372
Dim2
Dim1
marker
400
300
200
100
0
10
5
0
−5
−10
−15
−25
0
25
50

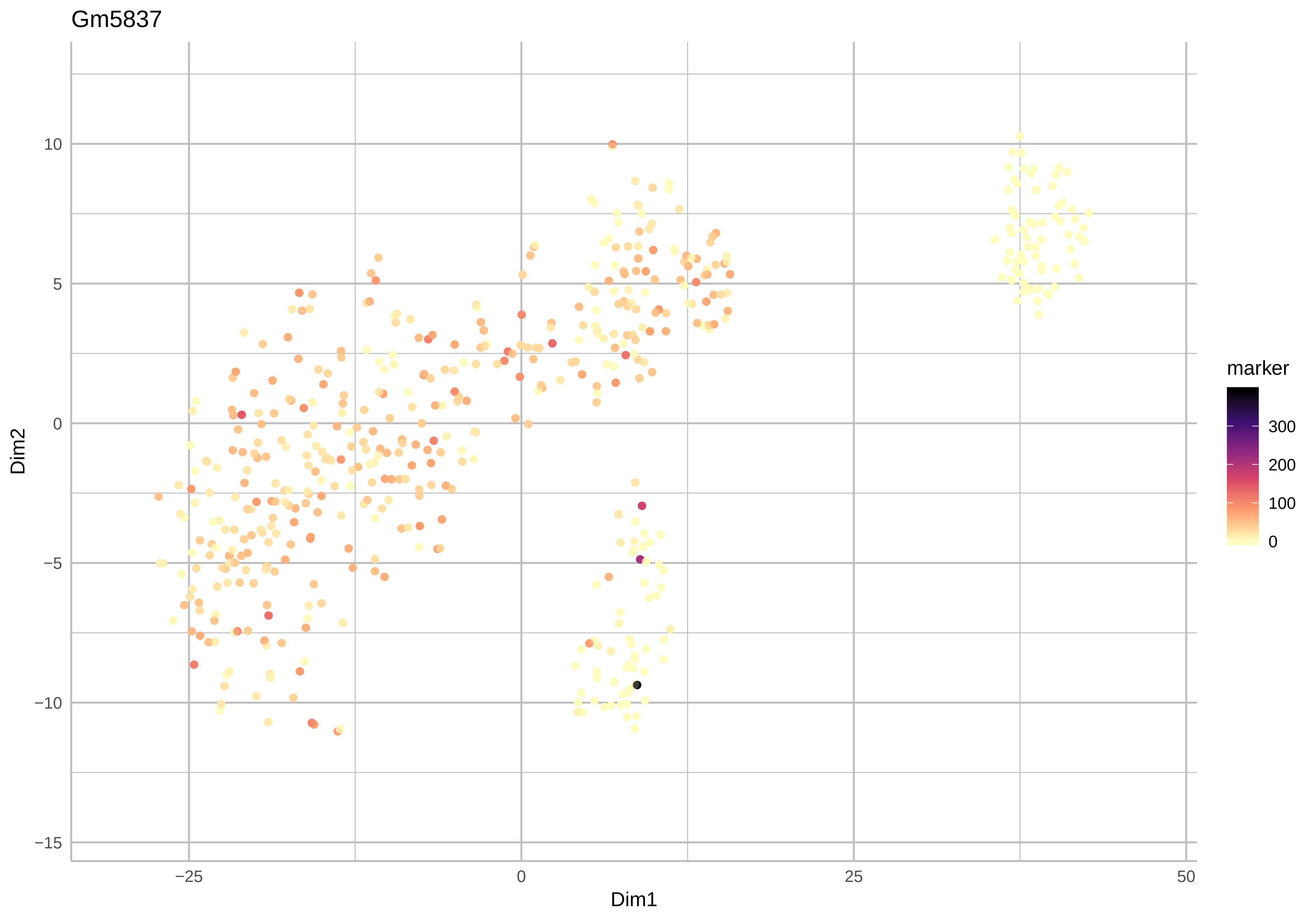

Gm5837
Dim2
Dim1
10
5
0
−5
−10
−15
−25
0
25
50
marker
300
200
100
0

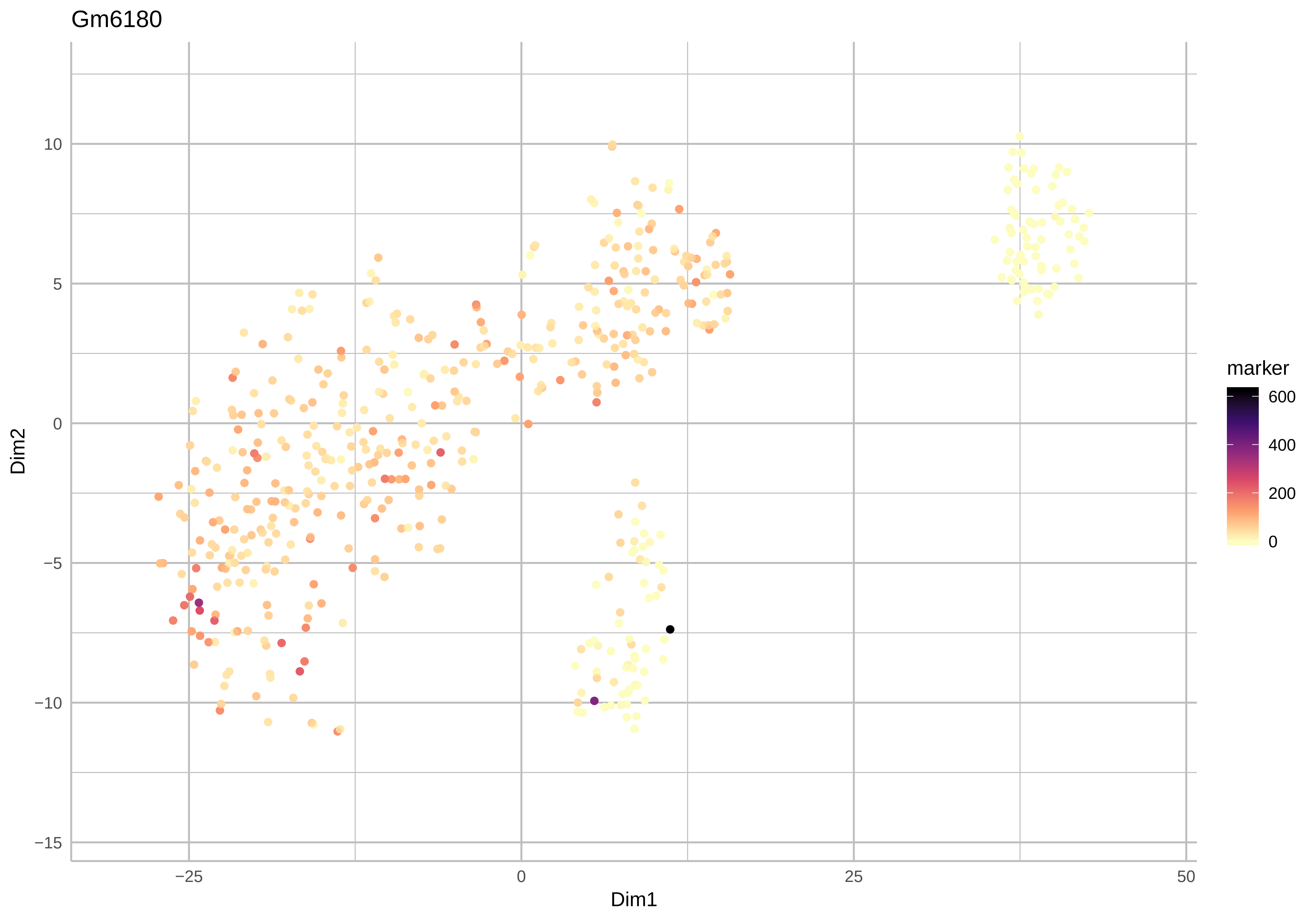

Gm6180
Dim2
Dim1
10
5
0
−5
−10
−15
−25
0
25
50
marker
600
400
200
0

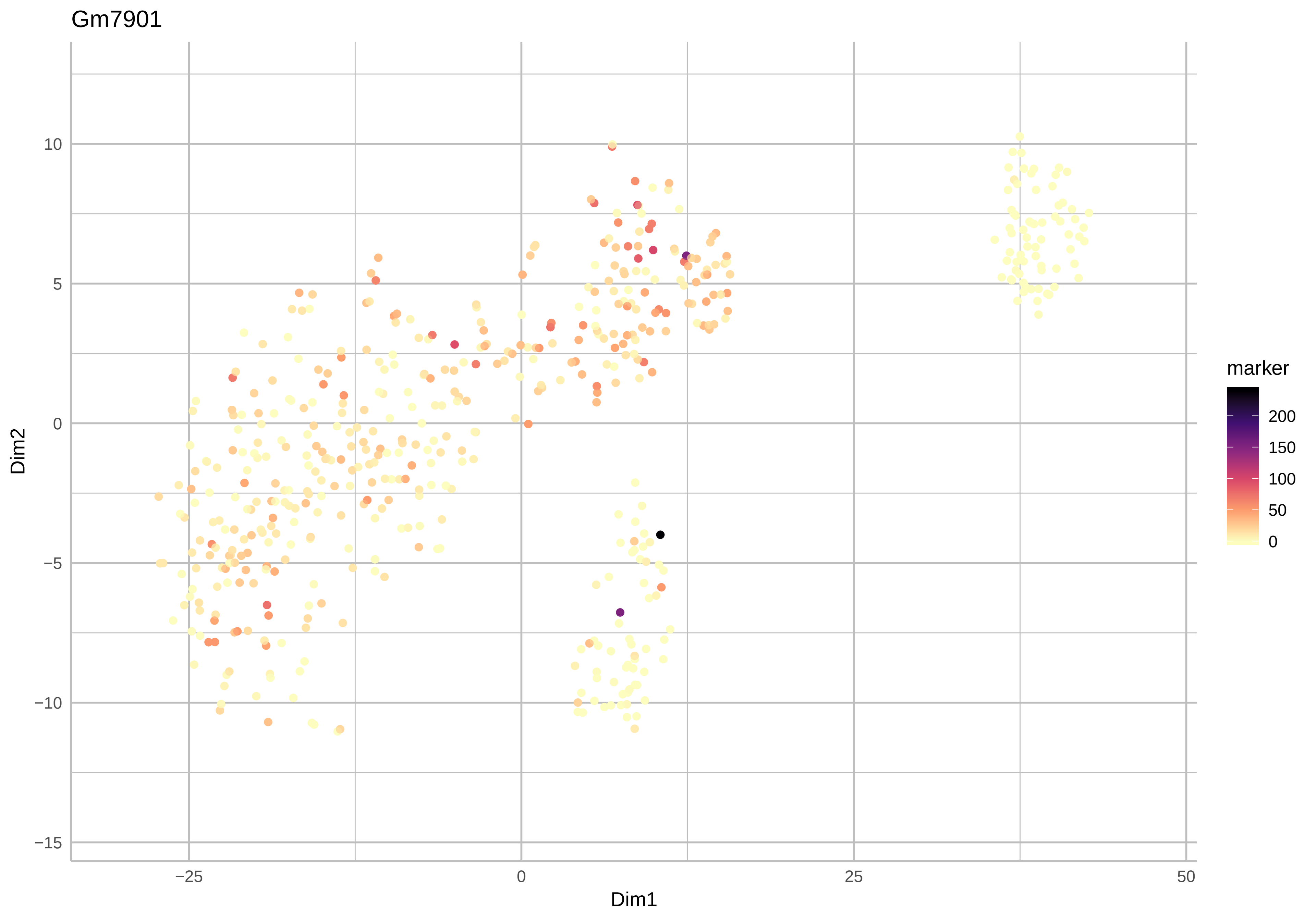
Gm7901
Dim2
Dim1
marker
10
5
0
−5
−10
−15
−25
0
25
50
200
150
100
50
0

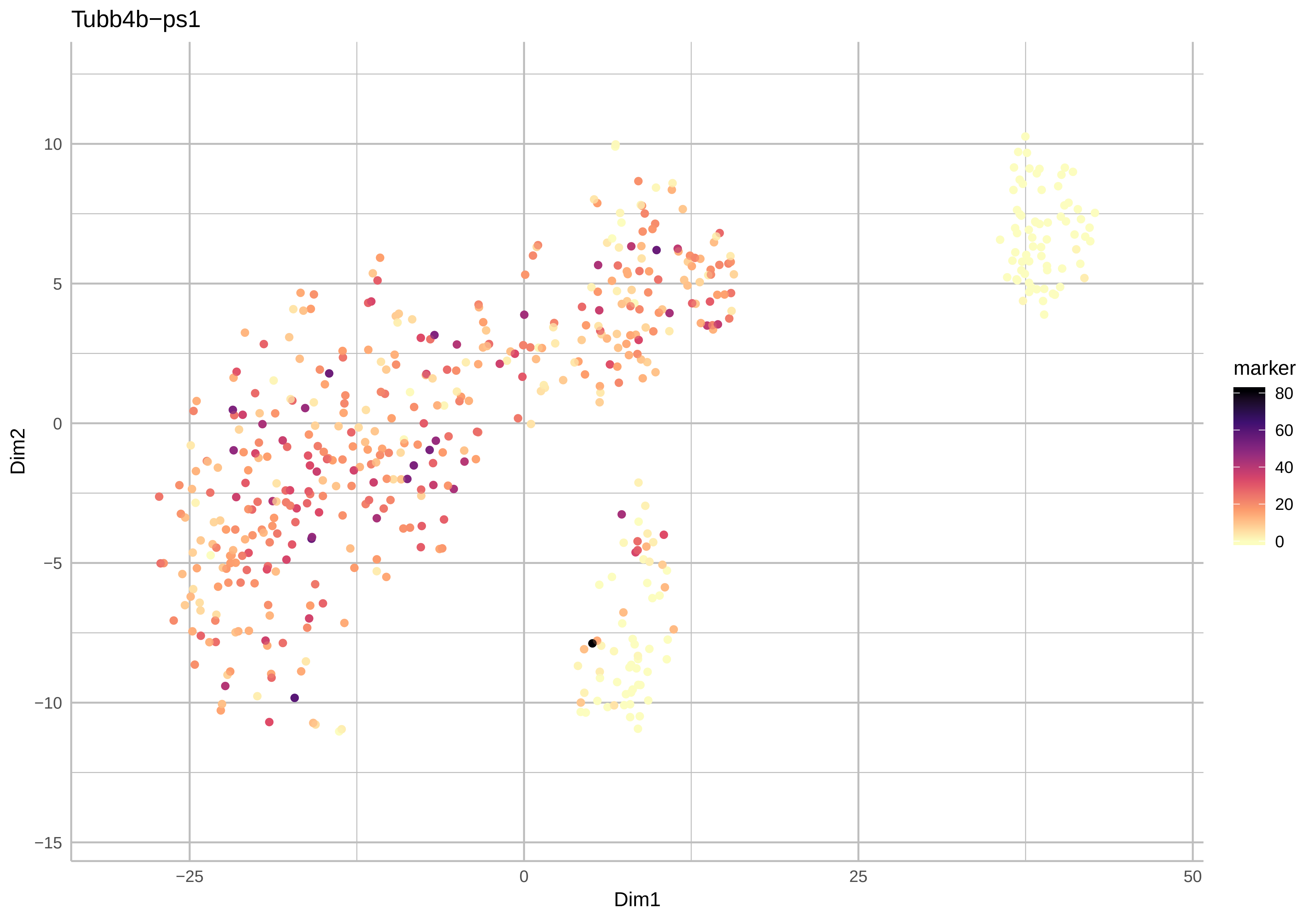
Tubb4b−ps1
Dim2
Dim1
marker
80
60
40
20
0
10
5
0
−5
−10
−15
−25
0
25
50

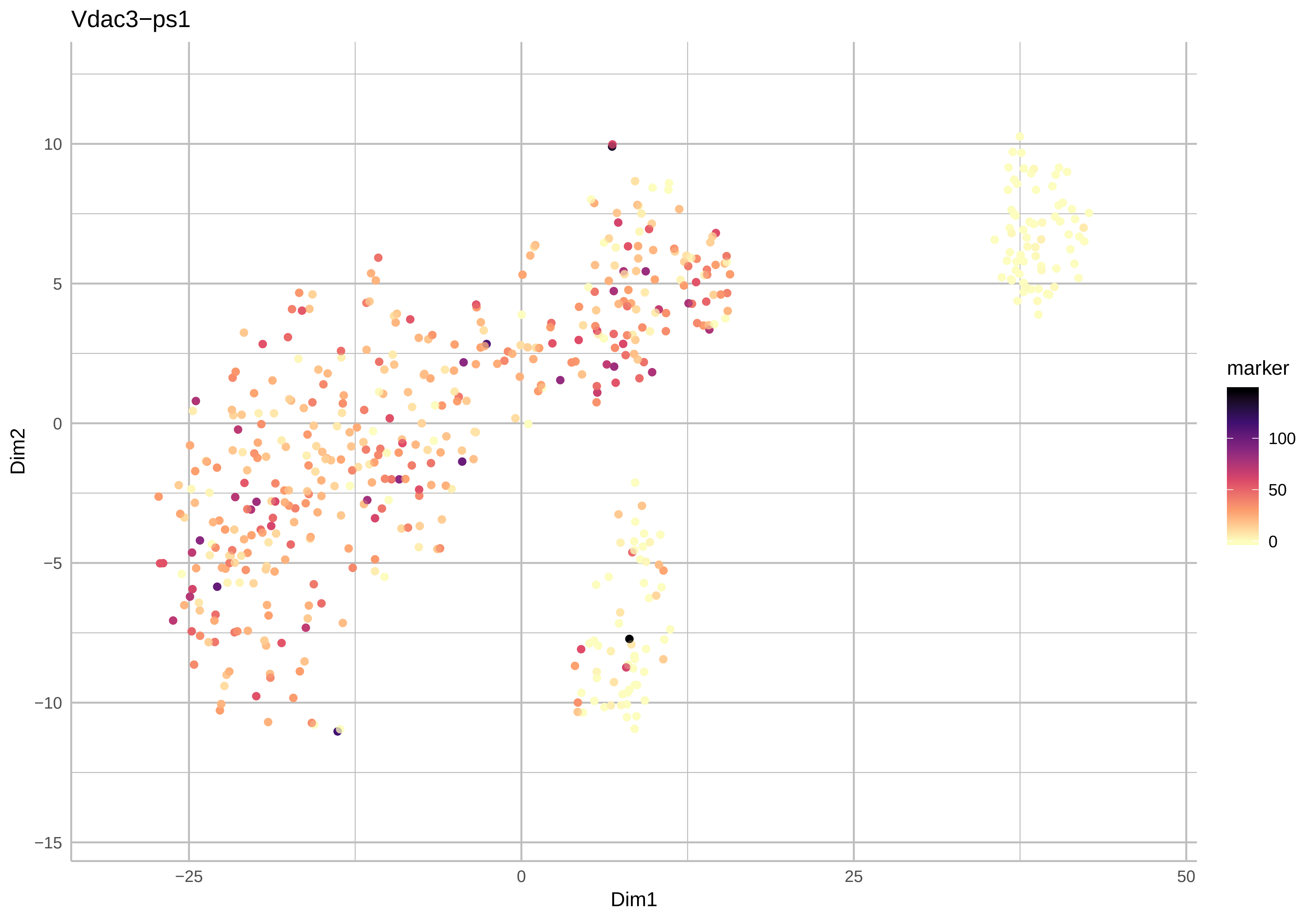

Vdac3−ps1
Dim2
10
5
0
−5
−10
−15
−25
0
25
50
Dim1
marker
100
50
0

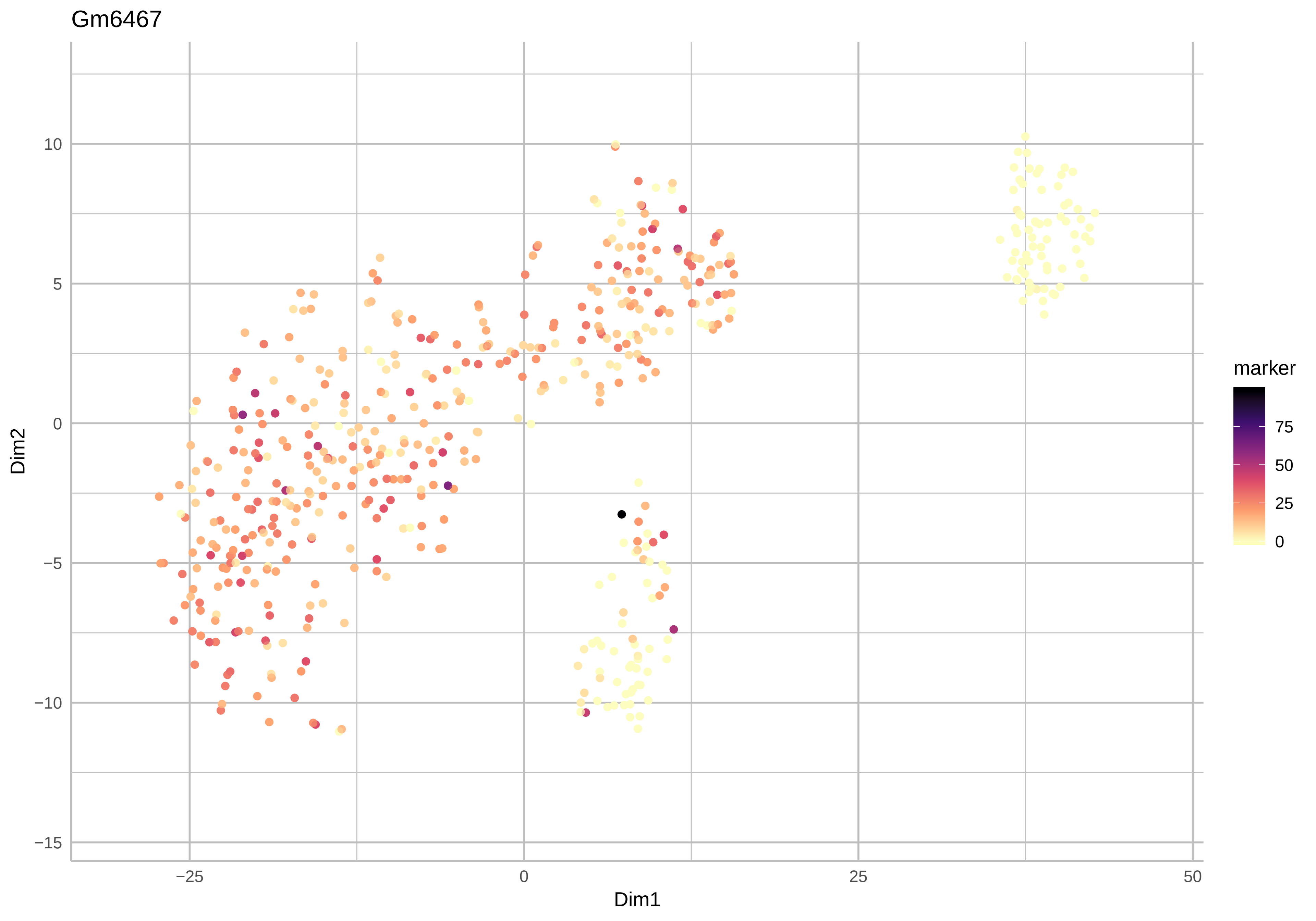
Gm6467
Dim1
Dim2
marker
0
25
50
75
−25
0
25
50
−15
−10
−5
0
5
10

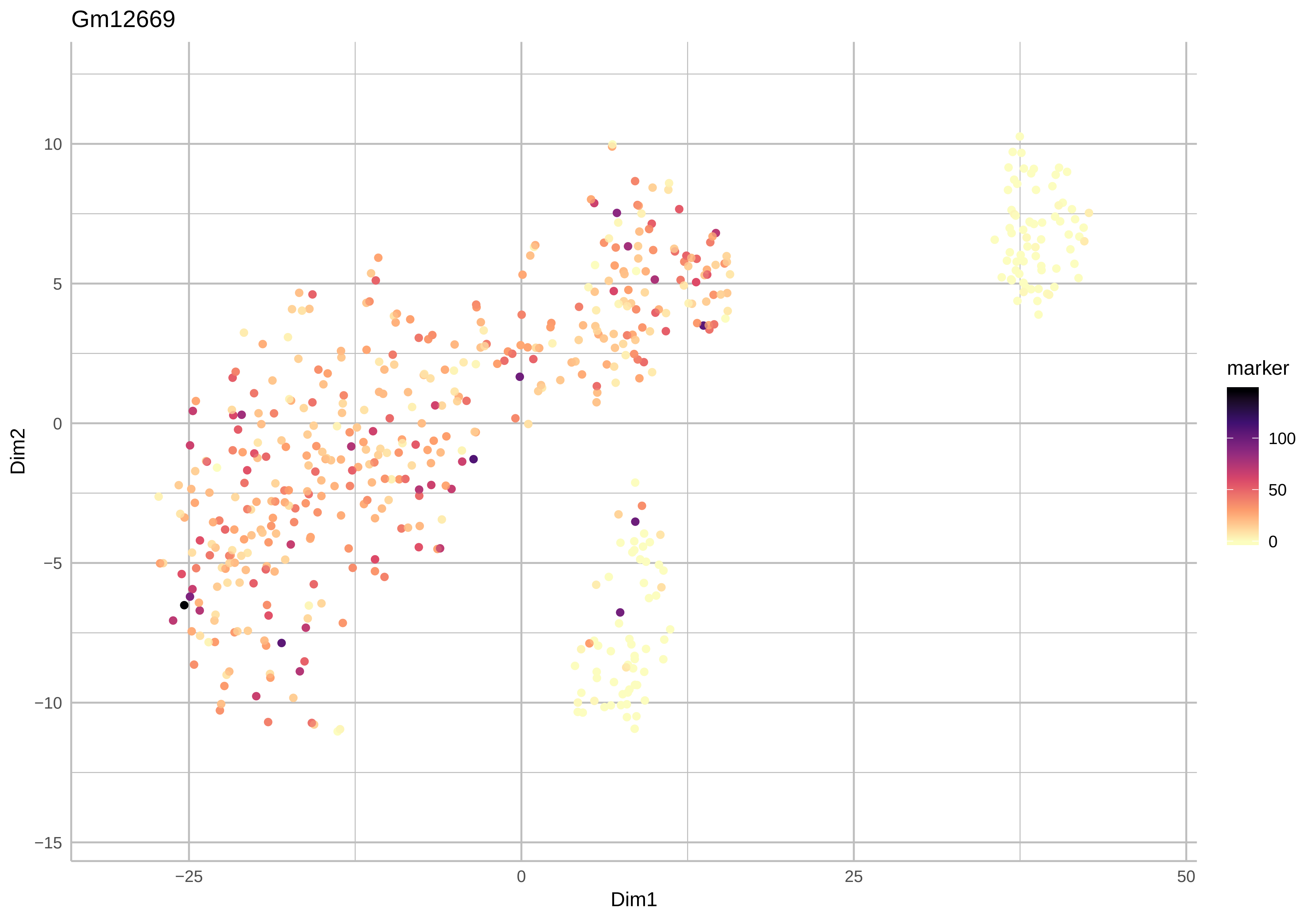
Gm12669
Dim2
Dim1
10
5
0
−5
−10
−15
−25
0
25
50
marker
100
50
0

Gm7293

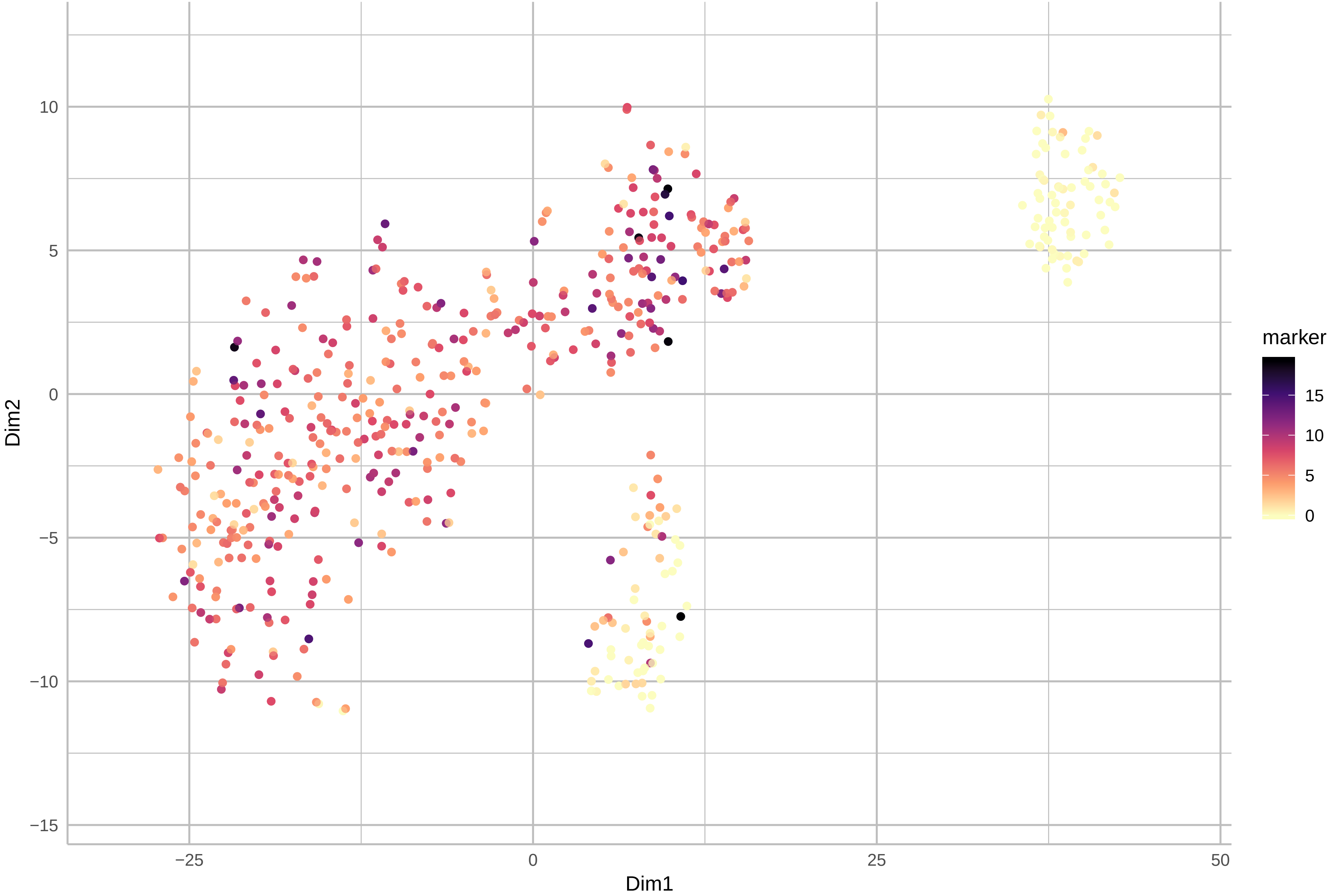

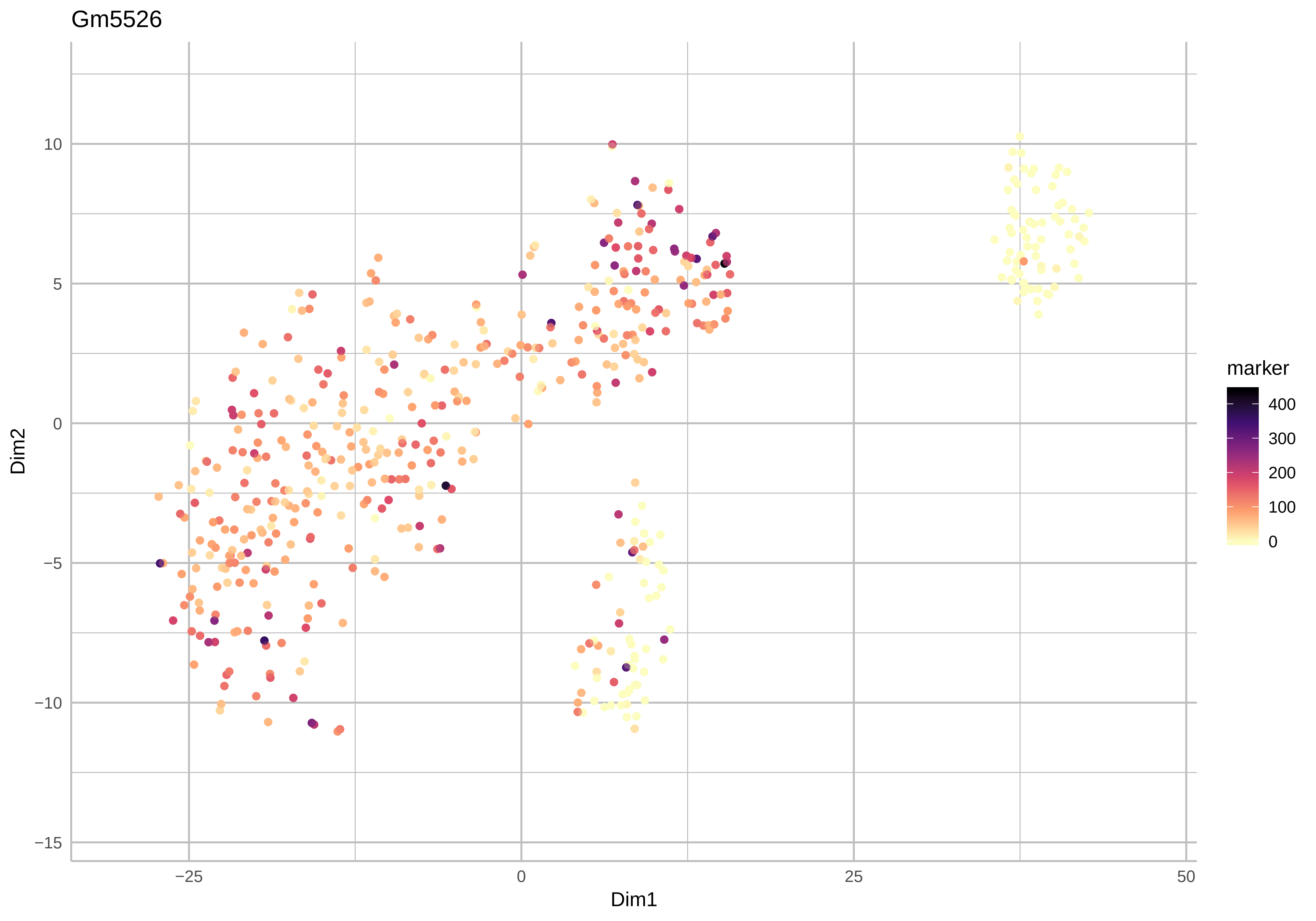
Gm5526
Dim2
Dim1
10
5
0
−5
−10
−15
−25
0
25
50
marker
400
300
200
100
0

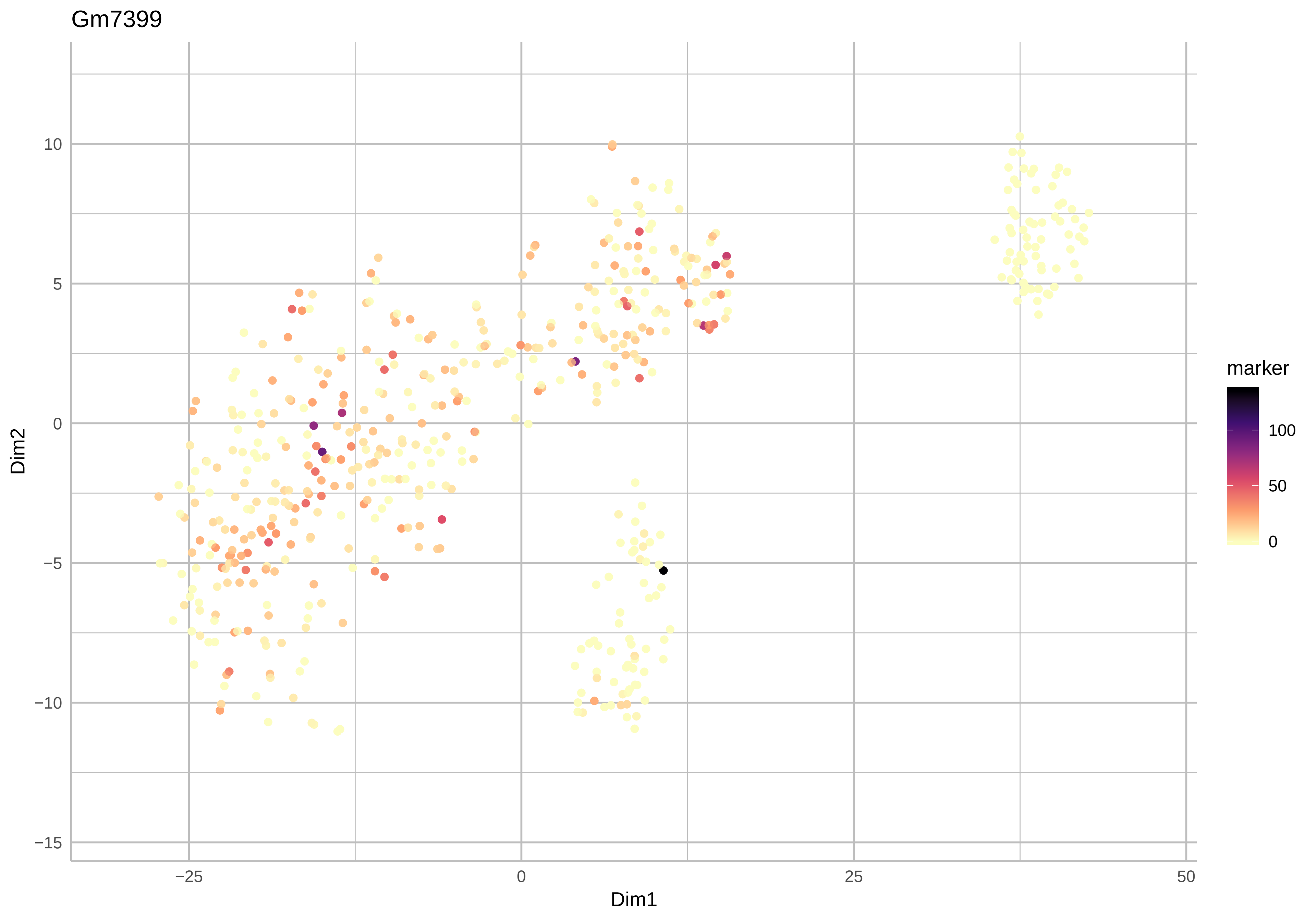
Gm7399
Dim2
10
5
0
−5
−10
−15
Dim1
−25
0
25
50
marker
100
50
0

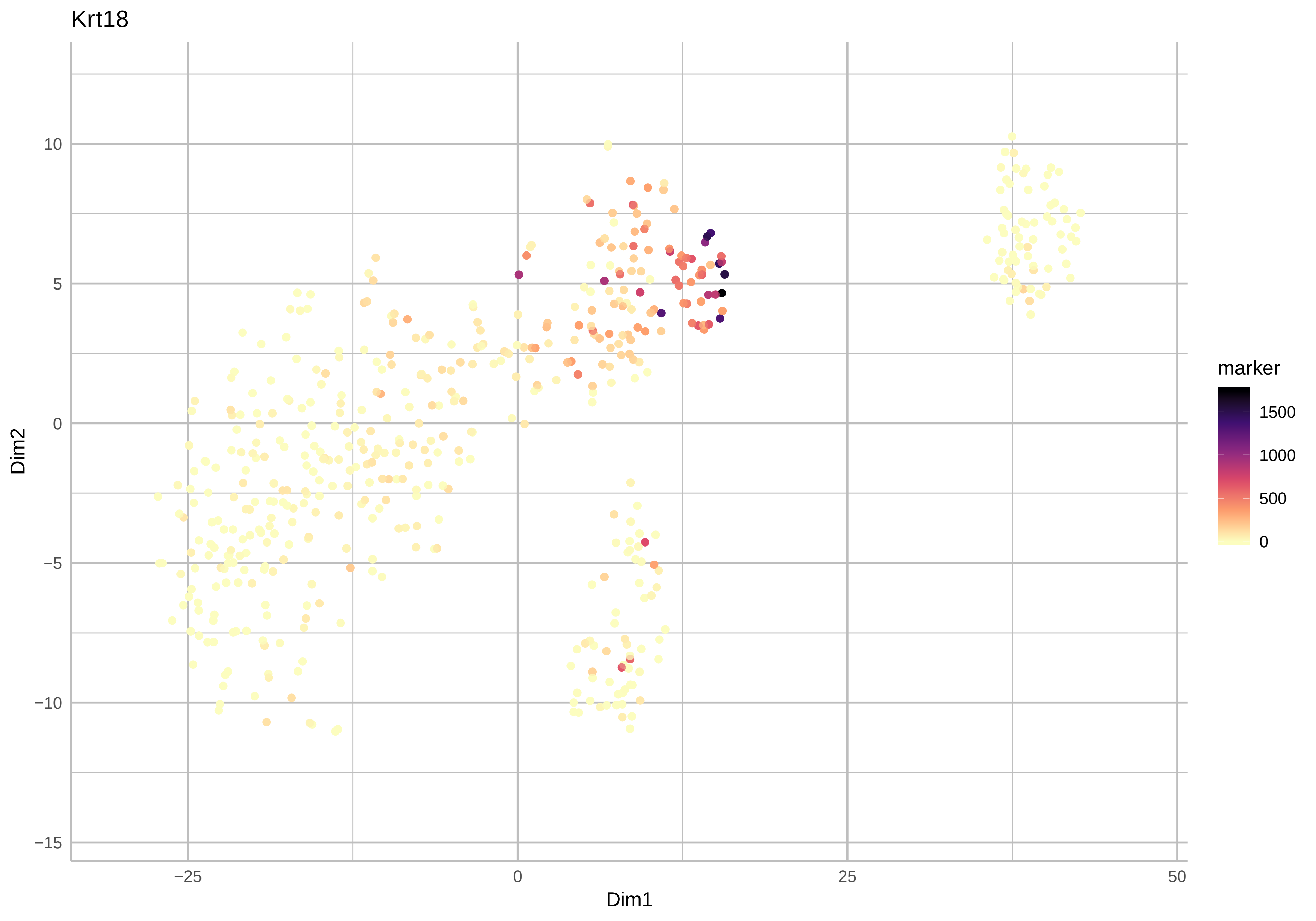
Krt18
Dim2
Dim1
10
5
0
−5
−10
−15
−25
0
25
50
marker
1500
1000
500
0

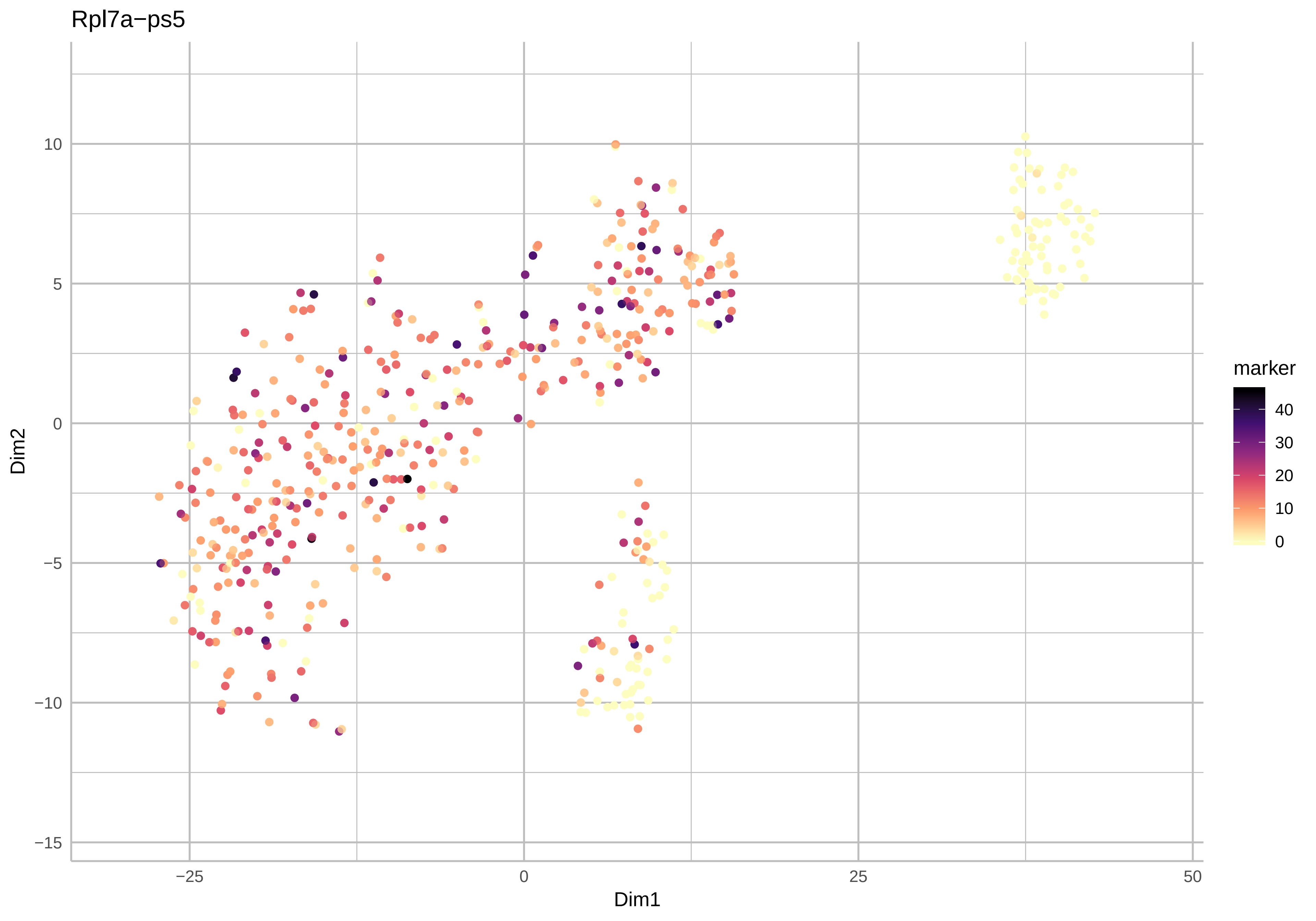

Rpl7a−ps5
Dim2
Dim1
10
5
0
−5
−10
−15
−25
0
25
50
marker
40
30
20
10
0

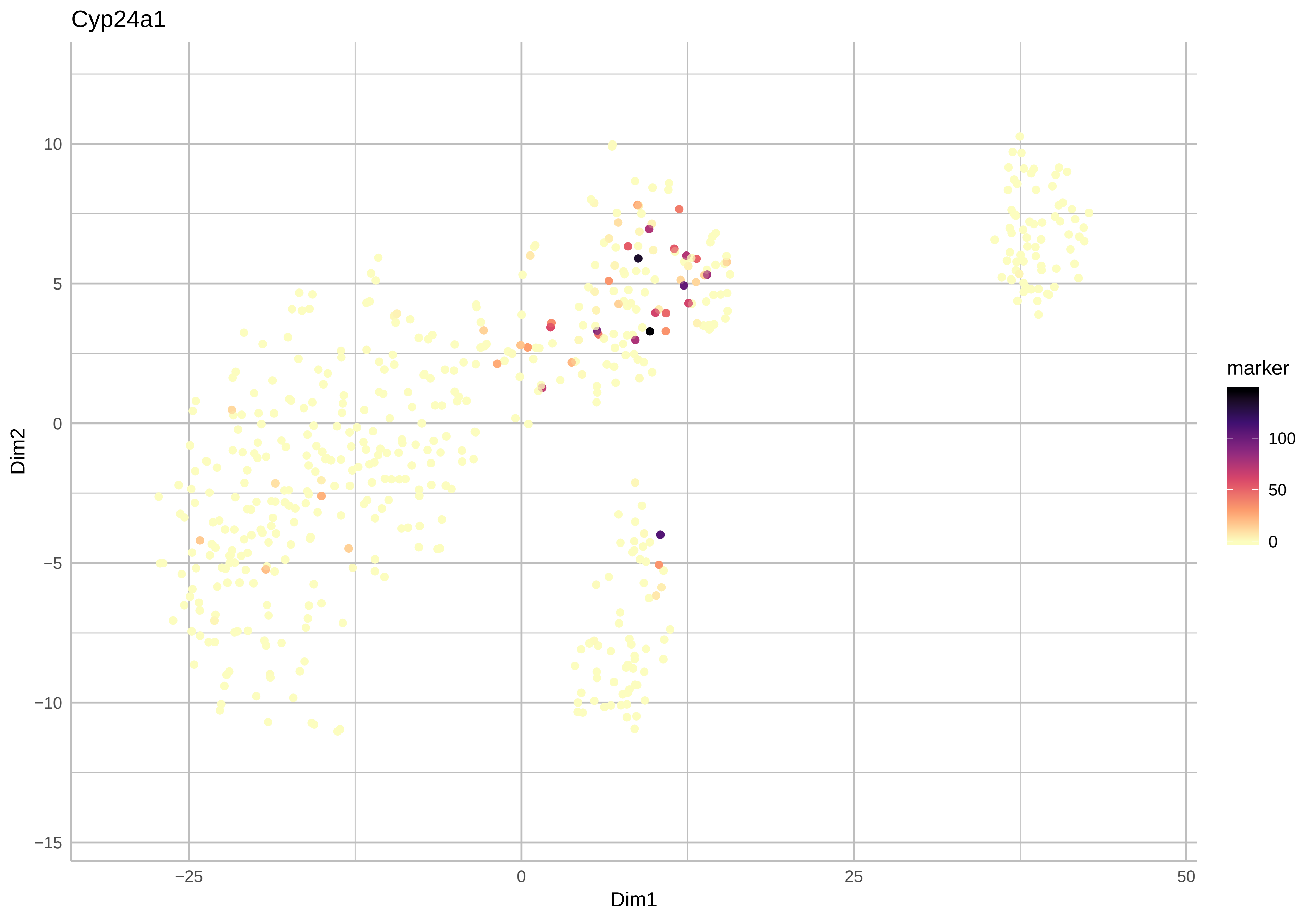
Cyp24a1
Dim2
Dim1
10
5
0
−5
−10
−15
−25
0
25
50
marker
100
50
0

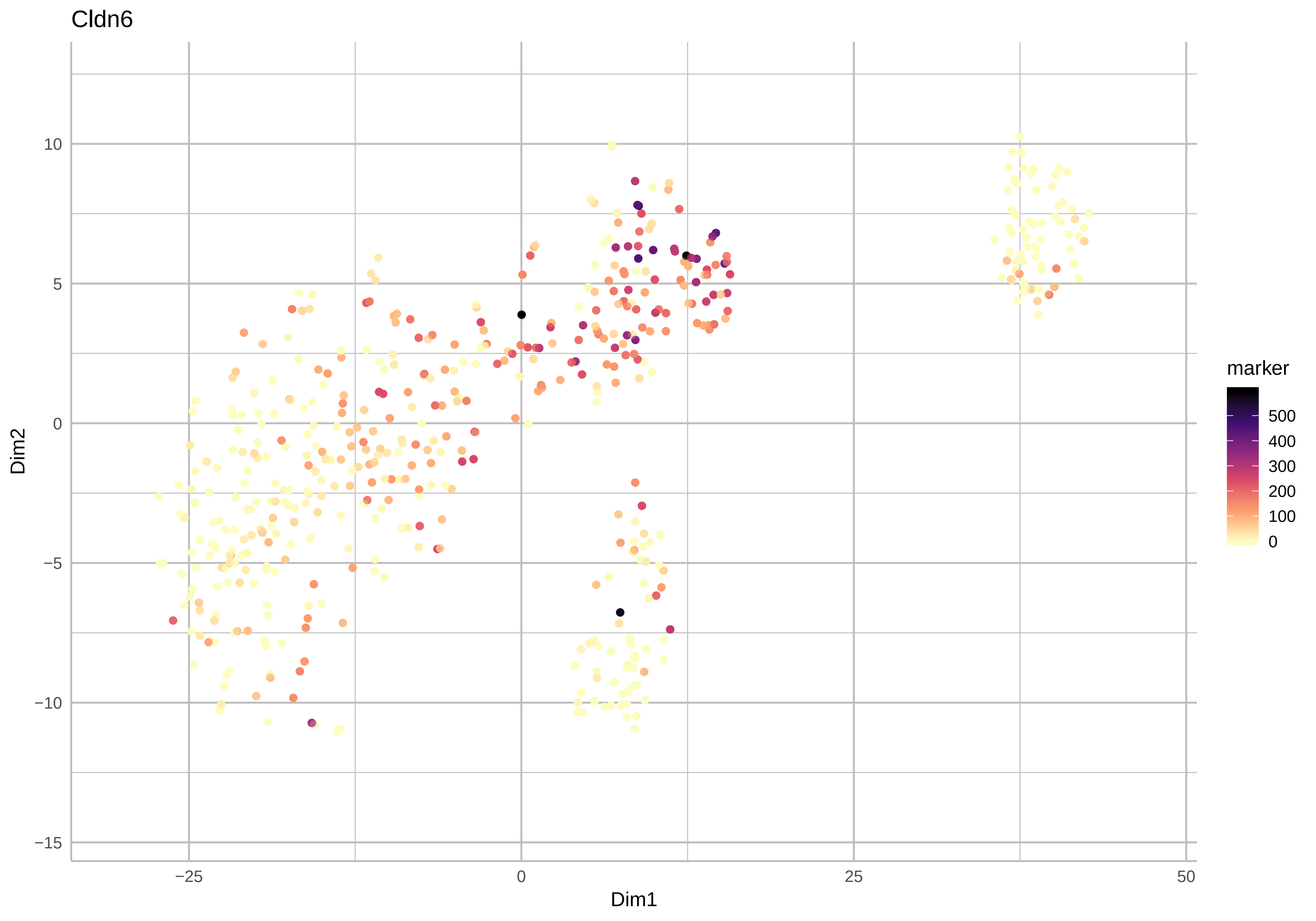

Cldn6
10
5
0
−5
−10
−15
Dim2
−25
0
25
50
Dim1
marker
500
400
300
200
100
0

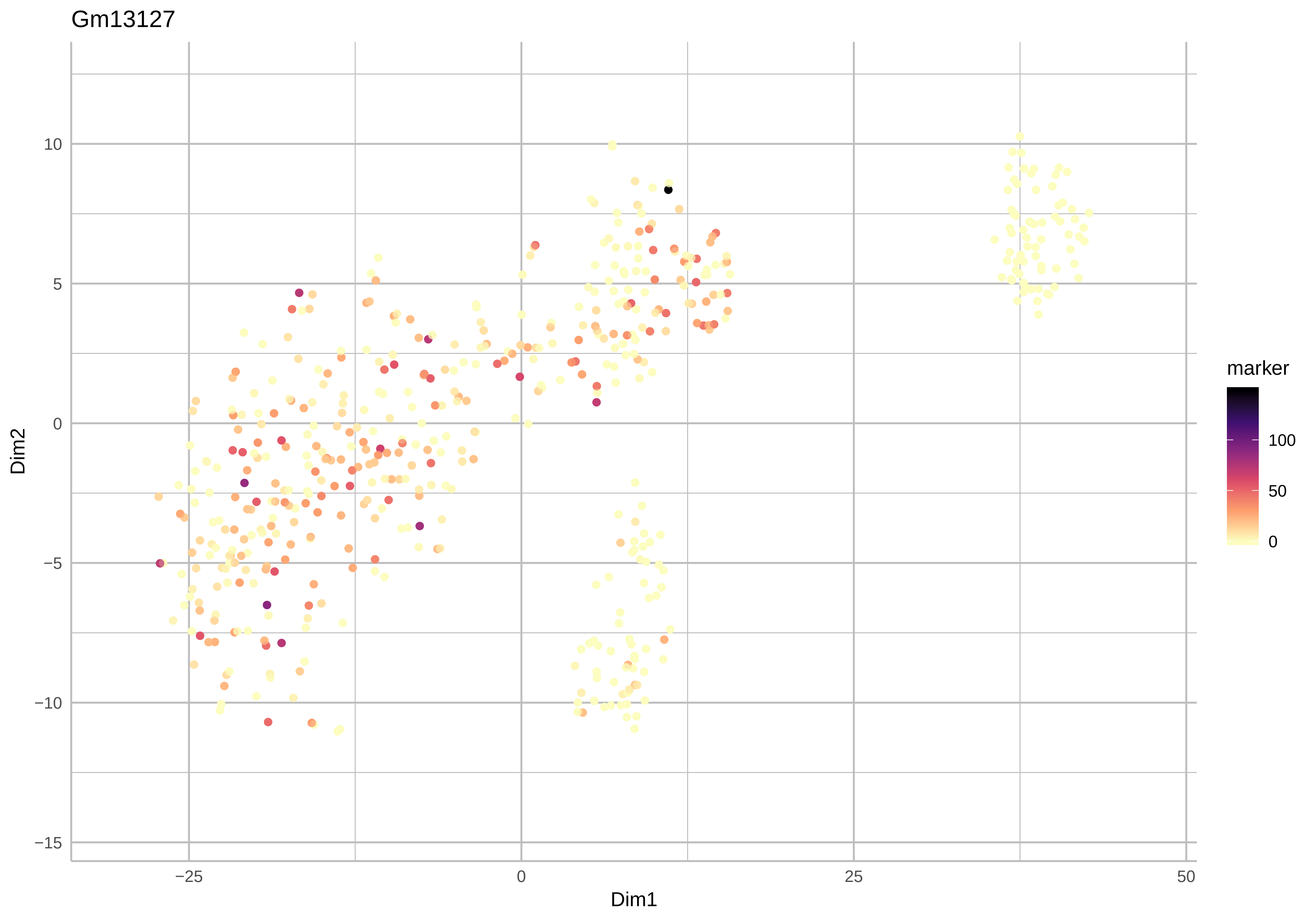

Gm13127
Dim2
Dim1
10
5
0
−5
−10
−15
−25
0
25
50
marker
100
50
0

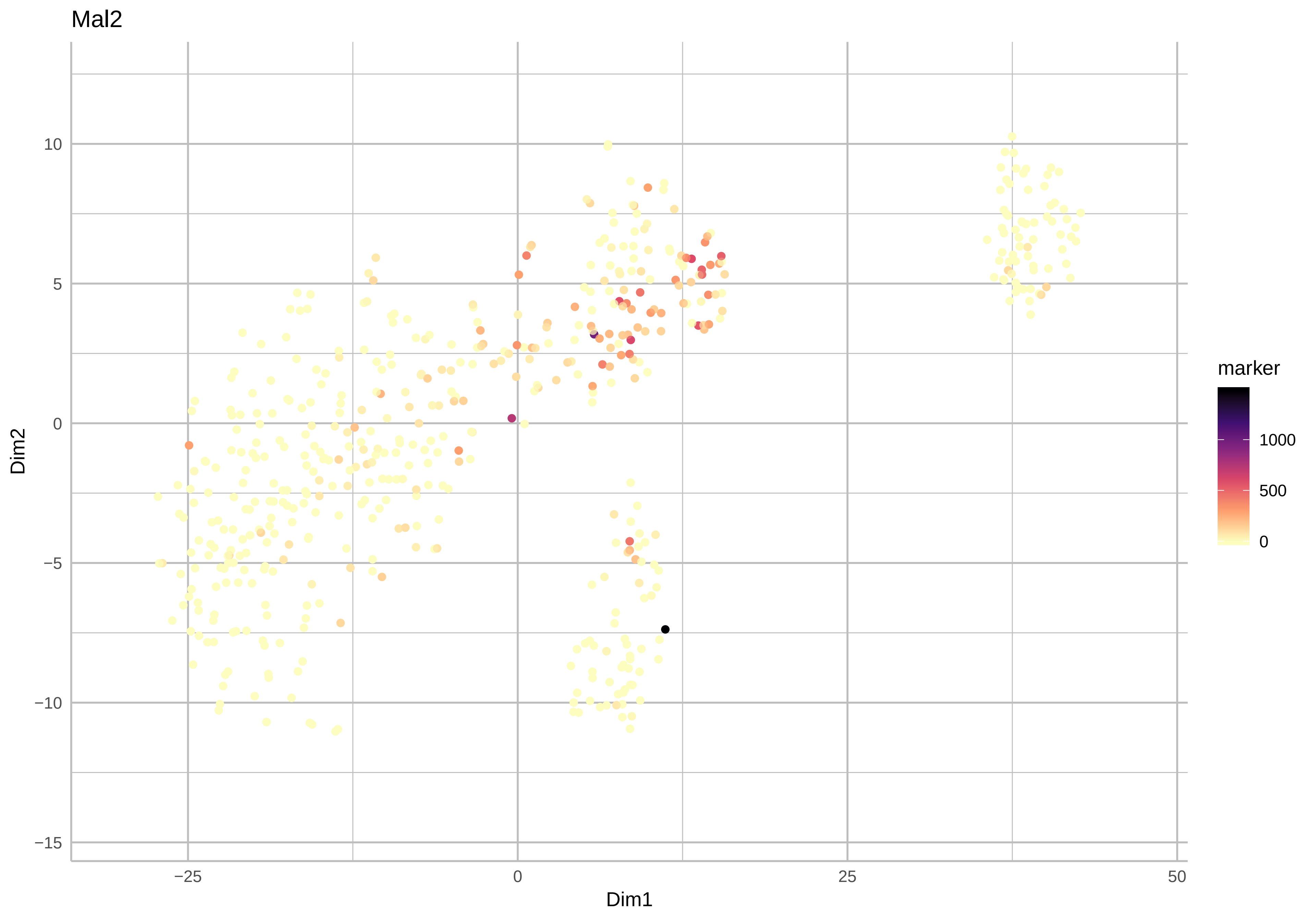

Mal2
Dim2
Dim1
10
5
0
−5
−10
−15
−25
0
25
50
marker
1000
500
0

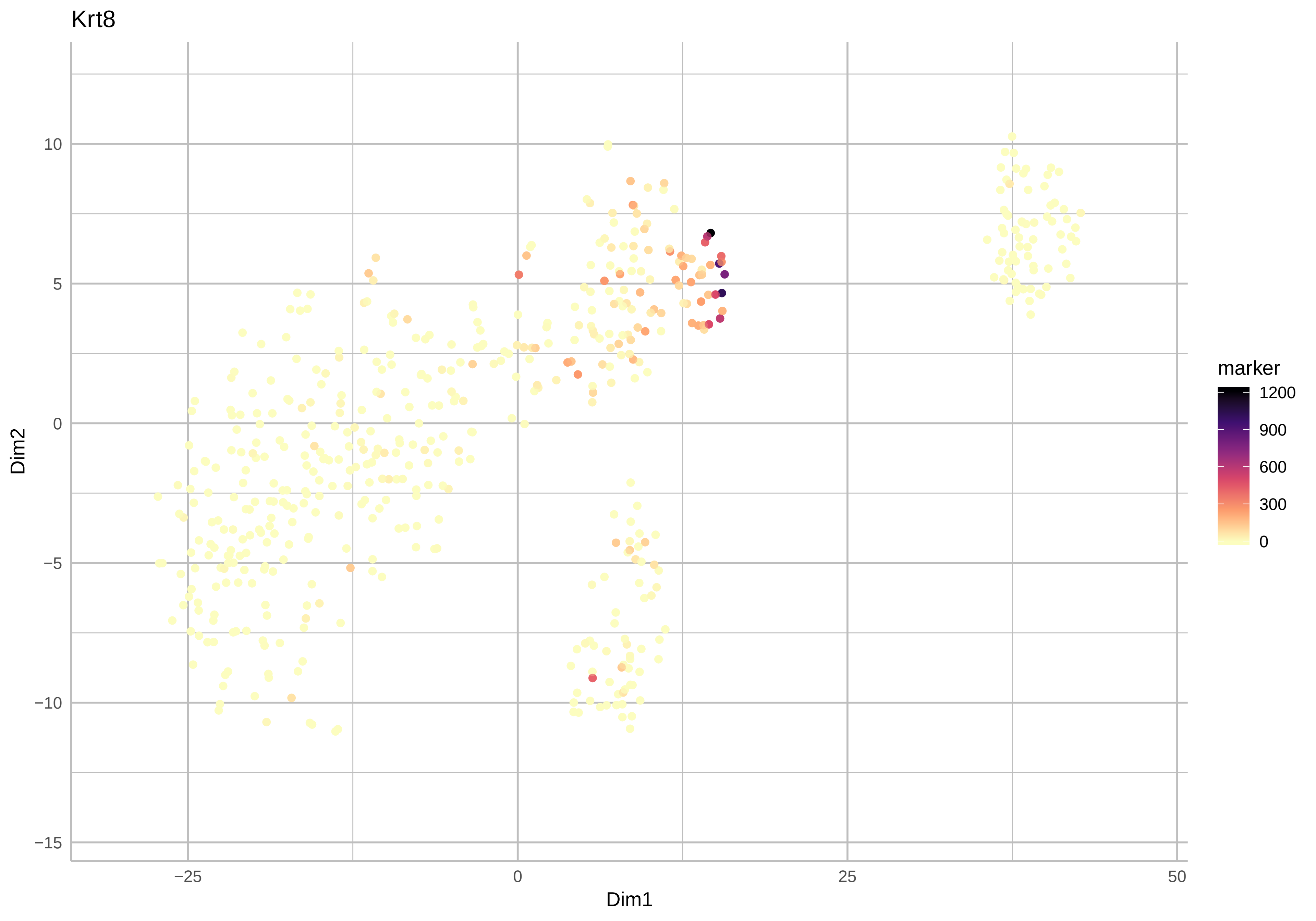

Krt8
Dim2
Dim1
10
5
0
−5
−10
−15
−25
0
25
50
marker
1200
900
600
300
0

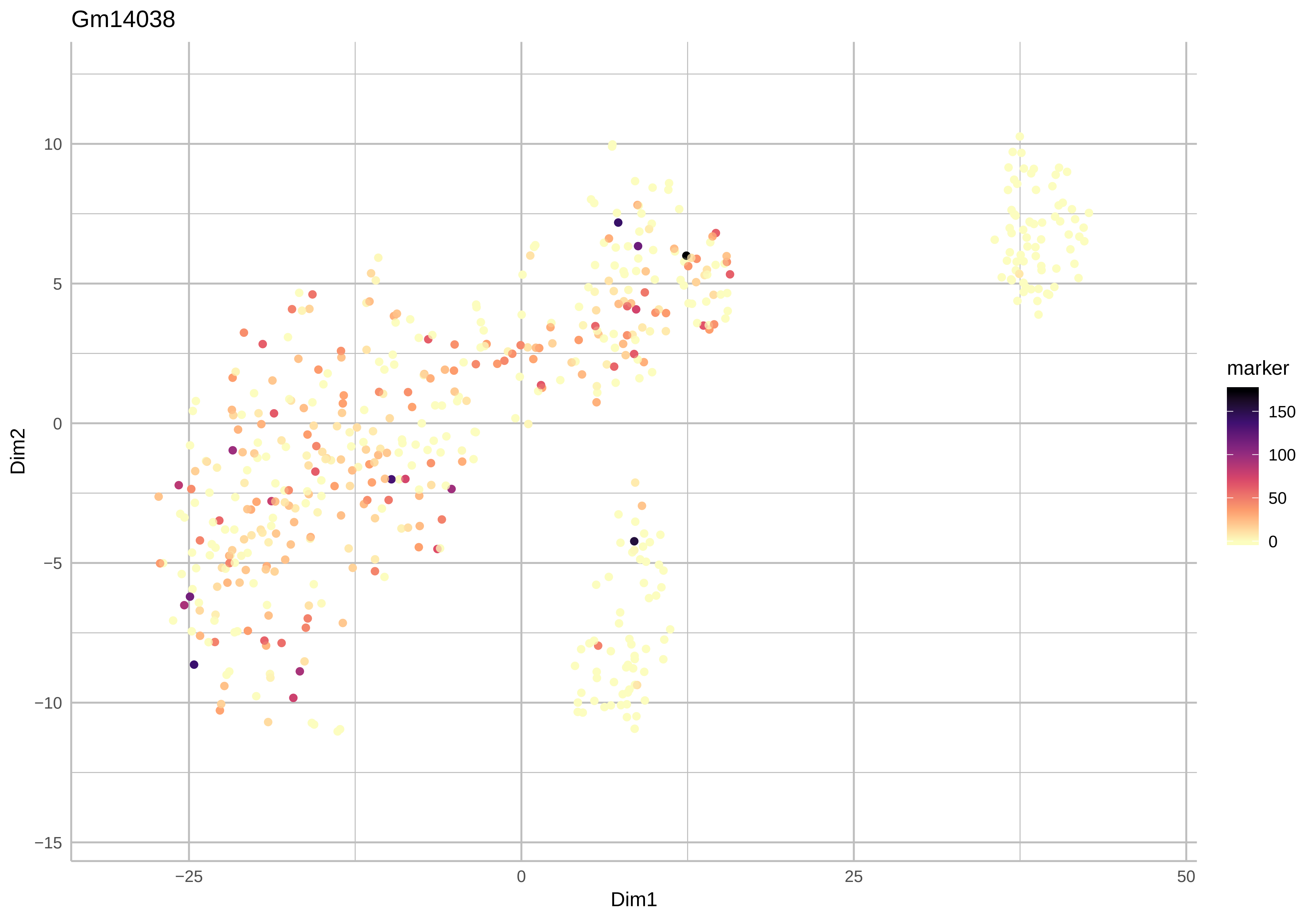

Gm14038
Dim2
10
5
0
−5
−10
−15
−25
0
25
50
Dim1
marker
150
100
50
0

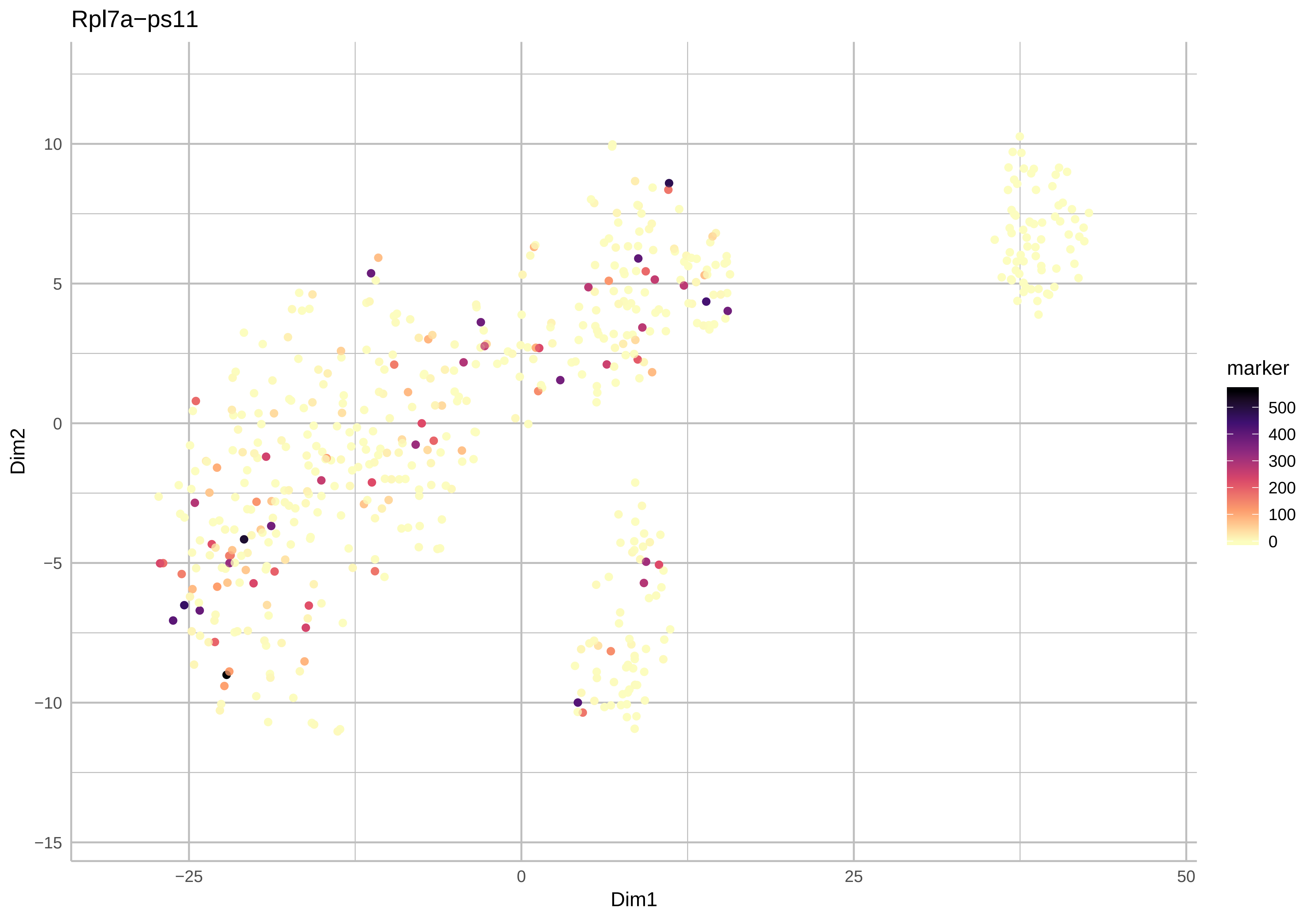

Rpl7a−ps11
Dim2
Dim1
10
5
0
−5
−10
−15
−25
0
25
50
marker
500
400
300
200
100
0

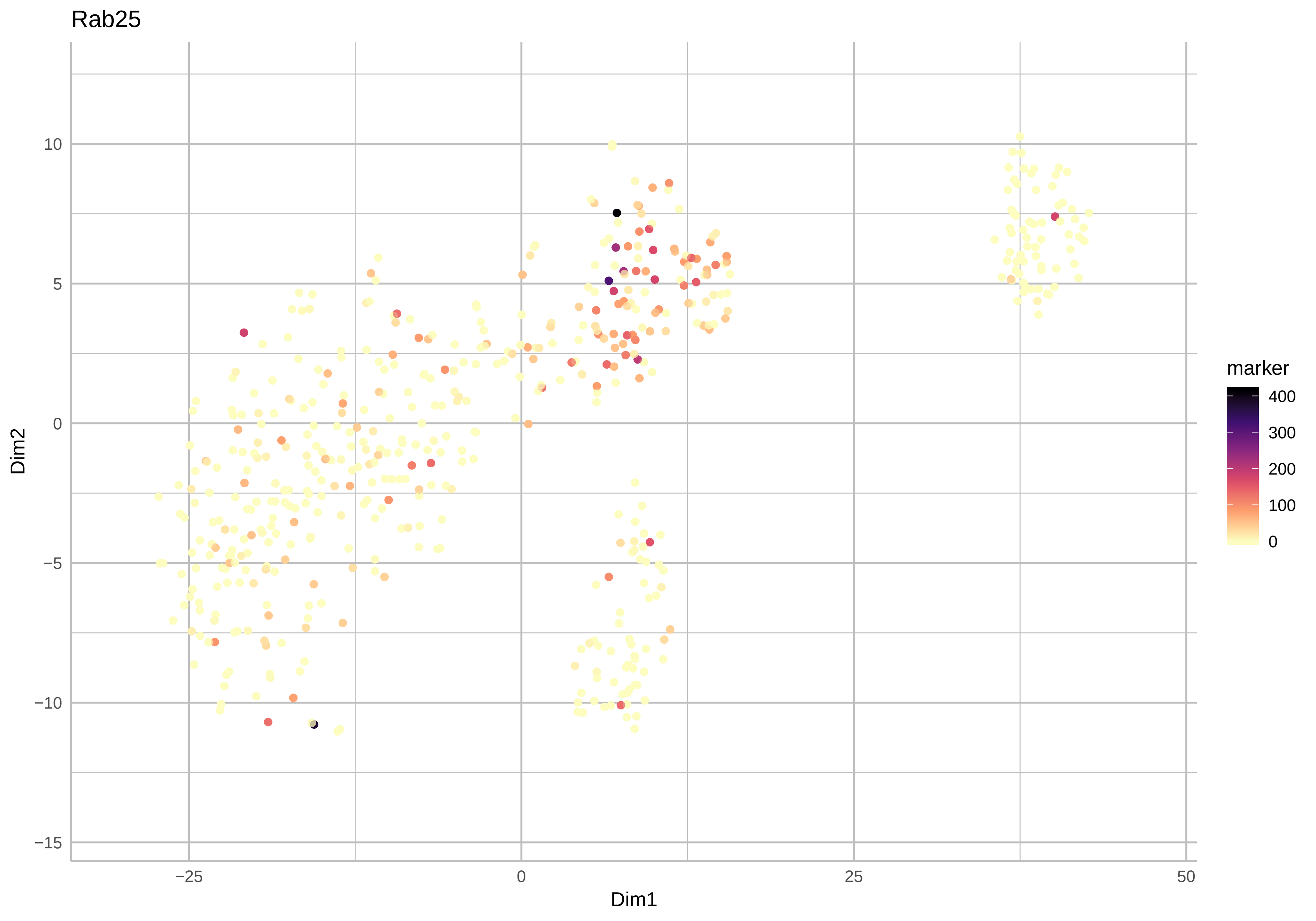

Rab25
Dim2
Dim1
10
5
0
−5
−10
−15
−25
0
25
50
marker
400
300
200
100
0

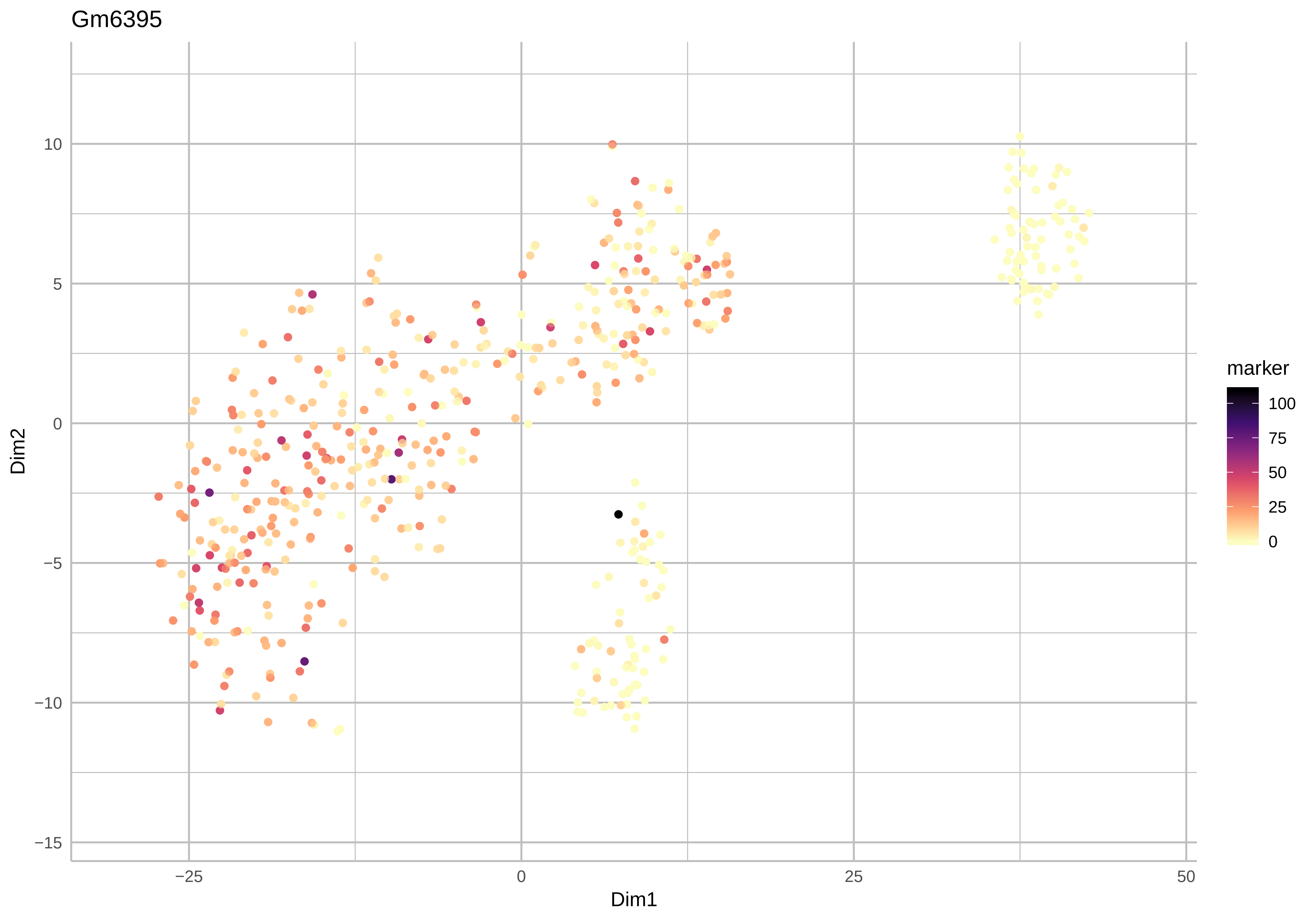
Gm6395
10
5
0
−5
−10
−15
−25
0
25
50
Dim1
Dim2
marker
100
75
50
25
0

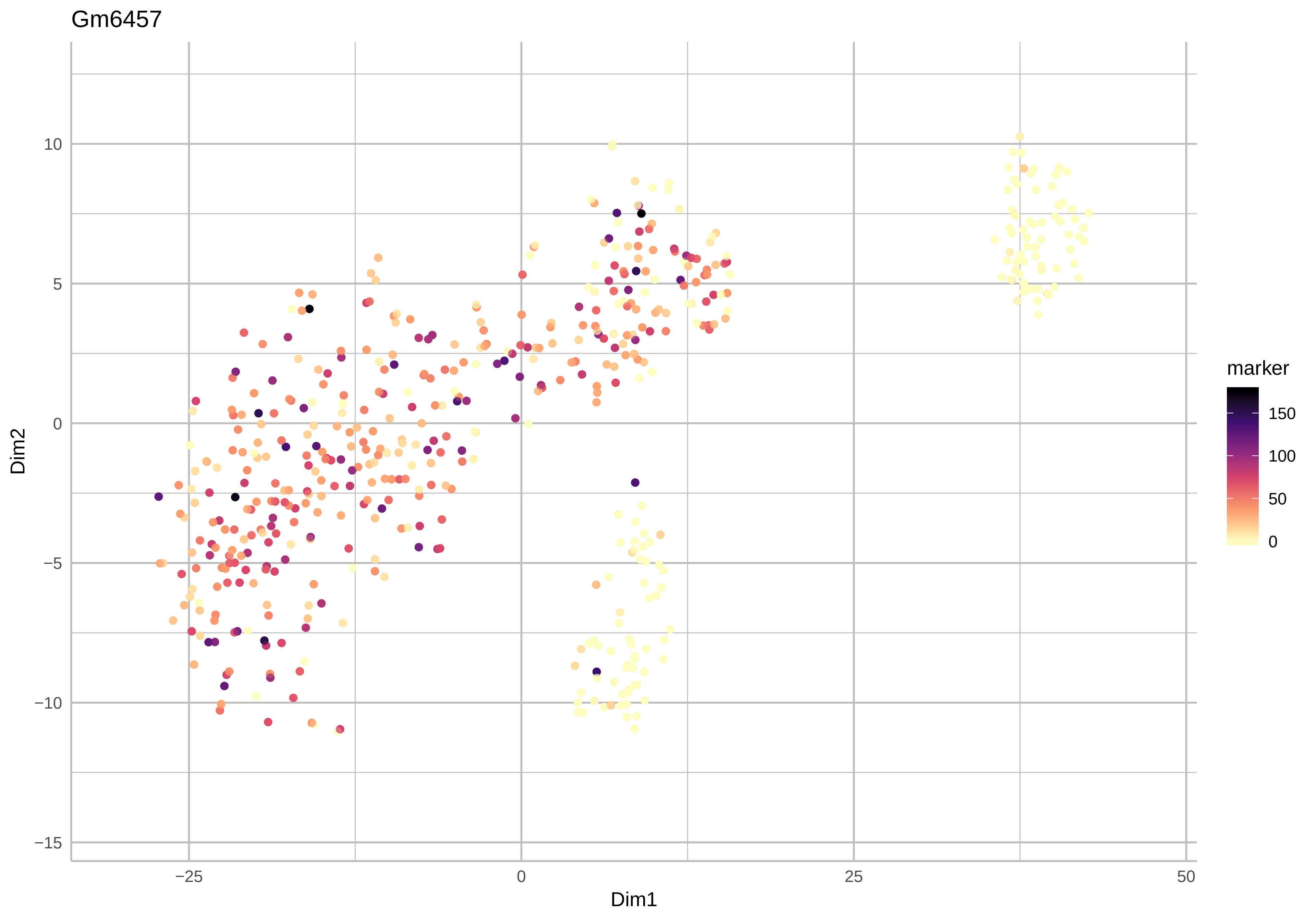
Gm6457
Dim2
Dim1
marker
150
100
50
0
10
5
0
−5
−10
−15
−25
0
25
50

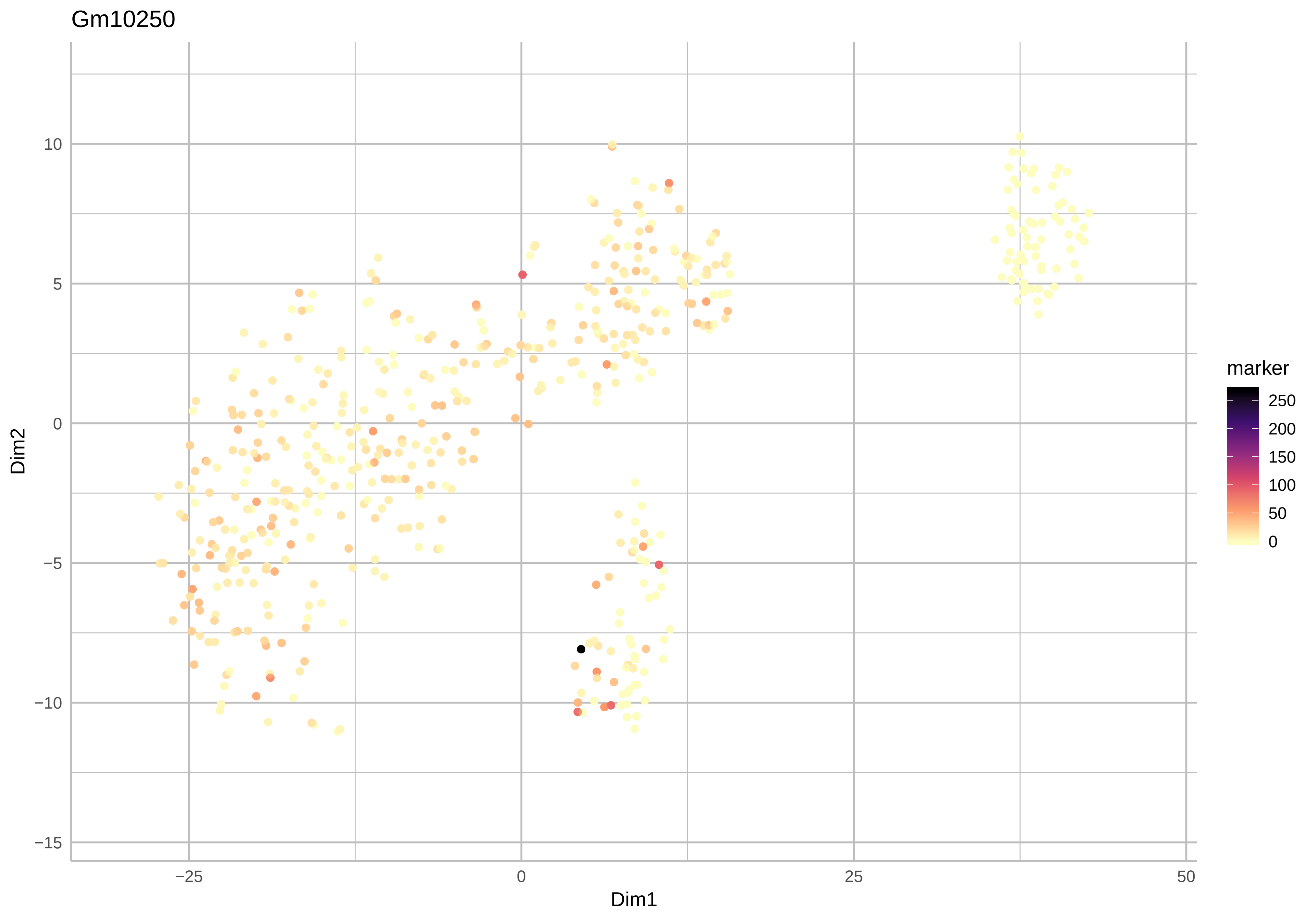

Gm10250
Dim2
Dim1
10
5
0
−5
−10
−15
−25
0
25
50
marker
250
200
150
100
50
0

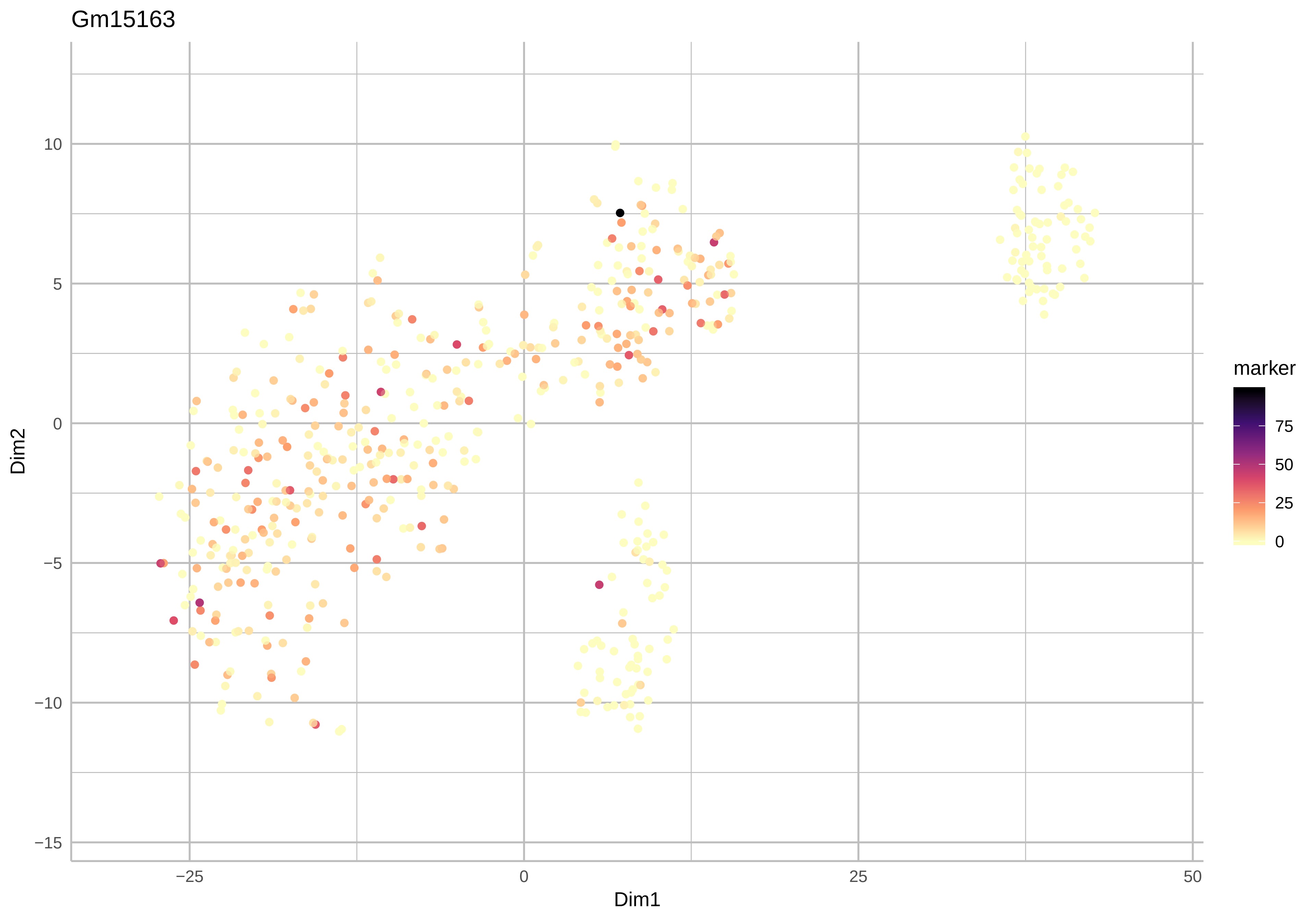
Gm15163
Dim2
Dim1
10
5
0
−5
−10
−15
−25
0
25
50
marker
75
50
25
0

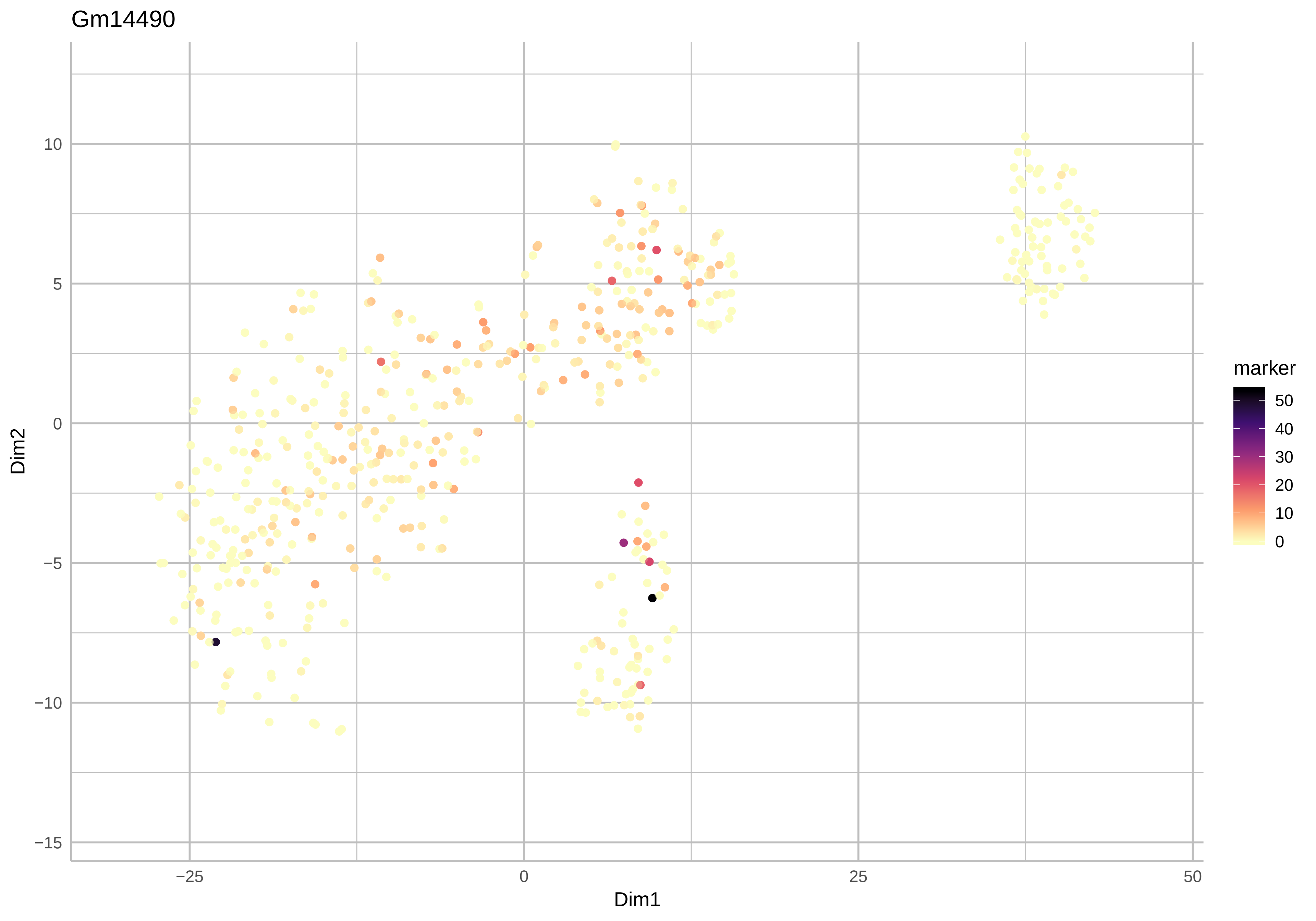

Gm14490
Dim2
Dim1
marker
50
40
30
20
10
0
10
5
0
−5
−10
−15
−25
0
25
50

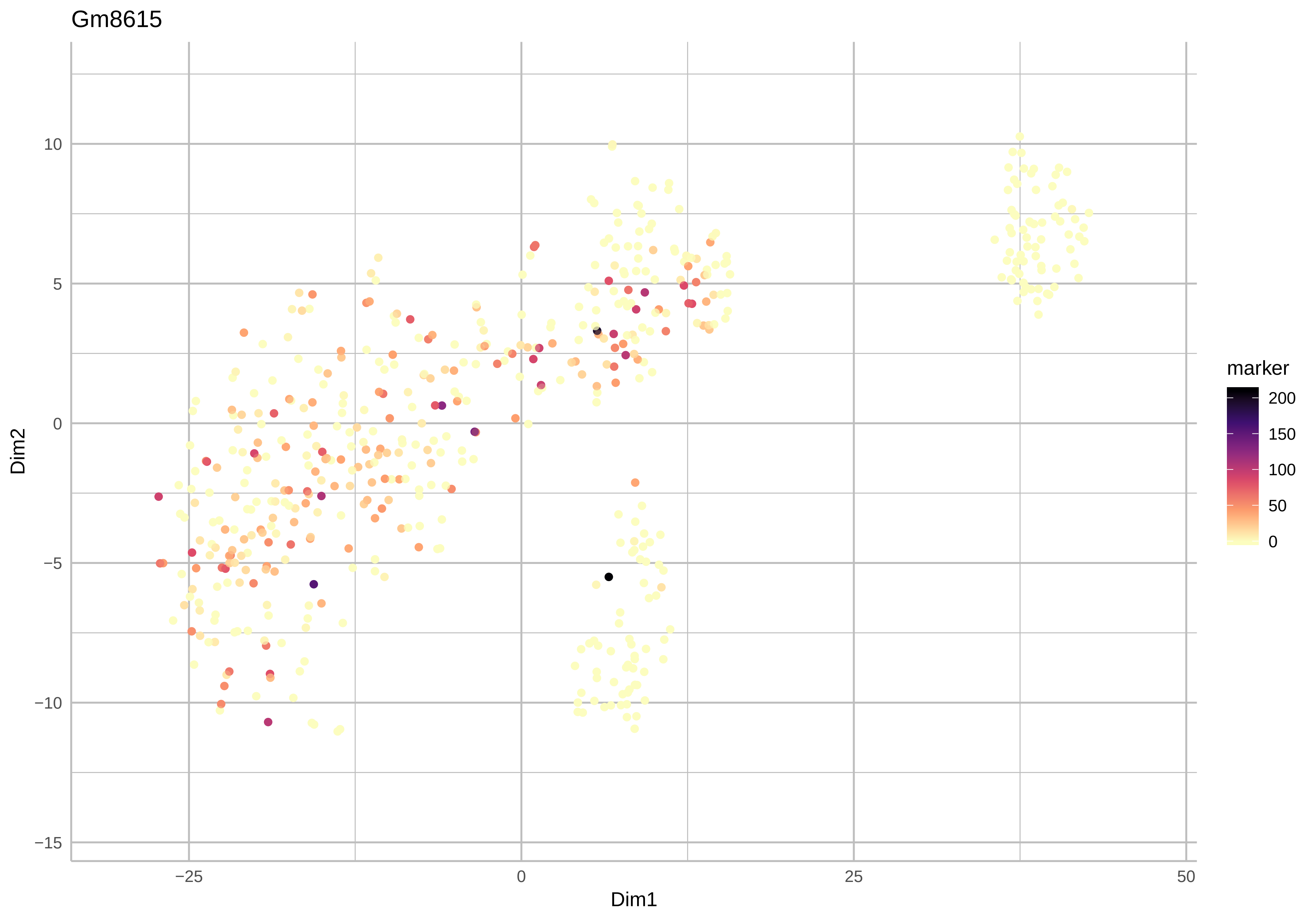

Gm8615
Dim2
Dim1
marker
200
150
100
50
0
10
5
0
−5
−10
−15
−25
0
25
50

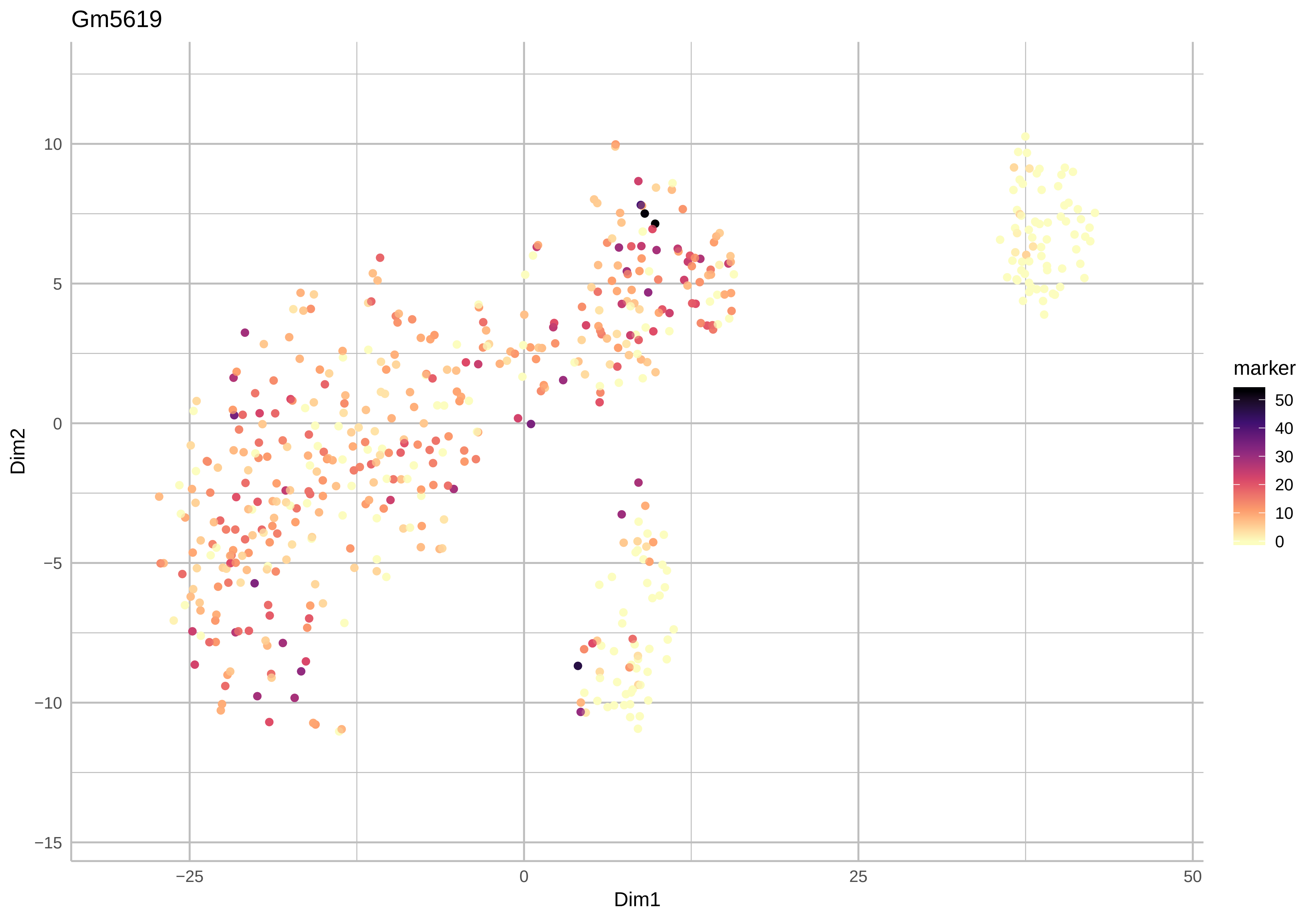

Gm5619
Dim2
Dim1
10
5
0
−5
−10
−15
−25
0
25
50
marker
50
40
30
20
10
0

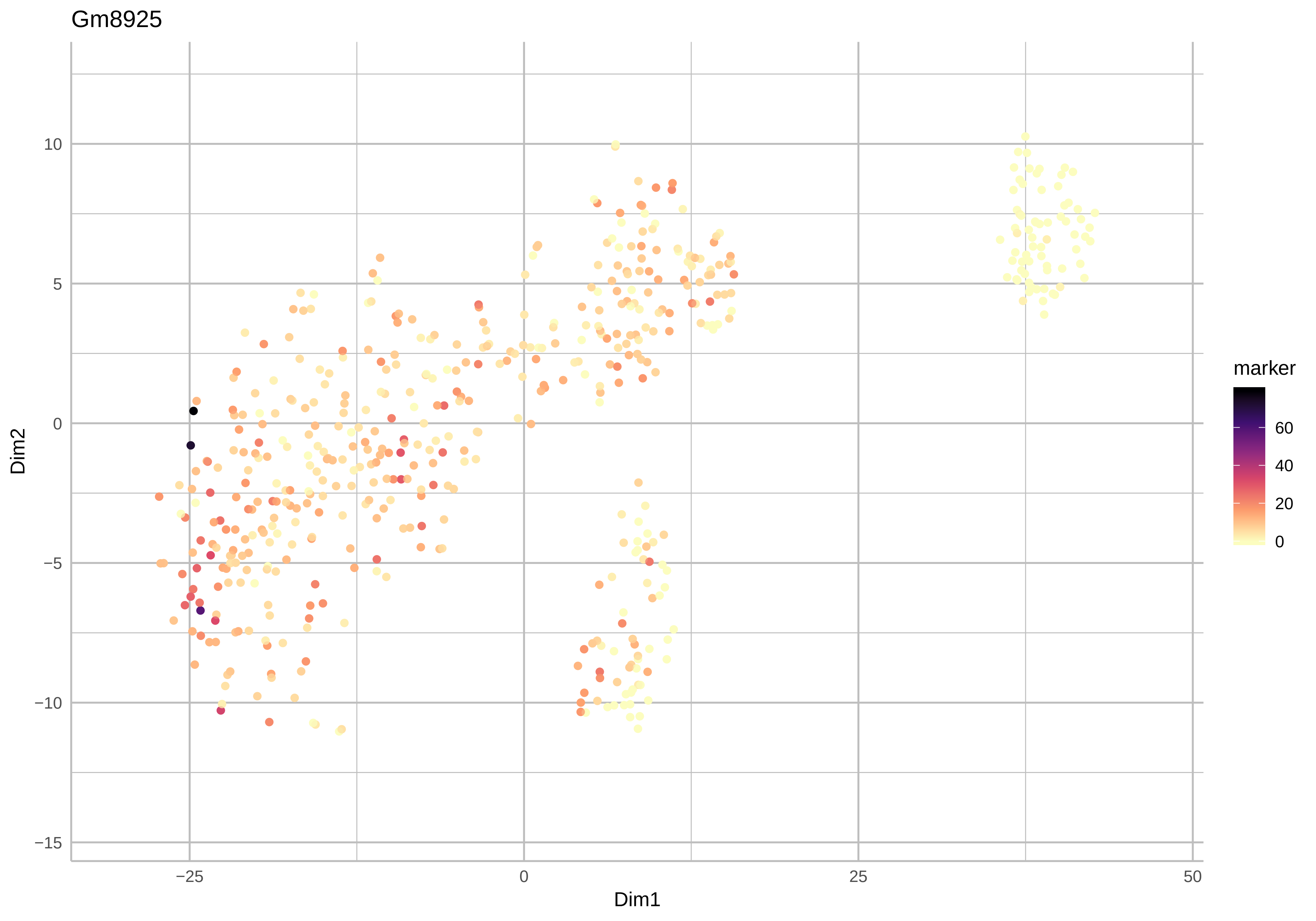

Gm8925
Dim2
Dim1
10
5
0
−5
−10
−15
−25
0
25
50
marker
60
40
20
0

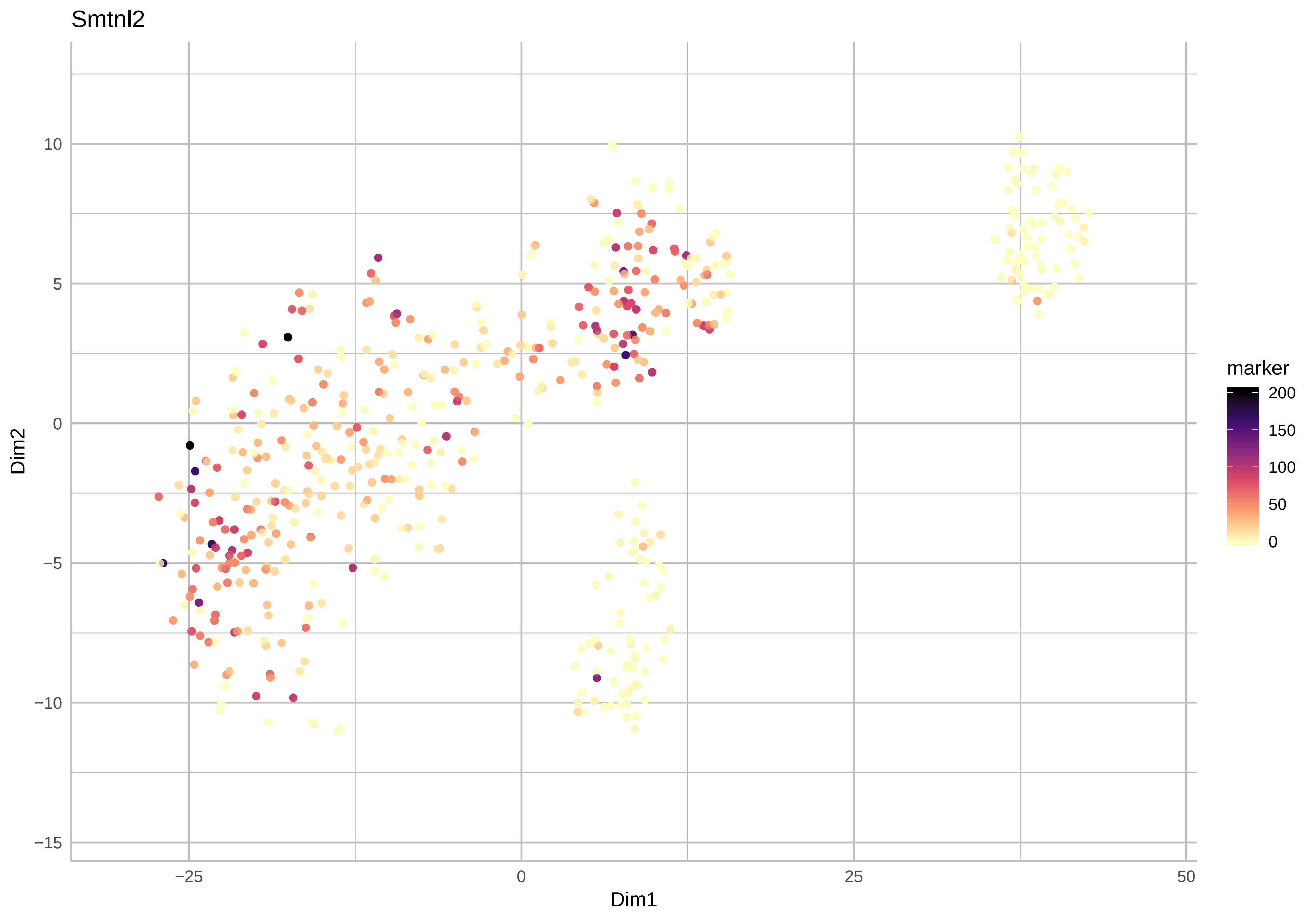
Smtnl2
10
5
0
−5
−10
−15
−25
0
25
50
Dim1
Dim2
marker
200
150
100
50
0

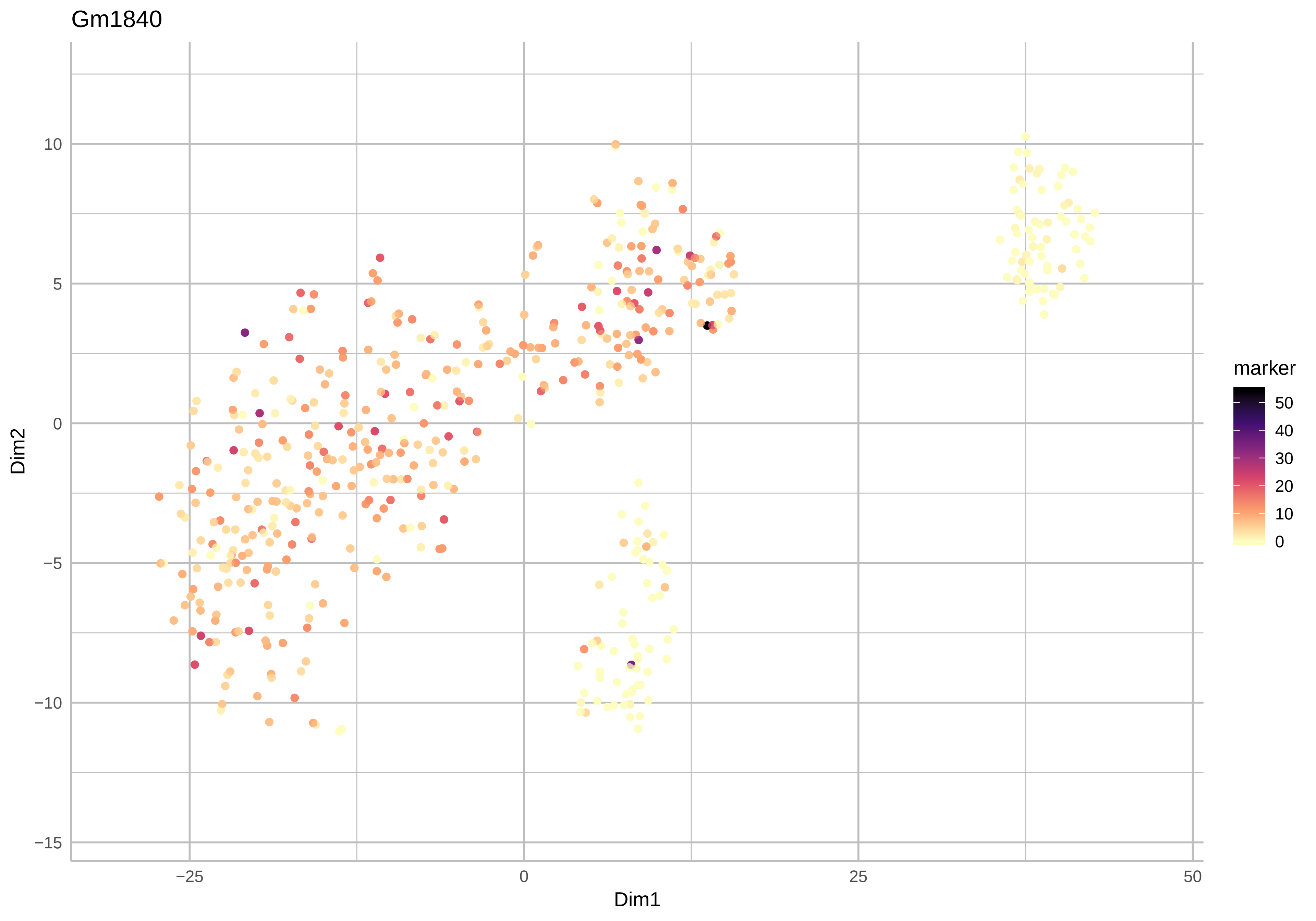
Gm1840
Dim2
Dim1
10
5
0
−5
−10
−15
−25
0
25
50
marker
50
40
30
20
10
0

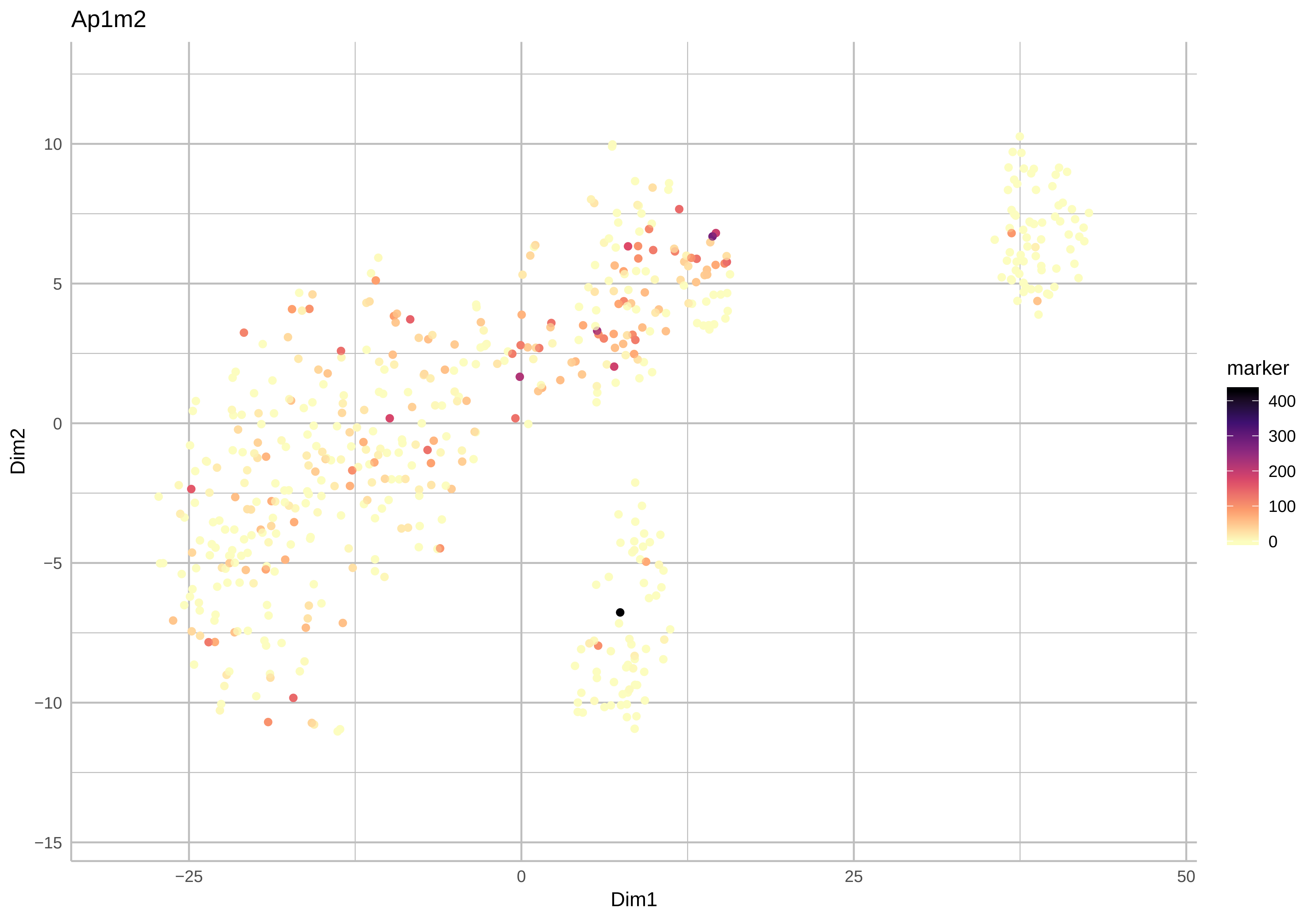

Ap1m2
Dim2
Dim1
10
5
0
−5
−10
−15
−25
0
25
50
marker
400
300
200
100
0

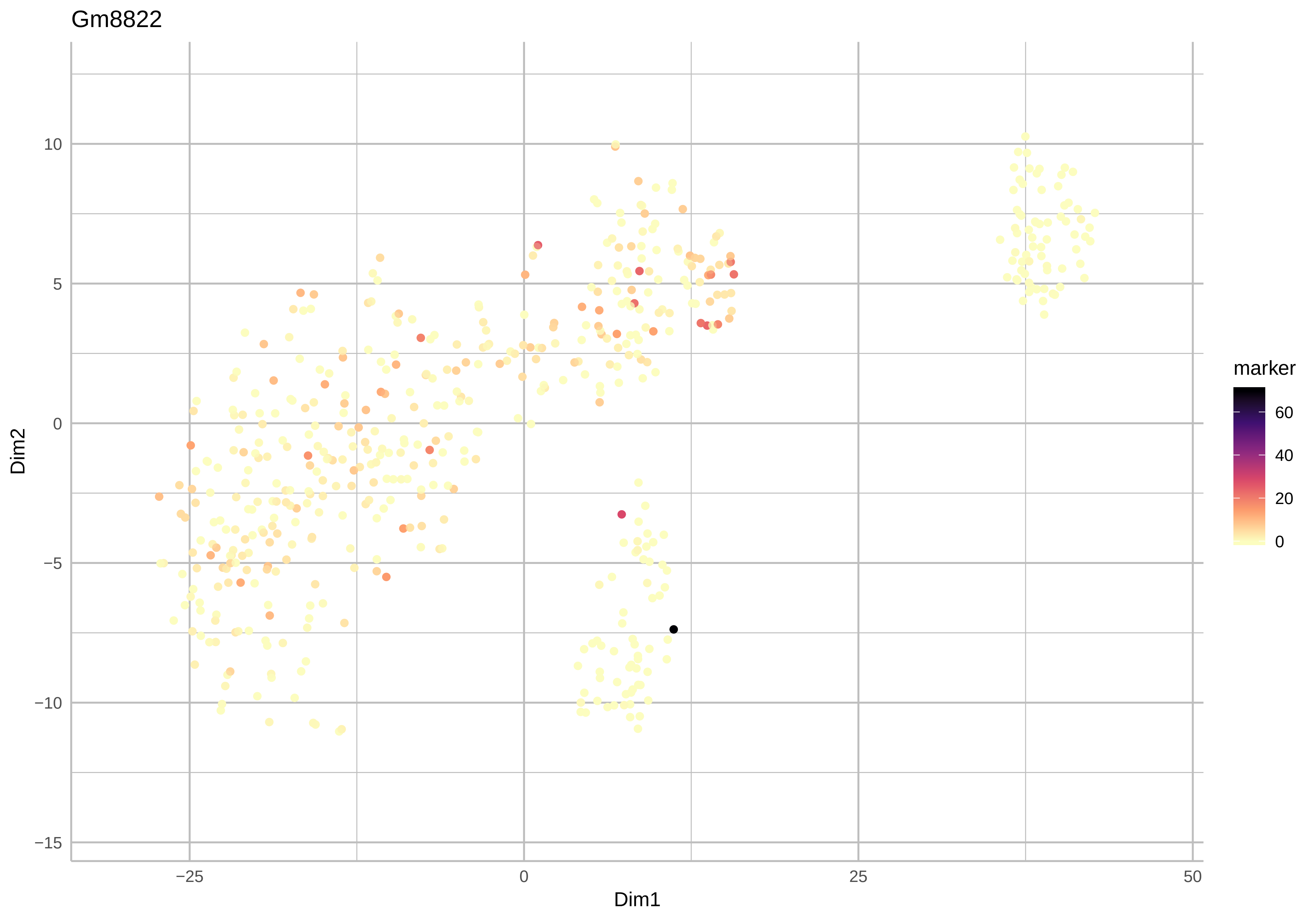

Gm8822
Dim2
Dim1
10
5
0
−5
−10
−15
−25
0
25
50
marker
60
40
20
0

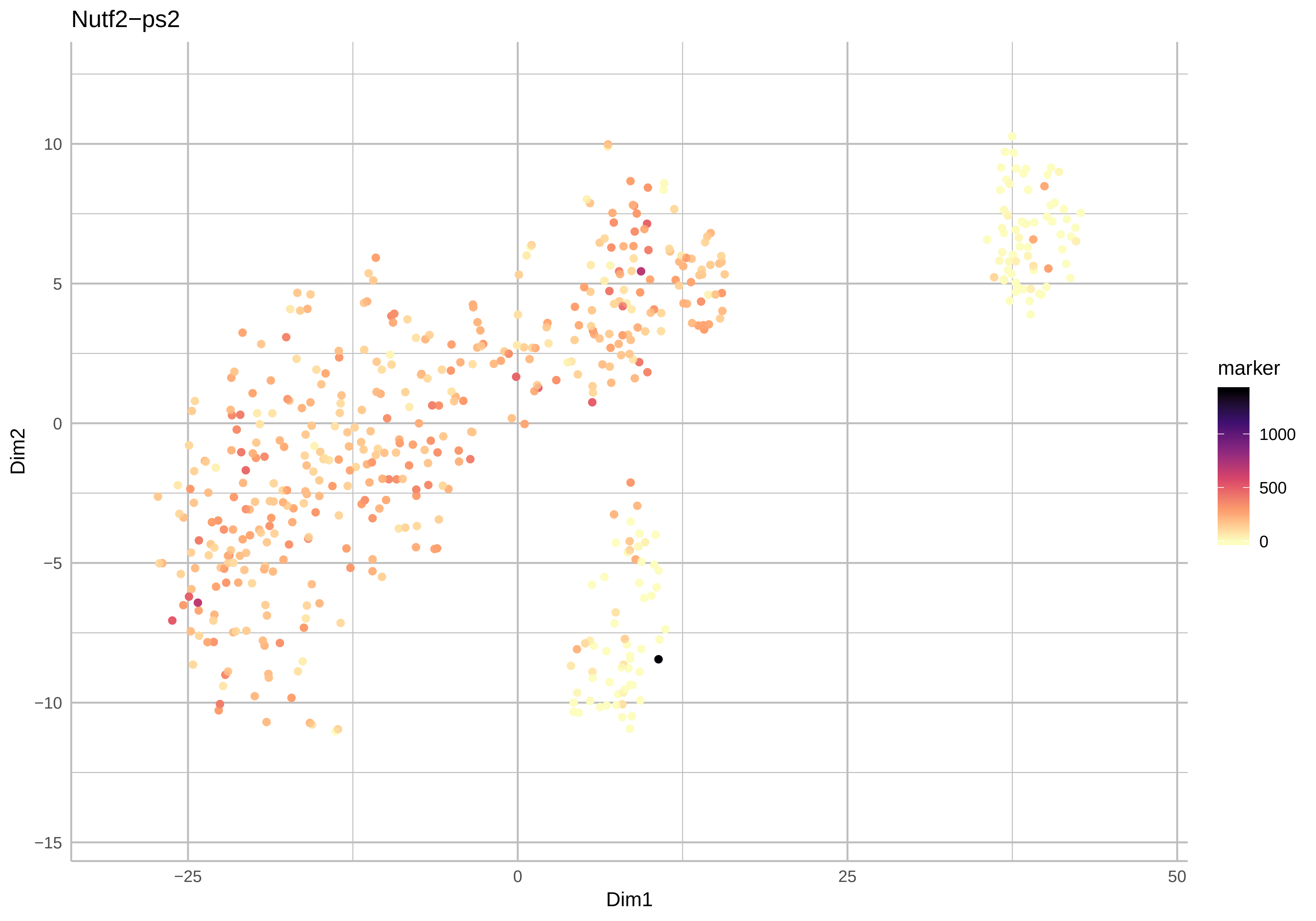

Nutf2−ps2
Dim2
Dim1
10
5
0
−5
−10
−15
−25
0
25
50
marker
1000
500
0

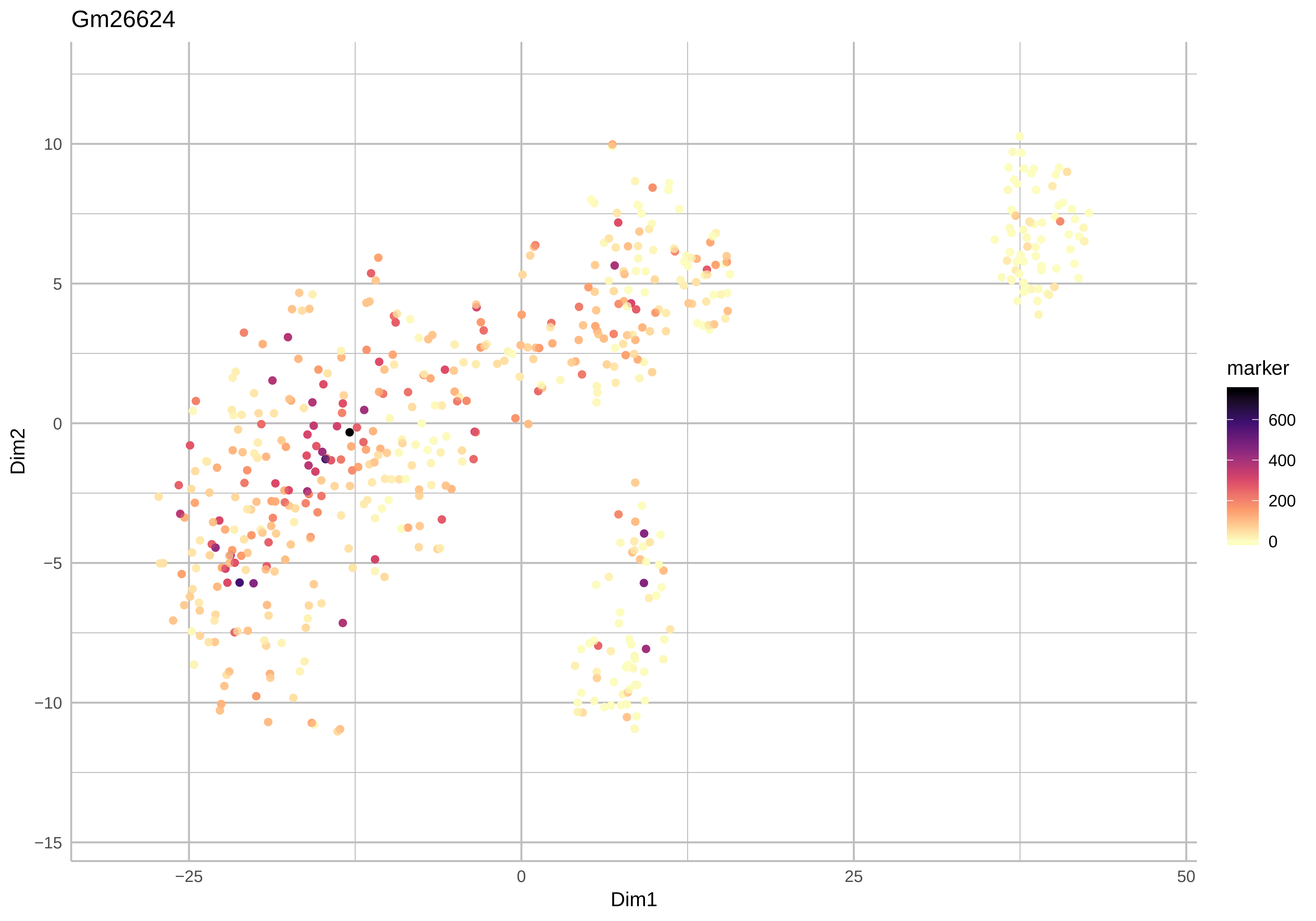
Gm26624
Dim2
Dim1
10
5
0
−5
−10
−15
−25
0
25
50
marker
600
400
200
0

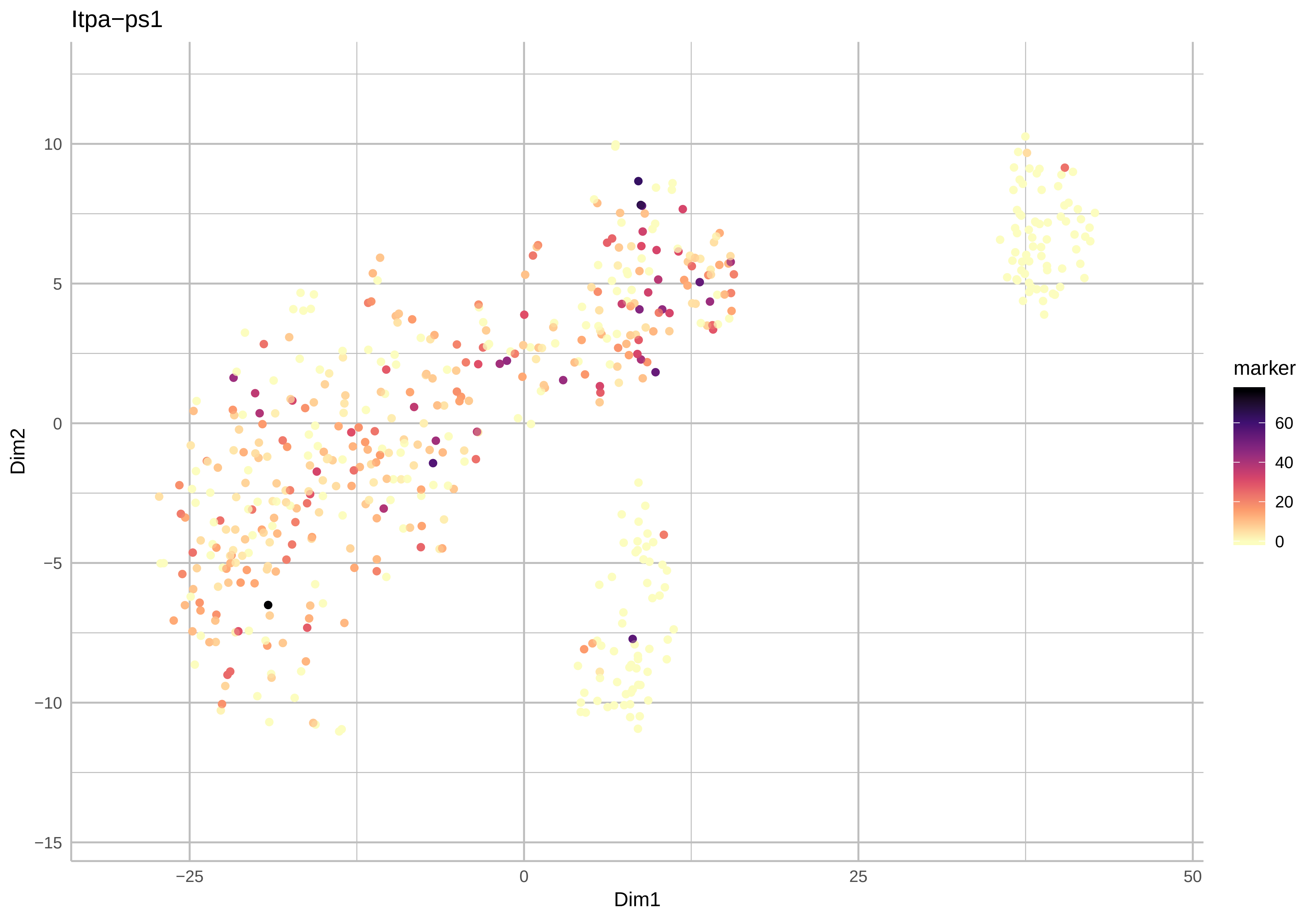

Itpa−ps1
10
5
0
−5
−10
−15
Dim2
−25
0
25
50
Dim1
marker
60
40
20
0

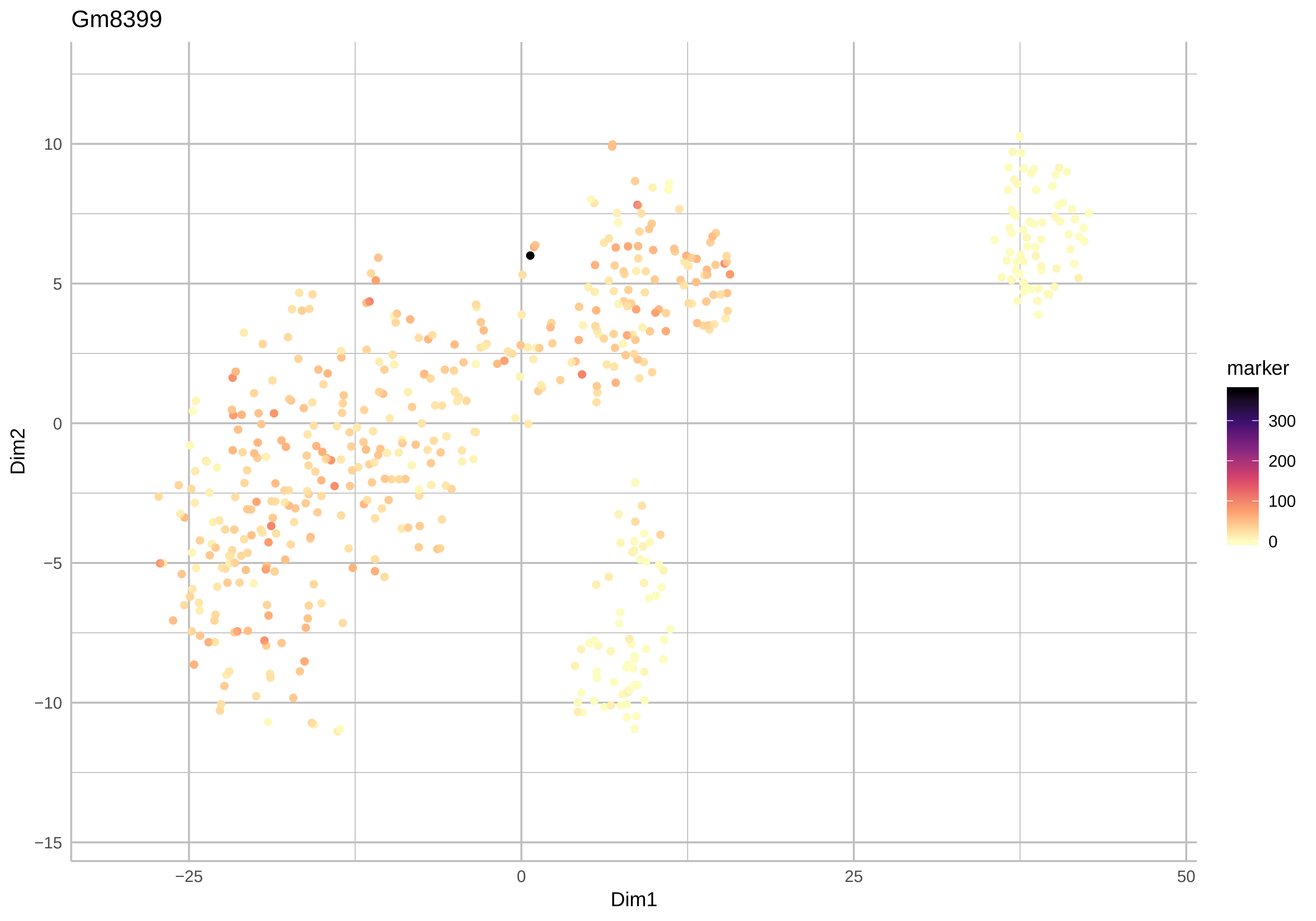
Gm8399
Dim2
Dim1
10
5
0
−5
−10
−15
−25
0
25
50
marker
300
200
100
0

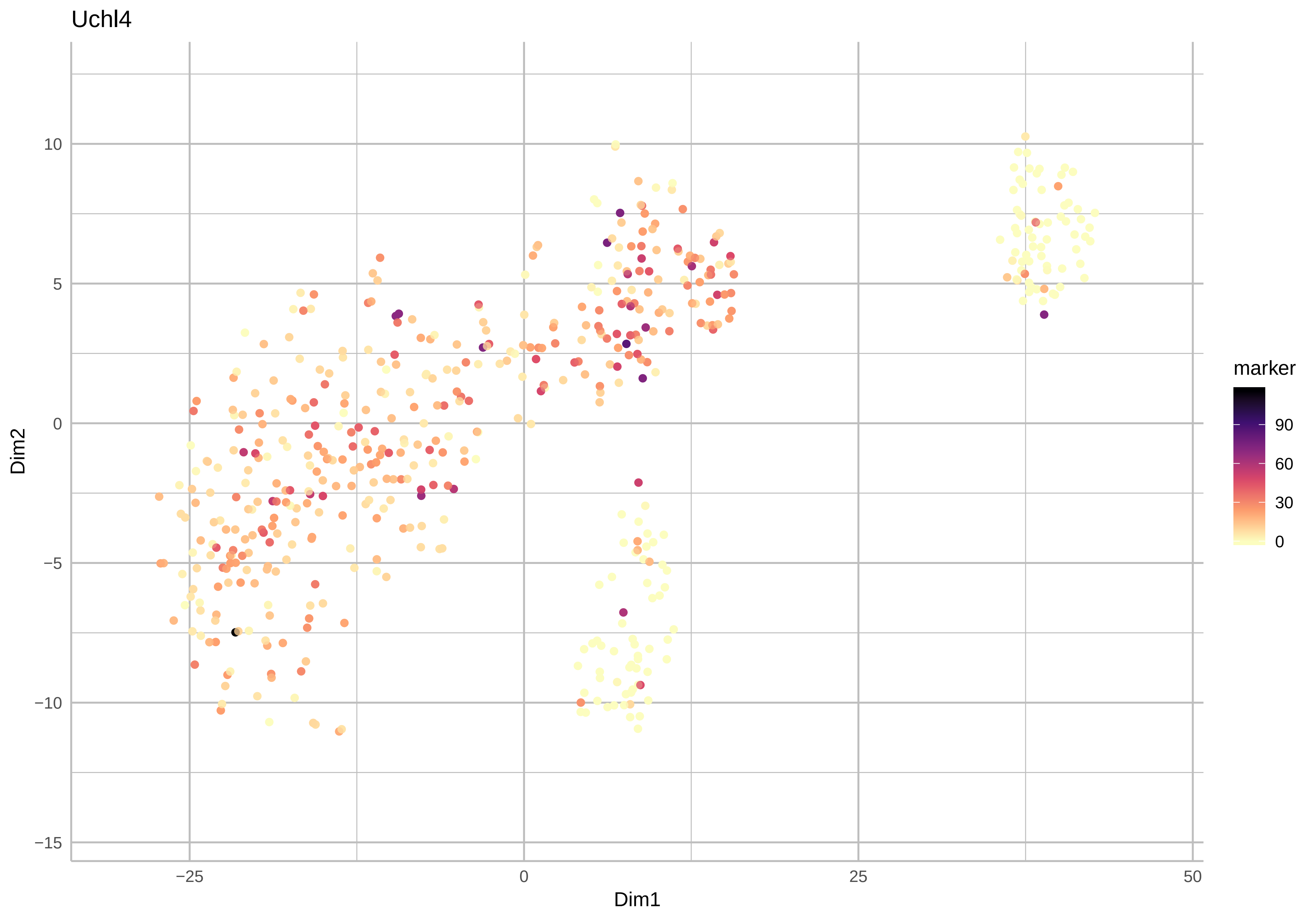

Uchl4
10
5
0
−5
−10
−15
Dim2
−25
0
25
50
Dim1
marker
90
60
30
0

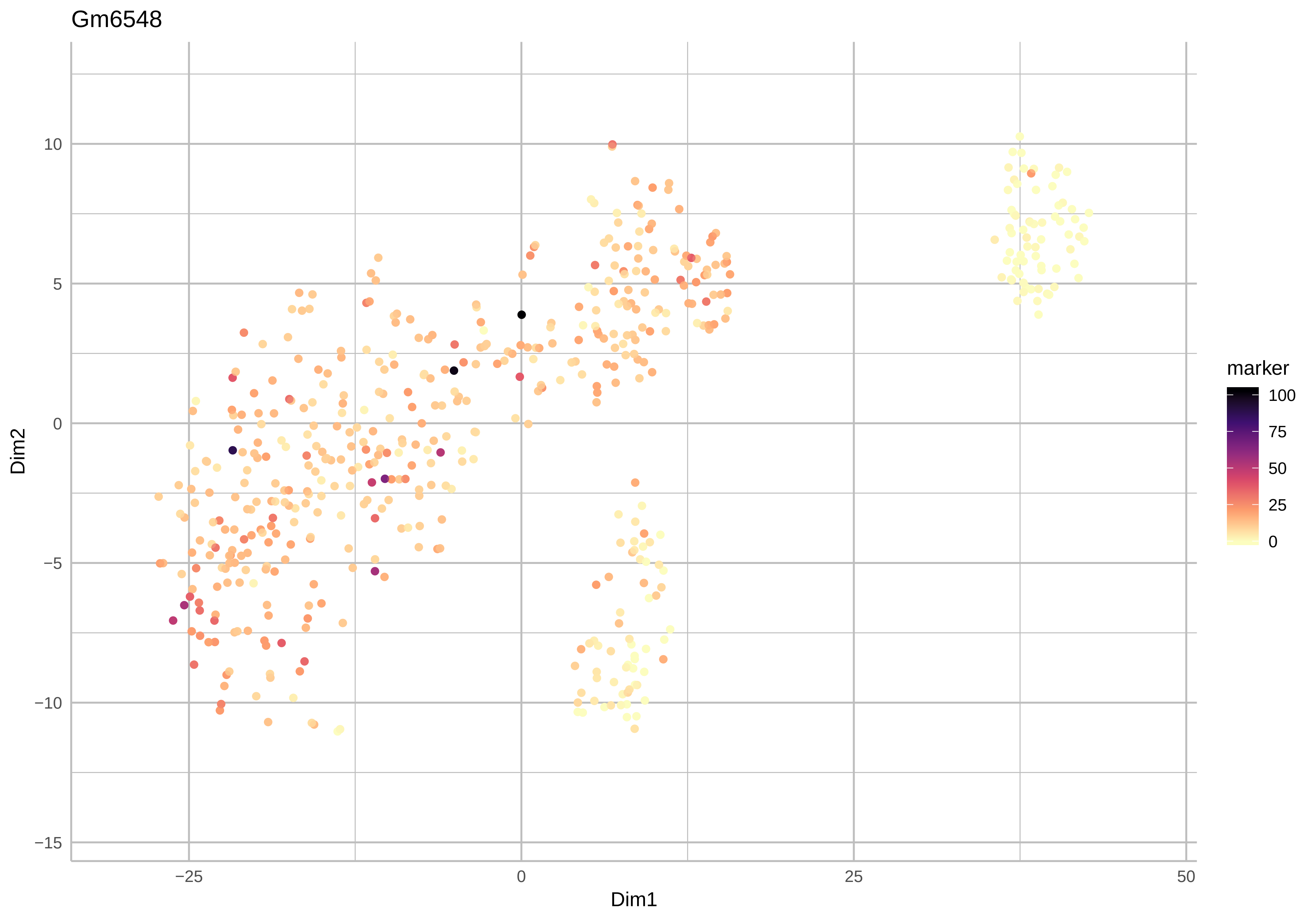

Gm6548
10
5
0
−5
−10
−15
−25
0
25
50
Dim1
Dim2
marker
100
75
50
25
0

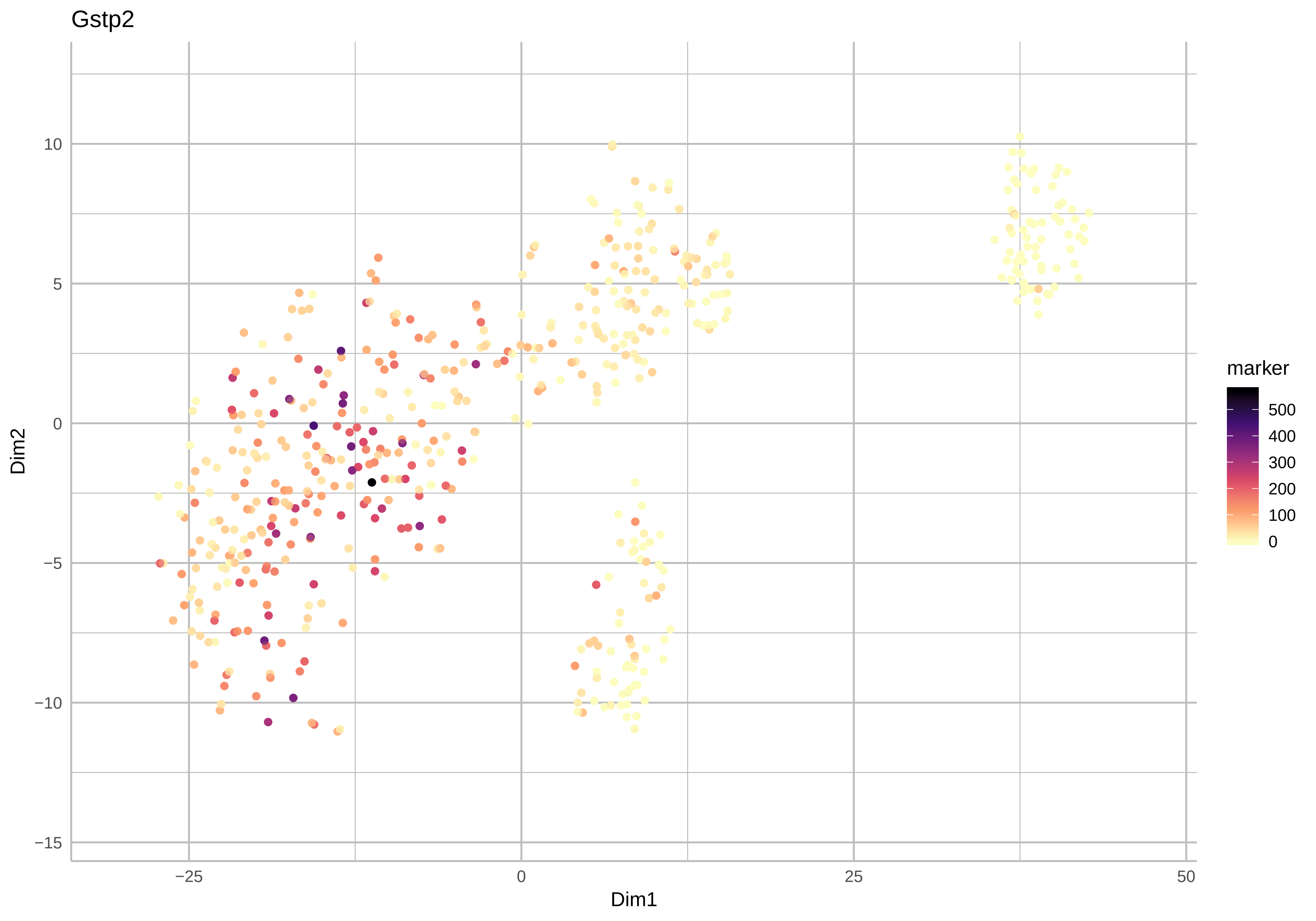

Gstp2
10
5
0
−5
−10
−15
Dim2
−25
0
25
50
Dim1
marker
500
400
300
200
100
0

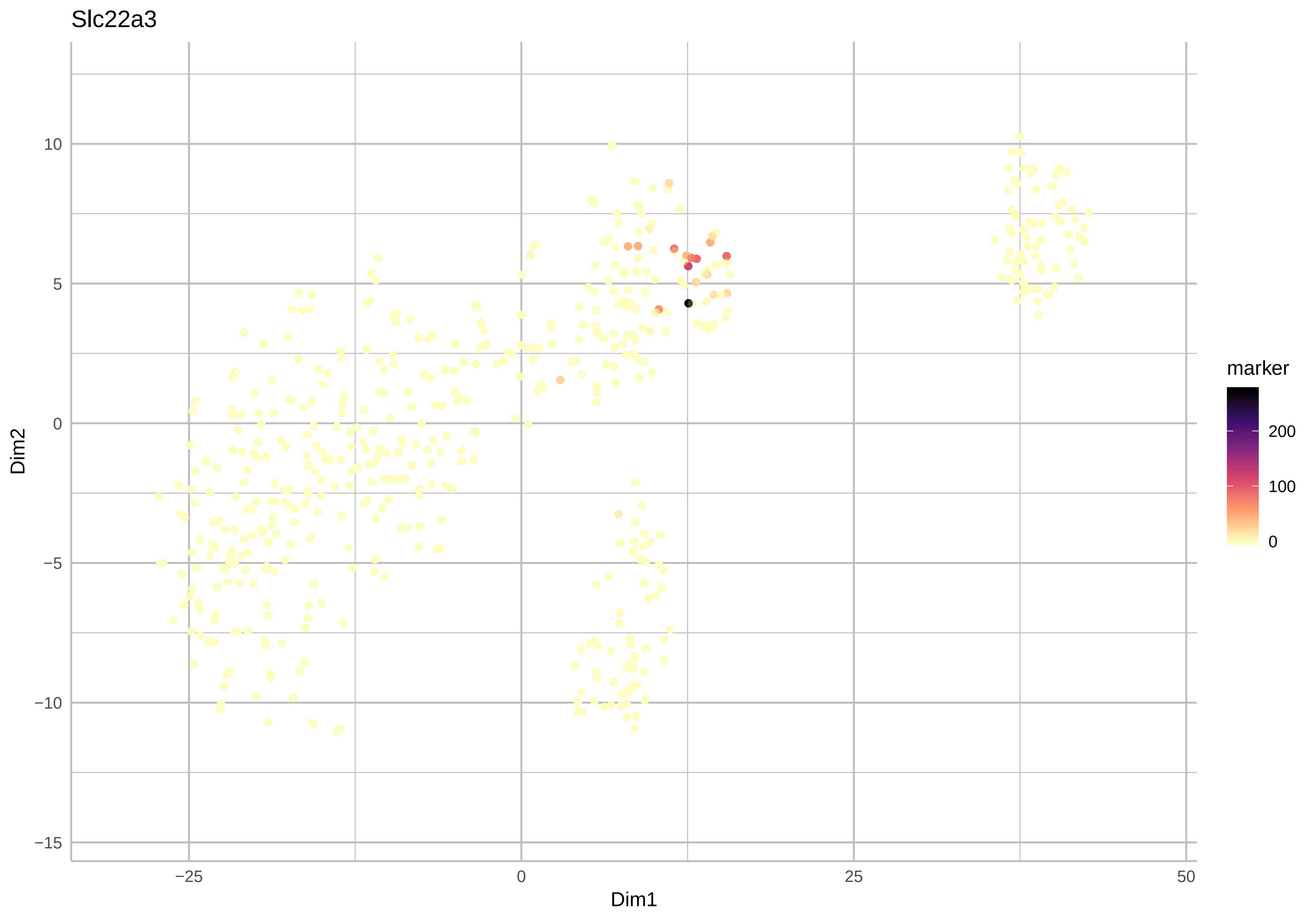

Slc22a3
Dim2
Dim1
10
5
0
−5
−10
−15
−25
0
25
50
marker
200
100
0

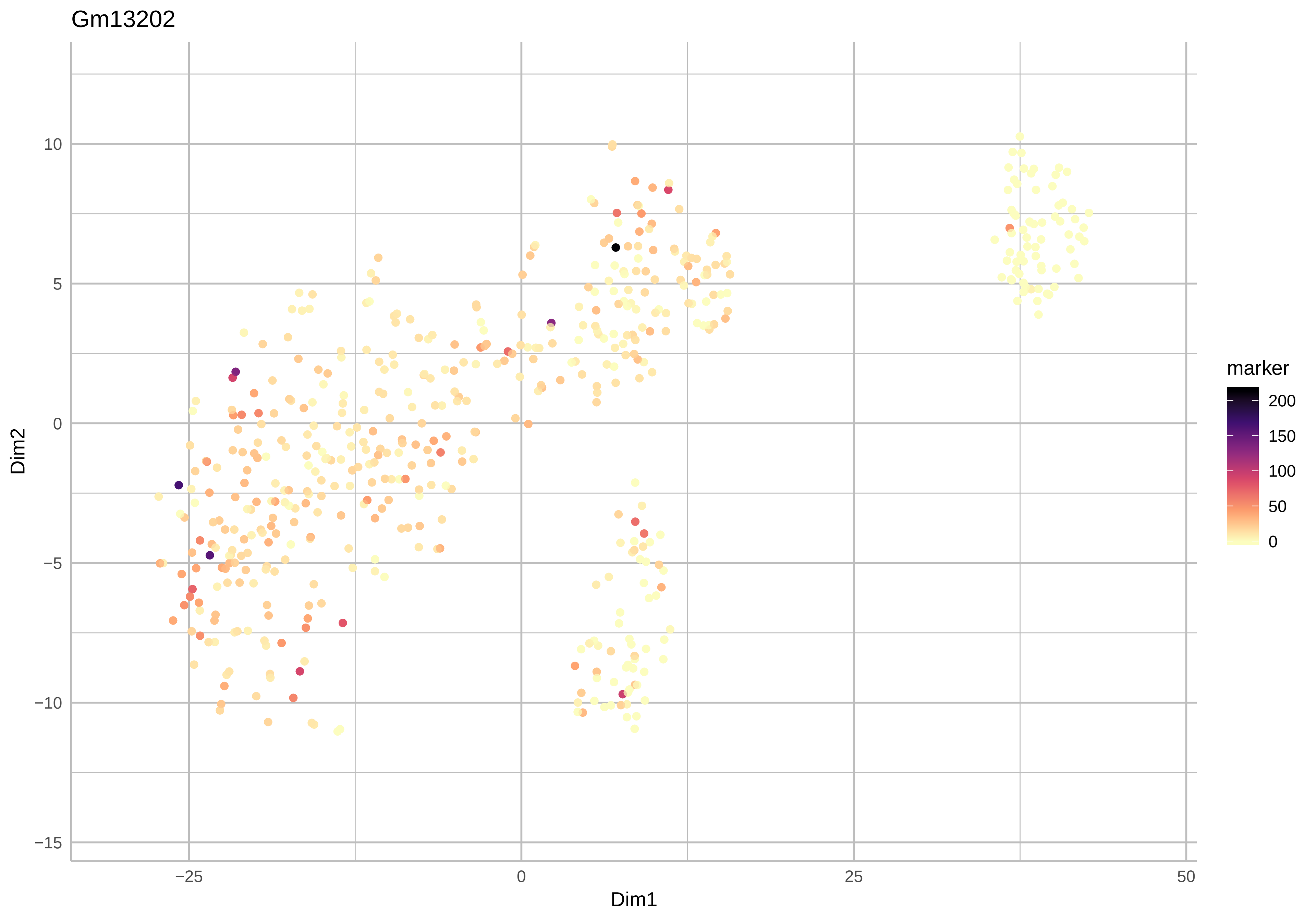
Gm13202
Dim2
Dim1
10
5
0
−5
−10
−15
−25
0
25
50
marker
200
150
100
50
0

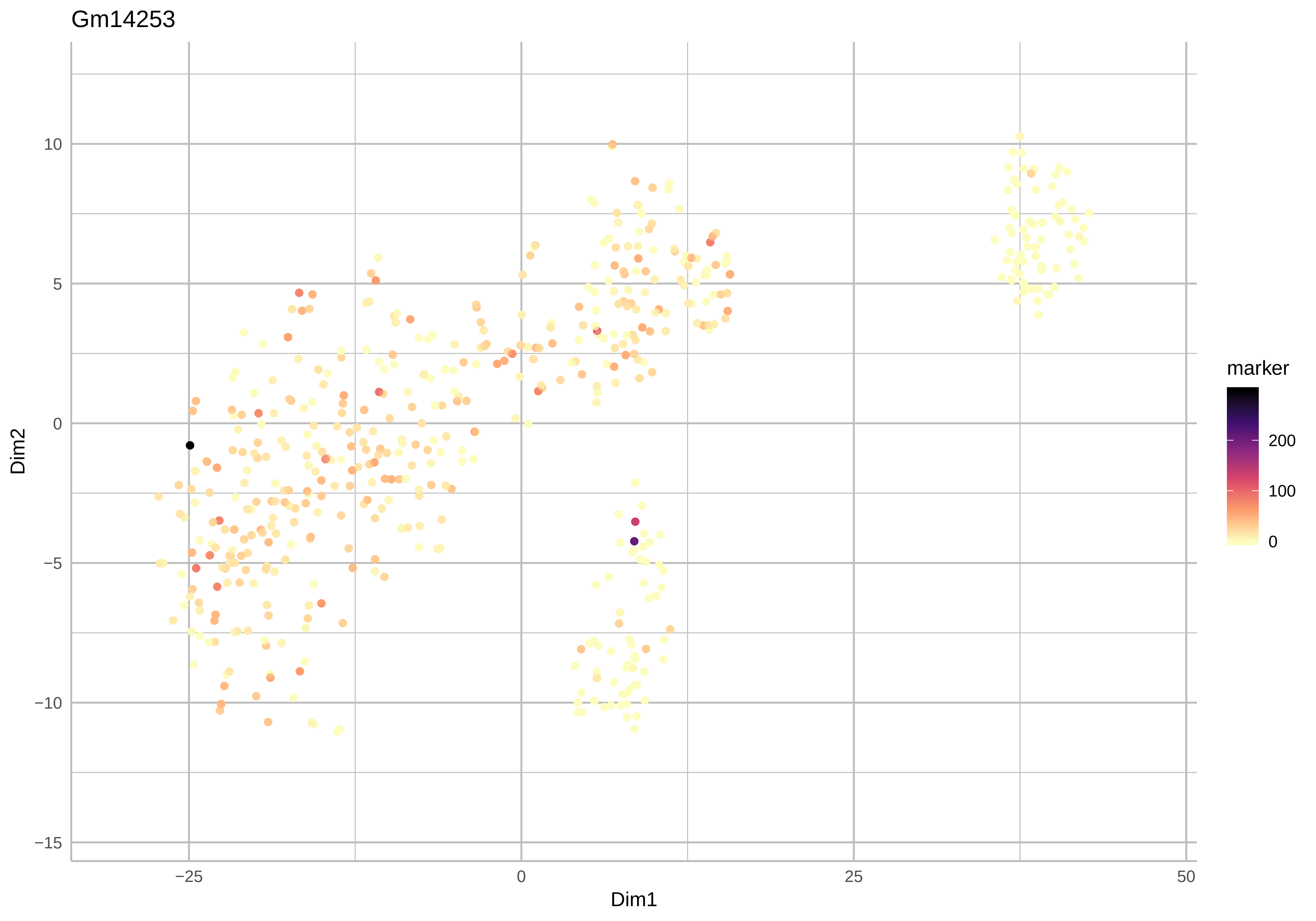

Gm14253
Dim2
Dim1
marker
200
100
0
10
5
0
−5
−10
−15
−25
0
25
50

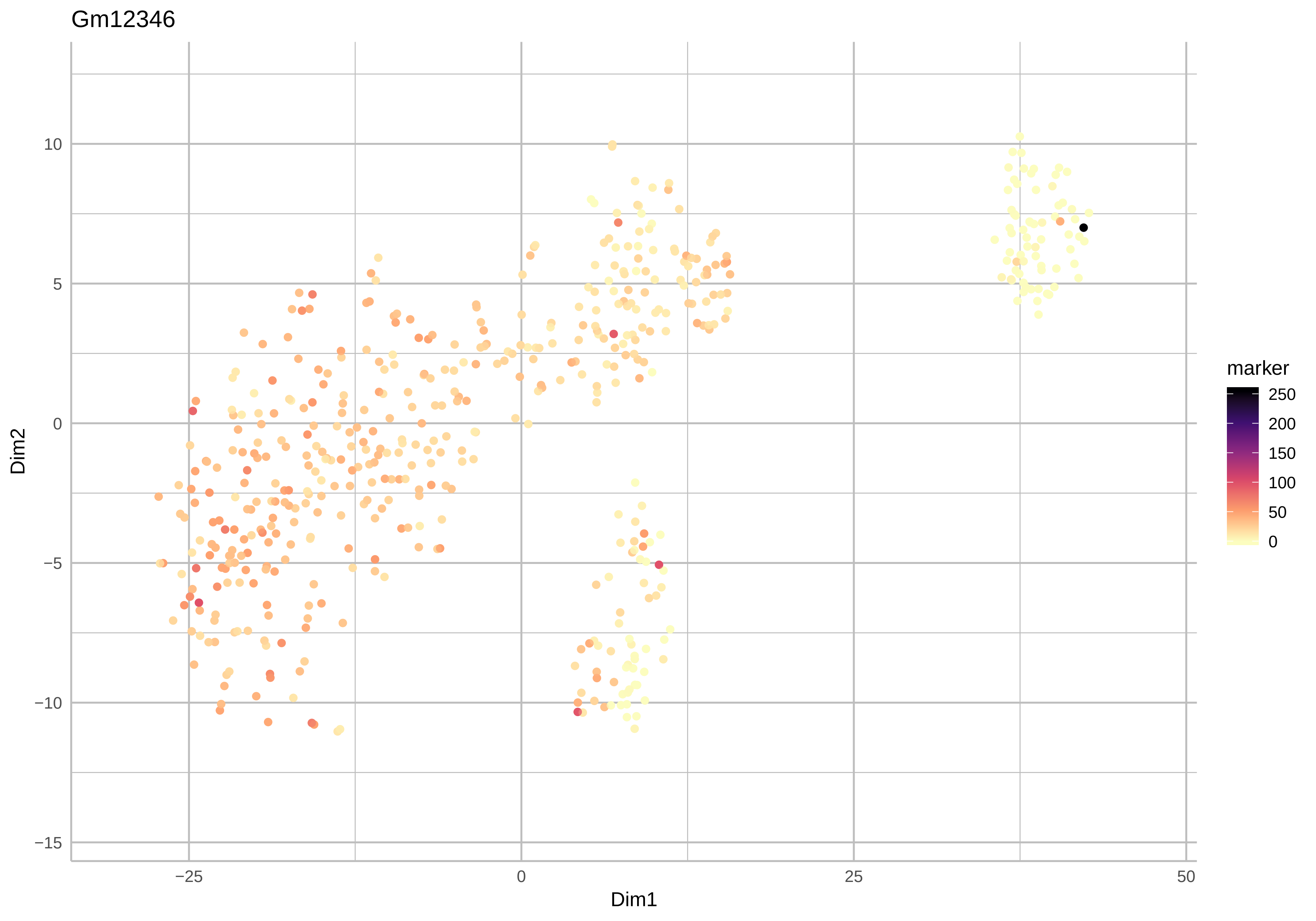

Gm12346
10
5
0
−5
−10
−15
Dim2
−25
0
25
50
Dim1
marker
250
200
150
100
50
0

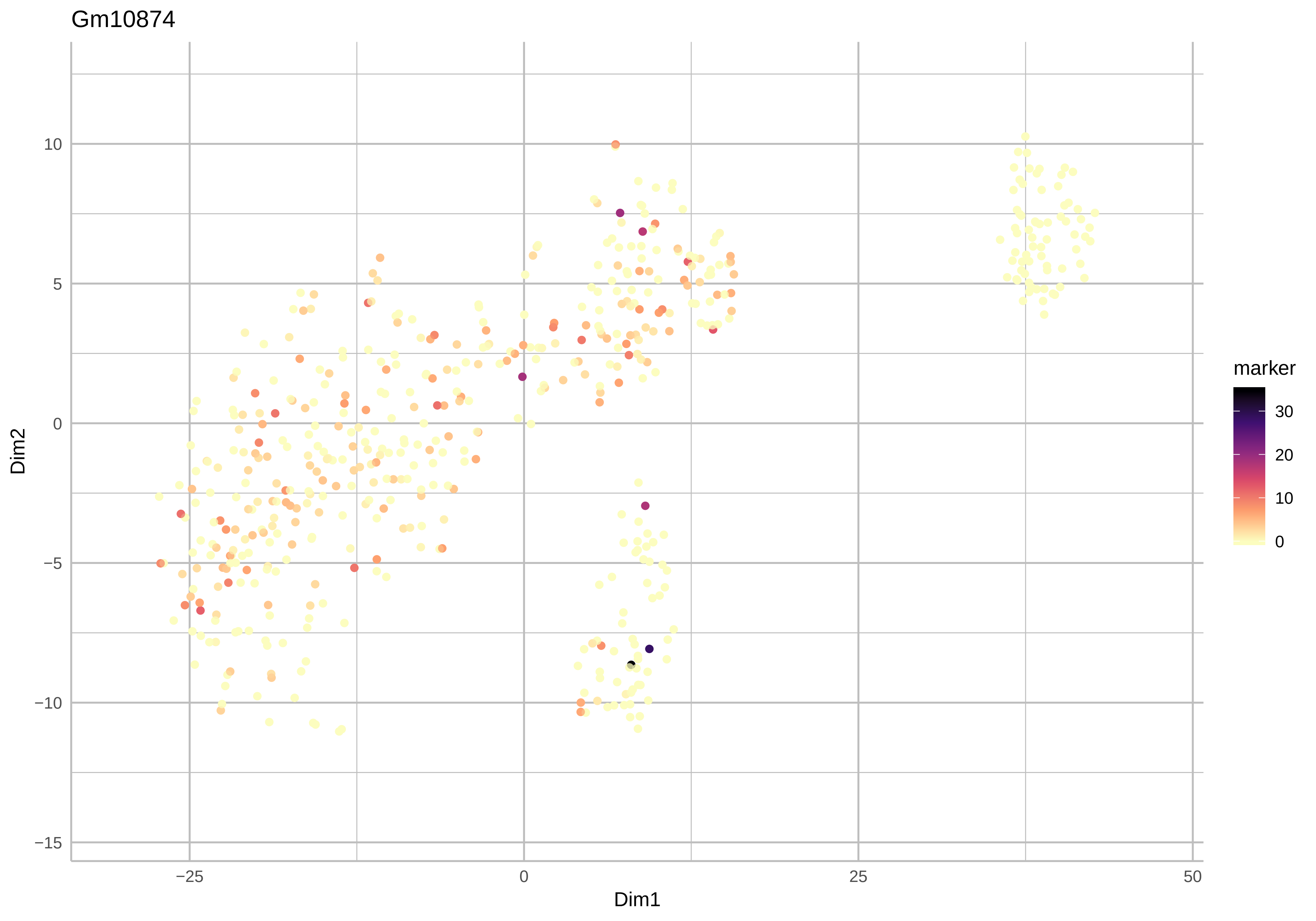

Gm10874
10
5
0
−5
−10
−15
Dim2
−25
0
25
50
Dim1
marker
30
20
10
0

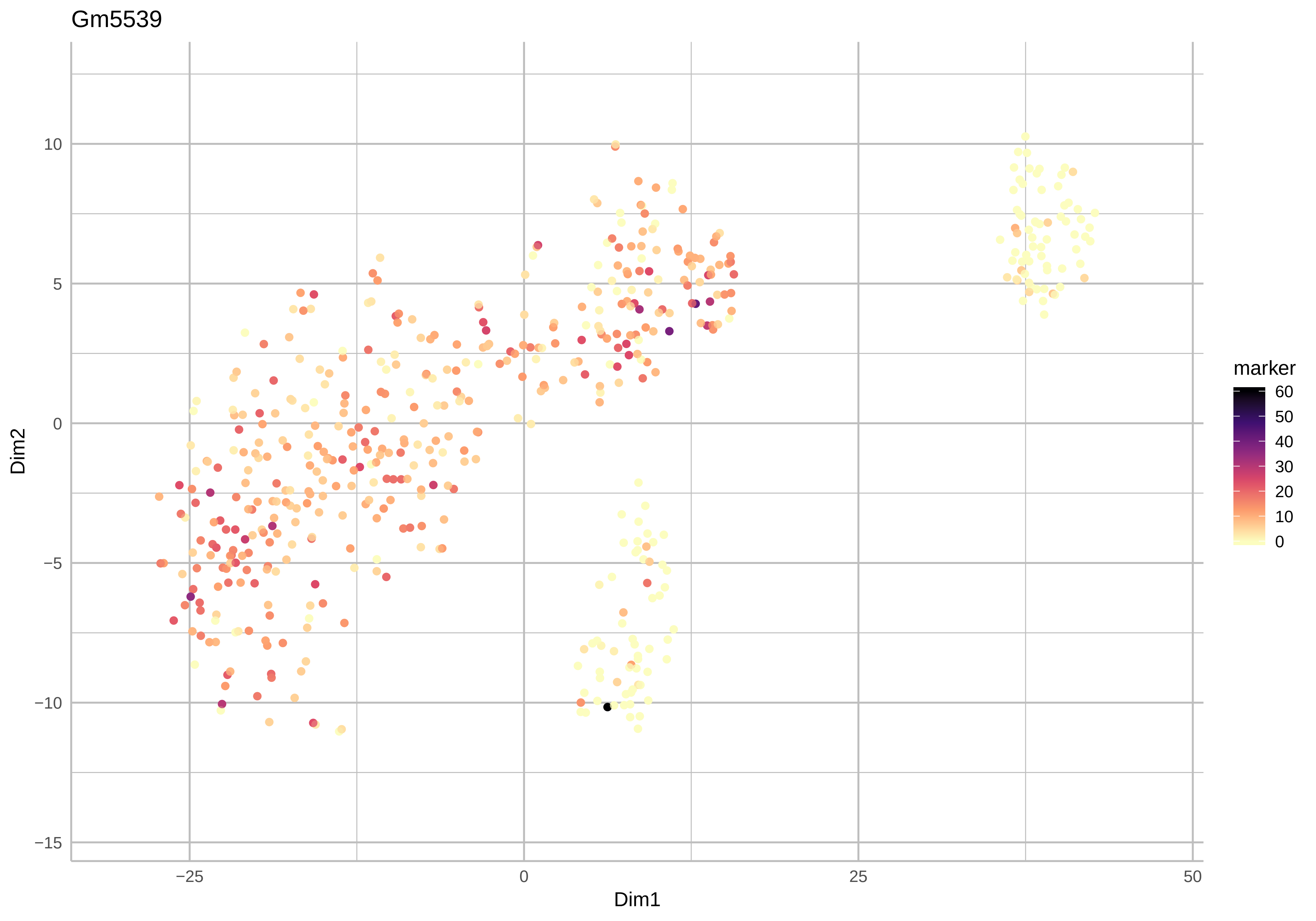

Gm5539
Dim2
Dim1
10
5
0
−5
−10
−15
−25
0
25
50
marker
60
50
40
30
20
10
0

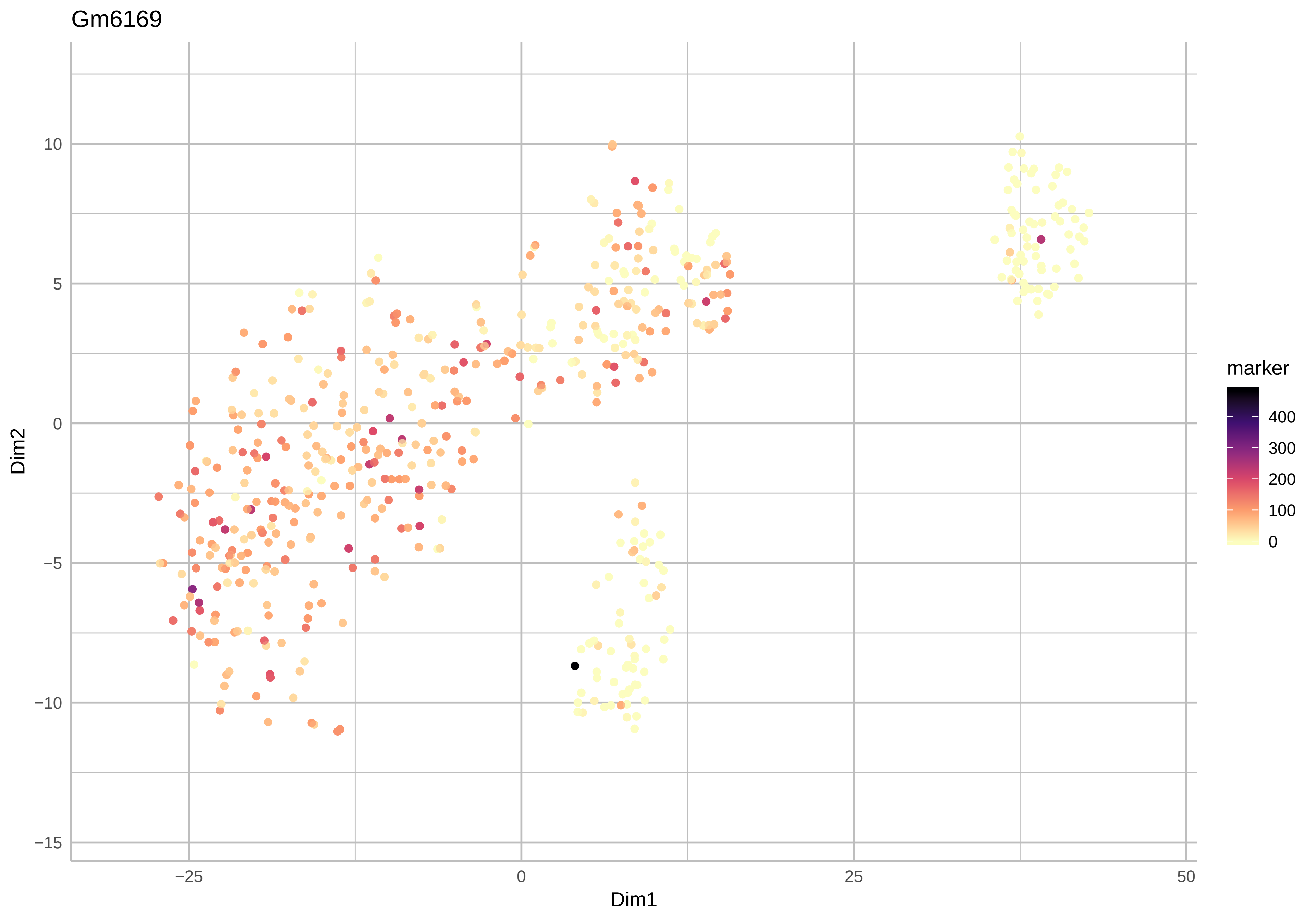

Gm6169
Dim2
Dim1
10
5
0
−5
−10
−15
−25
0
25
50
marker
400
300
200
100
0

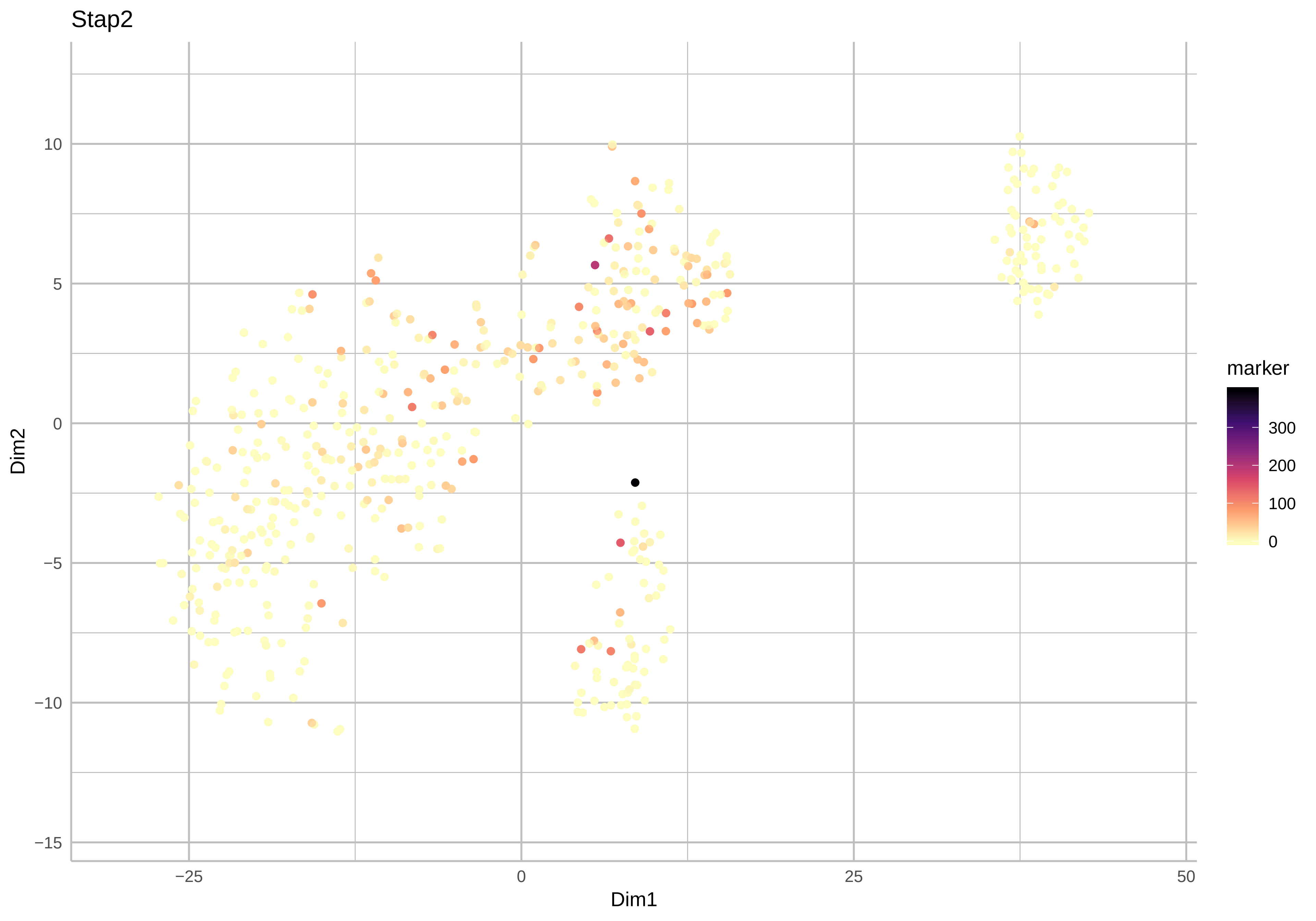

Stap2
Dim2
Dim1
10
5
0
−5
−10
−15
−25
0
25
50
marker
300
200
100
0

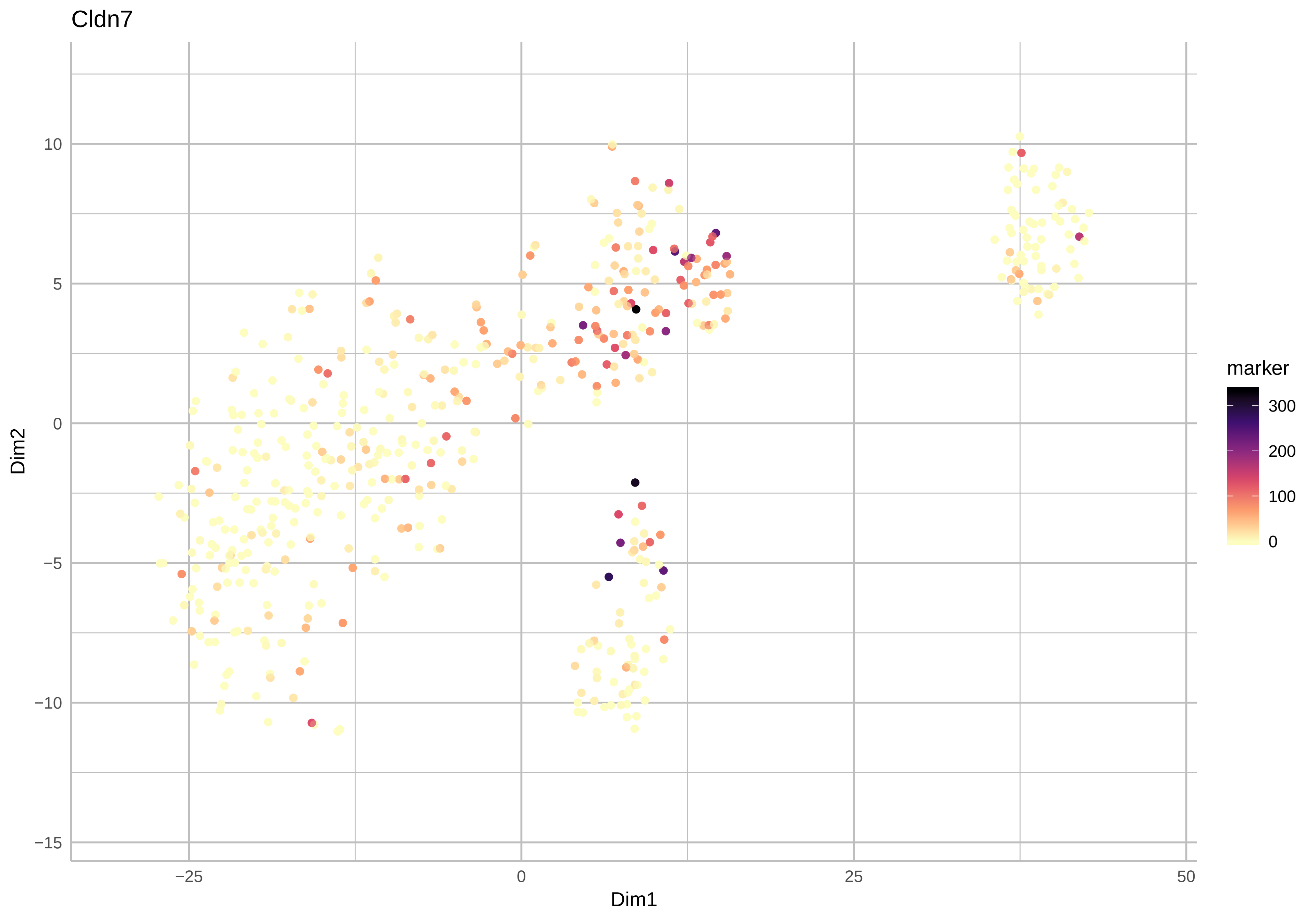
Cldn7
Dim2
Dim1
marker
300
200
100
0
10
5
0
−5
−10
−15
−25
0
25
50

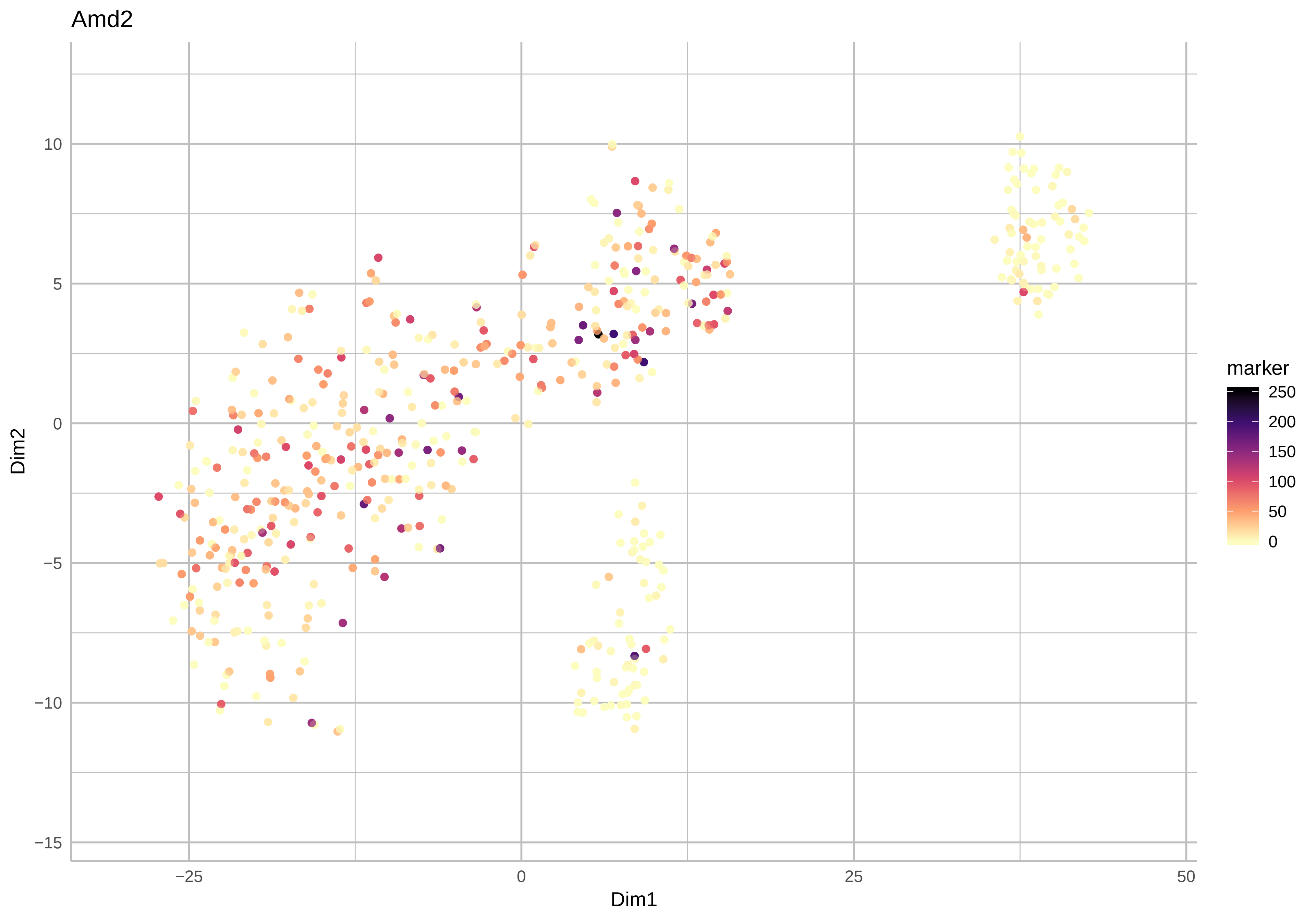

Amd2
Dim2
Dim1
10
5
0
−5
−10
−15
−25
0
25
50
marker
250
200
150
100
50
0

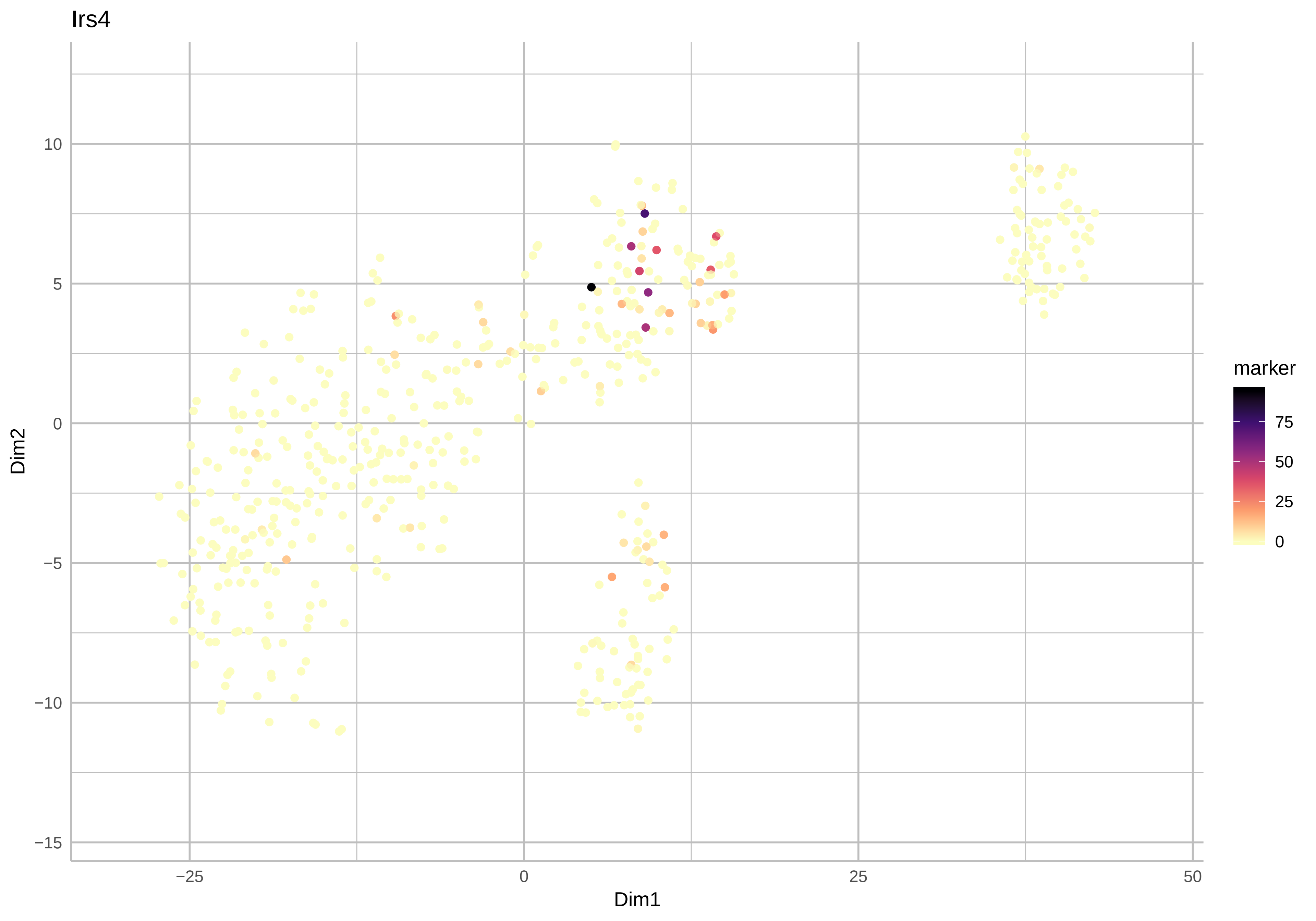
Irs4
Dim2
Dim1
10
5
0
−5
−10
−15
−25
0
25
50
marker
75
50
25
0

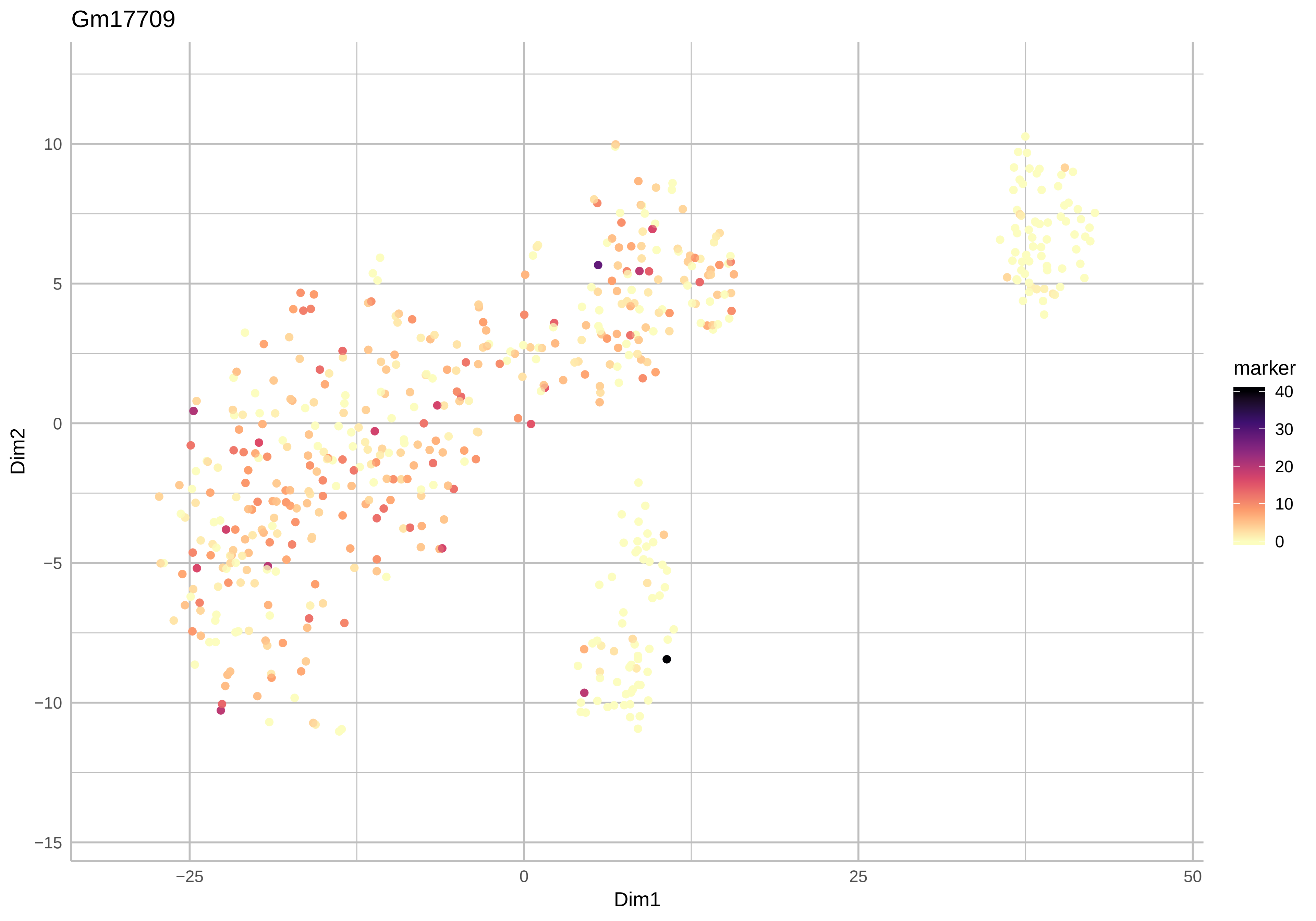
Gm17709
Dim2
10
5
0
−5
−10
−15
Dim1
−25
0
25
50
marker
40
30
20
10
0

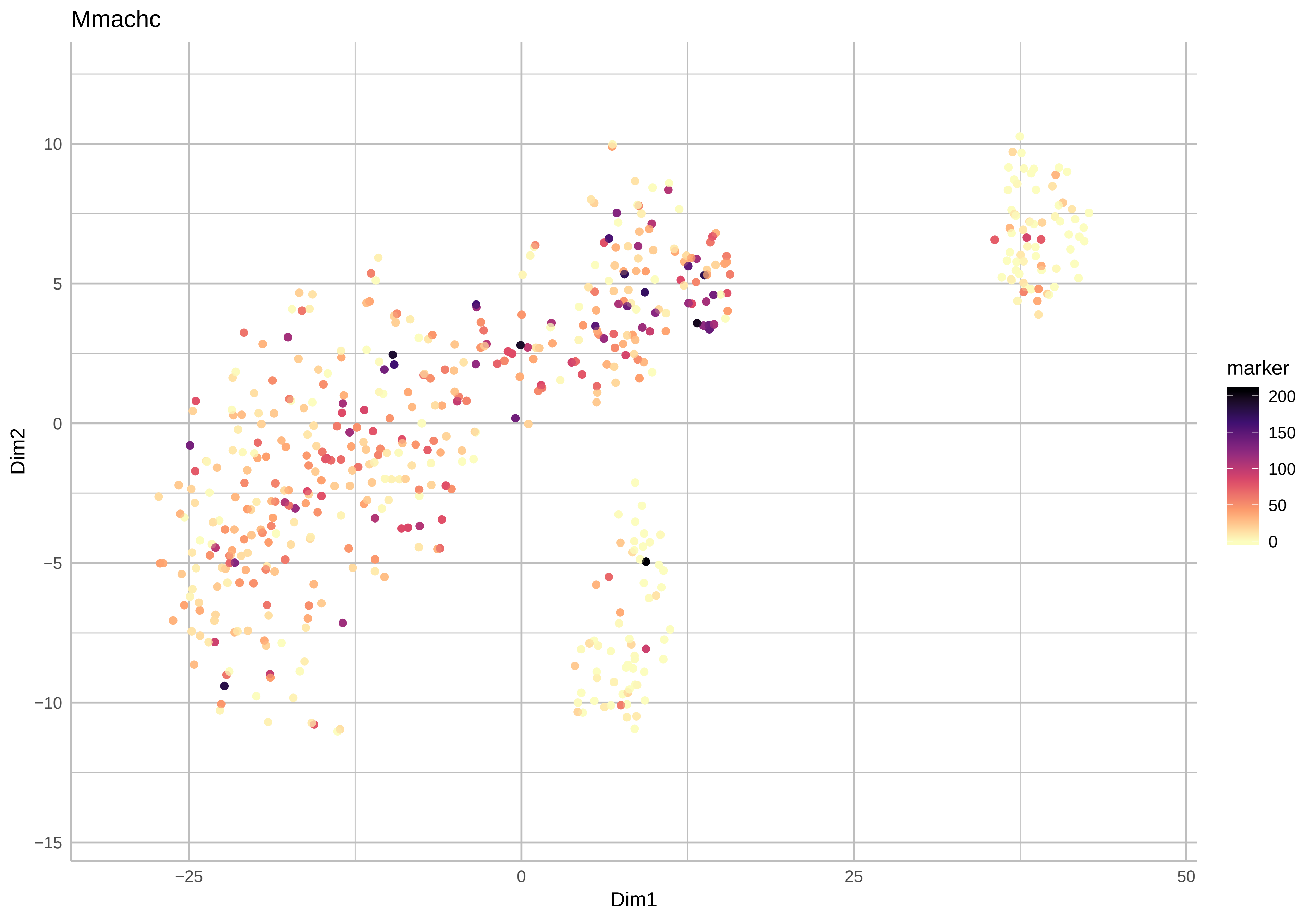

Mmachc
Dim2
Dim1
10
5
0
−5
−10
−15
−25
0
25
50
marker
200
150
100
50
0

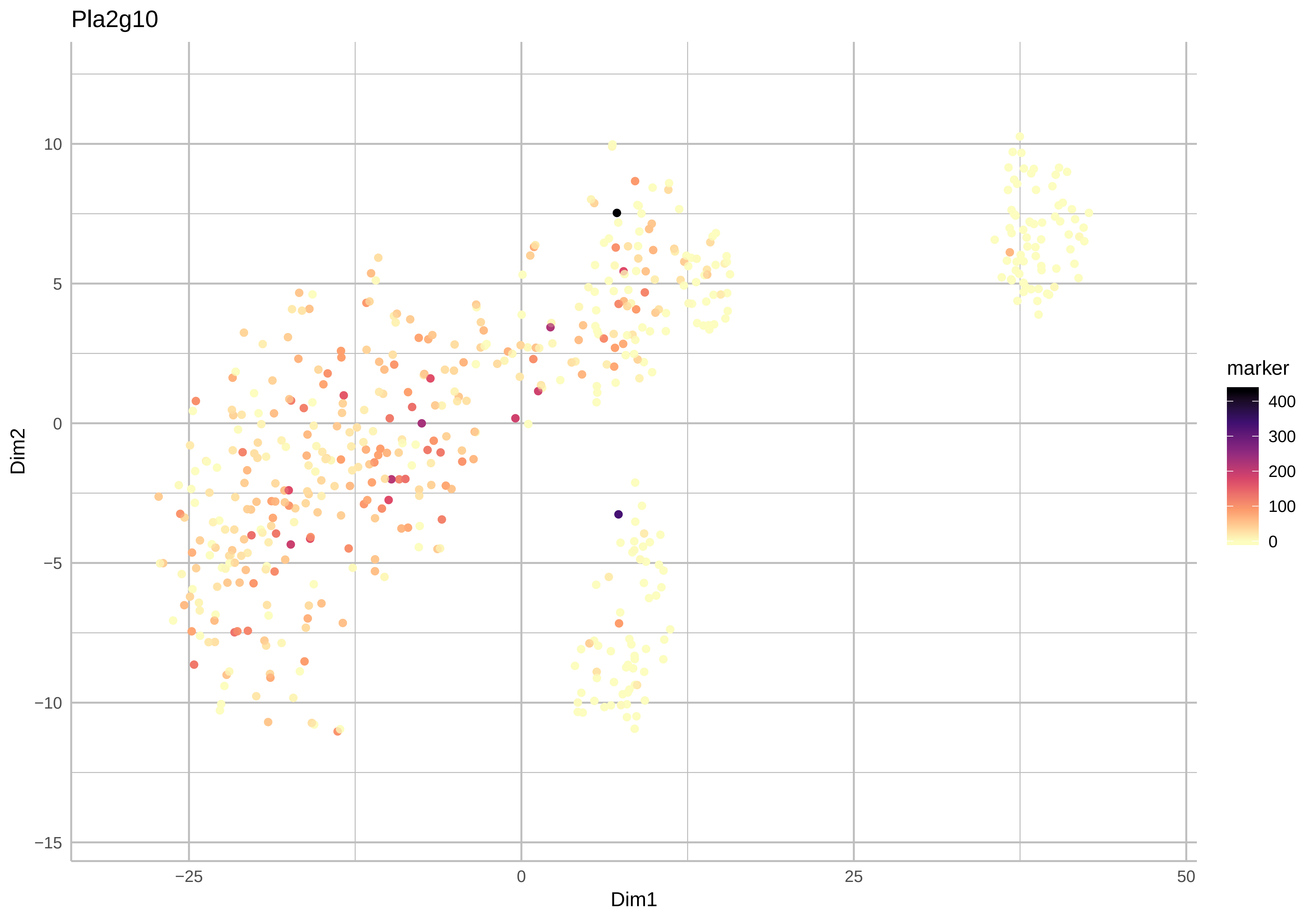

Pla2g10
Dim2
Dim1
marker
400
300
200
100
0
10
5
0
−5
−10
−15
−25
0
25
50

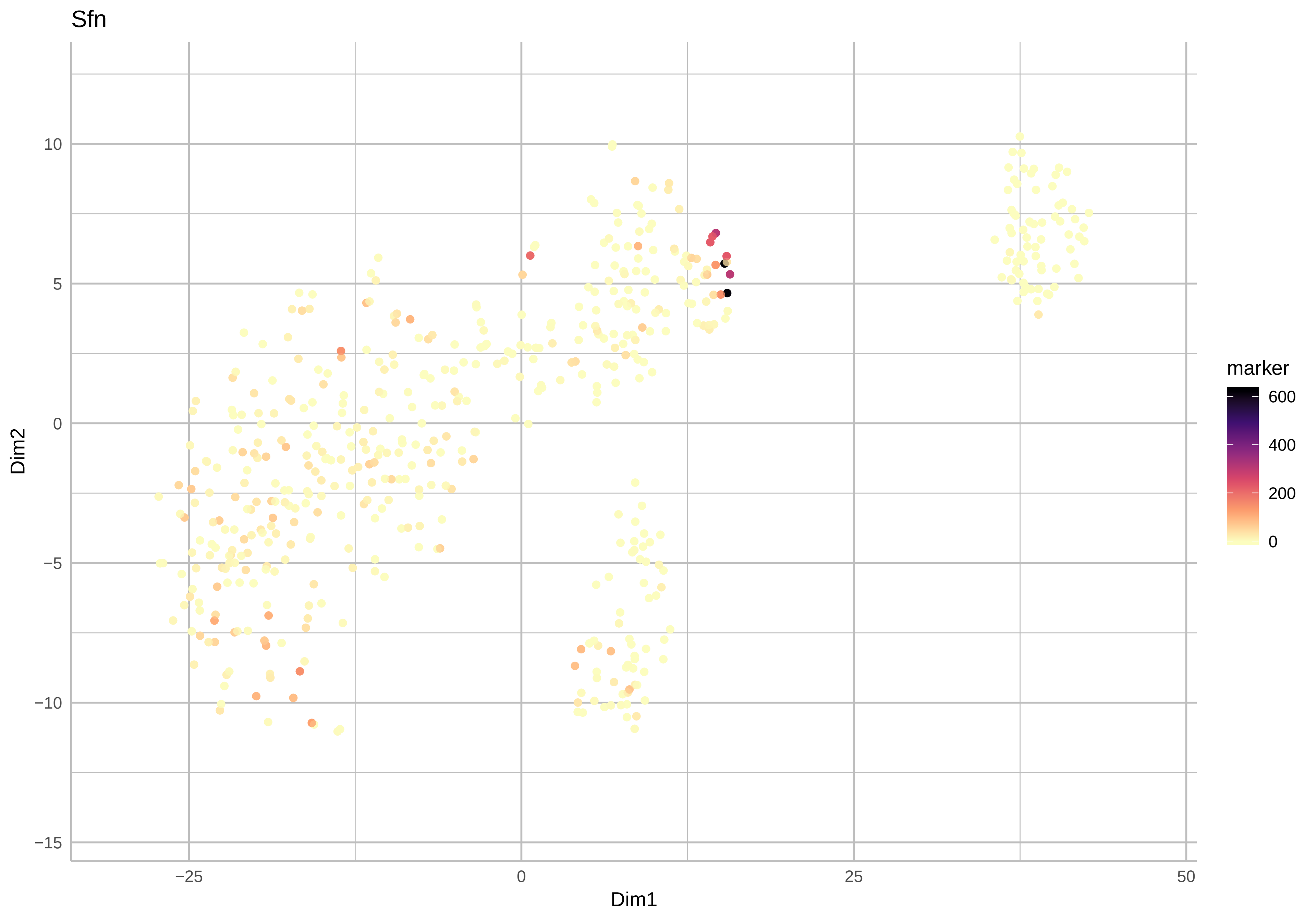

Sfn
10
5
0
−5
−10
−15
Dim2
−25
0
25
50
Dim1
marker
600
400
200
0

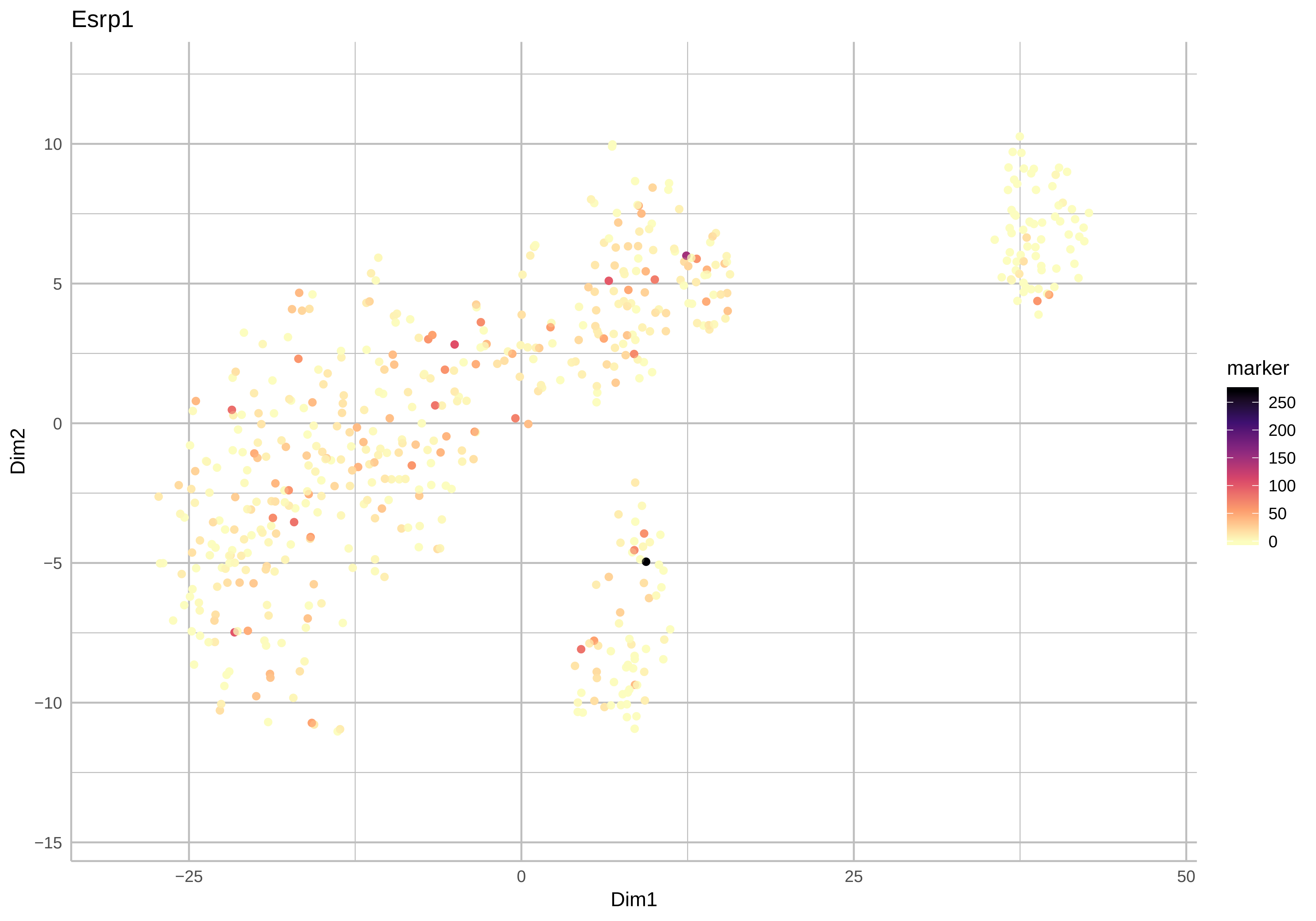

Esrp1
Dim2
Dim1
10
5
0
−5
−10
−15
−25
0
25
50
marker
250
200
150
100
50
0

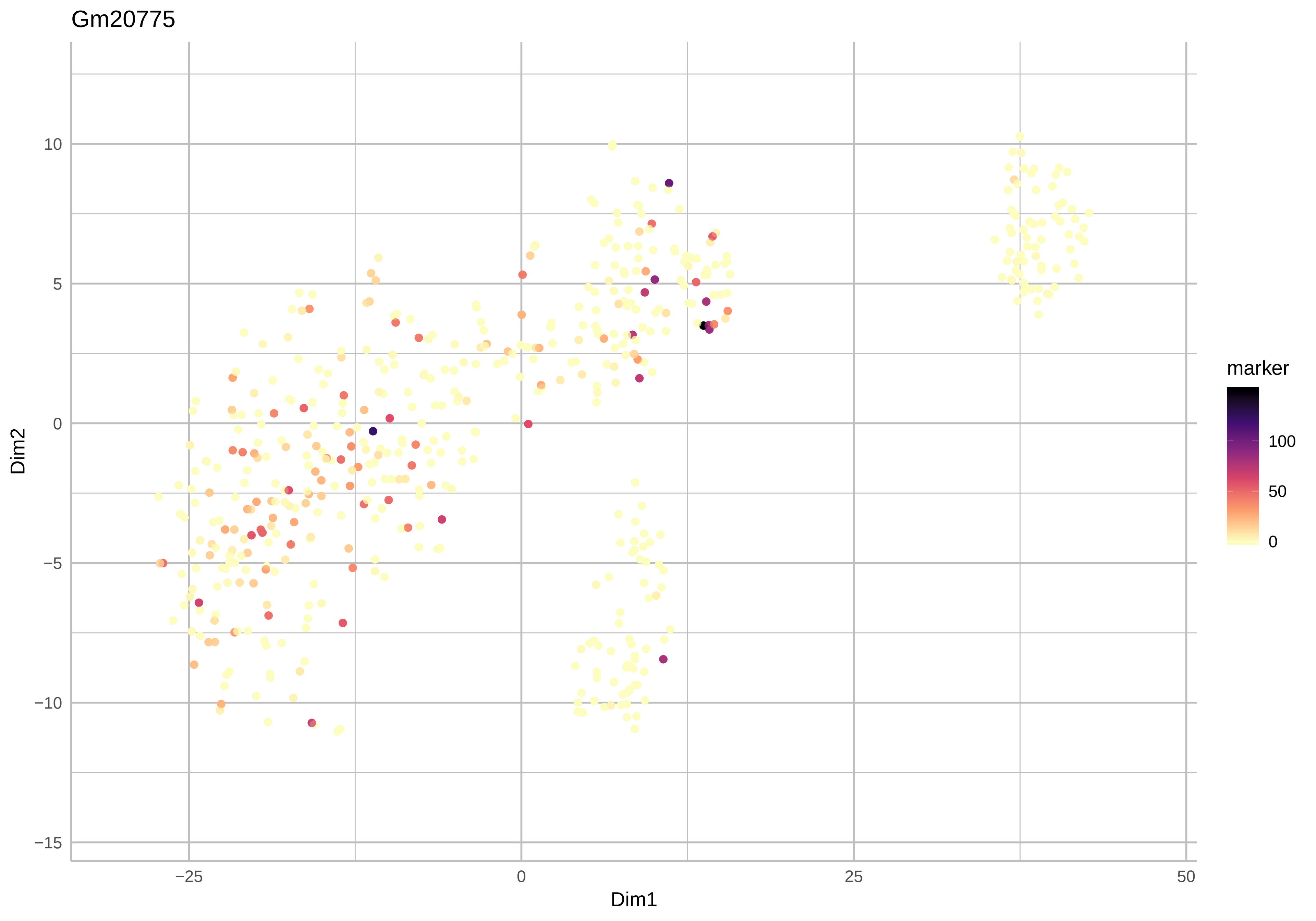

Gm20775
10
5
0
−5
−10
−15
Dim2
−25
0
25
50
Dim1
marker
100
50
0

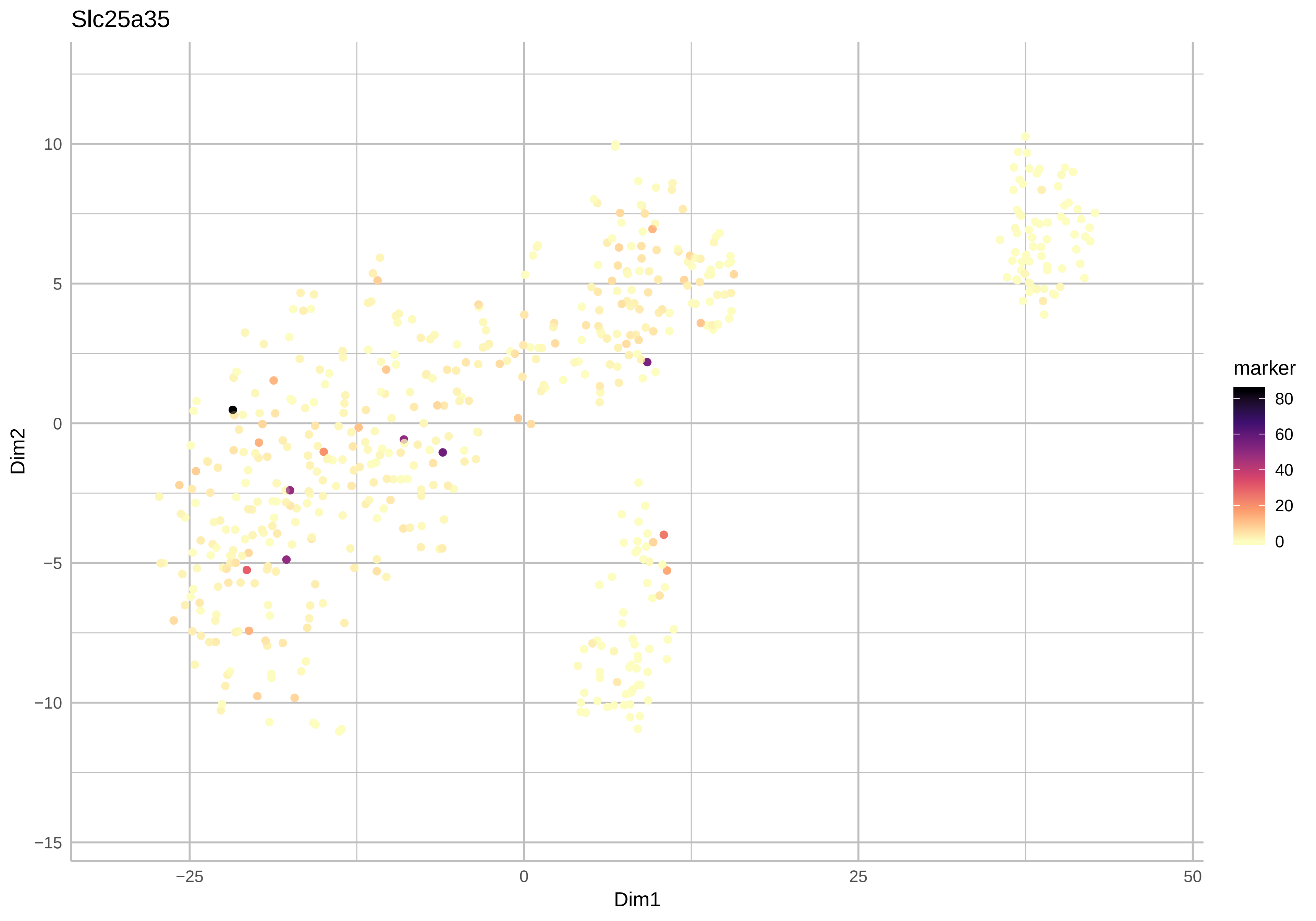
Slc25a35
Dim2
Dim1
marker
80
60
40
20
0
10
5
0
−5
−10
−15
−25
0
25
50

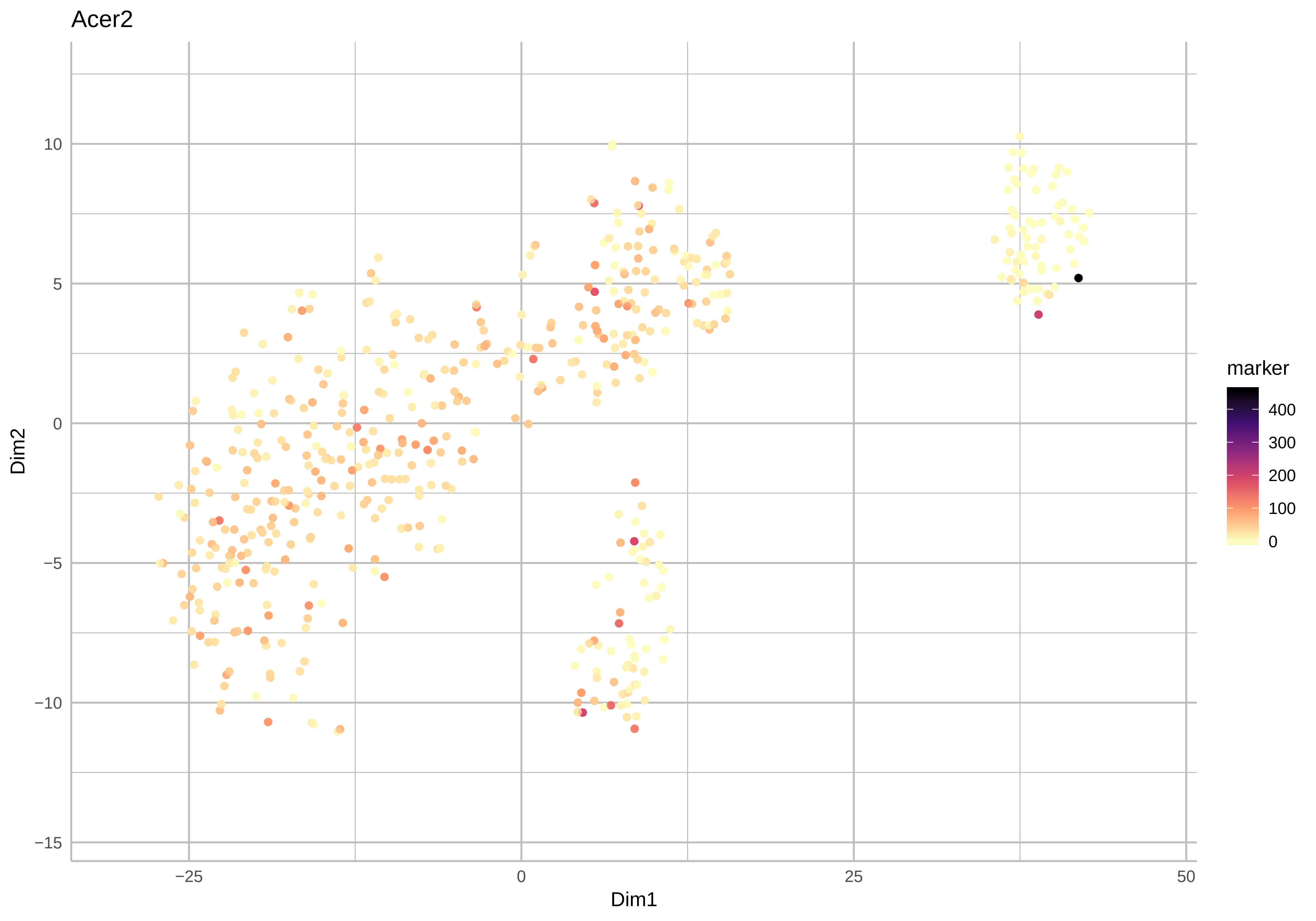
Acer2
10
5
0
−5
−10
−15
Dim2
−25
0
25
50
Dim1
marker
400
300
200
100
0

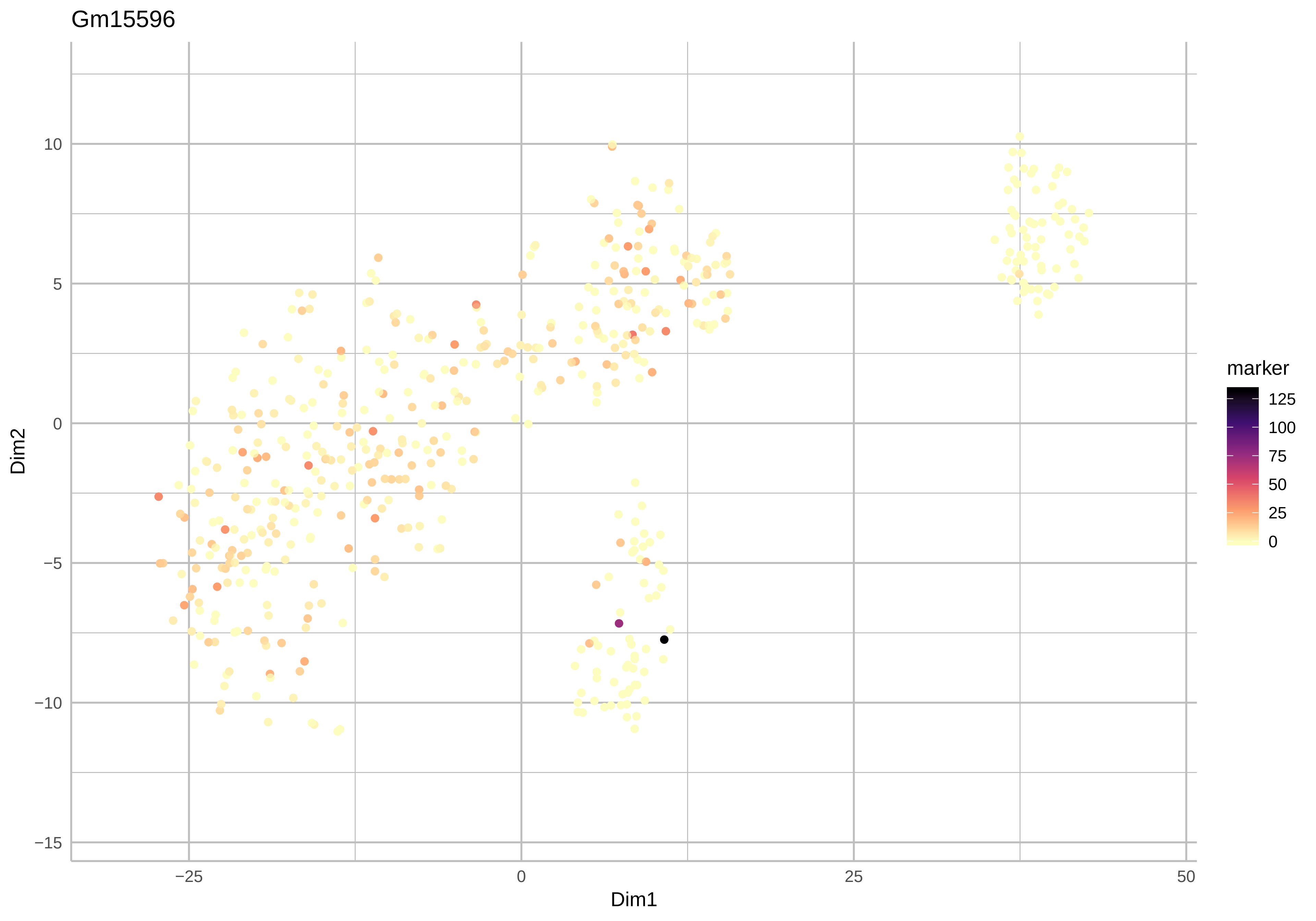

Gm15596
Dim2
Dim1
10
5
0
−5
−10
−15
−25
0
25
50
marker
125
100
75
50
25
0

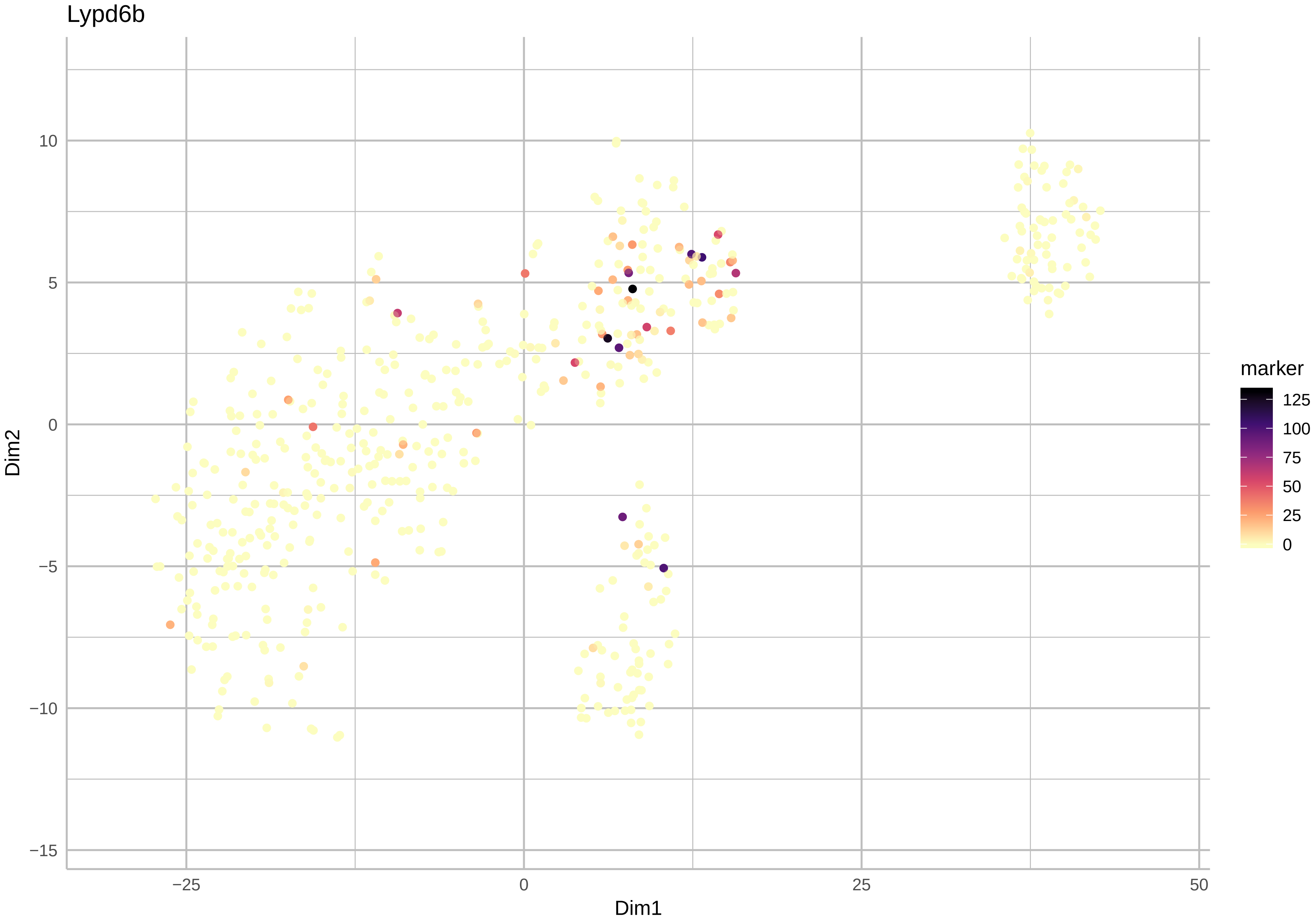

Lypd6b
10
5
0
−5
−10
−15
Dim2
−25
0
25
50
Dim1
marker
125
100
75
50
25
0

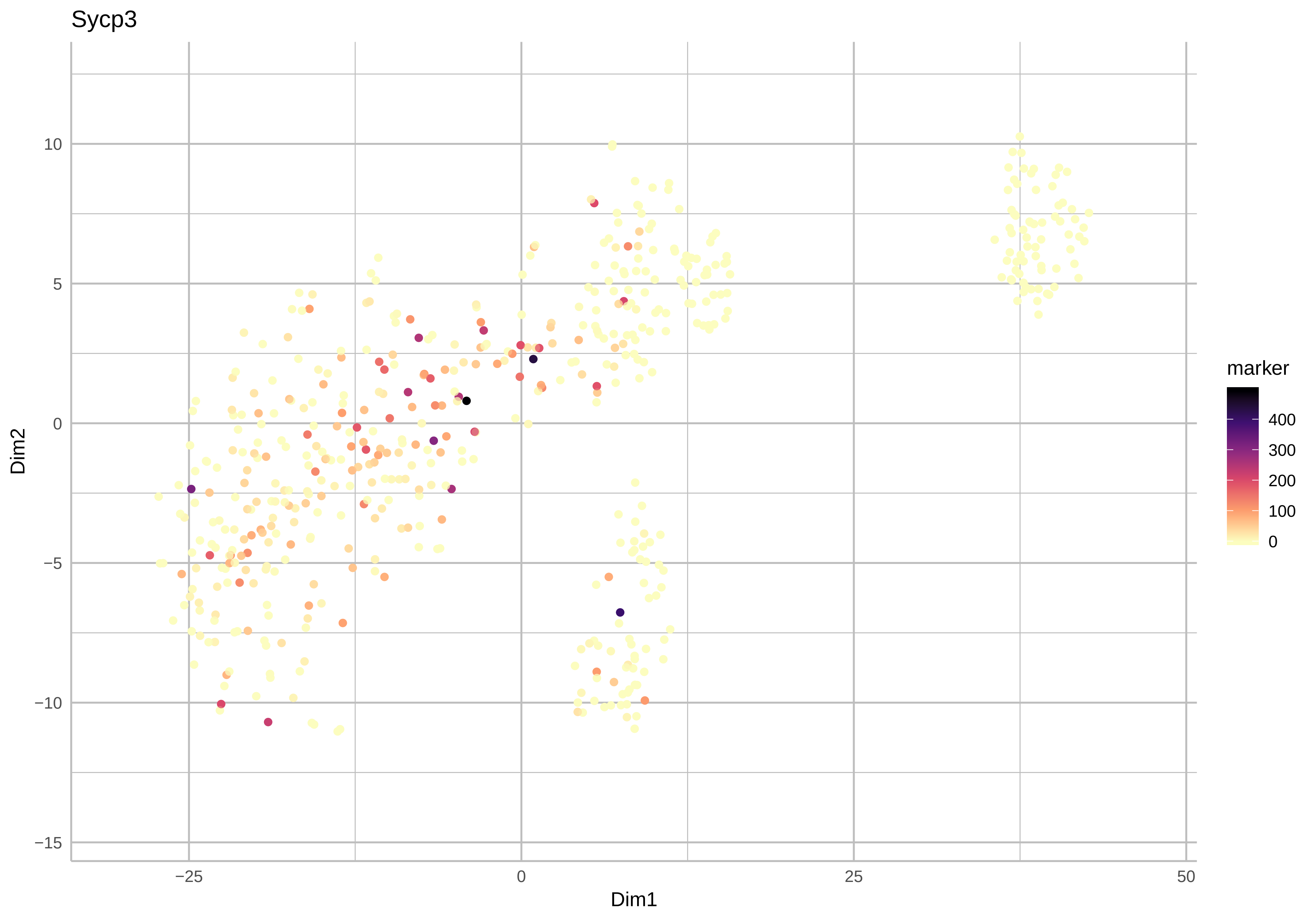
Sycp3
10
5
0
−5
−10
−15
Dim2
−25
0
25
50
Dim1
marker
400
300
200
100
0

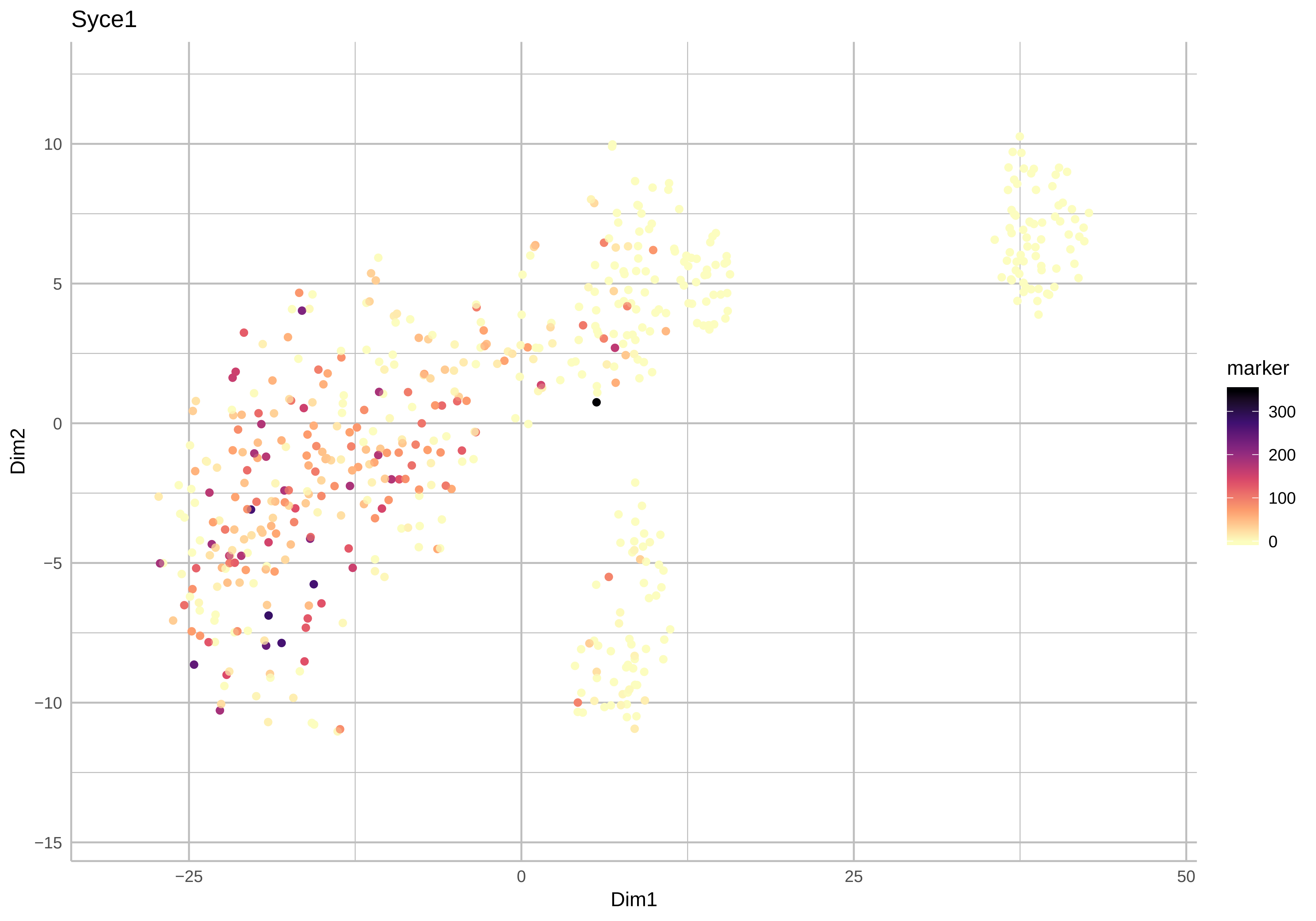

Syce1
10
5
0
−5
−10
−15
Dim2
−25
0
25
50
Dim1
marker
300
200
100
0

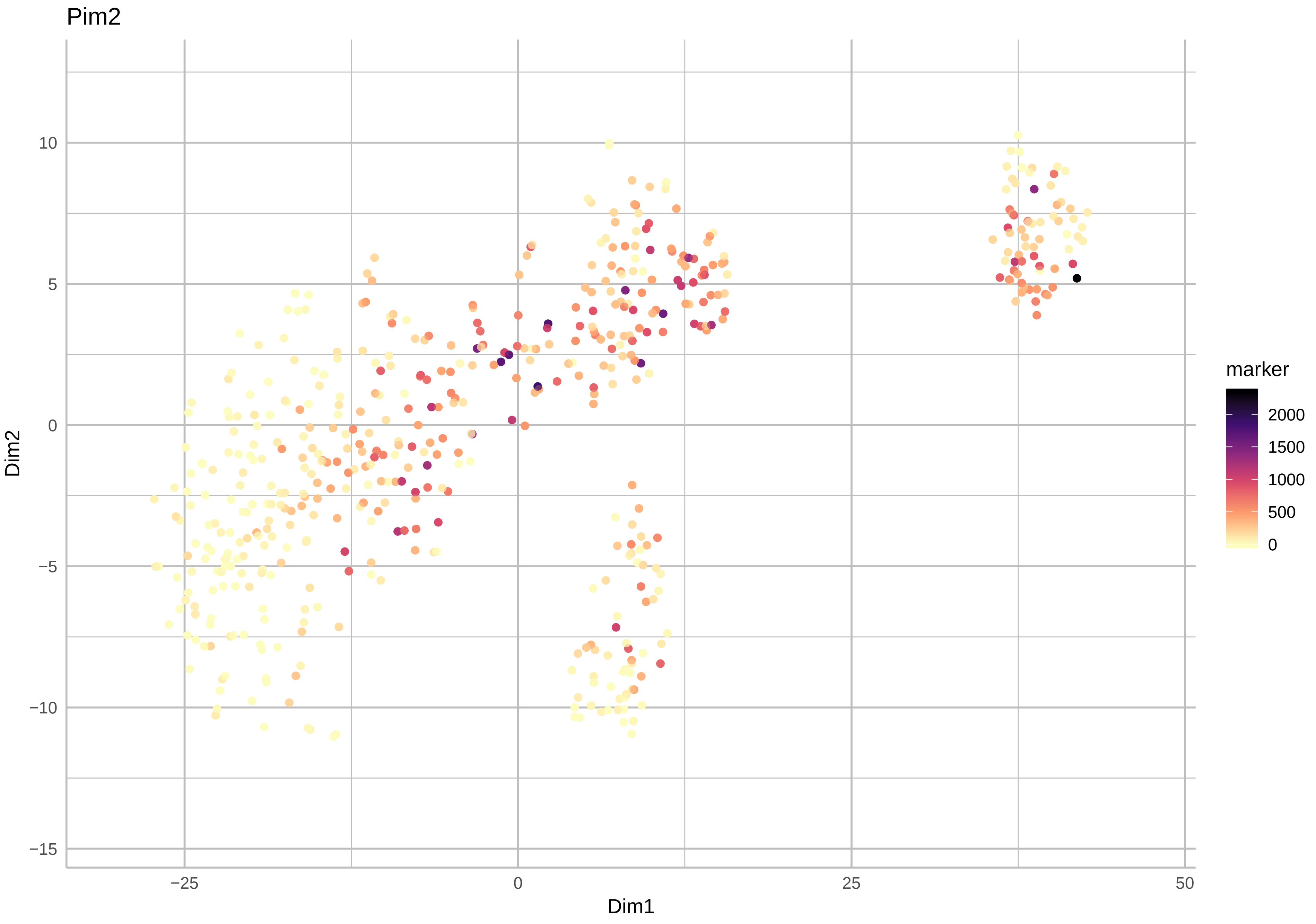
Pim2
Dim2
Dim1
marker
2000
1500
1000
500
0
10
5
0
−5
−10
−15
−25
0
25
50

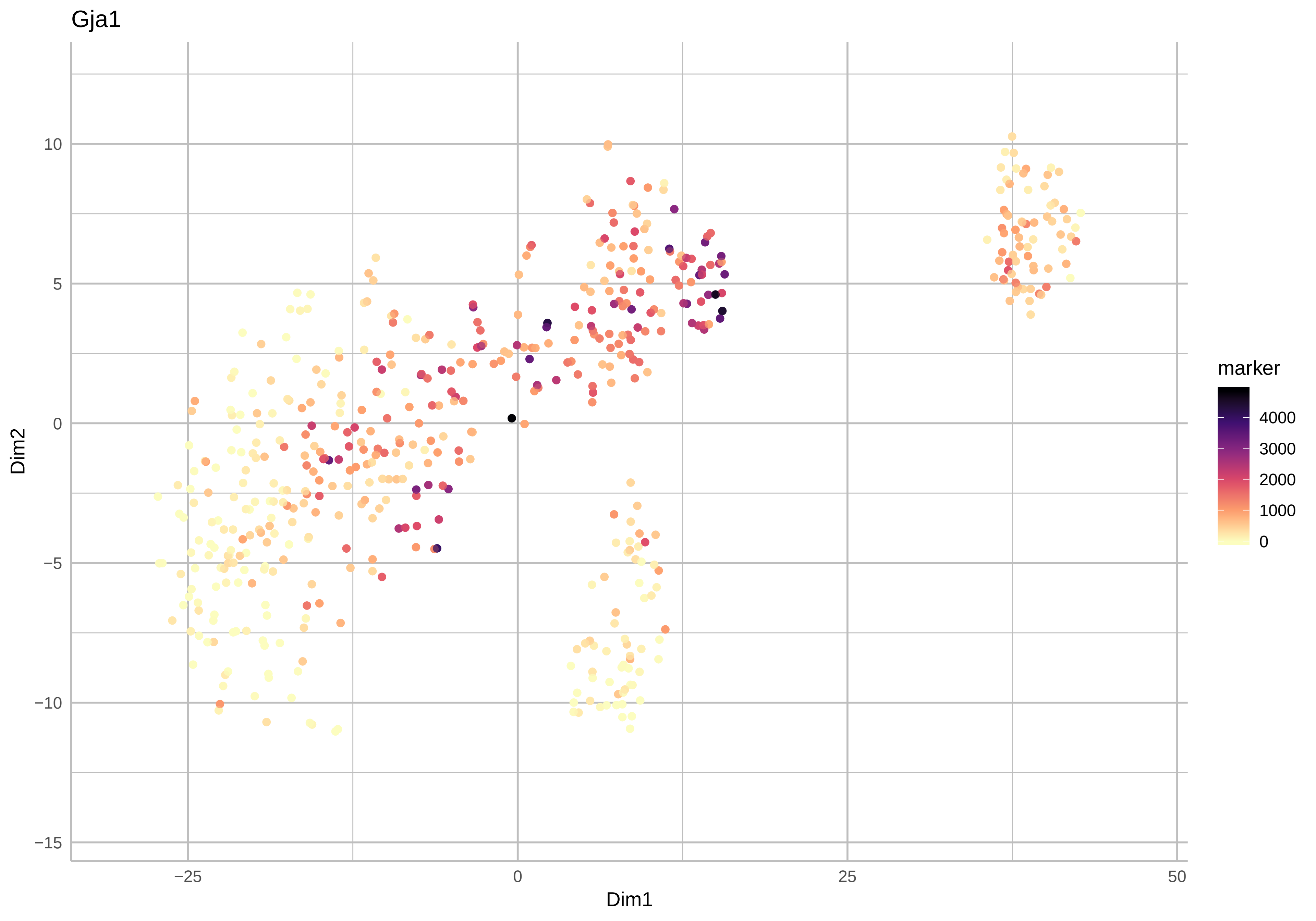

Gja1
Dim2
Dim1
10
5
0
−5
−10
−15
−25
0
25
50
marker
4000
3000
2000
1000
0

Car4

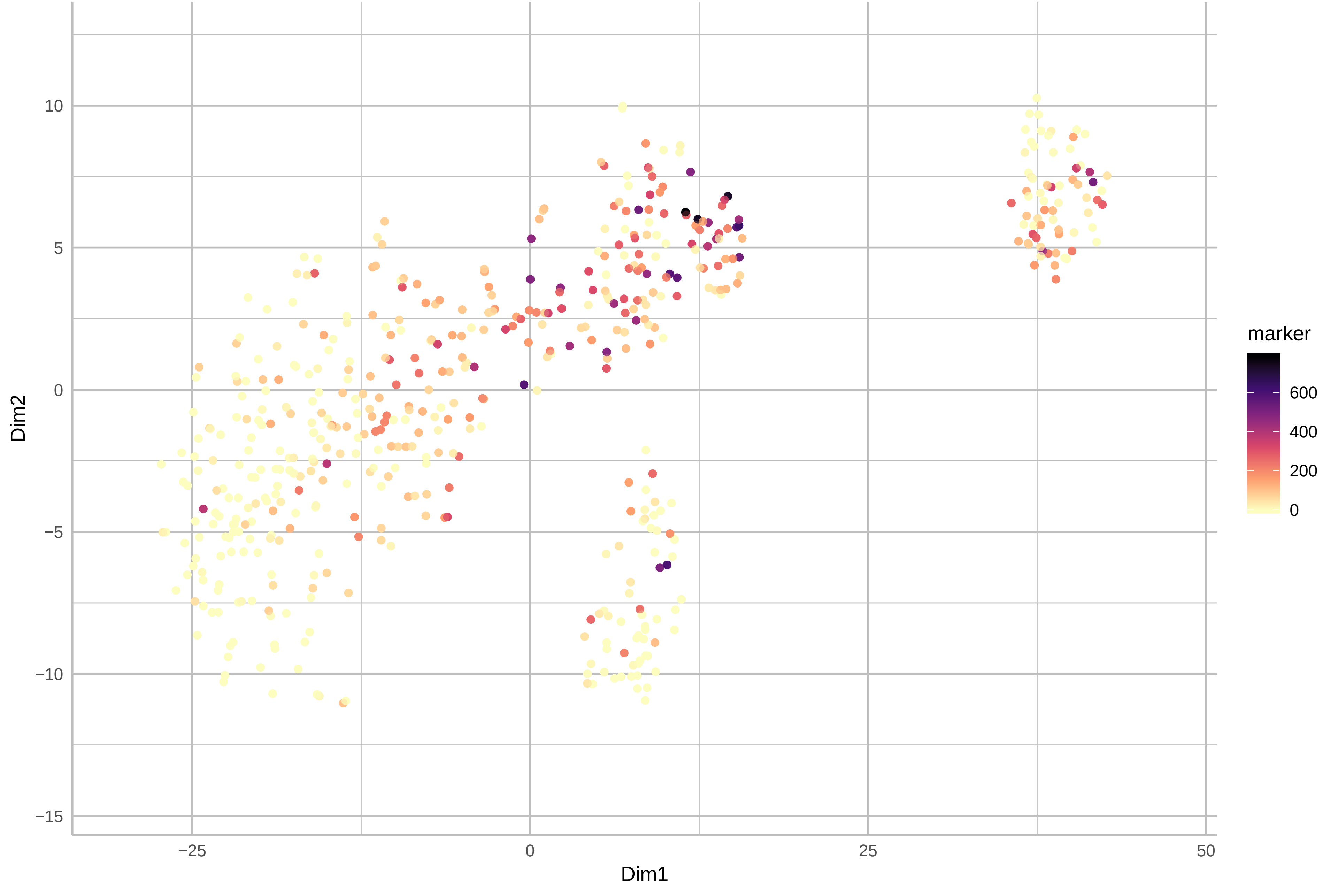

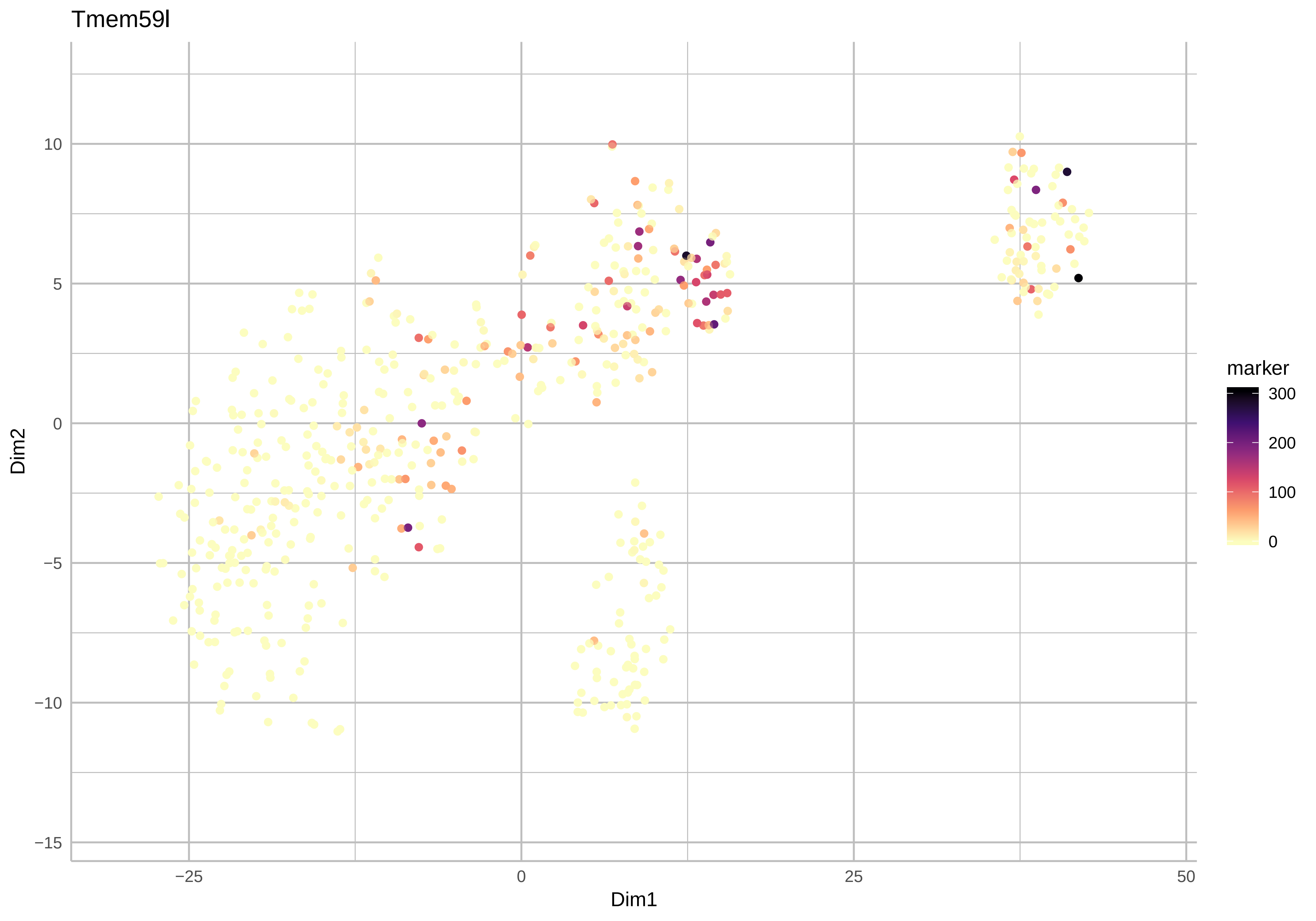

Tmem59l
10
5
0
−5
−10
−15
Dim2
−25
0
25
50
Dim1
marker
300
200
100
0

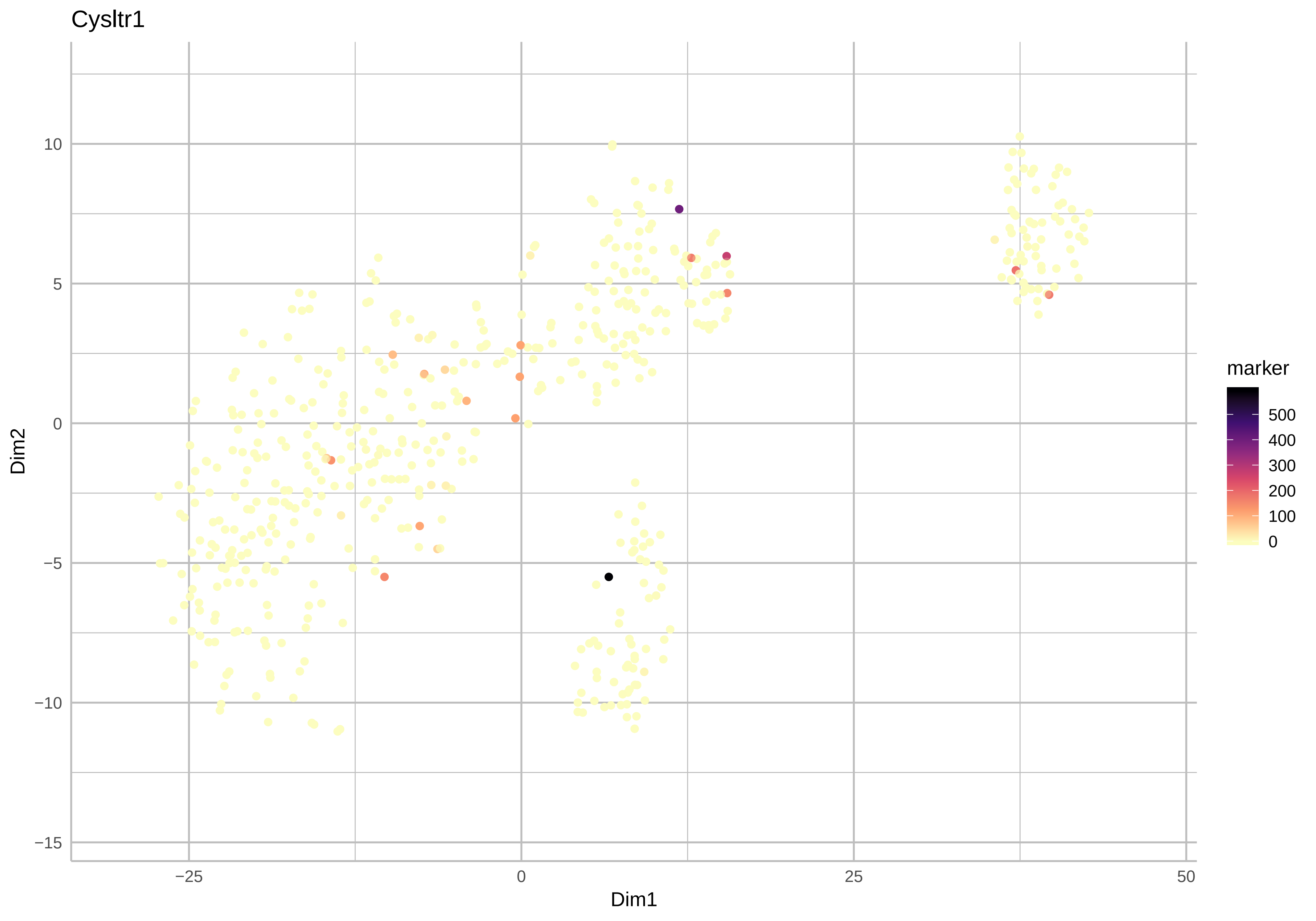

Cysltr1
Dim2
Dim1
10
5
0
−5
−10
−15
−25
0
25
50
marker
500
400
300
200
100
0

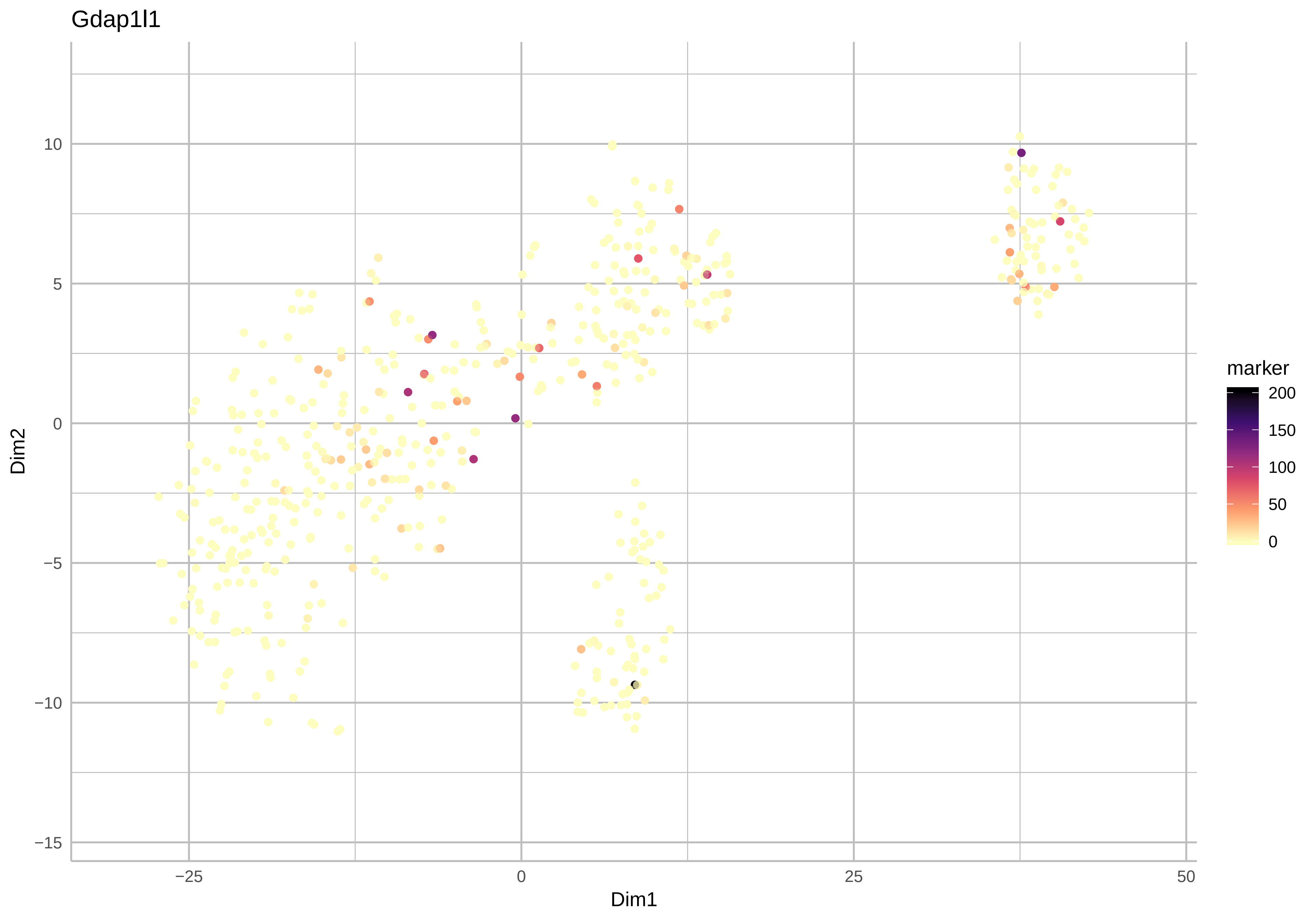

Gdap1l1
Dim2
Dim1
10
5
0
−5
−10
−15
−25
0
25
50
marker
200
150
100
50
0

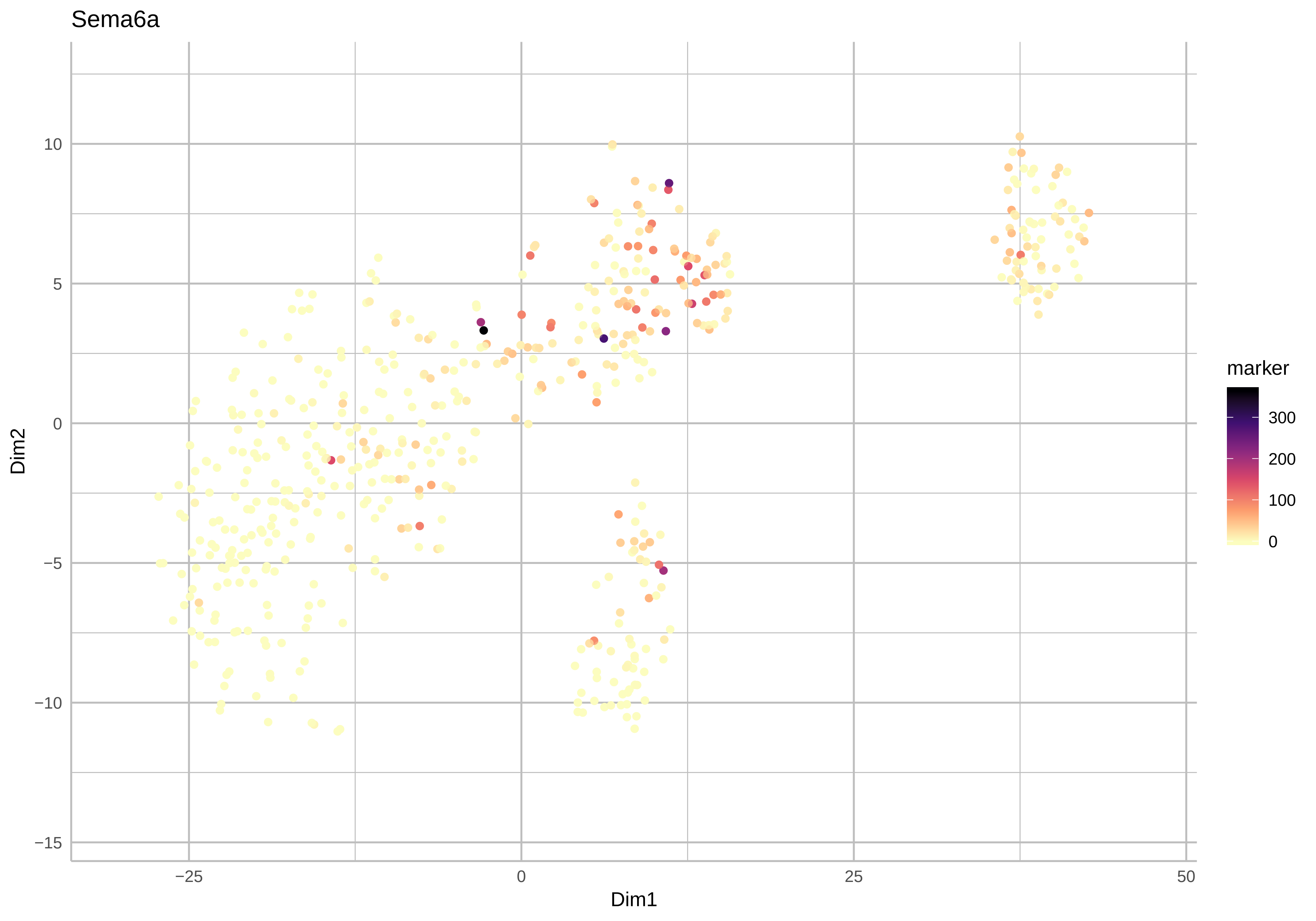

Sema6a
Dim2
Dim1
10
5
0
−5
−10
−15
−25
0
25
50
marker
300
200
100
0

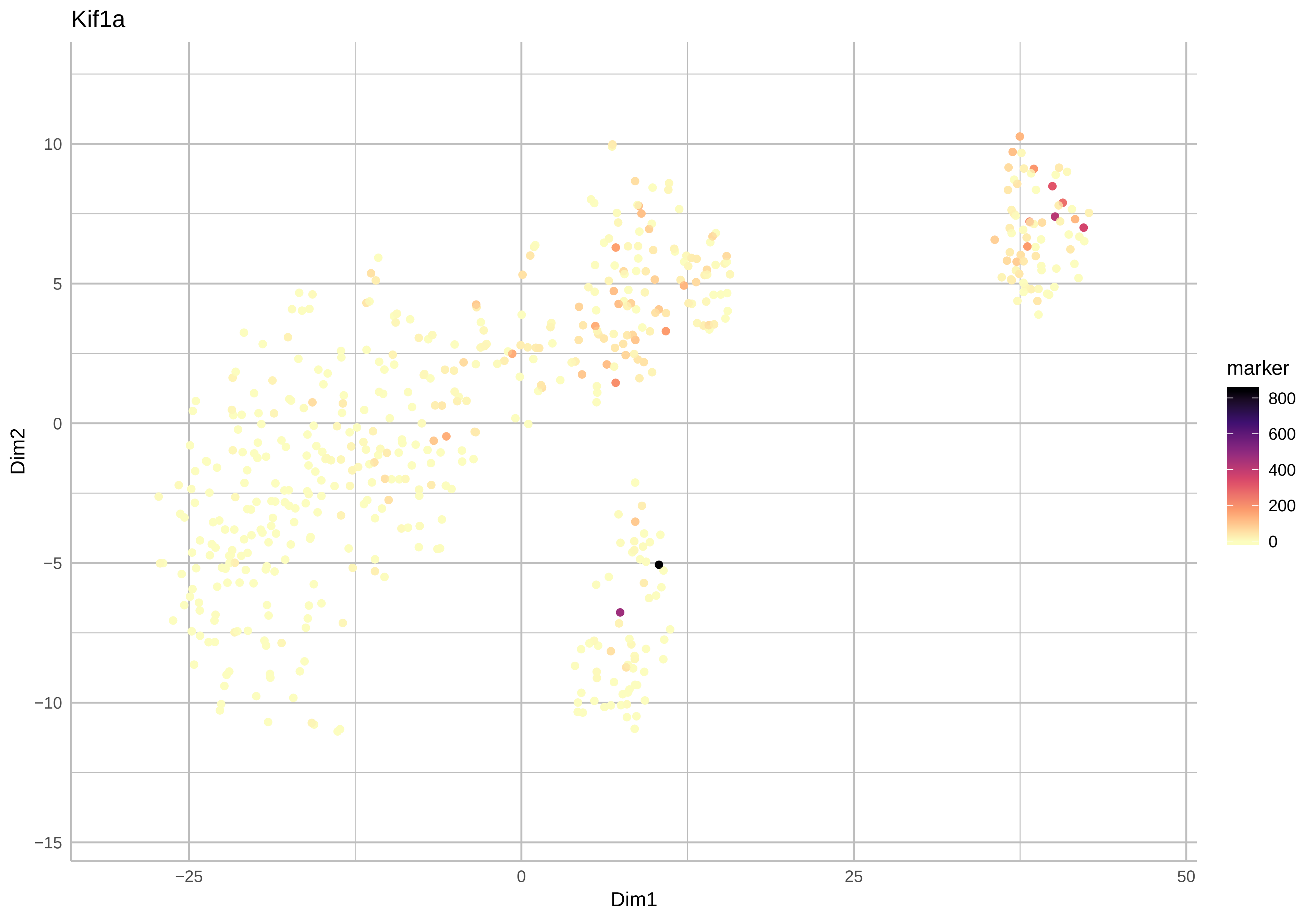
Kif1a
10
5
0
−5
−10
−15
Dim2
−25
0
25
50
Dim1
marker
800
600
400
200
0

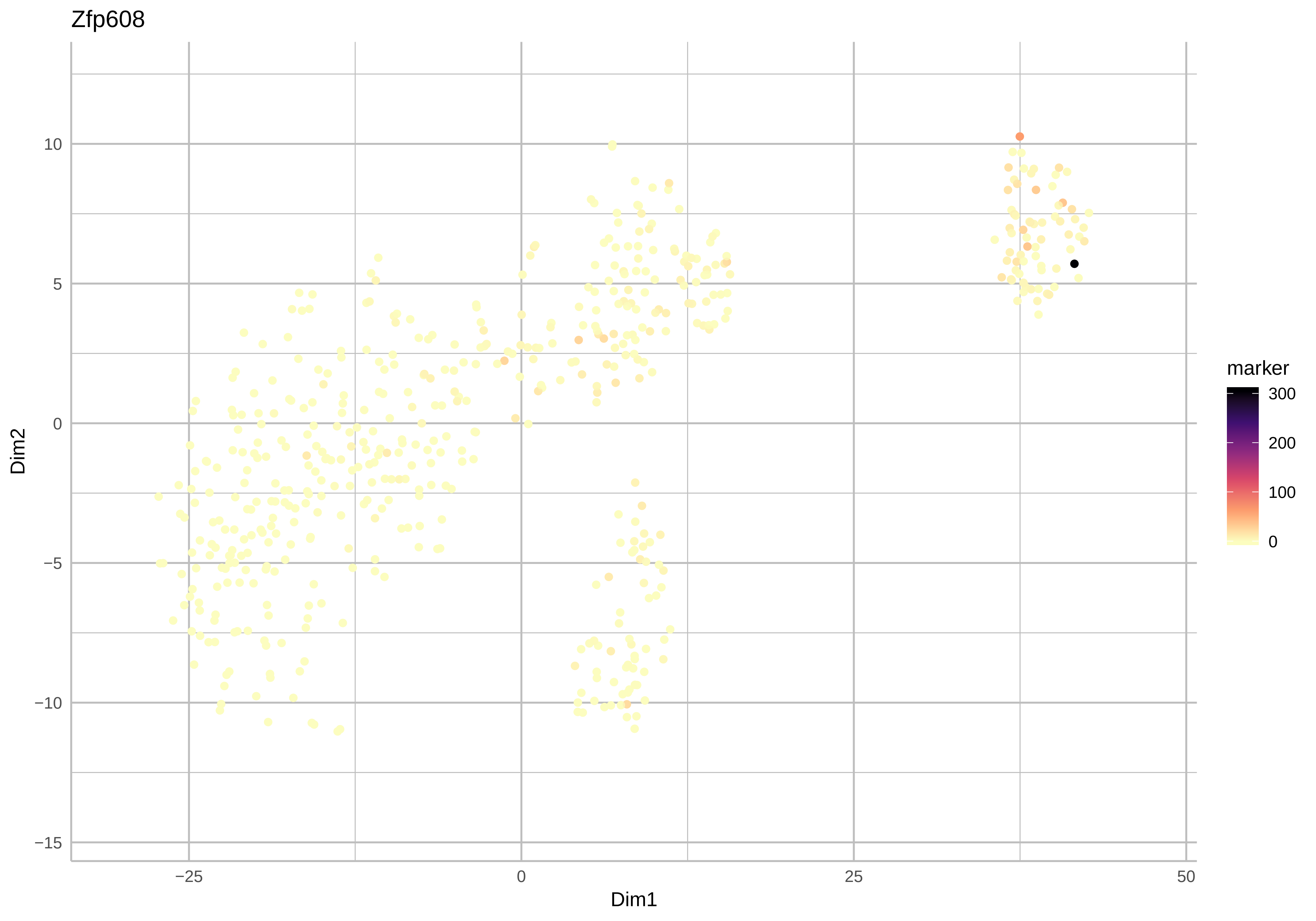

Zfp608
Dim2
Dim1
marker
300
200
100
0
10
5
0
−5
−10
−15
−25
0
25
50

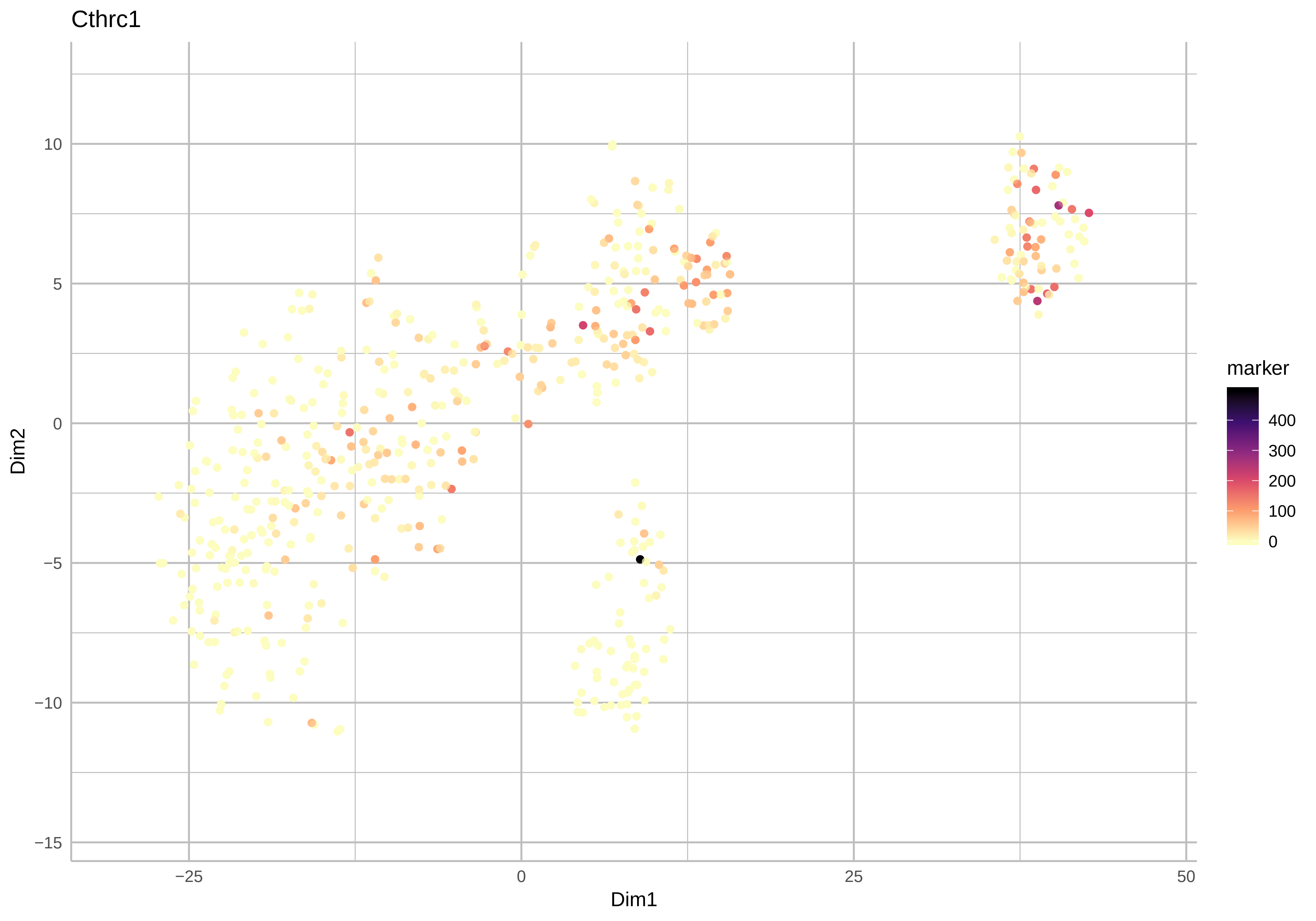
Cthrc1
Dim2
Dim1
10
5
0
−5
−10
−15
−25
0
25
50
marker
400
300
200
100
0

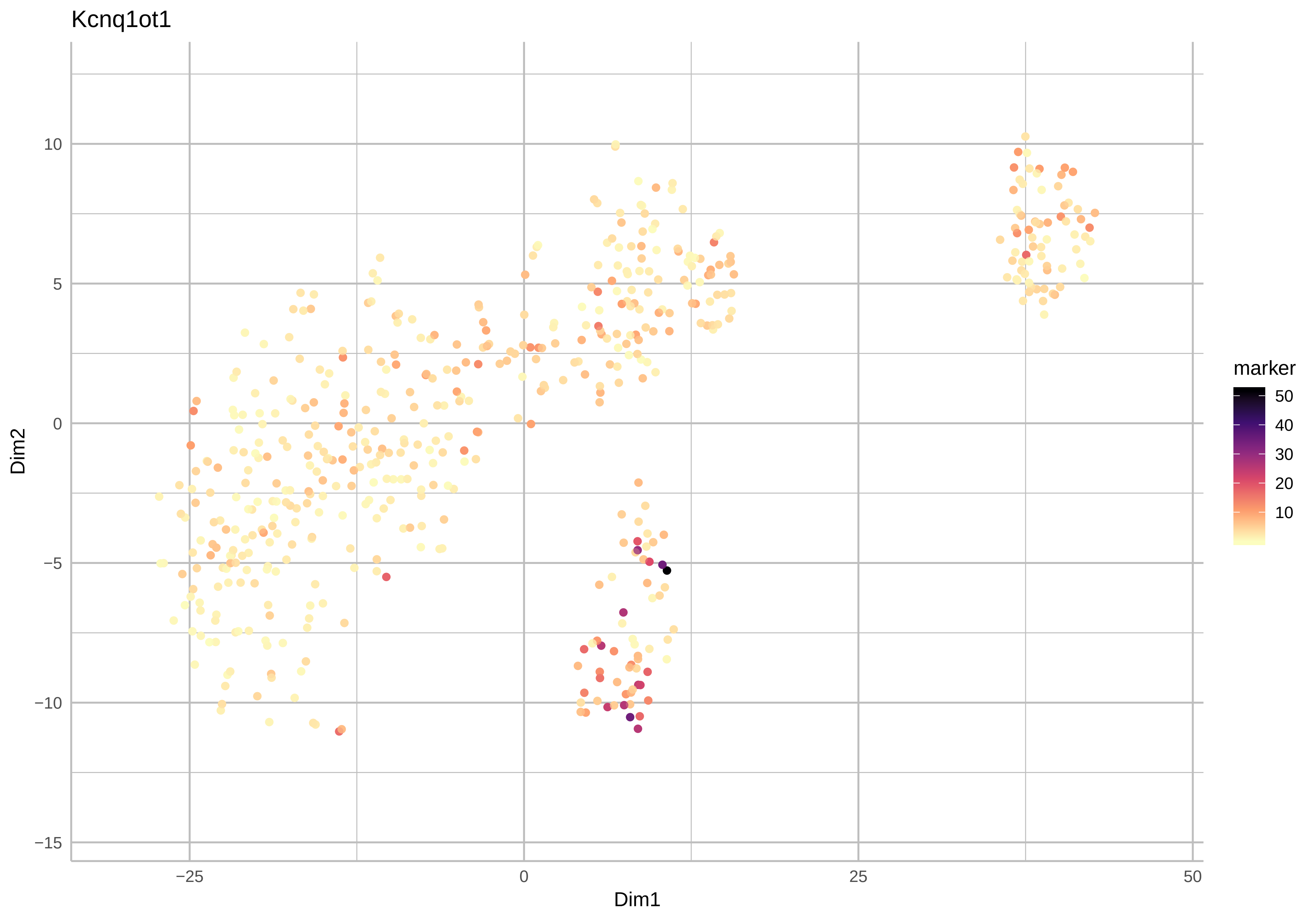

Kcnq1ot1
Dim2
Dim1
10
5
0
−5
−10
−15
−25
0
25
50
marker
50
40
30
20
10

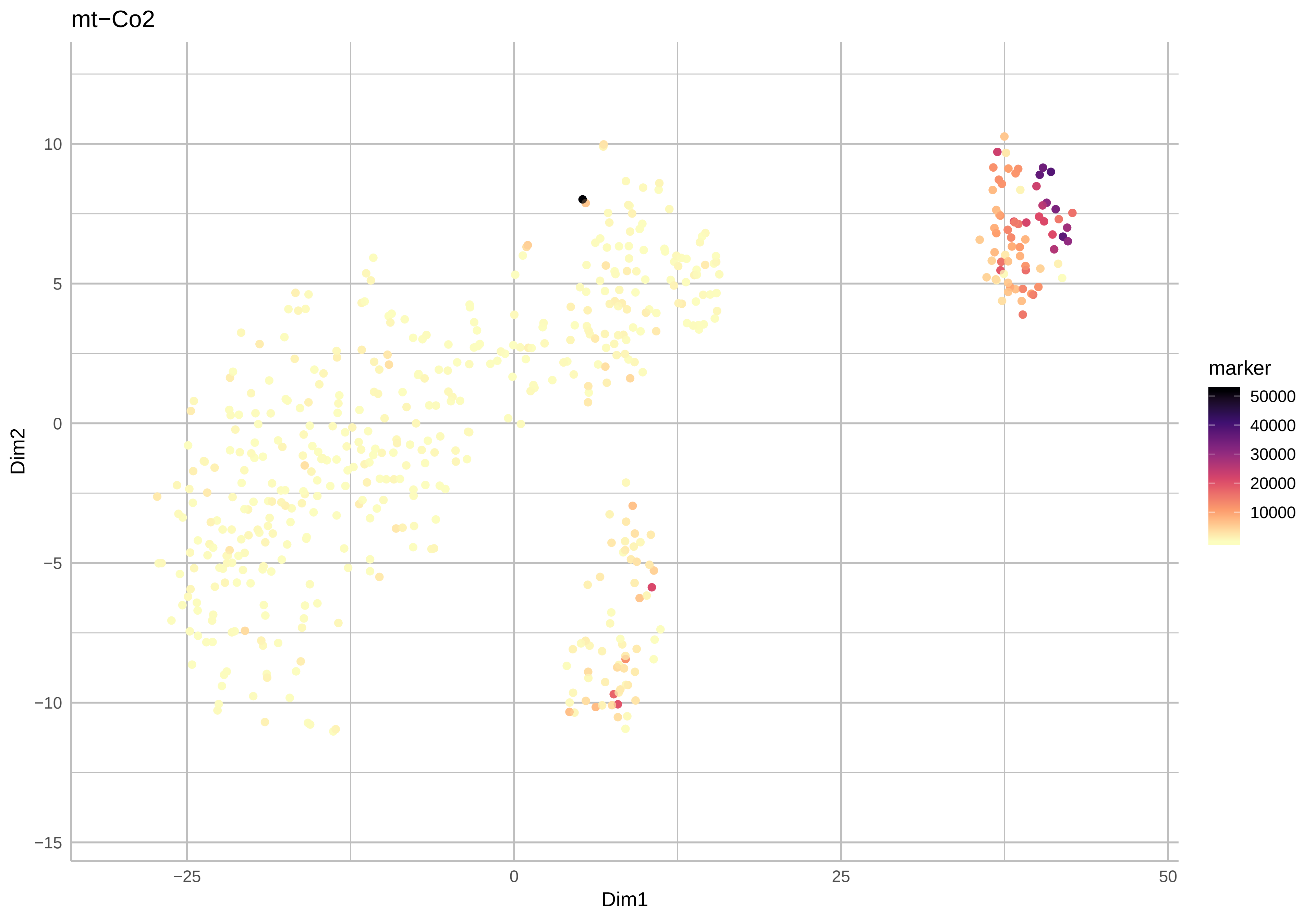
mt−Co2
Dim2
Dim1
10
5
0
−5
−10
−15
−25
0
25
50
marker
50000
40000
30000
20000
10000

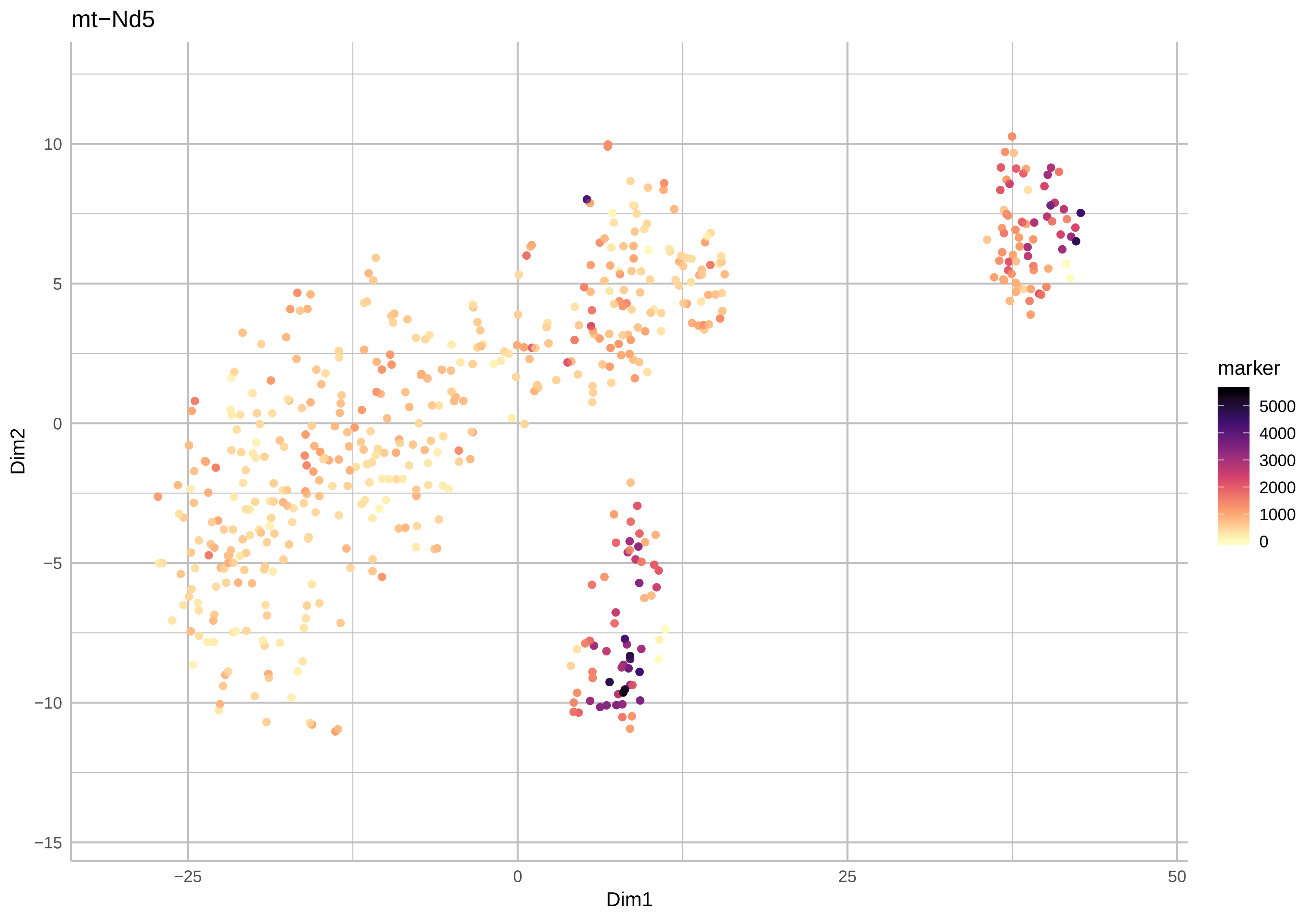
mt−Nd5
Dim2
Dim1
10
5
0
−5
−10
−15
−25
0
25
50
marker
5000
4000
3000
2000
1000
0

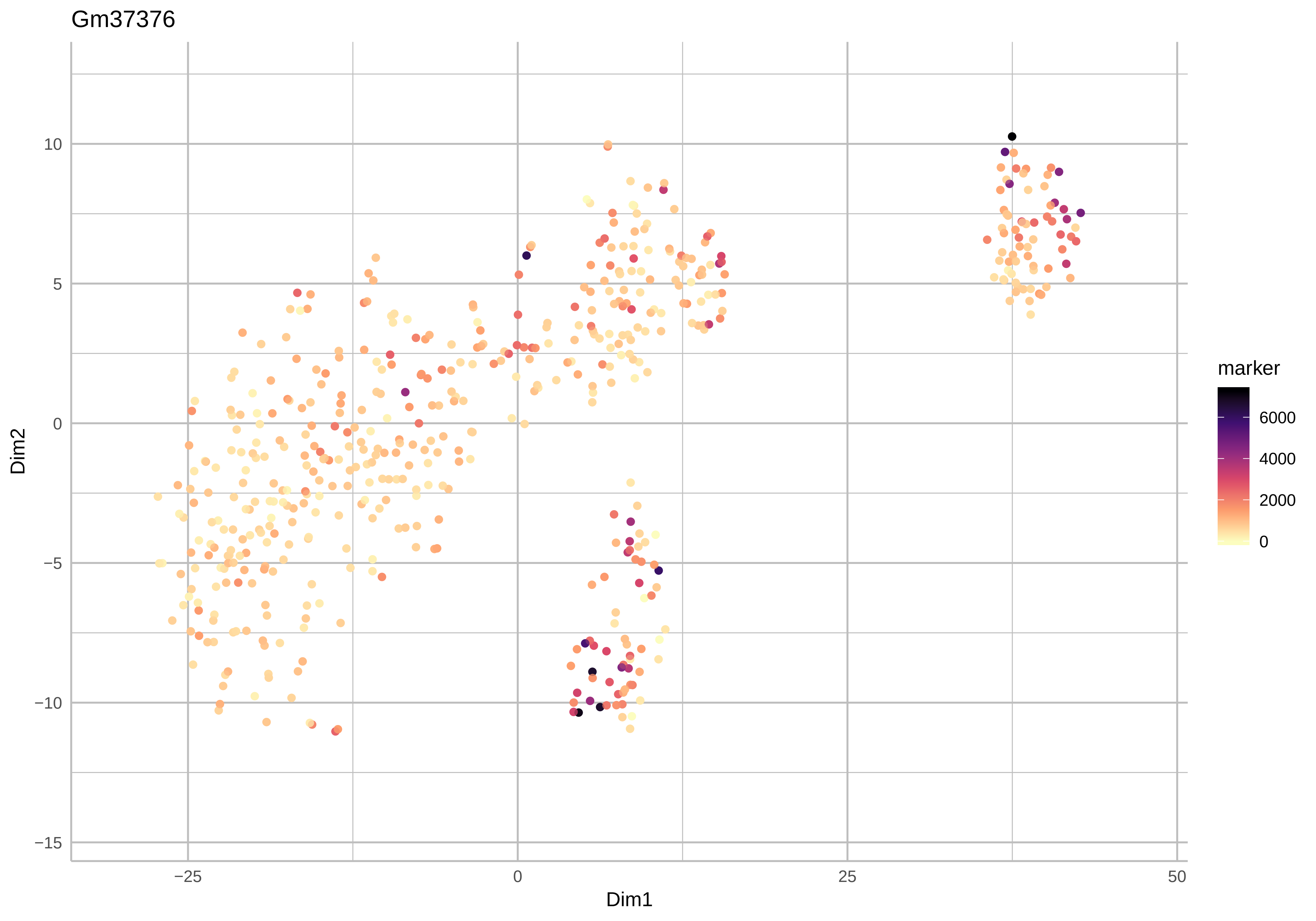

Gm37376
Dim2
Dim1
10
5
0
−5
−10
−15
−25
0
25
50
marker
6000
4000
2000
0

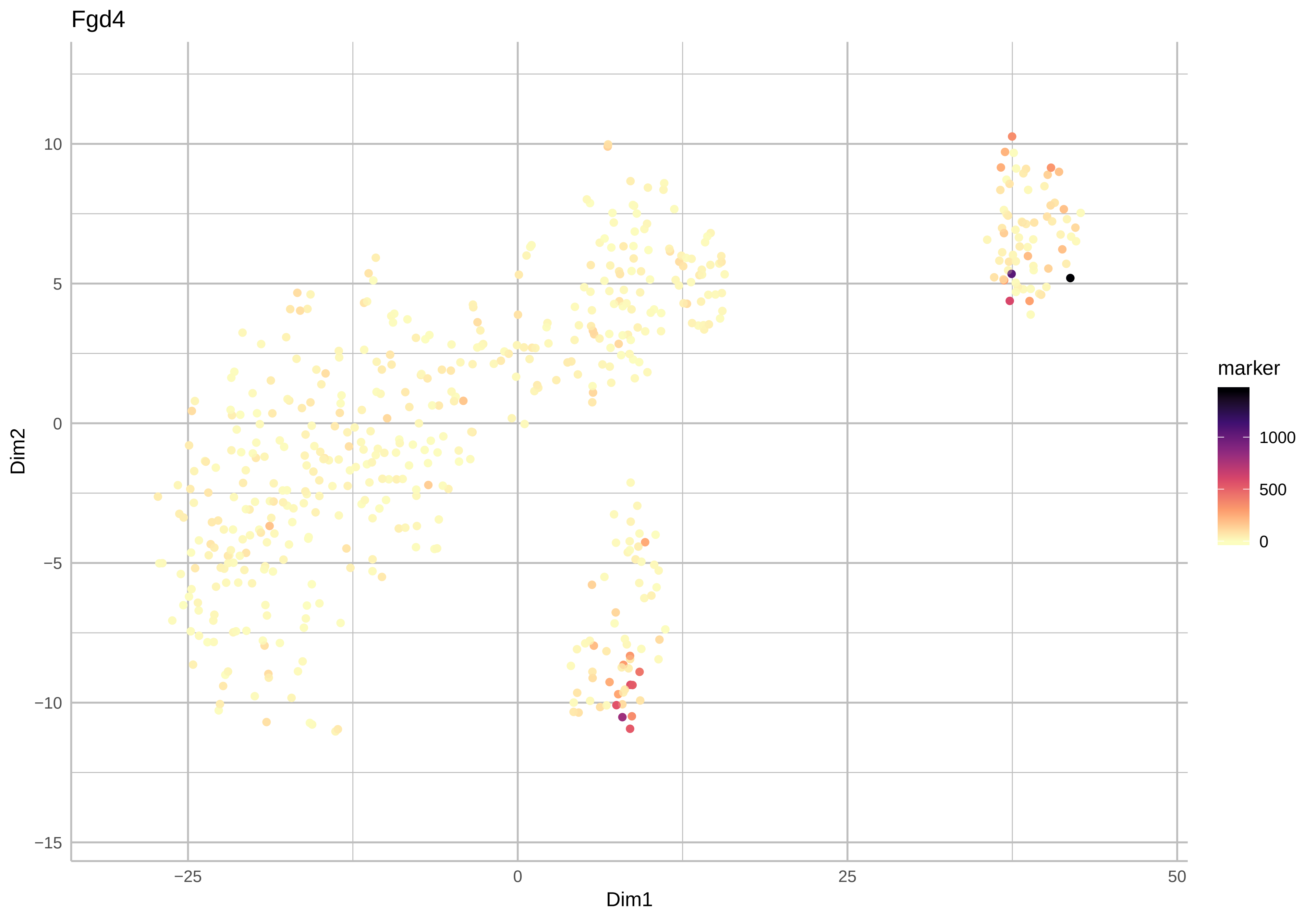
Fgd4
Dim2
Dim1
10
5
0
−5
−10
−15
−25
0
25
50
marker
1000
500
0

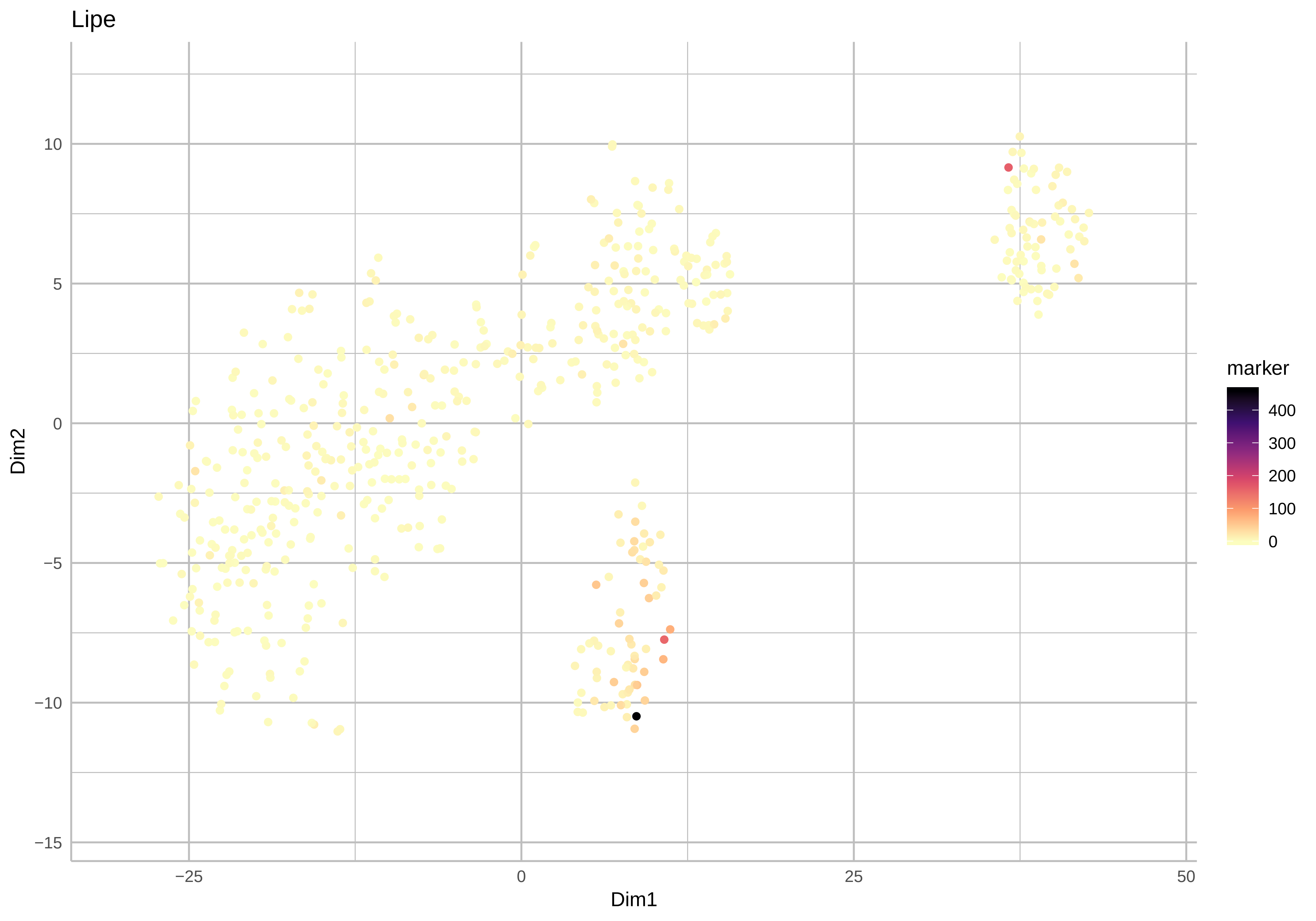
Lipe
Dim2
Dim1
10
5
0
−5
−10
−15
−25
0
25
50
marker
400
300
200
100
0

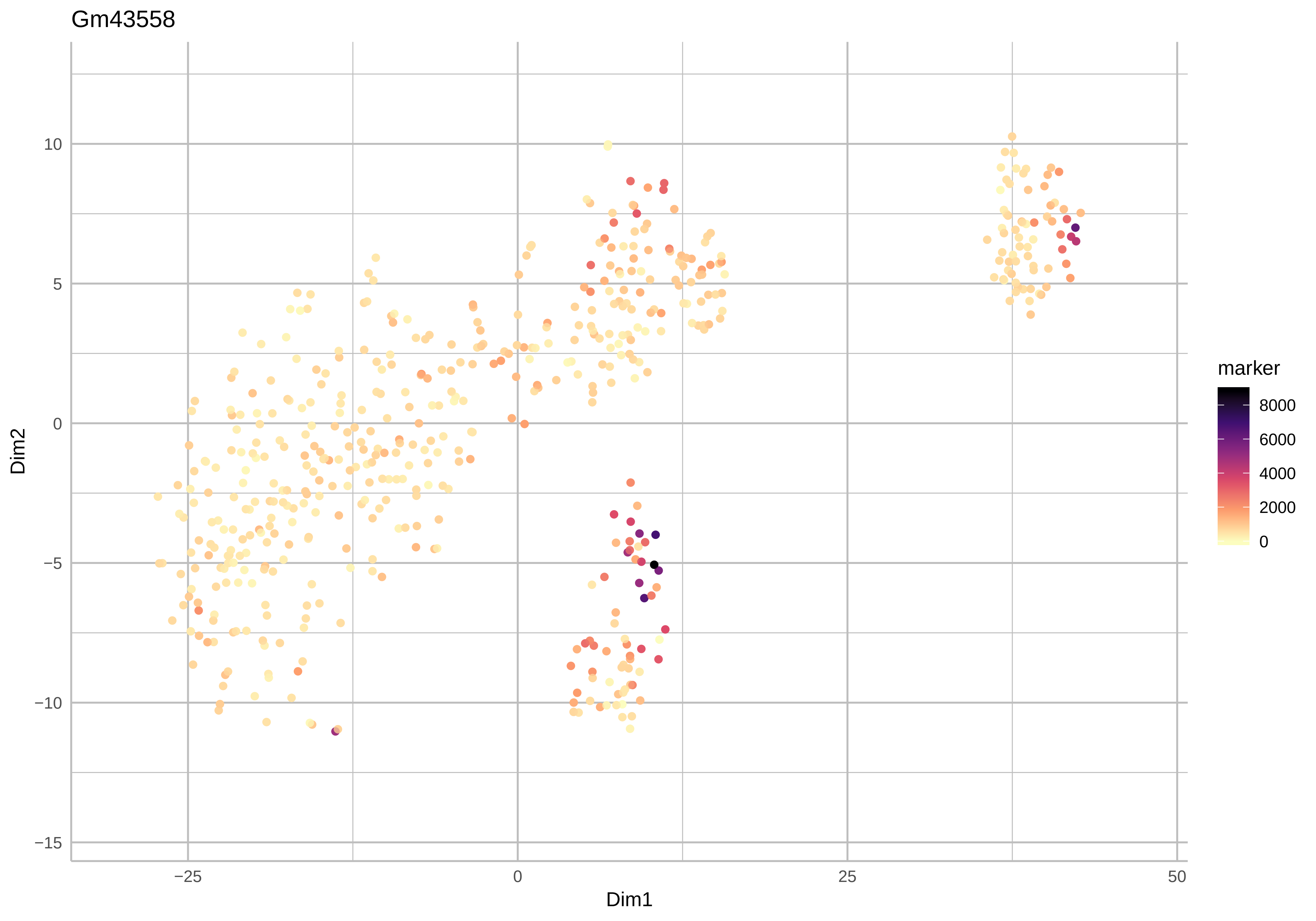

Gm43558
Dim2
Dim1
marker
8000
6000
4000
2000
0
10
5
0
−5
−10
−15
−25
0
25
50

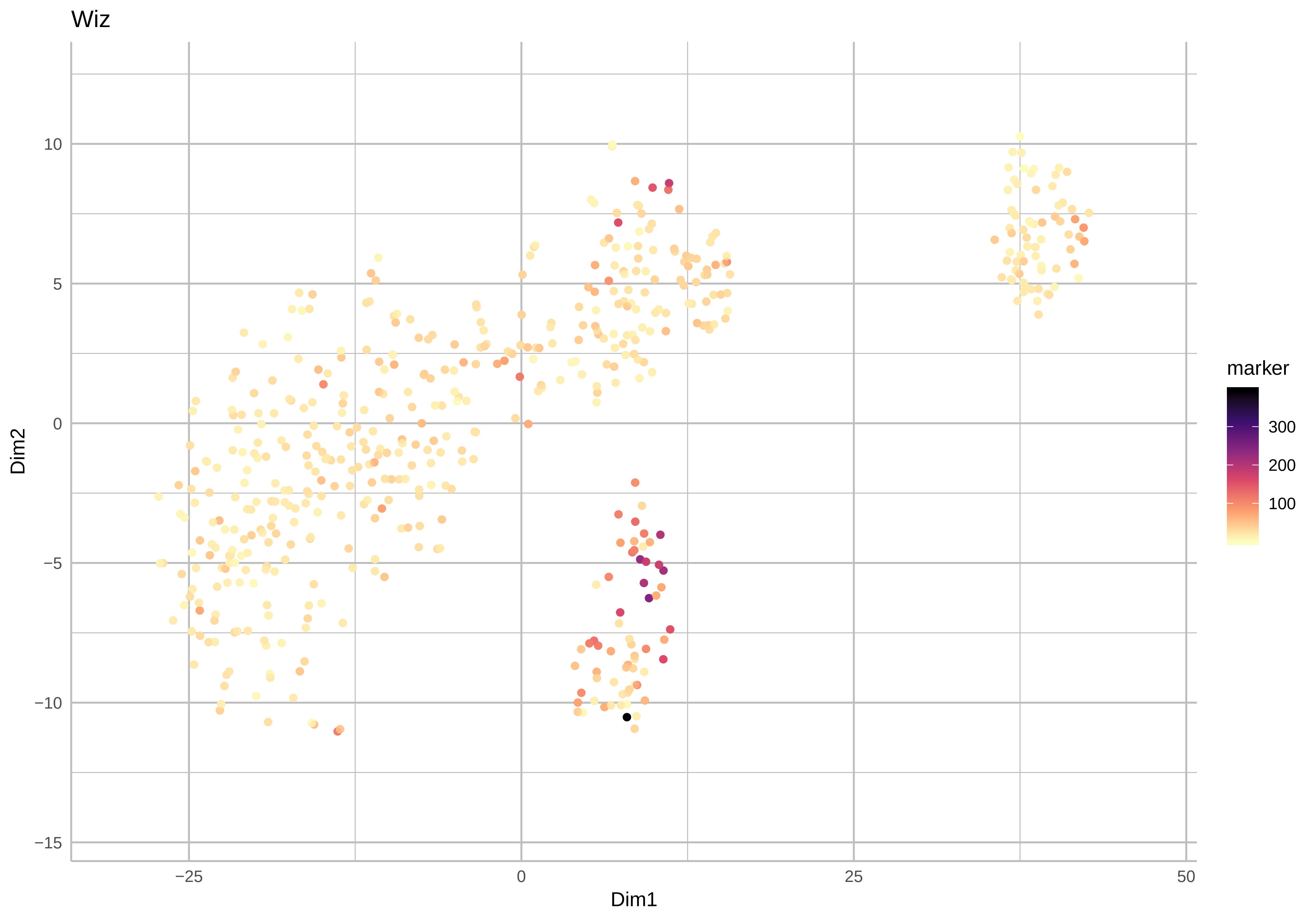
Wiz
10
5
0
−5
−10
−15
Dim2
−25
0
25
50
Dim1
marker
300
200
100

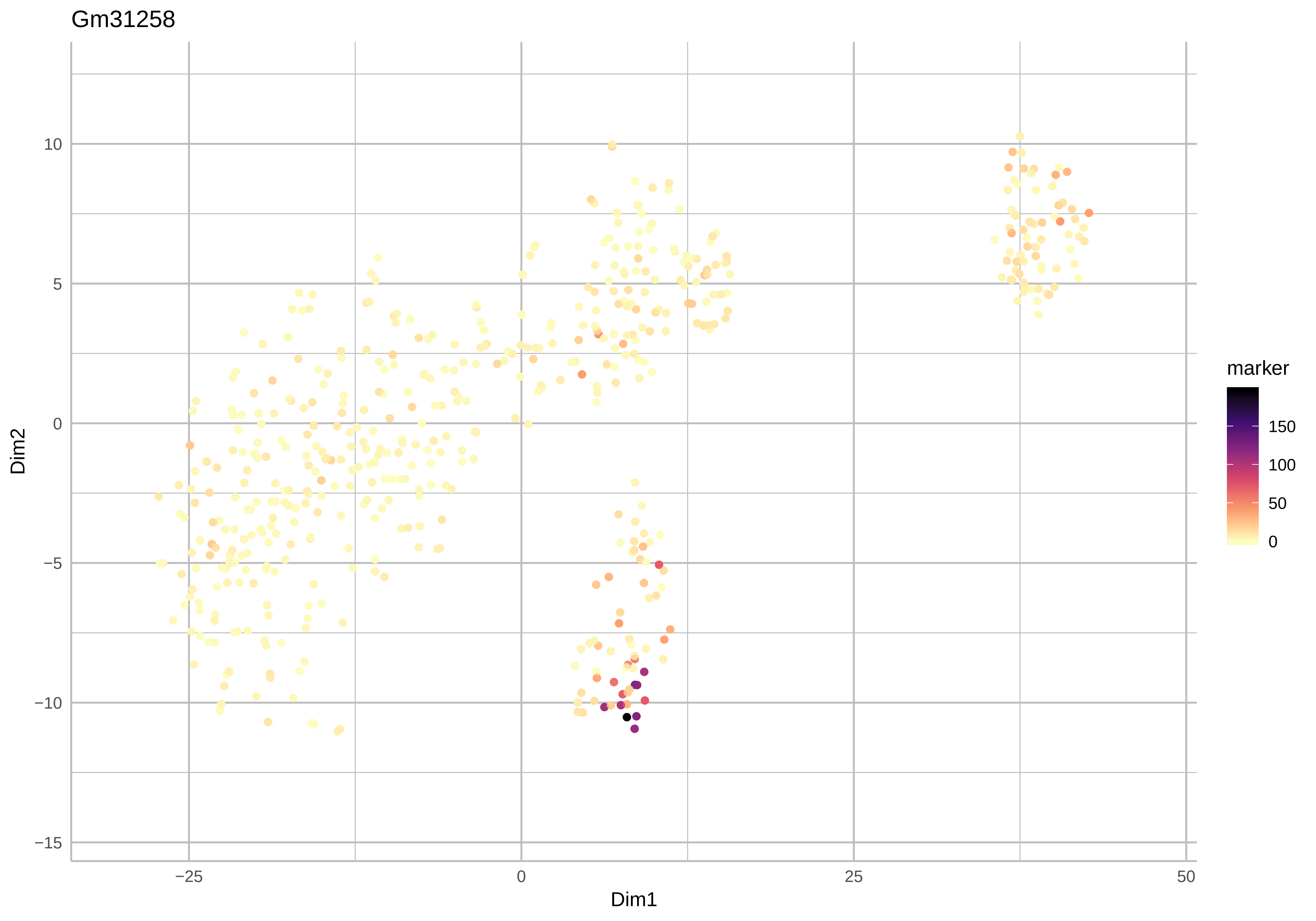
Gm31258
Dim2
Dim1
10
5
0
−5
−10
−15
−25
0
25
50
marker
150
100
50
0

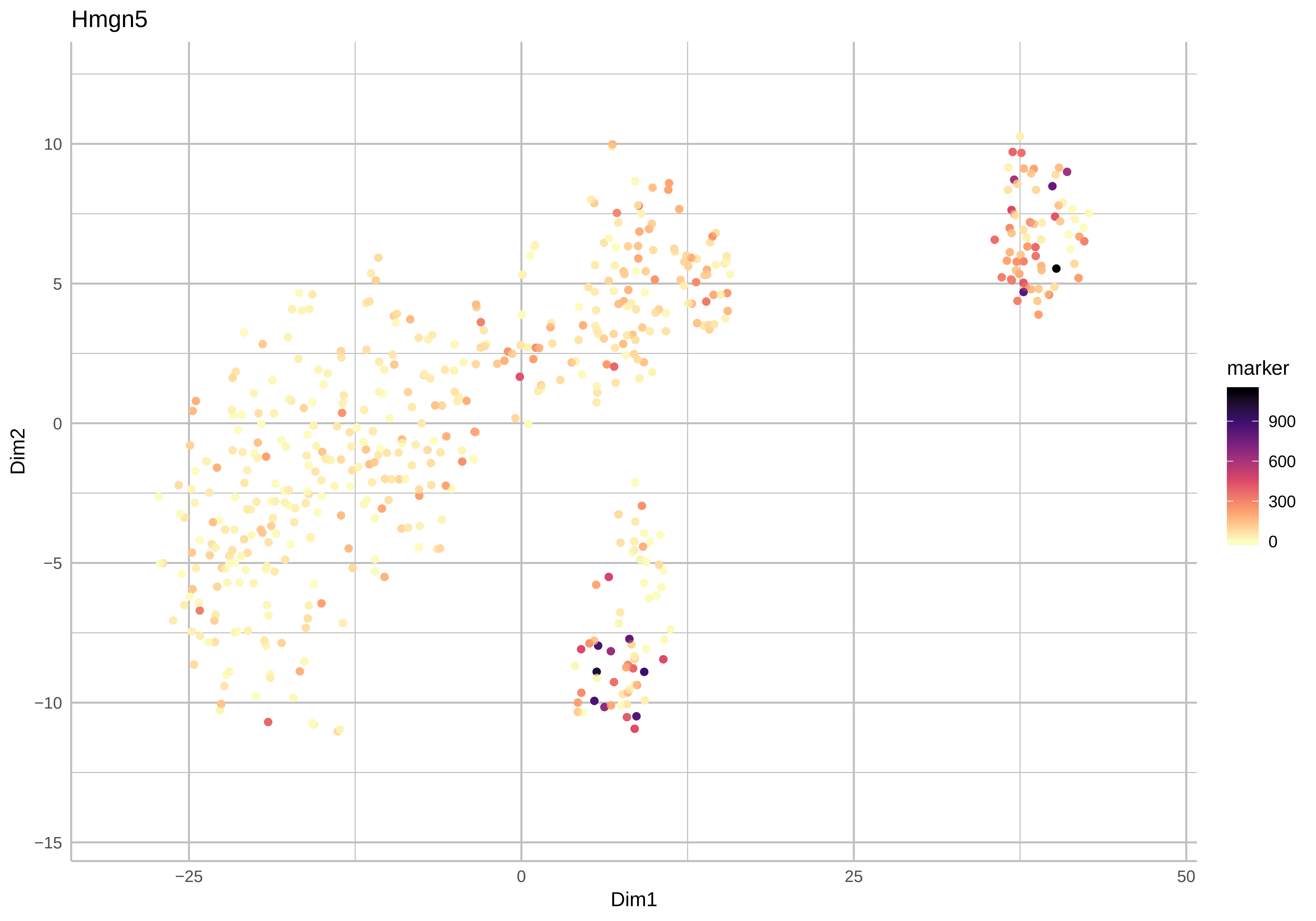
Hmgn5
Dim2
10
5
0
−5
−10
−15
Dim1
−25
0
25
50
marker
900
600
300
0

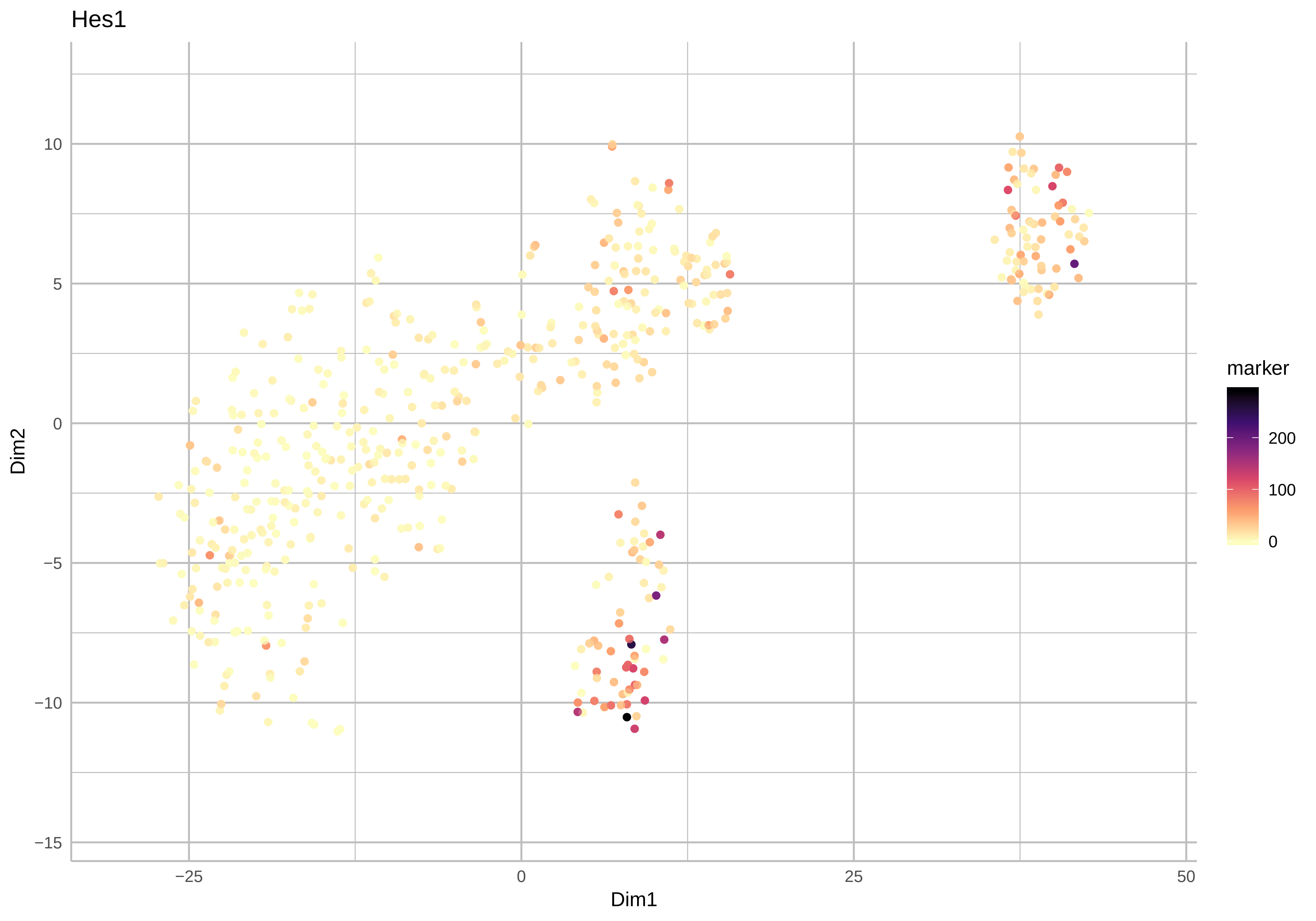

Hes1
Dim2
Dim1
10
5
0
−5
−10
−15
−25
0
25
50
marker
200
100
0

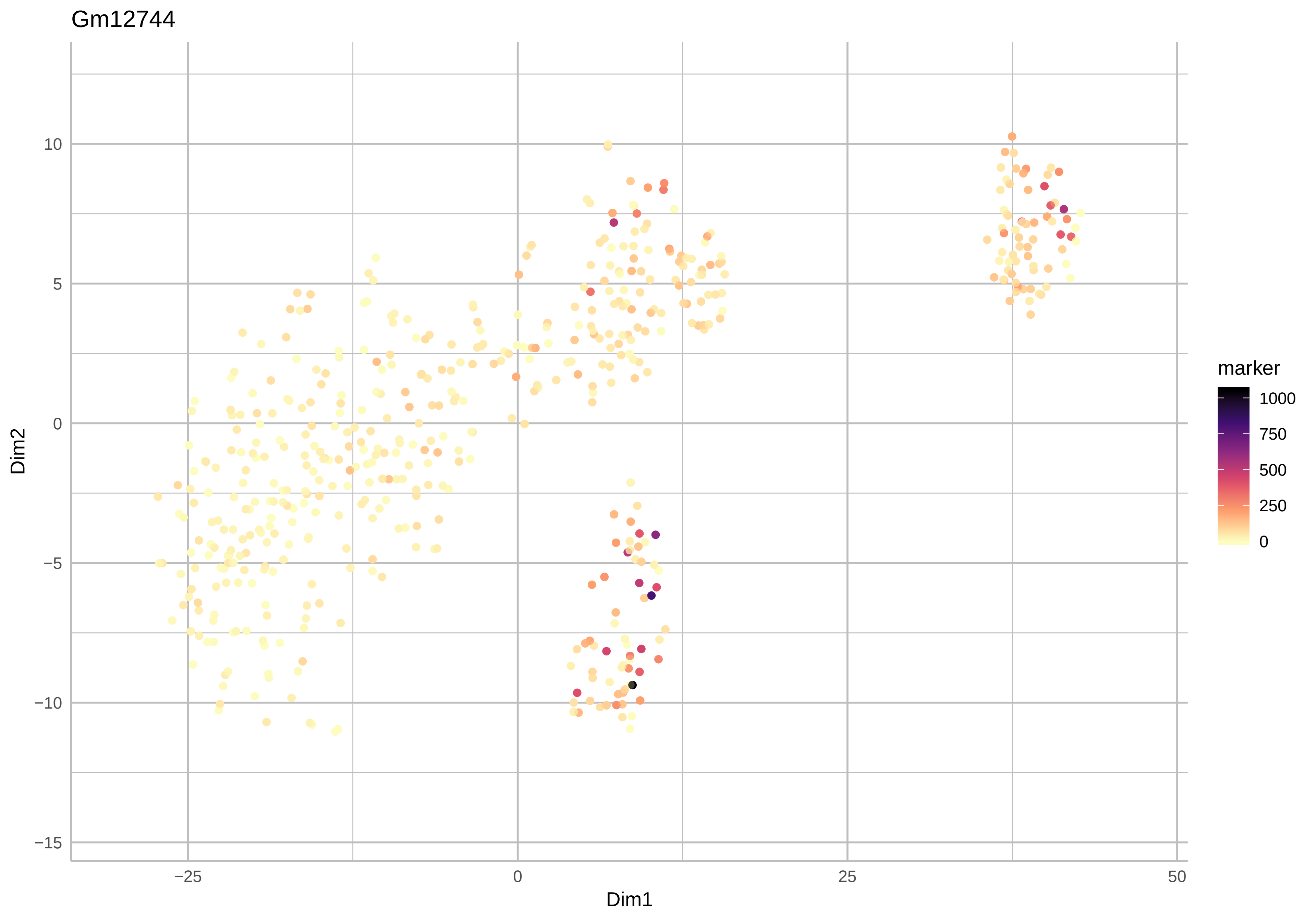

Gm12744
Dim2
Dim1
10
5
0
−5
−10
−15
−25
0
25
50
marker
1000
750
500
250
0

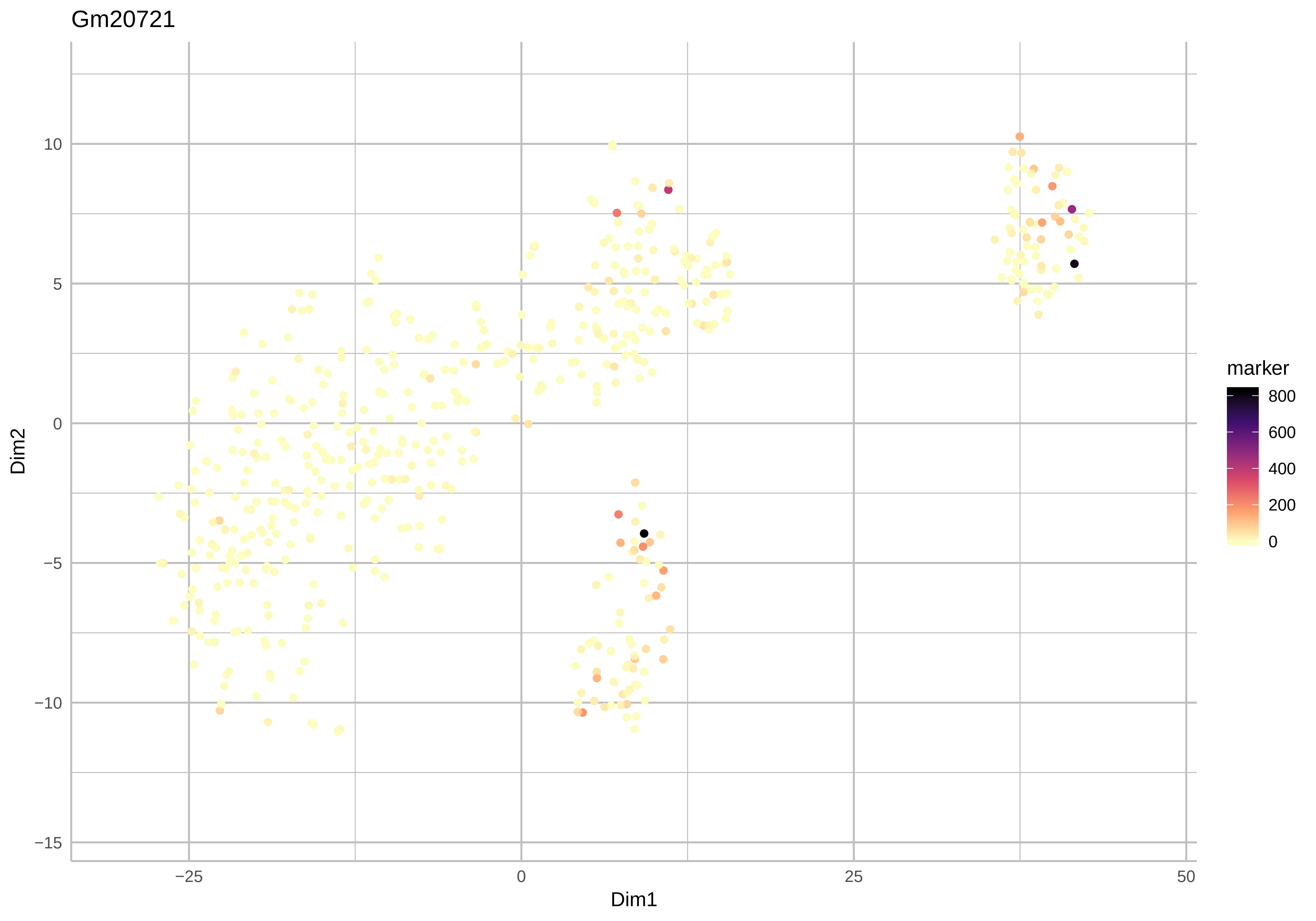

Gm20721
Dim2
Dim1
10
5
0
−5
−10
−15
−25
0
25
50
marker
800
600
400
200
0

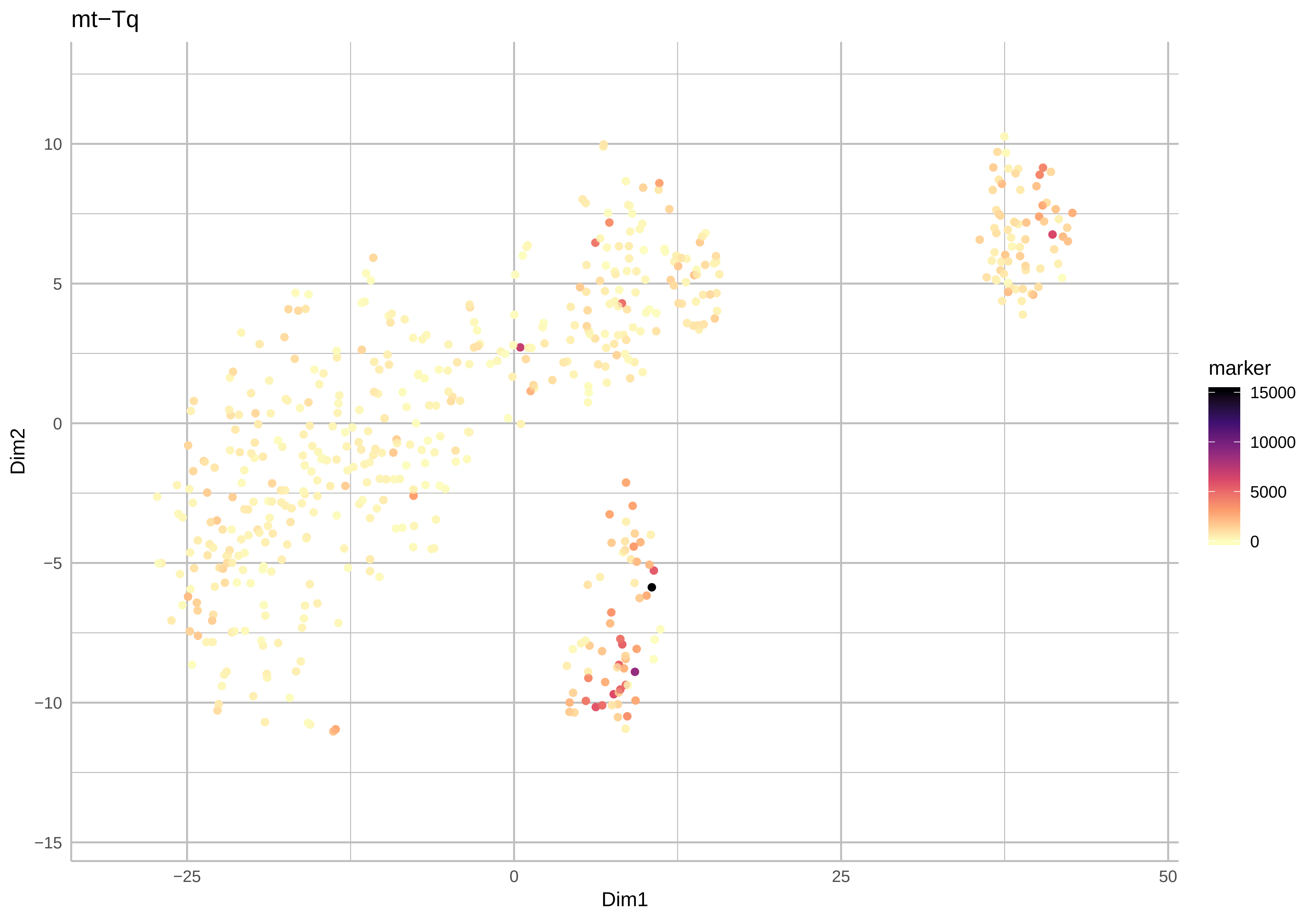

mt−Tq
Dim2
Dim1
marker
15000
10000
5000
0
10
5
0
−5
−10
−15
−25
0
25
50

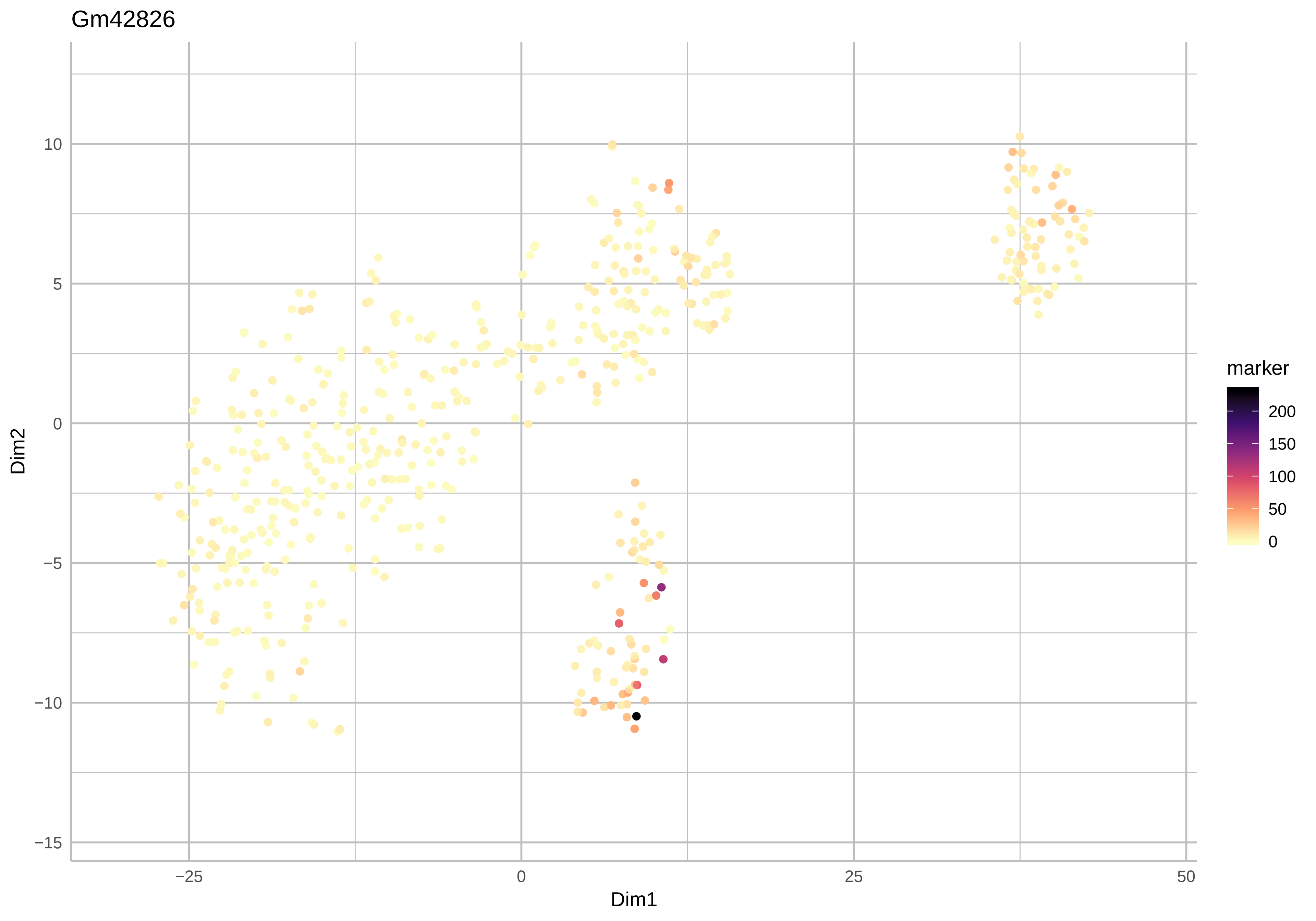
Gm42826
Dim2
Dim1
10
5
0
−5
−10
−15
−25
0
25
50
marker
200
150
100
50
0

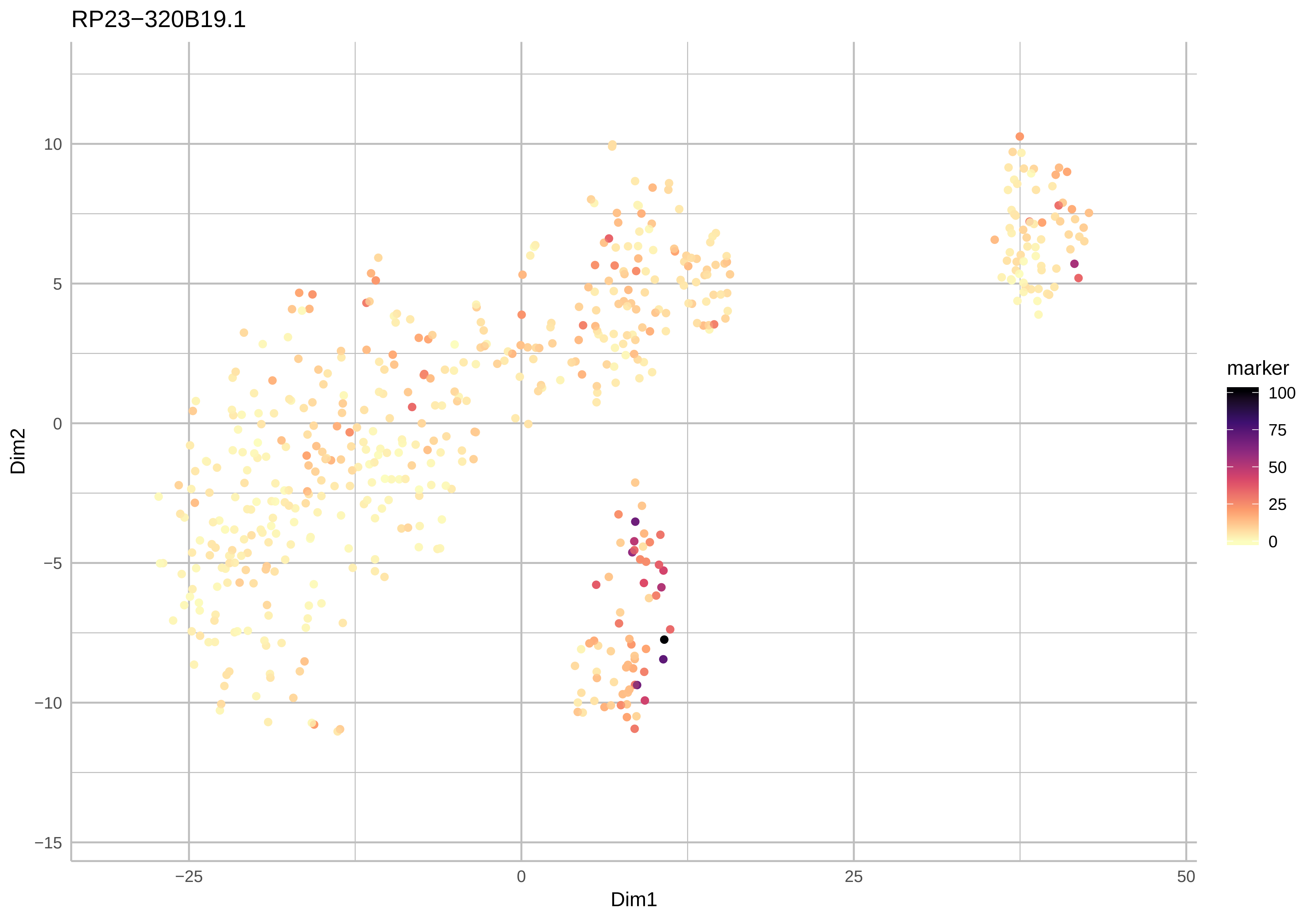
RP23−320B19.1
Dim2
10
5
0
−5
−10
−15
Dim1
−25
0
25
50
marker
100
75
50
25
0

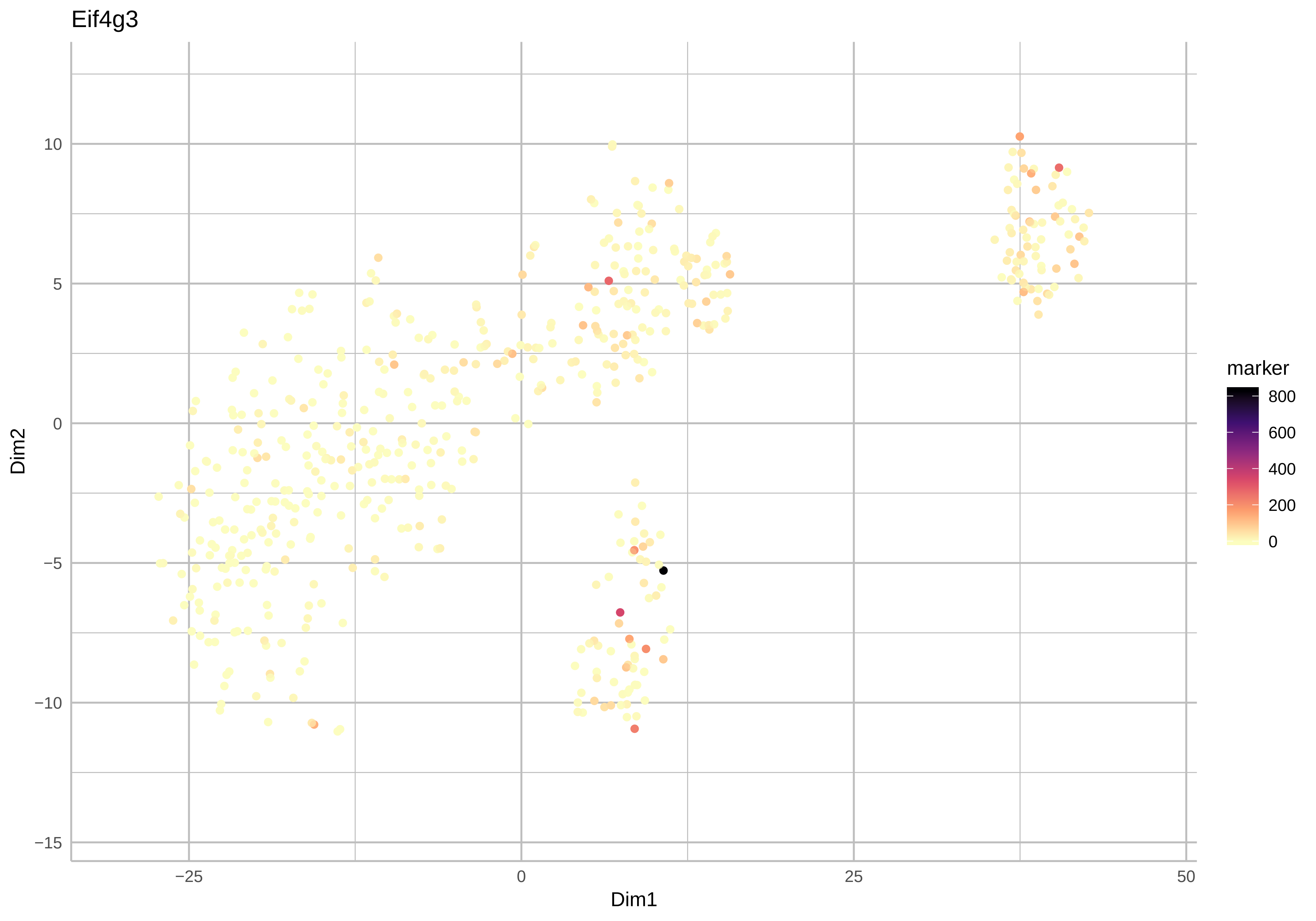

Eif4g3
Dim2
Dim1
10
5
0
−5
−10
−15
−25
0
25
50
marker
800
600
400
200
0

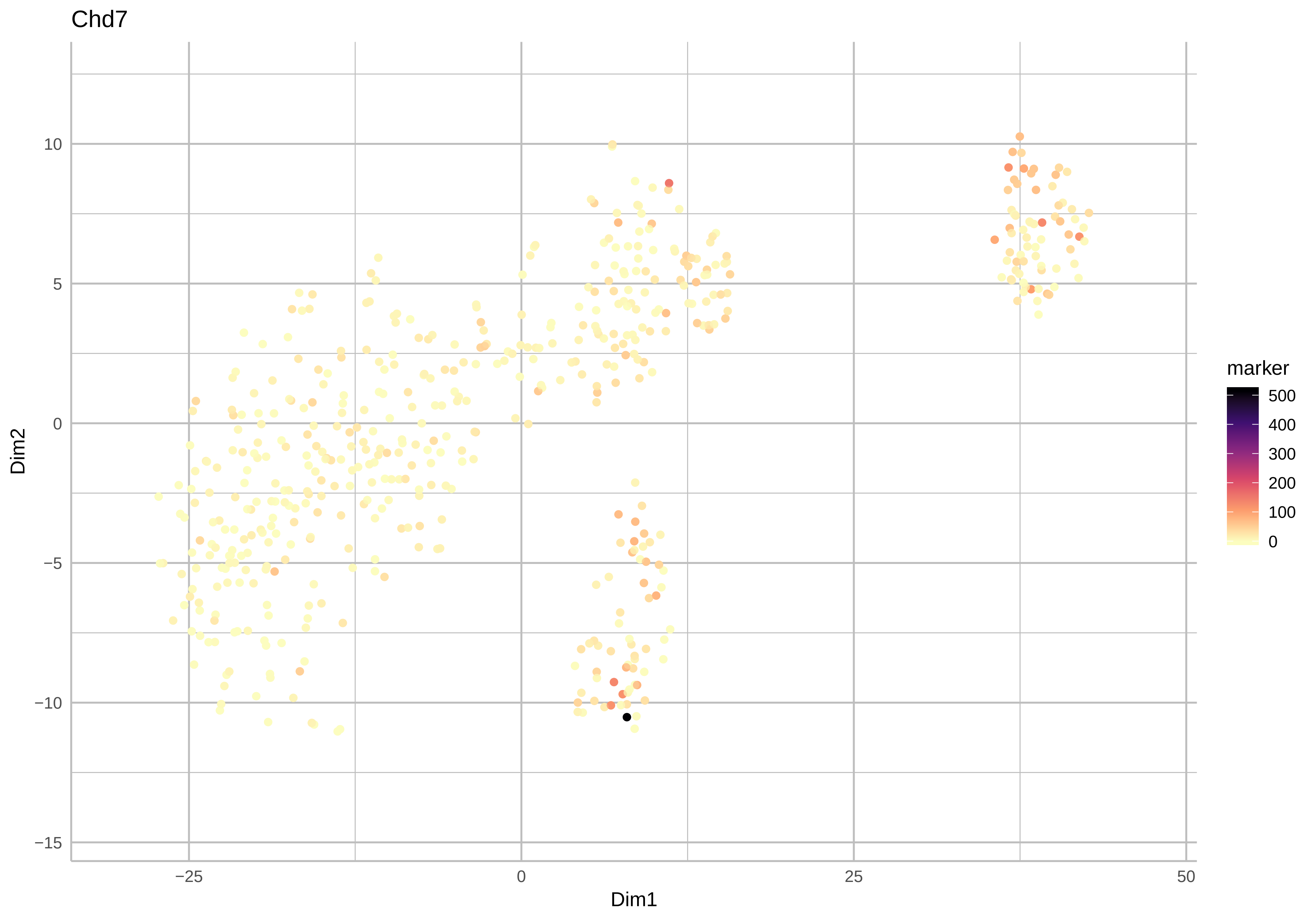
Chd7
Dim2
Dim1
10
5
0
−5
−10
−15
−25
0
25
50
marker
500
400
300
200
100
0

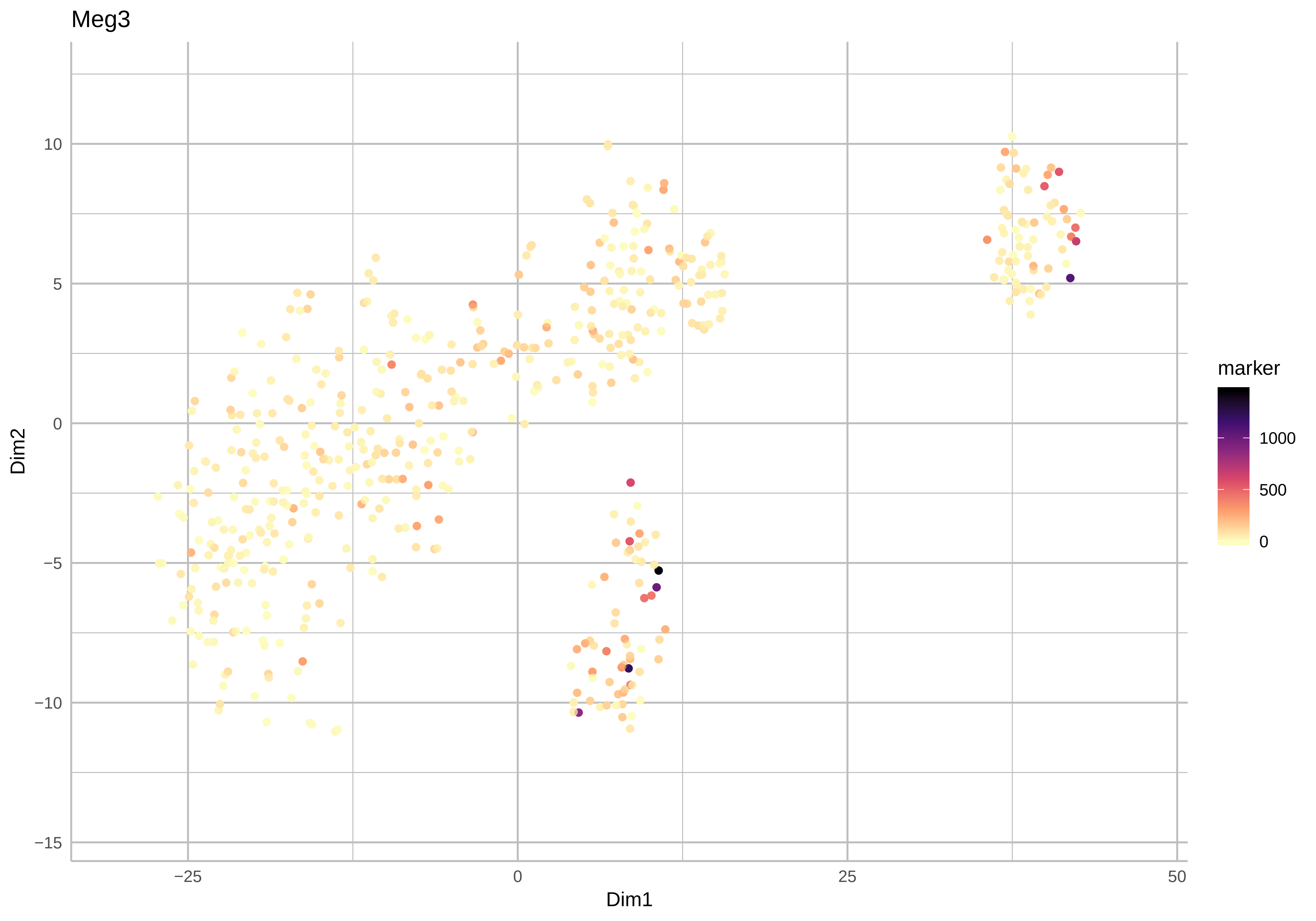
Meg3
Dim2
Dim1
10
5
0
−5
−10
−15
−25
0
25
50
marker
1000
500
0

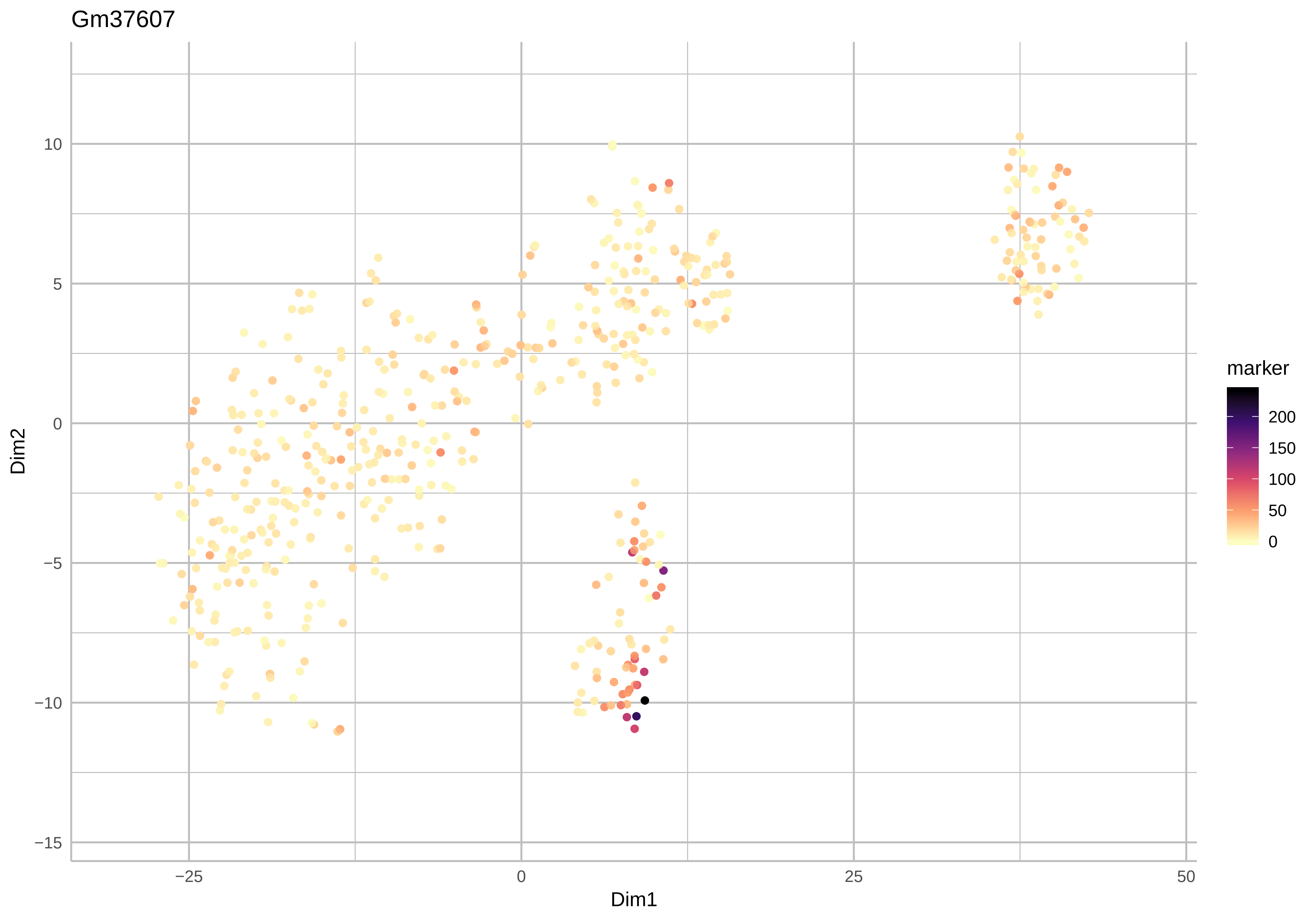

Gm37607
Dim2
Dim1
marker
200
150
100
50
0
10
5
0
−5
−10
−15
−25
0
25
50

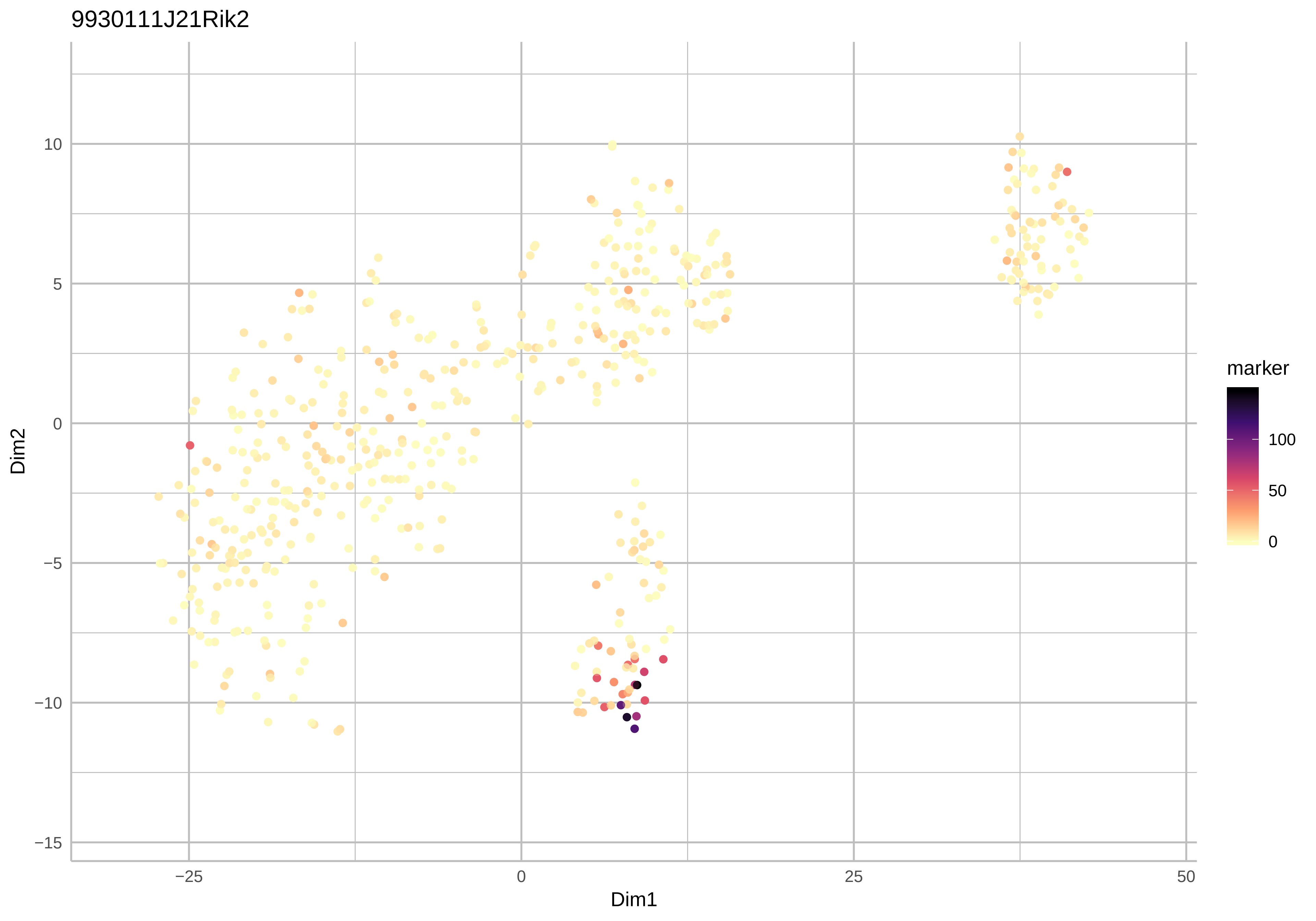

9930111J21Rik2
Dim2
Dim1
10
5
0
−5
−10
−15
−25
0
25
50
marker
100
50
0

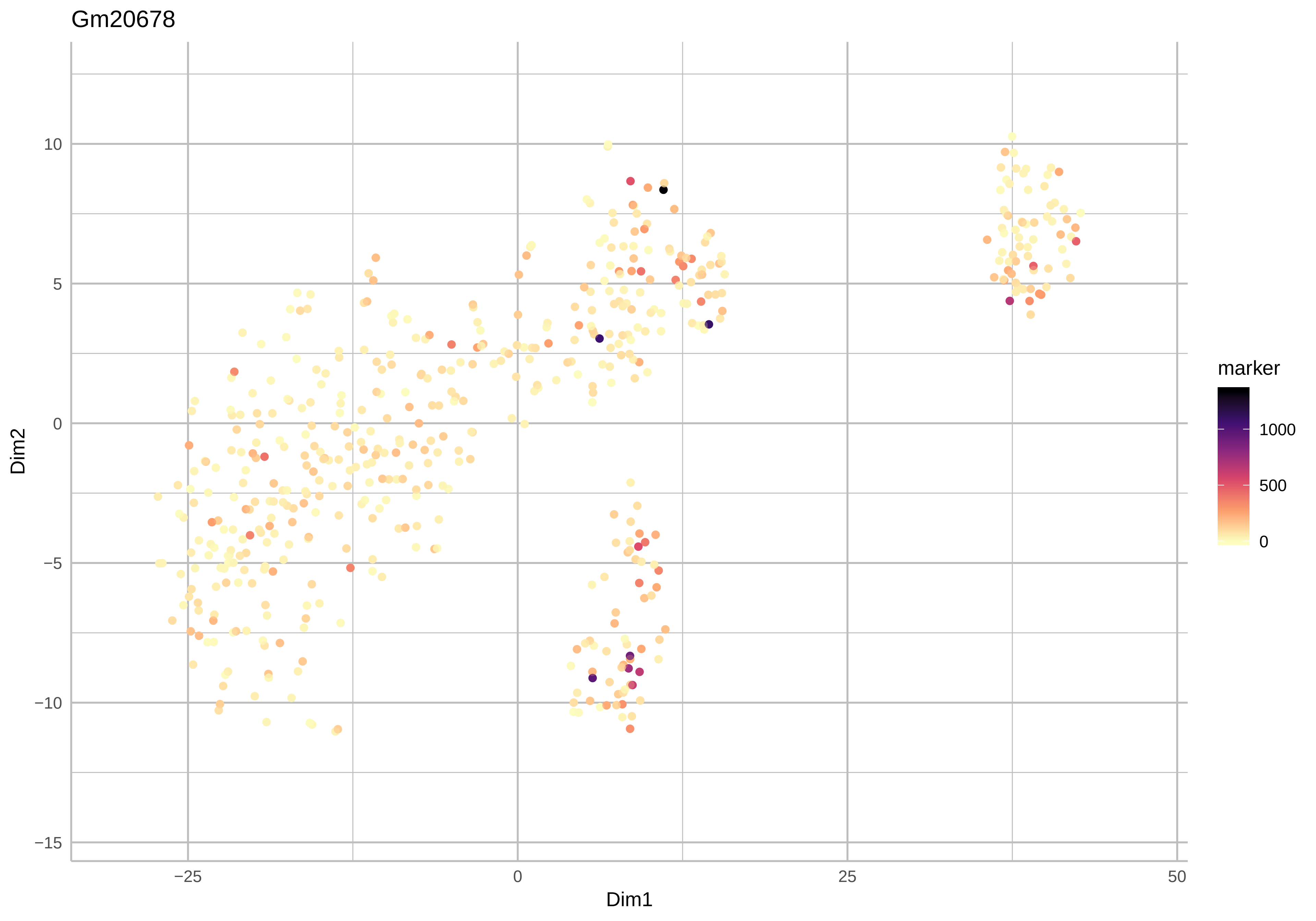

Gm20678
Dim2
Dim1
marker
10
5
0
−5
−10
−15
−25
0
25
50
1000
500
0

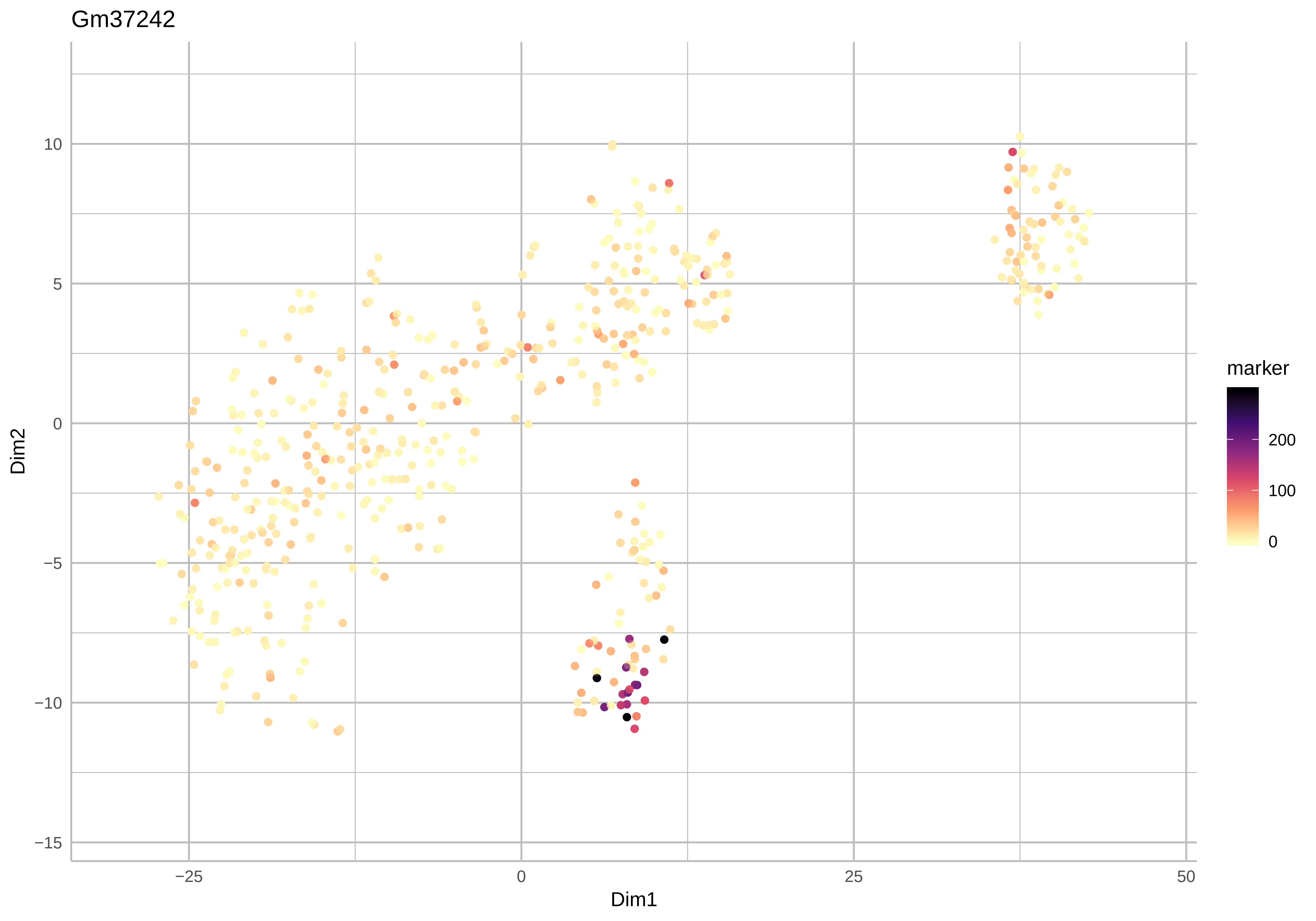

Gm37242
Dim2
Dim1
10
5
0
−5
−10
−15
−25
0
25
50
marker
200
100
0

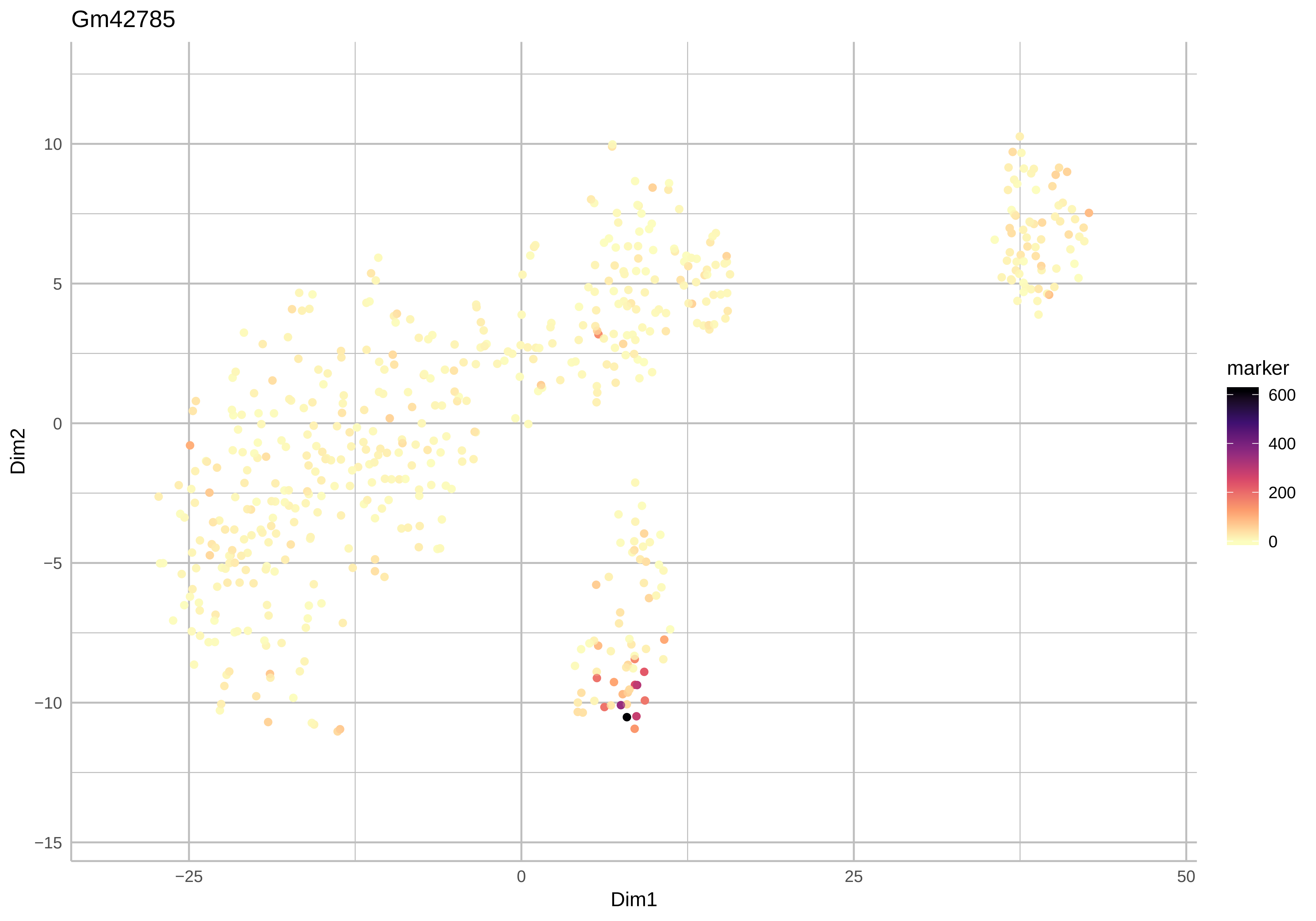
Gm42785
Dim2
10
5
0
−5
−10
−15
−25
0
25
50
Dim1
marker
600
400
200
0

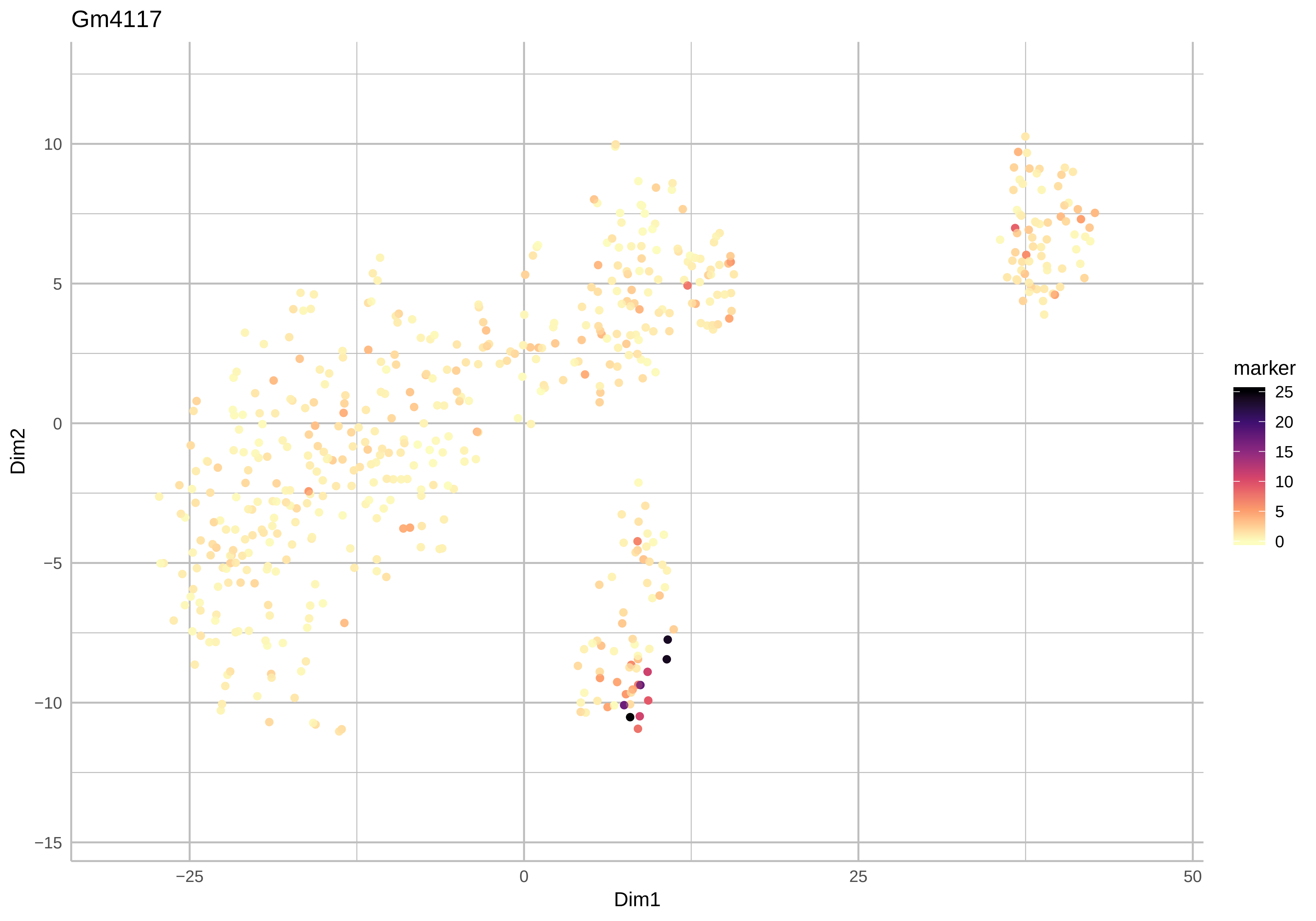
Gm4117
10
5
0
−5
−10
−15
Dim2
−25
0
25
50
Dim1
marker
25
20
15
10
5
0

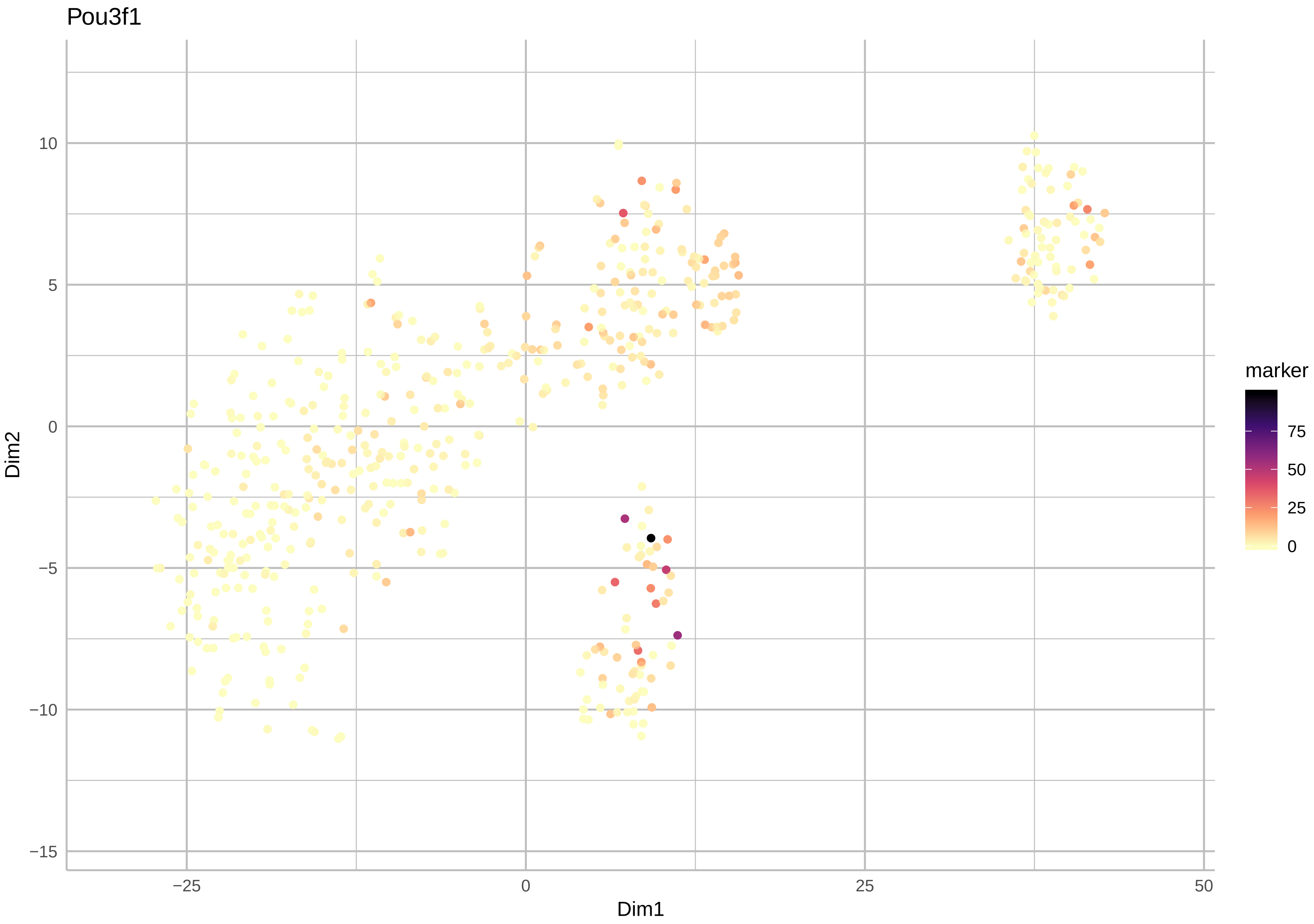
Pou3f1
Dim2
Dim1
10
5
0
−5
−10
−15
−25
0
25
50
marker
75
50
25
0

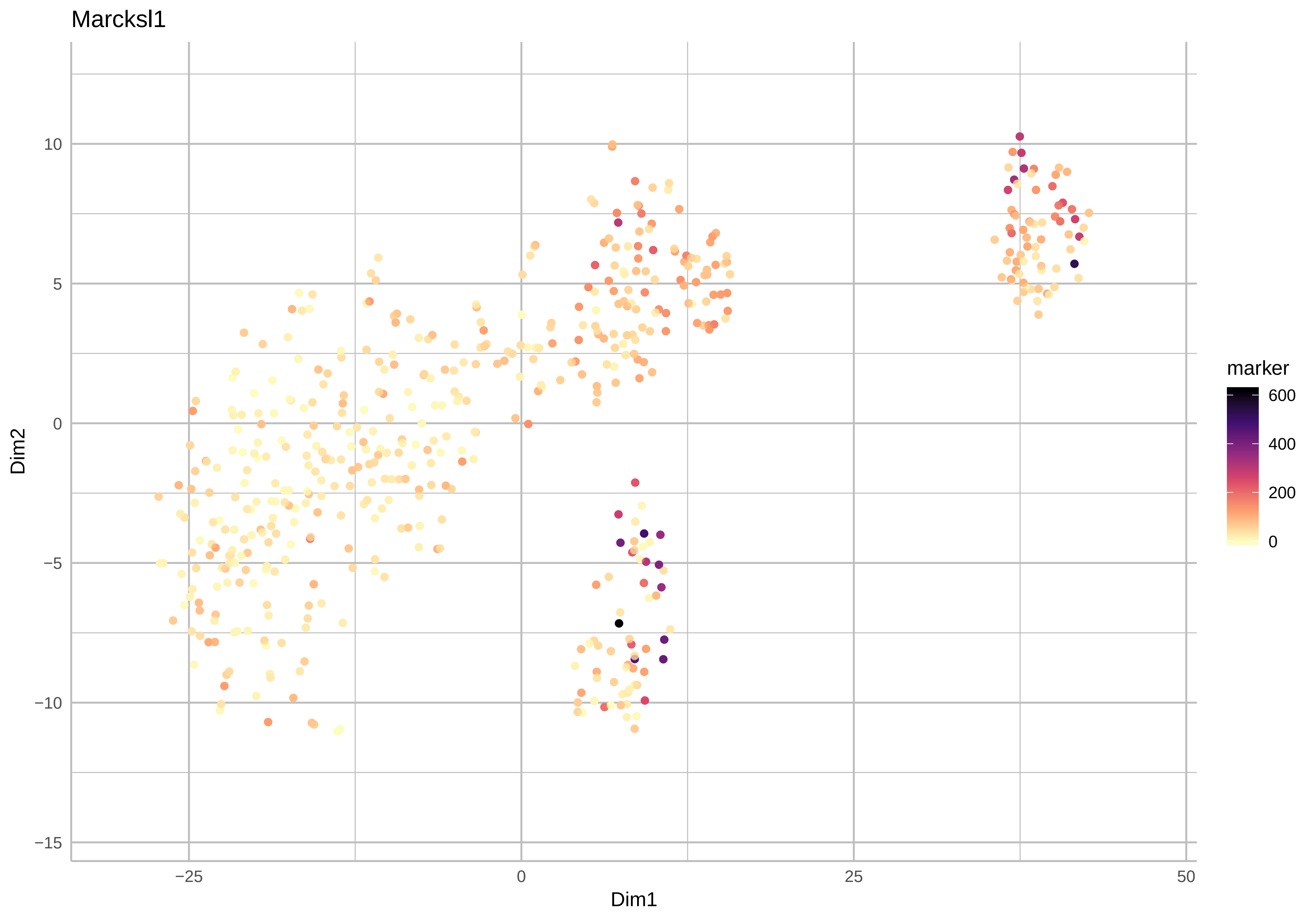
Marcksl1
Dim2
Dim1
10
5
0
−5
−10
−15
−25
0
25
50
marker
600
400
200
0

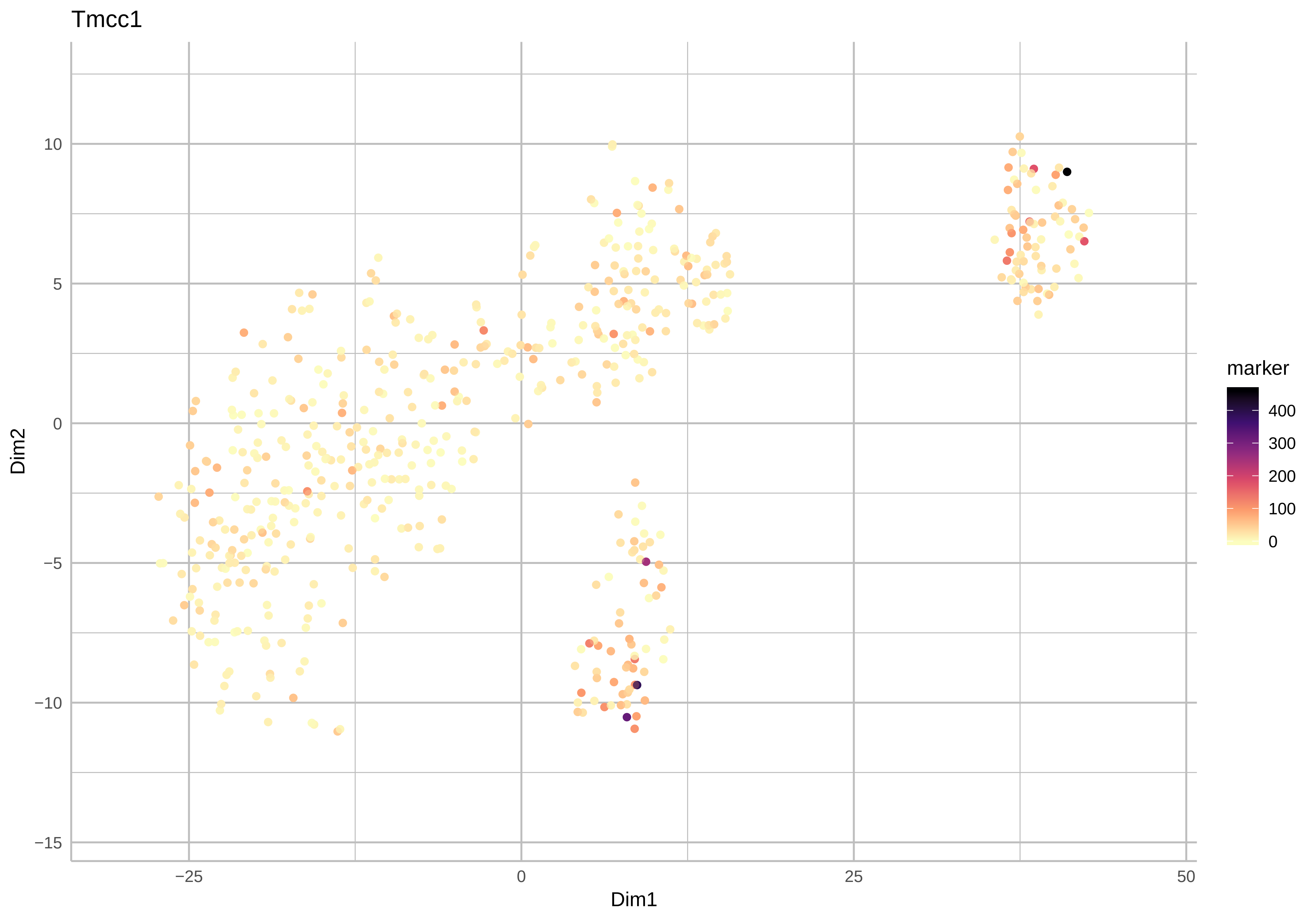

Tmcc1
Dim2
Dim1
10
5
0
−5
−10
−15
−25
0
25
50
marker
400
300
200
100
0

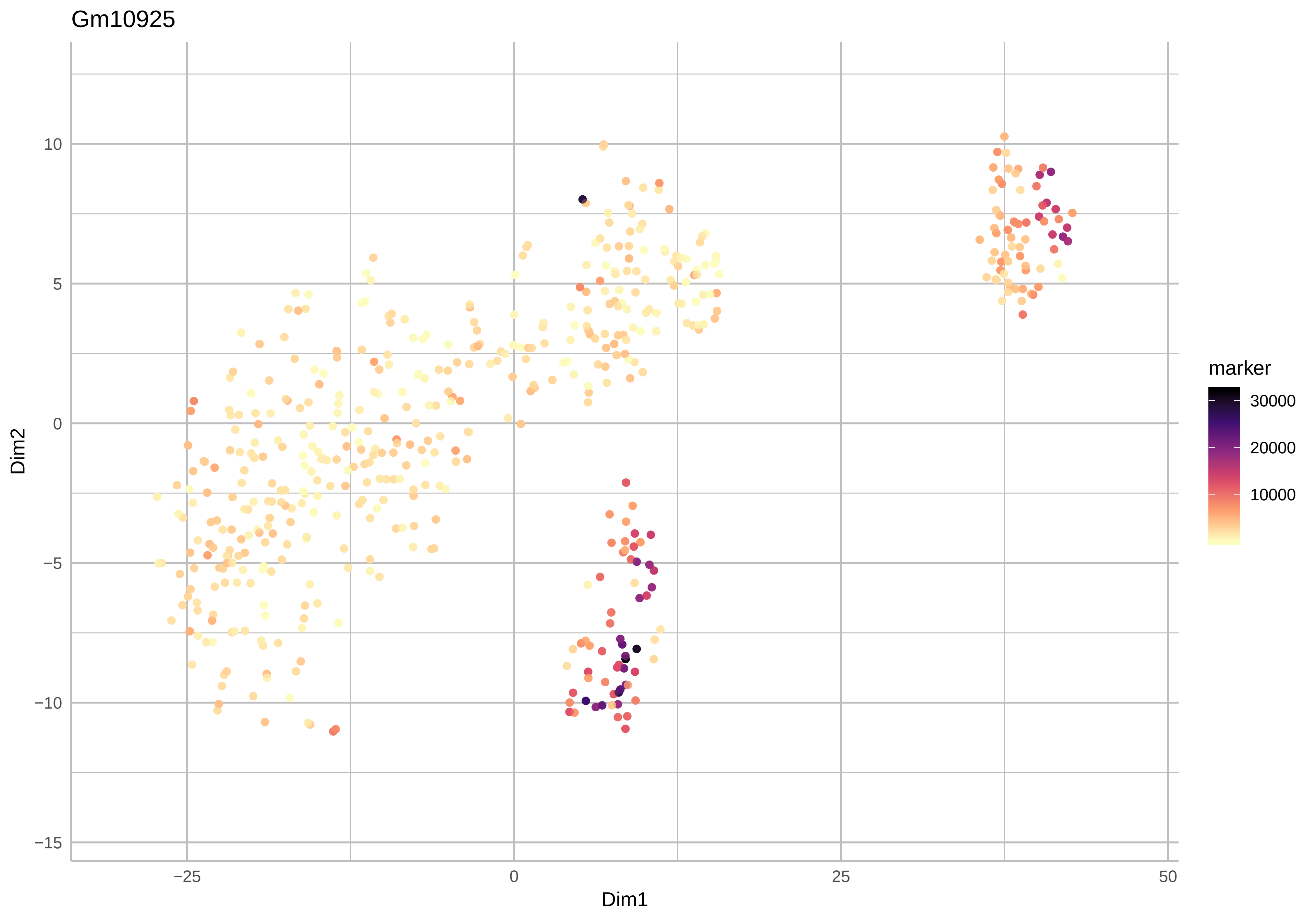
Gm10925
Dim2
Dim1
10
5
0
−5
−10
−15
−25
0
25
50
marker
30000
20000
10000

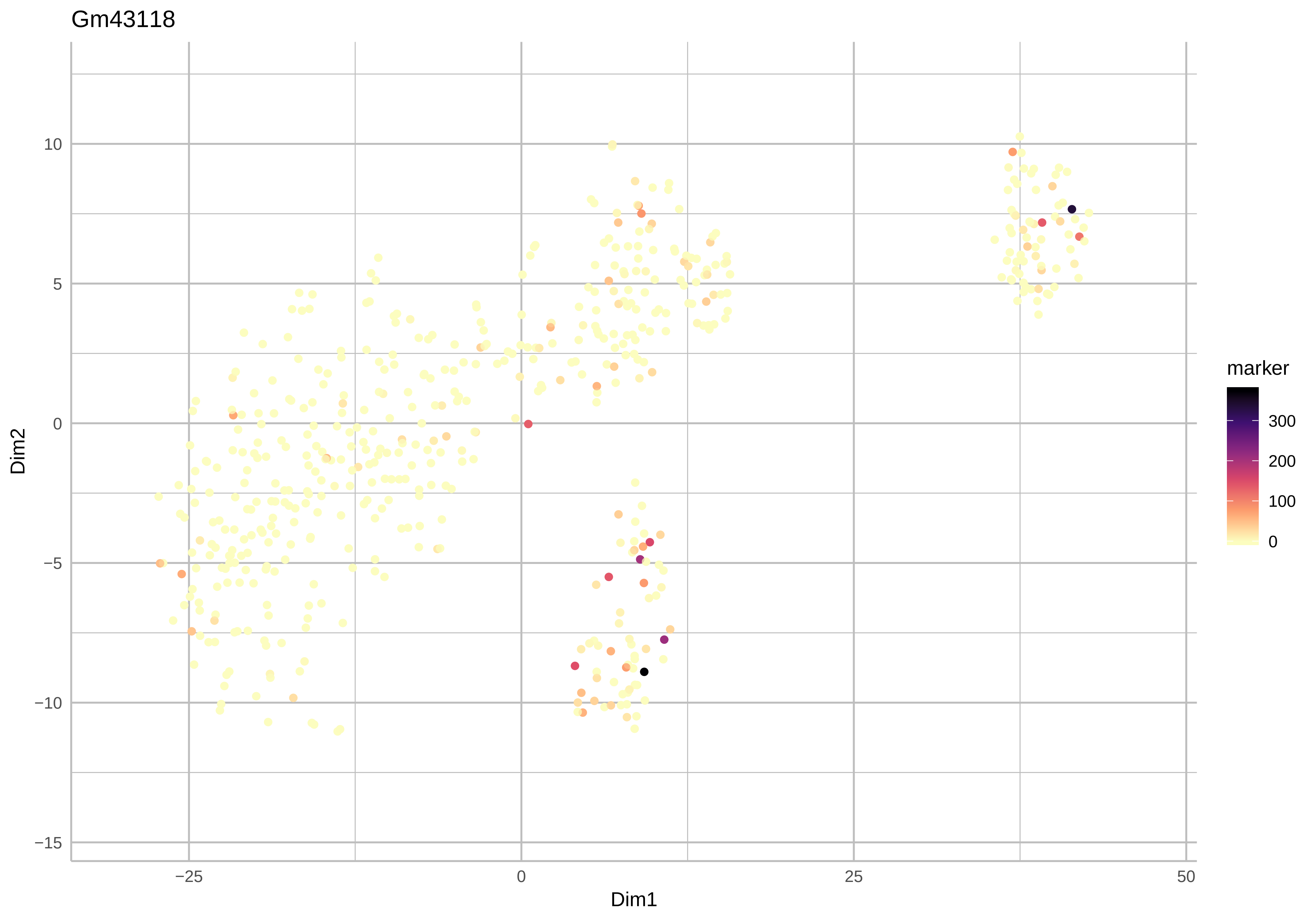
Gm43118
Dim1
Dim2
−25
0
25
50
10
5
0
−5
−10
−15
marker
300
200
100
0

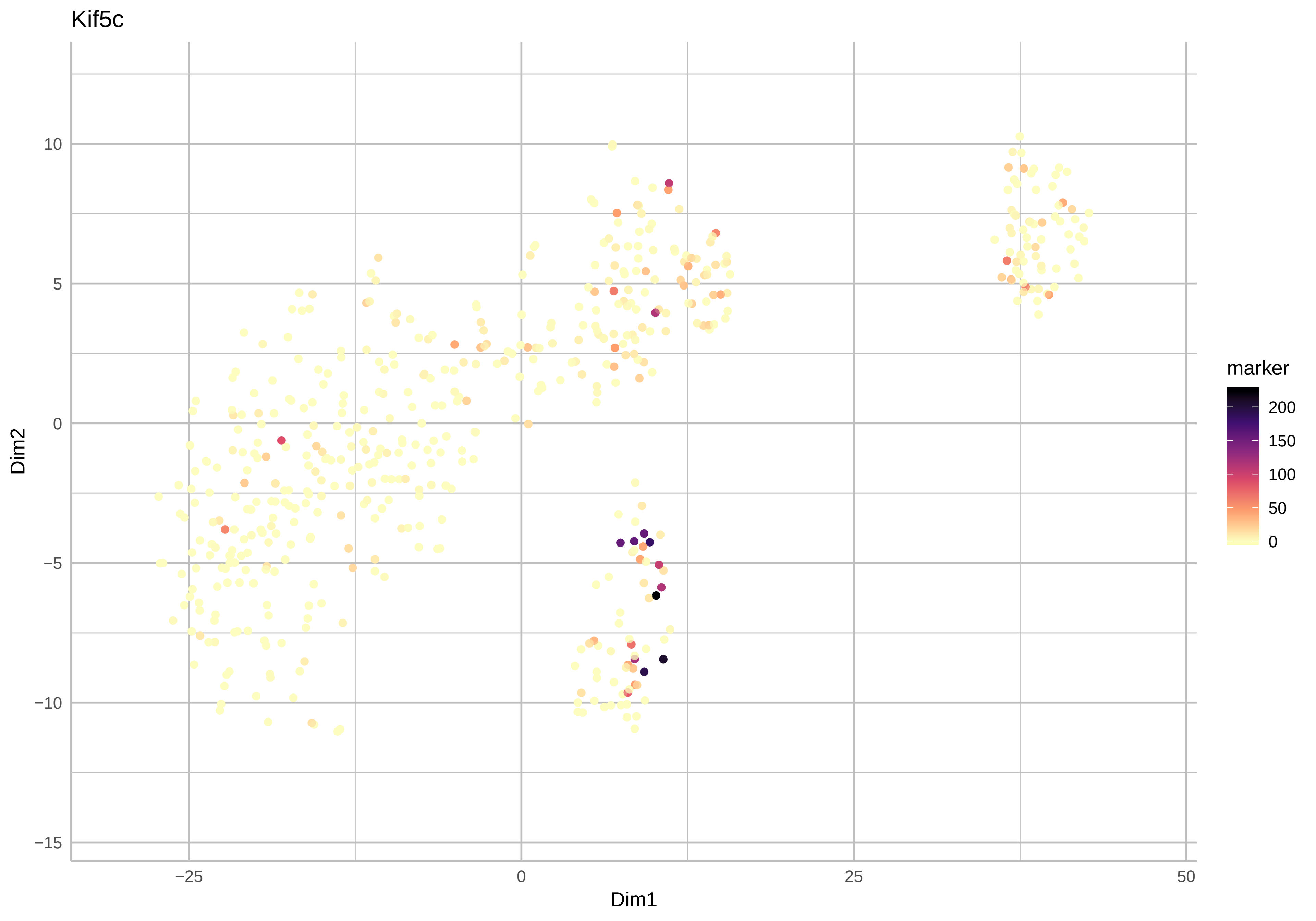

Kif5c
Dim2
Dim1
10
5
0
−5
−10
−15
−25
0
25
50
marker
200
150
100
50
0

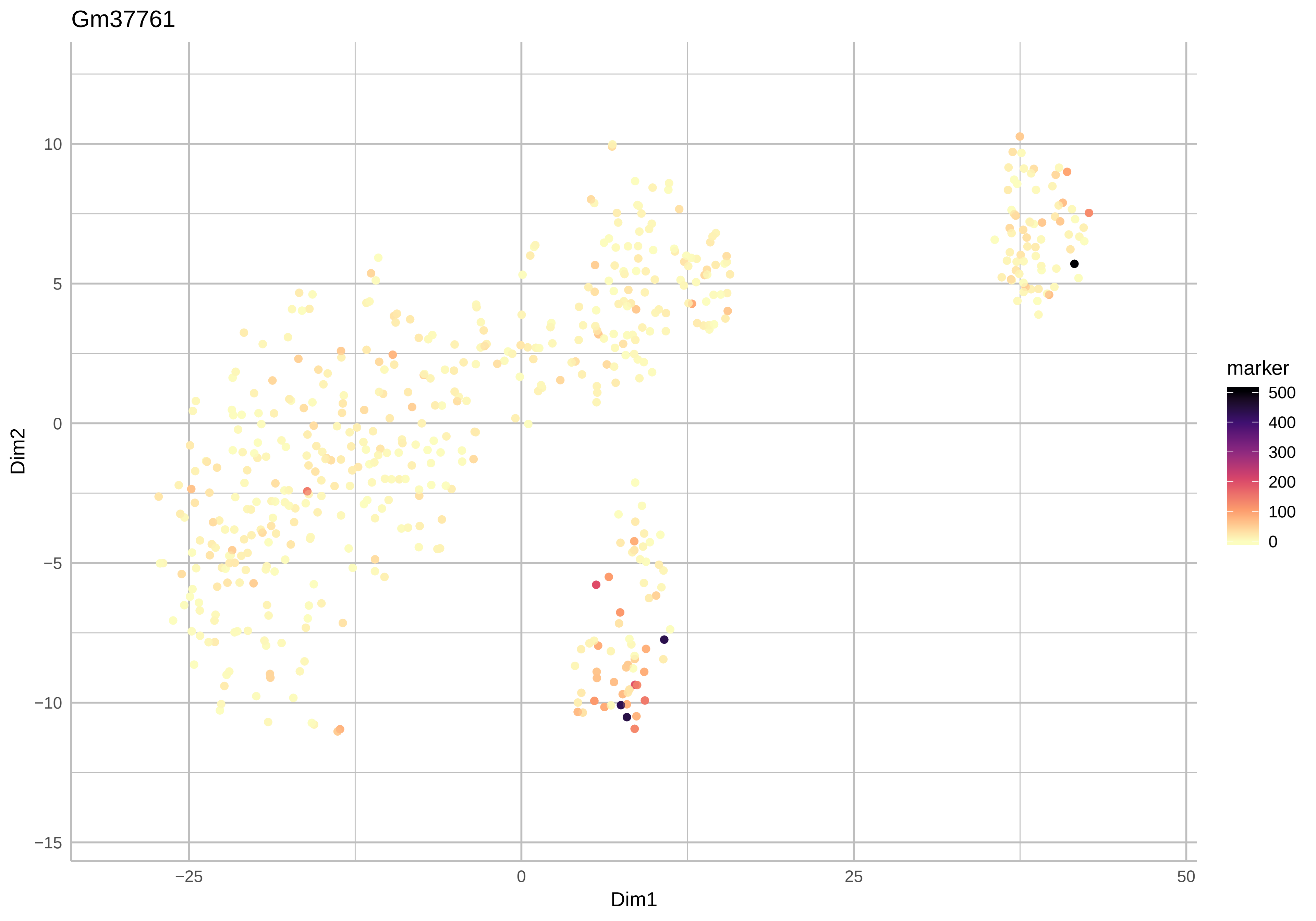

Gm37761
marker
500
400
300
200
100
0
Dim2
10
5
0
−5
−10
−15
−25
0
25
50
Dim1

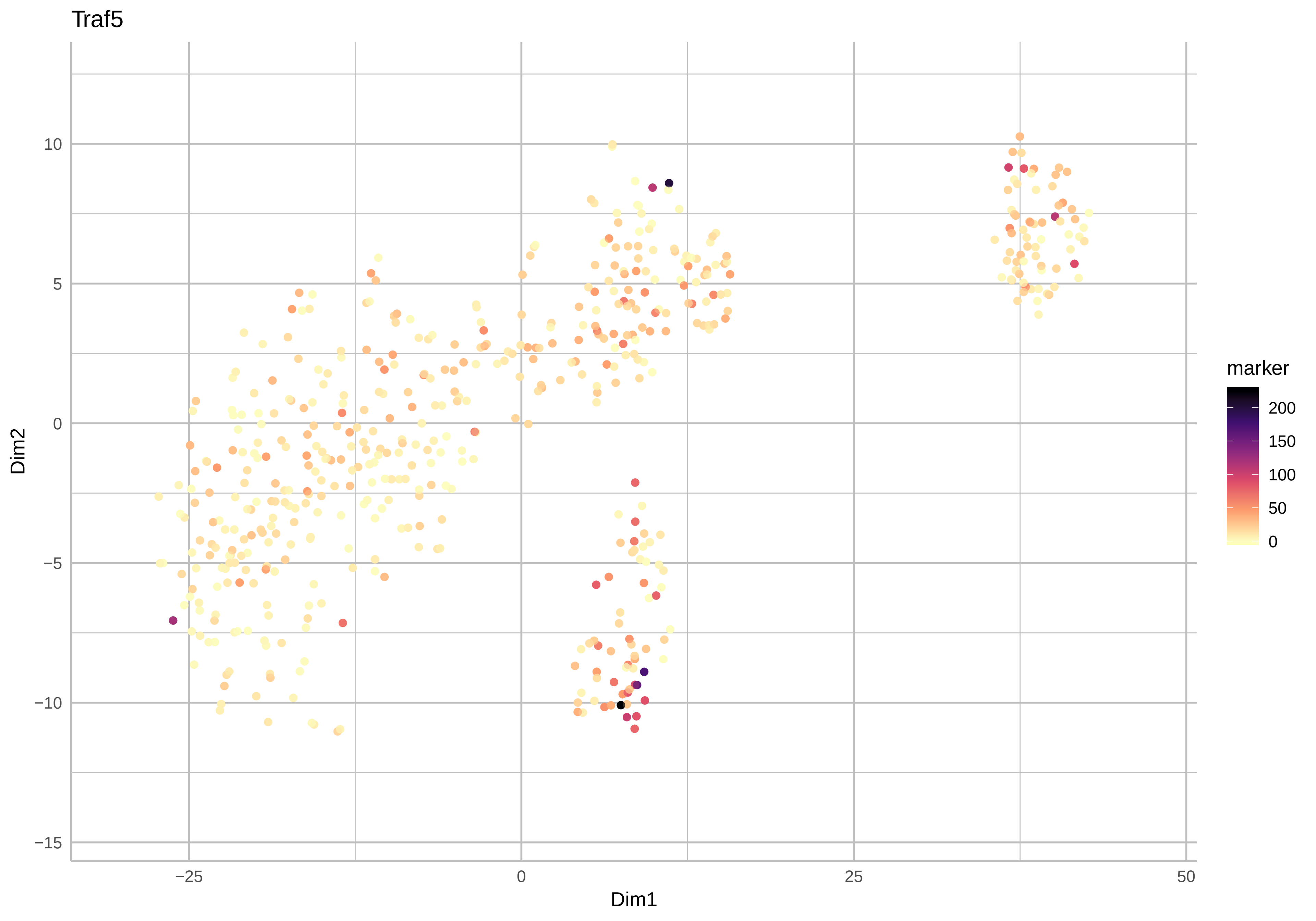

Traf5
Dim2
Dim1
10
5
0
−5
−10
−15
−25
0
25
50
marker
200
150
100
50
0

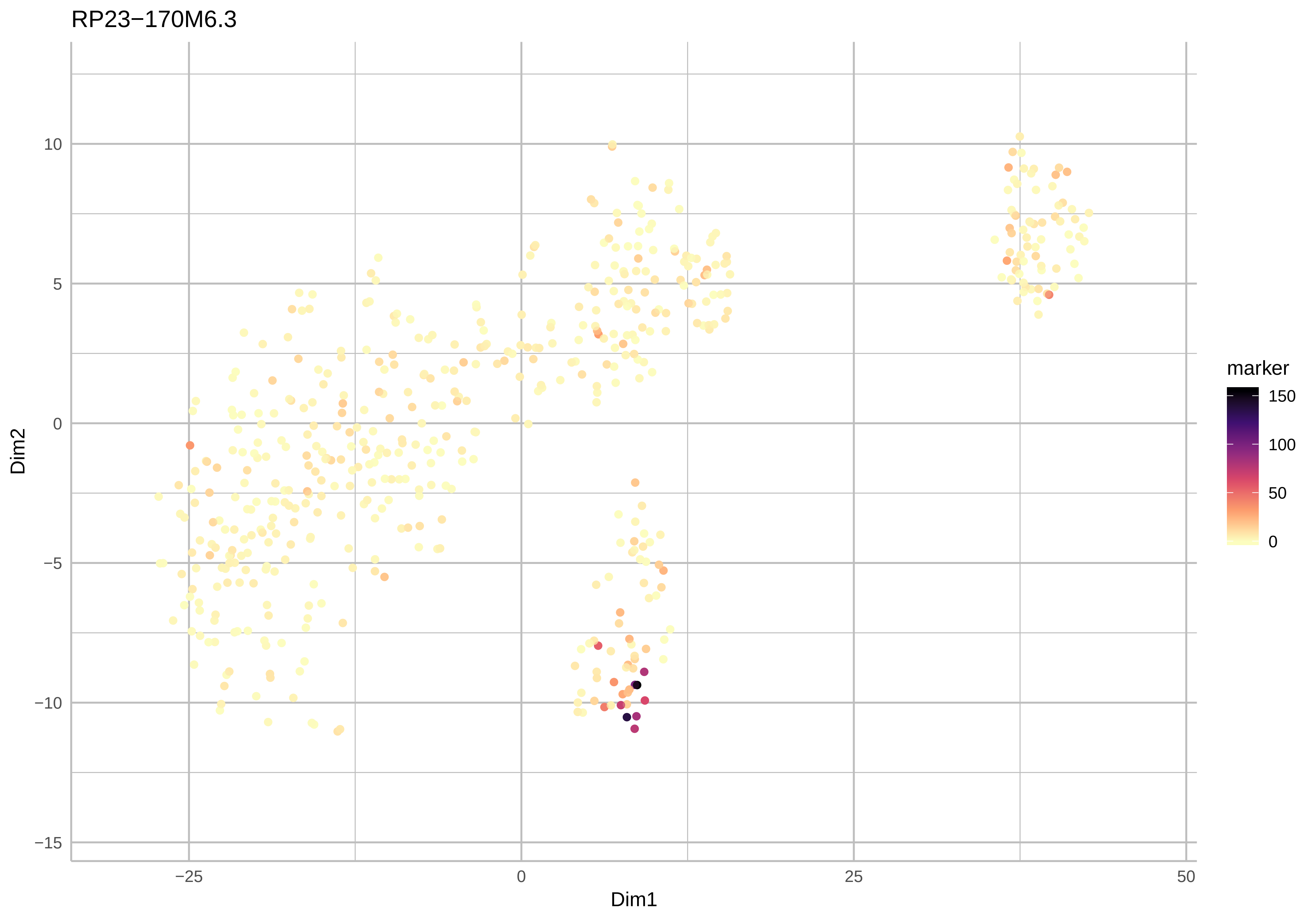
RP23−170M6.3
Dim2
Dim1
10
5
0
−5
−10
−15
−25
0
25
50
marker
150
100
50
0

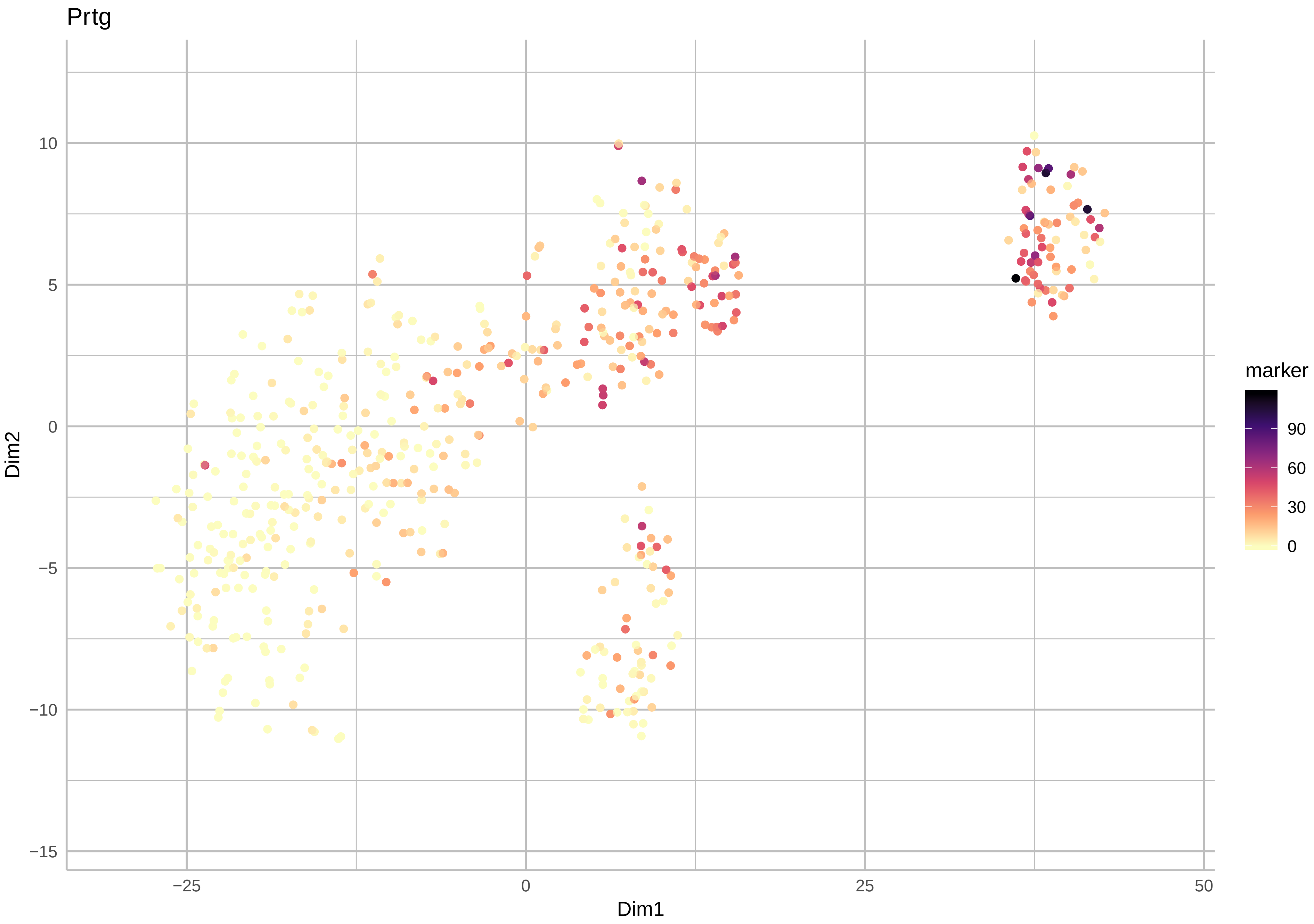

Prtg
10
5
0
−5
−10
−15
Dim2
−25
0
25
50
Dim1
marker
90
60
30
0

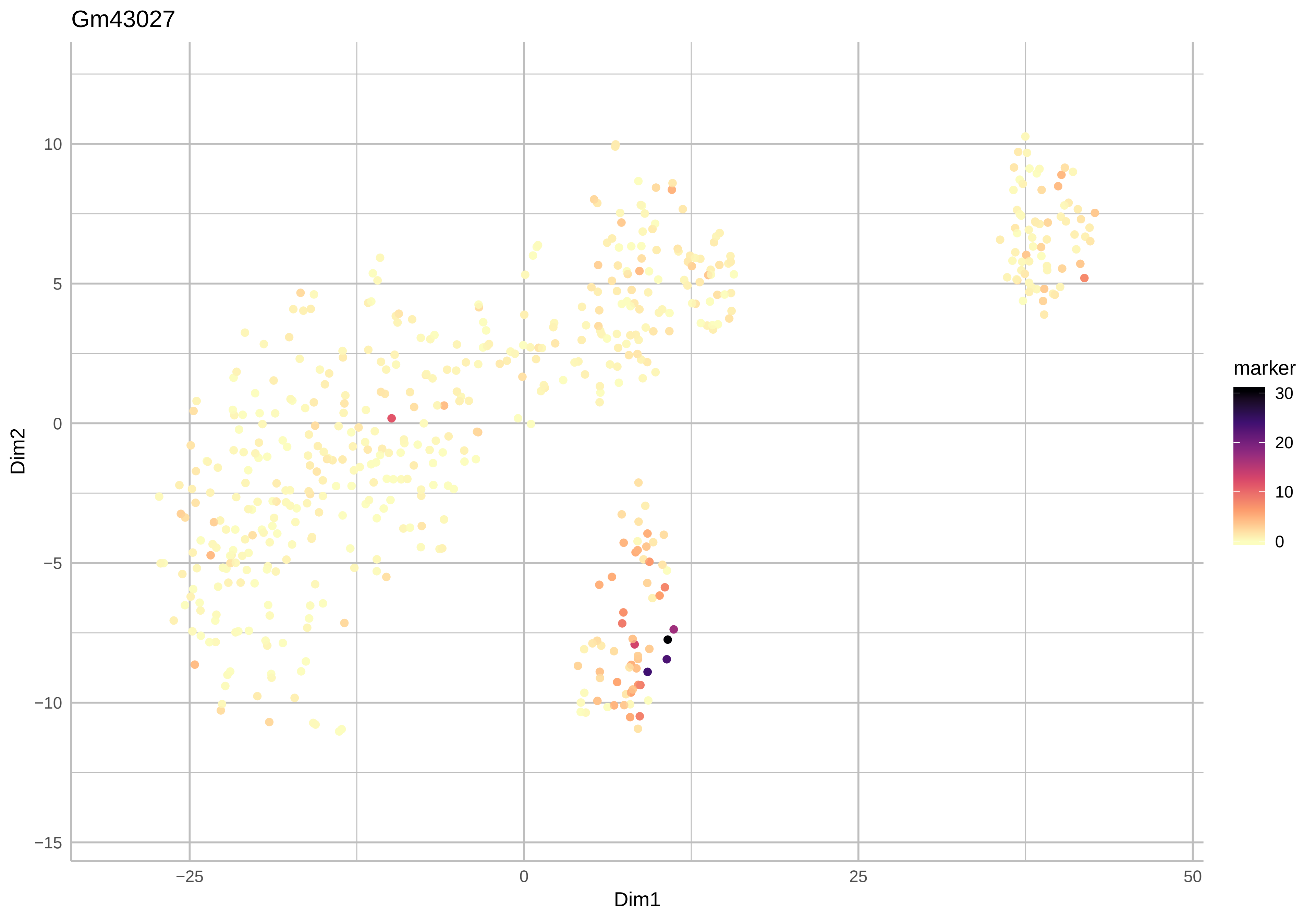
Gm43027
Dim2
Dim1
10
5
0
−5
−10
−15
−25
0
25
50
marker
30
20
10
0

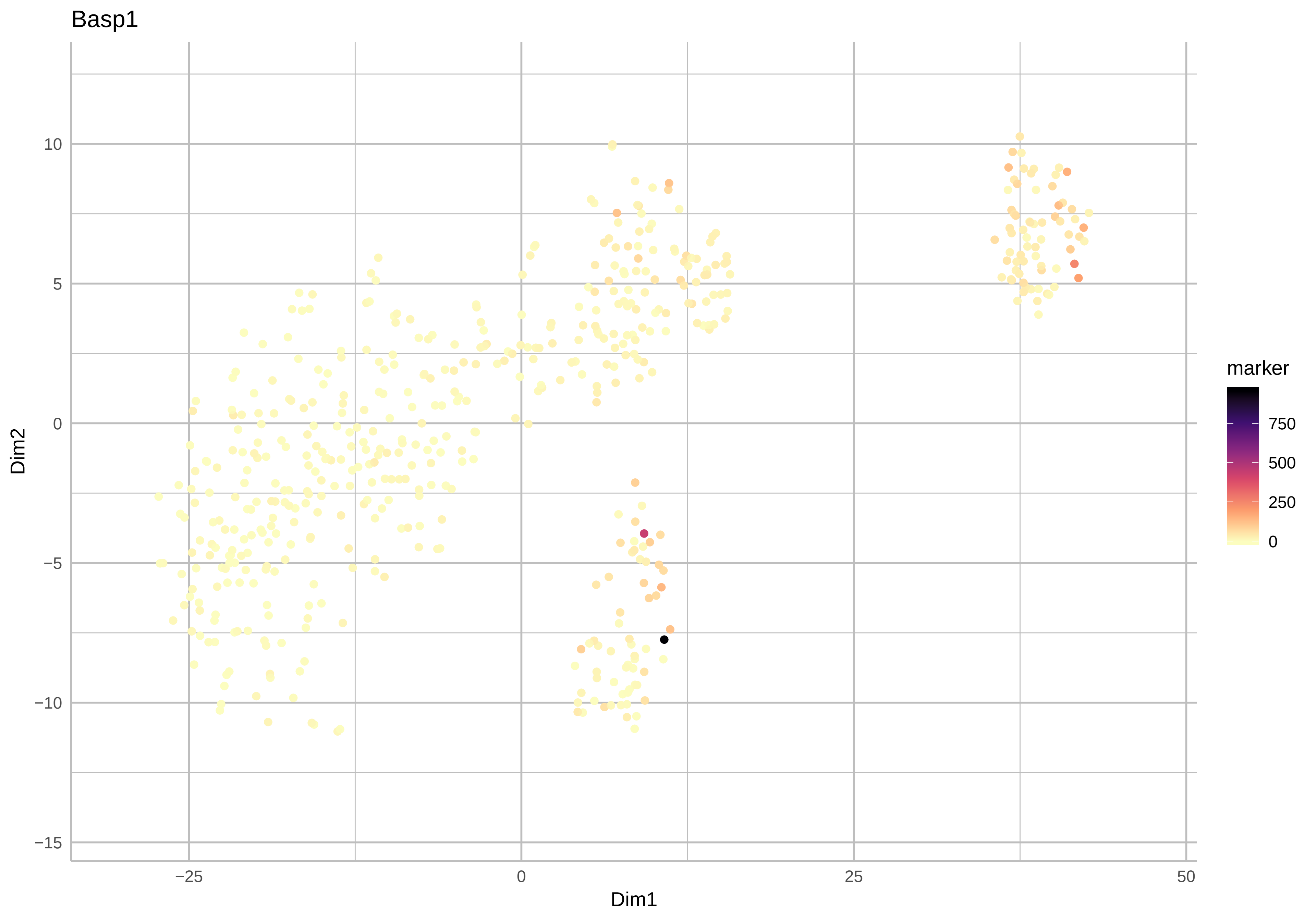

Basp1
10
5
0
−5
−10
−15
−25
0
25
50
Dim1
Dim2
marker
750
500
250
0

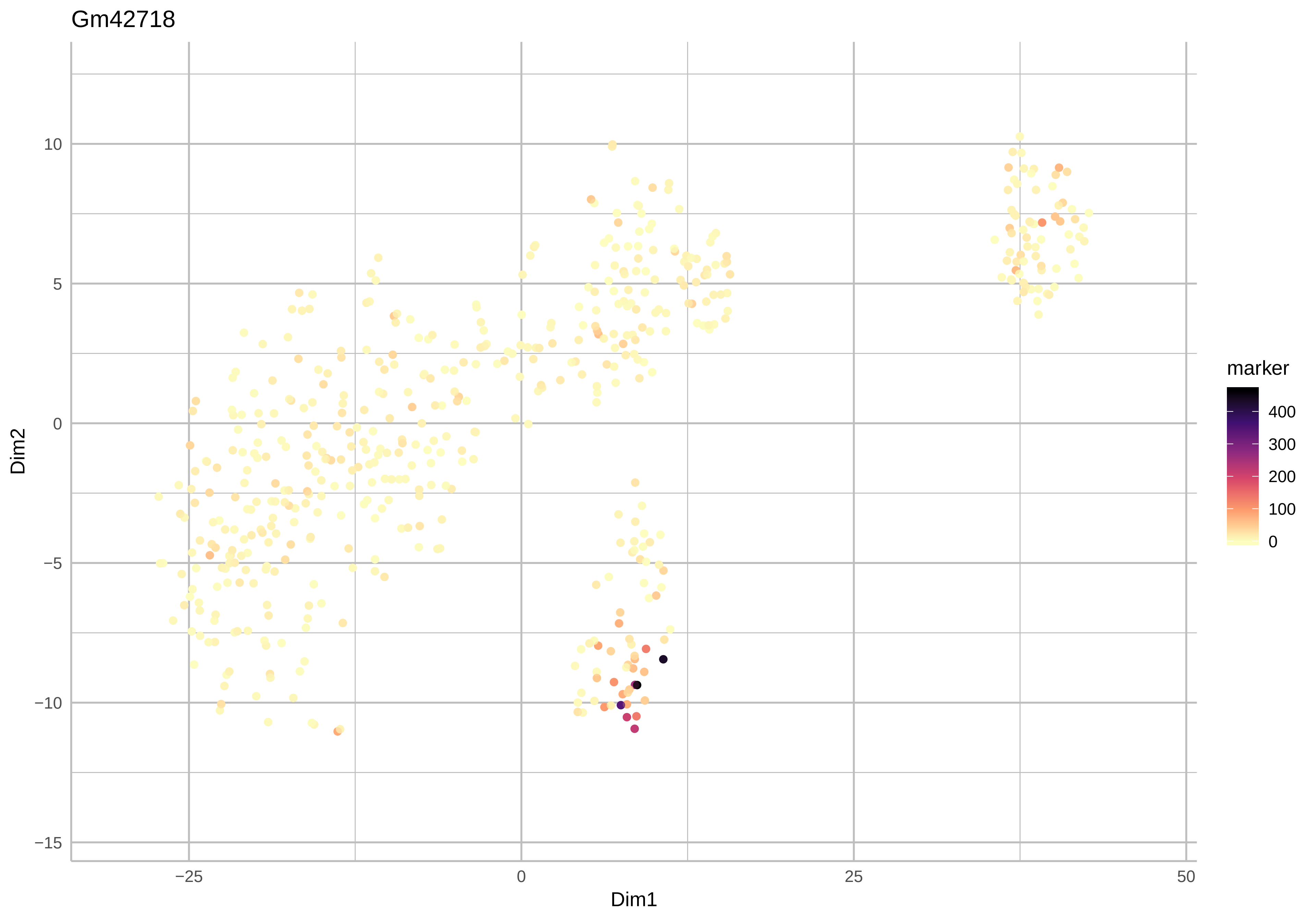

Gm42718
Dim2
10
5
0
−5
−10
−15
Dim1
−25
0
25
50
marker
400
300
200
100
0

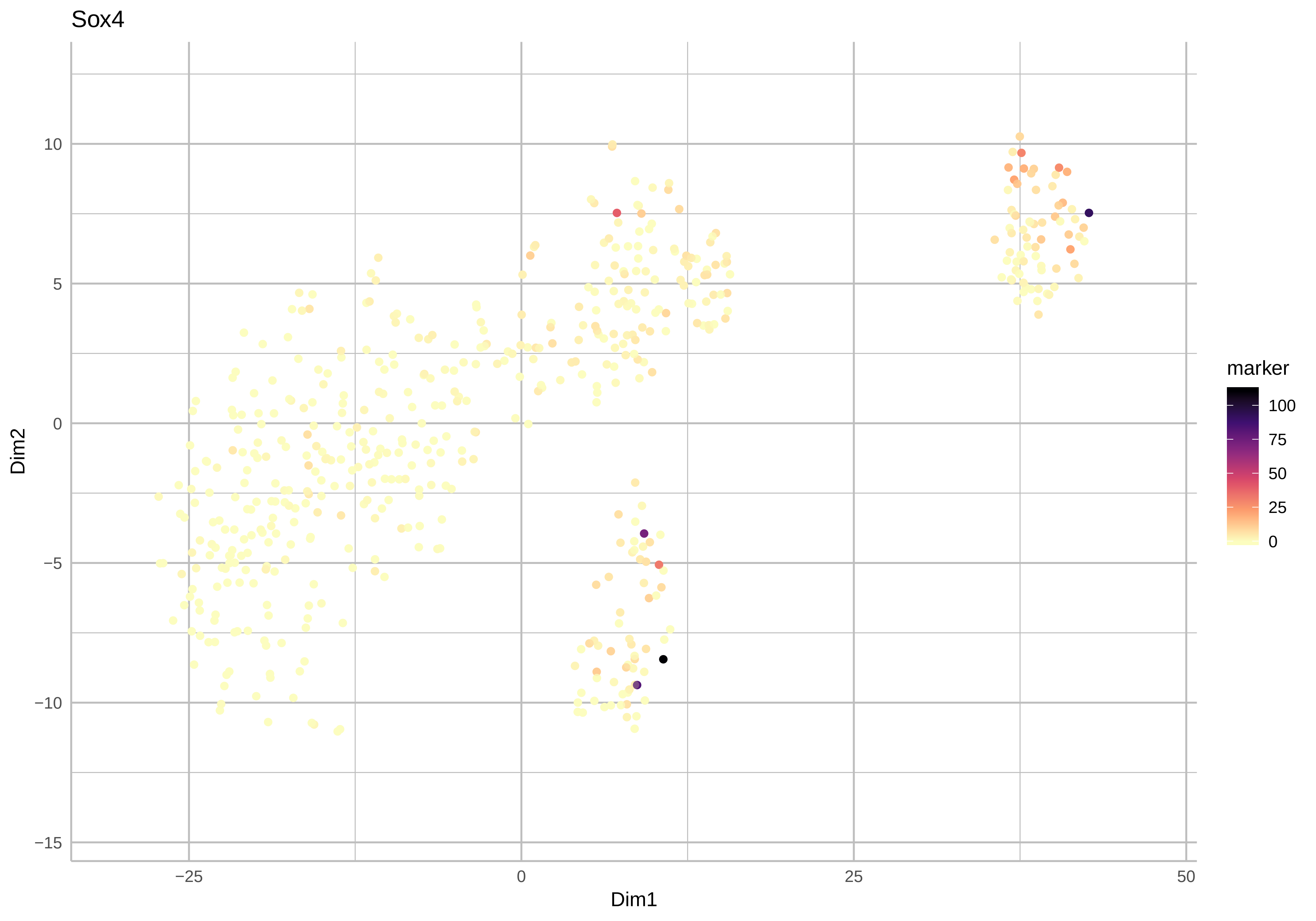
Sox4
Dim2
Dim1
10
5
0
−5
−10
−15
−25
0
25
50
marker
100
75
50
25
0

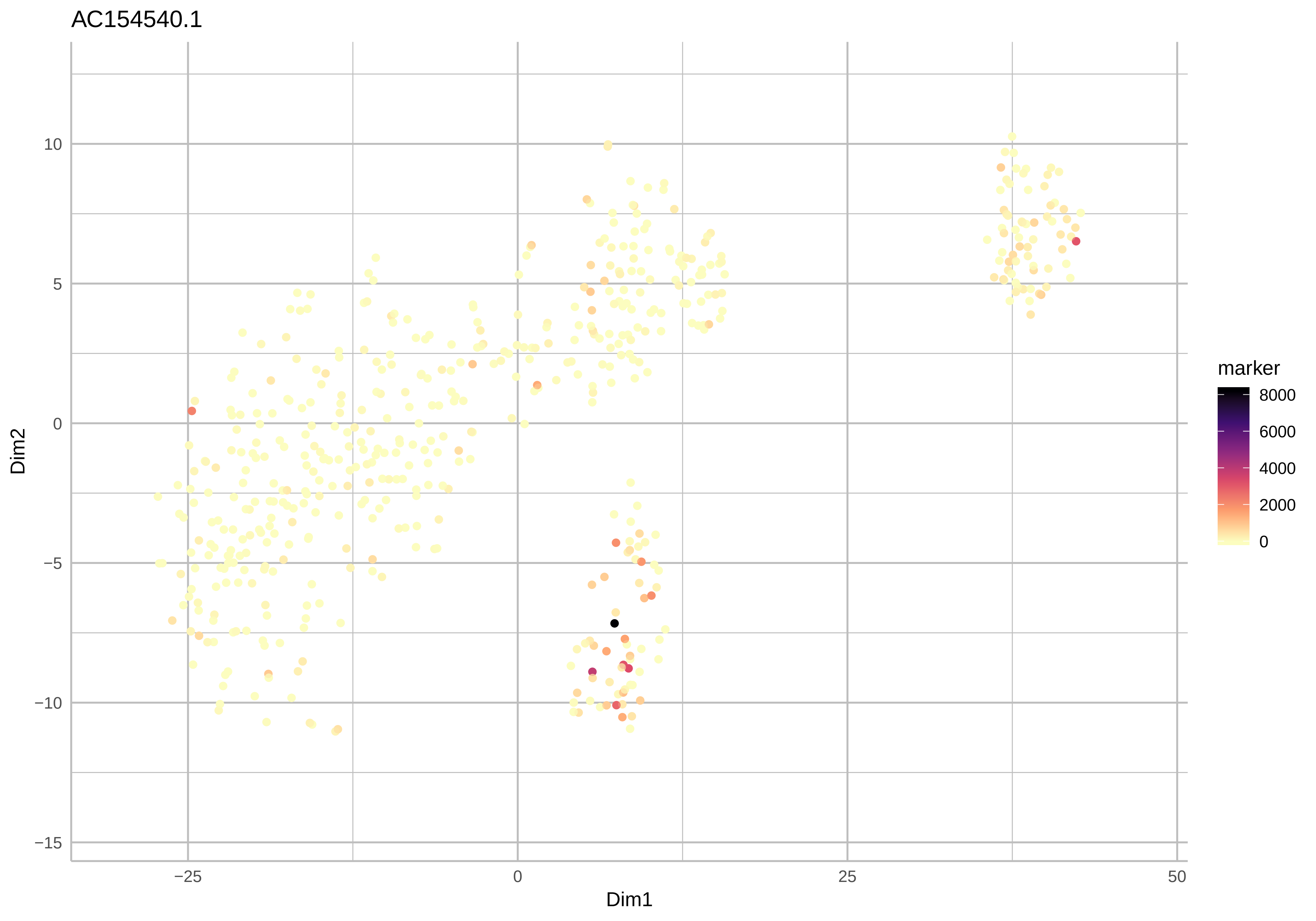
AC154540.1
Dim2
10
5
0
−5
−10
−15
−25
0
25
50
Dim1
marker
8000
6000
4000
2000
0

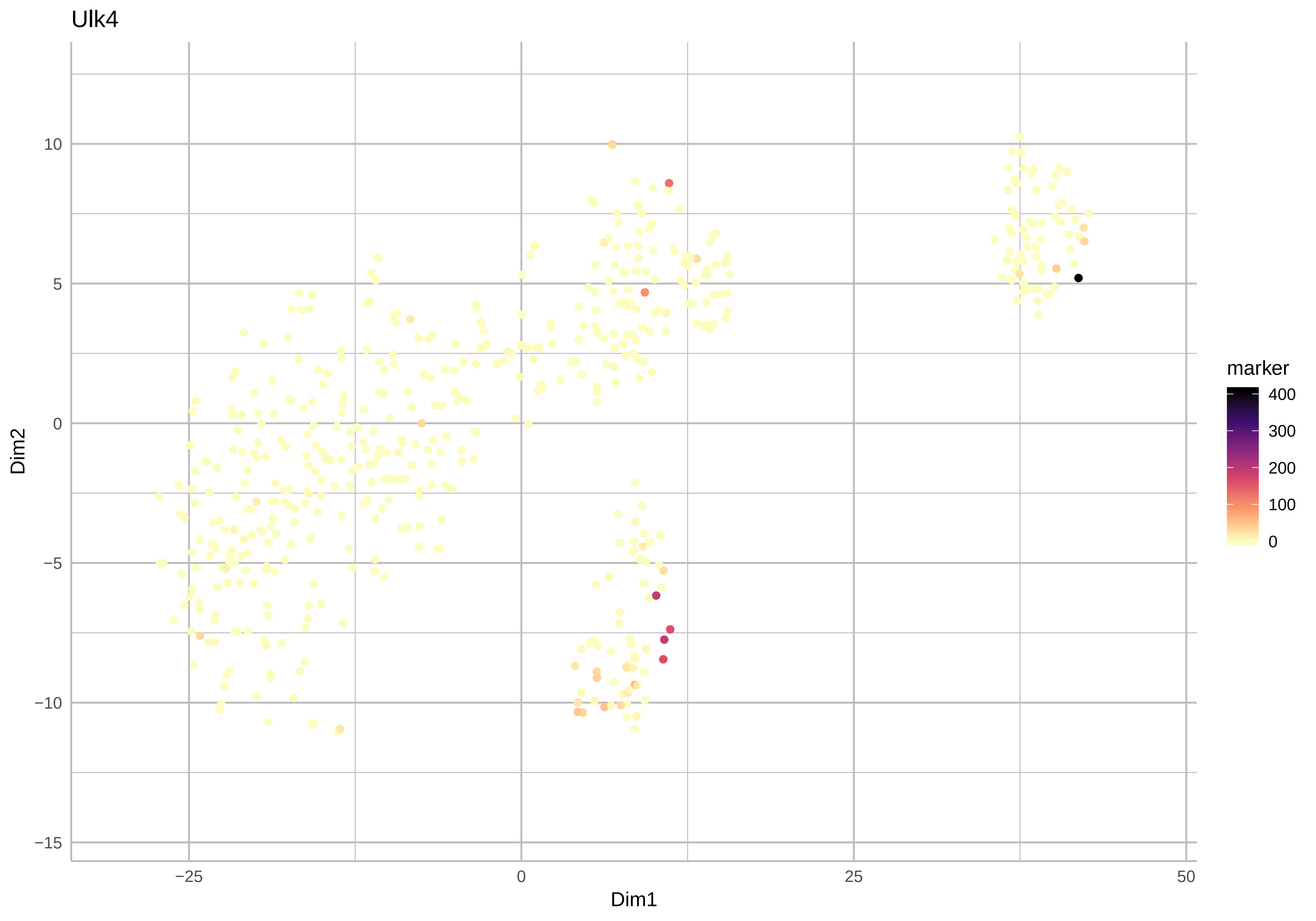
Ulk4
Dim2
10
5
0
−5
−10
−15
−25
0
25
50
Dim1
marker
400
300
200
100
0

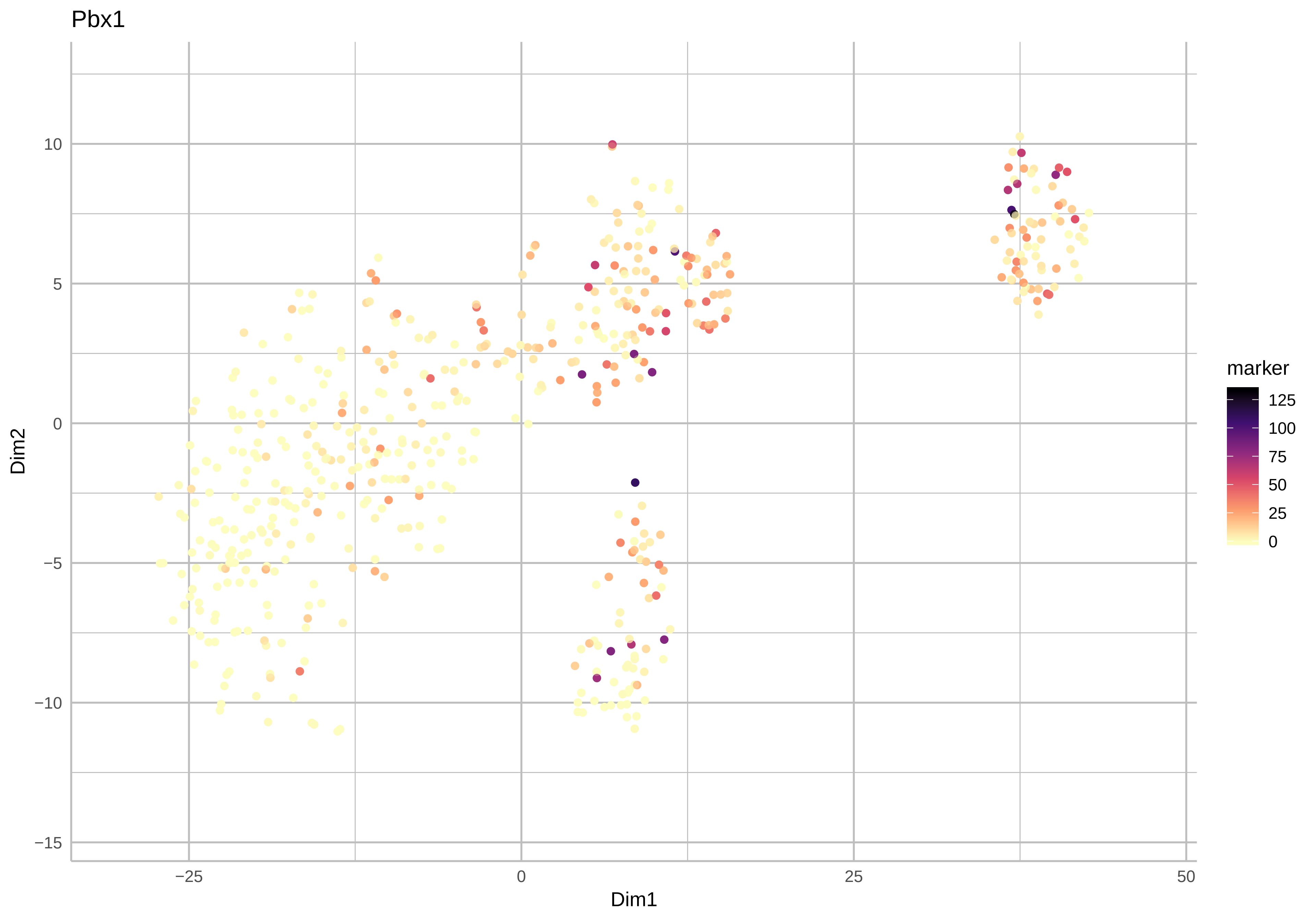

Pbx1
Dim2
Dim1
marker
125
100
75
50
25
0
10
5
0
−5
−10
−15
−25
0
25
50

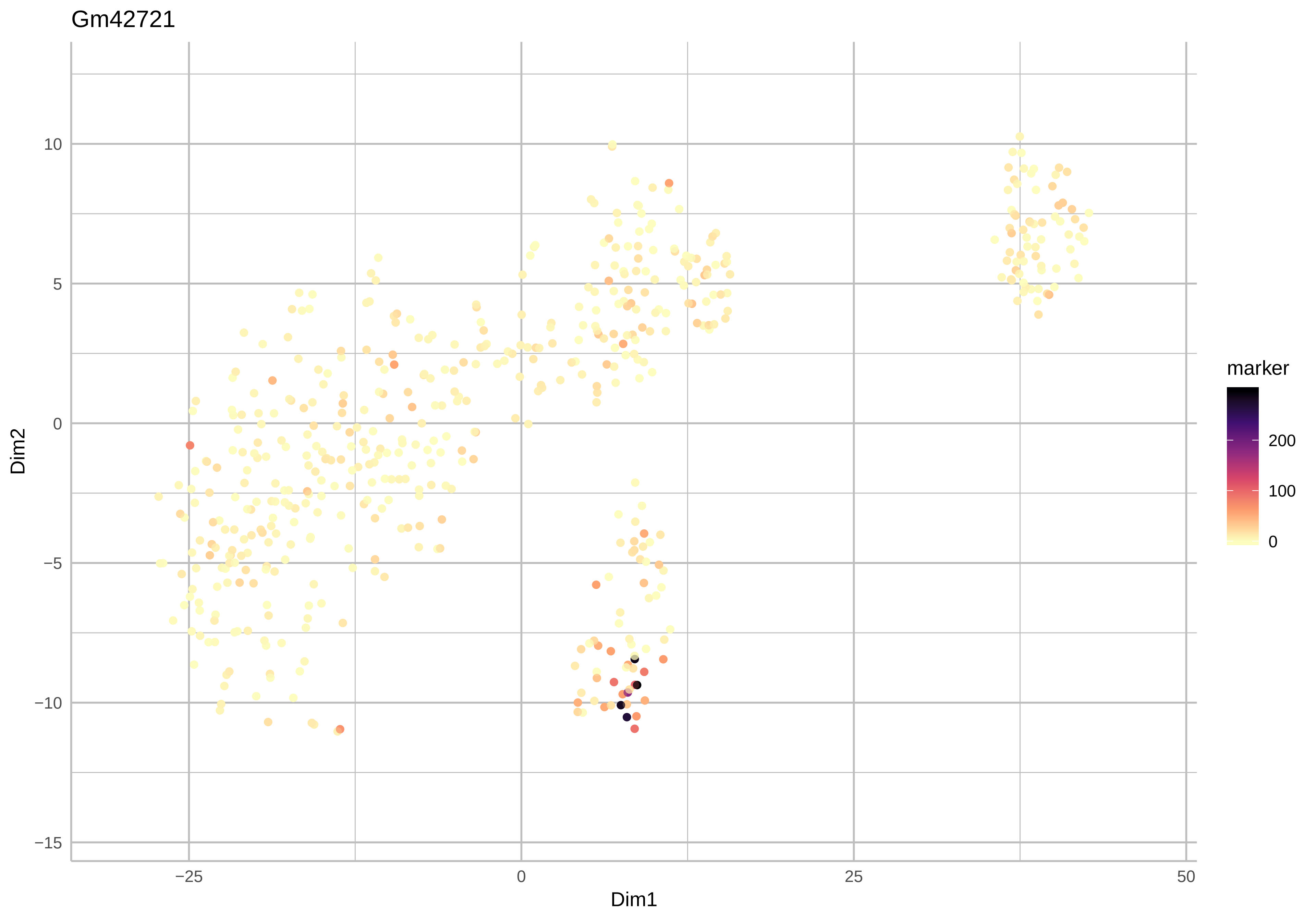
Gm42721
Dim2
Dim1
10
5
0
−5
−10
−15
−25
0
25
50
marker
200
100
0

Gm42566

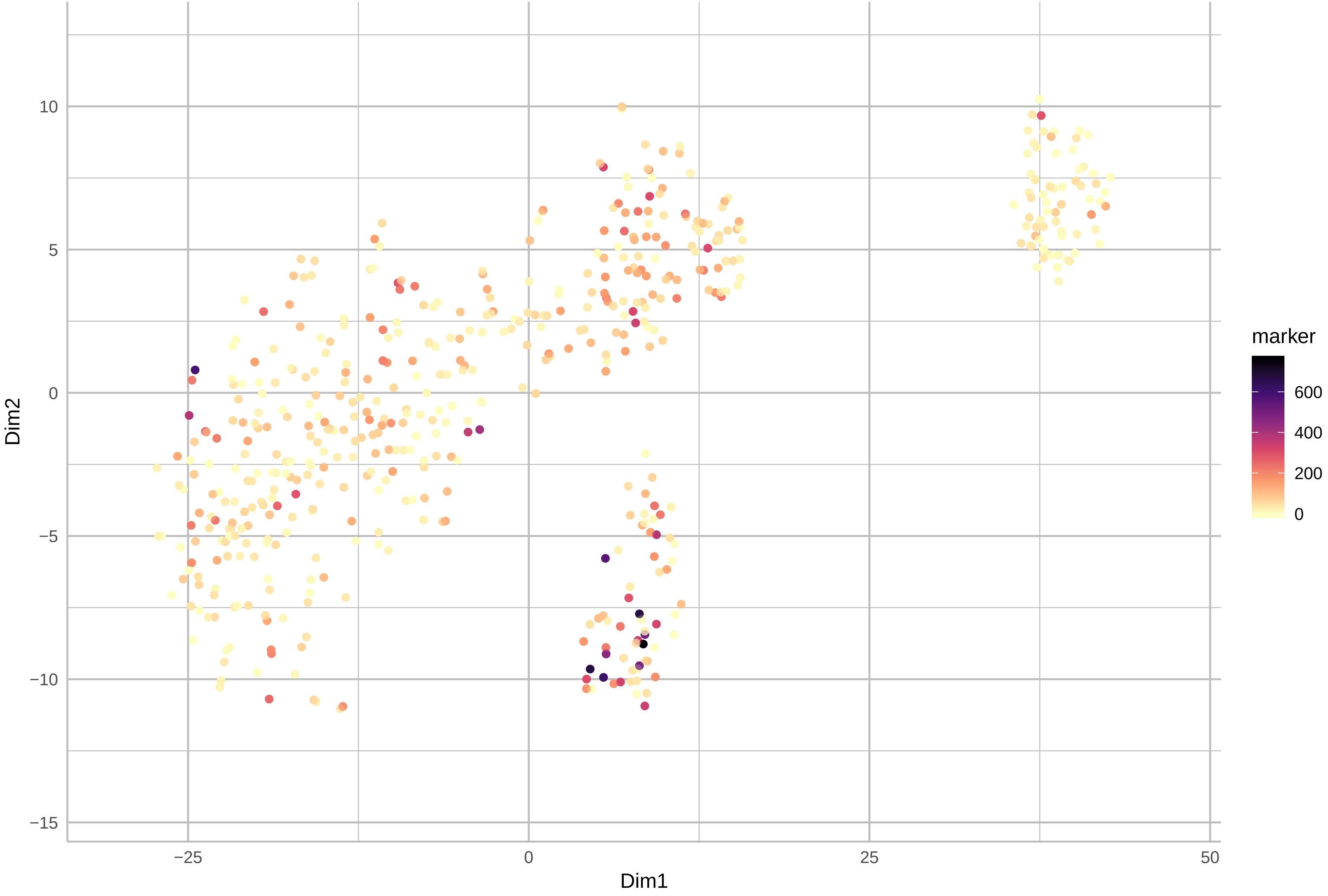

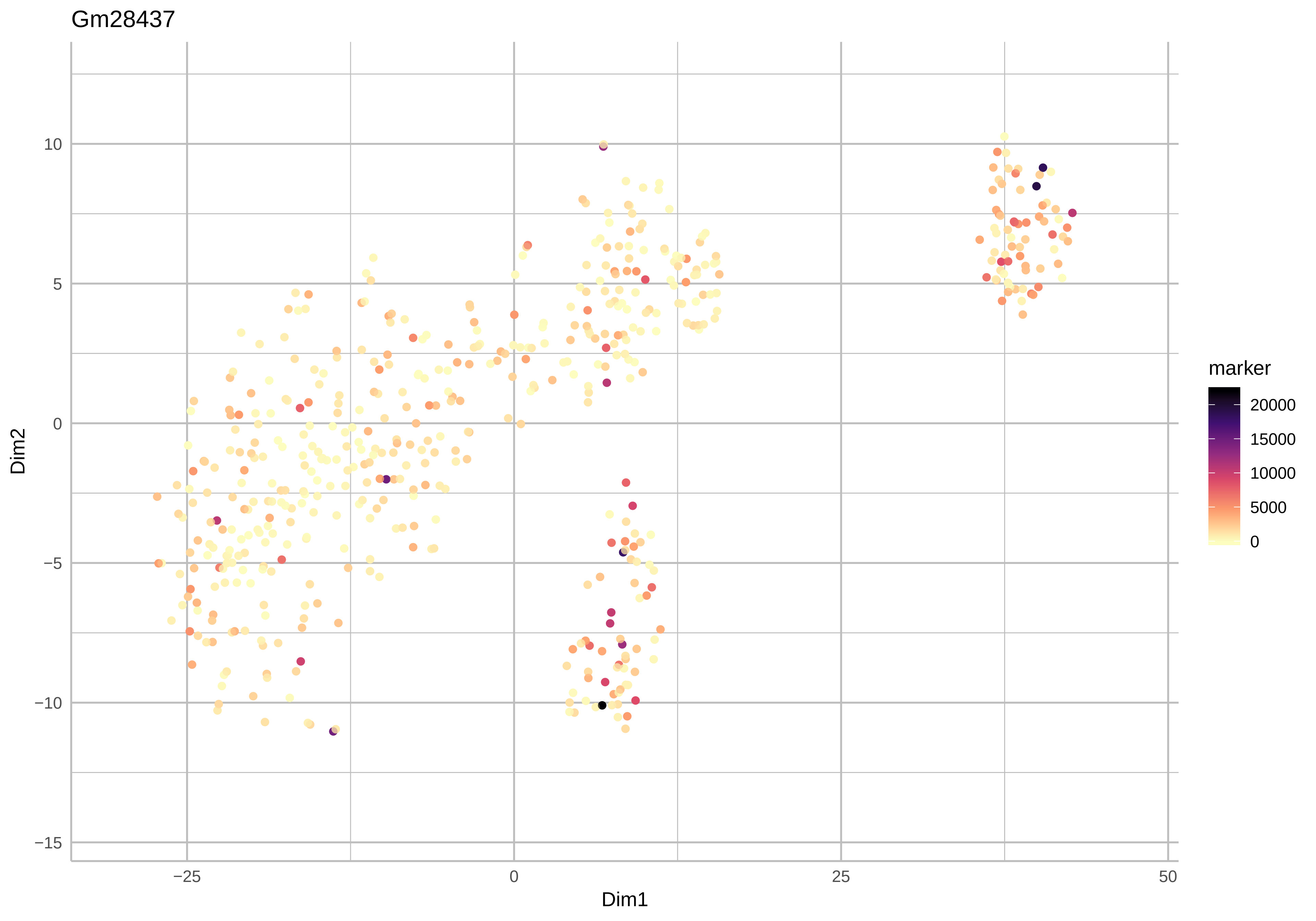

Gm28437
10
5
0
−5
−10
−15
Dim2
−25
0
25
50
Dim1
marker
20000
15000
10000
5000
0

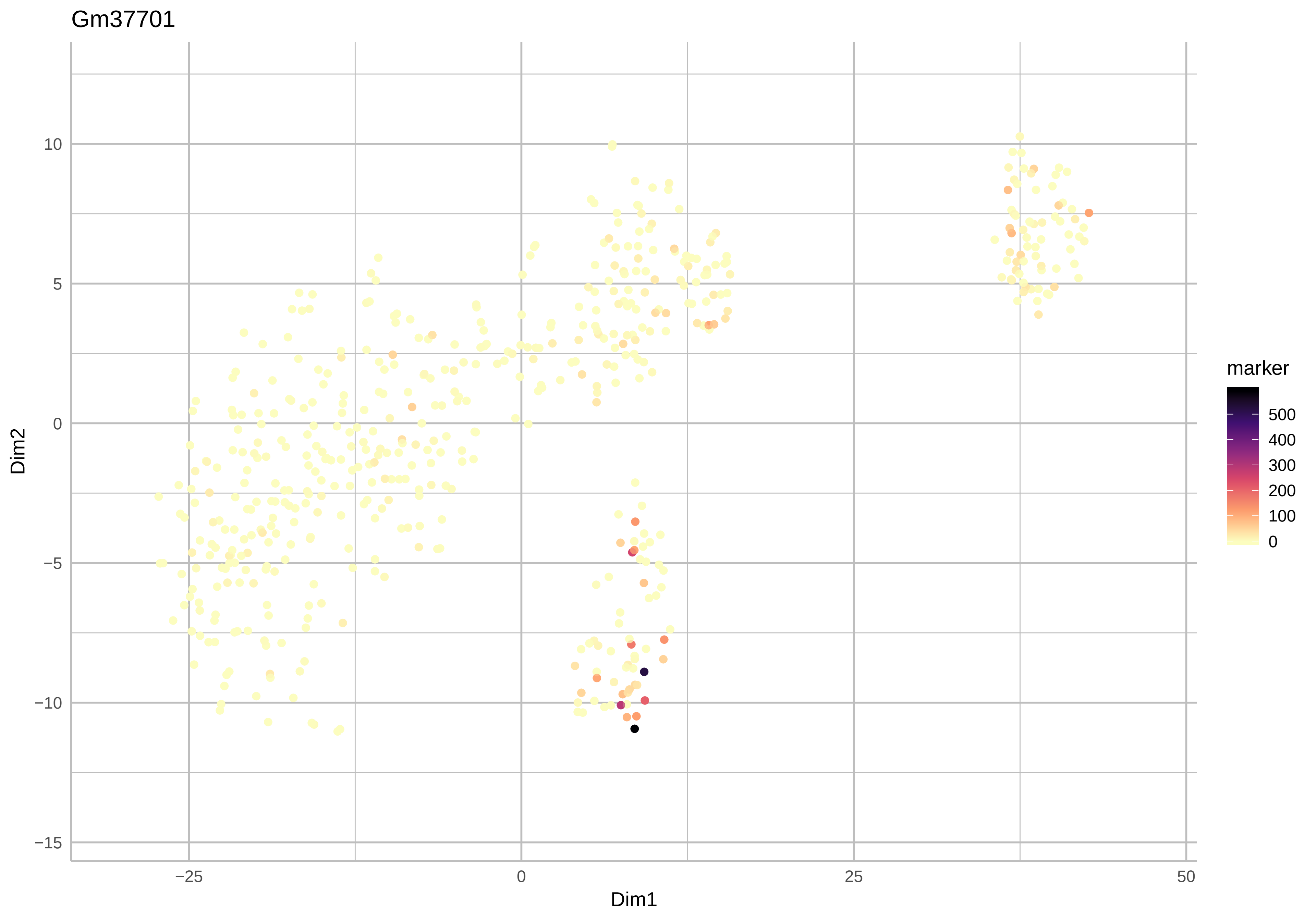

Gm37701
Dim2
Dim1
10
5
0
−5
−10
−15
−25
0
25
50
marker
500
400
300
200
100
0

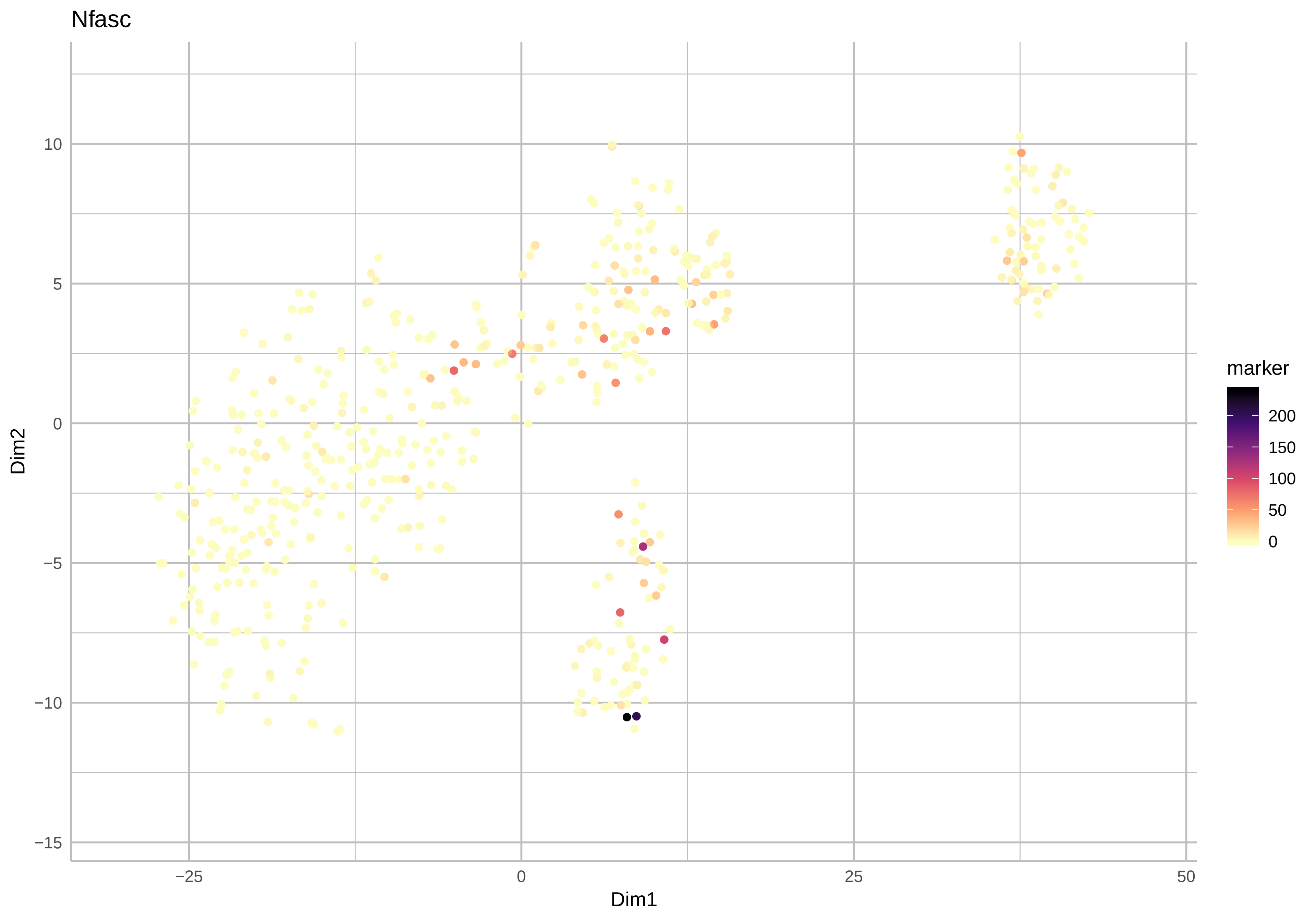
Nfasc
Dim2
Dim1
10
5
0
−5
−10
−15
−25
0
25
50
marker
200
150
100
50
0

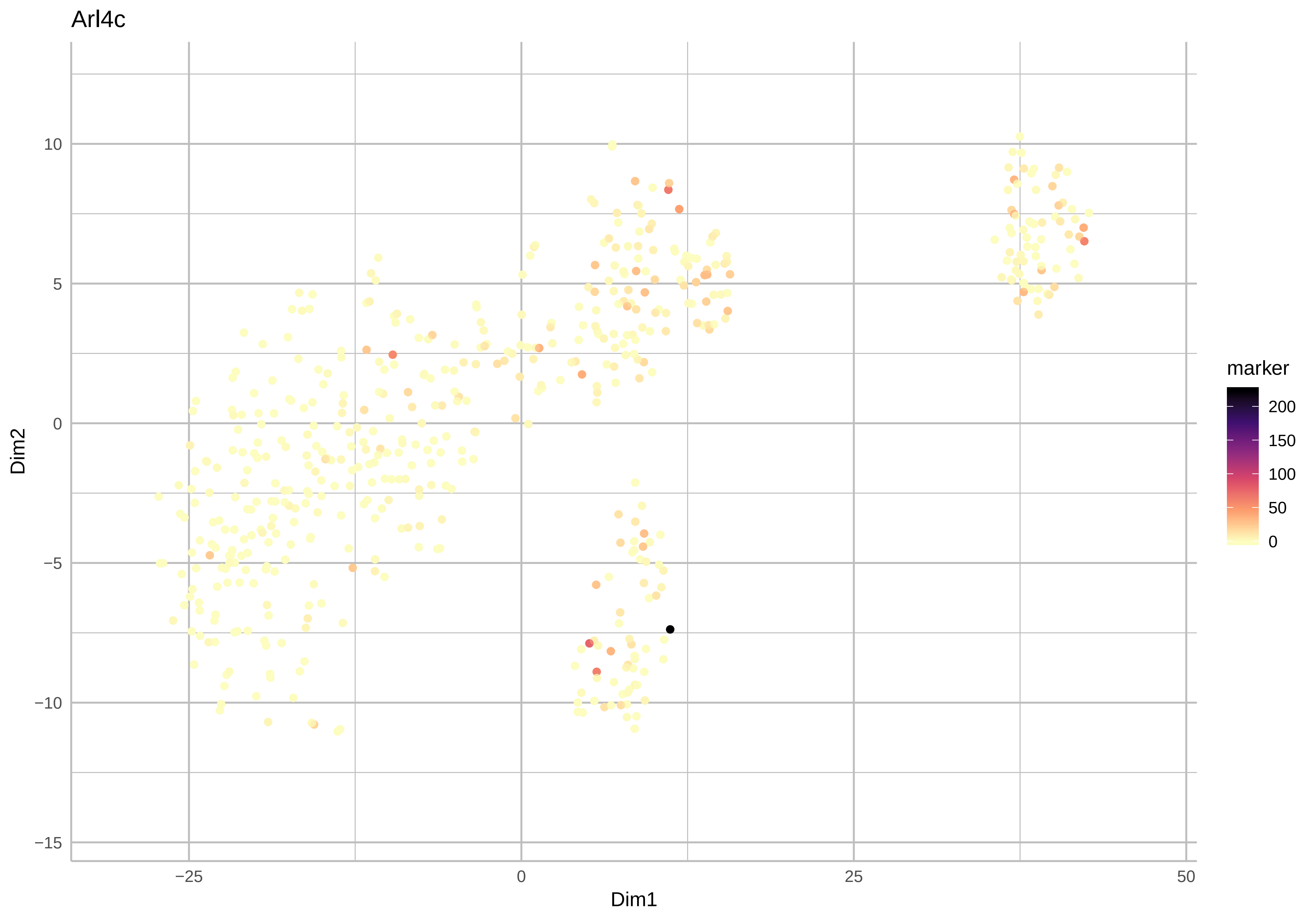
Arl4c
Dim2
Dim1
10
5
0
−5
−10
−15
−25
0
25
50
marker
200
150
100
50
0

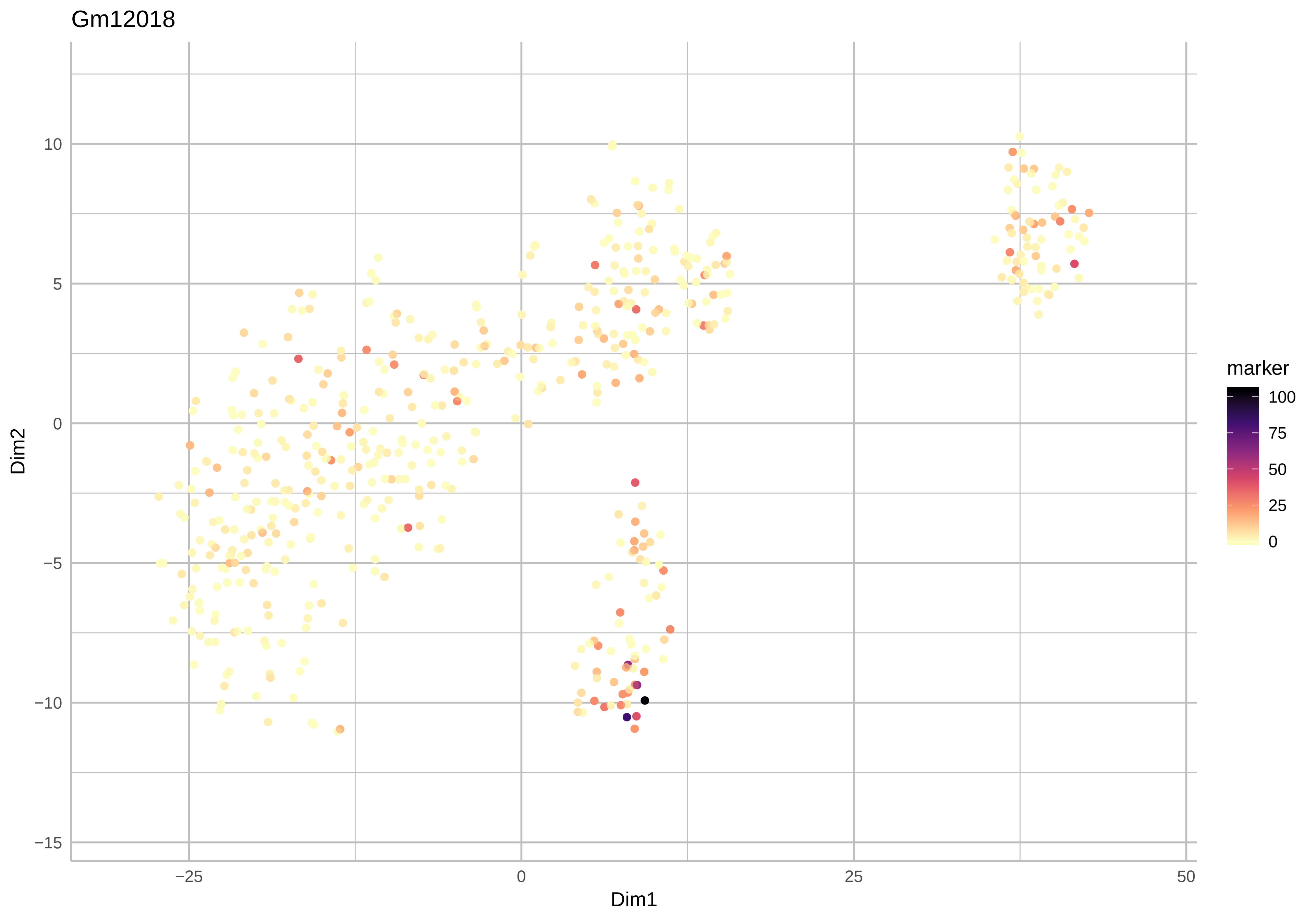

Gm12018
Dim2
Dim1
10
5
0
−5
−10
−15
−25
0
25
50
marker
100
75
50
25
0

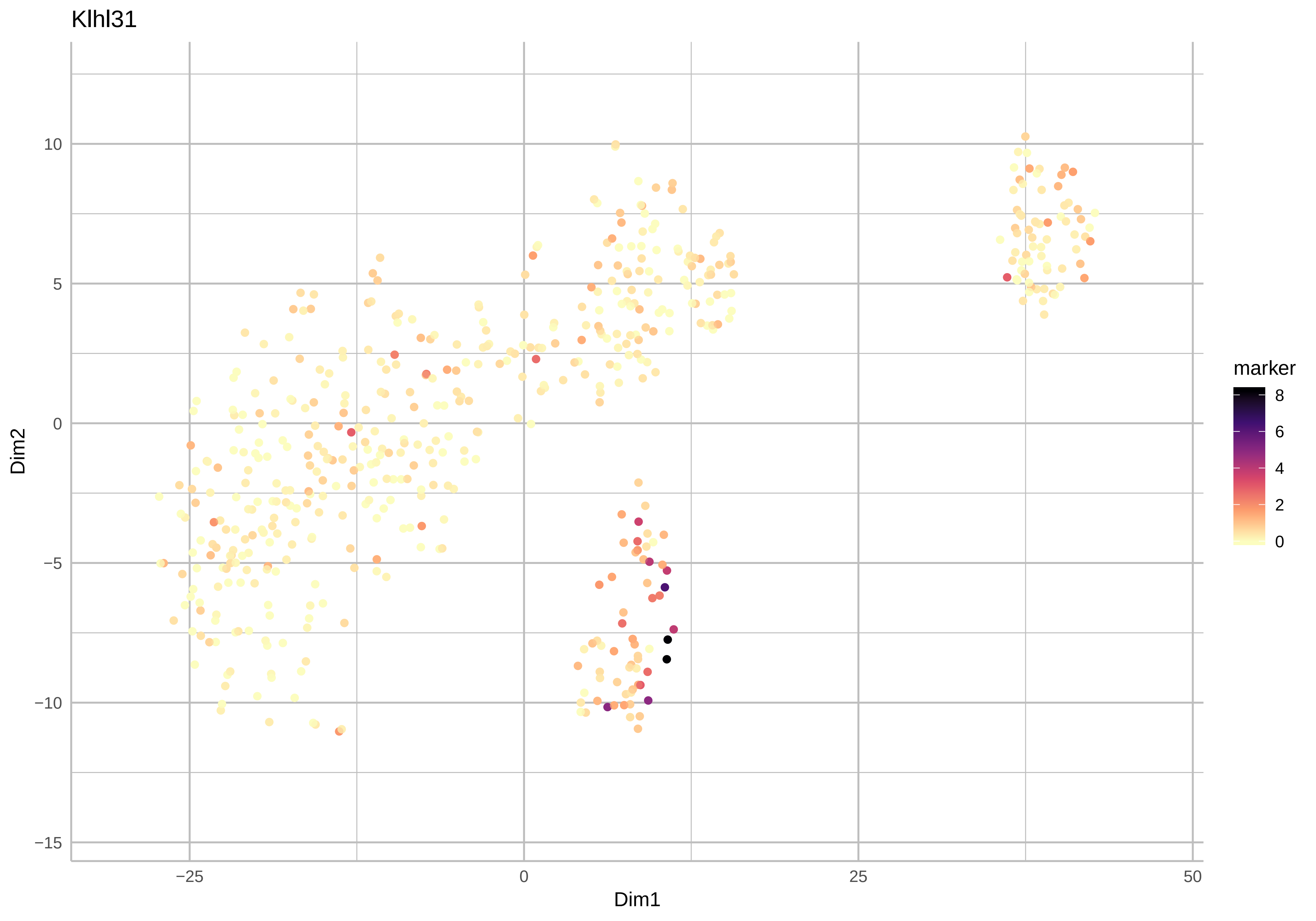

Klhl31
Dim2
10
5
0
−5
−10
−15
−25
0
25
50
Dim1
marker
8
6
4
2
0

Igfbp5

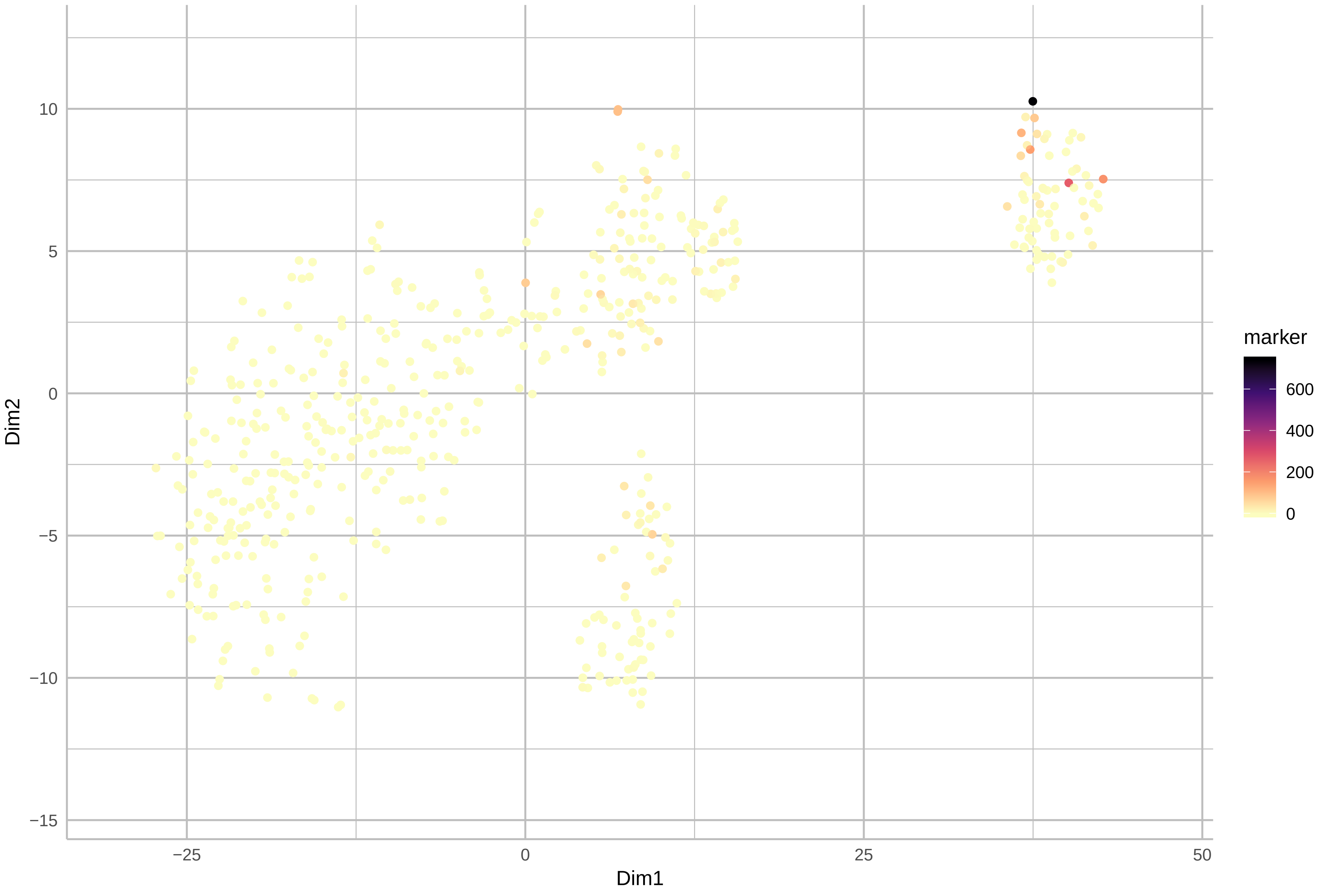

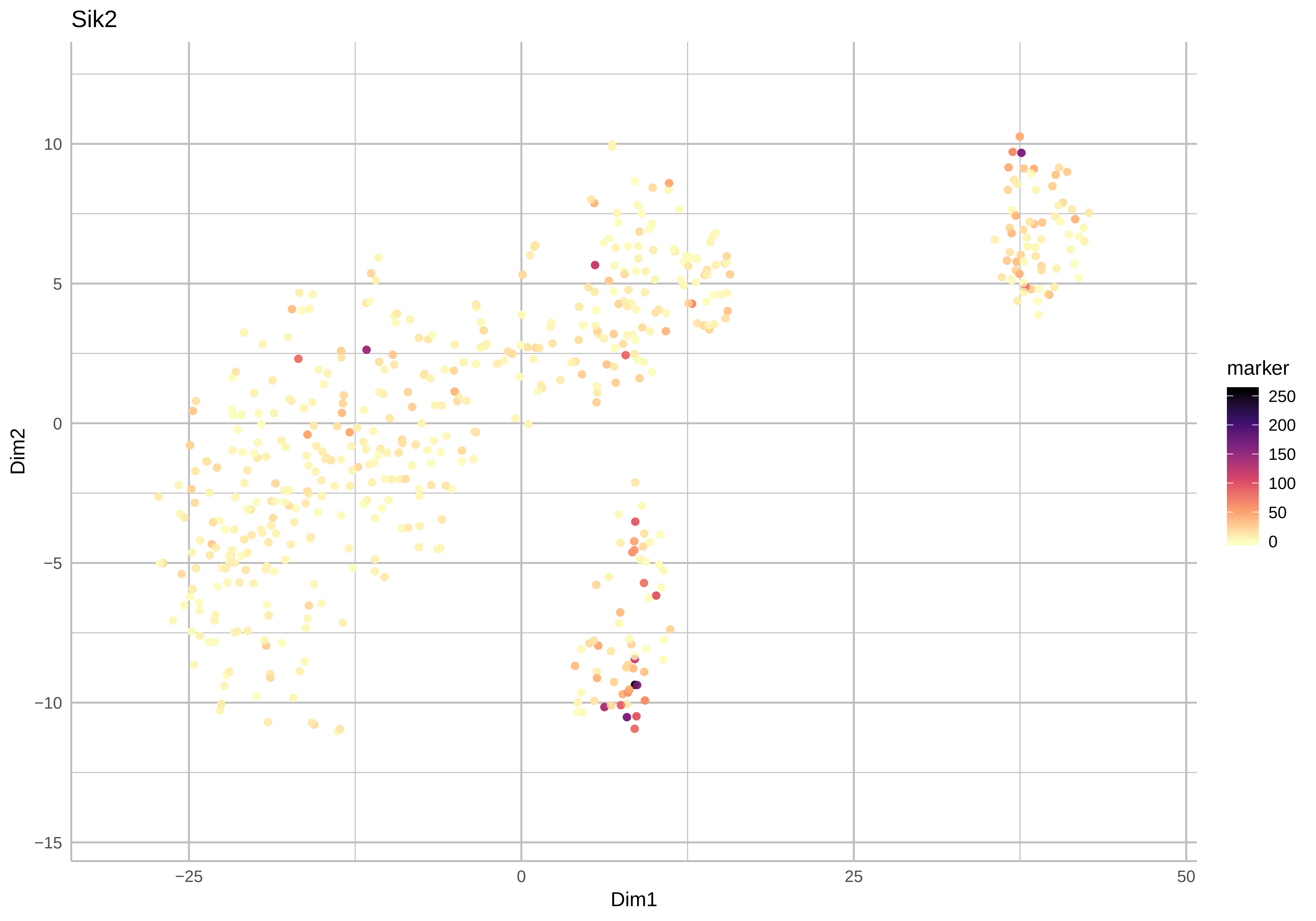

Sik2
Dim2
Dim1
10
5
0
−5
−10
−15
−25
0
25
50
marker
250
200
150
100
50
0

Cd24a

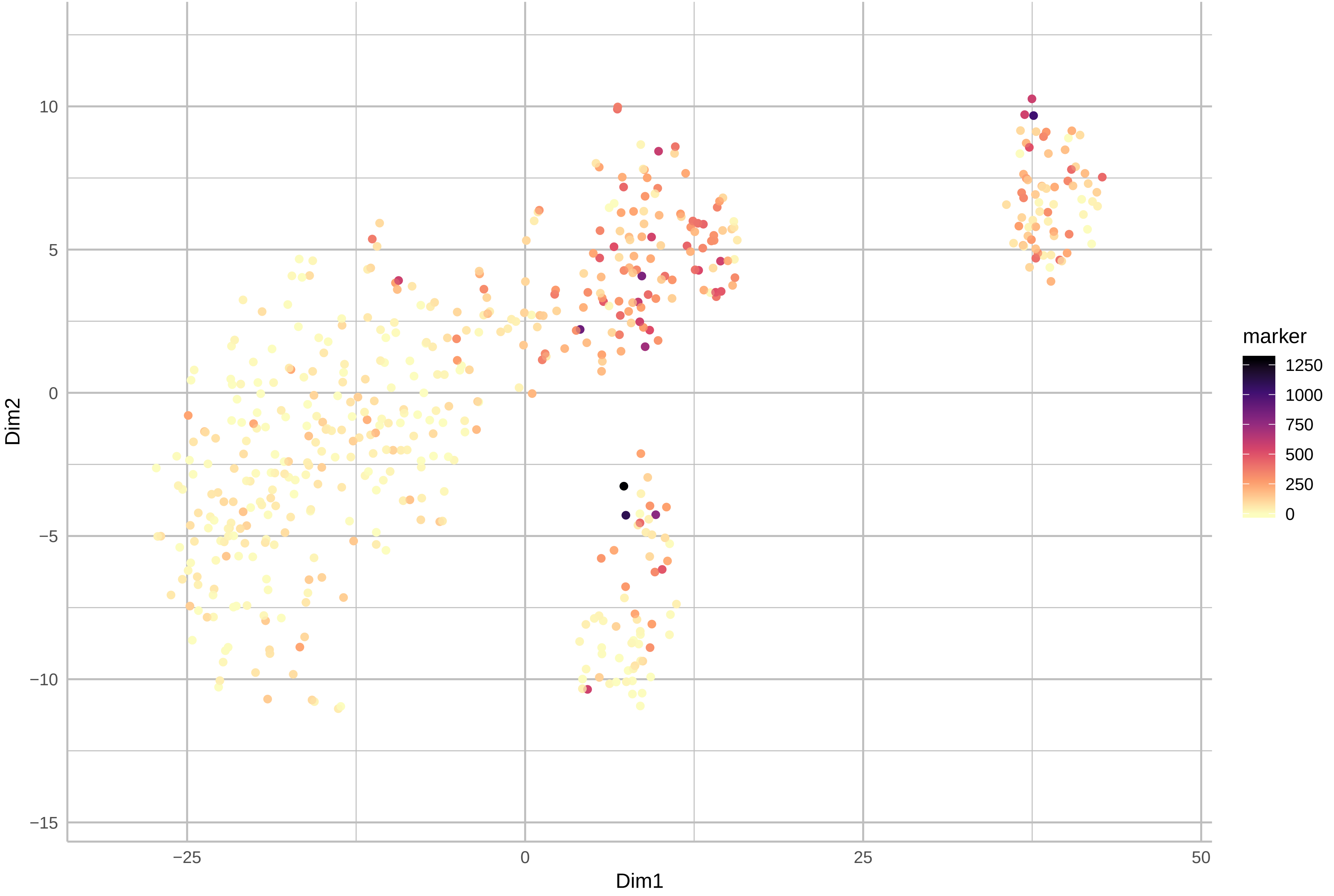

Gm42445

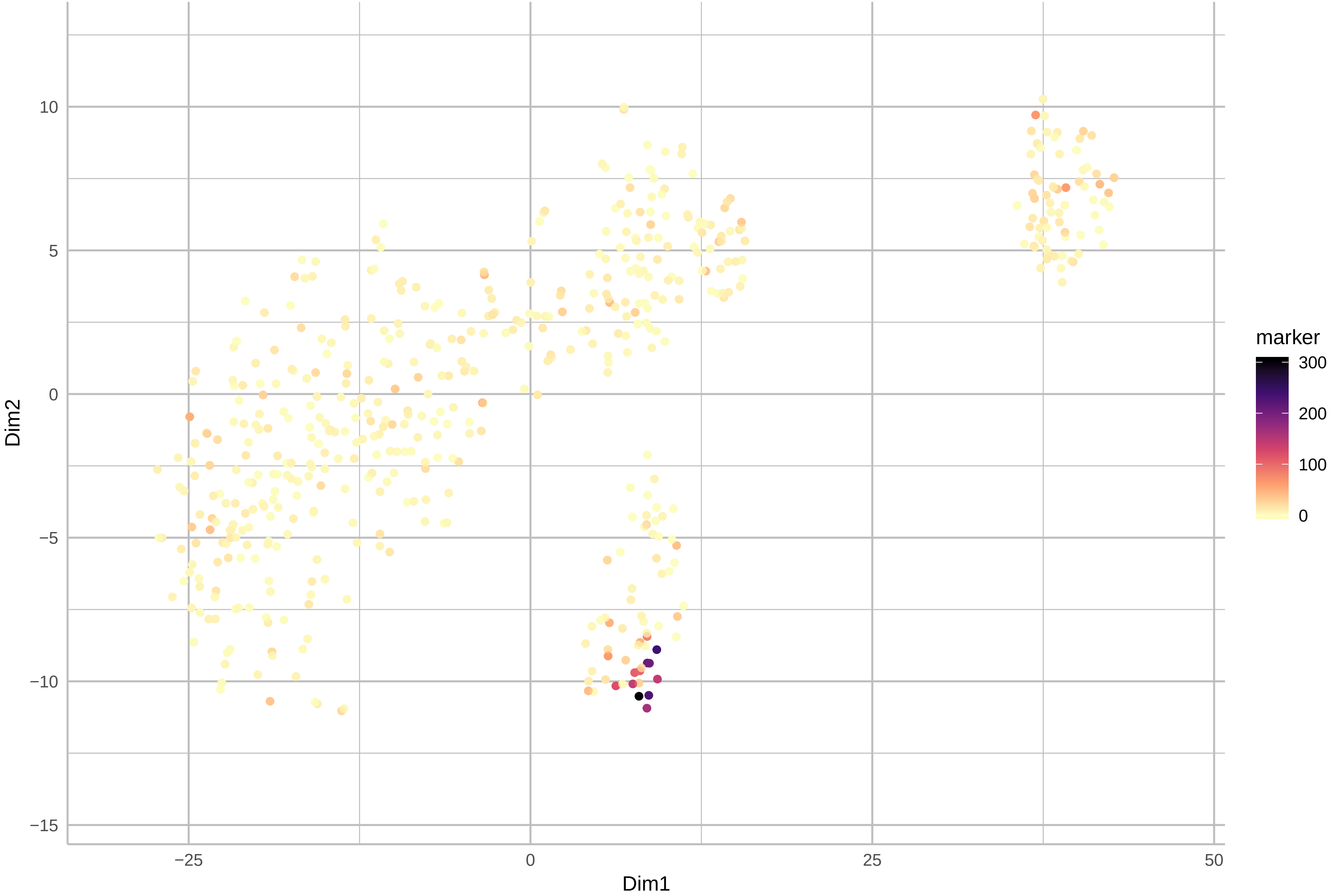

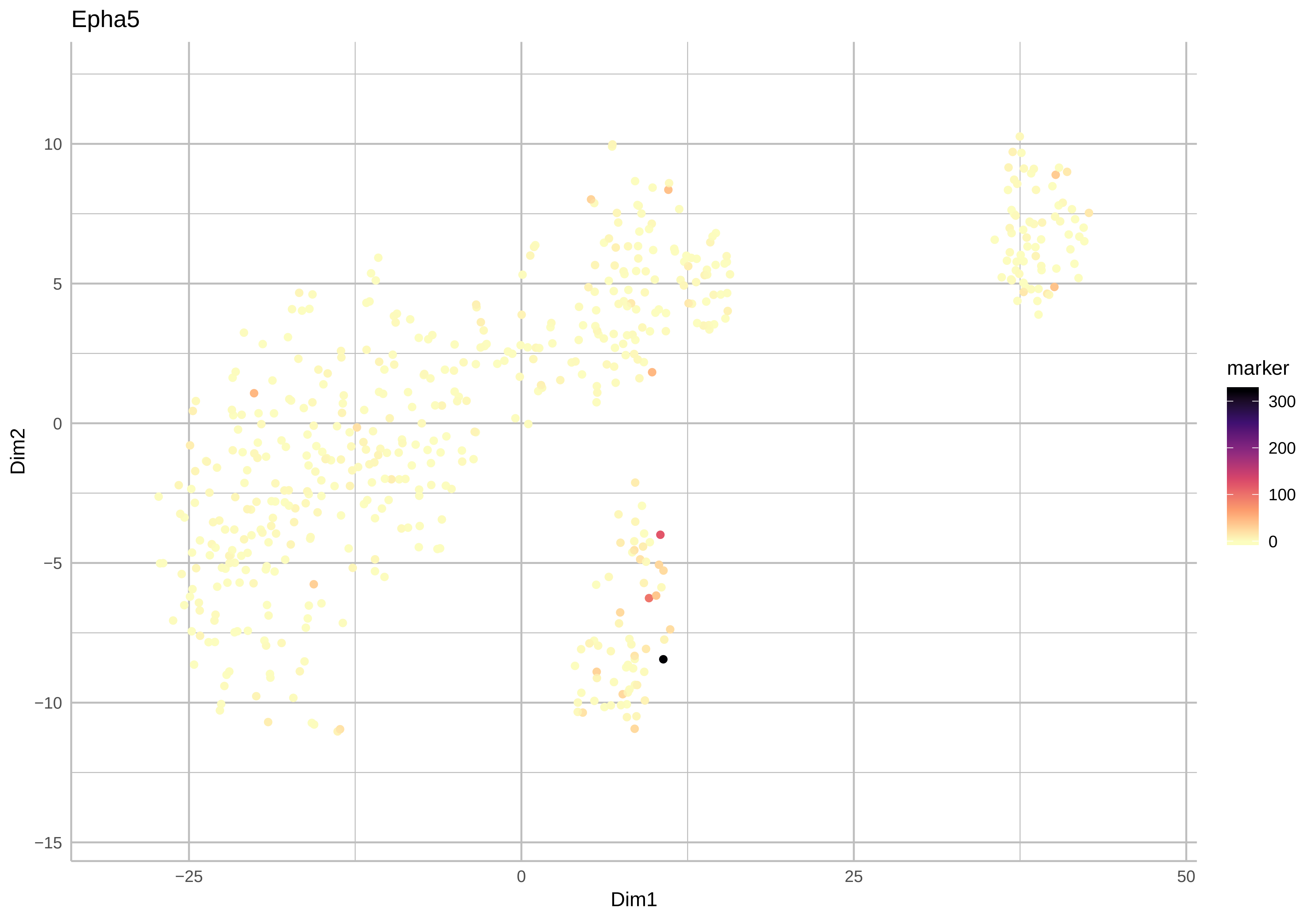
Epha5
Dim2
Dim1
marker
300
200
100
0
10
5
0
−5
−10
−15
−25
0
25
50

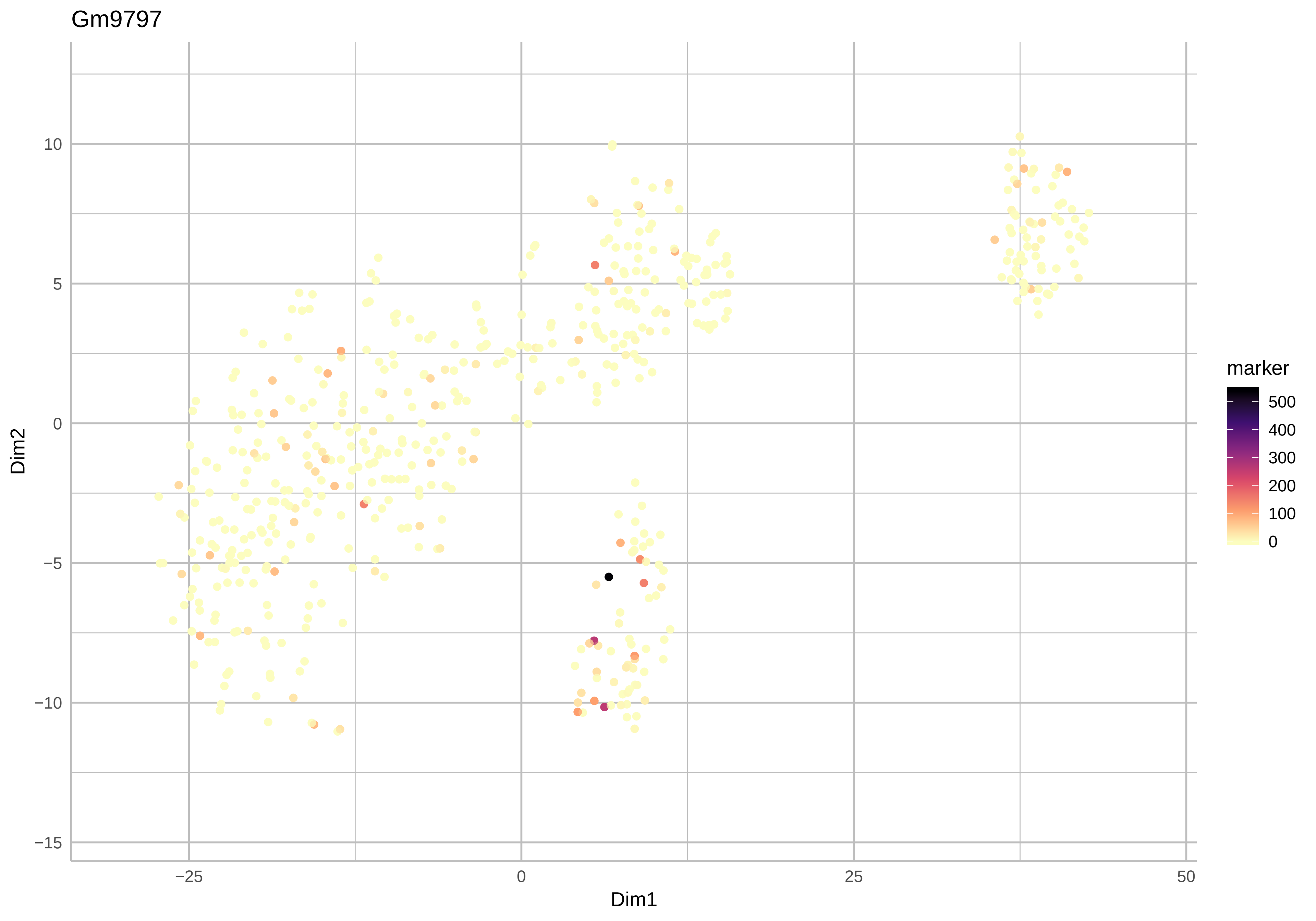

Gm9797
Dim2
Dim1
10
5
0
−5
−10
−15
−25
0
25
50
marker
500
400
300
200
100
0

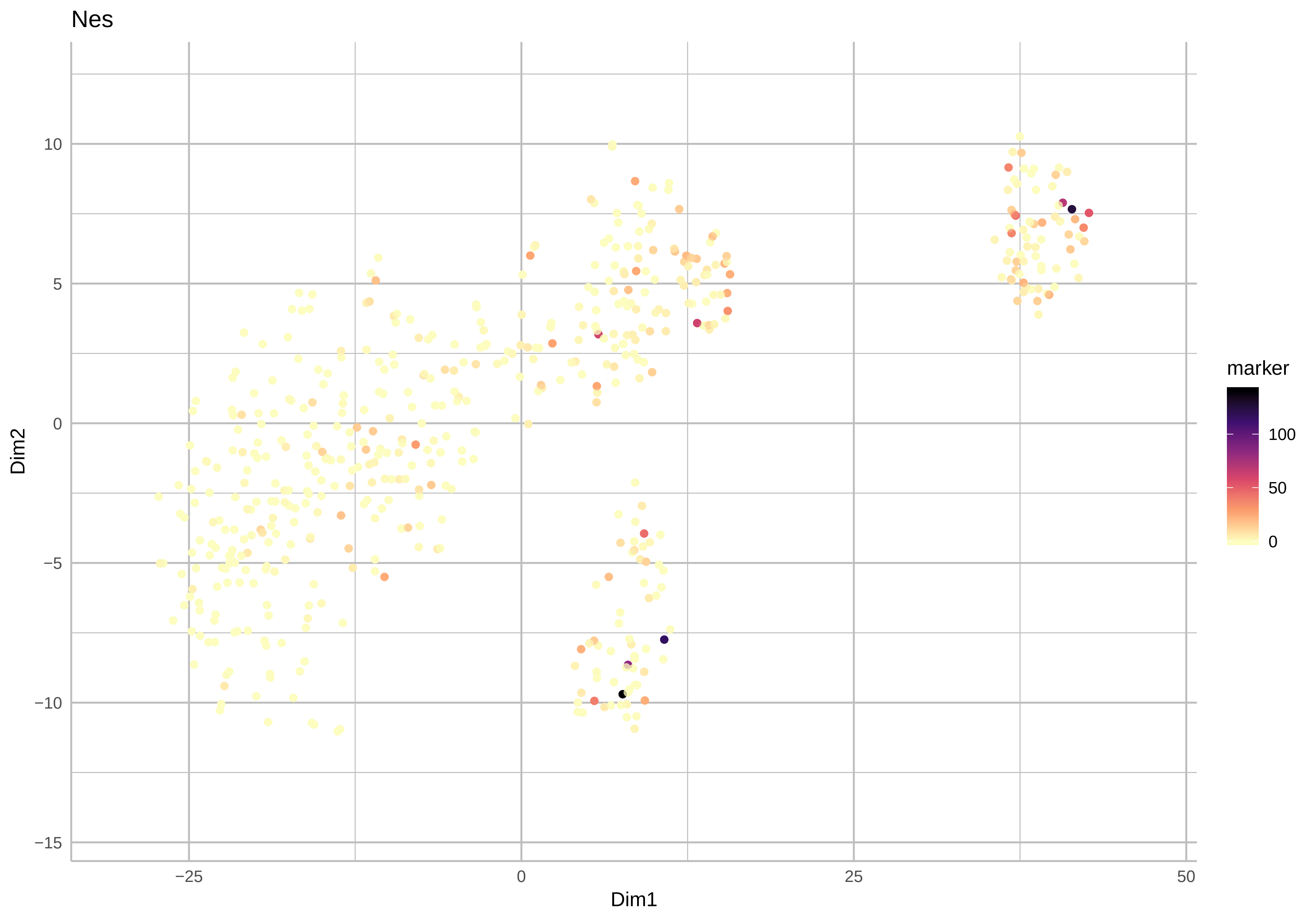
Nes
Dim2
Dim1
marker
100
50
0
10
5
0
−5
−10
−15
−25
0
25
50

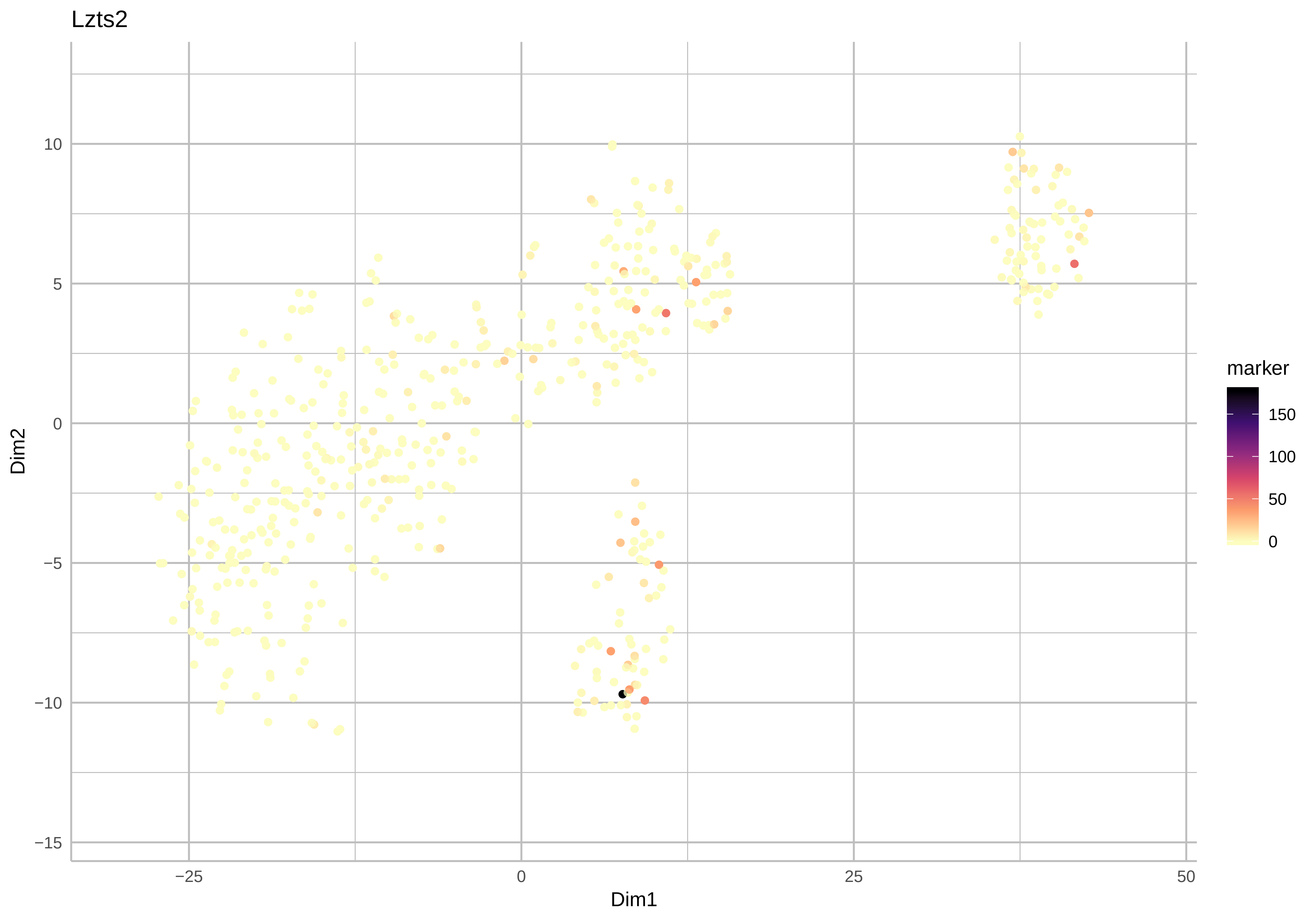

Lzts2
10
5
0
−5
−10
−15
Dim2
−25
0
25
50
Dim1
marker
150
100
50
0

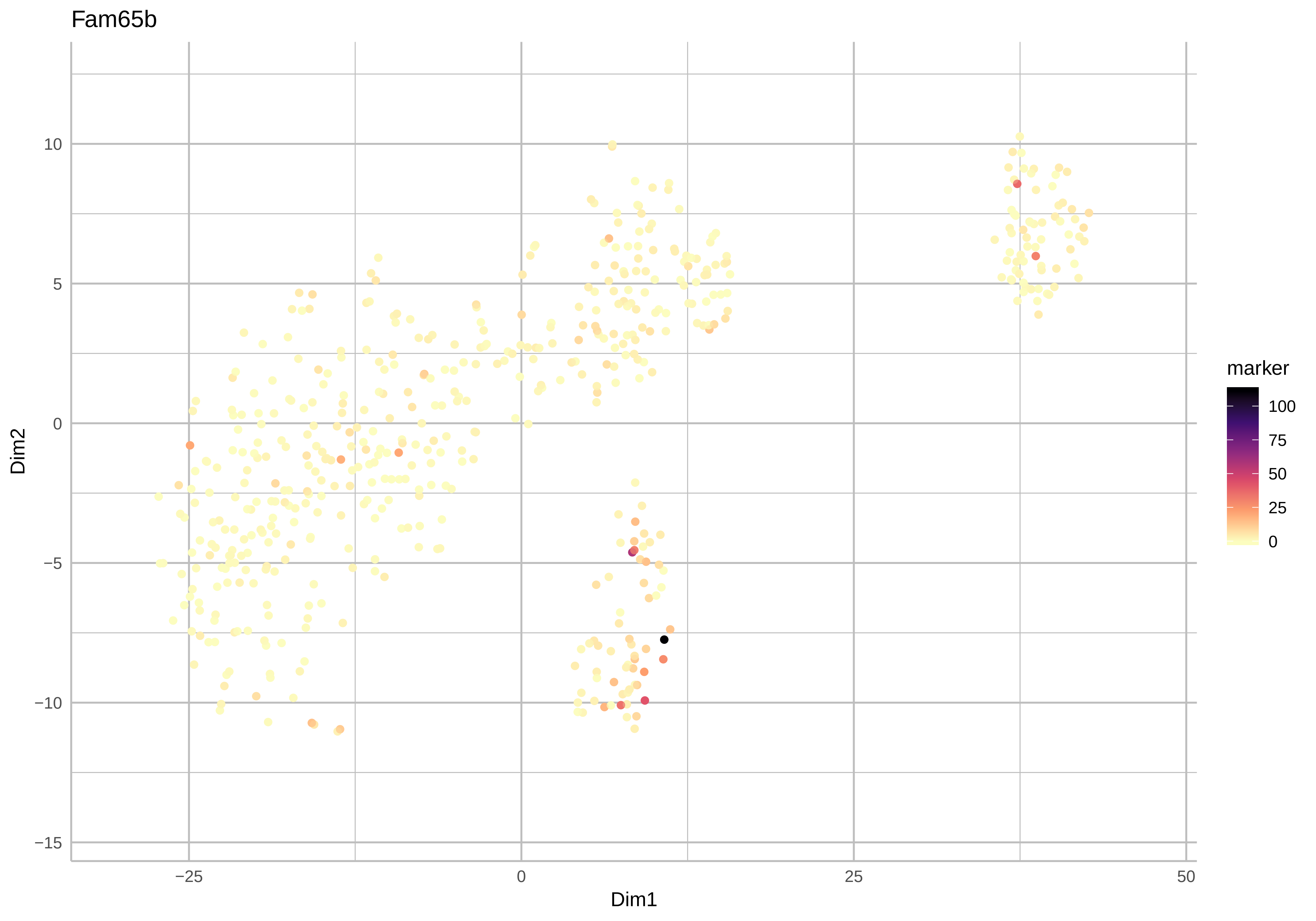

Fam65b
Dim2
Dim1
10
5
0
−5
−10
−15
−25
0
25
50
marker
100
75
50
25
0

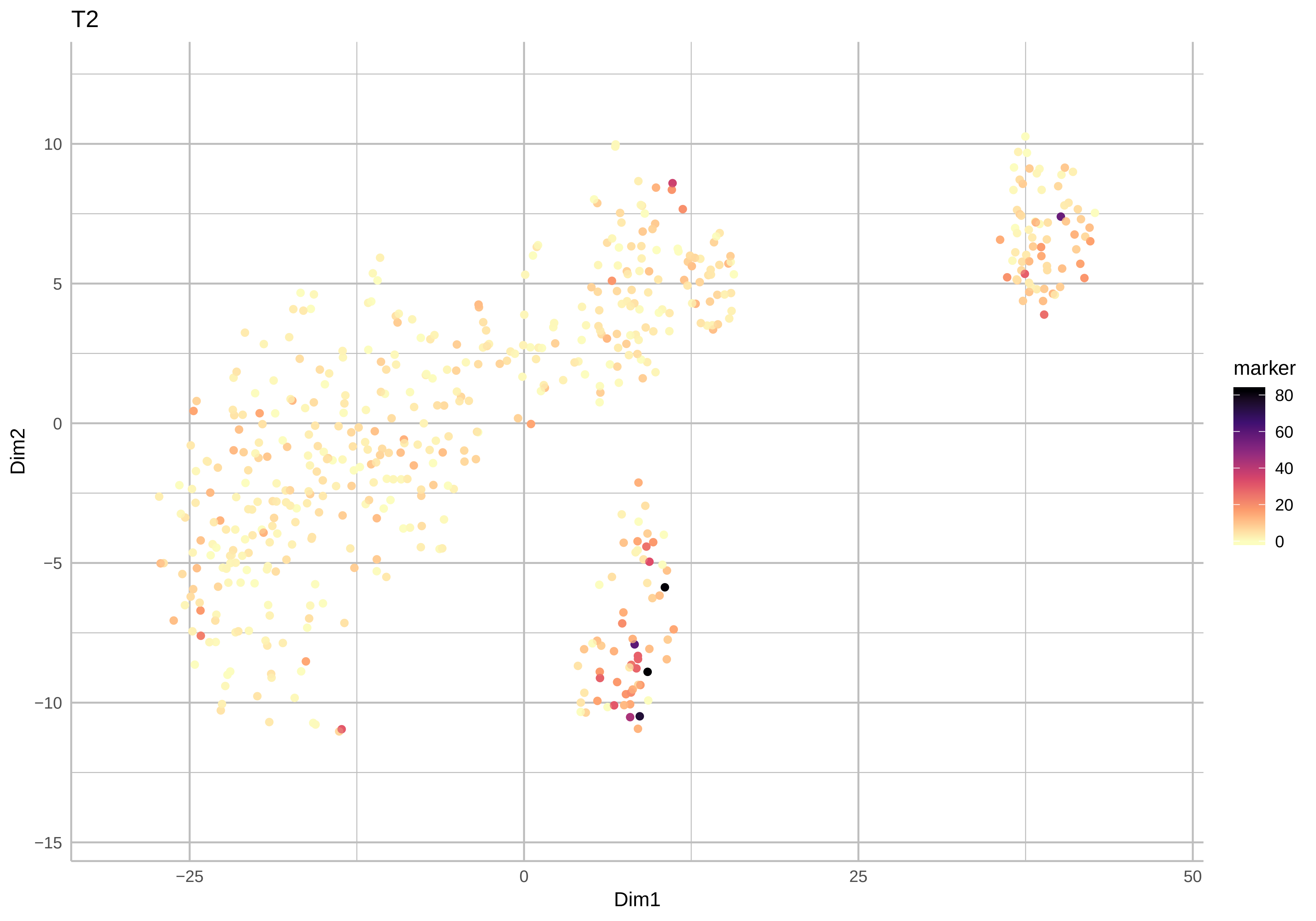

T2
Dim2
Dim1
10
5
0
−5
−10
−15
−25
0
25
50
marker
80
60
40
20
0

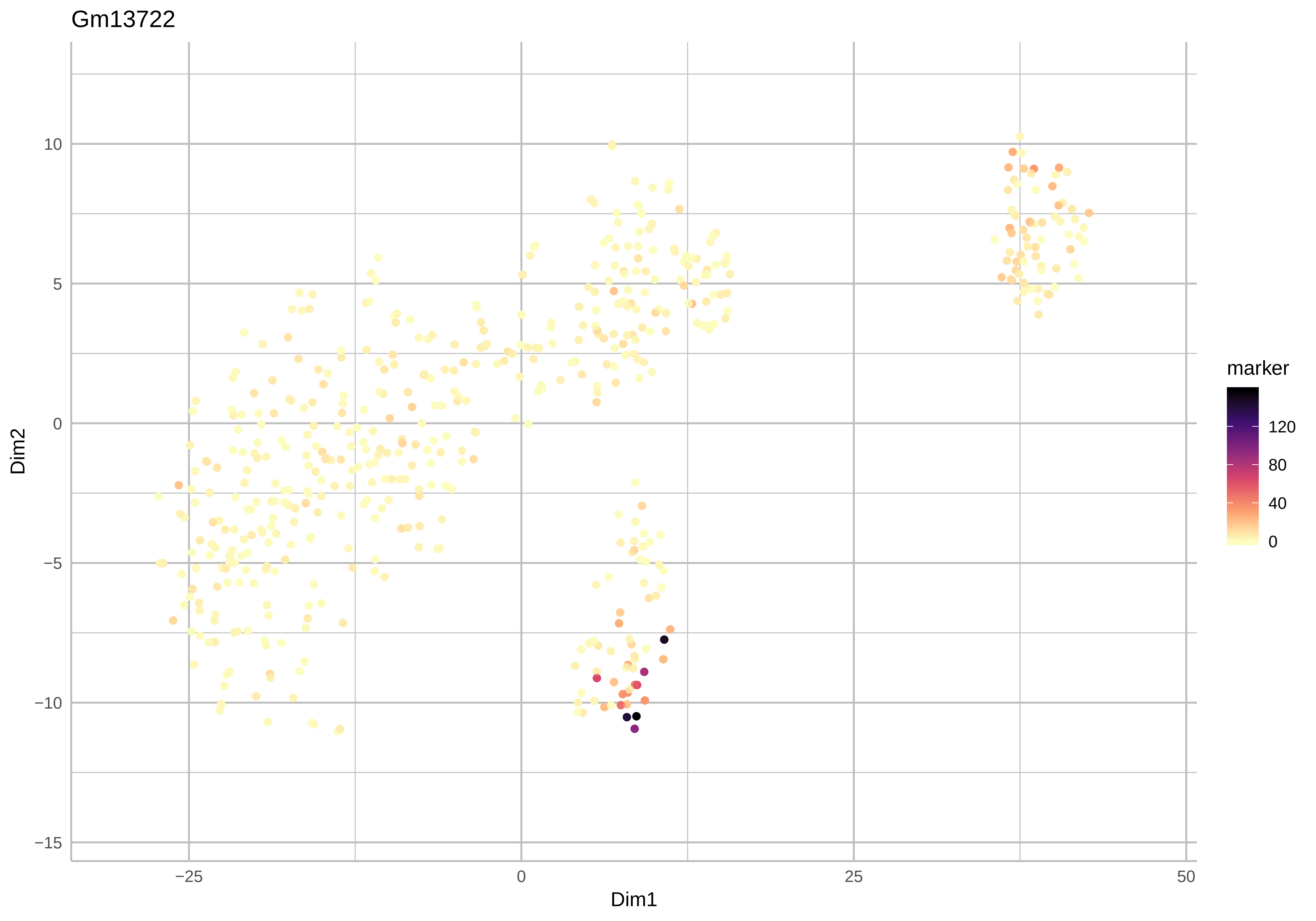
Gm13722
Dim2
Dim1
10
5
0
−5
−10
−15
−25
0
25
50
marker
120
80
40
0

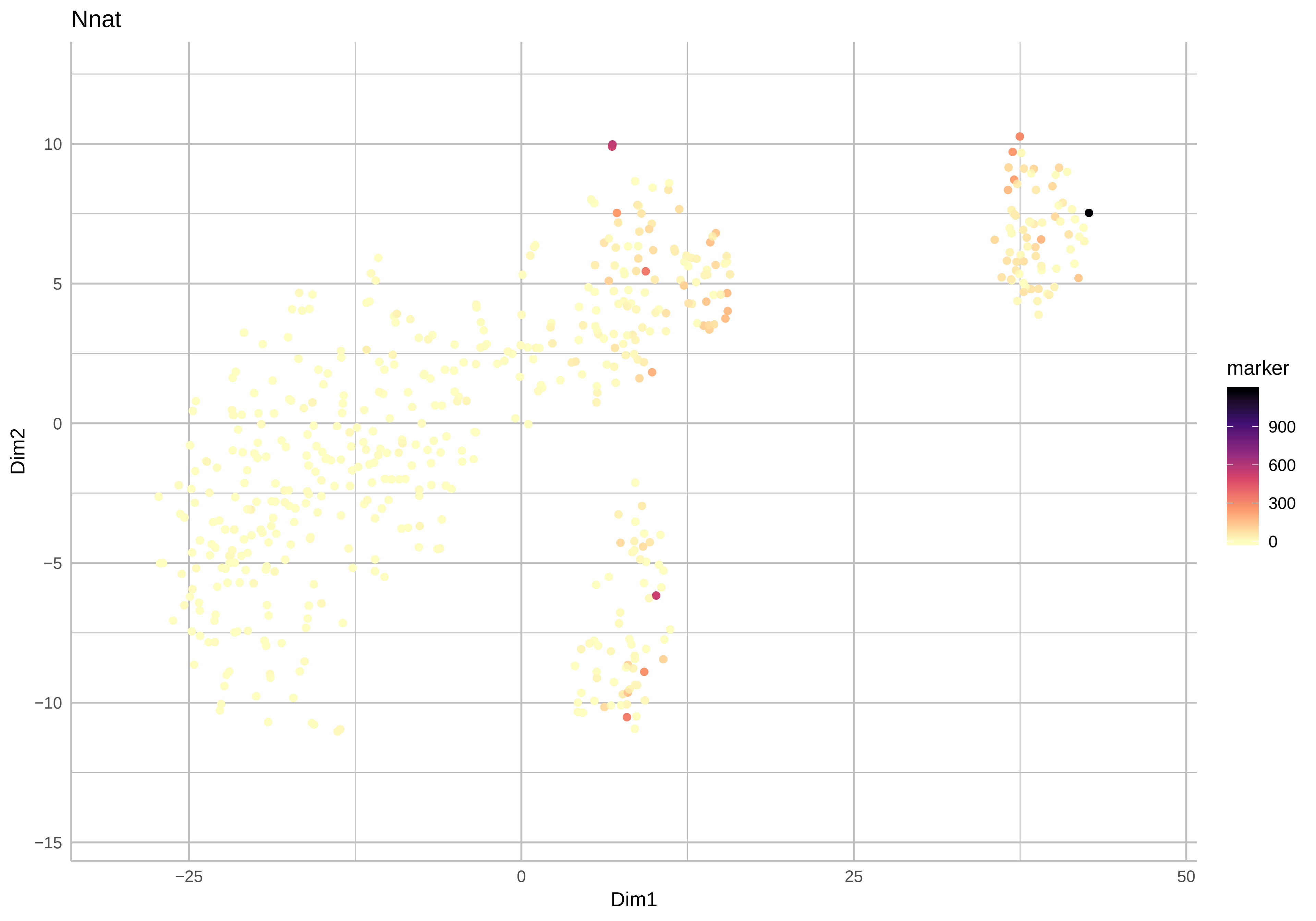
Nnat
Dim2
Dim1
10
5
0
−5
−10
−15
−25
0
25
50
marker
900
600
300
0

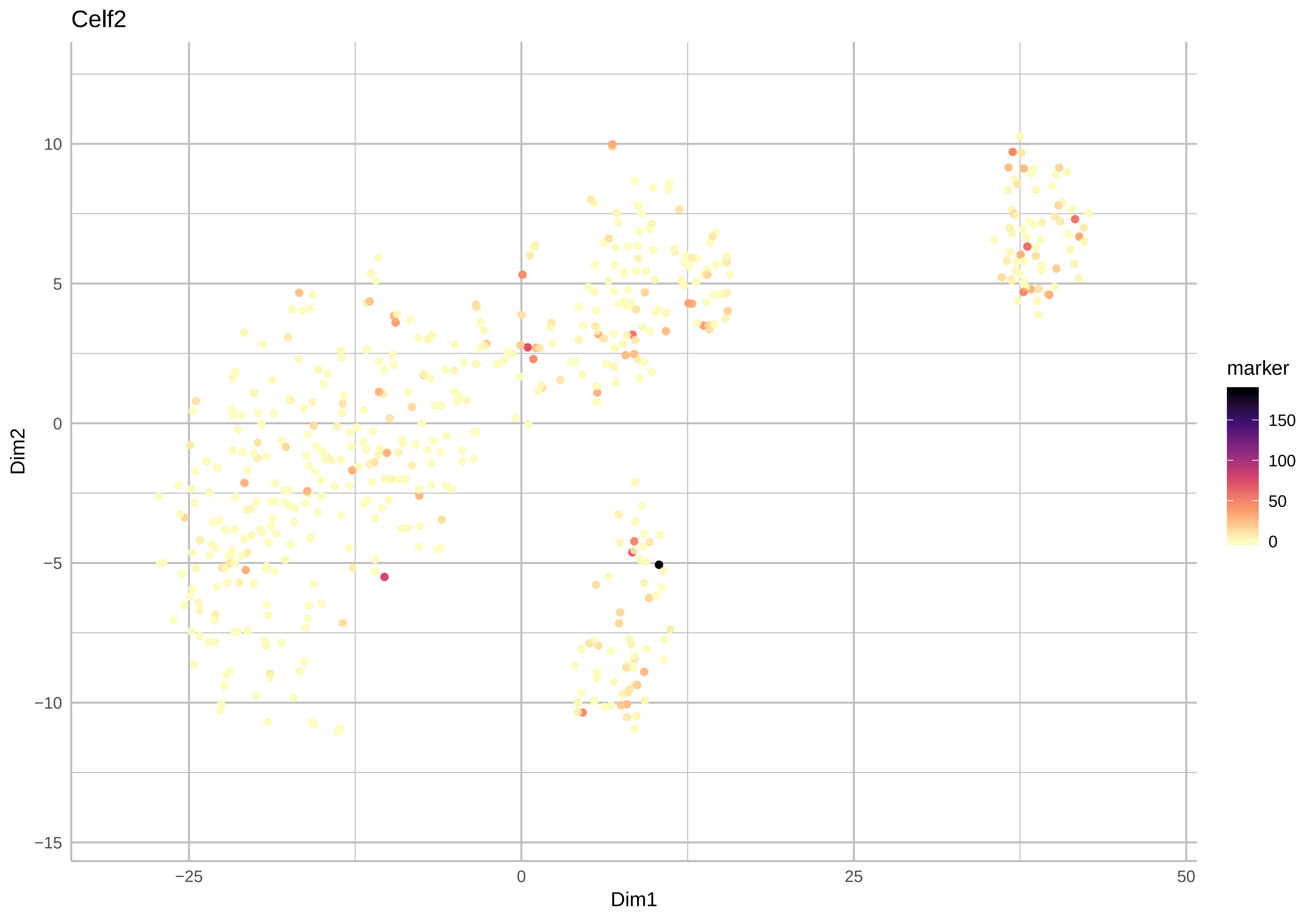
Celf2
Dim2
Dim1
marker
150
100
50
0
10
5
0
−5
−10
−15
−25
0
25
50

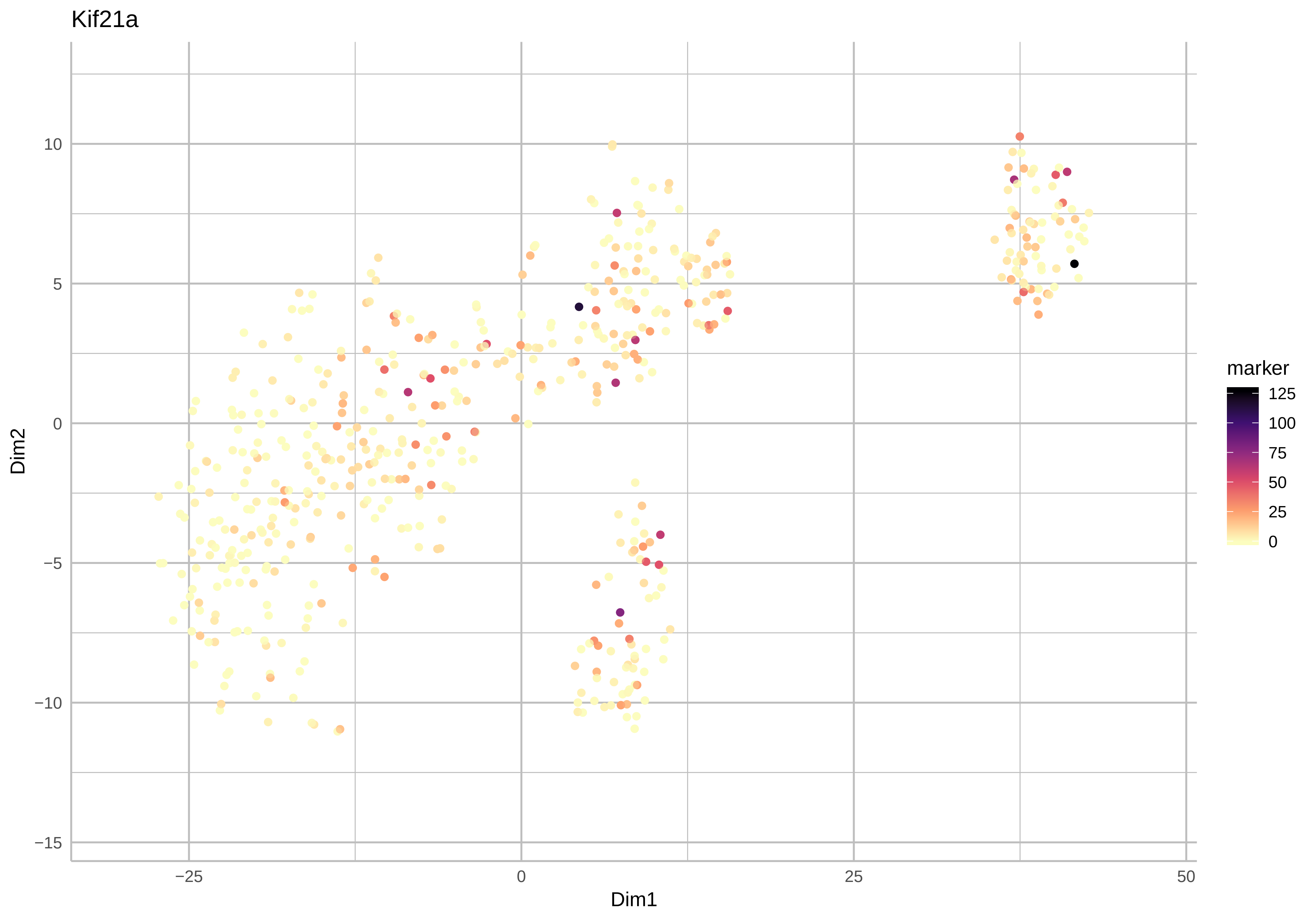
Kif21a
Dim2
Dim1
10
5
0
−5
−10
−15
−25
0
25
50
marker
125
100
75
50
25
0

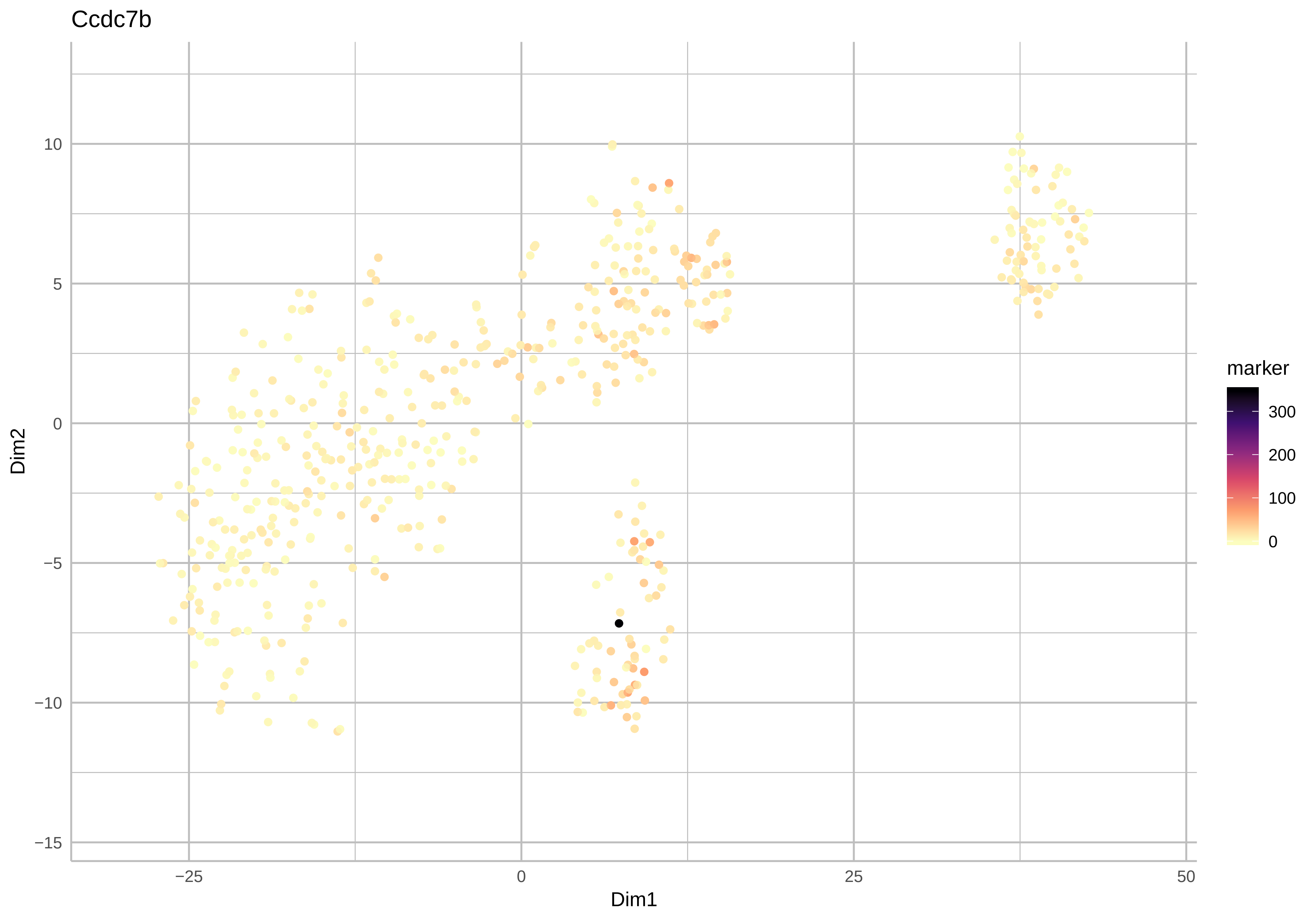
Ccdc7b
Dim2
Dim1
10
5
0
−5
−10
−15
−25
0
25
50
marker
300
200
100
0

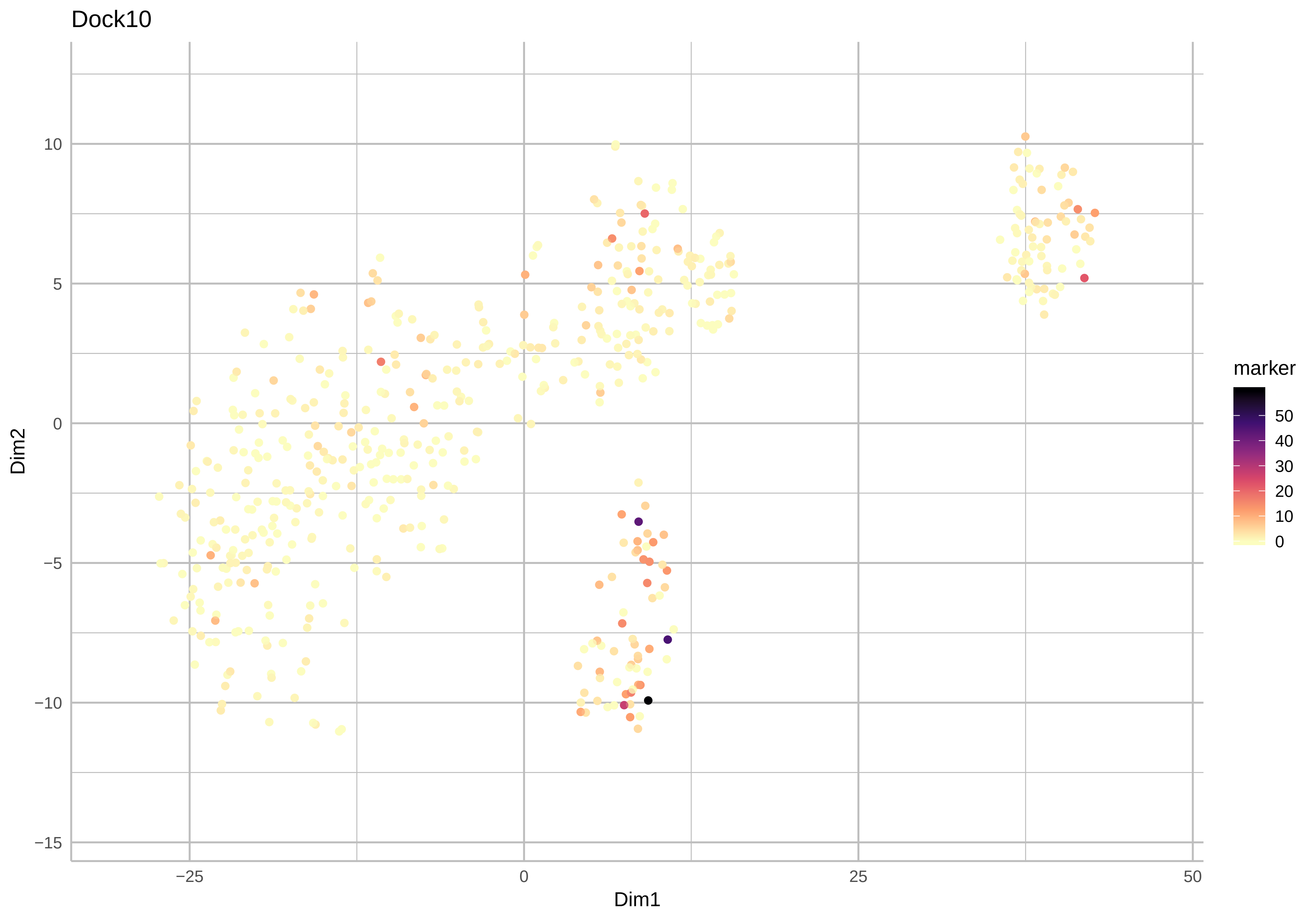
Dock10
Dim1
Dim2
marker
−25
0
25
50
10
5
0
−5
−10
−15
0
10
20
30
40
50

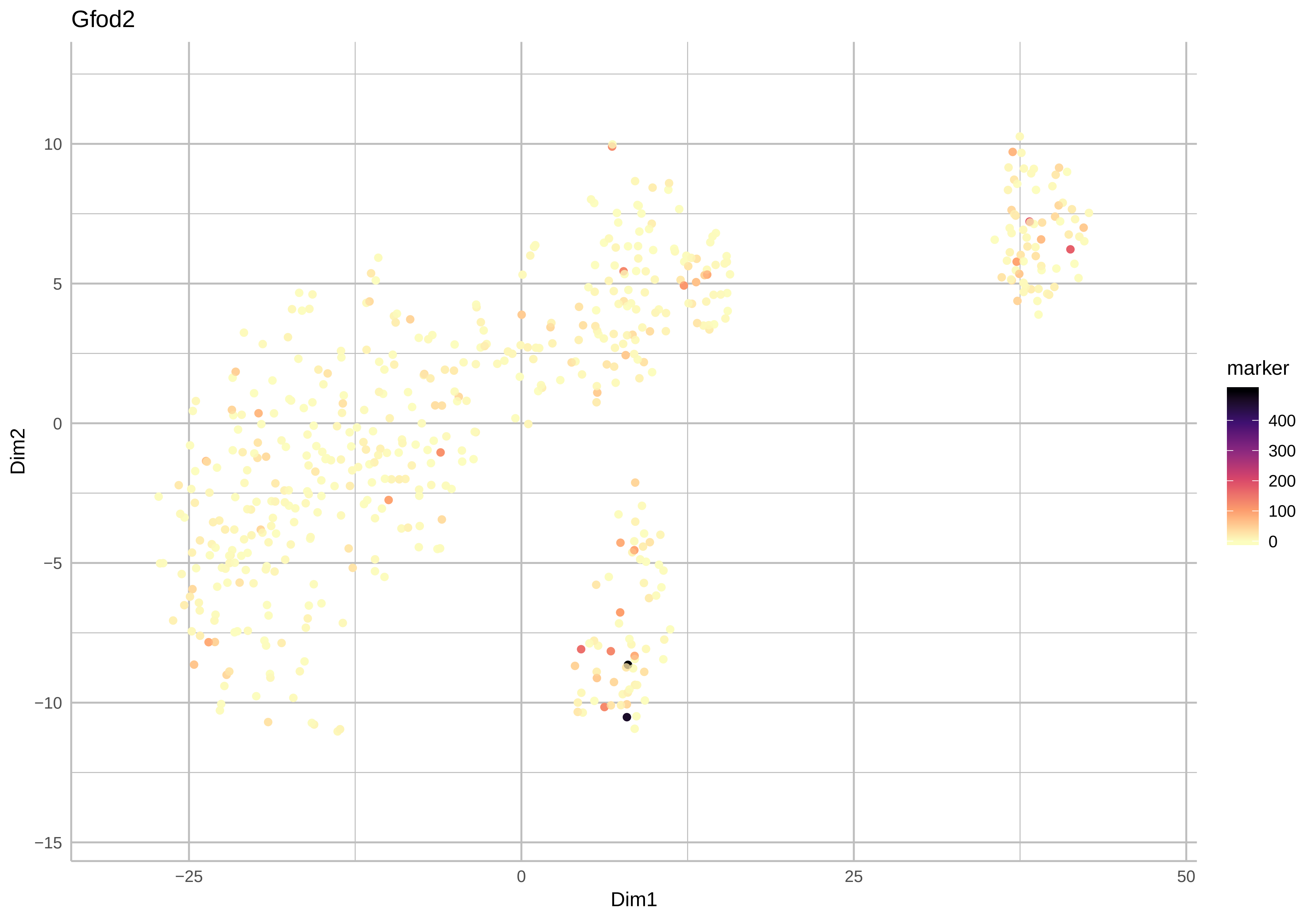
Gfod2
Dim2
Dim1
10
5
0
−5
−10
−15
−25
0
25
50
marker
400
300
200
100
0

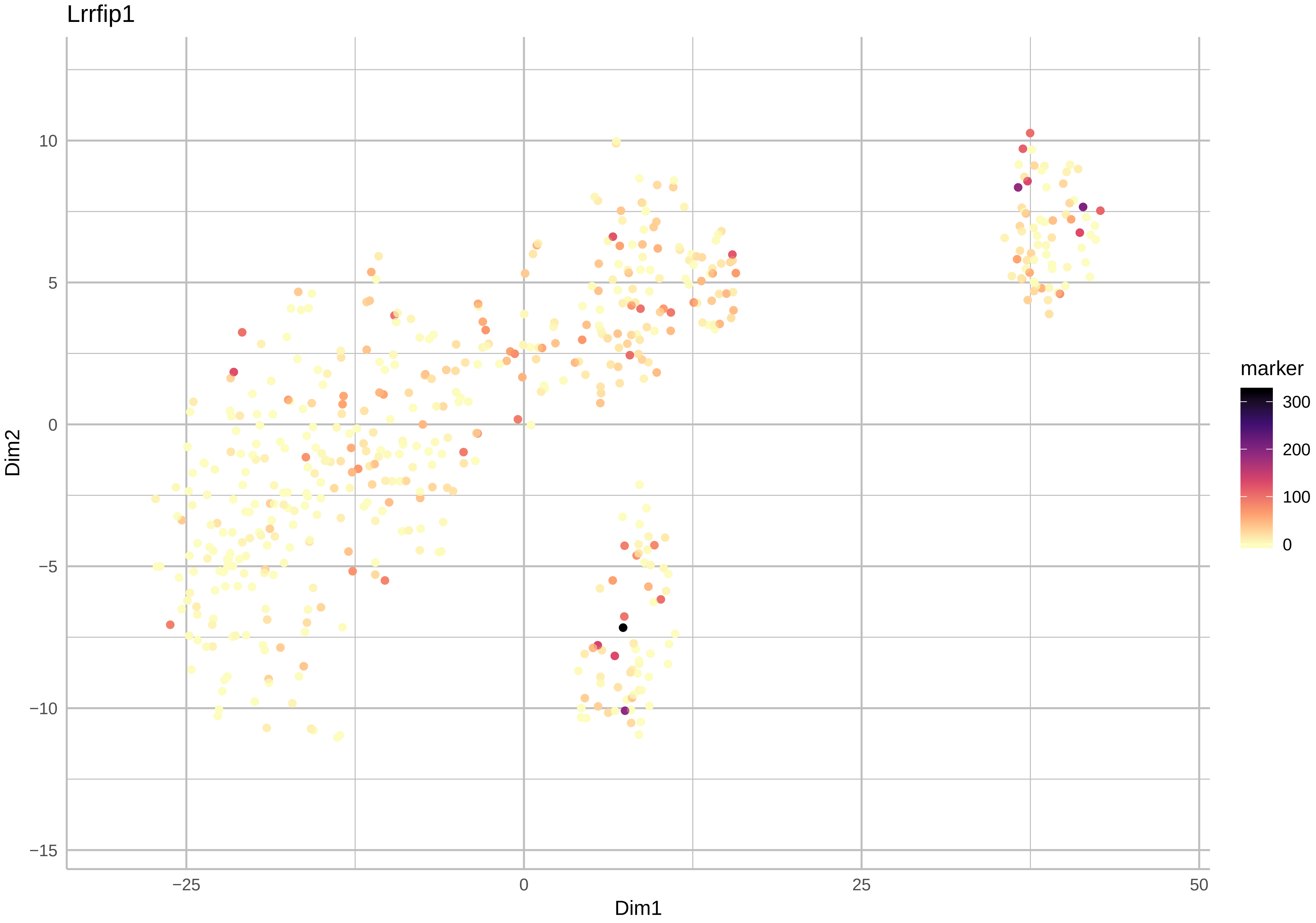
Lrrfip1
Dim2
Dim1
marker
300
200
100
0
10
5
0
−5
−10
−15
−25
0
25
50

Gm10660

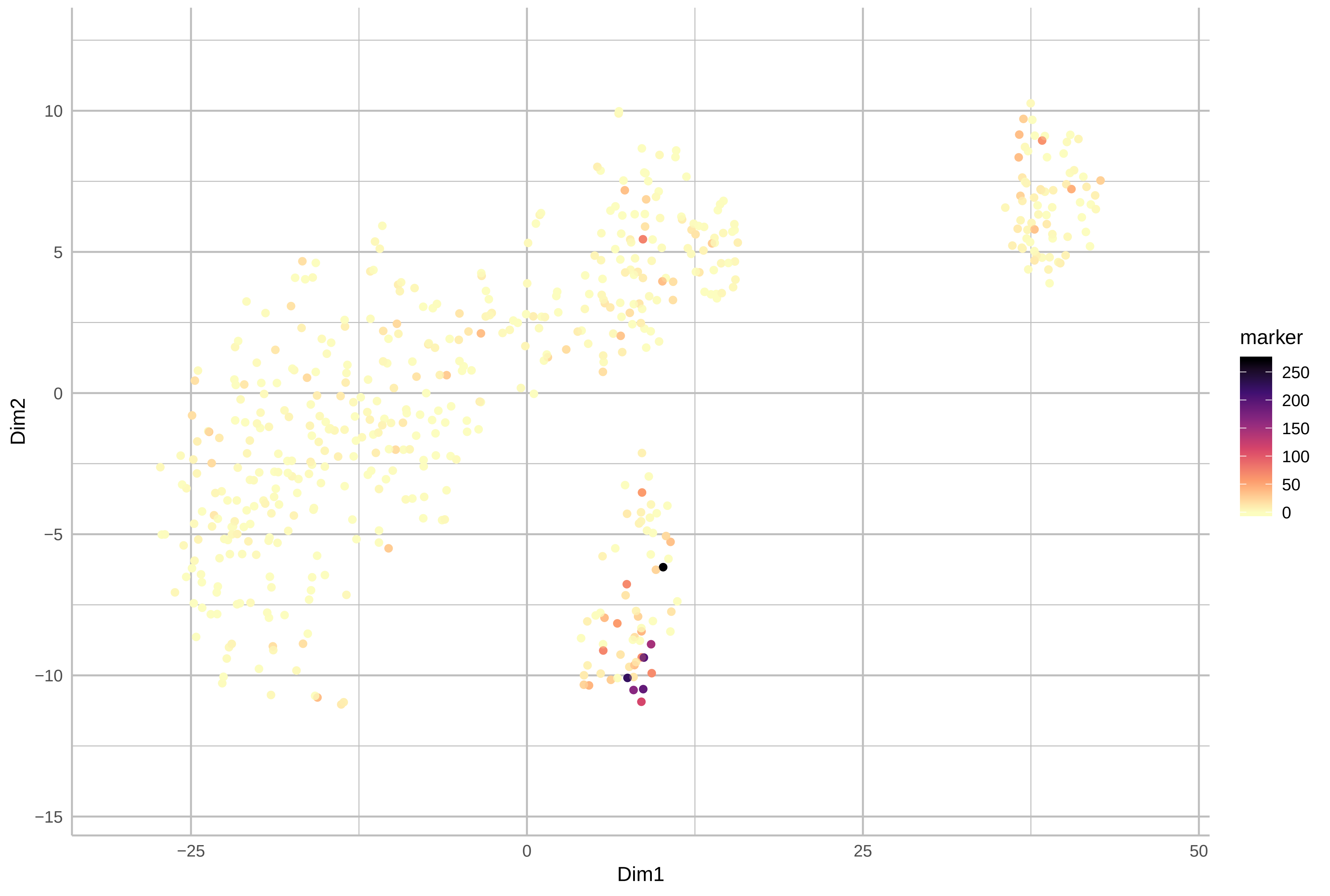

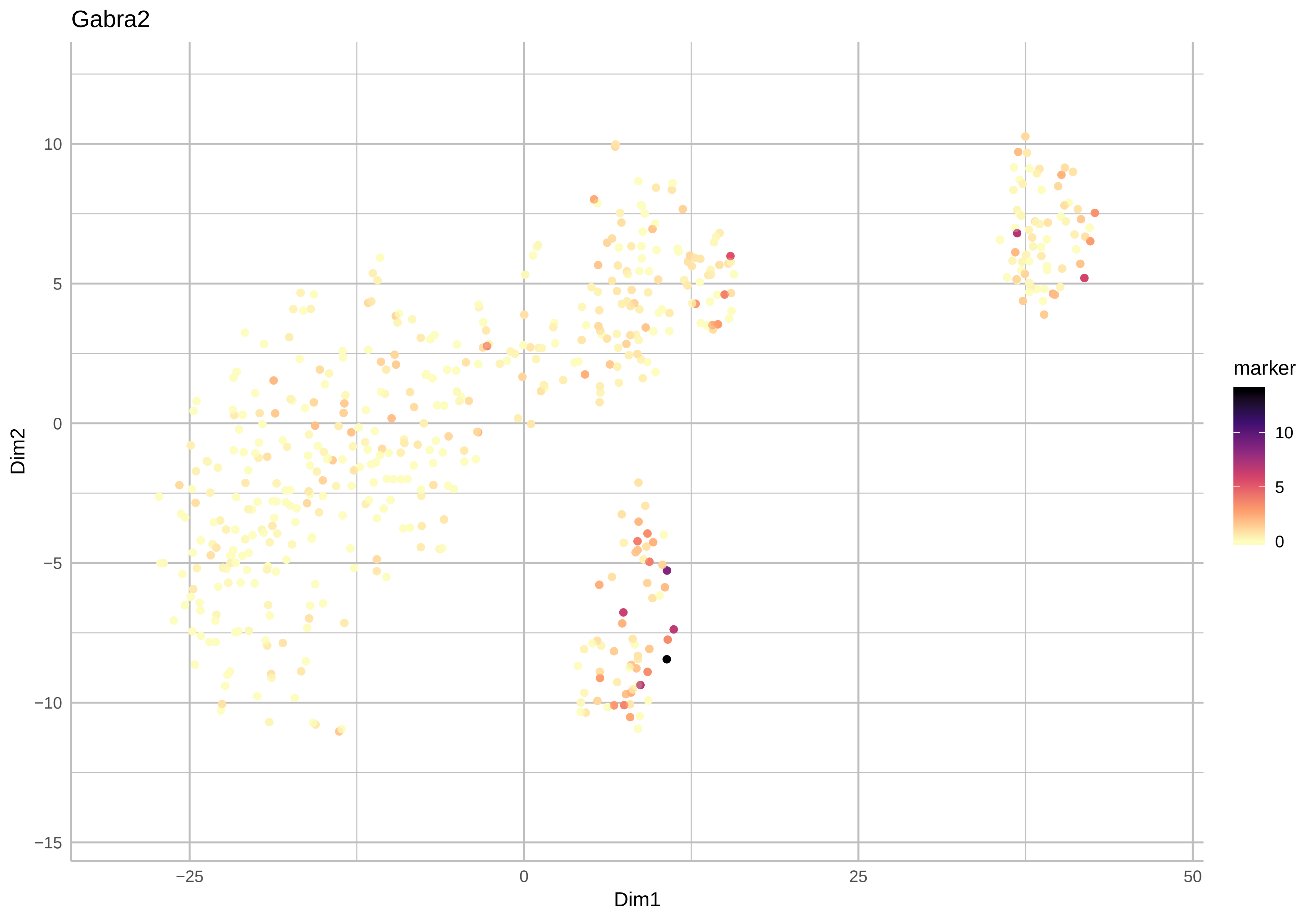

Gabra2
Dim2
Dim1
10
5
0
−5
−10
−15
−25
0
25
50
marker
10
5
0

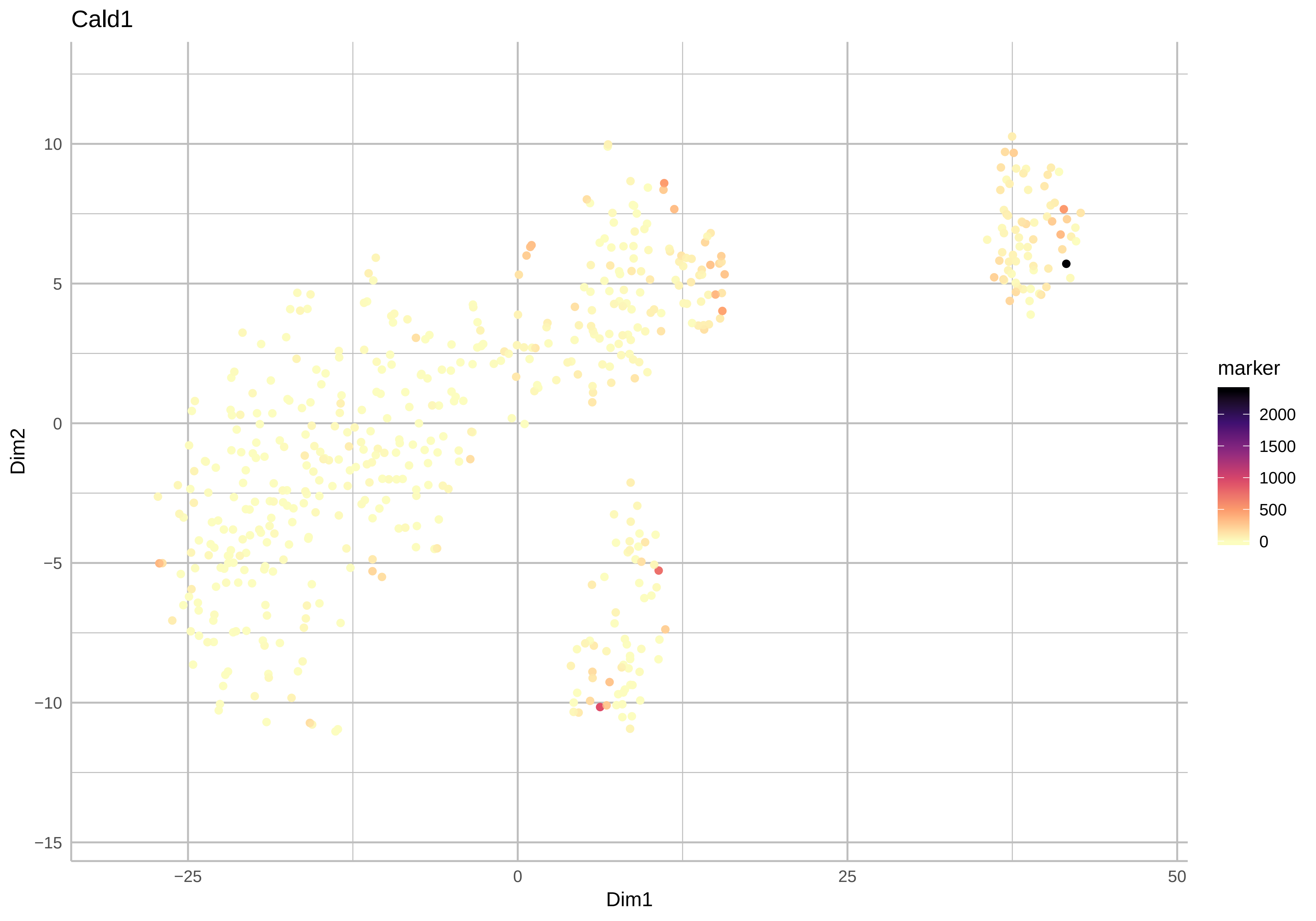
Cald1
10
5
0
−5
−10
−15
Dim2
−25
0
25
50
Dim1
marker
2000
1500
1000
500
0

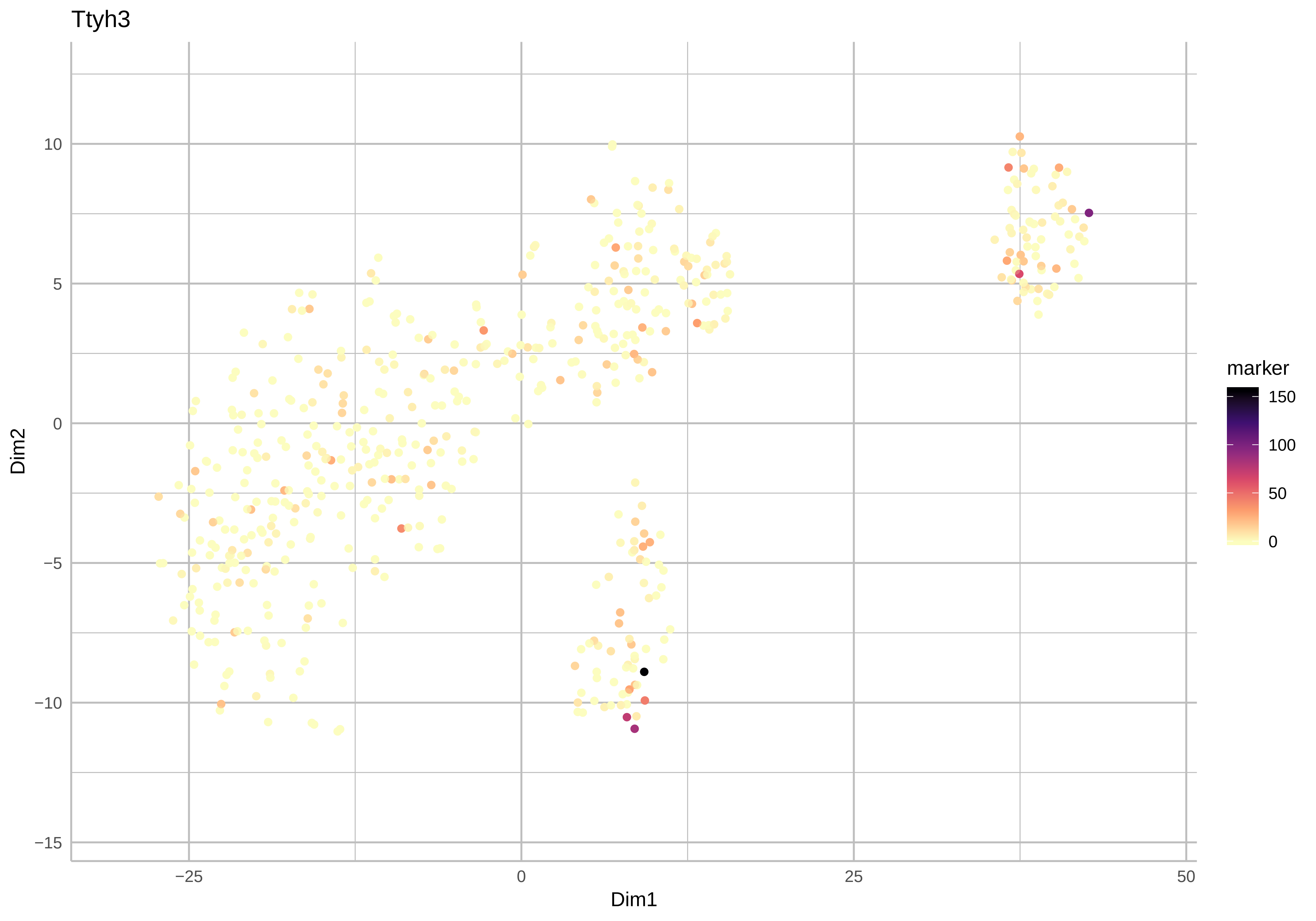
Ttyh3
Dim2
Dim1
10
5
0
−5
−10
−15
−25
0
25
50
marker
150
100
50
0

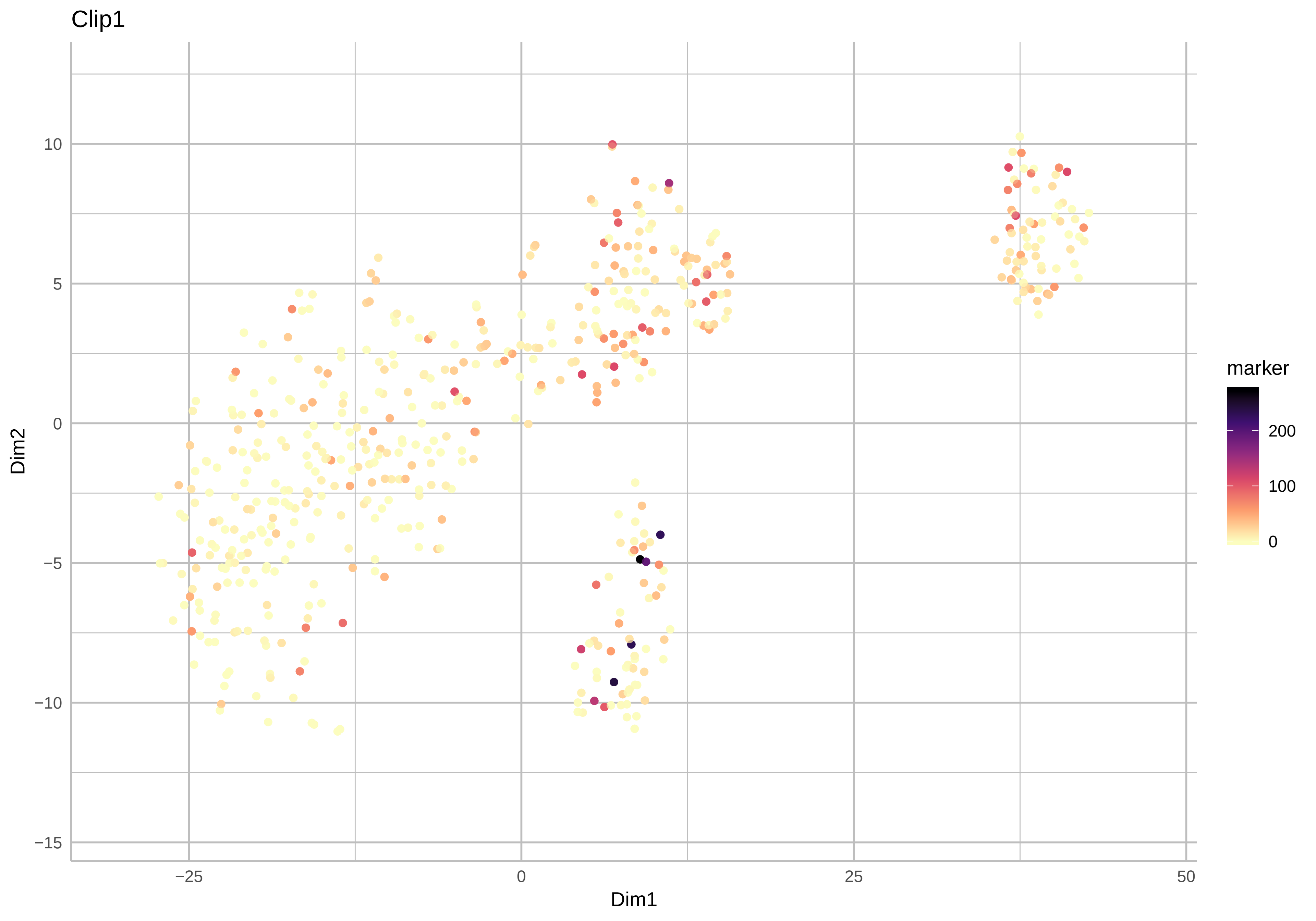
Clip1
Dim2
Dim1
marker
200
100
0
10
5
0
−5
−10
−15
−25
0
25
50

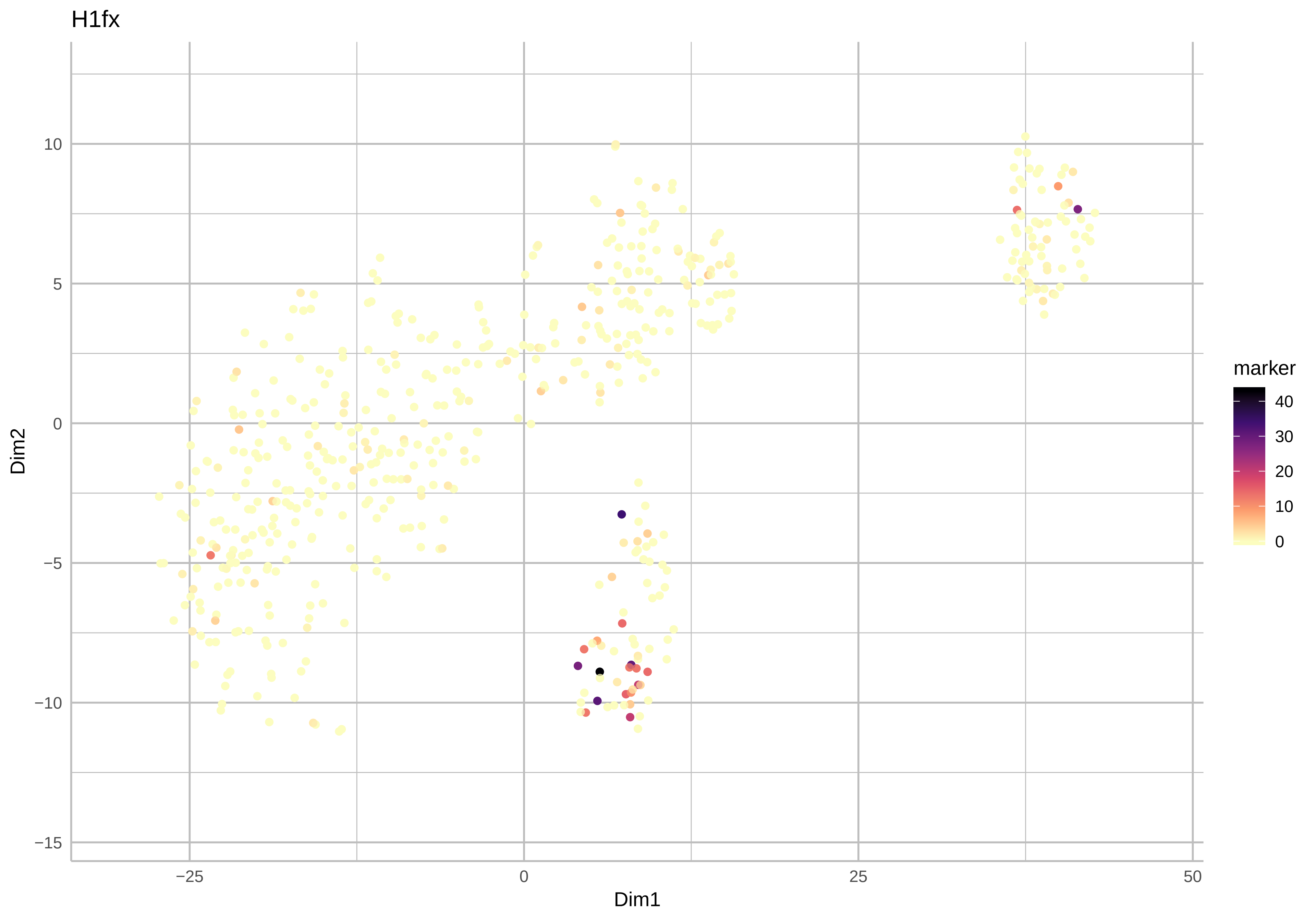
H1fx
Dim2
10
5
0
−5
−10
−15
Dim1
−25
0
25
50
marker
40
30
20
10
0

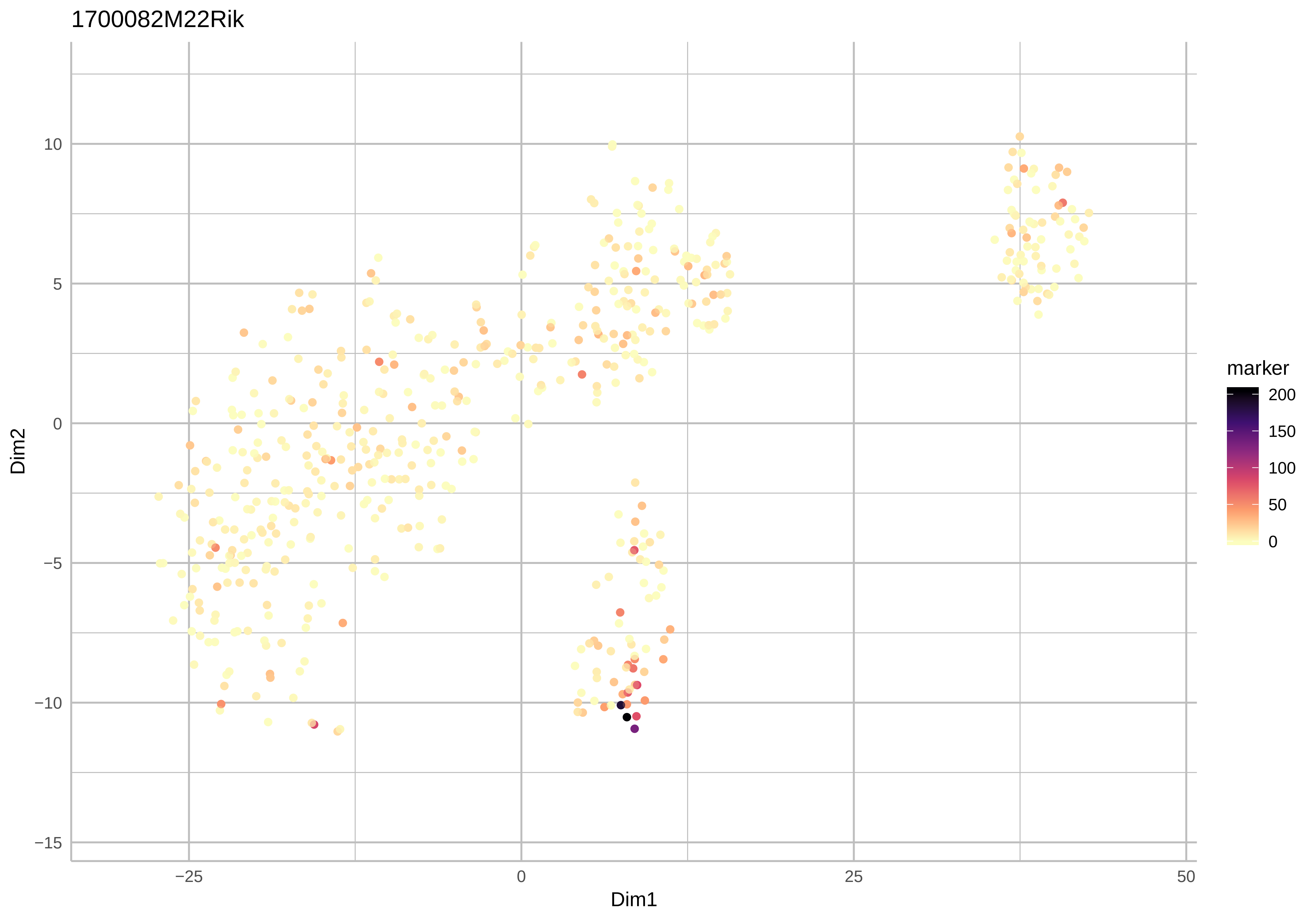
1700082M22Rik
Dim2
Dim1
10
5
0
−5
−10
−15
−25
0
25
50
marker
200
150
100
50
0

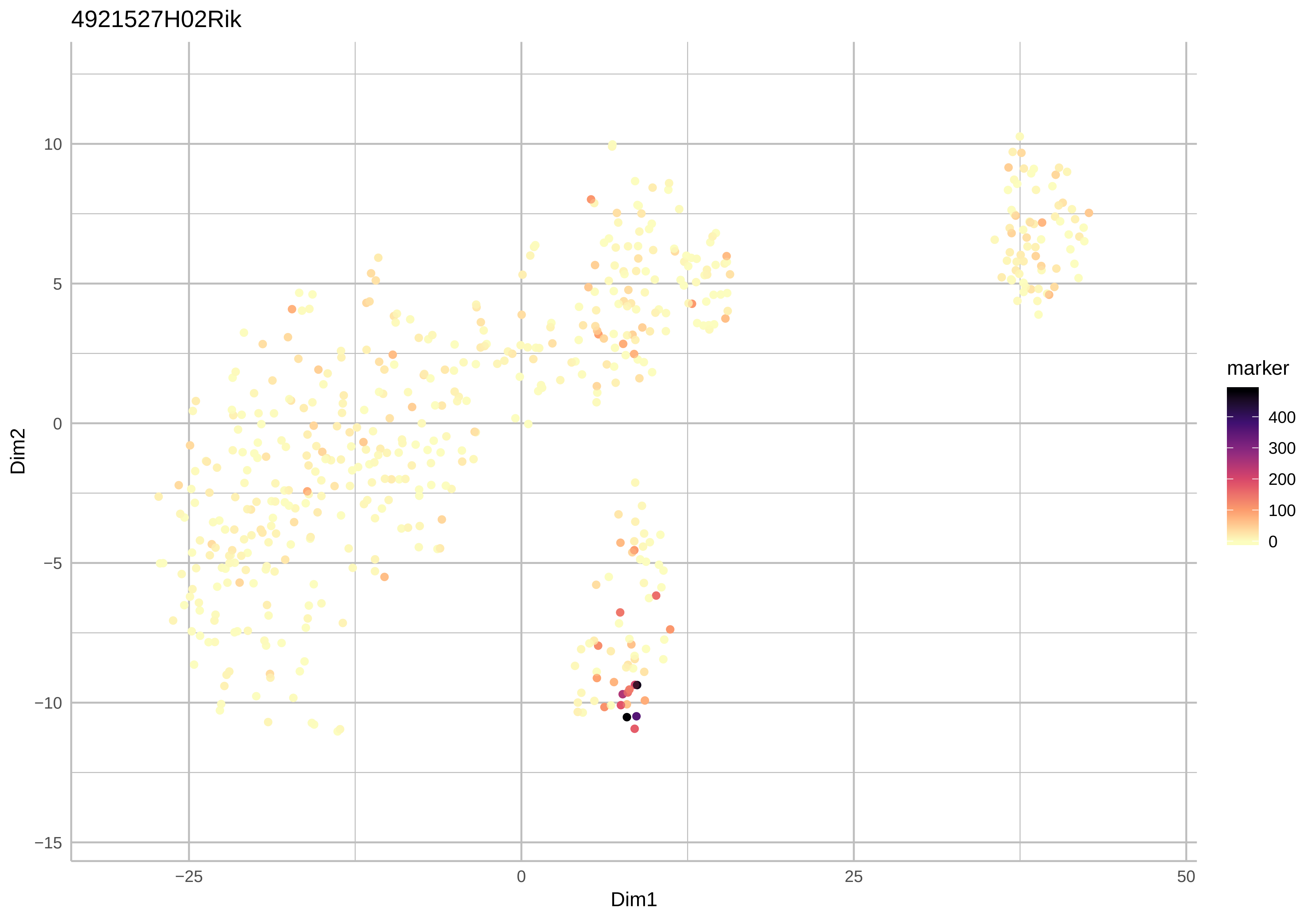
4921527H02Rik
Dim2
Dim1
10
5
0
−5
−10
−15
−25
0
25
50
marker
400
300
200
100
0

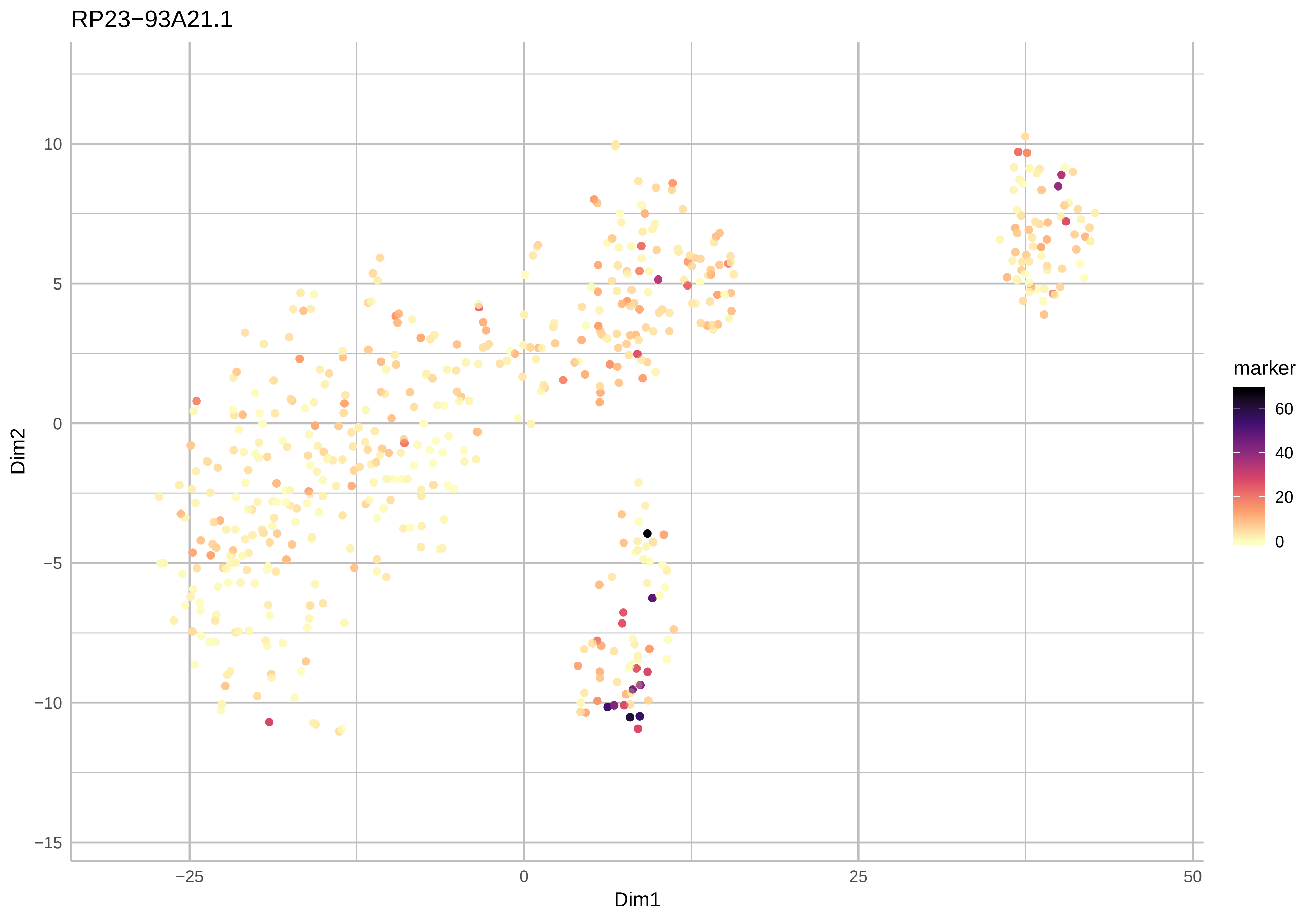
RP23−93A21.1
Dim2
Dim1
−25
0
25
50
10
5
0
−5
−10
−15
marker
60
40
20
0

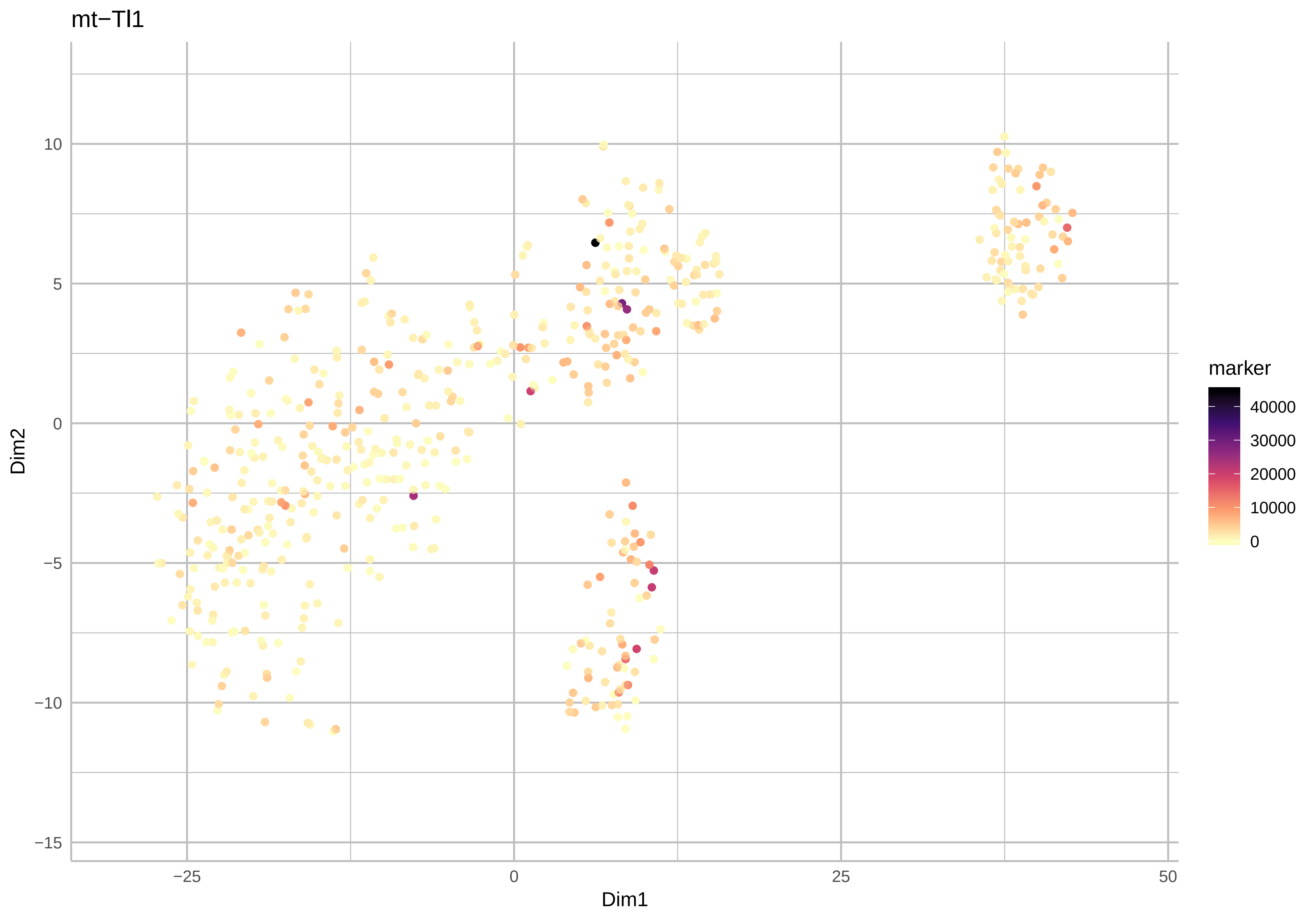
mt−Tl1
Dim2
Dim1
10
5
0
−5
−10
−15
−25
0
25
50
marker
40000
30000
20000
10000
0

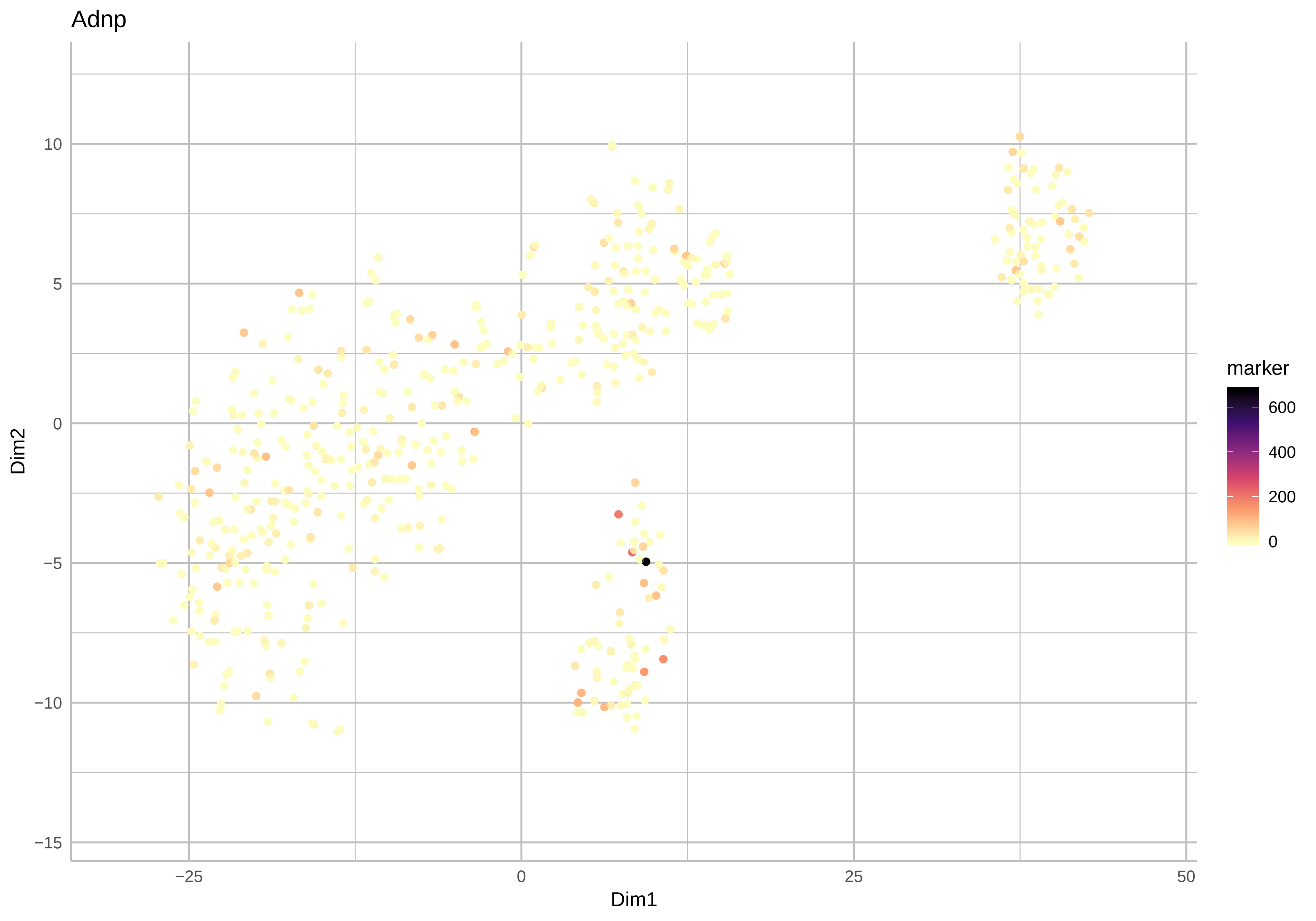

Adnp
Dim2
Dim1
10
5
0
−5
−10
−15
−25
0
25
50
marker
600
400
200
0

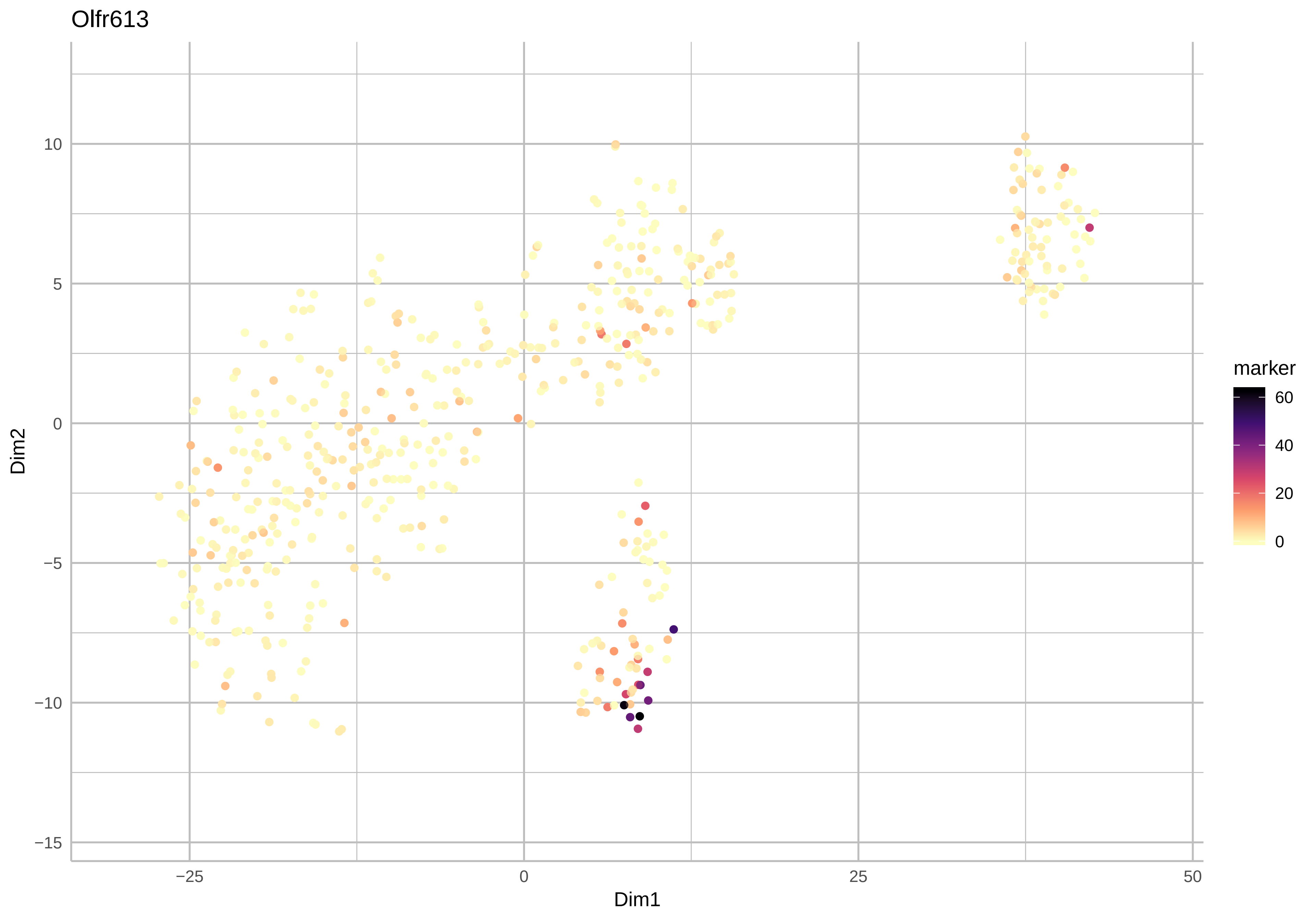

Olfr613
10
5
0
−5
−10
−15
Dim2
−25
0
25
50
Dim1
marker
60
40
20
0

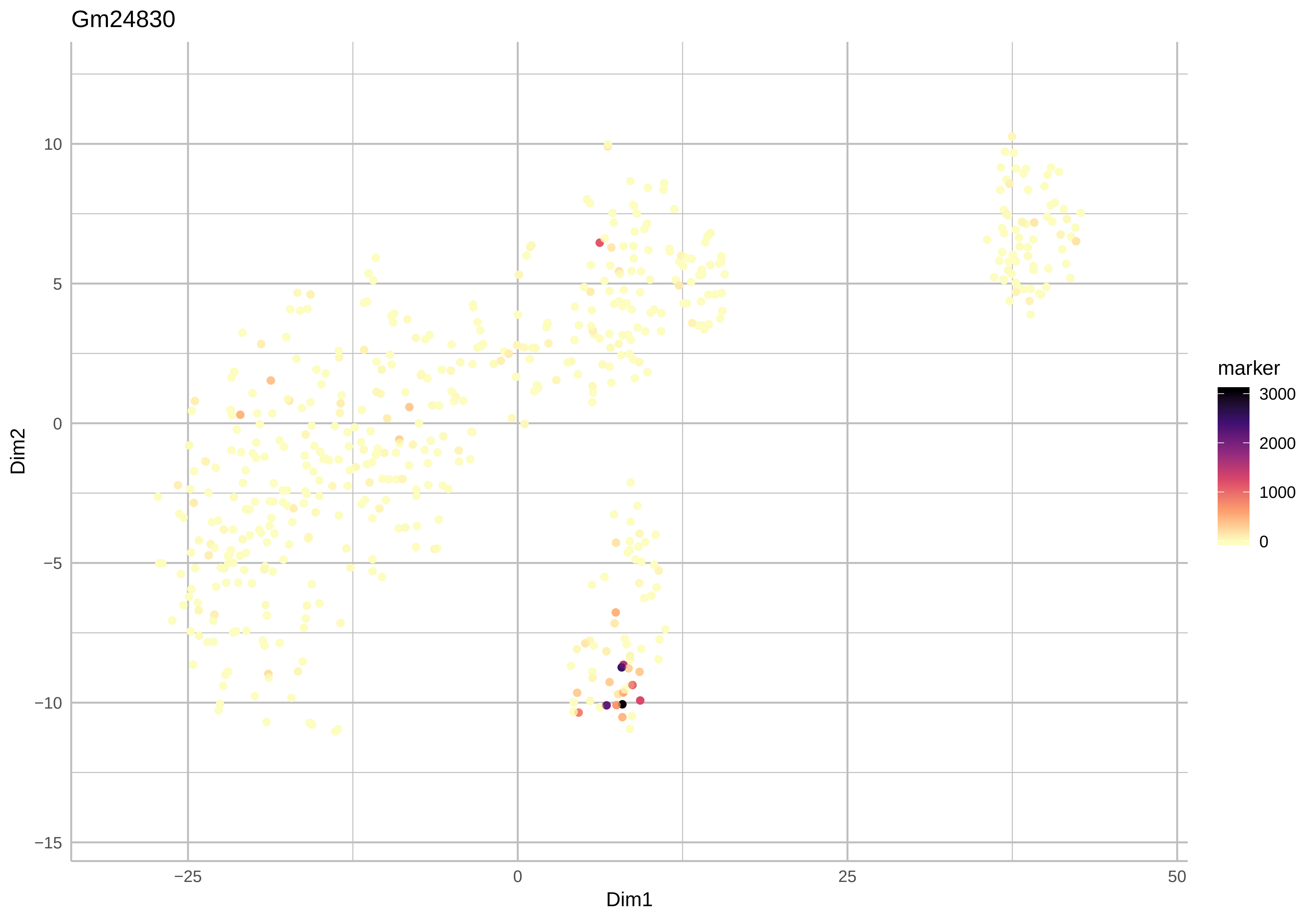
Gm24830
Dim2
Dim1
10
5
0
−5
−10
−15
−25
0
25
50
marker
3000
2000
1000
0

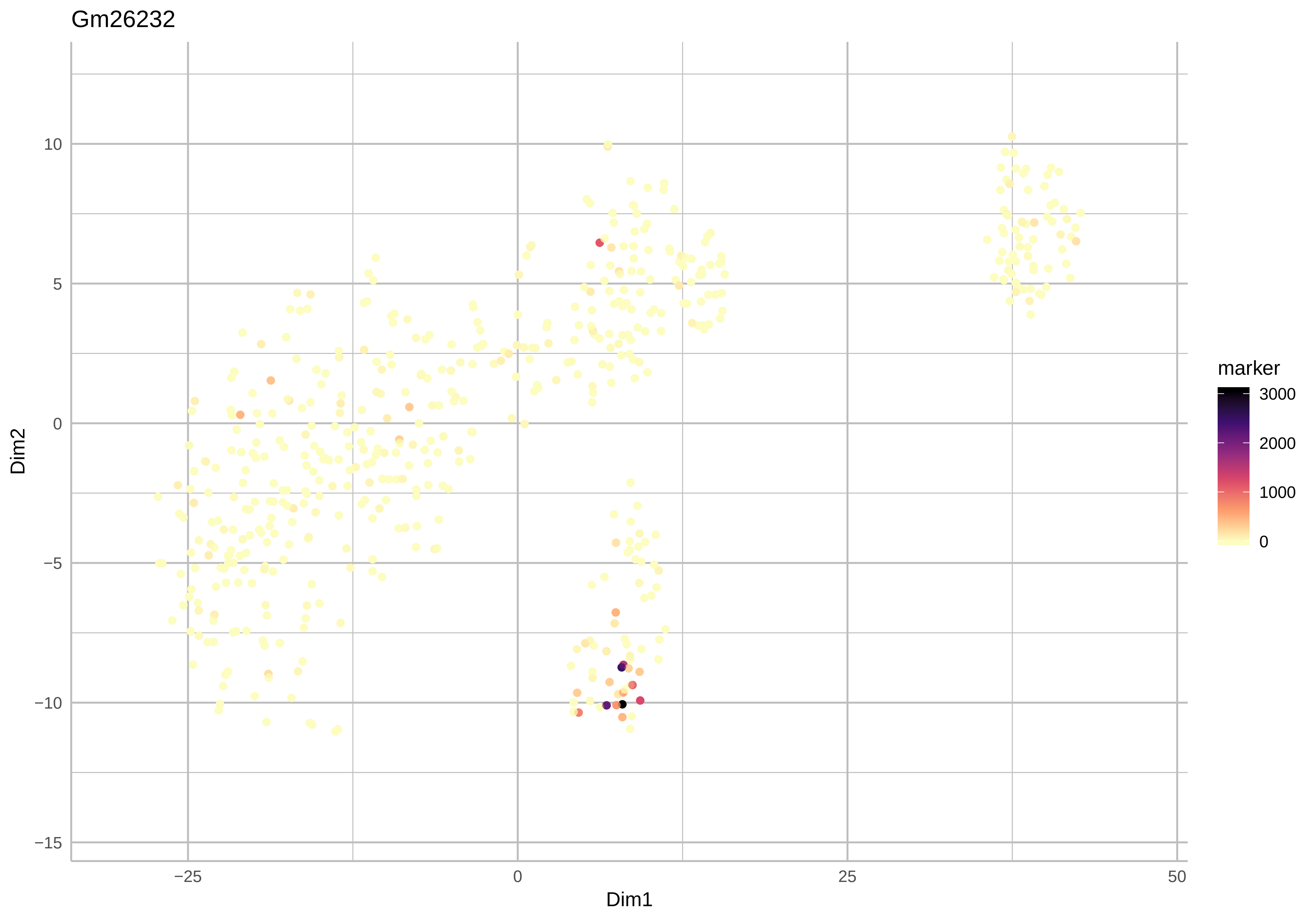

Gm26232
10
5
0
−5
−10
−15
−25
0
25
50
Dim1
Dim2
marker
3000
2000
1000
0

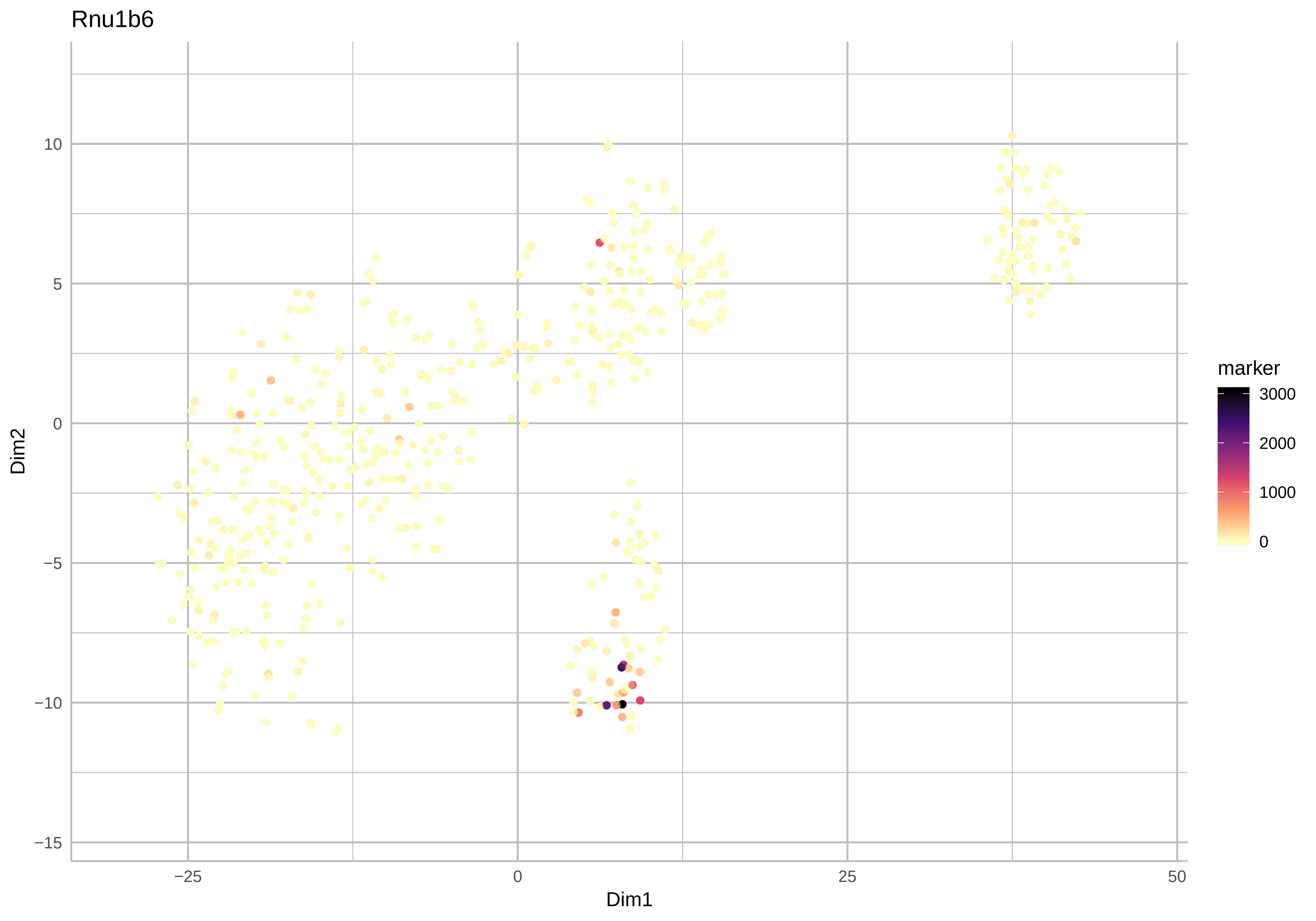
Rnu1b6
Dim2
Dim1
10
5
0
−5
−10
−15
−25
0
25
50
marker
3000
2000
1000
0

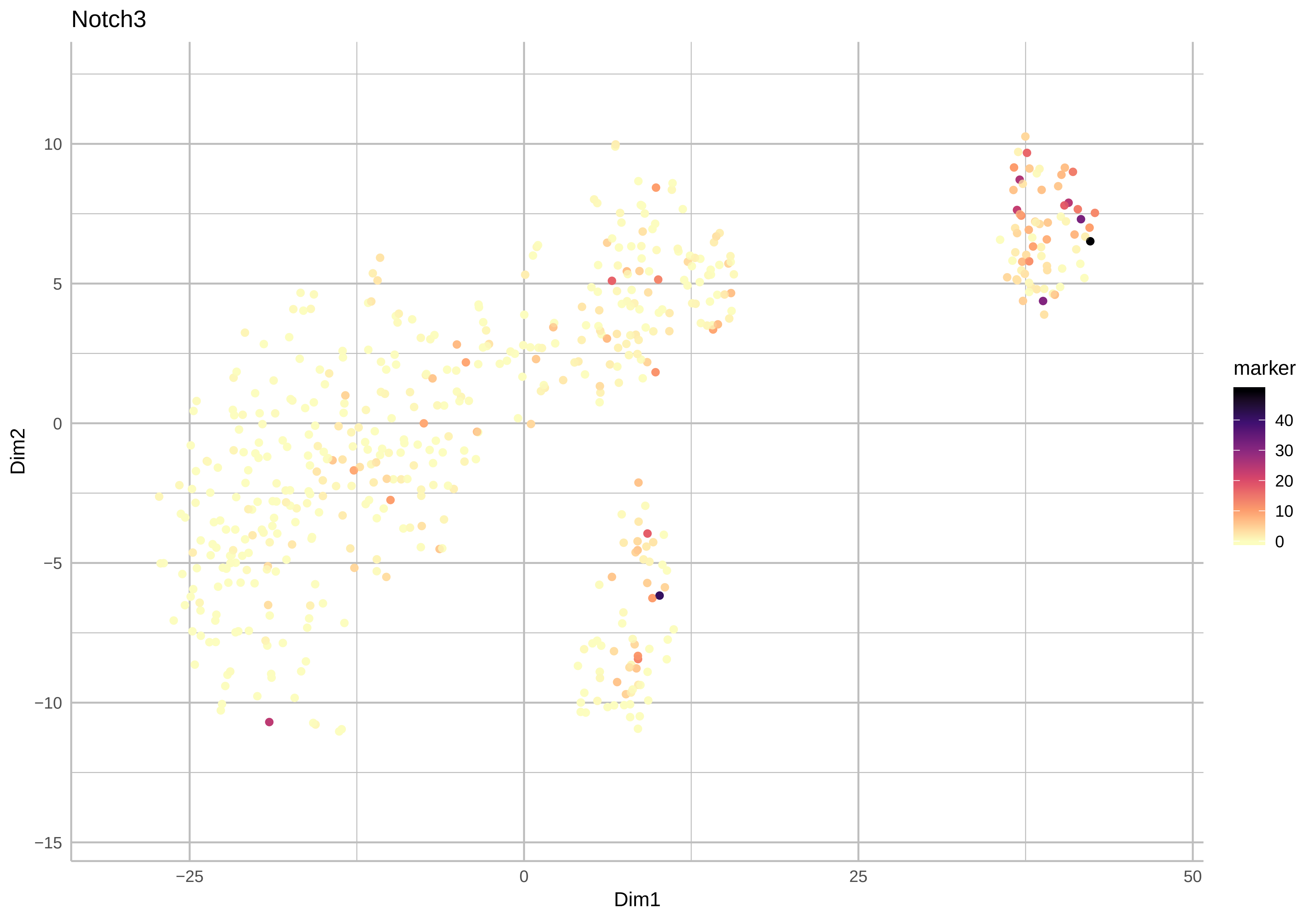

Notch3
Dim2
Dim1
10
5
0
−5
−10
−15
−25
0
25
50
marker
40
30
20
10
0

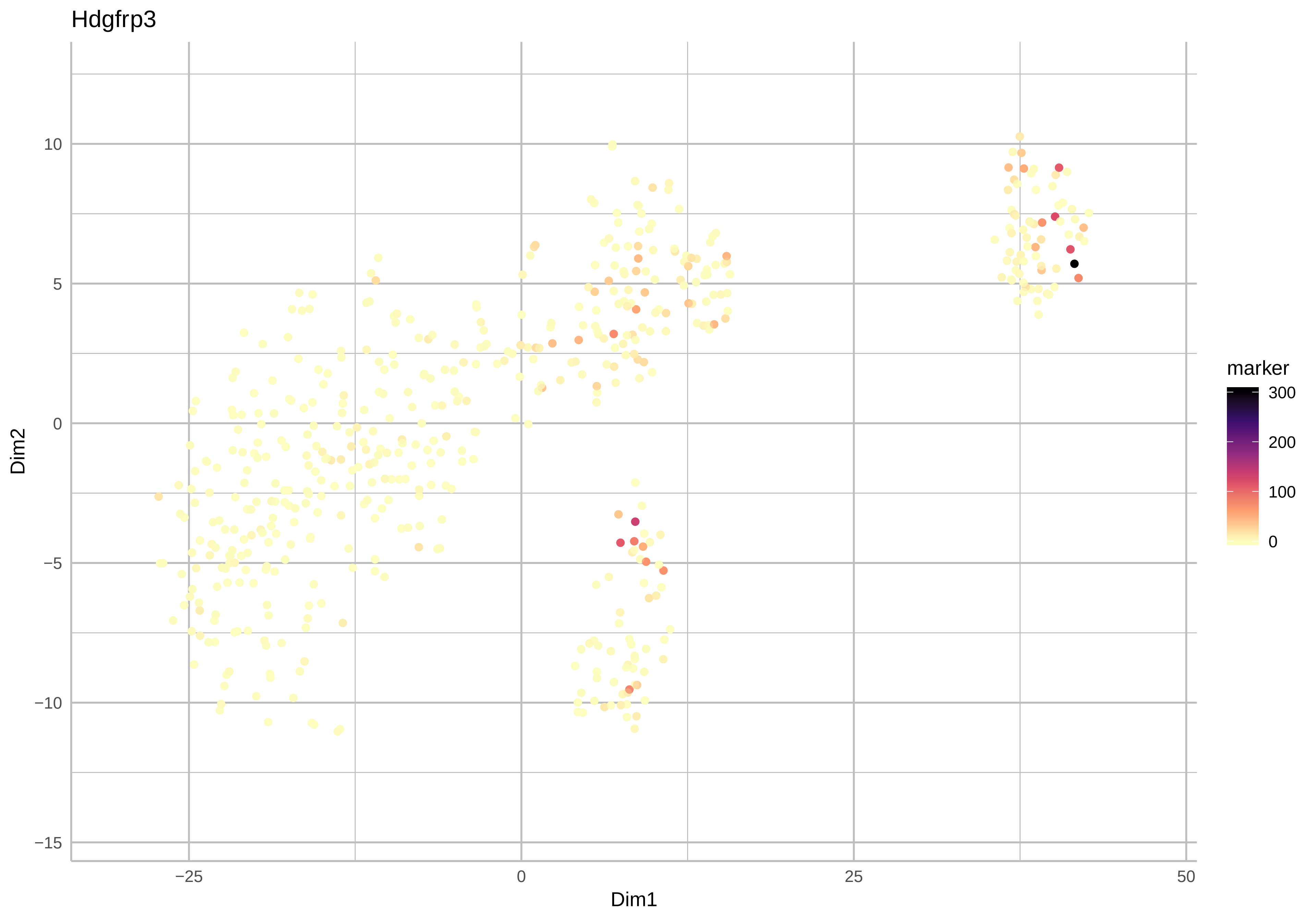
Hdgfrp3
Dim2
Dim1
10
5
0
−5
−10
−15
−25
0
25
50
marker
300
200
100
0

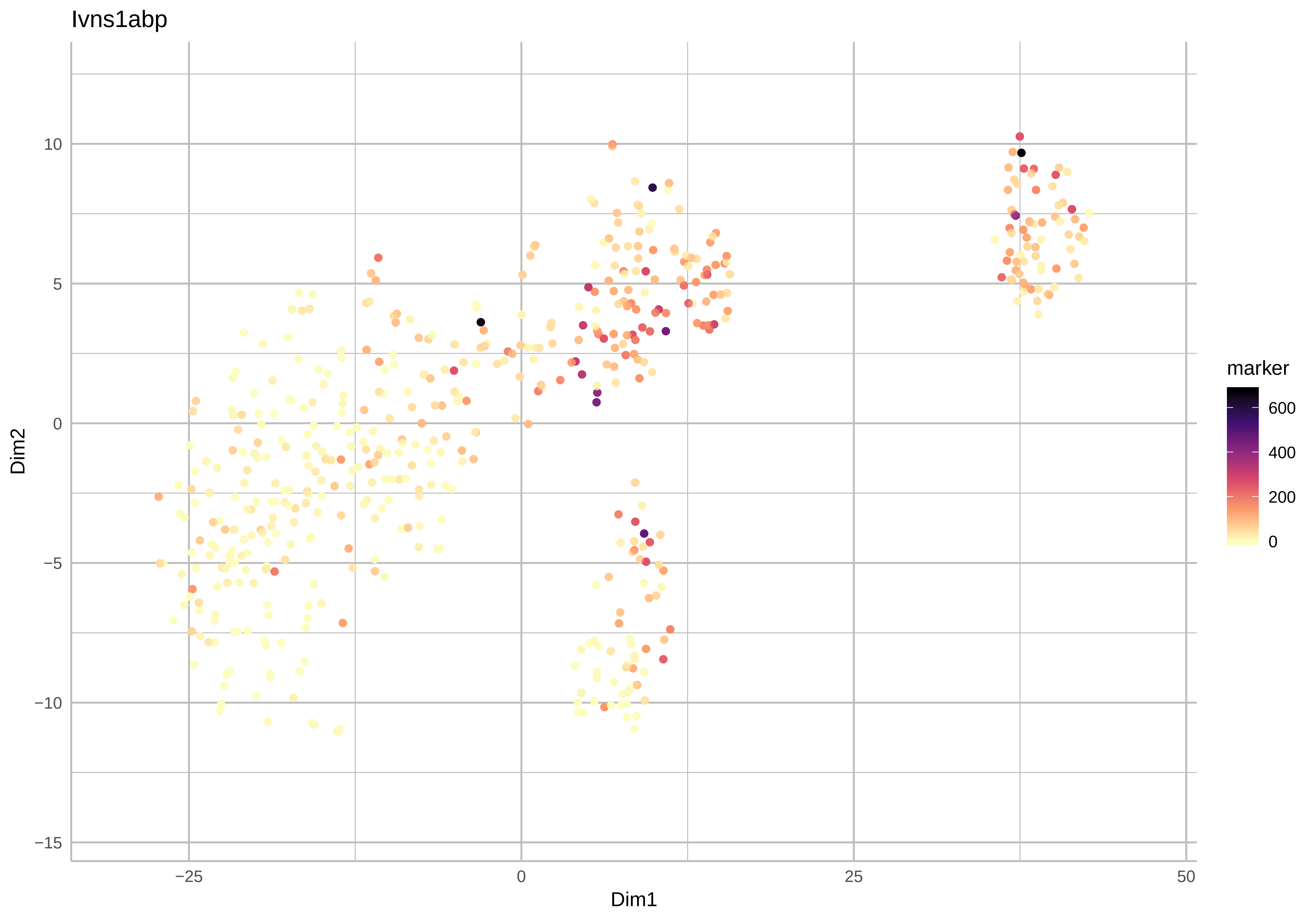

Ivns1abp
10
5
0
−5
−10
−15
−25
0
25
50
Dim1
Dim2
marker
600
400
200
0

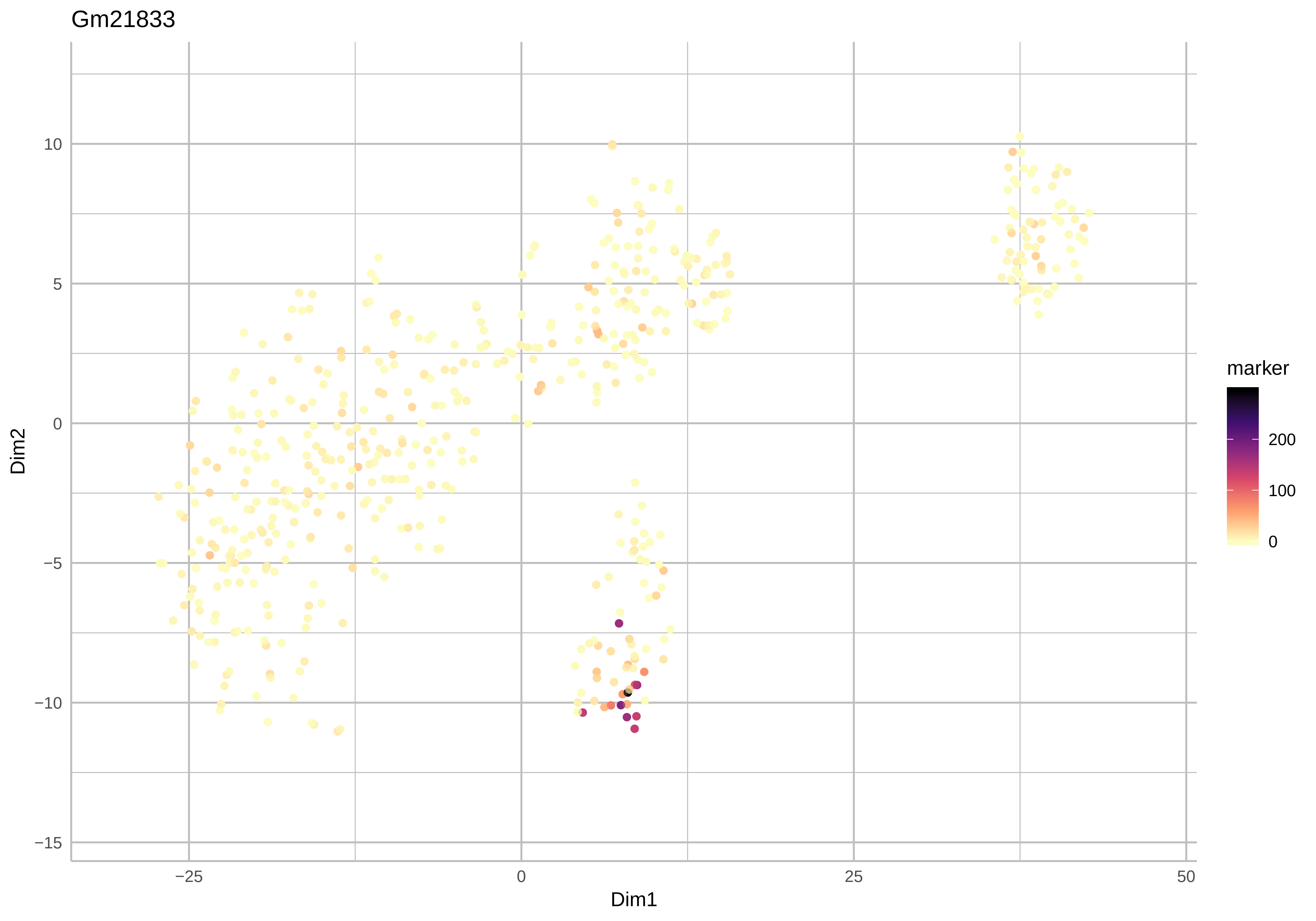
Gm21833
Dim2
10
5
0
−5
−10
−15
−25
0
25
50
Dim1
marker
200
100
0

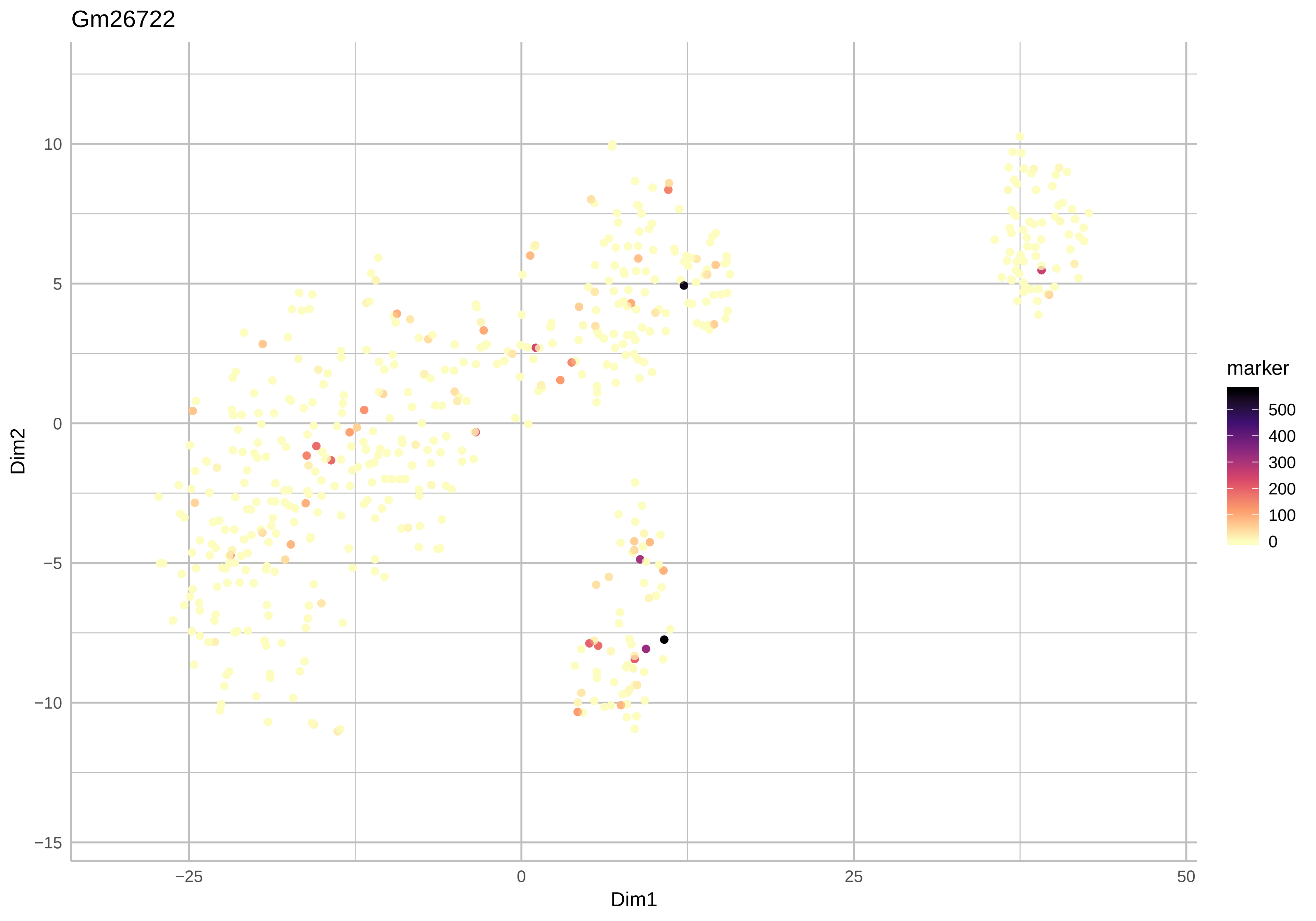

Gm26722
10
5
0
−5
−10
−15
−25
0
25
50
Dim1
Dim2
marker
500
400
300
200
100
0

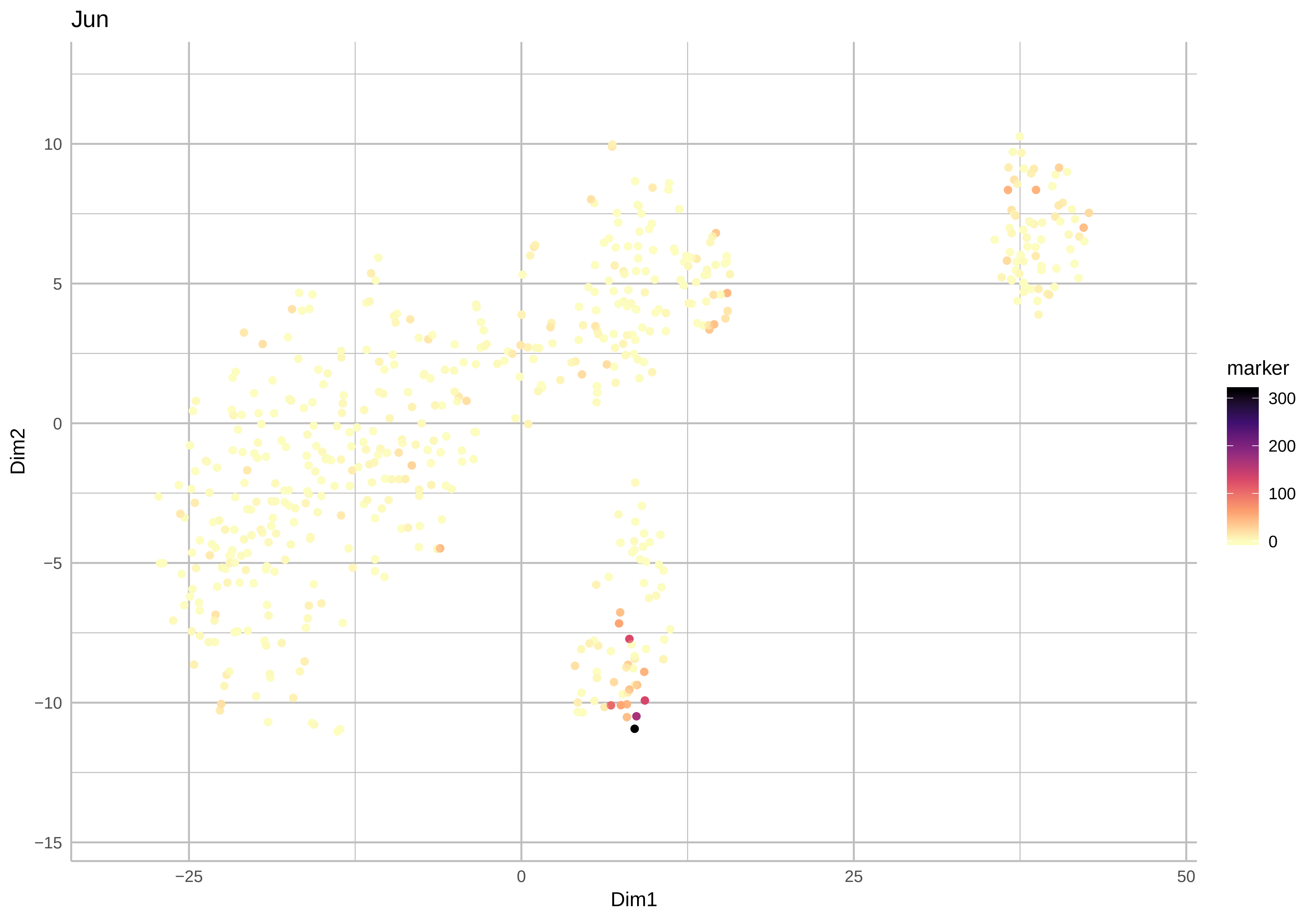

Jun
Dim2
Dim1
10
5
0
−5
−10
−15
−25
0
25
50
marker
300
200
100
0

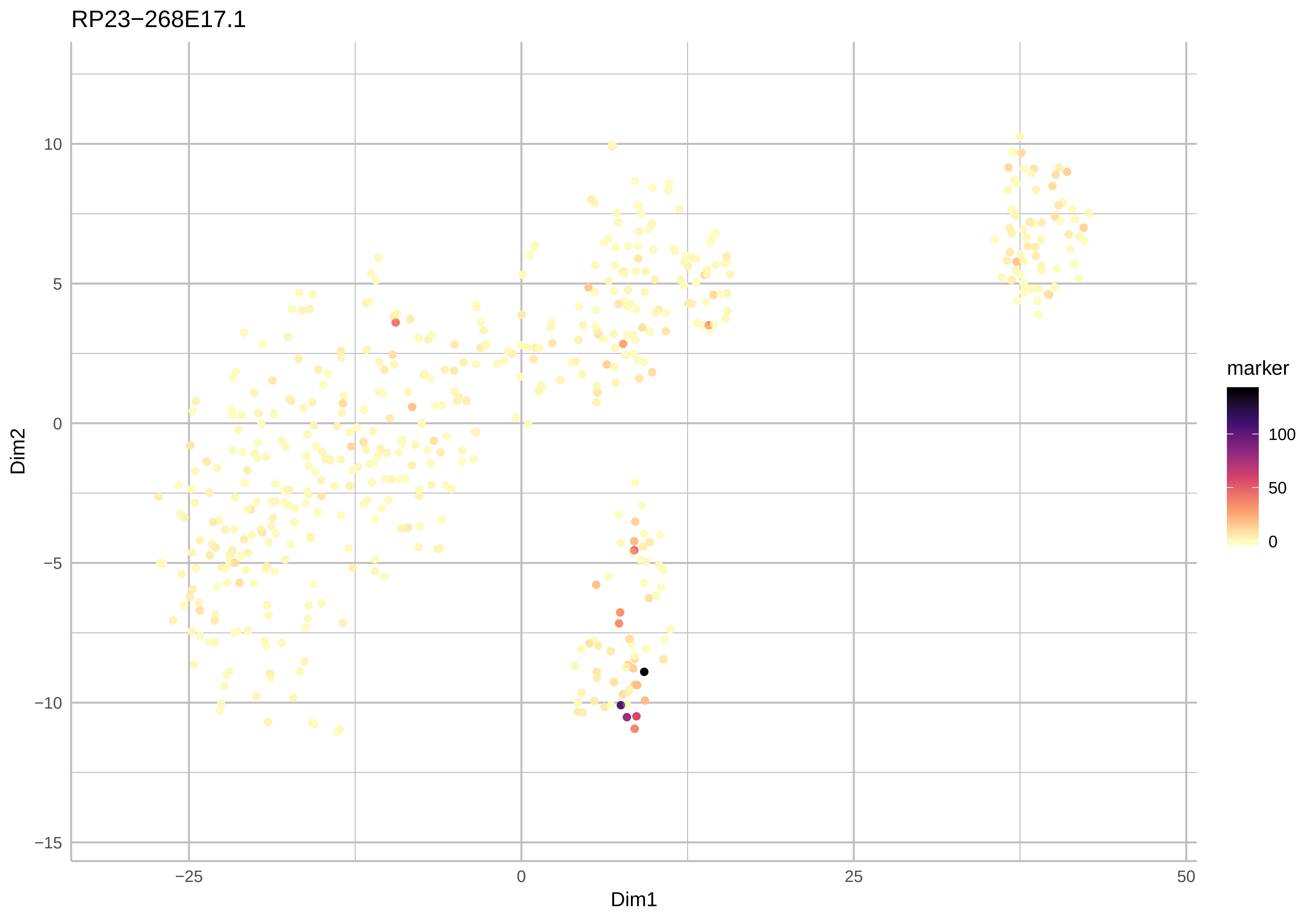

RP23−268E17.1
Dim2
Dim1
marker
100
50
0
10
5
0
−5
−10
−15
−25
0
25
50

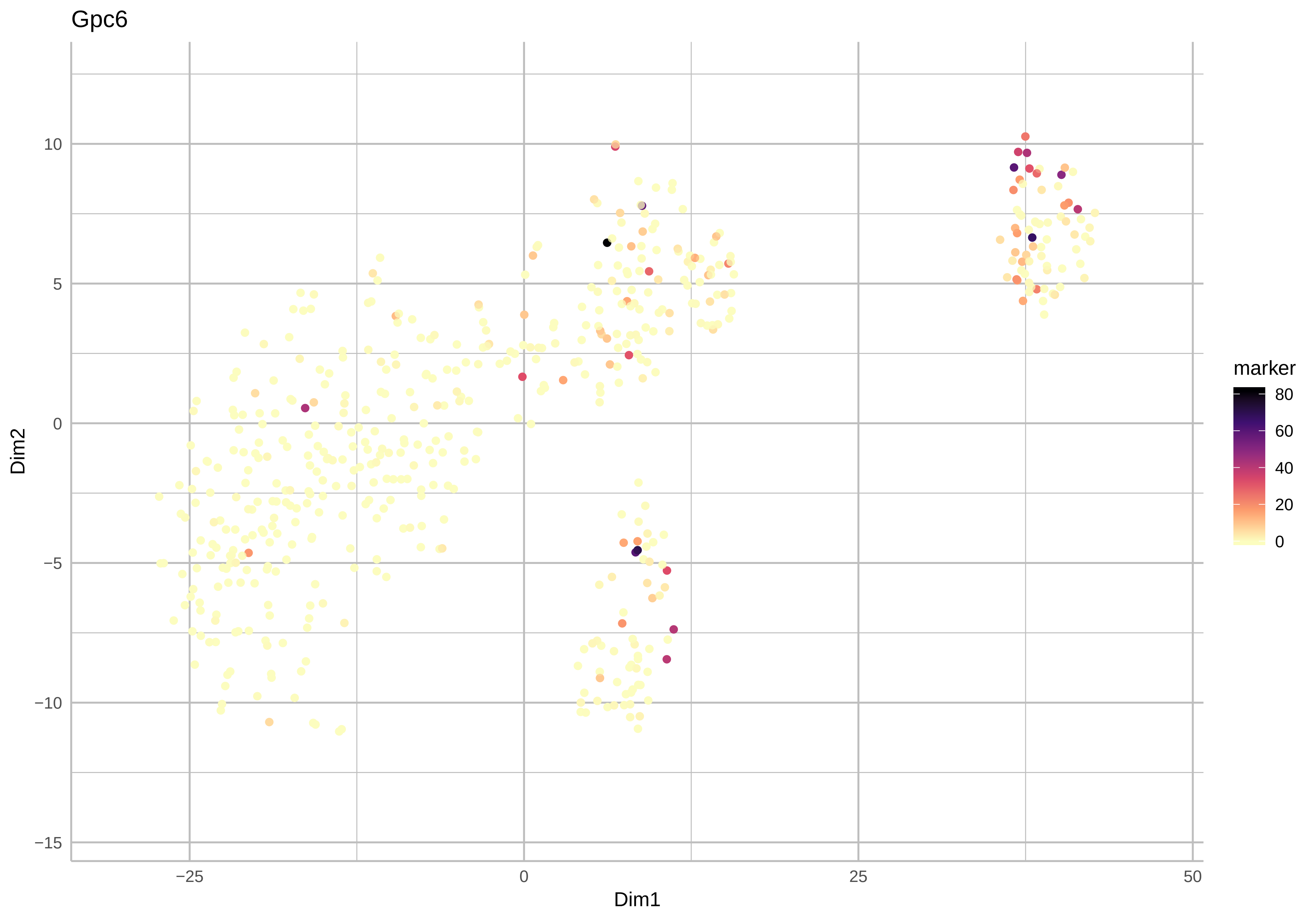

Gpc6
Dim2
Dim1
10
5
0
−5
−10
−15
−25
0
25
50
marker
80
60
40
20
0

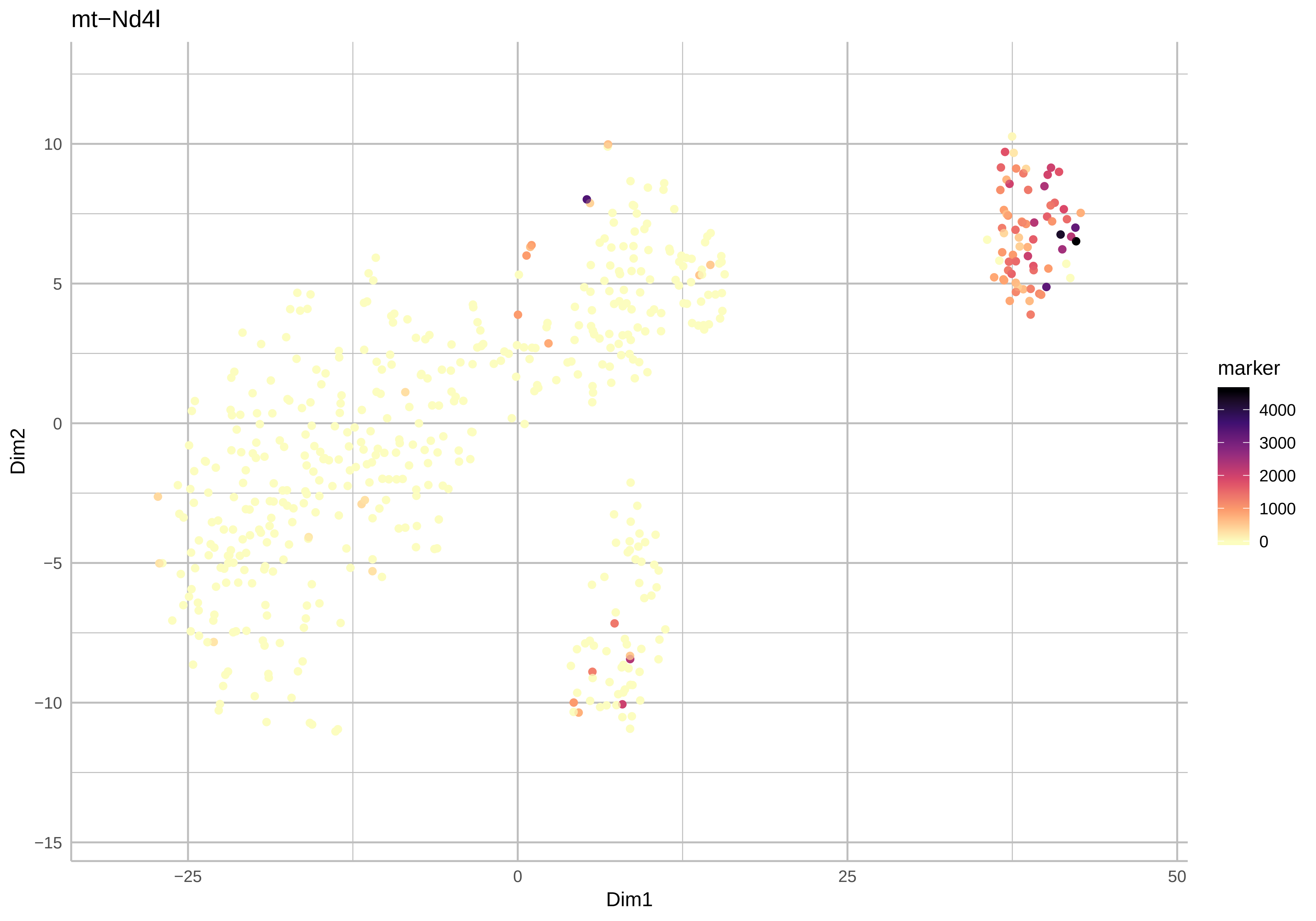
mt−Nd4l
Dim2
Dim1
10
5
0
−5
−10
−15
−25
0
25
50
marker
4000
3000
2000
1000
0

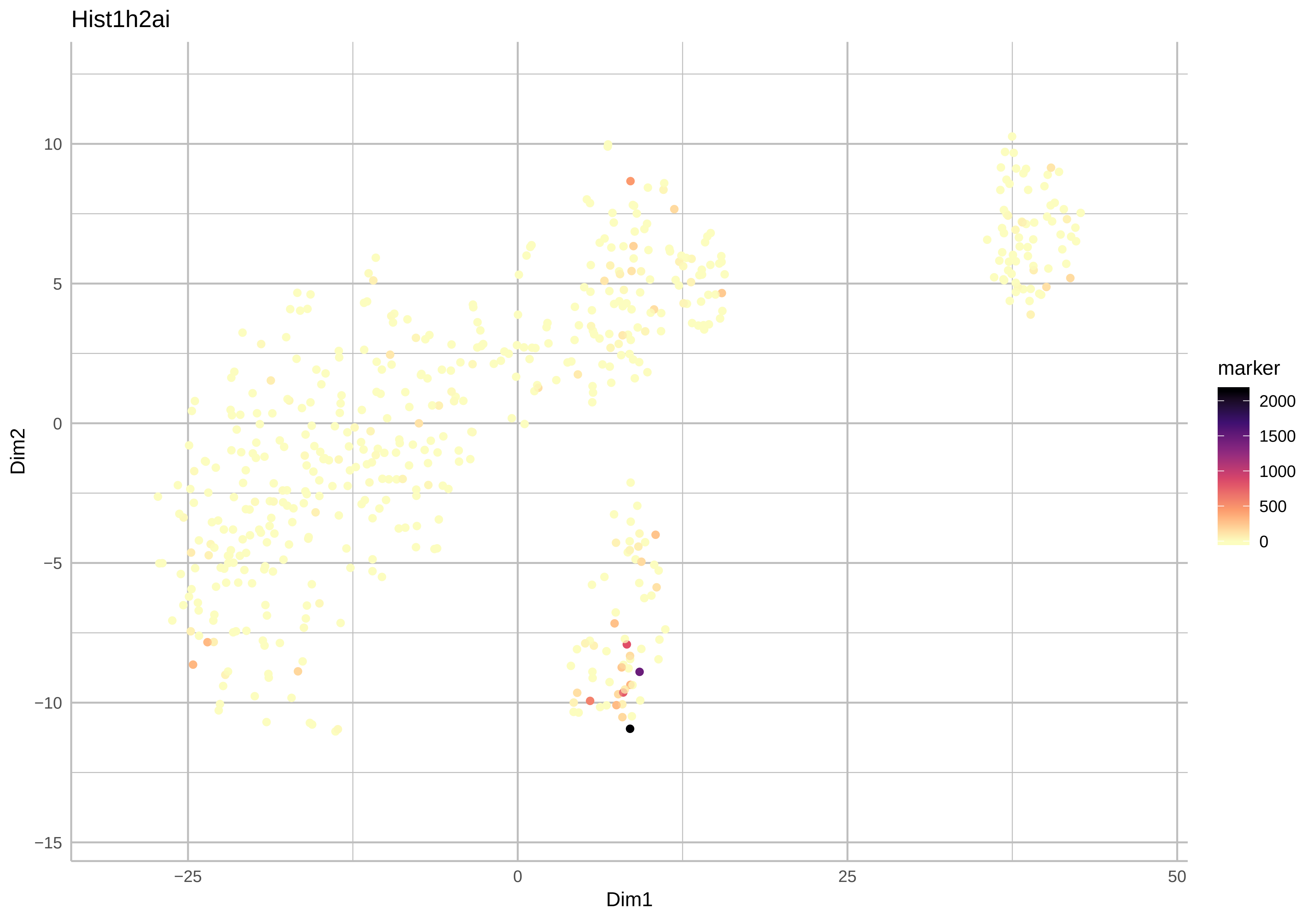

Hist1h2ai
Dim2
Dim1
10
5
0
−5
−10
−15
−25
0
25
50
marker
2000
1500
1000
500
0

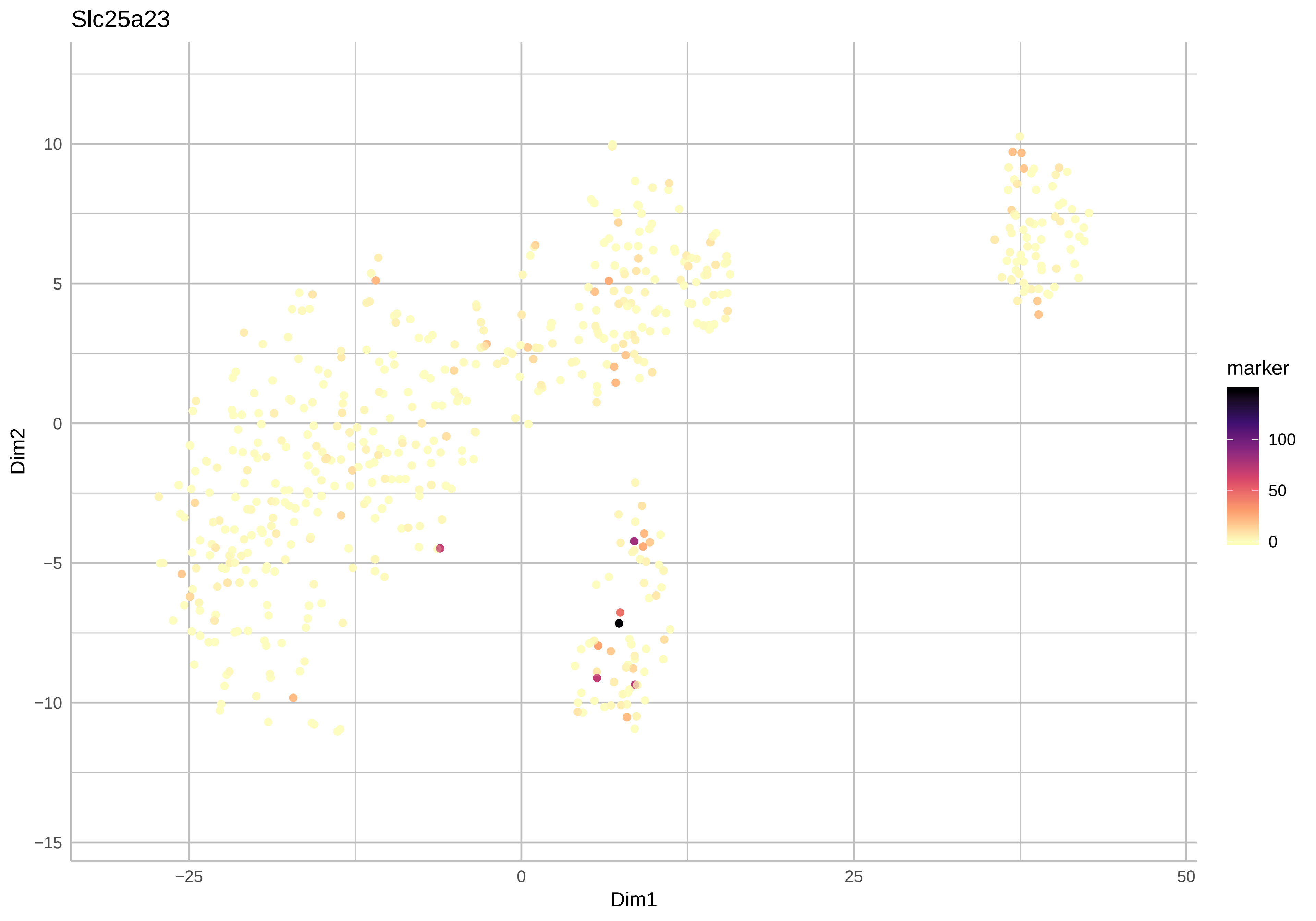
Slc25a23
Dim2
Dim1
10
5
0
−5
−10
−15
−25
0
25
50
marker
100
50
0

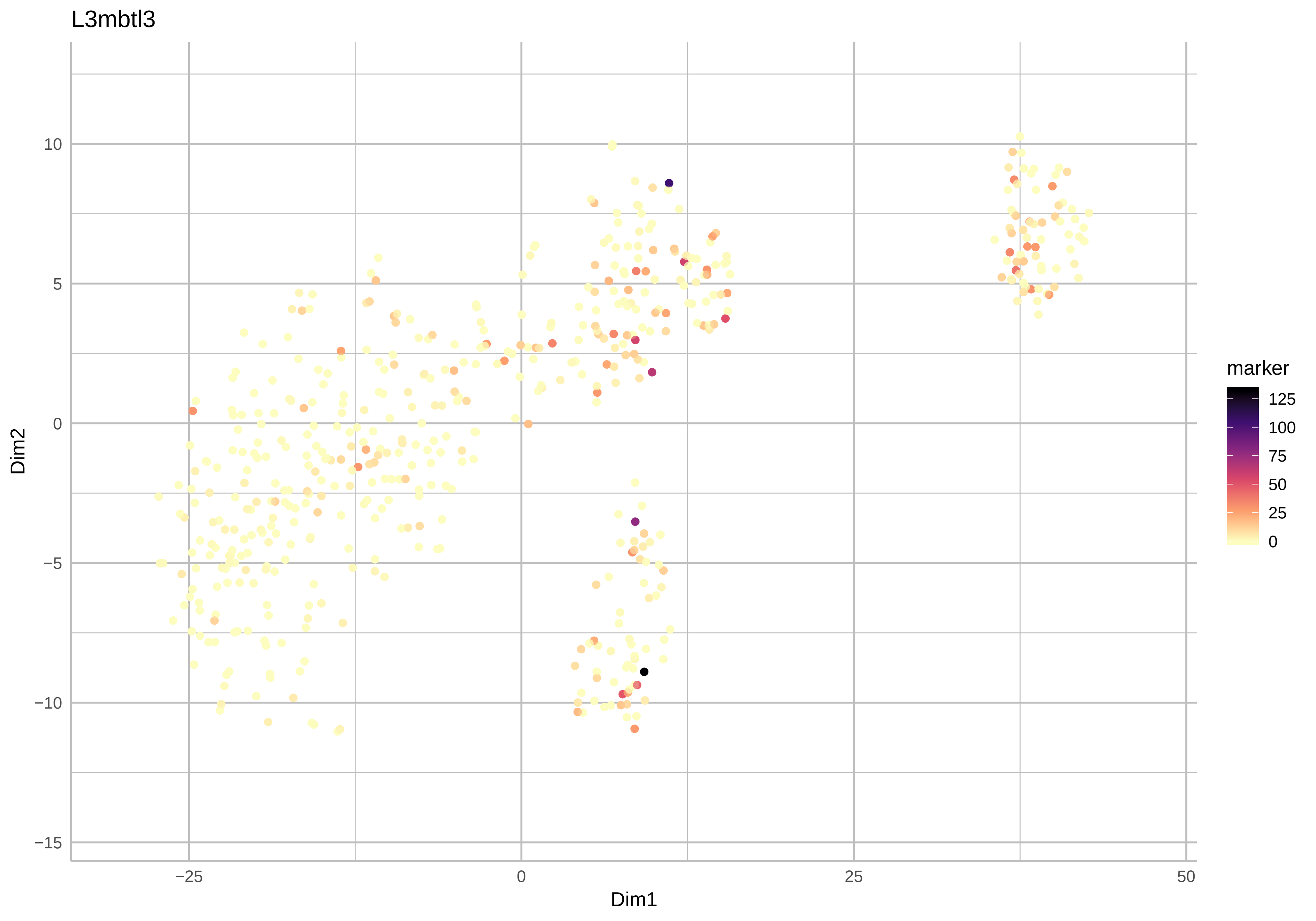

L3mbtl3
Dim2
Dim1
10
5
0
−5
−10
−15
−25
0
25
50
marker
125
100
75
50
25
0

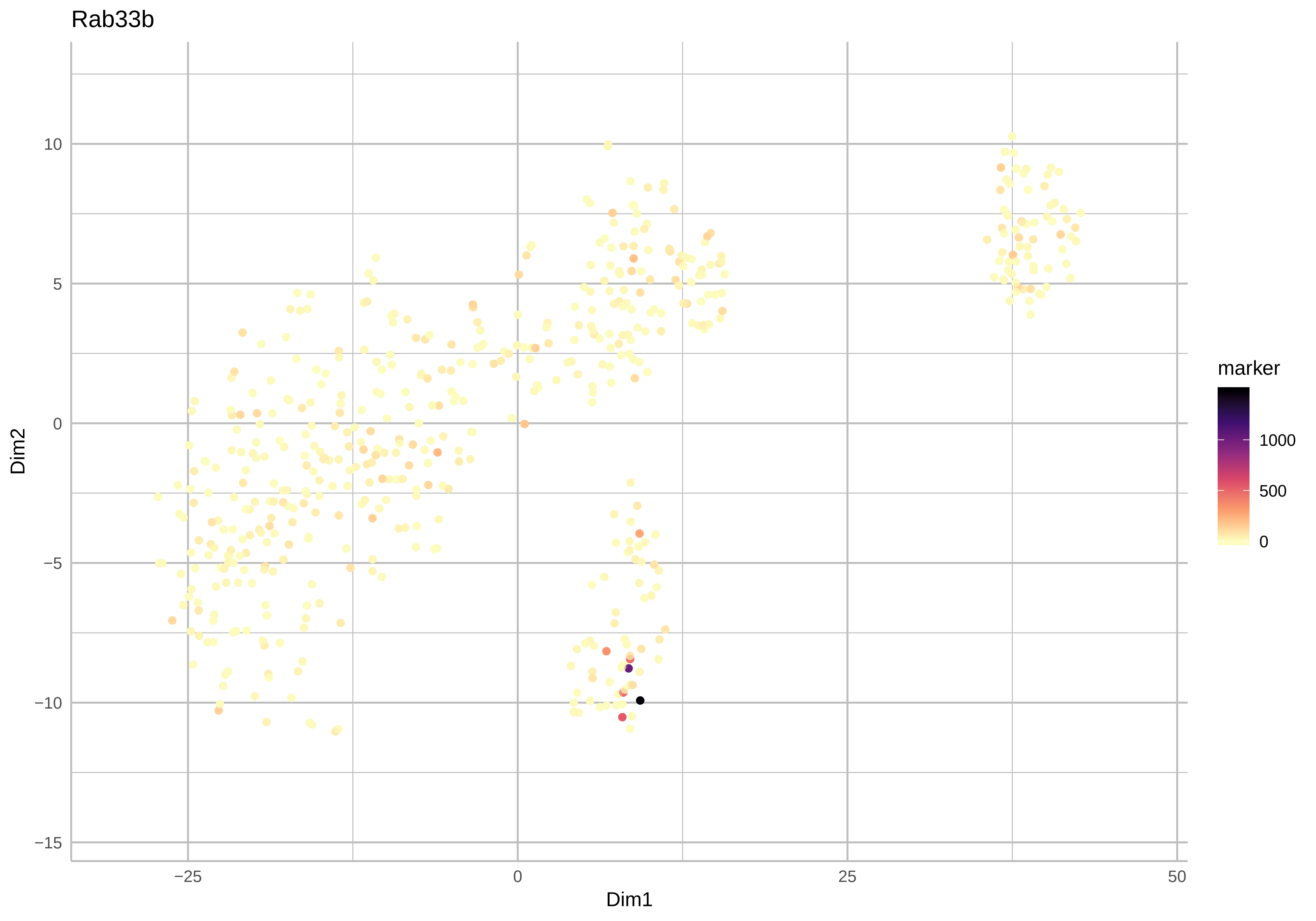

Rab33b
Dim2
Dim1
marker
1000
500
0
10
5
0
−5
−10
−15
−25
0
25
50

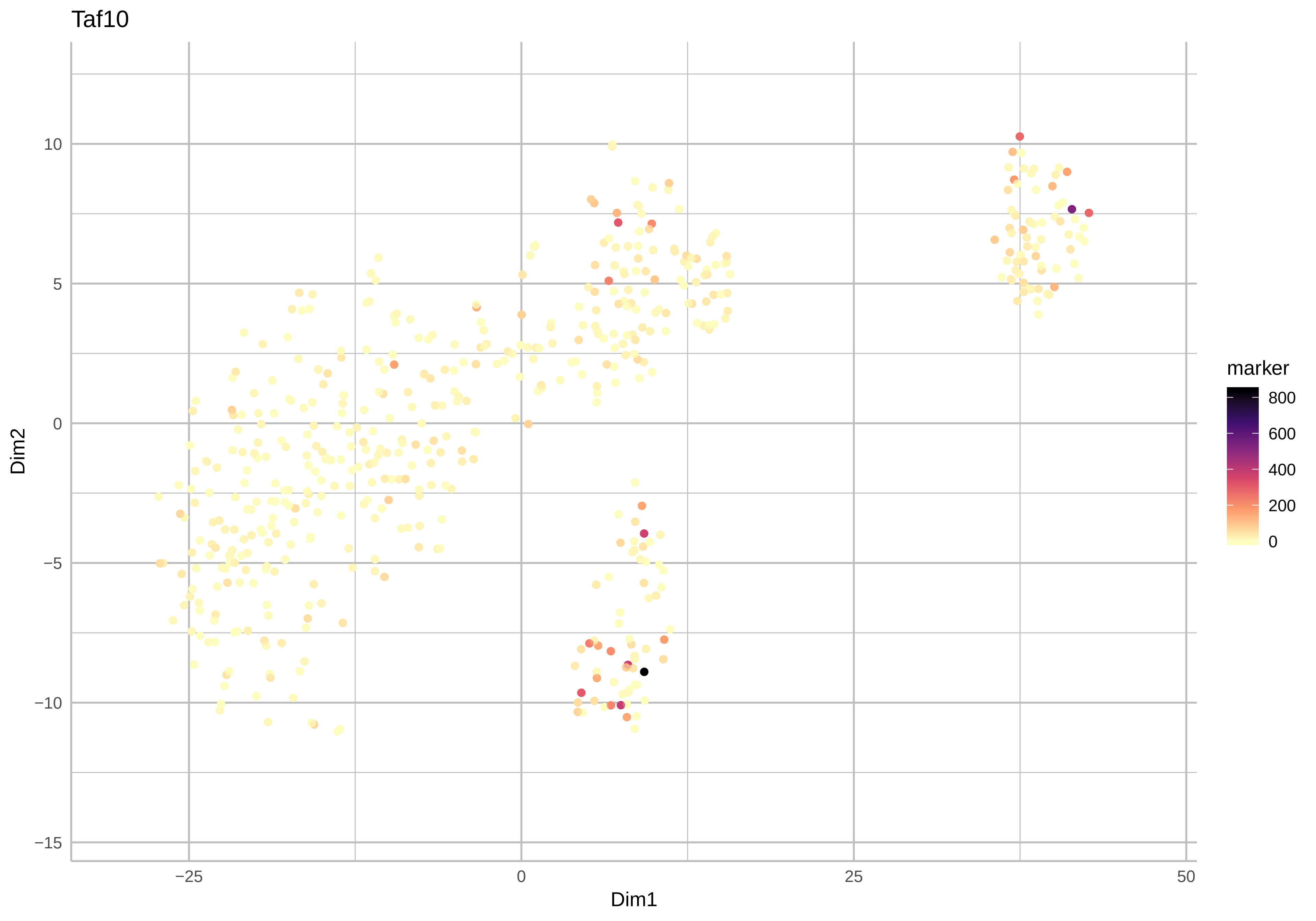

Taf10
Dim2
Dim1
−25
0
25
50
10
5
0
−5
−10
−15
marker
800
600
400
200
0

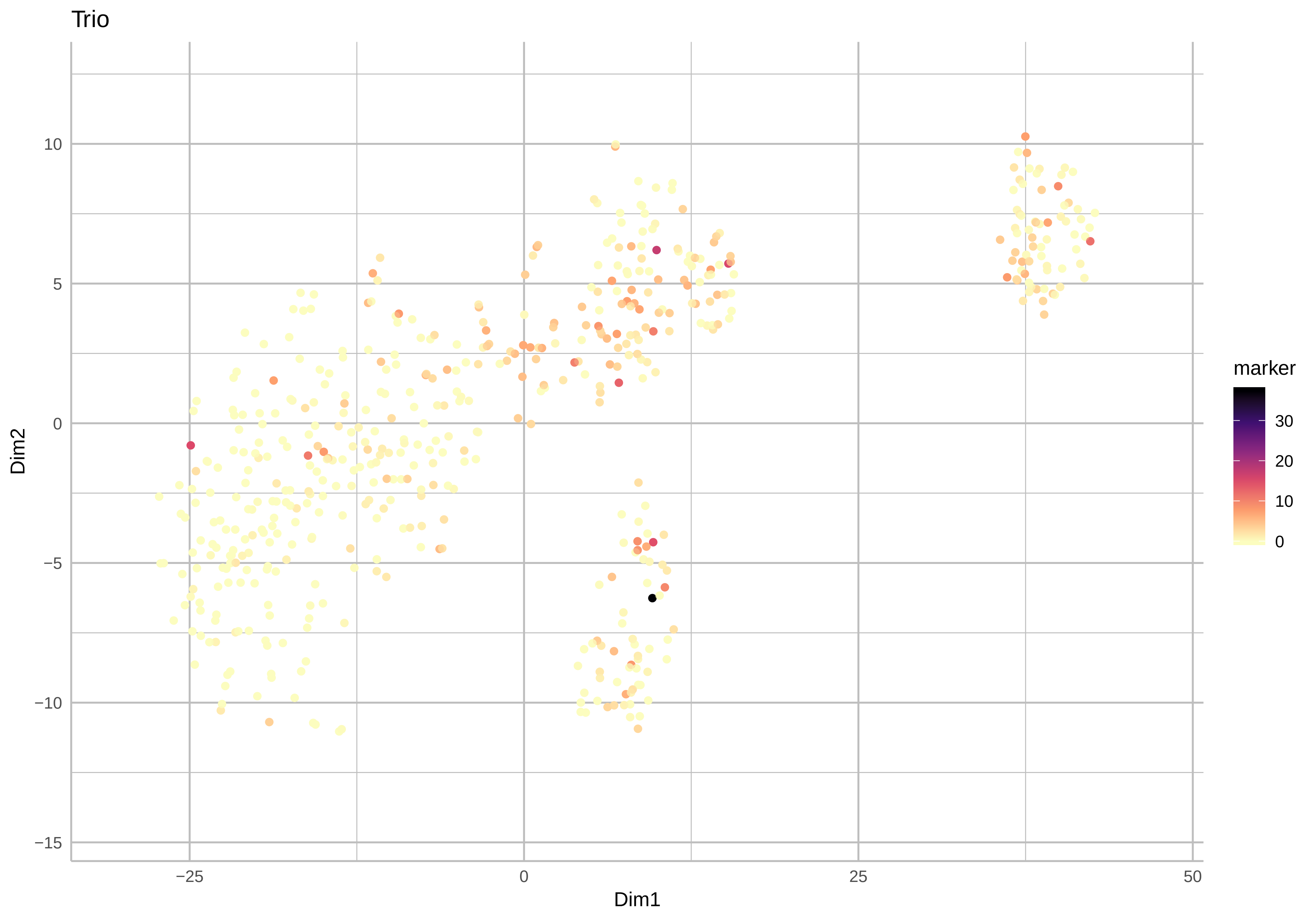

Trio
Dim2
Dim1
marker
30
20
10
0
10
5
0
−5
−10
−15
−25
0
25
50

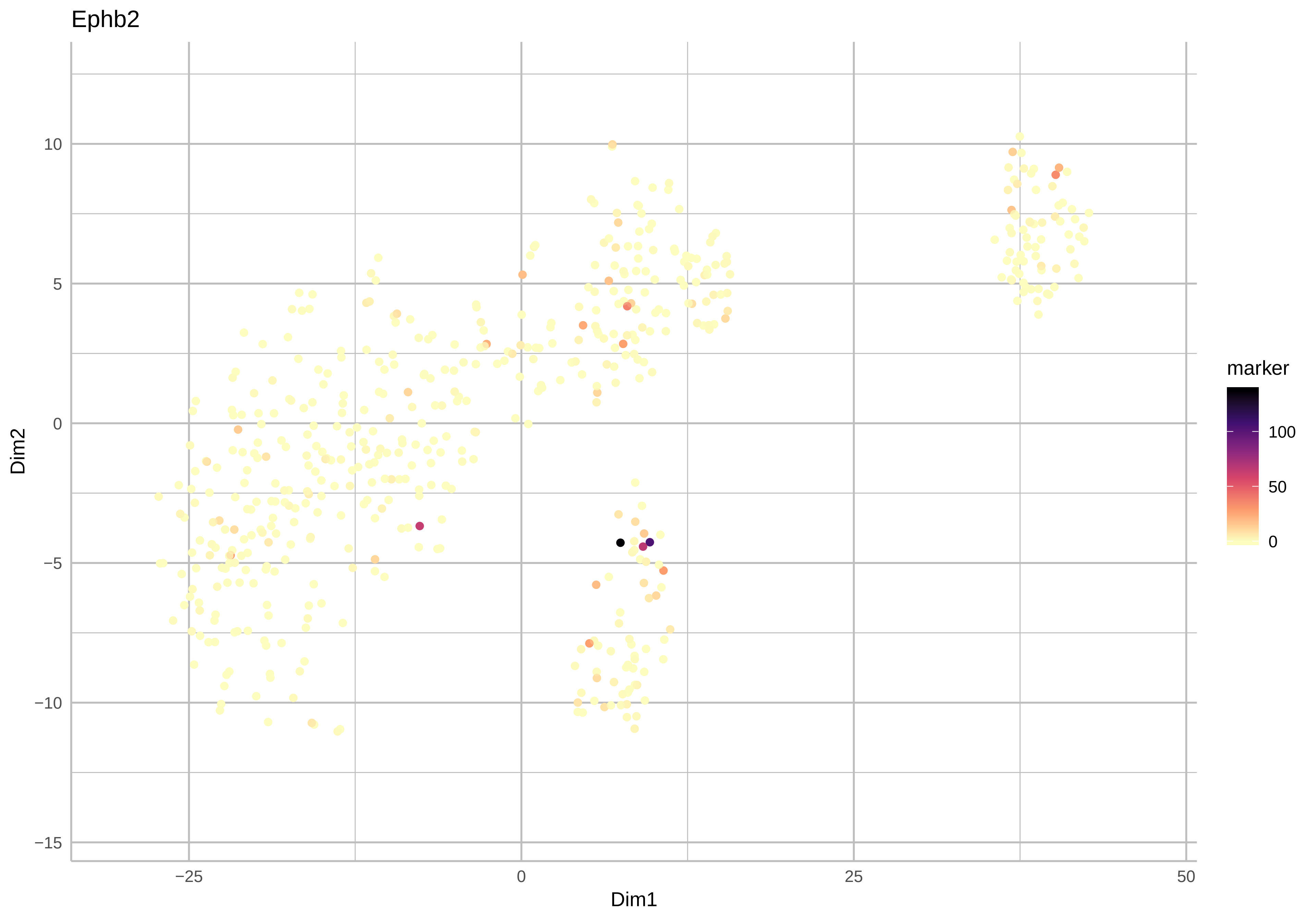
Ephb2
Dim2
Dim1
marker
10
5
0
−5
−10
−15
−25
0
25
50
100
50
0

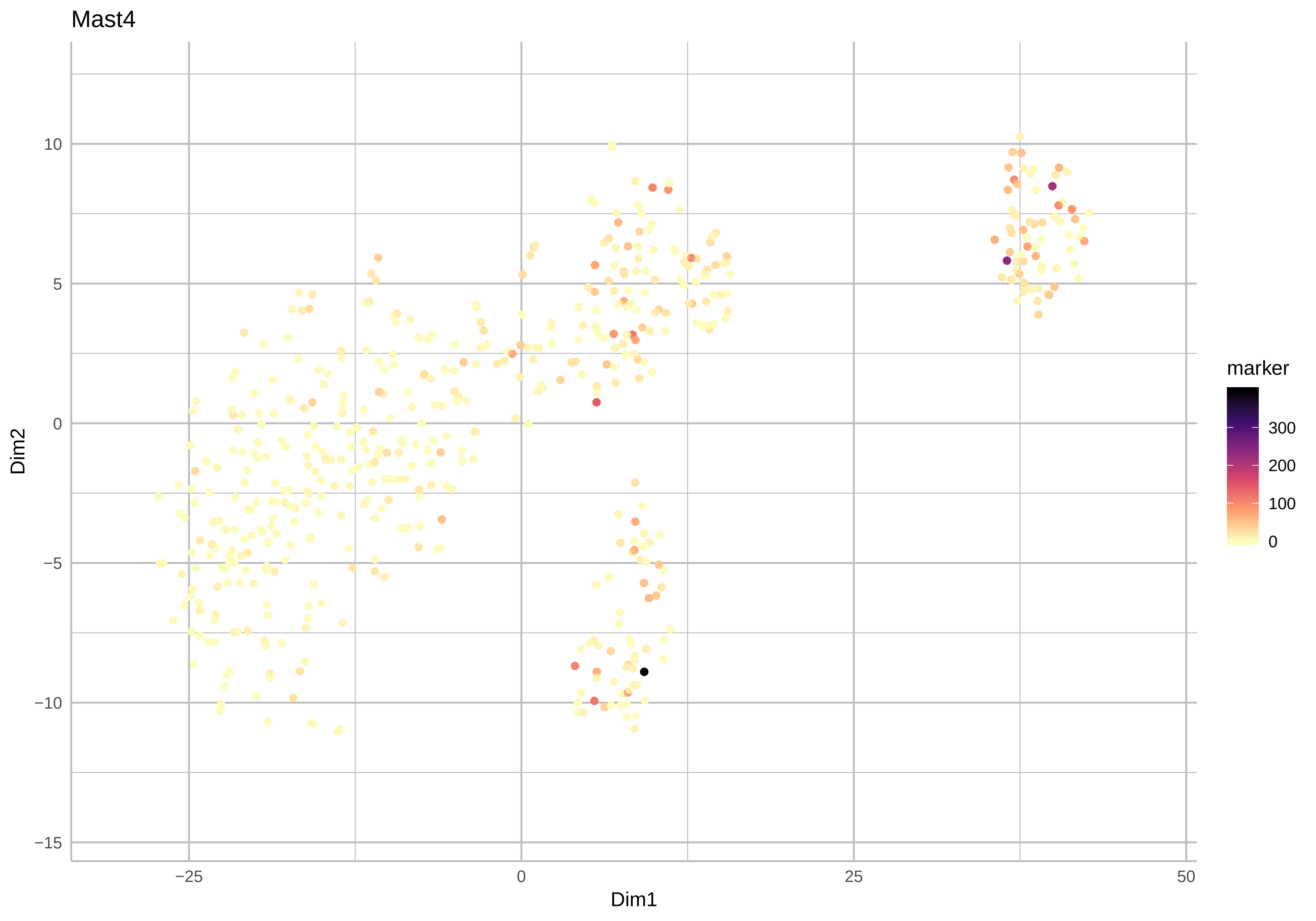

Mast4
10
5
0
−5
−10
−15
−25
0
25
50
Dim1
Dim2
marker
300
200
100
0

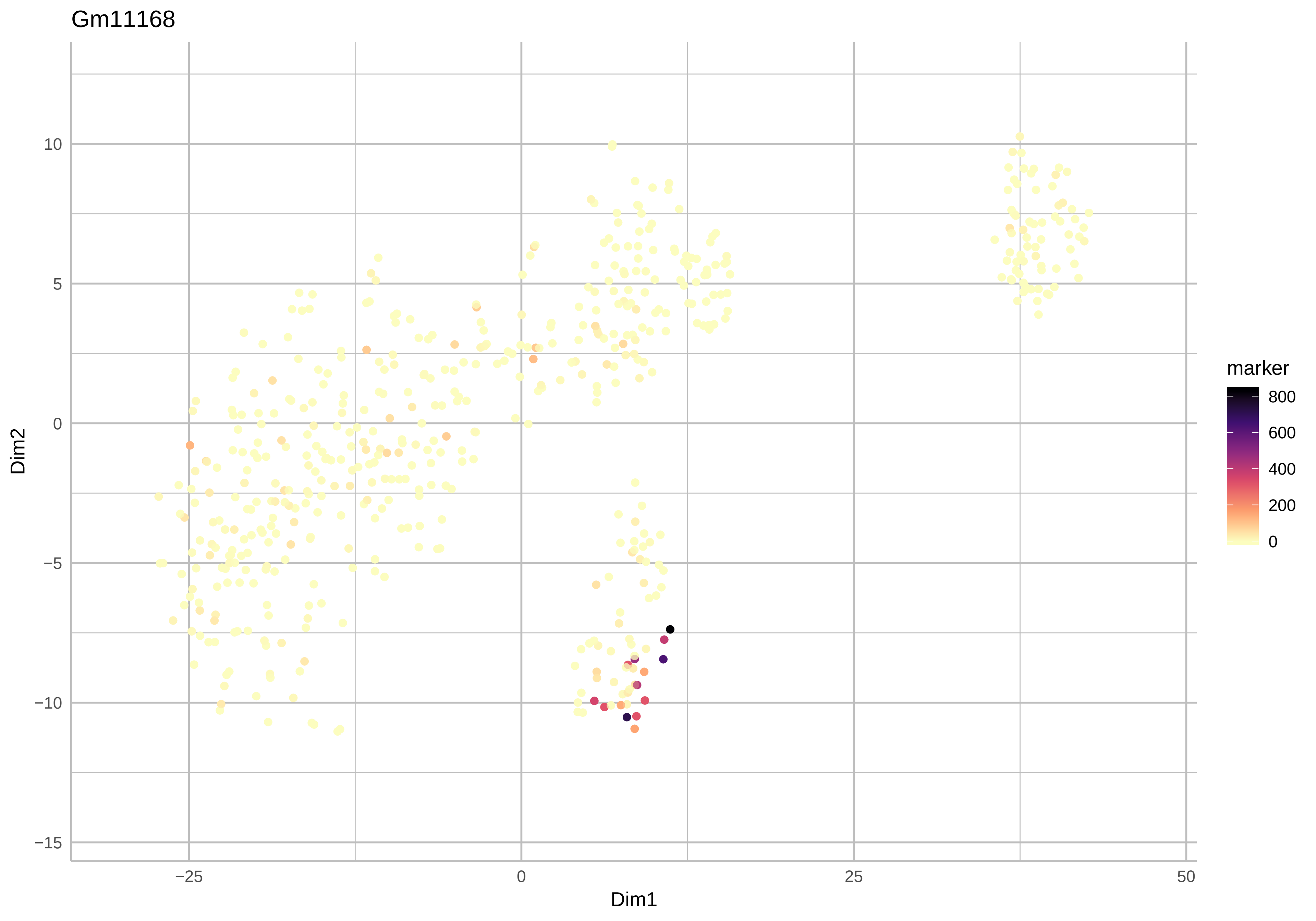

Gm11168
10
5
0
−5
−10
−15
Dim2
−25
0
25
50
Dim1
marker
800
600
400
200
0

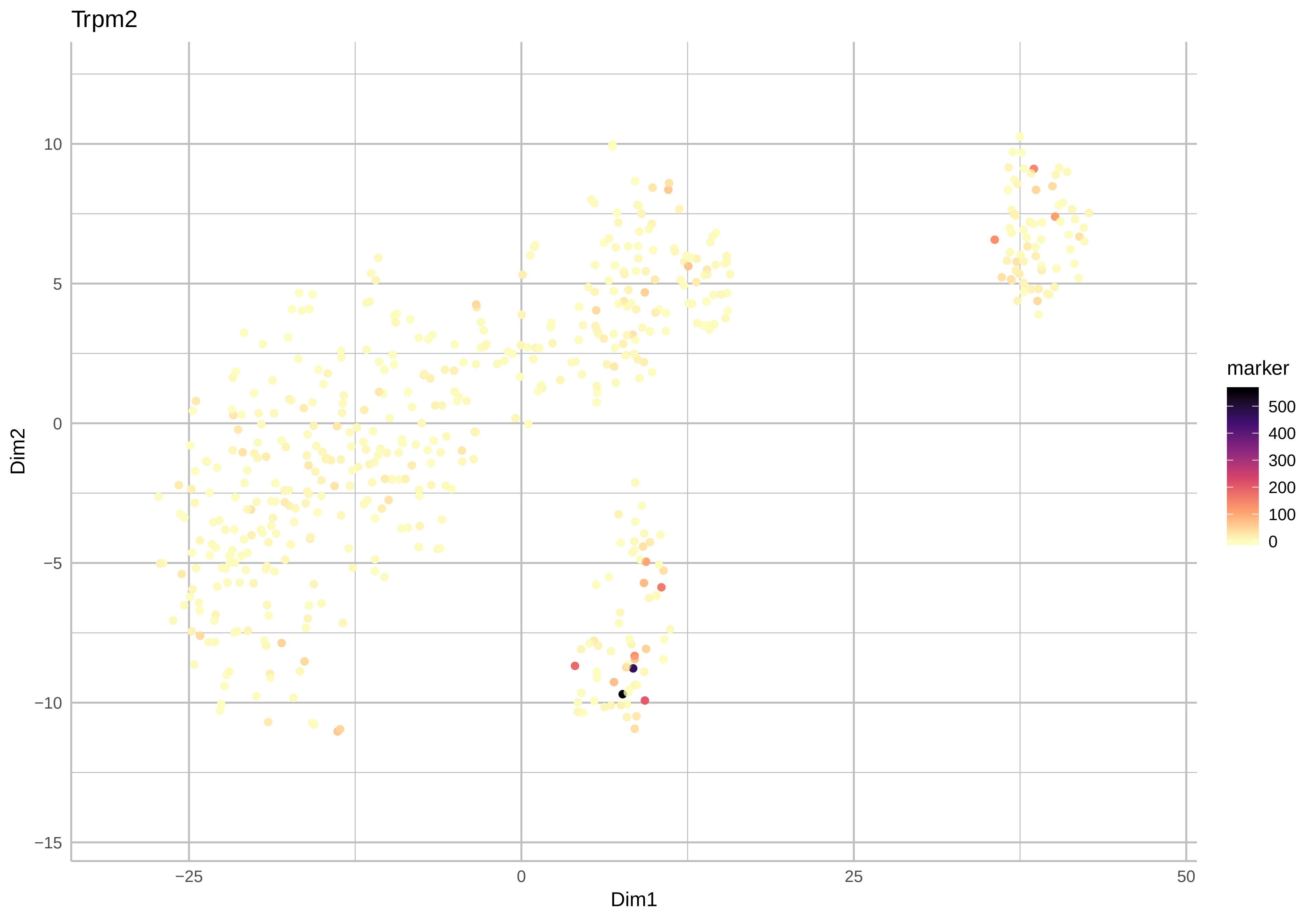
Trpm2
10
5
0
−5
−10
−15
−25
0
25
50
Dim1
Dim2
marker
500
400
300
200
100
0

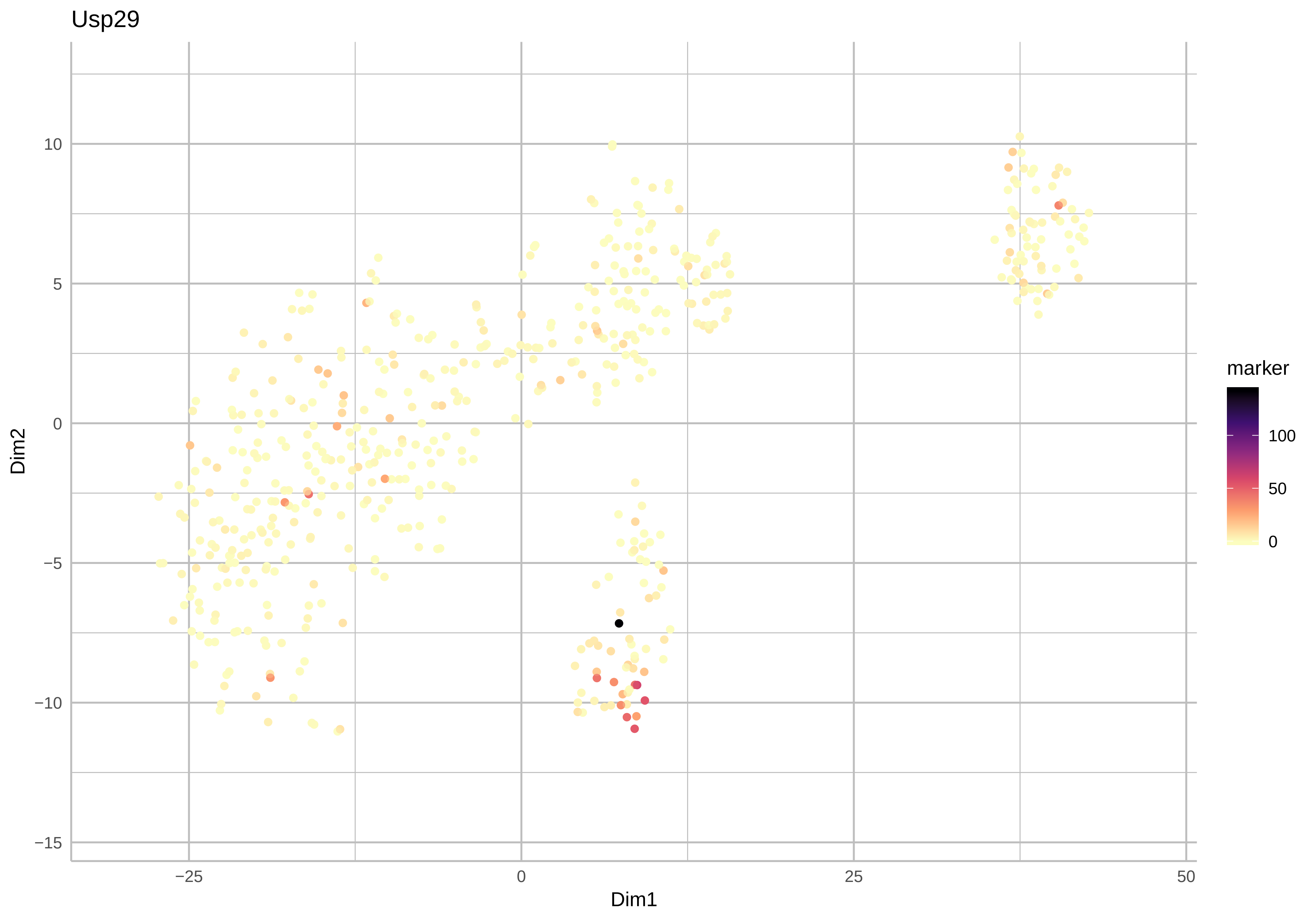

Usp29
Dim2
Dim1
marker
100
50
0
10
5
0
−5
−10
−15
−25
0
25
50

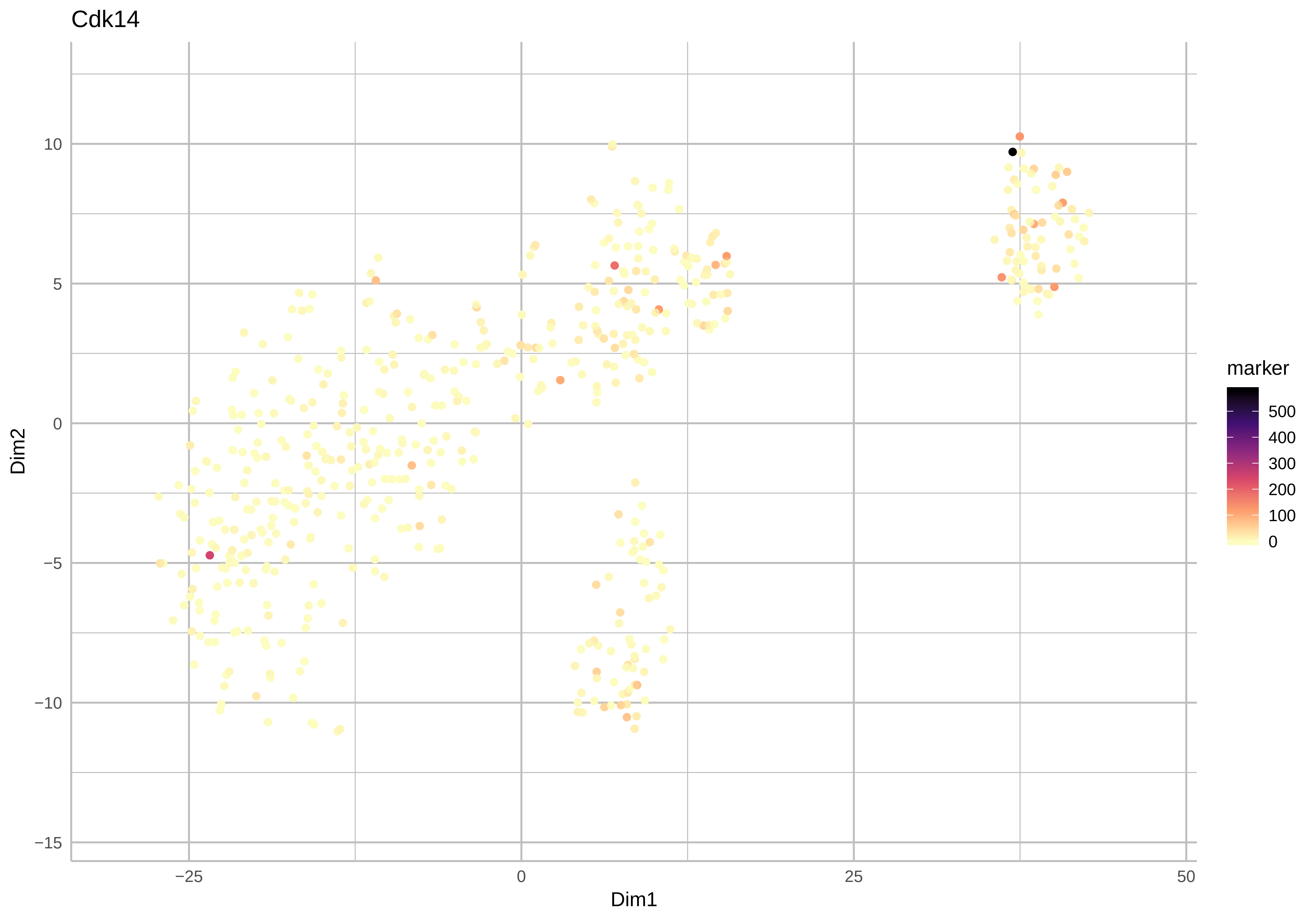

Cdk14
Dim2
Dim1
10
5
0
−5
−10
−15
−25
0
25
50
marker
500
400
300
200
100
0

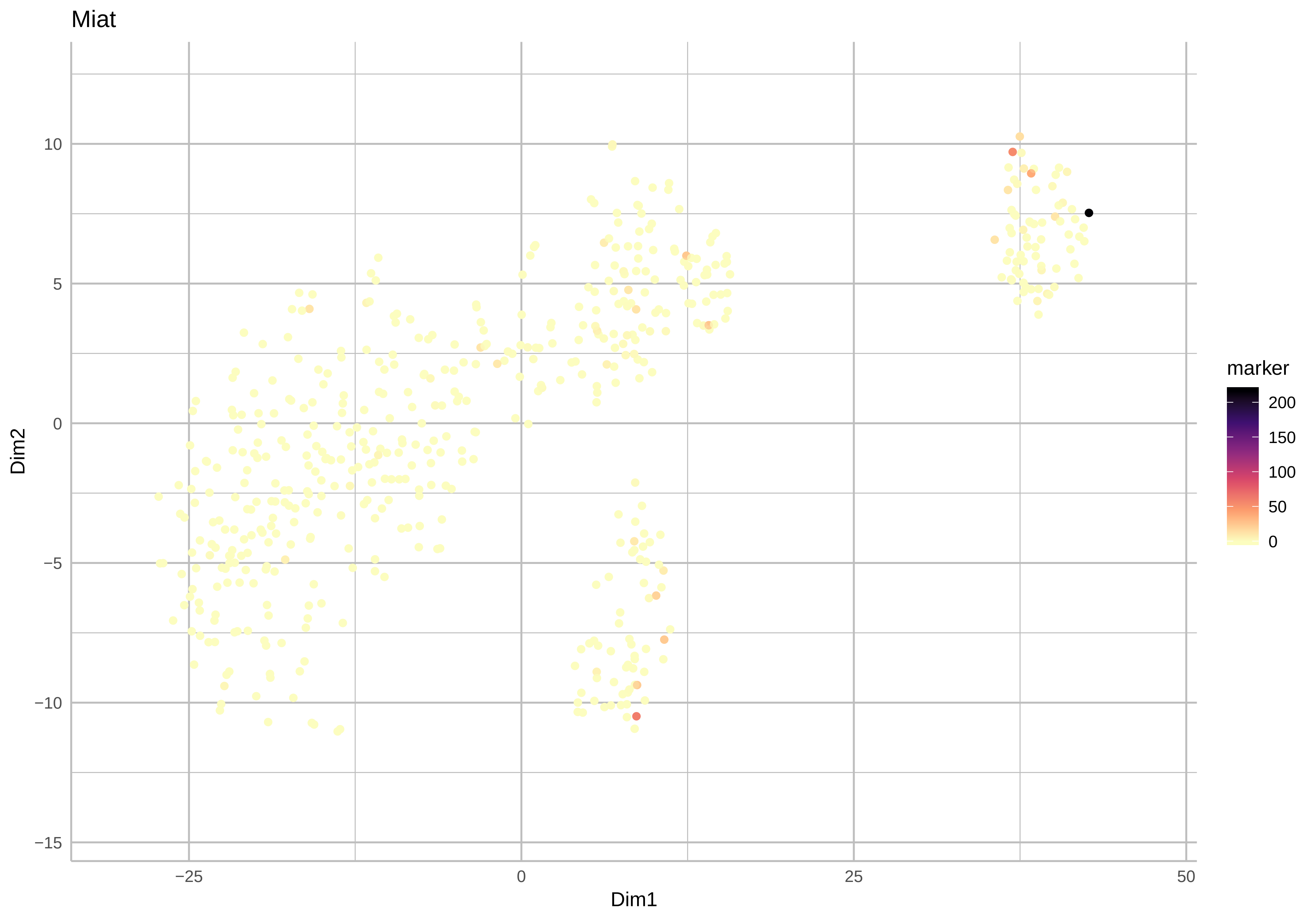

Miat
10
5
0
−5
−10
−15
Dim2
−25
0
25
50
Dim1
marker
200
150
100
50
0

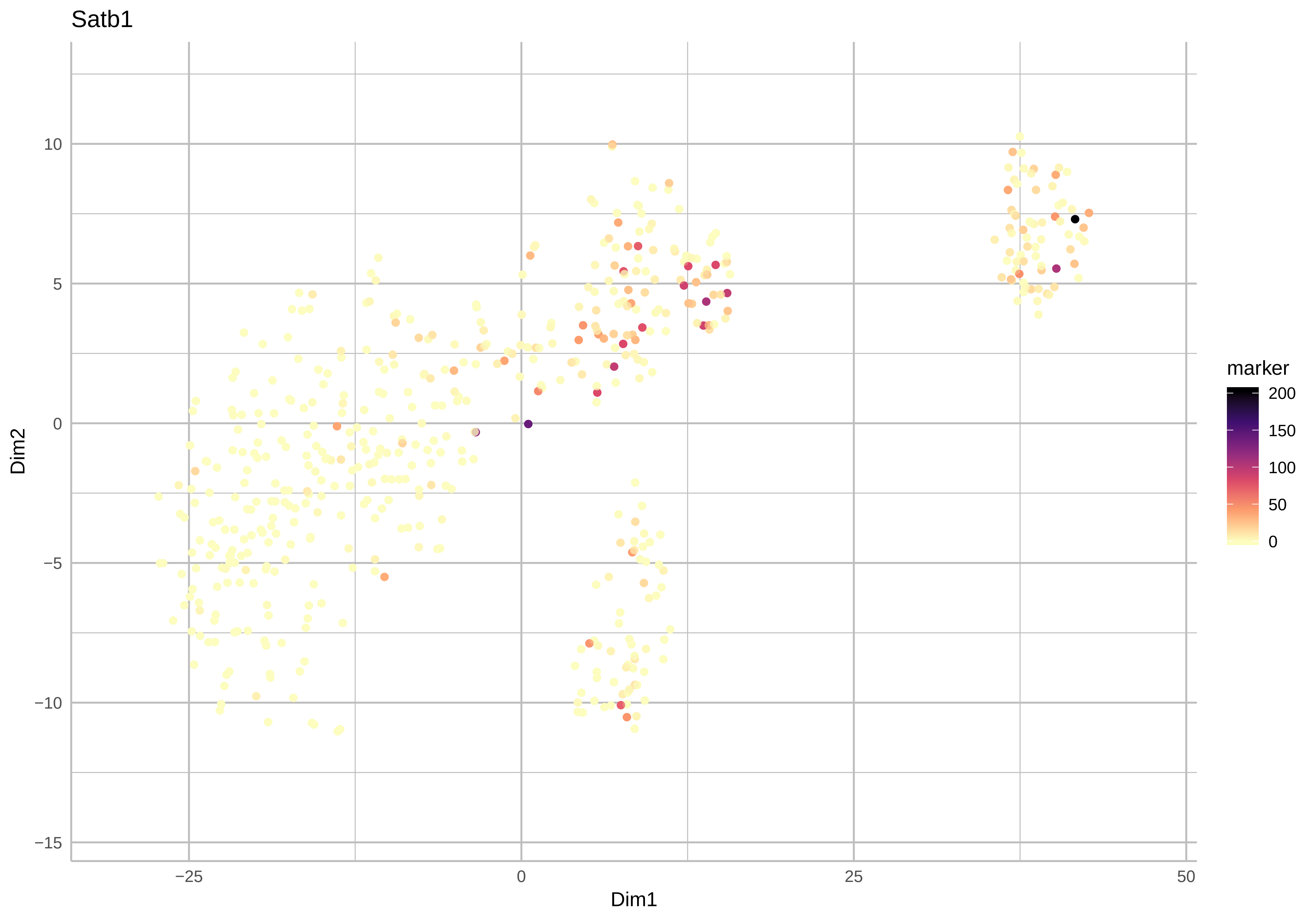
Satb1
Dim2
Dim1
10
5
0
−5
−10
−15
−25
0
25
50
marker
200
150
100
50
0

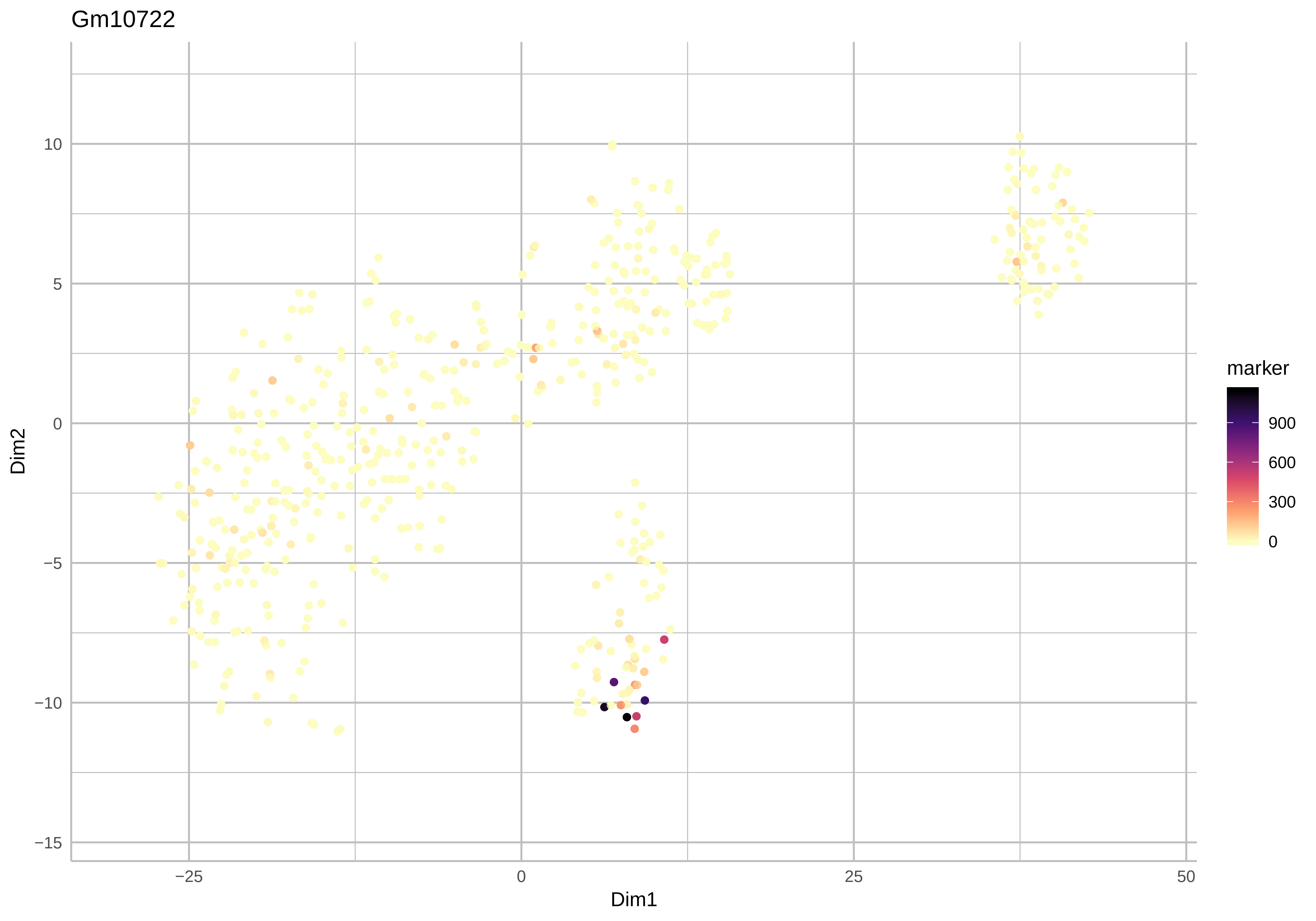

Gm10722
10
5
0
−5
−10
−15
−25
0
25
50
Dim1
Dim2
marker
900
600
300
0

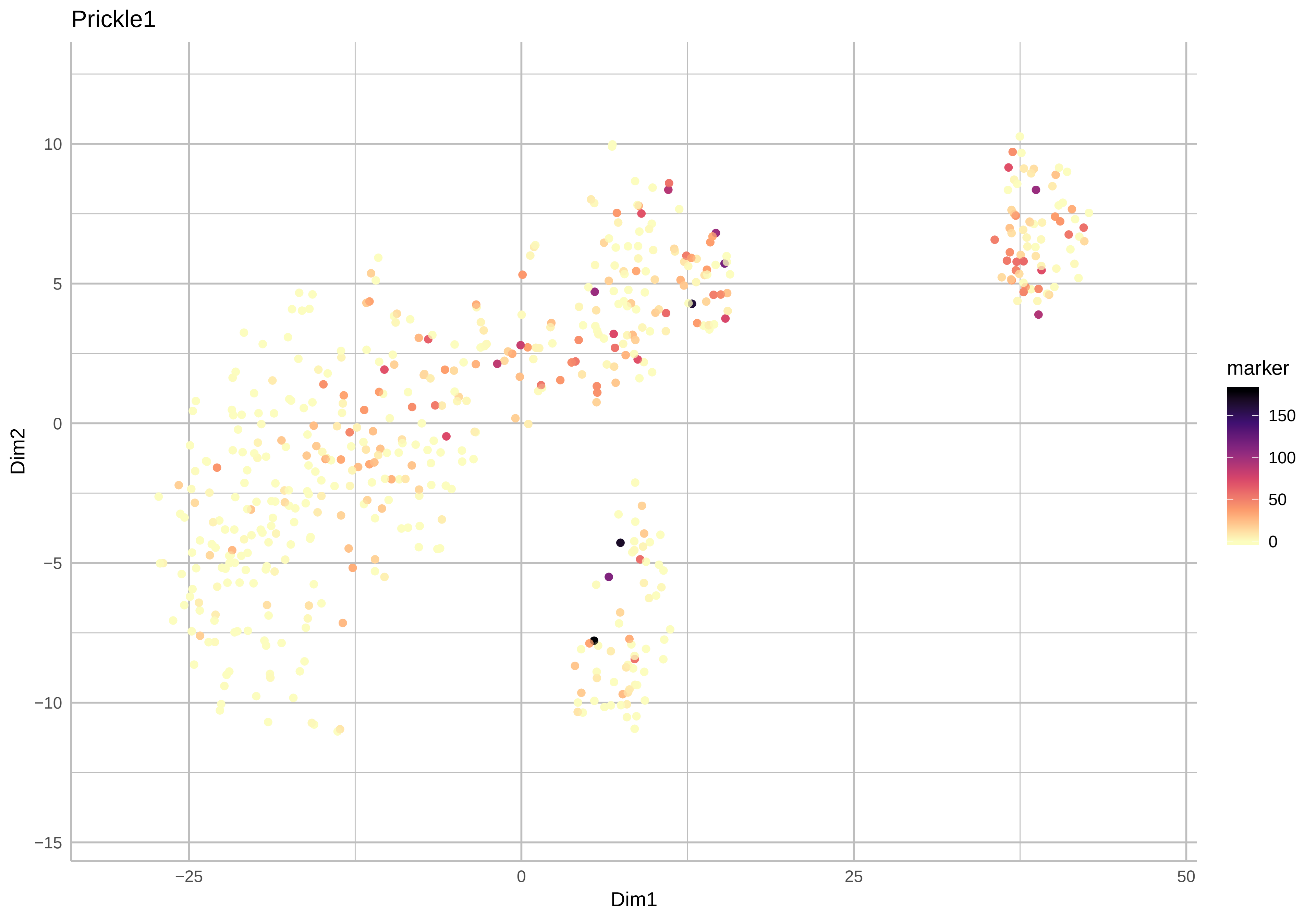

Prickle1
Dim2
Dim1
10
5
0
−5
−10
−15
−25
0
25
50
marker
150
100
50
0

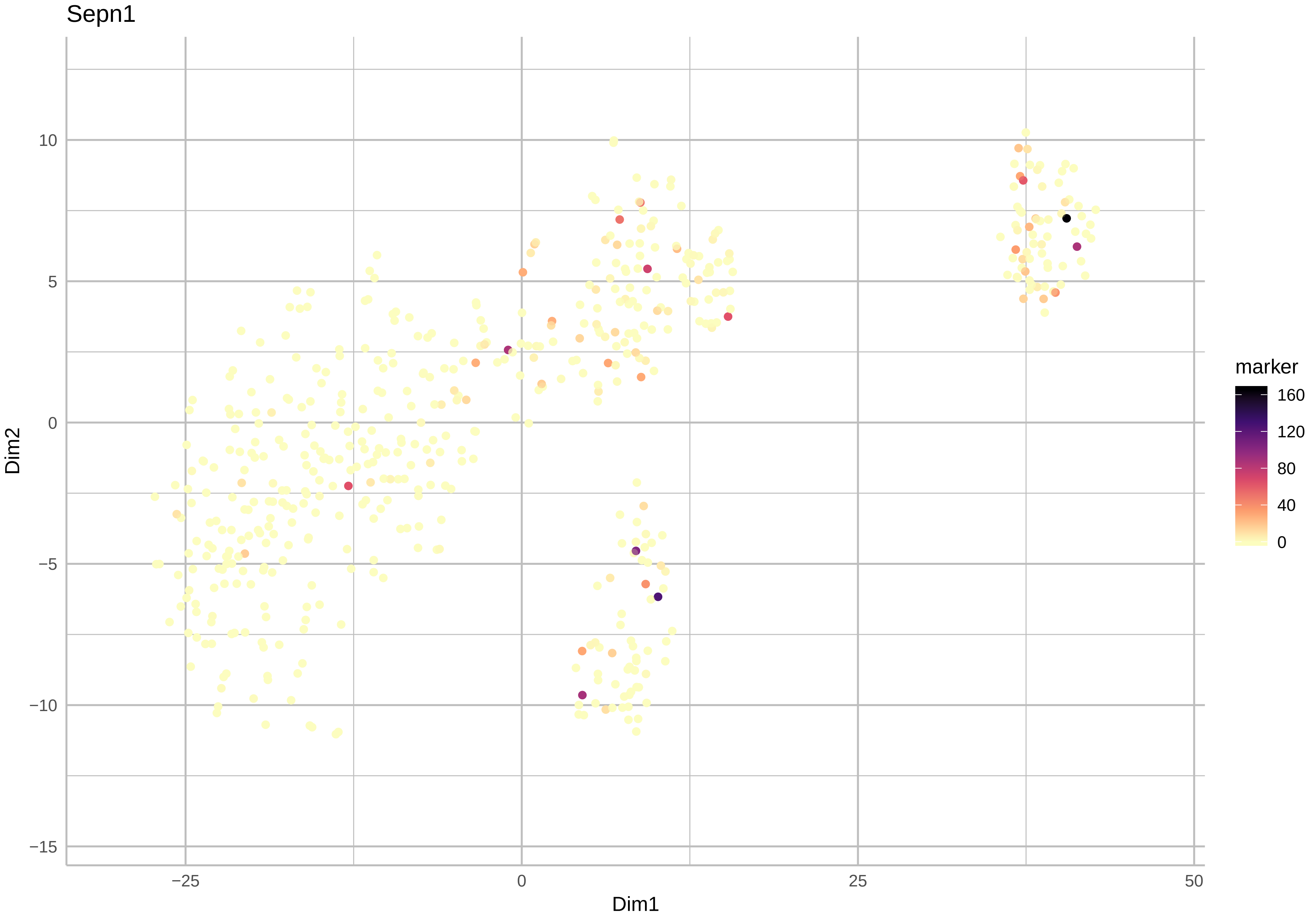
Sepn1
Dim2
Dim1
10
5
0
−5
−10
−15
−25
0
25
50
marker
160
120
80
40
0

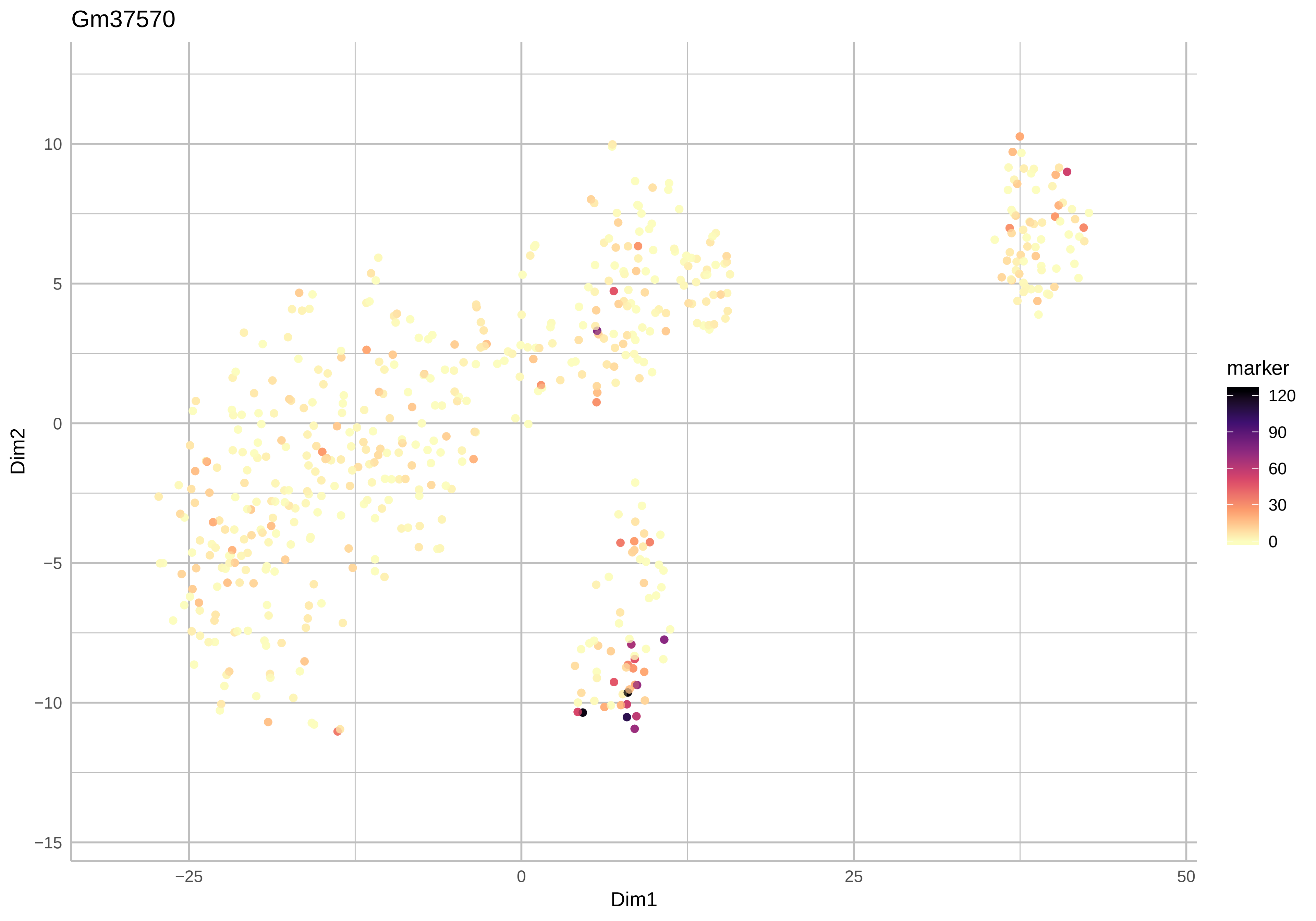
Gm37570
Dim2
Dim1
10
5
0
−5
−10
−15
−25
0
25
50
marker
120
90
60
30
0

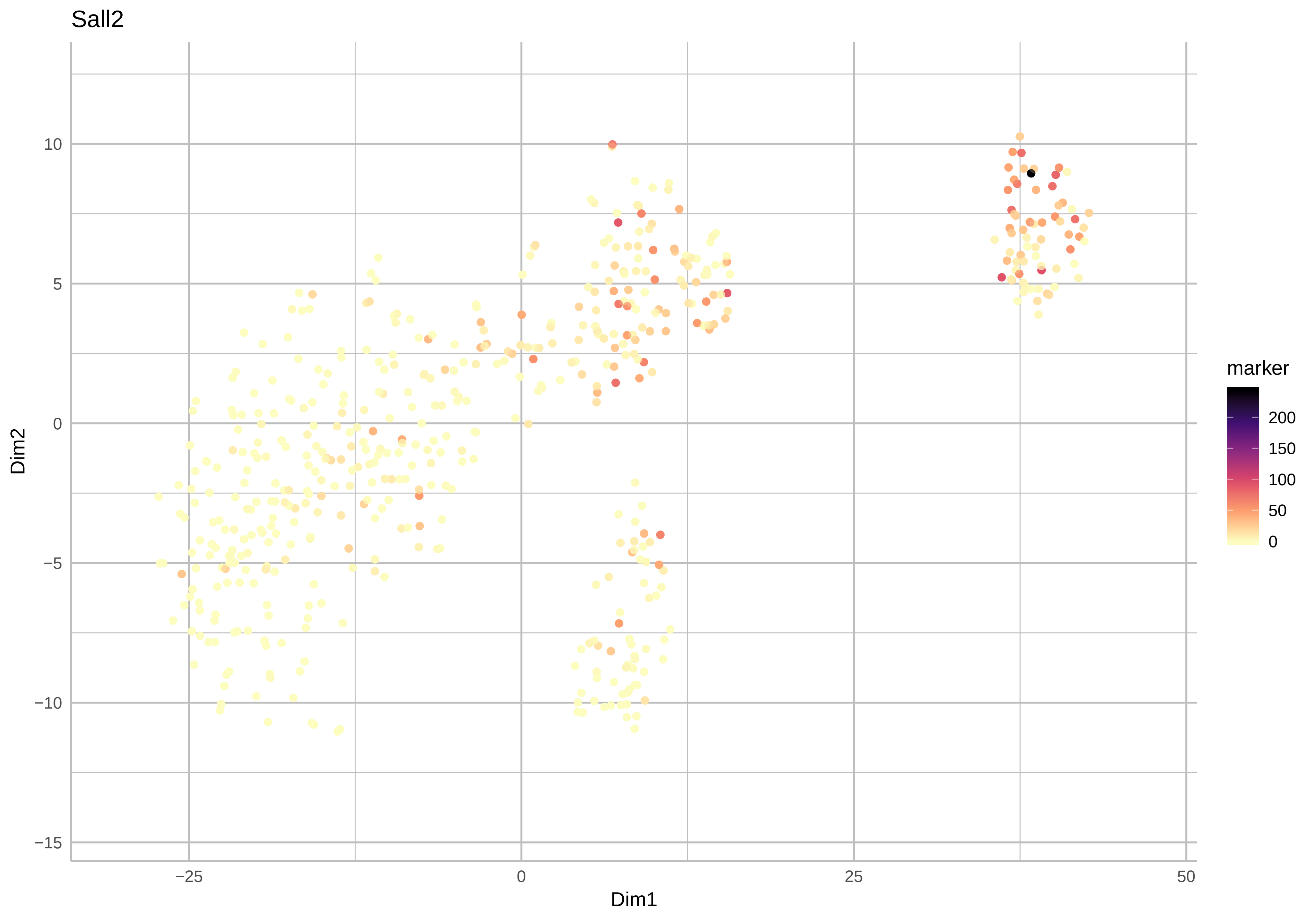

Sall2
Dim2
Dim1
10
5
0
−5
−10
−15
−25
0
25
50
marker
200
150
100
50
0

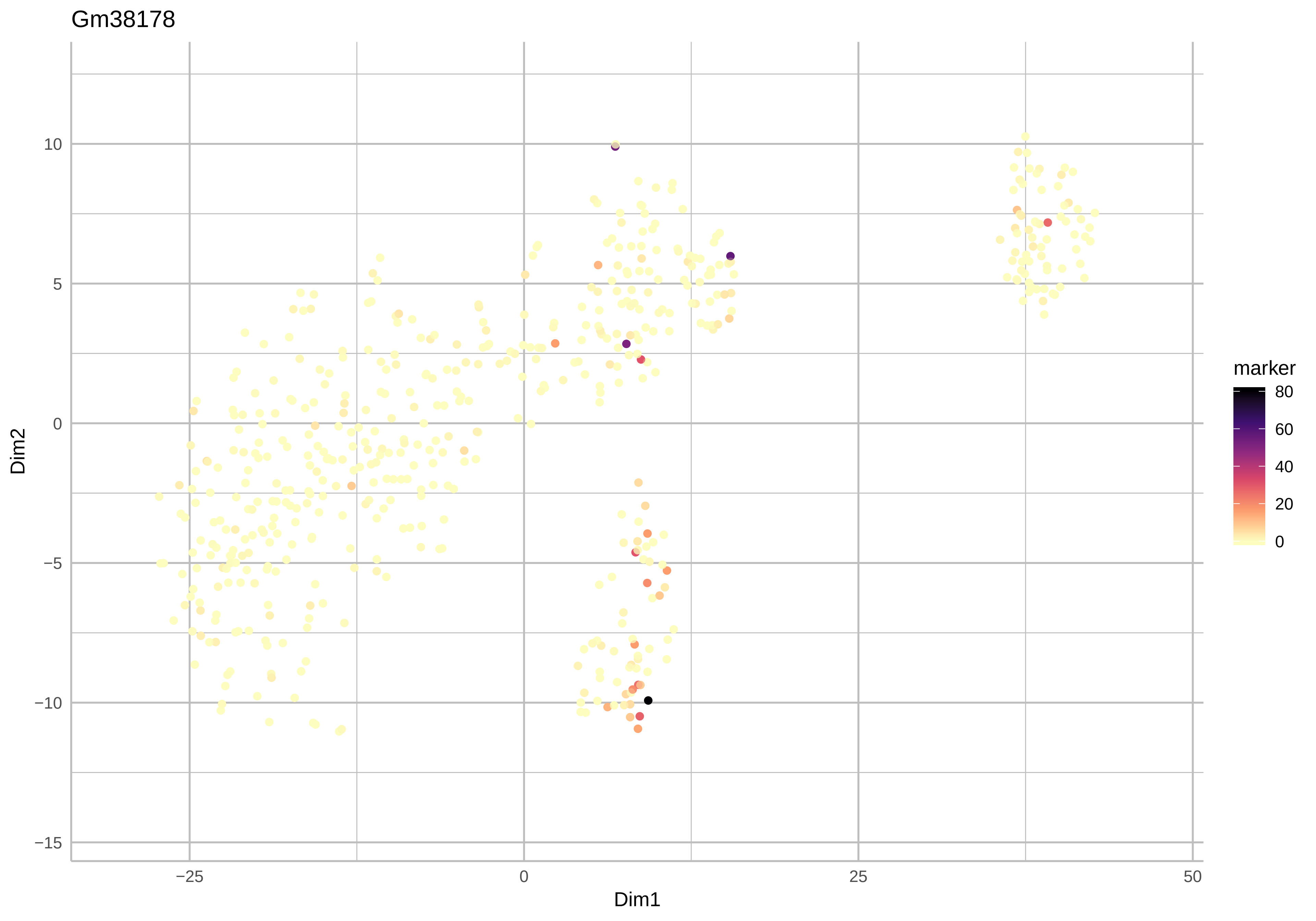

Gm38178
Dim2
Dim1
marker
80
60
40
20
0
10
5
0
−5
−10
−15
−25
0
25
50

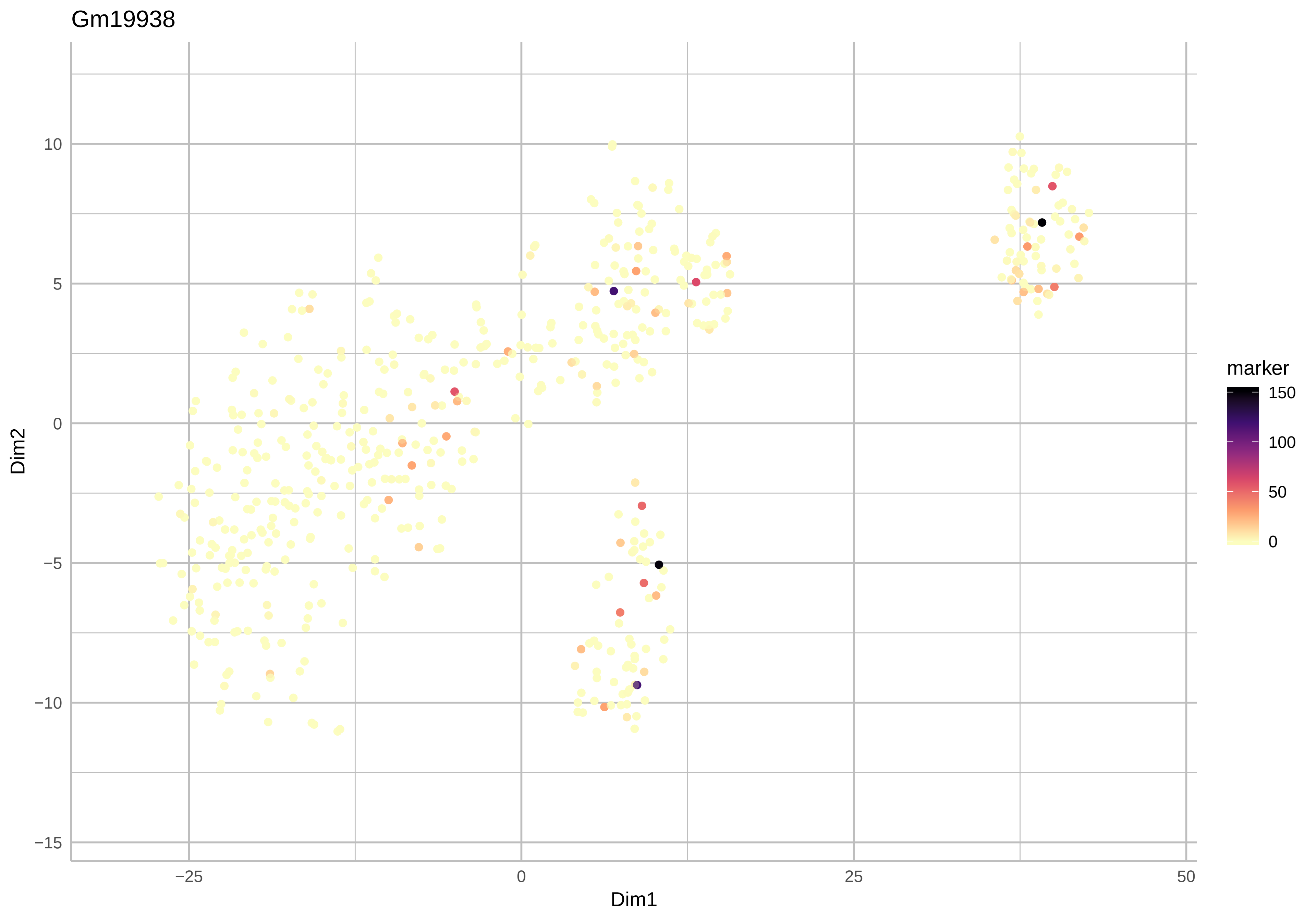

Gm19938
Dim2
Dim1
10
5
0
−5
−10
−15
−25
0
25
50
marker
150
100
50
0

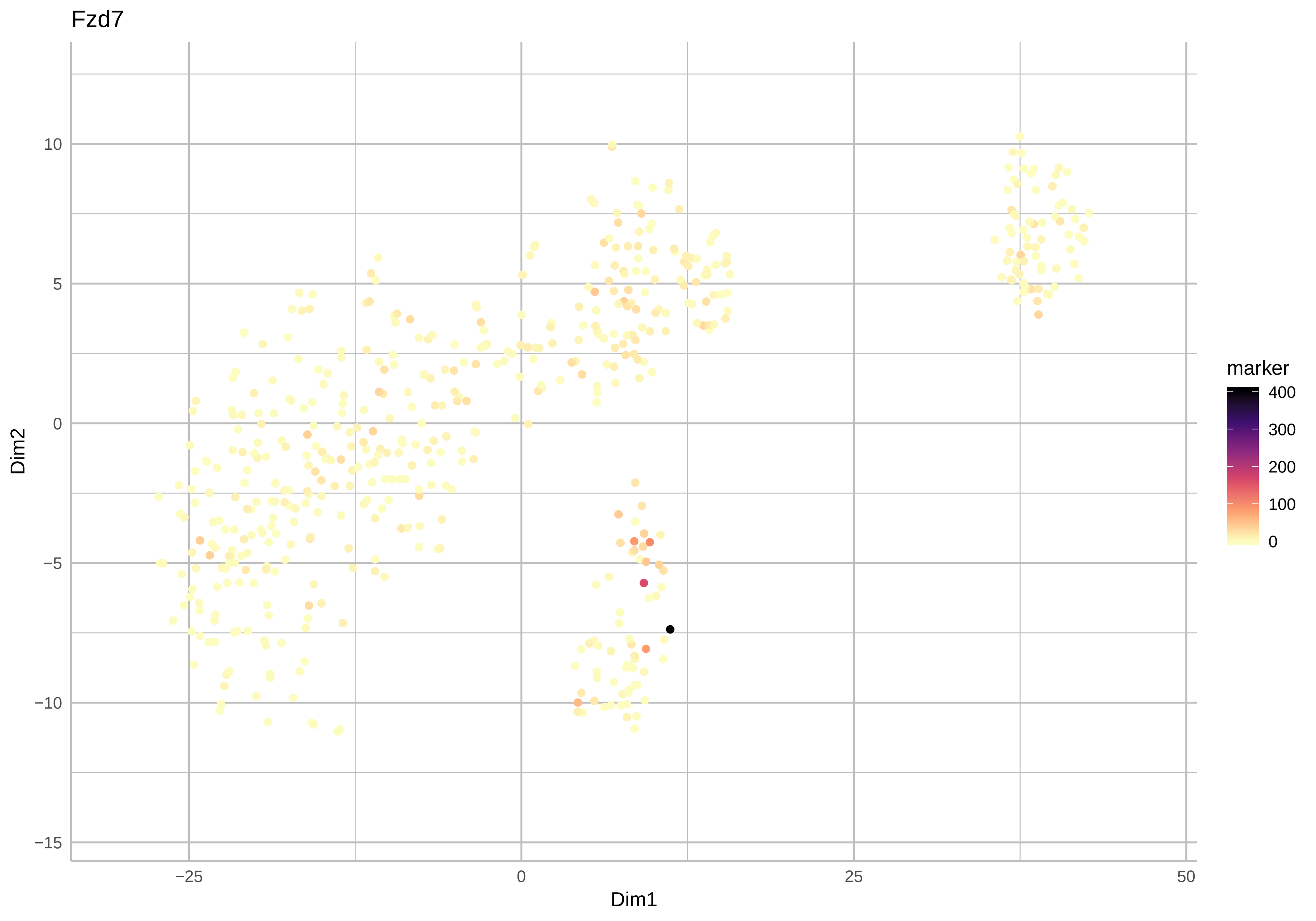

Fzd7
10
5
0
−5
−10
−15
Dim2
−25
0
25
50
Dim1
marker
400
300
200
100
0

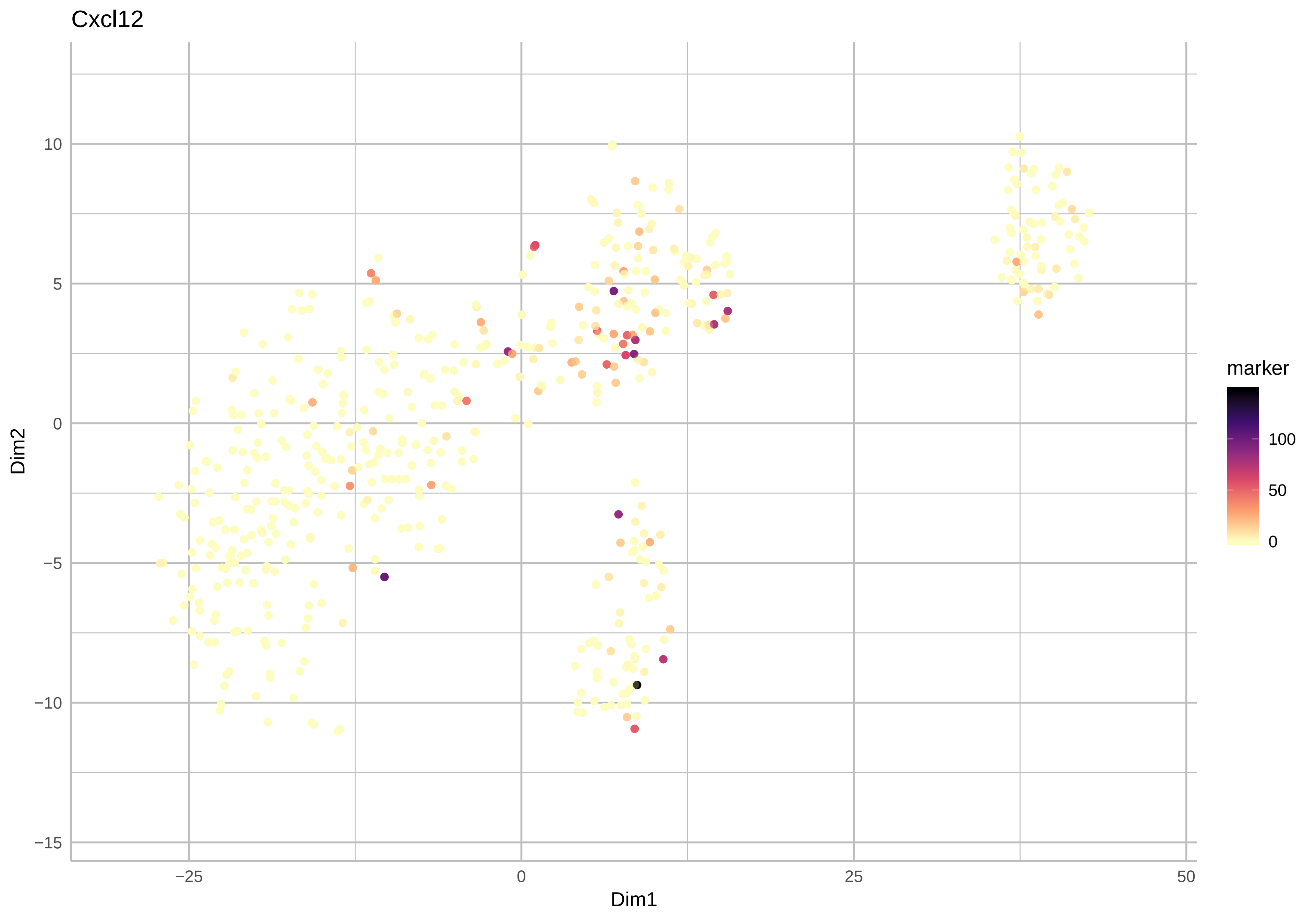

Cxcl12
Dim2
Dim1
10
5
0
−5
−10
−15
−25
0
25
50
marker
100
50
0

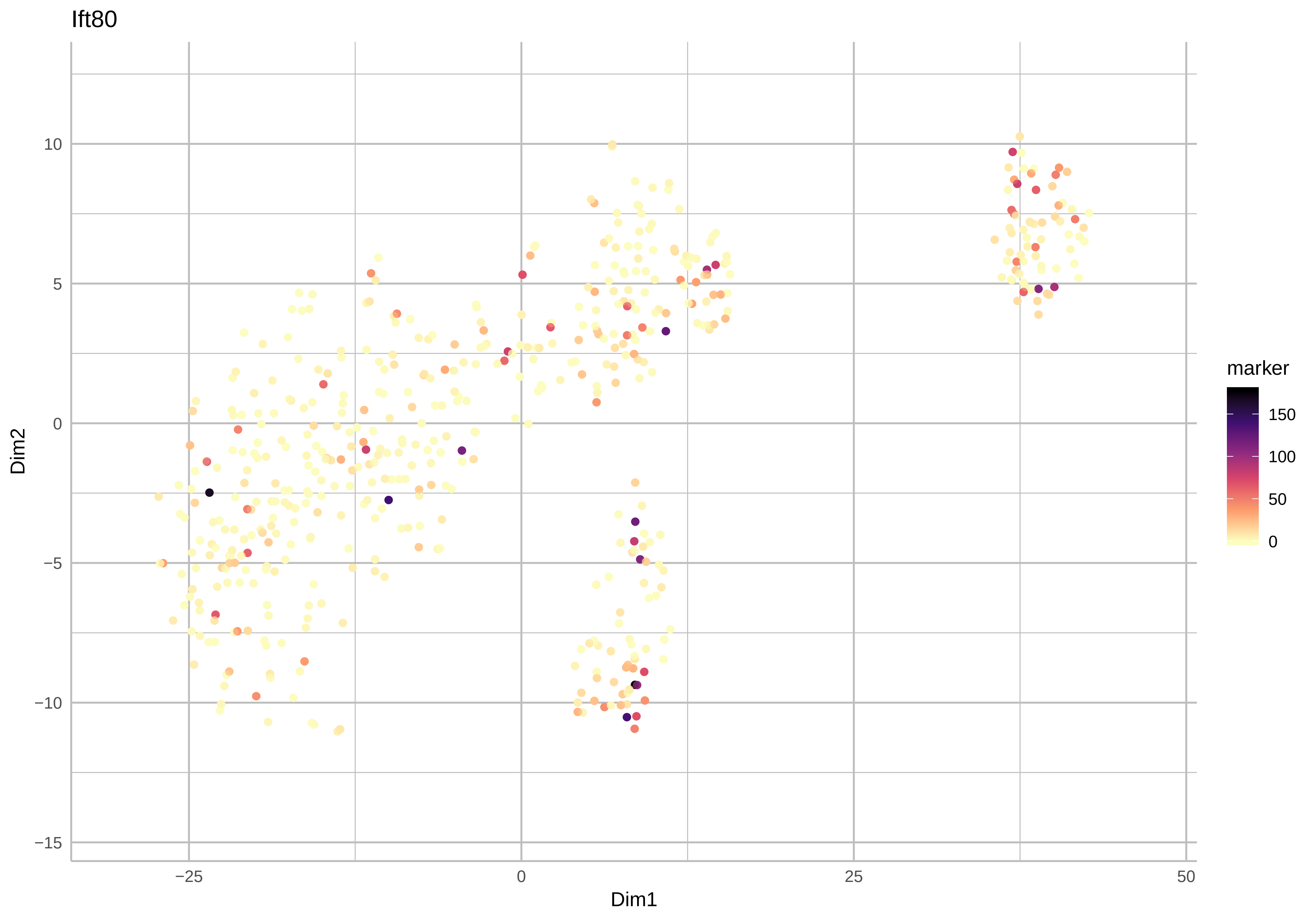
Ift80
Dim2
Dim1
marker
150
100
50
0
10
5
0
−5
−10
−15
−25
0
25
50

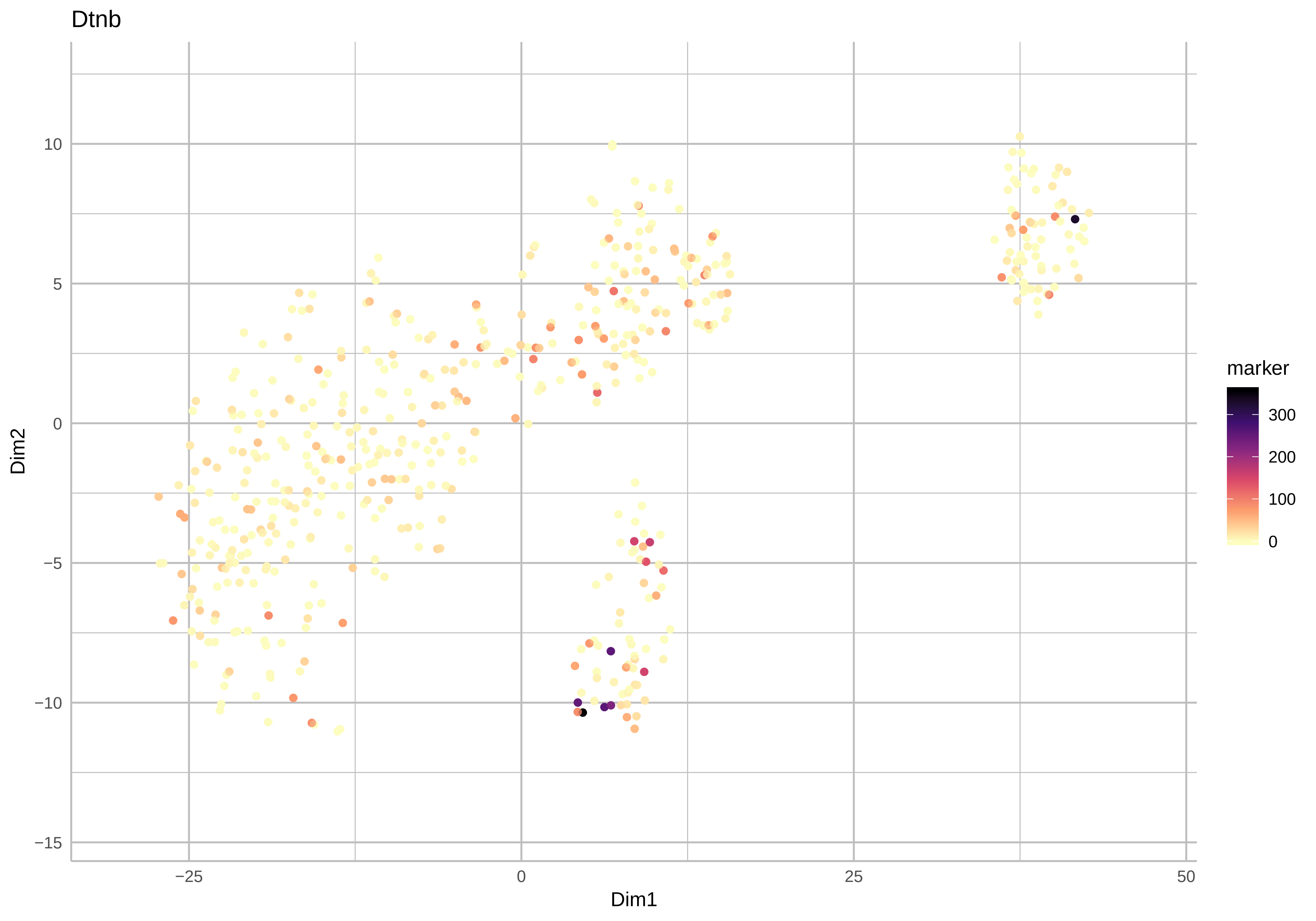
Dtnb
Dim2
Dim1
10
5
0
−5
−10
−15
−25
0
25
50
marker
300
200
100
0

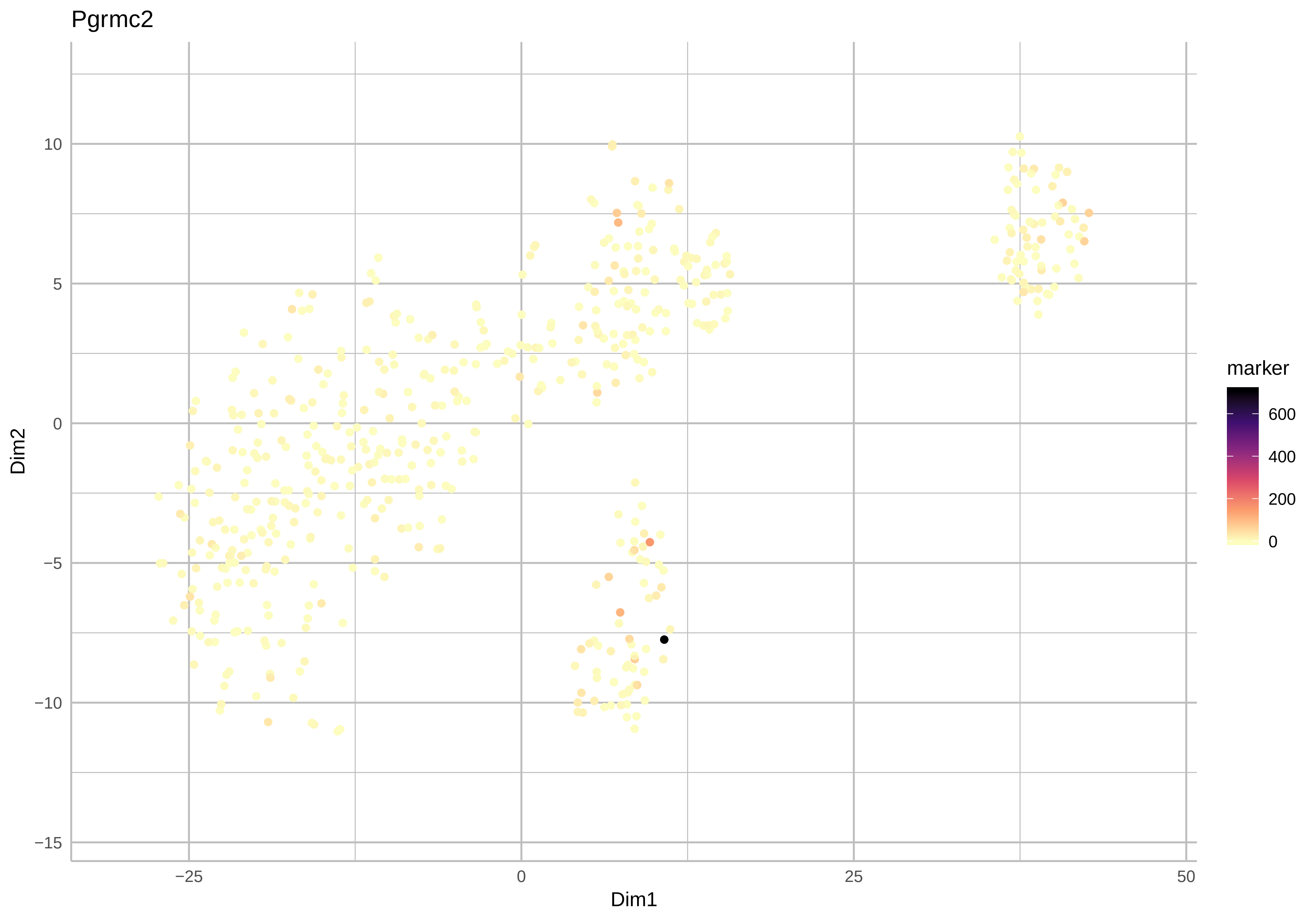
Pgrmc2
Dim2
Dim1
marker
600
400
200
0
10
5
0
−5
−10
−15
−25
0
25
50

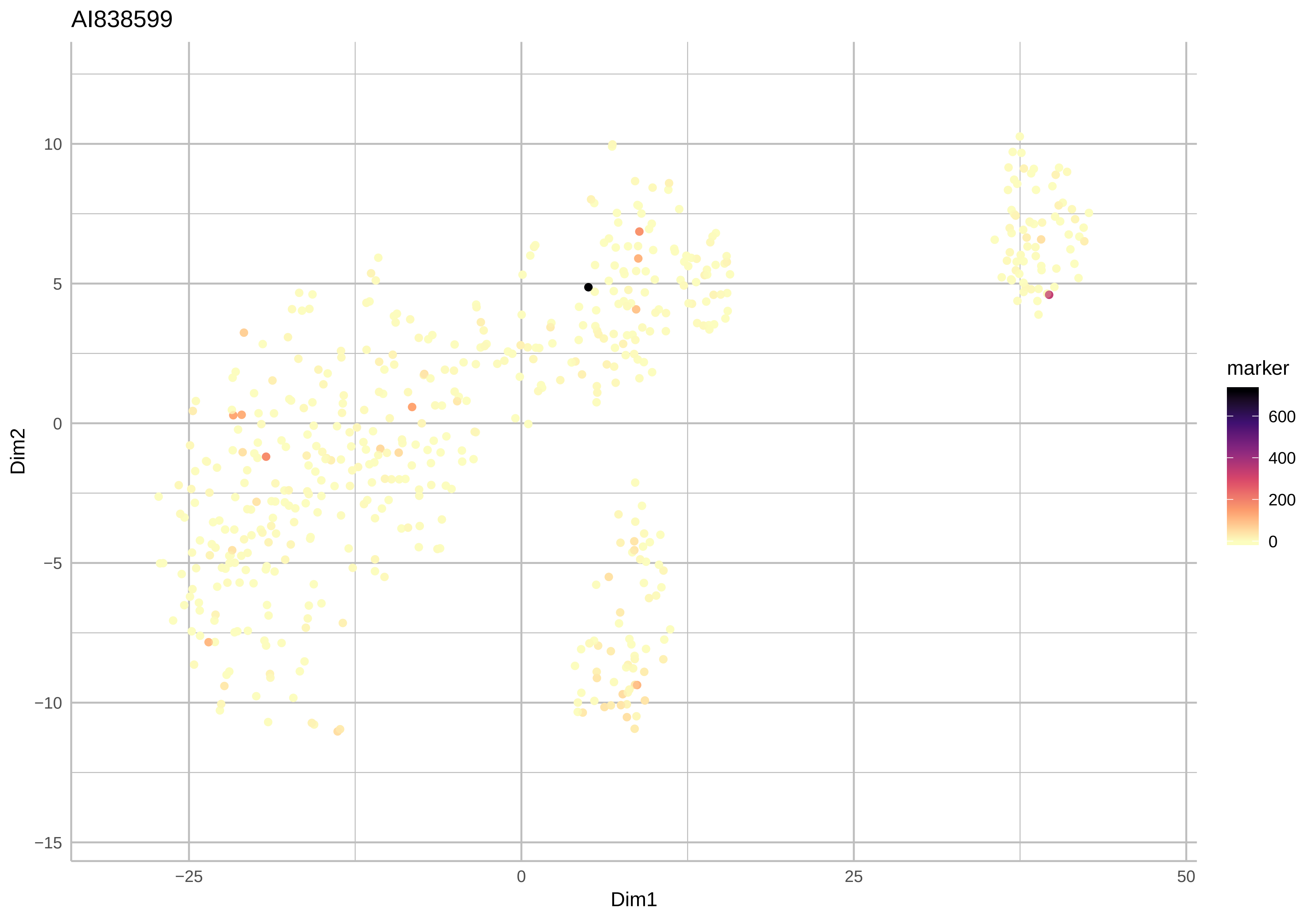

AI838599
Dim2
Dim1
10
5
0
−5
−10
−15
−25
0
25
50
marker
600
400
200
0

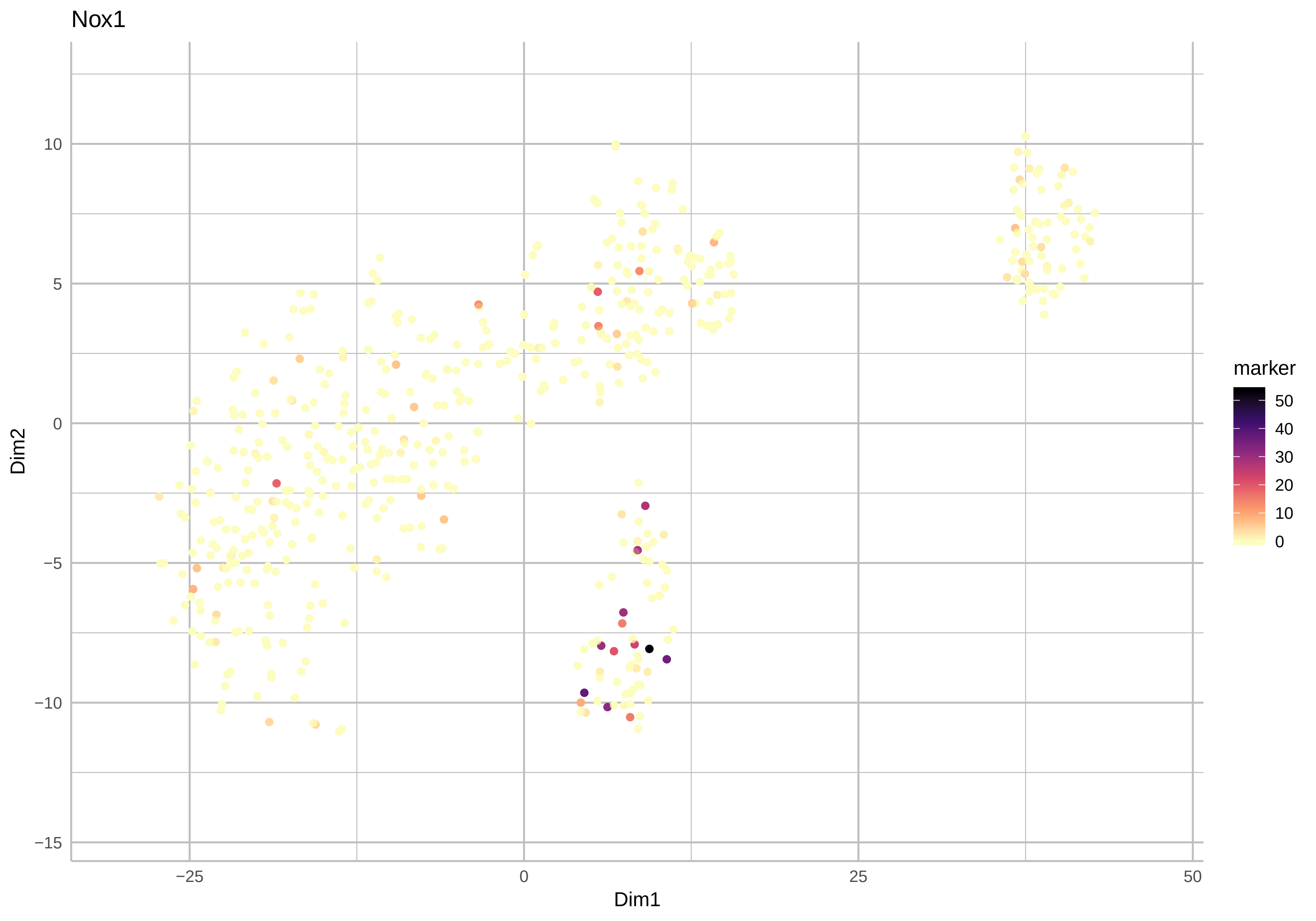

Nox1
Dim2
Dim1
10
5
0
−5
−10
−15
−25
0
25
50
marker
50
40
30
20
10
0

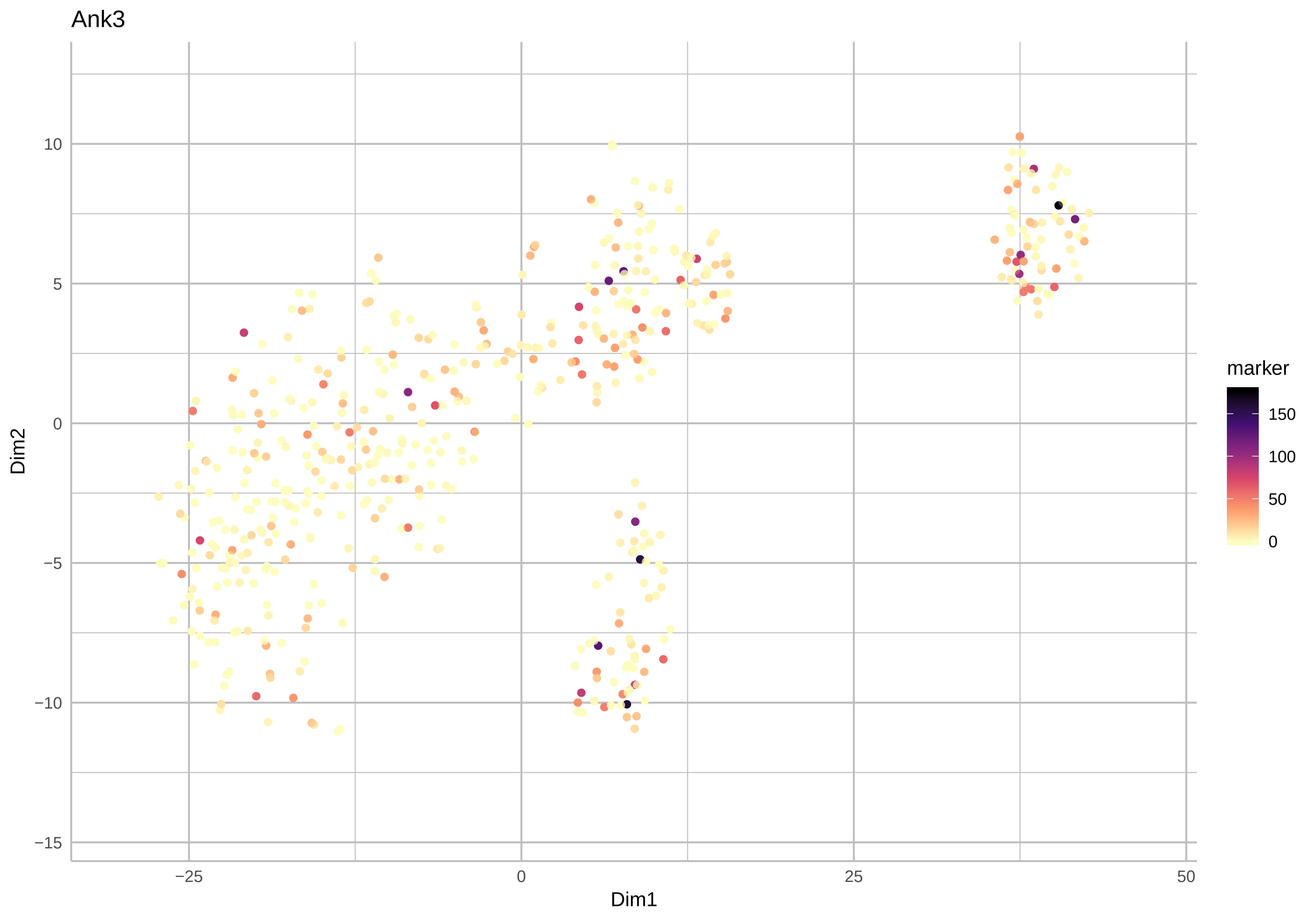
Ank3
Dim2
Dim1
−25
0
25
50
10
5
0
−5
−10
−15
marker
150
100
50
0

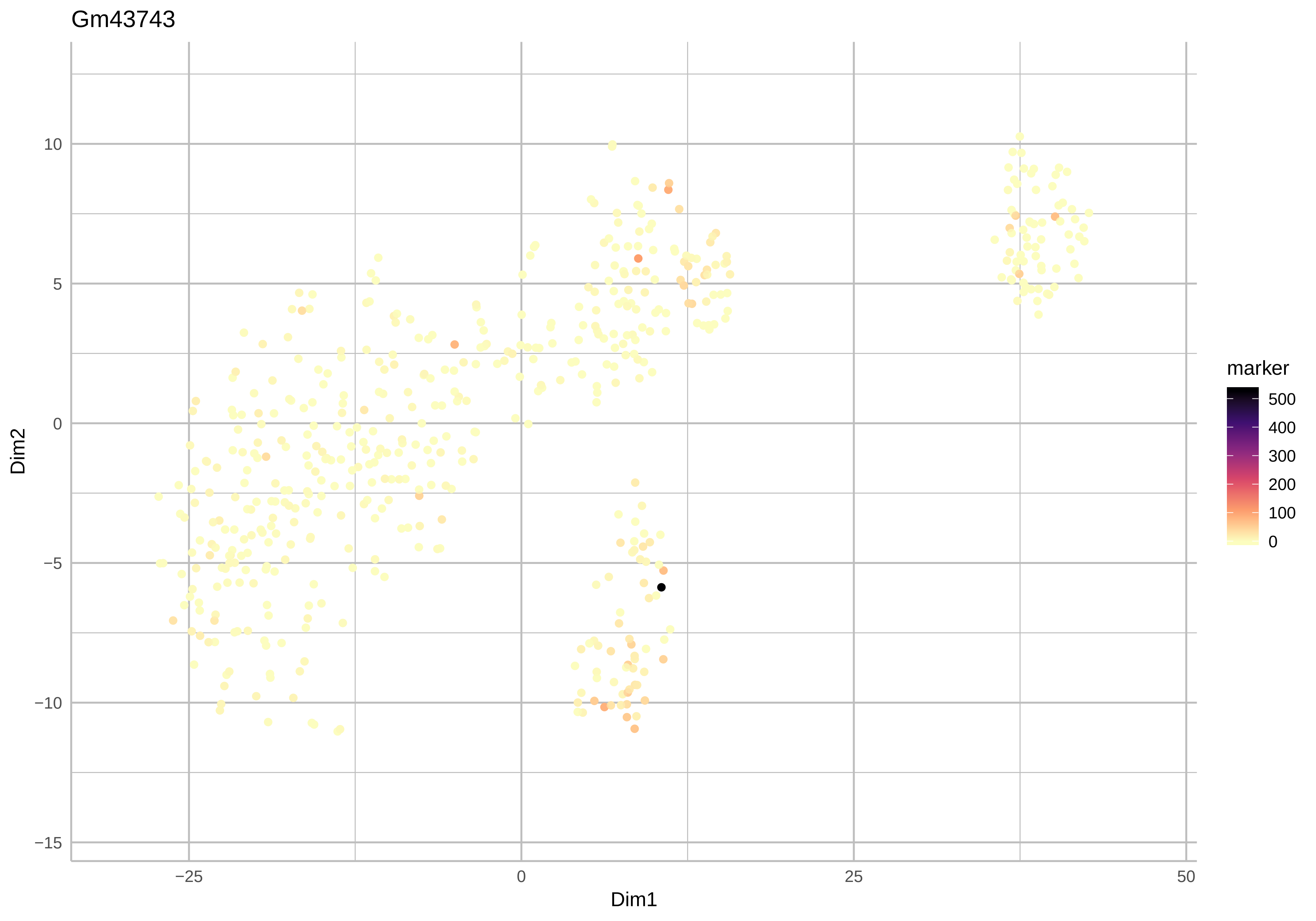
Gm43743
Dim2
Dim1
10
5
0
−5
−10
−15
−25
0
25
50
marker
500
400
300
200
100
0

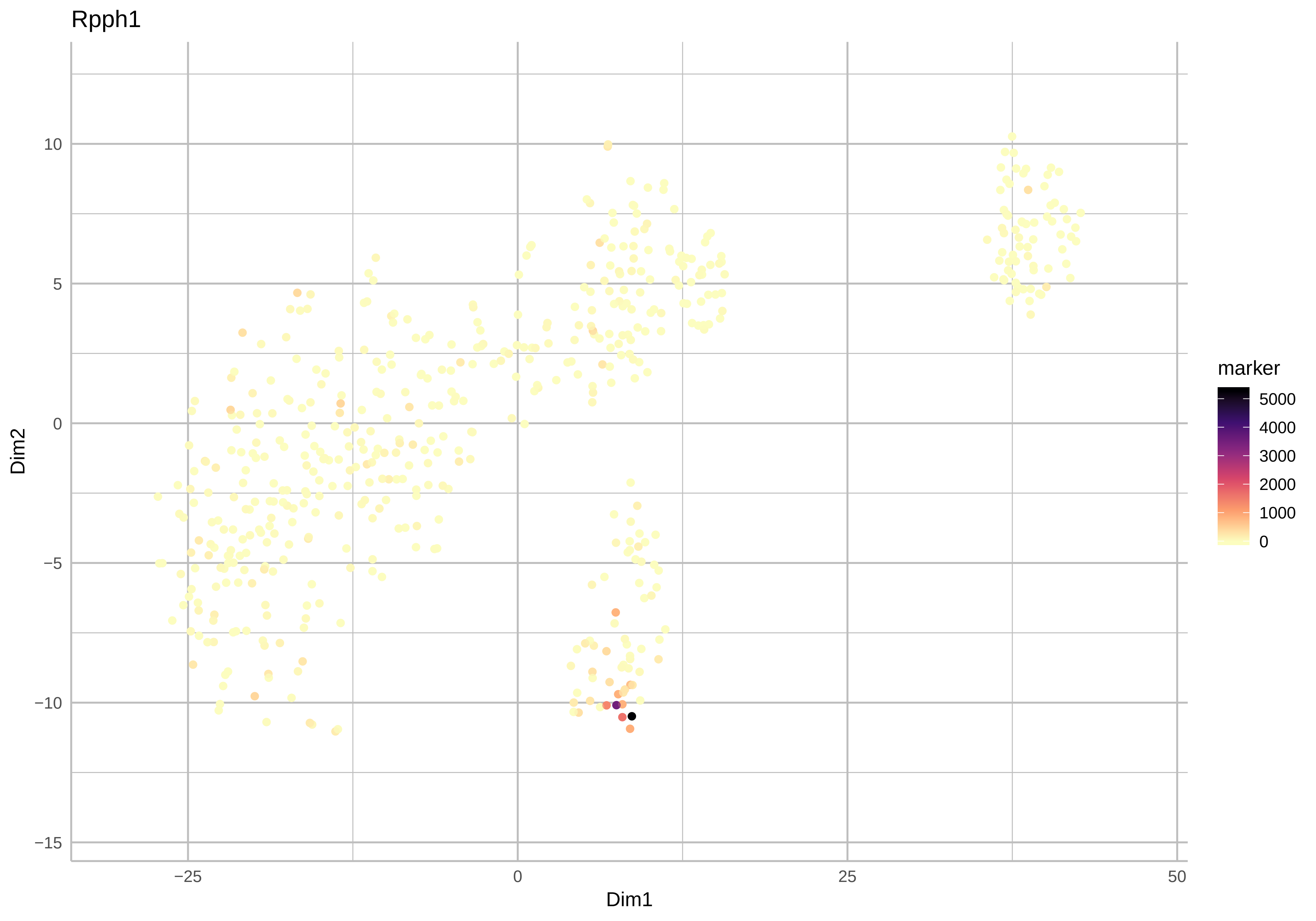
Rpph1
Dim2
Dim1
10
5
0
−5
−10
−15
−25
0
25
50
marker
5000
4000
3000
2000
1000
0

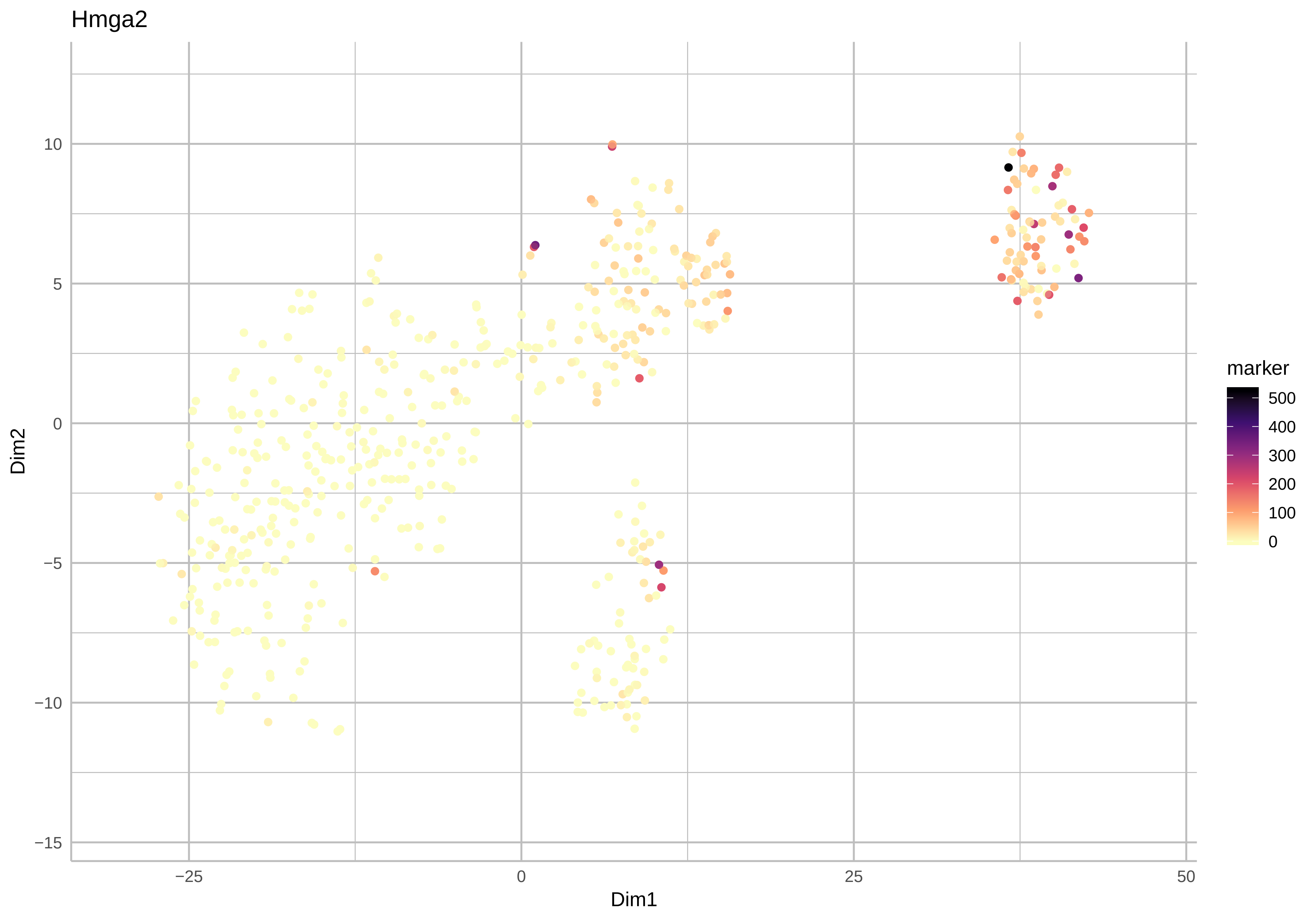

Hmga2
Dim2
Dim1
10
5
0
−5
−10
−15
−25
0
25
50
marker
500
400
300
200
100
0

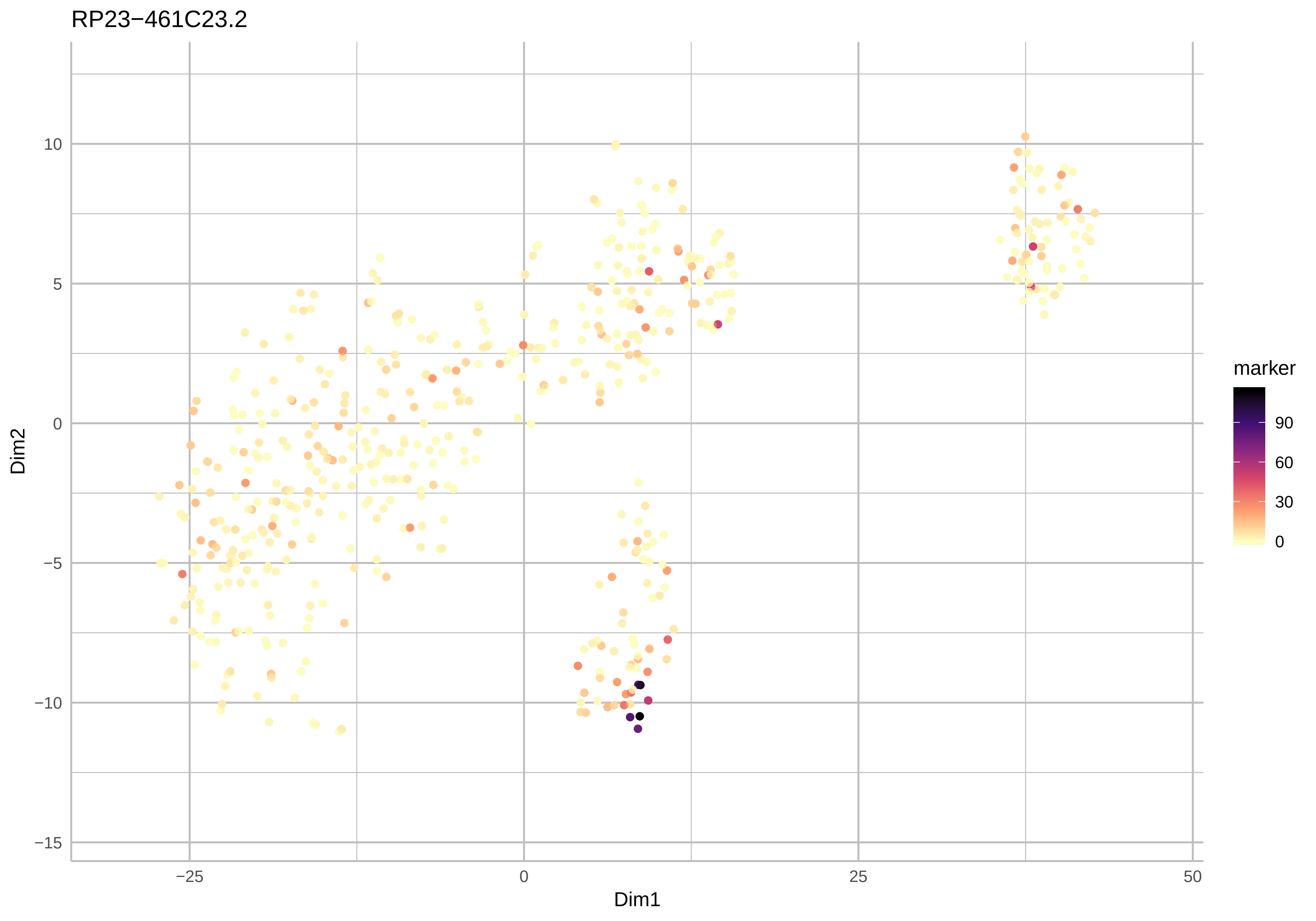
RP23−461C23.2
10
5
0
−5
−10
−15
−25
0
25
50
Dim1
Dim2
marker
90
60
30
0

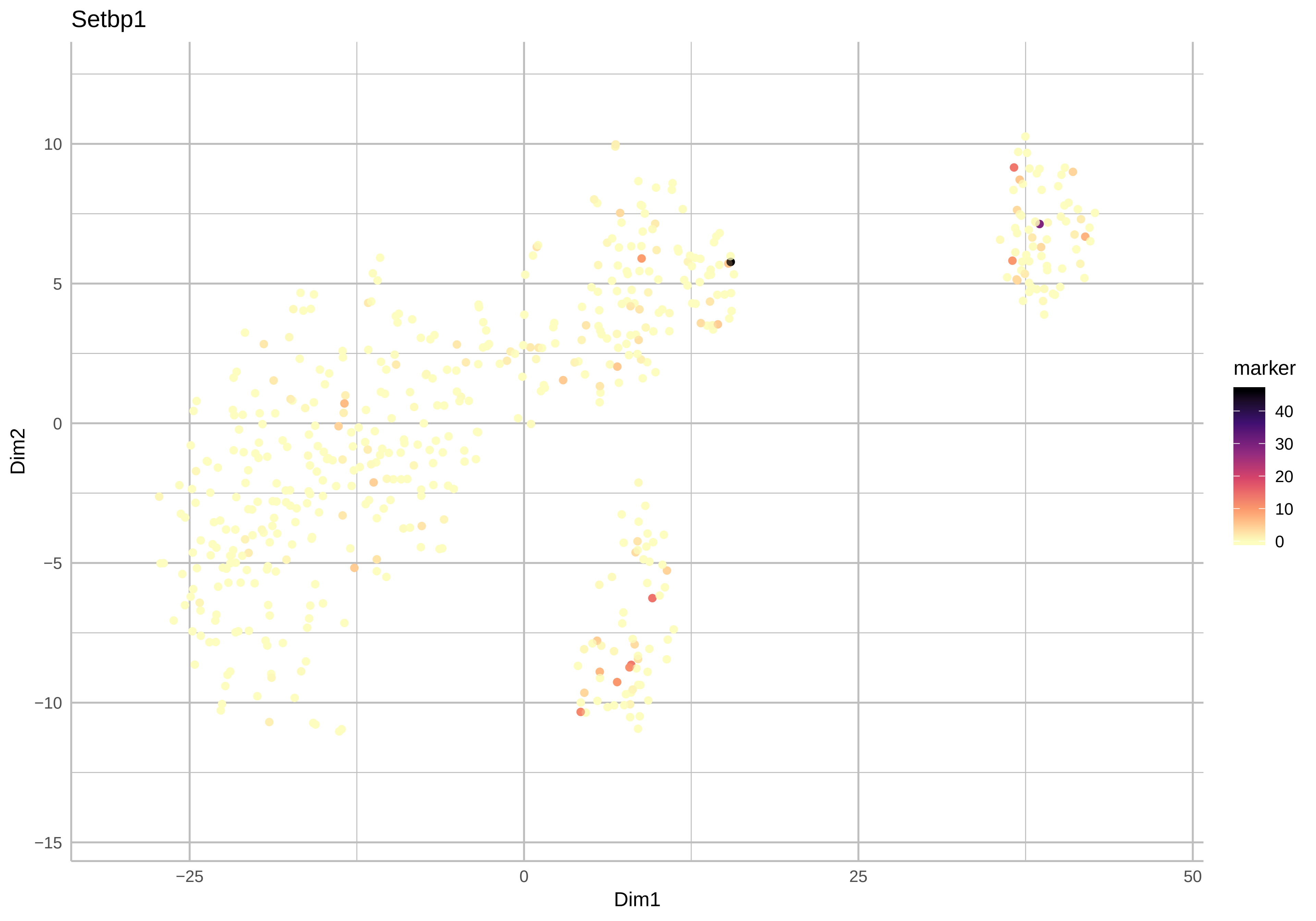
Setbp1
10
5
0
−5
−10
−15
−25
0
25
50
Dim1
Dim2
marker
40
30
20
10
0

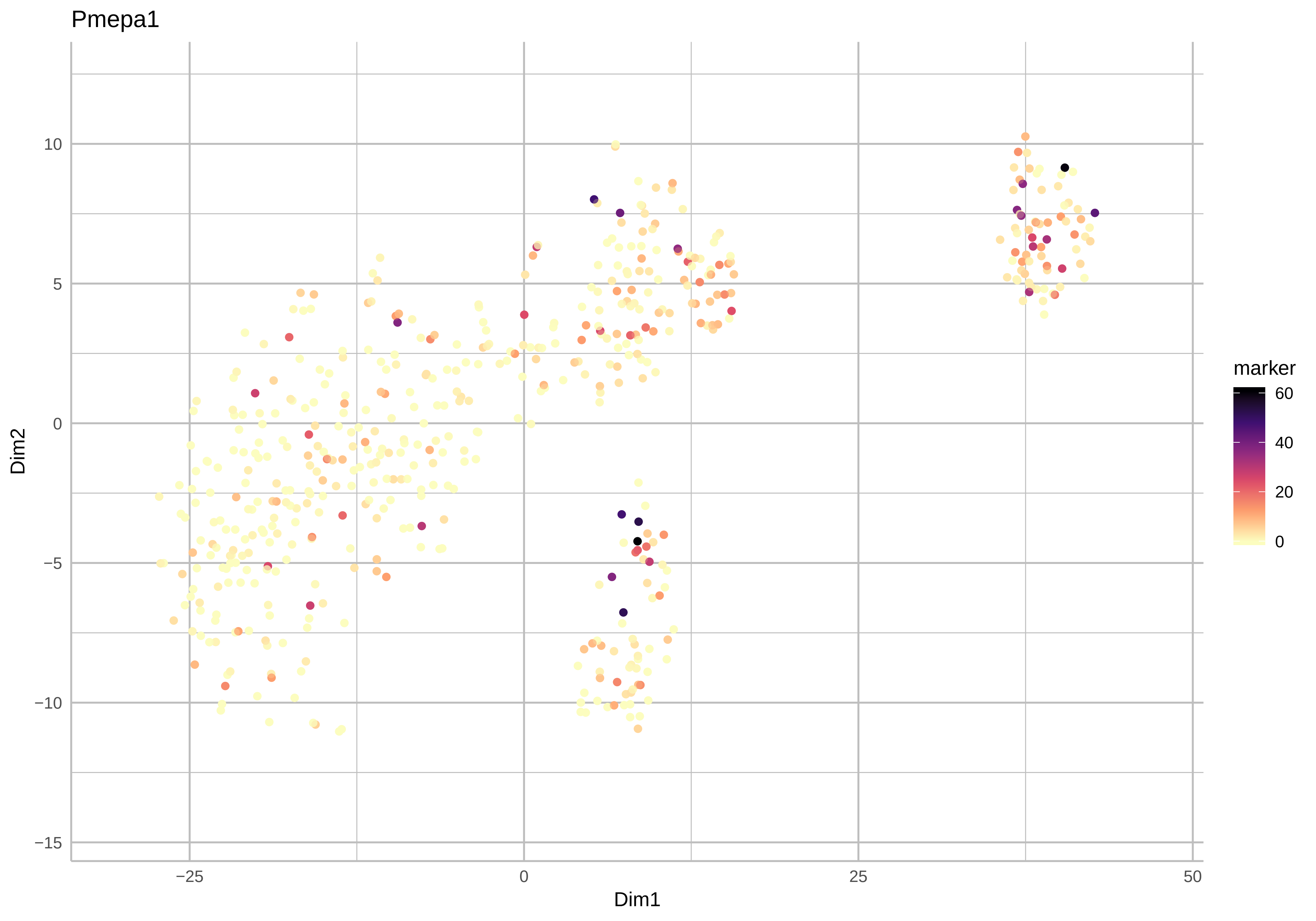

Pmepa1
Dim2
Dim1
10
5
0
−5
−10
−15
−25
0
25
50
marker
60
40
20
0

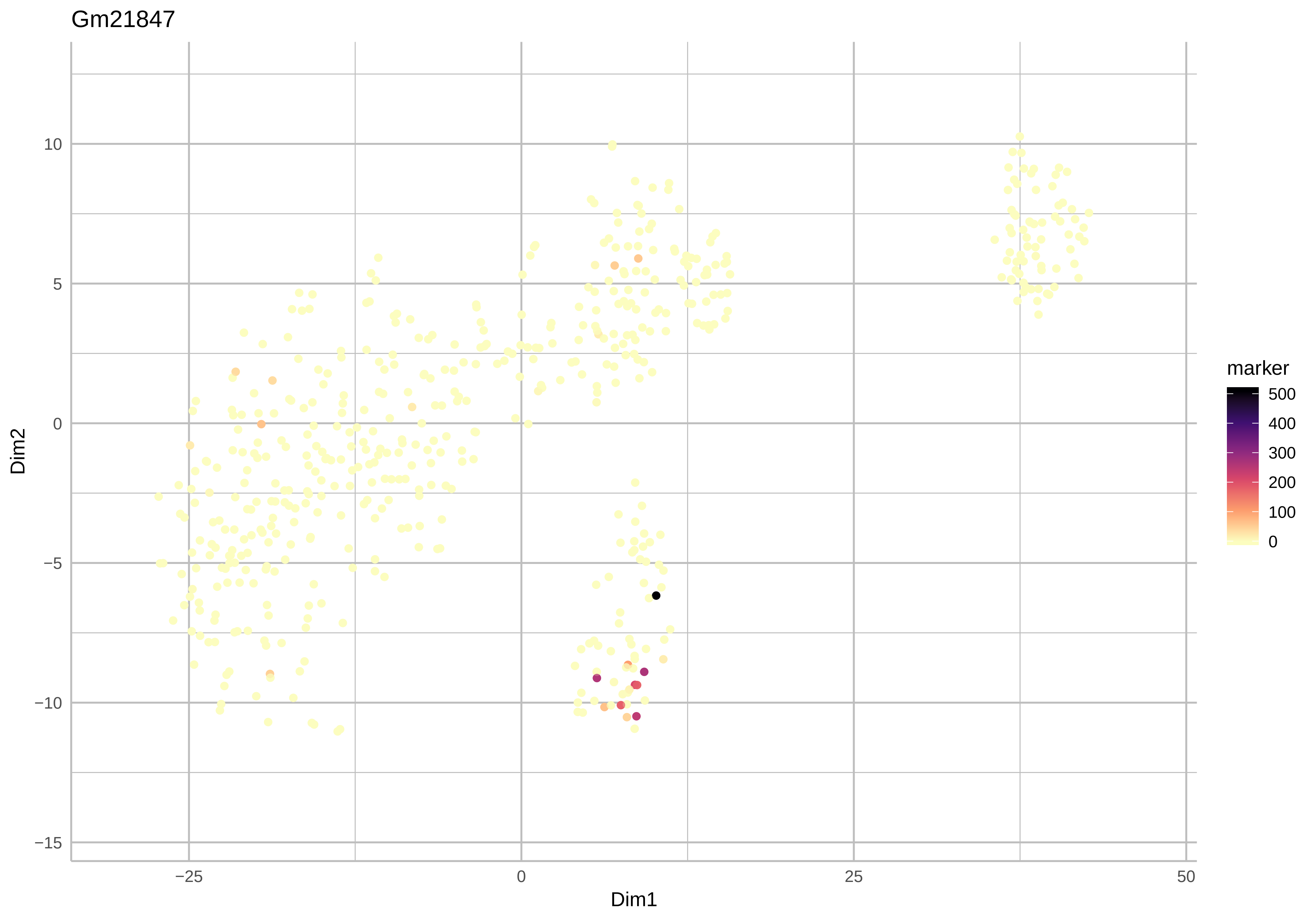
Gm21847
Dim2
Dim1
10
5
0
−5
−10
−15
−25
0
25
50
marker
500
400
300
200
100
0

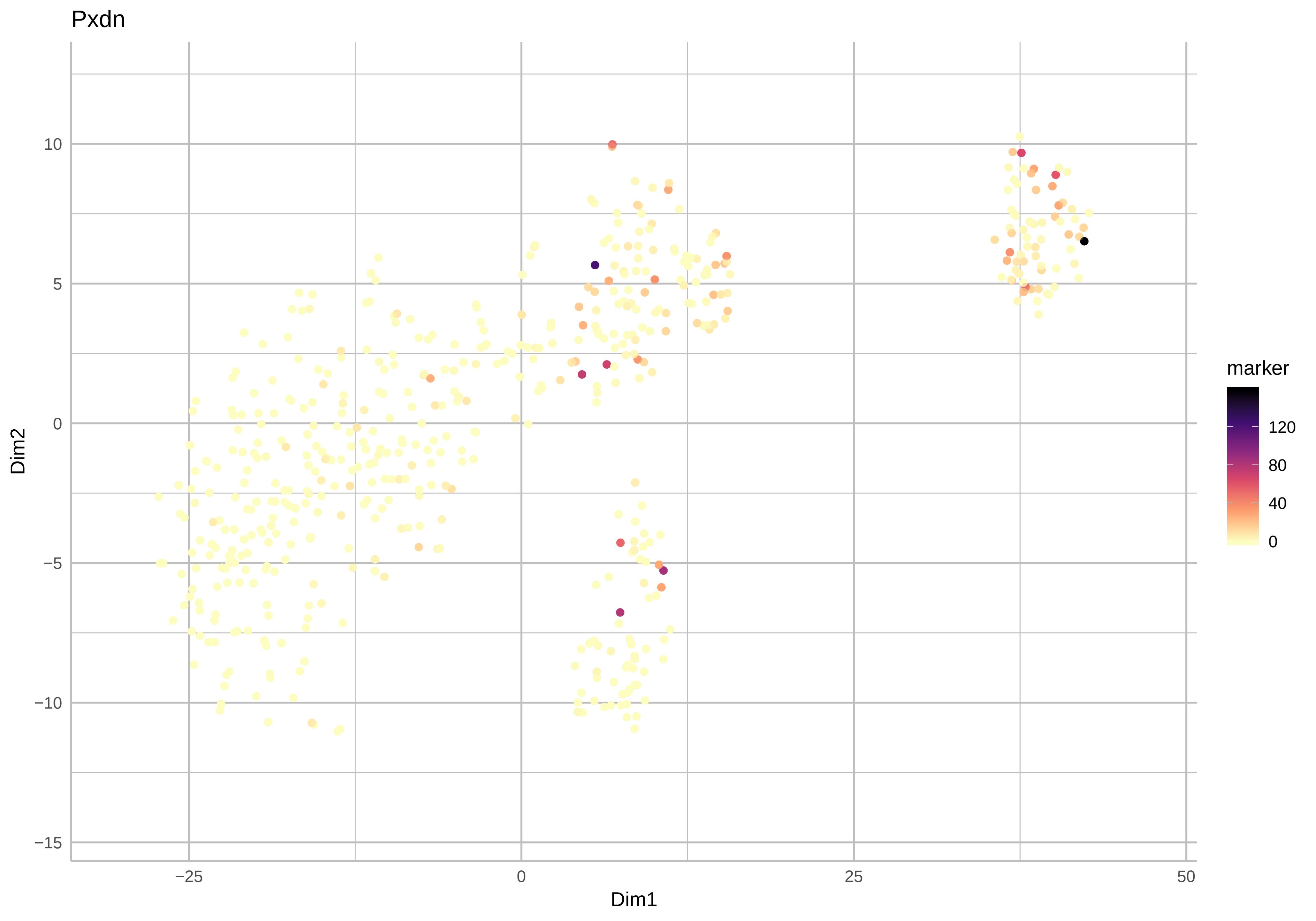
Pxdn
Dim2
Dim1
10
5
0
−5
−10
−15
−25
0
25
50
marker
120
80
40
0

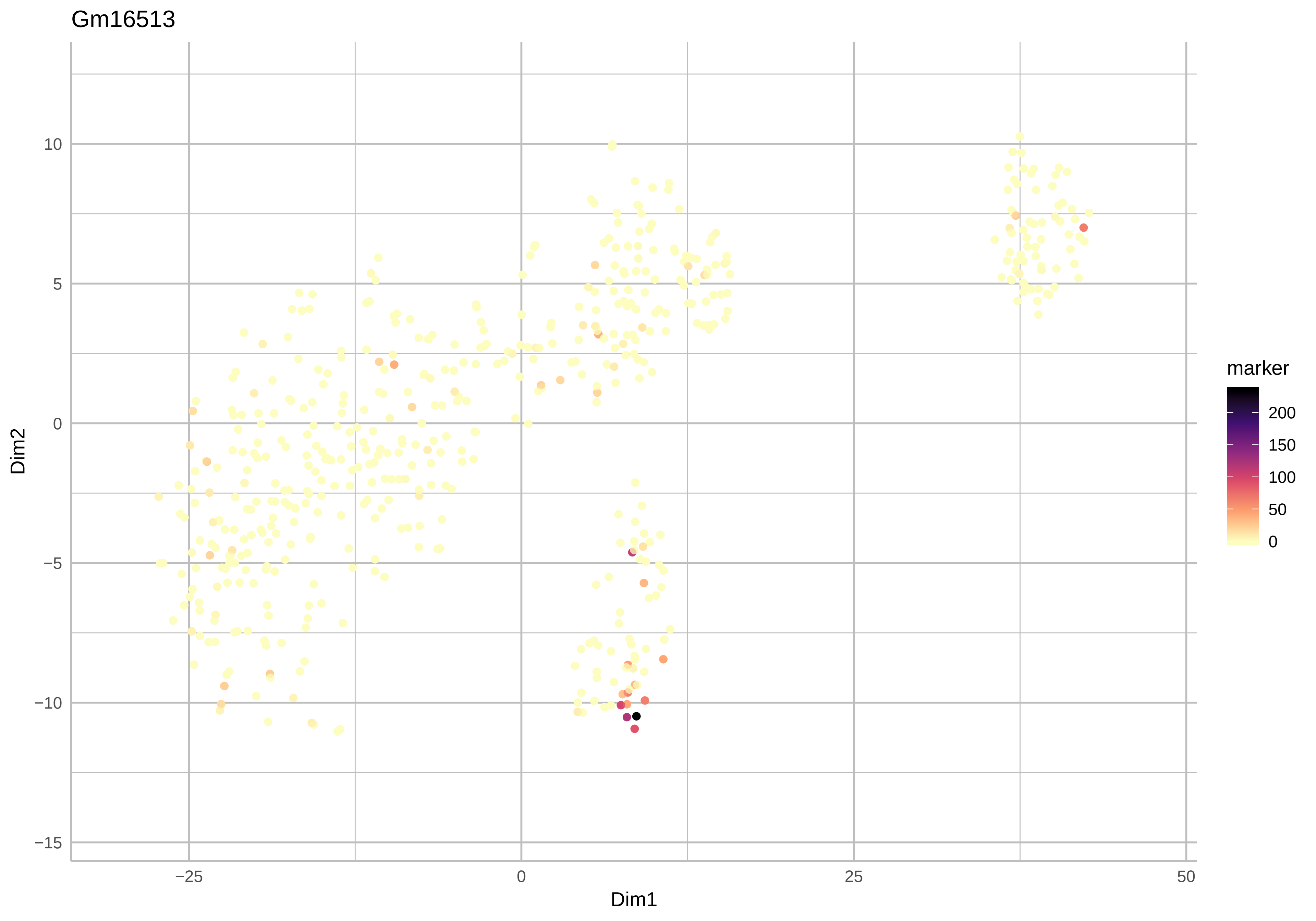

Gm16513
Dim2
Dim1
marker
200
150
100
50
0
10
5
0
−5
−10
−15
−25
0
25
50

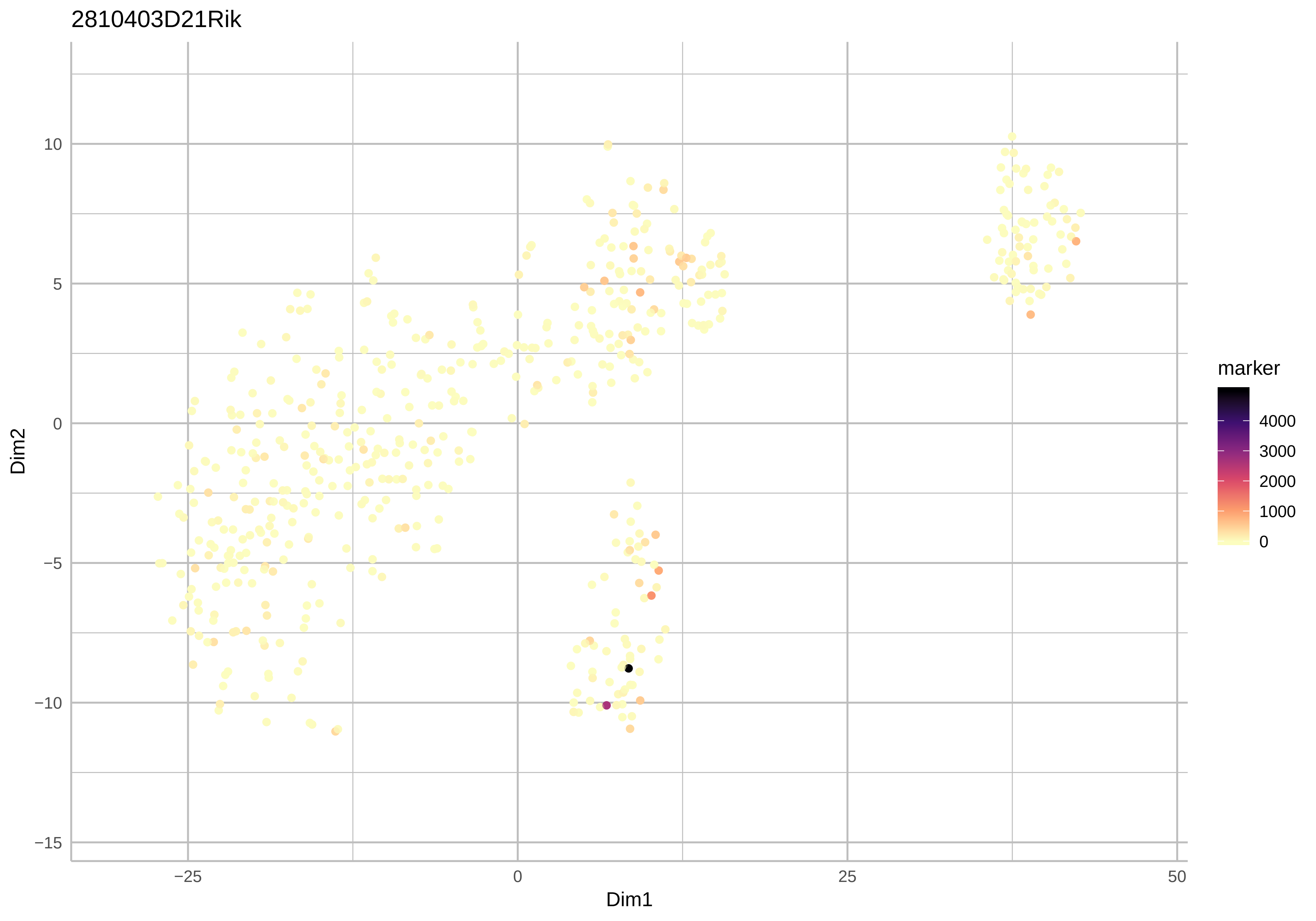

2810403D21Rik
Dim2
Dim1
10
5
0
−5
−10
−15
−25
0
25
50
marker
4000
3000
2000
1000
0

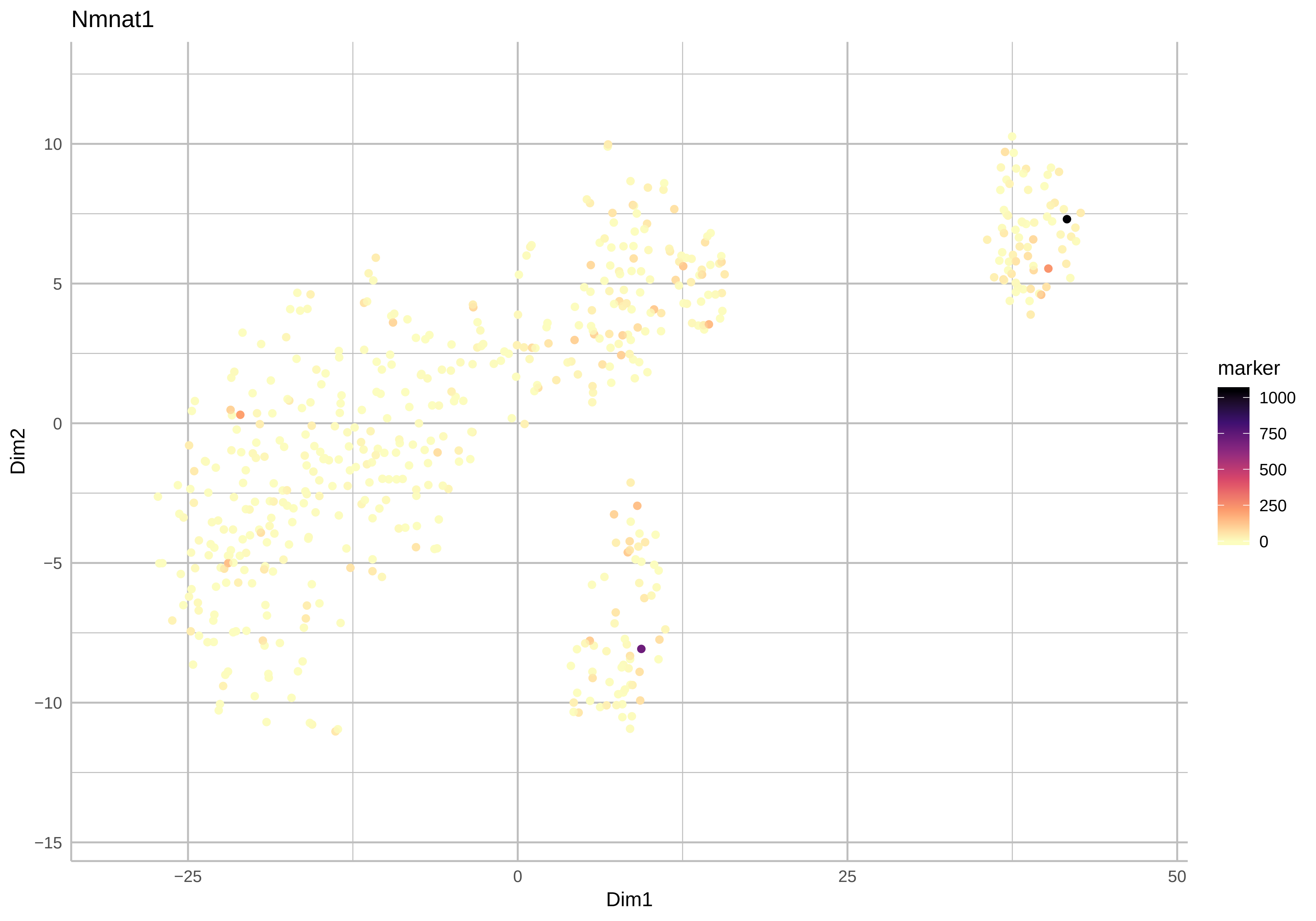

Nmnat1
Dim2
Dim1
marker
1000
750
500
250
0
10
5
0
−5
−10
−15
−25
0
25
50

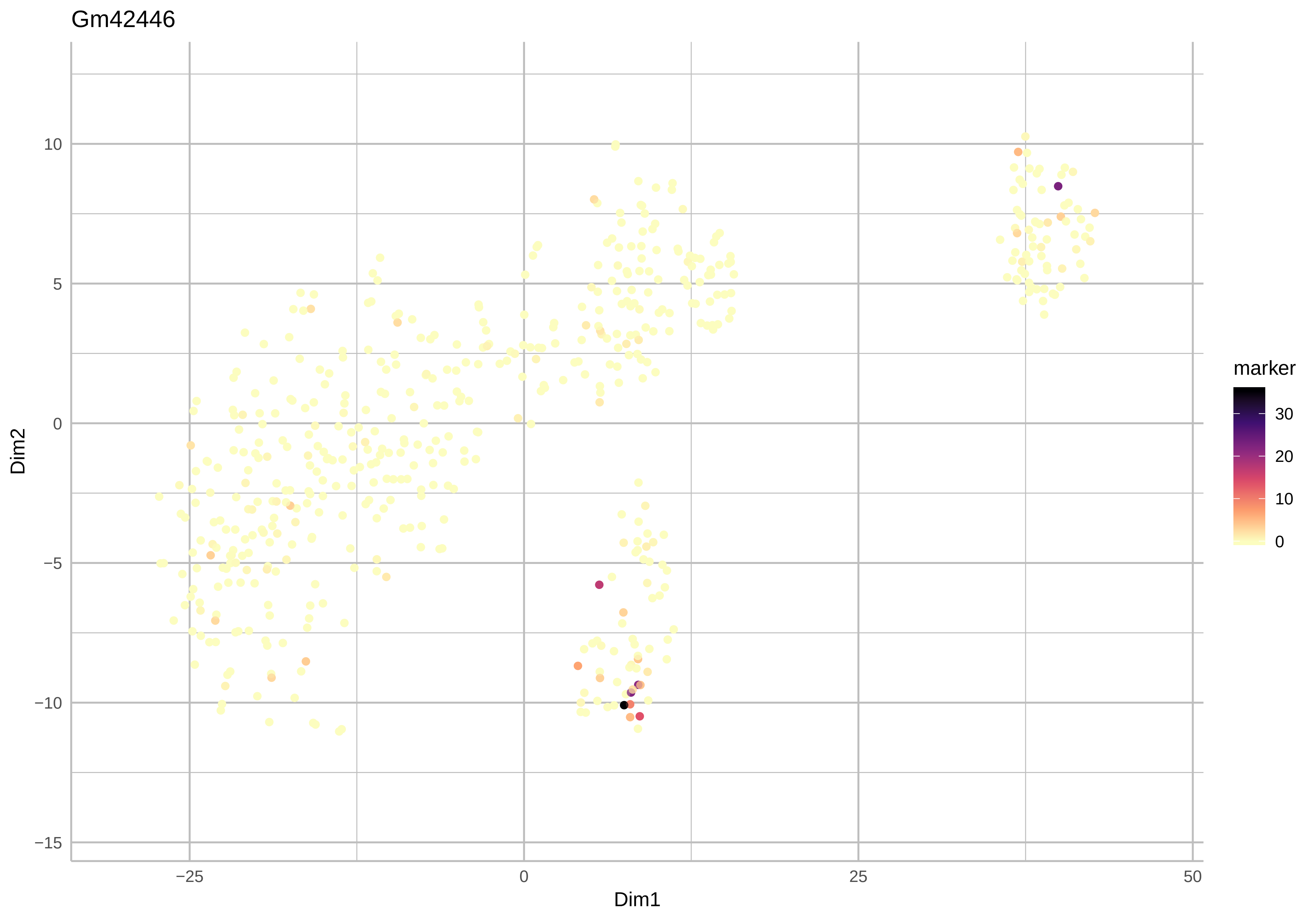

Gm42446
10
5
0
−5
−10
−15
−25
0
25
50
Dim1
Dim2
marker
30
20
10
0

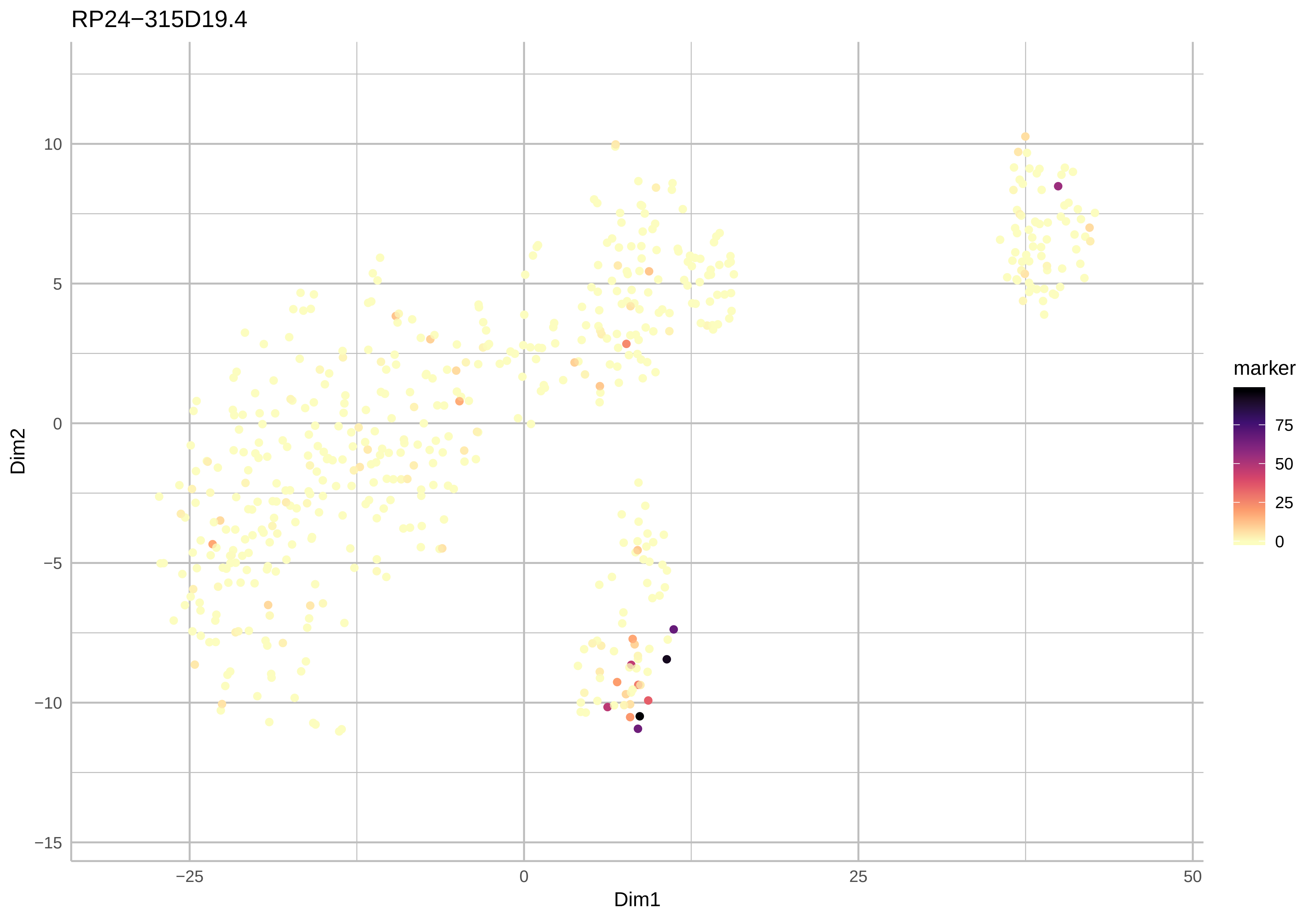

RP24−315D19.4
Dim2
Dim1
10
5
0
−5
−10
−15
−25
0
25
50
marker
75
50
25
0

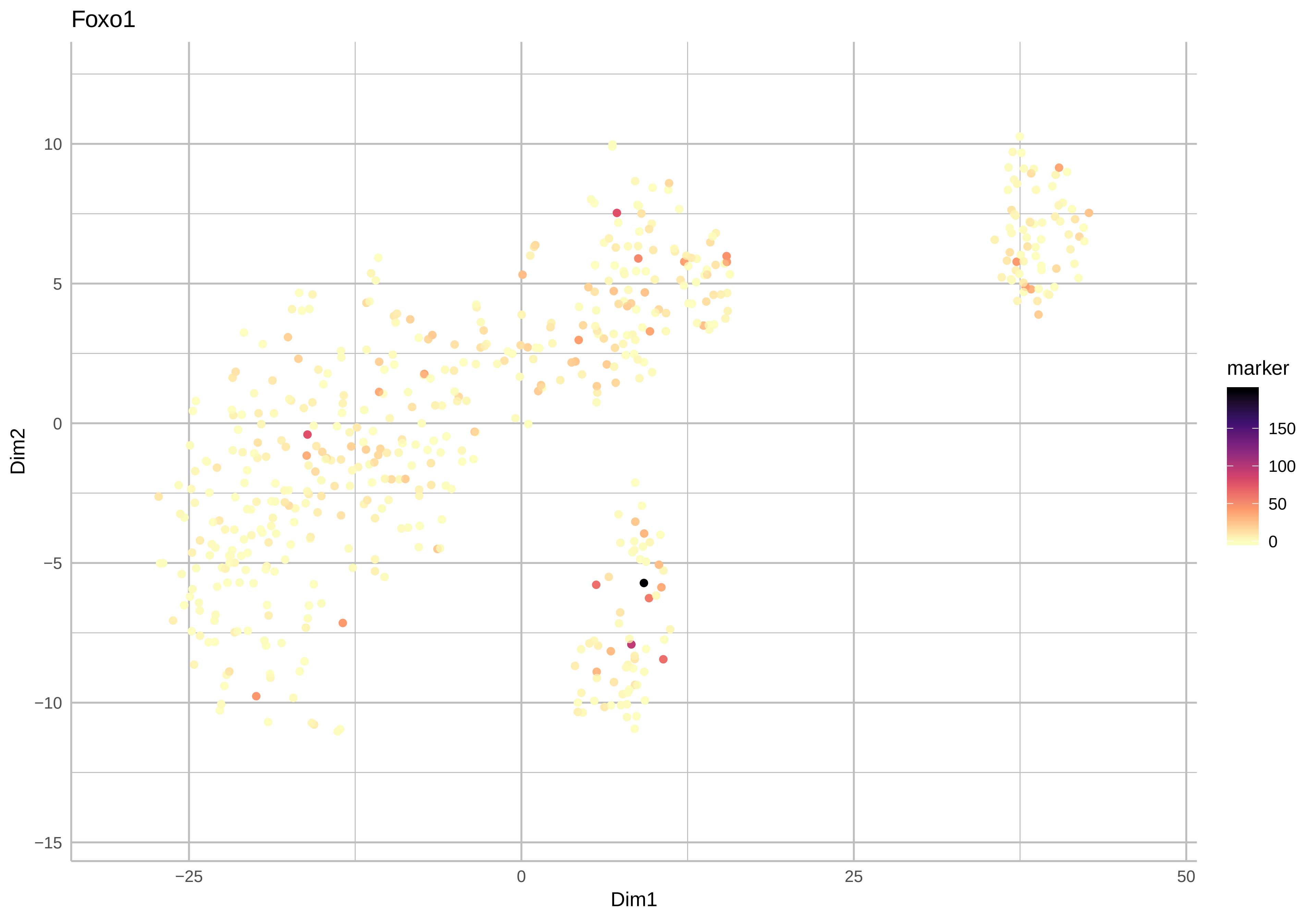
Foxo1
Dim2
Dim1
marker
150
100
50
0
10
5
0
−5
−10
−15
−25
0
25
50

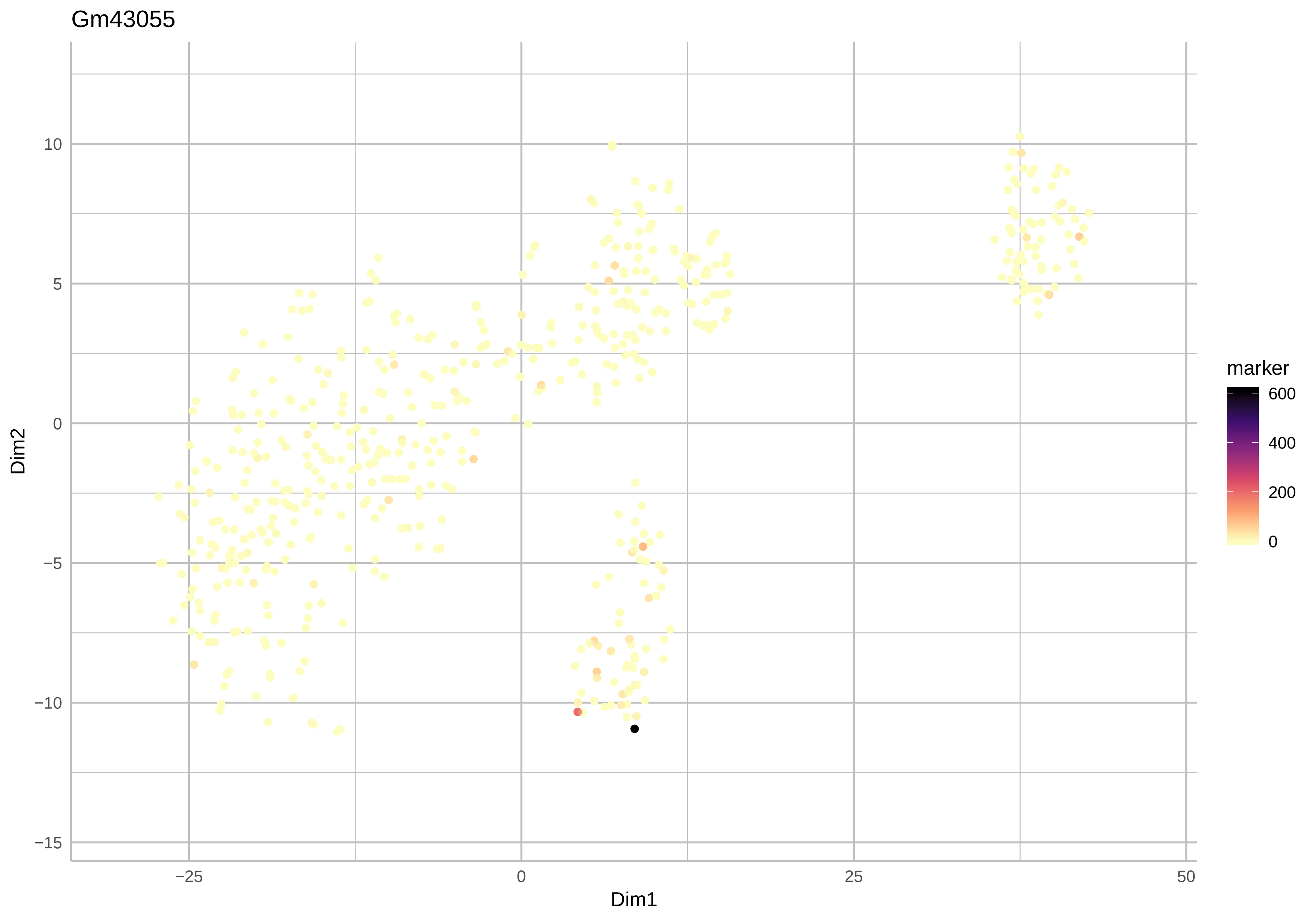
Gm43055
Dim2
Dim1
marker
600
400
200
0
10
5
0
−5
−10
−15
−25
0
25
50

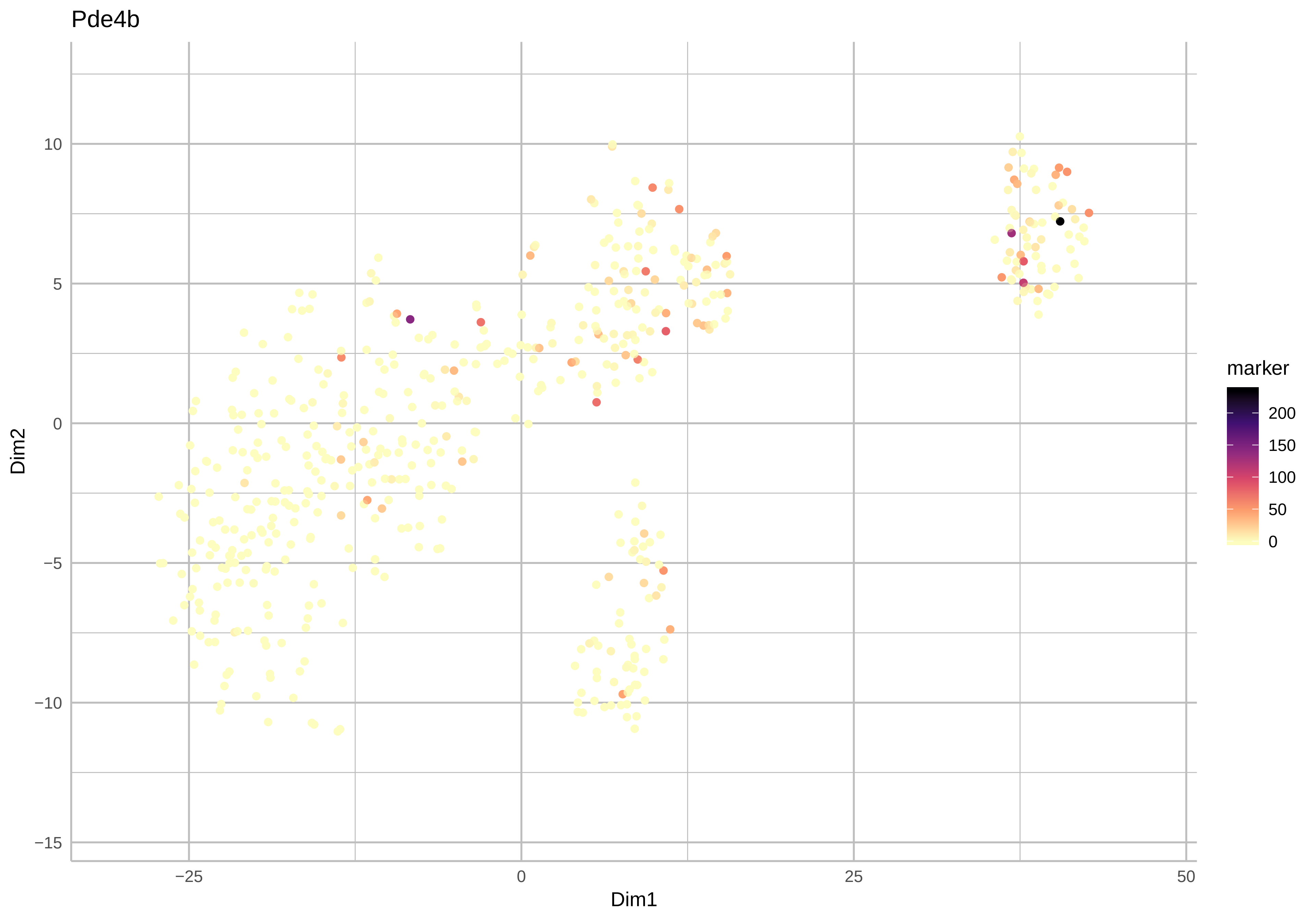

Pde4b
10
5
0
−5
−10
−15
−25
0
25
50
Dim1
Dim2
marker
200
150
100
50
0

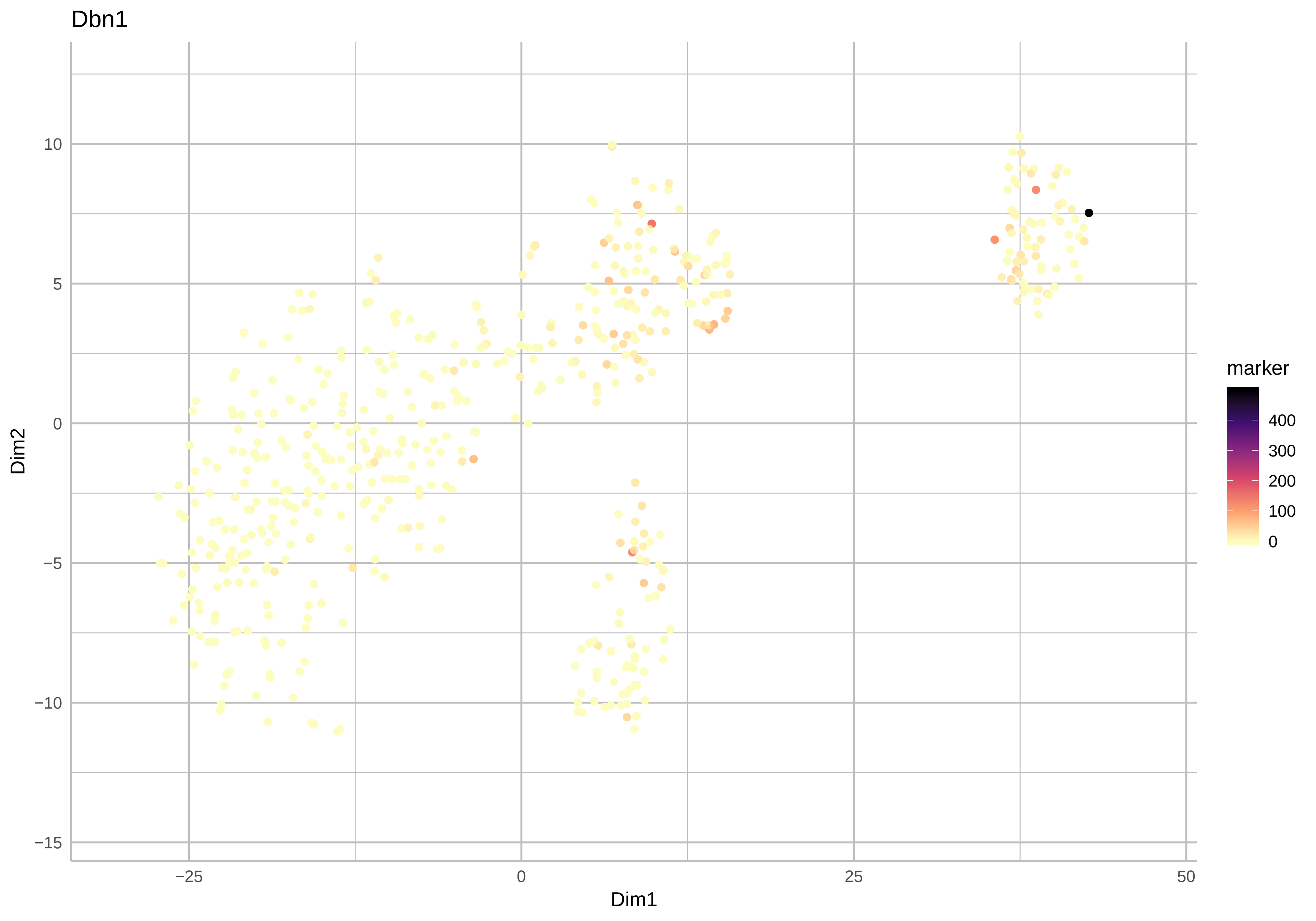

Dbn1
Dim2
Dim1
marker
400
300
200
100
0
10
5
0
−5
−10
−15
−25
0
25
50

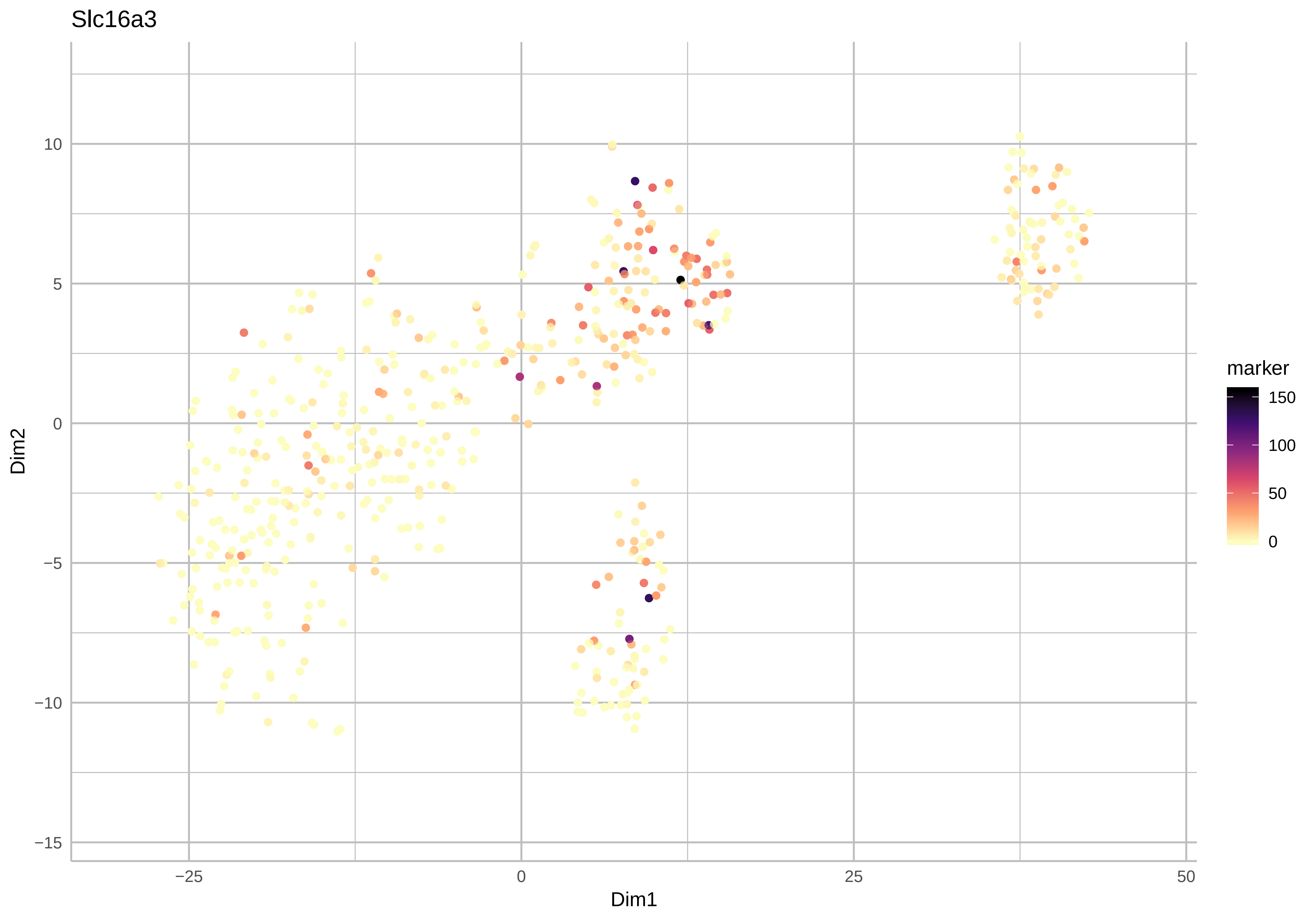
Slc16a3
Dim2
Dim1
10
5
0
−5
−10
−15
−25
0
25
50
marker
150
100
50
0

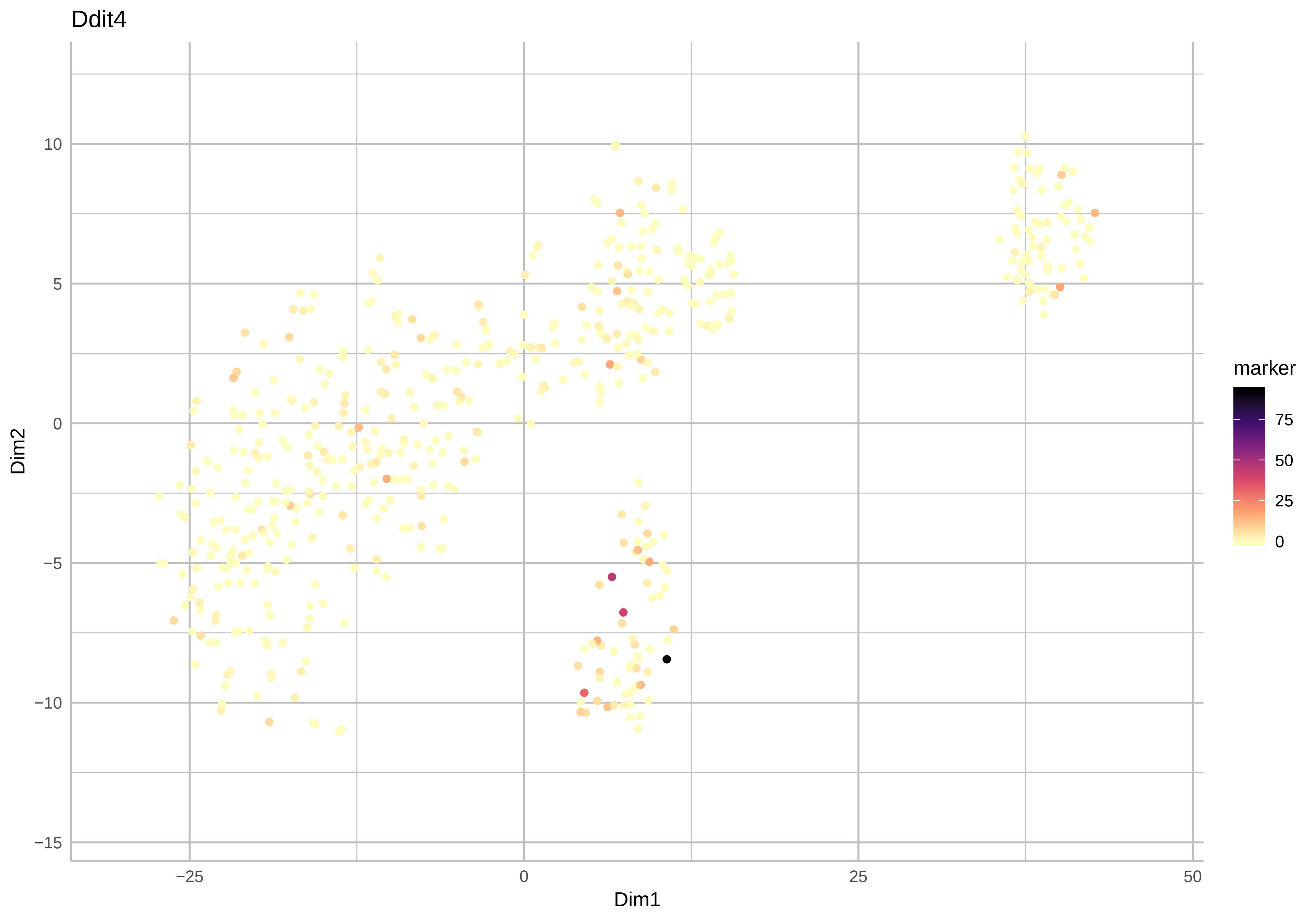
Ddit4
Dim2
10
5
0
−5
−10
−15
−25
0
25
50
Dim1
marker
75
50
25
0

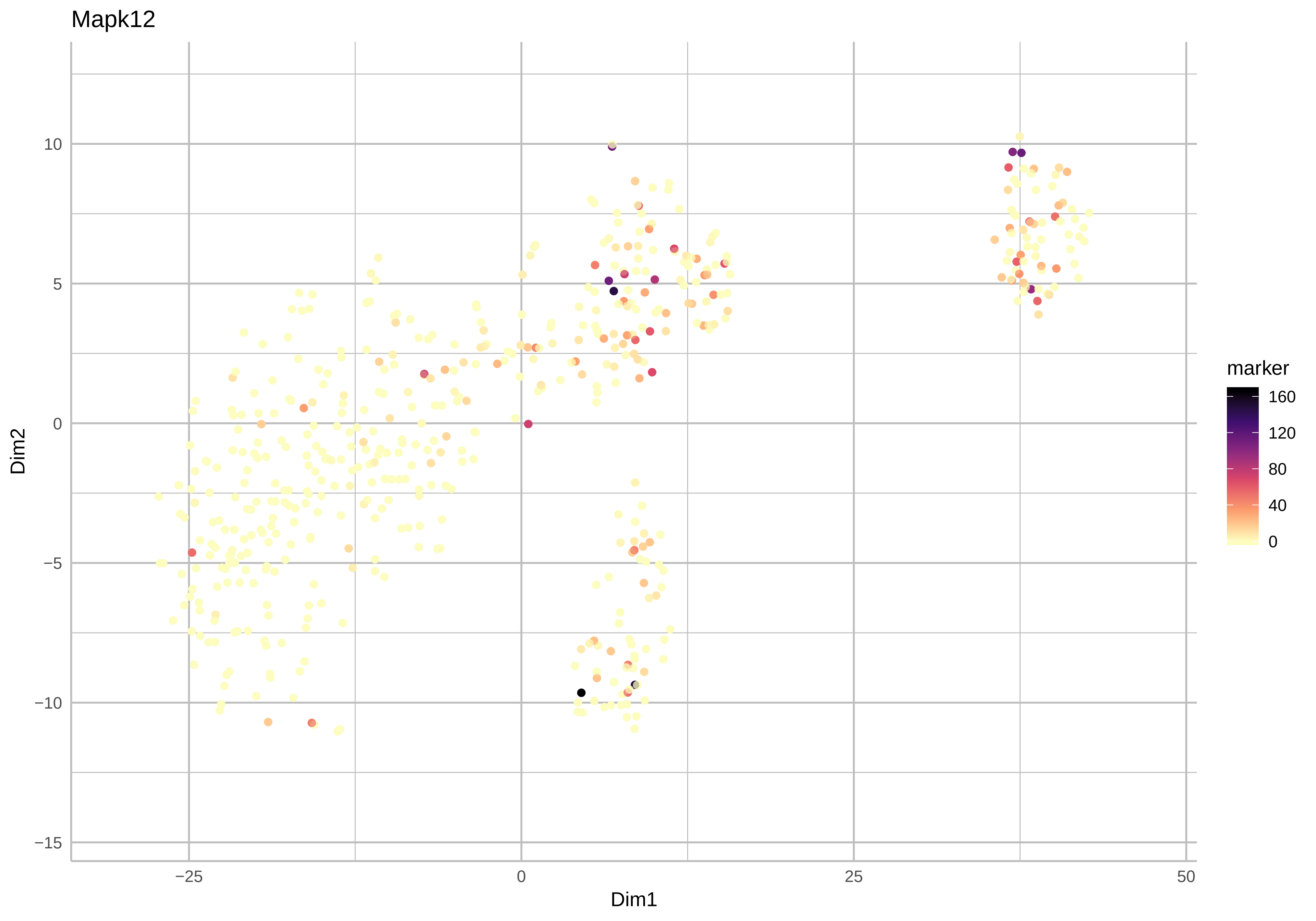
Mapk12
10
5
0
−5
−10
−15
−25
0
25
50
Dim1
Dim2
marker
160
120
80
40
0

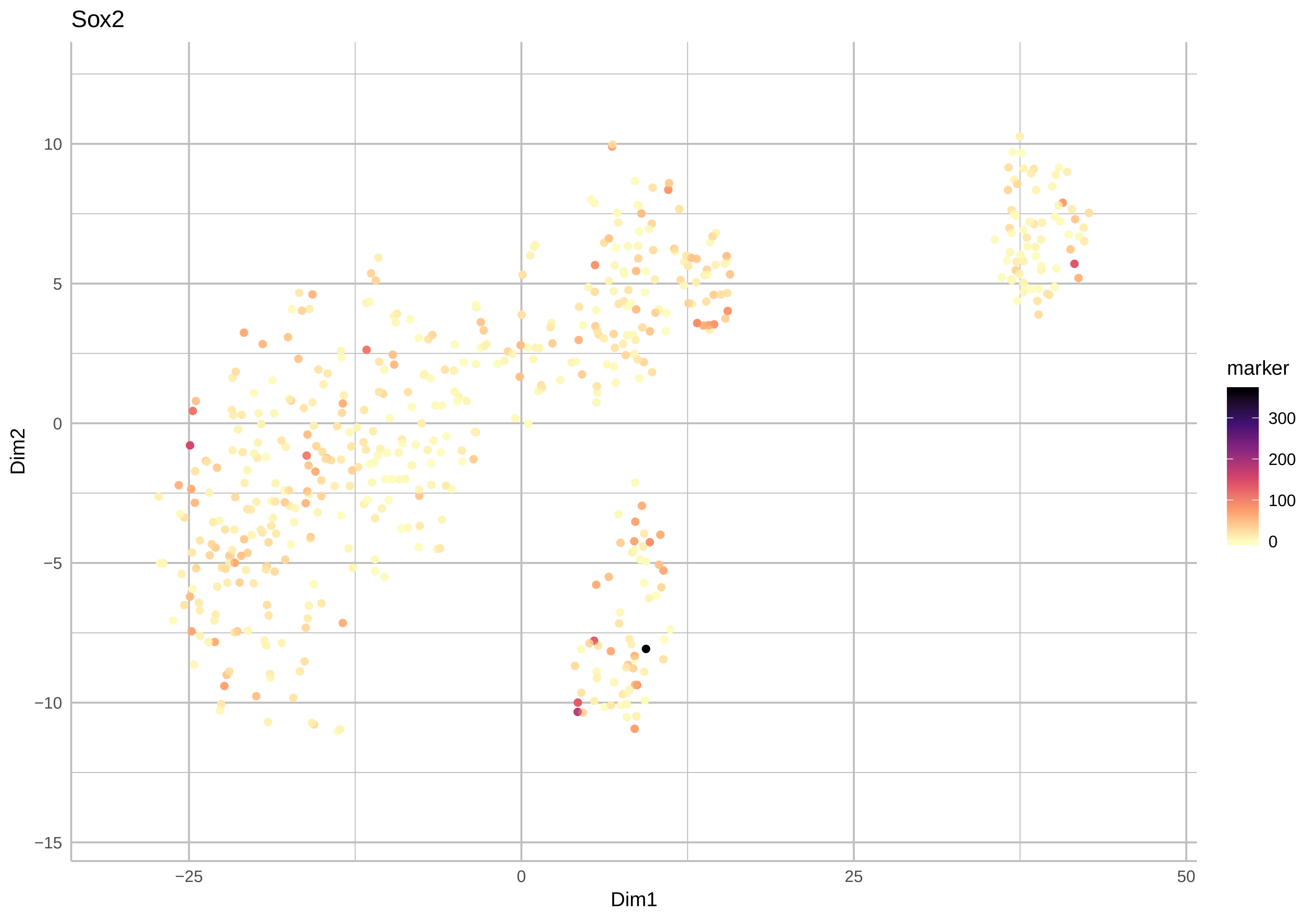

Sox2
Dim2
Dim1
10
5
0
−5
−10
−15
−25
0
25
50
marker
300
200
100
0

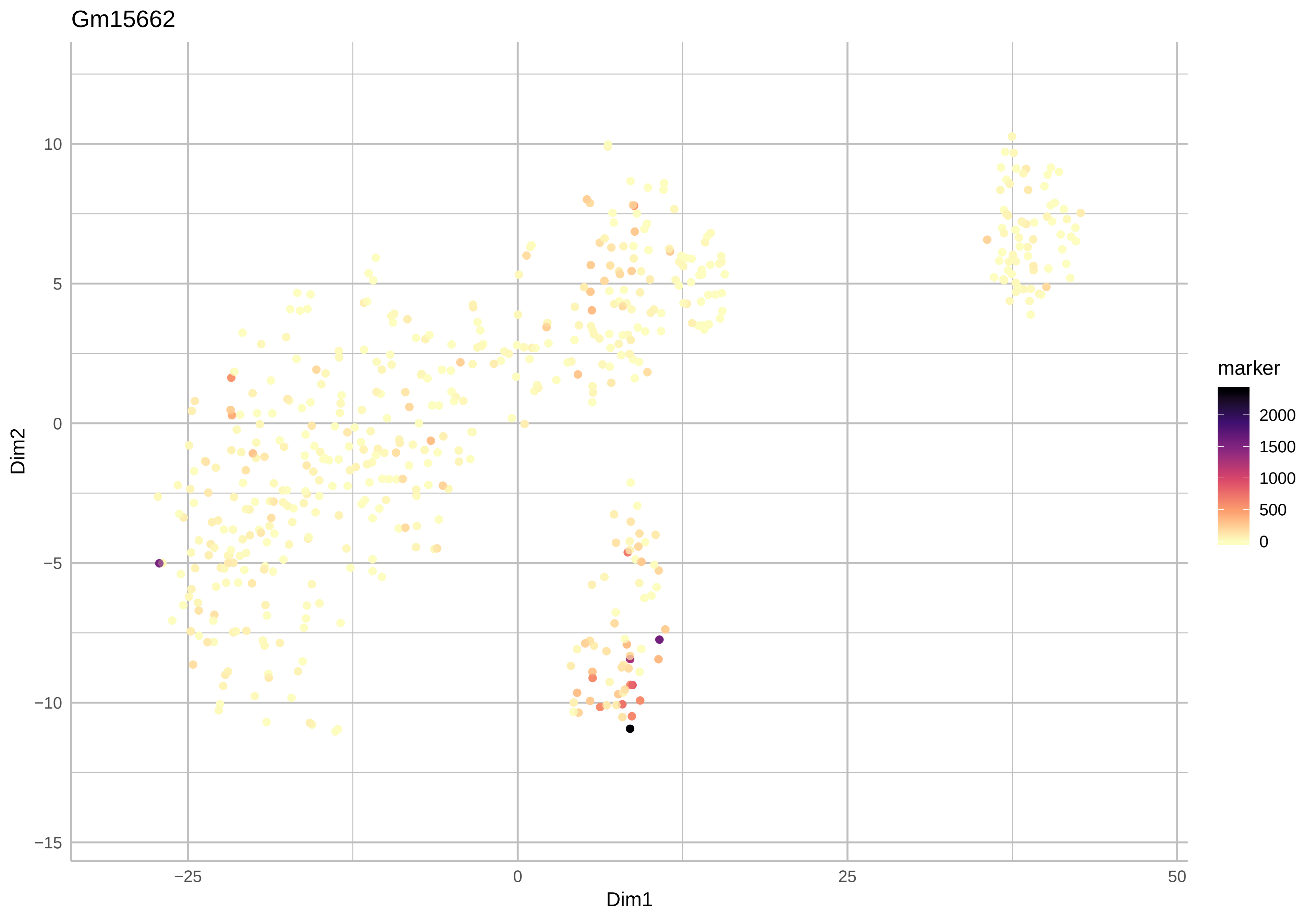
Gm15662
Dim2
Dim1
marker
2000
1500
1000
500
0
10
5
0
−5
−10
−15
−25
0
25
50

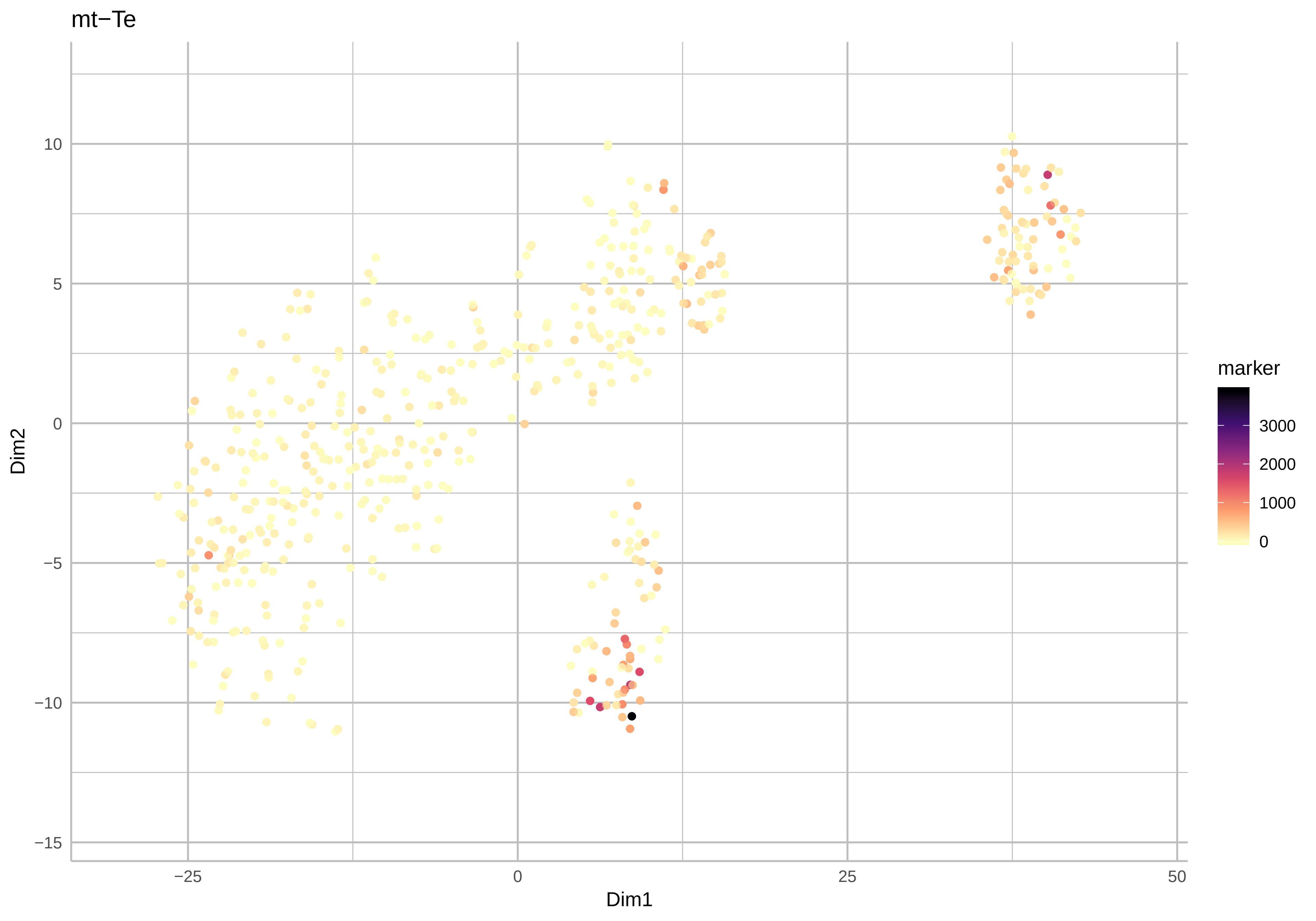
mt−Te
Dim2
Dim1
marker
3000
2000
1000
0
10
5
0
−5
−10
−15
−25
0
25
50

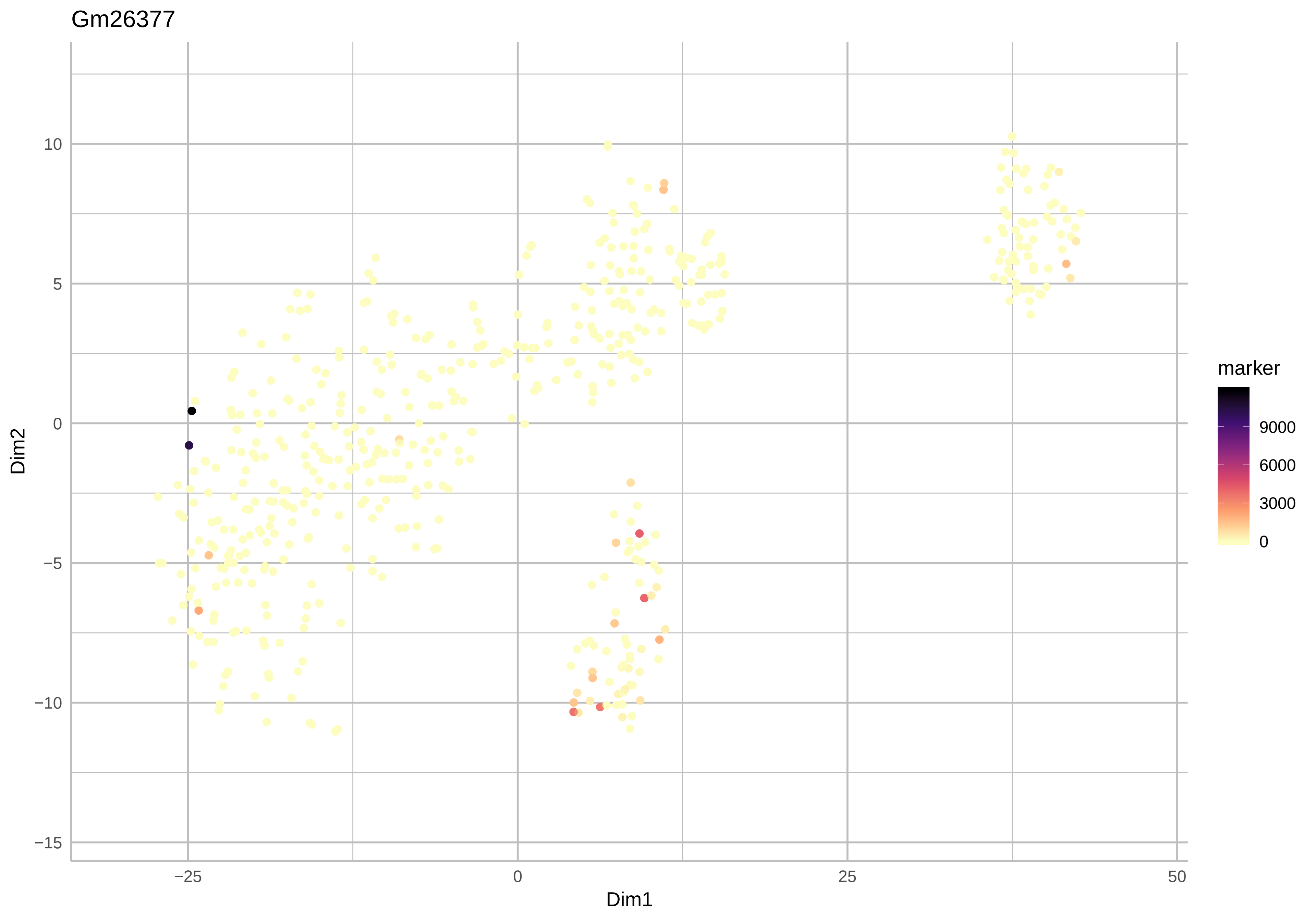
Gm26377
Dim2
Dim1
marker
9000
6000
3000
0
10
5
0
−5
−10
−15
−25
0
25
50

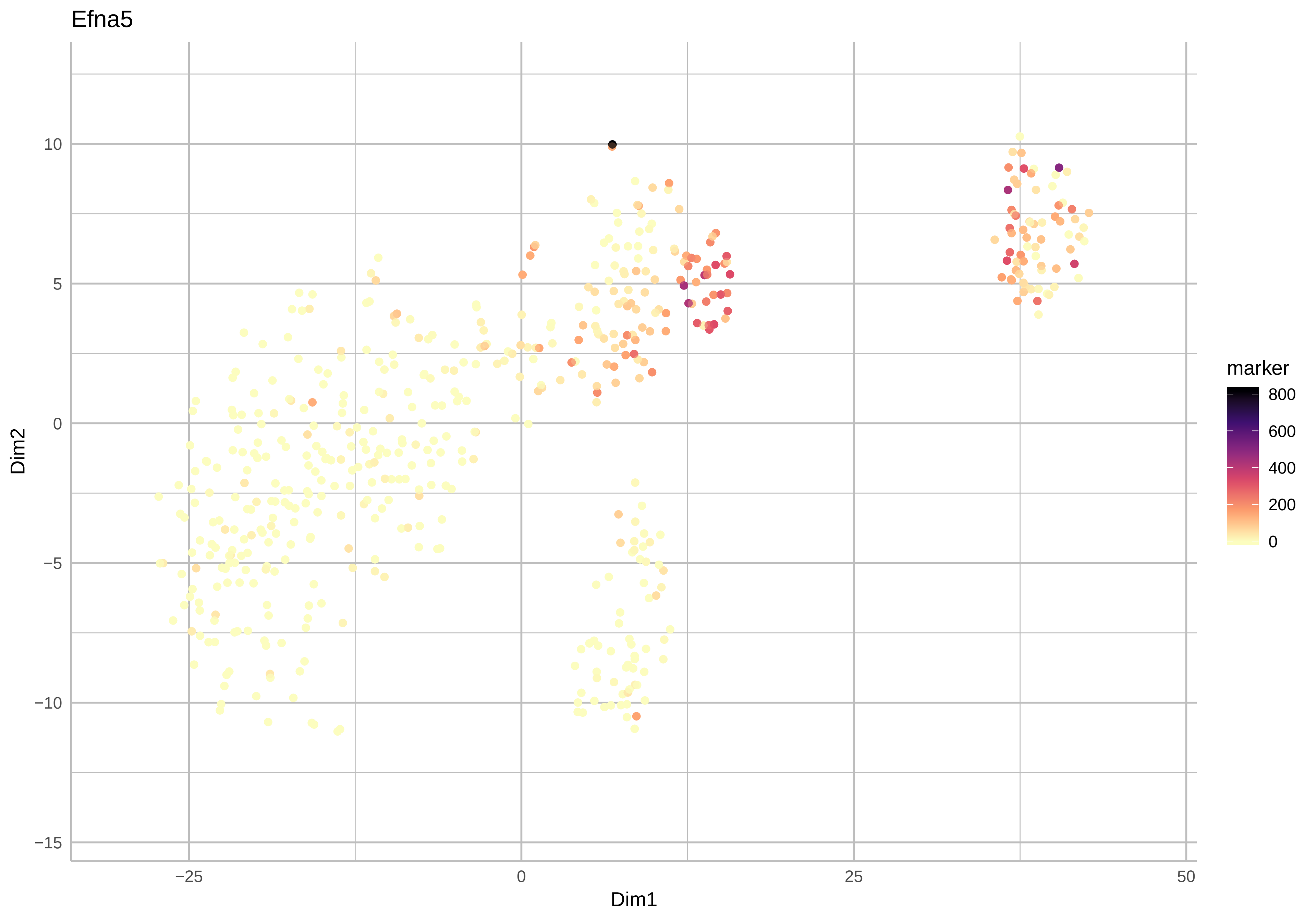
Efna5
Dim2
Dim1
10
5
0
−5
−10
−15
−25
0
25
50
marker
800
600
400
200
0

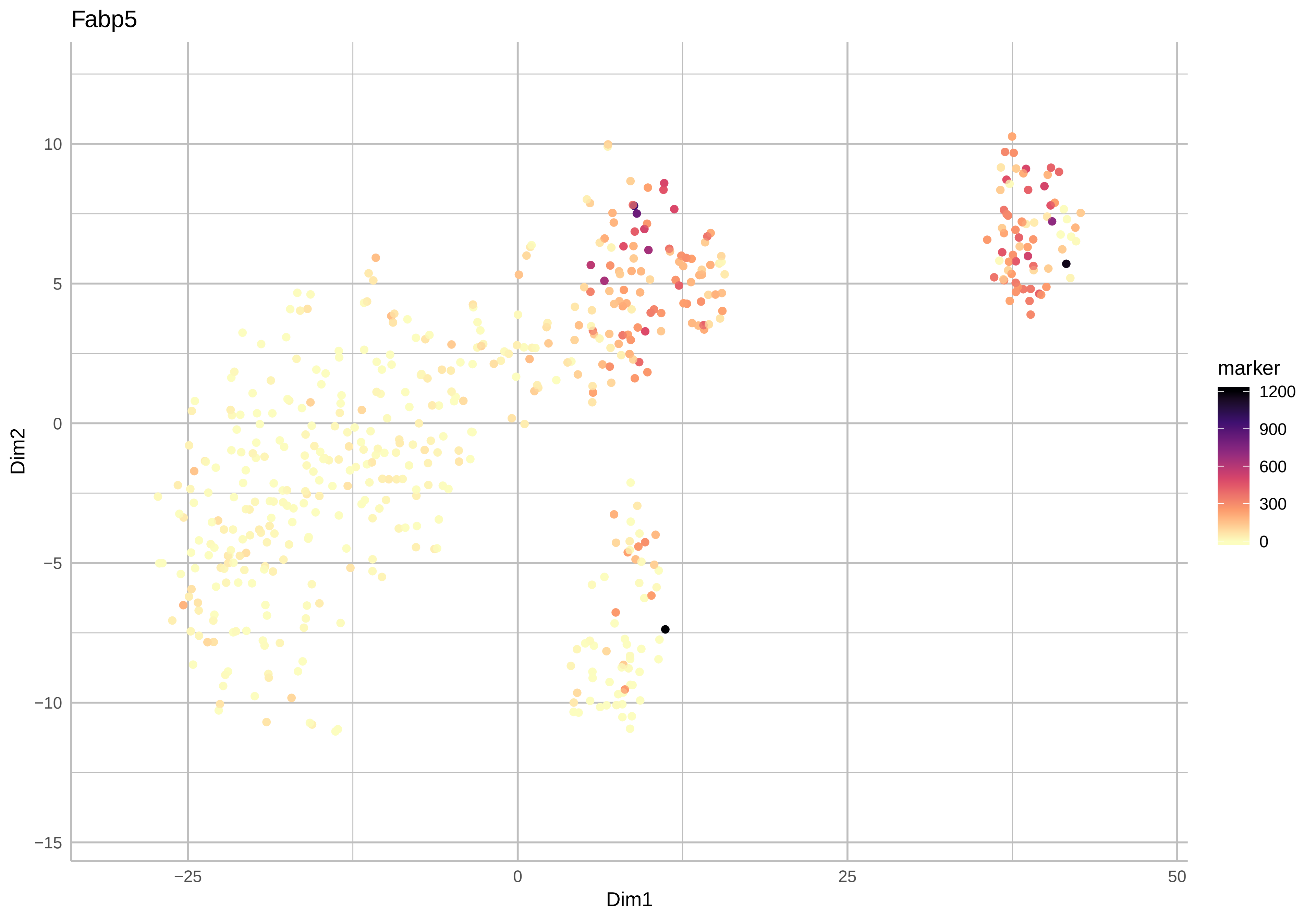
Fabp5
Dim2
Dim1
10
5
0
−5
−10
−15
−25
0
25
50
marker
1200
900
600
300
0

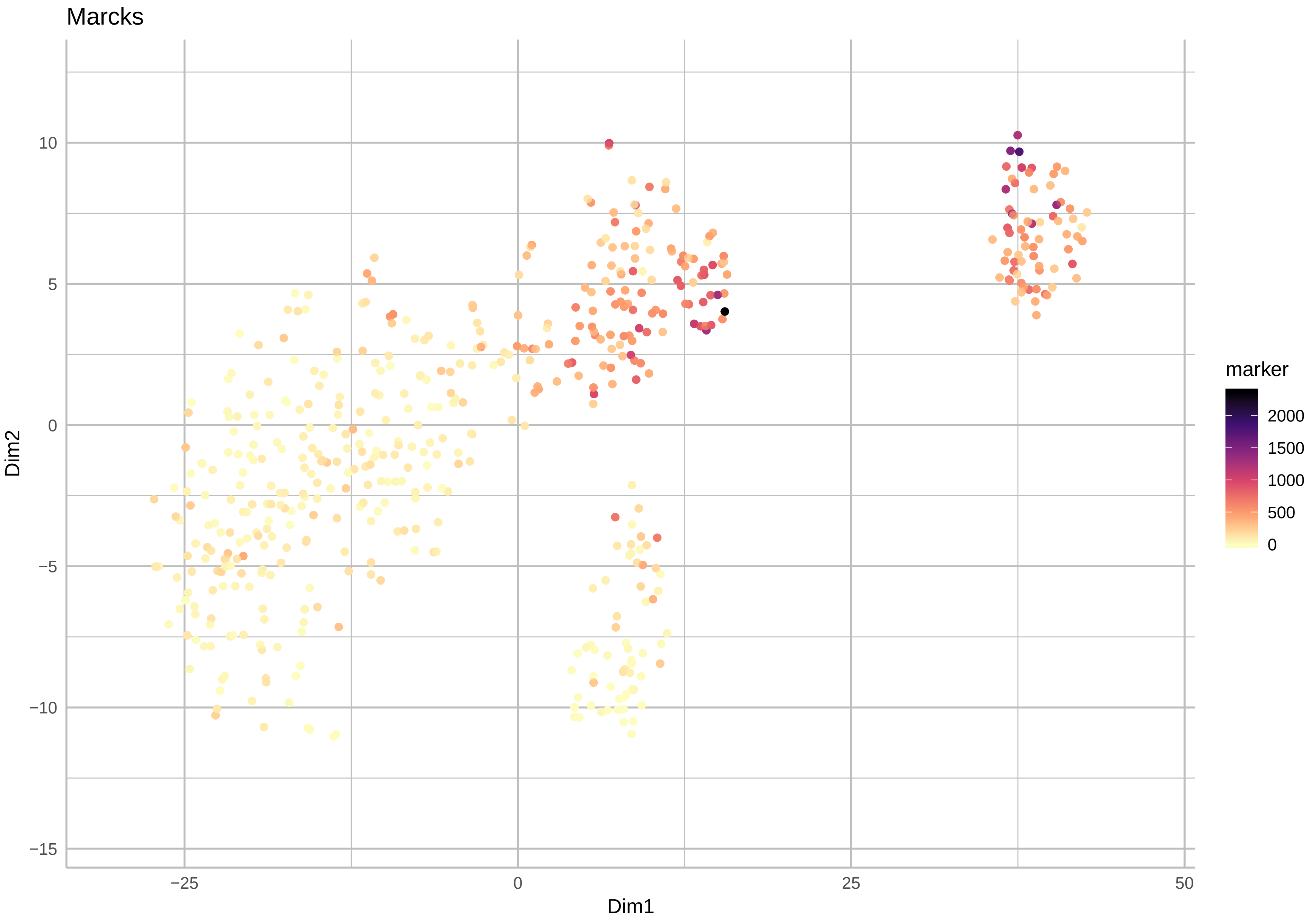
Marcks
Dim2
Dim1
10
5
0
−5
−10
−15
−25
0
25
50
marker
2000
1500
1000
500
0

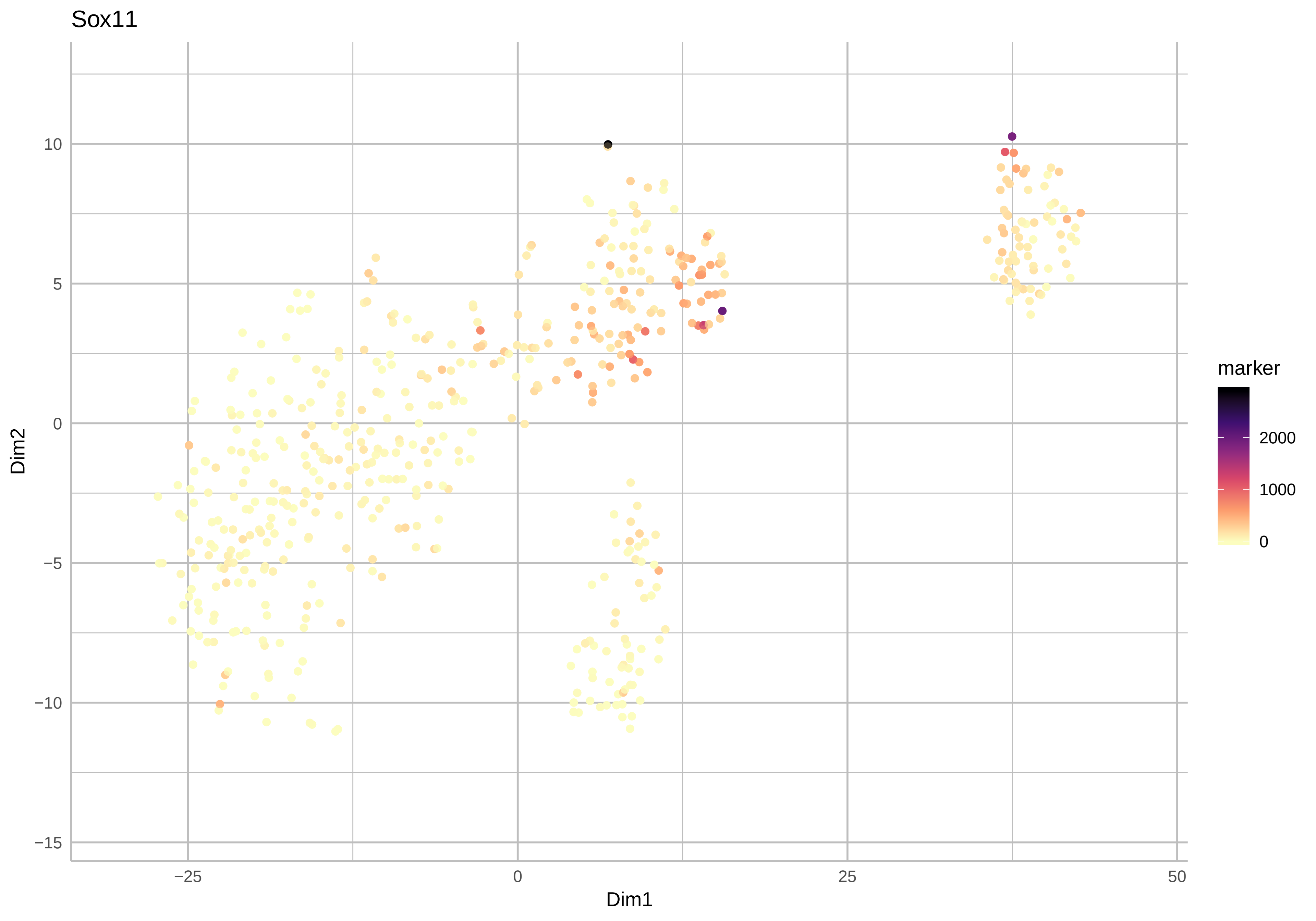

Sox11
Dim2
Dim1
10
5
0
−5
−10
−15
−25
0
25
50
marker
2000
1000
0

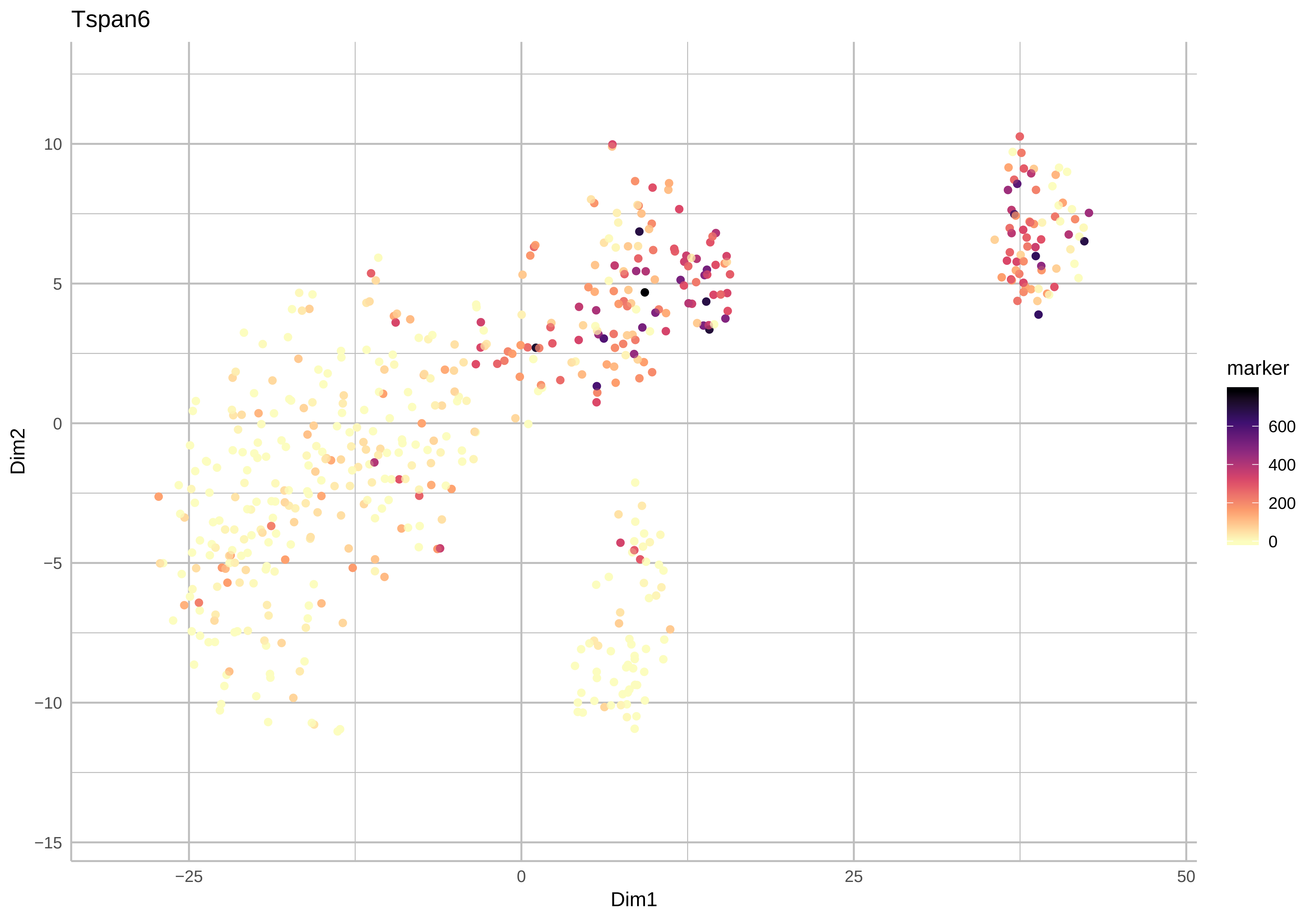

Tspan6
Dim2
Dim1
10
5
0
−5
−10
−15
−25
0
25
50
marker
600
400
200
0

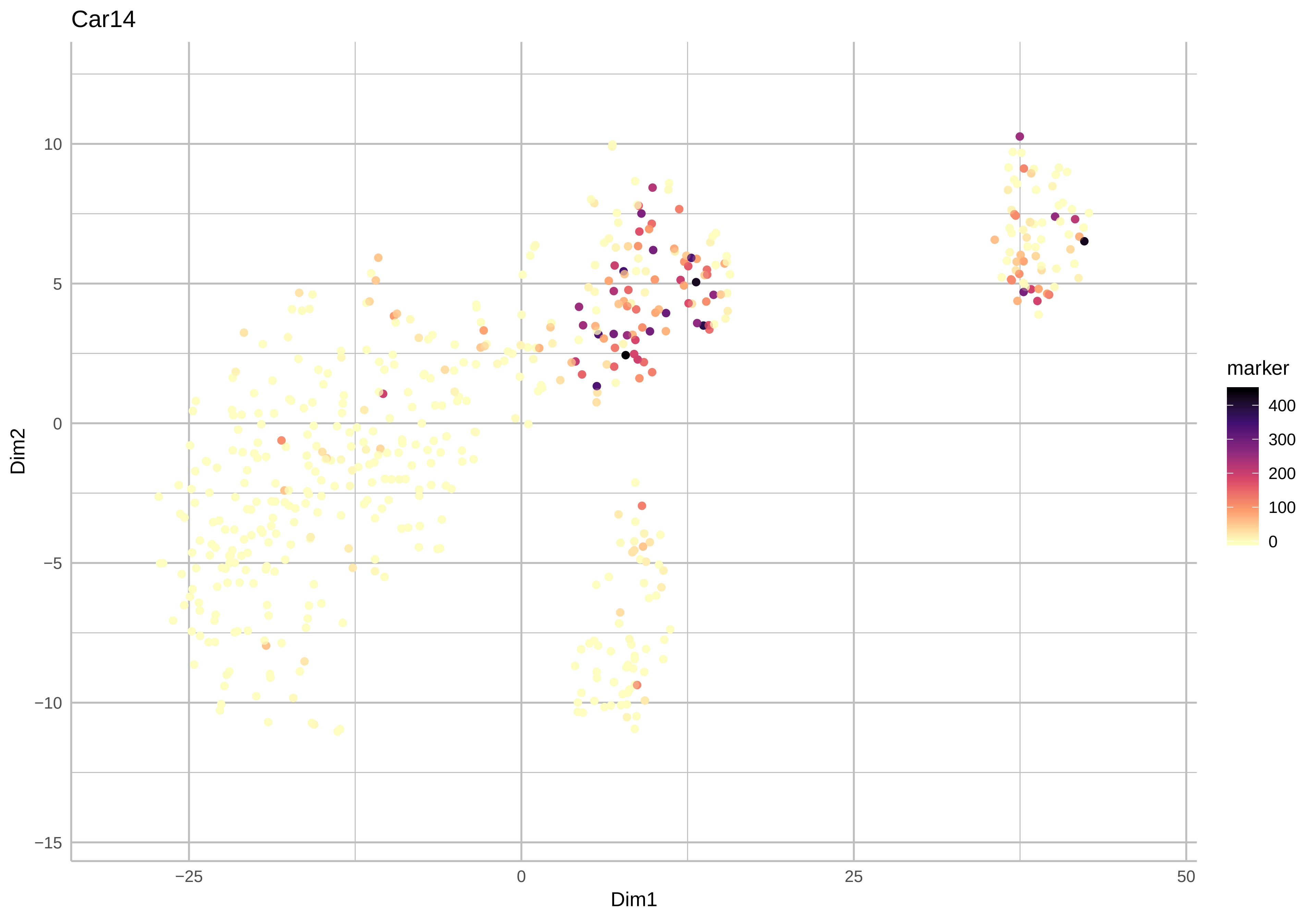

Car14
Dim2
Dim1
10
5
0
−5
−10
−15
−25
0
25
50
marker
400
300
200
100
0

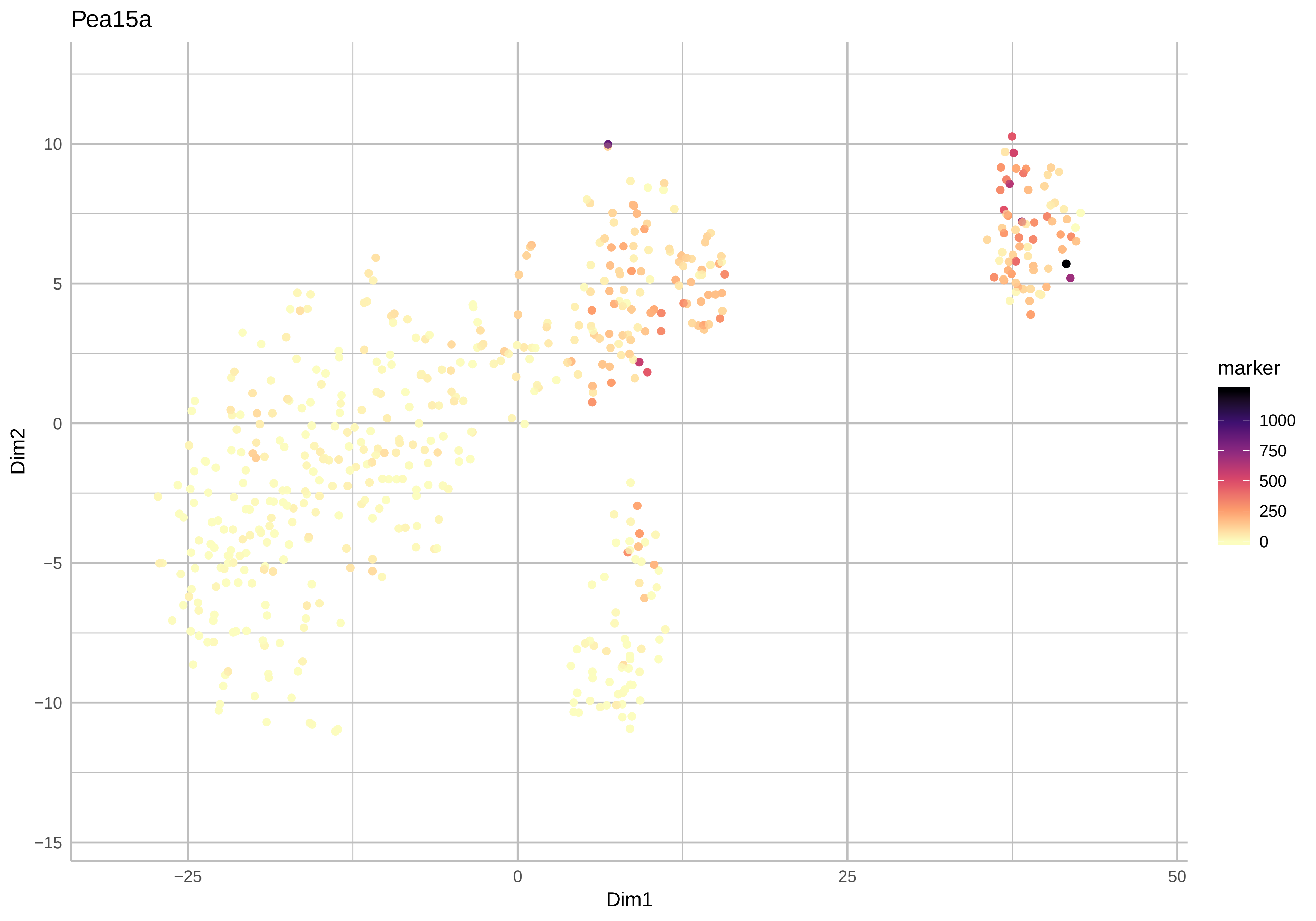

Pea15a
Dim2
10
5
0
−5
−10
−15
Dim1
−25
0
25
50
marker
1000
750
500
250
0

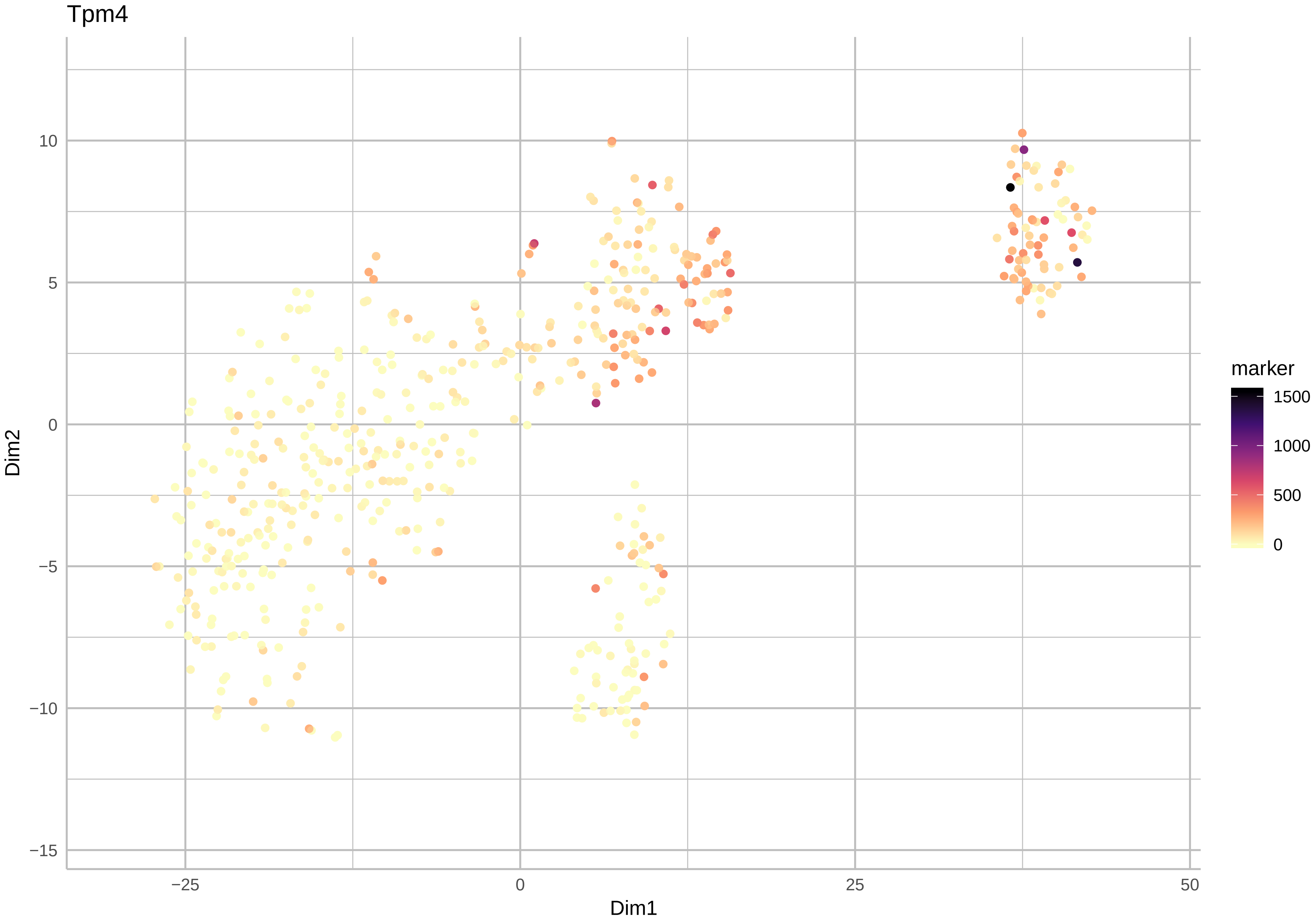

Tpm4
10
5
0
−5
−10
−15
Dim2
−25
0
25
50
Dim1
marker
1500
1000
500
0

Soat1

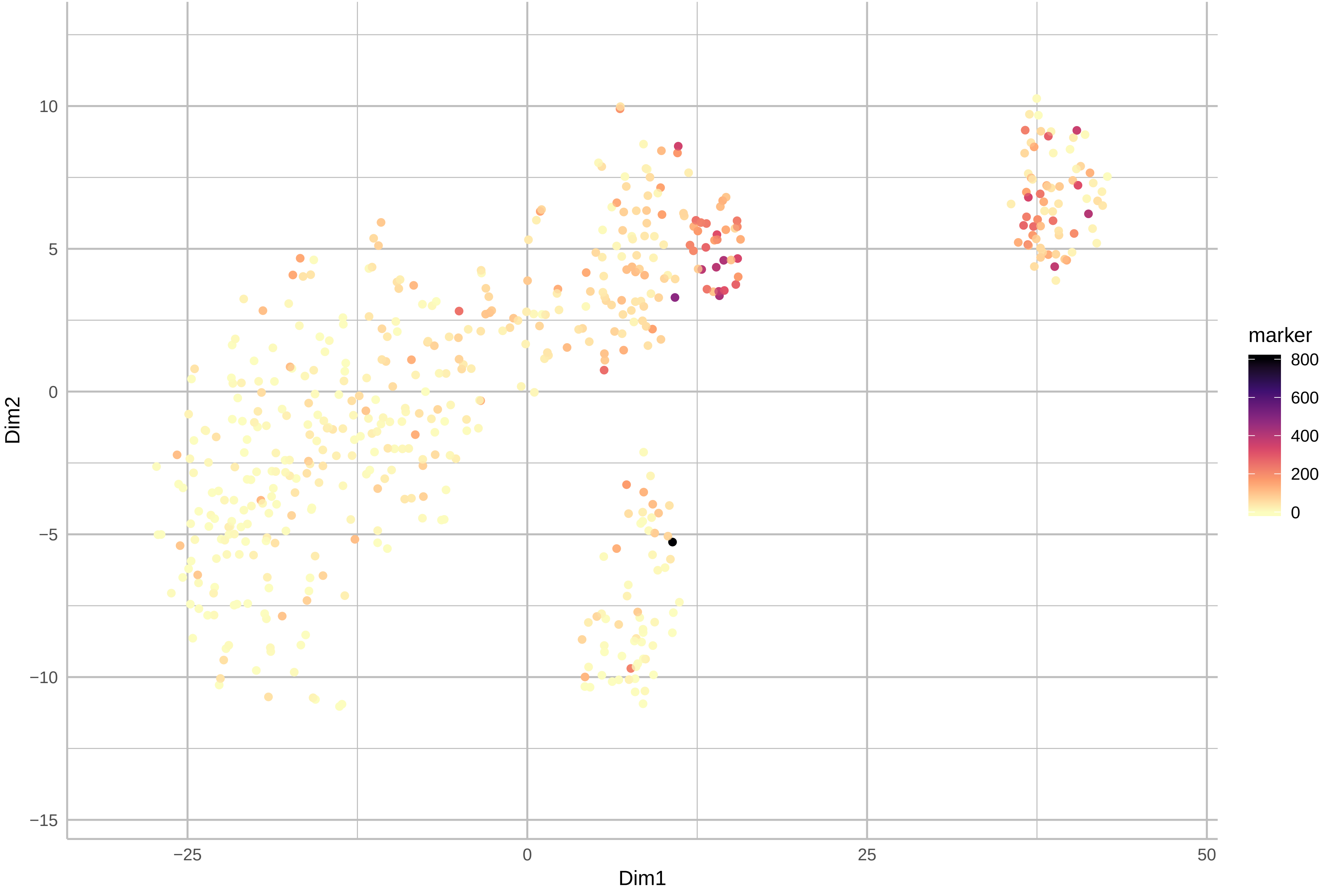

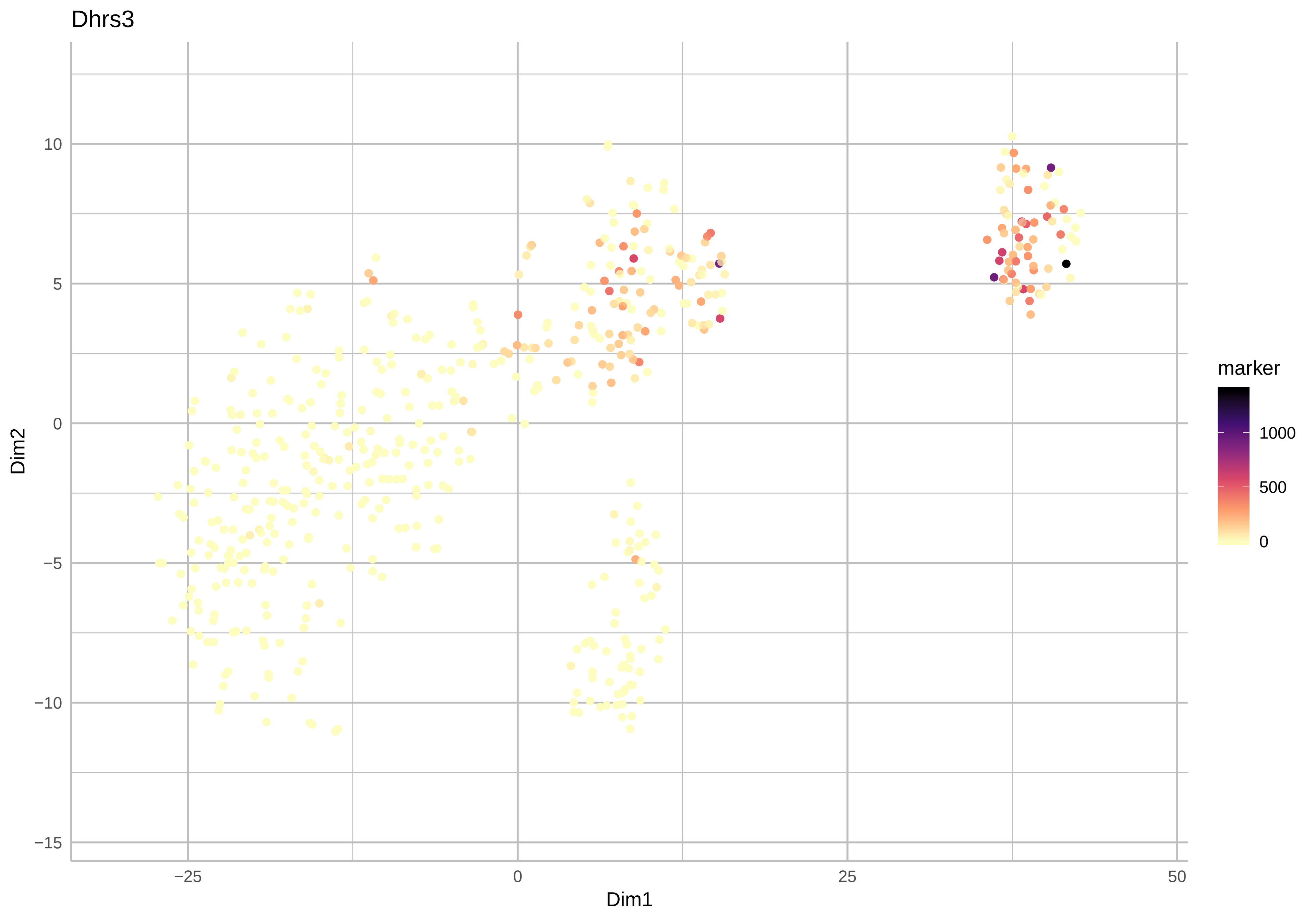

Dhrs3
Dim2
Dim1
10
5
0
−5
−10
−15
−25
0
25
50
marker
1000
500
0

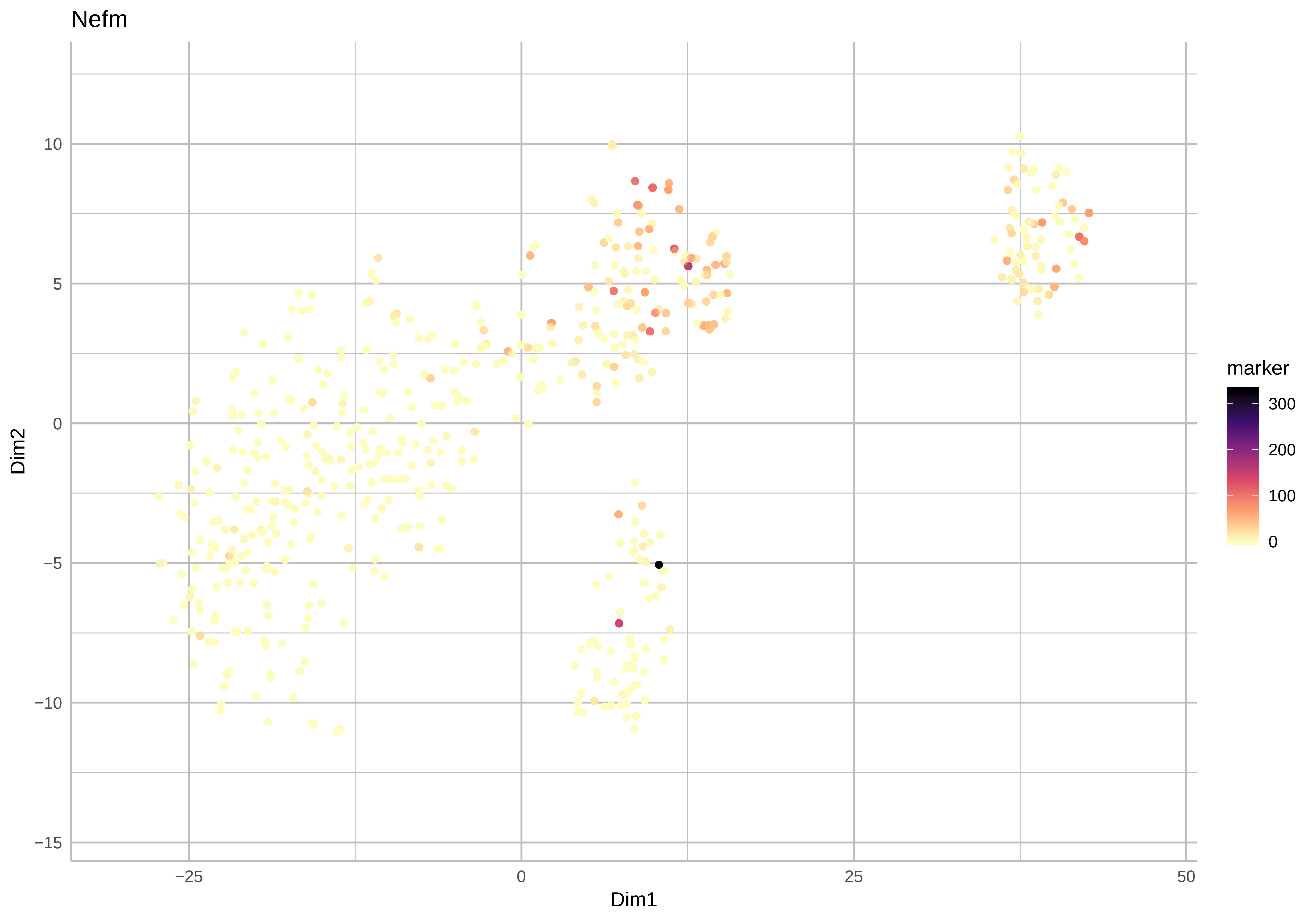

Nefm
10
5
0
−5
−10
−15
−25
0
25
50
Dim1
Dim2
marker
300
200
100
0

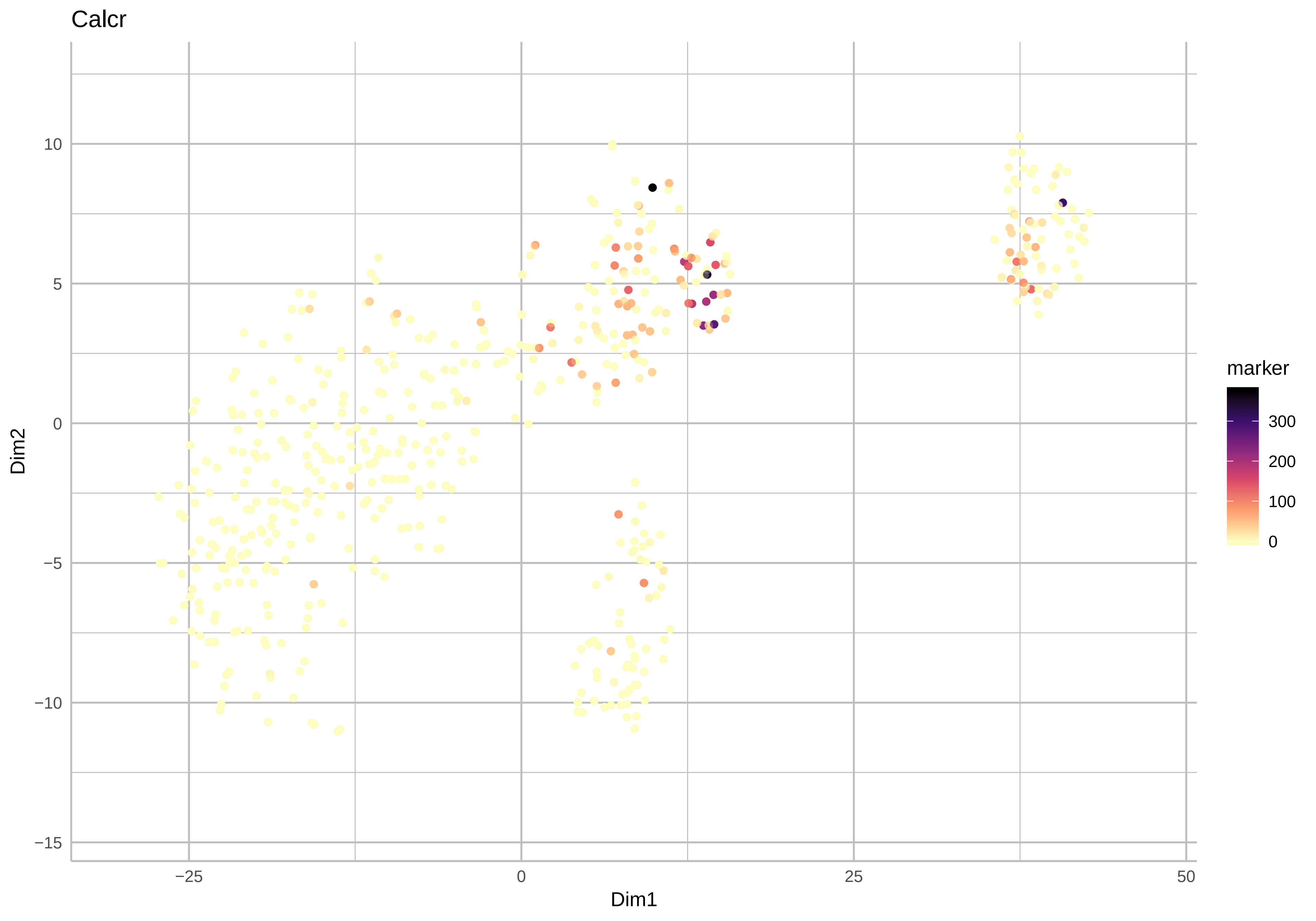

Calcr
10
5
0
−5
−10
−15
Dim2
−25
0
25
50
Dim1
marker
300
200
100
0

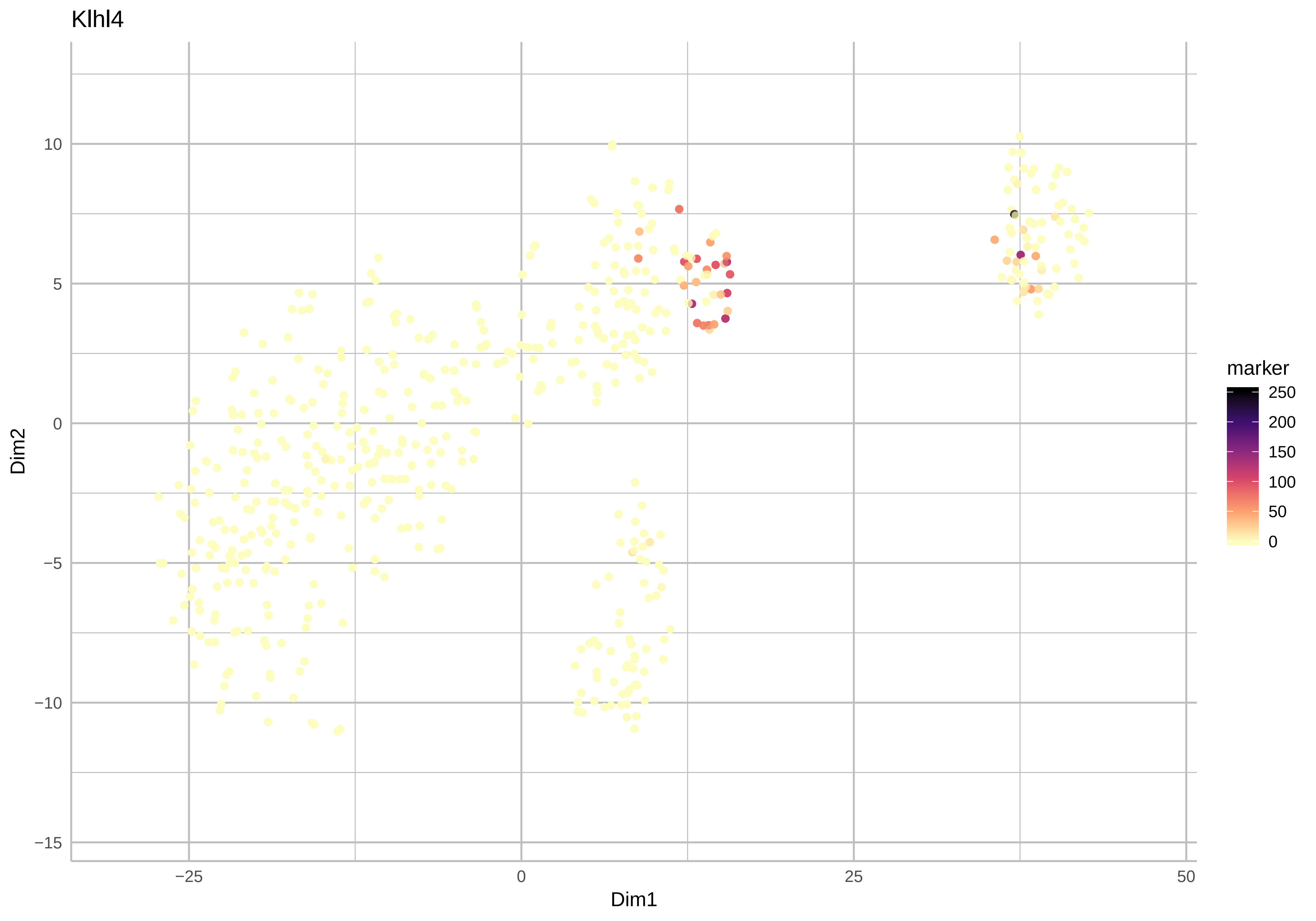
Klhl4
Dim2
Dim1
marker
250
200
150
100
50
0
10
5
0
−5
−10
−15
−25
0
25
50

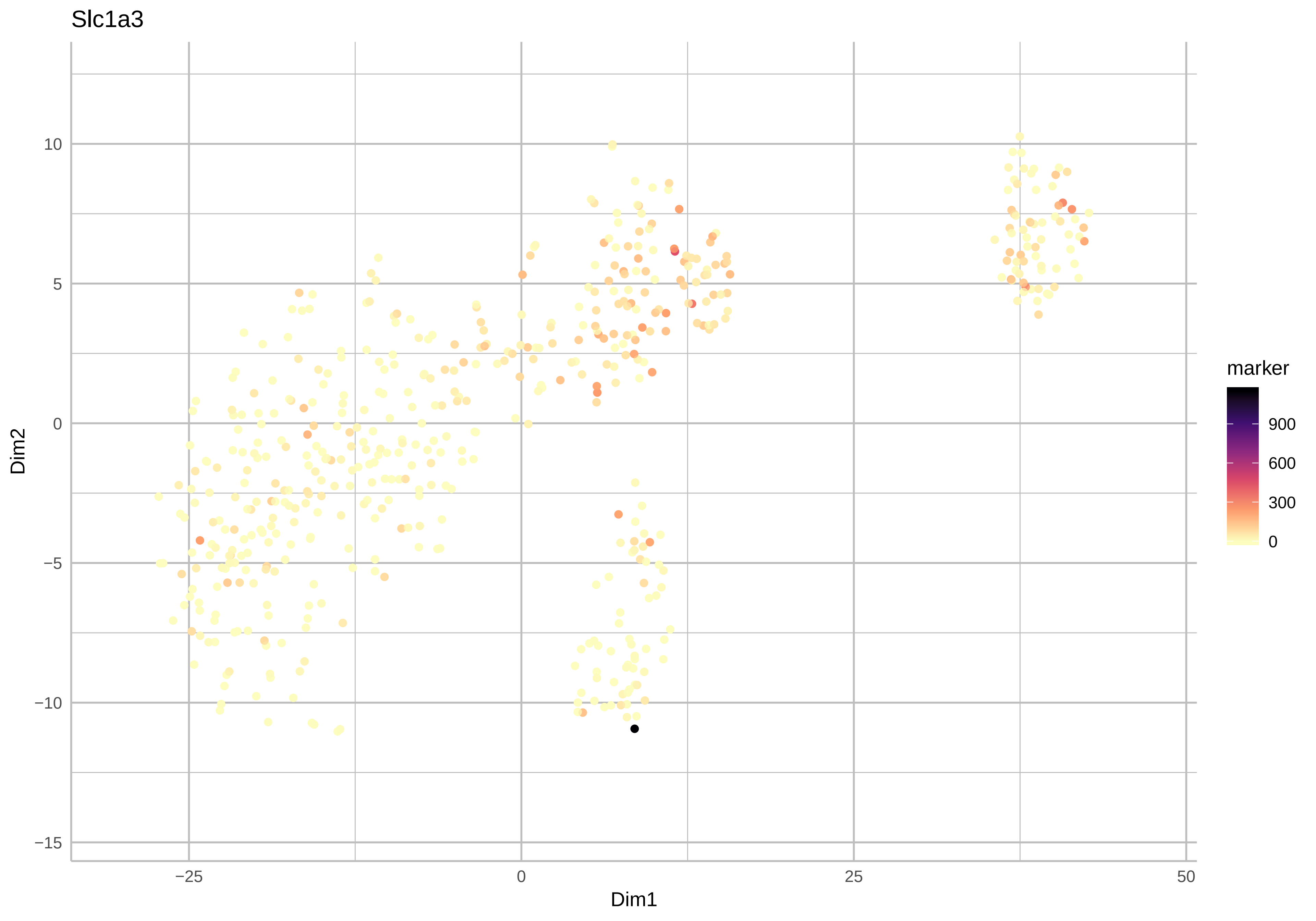
Slc1a3
Dim2
Dim1
10
5
0
−5
−10
−15
−25
0
25
50
marker
900
600
300
0

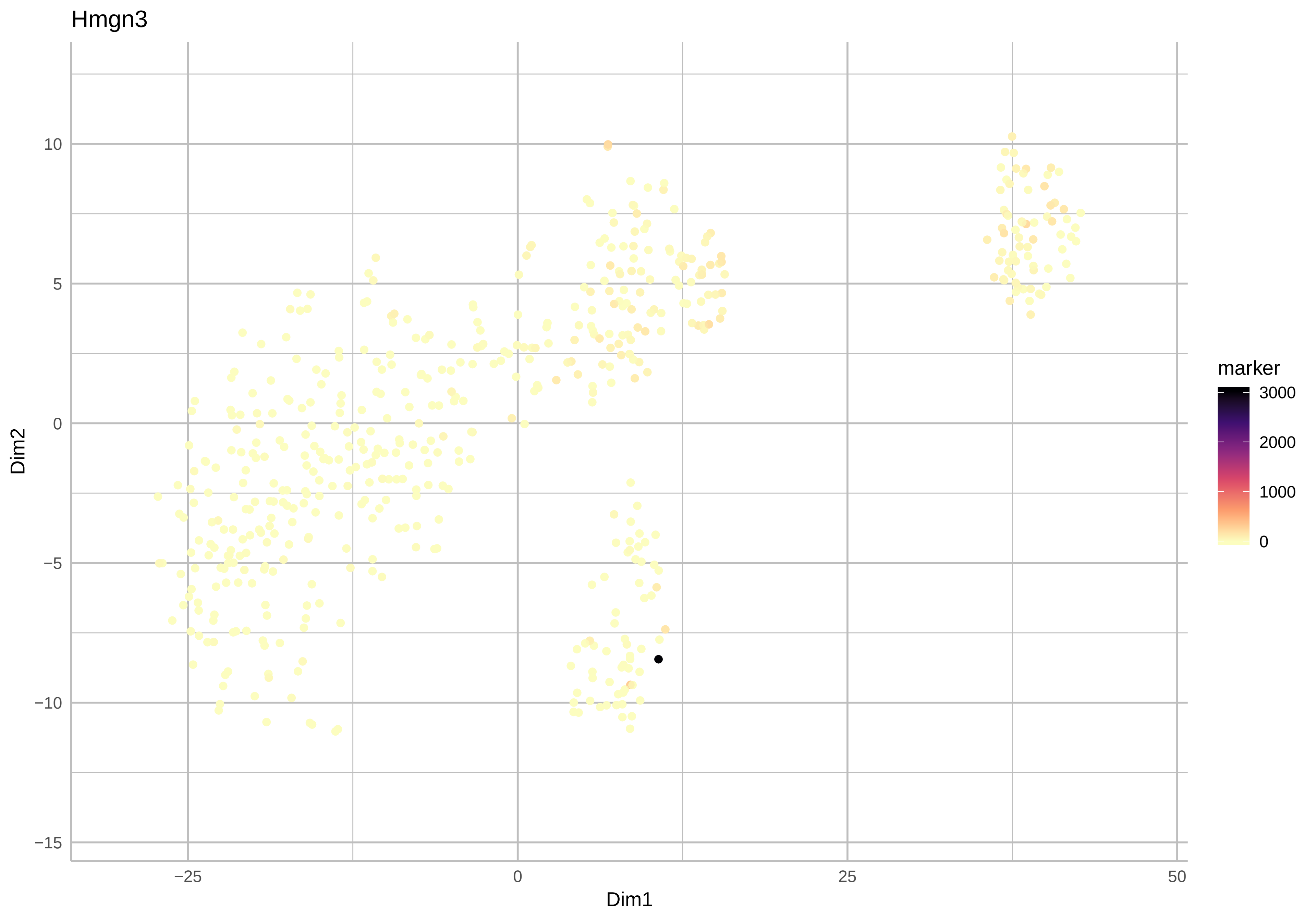
Hmgn3
Dim2
Dim1
10
5
0
−5
−10
−15
−25
0
25
50
marker
3000
2000
1000
0

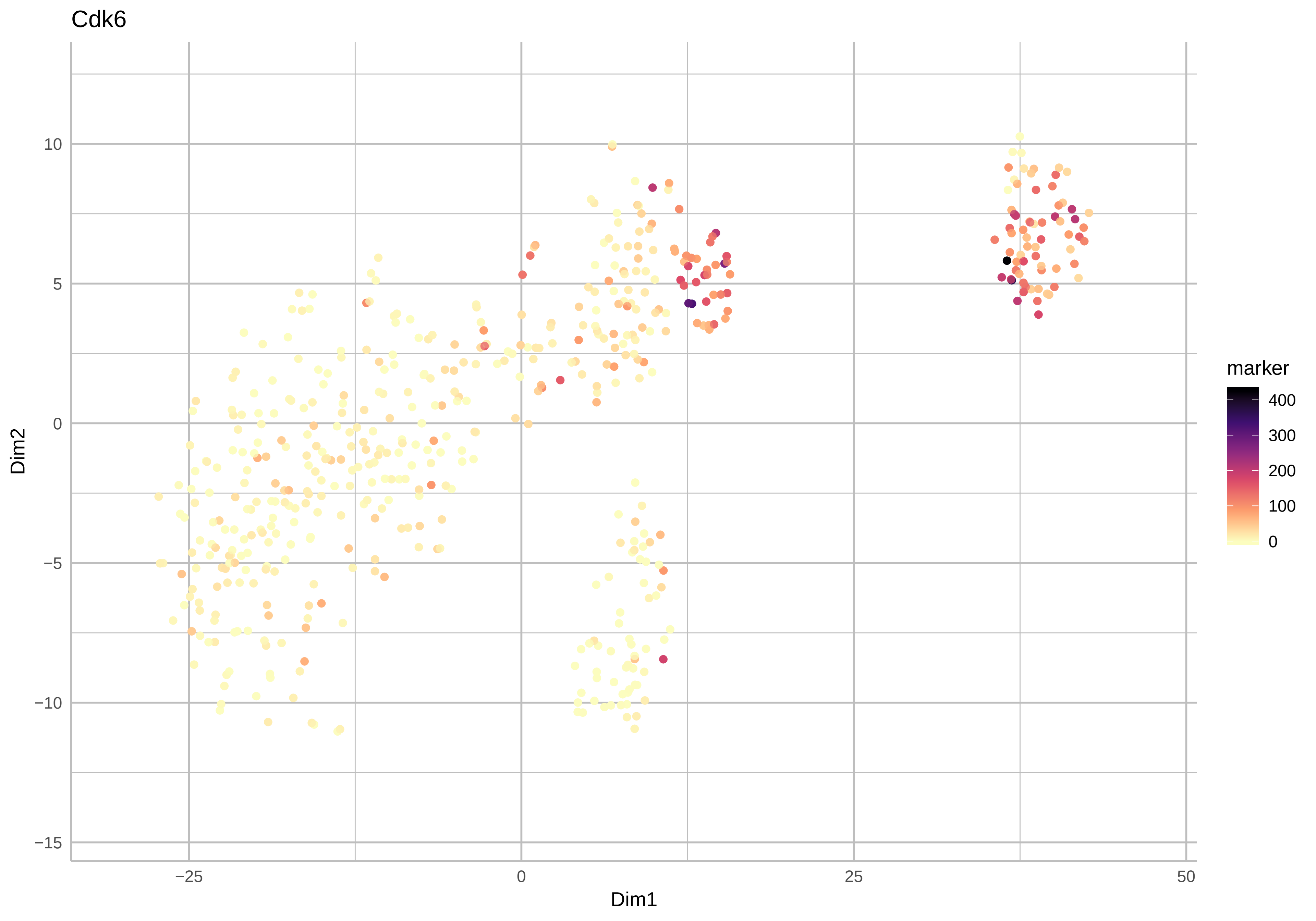
Cdk6
10
5
0
−5
−10
−15
−25
0
25
50
Dim1
Dim2
marker
400
300
200
100
0

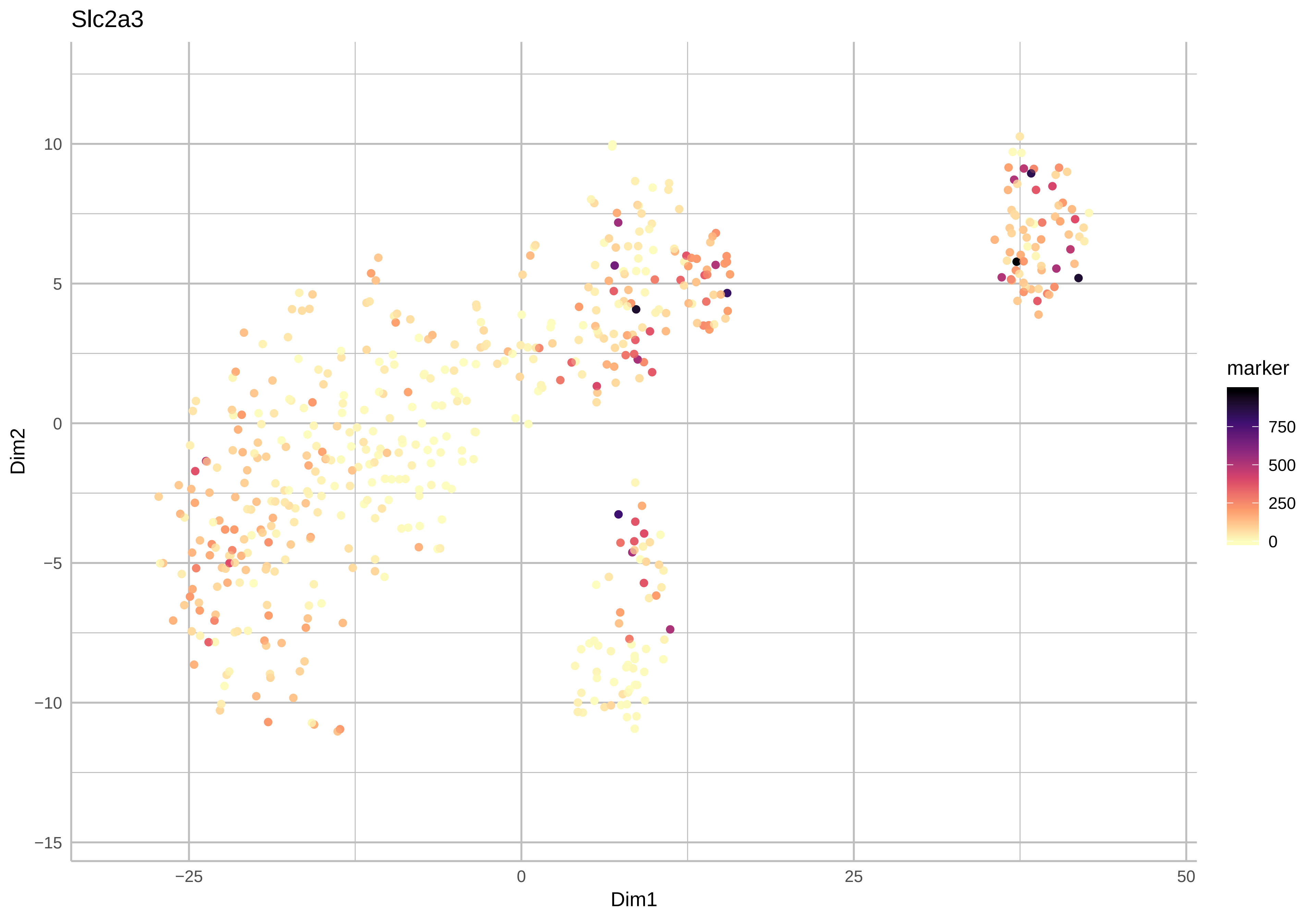
Slc2a3
Dim2
Dim1
10
5
0
−5
−10
−15
−25
0
25
50
marker
750
500
250
0

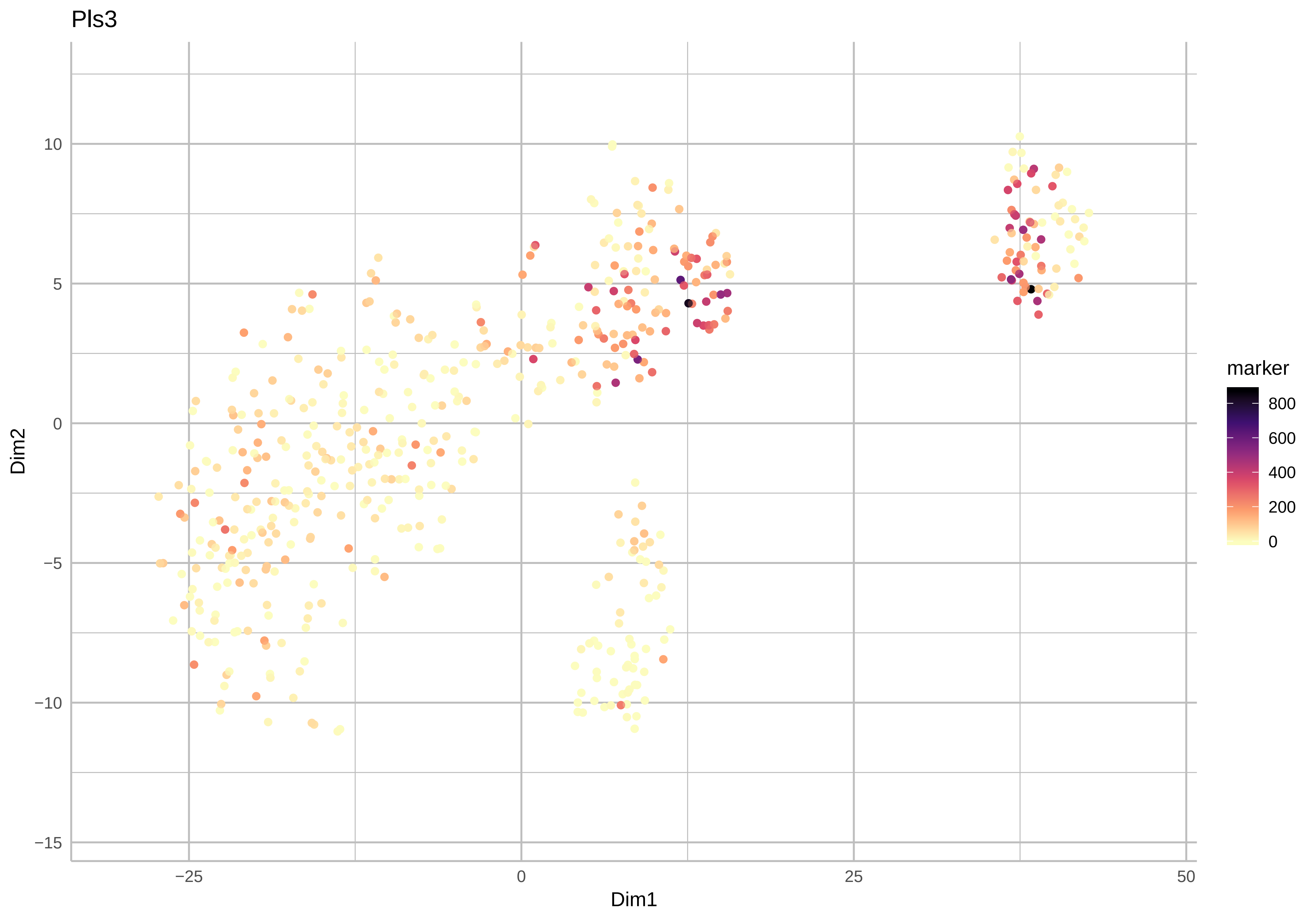

Pls3
Dim2
Dim1
10
5
0
−5
−10
−15
−25
0
25
50
marker
800
600
400
200
0

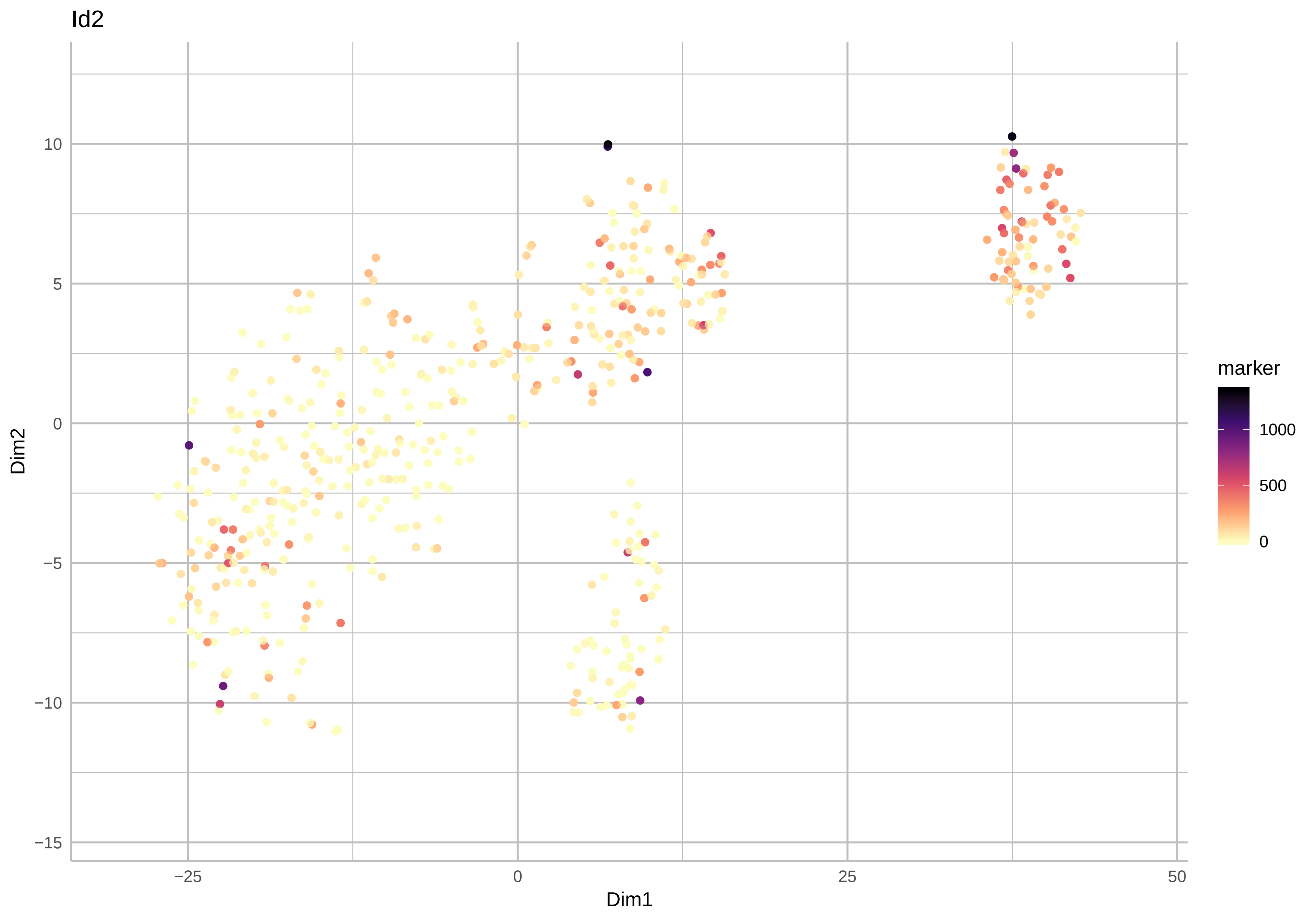
Id2
Dim2
Dim1
10
5
0
−5
−10
−15
−25
0
25
50
marker
1000
500
0

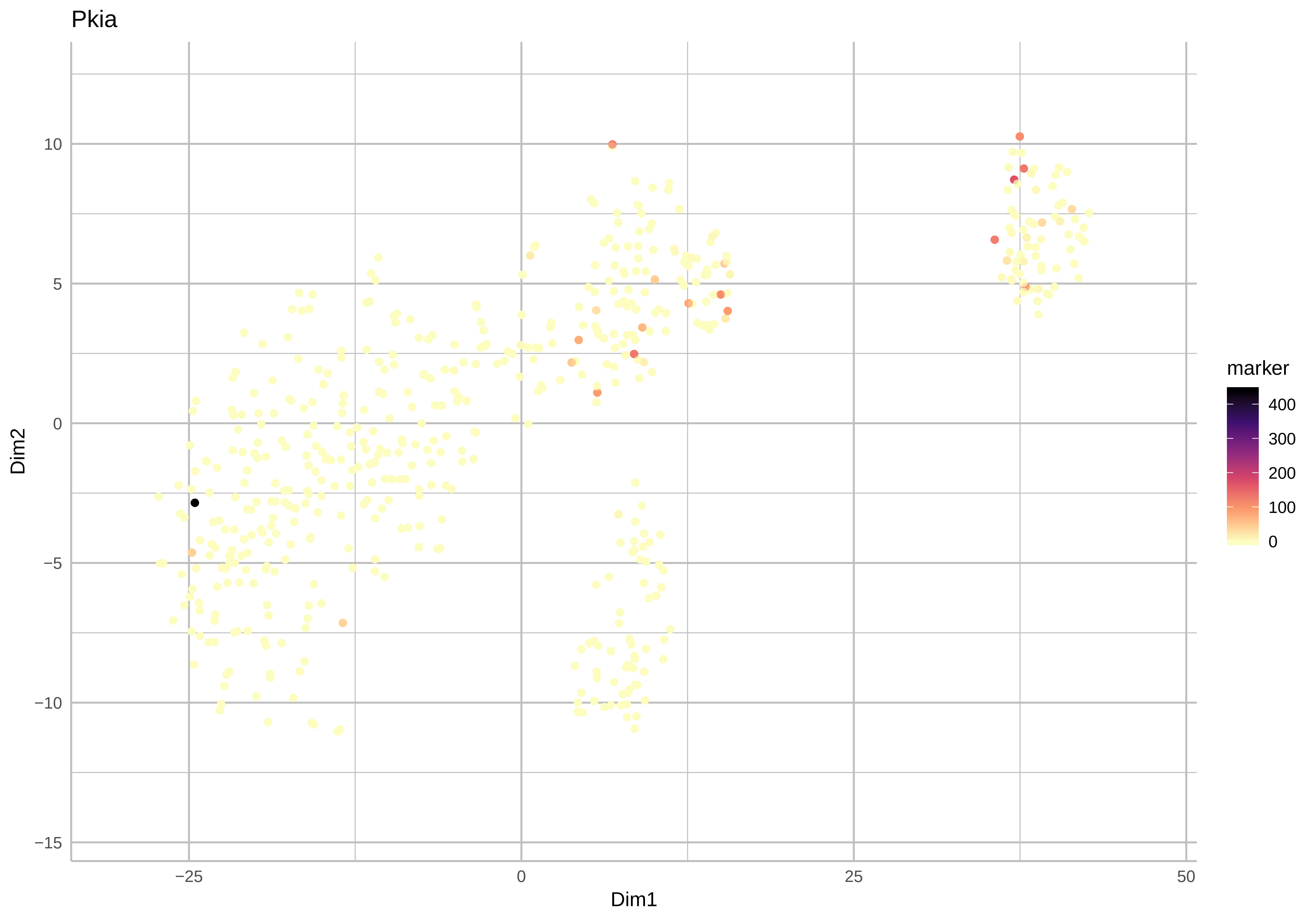

Pkia
Dim2
Dim1
10
5
0
−5
−10
−15
−25
0
25
50
marker
400
300
200
100
0

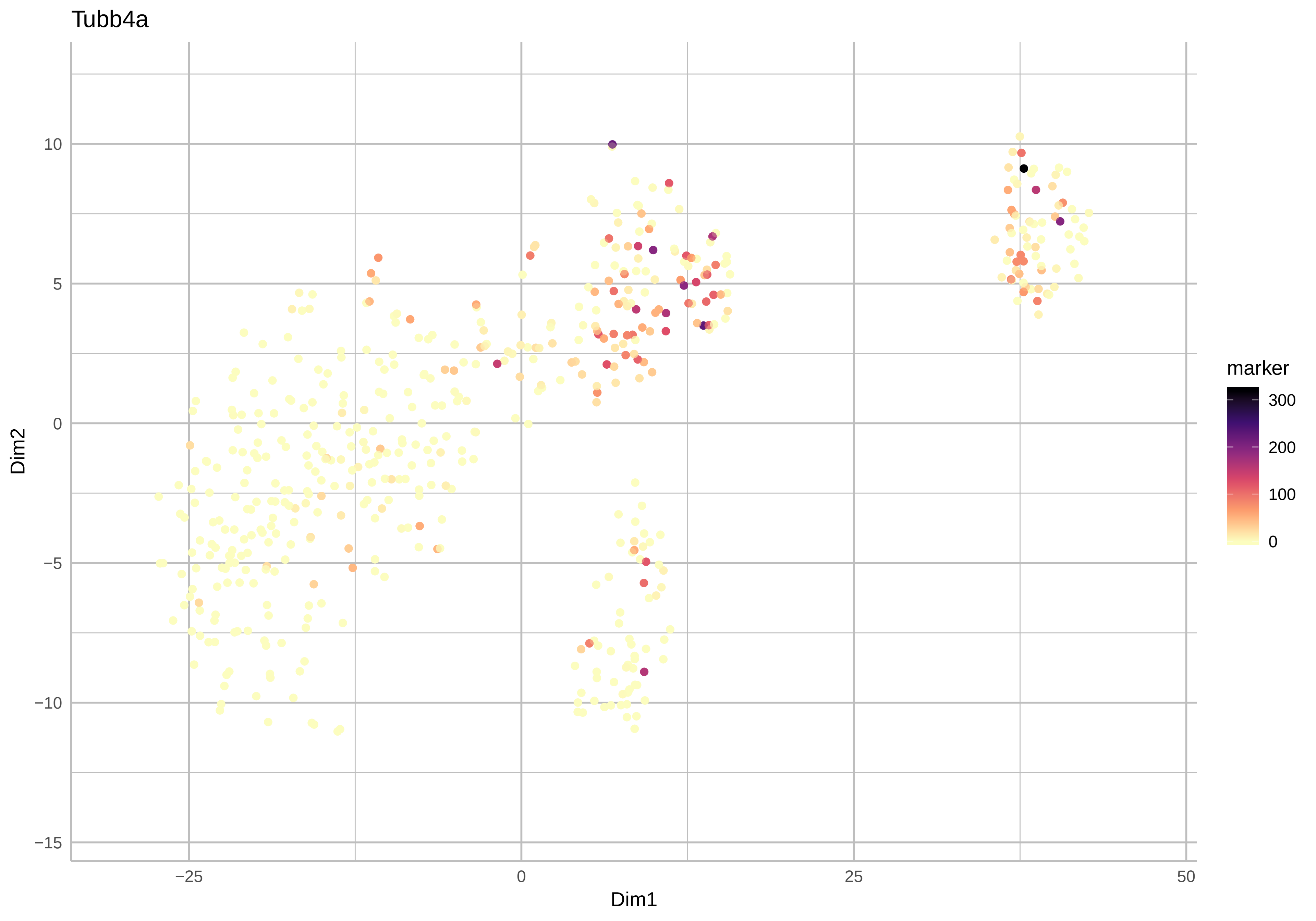

Tubb4a
Dim2
Dim1
10
5
0
−5
−10
−15
−25
0
25
50
marker
300
200
100
0

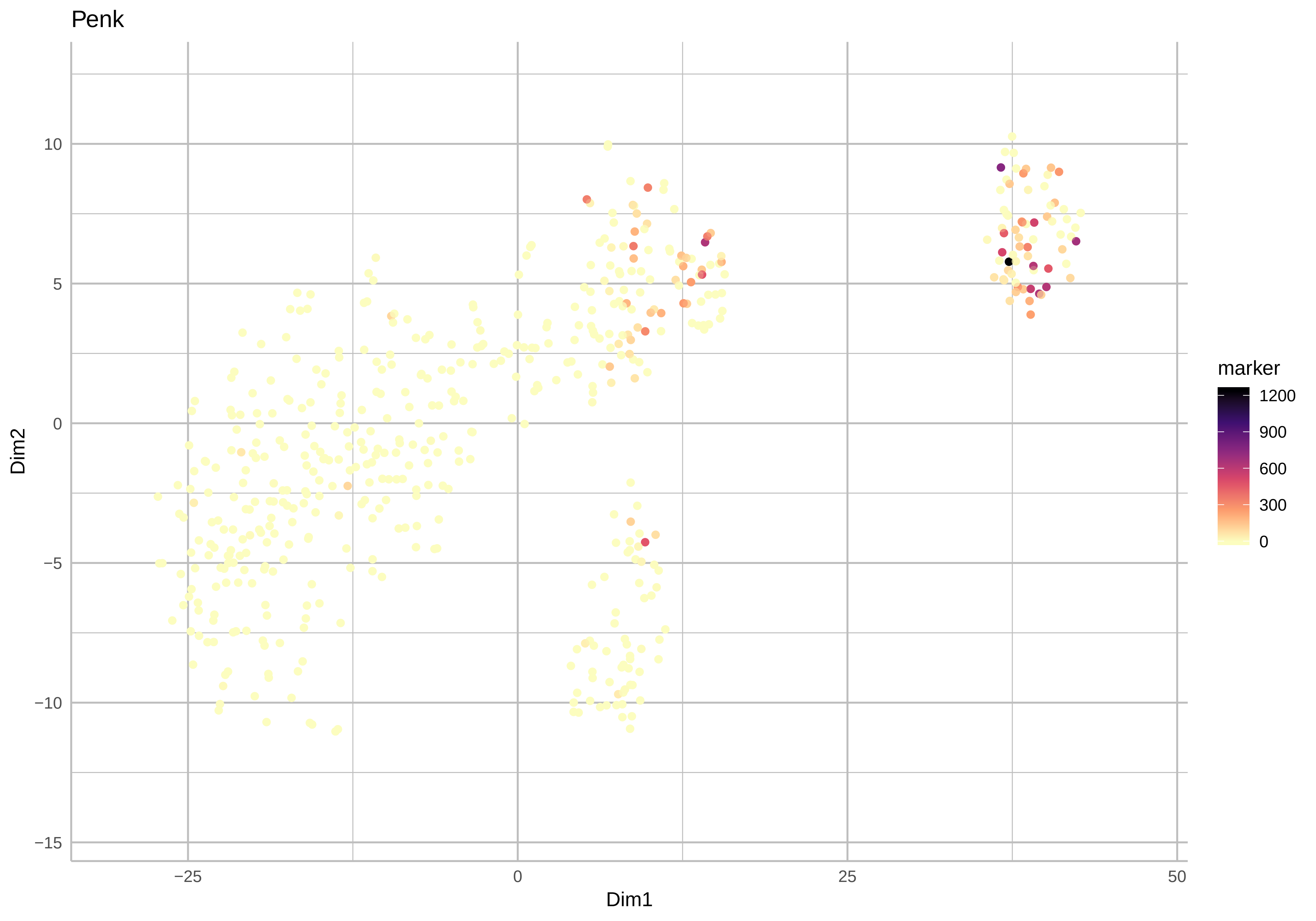

Penk
10
5
0
−5
−10
−15
Dim2
−25
0
25
50
Dim1
marker
1200
900
600
300
0

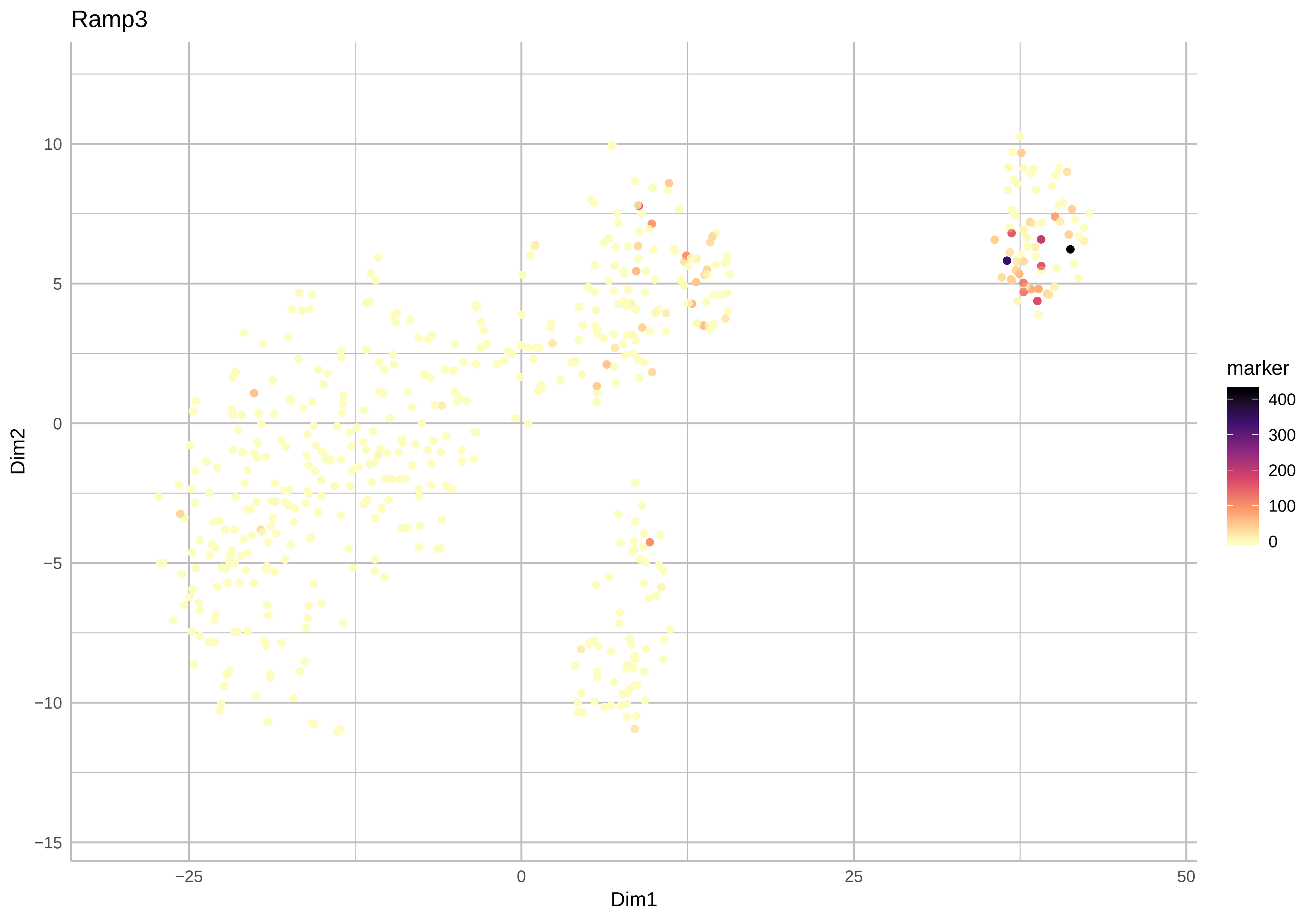
Ramp3
Dim2
Dim1
10
5
0
−5
−10
−15
−25
0
25
50
marker
400
300
200
100
0

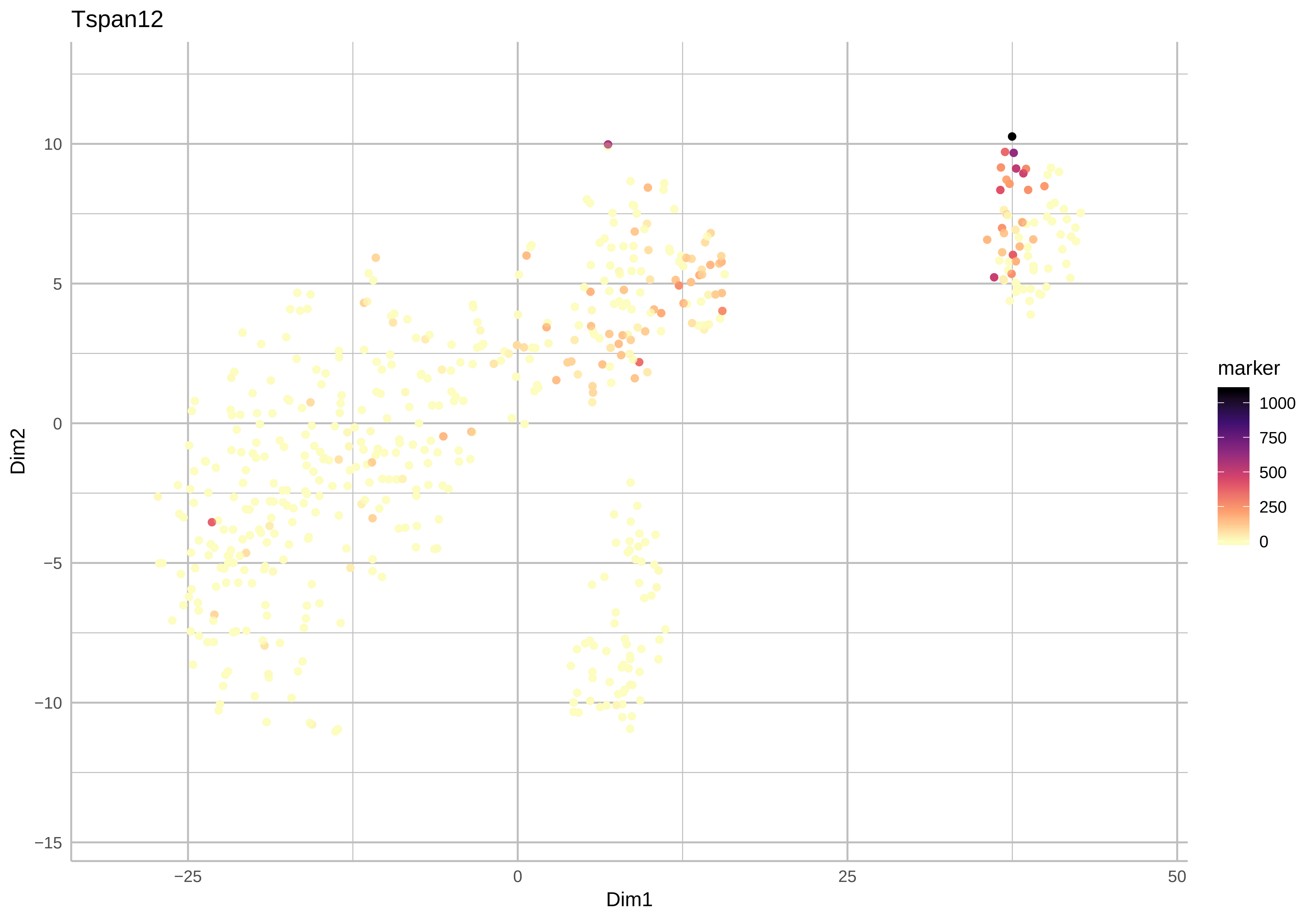

Tspan12
10
5
0
−5
−10
−15
Dim2
−25
0
25
50
Dim1
marker
1000
750
500
250
0

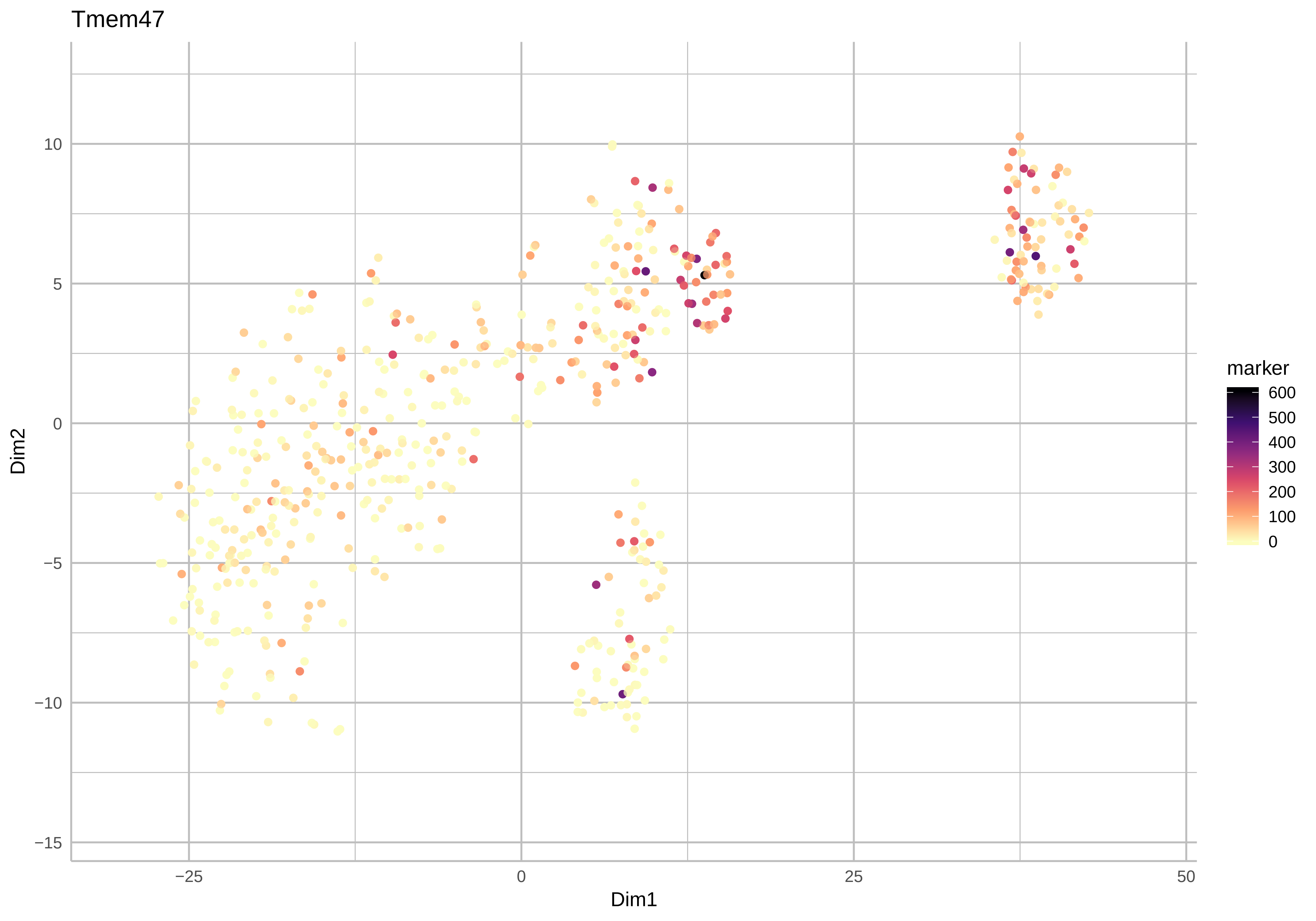
Tmem47
10
5
0
−5
−10
−15
Dim2
−25
0
25
50
Dim1
marker
600
500
400
300
200
100
0

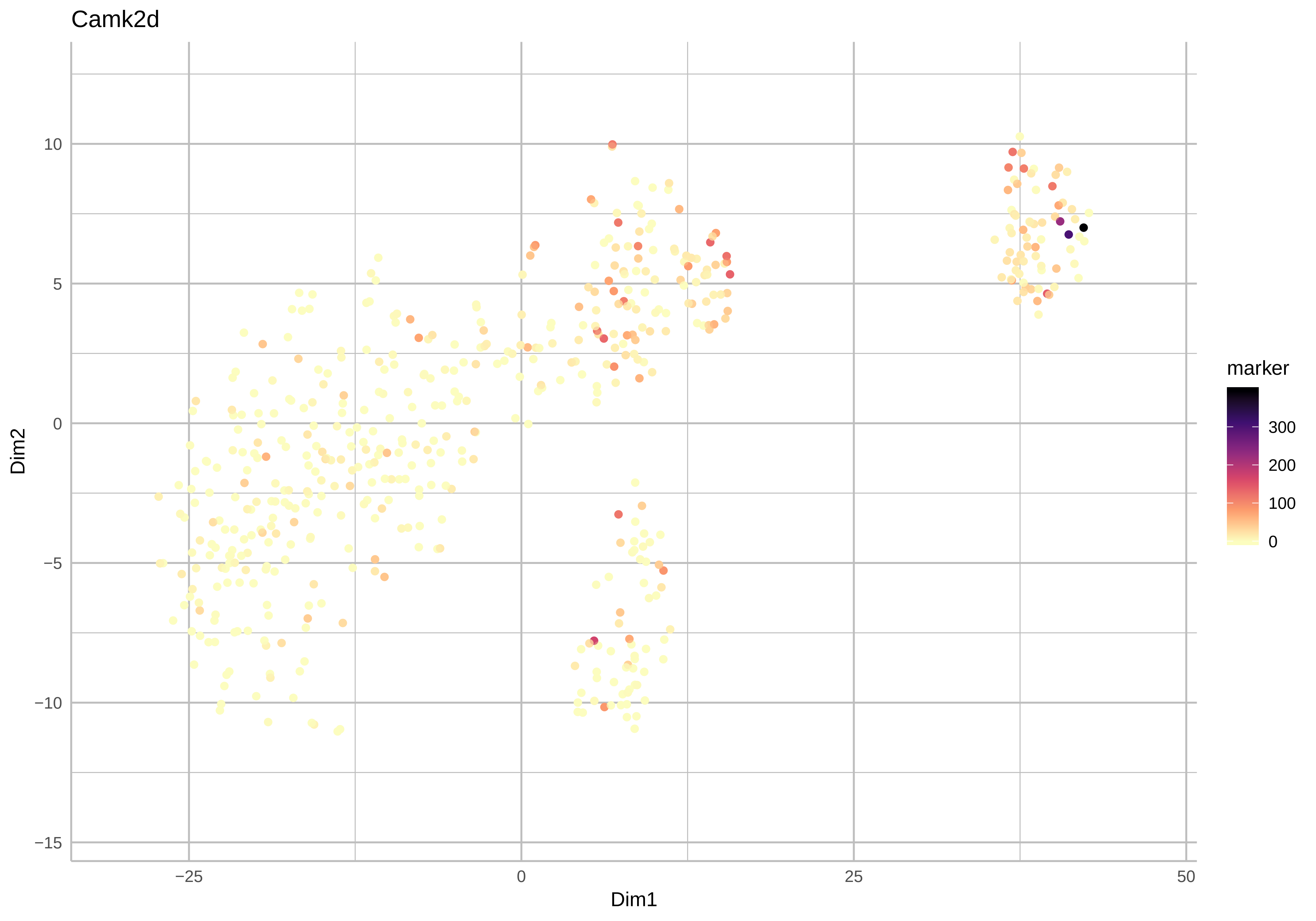
Camk2d
10
5
0
−5
−10
−15
−25
0
25
50
Dim1
Dim2
marker
300
200
100
0

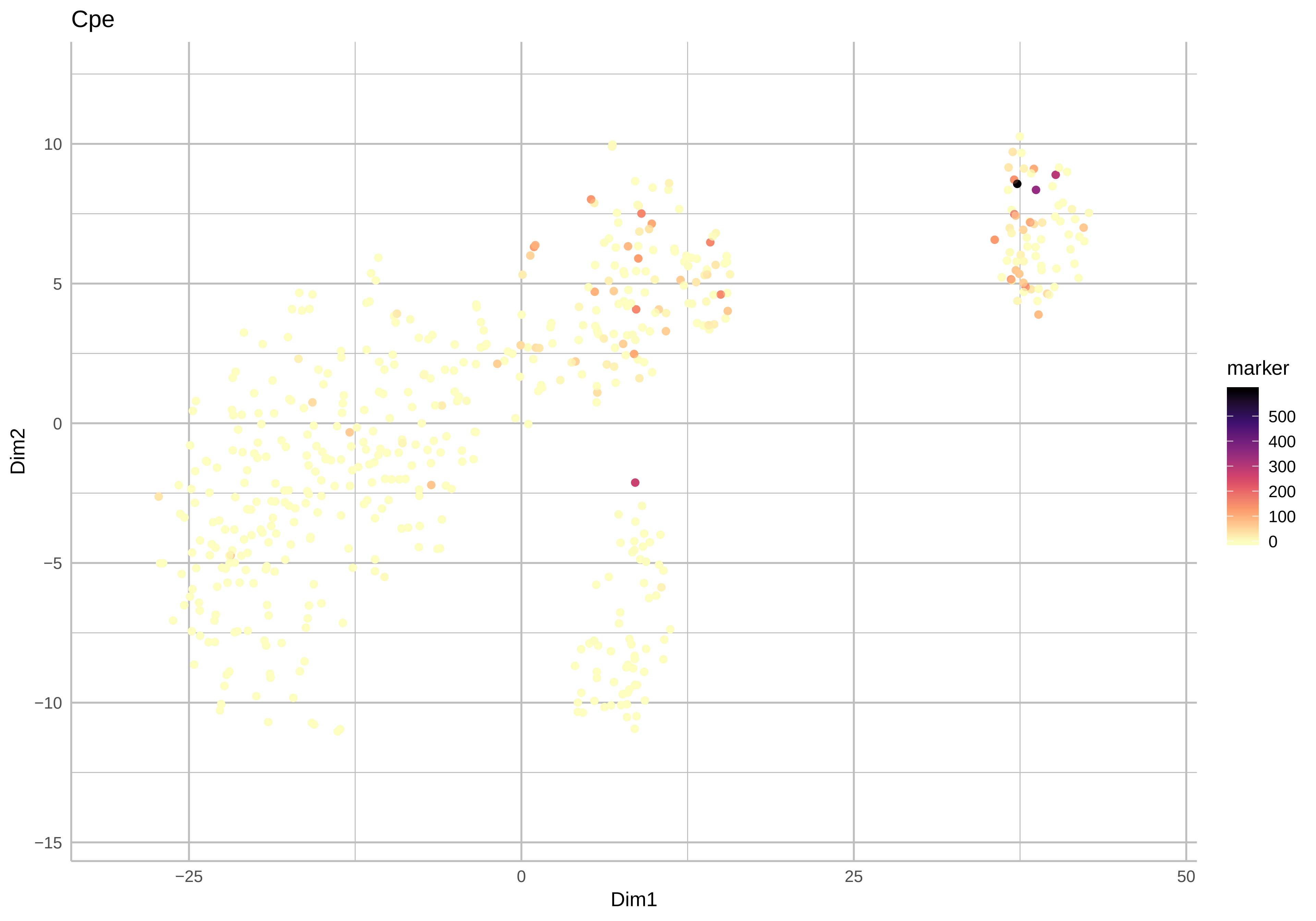
Cpe
Dim2
Dim1
10
5
0
−5
−10
−15
−25
0
25
50
marker
500
400
300
200
100
0

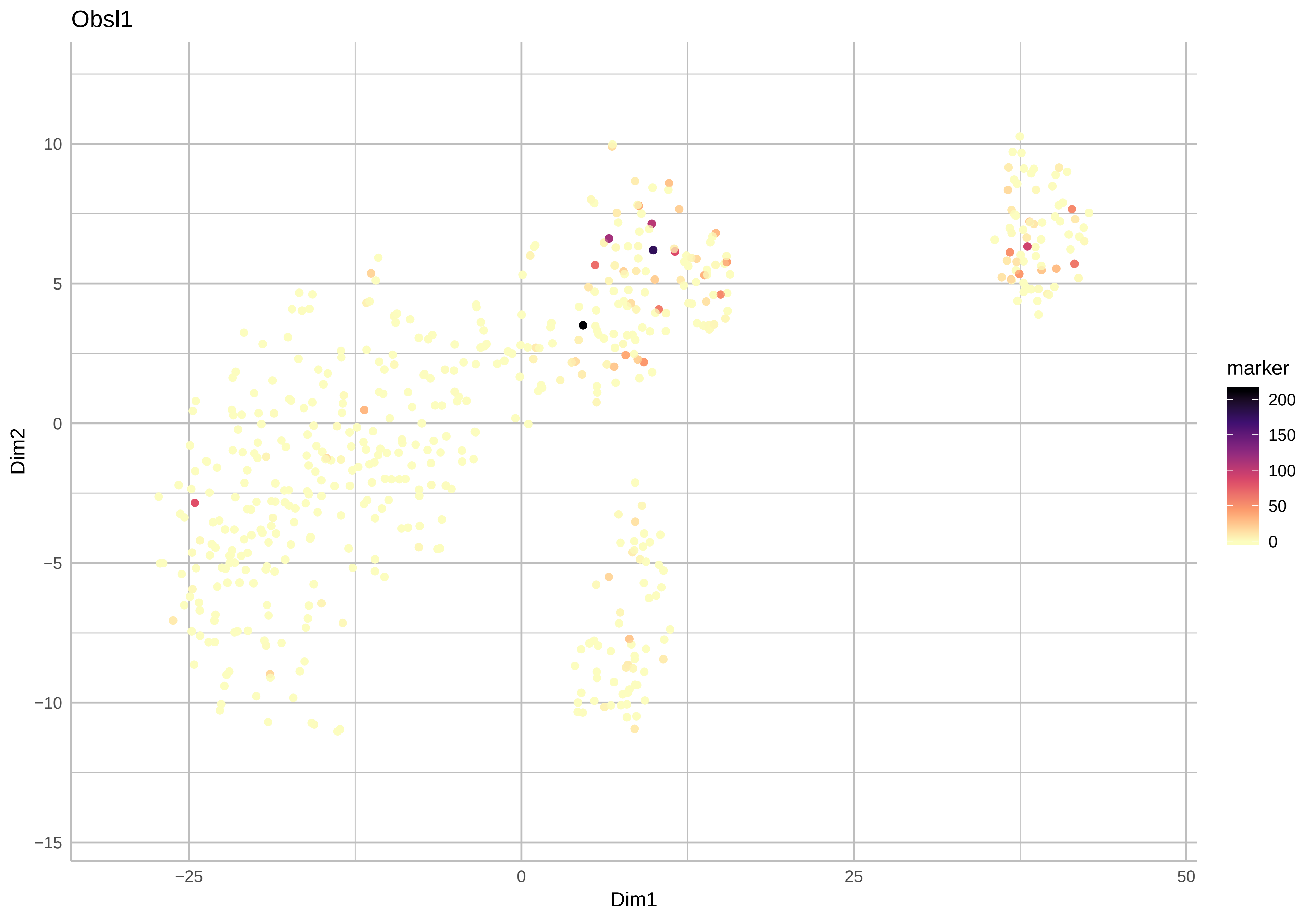
Obsl1
Dim2
Dim1
10
5
0
−5
−10
−15
−25
0
25
50
marker
200
150
100
50
0

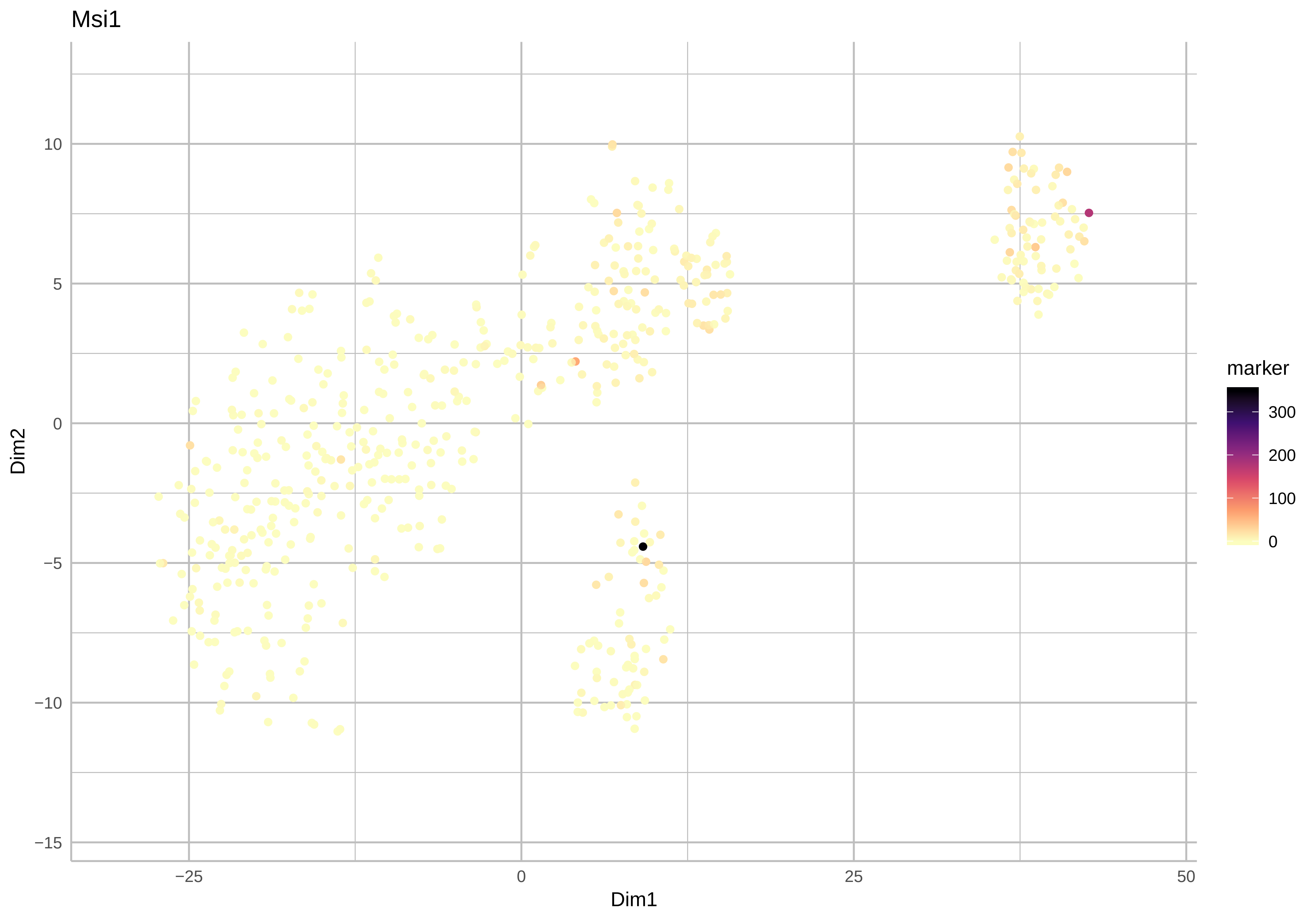
Msi1
Dim2
Dim1
10
5
0
−5
−10
−15
−25
0
25
50
marker
300
200
100
0

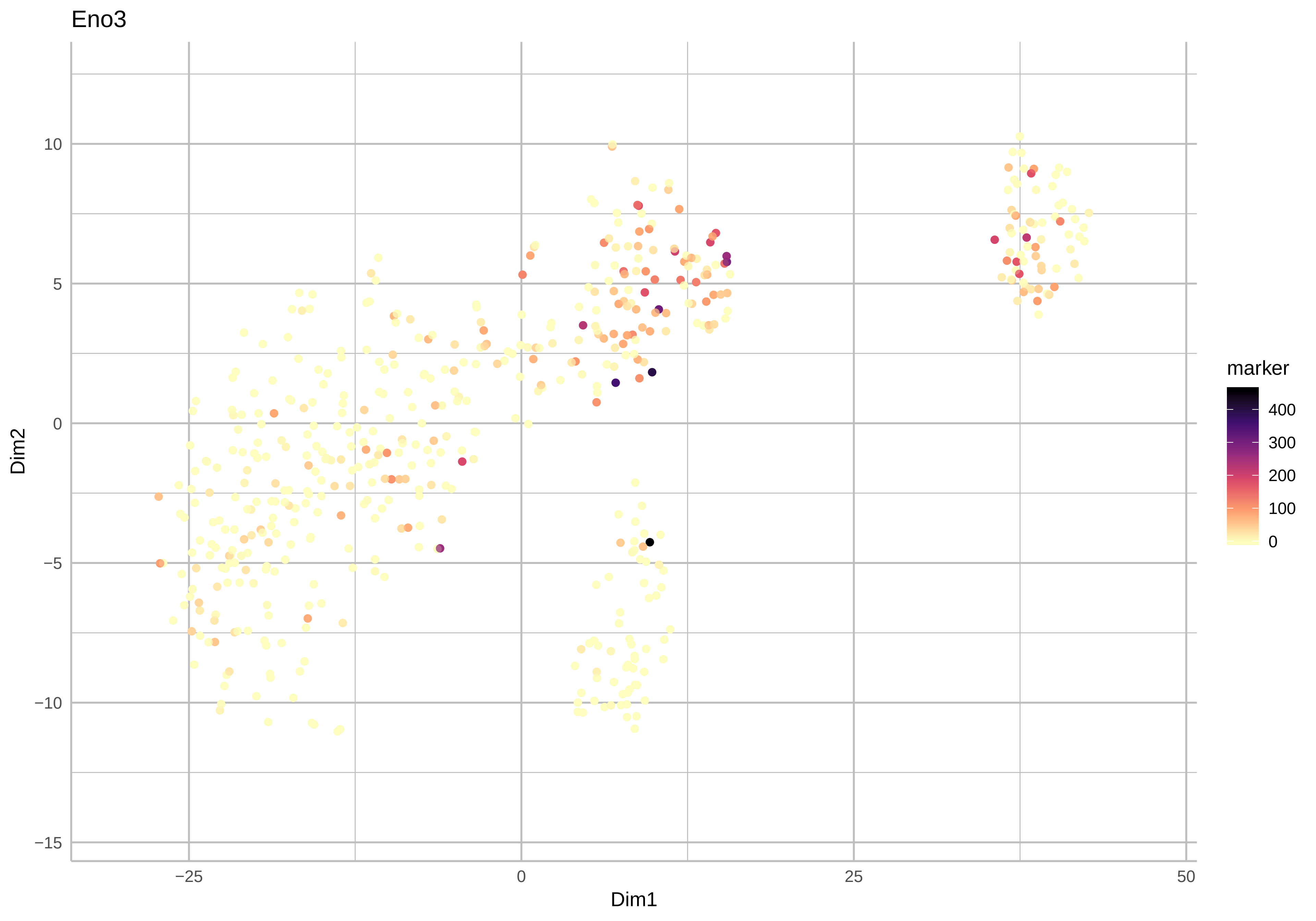
Eno3
Dim2
Dim1
10
5
0
−5
−10
−15
−25
0
25
50
marker
400
300
200
100
0

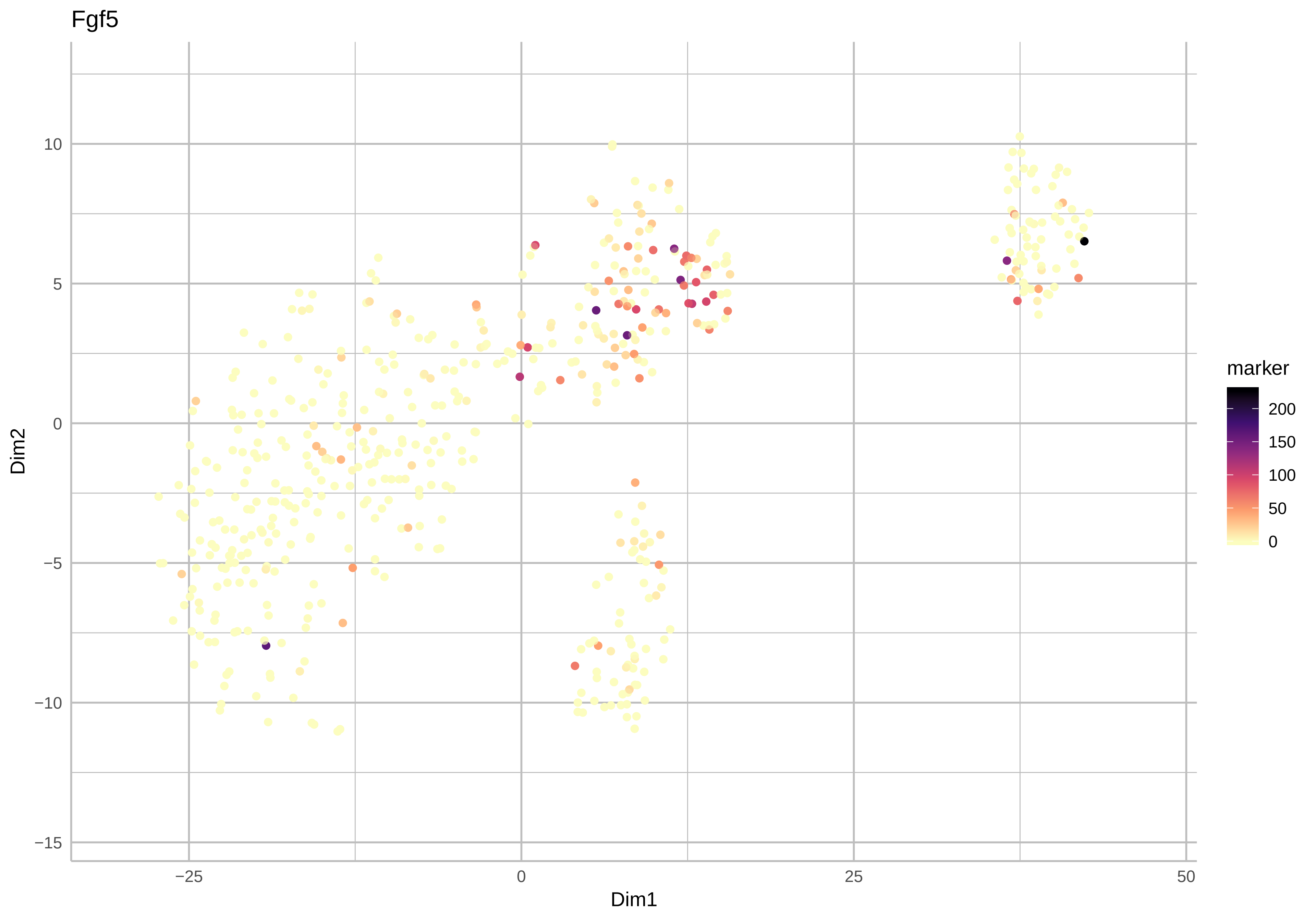

Fgf5
Dim2
Dim1
10
5
0
−5
−10
−15
−25
0
25
50
marker
200
150
100
50
0

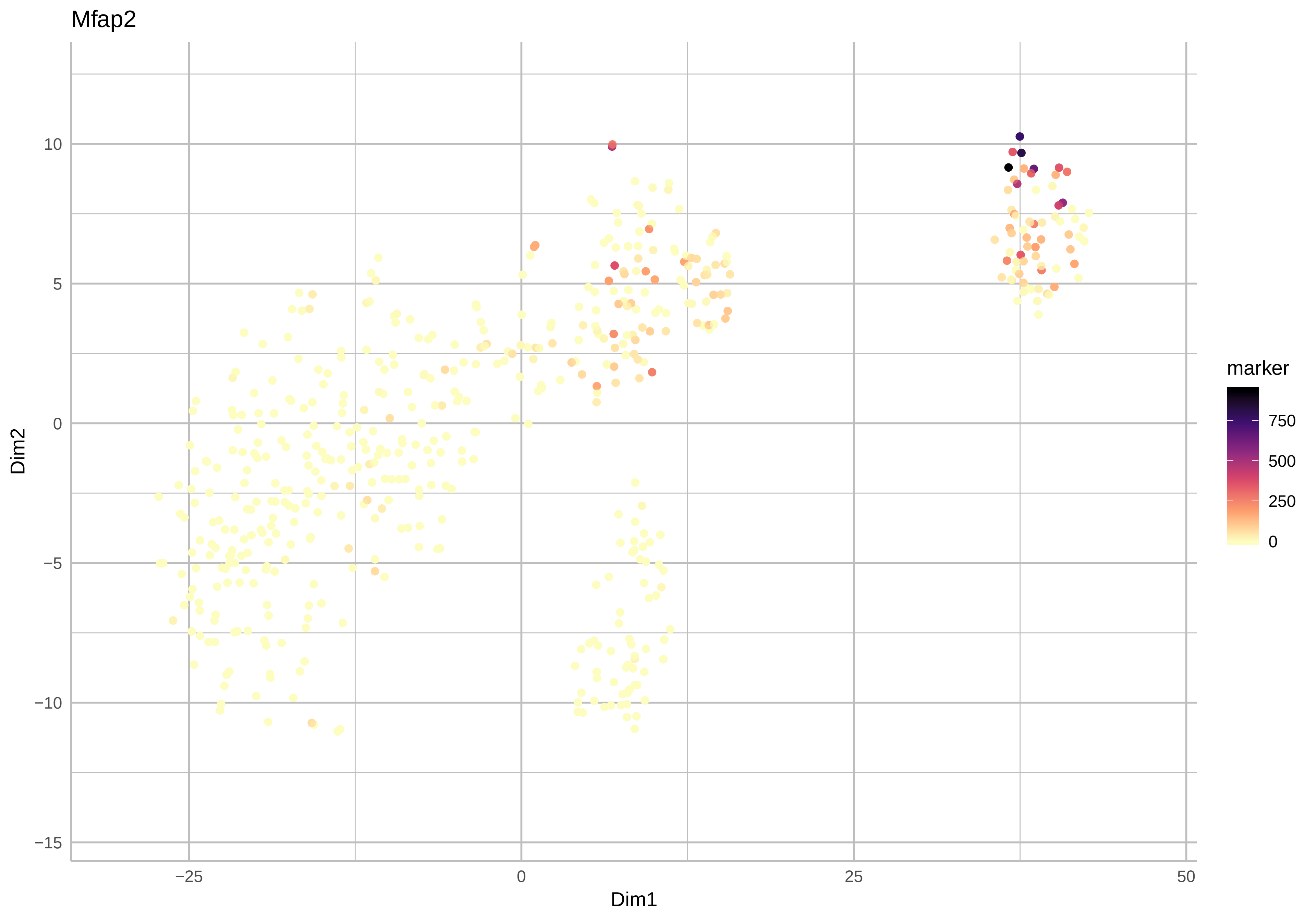
Mfap2
Dim2
Dim1
10
5
0
−5
−10
−15
−25
0
25
50
marker
750
500
250
0

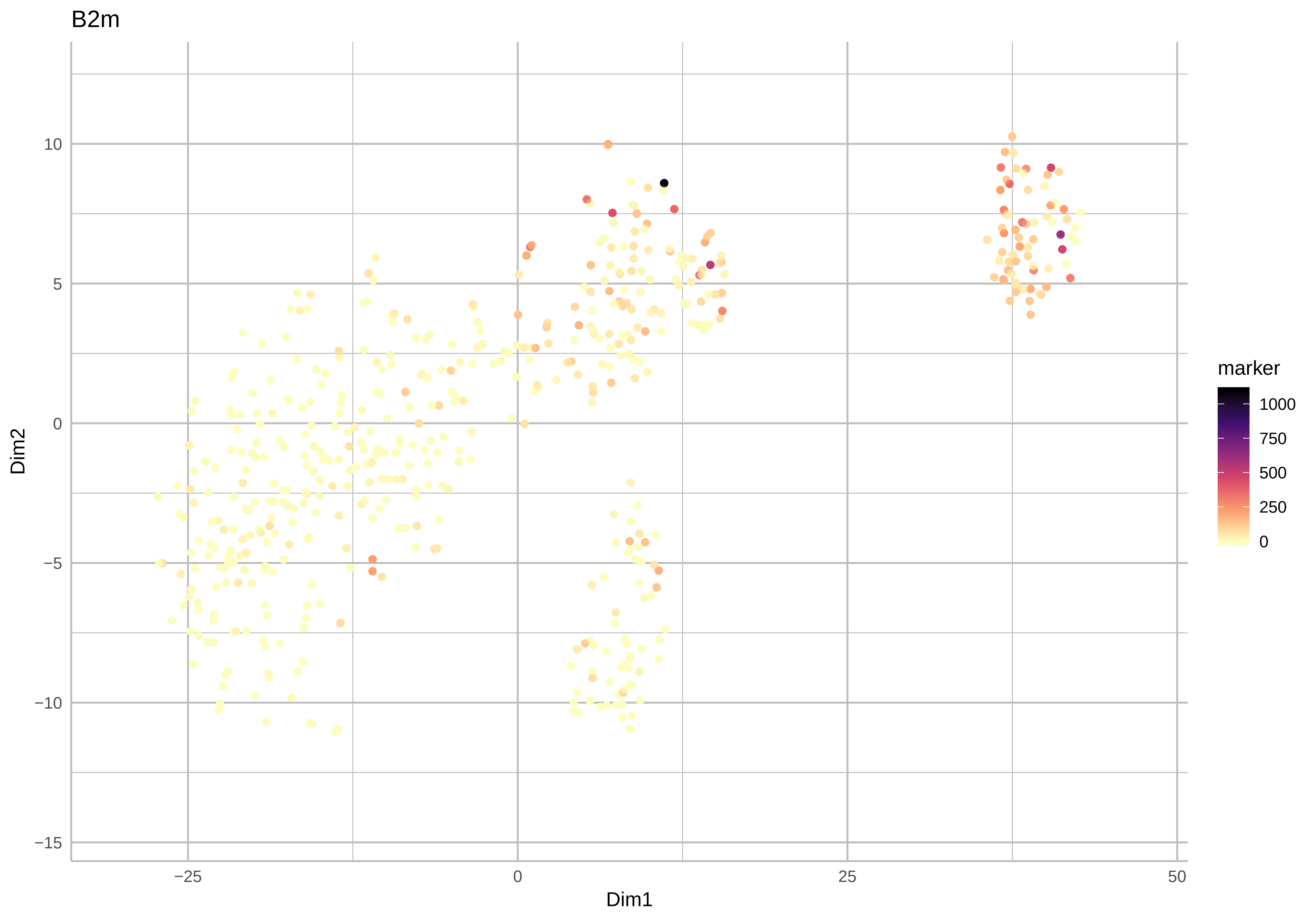
B2m
Dim2
Dim1
10
5
0
−5
−10
−15
−25
0
25
50
marker
1000
750
500
250
0

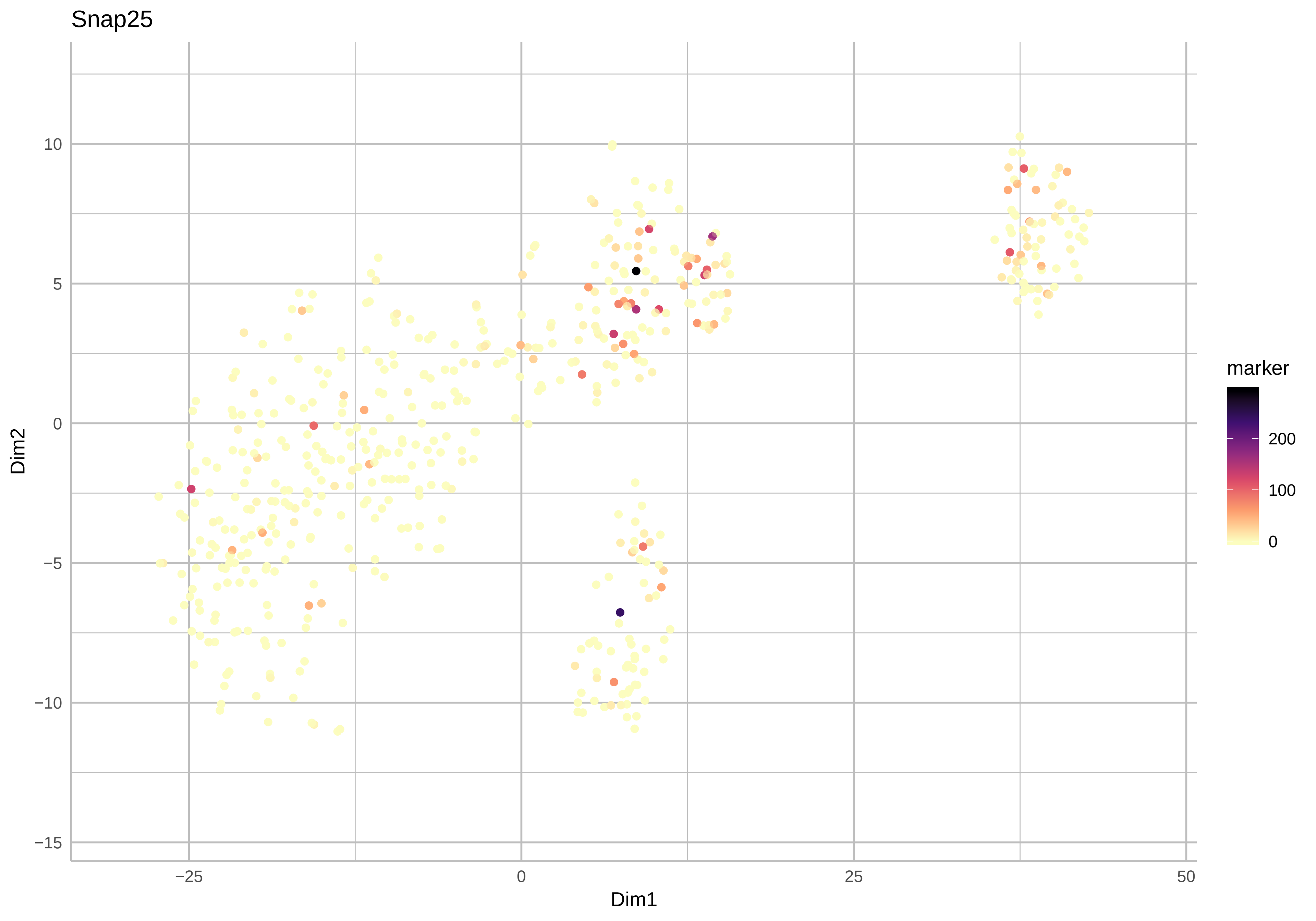

Snap25
10
5
0
−5
−10
−15
Dim2
−25
0
25
50
Dim1
marker
200
100
0

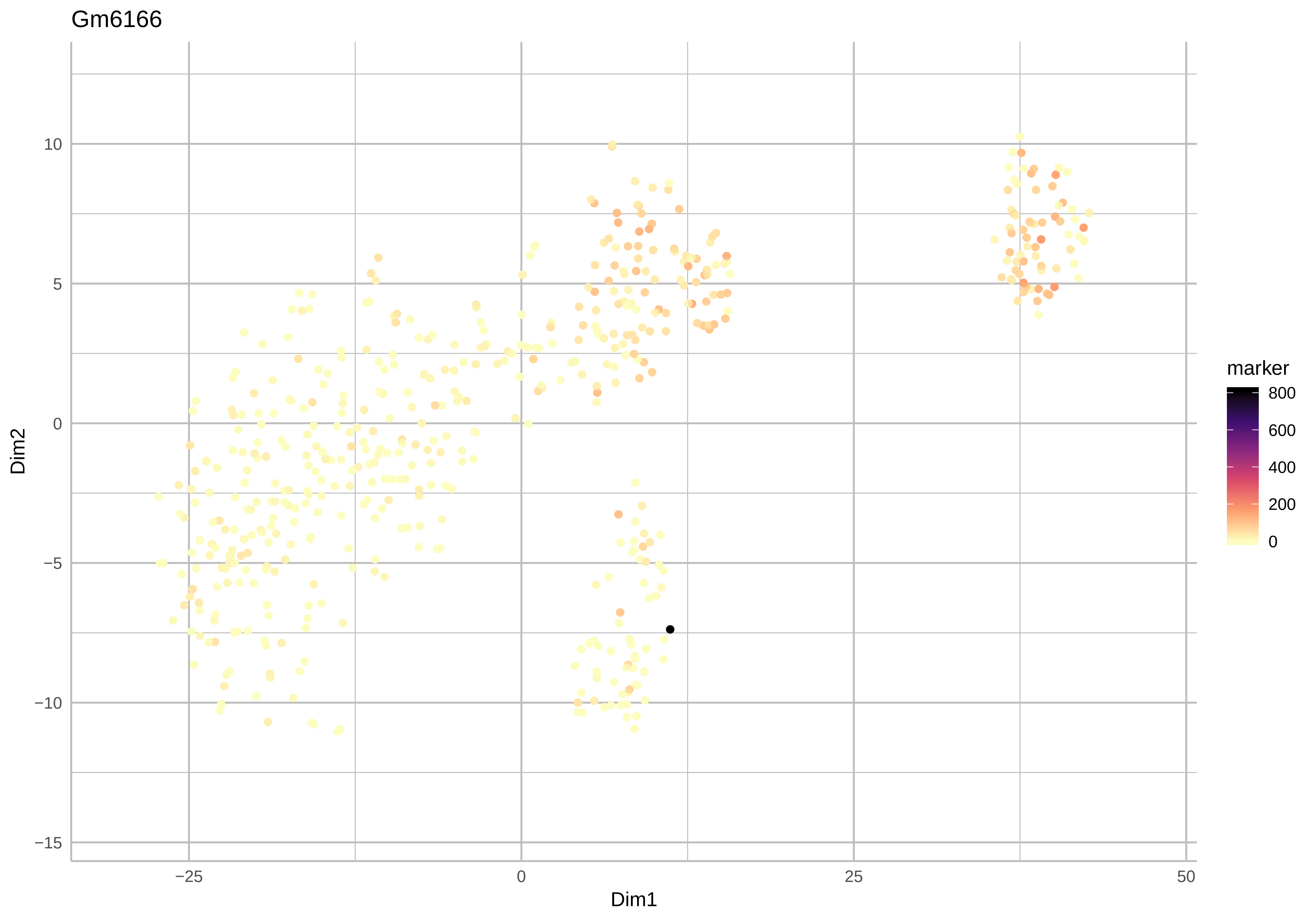

Gm6166
Dim2
Dim1
10
5
0
−5
−10
−15
−25
0
25
50
marker
800
600
400
200
0

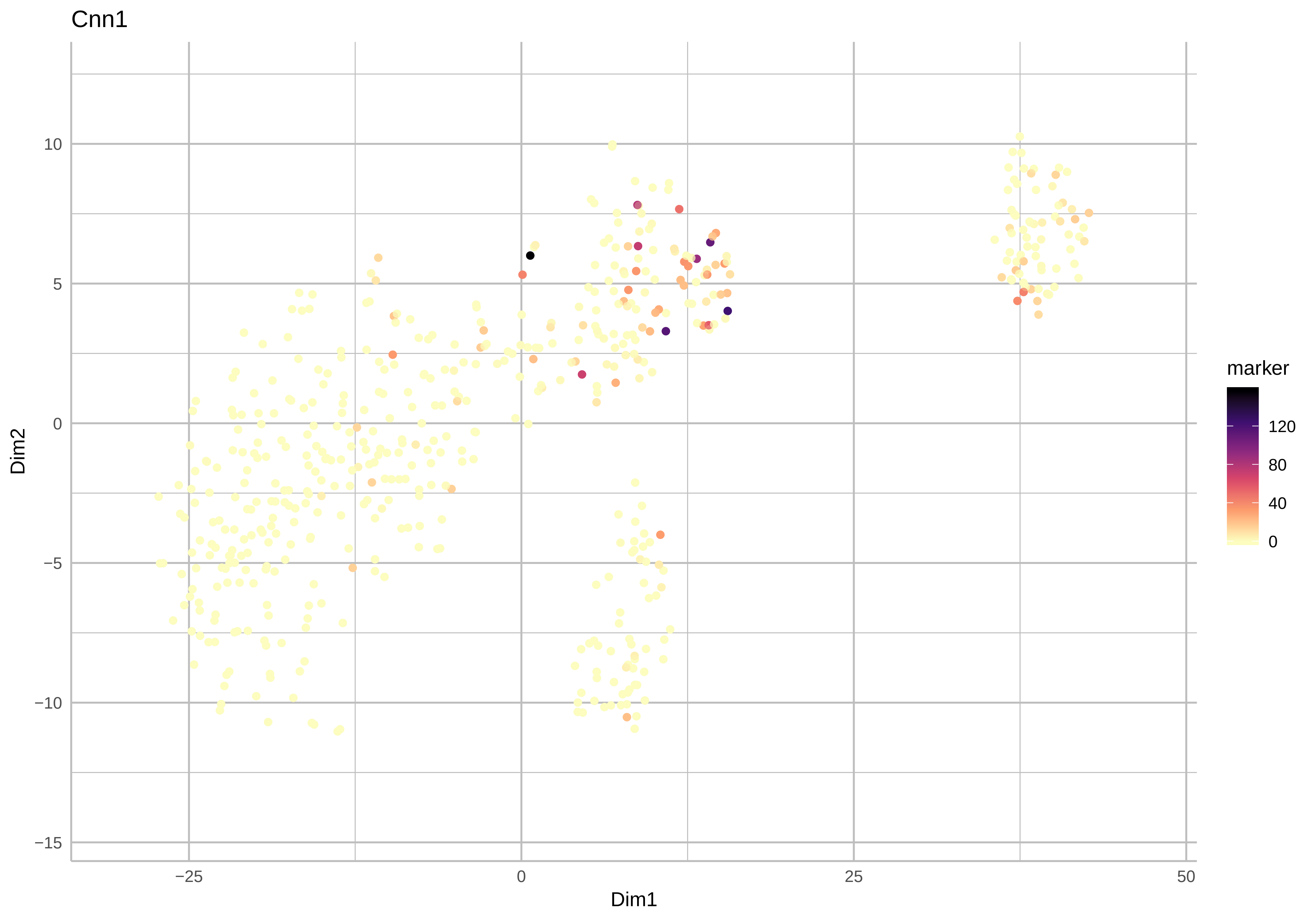
Cnn1
10
5
0
−5
−10
−15
Dim2
−25
0
25
50
Dim1
marker
120
80
40
0

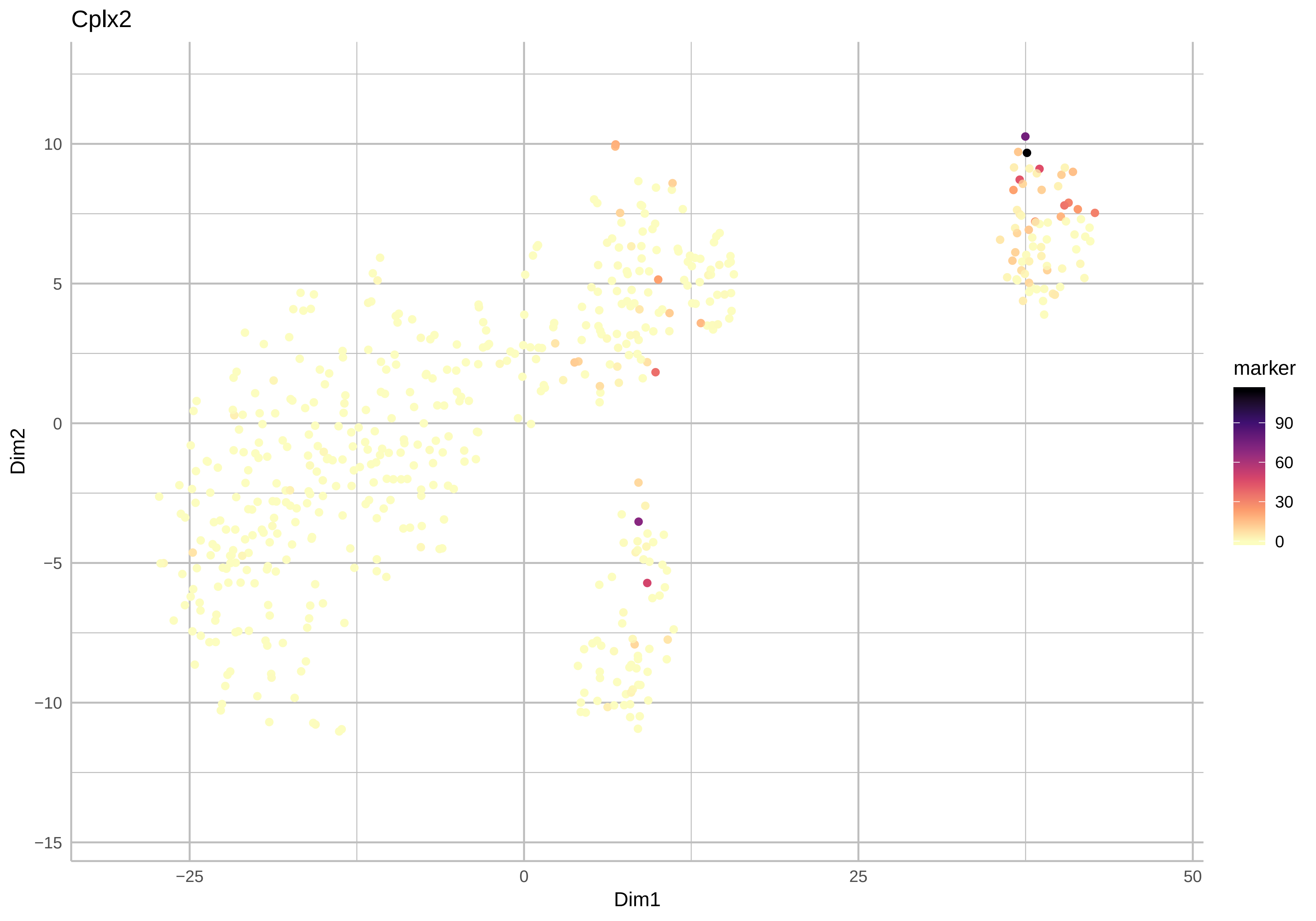

Cplx2
10
5
0
−5
−10
−15
−25
0
25
50
Dim1
Dim2
marker
90
60
30
0

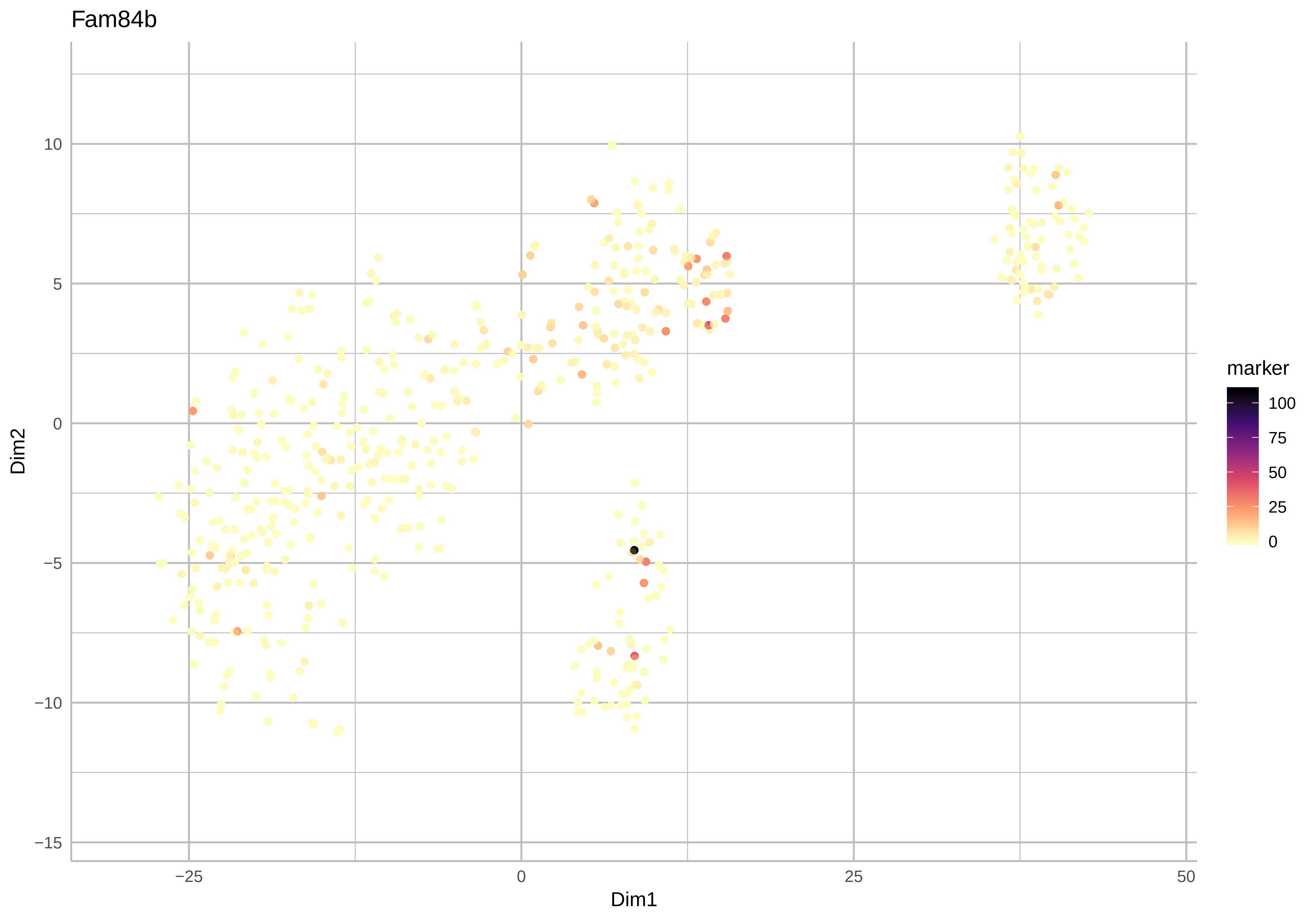

Fam84b
Dim2
Dim1
marker
100
75
50
25
0
10
5
0
−5
−10
−15
−25
0
25
50

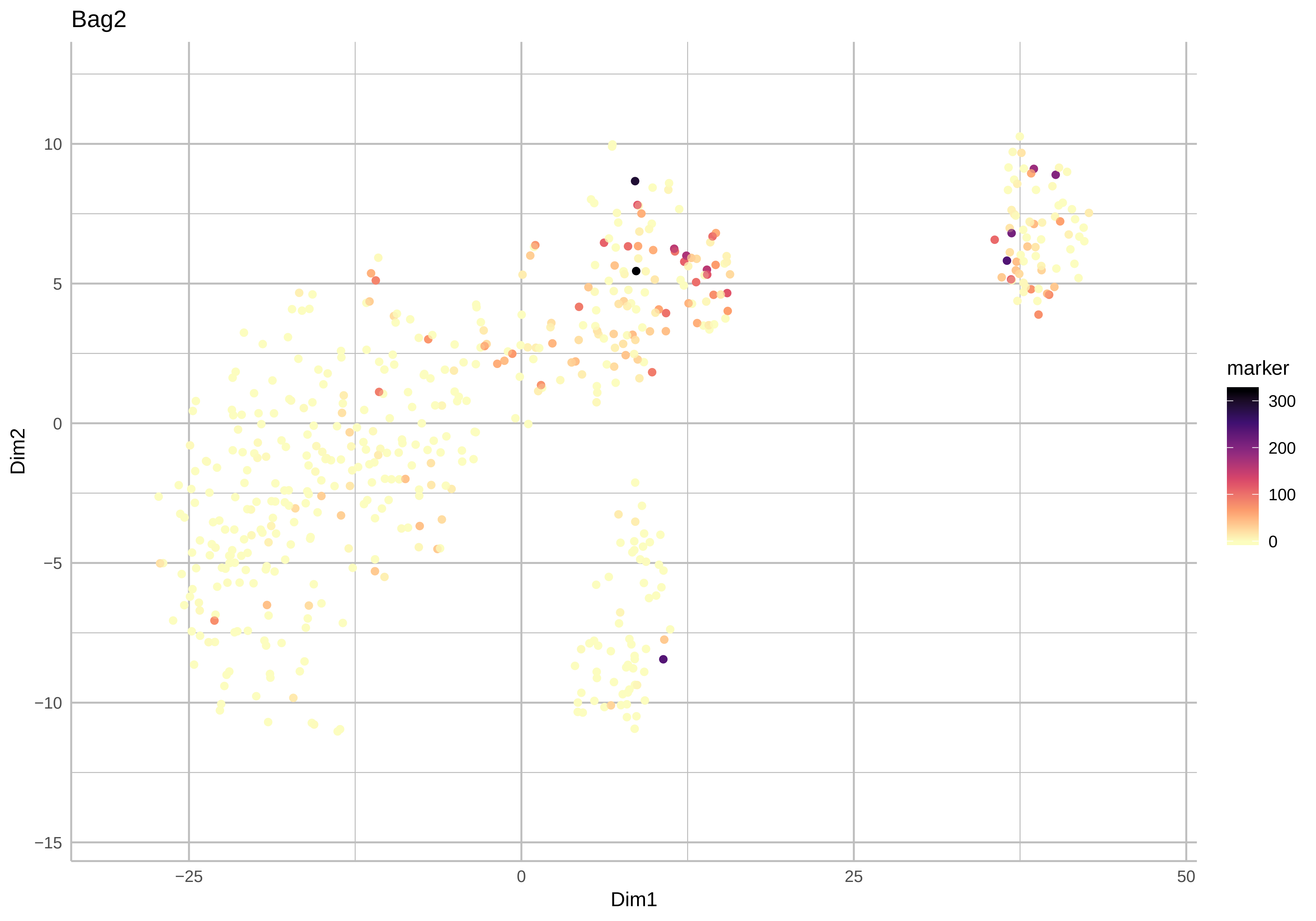
Bag2
Dim2
Dim1
10
5
0
−5
−10
−15
−25
0
25
50
marker
300
200
100
0

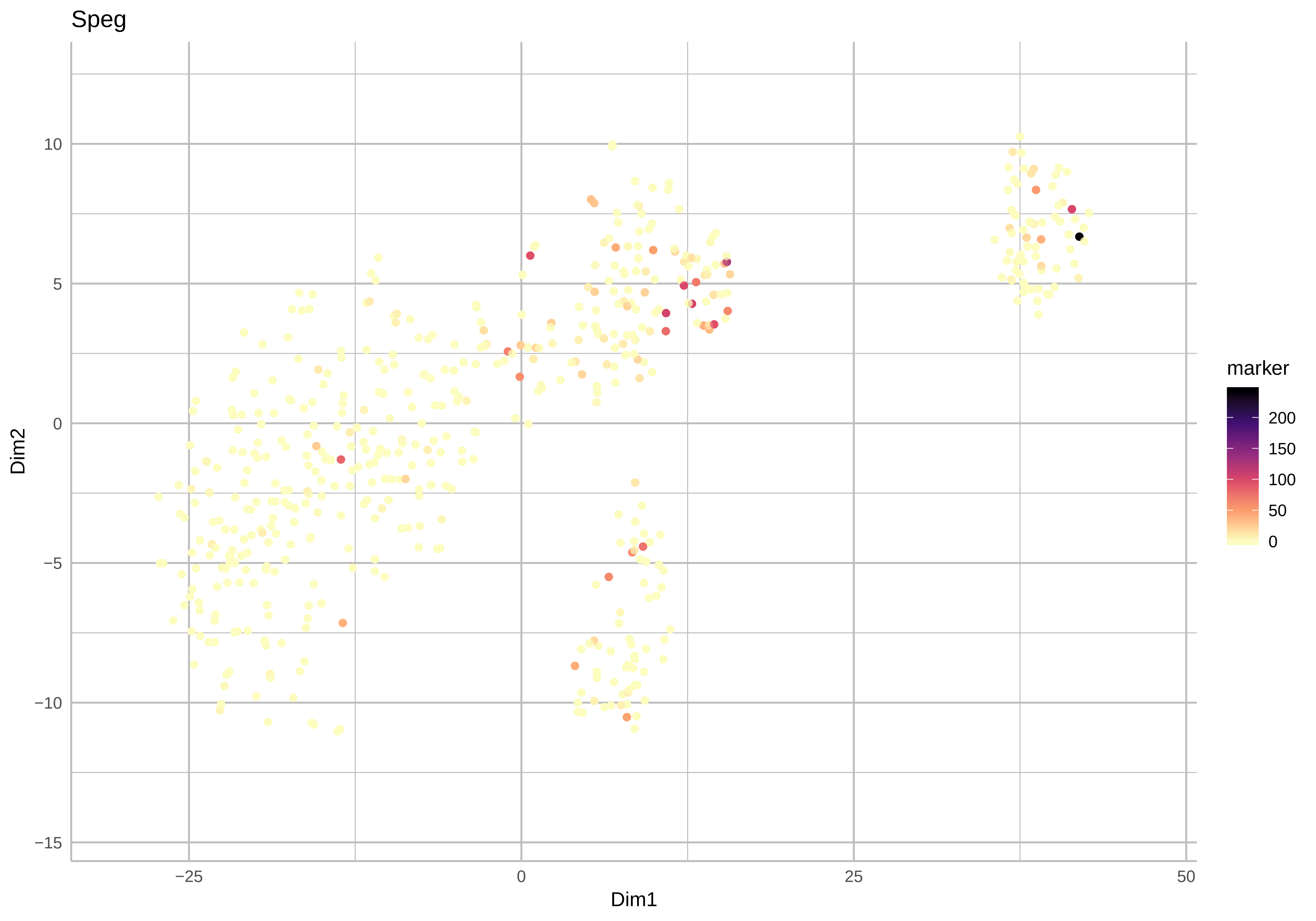
Speg
Dim2
Dim1
10
5
0
−5
−10
−15
−25
0
25
50
marker
200
150
100
50
0

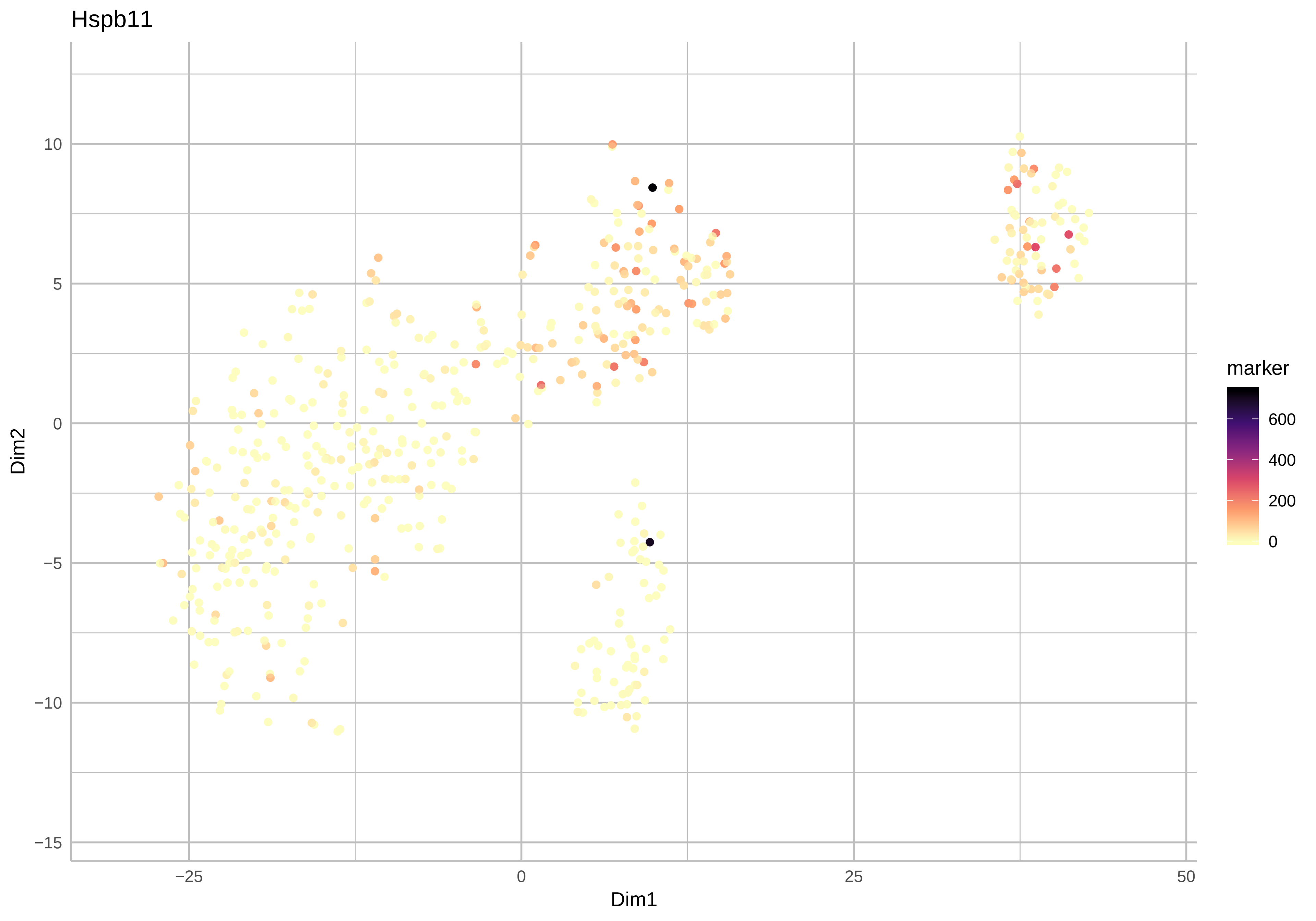
Hspb11
Dim2
Dim1
10
5
0
−5
−10
−15
−25
0
25
50
marker
600
400
200
0

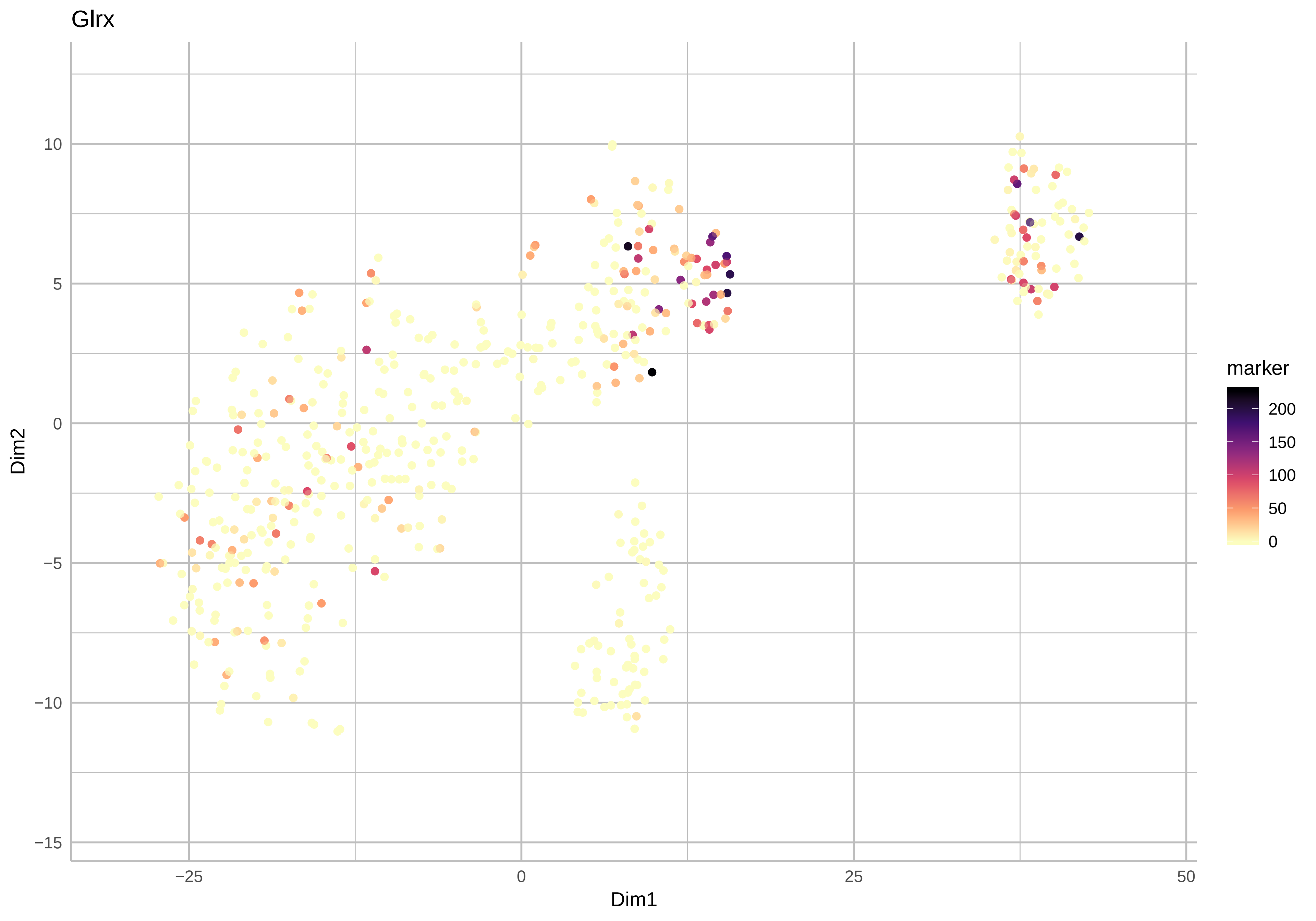
Glrx
Dim2
Dim1
10
5
0
−5
−10
−15
−25
0
25
50
marker
200
150
100
50
0

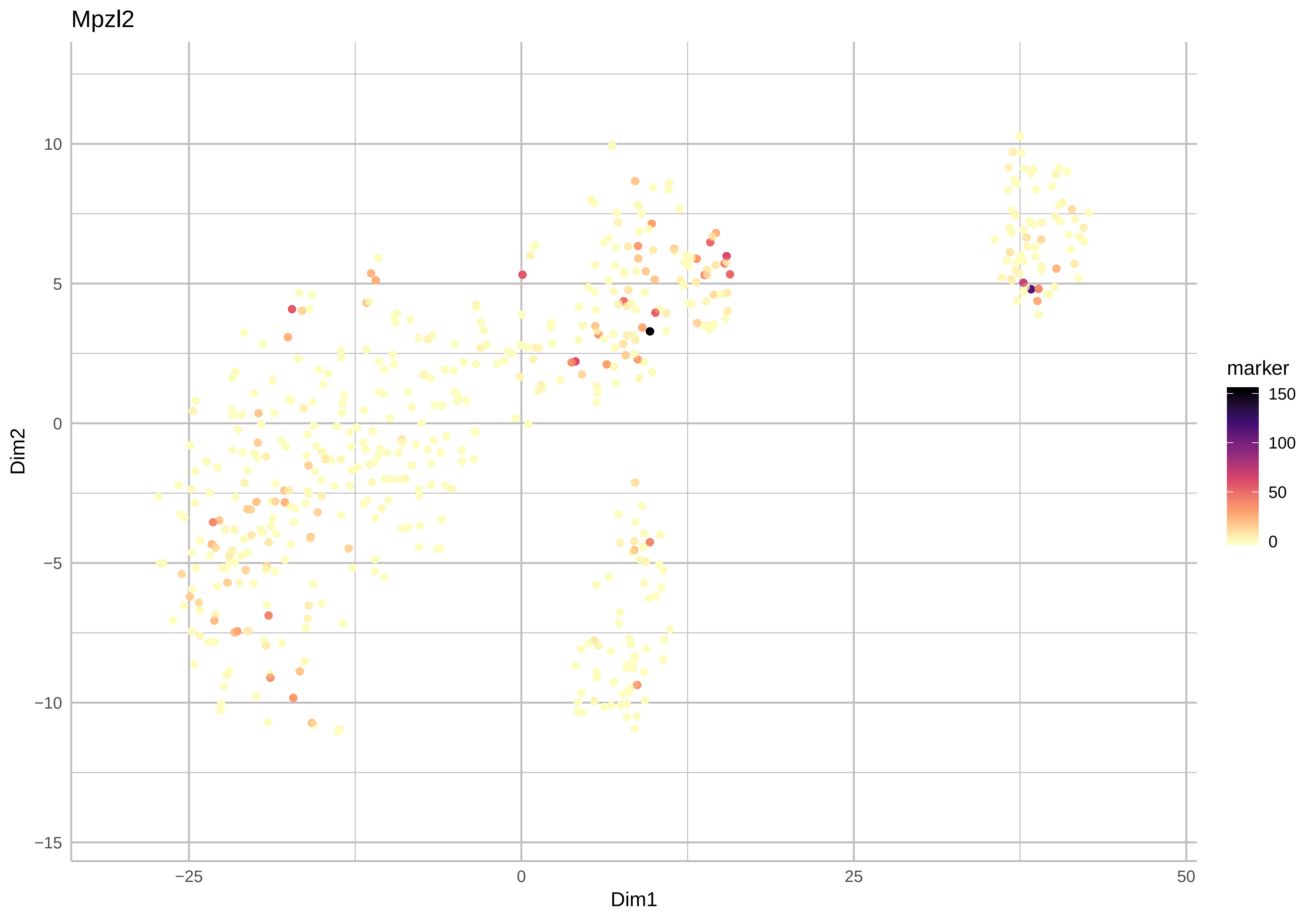

Mpzl2
10
5
0
−5
−10
−15
Dim2
−25
0
25
50
Dim1
marker
150
100
50
0

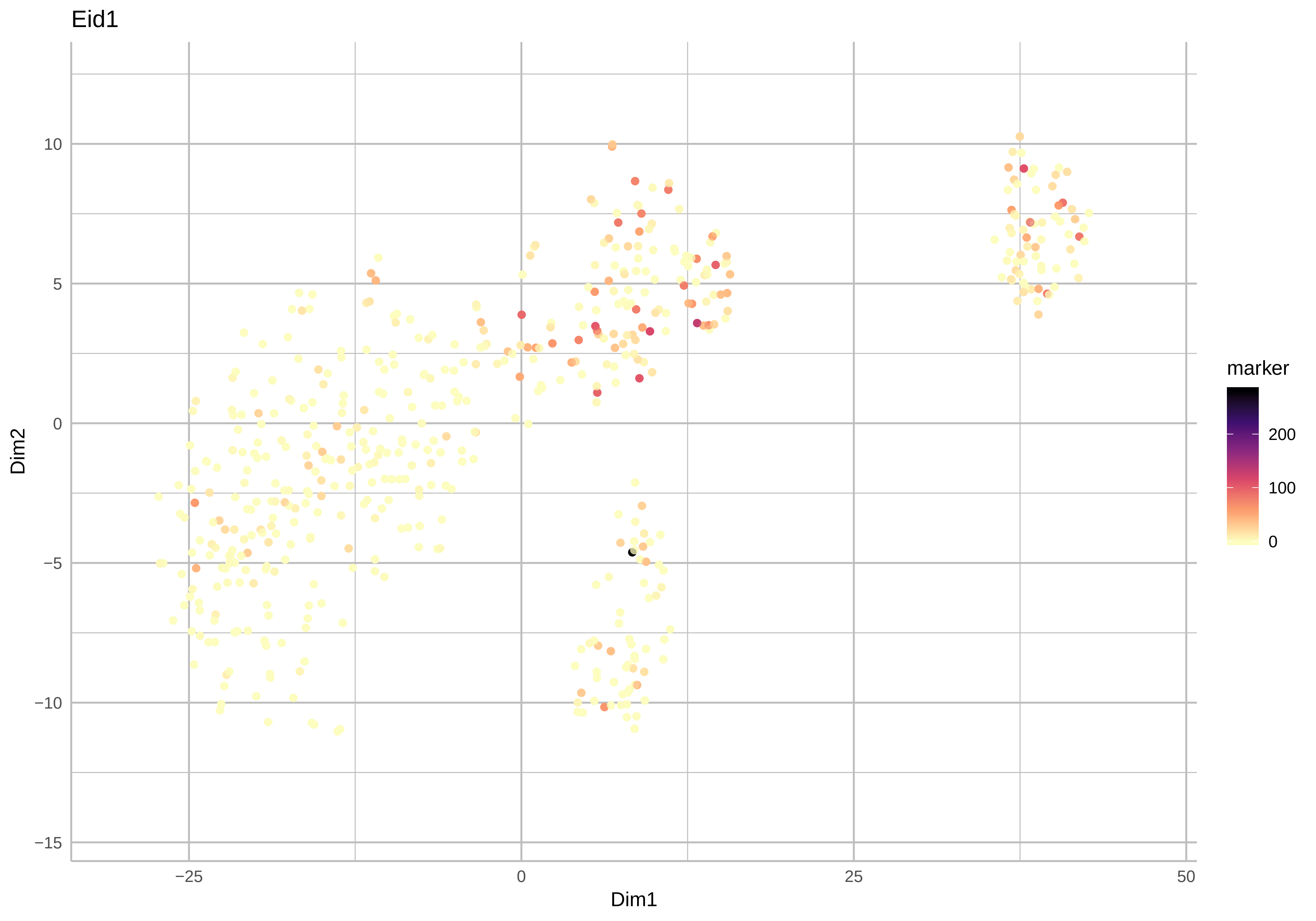

Eid1
Dim2
Dim1
10
5
0
−5
−10
−15
−25
0
25
50
marker
200
100
0

### Col5a2

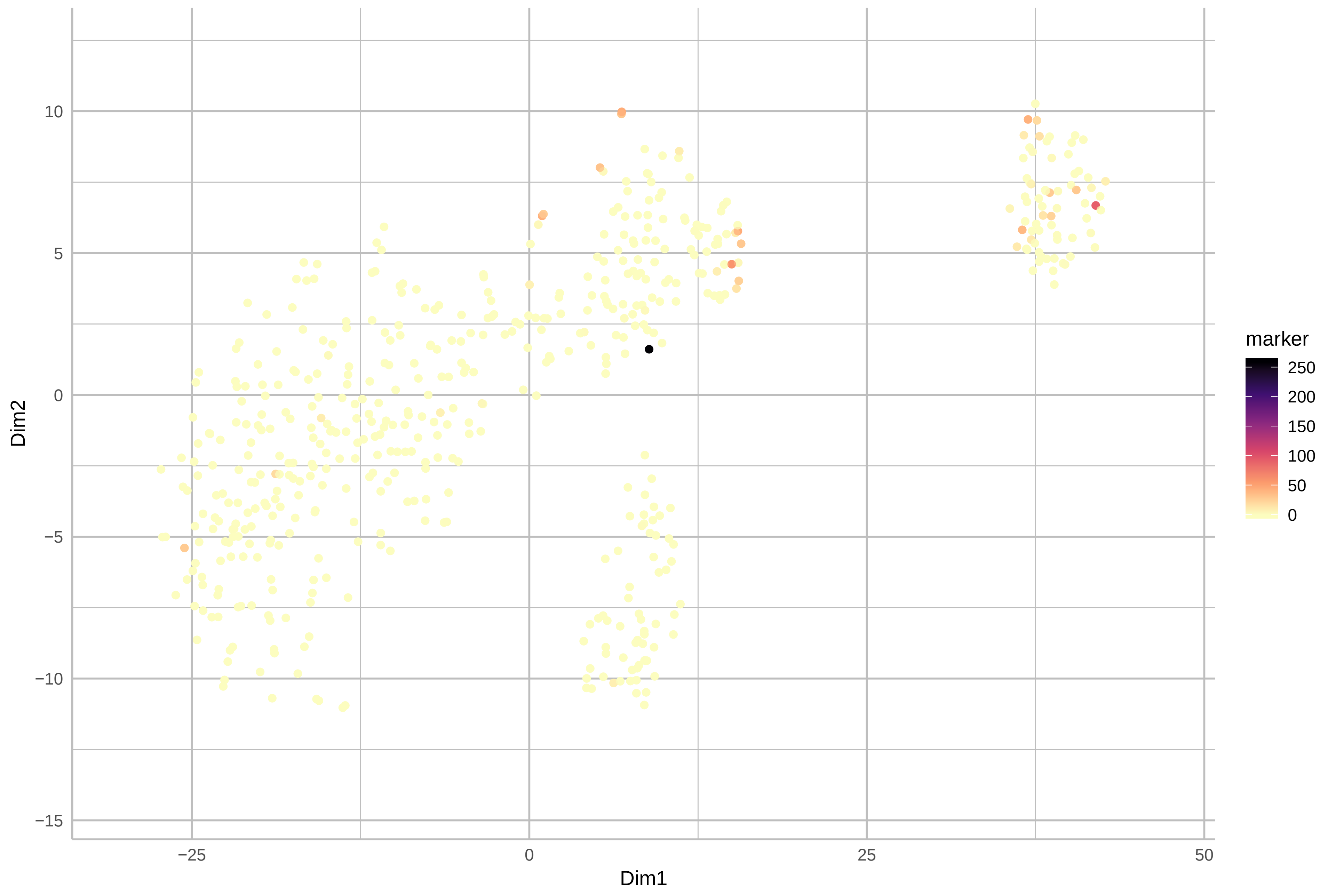

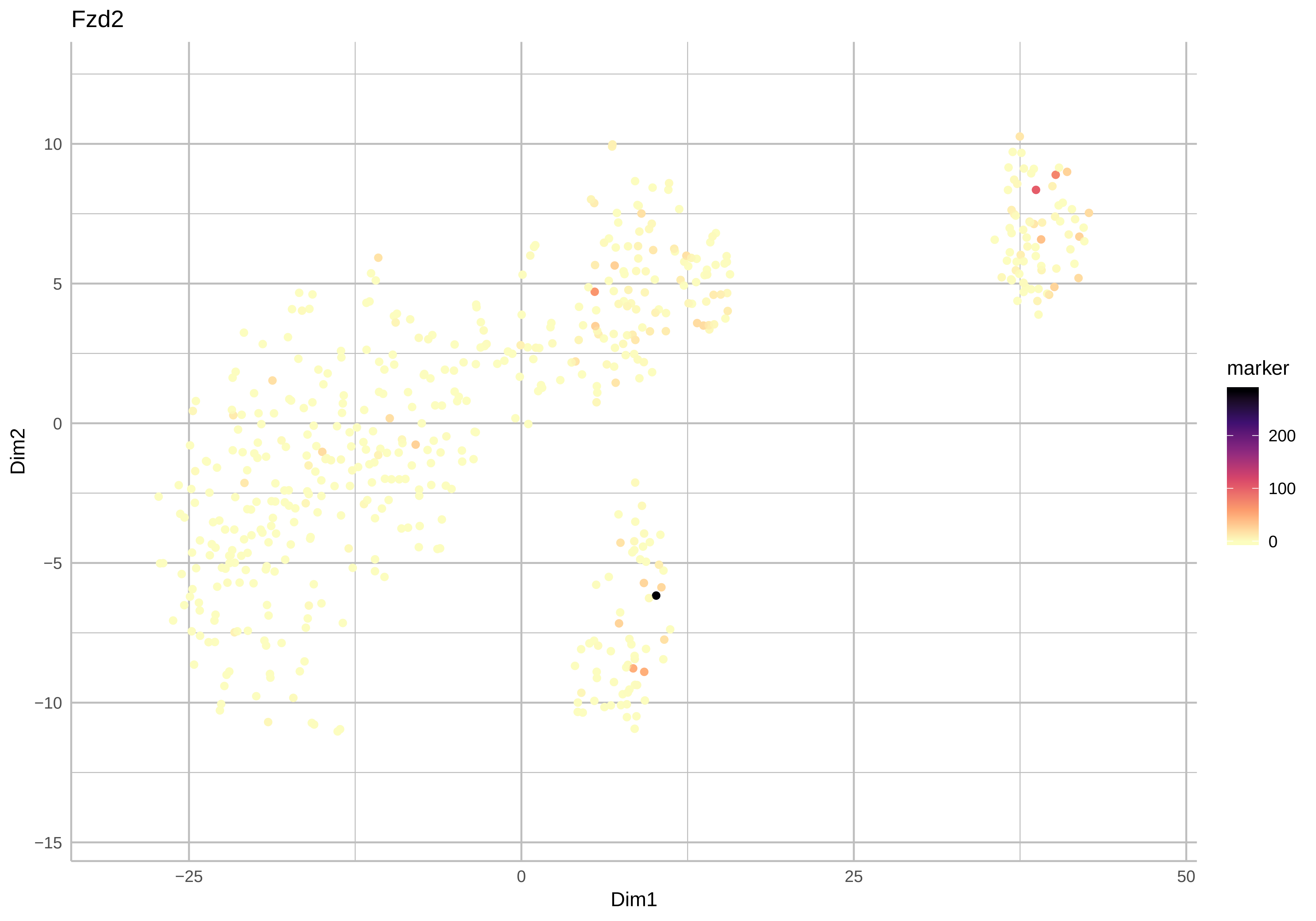
Fzd2
Dim2
Dim1
10
5
0
−5
−10
−15
−25
0
25
50
marker
200
100
0

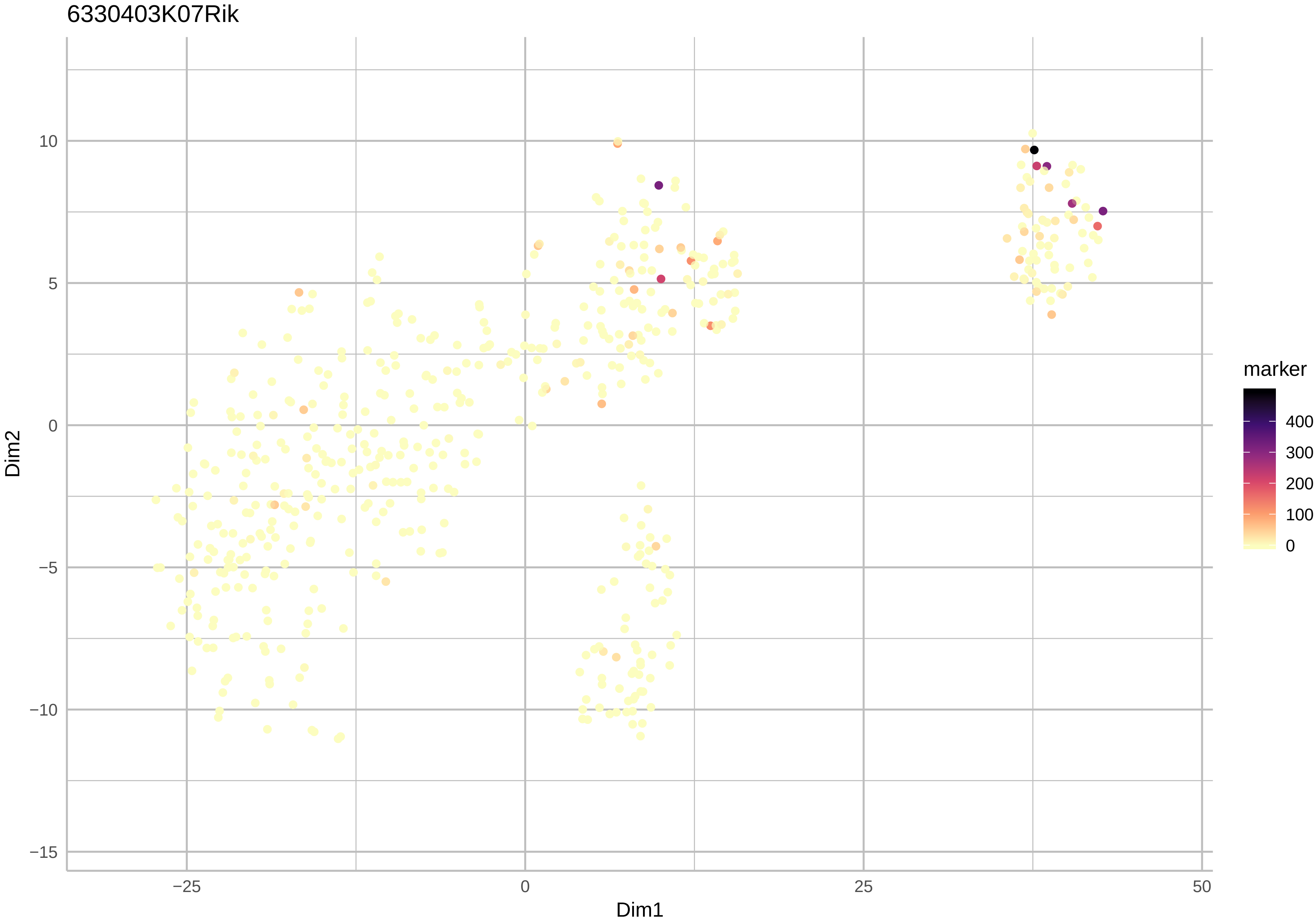

6330403K07Rik
Dim2
Dim1
10
5
0
−5
−10
−15
−25
0
25
50
marker
400
300
200
100
0

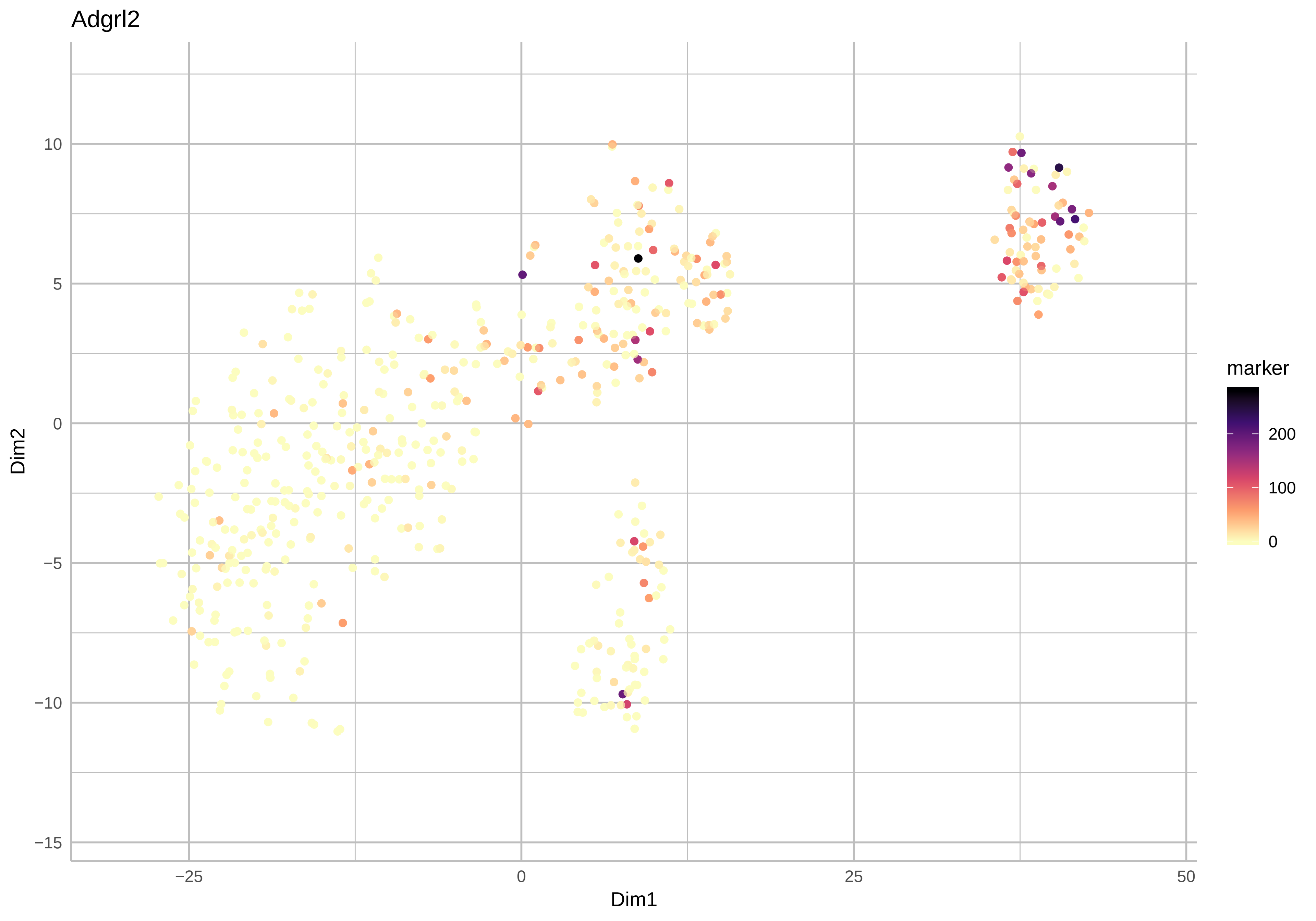
Adgrl2
10
5
0
−5
−10
−15
Dim2
−25
0
25
50
Dim1
marker
200
100
0

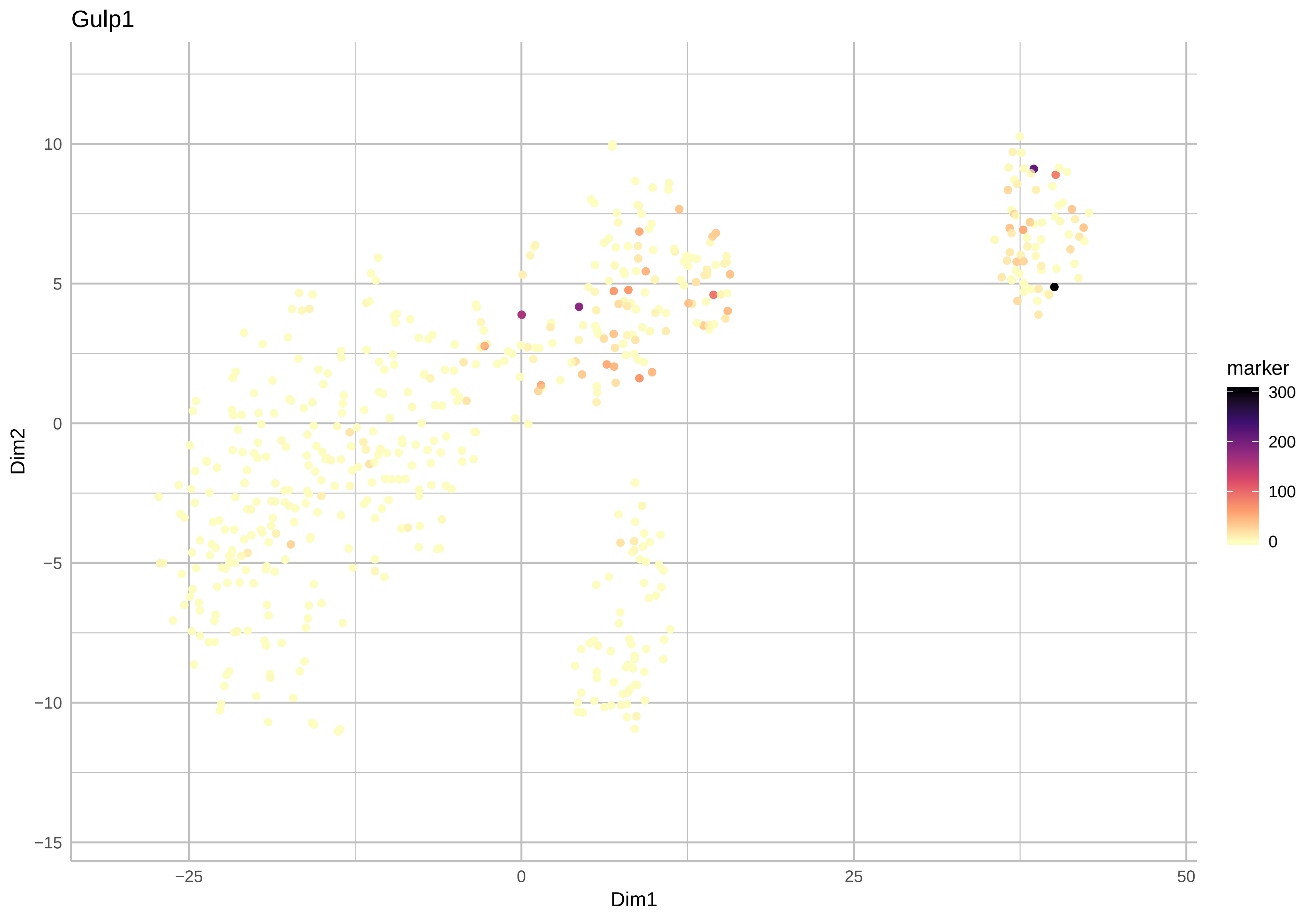

Gulp1
10
5
0
−5
−10
−15
−25
0
25
50
Dim1
Dim2
marker
300
200
100
0

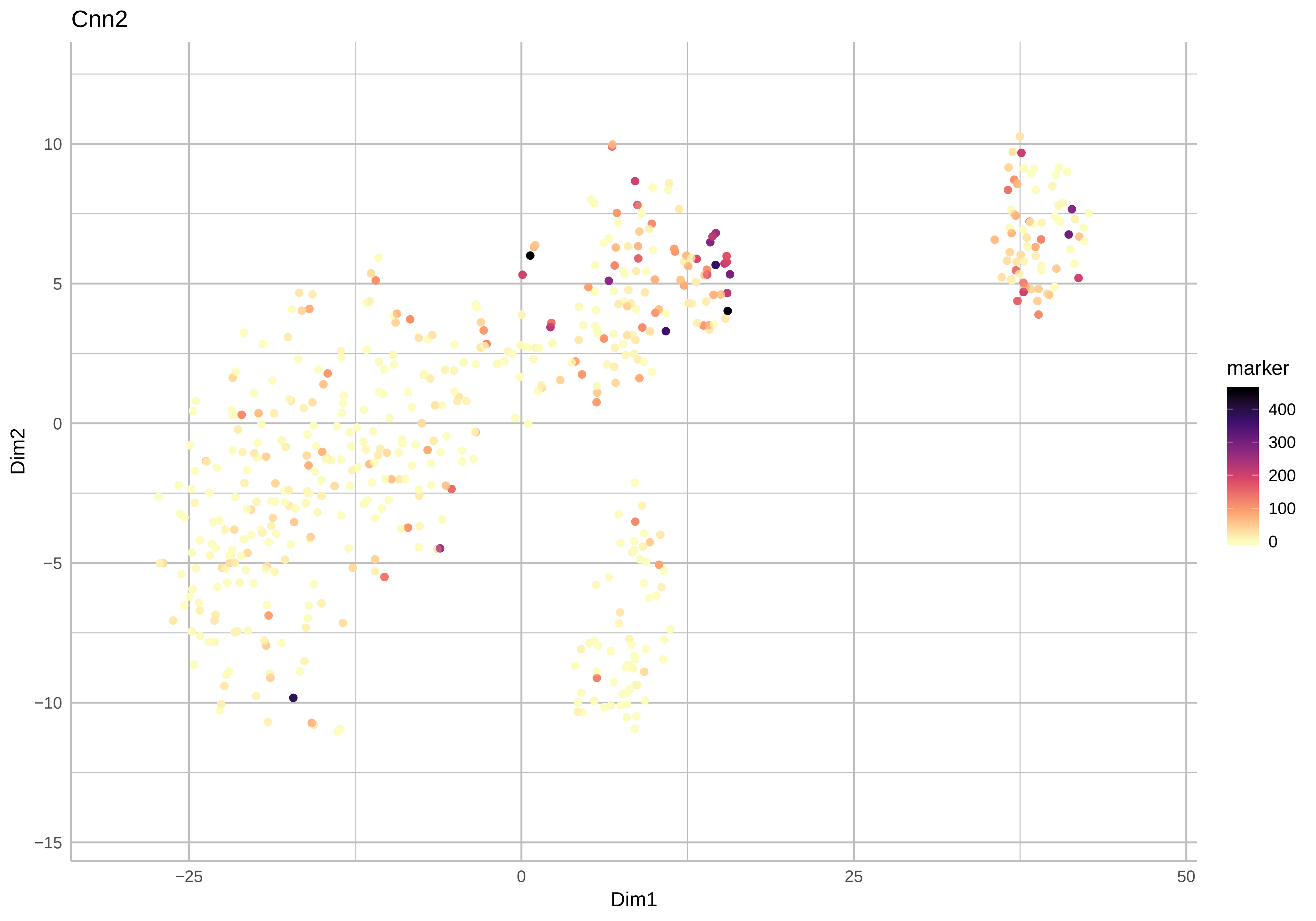

Cnn2
10
5
0
−5
−10
−15
Dim2
−25
0
25
50
Dim1
marker
400
300
200
100
0

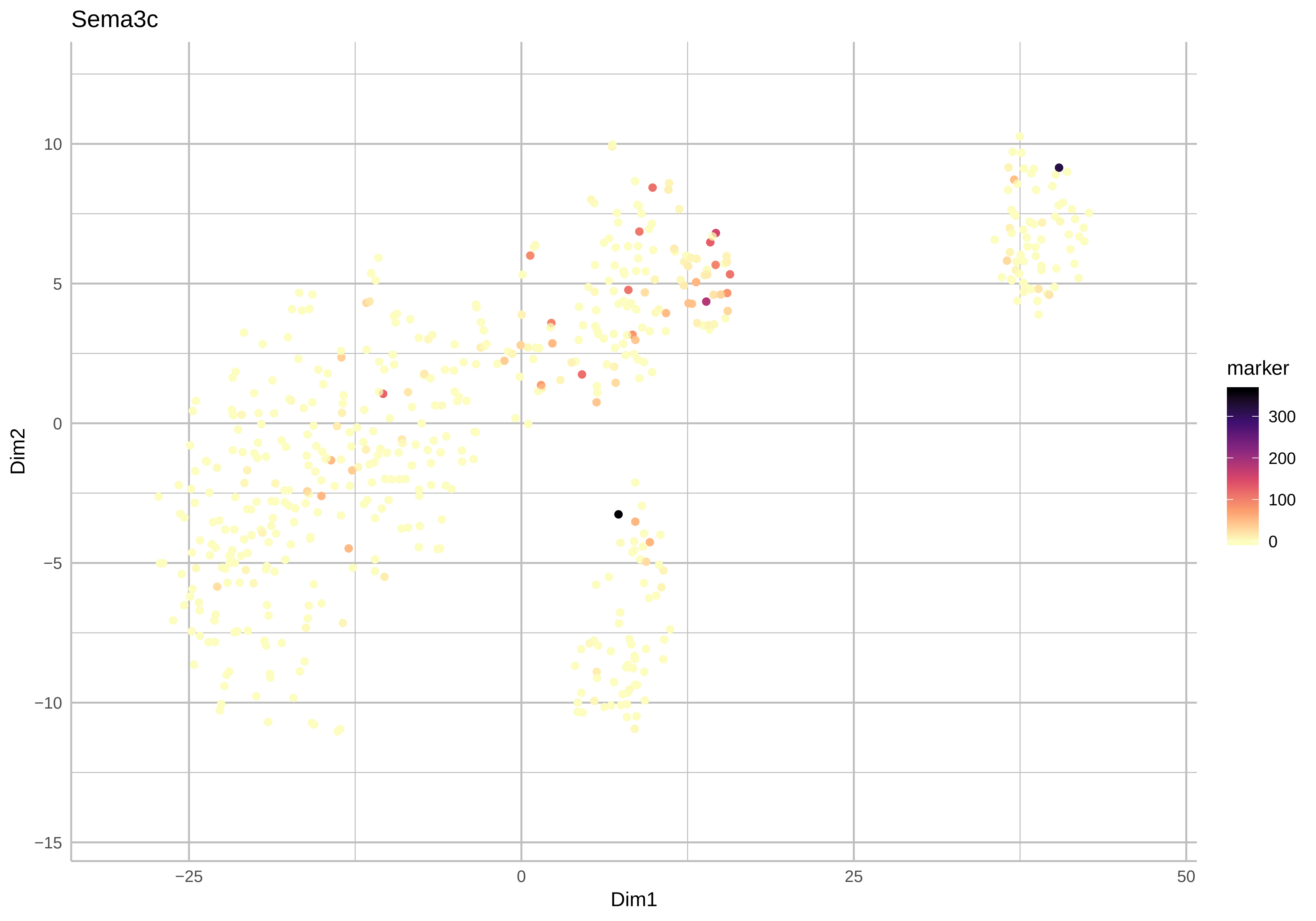

Sema3c
Dim2
Dim1
10
5
0
−5
−10
−15
−25
0
25
50
marker
300
200
100
0

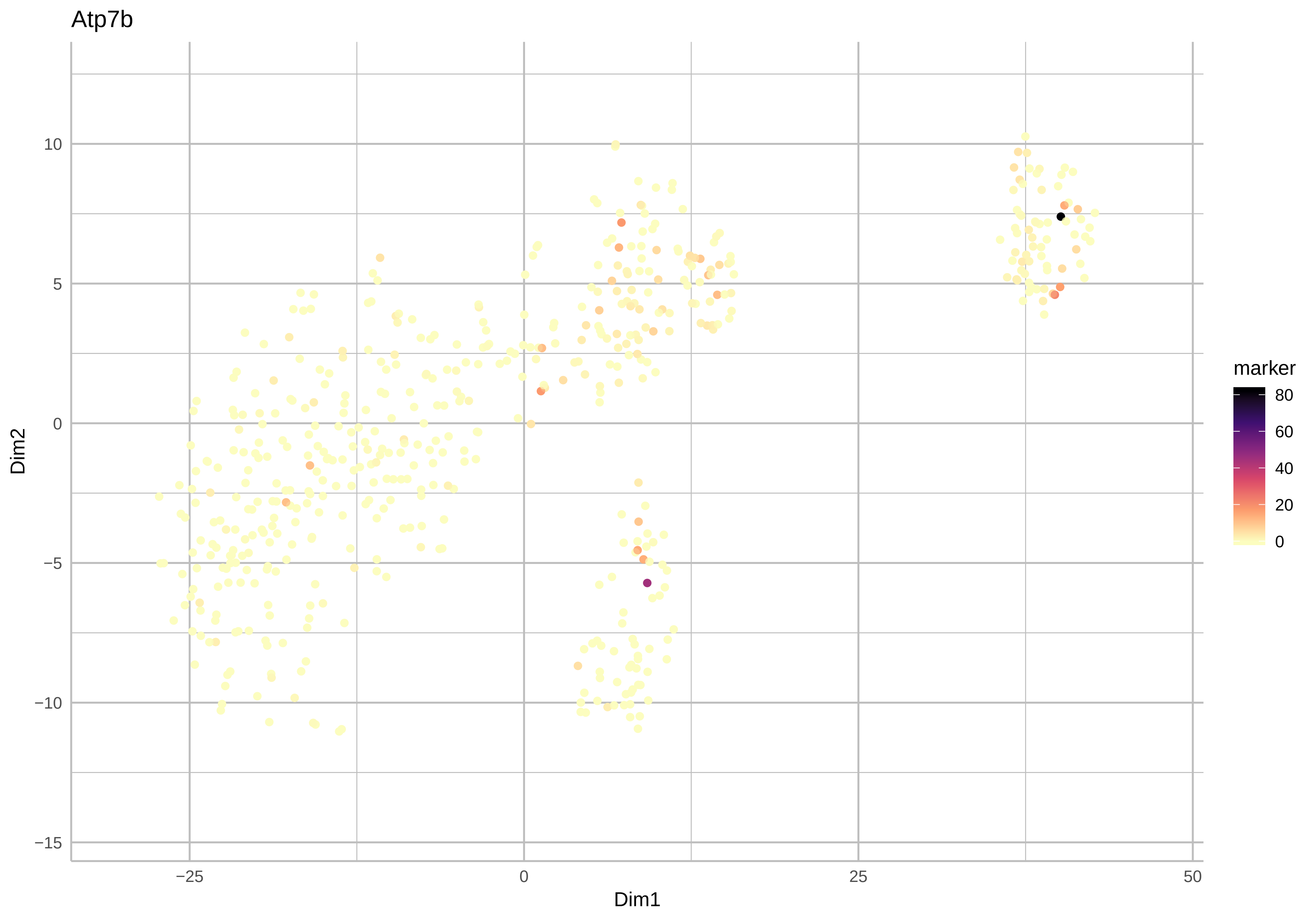

Atp7b
Dim2
Dim1
marker
80
60
40
20
0
10
5
0
−5
−10
−15
−25
0
25
50

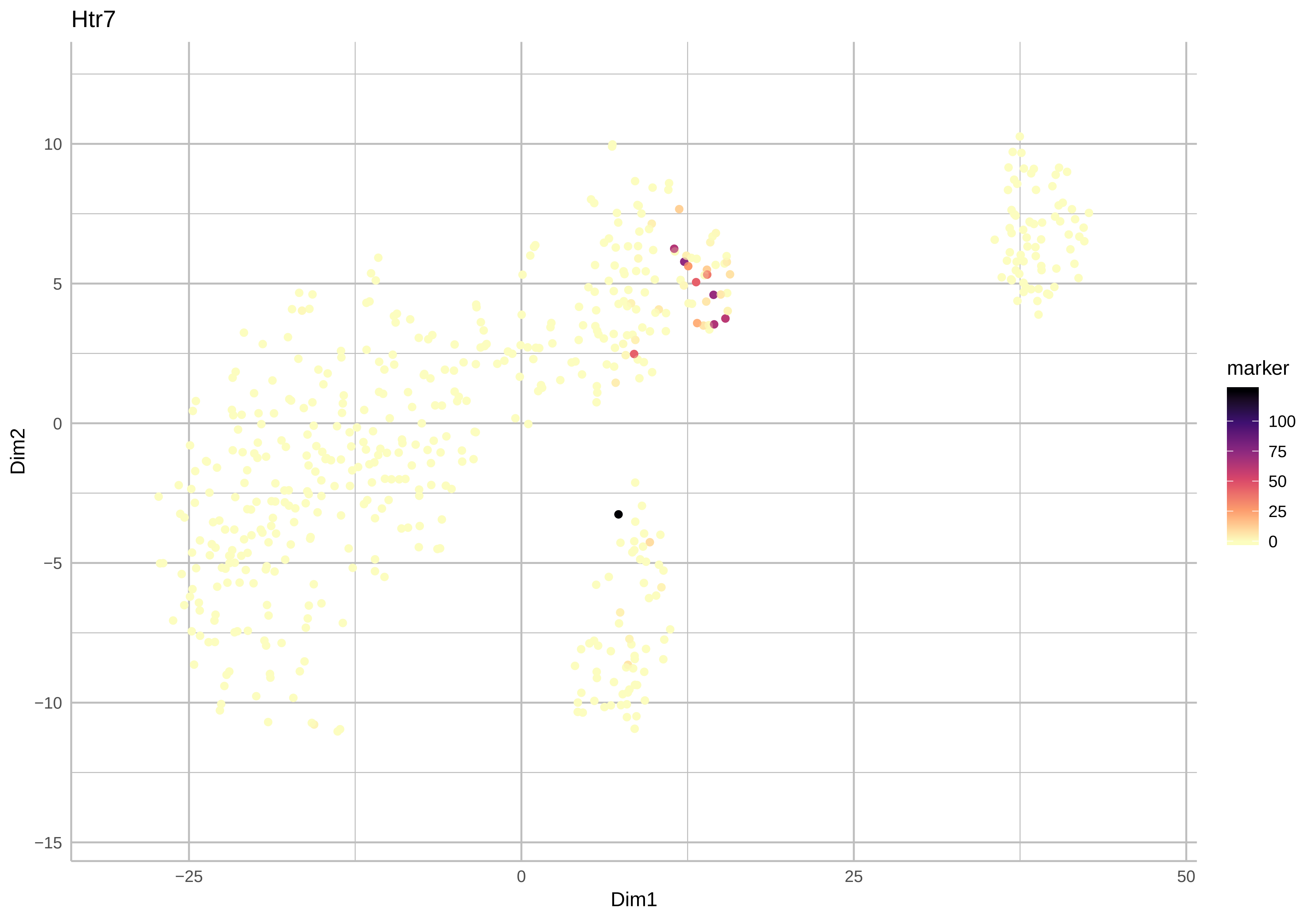

Htr7
10
5
0
−5
−10
−15
−25
0
25
50
Dim1
Dim2
marker
100
75
50
25
0

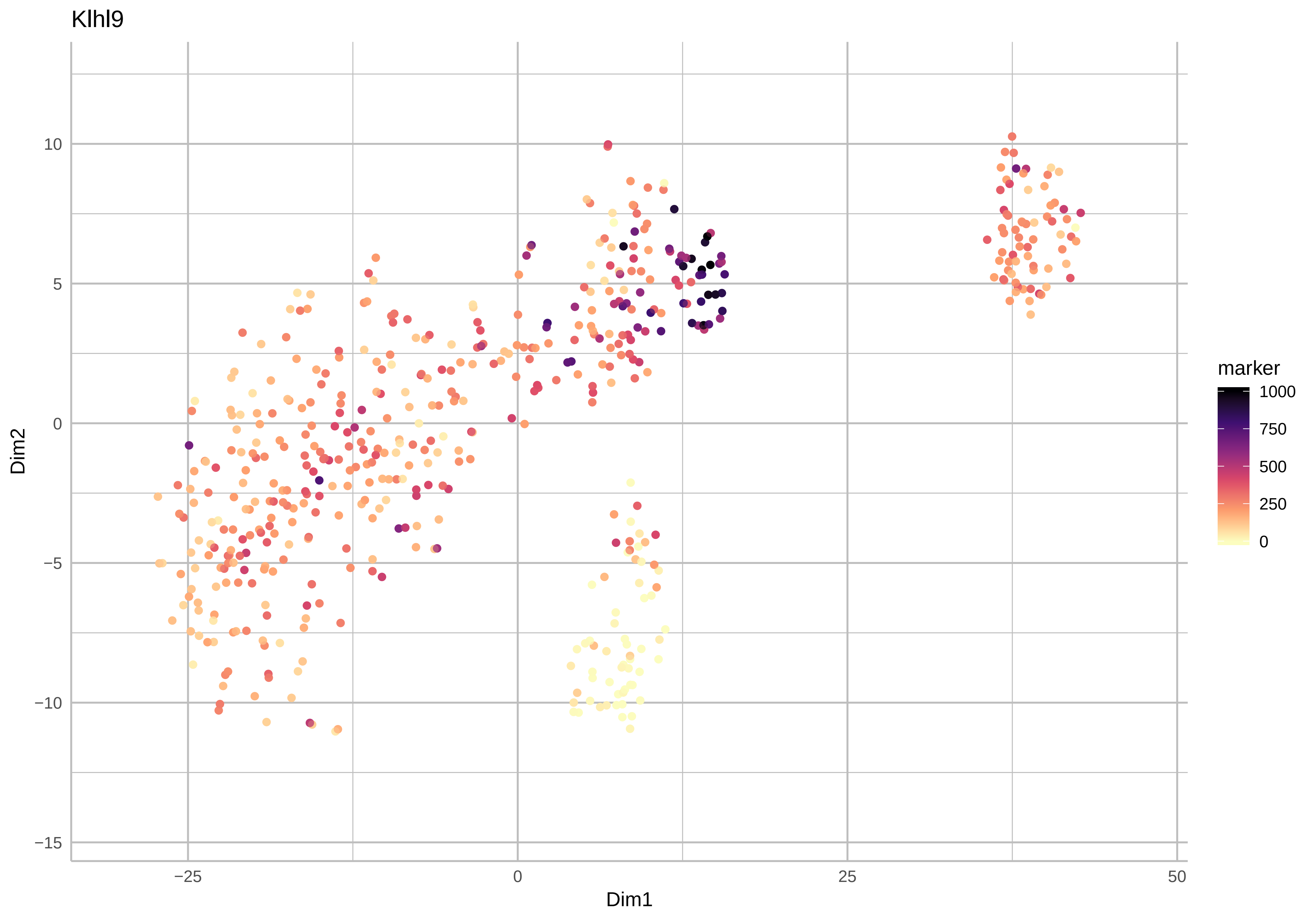
Klhl9
Dim2
10
5
0
−5
−10
−15
−25
0
25
50
Dim1
marker
1000
750
500
250
0

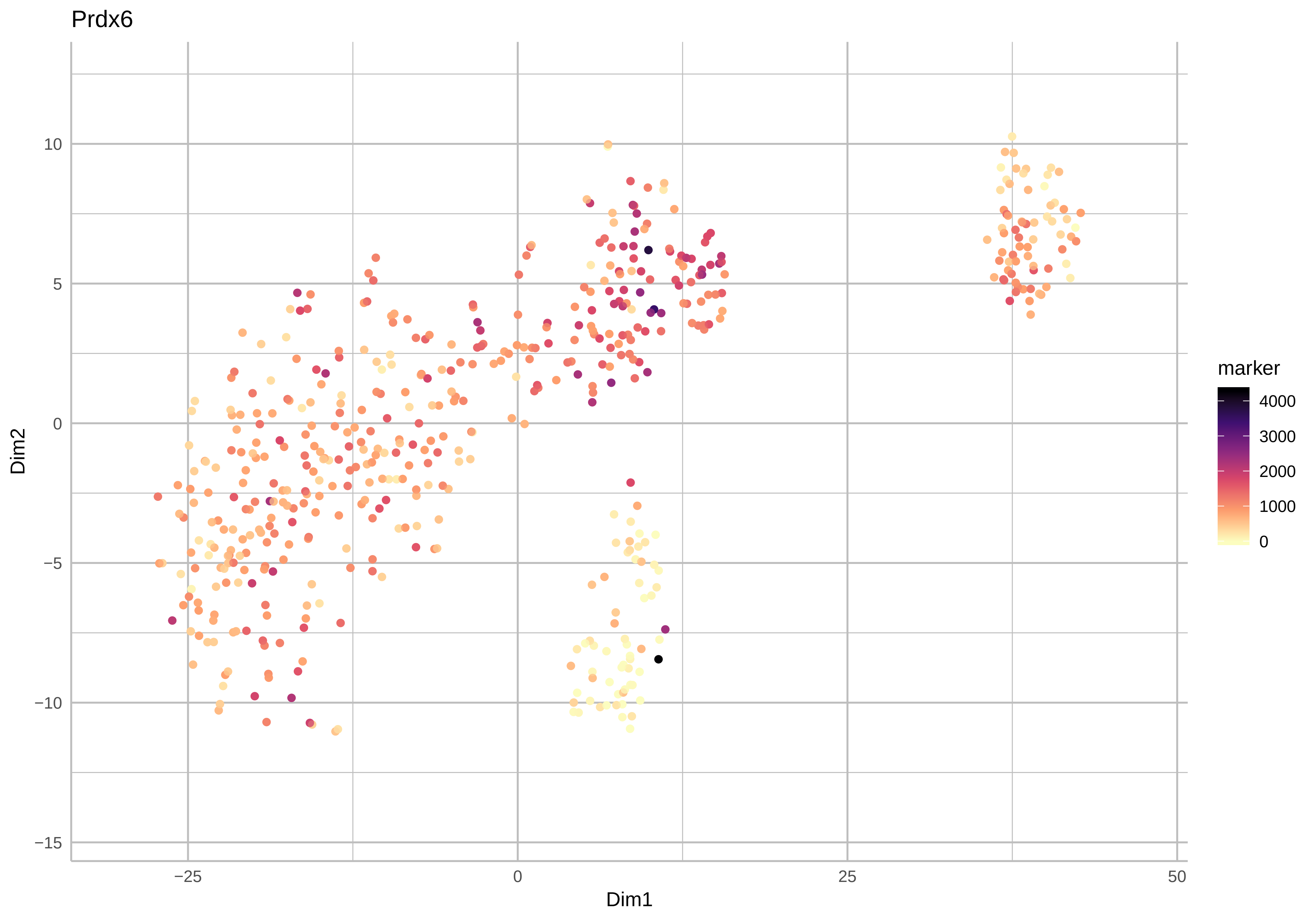

Prdx6
Dim2
Dim1
10
5
0
−5
−10
−15
−25
0
25
50
marker
4000
3000
2000
1000
0

Gmps

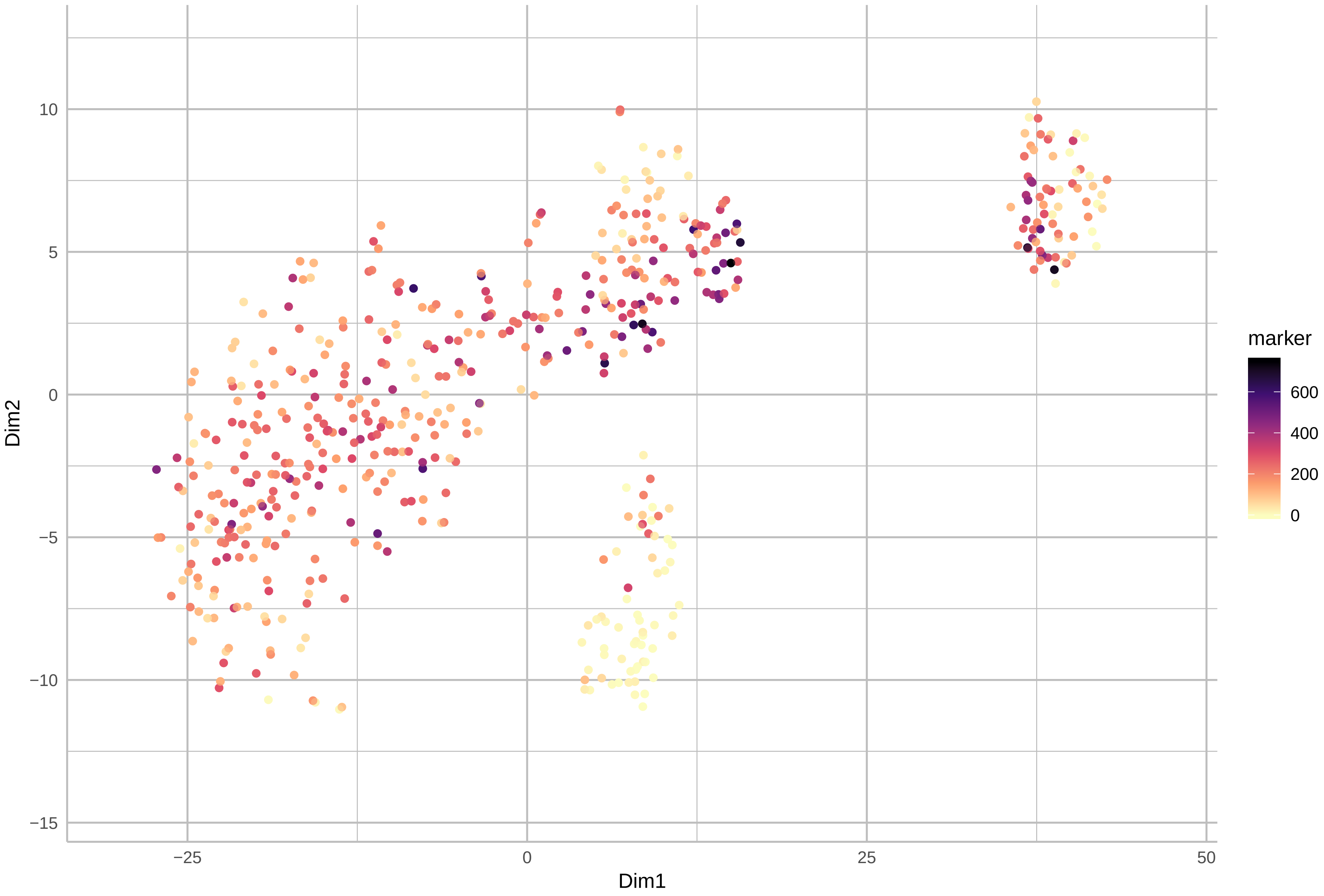

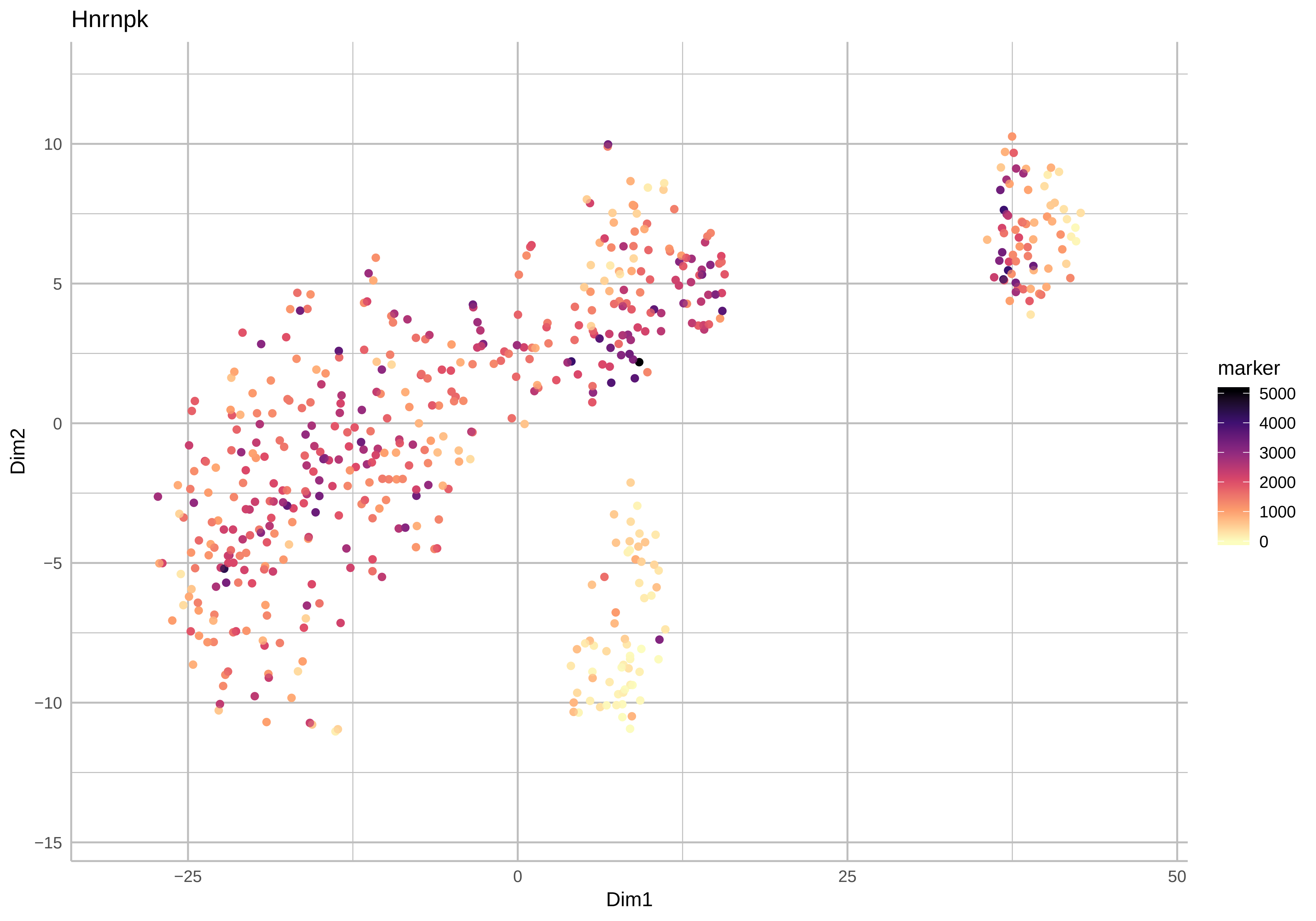

Hnrnpk
Dim2
Dim1
10
5
0
−5
−10
−15
−25
0
25
50
marker
5000
4000
3000
2000
1000
0

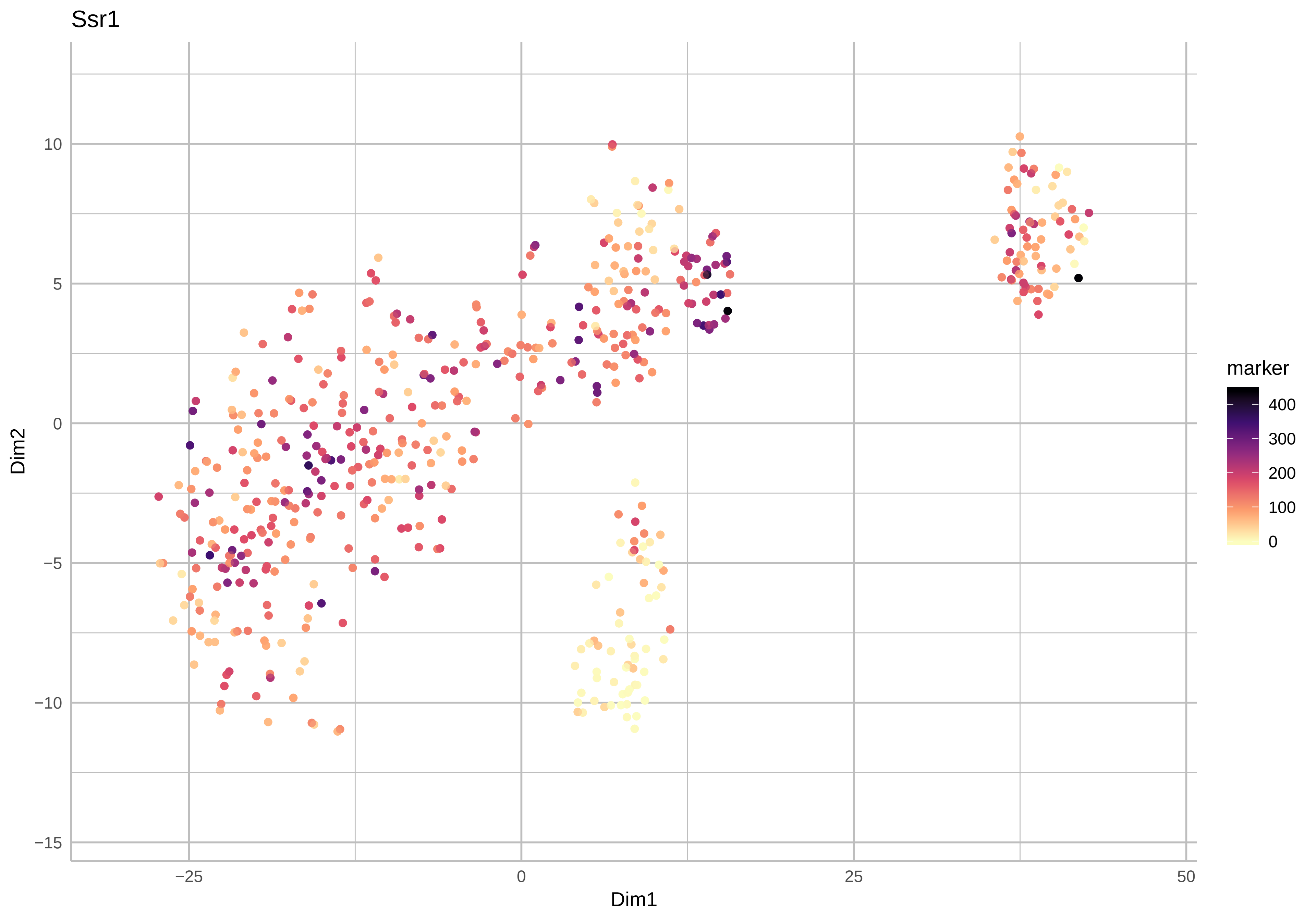

Ssr1
Dim2
Dim1
10
5
0
−5
−10
−15
−25
0
25
50
marker
400
300
200
100
0

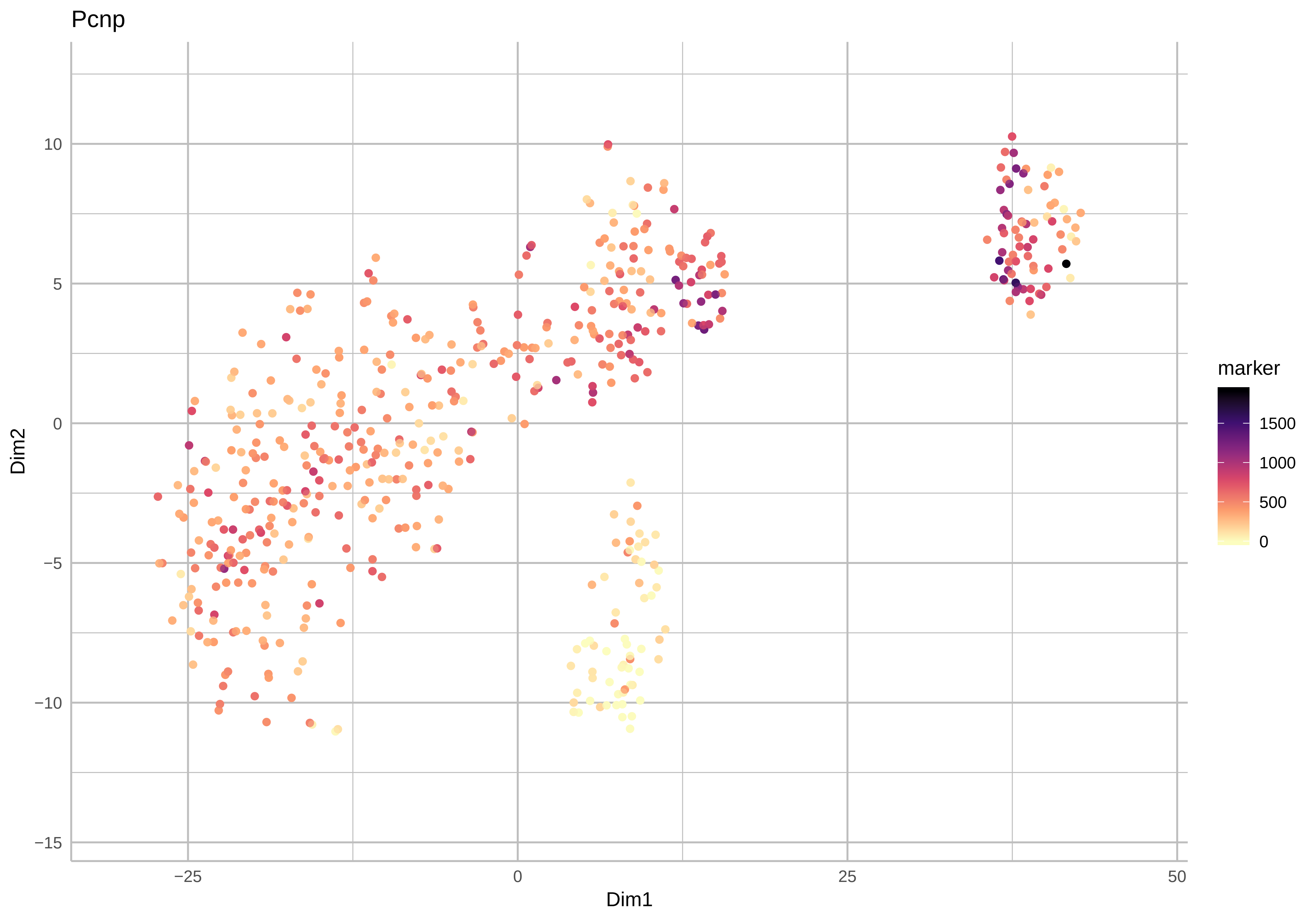

Pcnp
Dim2
Dim1
−25
0
25
50
10
5
0
−5
−10
−15
marker
1500
1000
500
0

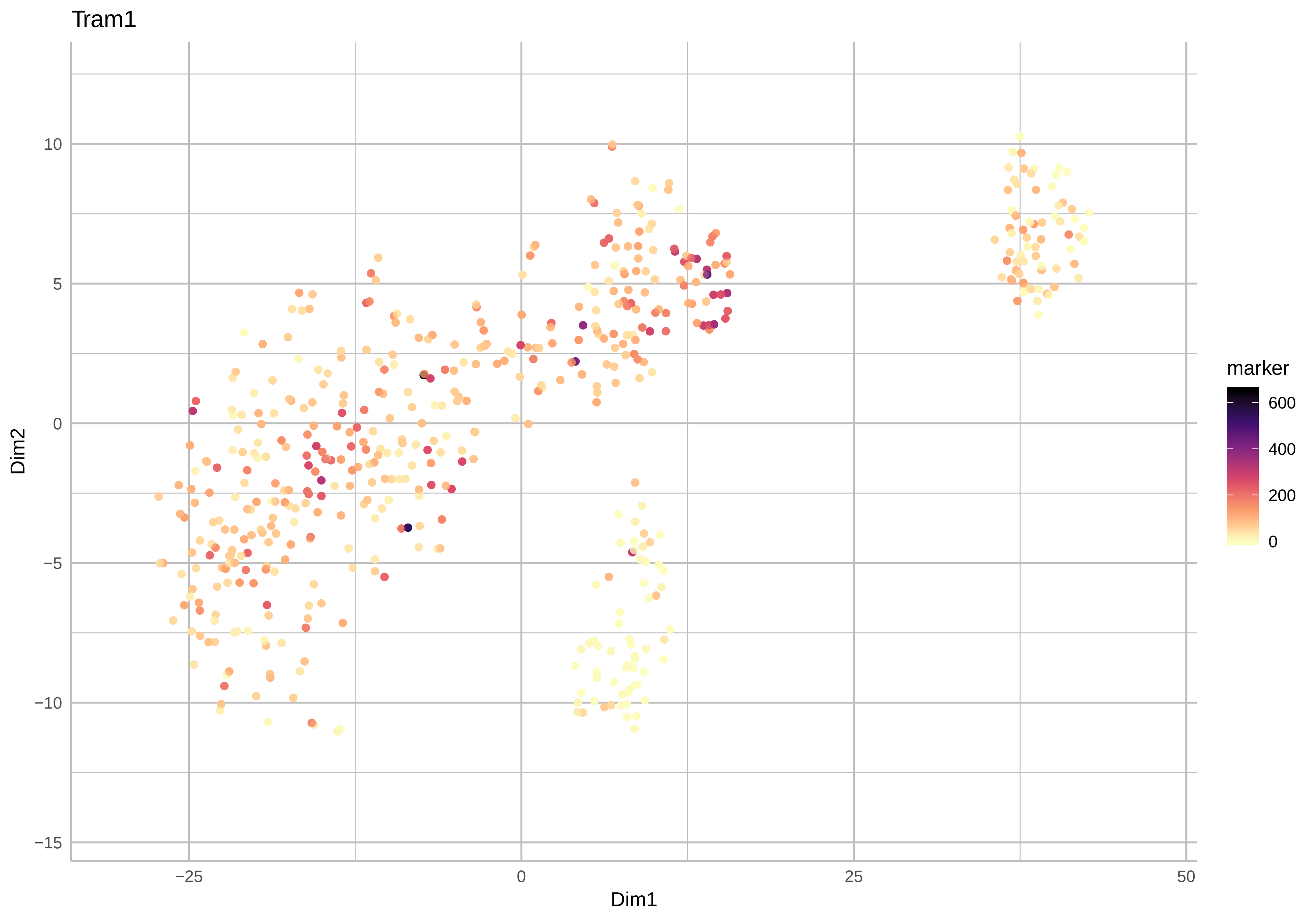

Tram1
Dim2
Dim1
10
5
0
−5
−10
−15
−25
0
25
50
marker
600
400
200
0

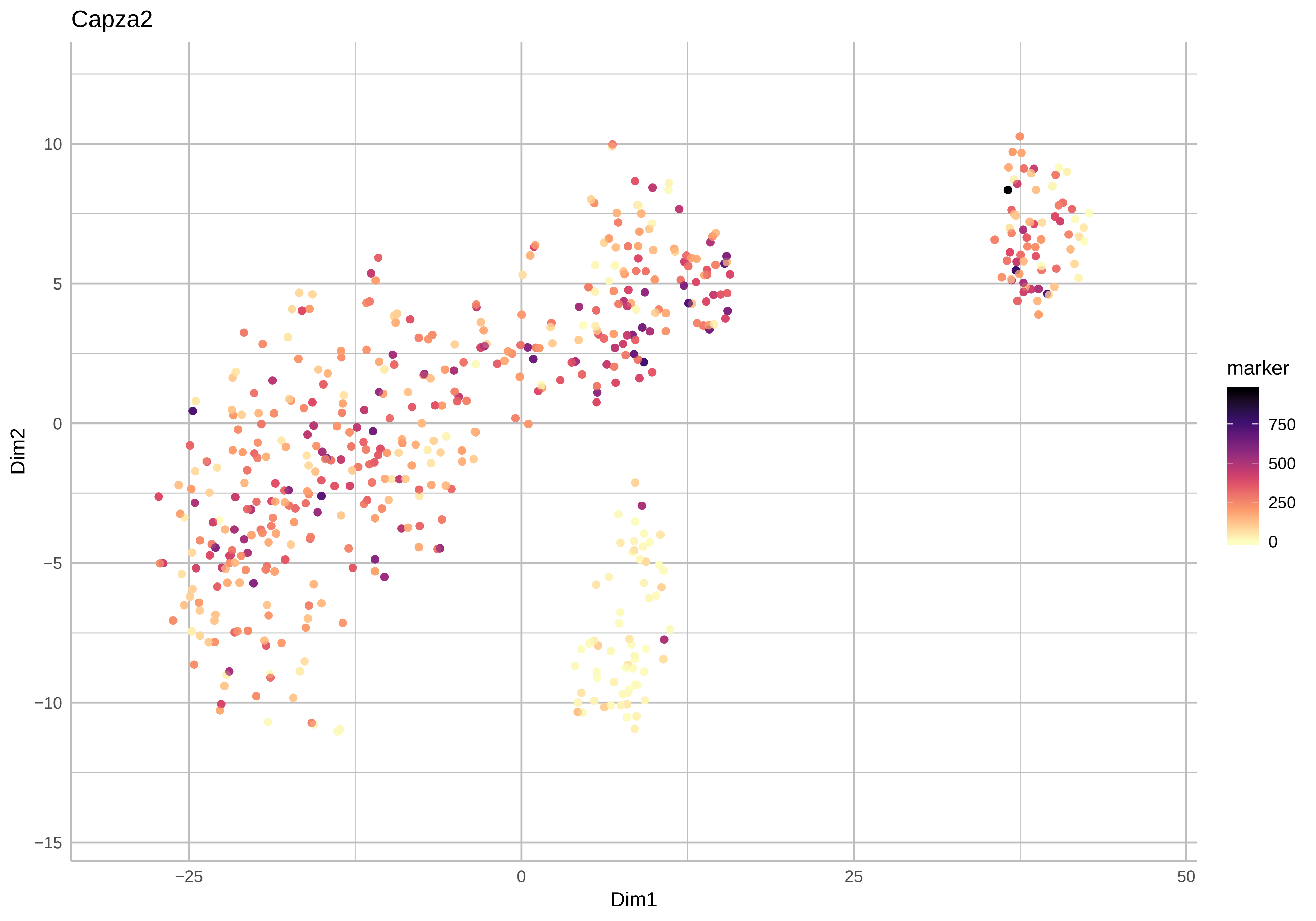

Capza2
10
5
0
−5
−10
−15
−25
0
25
50
Dim1
Dim2
marker
750
500
250
0

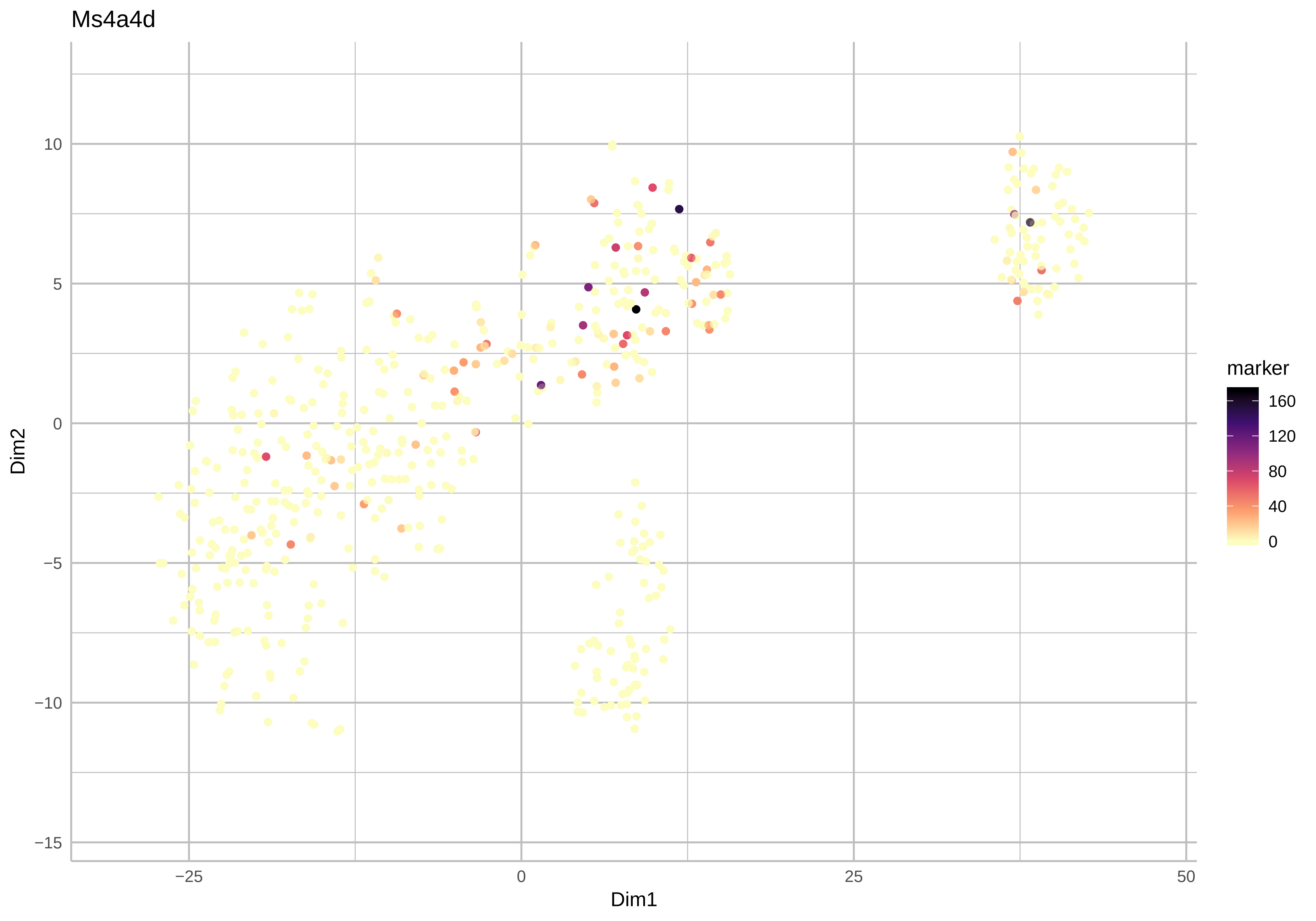
Ms4a4d
Dim2
Dim1
10
5
0
−5
−10
−15
−25
0
25
50
marker
160
120
80
40
0

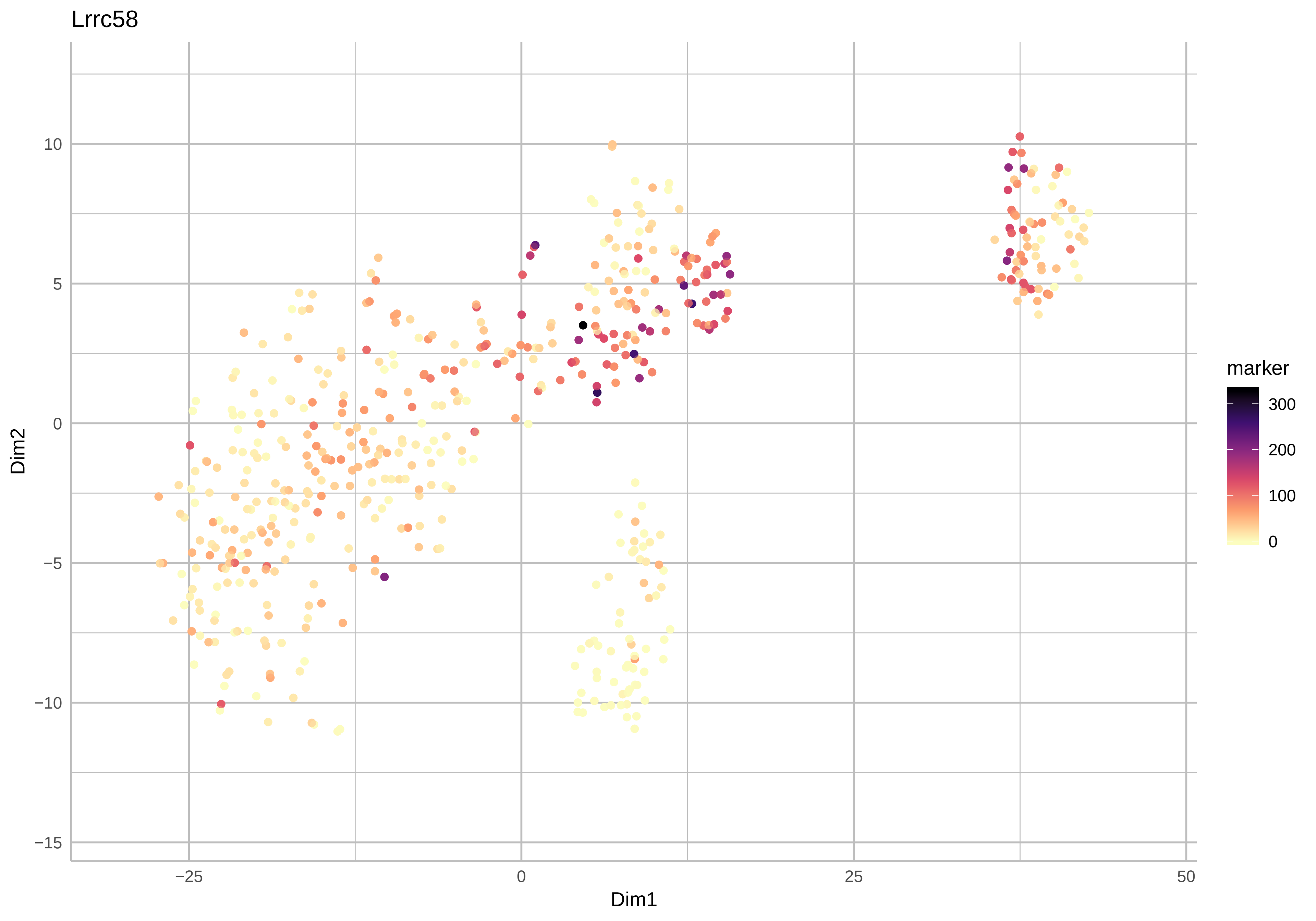

Lrrc58
Dim2
Dim1
10
5
0
−5
−10
−15
−25
0
25
50
marker
300
200
100
0

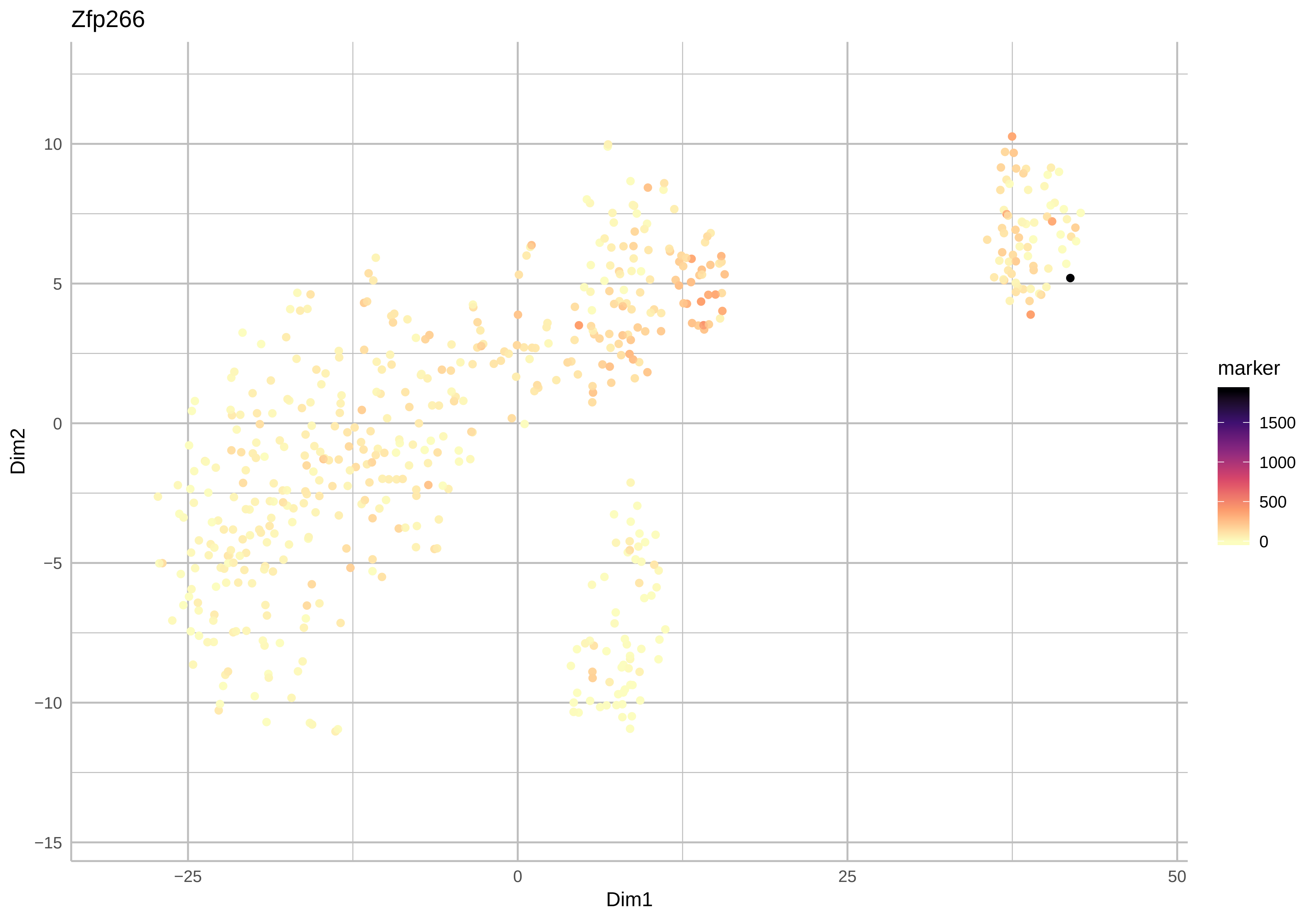

Zfp266
Dim2
Dim1
10
5
0
−5
−10
−15
−25
0
25
50
marker
1500
1000
500
0

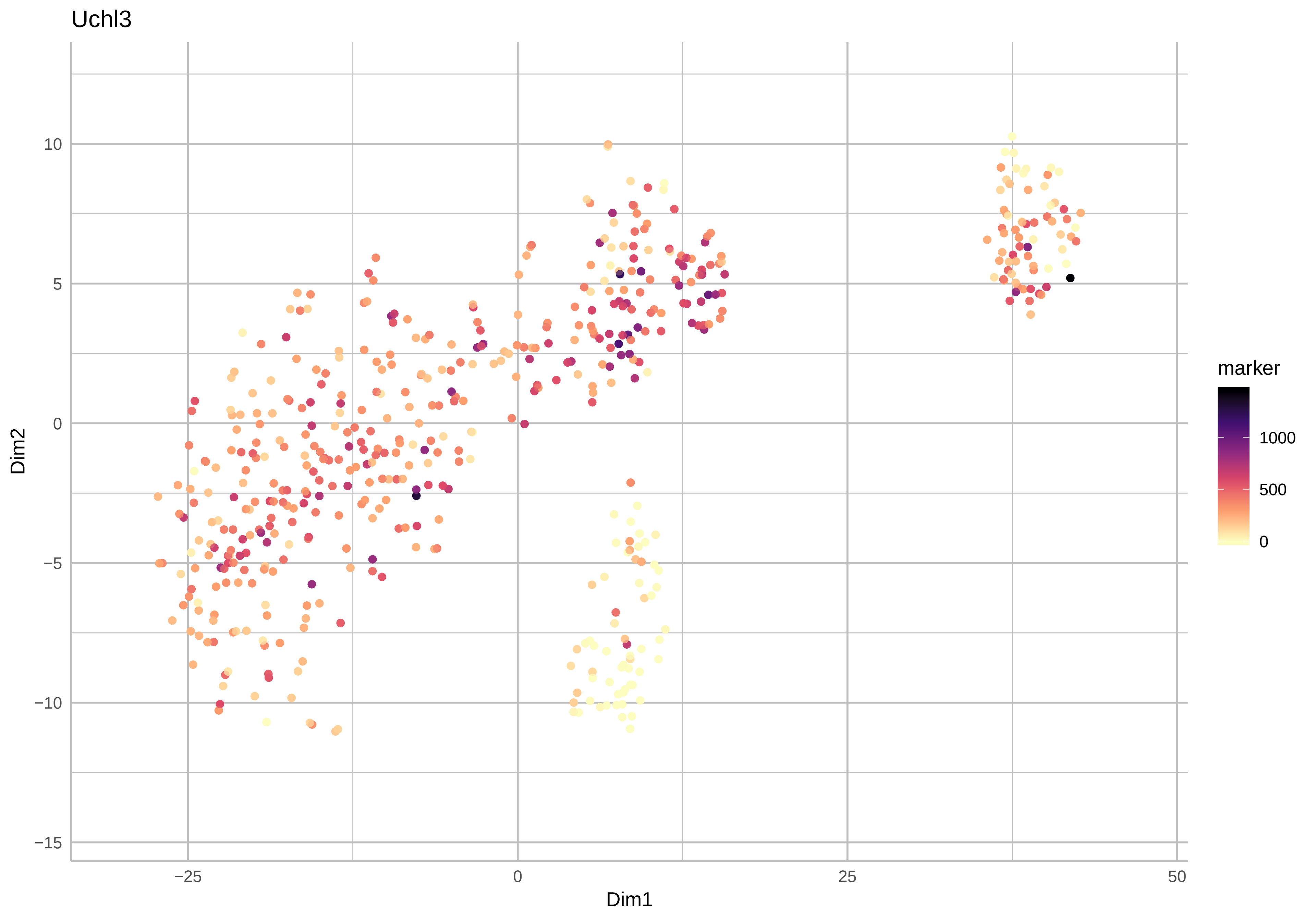
Uchl3
Dim2
Dim1
10
5
0
−5
−10
−15
−25
0
25
50
marker
1000
500
0

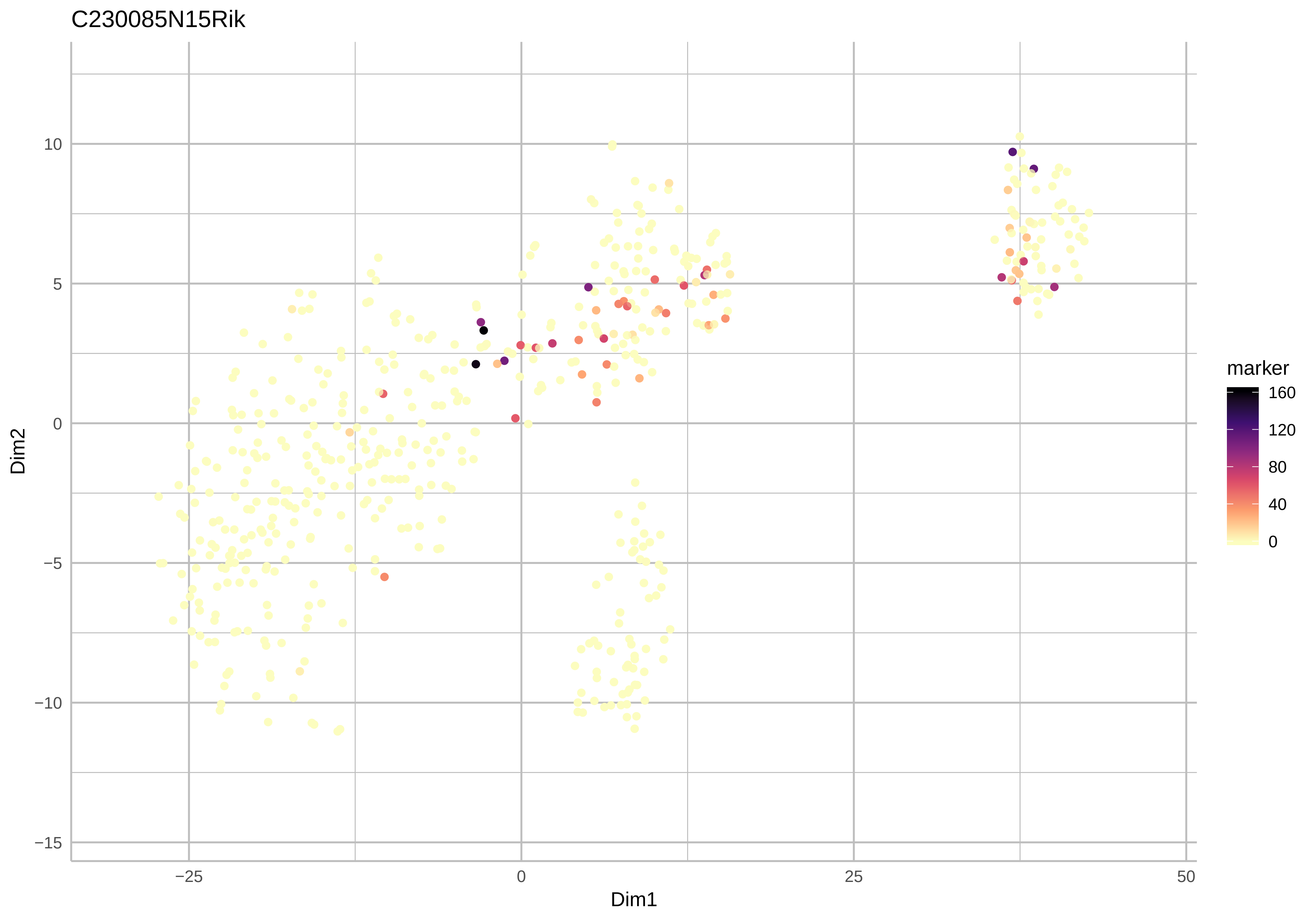
C230085N15Rik
Dim2
Dim1
10
5
0
−5
−10
−15
−25
0
25
50
marker
160
120
80
40
0

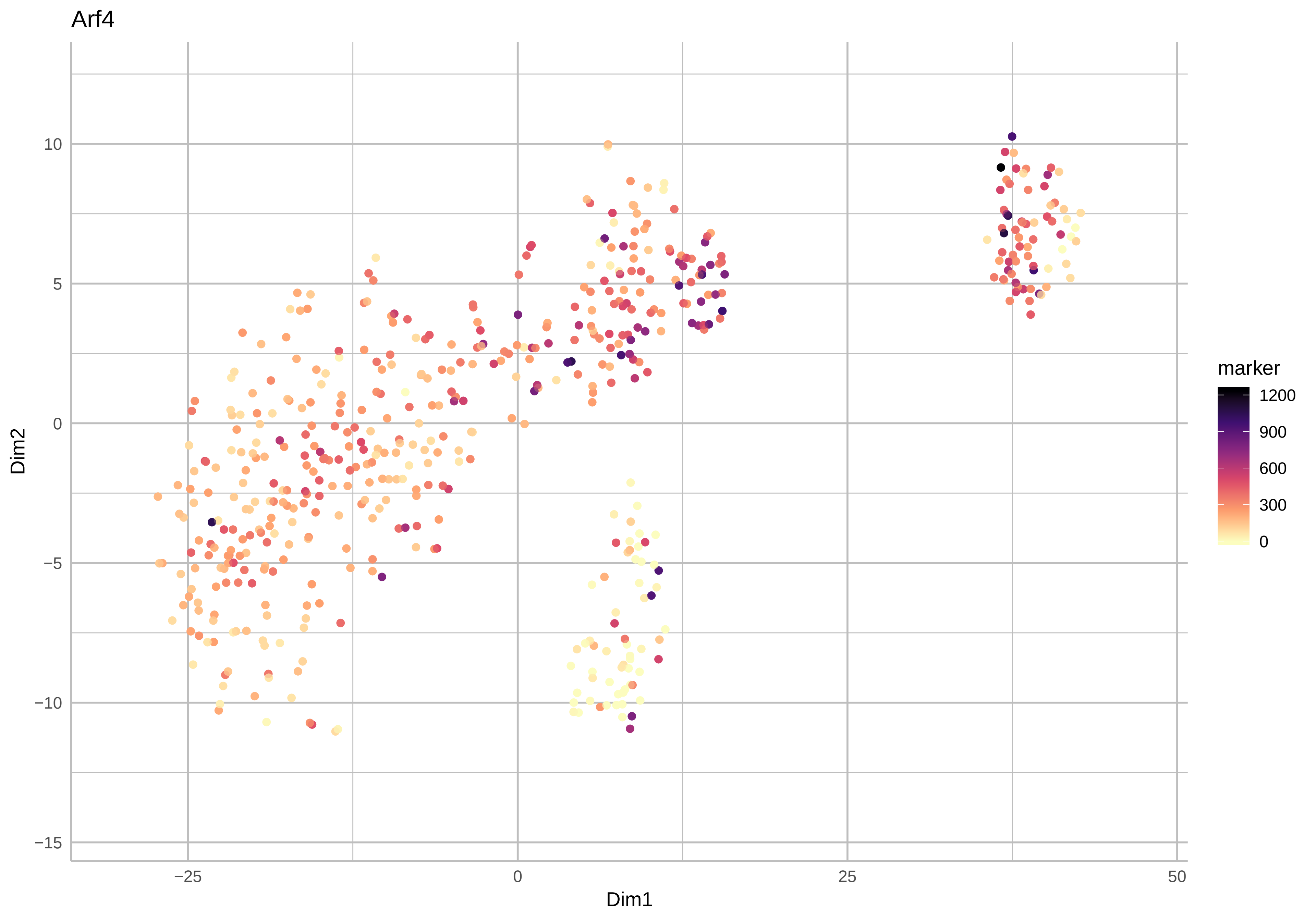
Arf4
Dim2
Dim1
10
5
0
−5
−10
−15
−25
0
25
50
marker
1200
900
600
300
0

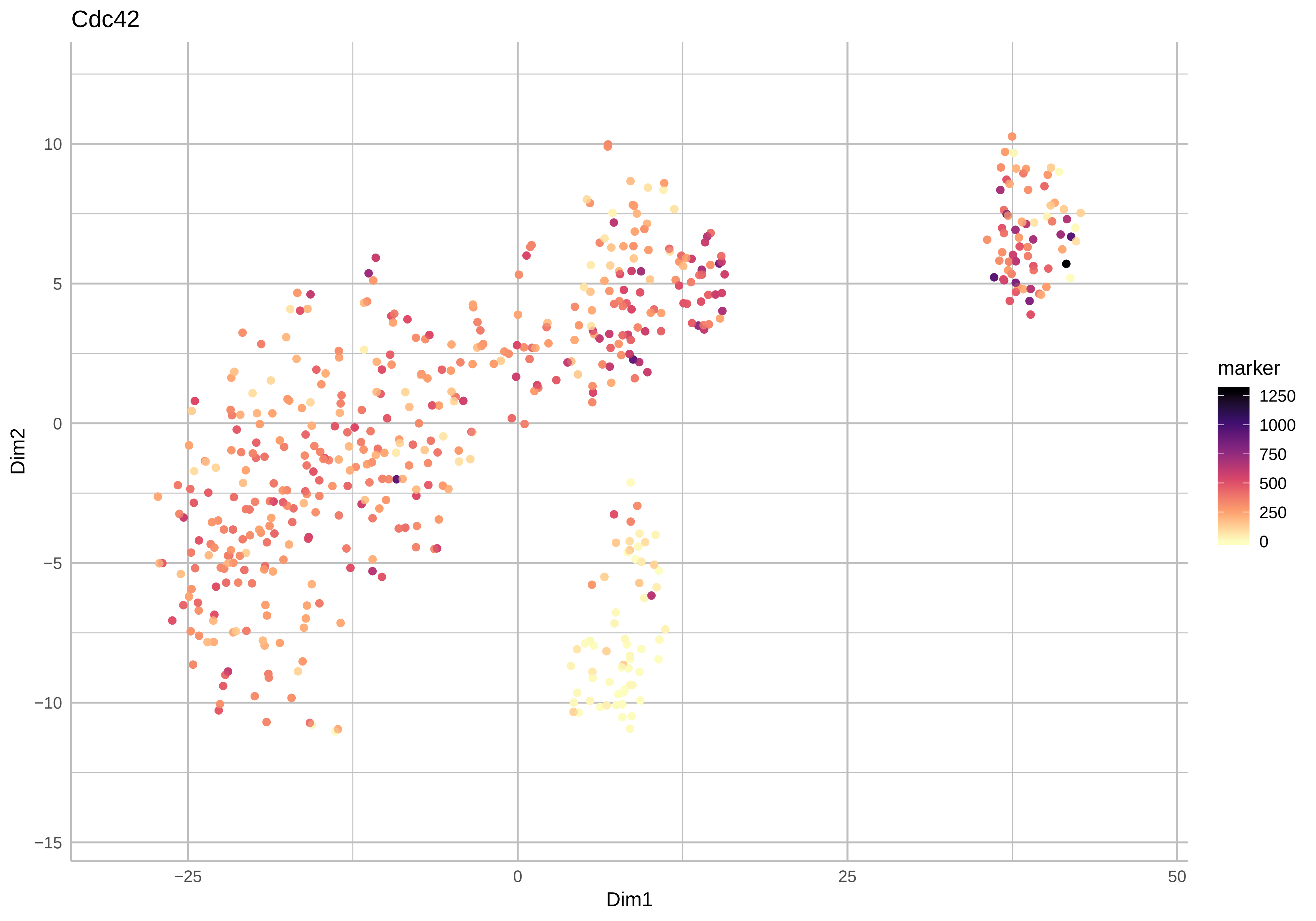

Cdc42
Dim2
Dim1
10
5
0
−5
−10
−15
−25
0
25
50
marker
1250
1000
750
500
250
0

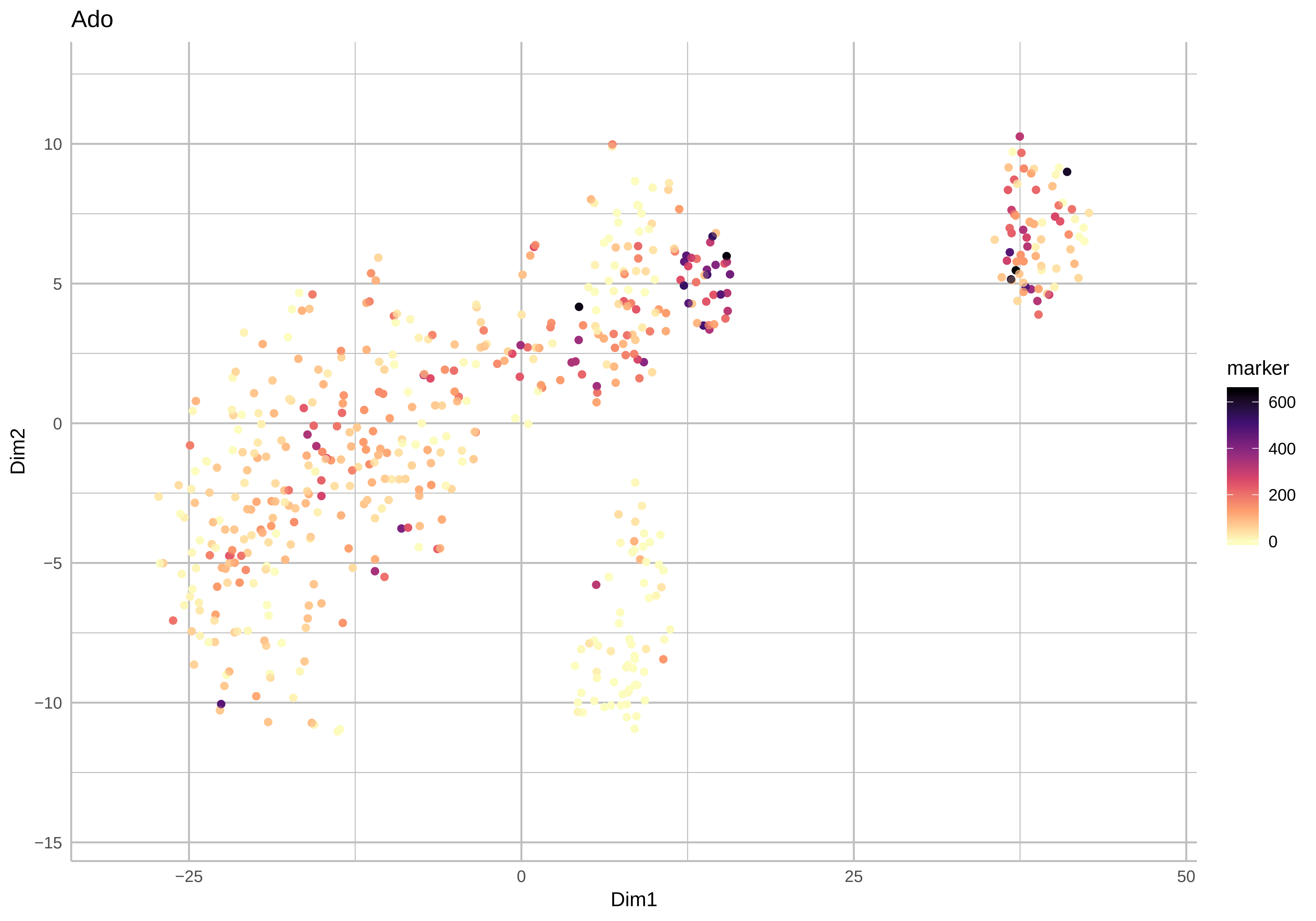

Ado
Dim2
10
5
0
−5
−10
−15
−25
0
25
50
Dim1
marker
600
400
200
0

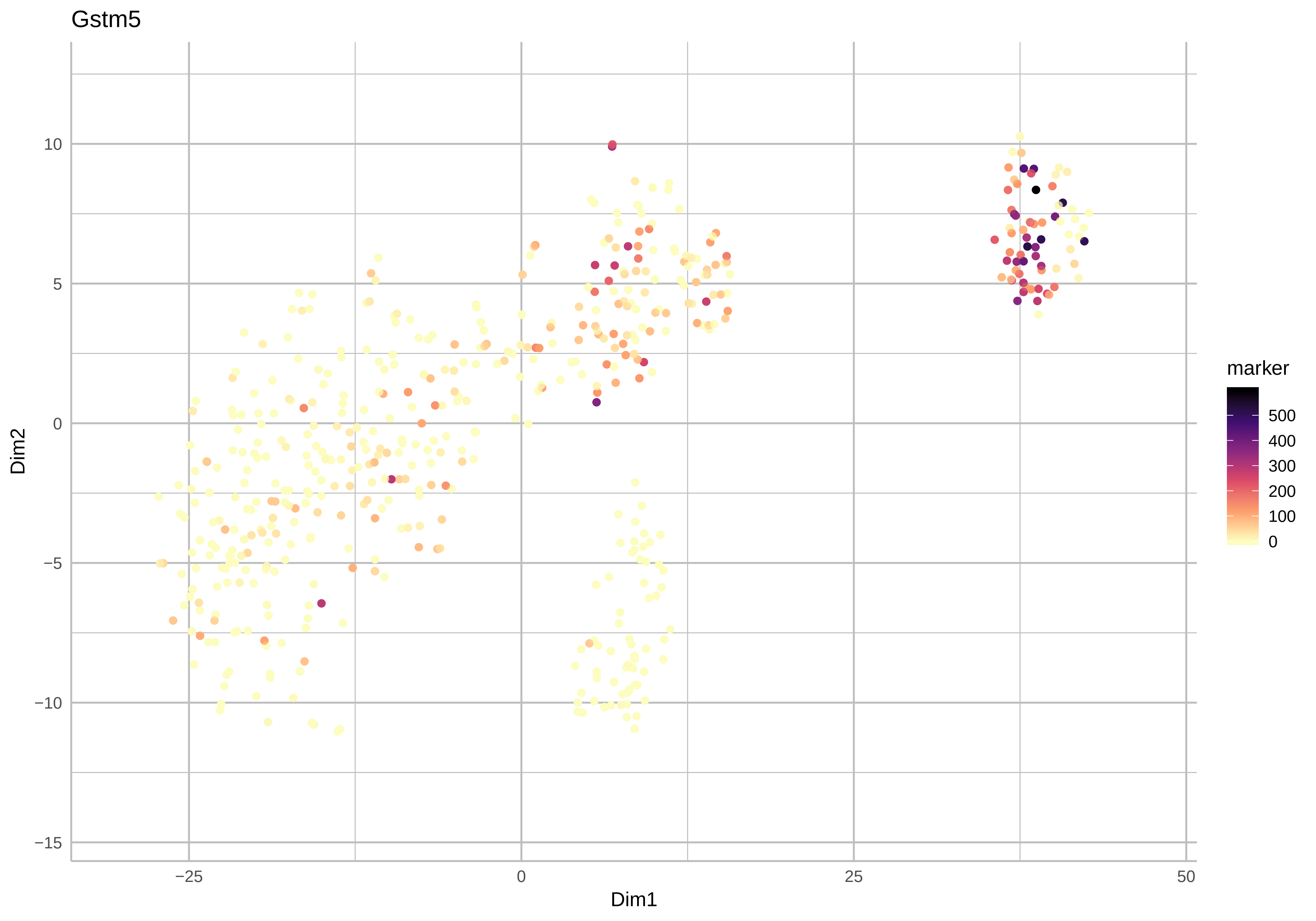

Gstm5
10
5
0
−5
−10
−15
Dim2
−25
0
25
50
Dim1
marker
500
400
300
200
100
0

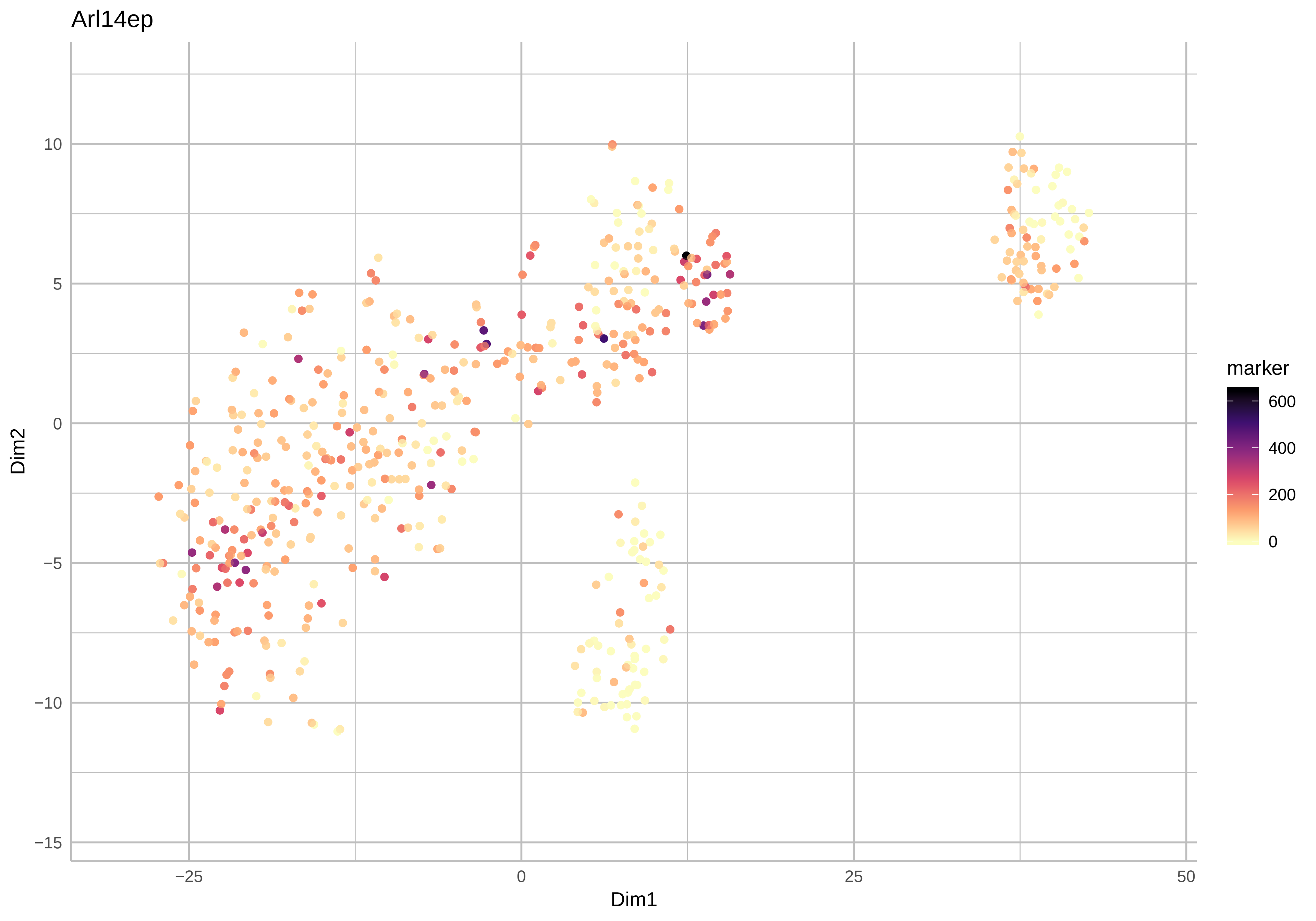

Arl14ep
10
5
0
−5
−10
−15
−25
0
25
50
Dim1
Dim2
marker
600
400
200
0

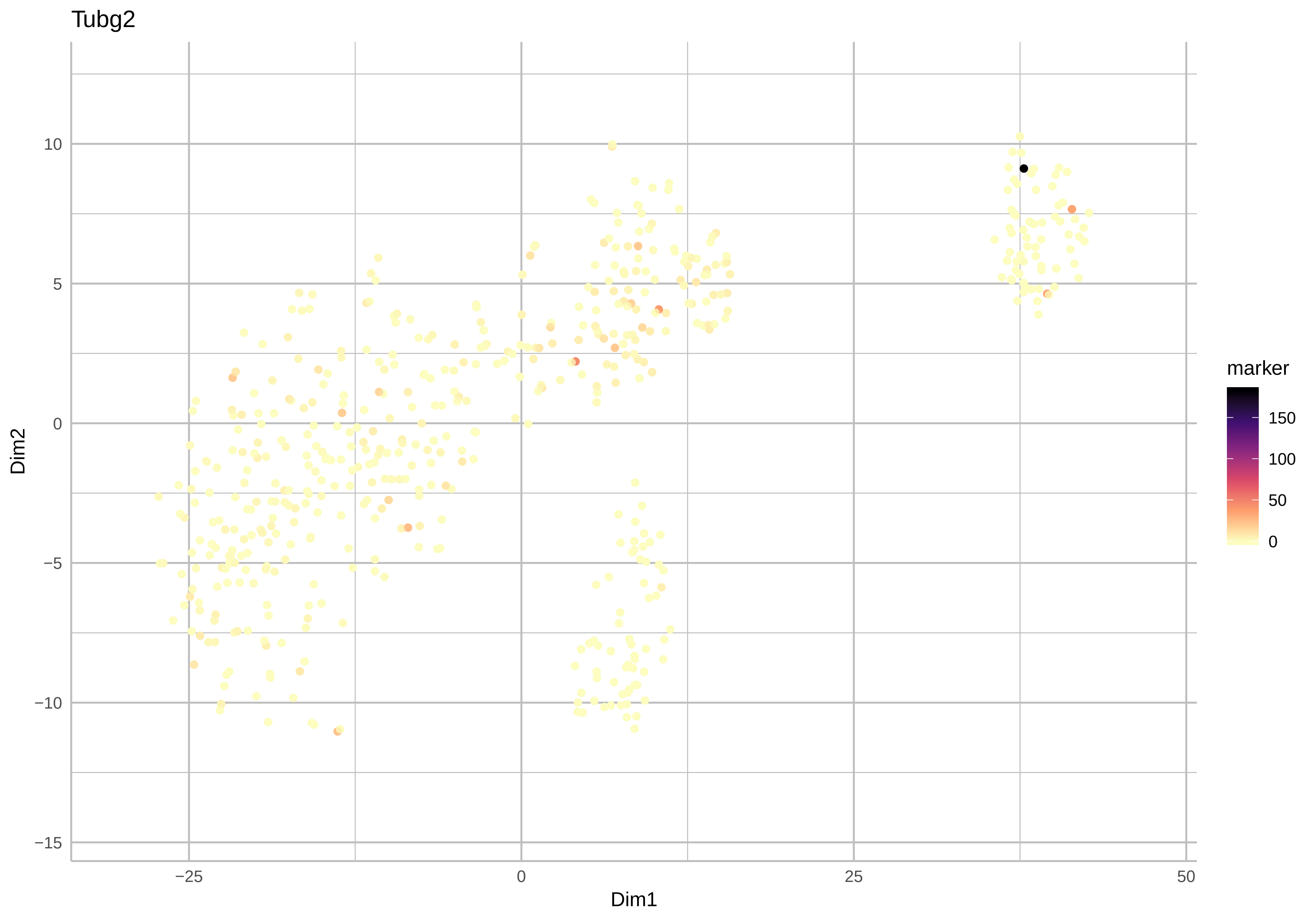
Tubg2
Dim2
Dim1
10
5
0
−5
−10
−15
−25
0
25
50
marker
150
100
50
0

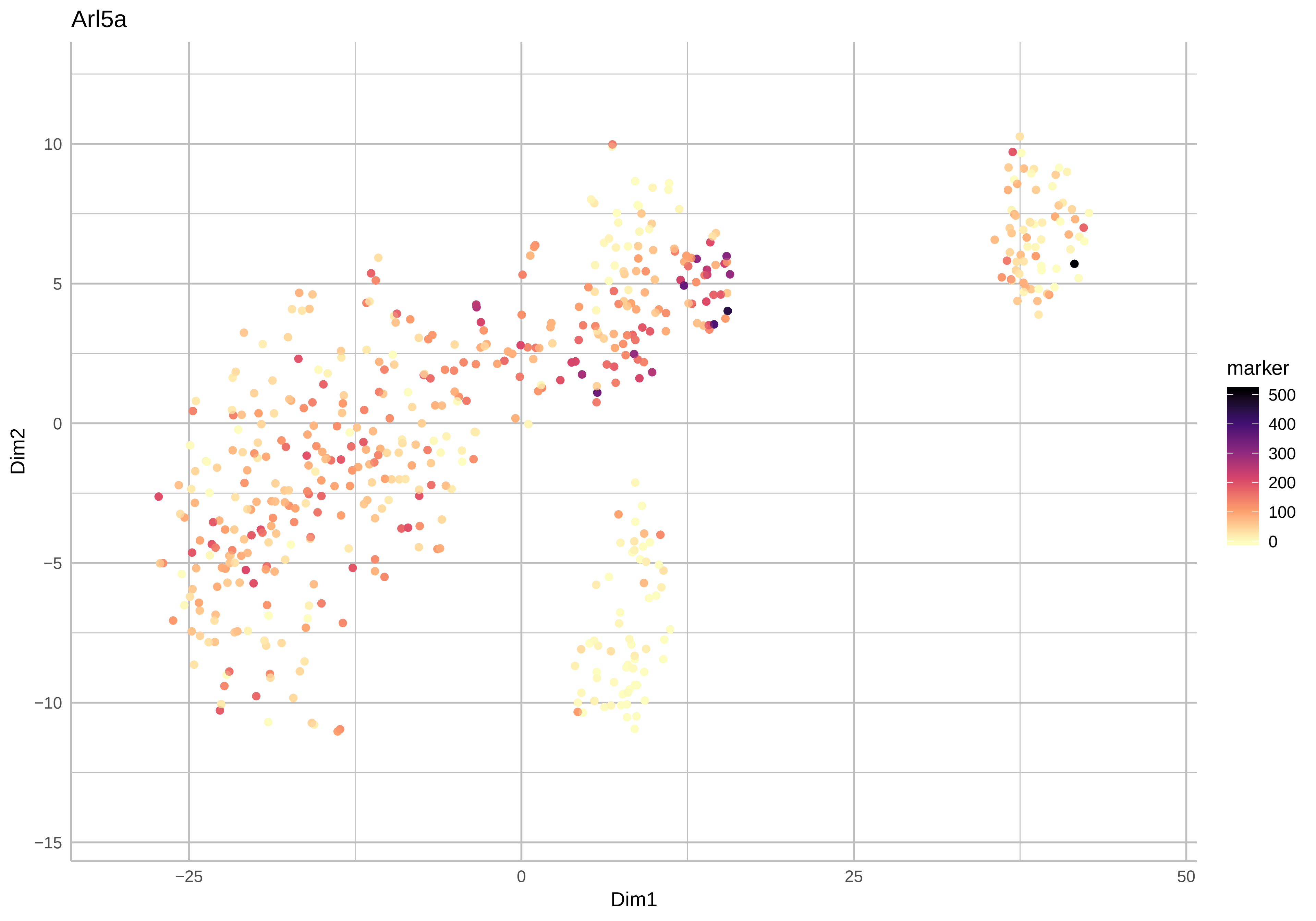
Arl5a
Dim2
Dim1
10
5
0
−5
−10
−15
−25
0
25
50
marker
500
400
300
200
100
0

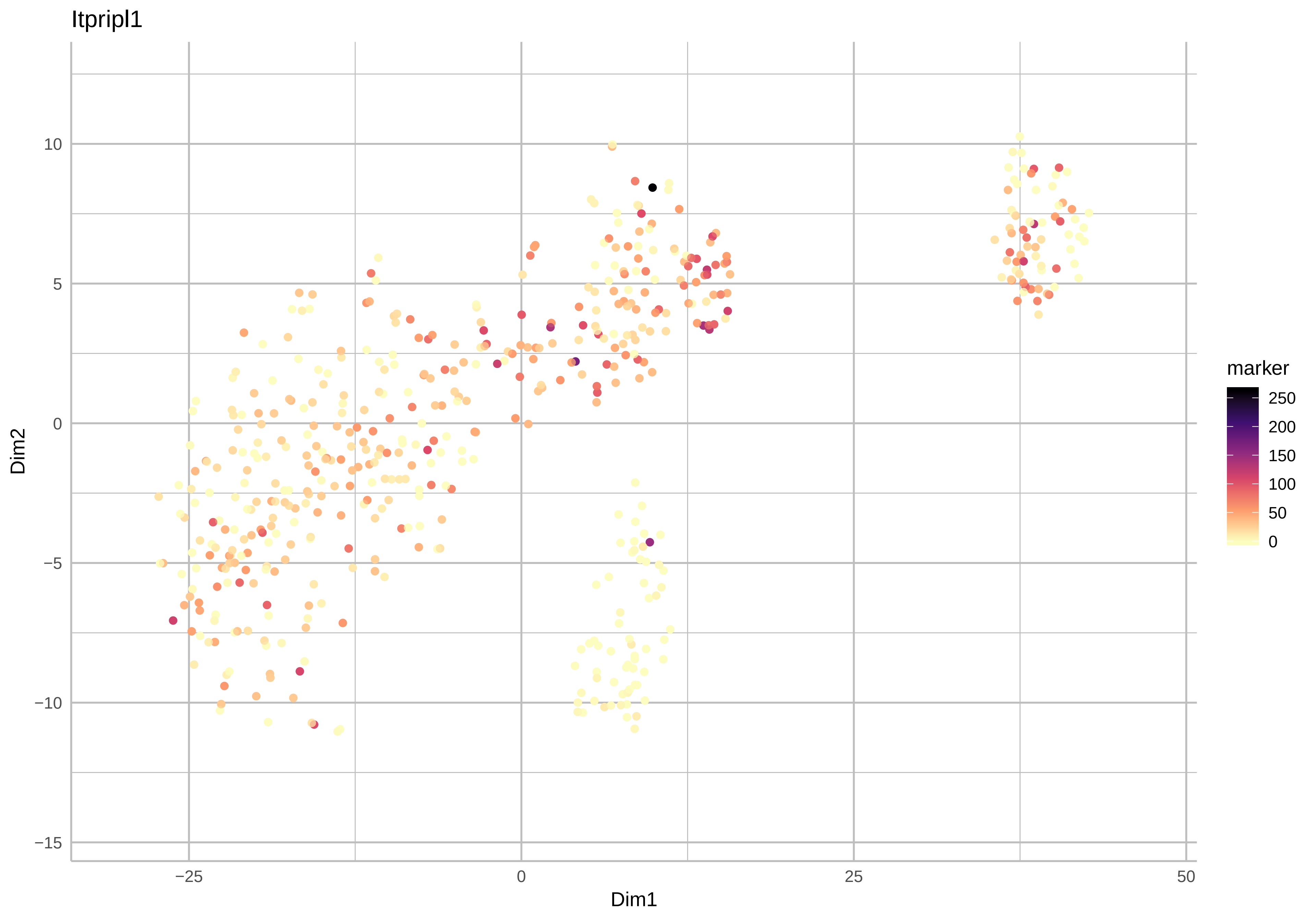

Itpripl1
Dim2
Dim1
marker
250
200
150
100
50
0
10
5
0
−5
−10
−15
−25
0
25
50

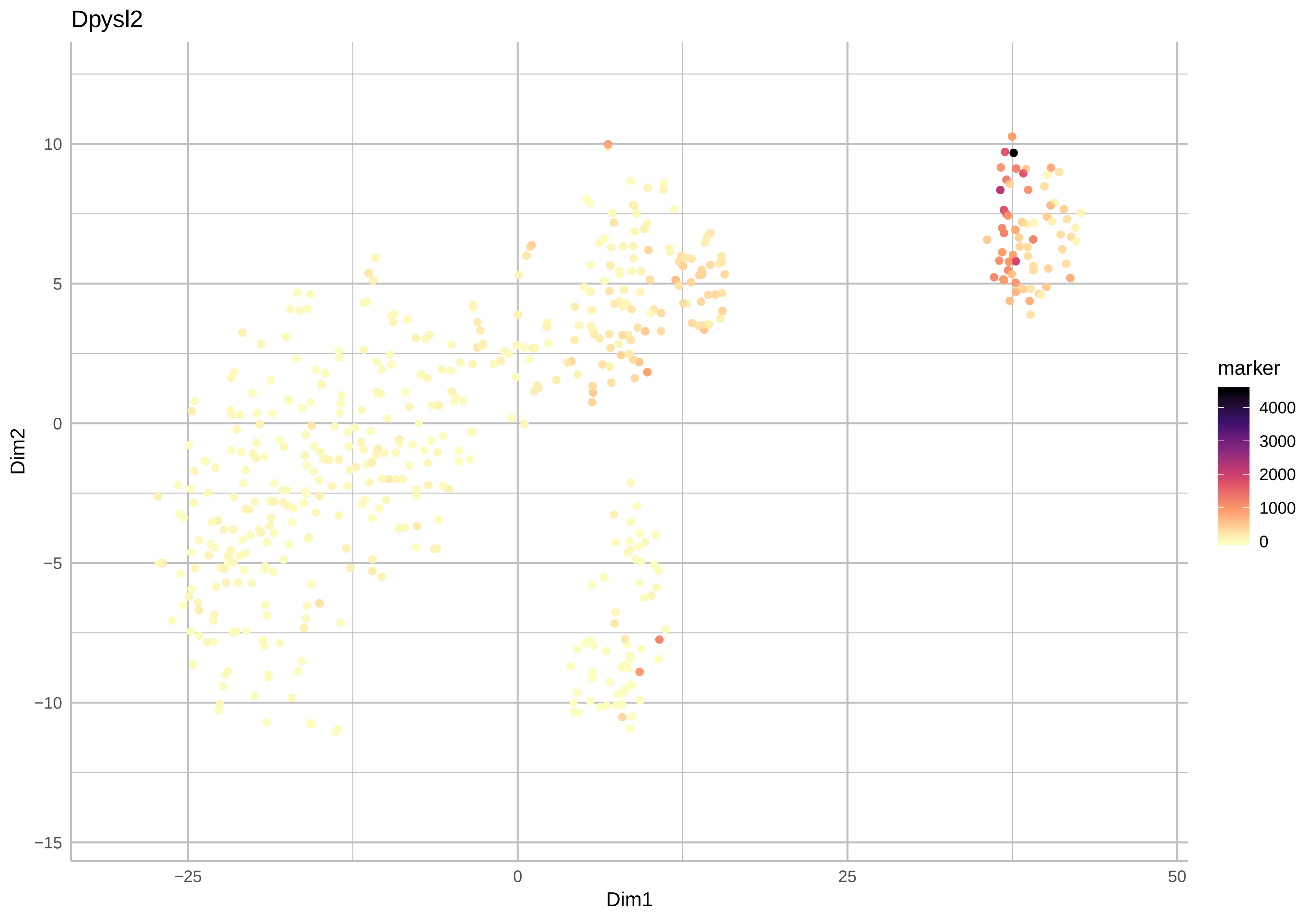

Dpysl2
Dim2
Dim1
10
5
0
−5
−10
−15
−25
0
25
50
marker
4000
3000
2000
1000
0

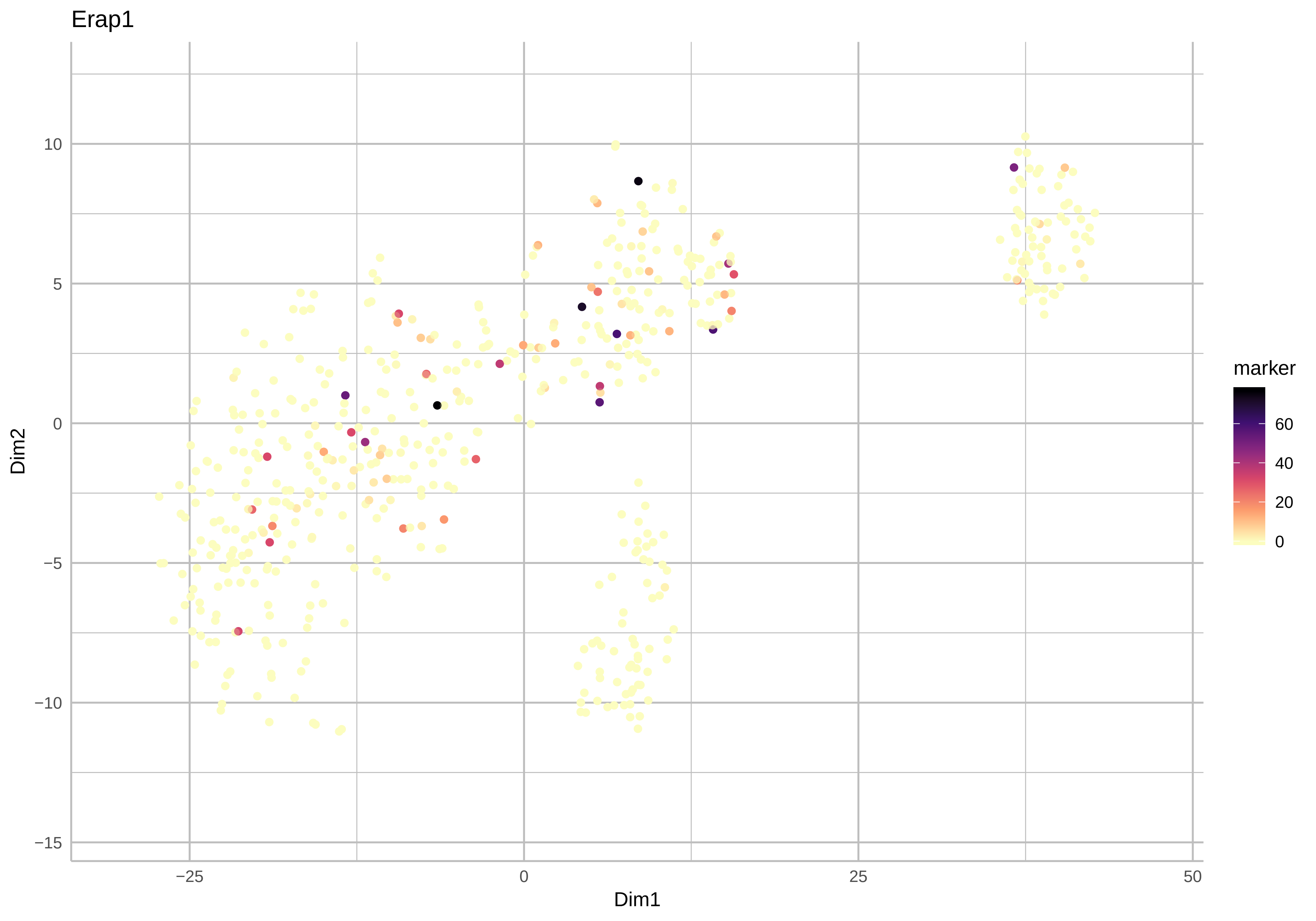

Erap1
Dim2
Dim1
10
5
0
−5
−10
−15
−25
0
25
50
marker
60
40
20
0

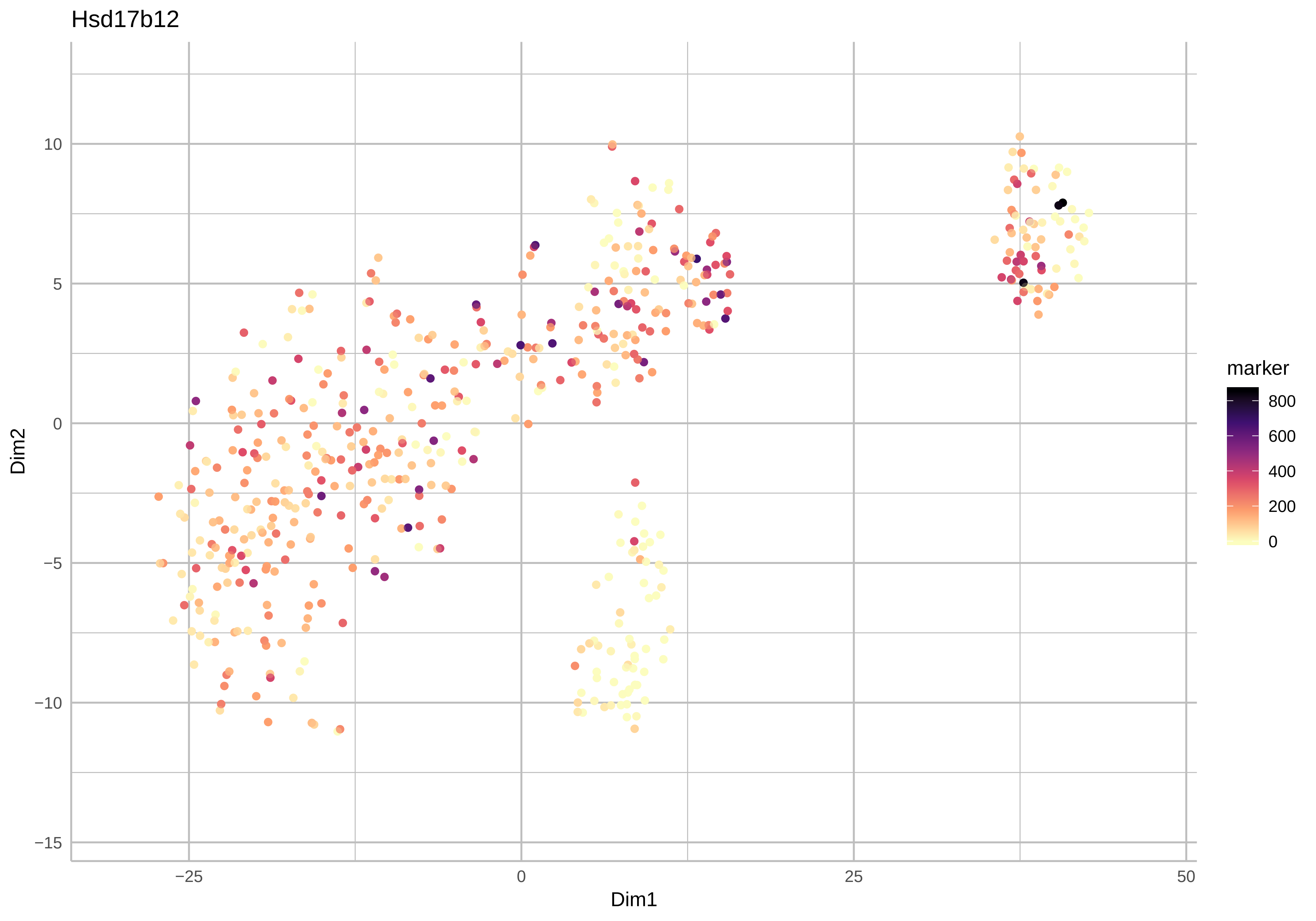

Hsd17b12
Dim2
Dim1
10
5
0
−5
−10
−15
−25
0
25
50
marker
800
600
400
200
0

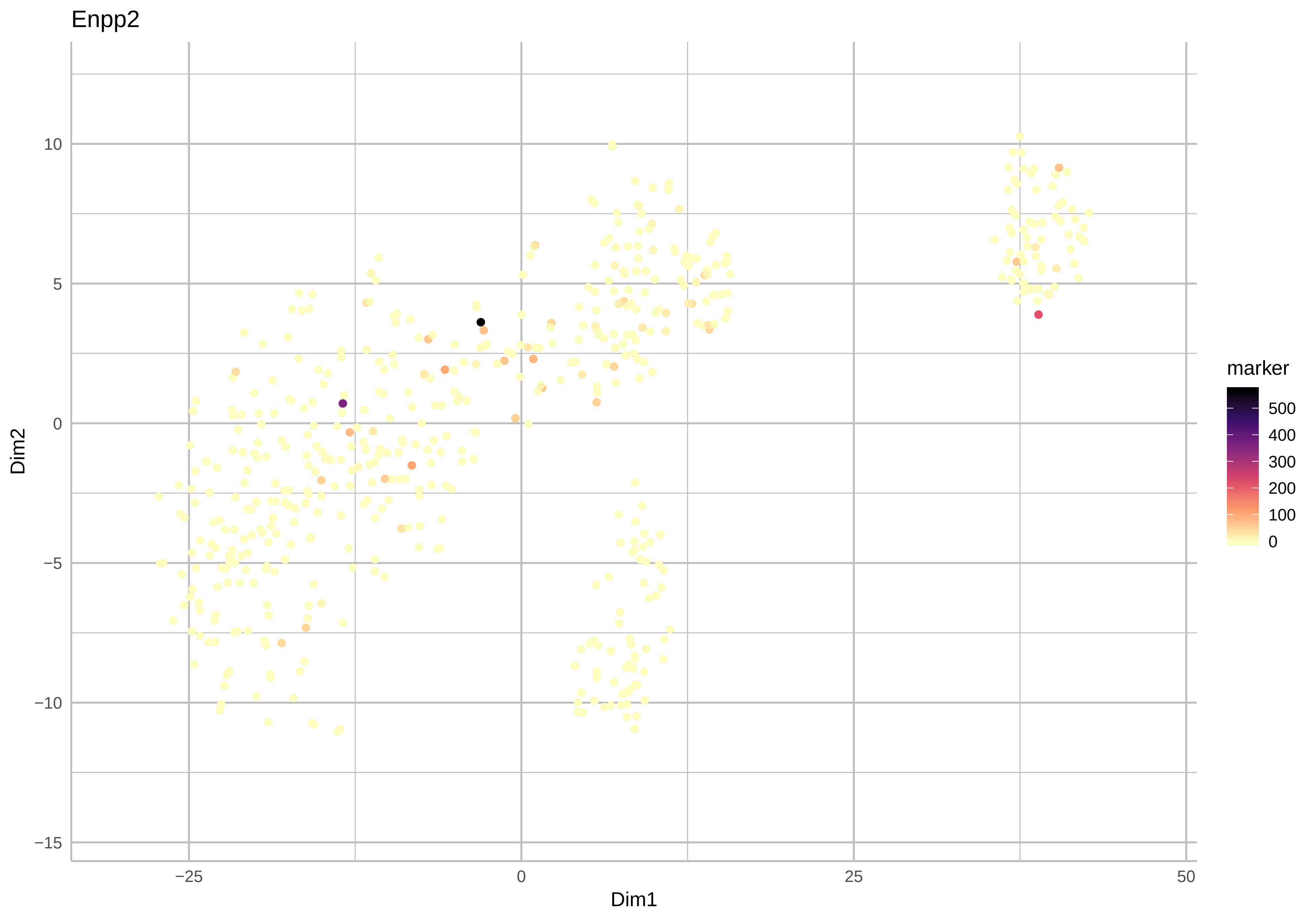
Enpp2
10
5
0
−5
−10
−15
Dim2
−25
0
25
50
Dim1
marker
500
400
300
200
100
0

Lrrc8a

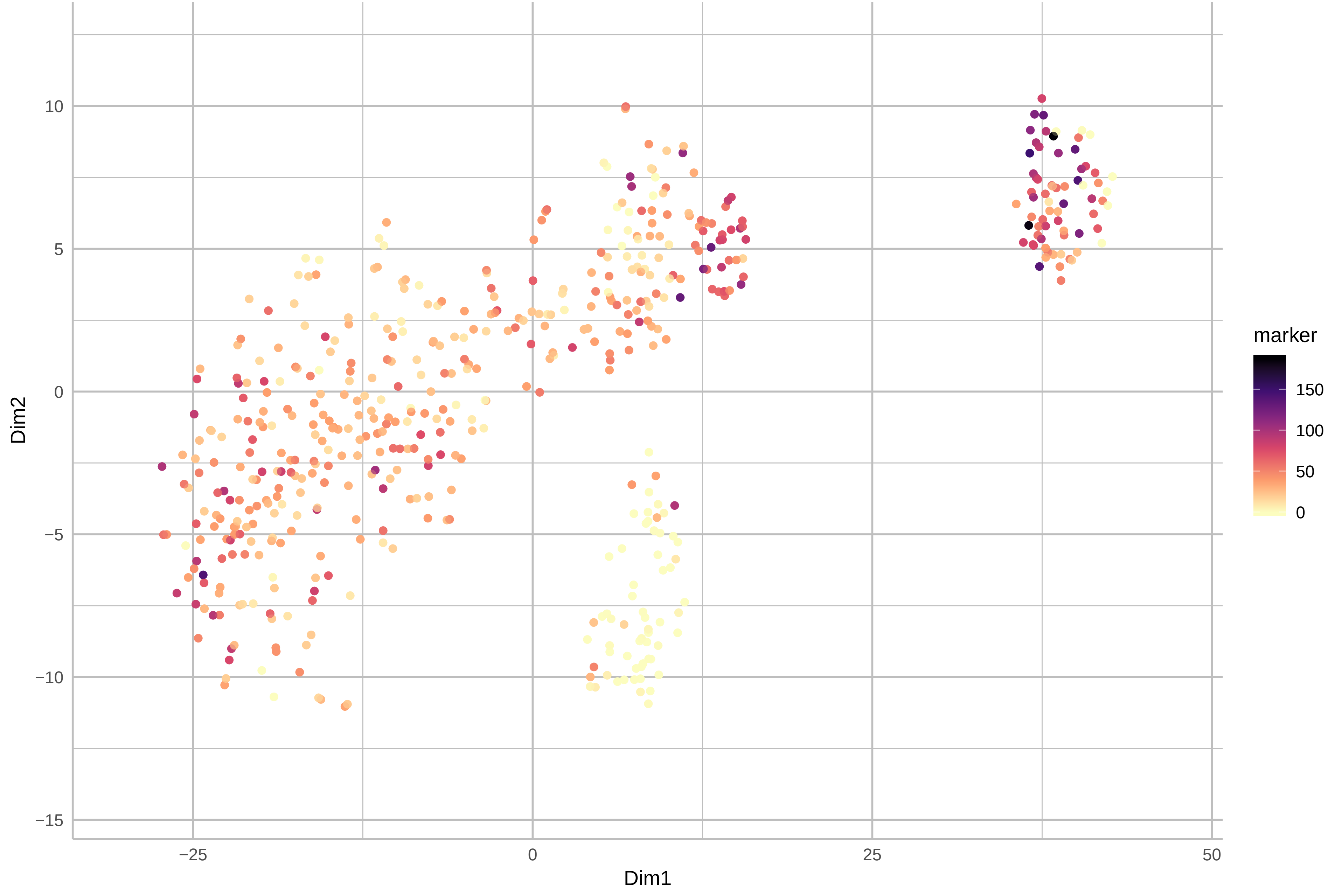

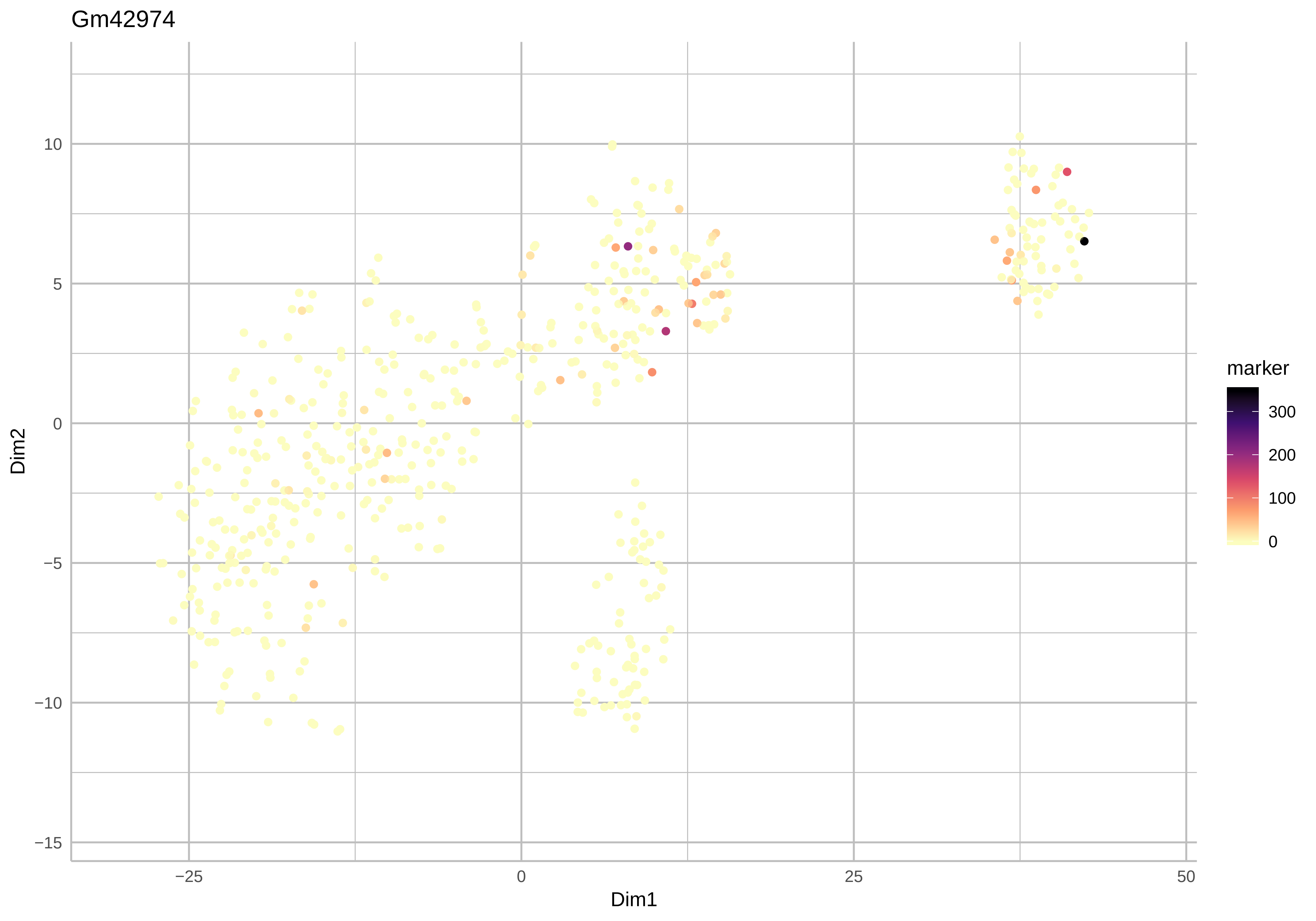

Gm42974
Dim2
Dim1
10
5
0
−5
−10
−15
−25
0
25
50
marker
300
200
100
0

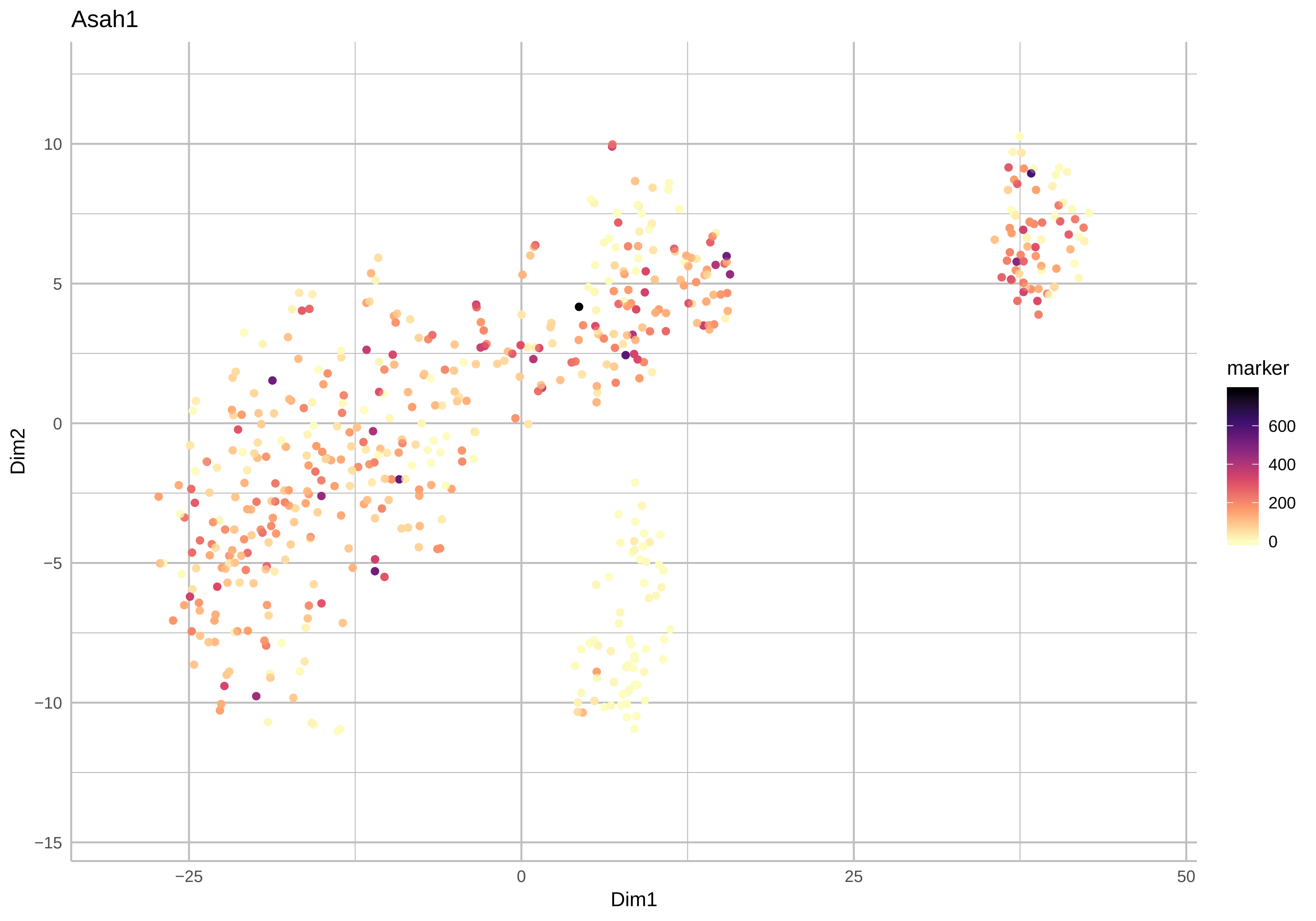
Asah1
Dim2
Dim1
marker
600
400
200
0
10
5
0
−5
−10
−15
−25
0
25
50

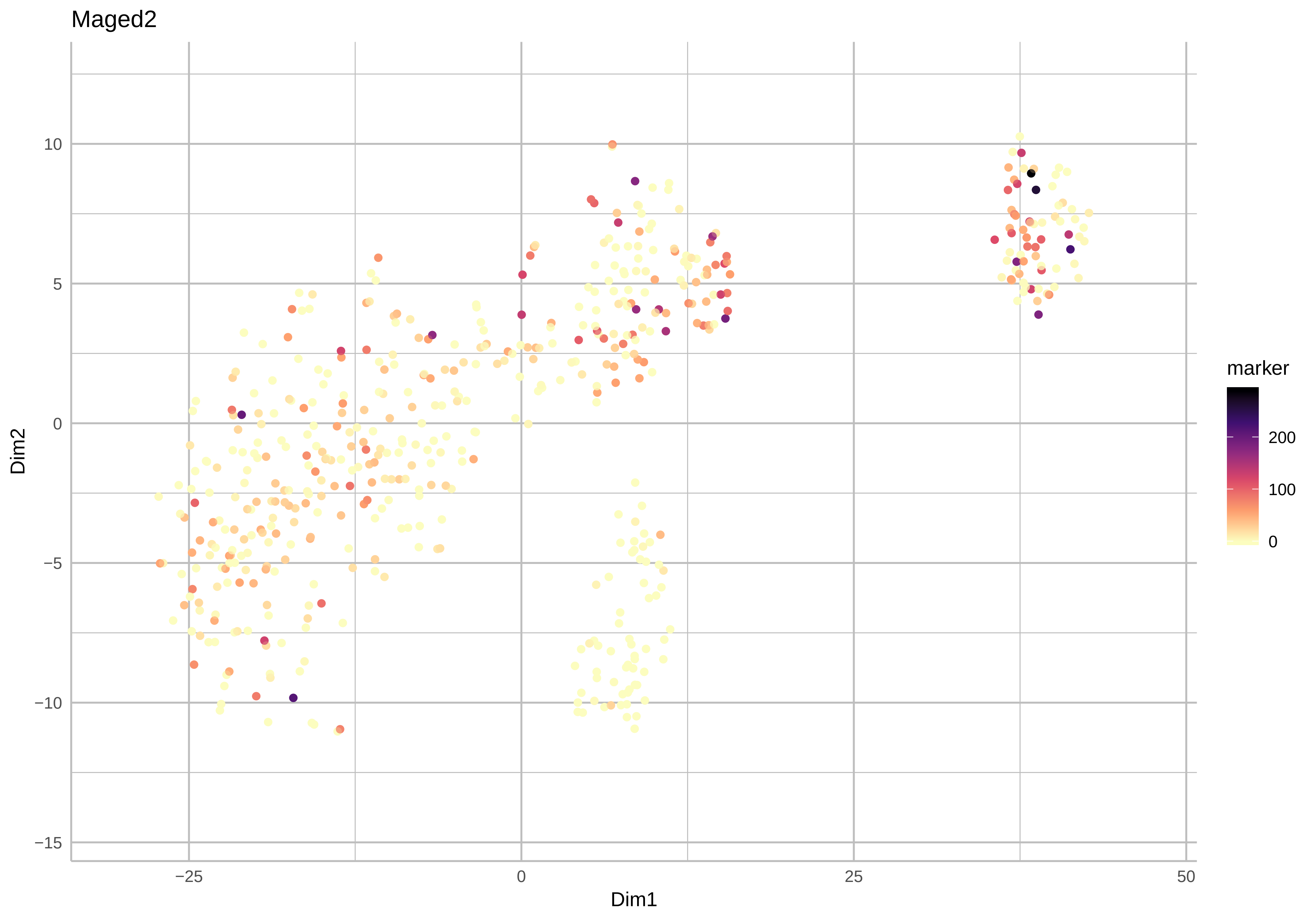
Maged2
Dim2
Dim1
10
5
0
−5
−10
−15
−25
0
25
50
marker
200
100
0

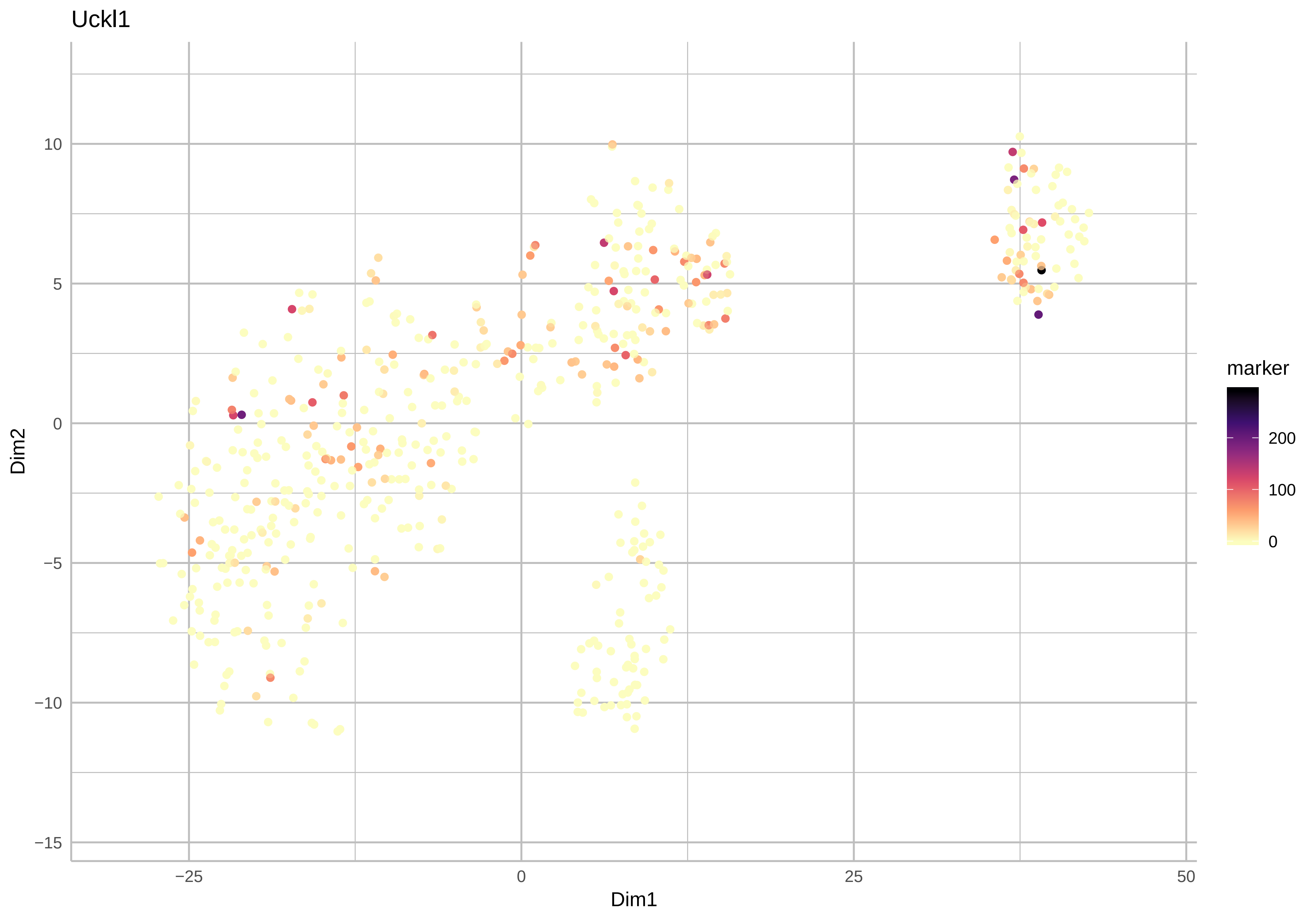
Uckl1
Dim2
Dim1
10
5
0
−5
−10
−15
−25
0
25
50
marker
200
100
0

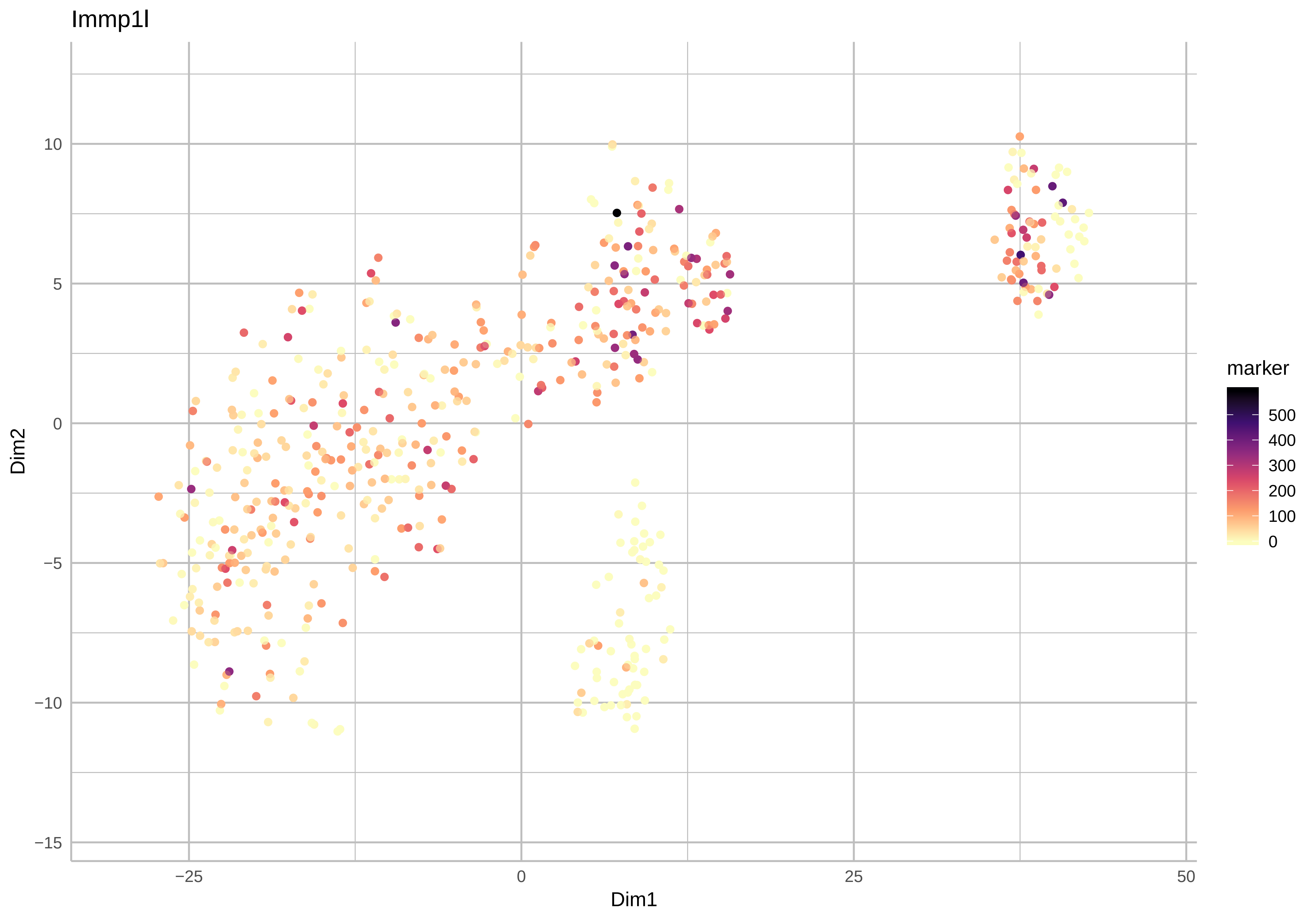

Immp1l
10
5
0
−5
−10
−15
Dim2
−25
0
25
50
Dim1
marker
500
400
300
200
100
0

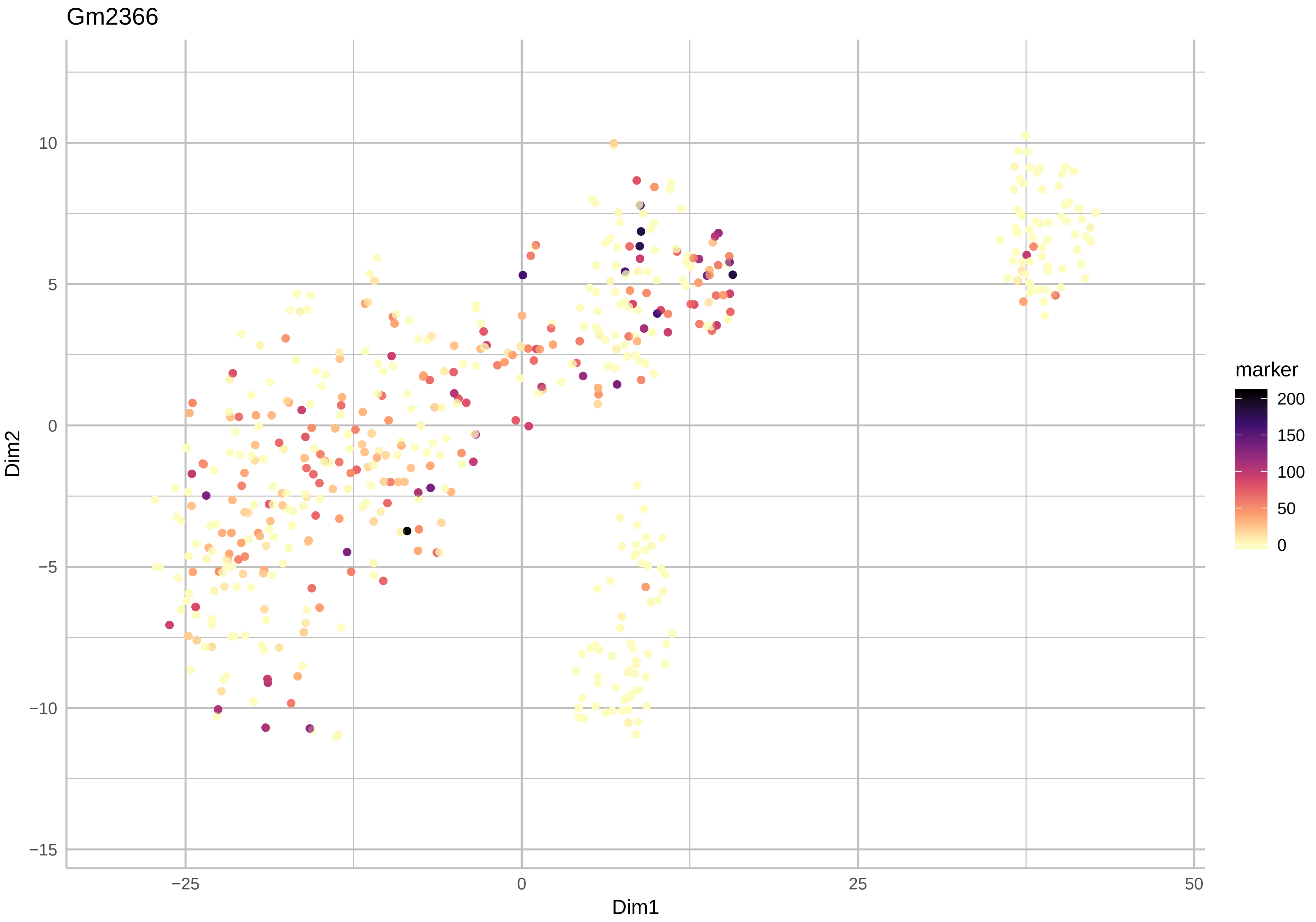
Gm2366
10
5
0
−5
−10
−15
Dim2
−25
0
25
50
Dim1
marker
200
150
100
50
0

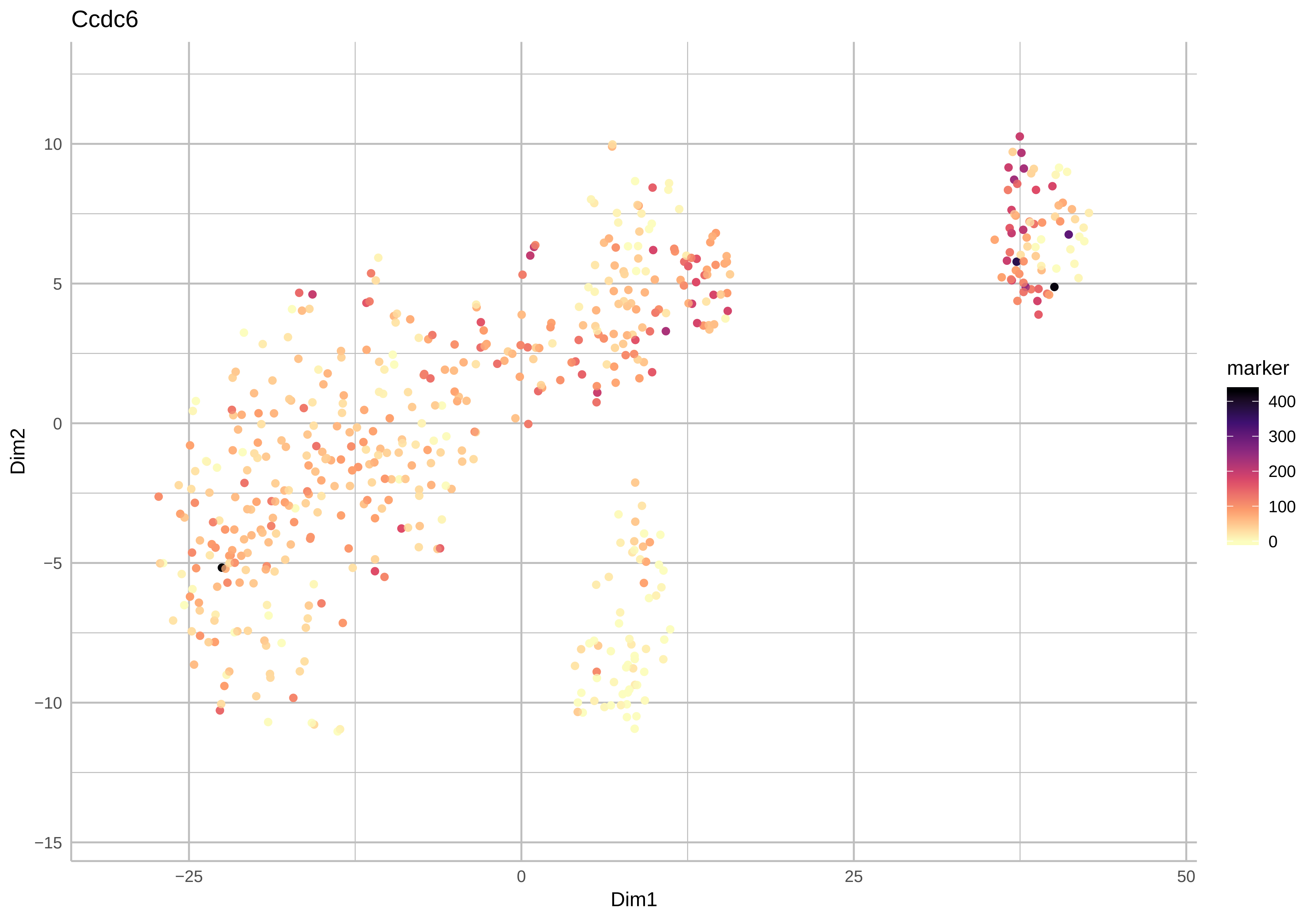

Ccdc6
Dim2
Dim1
marker
400
300
200
100
0
10
5
0
−5
−10
−15
−25
0
25
50

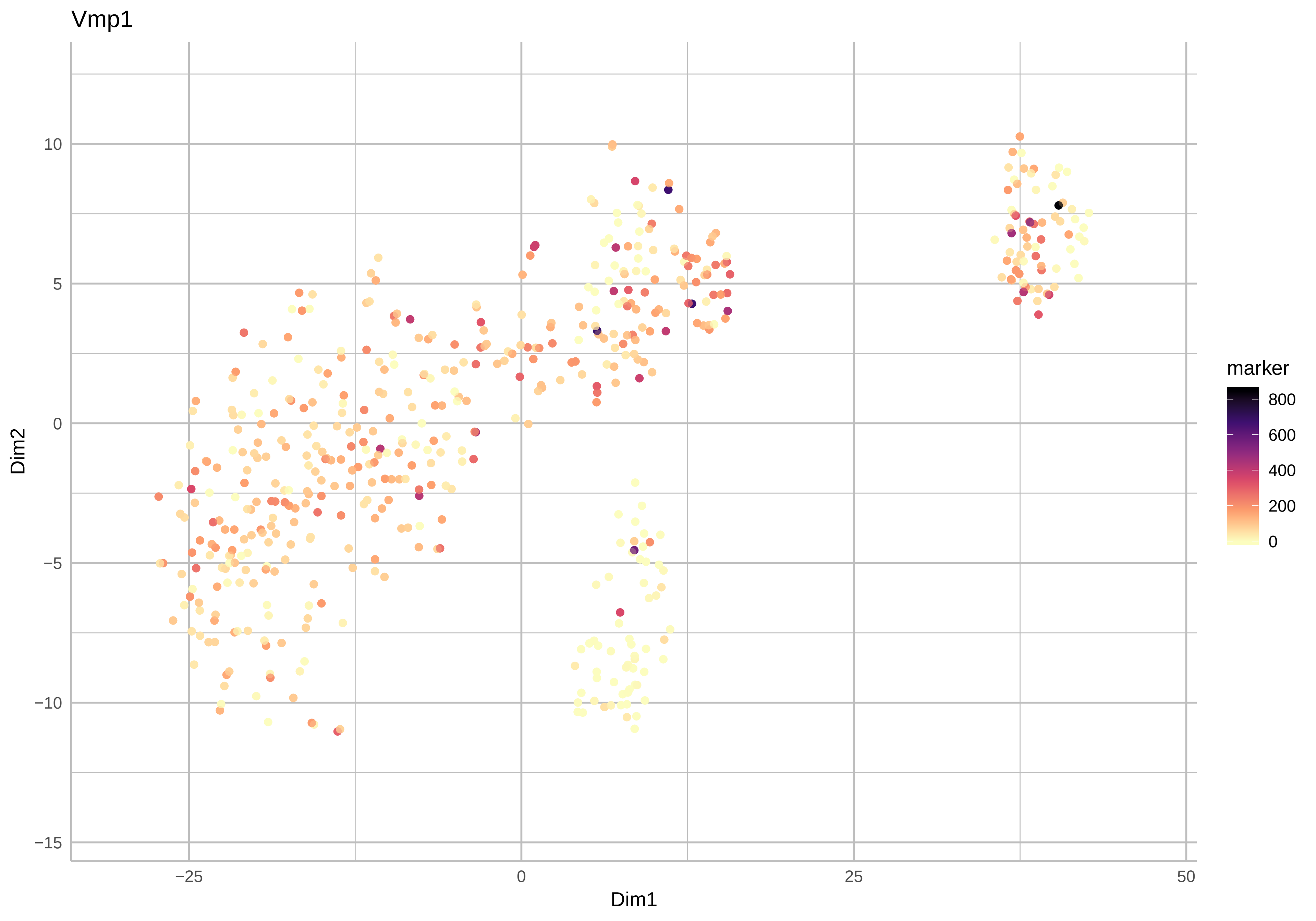

Vmp1
Dim2
Dim1
10
5
0
−5
−10
−15
−25
0
25
50
marker
800
600
400
200
0

Rab31

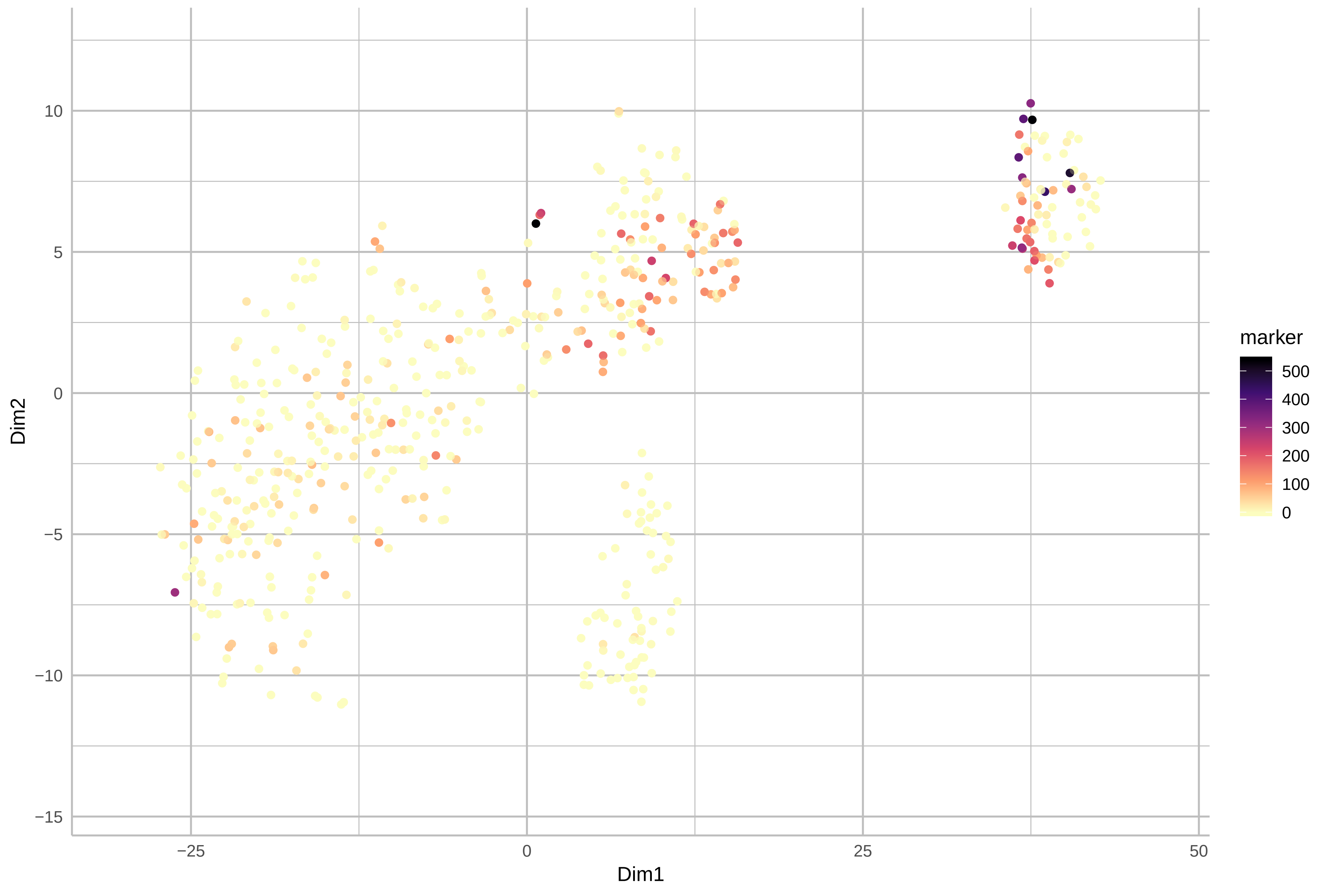

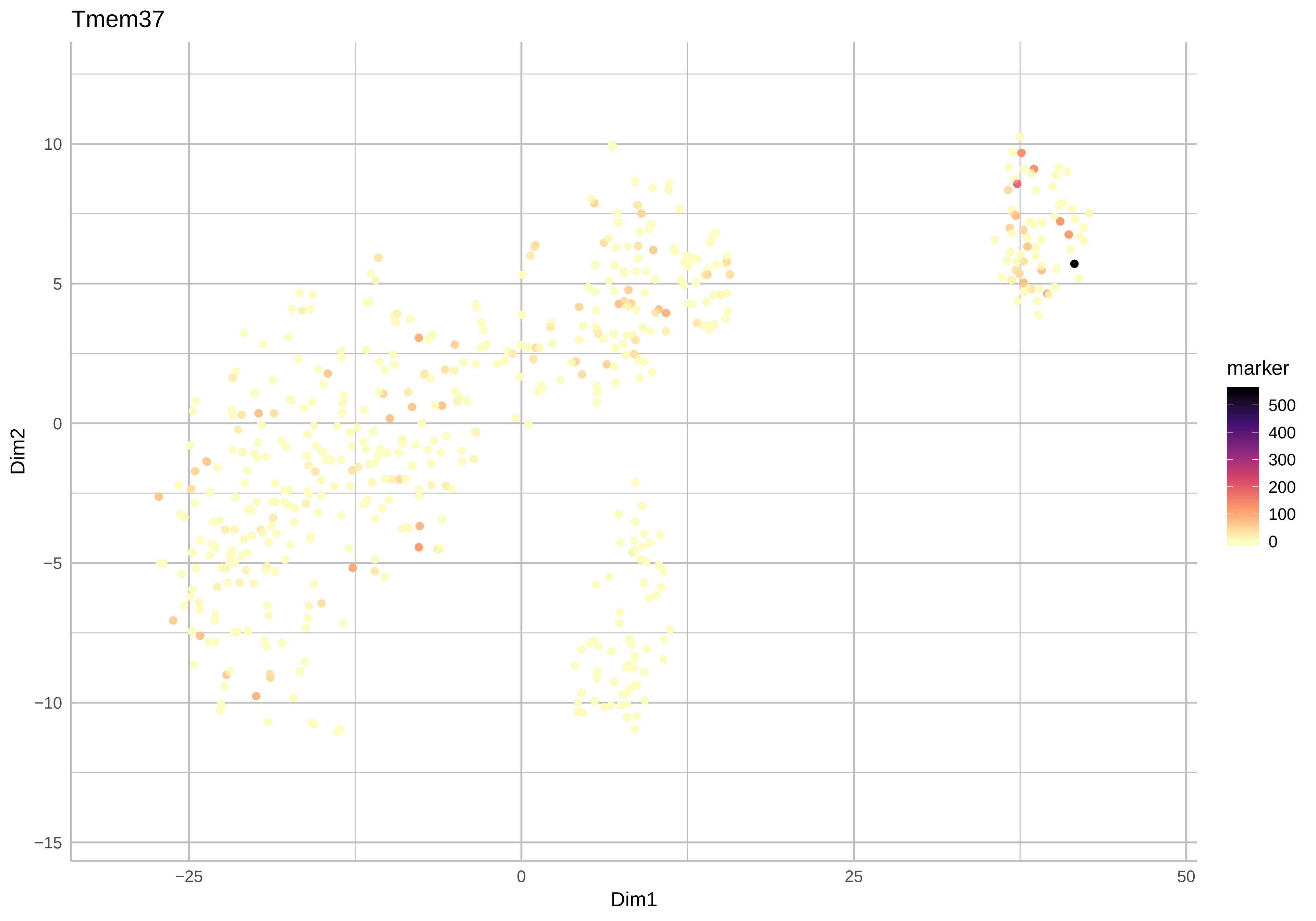

Tmem37
Dim2
Dim1
10
5
0
−5
−10
−15
−25
0
25
50
marker
500
400
300
200
100
0

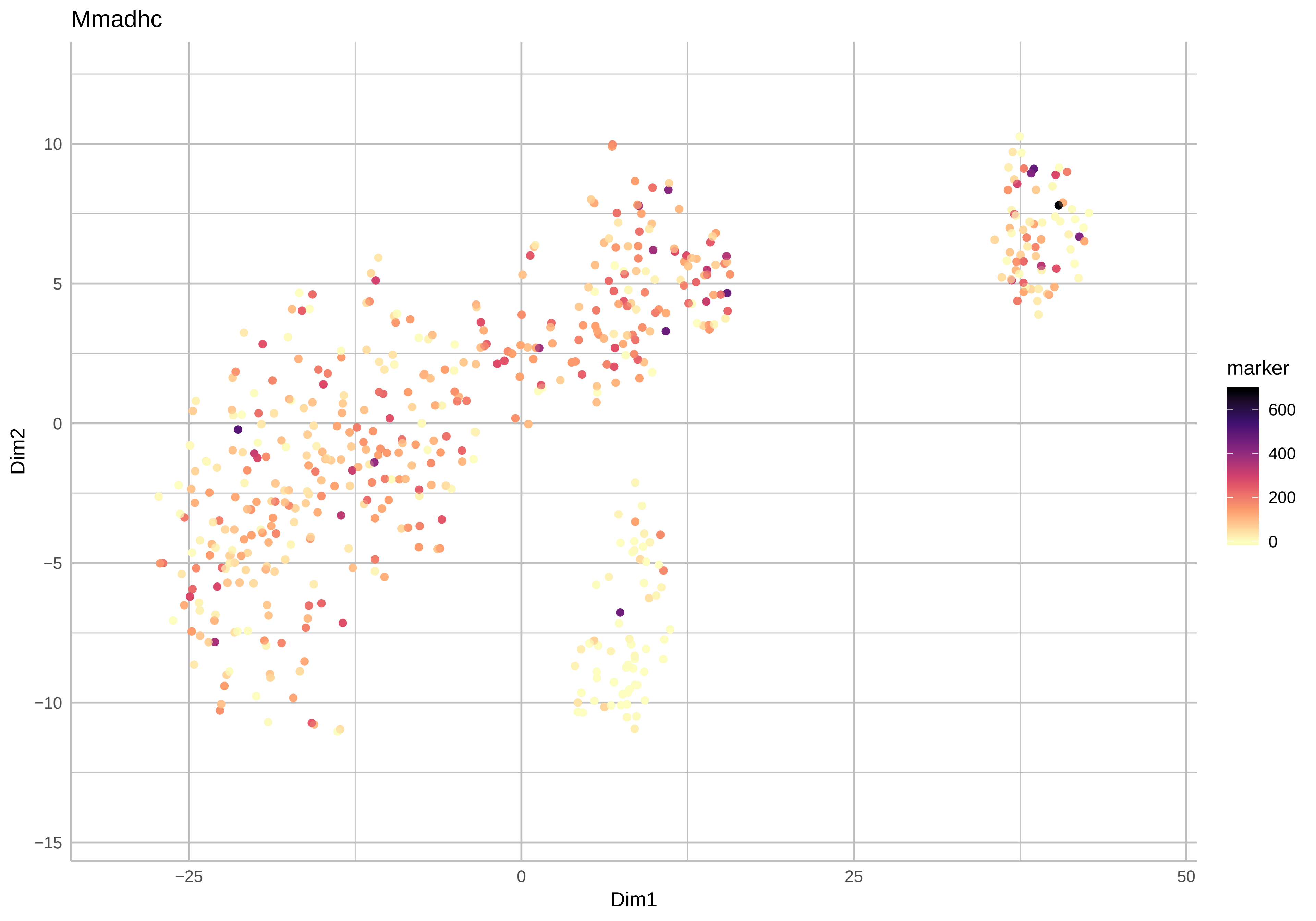

Mmadhc
Dim2
Dim1
10
5
0
−5
−10
−15
−25
0
25
50
marker
600
400
200
0

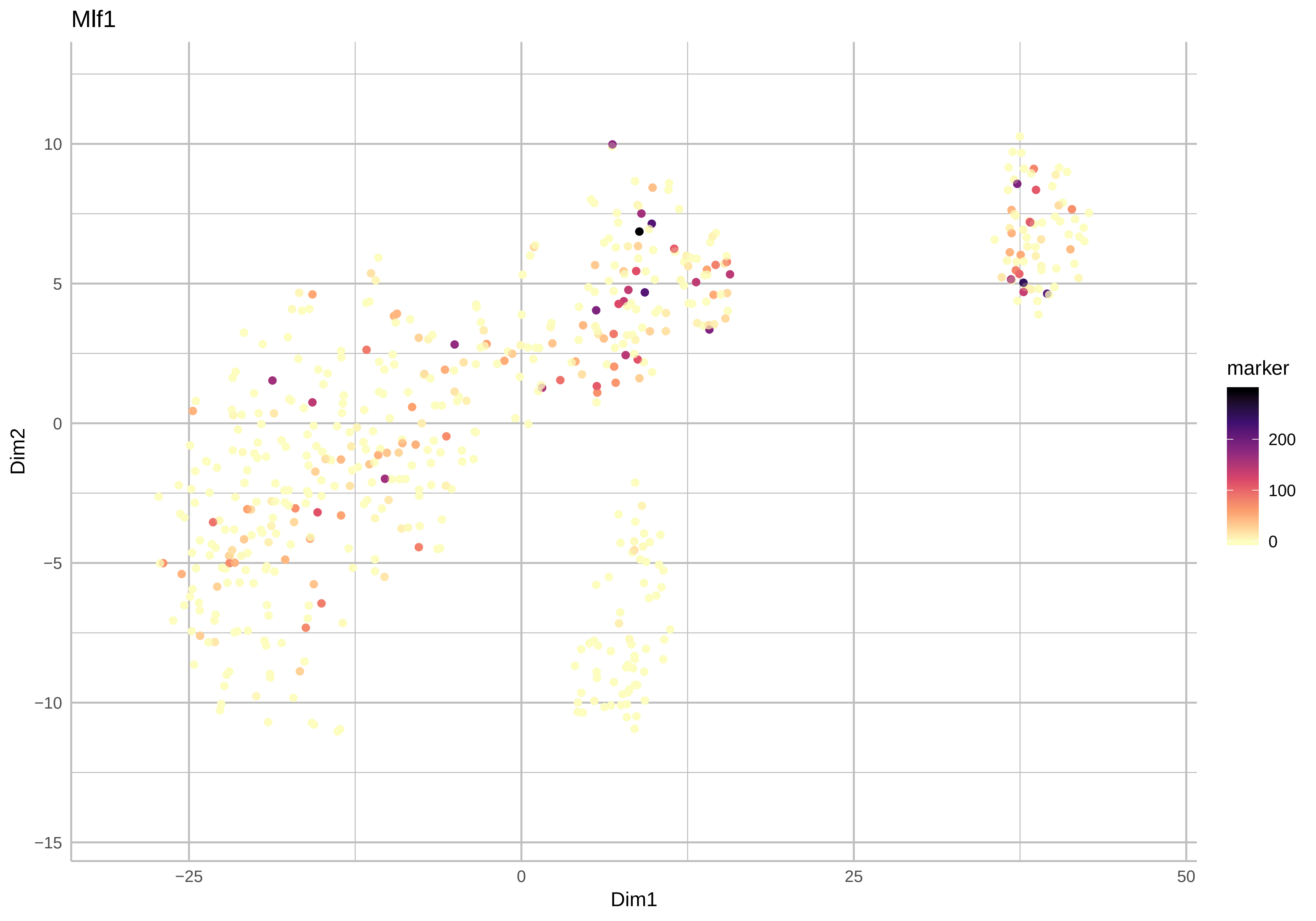

Mlf1
Dim2
Dim1
10
5
0
−5
−10
−15
−25
0
25
50
marker
200
100
0

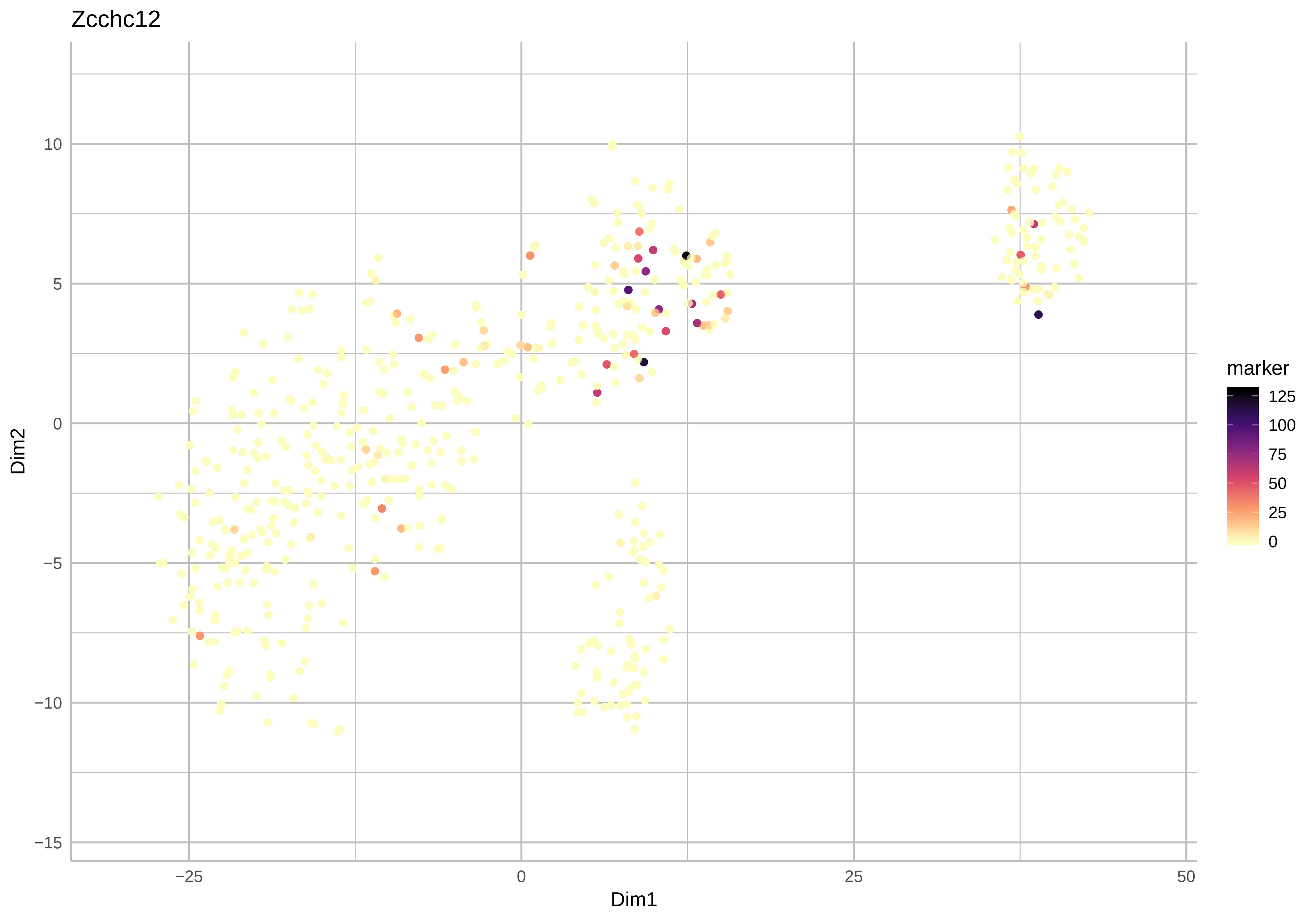

Zcchc12
Dim2
Dim1
10
5
0
−5
−10
−15
−25
0
25
50
marker
125
100
75
50
25
0

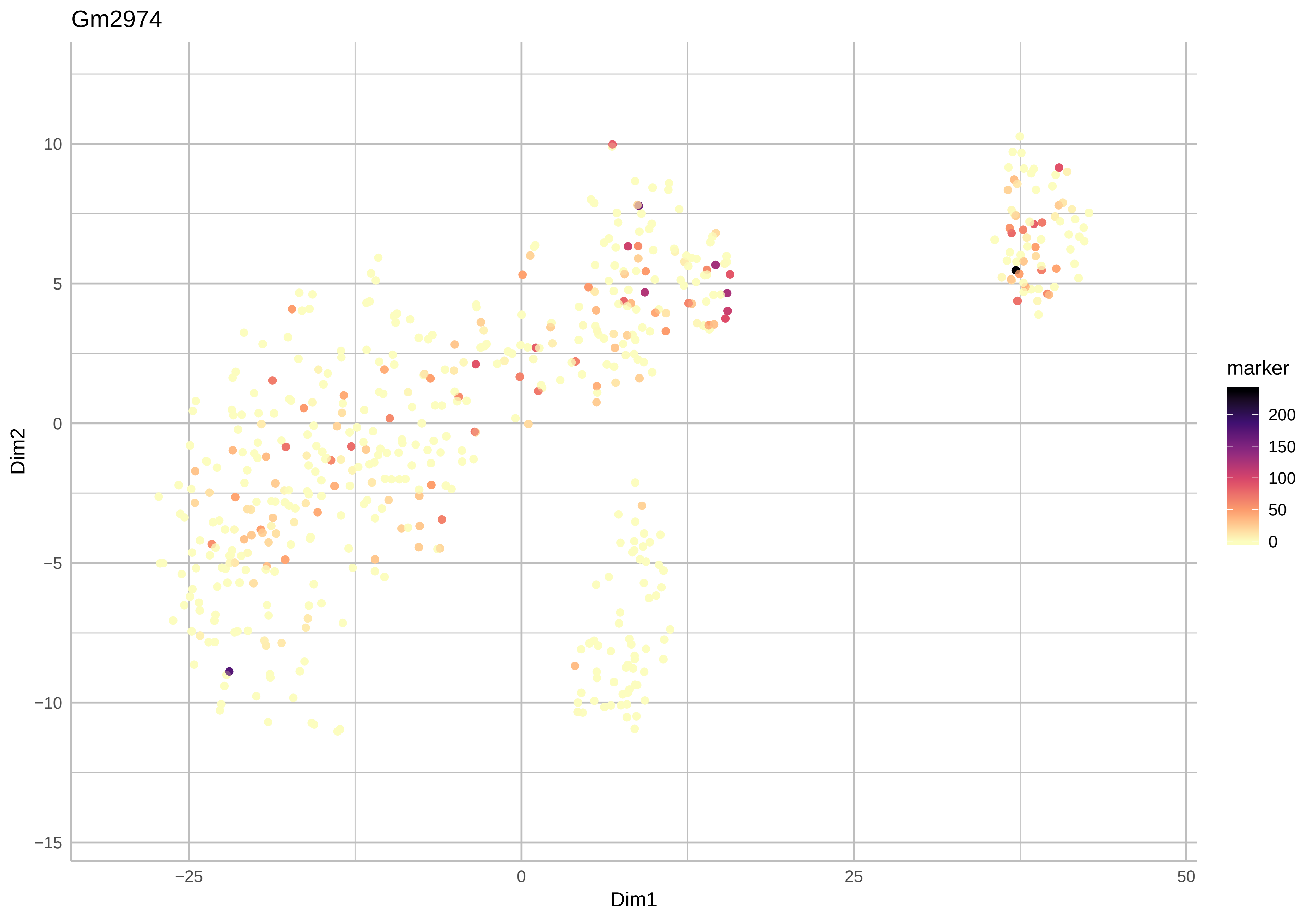

Gm2974
Dim2
Dim1
10
5
0
−5
−10
−15
−25
0
25
50
marker
200
150
100
50
0

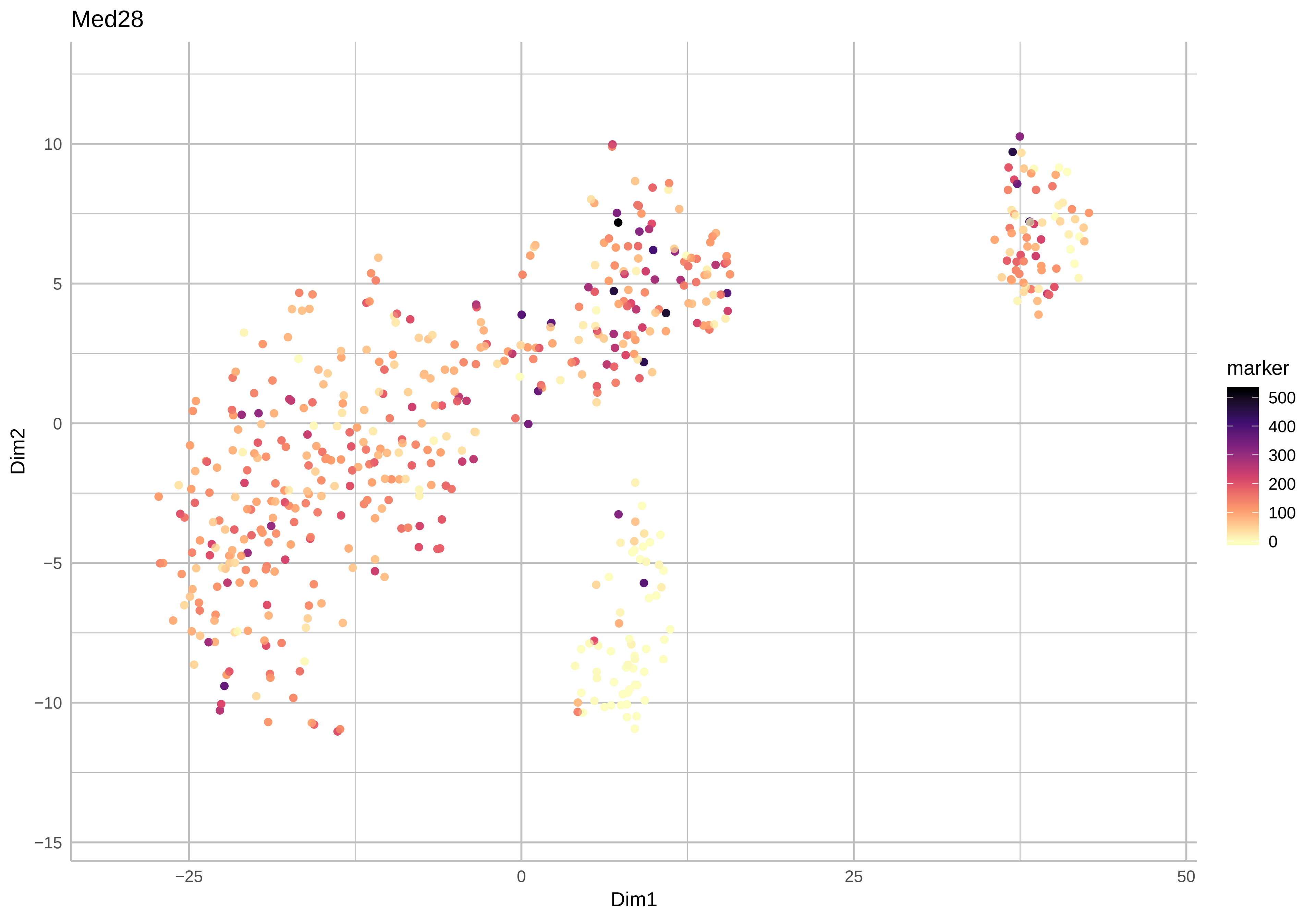

Med28
10
5
0
−5
−10
−15
Dim2
−25
0
25
50
Dim1
marker
500
400
300
200
100
0

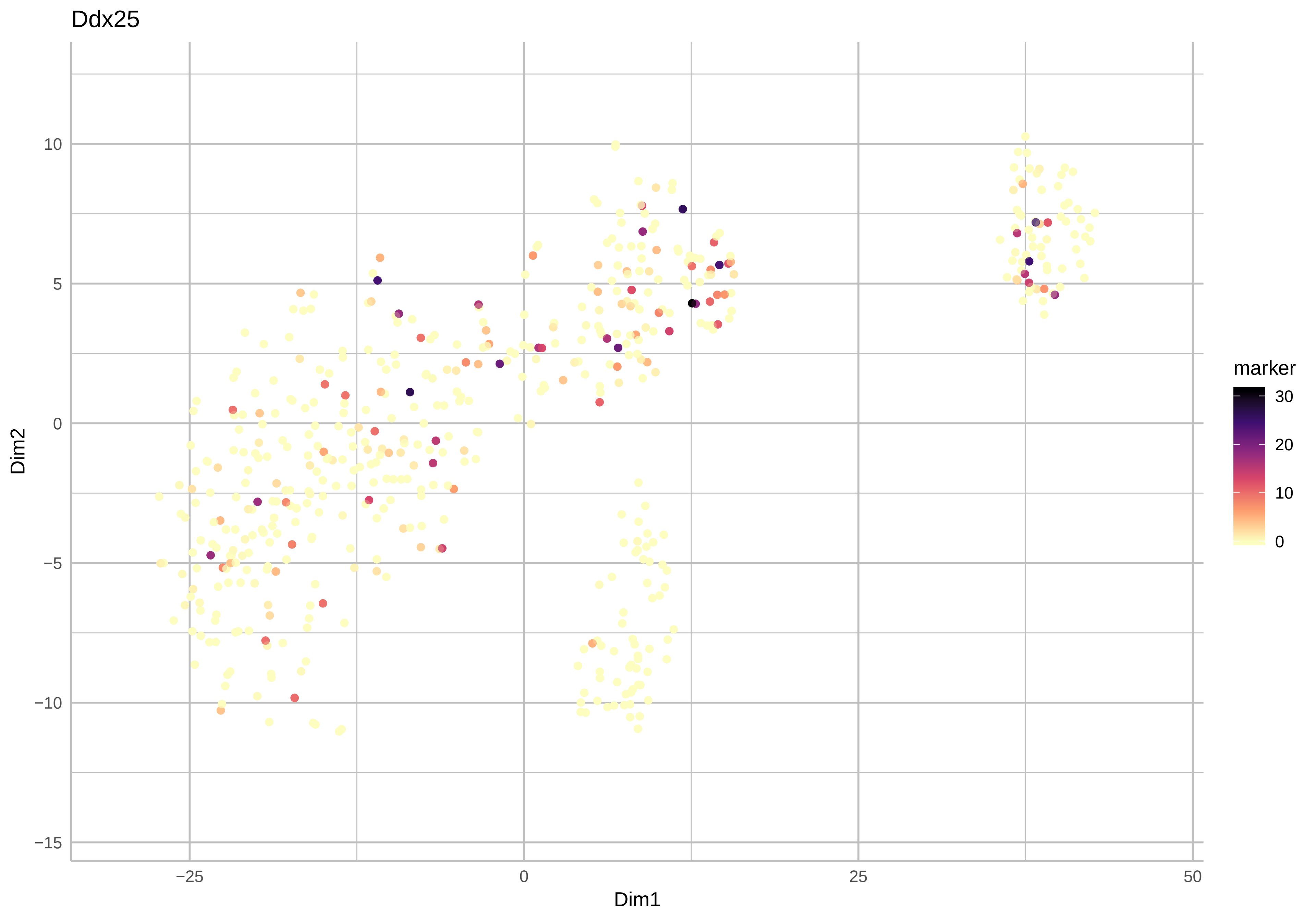
Ddx25
Dim2
Dim1
10
5
0
−5
−10
−15
−25
0
25
50
marker
30
20
10
0

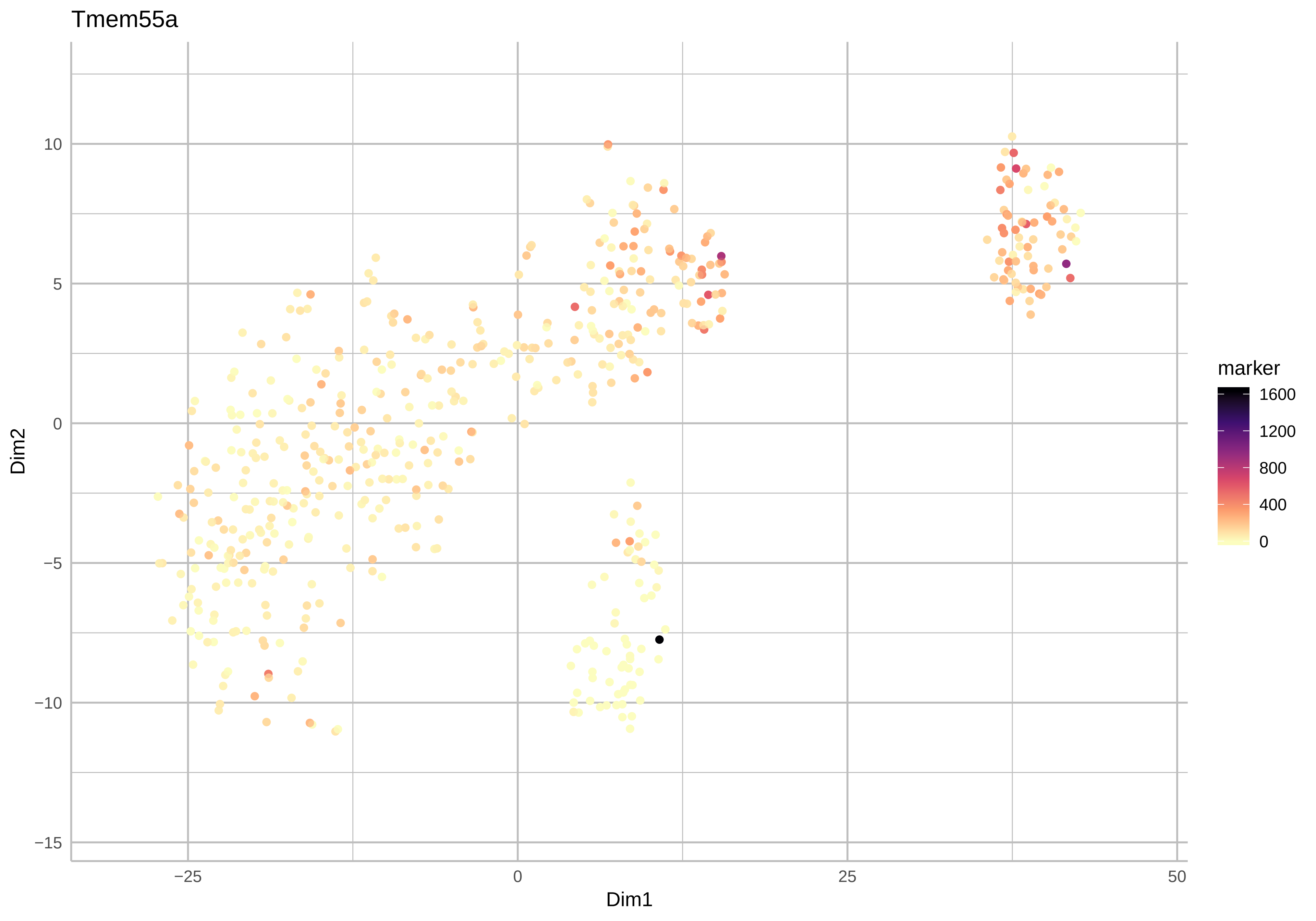
Tmem55a
Dim2
Dim1
10
5
0
−5
−10
−15
−25
0
25
50
marker
1600
1200
800
400
0

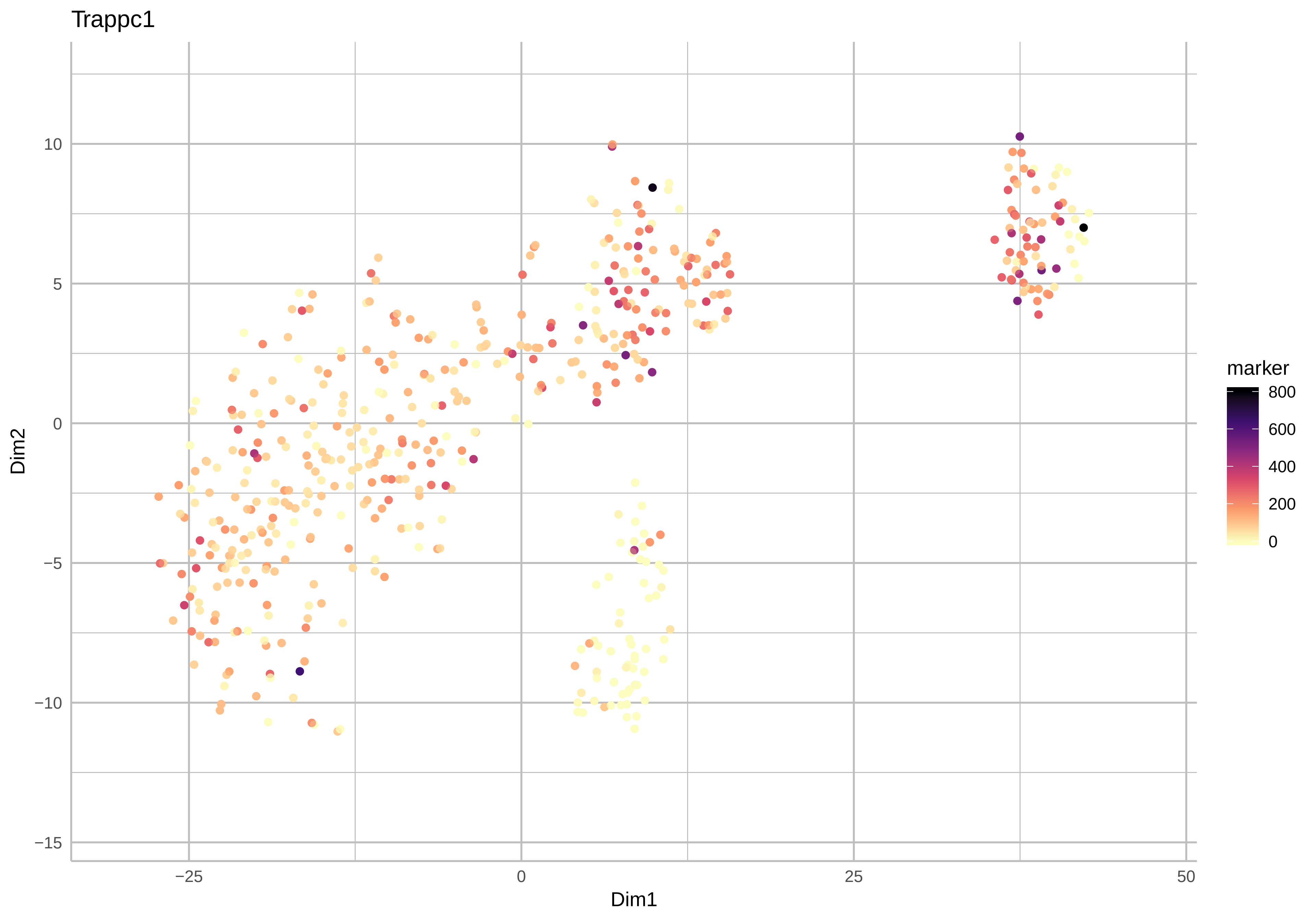

Trappc1
Dim2
Dim1
marker
800
600
400
200
0
10
5
0
−5
−10
−15
−25
0
25
50

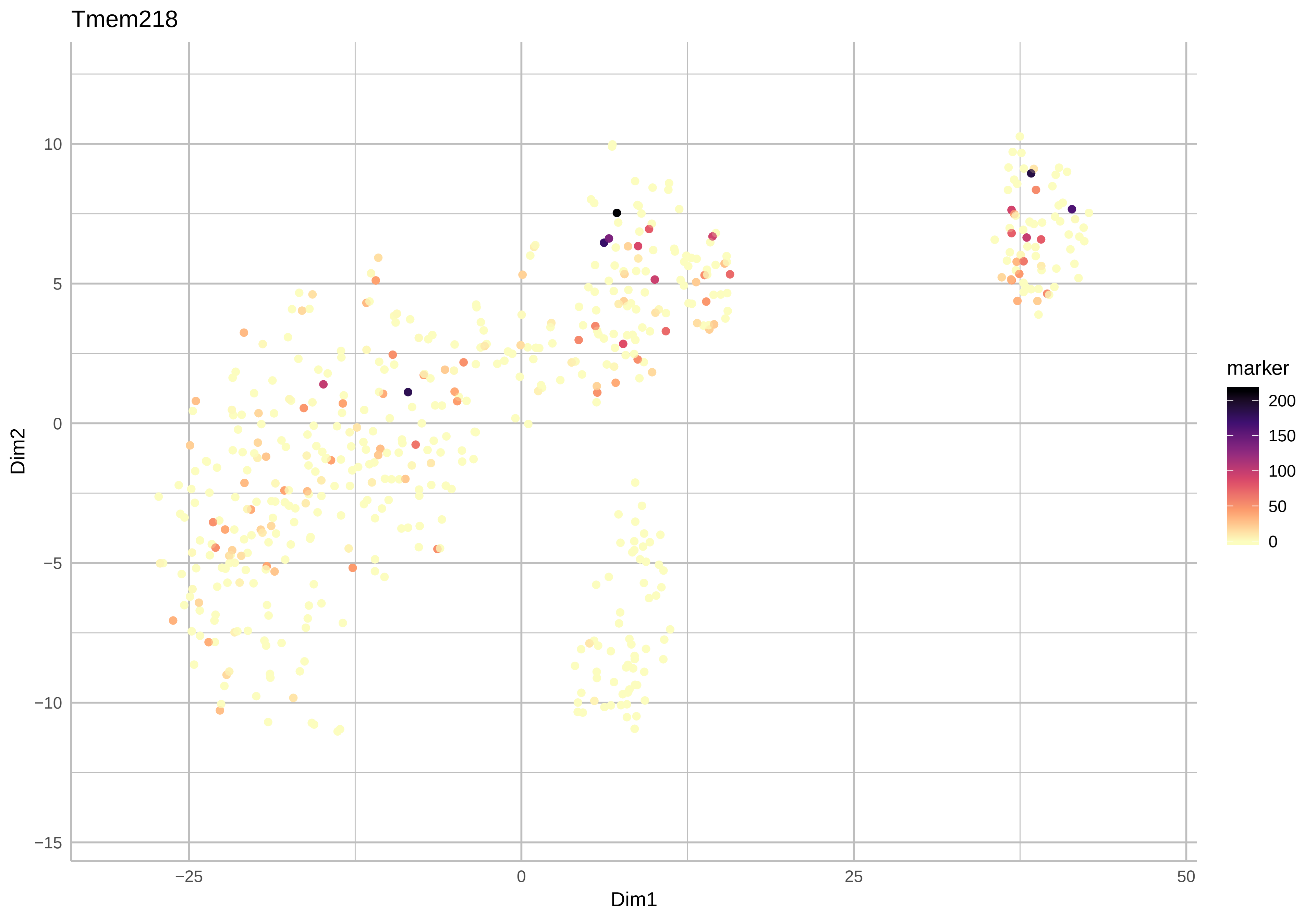
Tmem218
Dim2
Dim1
10
5
0
−5
−10
−15
−25
0
25
50
marker
200
150
100
50
0

Lrrc40

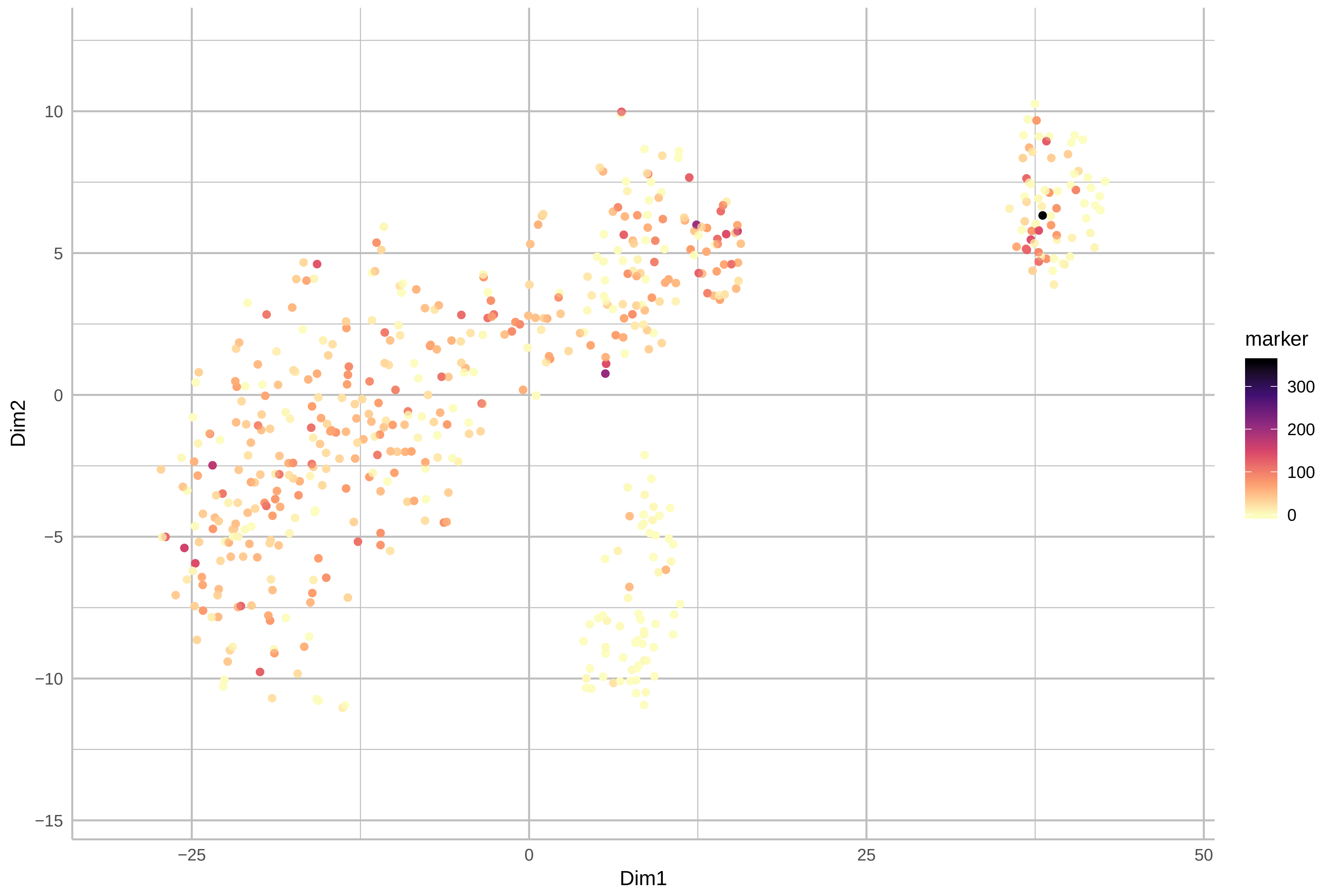

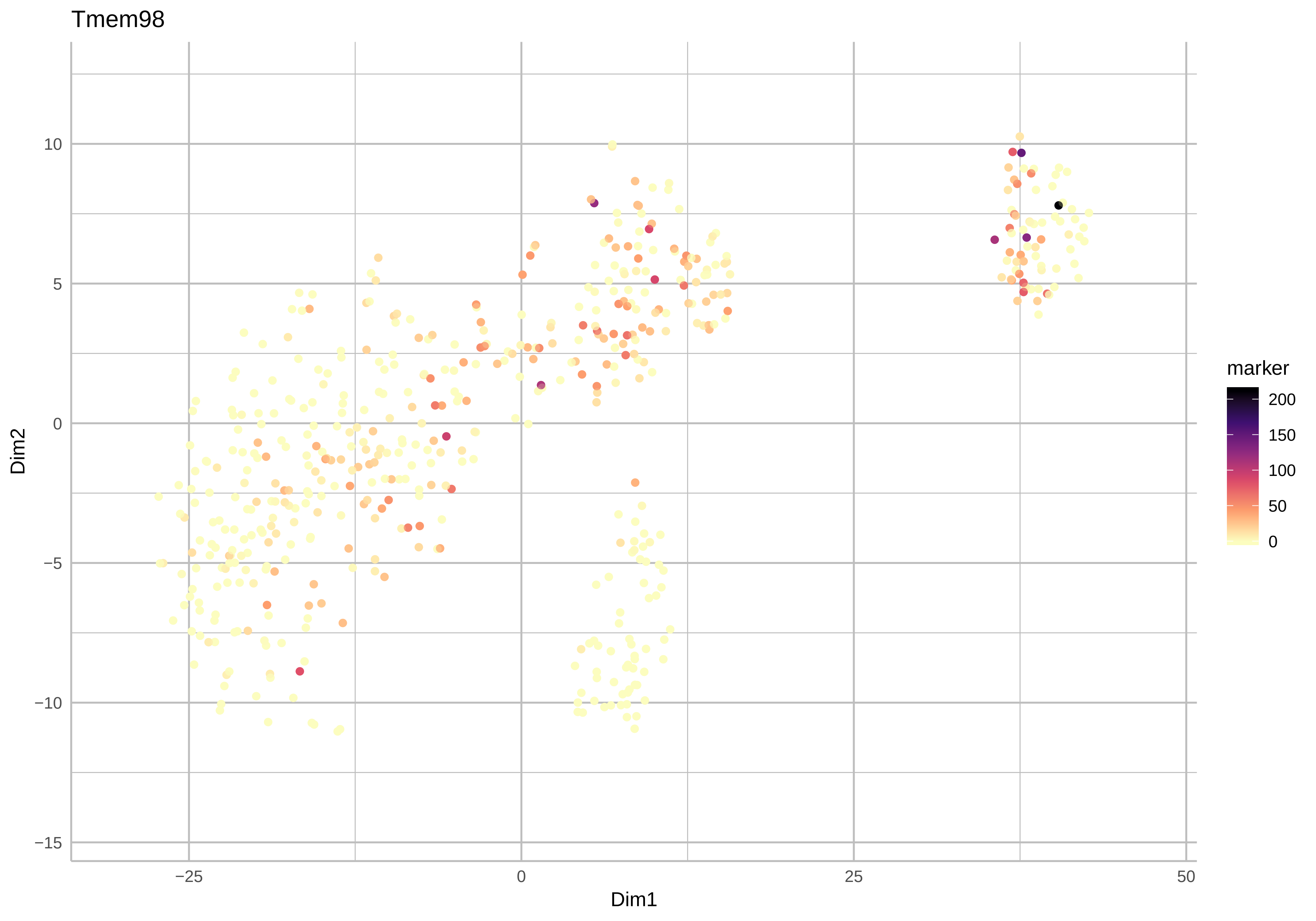

Tmem98
Dim2
Dim1
10
5
0
−5
−10
−15
−25
0
25
50
marker
200
150
100
50
0

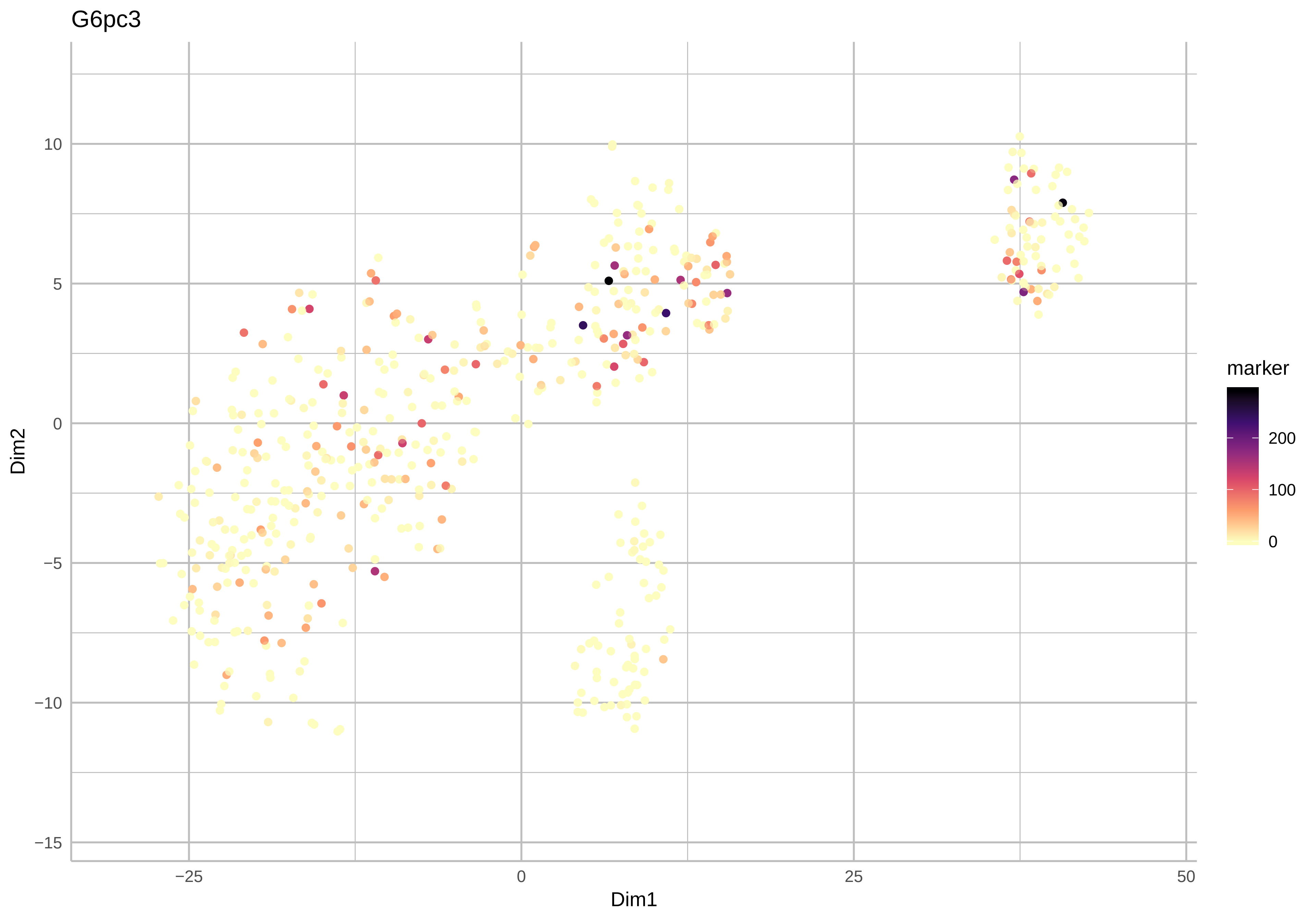

G6pc3
Dim2
Dim1
marker
200
100
0
10
5
0
−5
−10
−15
−25
0
25
50

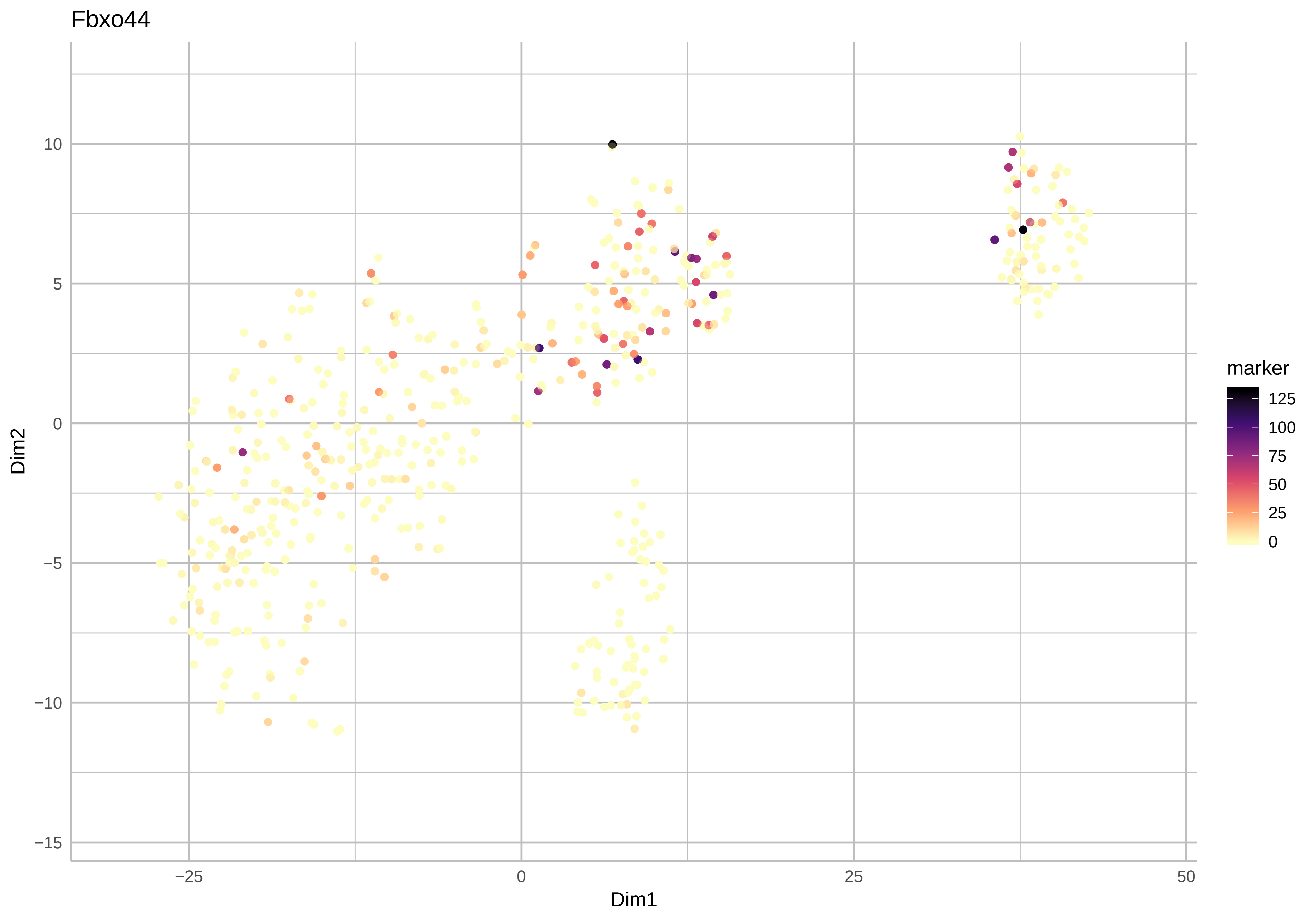

Fbxo44
Dim2
Dim1
10
5
0
−5
−10
−15
−25
0
25
50
marker
125
100
75
50
25
0

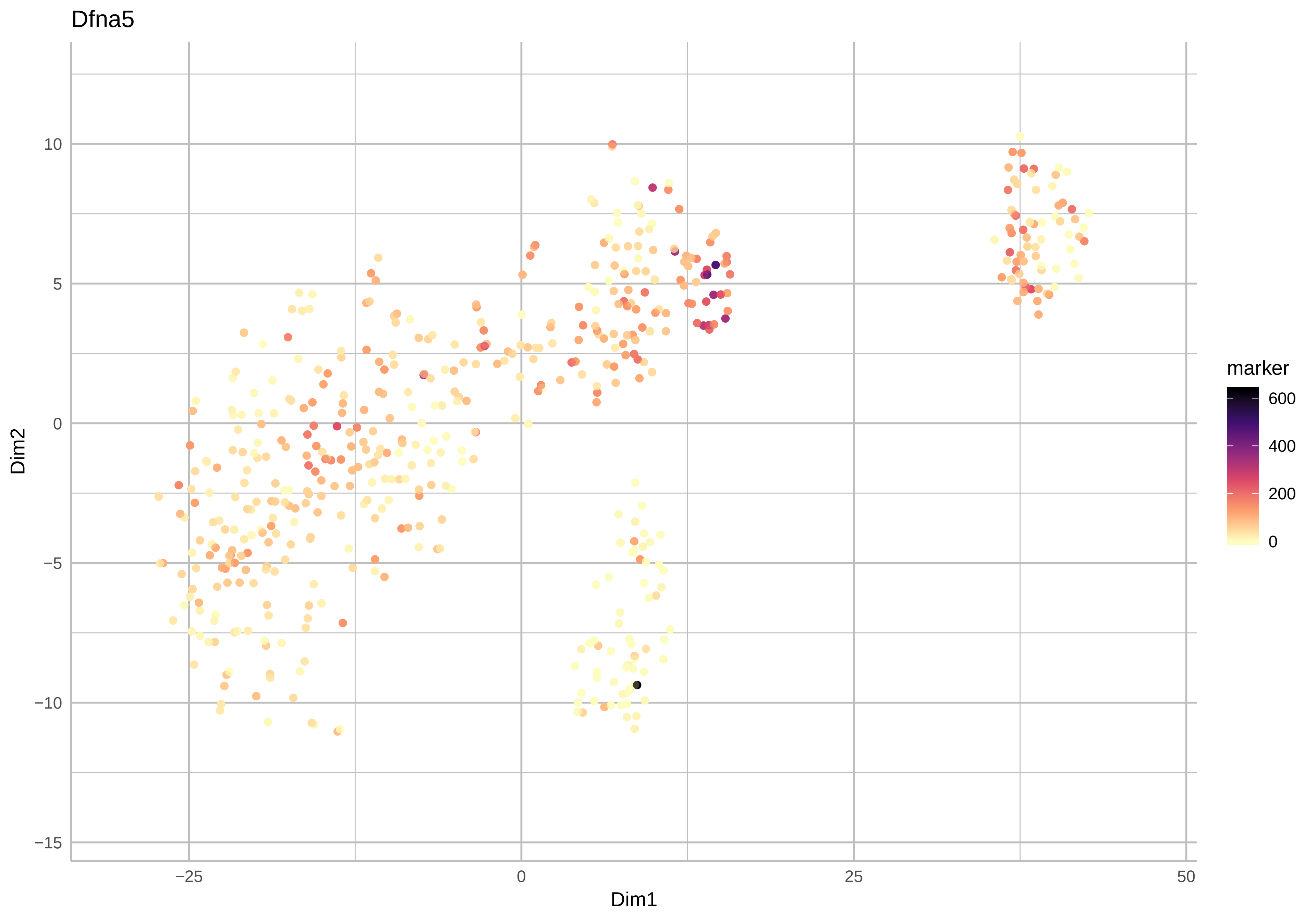

Dfna5
Dim2
Dim1
10
5
0
−5
−10
−15
−25
0
25
50
marker
600
400
200
0

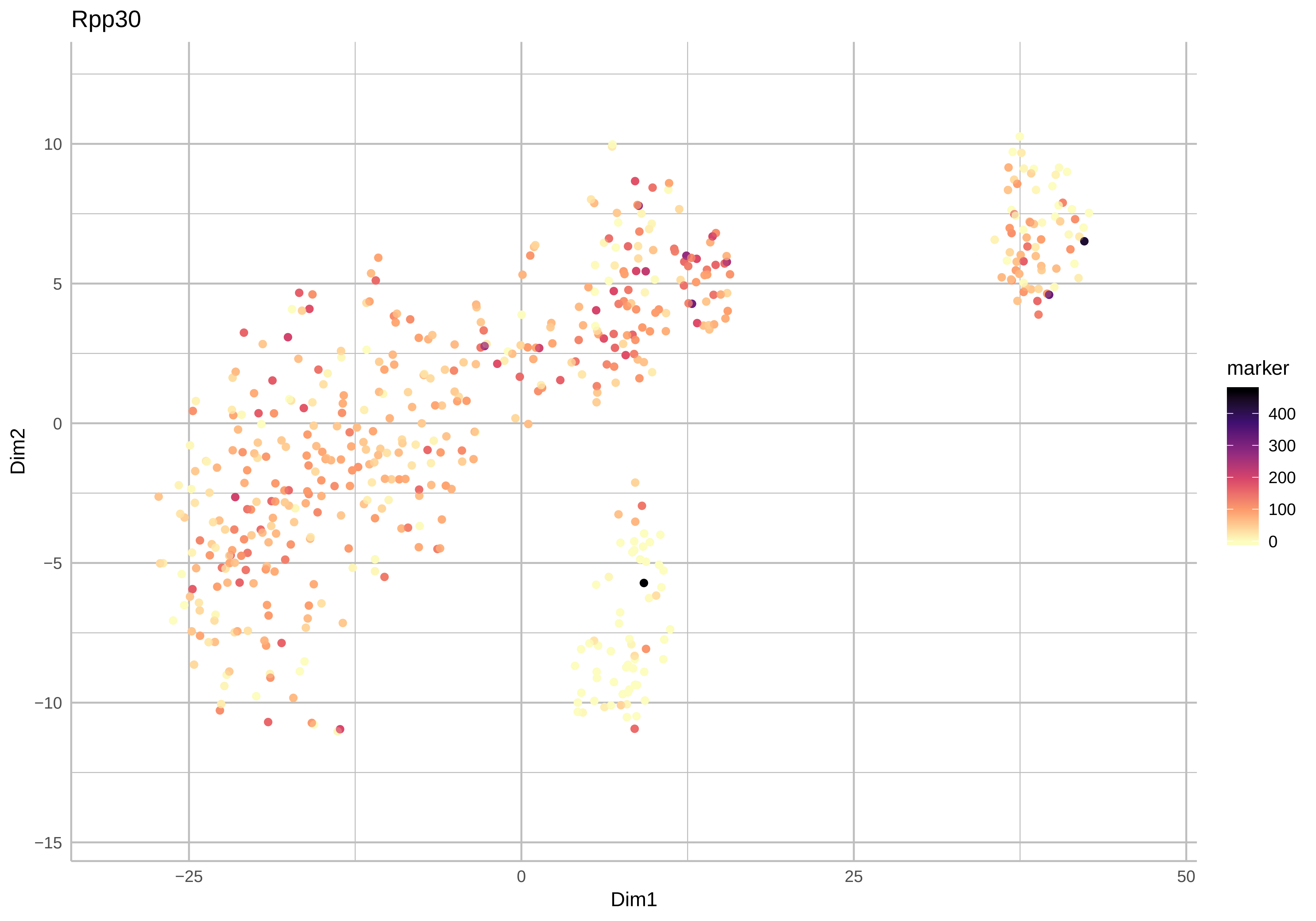

Rpp30
Dim2
Dim1
marker
400
300
200
100
0
10
5
0
−5
−10
−15
−25
0
25
50

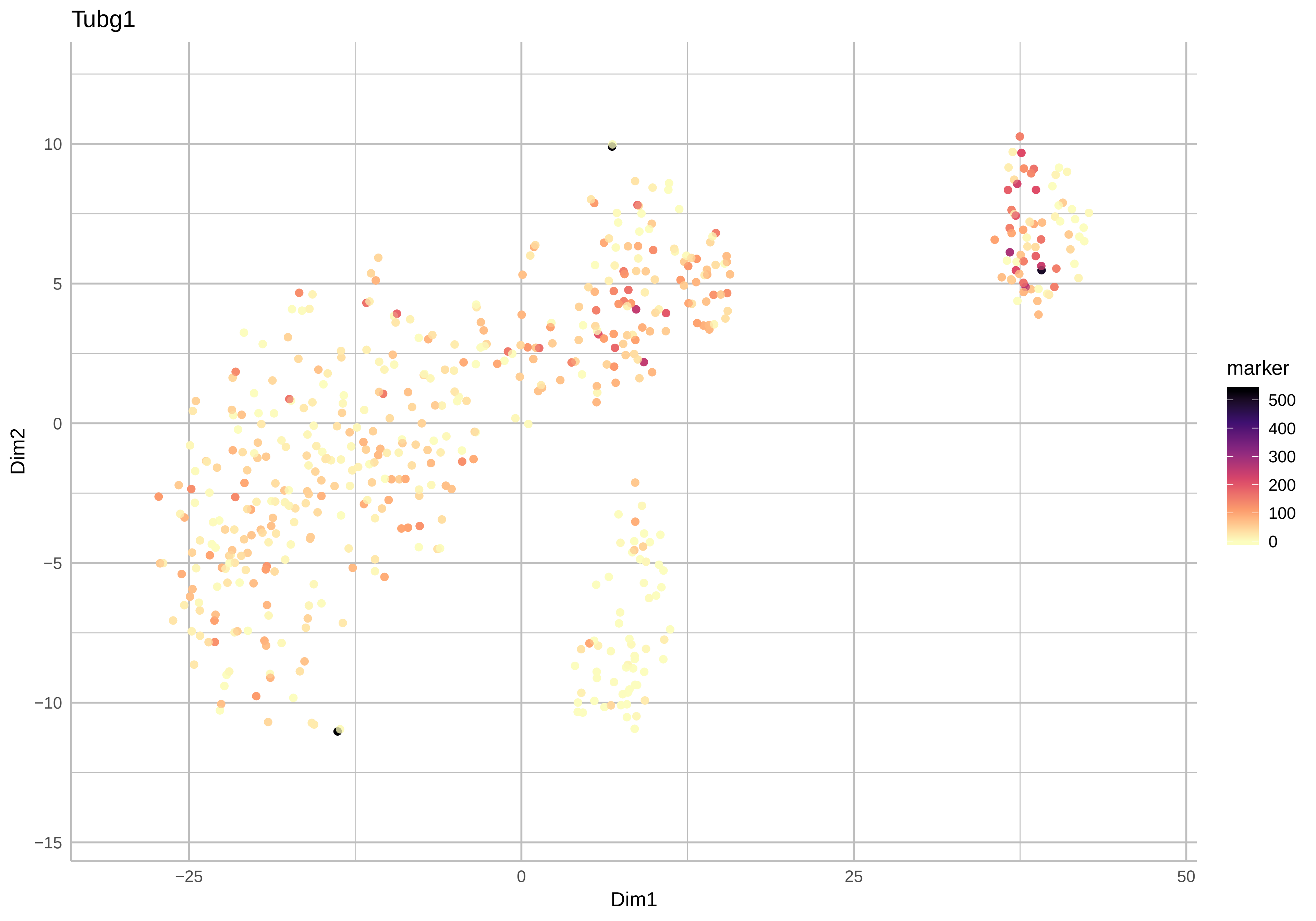

Tubg1
Dim2
Dim1
10
5
0
−5
−10
−15
−25
0
25
50
marker
500
400
300
200
100
0

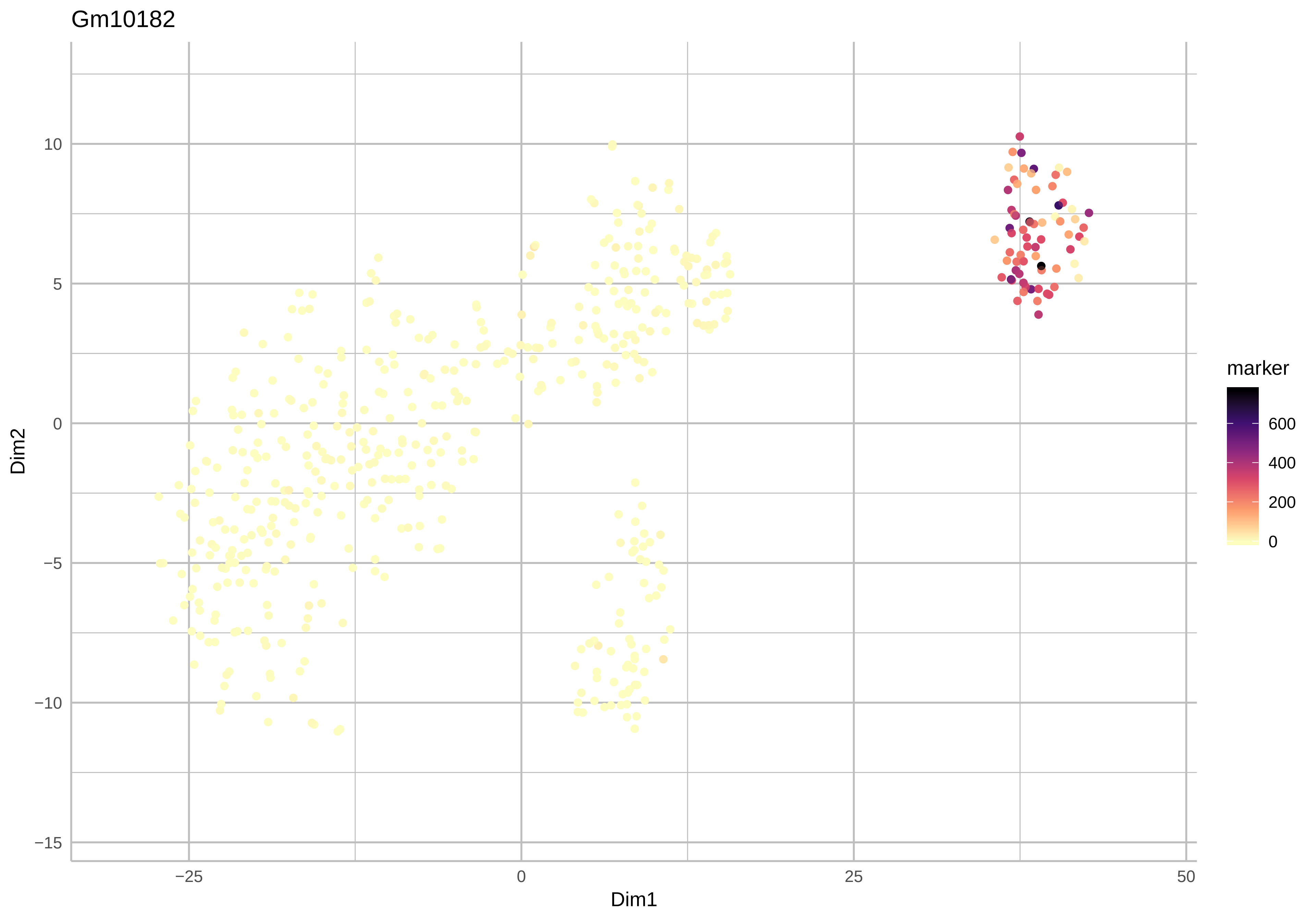

Gm10182
Dim2
Dim1
10
5
0
−5
−10
−15
−25
0
25
50
marker
600
400
200
0

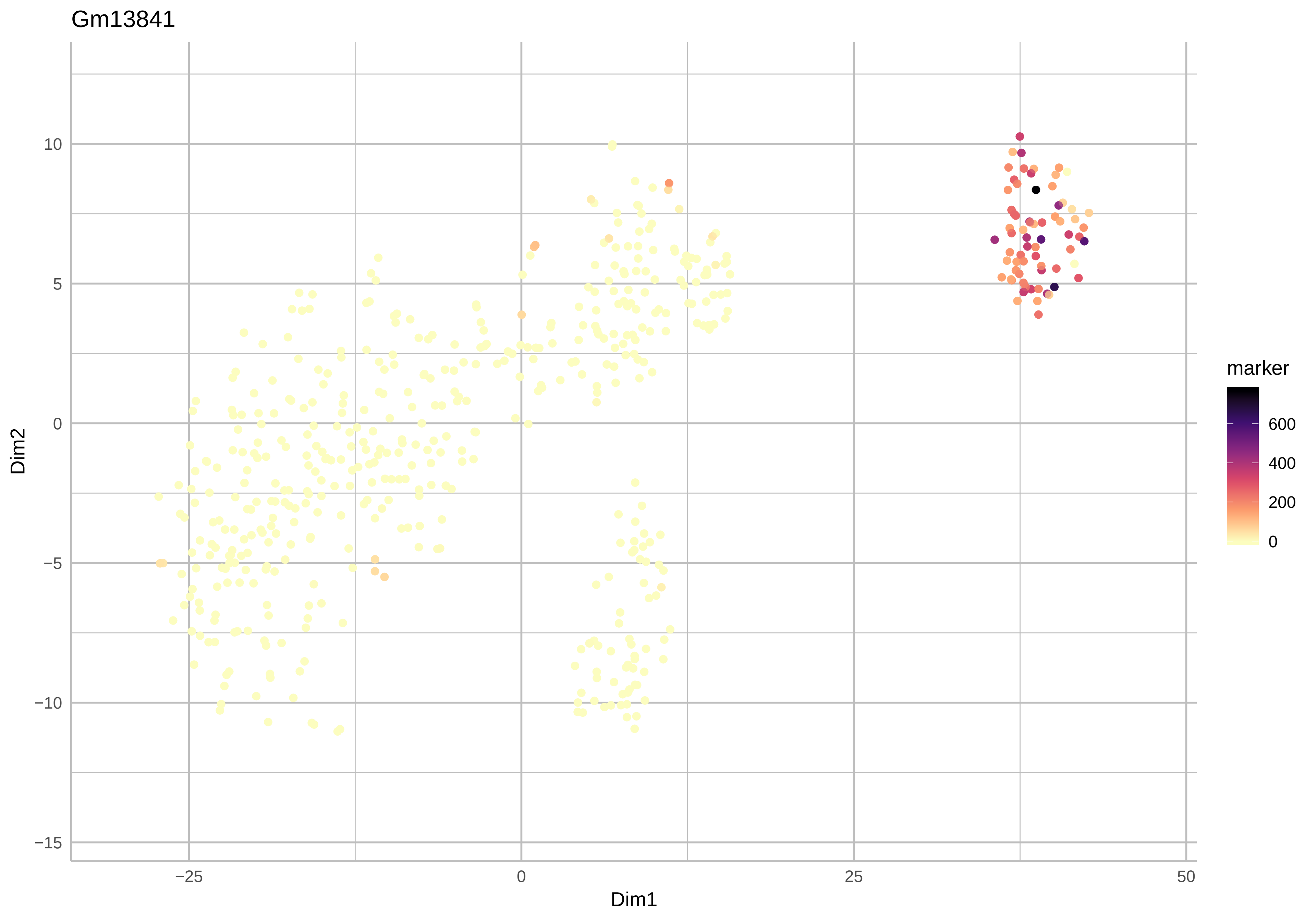

Gm13841
Dim2
Dim1
10
5
0
−5
−10
−15
−25
0
25
50
marker
600
400
200
0

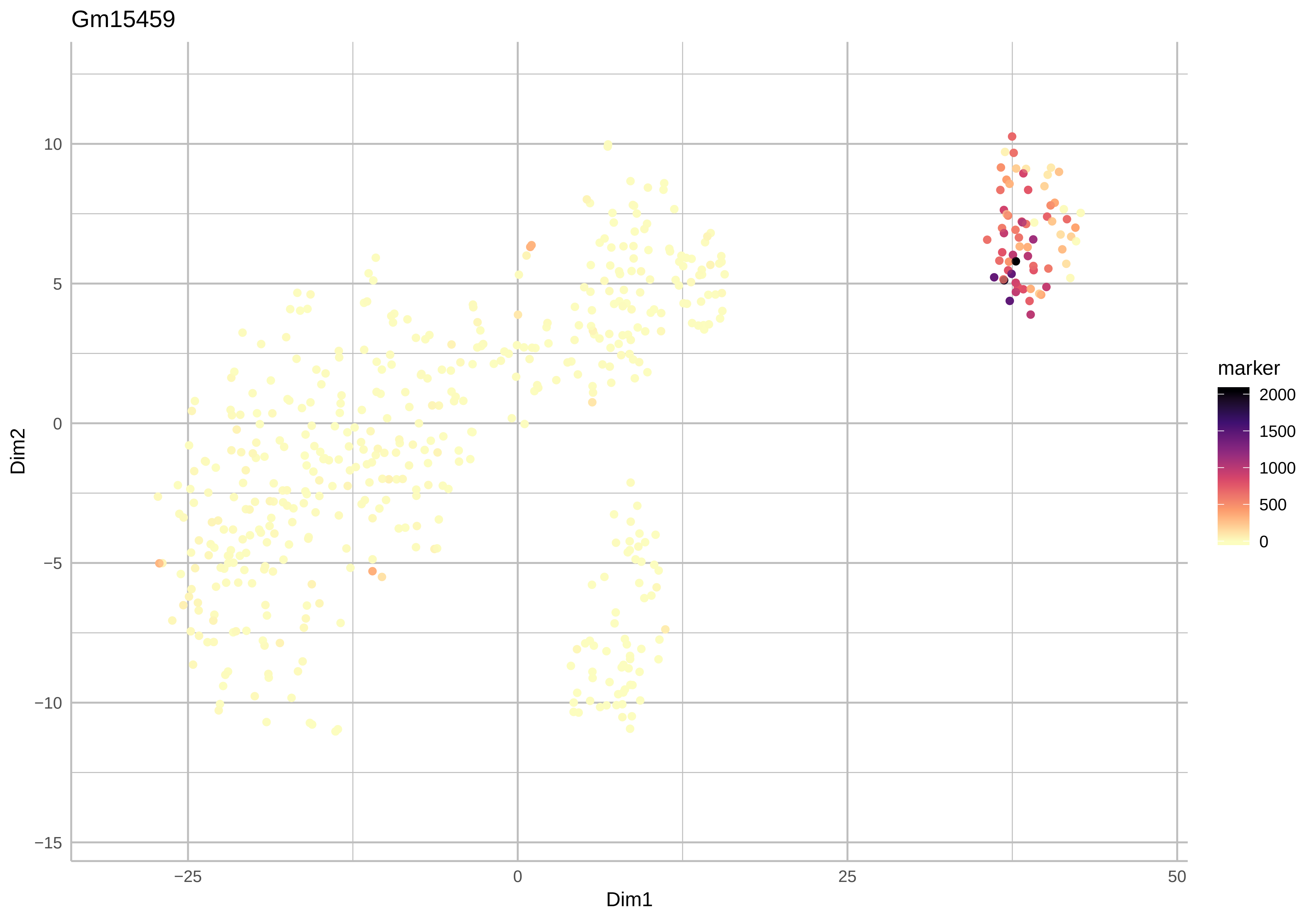
Gm15459
Dim2
Dim1
10
5
0
−5
−10
−15
−25
0
25
50
marker
2000
1500
1000
500
0

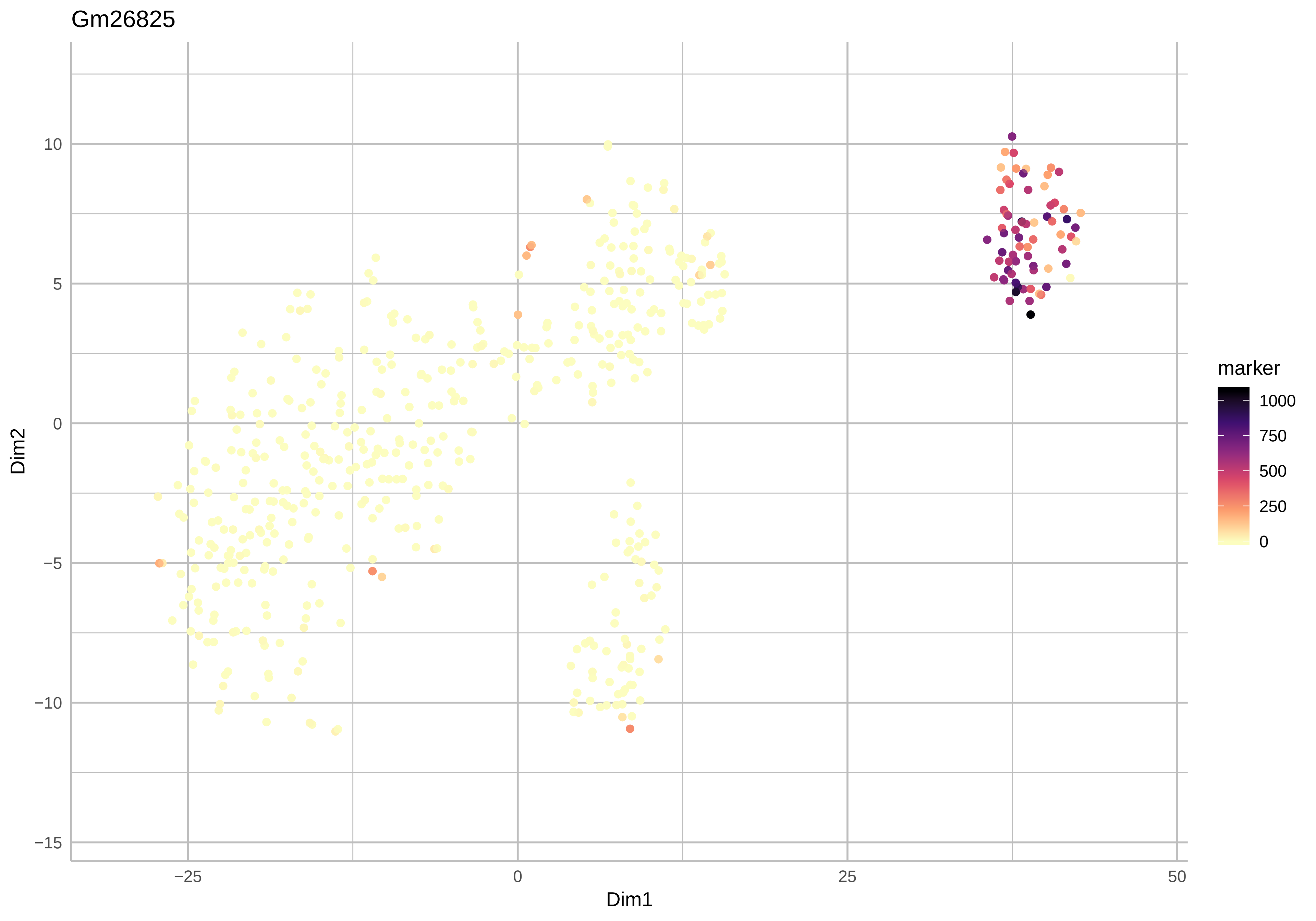

Gm26825
10
5
0
−5
−10
−15
−25
0
25
50
Dim1
Dim2
marker
1000
750
500
250
0

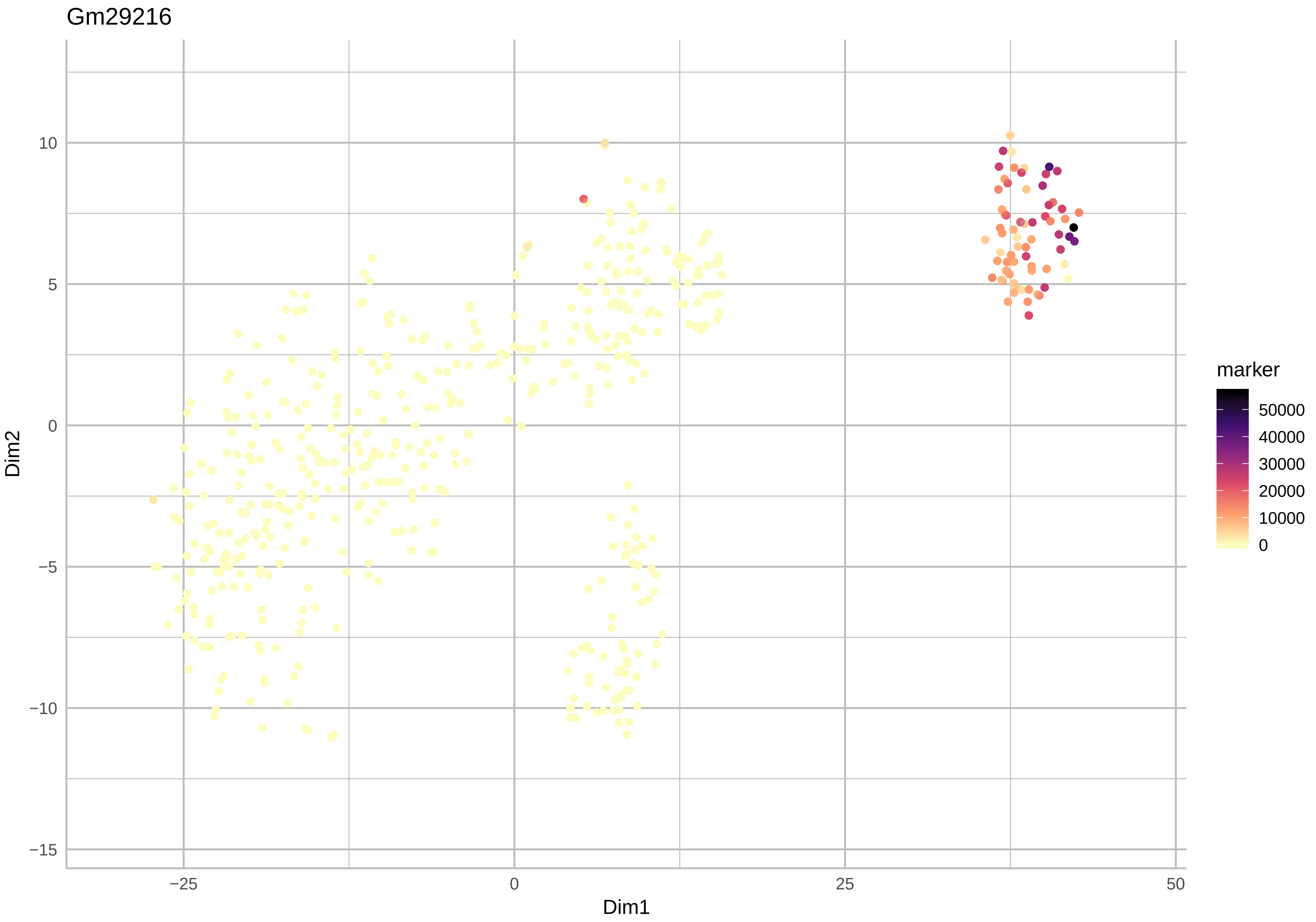
Gm29216
Dim2
Dim1
marker
50000
40000
30000
20000
10000
0
10
5
0
−5
−10
−15
−25
0
25
50

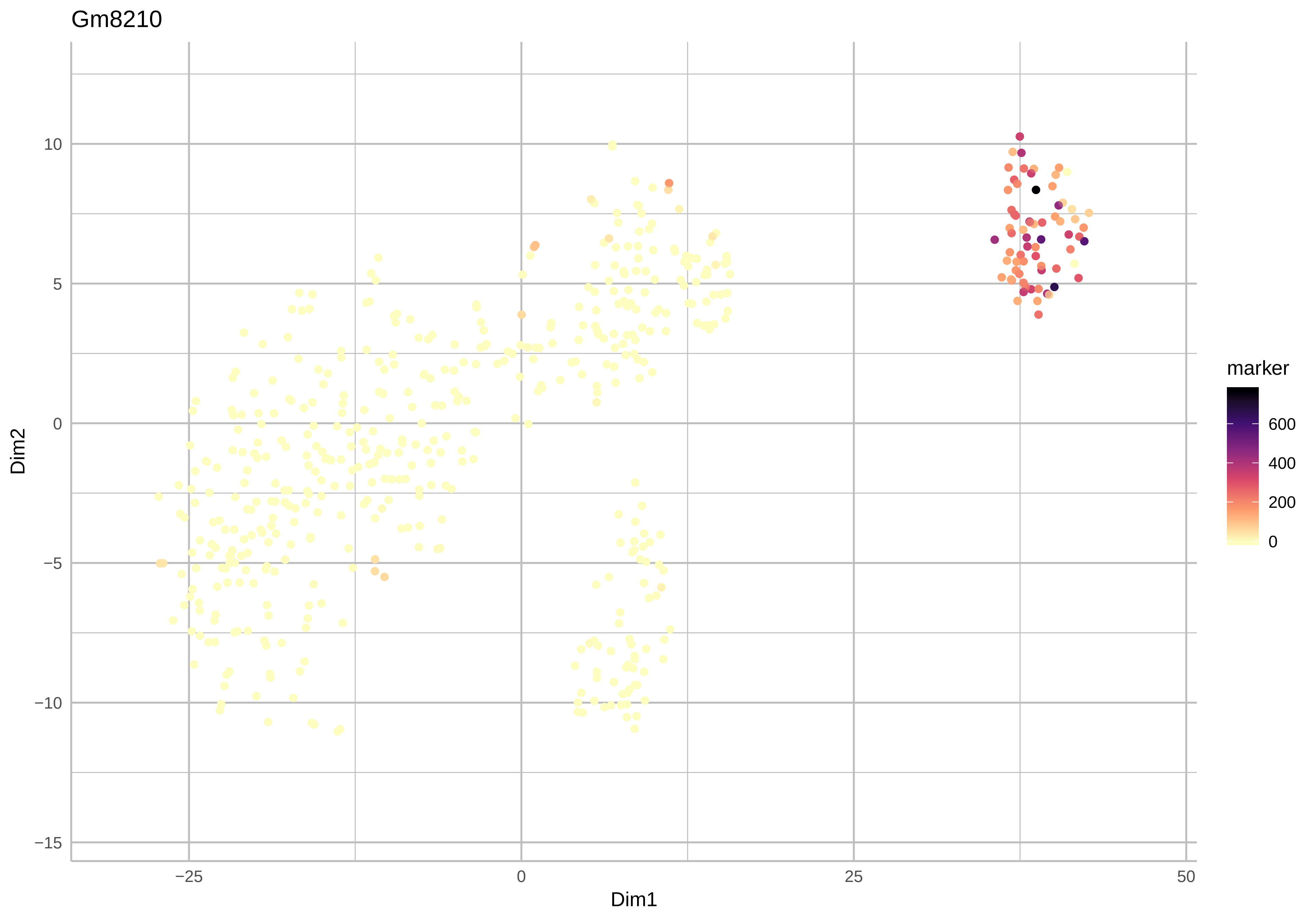

Gm8210
10
5
0
−5
−10
−15
Dim2
−25
0
25
50
Dim1
marker
600
400
200
0

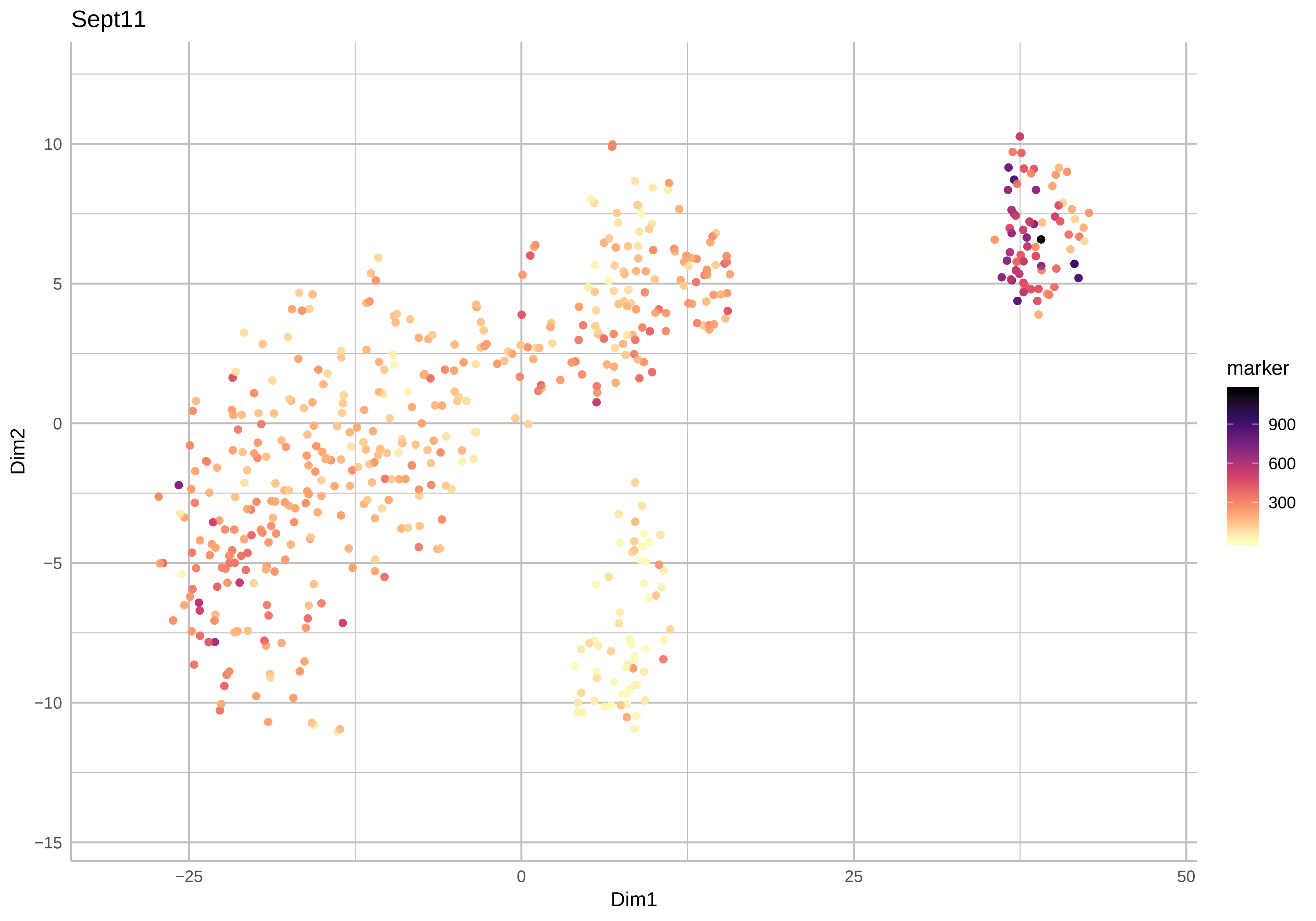
Sept11
Dim2
Dim1
marker
900
600
300
10
5
0
−5
−10
−15
−25
0
25
50

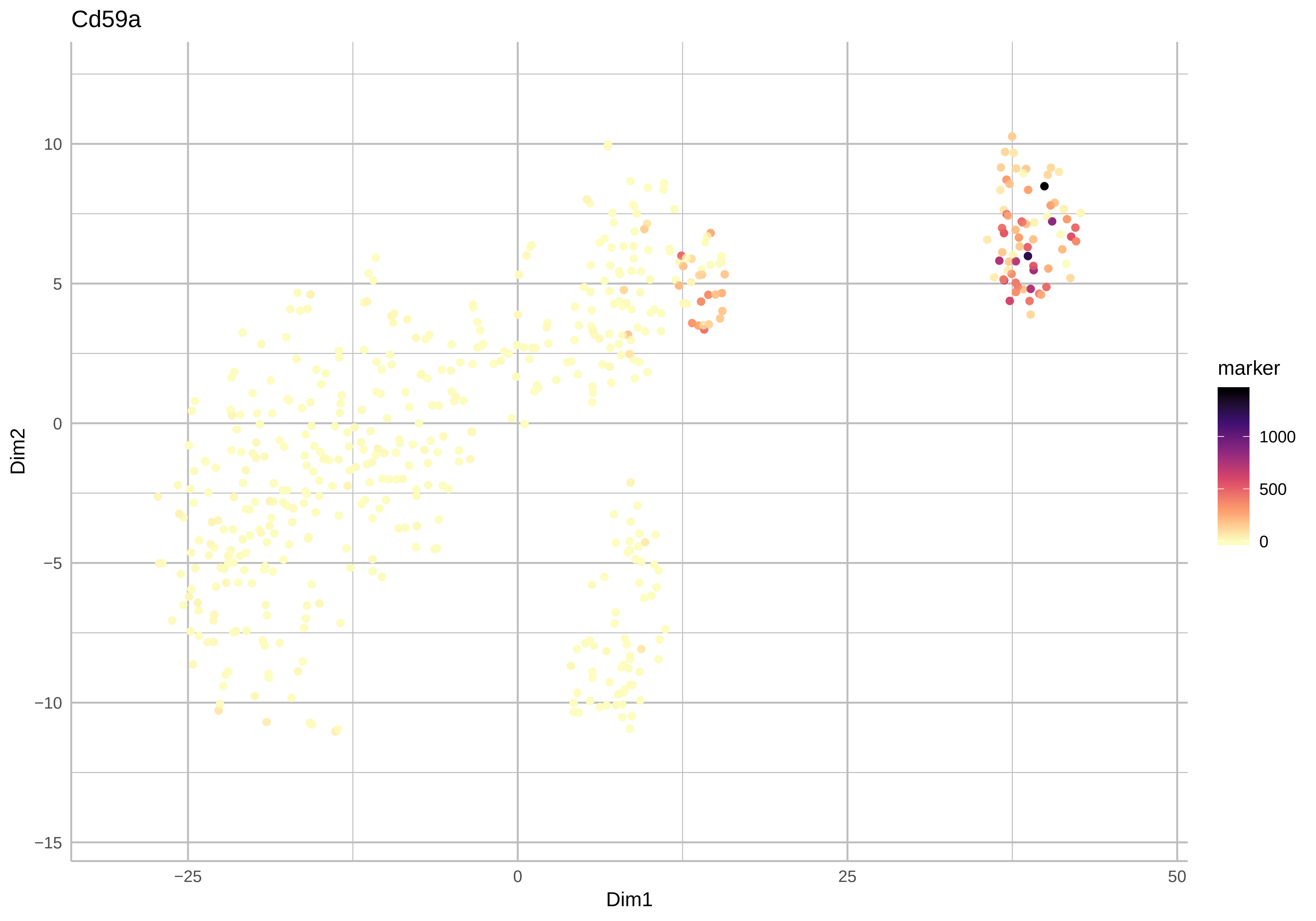

Cd59a
Dim2
Dim1
10
5
0
−5
−10
−15
−25
0
25
50
marker
1000
500
0

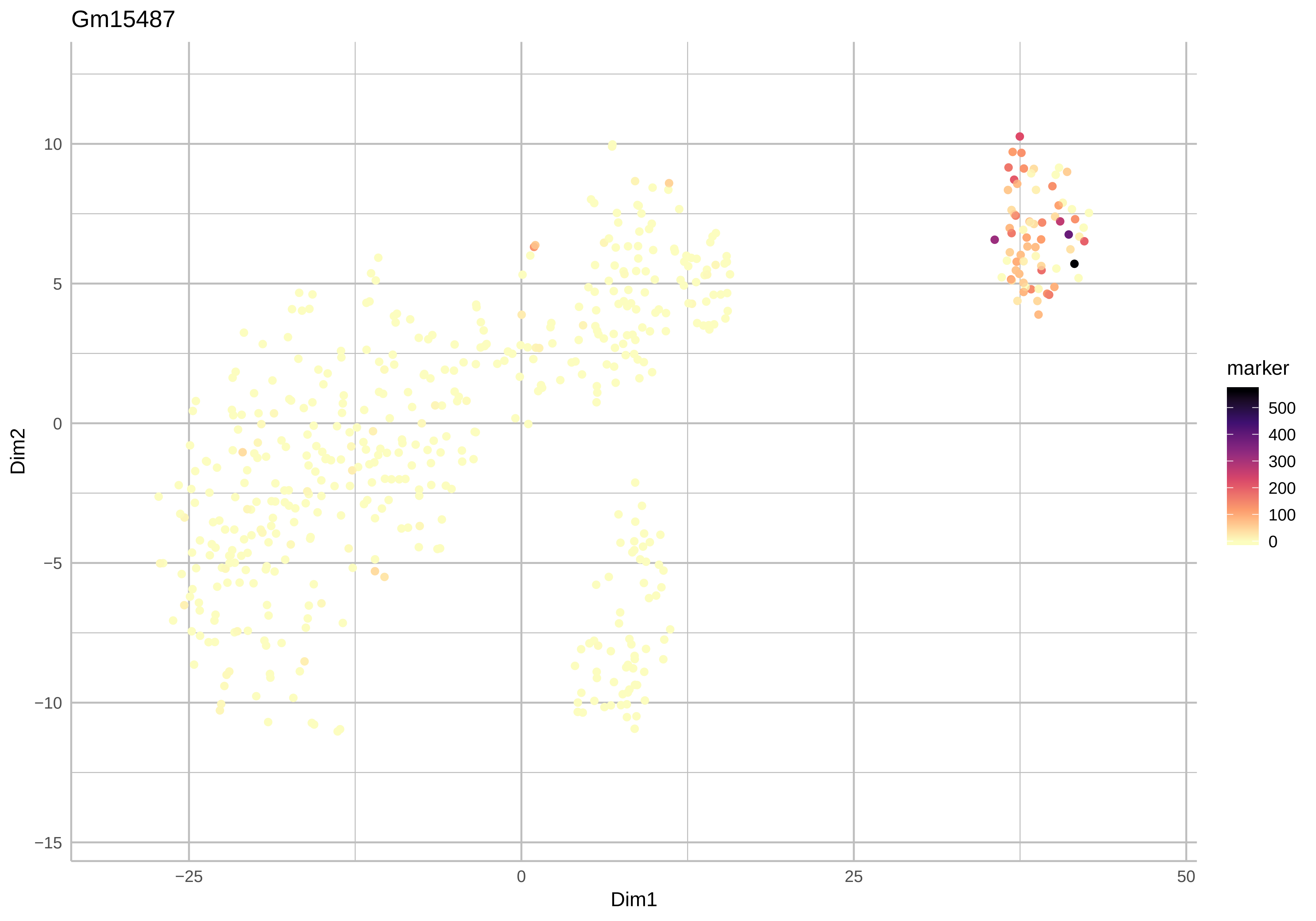
Gm15487
10
5
0
−5
−10
−15
Dim2
−25
0
25
50
Dim1
marker
500
400
300
200
100
0

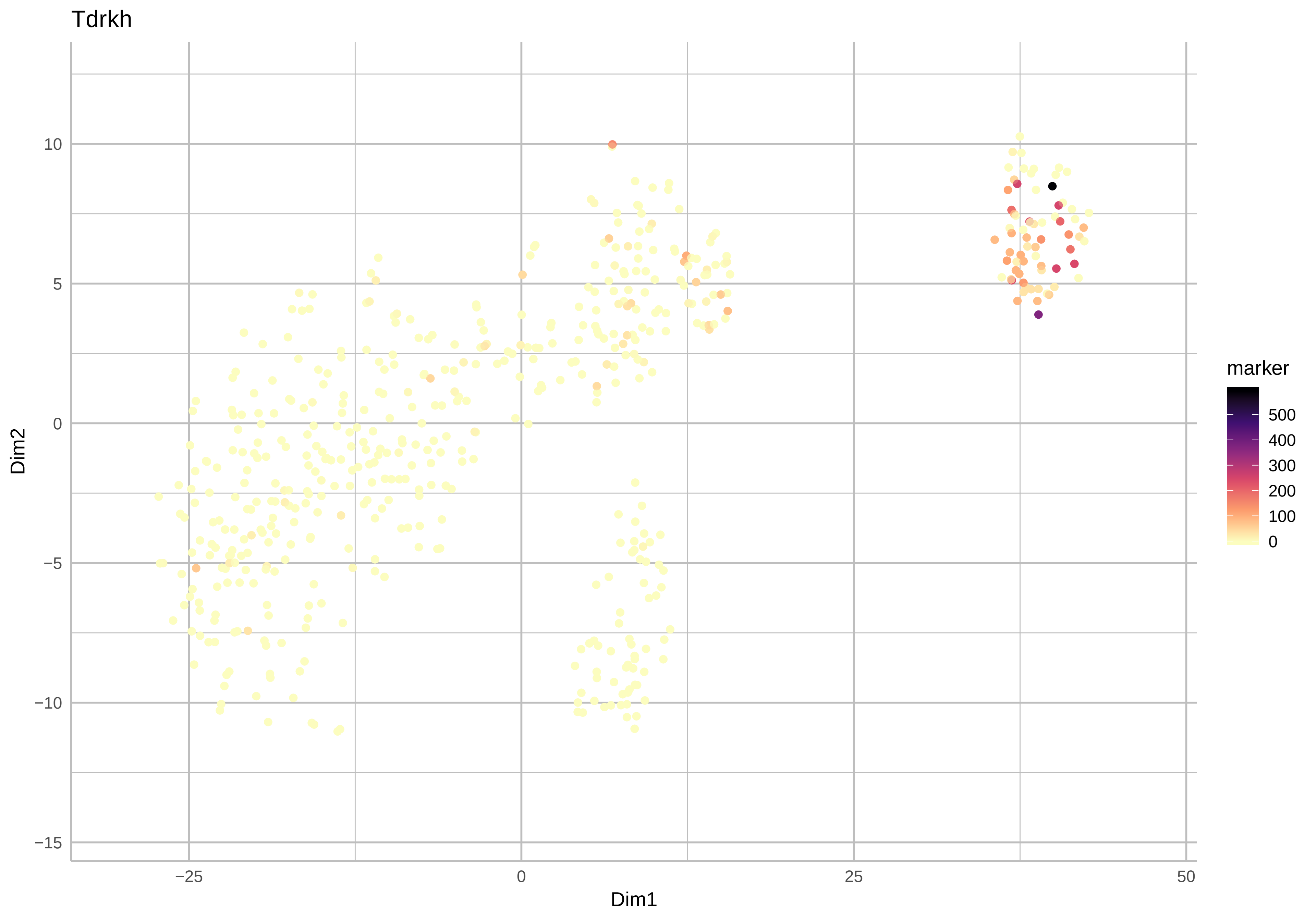

Tdrkh
10
5
0
−5
−10
−15
Dim2
−25
0
25
50
Dim1
marker
500
400
300
200
100
0

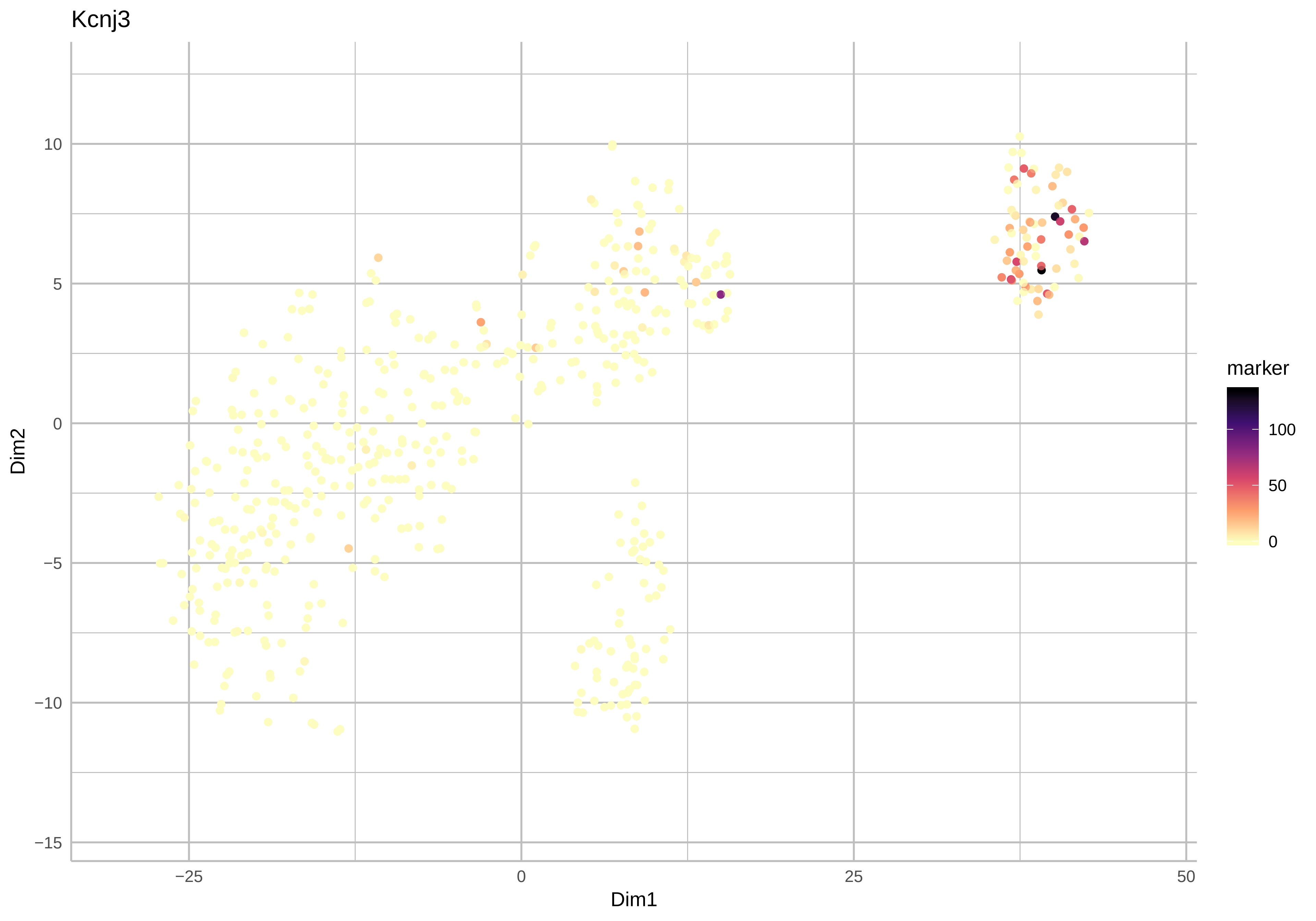
Kcnj3
Dim2
Dim1
marker
100
50
0
10
5
0
−5
−10
−15
−25
0
25
50

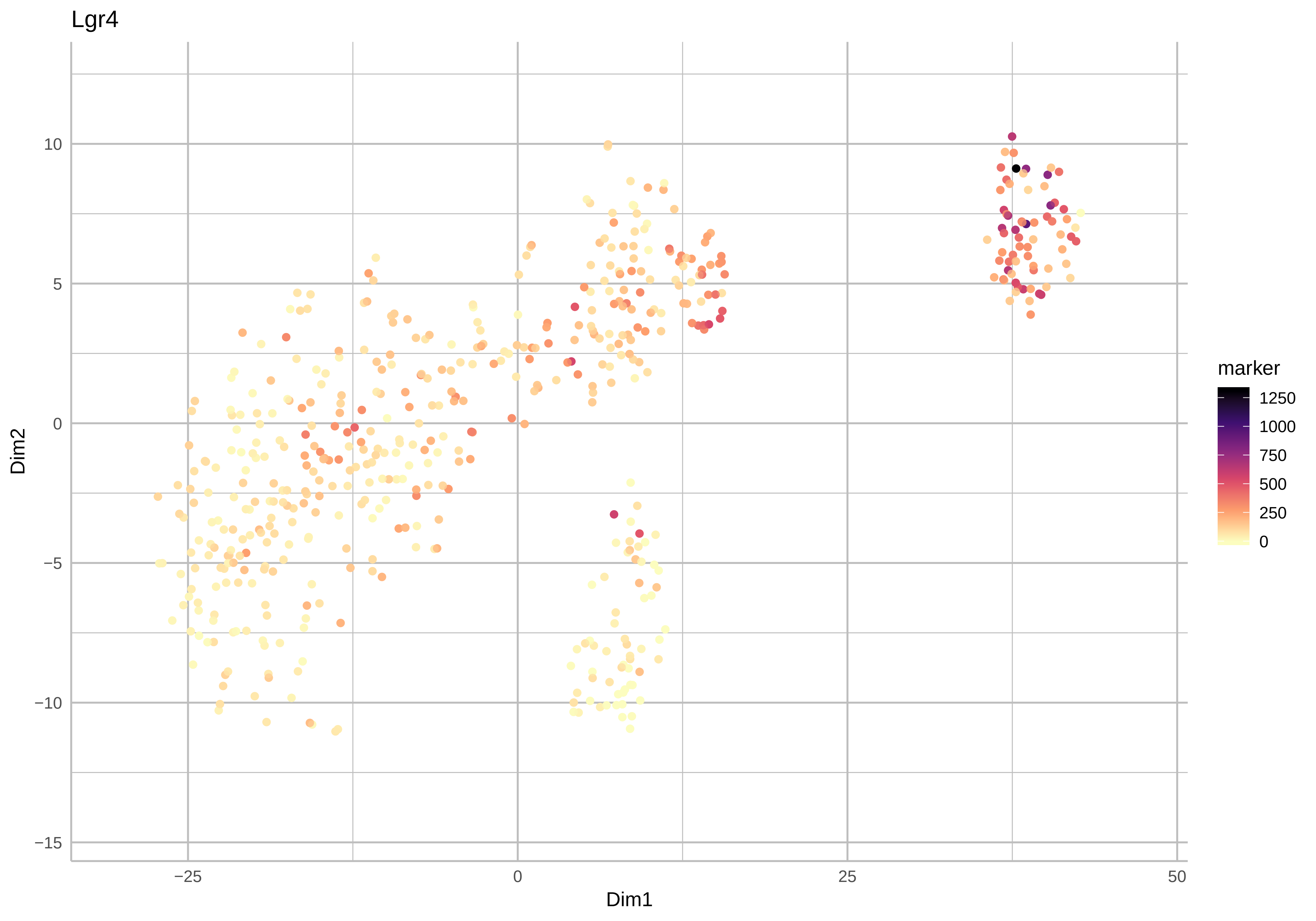
Lgr4
Dim2
Dim1
10
5
0
−5
−10
−15
−25
0
25
50
marker
1250
1000
750
500
250
0

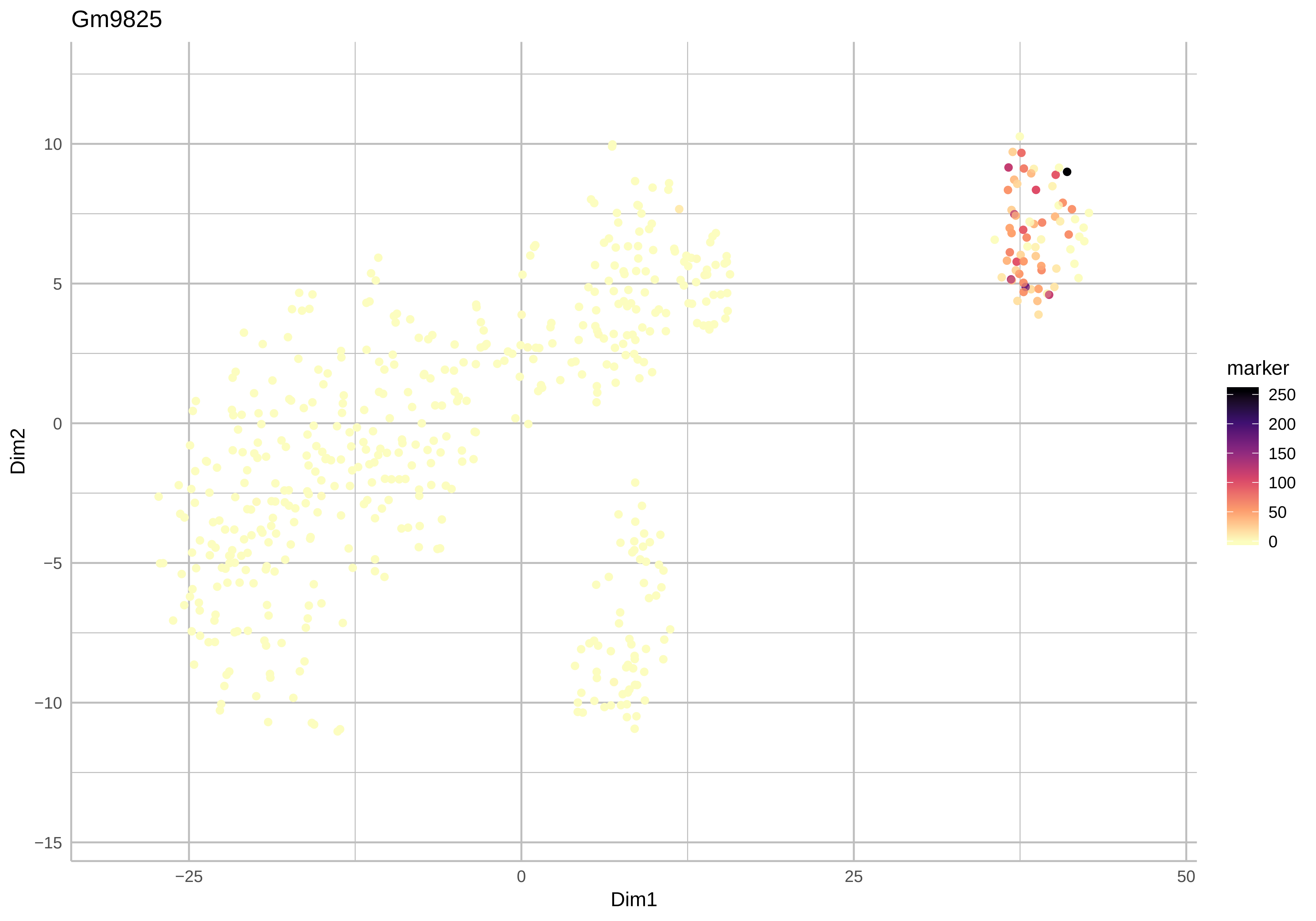

Gm9825
Dim2
Dim1
10
5
0
−5
−10
−15
−25
0
25
50
marker
250
200
150
100
50
0

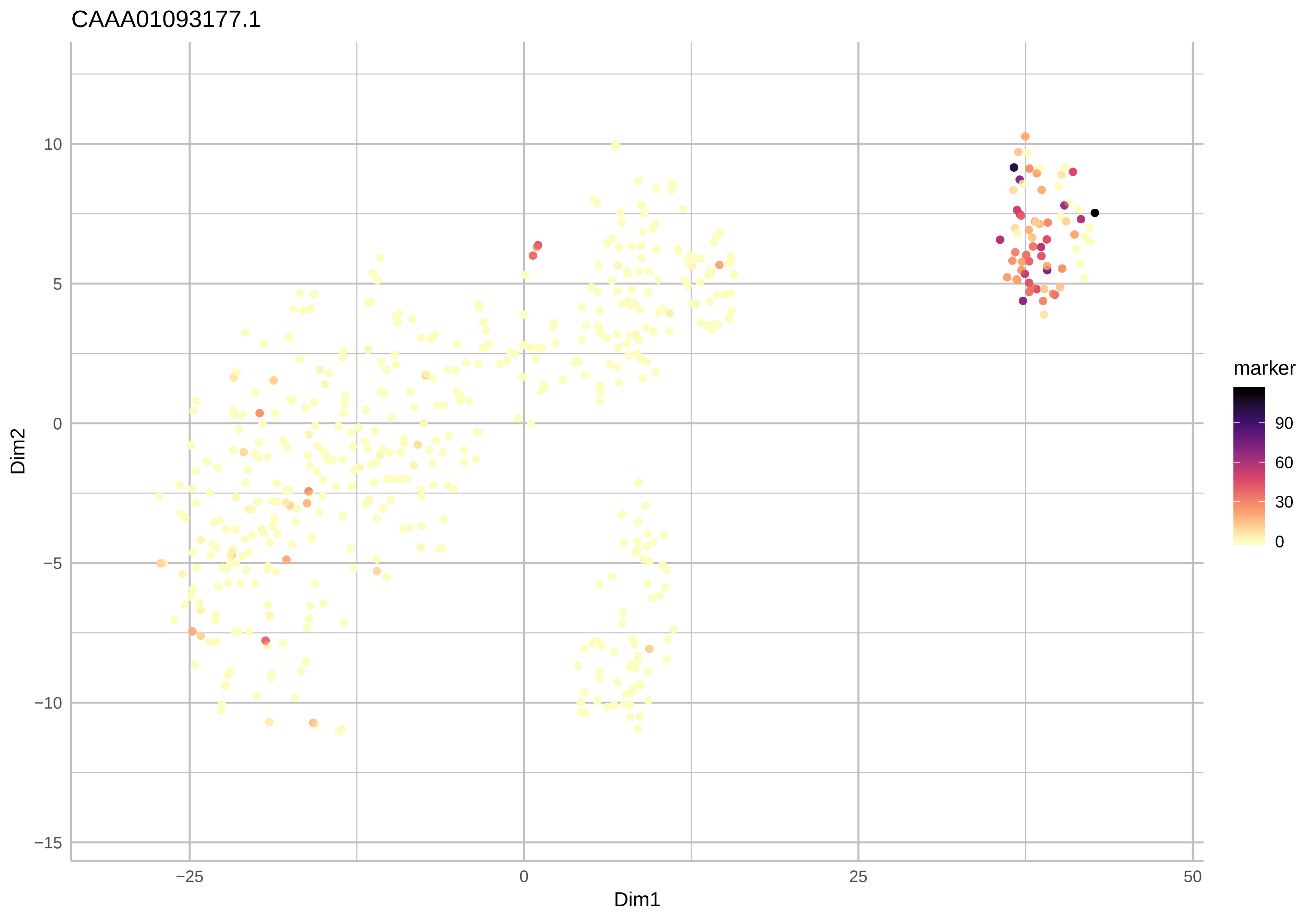
CAAA01093177.1
10
5
0
−5
−10
−15
−25
0
25
50
Dim1
Dim2
marker
90
60
30
0

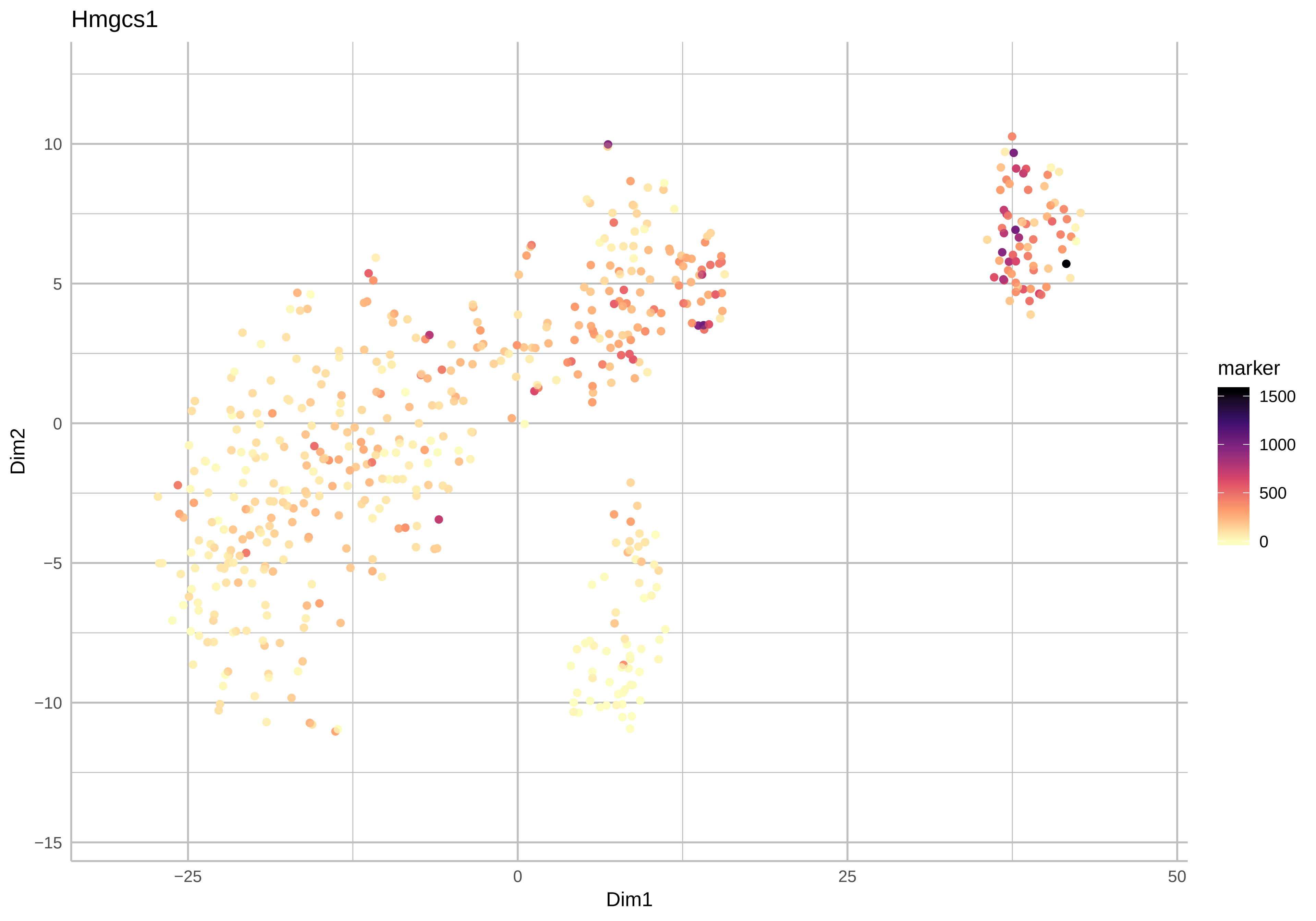

Hmgcs1
Dim2
Dim1
10
5
0
−5
−10
−15
−25
0
25
50
marker
1500
1000
500
0

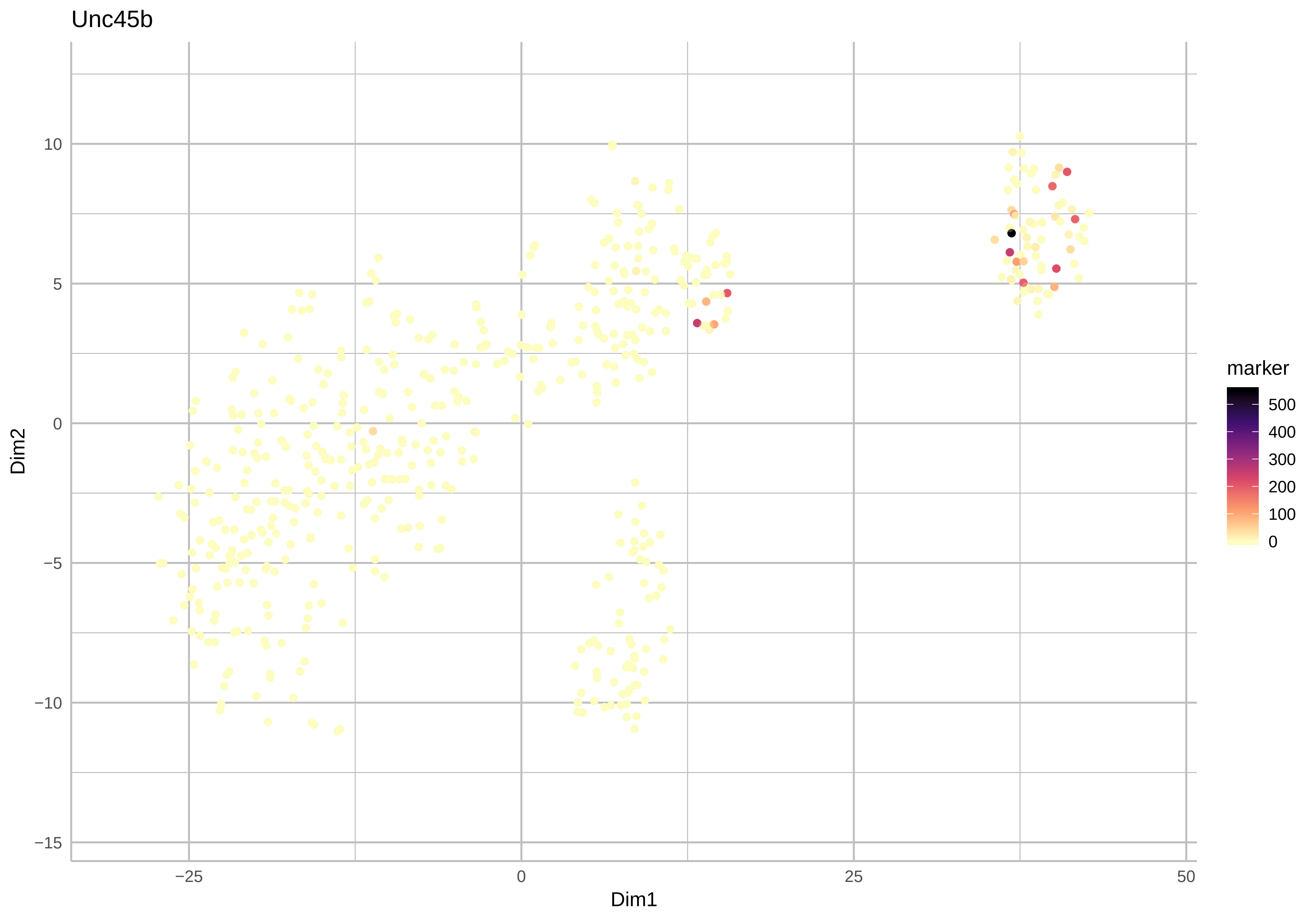

Unc45b
Dim2
Dim1
−25
0
25
50
10
5
0
−5
−10
−15
marker
500
400
300
200
100
0

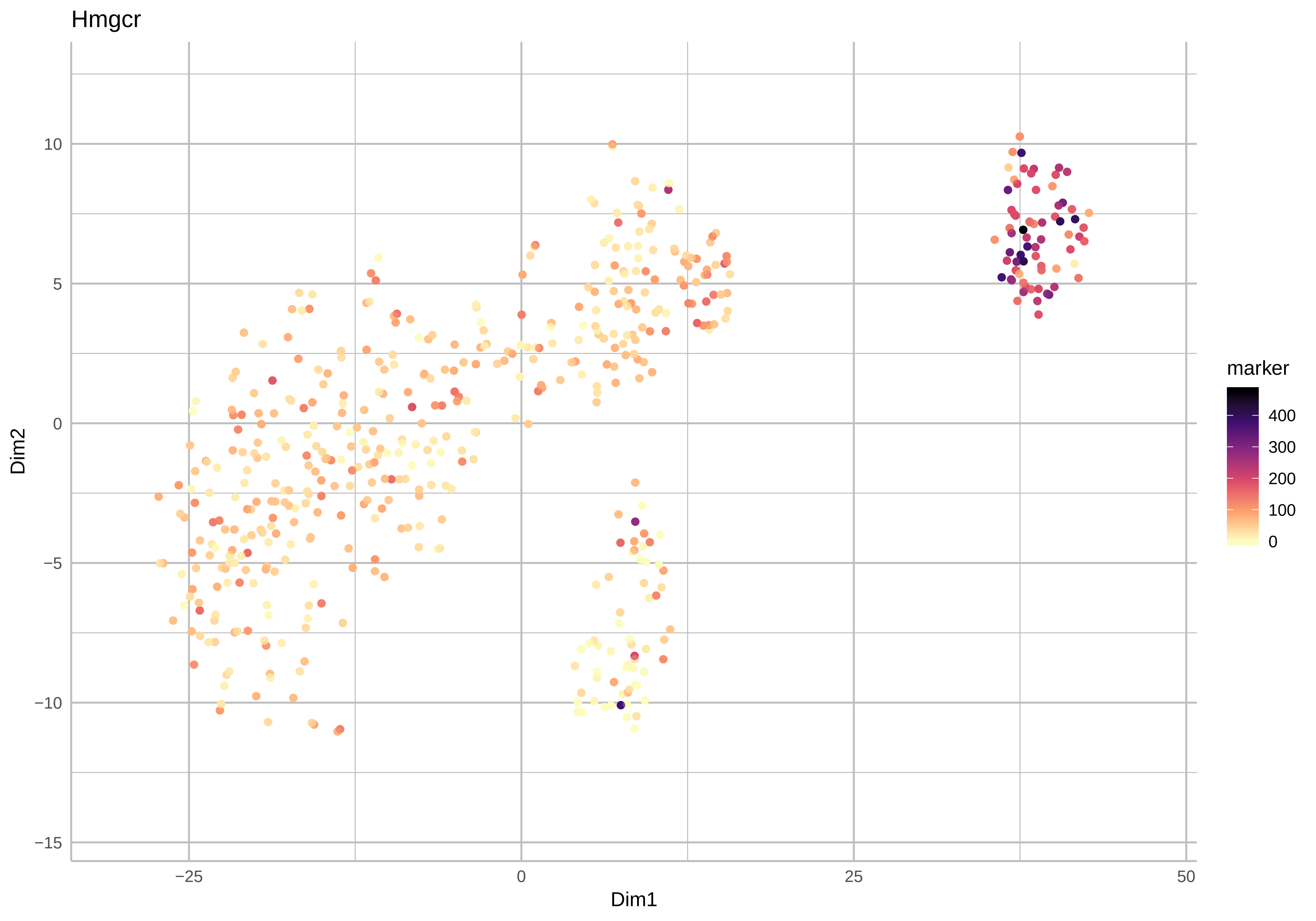

Hmgcr
Dim2
Dim1
10
5
0
−5
−10
−15
−25
0
25
50
marker
400
300
200
100
0

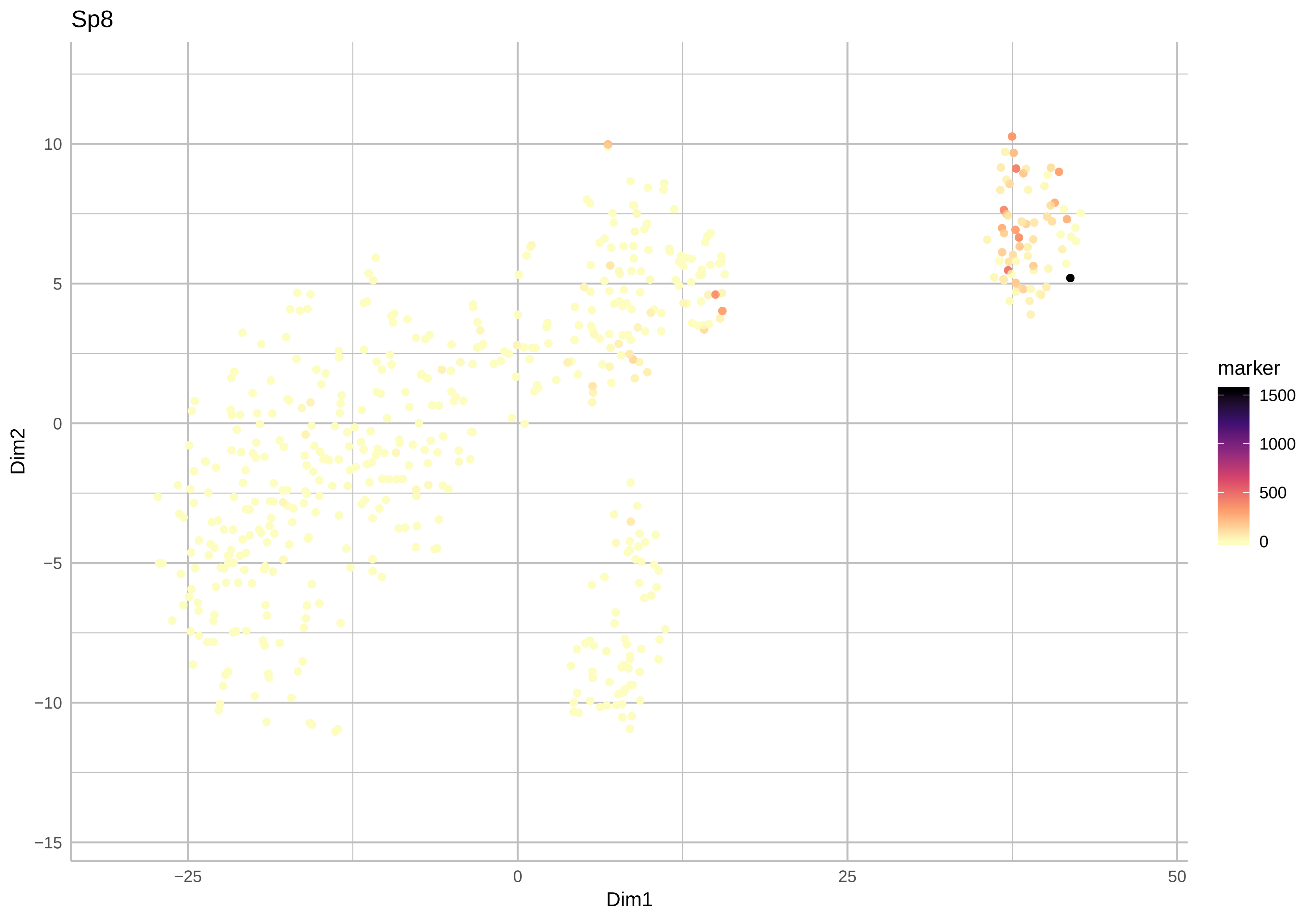
Sp8
10
5
0
−5
−10
−15
Dim2
−25
0
25
50
Dim1
marker
1500
1000
500
0

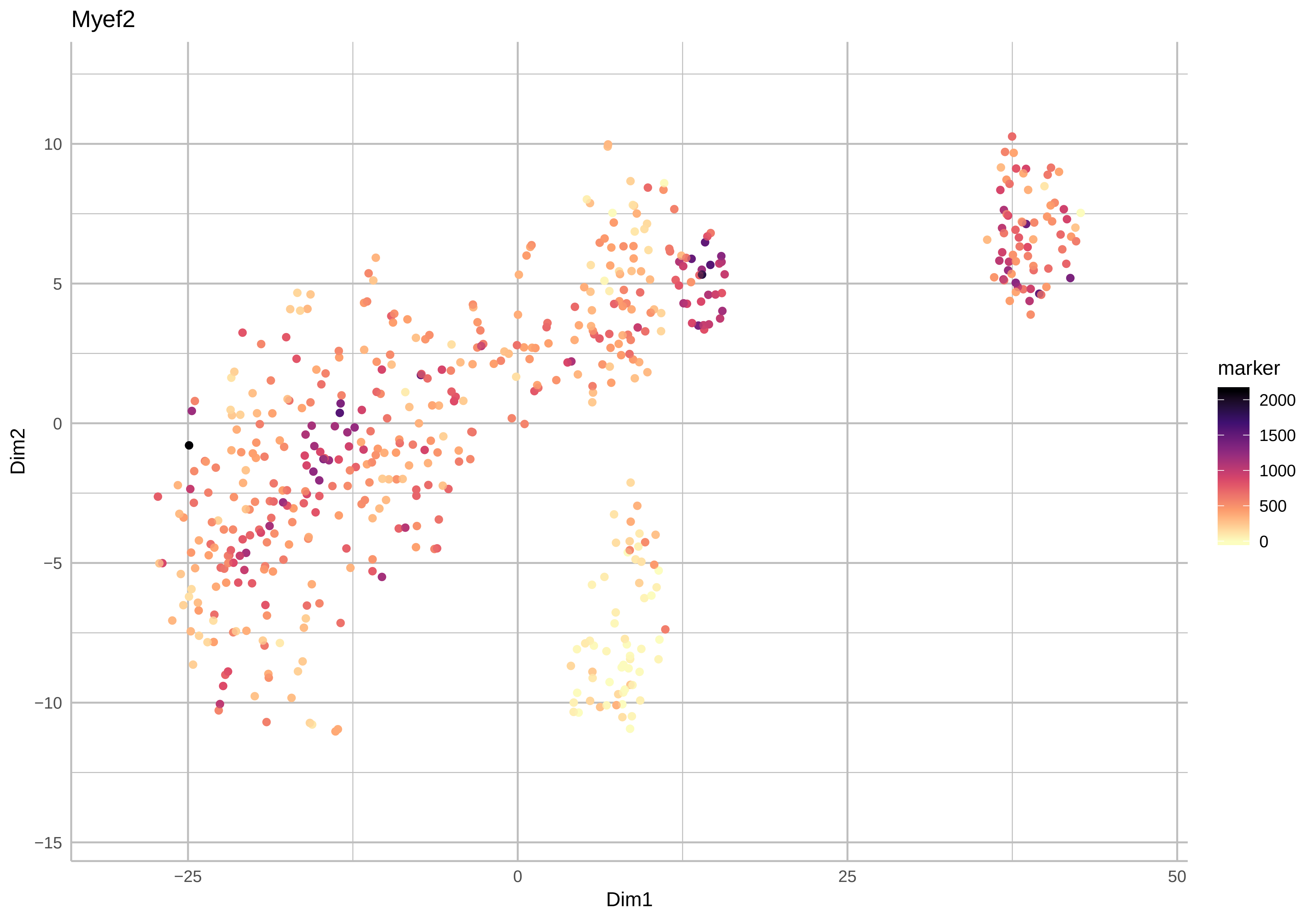

Myef2
Dim2
Dim1
10
5
0
−5
−10
−15
−25
0
25
50
marker
2000
1500
1000
500
0

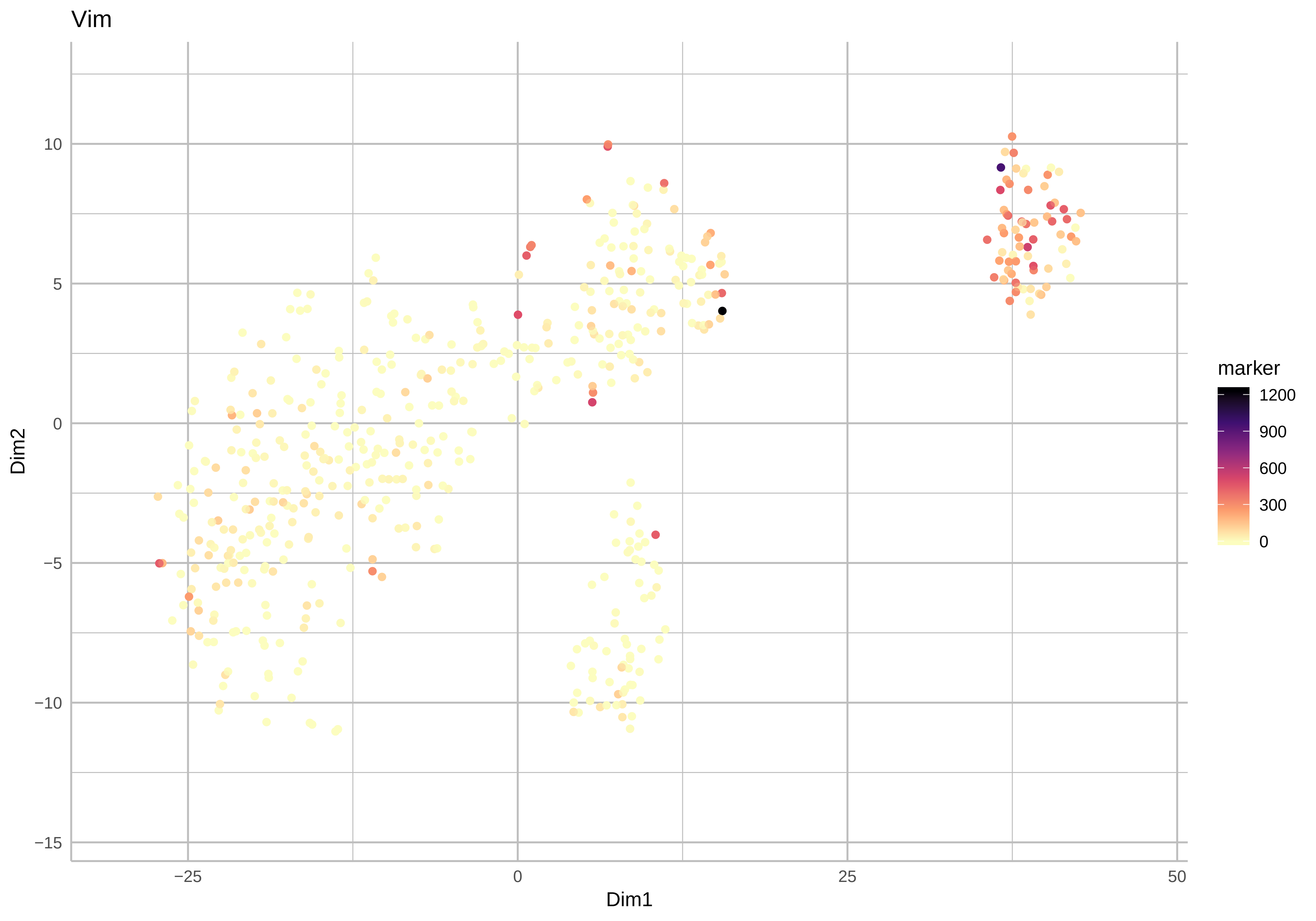
Vim
Dim2
Dim1
10
5
0
−5
−10
−15
−25
0
25
50
marker
1200
900
600
300
0

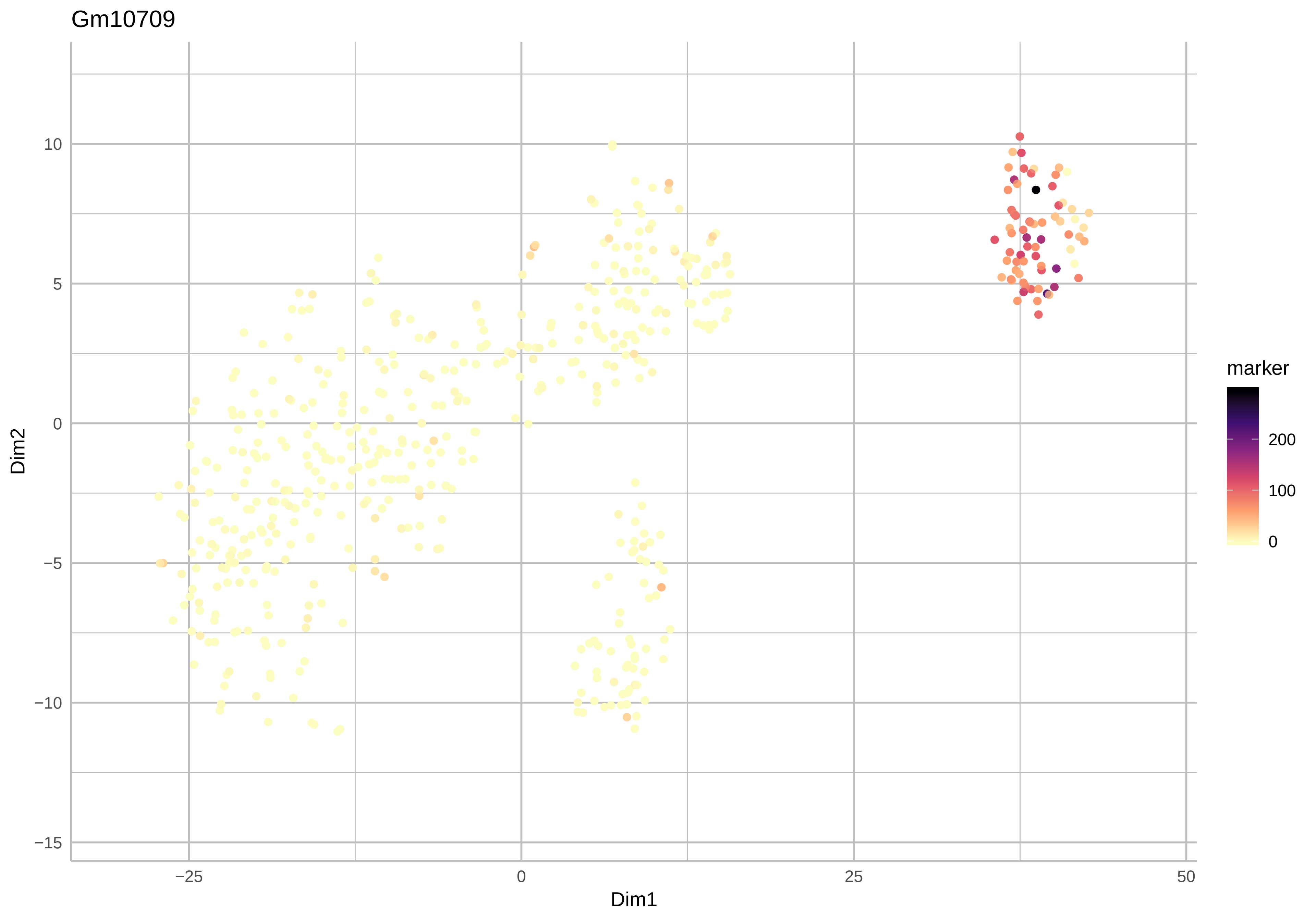

Gm10709
Dim2
Dim1
marker
200
100
0
10
5
0
−5
−10
−15
−25
0
25
50

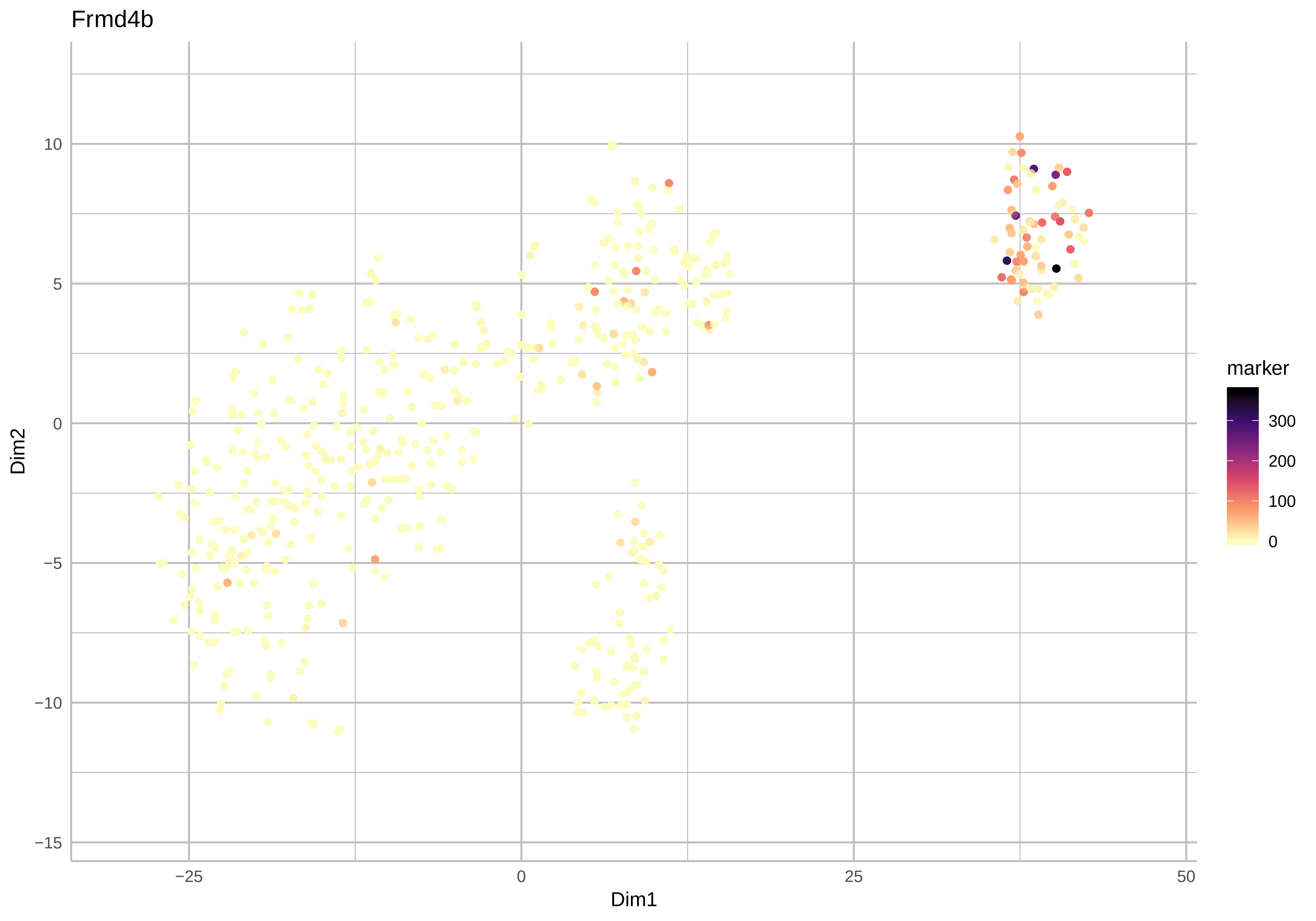
Frmd4b
Dim2
Dim1
10
5
0
−5
−10
−15
−25
0
25
50
marker
300
200
100
0

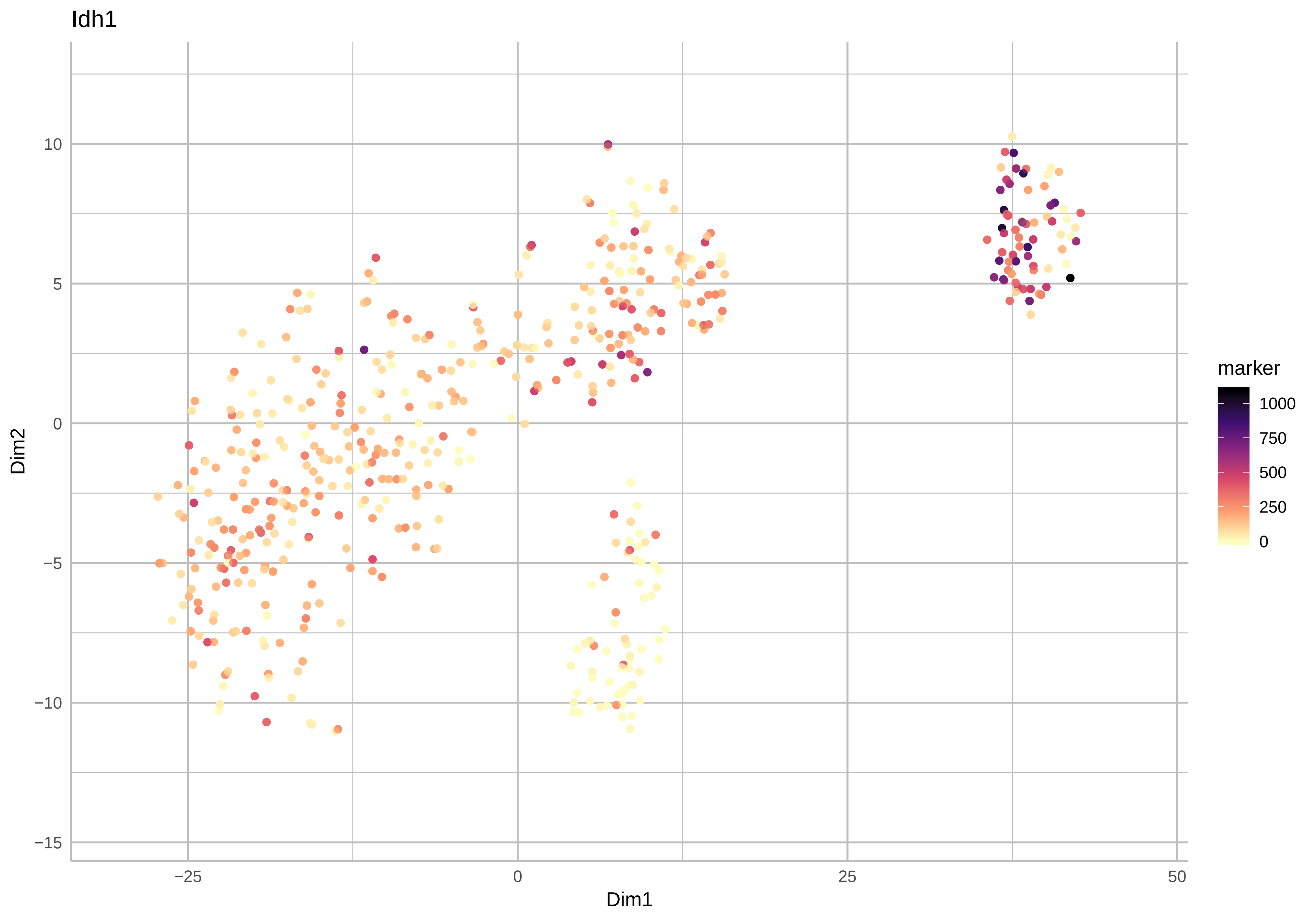

Idh1
Dim2
Dim1
10
5
0
−5
−10
−15
−25
0
25
50
marker
1000
750
500
250
0

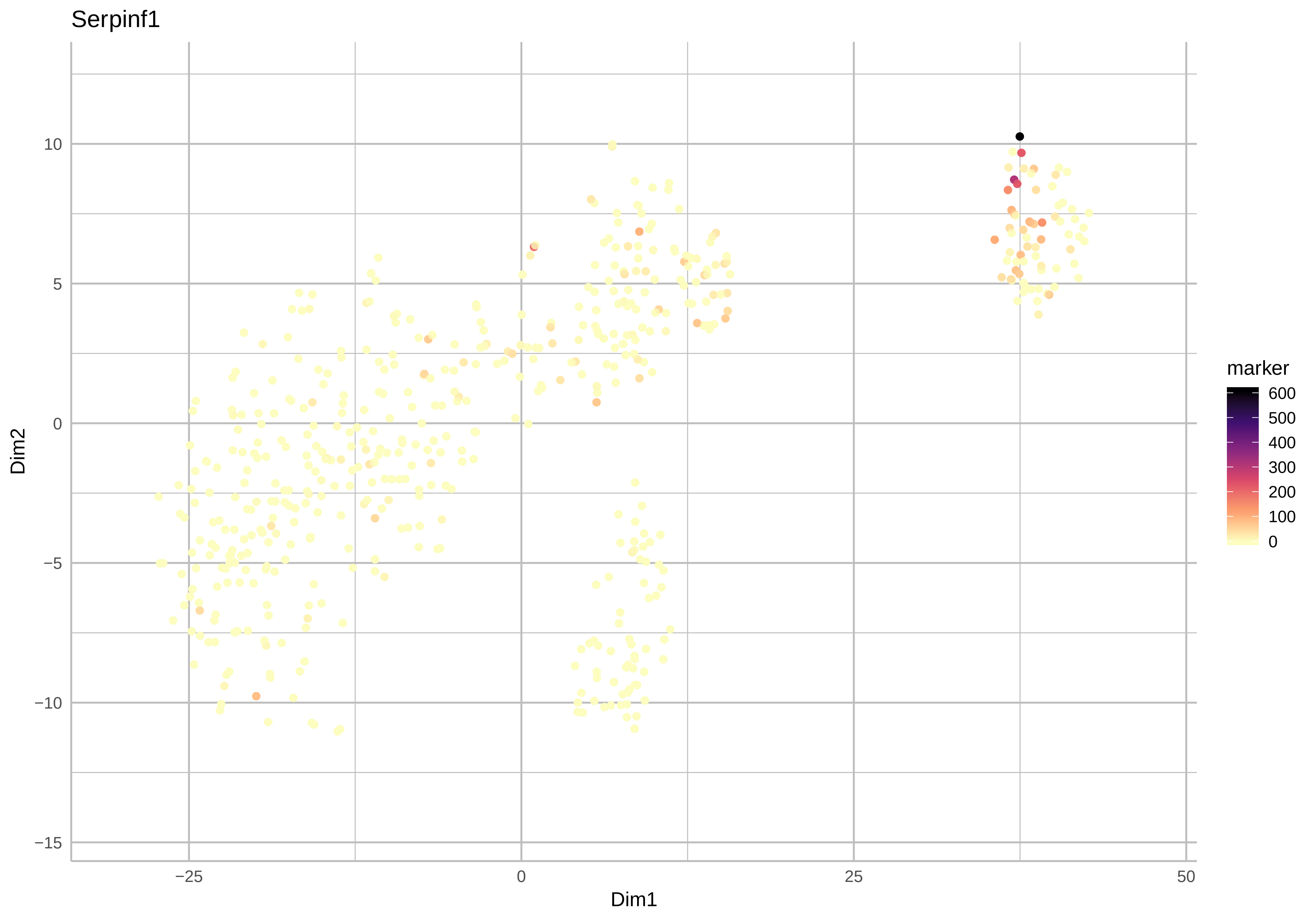

Serpinf1
Dim2
Dim1
10
5
0
−5
−10
−15
−25
0
25
50
marker
600
500
400
300
200
100
0

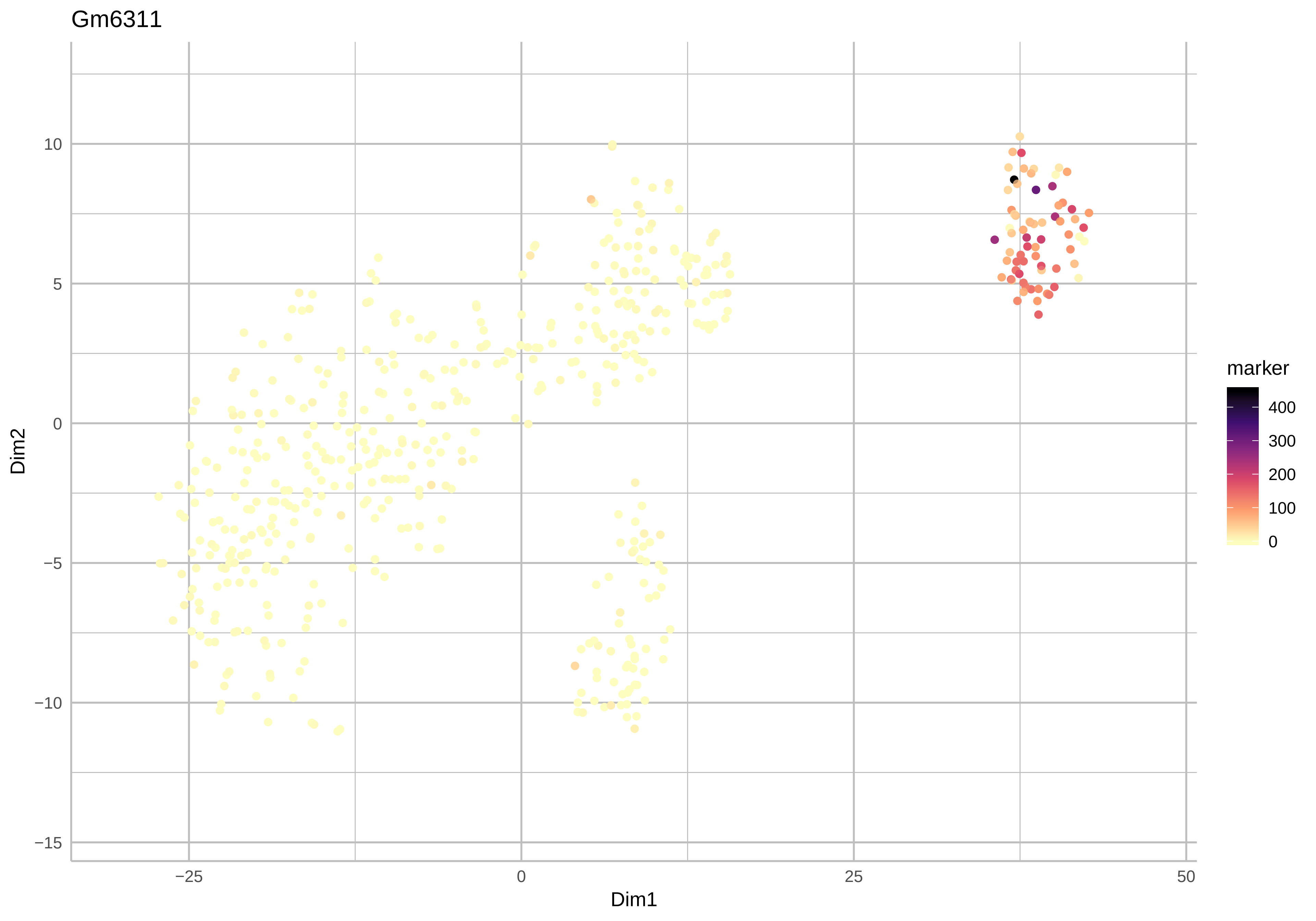
Gm6311
Dim1
Dim2
−25
0
25
50
10
5
0
−5
−10
−15
marker
400
300
200
100
0

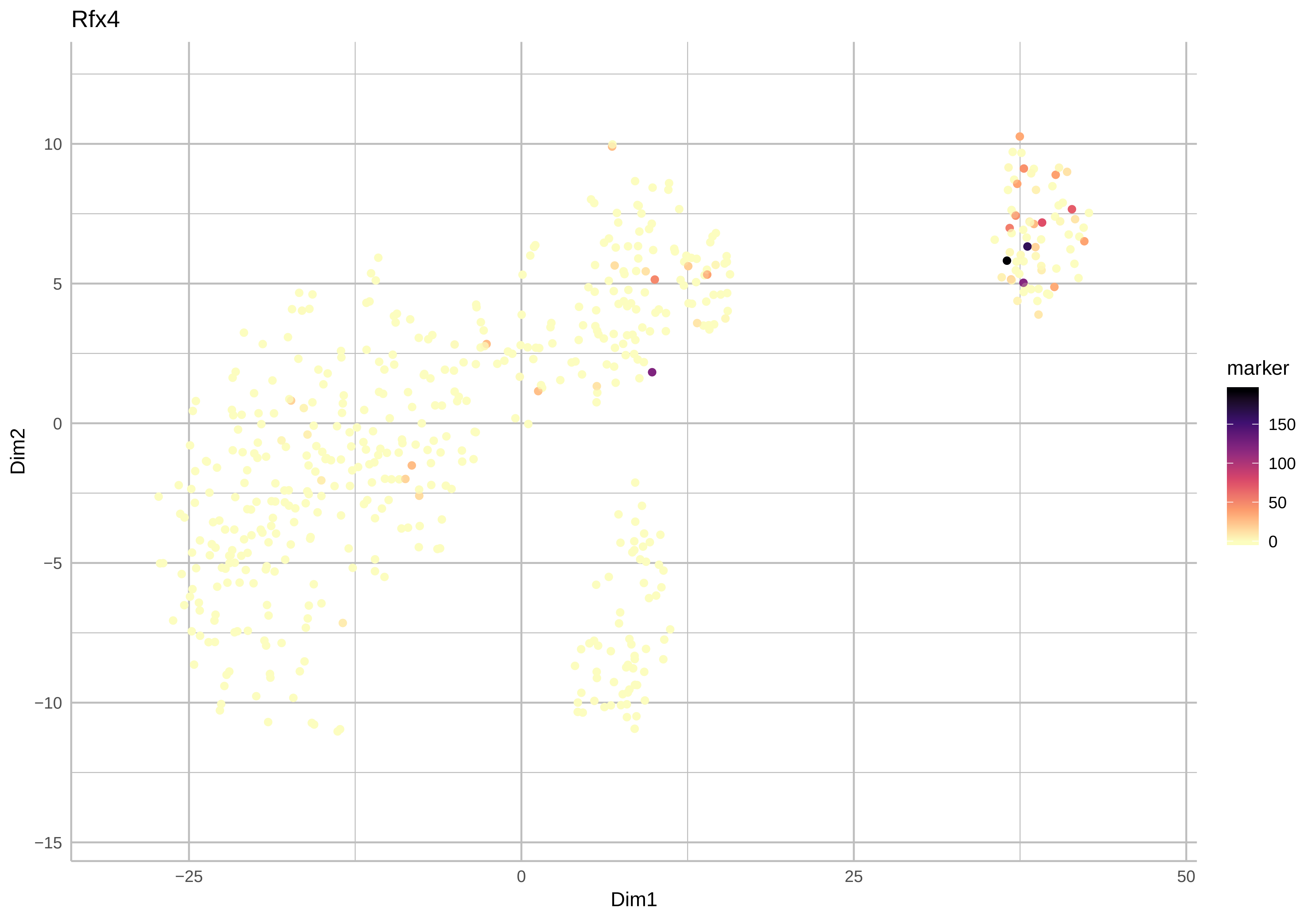
Rfx4
Dim2
Dim1
marker
150
100
50
0
10
5
0
−5
−10
−15
−25
0
25
50

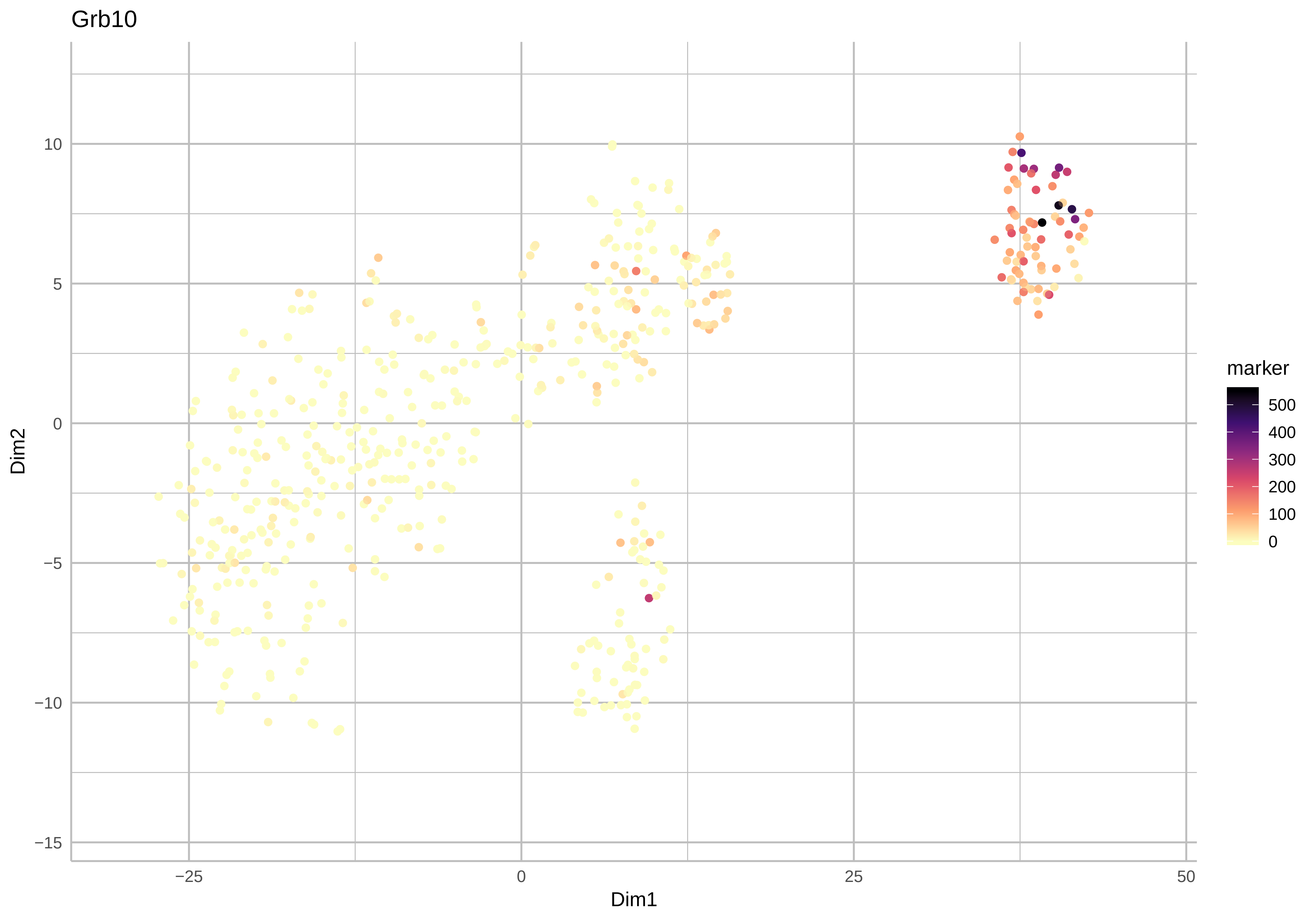

Grb10
10
5
0
−5
−10
−15
Dim2
−25
0
25
50
Dim1
marker
500
400
300
200
100
0

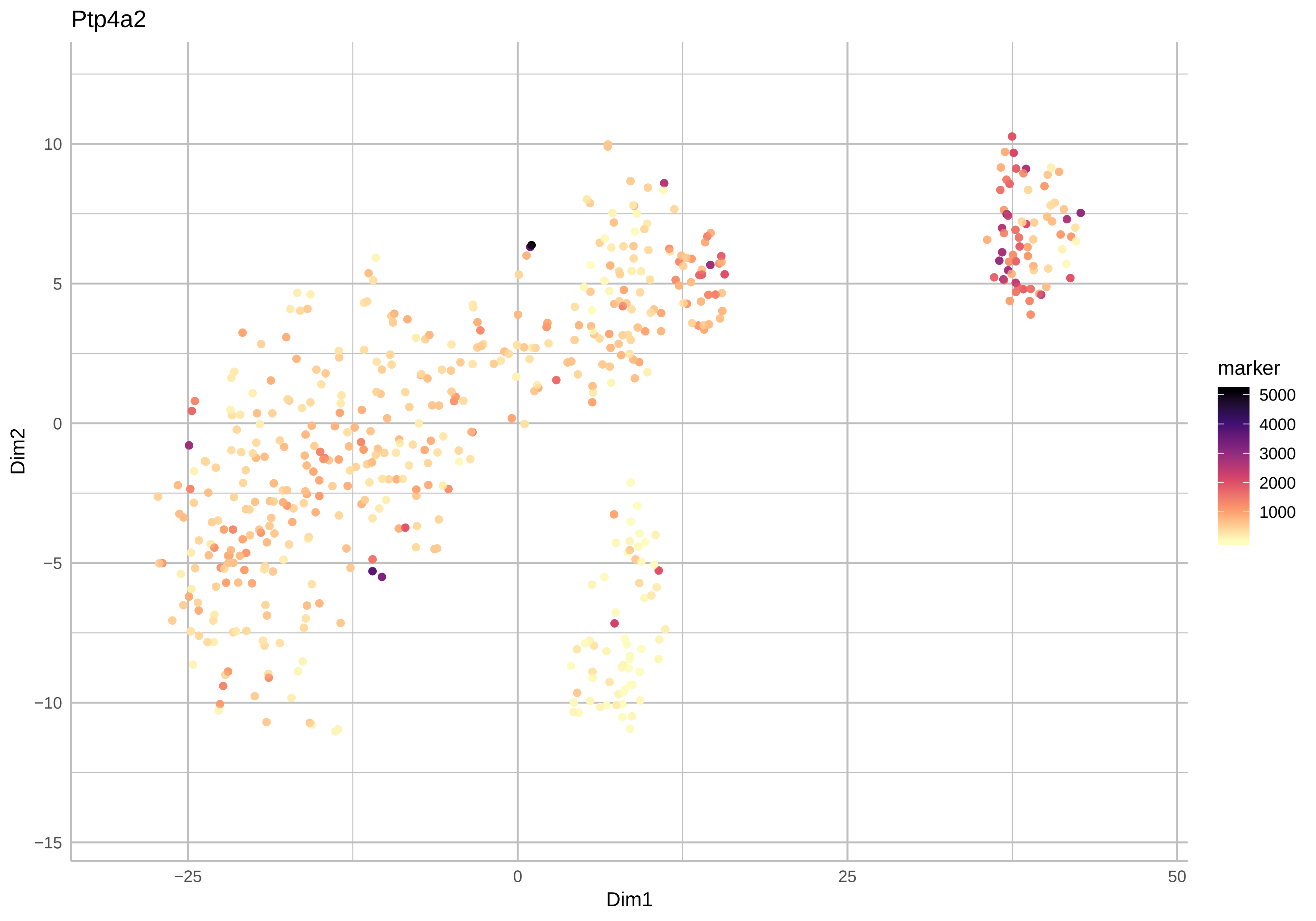

Ptp4a2
marker
5000
4000
3000
2000
1000
Dim2
10
5
0
−5
−10
−15
Dim1
−25
0
25
50

Gm4724

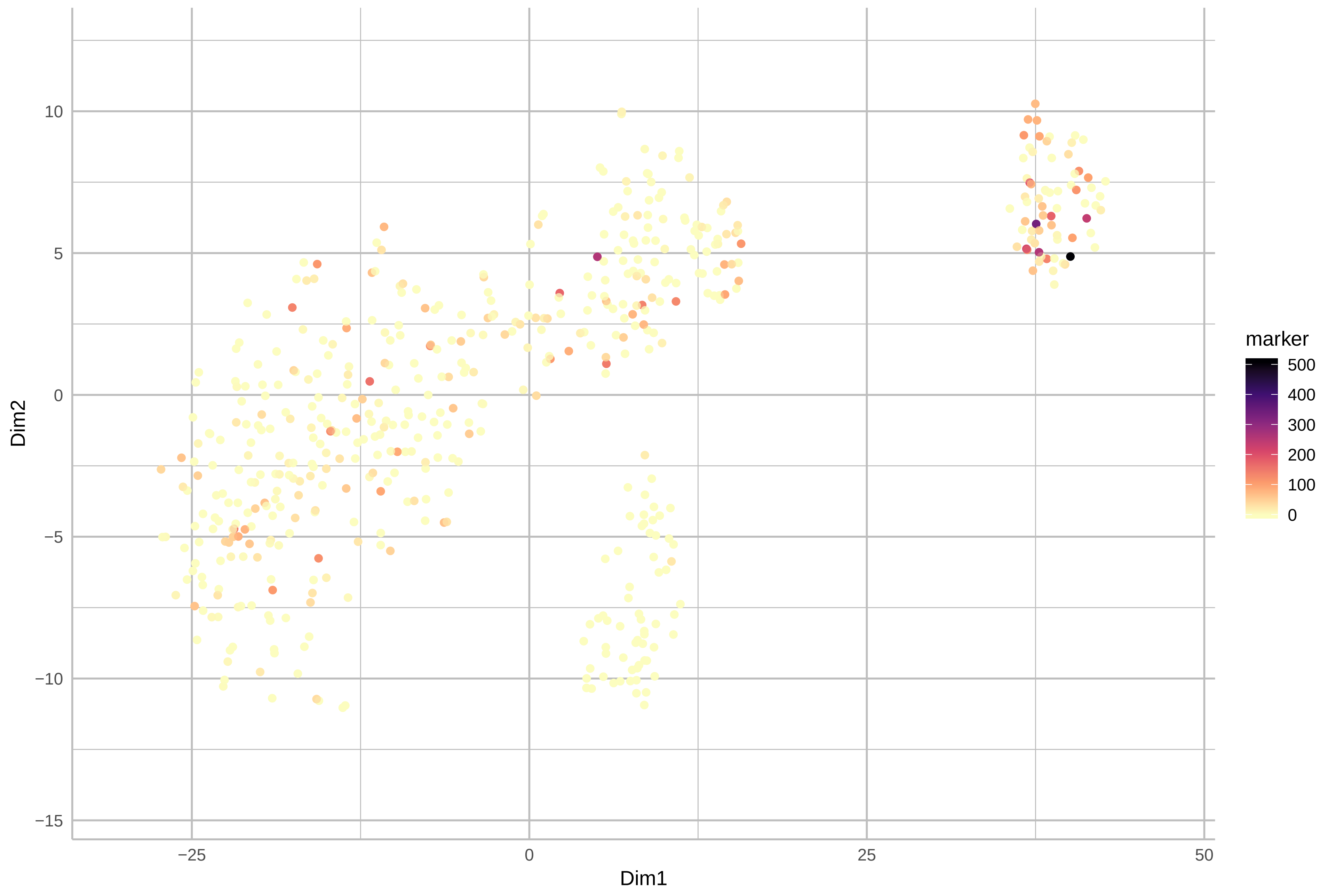

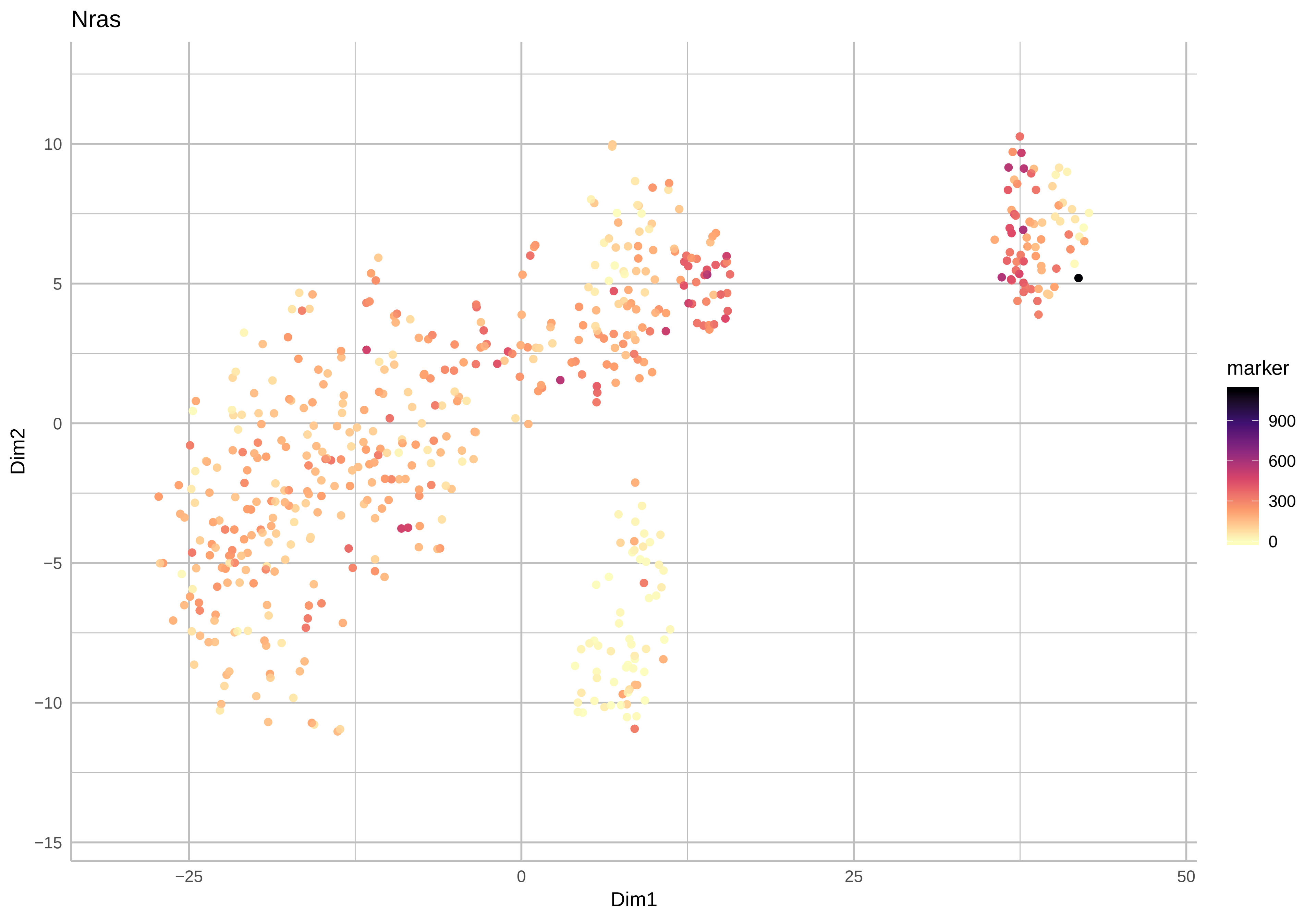
Nras
Dim2
Dim1
marker
900
600
300
0
10
5
0
−5
−10
−15
−25
0
25
50

Idi1

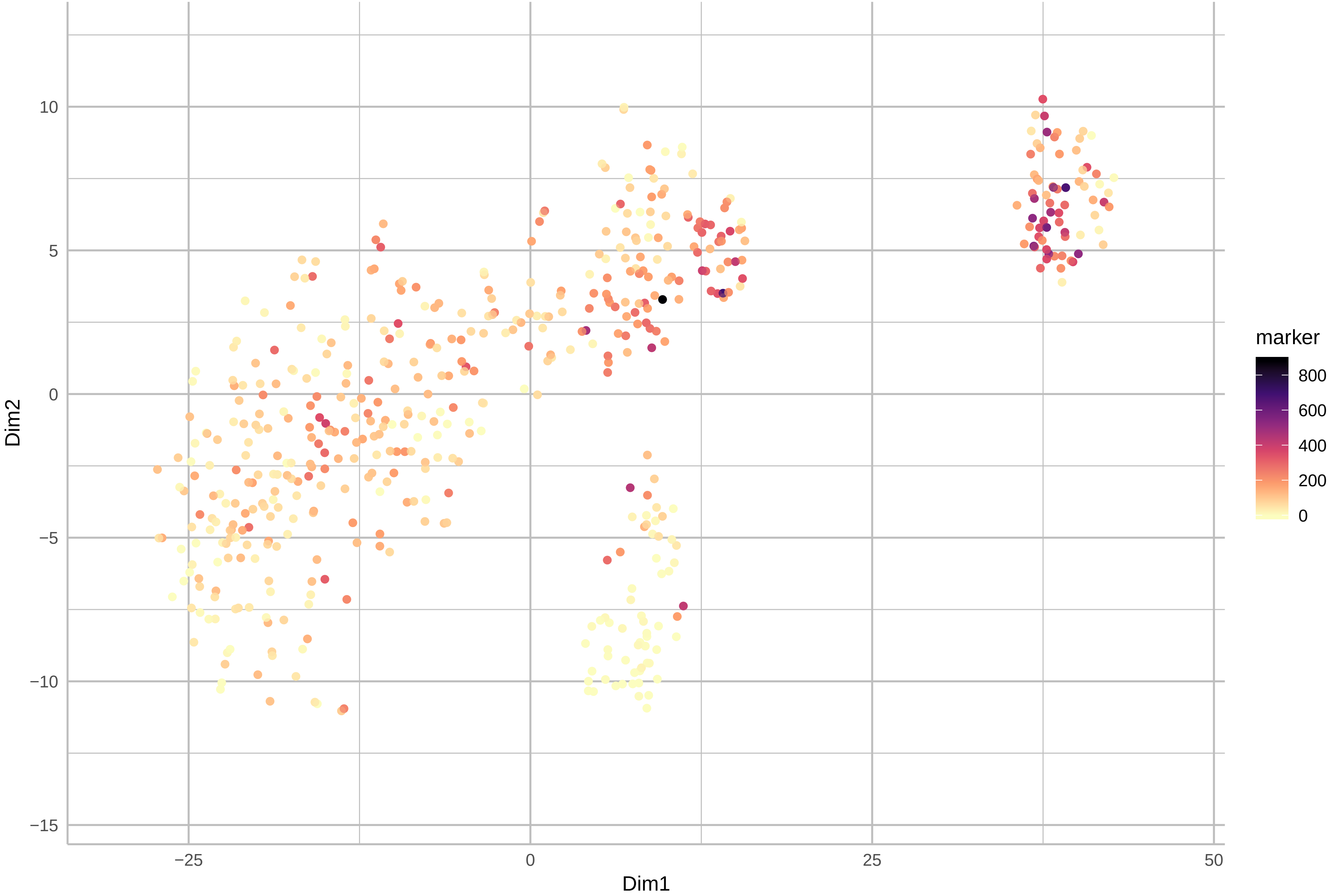

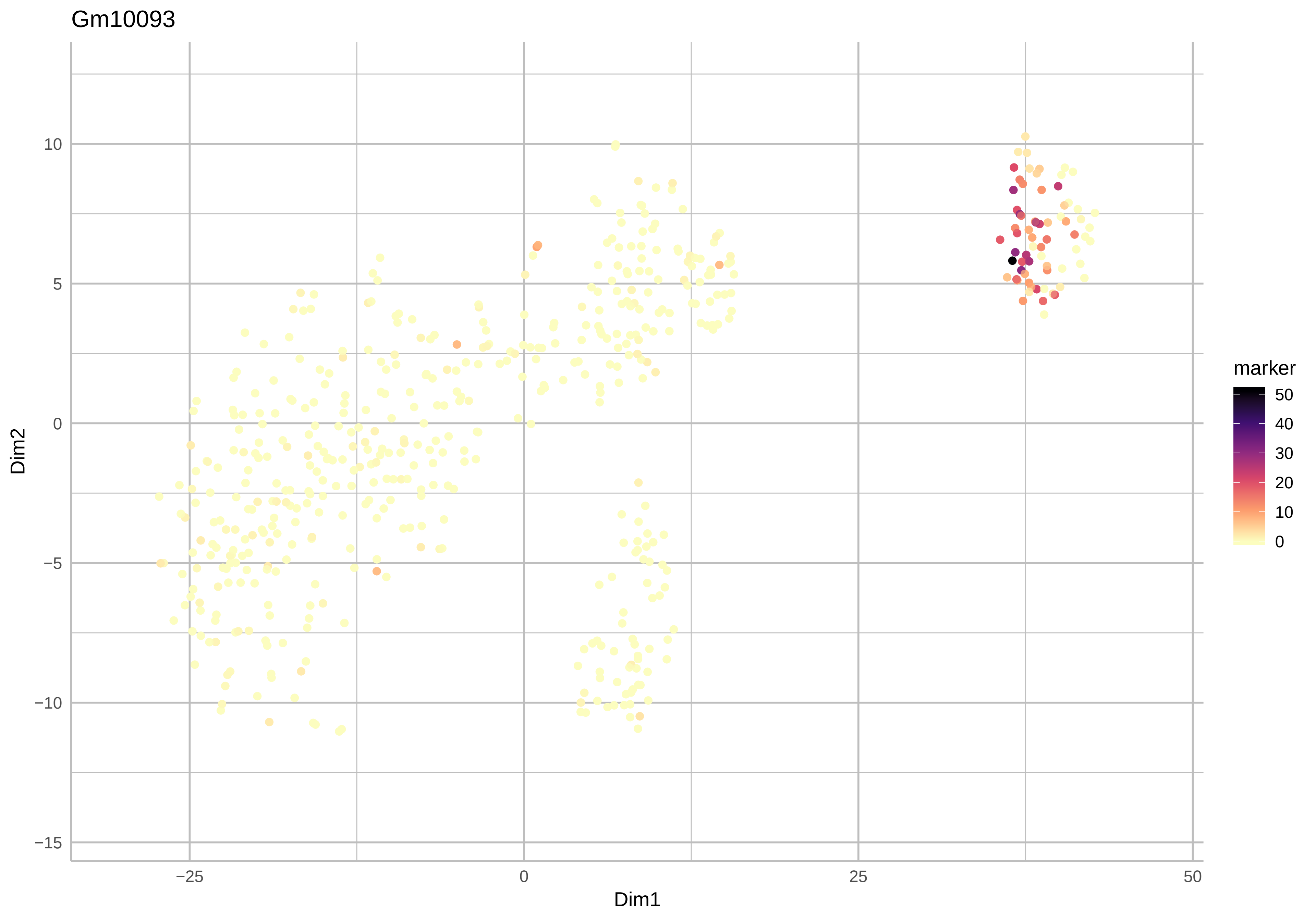
Gm10093
Dim2
10
5
0
−5
−10
−15
Dim1
−25
0
25
50
marker
50
40
30
20
10
0

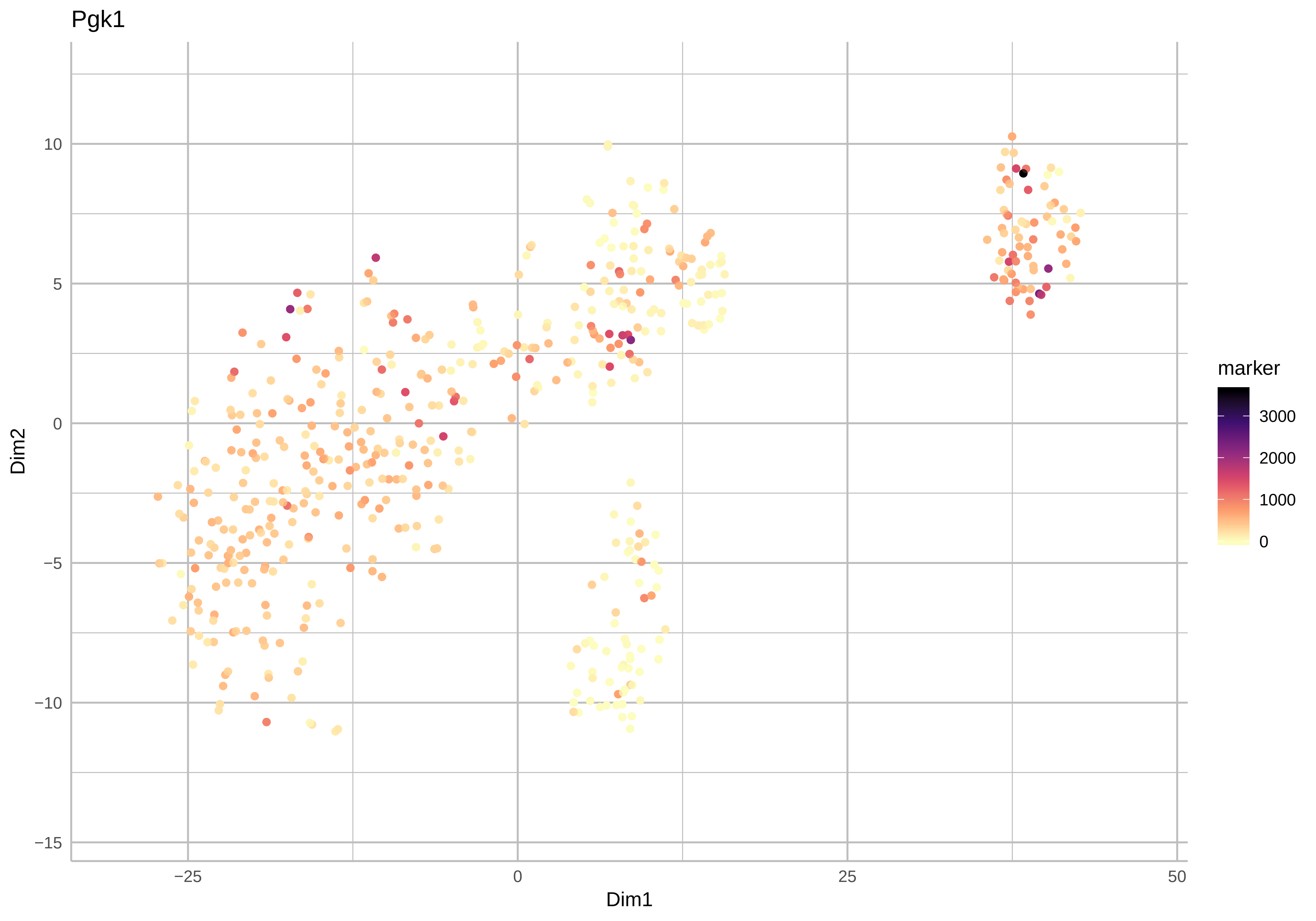

Pgk1
Dim2
10
5
0
−5
−10
−15
−25
0
25
50
Dim1
marker
3000
2000
1000
0

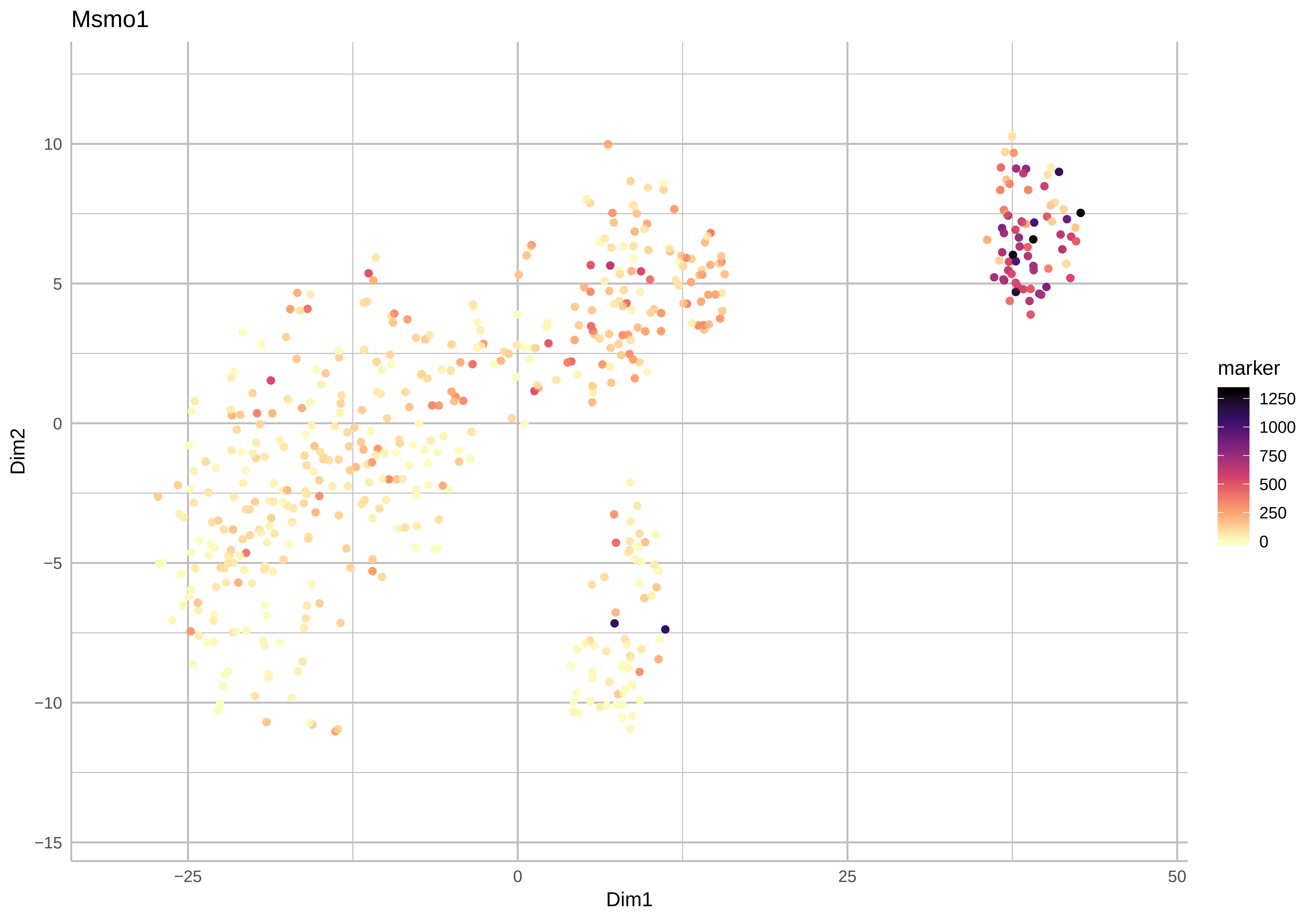

Msmo1
Dim1
Dim2
−25
0
25
50
10
5
0
−5
−10
−15
marker
1250
1000
750
500
250
0

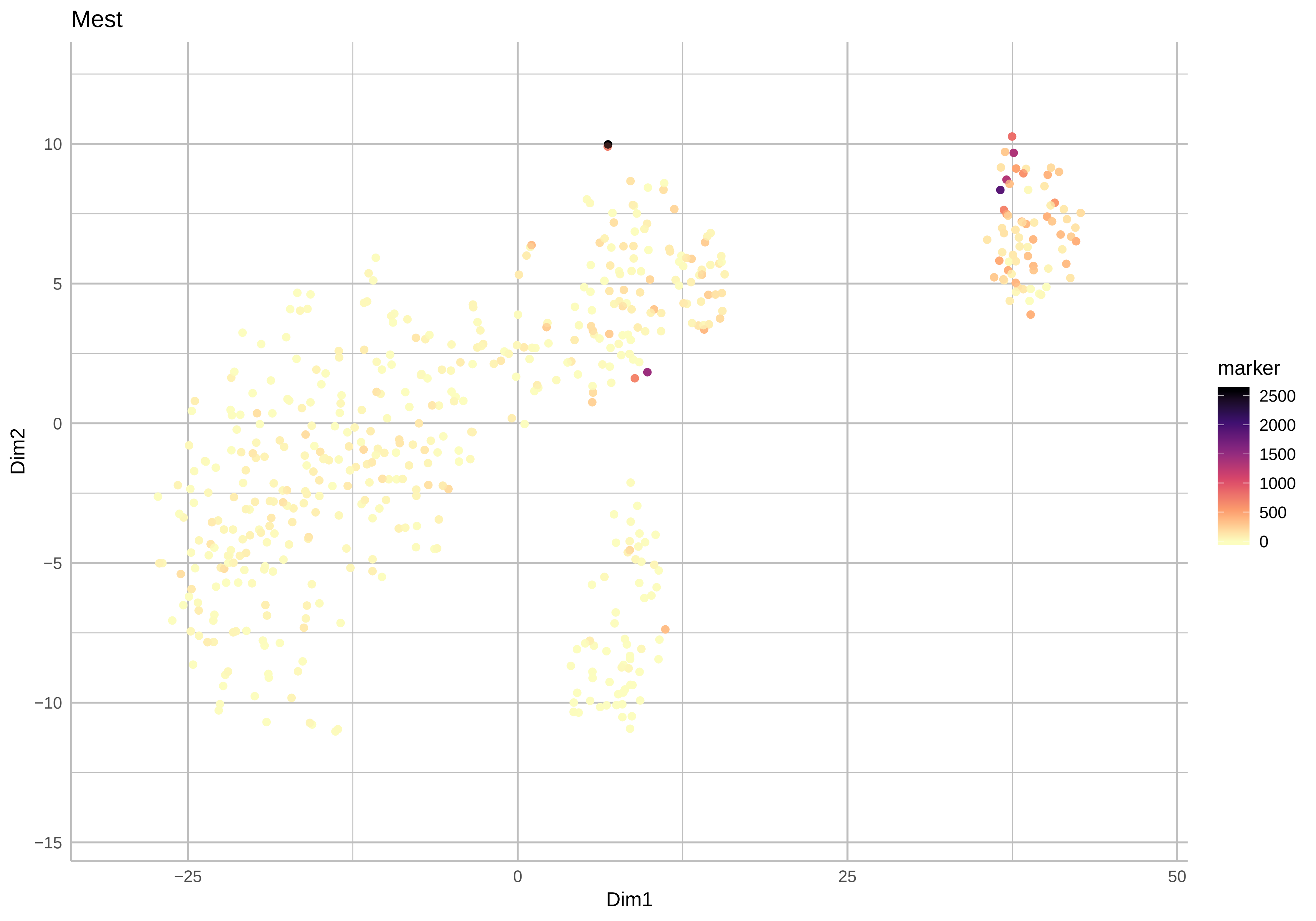
Mest
Dim2
Dim1
10
5
0
−5
−10
−15
−25
0
25
50
marker
2500
2000
1500
1000
500
0

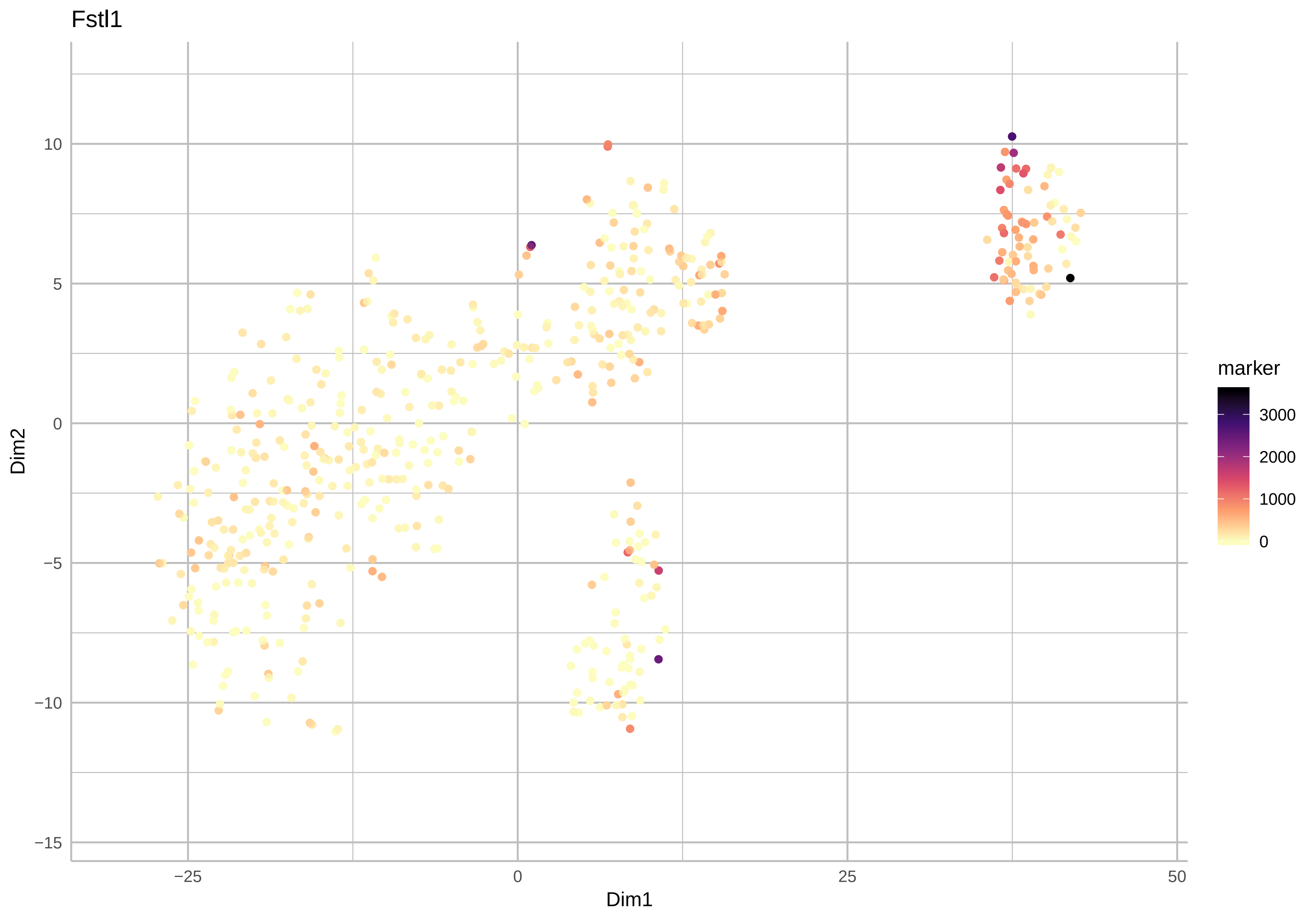

Fstl1
Dim2
Dim1
marker
3000
2000
1000
0
10
5
0
−5
−10
−15
−25
0
25
50

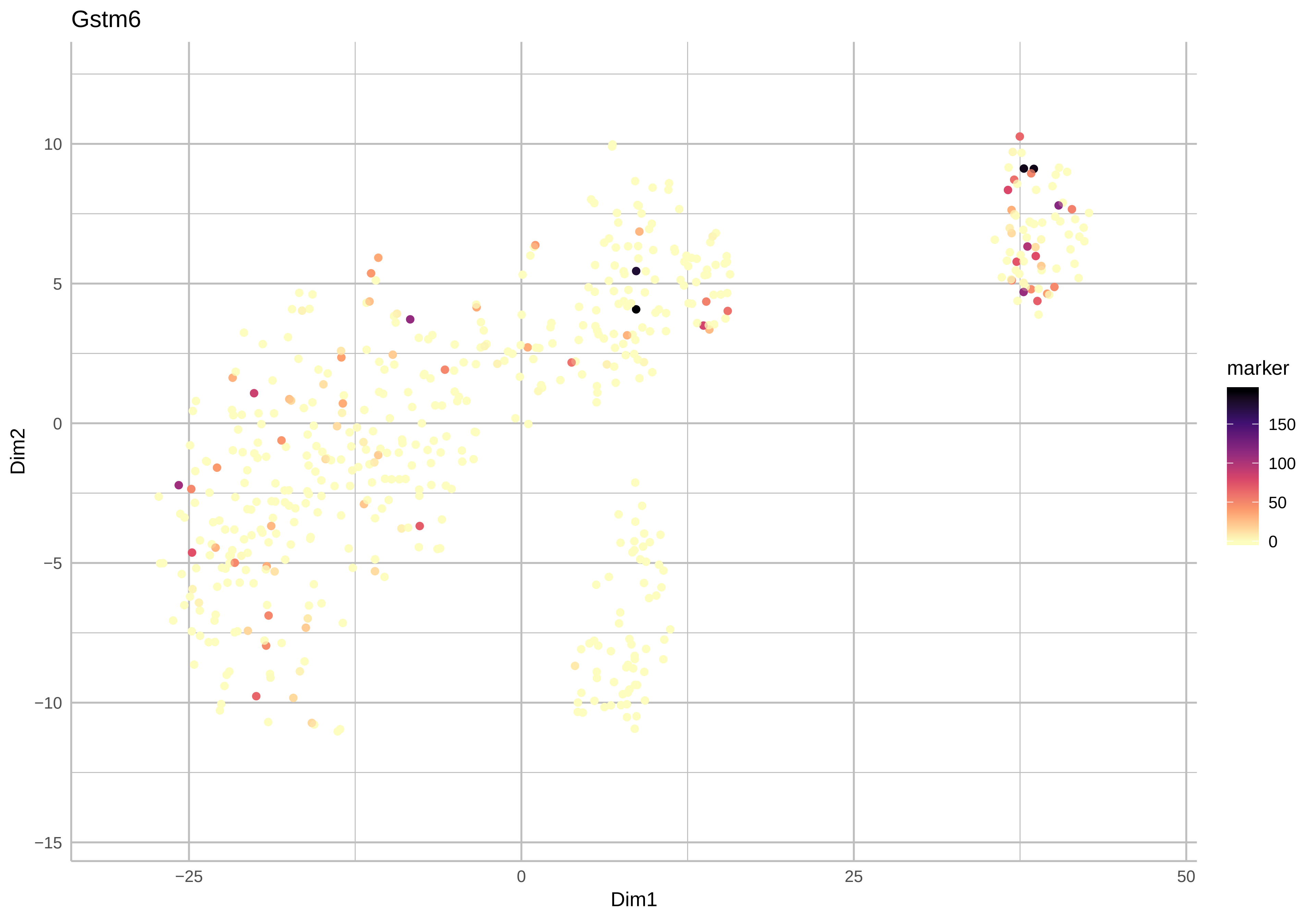

Gstm6
marker
150
100
50
0
10
5
0
−5
−10
−15
Dim2
−25
0
25
50
Dim1

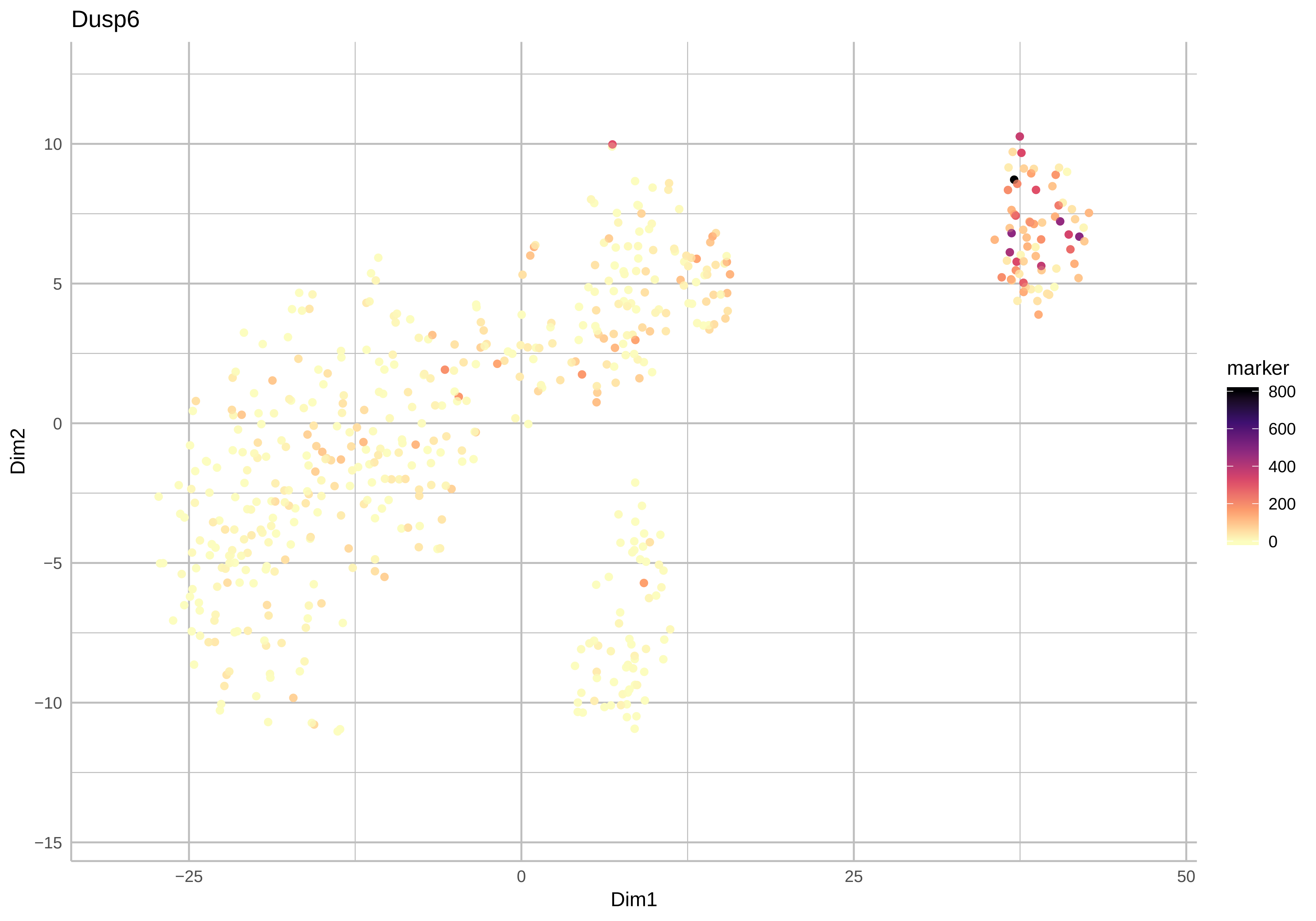

Dusp6
10
5
0
−5
−10
−15
Dim2
−25
0
25
50
Dim1
marker
800
600
400
200
0

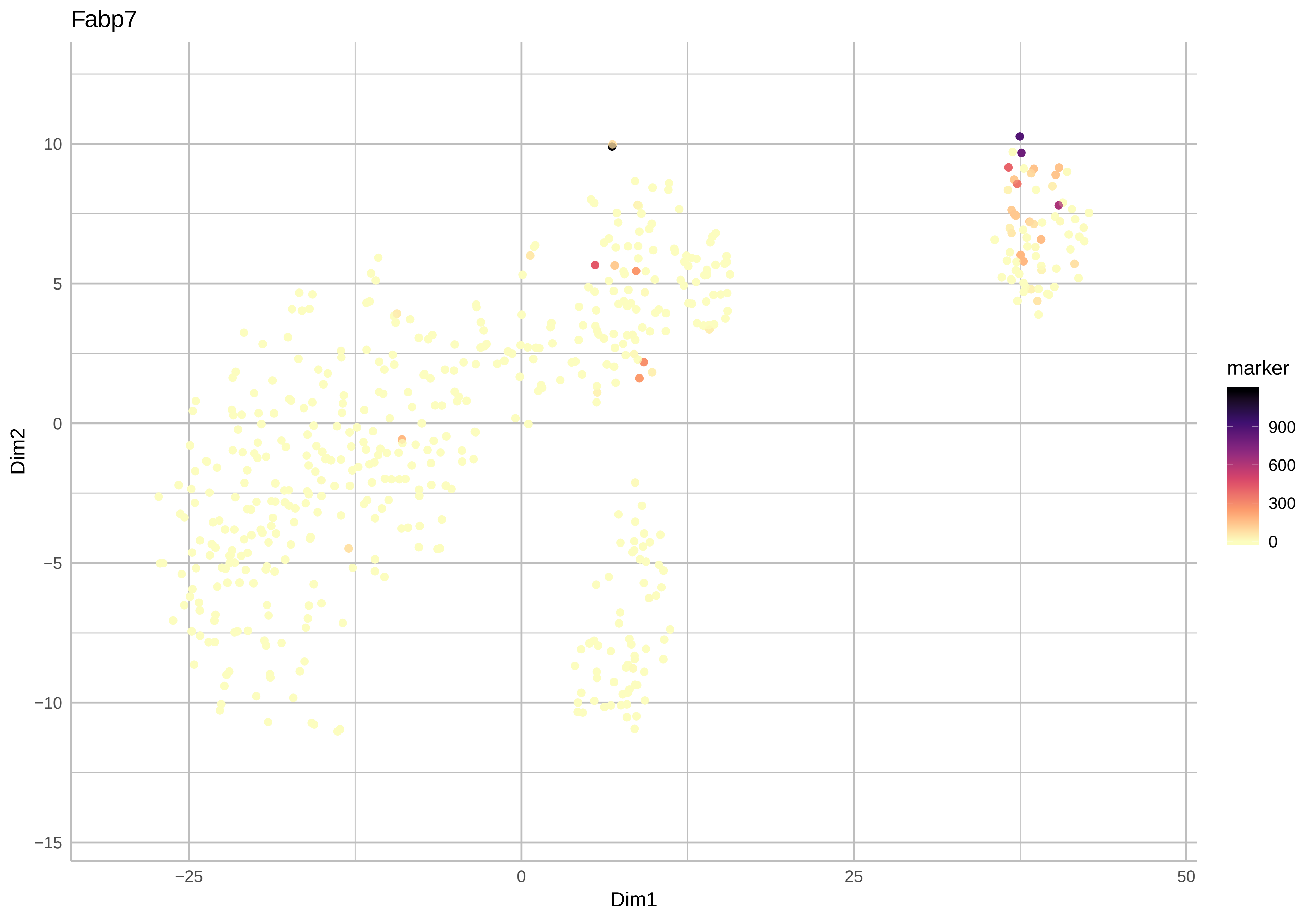

Fabp7
Dim2
Dim1
10
5
0
−5
−10
−15
−25
0
25
50
marker
900
600
300
0

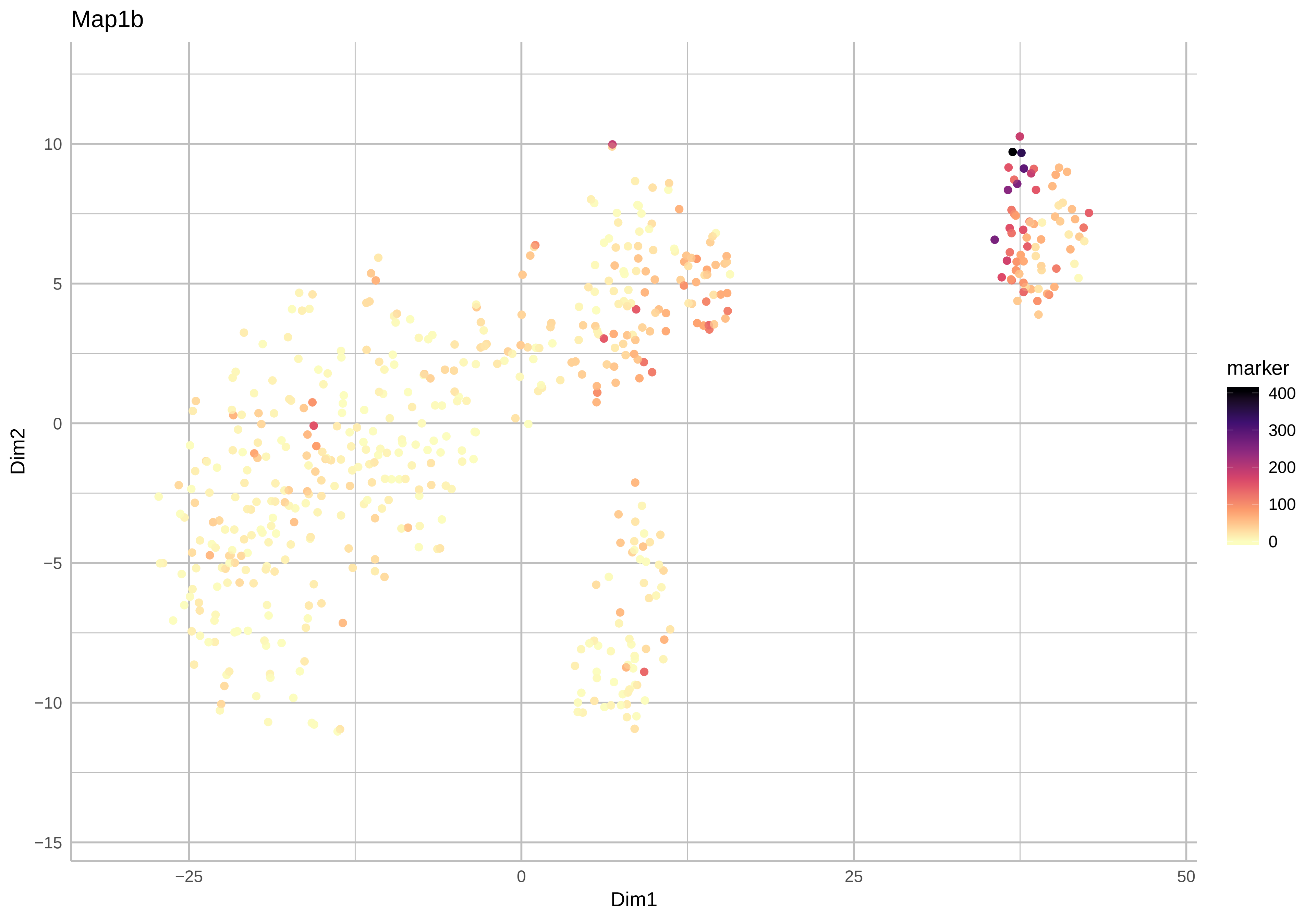

Map1b
10
5
0
−5
−10
−15
Dim2
−25
0
25
50
Dim1
marker
400
300
200
100
0

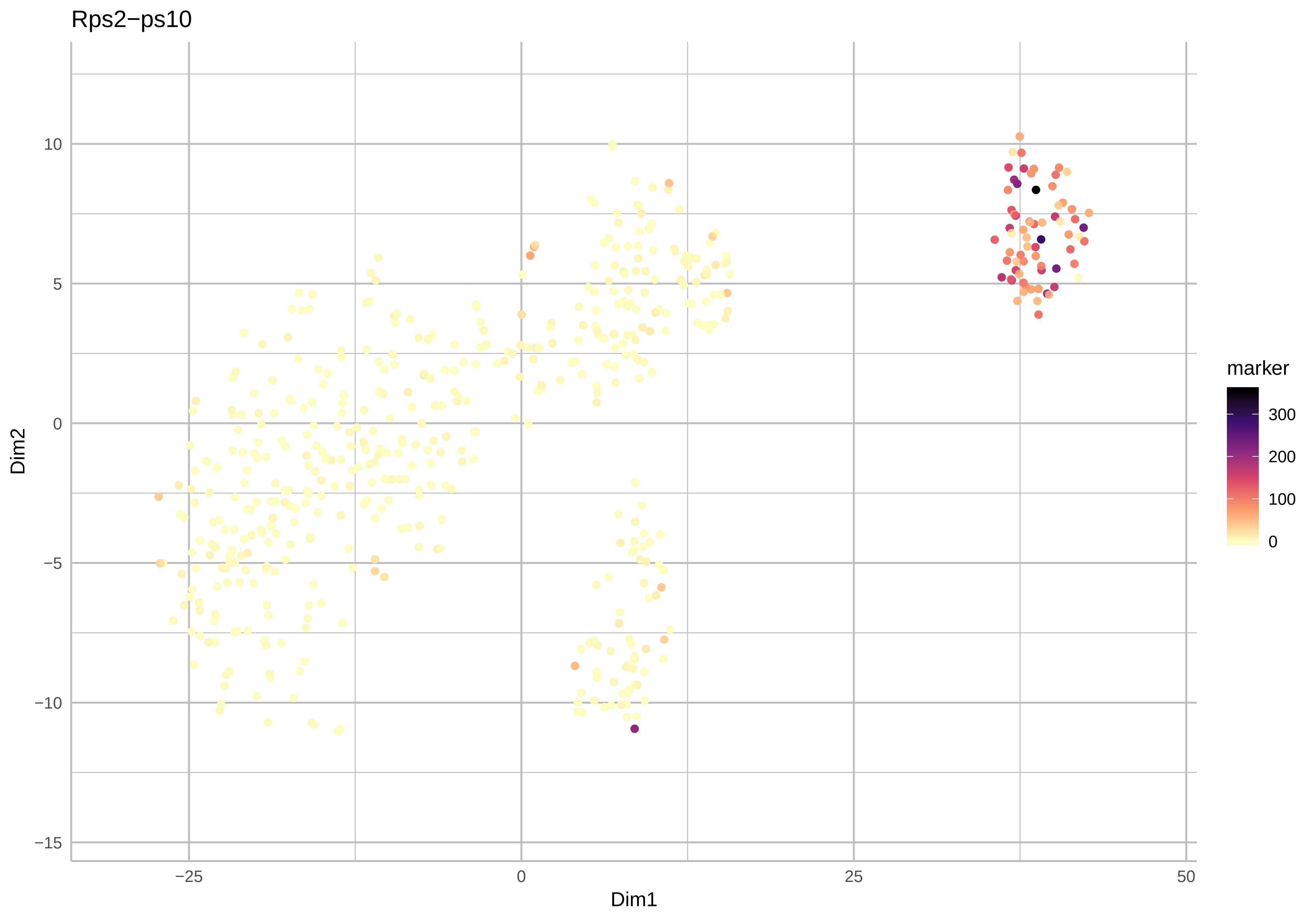
Rps2−ps10
Dim2
Dim1
10
5
0
−5
−10
−15
−25
0
25
50
marker
300
200
100
0

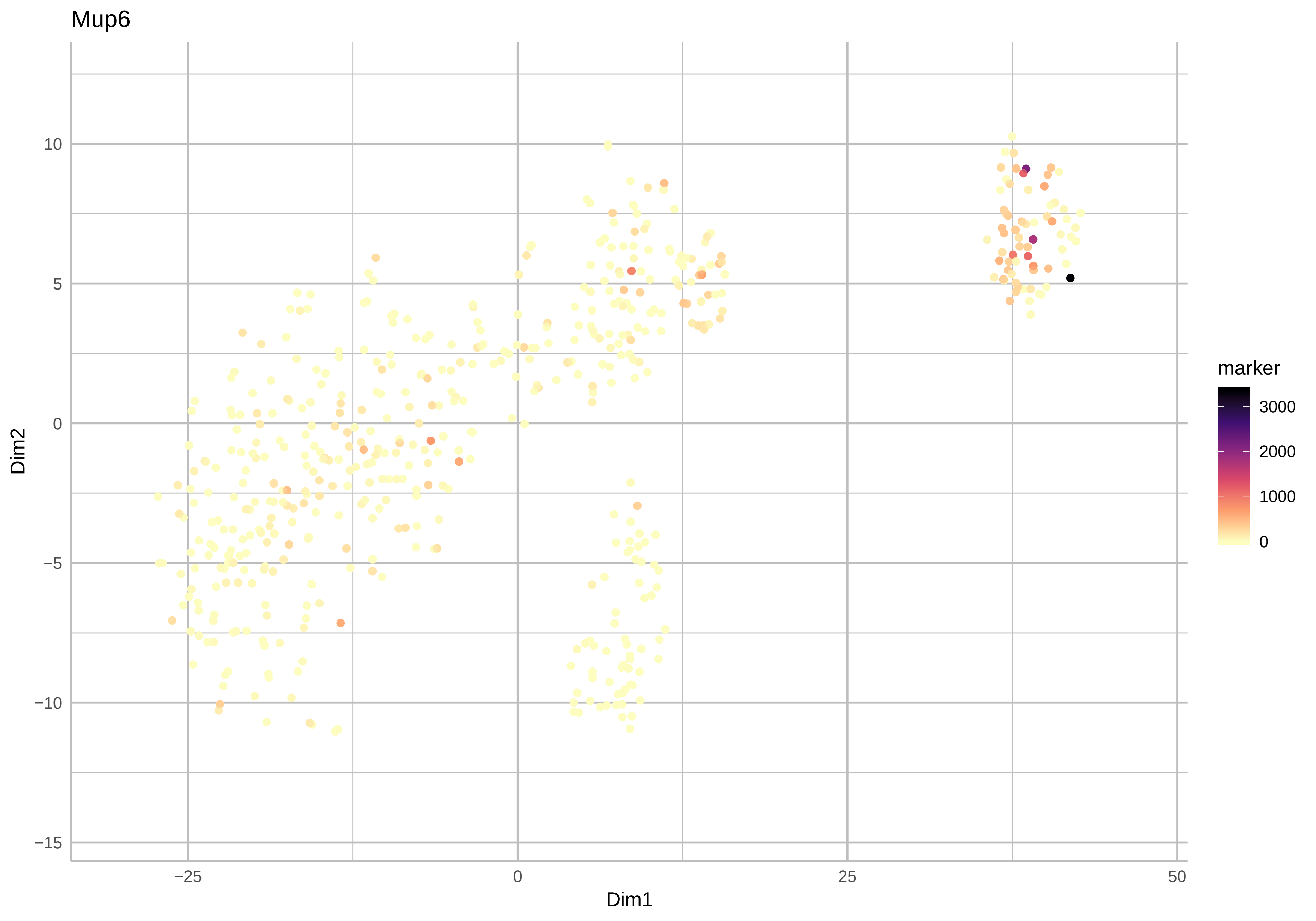

Mup6
Dim2
Dim1
10
5
0
−5
−10
−15
−25
0
25
50
marker
3000
2000
1000
0

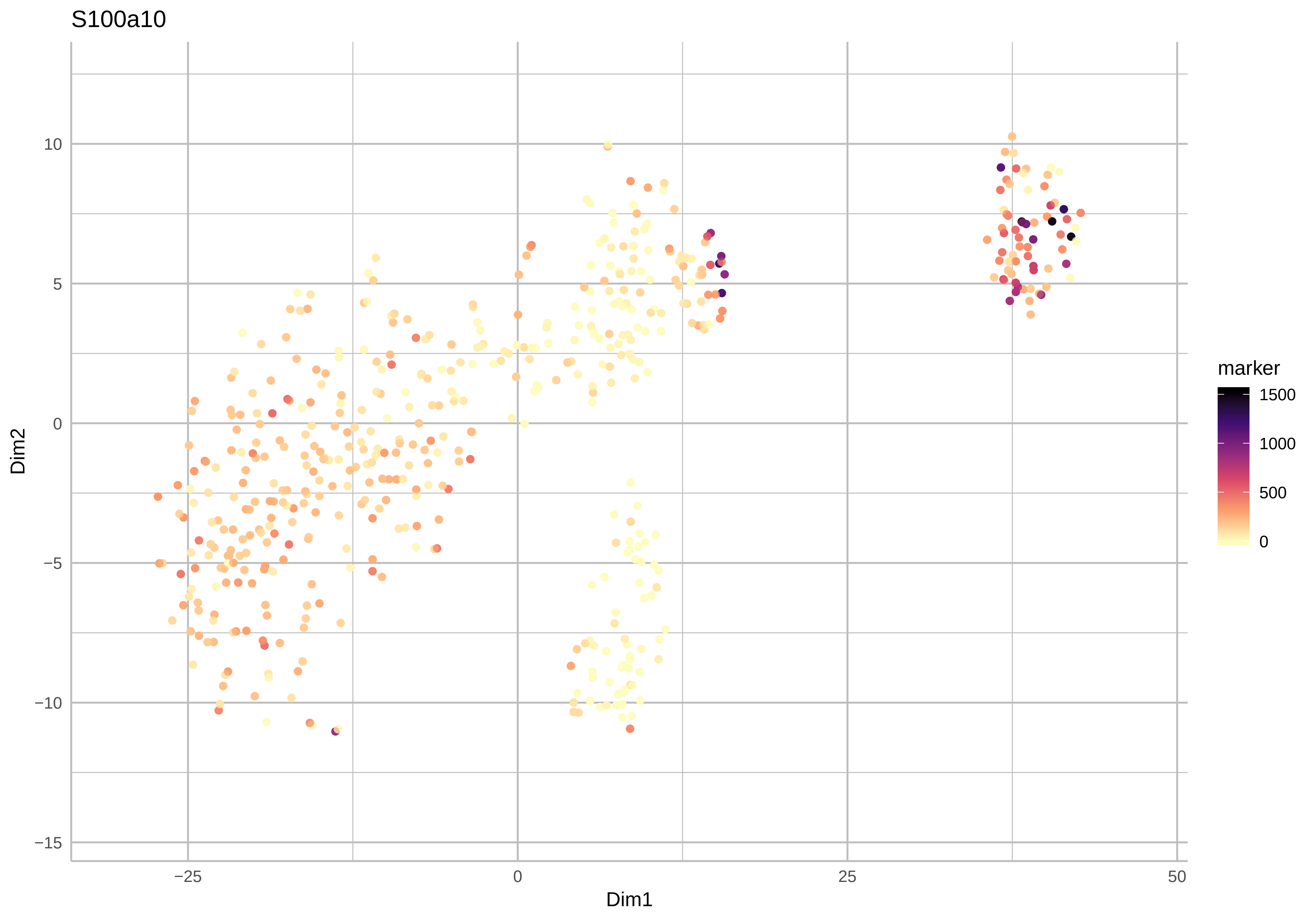

S100a10
Dim2
Dim1
10
5
0
−5
−10
−15
−25
0
25
50
marker
1500
1000
500
0

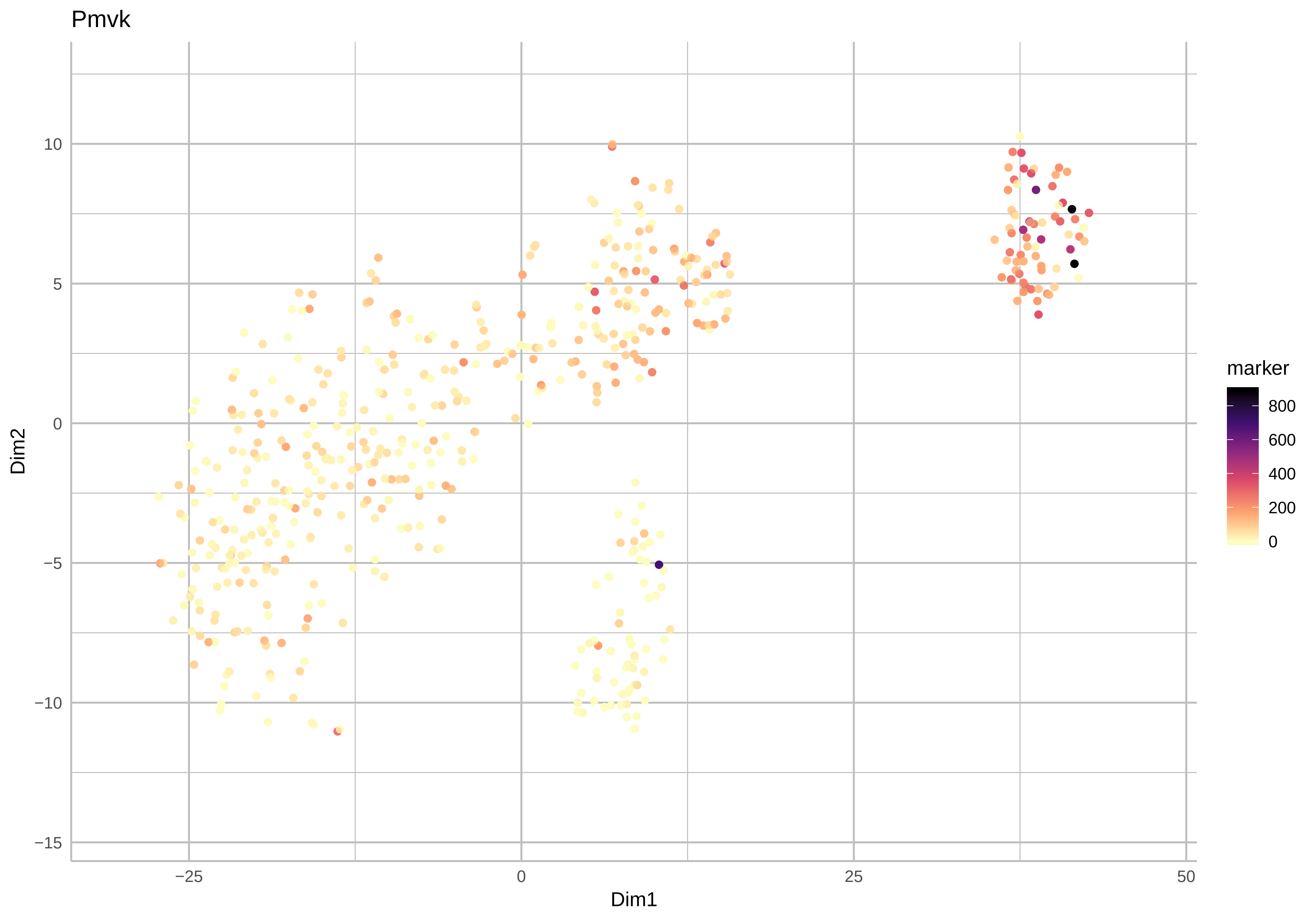

Pmvk
Dim2
Dim1
marker
800
600
400
200
0
10
5
0
−5
−10
−15
−25
0
25
50

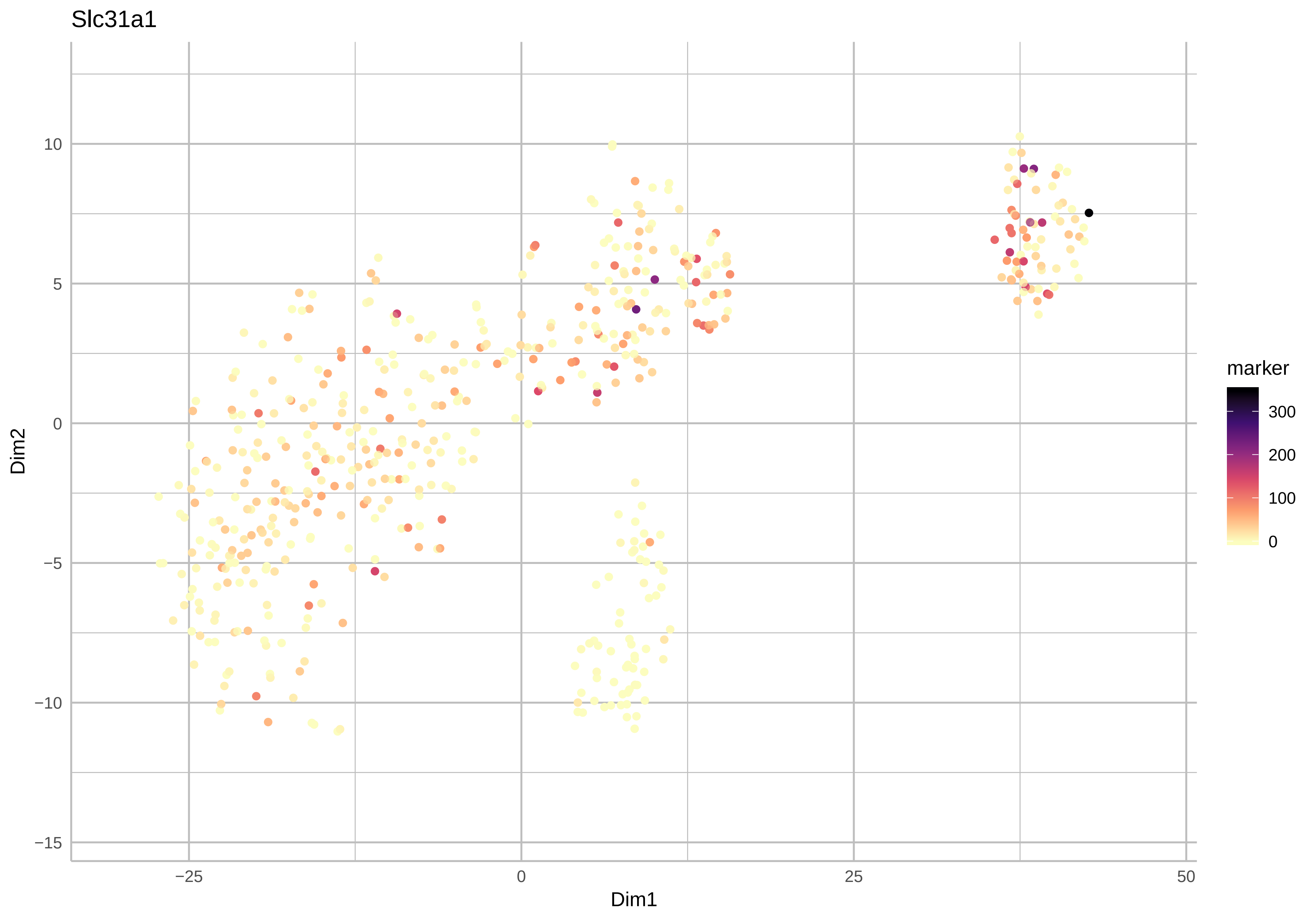

Slc31a1
Dim2
Dim1
10
5
0
−5
−10
−15
−25
0
25
50
marker
300
200
100
0

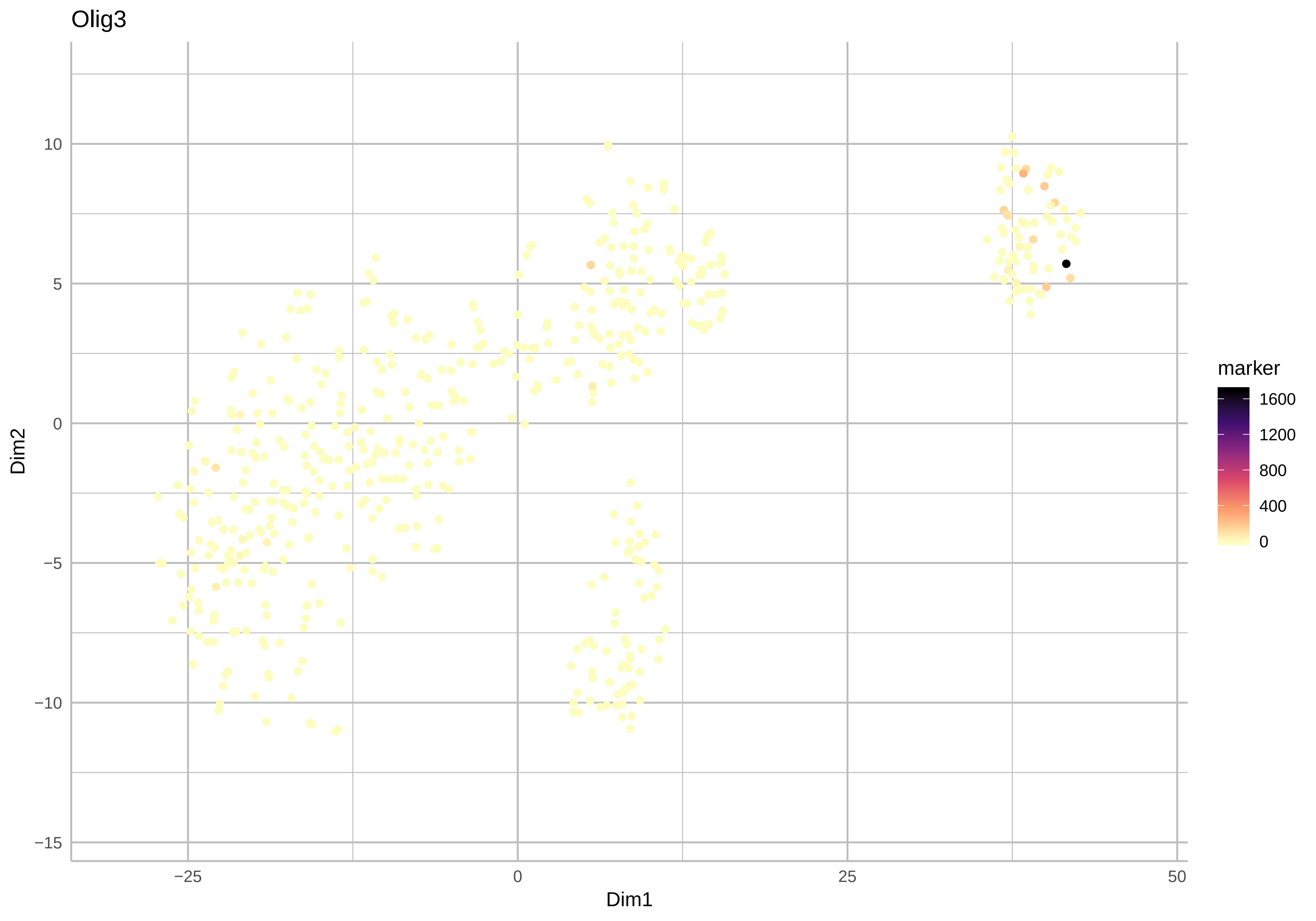
Olig3
Dim2
Dim1
10
5
0
−5
−10
−15
−25
0
25
50
marker
1600
1200
800
400
0

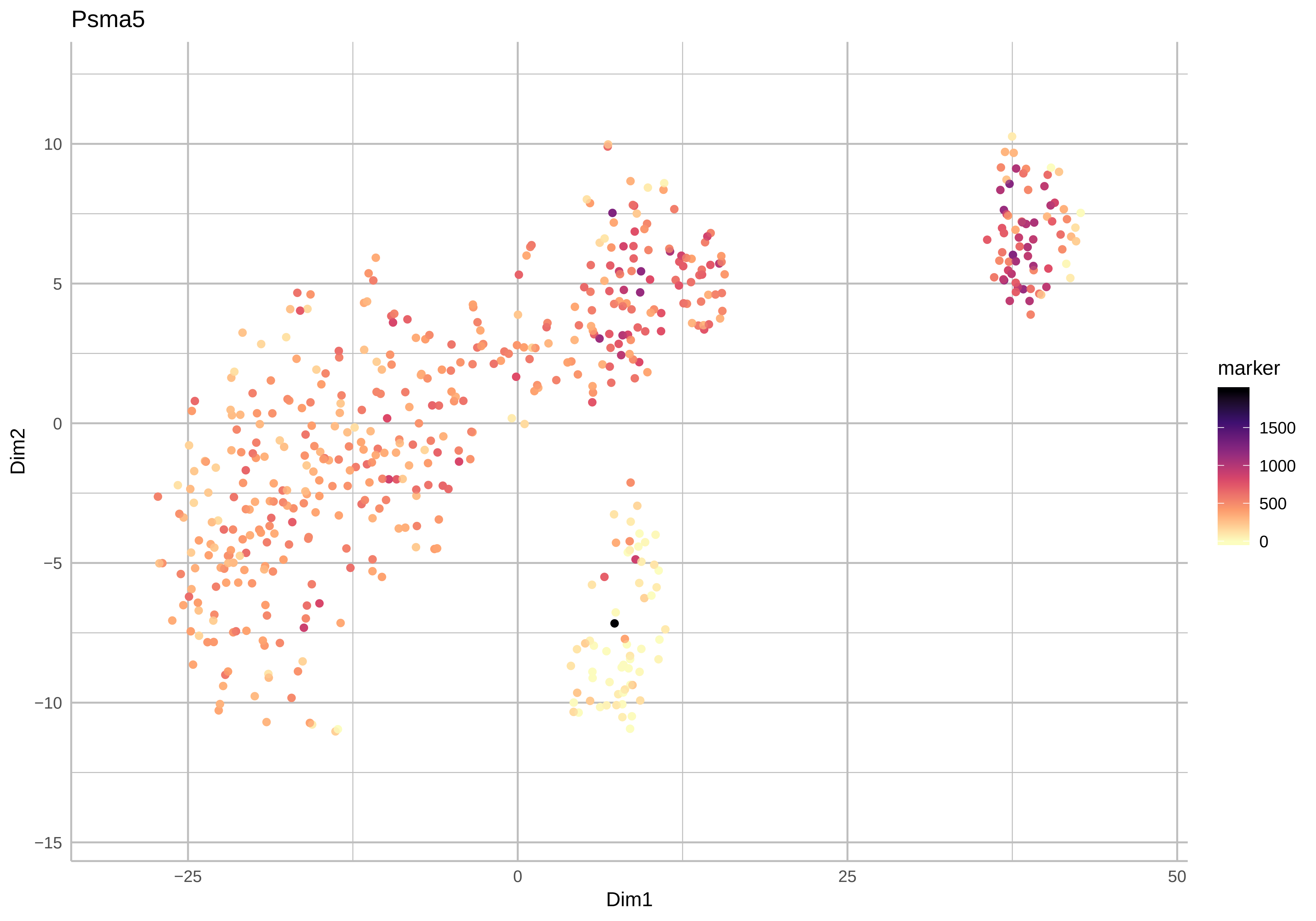

Psma5
Dim2
Dim1
10
5
0
−5
−10
−15
−25
0
25
50
marker
1500
1000
500
0

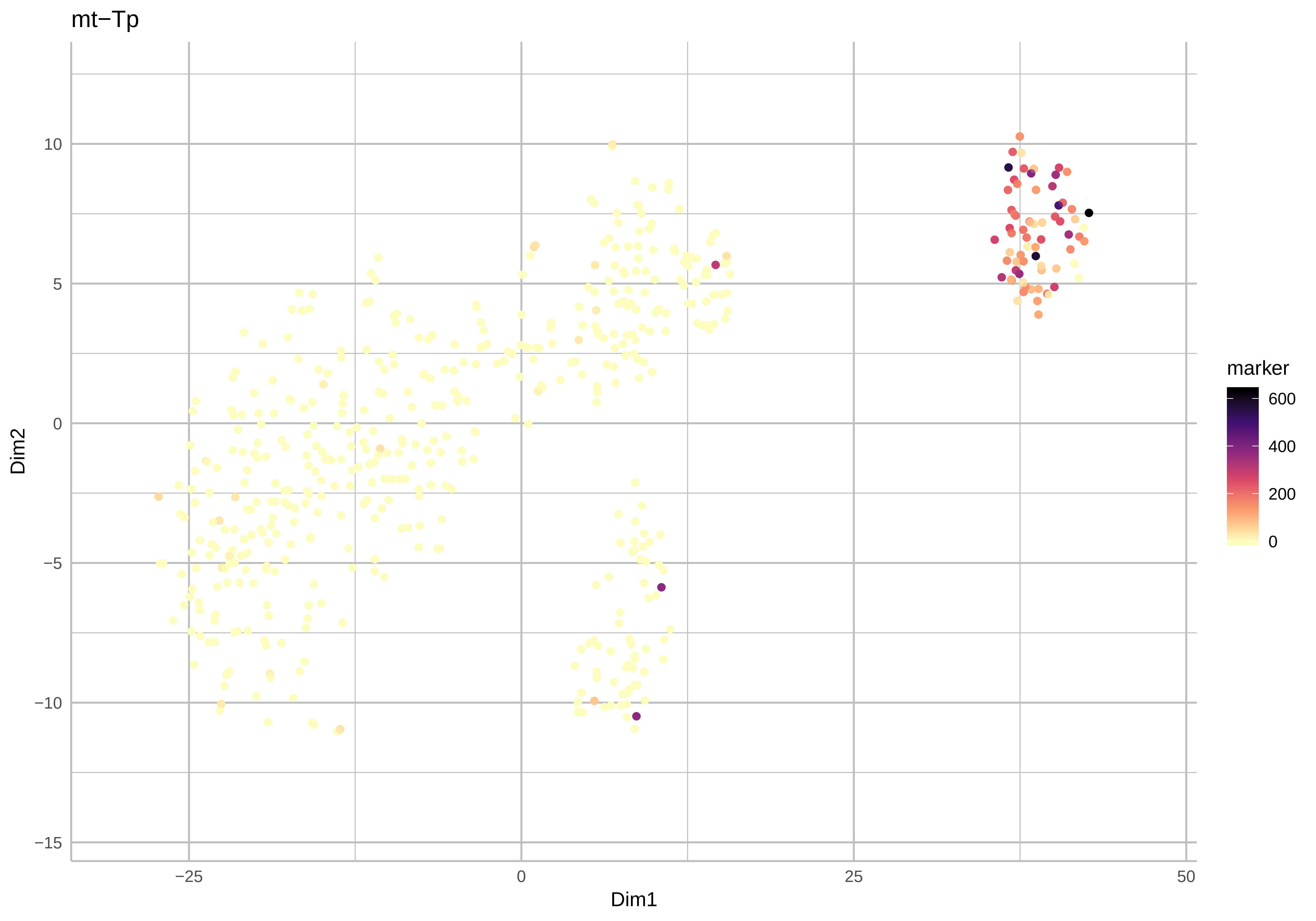
mt−Tp
Dim2
Dim1
10
5
0
−5
−10
−15
−25
0
25
50
marker
600
400
200
0

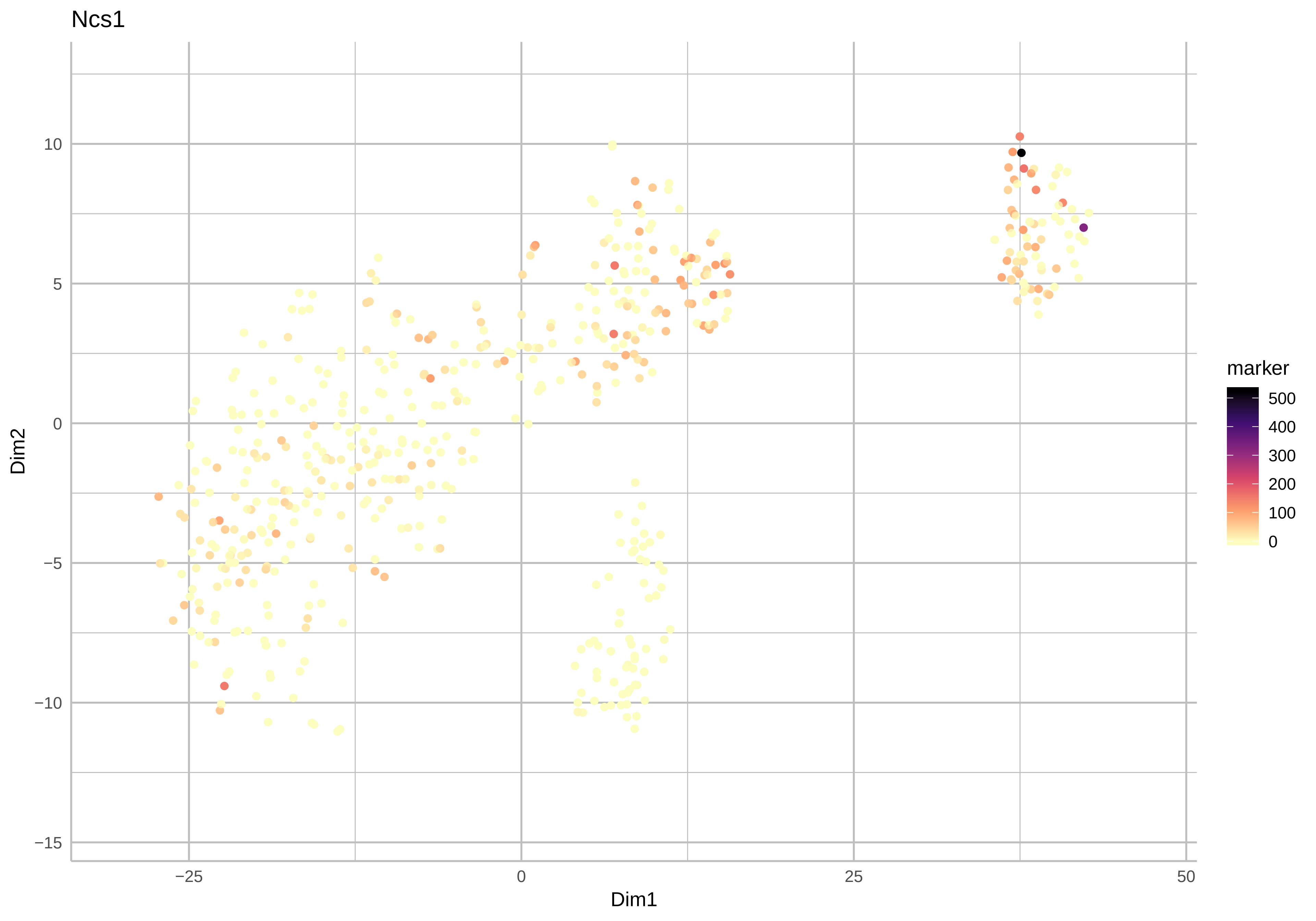
Ncs1
Dim2
Dim1
10
5
0
−5
−10
−15
−25
0
25
50
marker
500
400
300
200
100
0

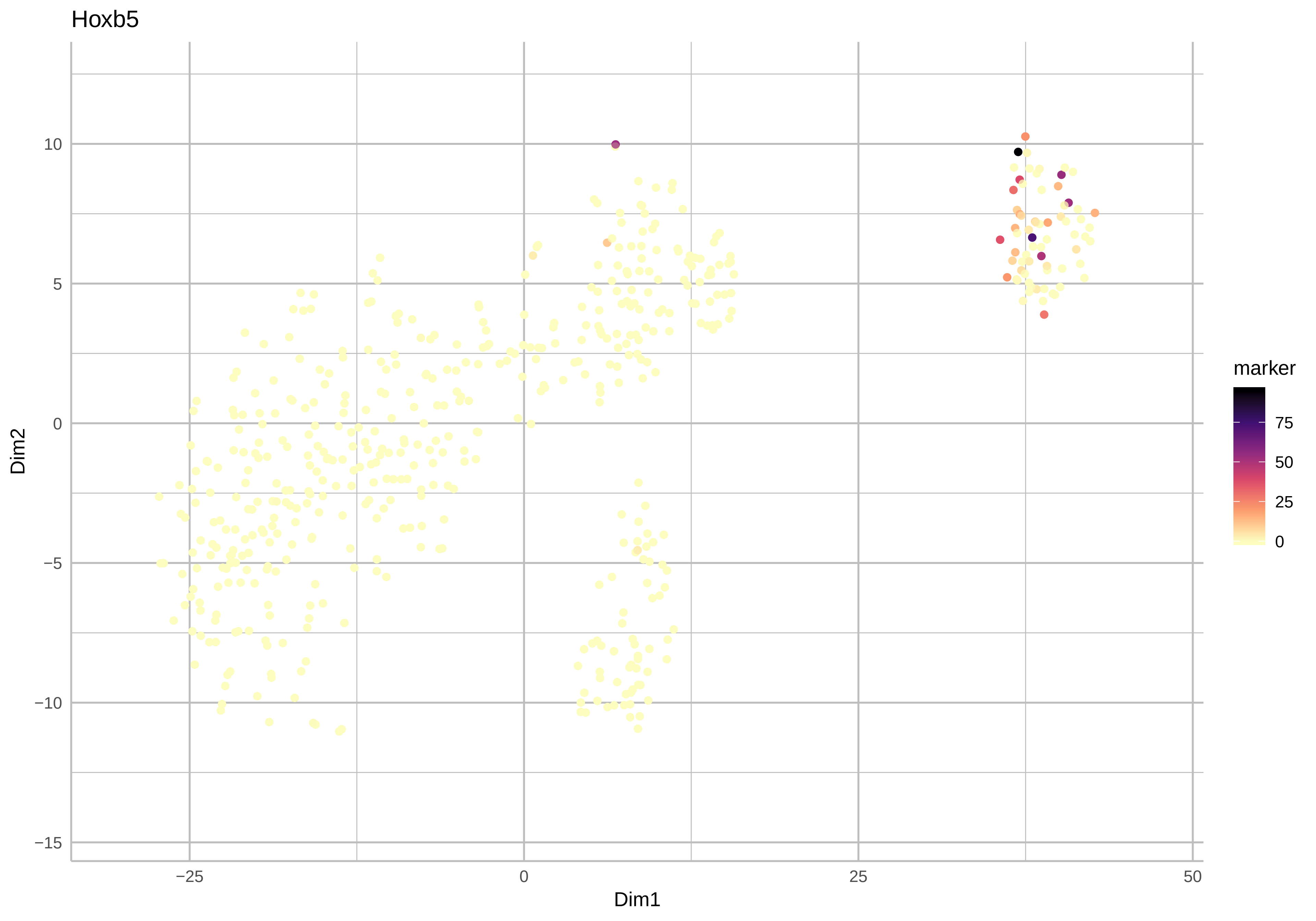

Hoxb5
10
5
0
−5
−10
−15
−25
0
25
50
Dim1
Dim2
marker
75
50
25
0

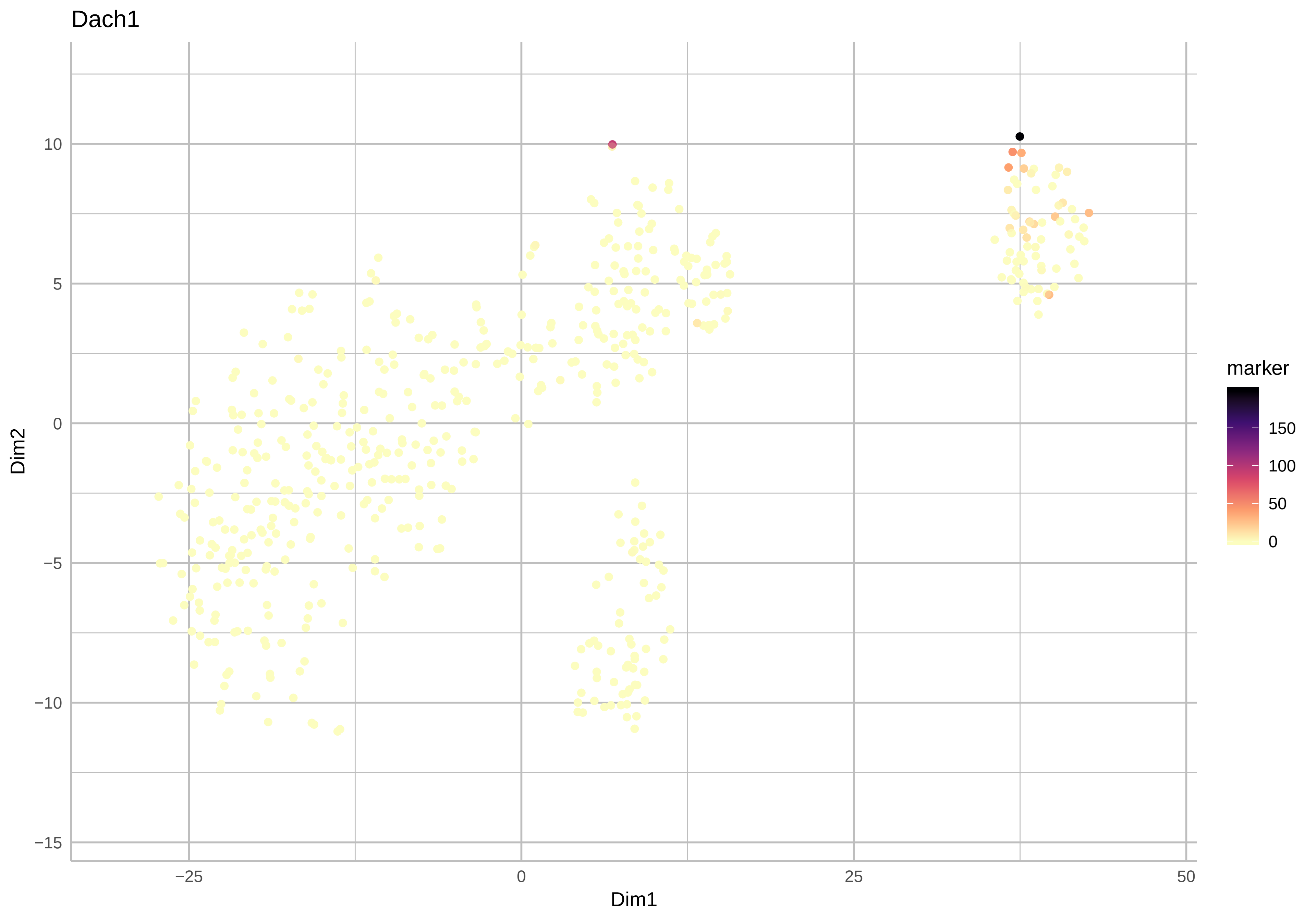
Dach1
Dim2
Dim1
10
5
0
−5
−10
−15
−25
0
25
50
marker
150
100
50
0

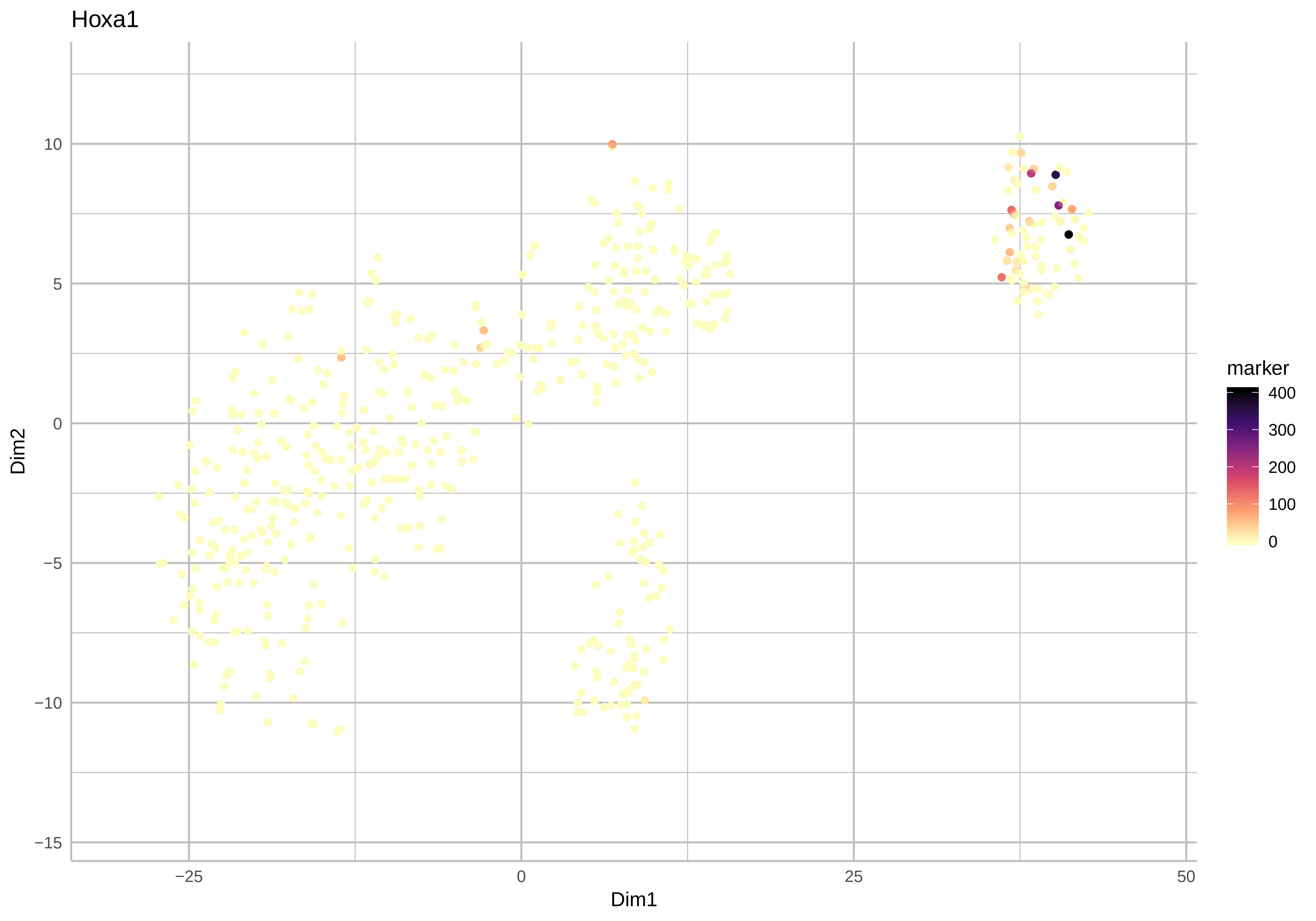

Hoxa1
10
5
0
−5
−10
−15
Dim2
−25
0
25
50
Dim1
marker
400
300
200
100
0

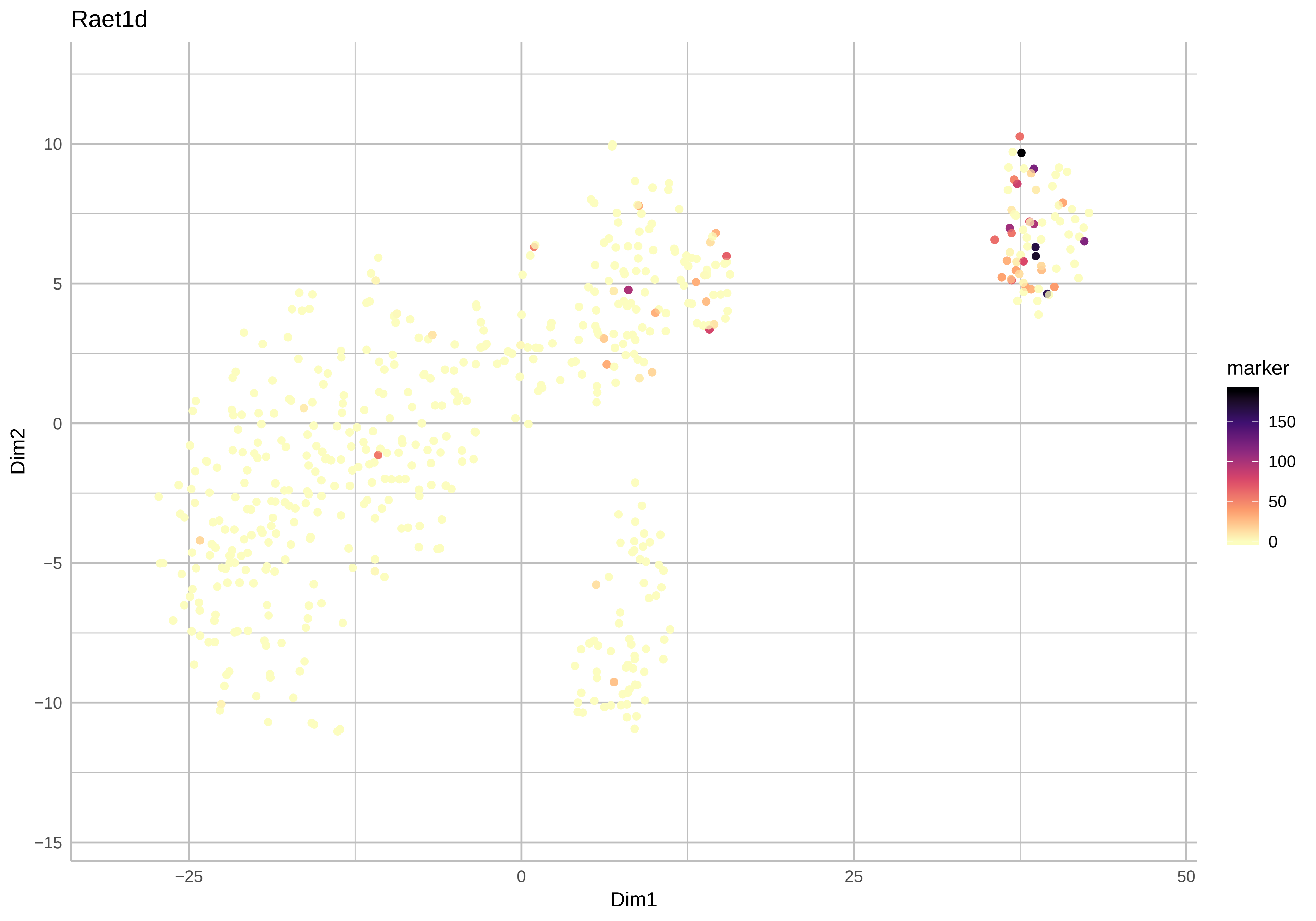

Raet1d
Dim2
Dim1
10
5
0
−5
−10
−15
−25
0
25
50
marker
150
100
50
0

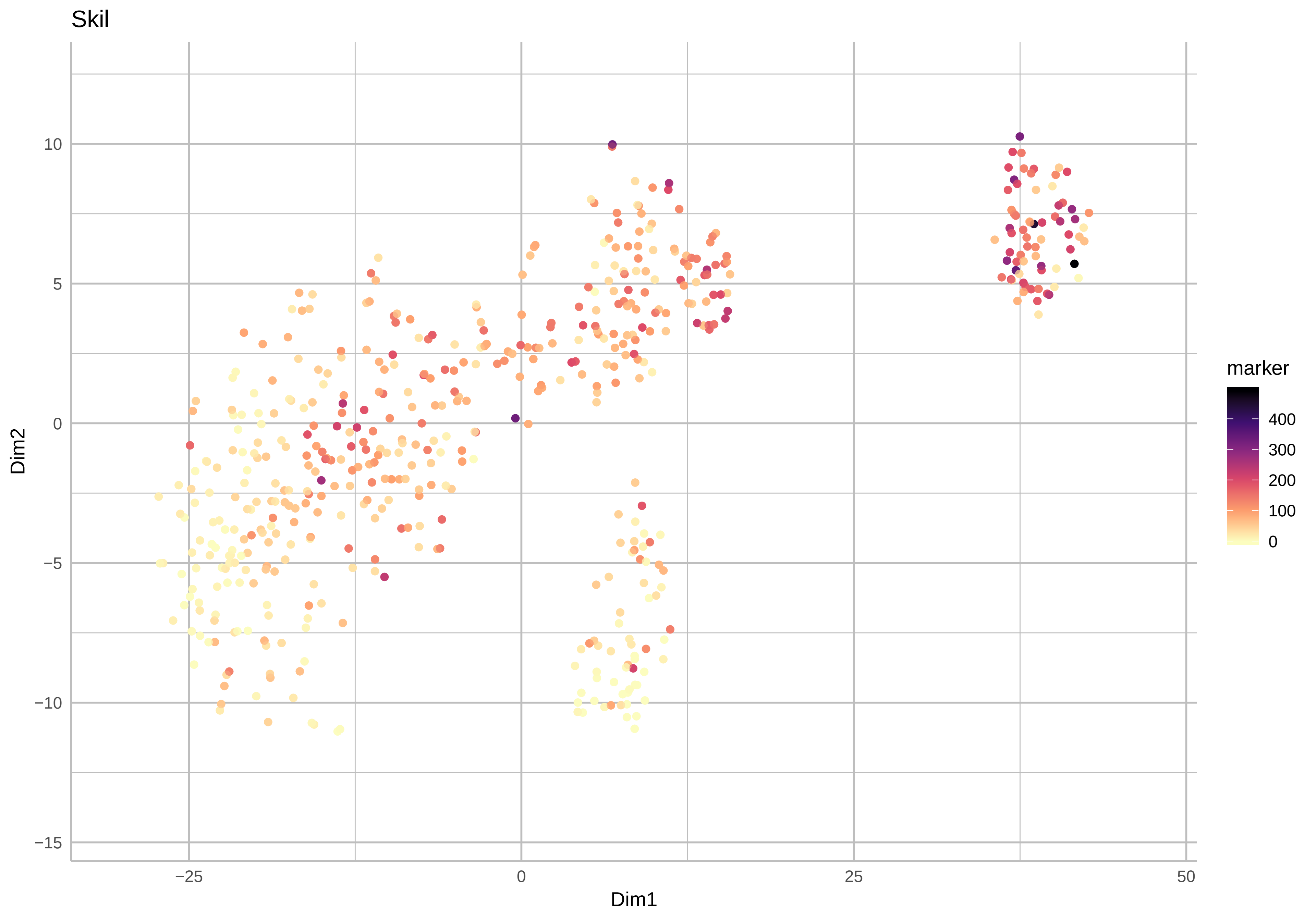

Skil
Dim2
Dim1
10
5
0
−5
−10
−15
−25
0
25
50
marker
400
300
200
100
0

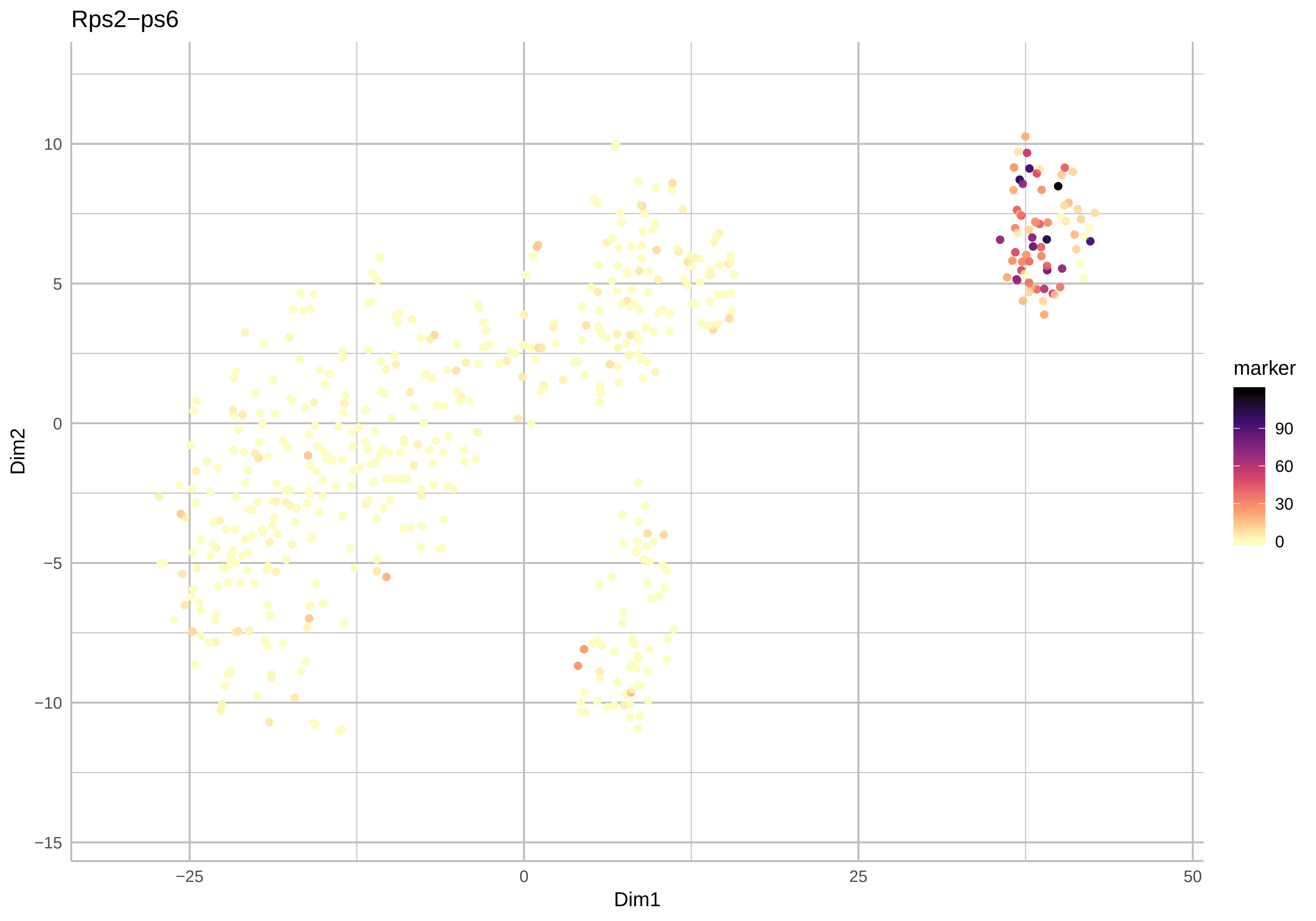
Rps2−ps6
Dim2
Dim1
−25
0
25
50
10
5
0
−5
−10
−15
marker
90
60
30
0

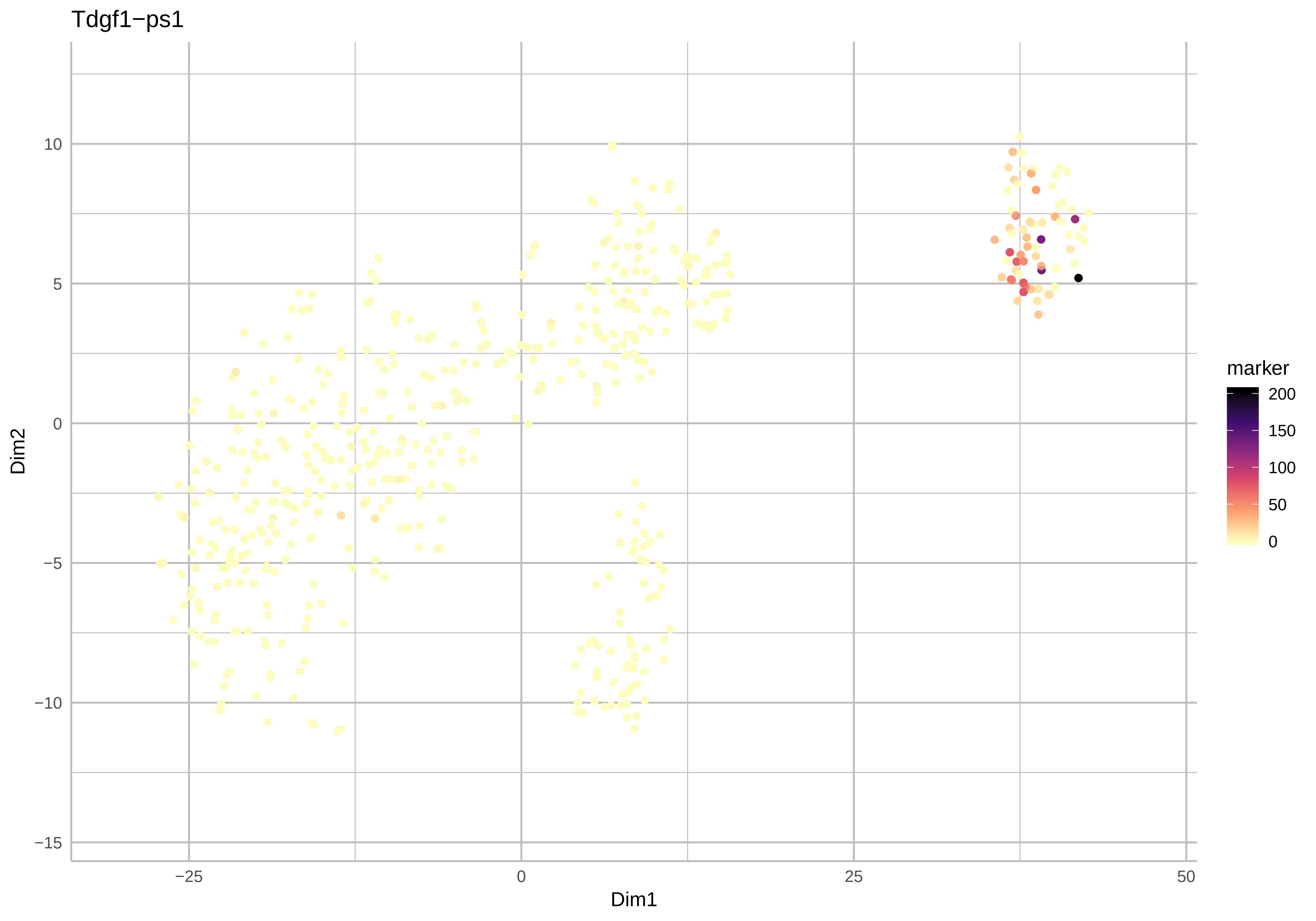
Tdgf1−ps1
Dim2
Dim1
10
5
0
−5
−10
−15
−25
0
25
50
marker
200
150
100
50
0

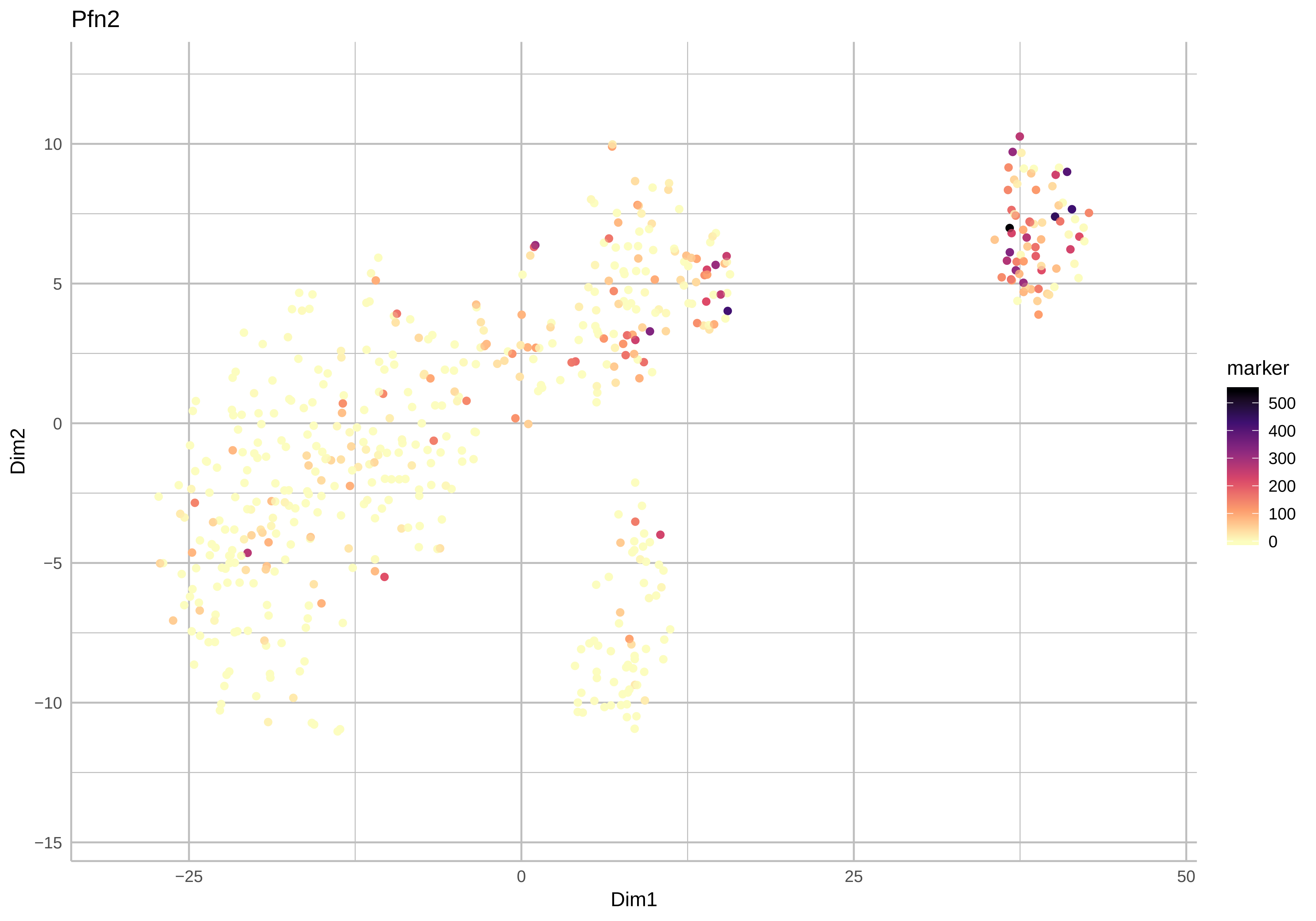

Pfn2
Dim2
Dim1
10
5
0
−5
−10
−15
−25
0
25
50
marker
500
400
300
200
100
0

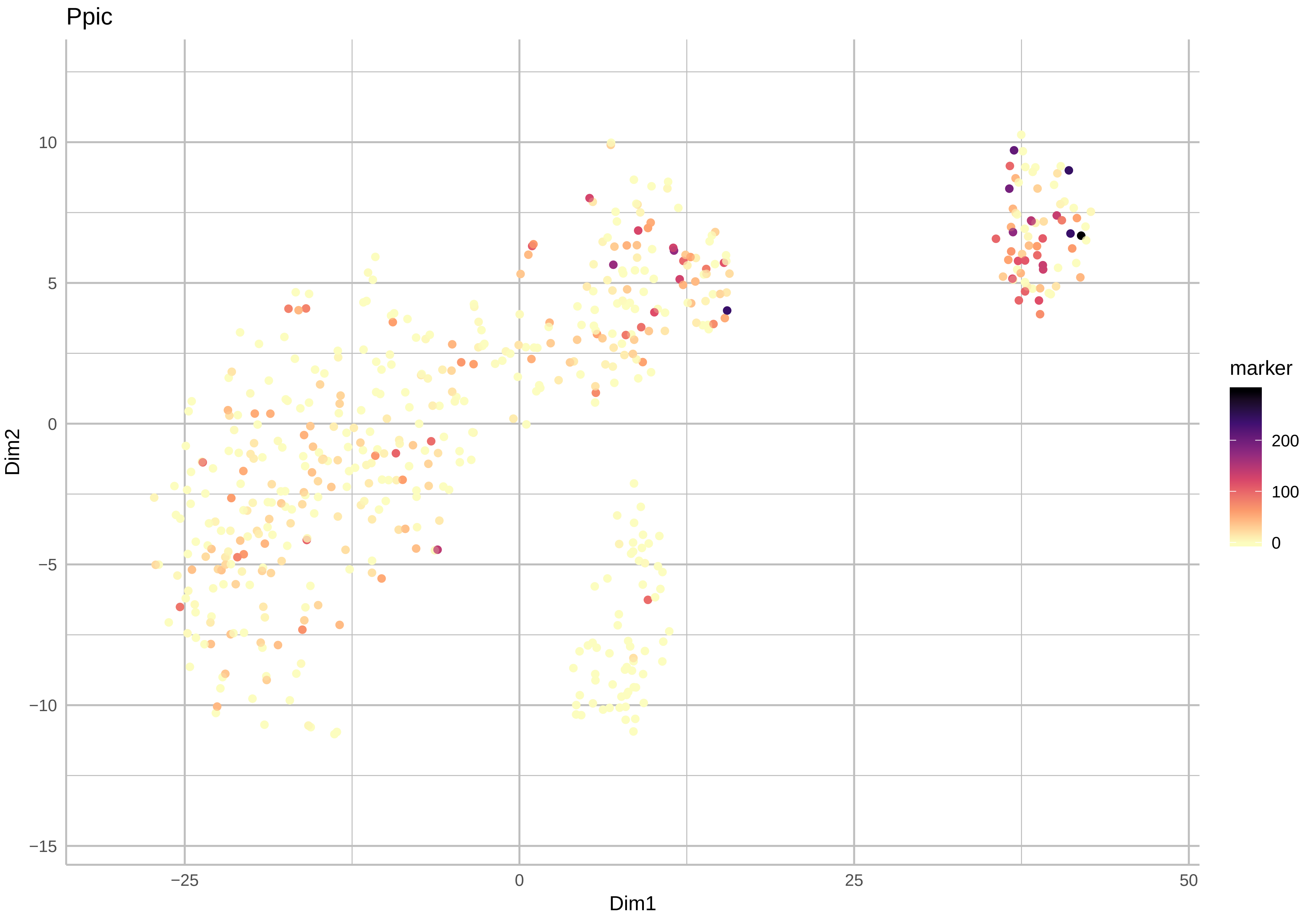

Ppic
10
5
0
−5
−10
−15
Dim2
−25
0
25
50
Dim1
marker
200
100
0

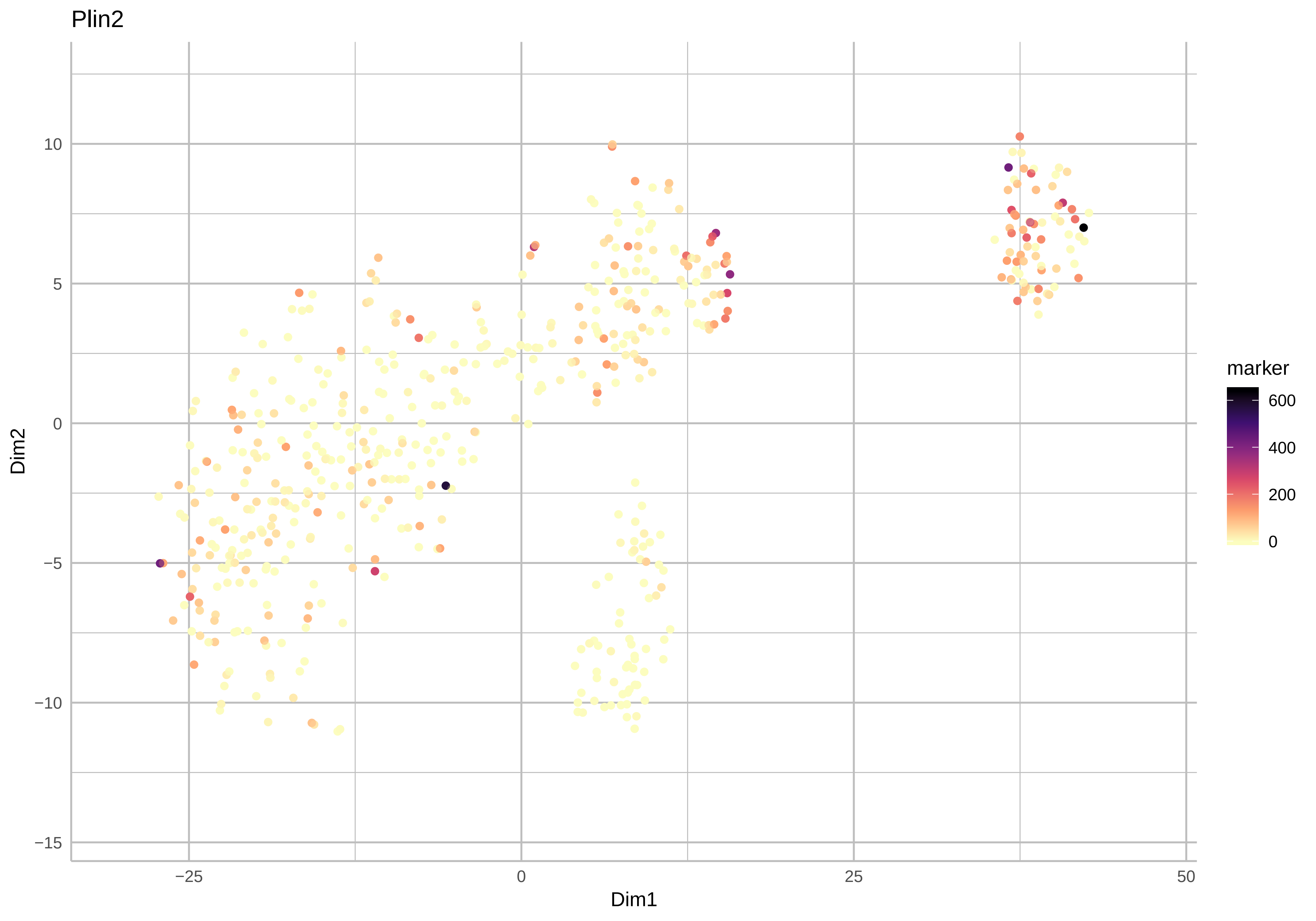
Plin2
10
5
0
−5
−10
−15
Dim2
−25
0
25
50
Dim1
marker
600
400
200
0

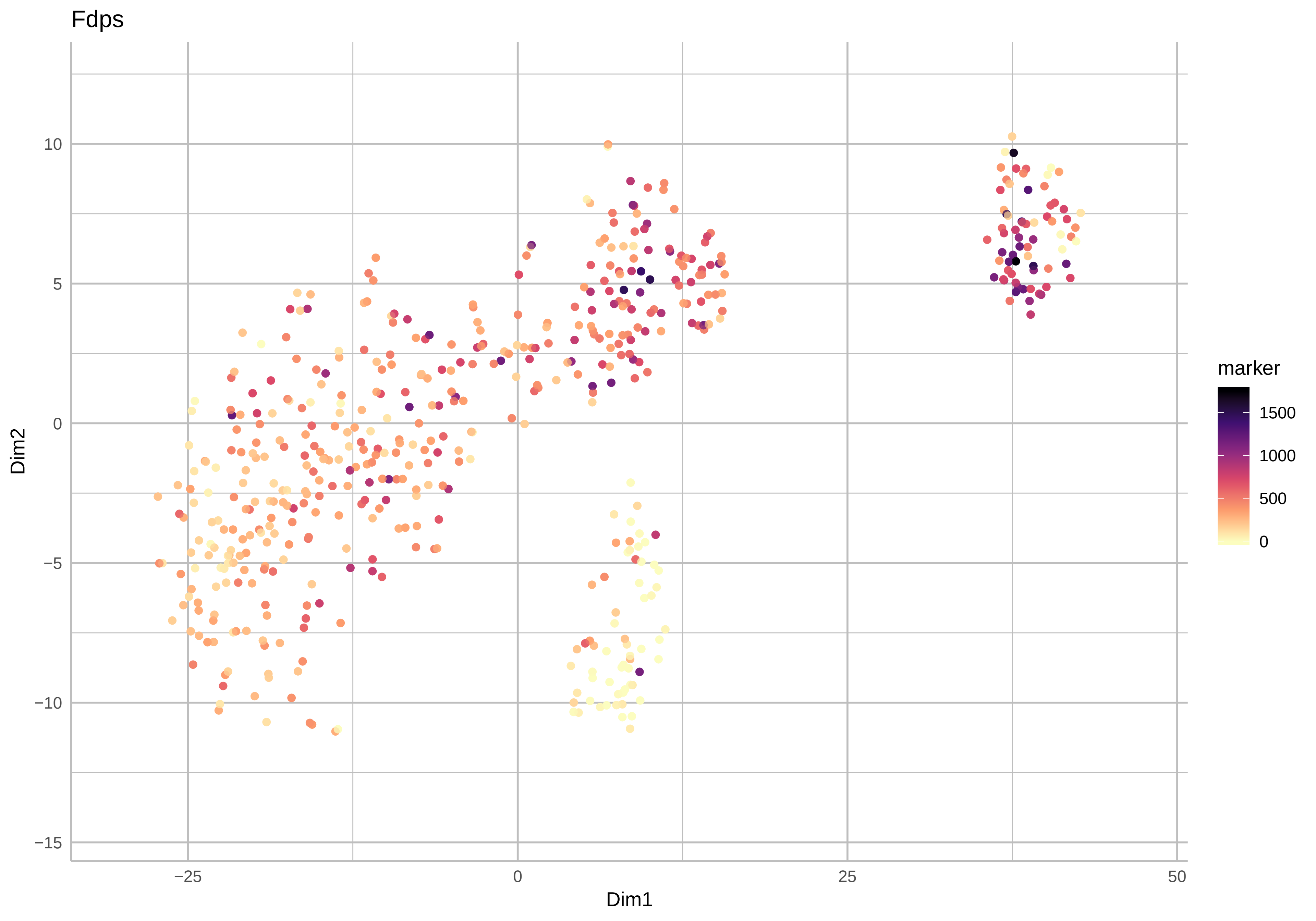

Fdps
10
5
0
−5
−10
−15
Dim2
−25
0
25
50
Dim1
marker
1500
1000
500
0

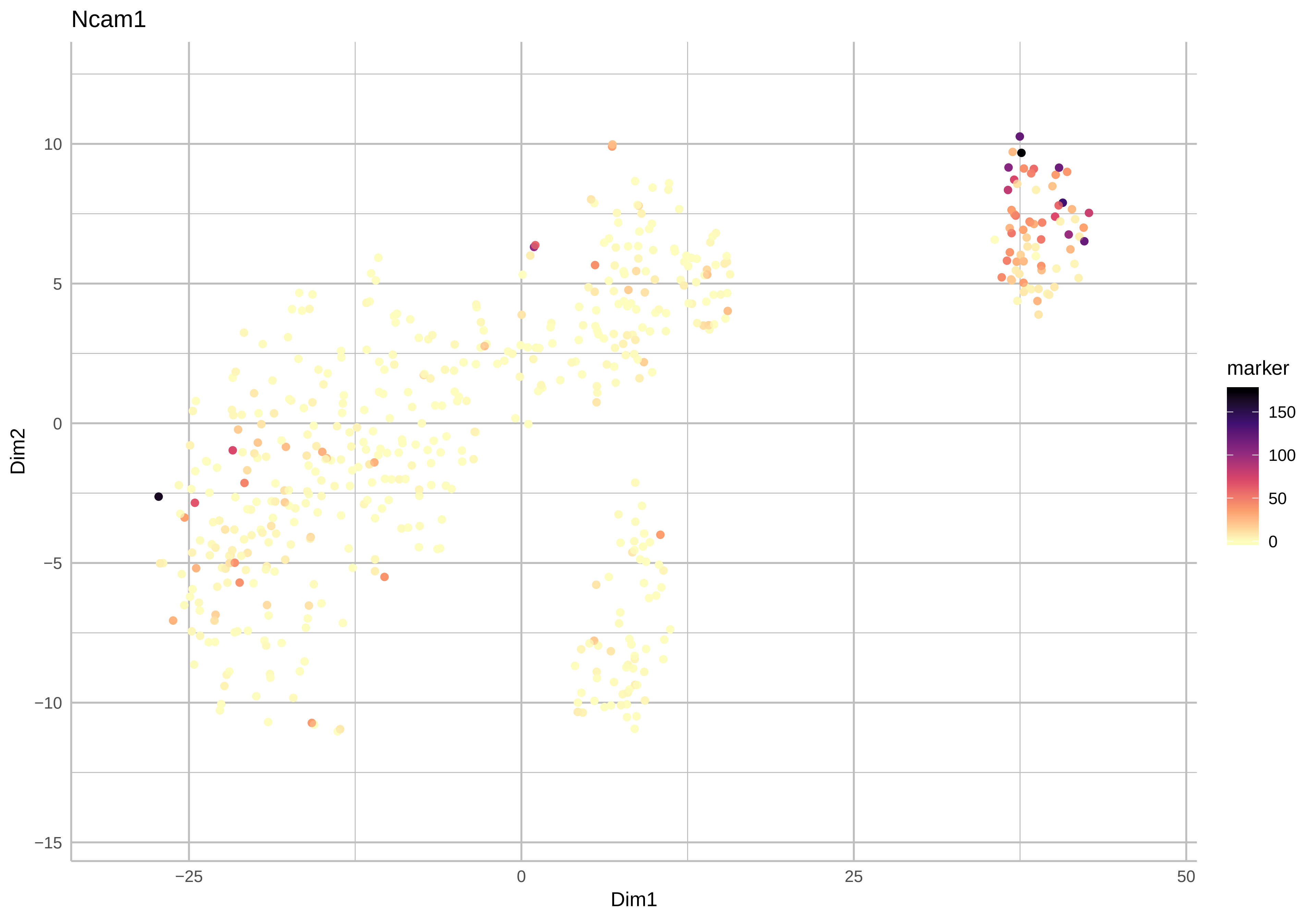
Ncam1
Dim2
Dim1
10
5
0
−5
−10
−15
−25
0
25
50
marker
150
100
50
0

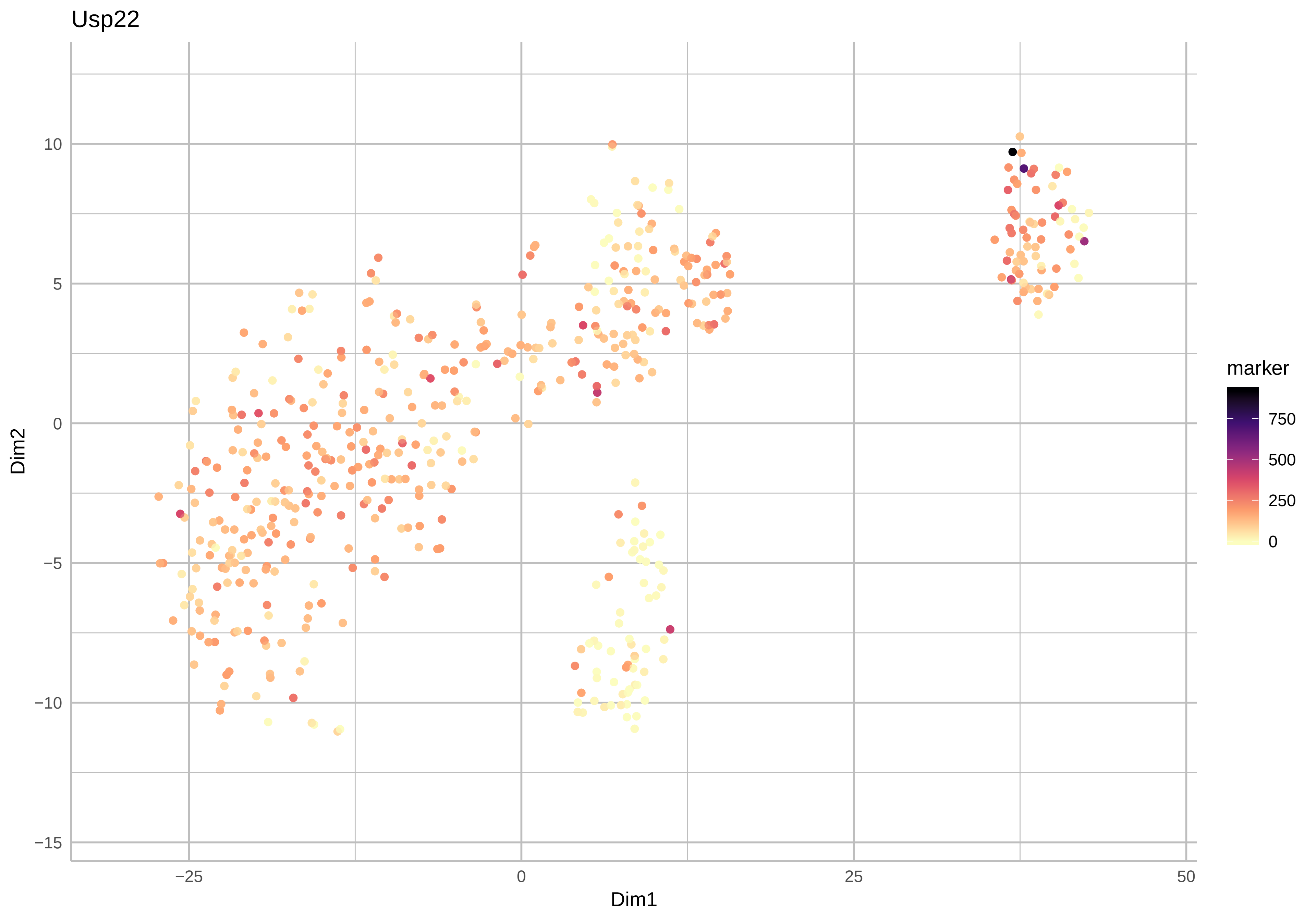

Usp22
Dim2
10
5
0
−5
−10
−15
−25
0
25
50
Dim1
marker
750
500
250
0

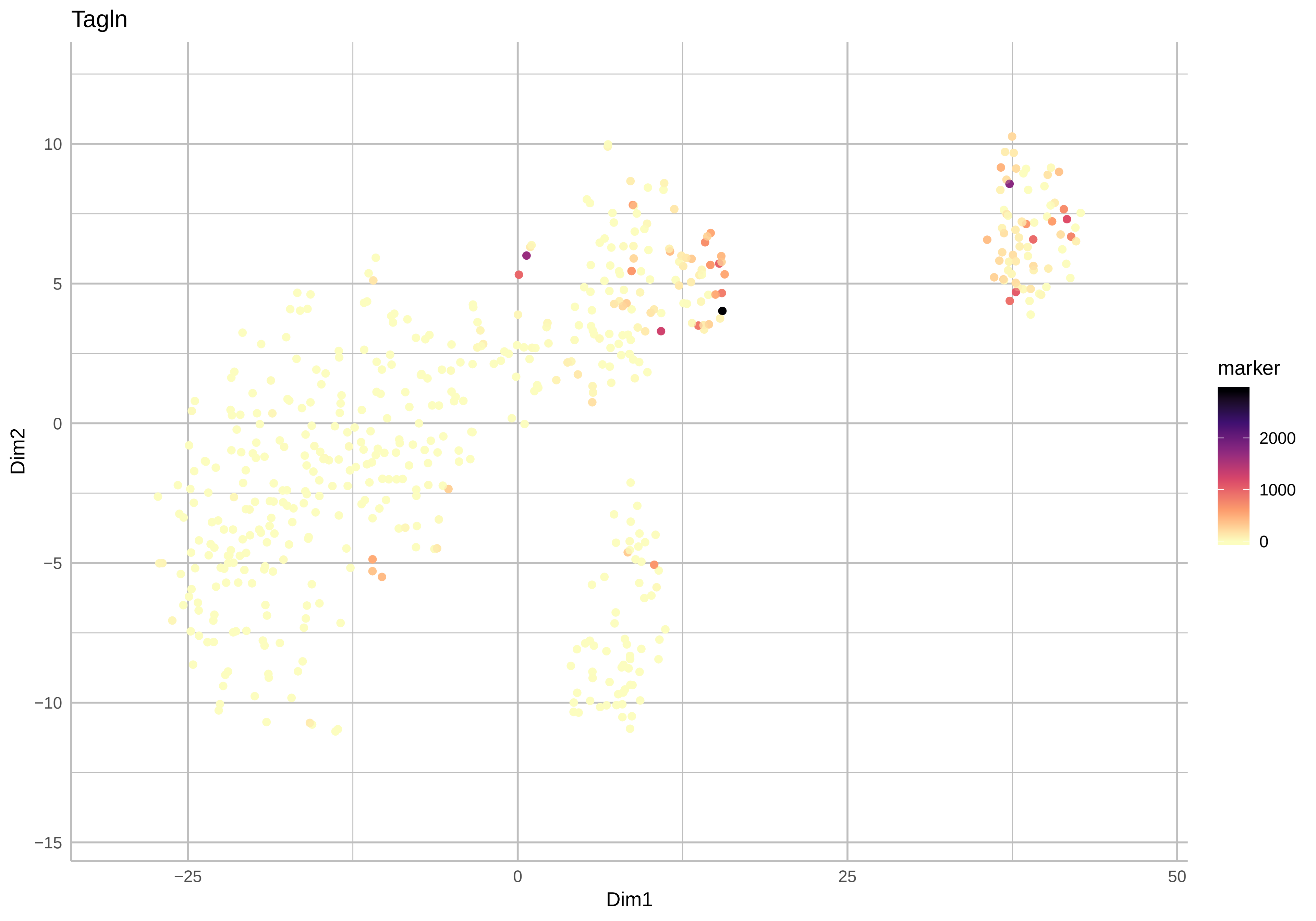

Tagln
Dim2
Dim1
marker
2000
1000
0
10
5
0
−5
−10
−15
−25
0
25
50

Hsd17b7

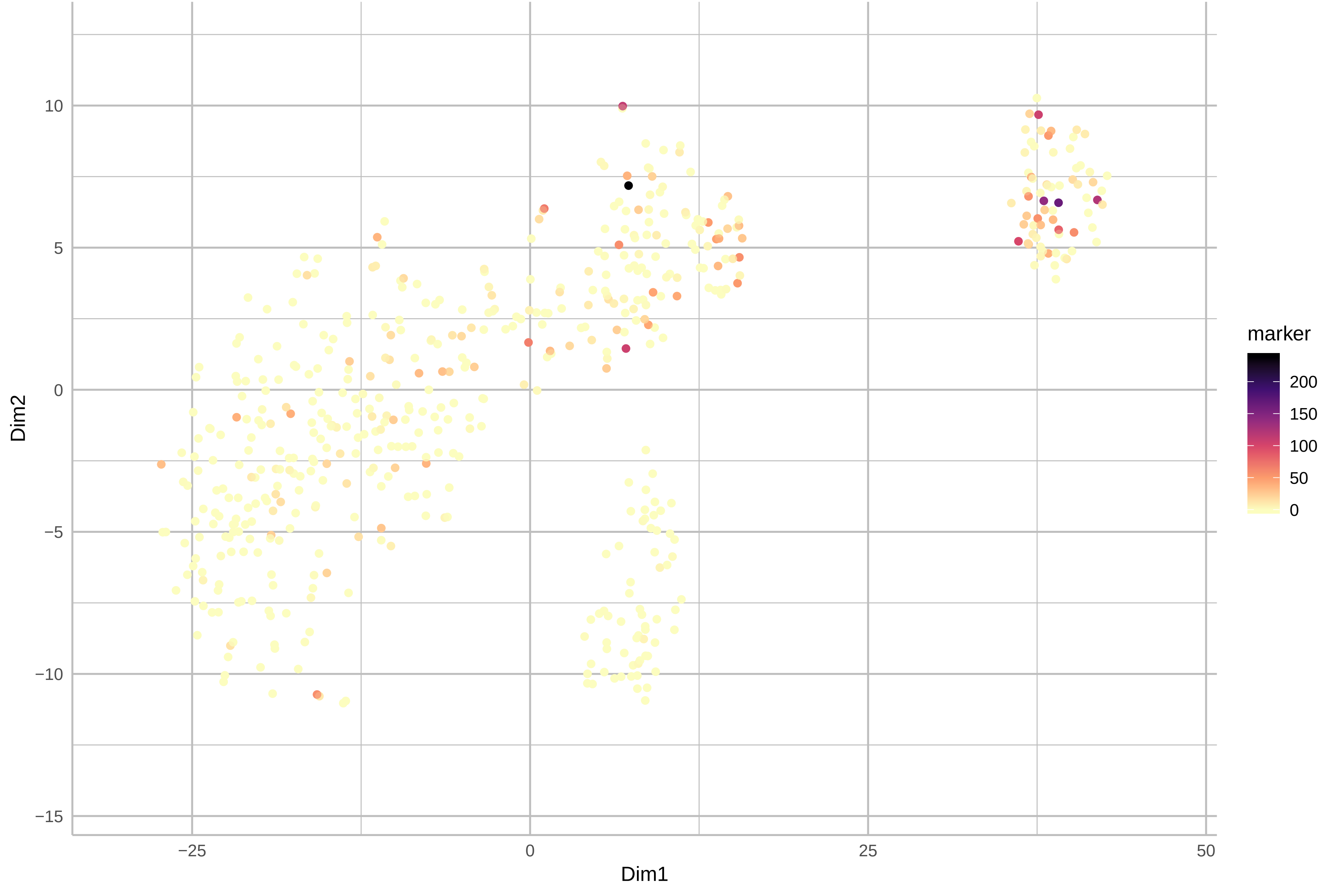

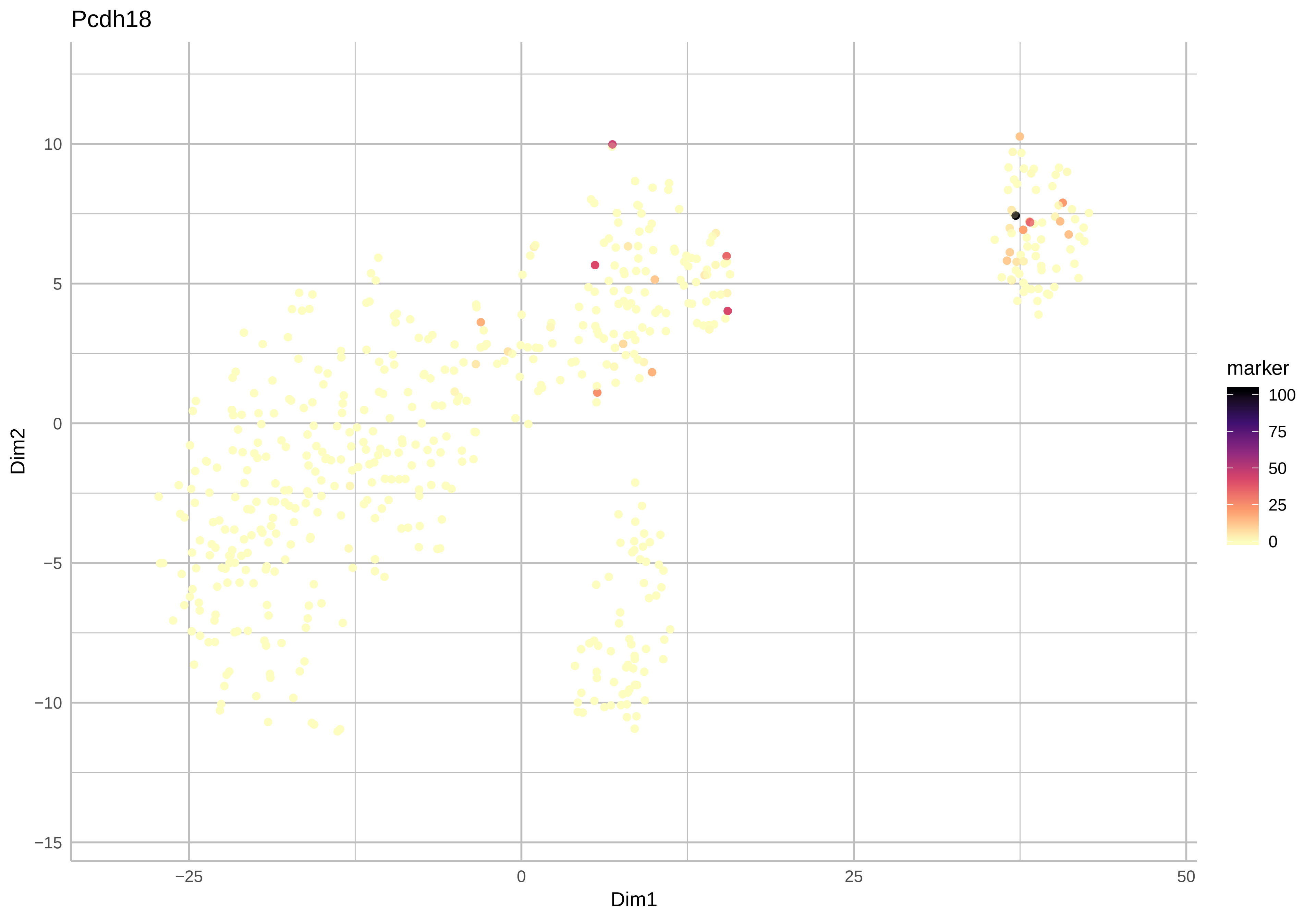

Pcdh18
10
5
0
−5
−10
−15
Dim2
−25
0
25
50
Dim1
marker
100
75
50
25
0

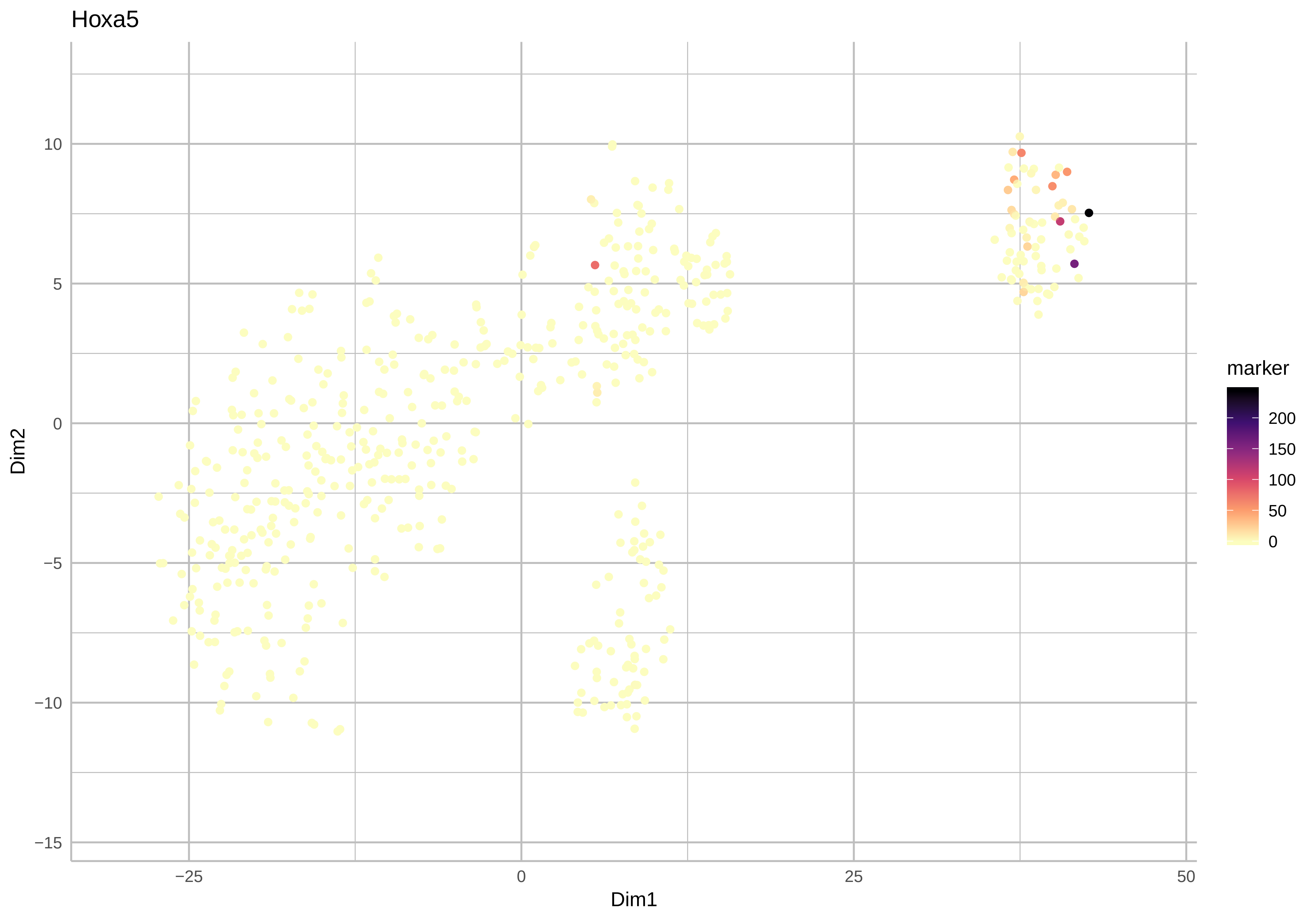

Hoxa5
Dim2
Dim1
10
5
0
−5
−10
−15
−25
0
25
50
marker
200
150
100
50
0

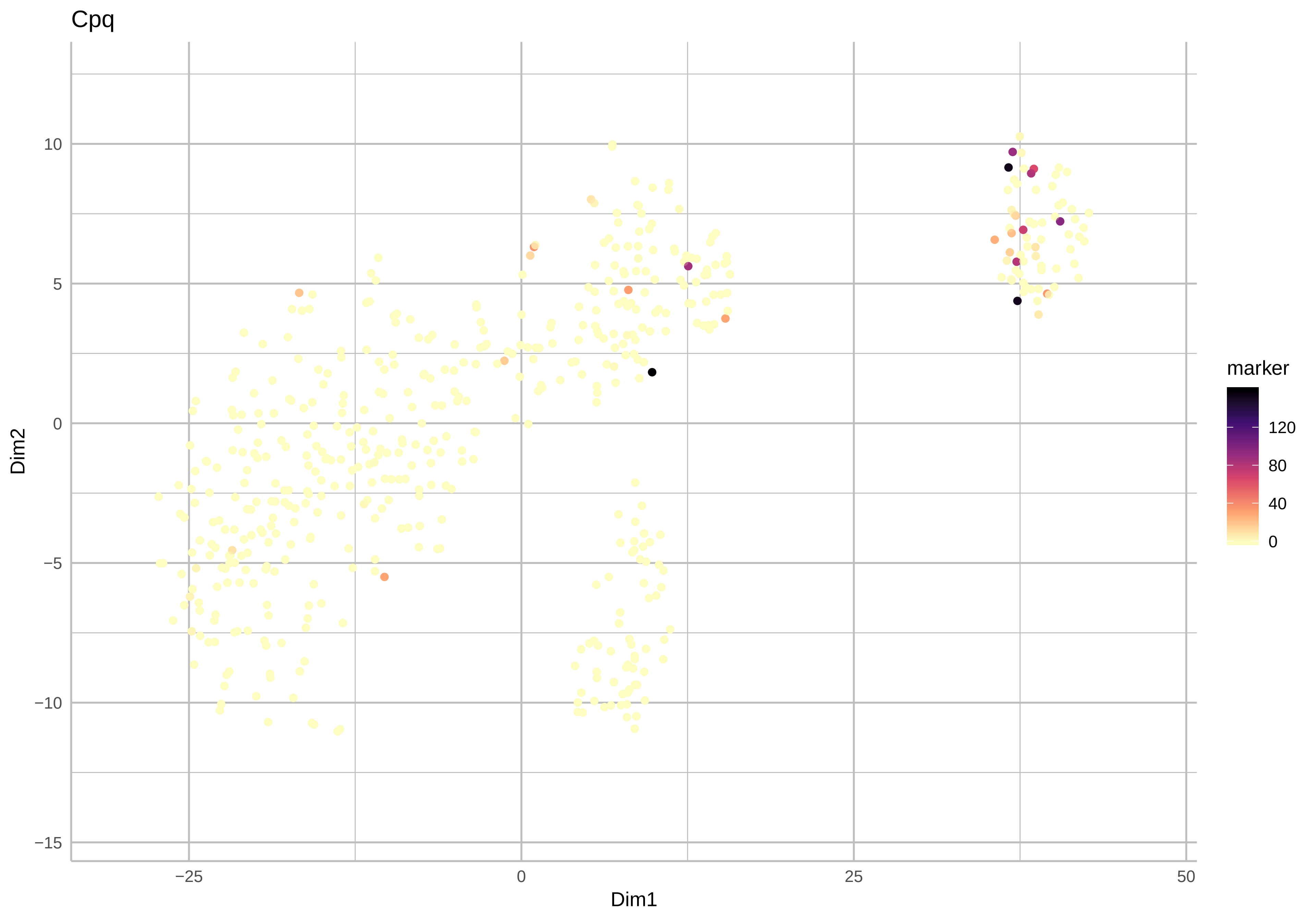
Cpq
Dim2
Dim1
10
5
0
−5
−10
−15
−25
0
25
50
marker
120
80
40
0

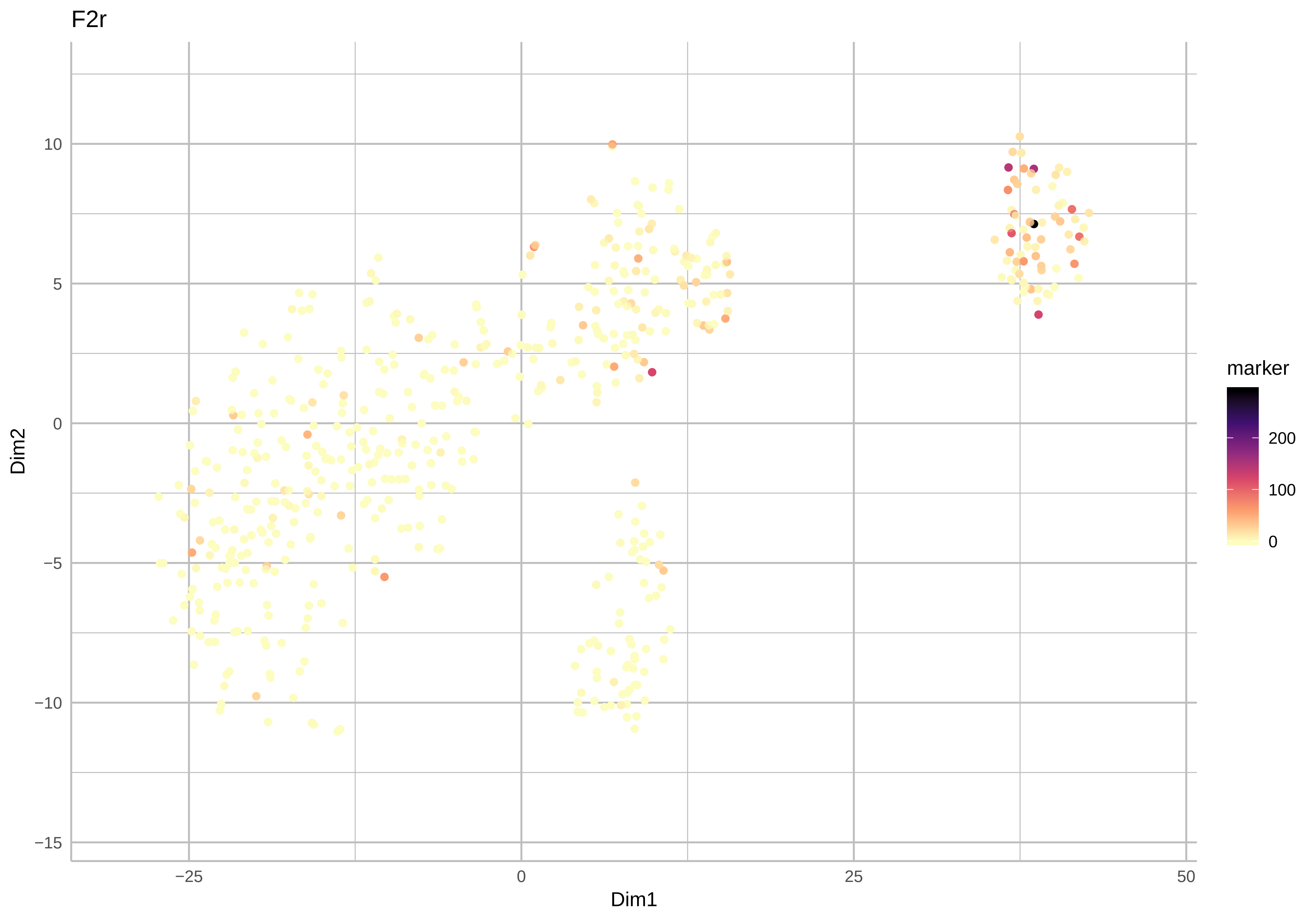

F2r
10
5
0
−5
−10
−15
Dim2
−25
0
25
50
Dim1
marker
200
100
0

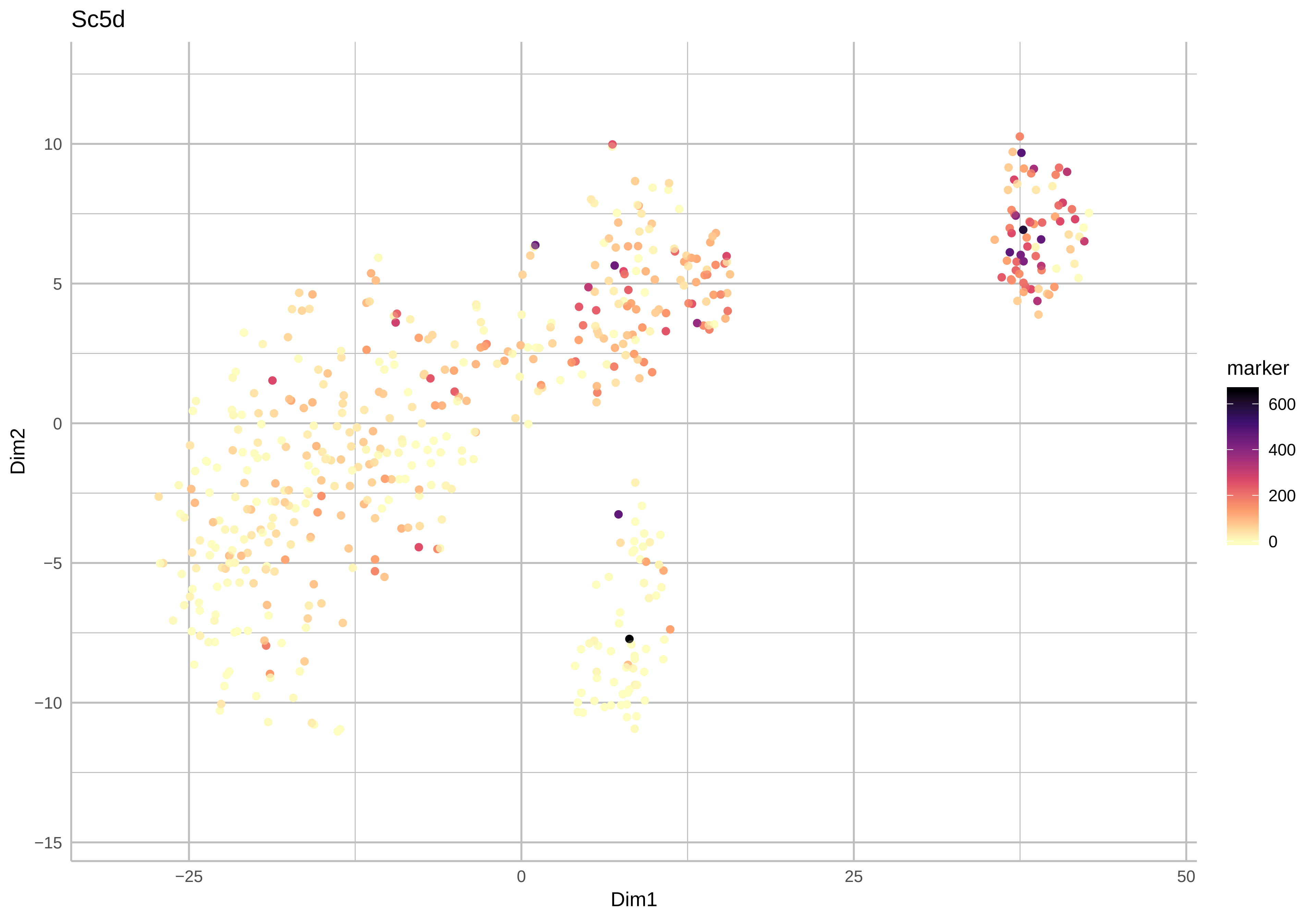
Sc5d
10
5
0
−5
−10
−15
Dim2
−25
0
25
50
Dim1
marker
600
400
200
0

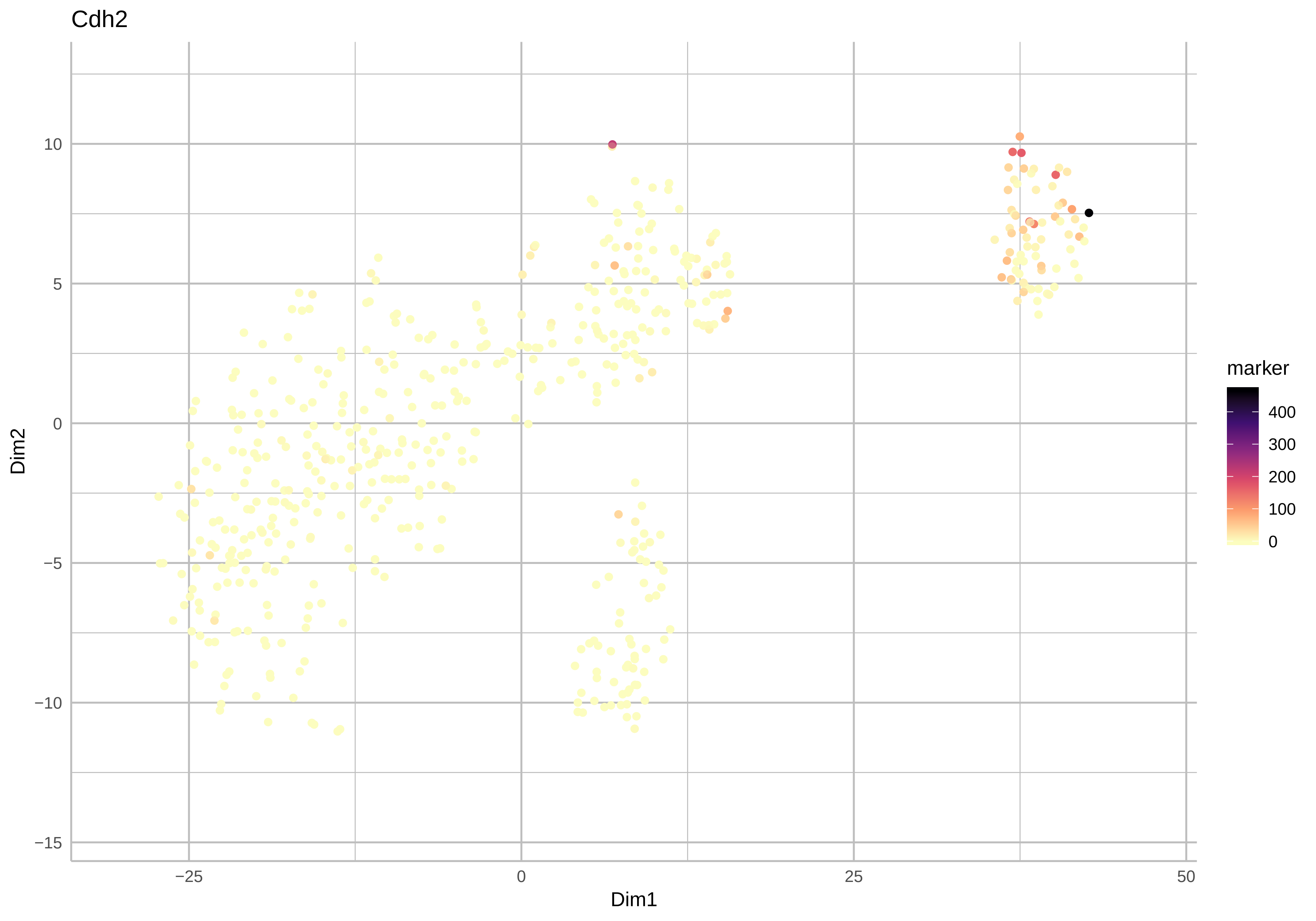

Cdh2
10
5
0
−5
−10
−15
−25
0
25
50
Dim1
Dim2
marker
400
300
200
100
0

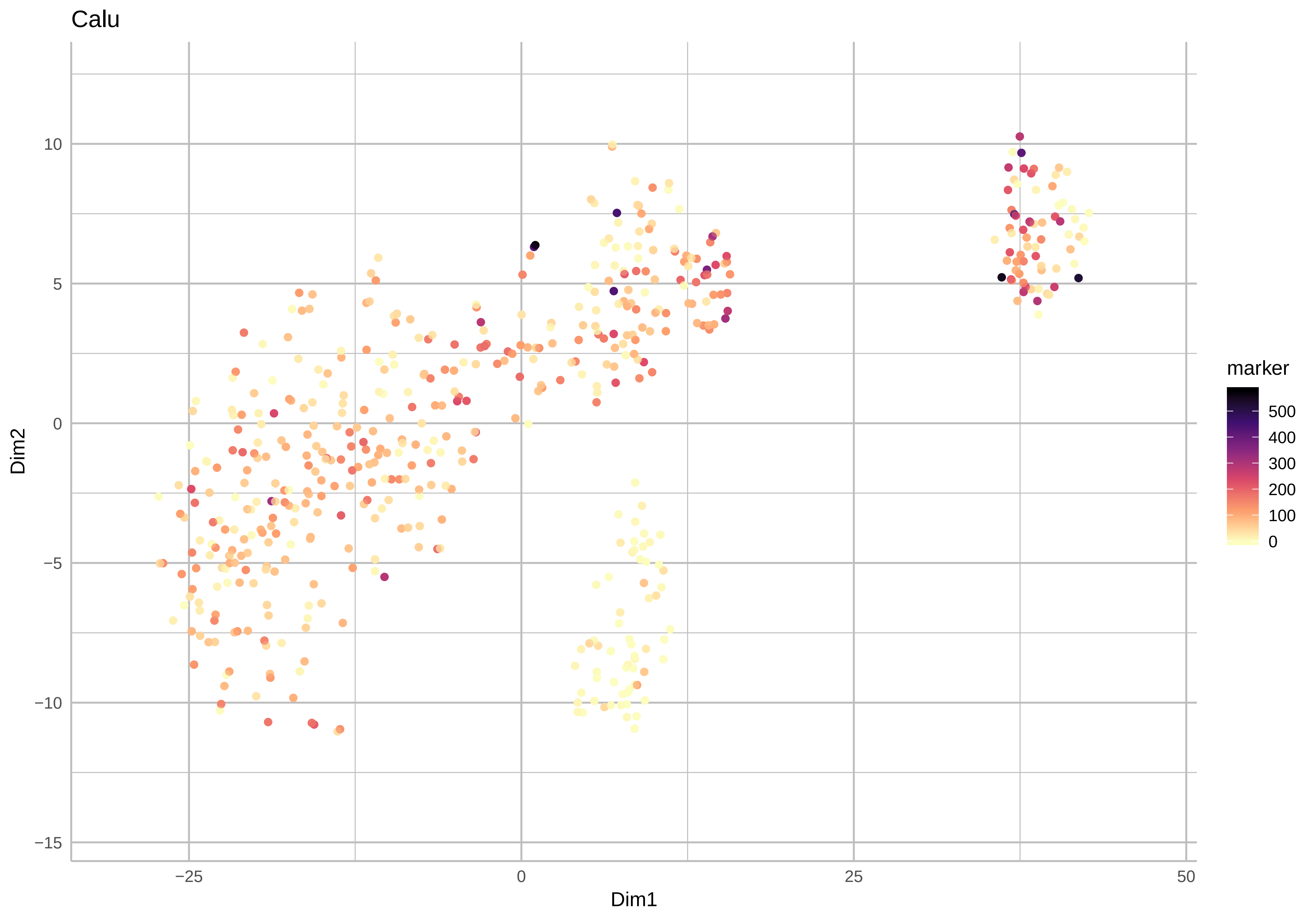

Calu
10
5
0
−5
−10
−15
Dim2
−25
0
25
50
Dim1
marker
500
400
300
200
100
0

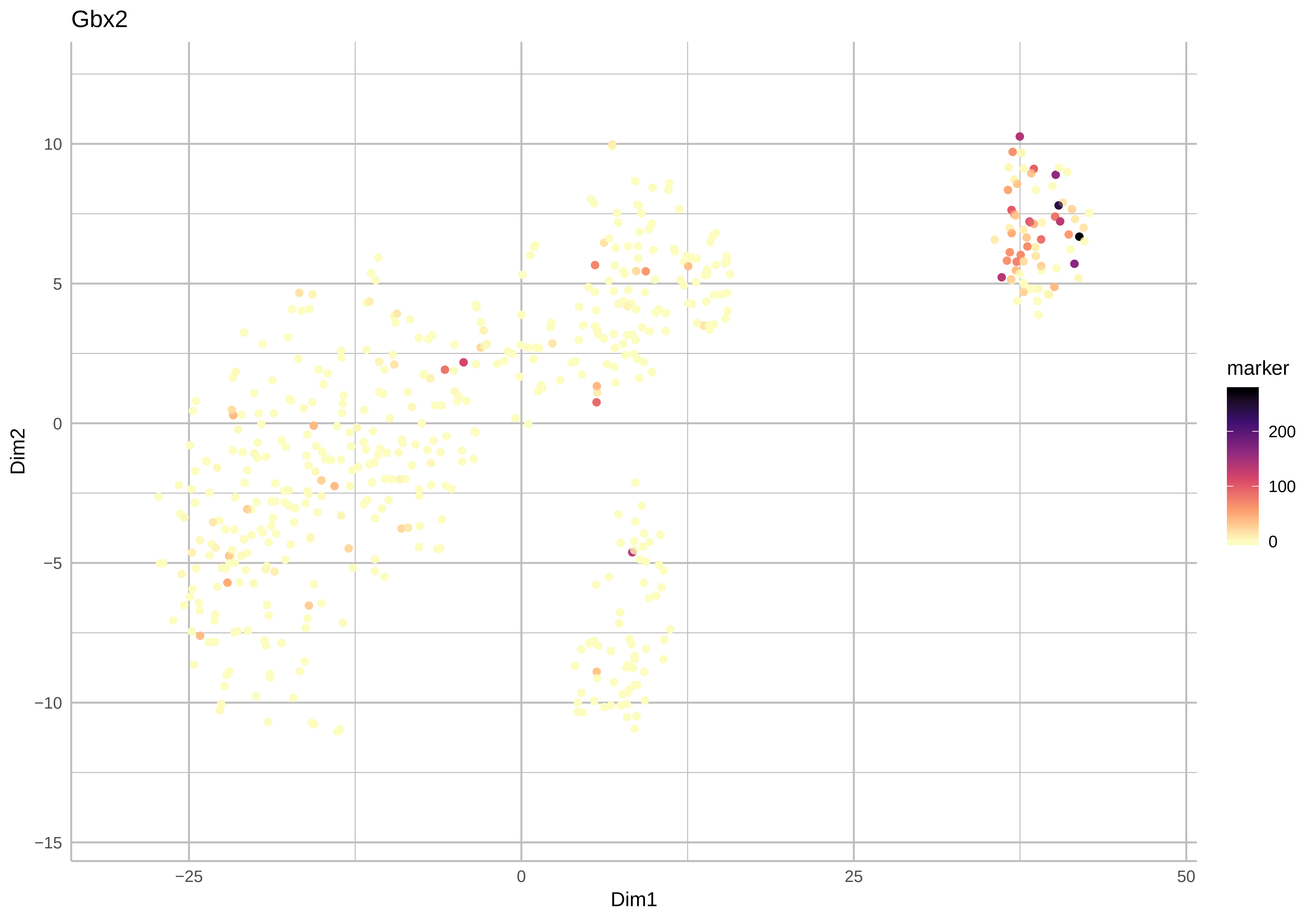

Gbx2
10
5
0
−5
−10
−15
−25
0
25
50
Dim1
Dim2
marker
200
100
0

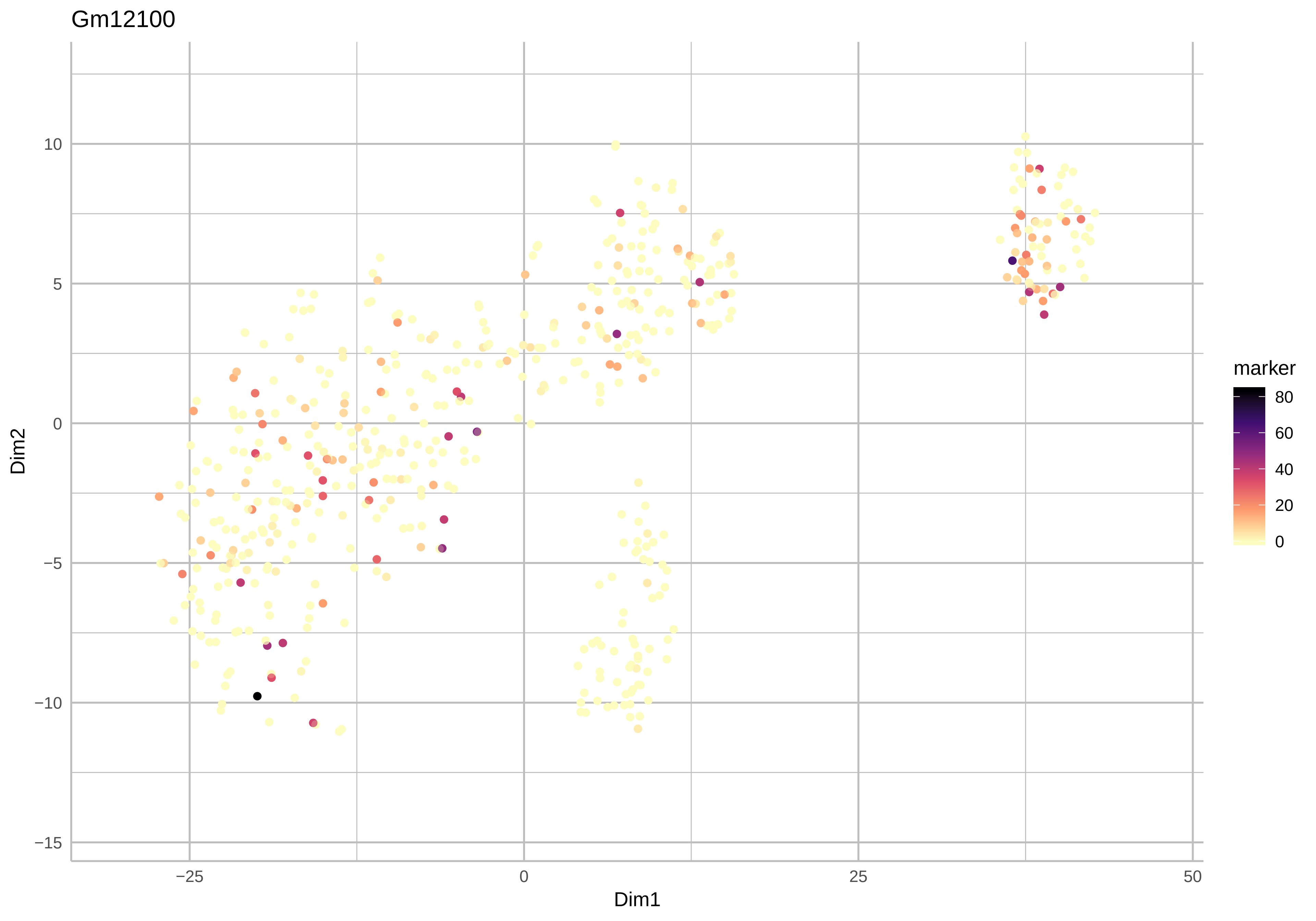

Gm12100
Dim2
Dim1
10
5
0
−5
−10
−15
−25
0
25
50
marker
80
60
40
20
0

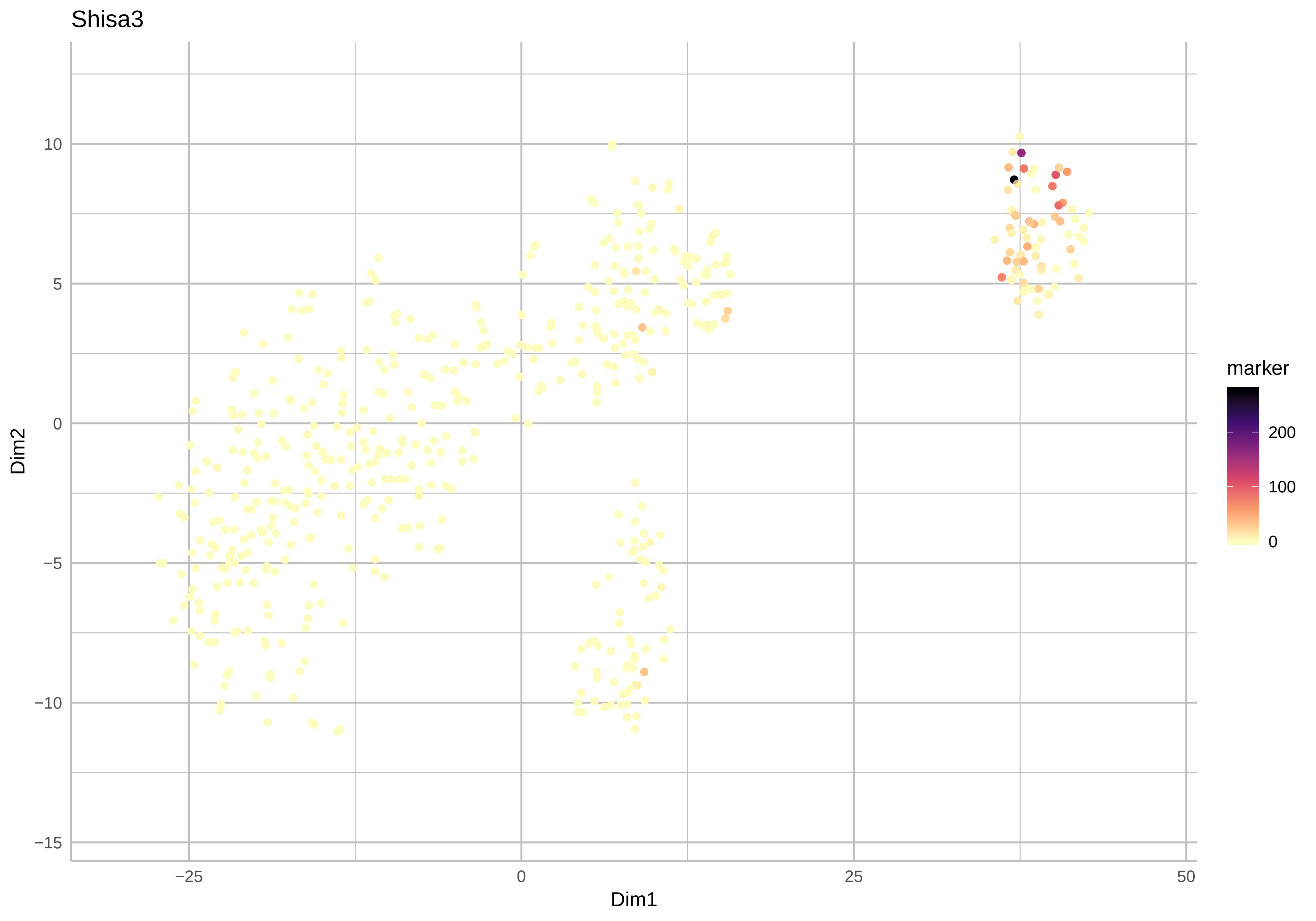
Shisa3
Dim2
Dim1
marker
200
100
0
10
5
0
−5
−10
−15
−25
0
25
50

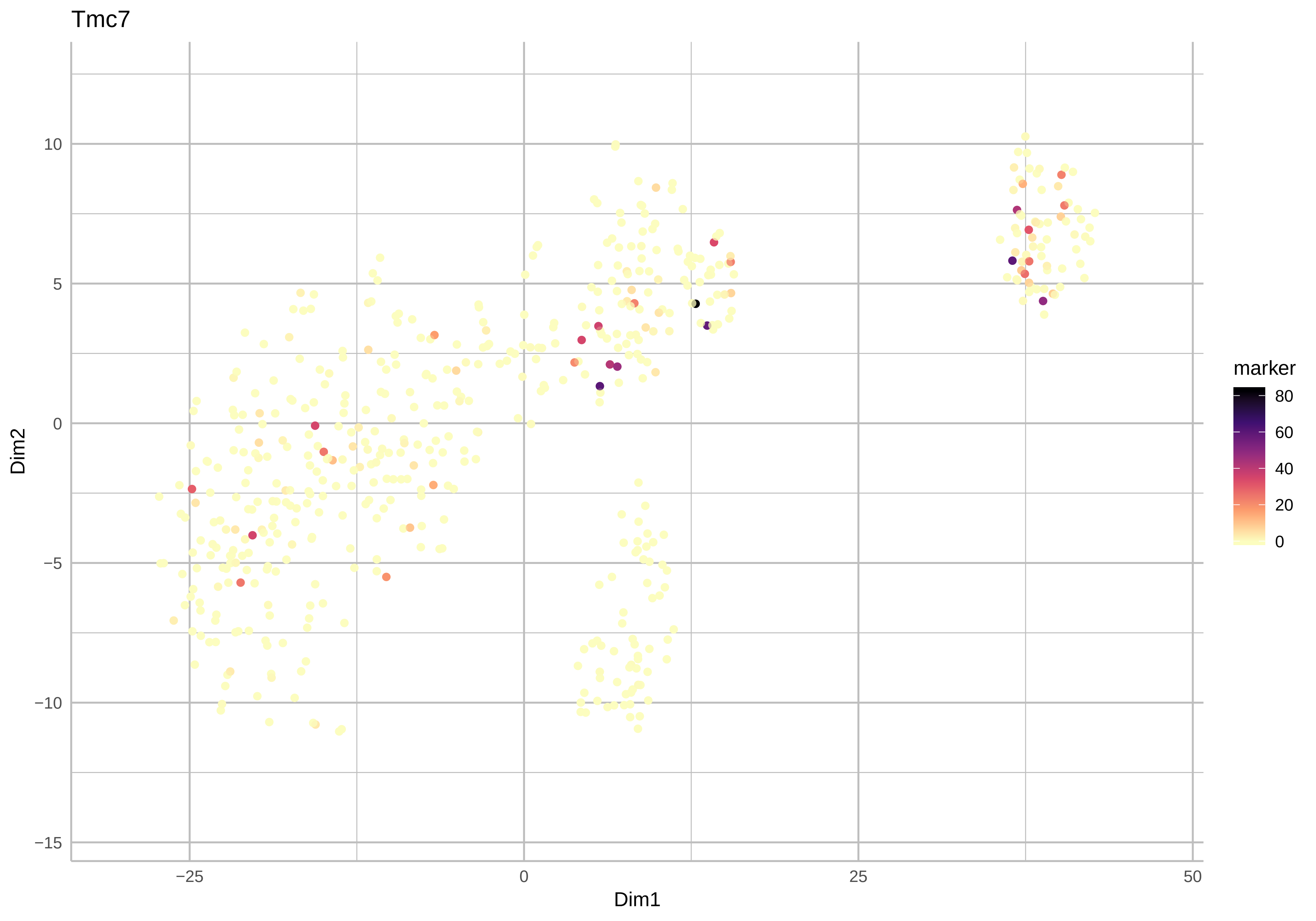

Tmc7
Dim2
Dim1
10
5
0
−5
−10
−15
−25
0
25
50
marker
80
60
40
20
0

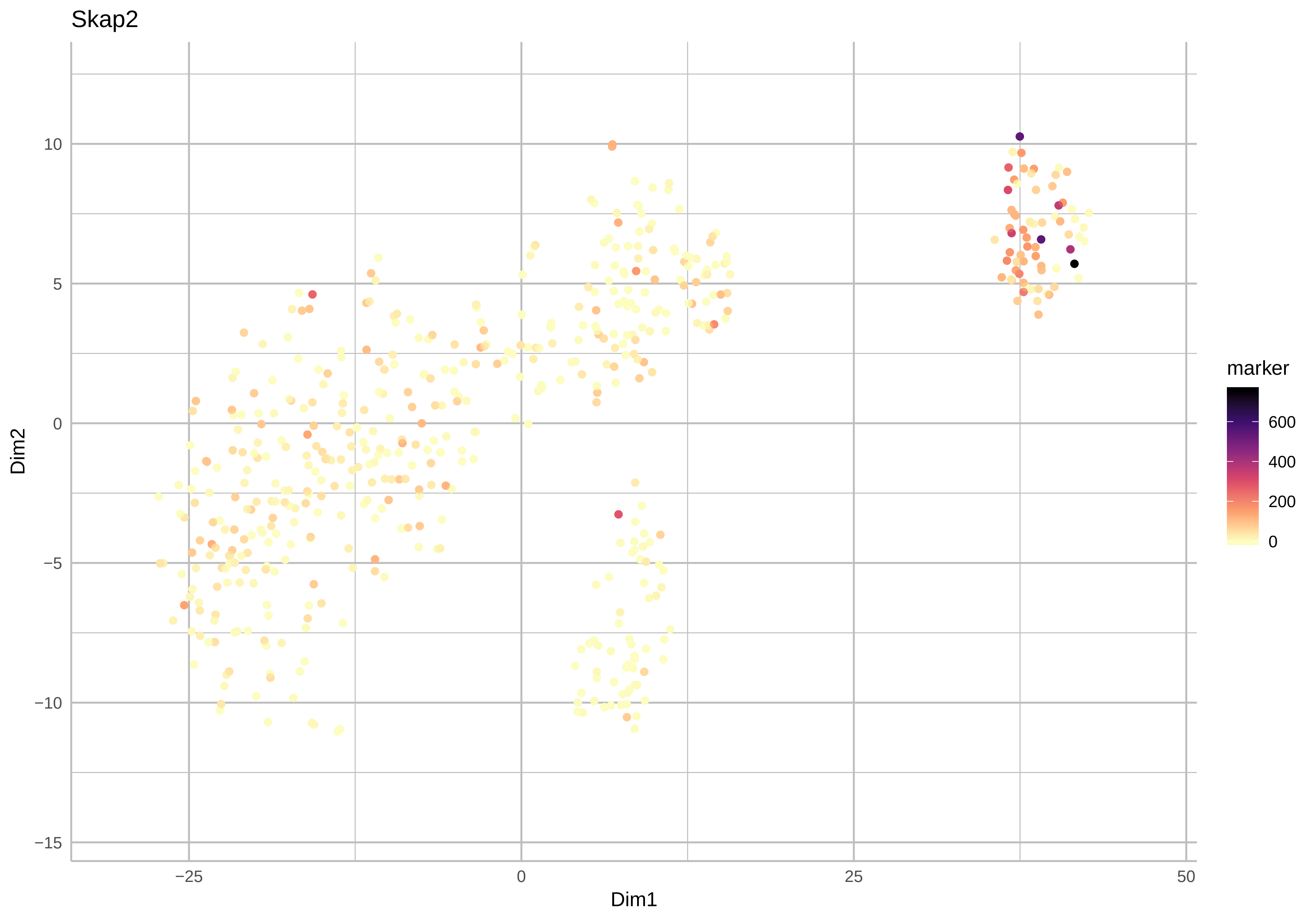

Skap2
Dim2
10
5
0
−5
−10
−15
Dim1
−25
0
25
50
marker
600
400
200
0

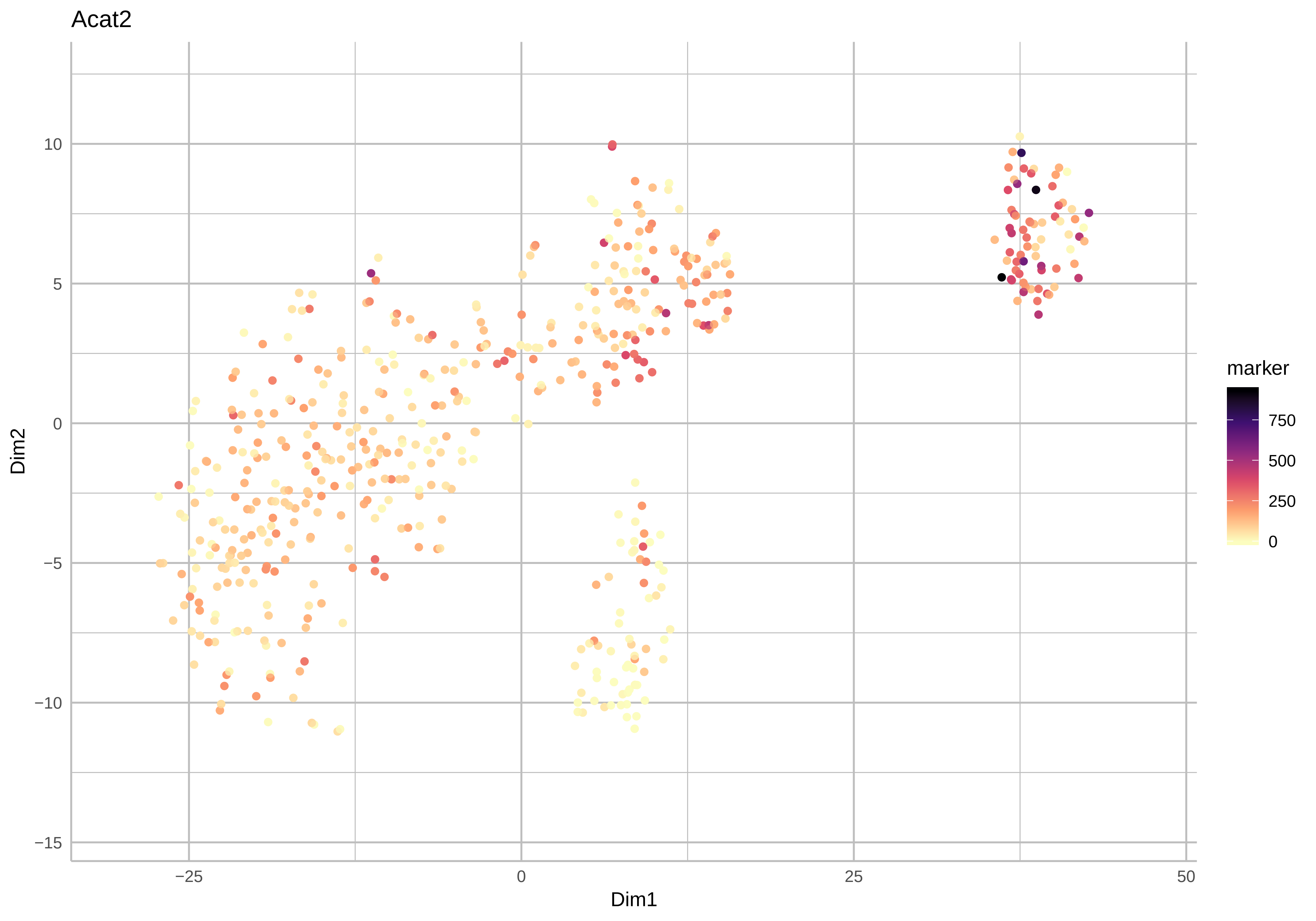
Acat2
Dim2
Dim1
10
5
0
−5
−10
−15
−25
0
25
50
marker
750
500
250
0

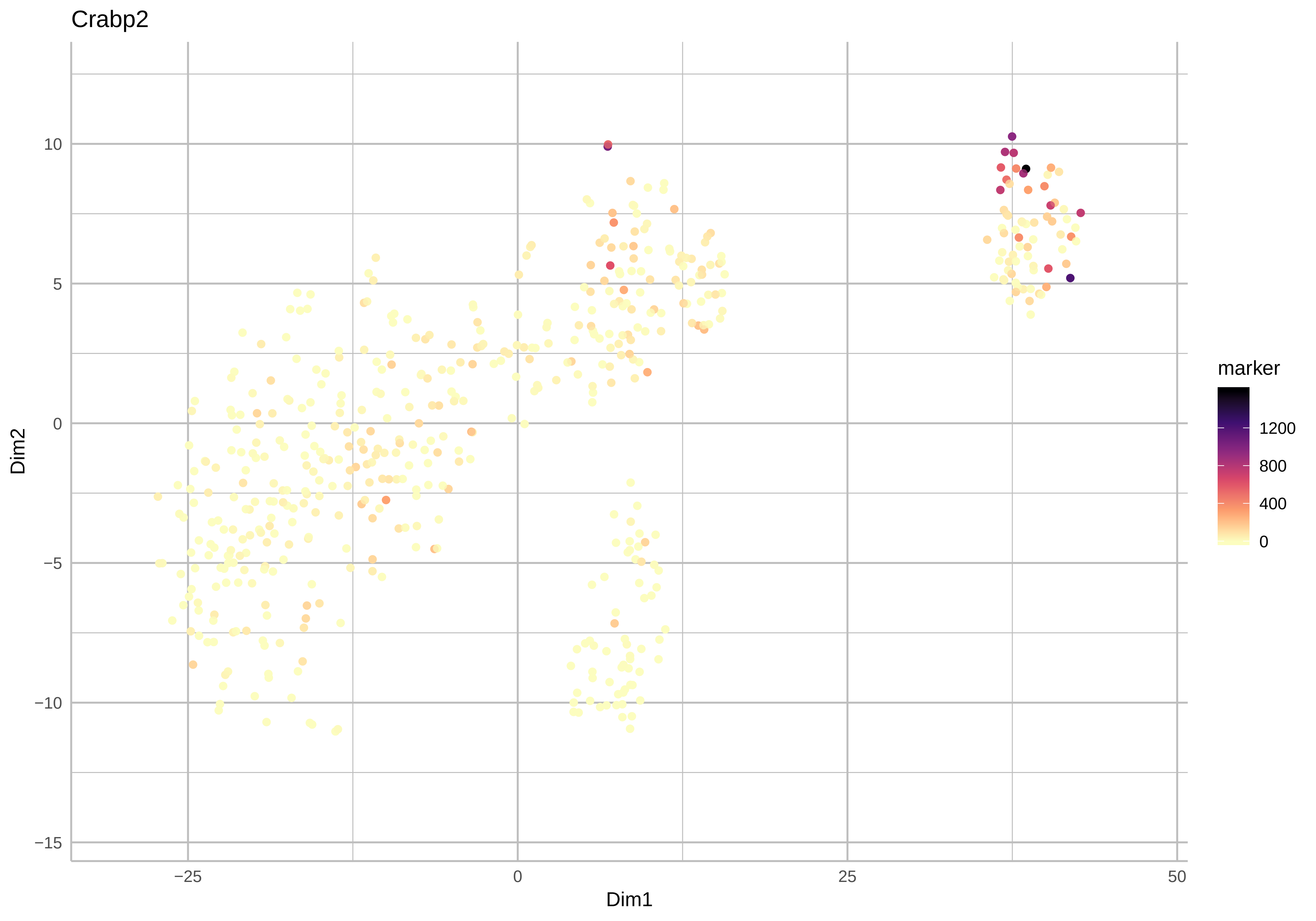

Crabp2
10
5
0
−5
−10
−15
Dim2
−25
0
25
50
Dim1
marker
1200
800
400
0

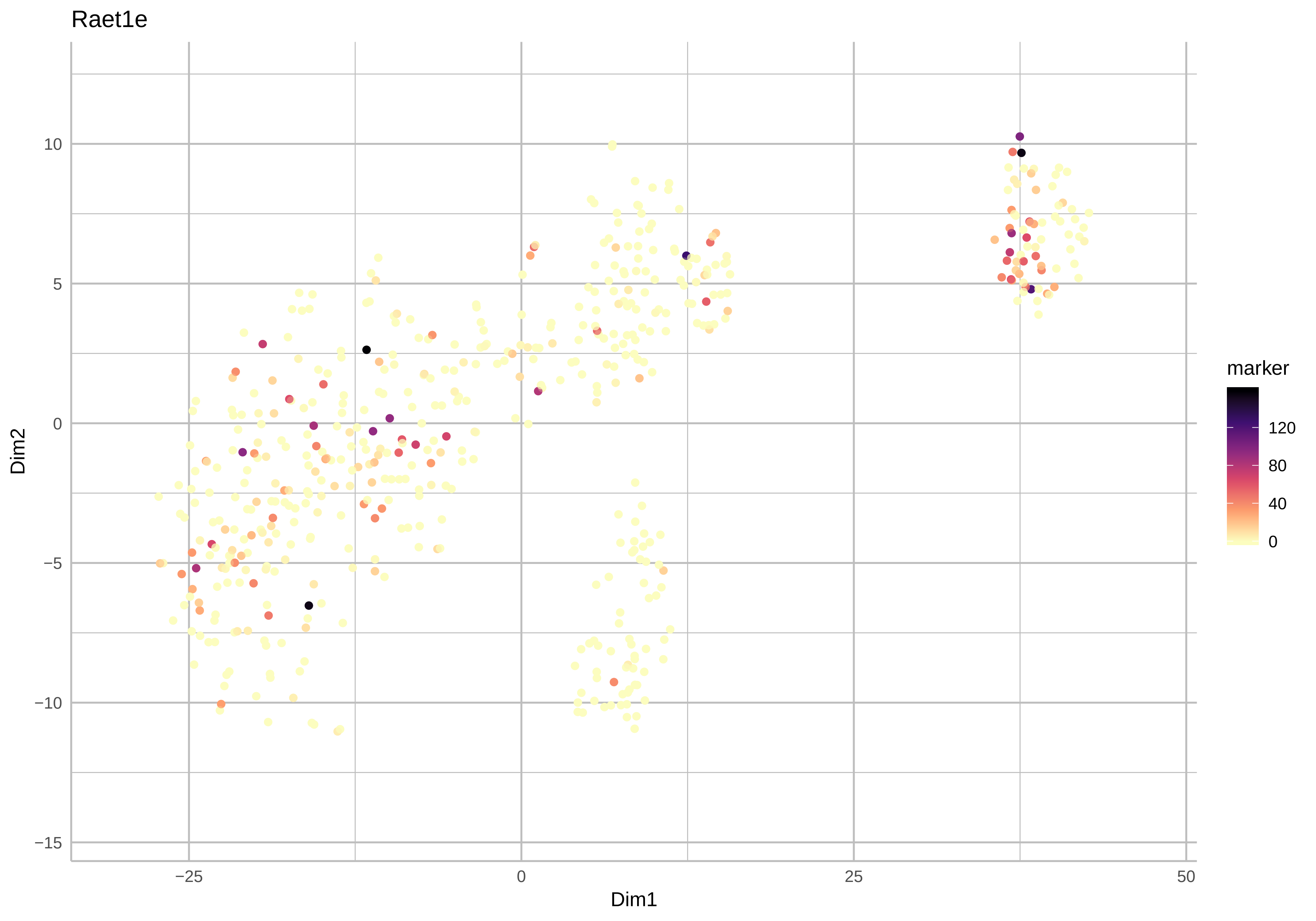
Raet1e
Dim1
Dim2
−25
0
25
50
10
5
0
−5
−10
−15
marker
120
80
40
0

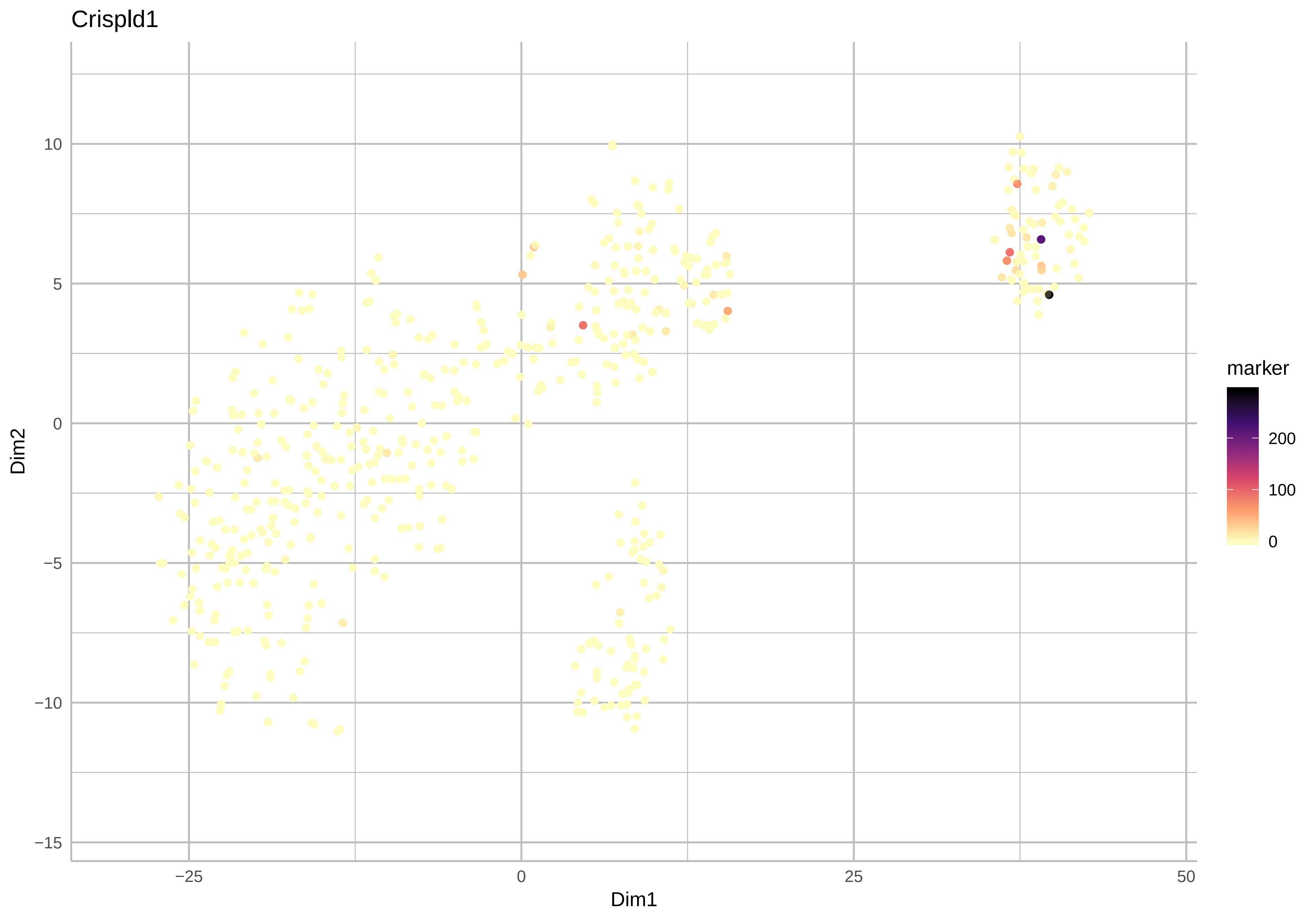
Crispld1
Dim2
Dim1
10
5
0
−5
−10
−15
−25
0
25
50
marker
200
100
0

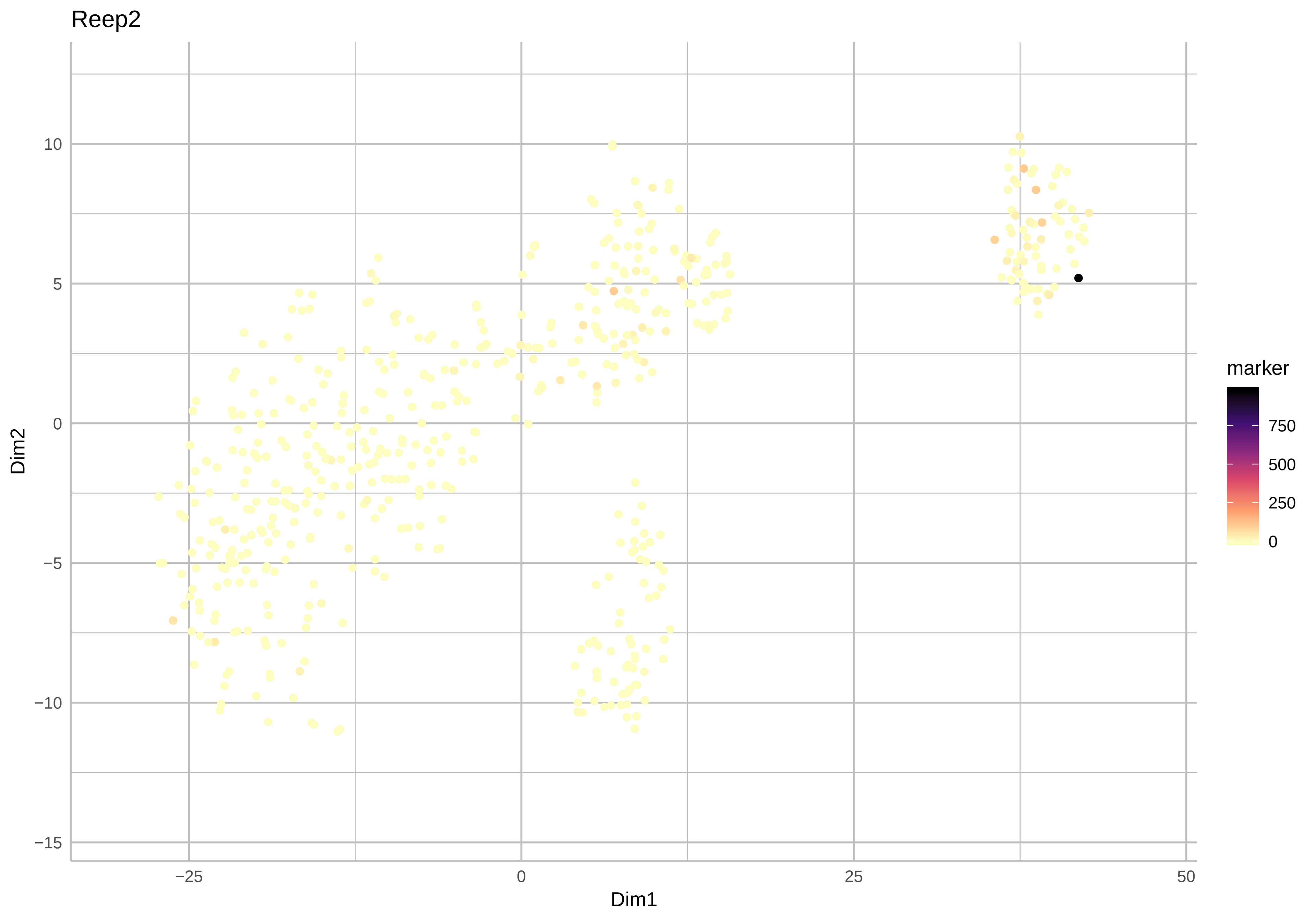

Reep2
10
5
0
−5
−10
−15
Dim2
−25
0
25
50
Dim1
marker
750
500
250
0

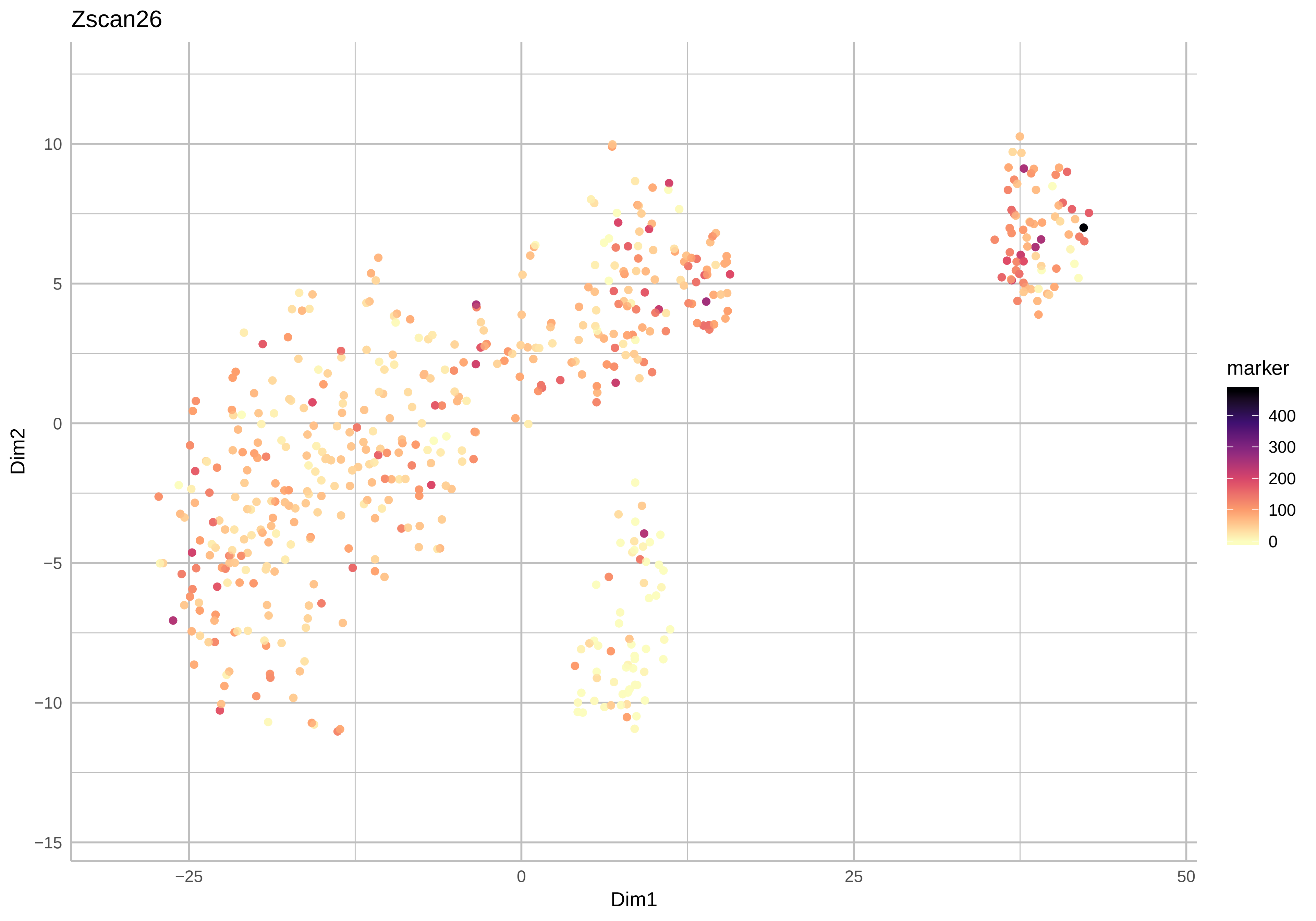

Zscan26
Dim2
Dim1
10
5
0
−5
−10
−15
−25
0
25
50
marker
400
300
200
100
0

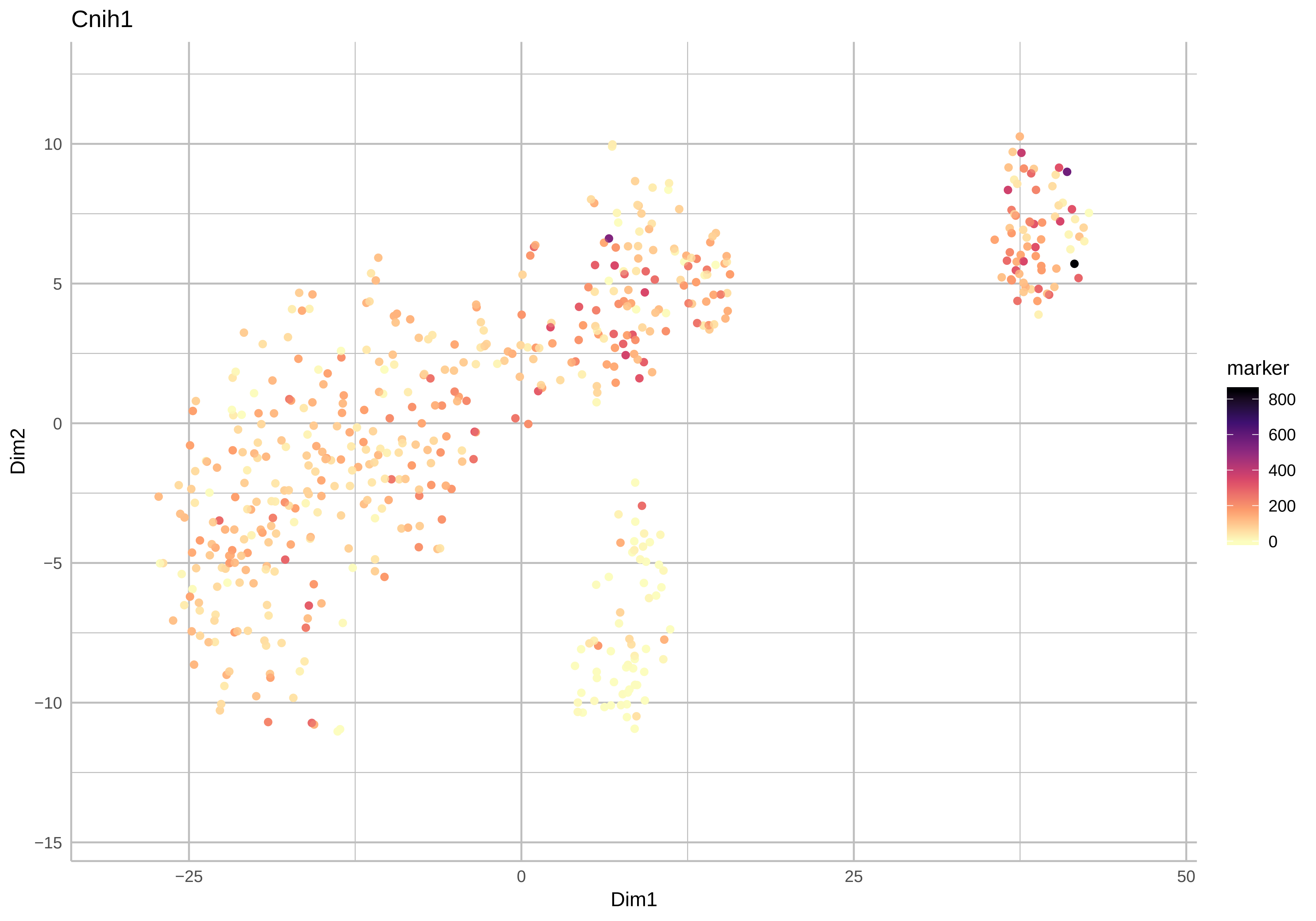

Cnih1
10
5
0
−5
−10
−15
Dim2
−25
0
25
50
Dim1
marker
800
600
400
200
0

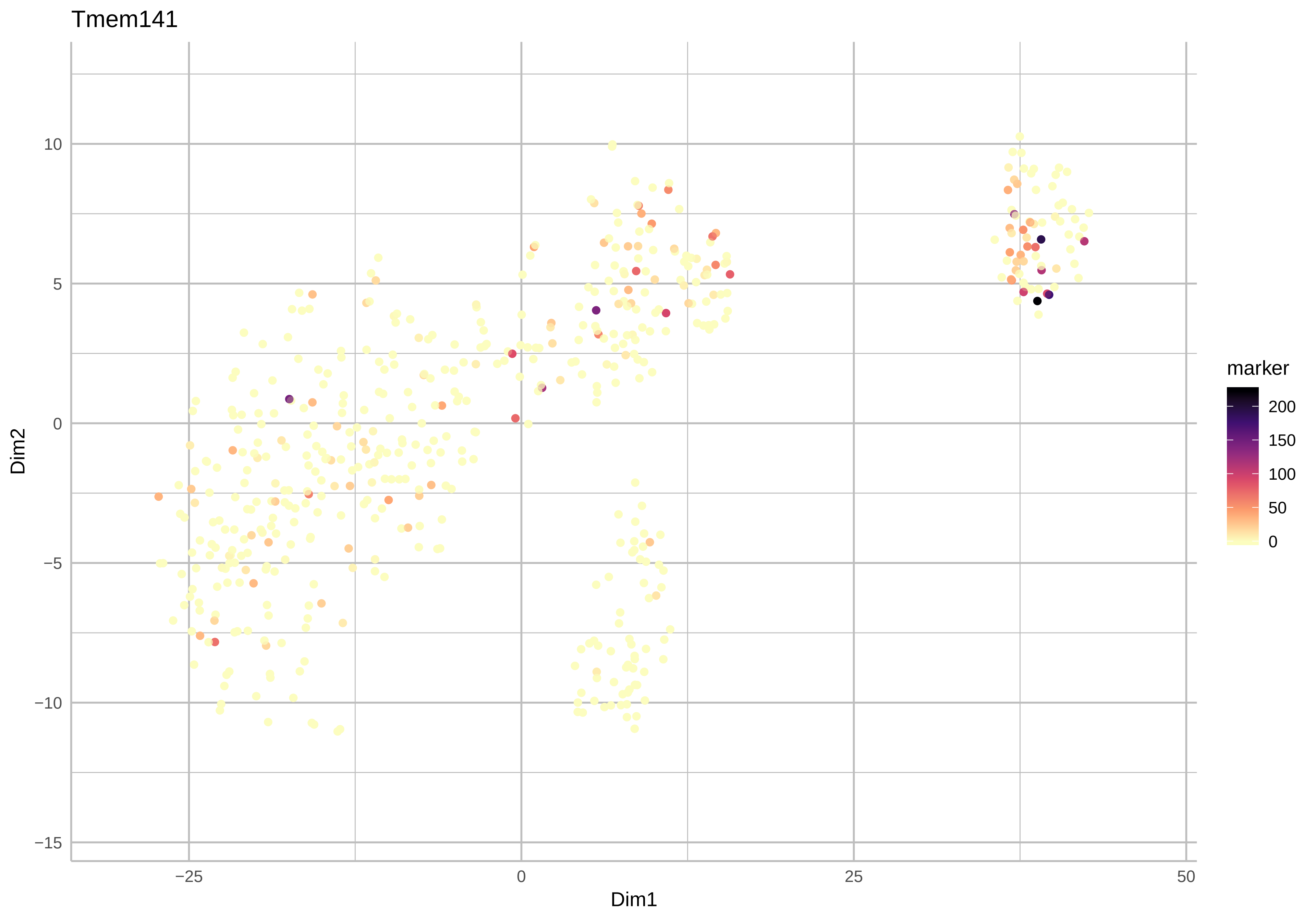

Tmem141
Dim2
10
5
0
−5
−10
−15
−25
0
25
50
Dim1
marker
200
150
100
50
0

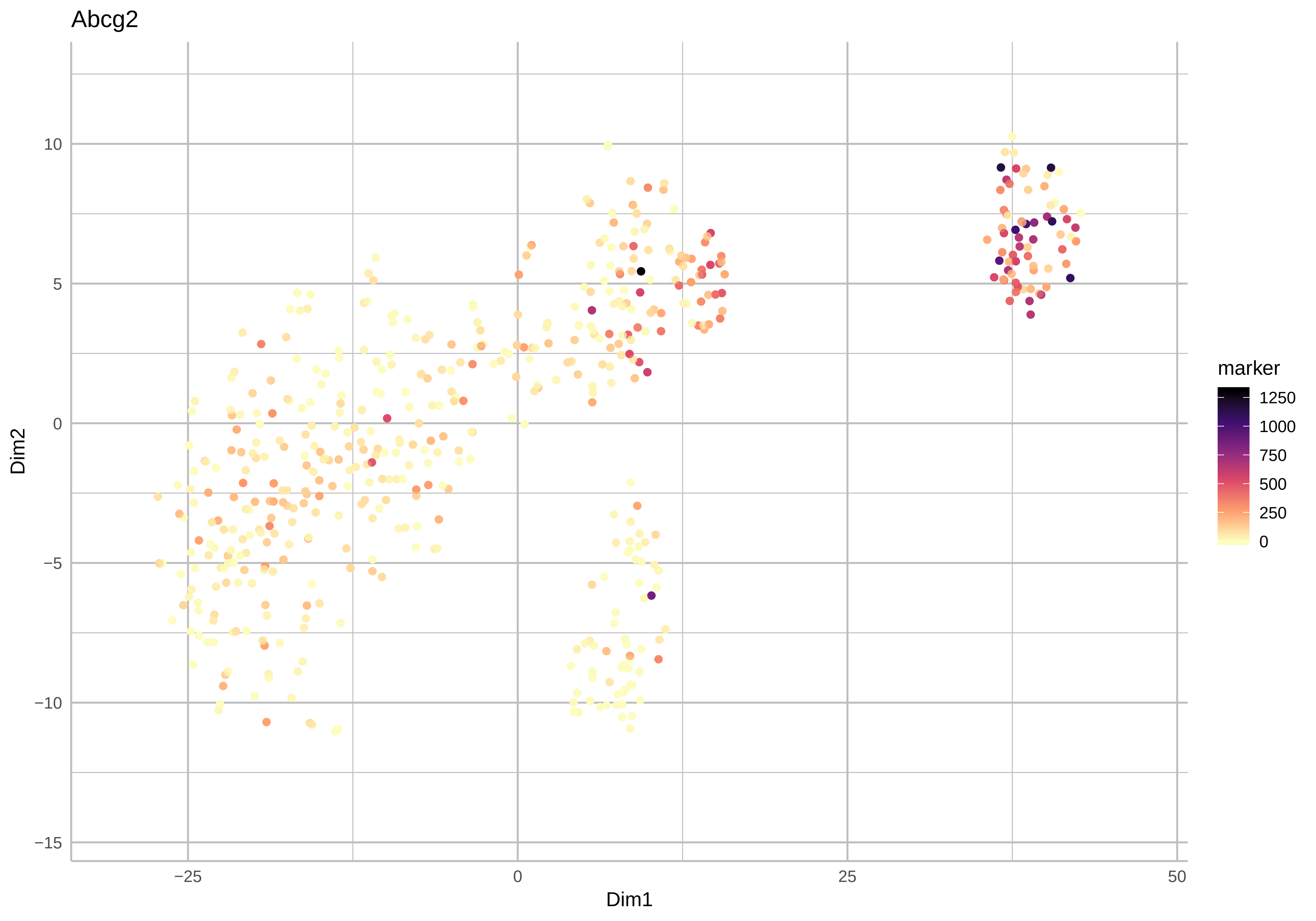
Abcg2
Dim2
Dim1
10
5
0
−5
−10
−15
−25
0
25
50
marker
1250
1000
750
500
250
0

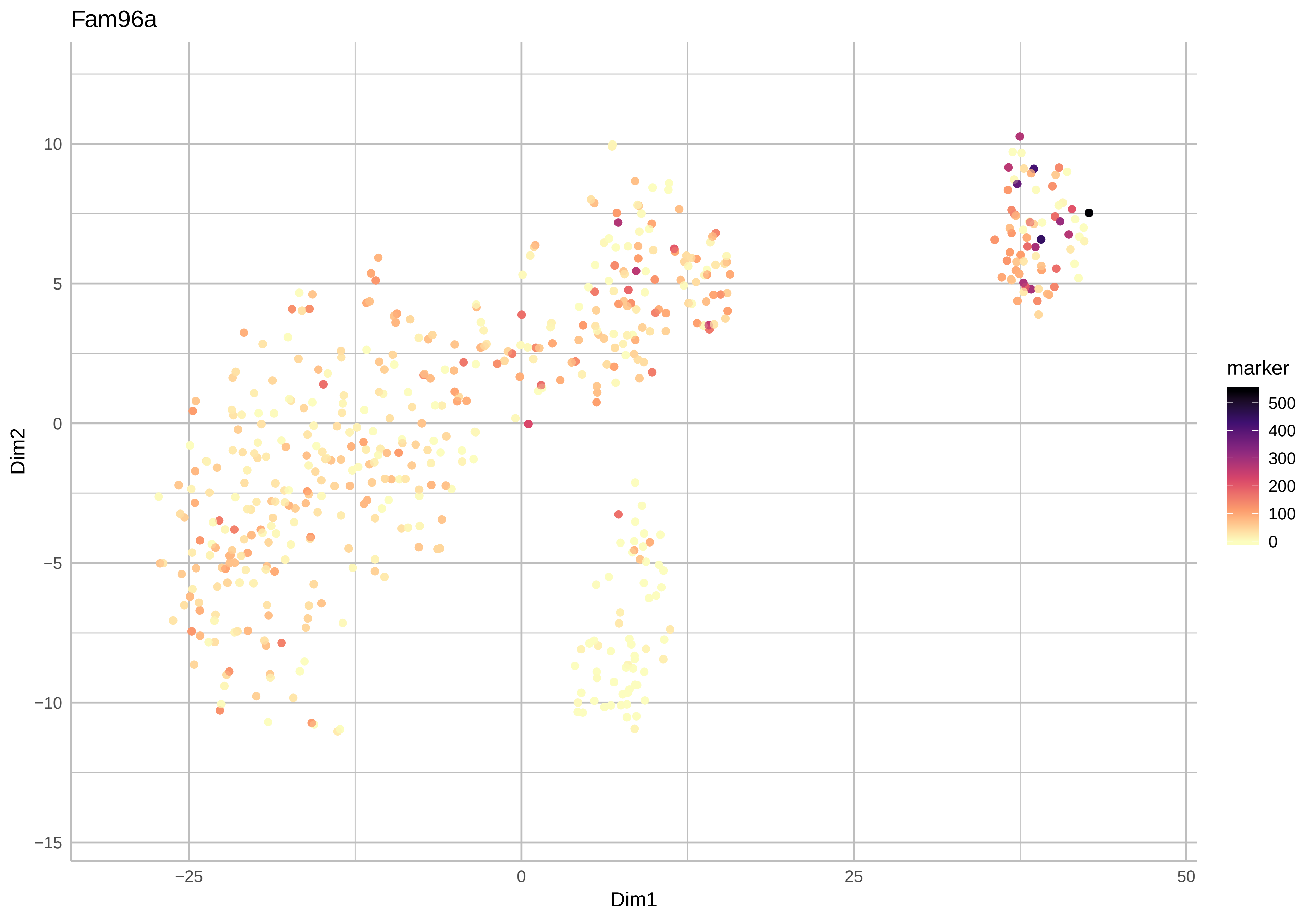

Fam96a
Dim2
Dim1
10
5
0
−5
−10
−15
−25
0
25
50
marker
500
400
300
200
100
0

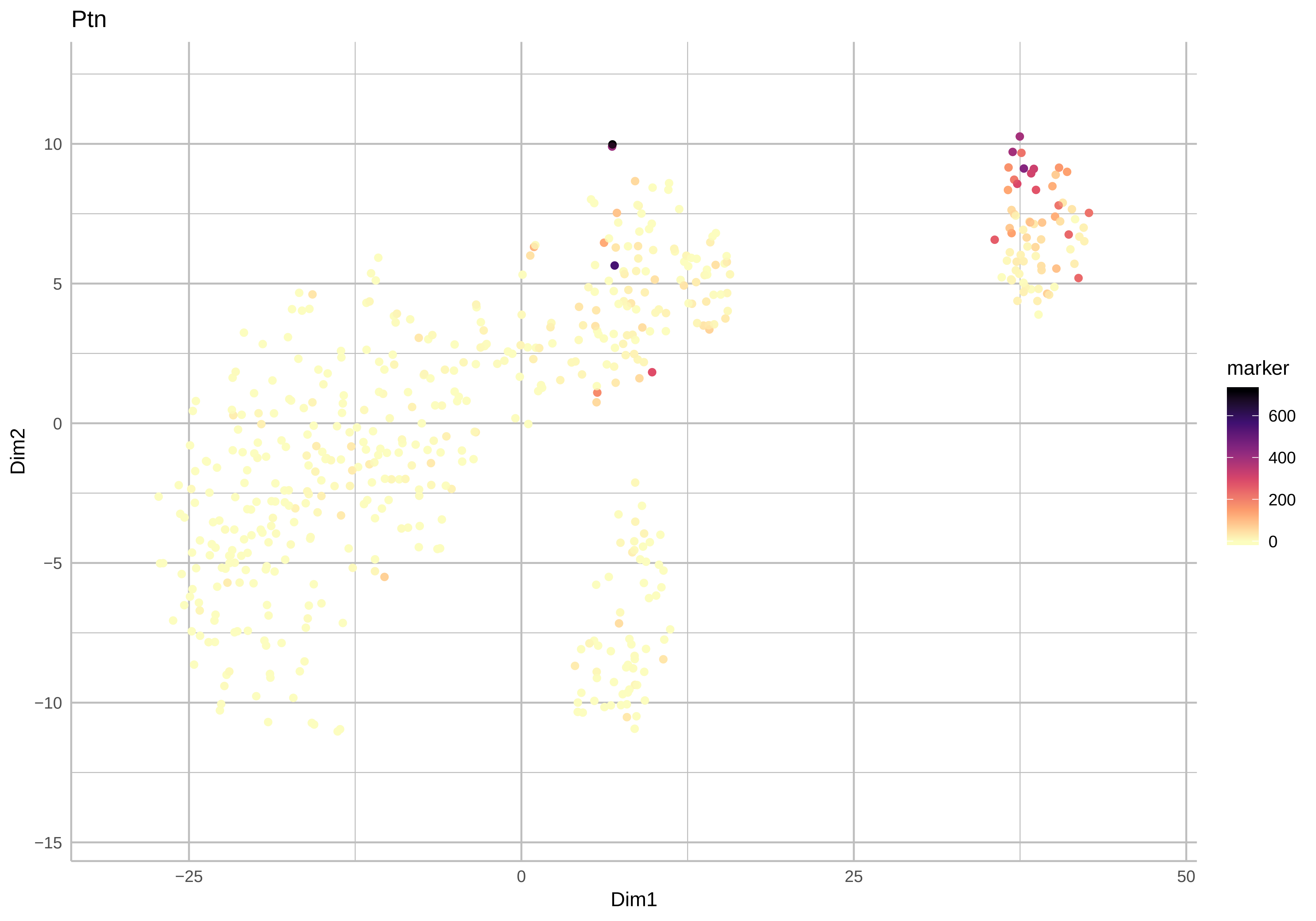
Ptn
Dim2
Dim1
10
5
0
−5
−10
−15
−25
0
25
50
marker
600
400
200
0

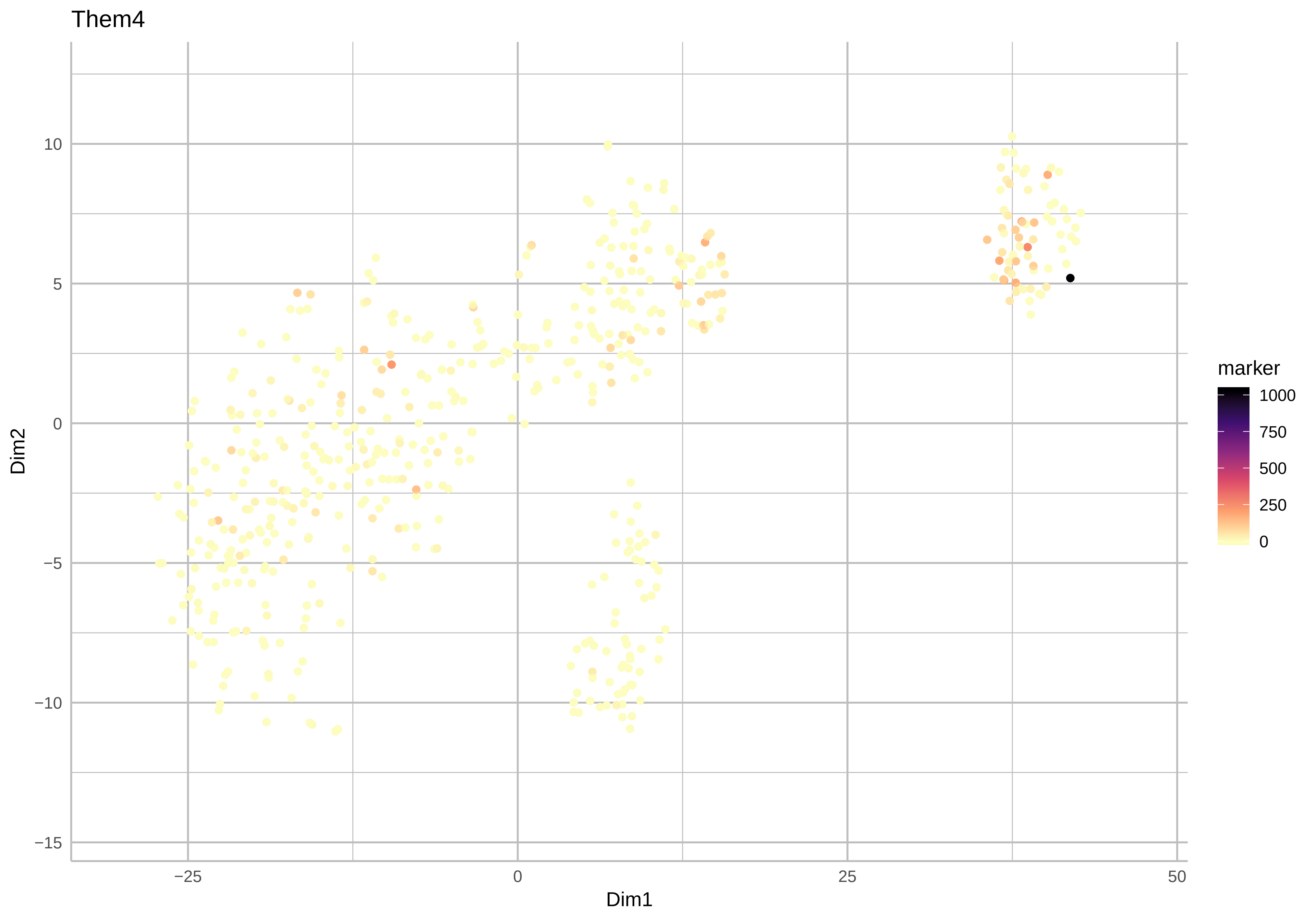
Them4
Dim2
Dim1
10
5
0
−5
−10
−15
−25
0
25
50
marker
1000
750
500
250
0

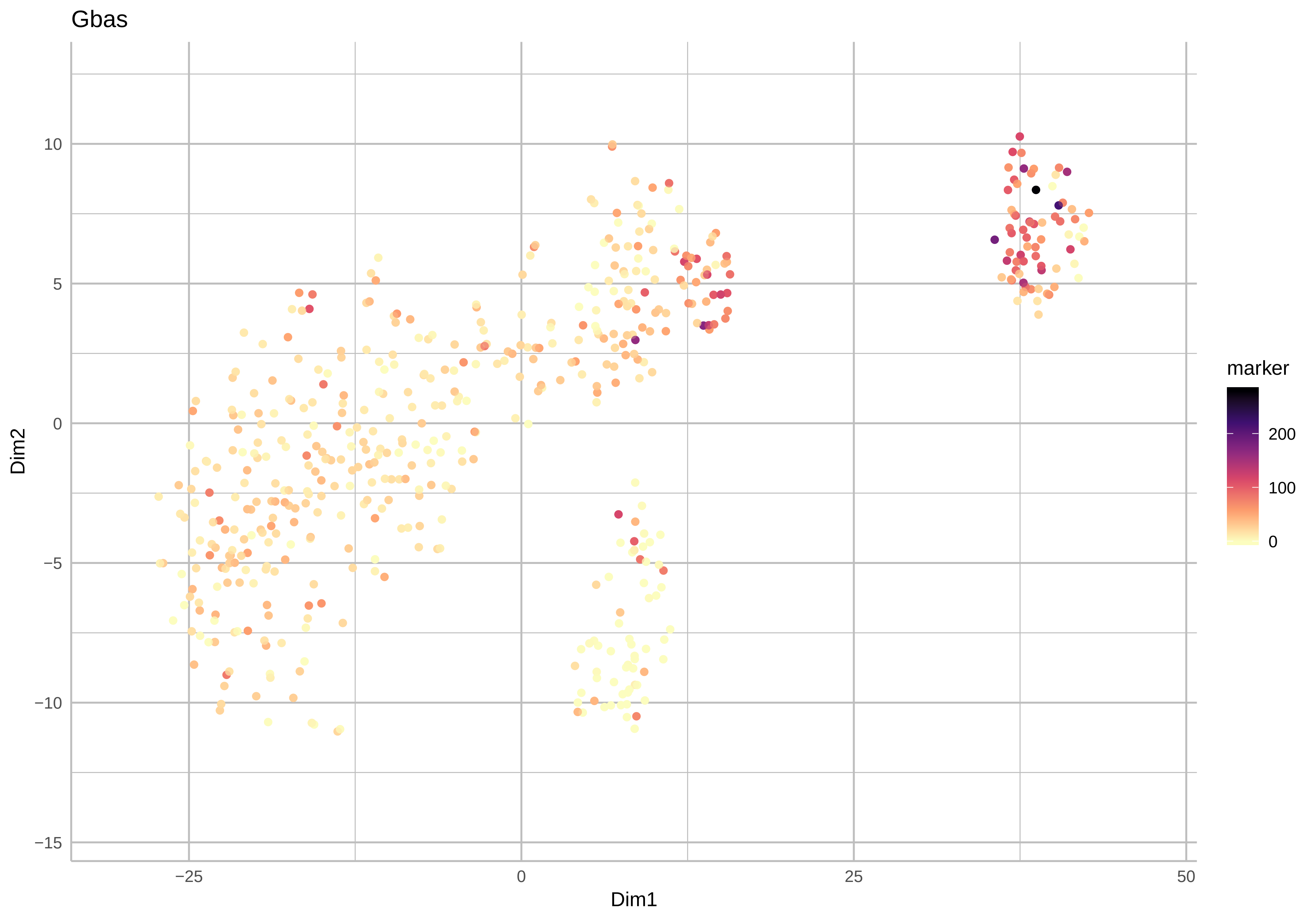
Gbas
Dim1
Dim2
marker
−25
0
25
50
10
5
0
−5
−10
−15
200
100
0

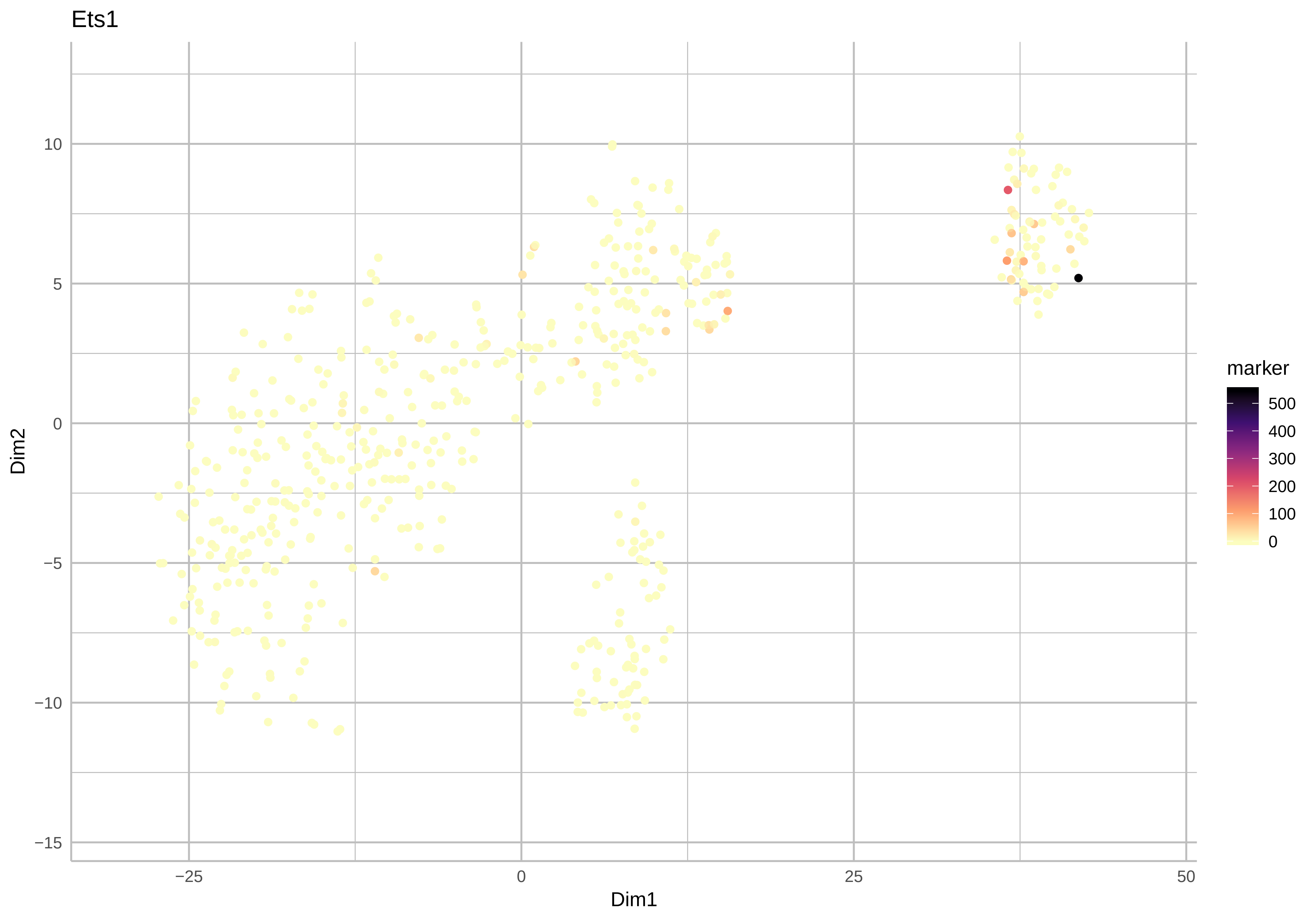
Ets1
10
5
0
−5
−10
−15
Dim2
−25
0
25
50
Dim1
marker
500
400
300
200
100
0

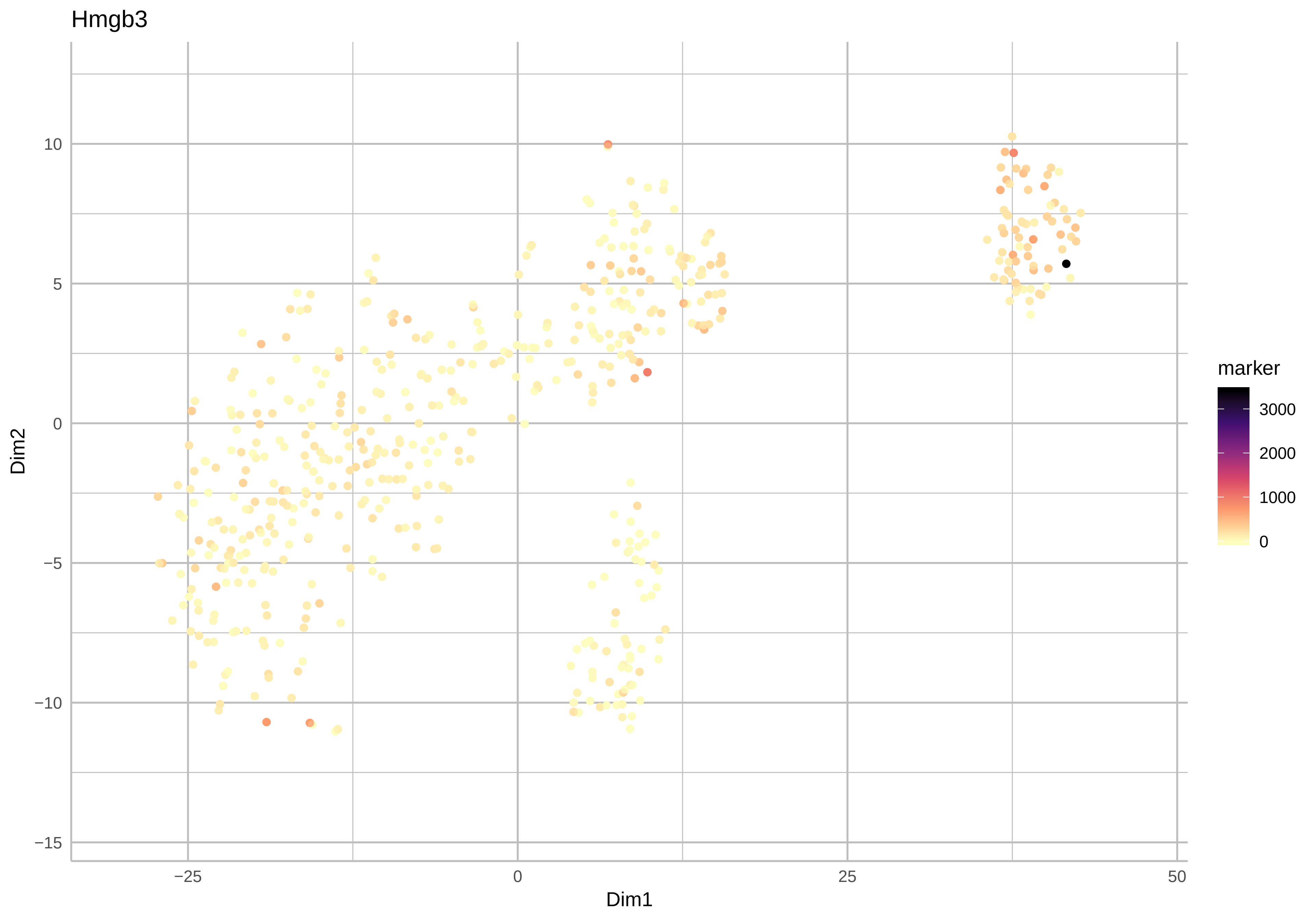
Hmgb3
Dim2
Dim1
10
5
0
−5
−10
−15
−25
0
25
50
marker
3000
2000
1000
0

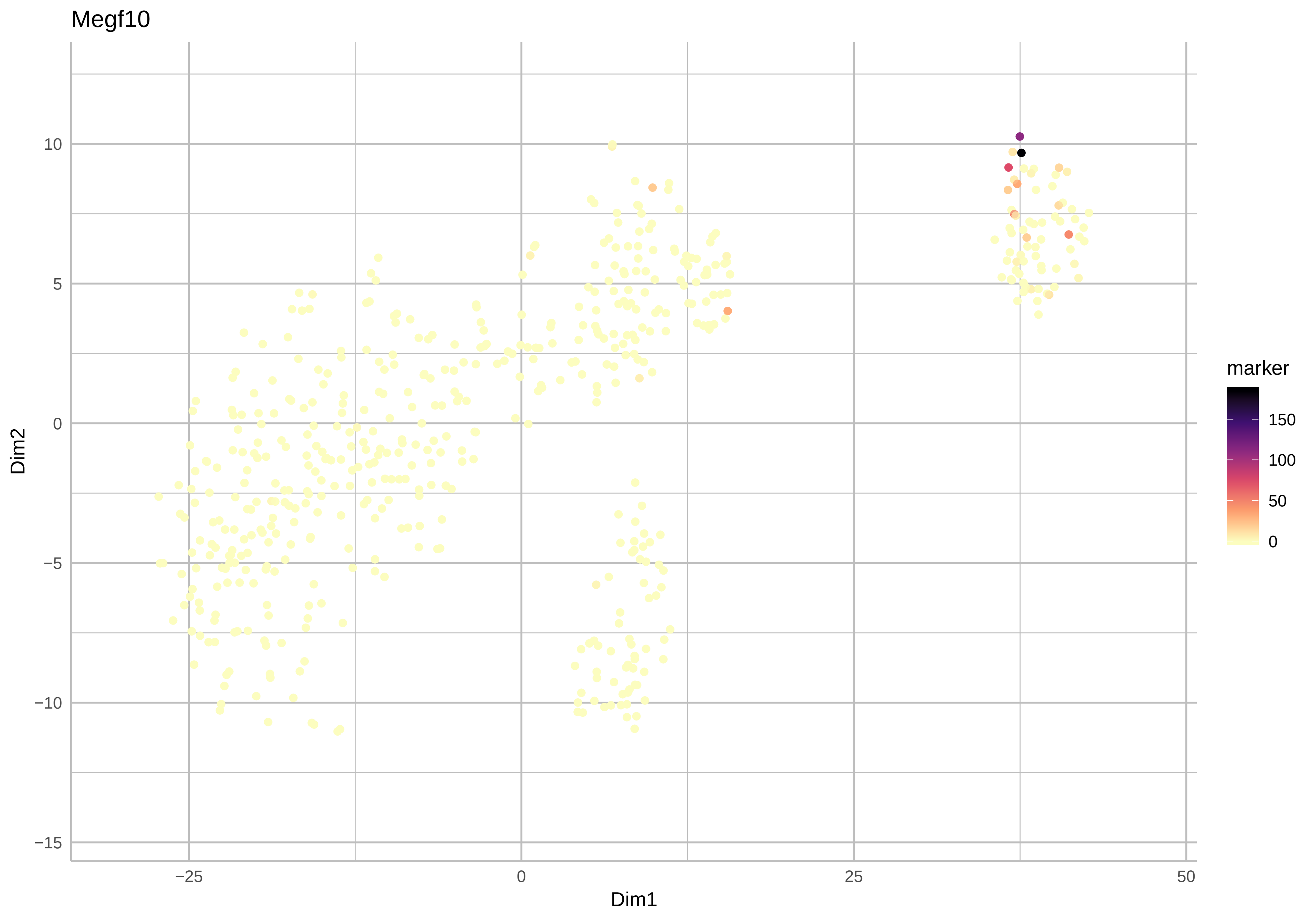

Megf10
10
5
0
−5
−10
−15
Dim2
−25
0
25
50
Dim1
marker
150
100
50
0

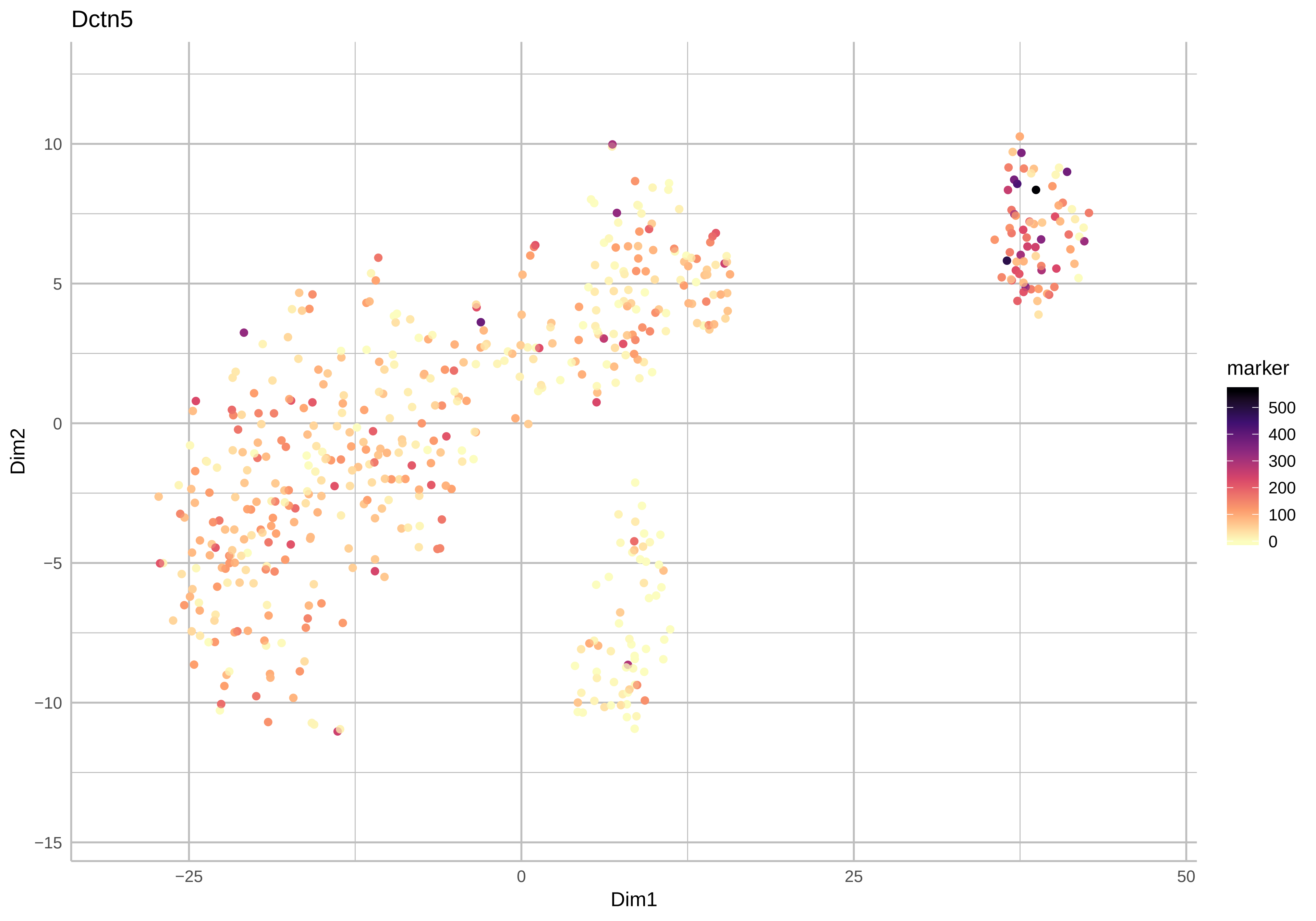
Dctn5
Dim2
10
5
0
−5
−10
−15
Dim1
−25
0
25
50
marker
500
400
300
200
100
0

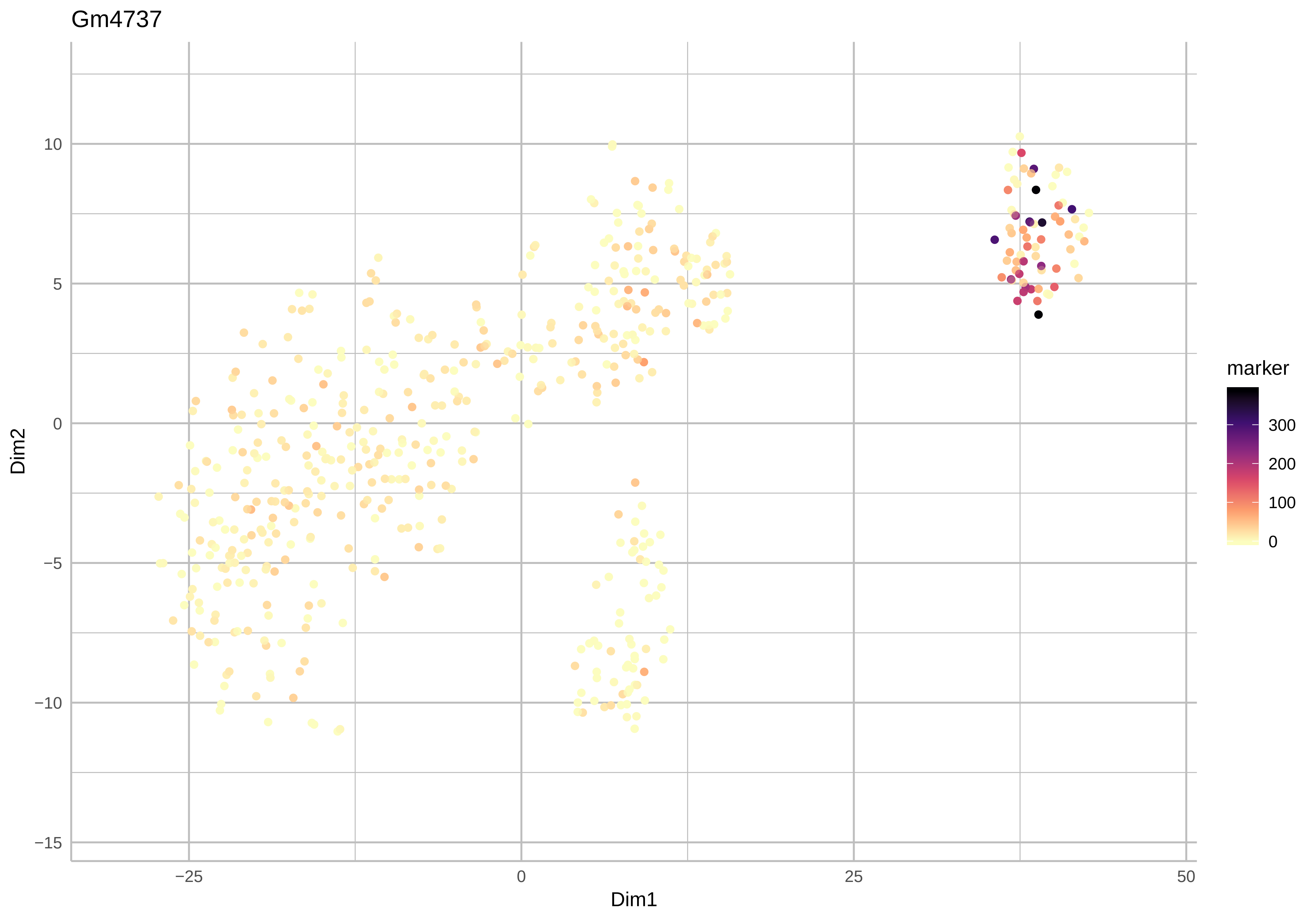
Gm4737
10
5
0
−5
−10
−15
−25
0
25
50
Dim1
Dim2
marker
300
200
100
0

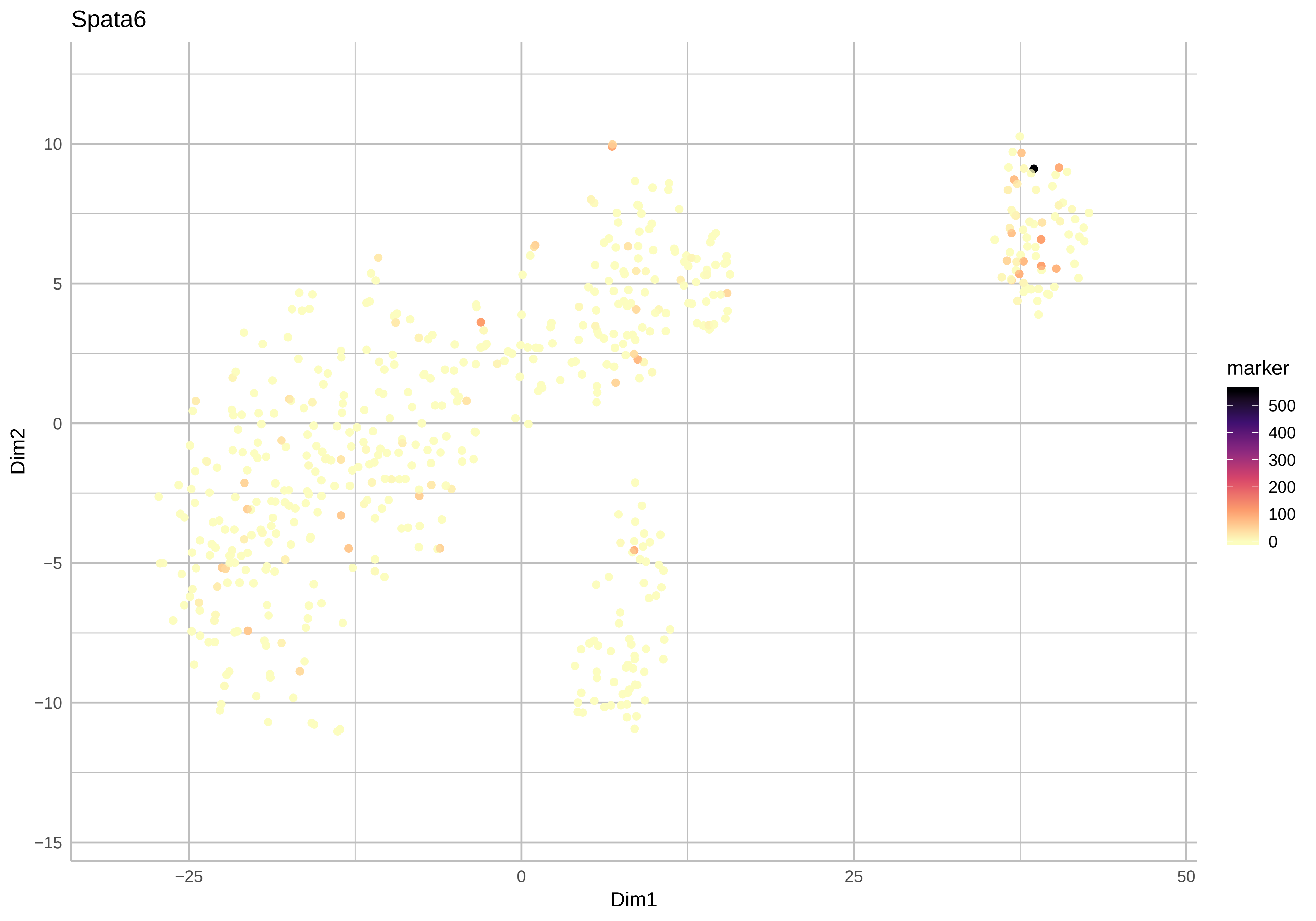
Spata6
Dim2
Dim1
10
5
0
−5
−10
−15
−25
0
25
50
marker
500
400
300
200
100
0

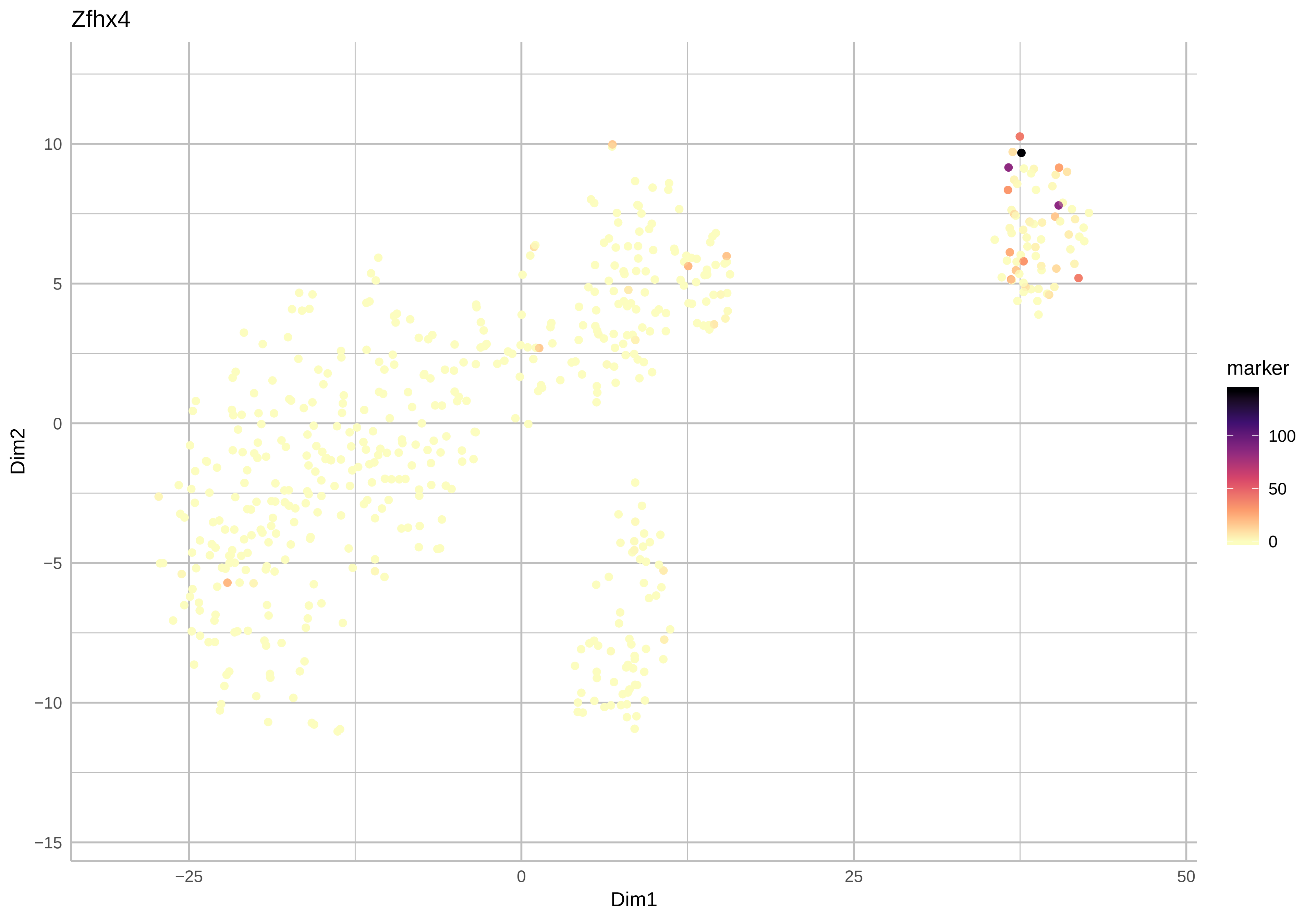
Zfhx4
Dim2
Dim1
10
5
0
−5
−10
−15
−25
0
25
50
marker
100
50
0

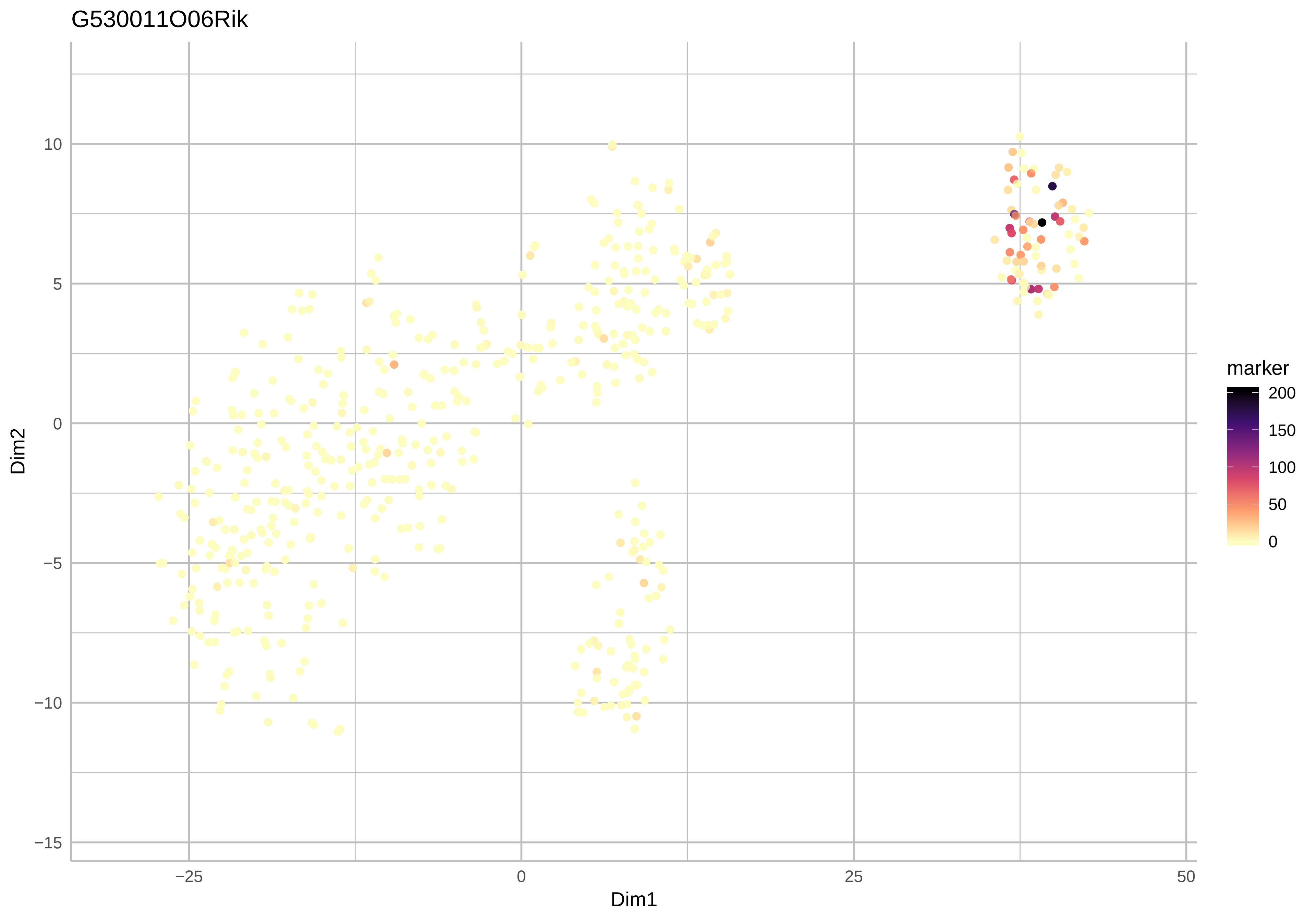
G530011O06Rik
Dim1
Dim2
−25
0
25
50
10
5
0
−5
−10
−15
marker
200
150
100
50
0

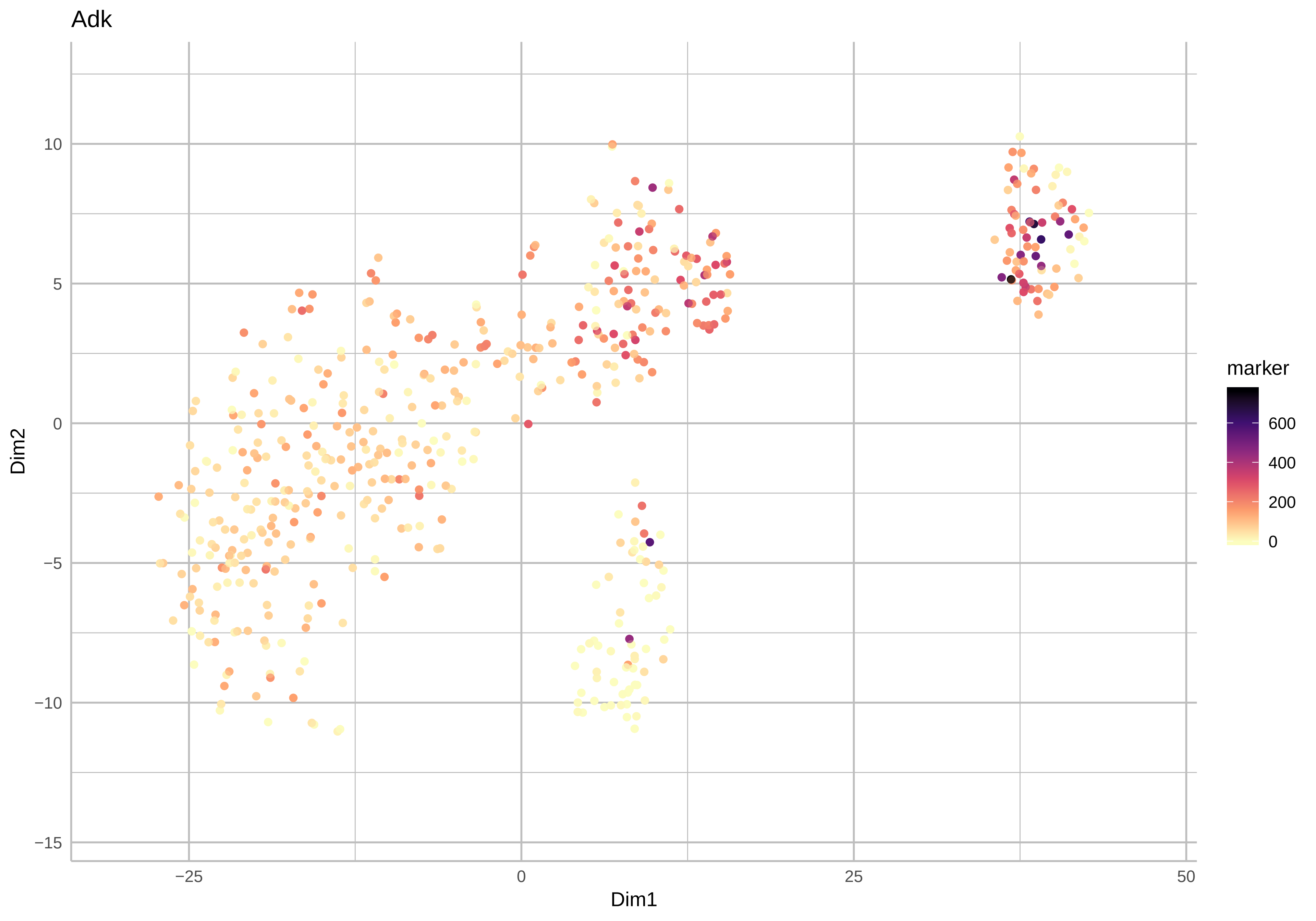

Adk
10
5
0
−5
−10
−15
Dim2
−25
0
25
50
Dim1
marker
600
400
200
0

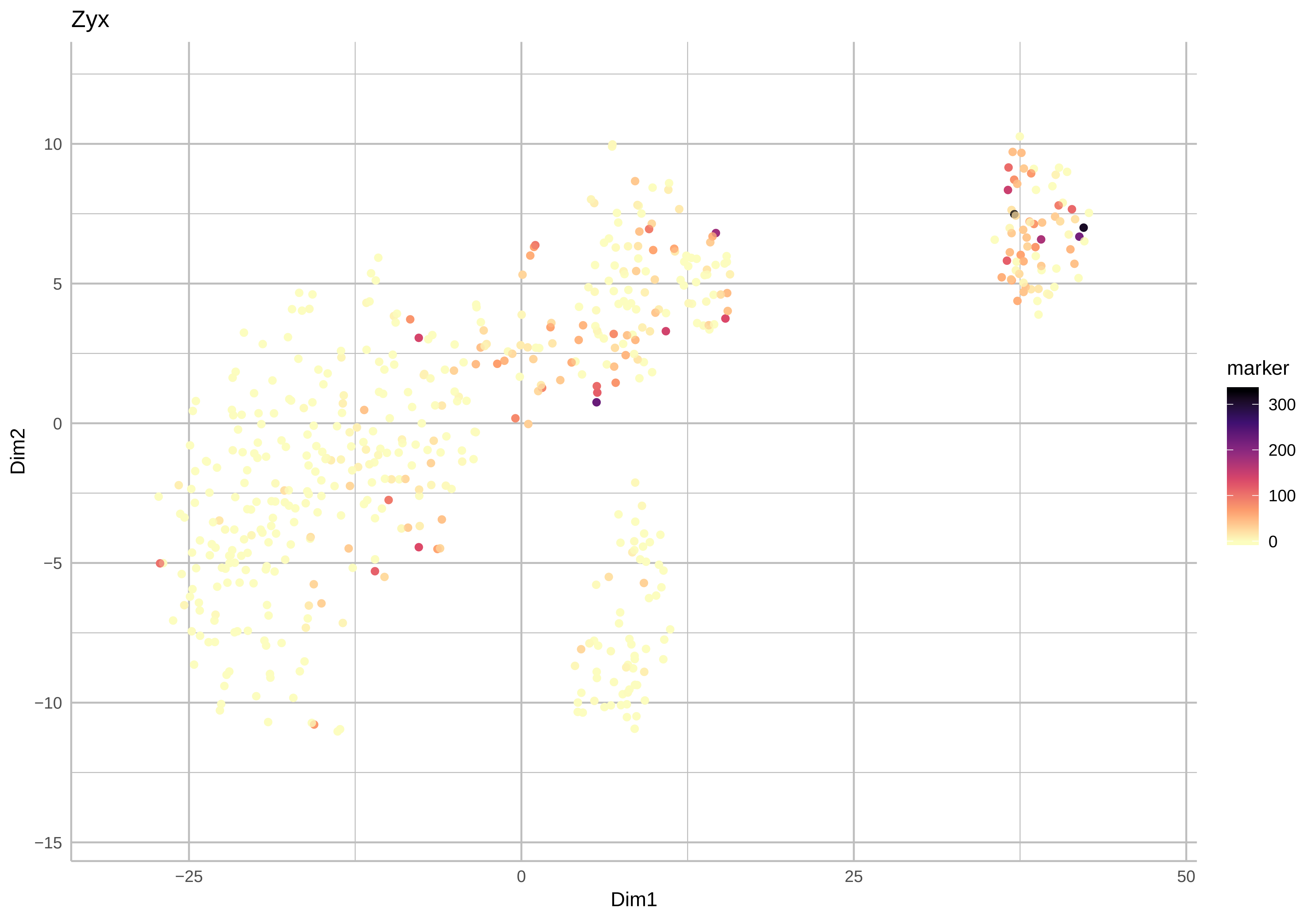
Zyx
Dim1
Dim2
−25
0
25
50
10
5
0
−5
−10
−15
marker
300
200
100
0

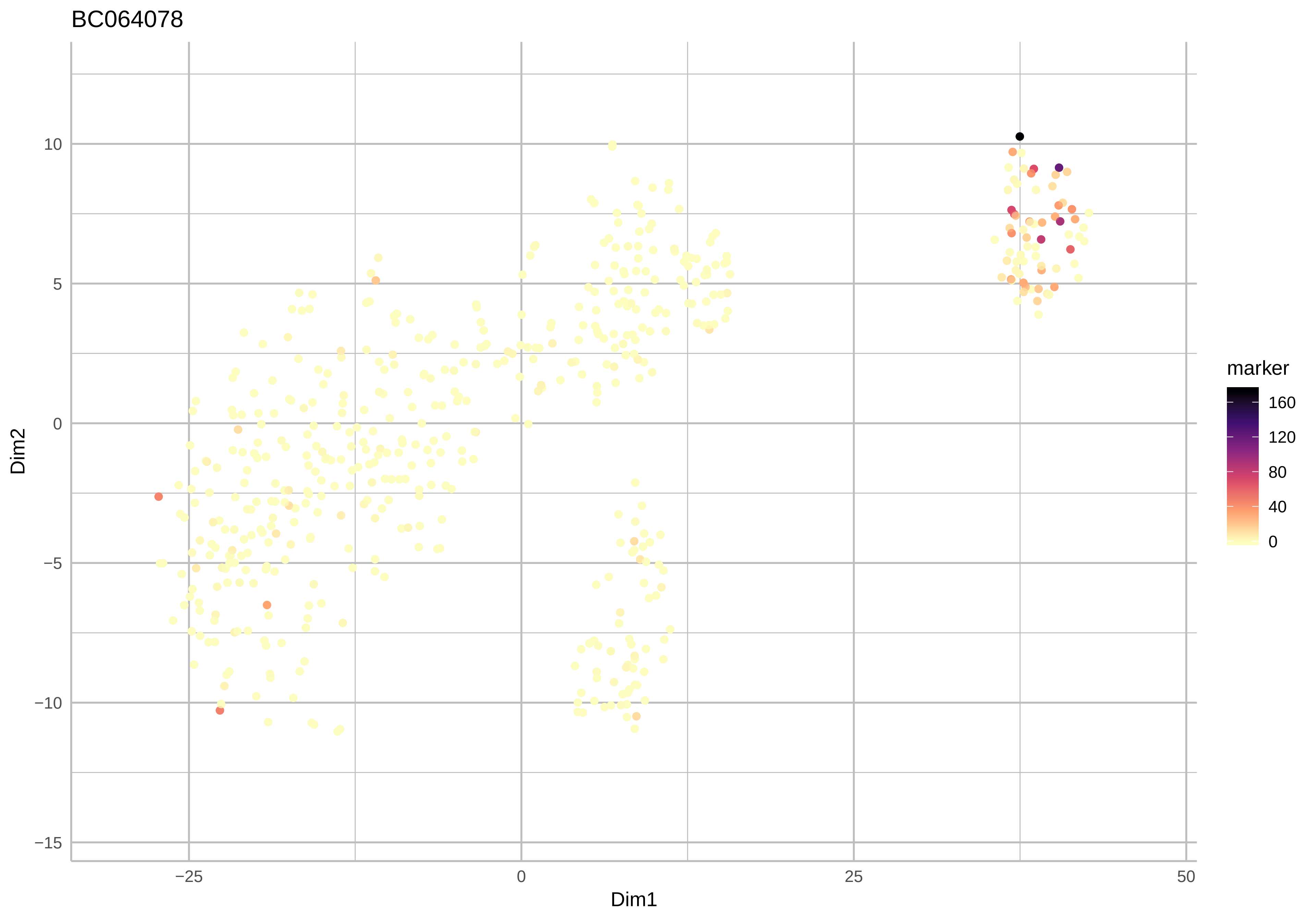

BC064078
Dim2
Dim1
10
5
0
−5
−10
−15
−25
0
25
50
marker
160
120
80
40
0

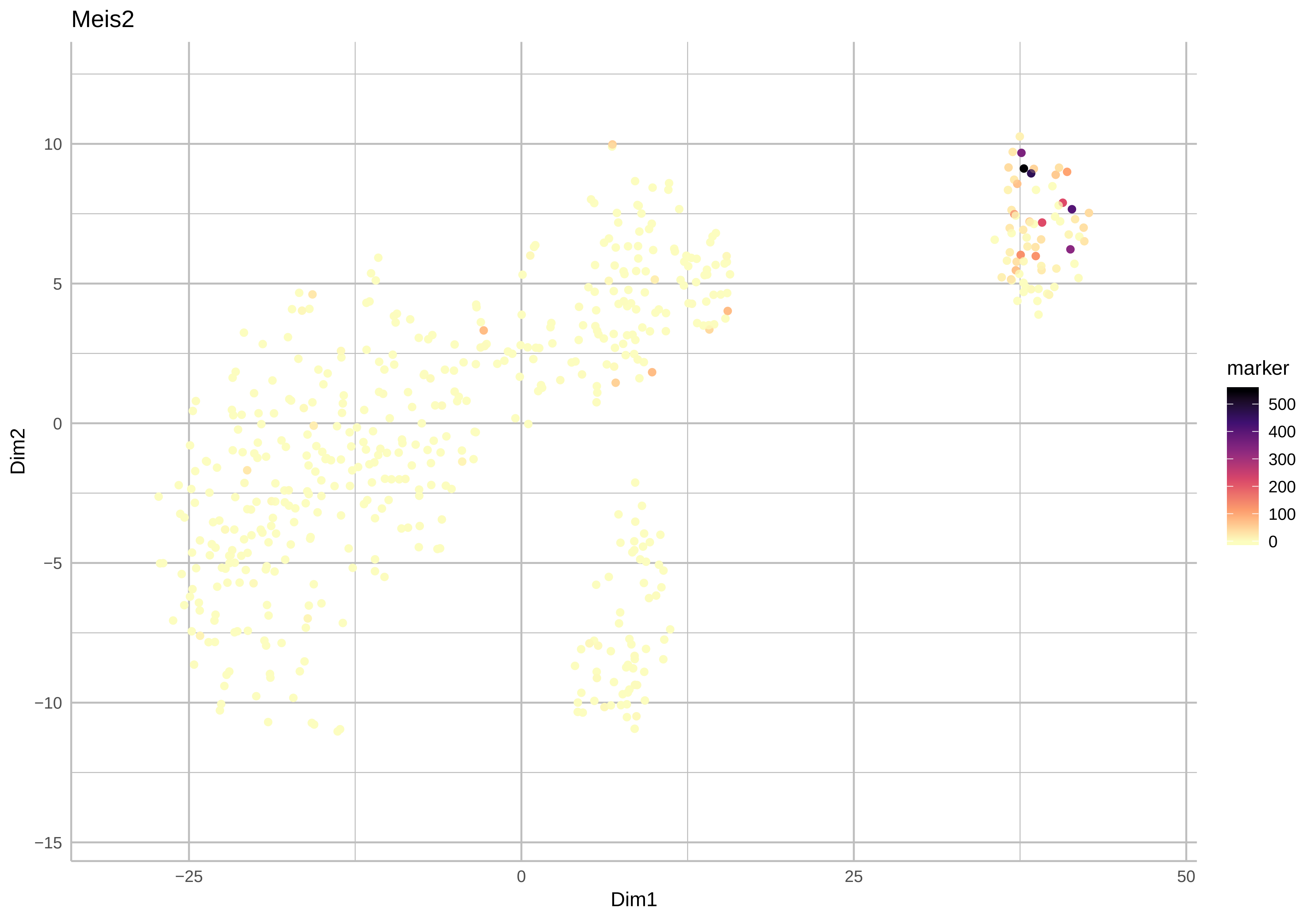
Meis2
Dim2
Dim1
10
5
0
−5
−10
−15
−25
0
25
50
marker
500
400
300
200
100
0

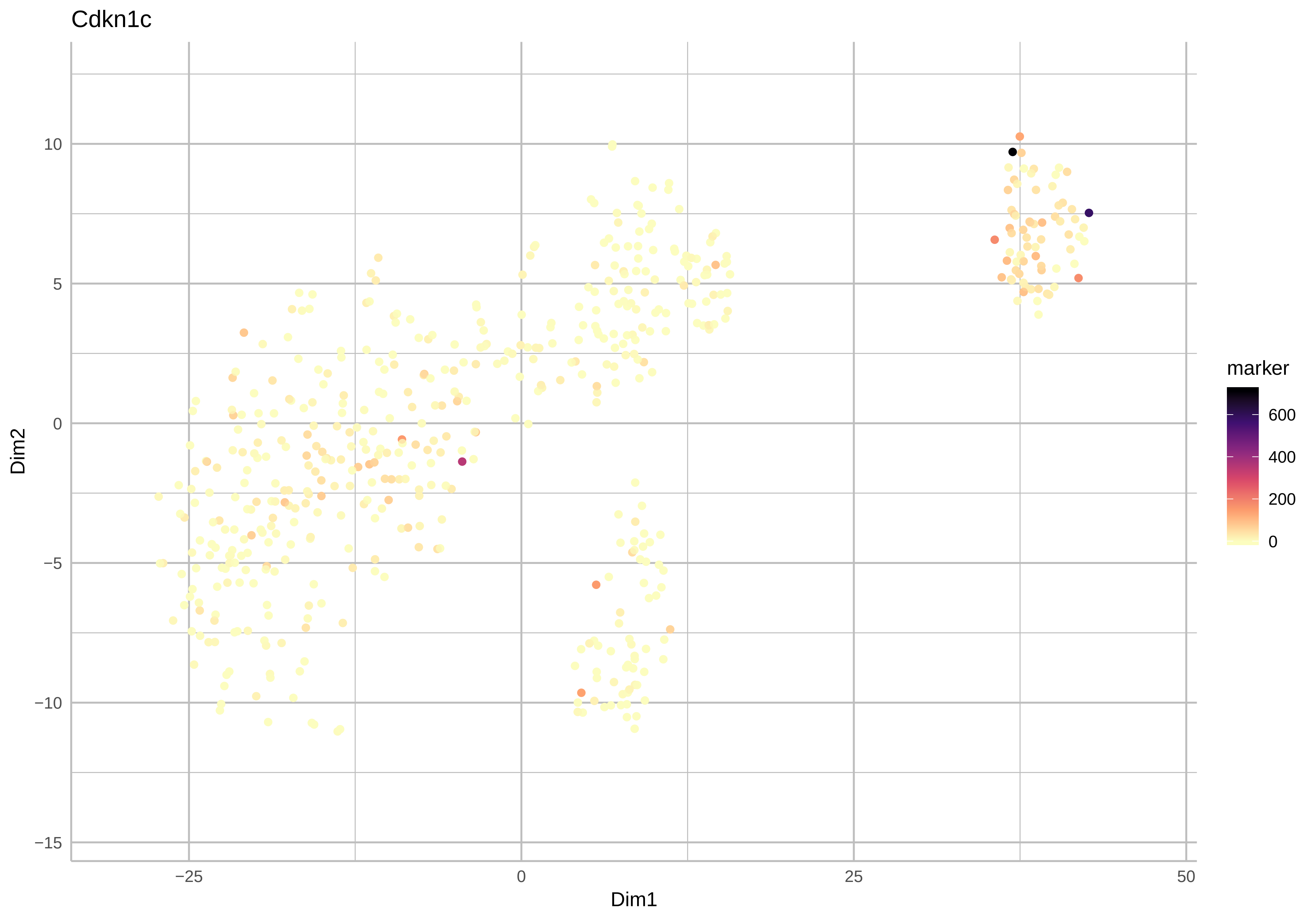

Cdkn1c
10
5
0
−5
−10
−15
Dim2
−25
0
25
50
Dim1
marker
600
400
200
0

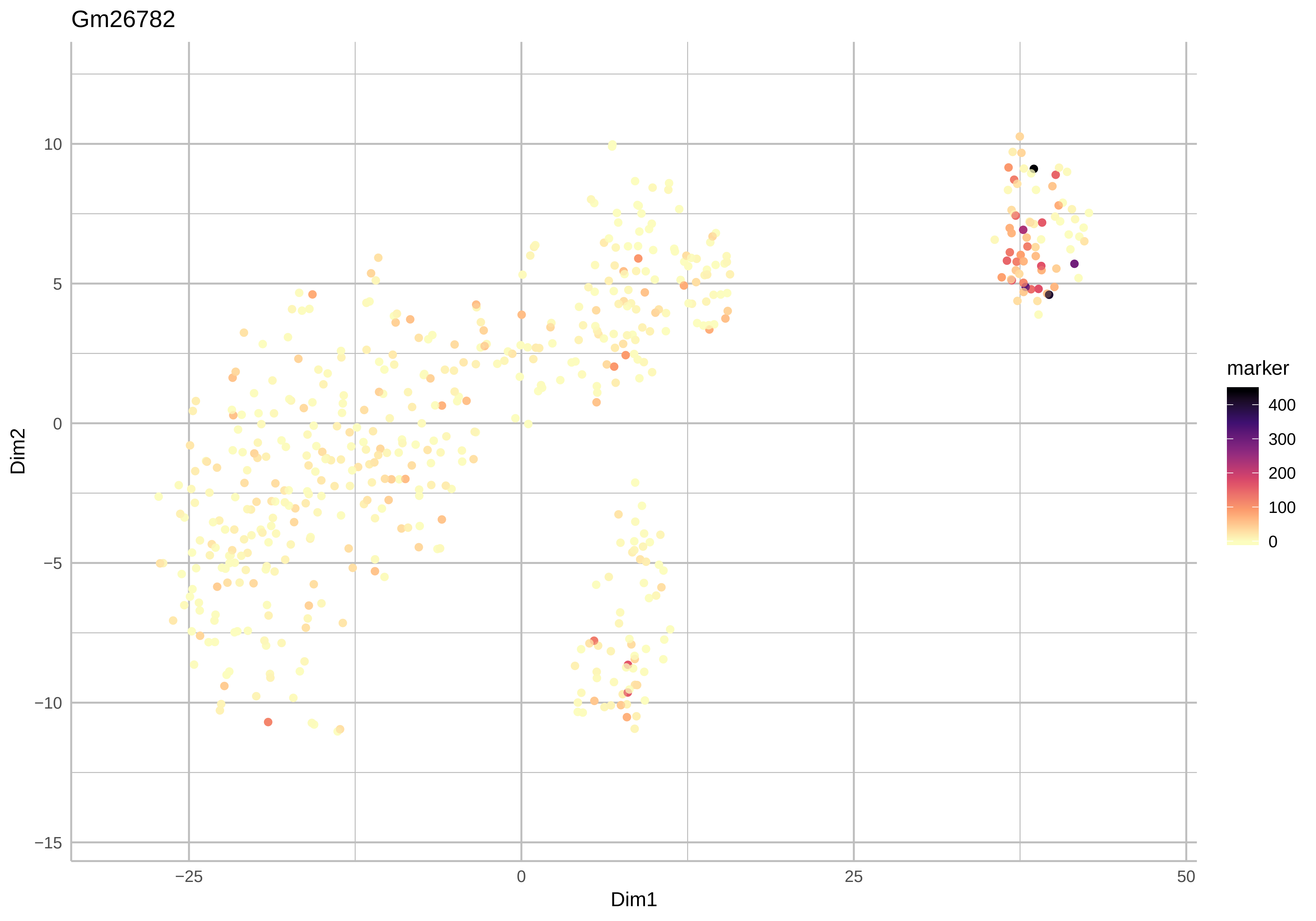

Gm26782
Dim2
Dim1
10
5
0
−5
−10
−15
−25
0
25
50
marker
400
300
200
100
0

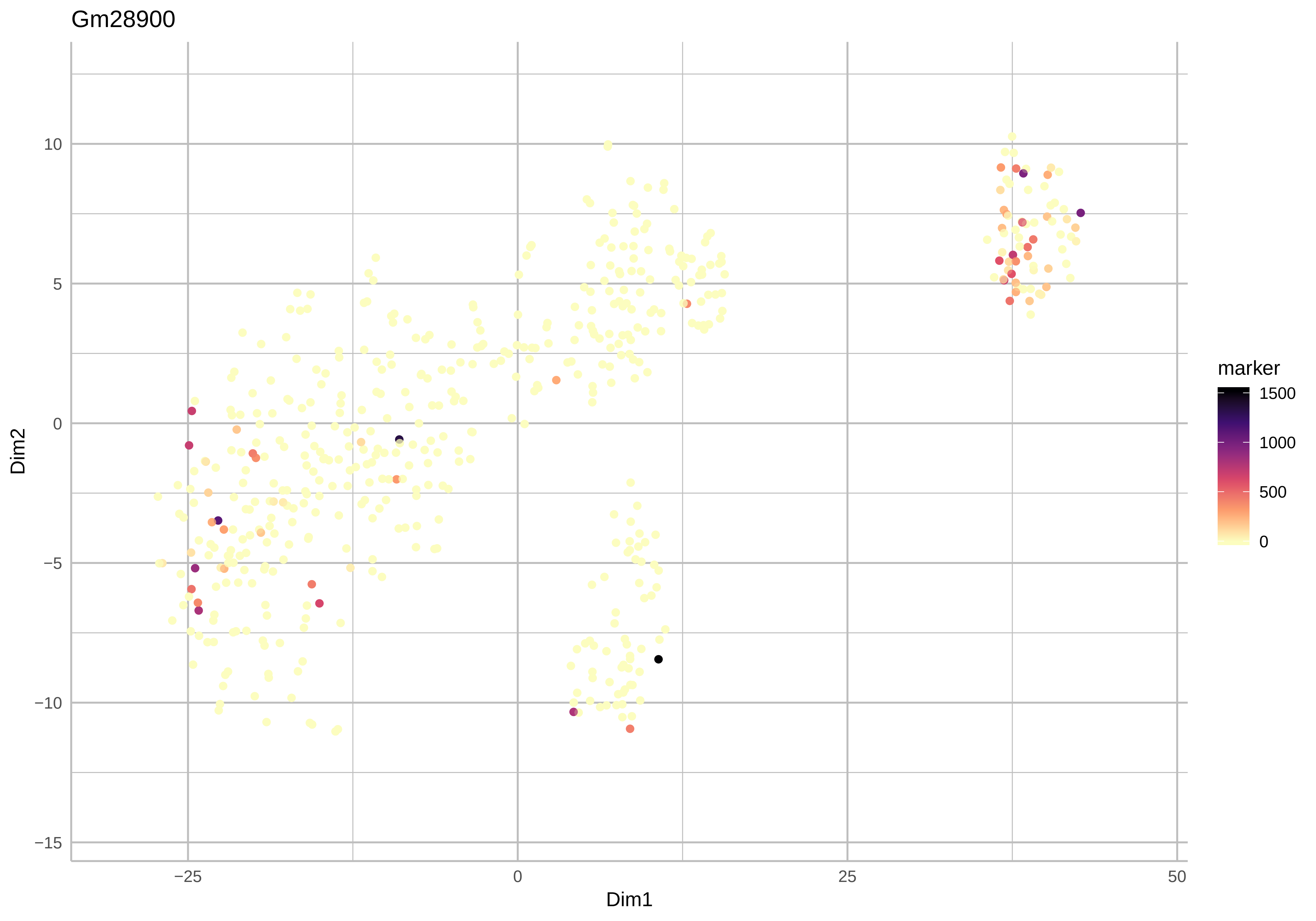
Gm28900
marker
1500
1000
500
0
Dim2
Dim1
10
5
0
−5
−10
−15
−25
0
25
50

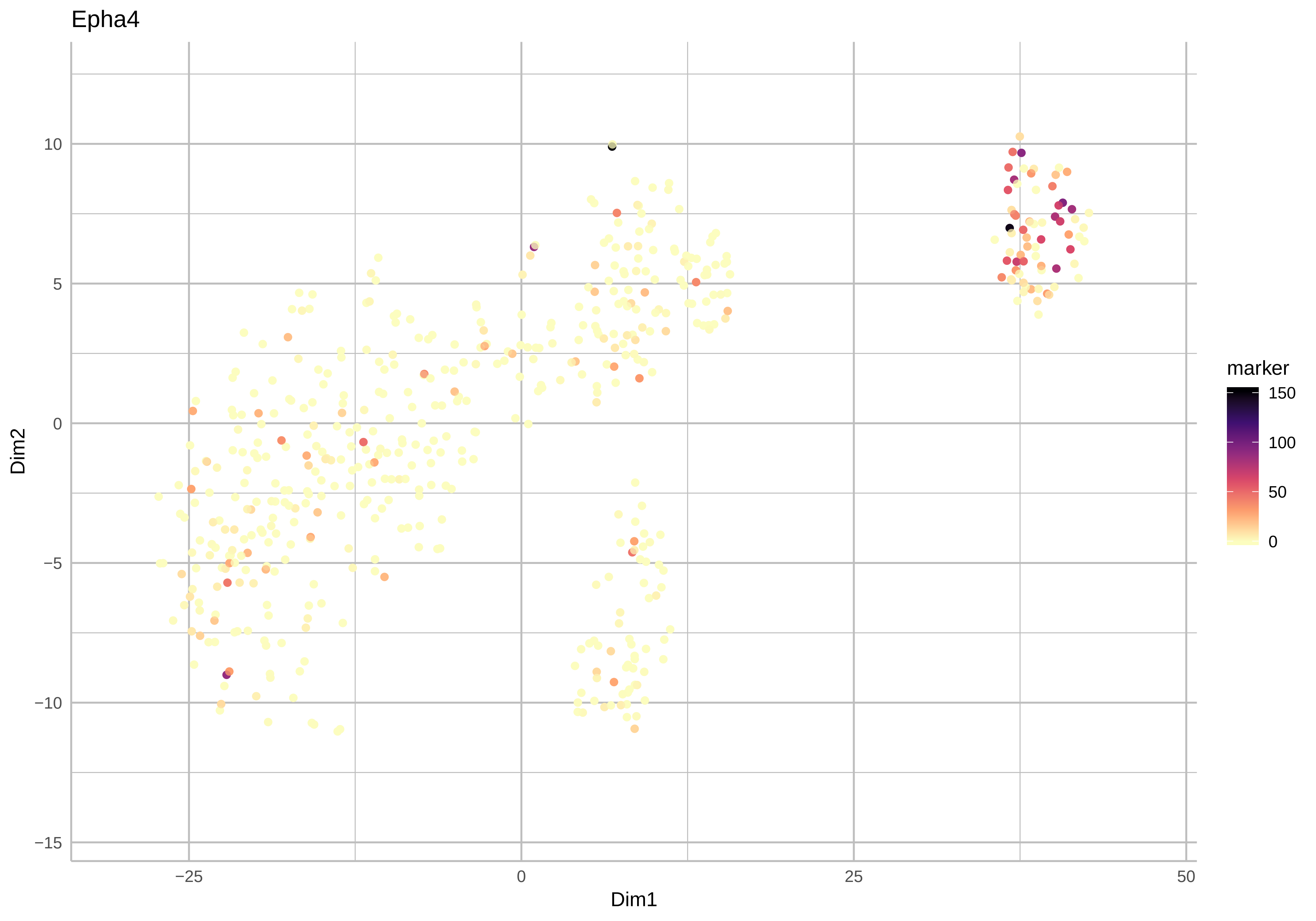

Epha4
Dim2
Dim1
marker
150
100
50
0
10
5
0
−5
−10
−15
−25
0
25
50

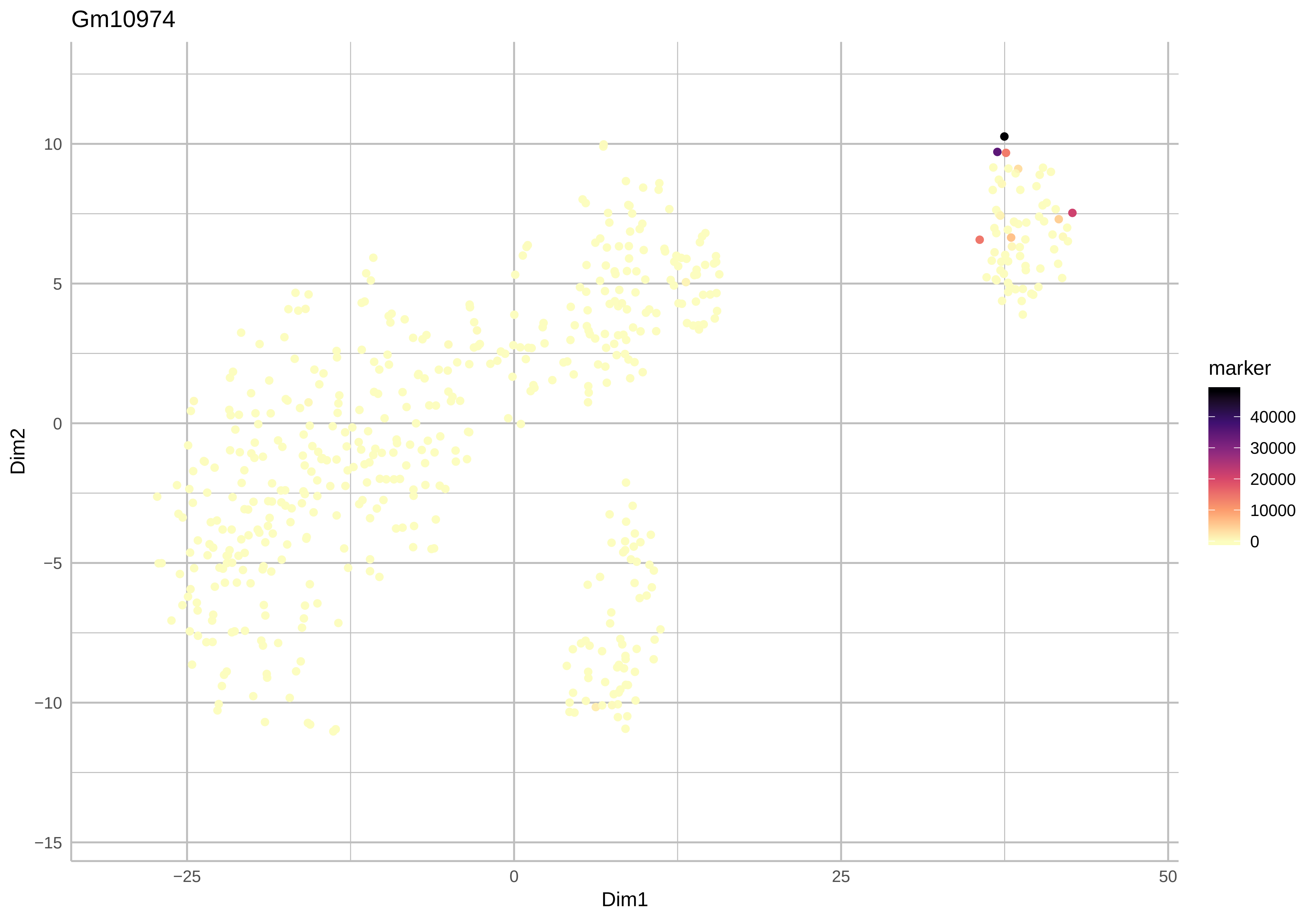

Gm10974
Dim2
Dim1
10
5
0
−5
−10
−15
−25
0
25
50
marker
40000
30000
20000
10000
0

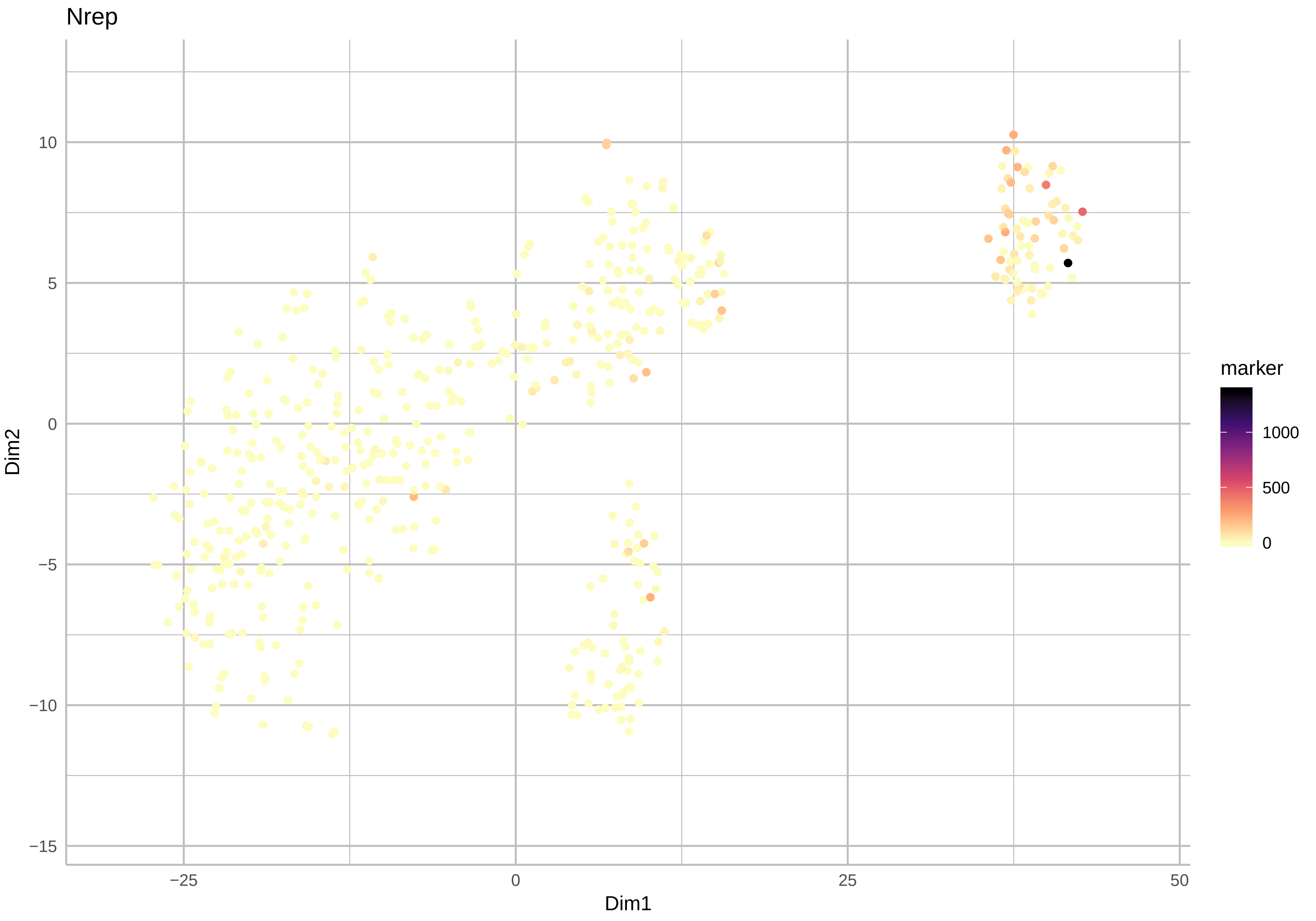

Nrep
10
5
0
−5
−10
−15
Dim2
−25
0
25
50
Dim1
marker
1000
500
0

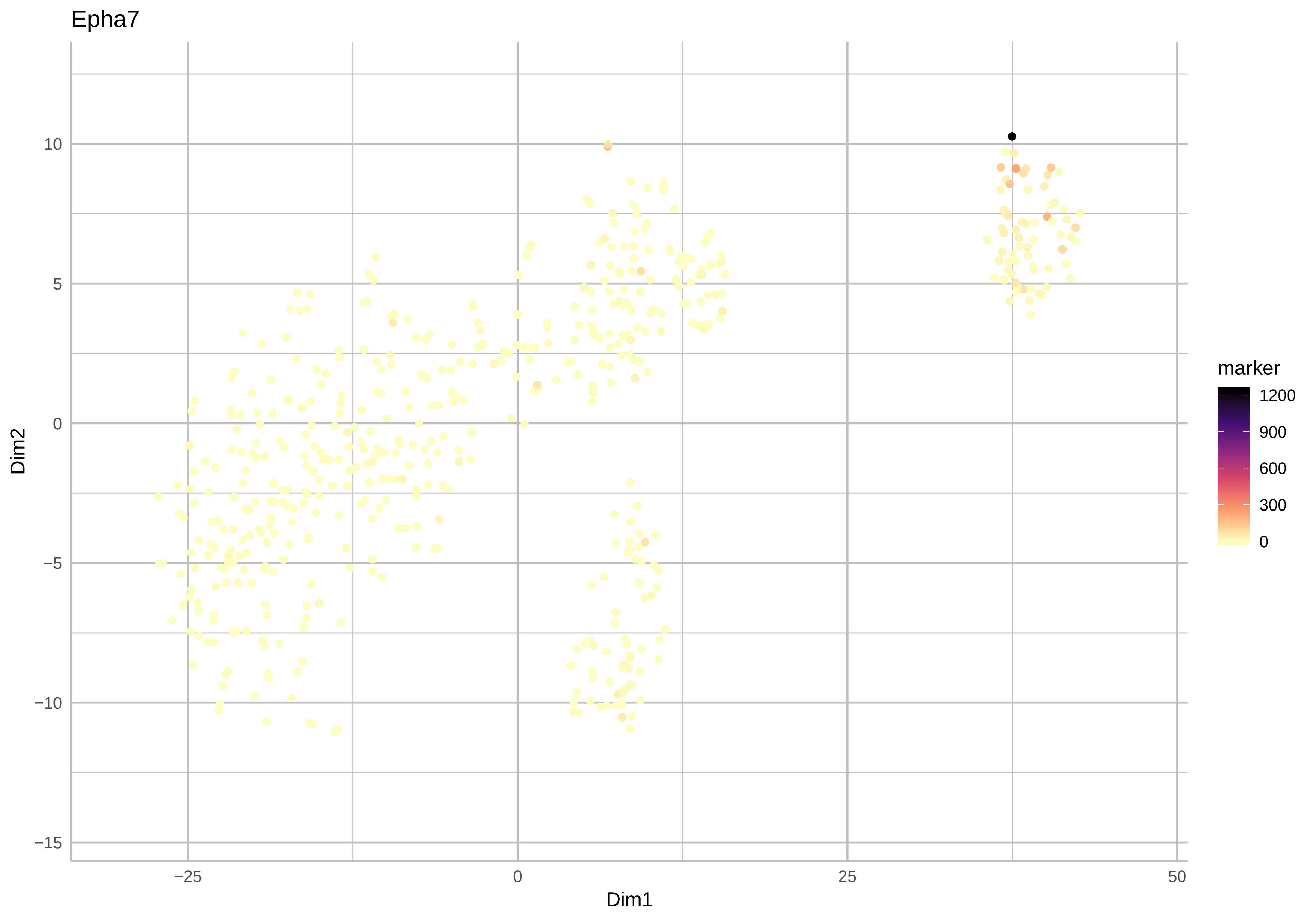

Epha7
10
5
0
−5
−10
−15
−25
0
25
50
Dim1
Dim2
marker
1200
900
600
300
0

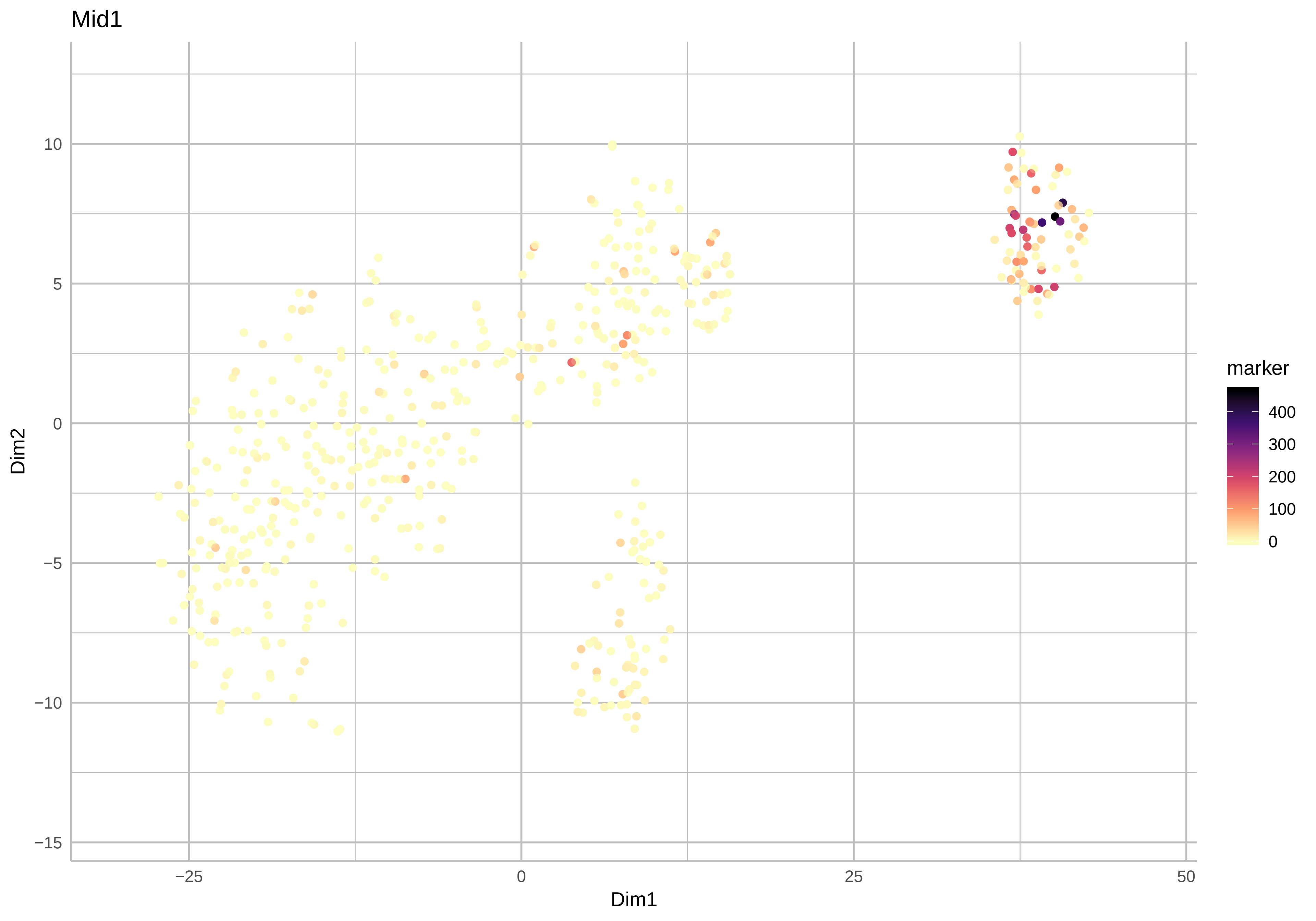

Mid1
Dim2
Dim1
10
5
0
−5
−10
−15
−25
0
25
50
marker
400
300
200
100
0

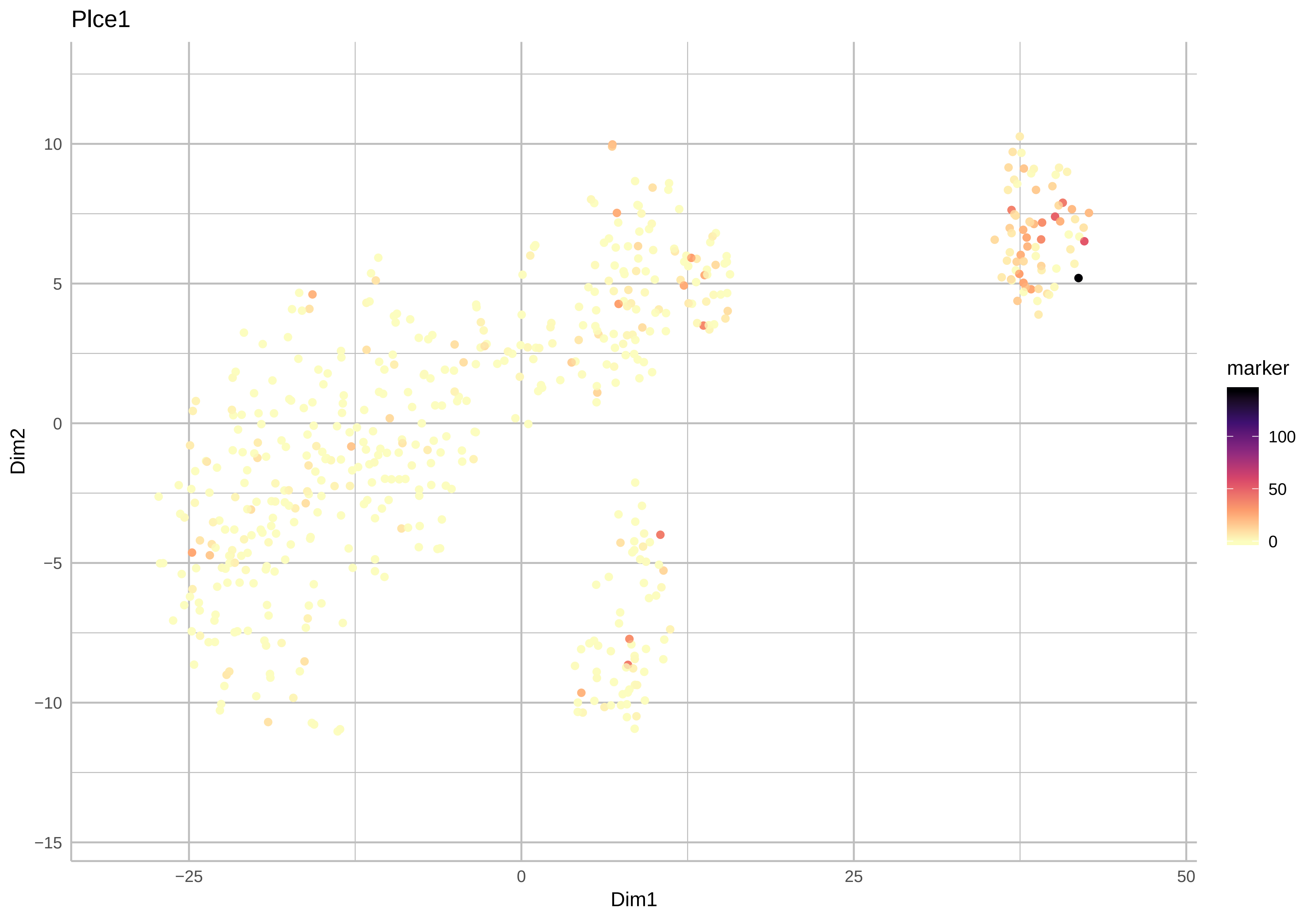
Plce1
Dim2
Dim1
10
5
0
−5
−10
−15
−25
0
25
50
marker
100
50
0

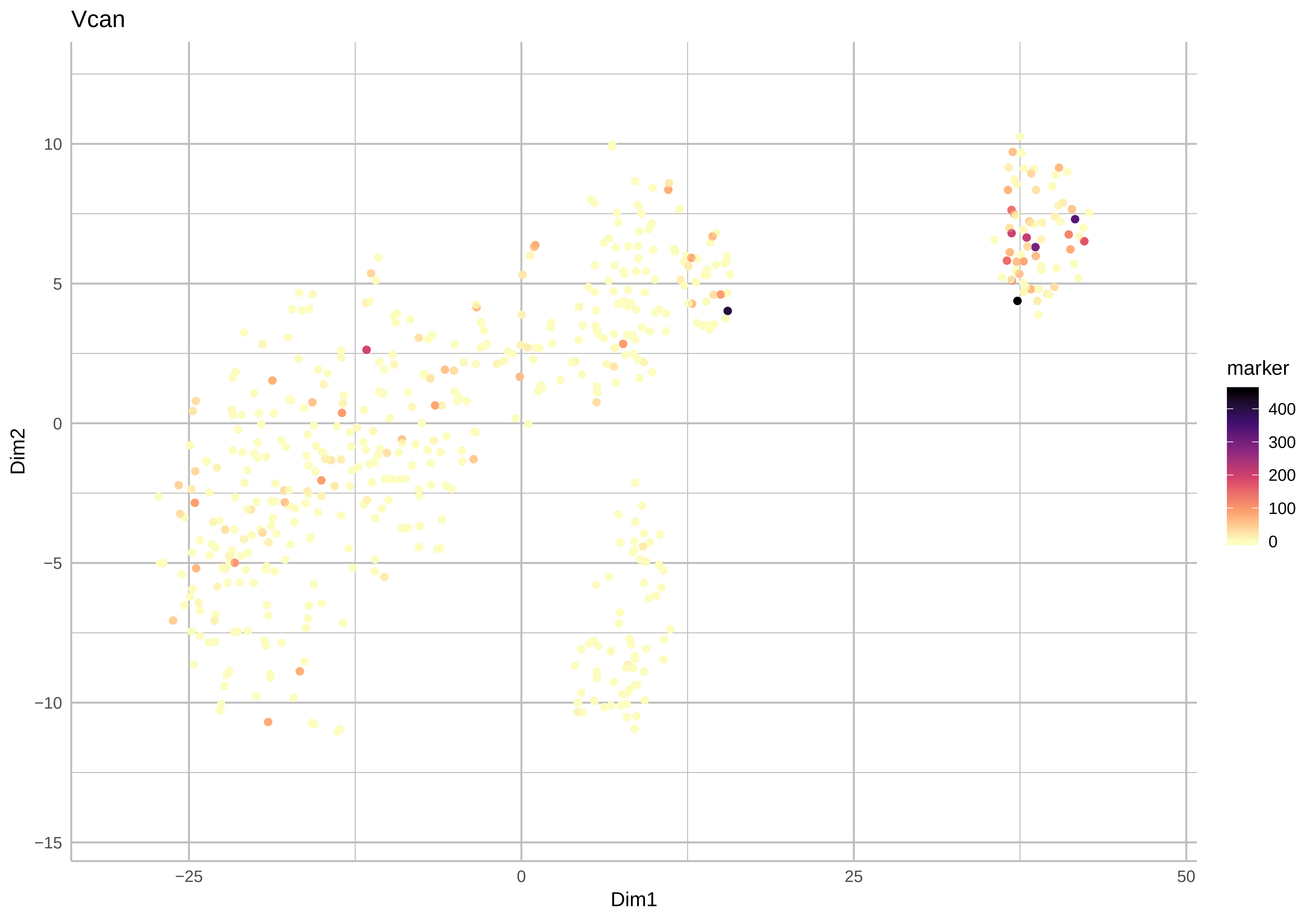
Vcan
Dim2
Dim1
10
5
0
−5
−10
−15
−25
0
25
50
marker
400
300
200
100
0

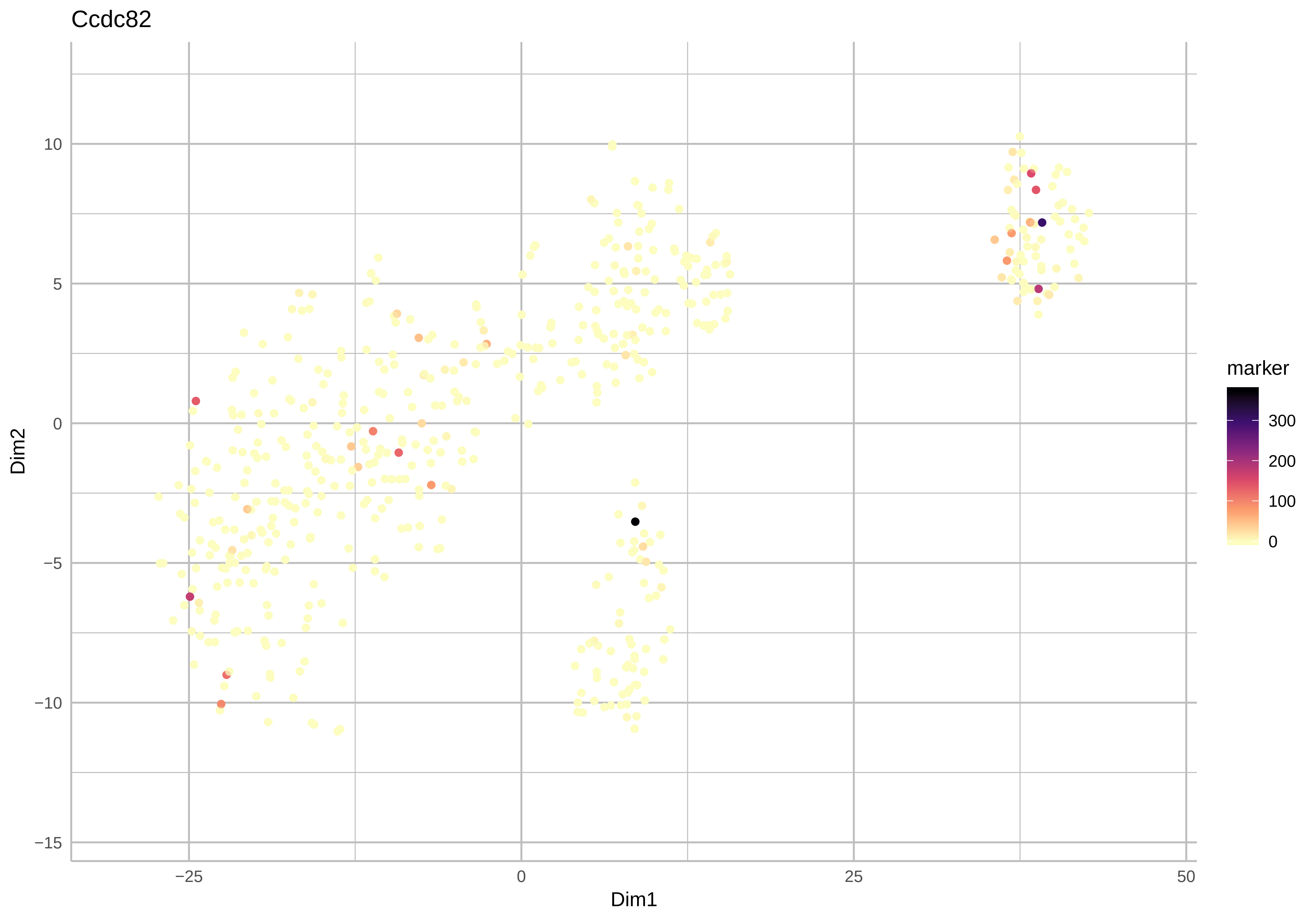

Ccdc82
Dim2
Dim1
10
5
0
−5
−10
−15
−25
0
25
50
marker
300
200
100
0

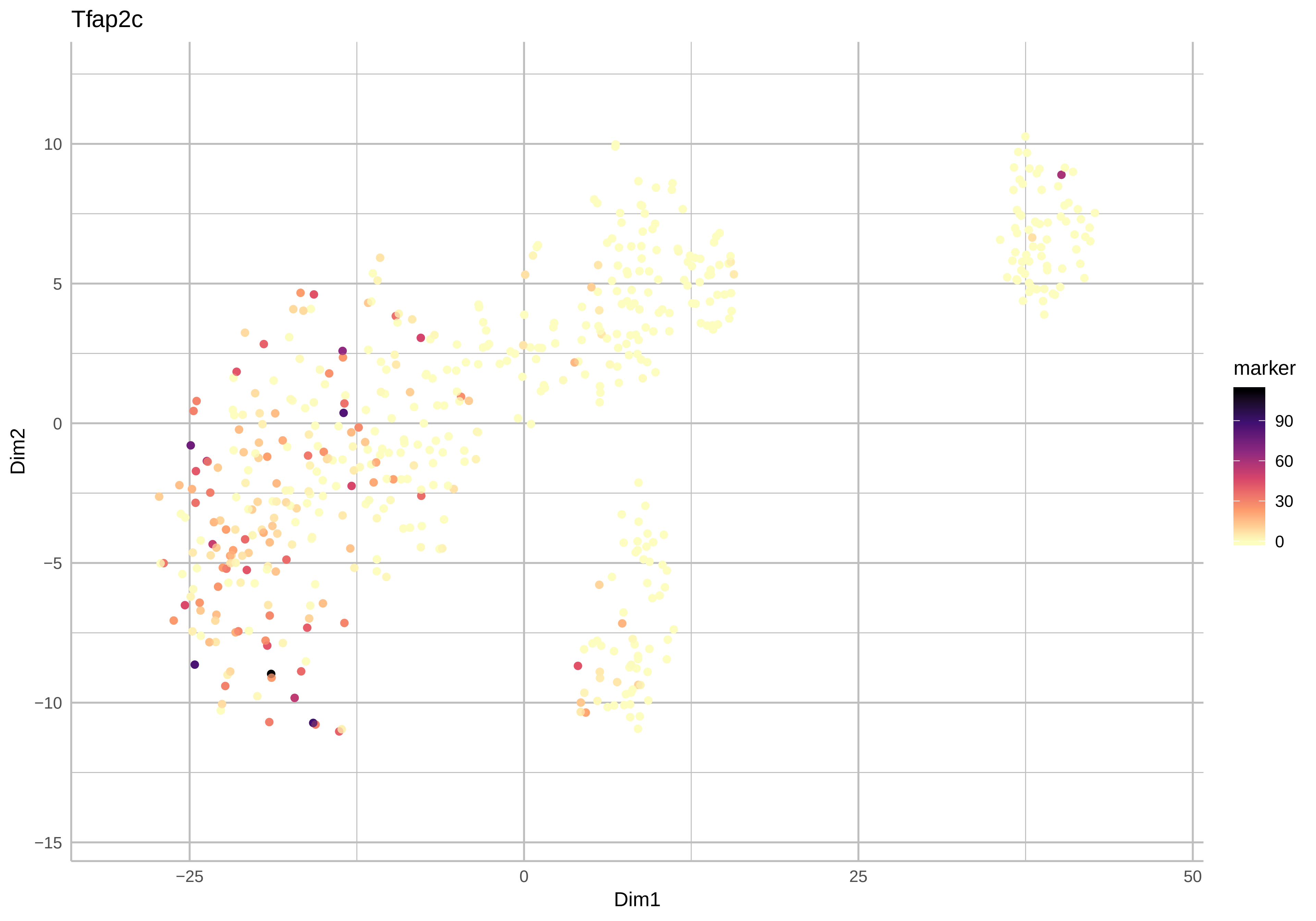
Tfap2c
Dim2
Dim1
10
5
0
−5
−10
−15
−25
0
25
50
marker
90
60
30
0

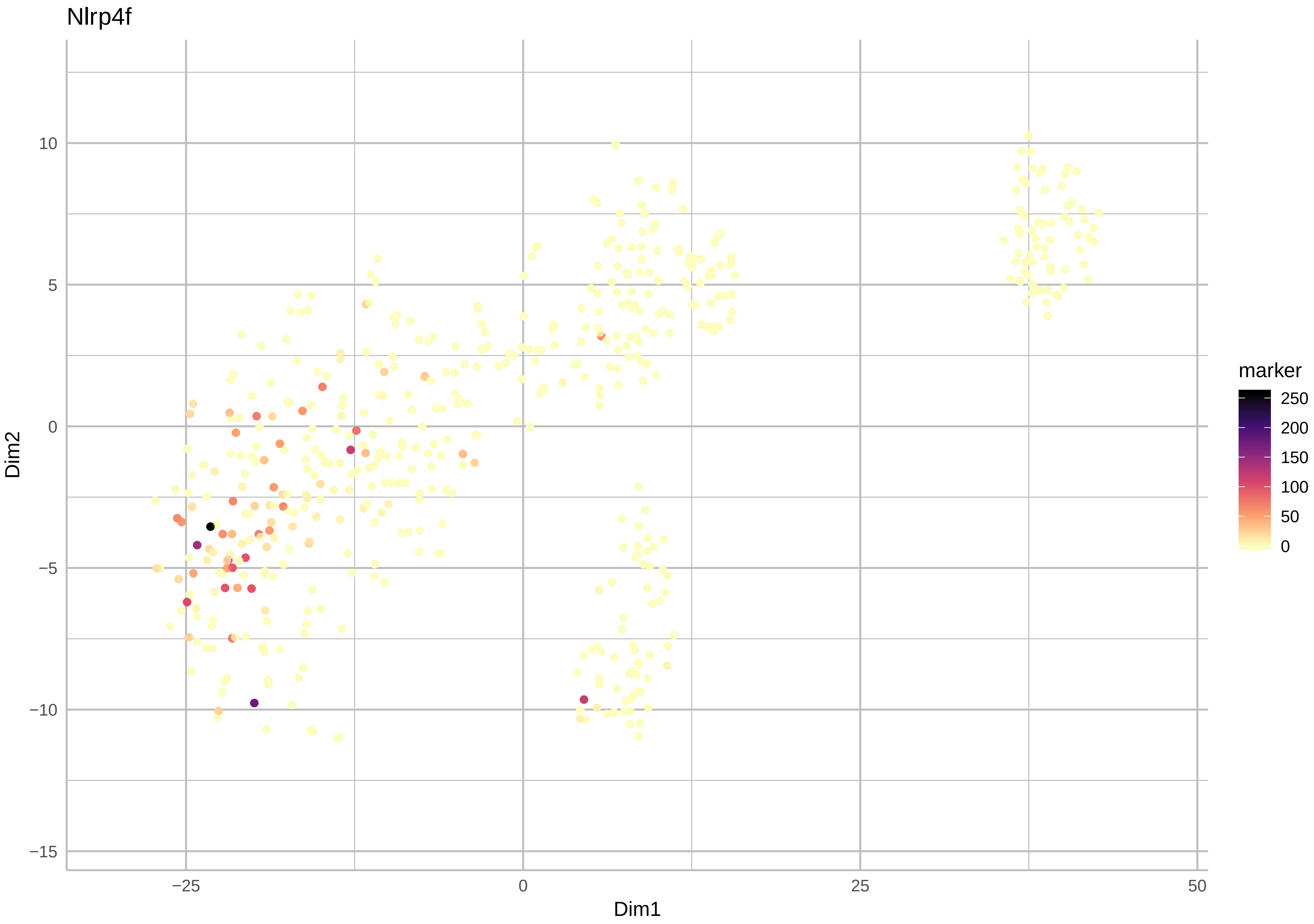
Nlrp4f
Dim2
Dim1
marker
250
200
150
100
50
0
10
5
0
−5
−10
−15
−25
0
25
50

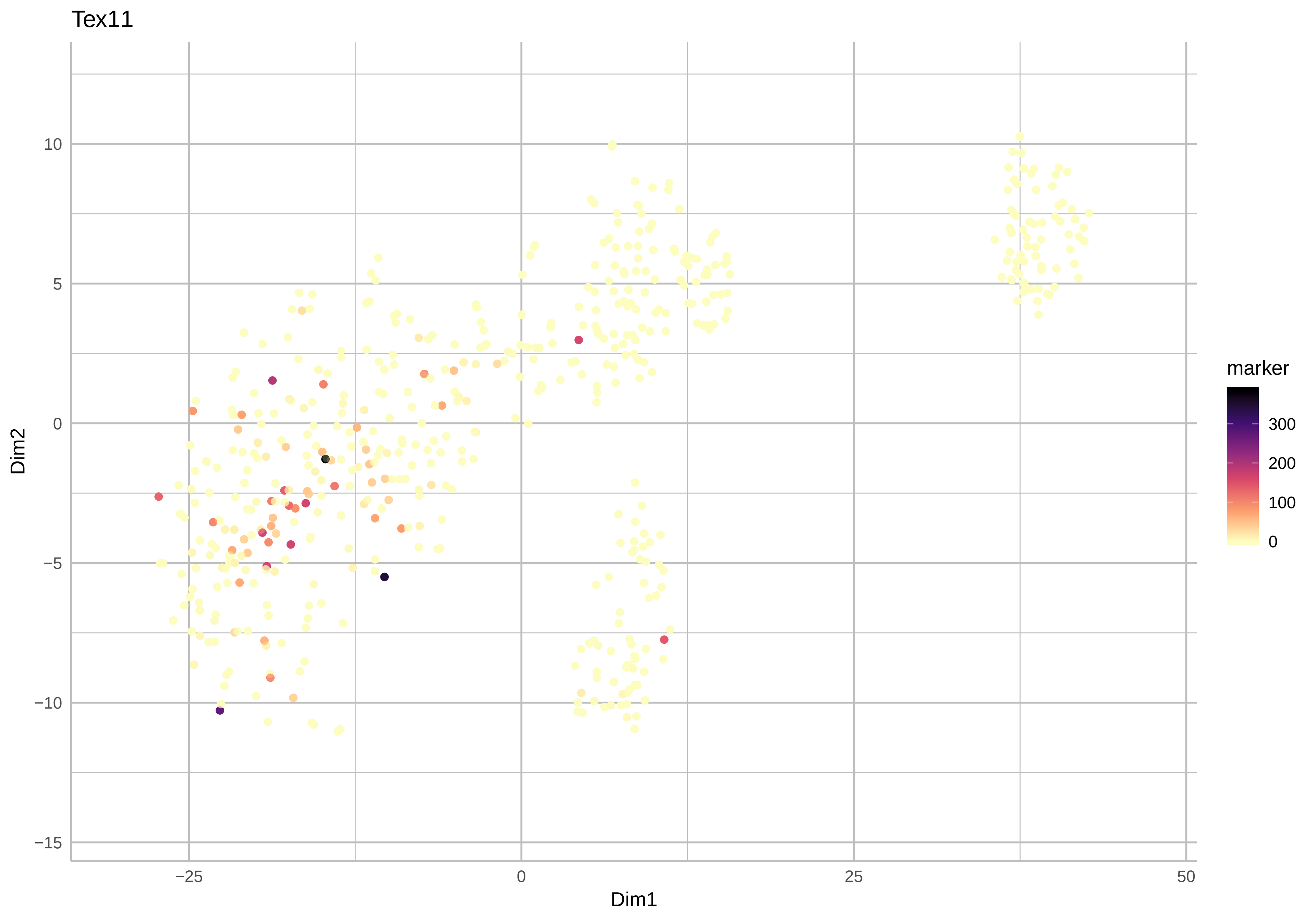

Tex11
Dim2
Dim1
10
5
0
−5
−10
−15
−25
0
25
50
marker
300
200
100
0

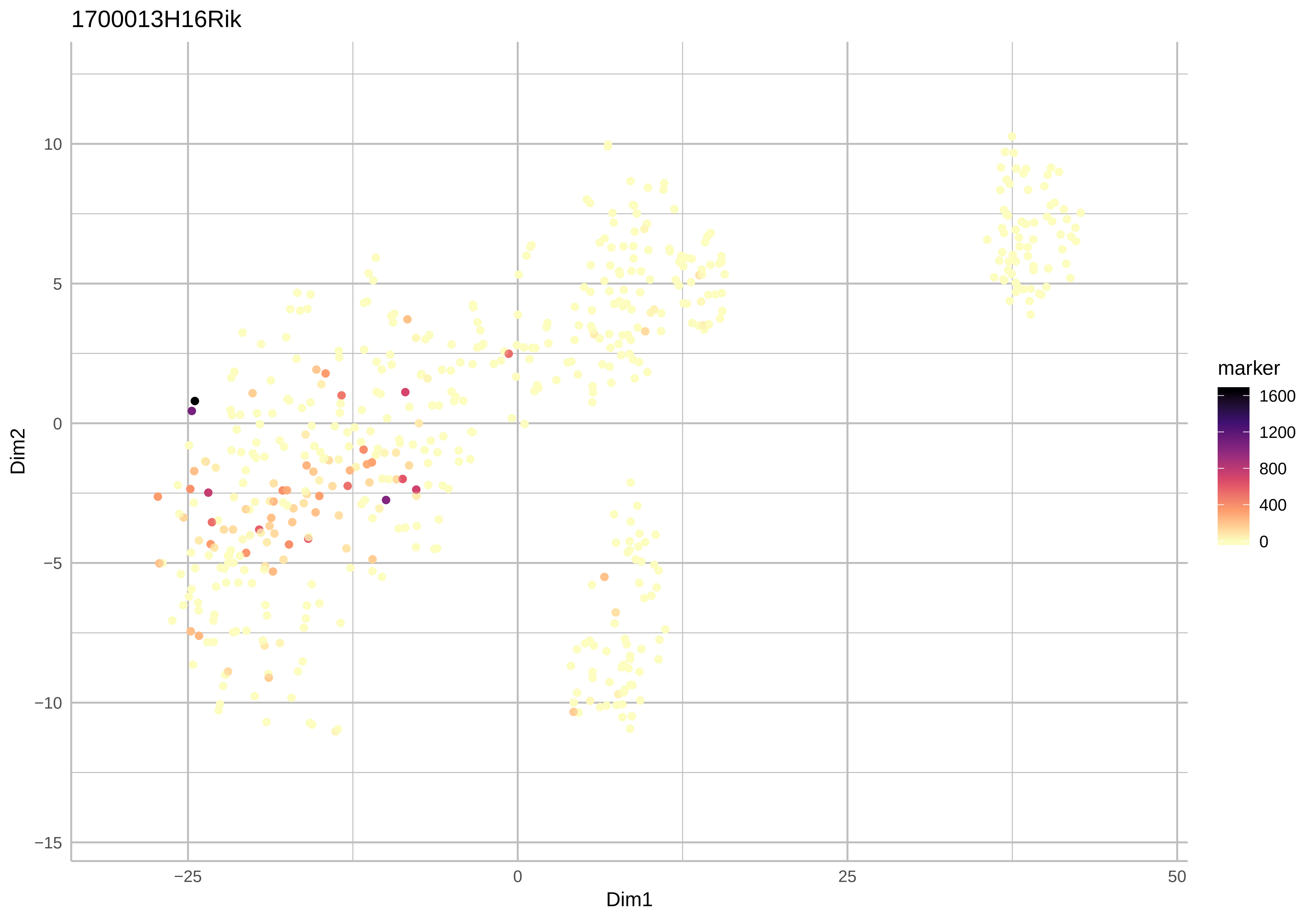
1700013H16Rik
Dim2
Dim1
marker
1600
1200
800
400
0
10
5
0
−5
−10
−15
−25
0
25
50

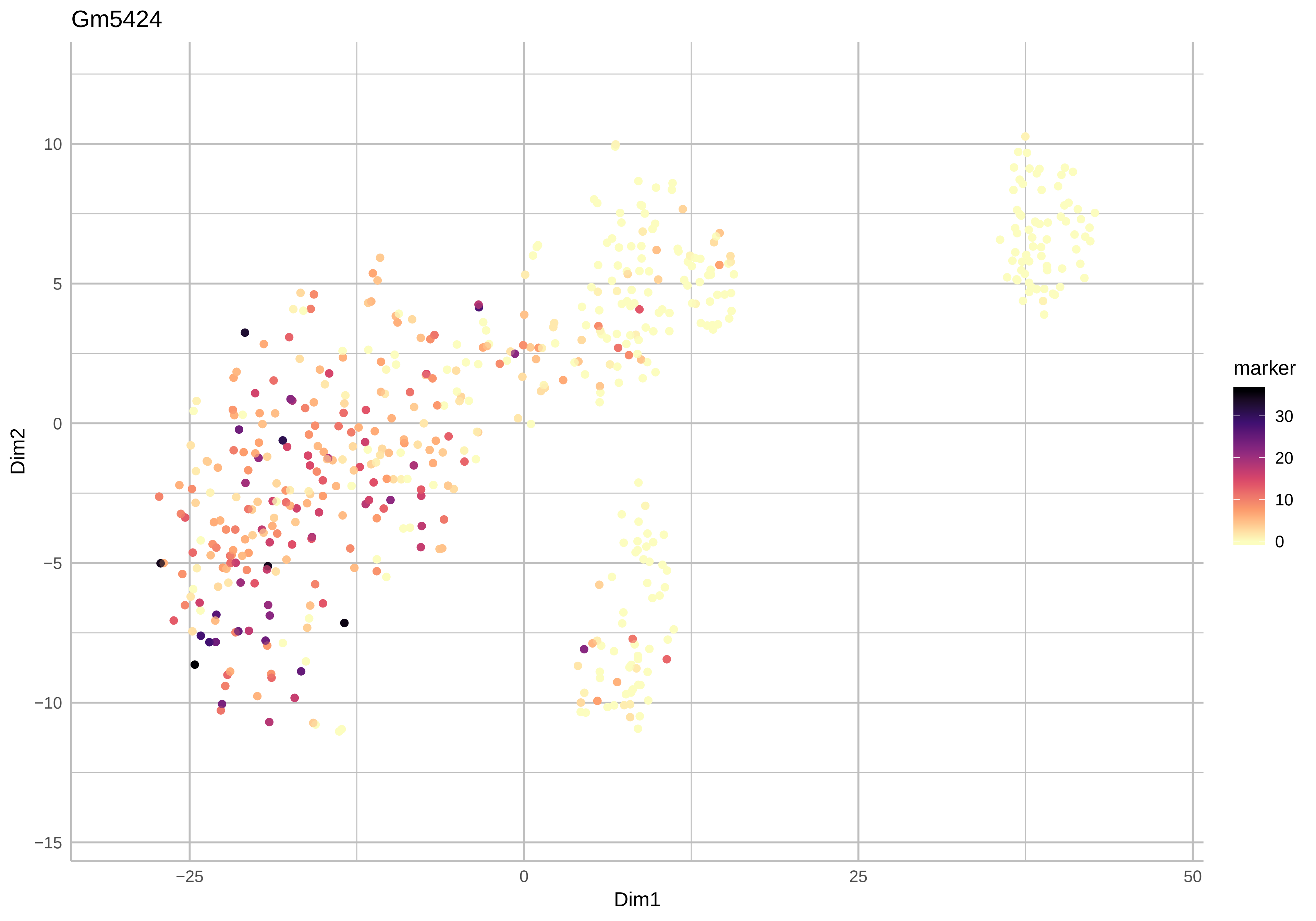

Gm5424
Dim2
Dim1
marker
30
20
10
0
10
5
0
−5
−10
−15
−25
0
25
50

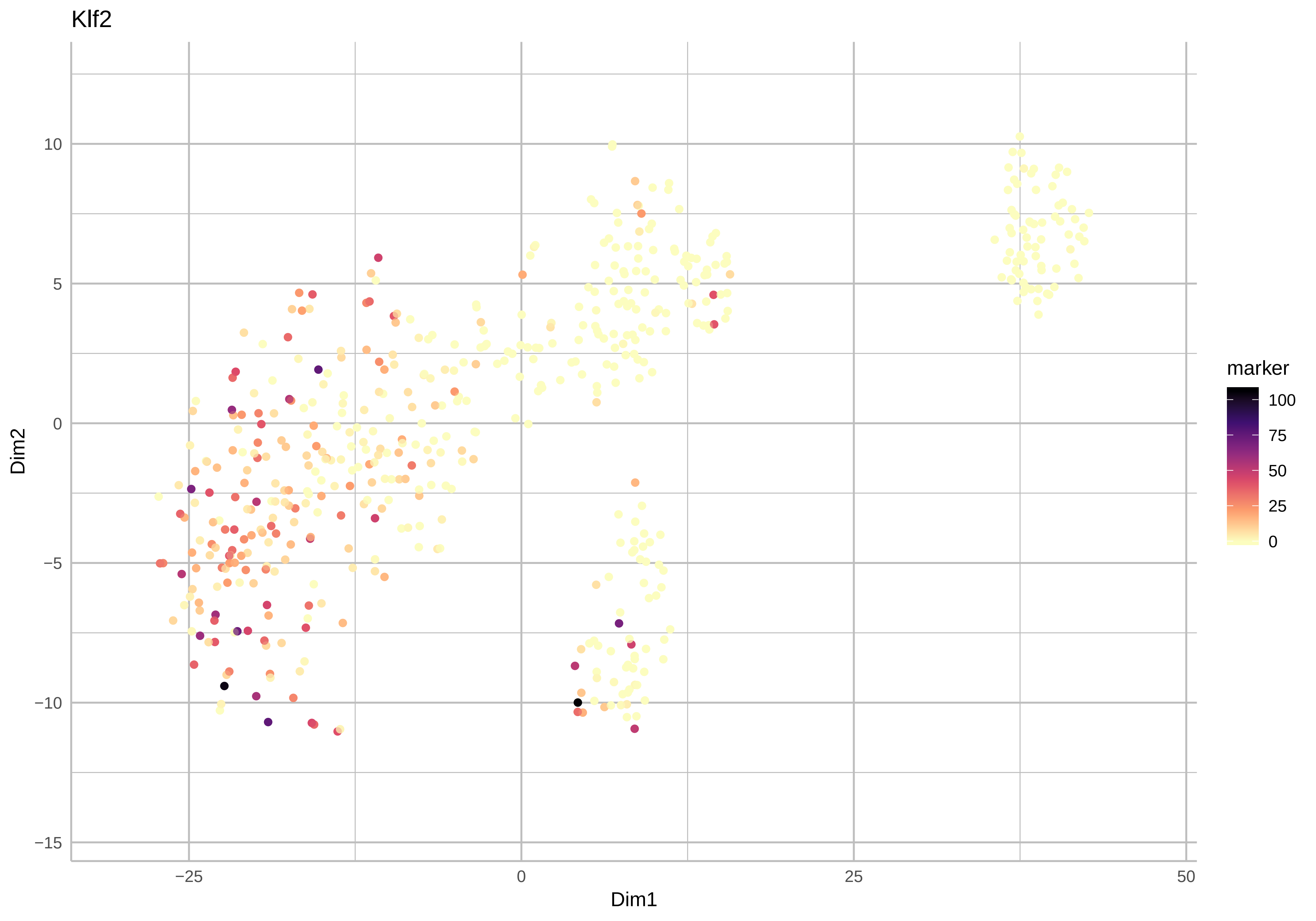
Klf2
10
5
0
−5
−10
−15
−25
0
25
50
Dim1
Dim2
marker
100
75
50
25
0

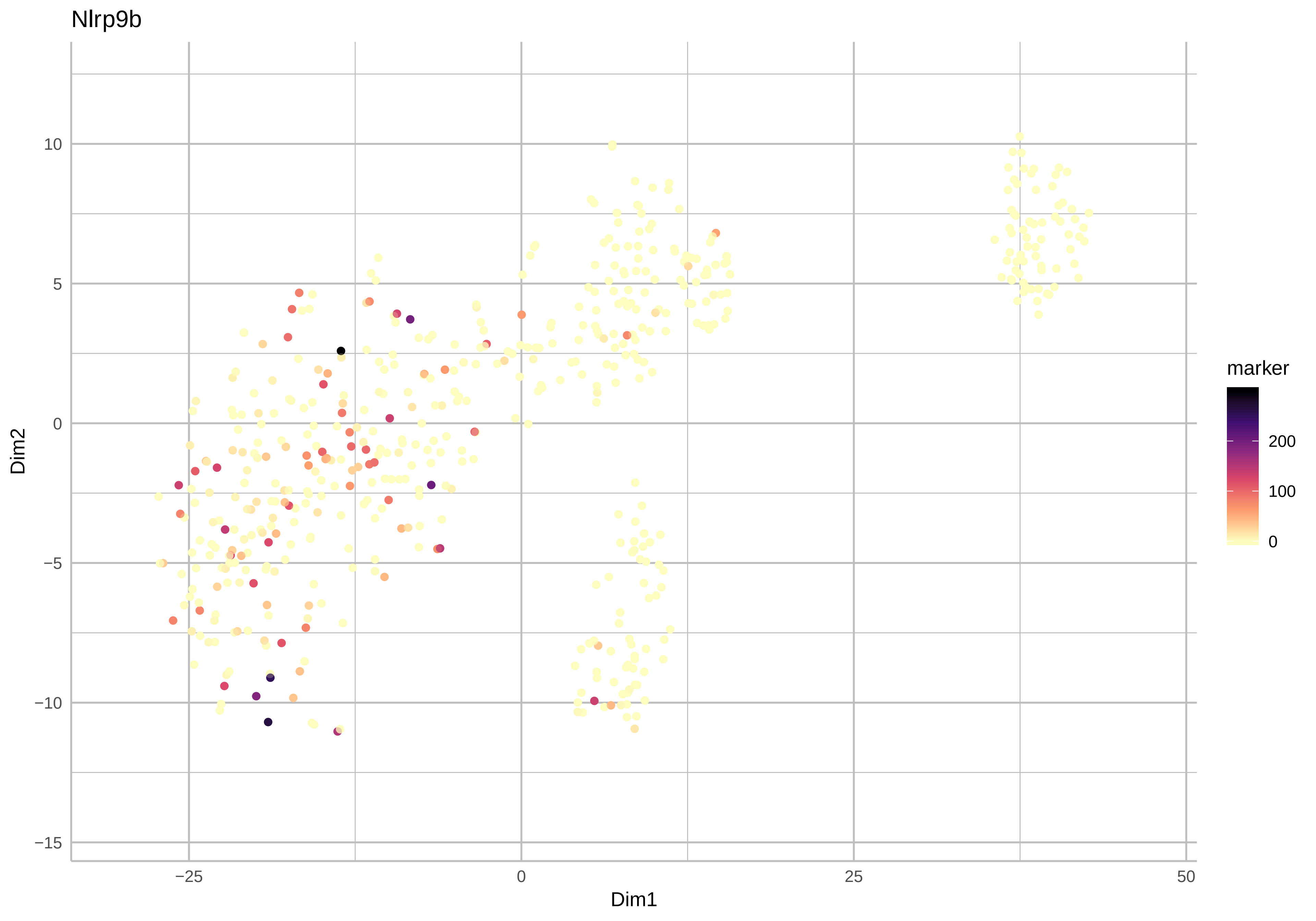
Nlrp9b
10
5
0
−5
−10
−15
−25
0
25
50
Dim1
Dim2
marker
200
100
0

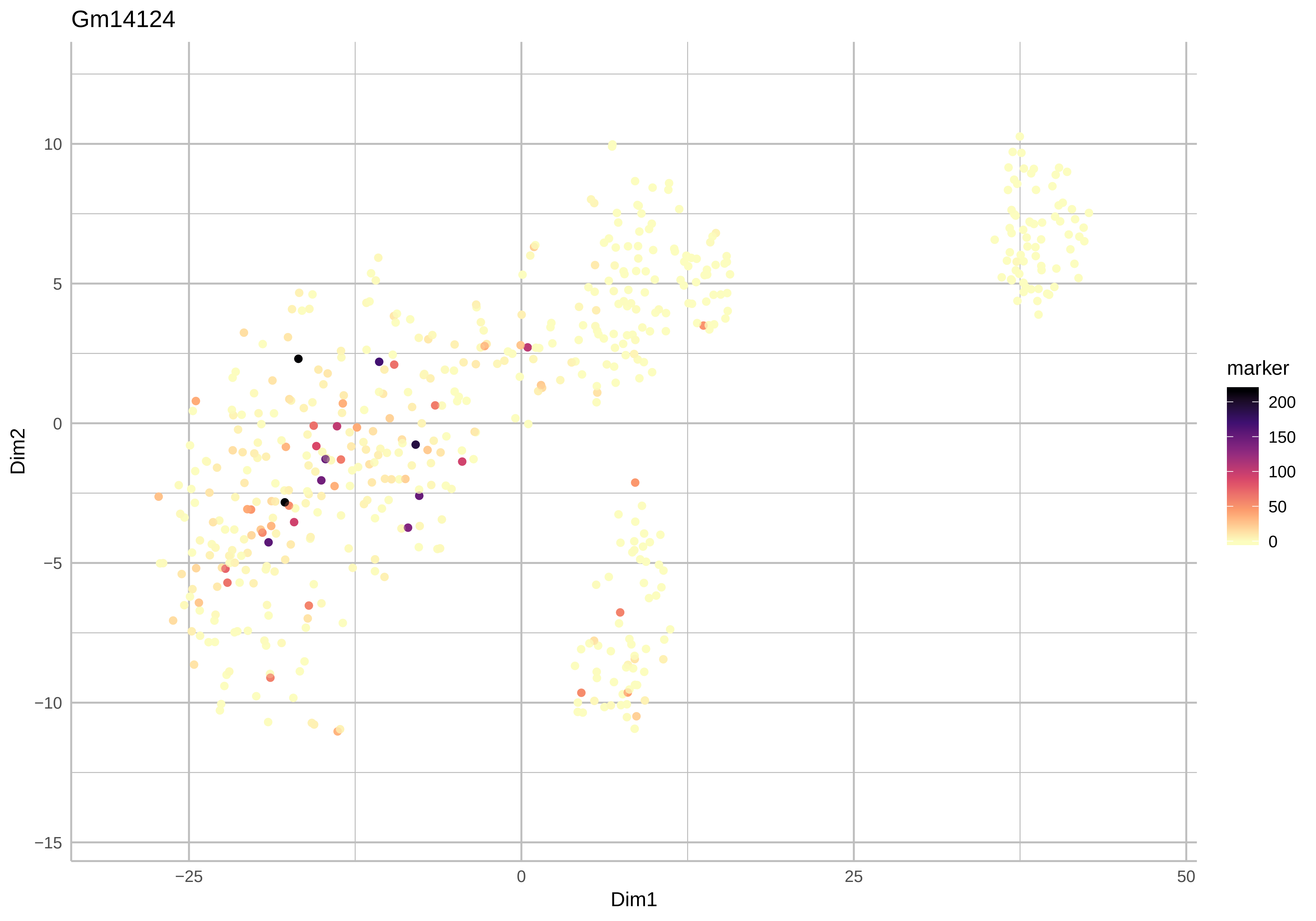
Gm14124
Dim2
Dim1
marker
200
150
100
50
0
10
5
0
−5
−10
−15
−25
0
25
50

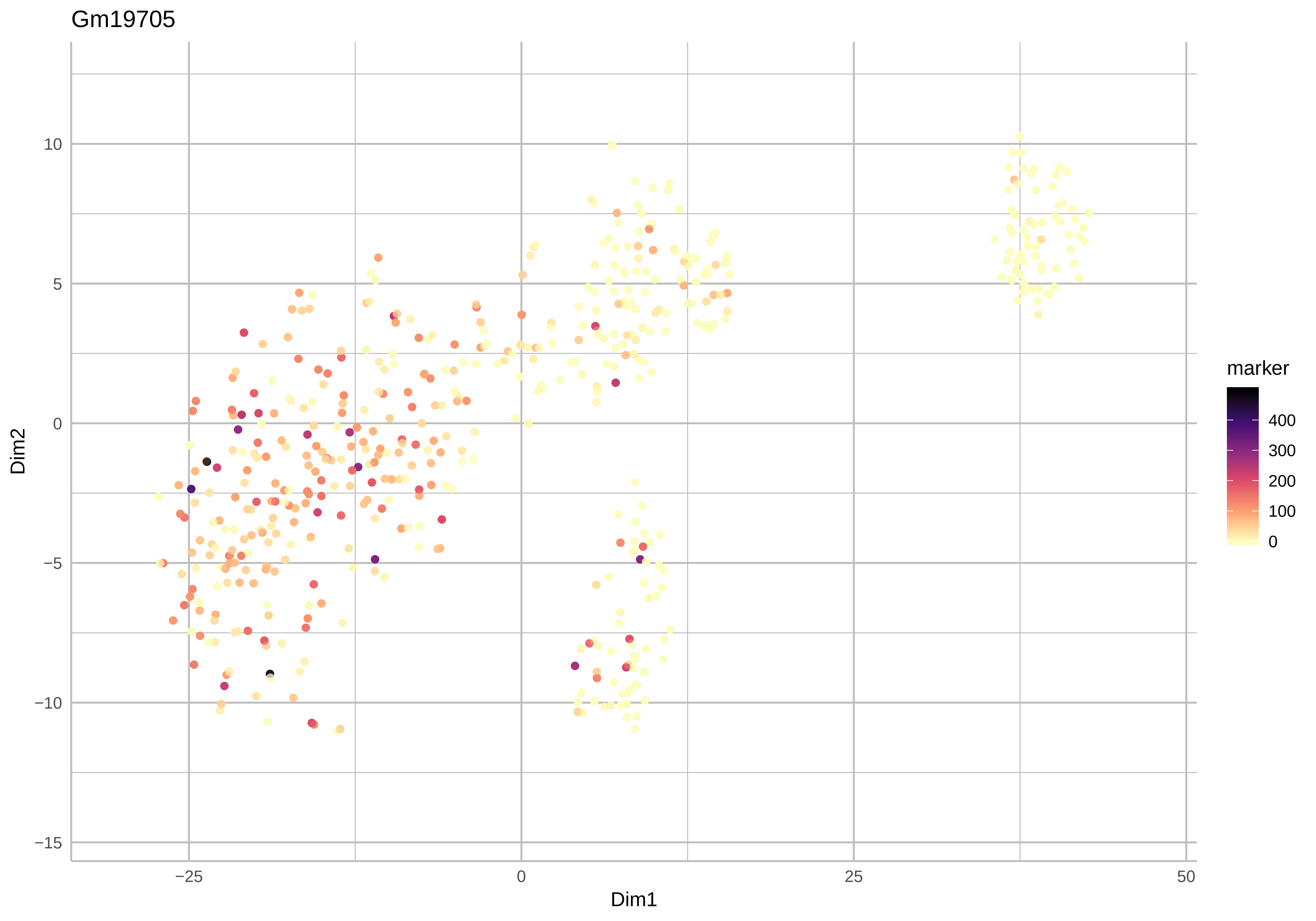

Gm19705
Dim2
Dim1
10
5
0
−5
−10
−15
−25
0
25
50
marker
400
300
200
100
0

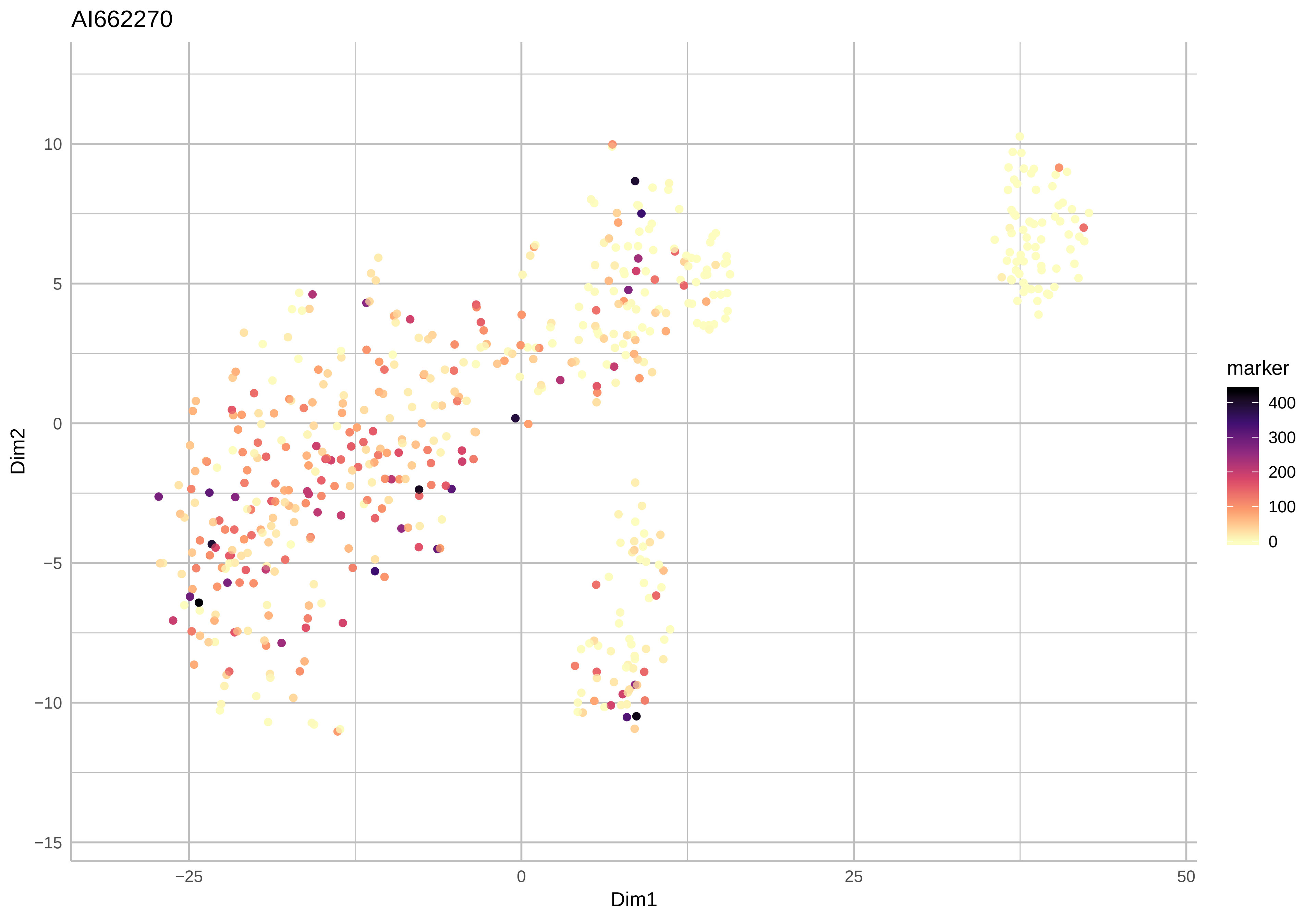
AI662270
Dim2
Dim1
10
5
0
−5
−10
−15
−25
0
25
50
marker
400
300
200
100
0

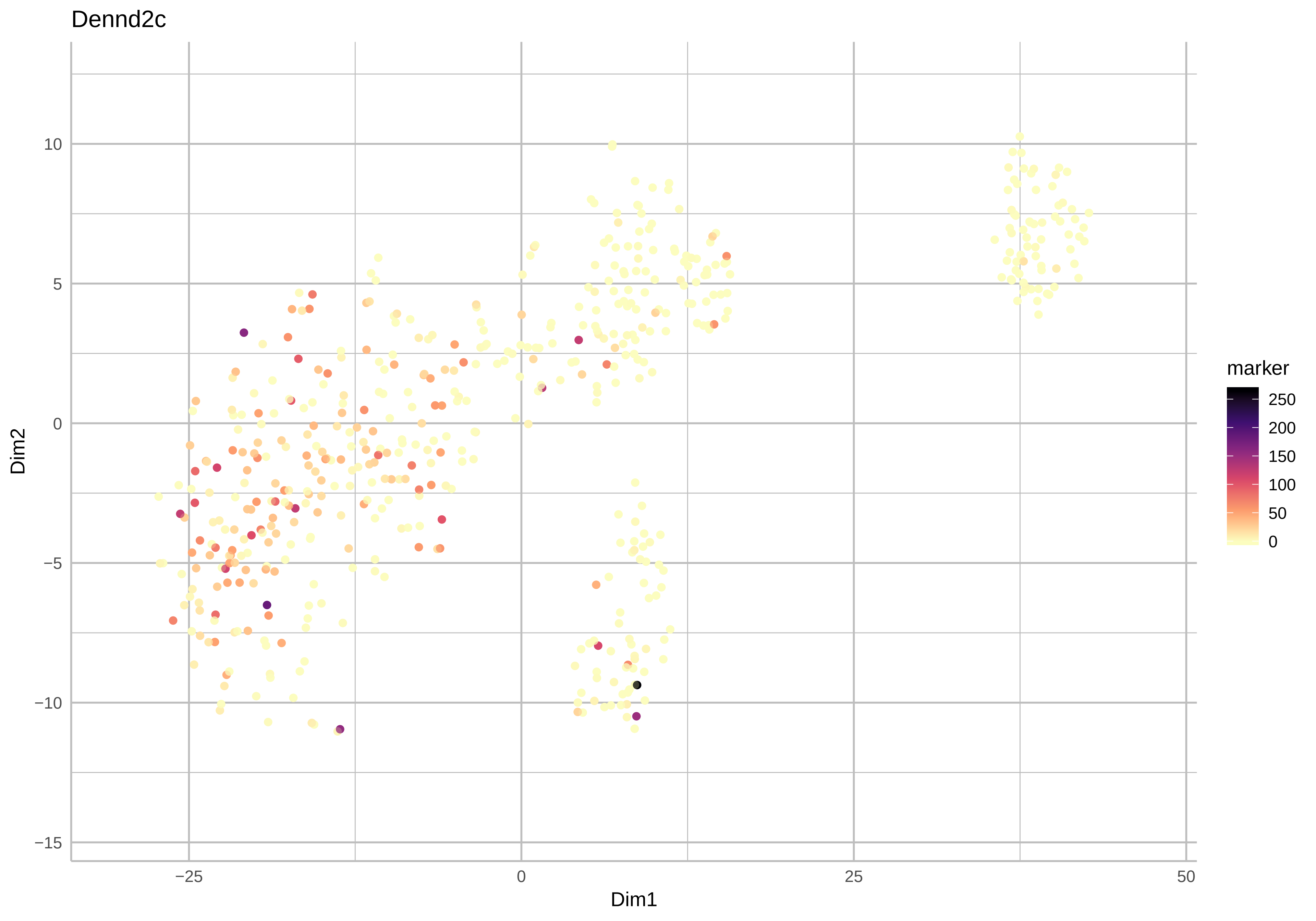

Dennd2c
10
5
0
−5
−10
−15
Dim2
−25
0
25
50
Dim1
marker
250
200
150
100
50
0

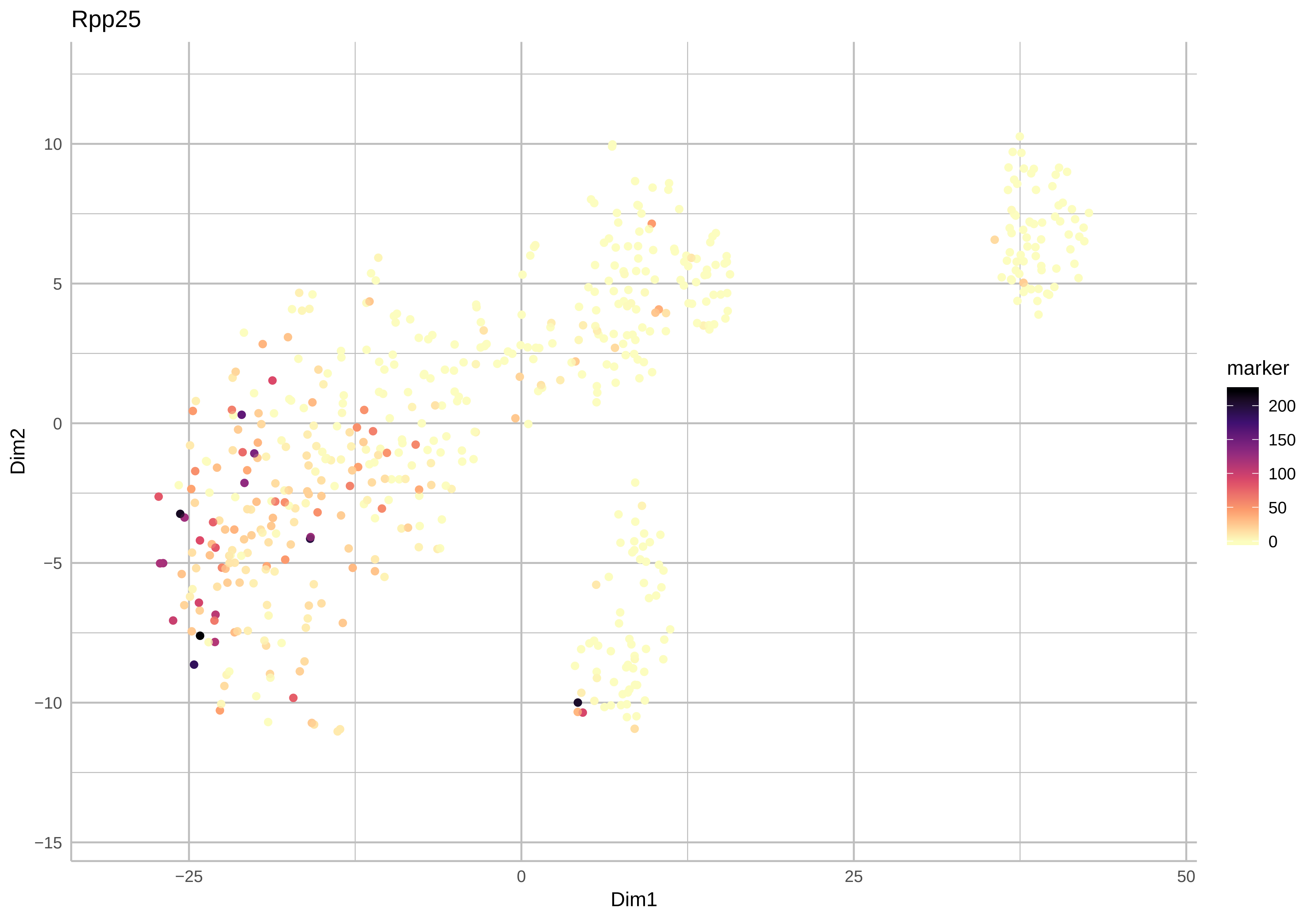

Rpp25
10
5
0
−5
−10
−15
Dim2
−25
0
25
50
Dim1
marker
200
150
100
50
0

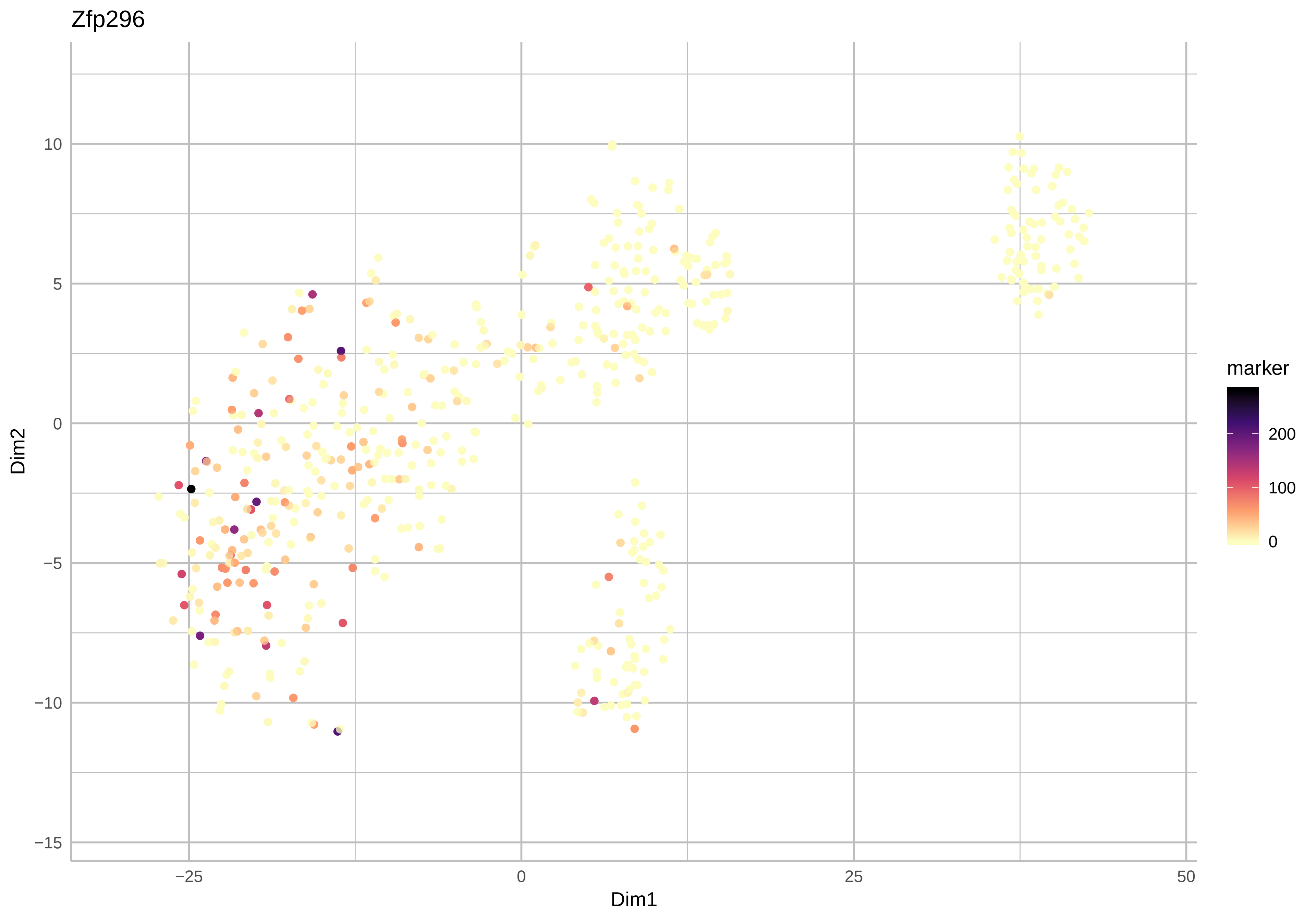

Zfp296
Dim2
10
5
0
−5
−10
−15
−25
0
25
50
Dim1
marker
200
100
0

Fgf4

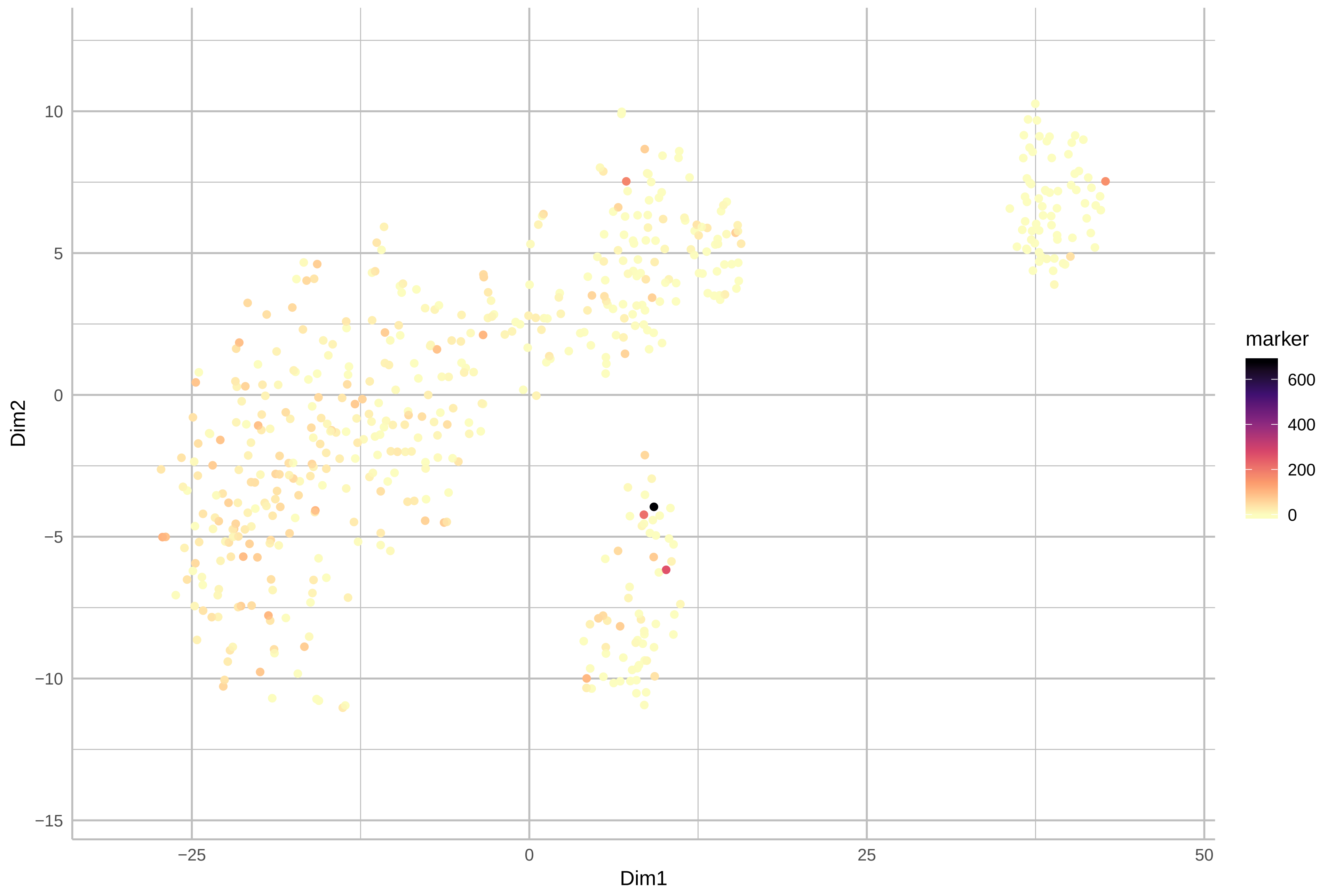

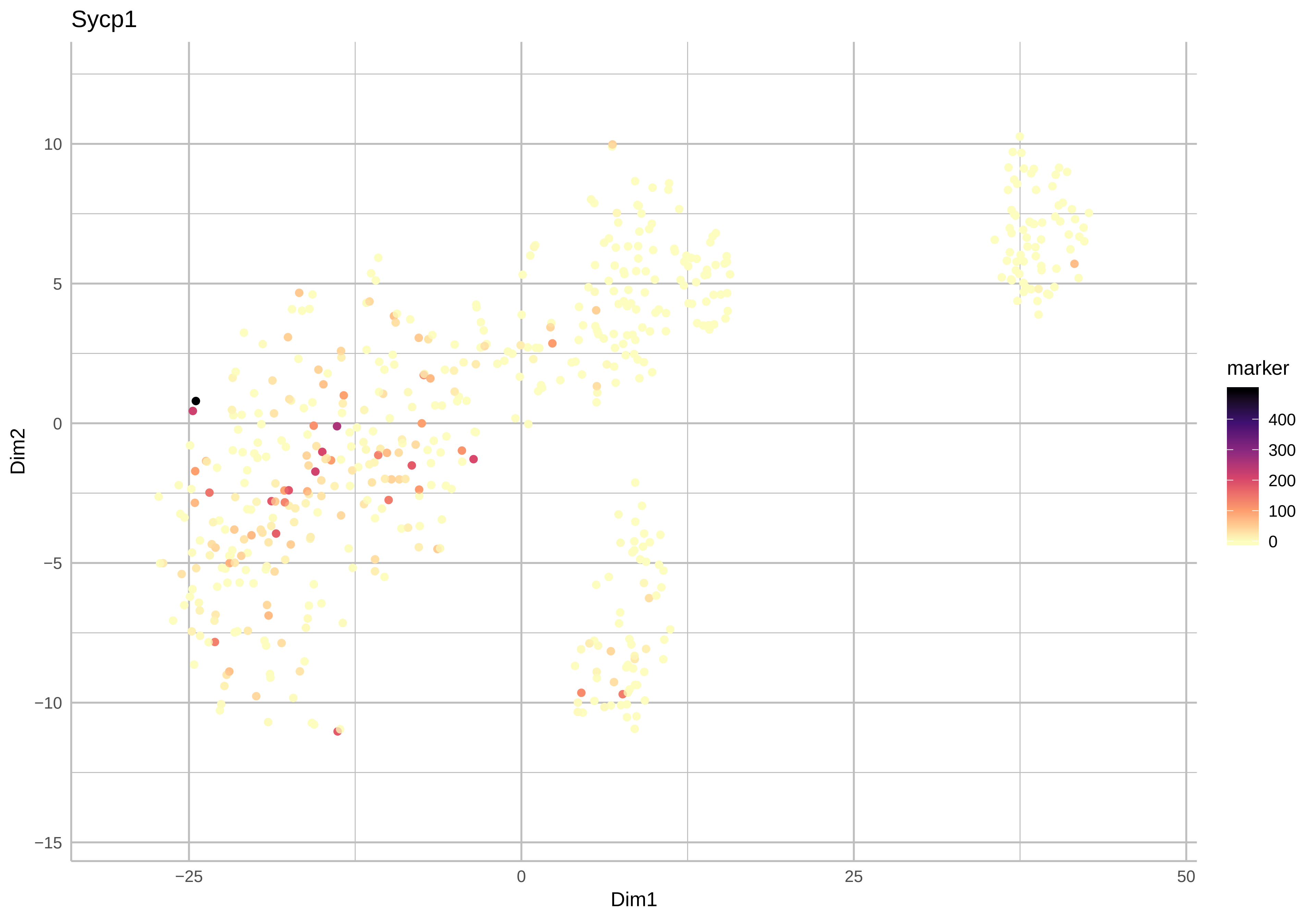

Sycp1
10
5
0
−5
−10
−15
Dim2
−25
0
25
50
Dim1
marker
400
300
200
100
0

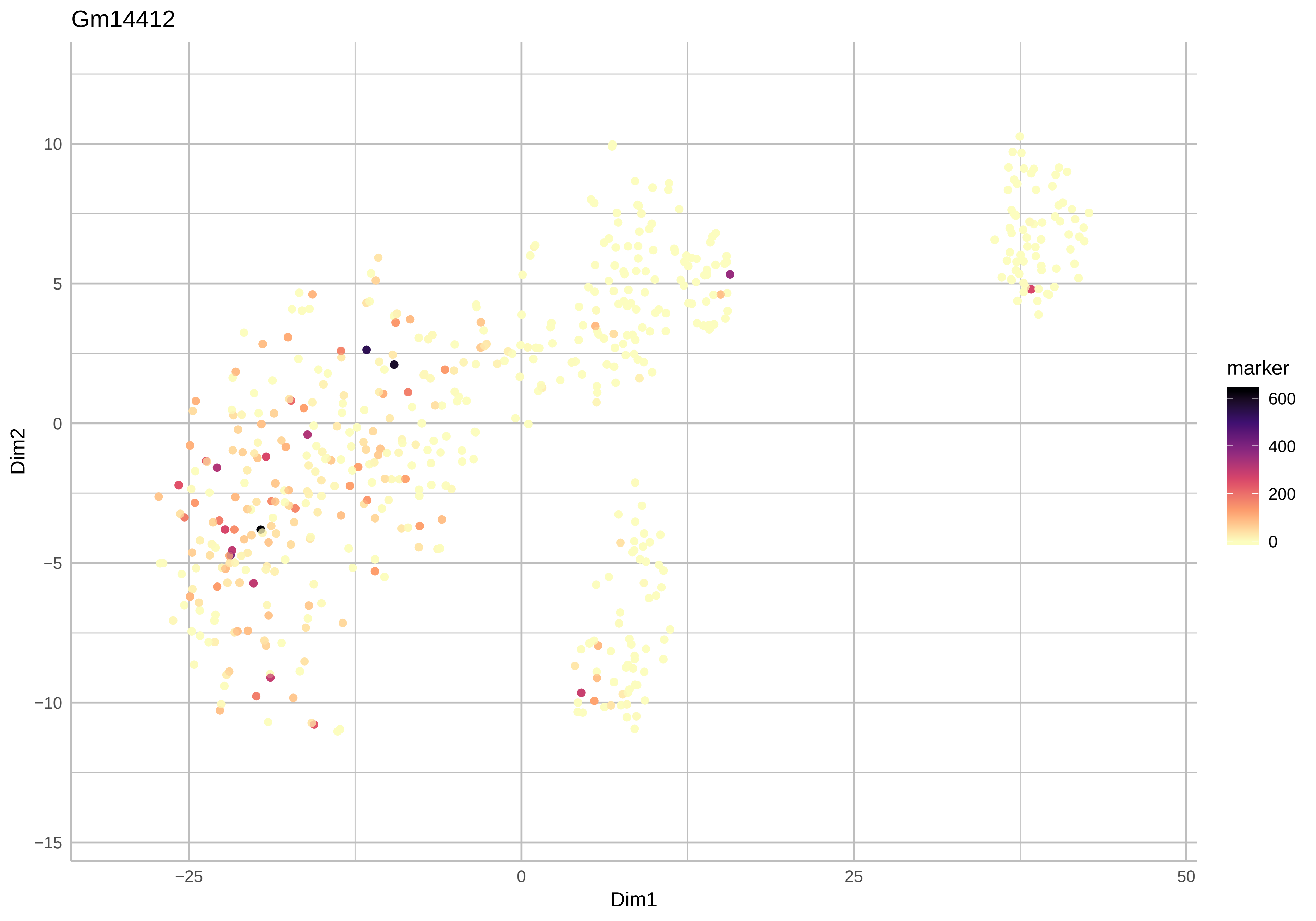

Gm14412
10
5
0
−5
−10
−15
Dim2
−25
0
25
50
Dim1
marker
600
400
200
0

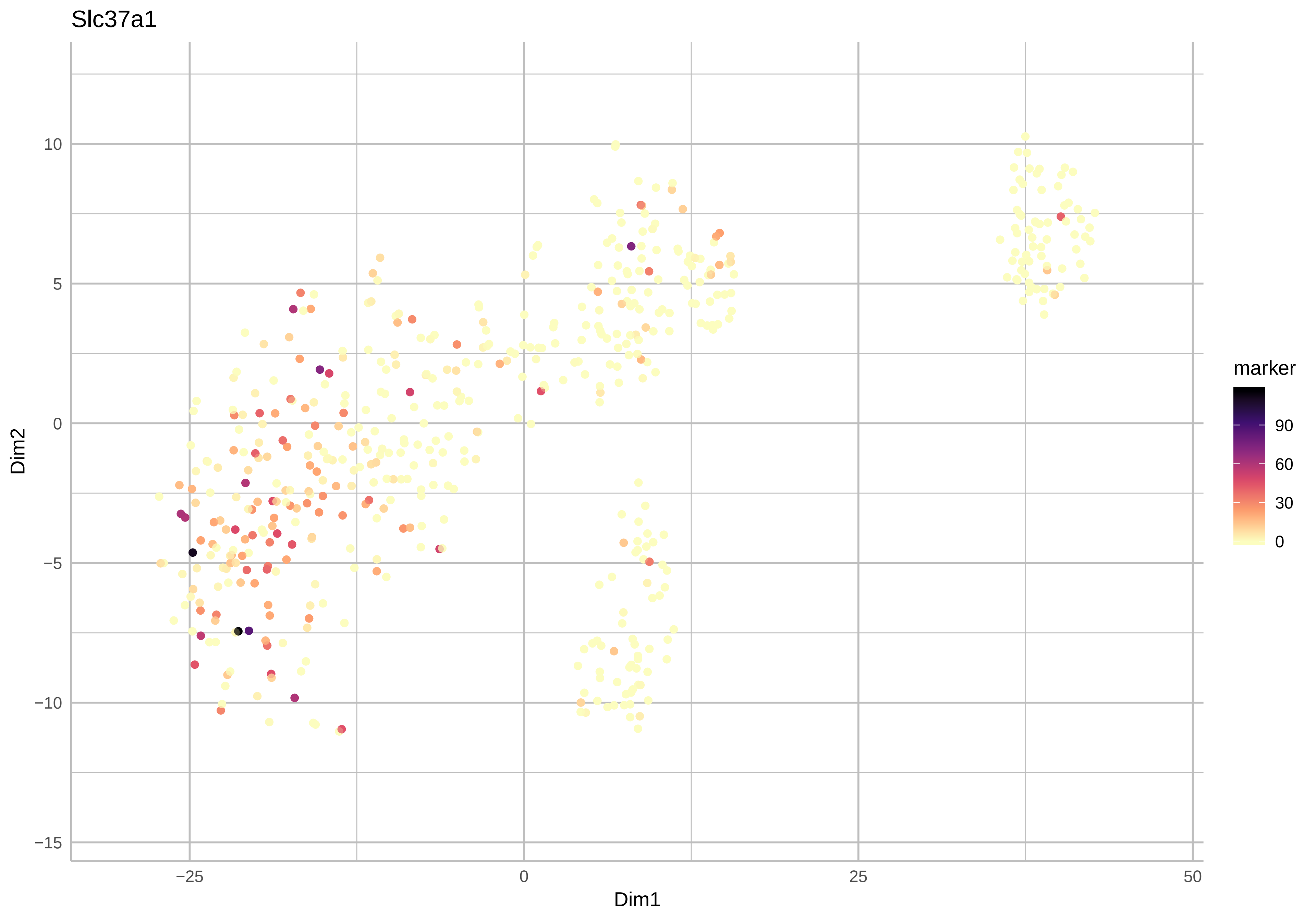
Slc37a1
Dim2
Dim1
marker
90
60
30
0
10
5
0
−5
−10
−15
−25
0
25
50

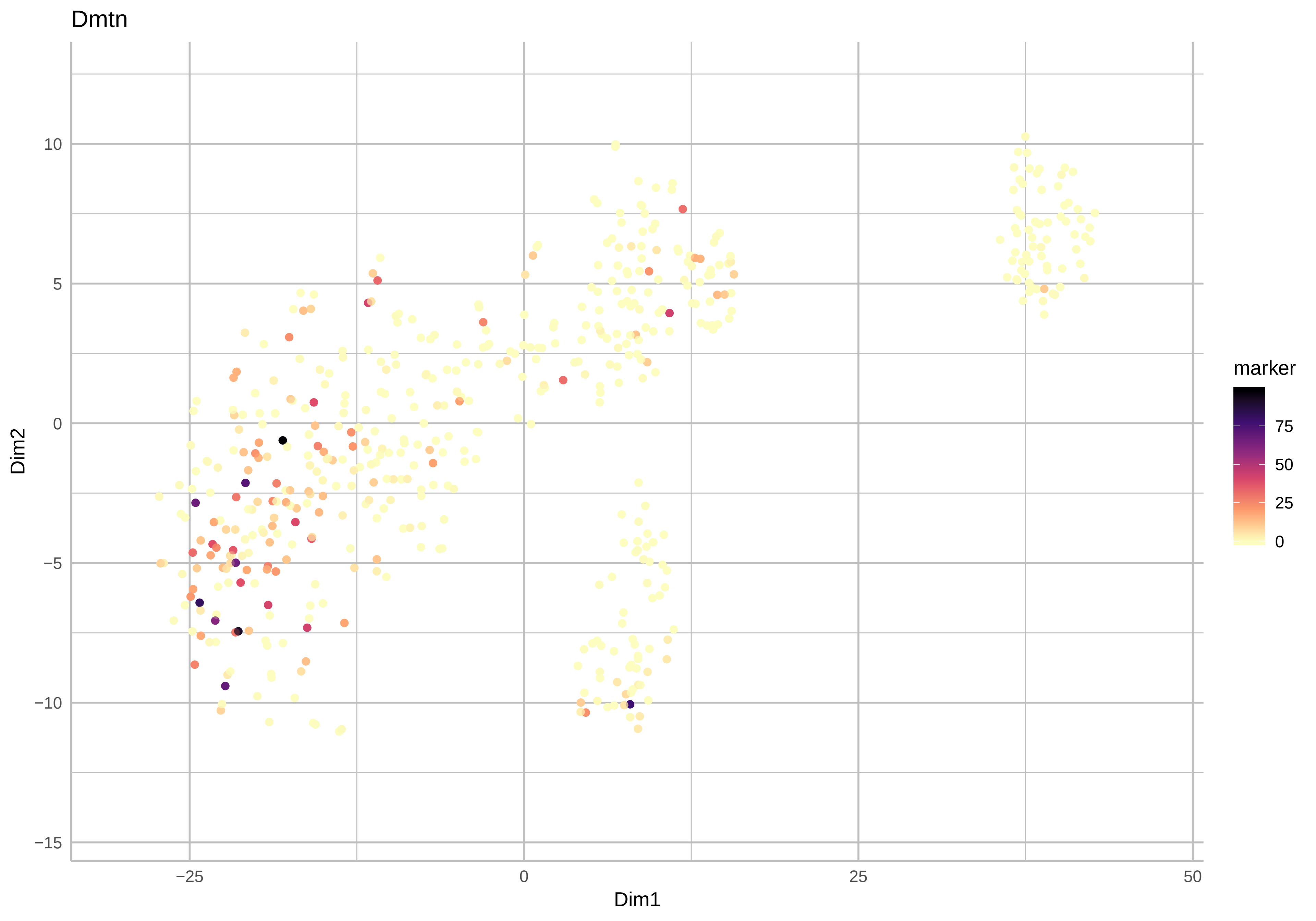

Dmtn
10
5
0
−5
−10
−15
−25
0
25
50
Dim1
Dim2
marker
75
50
25
0

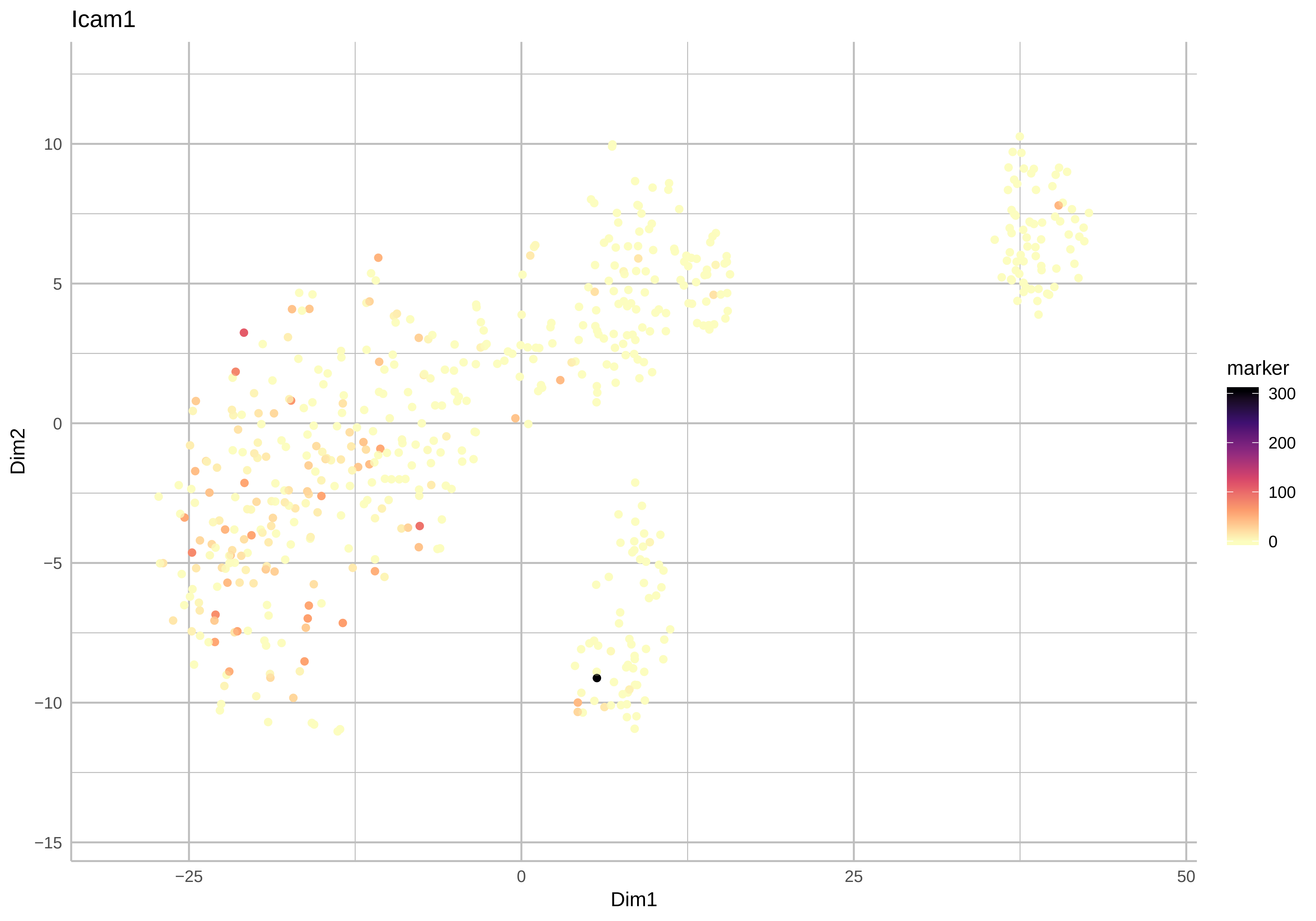

Icam1
Dim2
Dim1
10
5
0
−5
−10
−15
−25
0
25
50
marker
300
200
100
0

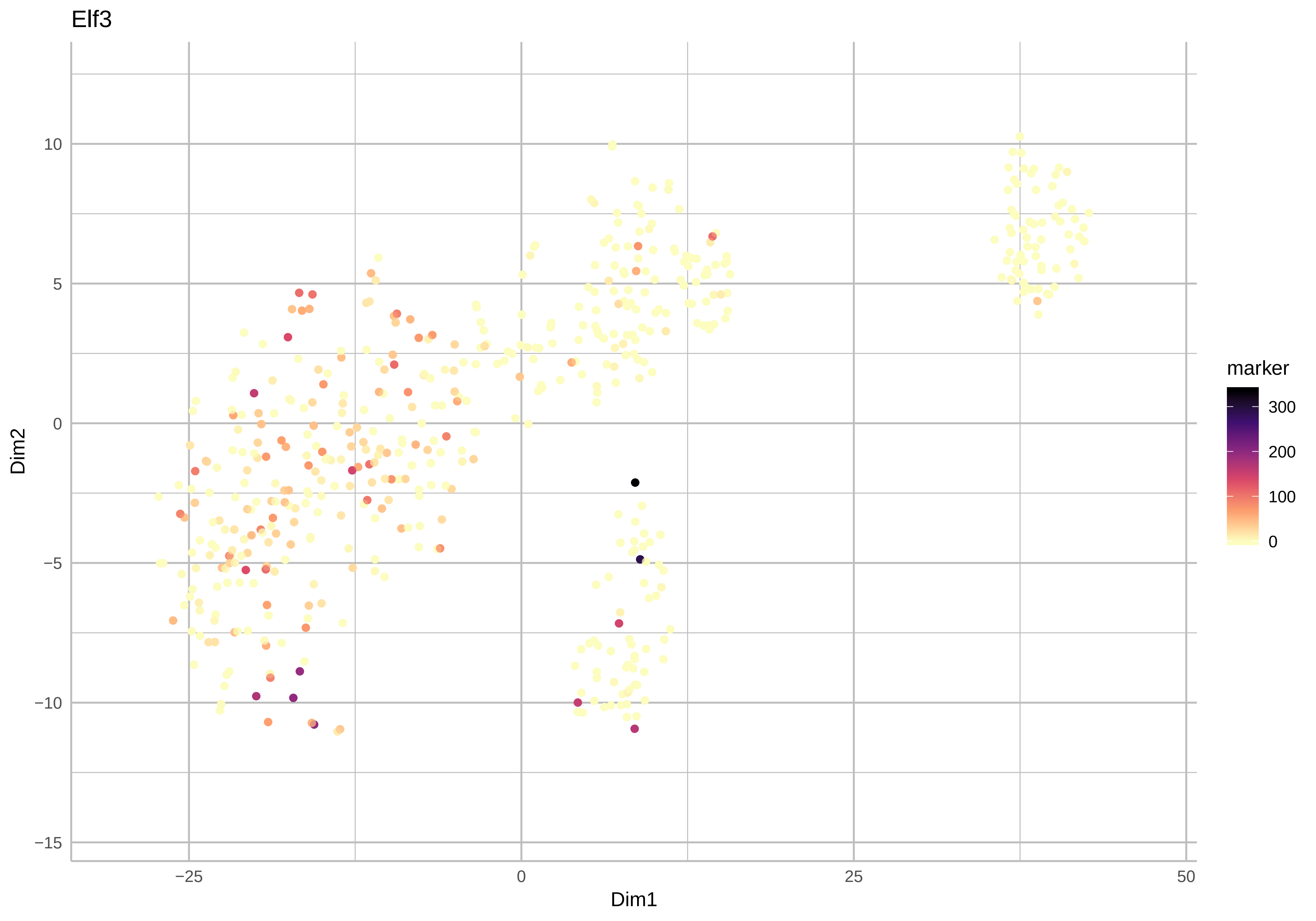

Elf3
Dim2
Dim1
marker
10
5
0
−5
−10
−15
−25
0
25
50
300
200
100
0

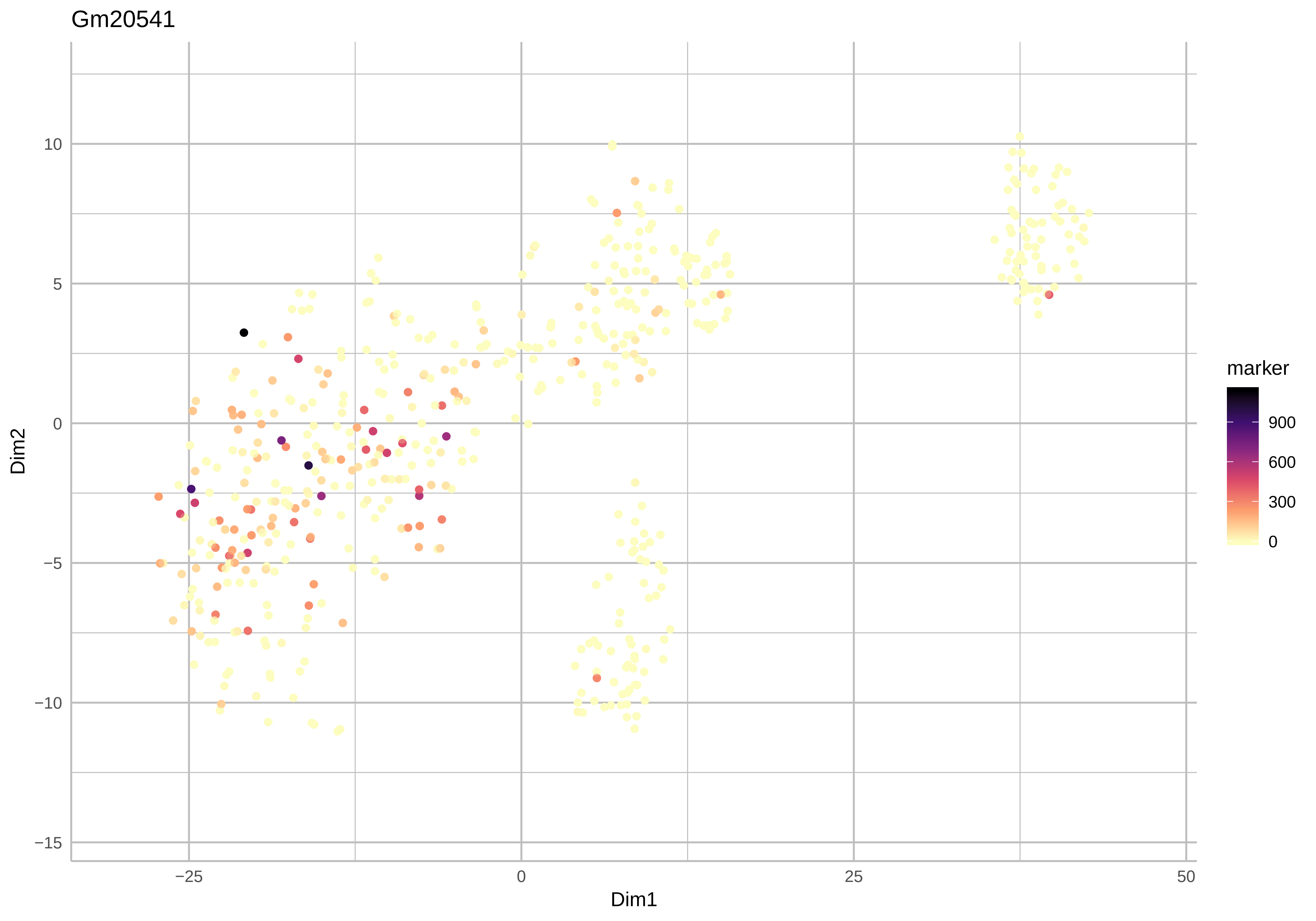
Gm20541
Dim2
Dim1
10
5
0
−5
−10
−15
−25
0
25
50
marker
900
600
300
0

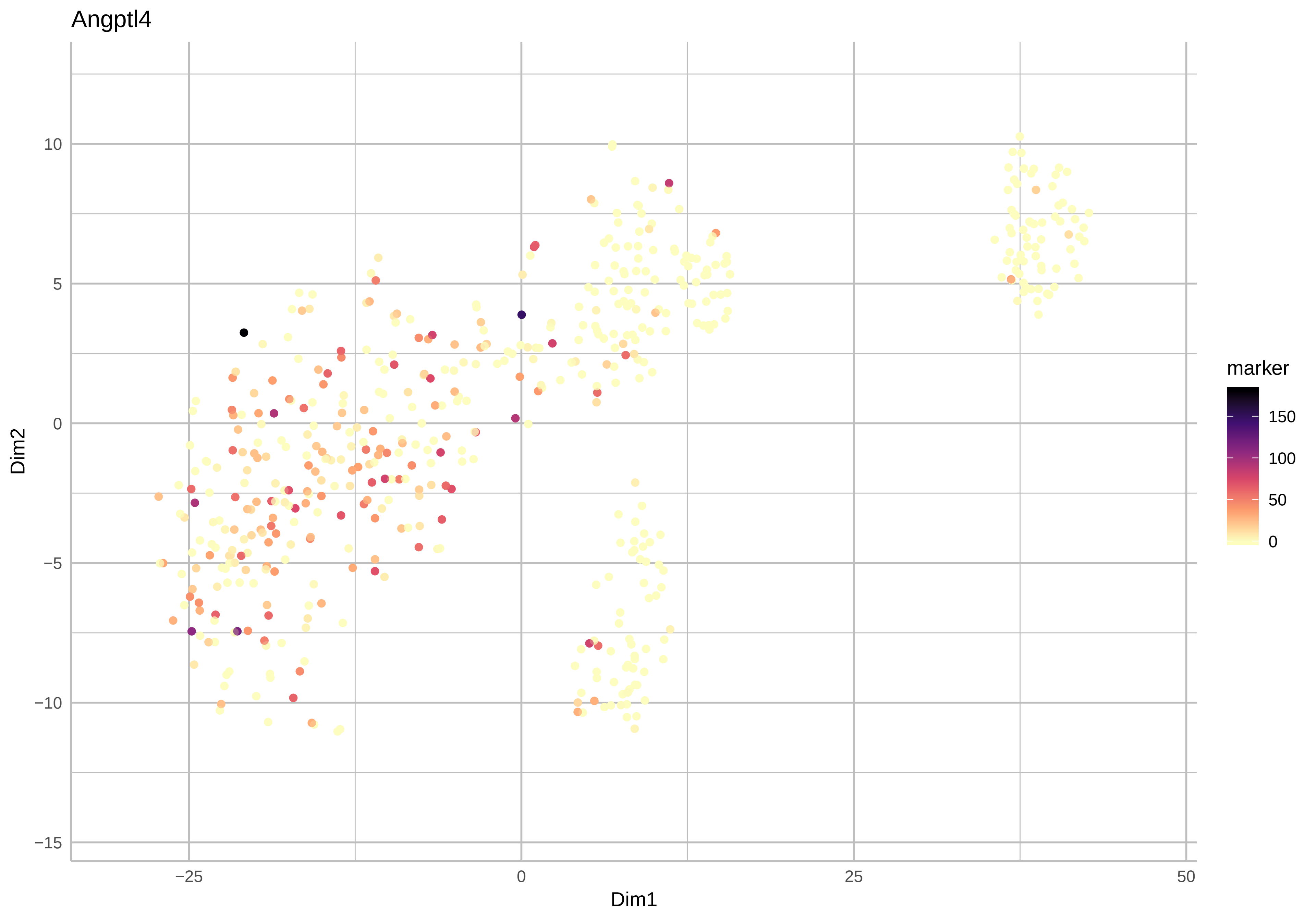
Angptl4
Dim2
Dim1
marker
150
100
50
0
10
5
0
−5
−10
−15
−25
0
25
50

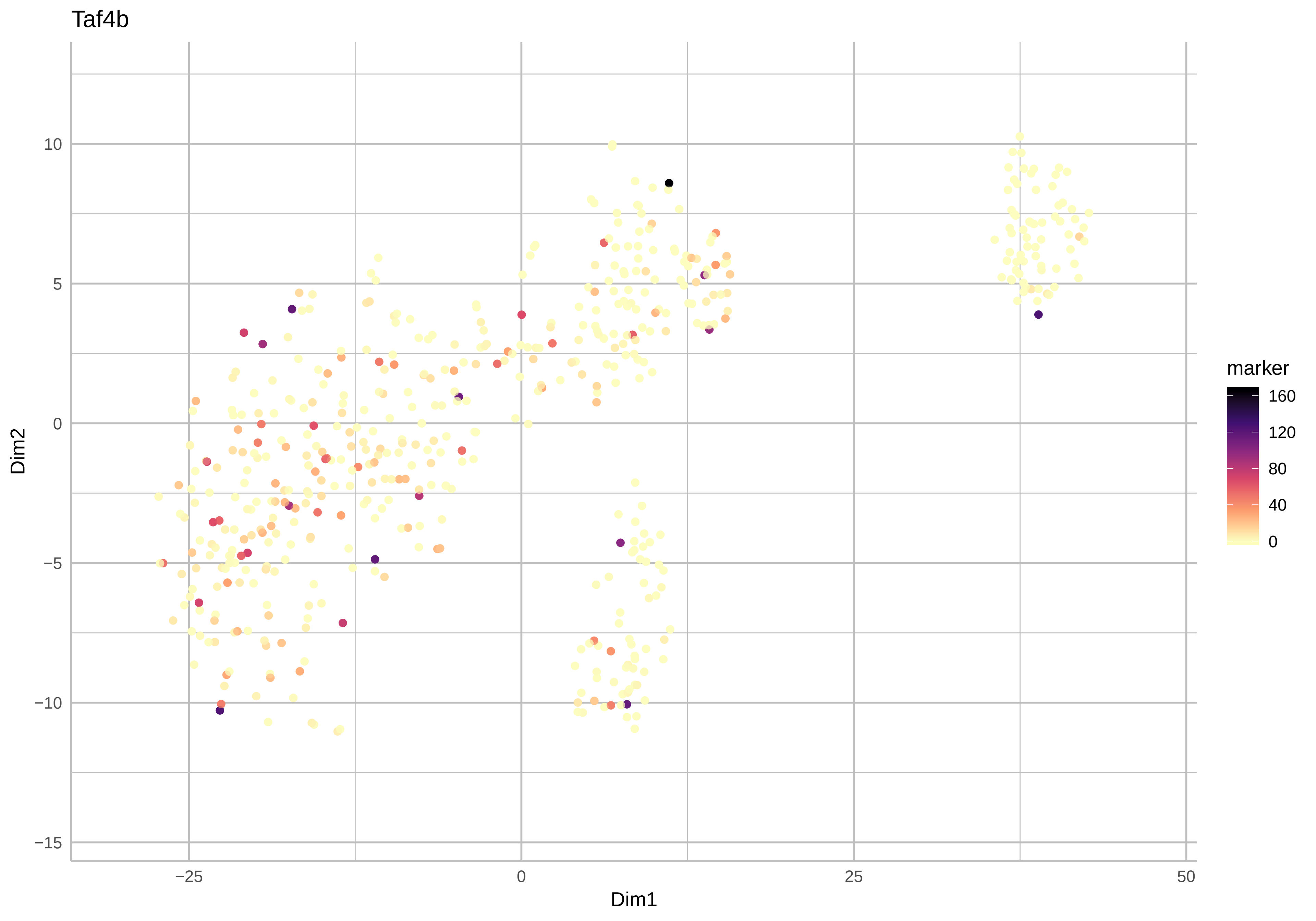

Taf4b
10
5
0
−5
−10
−15
Dim2
−25
0
25
50
Dim1
marker
160
120
80
40
0

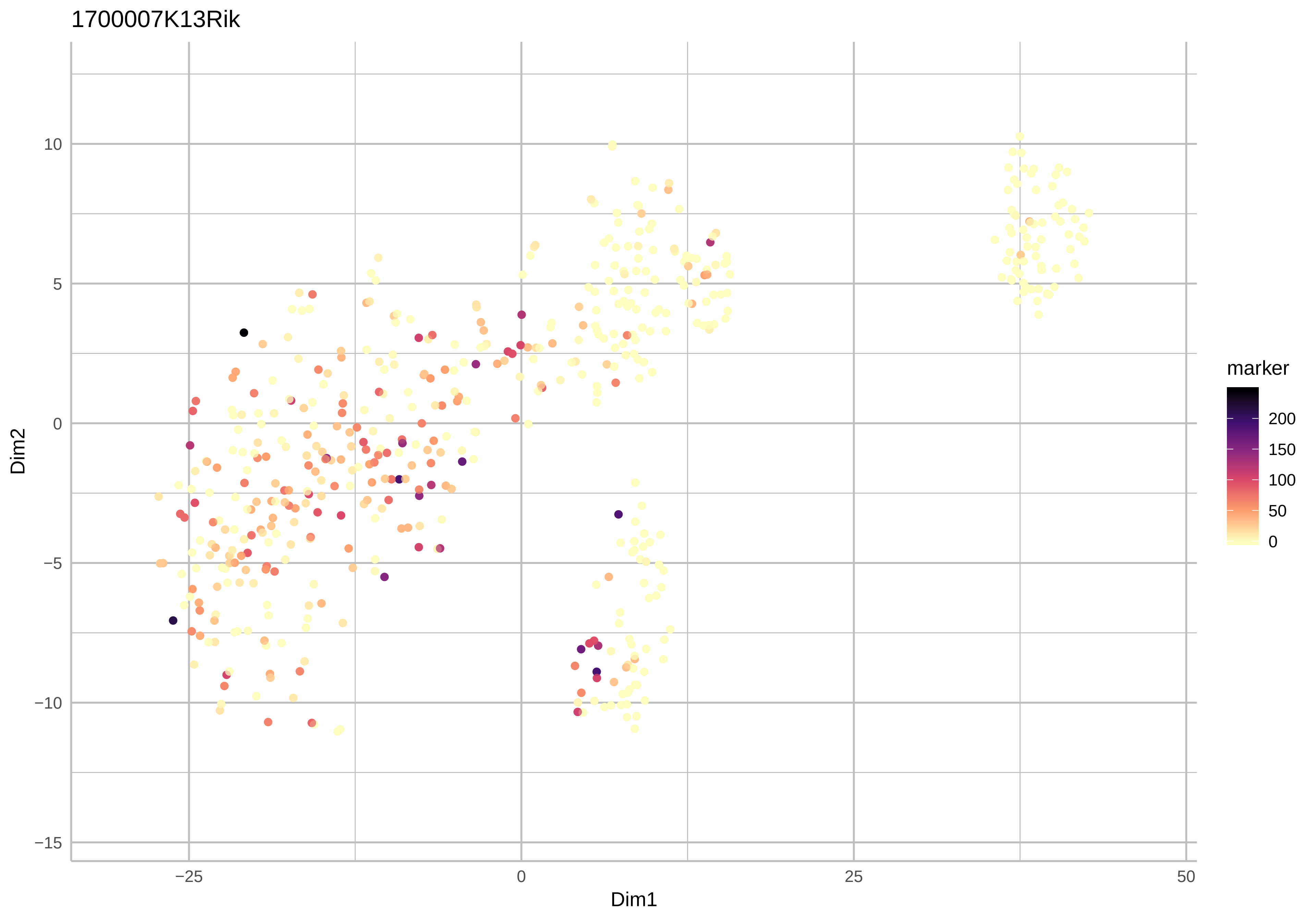

1700007K13Rik
Dim2
Dim1
10
5
0
−5
−10
−15
−25
0
25
50
marker
200
150
100
50
0

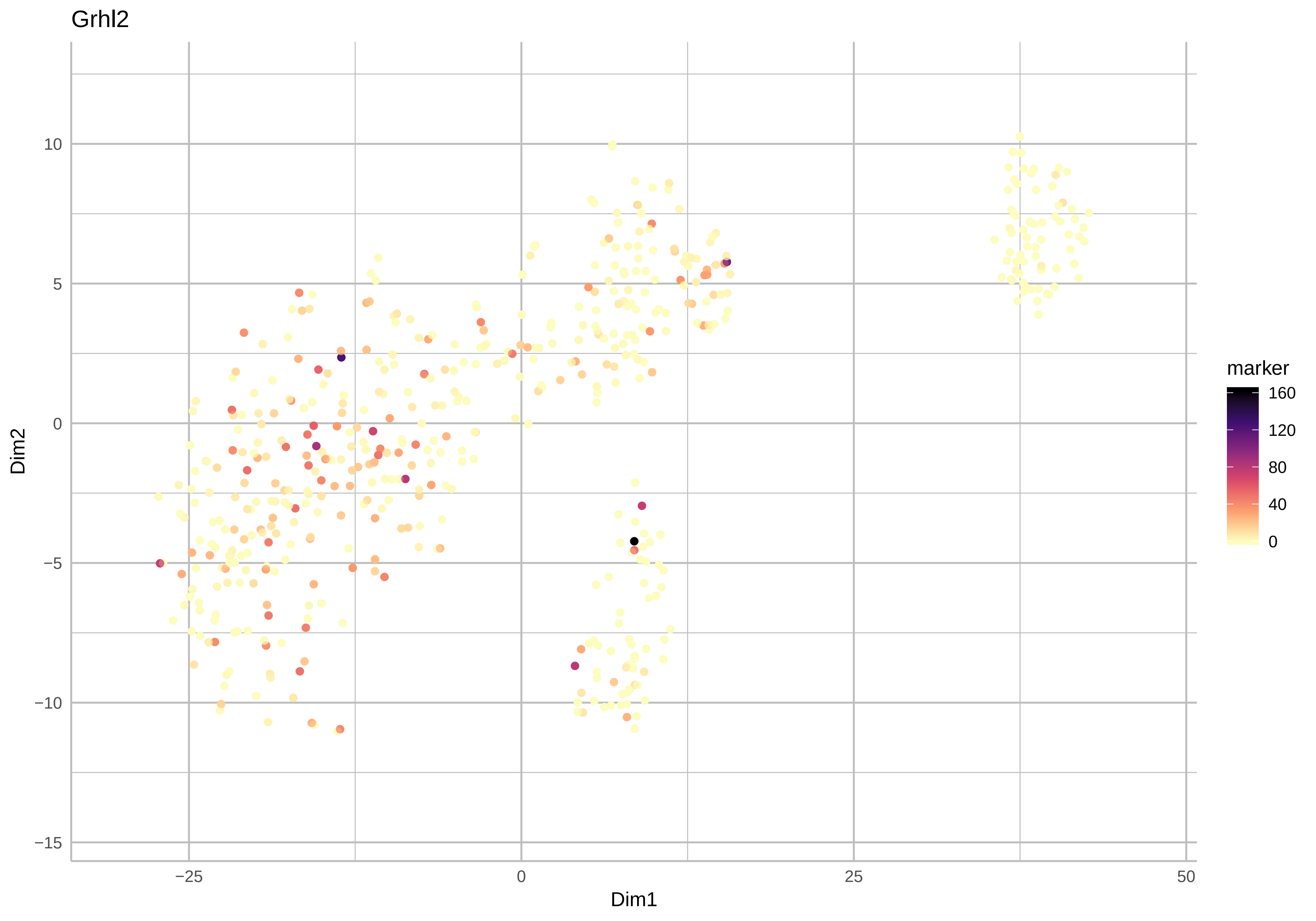

Grhl2
Dim2
Dim1
10
5
0
−5
−10
−15
−25
0
25
50
marker
160
120
80
40
0

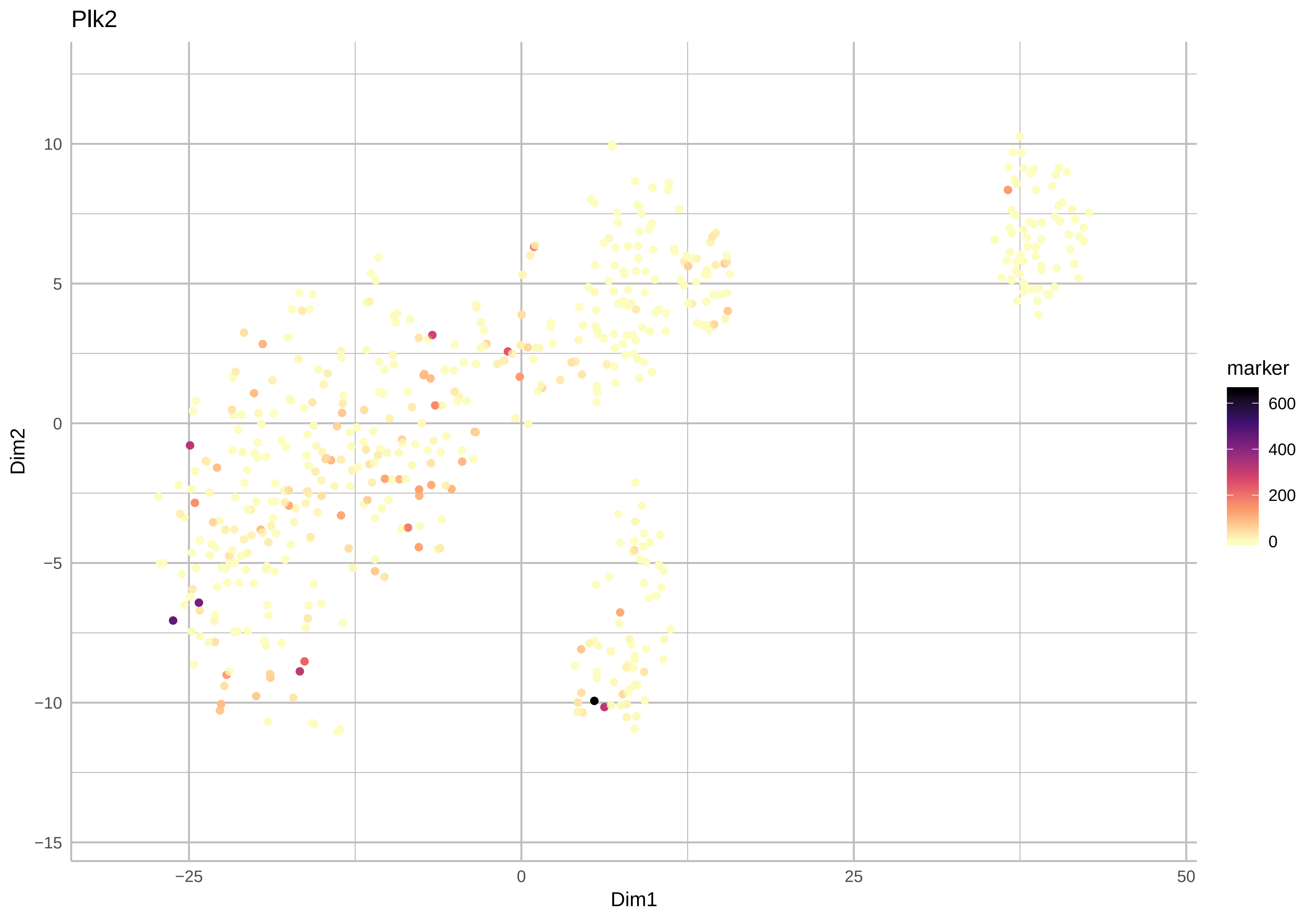
Plk2
Dim2
Dim1
10
5
0
−5
−10
−15
−25
0
25
50
marker
600
400
200
0

Fndc3c1

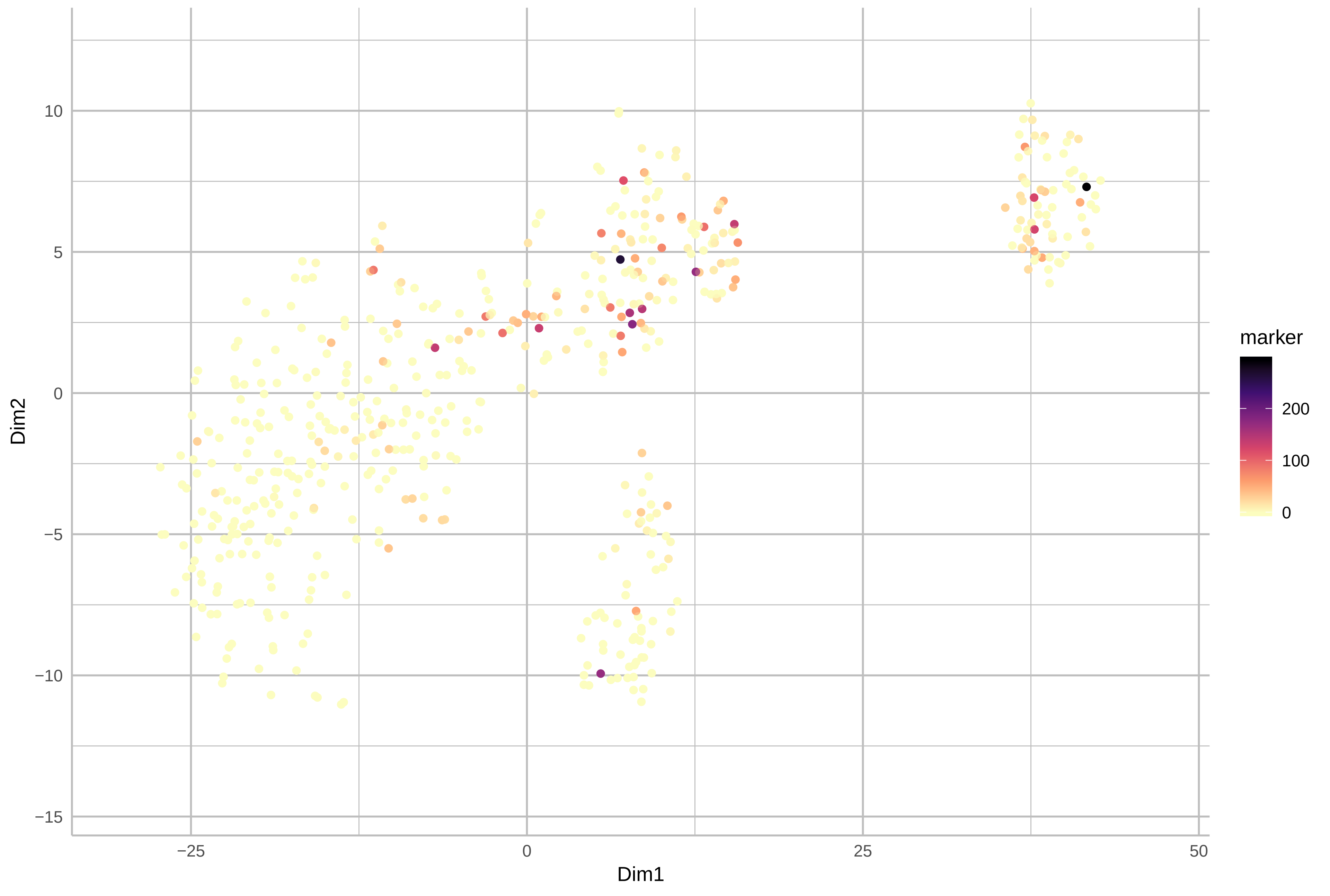

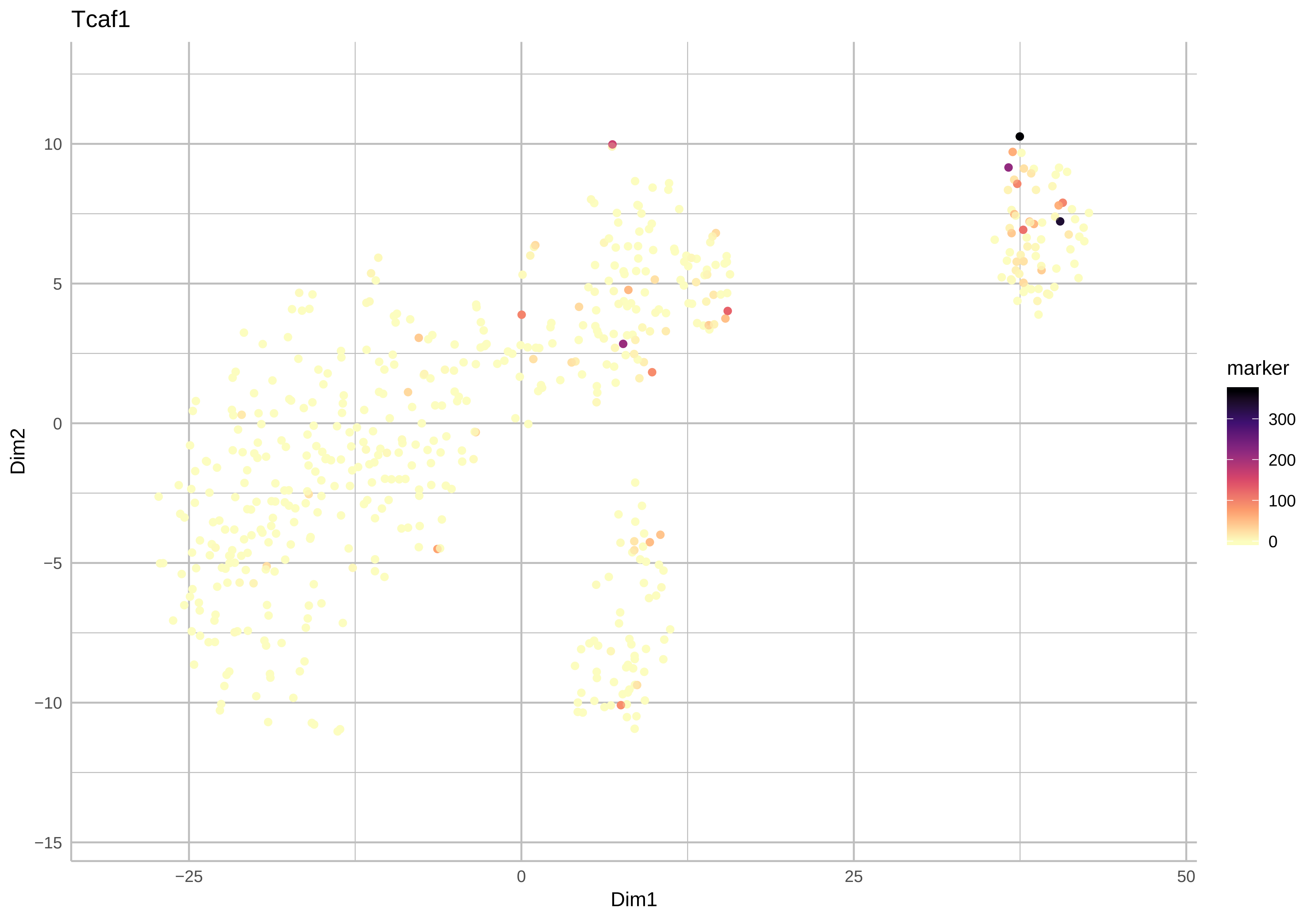

Tcaf1
Dim2
Dim1
10
5
0
−5
−10
−15
−25
0
25
50
marker
300
200
100
0

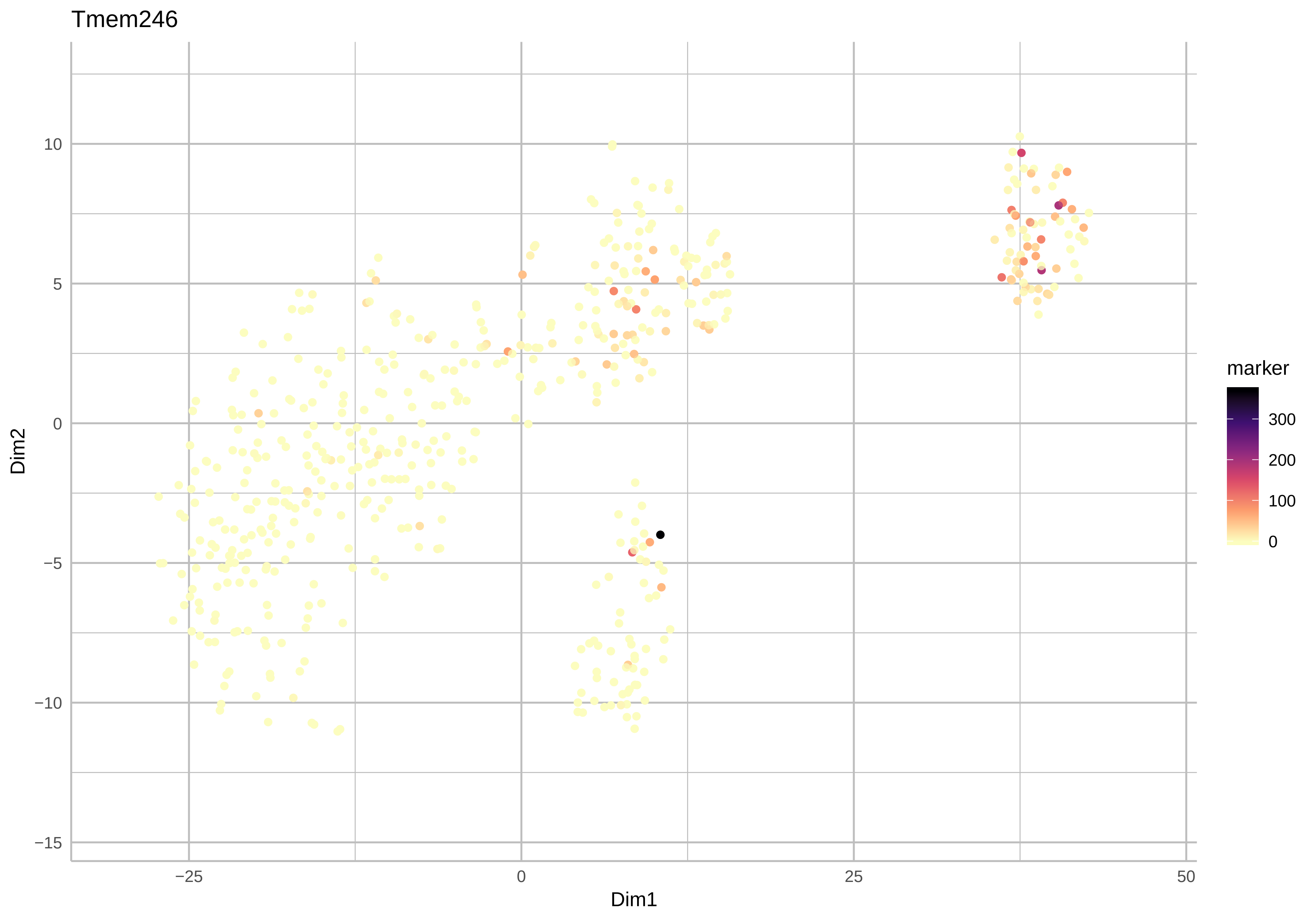
Tmem246
Dim2
10
5
0
−5
−10
−15
−25
0
25
50
Dim1
marker
300
200
100
0

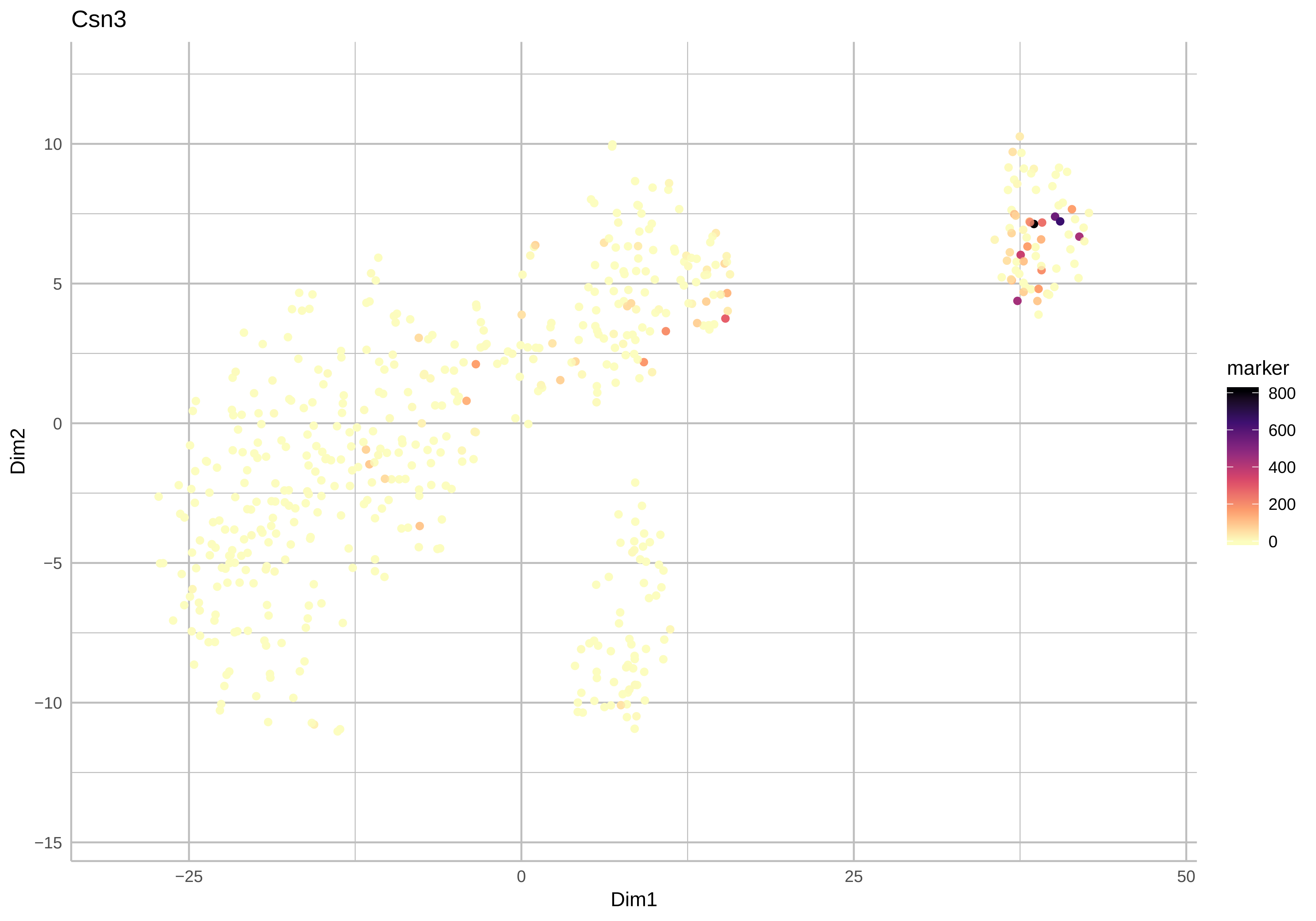
Csn3
Dim2
Dim1
10
5
0
−5
−10
−15
−25
0
25
50
marker
800
600
400
200
0

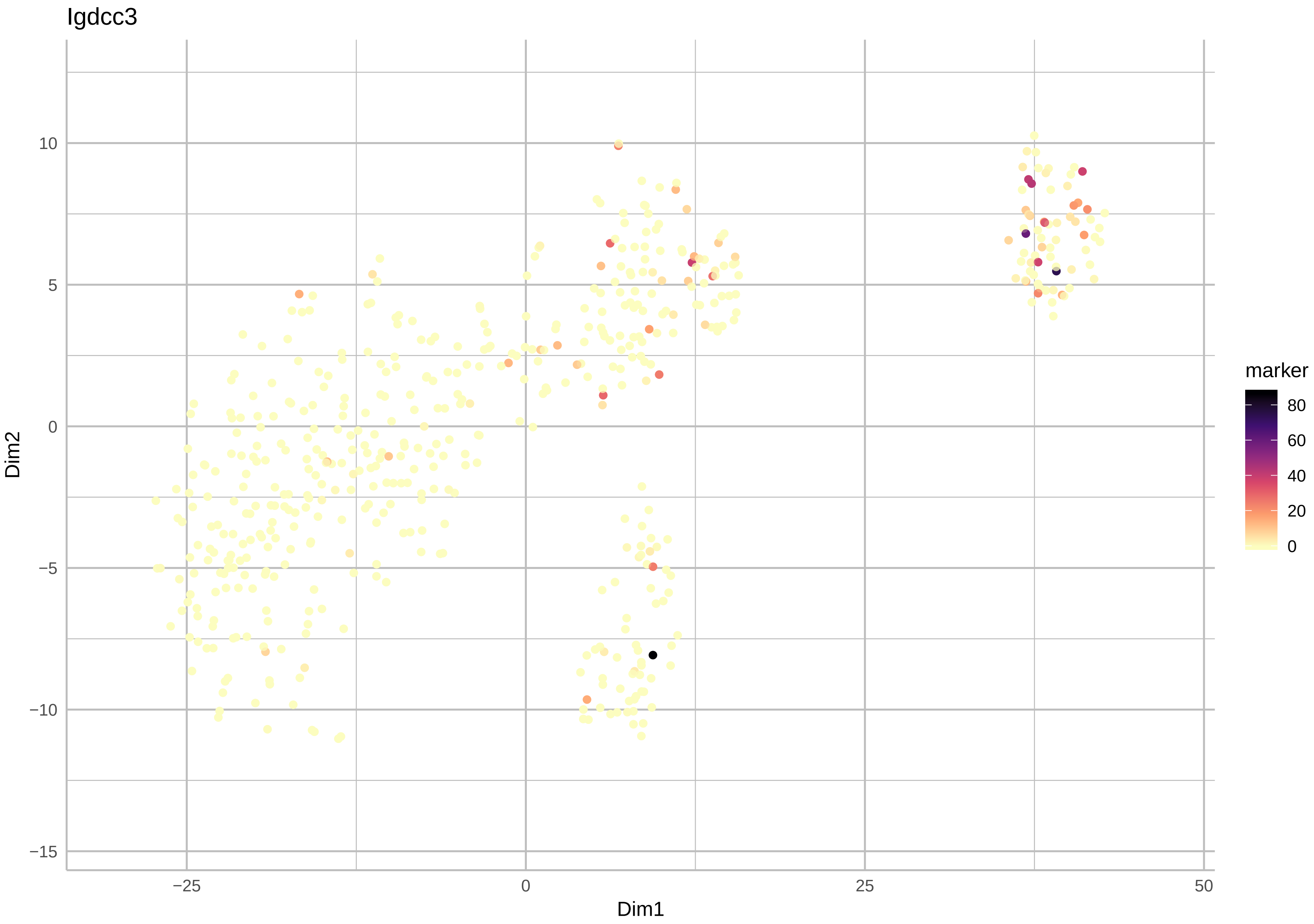

Igdcc3
Dim2
Dim1
marker
80
60
40
20
0
10
5
0
−5
−10
−15
−25
0
25
50

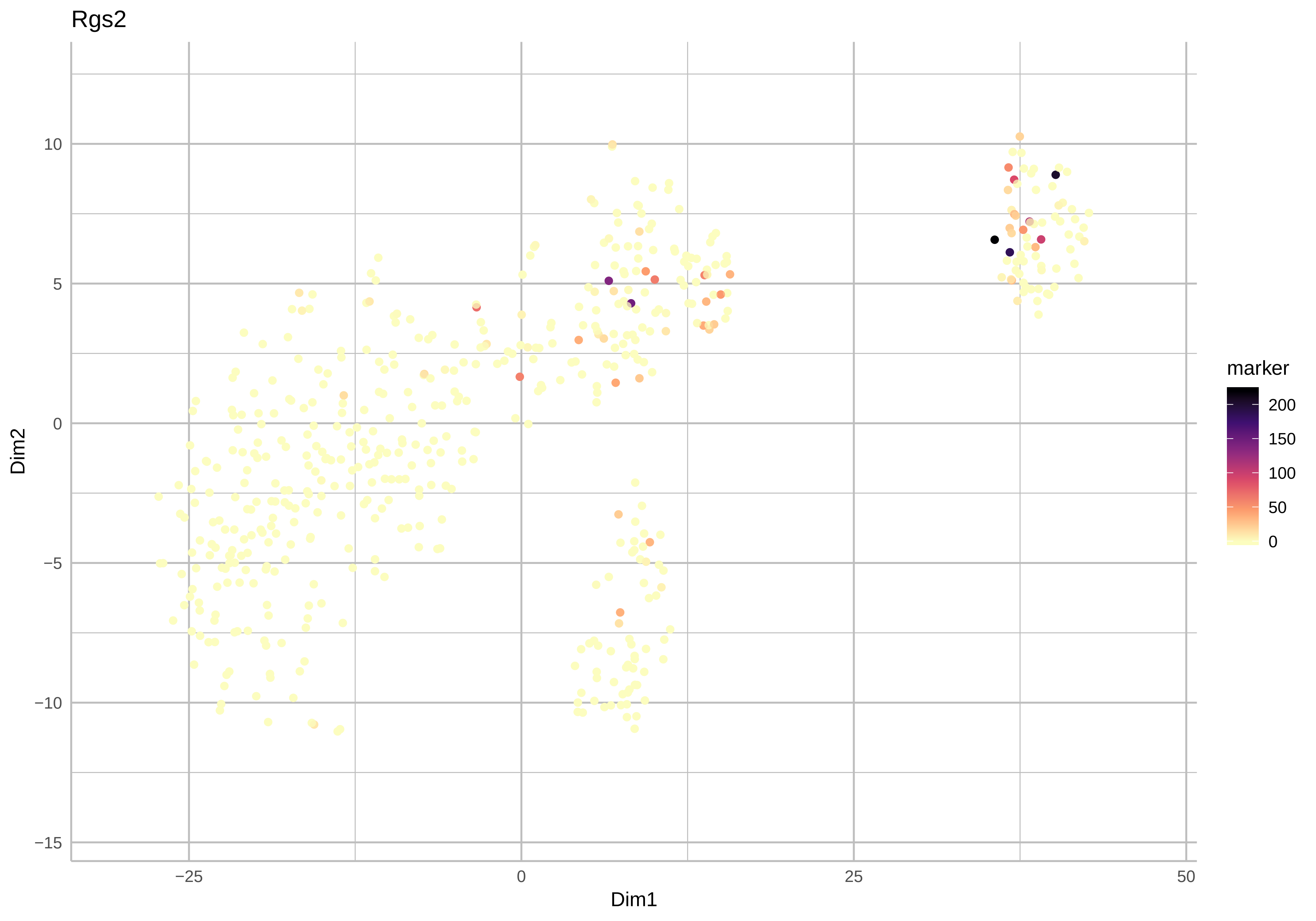

Rgs2
10
5
0
−5
−10
−15
Dim2
−25
0
25
50
Dim1
marker
200
150
100
50
0

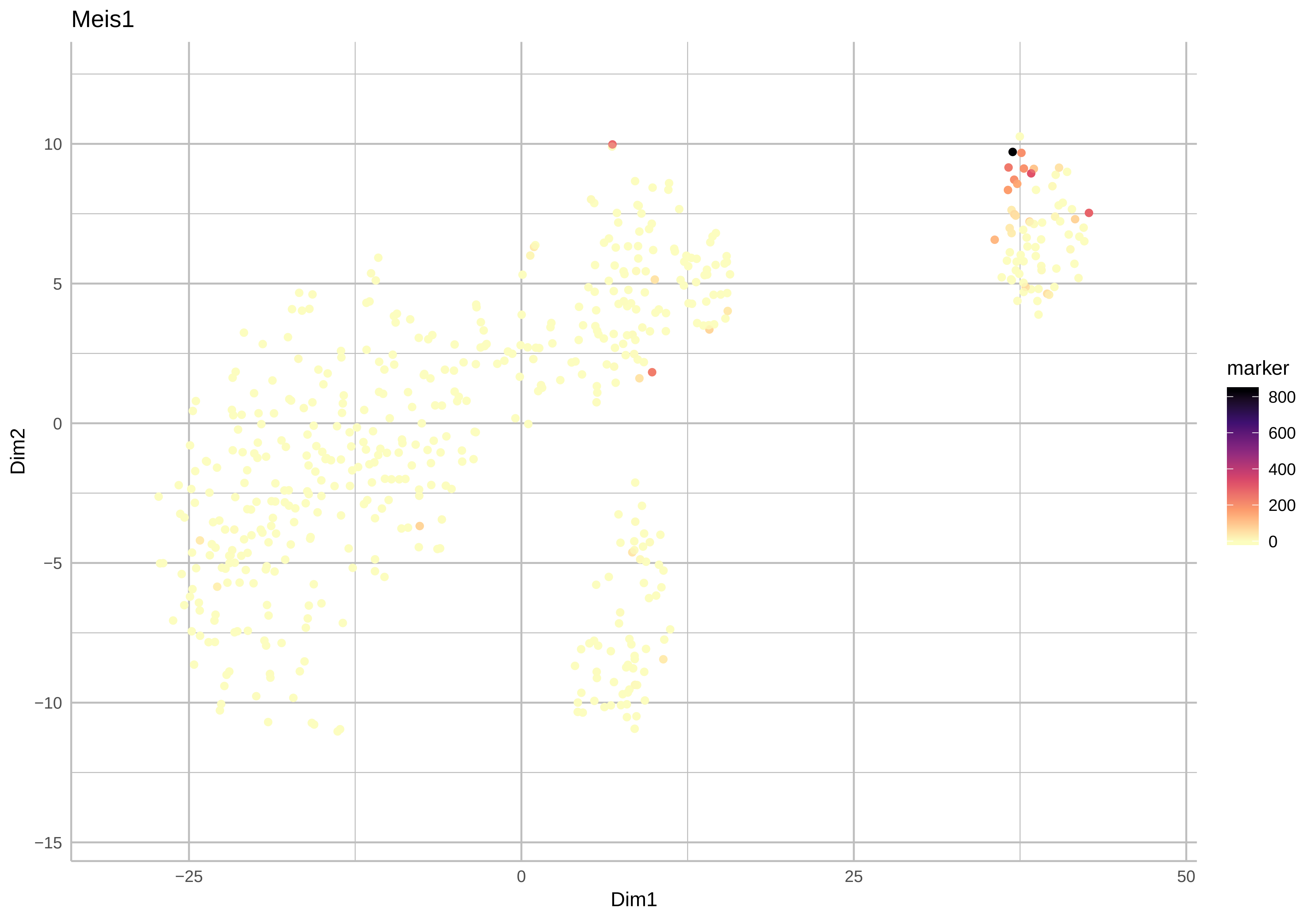
Meis1
Dim2
Dim1
10
5
0
−5
−10
−15
−25
0
25
50
marker
800
600
400
200
0

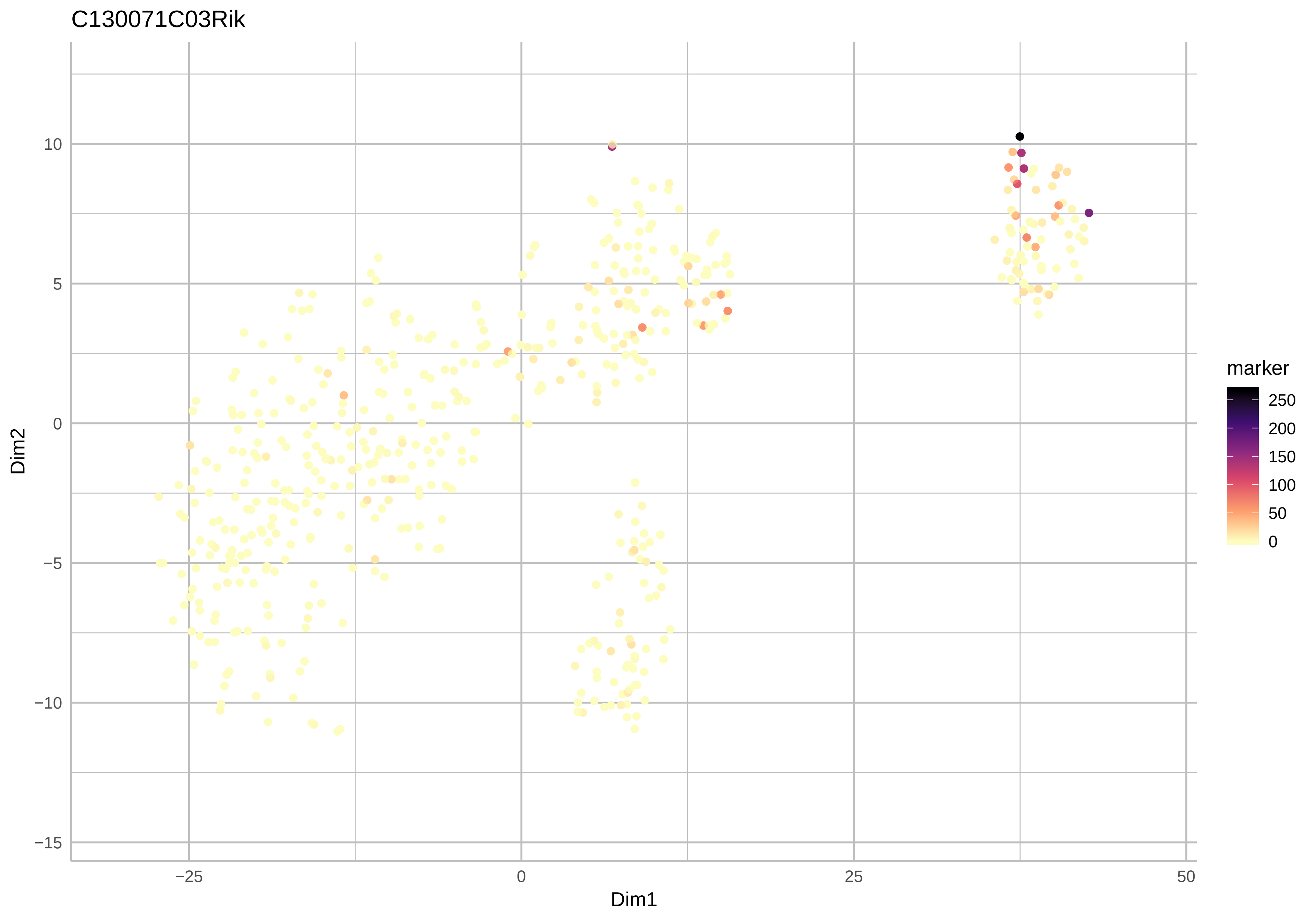
C130071C03Rik
Dim2
Dim1
10
5
0
−5
−10
−15
−25
0
25
50
marker
250
200
150
100
50
0

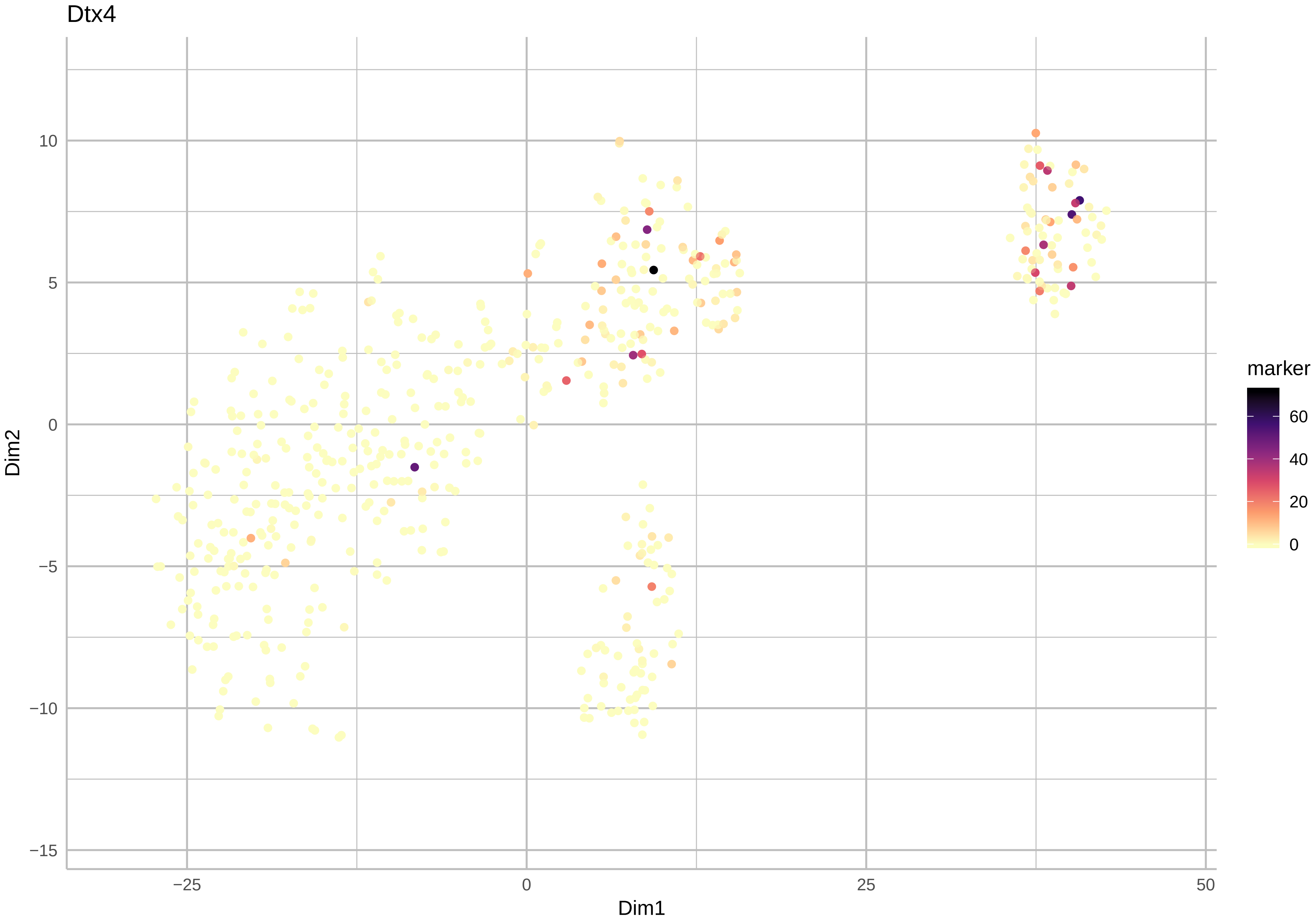

Dtx4
Dim2
Dim1
10
5
0
−5
−10
−15
−25
0
25
50
marker
60
40
20
0

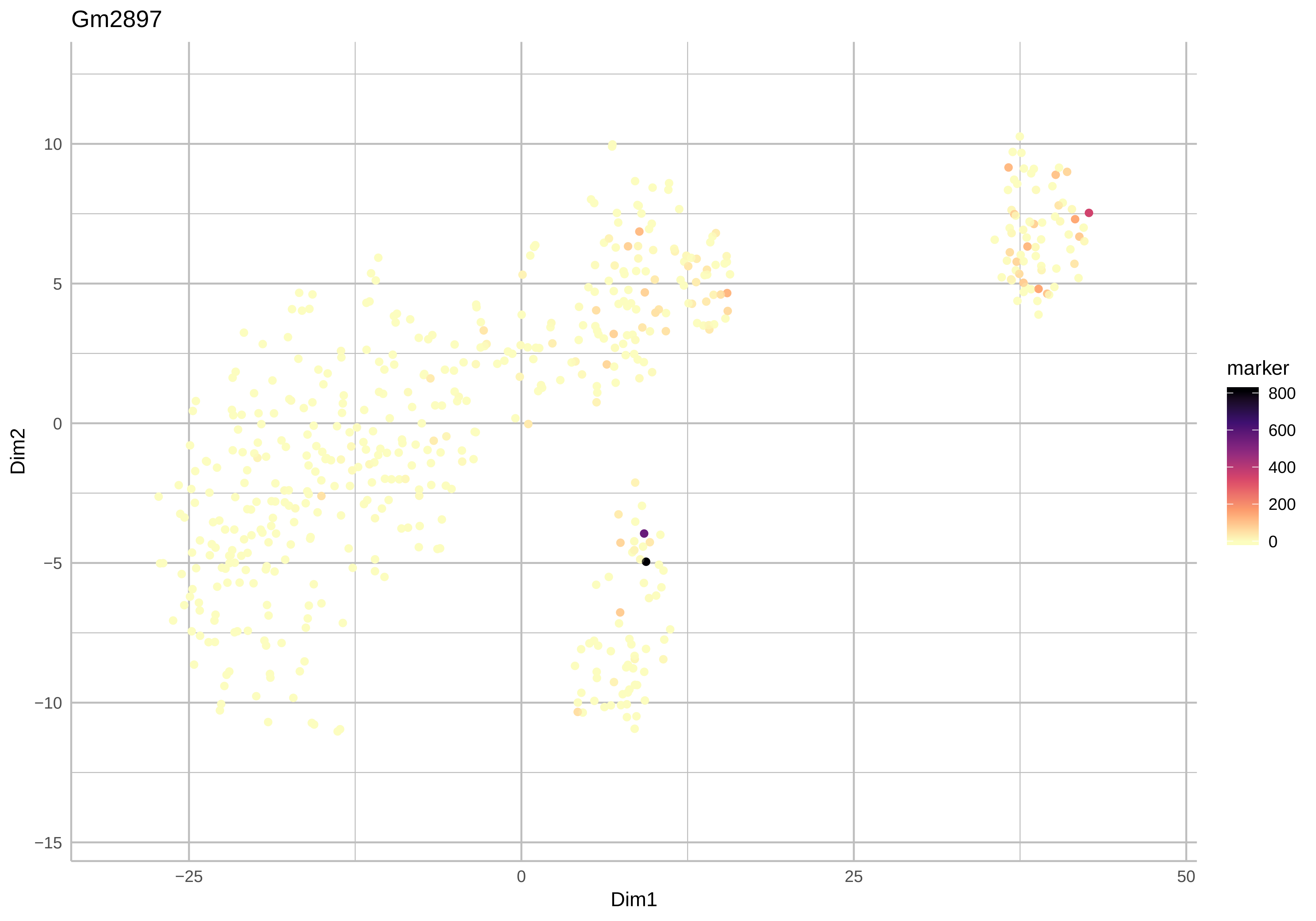
Gm2897
Dim2
Dim1
10
5
0
−5
−10
−15
−25
0
25
50
marker
800
600
400
200
0

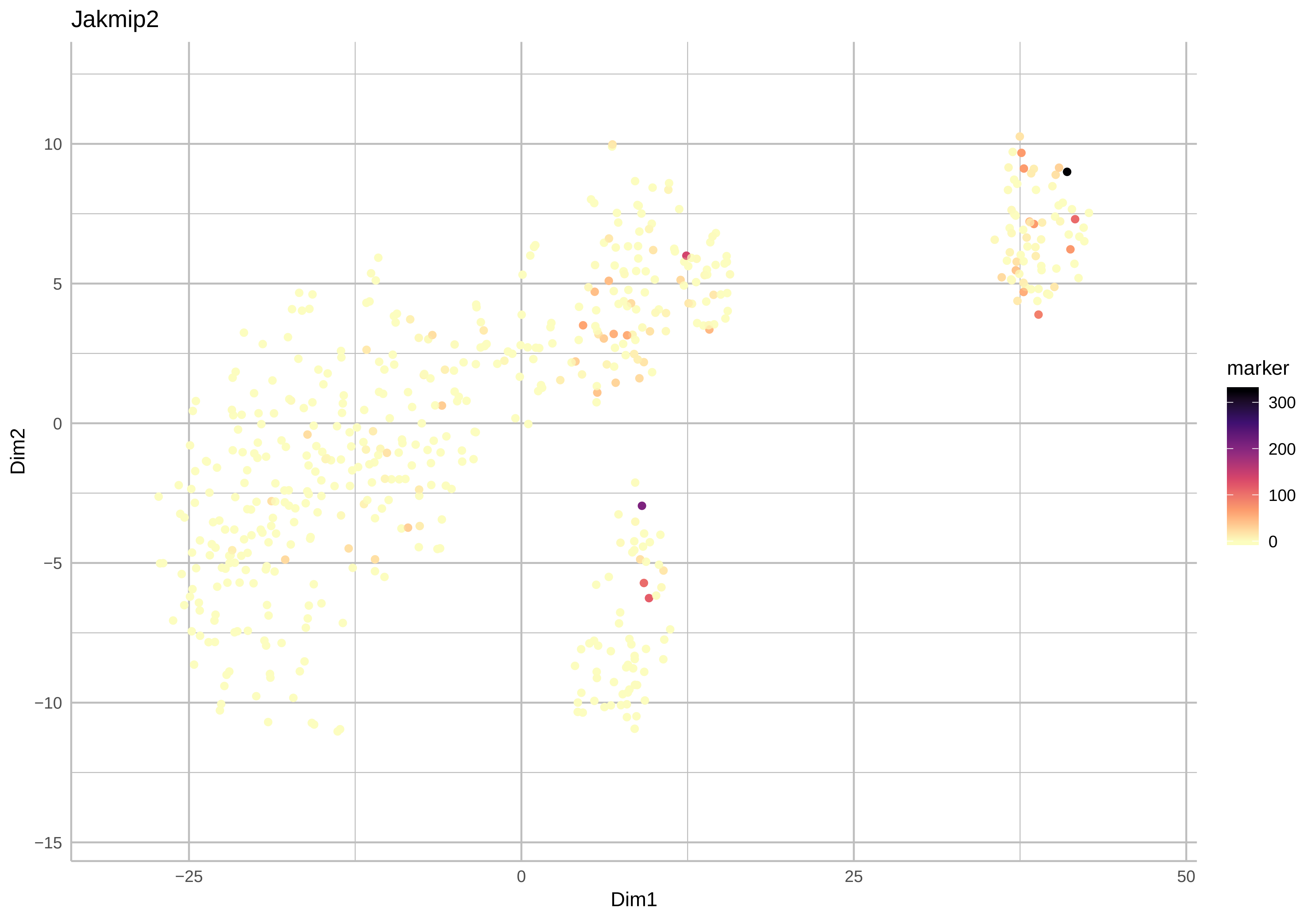

Jakmip2
Dim2
Dim1
10
5
0
−5
−10
−15
−25
0
25
50
marker
300
200
100
0

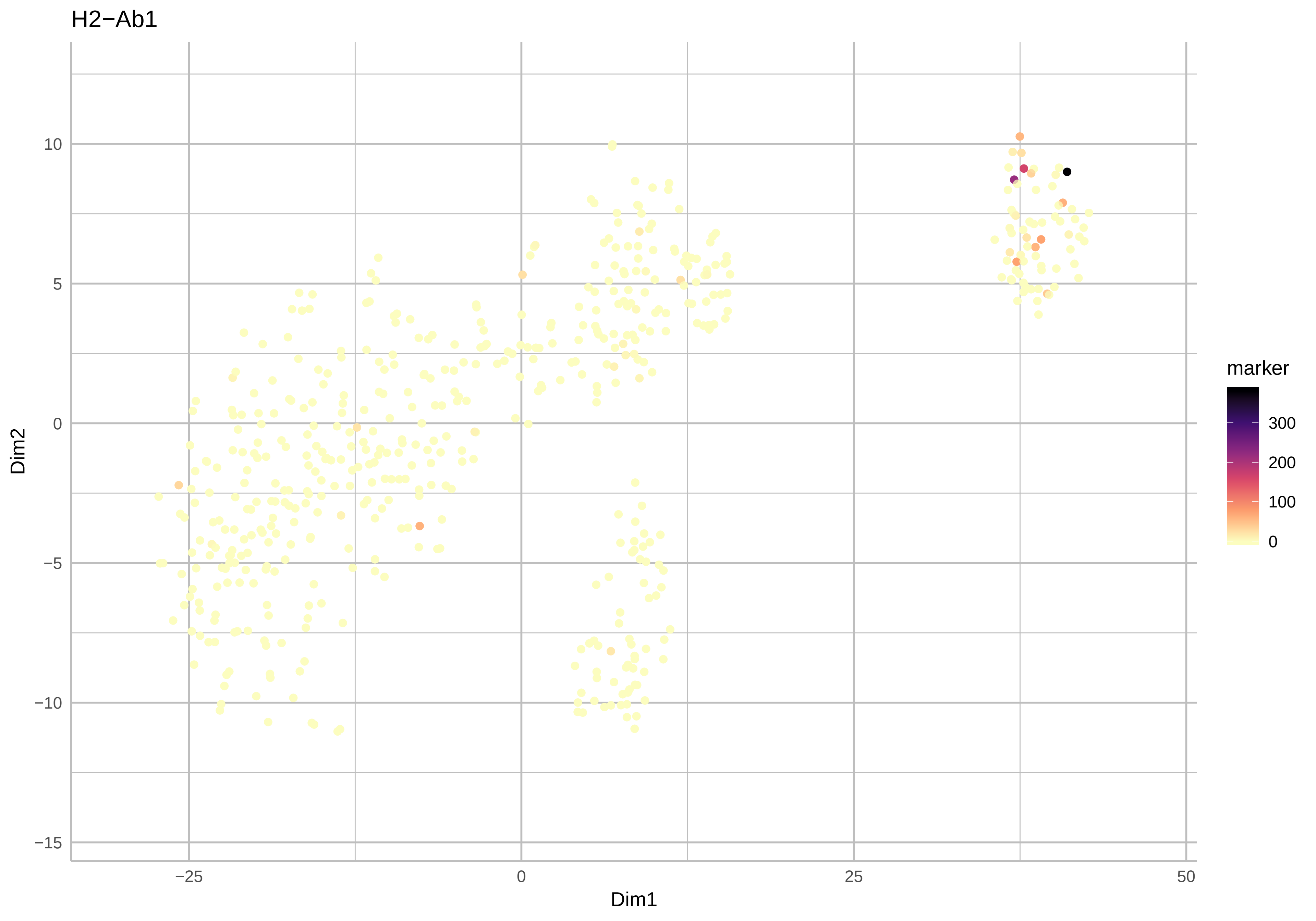
H2−Ab1
Dim2
Dim1
10
5
0
−5
−10
−15
−25
0
25
50
marker
300
200
100
0

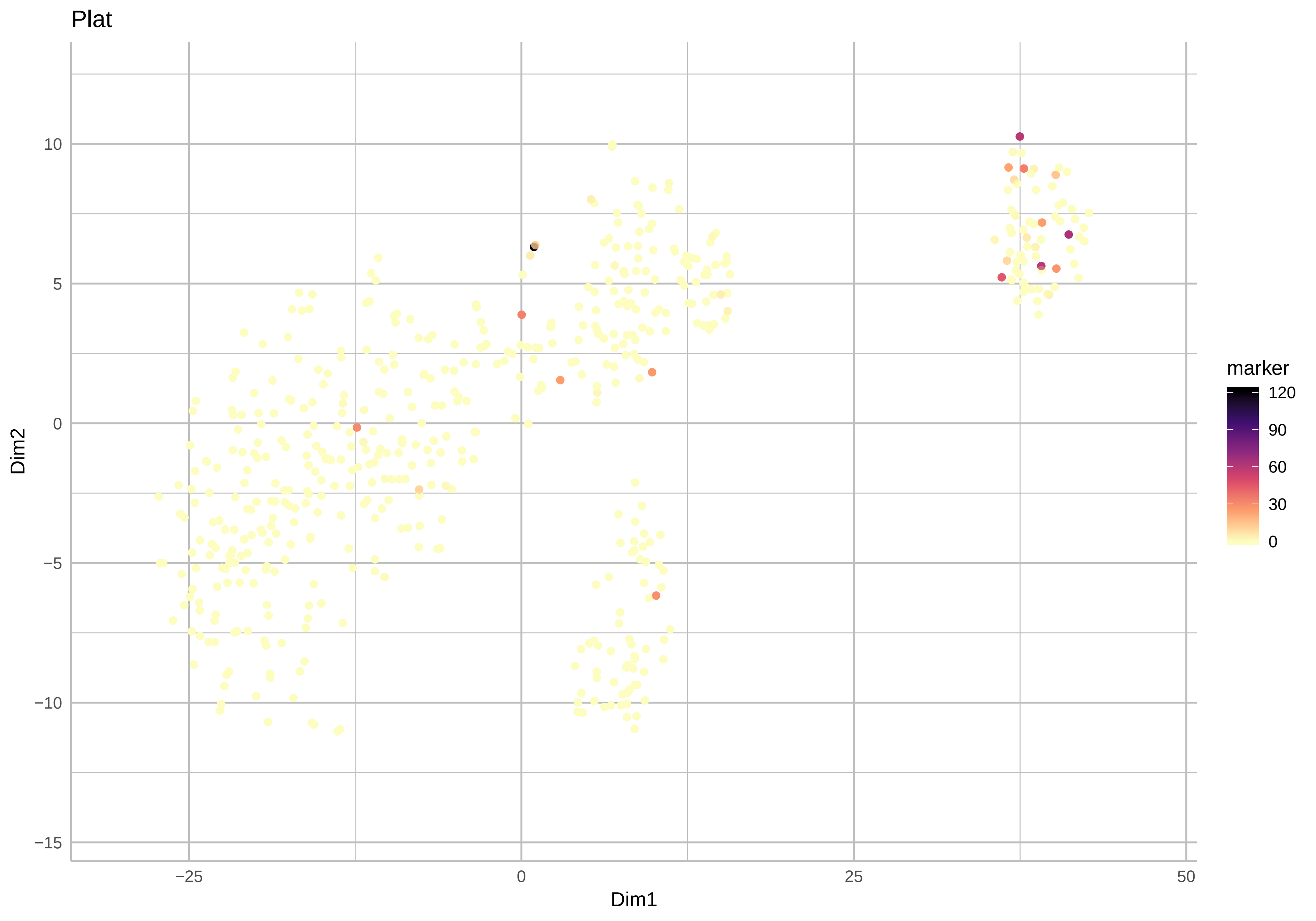

Plat
10
5
0
−5
−10
−15
Dim2
−25
0
25
50
Dim1
marker
120
90
60
30
0

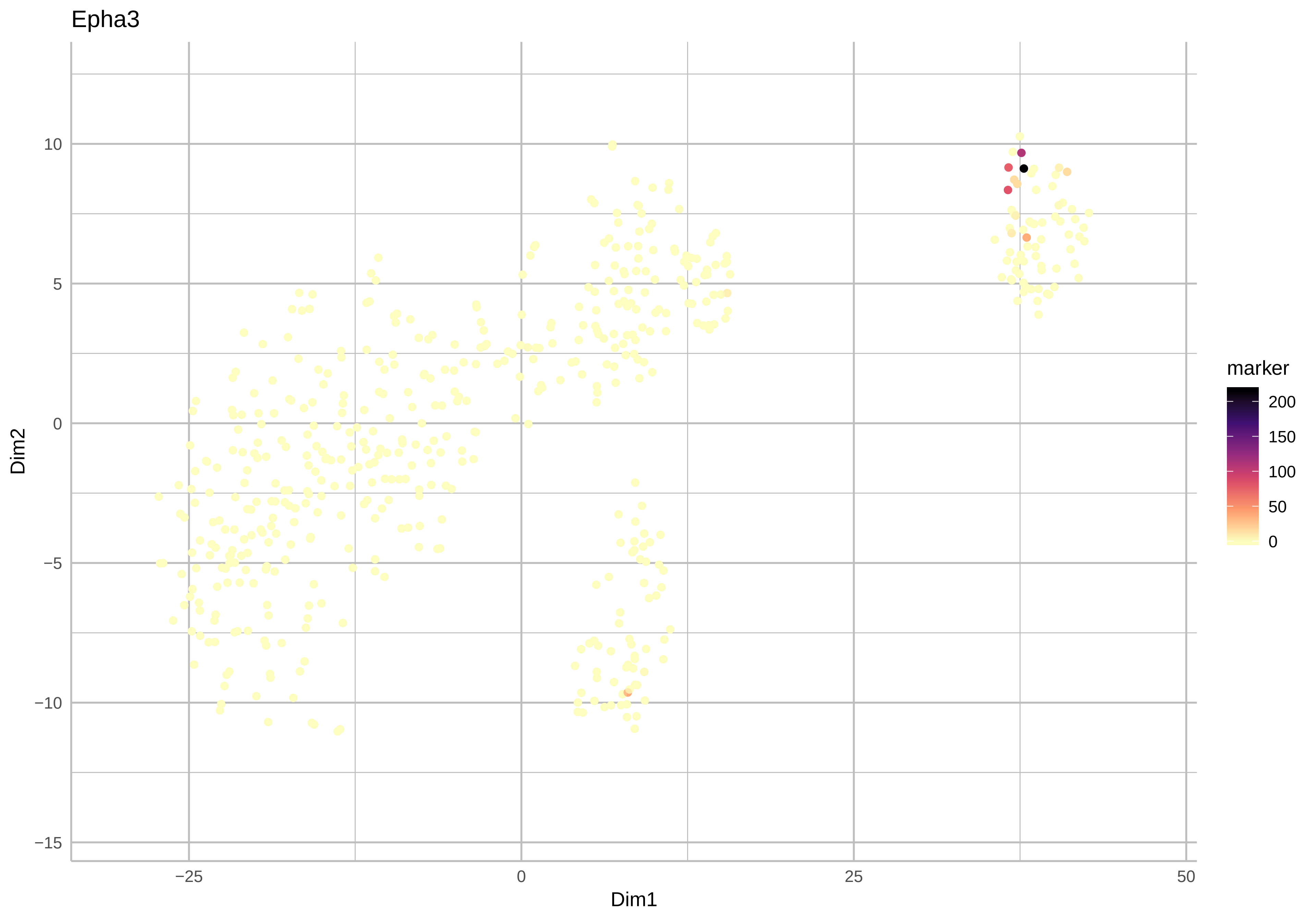

Epha3
10
5
0
−5
−10
−15
Dim2
−25
0
25
50
Dim1
marker
200
150
100
50
0

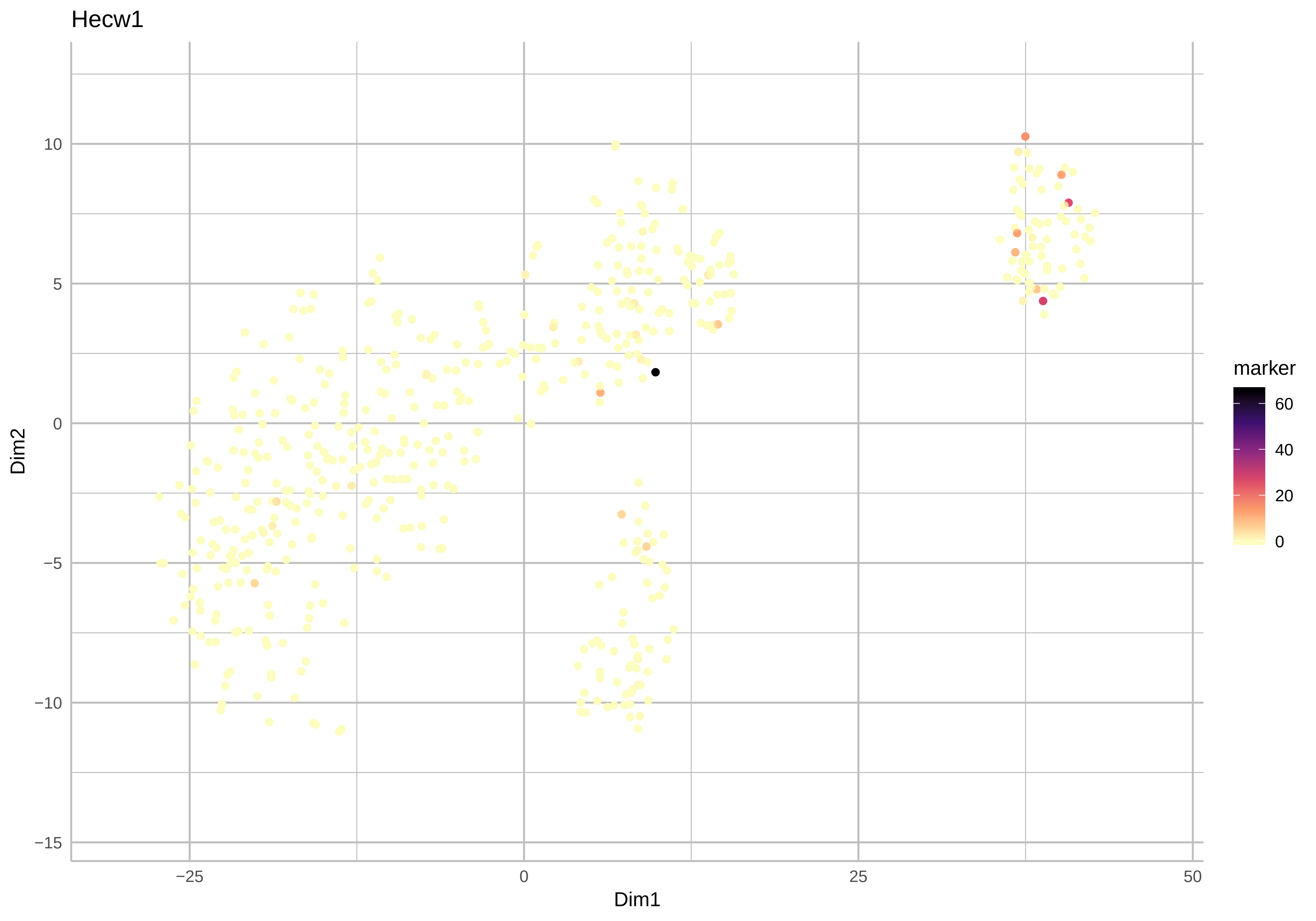

Hecw1
10
5
0
−5
−10
−15
Dim2
−25
0
25
50
Dim1
marker
60
40
20
0

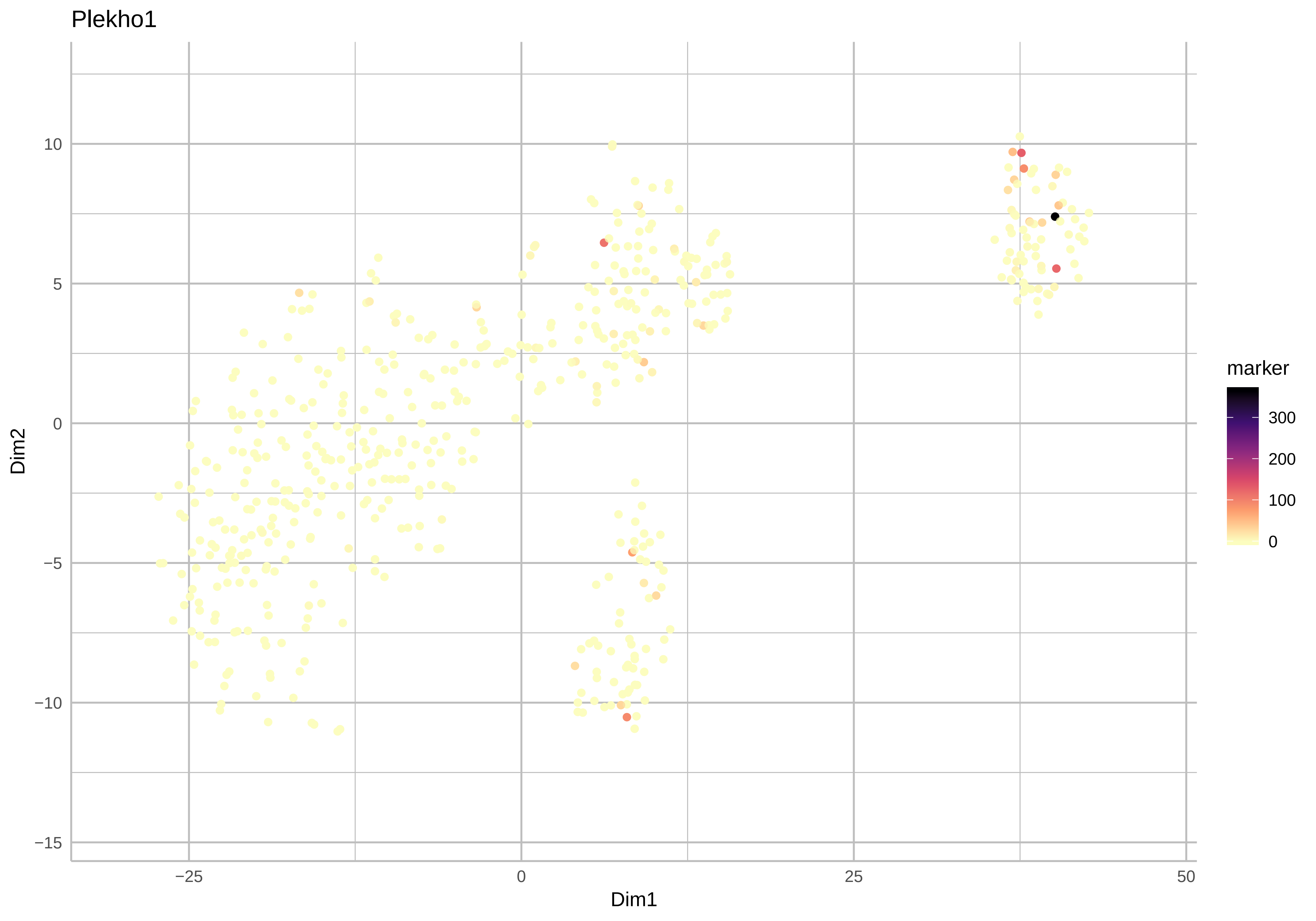

Plekho1
Dim2
Dim1
10
5
0
−5
−10
−15
−25
0
25
50
marker
300
200
100
0

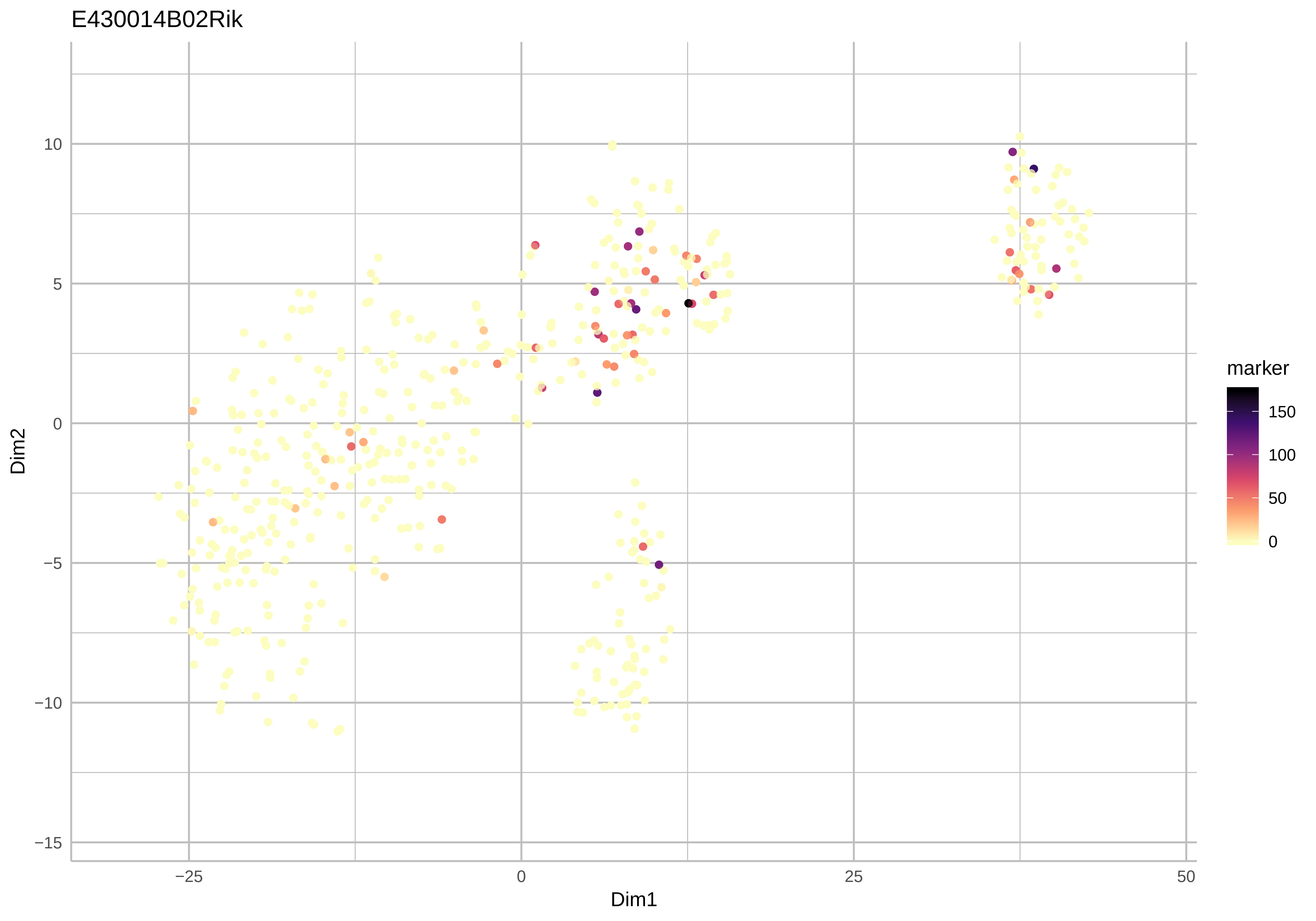

E430014B02Rik
Dim2
Dim1
10
5
0
−5
−10
−15
−25
0
25
50
marker
150
100
50
0

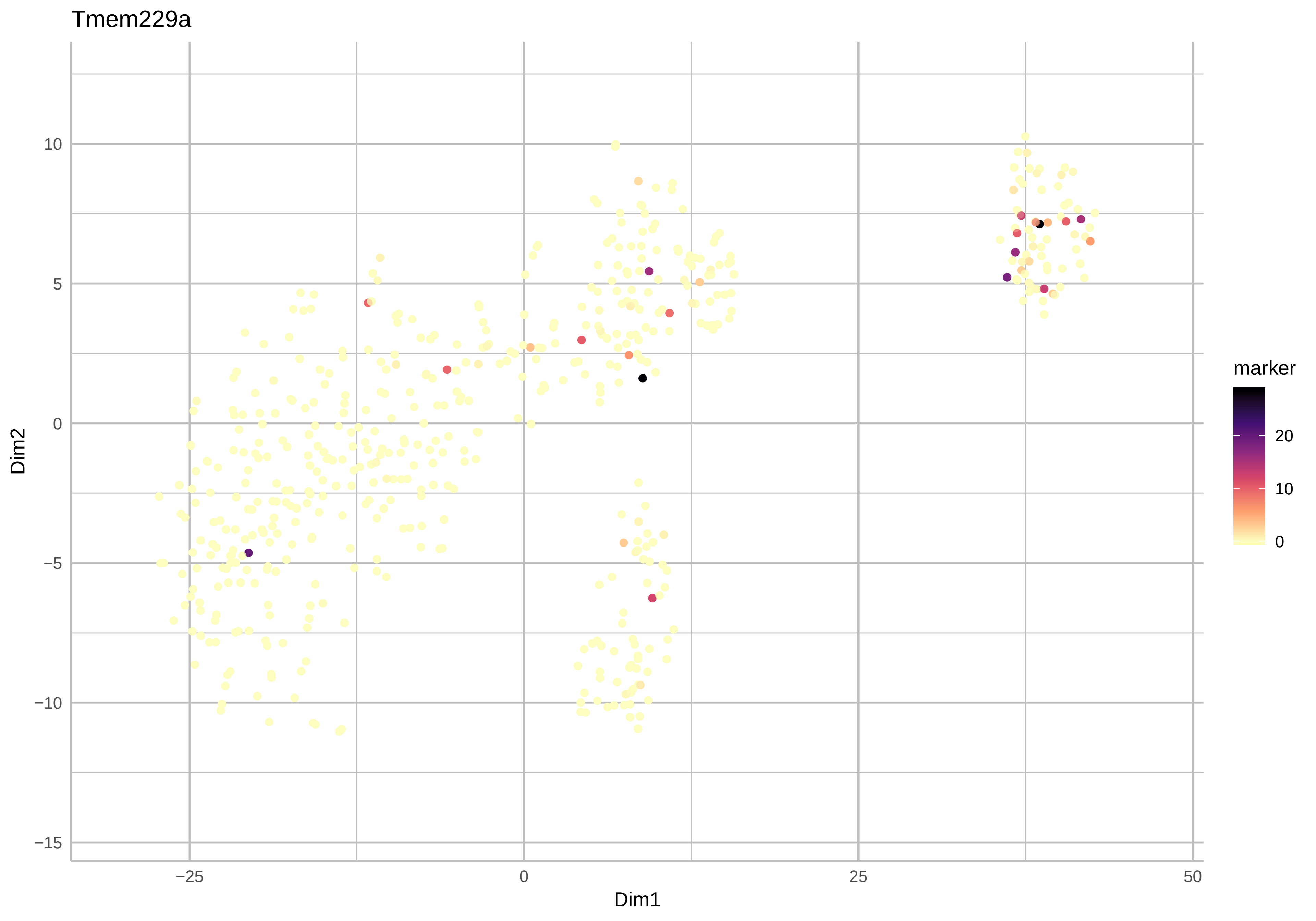

Tmem229a
10
5
0
−5
−10
−15
Dim2
−25
0
25
50
Dim1
marker
20
10
0

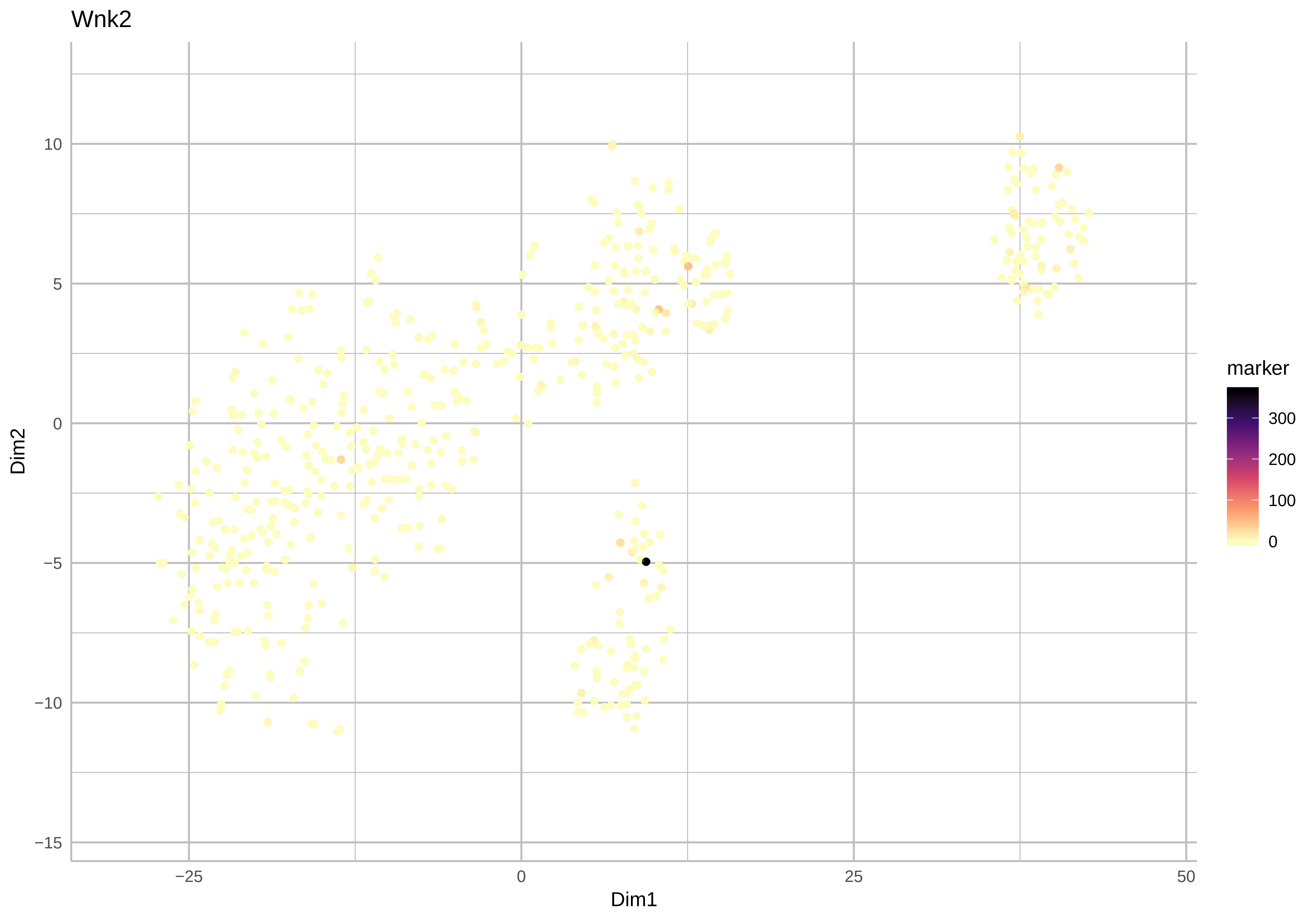

Wnk2
10
5
0
−5
−10
−15
Dim2
−25
0
25
50
Dim1
marker
300
200
100
0

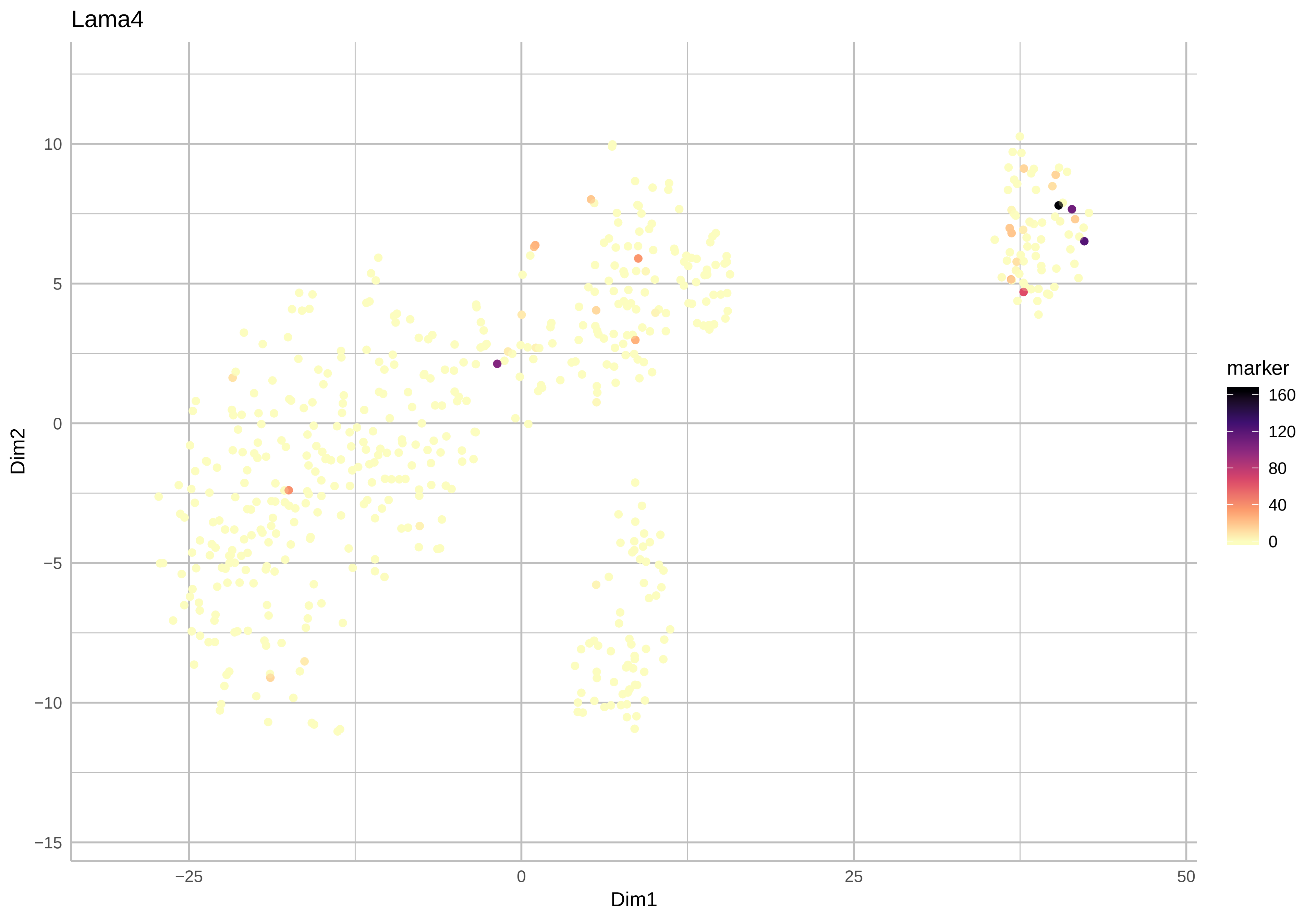
Lama4
10
5
0
−5
−10
−15
Dim2
−25
0
25
50
Dim1
marker
160
120
80
40
0

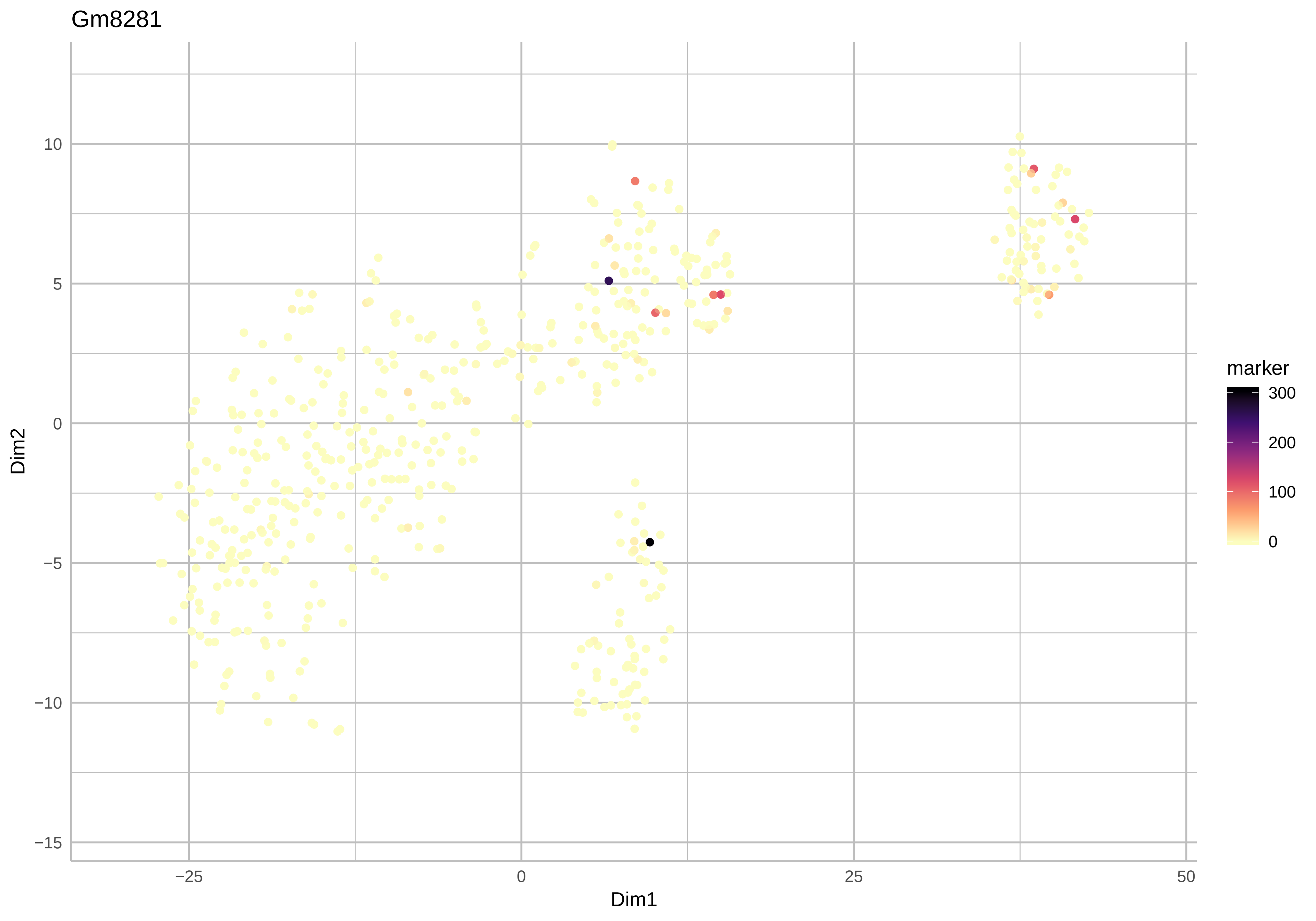

Gm8281
Dim2
Dim1
10
5
0
−5
−10
−15
−25
0
25
50
marker
300
200
100
0

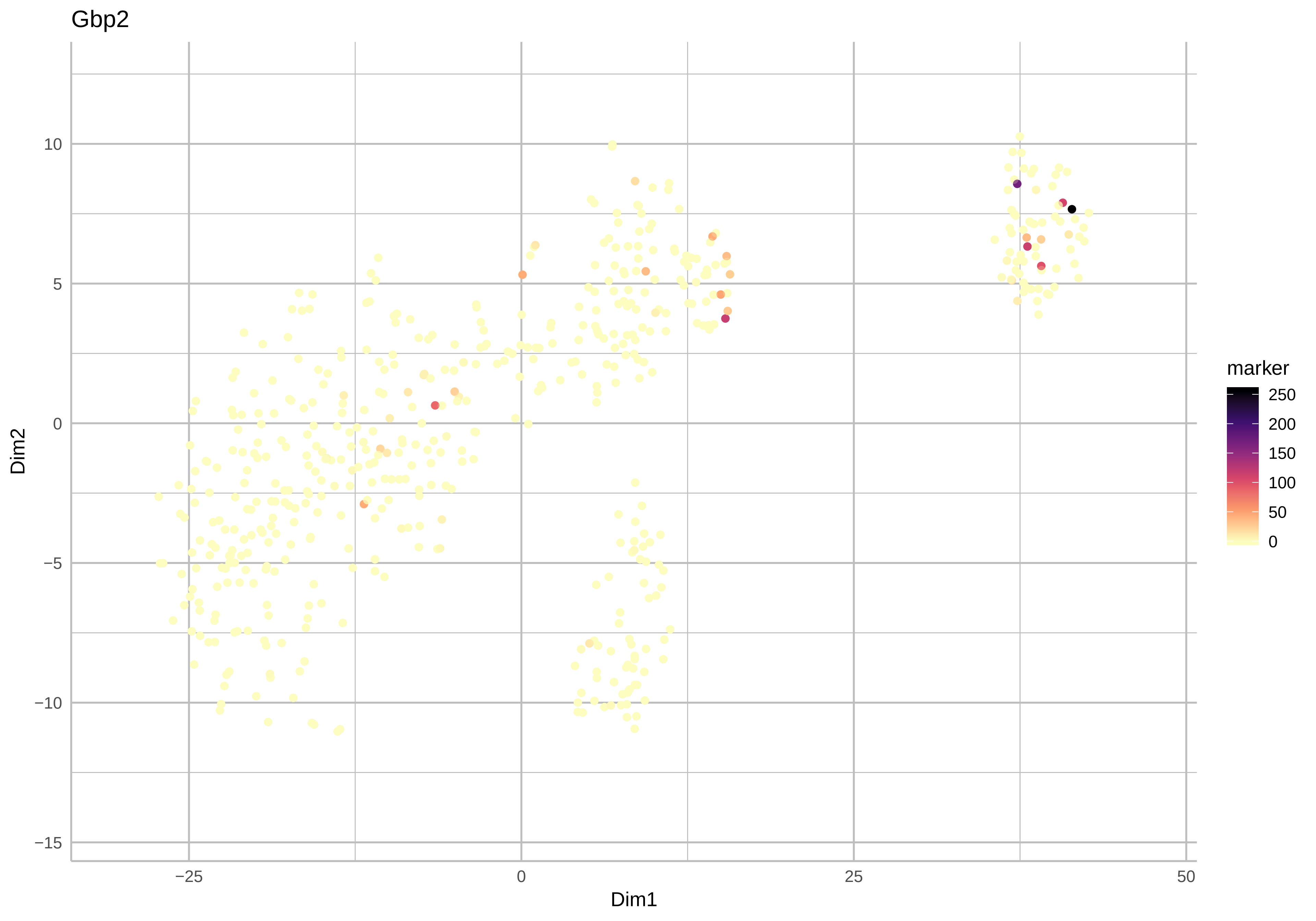
Gbp2
10
5
0
−5
−10
−15
Dim2
−25
0
25
50
Dim1
marker
250
200
150
100
50
0

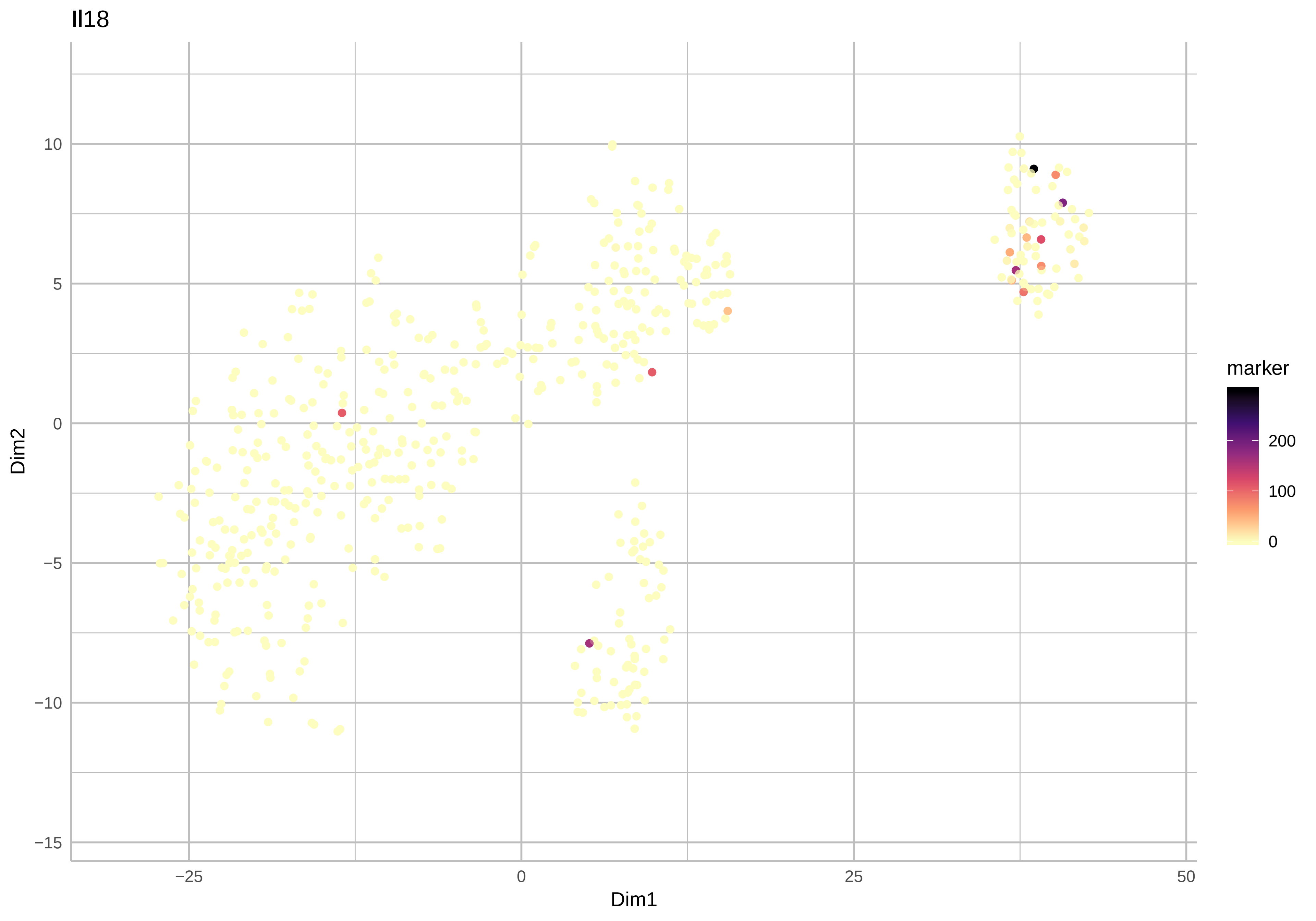

Il18
10
5
0
−5
−10
−15
Dim2
−25
0
25
50
Dim1
marker
200
100
0

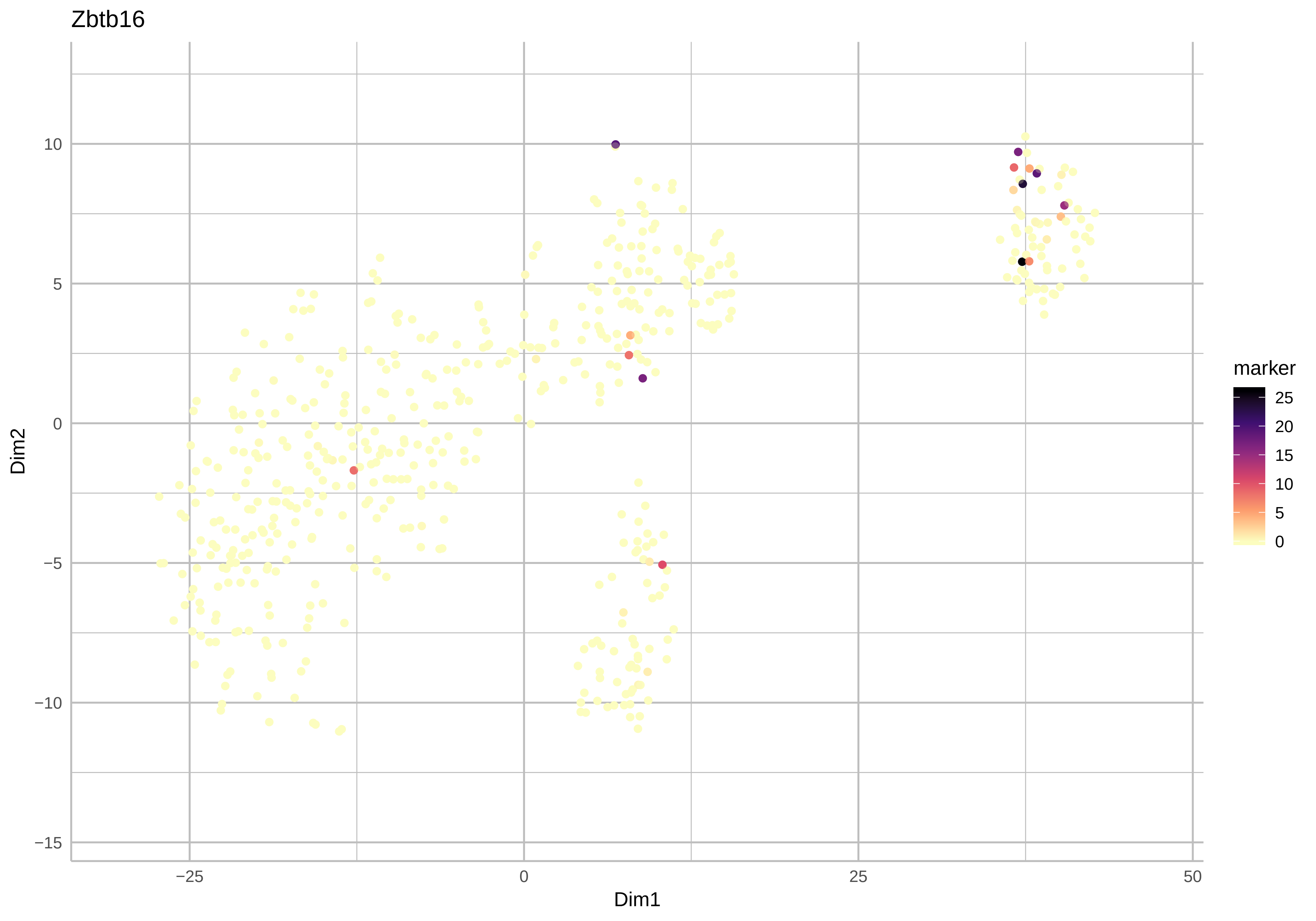

Zbtb16
Dim2
Dim1
10
5
0
−5
−10
−15
−25
0
25
50
marker
25
20
15
10
5
0

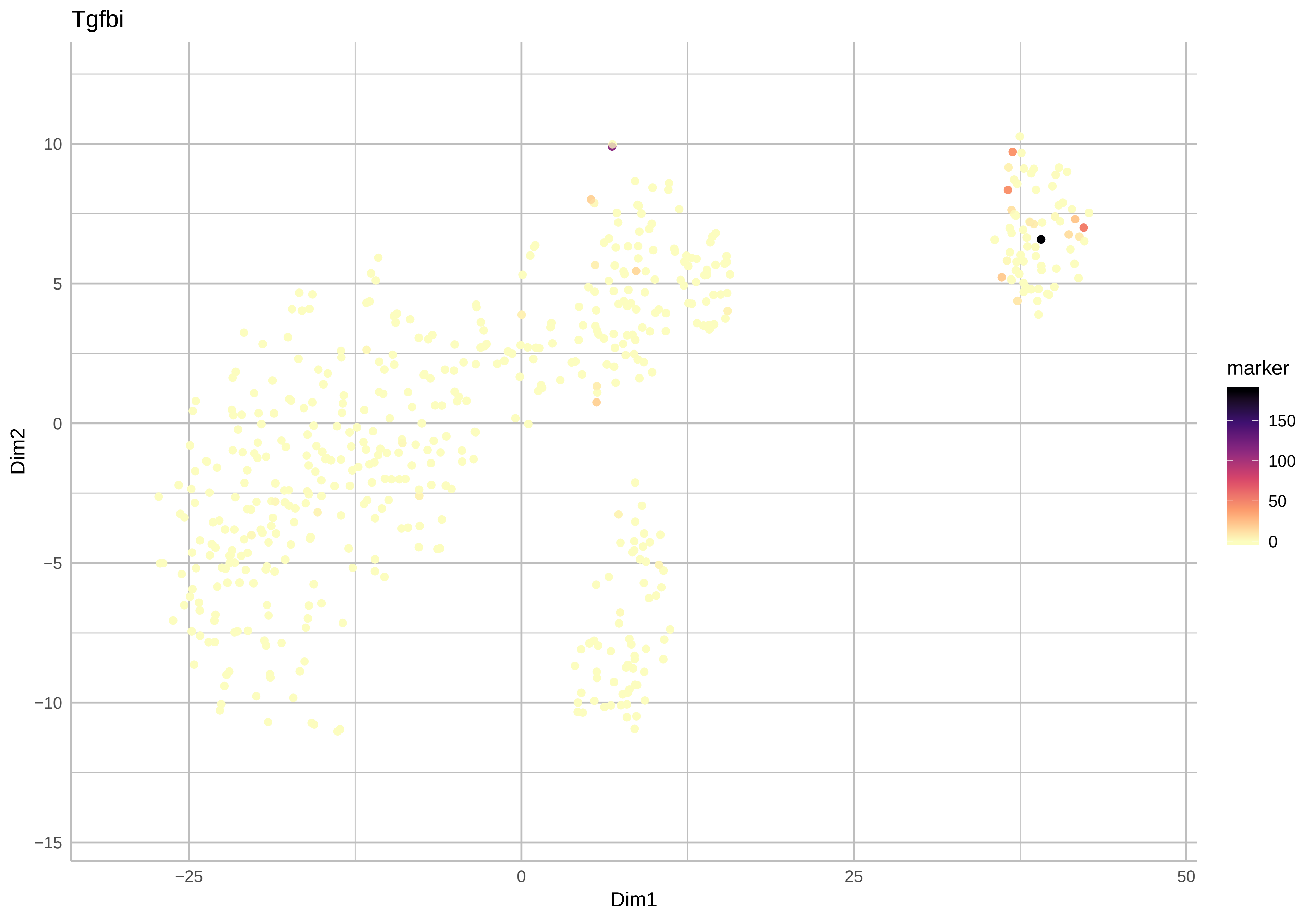

Tgfbi
Dim2
Dim1
marker
150
100
50
0
10
5
0
−5
−10
−15
−25
0
25
50

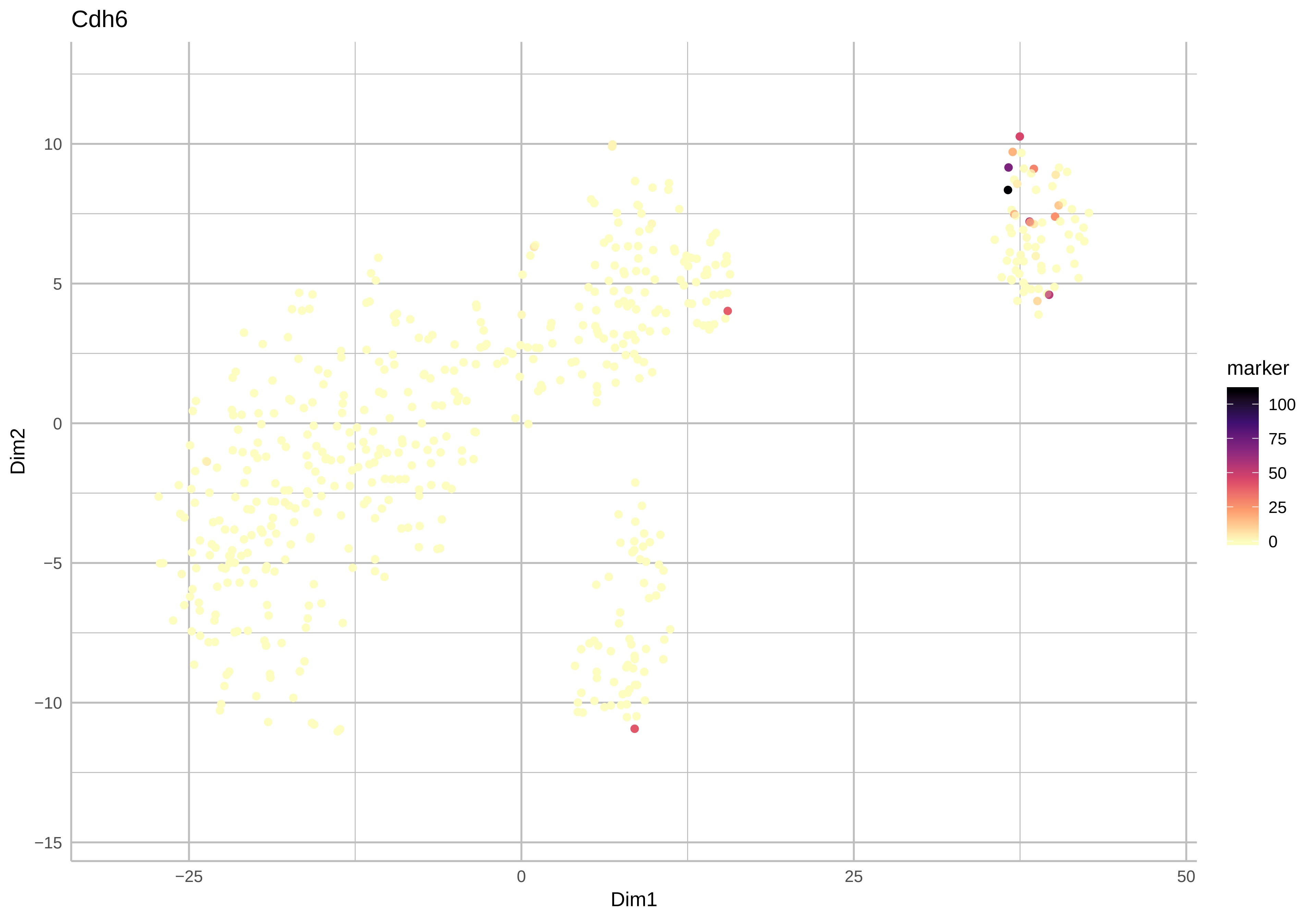

Cdh6
marker
100
75
50
25
0
Dim2
Dim1
10
5
0
−5
−10
−15
−25
0
25
50

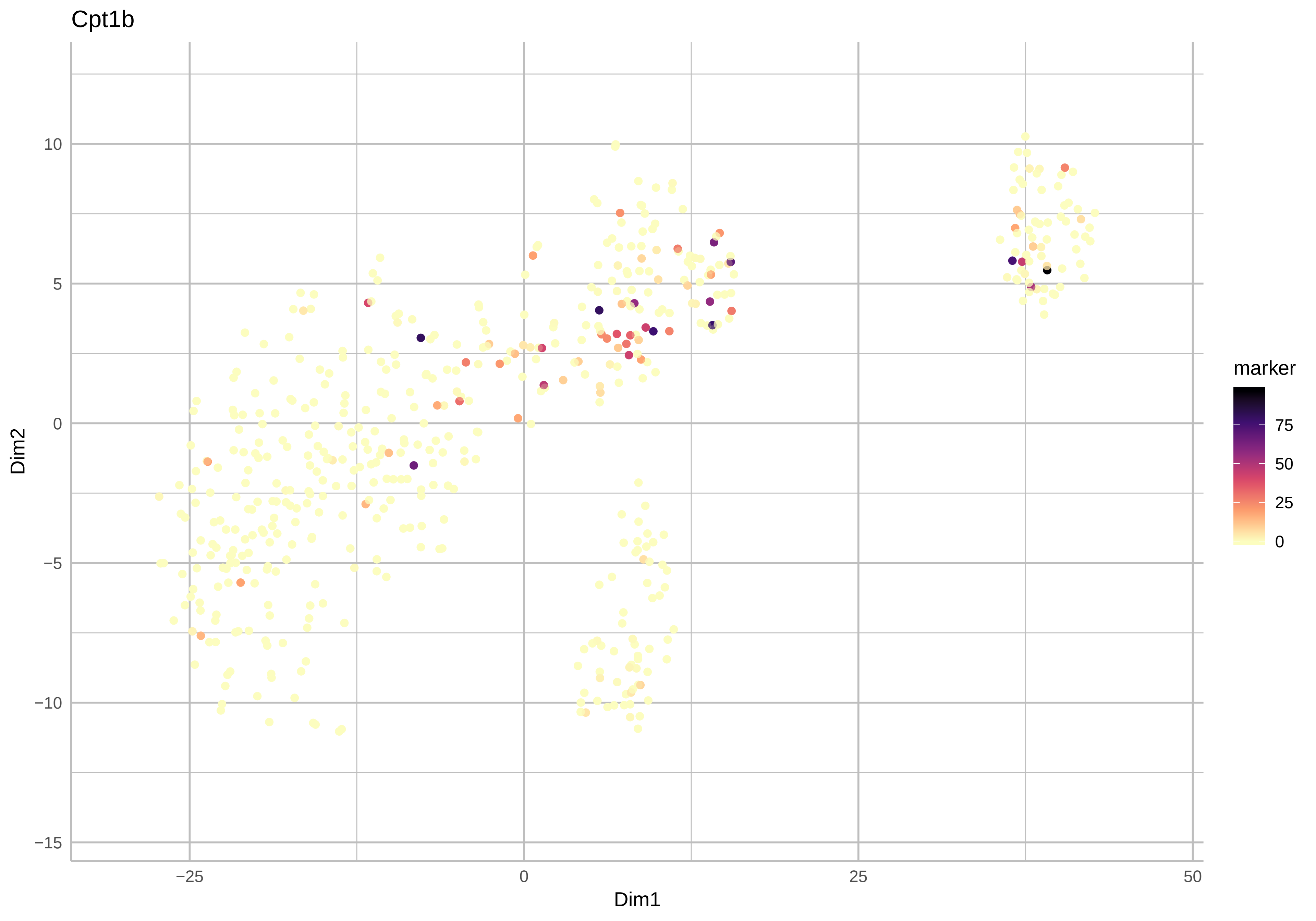

Cpt1b
Dim2
Dim1
10
5
0
−5
−10
−15
−25
0
25
50
marker
75
50
25
0

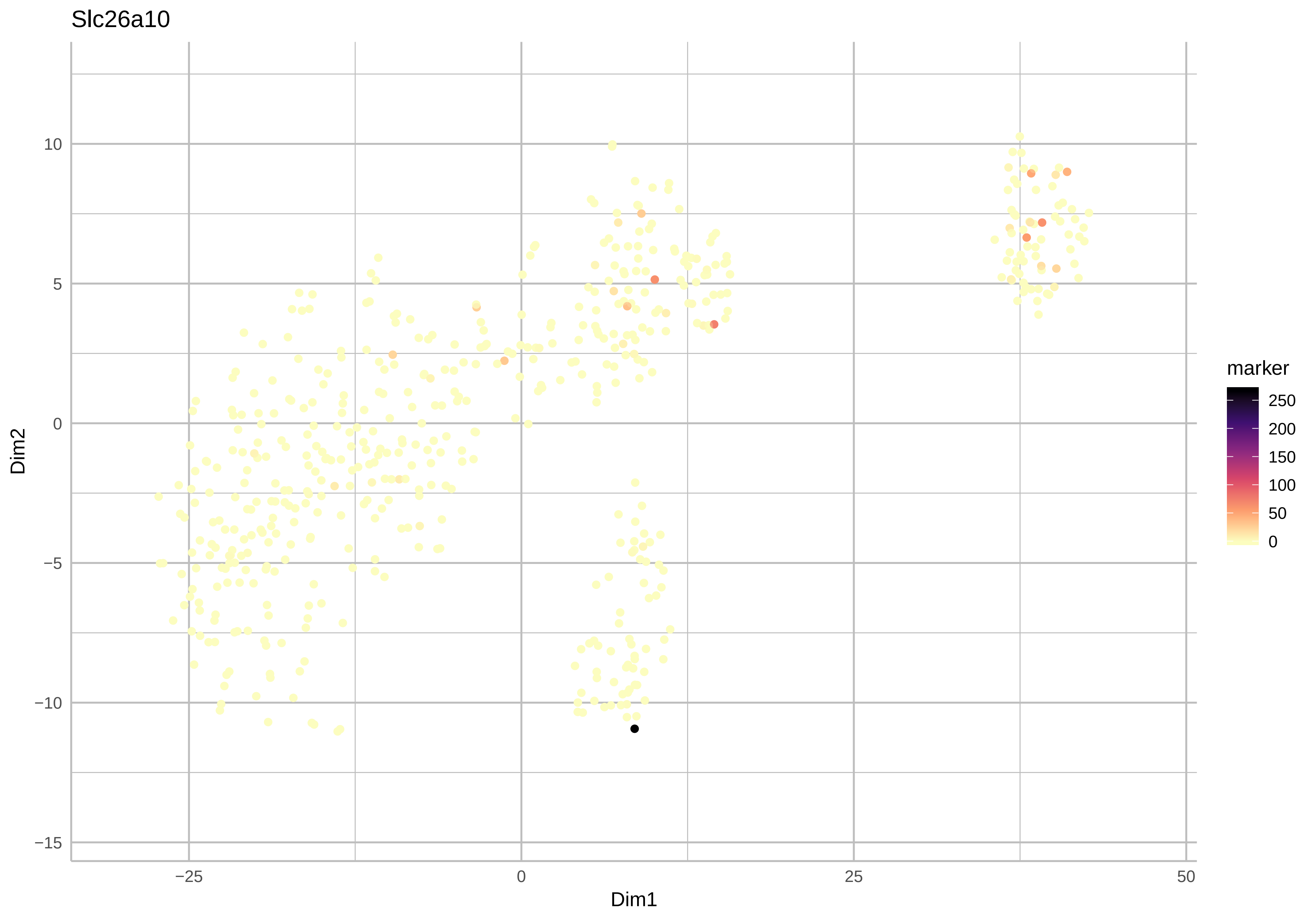
Slc26a10
Dim2
Dim1
10
5
0
−5
−10
−15
−25
0
25
50
marker
250
200
150
100
50
0

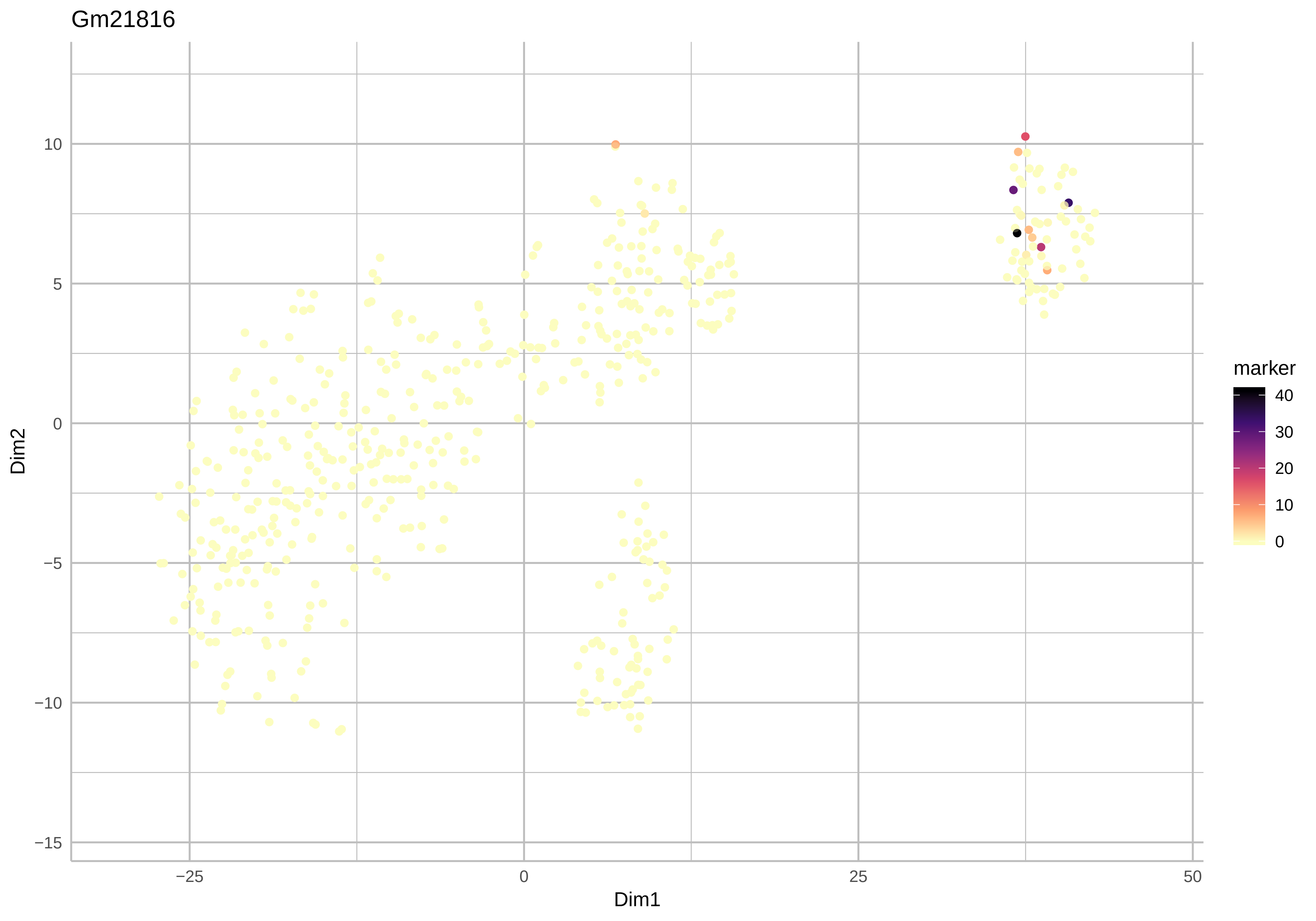

Gm21816
Dim2
Dim1
10
5
0
−5
−10
−15
−25
0
25
50
marker
40
30
20
10
0

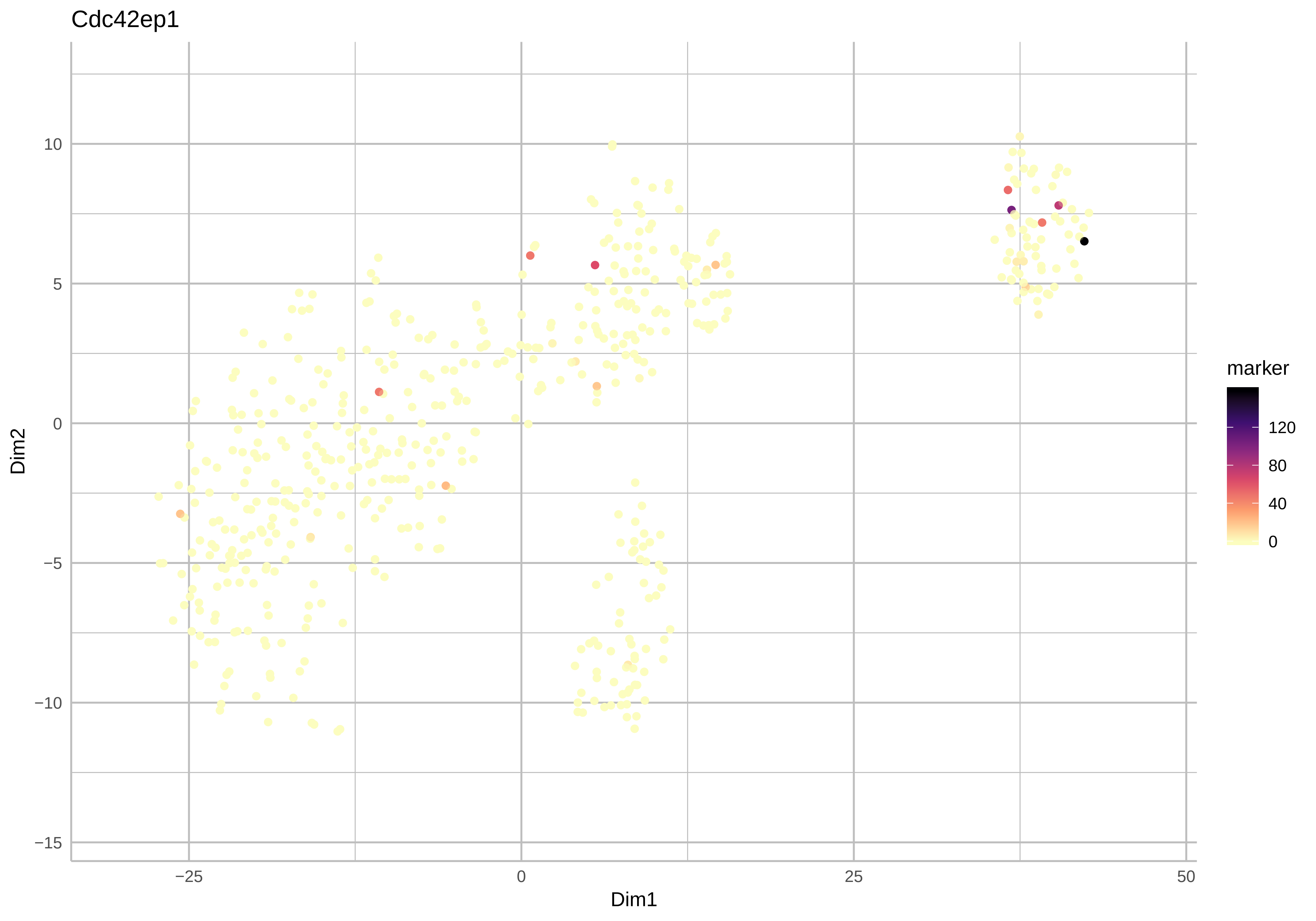

Cdc42ep1
10
5
0
−5
−10
−15
−25
0
25
50
Dim1
Dim2
marker
120
80
40
0

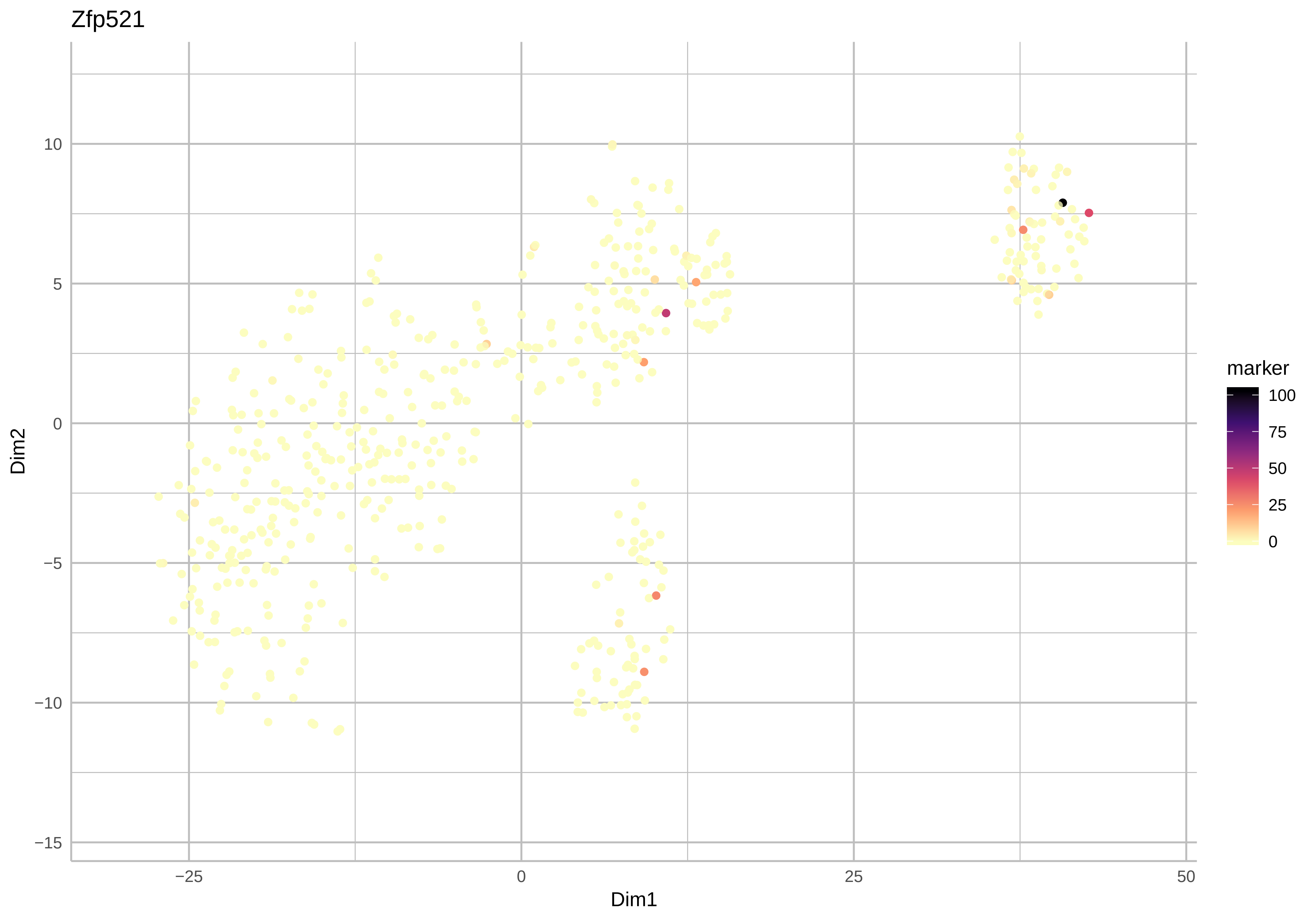
Zfp521
Dim2
Dim1
10
5
0
−5
−10
−15
−25
0
25
50
marker
100
75
50
25
0

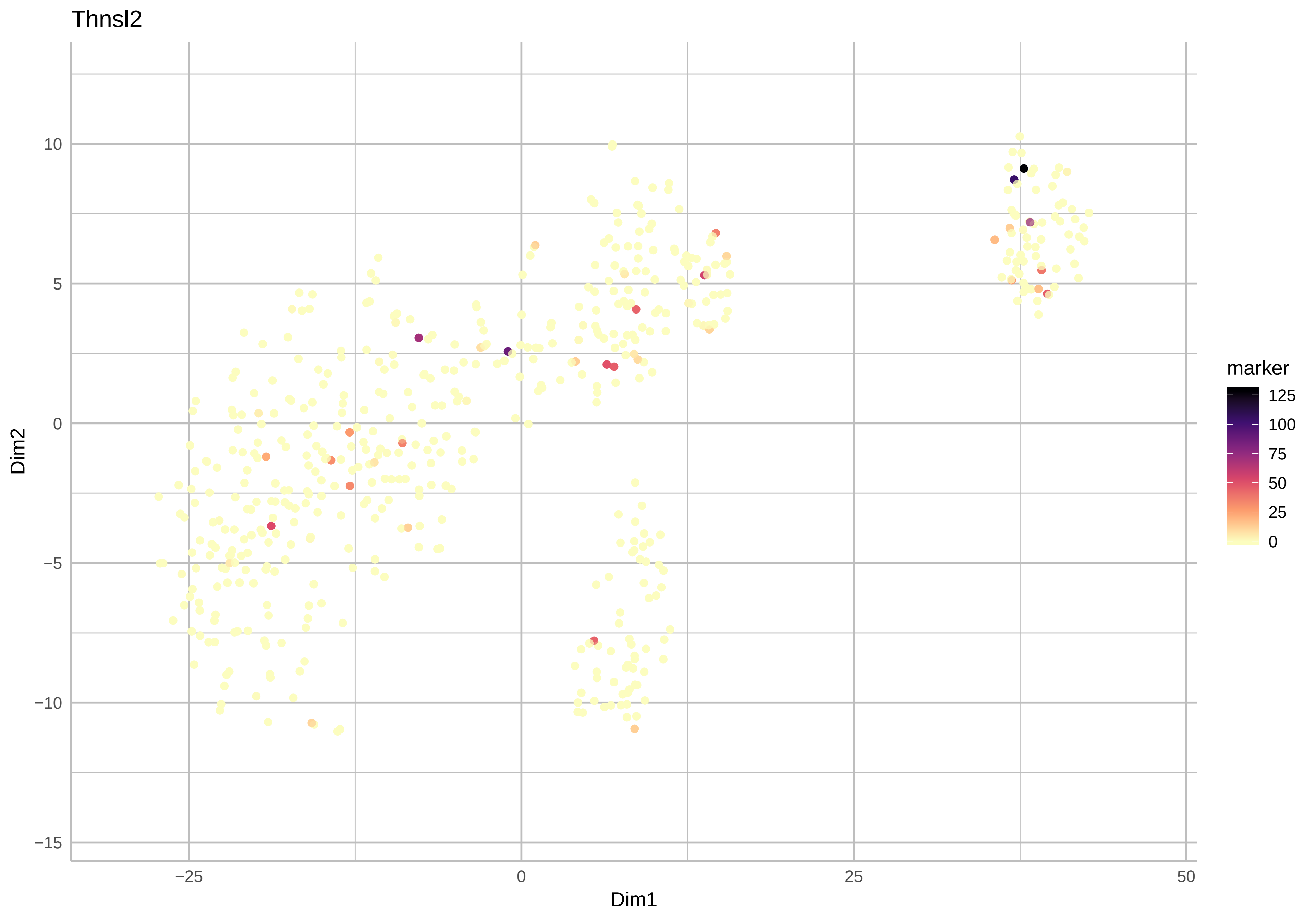
Thnsl2
Dim2
Dim1
marker
125
100
75
50
25
0
10
5
0
−5
−10
−15
−25
0
25
50

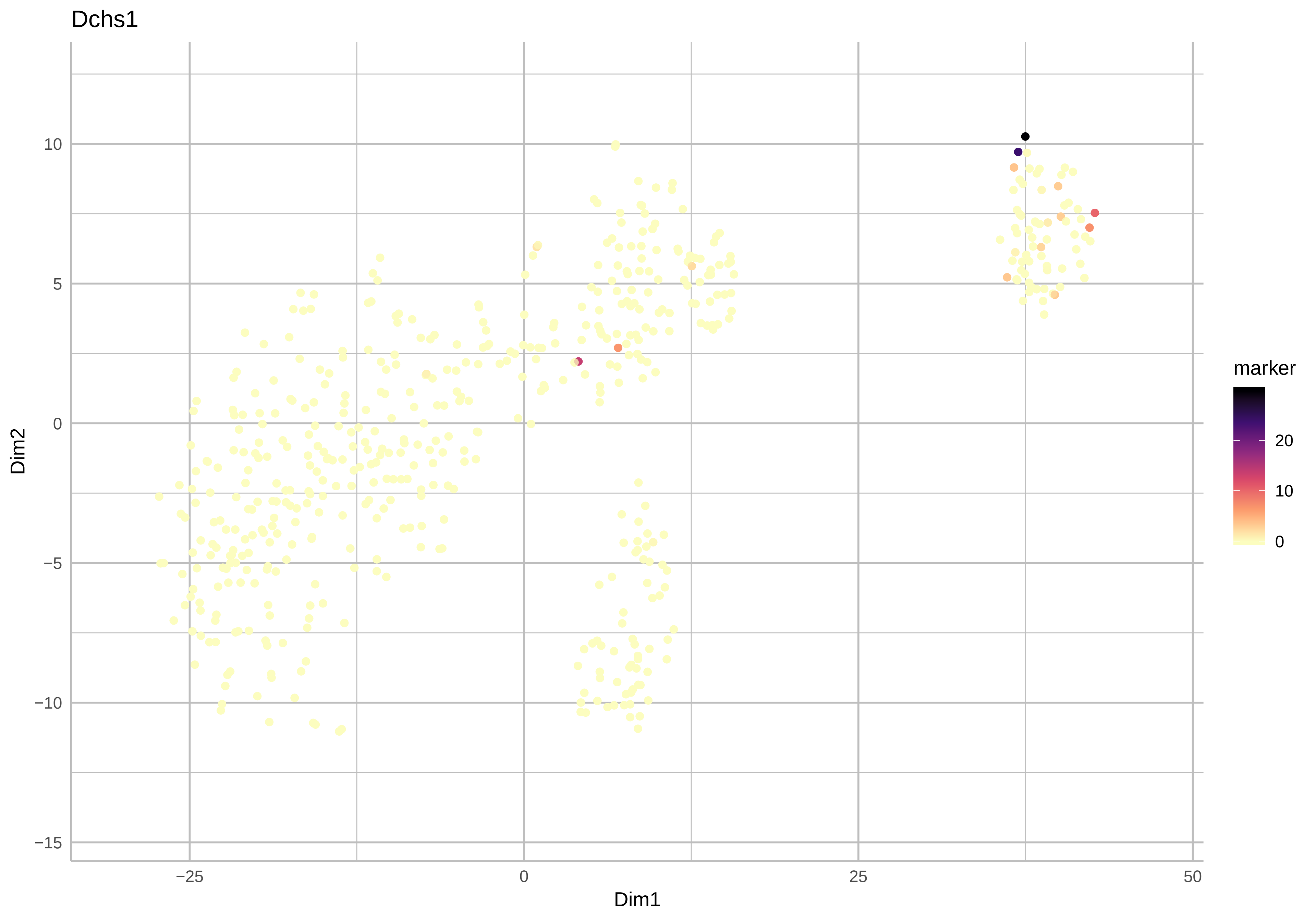

Dchs1
Dim2
Dim1
marker
20
10
0
10
5
0
−5
−10
−15
−25
0
25
50

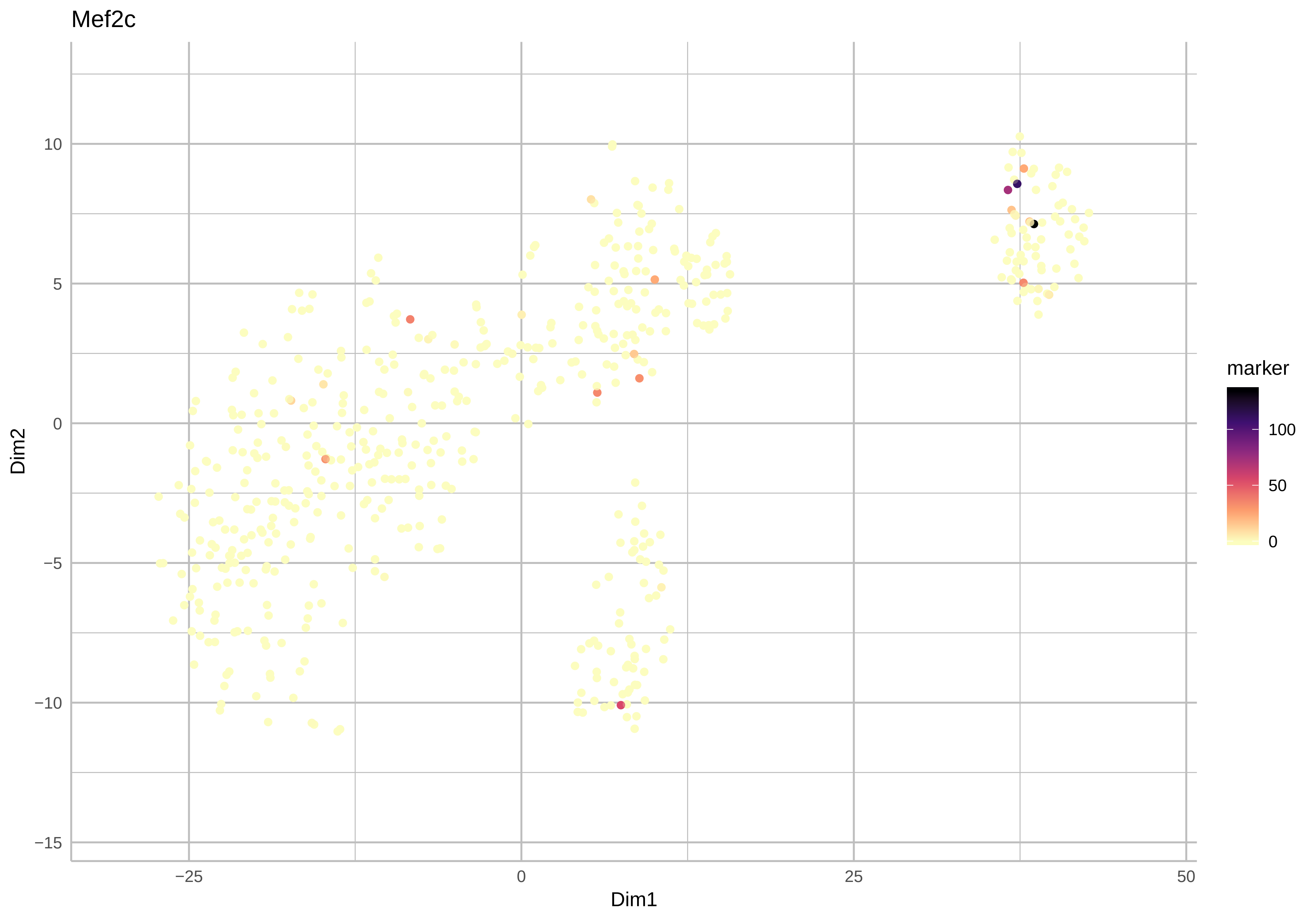

Mef2c
Dim2
Dim1
marker
100
50
0
10
5
0
−5
−10
−15
−25
0
25
50

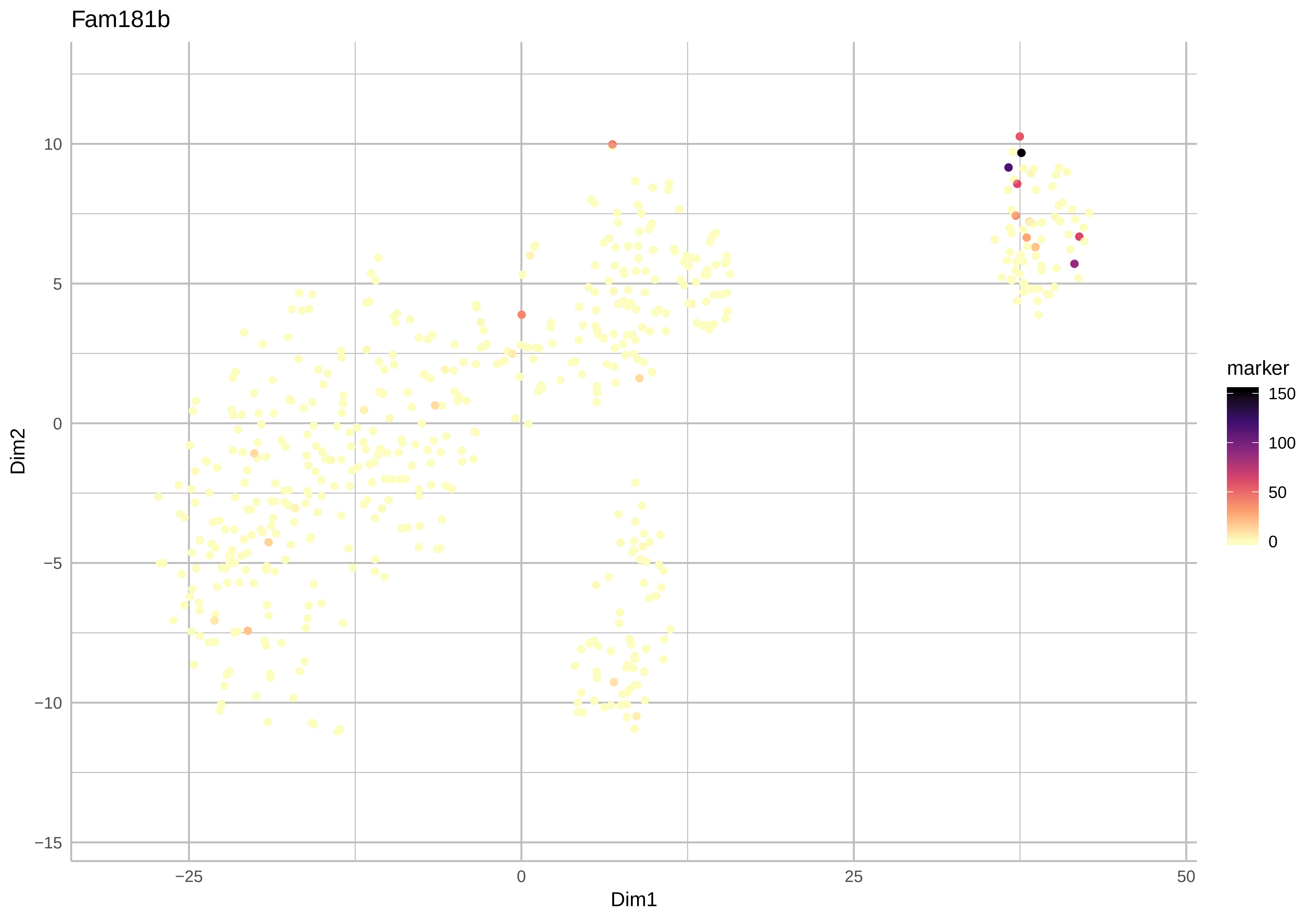

Fam181b
Dim2
Dim1
10
5
0
−5
−10
−15
−25
0
25
50
marker
150
100
50
0

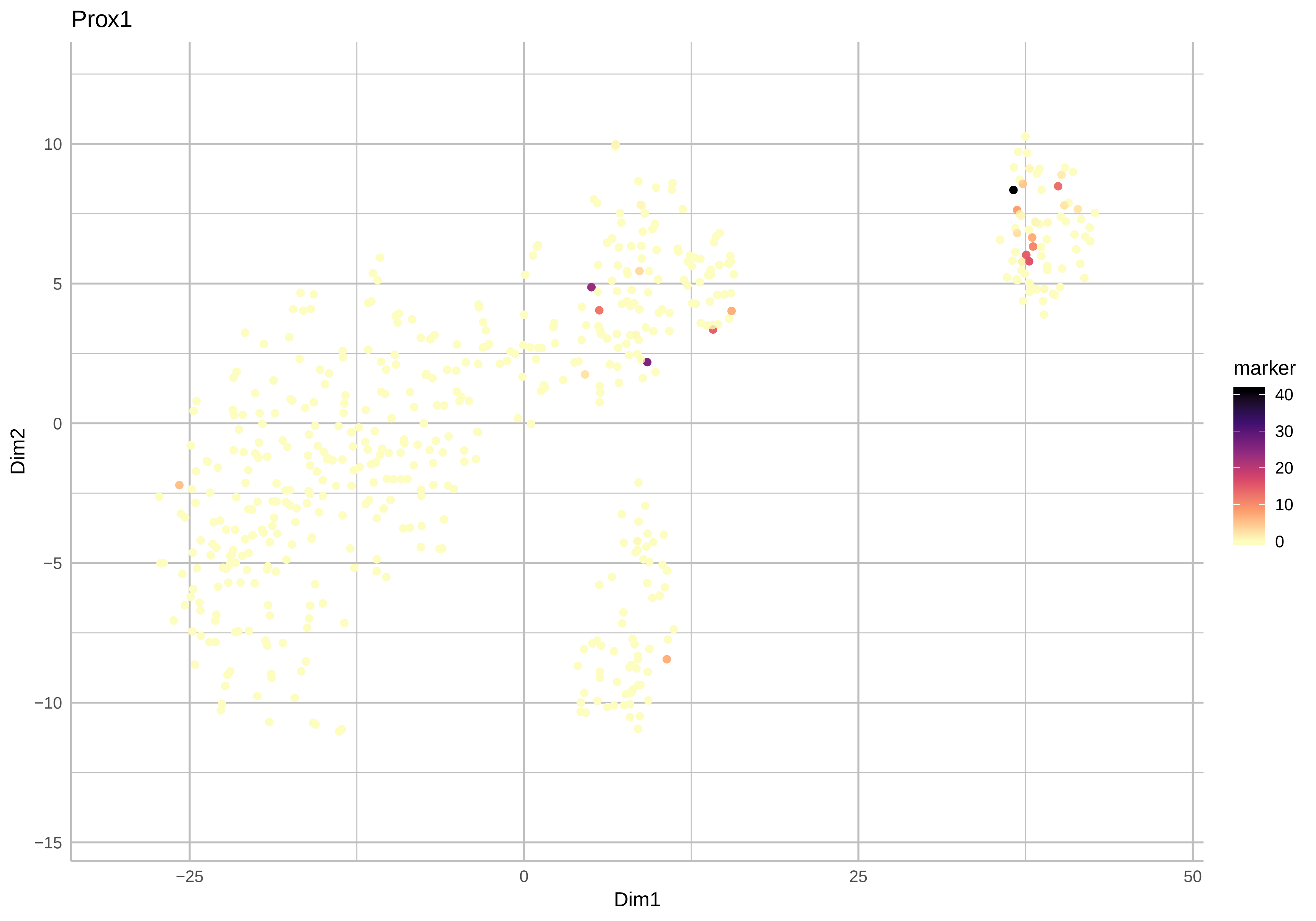
Prox1
Dim2
Dim1
10
5
0
−5
−10
−15
−25
0
25
50
marker
40
30
20
10
0

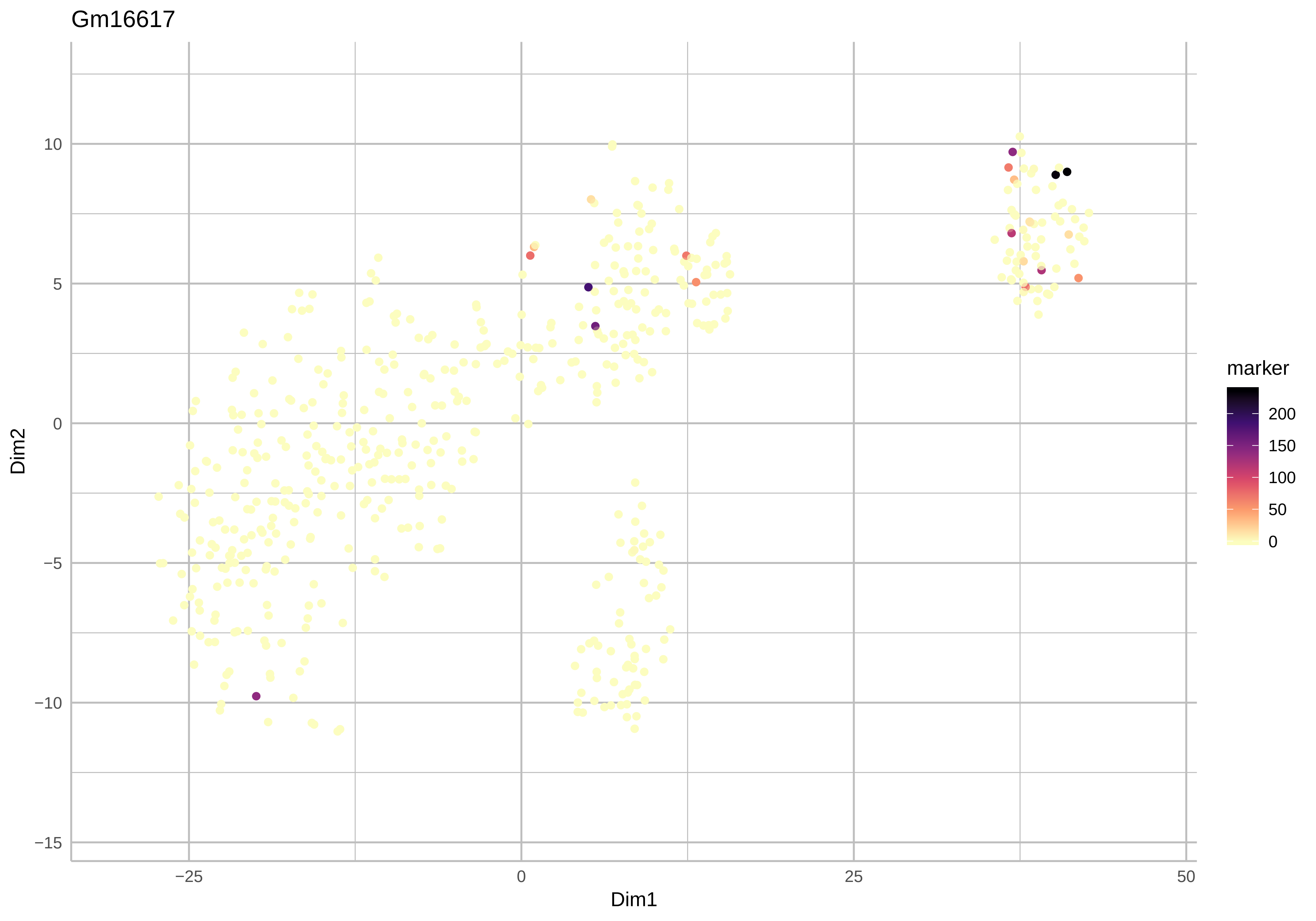

Gm16617
Dim2
Dim1
10
5
0
−5
−10
−15
−25
0
25
50
marker
200
150
100
50
0
