## Supplementary material for "Single-cell transcriptomics, scRNA-Seq and C1 CAGE discovered distinct phases of pluripotency during naïve-to-primed conversion in mice": 1_Mapping_and_preprocessing_for_Variant_Calling.sh

```
######   STAR Indexing ########

#!/bin/bash

STAR --runMode genomeGenerate --genomeDir ./b6_index --genomeFastaFiles UCSC_mm9.woYwRwMSMv5nSNP_b6_chr.fa --runThreadN 2

STAR --runMode genomeGenerate --genomeDir ./msm_index --genomeFastaFiles UCSC_mm9.woYwRwMSMv5nSNP_msm_chr.fa --runThreadN 2


#####  STAR Mapping  #########

##### 1 Example File ######

ID_array=(
'C10_CGAGGCTG-AGAGTAGA'
)

for ID in "${ID_array[@]}"

do

STAR --runThreadN 2 --genomeDir ./b6_index --readFilesIn /Volumes/ES_1d/1772-116-260_${ID}_L003_R1_001.fastq.gz /Volumes/ES_1d/1772-116-260_${ID}_L003_R2_001.fastq.gz --sjdbGTFfile /Users/STAR/genes.gtf --readFilesCommand gzcat --outFileNamePrefix /Volumes/ES_1d/ES_d1_b6_${ID}_

done


ID_array=(
'C10_CGAGGCTG-AGAGTAGA'
)

for ID in "${ID_array[@]}"

do

STAR --runThreadN 2 --genomeDir ./msm_index --readFilesIn /Volumes/ES_1d/1772-116-260_${ID}_L003_R1_001.fastq.gz /Volumes/ES_1d/1772-116-260_${ID}_L003_R2_001.fastq.gz --sjdbFileChrStartEnd /Volumes/SJ_out_tab/*SJ.out.tab --readFilesCommand gzcat --outFileNamePrefix /Volumes/SJ_out_tab/ES_d1_msm_${ID}_

done

######################################

#########       GATK variant detection for b6   #############


ID_array=(
'C10_CGAGGCTG-AGAGTAGA'
)

for ID in "${ID_array[@]}"
do

java -Xmx2g -jar picard.jar AddOrReplaceReadGroups I=ES_d0_b6_${ID}_Aligned.out.sam O=ES_d0_b6_${ID}_Aligned.out.bam SO=coordinate RGID=ES_d0_b6_${ID}_Aligned.out.bam RGLB=ES_d0_b6_${ID}_Aligned.out.bam RGPL=Illumina RGPU=run RGSM=ES_d0_b6_${ID}_Aligned.out.bam

java -Xmx2g -jar picard.jar MarkDuplicates I=ES_d0_b6_${ID}_Aligned.out.bam O=ES_d0_b6_${ID}_Aligned.out.rmdup.bam CREATE_INDEX=true METRICS_FILE=ES_d0_b6_${ID}_Aligned.out.rmdup.bam_metrics REMOVE_DUPLICATES=true  ASSUME_SORTED=true VALIDATION_STRINGENCY=LENIENT

java -Xmx4g -jar GenomeAnalysisTK.jar -T SplitNCigarReads -R UCSC_mm9.woYwRwMSMv5nSNP_b6_chr.fa -I ES_d0_b6_${ID}_Aligned.out.rmdup.bam -o /home/splitNCigarRead_bam/B6/${ID}.b6.rmDuplicates.split.bam -rf ReassignOneMappingQuality -RMQF 255 -RMQT 60 -U ALLOW_N_CIGAR_READS

done

#####  GATK variant detection for Msm  ############


ID_array=(
'C10_CGAGGCTG-AGAGTAGA'
)

for ID in "${ID_array[@]}"
do

java -Xmx2g -jar picard.jar AddOrReplaceReadGroups I=ES_d0_msm_${ID}_Aligned.out.sam O=ES_d0_msm_${ID}_Aligned.out.bam SO=coordinate RGID=ES_d0_msm_${ID}_Aligned.out.bam RGLB=ES_d0_msm_${ID}_Aligned.out.bam RGPL=Illumina RGPU=run RGSM=ES_d0_msm_${ID}_Aligned.out.bam


java -Xmx2g -jar picard.jar MarkDuplicates I=ES_d0_msm_${ID}_Aligned.out.bam O=ES_d0_msm_${ID}_Aligned.out.rmdup.bam CREATE_INDEX=true METRICS_FILE=ES_d0_msm_${ID}_Aligned.out.rmdup.bam_metrics REMOVE_DUPLICATES=true  ASSUME_SORTED=true VALIDATION_STRINGENCY=LENIENT


java -Xmx4g -jar GenomeAnalysisTK.jar -T SplitNCigarReads -R UCSC_mm9.woYwRwMSMv5nSNP_msm_chr.fa -I ES_d0_msm_${ID}_Aligned.out.rmdup.bam -o /home/splitNCigarRead_bam/MSM/${ID}.msm.rmDuplicates.split.bam -rf ReassignOneMappingQuality -RMQF 255 -RMQT 60 -U ALLOW_N_CIGAR_READS

done
```
