## Supplementary material for "Single-cell transcriptomics, scRNA-Seq and C1 CAGE discovered distinct phases of pluripotency during naïve-to-primed conversion in mice": 2_Variant_Calling_and_Allele_specific_read_counts_and_variant_classification.sh

```
#!/bin/bash


ID_array=(
'1772-116-259_A02'
'1772-116-259_A03'
'1772-116-259_A04'
'1772-116-259_A05'
'1772-116-259_A06'
'1772-116-259_A08'
'1772-116-259_A09'
'1772-116-259_A10'
'1772-116-259_B01'
'1772-116-259_B02'
'1772-116-259_B03'
'1772-116-259_B04'
'1772-116-259_B05'
'1772-116-259_B06'
'1772-116-259_B07'
'1772-116-259_B08'
'1772-116-259_B09'
'1772-116-259_B10'
'1772-116-259_B12'
'1772-116-259_C01'
'1772-116-259_C03'
'1772-116-259_C04'
'1772-116-259_C05'
'1772-116-259_C06'
'1772-116-259_C07'
'1772-116-259_C08'
'1772-116-259_C10'
'1772-116-259_C11'
'1772-116-259_C12'
'1772-116-259_D01'
'1772-116-259_D02'
'1772-116-259_D03'
'1772-116-259_D05'
'1772-116-259_D06'
'1772-116-259_D07'
'1772-116-259_D08'
'1772-116-259_D09'
'1772-116-259_D10'
'1772-116-259_D11'
'1772-116-259_D12'
'1772-116-259_E08'
'1772-116-259_E09'
'1772-116-259_E10'
'1772-116-259_E11'
'1772-116-259_F01'
'1772-116-259_F02'
'1772-116-259_F03'
'1772-116-259_F05'
'1772-116-259_F06'
'1772-116-259_F07'
'1772-116-259_F09'
'1772-116-259_F10'
'1772-116-259_F11'
'1772-116-259_F12'
'1772-116-259_G01'
'1772-116-259_G02'
'1772-116-259_G04'
'1772-116-259_G05'
'1772-116-259_G06'
'1772-116-259_G07'
'1772-116-259_G08'
'1772-116-259_G10'
'1772-116-259_G11'
'1772-116-259_G12'
'1772-116-259_H02'
'1772-116-259_H03'
'1772-116-259_H05'
'1772-116-259_H08'
'1772-116-259_H10'
'1772-116-259_H11'
'1772-116-259_H12'
'1772-116-260_A01'
'1772-116-260_A02'
'1772-116-260_A04'
'1772-116-260_A05'
'1772-116-260_A06'
'1772-116-260_A07'
'1772-116-260_A08'
'1772-116-260_A09'
'1772-116-260_A10'
'1772-116-260_A11'
'1772-116-260_A12'
'1772-116-260_B01'
'1772-116-260_B04'
'1772-116-260_B05'
'1772-116-260_B06'
'1772-116-260_B07'
'1772-116-260_B08'
'1772-116-260_B09'
'1772-116-260_B10'
'1772-116-260_B11'
'1772-116-260_B12'
'1772-116-260_C01'
'1772-116-260_C03'
'1772-116-260_C04'
'1772-116-260_C05'
'1772-116-260_C06'
'1772-116-260_C07'
'1772-116-260_C08'
'1772-116-260_C09'
'1772-116-260_C10'
'1772-116-260_C11'
'1772-116-260_C12'
'1772-116-260_D01'
'1772-116-260_D02'
'1772-116-260_D03'
'1772-116-260_D05'
'1772-116-260_D06'
'1772-116-260_D07'
'1772-116-260_D08'
'1772-116-260_D10'
'1772-116-260_D11'
'1772-116-260_D12'
'1772-116-260_E01'
'1772-116-260_E02'
'1772-116-260_E04'
'1772-116-260_E05'
'1772-116-260_E06'
'1772-116-260_E07'
'1772-116-260_E08'
'1772-116-260_E10'
'1772-116-260_E11'
'1772-116-260_E12'
'1772-116-260_F01'
'1772-116-260_F02'
'1772-116-260_F04'
'1772-116-260_F05'
'1772-116-260_F06'
'1772-116-260_F07'
'1772-116-260_F08'
'1772-116-260_F09'
'1772-116-260_F11'
'1772-116-260_F12'
'1772-116-260_G01'
'1772-116-260_G03'
'1772-116-260_G04'
'1772-116-260_G05'
'1772-116-260_G06'
'1772-116-260_G07'
'1772-116-260_G08'
'1772-116-260_G09'
'1772-116-260_G10'
'1772-116-260_G11'
'1772-116-260_G12'
'1772-116-260_H01'
'1772-116-260_H02'
'1772-116-260_H03'
'1772-116-260_H04'
'1772-116-260_H07'
'1772-116-260_H08'
'1772-116-260_H09'
'1772-116-260_H10'
'1772-116-260_H11'
'1772-116-260_H12'
'1772-116-261_A01'
'1772-116-261_A02'
'1772-116-261_A03'
'1772-116-261_A04'
'1772-116-261_A05'
'1772-116-261_A06'
'1772-116-261_A07'
'1772-116-261_A08'
'1772-116-261_A09'
'1772-116-261_A10'
'1772-116-261_A11'
'1772-116-261_B01'
'1772-116-261_B03'
'1772-116-261_B05'
'1772-116-261_B06'
'1772-116-261_B07'
'1772-116-261_B08'
'1772-116-261_B09'
'1772-116-261_B11'
'1772-116-261_B12'
'1772-116-261_C01'
'1772-116-261_C02'
'1772-116-261_C03'
'1772-116-261_C04'
'1772-116-261_C05'
'1772-116-261_C06'
'1772-116-261_C07'
'1772-116-261_C08'
'1772-116-261_C09'
'1772-116-261_C10'
'1772-116-261_C11'
'1772-116-261_D02'
'1772-116-261_D03'
'1772-116-261_D04'
'1772-116-261_D06'
'1772-116-261_D07'
'1772-116-261_D08'
'1772-116-261_D09'
'1772-116-261_D10'
'1772-116-261_D11'
'1772-116-261_D12'
'1772-116-261_E01'
'1772-116-261_E03'
'1772-116-261_E04'
'1772-116-261_E05'
'1772-116-261_E06'
'1772-116-261_E08'
'1772-116-261_E10'
'1772-116-261_E11'
'1772-116-261_F01'
'1772-116-261_F03'
'1772-116-261_F04'
'1772-116-261_F05'
'1772-116-261_F06'
'1772-116-261_F07'
'1772-116-261_F08'
'1772-116-261_F09'
'1772-116-261_F10'
'1772-116-261_F11'
'1772-116-261_F12'
'1772-116-261_G01'
'1772-116-261_G02'
'1772-116-261_G05'
'1772-116-261_G06'
'1772-116-261_G07'
'1772-116-261_G08'
'1772-116-261_G10'
'1772-116-261_G11'
'1772-116-261_H02'
'1772-116-261_H04'
'1772-116-261_H05'
'1772-116-261_H06'
'1772-116-261_H07'
'1772-116-261_H08'
'1772-116-261_H09'
'1772-116-261_H10'
'1772-116-261_H11'
'1772-116-262_A01'
'1772-116-262_A02'
'1772-116-262_A03'
'1772-116-262_A04'
'1772-116-262_A05'
'1772-116-262_A06'
'1772-116-262_A07'
'1772-116-262_A08'
'1772-116-262_B01'
'1772-116-262_B02'
'1772-116-262_B04'
'1772-116-262_B05'
'1772-116-262_B06'
'1772-116-262_B07'
'1772-116-262_B08'
'1772-116-262_B09'
'1772-116-262_B10'
'1772-116-262_B11'
'1772-116-262_B12'
'1772-116-262_C01'
'1772-116-262_C02'
'1772-116-262_C03'
'1772-116-262_C04'
'1772-116-262_C05'
'1772-116-262_C06'
'1772-116-262_C07'
'1772-116-262_C08'
'1772-116-262_C11'
'1772-116-262_C12'
'1772-116-262_D01'
'1772-116-262_D02'
'1772-116-262_D04'
'1772-116-262_D05'
'1772-116-262_D06'
'1772-116-262_D07'
'1772-116-262_D08'
'1772-116-262_D09'
'1772-116-262_D10'
'1772-116-262_D11'
'1772-116-262_D12'
'1772-116-262_E01'
'1772-116-262_E02'
'1772-116-262_E03'
'1772-116-262_E05'
'1772-116-262_E06'
'1772-116-262_E07'
'1772-116-262_E08'
'1772-116-262_E09'
'1772-116-262_E11'
'1772-116-262_E12'
'1772-116-262_F01'
'1772-116-262_F02'
'1772-116-262_F03'
'1772-116-262_F04'
'1772-116-262_F05'
'1772-116-262_F06'
'1772-116-262_F07'
'1772-116-262_F08'
'1772-116-262_F10'
'1772-116-262_F11'
'1772-116-262_F12'
'1772-116-262_G02'
'1772-116-262_G03'
'1772-116-262_G05'
'1772-116-262_G06'
'1772-116-262_G07'
'1772-116-262_G08'
'1772-116-262_G09'
'1772-116-262_G10'
'1772-116-262_G11'
'1772-116-262_G12'
'1772-116-262_H01'
'1772-116-262_H03'
'1772-116-262_H04'
'1772-116-262_H06'
'1772-116-262_H08'
'1772-116-262_H10'
'1772-116-262_H11'
'1772-116-262_H12'
'1772-116-263_A03'
'1772-116-263_A04'
'1772-116-263_A05'
'1772-116-263_A06'
'1772-116-263_A08'
'1772-116-263_A09'
'1772-116-263_A10'
'1772-116-263_A11'
'1772-116-263_B01'
'1772-116-263_B02'
'1772-116-263_B03'
'1772-116-263_B04'
'1772-116-263_B05'
'1772-116-263_B06'
'1772-116-263_B07'
'1772-116-263_B08'
'1772-116-263_B09'
'1772-116-263_C01'
'1772-116-263_C03'
'1772-116-263_C04'
'1772-116-263_C05'
'1772-116-263_C06'
'1772-116-263_C07'
'1772-116-263_C08'
'1772-116-263_C09'
'1772-116-263_C10'
'1772-116-263_C11'
'1772-116-263_D01'
'1772-116-263_D02'
'1772-116-263_D03'
'1772-116-263_D04'
'1772-116-263_D05'
'1772-116-263_D06'
'1772-116-263_D07'
'1772-116-263_D08'
'1772-116-263_D09'
'1772-116-263_D10'
'1772-116-263_D11'
'1772-116-263_E01'
'1772-116-263_E02'
'1772-116-263_E03'
'1772-116-263_E04'
'1772-116-263_E05'
'1772-116-263_E06'
'1772-116-263_E07'
'1772-116-263_E08'
'1772-116-263_E09'
'1772-116-263_E10'
'1772-116-263_E11'
'1772-116-263_F01'
'1772-116-263_F02'
'1772-116-263_F03'
'1772-116-263_F04'
'1772-116-263_F05'
'1772-116-263_F06'
'1772-116-263_F07'
'1772-116-263_F08'
'1772-116-263_F09'
'1772-116-263_F10'
'1772-116-263_F11'
'1772-116-263_G01'
'1772-116-263_G02'
'1772-116-263_G03'
'1772-116-263_G04'
'1772-116-263_G05'
'1772-116-263_G06'
'1772-116-263_G07'
'1772-116-263_G10'
'1772-116-263_G11'
'1772-116-263_H01'
'1772-116-263_H03'
'1772-116-263_H04'
'1772-116-263_H05'
'1772-116-263_H06'
'1772-116-263_H10'
'1772-116-263_H11'
'1772-123-289_A09'
'1772-123-289_B01'
'1772-123-289_B05'
'1772-123-289_B06'
'1772-123-289_C09'
'1772-123-289_C11'
'1772-123-289_D11'
'1772-123-289_F01'
'1772-123-289_F03'
'1772-123-289_F05'
'1772-123-289_F07'
'1772-123-289_F09'
'1772-123-291_A08'
'1772-123-291_A09'
'1772-123-291_A12'
'1772-123-291_B01'
'1772-123-291_B02'
'1772-123-291_B03'
'1772-123-291_B08'
'1772-123-291_B09'
'1772-123-291_B10'
'1772-123-291_B11'
'1772-123-291_C01'
'1772-123-291_C02'
'1772-123-291_C03'
'1772-123-291_C07'
'1772-123-291_D01'
'1772-123-291_D02'
'1772-123-291_D08'
'1772-123-291_D11'
'1772-123-291_E03'
'1772-123-291_E08'
'1772-123-291_E09'
'1772-123-291_F02'
'1772-123-291_F03'
'1772-123-291_F07'
'1772-123-291_F08'
'1772-123-291_F09'
'1772-123-291_F11'
'1772-123-291_G01'
'1772-123-291_G02'
'1772-123-291_G03'
'1772-123-291_G07'
'1772-123-291_H01'
'1772-123-291_H02'
'1772-123-291_H03'
'1772-123-291_H08'
'1772-123-291_H09'
'1772-151-313_A01'
'1772-151-313_A02'
'1772-151-313_A03'
'1772-151-313_A04'
'1772-151-313_A05'
'1772-151-313_A06'
'1772-151-313_A07'
'1772-151-313_A09'
'1772-151-313_A10'
'1772-151-313_A11'
'1772-151-313_A12'
'1772-151-313_B01'
'1772-151-313_B02'
'1772-151-313_B05'
'1772-151-313_B06'
'1772-151-313_B08'
'1772-151-313_B09'
'1772-151-313_B10'
'1772-151-313_B11'
'1772-151-313_B12'
'1772-151-313_C01'
'1772-151-313_C02'
'1772-151-313_C05'
'1772-151-313_C06'
'1772-151-313_C09'
'1772-151-313_C10'
'1772-151-313_C11'
'1772-151-313_C12'
'1772-151-313_D01'
'1772-151-313_D03'
'1772-151-313_D04'
'1772-151-313_D07'
'1772-151-313_D08'
'1772-151-313_D09'
'1772-151-313_D10'
'1772-151-313_D11'
'1772-151-313_D12'
'1772-151-313_E01'
'1772-151-313_E02'
'1772-151-313_E03'
'1772-151-313_E04'
'1772-151-313_E06'
'1772-151-313_E07'
'1772-151-313_E08'
'1772-151-313_E09'
'1772-151-313_E10'
'1772-151-313_E11'
'1772-151-313_E12'
'1772-151-313_F01'
'1772-151-313_F02'
'1772-151-313_F03'
'1772-151-313_F04'
'1772-151-313_F05'
'1772-151-313_F06'
'1772-151-313_F07'
'1772-151-313_F08'
'1772-151-313_F09'
'1772-151-313_F10'
'1772-151-313_G01'
'1772-151-313_G02'
'1772-151-313_G03'
'1772-151-313_G04'
'1772-151-313_G05'
'1772-151-313_G06'
'1772-151-313_G09'
'1772-151-313_G11'
'1772-151-313_G12'
'1772-151-313_H03'
'1772-151-313_H04'
'1772-151-313_H05'
'1772-151-313_H06'
'1772-151-313_H07'
'1772-151-313_H08'
'1772-151-313_H09'
'1772-151-313_H11'
'1772-151-313_H12'
)


##########    Merge b6 and msm bam files   ################


for ID in "${ID_array[@]}"

do


samtools sort -@ 8 -T /home/splitNCigarRead_bam/tmp/${ID}.b6 -o /home/splitNCigarRead_bam/MERGE/${ID}.b6.rmDuplicates.split.sort.bam /home/splitNCigarRead_bam/B6/${ID}.b6.rmDuplicates.split.bam


samtools sort -@ 8 -T /home/splitNCigarRead_bam/tmp/${ID}.msm -o /home/splitNCigarRead_bam/MERGE/${ID}.msm.rmDuplicates.split.sort.bam /home/splitNCigarRead_bam/MSM/${ID}.msm.rmDuplicates.split.bam


samtools merge /home/splitNCigarRead_bam/MERGE/${ID}.rmDuplicates.split.merge.bam /home/splitNCigarRead_bam/MERGE/${ID}.b6.rmDuplicates.split.sort.bam /home/splitNCigarRead_bam/MERGE/${ID}.msm.rmDuplicates.split.sort.bam


samtools sort -@ 8 -T /home/splitNCigarRead_bam/tmp/${ID}.merge -o /home/splitNCigarRead_bam/MERGE/${ID}.rmDuplicates.split.merge.sort.bam /home/splitNCigarRead_bam/MERGE/${ID}.rmDuplicates.split.merge.bam


samtools index -@ 8 /home/splitNCigarRead_bam/MERGE/${ID}.rmDuplicates.split.merge.sort.bam


rm /home/splitNCigarRead_bam/MERGE/${ID}.b6.rmDuplicates.split.sort.bam

rm /home/splitNCigarRead_bam/MERGE/${ID}.msm.rmDuplicates.split.sort.bam

rm /home/splitNCigarRead_bam/MERGE/${ID}.rmDuplicates.split.merge.bam


done


############    Detect variants on X Chromosome with GATK     ################


for ID in "${ID_array[@]}"

do


java -Xmx4g -jar GenomeAnalysisTK.jar -T HaplotypeCaller -R UCSC_mm9.woYwRwMSMv5nSNP_b6_chr.fa -I /home/splitNCigarRead_bam/MERGE/${ID}.rmDuplicates.split.merge.sort.bam -dontUseSoftClippedBases -stand_call_conf 20 -o /home/splitNCigarRead_bam/NC/${ID}.rmdup.split.merge_b6_msm.vcf -maxAltAlleles 15 -L chrX


vcftools --vcf /home/splitNCigarRead_bam/NC/${ID}.rmdup.split.merge_b6_msm.vcf --remove-indels --recode --recode-INFO-all --out /home/splitNCigarRead_bam/NC/${ID}.rmdup.split.merge_b6_msm_SNPs_only


java -Xmx4g -jar GenomeAnalysisTK.jar -T VariantFiltration -R UCSC_mm9.woYwRwMSMv5nSNP_b6_chr.fa -V /home/splitNCigarRead_bam/NC/${ID}.rmdup.split.merge_b6_msm_SNPs_only.recode.vcf -o /home/splitNCigarRead_bam/NC/${ID}.rmdup.split.merge_b6_msm_SNPs_only.recode.fil.vcf -window 35 -cluster 3


less /home/splitNCigarRead_bam/NC/${ID}.rmdup.split.merge_b6_msm_SNPs_only.recode.fil.vcf |grep "^#\|PASS" > /home/splitNCigarRead_bam/NC/${ID}.rmdup.split.merge_b6_msm_SNPs_only.recode.fil.PASS.vcf

less /home/splitNCigarRead_bam/NC/${ID}.rmdup.split.merge_b6_msm_SNPs_only.recode.fil.PASS.vcf |grep "^#\|chrX" > /home/splitNCigarRead_bam/NC/${ID}.rmdup.split.merge_b6_msm_SNPs_only.recode.fil.PASS.chrX.vcf


java -Xmx4g -jar /home/snpEff/snpEff.jar mm9 /home/splitNCigarRead_bam/NC/${ID}.rmdup.split.merge_b6_msm_SNPs_only.recode.fil.PASS.chrX.vcf > /home/splitNCigarRead_bam/NC/${ID}.rmdup.split.merge_b6_msm_SNPs_only.recode.fil.PASS.chrX.snpEff.vcf


java -Xmx4g -jar /home/snpEff/SnpSift.jar annotate /home/mm9dbsnp128.chrX.sort.vcf /home/splitNCigarRead_bam/NC/${ID}.rmdup.split.merge_b6_msm_SNPs_only.recode.fil.PASS.chrX.snpEff.vcf > /home/splitNCigarRead_bam/NC/${ID}.rmdup.split.merge_b6_msm_SNPs_only.recode.fil.PASS.chrX.snpEff.annotated.vcf


#####################   heterozygote SNP   #######################


less /home/splitNCigarRead_bam/NC/${ID}.rmdup.split.merge_b6_msm_SNPs_only.recode.fil.PASS.chrX.snpEff.annotated.vcf | java -jar ./snpEff/SnpSift.jar filter "(countHet() > 0)" > /home/splitNCigarRead_bam/NC/${ID}.rmdup.split.merge_b6_msm_SNPs_only.recode.fil.PASS.chrX.snpEff.annotated.hetero.vcf


#####################   Pipeline  for  b6  and  msm    #######################


###################    extract rs vcf files  annotated based on dbsnp128   #############


less /home/splitNCigarRead_bam/NC/${ID}.rmdup.split.merge_b6_msm_SNPs_only.recode.fil.PASS.chrX.snpEff.annotated.hetero.vcf | java -jar ./snpEff/SnpSift.jar filter "(exists ID) & ( ID =~ 'rs' )" > /home/splitNCigarRead_bam/NC/${ID}.rmdup.split.merge_b6_msm_SNPs_only.recode.fil.PASS.chrX.snpEff.annotated.hetero.rs.vcf


##################      extract coding variant    #################

less /home/splitNCigarRead_bam/NC/${ID}.rmdup.split.merge_b6_msm_SNPs_only.recode.fil.PASS.chrX.snpEff.annotated.hetero.rs.vcf |grep -v intergenic > /home/splitNCigarRead_bam/NC_re/${ID}.rmdup.split.merge_b6_msm_SNPs_only.recode.fil.PASS.chrX.snpEff.annotated.hetero.rs.no_intergenic.vcf


less /home/splitNCigarRead_bam/NC_re/${ID}.rmdup.split.merge_b6_msm_SNPs_only.recode.fil.PASS.chrX.snpEff.annotated.hetero.rs.no_intergenic.vcf | grep intron | grep -v synonymous |grep -v UTR | grep -v "^#" > /home/splitNCigarRead_bam/NC_re/${ID}.rmdup.split.merge_b6_msm_SNPs_only.recode.fil.PASS.chrX.snpEff.annotated.hetero.rs.no_intergenic.intron.no_UTR_no_synonymous_no_header.vcf


less  /home/splitNCigarRead_bam/NC_re/${ID}.rmdup.split.merge_b6_msm_SNPs_only.recode.fil.PASS.chrX.snpEff.annotated.hetero.rs.no_intergenic.intron.no_UTR_no_synonymous_no_header.vcf |cut -f 2 >  /home/splitNCigarRead_bam/NC_re/${ID}.rmdup.split.merge_b6_msm_SNPs_only.recode.fil.PASS.chrX.snpEff.annotated.hetero.rs.no_intergenic.intron.no_UTR_no_synonymous_no_header.position.txt


##################    filter out noncoding varinats       ###############


less  /home/splitNCigarRead_bam/NC_re/${ID}.rmdup.split.merge_b6_msm_SNPs_only.recode.fil.PASS.chrX.snpEff.annotated.hetero.rs.no_intergenic.vcf | grep -v "^#" |grep -vf  /home/splitNCigarRead_bam/NC_re/${ID}.rmdup.split.merge_b6_msm_SNPs_only.recode.fil.PASS.chrX.snpEff.annotated.hetero.rs.no_intergenic.intron.no_UTR_no_synonymous_no_header.position.txt >  /home/splitNCigarRead_bam/NC_re/${ID}.rmdup.split.merge_b6_msm_SNPs_only.recode.fil.PASS.chrX.snpEff.annotated.hetero.rs.no_intergenic.fil_noncoding.number.vcf


#################      get variant position     ###########################


less  /home/splitNCigarRead_bam/NC_re/${ID}.rmdup.split.merge_b6_msm_SNPs_only.recode.fil.PASS.chrX.snpEff.annotated.hetero.rs.no_intergenic.fil_noncoding.number.vcf | cut -f 1,2 >  /home/splitNCigarRead_bam/NC_re/${ID}.b6_position_coding.txt


less /home/splitNCigarRead_bam/NC/${ID}.rmdup.split.merge_b6_msm_SNPs_only.recode.fil.PASS.chrX.snpEff.annotated.hetero.rs.vcf |grep -v "^#" |cut -f 1,2 > /home/splitNCigarRead_bam/NC_re/${ID}.b6_position_noncoding.txt


#######################################################################


##################           MSM   processing           ##########################


##################     extract msm variants annotated based on MSMv4HQ.snp    ############


less /home/splitNCigarRead_bam/NC/${ID}.rmdup.split.merge_b6_msm_SNPs_only.recode.fil.PASS.chrX.snpEff.annotated.hetero.vcf | grep -v "^#" | cut -f 1,2,4,5 > /home/splitNCigarRead_bam/NC/${ID}.rmdup.split.merge_b6_msm_SNPs_only.recode.fil.PASS.chrX.snpEff.annotated.hetero.no_header.posi.bases.vcf


less /home/splitNCigarRead_bam/NC/${ID}.rmdup.split.merge_b6_msm_SNPs_only.recode.fil.PASS.chrX.snpEff.annotated.hetero.no_header.posi.bases.vcf |grep -wf MSMv4HQ.snp2.chrX_no_S_M_Y_W_R_K.from_mm10_to_mm9.log.mm9.base.txt > /home/splitNCigarRead_bam/NC/${ID}.rmdup.split.merge_b6_msm_SNPs_only.recode.fil.PASS.chrX.snpEff.annotated.hetero.no_header.posi.bases.fil_MSMv4HQ_snp2.vcf


##################      extract coding MSM variant    #################


less /home/splitNCigarRead_bam/NC_re/${ID}.rmdup.split.merge_b6_msm_SNPs_only.recode.fil.PASS.chrX.snpEff.annotated.hetero.no_header.posi.bases.fil_MSMv4HQ_snp2.vcf | grep -v intergenic > /home/splitNCigarRead_bam/NC_re/${ID}.rmdup.split.merge_b6_msm_SNPs_only.recode.fil.PASS.chrX.snpEff.annotated.hetero.no_header.posi.bases.fil_MSMv4HQ_snp2.no_intergenic.vcf


less /home/splitNCigarRead_bam/NC_re/${ID}.rmdup.split.merge_b6_msm_SNPs_only.recode.fil.PASS.chrX.snpEff.annotated.hetero.no_header.posi.bases.fil_MSMv4HQ_snp2.no_intergenic.vcf |grep intron | grep -v synonymous |grep -v UTR |grep -v "^#" > /home/splitNCigarRead_bam/NC_re/${ID}.rmdup.split.merge_b6_msm_SNPs_only.recode.fil.PASS.chrX.snpEff.annotated.hetero.no_header.posi.bases.fil_MSMv4HQ_snp2.no_intergenic.intron.no_UTR_no_synonymous_no_header.vcf


less /home/splitNCigarRead_bam/NC_re/${ID}.rmdup.split.merge_b6_msm_SNPs_only.recode.fil.PASS.chrX.snpEff.annotated.hetero.no_header.posi.bases.fil_MSMv4HQ_snp2.no_intergenic.intron.no_UTR_no_synonymous_no_header.vcf |cut -f 2 > /home/splitNCigarRead_bam/NC_re/${ID}.rmdup.split.merge_b6_msm_SNPs_only.recode.fil.PASS.chrX.snpEff.annotated.hetero.no_header.posi.bases.fil_MSMv4HQ_snp2.no_intergenic.intron.no_UTR_no_synonymous_no_header.position.txt


##################    filter out noncoding variants       ##############


less /home/splitNCigarRead_bam/NC_re/${ID}.rmdup.split.merge_b6_msm_SNPs_only.recode.fil.PASS.chrX.snpEff.annotated.hetero.no_header.posi.bases.fil_MSMv4HQ_snp2.no_intergenic.vcf |grep -v "^#" | grep -vf /home/splitNCigarRead_bam/NC_re/${ID}.rmdup.split.merge_b6_msm_SNPs_only.recode.fil.PASS.chrX.snpEff.annotated.hetero.no_header.posi.bases.fil_MSMv4HQ_snp2.no_intergenic.intron.no_UTR_no_synonymous_no_header.position.txt > /home/splitNCigarRead_bam/NC_re/${ID}.rmdup.split.merge_b6_msm_SNPs_only.recode.fil.PASS.chrX.snpEff.annotated.hetero.no_header.posi.bases.fil_MSMv4HQ_snp2.no_intergenic.fil_noncoding.number.vcf


####################    get variant position     #########################


less /home/splitNCigarRead_bam/NC_re/${ID}.rmdup.split.merge_b6_msm_SNPs_only.recode.fil.PASS.chrX.snpEff.annotated.hetero.no_header.posi.bases.fil_MSMv4HQ_snp2.no_intergenic.fil_noncoding.number.vcf | cut -f 1,2 > /home/splitNCigarRead_bam/NC_re/${ID}.msm.position_coding.txt


less /home/splitNCigarRead_bam/NC/${ID}.rmdup.split.merge_b6_msm_SNPs_only.recode.fil.PASS.chrX.snpEff.annotated.hetero.no_header.posi.bases.fil_MSMv4HQ_snp2.vcf | cut -f 1,2 > /home/splitNCigarRead_bam/NC_re/${ID}.msm.position_noncoding.txt


###################     bind B6 and MSM files       ################################################################


cat /home/splitNCigarRead_bam/NC_re/${ID}.b6_position_coding.txt /home/splitNCigarRead_bam/NC_re/${ID}.msm_position_coding.txt > /home/splitNCigarRead_bam/NC_re/${ID}.b6_msm_position_coding.txt

less /home/splitNCigarRead_bam/NC_re/${ID}.b6_msm_position_coding.txt |sort -k 1,1 -k 2,2n |uniq > /home/splitNCigarRead_bam/NC_re/${ID}.b6_msm_position_coding.sort.uniq.txt


cat /home/splitNCigarRead_bam/NC_re/${ID}.b6_position_noncoding.txt /home/splitNCigarRead_bam/NC_re/${ID}.msm_position_noncoding.txt > /home/splitNCigarRead_bam/NC_re/${ID}.b6_msm_position_noncoding.txt


less /home/splitNCigarRead_bam/NC_re/${ID}.b6_msm_position_noncoding.txt |sort -k 1,1 -k 2,2n |uniq > /home/splitNCigarRead_bam/NC_re/${ID}.b6_msm_position_noncoding.sort.uniq.txt


######################################################################################################################


done


#######     extract all variant positions detected by GATK on all single cell data    #######################


for ID in "${ID_array[@]}"

do

less /home/splitNCigarRead_bam/NC_re/${ID}.b6_msm_position_coding.sort.uniq.txt >> /home/splitNCigarRead_bam/NC_re/b6_msm_position_coding.sort.uniq.pre.txt

less /home/splitNCigarRead_bam/NC_re/${ID}.b6_msm_position_noncoding.sort.uniq.txt >> /home/splitNCigarRead_bam/NC_re/b6_msm_position_noncoding.sort.uniq.pre.txt

done


less /home/splitNCigarRead_bam/NC_re/b6_msm_position_coding.sort.uniq.pre.txt |sort -k 1,1 -k 2,2n |uniq > /home/splitNCigarRead_bam/NC_re/b6_msm_position_coding.sort.uniq.txt


less /home/splitNCigarRead_bam/NC_re/b6_msm_position_noncoding.sort.uniq.pre.txt |sort -k 1,1 -k 2,2n |uniq > /home/splitNCigarRead_bam/NC_re/b6_msm_position_noncoding.sort.uniq.txt


#######      Allele expression analysis      ################


for ID in "${ID_array[@]}"

do


	#########     position.txt contains the heterogyzous loci (chr + coordinate) returned by GATK and  I confirmed if the loci were registered in the dbSNP (Now I use snp128)     ############

	#########     I have confirmed variants detected by GATK are msm SNPs using database for msm.  ######################################### 

	#########       Using samtools to detect variants allelically for B6 and MSM                   #############


samtools mpileup -E -f UCSC_mm9.woYwRwMSMv5nSNP_b6_chr.fa -d 1000000 --position /home/splitNCigarRead_bam/NC_re/b6_msm_position_coding.sort.uniq.txt /home/splitNCigarRead_bam/MERGE/${ID}.rmDuplicates.split.merge.sort.bam    > /home/splitNCigarRead_bam/NC_re/${ID}.rmdup.split.merge_b6_msm.mpileup


perl pileup2base_no_strand.pl /home/splitNCigarRead_bam/NC_re/${ID}.rmdup.split.merge_b6_msm.mpileup 30 /home/splitNCigarRead_bam/NC_re/${ID}.rmdup.split.merge_b6_msm.perse30.txt


######  more_than_10


less /home/splitNCigarRead_bam/NC_re/${ID}.rmdup.split.merge_b6_msm.perse30.txt |grep -v ref |awk '{sum=($4+$5+$6+$7);if(sum > 9) print}' > /home/splitNCigarRead_bam/NC_re/${ID}.rmdup.split.merge_b6_msm.perse30.more_than_10.txt


less /home/splitNCigarRead_bam/NC_re/${ID}.rmdup.split.merge_b6_msm.perse30.txt |grep -v ref |awk '{sum=($4+$5+$6+$7);if(sum < 10) print}' > /home/splitNCigarRead_bam/NC_re/${ID}.rmdup.split.merge_b6_msm.perse30.less_than_10.txt


less /home/splitNCigarRead_bam/NC_re/${ID}.rmdup.split.merge_b6_msm.perse30.more_than_10.txt |grep -v ref |awk '{sum=($4+$5+$6+$7); print $0"\t"sum}' > /home/splitNCigarRead_bam/NC_re/${ID}.rmdup.split.merge_b6_msm.perse30.more_than_10.plus.sum.txt


less /home/splitNCigarRead_bam/NC_re/${ID}.rmdup.split.merge_b6_msm.perse30.more_than_10.plus.sum.txt |awk '{if($3 == "A") print $1"\t"$2"\t"$3"\t"$4"\t"$8}' > /home/splitNCigarRead_bam/NC_re/${ID}.rmdup.split.merge_b6_msm.perse30.more_than_10.plus.sum.A.txt


less /home/splitNCigarRead_bam/NC_re/${ID}.rmdup.split.merge_b6_msm.perse30.more_than_10.plus.sum.txt |awk '{if($3 == "T") print $1"\t"$2"\t"$3"\t"$5"\t"$8}' > /home/splitNCigarRead_bam/NC_re/${ID}.rmdup.split.merge_b6_msm.perse30.more_than_10.plus.sum.T.txt


less /home/splitNCigarRead_bam/NC_re/${ID}.rmdup.split.merge_b6_msm.perse30.more_than_10.plus.sum.txt |awk '{if($3 == "C") print $1"\t"$2"\t"$3"\t"$6"\t"$8}'  > /home/splitNCigarRead_bam/NC_re/${ID}.rmdup.split.merge_b6_msm.perse30.more_than_10.plus.sum.C.txt


less /home/splitNCigarRead_bam/NC_re/${ID}.rmdup.split.merge_b6_msm.perse30.more_than_10.plus.sum.txt |awk '{if($3 == "G") print $1"\t"$2"\t"$3"\t"$7"\t"$8}'  >  /home/splitNCigarRead_bam/NC_re/${ID}.rmdup.split.merge_b6_msm.perse30.more_than_10.plus.sum.G.txt


cat /home/splitNCigarRead_bam/NC_re/${ID}.rmdup.split.merge_b6_msm.perse30.more_than_10.plus.sum.A.txt /home/splitNCigarRead_bam/NC_re/${ID}.rmdup.split.merge_b6_msm.perse30.more_than_10.plus.sum.T.txt /home/splitNCigarRead_bam/NC_re/${ID}.rmdup.split.merge_b6_msm.perse30.more_than_10.plus.sum.C.txt /home/splitNCigarRead_bam/NC_re/${ID}.rmdup.split.merge_b6_msm.perse30.more_than_10.plus.sum.G.txt > /home/splitNCigarRead_bam/NC_re/${ID}.rmdup.split.merge_b6_msm.perse30.more_than_10.plus.sum.ATCG.txt


less /home/splitNCigarRead_bam/NC_re/${ID}.rmdup.split.merge_b6_msm.perse30.more_than_10.plus.sum.ATCG.txt |awk '{per=100*($4/$5); print $0"\t"per}' > /home/splitNCigarRead_bam/NC_re/${ID}.rmdup.split.merge_b6_msm.perse30.more_than_10.plus.sum.ATCG.percent.txt


less /home/splitNCigarRead_bam/NC_re/${ID}.rmdup.split.merge_b6_msm.perse30.more_than_10.plus.sum.ATCG.percent.txt |sort -k 1,1 -k 2,2n > /home/splitNCigarRead_bam/NC_re/${ID}.rmdup.split.merge_b6_msm.perse30.more_than_10.plus.sum.ATCG.percent.sort.txt


less /home/splitNCigarRead_bam/NC_re/${ID}.rmdup.split.merge_b6_msm.perse30.more_than_10.plus.sum.ATCG.percent.sort.txt |awk '{if($6>90) print $0"\t""b6"}' > /home/splitNCigarRead_bam/NC_re/${ID}.rmdup.split.merge_b6_msm.perse30.more_than_10.plus.sum.ATCG.percent.sort.b6.txt


less /home/splitNCigarRead_bam/NC_re/${ID}.rmdup.split.merge_b6_msm.perse30.more_than_10.plus.sum.ATCG.percent.sort.txt |awk '{if($6<10) print $0"\t""msm"}' > /home/splitNCigarRead_bam/NC_re/${ID}.rmdup.split.merge_b6_msm.perse30.more_than_10.plus.sum.ATCG.percent.sort.msm.txt


less /home/splitNCigarRead_bam/NC_re/${ID}.rmdup.split.merge_b6_msm.perse30.more_than_10.plus.sum.ATCG.percent.sort.txt |awk '{if($6<=90 && $6>=10) print $0"\t""biallele"}' > /home/splitNCigarRead_bam/NC_re/${ID}.rmdup.split.merge_b6_msm.perse30.more_than_10.plus.sum.ATCG.percent.sort.biallele.txt


cat /home/splitNCigarRead_bam/NC_re/${ID}.rmdup.split.merge_b6_msm.perse30.more_than_10.plus.sum.ATCG.percent.sort.b6.txt /home/splitNCigarRead_bam/NC_re/${ID}.rmdup.split.merge_b6_msm.perse30.more_than_10.plus.sum.ATCG.percent.sort.msm.txt /home/splitNCigarRead_bam/NC_re/${ID}.rmdup.split.merge_b6_msm.perse30.more_than_10.plus.sum.ATCG.percent.sort.biallele.txt > /home/splitNCigarRead_bam/NC_re/${ID}.rmdup.split.merge_b6_msm.perse30.more_than_10.plus.sum.ATCG.percent.sort.b6_msm_biallele.txt


less /home/splitNCigarRead_bam/NC_re/${ID}.rmdup.split.merge_b6_msm.perse30.more_than_10.plus.sum.ATCG.percent.sort.b6_msm_biallele.txt |sort -k 1,1 -k 2,2n > /home/splitNCigarRead_bam/NC_re/${ID}.rmdup.split.merge_b6_msm.perse30.more_than_10.plus.sum.ATCG.percent.sort.b6_msm_biallele.sort.txt


done
```
