## Supplementary material for "Single-cell transcriptomics, scRNA-Seq and C1 CAGE discovered distinct phases of pluripotency during naïve-to-primed conversion in mice": 3_Heatmap_for_allele_specific_expression.R

```
#  multimerge = function(mypath){
#  filenames=list.files(path=mypath, full.names=TRUE)
#  datalist = lapply(filenames, function(x){read.delim(file=x,header=T)})
#  Reduce(function(x,y) {merge(x,y,by = "variant", all = TRUE)}, datalist)}


# xci_pheatmap <- multimerge("/Users/pheatmap")

# dim(xci_pheatmap)

#  write.table(xci_pheatmap, "/Users/xci_pheatmap.txt",quote=F,col.names=T, append=F)

########################################################################################

# less /Users/xci_pheatmap.txt  |tr " " "\t" |cut -f 2- > /Users/xci_pheatmap.tab.txt

########################################################################################

xci_pheatmap2 <- read.table("/Users/xci_pheatmap.tab.txt", header=T, row.names = 1, check.names = FALSE)


dim(xci_pheatmap2)


cnt_na <- apply(xci_pheatmap2, 1, function(z) sum(is.na(z)))


xci_ptm2_255_fil <- xci_pheatmap2[cnt_na < 255,]


dim(xci_ptm2_255_fil)


xci_date <- xci_ptm2_255_fil[c(1:71,72:154,155:231,311:386,232:310,435:510,399:434,387:398)]

###################################################################################

#######  female_table_Preparation   ###############################################


library(ggplot2)

library(viridis)


meta <- read.csv("metadata_20190209230902.csv", header=TRUE, stringsAsFactors = F)

tpm <- read.table("geneTPM_M8_20190207033818_Kallisto.txt", header=T, row.names = 1, check.names = FALSE)
 
tpm <- tpm[, which(names(tpm) %in% meta$cell_id)]
 
meta <- meta[match(names(tpm), meta$cell_id),]

expr <- tpm[,!meta$discard]

meta <- meta[!meta$discard,]

meta$timePoint <- factor(meta$timePoint)

meta$timePoint <- factor(meta$timePoint, levels(meta$timePoint)[c(4:8,2,3,1)])

expr <- expr[,which(colSums(expr)>1)]

expr <- expr[which(rowSums(expr)>1),]

expr <- expr[which(rowSums(expr>0)>10),]

expressedGenes <- sapply(1:ncol(expr), function(x) length(which(expr[,x]>1)))

meta$nGenes <- expressedGenes

meta <- meta[which(meta$nGenes>50),]

expr <- expr[,which(names(expr) %in% meta$cell_id)]


expr <- expr[c(1:462,532:579)]

expr <- expr[c(1:462,463:498,499:510)]

colnames(xci_date) <- colnames(expr)


xci_date_no_MEF <- xci_date[, -which(colnames(xci_date) %in% c("1772-116-259_H11", "1772-116-261_C03", "1772-116-263_C06", "1772-116-263_H10", "1772-116-262_A02", "1772-116-262_C01", "1772-151-313_A05", "1772-151-313_A06", "1772-151-313_B11", "1772-151-313_C05", "1772-151-313_C10", "1772-151-313_D04", "1772-151-313_D07", "1772-151-313_D11", "1772-151-313_D12", "1772-151-313_E07", "1772-151-313_E08", "1772-151-313_E10", "1772-151-313_E11", "1772-151-313_E12", "1772-151-313_F07", "1772-151-313_F09", "1772-151-313_G05", "1772-151-313_H04", "1772-151-313_H05", "1772-151-313_H09", "1772-151-313_H11", "1772-123-291_A08", "1772-123-291_B10", "1772-123-291_C01", "1772-123-291_C02", "1772-123-291_C03", "1772-123-291_C07", "1772-123-291_E03", "1772-123-291_F08", "1772-123-291_F11", "1772-123-291_H01"))]


dim(xci_date_no_MEF)


annotation_col_cell_type_fin = data.frame(CellType = factor(rep(c("mES","Day1","Day2","Day3","Day4","MB#3P10","1EP7"),c(70,83,76,74,132,26,12))))


dim(annotation_col_cell_type_fin)


rownames(annotation_col_cell_type_fin) = colnames(xci_date_no_MEF)

CellType <- c("#F8766D","#E68613","#CD9600","#7CAE00","#00B8E7","#ED68ED","#FF68A1")


names(CellType) <- c("mES","Day1","Day2","Day3","Day4","MB#3P10","1EP7")


anno_colors_celltype <- list(CellType = CellType)


library(pheatmap)

pheatmap(xci_date_no_MEF, cluster_col = FALSE, cluster_rows=F, show_colnames = FALSE, show_rownames = T, color = colorRampPalette(c("navy", "green", "firebrick3"))(50), annotation_col = annotation_col_cell_type_fin, filename = "XCI_days_255_fil_fin_no_MEF.pdf", cellwidth = 0.8, cellheight = 5, fontsize = 7, annotation_colors = anno_colors_celltype, na_col = "#838B9B")


write.table(xci_date_no_MEF, "XCI_date.txt", quote=F, sep="\t", row.names=T, col.names=T)


######################    Pseudotime Heatmap   #######################


PT_F <- read.table("TSCAN_pseudotime_20190223042447.no_EpiSC.sort.txt", header = T, row.names = 1, check.name = FALSE)


pseudotimeIndex <- order(PT_F$Pseudotime, decreasing = F)


xci_date_no_MEF_pt <- xci_date_no_MEF[,pseudotimeIndex]


annotation_col_cell_type_fin_pt <- annotation_col_cell_type_fin[pseudotimeIndex,]


annotation_col_cell_type_fin_pt <- factor(annotation_col_cell_type_fin_pt, levels=c("mES","Day1","Day2","Day3","Day4","MB#3P10","1EP7"))


Annotation_pt <- data.frame(TimePoint = factor(annotation_col_cell_type_fin_pt, labels = levels(annotation_col_cell_type_fin_pt)))


rownames(Annotation_pt) <- names(xci_date_no_MEF_pt)


gg_color_hue <- function(n) {
    hues = seq(15, 375, length=n+1)
    hcl(h=hues, l=65, c=100)[1:n]
  }


col_pt <- gg_color_hue(16)[c(1:3,5,10,12,14)]


names(col_pt) <- levels(Annotation_pt$TimePoint)


ann_color_pt <- list(TimePoint = col_pt)


pheatmap(xci_date_no_MEF_pt, cluster_col = FALSE, cluster_rows=F, show_colnames = FALSE, show_rownames = T, color = colorRampPalette(c("navy", "green", "firebrick3"))(50), annotation_col = Annotation_pt, filename = "XCI_PT_255.pdf", cellwidth = 0.8, cellheight = 5, fontsize = 5, annotation_colors = ann_color_pt, na_col = "#838B9B")


####    Cluster pheatmap


PT_F_df <- as.data.frame(PT_F)


Cluster_F <- read.delim("tSNE_kMeans5_clusterData_kallisto_DGE_geneTPM_20190223045041.no_EpiSC.txt", header = T,  check.name = FALSE)


PT_Cluster_F <- cbind(PT_F_df,Cluster_F)


PT_Cluster_F_df <- as.data.frame(PT_Cluster_F)

PT_Cluster_F_df[order(PT_Cluster_F_df$Pseudotime),]


 Annotation_Cluster_pt <- data.frame(TimePoint = factor(annotation_col_cell_type_fin_pt, labels = levels(annotation_col_cell_type_fin_pt)), Cluster = factor(PT_Cluster_F_df[order(PT_Cluster_F_df$Pseudotime),]$Cluster))


rownames(Annotation_Cluster_pt) <- names(xci_date_no_MEF_pt)


#########################################


library(RColorBrewer)

col2 <- brewer.pal(4, "Dark2")


#########################################


col_Cluster <- col2

names(col_Cluster) <- levels(Annotation_Cluster_pt$Cluster)

ann_color_Cluster <- list(Cluster = col_Cluster)


#########################################

col_c_pt <- gg_color_hue(16)[c(1:3,5,10,12,14)]

names(col_c_pt) <- levels(Annotation_Cluster_pt$TimePoint)

ann_color_c_pt <- list(TimePoint = col_c_pt)

#########################################

annotation_colors = list(
  TimePoint = col_c_pt,
  Cluster = col_Cluster)


pheatmap(xci_date_no_MEF_pt, cluster_col = FALSE, cluster_rows=F, show_colnames = FALSE, show_rownames = T, annotation_col = Annotation_Cluster_pt, annotation_colors = annotation_colors, color = colorRampPalette(c("navy", "green", "firebrick3"))(50),  filename = "XCI_PT_255_190226_plus_Cluster.pdf", cellwidth = 0.8, cellheight = 5, fontsize = 5, na_col = "#838B9B")


write.table(xci_date_no_MEF_pt, "XCI_pseudotime.txt", quote=F, sep="\t", row.names=T, col.names=T)


}
 
main()
```
