## Supplementary material for "Single-cell transcriptomics, scRNA-Seq and C1 CAGE discovered distinct phases of pluripotency during naïve-to-primed conversion in mice": 4_XCI_definition.sh

```
#!/bin/bash

###########################


for ID in `seq 2 511`

do

i=$((ID-1))

less XCI_final_pt.tab.txt |cut -f 1,${ID} > XCI_final_pt.${i}.txt

done

for ID in `seq 1 510`

do

less XCI_final_pt.${ID}.txt |grep -v NA > XCI_final_pt.${ID}.no_NA.txt

done


############################


for ID in `seq 1 510`


do

less XCI_final_pt.${ID}.no_NA.txt |cut -f 2 |grep X > XCI_final_pt.${ID}.no_NA.single_cell_name.txt


less XCI_final_pt.${ID}.no_NA.txt |cut -f 2  |grep -v X |wc -l > XCI_define/XCI_final_pt.${ID}.no_NA.total_numbers.txt


less XCI_final_pt.${ID}.no_NA.txt |cut -f 2 |grep -v X |grep biallele |wc -l > XCI_final_pt.${ID}.no_NA.biallel_numers.txt


less XCI_final_pt.${ID}.no_NA.txt |cut -f 2 |grep -v X |grep b6 |wc -l > XCI_final_pt.${ID}.no_NA.b6_numbers.txt


less XCI_final_pt.${ID}.no_NA.txt |cut -f 2 |grep -v X |grep msm |wc -l > XCI_final_pt.${ID}.no_NA.msm_numbers.txt


paste XCI_final_pt.${ID}.no_NA.single_cell_name.txt XCI_final_pt.${ID}.no_NA.total_numbers.txt XCI_final_pt.${ID}.no_NA.biallel_numers.txt XCI_final_pt.${ID}.no_NA.b6_numbers.txt XCI_final_pt.${ID}.no_NA.msm_numbers.txt > XCI_final_pt.${ID}.no_NA.for_percentage.txt


less XCI_final_pt.${ID}.no_NA.for_percentage.txt | awk '{per=100*($3/$2); print per}' > XCI_final_pt.${ID}.no_NA.for_biallelic_percentage.txt

less XCI_final_pt.${ID}.no_NA.for_percentage.txt | awk '{per1=100*($4/$2); print per1}' > XCI_final_pt.${ID}.no_NA.for_b6_percentage.txt

less XCI_final_pt.${ID}.no_NA.for_percentage.txt | awk '{per2=100*($5/$2); print per2}' > XCI_final_pt.${ID}.no_NA.for_msm_percentage.txt


paste XCI_final_pt.${ID}.no_NA.single_cell_name.txt XCI_final_pt.${ID}.no_NA.for_biallelic_percentage.txt XCI_final_pt.${ID}.no_NA.for_b6_percentage.txt XCI_final_pt.${ID}.no_NA.for_msm_percentage.txt > XCI_final_pt.${ID}.no_NA.for_percentage_bi_b6_msm.txt


less XCI_final_pt.${ID}.no_NA.for_percentage.txt | awk '{if($2>=50) print $0"\t""Detected_Variants_50"; else if($2<50) print $0"\t""less_than_50"}' >> XCI_final_pt.no_NA.for_percentage_define_50.txt


less XCI_final_pt.${ID}.no_NA.for_percentage_bi_b6_msm.txt >> XCI_final_pt.no_NA.for_percentage_bi_b6_msm.txt


done


#####################


less XCI_final_pt.no_NA.for_percentage_bi_b6_msm.txt | awk '{if($4>72 || $3>72) print $0"\t""XCI"; else if($4>45 && $4<=72 || $3>45 && $3<=72) print $0"\t""XCI_Intermediate";  else print $0"\t""XC_Active"}' > XCI_final_pt.no_NA.for_percentage_bi_b6_msm_mono_allelic_ratio.retry.txt


less XCI_final_pt.no_NA.for_percentage_bi_b6_msm.txt | awk '{if($2>=11) print $1"\t""XC_Active"; else if($2>=0 && $2<11) print $1"\t""XCI"; else if($2==NULL) print $1"\t""Detected_Variants_0"}' > XCI_final_pt.no_NA.for_percentage_bi_b6_msm.2.retry.txt


less XCI_final_pt.no_NA.for_percentage_bi_b6_msm.2.retry.txt | cut -f 2 > XCI_final_pt.no_NA.for_percentage_bi_b6_msm.3.retry.txt


less XCI_final_pt.no_NA.for_percentage_bi_b6_msm_mono_allelic_ratio.retry.txt | cut -f 5 > XCI_final_pt.no_NA.for_percentage_bi_b6_msm_mono_allelic_ratio.2.retry.txt


less XCI_final_pt.no_NA.for_percentage_define_50.txt |cut -f 1,6 > XCI_final_pt.no_NA.for_percentage_define_50_2.txt


paste XCI_final_pt.no_NA.for_percentage_define_50_2.txt XCI_final_pt.no_NA.for_percentage_bi_b6_msm.3.retry.txt XCI_final_pt.no_NA.for_percentage_bi_b6_msm_mono_allelic_ratio.2.retry.txt > XCI_final_pt.no_NA.for_percentage_bi_b6_msm_XCI_define_file.retry.txt


less XCI_final_pt.no_NA.for_percentage_bi_b6_msm_XCI_define_file.retry.txt | awk '{if($3=="XC_Active" && $4=="XC_Active") print $1"\t""XC_Active"; else if($3=="XCI" && $4=="XCI") print $1"\t""XCI"; else if($2=="less_than_50") print $1"\t""No_definition"; else if($3=="Detected_Variants_0") print $1"\t""No_definition" ; else print $1"\t""XCI_Intermediate"}' > XCI_final_pt.no_NA.for_percentage_bi_b6_msm_XCI_define_file_defined.retry.txt


less tSNE_kMeans6_clusterData_kallisto_DGE_geneTPM_20170911.pseudotime.txt | cut -f 1-3 > tSNE_kMeans6_clusterData_kallisto_DGE_geneTPM_20170911.pseudotime.position.txt


less XCI_final_pt.no_NA.for_percentage_bi_b6_msm_XCI_define_file_defined.retry.txt | cut -f 2 > XCI_final_pt.no_NA.for_percentage_bi_b6_msm_XCI_define_file_defined.XCI.retry.txt


paste tSNE_kMeans6_clusterData_kallisto_DGE_geneTPM_20170911.pseudotime.position.txt XCI_final_pt.no_NA.for_percentage_bi_b6_msm_XCI_define_file_defined.XCI.retry.txt > XCI_final_pt.no_NA.for_percentage_bi_b6_msm_XCI_define_file_defined.TSNE.retry.txt


########################


rm XCI_final_pt.no_NA.for_percentage_bi_b6_msm_XCI_define_file_defined.XCI.retry.txt

rm tSNE_kMeans6_clusterData_kallisto_DGE_geneTPM_20170911.pseudotime.position.txt
```
