## Supplementary material for "Single-cell transcriptomics, scRNA-Seq and C1 CAGE discovered distinct phases of pluripotency during naïve-to-primed conversion in mice": 5_XCI_tSNE_plot.R

```
sne <- read.table("/Users/XCI_tSNE_NEW.txt", header = F, check.names = FALSE)


colnames(sne) <- c("Xchr_state","Dim1","Dim2","cell_id")


library(ggplot2)

col <- c("#000000","#7570B3","#66A61E", "#E7298A")

tsneRunPlot <- qplot(Dim1, Dim2, data=sne,
                     colour=Xchr_state) +
  scale_colour_manual(values=col,
                      name="",
                      labels=c("No_definition", "XCI","XCI_Intermediate", "XC_Active")) +
  theme(axis.ticks = element_blank()) +
  theme(panel.background = element_blank()) + theme(axis.line=element_line(color="grey")) +
  theme(panel.grid.minor=element_line(color="grey"), panel.grid.major=element_line(colour="grey")) +
  theme(text = element_text(family="Helvetica")) +
  scale_x_continuous(limits = c(min(sne$Dim1) + min(sne$Dim1)*0.1, max(sne$Dim1) + max(sne$Dim1)*0.1)) +
  scale_y_continuous(limits = c(min(sne$Dim2) + min(sne$Dim2)*0.3, max(sne$Dim2) + max(sne$Dim2)*0.2))

pdf(paste0("/Users/XCI_tSNE.pdf"), width = 12, height = 7, onefile = TRUE)

plot(tsneRunPlot)

dev.off()
```
