## Supplementary material for "Single-cell transcriptomics, scRNA-Seq and C1 CAGE discovered distinct phases of pluripotency during naïve-to-primed conversion in mice": C1_CAGE_DE_Tests.Rmd

````
---
title: "CTSS process 2"
author: "Jonathan Moody"
date: "05/11/2019"
output:
  html_document:
    keep_md: yes
    toc: true
    number_sections: true
    code_folding: hide
knit: (function(inputFile, encoding) {
  rmarkdown::render(inputFile, encoding = encoding, output_dir = "../reports") })
---

# Data loading

```{r}
library(data.table)
library(ggplot2)
library(cowplot)
library(RColorBrewer)
gg_color_hue <- function(n) {
  hues = seq(15, 375, length=n+1)
  hcl(h=hues, l=65, c=100)[1:n]
}
#options(width = 3000)
m <- readRDS('../data/CAGE_GENCODEvM21_mtx.Rds')

anno <- fread('/osc-fs_home/hon-chun/analysis/ILC2/resource/gene_model/GENCODE/FANTOM5_CAGE_clusters/annotation/GENCODEvM21//log/F5_CAGE_anno.GENCODEvM21.cage_cluster.info.tsv.gz')
setkey(anno, clusterID)
rownames(m) <- anno[rownames(m), clusterName]

write.table(m, '../data/CAGE_GENCODEvM1_mtx.tsv', col.names = T, row.names = T, sep='\t', quote=F)
system("gzip -f ../data/CAGE_GENCODEvM1_mtx.tsv")

# RNAseq meta
# meta <- fread('/home/jonathan/other_home/bottcher/Abe/mES2EpiSC/metadata_20180213150953.csv')

# CAGE meta
meta <- fread('/home/jonathan/other_home/bottcher/Abe/mES2EpiSC/metadata-C1CAGE_20180410112239.csv')
m <- m[, colnames(m) %in% meta[!is.na(Pseudotime), cell_id]] #discards one
setkey(meta, cell_id)
```

# Data exploration

```{r}
# Seurat object
library(Seurat)
meta_df <- as.data.frame(meta[colnames(m)])
rownames(meta_df) <- meta_df$cell_id
so <- CreateSeuratObject(m, min.cells = 3, project = "ES_C1", names.delim=NA, meta.data = as.data.frame(meta[cell_id %in% colnames(m)]))
so@meta.data <- meta_df

so <- NormalizeData(so, normalization.method = "LogNormalize", scale.factor = 10000)
so <- FindVariableFeatures(so, selection.method = "vst", nfeatures = 2000)
top10 <- head(VariableFeatures(so), 10)
plot1 <- VariableFeaturePlot(so)
plot2 <- LabelPoints(plot = plot1, points = top10, repel = TRUE)
plot2
so <- ScaleData(so, vars.to.regress = 'counts')
so <- RunPCA(so, features = VariableFeatures(object = so), verbose = FALSE)

#plot1 <- DimPlot(so, reduction = "pca")
plot2 <- ElbowPlot(so, ndims = 40)
plot2
#CombinePlots(plots = list(plot1, plot2))
DimPlot(so, reduction = "pca", group.by = 'timePoint')
so <- subset(so, PC_1<5)
DimPlot(so, reduction = "pca", group.by = 'timePoint')
FeaturePlot(so, reduction = "pca", features = 'Pseudotime')
VizDimLoadings(so, dims = 1:2, reduction = "pca")
saveRDS(so, '../generated/seurat_object.Rds')
```

# Marker genes

* Focusing on enhancers that are are 'marker genes' for a specific stage.

```{r, fig.height=6, fig.width=8, warning=FALSE, message=F}
#markers
so@meta.data$timePoint <- factor(so@meta.data$timePoint, levels = c("day0_mES", 'day1', 'day2', 'day3', 'day4', 'EpiSC'))
Idents(object = so) <- 'timePoint'
data.markers <- FindAllMarkers(so, only.pos = TRUE, min.pct = 0.1, logfc.threshold = 0.25)
data.markers <- as.data.table(data.markers)

top2 <- data.markers[, .SD[1:2], by=.(cluster)]
top5 <- data.markers[, .SD[1:5], by=.(cluster)]
DotPlot(so, features = unique(top5$gene))+coord_flip()
top5_e <- data.markers[grep('.E$', gene)]
top5_e <- top5_e[p_val_adj < 0.05]
DotPlot(so, features = unique(top5_e$gene))+coord_flip()
```

```{r, fig.width=10, fig.height=3}
#cols <- c("day0_mES" = "#d73027", "day1" = "#fc8d59", "day2" = "#fee090", "day3" = "#e0f3f8", "day4"="#91bfdb", "EpiSC"="#4575b4")
cols <-  gg_color_hue(16)[c(1:3,5,10,12)]
f <- function(gene_sel){
  df <- data.table(exp=so@assays$RNA@data[gene_sel, ], 
                   t=so@meta.data$Pseudotime, 
                   class=so@meta.data$timePoint)
  p1 <- (ggplot(df[exp > 0], aes(t, exp)) + geom_point(aes(color=class)) +stat_smooth())+ggtitle(gene_sel)+scale_color_manual(values = cols)+xlab('Pseudotime')+ylab('Expression (log)')
  df2 <- df[, .(cells_exp=.SD[exp>0, .N], .N), by=.(class)][, .(class, cells_exp, N, pc=round((cells_exp/N)*100))]
  p2 <- (ggplot(df2, aes(class, pc, fill=class)) +geom_bar(stat='identity') + ylab('% cells expressing')+ggtitle(gene_sel))+scale_fill_manual(values = cols)+xlab('Sample')
  print(plot_grid(p1, p2))
  return(1)
}
x <- top5_e[, f(gene), by=seq_len(nrow(top5_e))]
rm(x)
```
````
