## Supplementary material for "Single-cell transcriptomics, scRNA-Seq and C1 CAGE discovered distinct phases of pluripotency during naïve-to-primed conversion in mice": C1_CAGE_DE_Tests_2.Rmd

````
---
title: "C1 CAGE DE tests"
author: "Jonathan Moody"
date: "`r format(Sys.Date(), '%d %B, %Y')`"
output:
  html_document:
    keep_md: yes
    toc: true
    number_sections: true
    code_folding: hide
knit: (function(inputFile, encoding) {
  rmarkdown::render(inputFile, encoding = encoding, output_dir = "../reports") })
---
<style type="text/css">
.main-container {
  max-width: 1500px;
  margin-left: auto;
  margin-right: auto;
}
</style>

```{r setup, warning=FALSE, message=FALSE}
library(data.table)
library(ggplot2)
library(cowplot)
library(Seurat)
library(pheatmap)
library(RColorBrewer)
library(knitr)
options(width = 1500)
opts_chunk$set(tidy.opts=list(width.cutoff=1500))

gg_color_hue <- function(n) {
  hues = seq(15, 375, length=n+1)
  hcl(h=hues, l=65, c=100)[1:n]
}
col.gg <- gg_color_hue(16)[c(1:3,5,10,12,14,16)]
```

# Data loading

* Select features which are reasonably expressed / variable
```{r}
so <- readRDS('../generated/seurat_object.Rds')
so <- NormalizeData(so, normalization.method = "LogNormalize", scale.factor = 10000)

cells <- rowSums(as.matrix(so@assays$RNA@data) > 0)
so <- so[cells >= 5,]

anno <- fread('../annotation/bed/F5_CAGE_anno.GENCODEvM21.cage_cluster.annot.bed.gz')
setnames(anno, bed12)
anno <- anno[, .(chr, st, end, name, strand)]
setkey(anno, name)

so
```

* Add MB cluster data
```{r, warning=FALSE}
mbClusters <- fread('/home/bottcher/Abe/mES2EpiSC/C1CAGEresults/tSNE_kMeans6_clusterData_C1CAGE_d0-EpiSC_20180205152721.txt')
m <- as.matrix(mbClusters[,2:4])
rownames(m) <- mbClusters$V1
fix_cluster_n <- c(3,5,1,4,2,6)
m[, 'Cluster'] <- fix_cluster_n[m[, 'Cluster']]

table(MB_cluster_counts=m[, 'Cluster'])
table(After_filtering=m[rownames(m) %in% rownames(so@meta.data), 'Cluster'])

so[["tsne"]] <- CreateDimReducObject(embeddings = m[rownames(m) %in% rownames(so@meta.data), 1:2], key = "tsne_", assay = DefaultAssay(so))
so@meta.data$cluster <- m[rownames(so@meta.data), 3]
so$cluster <- as.factor(as.numeric(so$cluster))
Idents(so) <- 'cluster'

DimPlot(so, group.by = 'cluster', pt.size = 0.5)+scale_color_manual(values = brewer.pal(6, "Dark2"))
ggsave('../figs/C1_tsne_6clusters.pdf', width = 5, height = 3)
so <- so[, so$cluster != 6]
DimPlot(so, group.by = 'cluster', pt.size = 0.5)+scale_color_manual(values = brewer.pal(5, "Dark2"))
ggsave('../figs/C1_tsne_5clusters.pdf', width = 5, height = 3)
```

# DE Tests

* Cluster 3 has more differentially expressed promoters than the other clusters.
```{r, fig.height=6, fig.width=7, message=FALSE, eval=F}
res <- data.table(FindAllMarkers(so, only.pos = F, min.pct = 0.05, logfc.threshold = 0.25))
res_full <- res
res_full <- merge(res_full, anno, by.x='gene', by.y='name')
fwrite(res_full, '../data/DE_MBcluster_res.csv')
```

* Globally most DE promoters are downregulated in cluster 3
```{r, fig.width=10, fig.height=5}
res_full <- fread('../data/DE_MBcluster_res.csv')
res <- res_full[p_val_adj <= 0.05]
res <- res[order(p_val)][!duplicated(gene)][order(gene)]
fwrite(res, '../data/DE_MBcluster_res_sig.csv')
res[, table(`sig DE promoters per cluster`=as.numeric(cluster))]
res <- res[1:((floor(nrow(res)/10))*10)]

annotation_col <- data.frame(timePoint = factor(so@meta.data$timePoint, 
                             labels = c("day0_mES","day1","day2","day3","day4","EpiSC")),
                         cluster = as.factor(so@meta.data$cluster))

rownames(annotation_col) <- colnames(so@assays$RNA@data)

annotation_row <- data.frame(res$cluster, row.names = res$gene)

so <- ScaleData(so, features = res$gene)
mat <- (so@assays$RNA@scale.data)
mat <- mat[order(res$cluster, res$p_val), order(so@meta.data$cluster, so@meta.data$Pseudotime)]

ann.colors = list(timePoint = gg_color_hue(16)[c(1:3,5,10,12)],
                  cluster = brewer.pal(5, "Dark2"))
names(ann.colors$timePoint) <- c("day0_mES","day1","day2","day3","day4","EpiSC")
names(ann.colors$cluster) <- as.character(1:5)

x <- pheatmap(mat, 
        show_rownames = T, 
        show_colnames = F, 
        cluster_cols = F,
        cluster_rows = T,
        annotation_col = annotation_col, 
        #annotation_row = annotation_row,
        annotation_names_col = F,
        annotation_names_row = F,
        #cutree_rows= 10,
        kmeans_k = 10,
        main = NA,
        annotation_colors = ann.colors,
        filename = '../figs/DE_features_C1.pdf',
        width=10, height = 5
        )
df <- data.table(cluster = x$kmeans$cluster, feature=names(x$kmeans$cluster))
fwrite(df, '../generated/CAGE_heatmap_clusters.csv')
```

```{r, fig.width=10, fig.height=5}
grid::grid.newpage()
grid::grid.draw(x$gtable)
```

# Cluster 3 DE results

* Top 12 DE promoters in cluster 3
* Downregulated housekeeping genes
```{r}
res <- res_full[cluster == 3][p_val_adj <= 0.05]
res <- res[order(p_val)]

data.table(merge(res[1:12], anno, by.x='gene', by.y='name'))
topn <- res[1:12, gene]
DotPlot(so, features = unique(topn))+coord_flip()
pdf('../figs/DE_promoters_cluster3.pdf', height=5, width=7)
DotPlot(so, features = unique(topn))+coord_flip()
dev.off()
```

# Enhancers

* 12 enhancers show significant DE (p_val_adj <= 0.05) in cluster 3
* Two with increased expression
```{r}
merge(res[grepl('^e1@', gene)], anno, by.x='gene', by.y='name')
topn <- res[grepl('^e1@', gene), gene]
DotPlot(so, features = topn)+coord_flip()
pdf('../figs/DE_enhancers_cluster3.pdf', height=5, width=7)
DotPlot(so, features = topn)+coord_flip()
dev.off()
```

# Upregulated features in cluster 3

* Only 7 promoters show upregulation in cluster 3, of these two are enhancers also seen above.
* Enhancer ADDG10096077546.E is 135kb upstream of Eea1
* Enhancer ADDG12069361479.E is a mis-annotation and looks like the promotor of Rn7s2
```{r, fig.width=7, fig.height=4}
merge(res[avg_logFC > 0 & pct.1 > 0.8*pct.2], anno, by.x='gene', by.y='name')
topn <- res[!grepl('^e1@', gene) & avg_logFC > 0 & pct.1 > 0.8*pct.2, gene]
DotPlot(so, features = topn)+coord_flip()
pdf('../figs/DE_upregulated_cluster3.pdf', height=5, width=7)
DotPlot(so, features = topn)+coord_flip()
dev.off()
```

# day0 vs EpiSC

```{r}
Idents(so) <- so$timePoint
res <- data.table(FindMarkers(so, only.pos = F, min.pct = 0.05, logfc.threshold = 0.25, ident.1 = 'EpiSC', ident.2 = 'day0_mES'), keep.rownames = T)
setnames(res, 'rn', 'gene')
res <- merge(res, anno, by.x='gene', by.y='name')[order(p_val)]
fwrite(res, '../data/DE_day0_vs_EpiSC.csv')
```
````
