## Supplementary material for "Single-cell transcriptomics, scRNA-Seq and C1 CAGE discovered distinct phases of pluripotency during naïve-to-primed conversion in mice": C1_CAGE_Slingshot.Rmd

````
---
title: "CTSS process 3 - Pseudotime"
author: "Jonathan Moody"
date: "27/02/2019"
output:
  html_document:
    keep_md: yes
    toc: true
    number_sections: true
    code_folding: hide
knit: (function(inputFile, encoding) {
  rmarkdown::render(inputFile, encoding = encoding, output_dir = "../reports") })
---

# Data loading

* Select features which are reasonably expressed / variable
```{r, warning=FALSE, message=FALSE}
#library(reticulate)
#use_condaenv(condaenv="sc", conda="/home/jonathan/data/programs/miniconda_202002/bin/conda")
library(data.table)
library(ggplot2)
library(cowplot)
library(Seurat)
library(pheatmap)
library(RColorBrewer)
library(slingshot)
library(tradeSeq)
library(SingleCellExperiment)
```

```{r}
options(width = 3000)
so <- readRDS('../generated/seurat_object.Rds')
so <- NormalizeData(so, normalization.method = "LogNormalize", scale.factor = 10000)

cells <- rowSums(as.matrix(so@assays$RNA@data) > 0)
so <- so[cells >= 5,]

so <- FindVariableFeatures(so, selection.method = "vst", nfeatures = 10000)
x <- so@assays$RNA@meta.features
x$keep <- (x$vst.mean >= quantile(x$vst.mean, 0.2) & 
        x$vst.variance.standardized >= median(x$vst.variance.standardized)
        )

ggplot(x, aes(vst.mean, vst.variance.standardized, color=keep)) + geom_point() +scale_x_log10()

so <- so[x$keep == T, ]
so
```

* Add MB cluster data
```{r}
mbClusters <- fread('/home/bottcher/Abe/mES2EpiSC/C1CAGEresults/tSNE_kMeans6_clusterData_C1CAGE_d0-EpiSC_20180205152721.txt')
m <- as.matrix(mbClusters[,2:4])
rownames(m) <- mbClusters$V1
so[["tsne"]] <- CreateDimReducObject(embeddings = m[rownames(m) %in% rownames(so@meta.data), 1:2], key = "tsne_", assay = DefaultAssay(so))
so@meta.data$cluster <- m[rownames(so@meta.data), 3]
Idents(so) <- 'cluster'

DimPlot(so, reduction = "tsne", group.by = 'cluster', pt.size = 0.5)
FeaturePlot(so, features = 'Pseudotime', pt.size = 0.5)
DimPlot(so, reduction = "tsne", group.by = 'breaks', pt.size = 0.5)
```
#Pseudotime Bins

```{r}
so@meta.data$breaks <- cut(so@meta.data$Pseudotime, breaks = 10)
Idents(so) <- 'breaks'
av <- AverageExpression(so)[[1]]
x <- pheatmap(log1p(av), cluster_cols = F, cutree_rows = 5, show_rownames = F)
```

# Clustering

* Run PCA, define early and late clusters for easy DE test, compare with previous pseudotime
```{r}
so <- RunPCA(so, verbose = FALSE)
ElbowPlot(so, ndims = 30)
so <- FindNeighbors(so, dims = 1:10)
so <- FindClusters(so, algorithm = 4, resolution = c(1)) # 4 = leiden algorithm
so <- RunUMAP(so, dims = 1:10)

DimPlot(so, reduction = "pca", group.by = 'timePoint', pt.size = 0.5, label = T)
DimPlot(so, reduction = "pca", group.by = 'cluster', pt.size = 0.5, label = T)
DimPlot(so, reduction = "pca", group.by = 'RNA_snn_res.0.9', pt.size = 0.5, label = T)

DimPlot(so, reduction = "umap", group.by = 'timePoint', pt.size = 0.5, label = T)
DimPlot(so, reduction = "umap", group.by = 'cluster', pt.size = 0.5, label = T)
DimPlot(so, reduction = "umap", group.by = 'RNA_snn_res.0.9', pt.size = 0.5, label = T)
```

# Pseudotime curves with Slingshot

```{r}
cl <- so@meta.data$RNA_snn_res.0.9
lin <- getLineages(so@reductions$pca@cell.embeddings[, 1:8], clusterLabels = cl, start.clus = 2, end.clus=5)
crv <- getCurves(lin)
plotGeneCount(curve = crv, clusters = cl)
```

* Test possible values for K (number of knots in the model)

```{r, eval=F, fig.height=5, fig.width=10}
icMat <- evaluateK(counts = as.matrix(so@assays$RNA@counts), sds = crv, k=3:10, nGenes = 500, verbose=FALSE)
```

* Filter genes to model

```{r}
qt <- 0.8
k <- 7
x <- data.table(so@assays$RNA@meta.features, keep.rownames = T)
id <- x[vst.variance.standardized >= quantile(vst.variance.standardized, qt)]$rn
counts <- as.matrix(so@assays$RNA@counts)[id, ]
cat('All: ', nrow(counts), '\n')
cat('Enhancers: ', sum(grepl('.E$', rownames(counts))), '\n')
```

# Model fitting

* This fits the NB-GAM model for each gene with the lineage.

```{r, eval=F}
sce <- fitGAM(counts = counts,
              sds = crv,
              nknots = k
              )
saveRDS(sce, paste0('../generated/pseudo_model_',qt,'_',k,'.Rds'))
```

```{r}
sce <- readRDS(paste0('../generated/pseudo_model_',qt,'_',k,'.Rds'))
plotGeneCount(curve = crv, counts = counts, clusters = cl,
              models = sce)

f <- function(gene){
  return(plotGeneCount(crv, counts, gene = gene)+scale_color_gradient(low = "yellow", high = 'red', name='log2(count)')+ggtitle(gene))
}

f2 <- function(gene){
  return(plotSmoothers(sce, counts, gene = gene, lwd=0.5)+ggtitle(gene))
}
```

# associationTest

* The null hypothesis that all smoother coefficients are equal to each other. This can be interpreted as testing whether the smoothed gene expression is significantly changing along pseudotime.

```{r}
assoRes <- associationTest(sce)
res <- data.table(assoRes, keep.rownames = T)
res <- res[order(pvalue)]
```

## Top promoters
```{r, message=FALSE, warning=FALSE, fig.height=10, fig.width=12}
res[1:min(nrow(res), 9)]
p <- lapply(res[1:min(nrow(res), 9)]$rn, f)
CombinePlots(p, ncol = 3)
```

```{r, fig.height=10, fig.width=12}
p <- lapply(res[1:min(nrow(res), 9)]$rn, f2)
CombinePlots(p, ncol = 3)
```

## Enhancers
```{r, message=FALSE, warning=FALSE, fig.height=10, fig.width=12}
res[pvalue <= 0.1][grep('.E$', rn)][1:min(nrow(res), 9)]
p <- lapply(res[pvalue <= 0.1][grep('.E$', rn)][1:min(nrow(res), 9)]$rn, f)
CombinePlots(p)
```

```{r, message=FALSE, warning=FALSE, fig.height=10, fig.width=12}
p <- lapply(res[pvalue <= 0.1][grep('.E$', rn)][1:min(nrow(res), 9)]$rn, f2)
CombinePlots(p)
```

# startVsEndTest

* A Wald test that tests the null hypothesis that the expression at the starting point of the smoother (progenitor population) is identical to the expression at the end point of the smoother (differentiated population).

```{r, message=FALSE, warning=FALSE, fig.height=10, fig.width=12}
startRes <- startVsEndTest(sce)
res <- data.table(startRes, keep.rownames = T)
res <- res[order(pvalue)]
```

## Top promoters
```{r, message=FALSE, message=FALSE, warning=FALSE, fig.height=10, fig.width=12}
res[1:min(nrow(res), 9)]
p <- lapply(res[1:min(nrow(res), 9)]$rn, f)
CombinePlots(p)
```

```{r, message=FALSE, message=FALSE, warning=FALSE, fig.height=10, fig.width=12}
p <- lapply(res[1:min(nrow(res), 9)]$rn, f2)
CombinePlots(p)
```

## Enhancers
```{r, message=FALSE, message=FALSE, warning=FALSE, fig.height=10, fig.width=10}
res[pvalue <= 0.1][grep('.E$', rn)][1:min(nrow(res), 9)]
p <- lapply(res[pvalue <= 0.1][grep('.E$', rn)][1:min(nrow(res), 9)]$rn, f)
CombinePlots(p)
```

```{r, message=FALSE, message=FALSE, warning=FALSE, fig.height=10, fig.width=10}
p <- lapply(res[pvalue <= 0.1][grep('.E$', rn)][1:min(nrow(res), 9)]$rn, f2)
CombinePlots(p)
```

```{r}
colData(sce) <- cbind(colData(sce), so@meta.data)
reducedDims(sce) <- list(PCA=so@reductions$pca@cell.embeddings)
save(sce, startRes, assoRes, lin, file=paste0('../generated/pseudo_model_',qt,'_',k,'.Rdata'))
```
````
