## Supplementary material for "Single-cell transcriptomics, scRNA-Seq and C1 CAGE discovered distinct phases of pluripotency during naïve-to-primed conversion in mice": C1_CAGE_Slingshot_clustering.Rmd

````
---
title: "Pseudotime gene clustering"
author: "Jonathan Moody"
date: "`r format(Sys.Date(), '%d %B, %Y')`"
output:
  html_document:
    keep_md: yes
    toc: true
    number_sections: true
    code_folding: hide
knit: (function(inputFile, encoding) {
  rmarkdown::render(inputFile, encoding = encoding, output_dir = "../reports") })
---

```{r, warning=FALSE, message=FALSE}
library(data.table)
library(ggplot2)
library(cowplot)
library(pheatmap)
library(RColorBrewer)
library(slingshot)
library(tradeSeq)
library(SingleCellExperiment)
library(clusterExperiment)
f2 <- function(gene){
  return(plotSmoothers(sce, assay(sce, 'counts'), gene = gene, lwd=0.5)+ggtitle(gene))
} 
gg_color_hue <- function(n) {
  hues = seq(15, 375, length=n+1)
  hcl(h=hues, l=65, c=100)[1:n]
}
```

# Load data

* The slingshot pseudotime curve is based on the first pca plot, with 4 clusters.
* However cluster 4 here has a lot of overlap with cluster 3 in previous clustering by MB. (see comparison with MB_cluster)

```{r, fig.width=10, fig.height=5}
qt <- 0.8
k <- 7
nPointsClus <- 10
load(paste0('../generated/pseudo_model_',qt,'_',k,'.Rdata'))
```

```{r, warning=FALSE}
mbClusters <- fread('/home/bottcher/Abe/mES2EpiSC/C1CAGEresults/tSNE_kMeans6_clusterData_C1CAGE_d0-EpiSC_20180205152721.txt')
m <- as.matrix(mbClusters[,2:4])
rownames(m) <- mbClusters$V1
fix_cluster_n <- c(3,5,1,4,2,6)
m[, 'Cluster'] <- fix_cluster_n[m[, 'Cluster']]
colData(sce)$MB_cluster <- as.character(m[colnames(sce), 'Cluster'])
```

```{r, fig.height=4, fig.width=7}
crv <- getCurves(lin)
plotGeneCount(curve = crv, clusters = colData(sce)$RNA_snn_res.0.7, models = sce)

coldata <- as.data.table(colData(sce))
cols <-  gg_color_hue(16)[c(1:3,5,10,12)]

ggplot(coldata, aes(round(slingshot.pseudotime.curve1), fill=RNA_snn_res.0.7)) + geom_bar(position='stack') + xlab('Slingshot Pseudotime')+scale_color_manual(values = brewer.pal(5, "Dark2"))
ggsave('../figs/cells_clustering_1.pdf', height=4, width=7)

ggplot(coldata, aes(round(slingshot.pseudotime.curve1), fill=MB_cluster)) + geom_bar(position='stack') + xlab('Slingshot Pseudotime')+scale_fill_manual(values = brewer.pal(6, "Dark2"))
ggsave('../figs/cells_clustering_2.pdf', height=4, width=7)

ggplot(coldata, aes(round(slingshot.pseudotime.curve1), fill=timePoint)) + geom_bar(position='stack') + xlab('Slingshot Pseudotime')+scale_fill_manual(values = cols)
ggsave('../figs/cells_clustering_3.pdf', height=4, width=7)

```

# DE in MB clusters

* Promotors which were defined as DE in the previous cluster based analysis 
* Top 9 for each cluster plotted on the Slingshot pseudotime.

```{r}
cluster_de <- fread('../data/DE_MBcluster_res.csv')
cluster_de <- cluster_de[gene %in% rownames(sce)][!duplicated(gene)]
```

## MB Cluster 1 DE genes
```{r, fig.height=5, fig.width=12}
p <- lapply(cluster_de[cluster==1][1:9, gene], f2)
plot_grid(plotlist=p, ncol=3)
pdf('../figs/MB_custer1_DE_pseudotime.pdf', height=5, width=12)
plot_grid(plotlist=p, ncol=3)
dev.off()
```

## MB Cluster 2 DE genes
```{r, fig.height=5, fig.width=12}
p <- lapply(cluster_de[cluster==2][1:9, gene], f2)
plot_grid(plotlist=p, ncol=3)
pdf('../figs/MB_custer2_DE_pseudotime.pdf', height=5, width=12)
plot_grid(plotlist=p, ncol=3)
dev.off()
```

## MB Cluster 3 DE genes
```{r, fig.height=5, fig.width=12}
p <- lapply(cluster_de[cluster==3][1:9, gene], f2)
plot_grid(plotlist=p, ncol=3)
pdf('../figs/MB_custer3_DE_pseudotime.pdf', height=5, width=12)
plot_grid(plotlist=p, ncol=3)
dev.off()
```

## MB Cluster 4 DE genes
```{r, fig.height=5, fig.width=12}
p <- lapply(cluster_de[cluster==4][1:9, gene], f2)
plot_grid(plotlist=p, ncol=3)
pdf('../figs/MB_custer4_DE_pseudotime.pdf', height=5, width=12)
plot_grid(plotlist=p, ncol=3)
dev.off()
```

## MB Cluster 5 DE genes
```{r, fig.height=5, fig.width=12}
p <- lapply(cluster_de[cluster==5][1:9, gene], f2)
plot_grid(plotlist=p, ncol=3)
pdf('../figs/MB_custer5_DE_pseudotime.pdf', height=5, width=12)
plot_grid(plotlist=p, ncol=3)
dev.off()
```

# Gene clustering

* Cells are binned into 10 bins along the pseudotime.
* Expression is smoothed for each promoter across bins.
* Promotors are clustered into modules by thier variation in the pseudotime bins.
* Similar modules are merged.

```{r, eval=F}
genesUse <- union(rownames(assoRes[assoRes$pvalue < 0.05, ]),
               rownames(assoRes[startRes$pvalue < 0.05, ]))

clusPat <- clusterExpressionPatterns(sce, nPoints = nPointsClus, minSizes = 50,
                                     reduceMethod = "none", nReducedDims = nPointsClus,
                                     clusterFunction = "hierarchical01",
                                     subsampleArgs = list(clusterFunction='kmeans'),
                                     genes = genesUse,
                                     ncores=10)

save(sce, clusPat, startRes, assoRes, lin, file=paste0('../generated/pseudo_model_geneclus_',qt,'_',k,'.Rdata'))
```

```{r, fig.width=12, fig.height=6}
load(paste0('../generated/pseudo_model_geneclus_',qt,'_',k,'.Rdata'))
genesUse <- union(rownames(assoRes[assoRes$pvalue < 0.05, ]),
               rownames(assoRes[startRes$pvalue < 0.05, ]))
```

```{r, fig.width=7, fig.height=7, warning=FALSE}
clusPat$rsec <- mergeClusters(clusPat$rsec,mergeMethod="adjP",DEMethod="limma",cutoff=0.95,plot=T,plotInfo="mergeMethod") 
clusterLabels <- primaryCluster(clusPat$rsec)
cUniq <- sort(unique(clusterLabels))
cUniq <- cUniq[!cUniq == -1] # remove unclustered genes
```

# Plot Modules

* Expression in the smoothed pseudotime bins can be misleadings when the data is sparse.
* Enhancers from each bin are plotted below on the pseudotime axis, for comparison.

```{r, fig.height=5, fig.width=12}
anno_bed <- fread('../annotation/bed/F5_CAGE_anno.GENCODEvM21.cage_cluster.annot.bed.gz')
setnames(anno_bed, bed12)
anno_bed <- anno_bed[, .(chr, st, end, name, strand)]
anno <- fread('../annotation/log/F5_CAGE_anno.GENCODEvM21.cage_cluster.info.tsv.gz')
anno <- anno[, .(clusterName,  type, geneClassStr)]
anno <- merge(anno, anno_bed, by.x='clusterName', by.y='name')

cl <- data.table(clusterName = genesUse, cluster = clusterLabels)
cl <- cl[cluster != -1]
cl <- merge(cl, anno, by='clusterName')
cl <- cl[geneClassStr %in% c('enhancer_locus', 'protein_coding')]
setnames(cl, 'cluster', 'module')
fwrite(cl, '../data/Slingshot_clusters.csv')


cl[, table(type, module)]

x <- clusPat$yhatScaled
rownames(x) <- genesUse
colnames(x) <- 1:nPointsClus
x <- data.table(x, keep.rownames = T)
setnames(x, 'rn', 'clusterName')
x <- melt(x, id.vars = 'clusterName', variable.name = 'Pseudotime')
x <- merge(cl[module %in% 1:length(cUniq)], x, by='clusterName')

p <- (ggplot(x) 
  +geom_line(data=subset(x, type=='promoter'), aes(Pseudotime, value, color=type, group=clusterName))
  +geom_line(data=subset(x, type=='enhancer'), aes(Pseudotime, value, color=type, group=clusterName)) 
  +stat_smooth(data=x,  mapping = aes(x=as.numeric(Pseudotime), y=value, color='ClusterAverage'),  method='loess', span = 0.7)
  +scale_colour_manual(values = c('red','blue','grey'))
  +facet_wrap('module')
  +theme_bw()
  +ylab("Normalized expression")
  +xlab('Pseudotime bin')
  )
p
ggsave('../figs/feature_modules.pdf', height=5, width=12)
```
## Bin labels
```{r, fig.height=4, fig.width=7}
labs <- tradeSeq:::.getPredictRangeDf(colData(sce)$tradeSeq$dm,1,11)
coldata$pseudotime.bin <- cut(coldata$slingshot.pseudotime.curve1, breaks = labs$t1, include.lowest = T)
ggplot(coldata, aes(round(slingshot.pseudotime.curve1), fill=pseudotime.bin)) + geom_bar(position='stack') + xlab('Slingshot Pseudotime')
ggsave('../figs/cells_clustering_4.pdf', height=4, width=7)
table(bin=coldata$pseudotime.bin, MB_cluster=coldata$MB_cluster)
```


# Enhancers within each module

## Module 1
```{r, fig.height=10, fig.width=12}
p <- lapply(cl[module==1 & type=='enhancer']$clusterName, f2)
plot_grid(plotlist=p, ncol=3)
pdf('../figs/Module1_enhancers_pseudotime.pdf', height=10, width=12)
plot_grid(plotlist=p, ncol=3)
dev.off()
```

## Module 2
```{r, fig.height=10, fig.width=12}
p <- lapply(cl[module==2 & type=='enhancer']$clusterName, f2)
plot_grid(plotlist=p, ncol=3)
pdf('../figs/Module2_enhancers_pseudotime.pdf', height=10, width=12)
plot_grid(plotlist=p, ncol=3)
dev.off()
```

## Module 3
```{r, fig.height=6.5, fig.width=12}
p <- lapply(cl[module==3 & type=='enhancer']$clusterName, f2)
plot_grid(plotlist=p, ncol=3)
pdf('../figs/Module3_enhancers_pseudotime.pdf', height=6.5, width=12)
plot_grid(plotlist=p, ncol=3)
dev.off()
```

## Module 4
```{r, fig.height=12, fig.width=12}
p <- lapply(cl[module==4 & type=='enhancer']$clusterName, f2)
plot_grid(plotlist=p, ncol=3)
pdf('../figs/Module4_enhancers_pseudotime.pdf', height=12, width=12)
plot_grid(plotlist=p, ncol=3)
dev.off()
```

## Module 5
```{r, fig.height=4, fig.width=12}
p <- lapply(cl[module==5 & type=='enhancer']$clusterName, f2)
plot_grid(plotlist=p, ncol=3)
pdf('../figs/Module5_enhancers_pseudotime.pdf', height=4, width=12)
plot_grid(plotlist=p, ncol=3)
dev.off()
```
````
