## Supplementary material for "Single-cell transcriptomics, scRNA-Seq and C1 CAGE discovered distinct phases of pluripotency during naïve-to-primed conversion in mice": cellCycle_code.R

```
library(dplyr)
library(ggplot2)
library(edgeR)
library(gplots)
library(magrittr)
library(RColorBrewer)

###
# scRNA-seq data
# Kallisto geneCount_M8_20170908_Kallisto.txt includes 1EP7, MB#3P10, which older version geneCount_M8_20170808_Kallisto.txt did not
# orthologous mouse genes based on the cell cycle gene set from:
# "Identification of Genes Periodically Expressed in the Human Cell Cycle and Their Expression in Tumors"

meta <- read.csv("metadata_20190223045845.csv", header=TRUE, stringsAsFactors = F)
tpm <- read.table("geneCount_M8_20170908_Kallisto.txt", header=T, row.names = 1, check.names = FALSE)
tpm <- tpm[, which(names(tpm) %in% meta$cell_id)]
meta <- meta[match(names(tpm), meta$cell_id),]
expr <- tpm[,!meta$discard]
meta <- meta[!meta$discard,]
# the following must be TRUE
identical(meta$cell_id, names(expr))

cycleGenes <- read.table("mES2EpiSC_cellcyclegenes.txt", header=TRUE, sep = "\t", na.strings = c("", "NA"))
expressed <- rowSums(expr) > 0
expr <- expr[expressed,]

# makes list of 5 tables containing all the genes for each respective cell cycle phase
cycleGenes_list <- lapply(1:5,function(x){
  temp <- as.character(cycleGenes[,x])
  temp[temp!=""]
})

phaseScore <- sapply(cycleGenes_list, function(x){
  #### create table of each phase
  phase <- expr[rownames(expr) %in% unlist(x),]
  #### add average expression of all genes in the phase
  combined_matrix <- rbind(phase, average = apply(phase, 2, mean))
  #### use transpose to compute cor matrix
  cor_matrix <- cor(t(combined_matrix))
  #### take the numbers
  cor_vector <- cor_matrix[, dim(cor_matrix)[1]]
  #### restrict to correlation >= 0.3 
  phase_restricted <- phase[rownames(phase) %in% names(cor_vector[cor_vector >= 0.3]),]
  #### apply normalization to reads
  norm_factors <- calcNormFactors(phase_restricted, method = "TMM")
  phase_cpm <- cpm(phase_restricted, log = TRUE,
                   lib.size = colSums(expr) * norm_factors)
  #### output the phase specific scores (mean of normalized expression levels in the phase)
  apply(phase_cpm, 2, mean)
})

# normalization function
flexible_normalization <- function(data_in,by_row=TRUE){
  if(by_row){
    row_mean <- apply(data_in,1,mean)
    row_sd   <- apply(data_in,1,sd)
    output <- data_in
    for(i in 1:dim(data_in)[1]){
      output[i,] <- (data_in[i,] - row_mean[i])/row_sd[i]
    }
  }
  #### if by column
  if(!by_row){
    col_mean <- apply(data_in,2,mean)
    col_sd   <- apply(data_in,2,sd)
    output <- data_in
    for(i in 1:dim(data_in)[2]){
      output[,i] <- (data_in[,i] - col_mean[i])/col_sd[i]
    }
  }
  output
}

phaseScore_normed <- flexible_normalization(phaseScore, by_row=FALSE)
phaseScore_normed2 <- flexible_normalization(phaseScore_normed, by_row=TRUE)

# assign phase to cell
cellPhase <- apply(phaseScore_normed2, 1, function(x) colnames(cycleGenes)[which.max(x)])
assign_cellPhase <- data.frame(cellPhase); names(assign_cellPhase) <- "phase"
assign_cellPhase$phase <- factor(assign_cellPhase$phase, levels(assign_cellPhase$phase)[c(1,5,2,3,4)])

# save table with cell_id and assigned cell cycle phase for each cell_id
time <- paste(gsub("-|:", "", unlist(stringr::str_split(as.character(Sys.time()), " ", n = 2))), collapse = "")
write.csv(assign_cellPhase, file = paste0("cellCycle_metadata_", time, ".csv"), row.names = T)

#####

cellcycle <- read.csv("cellCycle_metadata_20190314160652.csv", header=TRUE, stringsAsFactors = F)
# start for cluster composition
sne <- fread("tSNE_kMeans5_clusterData_kallisto_DGE_geneTPM_20190223045041.txt")
sne <- as.data.frame(sne)
sne$Cluster <- factor(sne$Cluster)
sne$Cluster

sne <- sne[match(cellcycle$X, sne$cell_id),]
# the following must be TRUE
identical(sne$cell_id, cellcycle$X)
sne$phase <- factor(cellcycle$phase)
sne$phase <- factor(sne$phase, levels(sne$phase)[c(1,5,2,3,4)])


tsneCellCyclePlot <- qplot(Dim1, Dim2, data=sne, colour=phase) +
  scale_colour_brewer(palette = "Set1", name="") +
  theme(axis.ticks = element_blank()) + 
  theme(panel.background = element_blank()) + theme(axis.line=element_line(color="grey")) +
  theme(panel.grid.minor=element_line(color="grey"), panel.grid.major=element_line(colour="grey")) +
  theme(text = element_text(family="Helvetica")) +
  scale_x_continuous(limits = c(min(sne$Dim1) + min(sne$Dim1)*0.1, max(sne$Dim1) + max(sne$Dim1)*0.1)) +
  scale_y_continuous(limits = c(min(sne$Dim2) + min(sne$Dim2)*0.3, max(sne$Dim2) + max(sne$Dim2)*0.2))

time <- paste(gsub("-|:", "", unlist(stringr::str_split(as.character(Sys.time()), " ", n = 2))), collapse = "")
pdf(paste0("S3_tSNE_CellCycle_", time, ".pdf"), width = 12, height = 7, onefile = TRUE)
plot(tsneCellCyclePlot)
dev.off()


# save cluster composition as file
x1 <- subset(sne, Cluster==1)[,7]
x2 <- subset(sne, Cluster==2)[,7]
x3 <- subset(sne, Cluster==3)[,7]
x4 <- subset(sne, Cluster==4)[,7]
x5 <- subset(sne, Cluster==5)[,7]

cl1 <- as.data.frame(table(as.character(sne[match(x1, sne$cell_id),8]))); names(cl1) <- c("phase", "Cluster1")
cl2 <- as.data.frame(table(as.character(sne[match(x2, sne$cell_id),8]))); names(cl2) <- c("phase", "Cluster2")
cl3 <- as.data.frame(table(as.character(sne[match(x3, sne$cell_id),8]))); names(cl3) <- c("phase", "Cluster3")
cl4 <- as.data.frame(table(as.character(sne[match(x4, sne$cell_id),8]))); names(cl4) <- c("phase", "Cluster4")
cl5 <- as.data.frame(table(as.character(sne[match(x5, sne$cell_id),8]))); names(cl5) <- c("phase", "Cluster5")

allClusters <- merge(cl1, cl2, by = "phase", all = T)
allClusters <- merge(allClusters, cl3, by = "phase", all = T)
allClusters <- merge(allClusters, cl4, by = "phase", all = T)
allClusters <- merge(allClusters, cl5, by = "phase", all = T)
allClusters[is.na(allClusters)] <- 0
allClustersCycle <- allClusters

percentCycles <- data.frame(allClusters$phase,
                           allClusters$Cluster1/as.numeric(colSums(allClusters[-1]))[1]*100,
                           allClusters$Cluster2/as.numeric(colSums(allClusters[-1]))[2]*100,
                           allClusters$Cluster3/as.numeric(colSums(allClusters[-1]))[3]*100,
                           allClusters$Cluster4/as.numeric(colSums(allClusters[-1]))[4]*100,
                           allClusters$Cluster5/as.numeric(colSums(allClusters[-1]))[5]*100)
names(percentCycles) <- names(allClusters)

# save tSNE  and cellcycle data for reproducibility
time <- paste(gsub("-|:", "", unlist(stringr::str_split(as.character(Sys.time()), " ", n = 2))), collapse = "")
write.table(allClustersCycle, file = paste0("tSNE_kmeans5_cellcycleCounts_Kallisto_DGE_geneTPM_", time, ".txt"), row.names = FALSE, col.names = TRUE, sep = "\t", quote = FALSE) 
write.table(percentCycles, file = paste0("tSNE_kmeans5_cellcyclePercentage_Kallisto_DGE_geneTPM_", time, ".txt"), row.names = FALSE, col.names = TRUE, sep = "\t", quote = FALSE) 

# phase pie charts
blank_theme <- theme_minimal()+
  theme(
    axis.title.x = element_blank(),
    axis.title.y = element_blank(),
    panel.border = element_blank(),
    panel.grid=element_blank(),
    axis.ticks = element_blank(),
    plot.title=element_text(size=14, face="bold")
  )

percentCyclesLong <- melt(percentCycles); names(percentCyclesLong) <- c("phase", "Cluster", "Value")
percentCyclesLong$position <- cumsum(percentCyclesLong$Value) - percentCyclesLong$Value/2
percentCyclesLong$labels <- paste0(round(percentCyclesLong$Value, digits = 1), " %")
percentCyclesLong$phase <- factor(percentCyclesLong$phase, levels(percentCyclesLong$phase)[c(1,5,2,3,4)])

col <- brewer.pal(5, "Set1")
library(scales)
clusterPie <- ggplot(percentCyclesLong, aes(x = "", y = Value, fill = phase)) +
  geom_bar(aes(x = "", y = Value, fill = phase), width = 1, color = "black", stat = "identity") + 
  #geom_text(data = percentCyclesLong, aes(x = "", y = position, label = labels), size=2) +
  coord_polar(theta = "y") + 
  facet_grid(facets = .~Cluster, labeller = label_value) +
  scale_fill_manual(values = col) +
  blank_theme +
  theme(axis.text.x=element_blank())+
  theme(text = element_text(family="Helvetica"))

time <- paste(gsub("-|:", "", unlist(stringr::str_split(as.character(Sys.time()), " ", n = 2))), collapse = "")
pdf(paste0("S3_phase_pieCharts", time, ".pdf"), width = 12, height = 7, onefile = TRUE)
plot(clusterPie)
dev.off()  

# cluster pie charts
allClustersT <- as.data.frame(t(allClusters))
names(allClustersT) <- as.character(as.matrix(allClustersT[1,]))
allClustersT <- allClustersT[-1,]
allClustersT <- as.data.frame(apply(allClustersT, 2, function(x) as.numeric(as.character(x))))
allClustersT$Cluster <- names(allClusters)[-1]
percentT <- as.data.frame(sapply(1:5, function(x) allClustersT[,x]/as.numeric(colSums(allClustersT[,-6]))[x]*100))
names(percentT) <- names(allClustersT[-6])
percentT$Cluster <- allClustersT$Cluster

percentTLong <- melt(percentT); names(percentTLong) <- c("Cluster", "phase", "Value")
percentTLong$position <- cumsum(percentTLong$Value) - percentCyclesLong$Value/2
percentTLong$labels <- paste0(round(percentTLong$Value, digits = 1), " %")
percentTLong$phase <- factor(percentTLong$phase, levels(percentTLong$phase)[c(1,5,2,3,4)])

write.table(percentT, file = paste0("tSNE_percentClusterPerCycle_Kallisto_DGE_geneTPM_", time, ".txt"), row.names = FALSE, col.names = TRUE, sep = "\t", quote = FALSE) 

col <- brewer.pal(5, "Dark2")
library(scales)
phasePie <- ggplot(percentTLong, aes(x = "", y = Value, fill = Cluster)) +
  geom_bar(aes(x = "", y = Value, fill = Cluster), width = 1, color = "black", stat = "identity") + 
  #geom_text(data = percentTLong, aes(x = "", y = position, label = labels), size=2) +
  coord_polar(theta = "y") + 
  scale_fill_manual(values = col) +
  facet_grid(facets = .~phase, labeller = label_value) +
  blank_theme +
  theme(axis.text.x=element_blank()) +
  theme(text = element_text(family="Helvetica"))

time <- paste(gsub("-|:", "", unlist(stringr::str_split(as.character(Sys.time()), " ", n = 2))), collapse = "")
pdf(paste0("S3_cluster_pieCharts", time, ".pdf"), width = 12, height = 7, onefile = TRUE)
plot(phasePie)
dev.off()  
```
