## Supplementary material for "Single-cell transcriptomics, scRNA-Seq and C1 CAGE discovered distinct phases of pluripotency during naïve-to-primed conversion in mice": CTSS_process.Rmd

````
---
title: "ES C1 CAGE CTSS processing"
author: "Jonathan Moody"
output:
  html_document:
    keep_md: yes
    toc: true
    number_sections: true
    code_folding: hide
knit: (function(inputFile, encoding) {
  rmarkdown::render(inputFile, encoding = encoding, output_dir = "../generated") })
---

```{r setup, include=FALSE}
library(data.table)
library(ggplot2)
set.seed(2019)
```

```{bash, eval=F}
bash scripts/bed2ctss.sh &
```
  
```{r}
files <- system('ls ../data/ctss/*.bed', intern=T)
x <- lapply(files, fread)
rows <- sapply(x, nrow) # remove 1 row tables as some are empty
counts <- rbindlist(x[rows > 0])
setnames(counts, bed)
sums <- counts[, .(reads=sum(score), mito=.SD[chr=='chrM', sum(score)]), by=.(name)][order(name)]
sums[, mito_pc := (mito/reads)*100]
ggplot(sums, aes(reads)) + geom_histogram(bins=100) +geom_vline(xintercept = 100000, color='red')+geom_vline(xintercept = 629416, color='blue')
```

```{r}
anno <- fread('/osc-fs_home/hon-chun/analysis/ILC2/resource/gene_model/GENCODE/FANTOM5_CAGE_clusters/annotation/GENCODEvM21/bed/F5_CAGE_anno.GENCODEvM21.cage_cluster.coord.bed.gz') # mm10 is correct here
#anno <- fread('/osc-fs_home/hon-chun/resources/F5_DPI_annotation/mm9/F5_CAGE_Annotator/GENCODEvM1/bed/F5_CAGE_anno.GENCODEvM1.cage_cluster.coord.bed.gz')
setnames(anno, bed12)

setkey(counts, chr, st, end)
setkey(anno, chr, st, end)

ov <- foverlaps(counts, anno, nomatch = NULL)
setnames(ov, sub('^i.','ctss.',names(ov)))

ov <- ov[strand == '.' | ctss.strand == strand] # no antisense matches to promoters
sums <- merge(sums, ov[, .(f5_reads=sum(ctss.score), f5_feats=length(unique(name))), by=.(ctss.name)], by.x='name', by.y='ctss.name')
sums[, f5_pc := (f5_reads/reads)*100]

sums[, summary(f5_pc)]
fwrite(sums, '../data/metadata-C1CAGE_20180410112239_ctss_stats.csv')

m <- dcast.data.table(ov, formula = name ~ ctss.name, fun.aggregate = sum, value.var = 'ctss.score')
clusters <- m$name
m <- as.matrix(m[,-1])
rownames(m) <- clusters
all(colnames(m) == sums$name)
#m <- m[, colSums(m) > 10000]
#m <- t(t(m) / sums$reads * 10^6)
dim(m)
m[1:5,1:5]
saveRDS(m, '../data/CAGE_GENCODEvM21_mtx.Rds')
```
````
