## Supplementary material for "Single-cell transcriptomics, scRNA-Seq and C1 CAGE discovered distinct phases of pluripotency during naïve-to-primed conversion in mice": SCALE.Rmd

````
---
title: "ASE test"
author: "Jonathan Moody"
date: "04/11/2019"
output:
  html_document:
    keep_md: yes
    toc: true
    number_sections: true
    code_folding: hide
knit: (function(inputFile, encoding) {
  rmarkdown::render(inputFile, encoding = encoding, output_dir = "../reports") })
---

```{r}
library(data.table)
library(SCALE)
#options(width = 3000)
```
# Load data

* Select just protein coding genes
```{r}
alleleA <- readRDS('../generated/b6_matrix.Rds')
alleleB <- readRDS('../generated/msm_matrix.Rds')
spikein_input <- readRDS('../generated/spike_matrix.Rds')

spikein_input <-  spikein_input[, colnames(spikein_input) %in% c(colnames(alleleA), colnames(spikein_input)[1:2])]

anno <- fread('/osc-fs_home/hon-chun/resources/F5_DPI_annotation/mm9/F5_CAGE_Annotator/GENCODEvM1/log/F5_CAGE_anno.GENCODEvM1.transcript.info.tsv.gz')
anno <- anno[, .(geneID, geneName, geneClass)][!duplicated(geneID)]
setkey(anno, geneID)
rownames(alleleA) <- anno[rownames(alleleA), geneName]
rownames(alleleB) <- anno[rownames(alleleB), geneName]

#gid <- anno[geneClass=='protein_coding', geneName]
#alleleA <- alleleA[rownames(alleleA) %in% gid, ]
#alleleB <- alleleB[rownames(alleleB) %in% gid, ]

genename=rownames(alleleA)
sampname=colnames(alleleA)
head(colnames(alleleA))
head(rownames(alleleA))
rownames(spikein_input)
head(colnames(spikein_input))

```

# QC
```{r}
####################################################
####################################################
##########                                ##########
##########        quality control         ##########
##########                                ##########
####################################################
####################################################

N = apply(alleleA + alleleB, 2, sum)
lib.size = N/mean(N)
spikein_read=spikein_input[,-(1:2)]
# remove cells with extreme library size factors
lib.size.filter=(lib.size>0.5 & lib.size < 2.5)
alleleA=alleleA[,lib.size.filter]; alleleB=alleleB[,lib.size.filter]
N = apply(alleleA + alleleB, 2, sum)
lib.size = N/mean(N)
spikein_read=spikein_read[,lib.size.filter[colnames(spikein_read)]]

# remove cells with extreme ratios of spike-in versus endogenous reads
endo_readA=alleleA[,colnames(spikein_read)] # endogenous reads from allele A
endo_readB=alleleB[,colnames(spikein_read)] # endogenous reads from allele B
ratio=apply(spikein_read,2,sum)/apply(endo_readA+endo_readB,2,sum)
ratio.filter=(ratio >0.05 & ratio < 30)
alleleA=alleleA[,is.na(match(colnames(alleleA),colnames(spikein_read)[!ratio.filter]))]
alleleB=alleleB[,is.na(match(colnames(alleleB),colnames(spikein_read)[!ratio.filter]))]
N = apply(alleleA + alleleB, 2, sum)
lib.size = N/mean(N)
spikein_read=spikein_read[,ratio.filter]
spikein_input=cbind(spikein_input[,1:2],spikein_read)

```

# Heterogeneity
```{r}
####################################################
####################################################
##########                                ##########
##########          heterogeneity         ##########
##########                                ##########
####################################################
####################################################

# SCALE needs to be applied to a homogeneous population of cells.
# Below is sanity check on data heterogeneity using PCA and t-SNE.

read=alleleA+alleleB
lib.size.factor=matrix(ncol=ncol(read),nrow=nrow(read),apply(read,2,sum)/mean(apply(read,2,sum)),byrow=T)
read=read/lib.size.factor # adjust for sequencing depth
AR=alleleA/(alleleA+alleleB)

AR=AR[apply(read,1,sum)>10,] # remove silent, lowly expressed genes
read=read[apply(read,1,sum)>10,]

AR=as.matrix(AR[apply(read,1,sd)>quantile(apply(read,1,sd),0.9),]) # select for highly variable genes
read=as.matrix(read[apply(read,1,sd)>quantile(apply(read,1,sd),0.9),])

# SVD and t-SNE on total coverage
svd.read=svd(read)
plot(svd.read$v[,1], svd.read$v[,2], xlab='PC1',ylab='PC2')
scatterplot3d(svd.read$v[,1], svd.read$v[,2], svd.read$v[,3],
              xlab='PC1',ylab='PC2',zlab='PC3')
tsne_read=tsne(t(read),k=3,initial_dims=30)
plot(tsne_read[,1],tsne_read[,2],xlab='tSNE1',ylab='tSNE2')
scatterplot3d(tsne_read[,1],tsne_read[,2],tsne_read[,3],
              xlab='tSNE1',ylab='tSNE2',zlab='tSNE3')

# SVD and t-SNE on allelic ratio (AR) - Excludes all cells
# AR=AR[!apply(is.nan(AR),1,any),]
# svd.AR=svd(AR)
# plot(svd.AR$v[,1], svd.AR$v[,2], xlab='PC1',ylab='PC2')
# scatterplot3d(svd.AR$v[,1], svd.AR$v[,2], svd.AR$v[,3],
#               xlab='PC1',ylab='PC2',zlab='PC3')
# tsne_AR=tsne(t(AR),k=3,initial_dims=30)
# plot(tsne_AR[,1],tsne_AR[,2],xlab='tSNE1',ylab='tSNE2')
# scatterplot3d(tsne_AR[,1],tsne_AR[,2],tsne_AR[,3],
#               xlab='tSNE1',ylab='tSNE2',zlab='tSNE3')

```

# Technical variability

* Use spike ins to determine technical effects
```{r}
####################################################
####################################################
##########                                ##########
##########     technical variability      ##########
##########                                ##########
####################################################
####################################################

# Make sure the column names of spikein_input correspond to (a subset of)
# those of alleleA and alleleB.
abkt=tech_bias(spikein_input=spikein_input, alleleA = alleleA, 
               alleleB = alleleB, readlength = 50, pdf=FALSE)
```

# Gene classification

```{r}
gene_classify_mod <- function (alleleA, alleleB, epsilon = NULL) 
{
    if (is.null(epsilon)) {
        epsilon = 0.001
    }
    gene.category = rep(NA, nrow(alleleA))
    A.prop = B.prop = rep(NA, nrow(alleleA))
    results.list = vector(mode = "list", length = nrow(alleleA))
    n = ncol(alleleA)
    genename = rownames(alleleA)
    for (i in 1:nrow(alleleA)) {
        nA = alleleA[i, ]
        nB = alleleB[i, ]
        k = 4
        a = b = cura = curb = 1
        phi = rep(1, k)/k
        diff = 1
        numiters = 1
        while (diff > 1e-04 || numiters <= 30) {
            numiters = numiters + 1
            Zhat = Estep(a, b, phi, epsilon, n, nA, nB)
            curM = Mstep(Zhat, cura, n, nA, nB)
            curphi = curM$phihat
            cura = curb = curM$a.b
            diff = max(max(abs(phi - curphi)), max(abs(b - curb)))
            phi = curphi
            a = b = cura
            if (numiters >= 3000) 
                break
        }
        cell.post.filter = apply(Zhat, 1, max) > 0.8
        category = c("Off", "A", "B", "AB")
        results = table(category[apply(Zhat[cell.post.filter, 
            ], 1, which.max)])
        fwrite(data.table(class=category[apply(Zhat[cell.post.filter, 
            ], 1, which.max)], gene=genename[i], cell=colnames(alleleA)[cell.post.filter]), paste0('../generated/SCALE/',genename[i],'.csv'), col.names = F)
        results.list[[i]] = results
        if (setequal(names(results), "Off")) {
            gene.cate = "Silent"
        }
        else if (setequal(names(results), "A") | setequal(names(results), 
            c("A", "Off"))) {
            gene.cate = "MonoA"
        }
        else if (setequal(names(results), "B") | setequal(names(results), 
            c("B", "Off"))) {
            gene.cate = "MonoB"
        }
        else {
            gene.cate = "Biallelic"
        }
        if (!("Off" %in% names(results))) {
            results = c(results, Off = 0)
        }
        if (!("AB" %in% names(results))) {
            results = c(results, AB = 0)
        }
        if (!("A" %in% names(results))) {
            results = c(results, A = 0)
        }
        if (!("B" %in% names(results))) {
            results = c(results, B = 0)
        }
        A.prop[i] = round((results["A"] + results["AB"])/sum(results), 
            3)
        B.prop[i] = round((results["B"] + results["AB"])/sum(results), 
            3)
        if (A.prop[i] >= 0.05 & B.prop[i] >= 0.05 & A.prop[i] <= 
            0.95 & B.prop[i] <= 0.95) {
            gene.cate = "Biallelic.bursty"
        }
        gene.category[i] = gene.cate
        cat("Gene", i, ":", genename[i], ",", gene.cate, "\t")
        cat("A prop", A.prop[i], "B prop", B.prop[i], "\n")
    }
    gene.category[gene.category == "Biallelic"] = "Biallelic.nonbursty"
    gene.class.obj = list(gene.category = gene.category, A.prop = A.prop, 
        B.prop = B.prop, results.list = results.list)
    return(gene.class.obj)
}
```

```{r}
####################################################
####################################################
##########                                ##########
##########      gene classification       ##########
##########                                ##########
####################################################
####################################################

# Only first 200 genes are computed for demonstration.
#gene.class.obj=gene_classify_mod(alleleA = as.matrix(alleleA[1:20,]),
#                              alleleB = as.matrix(alleleB[1:20,]))
gene.class.obj=gene_classify_mod(alleleA = as.matrix(alleleA),
                             alleleB = as.matrix(alleleB))
A.prop=gene.class.obj$A.prop
B.prop=gene.class.obj$B.prop
gene.category=gene.class.obj$gene.category
results.list=gene.class.obj$results.list

# # Below is loading pre-computed gene classification results.
# data(gene.class.obj)
# A.prop=gene.class.obj$A.prop
# B.prop=gene.class.obj$B.prop
# gene.category=gene.class.obj$gene.category
# results.list=gene.class.obj$results.list

```

# Allelic bursting kinetics
```{r}
####################################################
####################################################
##########                                ##########
##########   allelic bursting kinetics    ##########
##########                                ##########
####################################################
####################################################

cellsize=rep(1,ncol(alleleA))  # cell size input
allelic.kinetics.obj=allelic_kinetics(alleleA = alleleA, 
                                      alleleB = alleleB, 
                                      abkt = abkt, 
                                      gene.category = gene.category, 
                                      cellsize = cellsize, pdf=TRUE)

# Below is loading pre-computed allelic kinetics.
# data(allelic.kinetics.obj)
bandwidth=allelic.kinetics.obj$bandwidth
konA=allelic.kinetics.obj$konA
koffA=allelic.kinetics.obj$koffA
sA=allelic.kinetics.obj$sA
konB=allelic.kinetics.obj$konB
koffB=allelic.kinetics.obj$koffB
sB=allelic.kinetics.obj$sB
sizeA=sA/koffA
sizeB=sB/koffB


####################################################
####################################################
##########                                ##########
########## differential bursting kinetics ##########
##########                                ##########
####################################################
####################################################
diff.allelic.obj=diff_allelic_bursting(alleleA = alleleA,
                                        alleleB = alleleB,
                                        cellsize = cellsize,
                                        gene.category = gene.category,
                                        abkt = abkt,
                                        allelic.kinetics.obj = allelic.kinetics.obj,
                                        mode = 'corrected')
pval.kon = diff.allelic.obj$pval.kon; pval.size = diff.allelic.obj$pval.size

# Below is loading pre-computed testing results.
# data(diff.allelic.obj)
# pval.kon = diff.allelic.obj$pval.kon; pval.size = diff.allelic.obj$pval.size

####################################################
####################################################
##########                                ##########
##########    non-independent bursting    ##########
##########                                ##########
####################################################
####################################################

non.ind.obj=non_ind_bursting(alleleA = alleleA, alleleB = alleleB,
                             gene.category = gene.category,
                             results.list = results.list)
pval.ind=non.ind.obj$pval.ind
non.ind.type=non.ind.obj$non.ind.type
```

# Save results
```{r}
####################################################
####################################################
##########                                ##########
##########            plot genes          ##########
##########                                ##########
####################################################
####################################################

i=which(genename=='Calm1')
allelic_plot(alleleA = alleleA, alleleB = alleleB,
             gene.class.obj = gene.class.obj,
             allelic.kinetics.obj = allelic.kinetics.obj,
             diff.allelic.obj = diff.allelic.obj,
             non.ind.obj = non.ind.obj, i= i)

####################################################
####################################################
##########                                ##########
##########     generate output tables     ##########
##########                                ##########
####################################################
####################################################

SCALE.output=output_table(alleleA=alleleA, alleleB=alleleB,
                          gene.class.obj = gene.class.obj,
                          allelic.kinetics.obj = allelic.kinetics.obj,
                          diff.allelic.obj = diff.allelic.obj,
                          non.ind.obj = non.ind.obj)
head(SCALE.output)
write.table(SCALE.output,file='../generated/SCALE.output.tab',col.names = T,row.names = F,quote = F,sep='\t')
```
````
