## Supplementary material for "Single-cell transcriptomics, scRNA-Seq and C1 CAGE discovered distinct phases of pluripotency during naïve-to-primed conversion in mice": SCALE_res.Rmd

````
---
title: "ASE test results"
author: "Jonathan Moody"
date: "04/11/2019"
output:
  html_document:
    keep_md: yes
    toc: true
    number_sections: true
    code_folding: hide
knit: (function(inputFile, encoding) {
  rmarkdown::render(inputFile, encoding = encoding, output_dir = "../reports") })
---
<style type="text/css">
.main-container {
  max-width: 3000px;
  margin-left: auto;
  margin-right: auto;
}
</style>
```{r}
library(data.table)
options(width = 3000)
res <- fread('../generated/SCALE.output.txt')

anno <- fread('/osc-fs_home/hon-chun/resources/F5_DPI_annotation/mm9/F5_CAGE_Annotator/GENCODEvM1/log/F5_CAGE_anno.GENCODEvM1.transcript.info.tsv.gz')
anno <- anno[, .(geneName, geneClass)]
anno <- anno[!duplicated(anno)]
res <- merge(res, anno[, .(geneName, geneClass)], by.x='genename', by.y='geneName')
```

# Enhancers
```{r}
res[grep('.E$', genename), table(gene.category)]
```

```{r}
df <- fread(cmd='cat ../generated/SCALE/*.csv', header=F)
setnames(df, c('cat','genename','cell'))
df2 <- dcast(df, genename~cell, value.var = 'cat')
df2 <- cbind(res[, 1:2, with=F], df2[, -1, with=F] )
fwrite(df2, '../generated/SCALE.output.cell.csv')
```
````
