## Supplementary material for "Single-cell transcriptomics, scRNA-Seq and C1 CAGE discovered distinct phases of pluripotency during naïve-to-primed conversion in mice": SCALE_res_rna.Rmd

````
---
title: "SCALE RNA results"
author: "Jonathan Moody"
date: "05/11/2019"
output:
  html_document:
    keep_md: yes
    toc: true
    number_sections: true
    code_folding: hide
knit: (function(inputFile, encoding) {
  rmarkdown::render(inputFile, encoding = encoding, output_dir = "../reports") })
---
<style type="text/css">
.main-container {
  max-width: 2000px;
  margin-left: auto;
  margin-right: auto;
}
</style>
```{r, message=FALSE, warning=FALSE}
suppressPackageStartupMessages(library(data.table))
suppressPackageStartupMessages(library(DT))
options(width = 2000)
res <- fread('../generated/SCALE.output_RNA.txt', na.strings = c('NA','-'))

anno <- fread('/osc-fs_home/hon-chun/resources/F5_DPI_annotation/mm9/F5_CAGE_Annotator/GENCODEvM1/log/F5_CAGE_anno.GENCODEvM1.transcript.info.tsv.gz')
anno <- anno[, .(geneName, geneClass)]
anno <- anno[!duplicated(anno)]
res <- merge(res, anno[, .(geneName, geneClass)], by.x='genename', by.y='geneName')

res[, gene.category := as.factor(gene.category)]
res[, geneClass := as.factor(geneClass)]
```

* konA/B = allele activation rate
* pvalue.kon = alleles have different activation rates
* SizeA/B = allele burst size
* pvalue.size = alleles have different burst sizes
* pval.ind = whether the two alleles burst independently
* “C” as coordinated bursting and “R” as repulsed bursting

# Enhancers

* Showing table for all expressed (not silent) enhancers
```{r}
res[grep('.E$', genename), table(gene.category)]

datatable(res[grepl('.E$', genename) & gene.category != 'Silent'], filter = 'top')
```

# Protein coding genes

* Protein coding gene categories:
* MonoA = b6 allele
* MonoB = msm allele
* Many more b6 than msm

```{r}
res[geneClass == 'protein_coding', table(gene.category)]
```

# All - results with pvalues

* Results from all gene categories where pvalues could be calculated (Must be biallelic bursty).
```{r}
res[(!is.na(pval.kon) | !is.na(pval.size) | !is.na(pval.ind)), table(geneClass)]
datatable(res[(!is.na(pval.kon) | !is.na(pval.size) | !is.na(pval.ind))], filter='top')
```

```{r}
df <- fread(cmd='cat ../generated/SCALE_RNA/*.csv', header=F)
setnames(df, c('cat','genename','cell'))
df2 <- dcast(df, genename~cell, value.var = 'cat')
df2 <- cbind(res[, 1:2, with=F], df2[, -1, with=F] )
fwrite(df2, '../generated/SCALE.output_RNA.cell.csv')
```
````
