## Supplementary material for "Single-cell transcriptomics, scRNA-Seq and C1 CAGE discovered distinct phases of pluripotency during naïve-to-primed conversion in mice": Supplementary_SourceCode.R

```
########## Workflow (excluding allelic expression and promotor and enhancer analysis) #########

# list of required R packages
library(data.table)
library(TSCAN)
library(magrittr)
library(ggplot2)
library(Rtsne)
library(stringr)
library(gplots)
library(viridisLite)
library(scde)
library(topGO)
library(AnnotationDbi)
library(org.Mm.eg.db)
library(Rgraphviz)
library(genefilter)
library(ALL)
library(pheatmap)
library(tidyr)
library(RColorBrewer)
library(CAGEr)
library(MultiAssayExperiment)
library(SummarizedExperiment)
library(vegan)

##########
# Tagdust filtering of raw FASTQ files

# tagdust.arch files contains the line:
# tagdust -1 R:N
# tagdust.fa file contains all base sequences and annotations of sequences to remove from the FASTQ files

#!bin/bash -e

mkdir -p tagdustOutput

TAGDUST=/bin/tagdust-2.13/src/tagdust
$TAGDUST | grep Copyright
BASEDIR=FASTQfiles/
  time (cd $BASEDIR/ ; ls ) |
  sed 's/_R._001\.fastq.gz//g' |
  uniq |
  parallel -j1 'tagdust-2.13/src/tagdust \
	-t 12 \
	-l tagdustOutput \
	-arch tagdust.arch \
	-o tagdustOutput/{} \
	-a tagdustOutput/{} \
	-ref tagdust.fa \
	FASTQfiles/{}_R1_001.fastq.gz FASTQfiles/{}_R2_001.fastq.gz' 

##########
# Kallisto expression quantification

# download and modify transcript vM8 FASTA file from GENCODE
# ftp://ftp.ebi.ac.uk/pub/databases/gencode/Gencode_mouse/release_M8/
#!bin/bash

awk 'FS="\t" {if($3=="transcript"){print $9}}' gencode.vM8.chr_patch_hapl_scaff.annotation.gtf |
  cut -d ';' -f 2,5 |
  cut -d '"' -f2,4 |
  sed 's/"/\t/g' > transcript2geneGTF.txt

grep ">" gencode.vM8.transcripts.fa |
  cut -d '|' -f1,6 |
  sed 's/|/\t/g' |
  sed 's/>//g' > transcript2geneFasta.txt


# download and modify gencode.vM8.transcripts.fa to only include Ensemble transcript IDs (>ENSMUST) for each transcript
#!bin/bash

kallisto index -i kallisto_M8 gencode.vM8.EnsembleTranscriptID.fa &&

mkdir -p kallistoOutput

BASEDIR=tagdustOutput/fastqOut
time (cd $BASEDIR ; ls) |
  sed 's/_R.\.fastq//g' |
  uniq |
  parallel -j2 'kallisto quant \
  -t 10 \
  -i kallisto_M8.idx \
  -o kallistoOutput/{}/ \
  tagdustOutput/fastqOut/{}_R1.fastq \
  tagdustOutput/fastqOut/{}_R2.fastq \
  -b 100 \
  --bias'

##########
# combine expression column of all single cell Kallisto output files into one table

#!bin/bash -e

# create output directories

mkdir -p finalFiles intermediateFiles qTable

(cd kallistoOutput/ ; ls ) |
  parallel -j1 'sed 1d kallistoOutput/{}/abundance.tsv > finalFiles/{}_tquant.txt' &&
  
for i in finalFiles/*_tquant.txt
do
  CELL_ID=$(basename $i _tquant.txt)
  HEADER_FILE="intermediateFiles/${CELL_ID}.header"
  # create header file with: gene_name TPM per cell
  echo -e "gene\t$CELL_ID" > $HEADER_FILE
  # append header file with expression values for each cell
  # column 4 contains count values and column 5 TPM values
  cat $i | cut -f 1,5 >> $HEADER_FILE
done &&
  
# make sure that only 1 particualar header file is chosen
cut -f1 intermediateFiles/$(cd kallistoOutput/ ; ls | head -n1).header |
paste -s > qTable/kallistoTPM
# Column 2 of each *.header file becomes a line of the qTable/kallistoTPM file
for FILE in intermediateFiles/*.header
do
cut -f2 $FILE |
  paste -s  >> qTable/kallistoTPM
done &&
  
rm -rf finalFiles intermediateFiles

##########
# transpose transcript expression table

#!usr/bin/Rscript

args = commandArgs(trailingOnly=TRUE)

expressionTable <- function(expressionTable, outputFileName) {
  z <- as.data.frame(fread(expressionTable, stringsAsFactors=F, header=T))
  cellNames <- z[,1]
  y <- z[,2:ncol(z)]
  Z <- t(y)
  x <- as.data.frame(Z)
  colnames(x) <- cellNames
  rownames(x) <- colnames(z)[-1]
  write.table(x, outputFileName, sep="\t", row.names=T, col.names=T, quote=F)
}

expressionTable(args[1], args[2])

#########

# The file transcript2gene.txt was created by parsing the gencode.vM8.chr_patch_hapl_scaff.annotation.gtf file
# It has 2 columns, with column one containing each transcript ID and column 2 having the corresponding gene symbols for each transcript
# metadata is a table that was manually curated and contains useful cell sample metadata
# the metadata table version used here includes a Pseudotime column for convenience as well

# load input files
meta <- read.csv("metadata_20190223045845.csv", header=TRUE, stringsAsFactors = F)
# transcriptTPM_M8_20190206_Kallisto.txt is output of previous step
expr <- read.table("transcriptTPM_M8_20190206_Kallisto.txt", header=T, row.names = 1, check.names = FALSE)
# links transcript IDs to gene names
tr2gene <- read.table("transcript2gene.txt", header=F, check.names = FALSE)
names(tr2gene) <-  c("transcriptID", "gene", "temp"); tr2gene <- tr2gene[,1:2]
tr2gene <- tr2gene[order(tr2gene$gene, decreasing = F),]
tr2gene$transcriptID <- as.character(tr2gene$transcriptID)
# sort transcript IDs and check that the order is the same in transcript expression table and gene reference table
test1 <- expr[order(rownames(expr), decreasing = T),]
test2 <- tr2gene[order(tr2gene$transcriptID, decreasing = T),]
# check that transcript IDs are identical in both tables
# the following must be TRUE
identical(rownames(test1), test2$transcriptID)
# add column with gene names to expression table
expr <- expr[match(tr2gene$transcriptID, rownames(expr)),]
expr$gene <- tr2gene$gene
# aggregate transcript expression to gene level
geneExpr <- aggregate(expr[,1:ncol(expr)-1], by=list(expr$gene), FUN=sum)
rownames(geneExpr) <- geneExpr$Group.1; geneExpr <- geneExpr[,-1]
# remove cells that are missing in Kallisto output
geneExpr <- geneExpr[, which(names(geneExpr) %in% meta$cell_id)]
meta <- meta[match(names(geneExpr), meta$cell_id),]
# reorder columns in expression table according to time points
expr <- geneExpr[,order(meta$Order, decreasing = F)]
# save gene expression table as file
time <- paste(gsub("-|:", "", unlist(stringr::str_split(as.character(Sys.time()), " ", n = 2))), collapse = "")
write.table(expr, file = paste0("geneTPM_M8_", time, "_Kallisto.txt"), append=FALSE, sep="\t", quote=FALSE, row.names=TRUE, col.names=TRUE)

##########
# TSCAN pseudotime sorting

# scRNA-seq data
meta <- read.csv("metadata_20190223045845.csv", header=TRUE, stringsAsFactors = F)
Tpm <- read.table("geneTPM_M8_20190207033818_Kallisto.txt", header=T, row.names = 1, check.names = FALSE)
# make sure meta data and expression table have same samples
meta <- meta[match(names(Tpm), meta$cell_id),]
# the following must be TRUE
identical(names(Tpm), meta$cell_id)

# preliminary DGE result between different time points
dge <- read.table("scde_d0-EpiSC_results_batch-effect-corrected_20190223041832.txt", header=T, row.names = 1, check.names = FALSE)
dge <- dge[which(dge$`p-adjusted`<0.01),]
genes <- which(row.names(Tpm) %in% row.names(dge))
markers <- Tpm[genes, which(meta$discard == FALSE)]

# return expression table without cells that failed quality checks
expr <- Tpm[, which(meta$discard == FALSE)]

# reorder factor levels of timePoint column
RNA <- c(2:6,8,9,1)
meta$timePoint <- factor(meta$timePoint)
meta$timePoint <- factor(meta$timePoint, levels(meta$timePoint)[RNA])
meta <- meta[which(meta$discard == FALSE),]

# filter cells and DGE genes based on minimum expression thresholds
tpm <- markers
tpm <- tpm[,which(colSums(tpm)>1)]
tpm <- tpm[which(rowSums(tpm)>1),]
tpm <- tpm[which(rowSums(tpm>0)>10),]
# the following must be TRUE
identical(names(tpm), meta$cell_id)

# start pseudotime sorting
prepro <- preprocess(tpm, takelog = TRUE, logbase = 2, pseudocount = 1)
exclust <-  exprmclust(prepro, modelNames = "VVV", reduce = T)
plotmclust(exclust, x = 1, y = 2, MSTorder = c(2,1,3,4), show_tree = T,
           show_cell_names = F, cell_name_size = 2, markerexpr = NULL)
scanOrder <- TSCANorder(exclust, MSTorder = c(2,1,3,4), orderonly = F, flip = F, listbranch = F)
timePoint <- meta$timePoint[match(scanOrder$sample_name, meta$cell_id)]
scanOrder <- cbind(scanOrder, timePoint)
# save pseudotime sorting as file
time <- paste(gsub("-|:", "", unlist(stringr::str_split(as.character(Sys.time()), " ", n = 2))), collapse = "")
write.table(scanOrder, file = paste0("TSCAN_pseudotime_", time, ".txt"), row.names = TRUE, col.names = TRUE, sep = "\t", quote = FALSE) 

###########
# Figure 1B

# set color scheme
gg_color_hue <- function(n) {
  hues = seq(15, 375, length=n+1)
  hcl(h=hues, l=65, c=100)[1:n]
}
col <- gg_color_hue(16)[c(1:3,5,10,12,14,16)]

# filter cells and all genes based on minimum expression thresholds
expr <- expr[,which(colSums(expr)>1)]
expr <- expr[which(rowSums(expr)>1),]
expr <- expr[which(rowSums(expr>0)>10),]
# the following must be TRUE
identical(names(expr), meta$cell_id)

expressedGenes <- sapply(1:ncol(expr), function(x) length(which(expr[,x]>1)))
meta$nGenes <- expressedGenes
meta <- meta[which(meta$nGenes>50),]

exprGenes <- expr[,which(names(expr) %in% meta$cell_id)]
# the following must be TRUE
identical(meta$cell_id, colnames(exprGenes))

geneNumberBox <- ggplot(meta, aes(x=timePoint, y=nGenes, fill = timePoint)) +
  geom_boxplot() + 
  xlab("") + 
  ylab("Number of expressed Genes")  +
  theme(axis.text.x = element_text(angle = 45, hjust = 0.4, vjust = 0.5)) + 
  #  scale_y_continuous(limits = c(0, max(meta$nGenes) + max(meta$nGenes)*0.2)) +
  scale_fill_manual(values=col, 
                    name="",
                    labels=c("mES - 1772-116-259", "Day 1 - 1772-116-260", "Day 2 - 1772-116-261", "Day 3 - 1772-116-263", "Day 4 - 1772-116-262, 1772-151-313", "EpiSC - 1772-116-268", "MB#3P10 - 1772-123-291", "1EP7 - 1772-123-289")) +
  scale_colour_manual(values=col, 
                      name="",
                      labels=c("mES - 1772-116-259", "Day 1 - 1772-116-260", "Day 2 - 1772-116-261", "Day 3 - 1772-116-263", "Day 4 - 1772-116-262, 1772-151-313", "EpiSC - 1772-116-268", "MB#3P10 - 1772-123-291", "1EP7 - 1772-123-289")) +
  theme(axis.ticks = element_blank()) + 
  theme(panel.background = element_blank()) + theme(axis.line=element_line(color="grey")) +
  theme(panel.grid.minor=element_line(color="grey"), panel.grid.major=element_line(colour="grey")) +
  theme(text = element_text(family="Helvetica"))

time <- paste(gsub("-|:", "", unlist(stringr::str_split(as.character(Sys.time()), " ", n = 2))), collapse = "")
pdf(paste0("1B_exressedGenesPerTimePoint_", time, ".pdf"), width = 12, height = 7, onefile = TRUE)
plot(geneNumberBox)
dev.off()

#Figure 1C

nearCell <- as.data.frame(cor(expr, method = "spearman"))
meta$nearestCellCorr <- sapply(1:ncol(nearCell), function(x) sort(nearCell[,x], decreasing = T)[2])

nnBox <- ggplot(meta, aes(x=timePoint, y=nearestCellCorr, fill = timePoint)) +
  geom_boxplot() +
  ylim(0,1) +
  xlab("") +
  ylab("Spearman correlation to nearest cell") +
  theme(axis.text.x = element_text(angle = 45, hjust = 0.4, vjust = 0.5)) + 
  scale_fill_manual(values=col, 
                    name="",
                    labels=c("mES", "Day 1", "Day 2", "Day 3", "Day 4", "EpiSC", "MB#3P10", "1EP7")) +
  scale_colour_manual(values=col, 
                      name="",
                      labels=c("mES", "Day 1", "Day 2", "Day 3", "Day 4", "EpiSC", "MB#3P10", "1EP7")) +
  theme(axis.ticks = element_blank()) + 
  theme(panel.background = element_blank()) + theme(axis.line=element_line(color="grey")) +
  theme(panel.grid.minor=element_line(color="grey"), panel.grid.major=element_line(colour="grey")) +
  theme(text = element_text(family="Helvetica"))

time <- paste(gsub("-|:", "", unlist(stringr::str_split(as.character(Sys.time()), " ", n = 2))), collapse = "")
pdf(paste0("1C_nearestneighbourCorrelation_", time, ".pdf"), width = 12, height = 7, onefile = TRUE)
plot(nnBox)
dev.off()

#Figure 1D
# marker distribution boxplot
marker.set <- c("Esrrb", "Nr0b1", "Dppa4", "Zfp42", "Pou5f1", "Sox2", "Sox4", "Cd24a","Dnmt3b")

plotMarkers<- function(counter) {
  
  gene <- as.numeric(expr[grep(paste(marker.set[counter], "$", sep = ""), row.names(expr)),])
  meta$marker <- log2(gene+1)
  
  ggplot(meta, aes(x = timePoint, y = marker, fill = timePoint)) + 
    geom_boxplot(varwidth = FALSE) +
    theme(axis.text.x = element_text(angle = 45, hjust = 0.4, vjust = 0.5)) + 
    scale_fill_manual(values=col, 
                      name="",
                      labels=c("mES", "Day 1", "Day 2", "Day 3", "Day 4", "EpiSC", "MB#3P10", "1EP7")) +
    scale_colour_manual(values=col, 
                        name="",
                        labels=c("mES", "Day 1", "Day 2", "Day 3", "Day 4", "EpiSC", "MB#3P10", "1EP7")) +
    theme(axis.ticks = element_blank()) + 
    theme(panel.background = element_blank()) + theme(axis.line=element_line(color="grey")) +
    theme(panel.grid.minor=element_line(color="grey"), panel.grid.major=element_line(colour="grey")) +
    theme(text = element_text(family="Helvetica")) +
    theme(axis.text.x = element_text(angle = 45, hjust = 0.4, vjust = 0.5)) + 
    xlab("") +
    ylab(paste0("log2(", marker.set[counter], ")")) + 
    guides(fill=FALSE)
}

time <- paste(gsub("-|:", "", unlist(stringr::str_split(as.character(Sys.time()), " ", n = 2))), collapse = "")
counter <- 1:length(marker.set)
pdf(paste0("1D_", time, ".pdf"), width = 10, height = 7, onefile = TRUE)
sapply(counter, function(x) plot(plotMarkers(x)))
dev.off()

###########
# PCA

# convert to log matrix
matExpr <- as.matrix(tpm)
logtpm <- log10(matExpr+1)
# run PCA 
base.pca <- prcomp(t(logtpm))
pca <- data.frame(base.pca$x[,1], base.pca$x[,2]); colnames(pca) <- c("PC1", "PC2")

# calculate component variance
eigen <- (base.pca$sdev)^2
variances <- eigen*100/sum(eigen)
variances[1:2]

# the following must be TRUE
identical(rownames(pca), meta$cell_id)
pca$timePoint <- meta$timePoint

# Figure 2B
# plot PC1 and PC2
pcaPlot <- qplot(PC1, PC2, data=pca, colour=timePoint,
                 xlab = paste('PCA1 ', round(variances[1], digits = 2), '%'), 
                 ylab = paste(paste('PCA2 ', round(variances[2], digits = 2), '%'))) +
  scale_colour_manual(values=col, 
                      name="",
                      labels=c("mES - 1772-116-259", "Day 1 - 1772-116-260", "Day 2 - 1772-116-261", "Day 3 - 1772-116-263", "Day 4 - 1772-116-262, 1772-151-313", "EpiSC - 1772-116-268", "MB#3P10 - 1772-123-291", "1EP7 - 1772-123-289")) +
  theme(axis.ticks = element_blank()) + 
  theme(panel.background = element_blank()) + theme(axis.line=element_line(color="grey")) +
  theme(panel.grid.minor=element_line(color="grey"), panel.grid.major=element_line(colour="grey")) +
  theme(text = element_text(family="Helvetica")) +
  scale_x_continuous(limits = c(min(pca$PC1) + min(pca$PC1)*0.1, max(pca$PC1) + max(pca$PC1)*0.1)) +
  scale_y_continuous(limits = c(min(pca$PC2) + min(pca$PC2)*0.1, max(pca$PC2) + max(pca$PC2)*0.1))

time <- paste(gsub("-|:", "", unlist(stringr::str_split(as.character(Sys.time()), " ", n = 2))), collapse = "")
pdf(paste0("2B_PCA_Kallisto_", time, ".pdf"), width = 12, height = 7, onefile = TRUE)
plot(pcaPlot)
dev.off()

pca$Run <- meta$Run
# save PCA data as file
time <- paste(gsub("-|:", "", unlist(stringr::str_split(as.character(Sys.time()), " ", n = 2))), collapse = "")
write.table(pca, file = paste0("pcaData_kallisto_geneTPM_", time, ".txt"), row.names = TRUE, col.names = TRUE, sep = "\t", quote = FALSE) 

##########
# tSNE and kMeans clustering

logtpmT <- t(logtpm)
# run tSNE 
tsne <- Rtsne(logtpmT)
Run <- as.character(sapply(rownames(logtpmT), function(x) str_split(x, "_", n=2)[[1]][[1]]))
Run <- factor(Run)
Run <- factor(Run, levels(Run)[c(1:3,5,4,9,6,8,7)])
sne <- data.frame(tsne$Y[,1], tsne$Y[,2], Run); colnames(sne) <- c("Dim1", "Dim2", "Run")

# Figure 2C/D
# the following must be TRUE

identical(colnames(logtpm), meta$cell_id)
sne$timePoint <- meta$timePoint

tsneRunPlot <- qplot(Dim1, Dim2, data=sne,
                     colour=timePoint) +
  scale_colour_manual(values=col, 
                      name="",
                      labels=c("mES", "Day 1", "Day 2", "Day 3", "Day 4", "EpiSC", "MB#3P10", "1EP7")) +
  theme(axis.ticks = element_blank()) + 
  theme(panel.background = element_blank()) + theme(axis.line=element_line(color="grey")) +
  theme(panel.grid.minor=element_line(color="grey"), panel.grid.major=element_line(colour="grey")) +
  theme(text = element_text(family="Helvetica")) +
  scale_x_continuous(limits = c(min(sne$Dim1) + min(sne$Dim1)*0.1, max(sne$Dim1) + max(sne$Dim1)*0.1)) +
  scale_y_continuous(limits = c(min(sne$Dim2) + min(sne$Dim2)*0.3, max(sne$Dim2) + max(sne$Dim2)*0.2))

time <- paste(gsub("-|:", "", unlist(stringr::str_split(as.character(Sys.time()), " ", n = 2))), collapse = "")
pdf(paste0("2C_tSNE_Kallisto_", time, ".pdf"), width = 12, height = 7, onefile = TRUE)
plot(tsneRunPlot)
dev.off()

# kMeans clustering
sne$Cluster <- factor(kmeans(sne[,1:2], centers = 5)[[1]])
# adjust cluster name to reflect pseudotime order
test <- as.numeric(sne$Cluster)
test <- gsub('5', 'd', test)
test <- gsub('3', 'f', test)
test <- gsub('4', '3', test)
test <- gsub('1', '4', test)
test <- gsub('d', '1', test)
test <- gsub('2', '5', test)
test <- gsub('f', '2', test)
test <- factor(as.numeric(test))
sne2 <- sne; sne2$Cluster <- test
sne$Cluster <- test

tsnekmeansPlot <- qplot(Dim1, Dim2, data=sne, colour=Cluster) +
  scale_colour_brewer(palette = "Dark2", name="") +
  theme(axis.ticks = element_blank()) + 
  theme(panel.background = element_blank()) + theme(axis.line=element_line(color="grey")) +
  theme(panel.grid.minor=element_line(color="grey"), panel.grid.major=element_line(colour="grey")) +
  theme(text = element_text(family="Helvetica")) +
  scale_x_continuous(limits = c(min(sne$Dim1) + min(sne$Dim1)*0.1, max(sne$Dim1) + max(sne$Dim1)*0.1)) +
  scale_y_continuous(limits = c(min(sne$Dim2) + min(sne$Dim2)*0.3, max(sne$Dim2) + max(sne$Dim2)*0.2))

time <- paste(gsub("-|:", "", unlist(stringr::str_split(as.character(Sys.time()), " ", n = 2))), collapse = "")
pdf(paste0("2D_kMeans5_Kallisto_", time, ".pdf"), width = 12, height = 7, onefile = TRUE)
plot(tsnekmeansPlot)
dev.off()

# save tSNE and kMeans data as file
sne$cell_id <- meta$cell_id
time <- paste(gsub("-|:", "", unlist(stringr::str_split(as.character(Sys.time()), " ", n = 2))), collapse = "")
write.table(sne, file = paste0("tSNE_kMeans5_clusterData_kallisto_DGE_geneTPM_", time, ".txt"), row.names = TRUE, col.names = TRUE, sep = "\t", quote = FALSE) 

# start for cluster composition
sne <- fread("tSNE_kMeans5_clusterData_kallisto_DGE_geneTPM_20190223045041.txt")
sne <- as.data.frame(sne)
sne$Cluster <- factor(sne$Cluster)
sne$Cluster

# save cluster composition as file
row.names(sne) <- sne$cell_id
x1 <- row.names(subset(sne, Cluster==1))
x2 <- row.names(subset(sne, Cluster==2))
x3 <- row.names(subset(sne, Cluster==3))
x4 <- row.names(subset(sne, Cluster==4))
x5 <- row.names(subset(sne, Cluster==5))

cl1 <- as.data.frame(table(as.character(meta[match(x1, meta$cell_id),129]))); names(cl1) <- c("timePoint", "Cluster1")
cl2 <- as.data.frame(table(as.character(meta[match(x2, meta$cell_id),129]))); names(cl2) <- c("timePoint", "Cluster2")
cl3 <- as.data.frame(table(as.character(meta[match(x3, meta$cell_id),129]))); names(cl3) <- c("timePoint", "Cluster3")
cl4 <- as.data.frame(table(as.character(meta[match(x4, meta$cell_id),129]))); names(cl4) <- c("timePoint", "Cluster4")
cl5 <- as.data.frame(table(as.character(meta[match(x5, meta$cell_id),129]))); names(cl5) <- c("timePoint", "Cluster5")

allClusters <- merge(cl1, cl2, by = "timePoint", all = T)
allClusters <- merge(allClusters, cl3, by = "timePoint", all = T)
allClusters <- merge(allClusters, cl4, by = "timePoint", all = T)
allClusters <- merge(allClusters, cl5, by = "timePoint", all = T)
allClusters[is.na(allClusters)] <- 0

percentTimes <- data.frame(allClusters$timePoint,
                           allClusters$Cluster1/as.numeric(colSums(allClusters[-1]))[1]*100,
                           allClusters$Cluster2/as.numeric(colSums(allClusters[-1]))[2]*100,
                           allClusters$Cluster3/as.numeric(colSums(allClusters[-1]))[3]*100,
                           allClusters$Cluster4/as.numeric(colSums(allClusters[-1]))[4]*100,
                           allClusters$Cluster5/as.numeric(colSums(allClusters[-1]))[5]*100)
                           #allClusters$Cluster6/as.numeric(colSums(allClusters[-1]))[6]*100)
names(percentTimes) <- names(allClusters)

# save tSNE  and cluster data for reproducibility
time <- paste(gsub("-|:", "", unlist(stringr::str_split(as.character(Sys.time()), " ", n = 2))), collapse = "")
write.table(sne, file = paste0("tSNE_kMeans5_clusterData_kallisto_DGE_geneTPM_", time, ".txt"), row.names = F, col.names = TRUE, sep = "\t", quote = FALSE) 
write.table(allClusters, file = paste0("tSNE_kmeans5_clusterCounts_Kallisto_DGE_geneTPM_", time, ".txt"), row.names = FALSE, col.names = TRUE, sep = "\t", quote = FALSE) 
write.table(percentTimes, file = paste0("tSNE_kmeans5_clusterPercentage_Kallisto_DGE_geneTPM_", time, ".txt"), row.names = FALSE, col.names = TRUE, sep = "\t", quote = FALSE) 

##########

# Figure 2E
sne$Pseudotime <- scanOrder[match(rownames(sne), rownames(scanOrder)), 3]

tsneDensityPlot <- ggplot(sne, aes(Dim1, Dim2, colour = Pseudotime)) +
  theme(axis.ticks = element_blank()) + 
  theme(panel.background = element_blank()) + 
  theme(axis.line=element_line(color="grey")) +
  theme(panel.grid.minor=element_line(color="grey"), panel.grid.major=element_line(colour="grey")) +
  stat_density2d(aes(fill = ..level.., alpha = ..level..), binwidth = 0.0001, geom = "polygon", position = "jitter") +
  scale_fill_continuous(low="white", high="black", name = "") + 
  guides(alpha = "none") +
  geom_point() +
  scale_color_gradientn(colours = plasma(6), guide = "colourbar", name= "") +
  scale_x_continuous(limits = c(min(sne$Dim1) + min(sne$Dim1)*0.1, max(sne$Dim1) + max(sne$Dim1)*0.1)) +
  scale_y_continuous(limits = c(min(sne$Dim2) + min(sne$Dim2)*0.3, max(sne$Dim2) + max(sne$Dim2)*0.2)) +
  theme(text = element_text(family="Helvetica")) 

time <- paste(gsub("-|:", "", unlist(stringr::str_split(as.character(Sys.time()), " ", n = 2))), collapse = "")
pdf(paste0("2E_tSNE_pseudotimeDensity_", time, ".pdf"), width = 10, height = 7, onefile = TRUE)
plot(tsneDensityPlot)
dev.off()

# Figure 2F
# add Psuedotime to meta data table
pseudoTime <- scanOrder[match(meta$cell_id, scanOrder$sample_name),3]
meta$Pseudotime <- pseudoTime

pseudotimeBox <- ggplot(meta, aes(x = timePoint, y = Pseudotime, fill = timePoint)) + 
  geom_boxplot() +
  coord_flip() + 
  scale_fill_manual(values=col, 
                    name="",
                    labels=c("mES", "Day 1", "Day 2", "Day 3", "Day 4", "EpiSC", "MB#3P10", "1EP7")) +
  scale_colour_manual(values=col, 
                      name="",
                      labels=c("mES", "Day 1", "Day 2", "Day 3", "Day 4", "EpiSC", "MB#3P10", "1EP7")) +
  theme(axis.ticks = element_blank()) + 
  theme(panel.background = element_blank()) + theme(axis.line=element_line(color="grey")) +
  theme(panel.grid.minor=element_line(color="grey"), panel.grid.major=element_line(colour="grey")) +
  theme(axis.title = element_text(size=14, family="Helvetica"),
        axis.text = element_text(size=8, family="Helvetica")) +
  scale_x_discrete(limits=rev(levels(meta$timePoint)), name="") + 
  guides(fill=FALSE)

time <- paste(gsub("-|:", "", unlist(stringr::str_split(as.character(Sys.time()), " ", n = 2))), collapse = "")
pdf(paste0("2E_pseudotime_boxplot_", time, ".pdf"), width = 10, height = 7, onefile = TRUE)
plot(pseudotimeBox)
dev.off()

###########
# update pseudotime in metadata 
# load pseudotime table
pseudo <- read.table("TSCAN_pseudotime_20190223042447.txt", header=T, row.names = 1, check.names = FALSE)
pseudo$cell_id <- as.character(pseudo$sample_name)
meta <- read.csv("metadata_intermediary_20190223045613.csv", header=TRUE, stringsAsFactors = F)
# update Pseudotime in meta data table
meta <- meta[,-131]
meta <- merge(meta, pseudo[,c(3,5)], by = 'cell_id', all = T)
meta<- meta[,c(1:130,132,131)]
# save updated metadata table in file
time <- paste(gsub("-|:", "", unlist(stringr::str_split(as.character(Sys.time()), " ", n = 2))), collapse = "")
# this saved metadata table is used from the beginning of the workflow description
write.csv(meta, paste0("metadata_", time, ".csv"), row.names = F)

##########
# SCDE differential gene expression on kMeans clusters

meta <- read.csv("metadata_20190223045845.csv", header=TRUE, stringsAsFactors = F)
counts <- read.table("geneCount_M8_20170908_Kallisto.txt", header=T, row.names = 1, check.names = FALSE)

counts <- counts[, which(names(counts) %in% meta$cell_id)]
# make sure meta data and expression table have same samples
meta <- meta[match(names(counts), meta$cell_id),]
# the following must be TRUE
identical(names(counts), meta$cell_id)

# remove failed cells
RNA <- c(2:6,8,1,9)
meta$timePoint <- factor(meta$timePoint)
meta$timePoint <- factor(meta$timePoint, levels(meta$timePoint)[RNA])
Red <- meta[which(meta$discard == FALSE),]
counts <- counts[,which(meta$discard == FALSE)]

# round to integers, in case if estimated count values
counts <- round(counts,0)
# remove non-expressed genes
counts <- counts[rowSums(counts)>0,]
nGenes <- length(counts[,1])

coverage <- colSums(counts)/nGenes

# filter out cells with low coverage
counts <- counts[,coverage>1]
Red <- Red[coverage>1,]

# each cpu core utilizes around 45Gb
n.cores <- 1

# data prep
cd <- clean.counts(counts, min.lib.size=1000, min.reads = 1, min.detected = 1)

# make group factor for DGE
#Red <- Red[order(Red$timePoint, decreasing = F),]
cdGroups <- Red[which(Red$cell_id %in% colnames(cd)),]
cd <- cd[,match(Red$cell_id, names(cd))]
# the following must be TRUE
identical(colnames(cd), cdGroups$cell_id)
groups <- cdGroups$timePoint
names(groups) <- colnames(cd)  
table(groups)

# convert all values to stored integers
cd <- as.data.frame(apply(cd, 2, function(x) {storage.mode(x) <- 'integer'; x}))

# fitting error models
# this step takes a quite long and is demanding on RAM
#scde.fitted.model <- scde.error.models(counts = cd, groups = groups, n.cores = n.cores, threshold.segmentation = TRUE, save.crossfit.plots = FALSE, save.model.plots = FALSE, verbose = 1)
#time <- paste(gsub("-|:", "", unlist(stringr::str_split(as.character(Sys.time()), " ", n = 2))), collapse = "")
#save(scde.fitted.model,file=paste0("scde_scRNA-seq_Kallisto_", time, ".RData"))

# to save time load the precalculated error models
load("scde_scRNA-seq_Kallisto_20170908.RData")
valid.cells <- scde.fitted.model$corr.a > 0
table(valid.cells)

scde.fitted.model <- scde.fitted.model[valid.cells, ]
# make sure that cd and scde.fitted.model contain exactly the same cells
scde.fitted.model <- scde.fitted.model[match(names(cd), rownames(scde.fitted.model)),]
# the following must be TRUE
identical(names(cd),  rownames(scde.fitted.model ))

scde.prior <- scde.expression.prior(models=scde.fitted.model, counts = cd, length.out = 400, show.plot = FALSE)

# prepare factor levels for SCDE DGE of kmeans clusters
c1 <- x1; names(c1) <- rep("Cluster1", length(c1))
c2 <- x2; names(c2) <- rep("Cluster2", length(c2))
c3 <- x3; names(c3) <- rep("Cluster3", length(c3))
c4 <- x4; names(c4) <- rep("Cluster4", length(c4))
c5 <- x5; names(c5) <- rep("Cluster5", length(c5))
#c6 <- x6; names(c6) <- rep("Cluster6", length(c6))
temp <- factor(names(c1))
names(temp) <- as.character(c1); c1 <- temp
temp <- factor(names(c2))
names(temp) <- as.character(c2); c2 <- temp
temp <- factor(names(c3))
names(temp) <- as.character(c3); c3 <- temp
temp <- factor(names(c4))
names(temp) <- as.character(c4); c4 <- temp
temp <- factor(names(c5))
names(temp) <- as.character(c5); c5 <- temp

# manually change here for individual cluster comparisons
pairList <- list(c1, c2, c3, c4, c5)

# make pairwise cluster comparisons
# restart loop by increasing start number by 1 compared to previous loop run until 4
start <- 4
for (i in start:length(pairList)) {
  pair <- unlist(list(pairList[[start]], pairList[[i+1]]))
  #pair <- unlist(list(c1, c2))
  # make sure only cells that are present in the SCDE cd matrix are used
  pair <- pair[names(pair) %in% colnames(cd)]
  
  index <- match(names(pair), names(groups))
  groupsDGE <- pair
  
  # run differential expression tests on all genes with batch correction
  # this can take several minutes 
  batch <- as.factor(ifelse(rbinom(nrow(scde.fitted.model[index,]), 1, 0.5) == 1, "batch1", "batch2"))
  ediff <- scde.expression.difference(scde.fitted.model[index,], cd[,index], scde.prior, groups = groupsDGE, batch = batch, n.randomizations  =  100, n.cores  =  n.cores, verbose  =  1)
  # show the top DGE genes
  head(ediff[[1]][order(ediff[[1]][,5], decreasing = TRUE),], 15)
  
  # 2-tailed p-value
  p.values <- 2*pnorm(abs(ediff[[1]][,5]),lower.tail=F) 
  # adjust to control for FDR
  p.values.adj <- 2*pnorm(abs(ediff[[1]][,6]),lower.tail=F) 
  significant.genes <- which(p.values.adj<0.05)
  length(significant.genes)
  
  # get fold changes
  # order by p-value
  ord <- order(p.values.adj[significant.genes]) 
  de <- cbind(ediff[[1]][significant.genes,1:3],p.values.adj[significant.genes])[ord,]
  colnames(de) <- c("Lower-bound","log2_FC","Upper-bound","p-adjusted")
  
  # get time stamp for output file name
  time <- paste(gsub("-|:", "", unlist(stringr::str_split(as.character(Sys.time()), " ", n = 2))), collapse = "")
  # make file name based on Cluster pair
  name <- paste0(levels(pair)[1], "-", levels(pair)[2])   
  # save DGE batch effect corrected results in file
  write.table(de, file = paste0("","scde_", name, "_results_ba
                                tch-effect-corrected_", time, ".txt"), row.names = TRUE, col.names = TRUE, sep = "\t", quote = FALSE) 
  
  # make table for non batch effect corrected DGE results
  # 2-tailed p-value
  p.values <- 2*pnorm(abs(ediff[[2]][,5]),lower.tail=F) 
  # adjust to control for FDR
  p.values.adj <- 2*pnorm(abs(ediff[[2]][,6]),lower.tail=F) 
  significant.genes <- which(p.values.adj<0.05)
  length(significant.genes)
  
  # get fold changes
  # order by p-value
  ord <- order(p.values.adj[significant.genes]) 
  de <- cbind(ediff[[2]][significant.genes,1:3],p.values.adj[significant.genes])[ord,]
  colnames(de) <- c("Lower-bound","log2_FC","Upper-bound","p-adjusted")
  # save DGE results in file
  write.table(de, file = paste0("", "scde_", name, "_results_", time, ".txt"), row.names = TRUE, col.names = TRUE, sep = "\t", quote = FALSE) 
}

###########
# cluster specific genes

# find marker genes
dge <- read.table("scde_Cluster1-Cluster2_results_batch-effect-corrected_20190223052340.txt", header=T, row.names = 1, check.names = FALSE)
mark1 <- dge[which(dge$`p-adjusted`<0.01),]
dge <- read.table("scde_Cluster1-Cluster3_results_batch-effect-corrected_20190223054004.txt", header=T, row.names = 1, check.names = FALSE)
mark2 <- dge[which(dge$`p-adjusted`<0.01),]
dge <- read.table("scde_Cluster1-Cluster4_results_batch-effect-corrected_20190223060011.txt", header=T, row.names = 1, check.names = FALSE)
mark3 <- dge[which(dge$`p-adjusted`<0.01),]
dge <- read.table("scde_Cluster1-Cluster5_results_batch-effect-corrected_20190223061739.txt", header=T, row.names = 1, check.names = FALSE)
mark4 <- dge[which(dge$`p-adjusted`<0.01),]
#dge <- read.table("February2019/scde_Cluster1-Cluster6_results_batch-effect-corrected_20190213125635.txt", header=T, row.names = 1, check.names = FALSE)
#mark5 <- dge[which(dge$`p-adjusted`<0.01),]
dge <- read.table("scde_Cluster2-Cluster3_results_batch-effect-corrected_20190223063415.txt", header=T, row.names = 1, check.names = FALSE)
mark6 <- dge[which(dge$`p-adjusted`<0.01),]
dge <- read.table("scde_Cluster2-Cluster4_results_batch-effect-corrected_20190223065248.txt", header=T, row.names = 1, check.names = FALSE)
mark7 <- dge[which(dge$`p-adjusted`<0.01),]
dge <- read.table("scde_Cluster2-Cluster5_results_batch-effect-corrected_20190223070917.txt", header=T, row.names = 1, check.names = FALSE)
mark8 <- dge[which(dge$`p-adjusted`<0.01),]
#dge <- read.table("February2019/scde_Cluster2-Cluster6_results_batch-effect-corrected_20190213145625.txt", header=T, row.names = 1, check.names = FALSE)
#mark9 <- dge[which(dge$`p-adjusted`<0.01),]
dge <- read.table("scde_Cluster3-Cluster4_results_batch-effect-corrected_20190223134622.txt", header=T, row.names = 1, check.names = FALSE)
mark10 <- dge[which(dge$`p-adjusted`<0.01),]
dge <- read.table("scde_Cluster3-Cluster5_results_batch-effect-corrected_20190223135804.txt", header=T, row.names = 1, check.names = FALSE)
mark11 <- dge[which(dge$`p-adjusted`<0.01),]
#dge <- read.table("February2019/scde_Cluster3-Cluster6_results_batch-effect-corrected_20190213160716.txt", header=T, row.names = 1, check.names = FALSE)
#mark12 <- dge[which(dge$`p-adjusted`<0.01),]
dge <- read.table("scde_Cluster4-Cluster5_results_batch-effect-corrected_20190223141759.txt", header=T, row.names = 1, check.names = FALSE)
mark13 <- dge[which(dge$`p-adjusted`<0.01),]

# combine all DE cluster genes in list
result.logFC1 <- unlist(list(mark1$log2_FC, mark2$log2_FC, mark6$log2_FC,
                            mark7$log2_FC, mark10$log2_FC, mark11$log2_FC, mark13$log2_FC))
markGenes1 <- unlist(list(row.names(mark1), row.names(mark2), row.names(mark6),
                         row.names(mark7), row.names(mark10), row.names(mark11), row.names(mark13)))

result.logFC2 <- unlist(list(mark3$log2_FC, mark4$log2_FC, mark8$log2_FC))
markGenes2 <- unlist(list(row.names(mark3), row.names(mark4), row.names(mark8)))

marksFC1 <-markGenes1[which(result.logFC1 > 2)]
marksFC1 <- c(marksFC1, markGenes1[which(result.logFC1 < -2)])
# apply stricter selection for cluster comparisons with very large number of DE genes
marksFC2 <-markGenes2[which(result.logFC2 > 5)]
marksFC2 <- c(marksFC2, markGenes2[which(result.logFC2 < -5)])

marker.set <- unlist(list(marksFC1, marksFC2))
marker.set <- unique(marker.set)
marker.set <- marker.set[!is.na(marker.set)]

# DE genes screening plots
exprSet <- expr
# the following must be TRUE
identical(sne$cell_id, meta$cell_id)

plotMarkers<- function(counter) {
  
  gene <- as.numeric(exprSet[grep(paste(marker.set[counter], "$", sep = ""), row.names(exprSet)),])
  sne$marker <- gene
  
  qplot(Dim1, Dim2, data=sne,
        colour=marker) +
    ggtitle(paste(marker.set[counter])) +
    theme(axis.ticks = element_blank()) + 
    theme(panel.background = element_blank()) + theme(axis.line=element_line(color="grey")) +
    theme(panel.grid.minor=element_line(color="grey"), panel.grid.major=element_line(colour="grey")) +
    geom_point(alpha = 0.4) +
    scale_colour_gradientn(colours = rev(magma(6)), guide = "colourbar") +
    theme(text = element_text(family="Helvetica")) +
    scale_x_continuous(limits = c(min(sne$Dim1) + min(sne$Dim1)*0.1, max(sne$Dim1) + max(sne$Dim1)*0.1)) +
    scale_y_continuous(limits = c(min(sne$Dim2) + min(sne$Dim2)*0.3, max(sne$Dim2) + max(sne$Dim2)*0.2))
}
time <- paste(gsub("-|:", "", unlist(stringr::str_split(as.character(Sys.time()), " ", n = 2))), collapse = "")
counter <- 1:length(marker.set)
pdf(paste0("tSNEheatscaleMarkersKallisto_", time, ".pdf"), width = 10, height = 7, onefile = TRUE)
sapply(counter, function(x) plot(plotMarkers(x)))
dev.off()

# manually selected genes
pluripotent <- c('Pou5f1', 'Sox2', 'Nanog')
naive <- c('Zfp42', 'Klf2', 'Klf4', 'Prdm14', 'Esrrb', 'Tfcp2l1', 'Nr0b1', 'Nr5a2', 'Dppa3', 'FoxD3', 'Blimp1',
           'Myc', 'Sall4', 'Tcf3', 'Stat3')
primed <- c('Otx2', 'Pou3f1', 'Fgf5', 'Zic3', 'Sox3')
xlinked <- c('Xist', 'Tsix', 'Rnf12', 'Pgk1', 'Zfx', 'Pdha1', 'Hprt', 'Phka')
signaling <- c('Fgfr2', 'Fgfr1', 'Smad1', 'Smad5', 'Smad8', 'Snail', 'Egr1', 'Smad2', 'Smad3', 'Nodal', 'Wnt3a', 'Wnt8a', 
               'Axin2', 'Lefty2', 'Eras', 'Neurog1', 'Pax6', 'Meis1', 'Foxb1', 'Socs3', 'Gp130', 'Lier', 'Jak1', 'Jak2', 
               'Tyk2', 'Shp2', 'Gab1', 'Gab2', 'Sos1', 'Sos2', 'Bmpr2', 'ActrIIa', 'ActrIIb', 'Alk3', 'Alk6', 'Alk2', 
               'Alk4', 'Alk5', 'Alk7', 'Gdf8')
te <- c('Gata3', 'Tead4', 'Elf5', 'Tfap2c', 'Hand1')
xen <- c('HNF4a', 'Foxa3', 'Dab2', 'Sox6', 'Sox17', 'Gata4', 'Gata6', 'Cdx1')

# custom gene list
up6 <- c("Gm26870", "Rn7sk", "Gm23935", "Gm24245", "Gm24270", "Gm10718", "Itga7", "Gm13577", "Rpph1", "Gm26205", "Ccdc36", "Gm22710", "Gm12918", "H1fx", "Plb1", "Gm21847", "Rnu1b6", "Gm10801")
down6 <- c("Gm15801", "Gm8783", "Dusp12", "Gm10131", "Sall4", "Hmgb2", "Snrpn", "Uhrf1", "Lin28a", "Ccnb2", "Cdca3", "Mki67", "Pou5f1", "Cdca8", "Ccna2", "Foxh1", "Cdc20", "Utf1", "Dnmt3b", "Ccnb1", "Eras", "Ccnf", "Cdca7", "Cdc7")
Rn7sk_related <- c("Larp7", "Mepce", "Brd4", "Cdk9", "Nelfa", "Cdkn2b", "Cdkn1b", "Cdkn1a", "Nelfb", "Nelfc", "Nelfe", "Supt4a", "Supt5")

allGenes <- c(naive,primed,pluripotent,xlinked,signaling,te,xen, up6,down6,Rn7sk_related, "Cdh1", "Cdh2")
marker.set <- unique(allGenes)

# custom genes screening plots
exprSet <- expr

plotMarkers<- function(counter) {
  
  gene <- as.numeric(exprSet[grep(paste(marker.set[counter], "$", sep = ""), row.names(exprSet)),])
  sne$marker <- gene
  
  qplot(Dim1, Dim2, data=sne,
        colour=marker) +
    ggtitle(paste(marker.set[counter])) +
    theme(axis.ticks = element_blank()) + 
    theme(panel.background = element_blank()) + theme(axis.line=element_line(color="grey")) +
    theme(panel.grid.minor=element_line(color="grey"), panel.grid.major=element_line(colour="grey")) +
    geom_point(alpha = 0.4) +
    scale_colour_gradientn(colours = rev(magma(6)), guide = "colourbar") +
    theme(text = element_text(family="Helvetica")) +
    scale_x_continuous(limits = c(min(sne$Dim1) + min(sne$Dim1)*0.1, max(sne$Dim1) + max(sne$Dim1)*0.1)) +
    scale_y_continuous(limits = c(min(sne$Dim2) + min(sne$Dim2)*0.3, max(sne$Dim2) + max(sne$Dim2)*0.2)) 
}
time <- paste(gsub("-|:", "", unlist(stringr::str_split(as.character(Sys.time()), " ", n = 2))), collapse = "")
counter <- 1:length(marker.set)
pdf(paste0("tSNEheatscaleCustomGenesKallisto_", time, ".pdf"), width = 10, height = 7, onefile = TRUE)
sapply(counter, function(x) plot(plotMarkers(x)))
dev.off()

# Figure 3B
# marker dynamics plot
exprSet <- expr
marker.set <- c('Nlrp4f', 'Rn7sk', 'Cd59a')

plotMarkers<- function(counter) {
  
  gene <- as.numeric(expr[grep(paste(marker.set[counter], "$", sep = ""), row.names(expr)),])
  meta$marker <- gene
  
  qplot(Pseudotime, marker, data=meta, colour=timePoint) +
    theme(axis.ticks = element_blank()) + 
    scale_fill_manual(values=col, 
                      name="",
                      labels=c("mES", "Day 1", "Day 2", "Day 3", "Day 4", "EpiSC", "MB#3P10", "1EP7")) +
    scale_colour_manual(values=col, 
                        name="",
                        labels=c("mES", "Day 1", "Day 2", "Day 3", "Day 4", "EpiSC", "MB#3P10", "1EP7")) +
    theme(axis.ticks = element_blank()) + 
    theme(panel.background = element_blank()) + theme(axis.line=element_line(color="grey")) +
    theme(panel.grid.minor=element_line(color="grey"), panel.grid.major=element_line(colour="grey")) +
    theme(text = element_text(family="Helvetica")) +
    ylab(paste0(marker.set[counter])) + 
    guides(fill=FALSE)
}

time <- paste(gsub("-|:", "", unlist(stringr::str_split(as.character(Sys.time()), " ", n = 2))), collapse = "")
counter <- 1:length(marker.set)
pdf(paste0("3B_", time, ".pdf"), width = 10, height = 7, onefile = TRUE)
sapply(counter, function(x) plot(plotMarkers(x)))
dev.off()

###########
# GO analysis
# this was not shown in the manuscript, but is included as an alternative to Enrichr
# loads function topDiffGenes
data(geneList)

meta <- read.csv("metadata_20190223045845.csv", header=TRUE, stringsAsFactors = F)
tpm <- read.table("geneTPM_M8_20190207033818_Kallisto.txt", header=T, row.names = 1, check.names = FALSE)
tpm <- tpm[, which(names(tpm) %in% meta$cell_id)]
meta <- meta[match(names(tpm), meta$cell_id),]
expr <- tpm[,!meta$discard]
meta <- meta[!meta$discard,]
# the following must be TRUE
identical(meta$cell_id, names(expr))

files <- list.files("")
files <- files[grep("-Cluster[0-9]_results_batch-effect-corrected_20190223", files)]

# the following will loop over all cluster DGE files and save the result for each cluster pair in a file
for(h in 1:length(files)) {
  dge <- read.table(paste0("", files[h]), header=T, row.names = 1, check.names = FALSE)
  mark <- dge[which(dge$`p-adjusted`<0.01),]
  
  name <- regexpr("_[A-z0-9]+-[A-z0-9]+[0-9]_", files[h]) %>%
    regmatches(files[h], .) %>% gsub("_", "", .)
  
  clu <- gsub("Cluster", "", name) %>% strsplit(., split = "-") %>% unlist
  
  pair <- unlist(list(eval(parse(text = paste0("c", clu[1]))), eval(parse(text = paste0("c", clu[2])))))
  index <- match(names(pair), names(expr))
  groups <- expr[,index]
  
  # make table with only selected DGE genes and cluster cells
  filter <- groups[rownames(groups) %in% rownames(mark),]
  filter <- filter[match(rownames(mark), rownames(filter)),]
  # the following must be TRUE
  identical(rownames(filter), rownames(mark))
  
  selProbes <- genefilter(filter, filterfun(pOverA(0.20, log2(100)), function(x) (IQR(x) > 0.25)))
  eset <- filter[selProbes, ]
  
  ###
  
  genes <- mark[rownames(mark) %in% rownames(eset),]
  # the following must be TRUE
  identical(rownames(genes), rownames(eset))
  
  anno <- AnnotationDbi::select(org.Mm.eg.db,
                                keys = rownames(genes),
                                columns = c("ENSEMBL", "REFSEQ", "SYMBOL", "GENENAME"),
                                keytype = "SYMBOL")
  
  gene_pVal <- as.numeric(genes$`p-adjusted`)
  names(gene_pVal) <- rownames(genes)
  
  reference <- "org.Mm.eg.db"
  
  onts <- c( "MF", "BP", "CC" )
  tidyGenes <- as.list(onts)
  names(tidyGenes) = onts
  linked <- as.list(onts)
  names(linked) = onts
  tab = as.list(onts)
  names(tab) = onts
  
  for(i in 1:3){
    # prepare data
    tgd <- new( "topGOdata", description = paste0("tSNE ", name, " DGE"), ontology=onts[i], 
                allGenes = gene_pVal, geneSel = topDiffGenes, nodeSize=5,
                annot=annFUN.org, mapping="org.Mm.eg.db", ID = "symbol" )
    
    # run tests
    resultFisher.elim <- runTest(tgd, algorithm = "elim", statistic = "Fisher" )
    resultFisher.classic <- runTest(tgd, algorithm = "classic", statistic = "Fisher" )
    resultKS.elim <- runTest(tgd, algorithm = "elim", statistic = "ks")
    resultKS.classic <- runTest(tgd, algorithm = "classic", statistic = "ks")
    
    if(length(score(resultKS.elim)) > 99) {
      # results table
      tab[[i]] <- GenTable( tgd, classicFisher = resultFisher.classic,
                            elimFisher = resultFisher.elim, 
                            classicKolmogorovSmirnov = resultKS.classic,
                            elimKolmogorovSmirnov = resultKS.elim,
                            orderBy = "elimKolmogorovSmirnov" , topNodes = 100)
      
      go <- tab[[i]]$GO.ID[tab[[i]]$`Rank in elimFisher`] 
      go.genes <- genesInTerm(tgd, whichGO = go) 
      
      linked[[i]] <- as.data.frame(do.call(rbind, lapply(1:length(go.genes),
                                                         function(x) cbind(names(go.genes)[x],
                                                                           go.genes[[x]]))))
      linked[[i]]$V3 <- onts[i]
      names(linked[[i]]) <- c("GO.ID", "gene", "GO.type")
      
      # make tidy gene list
      library(tidyr)
      temp <- plyr::ldply(go.genes, rbind); names(temp)[1] <- "GO.ID"
      tidyGenes[[i]] <- unite(temp, col = "genes", 2:ncol(temp), sep = ", ")
      tidyGenes[[i]]$genes <- gsub(pattern = ", NA", x = tidyGenes[[i]]$genes, replacement = "")
    }
    else {
      # results table
      tab[[i]] <- GenTable( tgd, classicFisher = resultFisher.classic,
                            elimFisher = resultFisher.elim, 
                            classicKolmogorovSmirnov = resultKS.classic,
                            elimKolmogorovSmirnov = resultKS.elim,
                            orderBy = "elimKolmogorovSmirnov" , topNodes = length(score(resultKS.elim))) 
      go <- tab[[i]]$GO.ID[tab[[i]]$`Rank in elimFisher`] 
      go.genes <- genesInTerm(tgd, whichGO = go) 
      
      linked[[i]] <- as.data.frame(do.call(rbind, lapply(1:length(go.genes),
                                                         function(x) cbind(names(go.genes)[x],
                                                                           go.genes[[x]]))))
      linked[[i]]$V3 <- onts[i]
      names(linked[[i]]) <- c("GO.ID", "gene", "GO.type")
      
      # make tidy gene list
      library(tidyr)
      temp <- plyr::ldply(go.genes, rbind); names(temp)[1] <- "GO.ID"
      tidyGenes[[i]] <- unite(temp, col = "genes", 2:ncol(temp), sep = ", ")
      tidyGenes[[i]]$genes <- gsub(pattern = ", NA", x = tidyGenes[[i]]$genes, replacement = "")
    }
  }
  
  # Molecular Function top scores
  MF <- tab[[1]]; MF$GO.type <- "MF"
  tidyMF <- tidyGenes[[1]][match(MF$GO.ID, tidyGenes[[1]]$GO.ID),]
  # the following must be TRUE
  identical(tidyMF$GO.ID, MF$GO.ID)
  MF$genes <- tidyMF$genes
  MF <- MF[,c(11,1:10,12)]
  
  # Biological Process top scores
  BP <- tab[[2]]; BP$GO.type <- "BP"
  tidyBP <- tidyGenes[[2]][match(BP$GO.ID, tidyGenes[[2]]$GO.ID),]
  # the following must be TRUE
  identical(tidyBP$GO.ID, BP$GO.ID)
  BP$genes <- tidyBP$genes
  BP <- BP[,c(11,1:10,12)]
  
  # Cellular Component top scores
  CC <- tab[[3]]; CC$GO.type <- "CC"
  tidyCC <- tidyGenes[[3]][match(CC$GO.ID, tidyGenes[[3]]$GO.ID),]
  # the following must be TRUE
  identical(tidyCC$GO.ID, CC$GO.ID)
  CC$genes <- tidyCC$genes
  CC <- CC[,c(11,1:10,12)]
  
  combinedGO <- rbind(MF, BP, CC)
  combinedGenes <- rbind(linked[[1]], linked[[2]], linked[[3]])
  
  time <- paste(gsub("-|:", "", unlist(stringr::str_split(as.character(Sys.time()), " ", n = 2))), collapse = "")
  write.table(combinedGO, file = paste0("", "tSNE_", name, "_GO_", time, ".txt"), row.names = TRUE, col.names = TRUE, sep = "\t", quote = FALSE)
  write.table(combinedGenes, file = paste0("", "tSNE_", name, "_longFormat_GO_genes_", time, ".txt"), row.names = TRUE, col.names = TRUE, sep = "\t", quote = FALSE)
  name
}

###########
# Supplementary Figures
pca <- fread("pcaData_kallisto_geneTPM_20190223043224.txt")
pca <- as.data.frame(pca); names(pca)[1] <- "cell_id"

pca$Cluster <- sne$Cluster
# the following must be TRUE
identical(sne$cell_id, pca$cell_id)

pcakmeansPlot <- qplot(PC1, PC2, data=pca, colour=Cluster, 
                       xlab = paste('PCA1 ', round(variances[1], digits = 2), '%'), 
                       ylab = paste(paste('PCA2 ', round(variances[2], digits = 2), '%'))) +
  scale_colour_brewer(palette = "Dark2", name="") +
  theme(axis.ticks = element_blank()) + 
  theme(panel.background = element_blank()) + theme(axis.line=element_line(color="grey")) +
  theme(panel.grid.minor=element_line(color="grey"), panel.grid.major=element_line(colour="grey")) +
  theme(text = element_text(family="Helvetica")) +
  scale_x_continuous(limits = c(min(pca$PC1) + min(pca$PC1)*0.1, max(pca$PC1) + max(pca$PC1)*0.1)) +
  scale_y_continuous(limits = c(min(pca$PC2) + min(pca$PC2)*0.1, max(pca$PC2) + max(pca$PC2)*0.1))

time <- paste(gsub("-|:", "", unlist(stringr::str_split(as.character(Sys.time()), " ", n = 2))), collapse = "")
pdf(paste0("S_PCA_kMeans_Kallisto_", time, ".pdf"), width = 12, height = 7, onefile = TRUE)
plot(pcakmeansPlot)
dev.off()

pca$Pseudotime <- meta$Pseudotime
# the following must be TRUE
identical(meta$cell_id, pca$cell_id)

pcaPseudotimePlot <- qplot(PC1, PC2, data=pca, colour=Pseudotime,
                           xlab = paste('PCA1 ', round(variances[1], digits = 2), '%'), 
                           ylab = paste(paste('PCA2 ', round(variances[2], digits = 2), '%'))) +
  theme(axis.ticks = element_blank()) + 
  theme(panel.background = element_blank()) + theme(axis.line=element_line(color="grey")) +
  theme(panel.grid.minor=element_line(color="grey"), panel.grid.major=element_line(colour="grey")) +
  theme(text = element_text(family="Helvetica")) +
  scale_x_continuous(limits = c(min(pca$PC1) + min(pca$PC1)*0.1, max(pca$PC1) + max(pca$PC1)*0.1)) +
  scale_y_continuous(limits = c(min(pca$PC2) + min(pca$PC2)*0.1, max(pca$PC2) + max(pca$PC2)*0.1)) +
  scale_color_gradientn(colours = plasma(6), guide = "colourbar", name= "") 

time <- paste(gsub("-|:", "", unlist(stringr::str_split(as.character(Sys.time()), " ", n = 2))), collapse = "")
pdf(paste0("S_PCA_pseudotime_Kallisto_", time, ".pdf"), width = 12, height = 7, onefile = TRUE)
plot(pcaPseudotimePlot)
dev.off()

############

sne <- as.data.frame(fread("tSNE_kMeans5_clusterData_kallisto_DGE_geneTPM_20190223045041.txt"))
meta <- as.data.frame(read.csv("metadata_20190223045845.csv", header=TRUE, stringsAsFactors = F))
tpm <- read.table("geneTPM_M8_20190207033818_Kallisto.txt", header=T, row.names = 1, check.names = FALSE)
tpm <- tpm[, which(names(tpm) %in% meta$cell_id)]
meta <- meta[match(names(tpm), meta$cell_id),]
expr <- tpm[,!meta$discard]

# make timePoint column a factor
RNA <- c(2:6,8,9,1)
meta$timePoint <- factor(meta$timePoint)
meta$timePoint <- factor(meta$timePoint, levels(meta$timePoint)[RNA])
meta <- meta[which(meta$discard == FALSE),]
# the following must be TRUE
identical(meta$cell_id, names(expr))

# scRNA-seq data
gg_color_hue <- function(n) {
  hues = seq(15, 375, length=n+1)
  hcl(h=hues, l=65, c=100)[1:n]
}
col <- gg_color_hue(16)[c(1:3,5,10,12,14,16)]

dge <- read.table("scde_d0-EpiSC_results_batch-effect-corrected_20190223041832.txt", header=T, row.names = 1, check.names = FALSE)
dge <- dge[which(dge$`p-adjusted`<0.01),]
genes <- which(row.names(expr) %in% row.names(dge))
markers <- expr[genes,]

# filter genes for heatmaps
markers <- markers[,which(colSums(markers)>1)]
markers <- markers[which(rowSums(markers)>1),]
markers <- markers[which(rowSums(markers>0)>10),]

meta <- meta[match(names(markers), meta$cell_id),]
# the following must be TRUE
identical(names(markers), meta$cell_id)
pseudotimeIndex <- order(meta$Pseudotime, decreasing = F)
markers <- markers[,pseudotimeIndex]

heat.vals <- log2(markers+1) - rowMeans(log2(markers+1))
timePoint <- meta$timePoint[match(names(heat.vals), meta$cell_id)]
levels(timePoint) <- c("mES", "Day 1", "Day 2", "Day 3", "Day 4", "EpiSC", "MB#3P10", "1EP7")
annotation <- data.frame(timePoint = factor(timePoint, labels = levels(timePoint)))
rownames(annotation) <- names(heat.vals)
names(col) <- levels(annotation$timePoint)
ann_color <- list(timePoint = col) 

# Figure heatmap mES vs EpiSC DE genes of pseudotime ordered cells
time <- paste(gsub("-|:", "", unlist(stringr::str_split(as.character(Sys.time()), " ", n = 2))), collapse = "")
pseudoHeat <- pheatmap(heat.vals, cluster_rows = T, cluster_cols = F, annotation = annotation, annotation_names_col = F,
                       kmeans_k = 20, treeheight_row = 70, annotation_colors = ann_color,
                       show_rownames = T, show_colnames = F, main = "mES vs EpiSC DE genes of kMeans cluster and pseudotime ordered cells", 
                       width = 14, height = 10, filename = paste0("S_d0-EpiSC_heatmap_14x10_", time, ".pdf"))

# get genes for kmeans gene clusters
clustGenes <- data.frame(as.integer(pseudoHeat$kmeans$cluster), names(pseudoHeat$kmeans$cluster))
names(clustGenes) <- c("cluster", "allGenes")
geneList <- data.frame(aggregate(clustGenes, by = list(clustGenes$cluster), FUN =paste))
geneList <- geneList[,c(1,3)]; names(geneList) <- c("cluster", "allGenes")
temp <- geneList$allGenes
geneList <- do.call(rbind, lapply(1:length(geneList$allGenes), function(x) cbind(geneList[x,], geneList$allGenes[[x]])))
names(geneList)[3] <- "gene"
# make tidy gene list
temp <- plyr::ldply(temp, rbind)
tidyGenes <- unite(temp, col = "genes", 1:ncol(temp), sep = ", ")
tidyGenes$genes <- gsub(pattern = ", NA", x = tidyGenes$genes, replacement = "")
write.table(tidyGenes, file = paste0("tscanHeatmap_geneClusters_d0-EpiSC_DGE_", time, ".txt"), row.names = TRUE, col.names = TRUE, sep = "\t", quote = FALSE) 
# long format gene list
geneList <- geneList[,c(1,3)]
write.table(geneList, file = paste0("tscanHeatmap_geneClusters_long_d0-EpiSC_DGE_", time, ".txt"), row.names = TRUE, col.names = TRUE, sep = "\t", quote = FALSE) 
# save cluster x cell order of row means corrected log2TPM values
clustCenters <- pseudoHeat$kmeans$centers; rownames(clustCenters) <- rownames(tidyGenes)
write.table(clustCenters, file = paste0("tscanHeatmap_clusterExpression_d0-EpiSC_DGE_", time, ".txt"), row.names = TRUE, col.names = TRUE, sep = "\t", quote = FALSE) 


# Figure heatmap X-linked genes scRNA-seq
# chromosome linked gene filtering
chr <- read.table("M8_GTF_geneInfo.txt", header=F, check.names = FALSE, stringsAsFactors = F)
# match with expression table genes
chr <- chr[match(rownames(expr), chr$V5),]
identical(rownames(expr), chr$V5)

# subset for gonosomes and autosomes
chrX <- expr[which(chr$V1 == "chrX"),]
chrY <- expr[which(chr$V1 == "chrY"),]
chrM <- expr[which(chr$V1 == "chrM"),]
chrA <- expr[which(chr$V1 != "chrX" & chr$V1 != "chrY" & chr$V1 != "chrM"),]

# filter genes for heatmaps
markers <- chrX
markers <- markers[,which(colSums(markers)>1)]
markers <- markers[which(rowSums(markers)>1),]
markers <- markers[which(rowSums(markers>0)>10),]

meta <- meta[match(names(markers), meta$cell_id),]
# the following must be TRUE
identical(names(markers), meta$cell_id)
pseudotimeIndex <- order(meta$Pseudotime, decreasing = F)
markers <- markers[,pseudotimeIndex]

heat.vals <- log2(markers+1) - rowMeans(log2(markers+1))
timePoint <- meta$timePoint[match(names(heat.vals), meta$cell_id)]
levels(timePoint) <- c("mES", "Day 1", "Day 2", "Day 3", "Day 4", "EpiSC", "MB#3P10", "1EP7")
annotation <- data.frame(timePoint = factor(timePoint, labels = levels(timePoint)))
rownames(annotation) <- names(heat.vals)
names(col) <- levels(annotation$timePoint)
ann_color <- list(timePoint = col) 

time <- paste(gsub("-|:", "", unlist(stringr::str_split(as.character(Sys.time()), " ", n = 2))), collapse = "")
pseudoHeat <- pheatmap(heat.vals, cluster_rows = T, cluster_cols = F, annotation = annotation, annotation_names_col = F,
                       kmeans_k = 20, treeheight_row = 70, annotation_colors = ann_color,
                       show_rownames = T, show_colnames = F, main = "X linked genes of tSNE cluster and pseudotime ordered cells", 
                       width = 14, height = 10, filename = paste0("S_X-genes_heatmap_14x10_", time, ".pdf"))

# get genes for kmeans gene clusters
clustGenes <- data.frame(as.integer(pseudoHeat$kmeans$cluster), names(pseudoHeat$kmeans$cluster))
names(clustGenes) <- c("cluster", "allGenes")
geneList <- data.frame(aggregate(clustGenes, by = list(clustGenes$cluster), FUN =paste))
geneList <- geneList[,c(1,3)]; names(geneList) <- c("cluster", "allGenes")
temp <- geneList$allGenes
geneList <- do.call(rbind, lapply(1:length(geneList$allGenes), function(x) cbind(geneList[x,], geneList$allGenes[[x]])))
names(geneList)[3] <- "gene"
# make tidy gene list
library(tidyr)
temp <- plyr::ldply(temp, rbind)
tidyGenes <- unite(temp, col = "genes", 1:ncol(temp), sep = ", ")
tidyGenes$genes <- gsub(pattern = ", NA", x = tidyGenes$genes, replacement = "")
write.table(tidyGenes, file = paste0("tscanHeatmap_geneClusters_X-genes_", time, ".txt"), row.names = TRUE, col.names = TRUE, sep = "\t", quote = FALSE) 
# long format gene list
geneList <- geneList[,c(1,3)]
write.table(geneList, file = paste0("tscanHeatmap_geneClusters_long_X-genes_", time, ".txt"), row.names = TRUE, col.names = TRUE, sep = "\t", quote = FALSE) 
# save cluster x cell order of row means corrected log2TPM values
clustCenters <- pseudoHeat$kmeans$centers; rownames(clustCenters) <- rownames(tidyGenes)
write.table(clustCenters, file = paste0("tscanHeatmap_clusterExpression_X-genes_", time, ".txt"), row.names = TRUE, col.names = TRUE, sep = "\t", quote = FALSE) 

# Figure autosomal genes
# filter genes for heatmaps
markers <- chrA
markers <- markers[,which(colSums(markers)>1)]
markers <- markers[which(rowSums(markers)>1),]
markers <- markers[which(rowSums(markers>0)>10),]

meta <- meta[match(names(markers), meta$cell_id),]
# the following must be TRUE
identical(names(markers), meta$cell_id)
pseudotimeIndex <- order(meta$Pseudotime, decreasing = F)
markers <- markers[,pseudotimeIndex]

heat.vals <- log2(markers+1) - rowMeans(log2(markers+1))
timePoint <- meta$timePoint[match(names(heat.vals), meta$cell_id)]
levels(timePoint) <- c("mES", "Day 1", "Day 2", "Day 3", "Day 4", "EpiSC", "MB#3P10", "1EP7")
annotation <- data.frame(timePoint = factor(timePoint, labels = levels(timePoint)))
rownames(annotation) <- names(heat.vals)
names(col) <- levels(annotation$timePoint)
ann_color <- list(timePoint = col) 

time <- paste(gsub("-|:", "", unlist(stringr::str_split(as.character(Sys.time()), " ", n = 2))), collapse = "")
pseudoHeat <- pheatmap(heat.vals, cluster_rows = T, cluster_cols = F, annotation = annotation, annotation_names_col = F,
                       kmeans_k = 20, treeheight_row = 70, annotation_colors = ann_color,
                       show_rownames = T, show_colnames = F, main = "Expressed autosomal genes of tSNE cluster and pseudotime ordered cells", 
                       width = 14, height = 10, filename = paste0("S_Autosomes_heatmap_14x10_", time, ".pdf"))

# get genes for kmeans gene clusters
clustGenes <- data.frame(as.integer(pseudoHeat$kmeans$cluster), names(pseudoHeat$kmeans$cluster))
names(clustGenes) <- c("cluster", "allGenes")
geneList <- data.frame(aggregate(clustGenes, by = list(clustGenes$cluster), FUN =paste))
geneList <- geneList[,c(1,3)]; names(geneList) <- c("cluster", "allGenes")
temp <- geneList$allGenes
geneList <- do.call(rbind, lapply(1:length(geneList$allGenes), function(x) cbind(geneList[x,], geneList$allGenes[[x]])))
names(geneList)[3] <- "gene"
# make tidy gene list
library(tidyr)
temp <- plyr::ldply(temp, rbind)
tidyGenes <- unite(temp, col = "genes", 1:ncol(temp), sep = ", ")
tidyGenes$genes <- gsub(pattern = ", NA", x = tidyGenes$genes, replacement = "")
write.table(tidyGenes, file = paste0("tscanHeatmap_geneClusters_Autosomes_", time, ".txt"), row.names = TRUE, col.names = TRUE, sep = "\t", quote = FALSE) 
# long format gene list
geneList <- geneList[,c(1,3)]
write.table(geneList, file = paste0("tscanHeatmap_geneClusters_long_Autosomes_", time, ".txt"), row.names = TRUE, col.names = TRUE, sep = "\t", quote = FALSE) 
# save cluster x cell order of row means corrected log2TPM values
clustCenters <- pseudoHeat$kmeans$centers; rownames(clustCenters) <- rownames(tidyGenes)
write.table(clustCenters, file = paste0("tscanHeatmap_clusterExpression_Autosomes_", time, ".txt"), row.names = TRUE, col.names = TRUE, sep = "\t", quote = FALSE) 


# Figure heatmap of all expressed genes
# filter genes for heatmaps
markers <- expr
markers <- markers[,which(colSums(markers)>1)]
markers <- markers[which(rowSums(markers)>1),]
markers <- markers[which(rowSums(markers>0)>10),]

meta <- meta[match(names(markers), meta$cell_id),]
# the following must be TRUE
identical(names(markers), meta$cell_id)
pseudotimeIndex <- order(meta$Pseudotime, decreasing = F)
markers <- markers[,pseudotimeIndex]

heat.vals <- log2(markers+1) - rowMeans(log2(markers+1))
timePoint <- meta$timePoint[match(names(heat.vals), meta$cell_id)]
levels(timePoint) <- c("mES", "Day 1", "Day 2", "Day 3", "Day 4", "EpiSC", "MB#3P10", "1EP7")
annotation <- data.frame(timePoint = factor(timePoint, labels = levels(timePoint)))
rownames(annotation) <- names(heat.vals)
names(col) <- levels(annotation$timePoint)
ann_color <- list(timePoint = col) 

time <- paste(gsub("-|:", "", unlist(stringr::str_split(as.character(Sys.time()), " ", n = 2))), collapse = "")
pseudoHeat <- pheatmap(heat.vals, cluster_rows = T, cluster_cols = F, annotation = annotation, annotation_names_col = F,
                       kmeans_k = 20, treeheight_row = 70, annotation_colors = ann_color,
                       show_rownames = T, show_colnames = F, main = "All expressed genes of pseudotime ordered cells", 
                       width = 14, height = 10, filename = paste0("S_AllGenes_heatmap_14x10_", time, ".pdf"))

# get genes for kmeans gene clusters
clustGenes <- data.frame(as.integer(pseudoHeat$kmeans$cluster), names(pseudoHeat$kmeans$cluster))
names(clustGenes) <- c("cluster", "allGenes")
geneList <- data.frame(aggregate(clustGenes, by = list(clustGenes$cluster), FUN =paste))
geneList <- geneList[,c(1,3)]; names(geneList) <- c("cluster", "allGenes")
temp <- geneList$allGenes
geneList <- do.call(rbind, lapply(1:length(geneList$allGenes), function(x) cbind(geneList[x,], geneList$allGenes[[x]])))
names(geneList)[3] <- "gene"
# make tidy gene list
library(tidyr)
temp <- plyr::ldply(temp, rbind)
tidyGenes <- unite(temp, col = "genes", 1:ncol(temp), sep = ", ")
tidyGenes$genes <- gsub(pattern = ", NA", x = tidyGenes$genes, replacement = "")
write.table(tidyGenes, file = paste0("tscanHeatmap_geneClusters_AllGenes_", time, ".txt"), row.names = TRUE, col.names = TRUE, sep = "\t", quote = FALSE) 
# long format gene list
geneList <- geneList[,c(1,3)]
write.table(geneList, file = paste0("tscanHeatmap_geneClusters_long_AllGenes_", time, ".txt"), row.names = TRUE, col.names = TRUE, sep = "\t", quote = FALSE) 
# save cluster x cell order of row means corrected log2TPM values
clustCenters <- pseudoHeat$kmeans$centers; rownames(clustCenters) <- rownames(tidyGenes)
write.table(clustCenters, file = paste0("tscanHeatmap_clusterExpression_AllGenes_", time, ".txt"), row.names = TRUE, col.names = TRUE, sep = "\t", quote = FALSE) 


# Figure 5C X/A
markers <- chrX
markers <- markers[,which(colSums(markers)>1)]
markers <- markers[which(rowSums(markers)>1),]
markers <- markers[which(rowSums(markers>0)>10),]

meta <- meta[match(names(markers), meta$cell_id),]
# the following must be TRUE
identical(names(markers), meta$cell_id)
meta$Xsum <- as.numeric(colSums(markers))

markers <- chrA
markers <- markers[,which(colSums(markers)>1)]
markers <- markers[which(rowSums(markers)>1),]
markers <- markers[which(rowSums(markers>0)>10),]

meta <- meta[match(names(markers), meta$cell_id),]
# the following must be TRUE
identical(names(markers), meta$cell_id)
meta$Asum <- as.numeric(colSums(markers))
meta$ratio <- meta$Xsum/meta$Asum
meta$ratio

ratioBox <- ggplot(meta, aes(x = timePoint, y = ratio, fill = timePoint)) + 
  geom_boxplot(outlier.shape = 1) + 
  theme(axis.ticks = element_blank()) + 
  theme(panel.background = element_blank()) + theme(axis.line=element_line(color="grey")) +
  theme(panel.grid.minor=element_line(color="grey"), panel.grid.major=element_line(colour="grey")) +
  theme(text = element_text(family="Helvetica")) +
  theme(axis.text.x = element_text(angle = 45, hjust = 0.4, vjust = 0.5)) + 
  scale_fill_manual(values=col, 
                    name="",
                    labels=c("mES", "Day 1", "Day 2", "Day 3", "Day 4", "EpiSC", "MB#3P10", "1EP7")) +
  scale_colour_manual(values=col, 
                      name="",
                      labels=c("mES", "Day 1", "Day 2", "Day 3", "Day 4", "EpiSC", "MB#3P10", "1EP7")) +
  xlab("") +
  ylab("chrX expressin sum/chrA expression sum") + 
  guides(fill=FALSE)

time <- paste(gsub("-|:", "", unlist(stringr::str_split(as.character(Sys.time()), " ", n = 2))), collapse = "")
pdf(paste0("5C_X_A_", time, ".pdf"), width = 10, height = 7, onefile = TRUE)
plot(ratioBox)
dev.off()

#####

# C1-CAGE data
# MOIRAI based expression table

library("CAGEr")
library("MultiAssayExperiment")
library("SummarizedExperiment")
library("magrittr")

bedFilesPaths <- function(runID, stamp)
  paste0("BED12files/"
         , runID
         , ".OP-WORKFLOW-CAGEscan-short-reads-v2.0."
         , timeStamp
         , "/CAGEscan_fragments") %>%
  list.files(., full.names = TRUE)

run_1772_144_108_bed <- bedFilesPaths("1772-144-108", "20160905162245")
run_1772_144_109_bed <- bedFilesPaths("1772-144-109", "20160905162341")
run_1772_146_168_bed <- bedFilesPaths("1772-146-168", "20160905162426")
run_1772_146_169_bed <- bedFilesPaths("1772-146-169", "20160905165237")
run_1772_146_170_bed <- bedFilesPaths("1772-146-170", "20160905165419")
run_1772_146_167_bed <- bedFilesPaths("1772-146-167", "20160905165526")
run_1772_146_171_bed <- bedFilesPaths("1772-146-171", "20160905165553")
run_1772_123_295_bed <- bedFilesPaths("1772-123-295", "20170309172257")
run_1772_123_296_bed <- bedFilesPaths("1772-123-296", "20170309172353")

# remove empty BED files from list
run_1772_144_108_bed <- run_1772_144_108_bed[-38]
run_1772_146_168_bed <- run_1772_146_168_bed[-c(8,20,44,56,68,80,88,92)]
# combine all path information
bedFilesPaths <- c(run_1772_144_108_bed, run_1772_144_109_bed, run_1772_146_168_bed, run_1772_146_169_bed, run_1772_146_170_bed, run_1772_146_167_bed, run_1772_146_171_bed, run_1772_123_295_bed, run_1772_123_296_bed)

ce <- DataFrame(inputFiles = bedFilesPaths
                , inputFilesType = "bed"
                , sampleLabels = bedFilesPaths %>%
                  basename %>%
                  sub(".bed", "", .) %>%
                  sub("^", "run", .) %>%
                  make.names)
rownames(ce) <- ce$sampleLabels
ce <- CAGEexp(colData = ce)
genomeName(ce) <- "BSgenome.Mmusculus.UCSC.mm10"

# load data in the CAGEexp object
getCTSS(ce, useMulticore = TRUE)

# save CAGEexp object
saveRDS(ce, "CAGEexp_mm10.rds")

###

# make combined annotation file
#!bin/bash
awk 'FS="\t" {if($3=="gene"){print $1,$4,$5,$7}}' gencode.vM8.chr_patch_hapl_scaff.annotation.gtf > temp1
awk 'FS="\t" {if($3=="gene"){print $1,$4,$5,$7,$9}}' gencode.vM8.chr_patch_hapl_scaff.annotation.gtf | cut -d ';' -f 2,4 | cut -d '"' -f2,4 | sed 's/"/\t/g' > temp2
paste temp1 temp2 > temp3
awk '{print $1,$2,$3,$4,$6,$5}' temp3 | sed 's/ \+/\t/g' | grep 'chr' - > temp4
awk '{$2 = $2 - 1; print}' temp4 | sed 's/ \+/\t/g' > gencode.vM8.feature.bed

# concatenate the different source bed files
# Fantom5 enhancer map and UCSC liftover tool was used to liftover mm9 to mm10

awk '{print $1,$2,$3,$6,$4,$5}' mm10_permissive_enhancers_phase_1_and_2.bed | sed 's/ \+/\t/g' > enhancers_temp.bed
awk 'BEGIN{FS=OFS="\t"} {$5 = "mm10_enhancer."NR; $6 = "enhancer"; print}' enhancers_temp.bed > mm10_enhancerFeature.bed
awk '{print $1,$2,$3,$6,$4,$5}' NAST_mm10.bed | sed 's/ \+/\t/g' > NAST_temp.bed
awk 'BEGIN{FS=OFS="\t"} {$5 = "mm10_nast."NR; $6 = "NAST"; print}' NAST_temp.bed > mm10_NASTFeature.bed
awk '{print $1,$2,$3,$6,$4,$5}' Mm_EPDnew_002_mm10.one.bed | sed 's/ \+/\t/g' > EPD_temp.bed
awk 'BEGIN{FS=OFS="\t"} {$5 = "mm10_epd."NR; $6 = "promoter"; print}' EPD_temp.bed > mm10_EPDFeature.bed
awk '{print $1,$2,$3,$6,$4,$5}' mm10_v3.CAGE_peaks_merged.bed | sed 's/ \+/\t/g' > F5_temp.bed
awk 'BEGIN{FS=OFS="\t"} {$5 = "mm10_fantom5."NR; $6 = "promoter"; print}' F5_temp.bed > mm10_F5Feature.bed
cat gencode.vM8.Feature.bed mm10_F5Feature.bed mm10_enhancerFeature.bed mm10_NASTFeature.bed mm10_EPDFeature.bed > concat.bed

# remove duplicate coordinate rows, but keep overlapping entries that are not congruent 
sort -k1,1 -k2,2n -k3,3n -u concat.bed > sorted.concat.bed
bedtools annotate -both -i sorted.concat.bed -files gencode.vM8.Feature.bed mm10_F5Feature.bed mm10_enhancerFeature.bed mm10_NASTFeature.bed mm10_EPDFeature.bed > mm10_sourceAnnot.bed

# clean intermediary files
rm -rf *temp* *Feature.bed refTSS_mm10.bed *concat.bed

###
# create the expression matrix
# load data
ce <- readRDS("CAGEexp_mm10.rds")
# define groups
ce$group <- ce$sampleLabels %>% sub(pat="_.*", repl="")  %>% factor

# expression per gene
annot <- read.delim("mm10_sourceAnnotation_20180129142414.bed", stringsAsFactors = TRUE, header = TRUE)
levels(annot$Strand) <- c("-", "*", "+")
summary(annot)
gr <- GRanges( seqnames = annot$Chromosome
               , ranges = IRanges( start = annot$Start + 1
                                   , end = annot$End)
               , strand = annot$Strand)
mcols(gr) <- annot[, -(1:4)]

gr$transcript_type <- gr$Feature

gr$gene_name <- gr$Feature
annotateCTSS(ce, gr)
CTSScoordinatesGR(ce)$Feature <- CTSScoordinatesGR(ce)$genes

gr$gene_name <- gr$Source
annotateCTSS(ce, gr)
CTSScoordinatesGR(ce)$Source <- CTSScoordinatesGR(ce)$genes

gr$gene_name <- gr$ID
annotateCTSS(ce, gr)

CTSScoordinatesGR(ce)

# feature count
hAnnot <- data.frame(c("",  "EPD", "NAST", "F5enhancer", "F5promoter", "Gencode"), c("No Match",  "EPD", "NAST", "F5enhancer", "F5promoter", "Gencode"), stringsAsFactors = F)

flatAnnots <- smallCAGEqc::hannot(decode(CTSScoordinatesGR(ce)$Source), hAnnot)

SourceSummary <- rowsum(CTSStagCountDf(ce), flatAnnots) %>% t %>% DataFrame

customScope <- function(libs) 
  list(libs = libs
       , columns = colnames(libs)
       , total = rowSums(libs))

smallCAGEqc::plotAnnot(SourceSummary %>% data.frame, "custom", GROUP = ce$group, customScope = customScope, TITLE = "")

# save gene level expression tables
time <- paste(gsub("-|:", "", unlist(stringr::str_split(as.character(Sys.time()), " ", n = 2))), collapse = "")

CTSStoGenes(ce)
data.table::fwrite(as.data.frame(assay(GeneExpSE(ce)))
                   , sep = "\t"
                   , row.names = TRUE
                   , paste0("geneExpressionTable_", time, ".tsv"))
###
# C1CAGE tSNE results have been obtained analogue to the method described for Fluidigm scRNA-seq
sne <- read.table("tSNE_kMeans6_clusterData_C1CAGE_d0-EpiSC_20180205152721.txt", header=T, row.names = 1, check.names = FALSE)
metaC1 <- read.csv("metadata-C1CAGE_20180410112239.csv", header=TRUE, stringsAsFactors = F)
count <- read.table("geneExpressionTable_20180130025537.tsv", header=T, row.names = 1, check.names = FALSE)
cells <- str_replace(names(count), "run", ""); cells <- str_replace_all(cells, "\\.", "-")
names(count) <- cells
metaC1 <- metaC1[which(metaC1$cell_id %in% names(count)),]
metaC1 <- metaC1[match(names(count), metaC1$cell_id),]
# the following must be TRUE
identical(metaC1$cell_id, colnames(count))

# make timePoint column a factor
# remove failed cells
meta <- metaC1[!metaC1$discard,]
meta$timePoint <- factor(meta$timePoint)
expr <- count[,!metaC1$discard]

meta <- meta[match(row.names(sne), meta$cell_id),]
# the following must be TRUE
identical(row.names(sne), meta$cell_id)

expr <- expr[,which(colSums(expr)>1)]
expr <- expr[which(rowSums(expr)>1),]
expr <- expr[which(rowSums(expr>0)>10),]

expr <- expr[,match(meta$cell_id, names(expr))]
# the following must be TRUE
identical(names(expr), meta$cell_id)

# color scheme
gg_color_hue <- function(n) {
  hues = seq(15, 375, length=n+1)
  hcl(h=hues, l=65, c=100)[1:n]
}
col <- gg_color_hue(16)[c(1:3,5,10,12)]

#Figure 4A
test <- as.numeric(sne$Cluster)
test <- gsub('3', 'd', test)
test <- gsub('5', 'f', test)
test <- gsub('1', '3', test)
test <- gsub('d', '1', test)
test <- gsub('2', '5', test)
test <- gsub('f', '2', test)
test <- factor(as.numeric(test))
sne$Cluster <- test
# manually put biased cell from cluster 6 into cluster 3
sne[rownames(sne) == '1772-146-170_D04', 'Cluster'] <- 4

tsnekmeansPlot <- qplot(Dim1, Dim2, data=sne,
                        colour=Cluster) +
  scale_colour_brewer(palette = "Dark2", name="") +
  theme(axis.ticks = element_blank()) + 
  theme(panel.background = element_blank()) + theme(axis.line=element_line(color="grey")) +
  theme(panel.grid.minor=element_line(color="grey"), panel.grid.major=element_line(colour="grey")) +
  scale_x_continuous(limits = c(min(sne$Dim1) + min(sne$Dim1)*0.1, max(sne$Dim1) + max(sne$Dim1)*0.1)) +
  scale_y_continuous(limits = c(min(sne$Dim2) + min(sne$Dim2)*0.3, max(sne$Dim2) + max(sne$Dim2)*0.2)) +
  theme(text = element_text(family="Helvetica")) 

time <- paste(gsub("-|:", "", unlist(stringr::str_split(as.character(Sys.time()), " ", n = 2))), collapse = "")
pdf(paste0("4A_tSNE_C1CAGE_d0-EpiSC_kMeans_", time ,".pdf"), width = 12, height = 7, onefile = T)
plot(tsnekmeansPlot)
dev.off()

# save updated kMeans data as file
sne$cell_id <- meta$cell_id
time <- paste(gsub("-|:", "", unlist(stringr::str_split(as.character(Sys.time()), " ", n = 2))), collapse = "")
write.table(sne, file = paste0("tSNE_kMeans6_clusterData_C1CAGE_d0-EpiSC_", time, ".txt"), row.names = TRUE, col.names = TRUE, sep = "\t", quote = FALSE) 

#Figure S7G
metaC1 <- read.csv("metadata-C1CAGE_20180410112239.csv", header=TRUE, stringsAsFactors = F)
count <- read.table("nastExpressionTable_20180215060957.tsv", header=T, row.names = 1, check.names = FALSE)
cells <- str_replace(names(count), "run", ""); cells <- str_replace_all(cells, "\\.", "-")
names(count) <- cells
metaC1 <- metaC1[which(metaC1$cell_id %in% names(count)),]
metaC1 <- metaC1[match(names(count), metaC1$cell_id),]
# the following mu be TRUEst
identical(metaC1$cell_id, colnames(count))

# make timePoint column a factor
# remove failed cells
meta <- metaC1[!metaC1$discard,]
meta$timePoint <- factor(meta$timePoint)
expr <- count[,!metaC1$discard]

meta <- meta[match(row.names(sne), meta$cell_id),]
# the following must be TRUE
identical(row.names(sne), meta$cell_id)

expr <- expr[,which(colSums(expr)>1)]
expr <- expr[which(rowSums(expr)>1),]
expr <- expr[which(rowSums(expr>0)>10),]

expr <- expr[,match(meta$cell_id, names(expr))]
# the following mu be TRUE
identical(names(expr), meta$cell_id)

sne$Pseudotime <- meta$Pseudotime
sne <- sne[order(sne$Cluster, sne$Pseudotime, decreasing = F),]
expr <- expr[,match(rownames(sne), names(expr))]

col2 <- brewer.pal(6, "Dark2")

heat.vals <- log2(expr+1) - rowMeans(log2(expr+1))
timePoint <- meta$timePoint[match(names(heat.vals), meta$cell_id)]
levels(timePoint) <- c("mES", "Day 1", "Day 2", "Day 3", "Day 4", "EpiSC")
cluster <- sne$Cluster[match(names(heat.vals), rownames(sne))]
annotation <- data.frame(timePoint = factor(timePoint, labels = levels(timePoint)), cluster = factor(cluster))
rownames(annotation) <- names(heat.vals)
names(col) <- levels(annotation$timePoint)
names(col2) <- levels(annotation$cluster)
ann_color <- list(timePoint = col, cluster = col2) 

time <- paste(gsub("-|:", "", unlist(stringr::str_split(as.character(Sys.time()), " ", n = 2))), collapse = "")
pseudoHeat <- pheatmap(heat.vals, cluster_rows = T, cluster_cols = F, annotation = annotation, annotation_names_col = F,
                       kmeans_k = 20, treeheight_row = 70, annotation_colors = ann_color,
                       show_rownames = T, show_colnames = F, main = "All expressed NASTs of tSNE cluster and pseudotime ordered cells", 
                       width = 14, height = 10, filename = paste0("S_All_NAST_heatmap_14x10_", time, ".pdf"))

# get genes for kmeans gene clusters
clustGenes <- data.frame(as.integer(pseudoHeat$kmeans$cluster), names(pseudoHeat$kmeans$cluster))
names(clustGenes) <- c("cluster", "allGenes")
geneList <- data.frame(aggregate(clustGenes, by = list(clustGenes$cluster), FUN =paste))
geneList <- geneList[,c(1,3)]; names(geneList) <- c("cluster", "allGenes")
temp <- geneList$allGenes
geneList <- do.call(rbind, lapply(1:length(geneList$allGenes), function(x) cbind(geneList[x,], geneList$allGenes[[x]])))
names(geneList)[3] <- "gene"
# make tidy gene list
temp <- plyr::ldply(temp, rbind); names(temp)[1] <- "cluster"
tidyGenes <- unite(temp, col = "genes", 2:ncol(temp), sep = ", ")
tidyGenes$genes <- gsub(pattern = ", NA", x = tidyGenes$genes, replacement = "")
write.table(tidyGenes, file = paste0("tscanHeatmap_geneClusters_AllGenes_NAST_", time, ".txt"), row.names = TRUE, col.names = TRUE, sep = "\t", quote = FALSE) 
# long format gene list
geneList <- geneList[,c(1,3)]
write.table(geneList, file = paste0("tscanHeatmap_geneClusters_long_AllGenes_NAST_", time, ".txt"), row.names = TRUE, col.names = TRUE, sep = "\t", quote = FALSE) 
# save cluster x cell order of row means corrected log2TPM values
clustCenters <- pseudoHeat$kmeans$centers; rownames(clustCenters) <- tidyGenes$cluster
write.table(clustCenters, file = paste0("tscanHeatmap_clusterExpression_AllGenes_NAST_", time, ".txt"), row.names = TRUE, col.names = TRUE, sep = "\t", quote = FALSE) 

# screening plots for C1-CAGE t-SNE clustering
# used for Figure S7
exprSet <- expr
allGenes <- c('Pou5f1', 'Nanog', 'Klf4', 'Esrrb', 'Zfp42', 'Fgf5', 'Sox4', 'Zeb2', 'H1fx', 'Rn7sk', 'Xist', 'Tsix',
              'Eif2s3y', 'Ddx3y', 'Vim', 'Nlrp4f', 'Cd59a', 'Cdh1', 'Cdh2', 'Itga7', 'Rpph1', 'Ccdc36', 'Fam166a',
              'Trp53', 'Ccnb2', 'Tmem263')

# custom gene list plot
plotMarkers <- function(counter) {
  
  gene <- as.numeric(exprSet[grep(paste(allGenes[counter], "$", sep = ""), row.names(exprSet)),][1,])
  sne$marker <- gene
  
  qplot(Dim1, Dim2, data=sne,
        colour=marker) +
    ggtitle(paste(allGenes[counter])) +
    theme(axis.ticks = element_blank()) + 
    theme(panel.background = element_blank()) + theme(axis.line=element_line(color="grey")) +
    theme(panel.grid.minor=element_line(color="grey"), panel.grid.major=element_line(colour="grey")) +
    geom_point(alpha = 0.4) +
    scale_colour_gradientn(colours = rev(magma(6)), guide = "colourbar") +
    scale_x_continuous(limits = c(min(sne$Dim1) + min(sne$Dim1)*0.1, max(sne$Dim1) + max(sne$Dim1)*0.1)) +
    scale_y_continuous(limits = c(min(sne$Dim2) + min(sne$Dim2)*0.3, max(sne$Dim2) + max(sne$Dim2)*0.2)) +
    theme(text = element_text(family="Helvetica"))  
}

time <- paste(gsub("-|:", "", unlist(stringr::str_split(as.character(Sys.time()), " ", n = 2))), collapse = "")
counter <- 1:length(allGenes)
pdf(paste0("tSNE_heatscaleCustomGenesC1CAGE_", time, ".pdf"), width = 10, height = 7, onefile = TRUE)
sapply(counter, function(x) plot(plotMarkers(x)))
dev.off()
```
